# Gene conversion is a key driver of diversity hotspots in *M. tuberculosis* antigens and virulence-associated loci

**DOI:** 10.64898/2026.02.26.708061

**Authors:** Maximillian G. Marin, Natalia Quinones-Olvera, Hu Jin, Michael A. Harris, Brendan M. Jeffrey, Alex Rosenthal, Kenan C. Murphy, Christopher Sassetti, Heng Li, Maha R. Farhat

## Abstract

Despite the long-held view of *Mycobacterium tuberculosis* (*Mtb*) as a genetically conserved pathogen, many genomic regions remain poorly resolved due to high sequence homology and repetitive content. Using complete genome assemblies generated from long-read sequencing of 151 globally representative clinical isolates, we comprehensively analyzed genome-wide patterns of genetic diversity and evolution across the *Mtb* genome. Our analysis uncovers pronounced diversity hotspots within paralogous regions generated by recurrent gene conversion between homologous genes. In many cases, these hotspots exhibit more than an order of magnitude greater genetic diversity than the rest of the *Mtb* genome, which is otherwise characterized by remarkably low variation. Mutations within these regions display clustered substitution patterns, excess paralog-matching variants, and distinct mutational spectra consistent with ongoing gene conversion. Our analysis identifies over 300 individual gene conversion events distributed throughout the *Mtb* phylogeny. These gene conversion events occur predominantly within gene families associated with virulence and host-pathogen interactions, including the PE, PPE, and ESX families. Several of the most pronounced diversity hotspots occur in antigens encoded within paralogous regions. Among these, the vaccine candidate PPE18 harbors mutations in validated epitope sequences and predicted alterations in HLA-II binding. Together, these findings demonstrate that gene conversion actively shapes antigenic and virulence-associated diversity in *Mtb*.

**Significance Statement:** *Mycobacterium tuberculosis*, the bacterium that causes tuberculosis, is widely considered a genetically conserved pathogen. However, repetitive regions comprising nearly 10% of its genome have historically been excluded from genomic analyses. Using complete genome assemblies from diverse clinical isolates, we show that gene conversion, a form of recombination between paralogous genes within the same genome, is a major driver of diversity in these previously unresolved regions. Gene conversion generates pronounced diversity hotspots that are often more than ten times more variable than the genomic background. Many of these hotspots occur in genes involved in virulence and host interaction. Notably, the vaccine antigen PPE18 undergoes recurrent gene conversion, highlighting a mechanism that can shape antigen diversity relevant to tuberculosis vaccine development.

## Introduction

The bacteria of the *Mycobacterium tuberculosis* (*Mtb*) complex (Mtbc) are the causative agent of tuberculosis (TB), a disease responsible for over 1.2 million deaths per year (1). Understanding the forces driving *Mtb* genome evolution has advanced the development of DNA-based diagnostics for antibiotic resistance and allowed for improved detection of disease transmission in TB endemic areas (2). As an obligate human pathogen with no known environmental reservoir, *Mycobacterium tuberculosis* exhibits a strictly clonal population structure with low levels of genetic diversity and a limited accessory genome (3–5). Genome-wide population genomic analyses have found no measurable evidence of ongoing horizontal gene transfer among circulating *Mtb* strains(6). Experimental mating studies have likewise failed to demonstrate interstrain genetic exchange, further supporting the species’ minimal accessory genome (7, 8). Consequently, genetic variation within the species is thought to arise through endogenous mutational processes, dominated by single-nucleotide substitutions and small insertions or deletions, alongside occasional structural variants(5).

Most genomic analyses of *Mtb* have relied on short-read sequencing, which can limit the accuracy of read alignment and variant detection in repetitive or highly homologous genomic regions(9–11). As a result, these repetitive and paralogous regions, comprising nearly 10% of the *Mtb* genome, are frequently excluded from genomic analyses (9–11). Notably, these excluded regions are enriched for the PE, PPE, and Esx gene families, which encode substrates of the ESX secretion systems (3, 5, 12). These families are of particular interest because they expanded in copy number and diversified during the emergence of *Mtb* as an obligate pathogen from environmental mycobacterial ancestors(3, 13–18). Members of these protein families are frequently secreted or localized to the cell surface and have been implicated in diverse aspects of the host–pathogen interface, including modulation of cell wall composition, nutrient acquisition during infection, tissue tropism, and host immune responses (17–22). Multiple comparative studies have shown that the PE, PPE, and ESX families are among the most sequence-variable genes in the Mtb genome, but these studies have been limited in scope or resolution by relying largely on Sanger sequencing of targeted loci or short-read whole-genome sequencing (23–25). Consequently, key questions have remained open, including the full extent of diversity across the species, the mechanisms generating it, and its functional consequences at the host-pathogen interface(5).

Recent advances in long-read sequencing now enable routine assembly of complete bacterial genomes (26–28), permitting confident resolution of repetitive and paralogous regions of the *Mtb* genome. Interrogating the mutational processes and evolutionary forces acting within these regions is therefore critical for understanding their function at the host-pathogen interface(5). The high sequence homology among paralogous loci creates the potential for gene conversion, a form of non-reciprocal intrachromosomal homologous recombination in which short tracts of sequence are copied from one paralogous locus to another (29–31). Gene conversion is a well-recognized evolutionary process among paralogous genes that can function both as a homogenizing force and as a diversity-generating mechanism through interlocus sequence transfer (29, 30, 32). Multiple earlier studies found strong evidence for gene conversion among specific ESX, PE, and PPE genes, although these were largely restricted to individual loci or limited by the resolution of short-read data (24, 25, 33). More recently, two studies demonstrated that long-read sequencing enables confident detection of individual gene-conversion events in clinical Mtb isolates, providing direct evidence that gene conversion contributes to genetic diversification and implicating it in antigenic variation (34, 35).

However, the genome-wide prevalence, evolutionary timing, and functional impact of gene conversion across globally diverse *Mtb* lineages have not been systematically evaluated. Here, we analyze the genome-wide diversity across all major lineages of human-adapted *M. tuberculosis* using complete genome assemblies generated from matched long- and short-read sequencing data. Our analysis reveals gene conversion as a pervasive mutational mechanism across paralogous loci, with recurrent events generating localized diversity hotspots in genes implicated in host-pathogen interaction, including several experimentally validated T cell antigens. Together, these findings reveal gene conversion as a pervasive force shaping species-level diversity at the host-pathogen interface.

## Results

### Hotspots of extreme genetic diversity are found within paralogous genomic regions

We curated a dataset of complete genome assemblies from 151 clinical isolates spanning the global diversity of human adapted *Mtb* (lineages 1–6 and 8, **Figure S1, Table S1**). All assemblies were generated using a hybrid *de novo* genome assembly approach that integrated both long- and short-read sequencing data to ensure high assembly accuracy and completeness (**Methods**). The final polished assemblies demonstrated the expected high similarity between *Mtb* genomes: average nucleotide identity (99.8–100%), genome size (4.38–4.44 Mb), and number of predicted coding sequences (4020–4135 CDSs) (**Table S2**).

Across this population of high-confidence *Mtb* genomes, we measured nucleotide diversity in non-overlapping 1-kb windows tiled across the genome (**Methods**). The median nucleotide diversity of the *Mtb* genome was low (0.2 substitutions/kb). However, a small subset of regions showed extreme diversity (Modified Z-score > 10, 12–83× the genome median; **Figure 1A**). These diversity hotspots were distributed across 37 distinct 1-kb genomic windows (2.3–15.5 substitutions/kb; **Table S3**). Notably, the hotspots were significantly enriched for genes belonging to three categories: PE/PPE (15 genes), ESX (4 genes), and REP13E12 repeats (8 genes) (Fisher’s exact test, Bonferroni-corrected, p_adj < 0.05; **Figure 1B**, **Table S4**). Members of these gene families are often characterized by both multi-copy paralogs distributed throughout the genome and locally repetitive sequences (15, 18), leading to their frequent exclusion from genomic analyses using short-read sequencing (10).

**Figure 1.**
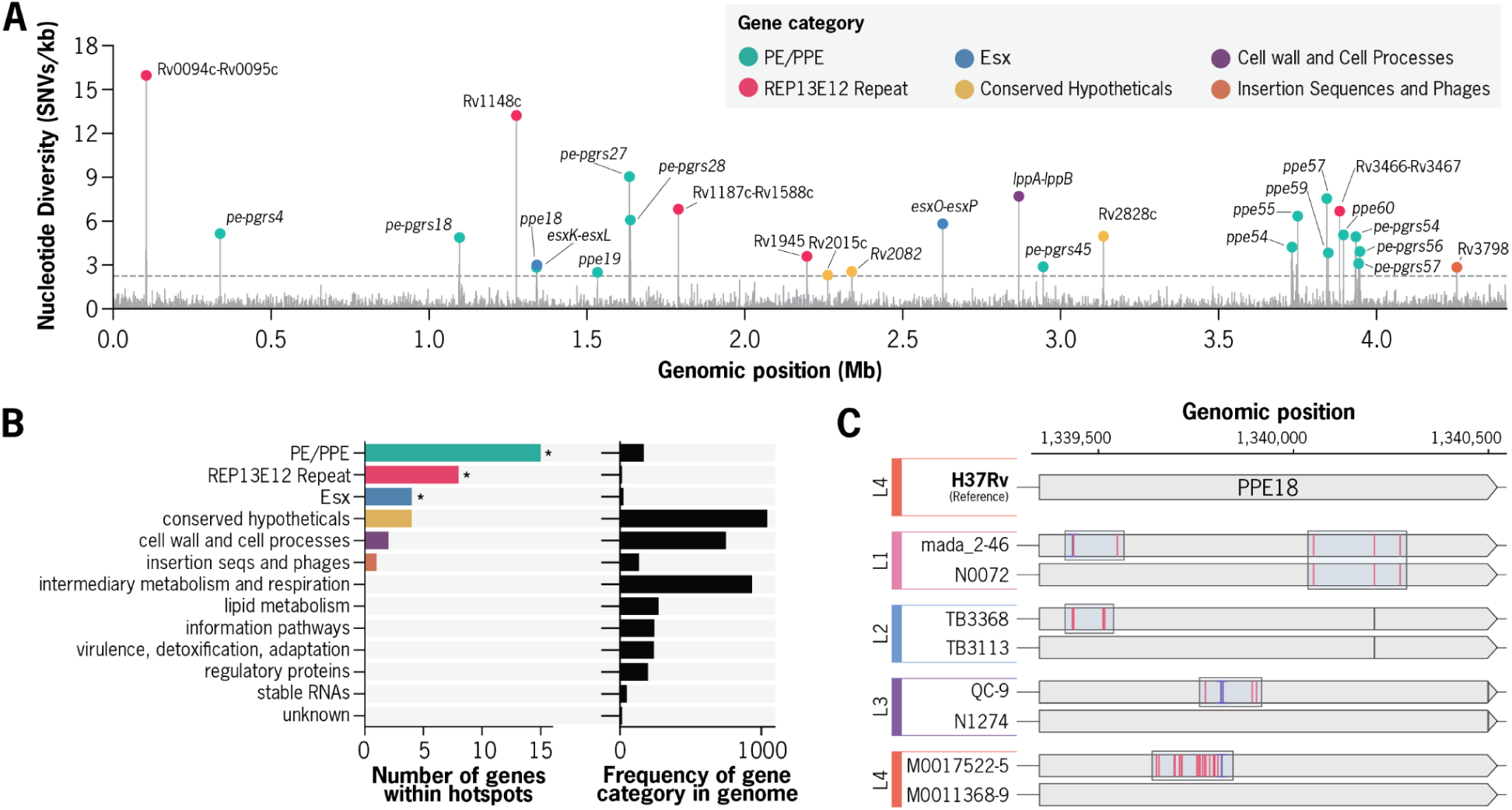
Characterization of nucleotide diversity hotspots within the *Mtb* genome. **a,** Genome-wide distribution of nucleotide diversity (π) across all non-overlapping 1-kb windows measured for complete Mycobacterium tuberculosis (Mtb) genome assemblies. A total of 37 1-kb windows were identified as diversity hotspots, with π values ranging from 2.3 to 15.5 substitutions per kb (up to 83-fold higher than the genome-wide median). Each diversity hotspot is colored based on the functional category of the associated gene(s). b, Functional classification of genes within diversity hotspots. **c,** Example of mutation patterns in the *ppe18* gene across representative *Mtb* isolates from distinct lineages. Red bars indicate substitution mutation positions relative to the H37Rv reference genome. Multiple dense tracts of substitutions are observed in different distinct of the *ppe18* gene across multiple *Mtb* genomes.

We hypothesized that elevated diversification in these hotspots may be related to some aspect of their non-unique sequence content, potentially through homology-mediated mutational processes. To evaluate this hypothesis, we systematically annotated multiple classes of non-unique sequence content across the *Mtb* genome, including multi-copy paralogous regions (PRs) as well as repetitive or low-complexity sequences (**Supplementary Results, Figure S2, Table S5**). Of the 37 diversity hotspots identified, 36 were specifically located within PRs; moreover, the majority (31/37) of hotspots were themselves paralogous to at least one other hotspot (**Figure S3**).

Because non-unique sequences can be more prone to assembly errors, we took two complementary approaches to confirm the accuracy of the variant calls underlying these hotspots. First, we manually inspected the assembly-to-reference and long-read alignments at all 37 hotspots across all genomes, confirming that the variant calls in these regions were well-supported and free of obvious structural artifacts (**Methods**, **Supplemental Results**). Second, we systematically screened every assembly for misassemblies using NucFlag, a reference-free assembly quality-control method (**Methods**). Across all 151 assemblies, only 29 of the 220,430 SNP calls (0.013%) fell within a NucFlag-flagged region, and these flagged regions were rare and predominantly structural (**Dataset S15**). Importantly, none of these misassembly-associated SNPs occurred within a diversity hotspot: of the 24,463 SNP calls located in hotspots, none overlapped with a flagged region, indicating the hotspots are not driven by assembly error (**Dataset S15**). During careful inspection of the data, we observed that mutations in these regions frequently occurred as clustered tracts of substitutions, as exemplified by the *ppe18* locus across a diverse panel of isolates (**Figure 1C, Supplemental Data**).

### Paralogous regions of the *Mtb* genome harbor distinct mutational patterns suggestive of recurrent gene conversion

Given the strong overlap between diversity hotspots and paralogous genomic regions, we hypothesized that elevated diversity in these regions is driven by gene conversion between paralogs rather than by rapid accumulation of independent point mutations. Frequent gene conversion would be expected to produce distinct mutational characteristics, including closely clustered substitutions that simultaneously match sequence variants found in other paralogs(36). To explore this hypothesis, we compared mutational characteristics of paralogous regions (PRs) to those of the rest of the *Mtb* genome. To enable phylogenetically informed analysis of substitution events across the evolutionary history of all 151 genomes, we reconstructed a genome-wide phylogeny and inferred ancestral states for all detected mutations (**Methods**).

We found that PRs harbor multiple distinct mutational characteristics consistent with recurrent mutation by gene conversion (**Figure S4**). First, substitutions in PRs exhibit significantly higher mutational density than those in non-PR regions (median inter-substitution distance of 8 bp versus 1016 bp, respectively; Mann-Whitney U Test, p < 2.2 × 10⁻¹⁶; **Figure S4A**).

Further, a subset (60/157 tested) of PRs exhibited a significant bias toward substitutions that exactly match variants found in other paralogs (hypergeometric test, Bonferroni-corrected, p_adj < 0.01; Figure 2B, **Dataset S4**). Notably, these PRs encompassed nearly all (33/37) previously identified diversity hotspots, including *ppe18*, *ppe60*, *pe-pgrs27*, *pe-pgrs28*, *esxO-esxP*, Rv0094c-Rv0095c, and Rv3466-Rv3467.

**Figure 2.**
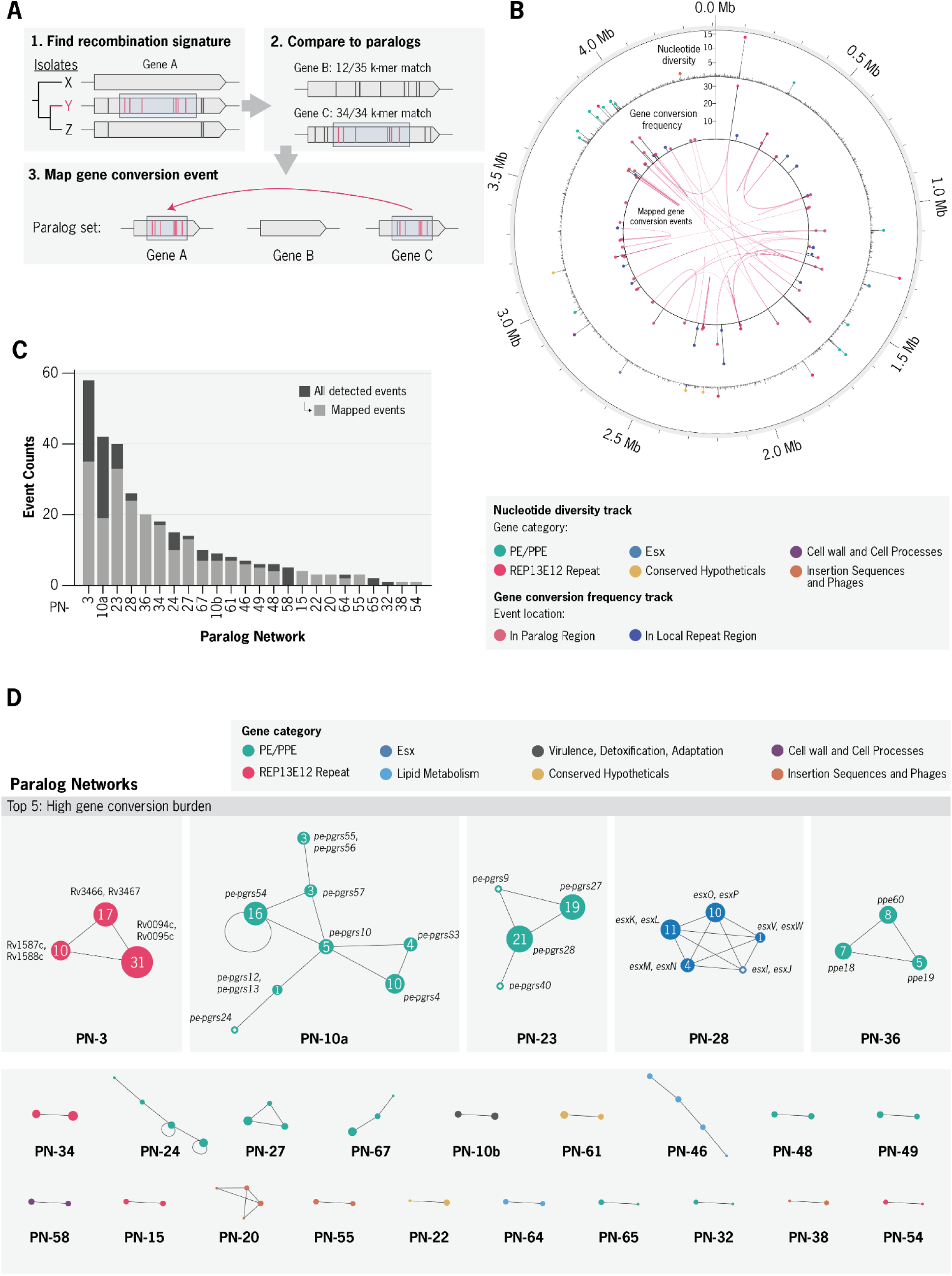
Systematic detection of gene conversion events throughout *Mtb* evolution. **a,** Overview of the computational pipeline to detect gene conversion (GC) events. **b,** Circular representation of the Mtb genome showing the genomic distribution of detected GCEs. Outer track: nucleotide diversity (π) per 1-kb window. Inner track: number of GCEs per genomic region. Red arcs represent donor–recipient pairs with at least one mapped GC event. **c,** Barplot showing the number of GCEs detected per paralog network. **d,** Network diagram of all recombining paralog networks. Each node represents a paralogous region (PR) with node size that reflects the total number of GCEs detected in that region. Edges represent homology between regions. Nodes are colored by the annotated gene category.

In parallel, mutational spectrum analysis revealed a significantly altered substitution profile in PRs relative to the rest of the genome (**Figure S4C–D**): PRs were depleted in C>T/G>A substitutions (p_adj = 9.6 × 10⁻⁵) and enriched in T>A/A>T substitutions (p_adj = 2.6 × 10⁻³; Mann-Whitney U test, Bonferroni-corrected; **Dataset S4**), suggesting the contribution of mutational processes beyond canonical point mutation.

Gene conversion relies on homologous recombination (HR) processes, which in mycobacteria predominantly proceeds through synthesis-dependent strand annealing (SDSA) (31, 37). To ensure that the repair pathways likely to mediate gene conversion are functional across the global *Mtb* population, we first confirmed the gene integrity of all annotated DNA repair enzymes. We predicted loss-of-function (LoF) mutations in the 151 isolate dataset across 59 annotated genes associated with 13 DNA repair processes (**Table S6**). None of the 13 known SDSA/HR genes had any predicted LoF mutations. Only two DNA repair genes, *ligB* (Rv3062) and *radA* (Rv3585), were found to each contain a single LoF mutation in two independent isolates. Both of these genes are not related to SDSA/HR, and are non-essential for *in vitro* growth (37–41). Overall, these findings support that SDSA/HR pathways are functional across the global diversity of *M. tuberculosis* lineages.

### Frequent gene conversion between a small subset of paralogs drives their exceptional diversification

We next developed a multi-step phylogenetic workflow to study the dynamics of gene conversion through the evolutionary history of *M. tuberculosis*. To detect individual events, we first used Gubbins (42) to identify sets of clustered mutations that co-occurred contemporaneously along the phylogeny. Each of these detected gene conversion events (GCEs) is associated with both a set of mutations (≥ 4 SNPs) and a specific branch of the phylogeny where the event is predicted to have occurred. For further validation, the associated recombinant sequence of each detected event was then compared to the sequence of all known paralogs using a k-mer based similarity metric (**Methods, Figure 2A**). All cases where a GCE’s mutations closely matched the reference sequence of known paralog(s) were further classified as mapped GCEs (mGCEs).

Using this framework, we detected 324 gene conversion events (GCEs) across the evolutionary history of our dataset (**Figure 2**). GCEs predominantly overlapped with PRs in the genome (91%, 295 of 324), while the remainder were associated with either local repeats or low-complexity sequences (**Figure S5A**). Overall, detected GCEs were characterized by short, SNP-dense recombination tracts with the median event having a length of 101 bp (IQR: 33–244 bp) and 6 SNPs (IQR: 5–10) (**Figure S6**). Additionally, a majority (74%, 217) of GCEs in PRs mapped to a paralog sequence in the reference genome (**Figure S5B**). Among the mapped GCEs, the genomic distances between inferred donor and acceptor paralogs spanned 1 kb to 2.14 Mb, supporting recombination between both adjacent sequences as well as distant regions of the chromosome (**Figure S6**). There was a strong positive relationship between the detected GCE frequency and nucleotide diversity across all genomic windows overlapping with a PR (Spearman ρ = 0.52, p = 1.4 × 10⁻³⁰; **Figure S7**). Consistent with this pattern, a majority of detected GCEs (64%, 208) also directly mutated regions previously identified as diversity hotspots.

GCEs were detected across 76 distinct genes and were significantly enriched among genes in three categories: PE/PPE (36 genes, 169 GCEs), ESX (7 genes, 36 GCEs), and REP13E12 repeats (10 genes, 87 GCEs) (Fisher’s exact test, Bonferroni-corrected, p_adj < 0.05; **Table 1**). To further characterize the distribution of GCEs across the *Mtb* genome, we constructed a genome-wide homology graph in which each PR is represented as a node and edges connect regions with detectable sequence homology (**Supplemental Methods**). This approach grouped the 200 PRs into 71 discrete paralog networks. GCEs were highly concentrated in a subset of paralog networks (Gini = 0.85; **Figure 2C-D**), with only 24 networks containing any detected GCE activity. We further classified the 24 GCE-containing networks into 9 low-, 10 intermediate-, and 5 high-burden networks (**Methods, Table 2**).

**Table 1.** Enrichment of gene-conversion-affected genes across gene categories. Enrichment was tested at the gene level using a one-sided Fisher’s exact test (genes with ≥1 detected GCE vs. all other annotated genes), with p-values Bonferroni-corrected across 13 categories. Categories with p_adj < 0.05 were considered enriched. The number of detected GCEs is shown for context and was not the unit of testing, as individual events may affect multiple genes.

| Gene Category | # of detected GCEs | # unique genes with GCE | Total annotated genes in H37Rv genome | Odds Ratio | P_adj (Bonferroni) |
| --- | --- | --- | --- | --- | --- |
| PE & PPE proteins | 169 | 36 | 168 | 26.4 | $6.1 \times 10^{-30}$ |
| REP13E12 Repeat Regions | 87 | 10 | 14 | 151.5 | $3.4 \times 10^{-14}$ |
| ESX proteins | 36 | 7 | 23 | 25.3 | $1.5 \times 10^{-6}$ |
| conserved hypotheticals | 19 | 7 | 1042 | 0.3 | 1.000 |
| lipid metabolism | 10 | 3 | 272 | 0.6 | 1.000 |
| insertion sequences | 7 | 5 | 133 | 2.1 | 1.000 |
| cell wall and cell processes | 6 | 3 | 749 | 0.2 | 1.000 |
| Virulence, detoxification, adaptation | 5 | 3 | 239 | 0.7 | 1.000 |
| unknown | 3 | 1 | 15 | 3.8 | 1.000 |
| intermediary metabolism and respiration | 1 | 1 | 936 | 0.04 | 1.000 |
| information pathways | 0 | 0 | 242 | - | 1.000 |
| regulatory proteins | 0 | 0 | 198 | - | 1.000 |
| stable RNAs | 0 | 0 | 48 | - | 1.000 |

**Table 2.**
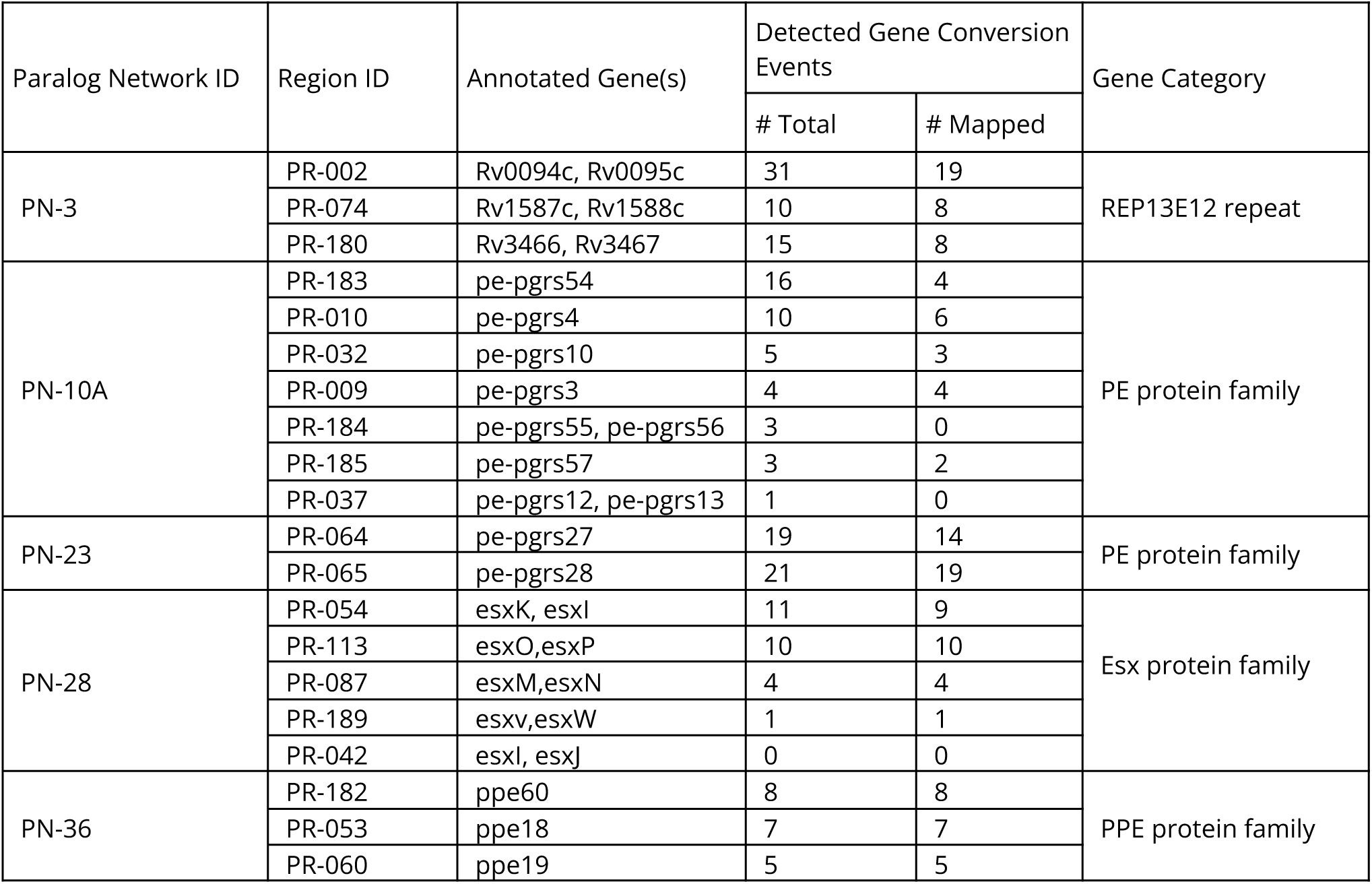
Top five paralog networks ranked by gene conversion burden. For each paralog network, all constituent paralogous regions are listed by unique ID and overlapping gene annotations. For each region, we report the total number of putative gene conversion events detected and the number of those events that mapped to a known paralog reference sequence. Paralog network burden was calculated as the total number of putative gene conversion events detected across all constituent regions.

The five high-burden paralog networks, designated PN-3, PN-10-A, PN-23, PN-28, and PN-36, collectively accounted for 55% of all detected GCEs (**Table 2, Methods**). Together, these networks exemplified distinct gene categories that are strongly enriched for gene conversion. PN-3 involved multiple REP13E12 repeat loci, while PN-10-A and PN-23 each comprised networks of PE-PGRS paralogs. PN-28 contained five distinct loci encoding pairs of ESX paralogs. Finally, PN-36 involved three PPE paralogs, *ppe18*, *ppe19*, and *ppe60* (**Figure 3**). Beyond these high-burden networks, gene conversion was also detected in genes implicated in additional pathways relevant to virulence and immunogenicity. These notably include toxin-antitoxin modules (*vapB25-vapC25* and *vapB31-vapC31*) and lipid metabolism genes (*ppsA*, *ppsB*, and *pks12*) (43–45) (**Figure S8**).

**Figure 3.**
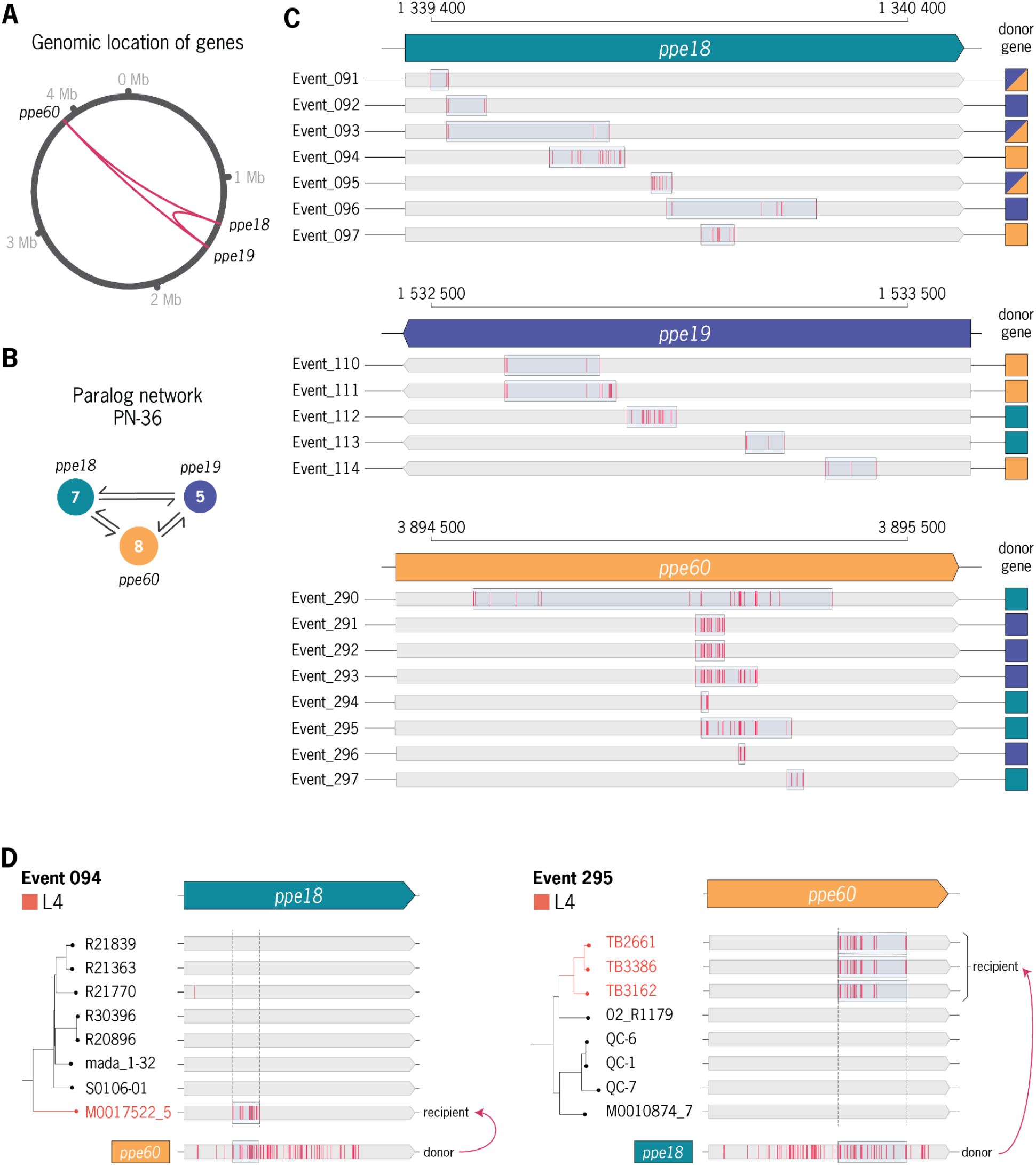
Detailed analysis of gene conversion signatures detected between ppe18, ppe19, ppe60 paralogs. **a,** Relative genomic location of PN-36 paralogs **b,** Paralog network graph labeled by number of detected GCEs per region. **c,** Overview of all gene conversion events detected per gene. Each event is labeled by the closest matching paralog(s) to the associated recombination tract. **d,** Phylogenetic context of mapped gene conversion events (Event-094 and Event-295) between *ppe18* and *ppe60*.

### General sequence characteristics do not explain detected gene conversion frequency

To determine what proportion of variance in gene conversion frequency between PRs can be explained by broad sequence features, we examined whether sequence divergence, genomic distance, paralog copy number, or GC content were predictive of the number of mapped gene conversion events (mGCEs) detected between paralog pairs. We modeled mGCE counts using a generalized linear model with a negative binomial distribution to account for overdispersion (**Methods**). We found that genomic distance, paralog copy number, and GC content were not significantly associated with gene conversion frequency (**Table S7-8**, **Figure S9**). However, higher sequence divergence between paralogs was associated with a reduction in the expected number of mGCEs (incidence rate ratio = 0.983 per additional SNP per kb; p = 2.9 × 10⁻¹⁰). This observation is consistent with the homology dependence of gene conversion, whereby increasing sequence divergence between paralogs reduces their ability to pair during homologous recombination (46, 47). Overall, gene conversion frequency within a region was not strongly predicted by the general sequence features evaluated (McFadden’s pseudo-R² = 0.14), suggesting that instead gene conversion activity varies in a locus-specific manner.

### Gene conversion is widespread throughout both recent and ancestral timescales of *Mtb* evolution

Having identified a subset of genomic loci as hotspots for repeated gene conversion, we next evaluated the phylogenetic distribution of GCEs. Across the dataset phylogeny, GCEs were broadly distributed, with events detected on both ancestral and terminal branches within every major lineage (**Figure 4**). Most GCEs (185, 57%) were inferred to have occurred on terminal branches, consistent with more recent evolution, while 139 (43%) occurred on internal branches, indicating that the associated mutations are shared across multiple descendant genomes (**Figure 4A**). In total, 32 GCEs affecting 20 unique genes were detected on the ancestral branches of lineages represented by two or more isolates (L1–L6). Several loci (e.g. *pe-pgrs27*, *ppe54*, and *ppe55*) underwent gene conversion convergently on the ancestral branches of multiple lineages (**Table S9**).

**Figure 4.**
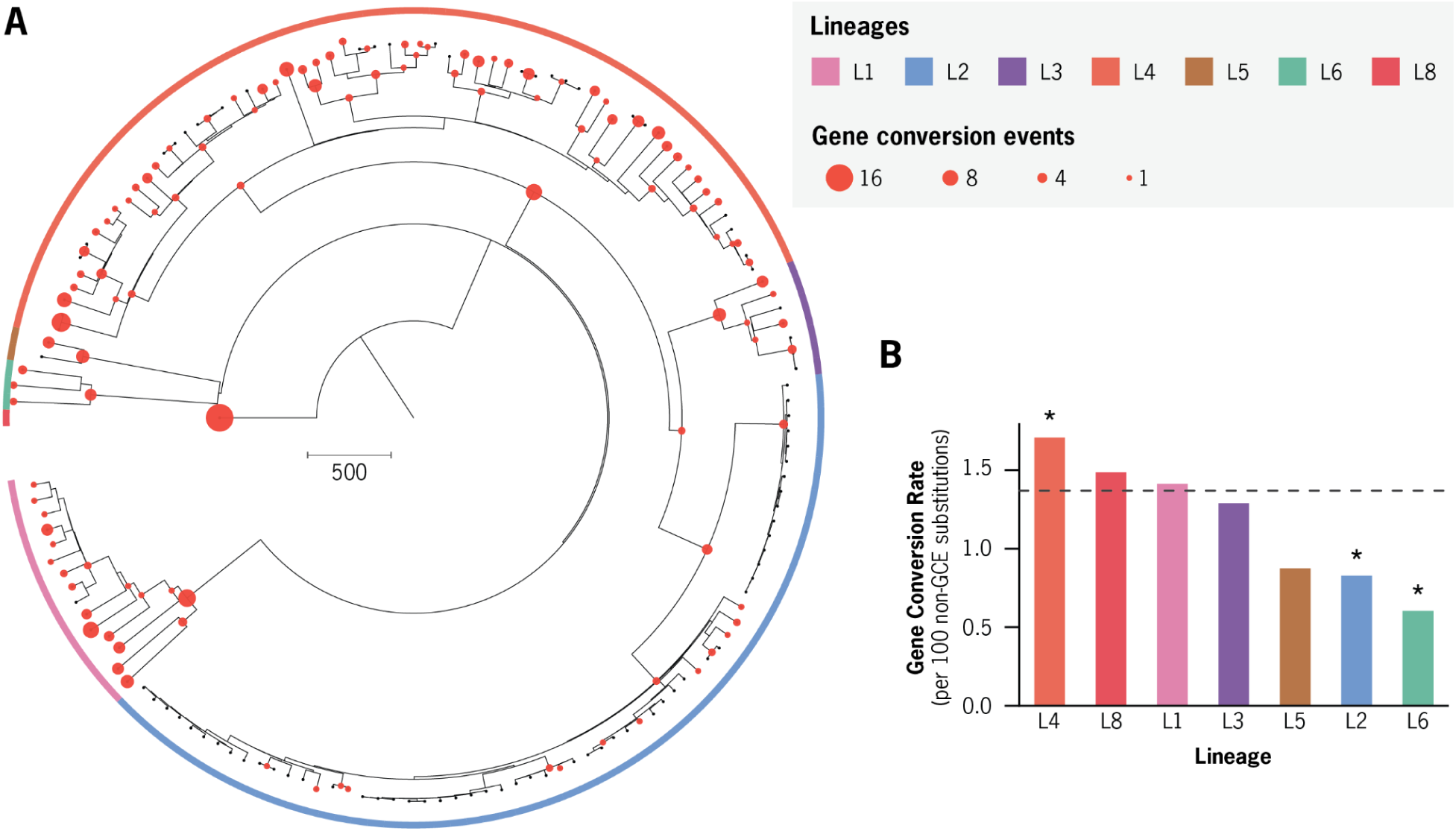
Signatures of gene conversion are widespread across the *Mtb* phylogeny. **a,** Maximum likelihood phylogeny of 151 complete *Mycobacterium tuberculosis* (Mtb) genomes, with branches colored by major phylogenetic lineage. Red circles indicate branches where mapped gene conversion (GC) events were confidently inferred. Circle size is proportional to the number of GC events detected per branch. **b,** Comparison of average GCE rates across all branches belonging to major Mtb lineages. The dashed horizontal line marks the phylogeny-wide average GCE rate.

To contextualize gene conversion within the broader evolutionary dynamics of Mtb, we quantified GCE frequency relative to the rate of recombination-independent (non-GCE) substitutions widely used for molecular clock estimates of Mtb evolutionary history(48, 49). The phylogeny-wide rate of GCEs was estimated to be 1.37 GCEs per 100 non-GCE substitutions. Accounting for the multiple substitutions introduced per event, GCEs contributed 0.124 substitutions for every non-GCE substitution accumulating along the branches of the phylogeny.

We next evaluated lineage-specific variation in gene conversion rates by modeling expected GCE frequencies using a Poisson distribution parameterized by the phylogeny-wide average rate (**Methods**). Of the seven lineages tested, Lineage 4 exhibited a significantly elevated rate of gene conversion, whereas Lineages 2 and 6 were characterized by significantly reduced rates (Benjamini–Hochberg FDR correction, q < 0.05; **Figure 4B**, **Table 3**). Overall, these results indicate that gene conversion has acted as a persistent source of genetic variation throughout the evolutionary history of *M. tuberculosis*, while also exhibiting significant rate variation between lineages.

**Table 3.** Comparison of detected gene conversion rates across phylogenetic branches of each *Mtb* lineage. For each lineage, detected gene conversion event (GCE) counts were compared to expectations under a two-sided Poisson test, with the expected count computed as the phylogeny-wide GCE rate (0.0137 GCEs per non-GCE SNP) scaled by the lineage’s total non-GCE SNP count. Branch-level statistics were summed across all branches assigned to each lineage. p-values were FDR-corrected (Benjamini–Hochberg) across the 7 tested lineages. Lineages with q < 0.05 were considered to have a significantly altered rate. The number of detected GCEs per 100 non-GCE SNPs is reported as the per-lineage rate. q, FDR-adjusted p-value.

| Lineage | Total Branches | # non-GCE SNPs | # GCE SNPs | # Detected GCEs | Detected GCE per 100 non-GCE SNPs | p-value | q-value (FDR Corrected) |
| --- | --- | --- | --- | --- | --- | --- | --- |
| 1 | 29 | 4458 | 578 | 63 | 1.41 | 0.839 | 0.899 |
| 2 | 121 | 3018 | 221 | 25 | 0.83 | 0.009 | 0.020 |
| 3 | 13 | 1551 | 133 | 20 | 1.29 | 0.899 | 0.899 |
| 4 | 123 | 9775 | 1534 | 167 | 1.71 | 0.006 | 0.020 |
| 5 | 3 | 1257 | 93 | 11 | 0.88 | 0.154 | 0.267 |
| 6 | 5 | 1984 | 111 | 12 | 0.61 | 0.002 | 0.013 |
| 8 | 1 | 1076 | 172 | 16 | 1.49 | 0.811 | 0.899 |

### Validation of individual gene conversion events in an independent dataset

We next asked whether signatures of gene conversion could be confidently recovered in a completely independent dataset of Mtb isolates. This was done in two stages: an initial screen of short-read WGS data for candidate gene conversion signatures, followed by complete long-read genome resequencing of selected isolates to evaluate the candidate events at base-level resolution. For this screen, we used a dataset of 937 clinical Mtb isolates (**Figure S10**) previously sequenced with short-read WGS. This cohort was chosen because archived genomic DNA remained available for many isolates, enabling subsequent long-read resequencing (PacBio HiFi). We then performed variant calling on these isolates and conservatively searched for gene conversion signatures using Gubbins, as in our primary analysis. Because short-read variant calling can be unreliable in paralogous genomic regions, we retained only the highest confidence SNP variants for detection of candidate gene conversion events (**Methods**).

Across these 937 clinical isolates, we identified 27 suspected gene conversion events (**Table S10**, **Figures S11–12**). Notably, 24 (89%) of these events occurred in genes that also harbored gene conversion events in our primary Mtb151 analysis. For each of the 27 events, we assessed whether archived genomic DNA from the corresponding isolates remained available in sufficient quantity for long-read resequencing. This was the case for 12 of the 27 events, which we advanced to PacBio HiFi WGS. For each event, we resequenced a target isolate (predicted to harbor the event’s mutations) and a phylogenetically matched control isolate (predicted to lack them), for a total of 22 isolates. Each isolate was then assembled into a complete genome using the same hybrid assembly approach as our primary dataset (**Methods**).

Using the complete genome assemblies of the 22 resequenced isolates, we independently detected gene conversion events and compared them to the events identified in the short-read screen. Across all 12 evaluated events, each SNP of the suspected gene conversion signature was independently called at the identical reference position and allele in the long-read assembly (86/86 SNPs; **Table S11**). Although every suspected event was confirmed, closer examination revealed that short-read WGS was substantially limited in its sensitivity to detect gene conversion. Within these same 12 confirmed events, short-read analysis recovered only 86 of the 199 SNPs resolved by the long-read assemblies (**Table S11**). Furthermore, analysis with the long-read assemblies detected 72 additional gene conversion events not found by the short-read screen, corresponding to a recall of 12/84 events (14.3%). Together, these results show that short-read WGS can confidently resolve fragments of genuine gene conversion events, but with far lower sensitivity than long-read sequencing.

### Gene conversion drives elevated diversity in a select group of *Mtb* antigens

The prevailing view is that *M. tuberculosis* antigens contain little to no sequence variation (10, 50), unlike many other viral and bacterial antigens that frequently exhibit diversifying selection (51). Because many known *M. tuberculosis* antigens are encoded within paralogous regions (PRs), they have historically been excluded or poorly resolved in short-read-based analyses. We therefore used our dataset of complete genomes to examine whether antigens encoded in paralogous regions (PRs) exhibit distinct patterns of sequence diversity and mutational burden relative to antigens located in unique genomic regions.

To facilitate this comparison, we processed *M. tuberculosis* CD4+ T cell epitope-mapping data from two large-scale peptide screening studies (52, 53); 18,371 unique peptides were assayed across 3,840 annotated *Mtb* proteins. From these, we classified 53 proteins as high-confidence T cell antigens containing a total of 226 experimentally validated epitope sequences **(Figure 5, Figure S13, Methods**). This list contains several well-characterized *Mtb* T cell antigens, including the IGRA diagnostic antigens (EsxA, EsxB) and nine additional antigens targeted by various vaccine candidates (**Table S12**). Of these 53 validated T cell antigens, 20 (38%) are encoded in PRs (**Figure S13**).

**Figure 5.**
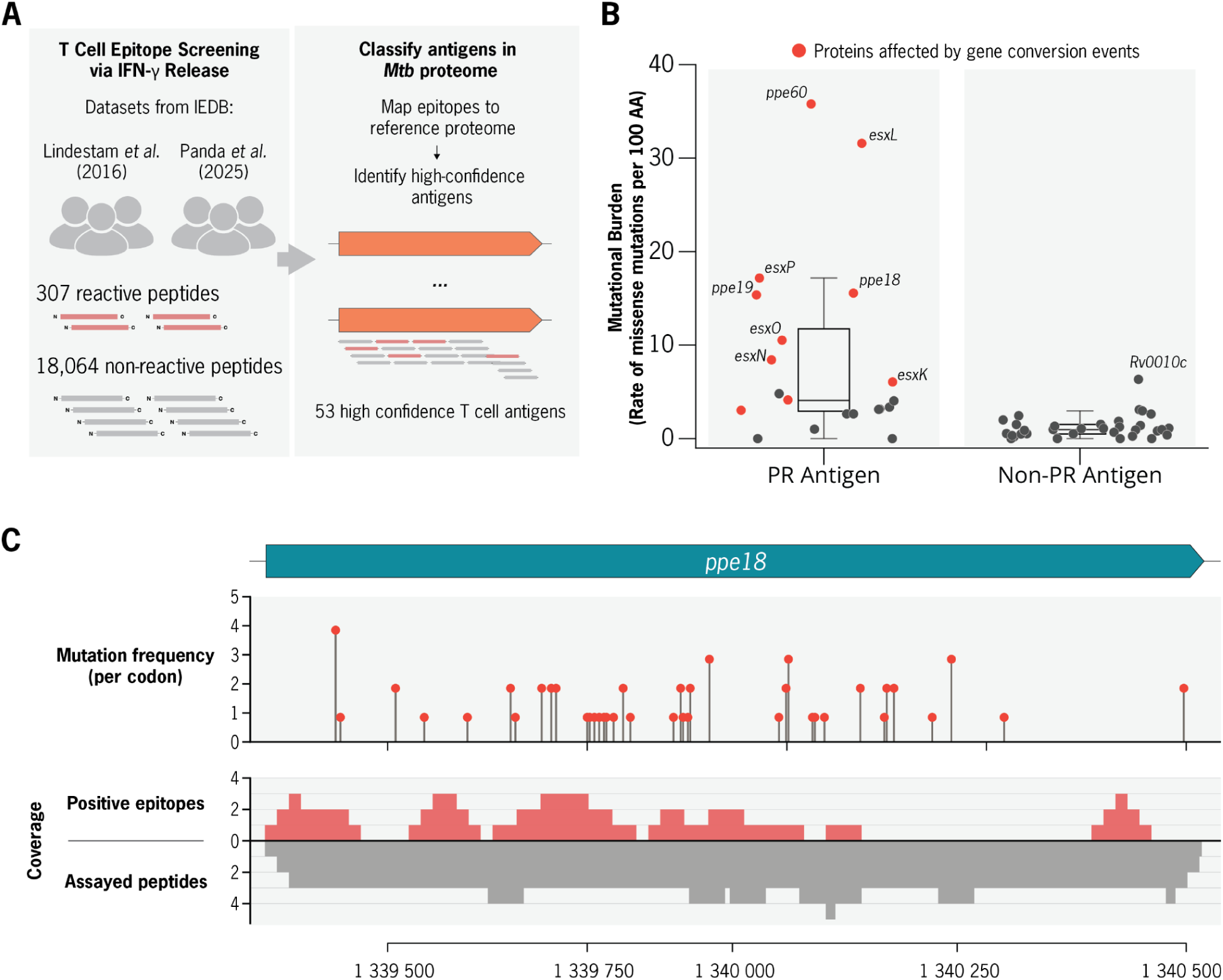
Evaluation of the mutational burden among validated T-cell antigens and epitopes. a, Overview of T-cell epitope data curation. Peptide reactivity data were obtained from two published CD4⁺ T-cell screening studies across TB patient cohorts, assaying 18,325 peptides spanning 3,840 Mtb proteins. **b,** Comparison of the mutational burden between antigens encoded within and outside paralogous regions. **c,** Visualization of non-synonymous mutation frequency and mapped epitope density within the PPE18 antigen. Top: frequency of non-synonymous mutations per codon across the PPE18 coding sequence. Bottom: mapped peptide screening data. The coverage of positive CD4⁺ T-cell epitopes (reactive in ≥1 patient) is shown above, while the overall coverage of assayed peptides per position is shown below in grey.

We next quantified length-normalized mutational burden (non-synonymous mutation events per 100 amino acids) across all annotated coding sequences in the *Mtb* genome (**Figure 5B**, **Figure S14**), comparing four protein categories: PR-associated antigens, unique antigens, PR-associated non-antigens, and unique non-antigens (**Table 4, Table S13**). We found that PR-antigens had the highest mutational burden (median = 4.1 mutations/100AA) and were significantly more diverse than unique antigens (median = 1.0 mutations/100AA; Mann-Whitney U test, Bonferroni-corrected p_adj = 7.8 × 10⁻⁵; **Table S13**). In contrast, unique antigens did not differ in mutational burden from unique non-antigens (p_adj = 1.000; **Table S13**). Additionally, PR-antigens had a higher mutational burden within validated epitope sequences than unique antigens (Mann-Whitney U test, p = 5.0 × 10⁻⁴; **Table S14**).

**Table 4.** Measured mutational burden compared across Antigens and non-antigens encoded in PR and non-PR regions of the *M. tuberculosis* genome. All annotated protein-coding genes in the H37Rv genome were classified based on overlap with the set of identified 53 high-confidence T cell antigens (Dataset S8, Methods) and previously identified paralogous regions of the genome, yielding four groups: antigens within paralogous regions (Ag-PR), antigens outside paralogous regions (Ag-Unq), non-antigens within paralogous regions (NonAg-PR), and non-antigens outside paralogous regions (NonAg-Unq). Normalized mutational burden is calculated as the number of detected non-synonymous mutation events per 100 coding residues; median and interquartile range are reported across all CDSs within each group.

| Antigen Classification | Genomic context | # of CDSs | **Normalized Mutational Burden**<br>(Detected non-synonymous mutation events per 100 coding residues) |  |
| --- | --- | --- | --- | --- |
|  |  |  | Median | Interquartile Range |
| Antigen | PR | 20 | 4.08 | 2.93 - 11.71 |
| Antigen | Non-PR | 33 | 0.96 | 0.49 - 1.51 |
| Non-Antigen | PR | 204 | 1.33 | 0.60 - 2.32 |
| Non-Antigen | Non-PR | 3584 | 0.76 | 0.39 - 1.23 |

Of the top ten antigens ranked by mutational burden, nine are encoded by paralogous genes belonging exclusively to two paralog networks (PN-28, PN-36) associated with frequent gene conversion (**Figure 5**, **Table 2)**. To explore this further, we next examined whether GCEs preferentially affected epitope sequences encoded within PRs. Among paralogous regions, gene conversion events were significantly enriched for those introducing at least one non-synonymous mutation into a validated T cell epitope: 25 of the 295 in-PR GCEs (8.5%) changed an epitope’s amino acid sequence, whereas epitope-coding sequences comprise only 1.71% of PR sequence by length (5.0-fold enrichment; hypergeometric test, p = 9.7 × 10⁻¹¹; **Methods**).

### Recurrent gene conversion generates sequence diversity across the vaccine antigen PPE18

Among the antigens characterized by both high mutational burden and frequent gene conversion, PPE18 stood out due to its clinical relevance as a potent antigen and an important vaccine target. Within PPE18, 44 of 61 non-synonymous mutations occurred within validated T cell epitope sequences. Modeling non-synonymous mutation rates across PPE18, epitope regions exhibited a 1.73-fold higher non-synonymous mutation rate than non-epitope regions (95% CI: 0.99–3.03; p = 0.055, Poisson regression), indicating a trend toward elevated epitope diversity that did not reach statistical significance. Using T cell response-frequency data available for a subset of PPE18 peptides (n = 50), we evaluated whether non-synonymous mutation burden was associated with the number of responding donors (n = 63 individuals with latent TB, **Methods**). T cell responsiveness was not associated with increased non-synonymous mutation burden (rate ratio = 1.14, 95% CI: 0.92–1.42; p = 0.223; negative binomial regression).

To further explore the impact of gene conversion mutations on PPE18 antigenicity, we performed *in silico* predictions of HLA-II binding using netMHCpanII-EL(54) (**Methods**). Binding predictions were generated for all possible 15-mer peptides within PPE18, both before and after mutation by each detected GCE, across a panel of 27 common HLA-II alleles (**Figure S15, Methods**). Of the seven GCEs detected in *ppe18*, five are predicted to change peptide binding for at least one HLA-II allele **(Table S14**). Across evaluated GCEs, both gains and losses in HLA-II binding were predicted, with gains occurring more often than losses (45 of 66 predicted binding changes were gains; binomial test, p = 0.004). Notable among these, Event-094 introduced 17 missense mutations in PPE18 and yielded 41 predicted peptide-HLA binding changes (26 gains and 15 losses). To assess population-specific patterns, we re-evaluated Lineage 4 GCEs against the seven most frequent HLA-II alleles in European populations, yielding results consistent with predictions based on the 27 global alleles (**Methods, Supplementary Results**).

## Discussion

In this work, we analyzed genome-wide sequence diversity across a diverse collection of complete *M. tuberculosis* genomes and found gene conversion to be a pervasive and active mutational process shaping species evolution. Long-read genome sequencing enabled resolution of variation within repetitive and paralogous regions that were largely inaccessible to short-read–based analyses. Across the evolutionary history of our analyzed isolates, we identified hundreds of gene conversion events within paralogous loci throughout the *M. tuberculosis* genome. Our analysis revealed that recurrent gene conversion at a subset of paralogous loci drives the formation of pronounced diversity hotspots throughout the *Mtb* genome. These findings demonstrate that paralogous regions of the *Mtb* genome follow evolutionary dynamics distinct from the rest of the chromosome, with important implications for studies of selection, adaptation, host–pathogen interactions, and vaccine design.

Closer examination of the genomic and phylogenetic distribution of gene conversion revealed marked concentration within a limited subset of paralog networks (**Figure 2**). Gene conversion events were significantly enriched within the PE, PPE, and ESX gene families, with notable events also detected in other virulence-associated systems, including toxin–antitoxin modules and PDIM-associated lipid biosynthesis genes. The most prominent hotspots involved PE-PGRS and PPE paralog sets (*pe-pgrs17*, *pe-pgrs18*, *ppe18*, *ppe19*, and *ppe60*), as well as ESX paralog pairs (*esxK-L*, *esxO-P*, *esxM-N*). REP13E12 repeat regions, which contain a putative prophage attachment site but lack defined biological function, also ranked among the most frequently recombined loci. Together, these findings indicate that recurrent gene conversion promotes diversification within specific paralogous loci, with many of the affected genes linked to host–pathogen interaction and virulence.

These findings also have important implications for the interpretation of selection in loci affected by gene conversion. Repeated gene conversion events can generate clustered tracts of substitutions that mimic convergent evolution, particularly when the same donor paralog serves as a template in independent recombination events. Although such mutations are technically homoplastic, their interpretation is complicated by the fact that they arise through templated copying from donor paralogs rather than independent *de novo* point mutations. More broadly, gene conversion violates the assumption of independent mutations underlying selection metrics such as dN/dS, because it can introduce linked synonymous and non-synonymous substitutions simultaneously within a single conversion tract(55). As a result, dN/dS values in these regions cannot be reliably interpreted as evidence for or against selection. Thus, strong conclusions about adaptive evolution in paralogous loci should not be drawn without explicitly accounting for gene conversion.

Previous population genomic analyses have suggested that *M. tuberculosis* antigens are generally conserved and under purifying selection (10, 50). However, these conclusions were largely drawn from short-read–based analyses that excluded antigens encoded within paralogous regions. Consistent with prior work, our analysis found that antigens located in unique genomic regions exhibit low mutational burden. In contrast, our analysis identified several PR-encoded antigens that undergo recurrent gene conversion and are among the proteins with the highest mutational burdens genome-wide. Because gene conversion introduces clustered, linked substitutions, we did not apply conventional metrics such as dN/dS to these loci, as their underlying assumptions are violated. Although mutational burden does not directly measure selection, our analysis identifies specific PR-encoded antigens that harbor substantially more non-synonymous substitutions than other antigens, in some cases differing by an order of magnitude. Together, these findings support a role for gene conversion in localized antigenic diversification within an otherwise low-diversity pathogen genome.

Among the PR-encoded antigens characterized by a high mutational burden, PPE18 stood out as particularly clinically relevant (**Figure 3**). PPE18 has long been recognized as a potent T cell antigen and is a component of multiple vaccine candidates currently in clinical development, including M72/AS01E and BNT164(52, 53, 56–58). Inspecting the distribution of mutations across PPE18, we found that its validated T cell epitopes are clearly not spared from mutation: a majority of non-synonymous mutations (44 of 61) fell within epitope sequences, and all seven gene conversion events detected in PPE18 generated a mutation in at least one epitope. However, this variation was not significantly concentrated in epitopes, showing only a subtle trend toward a higher mutation rate in epitope regions (1.73-fold; p = 0.055, Poisson regression). Even so, the substantial share of this variation within validated epitopes of a vaccine antigen makes its immunological impact a key concern for vaccine development. To begin to address this, we applied *in silico* binding prediction methods, which predicted that several gene conversion events may alter HLA-II binding among common human alleles. It is important to note that a T cell response depends not only on peptide presentation but also on T cell receptor recognition, which a mutation can disrupt independently of any effect on presentation. More broadly, *in silico* analyses alone cannot establish the immunological consequences of these variants. Determining whether gene conversion in PPE18 influences T cell responses and vaccine efficacy will therefore require future immunological and clinical studies designed to account for both pathogen and host diversity.

Our identification of gene conversion as an active mutational process is further supported by mutational characteristics that align closely with established mechanisms of homologous recombination. For example, the tract lengths of detected events (∼101 bp) are consistent with RecA-mediated strand invasion and synthesis-dependent strand annealing, the homologous recombination pathway described in mycobacteria(37, 59). In addition, substitution patterns within paralogous regions reveal a distinct enrichment of T>C transitions relative to the genomic background. This bias is consistent with the observation in many species that mismatch repair dynamics during recombination-associated DNA synthesis differ from those operating during DNA replication(36, 60). Notably, the enrichment of T>C substitutions parallels mutation spectra observed in nucS-deficient mycobacterial strains, further supporting the possibility that NucS-mediated mismatch repair is reduced during homologous recombination in mycobacteria(61–63). Over evolutionary time, such recombination-associated repair asymmetries could contribute to reshaping nucleotide composition within paralogous genomic regions.

This study has limitations. Our detection framework identifies gene conversion events based on clustered substitutions and signatures of rapid sequence change; consequently, events occurring between nearly identical paralogs are likely underdetected. In addition, older gene conversion events may be progressively eroded by subsequent mutation, reducing the clustering signal required for detection and complicating inference of the original donor paralog. Reliance on a single reference genome may also introduce subtle biases in event localization and paralog assignment. Although the dataset spans major *Mtb* lineages, certain geographic regions and sub-lineages remain underrepresented. Larger collections of complete genomes and reference-independent approaches will be needed to refine estimates of gene conversion prevalence and evolutionary timing.

In summary, we identify gene conversion as a central endogenous mechanism shaping genetic diversity in *M. tuberculosis*. This work reframes assumptions about *Mtb* genomic stability and highlights the need for recombination-aware evolutionary analyses and lineage-informed vaccine design. Future studies integrating broader sampling, longitudinal data, and functional immunological assays will further clarify the role of gene conversion in pathogen evolution and host–pathogen interactions.

## Methods

### Dataset of clinical *Mtb* isolates with long- and short-read WGS

Our primary dataset consisted of 151 complete *M. tuberculosis* genome assemblies (referred to as Mtb151), all previously generated using the same core hybrid (long- and short-read) assembly approach(64). The raw long-read and short-read sequencing data used to generate these assemblies were drawn from multiple published datasets(11, 64–69).

Each clinical isolate in the dataset had matched long-read and short-read WGS data. Short-read data were paired-end Illumina in all cases, while the long-read platform and chemistry varied across the contributing datasets. The long-read data spanned three types: PacBio CLR subreads (N = 48), Oxford Nanopore (ONT) R9.4.1 reads (N = 95), and PacBio CCS/HiFi reads (N = 8). Long-read and short-read data had median genome-wide sequencing depths of 149x (IQR: 100–230x) and 56x (IQR: 47–100x), respectively, relative to the H37Rv reference genome.

The core hybrid assembly approach was consistent across all isolates. First, *de novo* long-read assembly with iterative long-read polishing was performed using Flye (v2.6 or v2.9)(70). Second, short-read polishing of the resulting *de novo* assembly was performed with Pilon (v1.23) to correct small base-level errors using the matched Illumina reads(71). For ONT (R9.4.1) based assemblies, whose higher base-calling error rate typically requires additional correction, a Medaka (v1.5.0) long-read polishing step and a PolyPolish (v0.5.0) short-read polishing step were added(27, 72). Because the exact tool combination was tailored to the needs of each long-read data type, three variants of this pipeline were used, each described in full in the **Supplemental Methods**. All assemblies were annotated with Bakta (v4.8)(73). The dataset label, number of isolates contributed, long-read sequencing platform and read type, assembly pipeline, and source publication for each contributing dataset are summarized in **Table S1**. The per-isolate metadata and sequencing run accessions are provided in **Dataset S1**.

### Nucleotide diversity analysis and diversity hotspot identification

All substitution variant calls from each genome (**Supplemental Methods**) were merged using the bcftools merge (v1.20) to produce a single VCF file of all SNPs across the population. vcftools (v0.1.16) was then used to calculate nucleotide diversity (π) from this SNP callset across all 1-kb non-overlapping windows of the H37Rv genome (`--window-pi 1000`). The nucleotide diversity (π) metric was calculated as the average number of substitutions that exist between all pairwise comparisons of genomes within the specific window of the genome being evaluated (75). All variant-merging and nucleotide-diversity calculation steps are implemented in the Mtb.WGA.Core.V3.smk Snakemake workflow available in the manuscript repository.

To identify outlier regions of elevated diversity, we used a robust statistical method based on the Modified Z-Score(76). For each 1-kb window, the Modified Z Score (*Z_i_*) was calculated as:

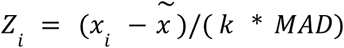

Where x*_i_* is the nucleotide diversity of the window, *x̃* is the median nucleotide diversity across all windows, *MAD* is the median absolute deviation of the distribution, *k*=1.4826 is a scaling constant that makes the statistic comparable to a standard z-score under normality assumptions (76). Windows with Zᵢ > 10 were classified as diversity hotspots. This approach is robust to outliers and does not assume a normal distribution of values, making it well-suited for the highly skewed distribution of nucleotide diversity scores. All population-wide nucleotide diversity stats calculated across all 1-kb windows are available in **Dataset S3**.

### Phylogeny inference

Genetic variants relative to the H37Rv reference genome were inferred for each complete genome assembly using `minimap2` alignment to the reference (**Supplemental Methods**). A concatenated SNP alignment was generated from the merged variant calls using `bcftools mpileup` (v1.20) [(76)], retaining only SNP positions with ≤ 10% missing or ambiguous calls across the dataset. From this SNP alignment, an approximately-maximum-likelihood phylogeny was inferred using FastTree (v2.1.10) under the generalized time-reversible (GTR) nucleotide substitution model (`FastTree -nt -gtr`)(77). This phylogeny was used as the starting tree for all downstream Gubbins analyses.

### Detection of putative gene conversion event signatures

Putative gene conversion events (GCEs) were identified using Gubbins (v3.2.1). Gubbins was run in extensive-search mode (`--extensive-search`) with a minimum of 4 SNPs required to call a recombination block (`--min-snps 4`), a minimum window size of 25 bp, and a maximum window size of 1000 bp (`--min-window-size 25 --max-window-size 1000`), using the initial dataset phylogeny inferred by FastTree as the starting tree. Each detected recombination tract defines a putative GCE, comprising a set of clustered SNPs and the phylogenetic branch on which they occurred. Per-event details for all 324 detected GCEs, including their phylogenetic assignments, are provided in Dataset S5, and the Gubbins final output phylogeny is provided in Newick format in Dataset S10. The full recombination-detection pipeline is available in the manuscript GitHub repository (`Snakemake_Workflows/2_CoreAnalysis_SMK/Mtb.WGA.Core.V3.smk`). The functions used to reformat and process the Gubbins results are provided in `gcutils/gubbinsfuncs.py`.

Detailed descriptions of the downstream analyses of these detected GCEs are provided in the **Supplemental Methods**. These include: (1) mapping of the recombination tract associated with each putative GCE to known paralog sequences ("Mapping of gene conversion events to donor paralogs"); (2) analysis of the genomic distribution of GCEs ("Paralog Network Construction and Gene Conversion Event Distribution Analysis"); and (3) analysis of the phylogenetic distribution of GCEs ("Evaluation of gene conversion frequency and lineage-specific rate variation across phylogeny").

## Acknowledgements

We thank the members of the Farhat lab for helpful discussions and comments on the research project and manuscript. We acknowledge the International Science and Technology Center for their support in establishing the TB Portal agreement with Georgia and CRDF Global for their support in establishing the TB Portal agreements with Azerbaijan and Moldova. Portions of this research were conducted on the O2 High Performance Compute Cluster, supported by the Research Computing Group at Harvard Medical School.

## Data, Materials, and Software Availability

All SRA/ENA run accessions and associated metadata for all *M. tuberculosis* isolates used in this study can be found in Supplemental Data 1. All supplemental materials, genome assemblies, and additional supporting data are available at https://doi.org/10.5281/zenodo.18346123. Code for data processing and analysis is available from the following GitHub repository, https://github.com/maxgmarin/mtb-geneconv-manuscript/. The Snakemake workflow engine (v7.32.4) was used for sequencing data processing(78).

