## Supplemental Data File 11 for "Gene conversion is a key driver of diversity hotspots in *M. tuberculosis* antigens and virulence-associated loci"

### SI-11 — Diversity Hotspot Visualizations

#### Overview

This file contains one visualization per identified diversity hotspot region defined in **Table S2**. Each visualization shows all variants detected across every genome in the dataset, placed in genomic and phylogenetic context. The goal is to transparently display the complete variant pattern for each region of interest.

#### What is shown

Each page corresponds to a single diversity hotspot region (Table S2). Within each page:

- Gene annotations and genomic coordinates: annotated genes in the region are drawn relative to H37Rv *[NC_000962.3]* genomic coordinates.
- Per-genome variant calls: variants identified from the genome assembly alignment are shown for each isolate in the dataset. Variants shown in each alignment are colored by type (SNP, Insertion, Deletion) following the key below.
- Dataset phylogeny: the phylogeny is displayed to the left, with each tip aligned horizontally to that isolate's alignment row, providing phylogenetic context for the variant patterns. Branches that belong to a known MTBC lineage are colored according to the shown key below.

#### Figure key

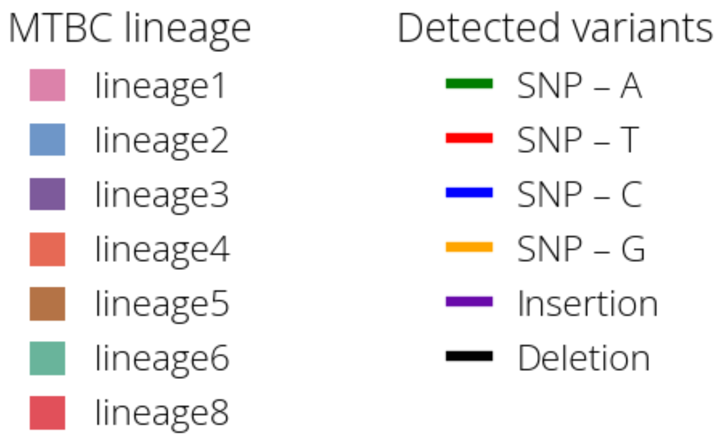

**Diversity Hotspot View - 01:**
**Genomic range shown: NC_000962.3:102000-107000**
**Gene(s) of interest: Rv0094c,Rv0095c**

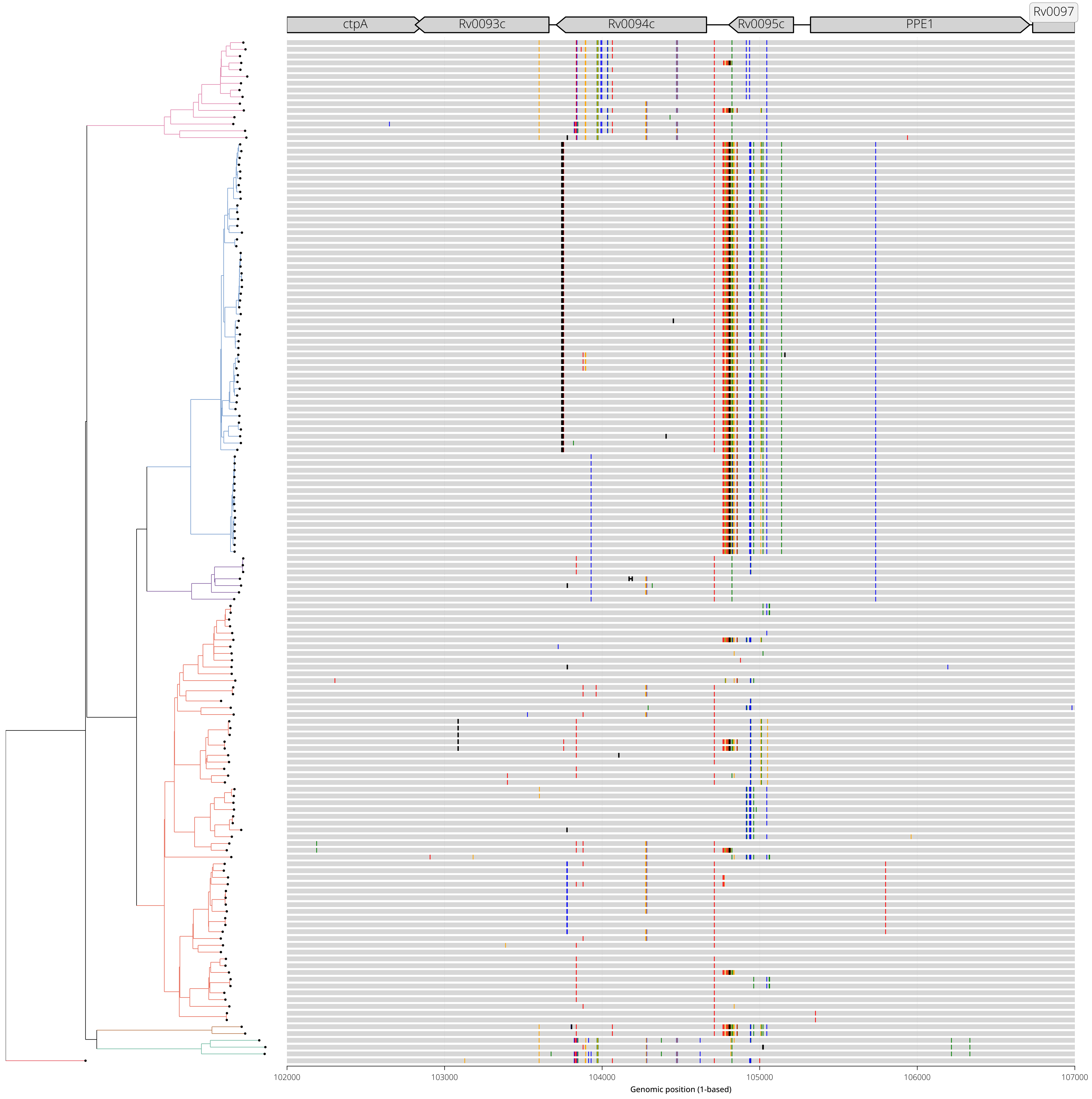

**Diversity Hotspot View - 02:**
**Genomic range shown: NC_000962.3:336000-341000**
**Gene(s) of interest: PE_PGRS4**

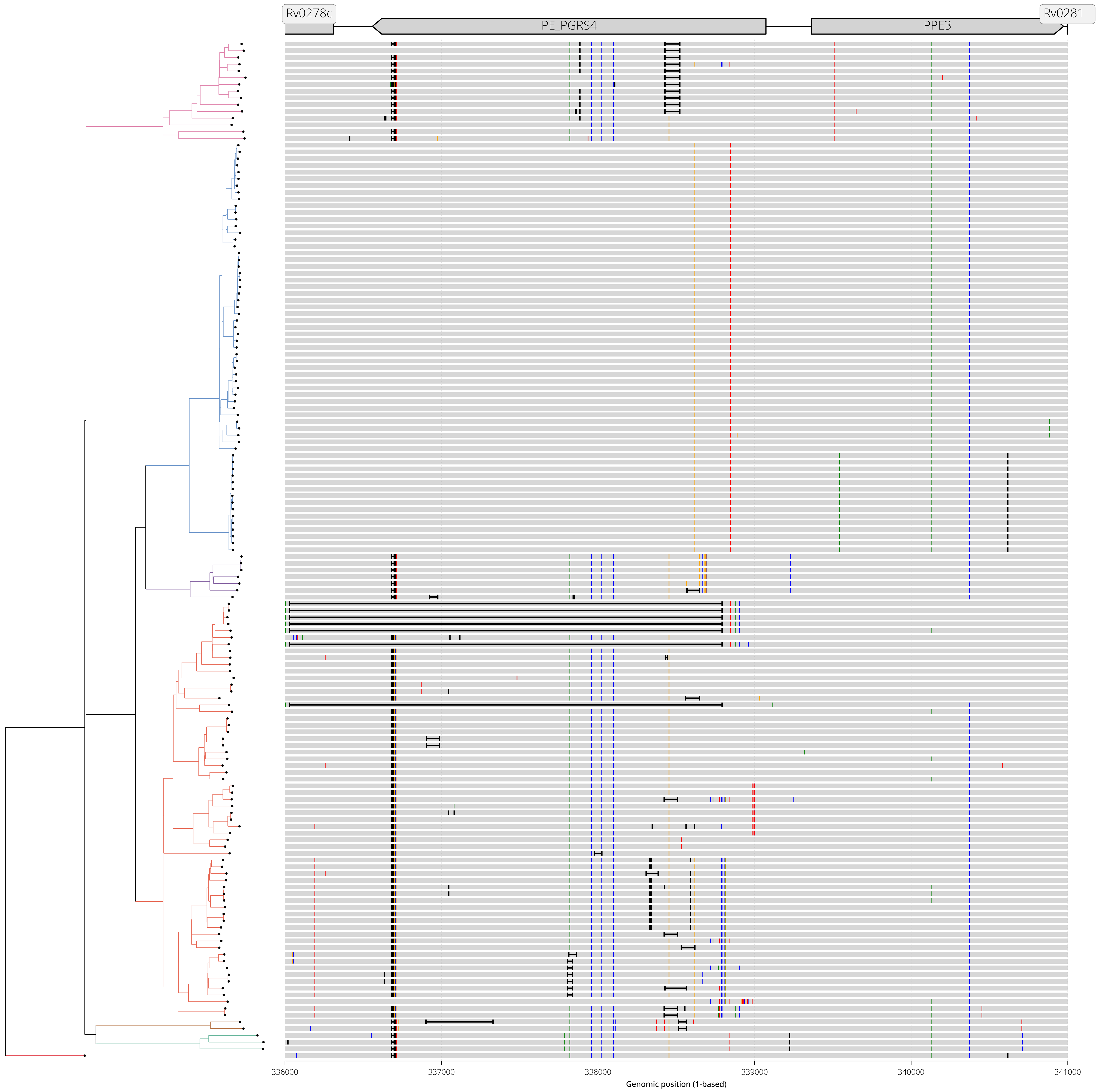

**Diversity Hotspot View - 03:**
**Genomic range shown: NC_000962.3:1095000-1098000**
**Gene(s) of interest: PE_PGRS18,mprA**

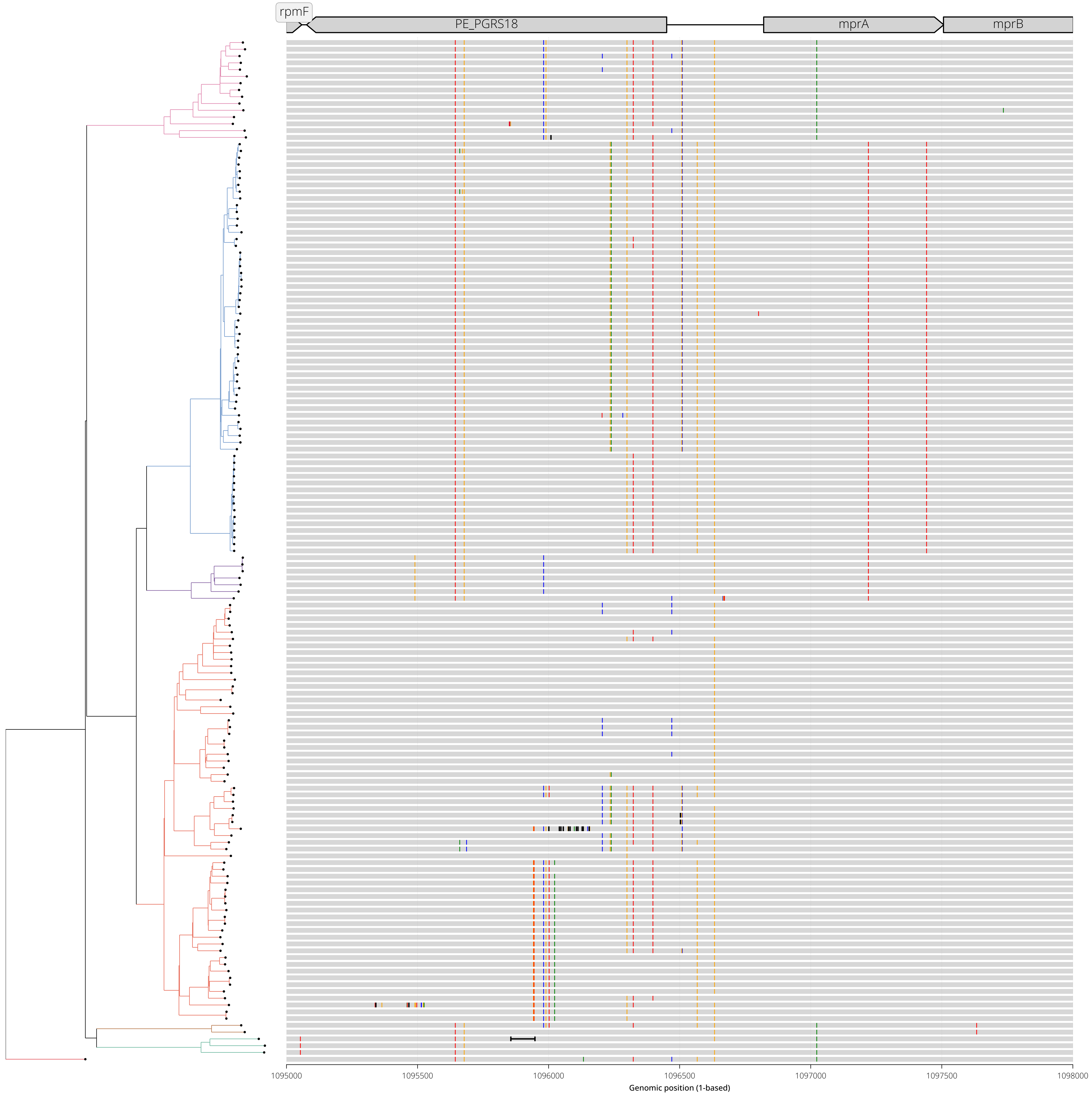

**Diversity Hotspot View - 04:**
**Genomic range shown: NC_000962.3:1274000-1279000**
**Gene(s) of interest: Rv1148c**

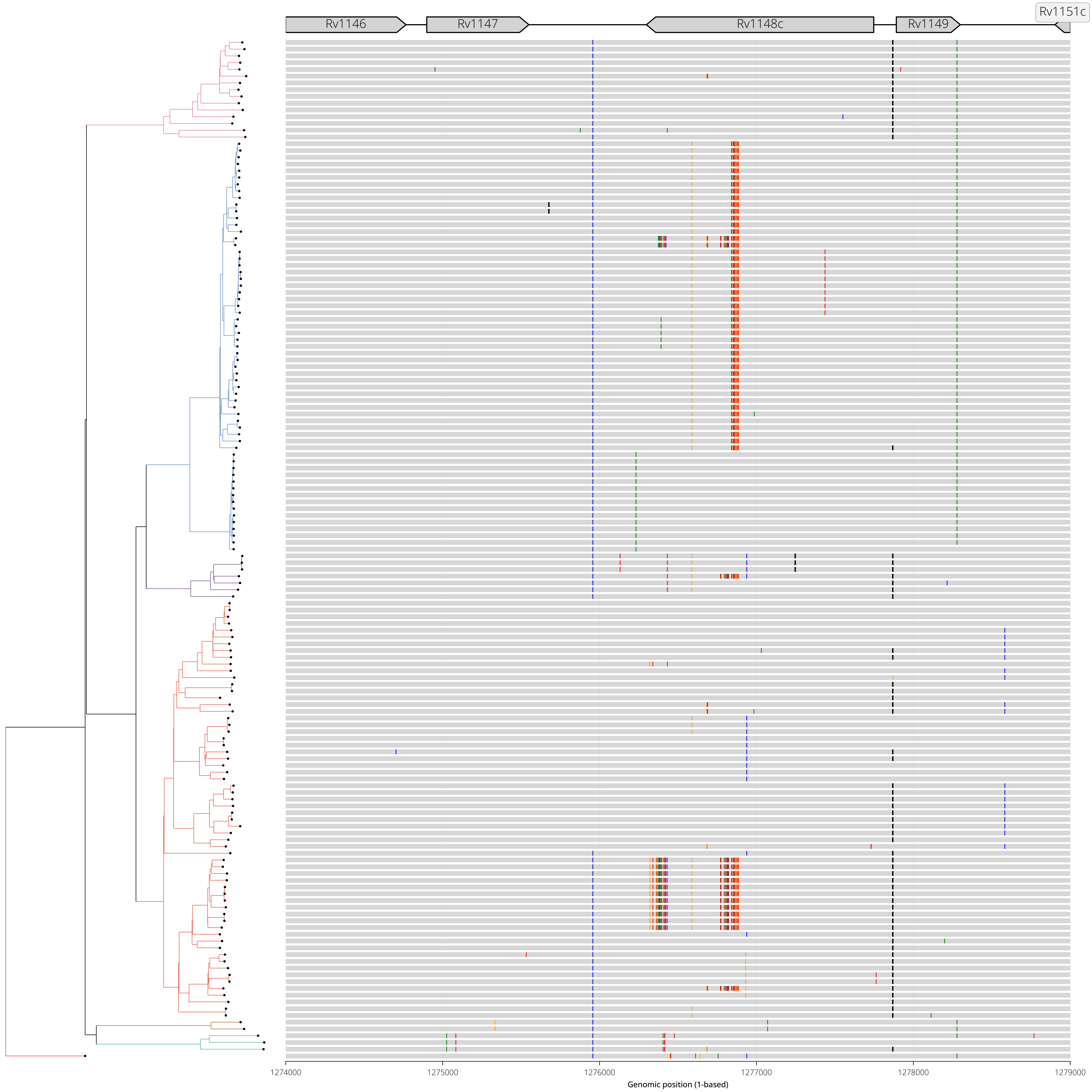

**Diversity Hotspot View - 05:**
**Genomic range shown: NC_000962.3:1338000-1343000**
**Gene(s) of interest: PPE18,esxK**

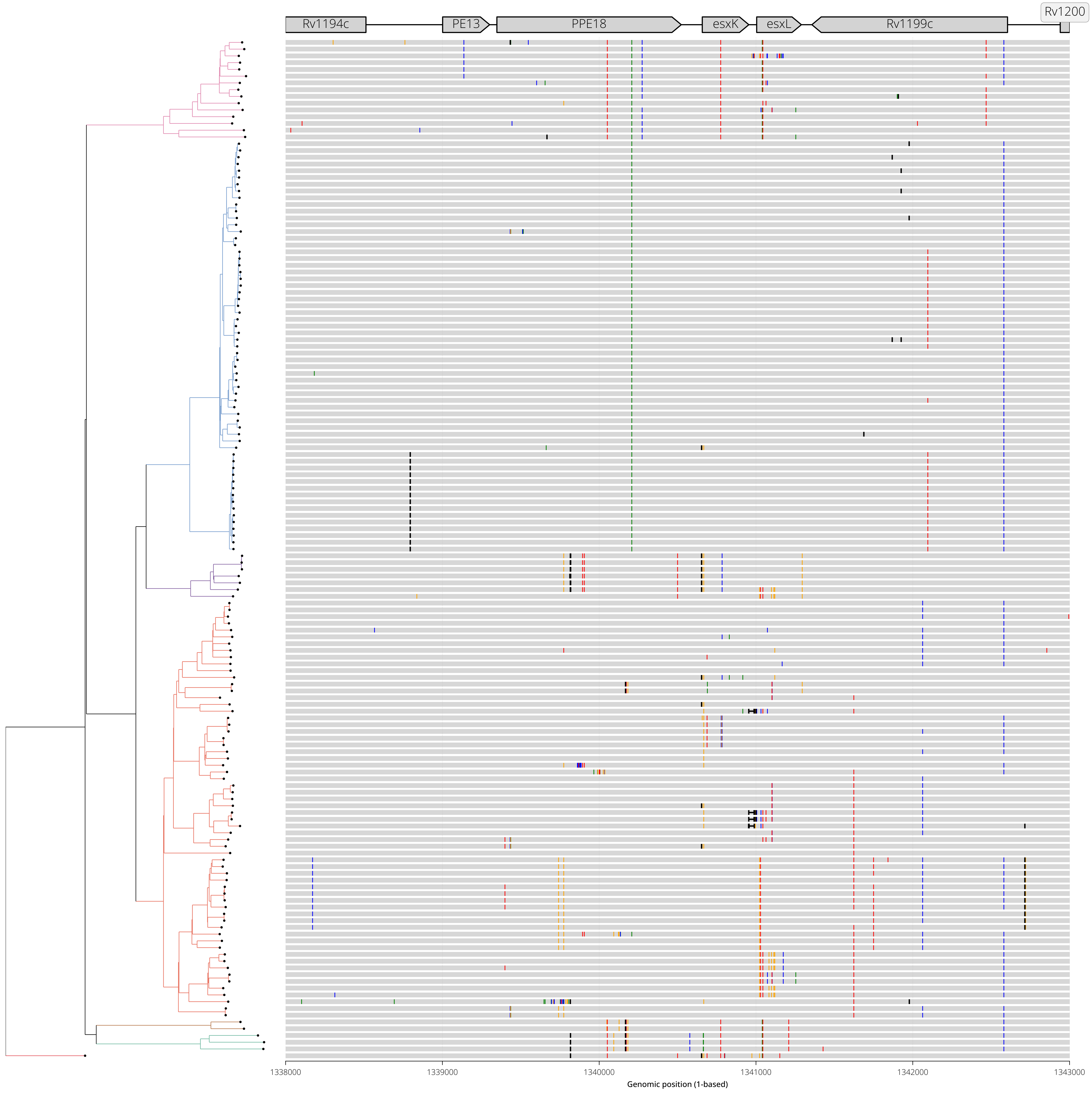

**Diversity Hotspot View - 06:**
**Genomic range shown: NC_000962.3:1339000-1344000**
**Gene(s) of interest: esxL,Rv1199c**

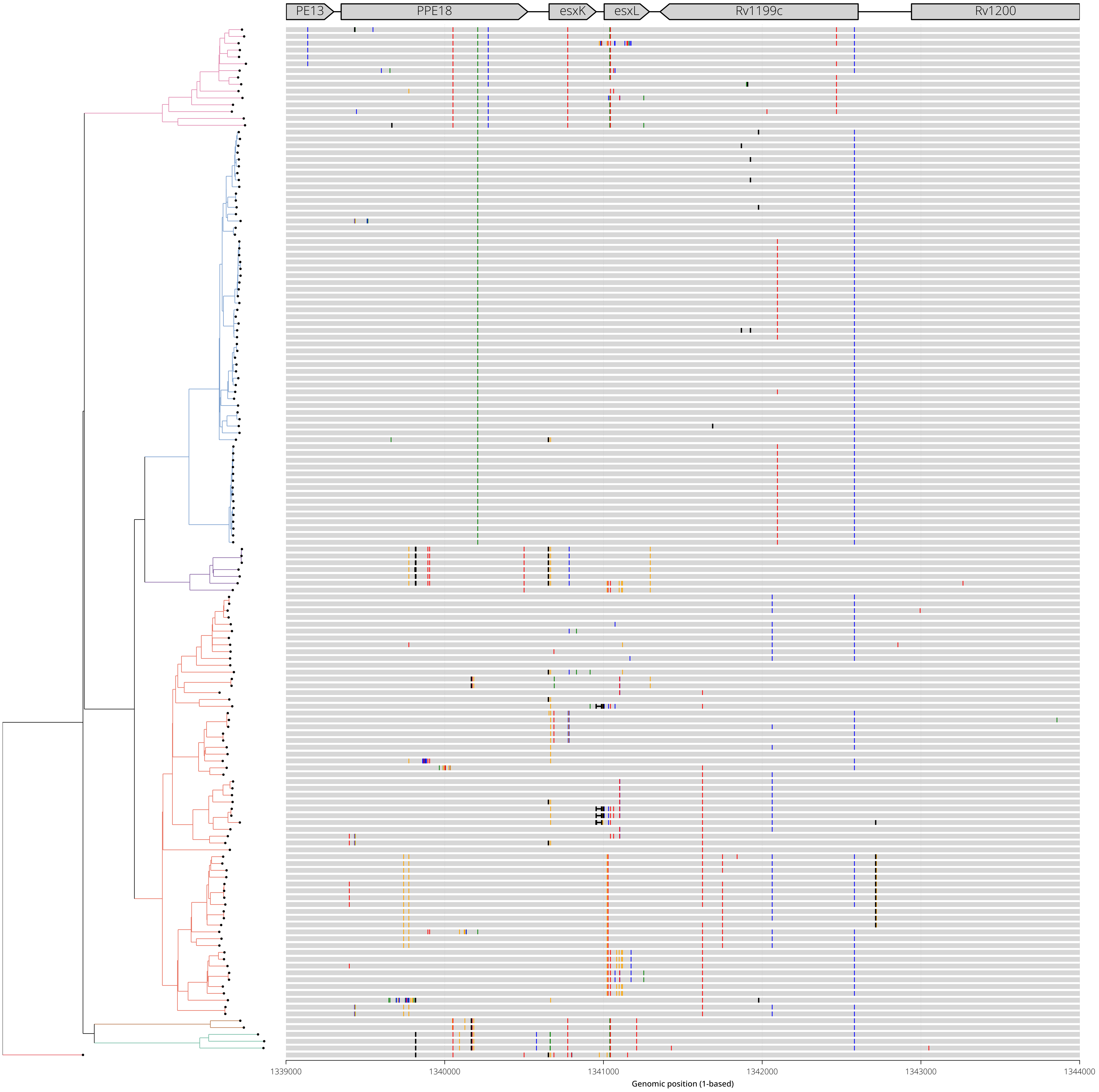

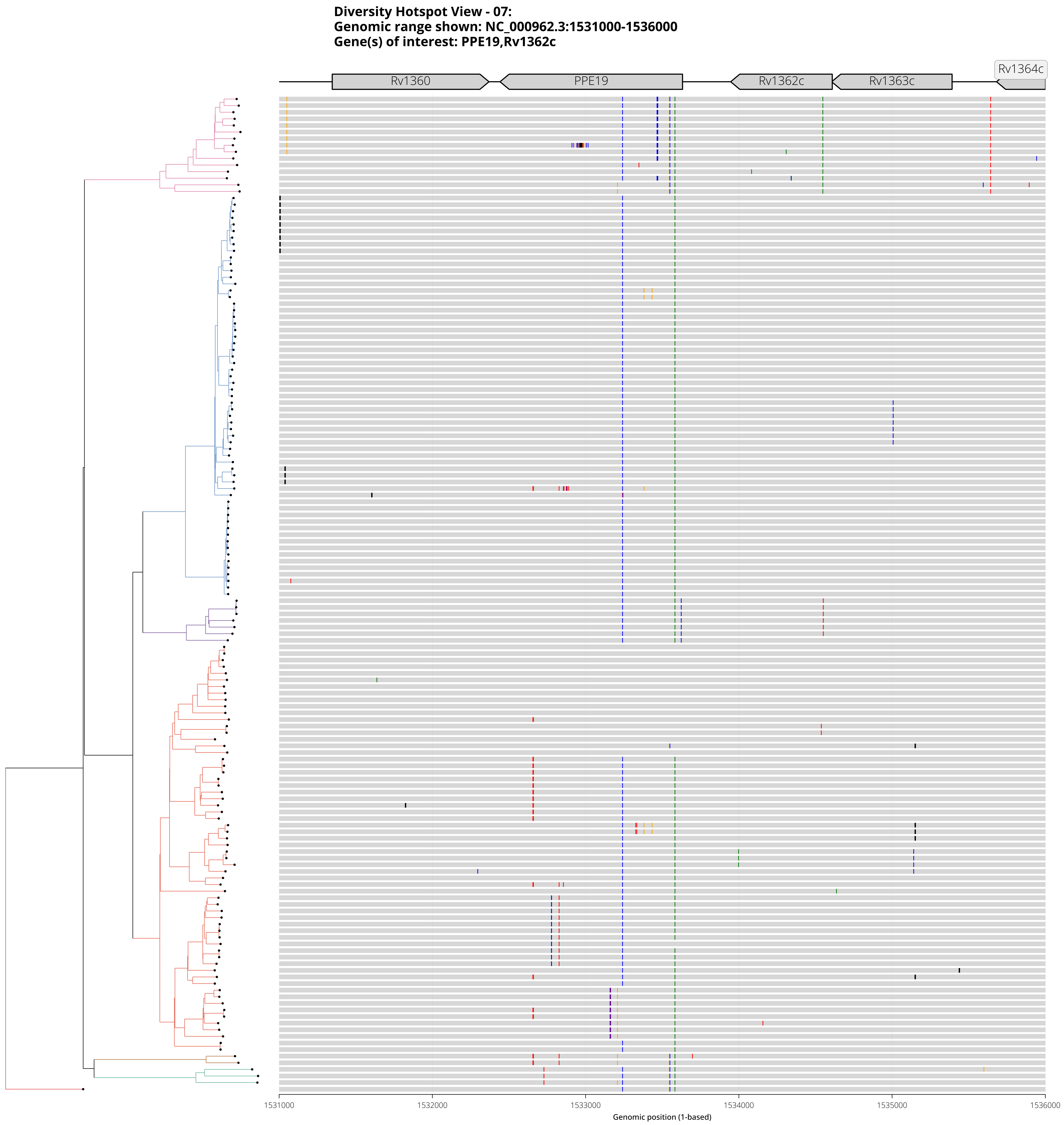
Diversity Hotspot View - 07:
Genomic range shown: NC_000962.3:1531000-1536000
Gene(s) of interest: PPE19,Rv1362c
Rv1360
PPE19
Rv1362c
Rv1363c
Rv1364c
1531000
1532000
1533000
1534000
1535000
1536000
Genomic position (1-based)

**Diversity Hotspot View - 08:**
**Genomic range shown: NC_000962.3:1631000-1638000**
**Gene(s) of interest: PE_PGRS27**

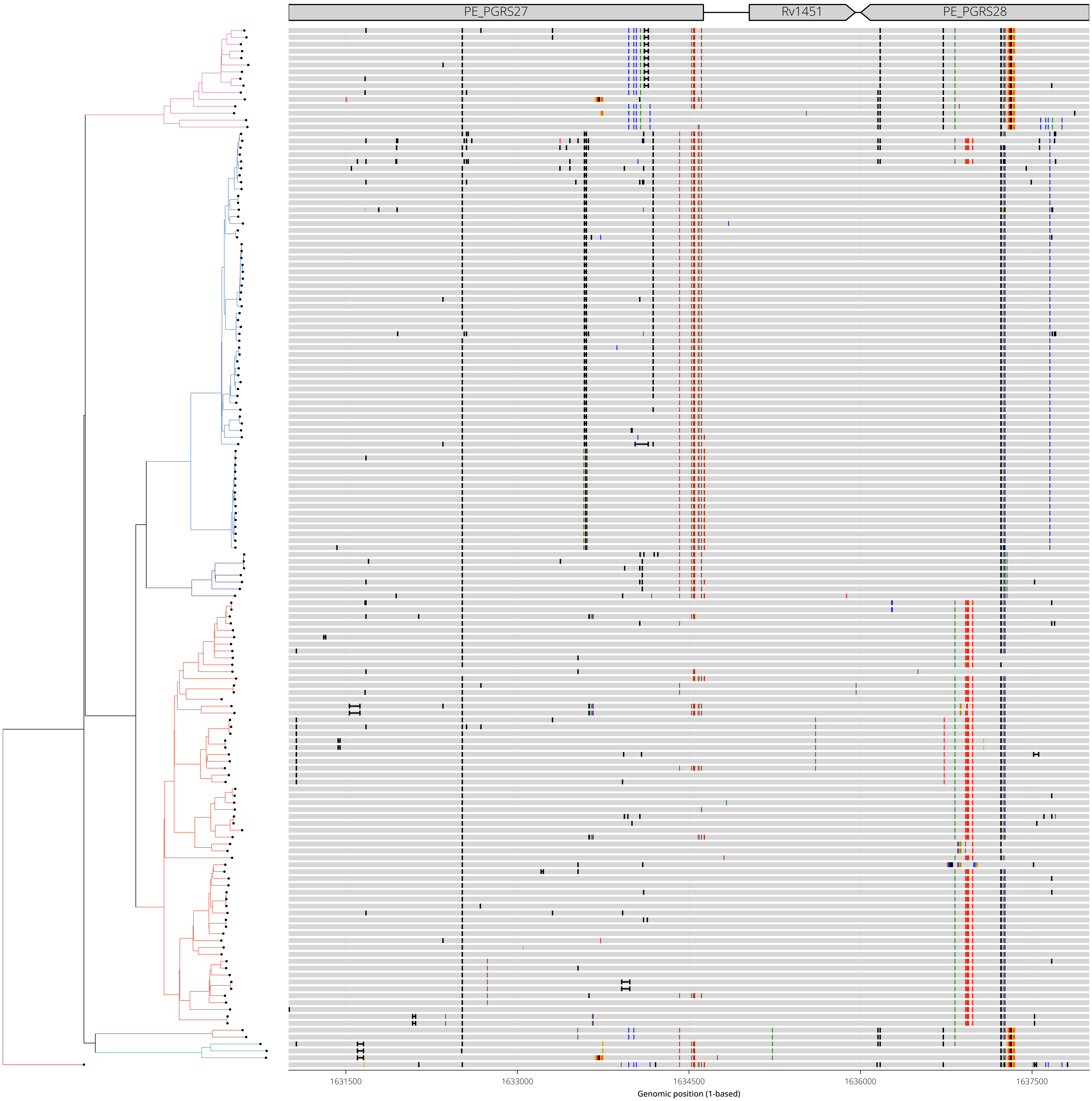

**Diversity Hotspot View - 09:**
**Genomic range shown: NC_000962.3:1634000-1641000**
**Gene(s) of interest: PE_PGRS28**

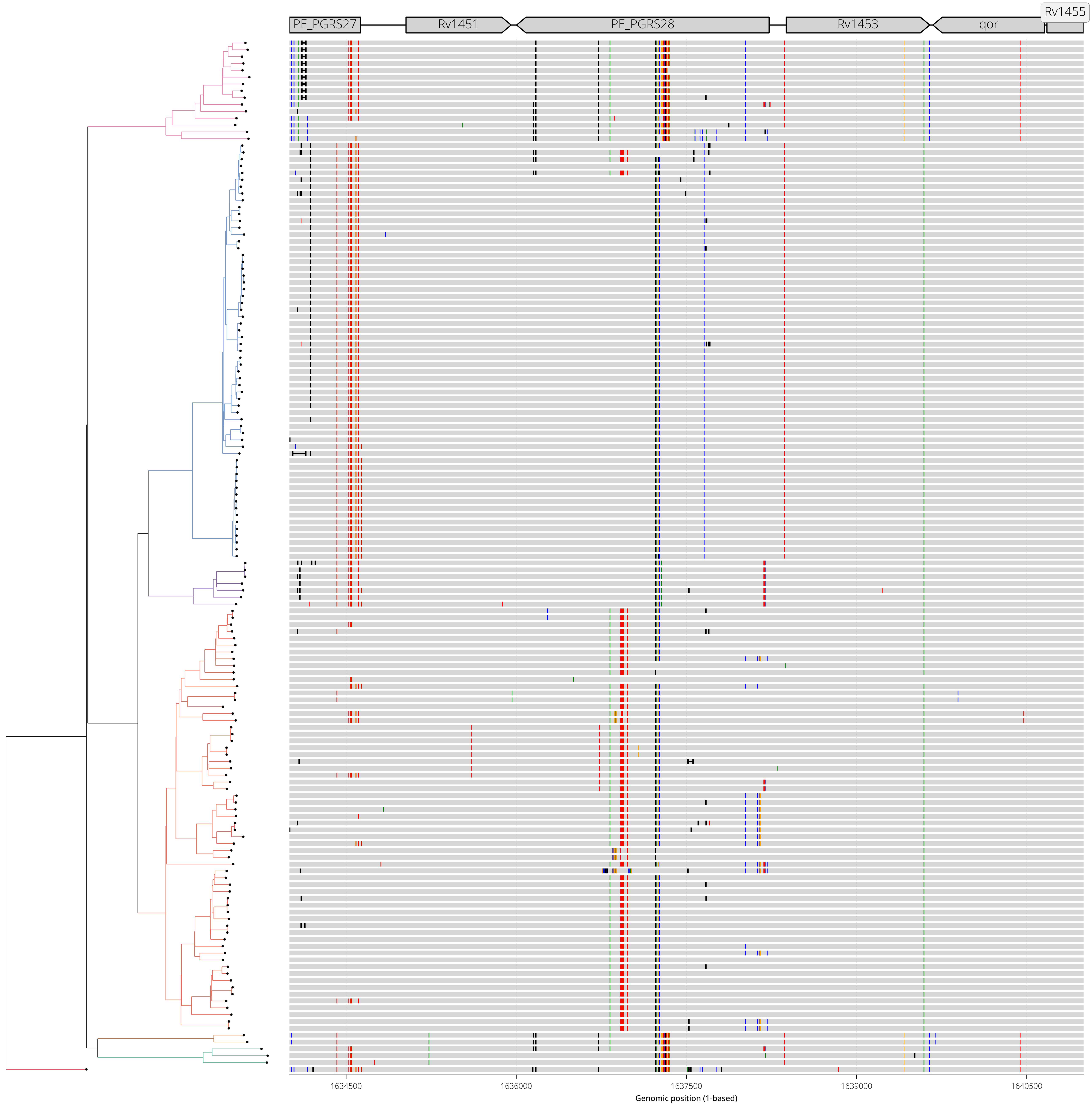

**Diversity Hotspot View - 10:**
**Genomic range shown: NC_000962.3:1787000-1792000**
**Gene(s) of interest: Rv1587c,Rv1588c**

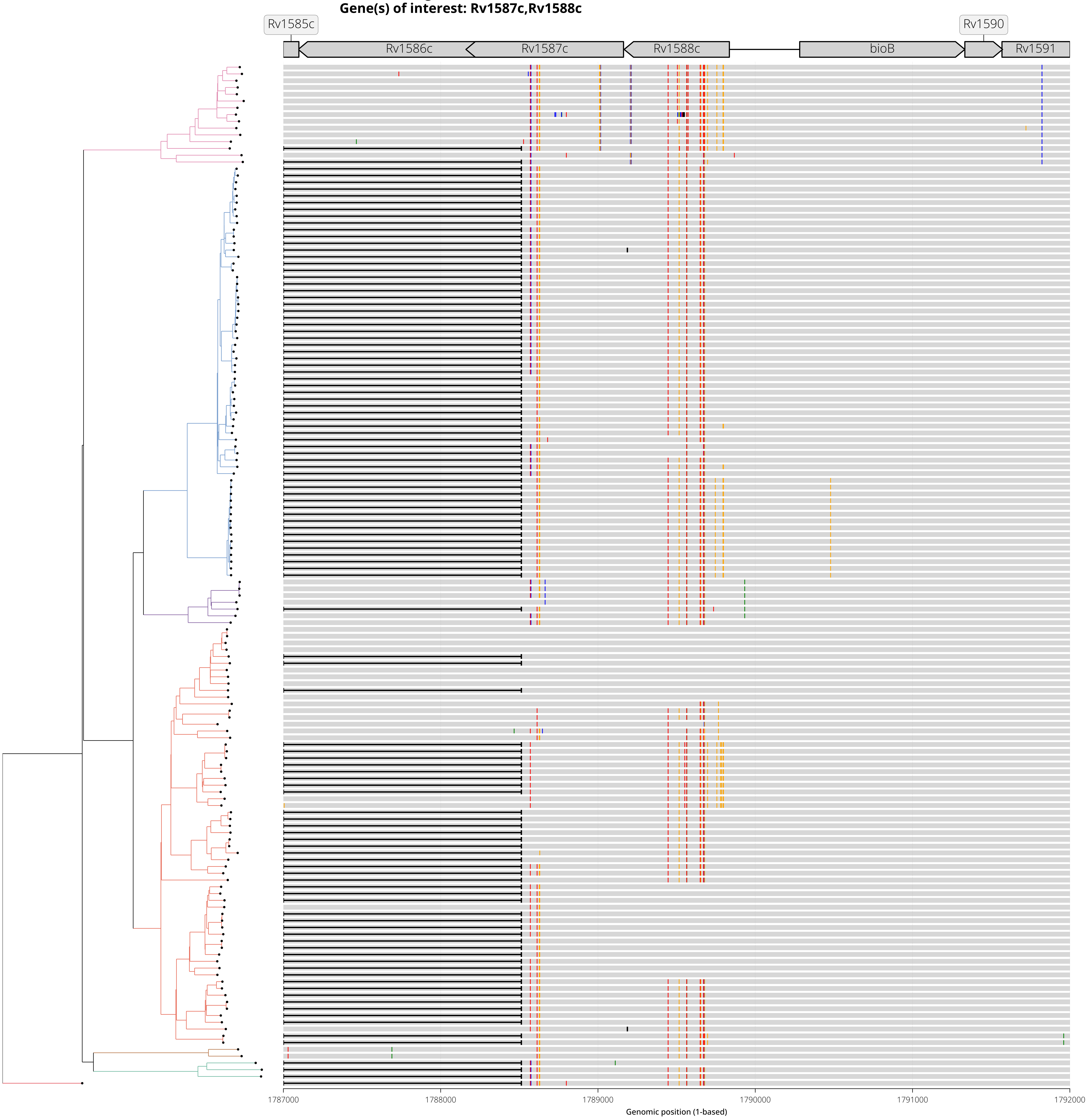

**Diversity Hotspot View - 11:**
**Genomic range shown: NC_000962.3:2194000-2199000**
**Gene(s) of interest: Rv1945**

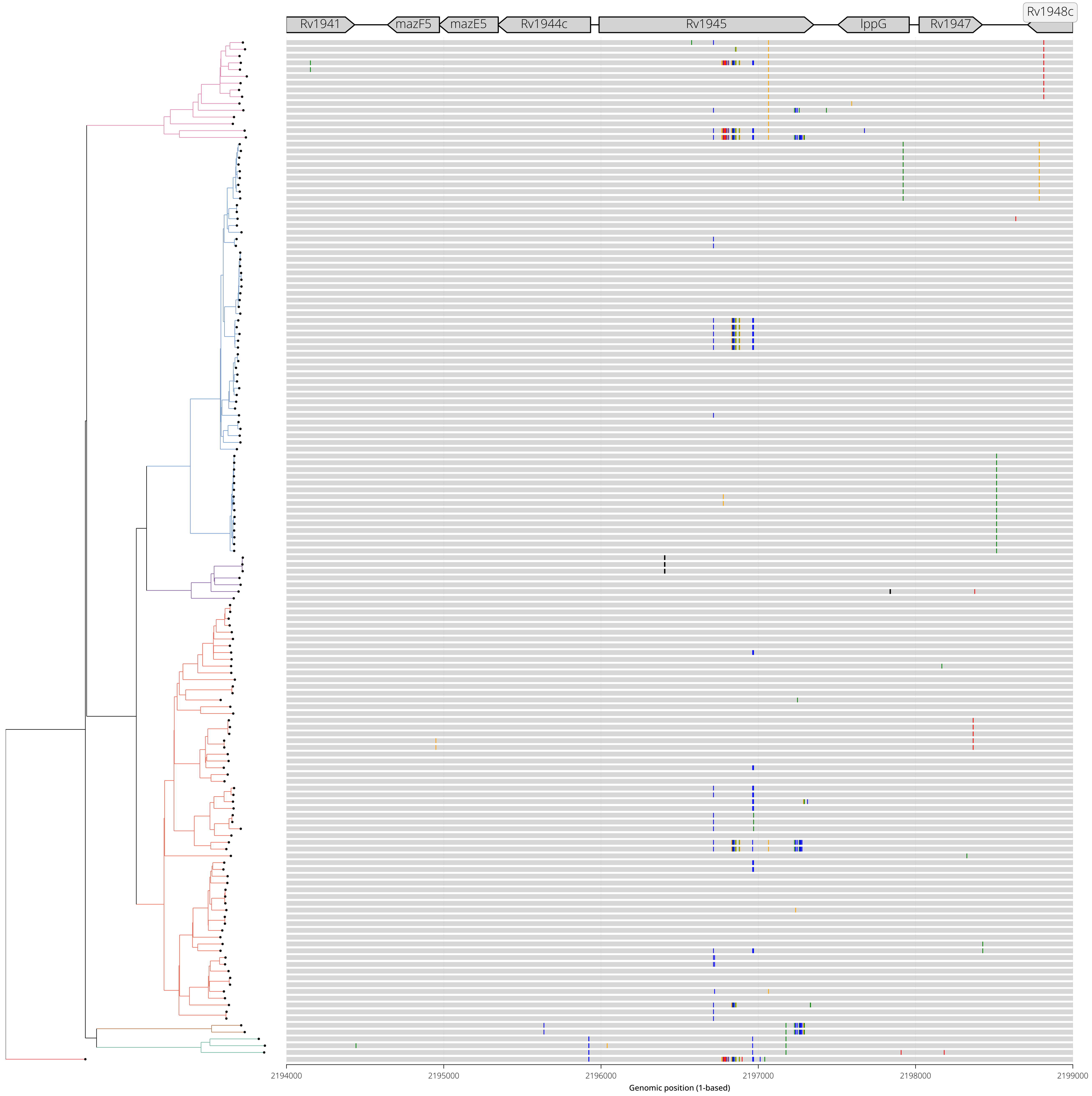

**Diversity Hotspot View - 12:**
**Genomic range shown: NC_000962.3:2260000-2265000**
**Gene(s) of interest: Rv2015c**

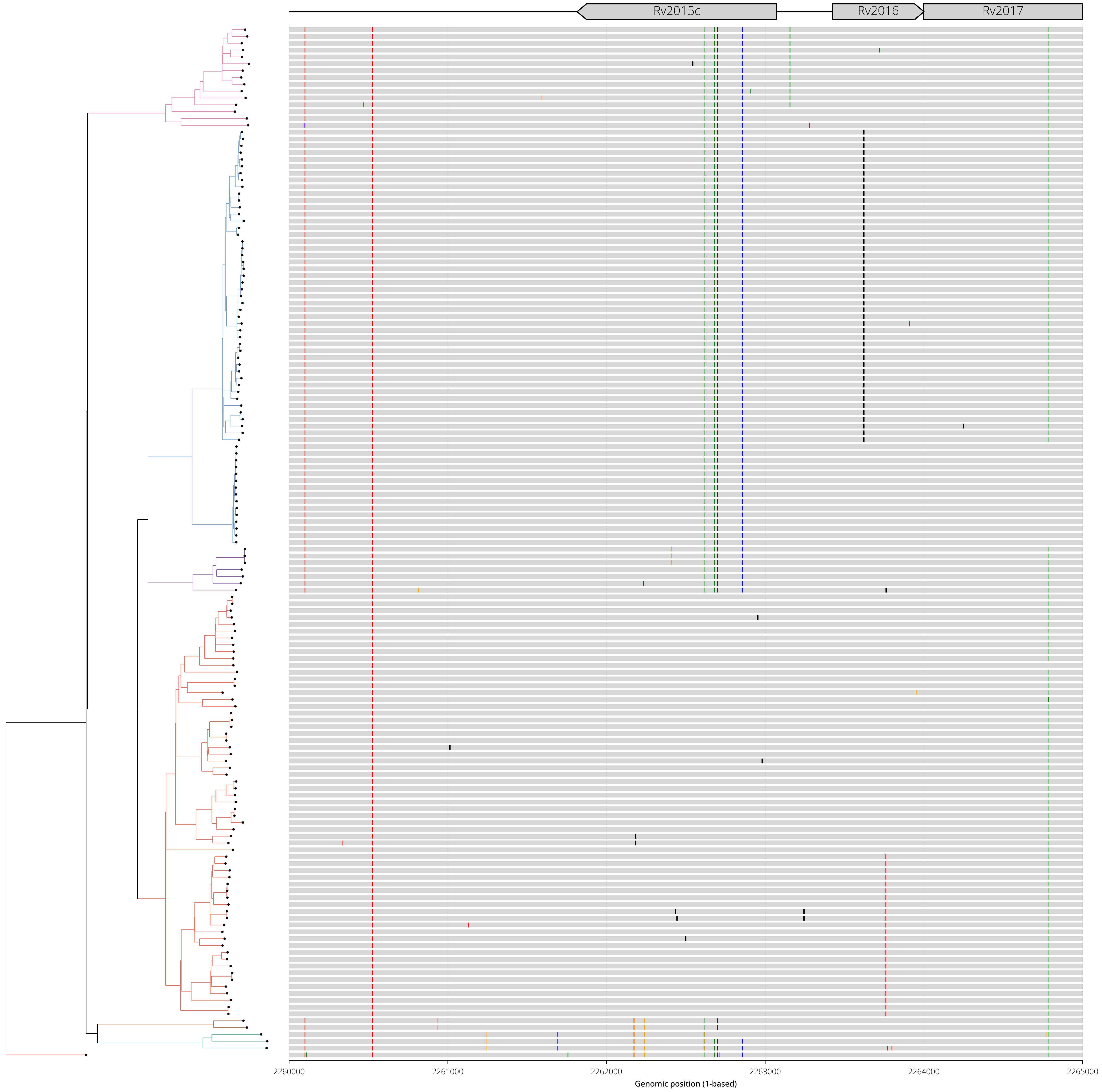

**Diversity Hotspot View - 13:**
**Genomic range shown: NC_000962.3:2336000-2341000**
**Gene(s) of interest: Rv2081c,Rv2082**

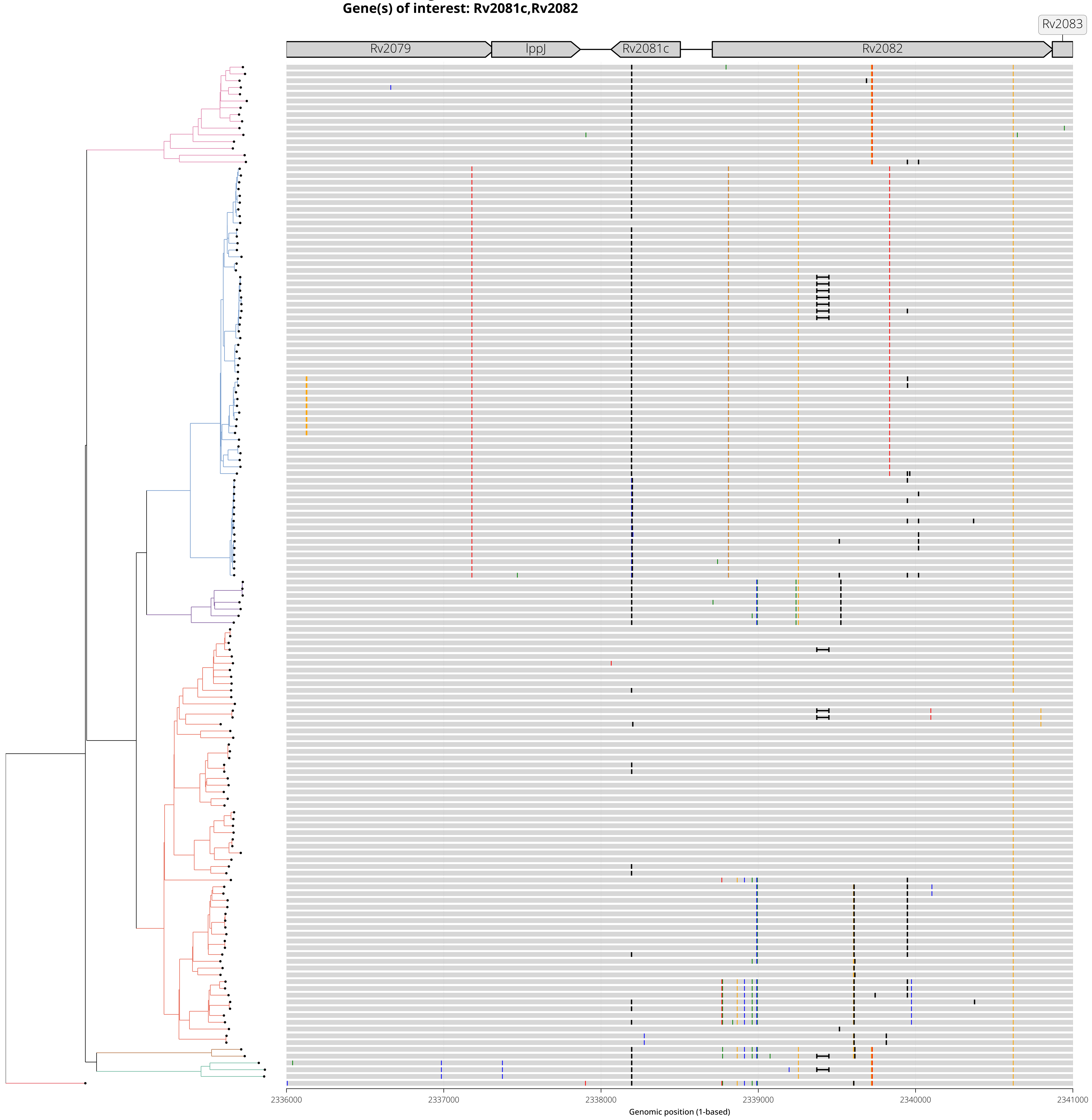

**Diversity Hotspot View - 14:**
**Genomic range shown: NC_000962.3:2624000-2629000**
**Gene(s) of interest: esxO,esxP,Rv2348c**

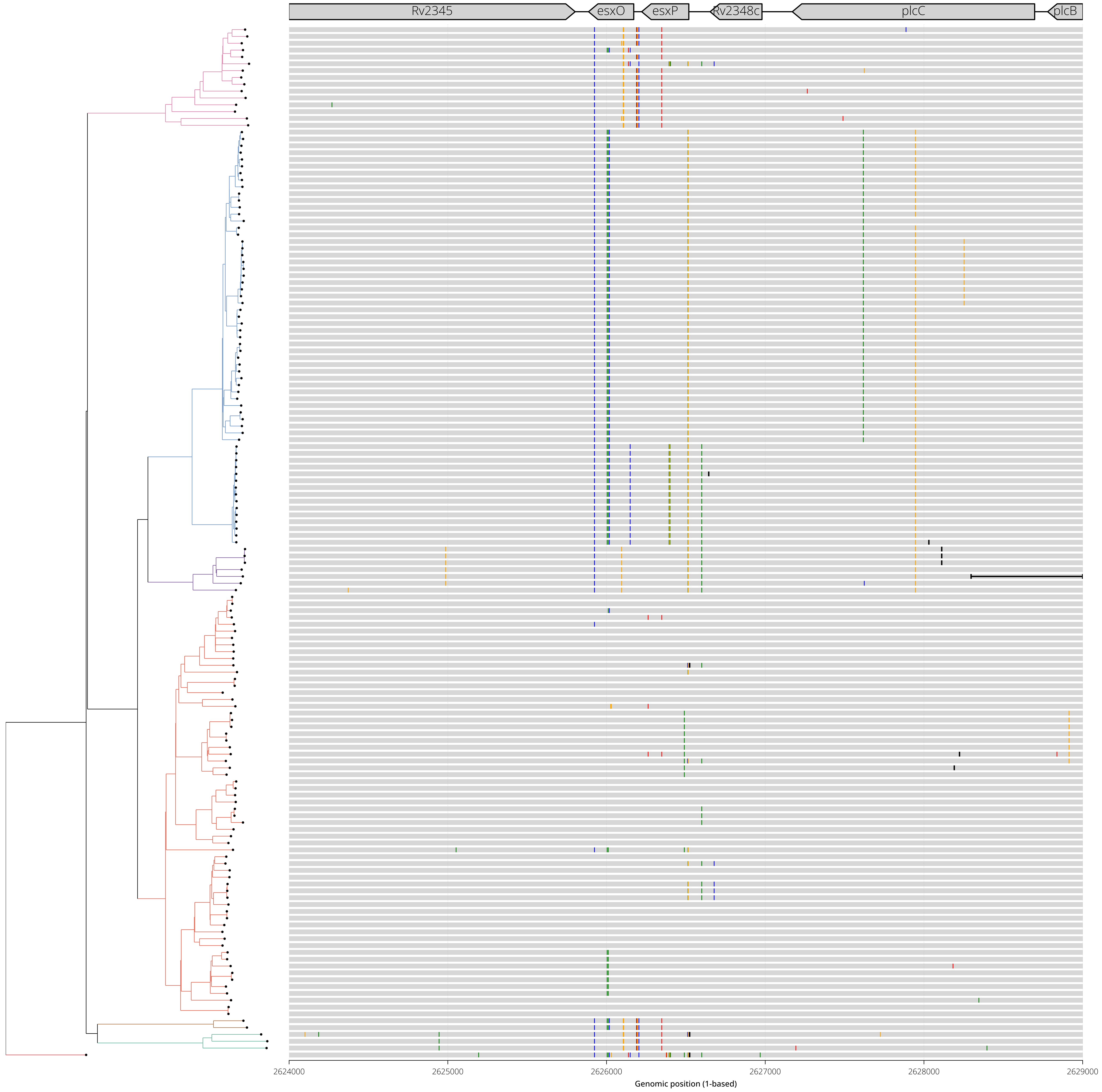

**Diversity Hotspot View - 15:**
**Genomic range shown: NC_000962.3:2865000-2870000**
**Gene(s) of interest: lppA,lppB,vapB18**

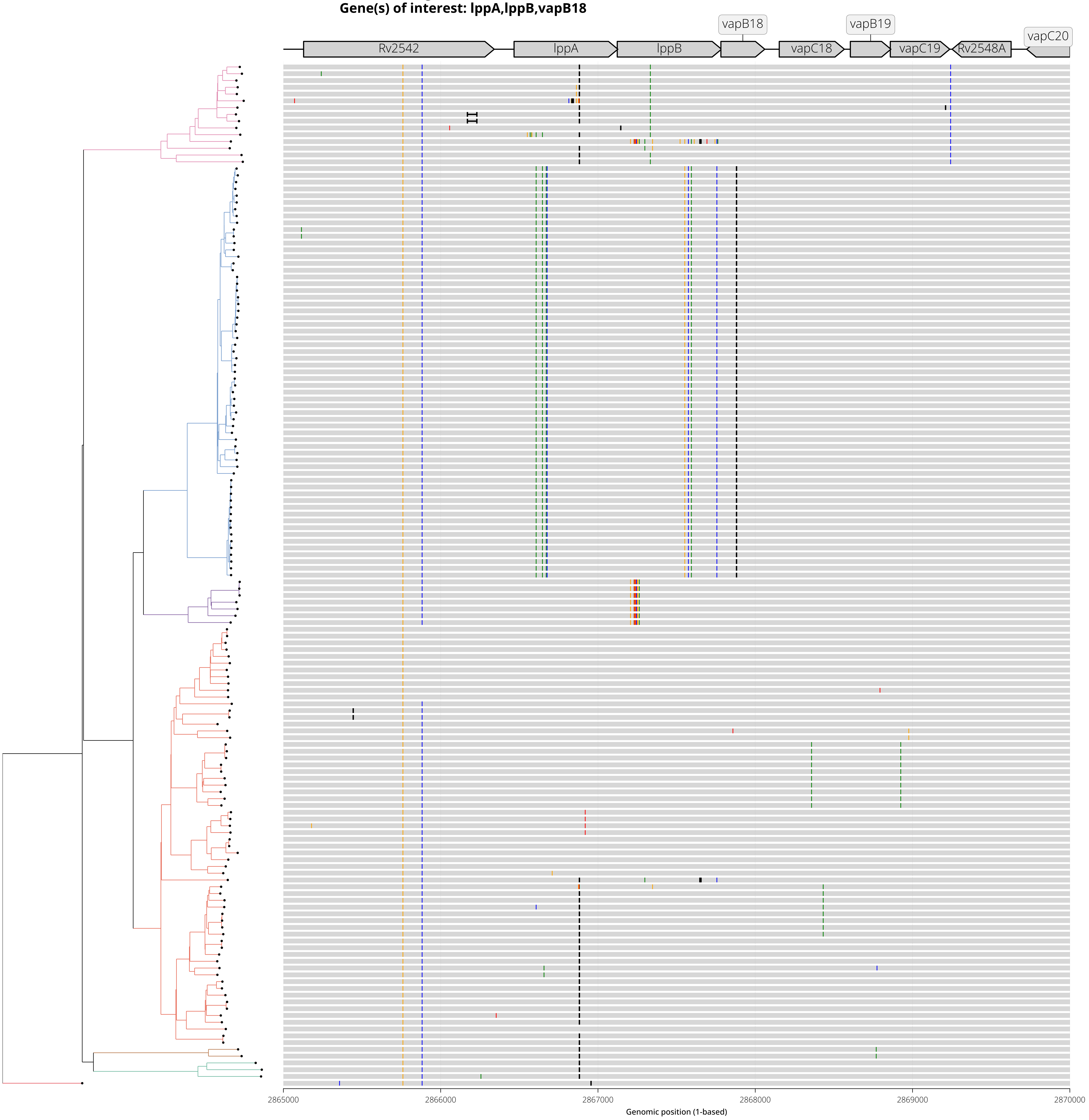

**Diversity Hotspot View - 16:**
**Genomic range shown: NC_000962.3:2942000-2947000**
**Gene(s) of interest: PE_PGRS45**

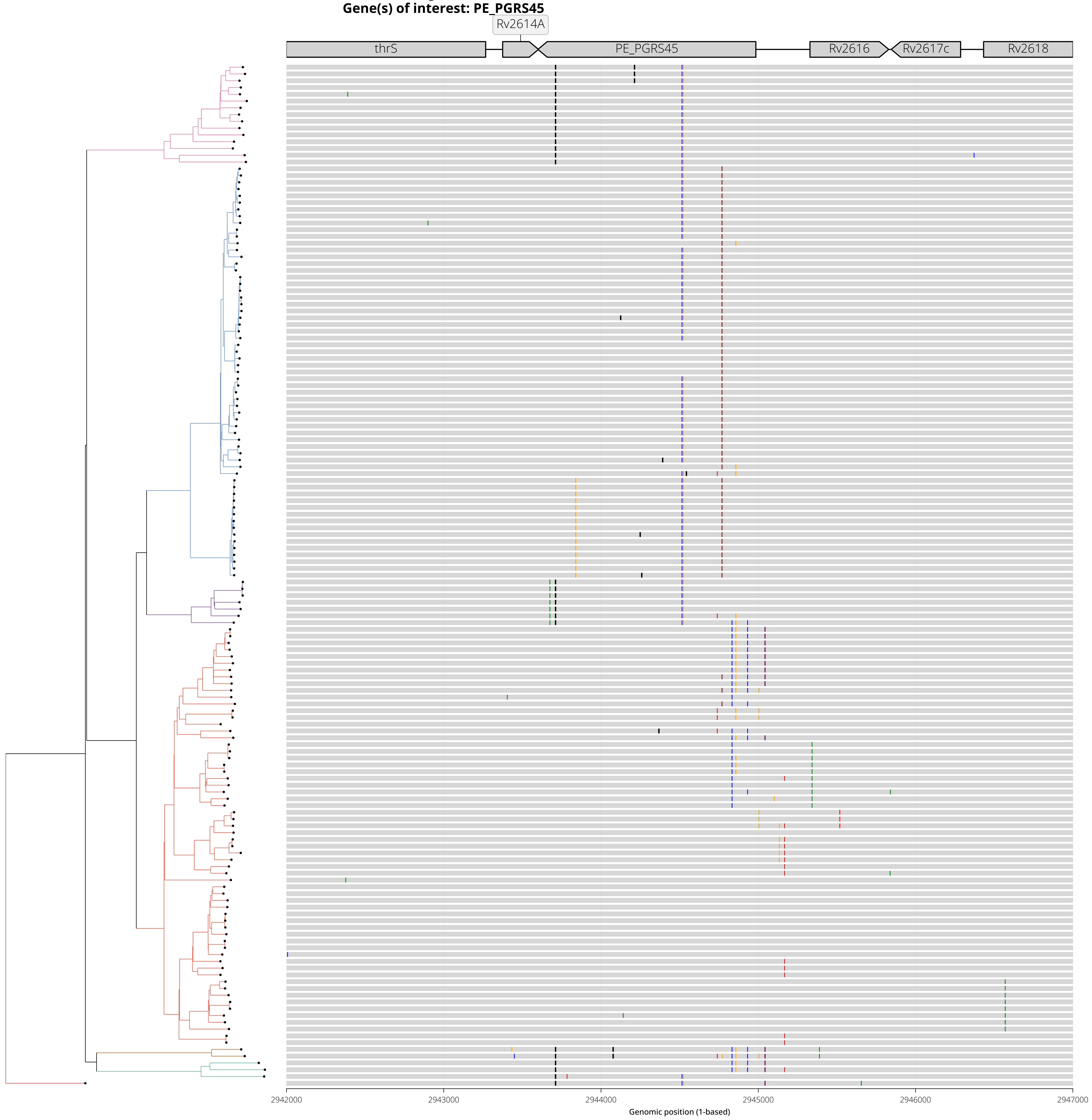

**Diversity Hotspot View - 17:**
**Genomic range shown: NC_000962.3:3133000-3138000**
**Gene(s) of interest: Rv2827c,Rv2828c**

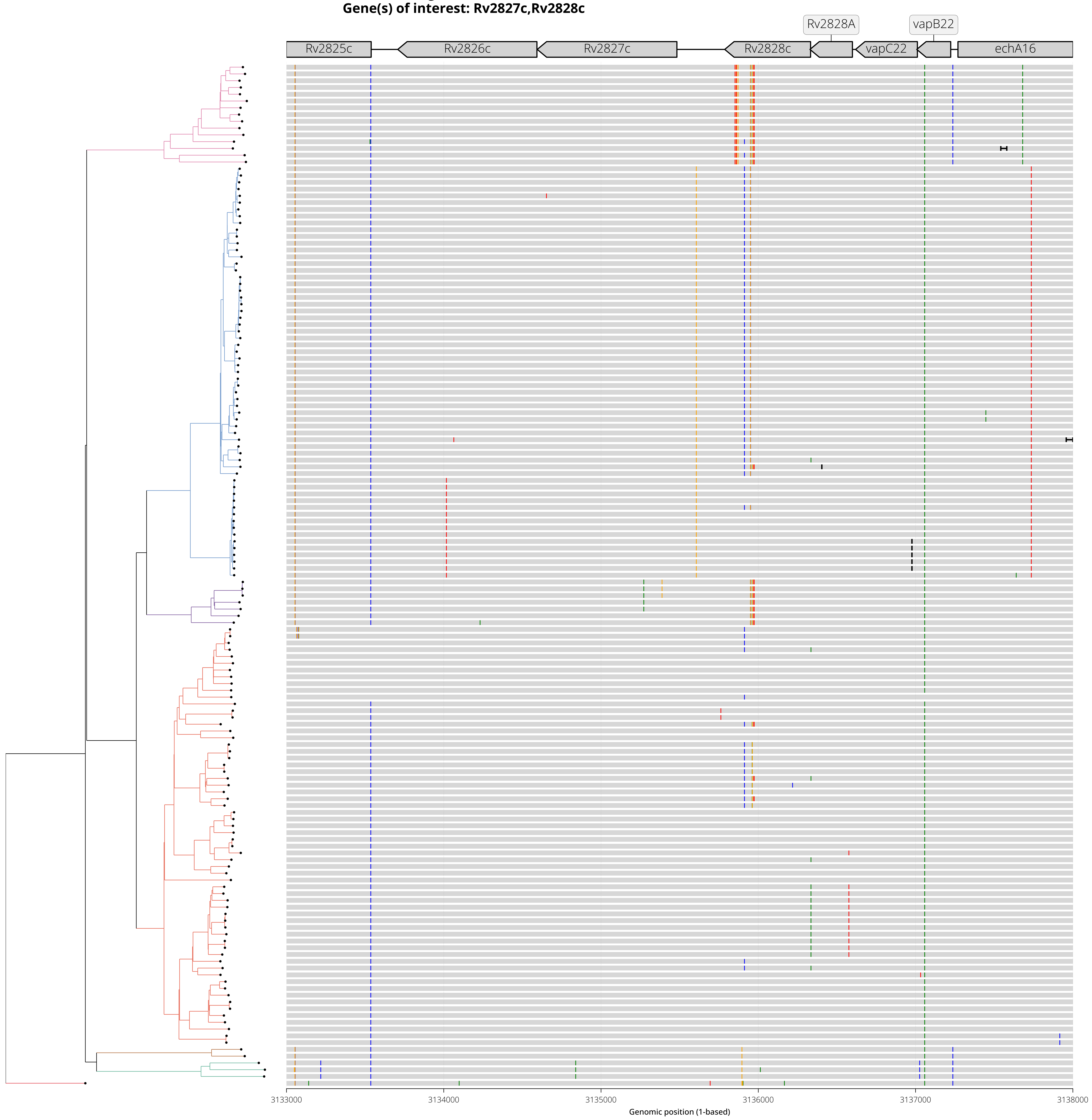

**Diversity Hotspot View - 18:**
**Genomic range shown: NC_000962.3:3728000-3737000**
**Gene(s) of interest: PPE54**

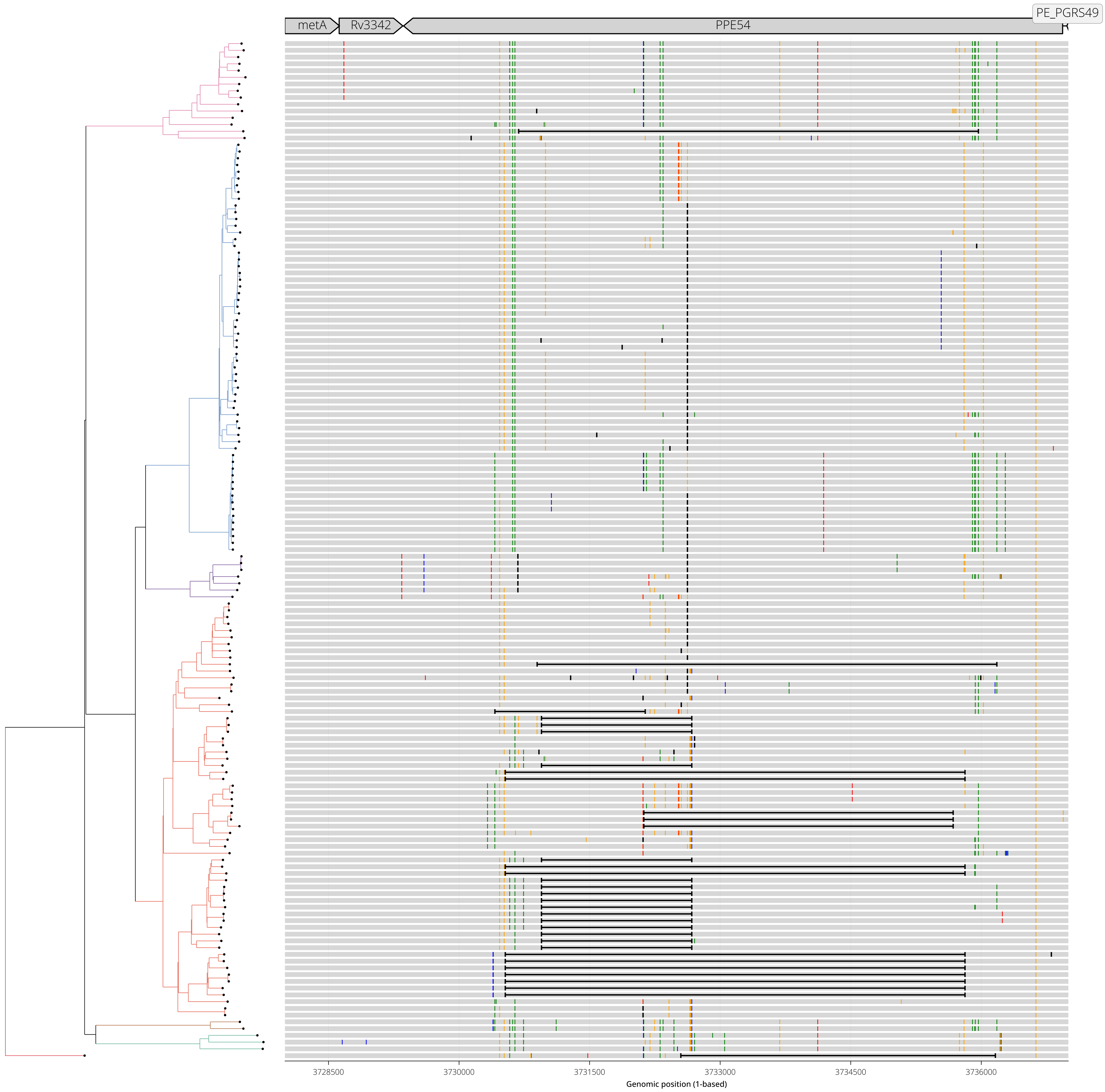

**Diversity Hotspot View - 19:**
**Genomic range shown: NC_000962.3:3746000-3755000**
**Gene(s) of interest: PPE55**

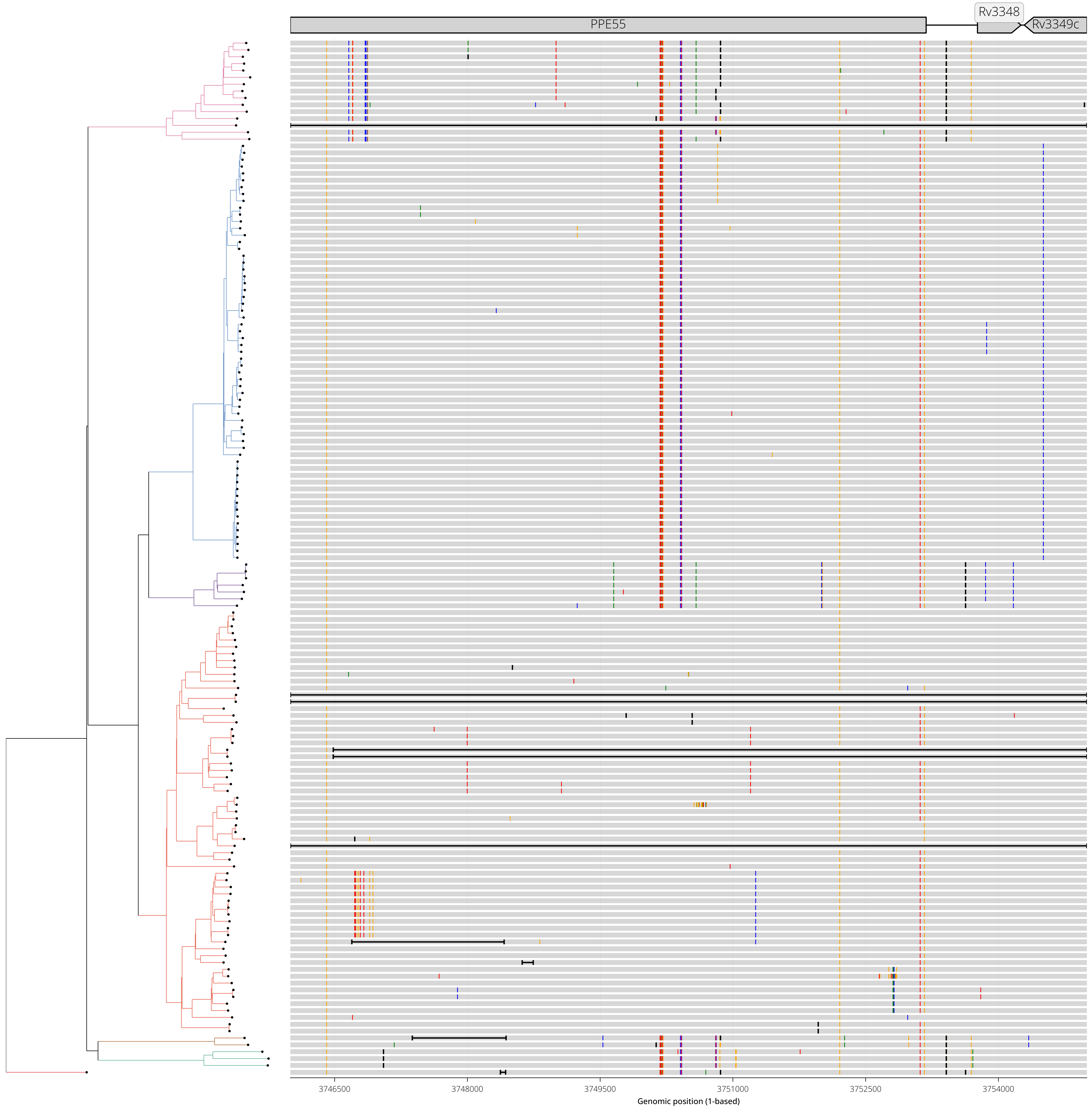

**Diversity Hotspot View - 20:**
**Genomic range shown: NC_000962.3:3841000-3844000**
**Gene(s) of interest: Rv3424c,PPE57**

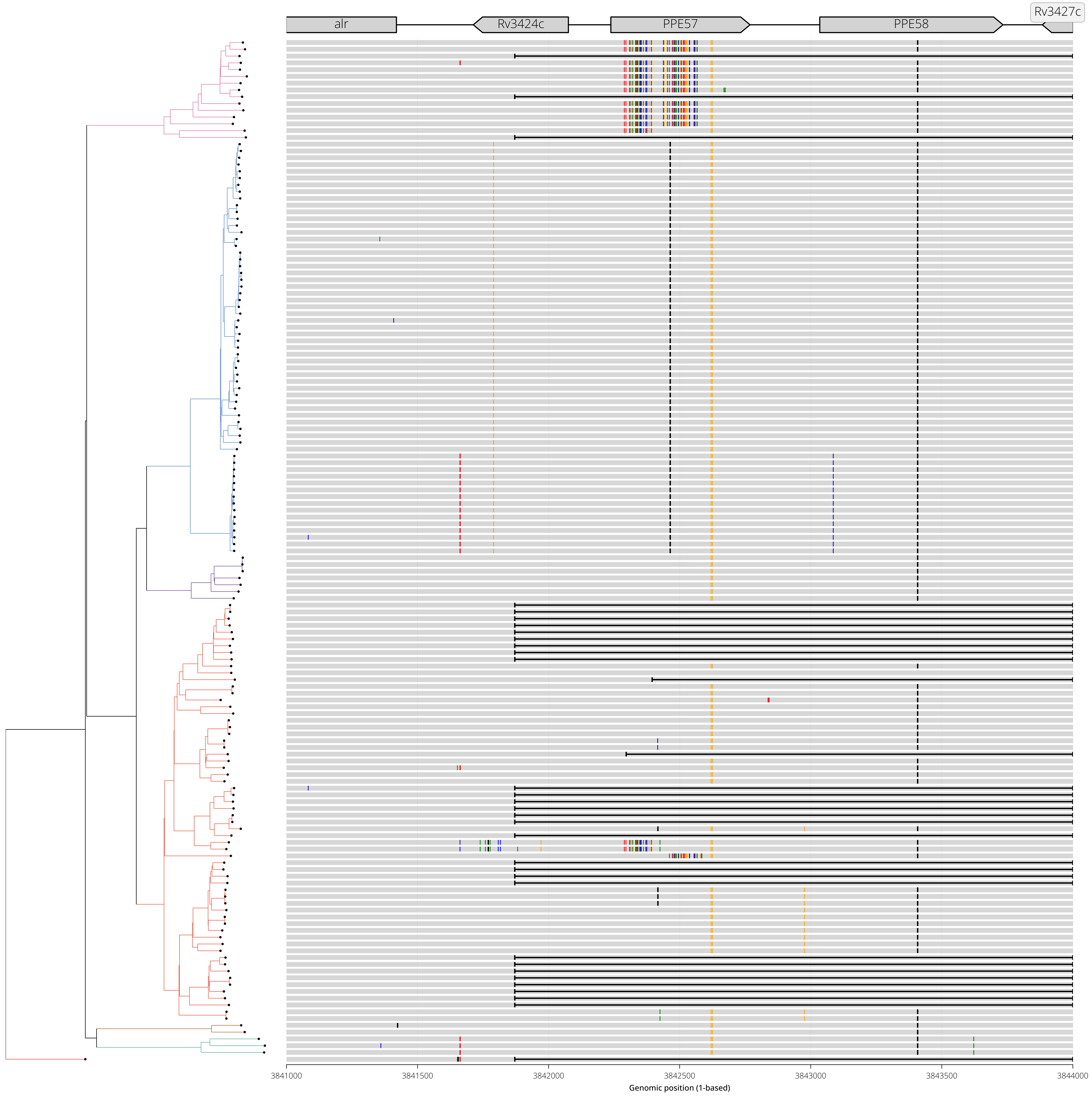

**Diversity Hotspot View - 21:**
**Genomic range shown: NC_000962.3:3845000-3850000**
**Gene(s) of interest: PPE59,Rv3430c**

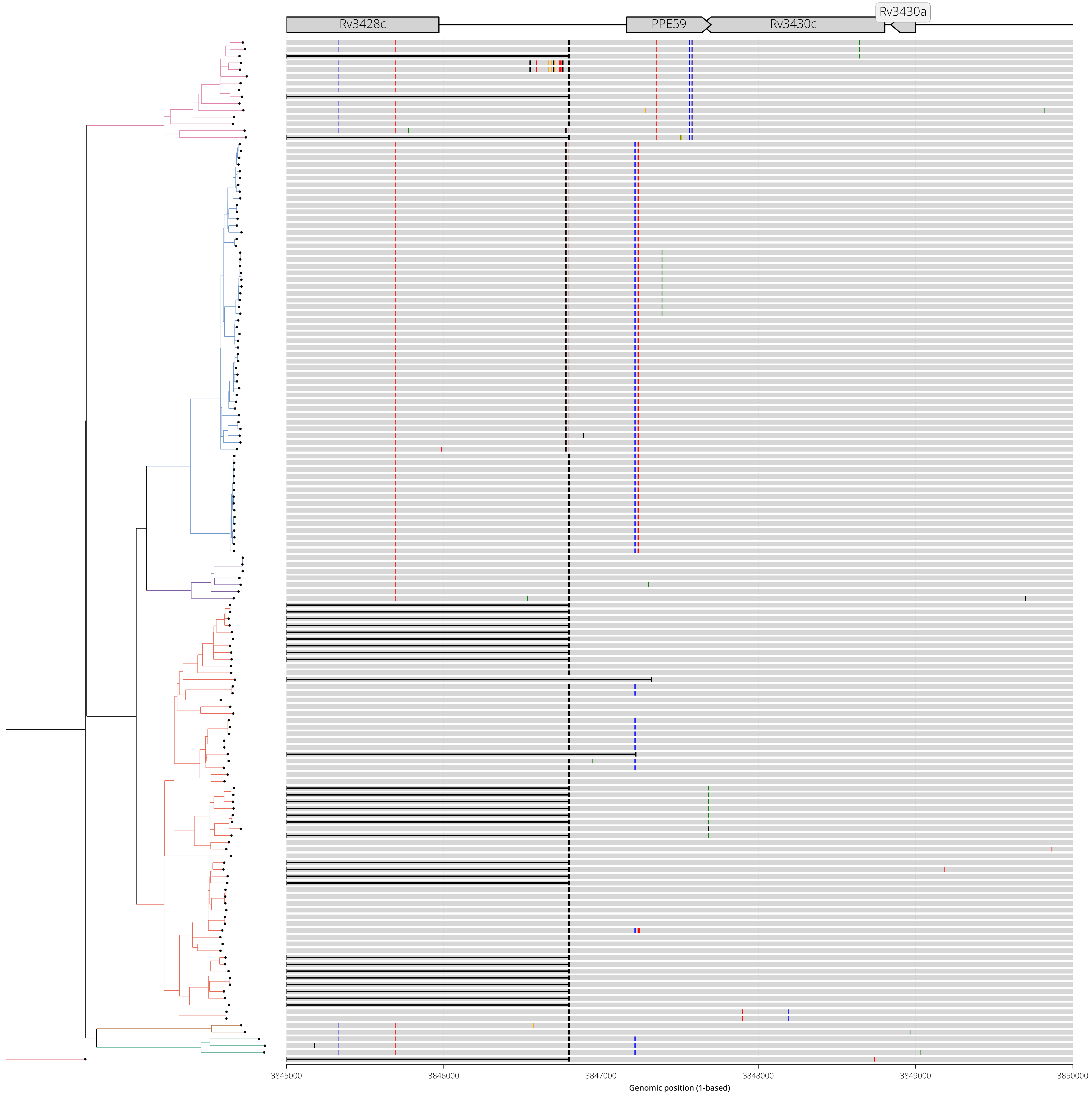

**Diversity Hotspot View - 22:**
**Genomic range shown: NC_000962.3:3881000-3886000**
**Gene(s) of interest: rmlC,Rv3466,Rv3467**

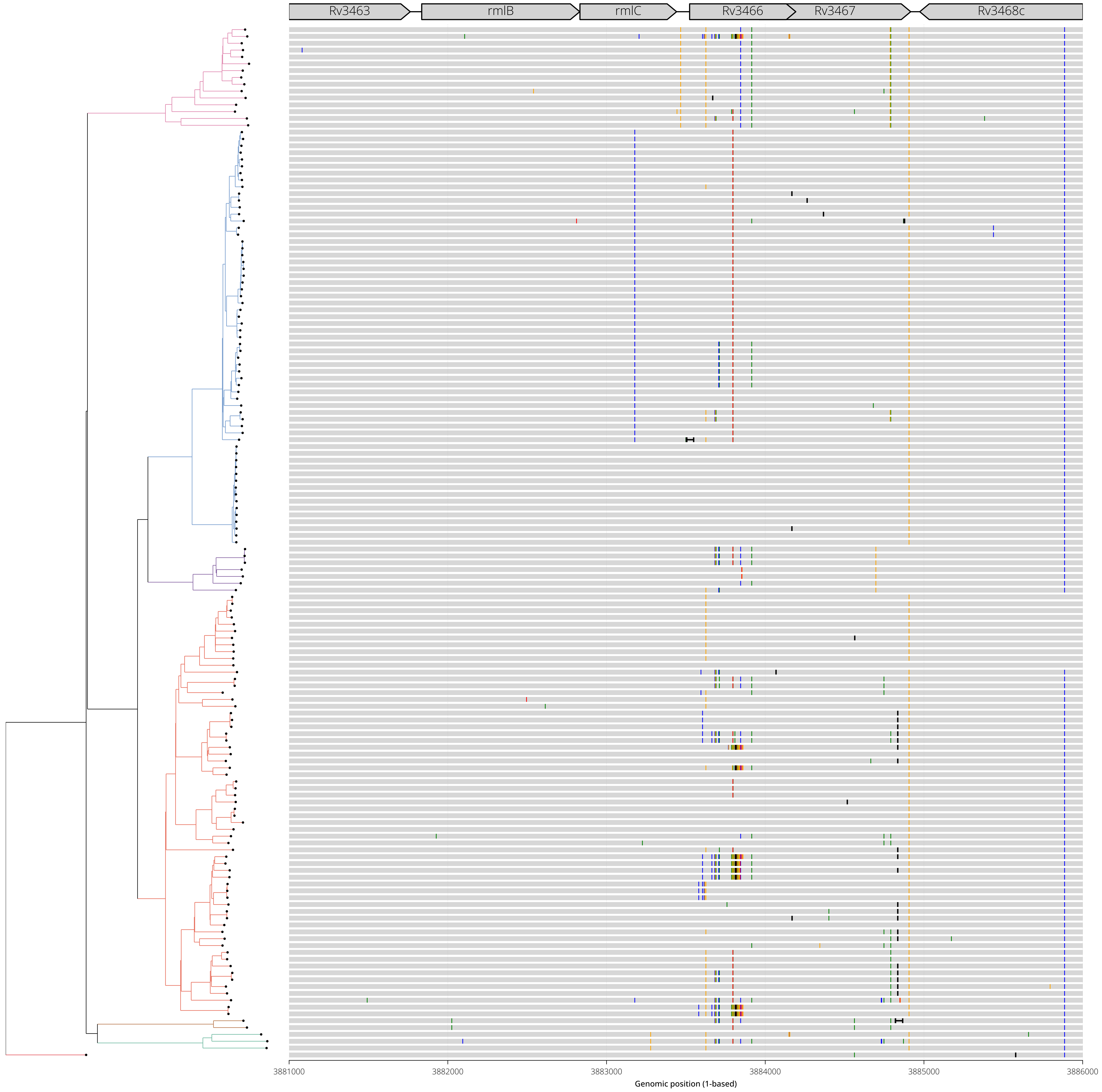

**Diversity Hotspot View - 23:**
**Genomic range shown: NC_000962.3:3893000-3898000**
**Gene(s) of interest: PPE60,Rv3479**

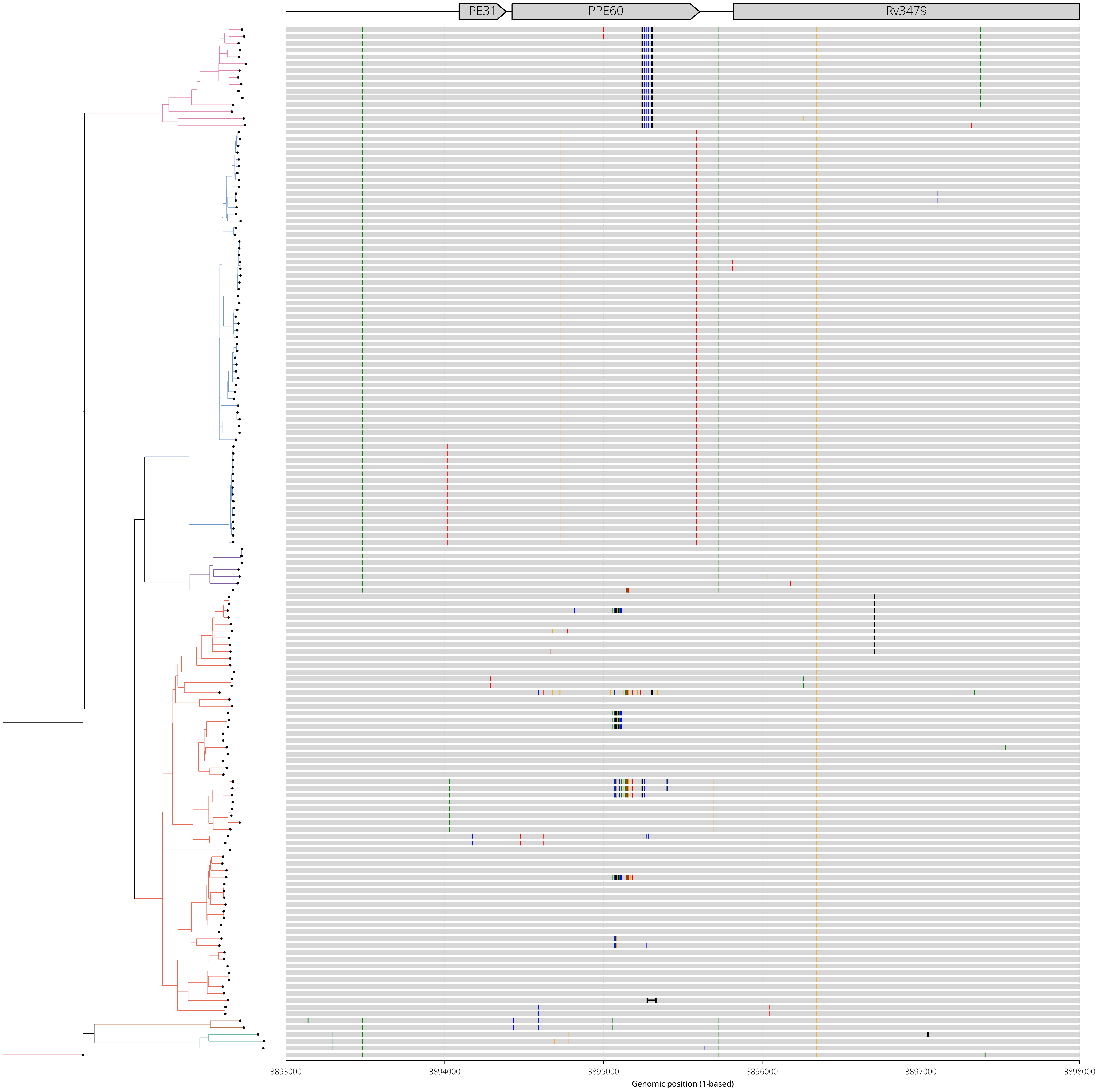

**Diversity Hotspot View - 24:**
**Genomic range shown: NC_000962.3:3931000-3938000**
**Gene(s) of interest: PE_PGRS54**

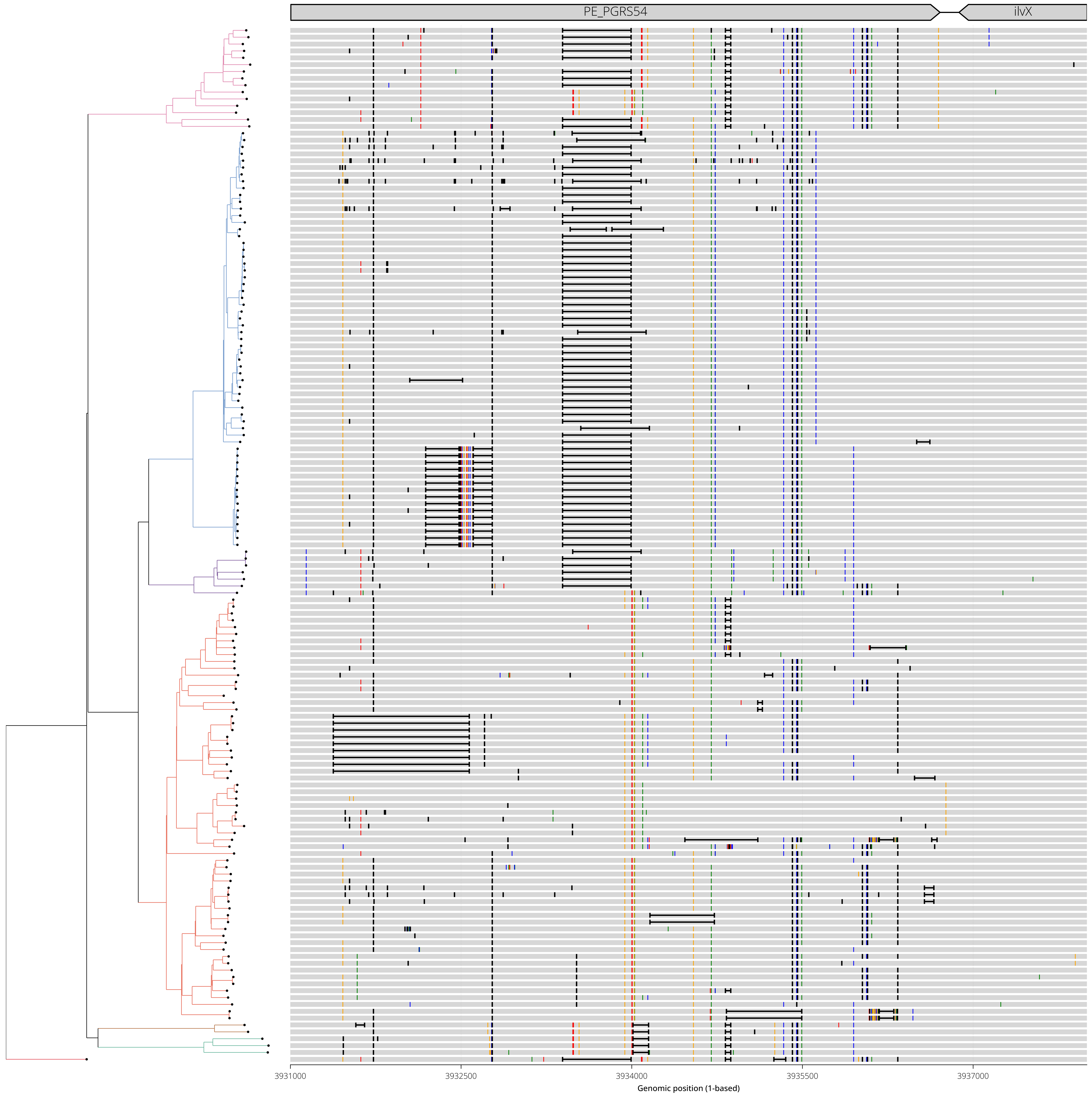

**Diversity Hotspot View - 25:**
**Genomic range shown: NC_000962.3:3940000-3947000**
**Gene(s) of interest: PE_PGRS56**

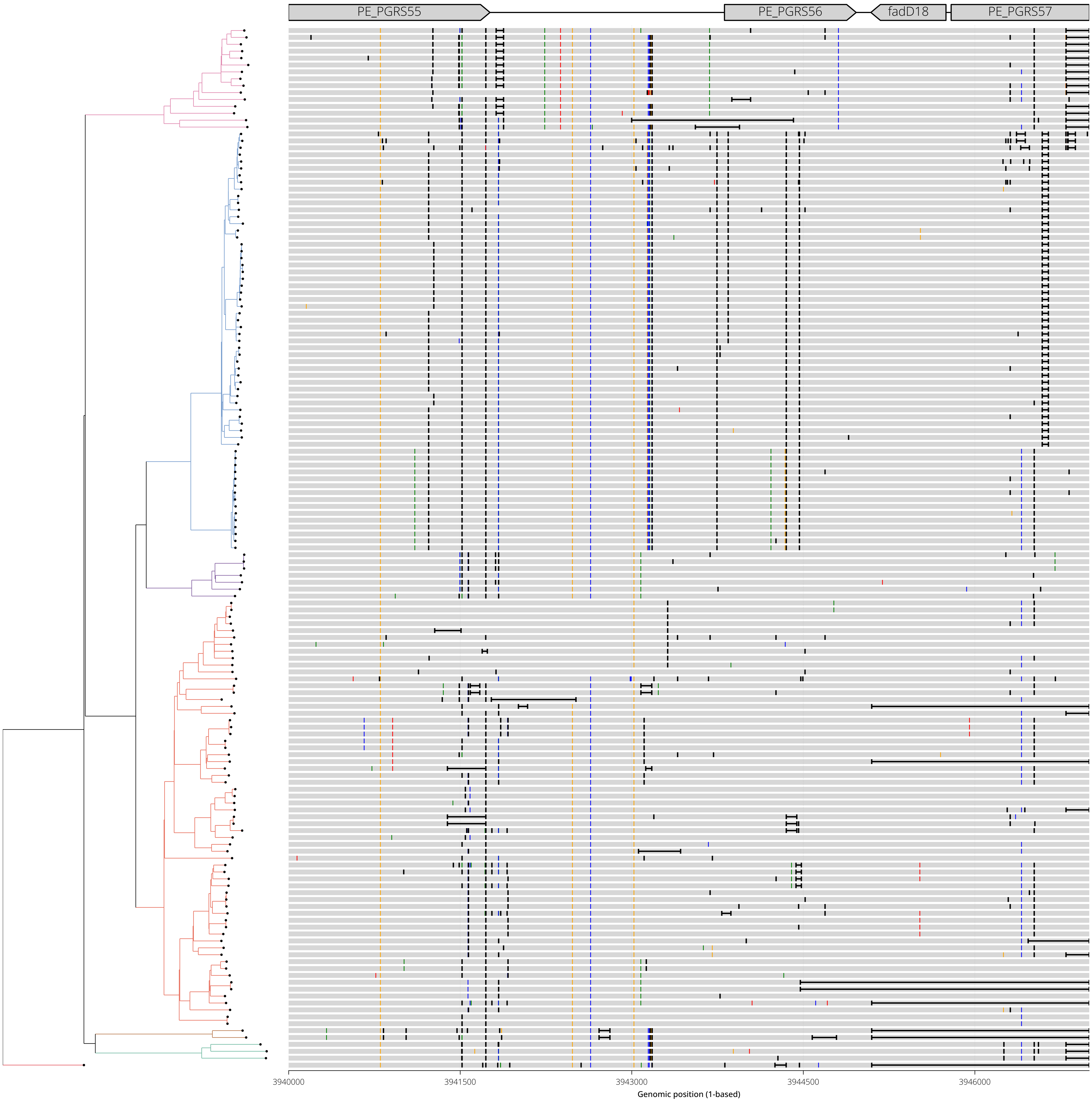

**Diversity Hotspot View - 26:**
**Genomic range shown: NC_000962.3:3944000-3951000**
**Gene(s) of interest: PE_PGRS57**

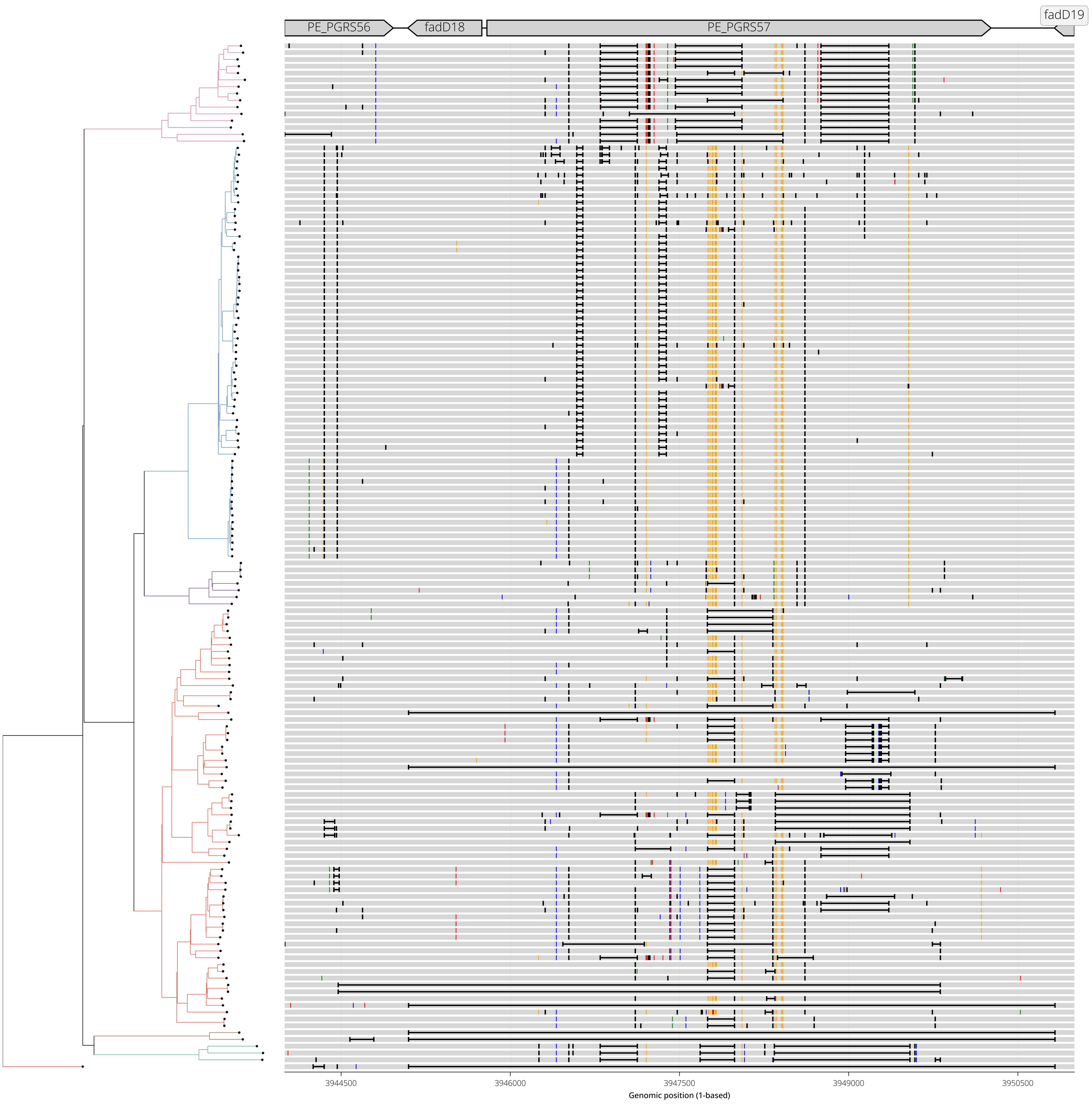

**Diversity Hotspot View - 27:**
**Genomic range shown: NC_000962.3:4252000-4257000**
**Gene(s) of interest: Rv3798,accD4**

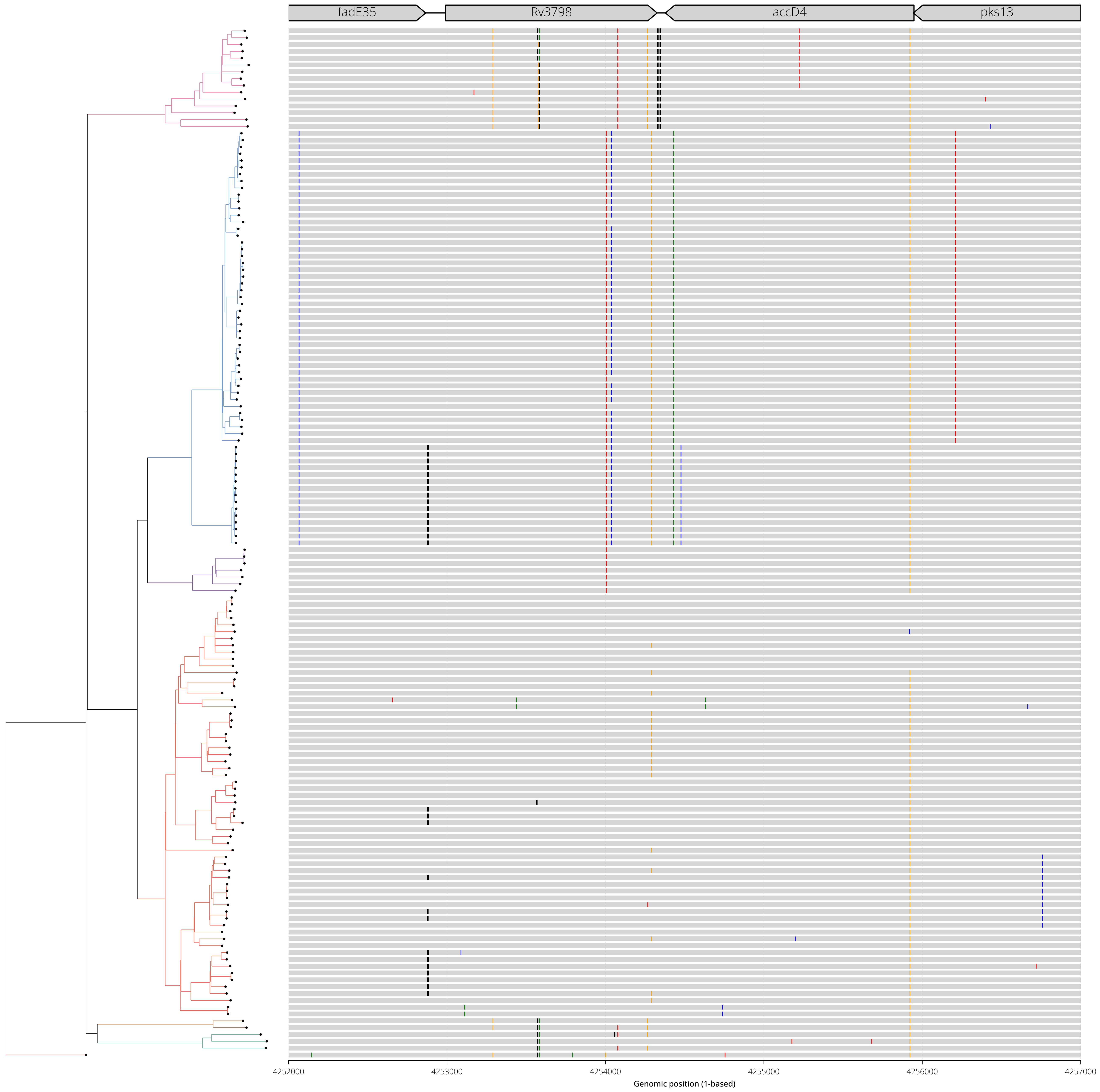
