## Supplemental Data File 12 for "Gene conversion is a key driver of diversity hotspots in *M. tuberculosis* antigens and virulence-associated loci"

### SI-12 — Per-paralogous-region visualization of gene conversion events in genomic and phylogenetic context

#### Overview

This file contains visualizations for each paralogous region identified to contain putative gene conversion events in our primary analysis. The purpose is to transparently show all detected events and where their associated mutations occurred both along the genome and across the phylogeny. All gene annotations and coordinates shown are relative to the H37Rv reference [NC_000962.3].

For each paralogous region, two complementary visualizations are generated:
View A (Genomic-context view) and View B (Phylogenetic-context view) All supporting raw data for these visualizations can be found in Supplemental Data files 5 and 6.

#### View A — Genomic-context view

A genome-browser-style view of the target region. From top to bottom:

- Paralogous region alignments: the span of each paralogous region aligned to the target region, with one alignment per row. Mismatches, insertions, and deletions between the paralog and the target region are shown using the coloring scheme defined in the "**Detected variants**" key below.
- Gene annotations: H37Rv annotated genes for the target region.
- Gene conversion events: each putative gene conversion event is drawn as a range with a red highlight. The substitutions associated with each event are drawn as vertical lines within the event box, colored by the mutant allele following the key below.

##### Figure key — View A (genomic context)

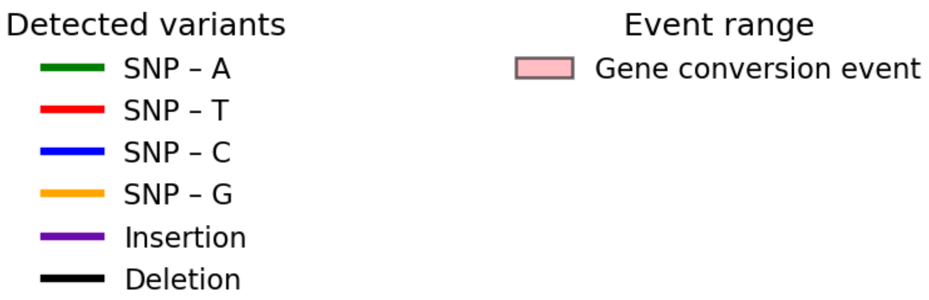

#### View B — Phylogenetic-context view

The target region is shown with detected variants per isolate placed in a phylogenetic context.

From top to bottom:

- Gene annotations: H37Rv annotated genes for the target region.
- Per-genome variant calls: the alignment of each genome in the dataset to the reference, with variants colored by the mutant allele following the key below.
- Dataset phylogeny: the phylogeny is displayed to the left, with each tip aligned horizontally to that genome's variant row.
- Gene conversion events: each event is drawn as a box spanning the genomic region it covers, overlaid on the genome alignments and aligned to the phylogeny branch on which it is inferred to have occurred. Each box is labeled with its unique event ID.

Together these tracks communicate for a given region which variants exist, where they occur along the genome, and where they fall in the phylogeny.

**Figure key — View B (phylogenetic context)**

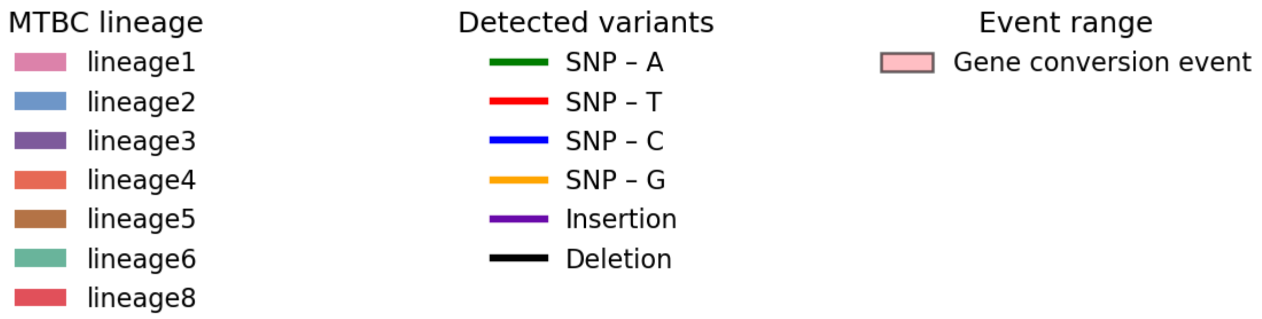

RegionID: PR_HmRegion_002 | Paralog Network ID: PR_Set_3
Genes: Rv0094c,Rv0095c | NC_000962.3:102905-105930
Mapped GCEs: 19 | Putative GCEs: 31

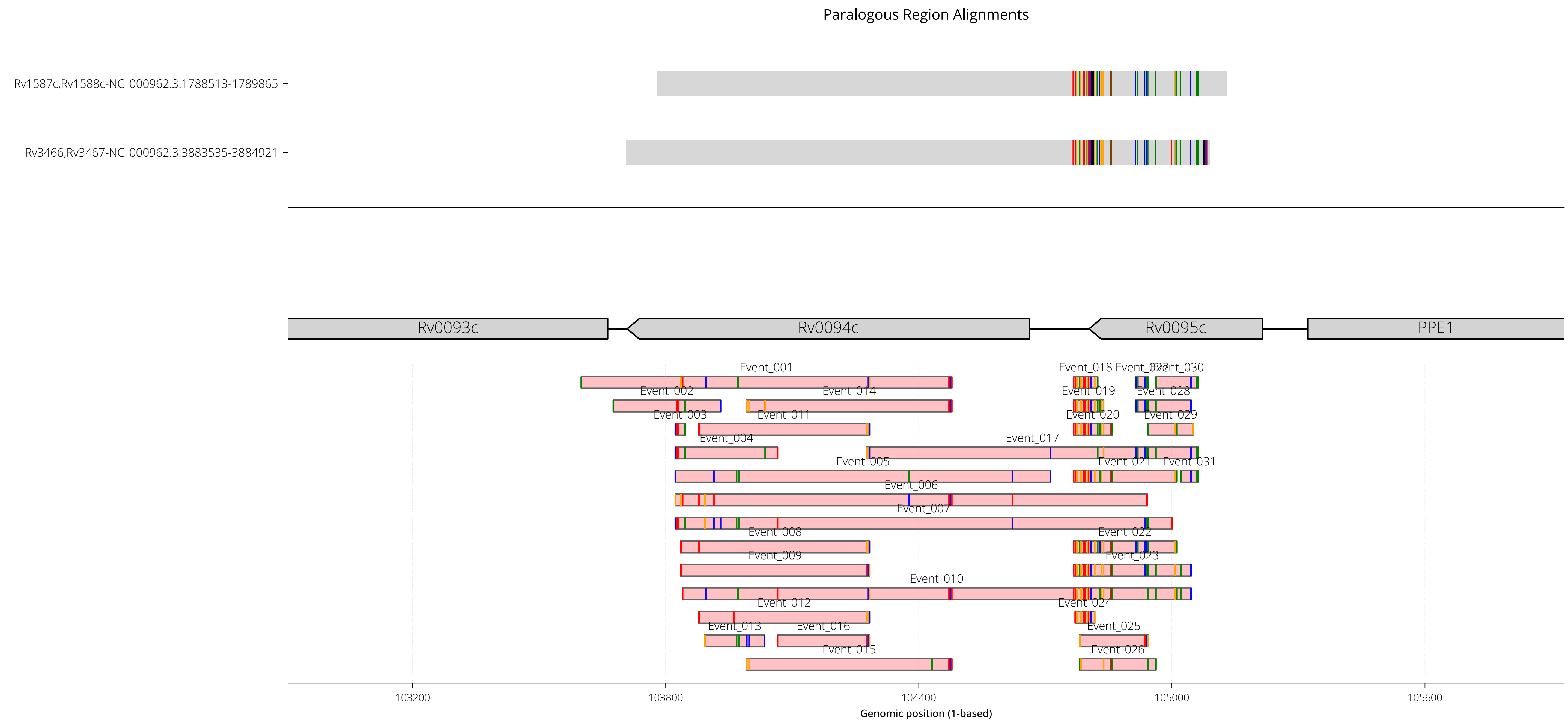

RegionID: PR_HmRegion_002 | Paralog Network ID: PR_Set_3
Genes: Rv0094c,Rv0095c | NC_000962.3:102905-105930
Mapped GCEs: 19 | Putative GCEs: 31

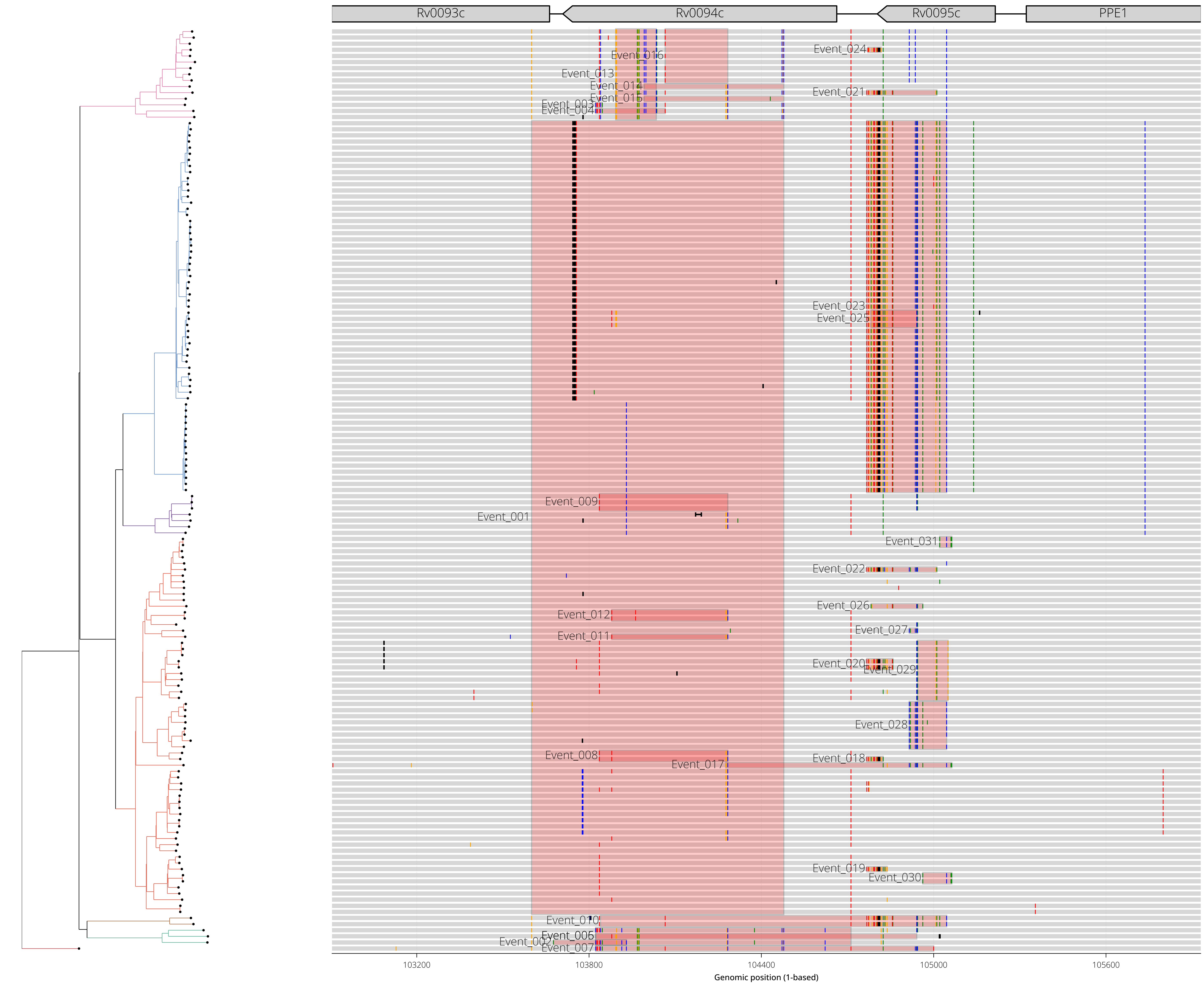

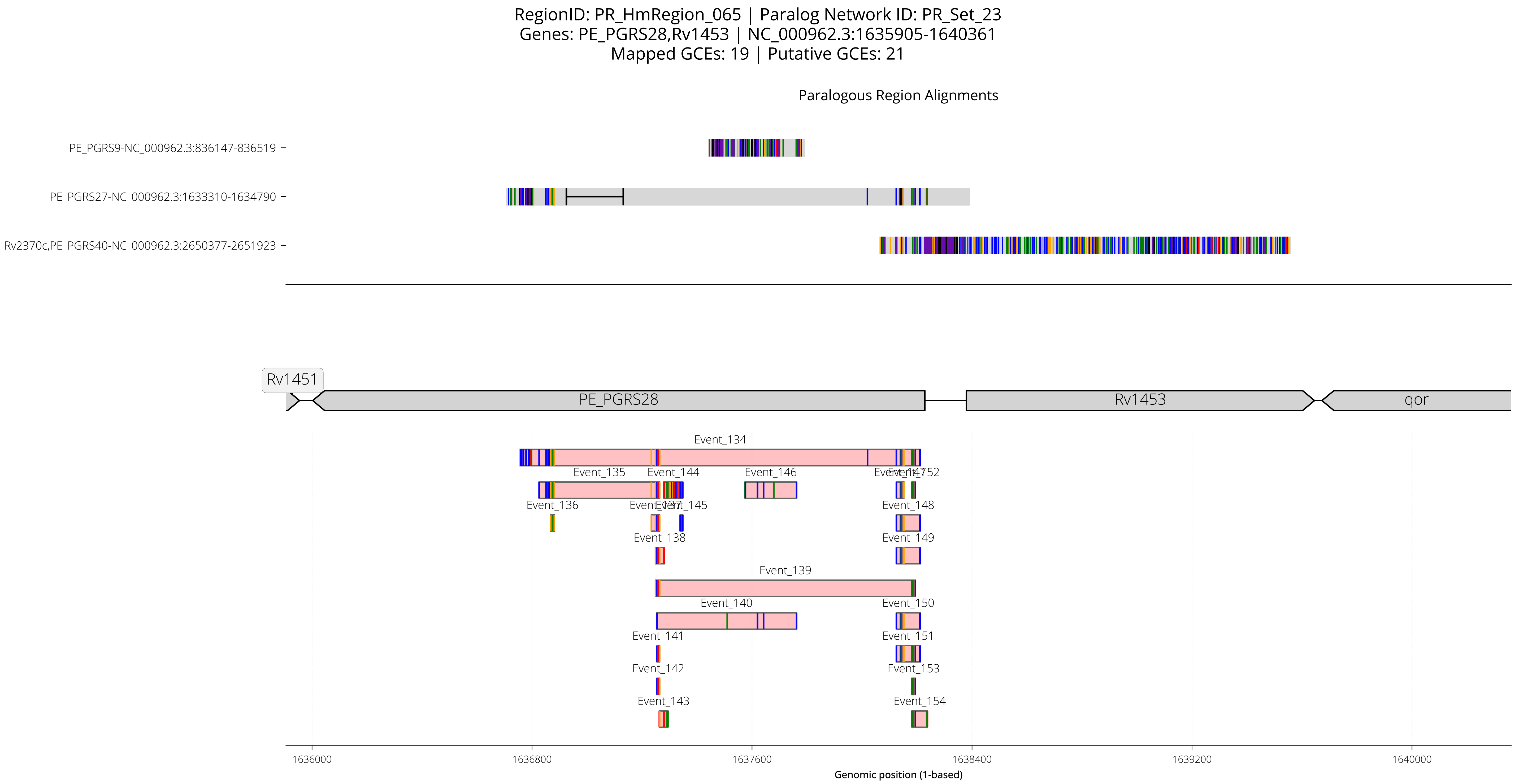

RegionID: PR_HmRegion_065 | Paralog Network ID: PR_Set_23
Genes: PE_PGRS28,Rv1453 | NC_000962.3:1635905-1640361
Mapped GCEs: 19 | Putative GCEs: 21
Paralogous Region Alignments
PE_PGRS9-NC_000962.3:836147-836519
PE_PGRS27-NC_000962.3:1633310-1634790
Rv2370c,PE_PGRS40-NC_000962.3:2650377-2651923
Rv1451
PE_PGRS28
Rv1453
qor
Event_134
Event_135
Event_144
Event_146
Event_147
Event_152
Event_136
Event_137
Event_145
Event_148
Event_138
Event_149
Event_139
Event_140
Event_150
Event_141
Event_151
Event_142
Event_153
Event_143
Event_154
1636000
1636800
1637600
1638400
1639200
1640000
Genomic position (1-based)

RegionID: PR_HmRegion_065 | Paralog Network ID: PR_Set_23
Genes: PE_PGRS28,Rv1453 | NC_000962.3:1635905-1640361
Mapped GCEs: 19 | Putative GCEs: 21

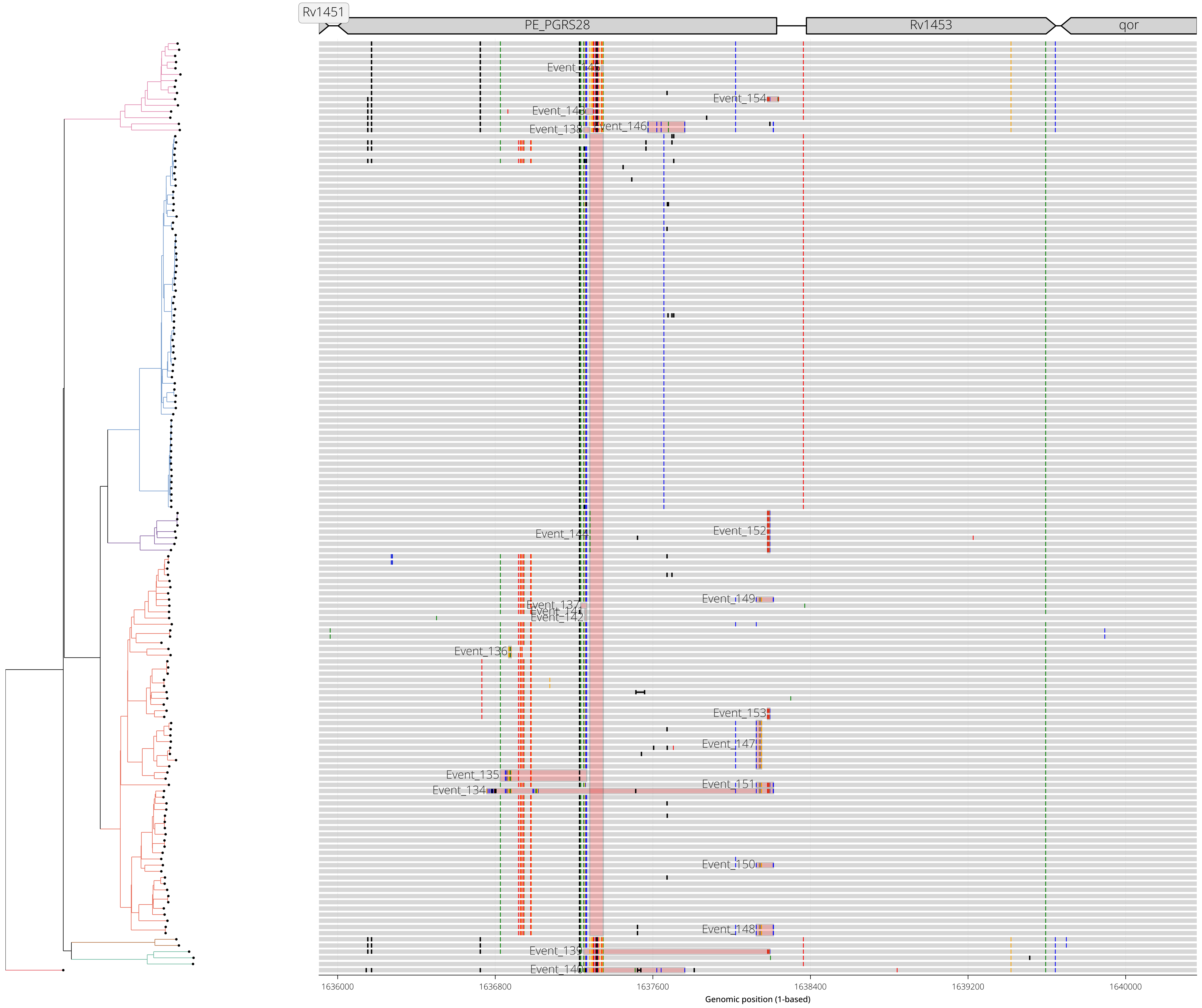

RegionID: PR_HmRegion_064 | Paralog Network ID: PR_Set_23
Genes: PE_PGRS27 | NC_000962.3:1632510-1635590
Mapped GCEs: 14 | Putative GCEs: 19

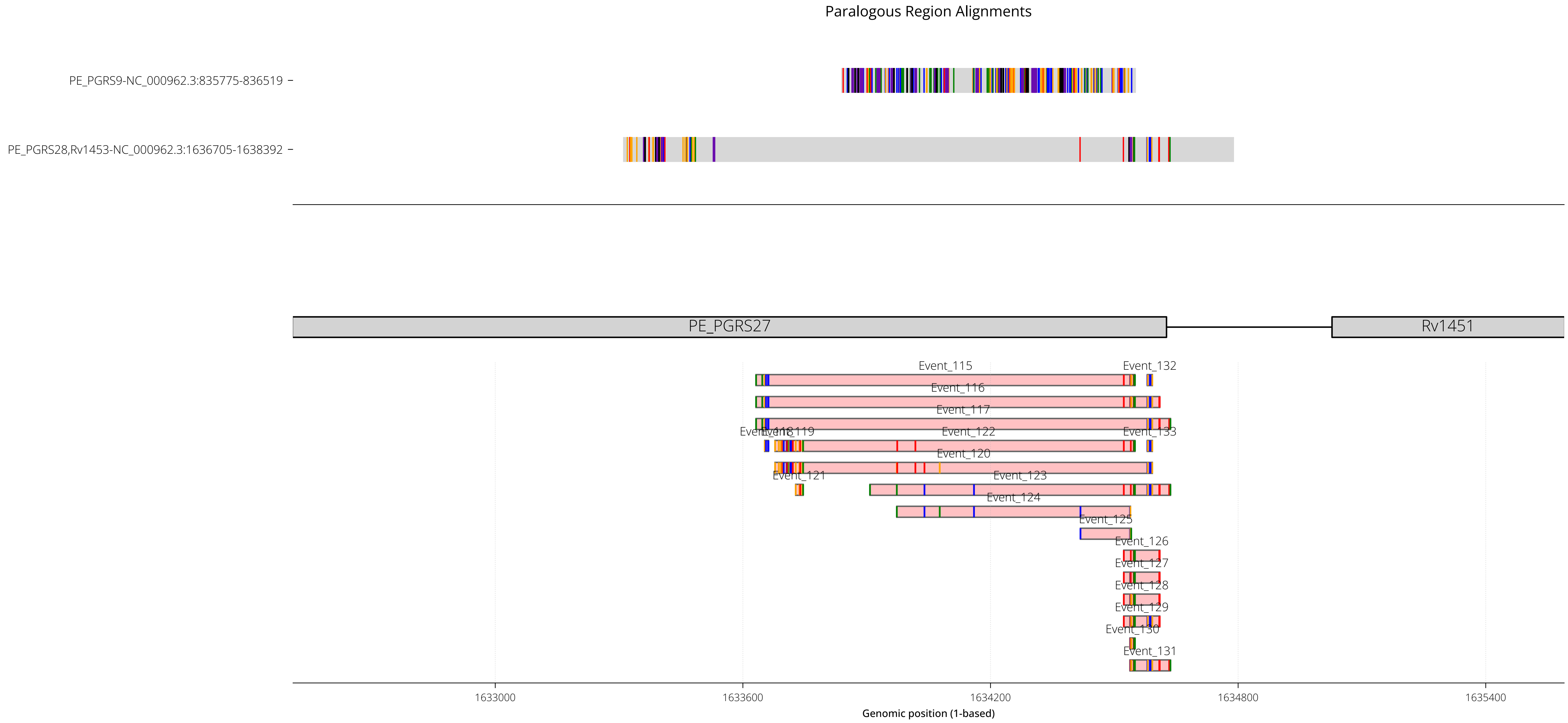

RegionID: PR_HmRegion_064 | Paralog Network ID: PR_Set_23
Genes: PE_PGRS27 | NC_000962.3:1632510-1635590
Mapped GCEs: 14 | Putative GCEs: 19

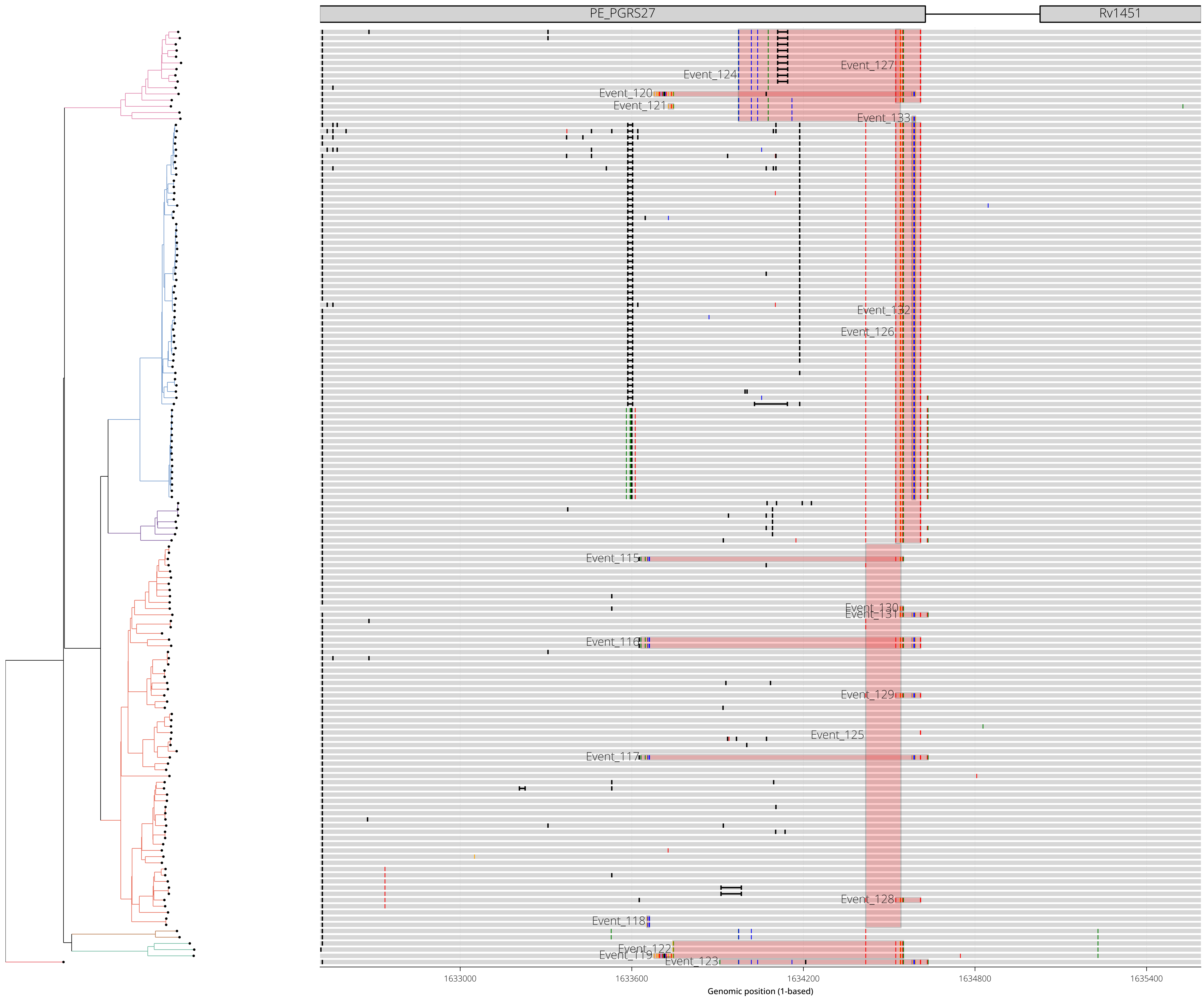

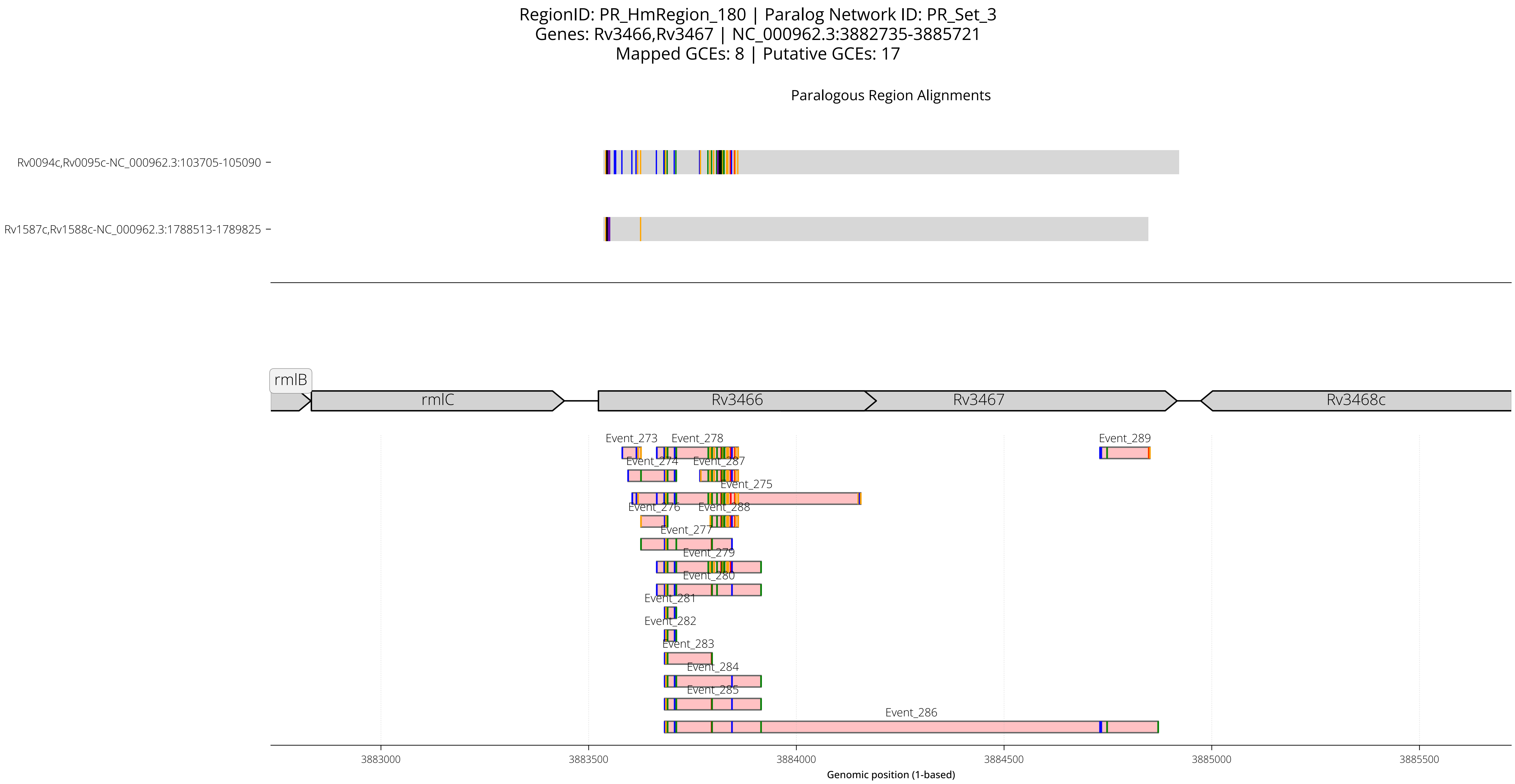

RegionID: PR_HmRegion_180 | Paralog Network ID: PR_Set_3
Genes: Rv3466,Rv3467 | NC_000962.3:3882735-3885721
Mapped GCEs: 8 | Putative GCEs: 17
Paralogous Region Alignments
Rv0094c,Rv0095c-NC_000962.3:103705-105090
Rv1587c,Rv1588c-NC_000962.3:1788513-1789825
rmlB
rmlC
Rv3466
Rv3467
Rv3468c
Event_273
Event_278
Event_289
Event_274
Event_287
Event_275
Event_276
Event_288
Event_277
Event_279
Event_280
Event_281
Event_282
Event_283
Event_284
Event_285
Event_286
3883000
3883500
3884000
3884500
3885000
3885500
Genomic position (1-based)

RegionID: PR_HmRegion_180 | Paralog Network ID: PR_Set_3
Genes: Rv3466,Rv3467 | NC_000962.3:3882735-3885721
Mapped GCEs: 8 | Putative GCEs: 17

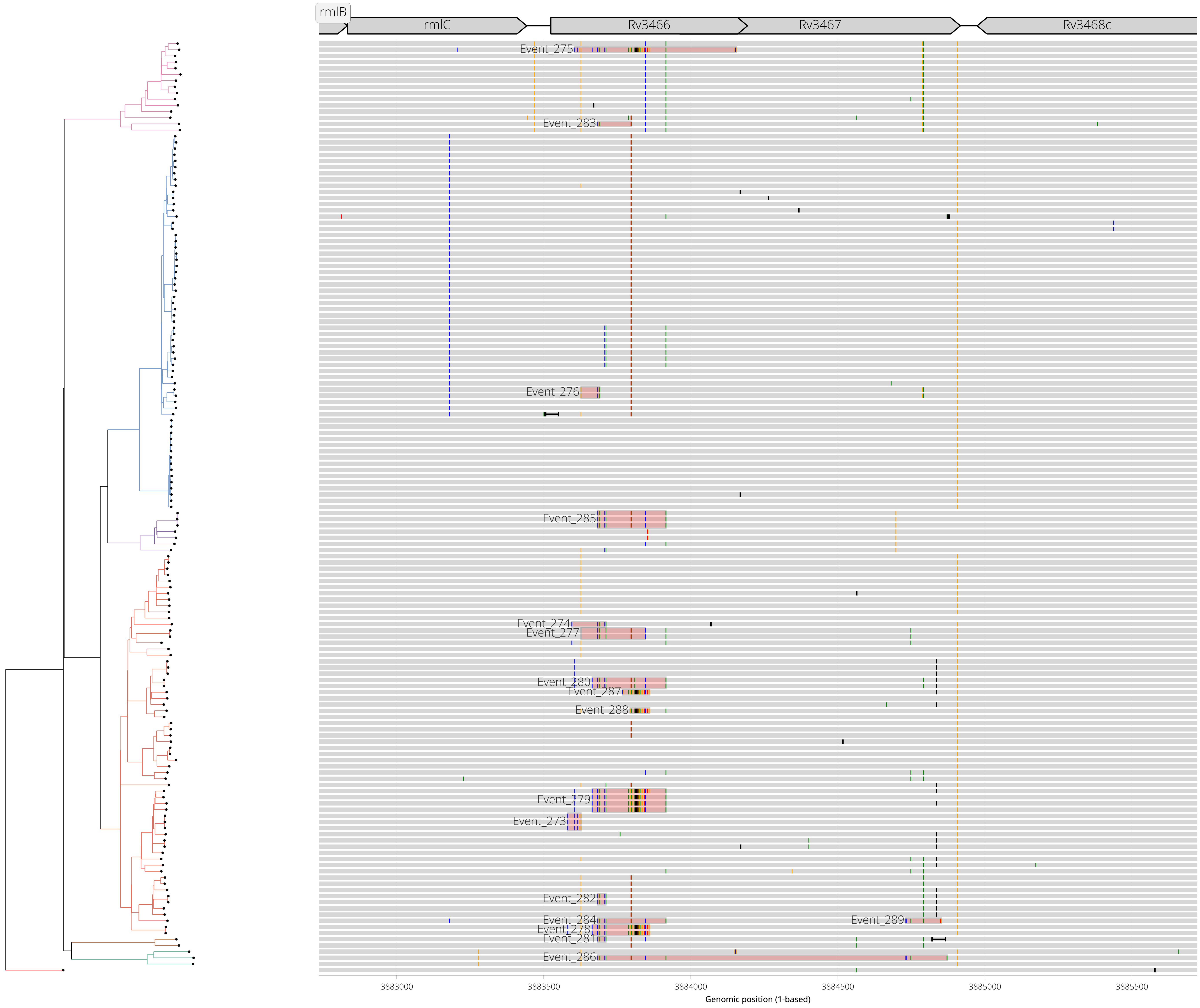

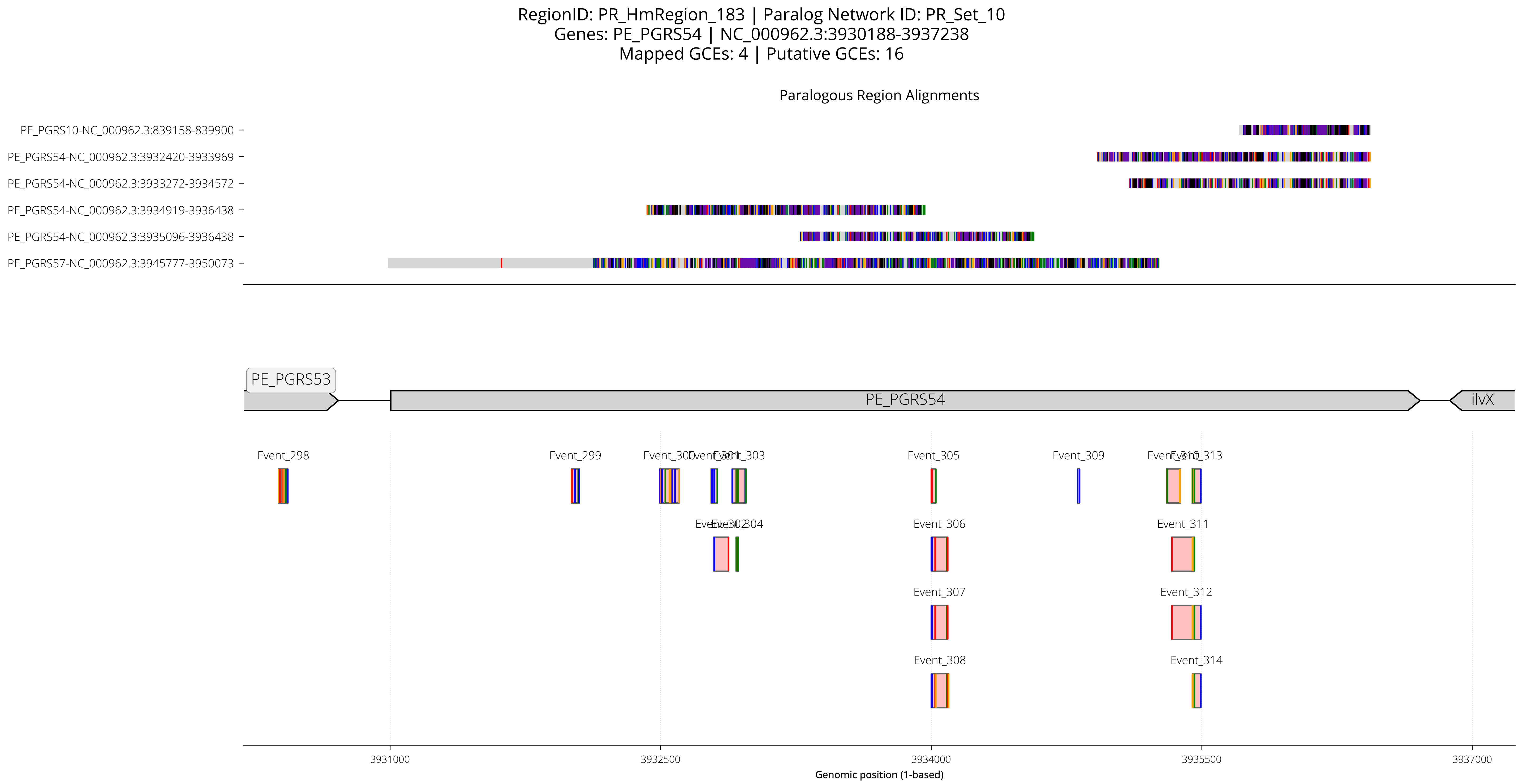
RegionID: PR_HmRegion_183 | Paralog Network ID: PR_Set_10
Genes: PE_PGRS54 | NC_000962.3:3930188-3937238
Mapped GCEs: 4 | Putative GCEs: 16
Paralogous Region Alignments
PE_PGRS10-NC_000962.3:839158-839900
PE_PGRS54-NC_000962.3:3932420-3933969
PE_PGRS54-NC_000962.3:3933272-3934572
PE_PGRS54-NC_000962.3:3934919-3936438
PE_PGRS54-NC_000962.3:3935096-3936438
PE_PGRS57-NC_000962.3:3945777-3950073
PE_PGRS53
PE_PGRS54
ilvX
Event_298
Event_299
Event_305
Event_309
Event_306
Event_311
Event_307
Event_312
Event_308
Event_314
3931000
3932500
3934000
3935500
3937000
Genomic position (1-based)

RegionID: PR_HmRegion_183 | Paralog Network ID: PR_Set_10
Genes: PE_PGRS54 | NC_000962.3:3930188-3937238
Mapped GCEs: 4 | Putative GCEs: 16

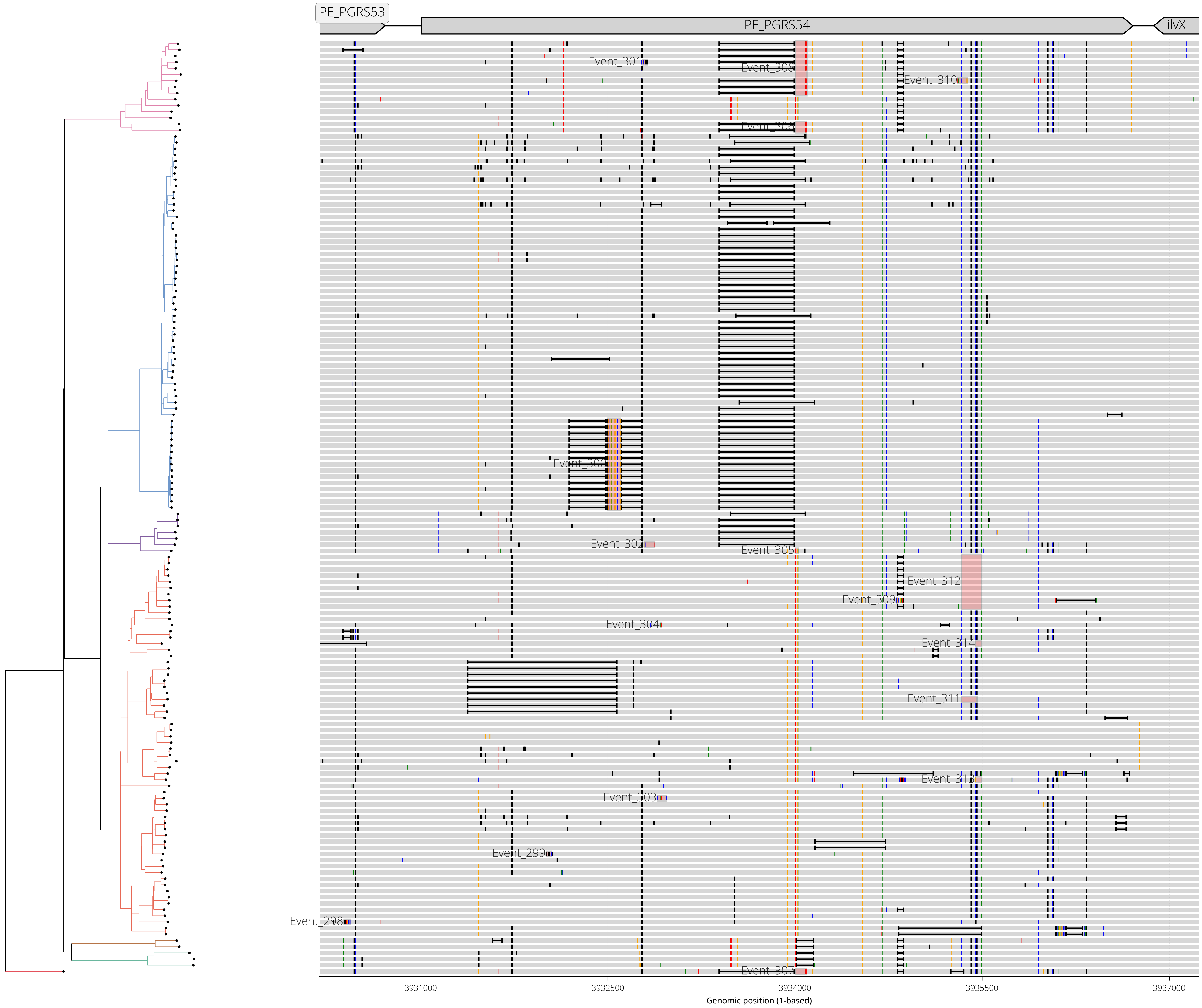

RegionID: PR_HmRegion_094 | Paralog Network ID: PR_Set_34
Genes: Rv1944c,Rv1945 | NC_000962.3:2195056-2198160
Mapped GCEs: 11 | Putative GCEs: 11

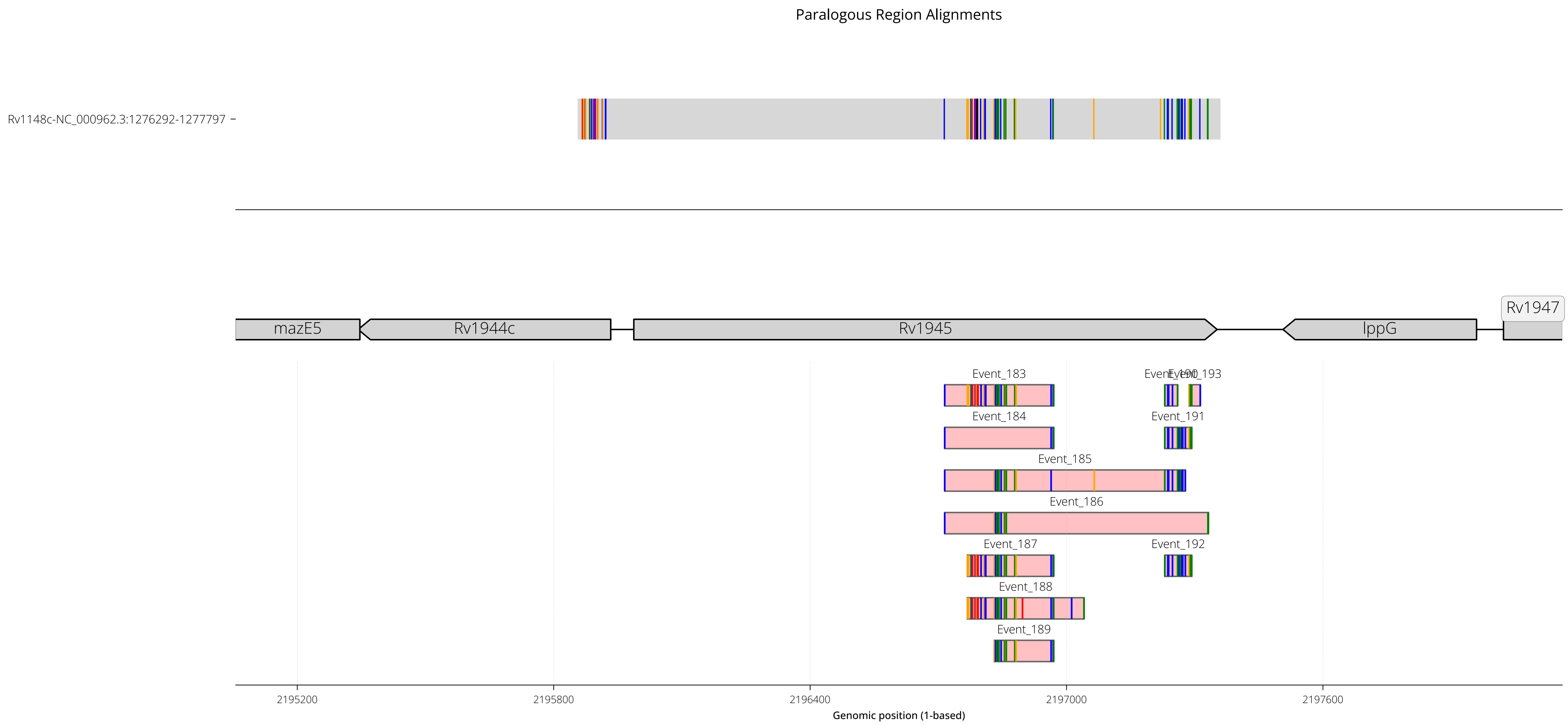

RegionID: PR_HmRegion_094 | Paralog Network ID: PR_Set_34
Genes: Rv1944c,Rv1945 | NC_000962.3:2195056-2198160
Mapped GCEs: 11 | Putative GCEs: 11

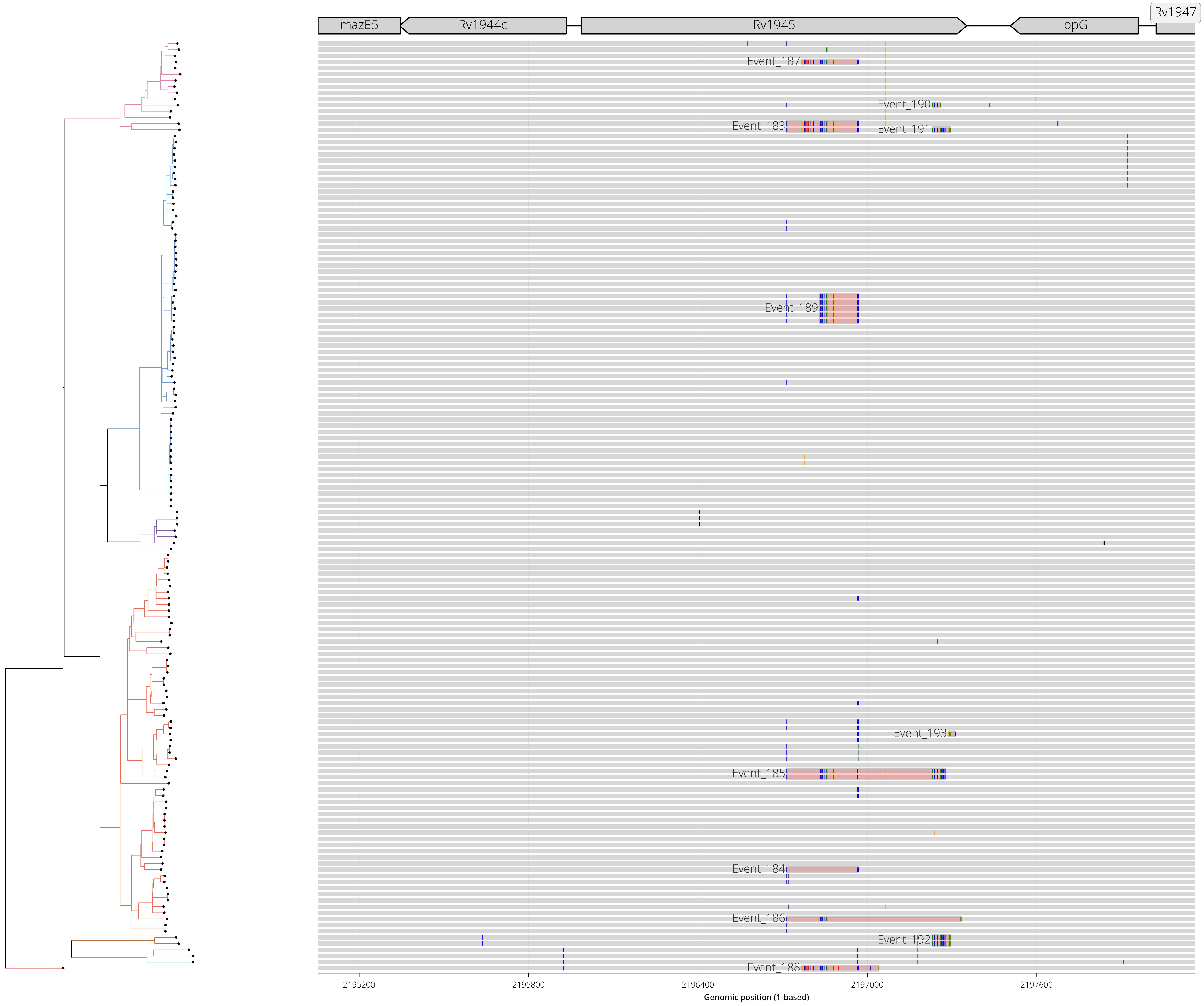

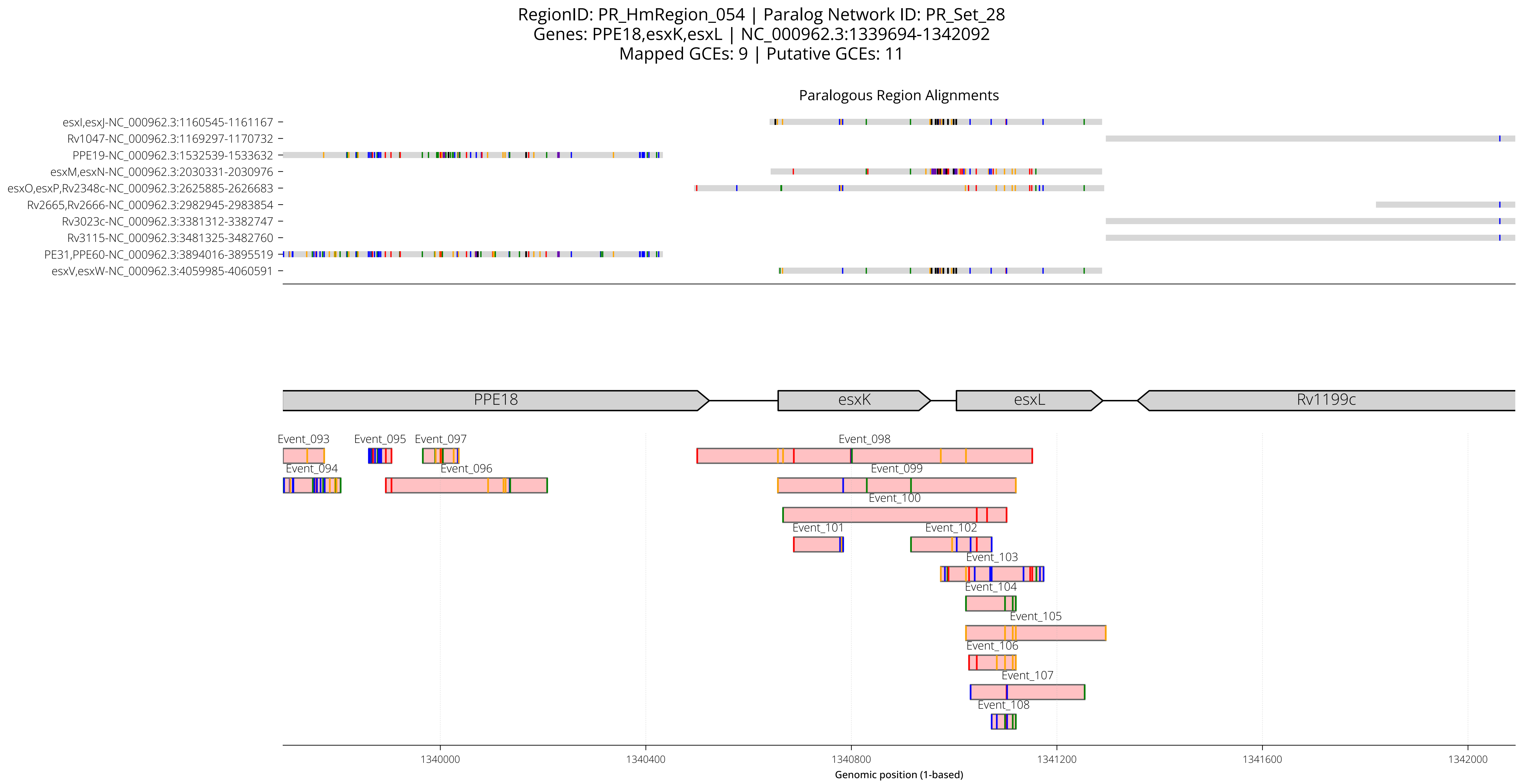

RegionID: PR_HmRegion_054 | Paralog Network ID: PR_Set_28
Genes: PPE18,esxK,esxL | NC_000962.3:1339694-1342092
Mapped GCEs: 9 | Putative GCEs: 11
Paralogous Region Alignments
esxI,esxJ-NC_000962.3:1160545-1161167
Rv1047-NC_000962.3:1169297-1170732
PPE19-NC_000962.3:1532539-1533632
esxM,esxN-NC_000962.3:2030331-2030976
esxO,esxP,Rv2348c-NC_000962.3:2625885-2626683
Rv2665,Rv2666-NC_000962.3:2982945-2983854
Rv3023c-NC_000962.3:3381312-3382747
Rv3115-NC_000962.3:3481325-3482760
PE31,PPE60-NC_000962.3:3894016-3895519
esxV,esxW-NC_000962.3:4059985-4060591
PPE18
esxK
esxL
Rv1199c
Event_093
Event_094
Event_095
Event_096
Event_097
Event_098
Event_099
Event_100
Event_101
Event_102
Event_103
Event_104
Event_105
Event_106
Event_107
Event_108
1340000
1340400
1340800
1341200
1341600
1342000
Genomic position (1-based)

RegionID: PR_HmRegion_054 | Paralog Network ID: PR_Set_28
Genes: PPE18,esxK,esxL | NC_000962.3:1339694-1342092
Mapped GCEs: 9 | Putative GCEs: 11

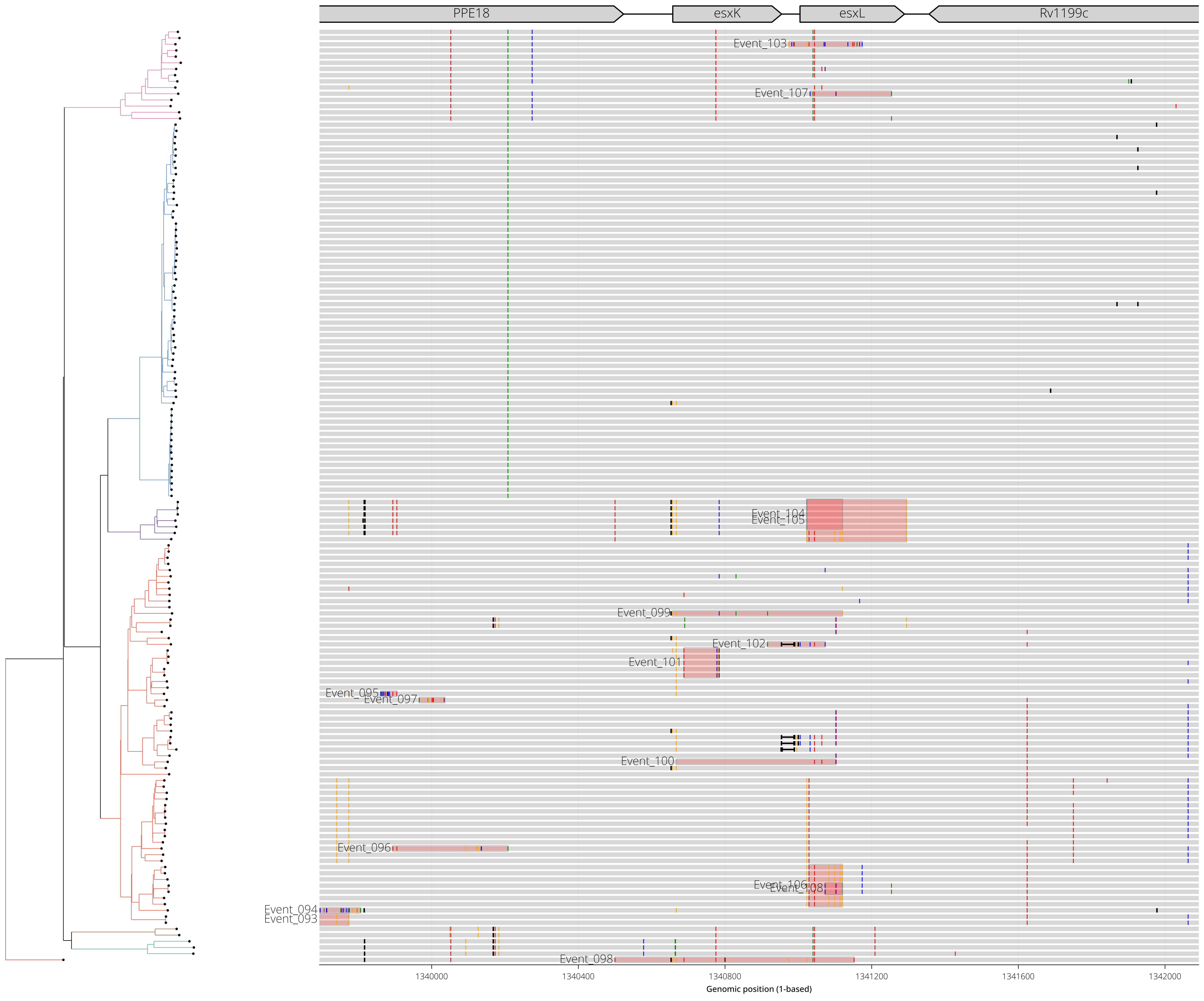

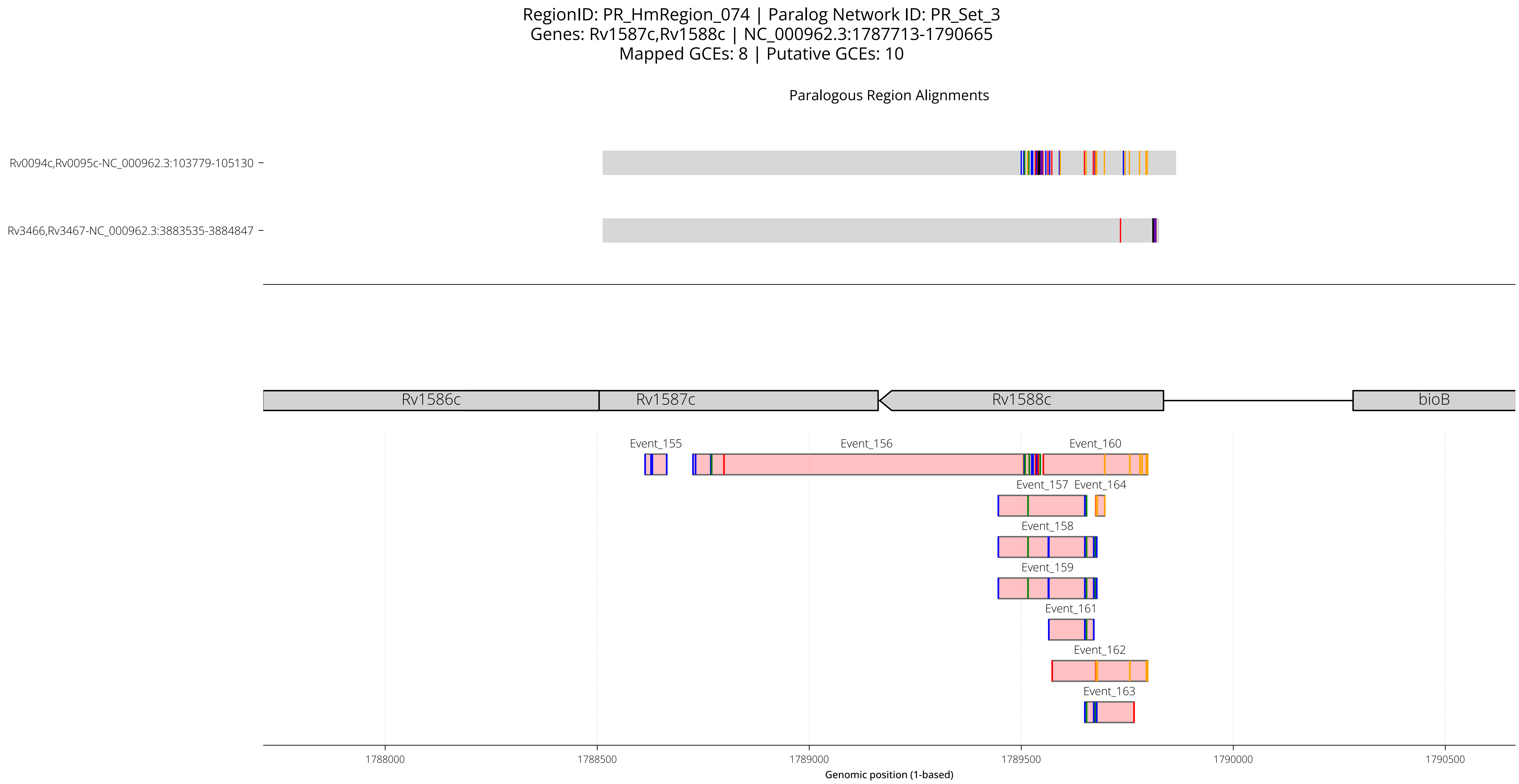
RegionID: PR_HmRegion_074 | Paralog Network ID: PR_Set_3
Genes: Rv1587c,Rv1588c | NC_000962.3:1787713-1790665
Mapped GCEs: 8 | Putative GCEs: 10
Paralogous Region Alignments
Rv0094c,Rv0095c-NC_000962.3:103779-105130
Rv3466,Rv3467-NC_000962.3:3883535-3884847
Rv1586c
Rv1587c
Rv1588c
bioB
Event_155
Event_156
Event_160
Event_157
Event_164
Event_158
Event_159
Event_161
Event_162
Event_163
1788000
1788500
1789000
1789500
1790000
1790500
Genomic position (1-based)

RegionID: PR_HmRegion_074 | Paralog Network ID: PR_Set_3
Genes: Rv1587c,Rv1588c | NC_000962.3:1787713-1790665
Mapped GCEs: 8 | Putative GCEs: 10

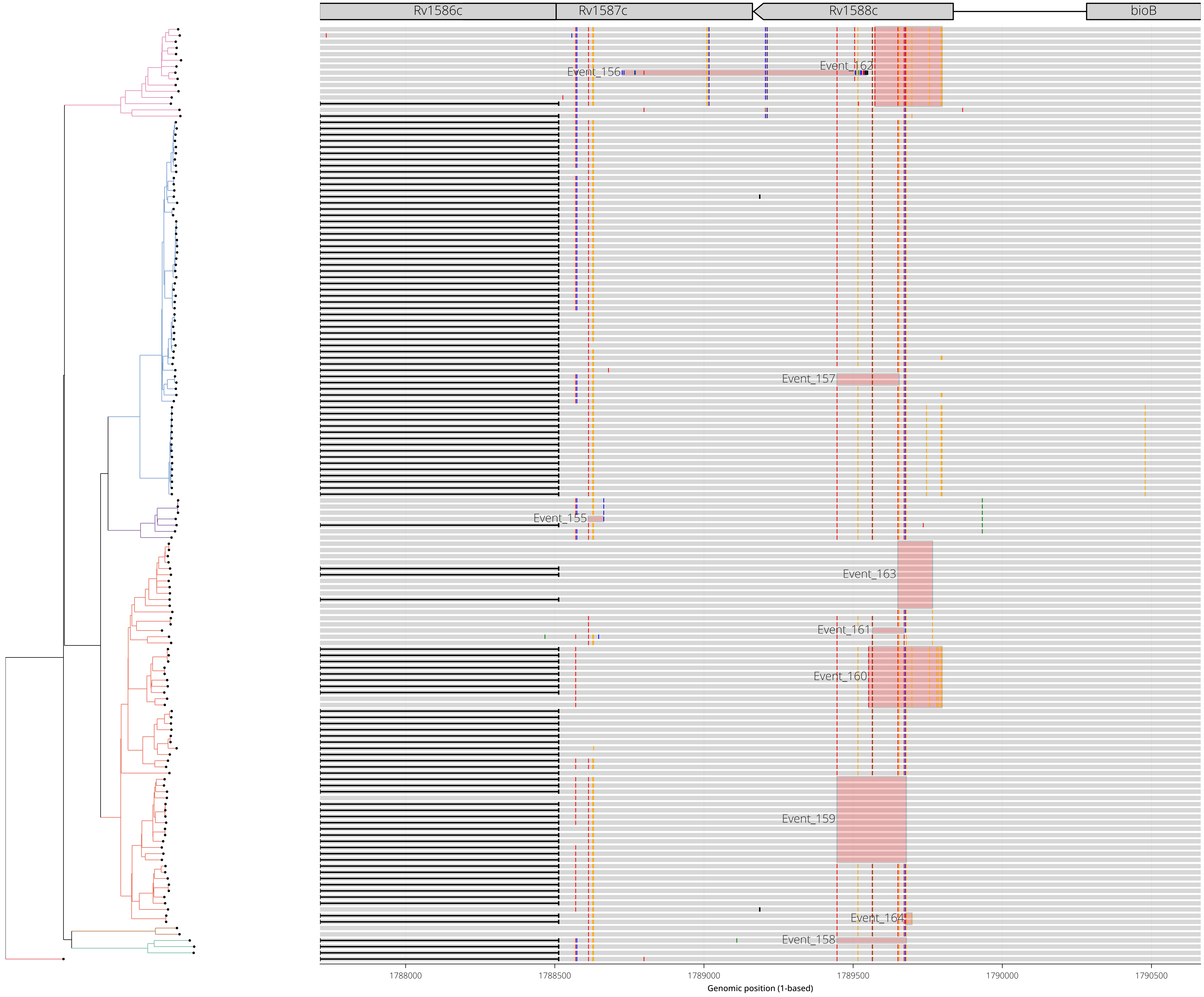

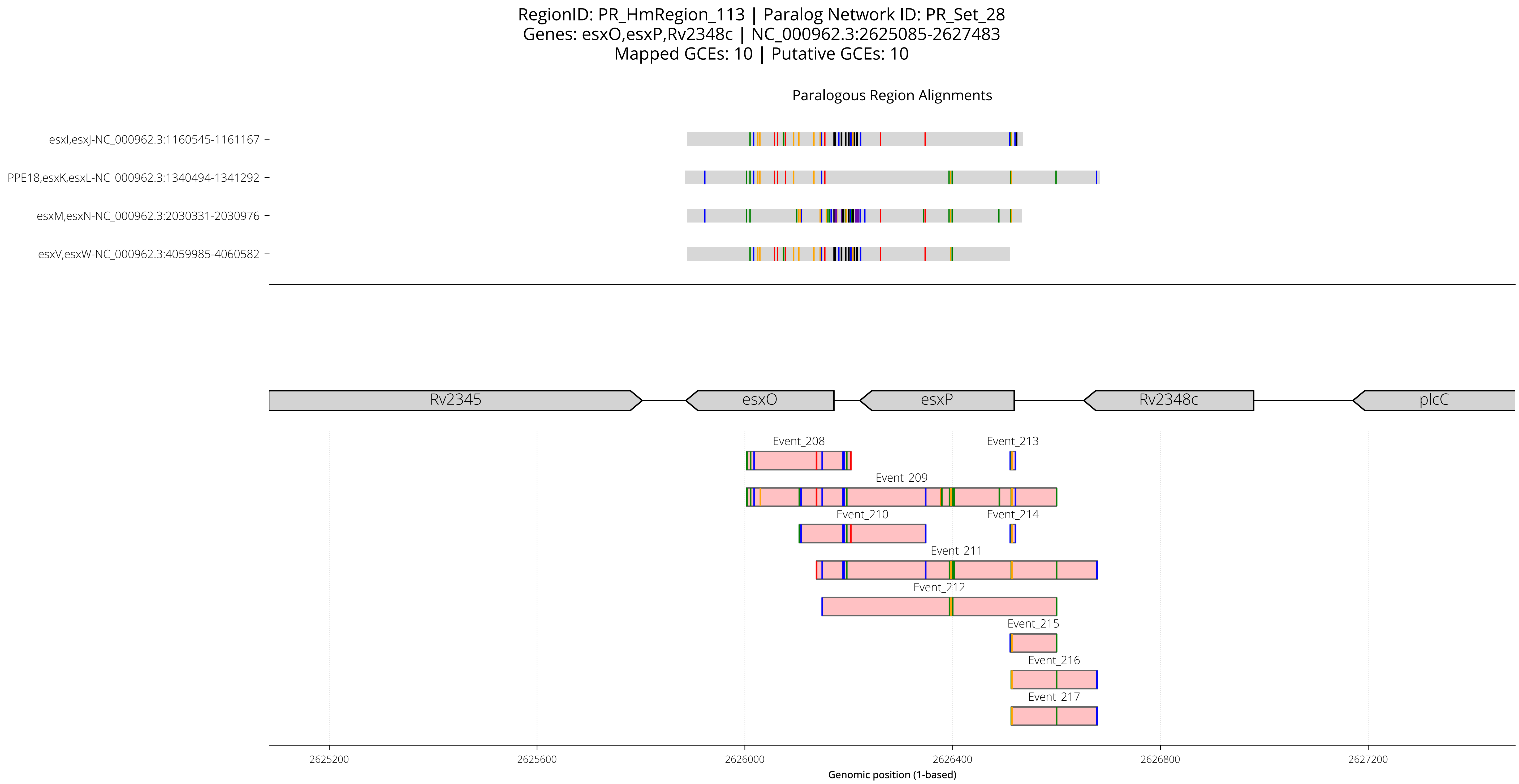
RegionID: PR_HmRegion_113 | Paralog Network ID: PR_Set_28
Genes: esxO,esxP,Rv2348c | NC_000962.3:2625085-2627483
Mapped GCEs: 10 | Putative GCEs: 10
Paralogous Region Alignments
esxI,esxJ-NC_000962.3:1160545-1161167
PPE18,esxK,esxL-NC_000962.3:1340494-1341292
esxM,esxN-NC_000962.3:2030331-2030976
esxV,esxW-NC_000962.3:4059985-4060582
Rv2345
esxO
esxP
Rv2348c
plcC
Event_208
Event_213
Event_209
Event_210
Event_214
Event_211
Event_212
Event_215
Event_216
Event_217
2625200
2625600
2626000
2626400
2626800
2627200
Genomic position (1-based)

RegionID: PR_HmRegion_113 | Paralog Network ID: PR_Set_28
Genes: esxO,esxP,Rv2348c | NC_000962.3:2625085-2627483
Mapped GCEs: 10 | Putative GCEs: 10

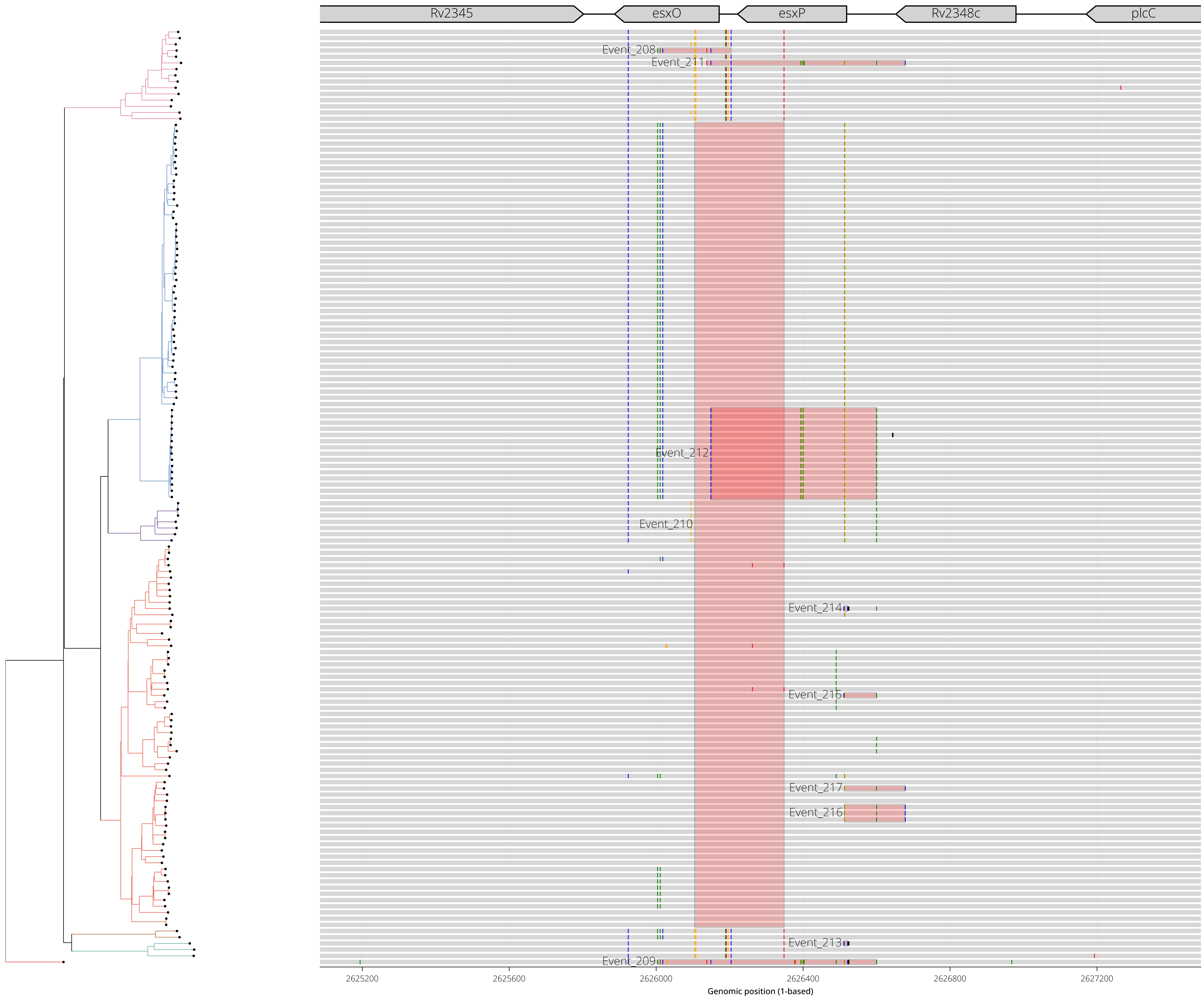

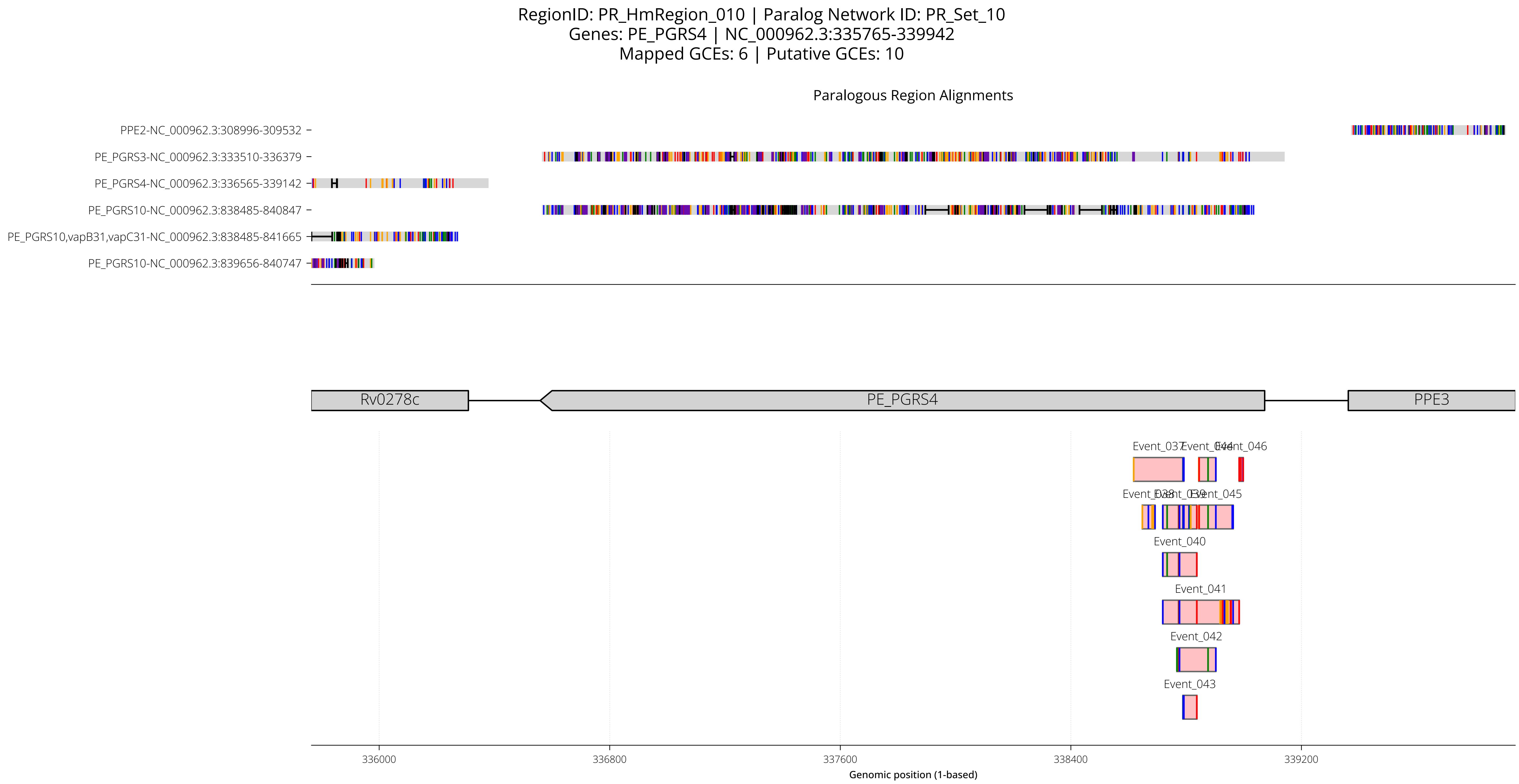

RegionID: PR_HmRegion_010 | Paralog Network ID: PR_Set_10
Genes: PE_PGRS4 | NC_000962.3:335765-339942
Mapped GCEs: 6 | Putative GCEs: 10
Paralogous Region Alignments
PPE2-NC_000962.3:308996-309532
PE_PGRS3-NC_000962.3:333510-336379
PE_PGRS4-NC_000962.3:336565-339142
PE_PGRS10-NC_000962.3:838485-840847
PE_PGRS10,vapB31,vapC31-NC_000962.3:838485-841665
PE_PGRS10-NC_000962.3:839656-840747
Rv0278c
PE_PGRS4
PPE3
Event_037
Event_046
Event_040
Event_041
Event_042
Event_043
336000
336800
337600
338400
339200
Genomic position (1-based)

RegionID: PR_HmRegion_010 | Paralog Network ID: PR_Set_10
Genes: PE_PGRS4 | NC_000962.3:335765-339942
Mapped GCEs: 6 | Putative GCEs: 10

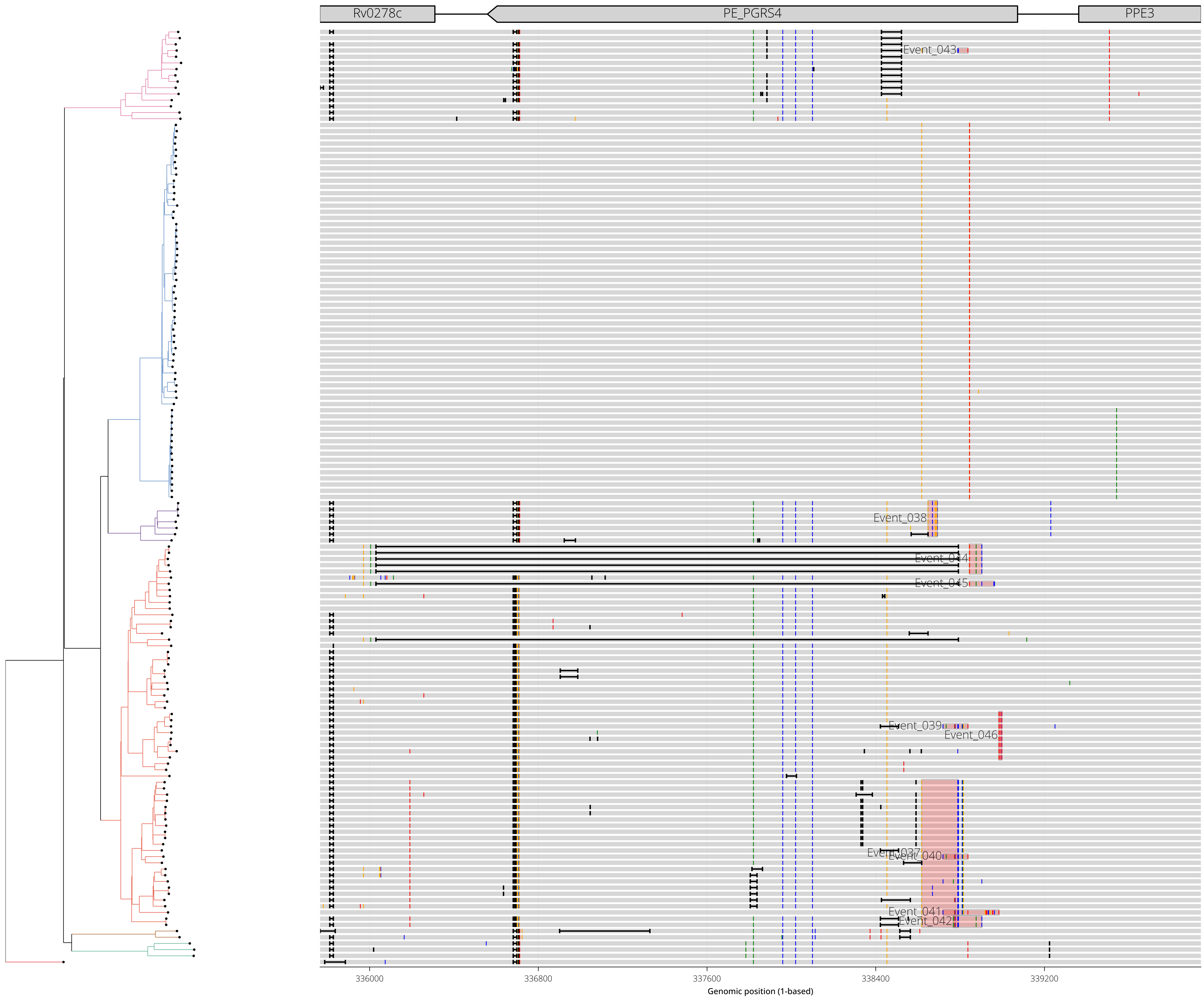

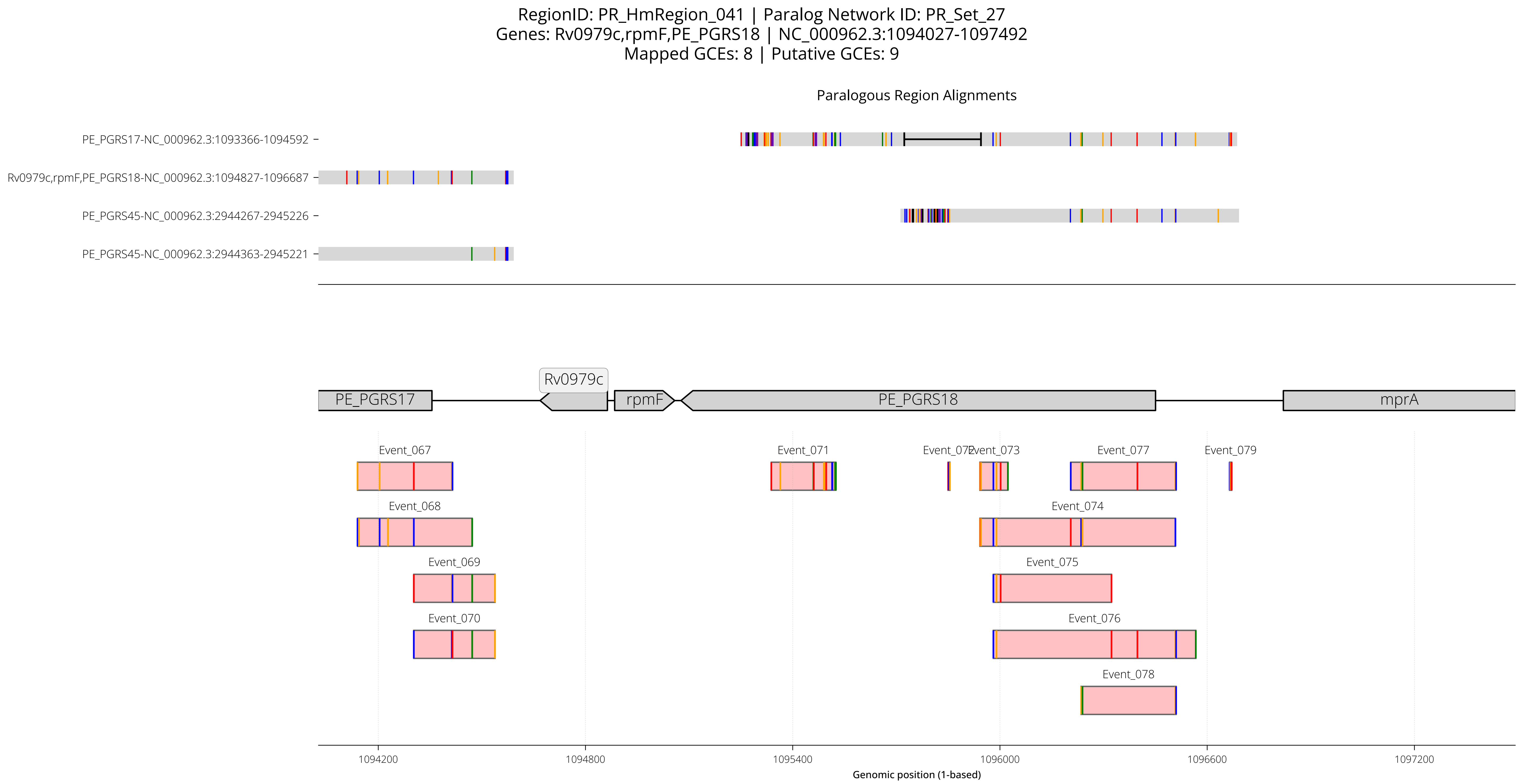

RegionID: PR_HmRegion_041 | Paralog Network ID: PR_Set_27
Genes: Rv0979c,rpmF,PE_PGRS18 | NC_000962.3:1094027-1097492
Mapped GCEs: 8 | Putative GCEs: 9
Paralogous Region Alignments
PE_PGRS17-NC_000962.3:1093366-1094592
Rv0979c,rpmF,PE_PGRS18-NC_000962.3:1094827-1096687
PE_PGRS45-NC_000962.3:2944267-2945226
PE_PGRS45-NC_000962.3:2944363-2945221
Rv0979c
PE_PGRS17
rpmF
PE_PGRS18
mprA
Event_067
Event_068
Event_069
Event_070
Event_071
Event_072
Event_073
Event_074
Event_075
Event_076
Event_077
Event_078
Event_079
1094200
1094800
1095400
1096000
1096600
1097200
Genomic position (1-based)

RegionID: PR_HmRegion_041 | Paralog Network ID: PR_Set_27
Genes: Rv0979c,rpmF,PE_PGRS18 | NC_000962.3:1094027-1097492
Mapped GCEs: 8 | Putative GCEs: 9

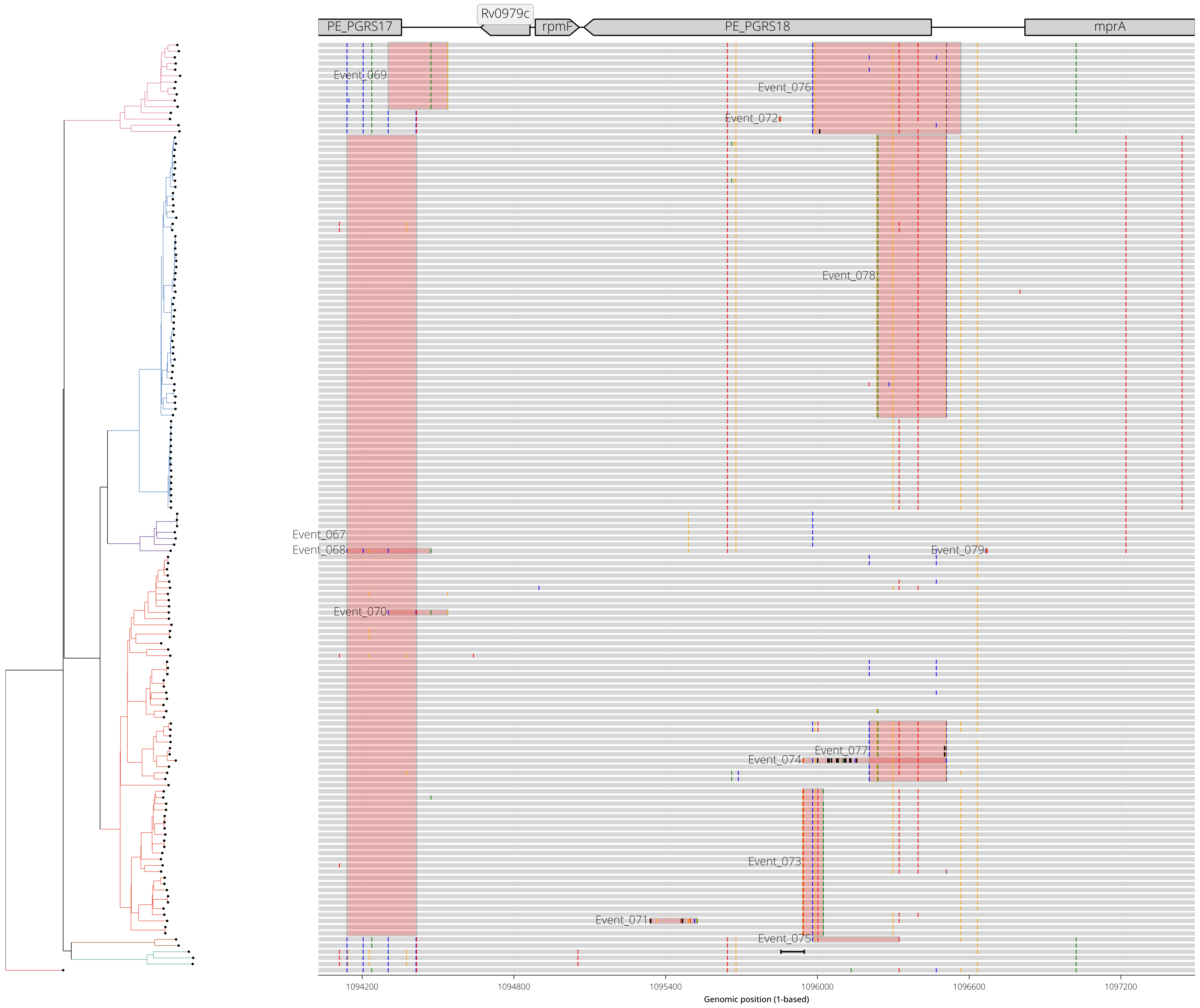

RegionID: PR_HmRegion_171 | Paralog Network ID: PR_Set_24
Genes: PPE55 | NC_000962.3:3742909-3753978
Mapped GCEs: 6 | Putative GCEs: 8

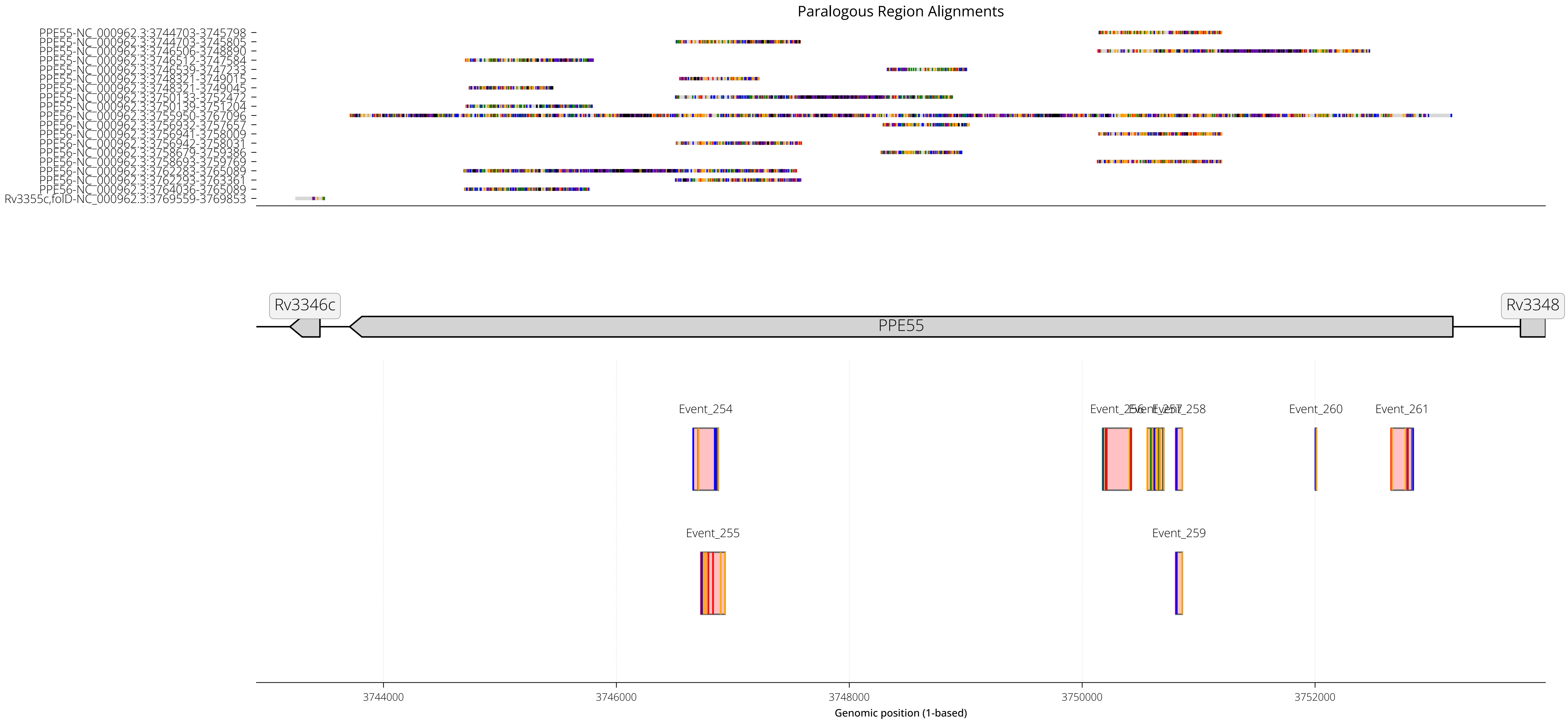

RegionID: PR_HmRegion_171 | Paralog Network ID: PR_Set_24
Genes: PPE55 | NC_000962.3:3742909-3753978
Mapped GCEs: 6 | Putative GCEs: 8

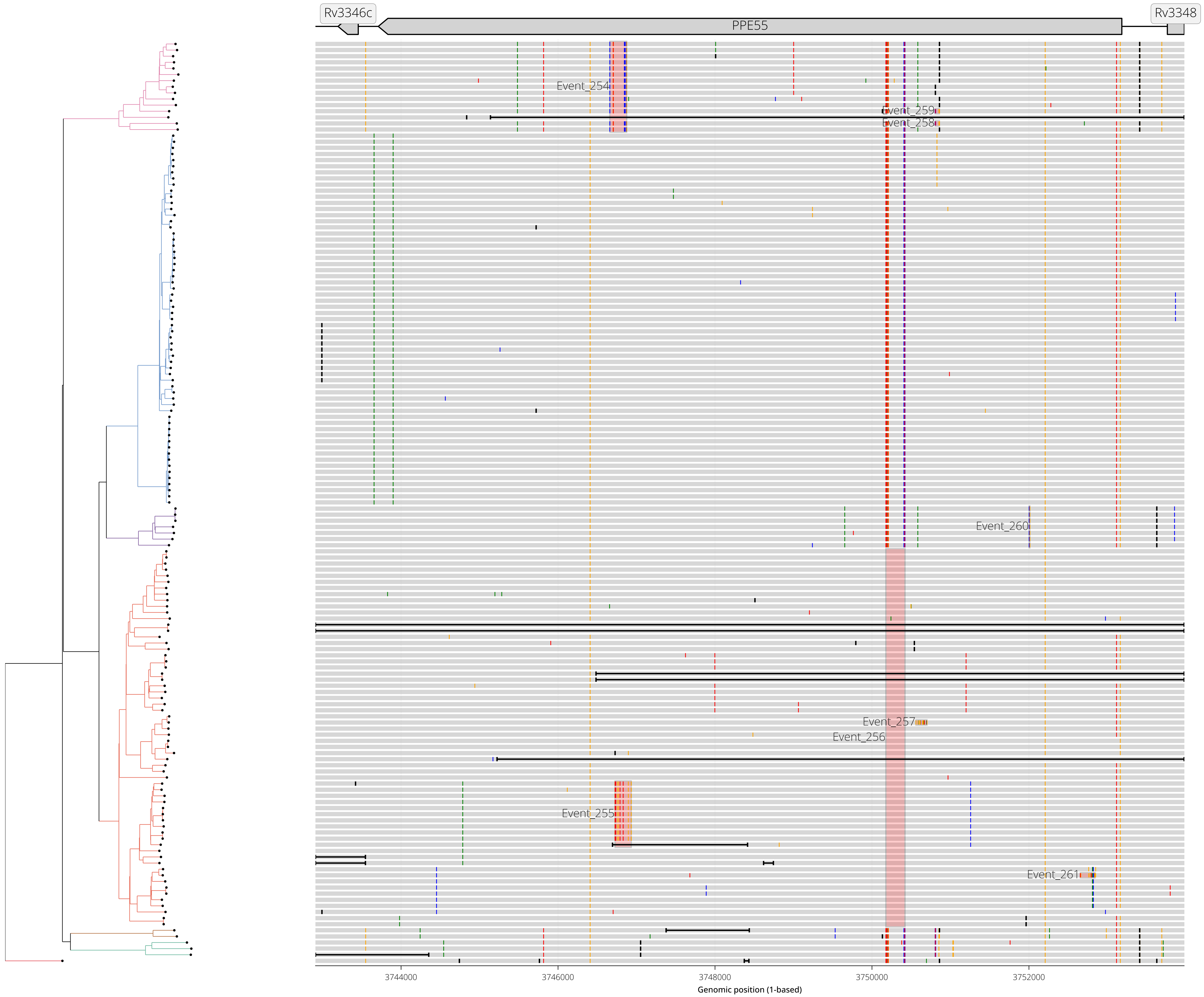

RegionID: PR_HmRegion_168 | Paralog Network ID: PR_Set_67
Genes: PPE54 | NC_000962.3:3734835-3737114
Mapped GCEs: 5 | Putative GCEs: 8

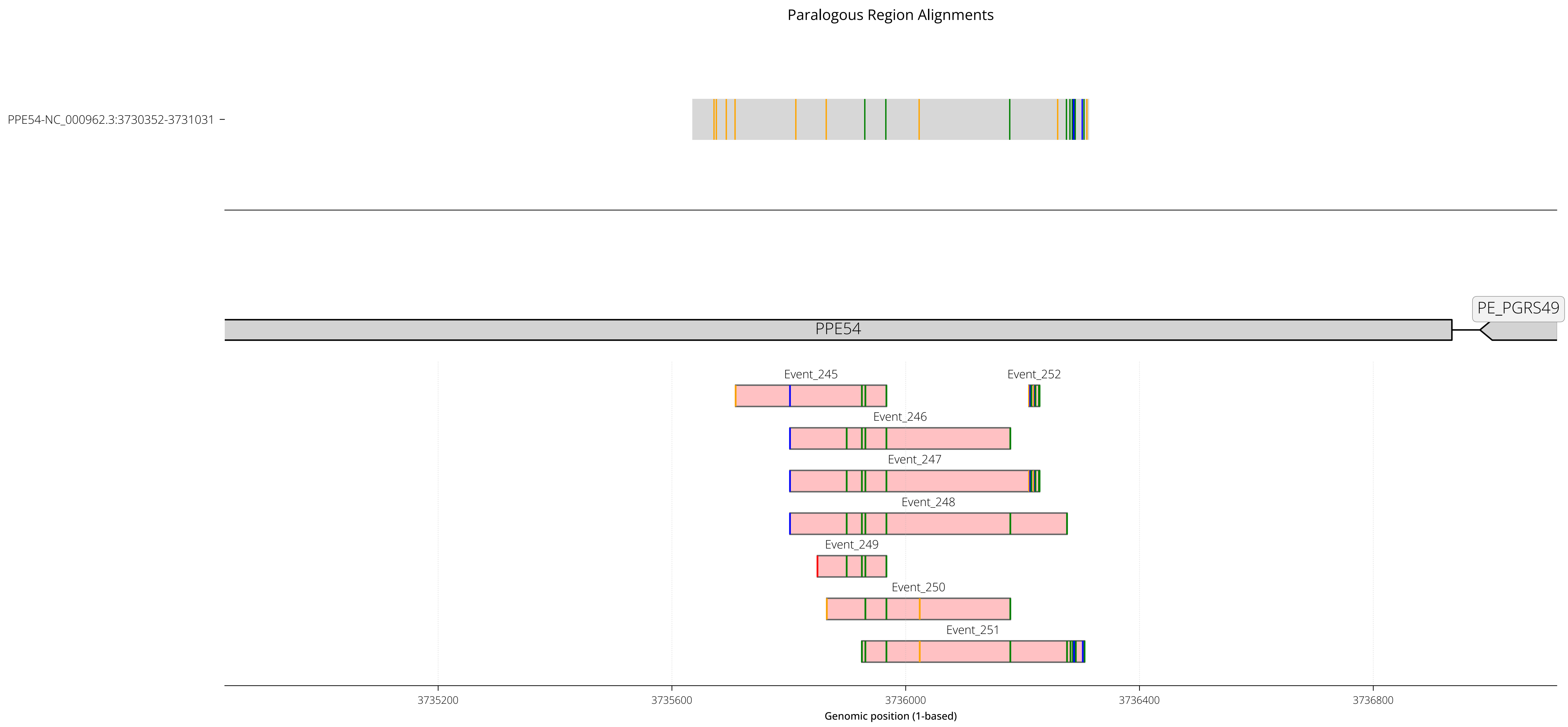

RegionID: PR_HmRegion_168 | Paralog Network ID: PR_Set_67
Genes: PPE54 | NC_000962.3:3734835-3737114
Mapped GCEs: 5 | Putative GCEs: 8

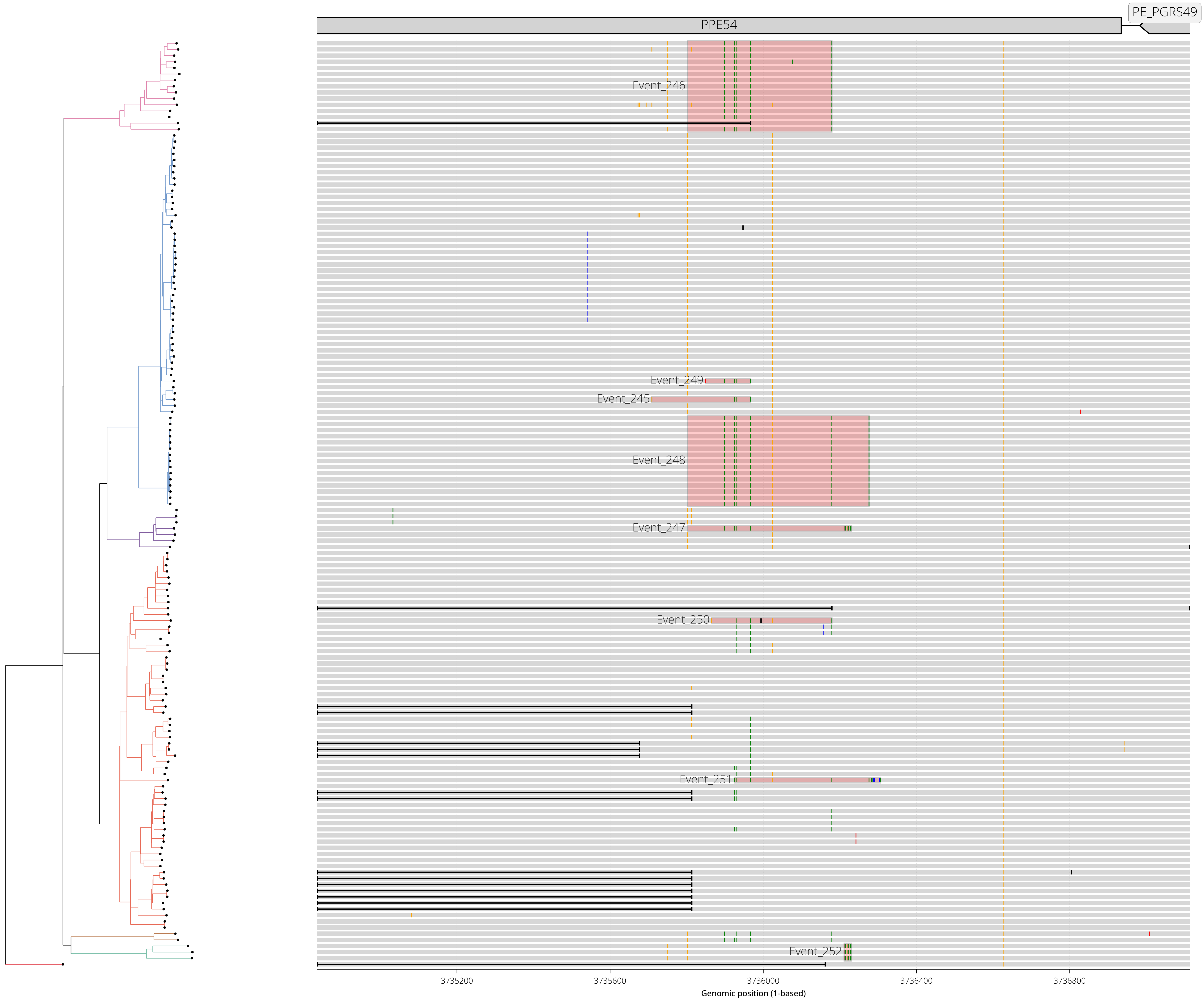

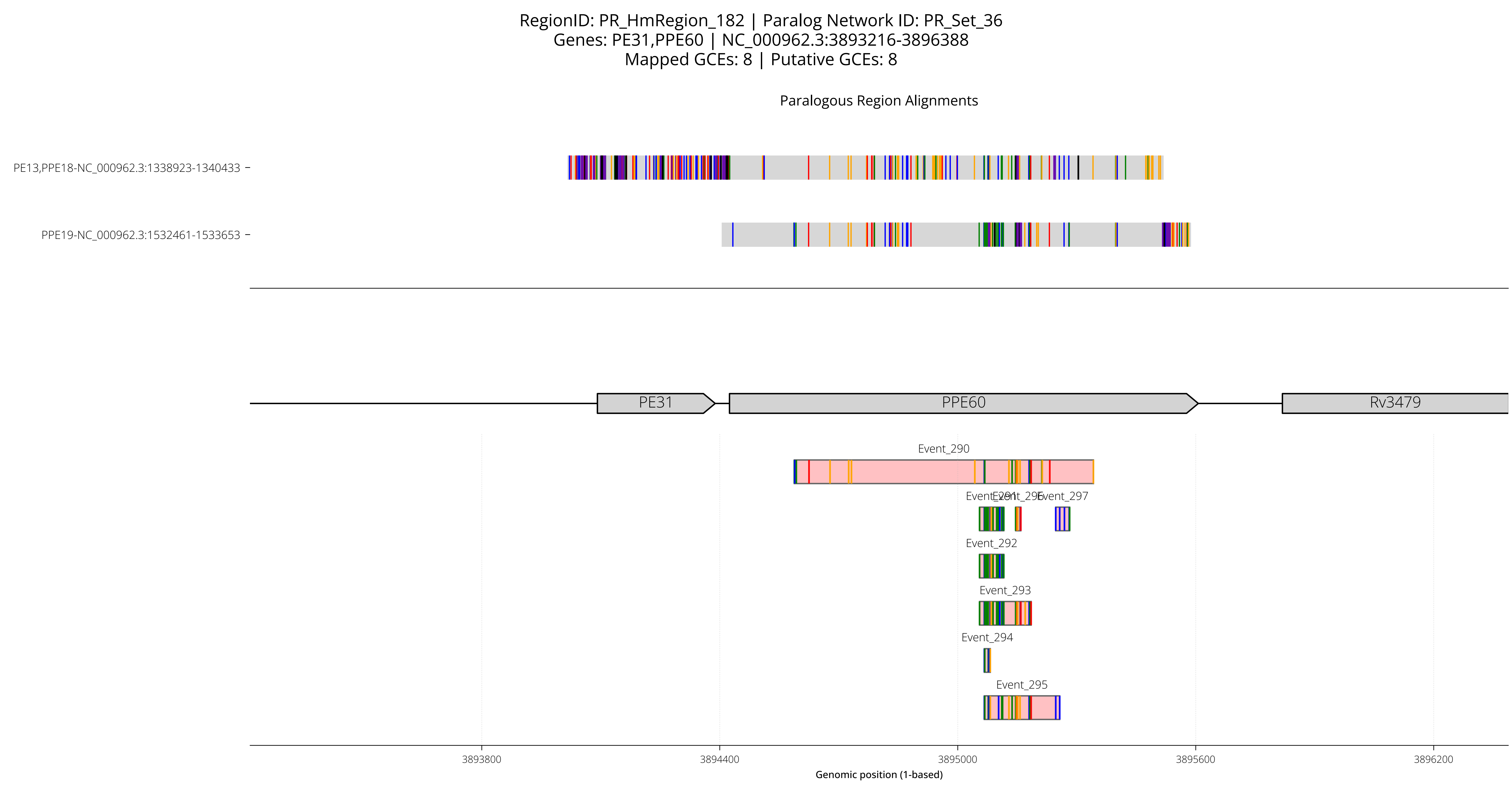
RegionID: PR_HmRegion_182 | Paralog Network ID: PR_Set_36
Genes: PE31,PPE60 | NC_000962.3:3893216-3896388
Mapped GCEs: 8 | Putative GCEs: 8
Paralogous Region Alignments
PE13,PPE18-NC_000962.3:1338923-1340433
PPE19-NC_000962.3:1532461-1533653
PE31
PPE60
Rv3479
Event_290
Event_291
Event_296
Event_297
Event_292
Event_293
Event_294
Event_295
3893800
3894400
3895000
3895600
3896200
Genomic position (1-based)

RegionID: PR_HmRegion_182 | Paralog Network ID: PR_Set_36
Genes: PE31,PPE60 | NC_000962.3:3893216-3896388
Mapped GCEs: 8 | Putative GCEs: 8

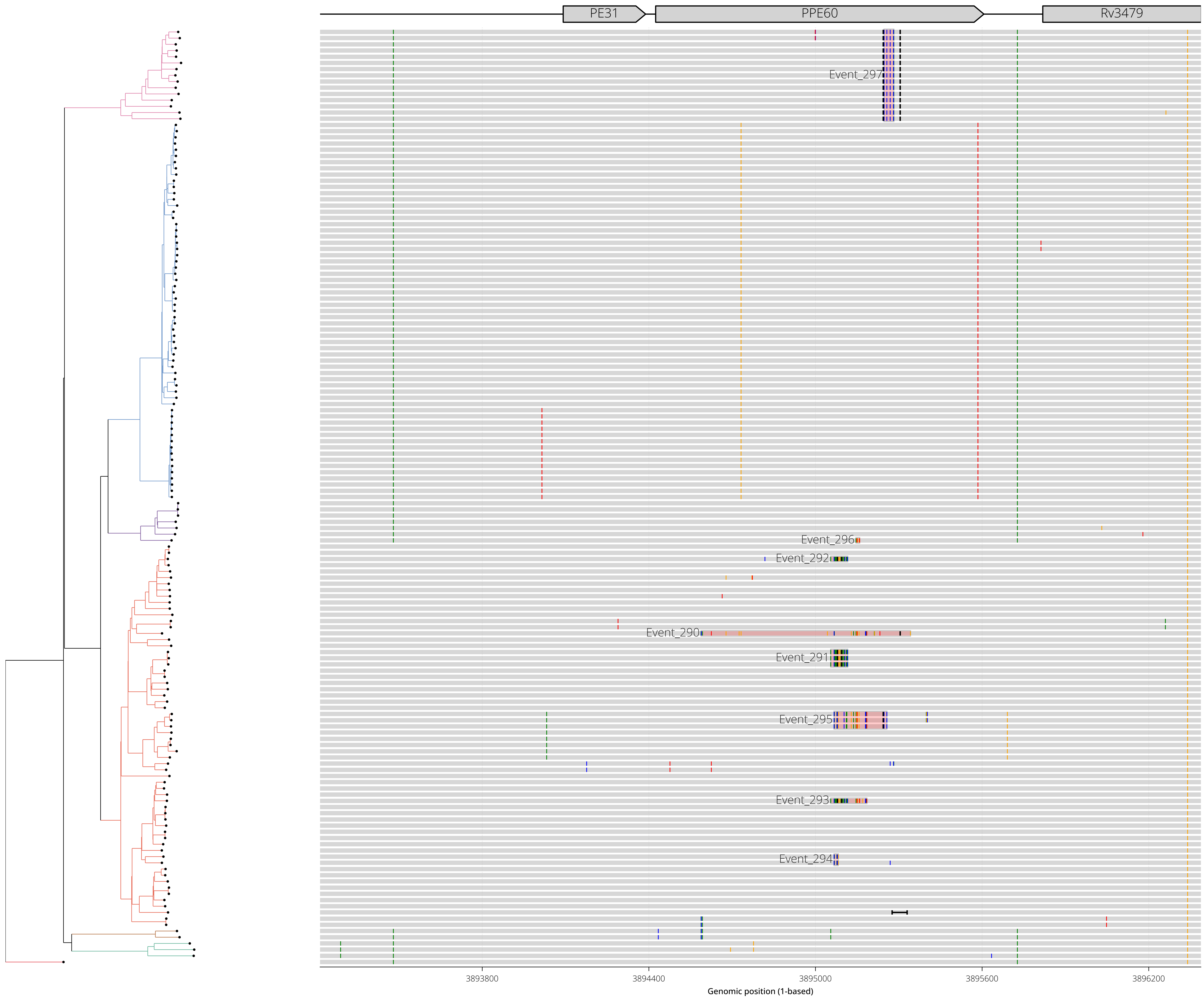

RegionID: PR_HmRegion_053 | Paralog Network ID: PR_Set_36
Genes: PE13,PPE18 | NC_000962.3:1338123-1341233
Mapped GCEs: 7 | Putative GCEs: 7

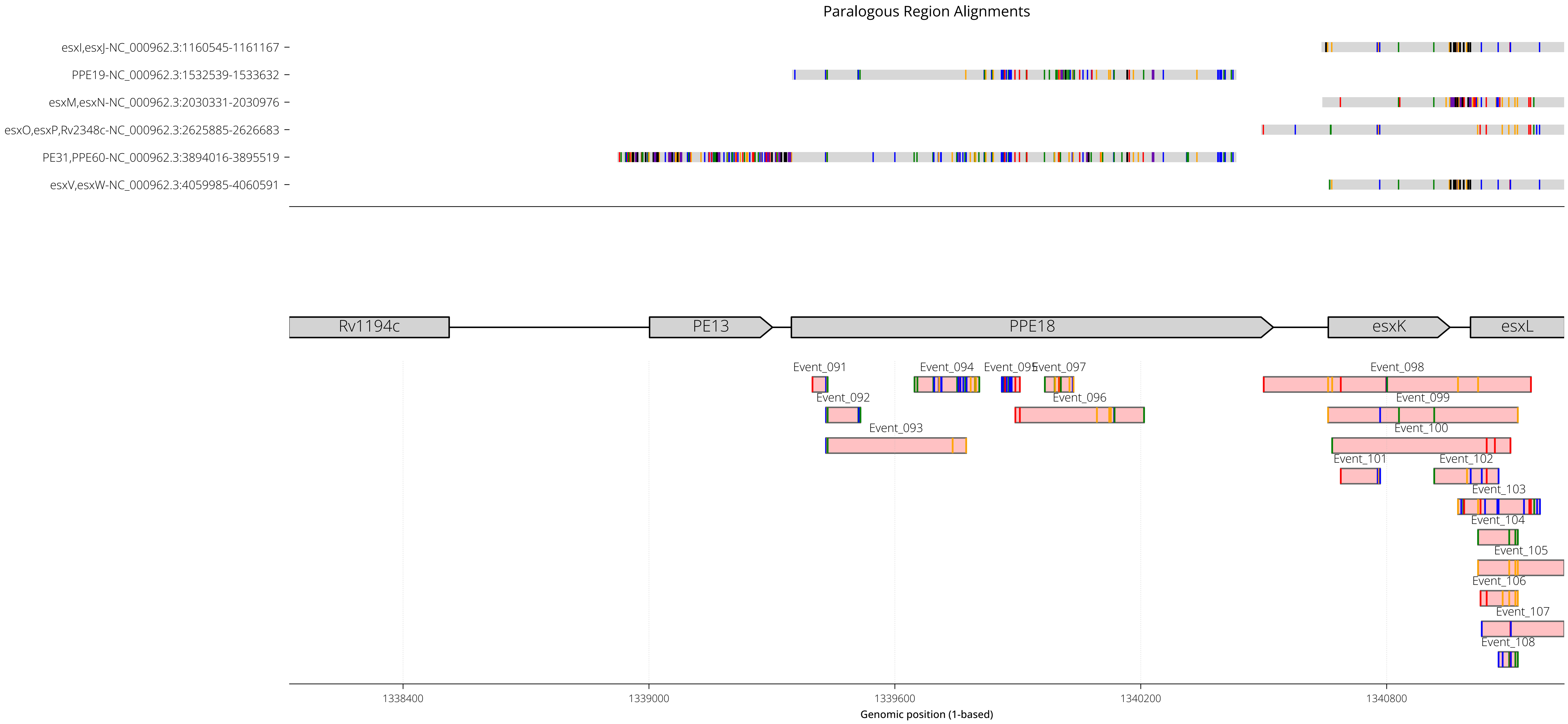

RegionID: PR_HmRegion_053 | Paralog Network ID: PR_Set_36
Genes: PE13,PPE18 | NC_000962.3:1338123-1341233
Mapped GCEs: 7 | Putative GCEs: 7

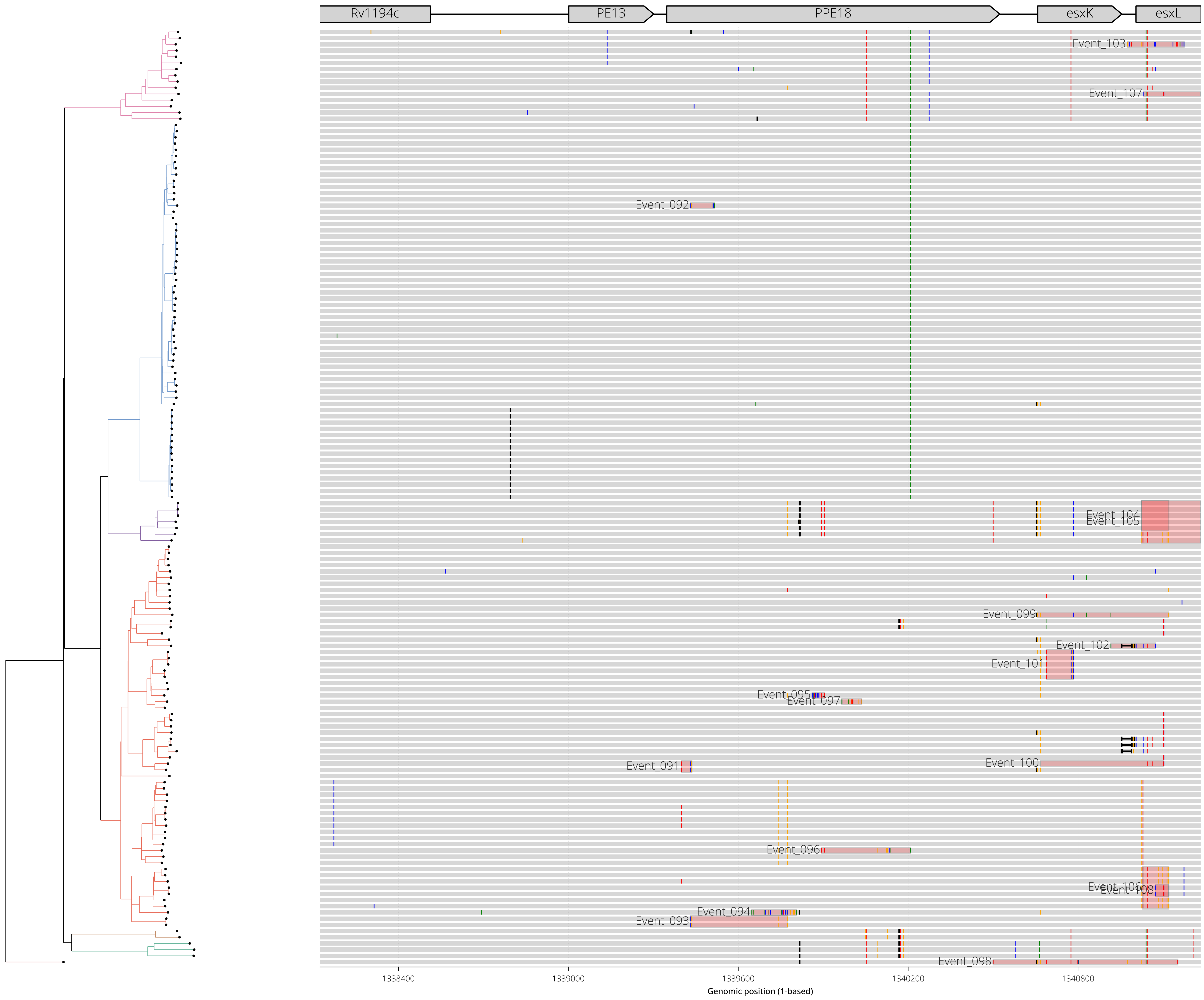

RegionID: PR_HmRegion_050 | Paralog Network ID: PR_Set_34
Genes: Rv1148c | NC_000962.3:1275492-1278597
Mapped GCEs: 6 | Putative GCEs: 7

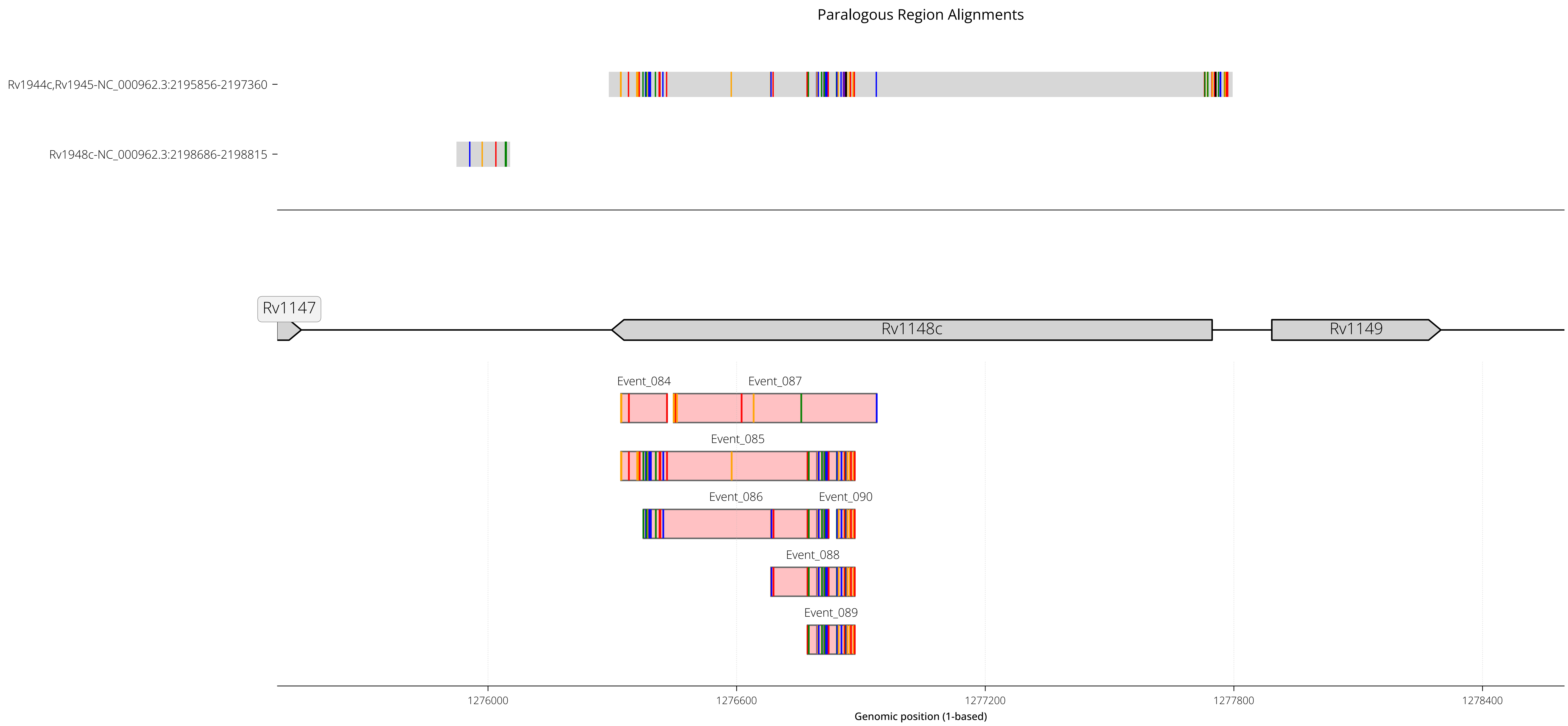

RegionID: PR_HmRegion_050 | Paralog Network ID: PR_Set_34
Genes: Rv1148c | NC_000962.3:1275492-1278597
Mapped GCEs: 6 | Putative GCEs: 7

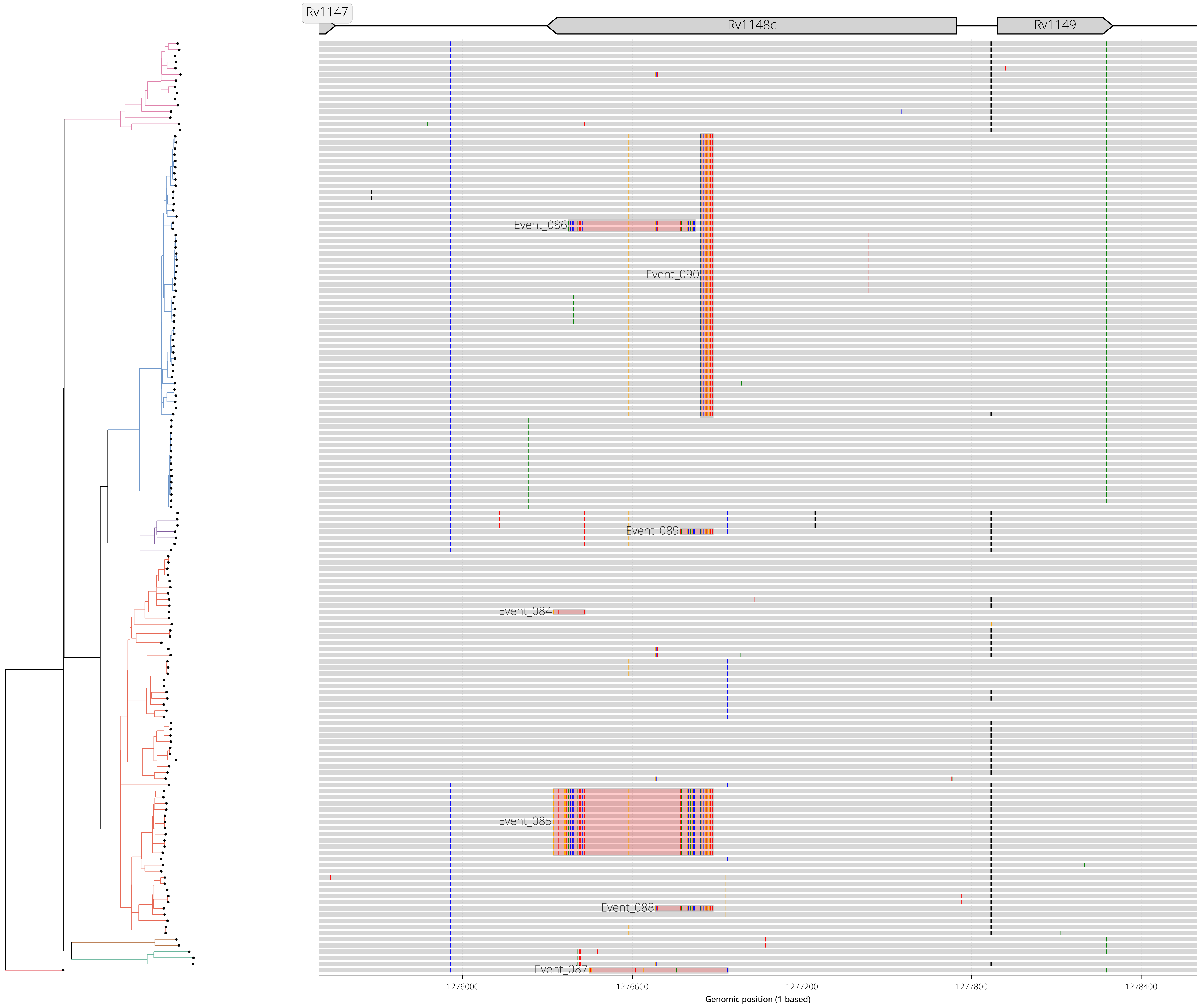

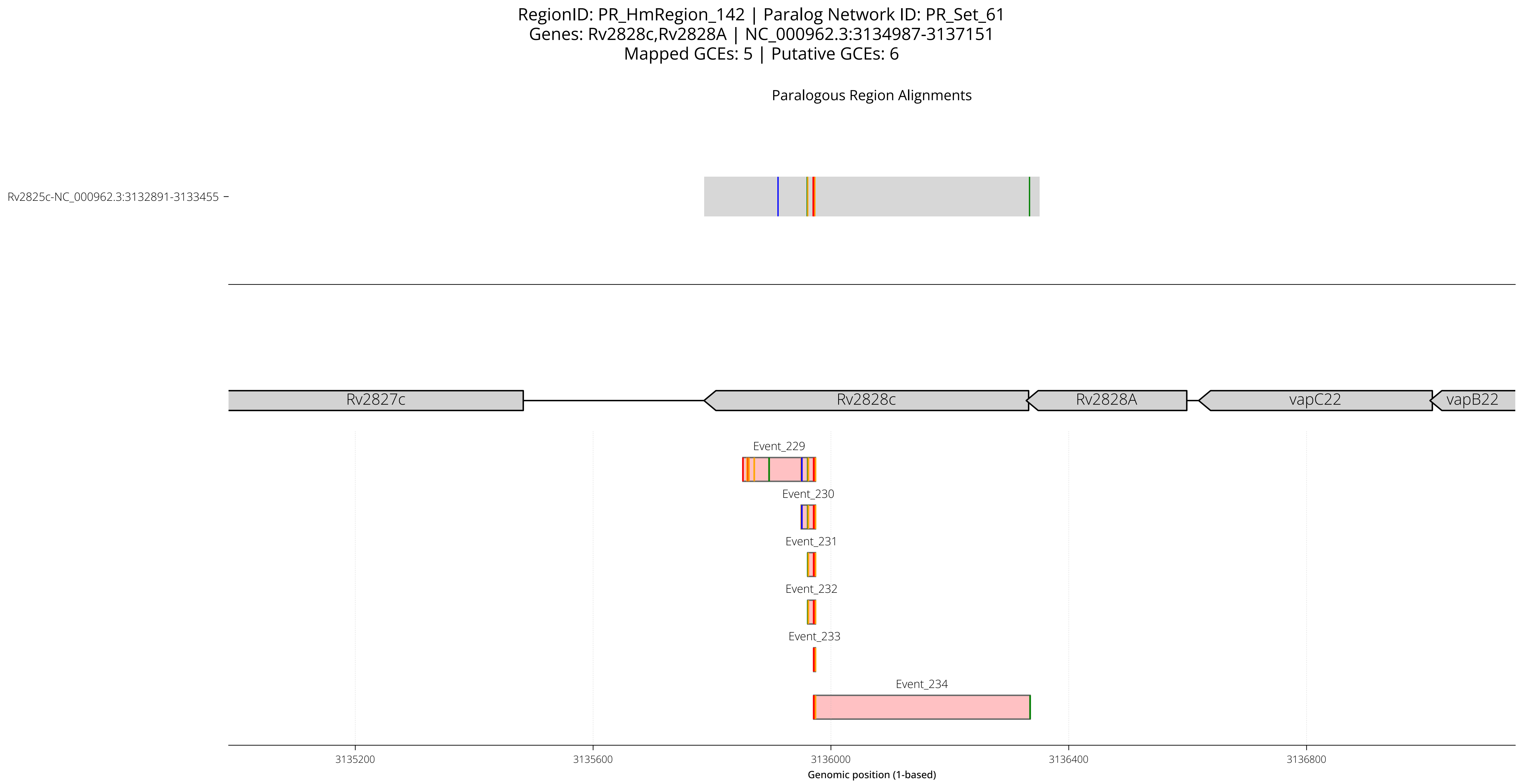
RegionID: PR_HmRegion_142 | Paralog Network ID: PR_Set_61
Genes: Rv2828c,Rv2828A | NC_000962.3:3134987-3137151
Mapped GCEs: 5 | Putative GCEs: 6
Paralogous Region Alignments
Rv2825c-NC_000962.3:3132891-3133455
Rv2827c
Rv2828c
Rv2828A
vapC22
vapB22
Event_229
Event_230
Event_231
Event_232
Event_233
Event_234
3135200
3135600
3136000
3136400
3136800
Genomic position (1-based)

RegionID: PR_HmRegion_142 | Paralog Network ID: PR_Set_61
Genes: Rv2828c,Rv2828A | NC_000962.3:3134987-3137151
Mapped GCEs: 5 | Putative GCEs: 6

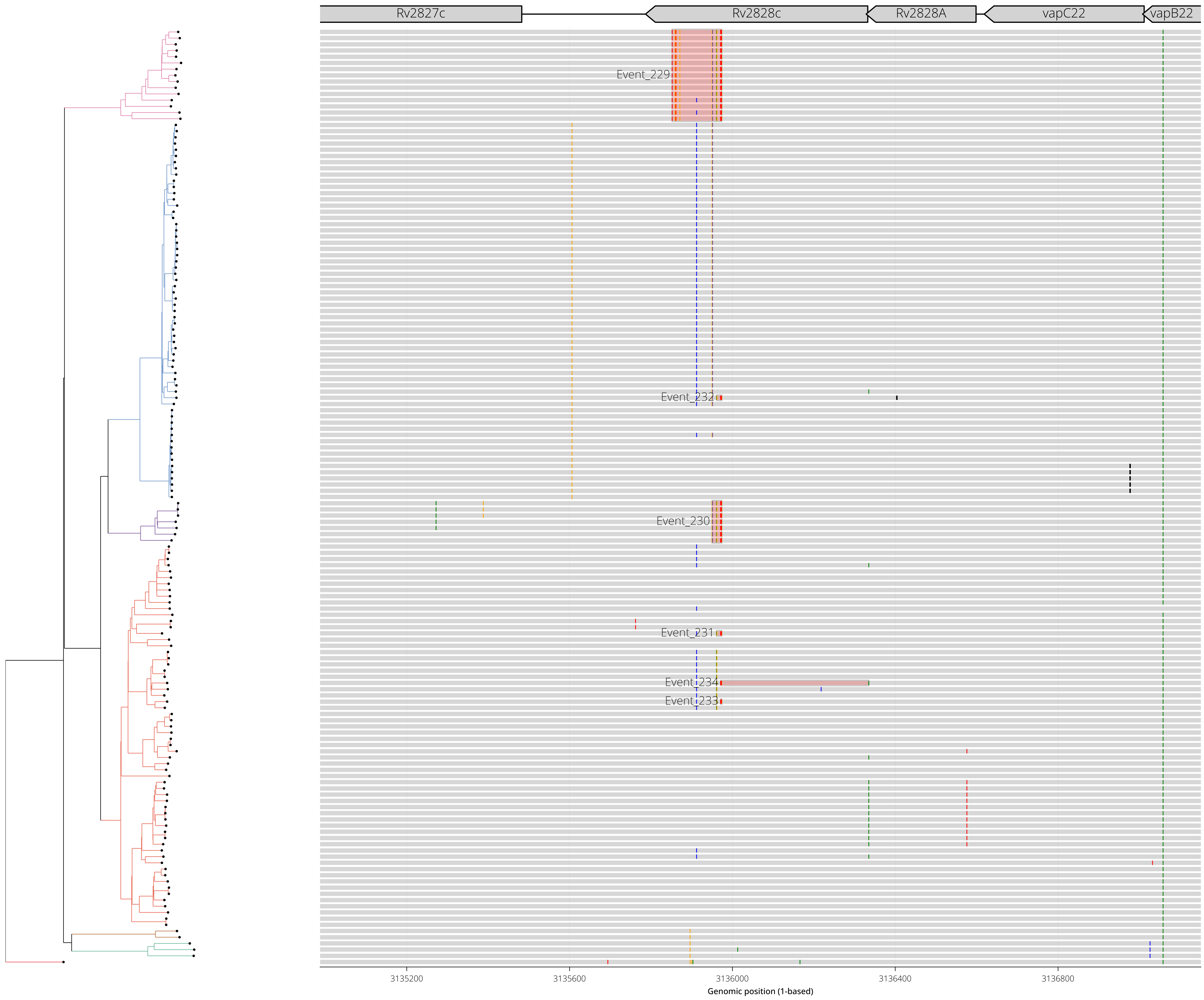

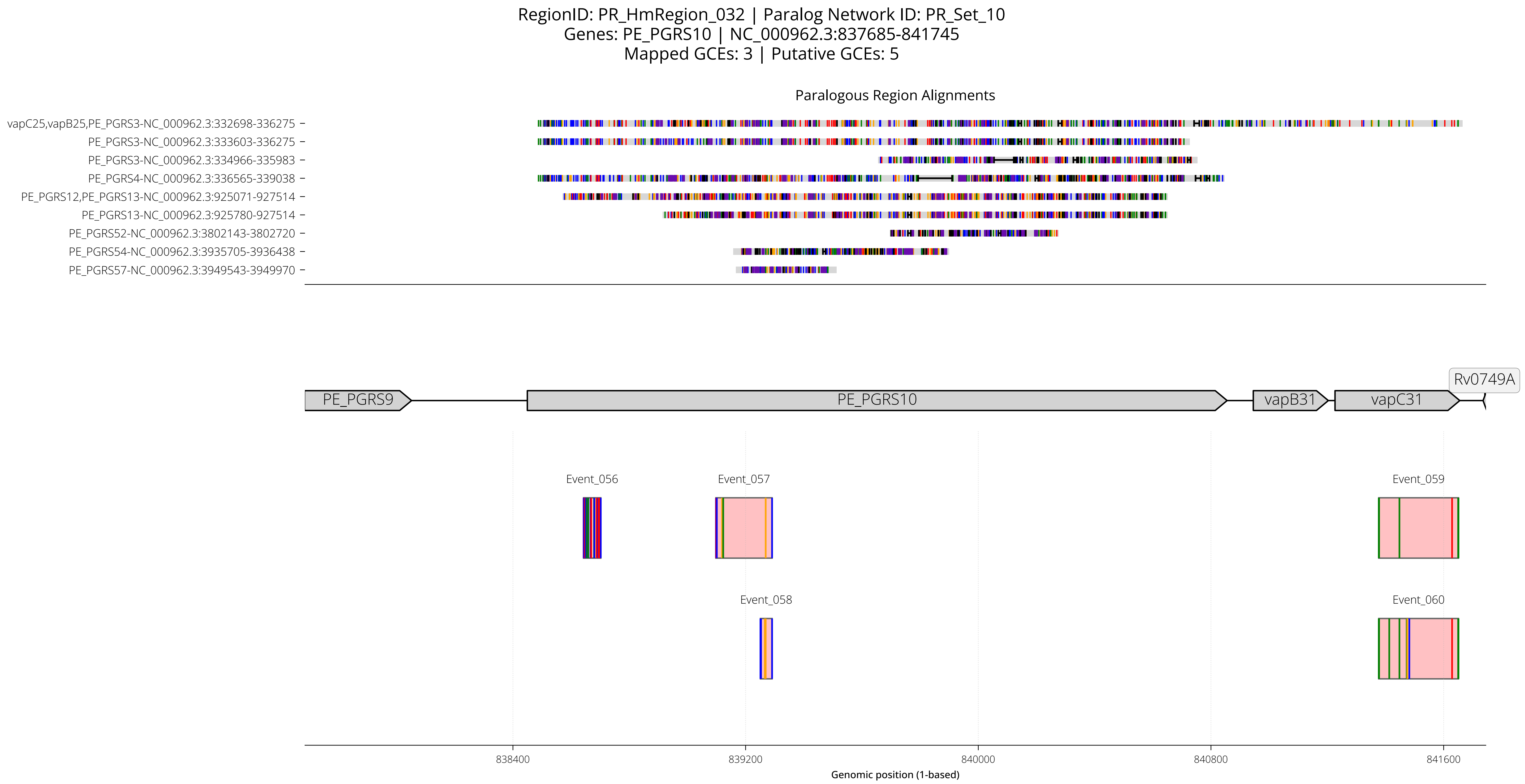
RegionID: PR_HmRegion_032 | Paralog Network ID: PR_Set_10
Genes: PE_PGRS10 | NC_000962.3:837685-841745
Mapped GCEs: 3 | Putative GCEs: 5
Paralogous Region Alignments
vapC25,vapB25,PE_PGRS3-NC_000962.3:332698-336275
PE_PGRS3-NC_000962.3:333603-336275
PE_PGRS3-NC_000962.3:334966-335983
PE_PGRS4-NC_000962.3:336565-339038
PE_PGRS12,PE_PGRS13-NC_000962.3:925071-927514
PE_PGRS13-NC_000962.3:925780-927514
PE_PGRS52-NC_000962.3:3802143-3802720
PE_PGRS54-NC_000962.3:3935705-3936438
PE_PGRS57-NC_000962.3:3949543-3949970
Rv0749A
PE_PGRS9
PE_PGRS10
vapB31
vapC31
Event_056
Event_057
Event_059
Event_058
Event_060
838400
839200
840000
840800
841600
Genomic position (1-based)

RegionID: PR_HmRegion_032 | Paralog Network ID: PR_Set_10
Genes: PE_PGRS10 | NC_000962.3:837685-841745
Mapped GCEs: 3 | Putative GCEs: 5

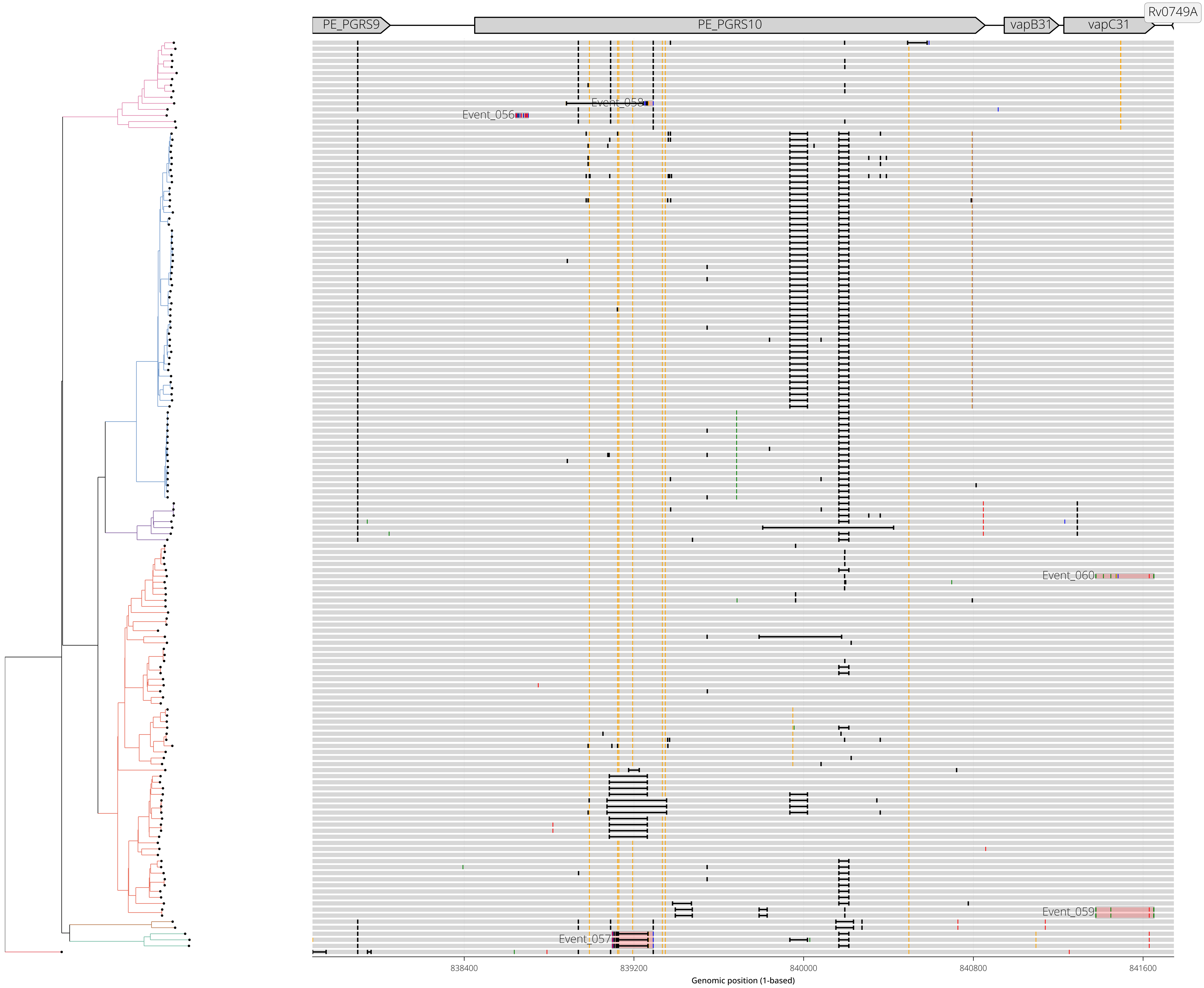

RegionID: PR_HmRegion_032_B | Paralog Network ID: PR_Set_10_B
Genes: vapB31,vapC31 | NC_000962.3:840146-842465
Mapped GCEs: 3 | Putative GCEs: 5

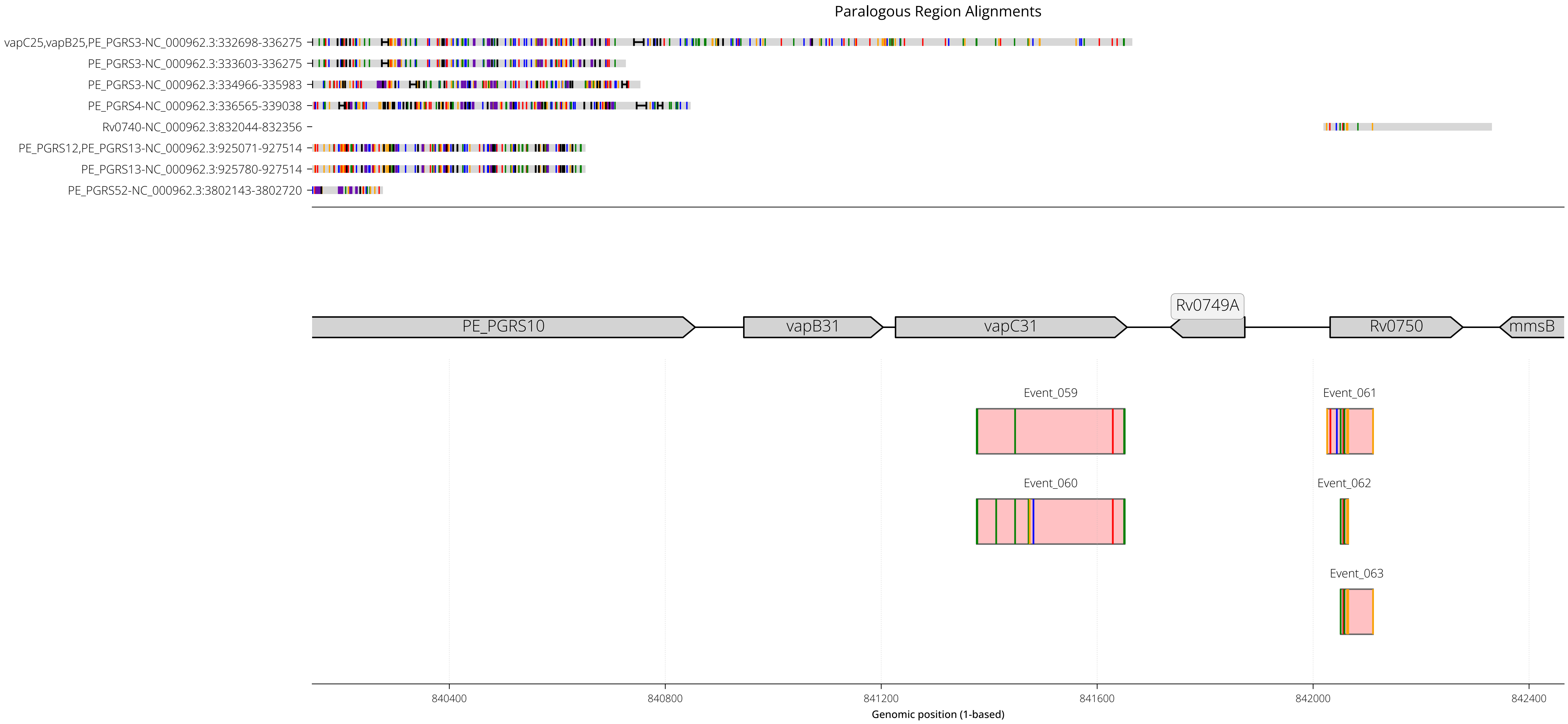

RegionID: PR_HmRegion_032_B | Paralog Network ID: PR_Set_10_B
Genes: vapB31,vapC31 | NC_000962.3:840146-842465
Mapped GCEs: 3 | Putative GCEs: 5

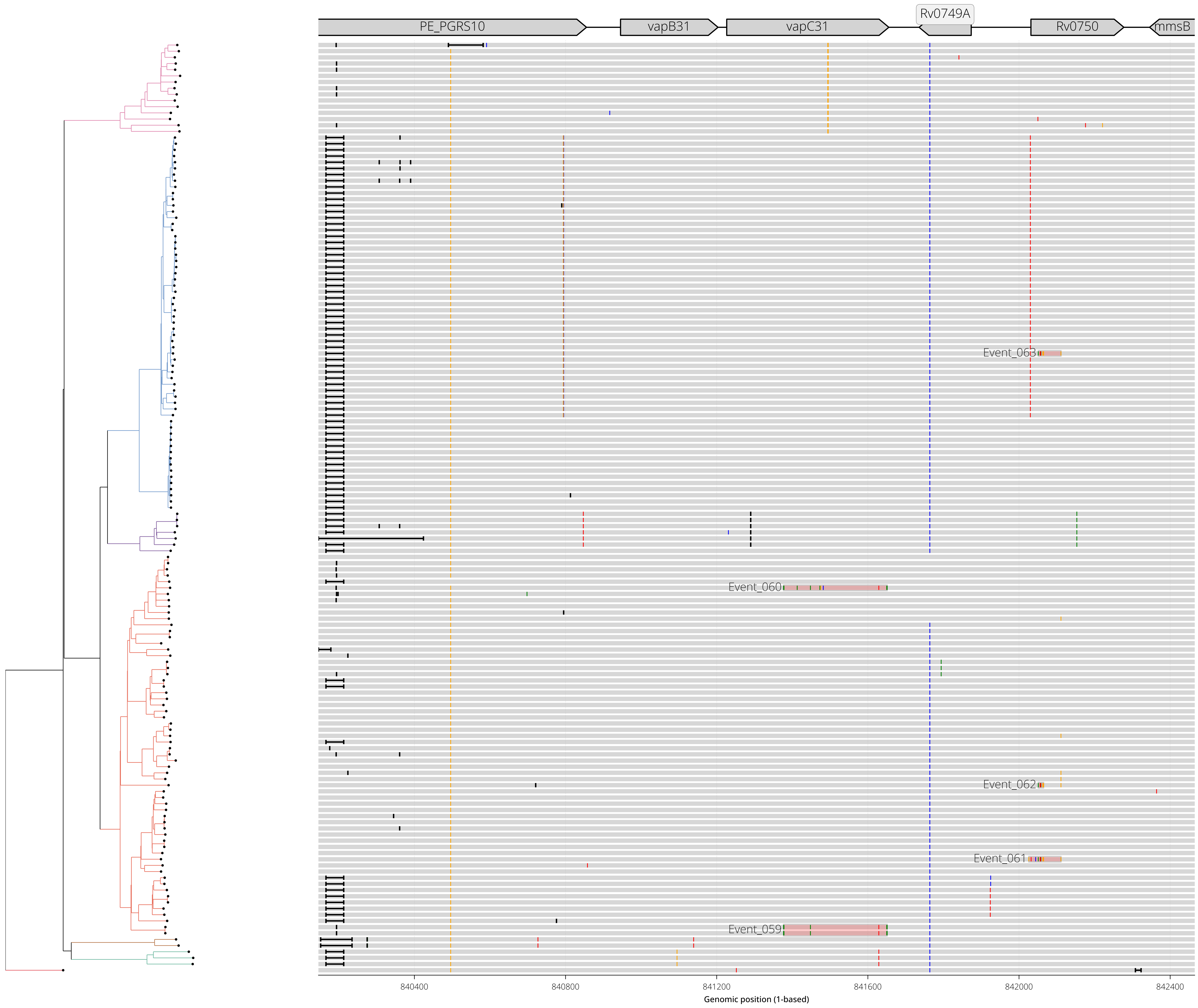

RegionID: PR_HmRegion_060 | Paralog Network ID: PR_Set_36
Genes: PPE19 | NC_000962.3:1531661-1534453
Mapped GCEs: 5 | Putative GCEs: 5

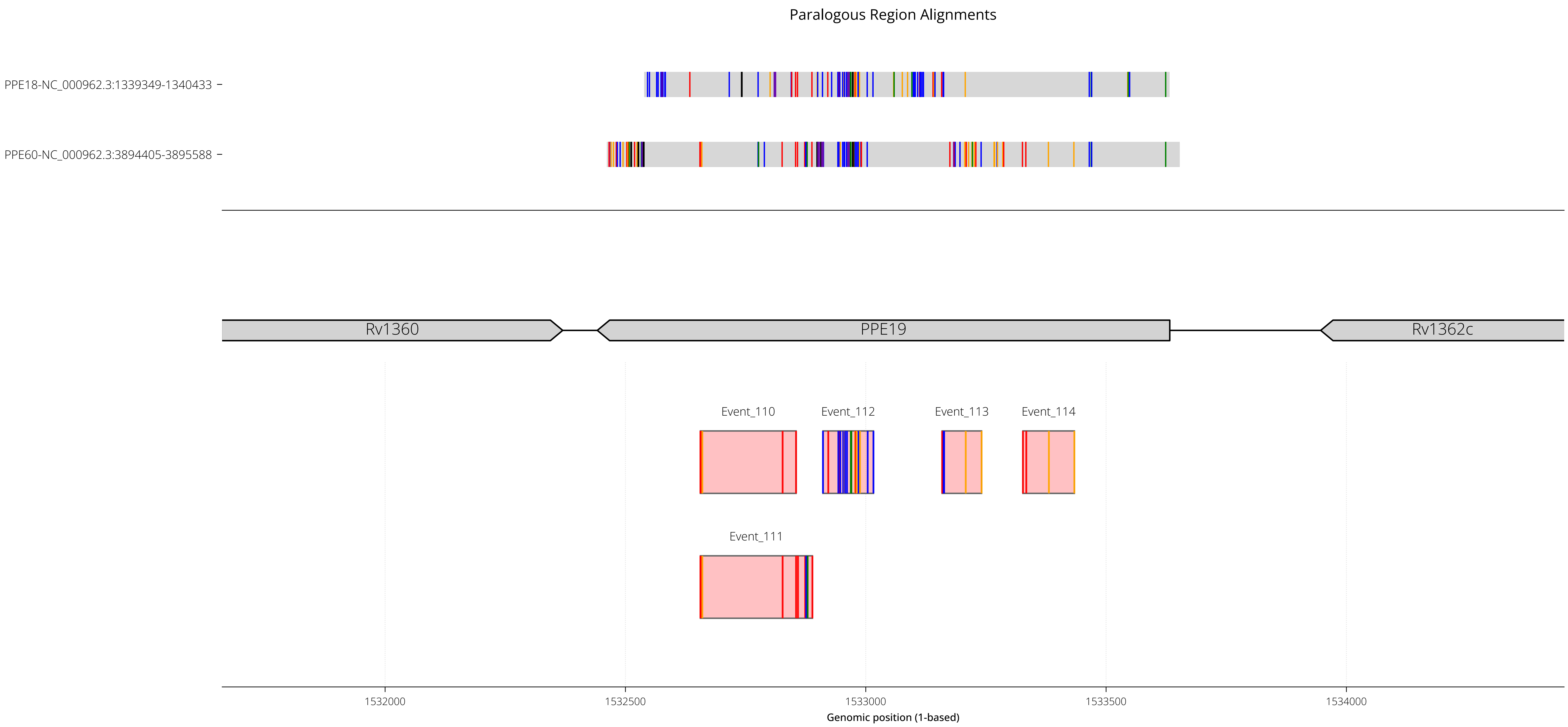

RegionID: PR_HmRegion_060 | Paralog Network ID: PR_Set_36
Genes: PPE19 | NC_000962.3:1531661-1534453
Mapped GCEs: 5 | Putative GCEs: 5

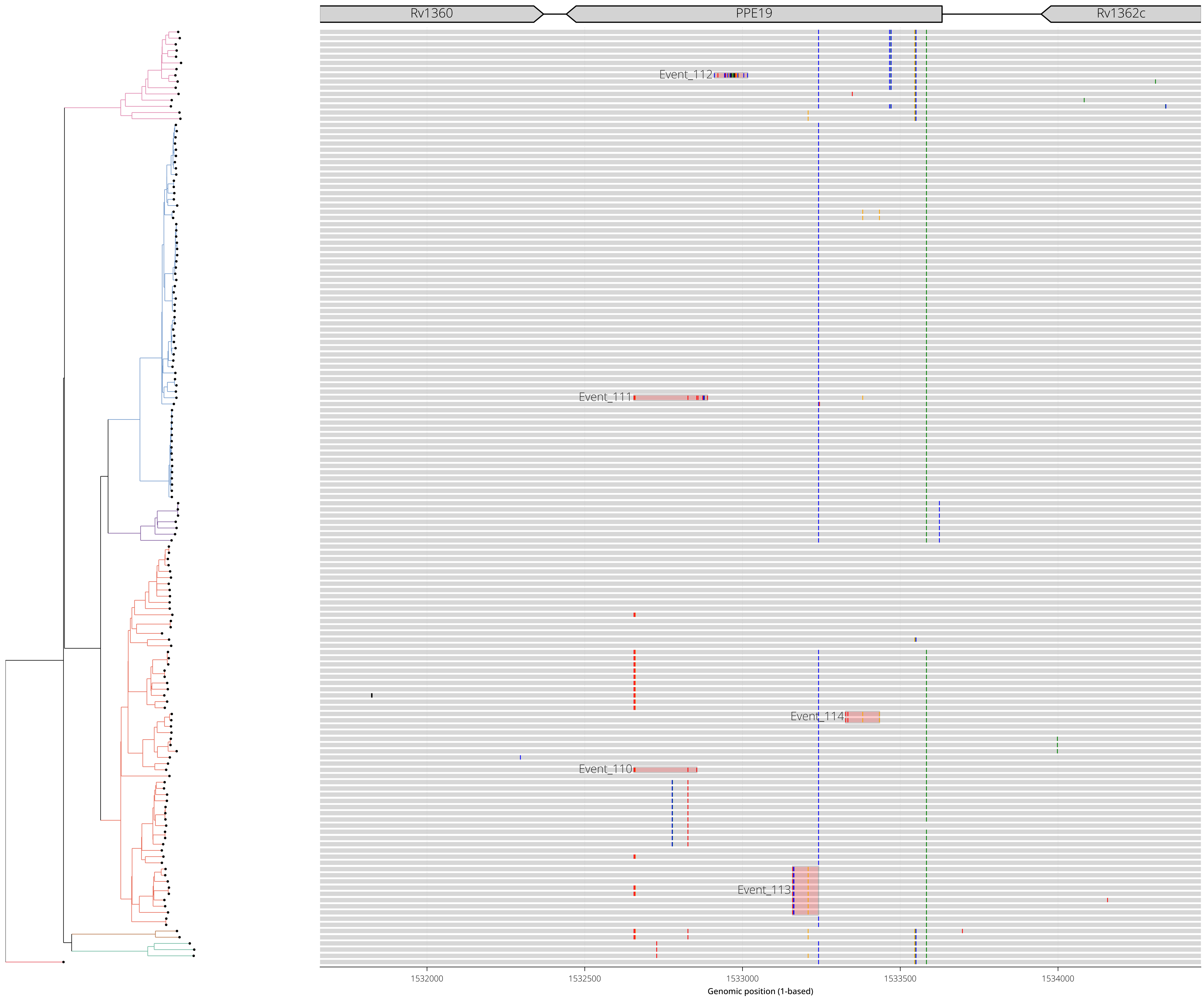

RegionID: PR_HmRegion_173 | Paralog Network ID: PR_Set_24
Genes: PPE56 | NC_000962.3:3755150-3767919
Mapped GCEs: 4 | Putative GCEs: 5

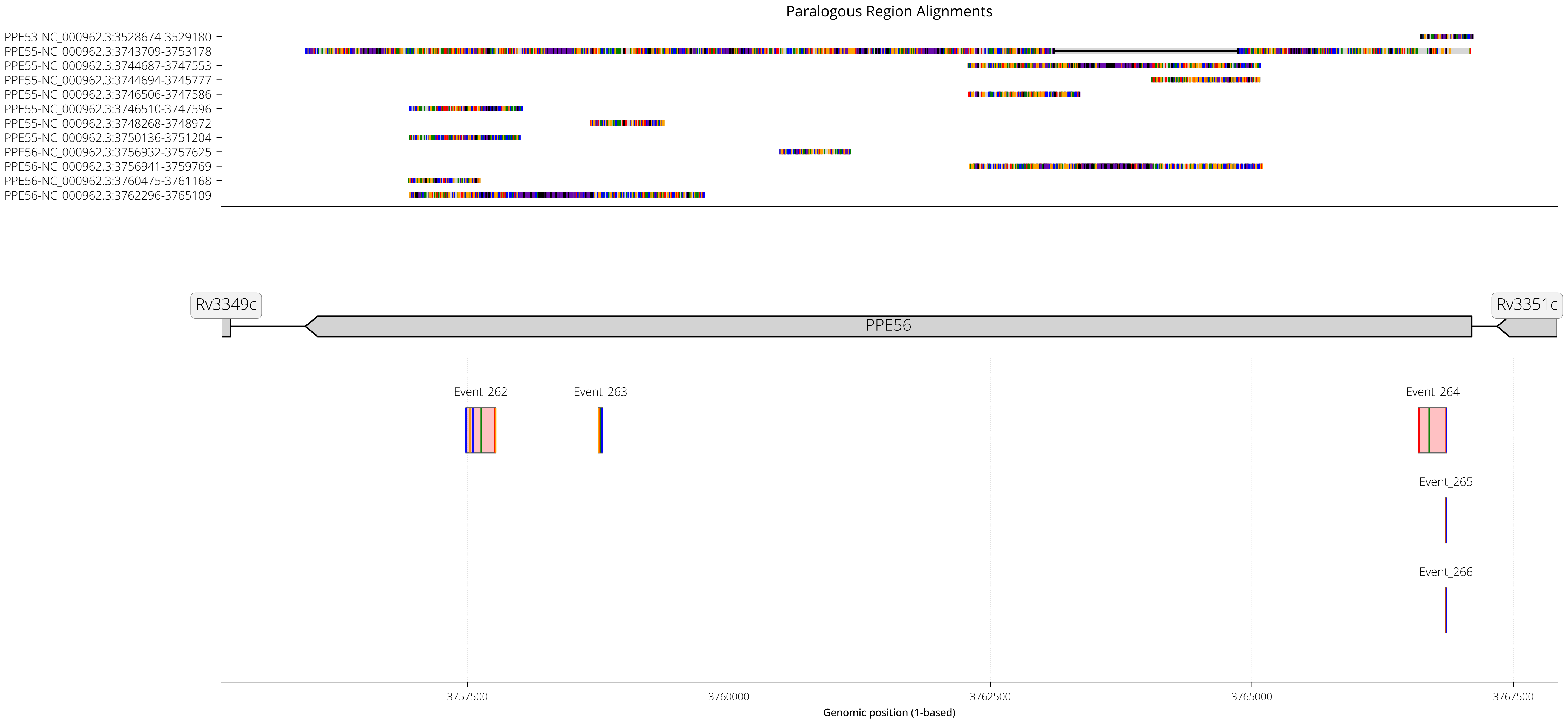

RegionID: PR_HmRegion_173 | Paralog Network ID: PR_Set_24
Genes: PPE56 | NC_000962.3:3755150-3767919
Mapped GCEs: 4 | Putative GCEs: 5

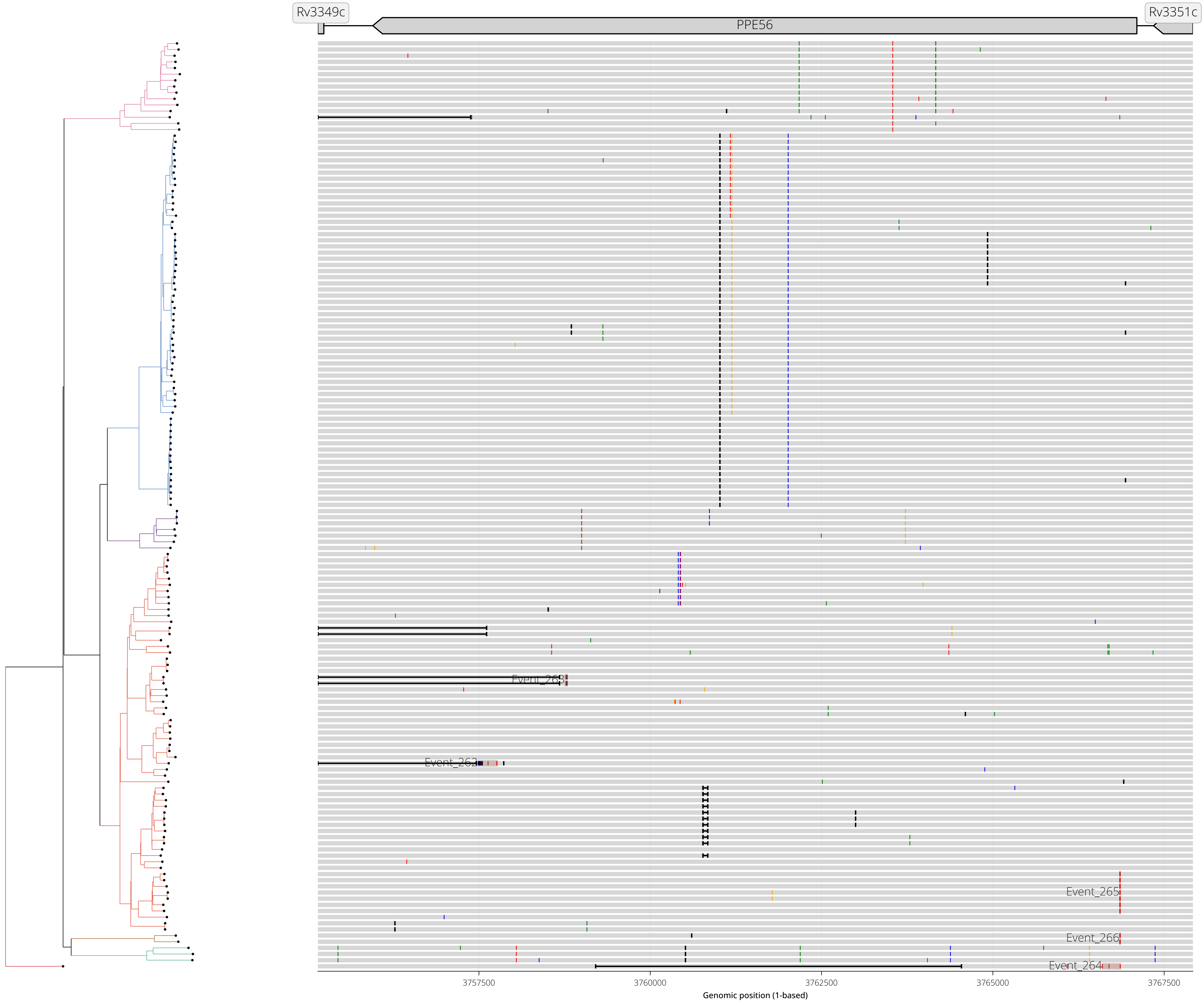

RegionID: PR_HmRegion_040 | Paralog Network ID: PR_Set_27
Genes: PE_PGRS17 | NC_000962.3:1092395-1095392
Mapped GCEs: 4 | Putative GCEs: 4

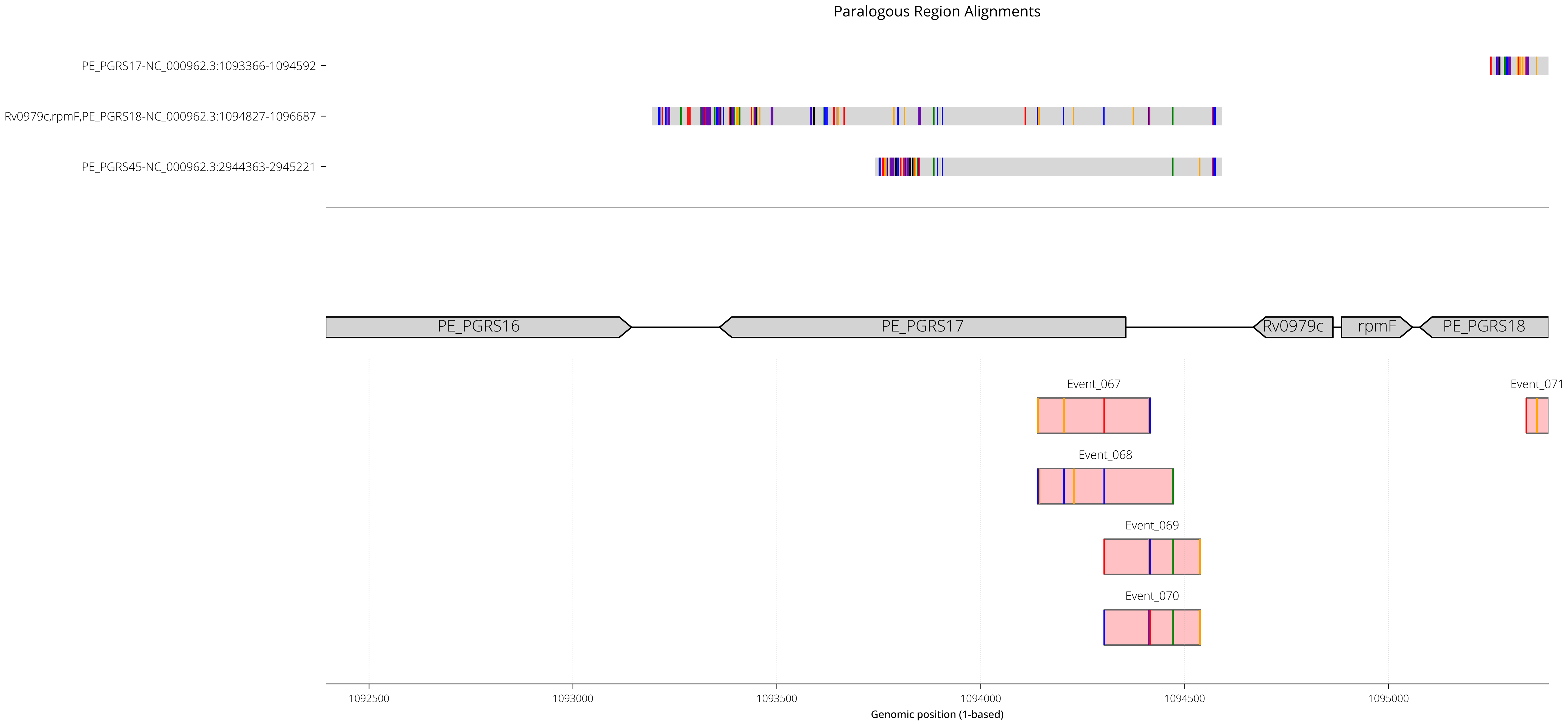

RegionID: PR_HmRegion_040 | Paralog Network ID: PR_Set_27
Genes: PE_PGRS17 | NC_000962.3:1092395-1095392
Mapped GCEs: 4 | Putative GCEs: 4

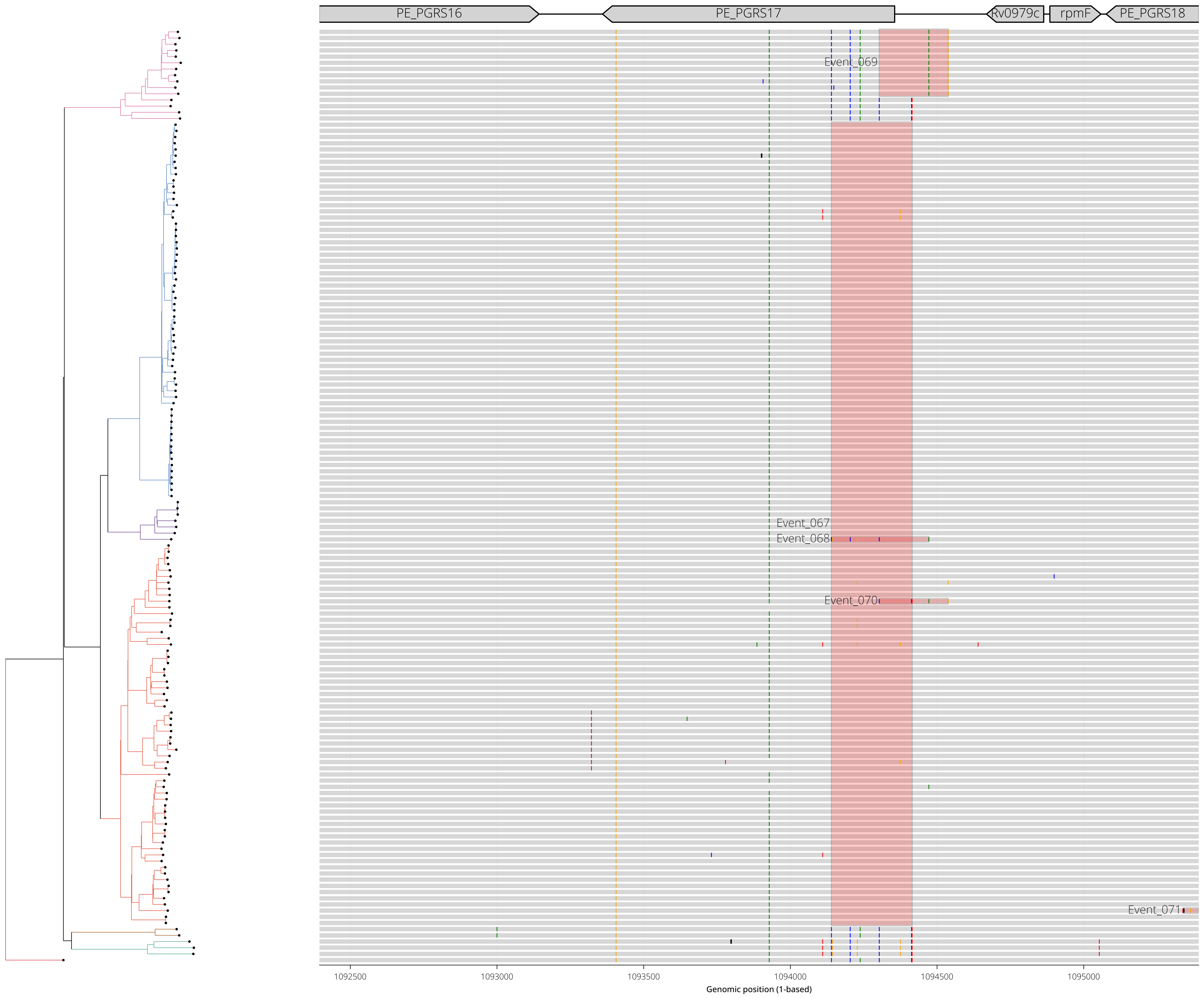

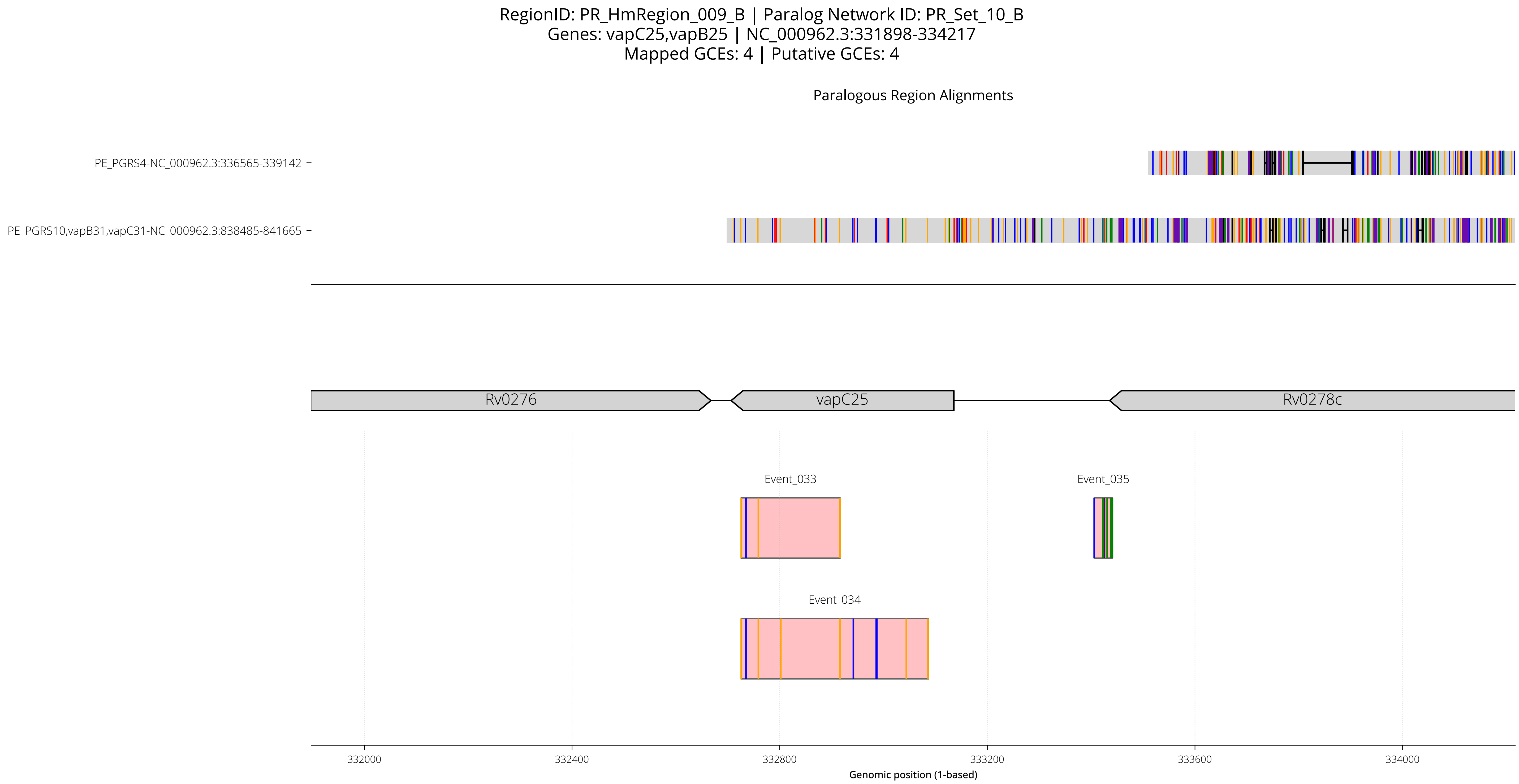
RegionID: PR_HmRegion_009_B | Paralog Network ID: PR_Set_10_B
Genes: vapC25,vapB25 | NC_000962.3:331898-334217
Mapped GCEs: 4 | Putative GCEs: 4
Paralogous Region Alignments
PE_PGRS4-NC_000962.3:336565-339142
PE_PGRS10,vapB31,vapC31-NC_000962.3:838485-841665
Rv0276
vapC25
Rv0278c
Event_033
Event_035
Event_034
332000
332400
332800
333200
333600
334000
Genomic position (1-based)

RegionID: PR_HmRegion_009_B | Paralog Network ID: PR_Set_10_B
Genes: vapC25,vapB25 | NC_000962.3:331898-334217
Mapped GCEs: 4 | Putative GCEs: 4

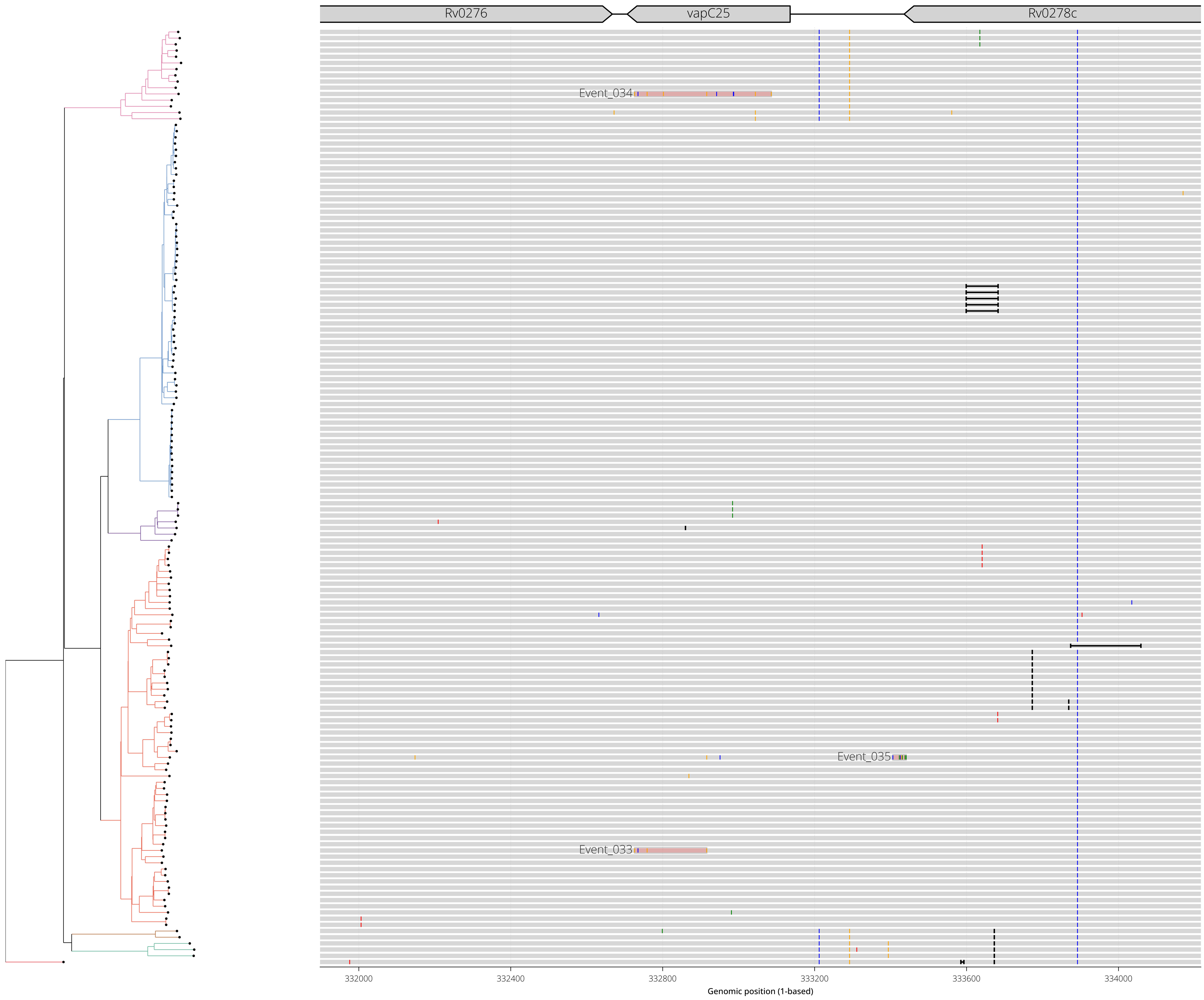

RegionID: PR_HmRegion_086 | Paralog Network ID: PR_Set_49
Genes: PPE27 | NC_000962.3:2027625-2030042
Mapped GCEs: 3 | Putative GCEs: 4

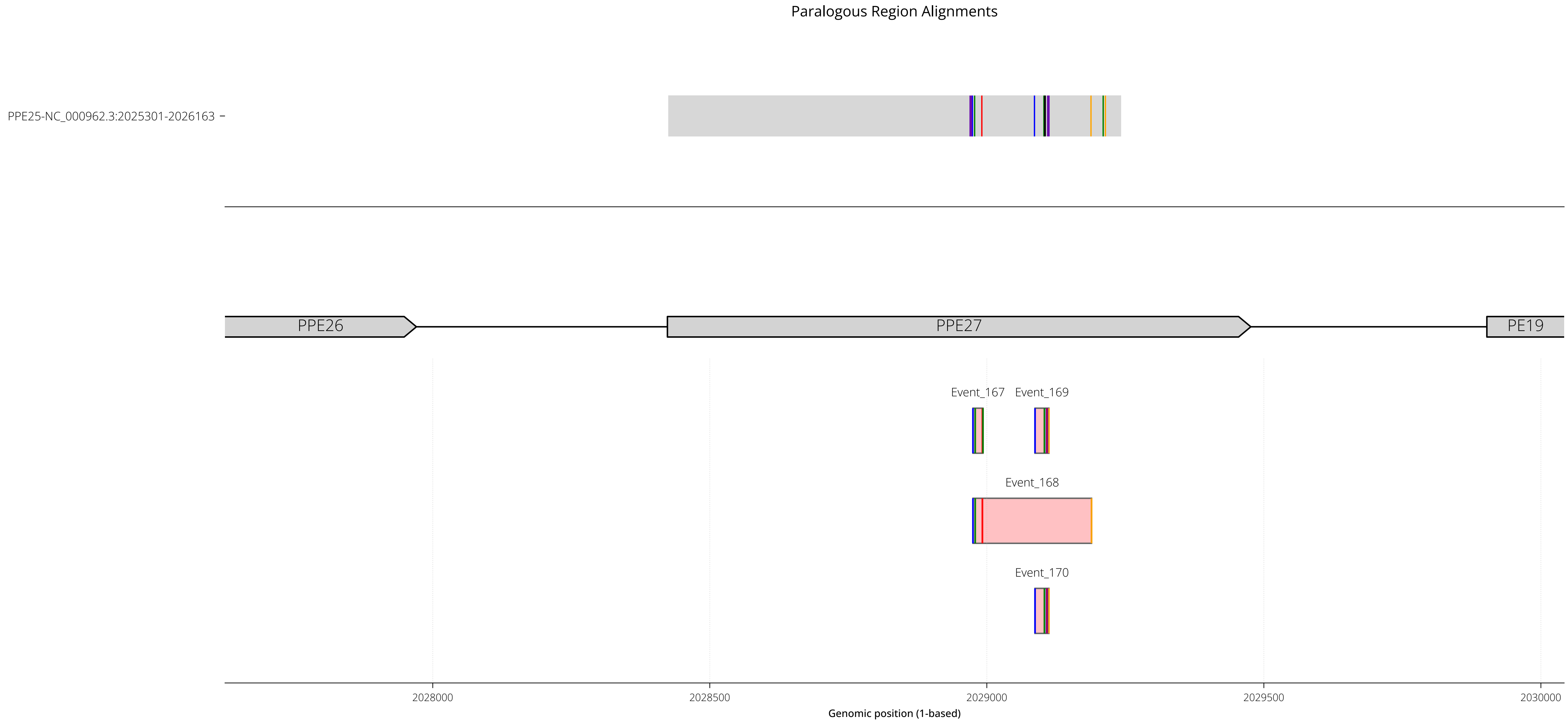

RegionID: PR_HmRegion_086 | Paralog Network ID: PR_Set_49
Genes: PPE27 | NC_000962.3:2027625-2030042
Mapped GCEs: 3 | Putative GCEs: 4

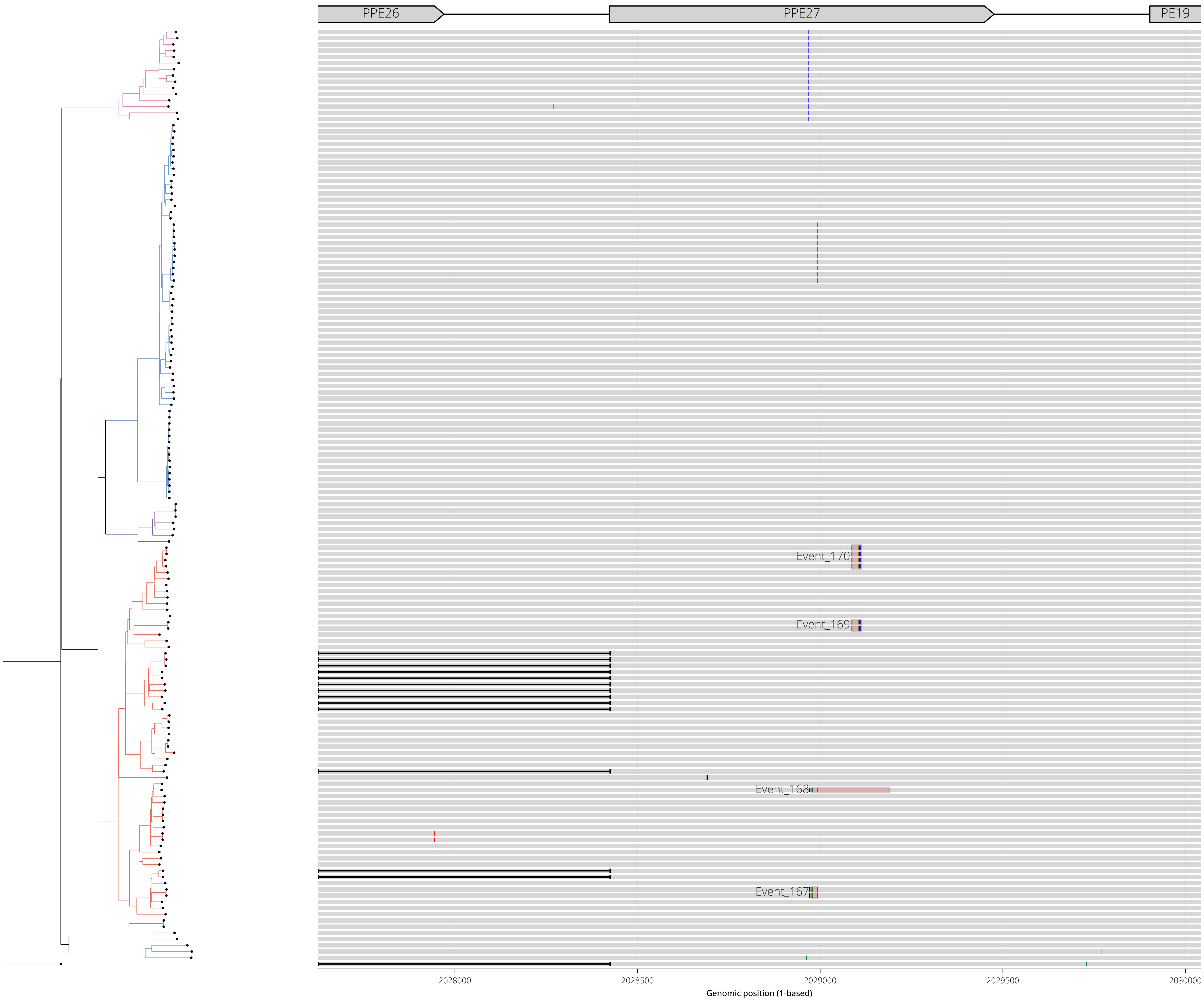

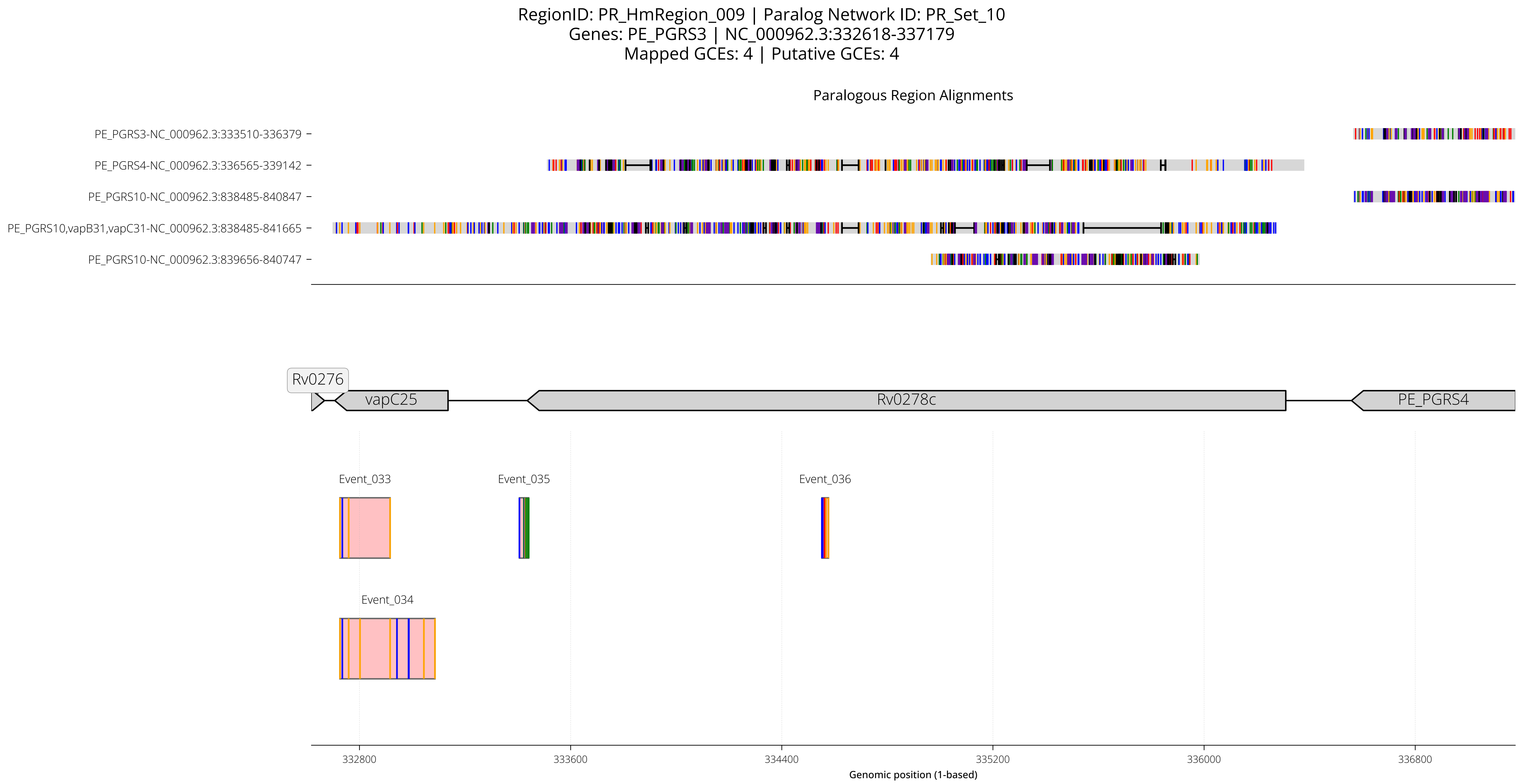

RegionID: PR_HmRegion_009 | Paralog Network ID: PR_Set_10
Genes: PE_PGRS3 | NC_000962.3:332618-337179
Mapped GCEs: 4 | Putative GCEs: 4
Paralogous Region Alignments
PE_PGRS3-NC_000962.3:333510-336379
PE_PGRS4-NC_000962.3:336565-339142
PE_PGRS10-NC_000962.3:838485-840847
PE_PGRS10,vapB31,vapC31-NC_000962.3:838485-841665
PE_PGRS10-NC_000962.3:839656-840747
Rv0276
vapC25
Rv0278c
PE_PGRS4
Event_033
Event_035
Event_036
Event_034
332800
333600
334400
335200
336000
336800
Genomic position (1-based)

RegionID: PR_HmRegion_009 | Paralog Network ID: PR_Set_10
Genes: PE_PGRS3 | NC_000962.3:332618-337179
Mapped GCEs: 4 | Putative GCEs: 4

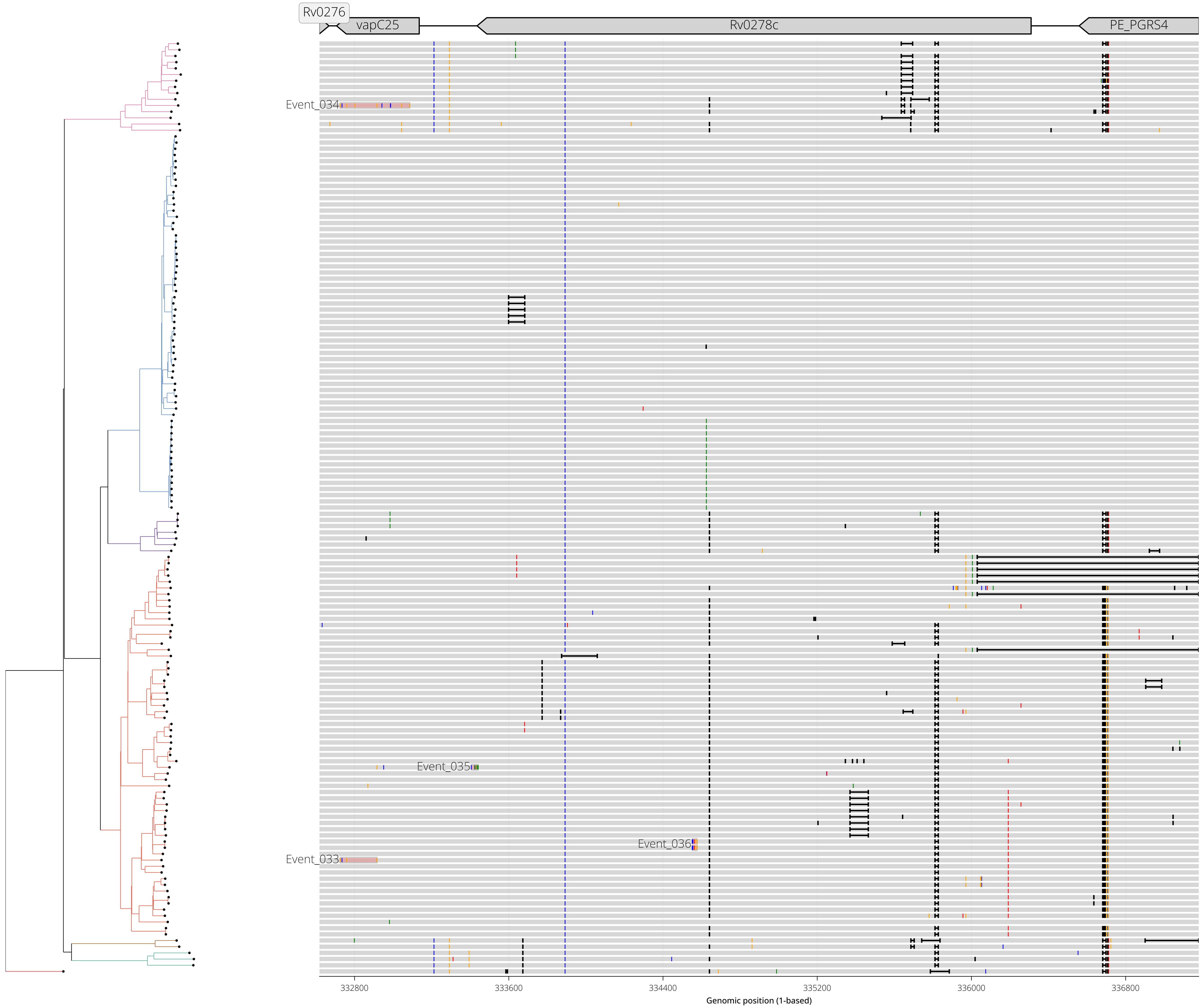

RegionID: PR_HmRegion_087 | Paralog Network ID: PR_Set_28
Genes: esxM,esxN | NC_000962.3:2029531-2031783
Mapped GCEs: 4 | Putative GCEs: 4

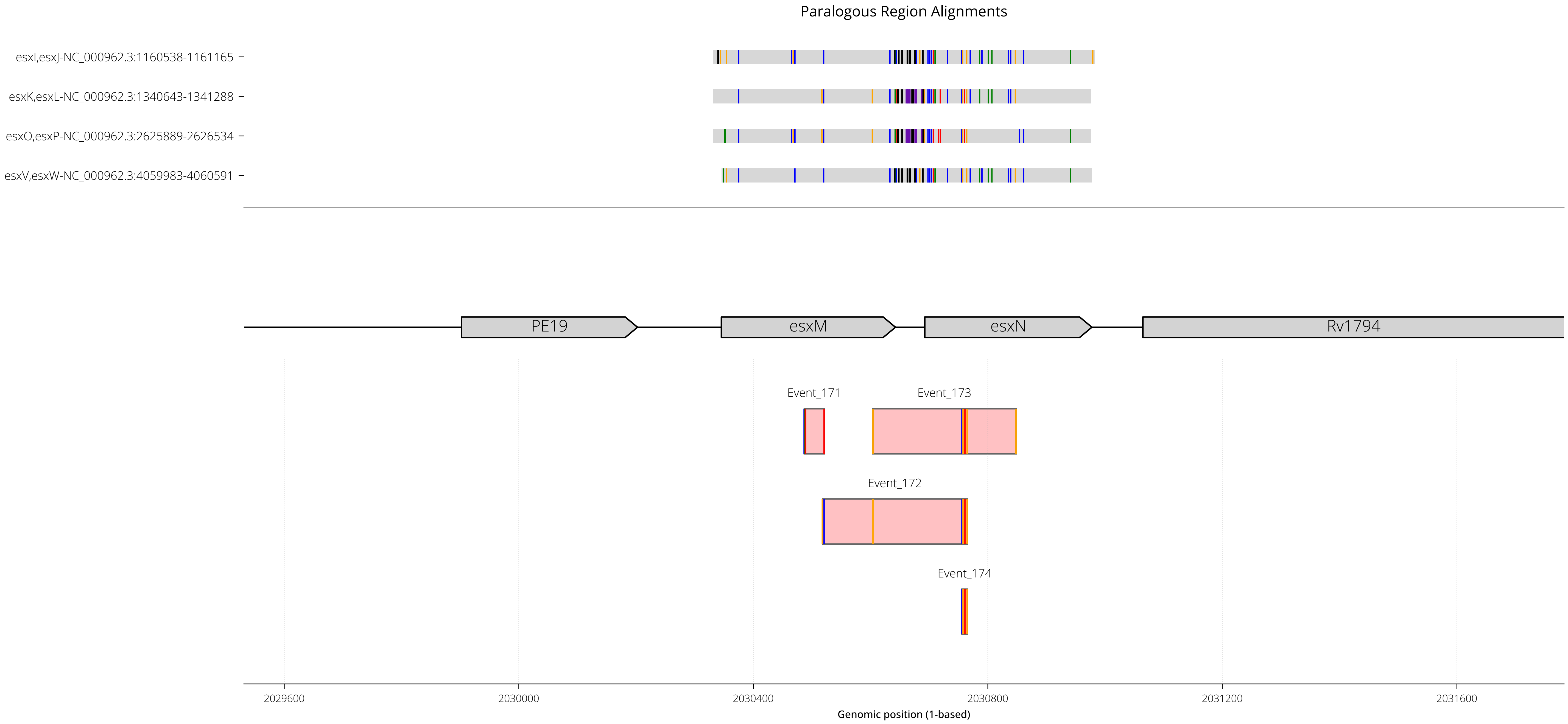

RegionID: PR_HmRegion_087 | Paralog Network ID: PR_Set_28
Genes: esxM,esxN | NC_000962.3:2029531-2031783
Mapped GCEs: 4 | Putative GCEs: 4

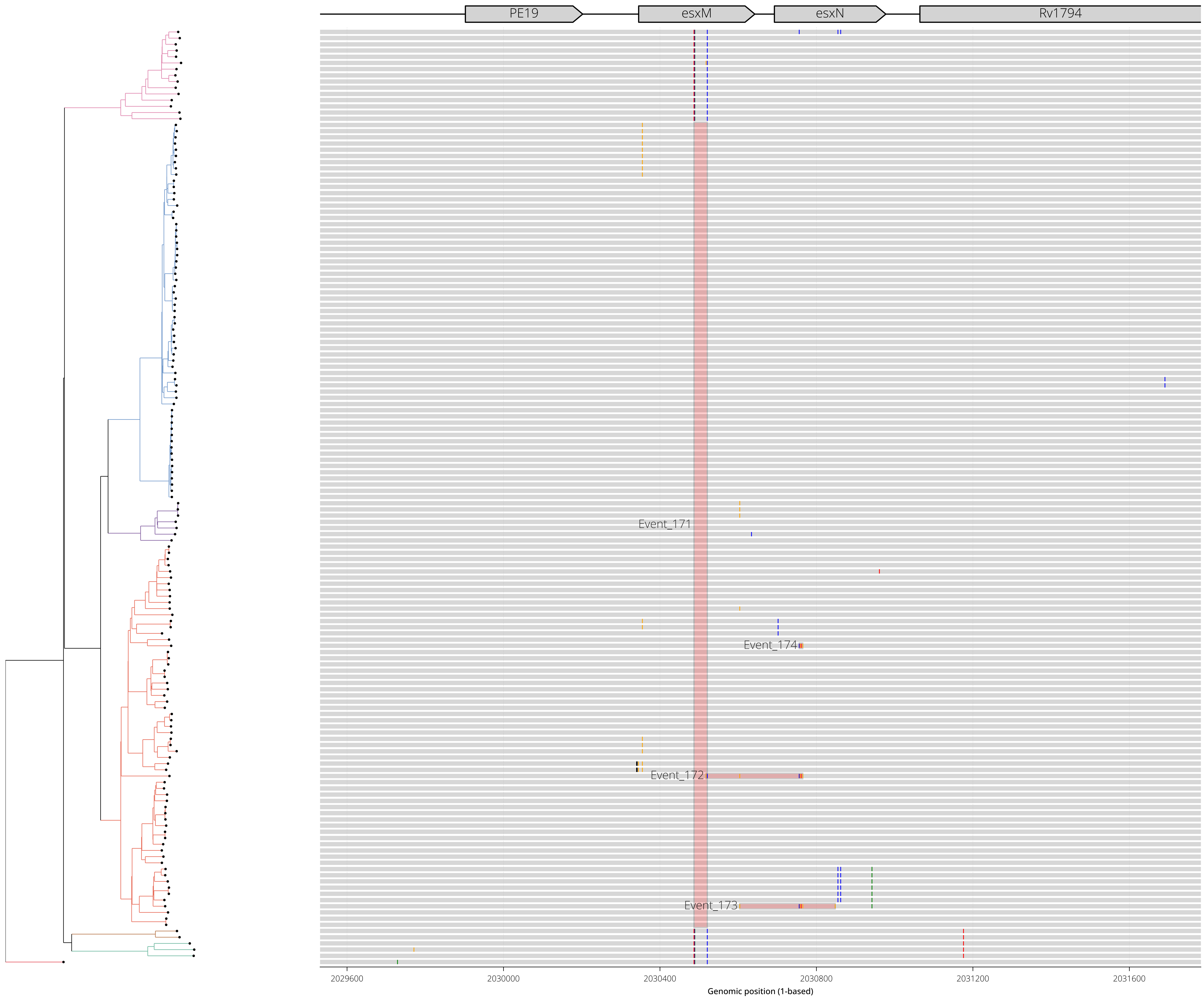

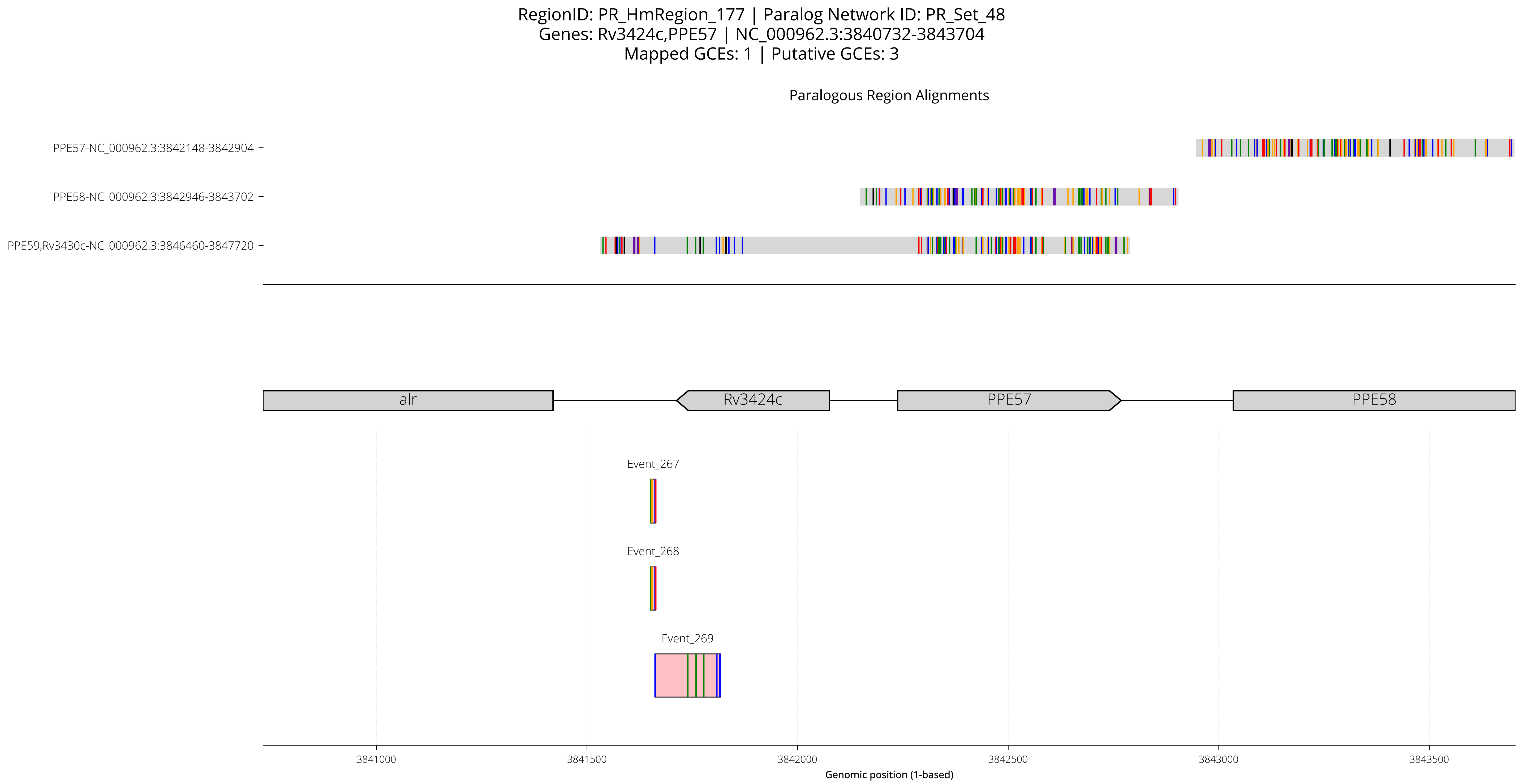
RegionID: PR_HmRegion_177 | Paralog Network ID: PR_Set_48
Genes: Rv3424c,PPE57 | NC_000962.3:3840732-3843704
Mapped GCEs: 1 | Putative GCEs: 3
Paralogous Region Alignments
PPE57-NC_000962.3:3842148-3842904
PPE58-NC_000962.3:3842946-3843702
PPE59,Rv3430c-NC_000962.3:3846460-3847720
alr
Rv3424c
PPE57
PPE58
Event_267
Event_268
Event_269
3841000
3841500
3842000
3842500
3843000
3843500
Genomic position (1-based)

RegionID: PR_HmRegion_177 | Paralog Network ID: PR_Set_48
Genes: Rv3424c,PPE57 | NC_000962.3:3840732-3843704
Mapped GCEs: 1 | Putative GCEs: 3

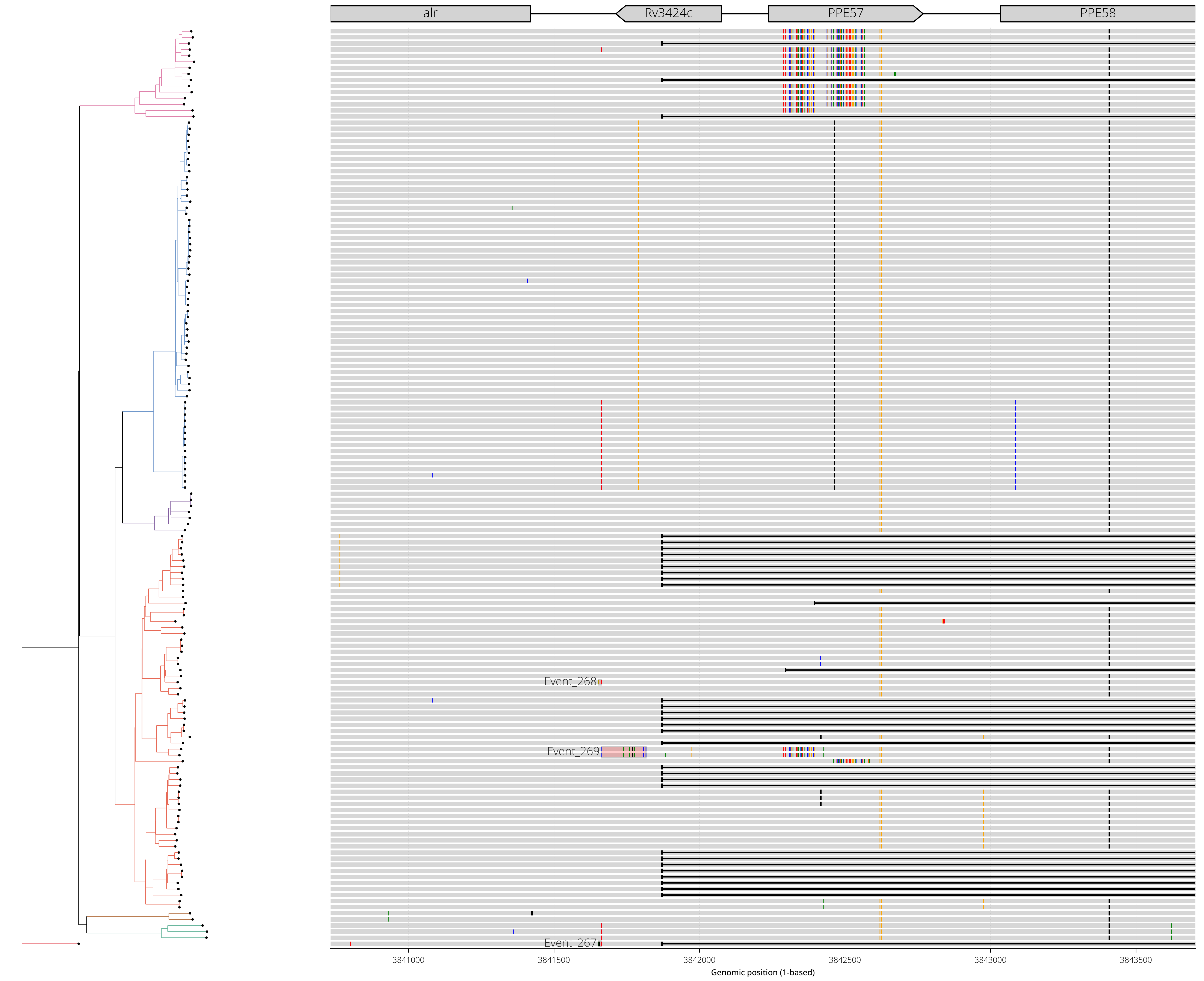

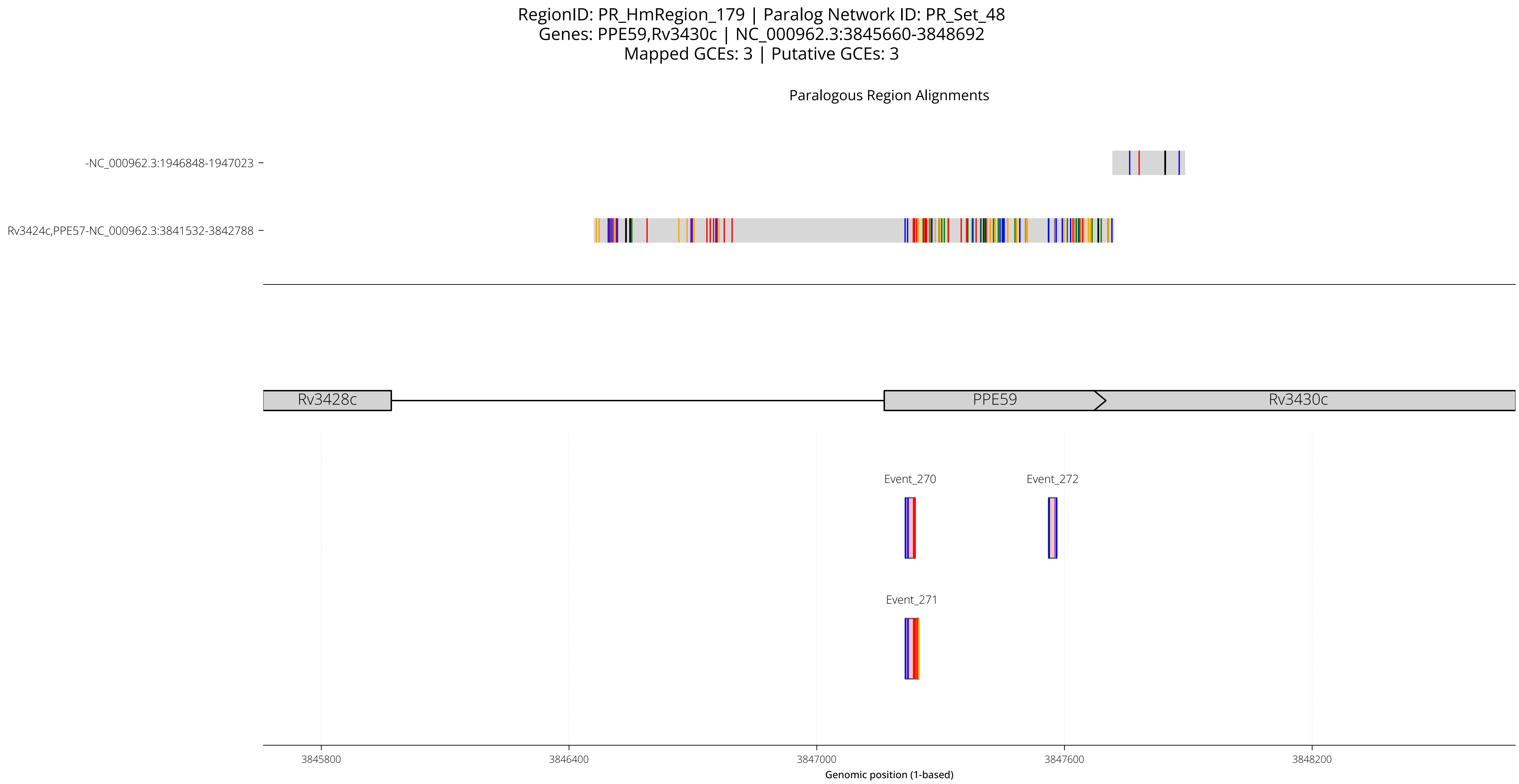
RegionID: PR_HmRegion_179 | Paralog Network ID: PR_Set_48
Genes: PPE59,Rv3430c | NC_000962.3:3845660-3848692
Mapped GCEs: 3 | Putative GCEs: 3
Paralogous Region Alignments
-NC_000962.3:1946848-1947023
Rv3424c,PPE57-NC_000962.3:3841532-3842788
Rv3428c
PPE59
Rv3430c
Event_270
Event_272
Event_271
3845800
3846400
3847000
3847600
3848200
Genomic position (1-based)

RegionID: PR_HmRegion_179 | Paralog Network ID: PR_Set_48
Genes: PPE59,Rv3430c | NC_000962.3:3845660-3848692
Mapped GCEs: 3 | Putative GCEs: 3

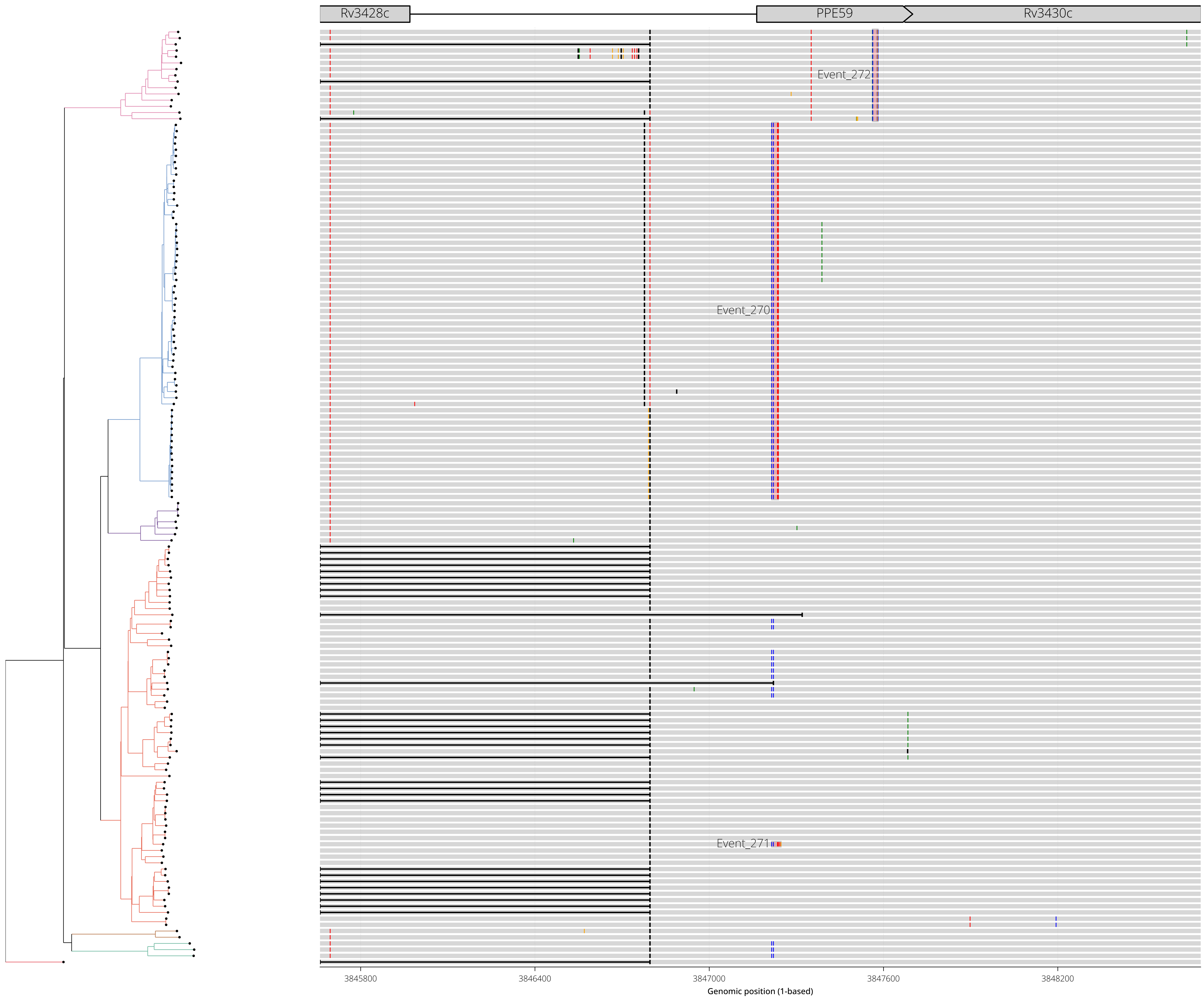

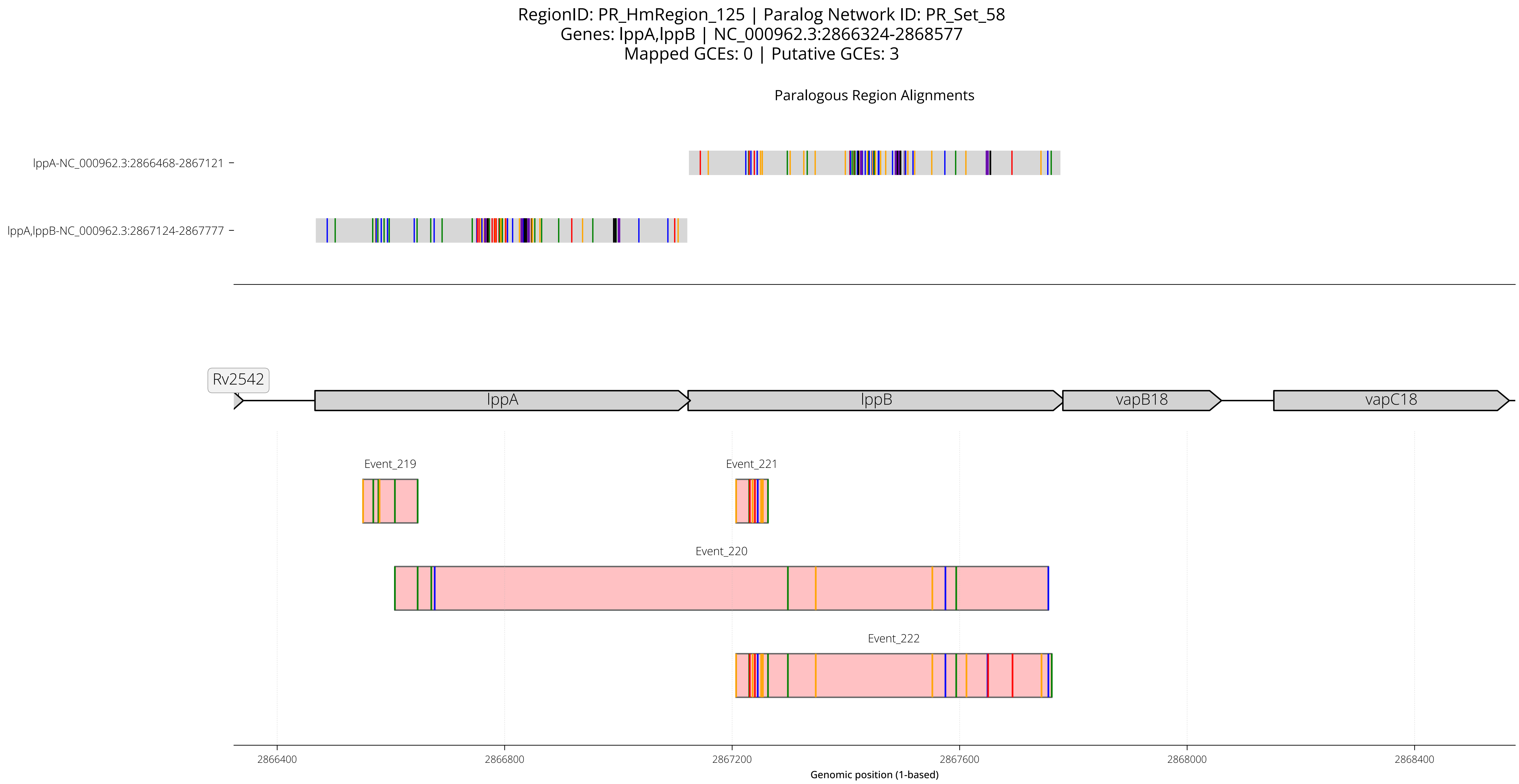
RegionID: PR_HmRegion_125 | Paralog Network ID: PR_Set_58
Genes: lppA,lppB | NC_000962.3:2866324-2868577
Mapped GCEs: 0 | Putative GCEs: 3
Paralogous Region Alignments
lppA-NC_000962.3:2866468-2867121
lppA,lppB-NC_000962.3:2867124-2867777
Rv2542
lppA
lppB
vapB18
vapC18
Event_219
Event_221
Event_220
Event_222
2866400
2866800
2867200
2867600
2868000
2868400
Genomic position (1-based)

RegionID: PR_HmRegion_125 | Paralog Network ID: PR_Set_58
Genes: lppA,lppB | NC_000962.3:2866324-2868577
Mapped GCEs: 0 | Putative GCEs: 3

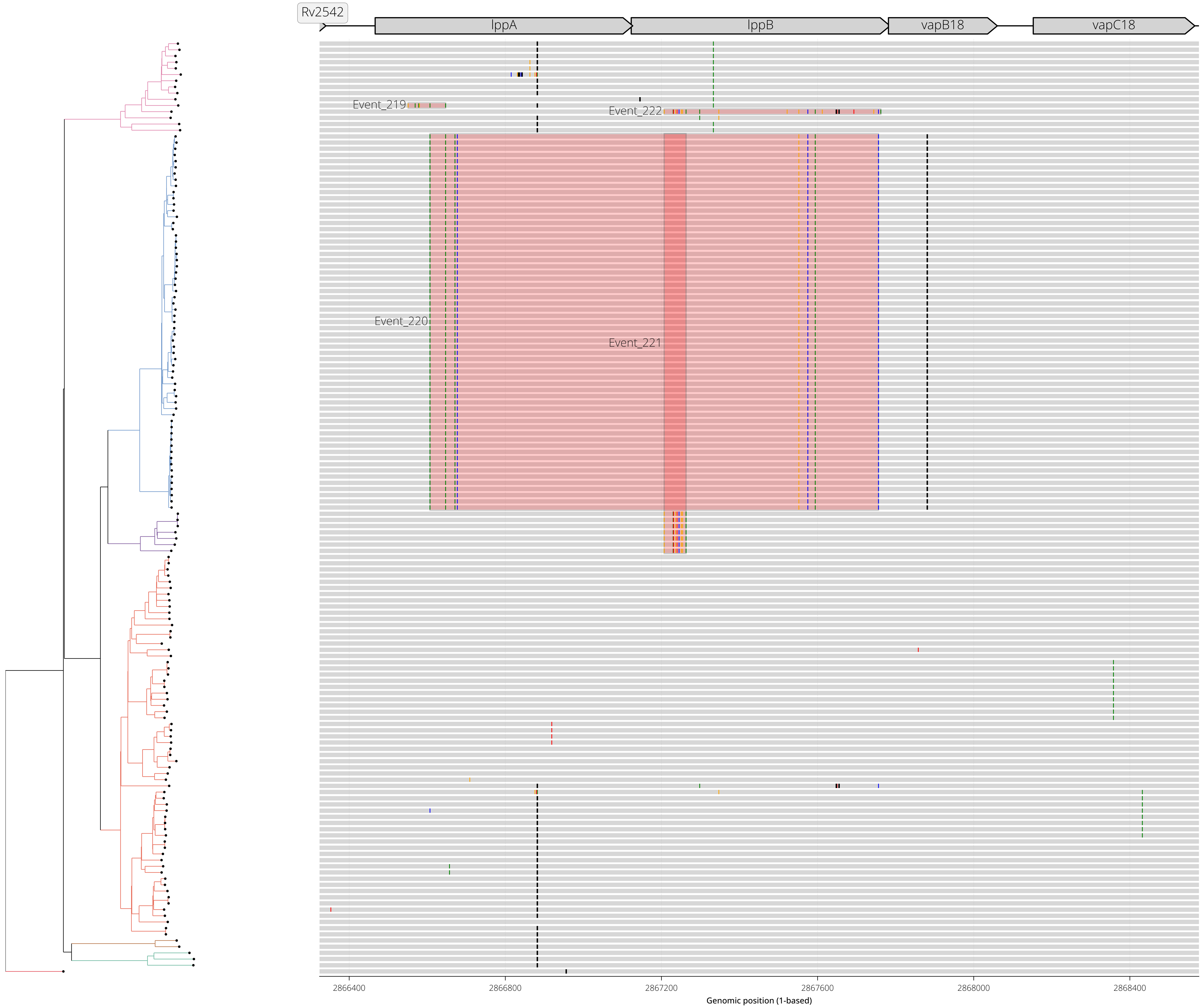

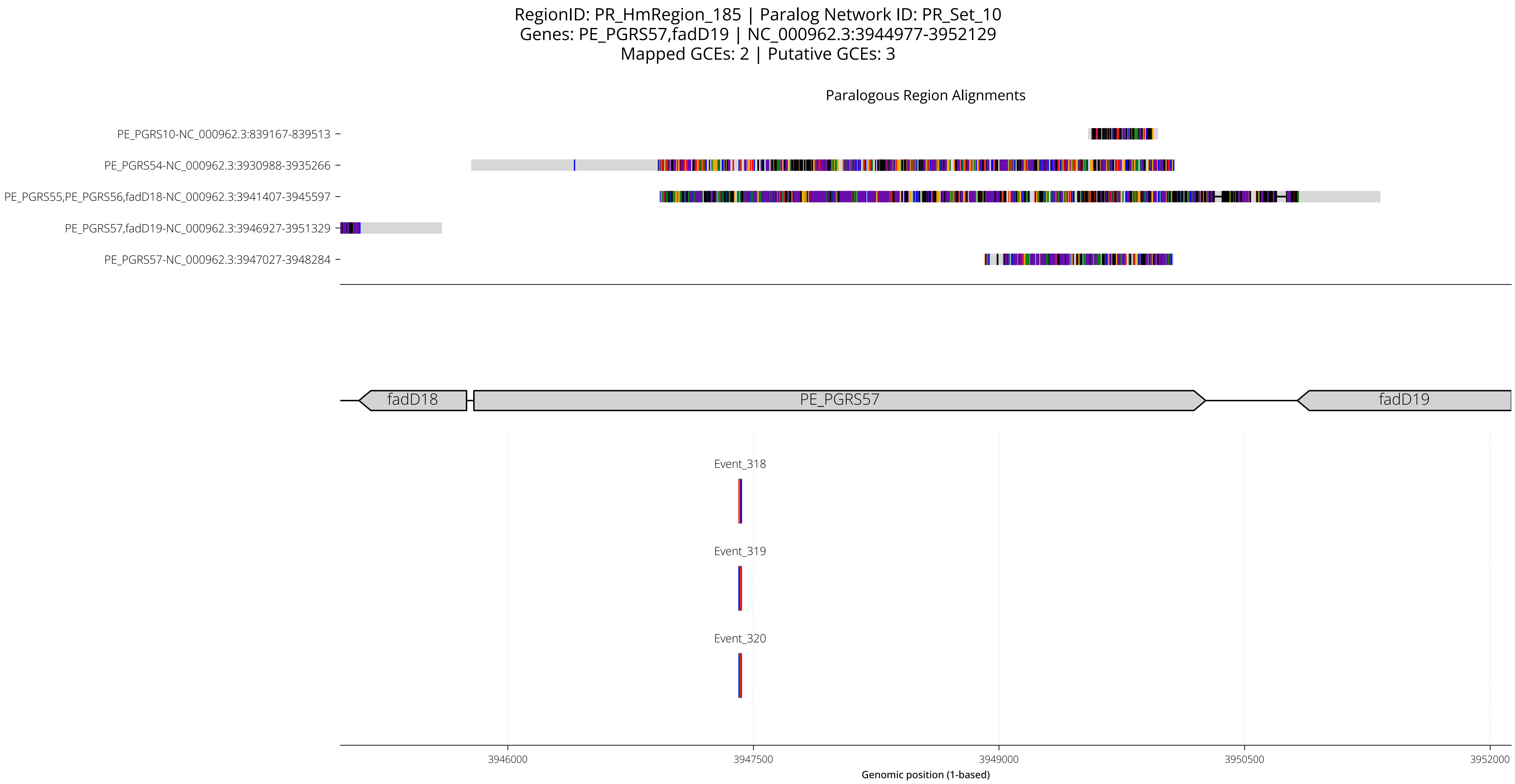
RegionID: PR_HmRegion_185 | Paralog Network ID: PR_Set_10
Genes: PE_PGRS57,fadD19 | NC_000962.3:3944977-3952129
Mapped GCEs: 2 | Putative GCEs: 3
Paralogous Region Alignments
PE_PGRS10-NC_000962.3:839167-839513
PE_PGRS54-NC_000962.3:3930988-3935266
PE_PGRS55,PE_PGRS56,fadD18-NC_000962.3:3941407-3945597
PE_PGRS57,fadD19-NC_000962.3:3946927-3951329
PE_PGRS57-NC_000962.3:3947027-3948284
fadD18
PE_PGRS57
fadD19
Event_318
Event_319
Event_320
3946000
3947500
3949000
3950500
3952000
Genomic position (1-based)

RegionID: PR_HmRegion_185 | Paralog Network ID: PR_Set_10
Genes: PE_PGRS57,fadD19 | NC_000962.3:3944977-3952129
Mapped GCEs: 2 | Putative GCEs: 3

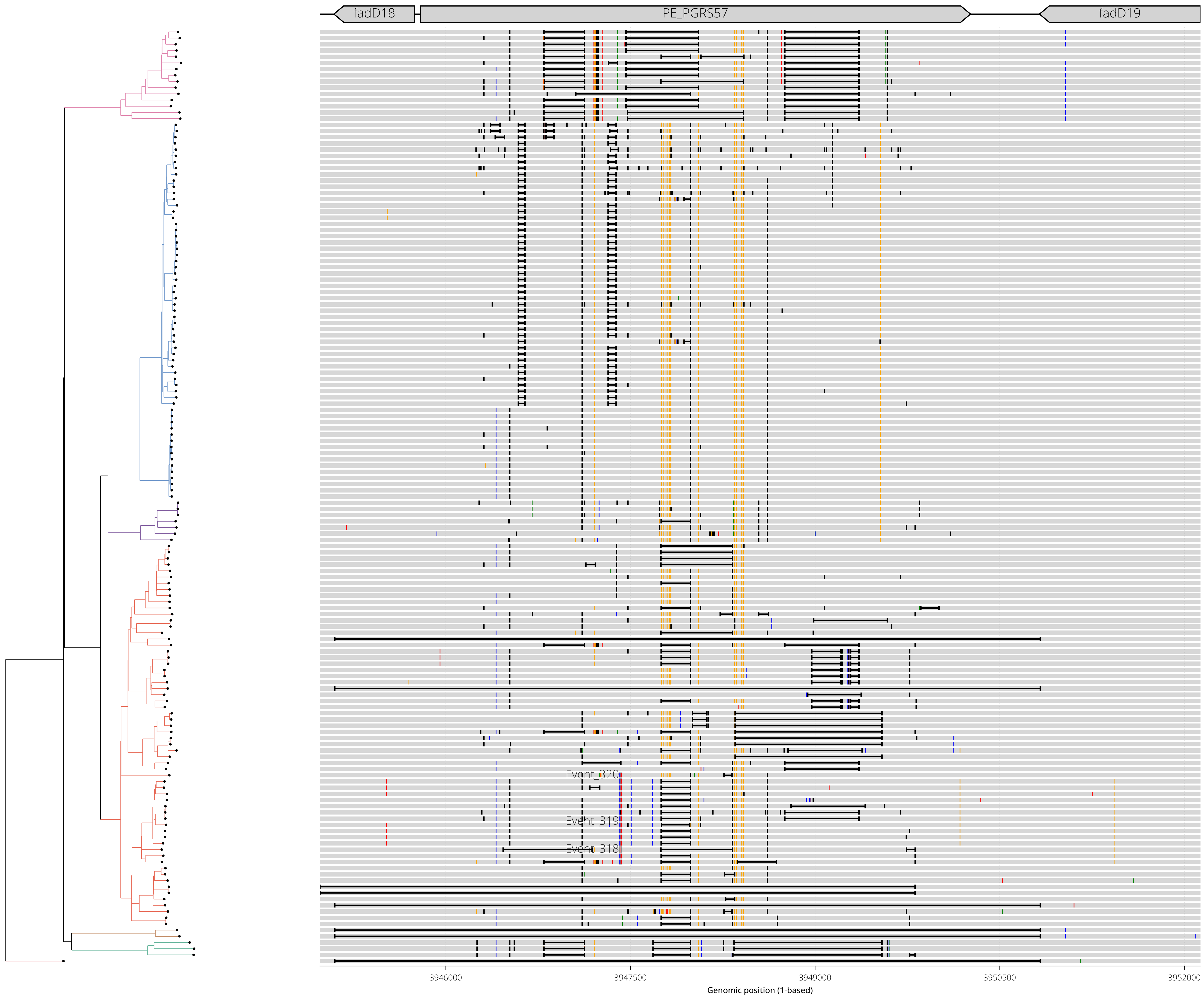

RegionID: PR_HmRegion_033 | Paralog Network ID: PR_Set_22
Genes: Rv0750 | NC_000962.3:841219-843131
Mapped GCEs: 3 | Putative GCEs: 3

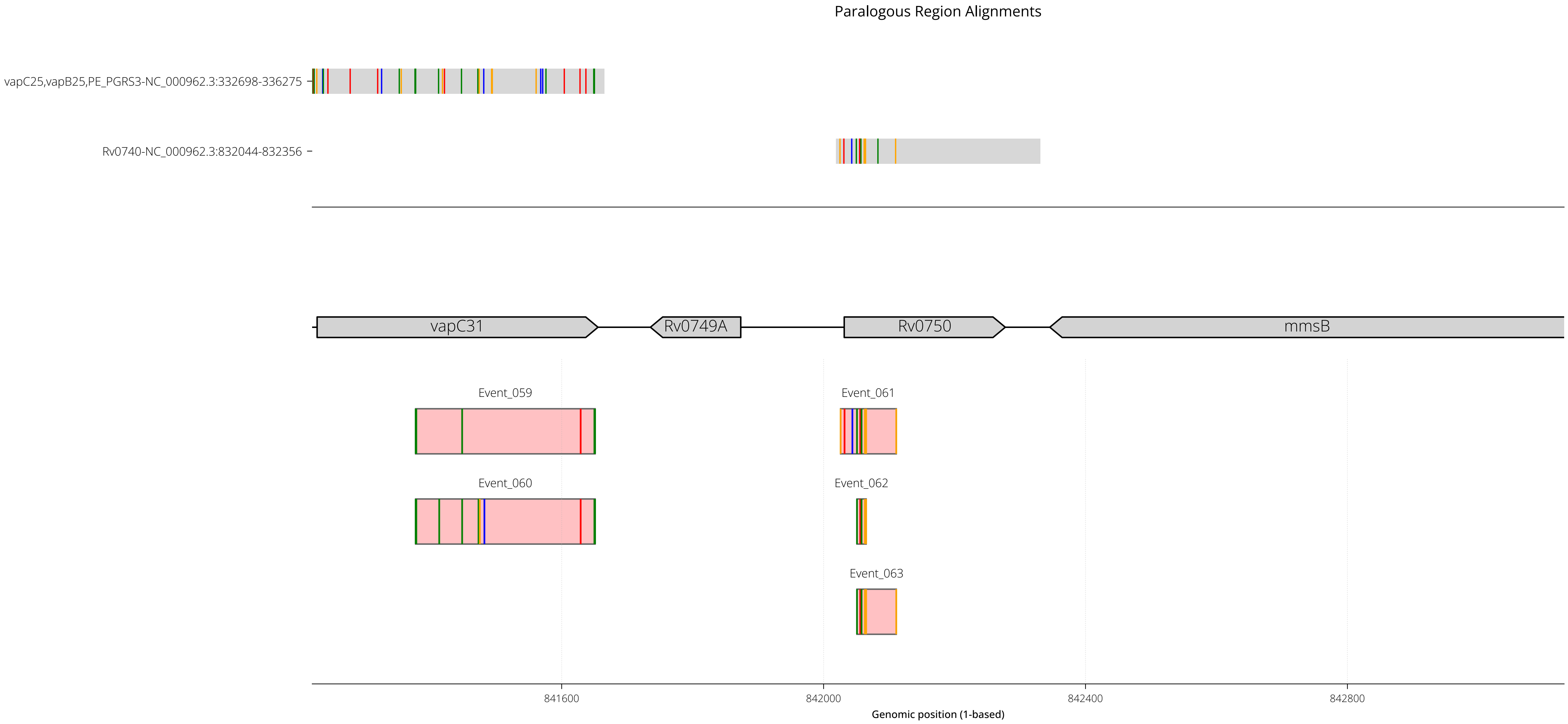

RegionID: PR_HmRegion_033 | Paralog Network ID: PR_Set_22
Genes: Rv0750 | NC_000962.3:841219-843131
Mapped GCEs: 3 | Putative GCEs: 3

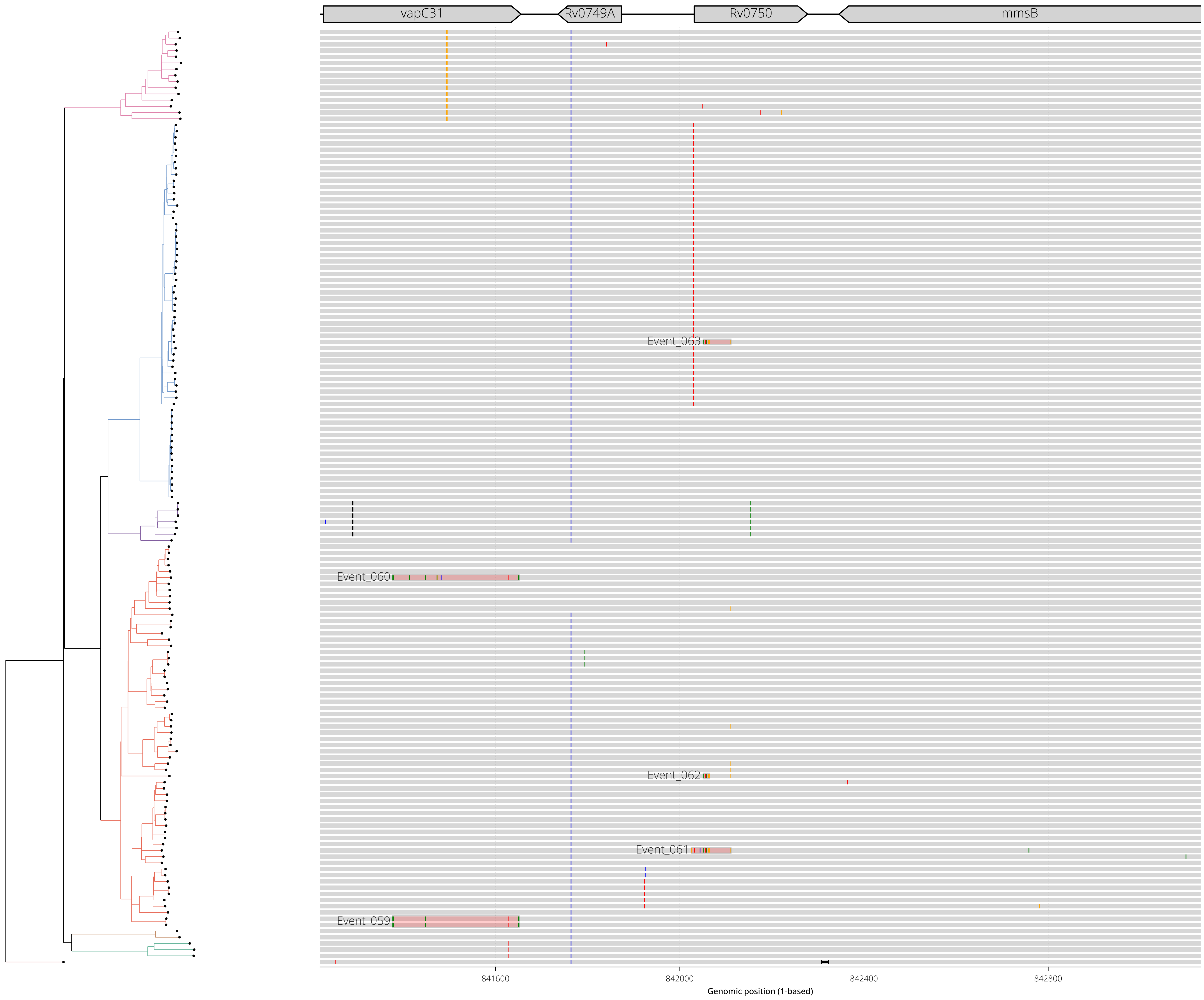

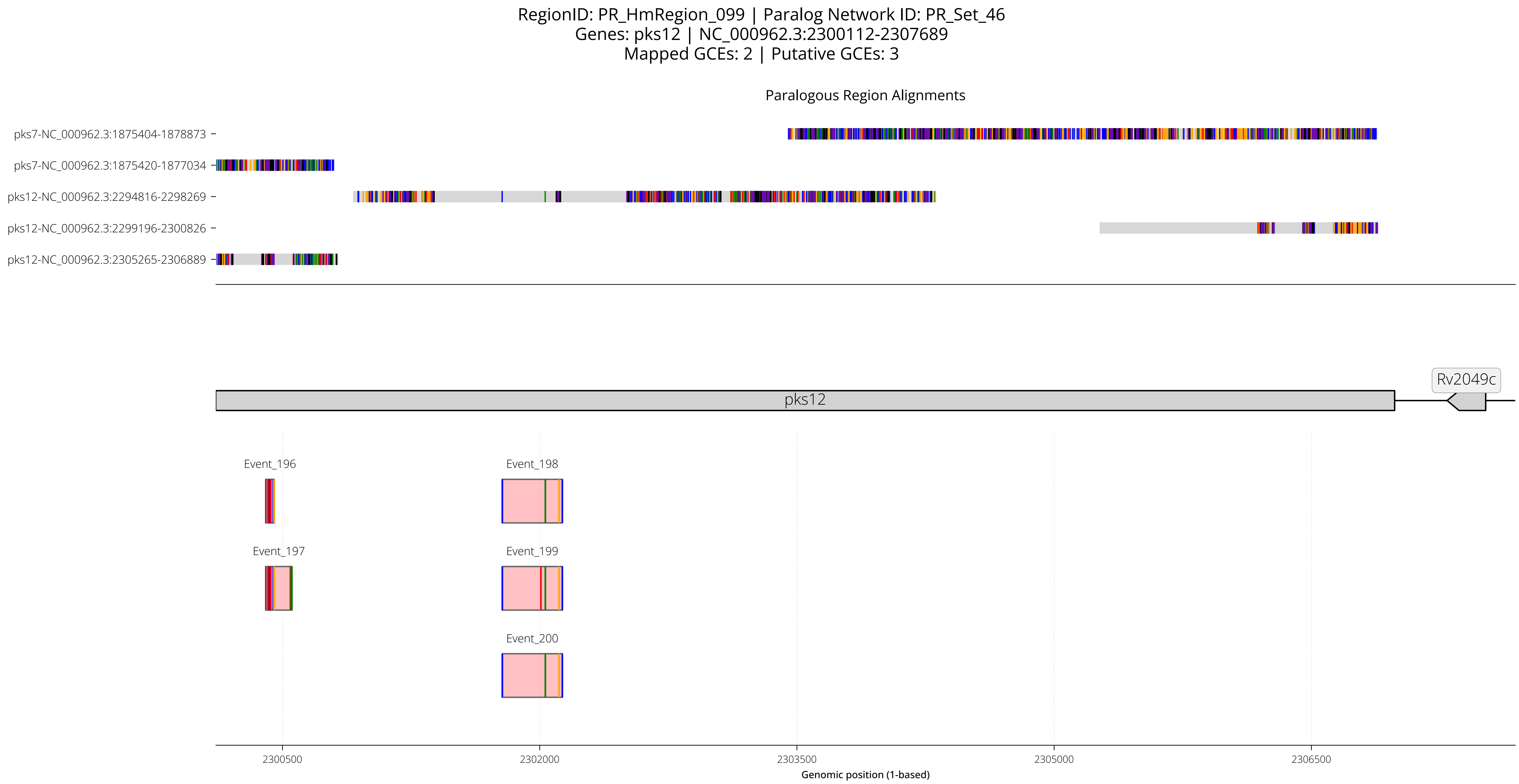
RegionID: PR_HmRegion_099 | Paralog Network ID: PR_Set_46
Genes: pks12 | NC_000962.3:2300112-2307689
Mapped GCEs: 2 | Putative GCEs: 3
Paralogous Region Alignments
pks7-NC_000962.3:1875404-1878873
pks7-NC_000962.3:1875420-1877034
pks12-NC_000962.3:2294816-2298269
pks12-NC_000962.3:2299196-2300826
pks12-NC_000962.3:2305265-2306889
Rv2049c
pks12
Event_196
Event_198
Event_197
Event_199
Event_200
2300500
2302000
2303500
2305000
2306500
Genomic position (1-based)

RegionID: PR_HmRegion_099 | Paralog Network ID: PR_Set_46
Genes: pks12 | NC_000962.3:2300112-2307689
Mapped GCEs: 2 | Putative GCEs: 3

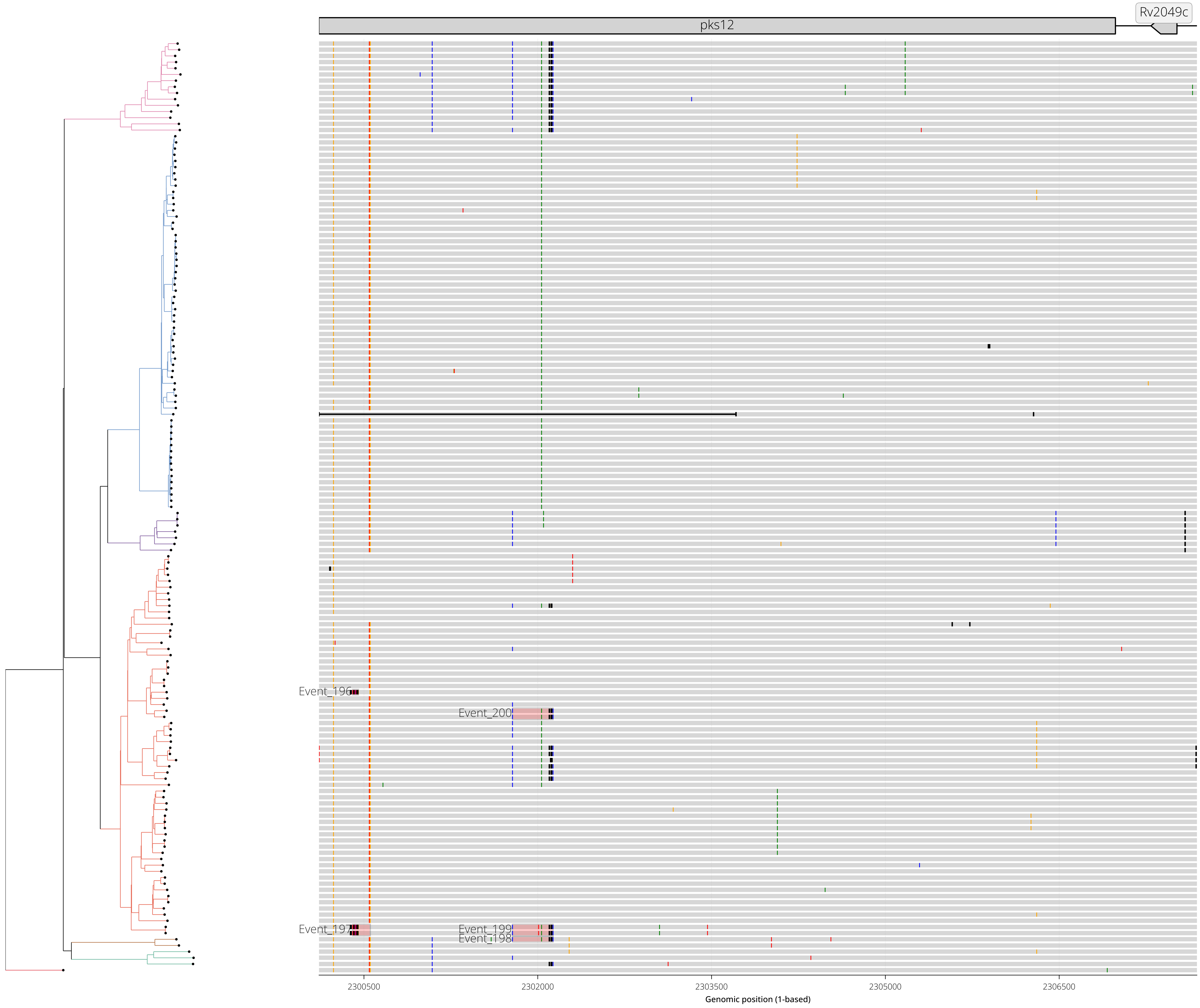

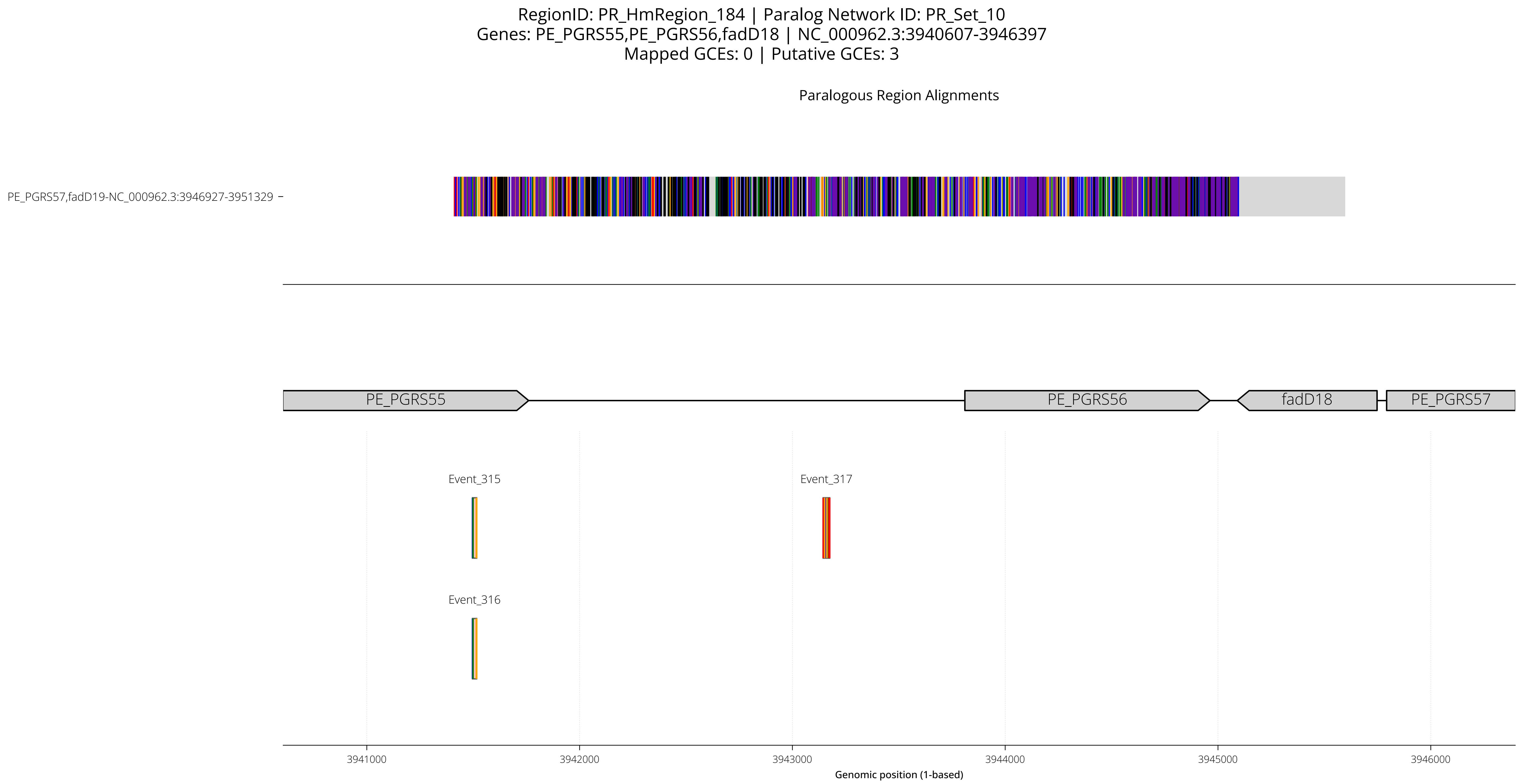
RegionID: PR_HmRegion_184 | Paralog Network ID: PR_Set_10
Genes: PE_PGRS55,PE_PGRS56,fadD18 | NC_000962.3:3940607-3946397
Mapped GCEs: 0 | Putative GCEs: 3
Paralogous Region Alignments
PE_PGRS57,fadD19-NC_000962.3:3946927-3951329
PE_PGRS55
PE_PGRS56
fadD18
PE_PGRS57
Event_315
Event_317
Event_316
3941000
3942000
3943000
3944000
3945000
3946000
Genomic position (1-based)

RegionID: PR_HmRegion_184 | Paralog Network ID: PR_Set_10
Genes: PE_PGRS55,PE_PGRS56,fadD18 | NC_000962.3:3940607-3946397
Mapped GCEs: 0 | Putative GCEs: 3

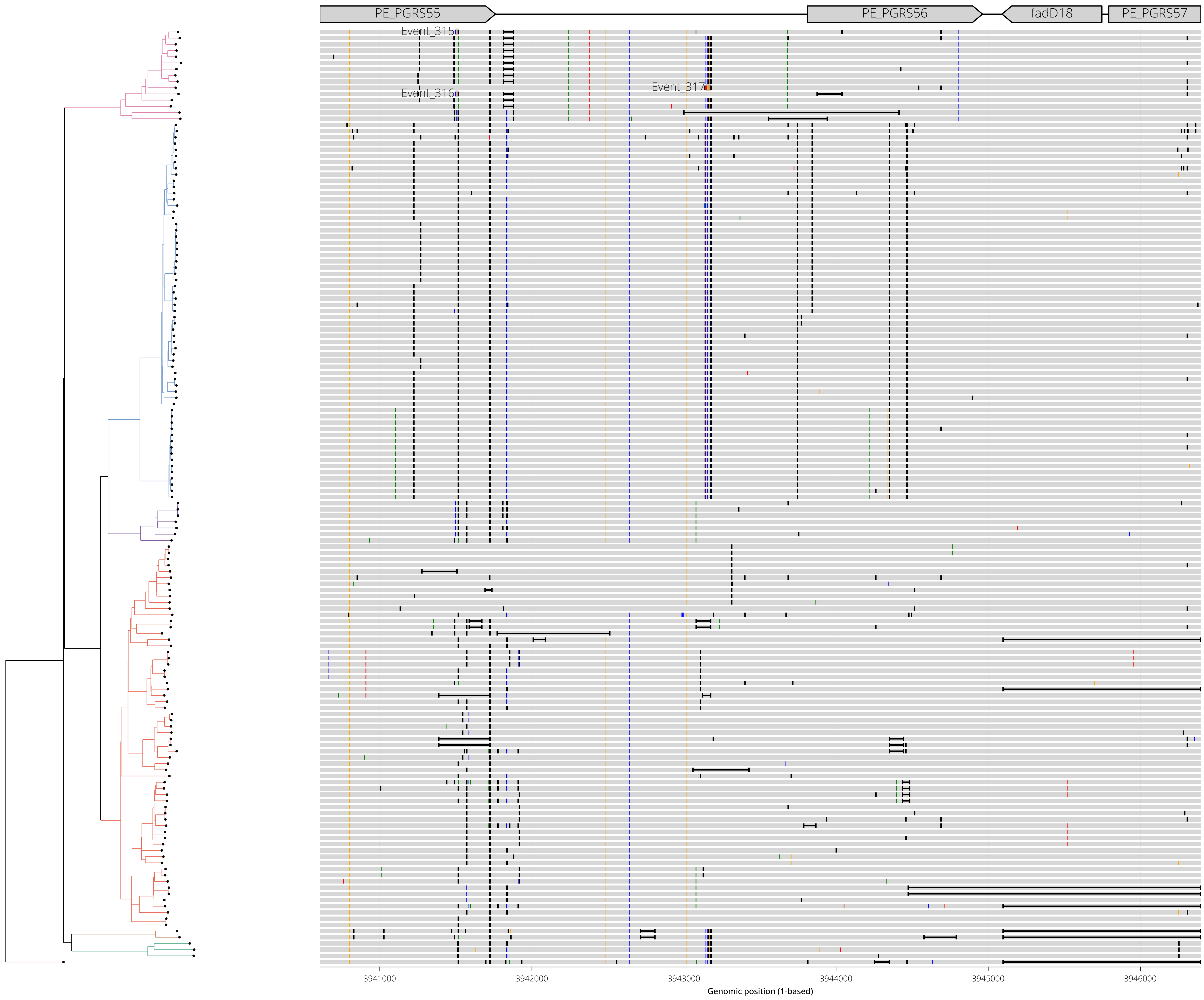

RegionID: PR_HmRegion_098 | Paralog Network ID: PR_Set_46
Genes: pks12 | NC_000962.3:2298396-2301626
Mapped GCEs: 2 | Putative GCEs: 2

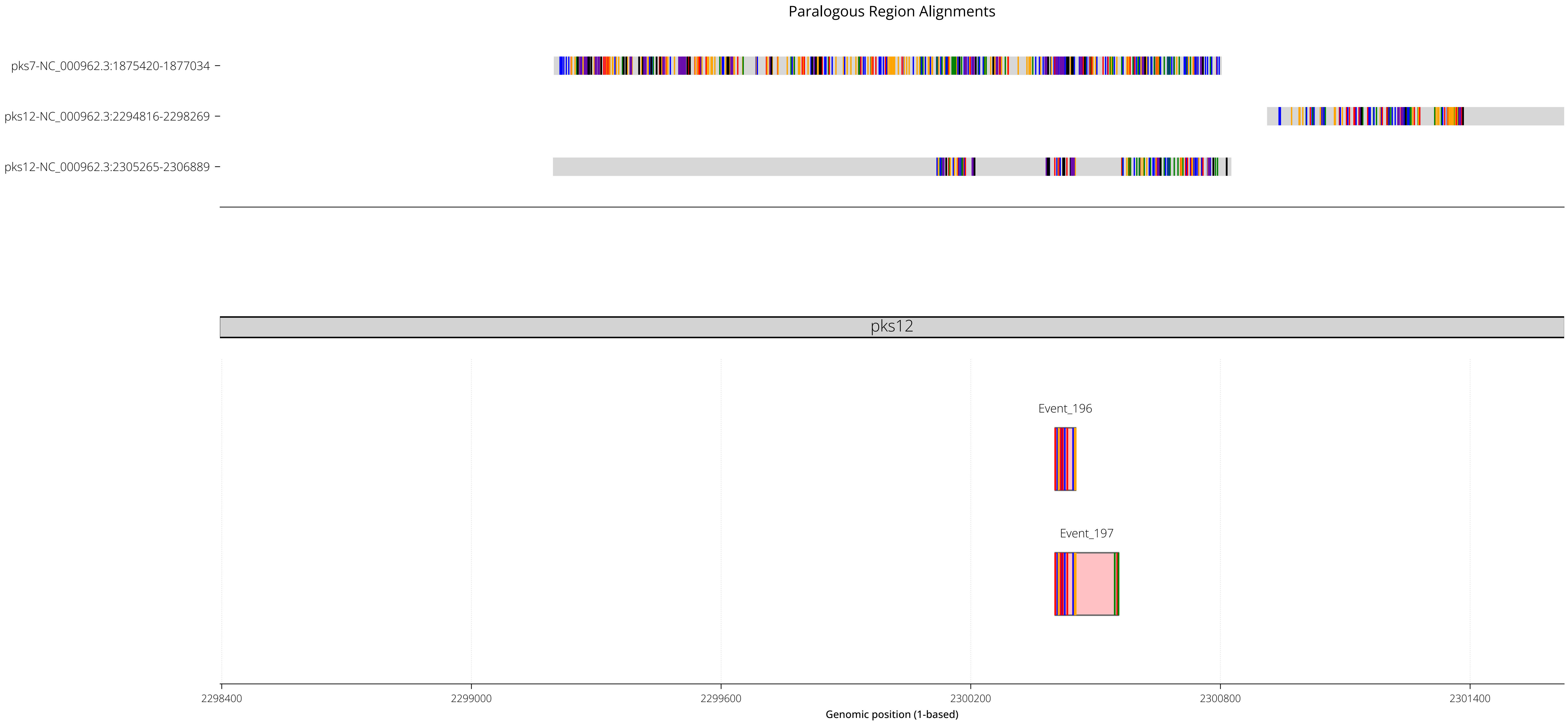

RegionID: PR_HmRegion_098 | Paralog Network ID: PR_Set_46
Genes: pks12 | NC_000962.3:2298396-2301626
Mapped GCEs: 2 | Putative GCEs: 2

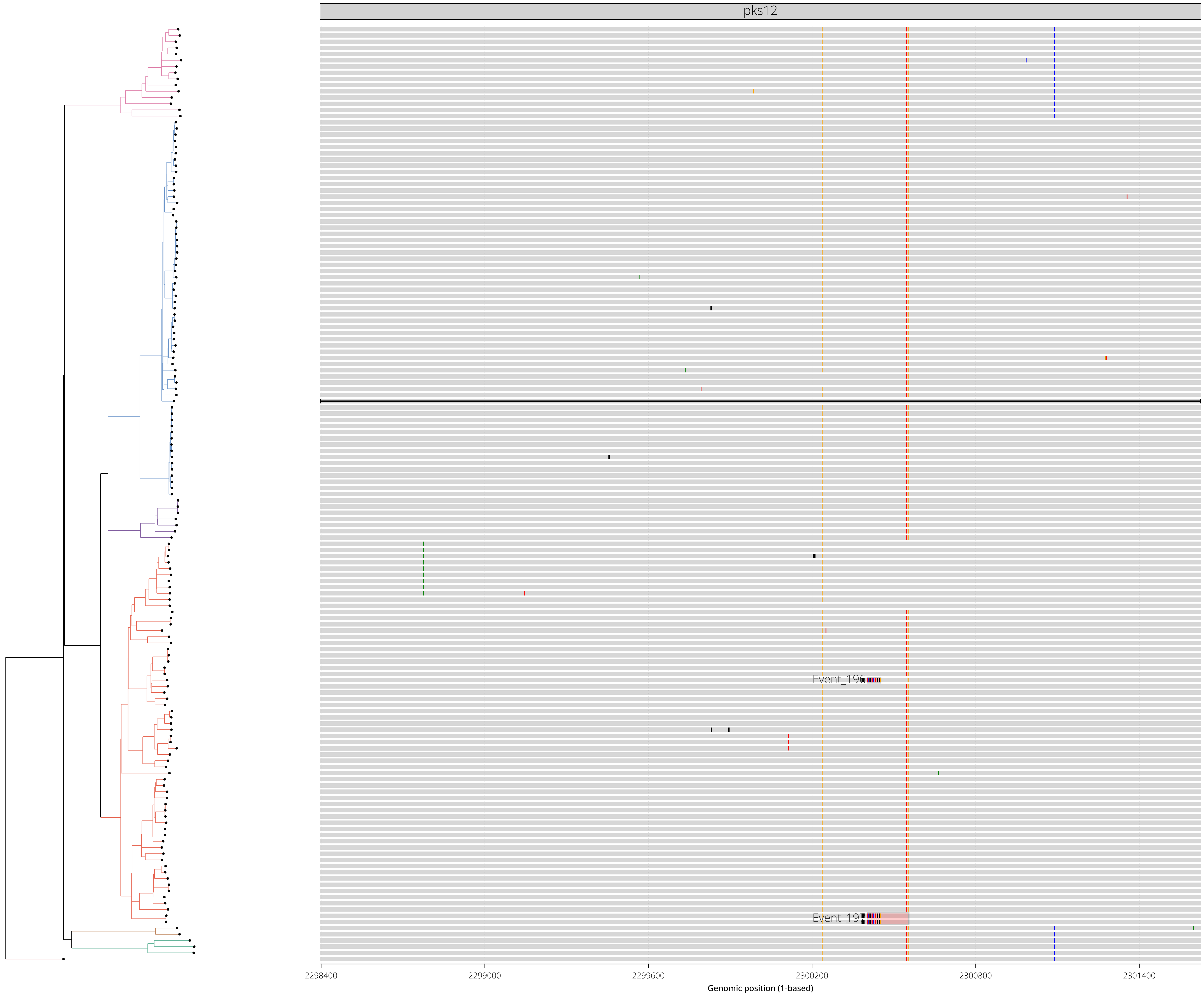

RegionID: PR_HmRegion_097 | Paralog Network ID: PR_Set_46
Genes: pks12 | NC_000962.3:2294016-2299069
Mapped GCEs: 2 | Putative GCEs: 2

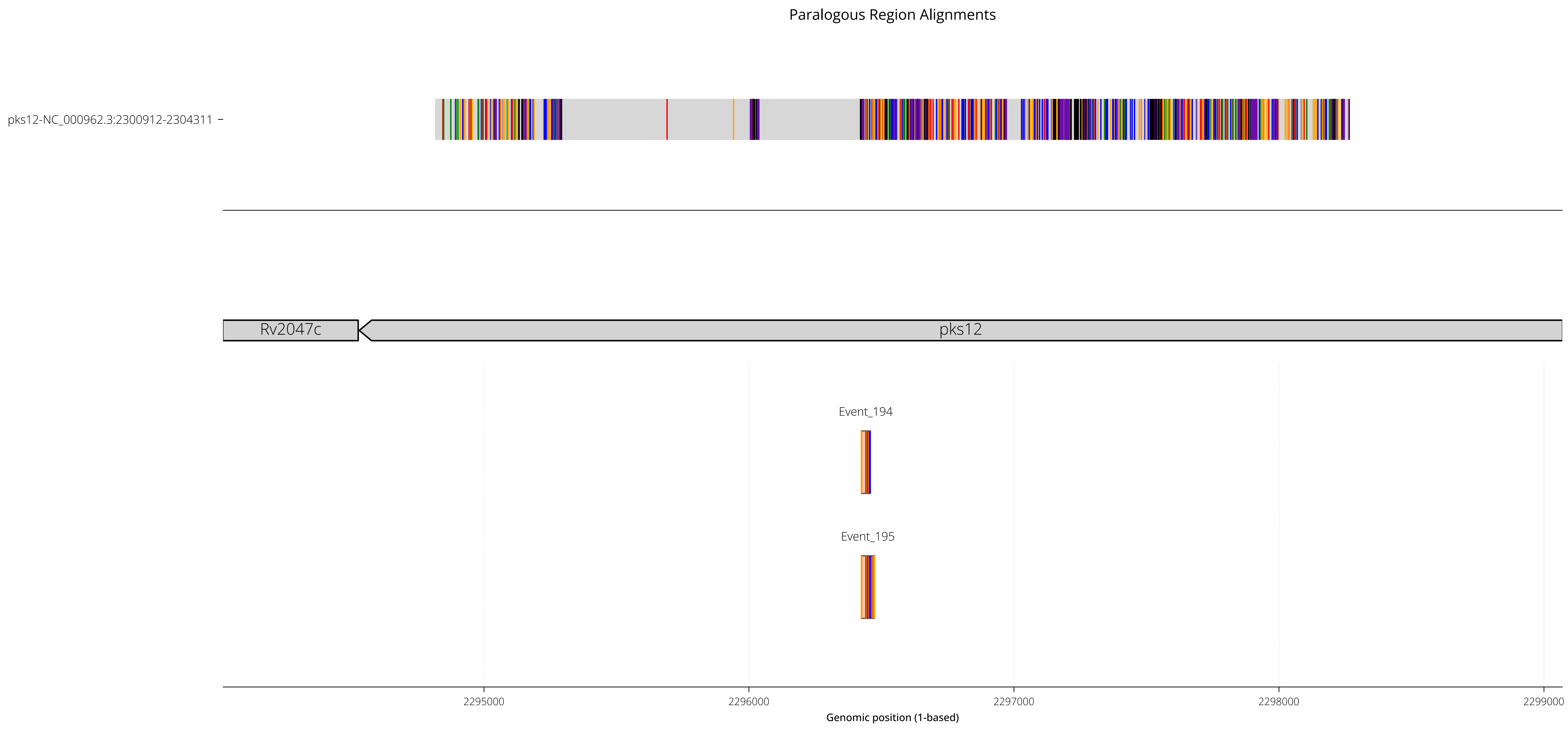

RegionID: PR_HmRegion_097 | Paralog Network ID: PR_Set_46
Genes: pks12 | NC_000962.3:2294016-2299069
Mapped GCEs: 2 | Putative GCEs: 2

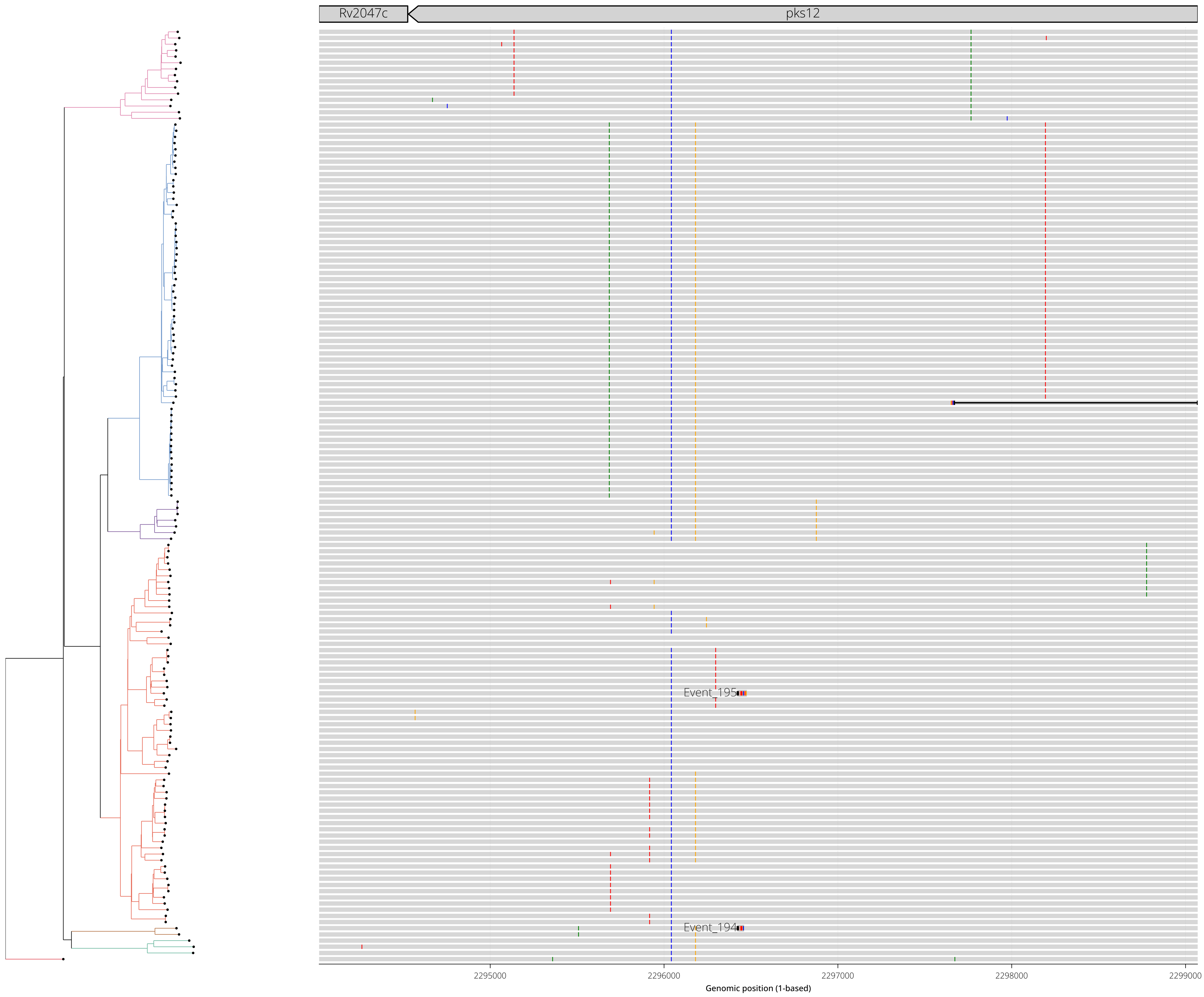

RegionID: PR_HmRegion_036 | Paralog Network ID: PR_Set_20
Genes: Rv0828c,Rv0829 | NC_000962.3:920779-922693
Mapped GCEs: 2 | Putative GCEs: 2

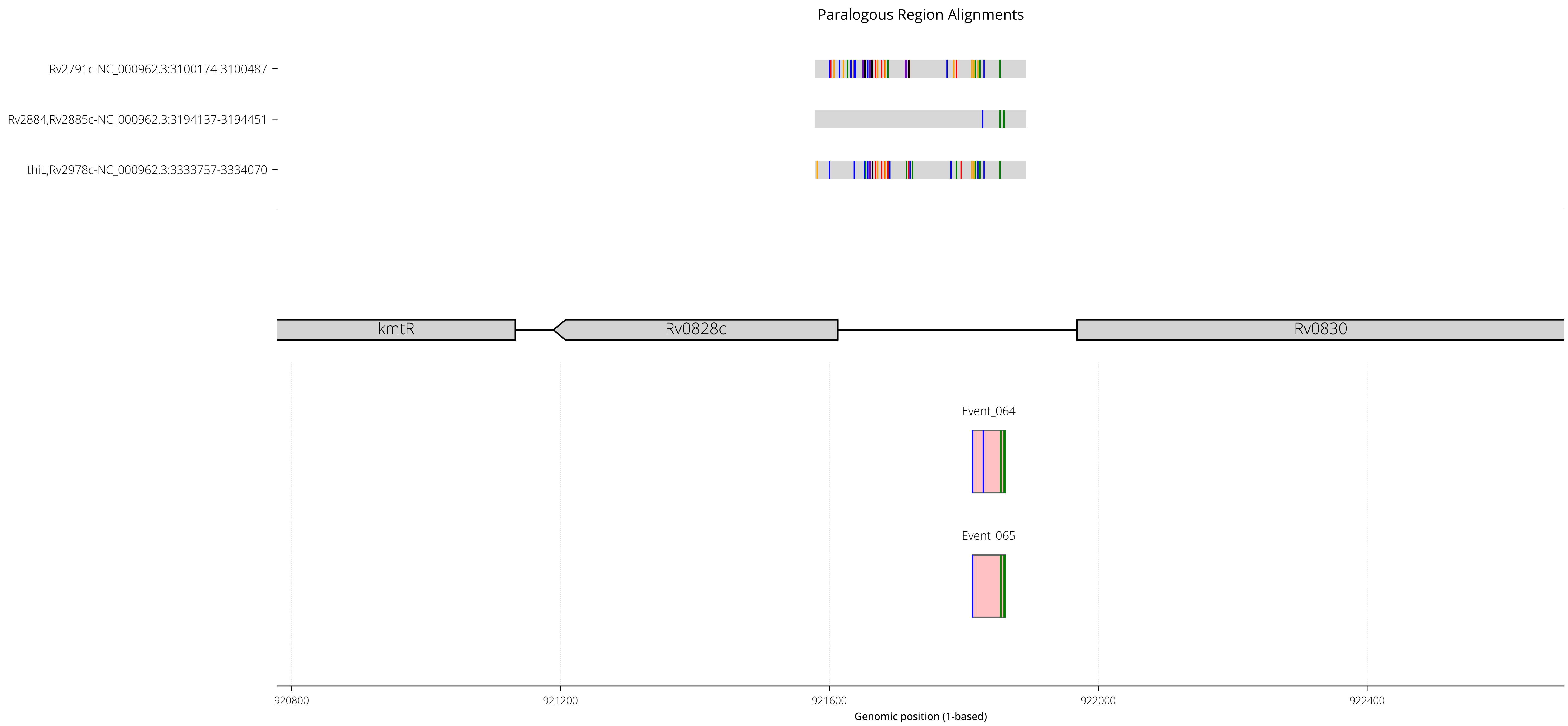

RegionID: PR_HmRegion_036 | Paralog Network ID: PR_Set_20
Genes: Rv0828c,Rv0829 | NC_000962.3:920779-922693
Mapped GCEs: 2 | Putative GCEs: 2

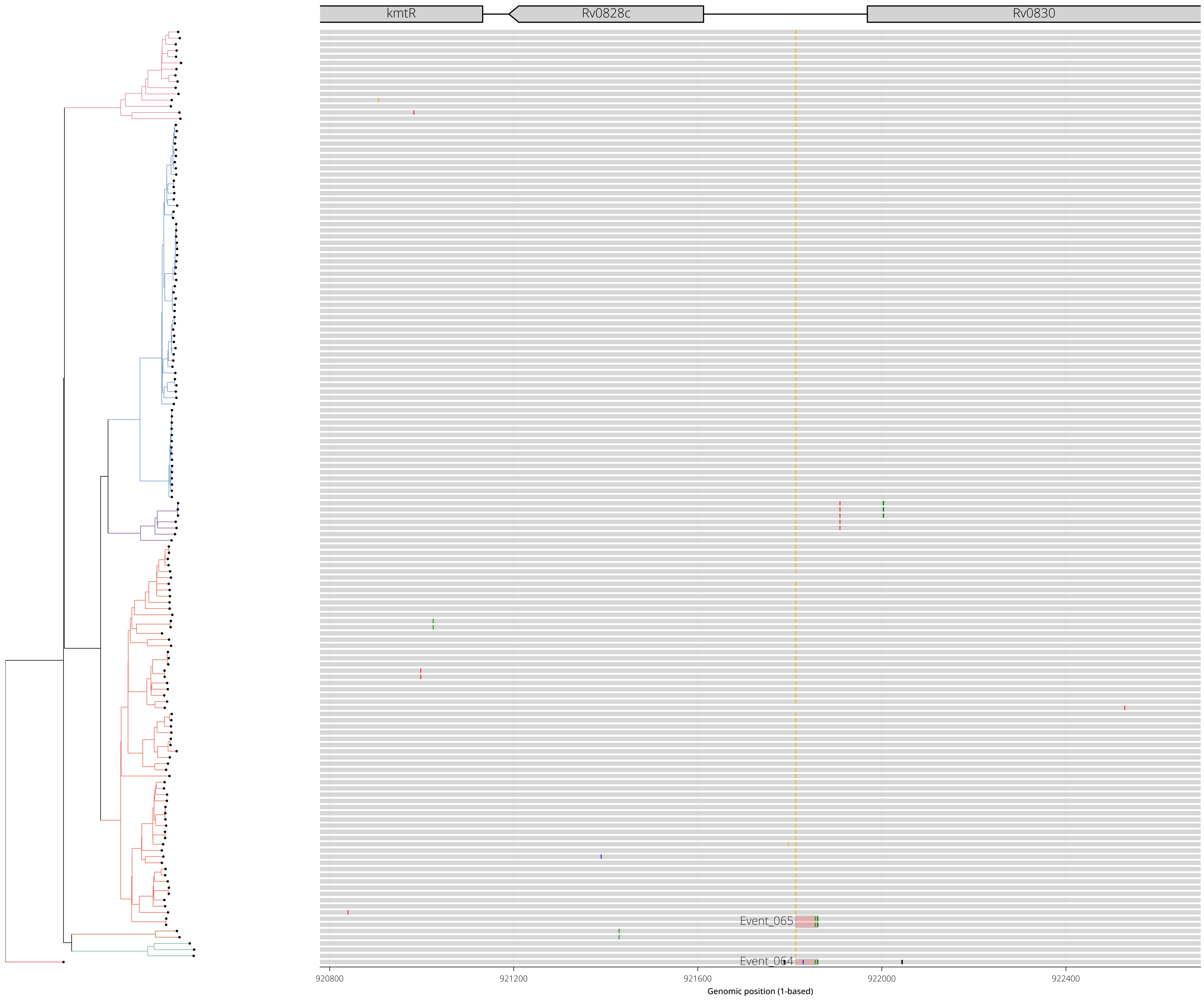

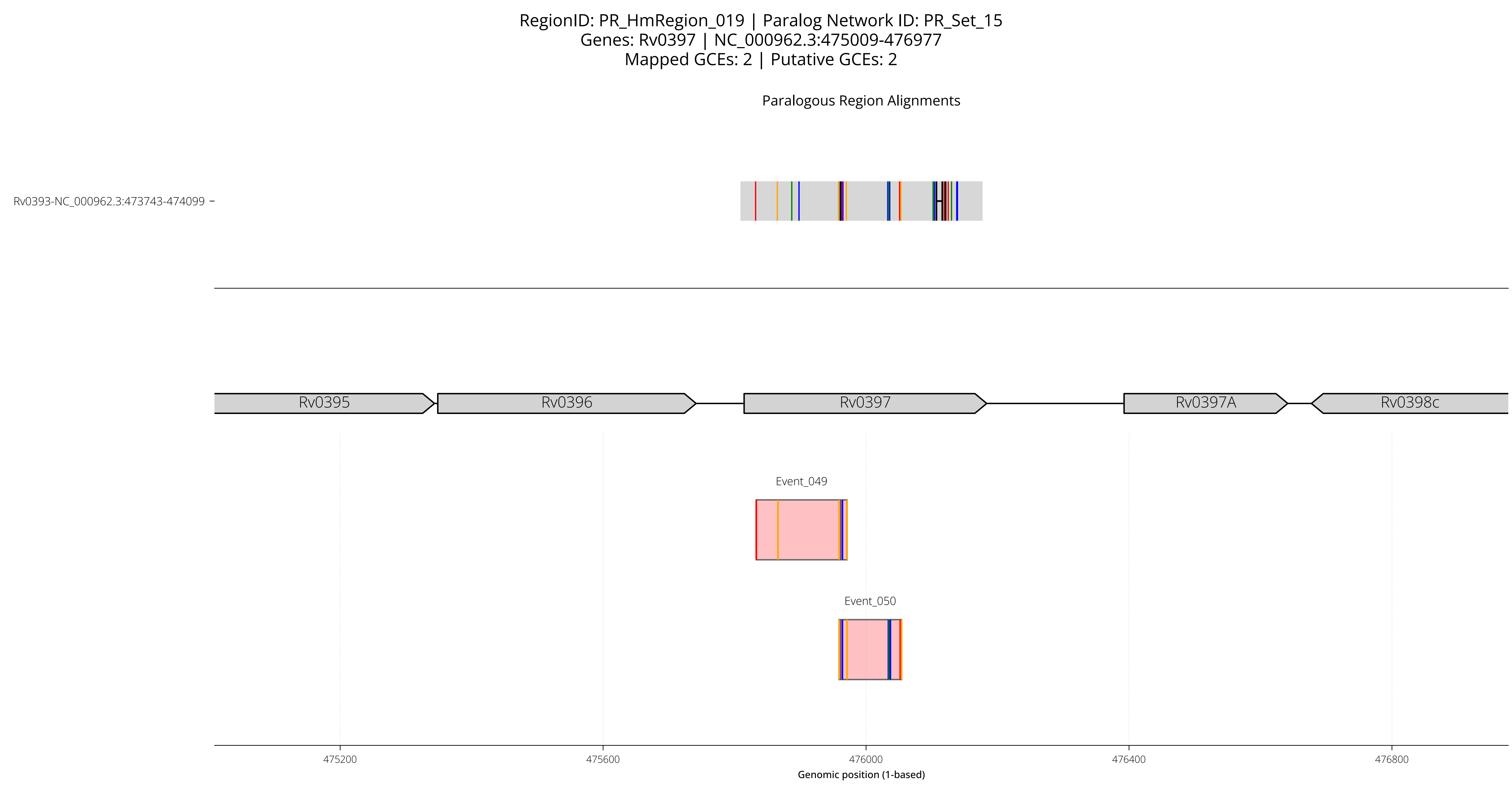

RegionID: PR_HmRegion_019 | Paralog Network ID: PR_Set_15
Genes: Rv0397 | NC_000962.3:475009-476977
Mapped GCEs: 2 | Putative GCEs: 2
Paralogous Region Alignments
Rv0393-NC_000962.3:473743-474099
Rv0395
Rv0396
Rv0397
Rv0397A
Rv0398c
Event_049
Event_050
475200
475600
476000
476400
476800
Genomic position (1-based)

RegionID: PR_HmRegion_019 | Paralog Network ID: PR_Set_15
Genes: Rv0397 | NC_000962.3:475009-476977
Mapped GCEs: 2 | Putative GCEs: 2

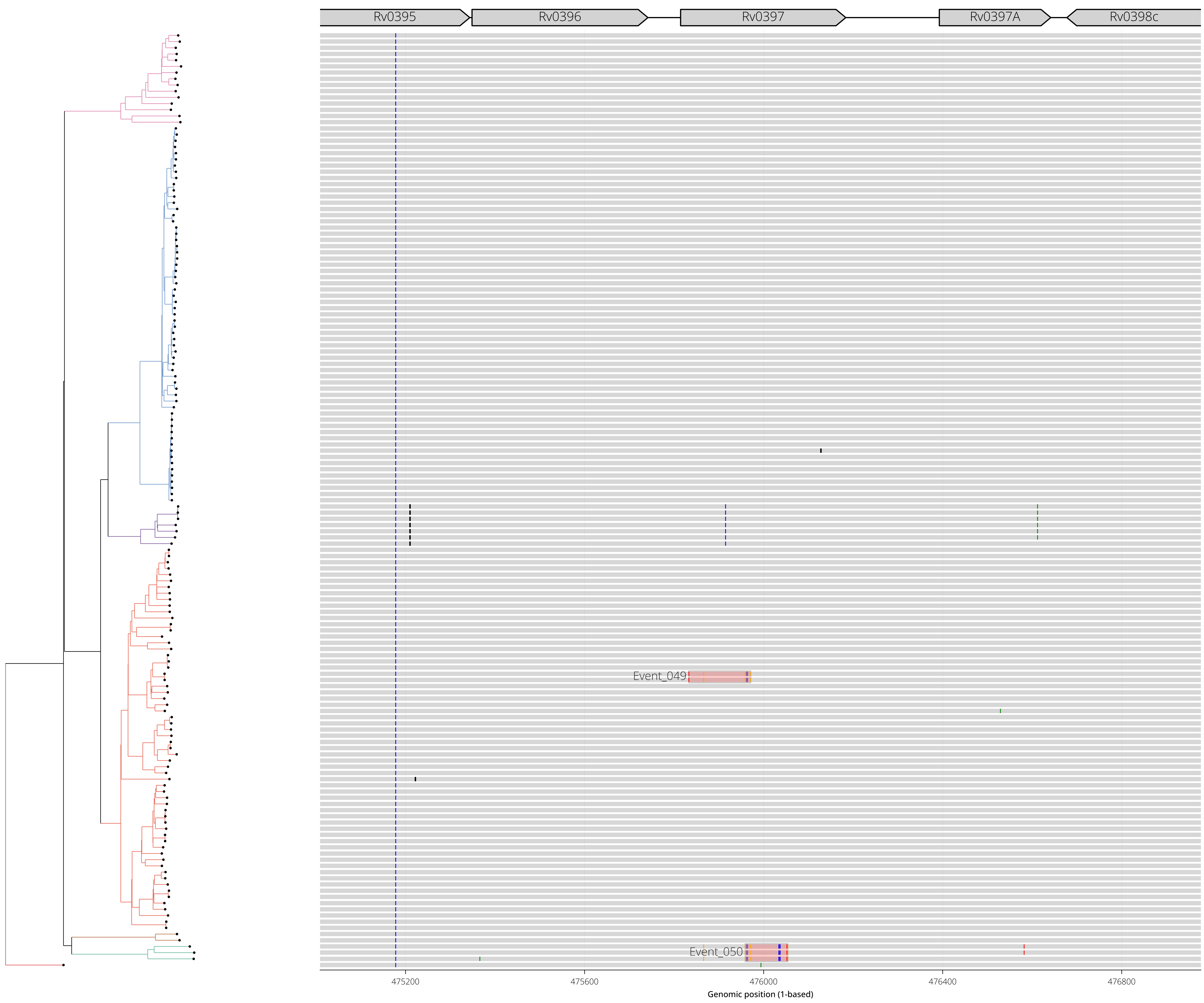

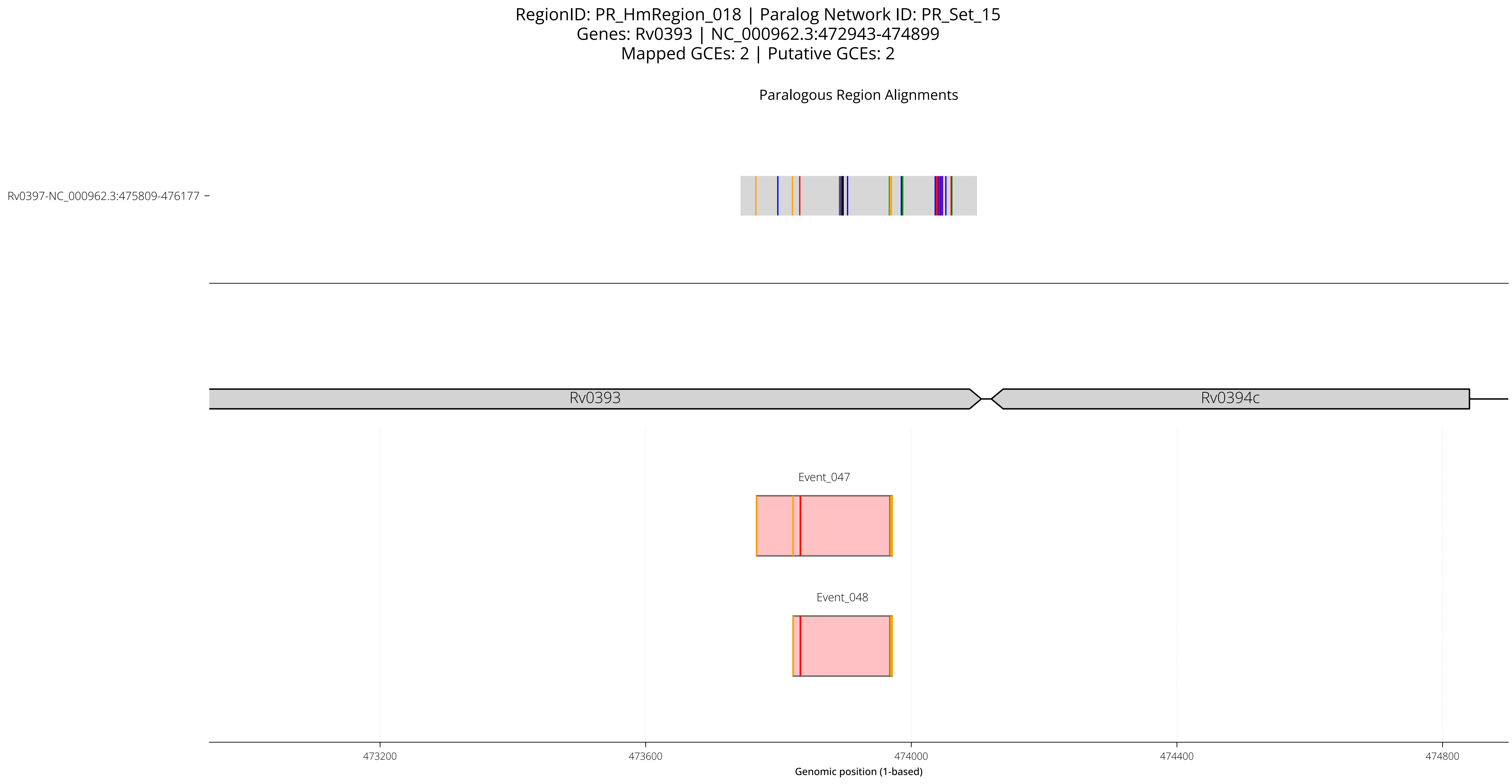

RegionID: PR_HmRegion_018 | Paralog Network ID: PR_Set_15
Genes: Rv0393 | NC_000962.3:472943-474899
Mapped GCEs: 2 | Putative GCEs: 2
Paralogous Region Alignments
Rv0397-NC_000962.3:475809-476177
Rv0393
Rv0394c
Event_047
Event_048
473200
473600
474000
474400
474800
Genomic position (1-based)

RegionID: PR_HmRegion_018 | Paralog Network ID: PR_Set_15
Genes: Rv0393 | NC_000962.3:472943-474899
Mapped GCEs: 2 | Putative GCEs: 2

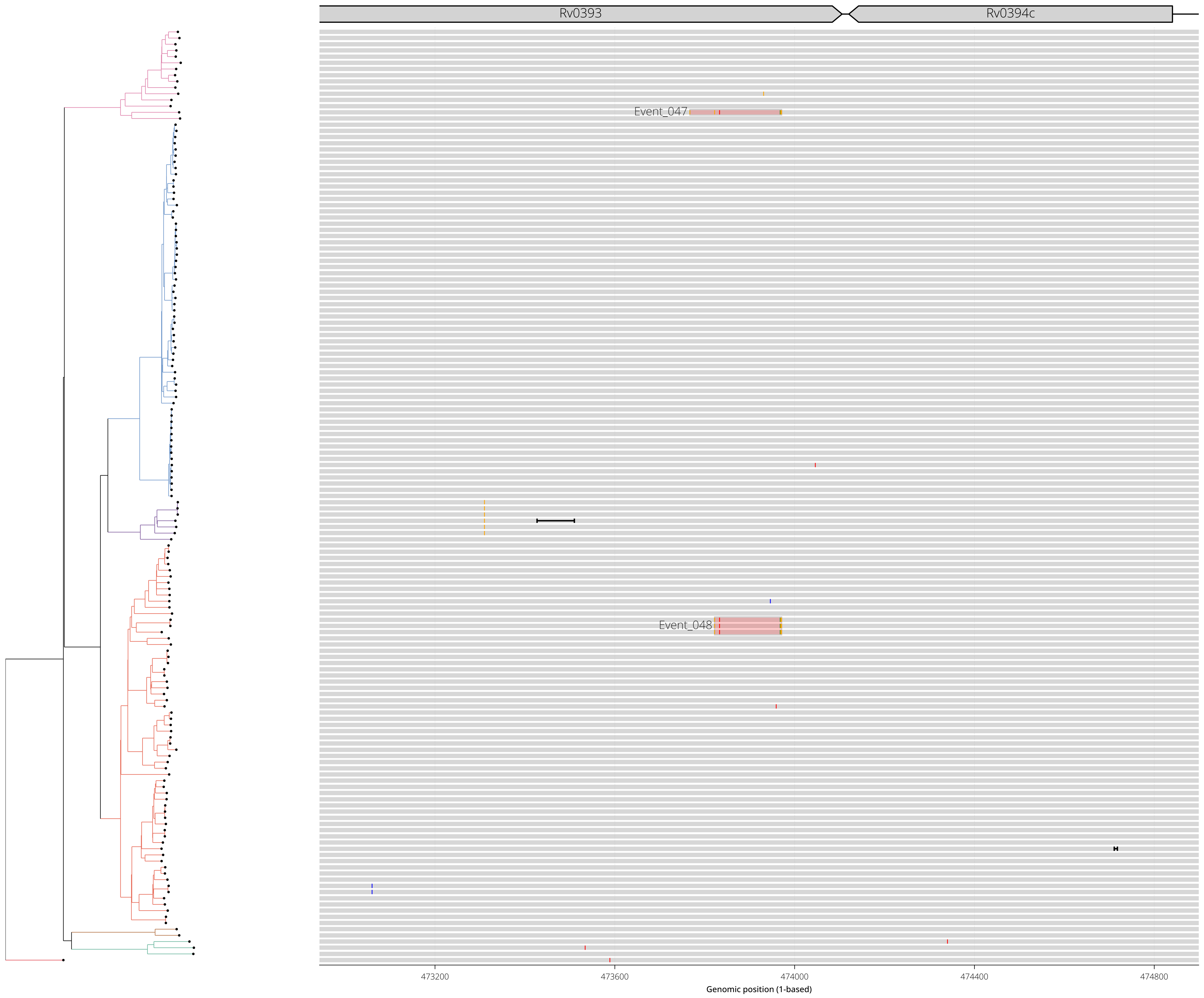

RegionID: PR_HmRegion_085 | Paralog Network ID: PR_Set_49
Genes: PPE25 | NC_000962.3:2024501-2026963
Mapped GCEs: 2 | Putative GCEs: 2

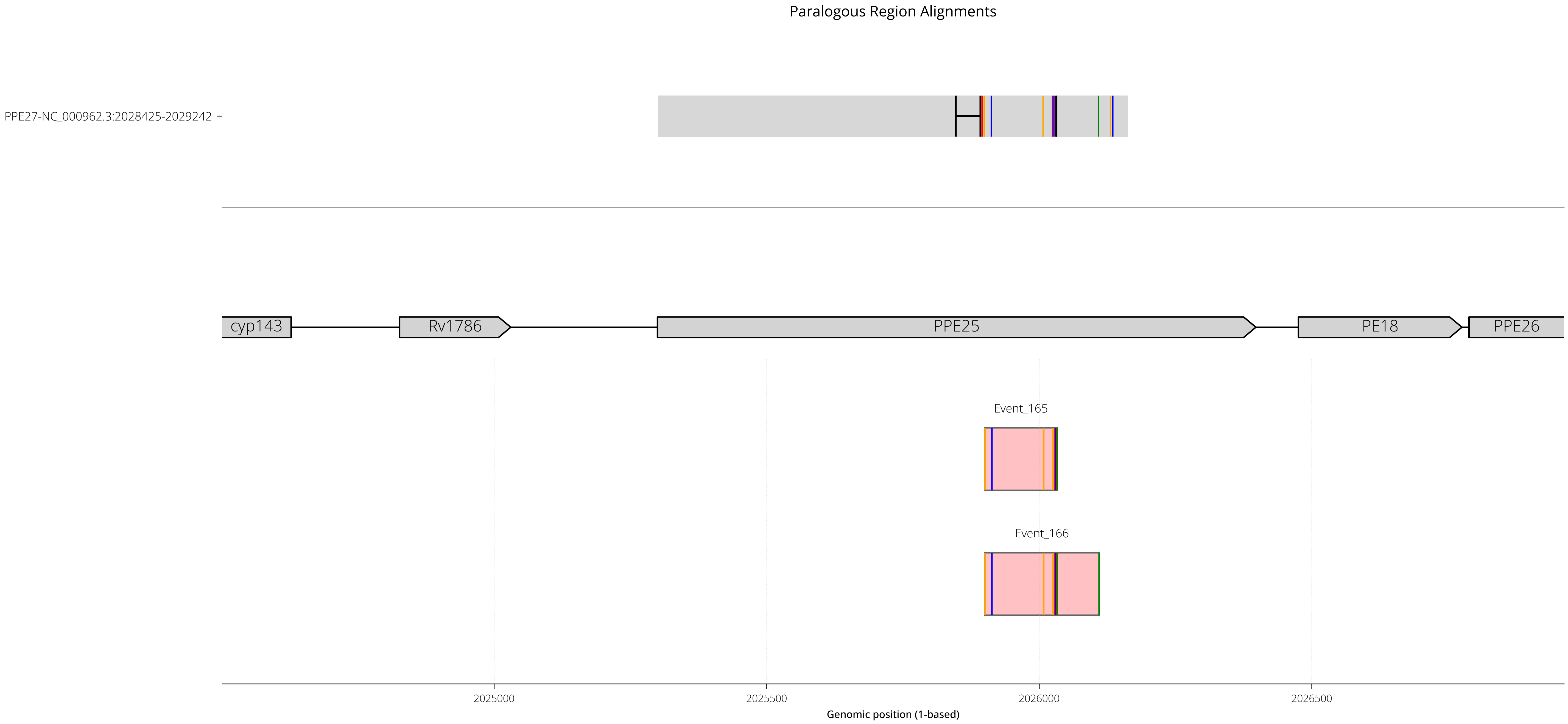

RegionID: PR_HmRegion_085 | Paralog Network ID: PR_Set_49
Genes: PPE25 | NC_000962.3:2024501-2026963
Mapped GCEs: 2 | Putative GCEs: 2

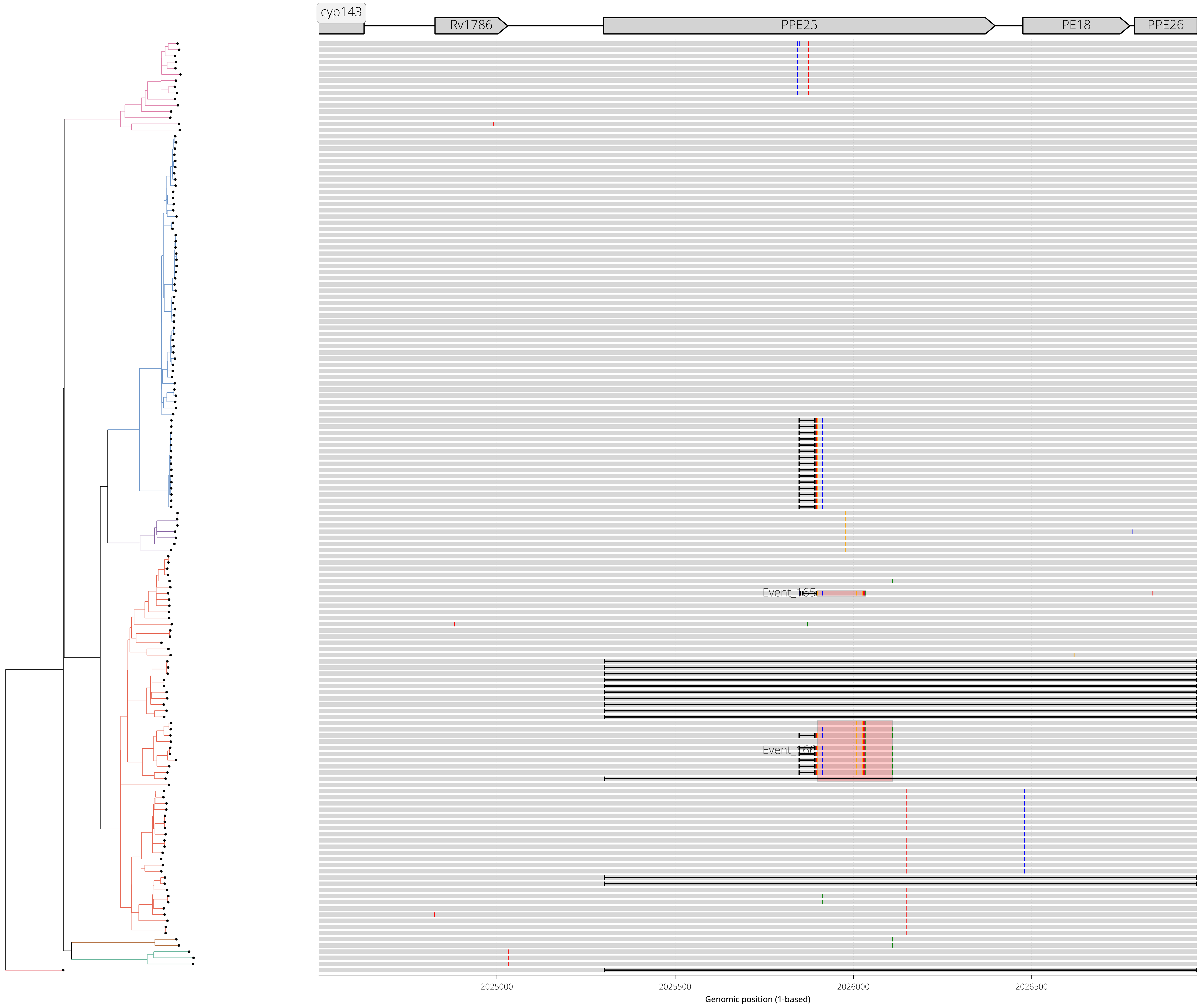

RegionID: PR_HmRegion_107 | Paralog Network ID: PR_Set_55
Genes: pknL,Rv2177c | NC_000962.3:2438339-2440748
Mapped GCEs: 2 | Putative GCEs: 2

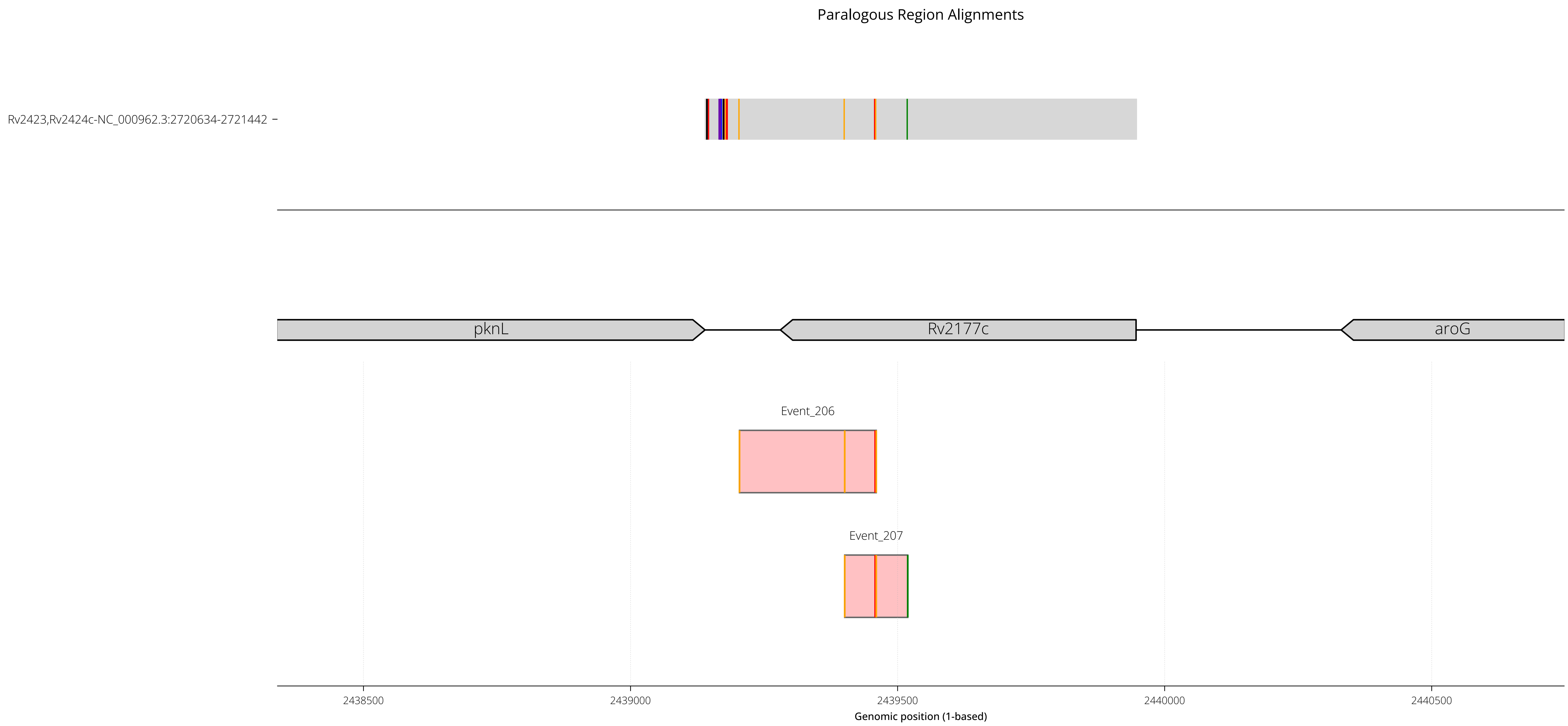

RegionID: PR_HmRegion_107 | Paralog Network ID: PR_Set_55
Genes: pknL,Rv2177c | NC_000962.3:2438339-2440748
Mapped GCEs: 2 | Putative GCEs: 2

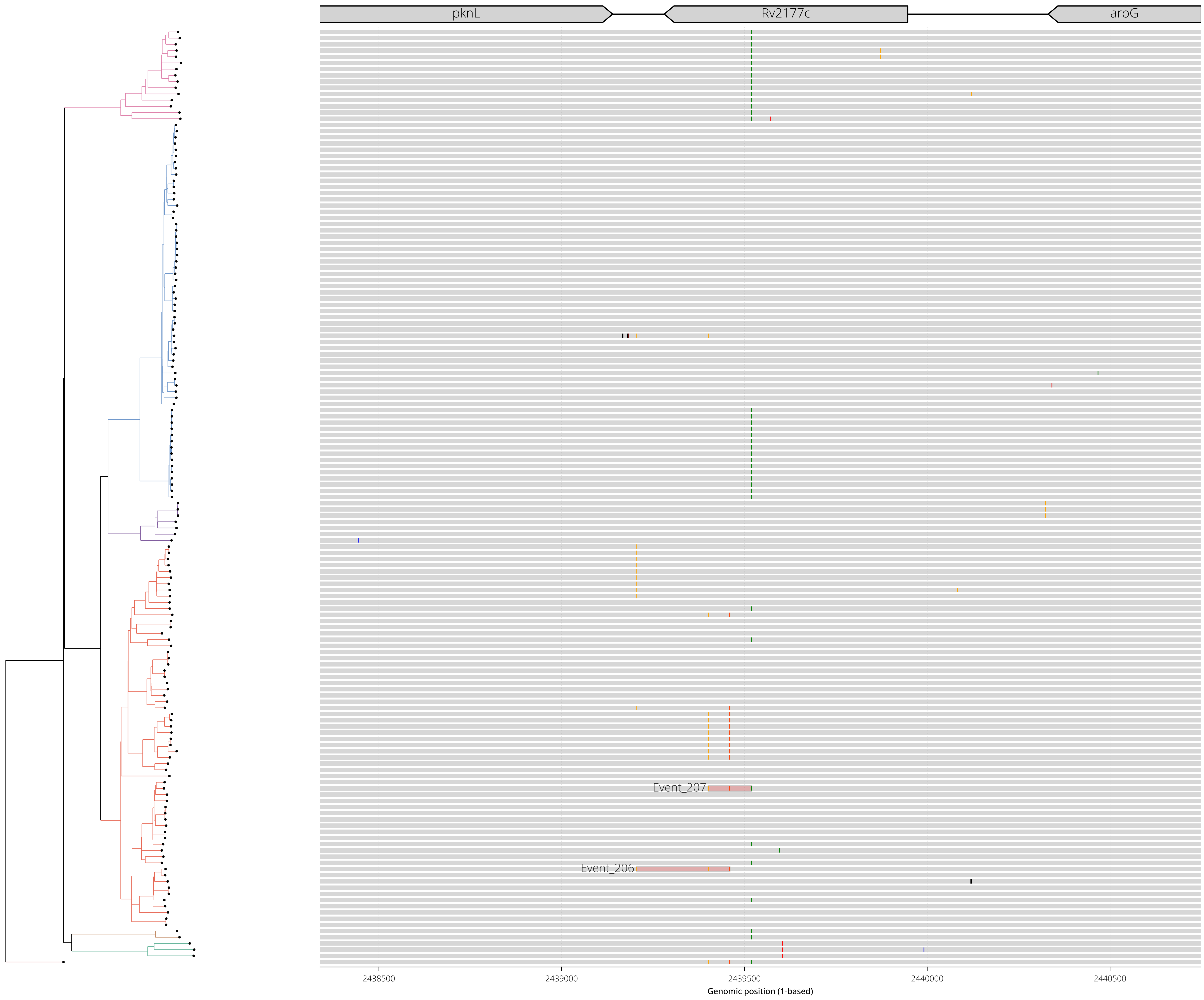

RegionID: PR_HmRegion_124 | Paralog Network ID: PR_Set_58
Genes: lppA | NC_000962.3:2865668-2867921
Mapped GCEs: 0 | Putative GCEs: 2

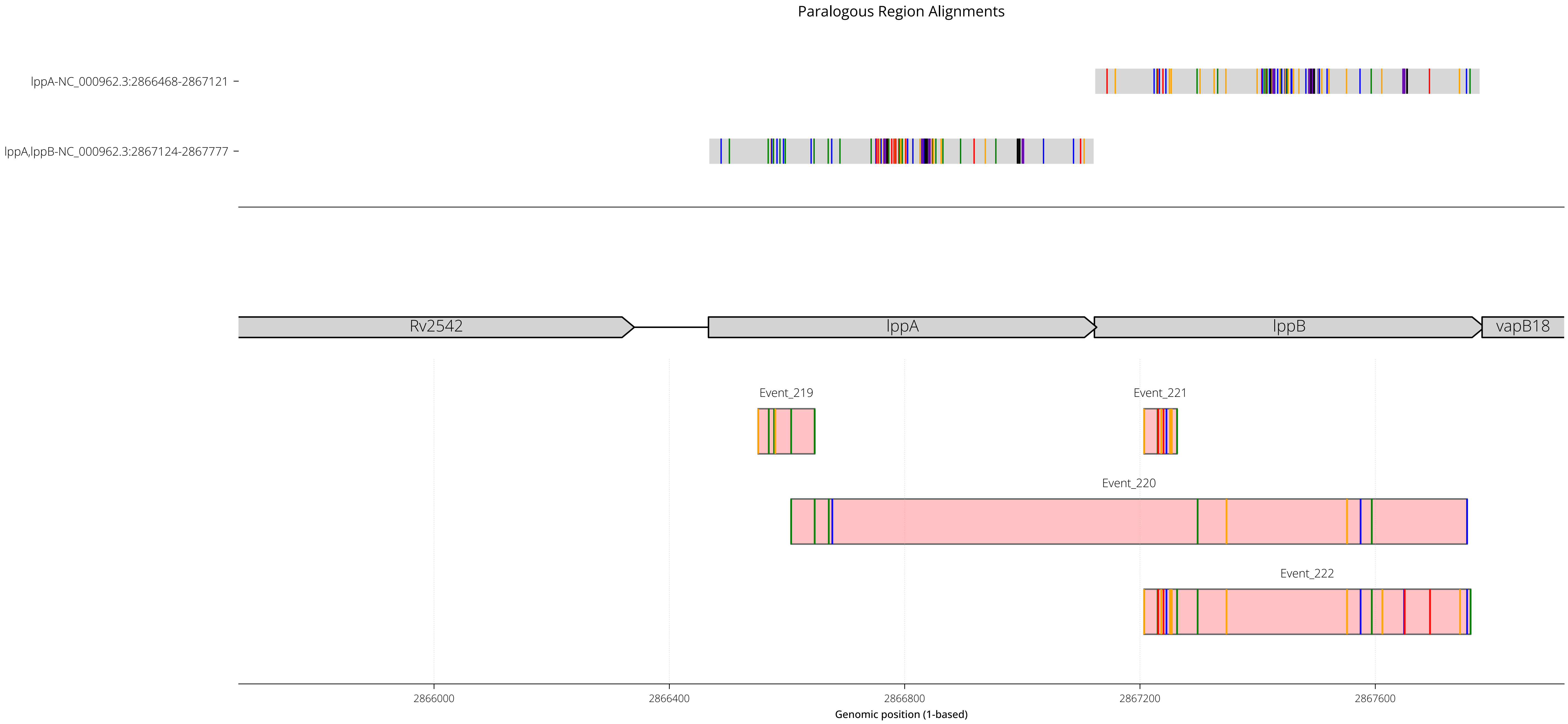

RegionID: PR_HmRegion_124 | Paralog Network ID: PR_Set_58
Genes: lppA | NC_000962.3:2865668-2867921
Mapped GCEs: 0 | Putative GCEs: 2

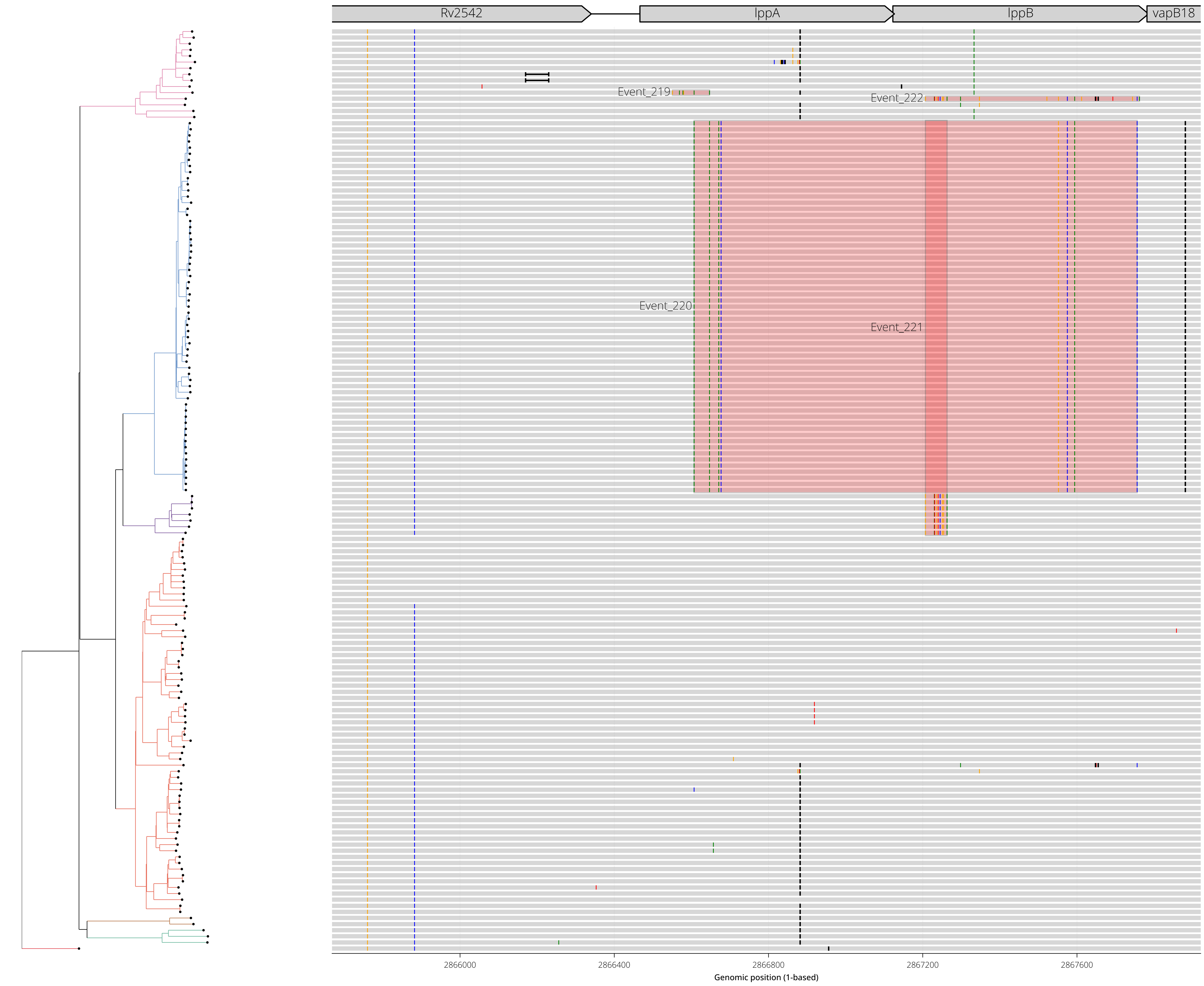

RegionID: PR_HmRegion_141 | Paralog Network ID: PR_Set_61
Genes: Rv2825c | NC_000962.3:3132091-3134255
Mapped GCEs: 2 | Putative GCEs: 2

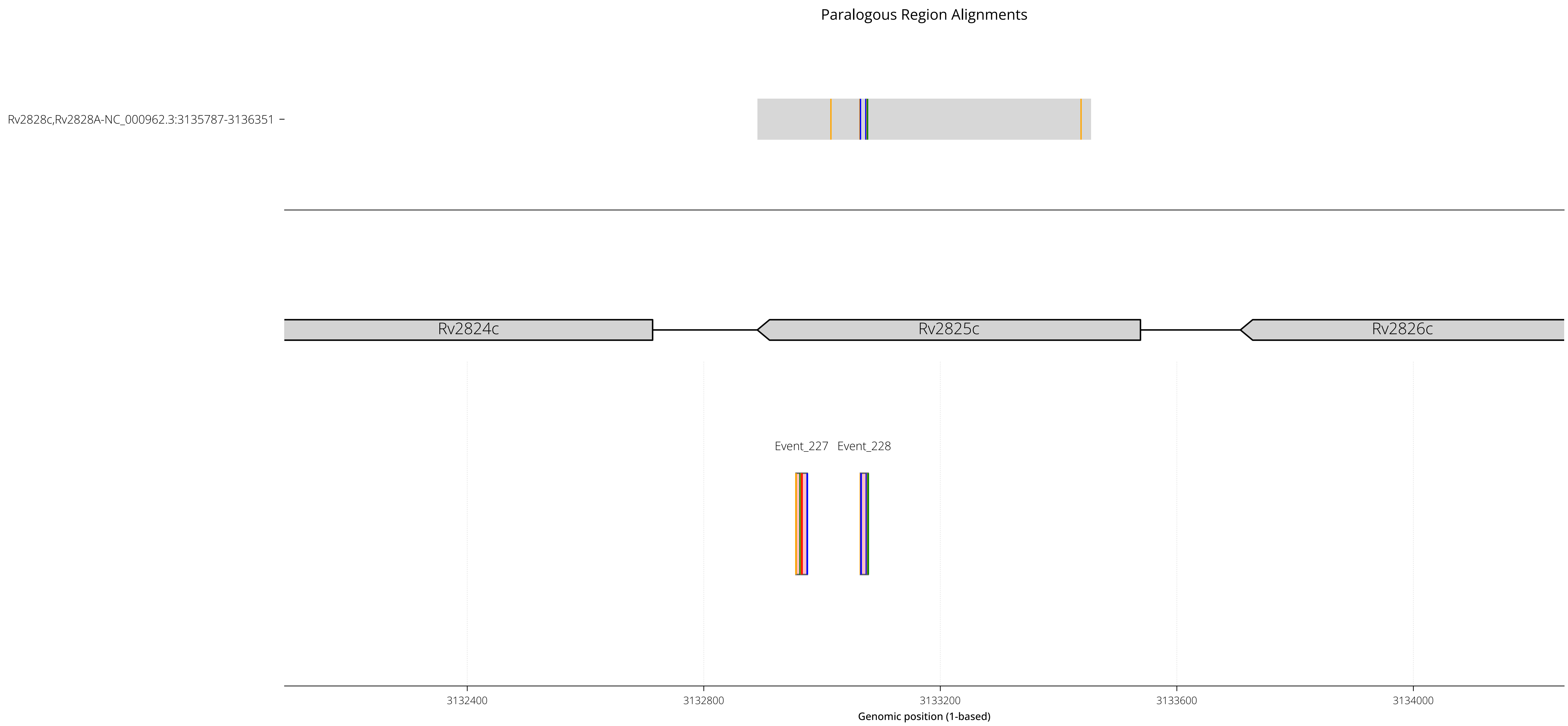

RegionID: PR_HmRegion_141 | Paralog Network ID: PR_Set_61
Genes: Rv2825c | NC_000962.3:3132091-3134255
Mapped GCEs: 2 | Putative GCEs: 2

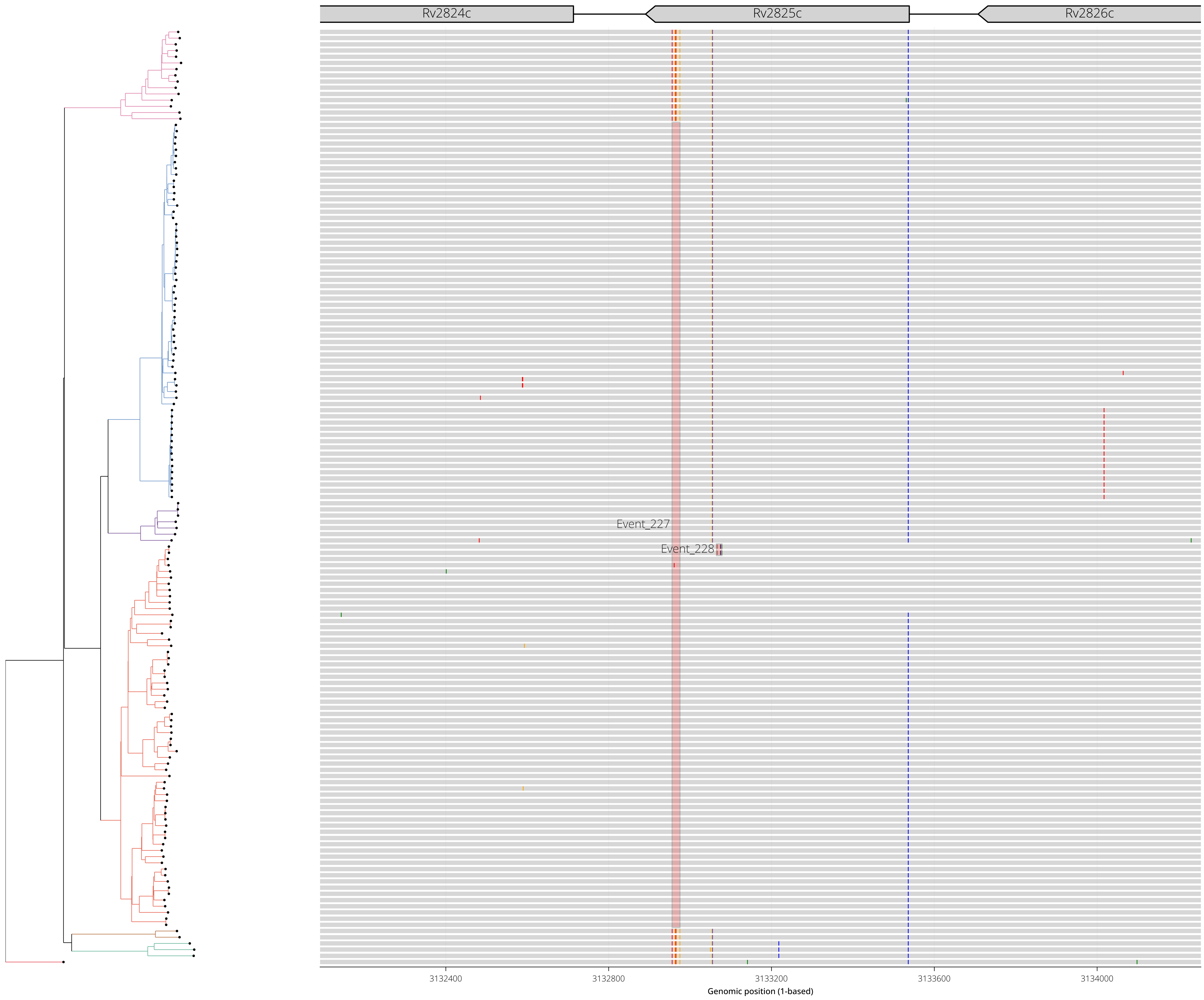

RegionID: PR_HmRegion_155 | Paralog Network ID: PR_Set_65
Genes: PPE46,PE27A | NC_000962.3:3376126-3379250
Mapped GCEs: 0 | Putative GCEs: 2

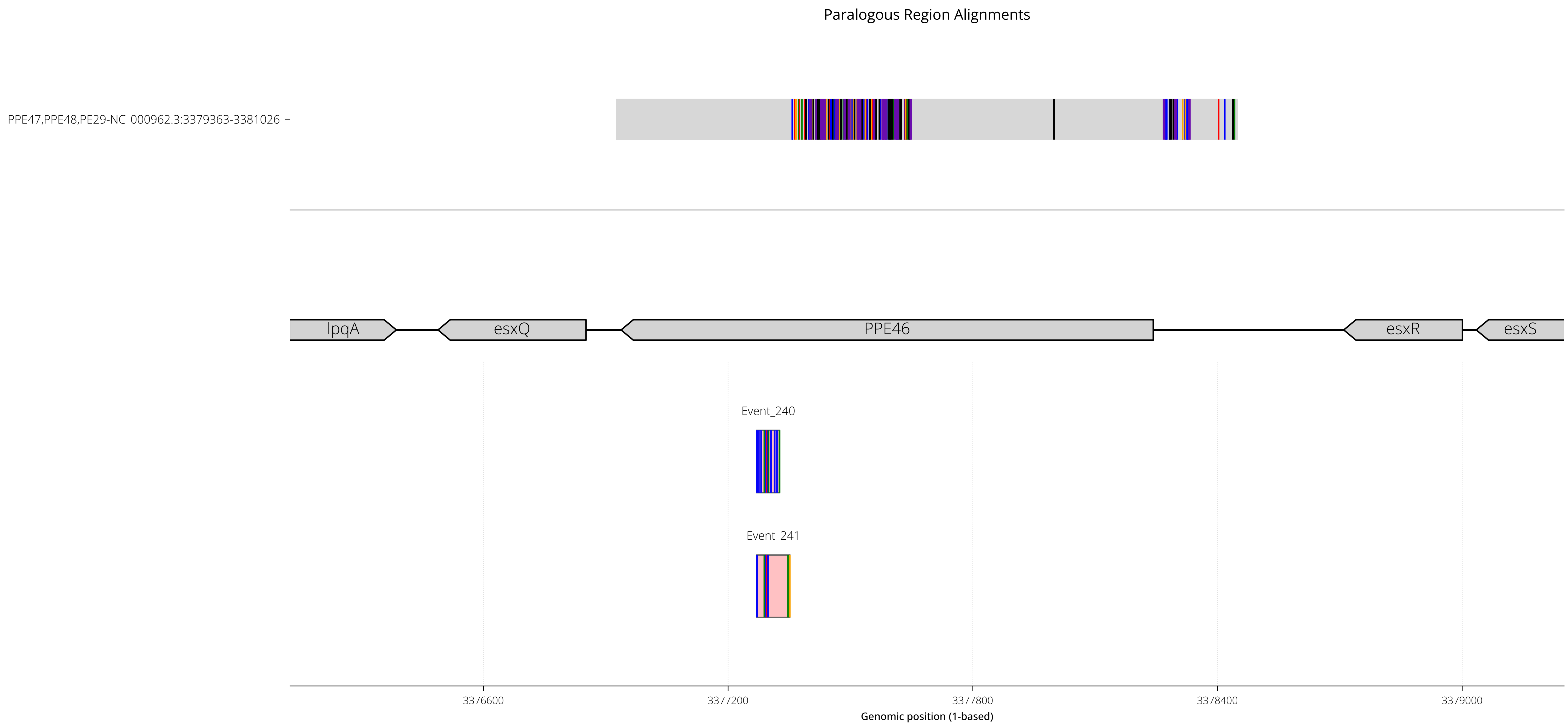

RegionID: PR_HmRegion_155 | Paralog Network ID: PR_Set_65
Genes: PPE46,PE27A | NC_000962.3:3376126-3379250
Mapped GCEs: 0 | Putative GCEs: 2

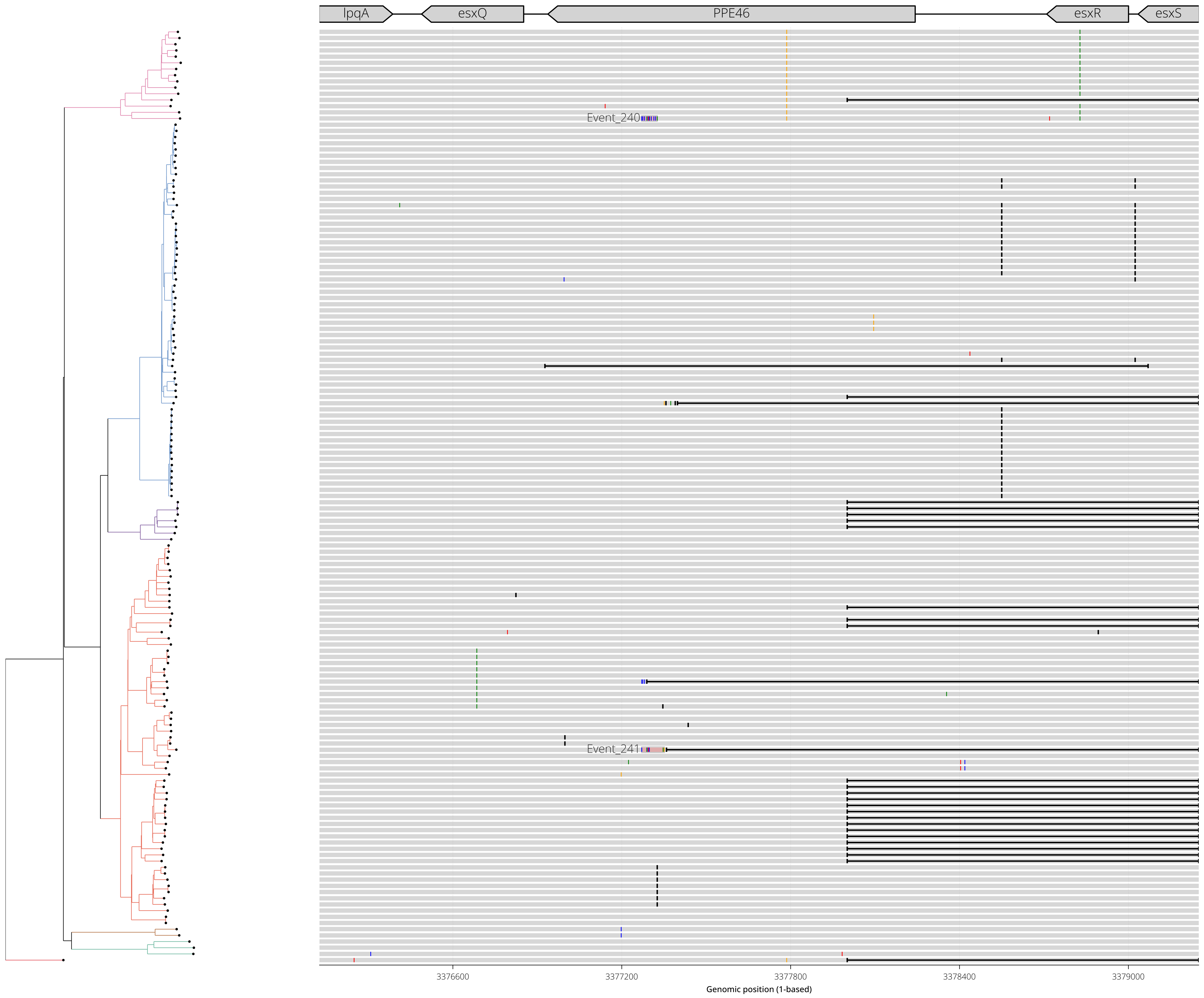

RegionID: PR_HmRegion_166 | Paralog Network ID: PR_Set_67
Genes: PPE54 | NC_000962.3:3729550-3731864
Mapped GCEs: 2 | Putative GCEs: 2

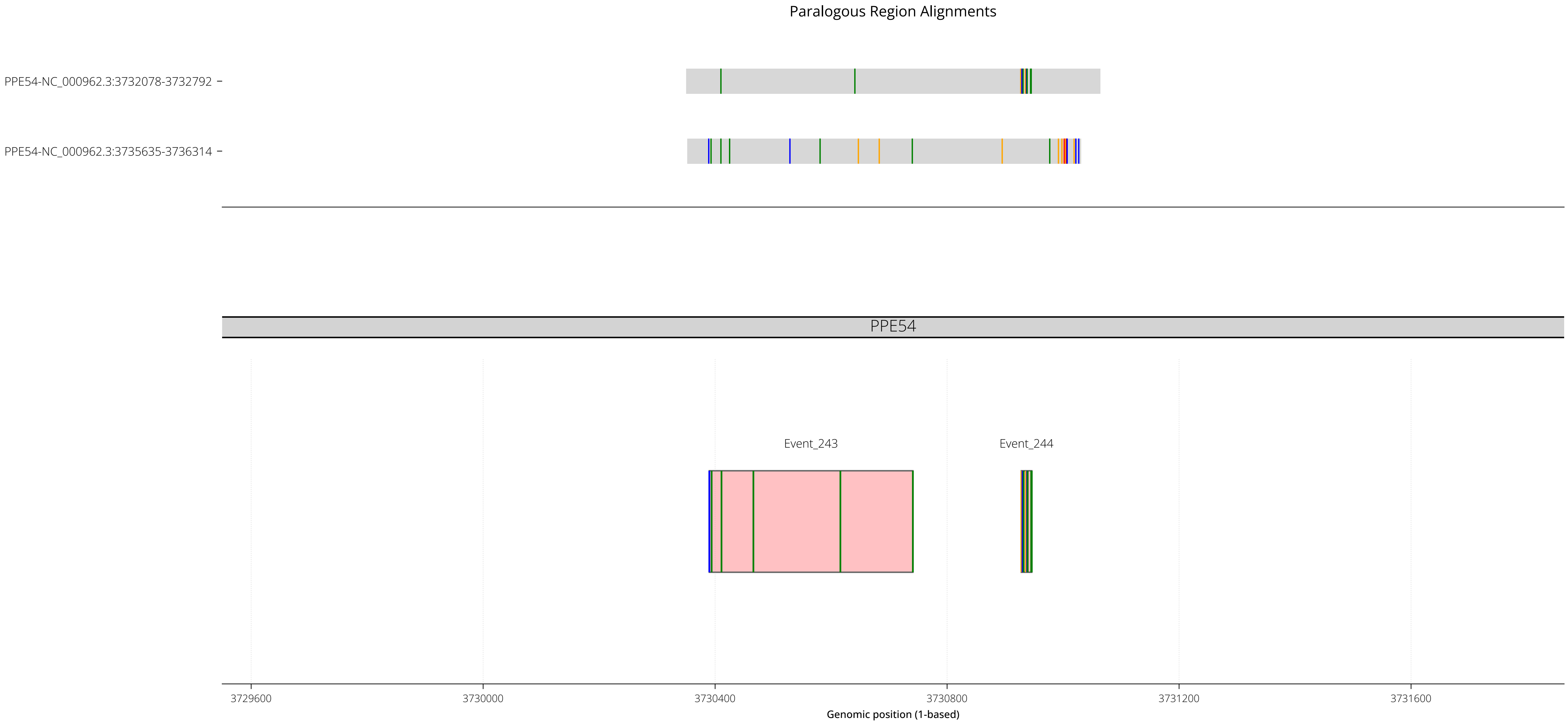

RegionID: PR_HmRegion_166 | Paralog Network ID: PR_Set_67
Genes: PPE54 | NC_000962.3:3729550-3731864
Mapped GCEs: 2 | Putative GCEs: 2

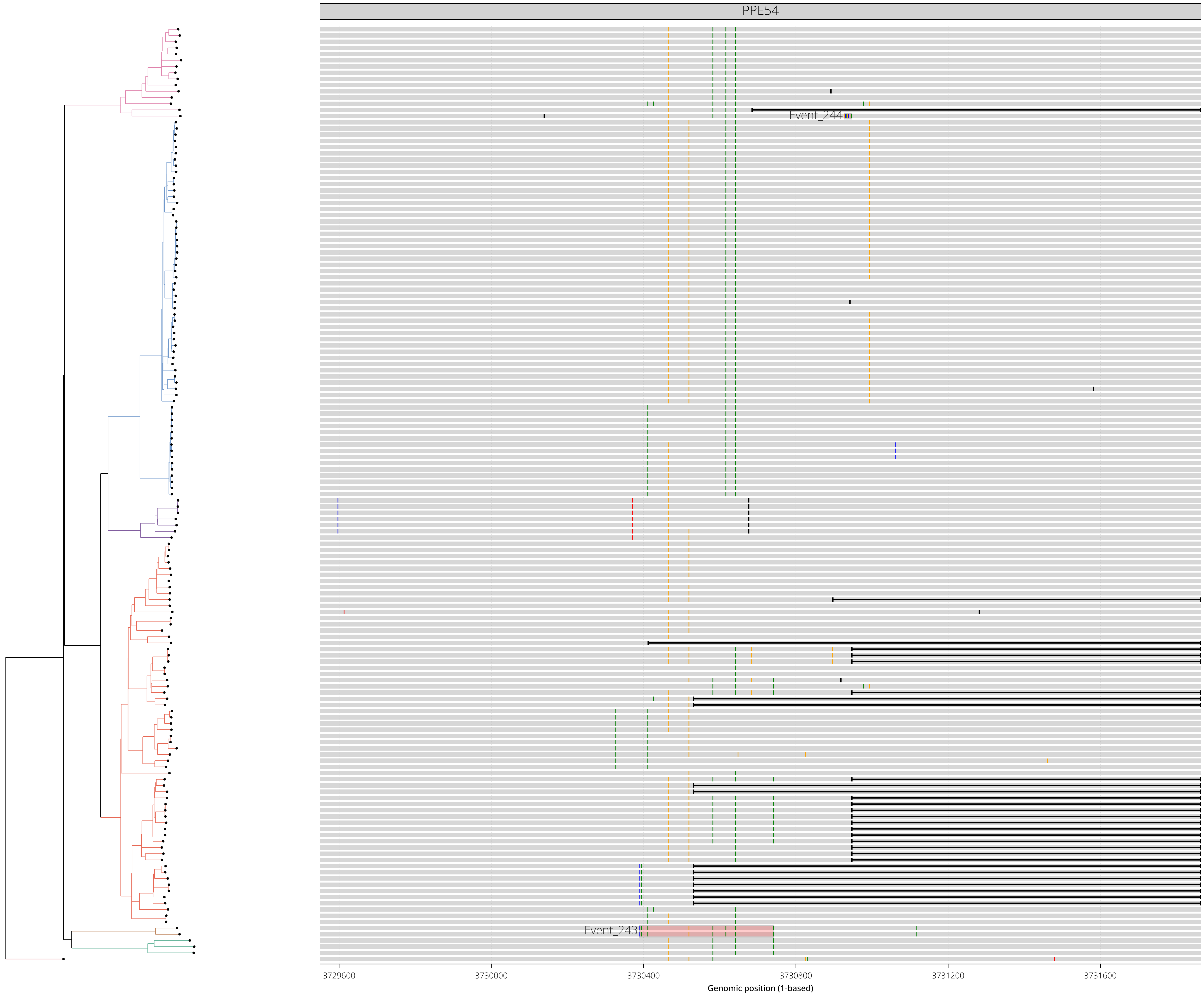

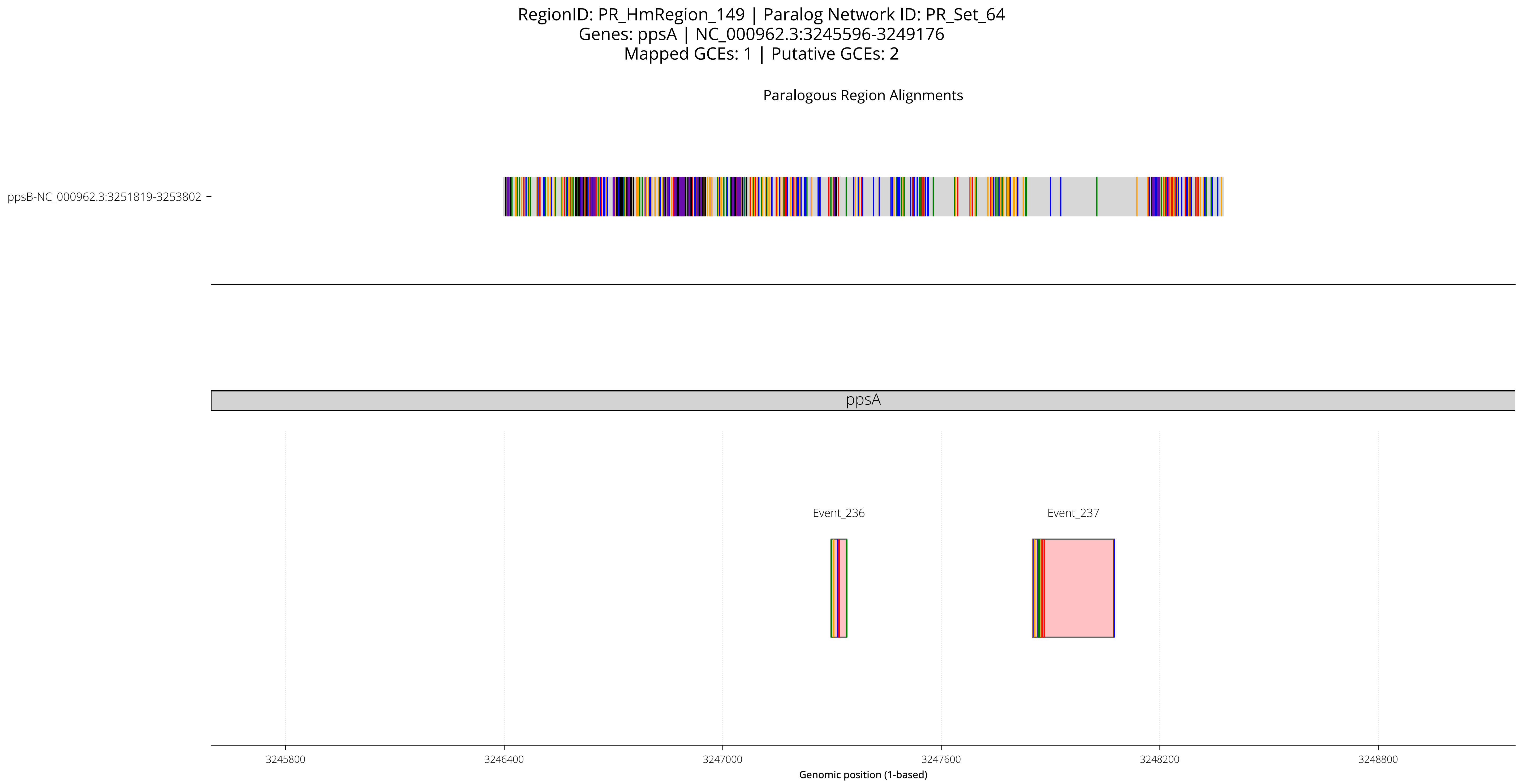

RegionID: PR_HmRegion_149 | Paralog Network ID: PR_Set_64
Genes: ppsA | NC_000962.3:3245596-3249176
Mapped GCEs: 1 | Putative GCEs: 2
Paralogous Region Alignments
ppsB-NC_000962.3:3251819-3253802
ppsA
Event_236
Event_237
3245800
3246400
3247000
3247600
3248200
3248800
Genomic position (1-based)

RegionID: PR_HmRegion_149 | Paralog Network ID: PR_Set_64
Genes: ppsA | NC_000962.3:3245596-3249176
Mapped GCEs: 1 | Putative GCEs: 2

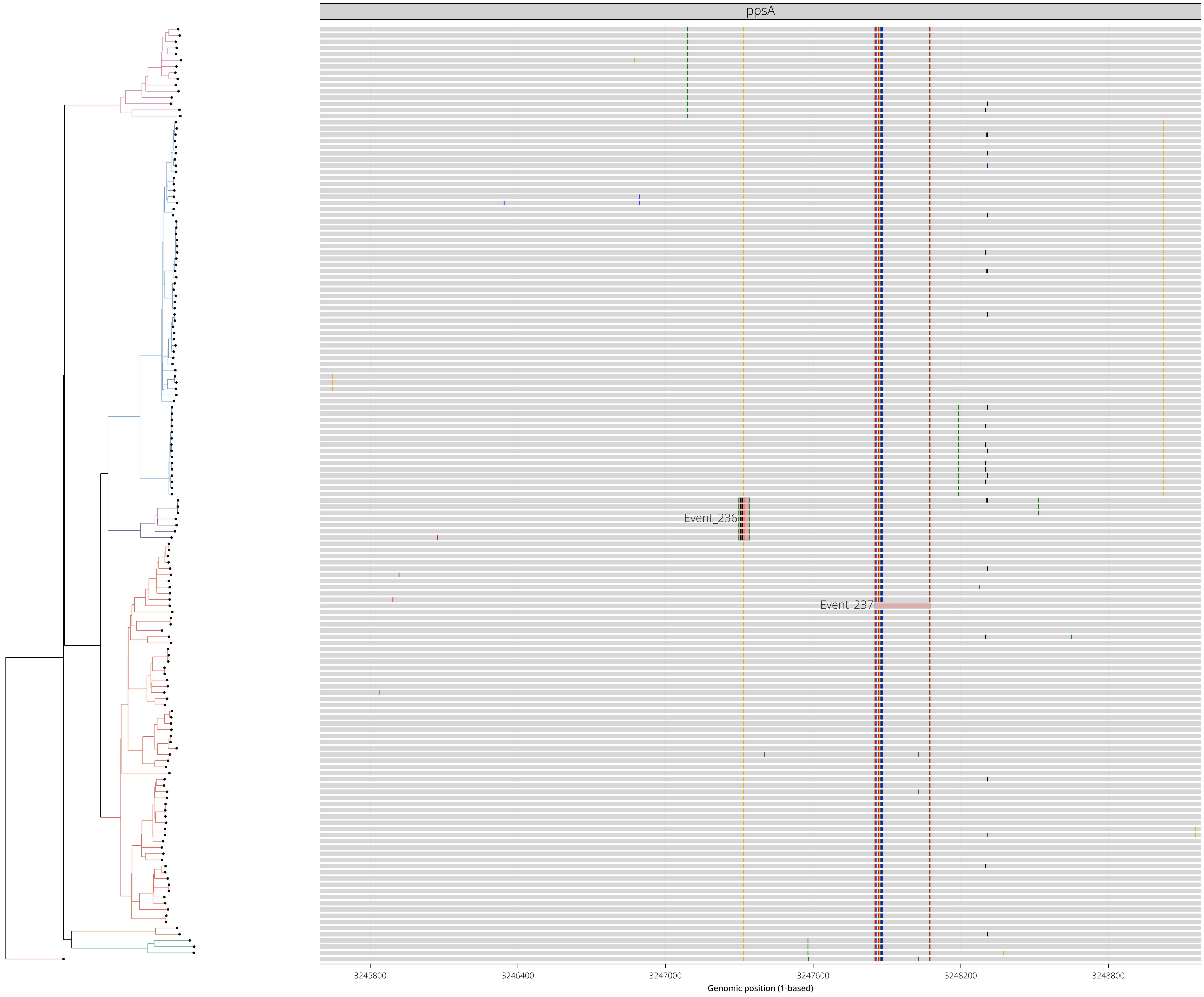

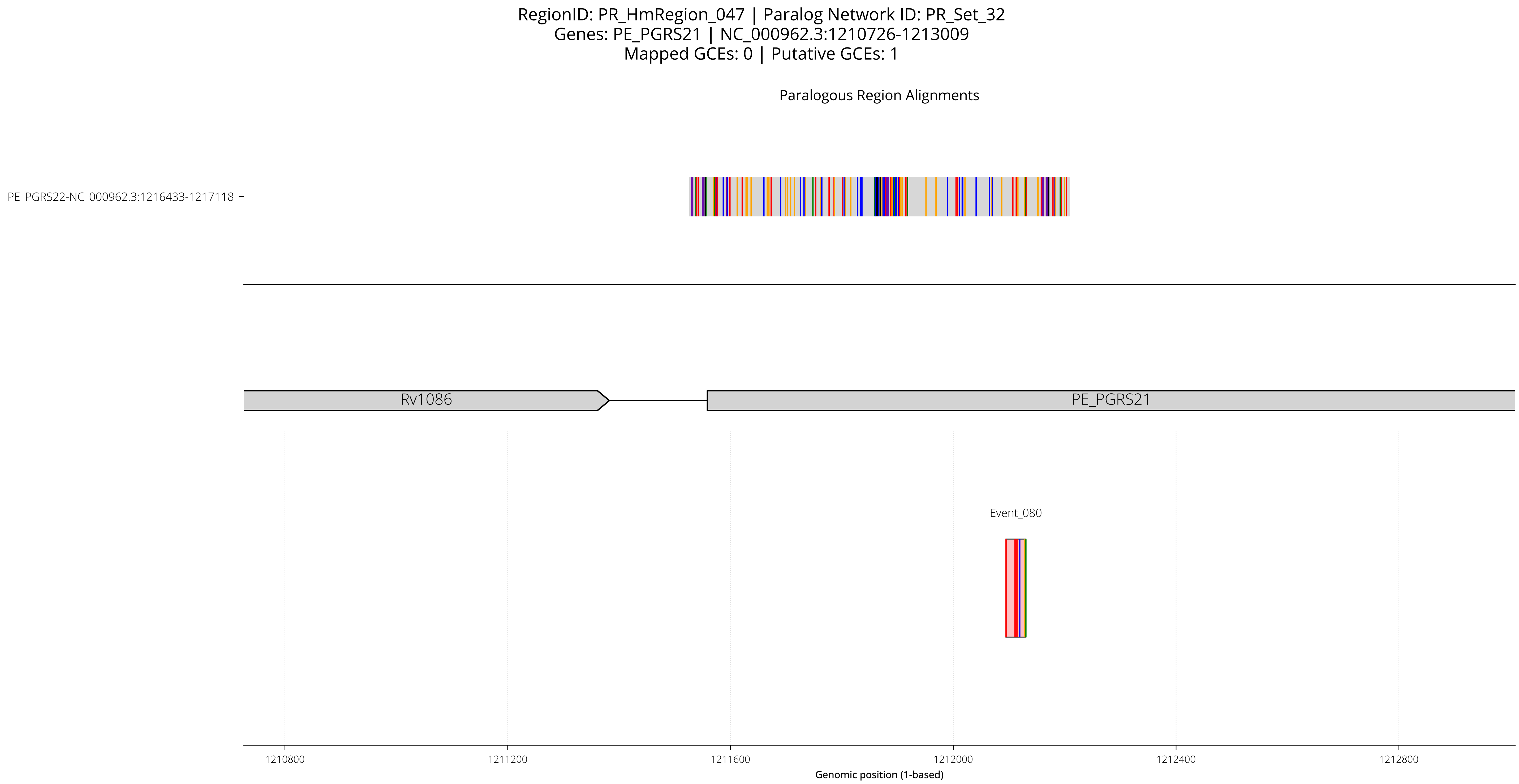

RegionID: PR_HmRegion_047 | Paralog Network ID: PR_Set_32
Genes: PE_PGRS21 | NC_000962.3:1210726-1213009
Mapped GCEs: 0 | Putative GCEs: 1
Paralogous Region Alignments
PE_PGRS22-NC_000962.3:1216433-1217118
Rv1086
PE_PGRS21
Event_080
1210800
1211200
1211600
1212000
1212400
1212800
Genomic position (1-based)

RegionID: PR_HmRegion_047 | Paralog Network ID: PR_Set_32
Genes: PE_PGRS21 | NC_000962.3:1210726-1213009
Mapped GCEs: 0 | Putative GCEs: 1

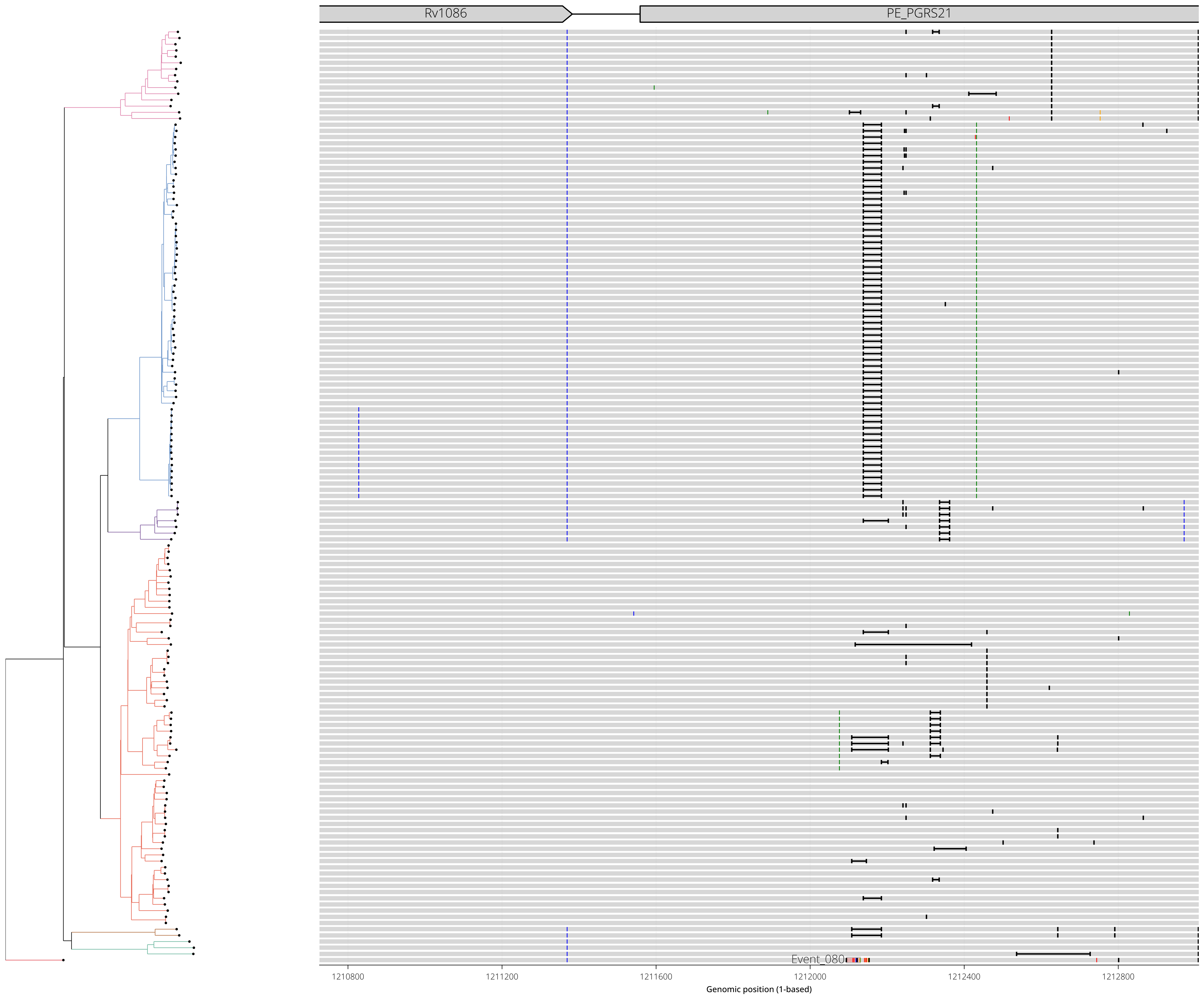

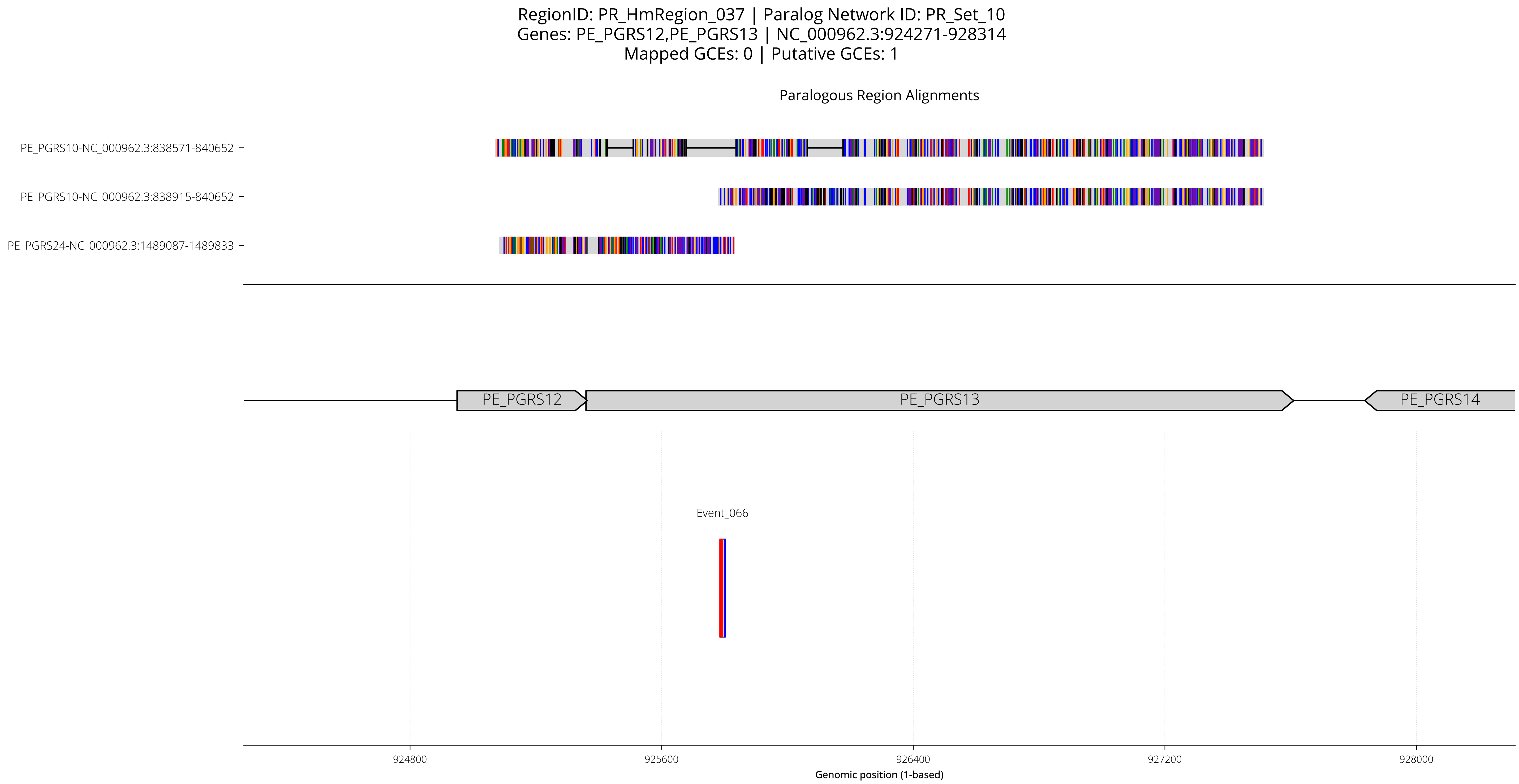
RegionID: PR_HmRegion_037 | Paralog Network ID: PR_Set_10
Genes: PE_PGRS12,PE_PGRS13 | NC_000962.3:924271-928314
Mapped GCEs: 0 | Putative GCEs: 1
Paralogous Region Alignments
PE_PGRS10-NC_000962.3:838571-840652
PE_PGRS10-NC_000962.3:838915-840652
PE_PGRS24-NC_000962.3:1489087-1489833
PE_PGRS12
PE_PGRS13
PE_PGRS14
Event_066
924800
925600
926400
927200
928000
Genomic position (1-based)

RegionID: PR_HmRegion_037 | Paralog Network ID: PR_Set_10
Genes: PE_PGRS12,PE_PGRS13 | NC_000962.3:924271-928314
Mapped GCEs: 0 | Putative GCEs: 1

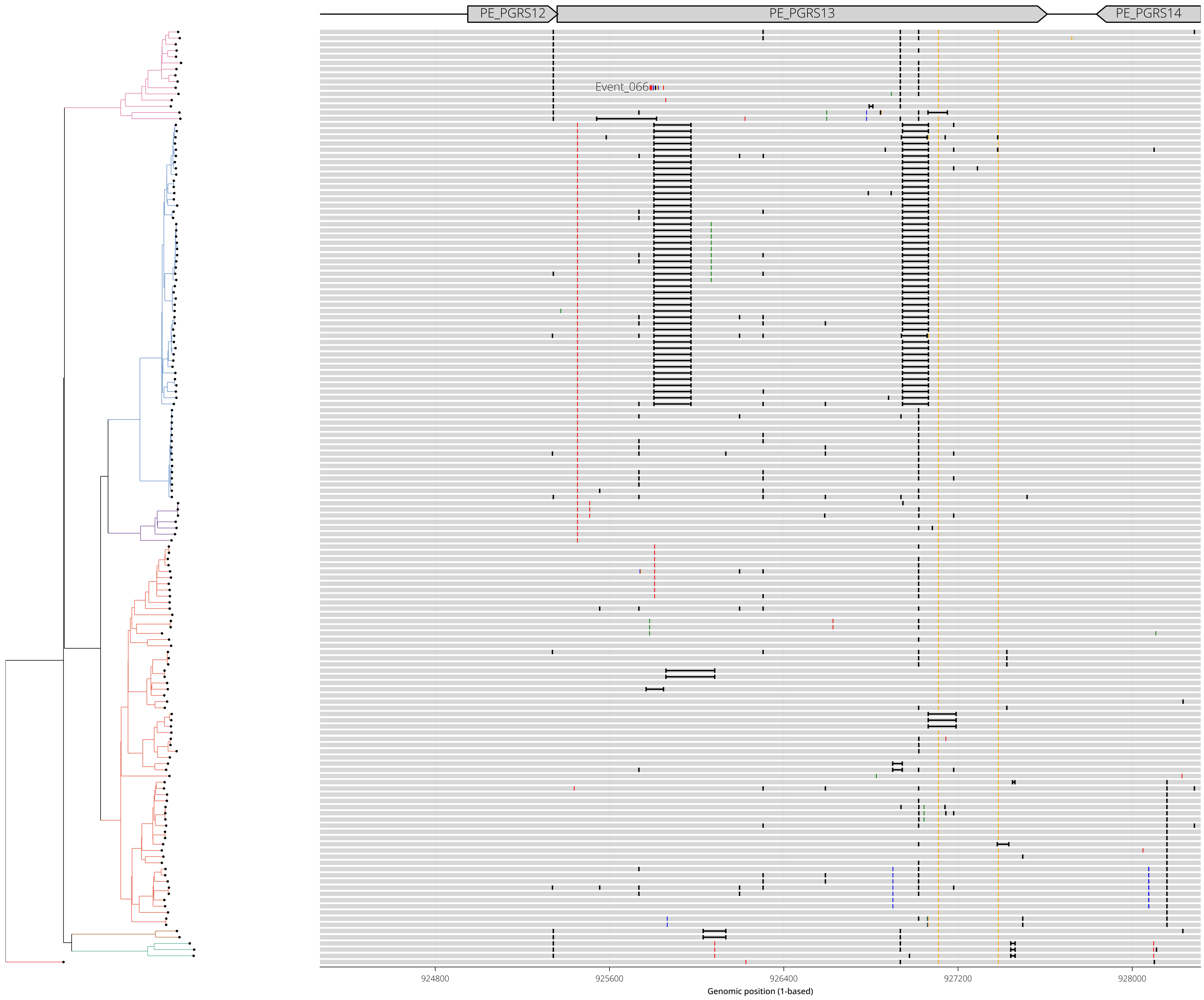

RegionID: PR_HmRegion_119 | Paralog Network ID: PR_Set_55
Genes: Rv2423,Rv2424c | NC_000962.3:2719834-2722242
Mapped GCEs: 1 | Putative GCEs: 1

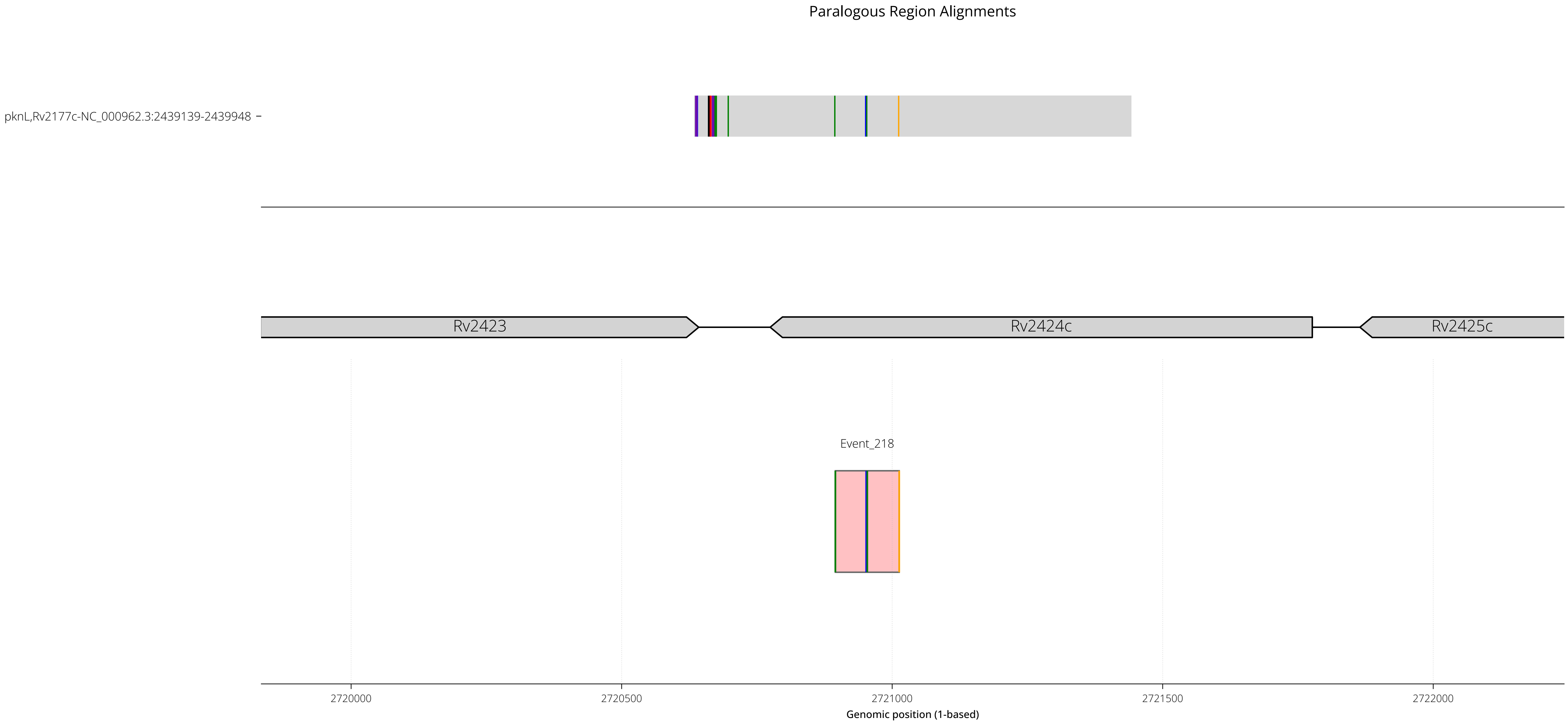

RegionID: PR_HmRegion_119 | Paralog Network ID: PR_Set_55
Genes: Rv2423,Rv2424c | NC_000962.3:2719834-2722242
Mapped GCEs: 1 | Putative GCEs: 1

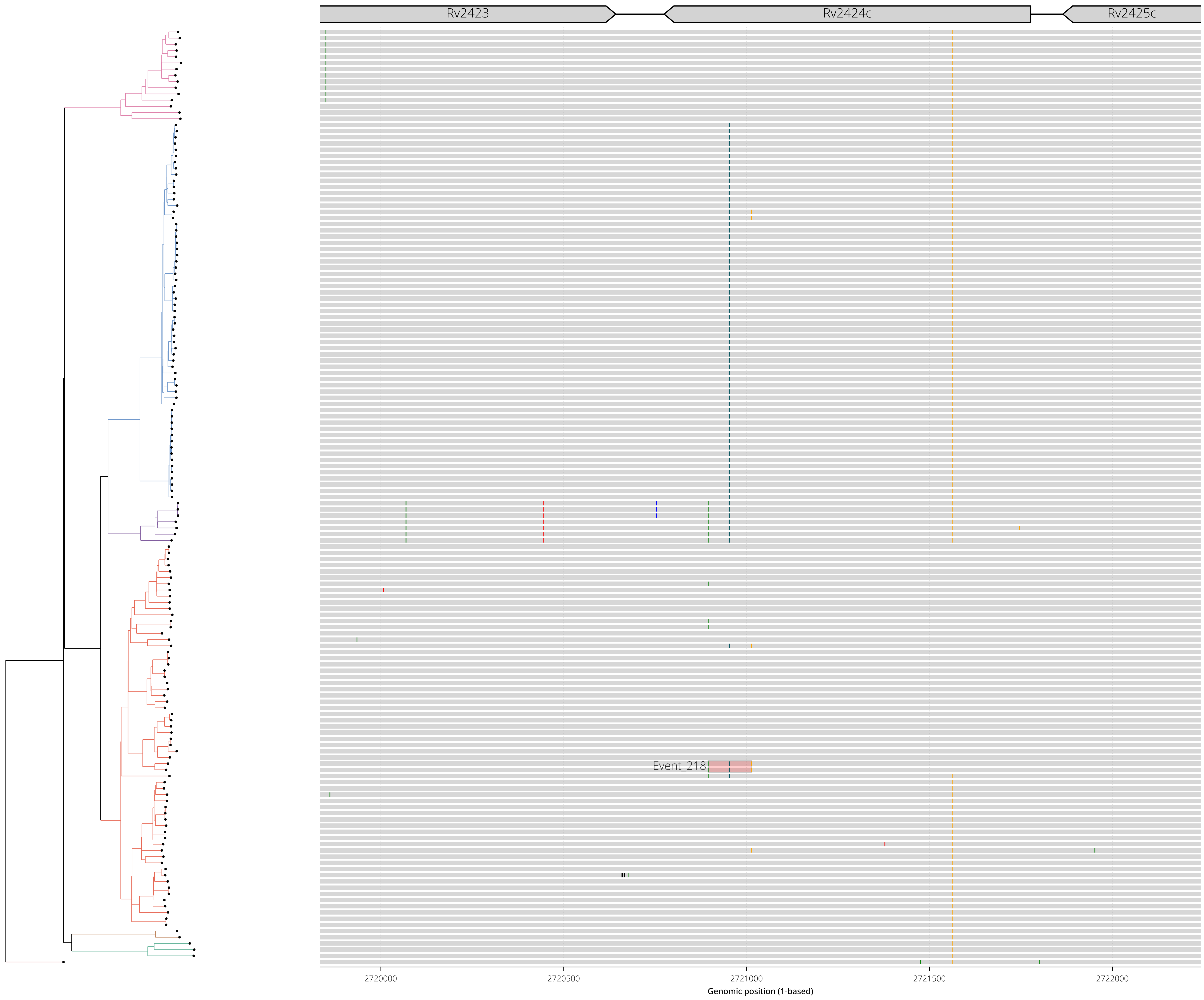

RegionID: PR_HmRegion_150 | Paralog Network ID: PR_Set_64
Genes: ppsB | NC_000962.3:3251019-3254602
Mapped GCEs: 1 | Putative GCEs: 1

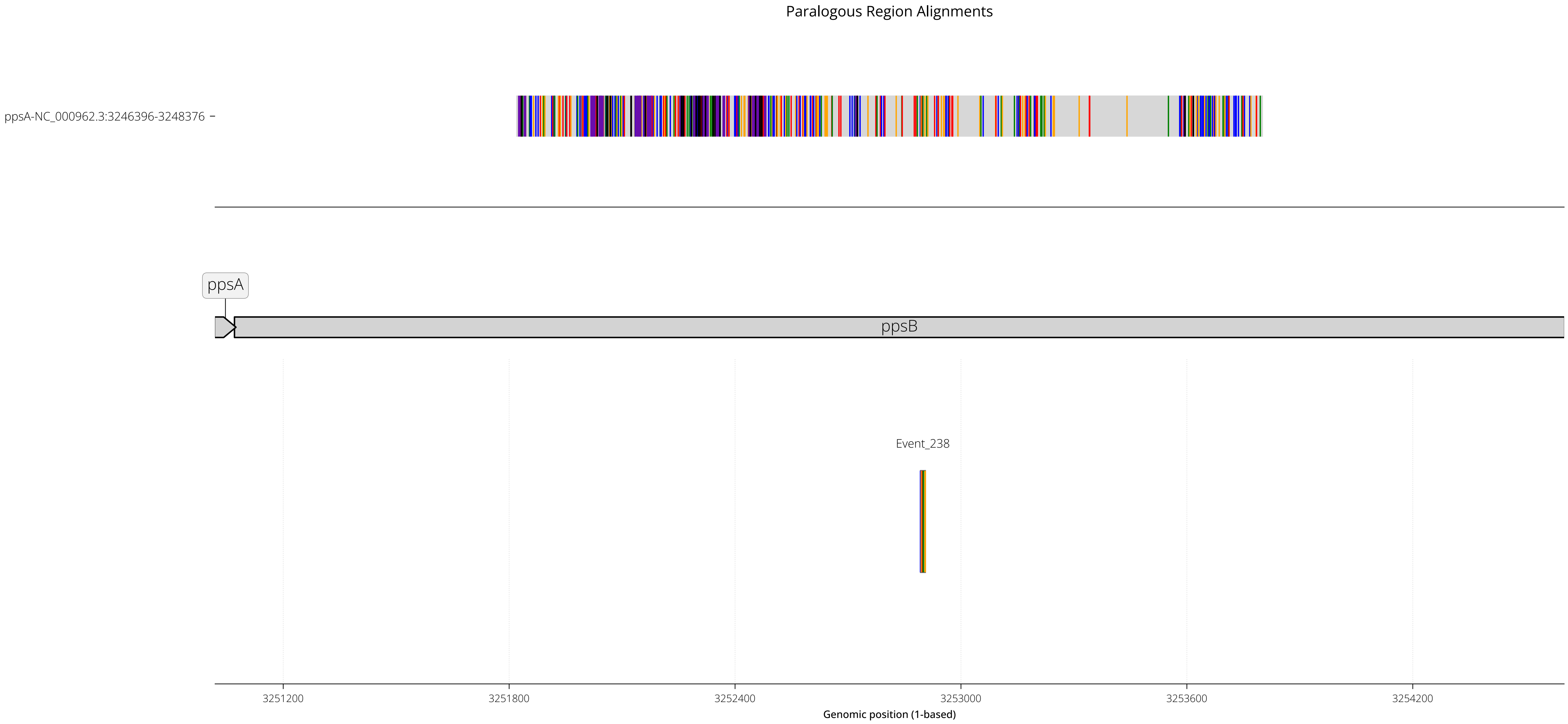

RegionID: PR_HmRegion_150 | Paralog Network ID: PR_Set_64
Genes: ppsB | NC_000962.3:3251019-3254602
Mapped GCEs: 1 | Putative GCEs: 1

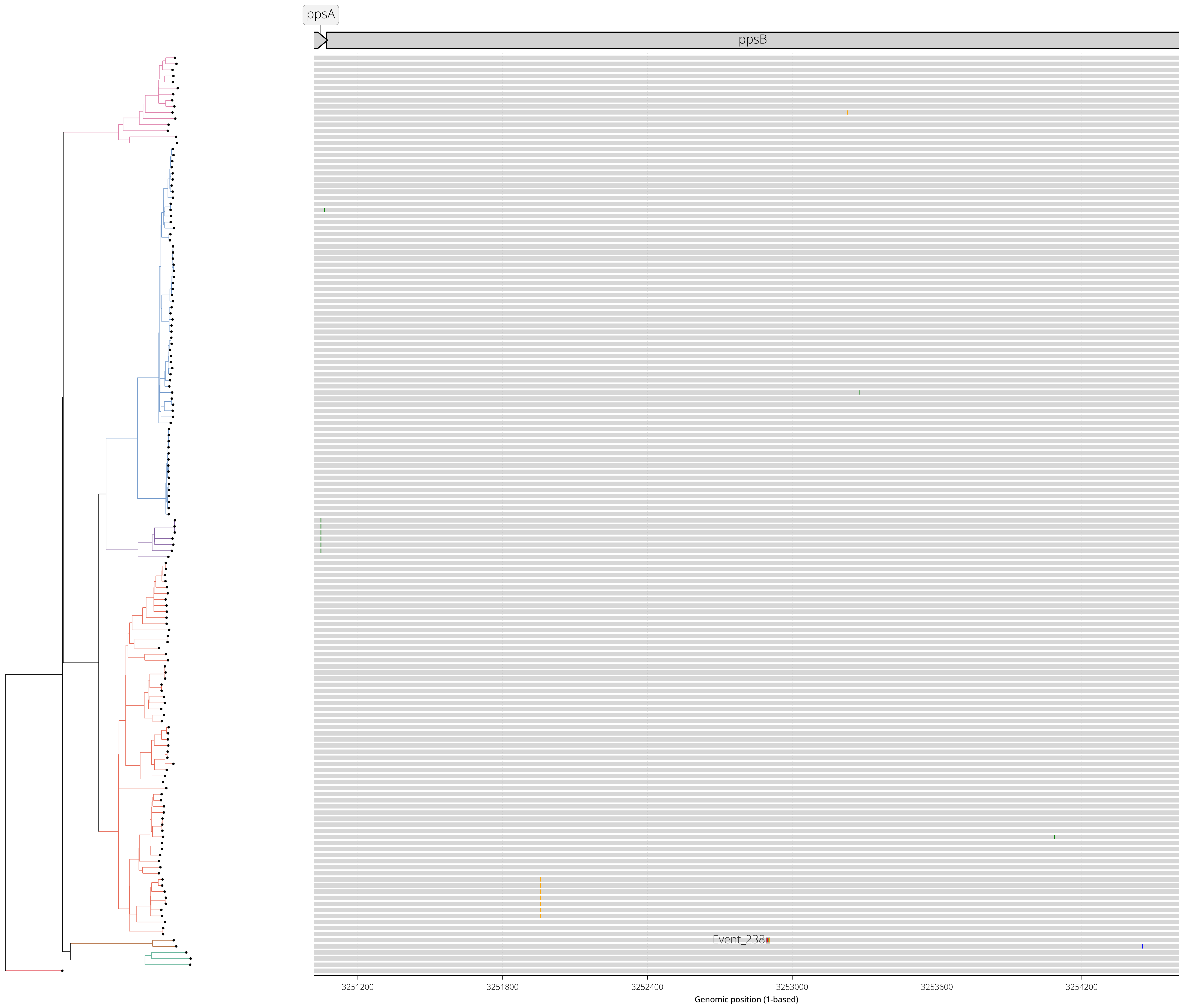

RegionID: PR_HmRegion_147 | Paralog Network ID: PR_Set_20
Genes: Rv2884,Rv2885c,Rv2886c | NC_000962.3:3193337-3197002
Mapped GCEs: 1 | Putative GCEs: 1

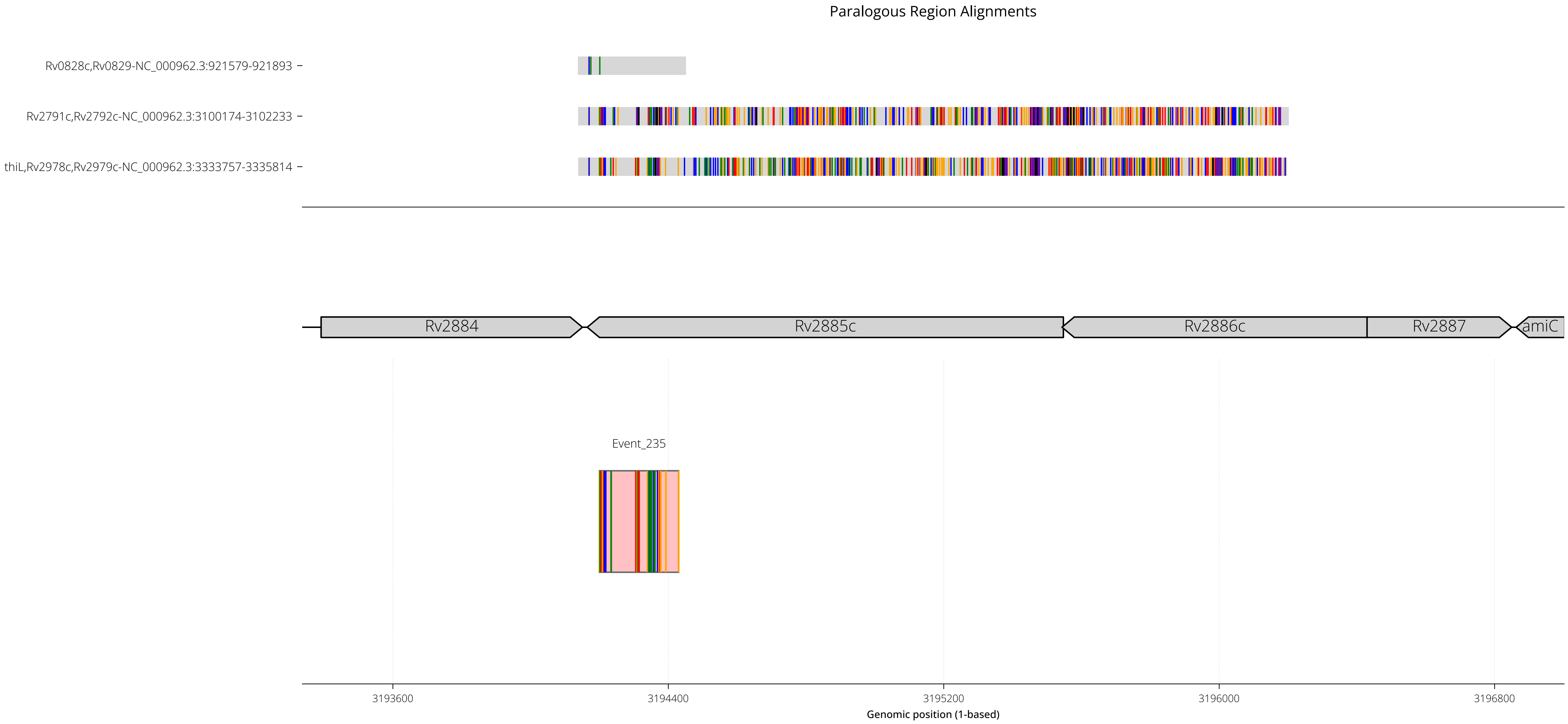

RegionID: PR_HmRegion_147 | Paralog Network ID: PR_Set_20
Genes: Rv2884,Rv2885c,Rv2886c | NC_000962.3:3193337-3197002
Mapped GCEs: 1 | Putative GCEs: 1

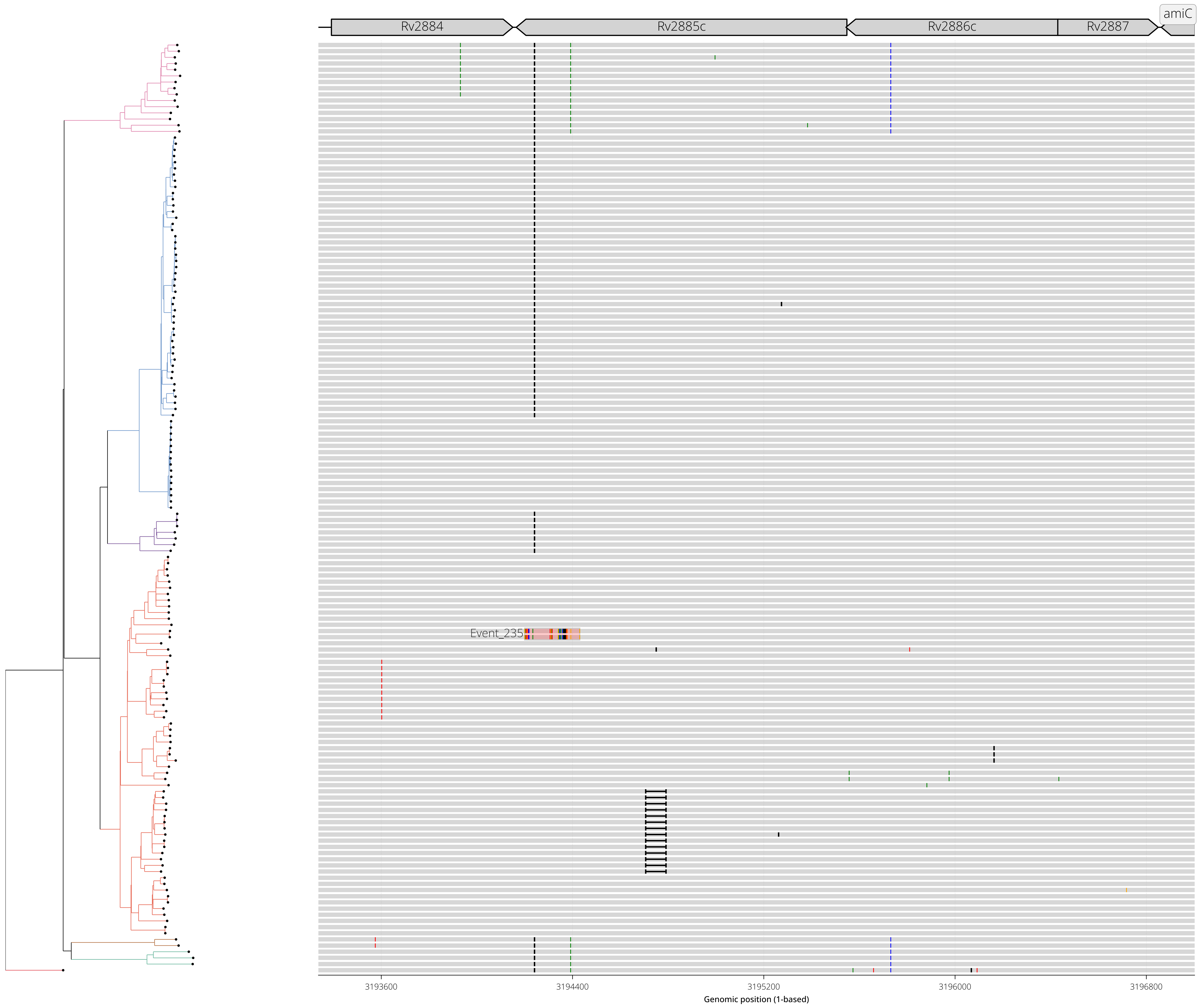

RegionID: PR_HmRegion_127 | Paralog Network ID: PR_Set_27
Genes: PE_PGRS45 | NC_000962.3:2943467-2946026
Mapped GCEs: 1 | Putative GCEs: 1

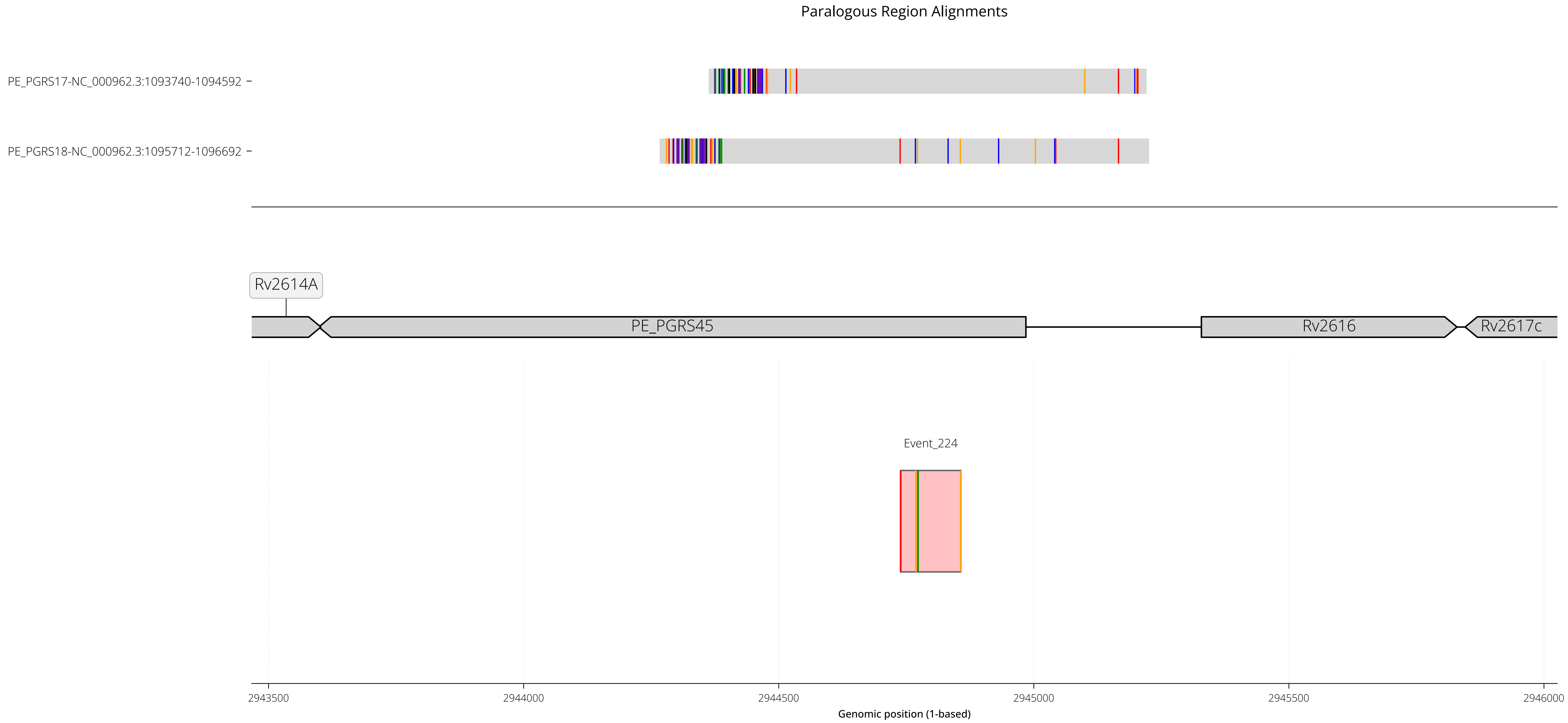

RegionID: PR_HmRegion_127 | Paralog Network ID: PR_Set_27
Genes: PE_PGRS45 | NC_000962.3:2943467-2946026
Mapped GCEs: 1 | Putative GCEs: 1

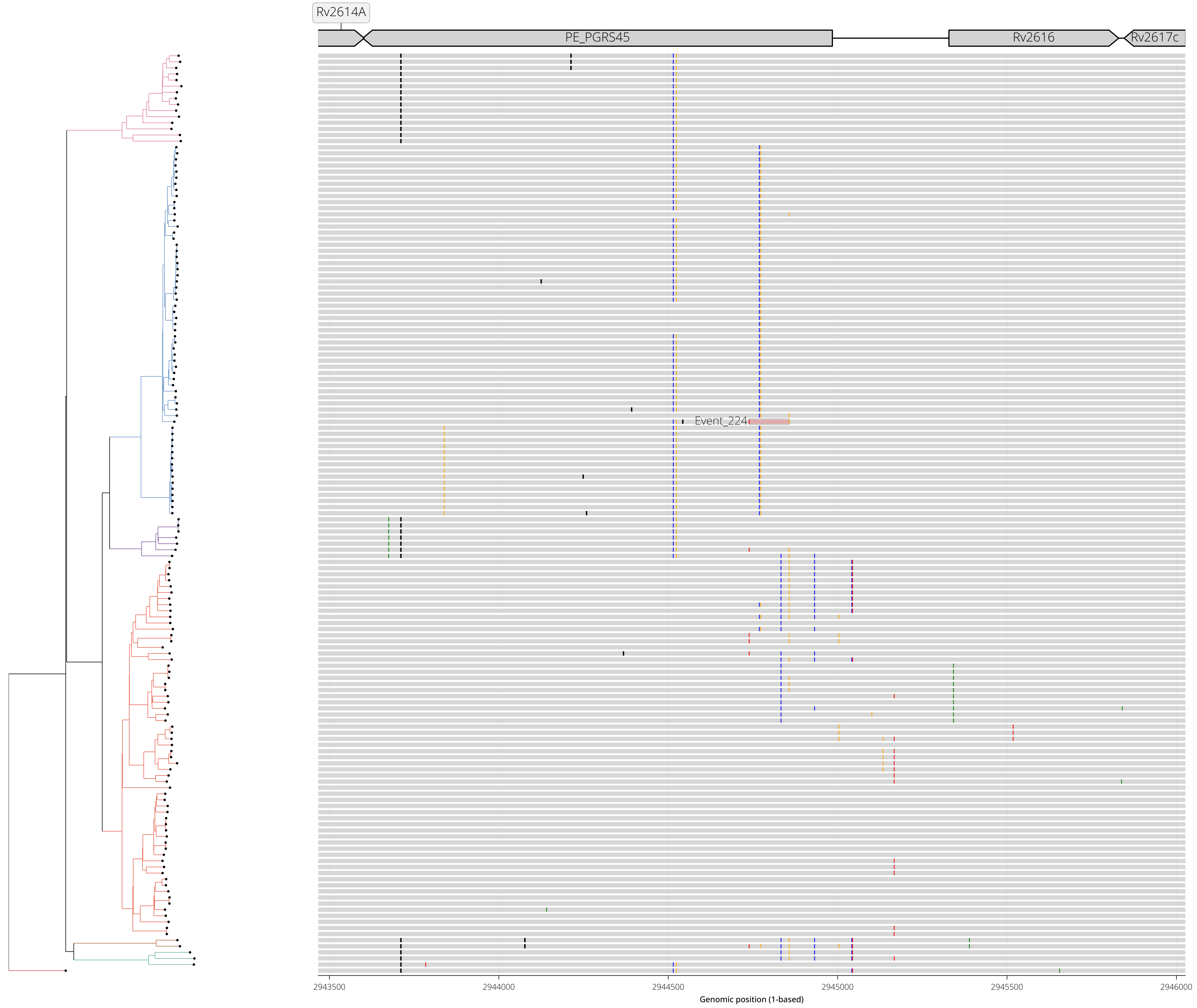

RegionID: PR_HmRegion_140 | Paralog Network ID: PR_Set_24
Genes: Rv2814c,Rv2815c | NC_000962.3:3118383-3124377
Mapped GCEs: 0 | Putative GCEs: 1

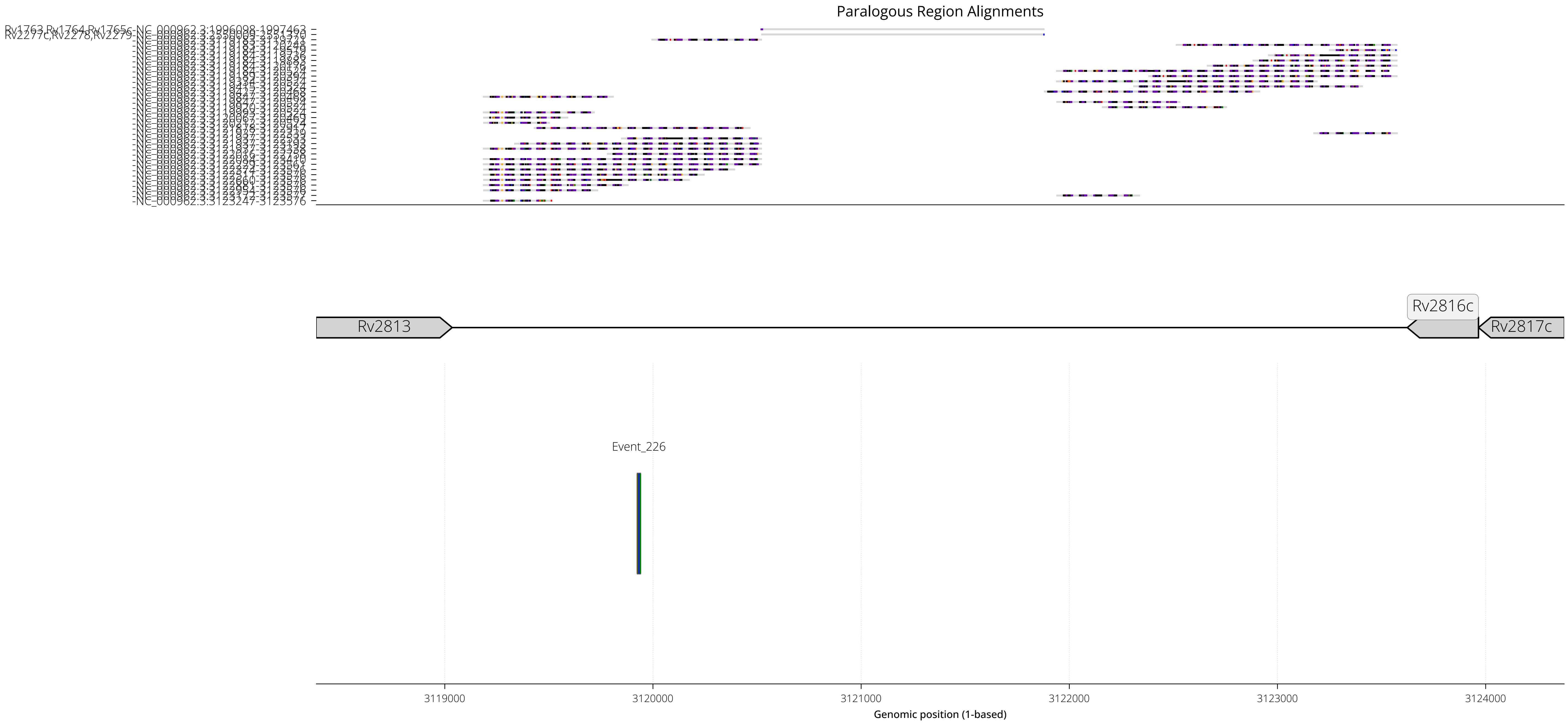

RegionID: PR_HmRegion_140 | Paralog Network ID: PR_Set_24
Genes: Rv2814c,Rv2815c | NC_000962.3:3118383-3124377
Mapped GCEs: 0 | Putative GCEs: 1

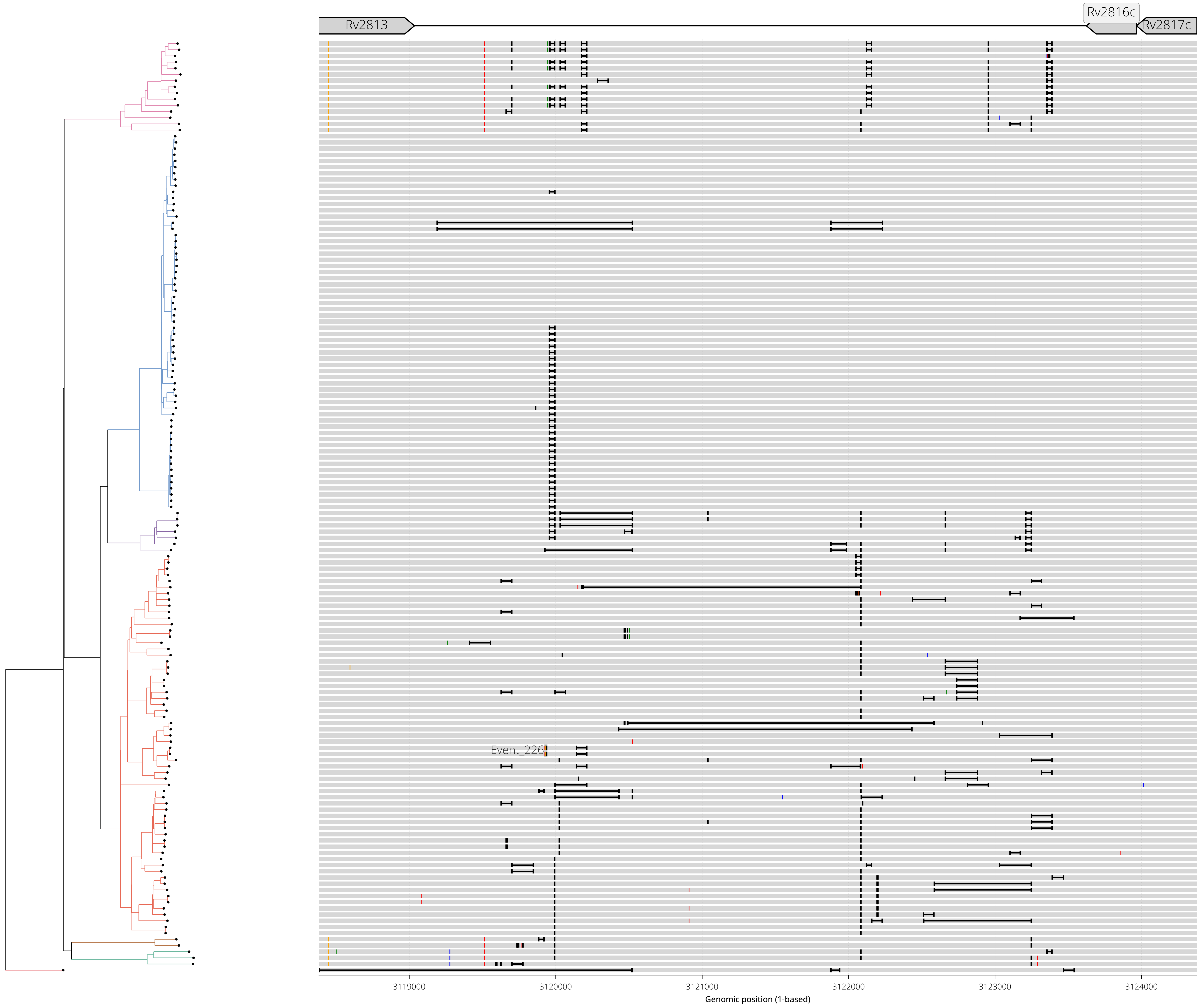

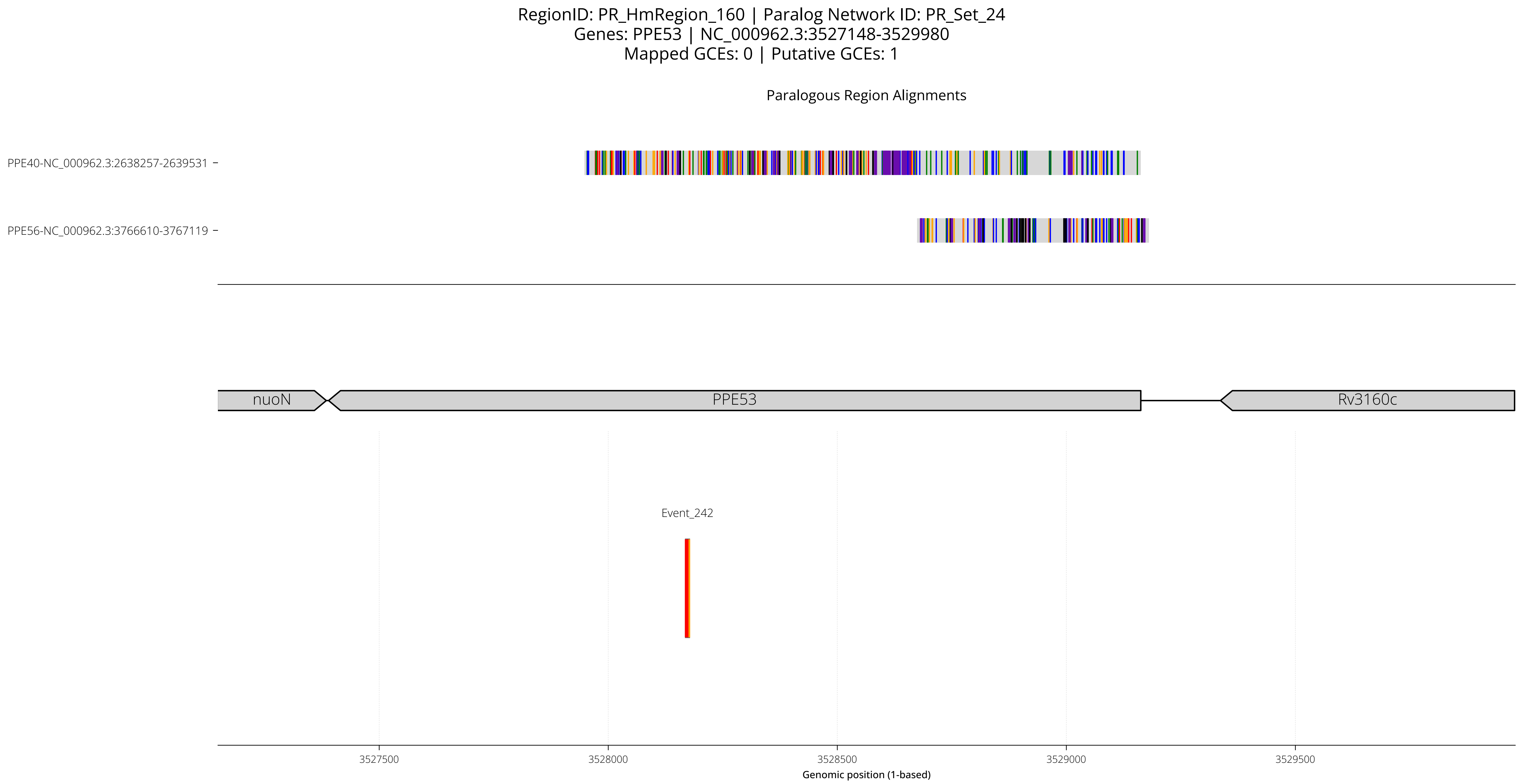
RegionID: PR_HmRegion_160 | Paralog Network ID: PR_Set_24
Genes: PPE53 | NC_000962.3:3527148-3529980
Mapped GCEs: 0 | Putative GCEs: 1
Paralogous Region Alignments
PPE40-NC_000962.3:2638257-2639531
PPE56-NC_000962.3:3766610-3767119
nuoN
PPE53
Rv3160c
Event_242
3527500
3528000
3528500
3529000
3529500
Genomic position (1-based)

RegionID: PR_HmRegion_160 | Paralog Network ID: PR_Set_24
Genes: PPE53 | NC_000962.3:3527148-3529980
Mapped GCEs: 0 | Putative GCEs: 1

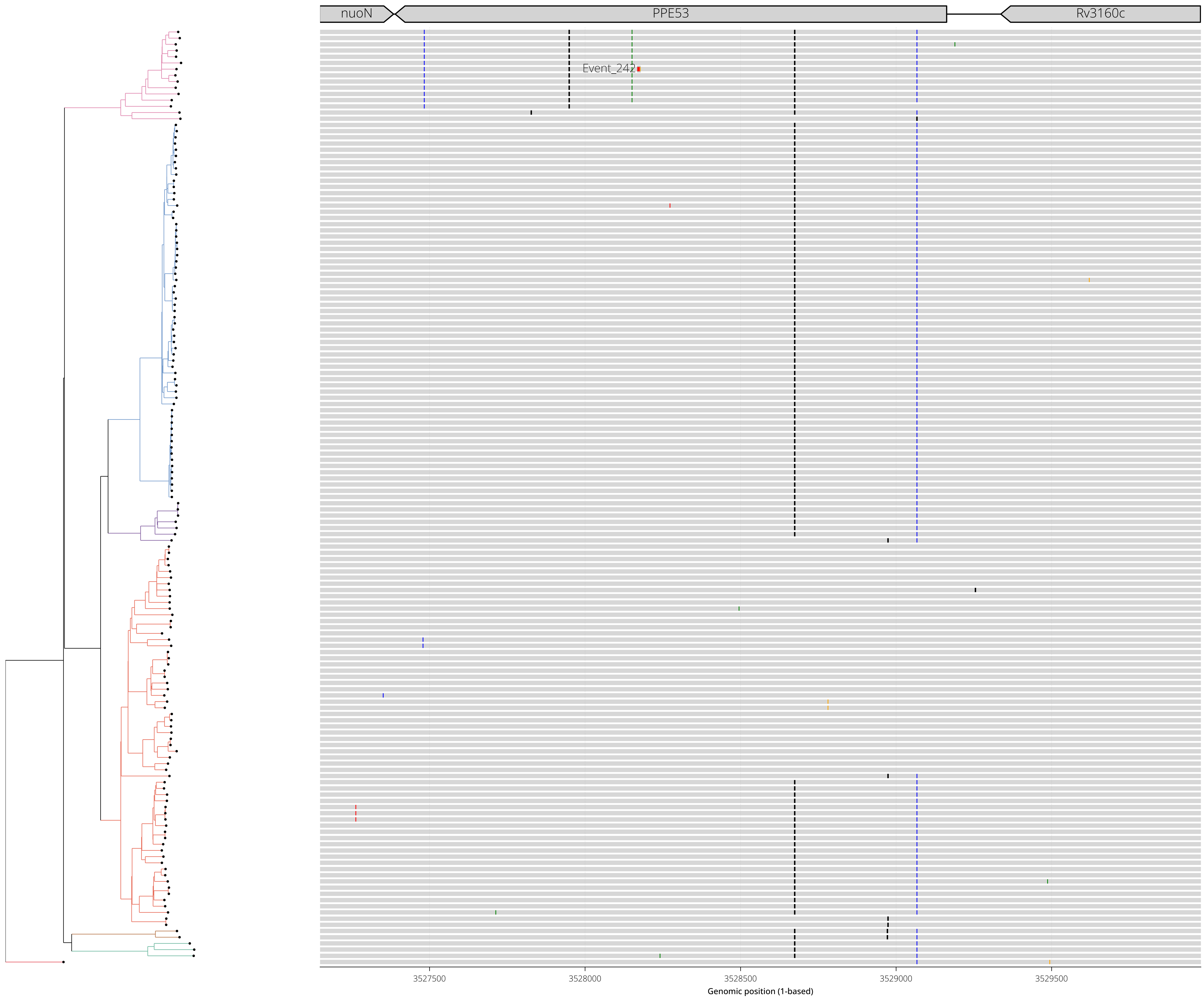

RegionID: PR_HmRegion_189 | Paralog Network ID: PR_Set_28
Genes: esxV,esxW | NC_000962.3:4059183-4061391
Mapped GCEs: 1 | Putative GCEs: 1

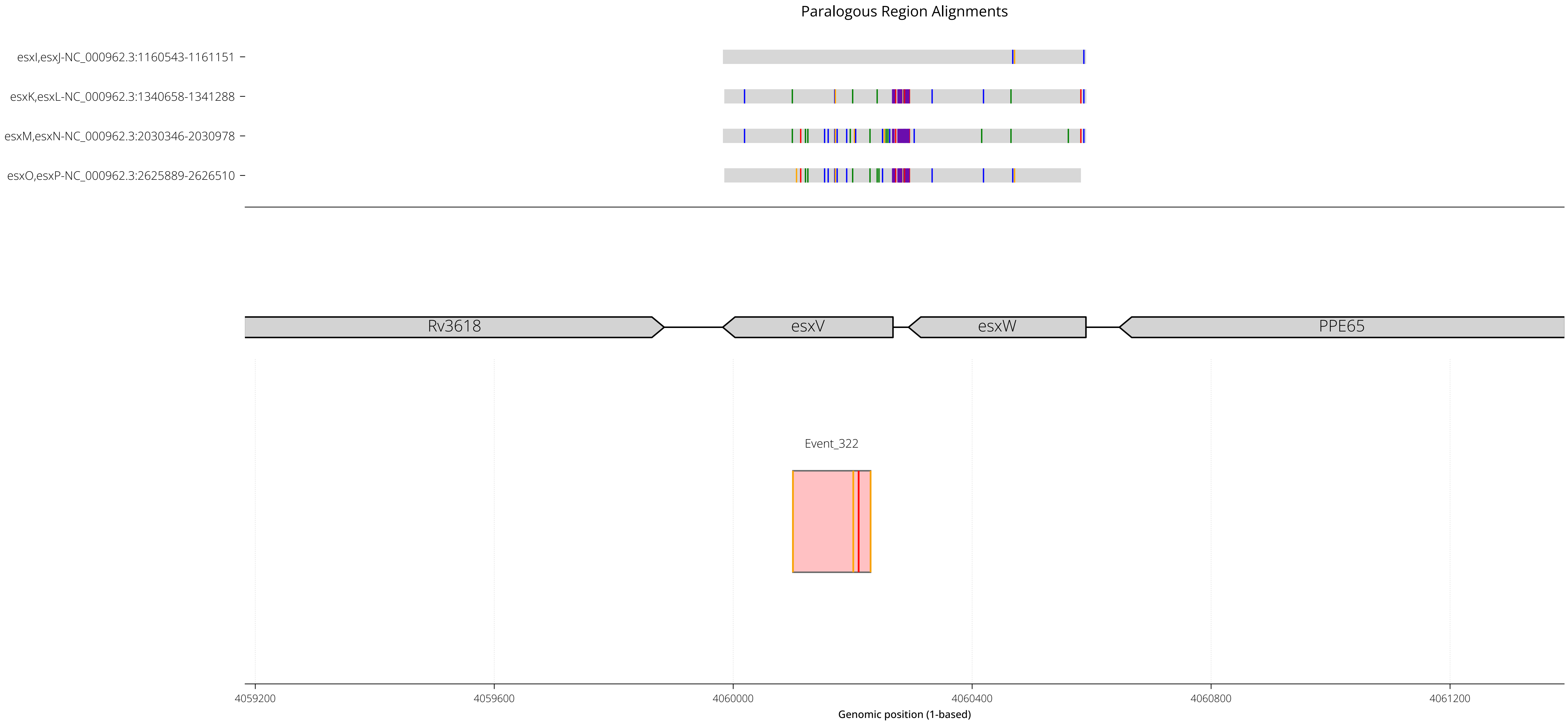

RegionID: PR_HmRegion_189 | Paralog Network ID: PR_Set_28
Genes: esxV,esxW | NC_000962.3:4059183-4061391
Mapped GCEs: 1 | Putative GCEs: 1

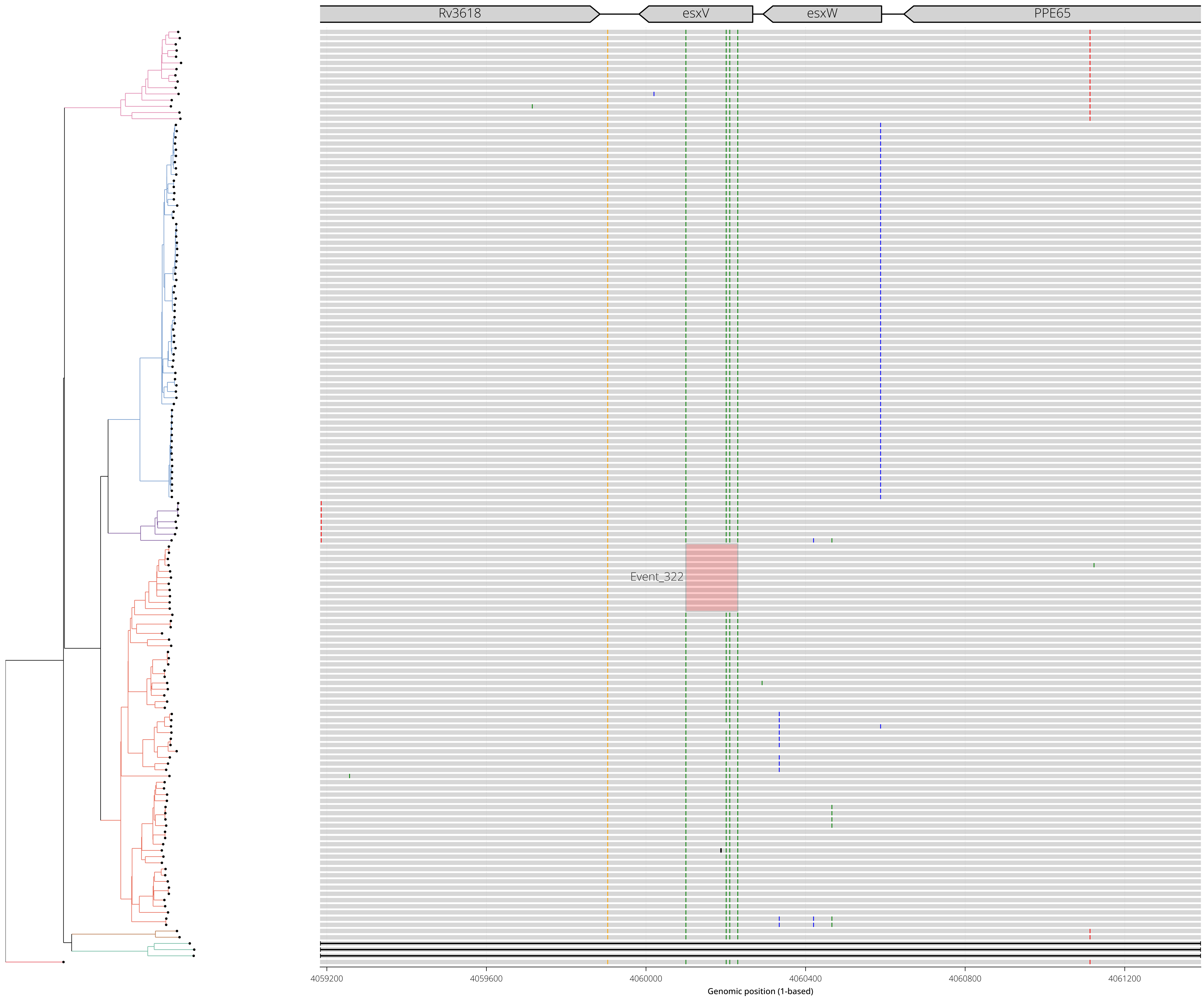

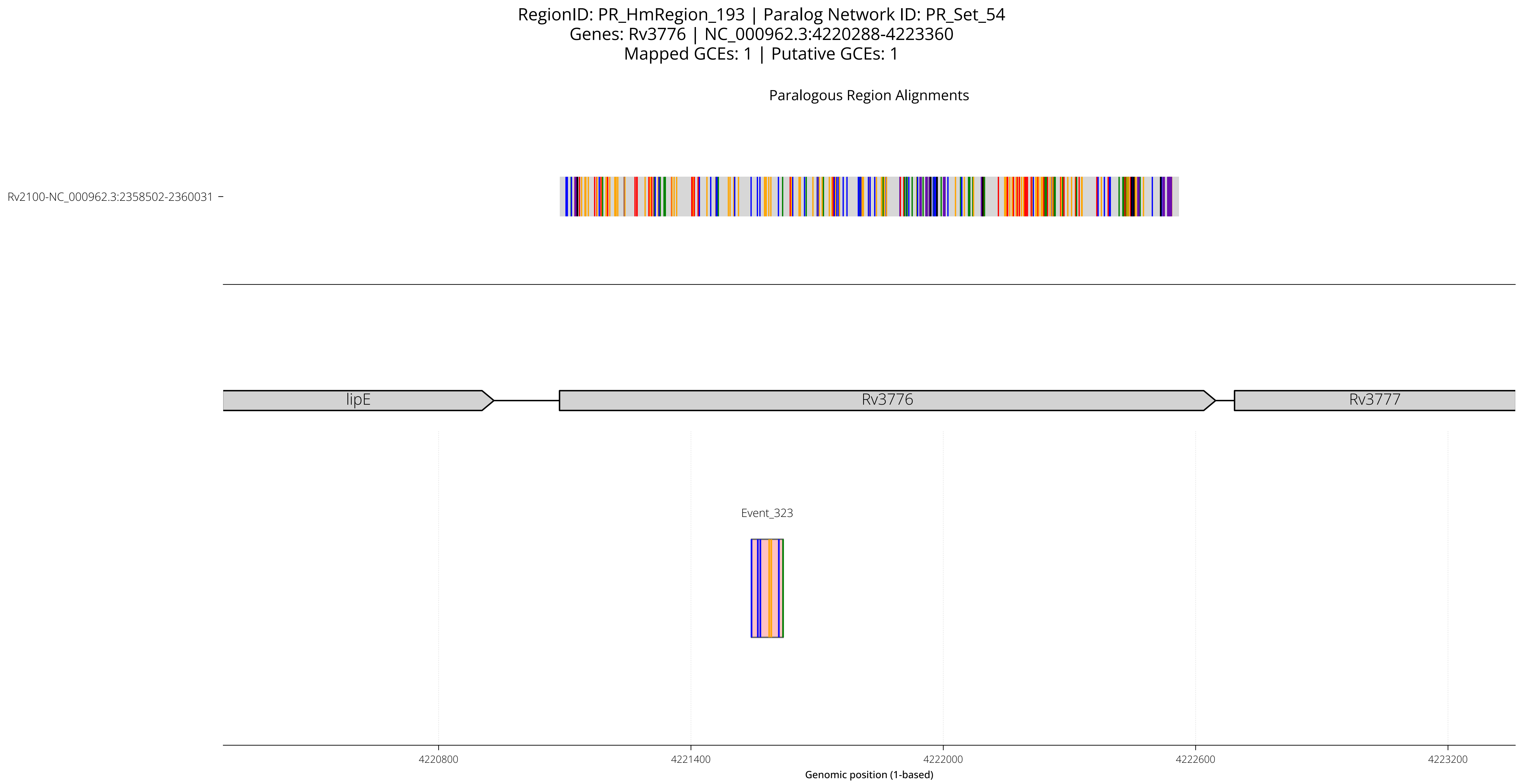

RegionID: PR_HmRegion_193 | Paralog Network ID: PR_Set_54
Genes: Rv3776 | NC_000962.3:4220288-4223360
Mapped GCEs: 1 | Putative GCEs: 1
Paralogous Region Alignments
Rv2100-NC_000962.3:2358502-2360031
lipE
Rv3776
Rv3777
Event_323
4220800
4221400
4222000
4222600
4223200
Genomic position (1-based)

RegionID: PR_HmRegion_193 | Paralog Network ID: PR_Set_54
Genes: Rv3776 | NC_000962.3:4220288-4223360
Mapped GCEs: 1 | Putative GCEs: 1

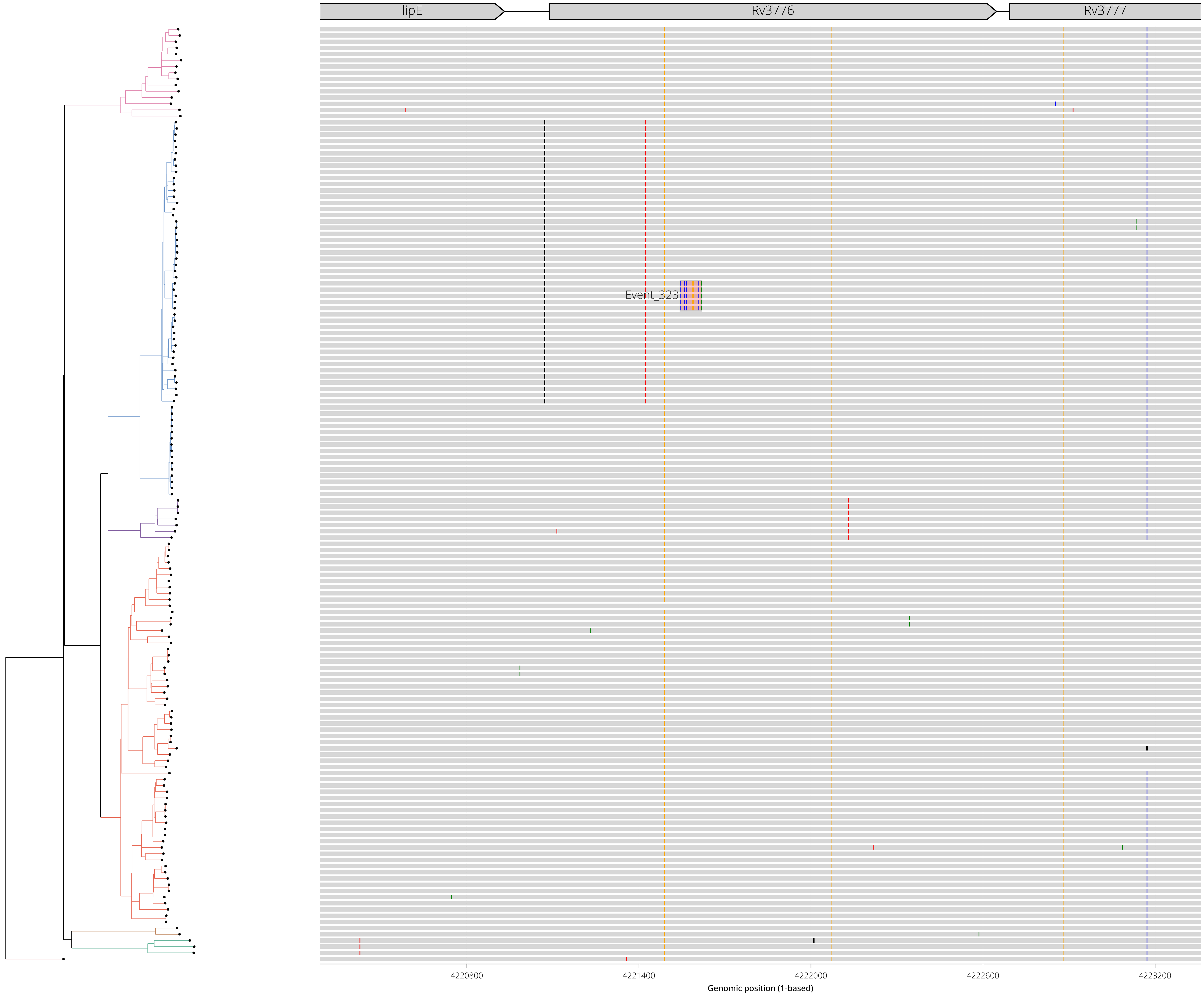

RegionID: PR_HmRegion_194 | Paralog Network ID: PR_Set_38
Genes: fadE35,Rv3798 | NC_000962.3:4252053-4255167
Mapped GCEs: 1 | Putative GCEs: 1

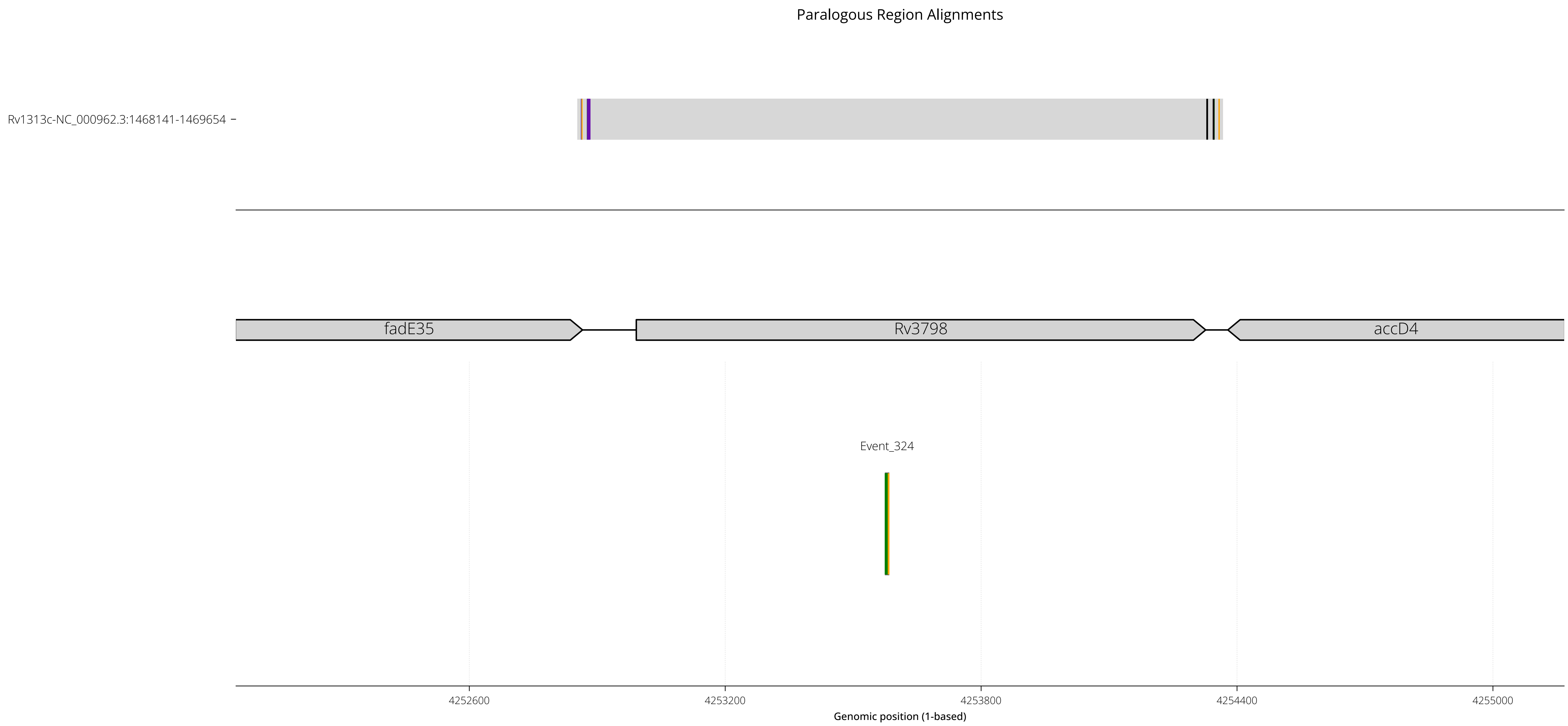

RegionID: PR_HmRegion_194 | Paralog Network ID: PR_Set_38
Genes: fadE35,Rv3798 | NC_000962.3:4252053-4255167
Mapped GCEs: 1 | Putative GCEs: 1

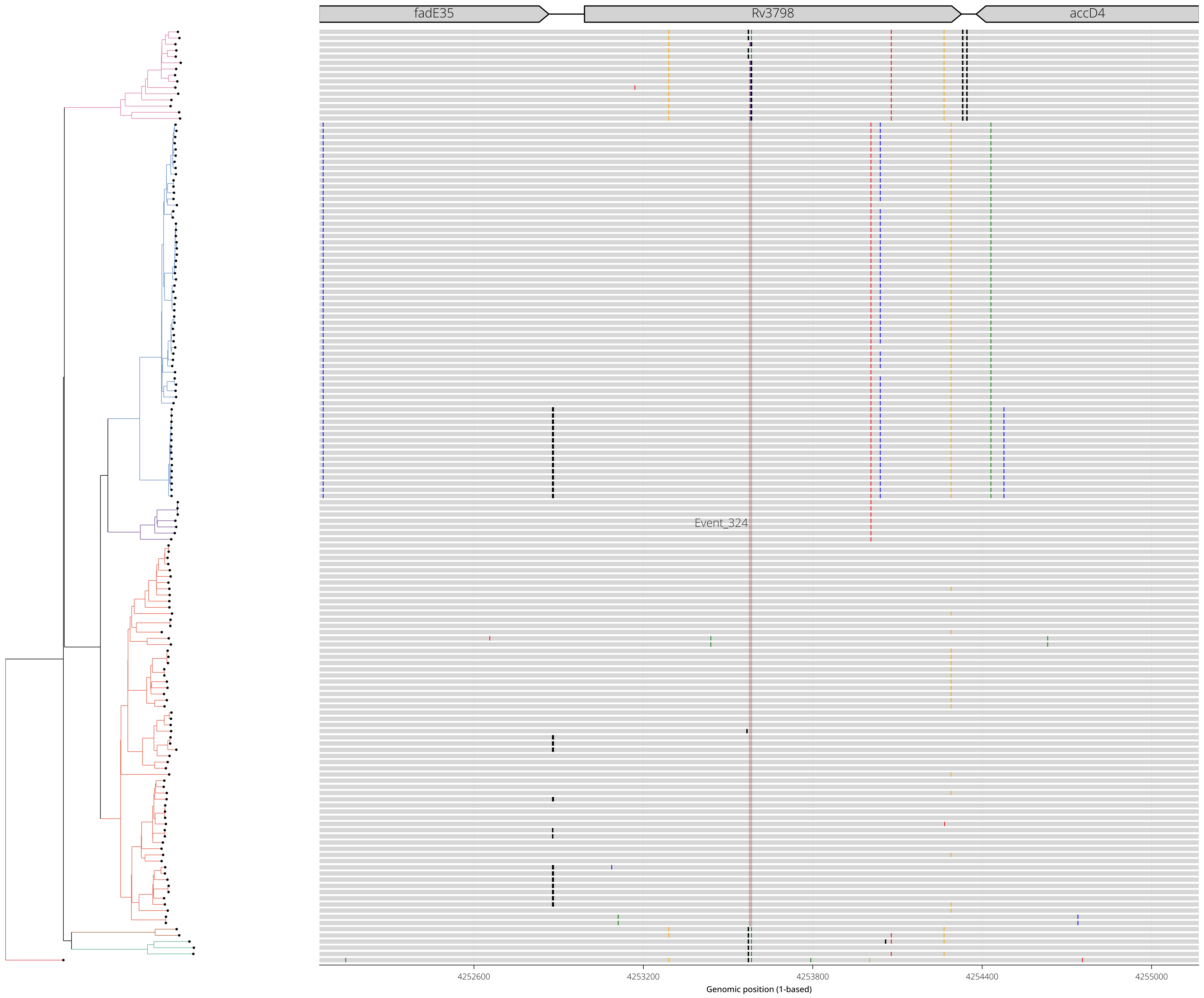
