## Supplemental Data File 13 for "Gene conversion is a key driver of diversity hotspots in *M. tuberculosis* antigens and virulence-associated loci"

### SI-13 — Per-event visualization of individual gene conversion events

#### Overview

This file contains visualizations for each putative gene conversion event detected across our primary analysis of 151 *M. tuberculosis* genomes (N = 324 events, **Supplemental Data File 5**). For each event, five complementary visualizations are generated. The purpose is to transparently document every detected event individually: the associated variants, the genomic location of all known paralogous sequences, the supporting k-mer evidence for accessing paralog similarity, and event placement in the phylogeny. All gene annotations and coordinates shown are relative to the H37Rv reference [NC_000962.3].

Five visualizations are generated per event:

- View A: Event versus reference paralog sequences: the event and its variants relative to the aligned paralogous regions from the H37Rv reference.
- View B: Genomic locations of paralogs relative to the event: a circular genome view linking the event to each paralogous regions' location in the H37Rv reference.
- View C: K-mer analysis and paralog mapping: per-paralog visualization of sequence similarity to the target event.
- View D: Sub-phylogeny view: the event placed in the local subtree containing it (zoomed in).
- View E: Full-phylogeny view: the same event placed in the complete dataset phylogeny (zoomed out).

Note that View's D and E show the same data as in SD-12's visualization's except only a single gene conversion event is highlighted at a time. All supporting raw data for these visualizations can be found in Supplemental Data files 5 and 6.

#### What is shown

#### View A — Event versus reference paralog sequences

A genome-browser-style view comparing the event to its candidate paralog sequences. From top to bottom:

- Paralog variant alignments: each paralogous region aligned to the affected region, with one alignment per row and its alignment span shown relative to the target region. Variants between the paralog and the region follow the same key used in SI-12.
- Gene annotations: H37Rv annotated genes for the region.
- Gene conversion event: the genomic span of this individual event, drawn as a range with a red highlight. Variants within the event box follow the variant key below.

**Figure key — View A:**

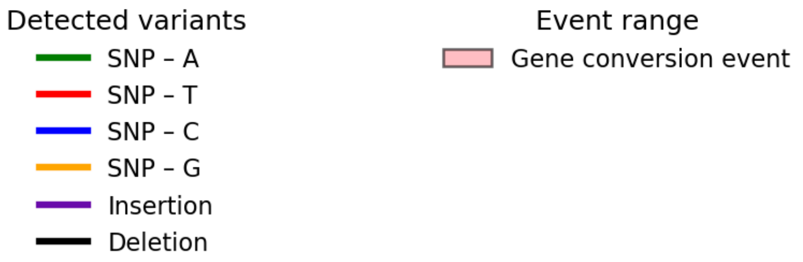


#### View B — Genomic locations of paralogs relative to the event

A circular view of the H37Rv genome showing the genomic location of the event and each of its candidate paralogs. Links are drawn from the event location (labeled with its event ID) to the reference location of each listed paralog.

**Figure key — View B:**

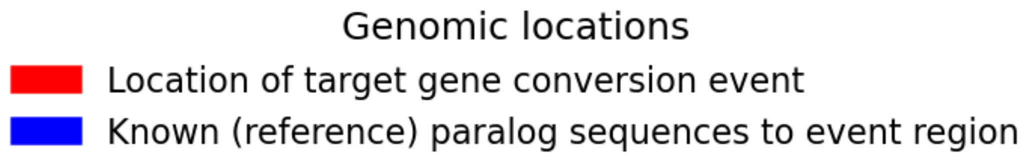

#### View C — K-mer analysis and paralog mapping

The top panel shows gene annotations with the substitutions associated with the event overlaid. Below, each candidate paralog is given an individual track showing how the k-mer profile of the event (the recombinant, gene-conversion-mutated tract) compares to that paralog sequence. The coloring indicates whether the recombinant tract's k-mer profile matches the paralog sequence, providing the evidence used to identify the most likely donor.

**Figure key — View C:**

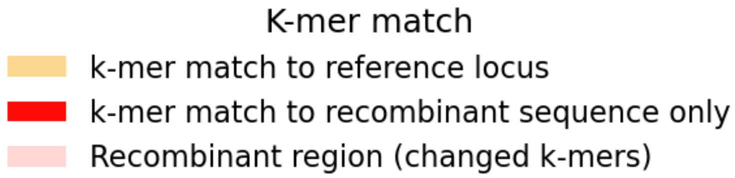

##### Views D and E — Phylogenetic-context views (sub-phylogeny and full phylogeny)

Both views place the single event of interest in phylogenetic context, using the same layout. From top to bottom:

- Gene annotations: H37Rv annotated genes for the region.
- Per-genome variant calls: the alignment of each shown genome to the reference, with variants colored by the mutant allele following the key below.
- Phylogeny: the phylogeny is displayed to the left, with each tip aligned horizontally to that genome's variant row. Branches are colored by MTBC lineage.
- Gene conversion event: the event is drawn as a box spanning the genomic region it covers, overlaid on the genome alignments and aligned to the phylogeny branch on which it is inferred to have occurred. The box is labeled with the event ID.

The two views differ only in phylogenetic scope. View D (sub-phylogeny view) is restricted to the local subtree starting at the ancestral branch immediately before the event is inferred to have occurred, giving a zoomed-in view of the event and only the isolates carrying the recombination. View E (full-phylogeny view) shows the complete phylogeny of all 151 genomes without zooming in, placing the event in the context of the entire dataset.

##### Figure key — Views D and E

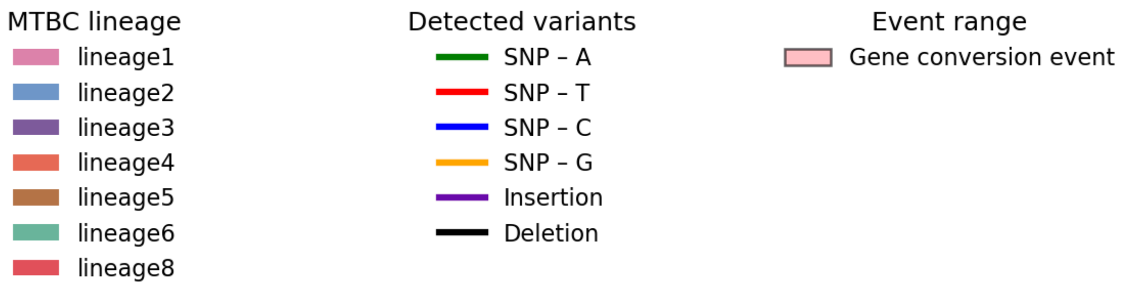

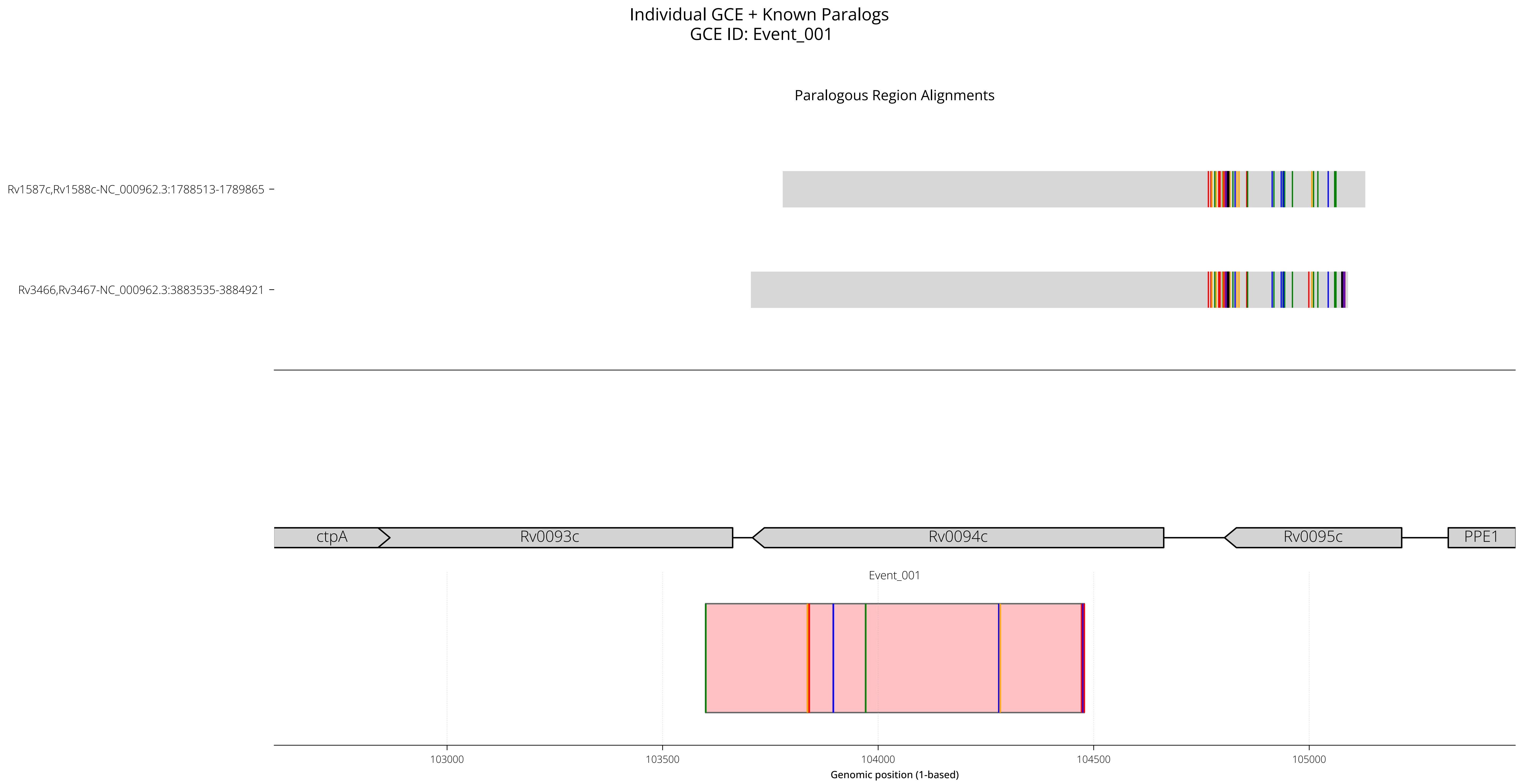
Individual GCE + Known Paralogs
GCE ID: Event_001
Paralogous Region Alignments
Rv1587c,Rv1588c-NC_000962.3:1788513-1789865
Rv3466,Rv3467-NC_000962.3:3883535-3884921
ctpA
Rv0093c
Rv0094c
Rv0095c
PPE1
Event_001
103000
103500
104000
104500
105000
Genomic position (1-based)

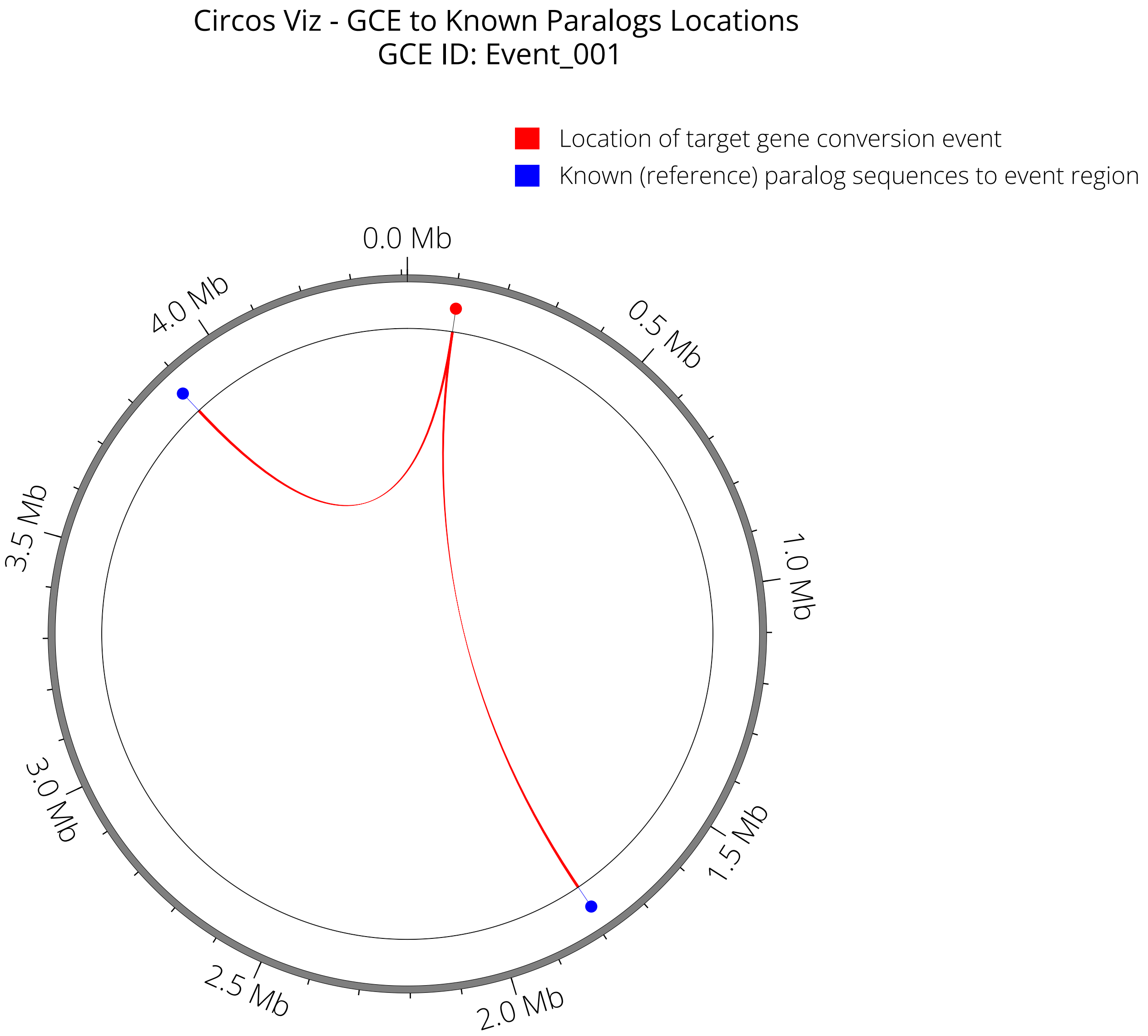
Circos Viz - GCE to Known Paralogs Locations
GCE ID: Event_001
Location of target gene conversion event
Known (reference) paralog sequences to event region
0.0 Mb
0.5 Mb
1.0 Mb
1.5 Mb
2.0 Mb
2.5 Mb
3.0 Mb
3.5 Mb
4.0 Mb

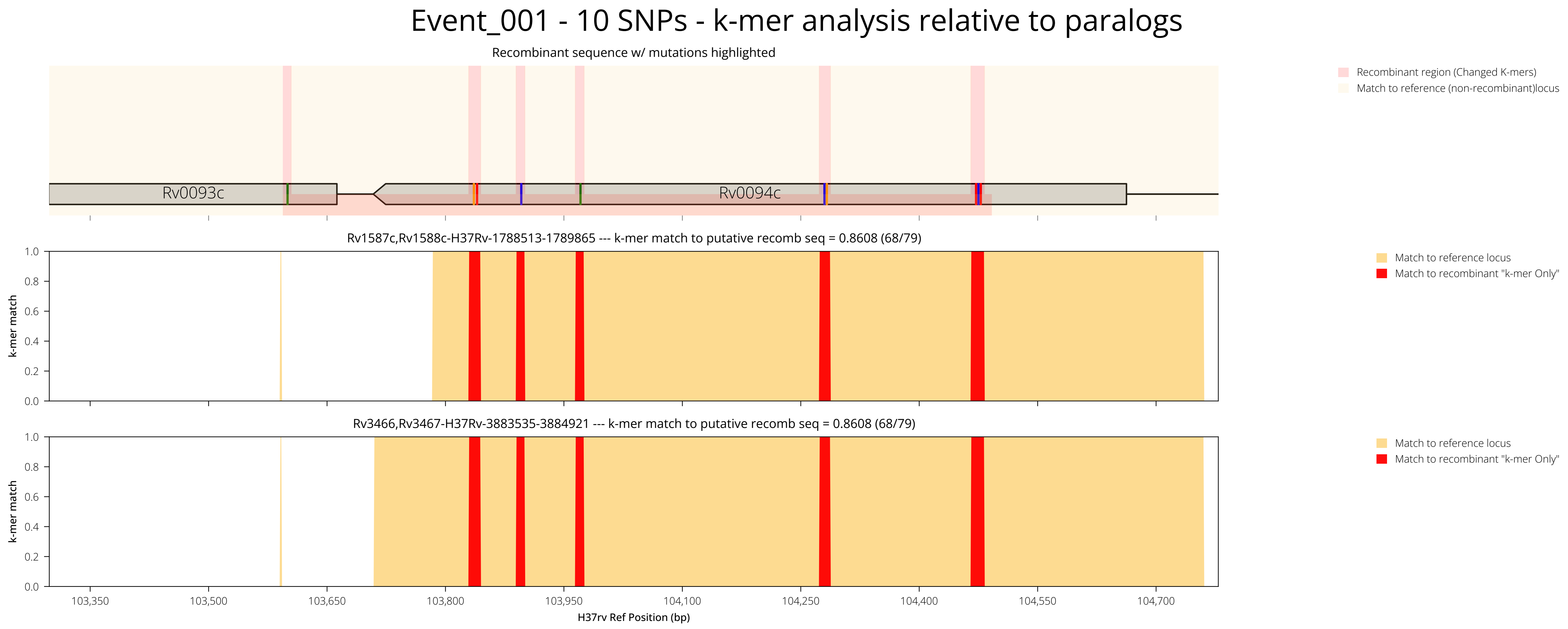

Event_001 - 10 SNPs - k-mer analysis relative to paralogs
Recombinant sequence w/ mutations highlighted
Recombinant region (Changed K-mers)
Match to reference (non-recombinant)locus
Rv0093c
Rv0094c
Rv1587c,Rv1588c-H37Rv-1788513-1789865 --- k-mer match to putative recomb seq = 0.8608 (68/79)
Match to reference locus
Match to recombinant "k-mer Only"
k-mer match
1.0
0.8
0.6
0.4
0.2
0.0
Rv3466,Rv3467-H37Rv-3883535-3884921 --- k-mer match to putative recomb seq = 0.8608 (68/79)
Match to reference locus
Match to recombinant "k-mer Only"
k-mer match
1.0
0.8
0.6
0.4
0.2
0.0
103,350
103,500
103,650
103,800
103,950
104,100
104,250
104,400
104,550
104,700
H37rv Ref Position (bp)

Sub-phylogeny Viz + Paralog Alignments
GCE ID: Event_001

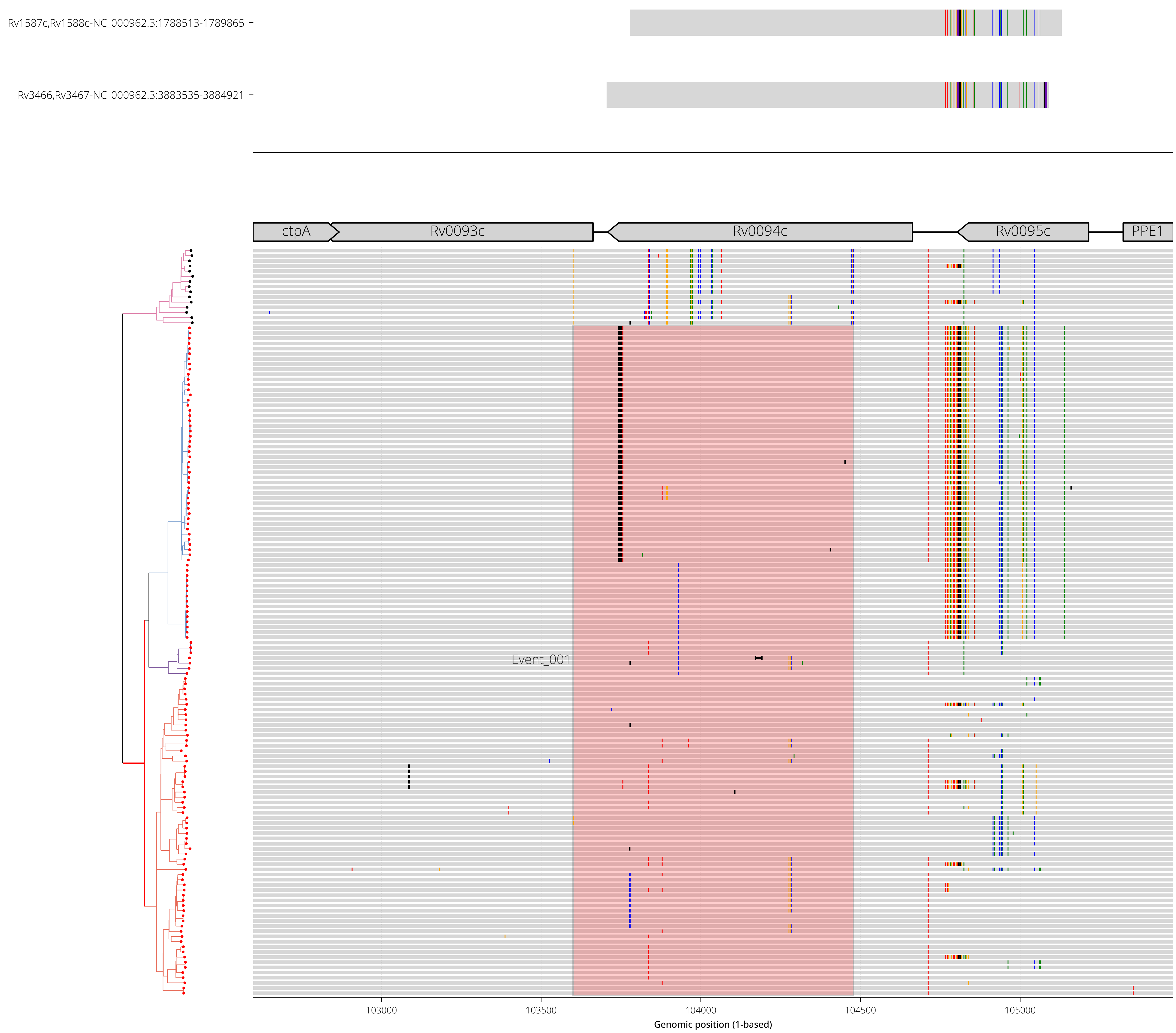

Full Phylogeny Viz + Paralog Alignments
GCE ID: Event_001

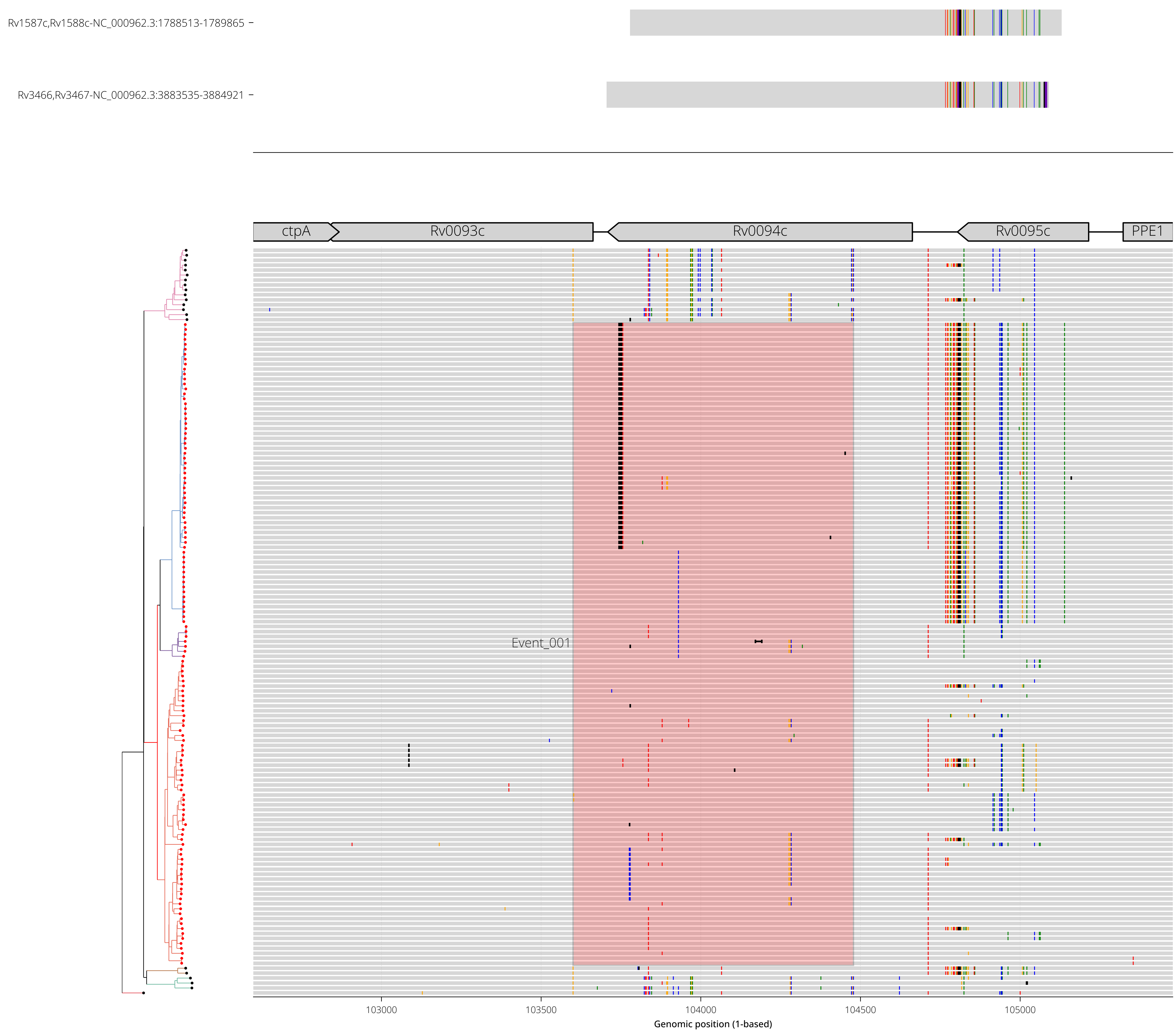

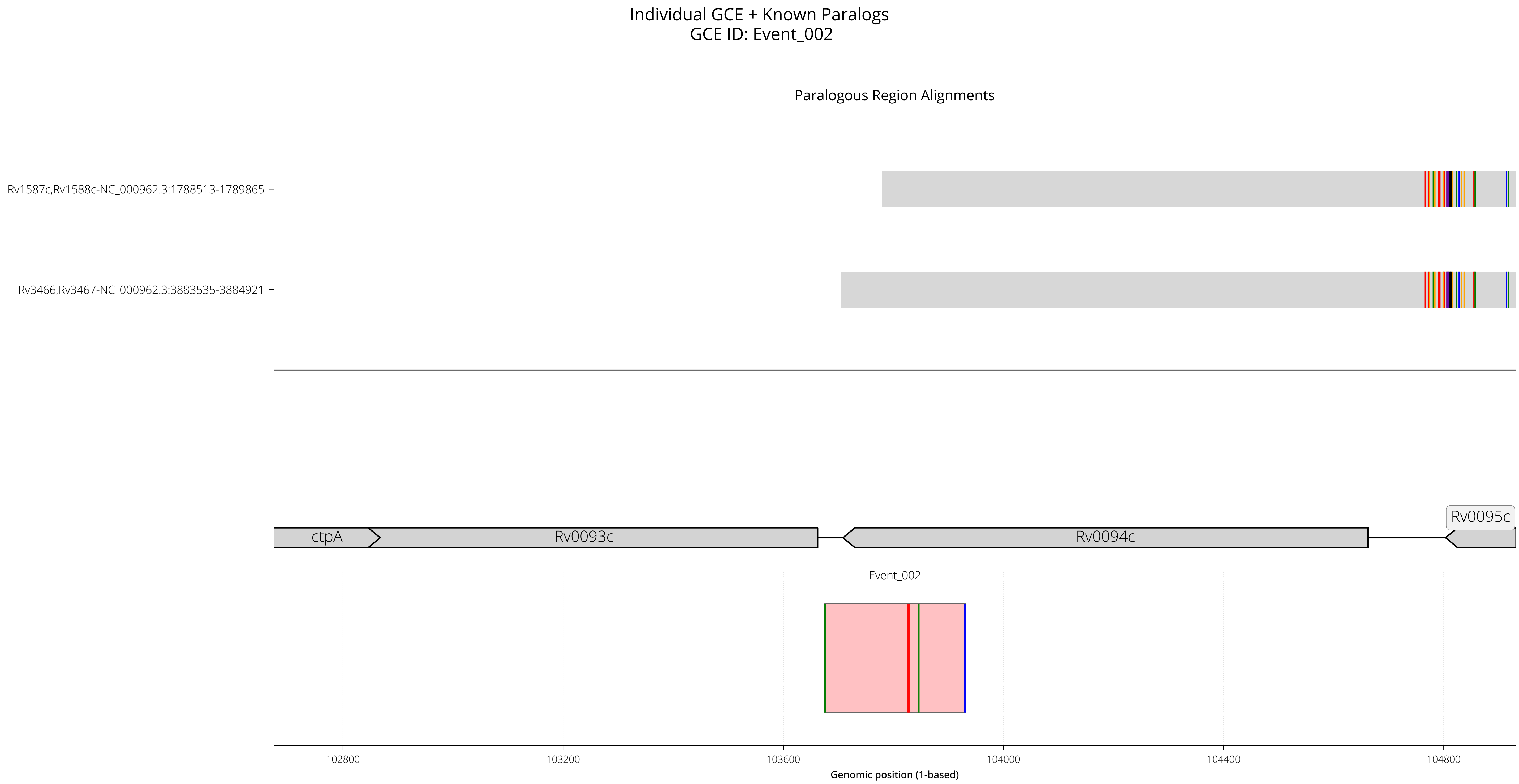

Individual GCE + Known Paralogs
GCE ID: Event_002
Paralogous Region Alignments
Rv1587c,Rv1588c-NC_000962.3:1788513-1789865
Rv3466,Rv3467-NC_000962.3:3883535-3884921
ctpA
Rv0093c
Rv0094c
Rv0095c
Event_002
102800
103200
103600
104000
104400
104800
Genomic position (1-based)

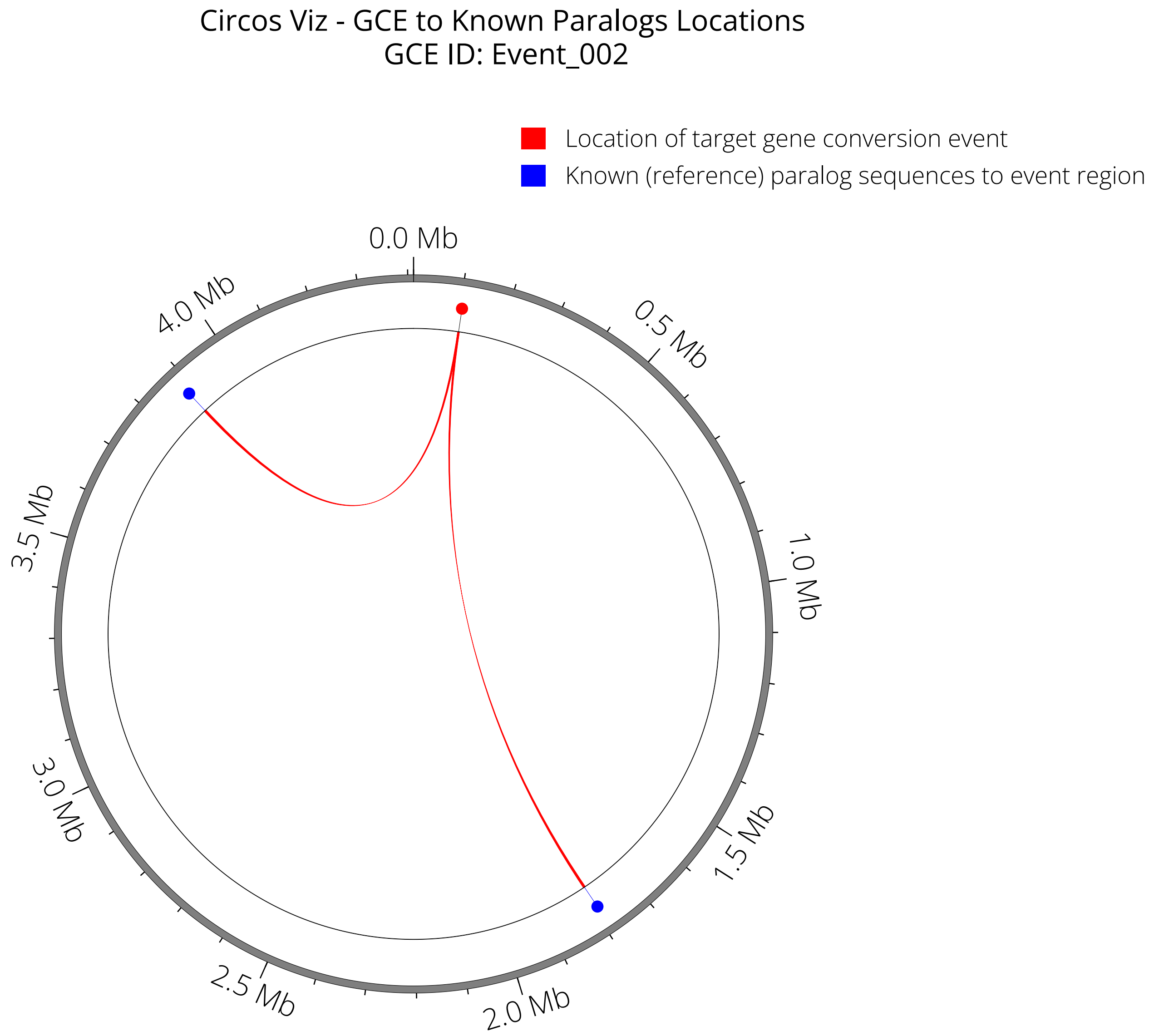
Circos Viz - GCE to Known Paralogs Locations
GCE ID: Event_002
Location of target gene conversion event
Known (reference) paralog sequences to event region
0.0 Mb
0.5 Mb
1.0 Mb
1.5 Mb
2.0 Mb
2.5 Mb
3.0 Mb
3.5 Mb
4.0 Mb

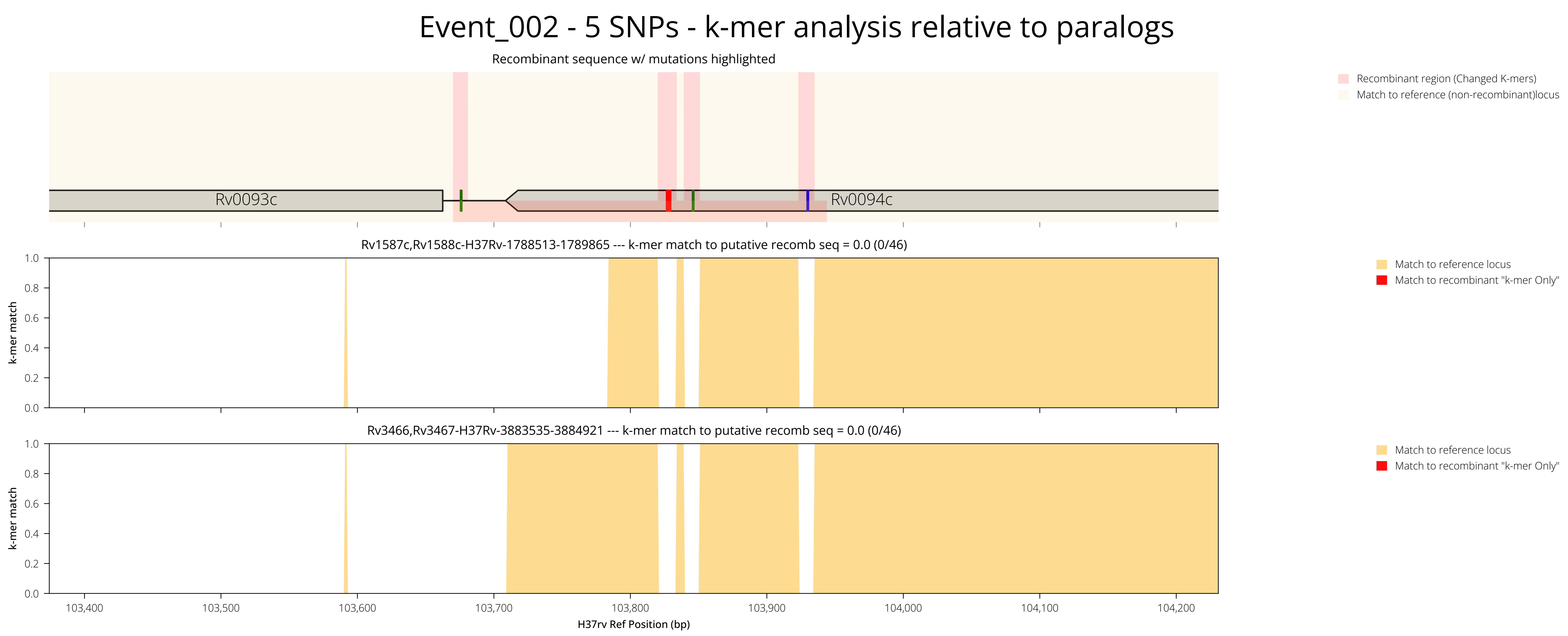

Event_002 - 5 SNPs - k-mer analysis relative to paralogs
Recombinant sequence w/ mutations highlighted
Rv0093c
Rv0094c
Recombinant region (Changed K-mers)
Match to reference (non-recombinant)locus
Rv1587c,Rv1588c-H37Rv-1788513-1789865 --- k-mer match to putative recomb seq = 0.0 (0/46)
k-mer match
1.0
0.8
0.6
0.4
0.2
0.0
Match to reference locus
Match to recombinant "k-mer Only"
Rv3466,Rv3467-H37Rv-3883535-3884921 --- k-mer match to putative recomb seq = 0.0 (0/46)
k-mer match
1.0
0.8
0.6
0.4
0.2
0.0
Match to reference locus
Match to recombinant "k-mer Only"
103,400
103,500
103,600
103,700
103,800
103,900
104,000
104,100
104,200
H37rv Ref Position (bp)

Sub-phylogeny Viz + Paralog Alignments
GCE ID: Event_002

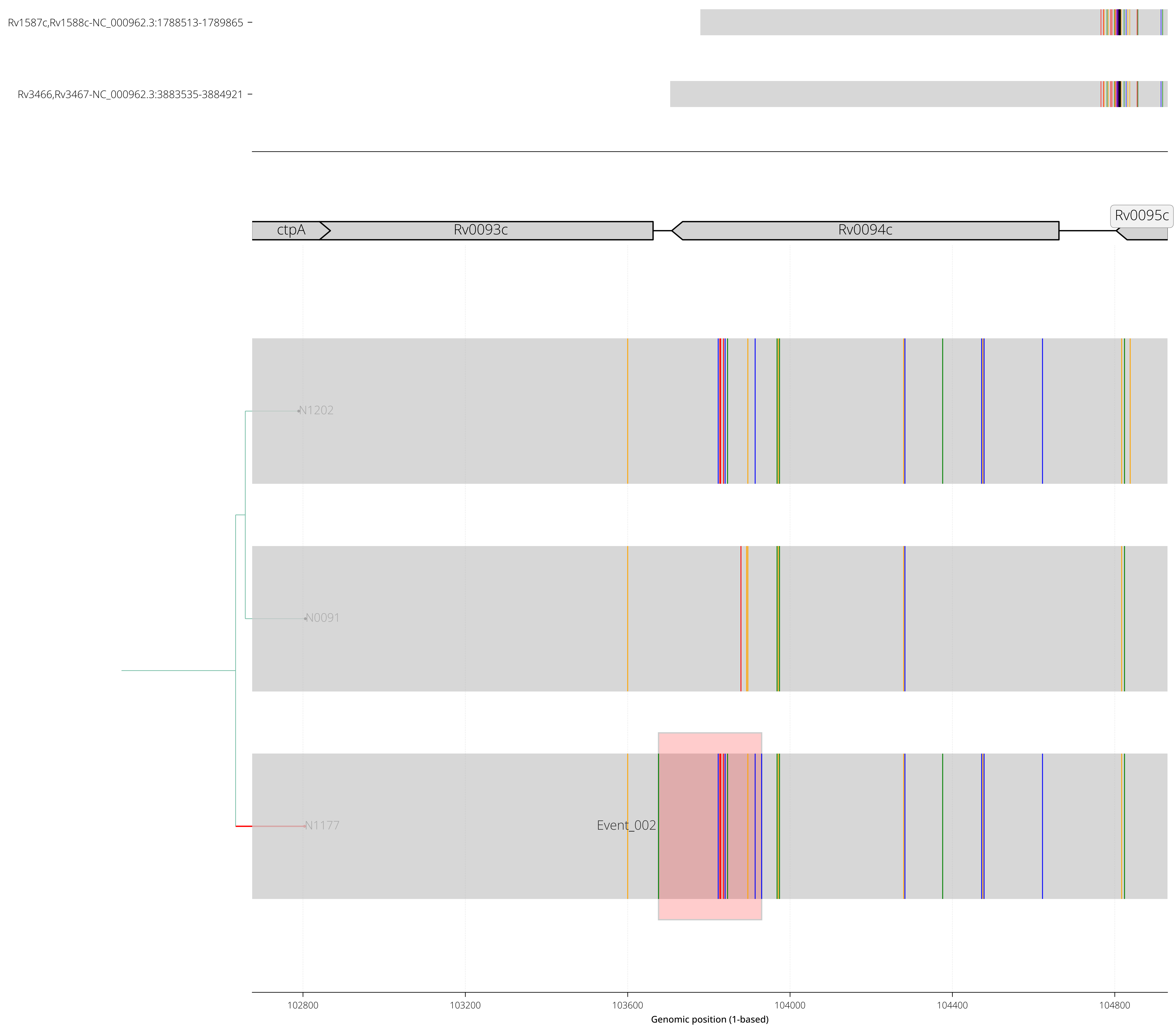

Full Phylogeny Viz + Paralog Alignments
GCE ID: Event_002

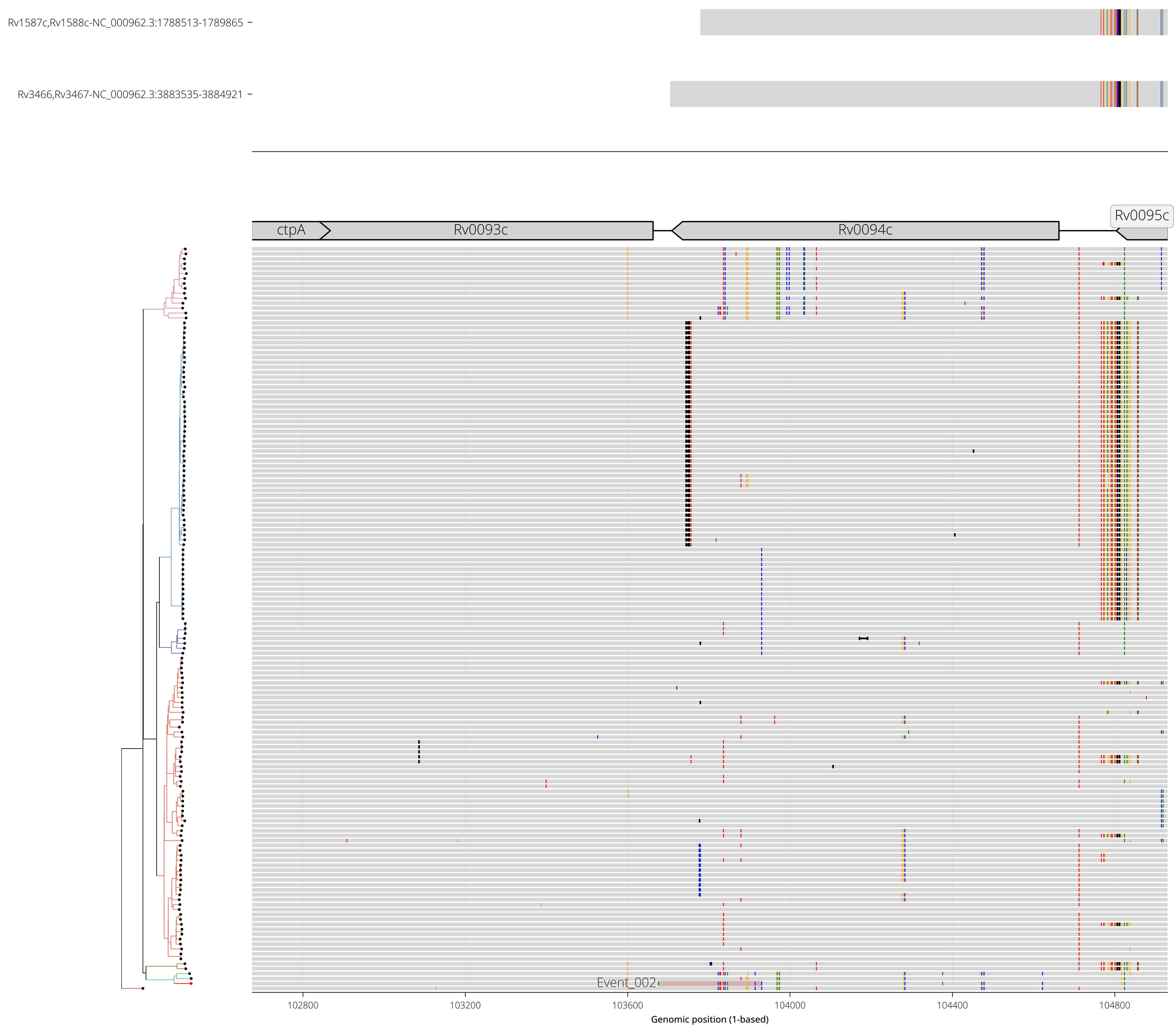

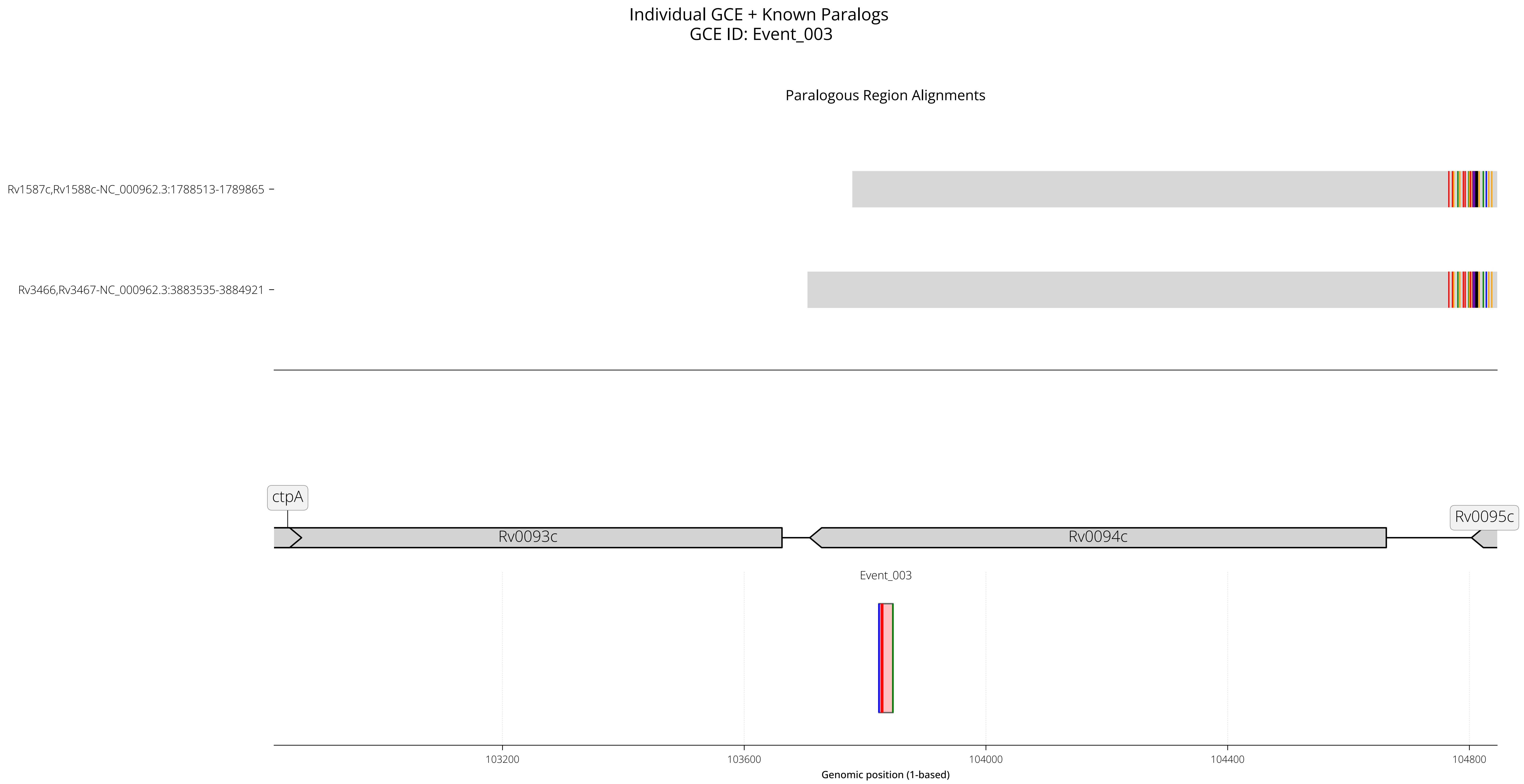
Individual GCE + Known Paralogs
GCE ID: Event_003
Paralogous Region Alignments
Rv1587c,Rv1588c-NC_000962.3:1788513-1789865
Rv3466,Rv3467-NC_000962.3:3883535-3884921
ctpA
Rv0093c
Rv0094c
Rv0095c
Event_003
103200
103600
104000
104400
104800
Genomic position (1-based)

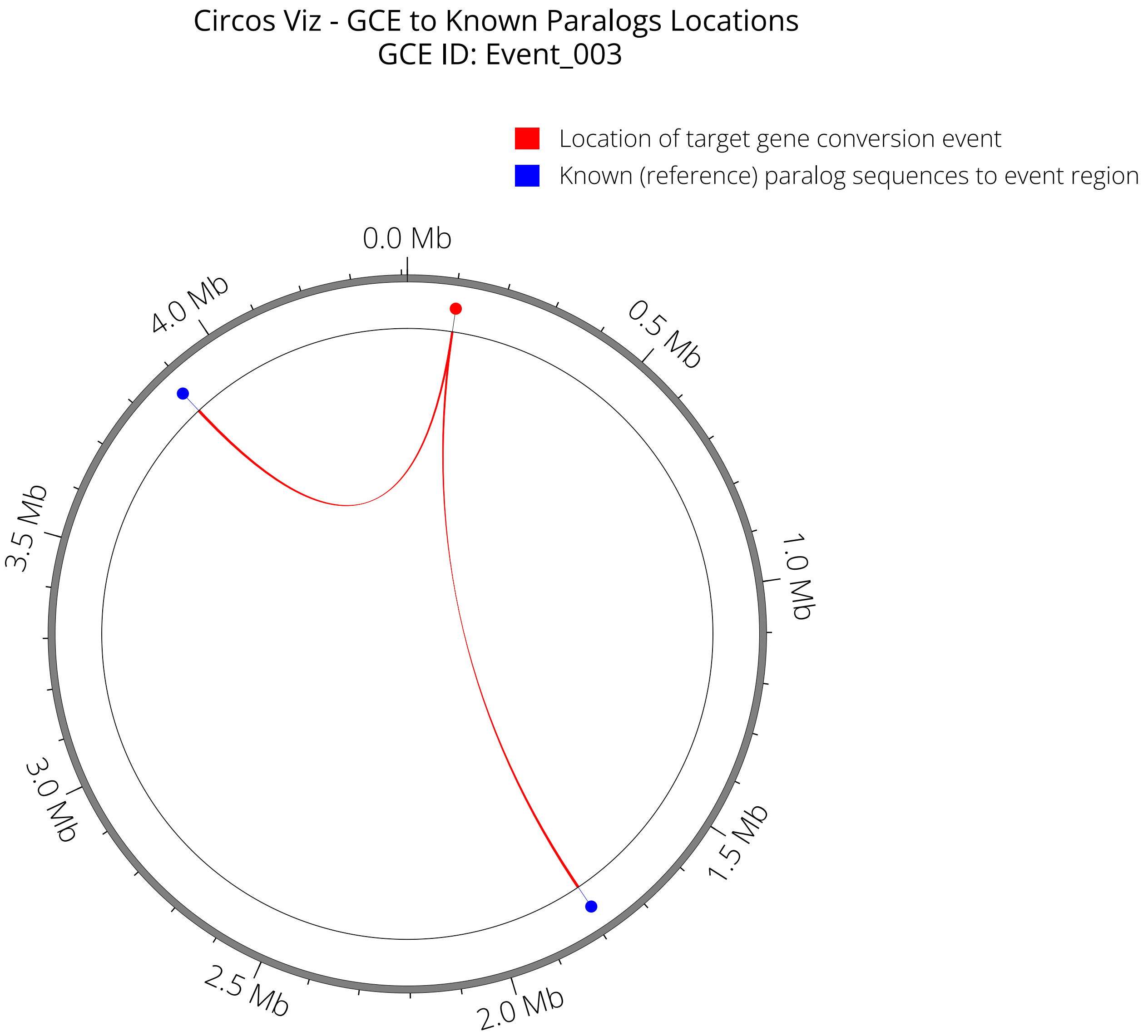
Circos Viz - GCE to Known Paralogs Locations
GCE ID: Event_003
Location of target gene conversion event
Known (reference) paralog sequences to event region
0.0 Mb
0.5 Mb
1.0 Mb
1.5 Mb
2.0 Mb
2.5 Mb
3.0 Mb
3.5 Mb
4.0 Mb

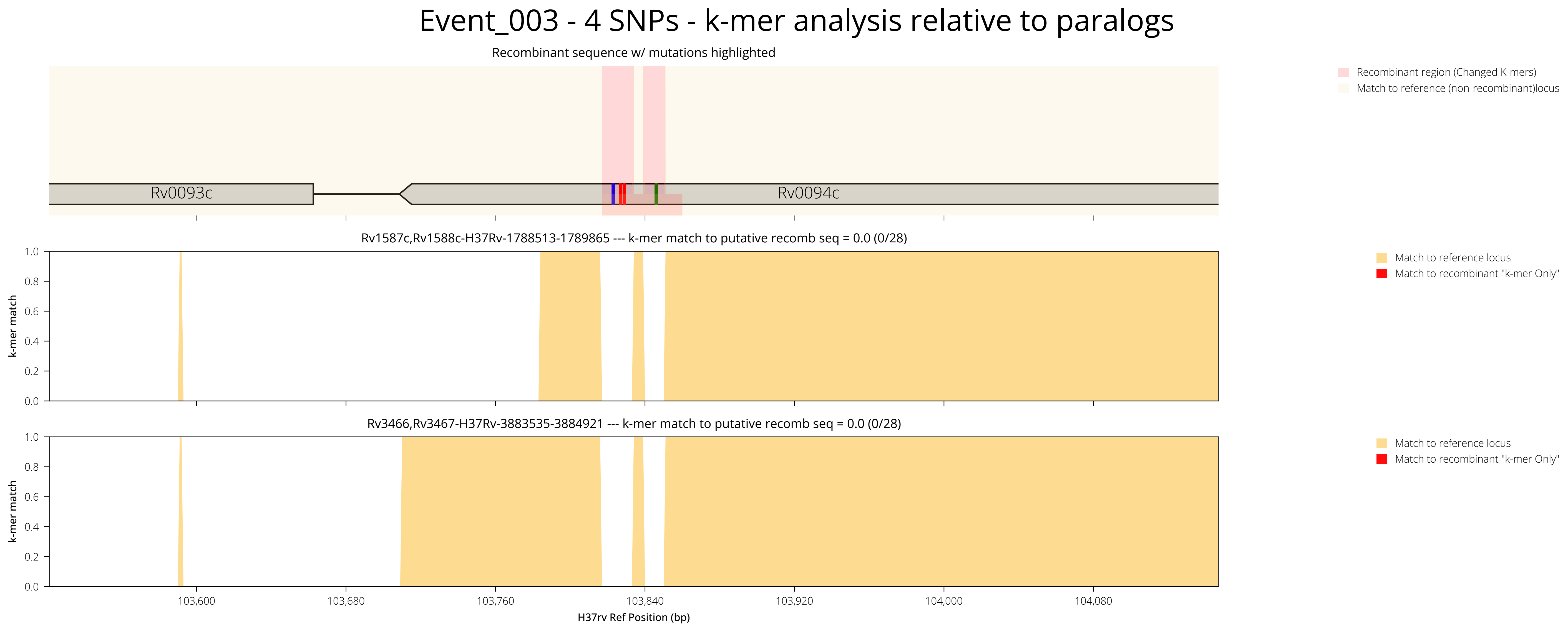

Event_003 - 4 SNPs - k-mer analysis relative to paralogs
Recombinant sequence w/ mutations highlighted
Recombinant region (Changed K-mers)
Match to reference (non-recombinant)locus
Rv0093c
Rv0094c
Rv1587c,Rv1588c-H37Rv-1788513-1789865 --- k-mer match to putative recomb seq = 0.0 (0/28)
Match to reference locus
Match to recombinant "k-mer Only"
k-mer match
1.0
0.8
0.6
0.4
0.2
0.0
Rv3466,Rv3467-H37Rv-3883535-3884921 --- k-mer match to putative recomb seq = 0.0 (0/28)
Match to reference locus
Match to recombinant "k-mer Only"
k-mer match
1.0
0.8
0.6
0.4
0.2
0.0
103,600
103,680
103,760
103,840
103,920
104,000
104,080
H37rv Ref Position (bp)

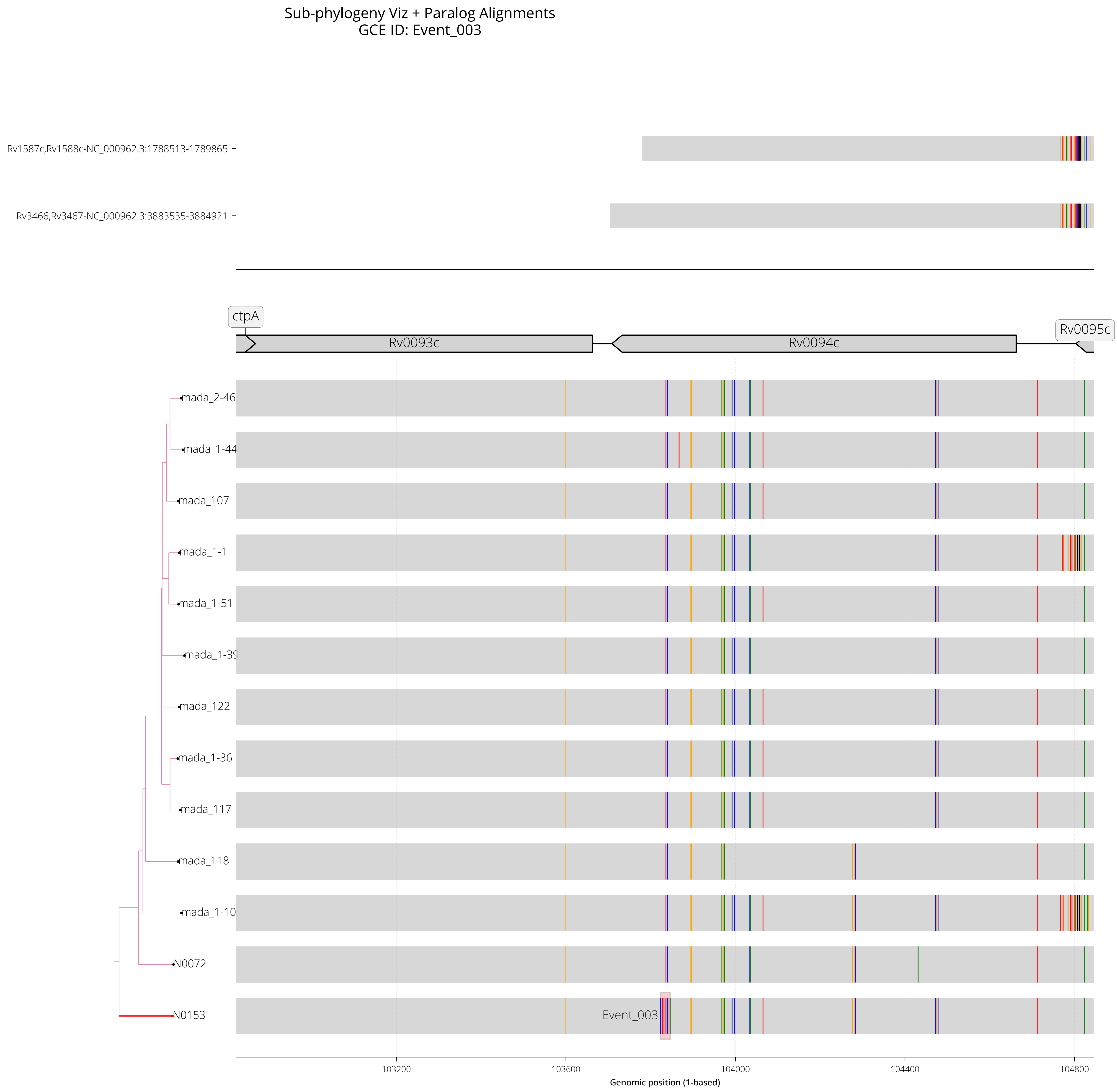

Sub-phylogeny Viz + Paralog Alignments
GCE ID: Event_003
Rv1587c,Rv1588c-NC_000962.3:1788513-1789865
Rv3466,Rv3467-NC_000962.3:3883535-3884921
ctpA
Rv0093c
Rv0094c
Rv0095c
mada_2-46
mada_1-44
mada_107
mada_1-1
mada_1-51
mada_1-39
mada_122
mada_1-36
mada_117
mada_118
mada_1-10
N0072
N0153
Event_003
103200
103600
104000
104400
104800
Genomic position (1-based)

Full Phylogeny Viz + Paralog Alignments
GCE ID: Event_003

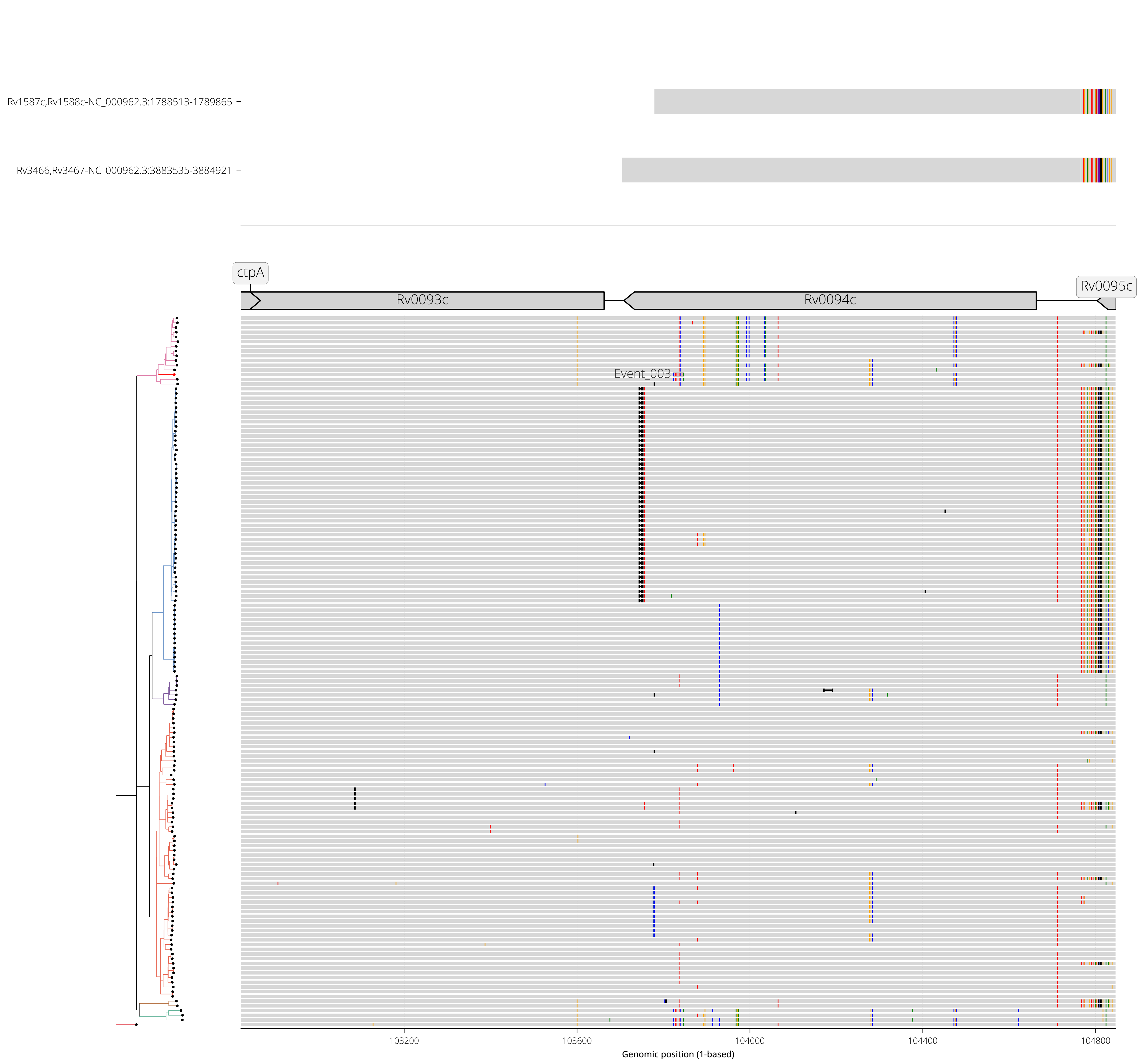

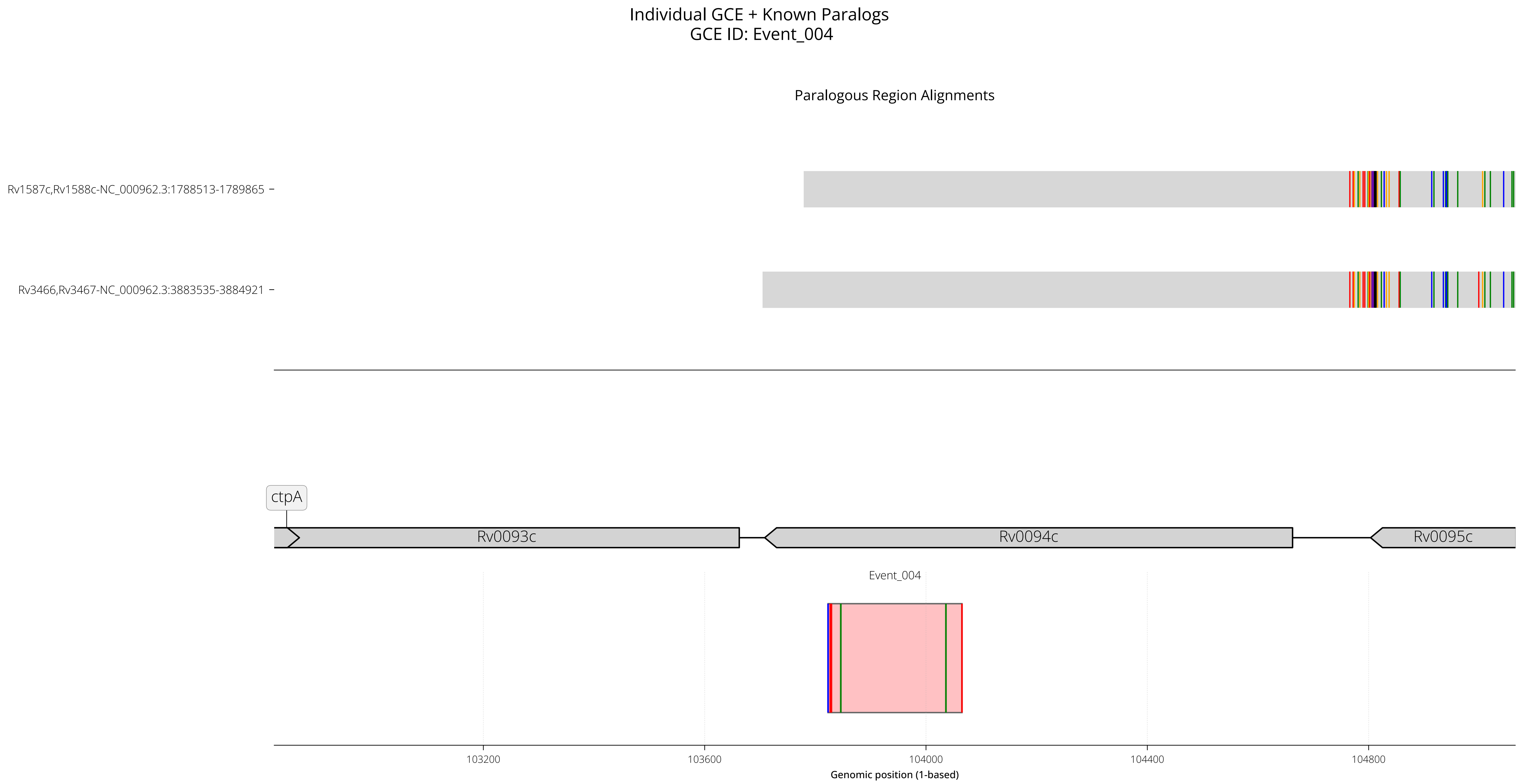
Individual GCE + Known Paralogs
GCE ID: Event_004
Paralogous Region Alignments
Rv1587c,Rv1588c-NC_000962.3:1788513-1789865
Rv3466,Rv3467-NC_000962.3:3883535-3884921
ctpA
Rv0093c
Rv0094c
Rv0095c
Event_004
103200
103600
104000
104400
104800
Genomic position (1-based)

Circos Viz - GCE to Known Paralogs Locations
GCE ID: Event_004

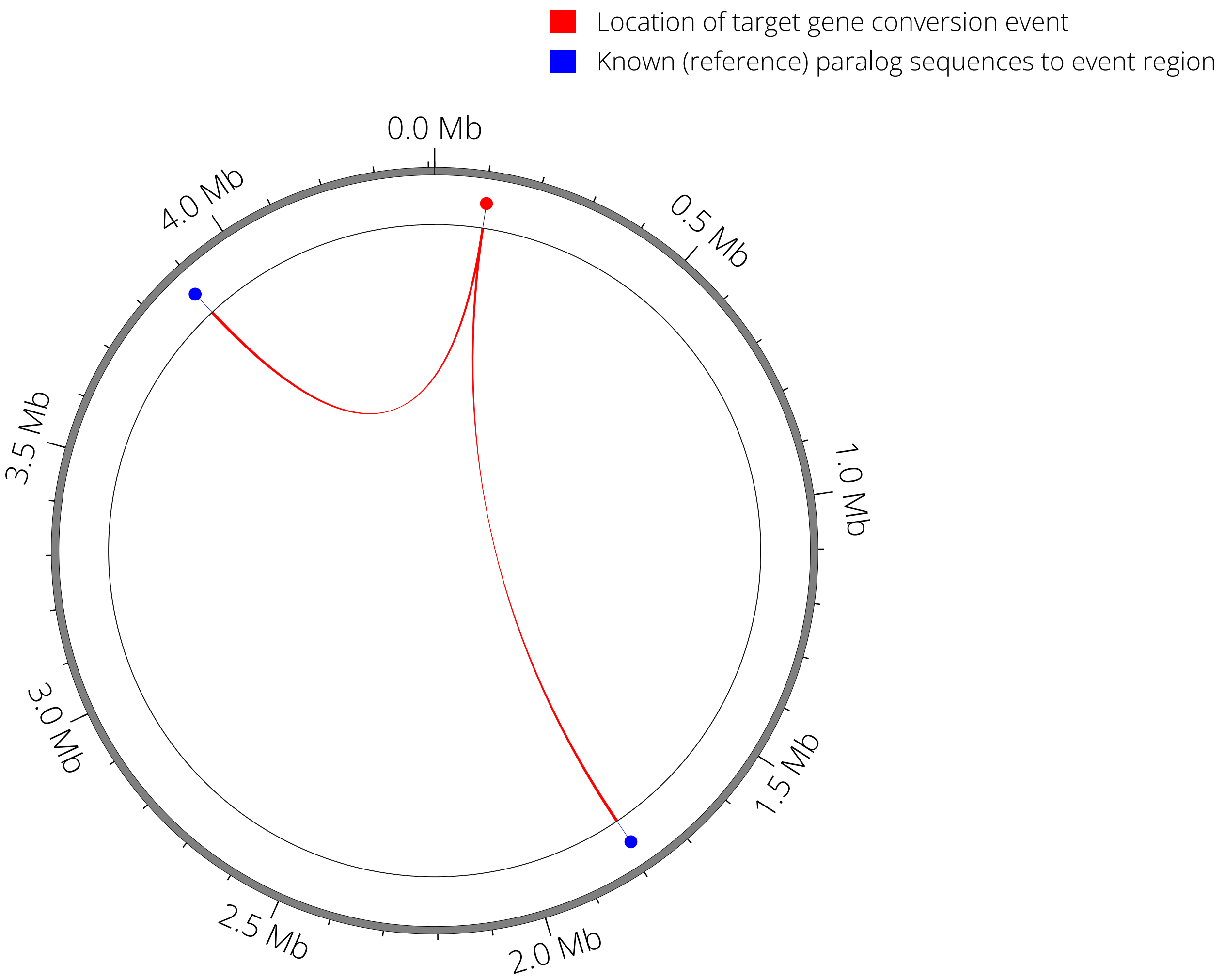

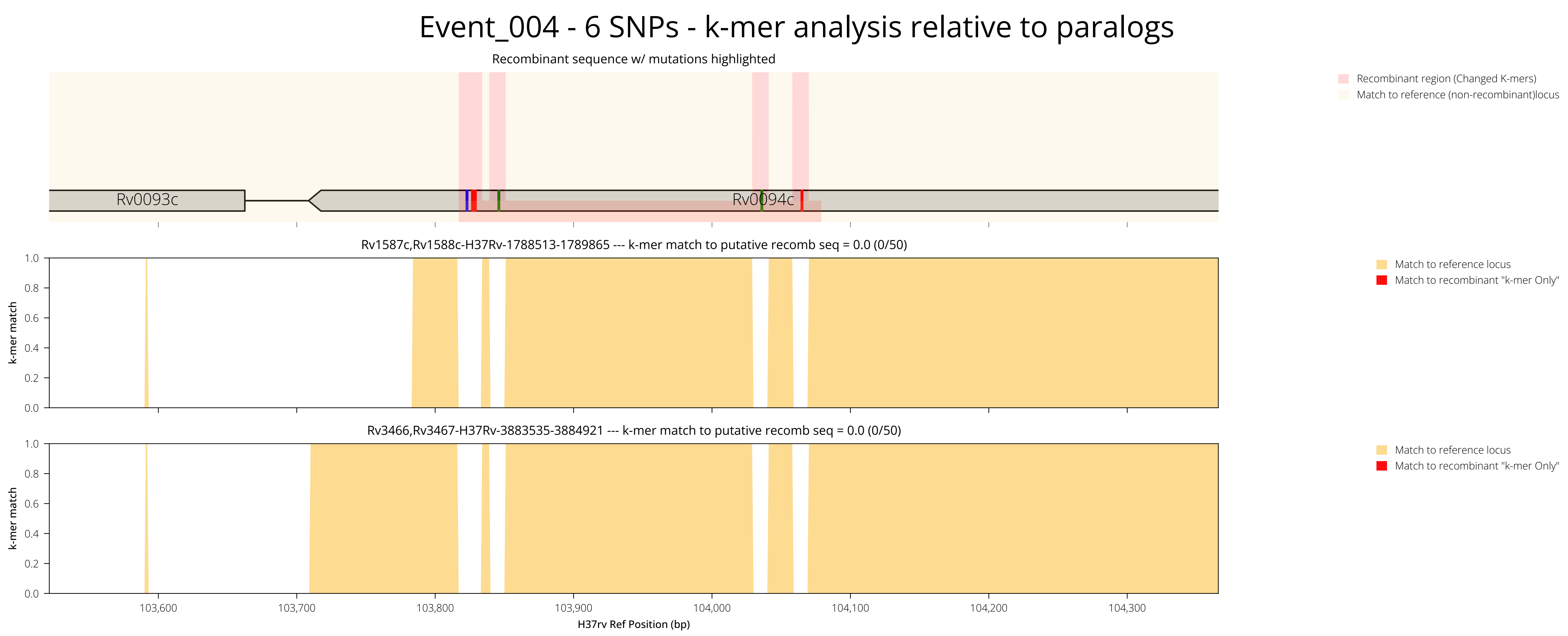
Event_004 - 6 SNPs - k-mer analysis relative to paralogs
Recombinant sequence w/ mutations highlighted
Recombinant region (Changed K-mers)
Match to reference (non-recombinant)locus
Rv0093c
Rv0094c
Rv1587c,Rv1588c-H37Rv-1788513-1789865 --- k-mer match to putative recomb seq = 0.0 (0/50)
Match to reference locus
Match to recombinant "k-mer Only"
k-mer match
1.0
0.8
0.6
0.4
0.2
0.0
Rv3466,Rv3467-H37Rv-3883535-3884921 --- k-mer match to putative recomb seq = 0.0 (0/50)
Match to reference locus
Match to recombinant "k-mer Only"
k-mer match
103,600
103,700
103,800
103,900
104,000
104,100
104,200
104,300
H37rv Ref Position (bp)

Sub-phylogeny Viz + Paralog Alignments
GCE ID: Event_004

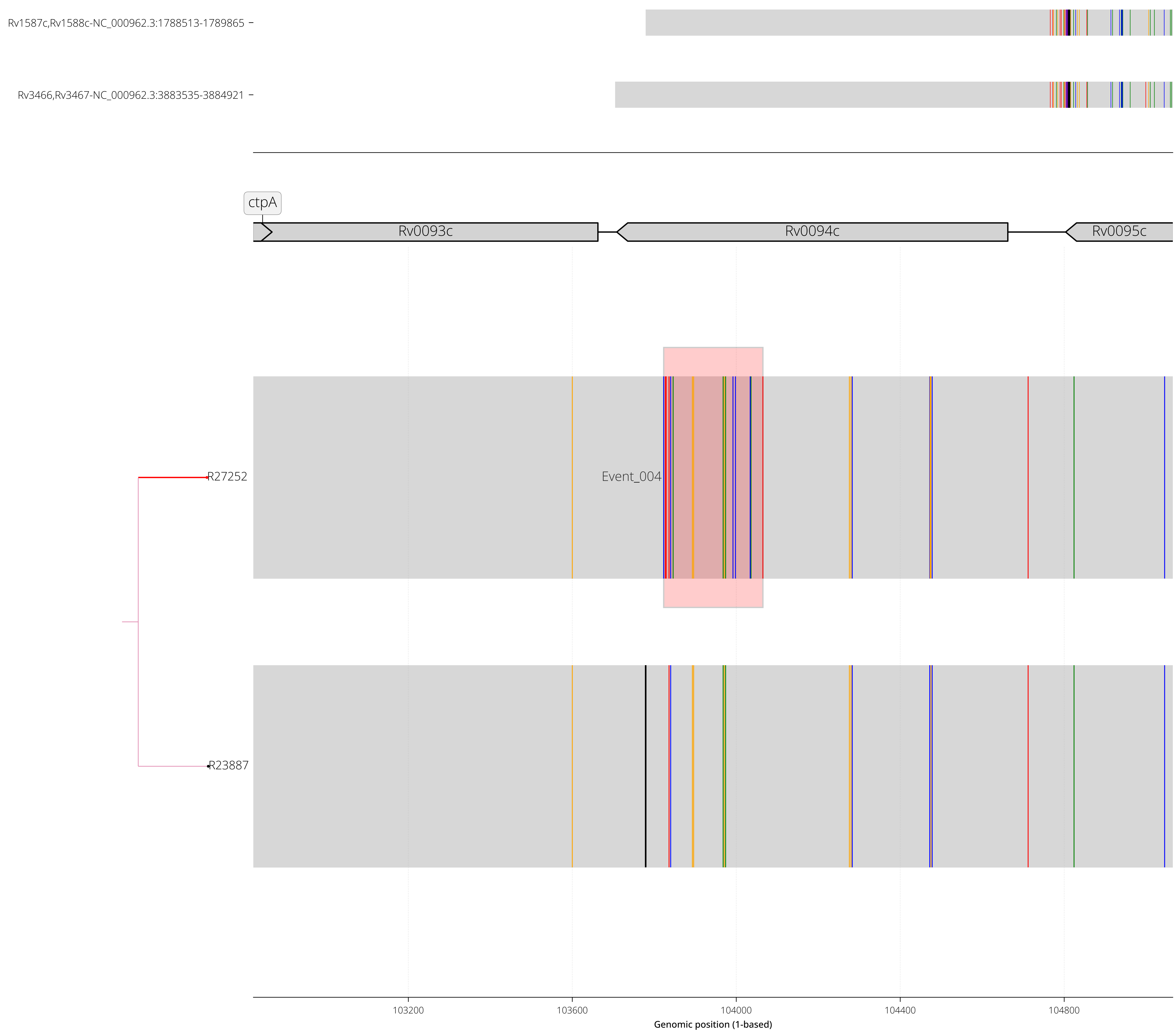

Full Phylogeny Viz + Paralog Alignments
GCE ID: Event_004

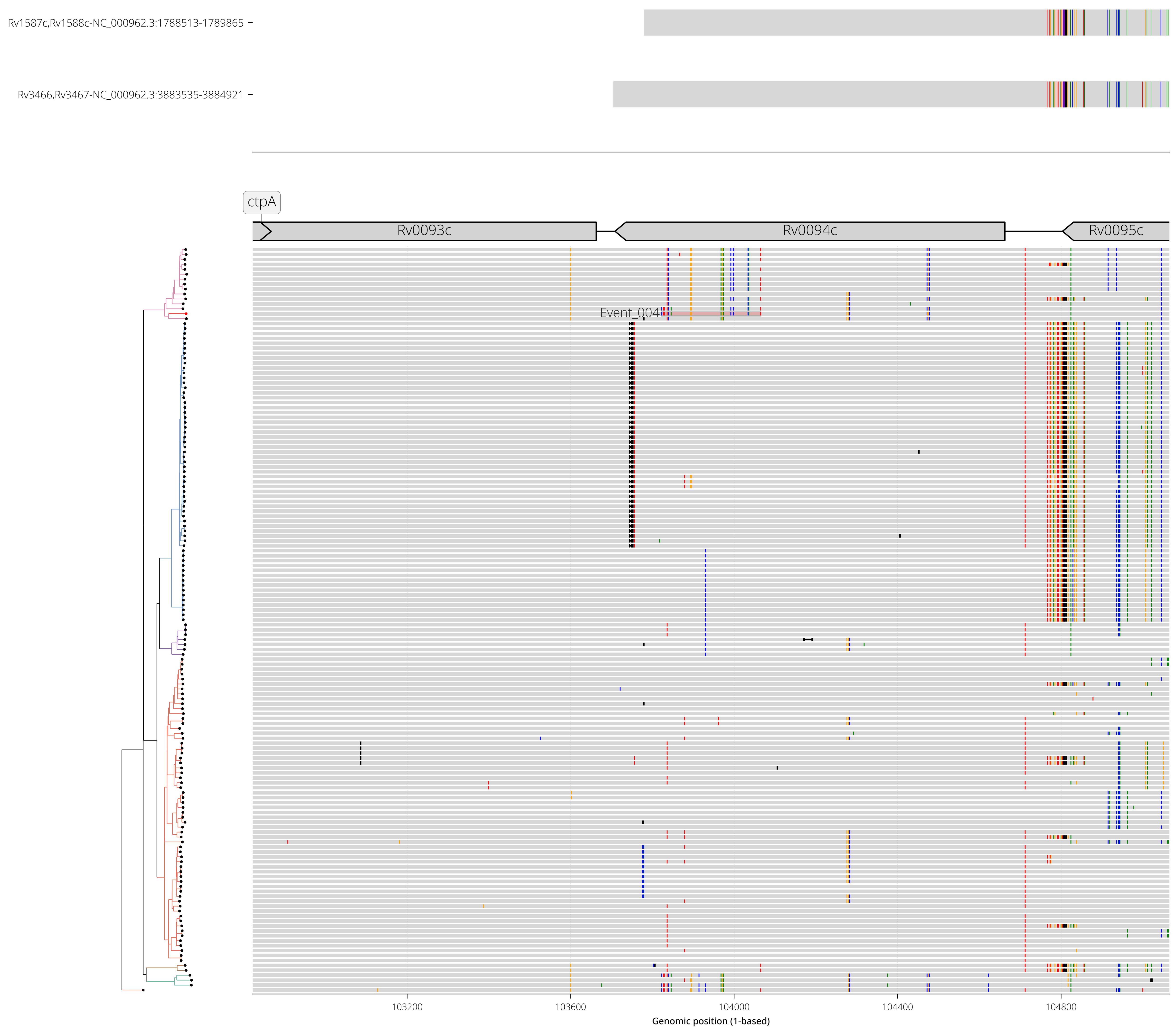

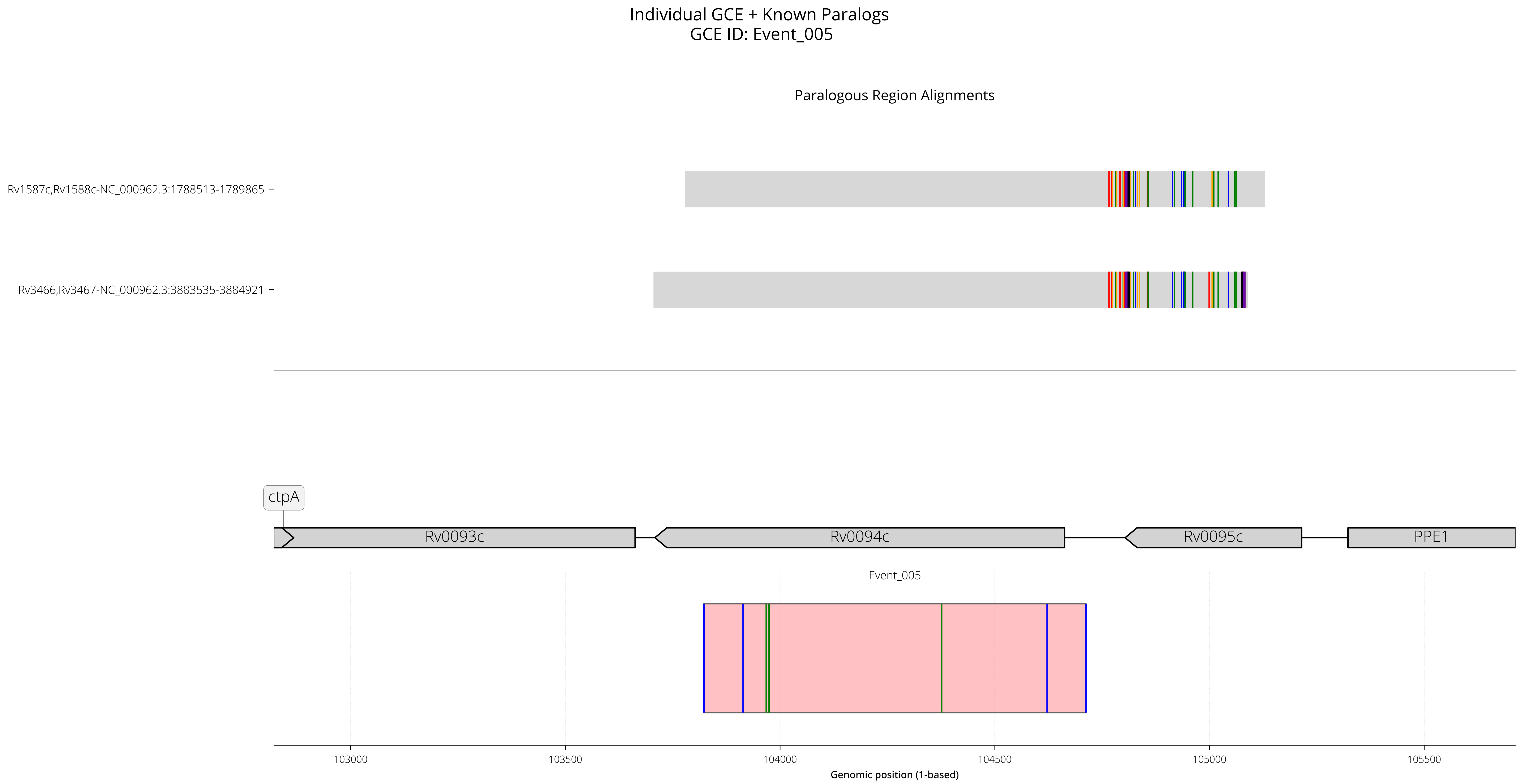
Individual GCE + Known Paralogs
GCE ID: Event_005
Paralogous Region Alignments
Rv1587c,Rv1588c-NC_000962.3:1788513-1789865
Rv3466,Rv3467-NC_000962.3:3883535-3884921
ctpA
Rv0093c
Rv0094c
Rv0095c
PPE1
Event_005
103000
103500
104000
104500
105000
105500
Genomic position (1-based)

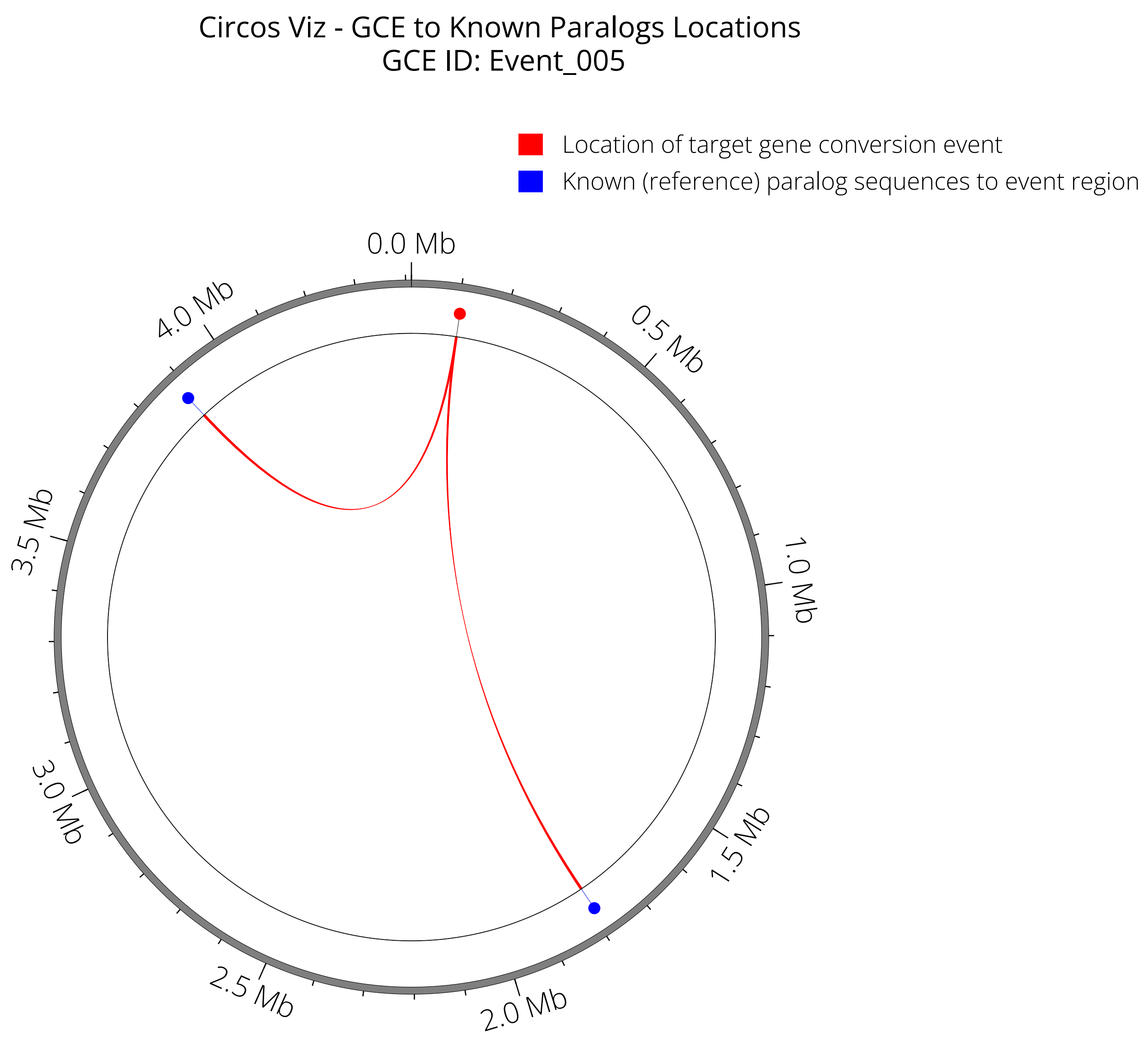

Circos Viz - GCE to Known Paralogs Locations
GCE ID: Event_005
Location of target gene conversion event
Known (reference) paralog sequences to event region
0.0 Mb
0.5 Mb
1.0 Mb
1.5 Mb
2.0 Mb
2.5 Mb
3.0 Mb
3.5 Mb
4.0 Mb

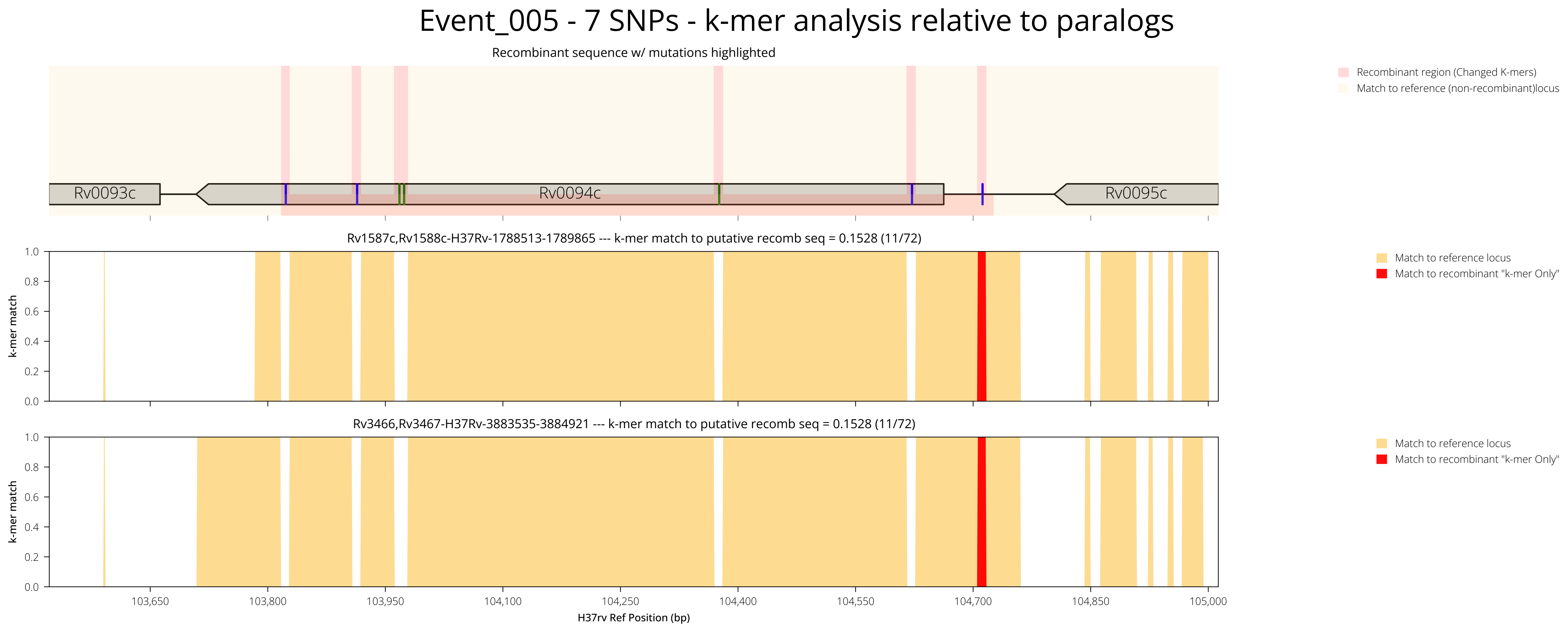
Event_005 - 7 SNPs - k-mer analysis relative to paralogs
Recombinant sequence w/ mutations highlighted
Recombinant region (Changed K-mers)
Match to reference (non-recombinant)locus
Rv0093c
Rv0094c
Rv0095c
Rv1587c,Rv1588c-H37Rv-1788513-1789865 --- k-mer match to putative recomb seq = 0.1528 (11/72)
Match to reference locus
Match to recombinant "k-mer Only"
Rv3466,Rv3467-H37Rv-3883535-3884921 --- k-mer match to putative recomb seq = 0.1528 (11/72)
Match to reference locus
Match to recombinant "k-mer Only"
k-mer match
1.0
0.8
0.6
0.4
0.2
0.0
103,650
103,800
103,950
104,100
104,250
104,400
104,550
104,700
104,850
105,000
H37rv Ref Position (bp)

Sub-phylogeny Viz + Paralog Alignments
GCE ID: Event_005

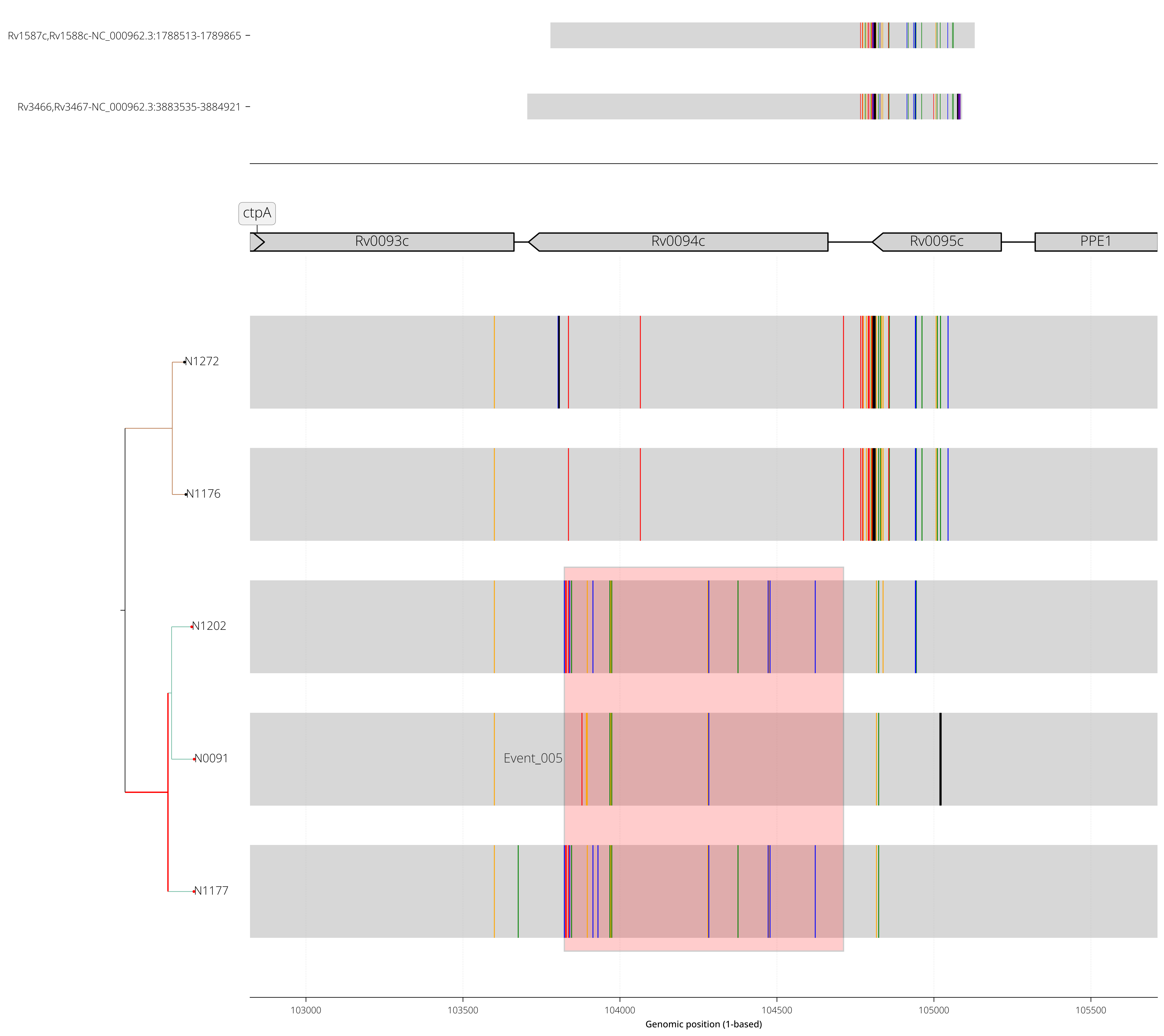

Full Phylogeny Viz + Paralog Alignments
GCE ID: Event_005

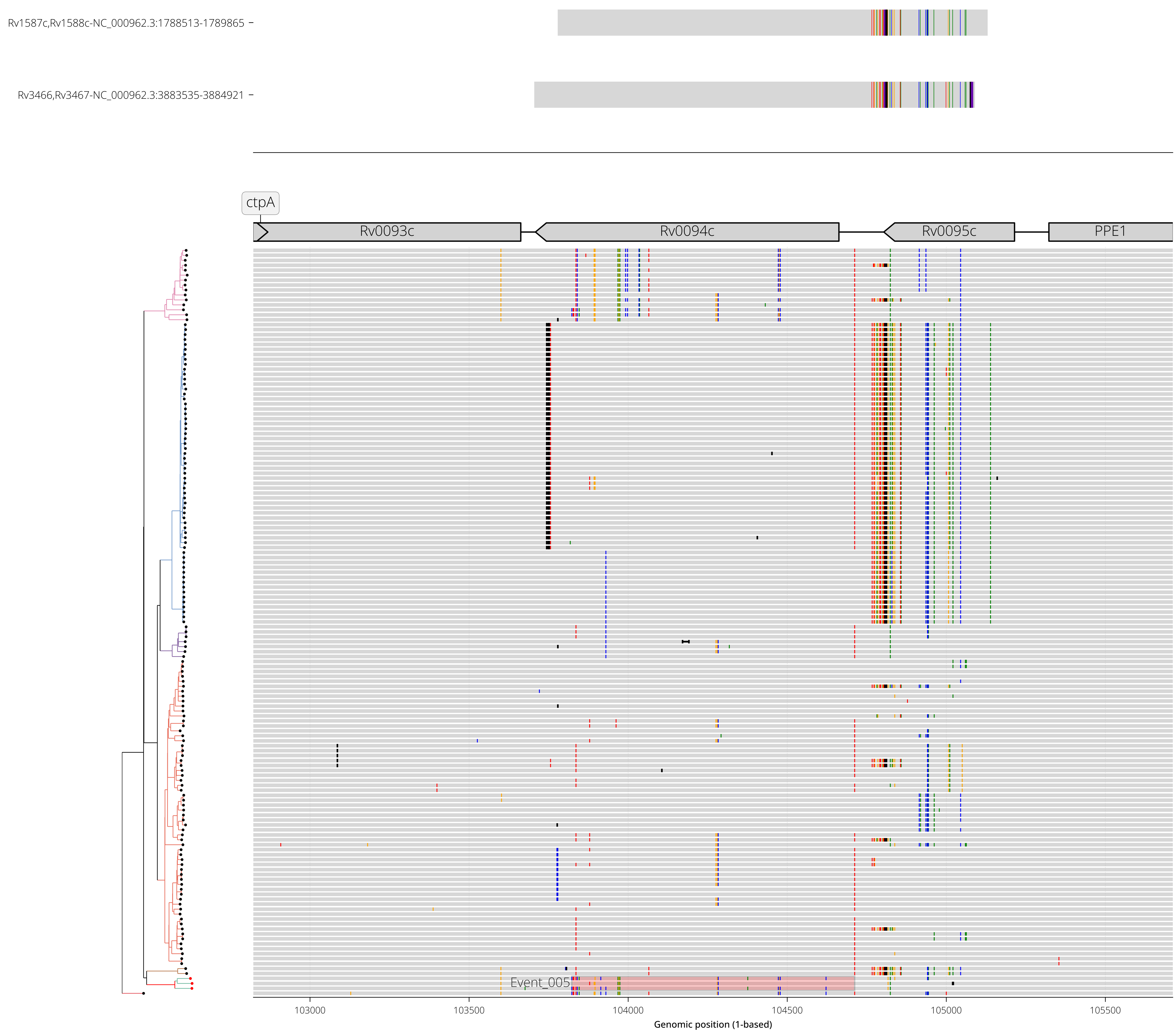

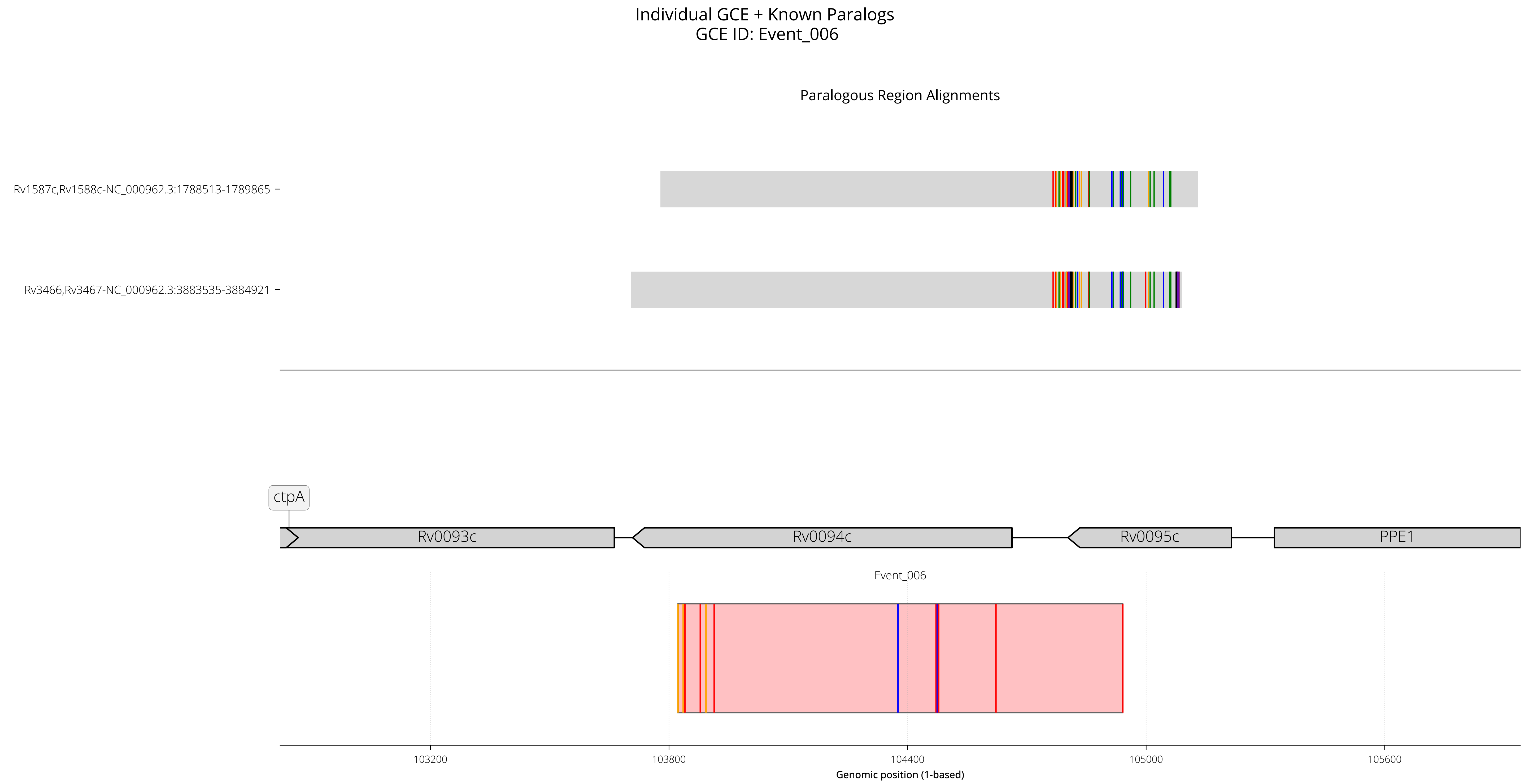

Individual GCE + Known Paralogs
GCE ID: Event_006
Paralogous Region Alignments
Rv1587c,Rv1588c-NC_000962.3:1788513-1789865
Rv3466,Rv3467-NC_000962.3:3883535-3884921
ctpA
Rv0093c
Rv0094c
Rv0095c
PPE1
Event_006
103200
103800
104400
105000
105600
Genomic position (1-based)

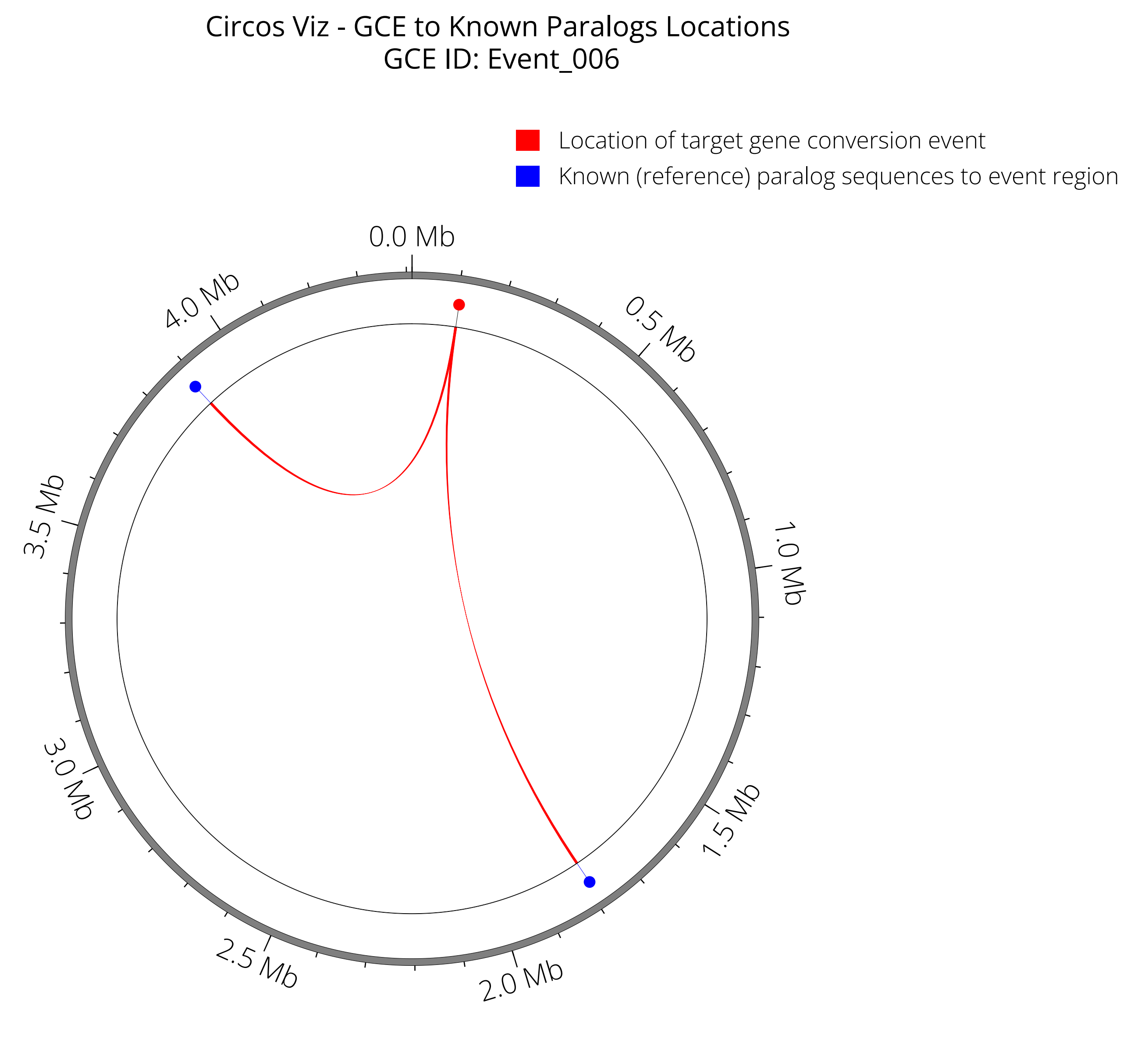
Circos Viz - GCE to Known Paralogs Locations
GCE ID: Event_006
Location of target gene conversion event
Known (reference) paralog sequences to event region
0.0 Mb
0.5 Mb
1.0 Mb
1.5 Mb
2.0 Mb
2.5 Mb
3.0 Mb
3.5 Mb
4.0 Mb

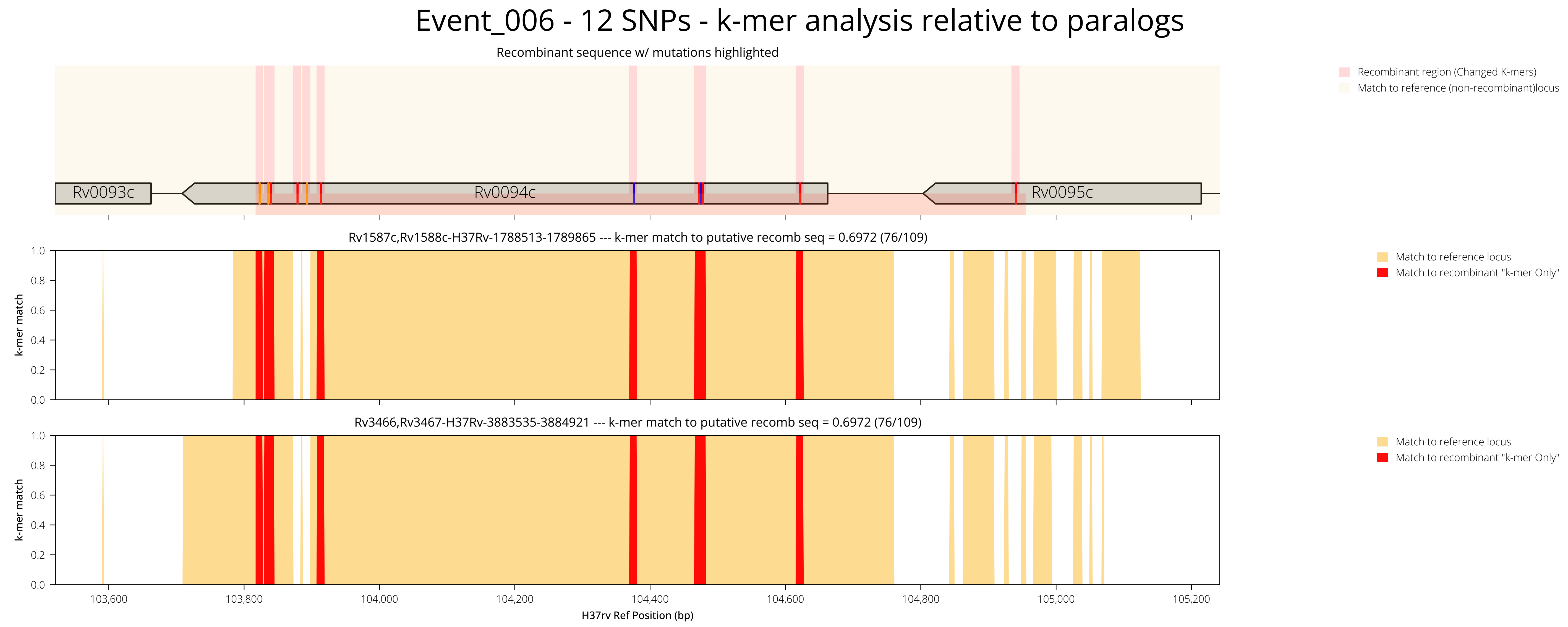

Event_006 - 12 SNPs - k-mer analysis relative to paralogs
Recombinant sequence w/ mutations highlighted
Rv0093c
Rv0094c
Rv0095c
Recombinant region (Changed K-mers)
Match to reference (non-recombinant)locus
Rv1587c,Rv1588c-H37Rv-1788513-1789865 --- k-mer match to putative recomb seq = 0.6972 (76/109)
Match to reference locus
Match to recombinant "k-mer Only"
k-mer match
1.0
0.8
0.6
0.4
0.2
0.0
Rv3466,Rv3467-H37Rv-3883535-3884921 --- k-mer match to putative recomb seq = 0.6972 (76/109)
Match to reference locus
Match to recombinant "k-mer Only"
k-mer match
103,600
103,800
104,000
104,200
104,400
104,600
104,800
105,000
105,200
H37rv Ref Position (bp)

Sub-phylogeny Viz + Paralog Alignments
GCE ID: Event_006

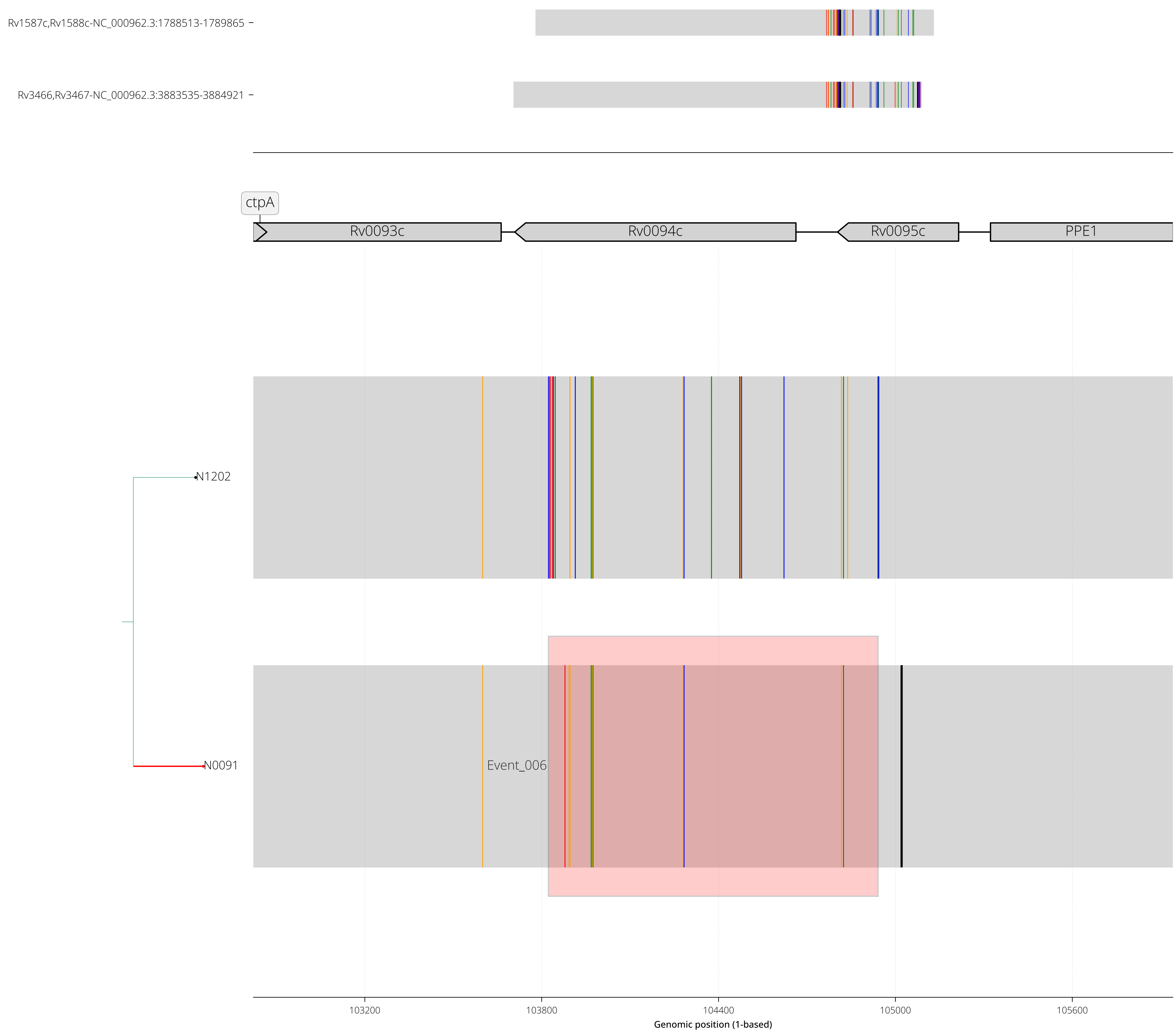

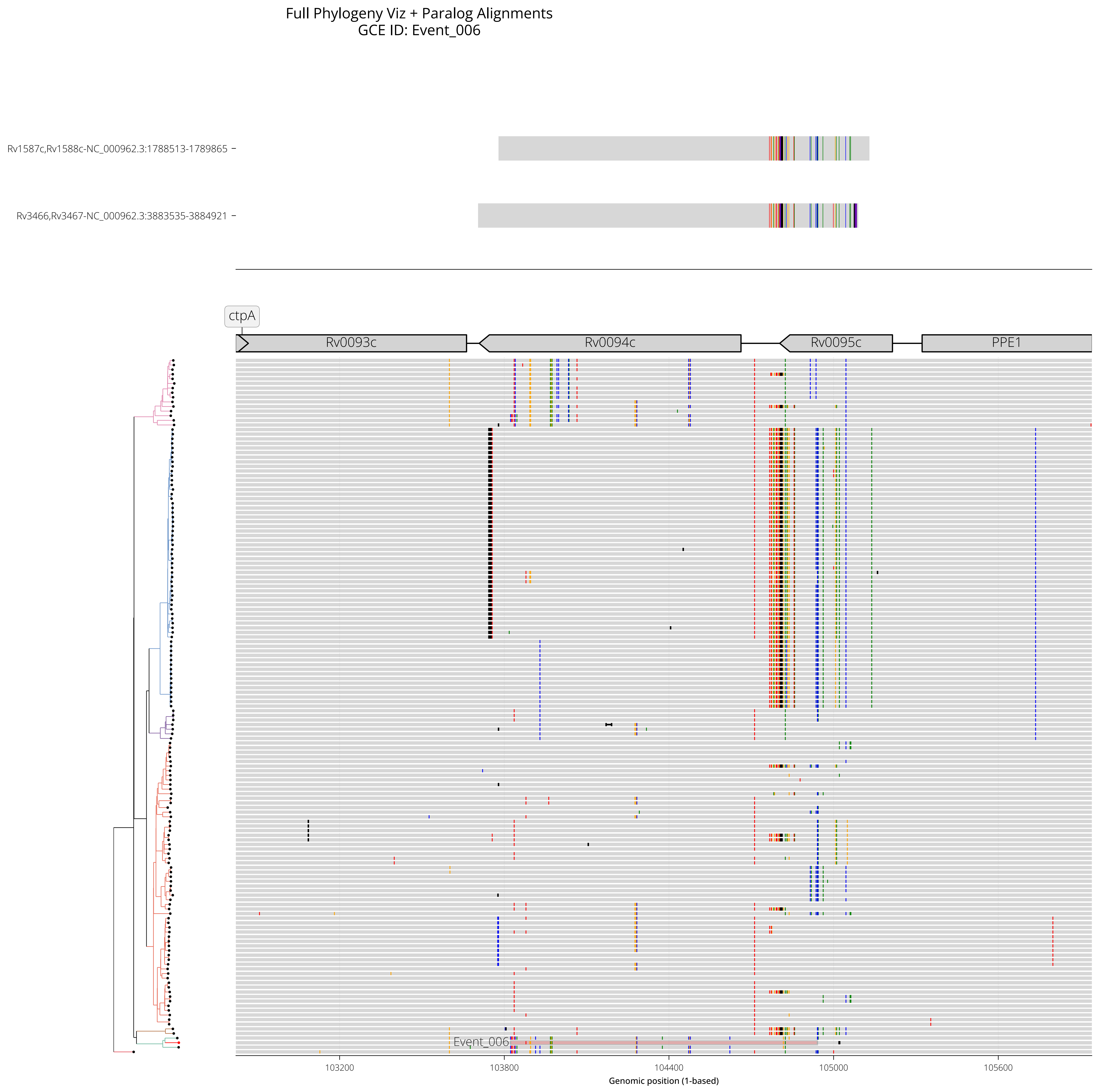

Full Phylogeny Viz + Paralog Alignments
GCE ID: Event_006
Rv1587c,Rv1588c-NC_000962.3:1788513-1789865
Rv3466,Rv3467-NC_000962.3:3883535-3884921
ctpA
Rv0093c
Rv0094c
Rv0095c
PPE1
Event_006
103200
103800
104400
105000
105600
Genomic position (1-based)

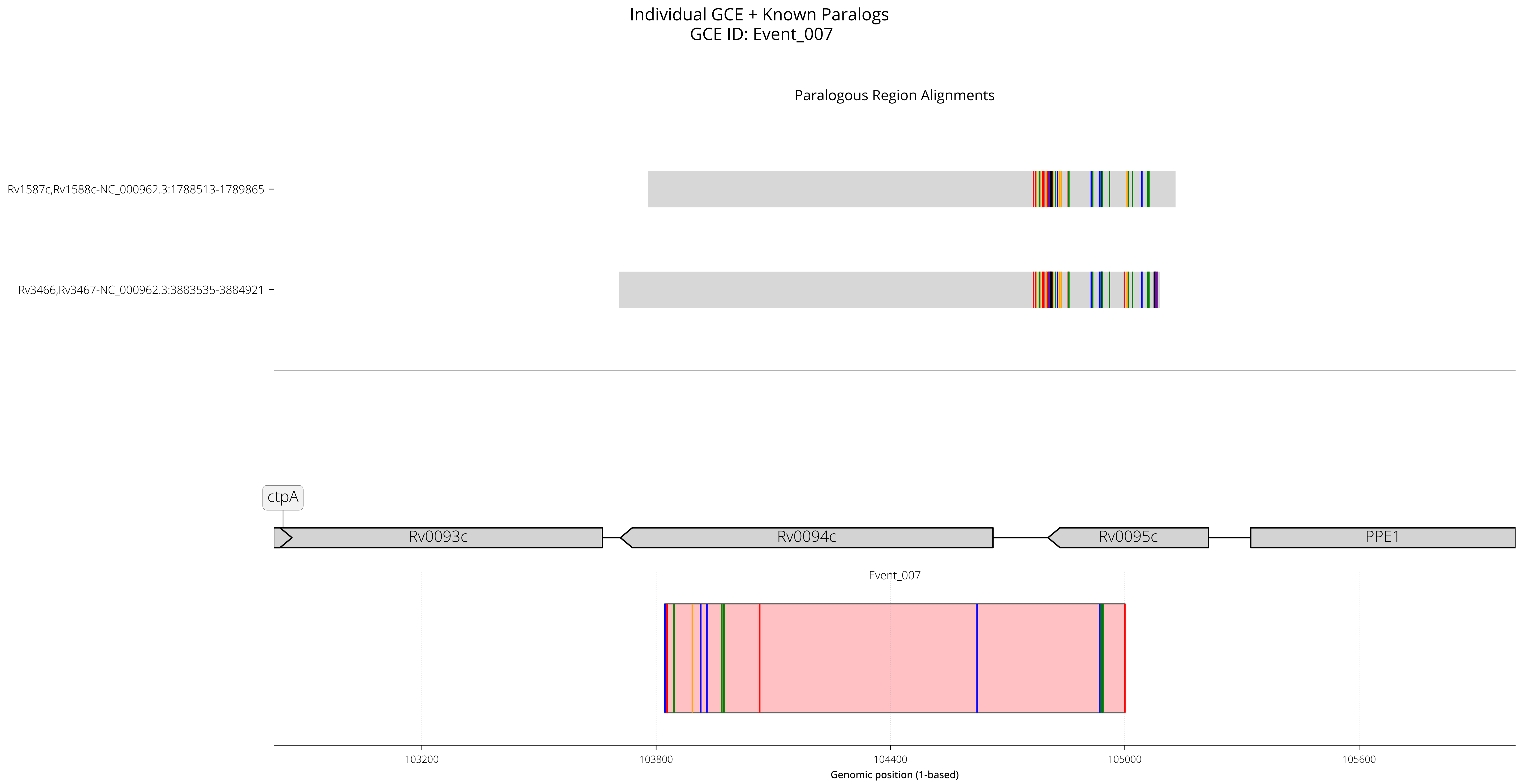
Individual GCE + Known Paralogs
GCE ID: Event_007
Paralogous Region Alignments
Rv1587c,Rv1588c-NC_000962.3:1788513-1789865
Rv3466,Rv3467-NC_000962.3:3883535-3884921
ctpA
Rv0093c
Rv0094c
Rv0095c
PPE1
Event_007
103200
103800
104400
105000
105600
Genomic position (1-based)

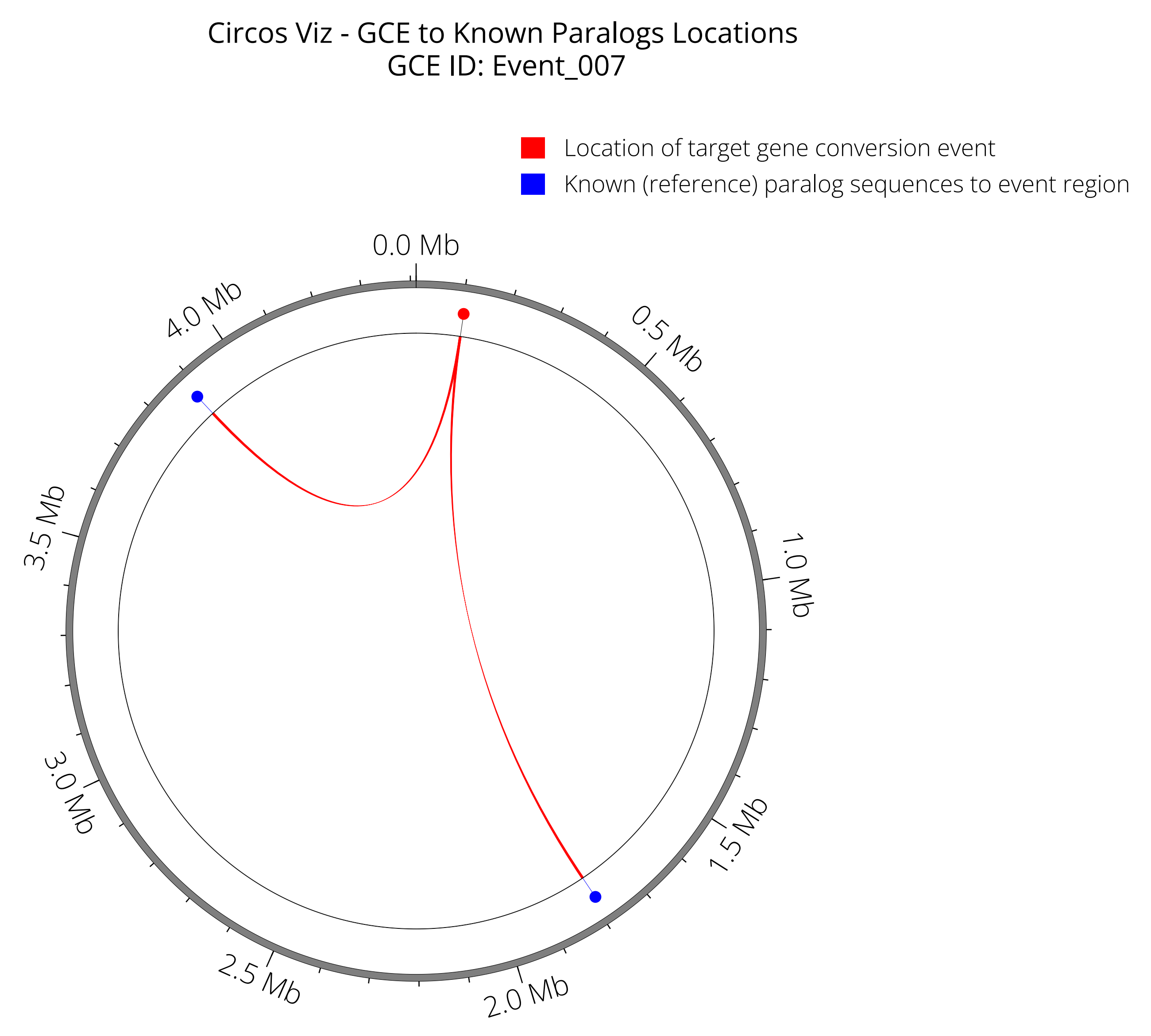

Circos Viz - GCE to Known Paralogs Locations
GCE ID: Event_007
Location of target gene conversion event
Known (reference) paralog sequences to event region
0.0 Mb
0.5 Mb
1.0 Mb
1.5 Mb
2.0 Mb
2.5 Mb
3.0 Mb
3.5 Mb
4.0 Mb

### Event_007 - 16 SNPs - k-mer analysis relative to paralogs

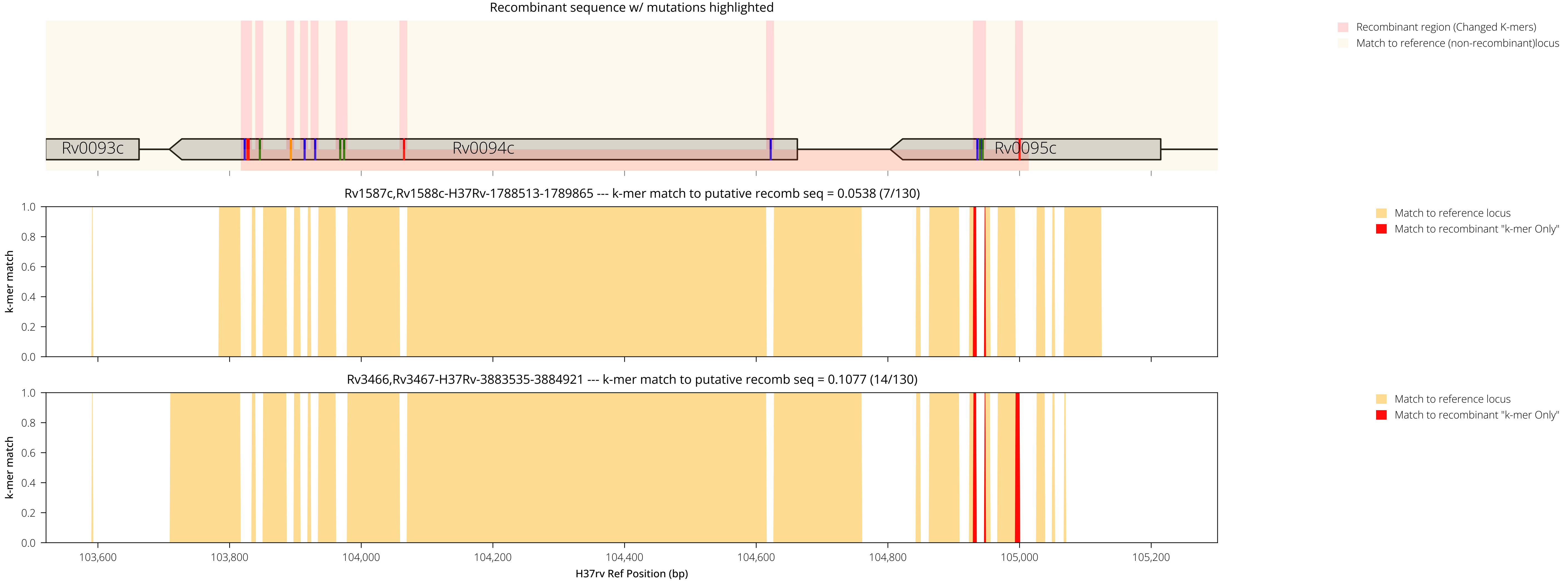

Sub-phylogeny Viz + Paralog Alignments
GCE ID: Event_007

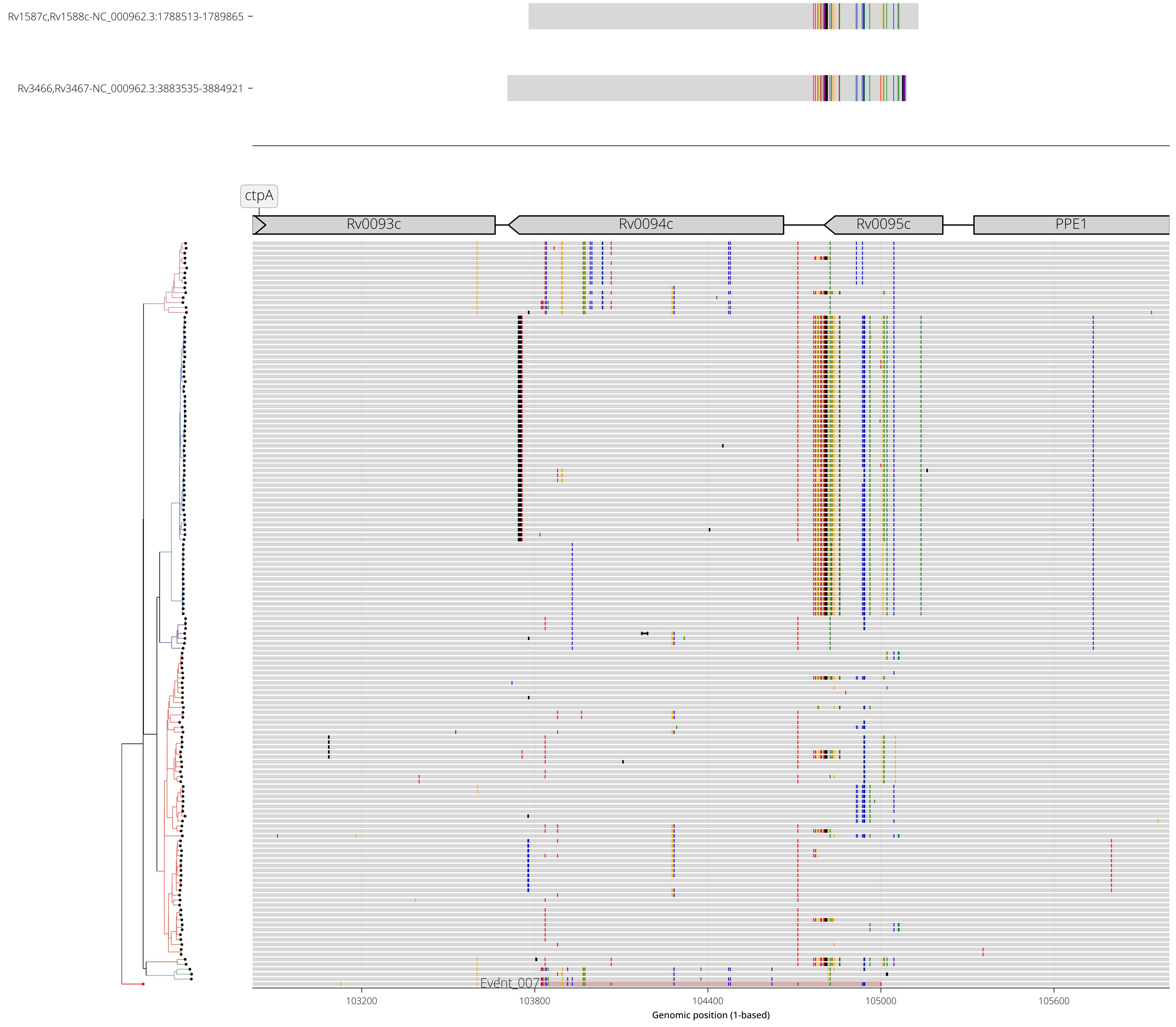

Full Phylogeny Viz + Paralog Alignments
GCE ID: Event_007

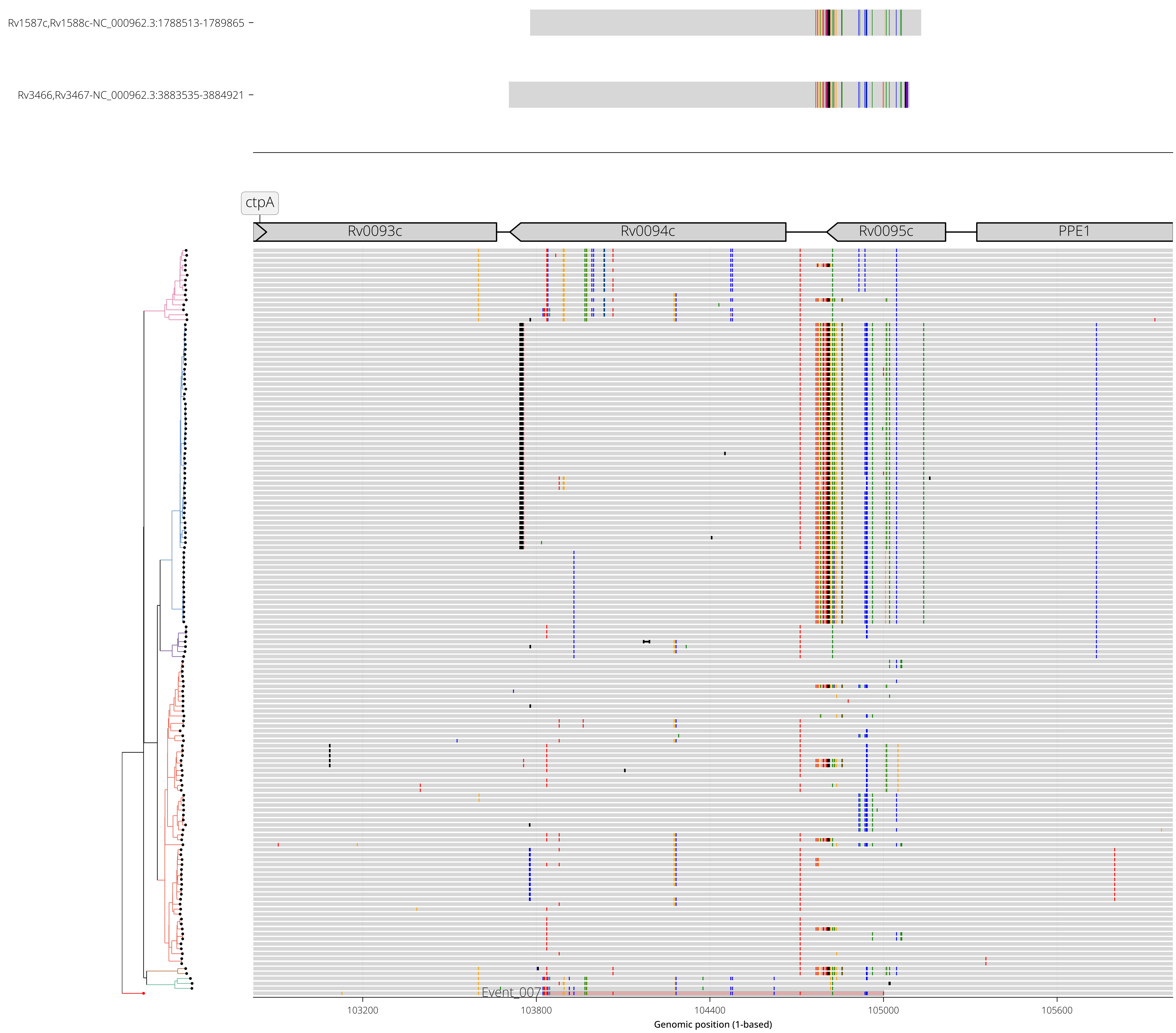

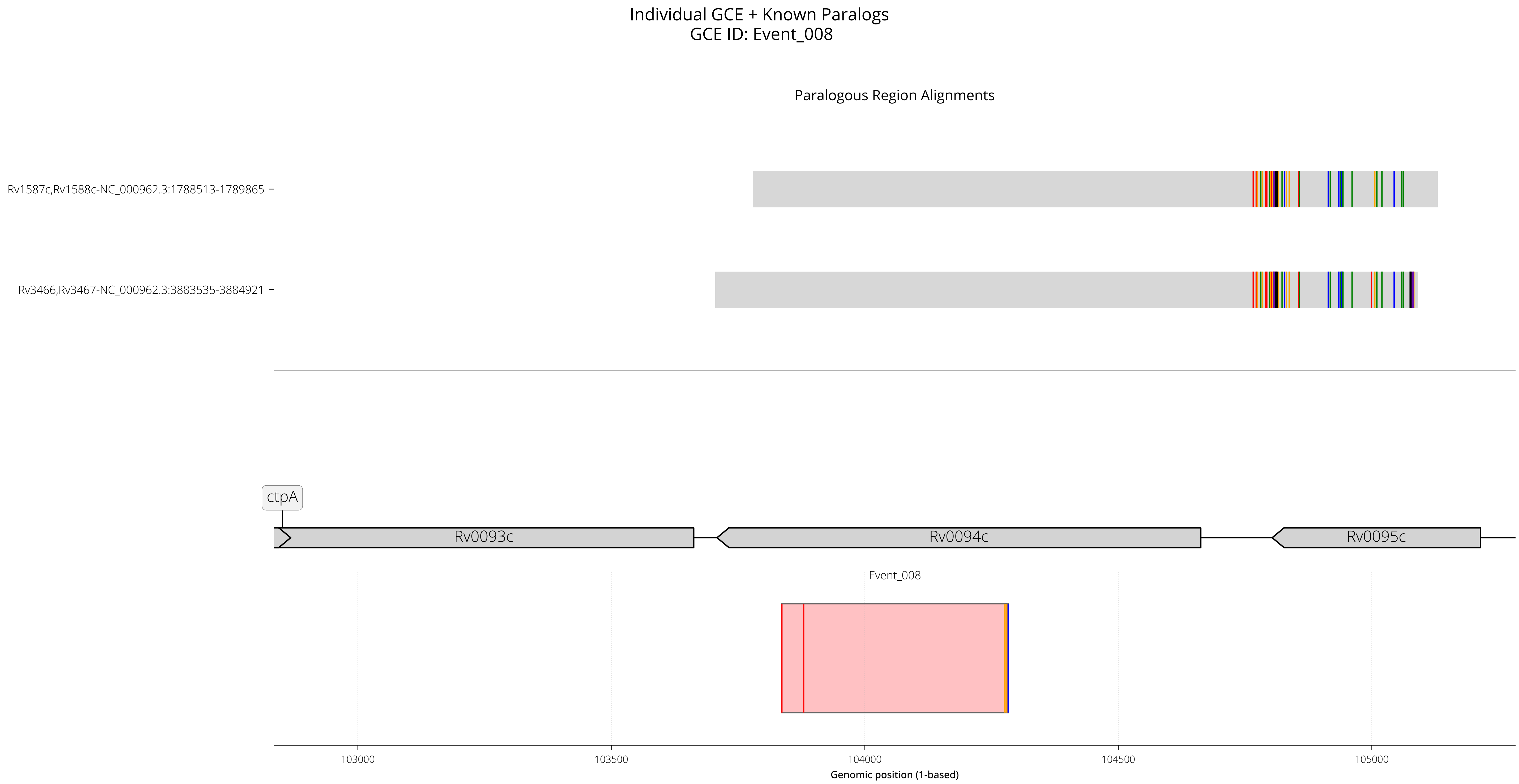

Individual GCE + Known Paralogs
GCE ID: Event_008
Paralogous Region Alignments
Rv1587c,Rv1588c-NC_000962.3:1788513-1789865
Rv3466,Rv3467-NC_000962.3:3883535-3884921
ctpA
Rv0093c
Rv0094c
Rv0095c
Event_008
103000
103500
104000
104500
105000
Genomic position (1-based)

Circos Viz - GCE to Known Paralogs Locations
GCE ID: Event_008

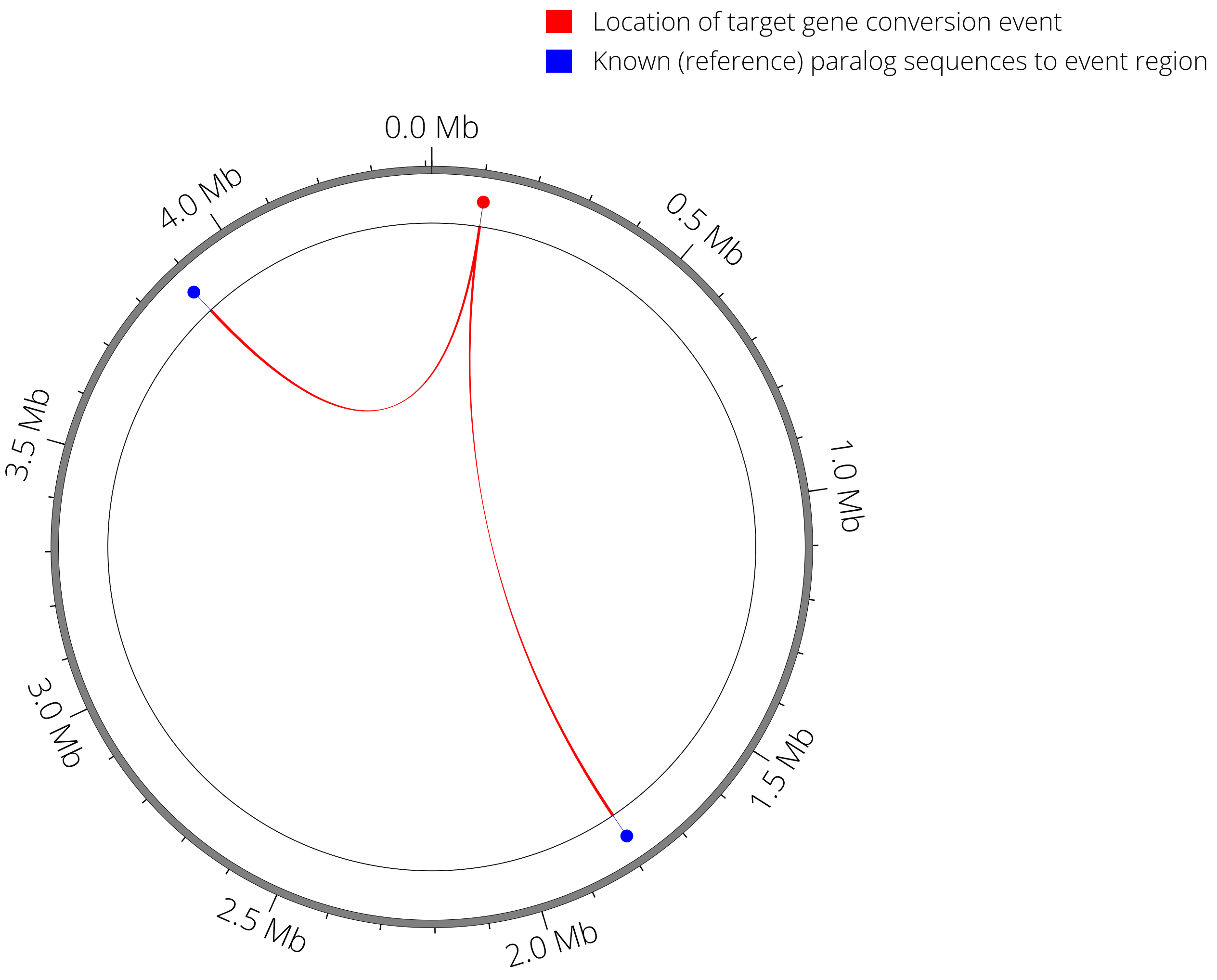

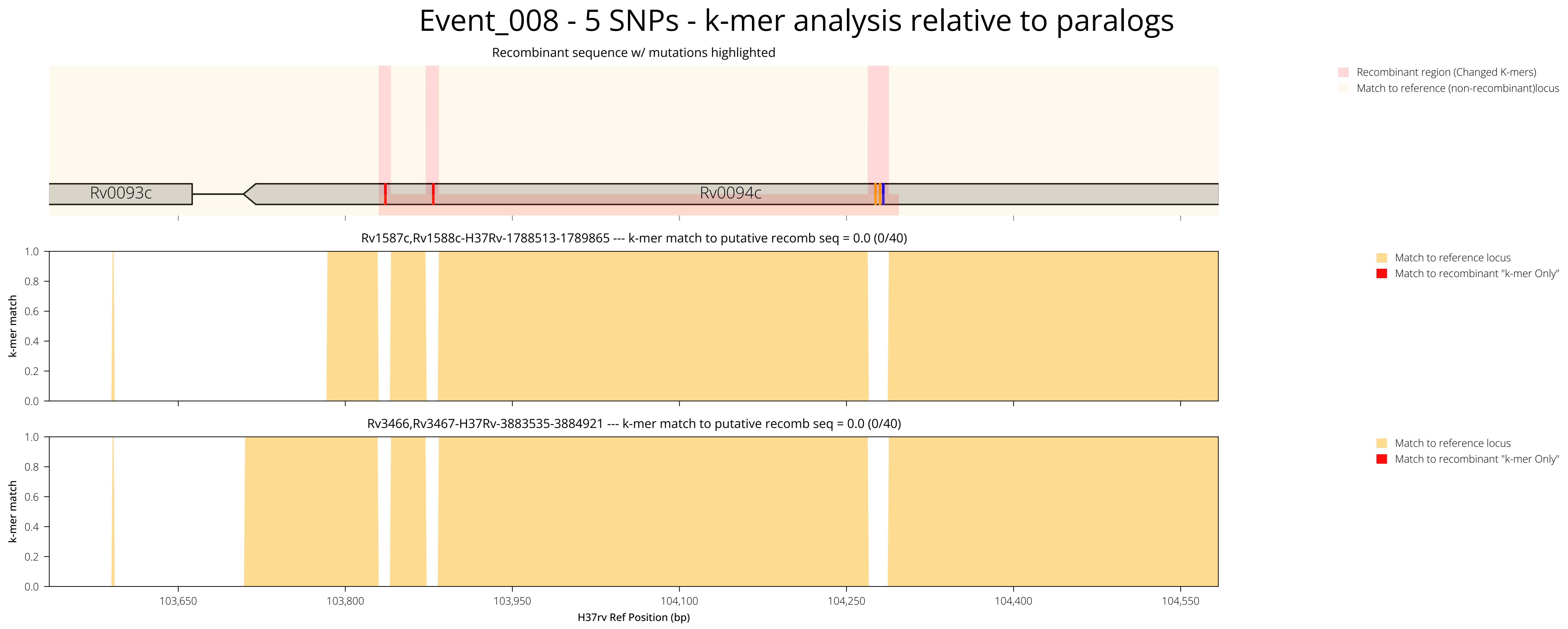

Event_008 - 5 SNPs - k-mer analysis relative to paralogs
Recombinant sequence w/ mutations highlighted
Rv0093c
Rv0094c
Recombinant region (Changed K-mers)
Match to reference (non-recombinant)locus
Rv1587c,Rv1588c-H37Rv-1788513-1789865 --- k-mer match to putative recomb seq = 0.0 (0/40)
Match to reference locus
Match to recombinant "k-mer Only"
Rv3466,Rv3467-H37Rv-3883535-3884921 --- k-mer match to putative recomb seq = 0.0 (0/40)
Match to reference locus
Match to recombinant "k-mer Only"
k-mer match
1.0
0.8
0.6
0.4
0.2
0.0
103,650
103,800
103,950
104,100
104,250
104,400
104,550
H37rv Ref Position (bp)

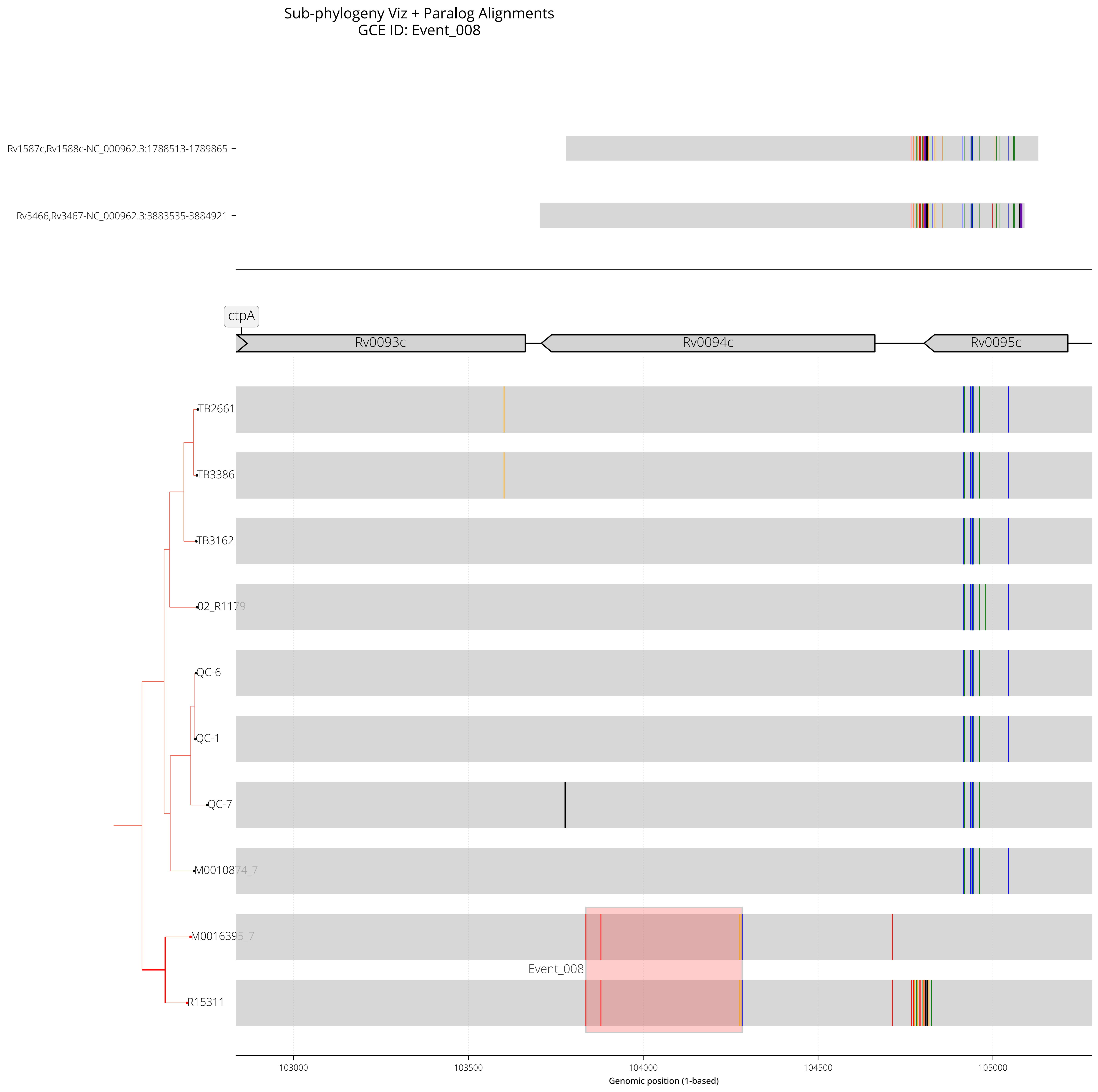
Sub-phylogeny Viz + Paralog Alignments
GCE ID: Event_008
Rv1587c,Rv1588c-NC_000962.3:1788513-1789865
Rv3466,Rv3467-NC_000962.3:3883535-3884921
ctpA
Rv0093c
Rv0094c
Rv0095c
TB2661
TB3386
TB3162
02_R1179
QC-6
QC-1
QC-7
M0010874_7
M0016395_7
Event_008
R15311
103000
103500
104000
104500
105000
Genomic position (1-based)

Full Phylogeny Viz + Paralog Alignments
GCE ID: Event_008

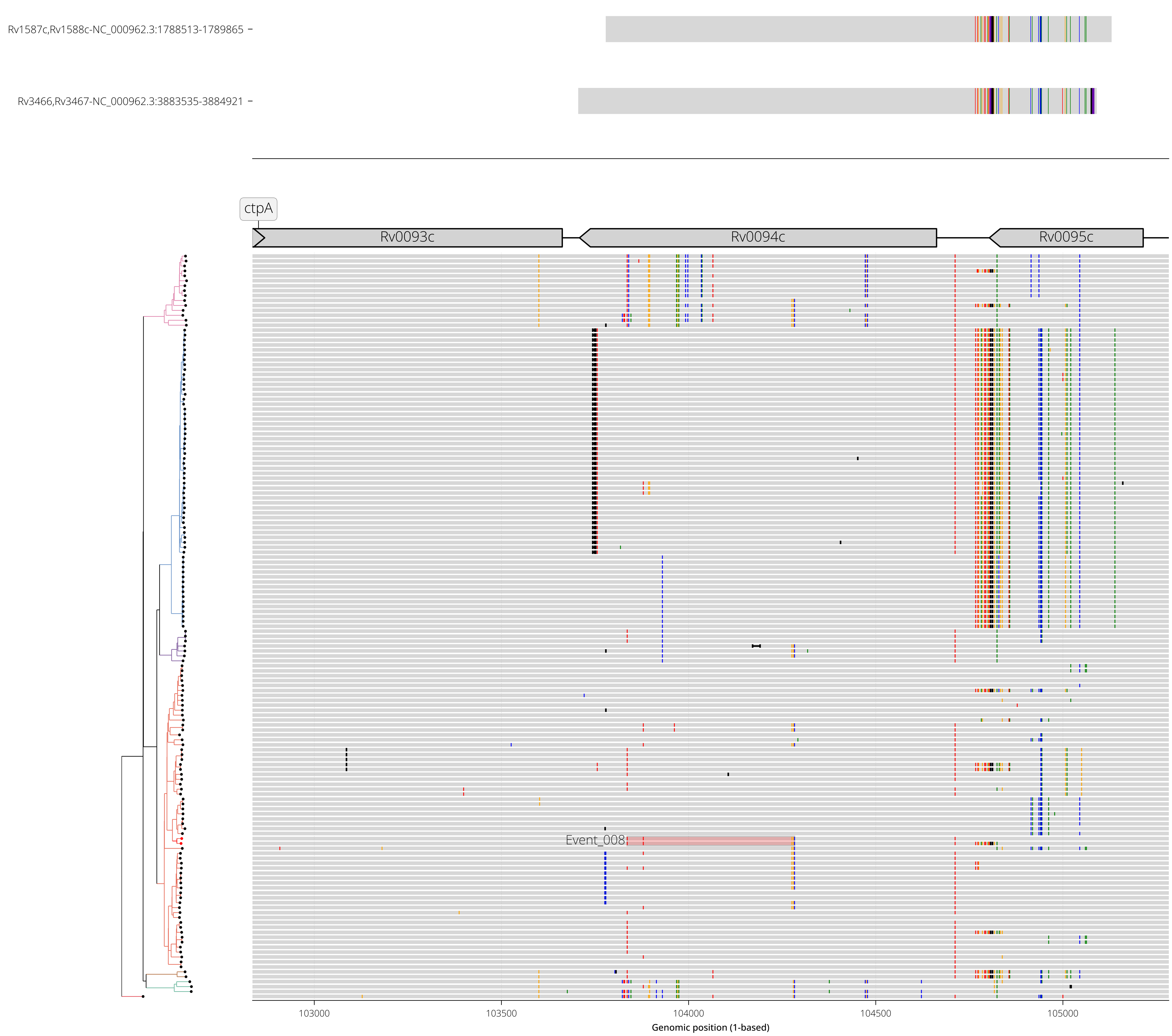

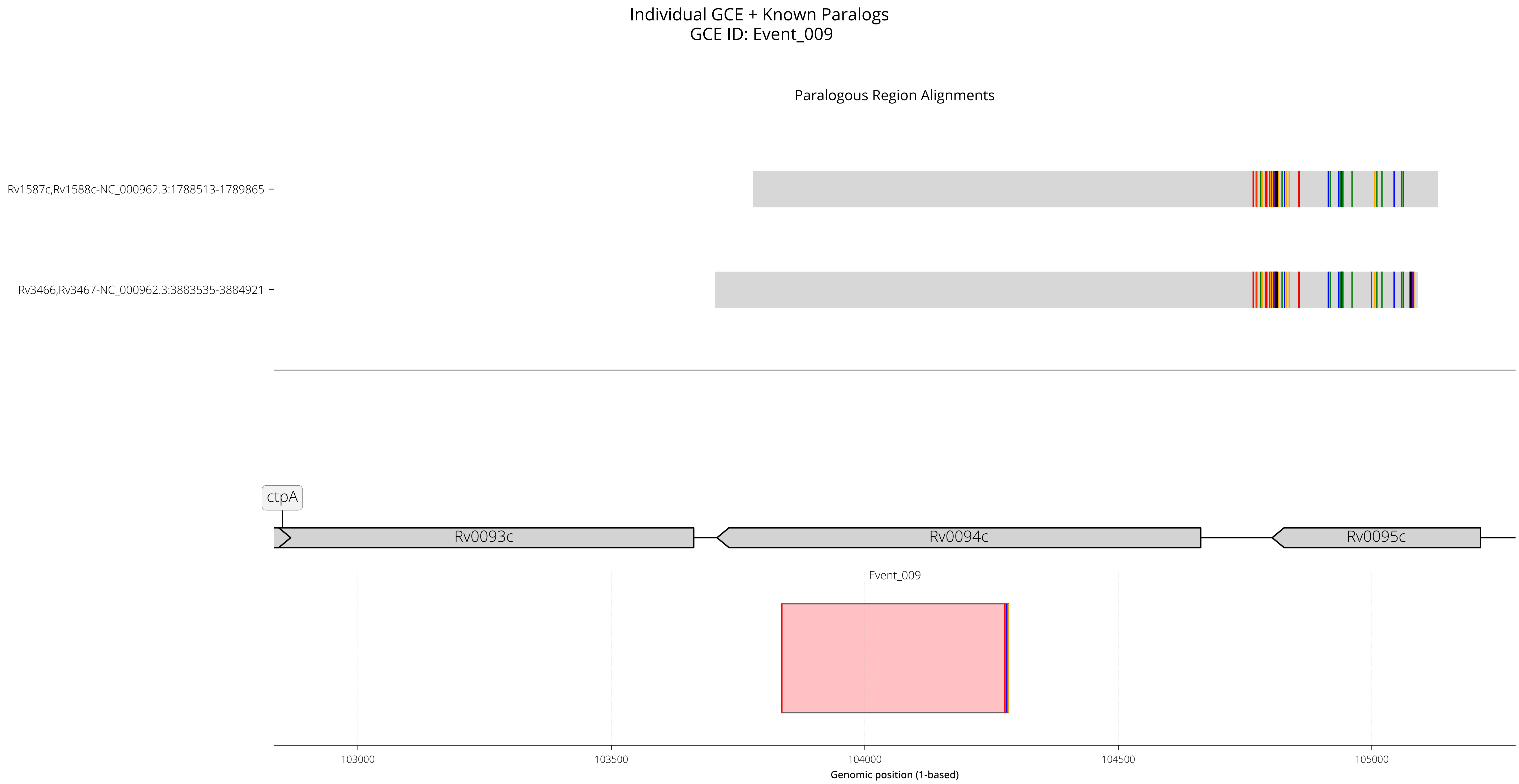

Individual GCE + Known Paralogs
GCE ID: Event_009
Paralogous Region Alignments
Rv1587c,Rv1588c-NC_000962.3:1788513-1789865
Rv3466,Rv3467-NC_000962.3:3883535-3884921
ctpA
Rv0093c
Rv0094c
Rv0095c
Event_009
103000
103500
104000
104500
105000
Genomic position (1-based)

Circos Viz - GCE to Known Paralogs Locations
GCE ID: Event_009

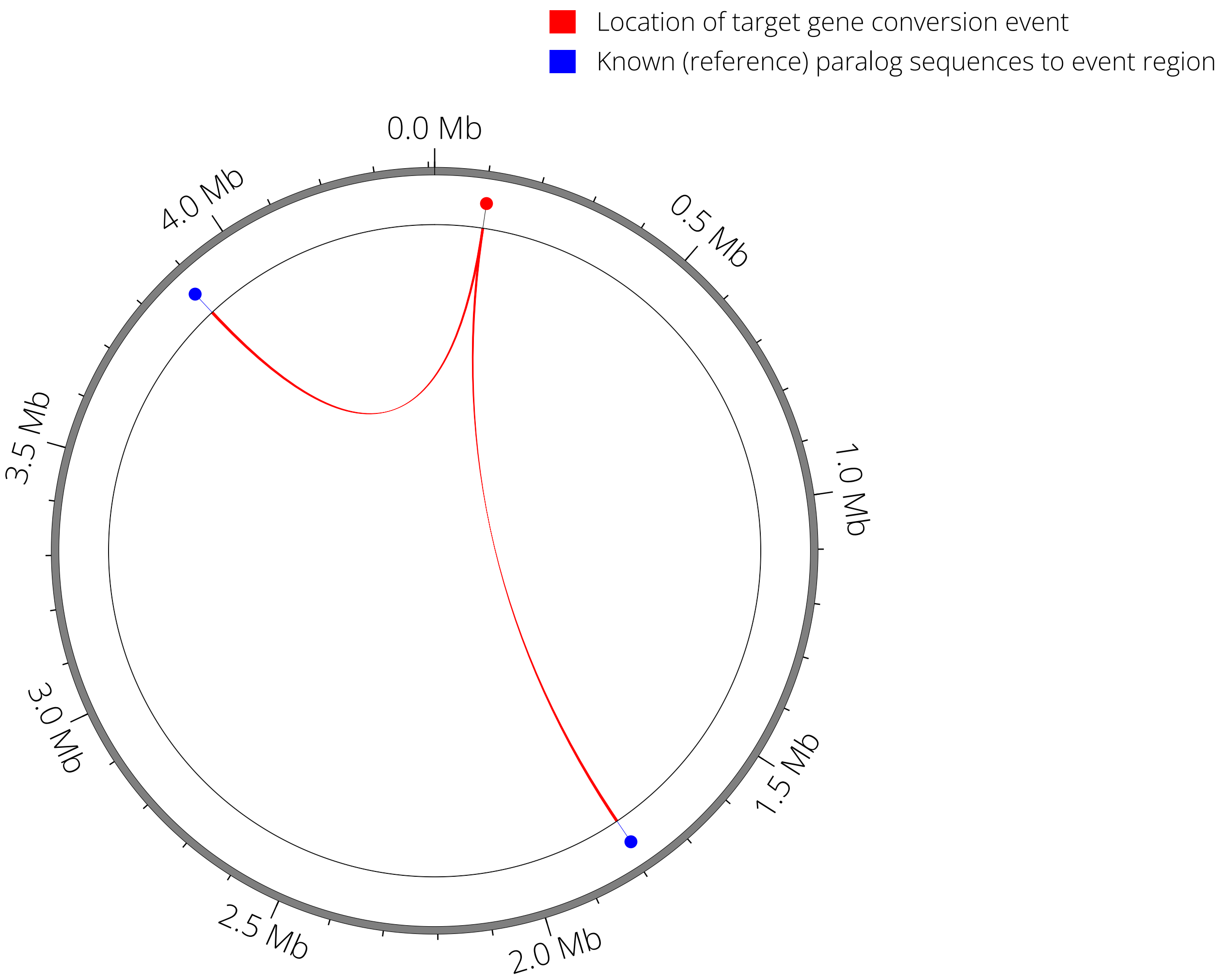

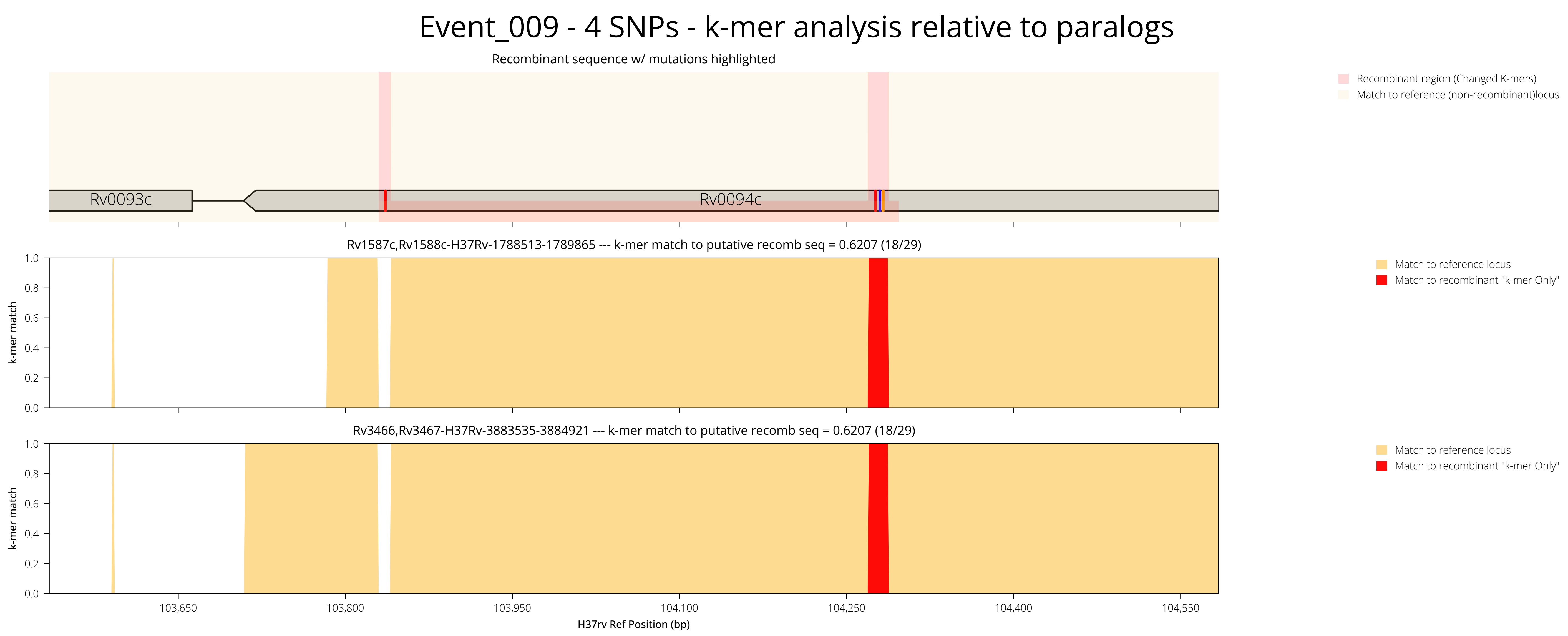
Event_009 - 4 SNPs - k-mer analysis relative to paralogs
Recombinant sequence w/ mutations highlighted
Recombinant region (Changed K-mers)
Match to reference (non-recombinant)locus
Rv0093c
Rv0094c
Rv1587c,Rv1588c-H37Rv-1788513-1789865 --- k-mer match to putative recomb seq = 0.6207 (18/29)
Match to reference locus
Match to recombinant "k-mer Only"
k-mer match
1.0
0.8
0.6
0.4
0.2
0.0
Rv3466,Rv3467-H37Rv-3883535-3884921 --- k-mer match to putative recomb seq = 0.6207 (18/29)
Match to reference locus
Match to recombinant "k-mer Only"
k-mer match
103,650
103,800
103,950
104,100
104,250
104,400
104,550
H37rv Ref Position (bp)

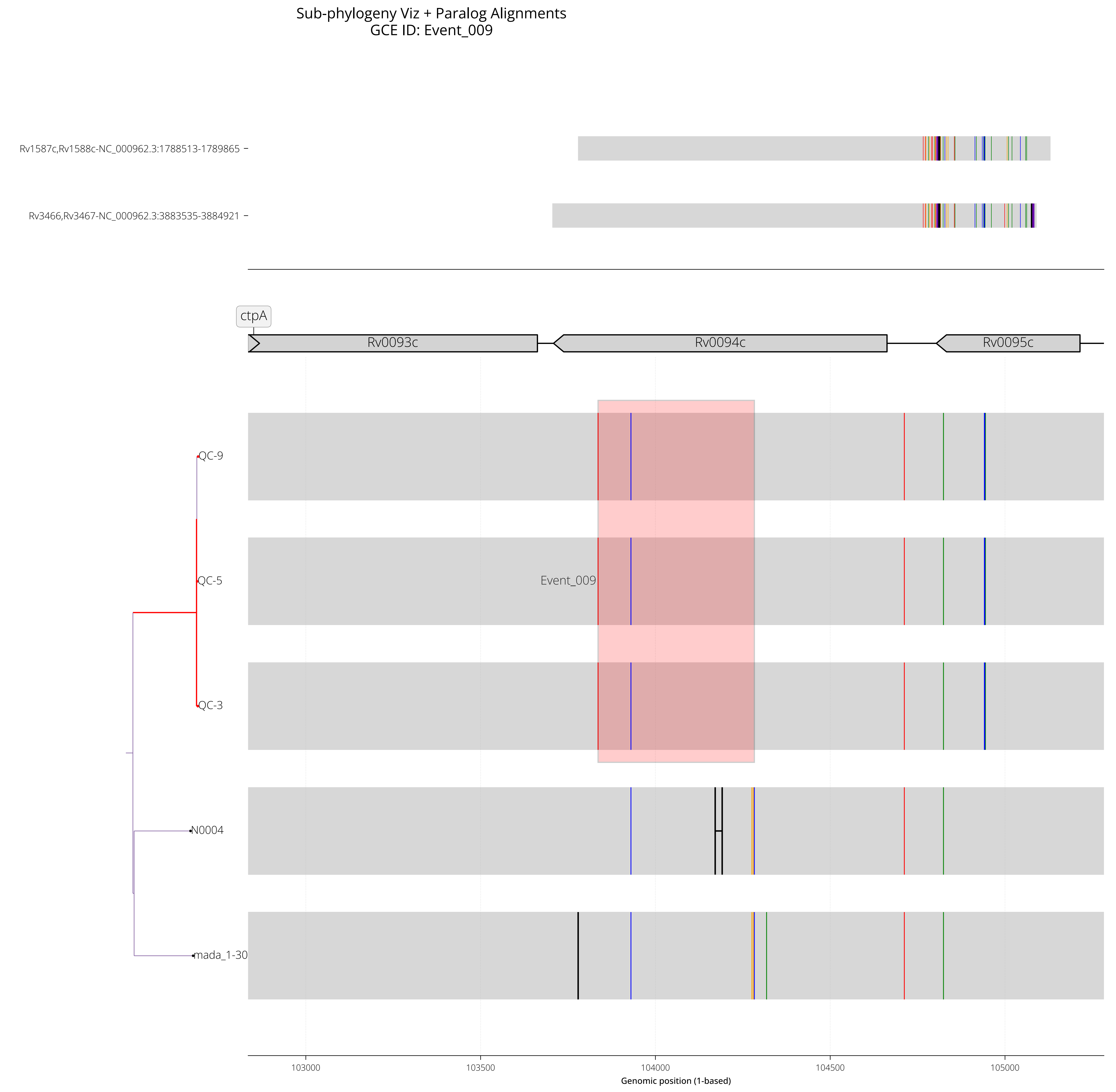
Sub-phylogeny Viz + Paralog Alignments
GCE ID: Event_009
Rv1587c,Rv1588c-NC_000962.3:1788513-1789865
Rv3466,Rv3467-NC_000962.3:3883535-3884921
ctpA
Rv0093c
Rv0094c
Rv0095c
QC-9
QC-5
QC-3
N0004
mada_1-30
Event_009
103000
103500
104000
104500
105000
Genomic position (1-based)

Full Phylogeny Viz + Paralog Alignments
GCE ID: Event_009

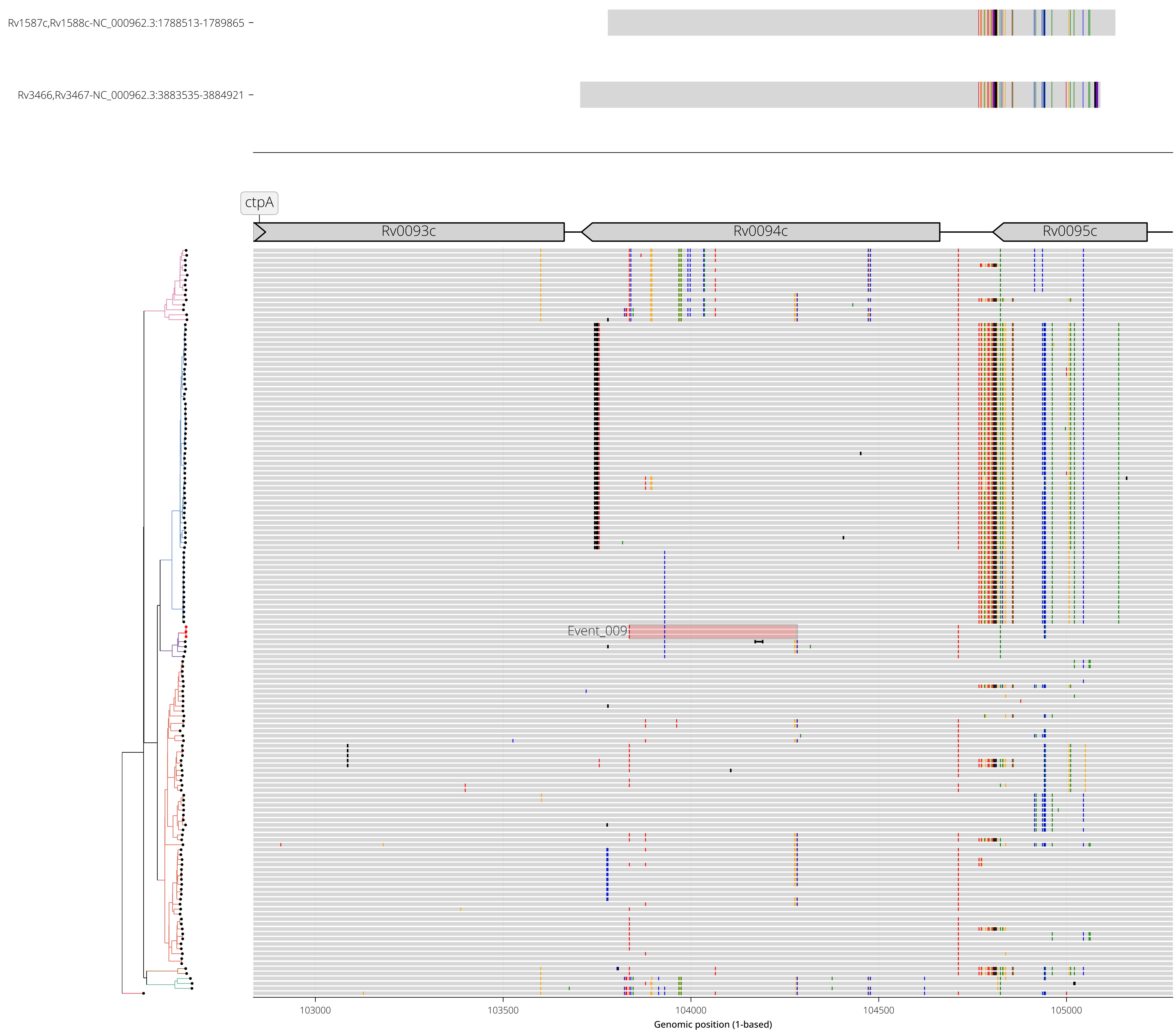

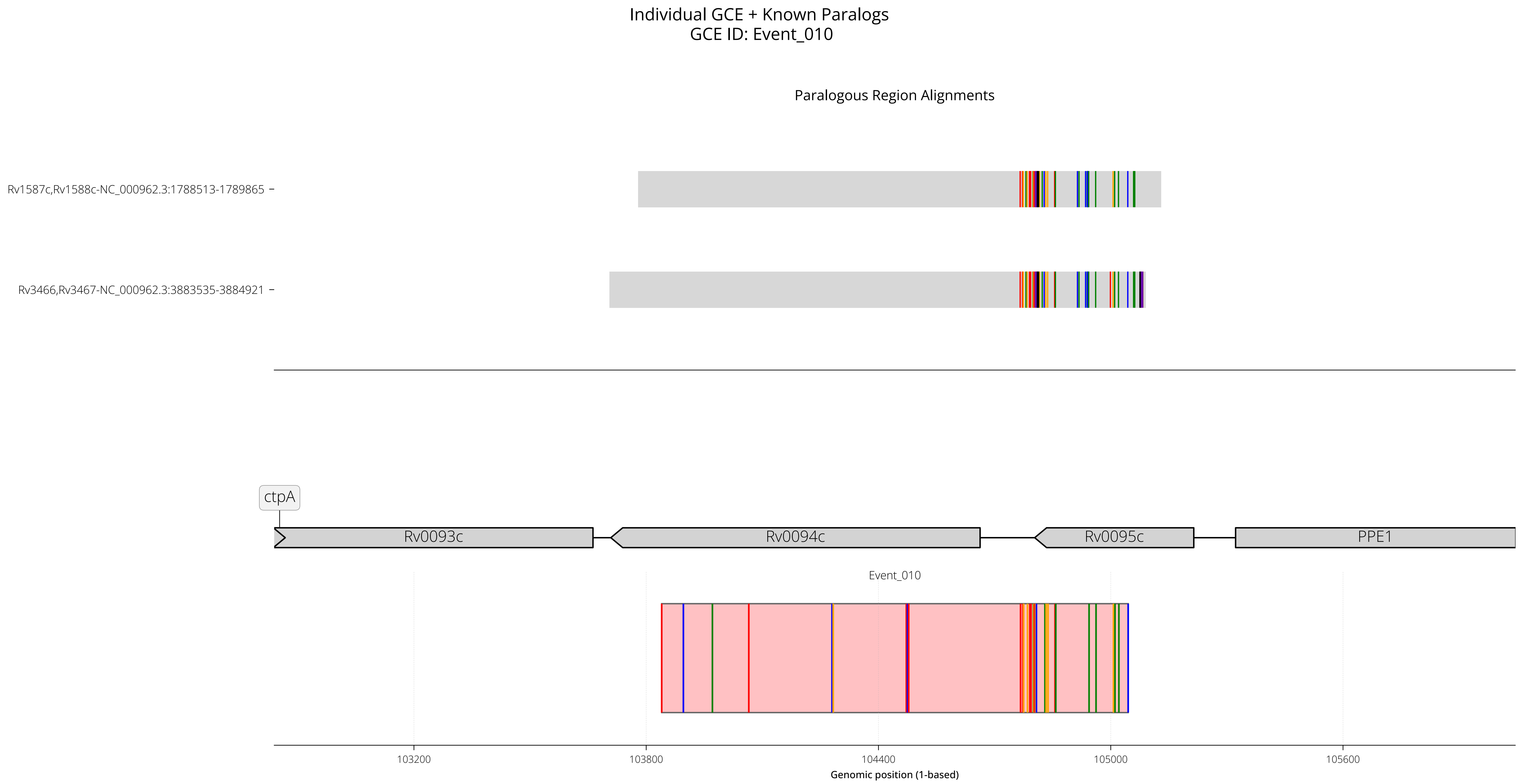
Individual GCE + Known Paralogs
GCE ID: Event_010
Paralogous Region Alignments
Rv1587c,Rv1588c-NC_000962.3:1788513-1789865
Rv3466,Rv3467-NC_000962.3:3883535-3884921
ctpA
Rv0093c
Rv0094c
Rv0095c
PPE1
Event_010
103200
103800
104400
105000
105600
Genomic position (1-based)

Circos Viz - GCE to Known Paralogs Locations
GCE ID: Event_010

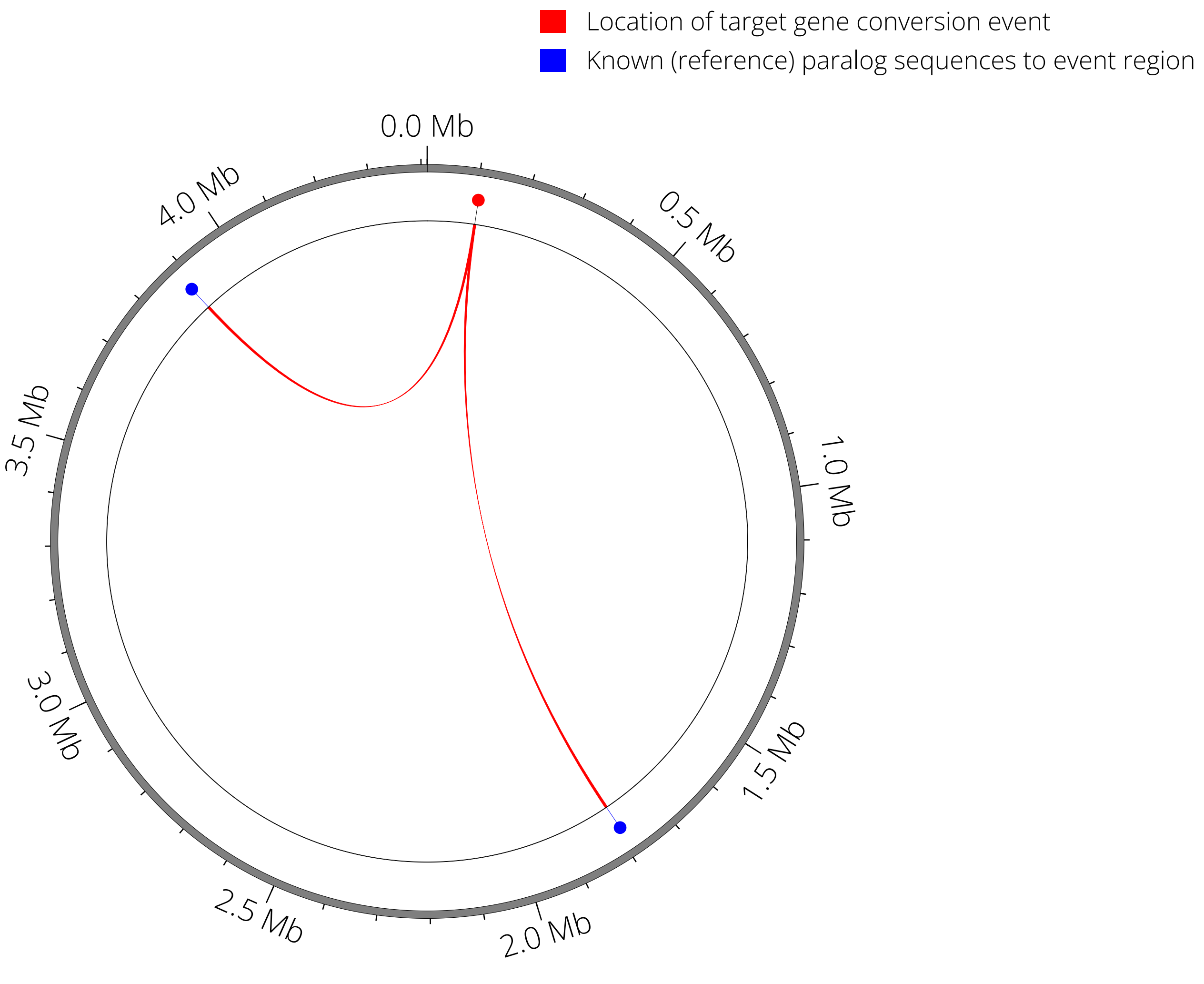

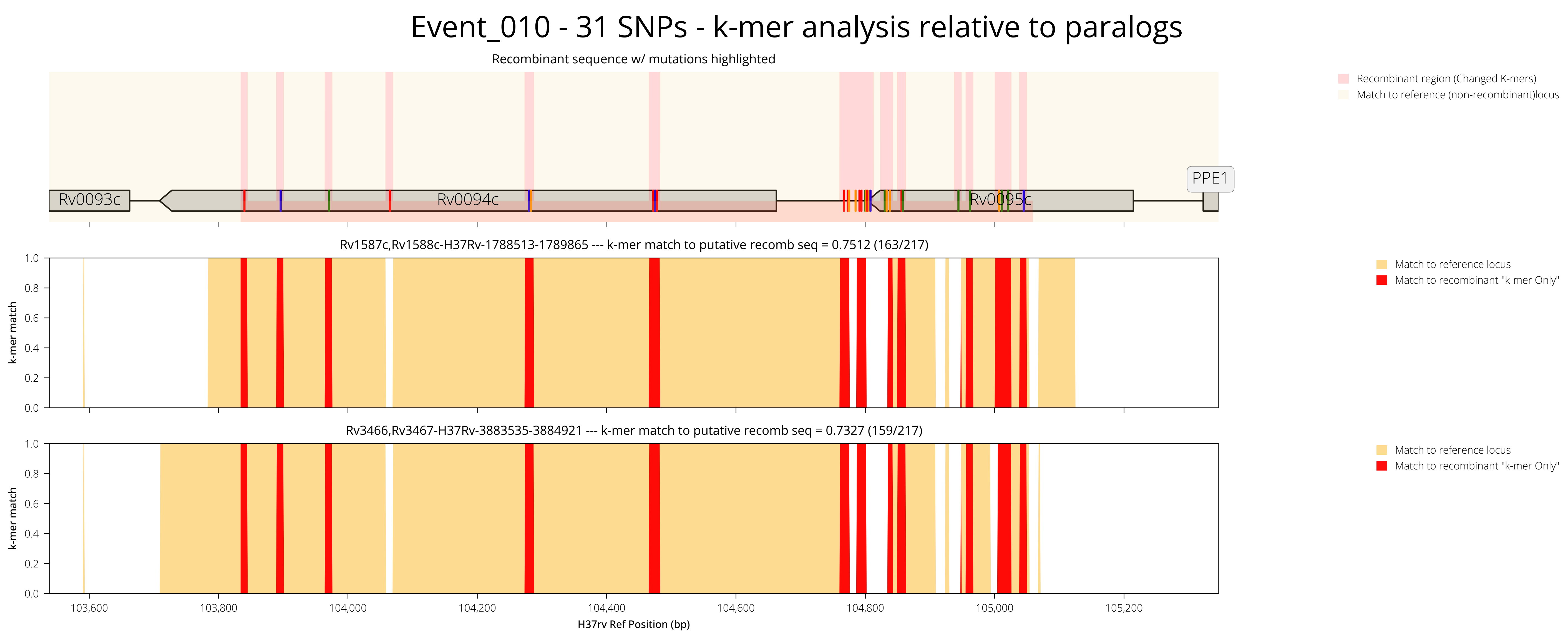
Event_010 - 31 SNPs - k-mer analysis relative to paralogs
Recombinant sequence w/ mutations highlighted
Recombinant region (Changed K-mers)
Match to reference (non-recombinant)locus
Rv0093c
Rv0094c
Rv0095c
PPE1
Rv1587c,Rv1588c-H37Rv-1788513-1789865 --- k-mer match to putative recomb seq = 0.7512 (163/217)
k-mer match
Match to reference locus
Match to recombinant "k-mer Only"
Rv3466,Rv3467-H37Rv-3883535-3884921 --- k-mer match to putative recomb seq = 0.7327 (159/217)
k-mer match
Match to reference locus
Match to recombinant "k-mer Only"
H37rv Ref Position (bp)

Sub-phylogeny Viz + Paralog Alignments
GCE ID: Event_010

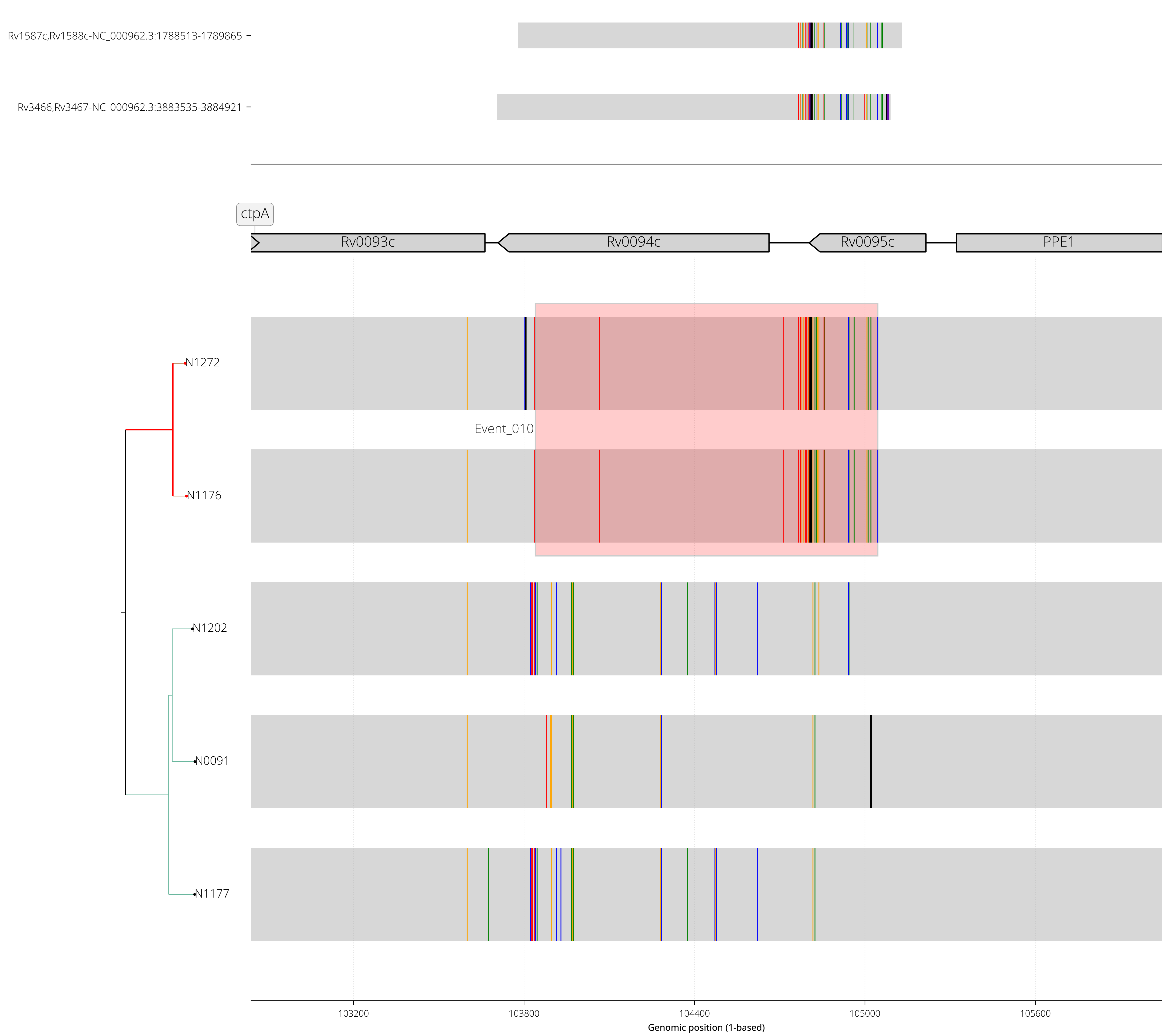

Full Phylogeny Viz + Paralog Alignments
GCE ID: Event_010

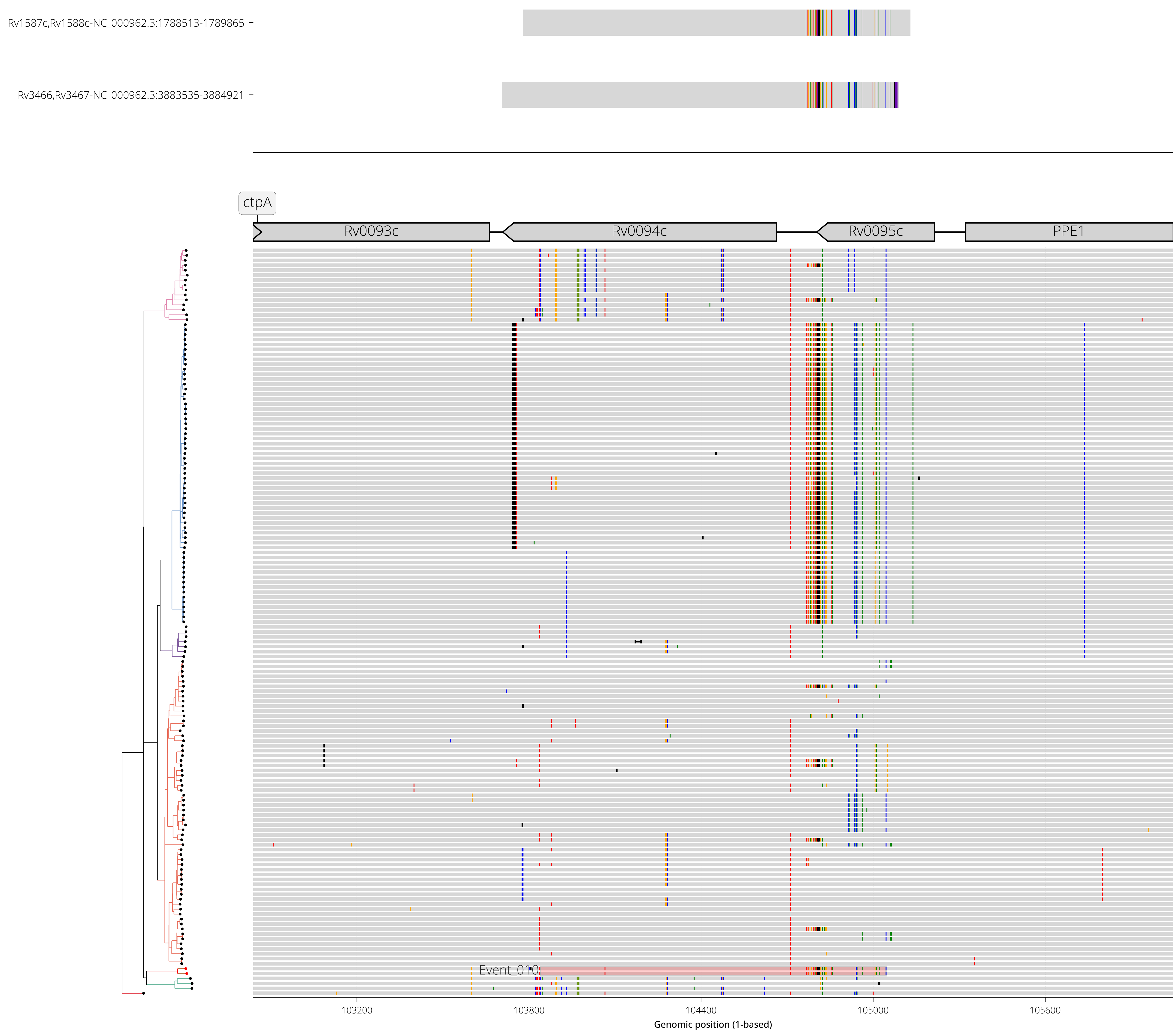

Individual GCE + Known Paralogs
GCE ID: Event_011

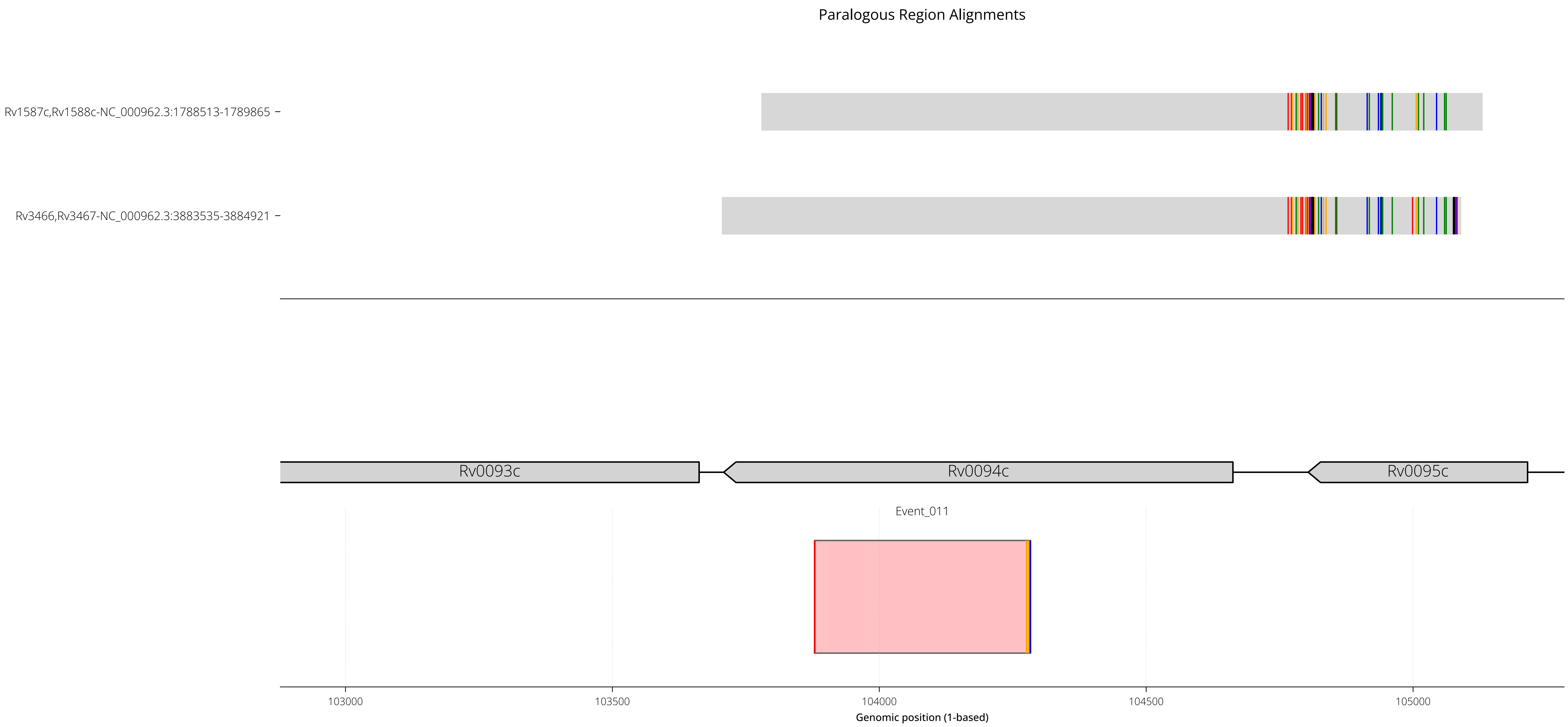

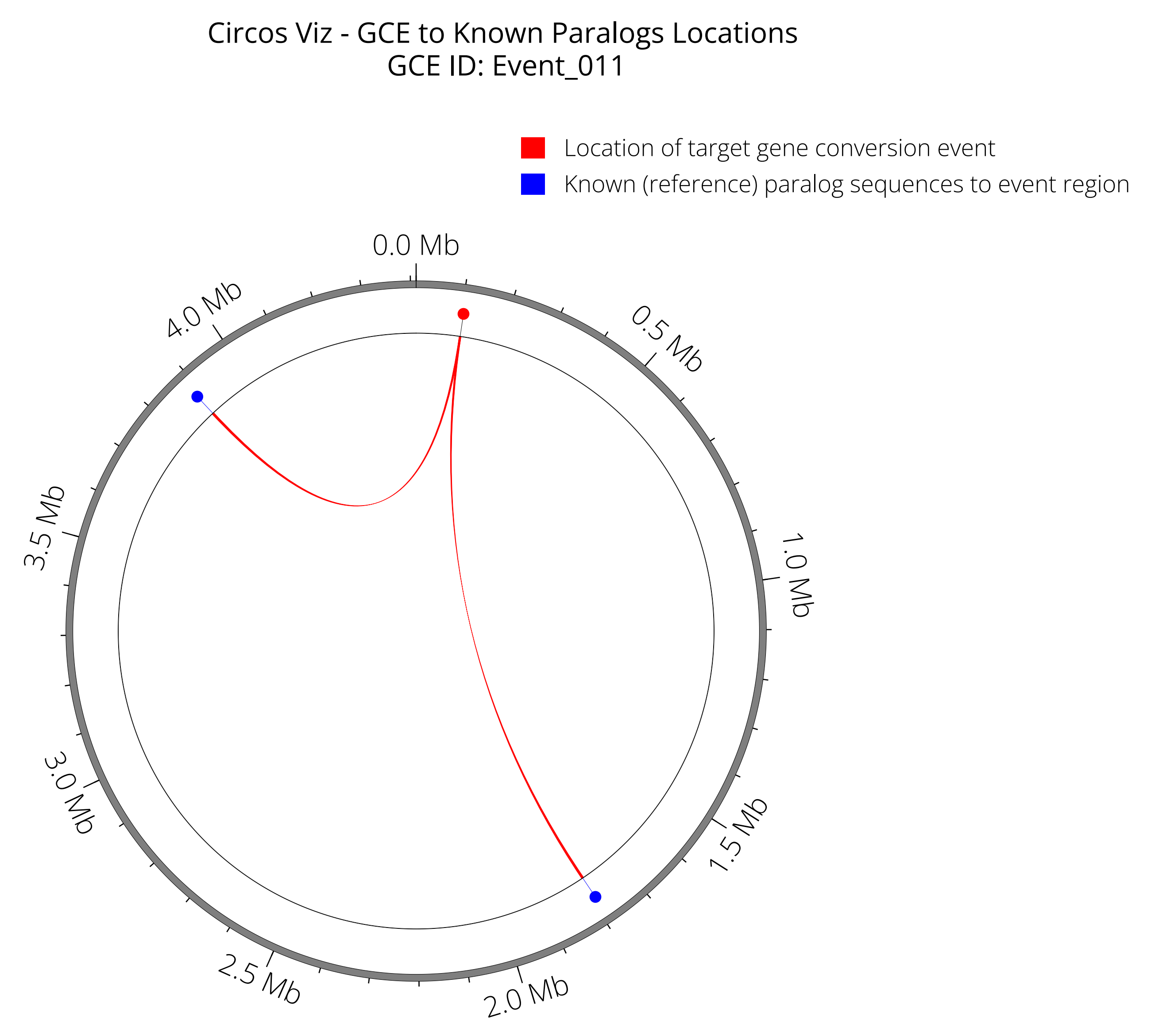
Circos Viz - GCE to Known Paralogs Locations
GCE ID: Event_011
Location of target gene conversion event
Known (reference) paralog sequences to event region
0.0 Mb
0.5 Mb
1.0 Mb
1.5 Mb
2.0 Mb
2.5 Mb
3.0 Mb
3.5 Mb
4.0 Mb

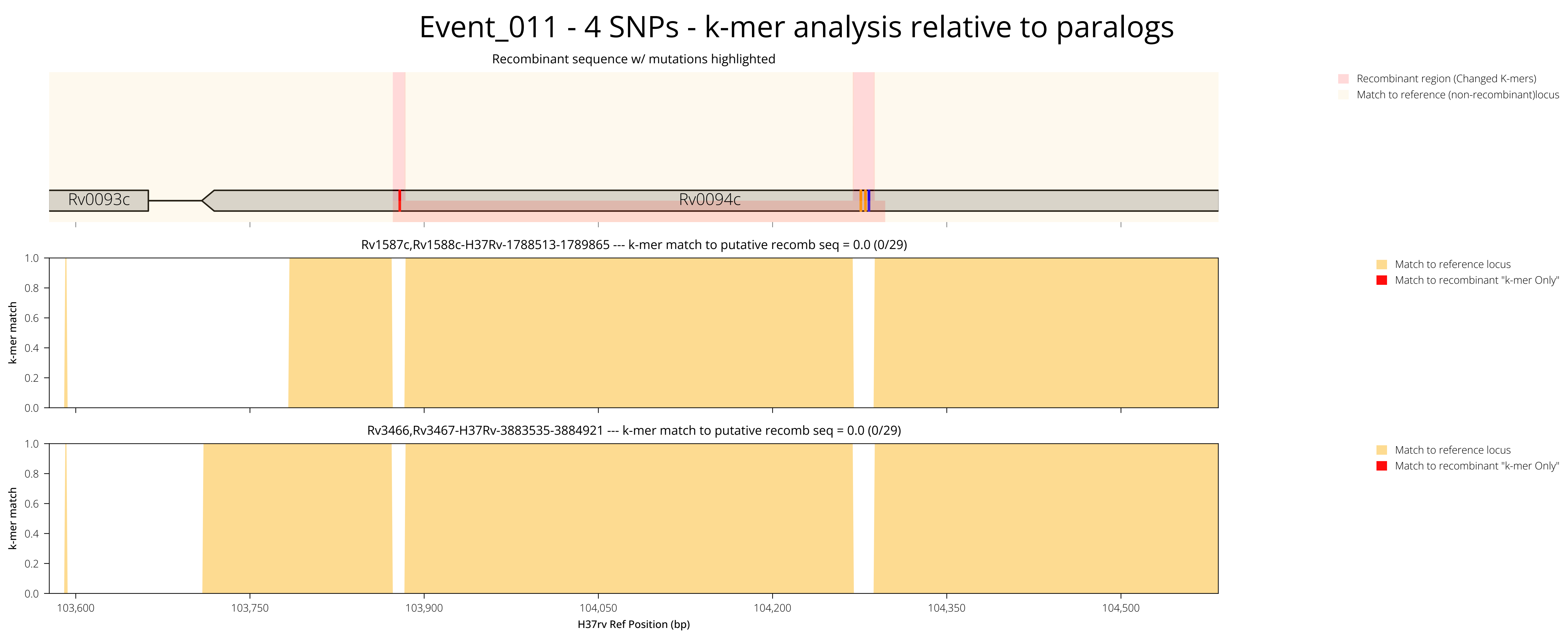

Event_011 - 4 SNPs - k-mer analysis relative to paralogs
Recombinant sequence w/ mutations highlighted
Recombinant region (Changed K-mers)
Match to reference (non-recombinant)locus
Rv0093c
Rv0094c
Rv1587c,Rv1588c-H37Rv-1788513-1789865 --- k-mer match to putative recomb seq = 0.0 (0/29)
k-mer match
1.0
0.8
0.6
0.4
0.2
0.0
Match to reference locus
Match to recombinant "k-mer Only"
Rv3466,Rv3467-H37Rv-3883535-3884921 --- k-mer match to putative recomb seq = 0.0 (0/29)
k-mer match
1.0
0.8
0.6
0.4
0.2
0.0
Match to reference locus
Match to recombinant "k-mer Only"
103,600
103,750
103,900
104,050
104,200
104,350
104,500
H37rv Ref Position (bp)

Sub-phylogeny Viz + Paralog Alignments
GCE ID: Event_011

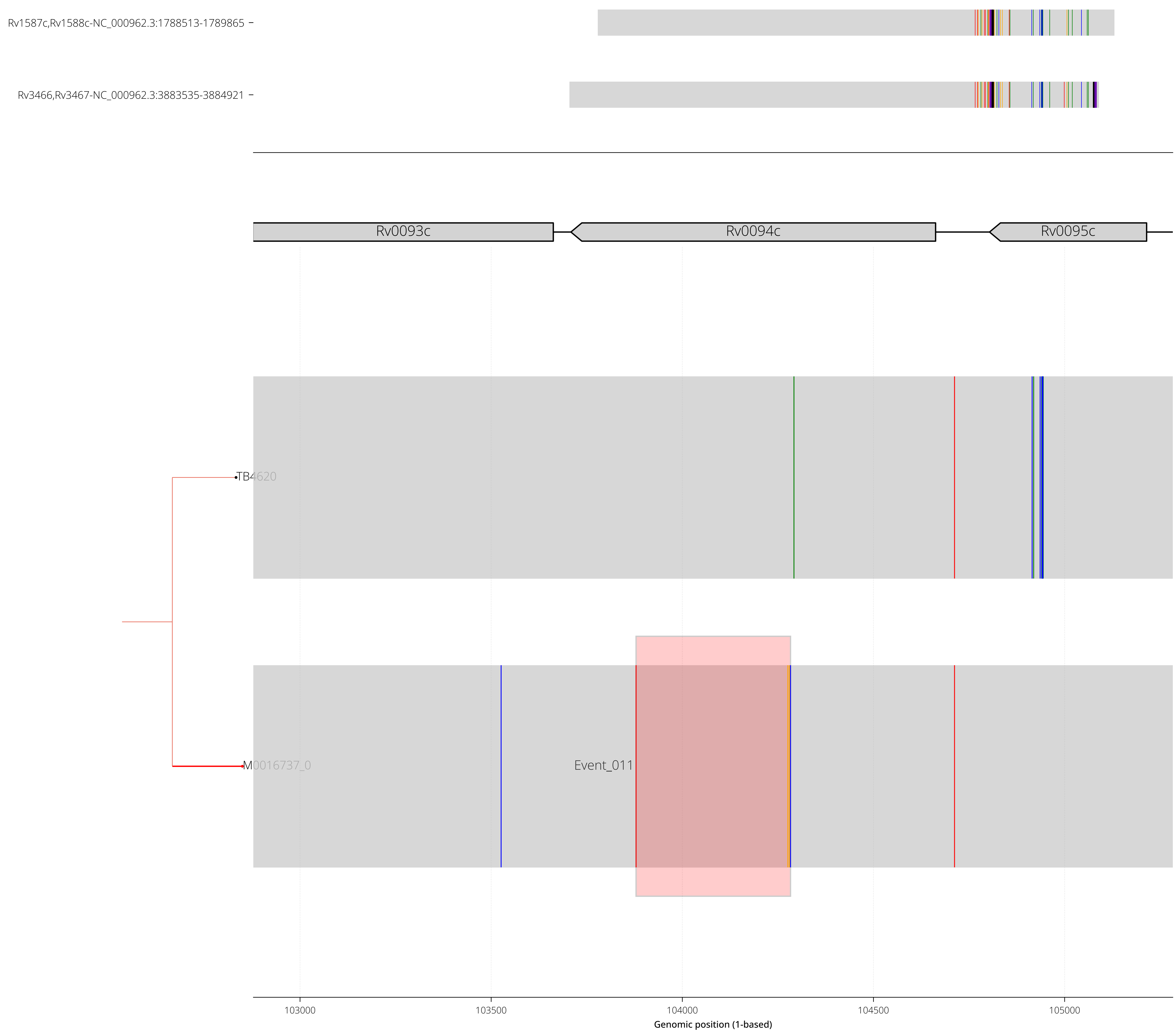

Full Phylogeny Viz + Paralog Alignments
GCE ID: Event_011

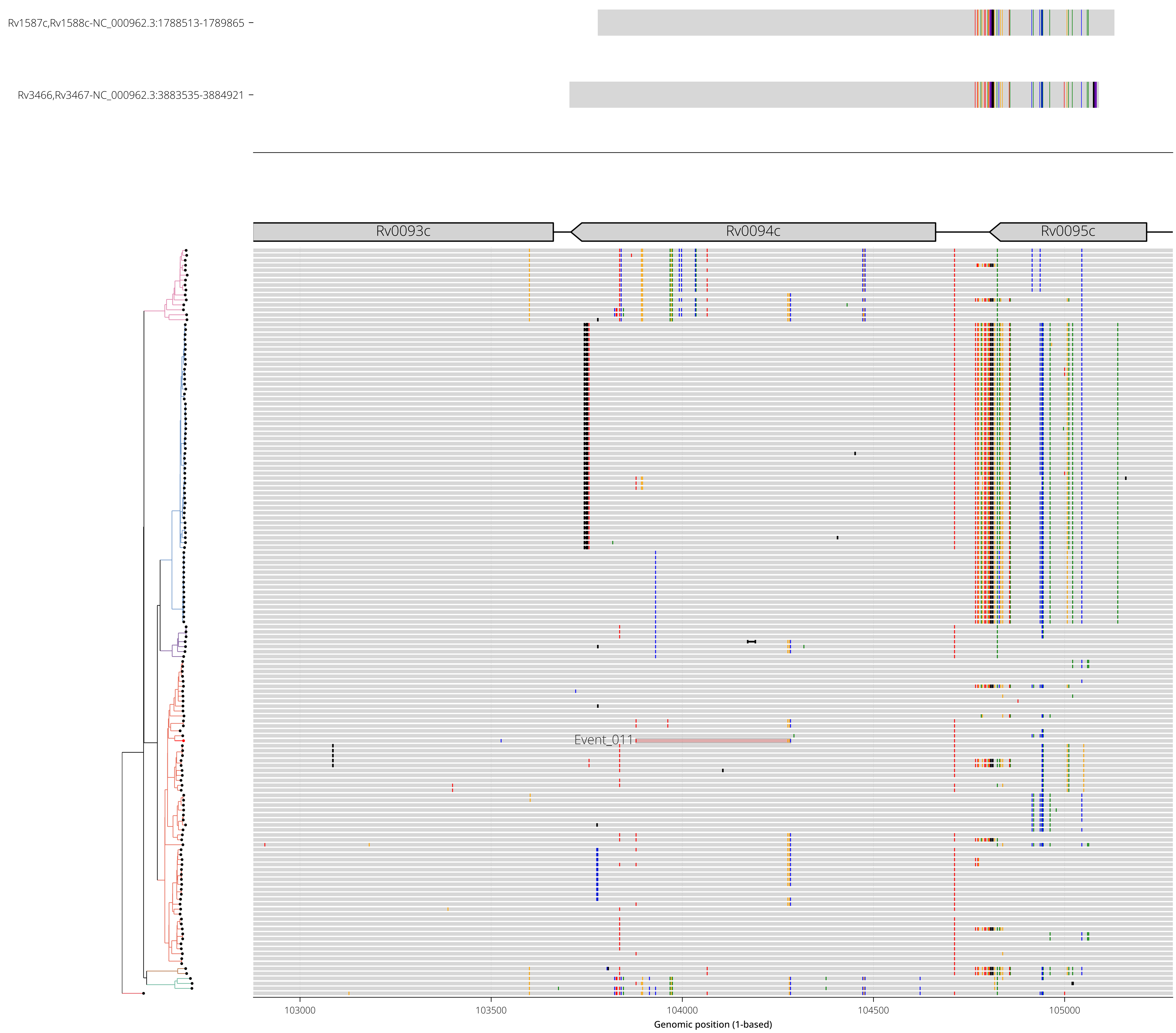

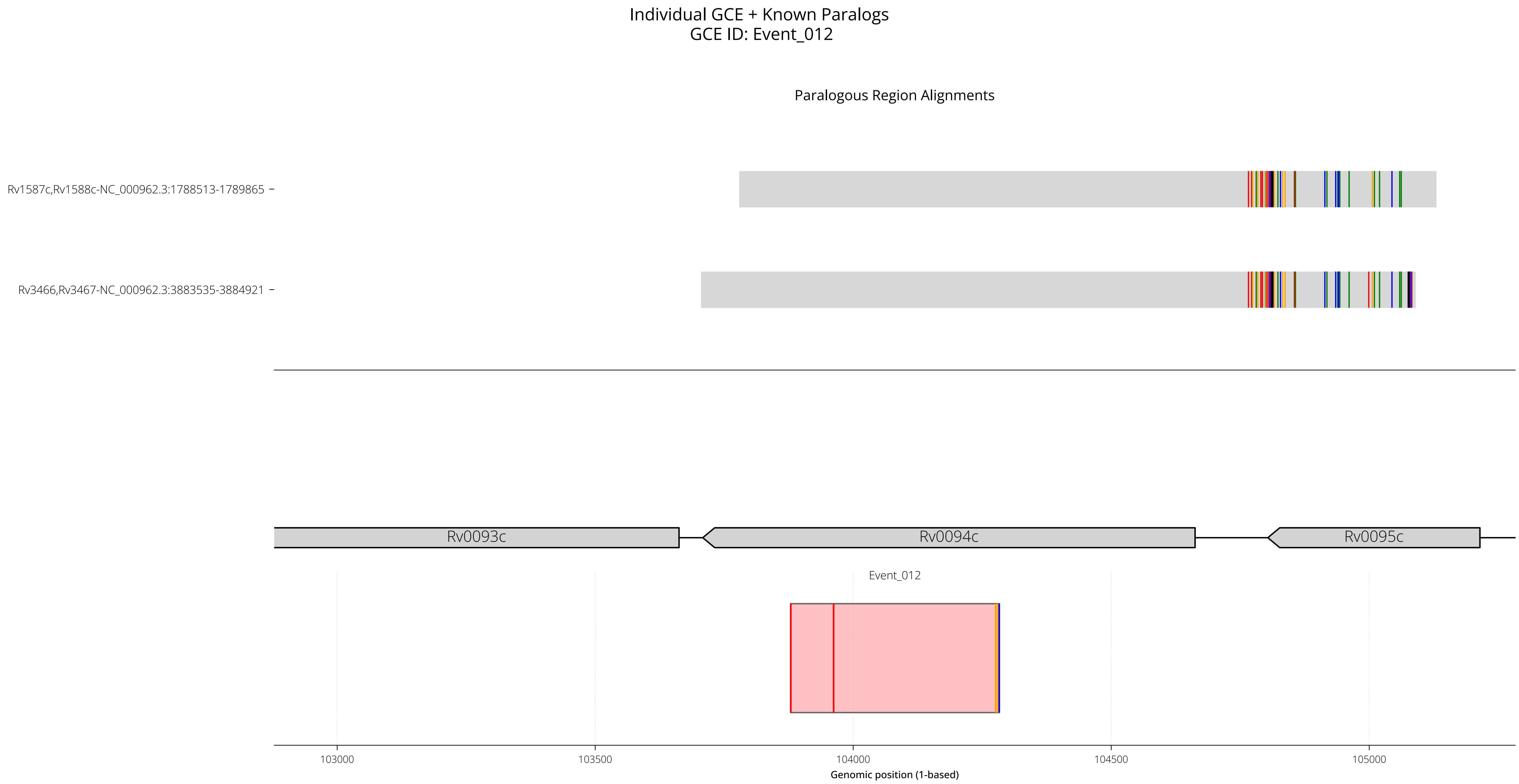

Individual GCE + Known Paralogs
GCE ID: Event_012
Paralogous Region Alignments
Rv1587c,Rv1588c-NC_000962.3:1788513-1789865
Rv3466,Rv3467-NC_000962.3:3883535-3884921
Rv0093c
Rv0094c
Rv0095c
Event_012
103000
103500
104000
104500
105000
Genomic position (1-based)

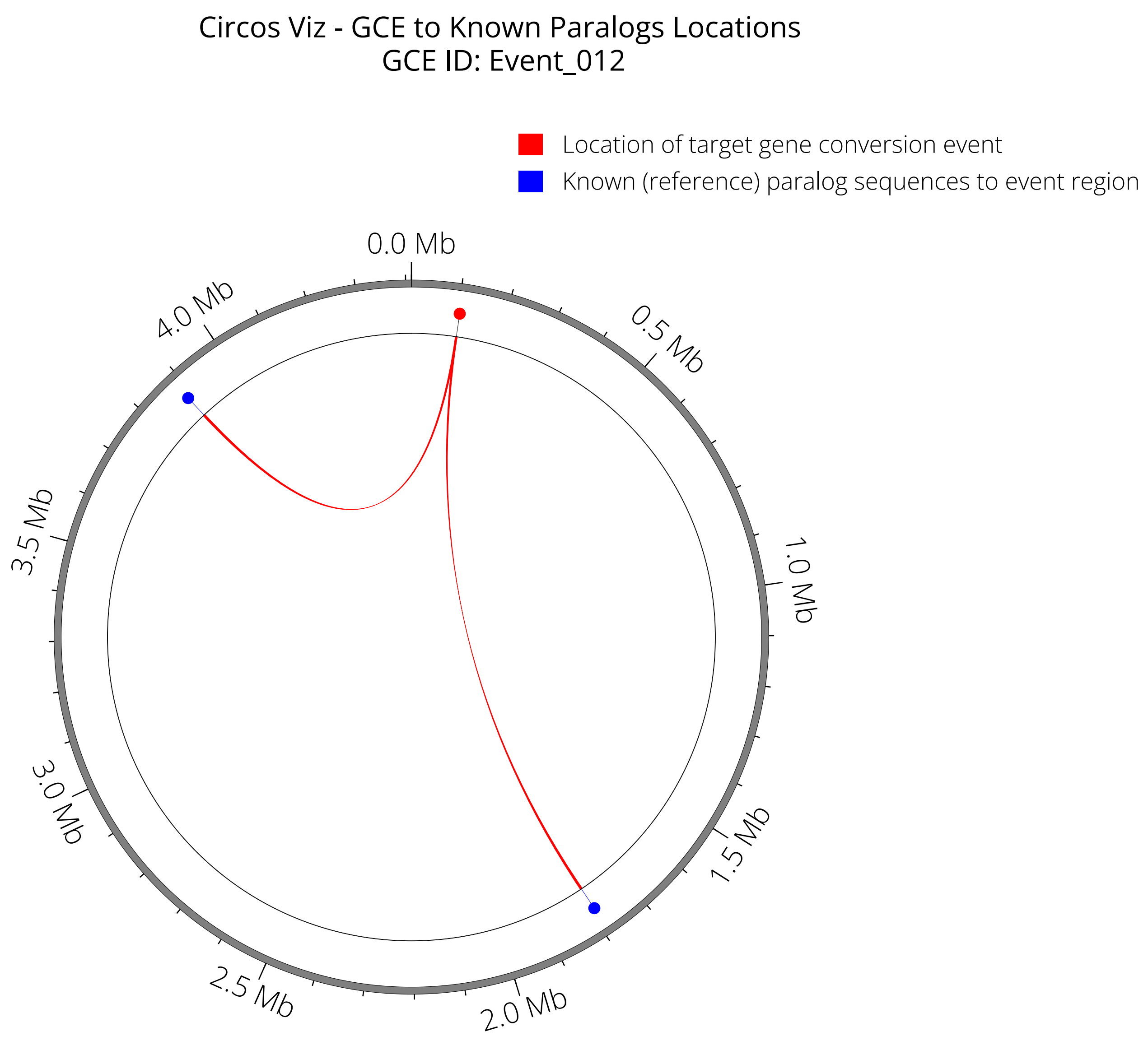

Circos Viz - GCE to Known Paralogs Locations
GCE ID: Event_012
Location of target gene conversion event
Known (reference) paralog sequences to event region
0.0 Mb
0.5 Mb
1.0 Mb
1.5 Mb
2.0 Mb
2.5 Mb
3.0 Mb
3.5 Mb
4.0 Mb

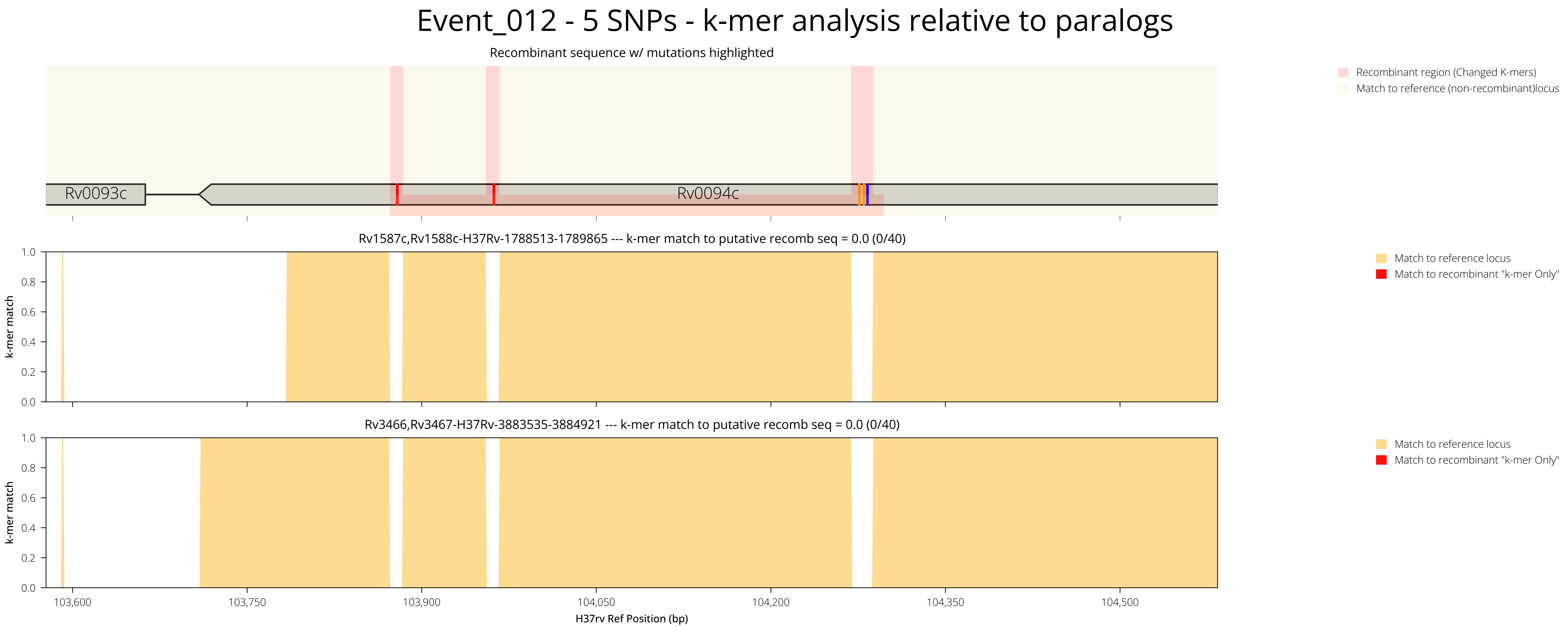

Event_012 - 5 SNPs - k-mer analysis relative to paralogs
Recombinant sequence w/ mutations highlighted
Rv0093c
Rv0094c
Recombinant region (Changed K-mers)
Match to reference (non-recombinant)locus
Rv1587c,Rv1588c-H37Rv-1788513-1789865 --- k-mer match to putative recomb seq = 0.0 (0/40)
Match to reference locus
Match to recombinant "k-mer Only"
Rv3466,Rv3467-H37Rv-3883535-3884921 --- k-mer match to putative recomb seq = 0.0 (0/40)
Match to reference locus
Match to recombinant "k-mer Only"
k-mer match
1.0
0.8
0.6
0.4
0.2
0.0
103,600
103,750
103,900
104,050
104,200
104,350
104,500
H37rv Ref Position (bp)

Sub-phylogeny Viz + Paralog Alignments
GCE ID: Event_012

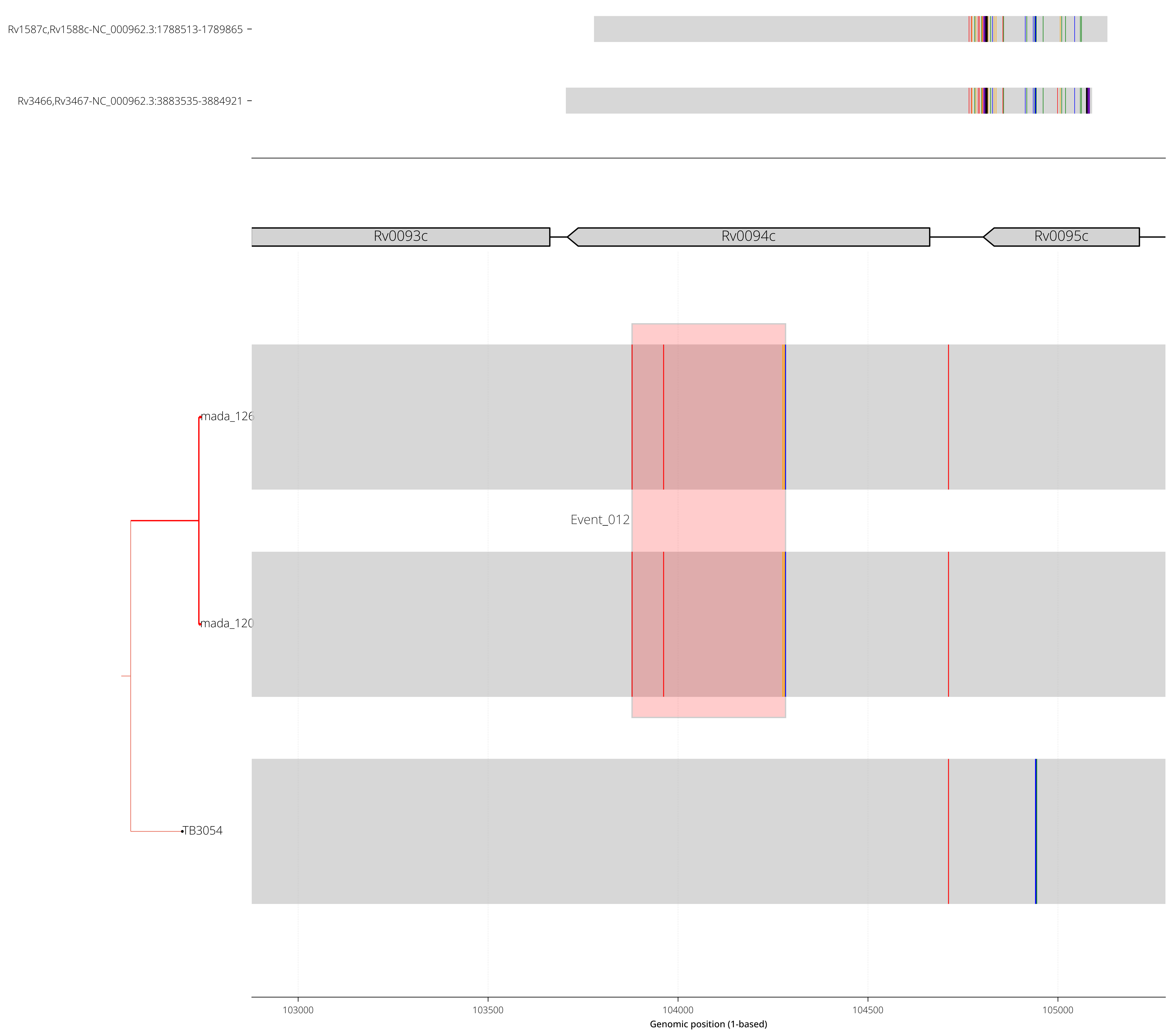

Full Phylogeny Viz + Paralog Alignments
GCE ID: Event_012

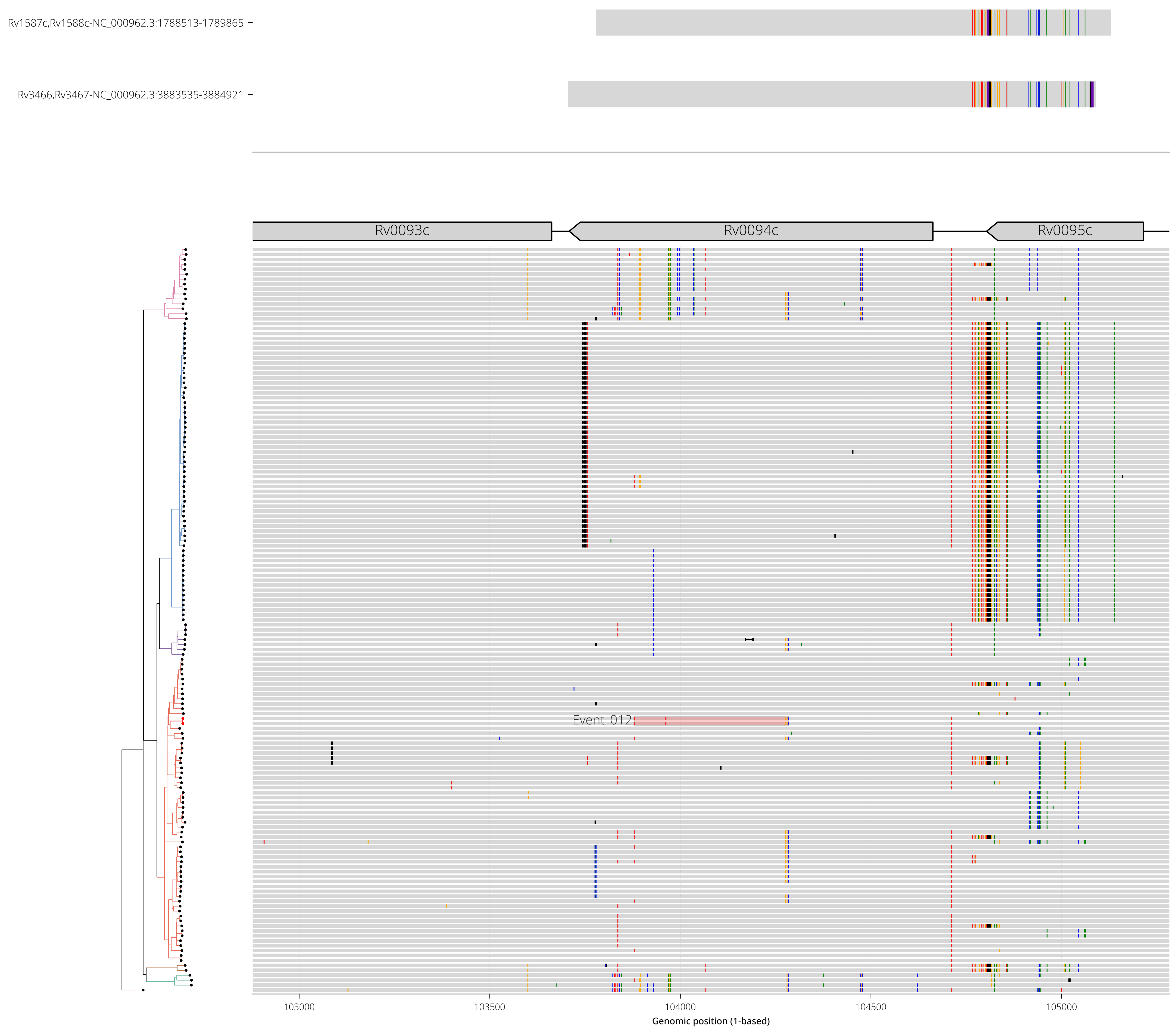

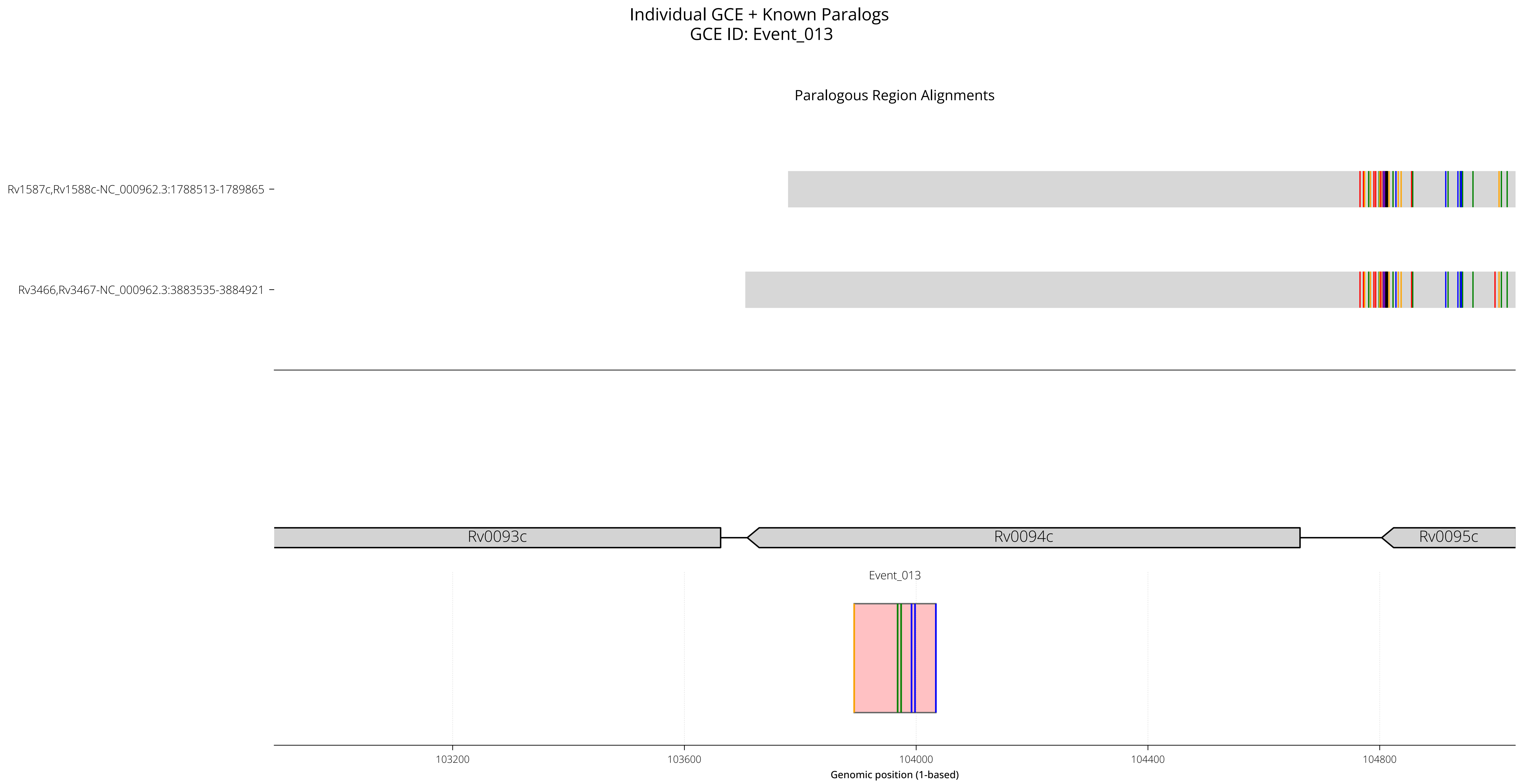
Individual GCE + Known Paralogs
GCE ID: Event_013
Paralogous Region Alignments
Rv1587c,Rv1588c-NC_000962.3:1788513-1789865
Rv3466,Rv3467-NC_000962.3:3883535-3884921
Rv0093c
Rv0094c
Rv0095c
Event_013
103200
103600
104000
104400
104800
Genomic position (1-based)

Circos Viz - GCE to Known Paralogs Locations
GCE ID: Event_013

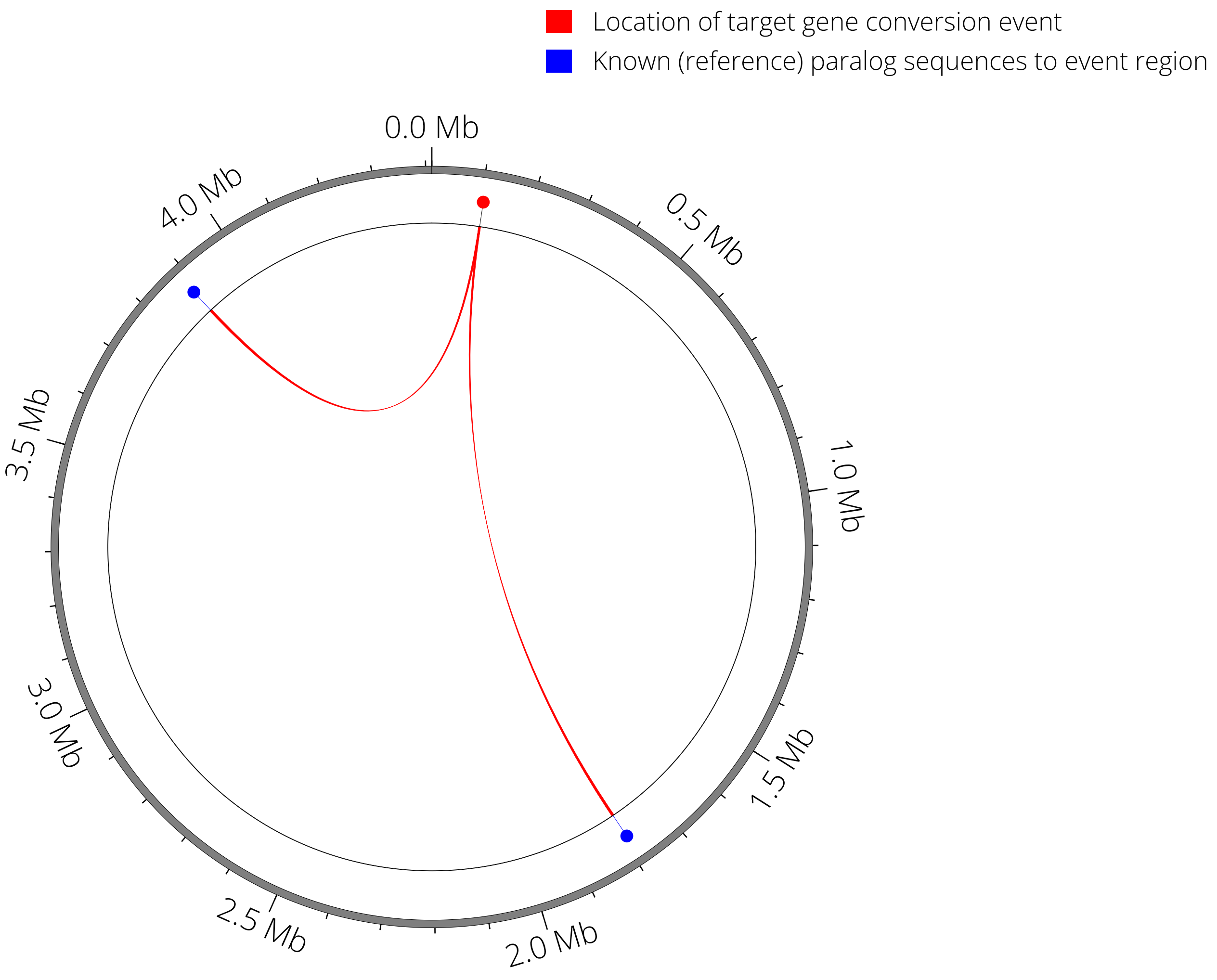

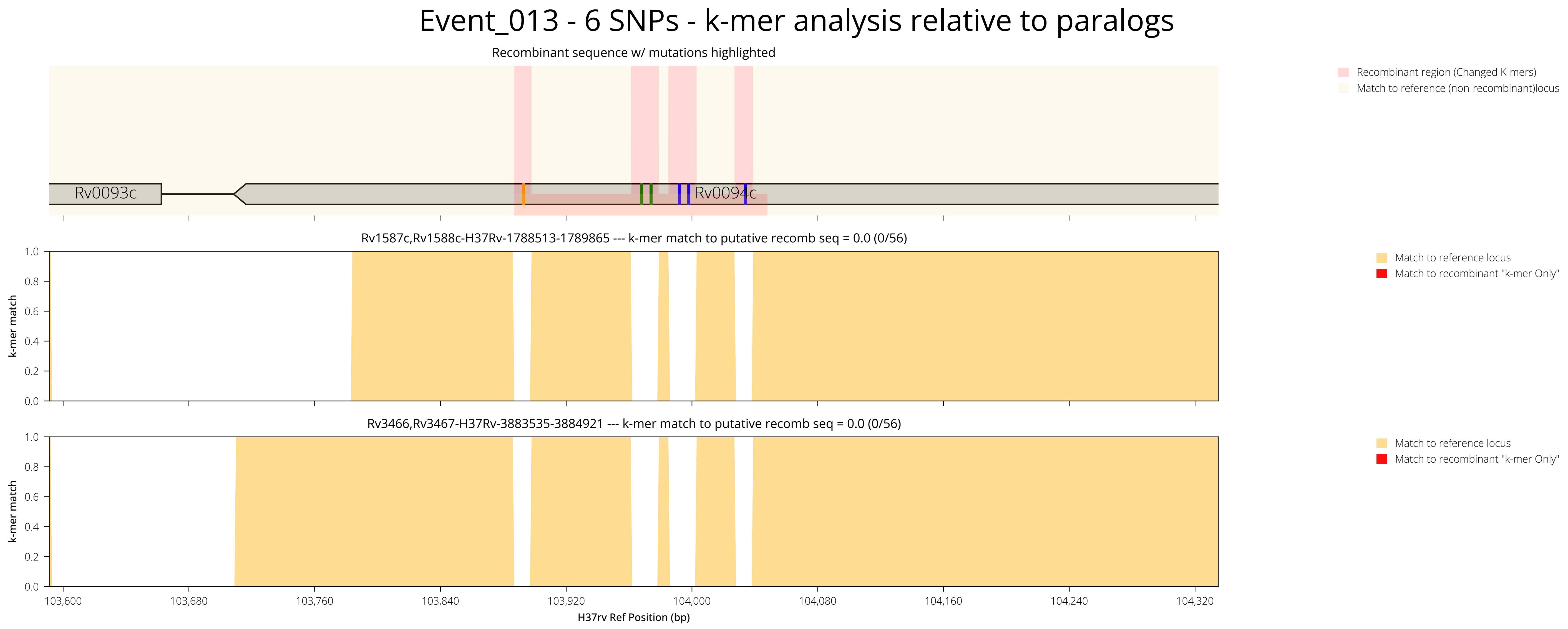
Event_013 - 6 SNPs - k-mer analysis relative to paralogs
Recombinant sequence w/ mutations highlighted
Rv0093c
Rv0094c
Recombinant region (Changed K-mers)
Match to reference (non-recombinant)locus
Rv1587c,Rv1588c-H37Rv-1788513-1789865 --- k-mer match to putative recomb seq = 0.0 (0/56)
k-mer match
Match to reference locus
Match to recombinant "k-mer Only"
Rv3466,Rv3467-H37Rv-3883535-3884921 --- k-mer match to putative recomb seq = 0.0 (0/56)
k-mer match
Match to reference locus
Match to recombinant "k-mer Only"
1.0
0.8
0.6
0.4
0.2
0.0
103,600
103,680
103,760
103,840
103,920
104,000
104,080
104,160
104,240
104,320
H37rv Ref Position (bp)

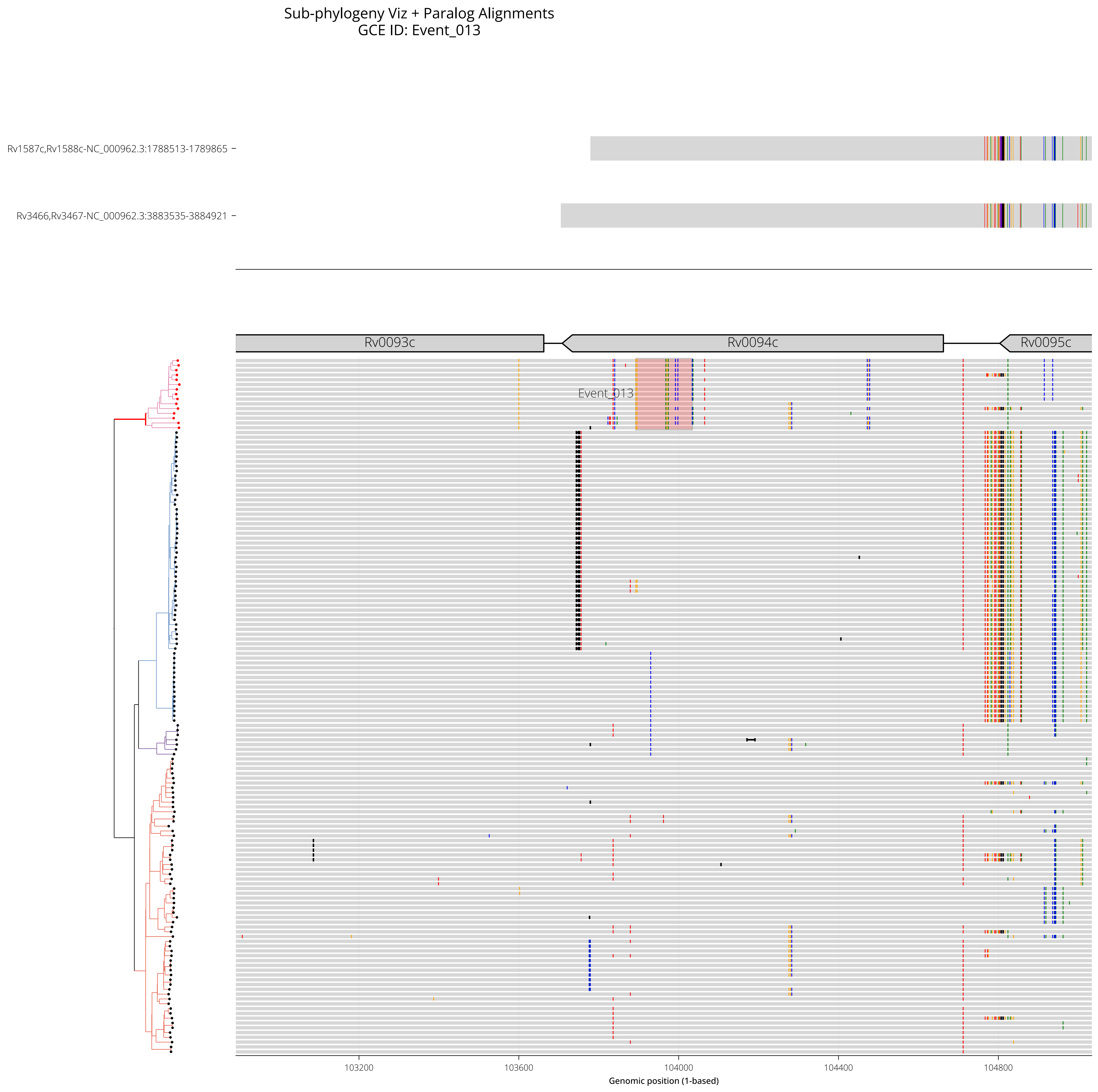
Sub-phylogeny Viz + Paralog Alignments
GCE ID: Event_013
Rv1587c,Rv1588c-NC_000962.3:1788513-1789865
Rv3466,Rv3467-NC_000962.3:3883535-3884921
Rv0093c
Rv0094c
Rv0095c
Event_013
103200
103600
104000
104400
104800
Genomic position (1-based)

Full Phylogeny Viz + Paralog Alignments
GCE ID: Event_013

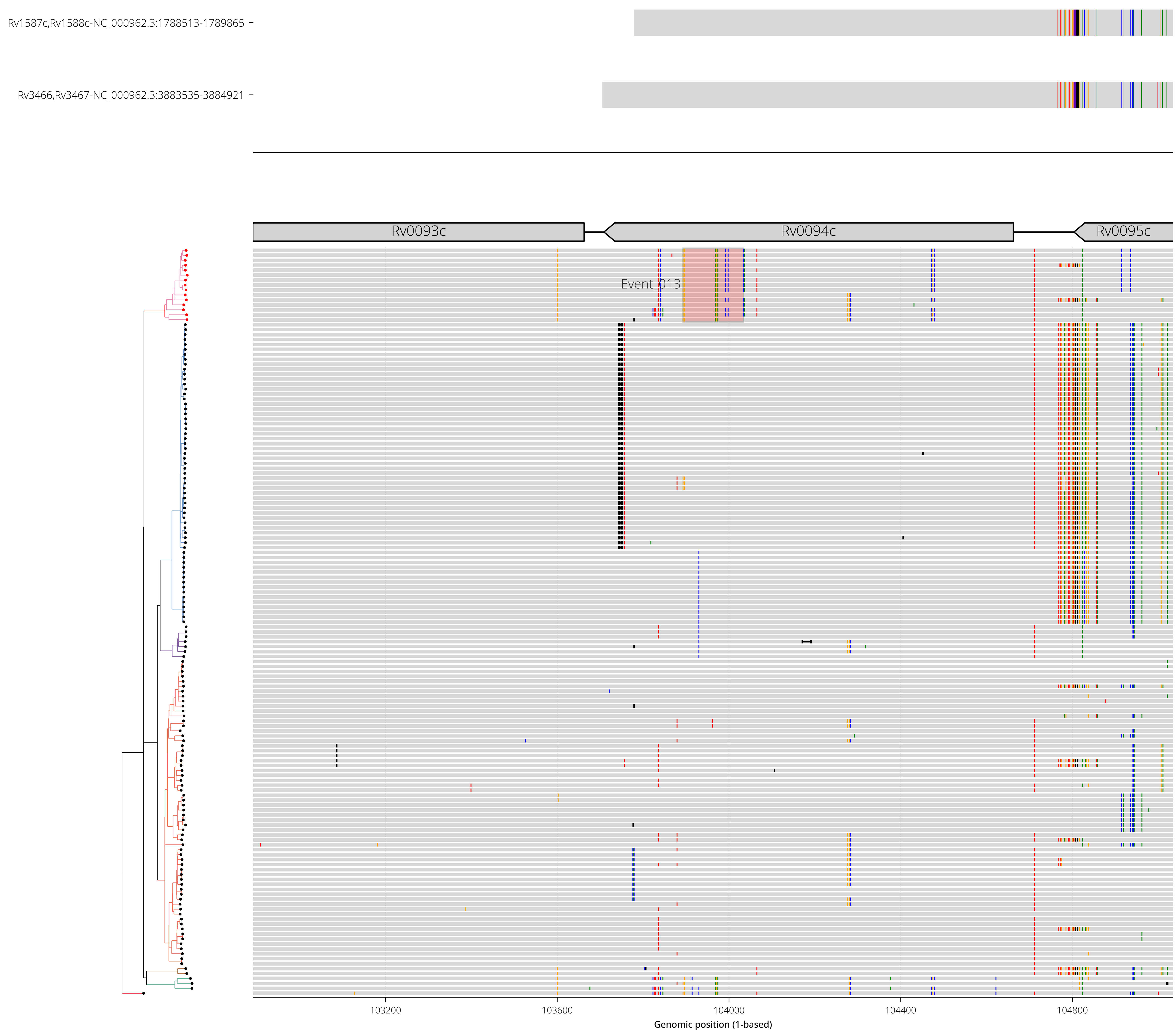

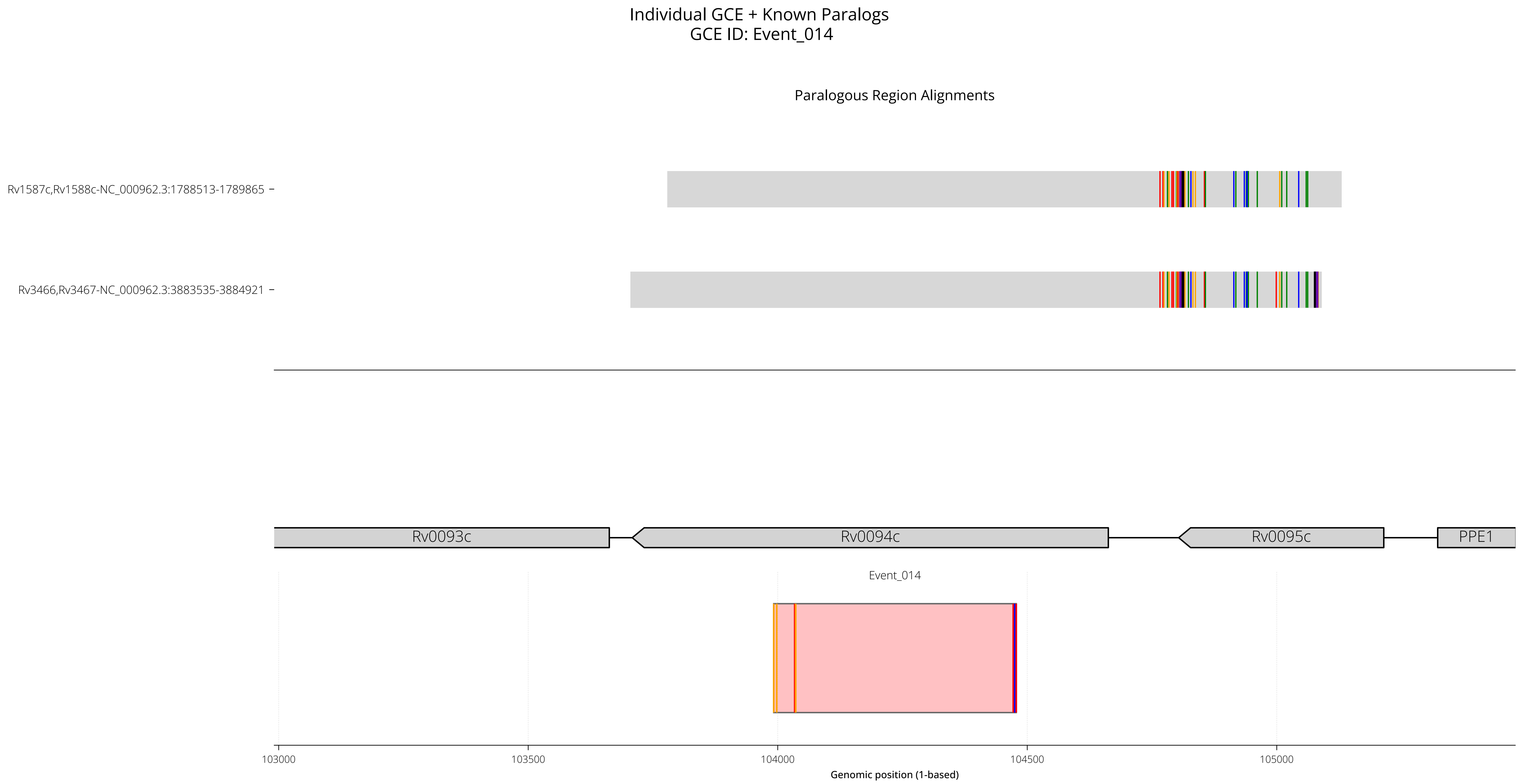
Individual GCE + Known Paralogs
GCE ID: Event_014
Paralogous Region Alignments
Rv1587c,Rv1588c-NC_000962.3:1788513-1789865
Rv3466,Rv3467-NC_000962.3:3883535-3884921
Rv0093c
Rv0094c
Rv0095c
PPE1
Event_014
103000
103500
104000
104500
105000
Genomic position (1-based)

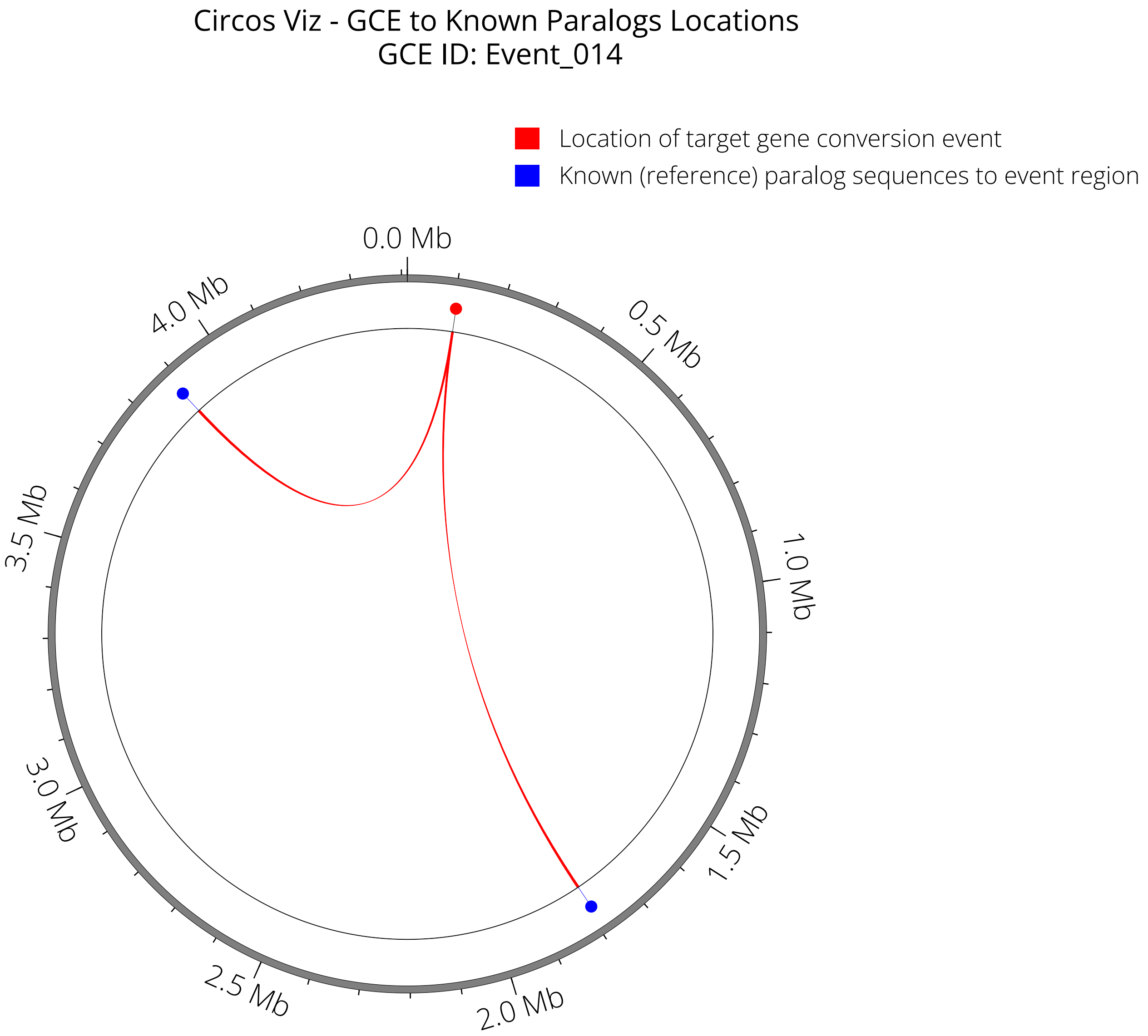
Circos Viz - GCE to Known Paralogs Locations
GCE ID: Event_014
Location of target gene conversion event
Known (reference) paralog sequences to event region
0.0 Mb
0.5 Mb
1.0 Mb
1.5 Mb
2.0 Mb
2.5 Mb
3.0 Mb
3.5 Mb
4.0 Mb

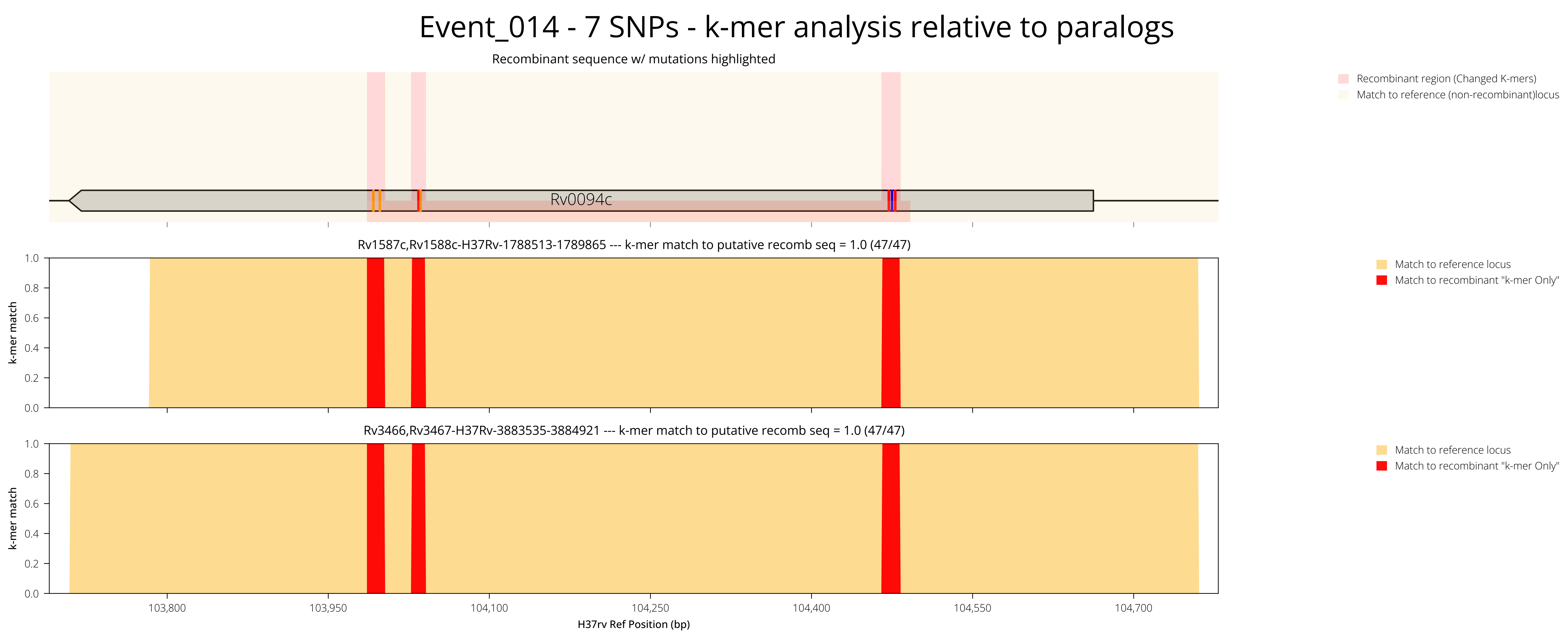

Event_014 - 7 SNPs - k-mer analysis relative to paralogs
Recombinant sequence w/ mutations highlighted
Recombinant region (Changed K-mers)
Match to reference (non-recombinant)locus
Rv0094c
Rv1587c,Rv1588c-H37Rv-1788513-1789865 --- k-mer match to putative recomb seq = 1.0 (47/47)
Match to reference locus
Match to recombinant "k-mer Only"
k-mer match
1.0
0.8
0.6
0.4
0.2
0.0
Rv3466,Rv3467-H37Rv-3883535-3884921 --- k-mer match to putative recomb seq = 1.0 (47/47)
Match to reference locus
Match to recombinant "k-mer Only"
k-mer match
1.0
0.8
0.6
0.4
0.2
0.0
103,800
103,950
104,100
104,250
104,400
104,550
104,700
H37rv Ref Position (bp)

Sub-phylogeny Viz + Paralog Alignments
GCE ID: Event_014

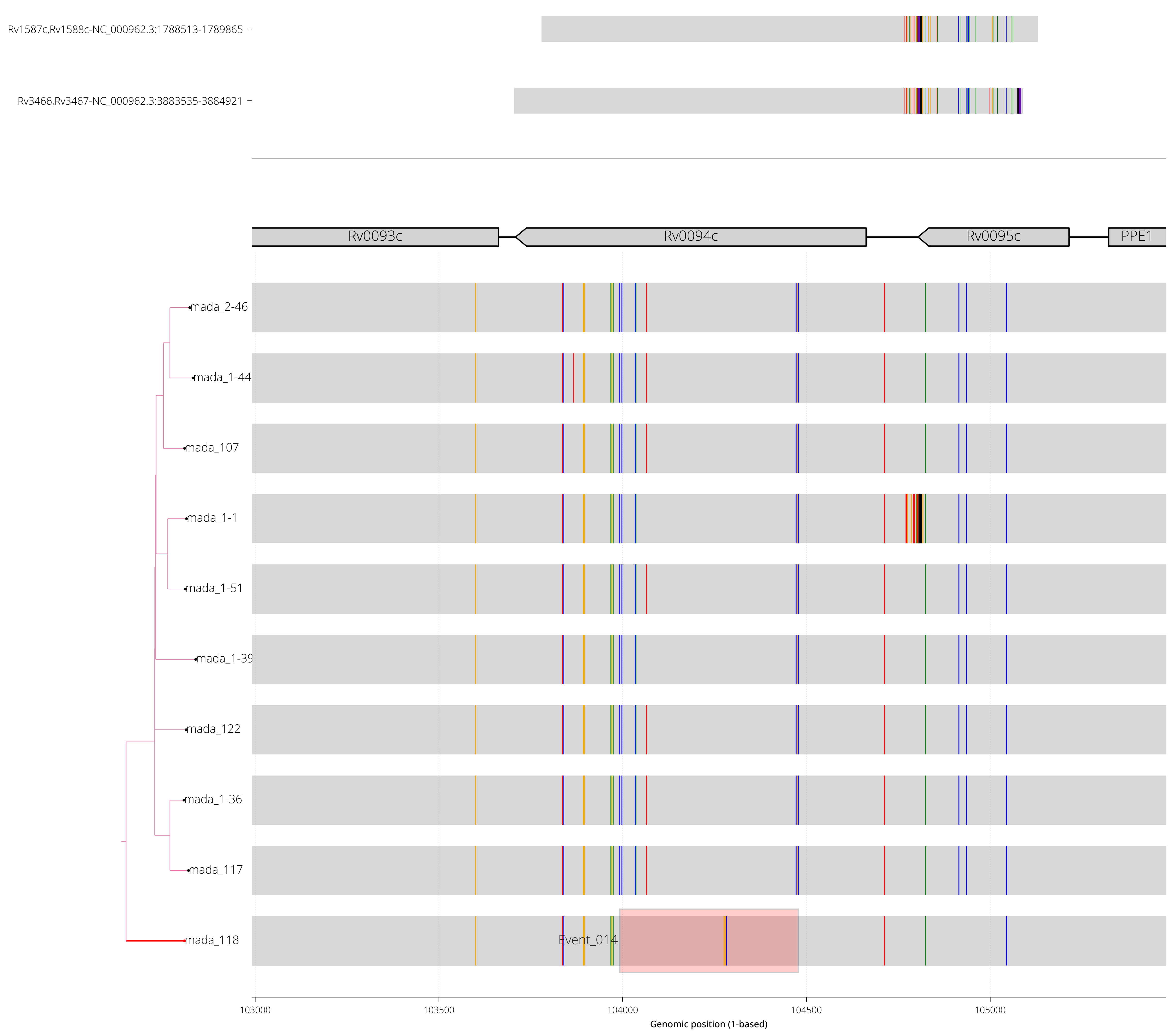

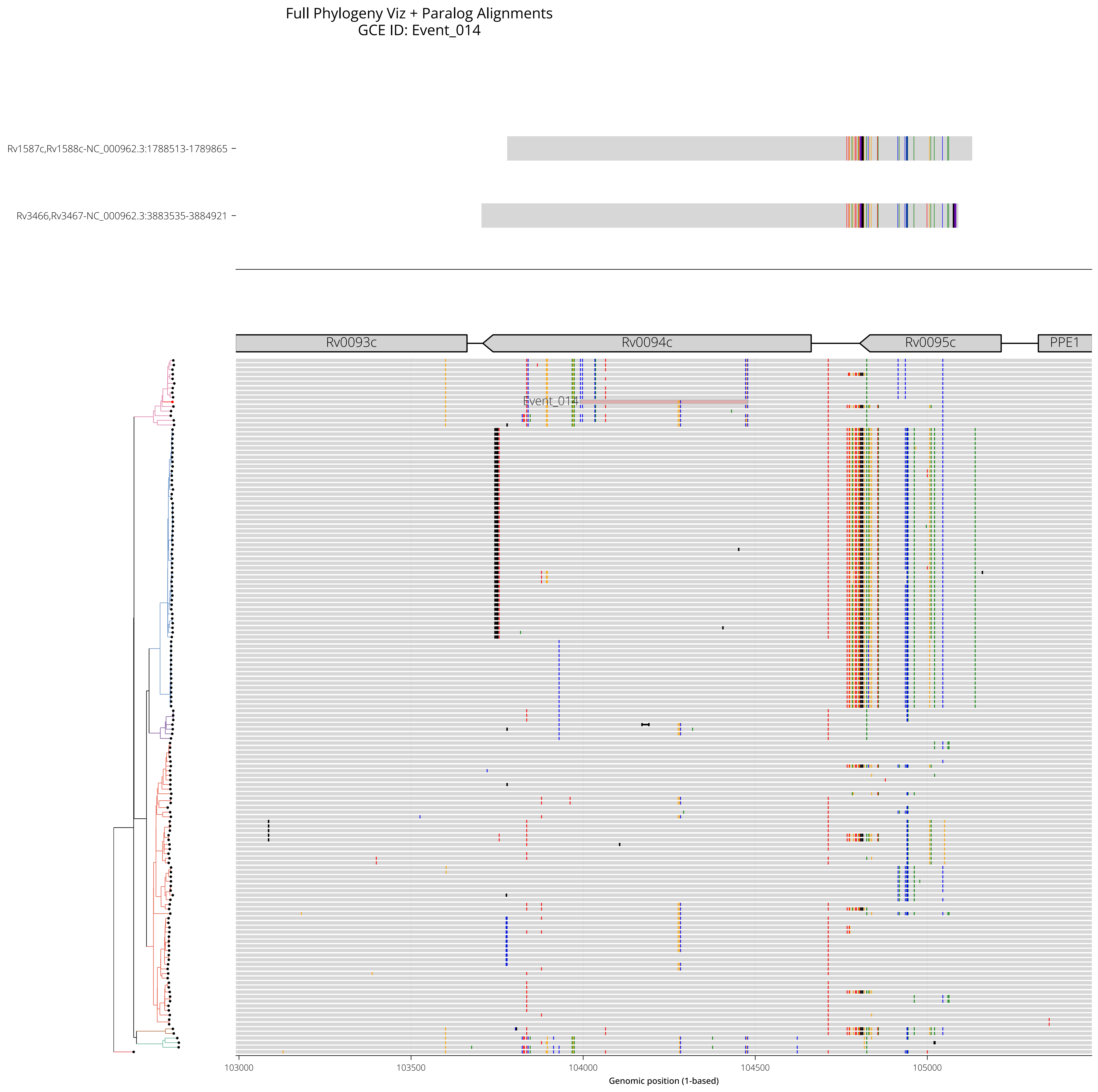
Full Phylogeny Viz + Paralog Alignments
GCE ID: Event_014
Rv1587c,Rv1588c-NC_000962.3:1788513-1789865
Rv3466,Rv3467-NC_000962.3:3883535-3884921
Rv0093c
Rv0094c
Rv0095c
PPE1
Event_014
103000
103500
104000
104500
105000
Genomic position (1-based)

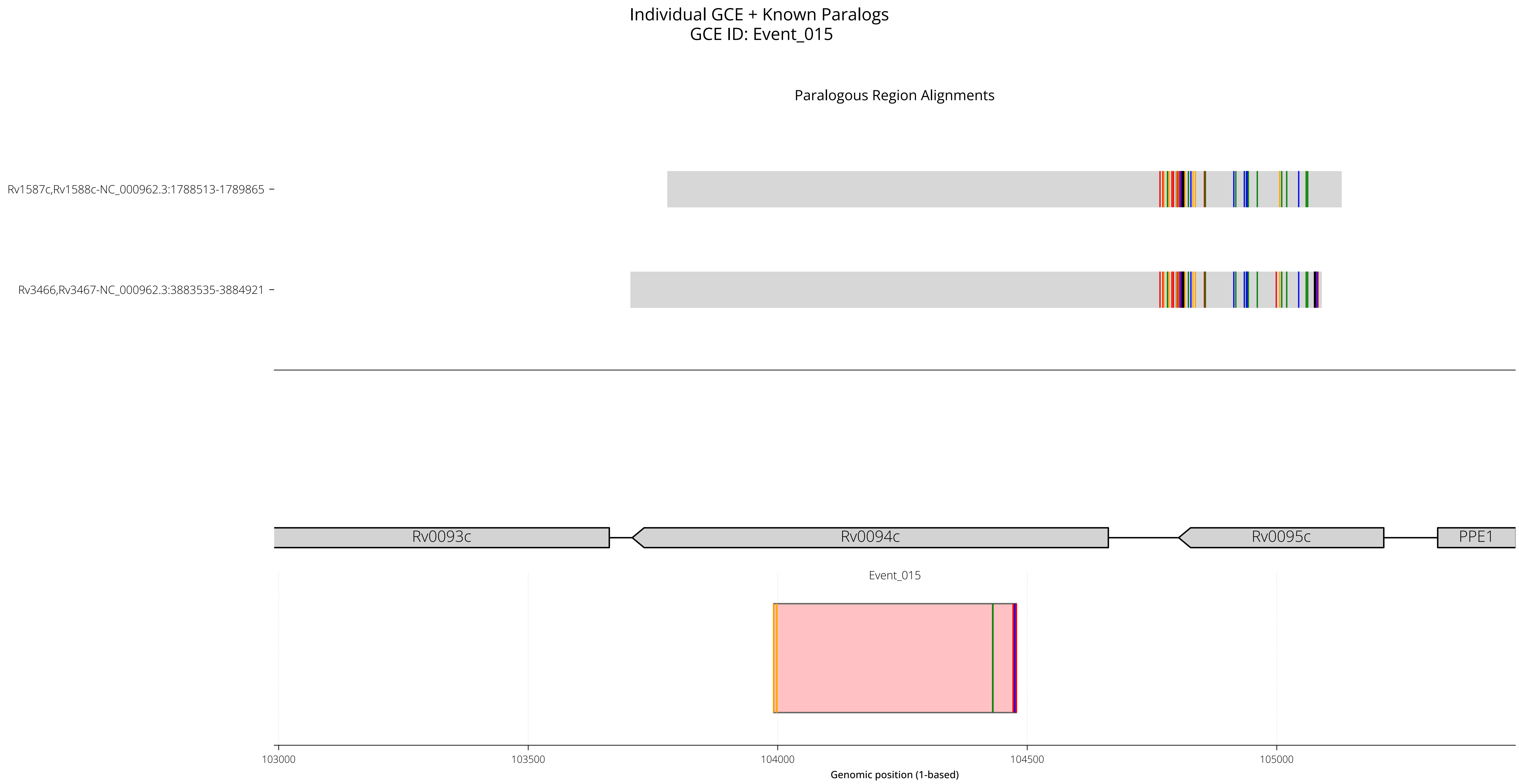
Individual GCE + Known Paralogs
GCE ID: Event_015
Paralogous Region Alignments
Rv1587c,Rv1588c-NC_000962.3:1788513-1789865
Rv3466,Rv3467-NC_000962.3:3883535-3884921
Rv0093c
Rv0094c
Rv0095c
PPE1
Event_015
103000
103500
104000
104500
105000
Genomic position (1-based)

Circos Viz - GCE to Known Paralogs Locations
GCE ID: Event_015

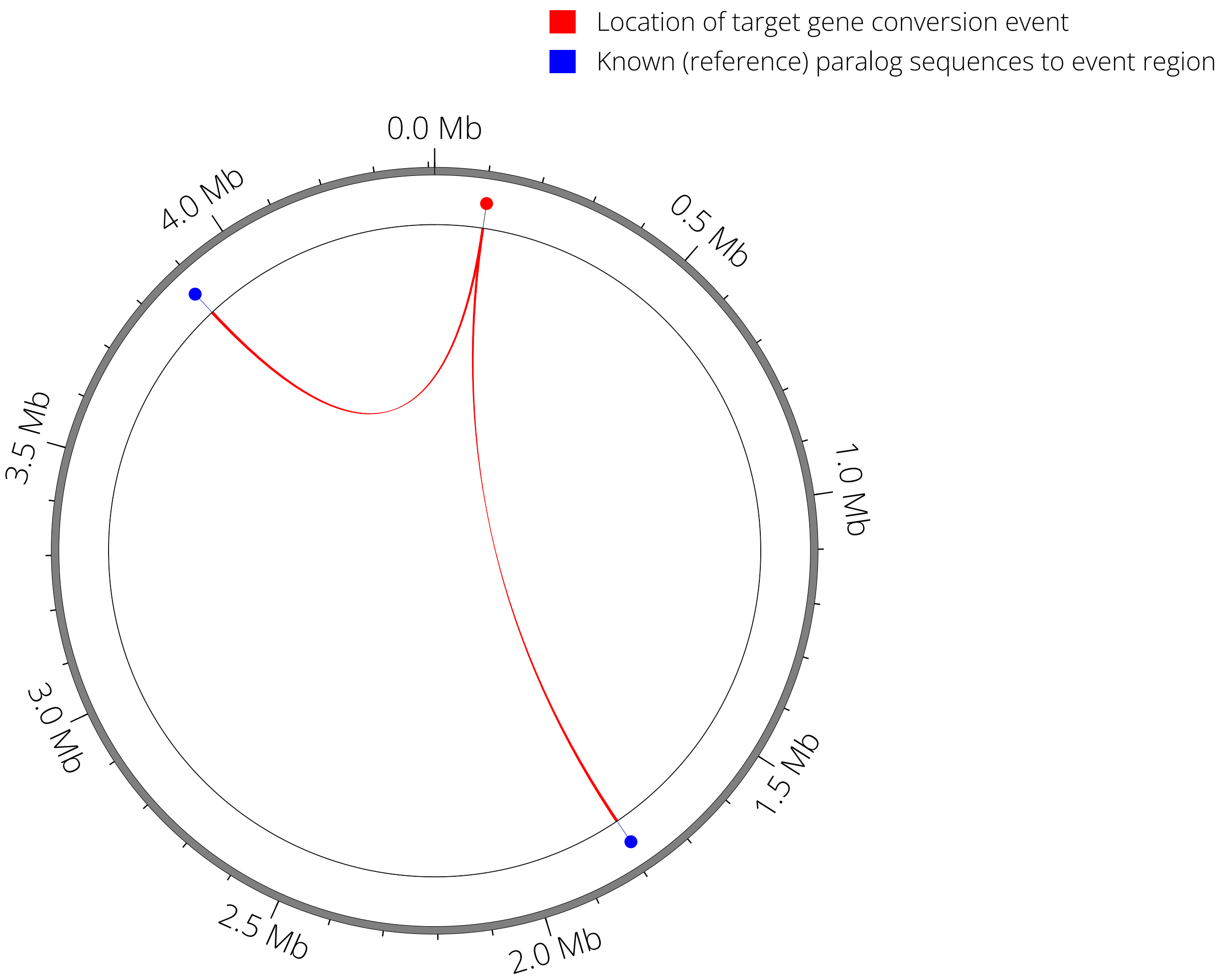

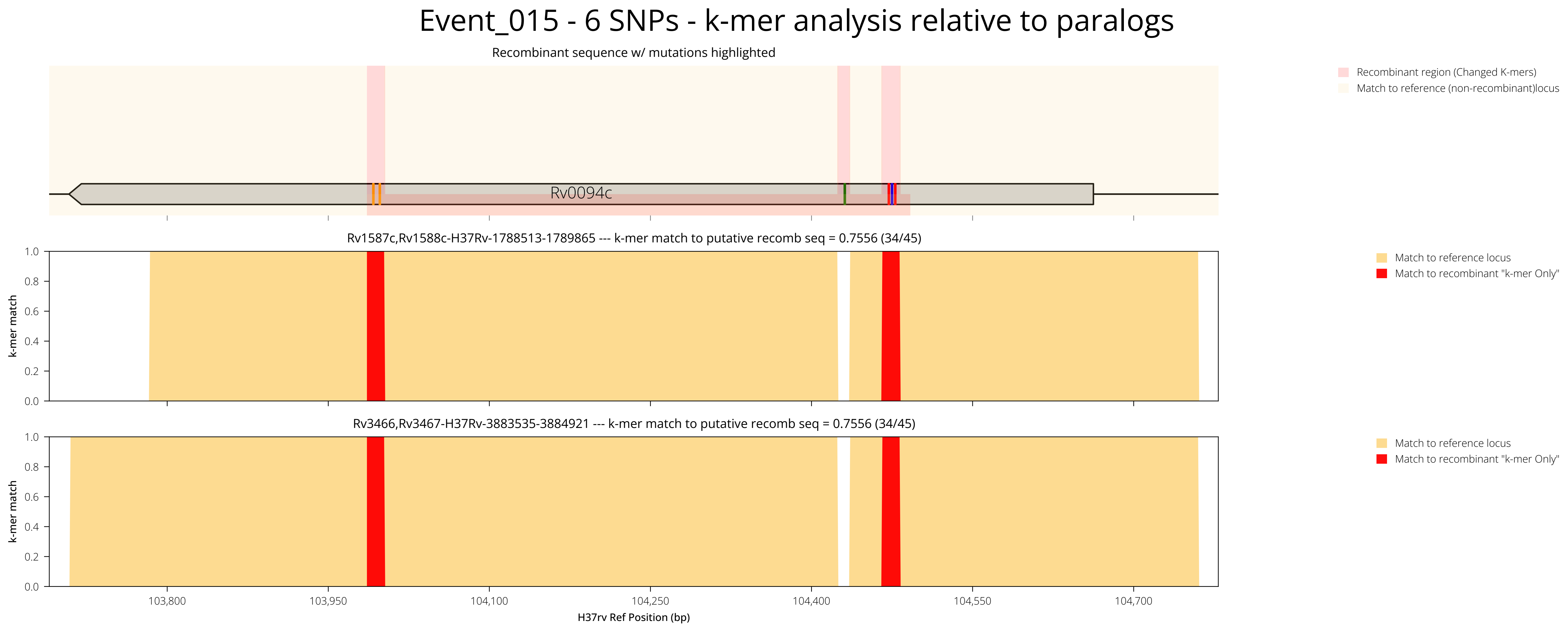

Event_015 - 6 SNPs - k-mer analysis relative to paralogs
Recombinant sequence w/ mutations highlighted
Recombinant region (Changed K-mers)
Match to reference (non-recombinant)locus
Rv0094c
Rv1587c,Rv1588c-H37Rv-1788513-1789865 --- k-mer match to putative recomb seq = 0.7556 (34/45)
Match to reference locus
Match to recombinant "k-mer Only"
k-mer match
1.0
0.8
0.6
0.4
0.2
0.0
Rv3466,Rv3467-H37Rv-3883535-3884921 --- k-mer match to putative recomb seq = 0.7556 (34/45)
Match to reference locus
Match to recombinant "k-mer Only"
k-mer match
103,800
103,950
104,100
104,250
104,400
104,550
104,700
H37rv Ref Position (bp)

Sub-phylogeny Viz + Paralog Alignments
GCE ID: Event_015

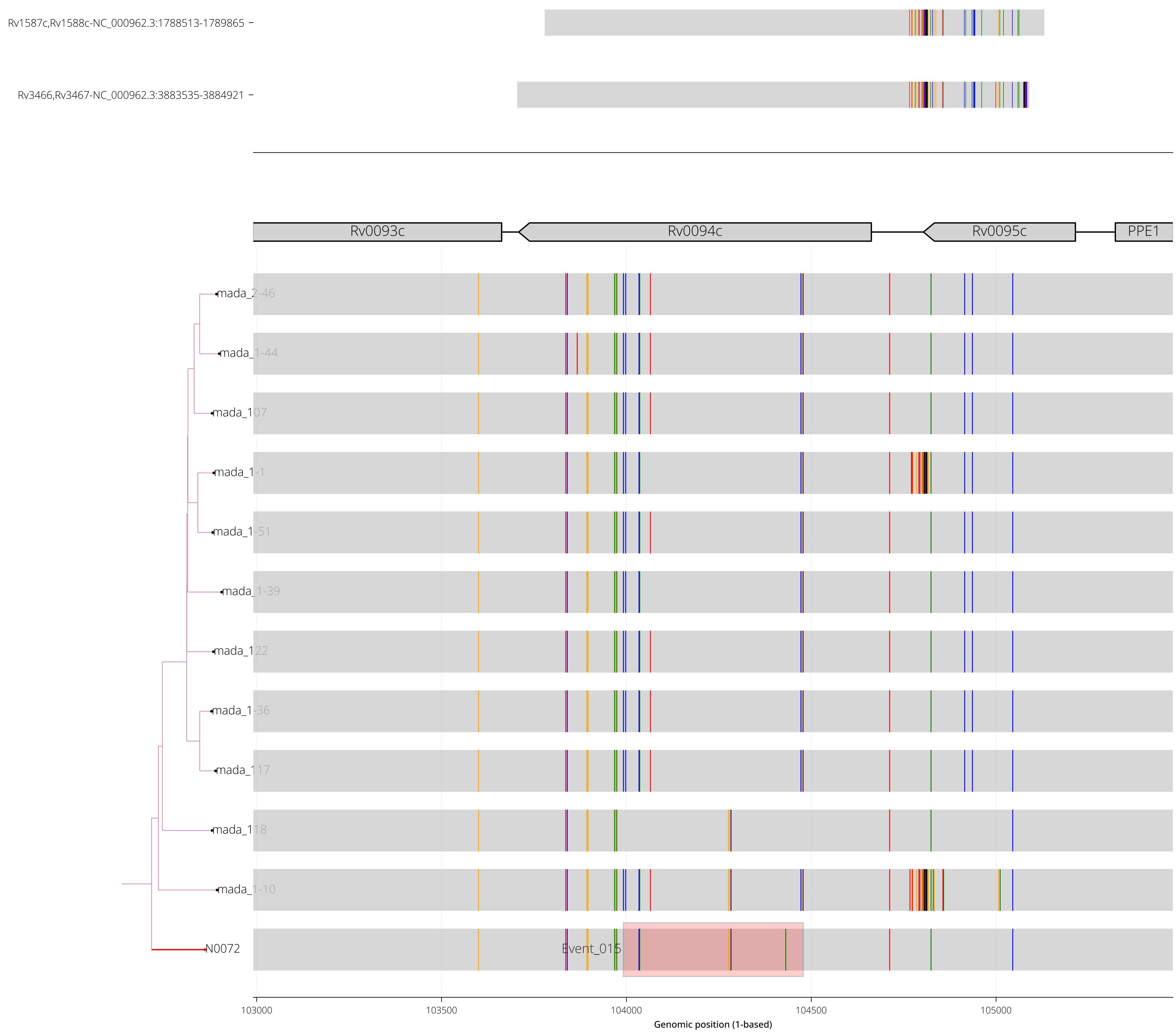

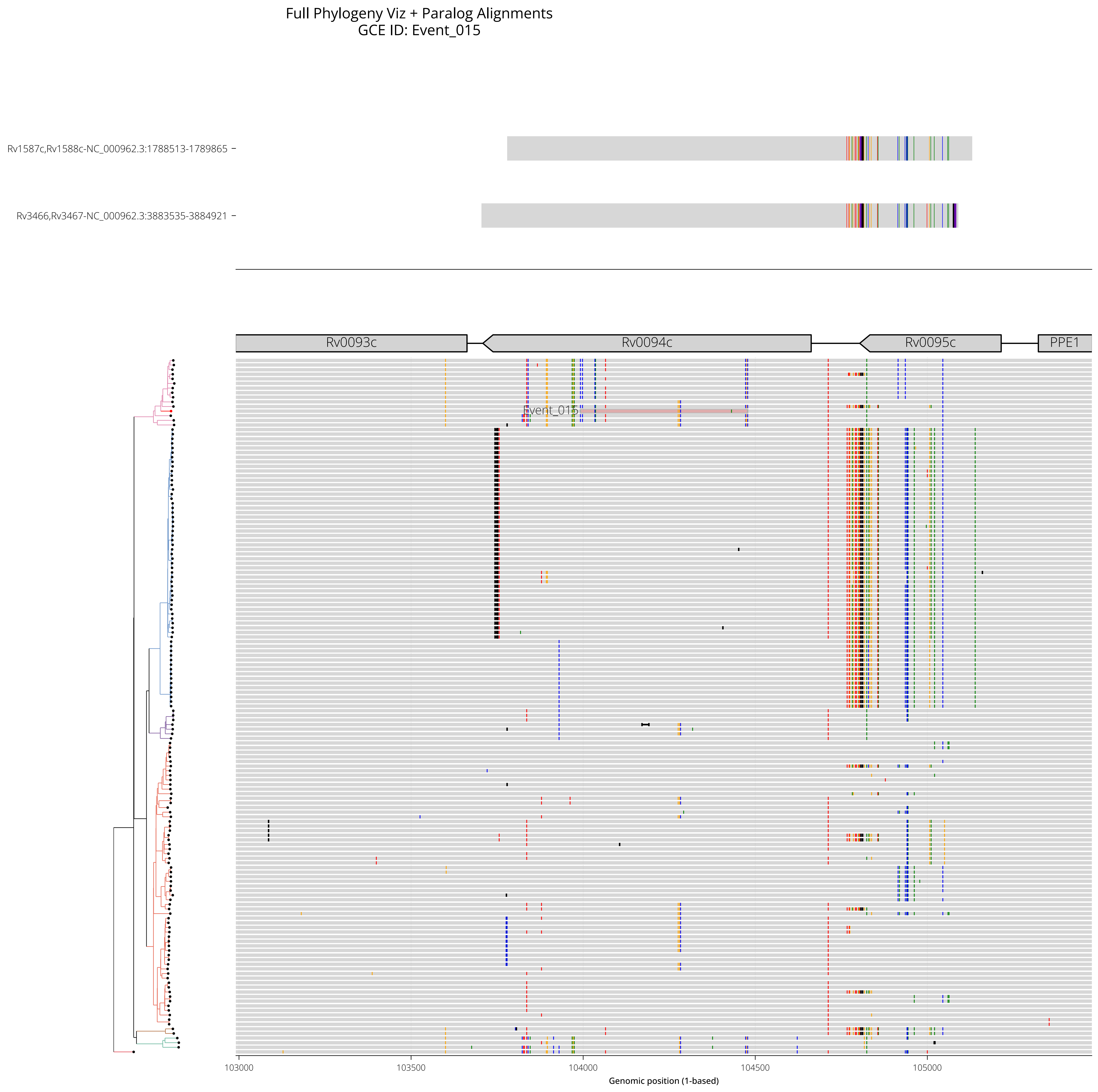

Full Phylogeny Viz + Paralog Alignments
GCE ID: Event_015
Rv1587c,Rv1588c-NC_000962.3:1788513-1789865
Rv3466,Rv3467-NC_000962.3:3883535-3884921
Rv0093c
Rv0094c
Rv0095c
PPE1
Event_015
103000
103500
104000
104500
105000
Genomic position (1-based)

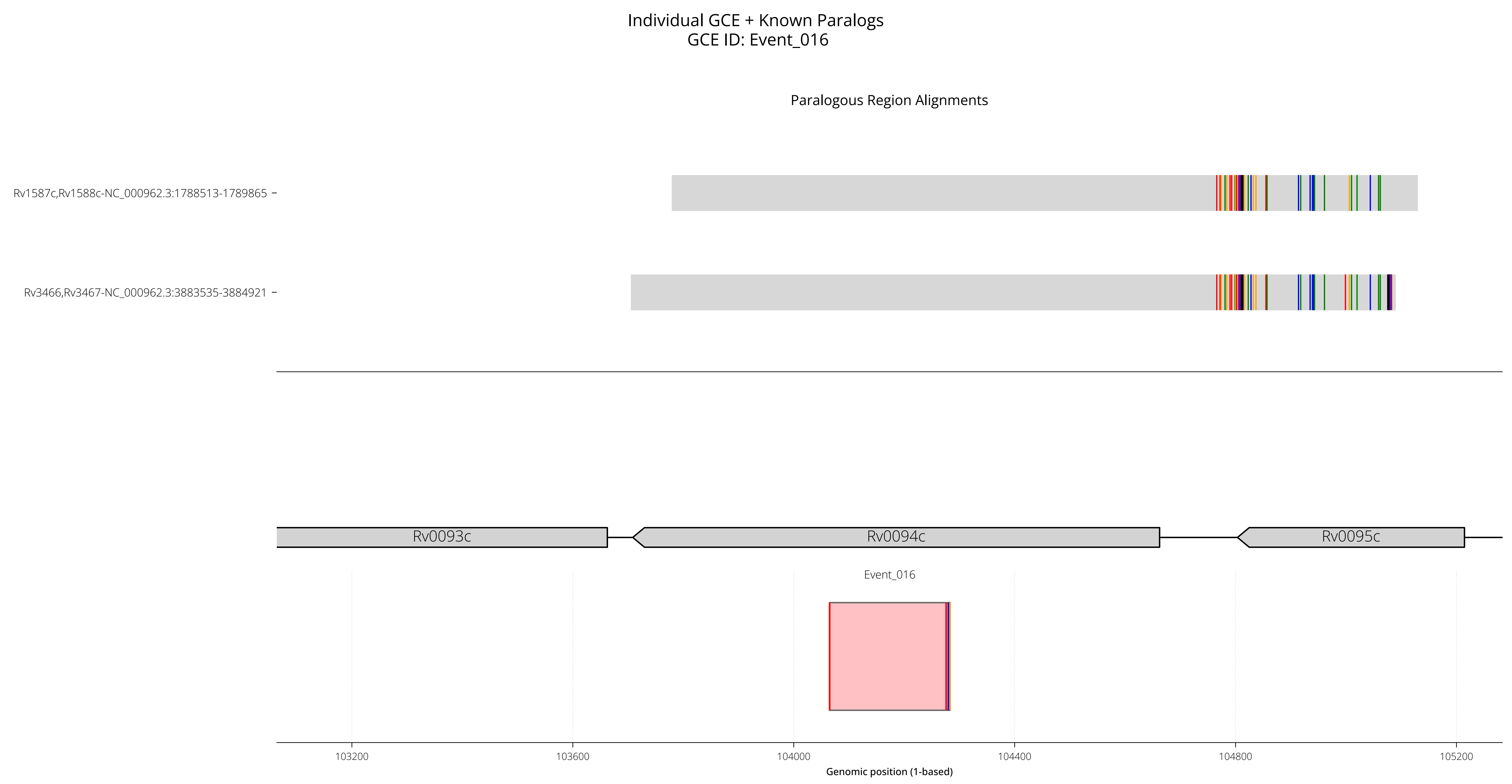
Individual GCE + Known Paralogs
GCE ID: Event_016
Paralogous Region Alignments
Rv1587c,Rv1588c-NC_000962.3:1788513-1789865
Rv3466,Rv3467-NC_000962.3:3883535-3884921
Rv0093c
Rv0094c
Rv0095c
Event_016
103200
103600
104000
104400
104800
105200
Genomic position (1-based)

Circos Viz - GCE to Known Paralogs Locations
GCE ID: Event_016

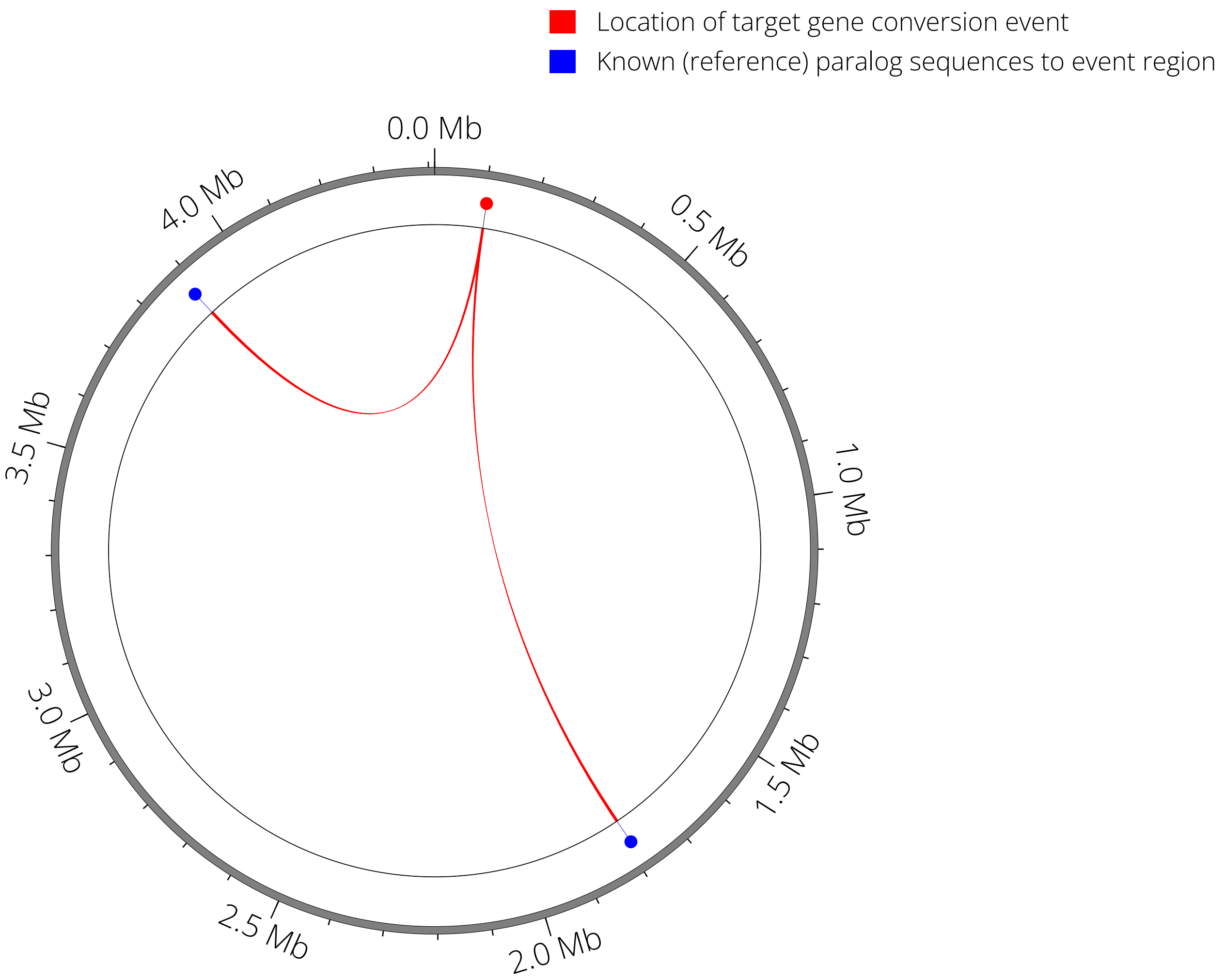

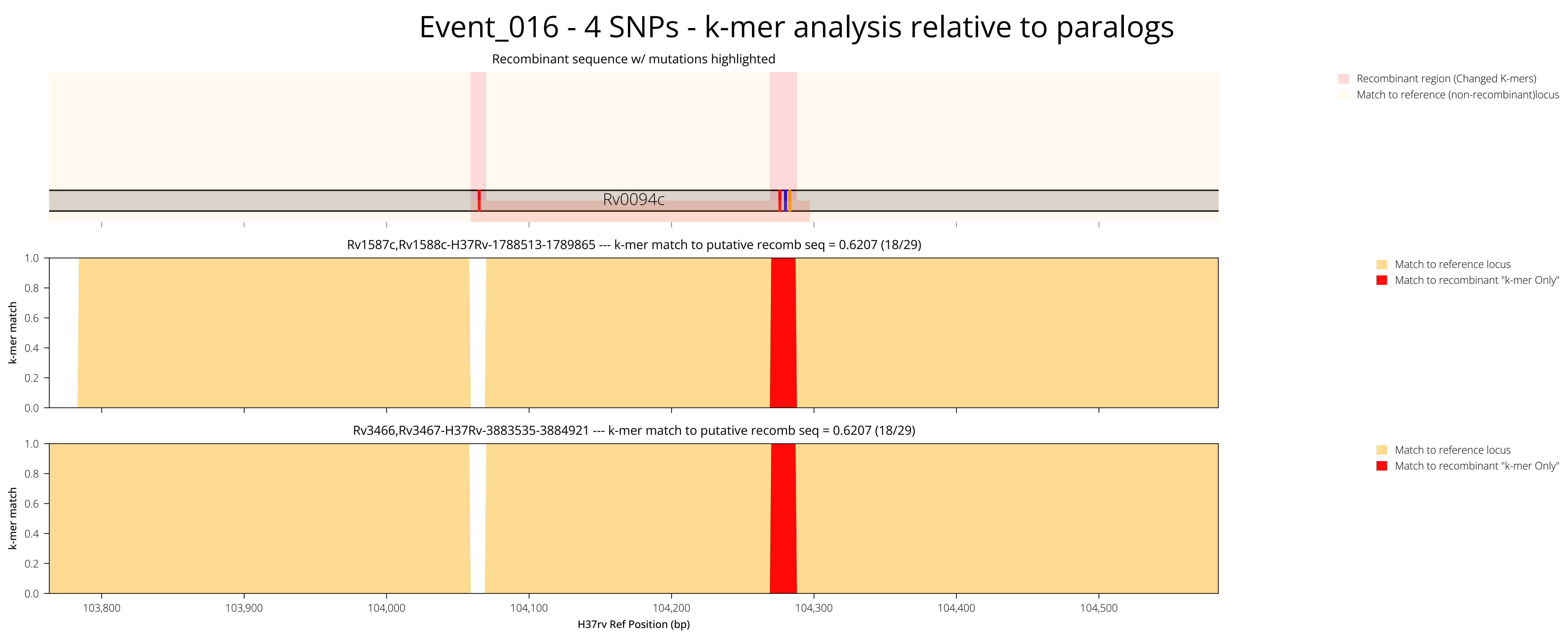

Event_016 - 4 SNPs - k-mer analysis relative to paralogs
Recombinant sequence w/ mutations highlighted
Rv0094c
Recombinant region (Changed K-mers)
Match to reference (non-recombinant)locus
Rv1587c,Rv1588c-H37Rv-1788513-1789865 --- k-mer match to putative recomb seq = 0.6207 (18/29)
Match to reference locus
Match to recombinant "k-mer Only"
Rv3466,Rv3467-H37Rv-3883535-3884921 --- k-mer match to putative recomb seq = 0.6207 (18/29)
Match to reference locus
Match to recombinant "k-mer Only"
k-mer match
1.0
0.8
0.6
0.4
0.2
0.0
103,800
103,900
104,000
104,100
104,200
104,300
104,400
104,500
H37rv Ref Position (bp)

Sub-phylogeny Viz + Paralog Alignments
GCE ID: Event_016

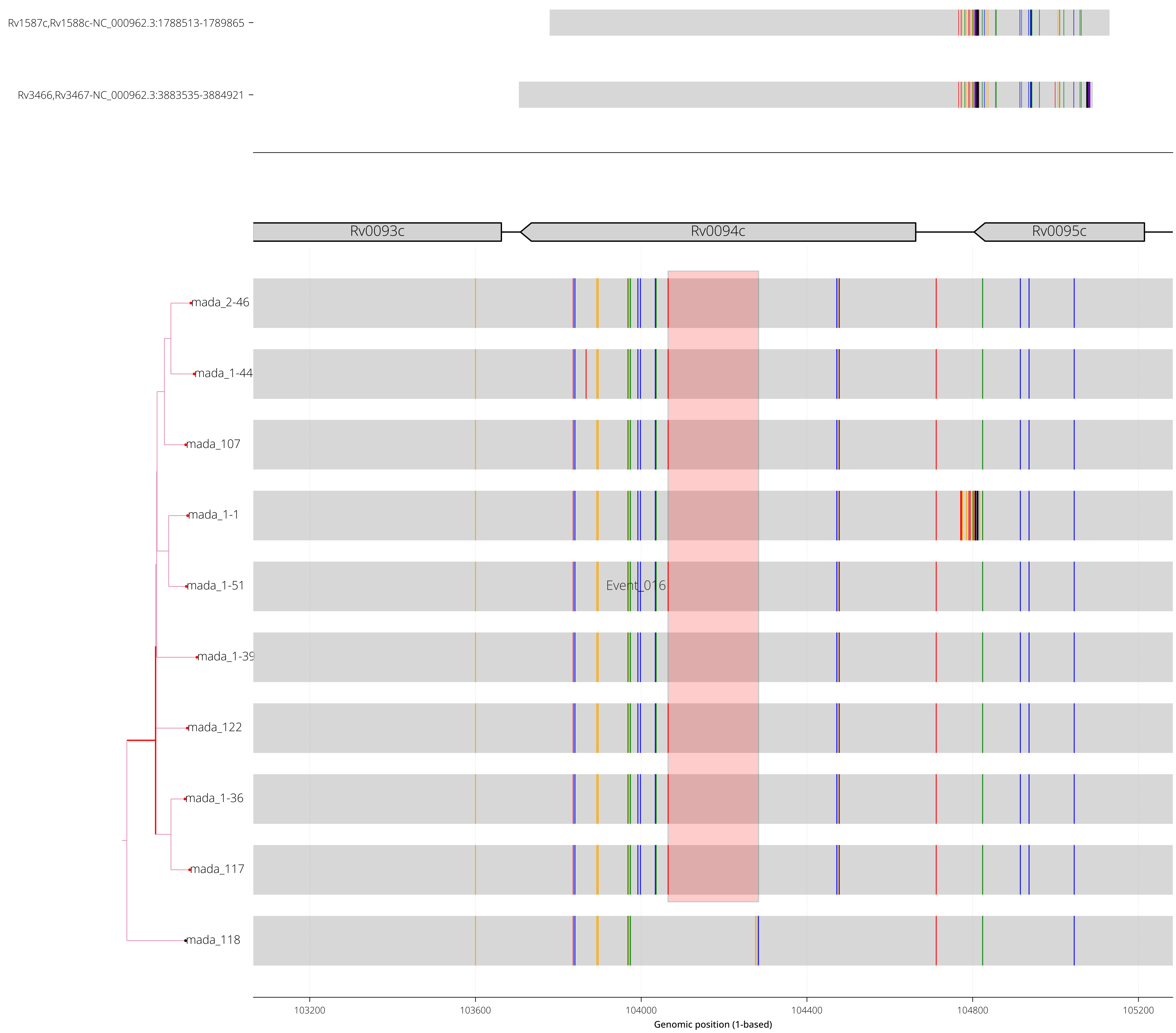

### Full Phylogeny Viz + Paralog Alignments
GCE ID: Event_016

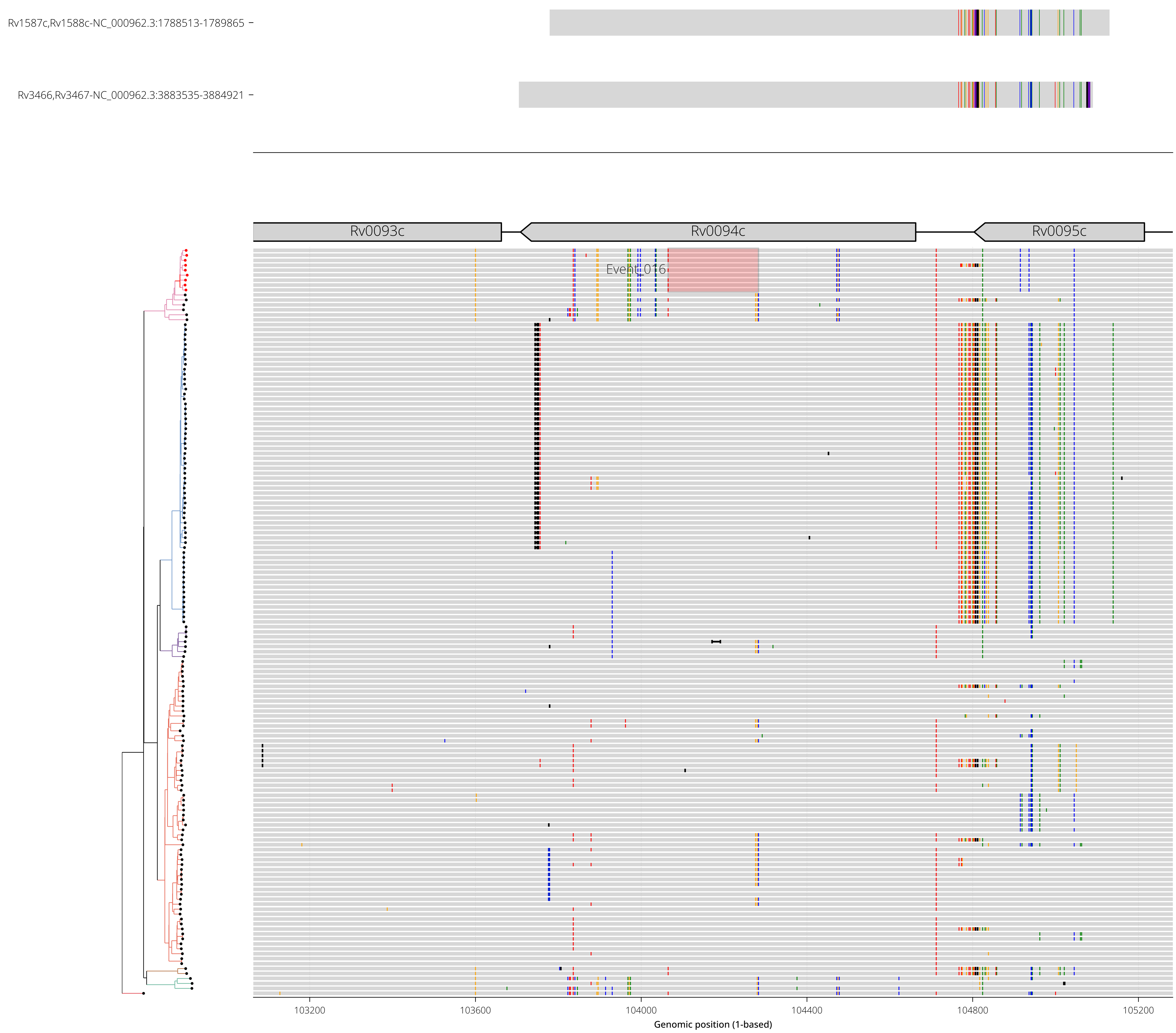

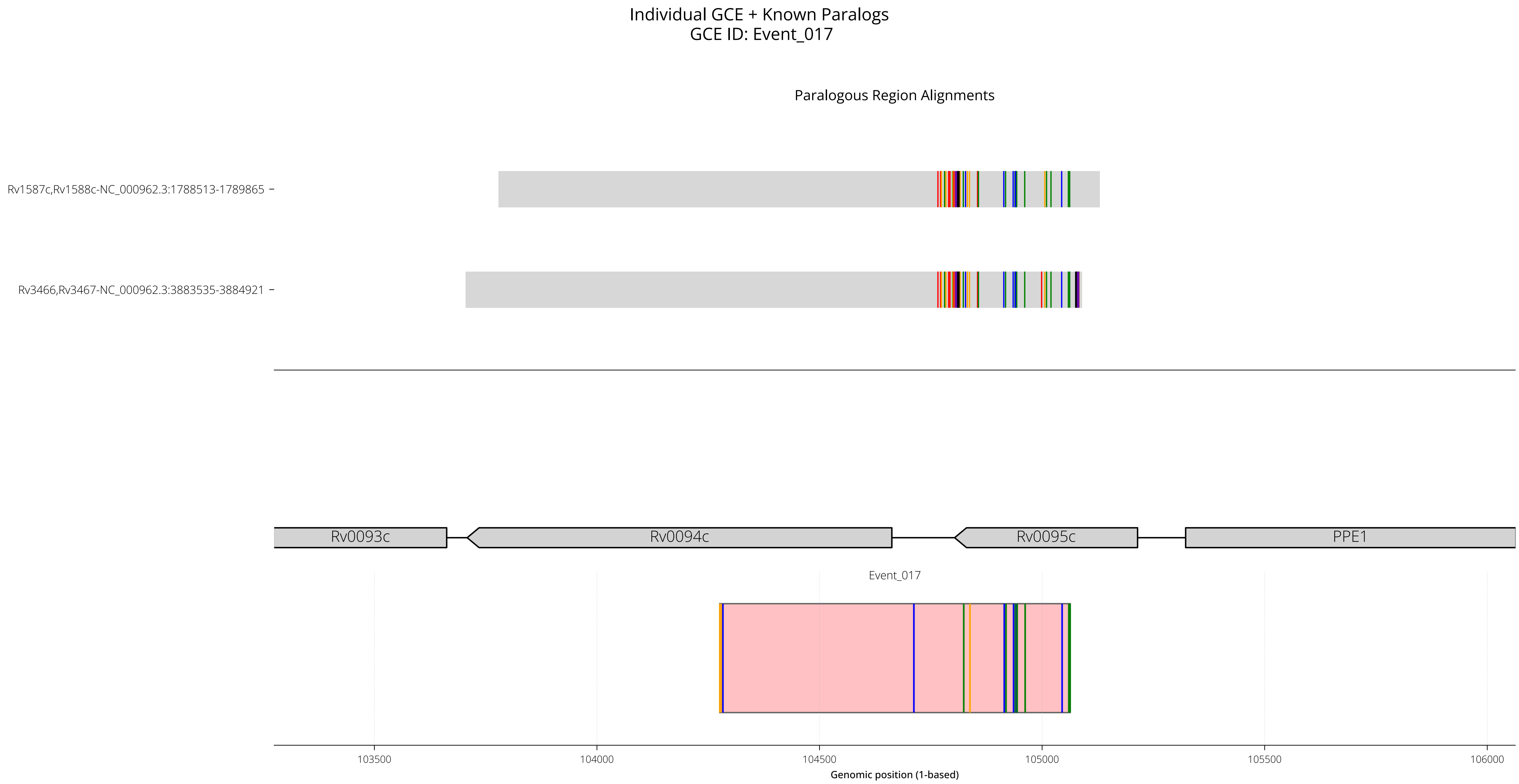

Individual GCE + Known Paralogs
GCE ID: Event_017
Paralogous Region Alignments
Rv1587c,Rv1588c-NC_000962.3:1788513-1789865
Rv3466,Rv3467-NC_000962.3:3883535-3884921
Rv0093c
Rv0094c
Rv0095c
PPE1
Event_017
103500
104000
104500
105000
105500
106000
Genomic position (1-based)

Circos Viz - GCE to Known Paralogs Locations
GCE ID: Event_017

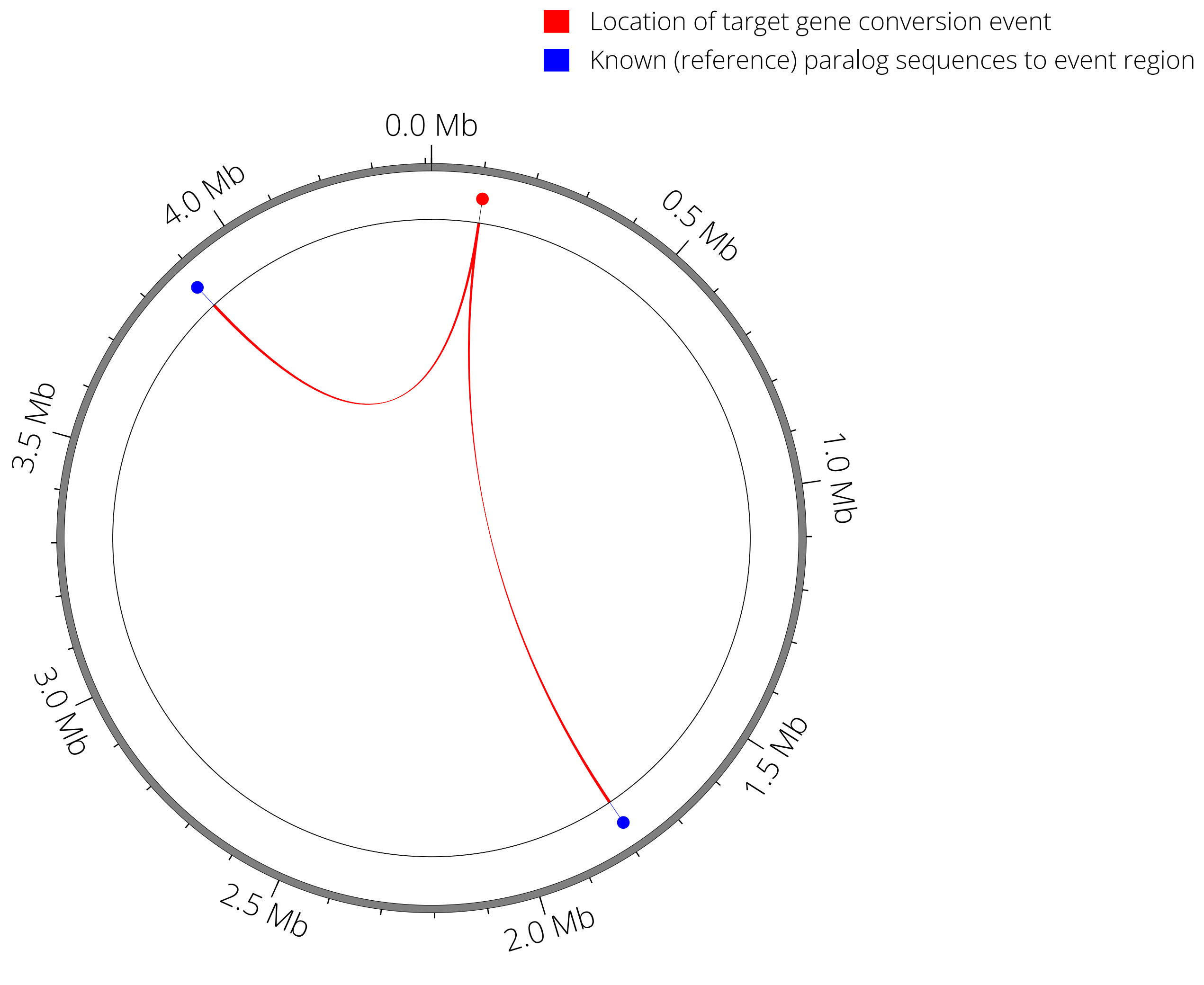

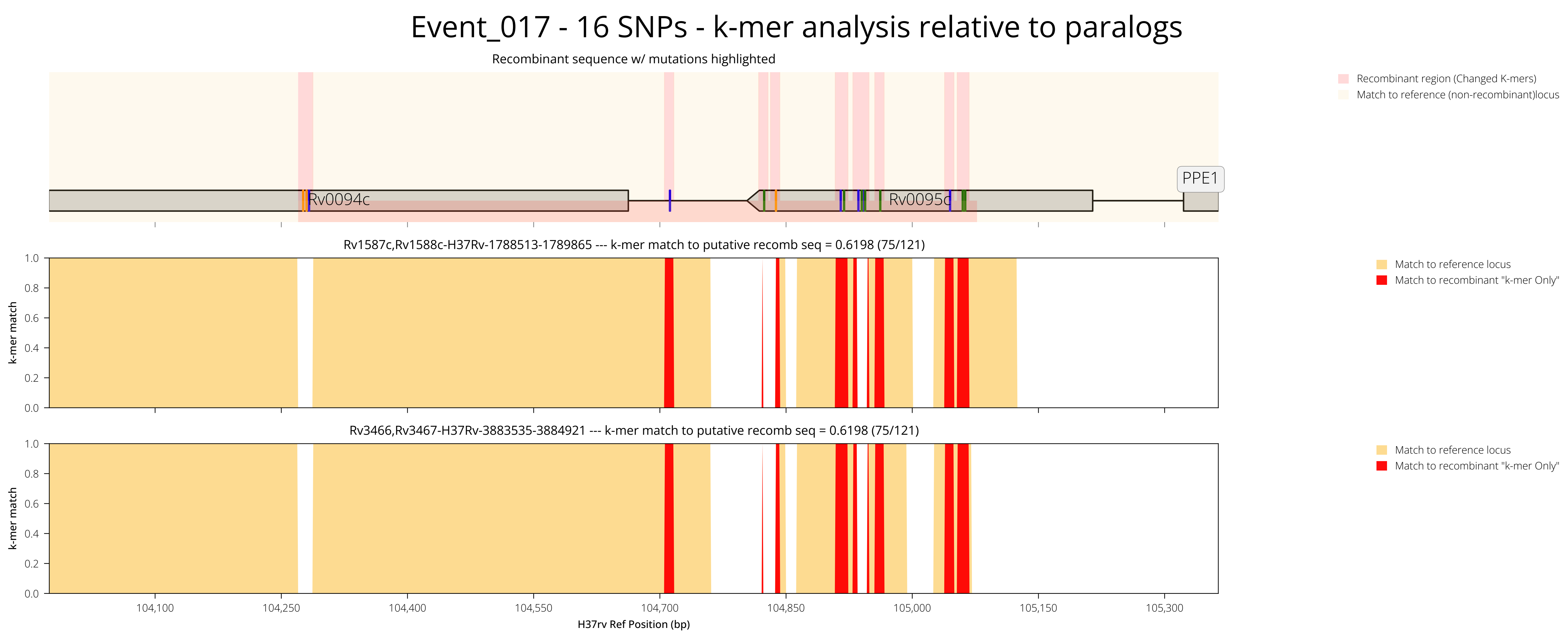

Event_017 - 16 SNPs - k-mer analysis relative to paralogs
Recombinant sequence w/ mutations highlighted
Rv0094c
Rv0095c
PPE1
Recombinant region (Changed K-mers)
Match to reference (non-recombinant)locus
Rv1587c,Rv1588c-H37Rv-1788513-1789865 --- k-mer match to putative recomb seq = 0.6198 (75/121)
Rv3466,Rv3467-H37Rv-3883535-3884921 --- k-mer match to putative recomb seq = 0.6198 (75/121)
k-mer match
Match to reference locus
Match to recombinant "k-mer Only"
1.0
0.8
0.6
0.4
0.2
0.0
104,100
104,250
104,400
104,550
104,700
104,850
105,000
105,150
105,300
H37rv Ref Position (bp)

Sub-phylogeny Viz + Paralog Alignments
GCE ID: Event_017

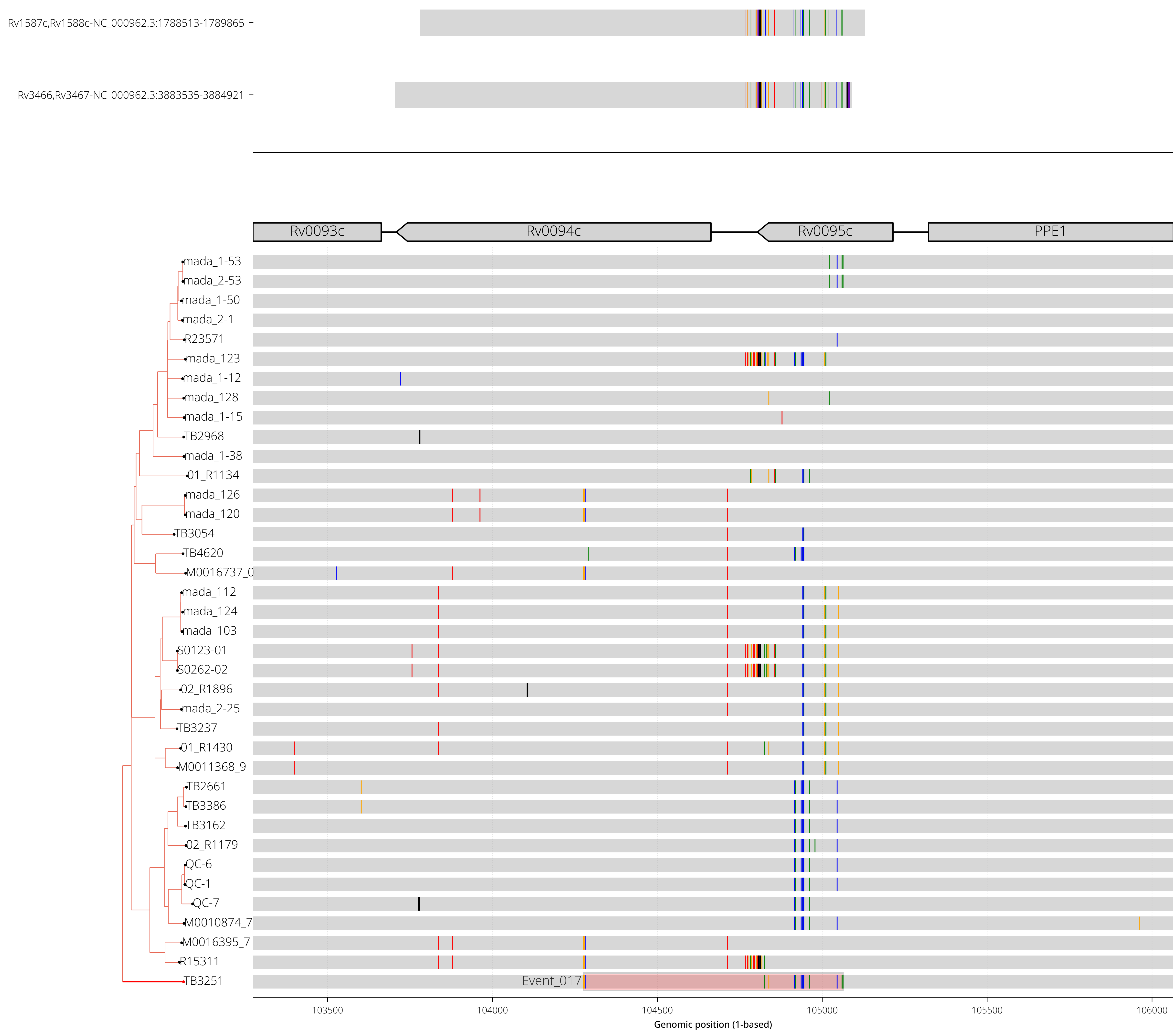

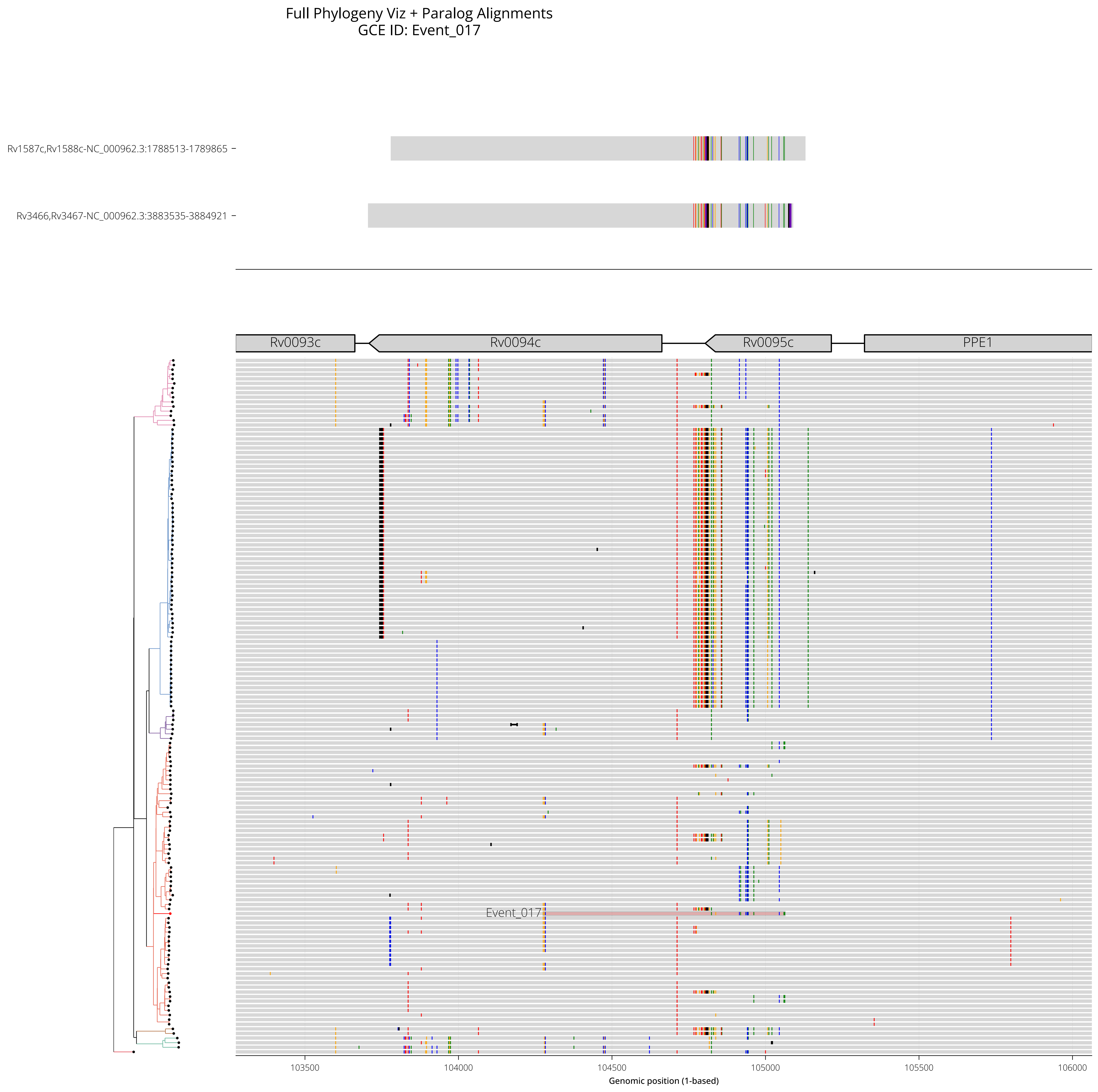

Full Phylogeny Viz + Paralog Alignments
GCE ID: Event_017
Rv1587c,Rv1588c-NC_000962.3:1788513-1789865
Rv3466,Rv3467-NC_000962.3:3883535-3884921
Rv0093c
Rv0094c
Rv0095c
PPE1
Event_017
103500
104000
104500
105000
105500
106000
Genomic position (1-based)

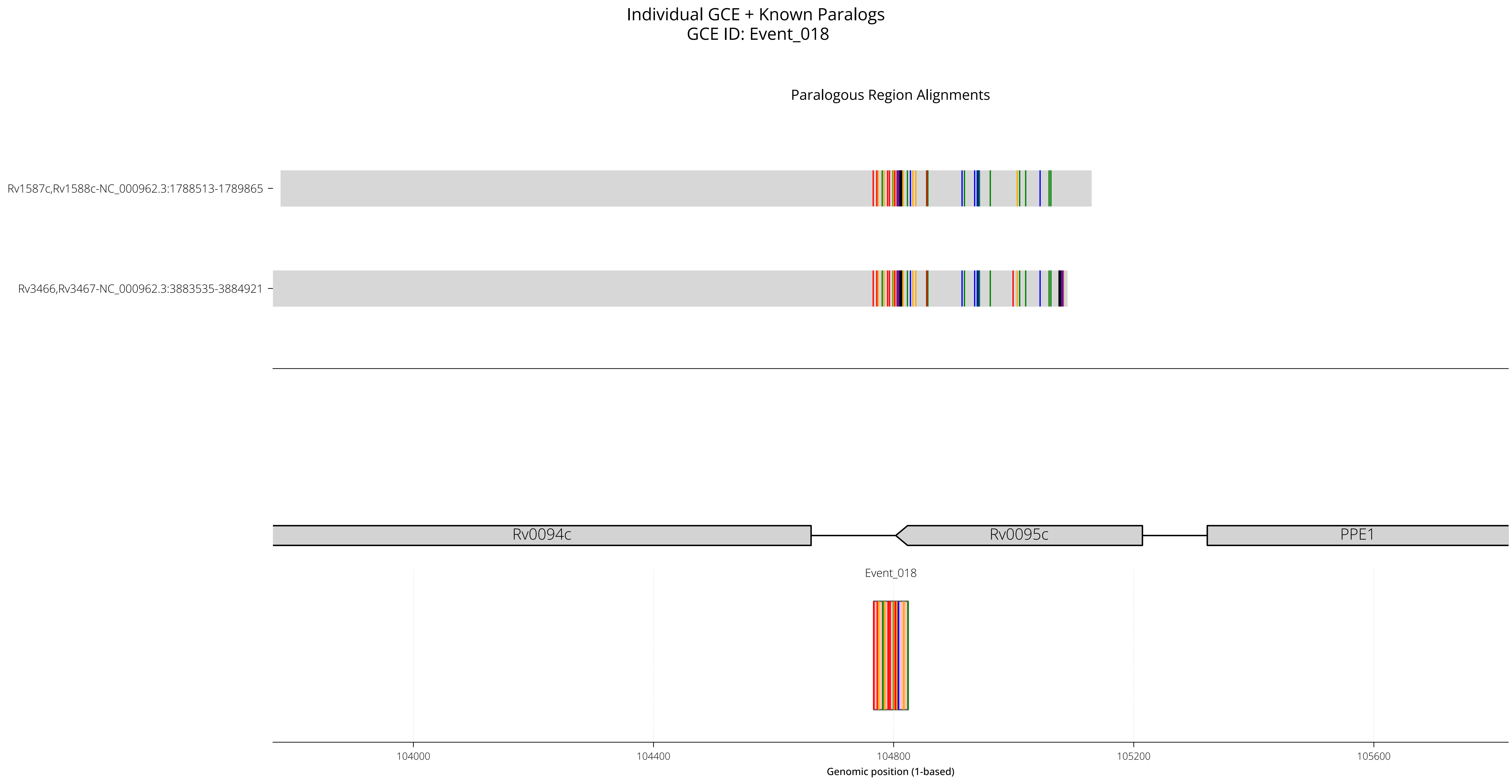
Individual GCE + Known Paralogs
GCE ID: Event_018
Paralogous Region Alignments
Rv1587c,Rv1588c-NC_000962.3:1788513-1789865
Rv3466,Rv3467-NC_000962.3:3883535-3884921
Rv0094c
Rv0095c
PPE1
Event_018
104000
104400
104800
105200
105600
Genomic position (1-based)

Circos Viz - GCE to Known Paralogs Locations
GCE ID: Event_018

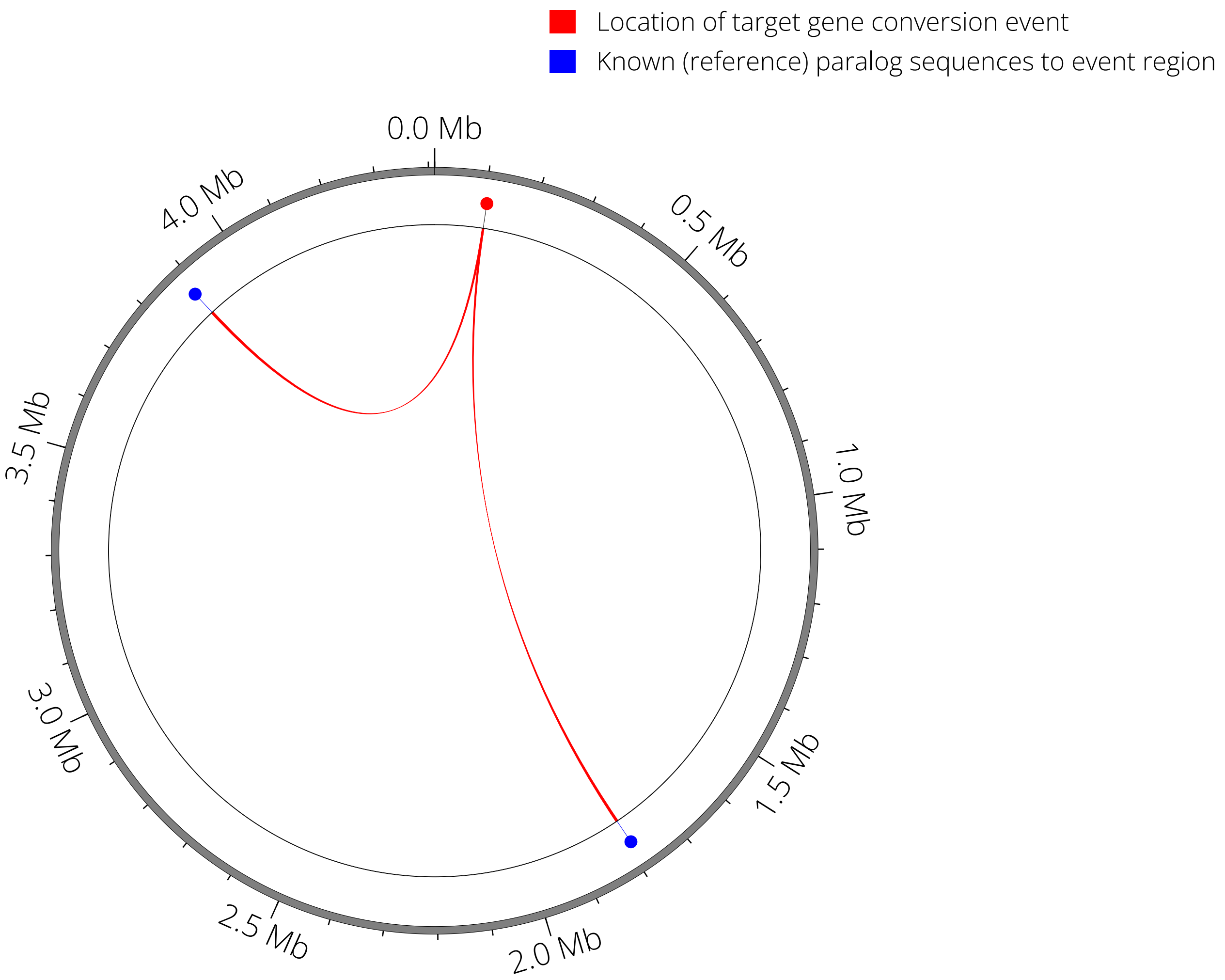

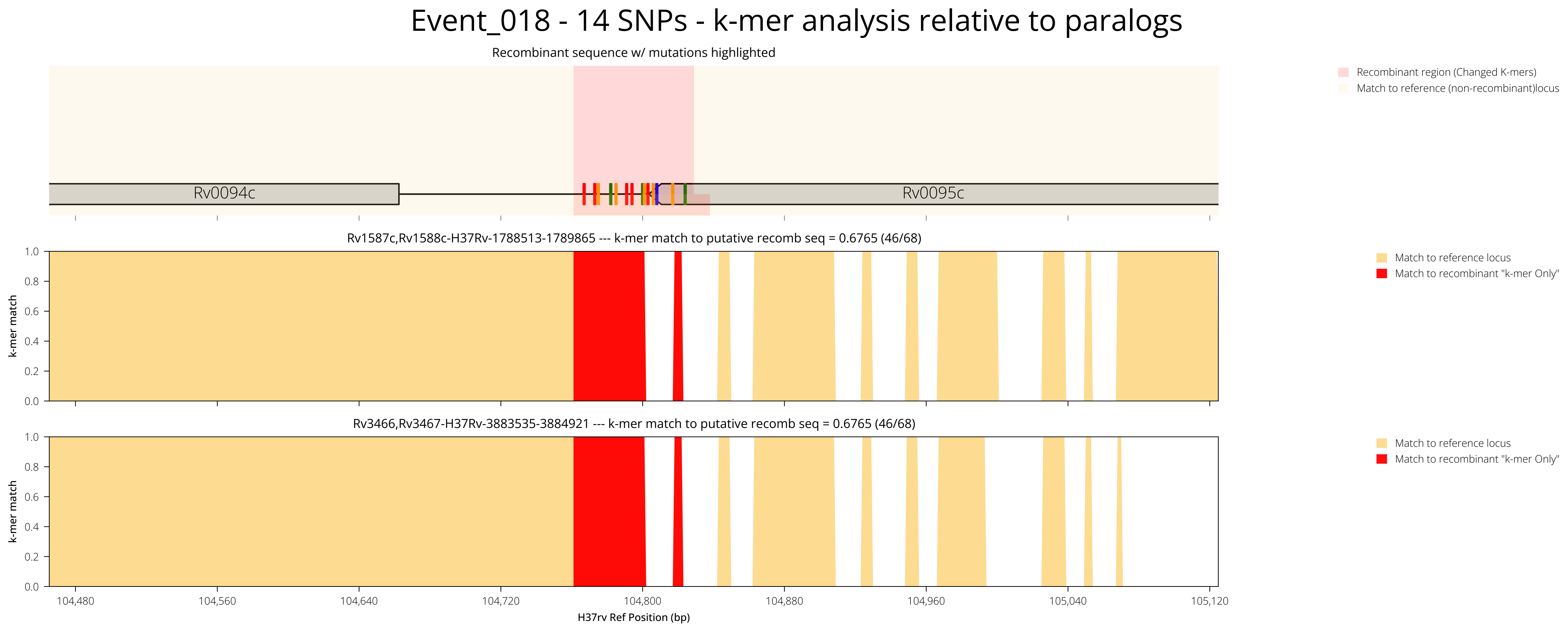

Event_018 - 14 SNPs - k-mer analysis relative to paralogs
Recombinant sequence w/ mutations highlighted
Rv0094c
Rv0095c
Recombinant region (Changed K-mers)
Match to reference (non-recombinant)locus
Rv1587c,Rv1588c-H37Rv-1788513-1789865 --- k-mer match to putative recomb seq = 0.6765 (46/68)
Match to reference locus
Match to recombinant "k-mer Only"
Rv3466,Rv3467-H37Rv-3883535-3884921 --- k-mer match to putative recomb seq = 0.6765 (46/68)
Match to reference locus
Match to recombinant "k-mer Only"
k-mer match
1.0
0.8
0.6
0.4
0.2
0.0
104,480
104,560
104,640
104,720
104,800
104,880
104,960
105,040
105,120
H37rv Ref Position (bp)

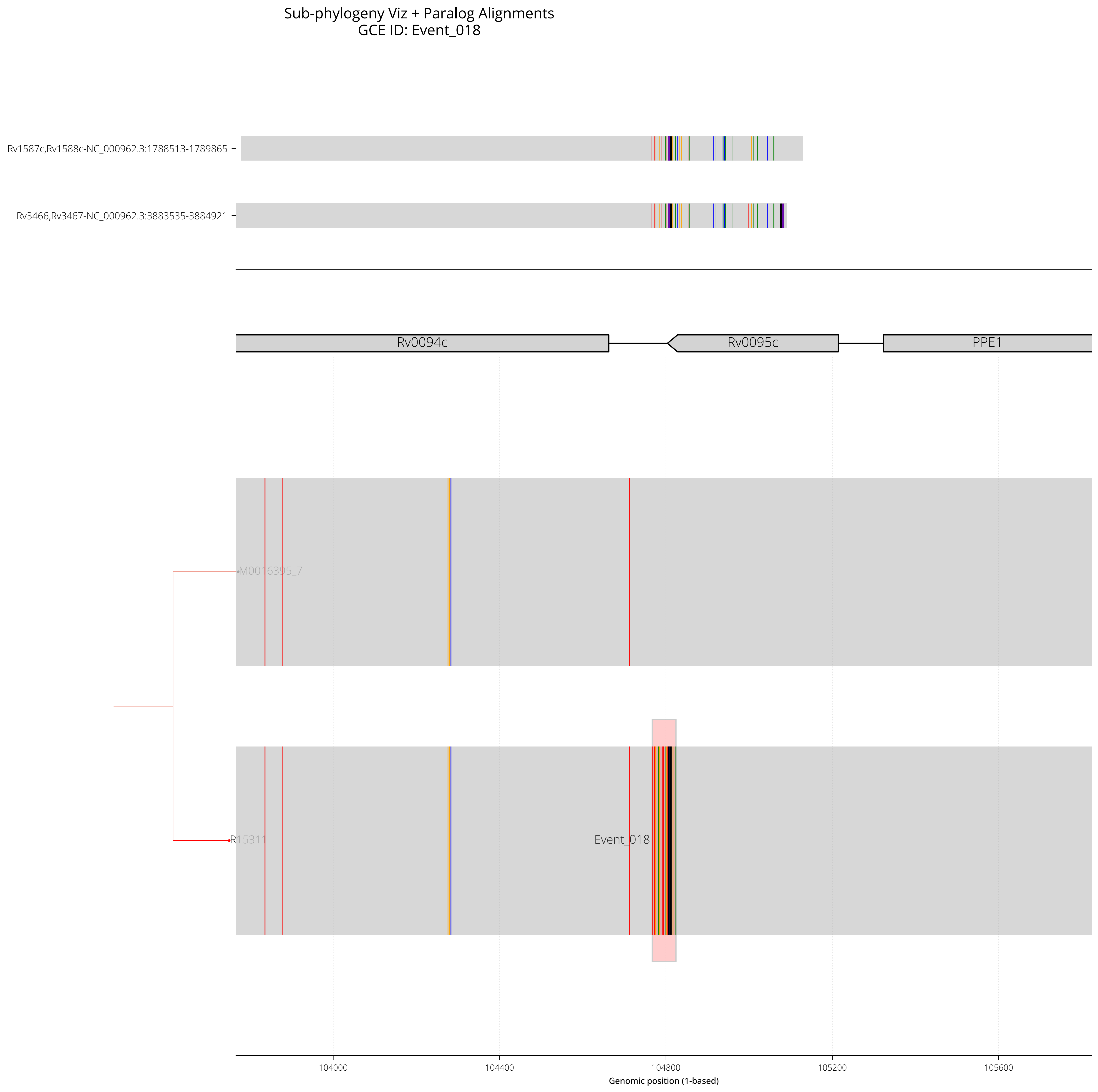
Sub-phylogeny Viz + Paralog Alignments
GCE ID: Event_018
Rv1587c,Rv1588c-NC_000962.3:1788513-1789865
Rv3466,Rv3467-NC_000962.3:3883535-3884921
Rv0094c
Rv0095c
PPE1
M0016395_7
R15311
Event_018
104000
104400
104800
105200
105600
Genomic position (1-based)

### Full Phylogeny Viz + Paralog Alignments
GCE ID: Event_018

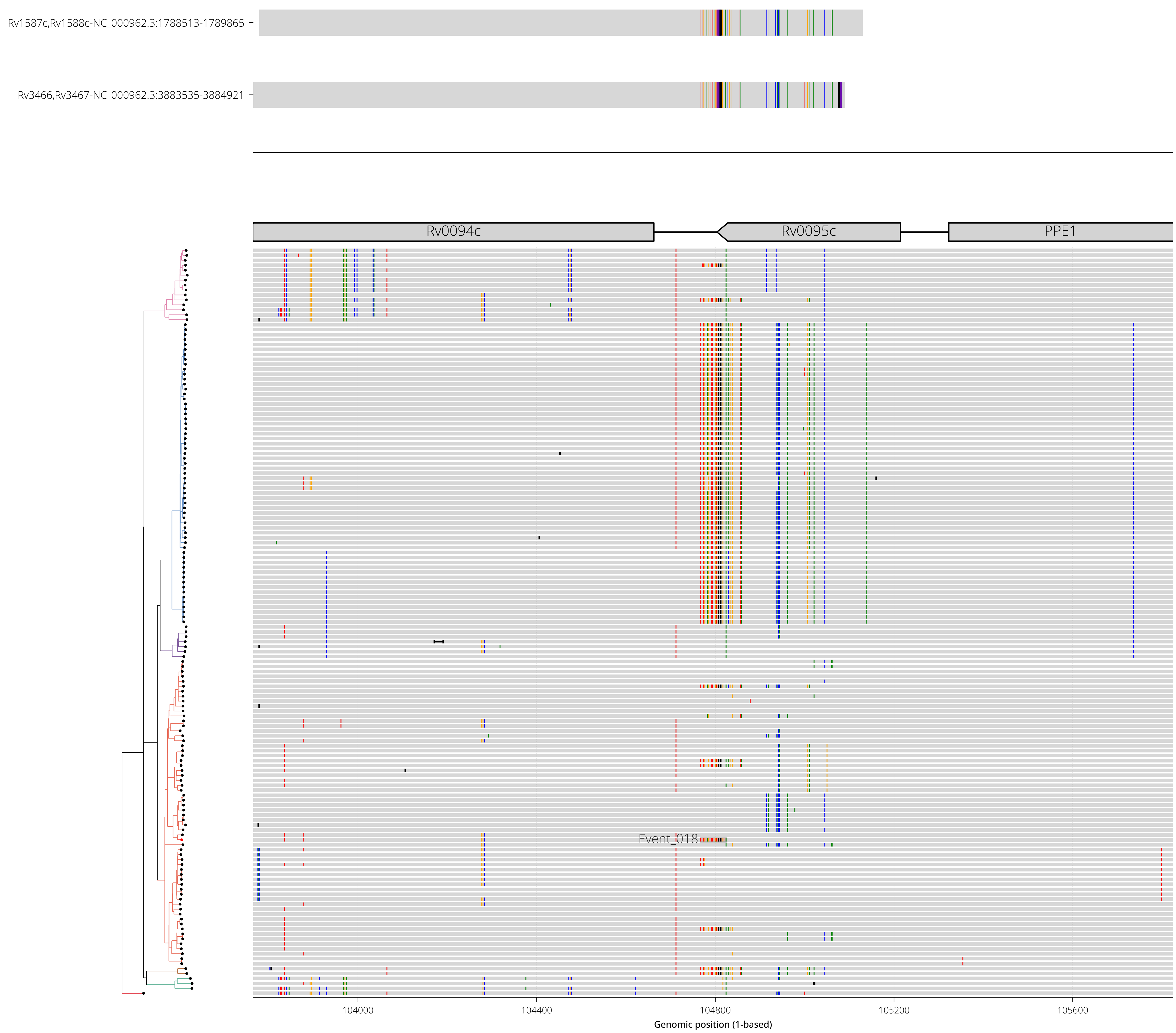

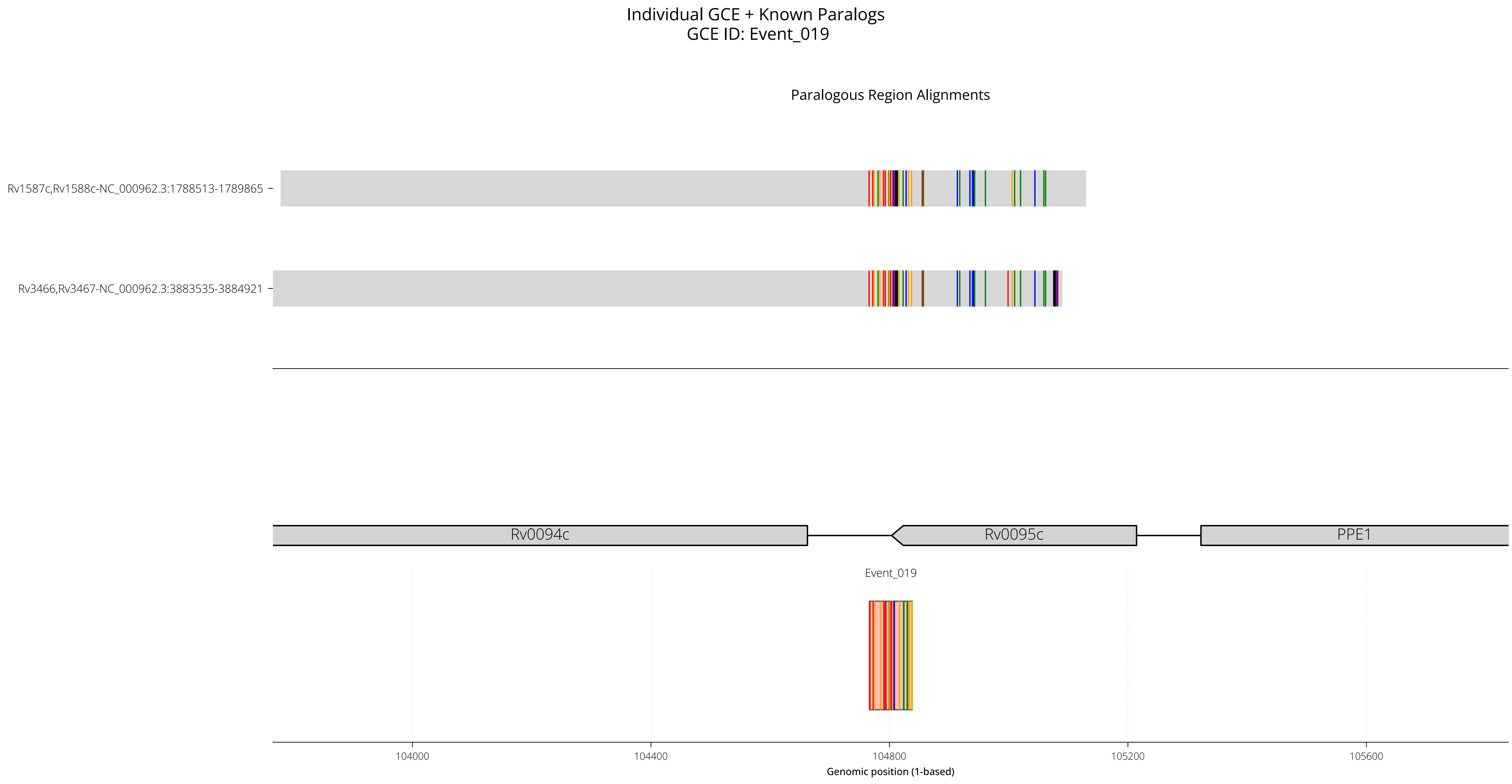
Individual GCE + Known Paralogs
GCE ID: Event_019
Paralogous Region Alignments
Rv1587c,Rv1588c-NC_000962.3:1788513-1789865
Rv3466,Rv3467-NC_000962.3:3883535-3884921
Rv0094c
Rv0095c
PPE1
Event_019
104000
104400
104800
105200
105600
Genomic position (1-based)

Circos Viz - GCE to Known Paralogs Locations
GCE ID: Event_019

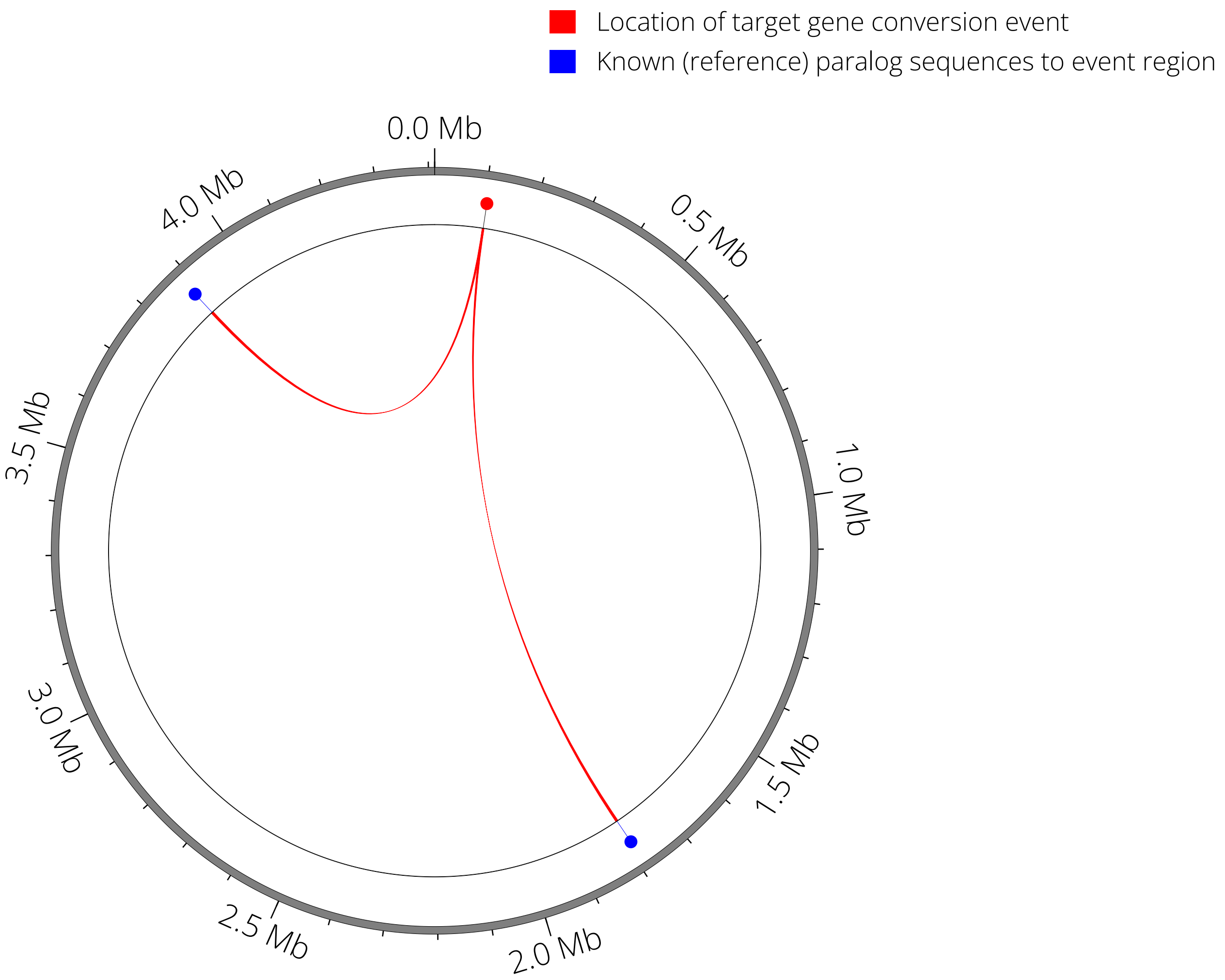

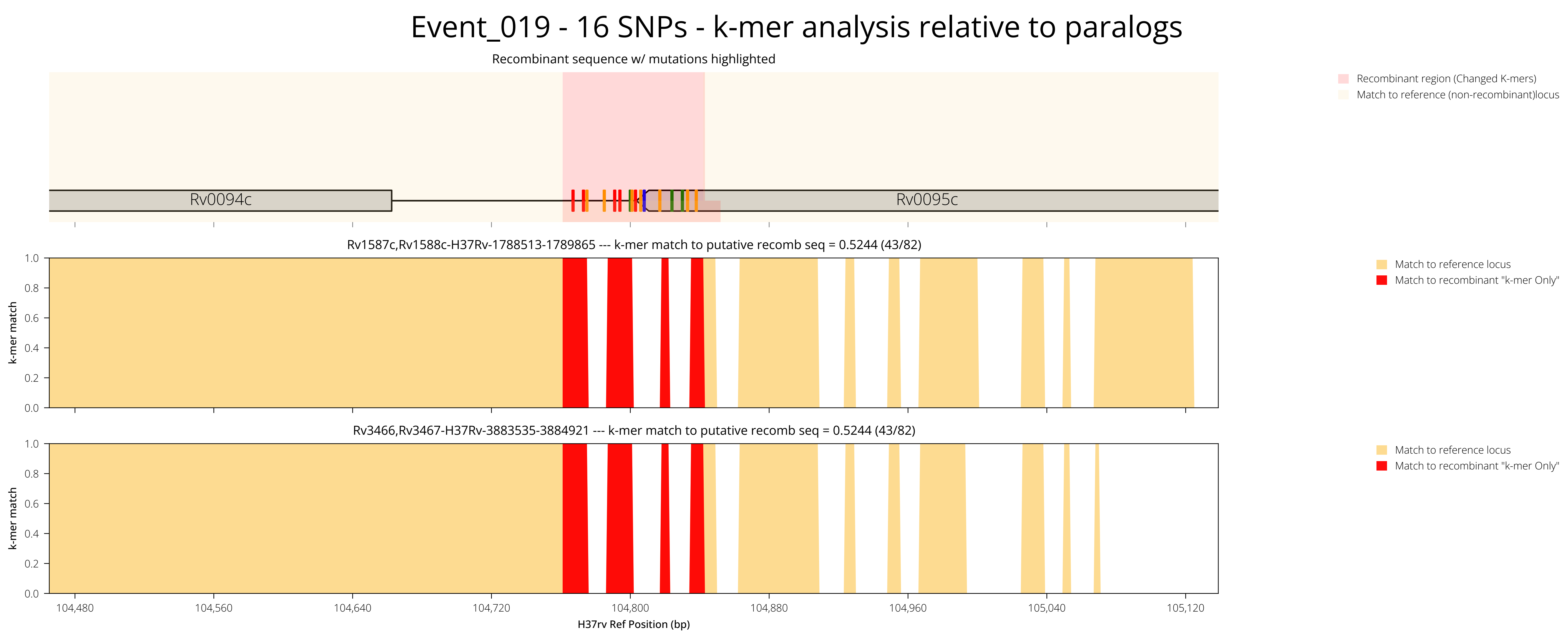

Event_019 - 16 SNPs - k-mer analysis relative to paralogs
Recombinant sequence w/ mutations highlighted
Recombinant region (Changed K-mers)
Match to reference (non-recombinant)locus
Rv0094c
Rv0095c
Rv1587c,Rv1588c-H37Rv-1788513-1789865 --- k-mer match to putative recomb seq = 0.5244 (43/82)
Match to reference locus
Match to recombinant "k-mer Only"
k-mer match
Rv3466,Rv3467-H37Rv-3883535-3884921 --- k-mer match to putative recomb seq = 0.5244 (43/82)
Match to reference locus
Match to recombinant "k-mer Only"
k-mer match
104,480
104,560
104,640
104,720
104,800
104,880
104,960
105,040
105,120
H37rv Ref Position (bp)

Sub-phylogeny Viz + Paralog Alignments
GCE ID: Event_019

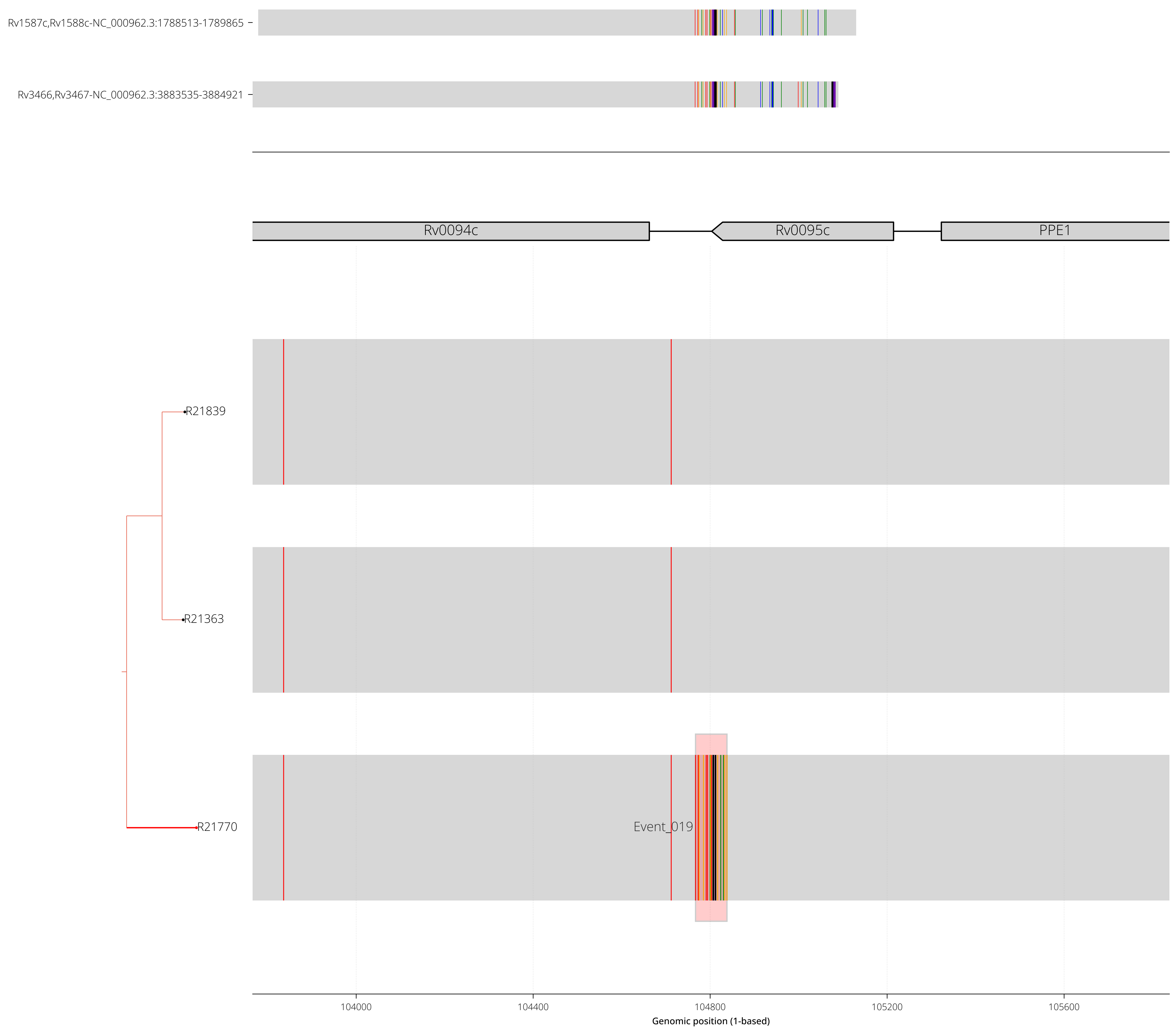

Full Phylogeny Viz + Paralog Alignments
GCE ID: Event_019

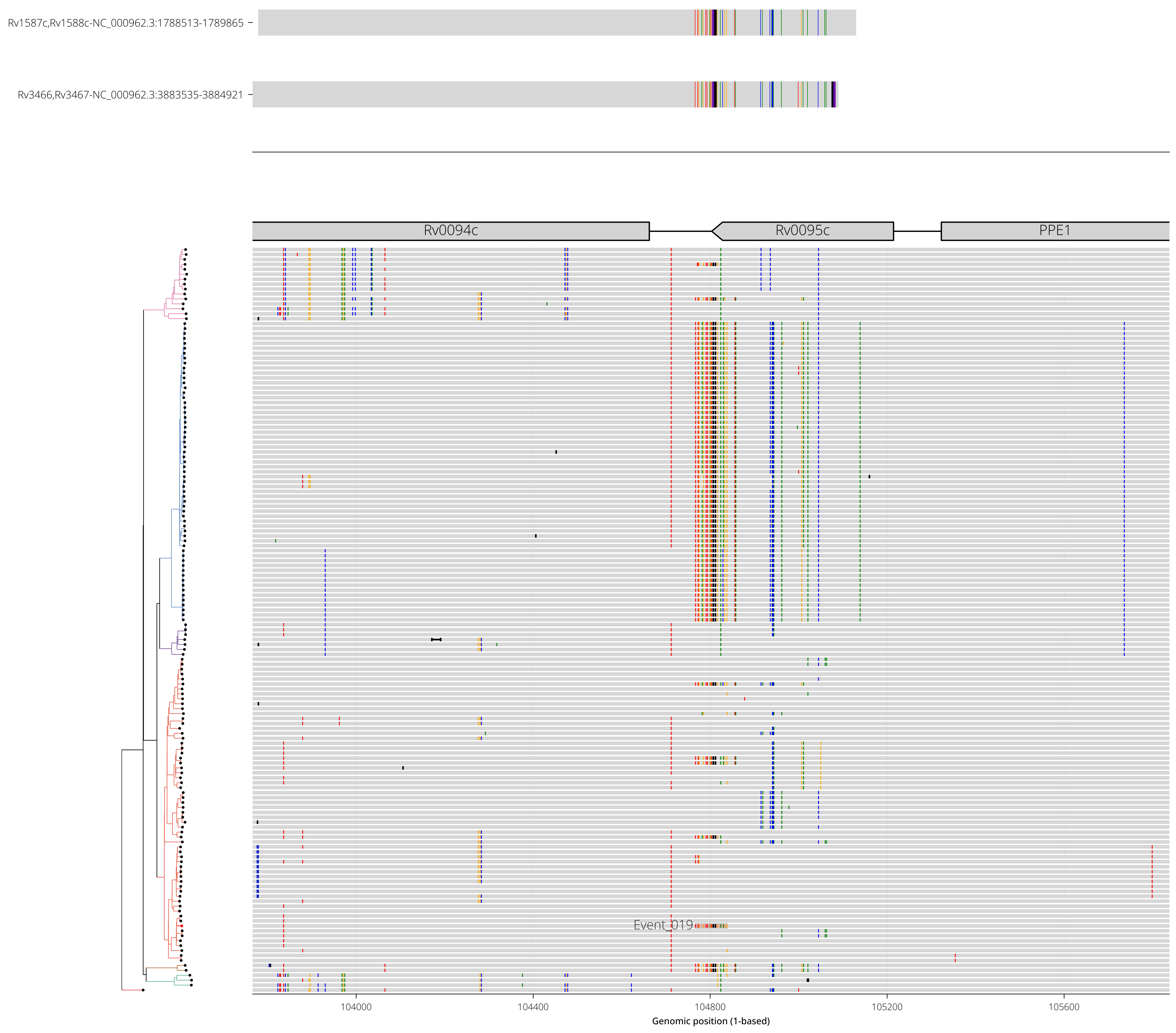

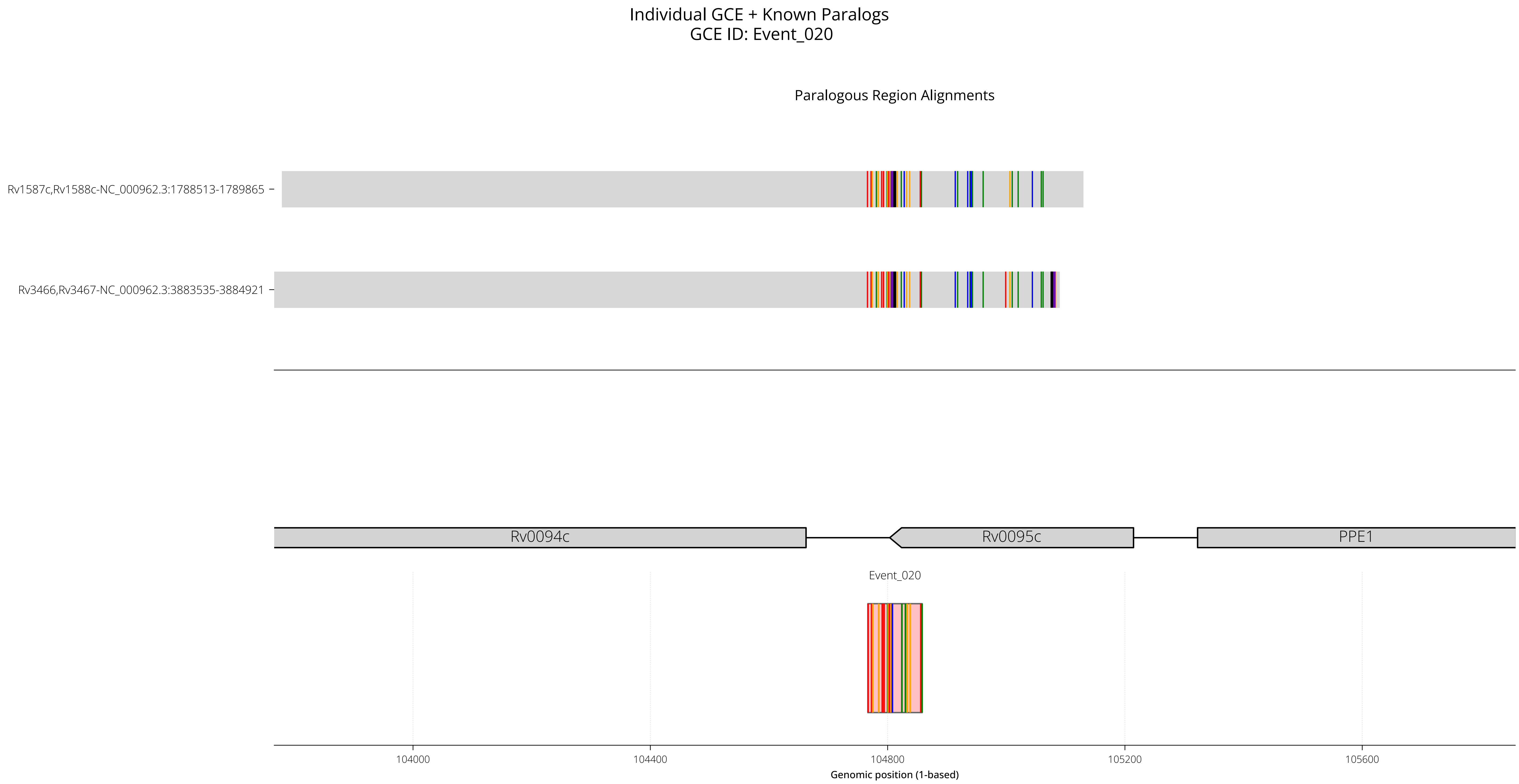
Individual GCE + Known Paralogs
GCE ID: Event_020
Paralogous Region Alignments
Rv1587c,Rv1588c-NC_000962.3:1788513-1789865
Rv3466,Rv3467-NC_000962.3:3883535-3884921
Rv0094c
Rv0095c
PPE1
Event_020
104000
104400
104800
105200
105600
Genomic position (1-based)

Circos Viz - GCE to Known Paralogs Locations
GCE ID: Event_020

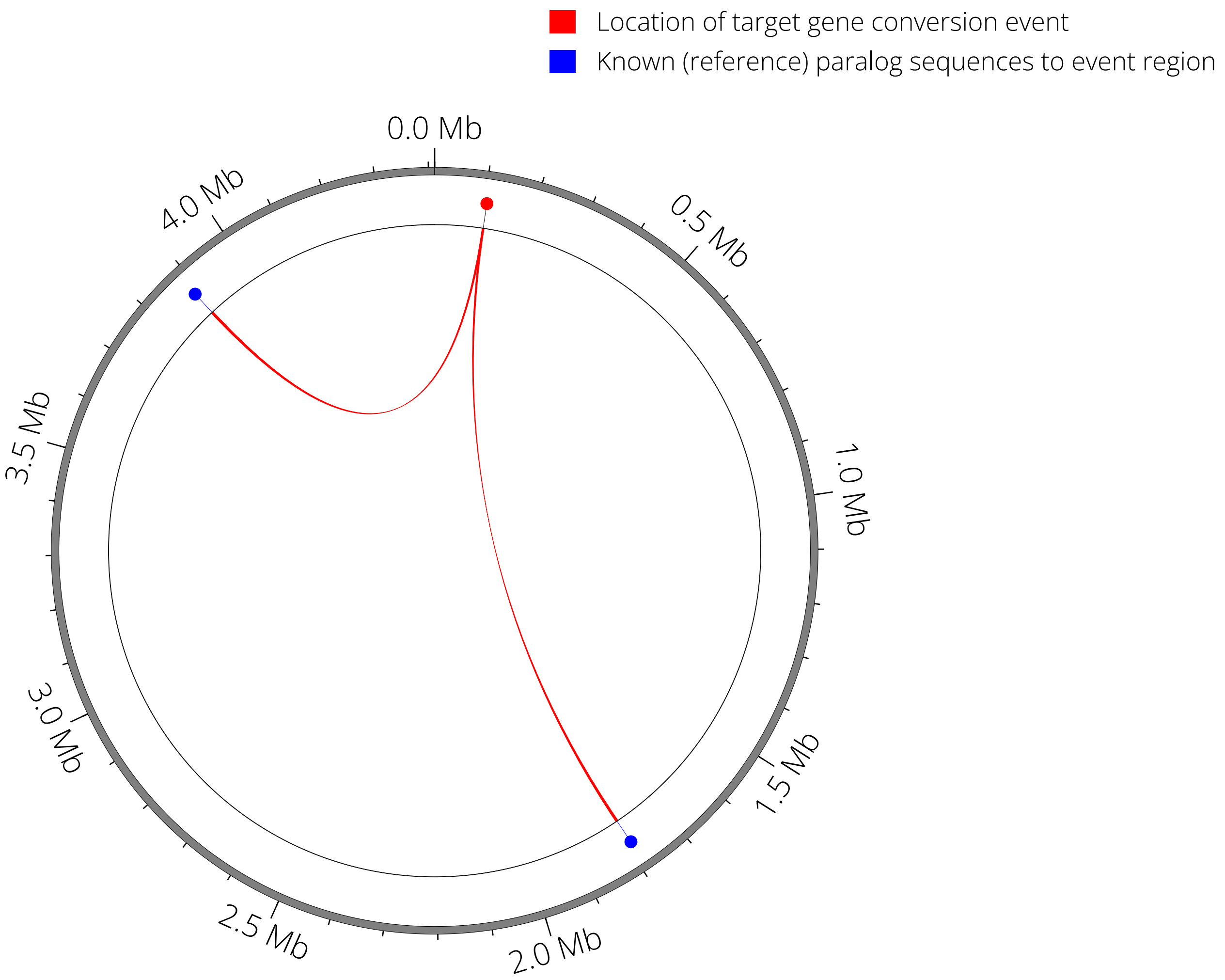

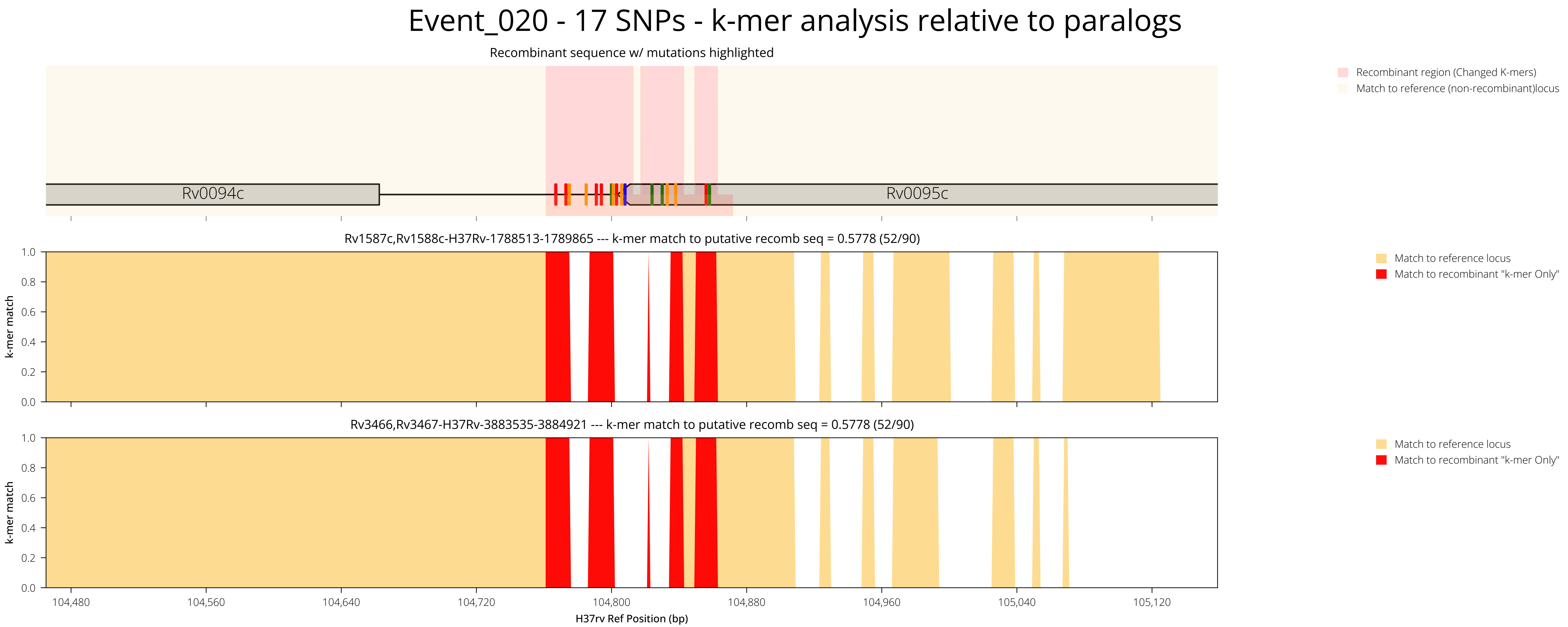

Event_020 - 17 SNPs - k-mer analysis relative to paralogs
Recombinant sequence w/ mutations highlighted
Recombinant region (Changed K-mers)
Match to reference (non-recombinant)locus
Rv0094c
Rv0095c
Rv1587c,Rv1588c-H37Rv-1788513-1789865 --- k-mer match to putative recomb seq = 0.5778 (52/90)
Match to reference locus
Match to recombinant "k-mer Only"
k-mer match
1.0
0.8
0.6
0.4
0.2
0.0
Rv3466,Rv3467-H37Rv-3883535-3884921 --- k-mer match to putative recomb seq = 0.5778 (52/90)
Match to reference locus
Match to recombinant "k-mer Only"
k-mer match
1.0
0.8
0.6
0.4
0.2
0.0
104,480
104,560
104,640
104,720
104,800
104,880
104,960
105,040
105,120
H37rv Ref Position (bp)

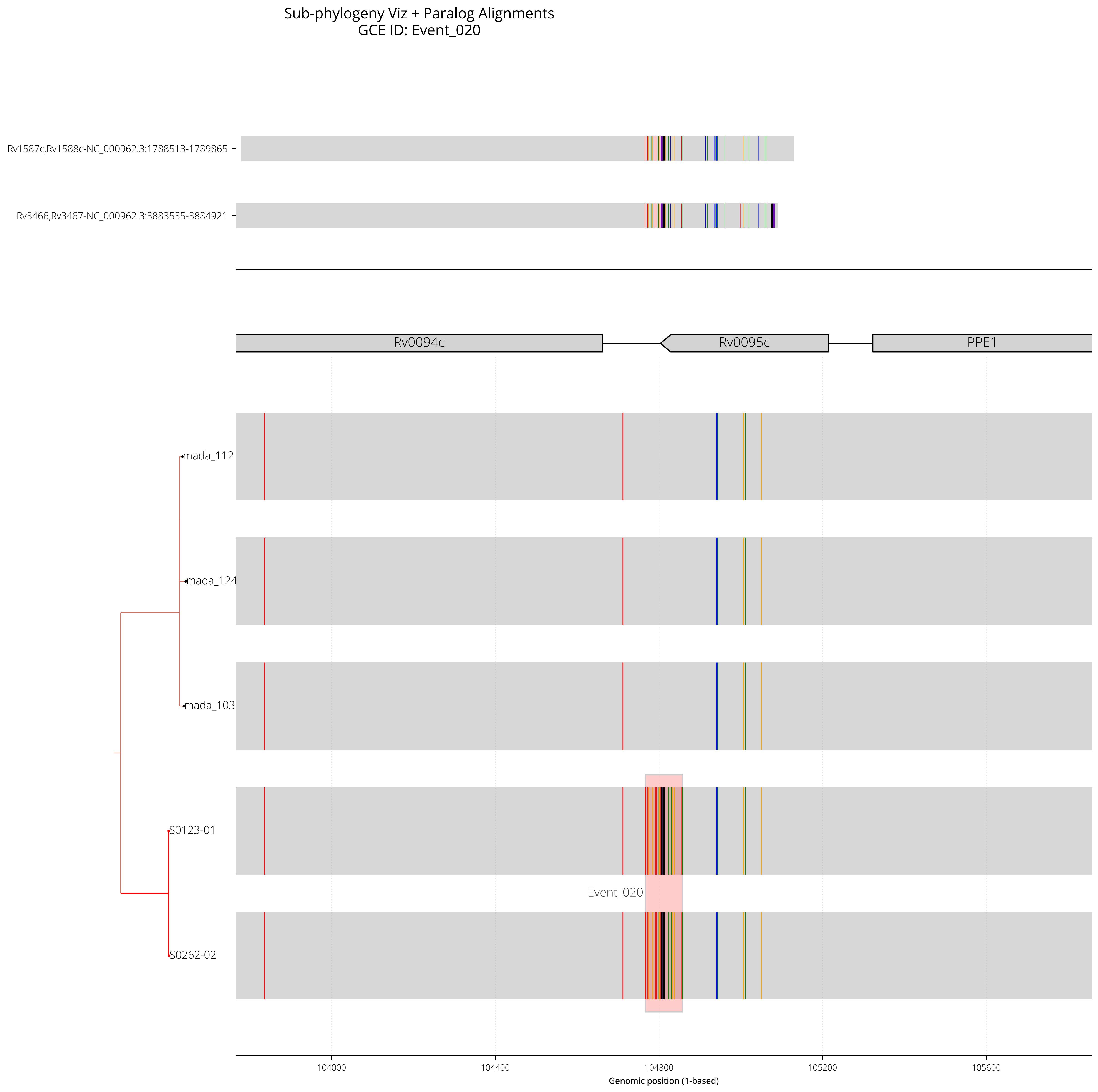

Sub-phylogeny Viz + Paralog Alignments
GCE ID: Event_020
Rv1587c,Rv1588c-NC_000962.3:1788513-1789865
Rv3466,Rv3467-NC_000962.3:3883535-3884921
Rv0094c
Rv0095c
PPE1
mada_112
mada_124
mada_103
S0123-01
S0262-02
Event_020
104000
104400
104800
105200
105600
Genomic position (1-based)

Full Phylogeny Viz + Paralog Alignments
GCE ID: Event_020

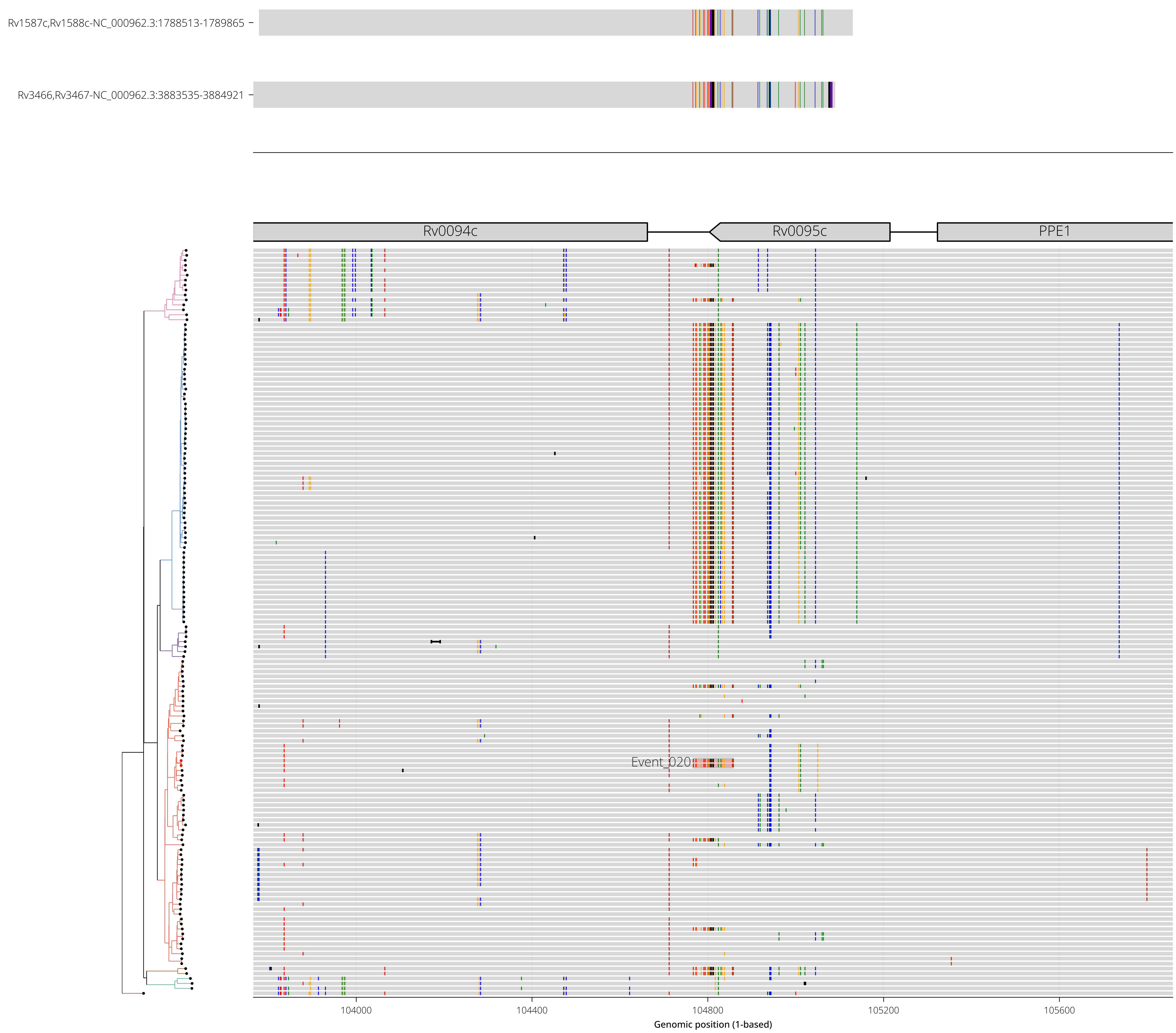

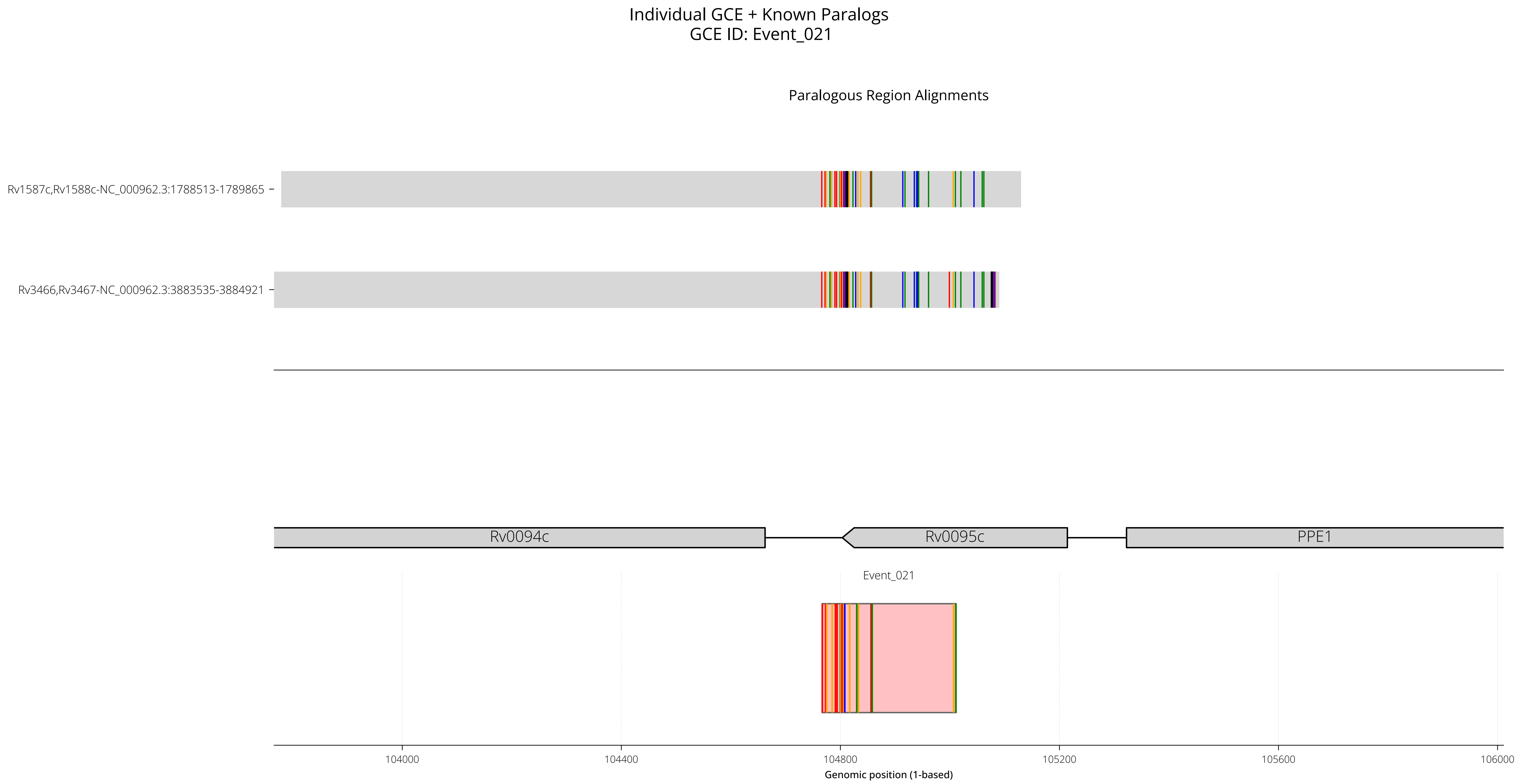

Individual GCE + Known Paralogs
GCE ID: Event_021
Paralogous Region Alignments
Rv1587c,Rv1588c-NC_000962.3:1788513-1789865
Rv3466,Rv3467-NC_000962.3:3883535-3884921
Rv0094c
Rv0095c
PPE1
Event_021
104000
104400
104800
105200
105600
106000
Genomic position (1-based)

### Circos Viz - GCE to Known Paralogs Locations
GCE ID: Event_021

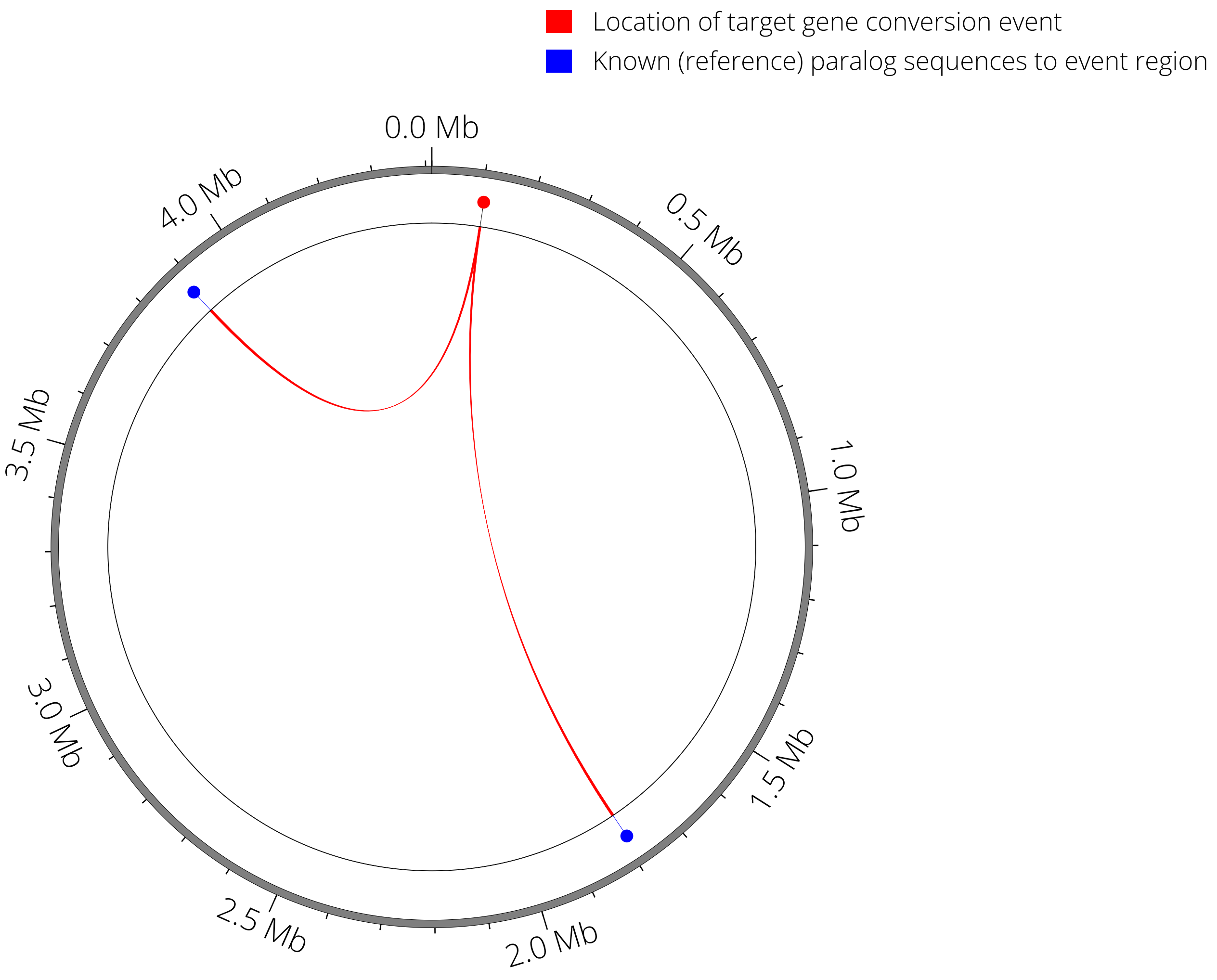

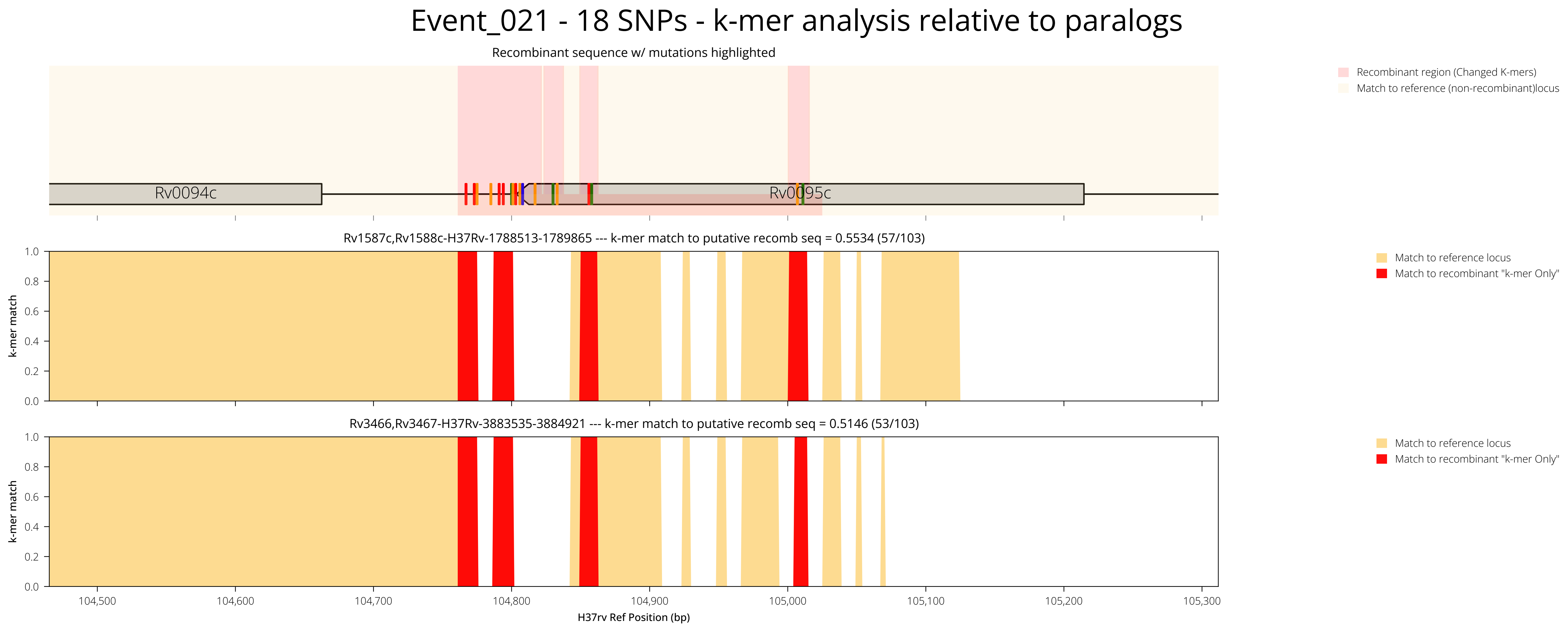

Event_021 - 18 SNPs - k-mer analysis relative to paralogs
Recombinant sequence w/ mutations highlighted
Rv0094c
Rv0095c
Recombinant region (Changed K-mers)
Match to reference (non-recombinant)locus
Rv1587c,Rv1588c-H37Rv-1788513-1789865 --- k-mer match to putative recomb seq = 0.5534 (57/103)
Match to reference locus
Match to recombinant "k-mer Only"
k-mer match
1.0
0.8
0.6
0.4
0.2
0.0
Rv3466,Rv3467-H37Rv-3883535-3884921 --- k-mer match to putative recomb seq = 0.5146 (53/103)
Match to reference locus
Match to recombinant "k-mer Only"
k-mer match
1.0
0.8
0.6
0.4
0.2
0.0
104,500
104,600
104,700
104,800
104,900
105,000
105,100
105,200
105,300
H37rv Ref Position (bp)

Sub-phylogeny Viz + Paralog Alignments
GCE ID: Event_021

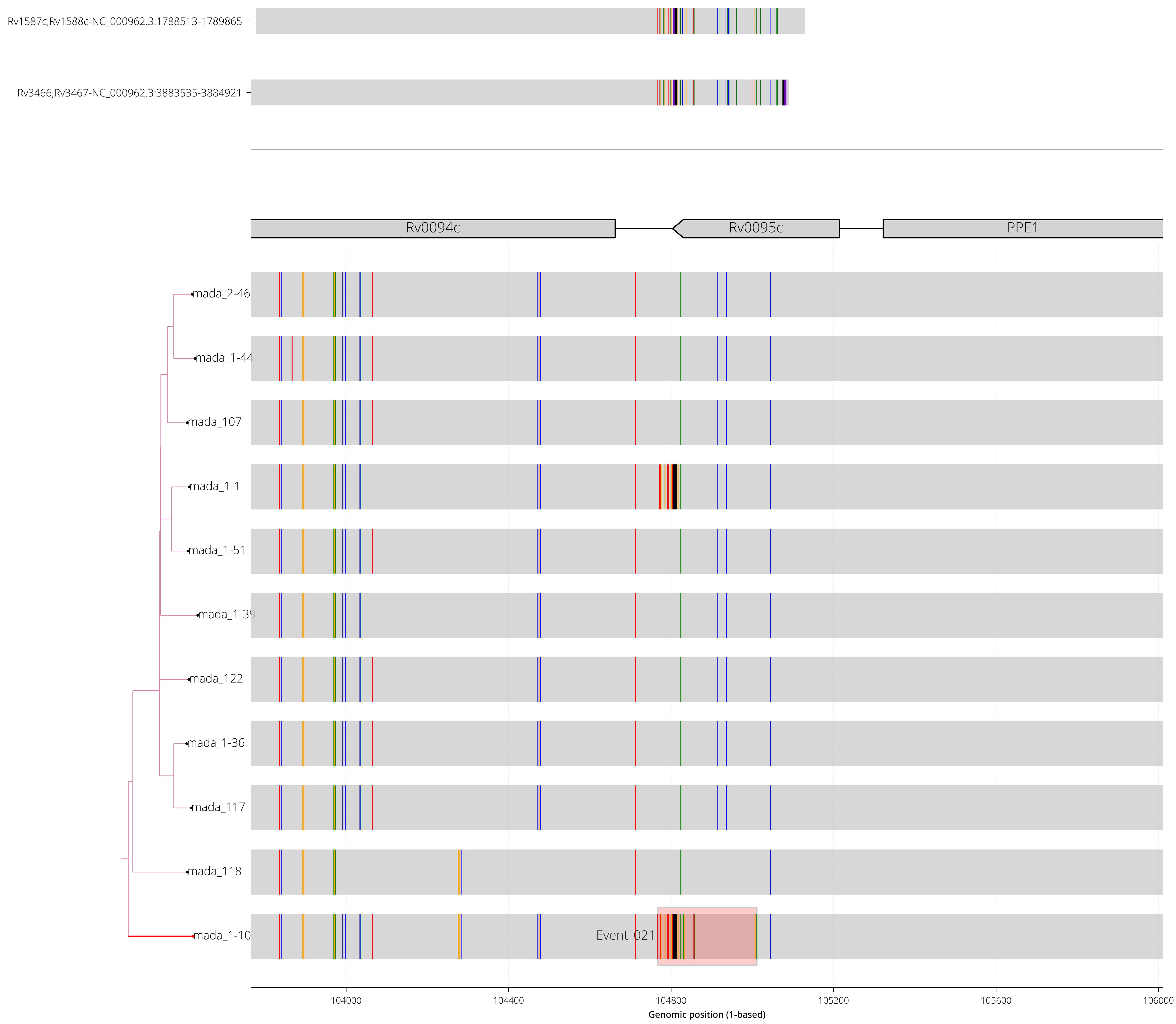

Full Phylogeny Viz + Paralog Alignments
GCE ID: Event_021

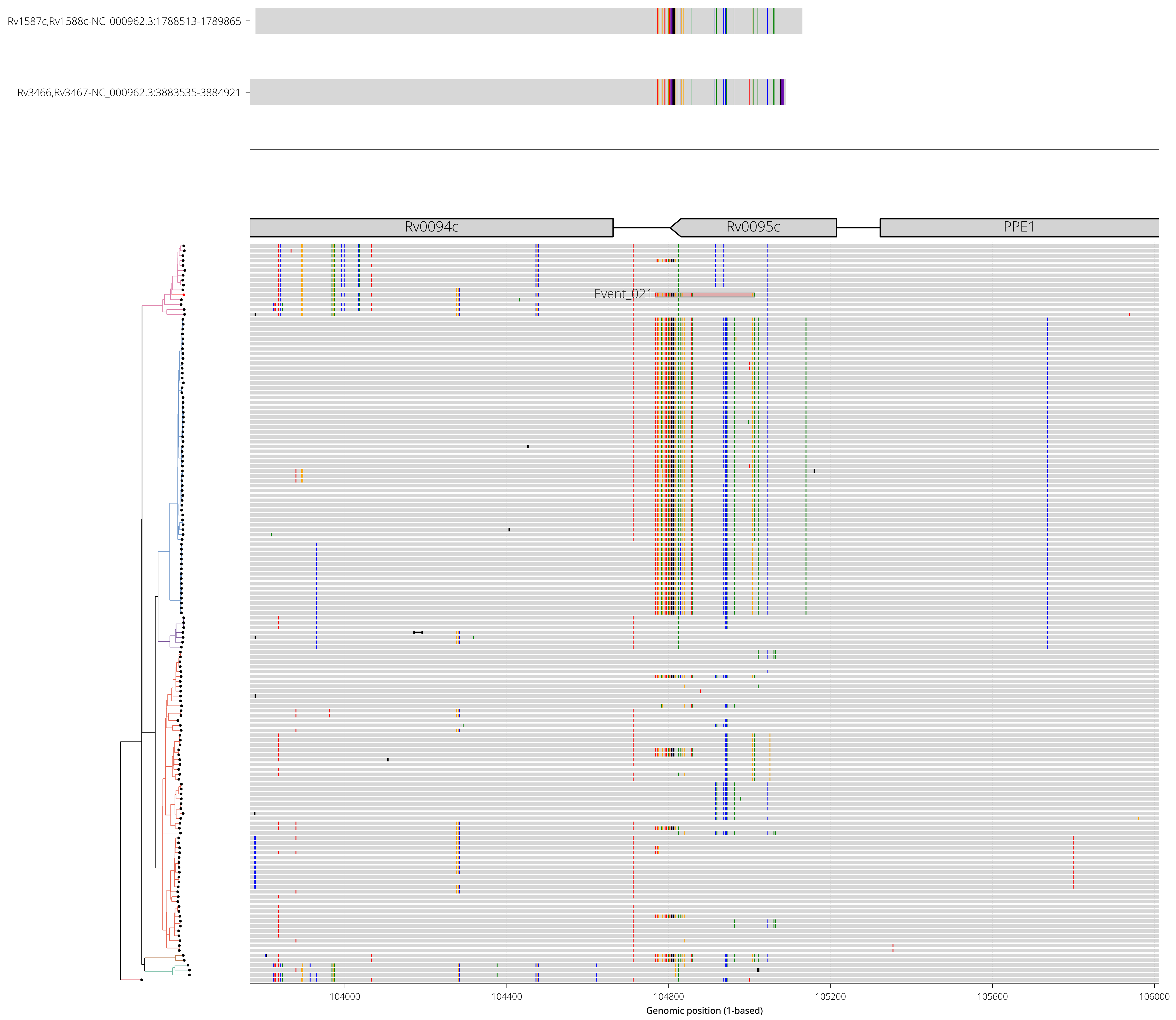

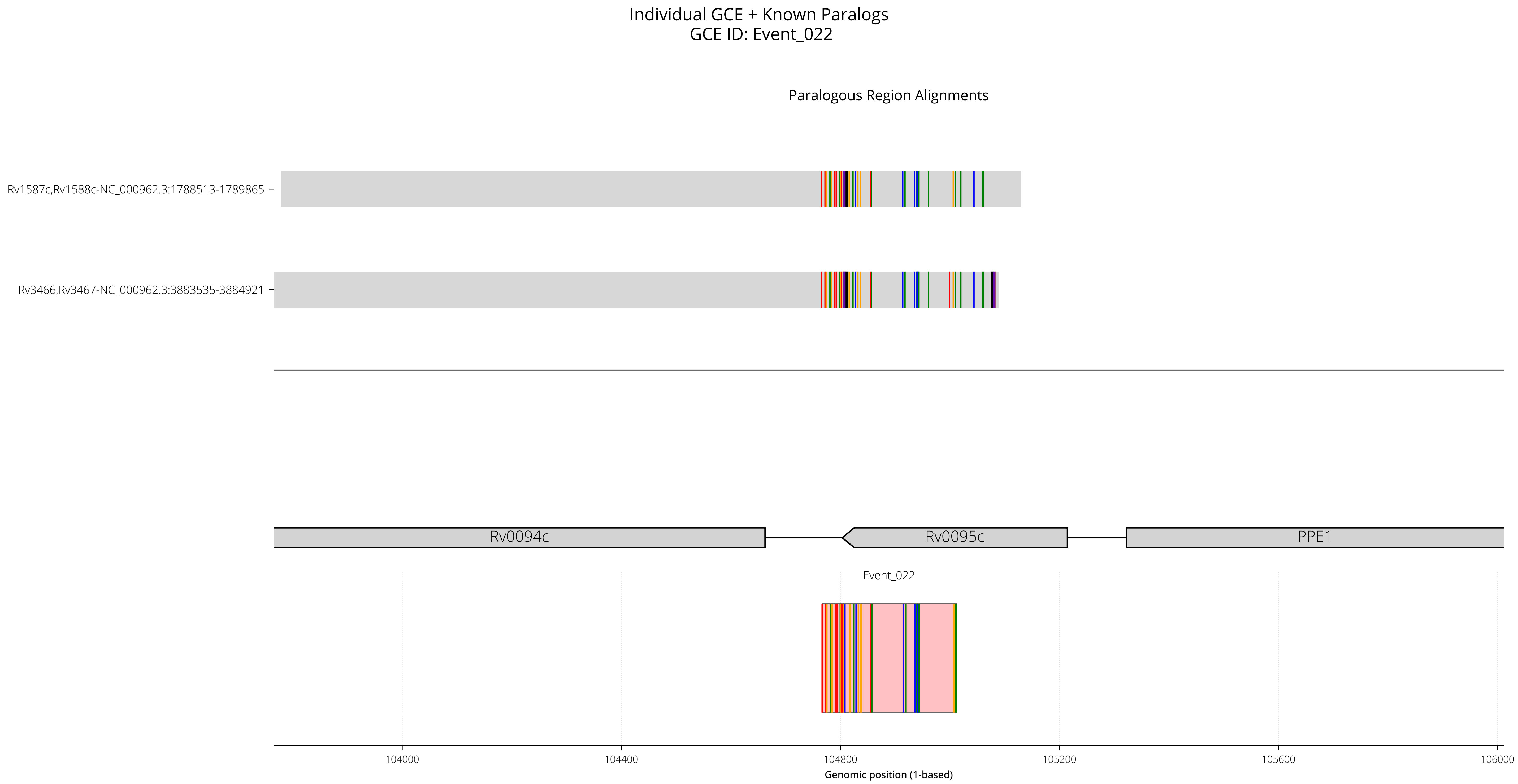
Individual GCE + Known Paralogs
GCE ID: Event_022
Paralogous Region Alignments
Rv1587c,Rv1588c-NC_000962.3:1788513-1789865
Rv3466,Rv3467-NC_000962.3:3883535-3884921
Rv0094c
Rv0095c
PPE1
Event_022
104000
104400
104800
105200
105600
106000
Genomic position (1-based)

Circos Viz - GCE to Known Paralogs Locations
GCE ID: Event_022

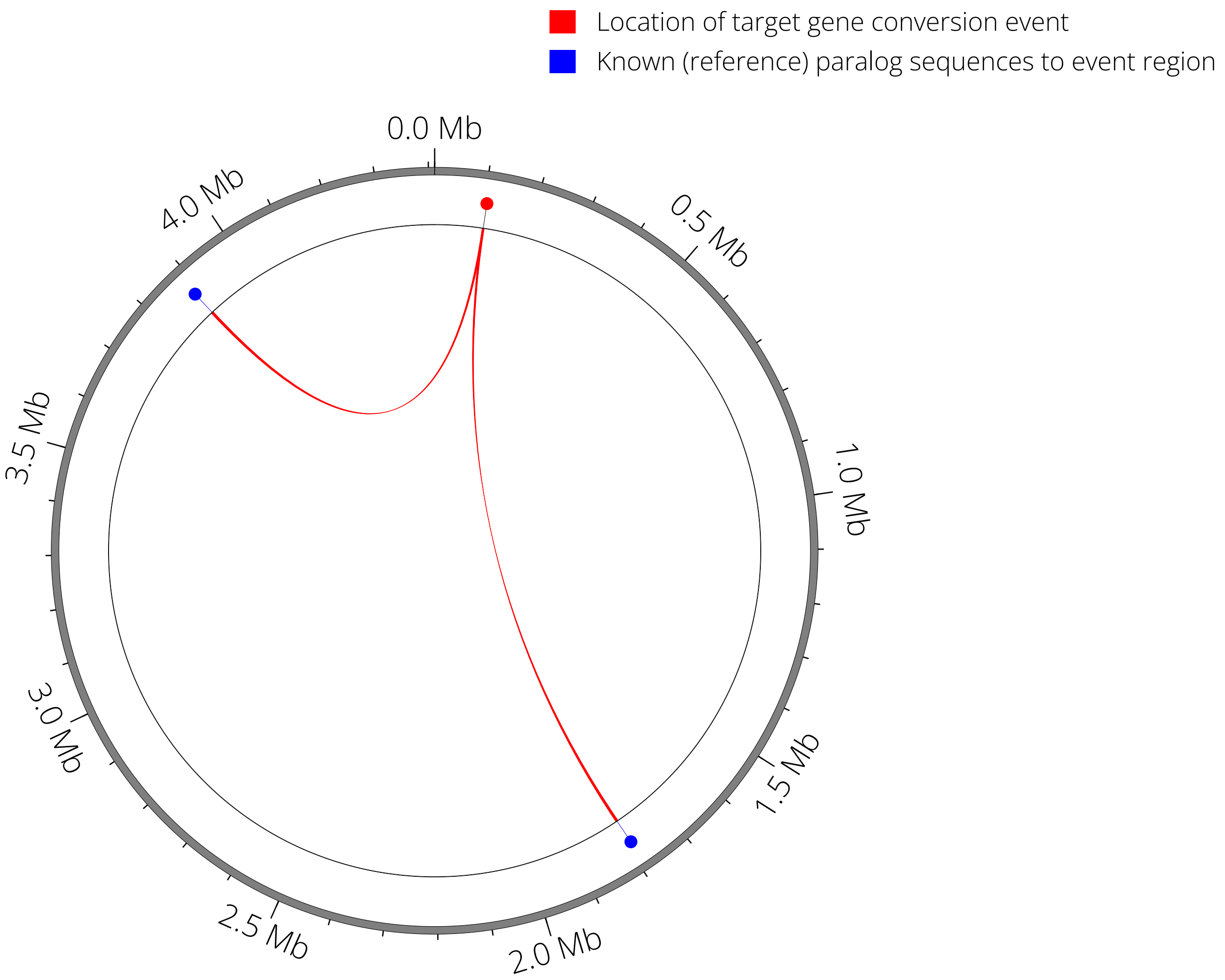

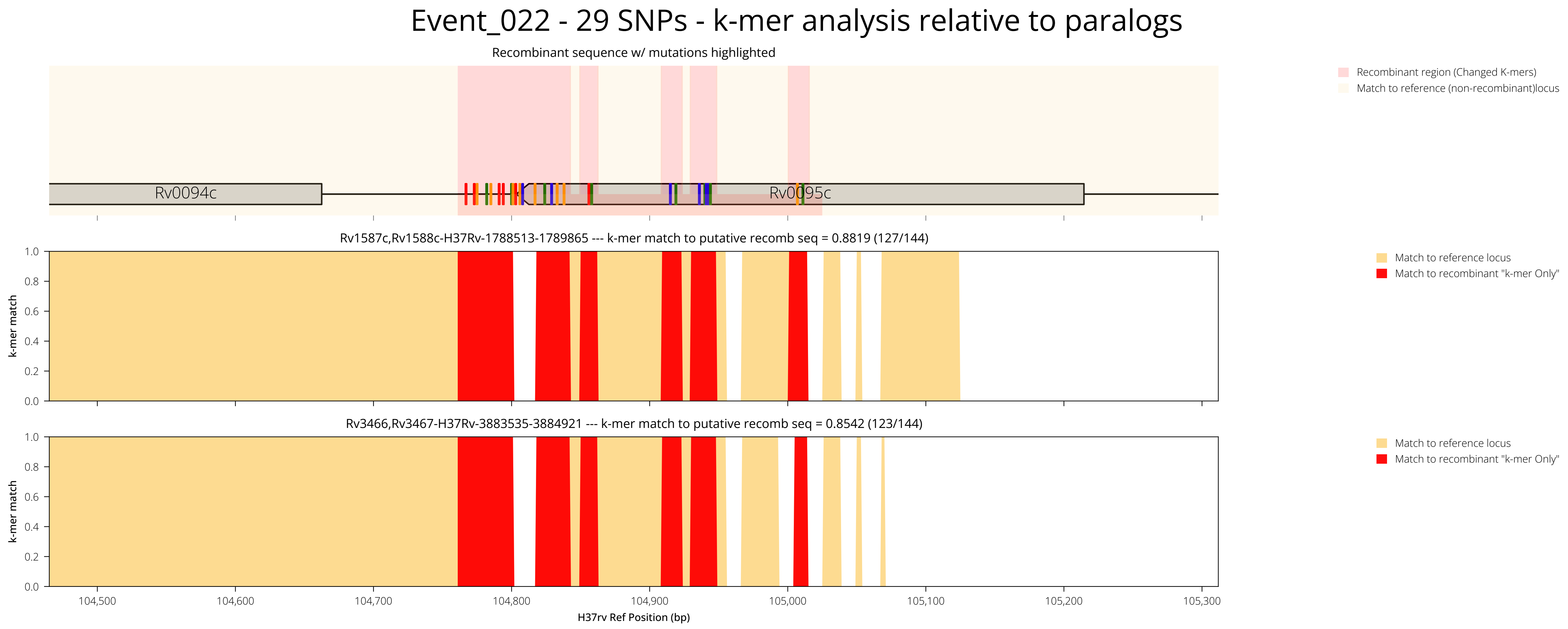

Event_022 - 29 SNPs - k-mer analysis relative to paralogs
Recombinant sequence w/ mutations highlighted
Rv0094c
Rv0095c
Recombinant region (Changed K-mers)
Match to reference (non-recombinant)locus
Rv1587c,Rv1588c-H37Rv-1788513-1789865 --- k-mer match to putative recomb seq = 0.8819 (127/144)
k-mer match
Match to reference locus
Match to recombinant "k-mer Only"
Rv3466,Rv3467-H37Rv-3883535-3884921 --- k-mer match to putative recomb seq = 0.8542 (123/144)
k-mer match
Match to reference locus
Match to recombinant "k-mer Only"
104,500
104,600
104,700
104,800
104,900
105,000
105,100
105,200
105,300
H37rv Ref Position (bp)

Sub-phylogeny Viz + Paralog Alignments
GCE ID: Event_022

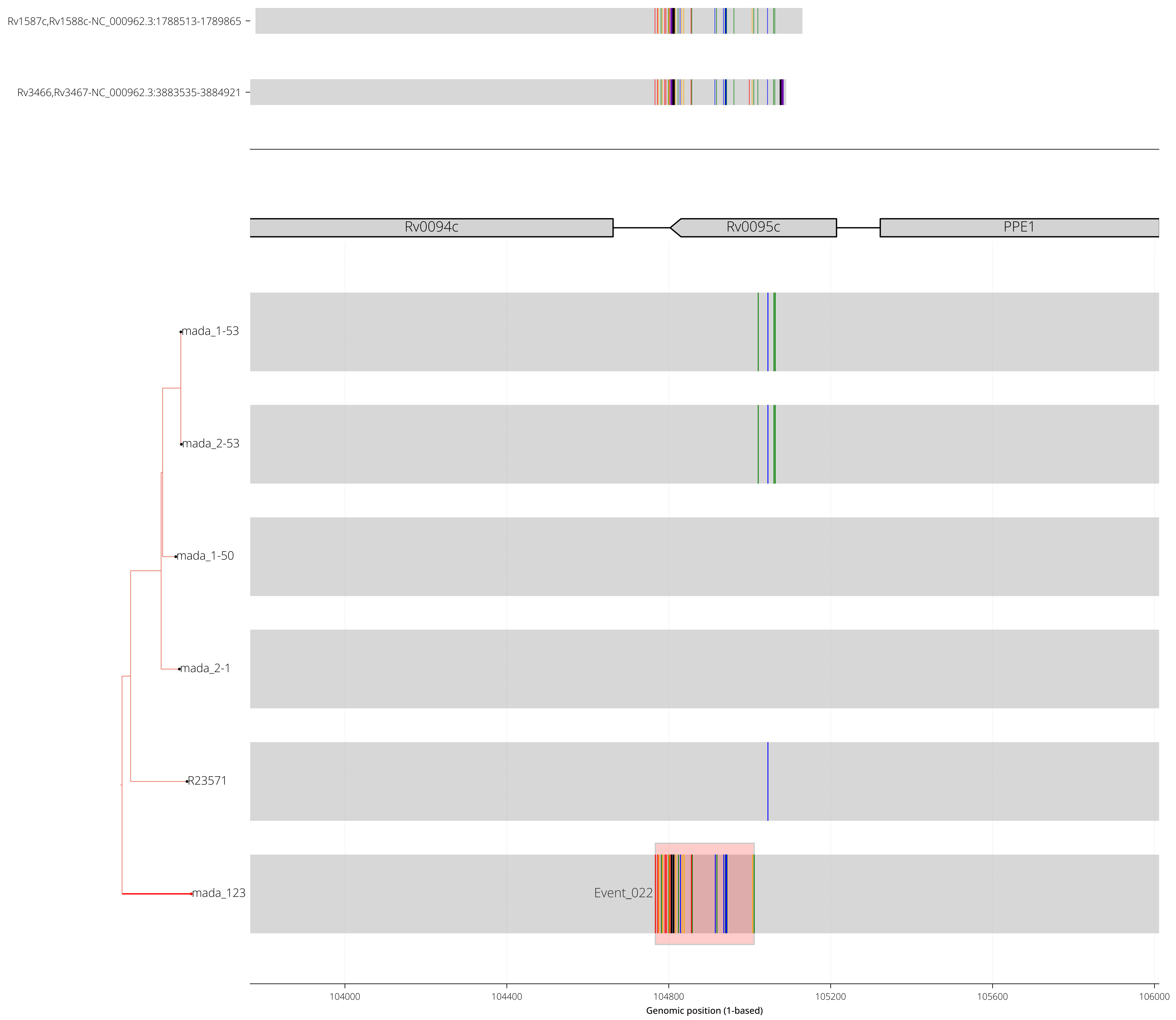

Full Phylogeny Viz + Paralog Alignments
GCE ID: Event_022

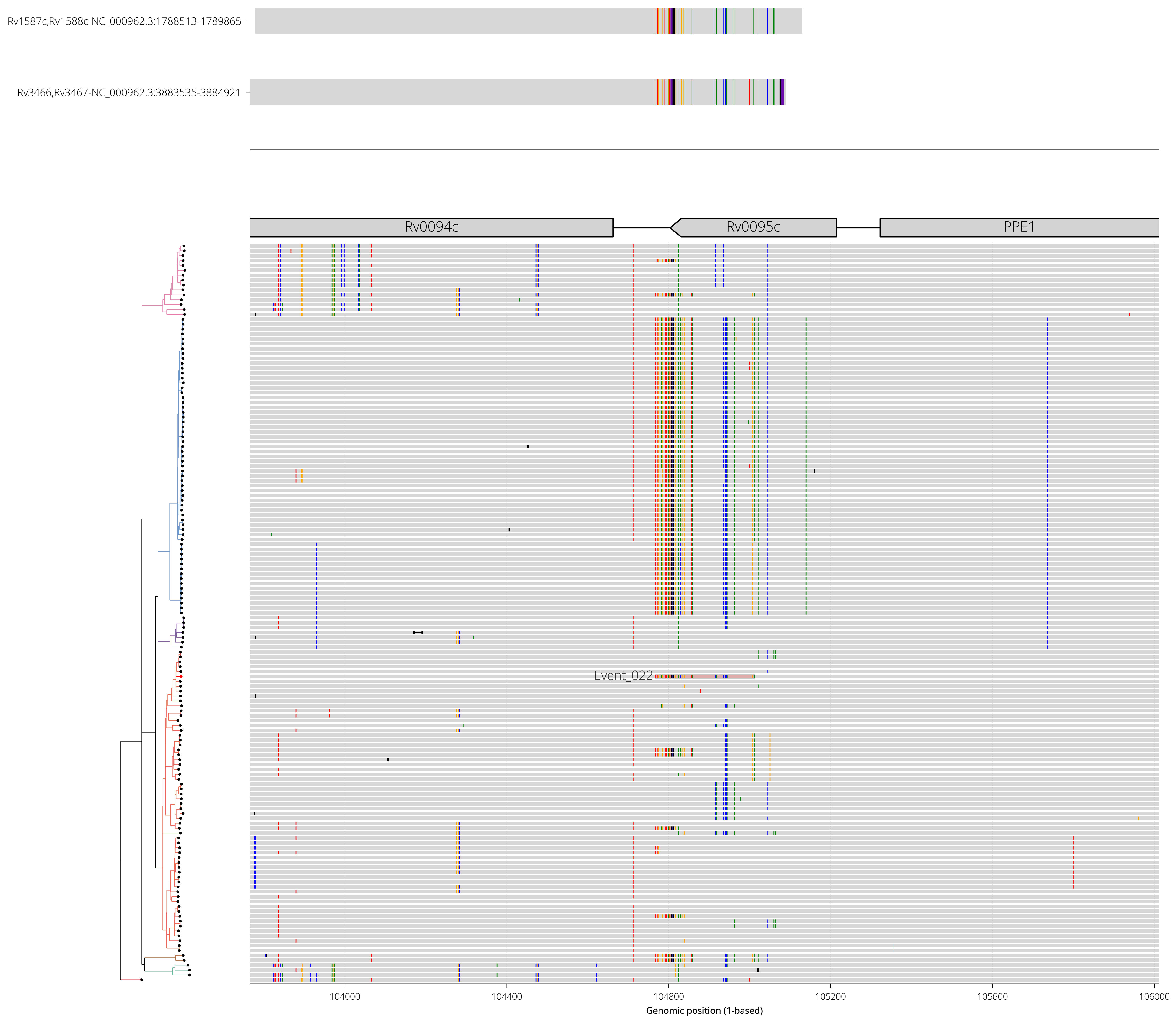

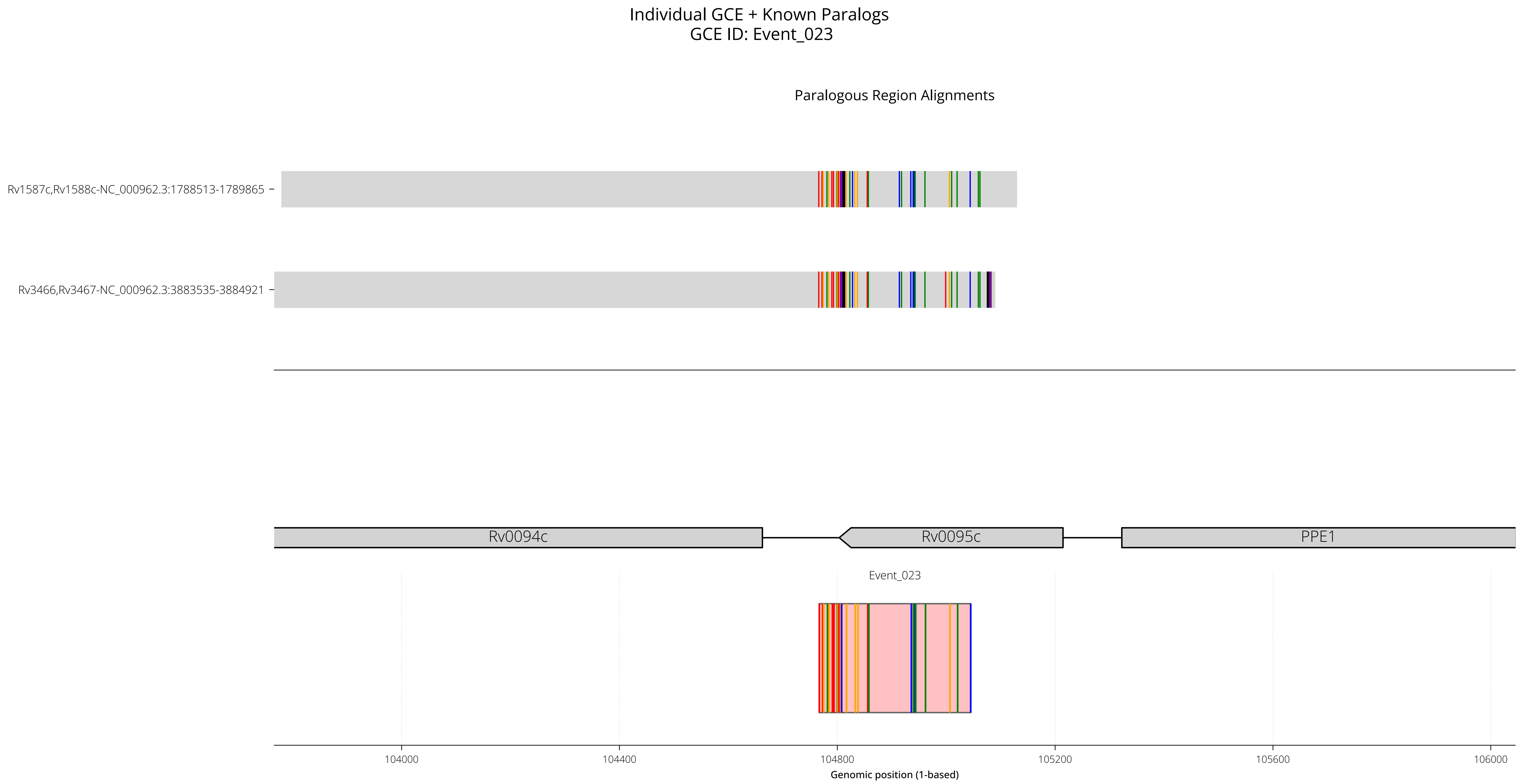

Individual GCE + Known Paralogs
GCE ID: Event_023
Paralogous Region Alignments
Rv1587c,Rv1588c-NC_000962.3:1788513-1789865
Rv3466,Rv3467-NC_000962.3:3883535-3884921
Rv0094c
Rv0095c
PPE1
Event_023
104000
104400
104800
105200
105600
106000
Genomic position (1-based)

Circos Viz - GCE to Known Paralogs Locations
GCE ID: Event_023

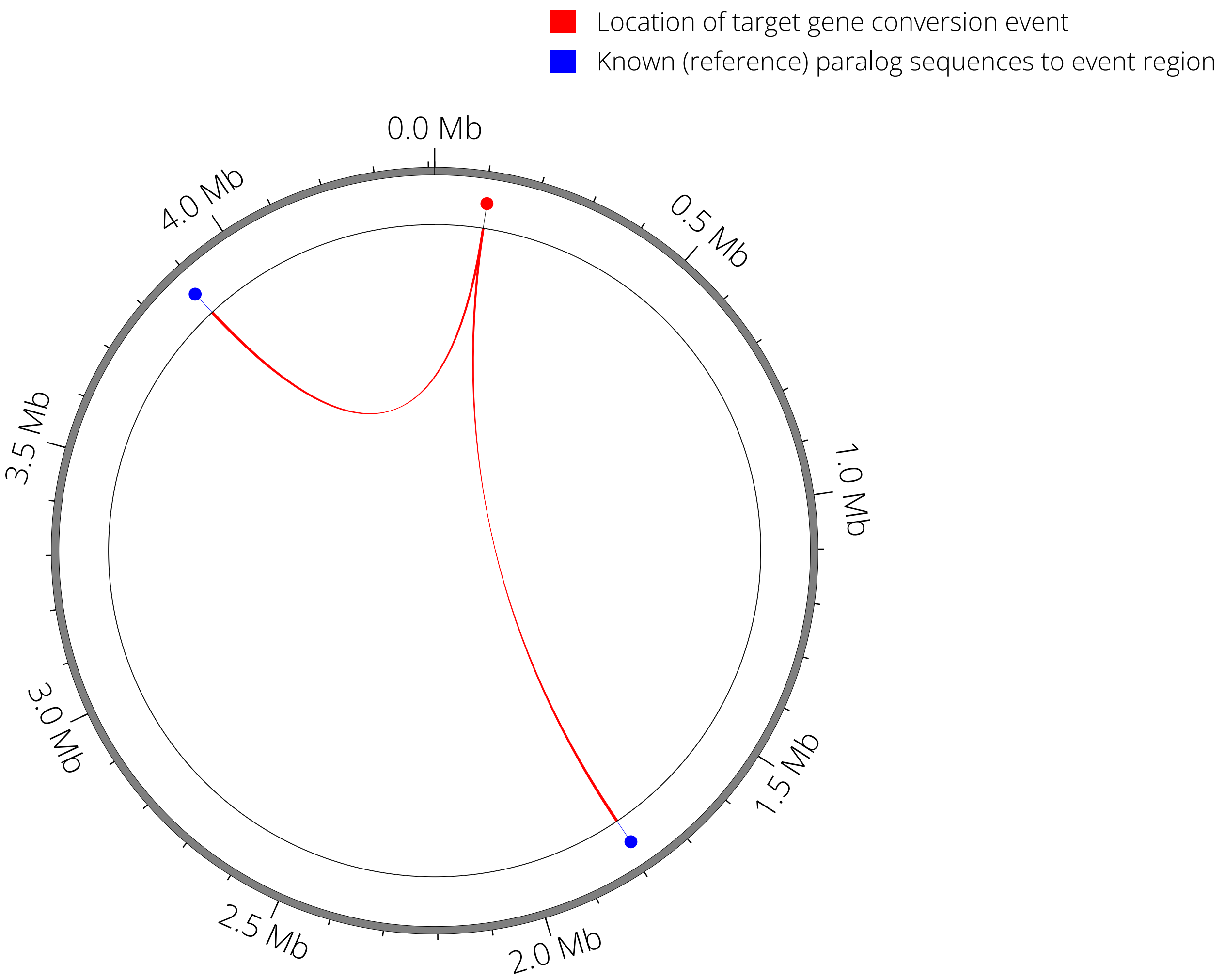

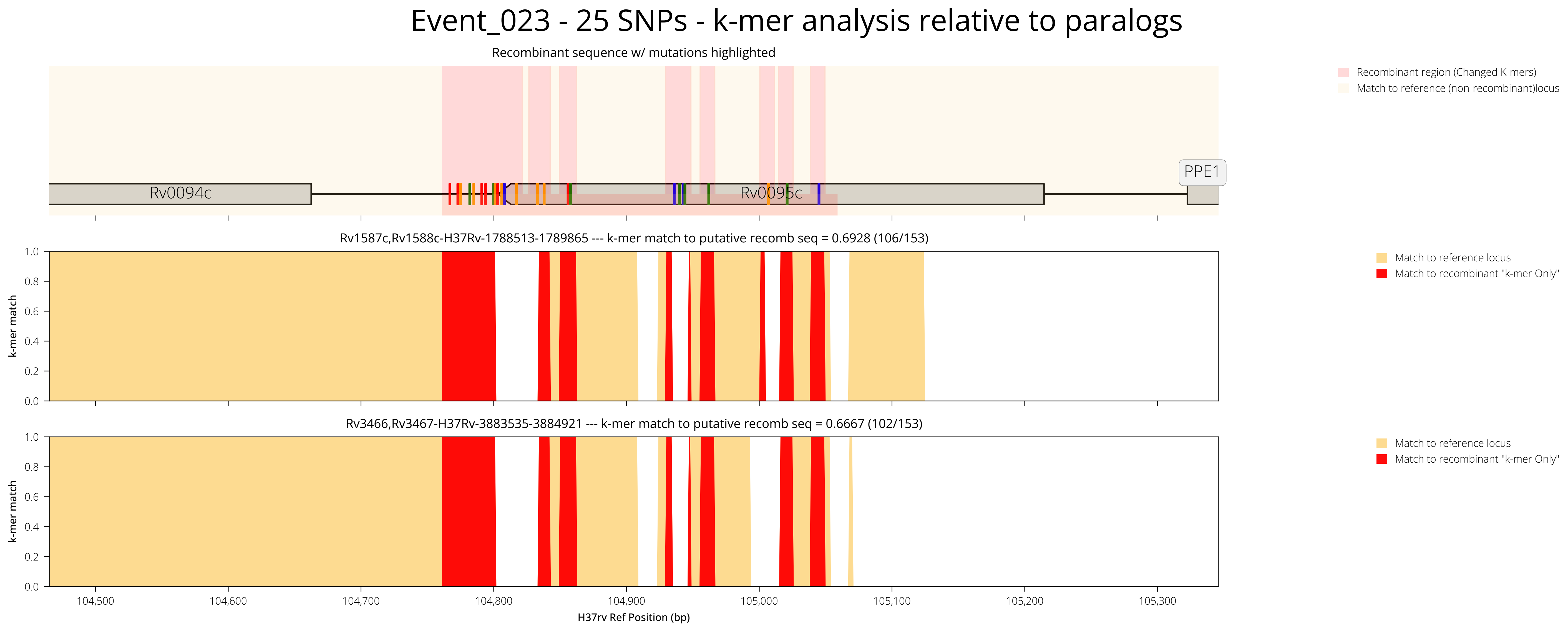

Event_023 - 25 SNPs - k-mer analysis relative to paralogs
Recombinant sequence w/ mutations highlighted
Recombinant region (Changed K-mers)
Match to reference (non-recombinant)locus
Rv0094c
Rv0095c
PPE1
Rv1587c,Rv1588c-H37Rv-1788513-1789865 --- k-mer match to putative recomb seq = 0.6928 (106/153)
Match to reference locus
Match to recombinant "k-mer Only"
k-mer match
1.0
0.8
0.6
0.4
0.2
0.0
Rv3466,Rv3467-H37Rv-3883535-3884921 --- k-mer match to putative recomb seq = 0.6667 (102/153)
Match to reference locus
Match to recombinant "k-mer Only"
k-mer match
104,500
104,600
104,700
104,800
104,900
105,000
105,100
105,200
105,300
H37rv Ref Position (bp)

Sub-phylogeny Viz + Paralog Alignments
GCE ID: Event_023

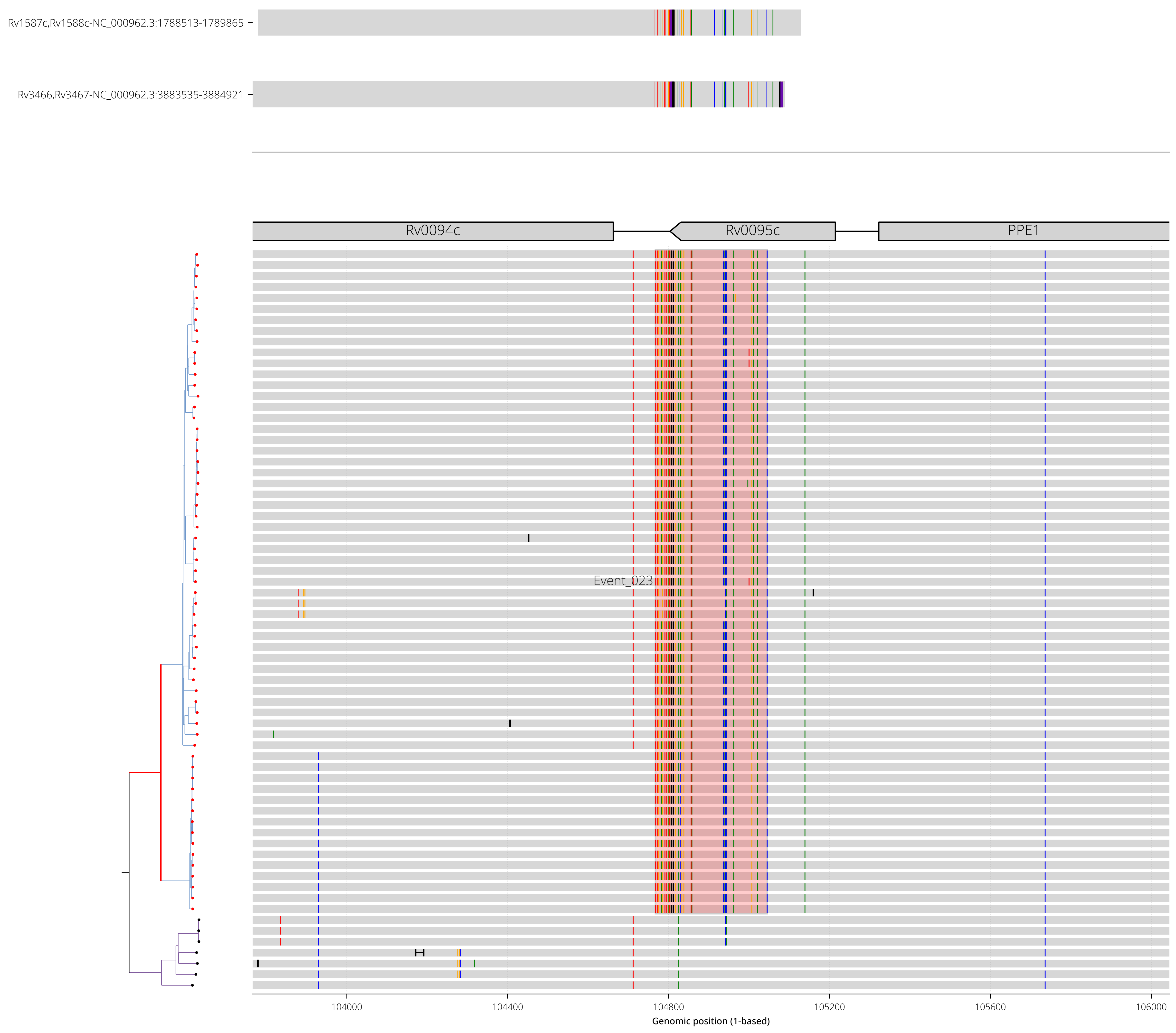

Full Phylogeny Viz + Paralog Alignments
GCE ID: Event_023

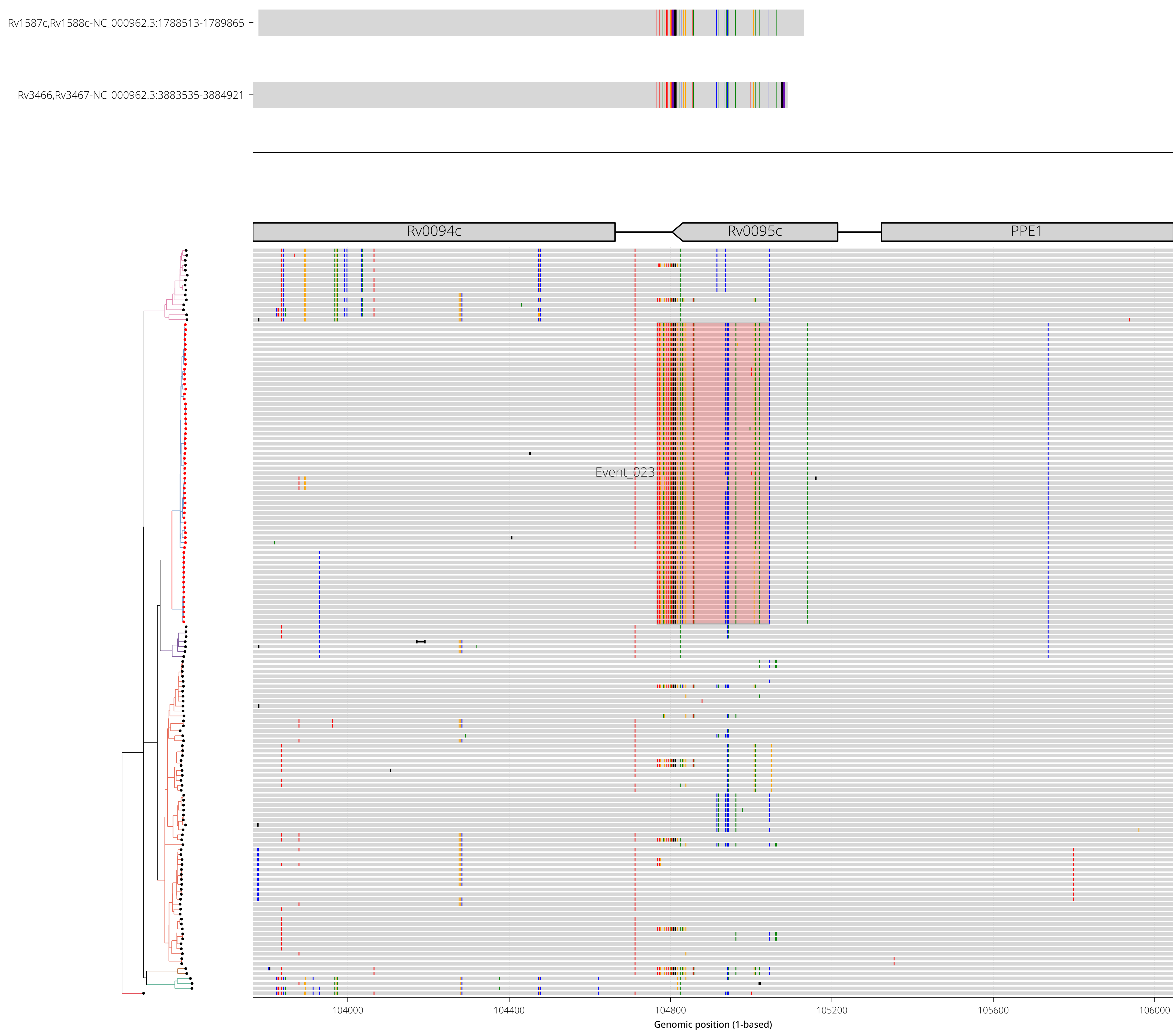

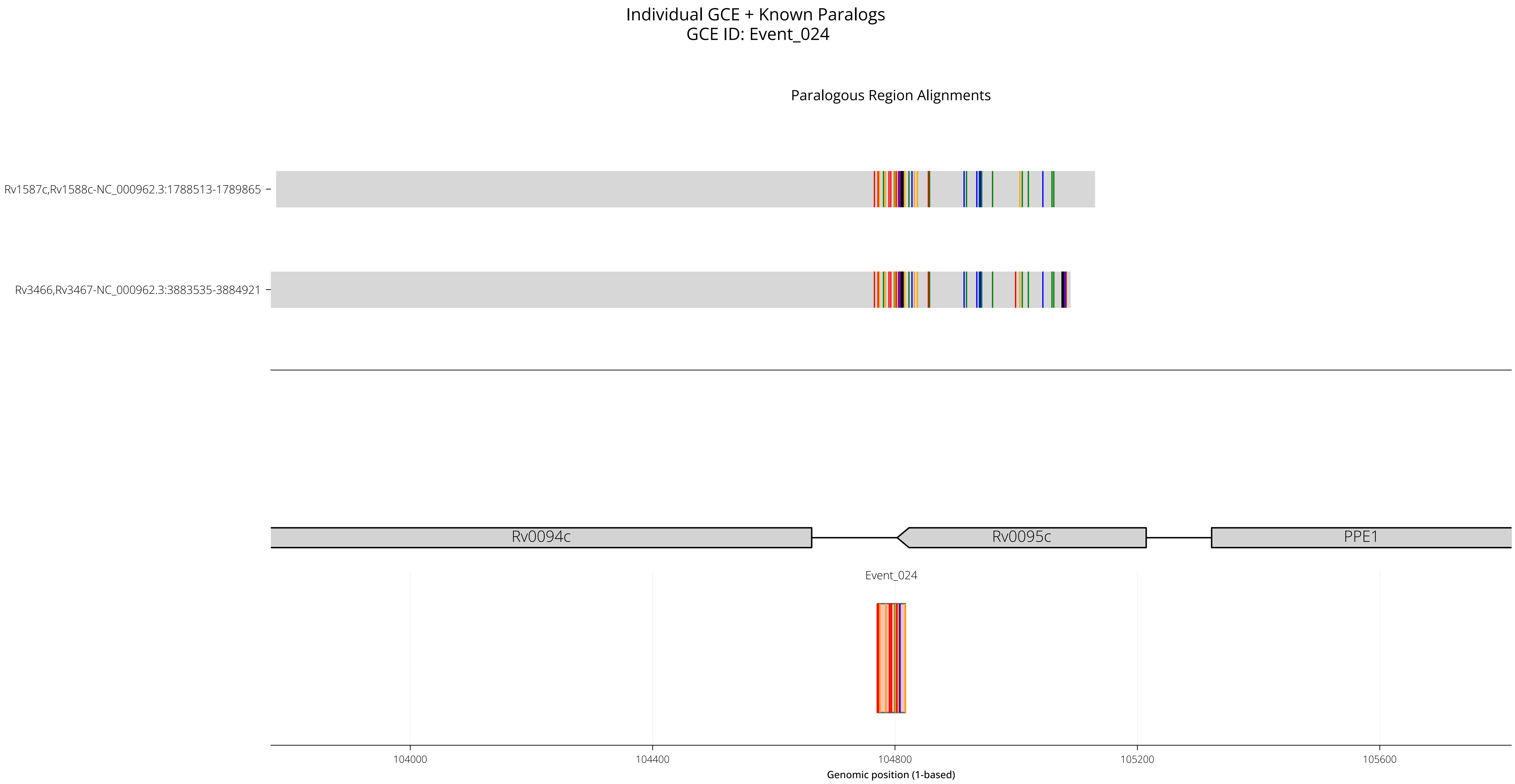
Individual GCE + Known Paralogs
GCE ID: Event_024
Paralogous Region Alignments
Rv1587c,Rv1588c-NC_000962.3:1788513-1789865
Rv3466,Rv3467-NC_000962.3:3883535-3884921
Rv0094c
Rv0095c
PPE1
Event_024
104000
104400
104800
105200
105600
Genomic position (1-based)

Circos Viz - GCE to Known Paralogs Locations
GCE ID: Event_024

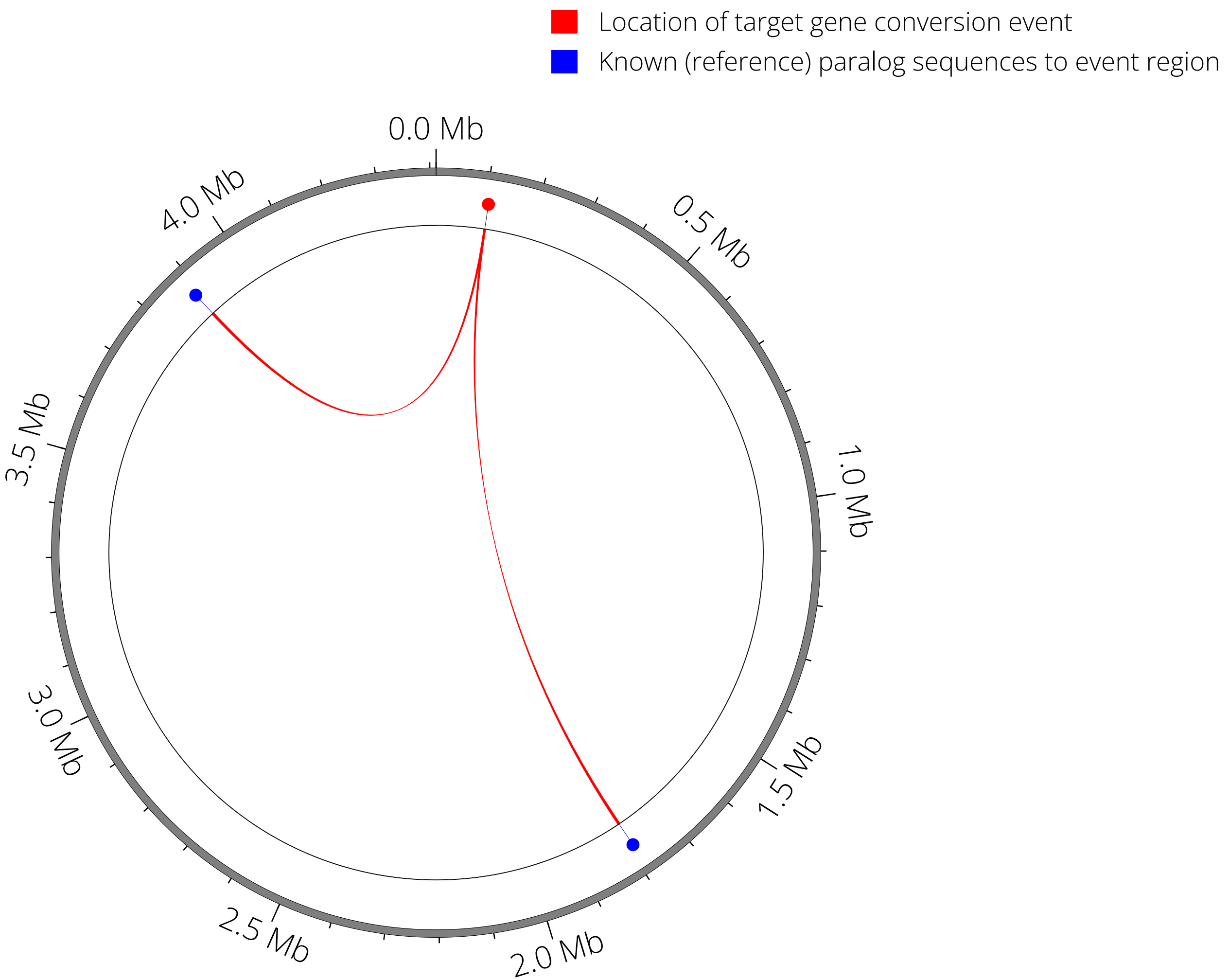

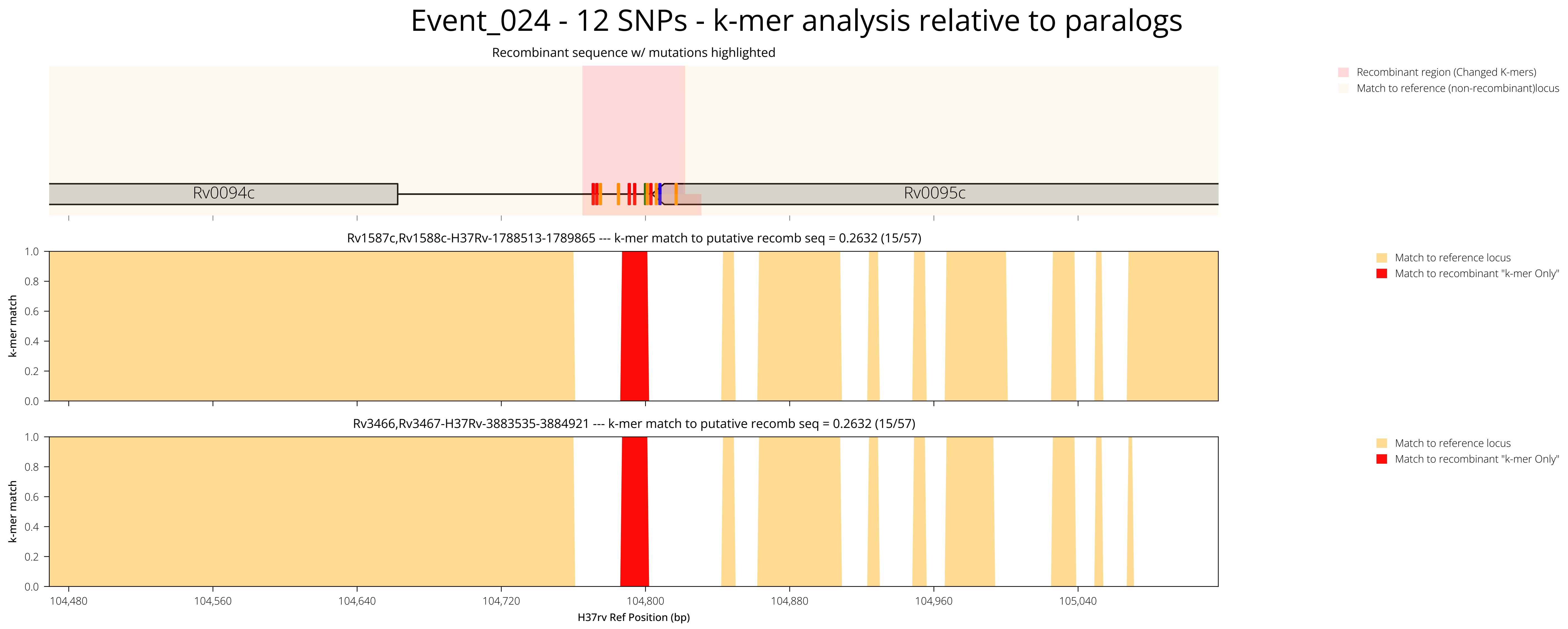

Event_024 - 12 SNPs - k-mer analysis relative to paralogs
Recombinant sequence w/ mutations highlighted
Rv0094c
Rv0095c
Recombinant region (Changed K-mers)
Match to reference (non-recombinant)locus
Rv1587c,Rv1588c-H37Rv-1788513-1789865 --- k-mer match to putative recomb seq = 0.2632 (15/57)
Match to reference locus
Match to recombinant "k-mer Only"
k-mer match
1.0
0.8
0.6
0.4
0.2
0.0
Rv3466,Rv3467-H37Rv-3883535-3884921 --- k-mer match to putative recomb seq = 0.2632 (15/57)
Match to reference locus
Match to recombinant "k-mer Only"
k-mer match
104,480
104,560
104,640
104,720
104,800
104,880
104,960
105,040
H37rv Ref Position (bp)

Sub-phylogeny Viz + Paralog Alignments
GCE ID: Event_024

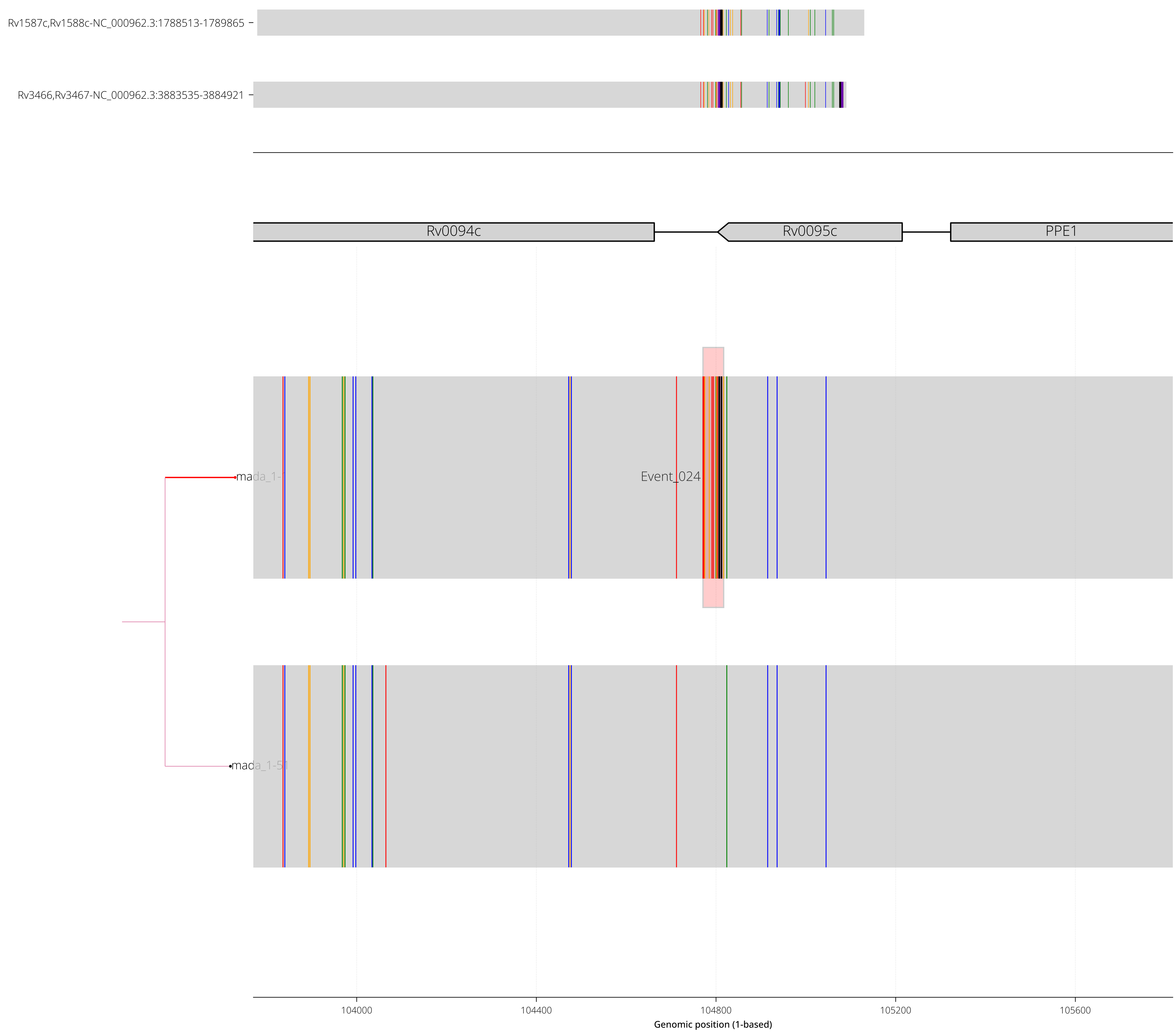

Full Phylogeny Viz + Paralog Alignments
GCE ID: Event_024

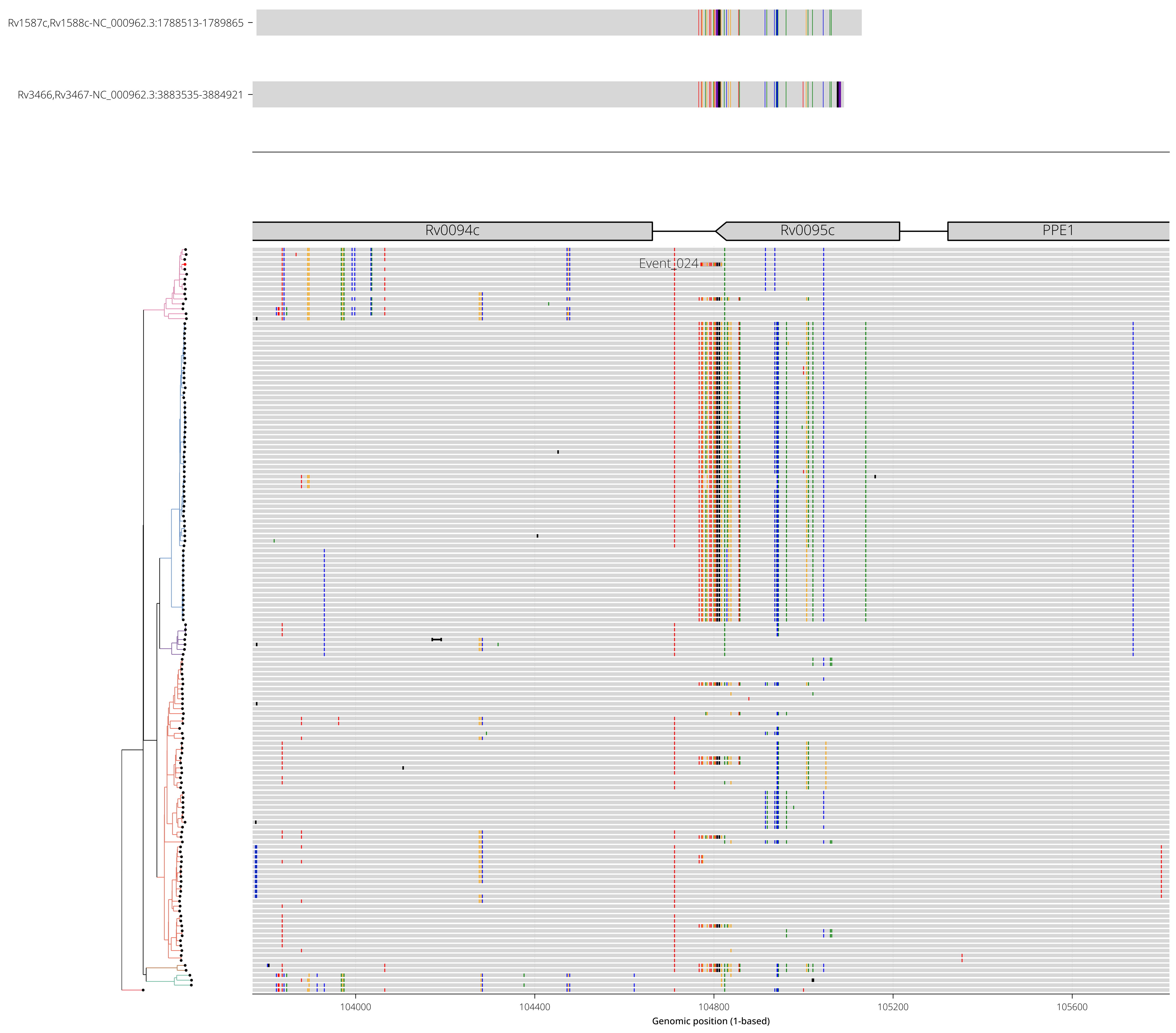

Individual GCE + Known Paralogs
GCE ID: Event_025

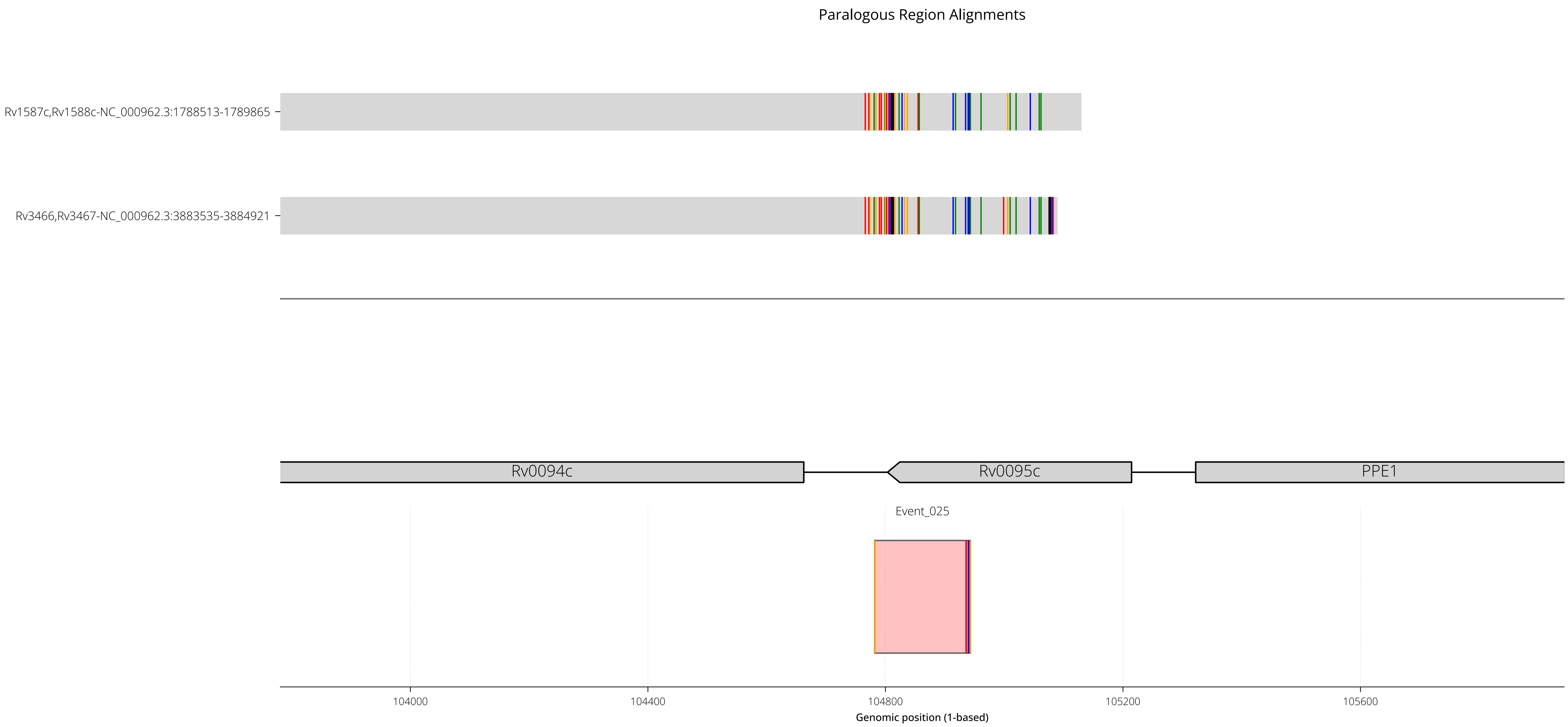

Circos Viz - GCE to Known Paralogs Locations
GCE ID: Event_025

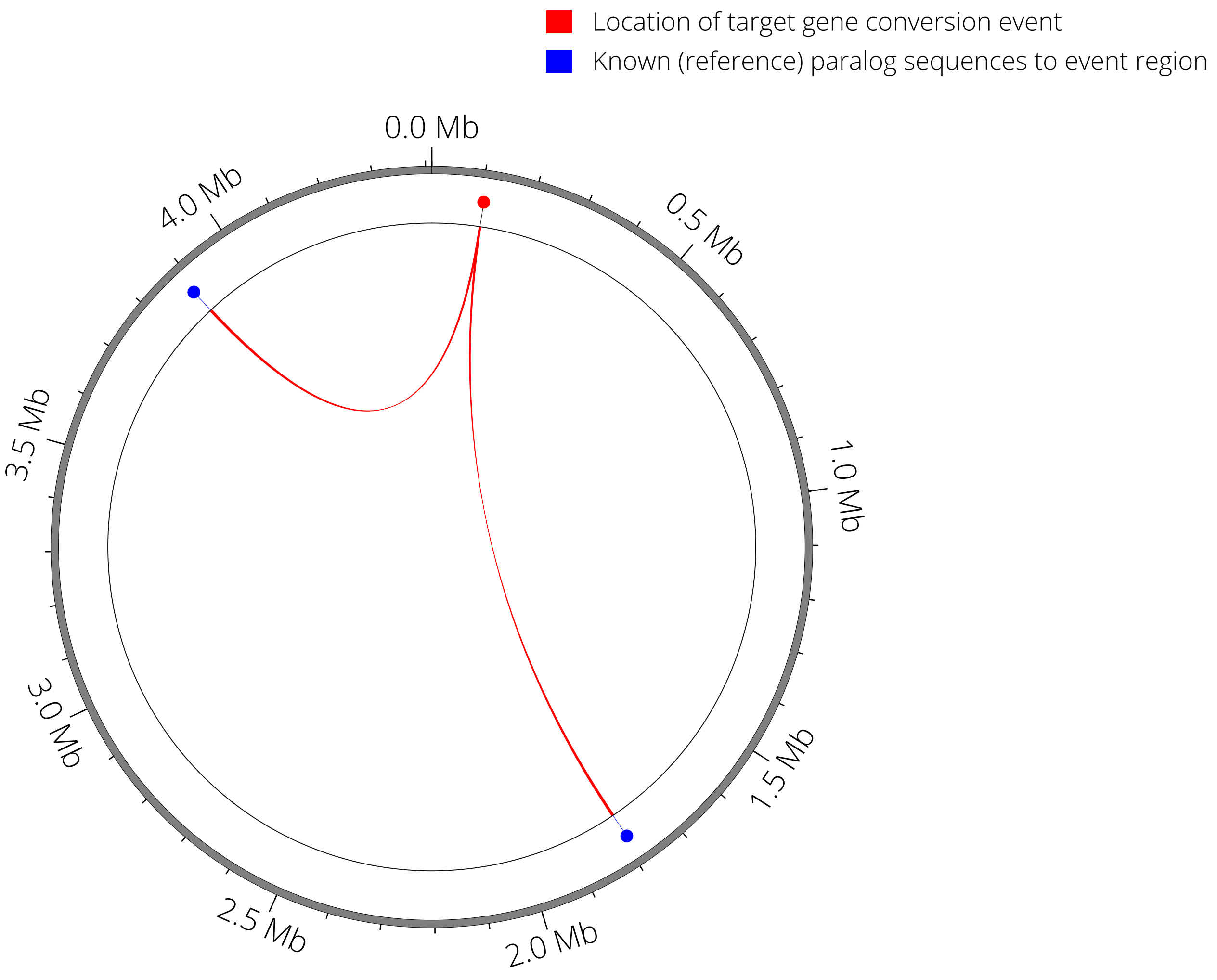

### Event_025 - 4 SNPs - k-mer analysis relative to paralogs

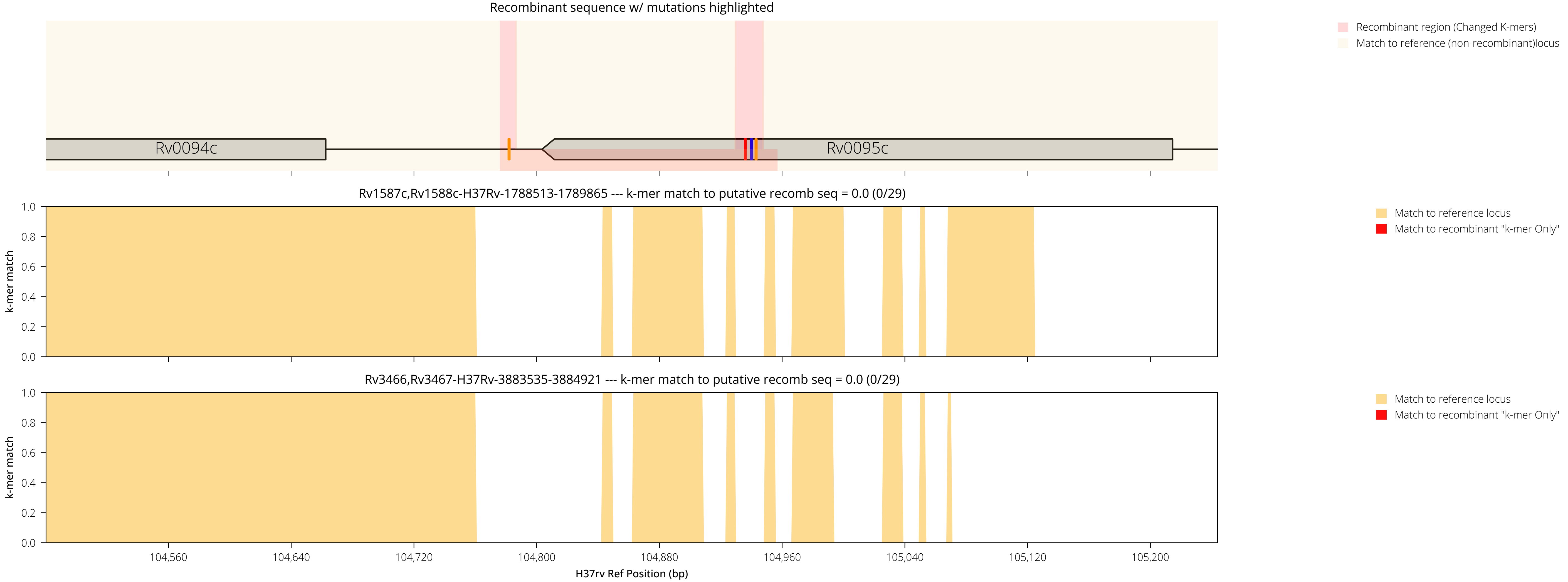

Sub-phylogeny Viz + Paralog Alignments
GCE ID: Event_025

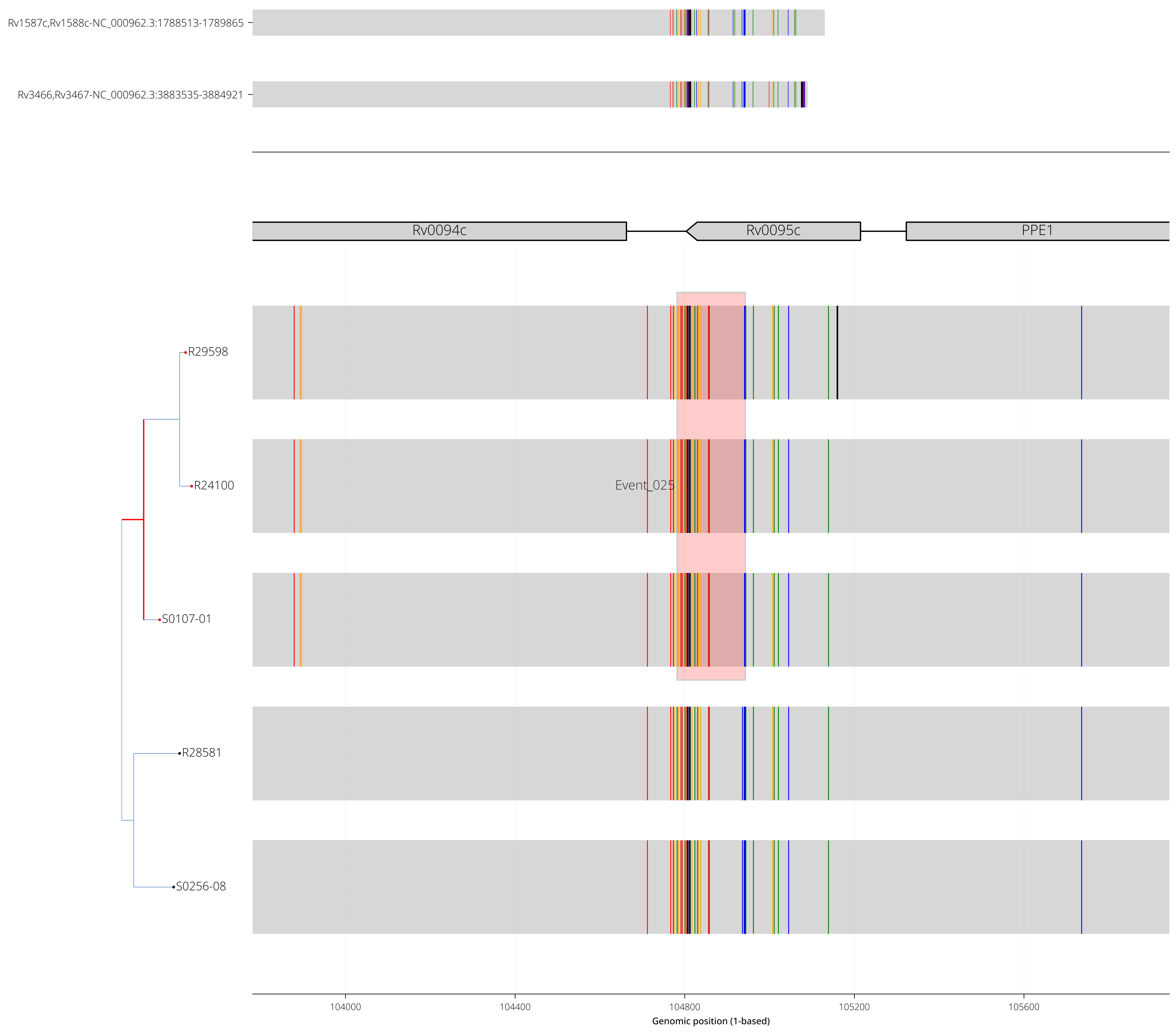

Full Phylogeny Viz + Paralog Alignments
GCE ID: Event_025

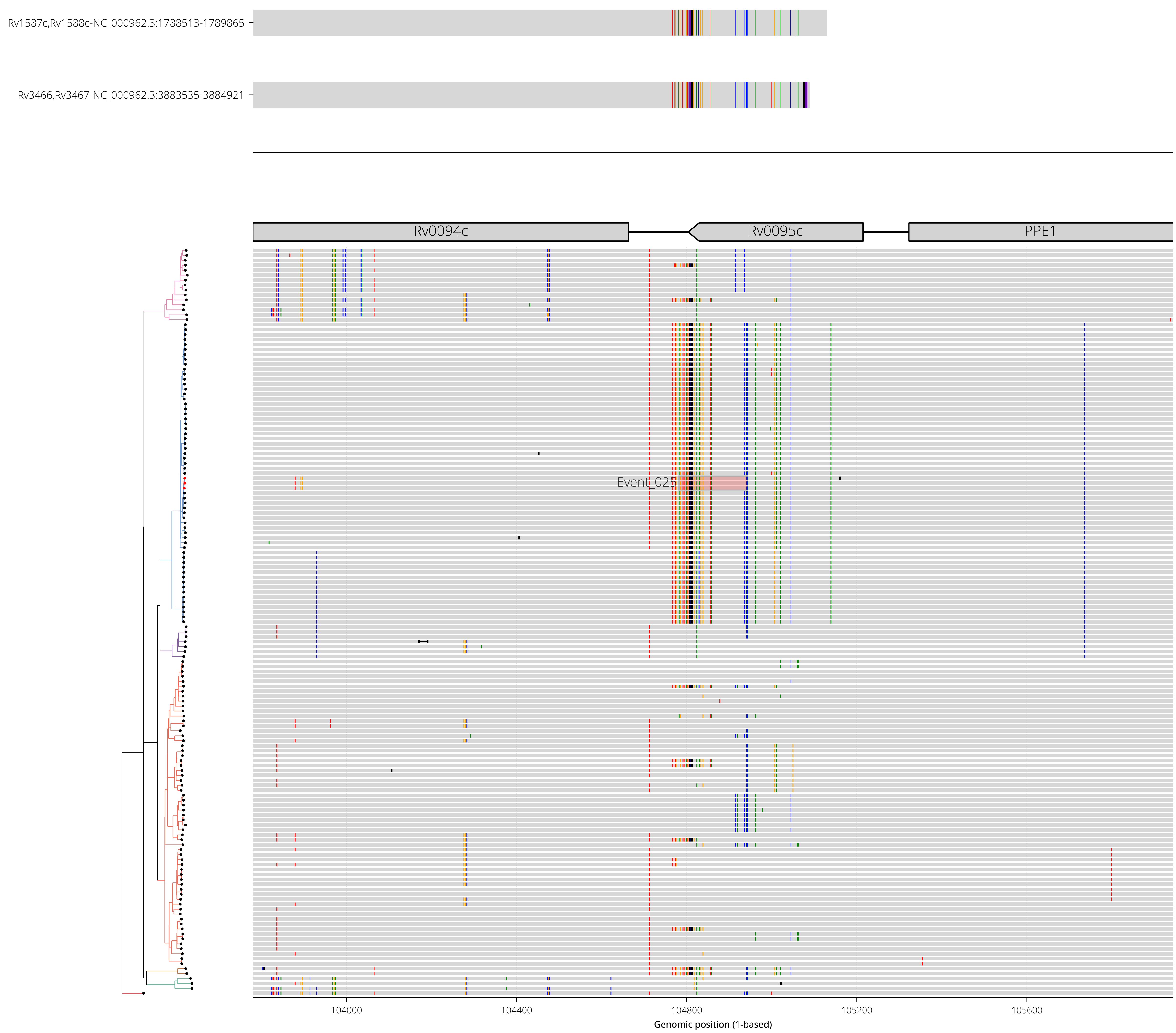

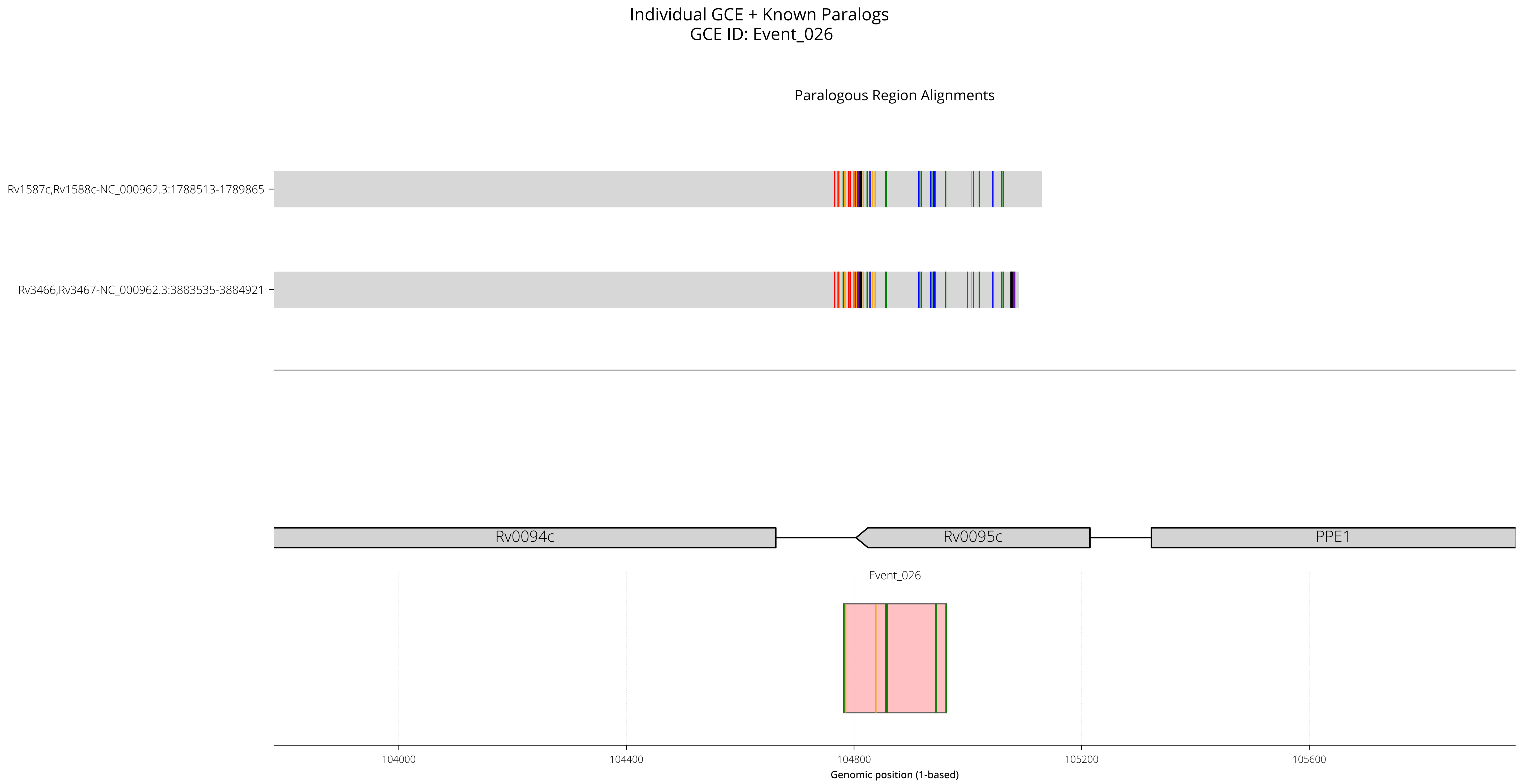

Individual GCE + Known Paralogs
GCE ID: Event_026
Paralogous Region Alignments
Rv1587c,Rv1588c-NC_000962.3:1788513-1789865
Rv3466,Rv3467-NC_000962.3:3883535-3884921
Rv0094c
Rv0095c
PPE1
Event_026
104000
104400
104800
105200
105600
Genomic position (1-based)

Circos Viz - GCE to Known Paralogs Locations
GCE ID: Event_026

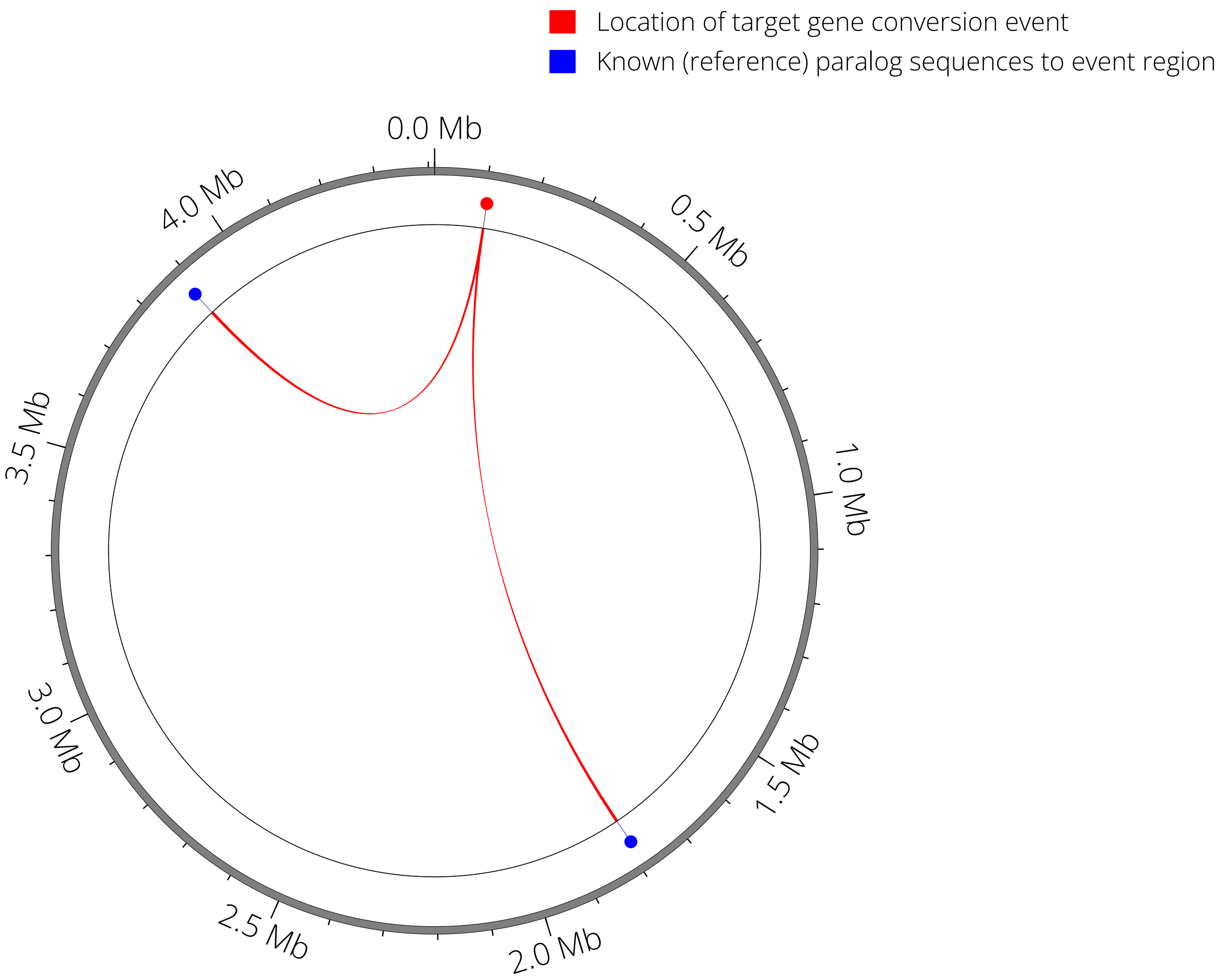

### Event_026 - 7 SNPs - k-mer analysis relative to paralogs

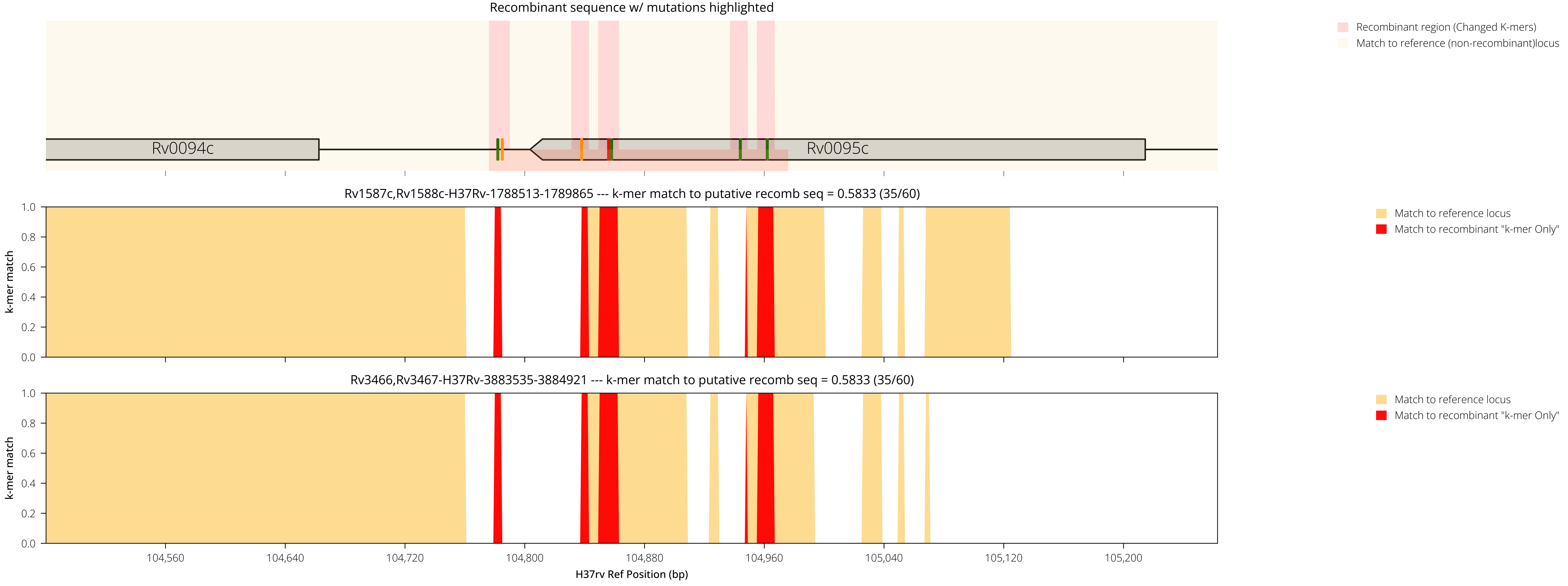

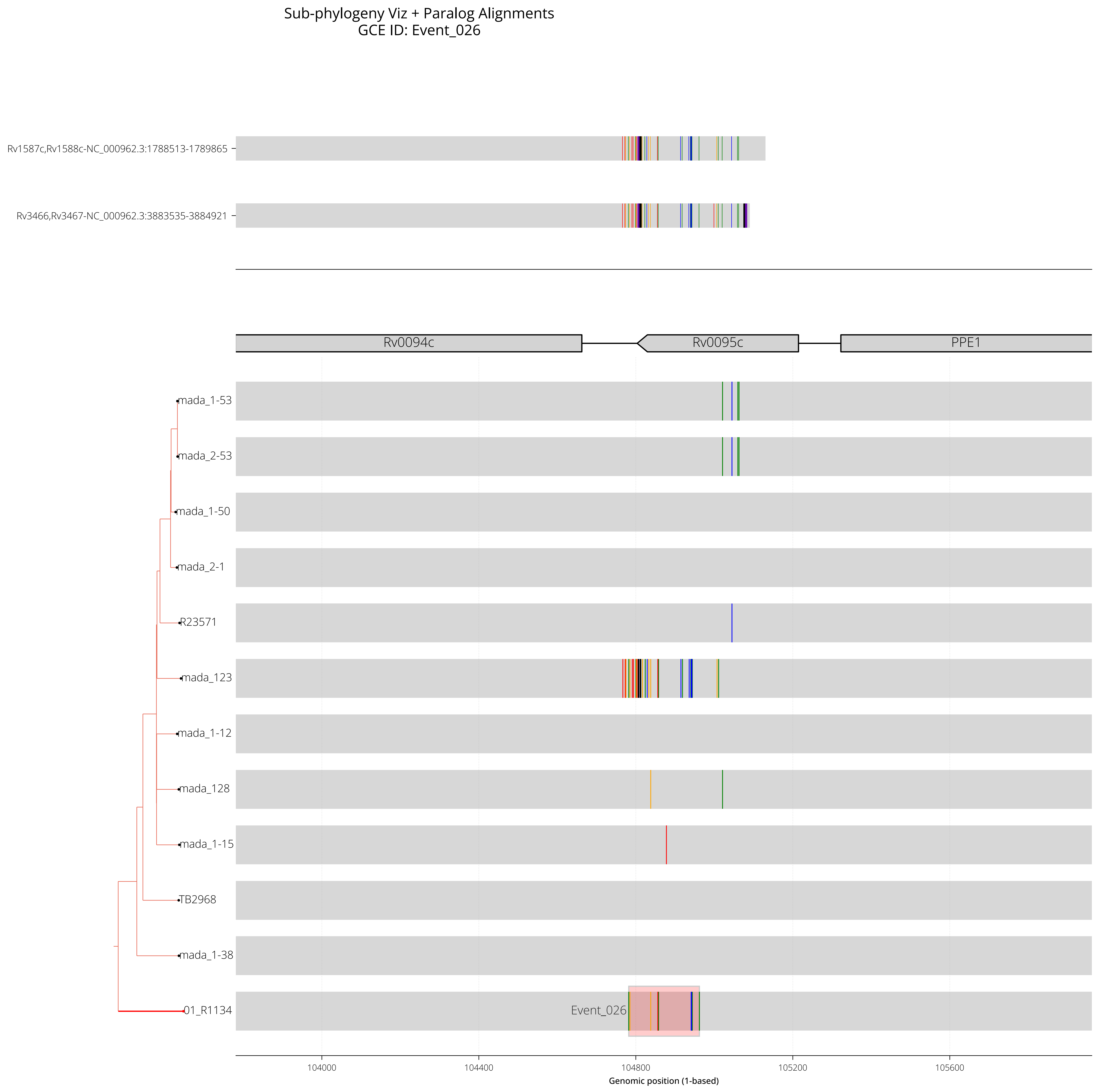
Sub-phylogeny Viz + Paralog Alignments
GCE ID: Event_026
Rv1587c,Rv1588c-NC_000962.3:1788513-1789865
Rv3466,Rv3467-NC_000962.3:3883535-3884921
Rv0094c
Rv0095c
PPE1
mada_1-53
mada_2-53
mada_1-50
mada_2-1
R23571
mada_123
mada_1-12
mada_128
mada_1-15
TB2968
mada_1-38
01_R1134
Event_026
104000
104400
104800
105200
105600
Genomic position (1-based)

Full Phylogeny Viz + Paralog Alignments
GCE ID: Event_026

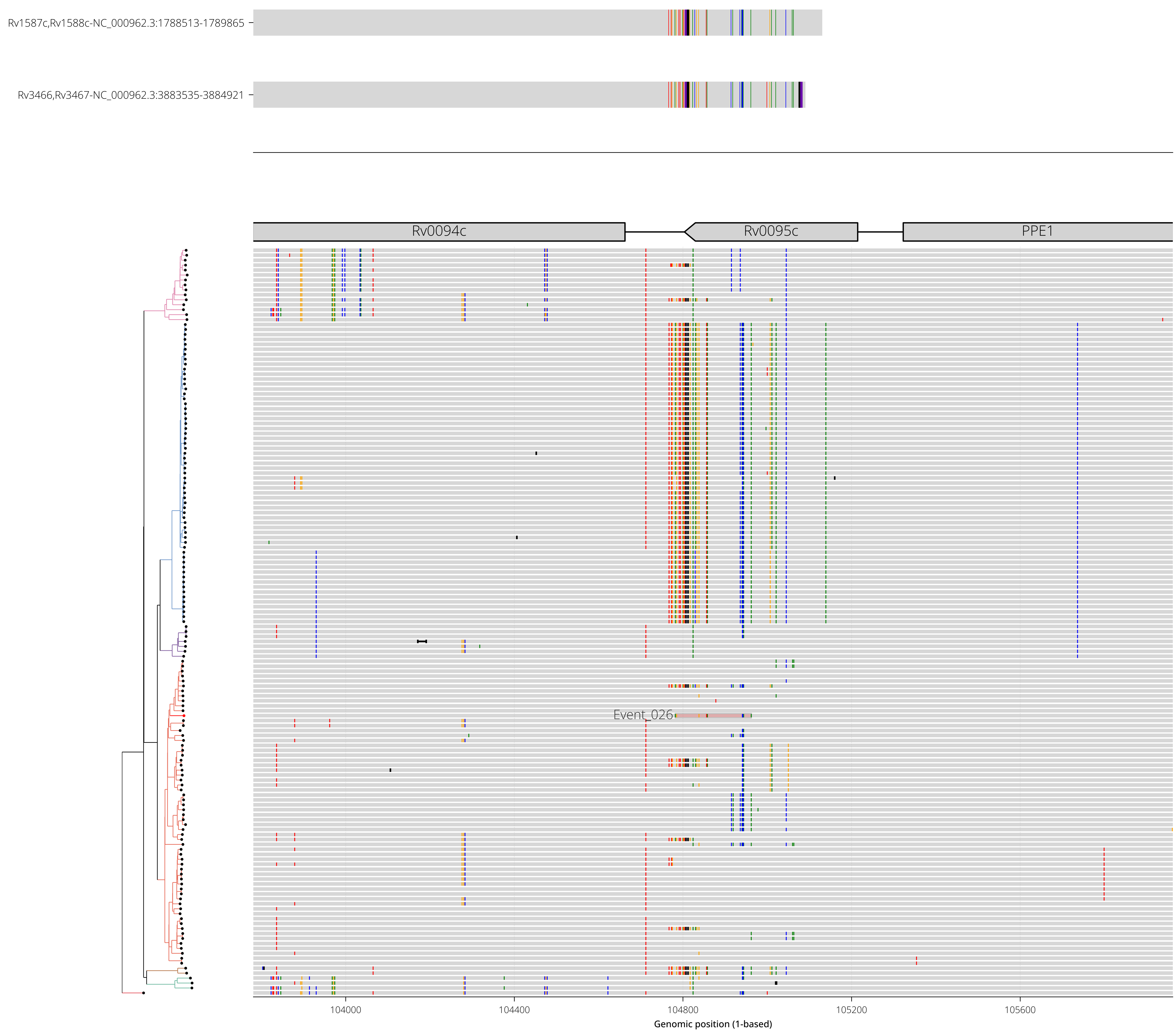

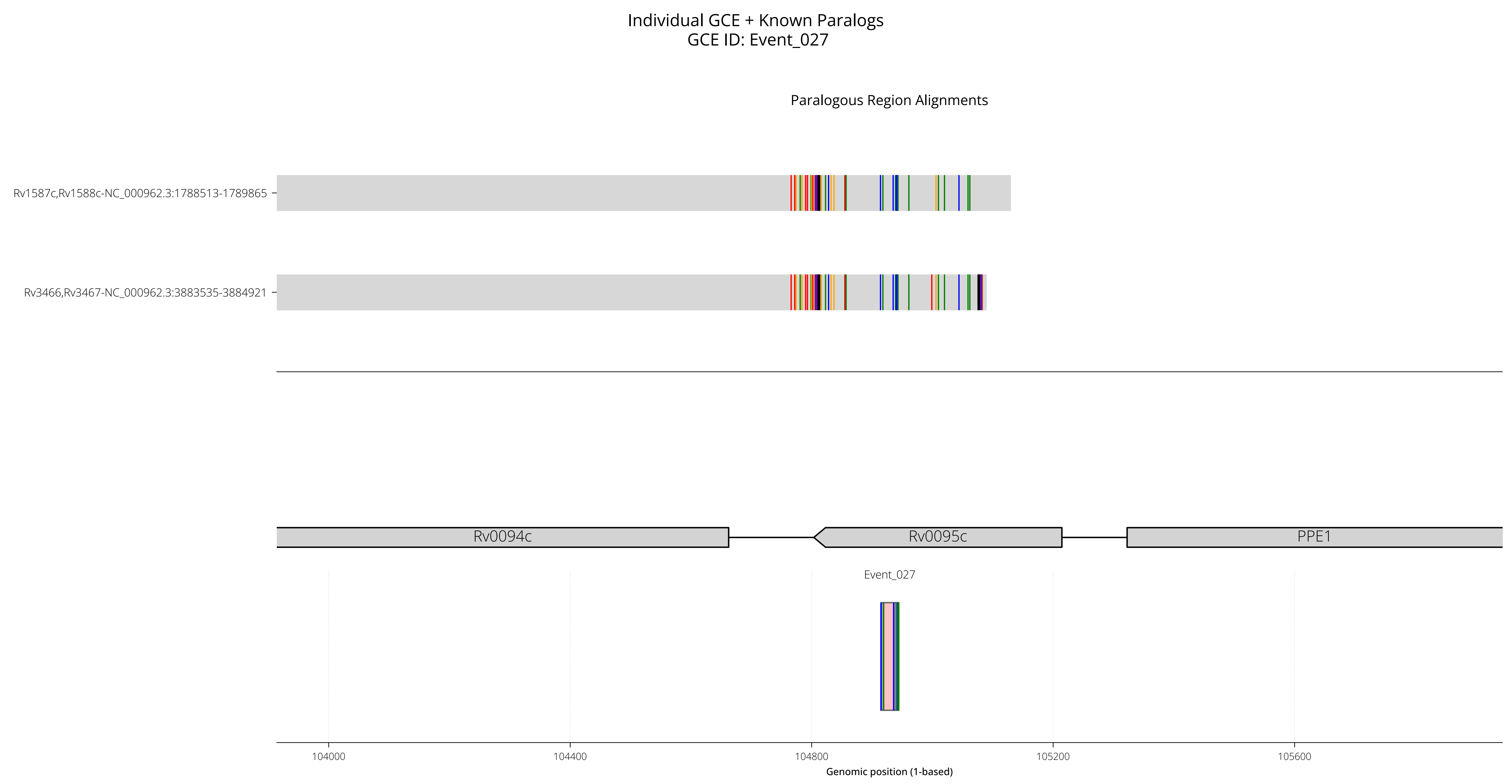

Individual GCE + Known Paralogs
GCE ID: Event_027
Paralogous Region Alignments
Rv1587c,Rv1588c-NC_000962.3:1788513-1789865
Rv3466,Rv3467-NC_000962.3:3883535-3884921
Rv0094c
Rv0095c
PPE1
Event_027
104000
104400
104800
105200
105600
Genomic position (1-based)

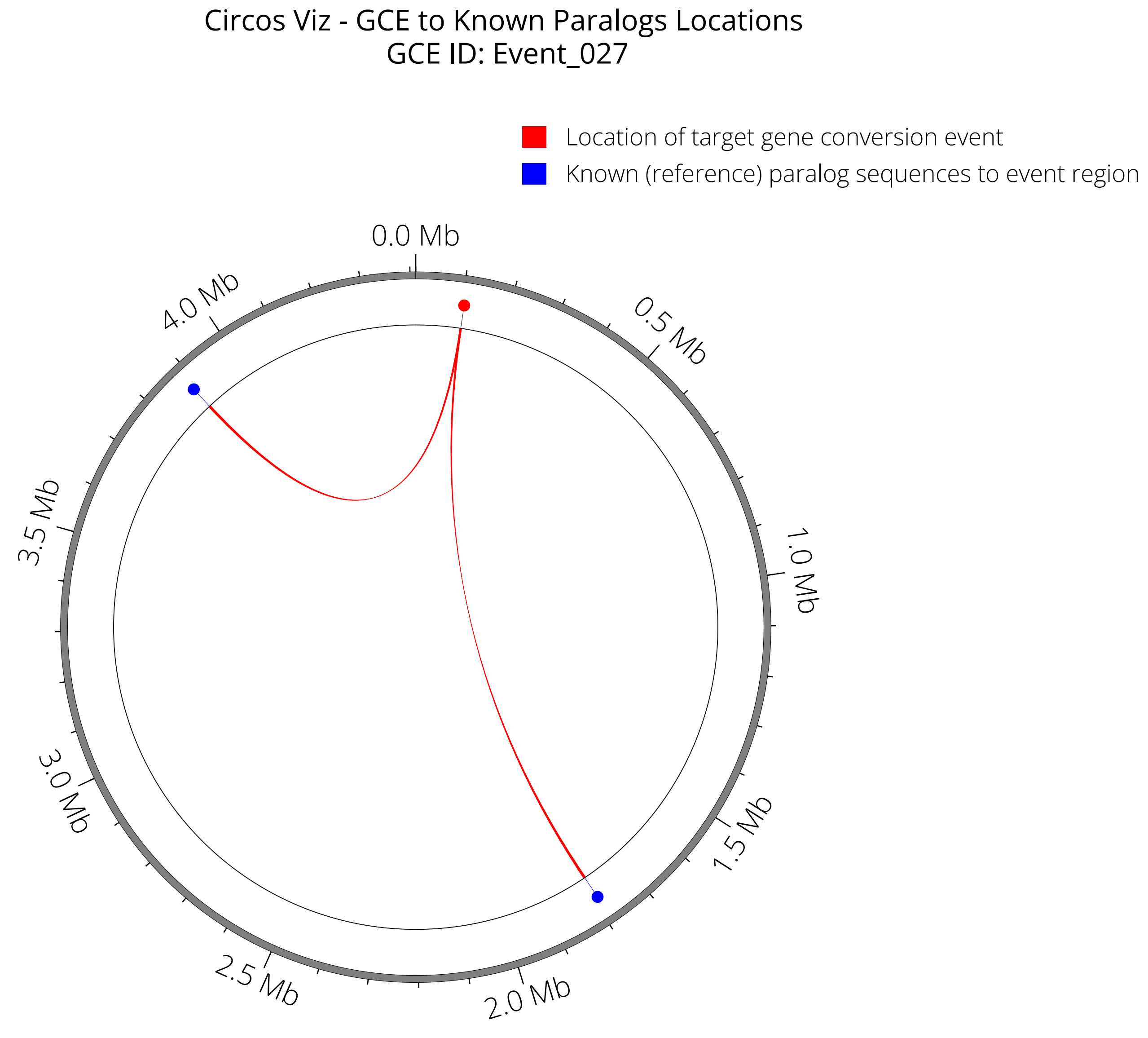
Circos Viz - GCE to Known Paralogs Locations
GCE ID: Event_027
Location of target gene conversion event
Known (reference) paralog sequences to event region
0.0 Mb
0.5 Mb
1.0 Mb
1.5 Mb
2.0 Mb
2.5 Mb
3.0 Mb
3.5 Mb
4.0 Mb

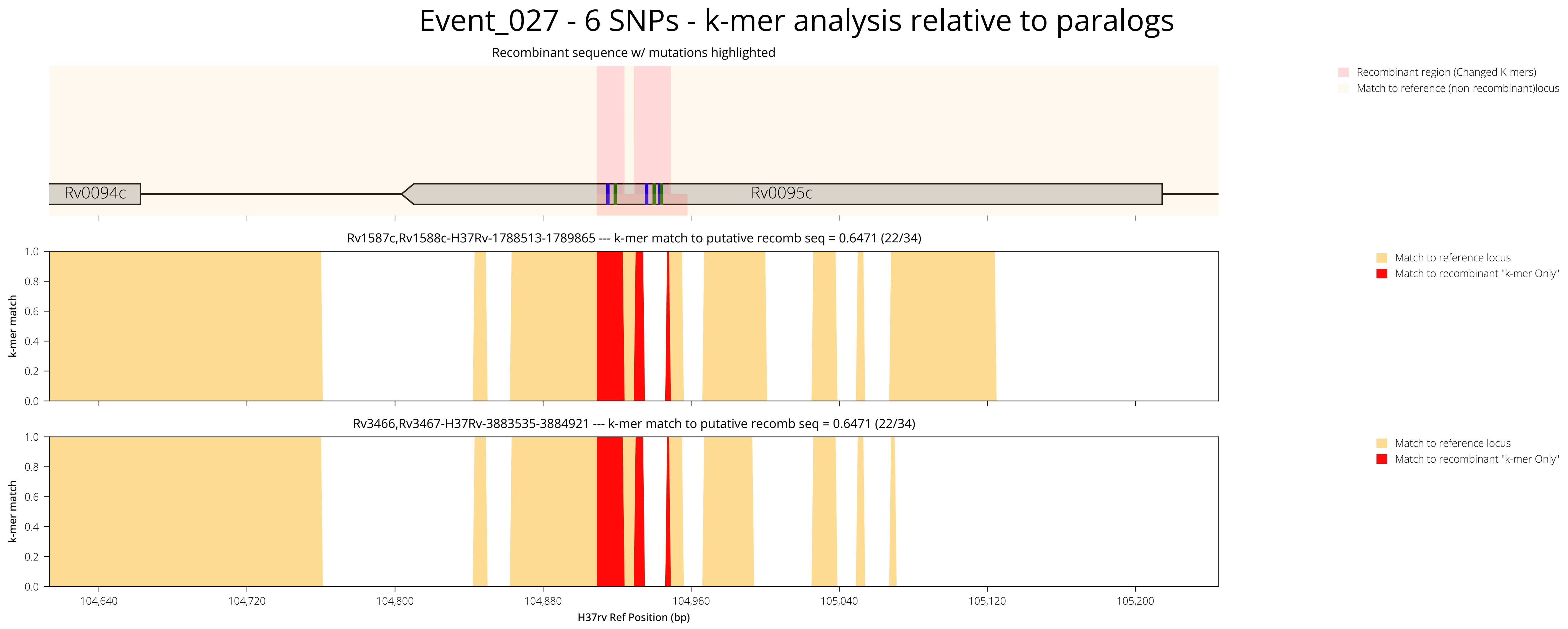

Event_027 - 6 SNPs - k-mer analysis relative to paralogs
Recombinant sequence w/ mutations highlighted
Recombinant region (Changed K-mers)
Match to reference (non-recombinant)locus
Rv0094c
Rv0095c
Rv1587c,Rv1588c-H37Rv-1788513-1789865 --- k-mer match to putative recomb seq = 0.6471 (22/34)
Match to reference locus
Match to recombinant "k-mer Only"
k-mer match
1.0
0.8
0.6
0.4
0.2
0.0
Rv3466,Rv3467-H37Rv-3883535-3884921 --- k-mer match to putative recomb seq = 0.6471 (22/34)
Match to reference locus
Match to recombinant "k-mer Only"
k-mer match
104,640
104,720
104,800
104,880
104,960
105,040
105,120
105,200
H37rv Ref Position (bp)

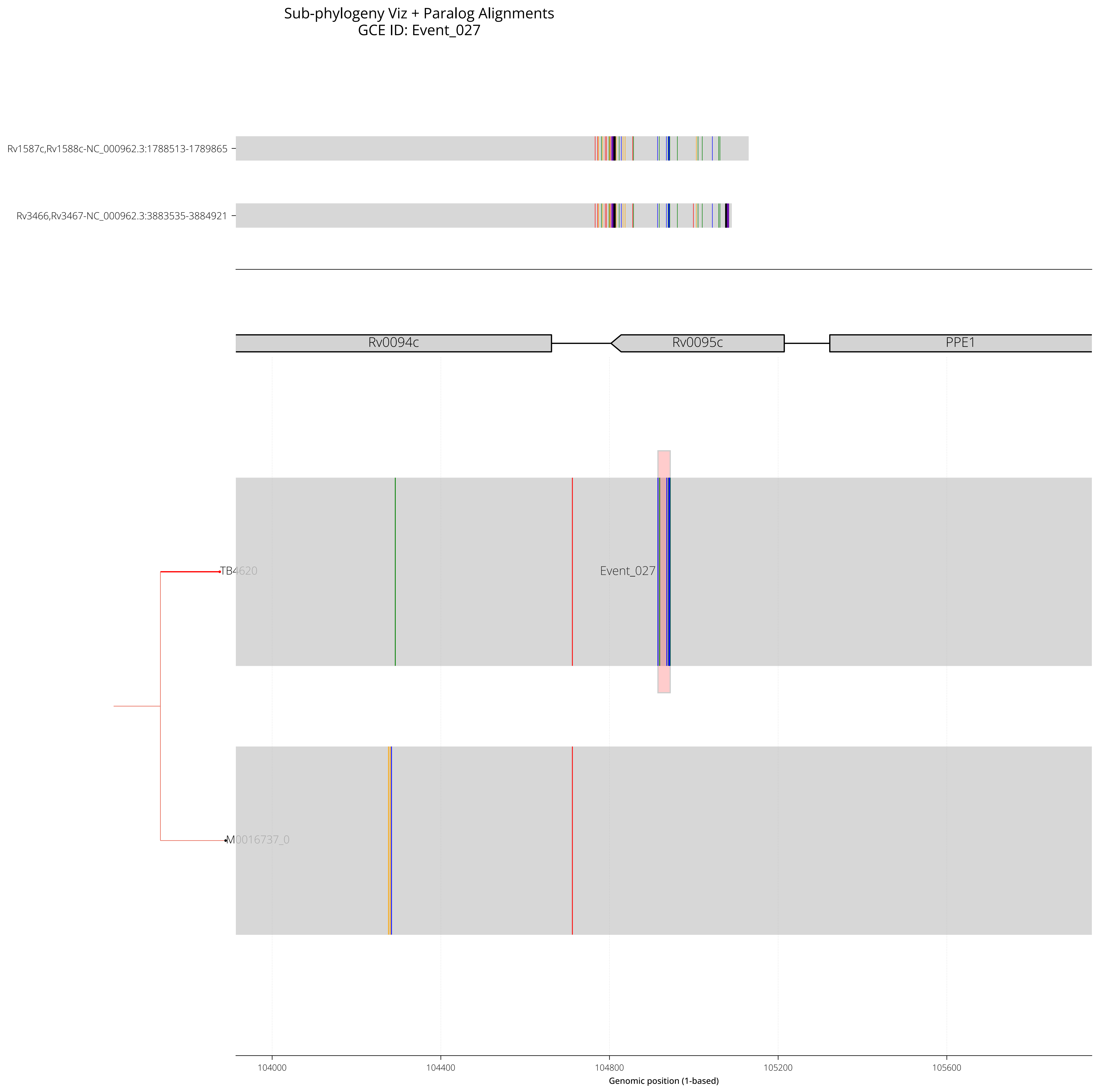

Sub-phylogeny Viz + Paralog Alignments
GCE ID: Event_027
Rv1587c,Rv1588c-NC_000962.3:1788513-1789865
Rv3466,Rv3467-NC_000962.3:3883535-3884921
Rv0094c
Rv0095c
PPE1
TB4620
Event_027
M0016737_0
104000
104400
104800
105200
105600
Genomic position (1-based)

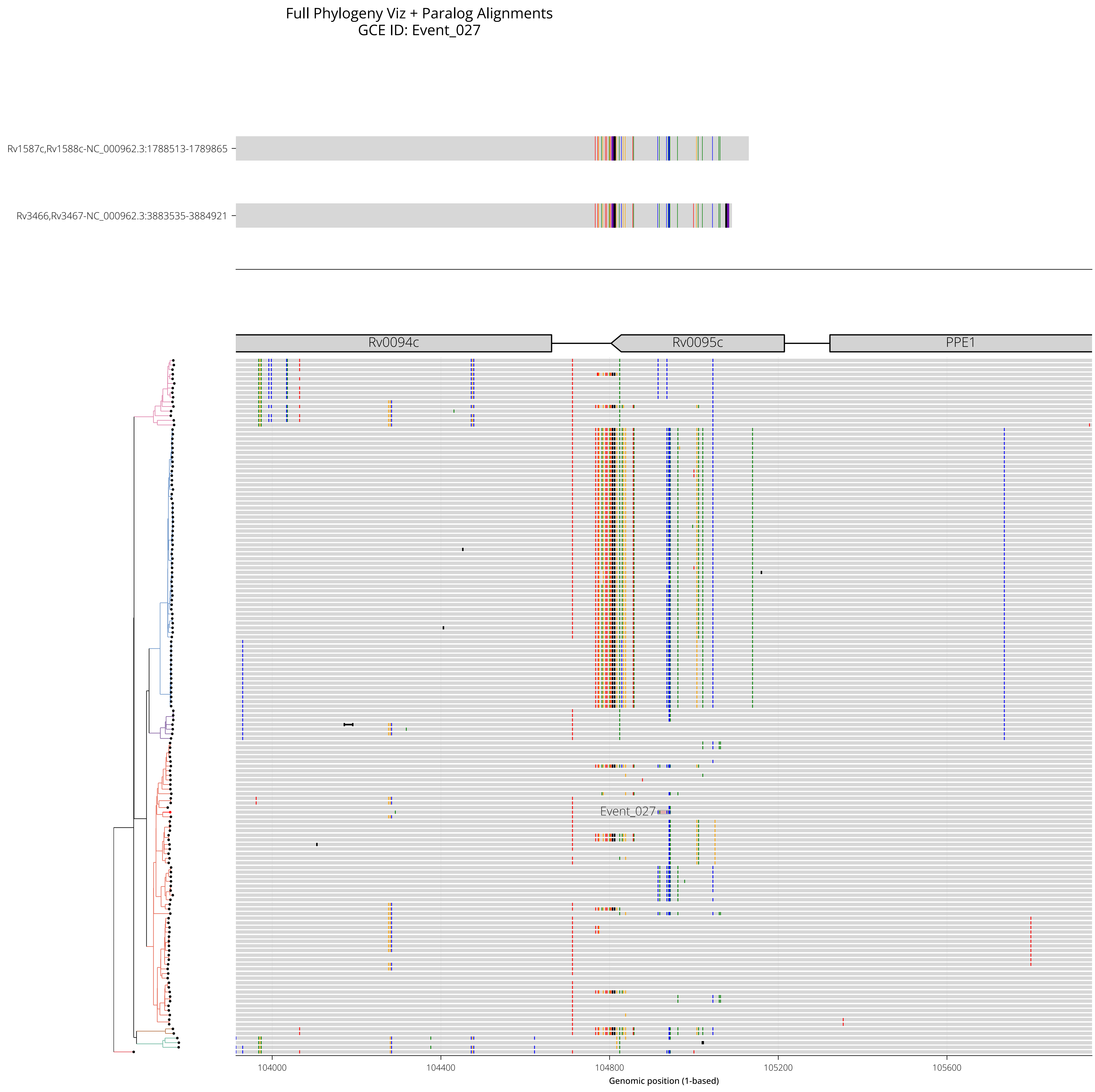

Full Phylogeny Viz + Paralog Alignments
GCE ID: Event_027
Rv1587c,Rv1588c-NC_000962.3:1788513-1789865
Rv3466,Rv3467-NC_000962.3:3883535-3884921
Rv0094c
Rv0095c
PPE1
Event_027
104000
104400
104800
105200
105600
Genomic position (1-based)

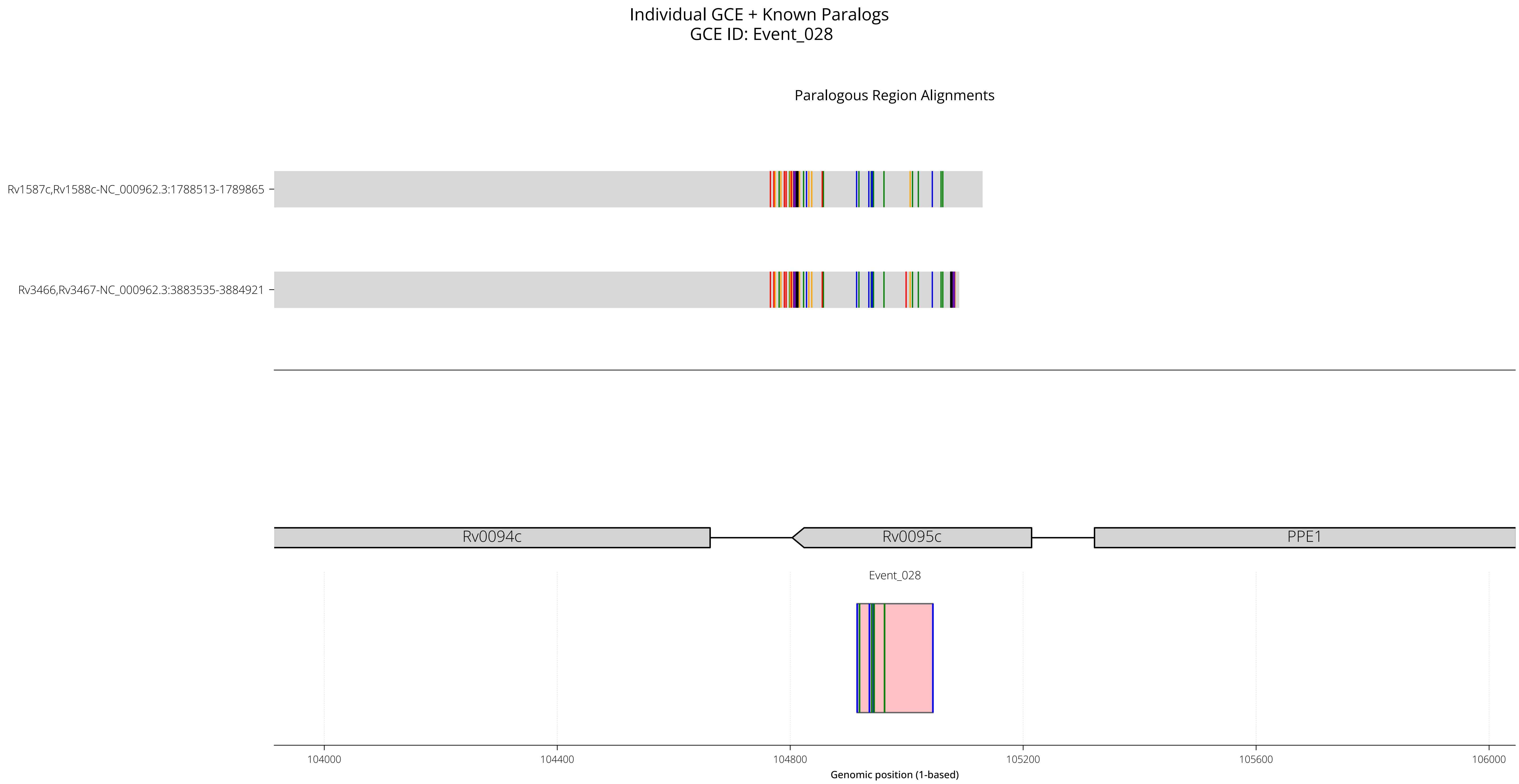

Individual GCE + Known Paralogs
GCE ID: Event_028
Paralogous Region Alignments
Rv1587c,Rv1588c-NC_000962.3:1788513-1789865
Rv3466,Rv3467-NC_000962.3:3883535-3884921
Rv0094c
Rv0095c
PPE1
Event_028
104000
104400
104800
105200
105600
106000
Genomic position (1-based)

Circos Viz - GCE to Known Paralogs Locations
GCE ID: Event_028

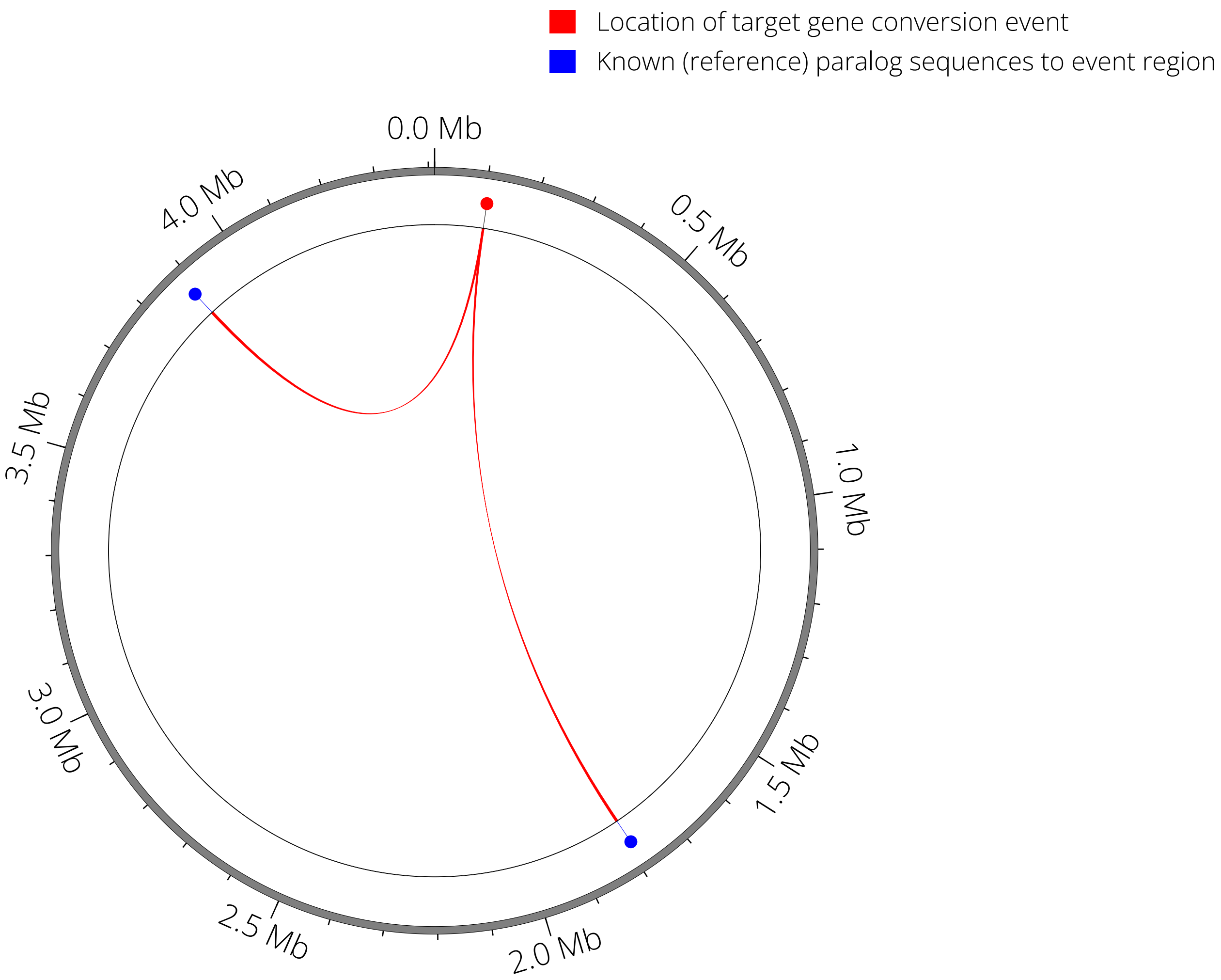

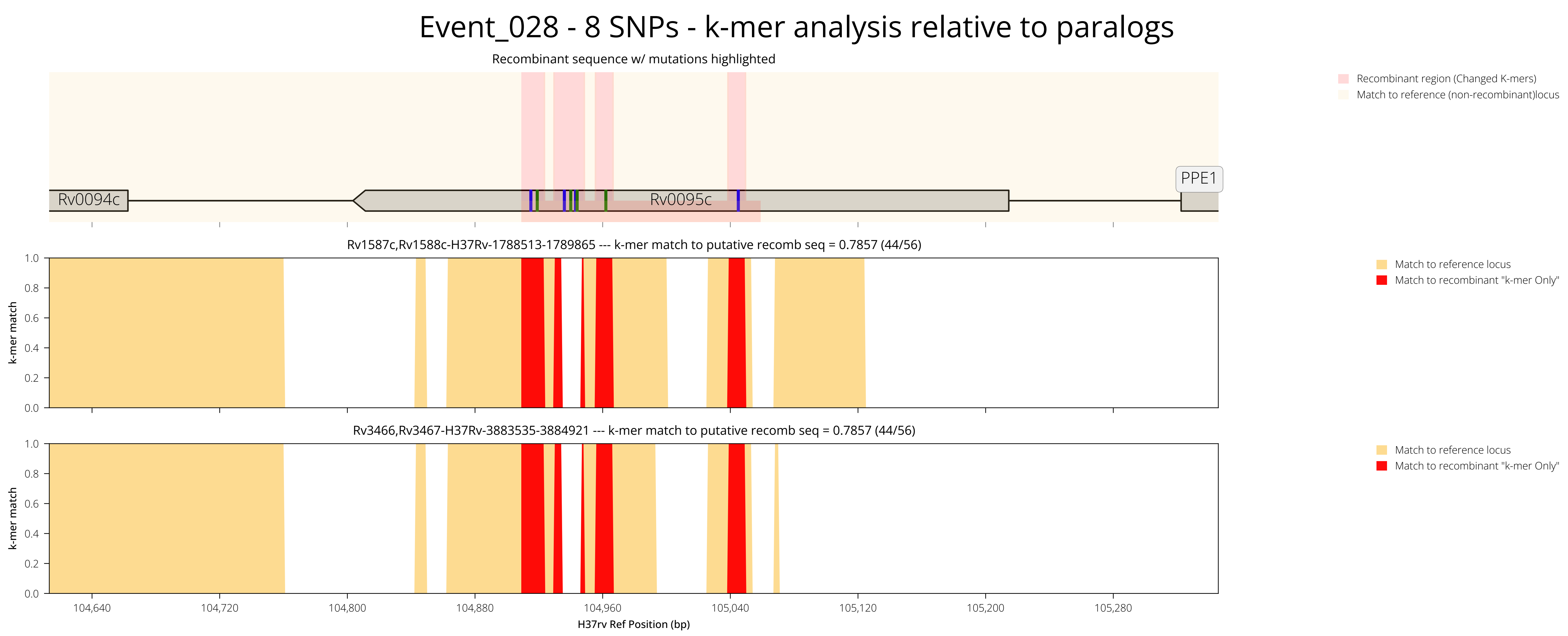

Event_028 - 8 SNPs - k-mer analysis relative to paralogs
Recombinant sequence w/ mutations highlighted
Recombinant region (Changed K-mers)
Match to reference (non-recombinant)locus
Rv0094c
Rv0095c
PPE1
Rv1587c,Rv1588c-H37Rv-1788513-1789865 --- k-mer match to putative recomb seq = 0.7857 (44/56)
Match to reference locus
Match to recombinant "k-mer Only"
k-mer match
Rv3466,Rv3467-H37Rv-3883535-3884921 --- k-mer match to putative recomb seq = 0.7857 (44/56)
Match to reference locus
Match to recombinant "k-mer Only"
k-mer match
104,640
104,720
104,800
104,880
104,960
105,040
105,120
105,200
105,280
H37rv Ref Position (bp)

Sub-phylogeny Viz + Paralog Alignments
GCE ID: Event_028

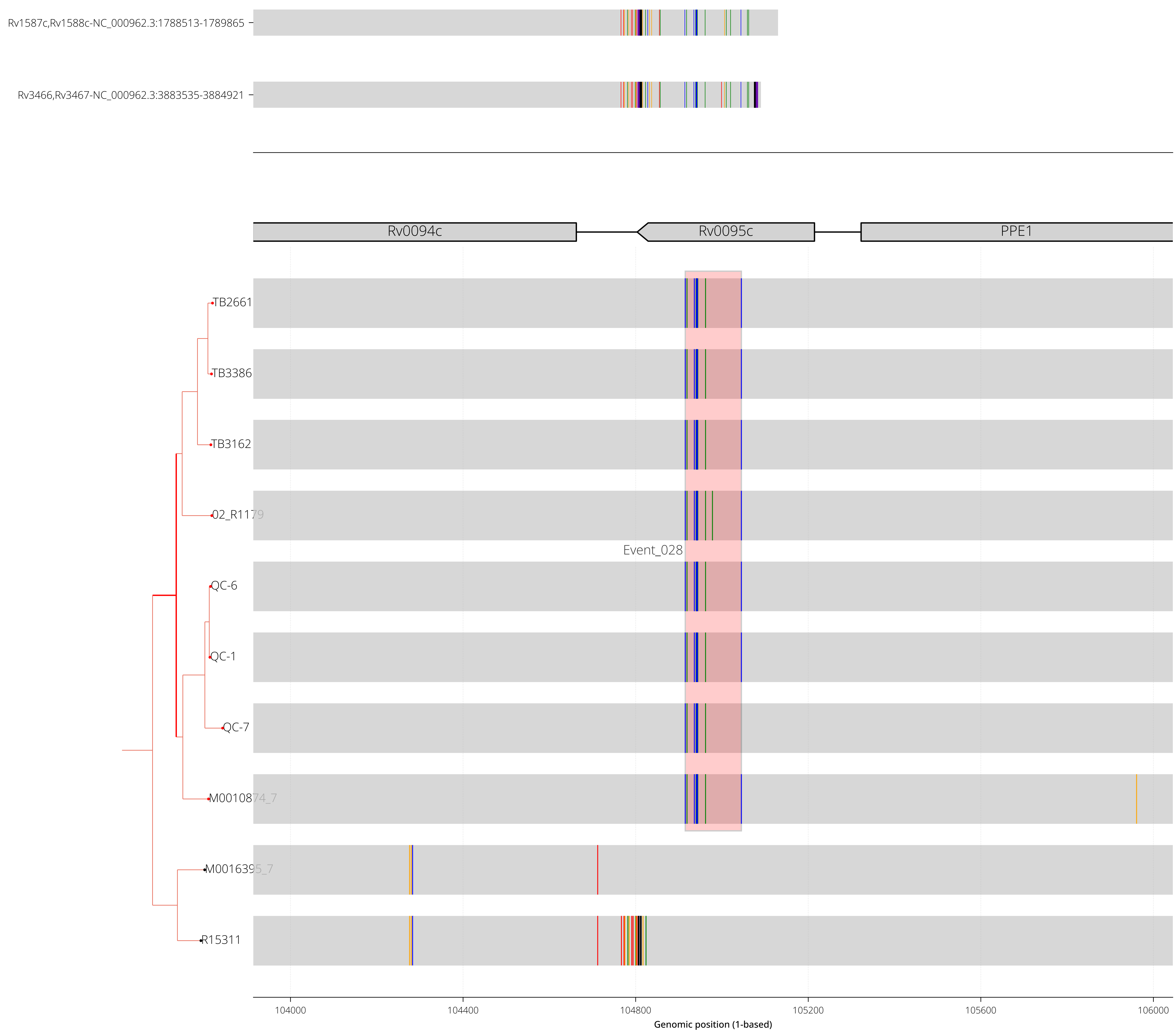

### Full Phylogeny Viz + Paralog Alignments
GCE ID: Event_028

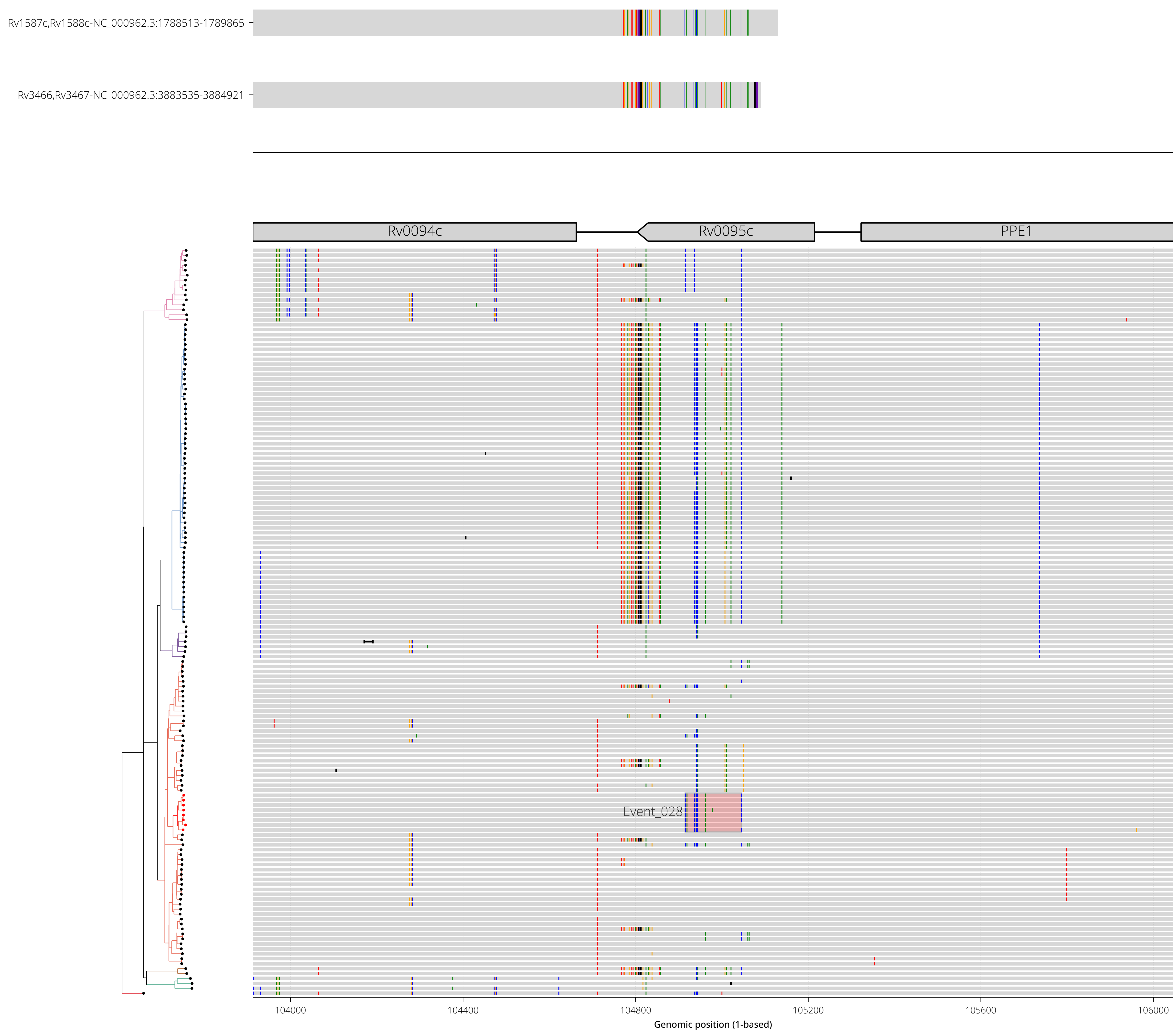

Individual GCE + Known Paralogs
GCE ID: Event_029

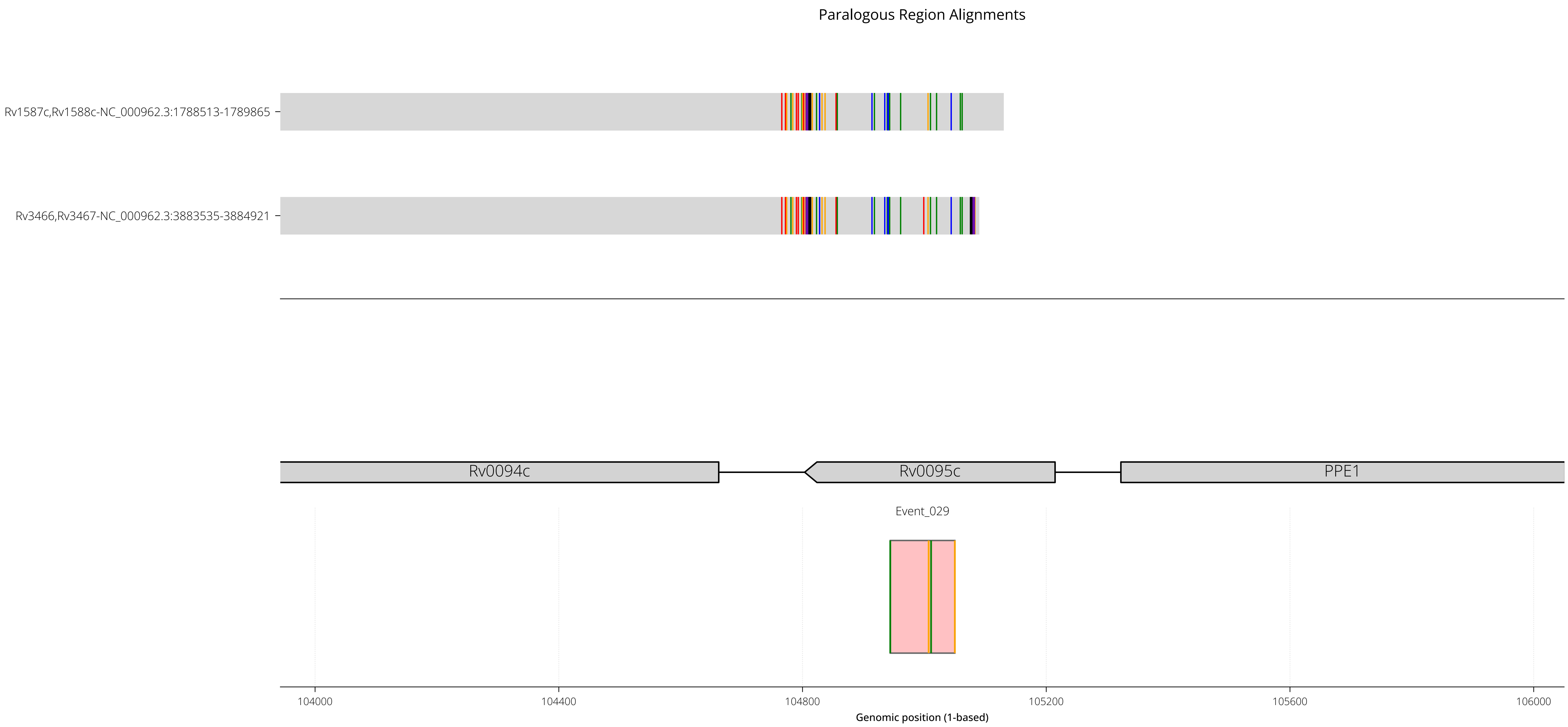

Circos Viz - GCE to Known Paralogs Locations
GCE ID: Event_029

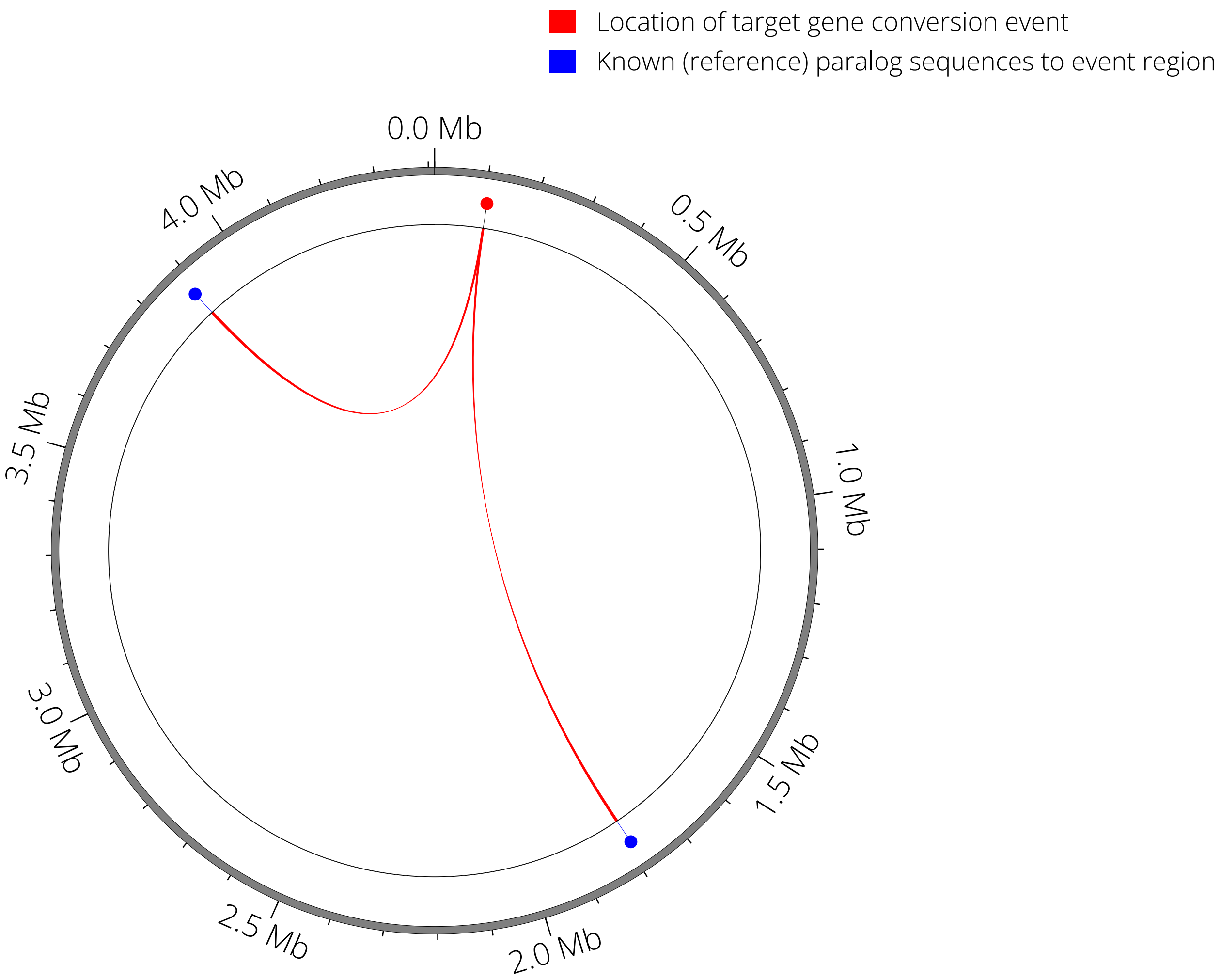

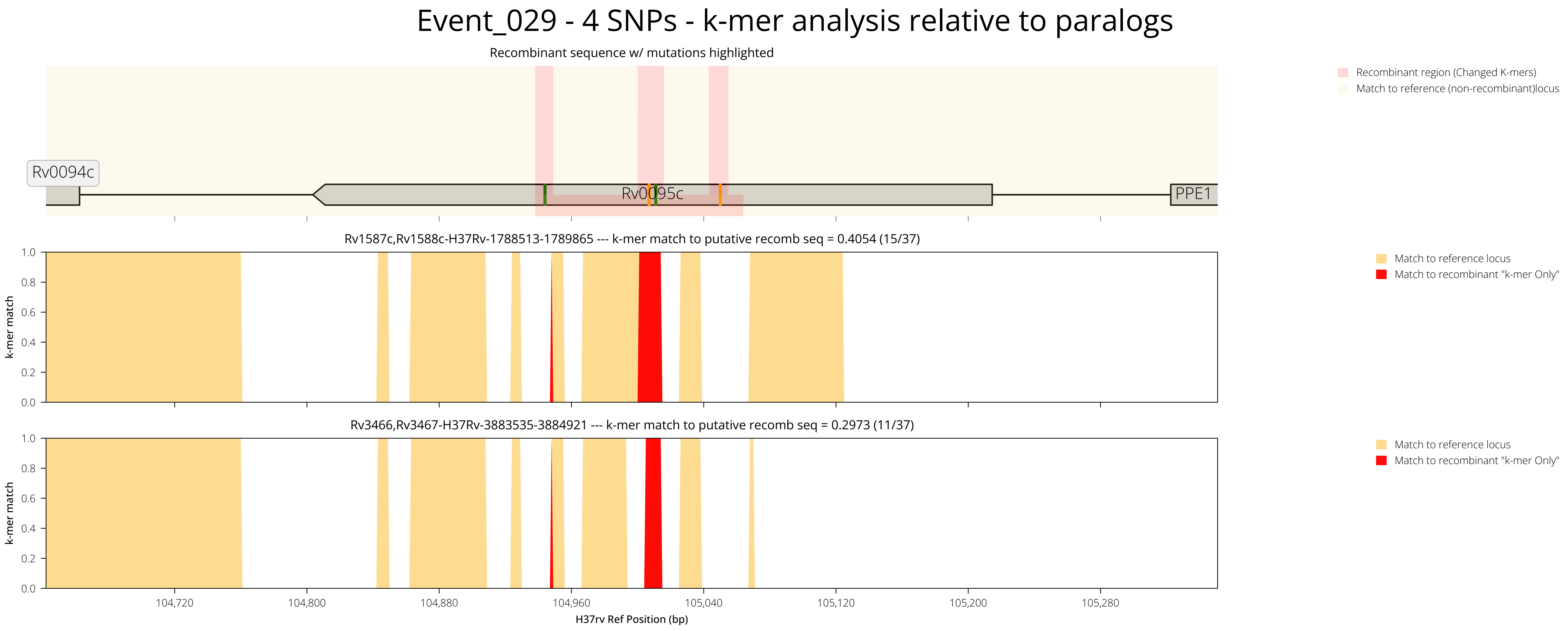
Event_029 - 4 SNPs - k-mer analysis relative to paralogs
Recombinant sequence w/ mutations highlighted
Recombinant region (Changed K-mers)
Match to reference (non-recombinant)locus
Rv0094c
Rv0095c
PPE1
Rv1587c,Rv1588c-H37Rv-1788513-1789865 --- k-mer match to putative recomb seq = 0.4054 (15/37)
Match to reference locus
Match to recombinant "k-mer Only"
Rv3466,Rv3467-H37Rv-3883535-3884921 --- k-mer match to putative recomb seq = 0.2973 (11/37)
Match to reference locus
Match to recombinant "k-mer Only"
k-mer match
1.0
0.8
0.6
0.4
0.2
0.0
104,720
104,800
104,880
104,960
105,040
105,120
105,200
105,280
H37rv Ref Position (bp)

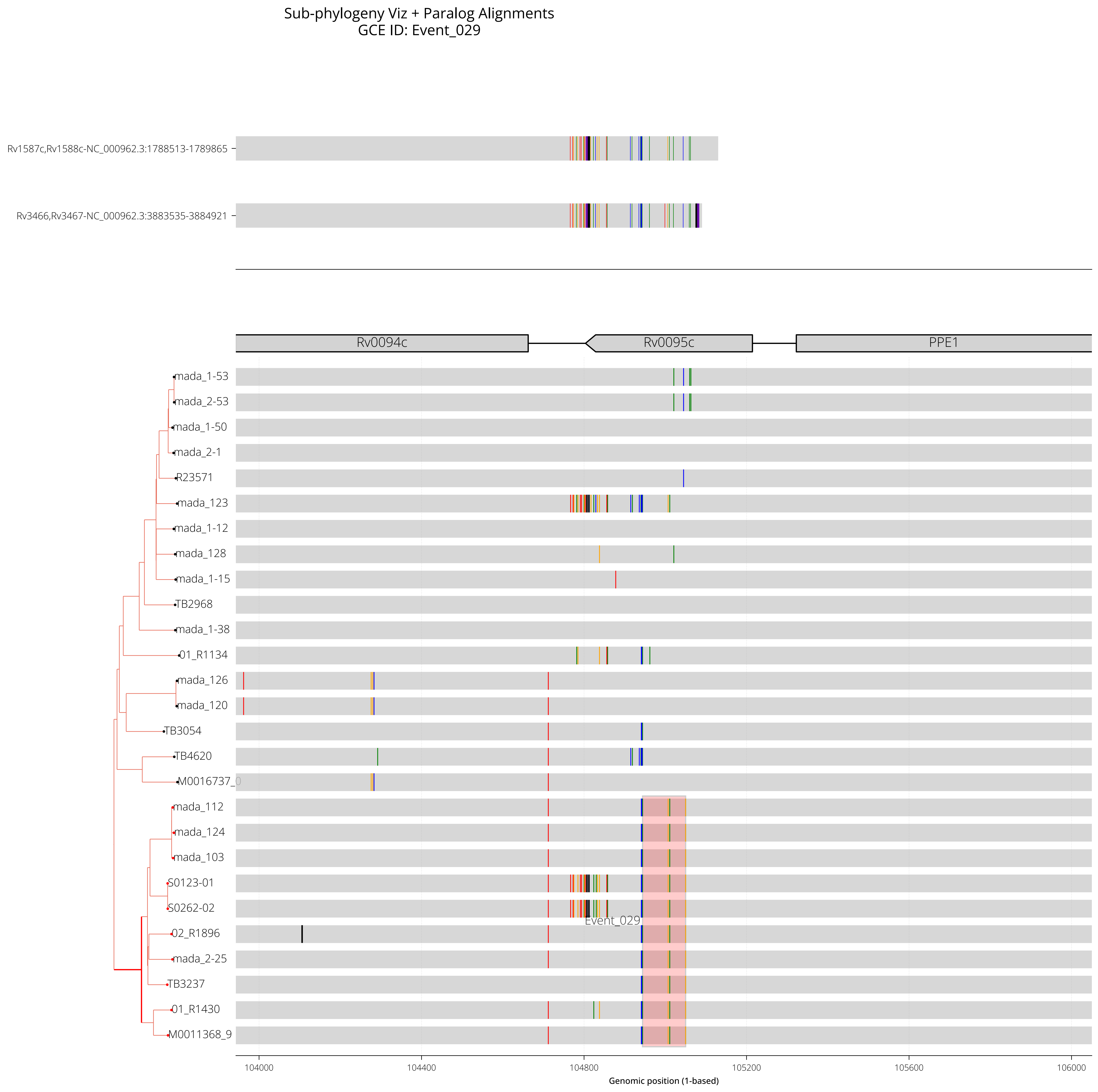

Sub-phylogeny Viz + Paralog Alignments
GCE ID: Event_029
Rv1587c,Rv1588c-NC_000962.3:1788513-1789865
Rv3466,Rv3467-NC_000962.3:3883535-3884921
Rv0094c
Rv0095c
PPE1
mada_1-53
mada_2-53
mada_1-50
mada_2-1
R23571
mada_123
mada_1-12
mada_128
mada_1-15
TB2968
mada_1-38
01_R1134
mada_126
mada_120
TB3054
TB4620
M0016737_0
mada_112
mada_124
mada_103
S0123-01
S0262-02
02_R1896
mada_2-25
TB3237
01_R1430
M0011368_9
Event_029
104000
104400
104800
105200
105600
106000
Genomic position (1-based)

Full Phylogeny Viz + Paralog Alignments
GCE ID: Event_029

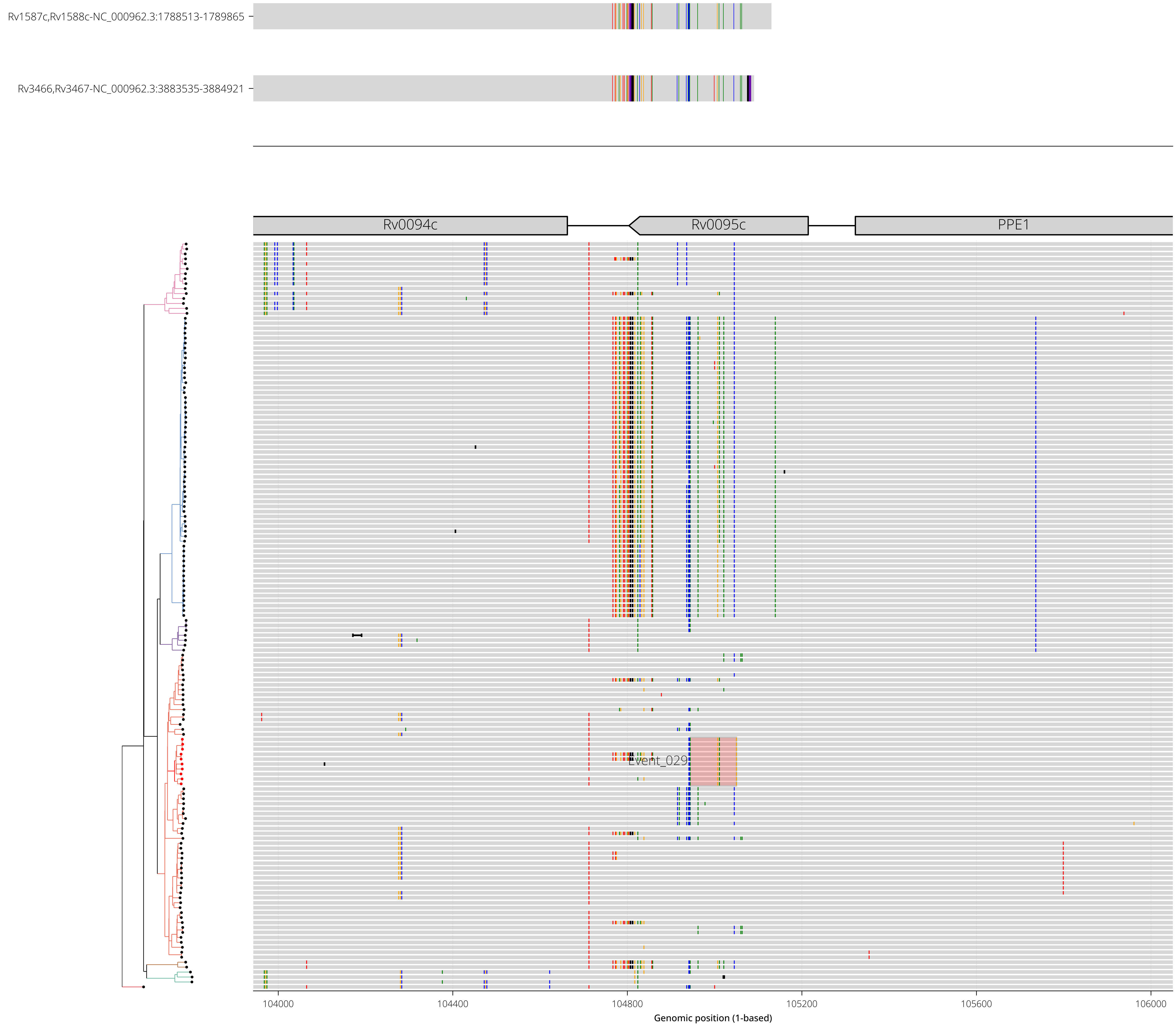

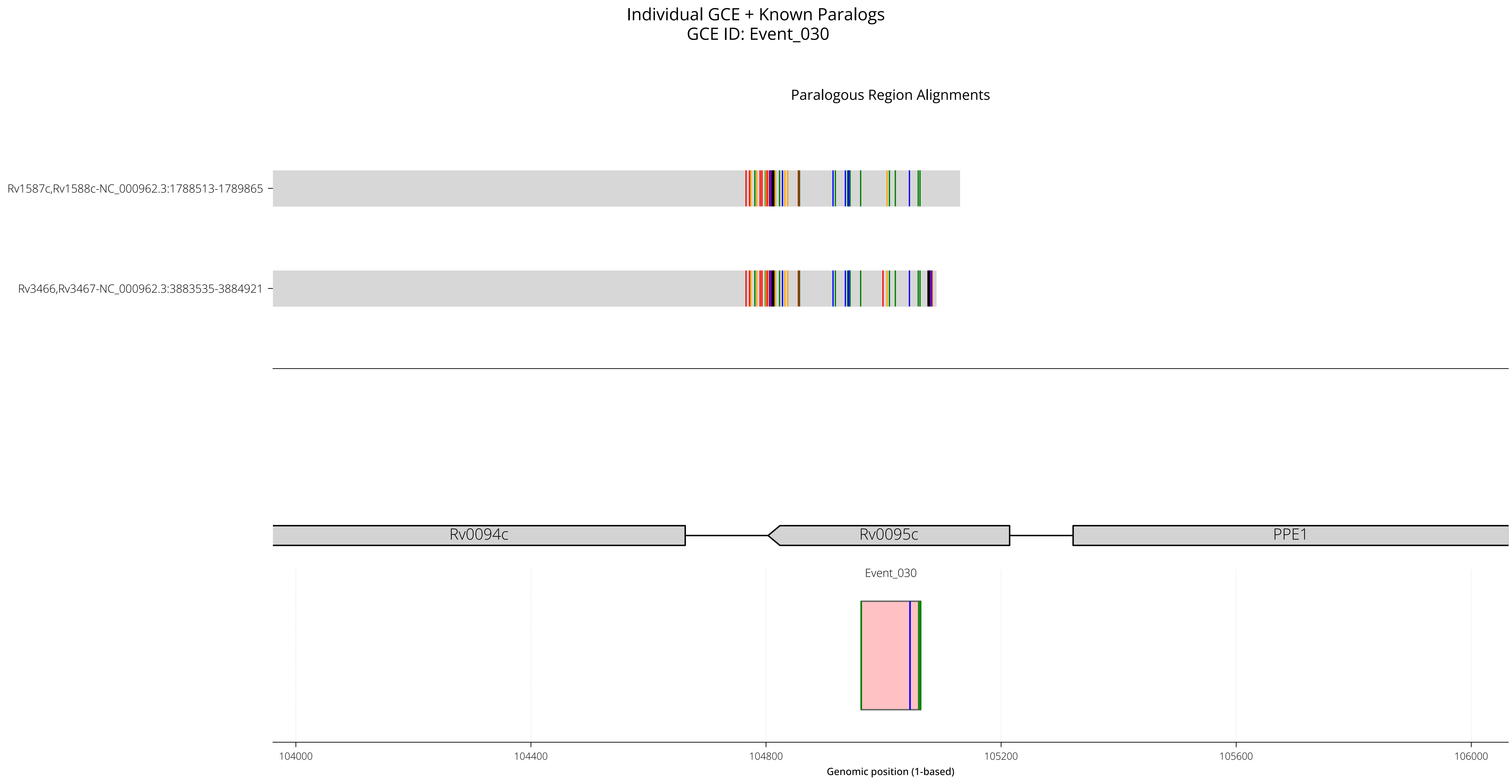
Individual GCE + Known Paralogs
GCE ID: Event_030
Paralogous Region Alignments
Rv1587c,Rv1588c-NC_000962.3:1788513-1789865
Rv3466,Rv3467-NC_000962.3:3883535-3884921
Rv0094c
Rv0095c
PPE1
Event_030
104000
104400
104800
105200
105600
106000
Genomic position (1-based)

Circos Viz - GCE to Known Paralogs Locations
GCE ID: Event_030

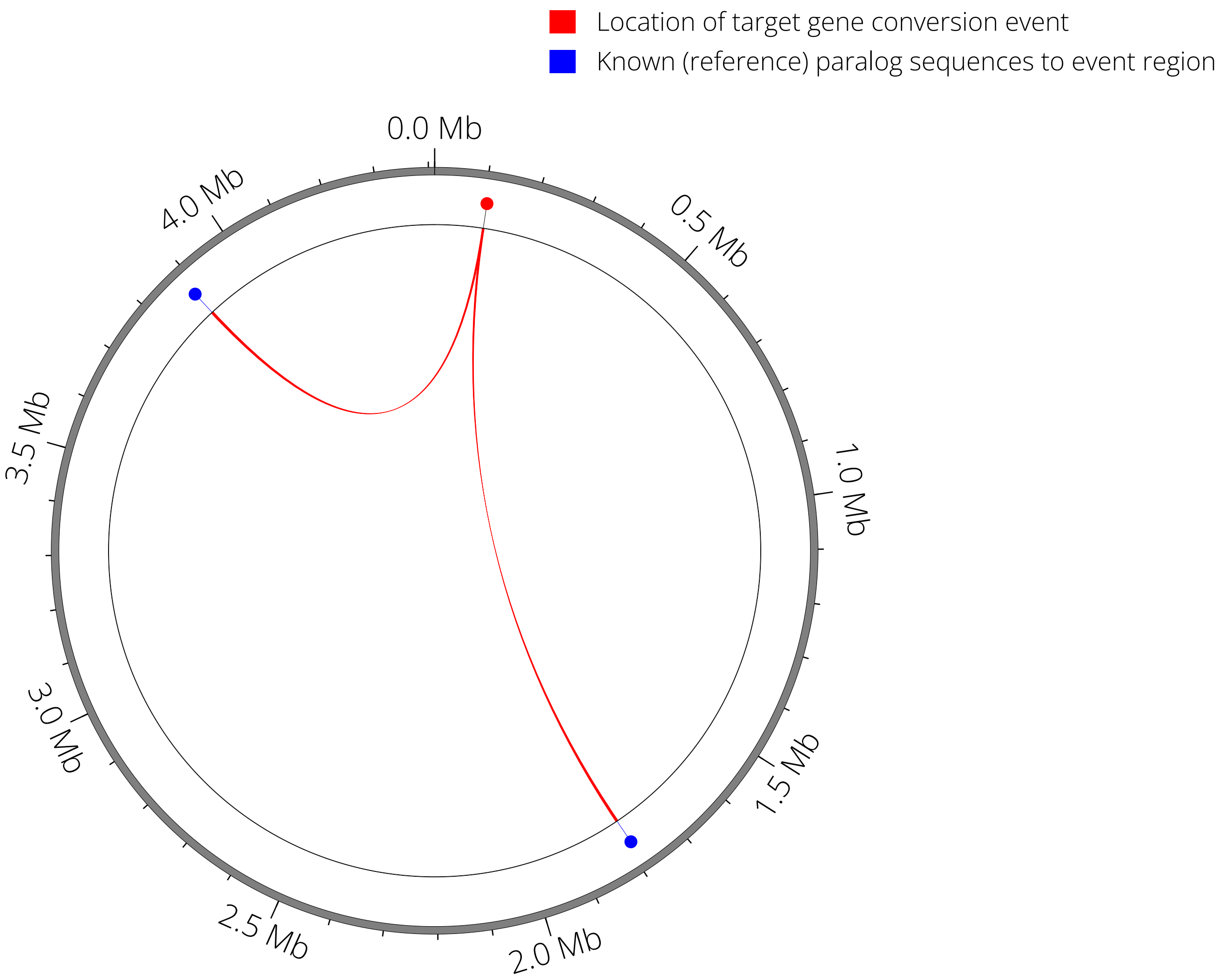

### Event_030 - 4 SNPs - k-mer analysis relative to paralogs

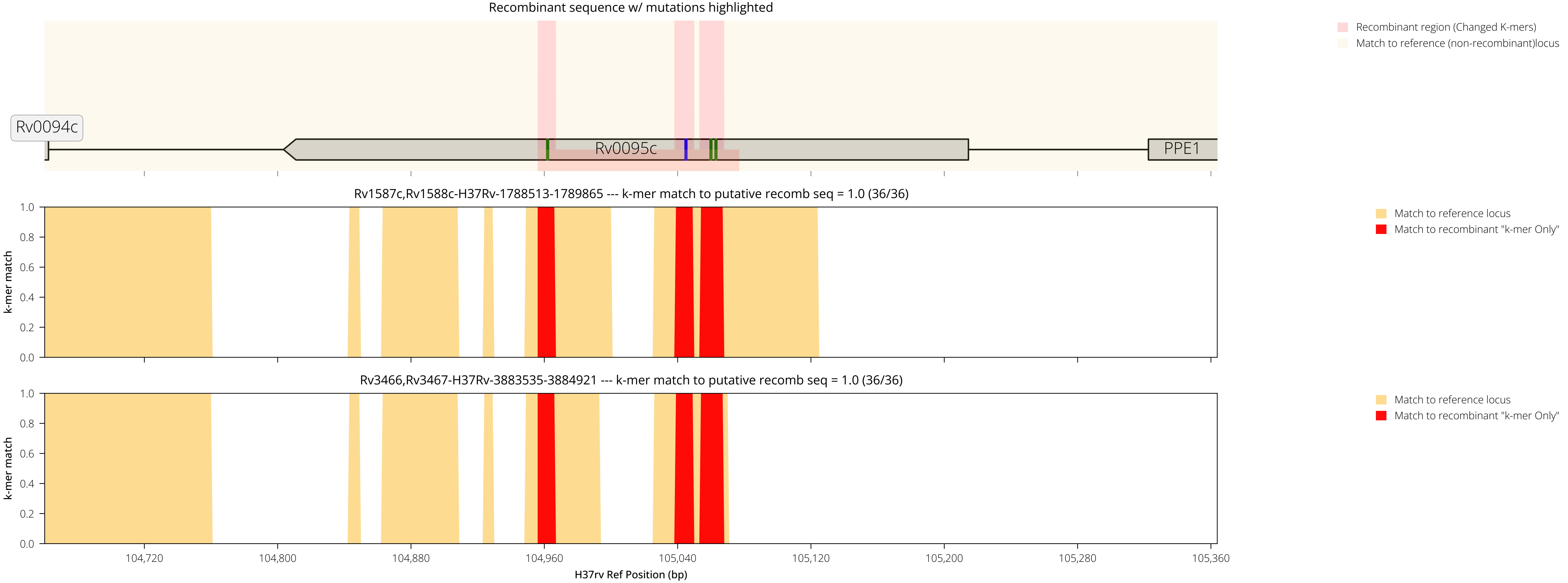

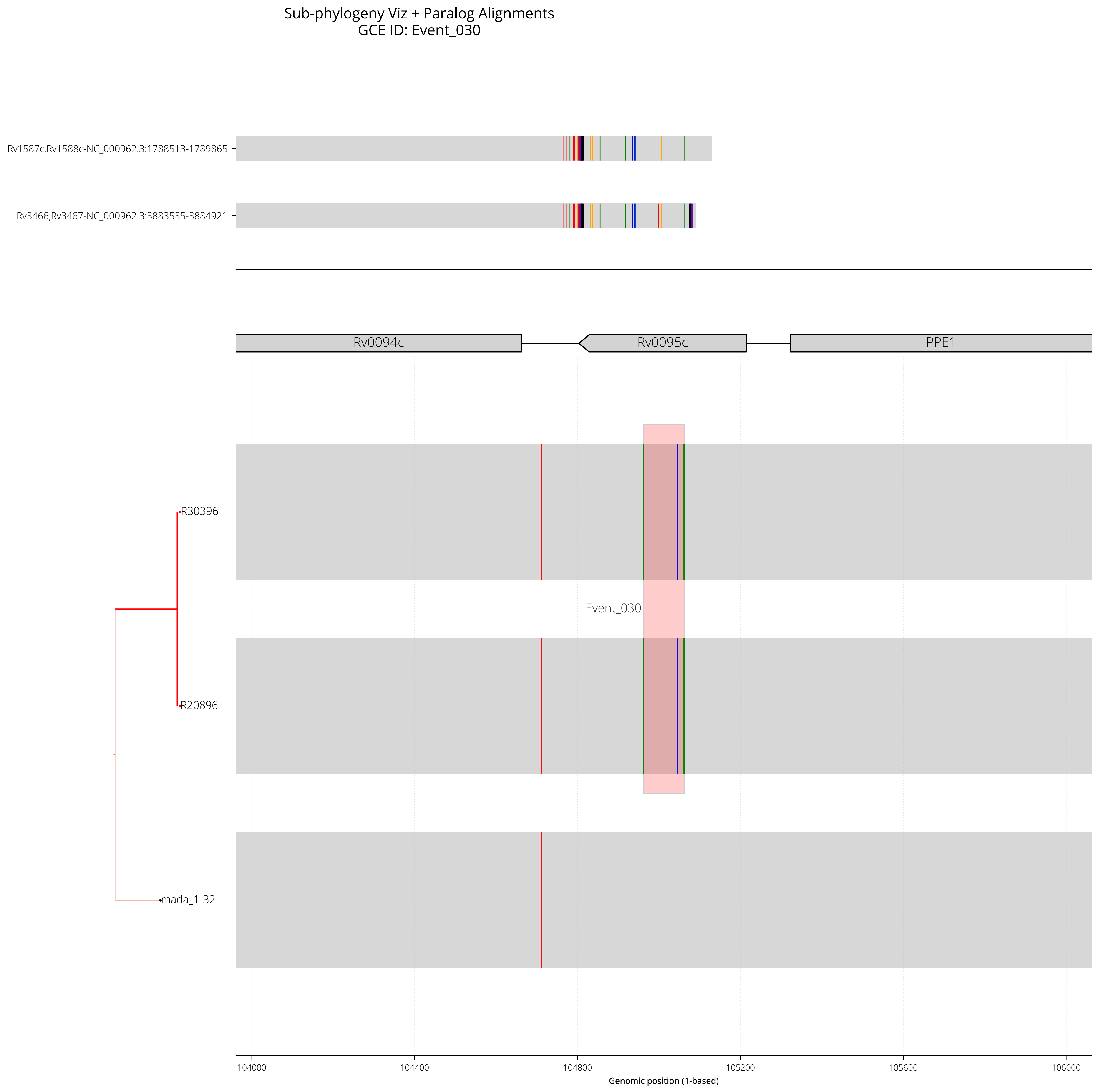
Sub-phylogeny Viz + Paralog Alignments
GCE ID: Event_030
Rv1587c,Rv1588c-NC_000962.3:1788513-1789865
Rv3466,Rv3467-NC_000962.3:3883535-3884921
Rv0094c
Rv0095c
PPE1
R30396
Event_030
R20896
mada_1-32
104000
104400
104800
105200
105600
106000
Genomic position (1-based)

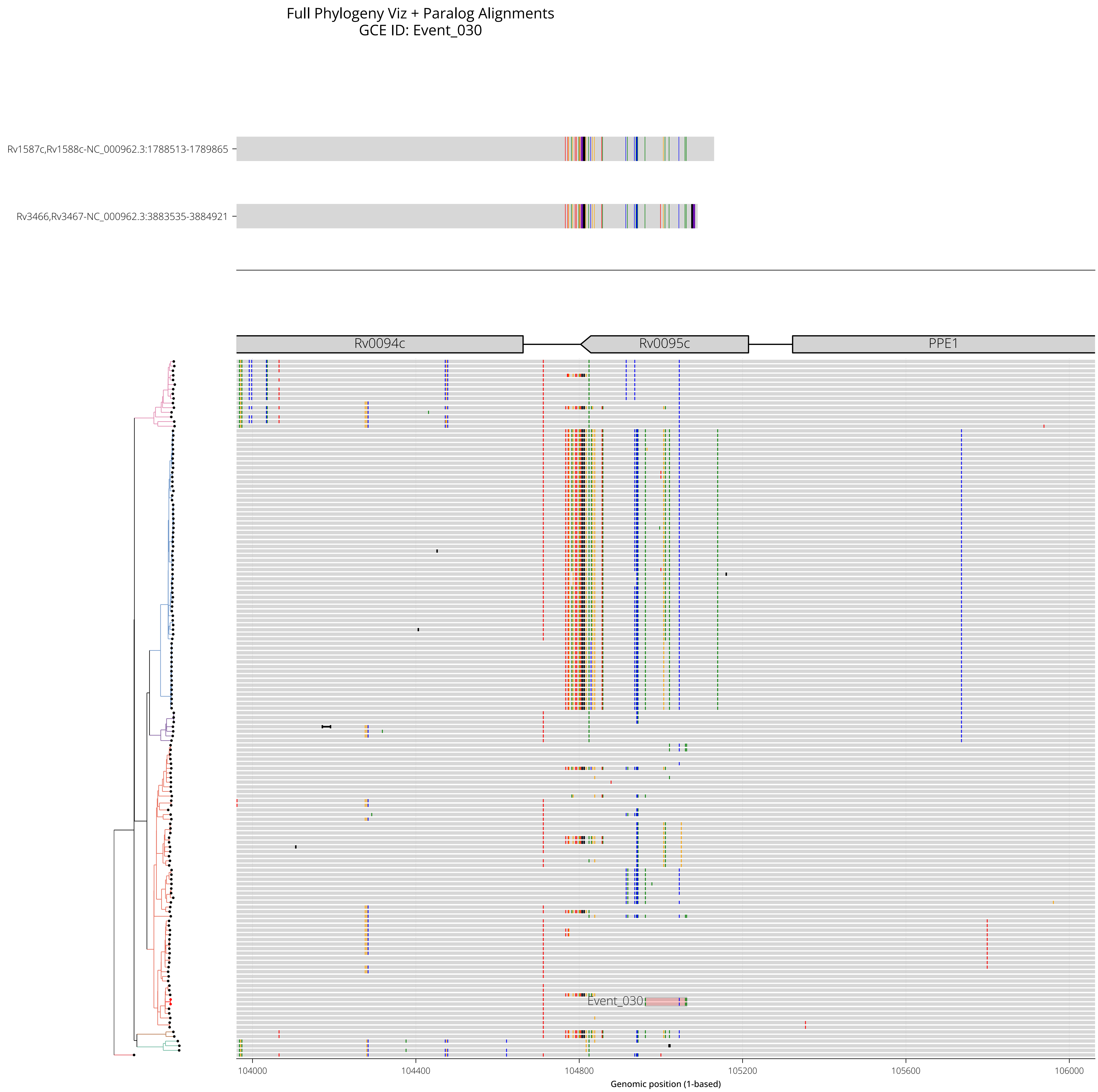

Full Phylogeny Viz + Paralog Alignments
GCE ID: Event_030
Rv1587c,Rv1588c-NC_000962.3:1788513-1789865
Rv3466,Rv3467-NC_000962.3:3883535-3884921
Rv0094c
Rv0095c
PPE1
Event_030
104000
104400
104800
105200
105600
106000
Genomic position (1-based)

Individual GCE + Known Paralogs
GCE ID: Event_031

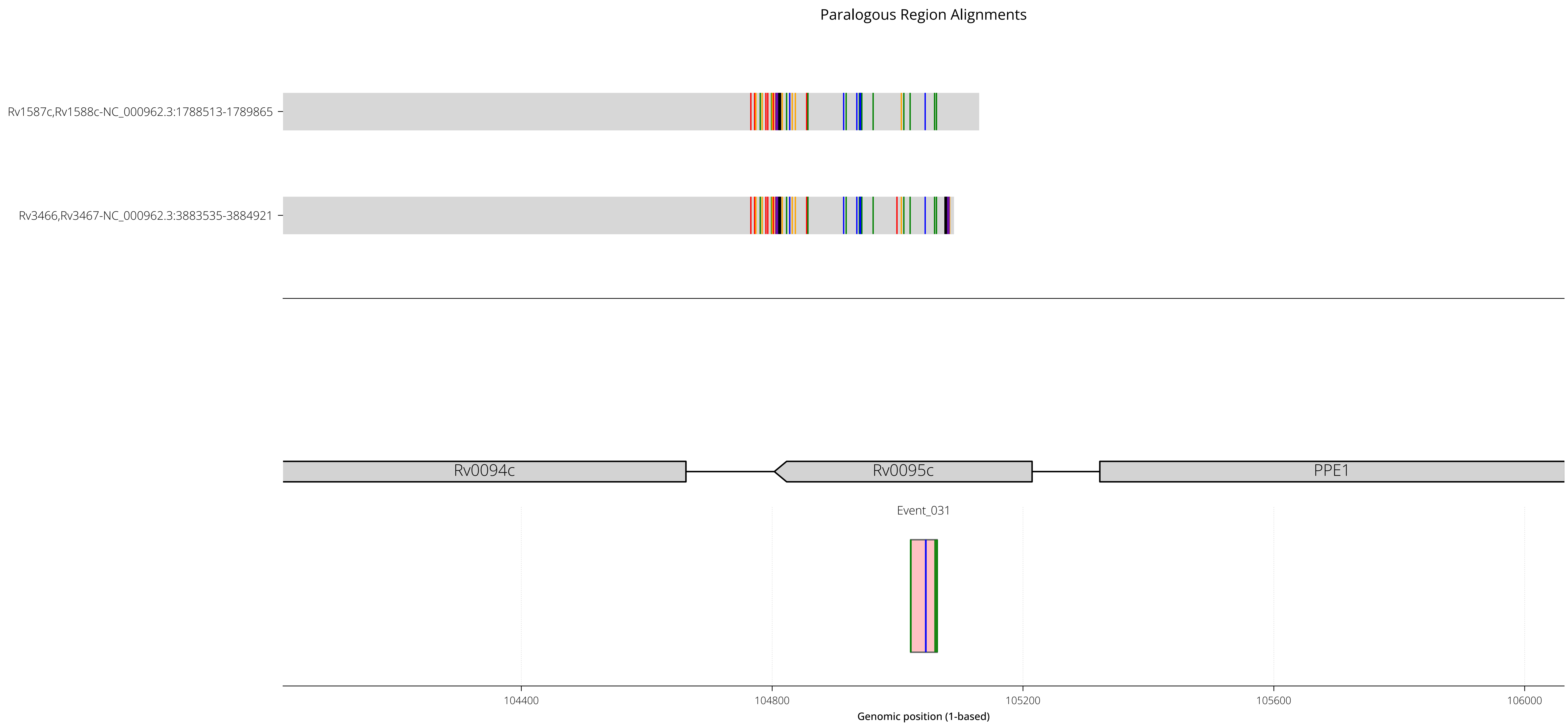

Circos Viz - GCE to Known Paralogs Locations
GCE ID: Event_031

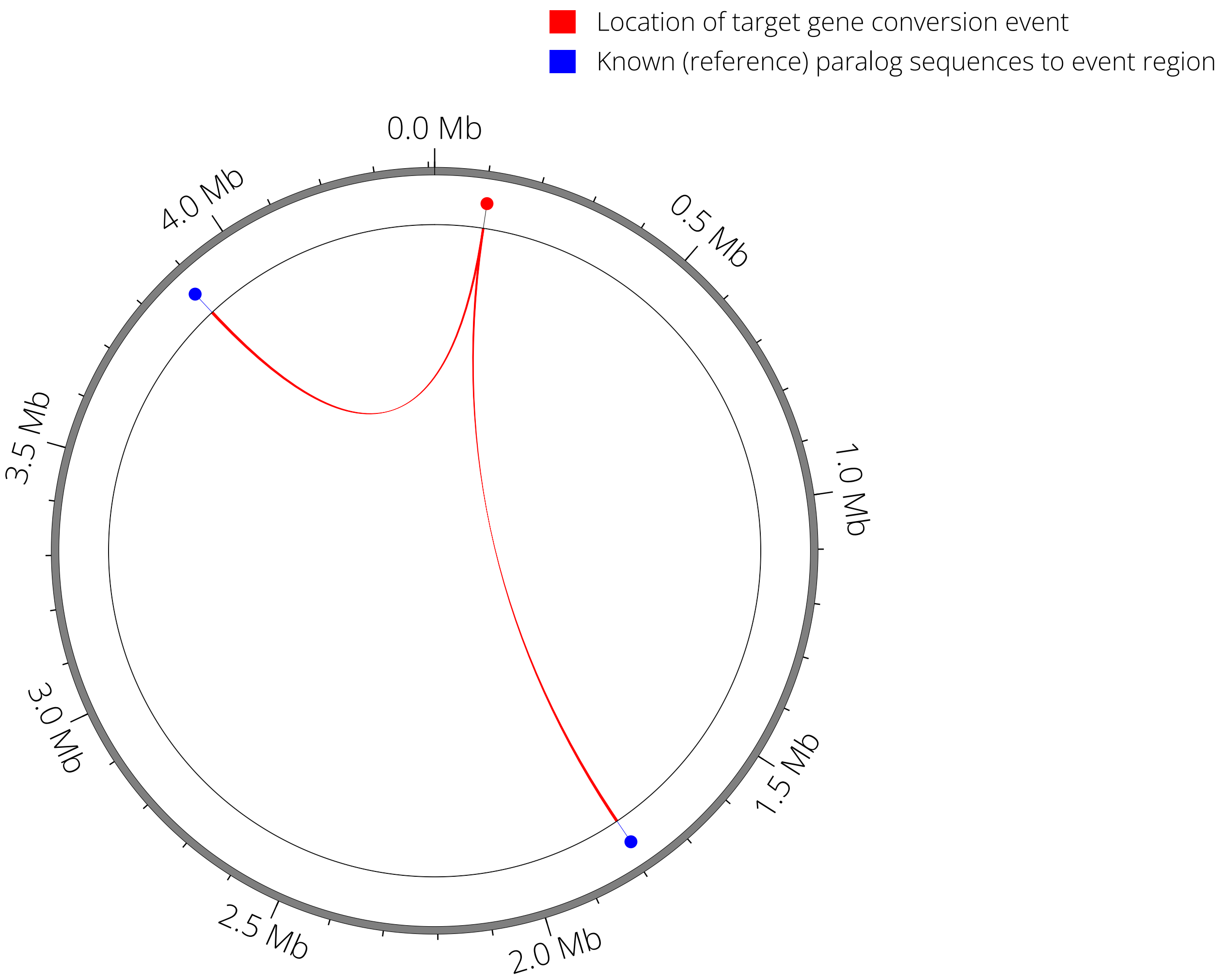

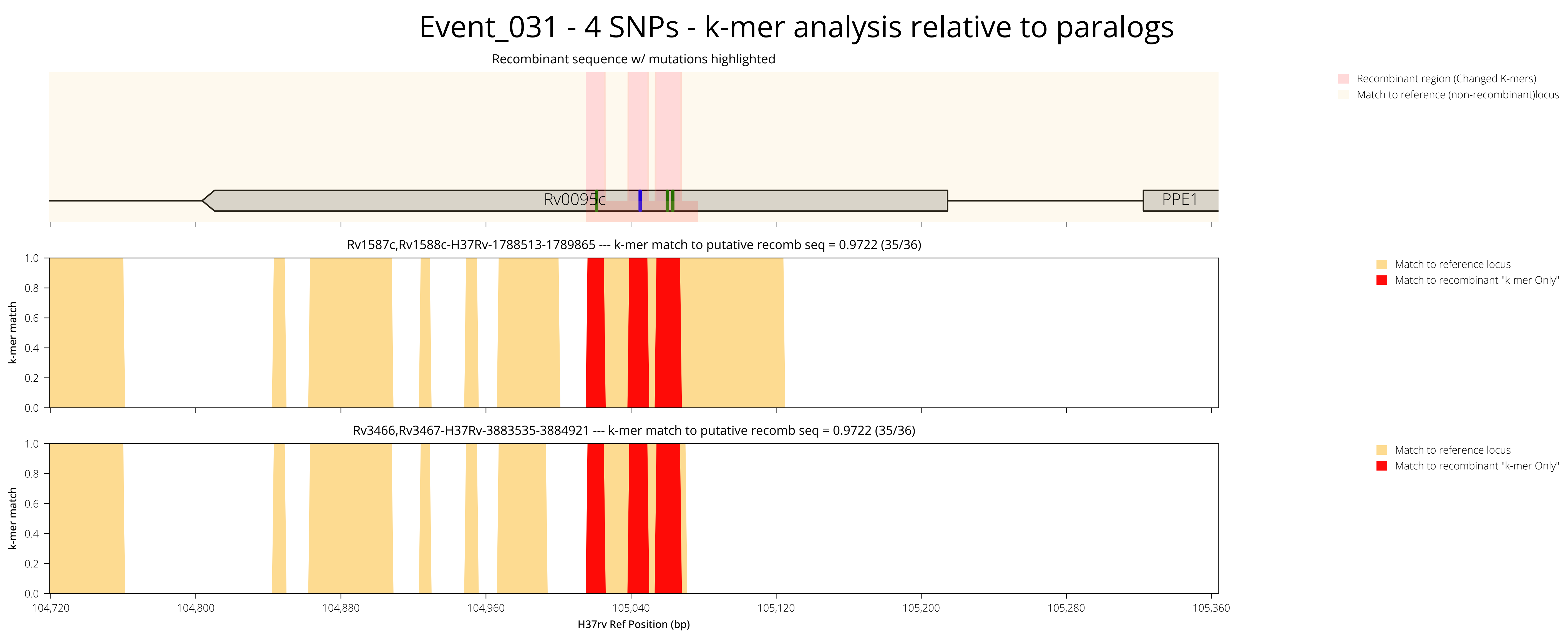
Event_031 - 4 SNPs - k-mer analysis relative to paralogs
Recombinant sequence w/ mutations highlighted
Recombinant region (Changed K-mers)
Match to reference (non-recombinant)locus
Rv0095c
PPE1
Rv1587c,Rv1588c-H37Rv-1788513-1789865 --- k-mer match to putative recomb seq = 0.9722 (35/36)
Match to reference locus
Match to recombinant "k-mer Only"
k-mer match
Rv3466,Rv3467-H37Rv-3883535-3884921 --- k-mer match to putative recomb seq = 0.9722 (35/36)
Match to reference locus
Match to recombinant "k-mer Only"
k-mer match
104,720
104,800
104,880
104,960
105,040
105,120
105,200
105,280
105,360
H37rv Ref Position (bp)

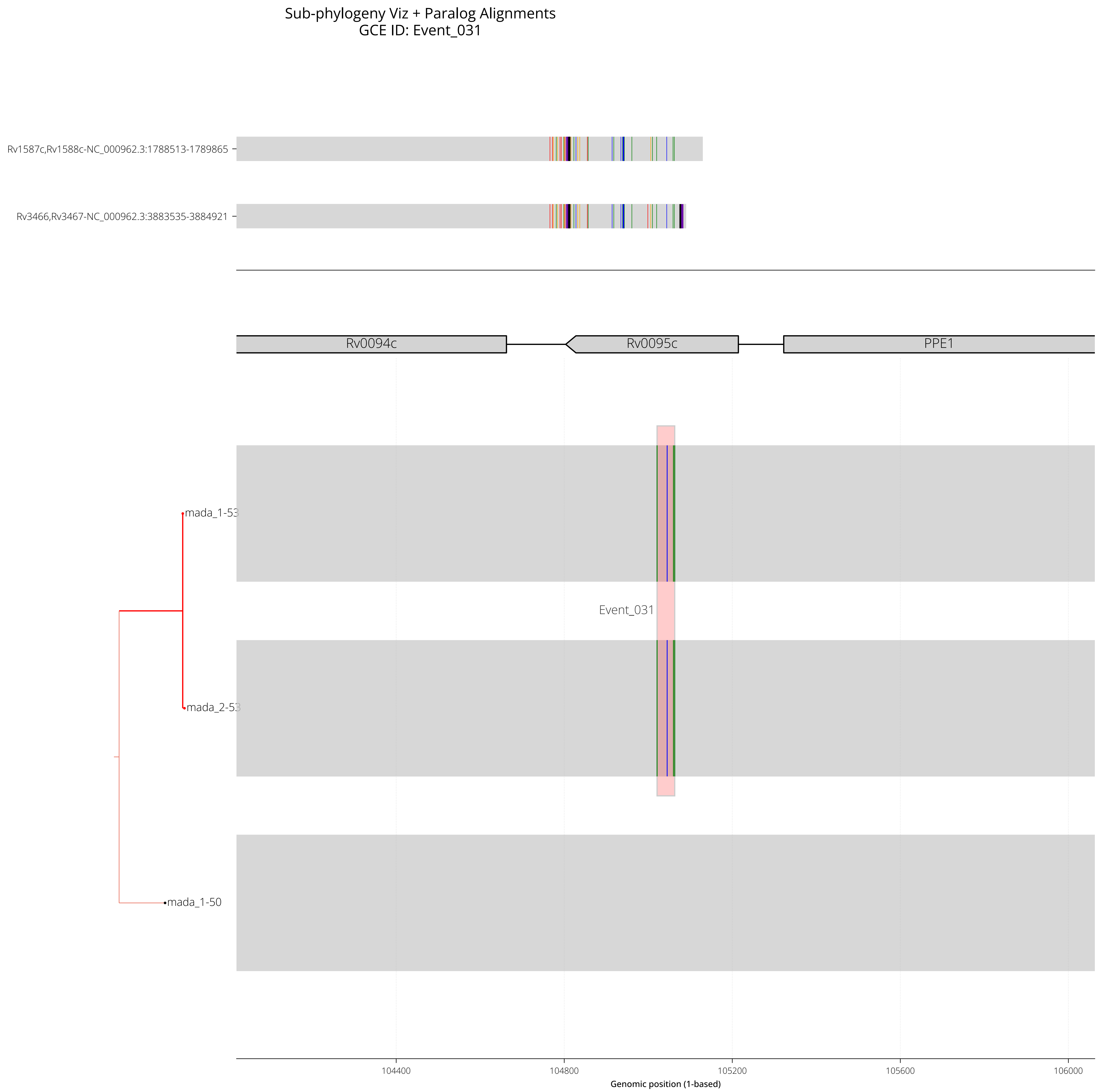

Sub-phylogeny Viz + Paralog Alignments
GCE ID: Event_031
Rv1587c,Rv1588c-NC_000962.3:1788513-1789865
Rv3466,Rv3467-NC_000962.3:3883535-3884921
Rv0094c
Rv0095c
PPE1
mada_1-53
mada_2-53
mada_1-50
Event_031
104400
104800
105200
105600
106000
Genomic position (1-based)

Full Phylogeny Viz + Paralog Alignments
GCE ID: Event_031

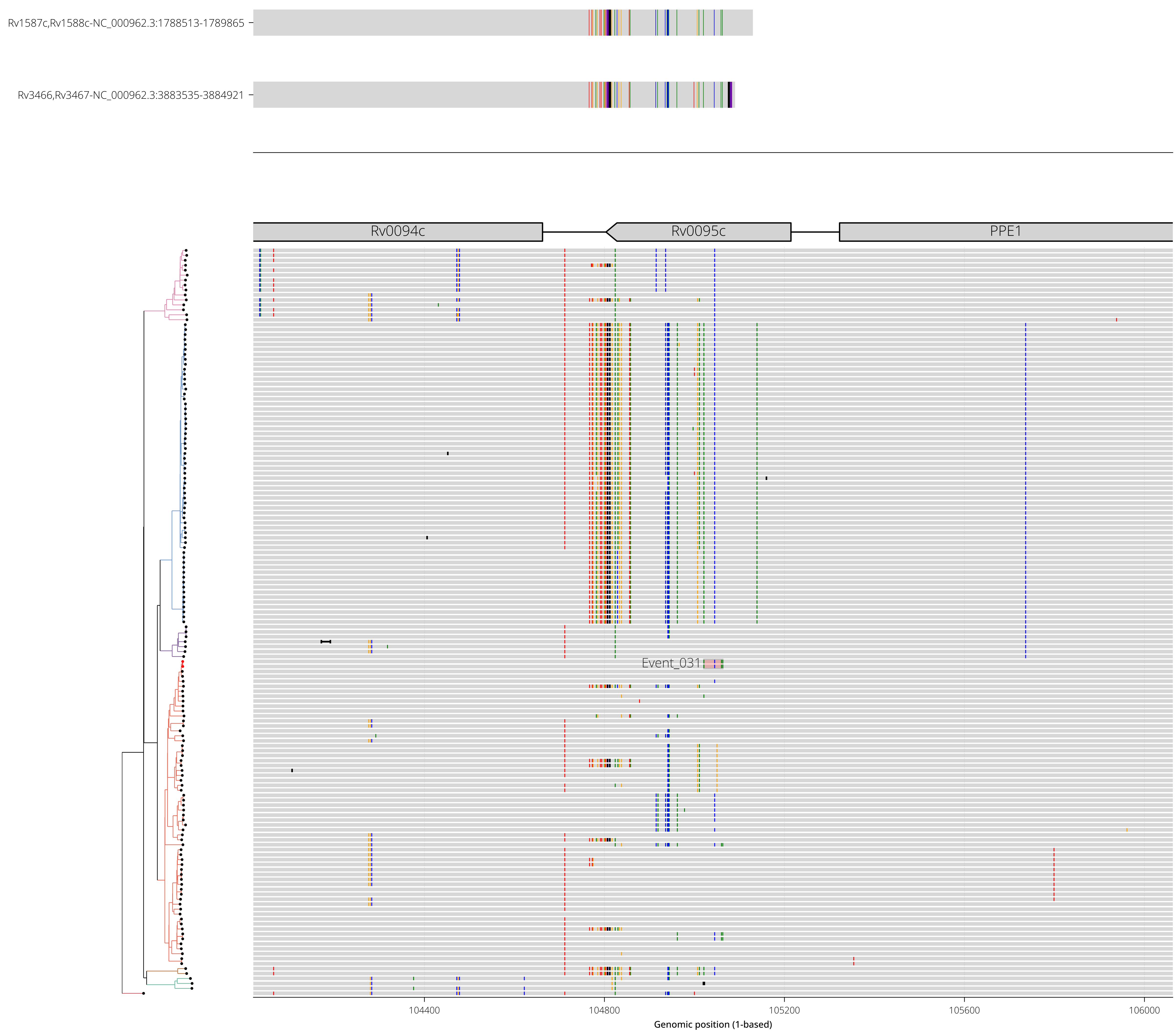

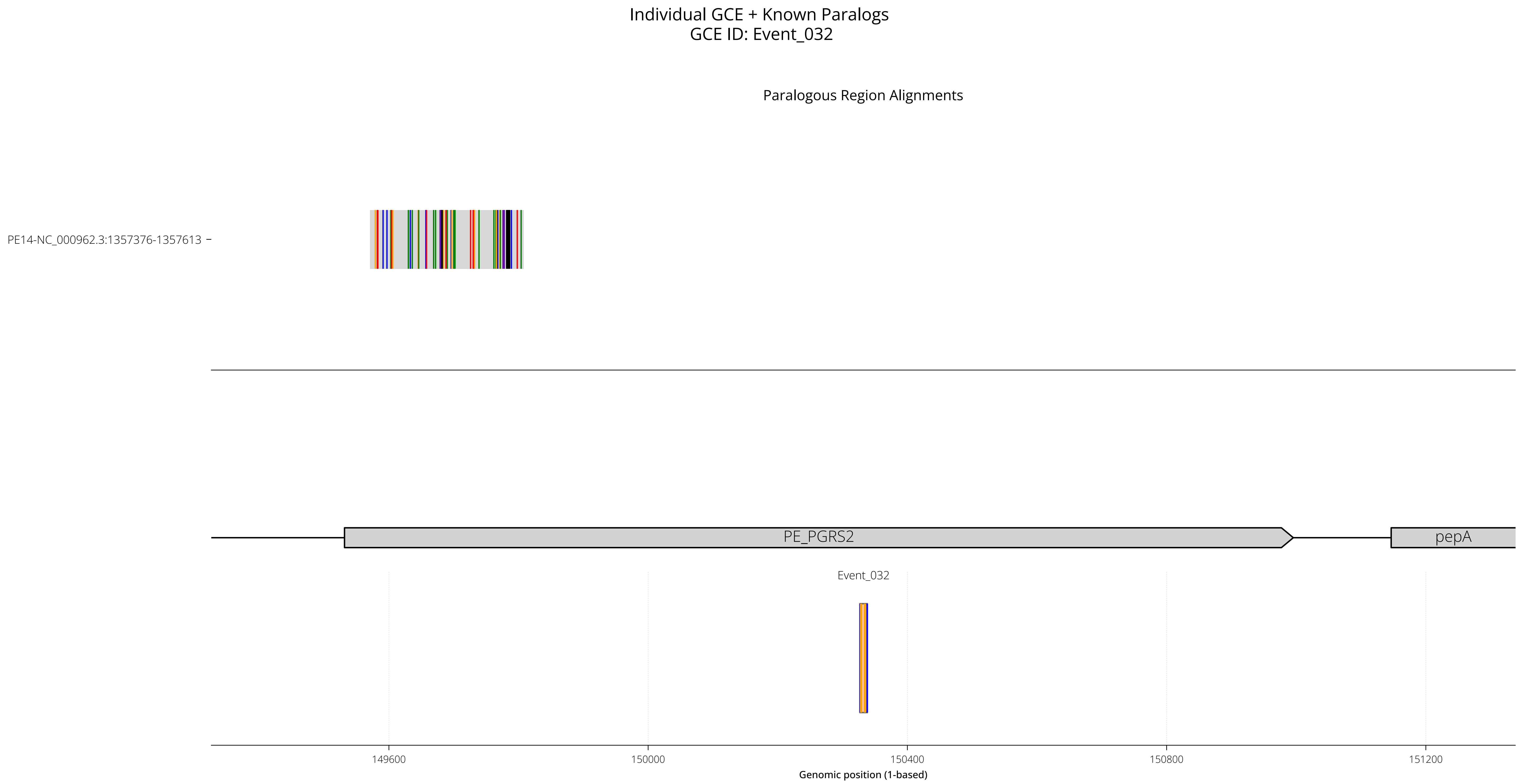

Individual GCE + Known Paralogs
GCE ID: Event_032
Paralogous Region Alignments
PE14-NC_000962.3:1357376-1357613
PE_PGRS2
pepA
Event_032
149600
150000
150400
150800
151200
Genomic position (1-based)

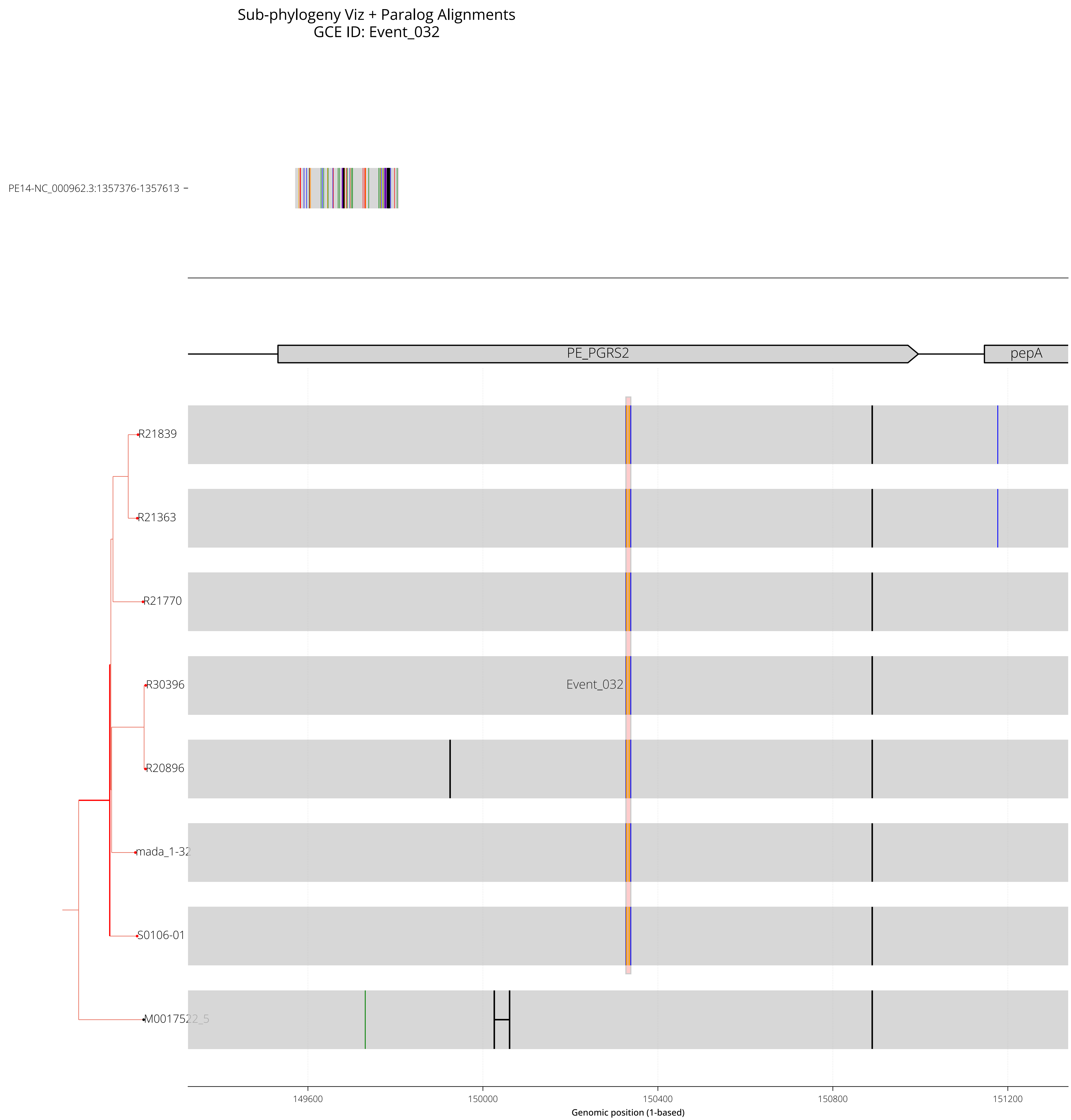

Sub-phylogeny Viz + Paralog Alignments
GCE ID: Event_032
PE14-NC_000962.3:1357376-1357613
PE_PGRS2
pepA
R21839
R21363
R21770
R30396
R20896
mada_1-32
S0106-01
M0017522_5
Event_032
149600
150000
150400
150800
151200
Genomic position (1-based)

Full Phylogeny Viz + Paralog Alignments
GCE ID: Event_032

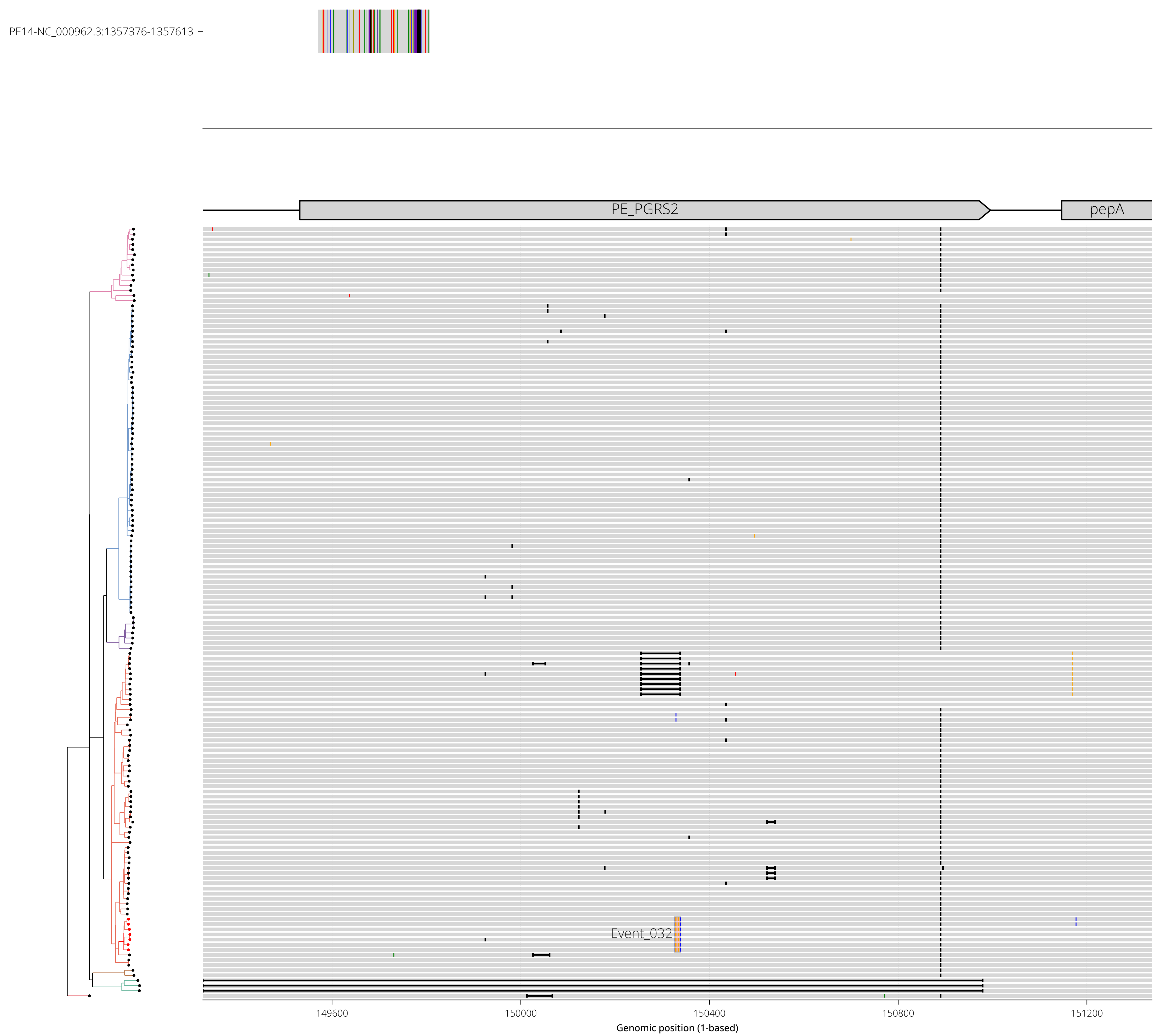

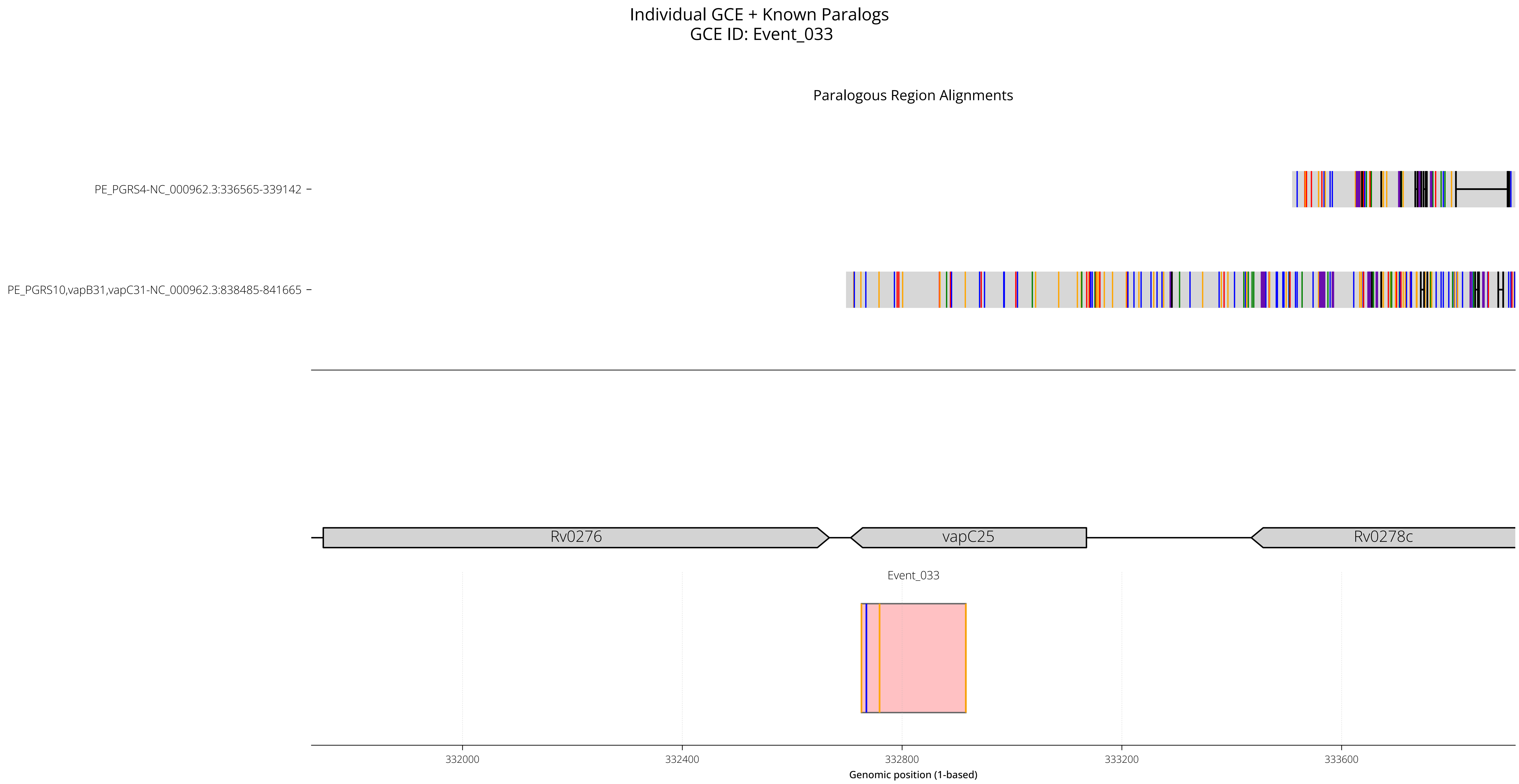

Individual GCE + Known Paralogs
GCE ID: Event_033
Paralogous Region Alignments
PE_PGRS4-NC_000962.3:336565-339142
PE_PGRS10,vapB31,vapC31-NC_000962.3:838485-841665
Rv0276
vapC25
Rv0278c
Event_033
332000
332400
332800
333200
333600
Genomic position (1-based)

Circos Viz - GCE to Known Paralogs Locations
GCE ID: Event_033

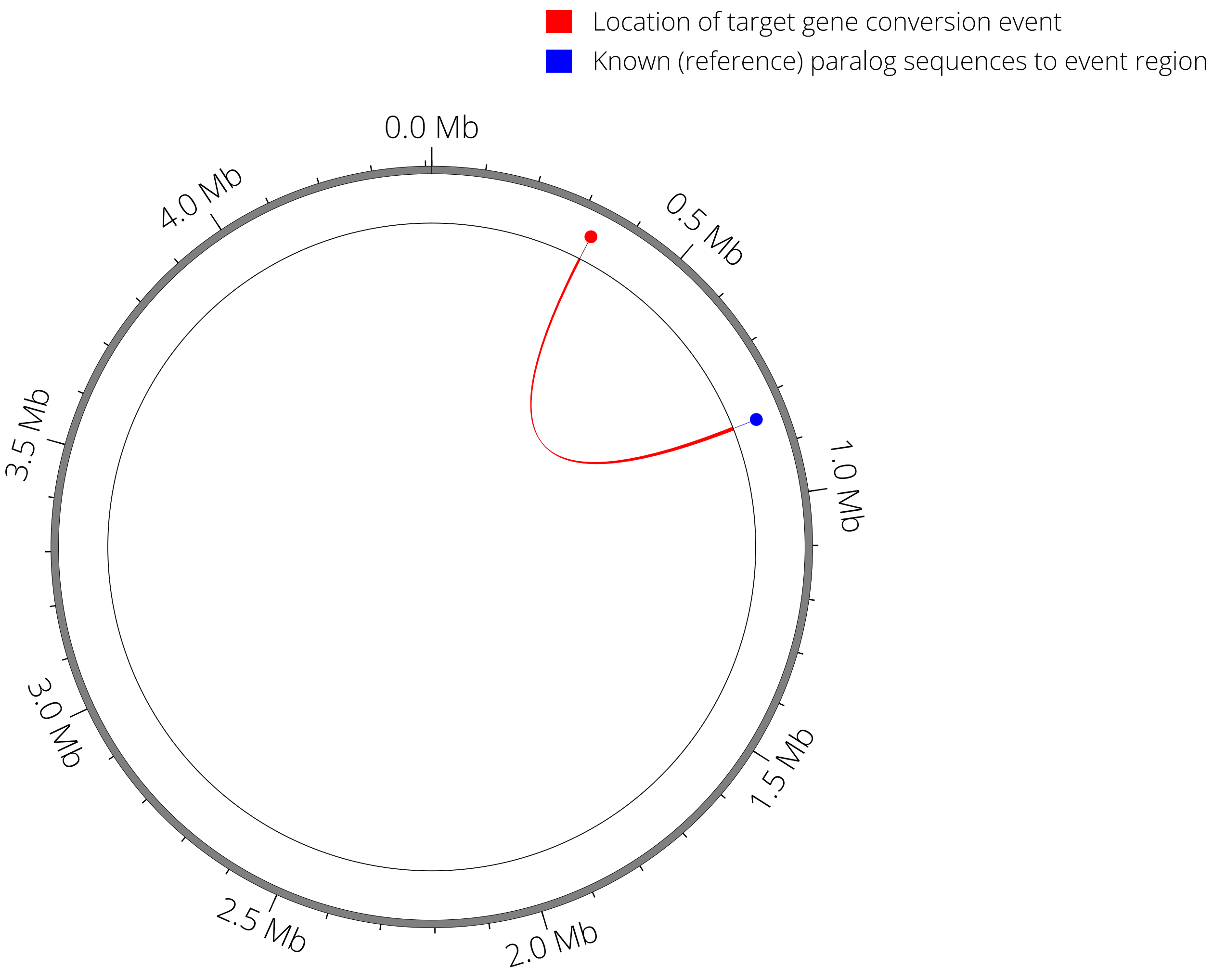

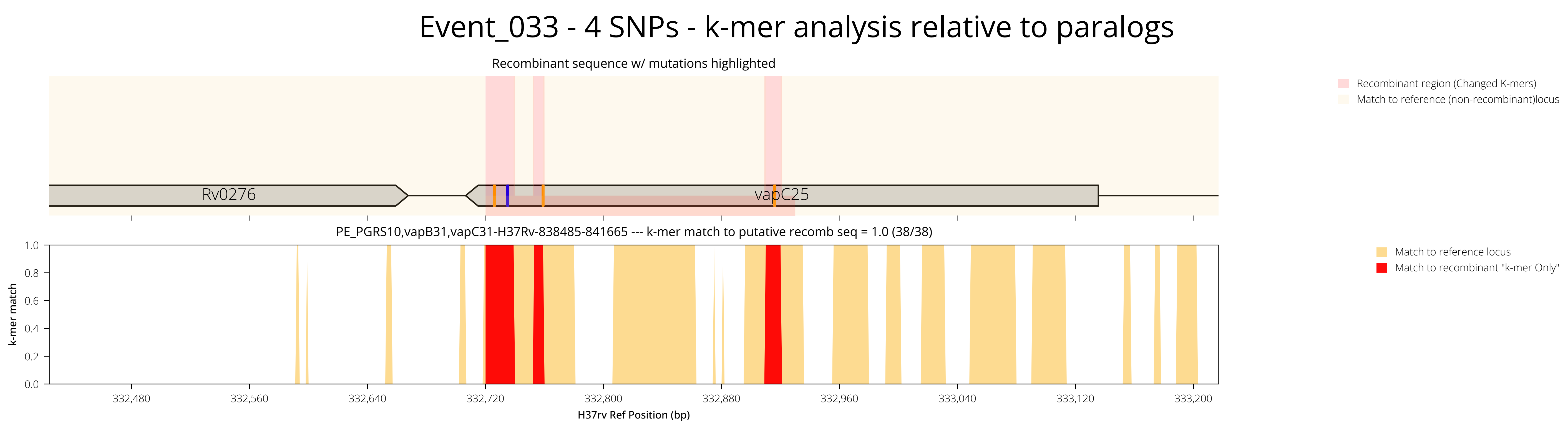

Event_033 - 4 SNPs - k-mer analysis relative to paralogs
Recombinant sequence w/ mutations highlighted
Rv0276
vapC25
Recombinant region (Changed K-mers)
Match to reference (non-recombinant)locus
PE_PGRS10,vapB31,vapC31-H37Rv-838485-841665 --- k-mer match to putative recomb seq = 1.0 (38/38)
Match to reference locus
Match to recombinant "k-mer Only"
k-mer match
1.0
0.8
0.6
0.4
0.2
0.0
332,480
332,560
332,640
332,720
332,800
332,880
332,960
333,040
333,120
333,200
H37rv Ref Position (bp)

### Sub-phylogeny Viz + Paralog Alignments
GCE ID: Event_033

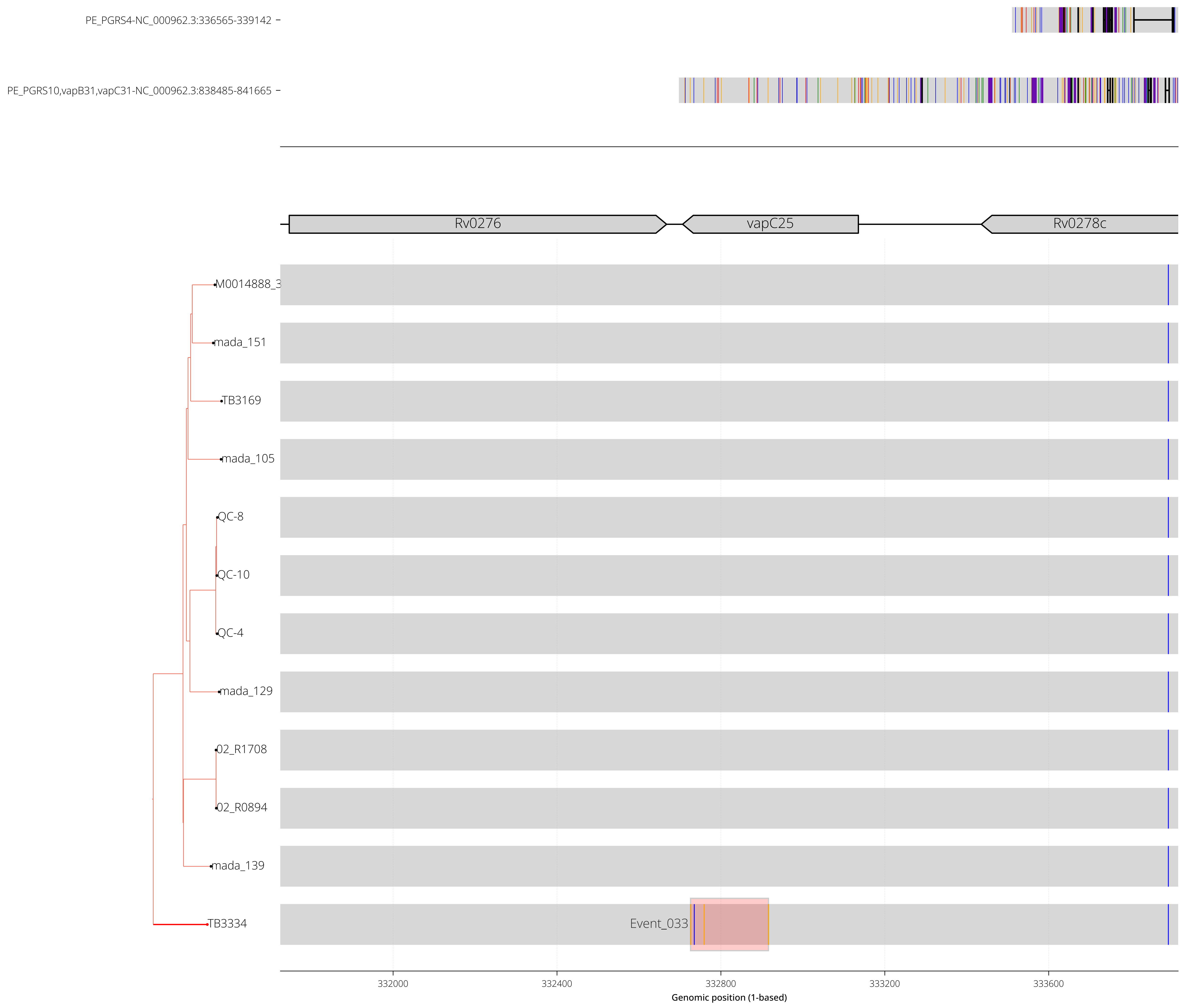

Full Phylogeny Viz + Paralog Alignments
GCE ID: Event_033

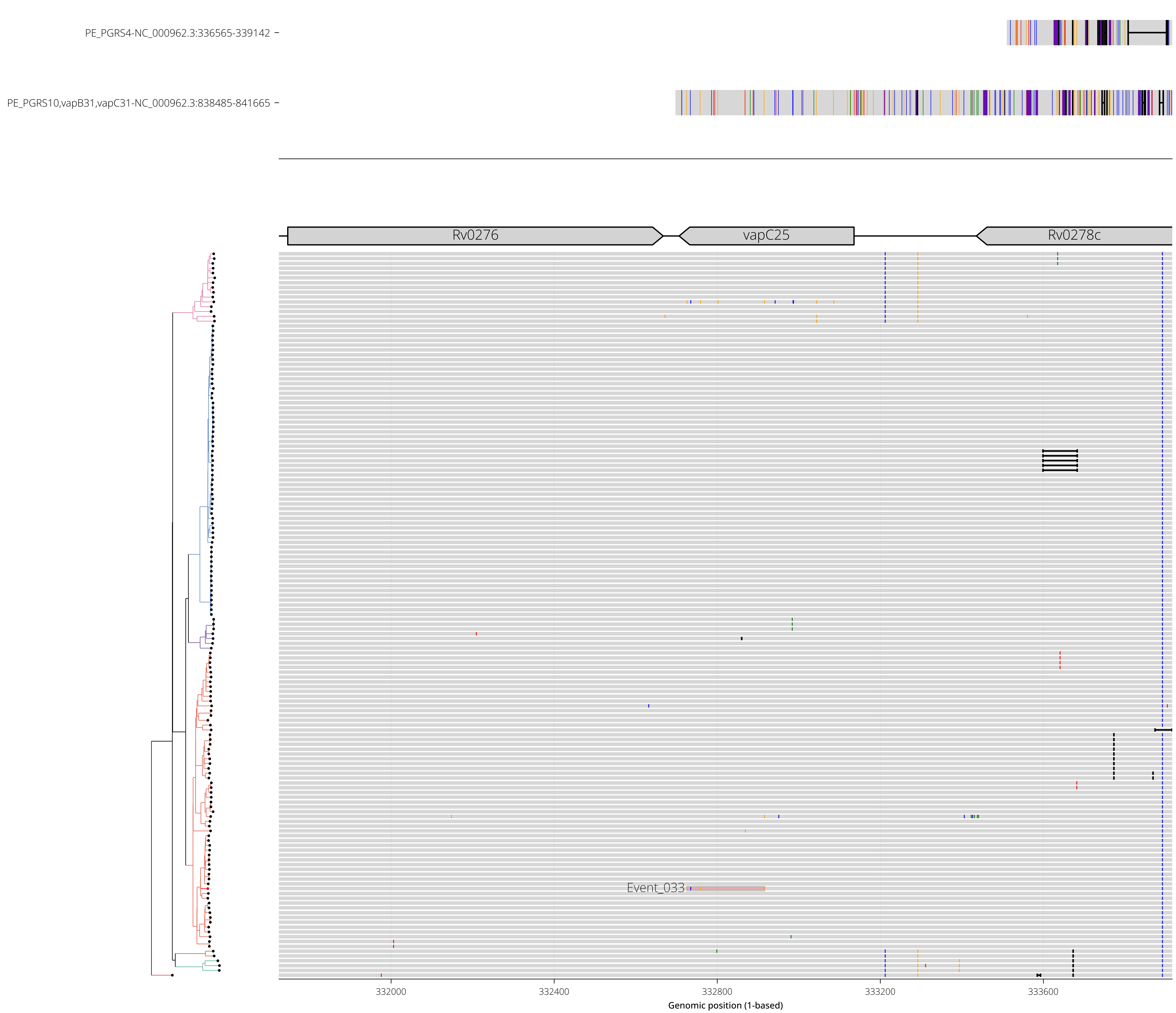

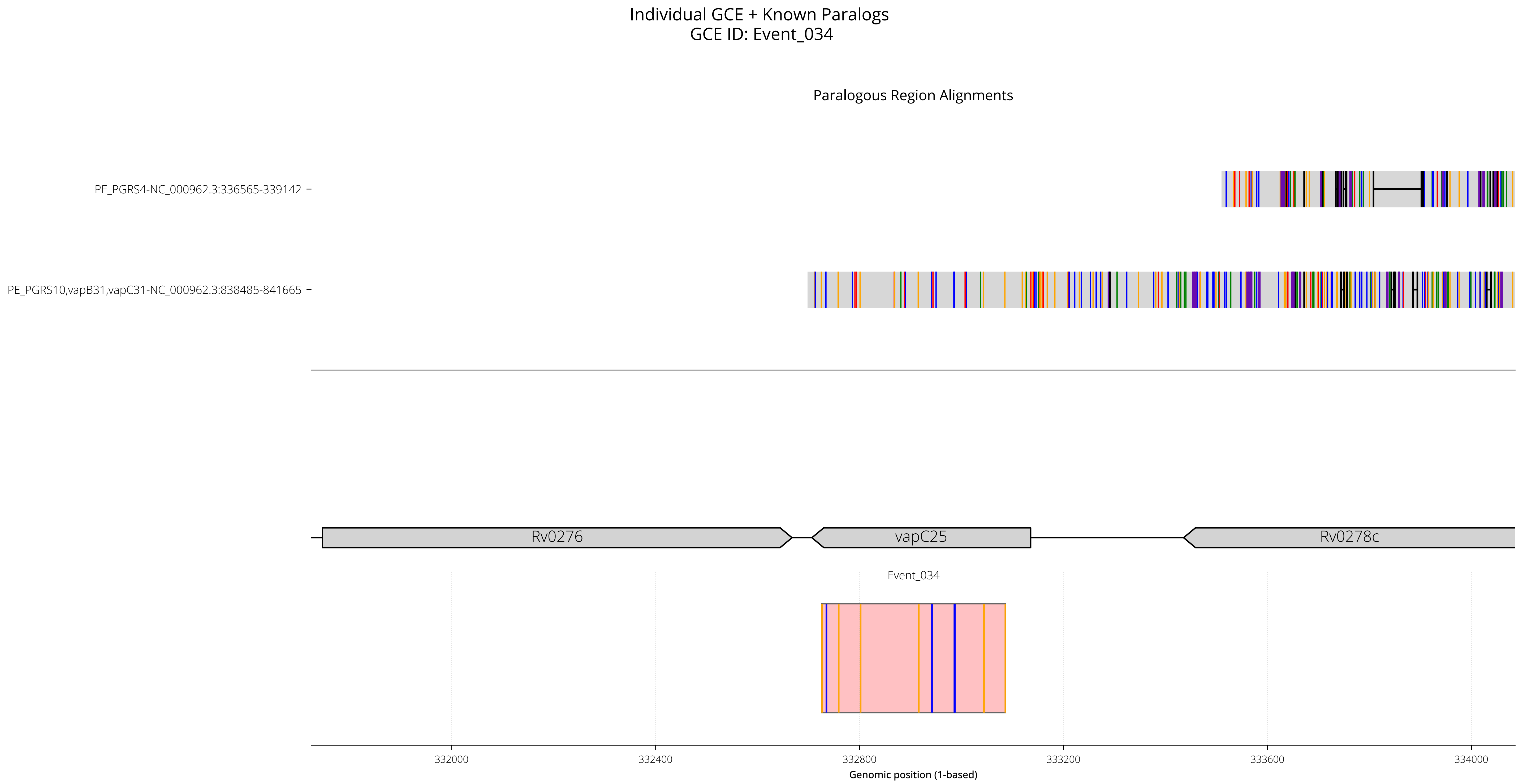

Individual GCE + Known Paralogs
GCE ID: Event_034
Paralogous Region Alignments
PE_PGRS4-NC_000962.3:336565-339142
PE_PGRS10,vapB31,vapC31-NC_000962.3:838485-841665
Rv0276
vapC25
Rv0278c
Event_034
332000
332400
332800
333200
333600
334000
Genomic position (1-based)

Circos Viz - GCE to Known Paralogs Locations
GCE ID: Event_034

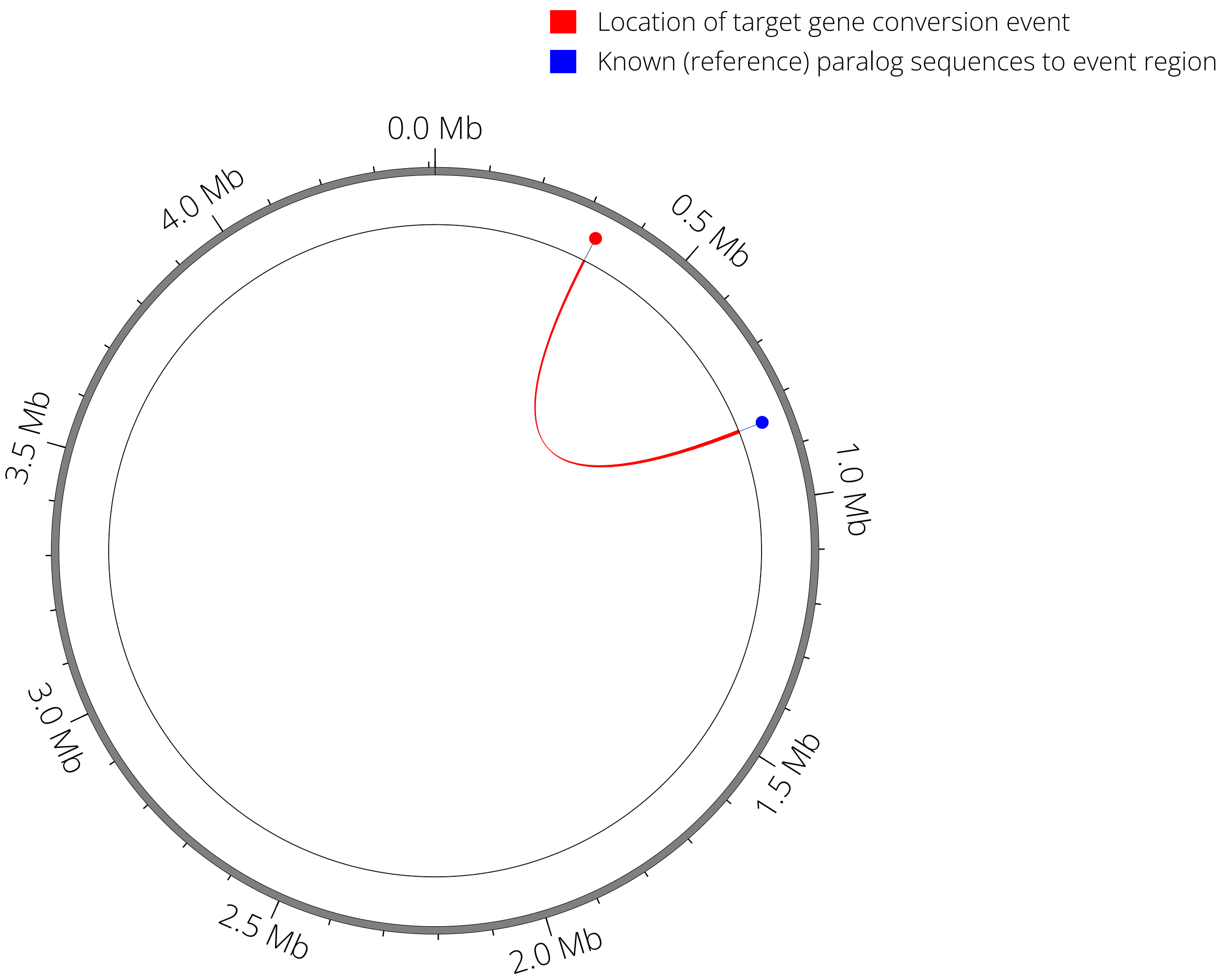

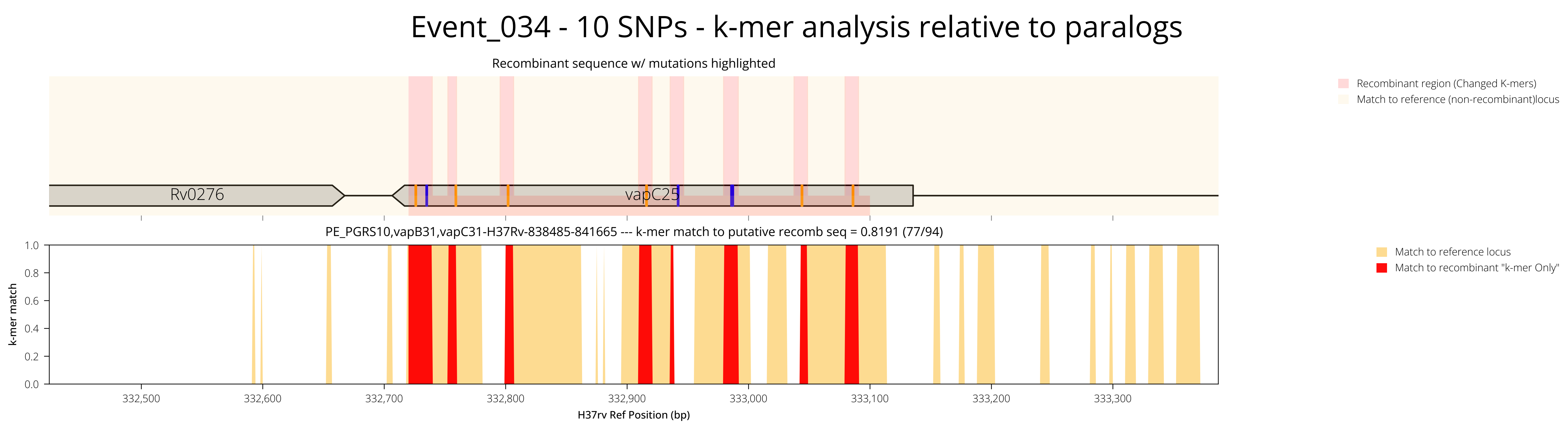

Event_034 - 10 SNPs - k-mer analysis relative to paralogs
Recombinant sequence w/ mutations highlighted
Recombinant region (Changed K-mers)
Match to reference (non-recombinant)locus
Rv0276
vapC25
PE_PGRS10,vapB31,vapC31-H37Rv-838485-841665 --- k-mer match to putative recomb seq = 0.8191 (77/94)
Match to reference locus
Match to recombinant "k-mer Only"
k-mer match
1.0
0.8
0.6
0.4
0.2
0.0
332,500
332,600
332,700
332,800
332,900
333,000
333,100
333,200
333,300
H37rv Ref Position (bp)

Sub-phylogeny Viz + Paralog Alignments
GCE ID: Event_034

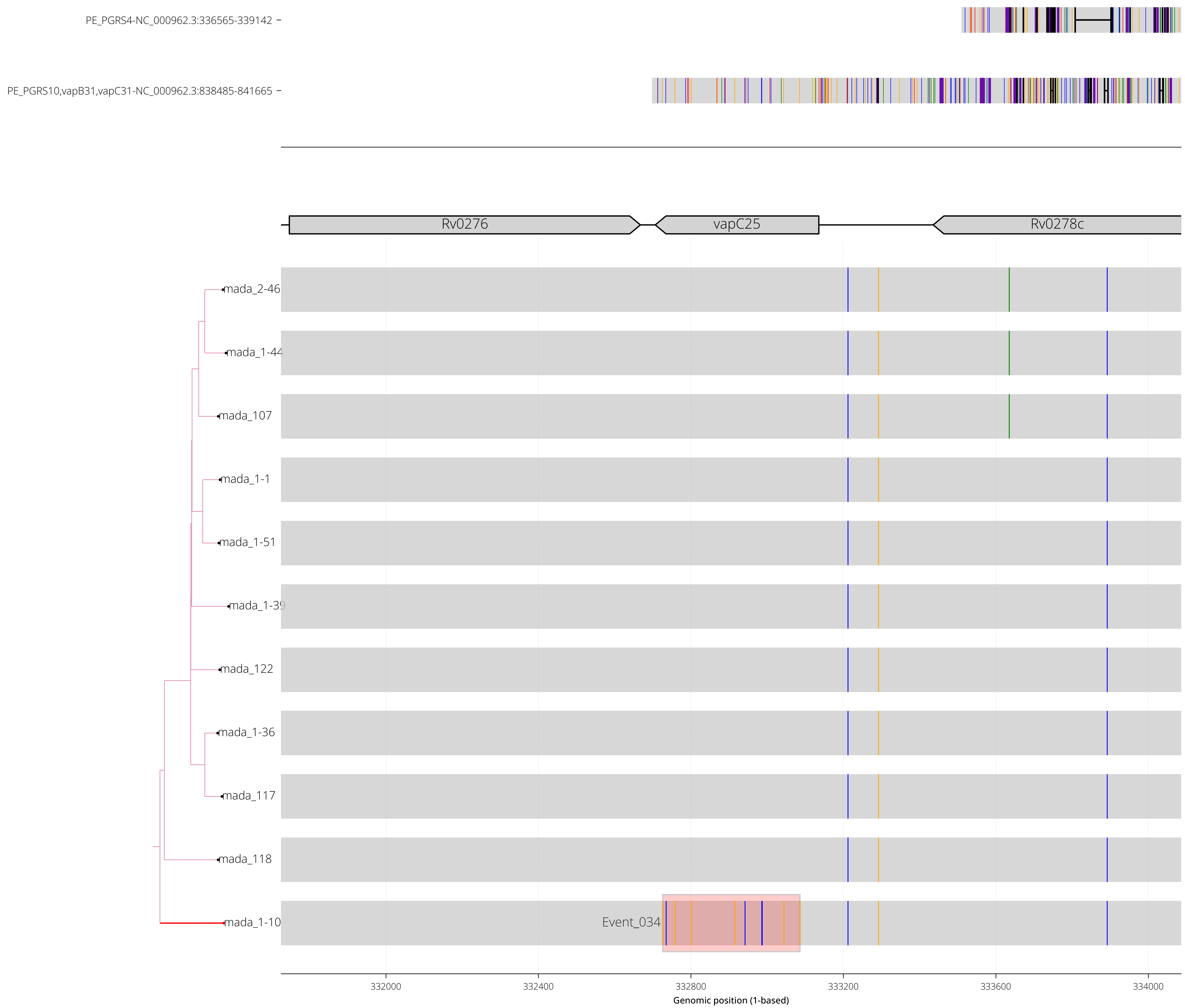

Full Phylogeny Viz + Paralog Alignments
GCE ID: Event_034

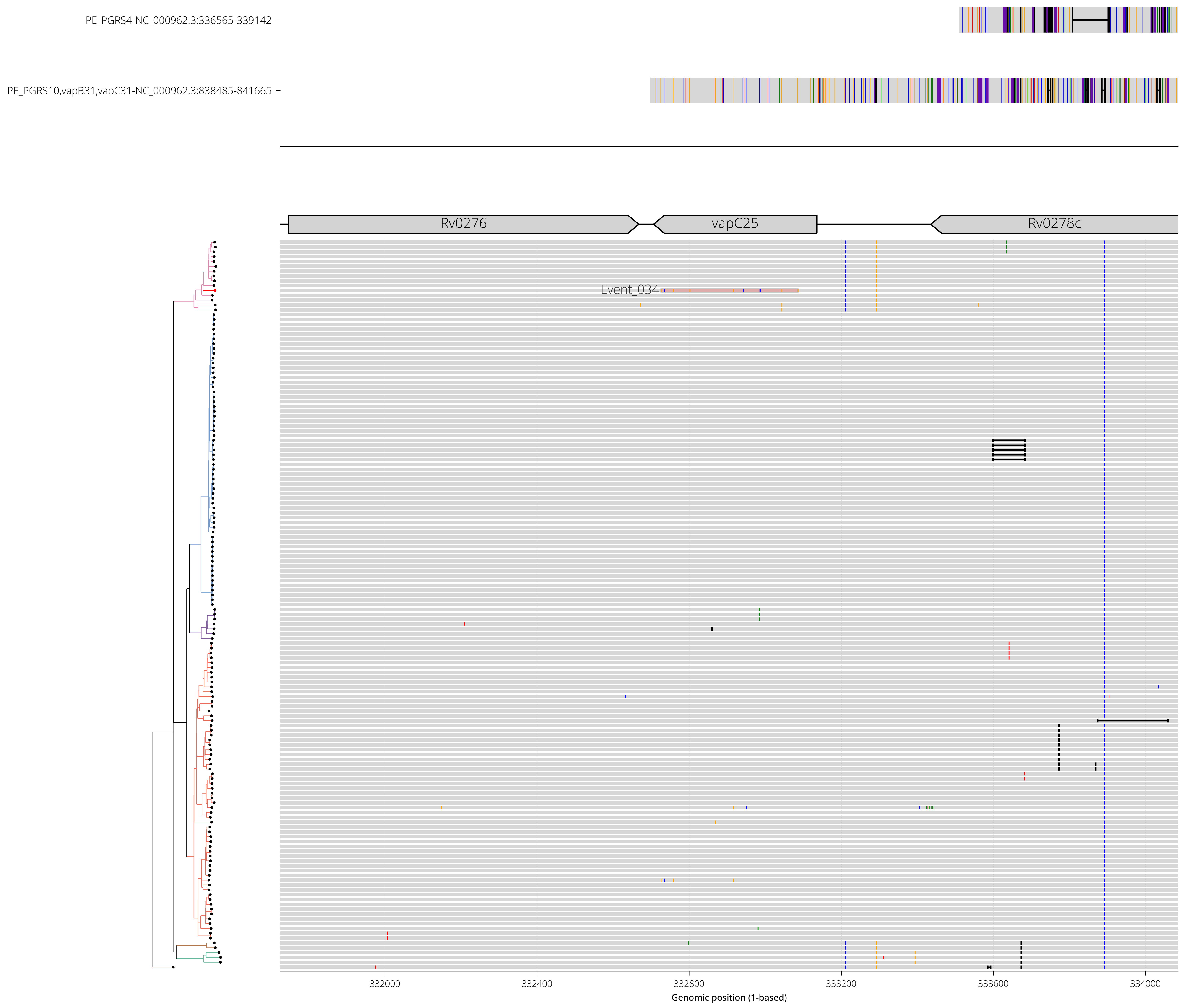

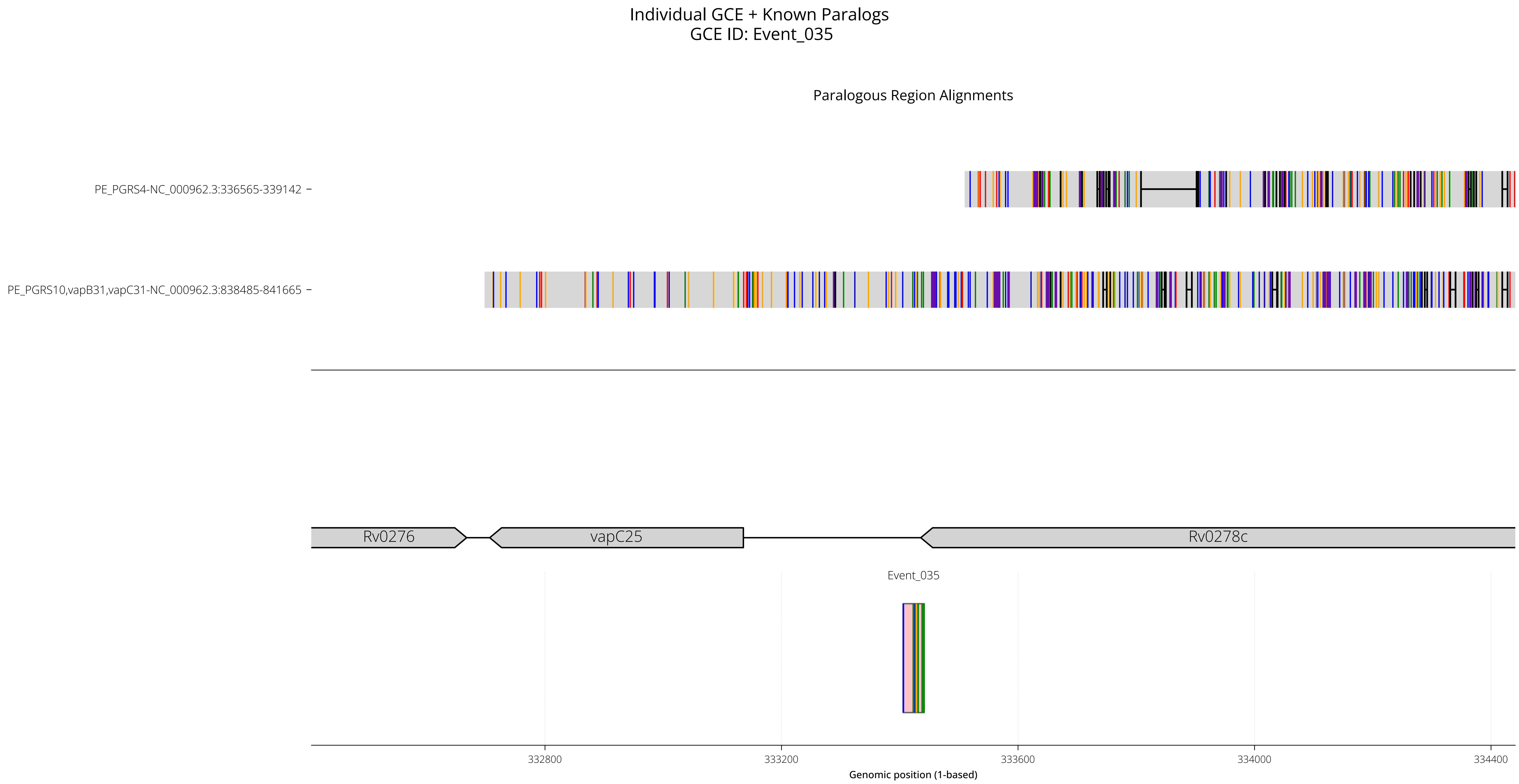

Individual GCE + Known Paralogs
GCE ID: Event_035
Paralogous Region Alignments
PE_PGRS4-NC_000962.3:336565-339142
PE_PGRS10,vapB31,vapC31-NC_000962.3:838485-841665
Rv0276
vapC25
Rv0278c
Event_035
332800
333200
333600
334000
334400
Genomic position (1-based)

Circos Viz - GCE to Known Paralogs Locations
GCE ID: Event_035

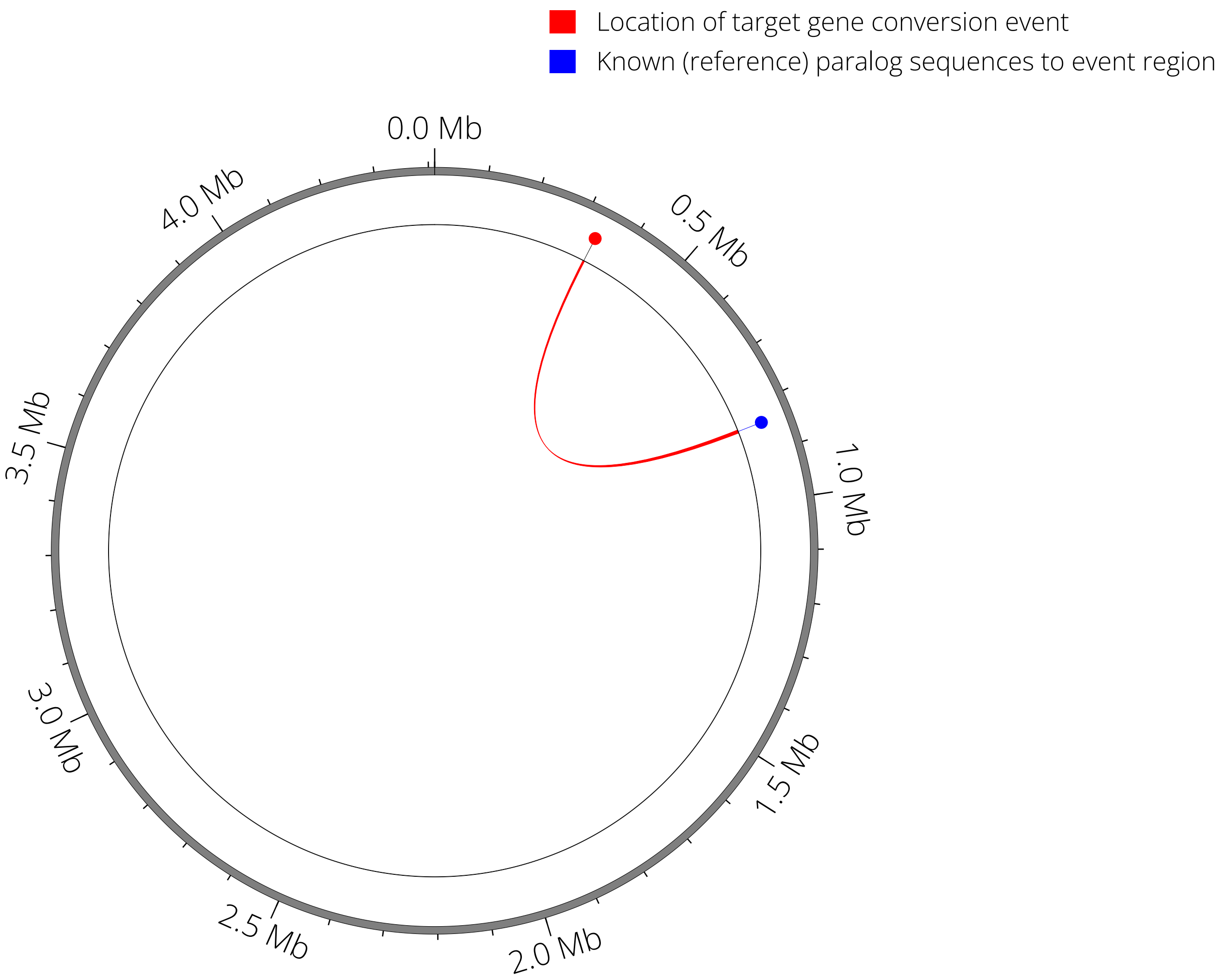

### Event_035 - 7 SNPs - k-mer analysis relative to paralogs

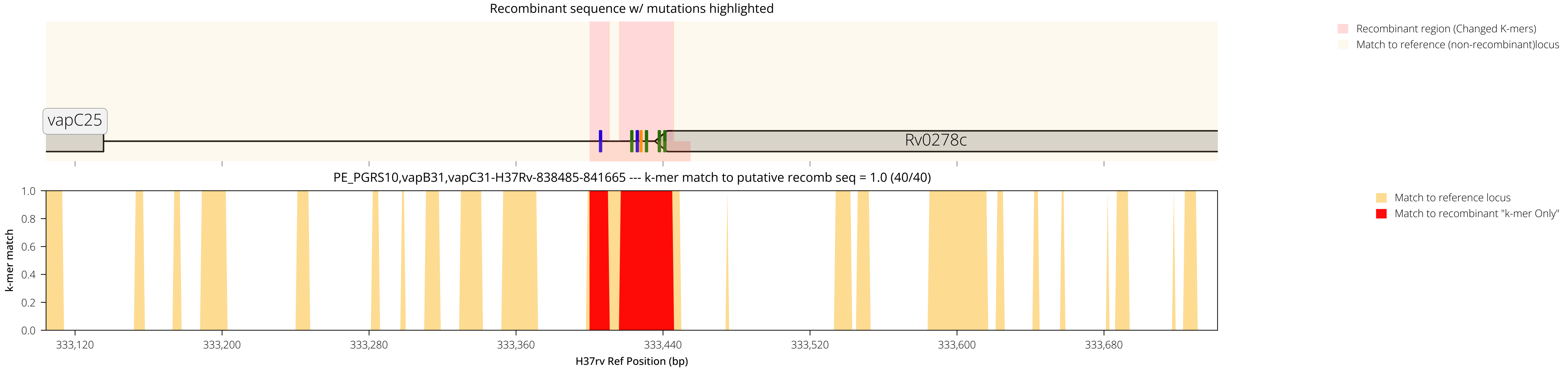

Sub-phylogeny Viz + Paralog Alignments
GCE ID: Event_035

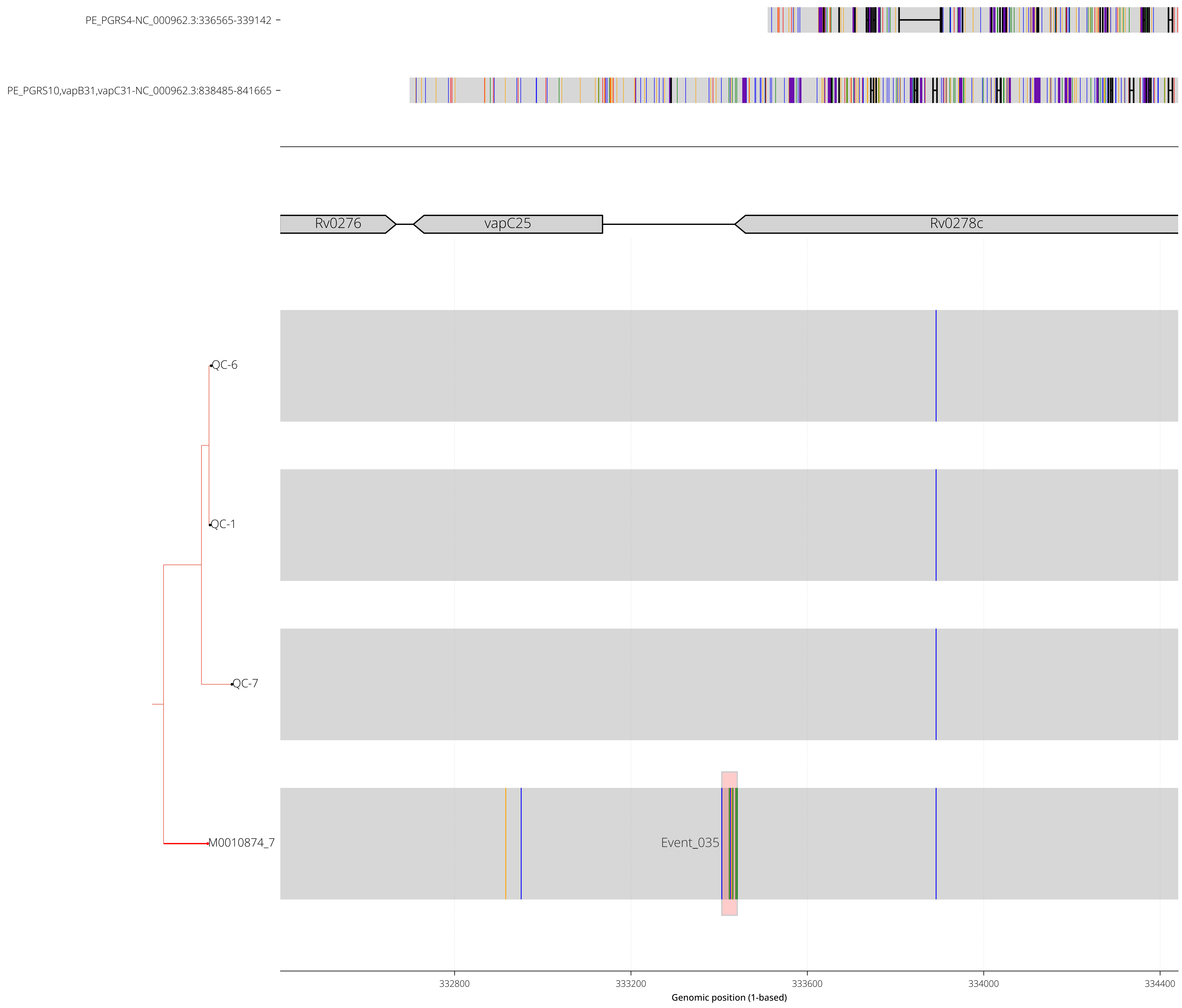

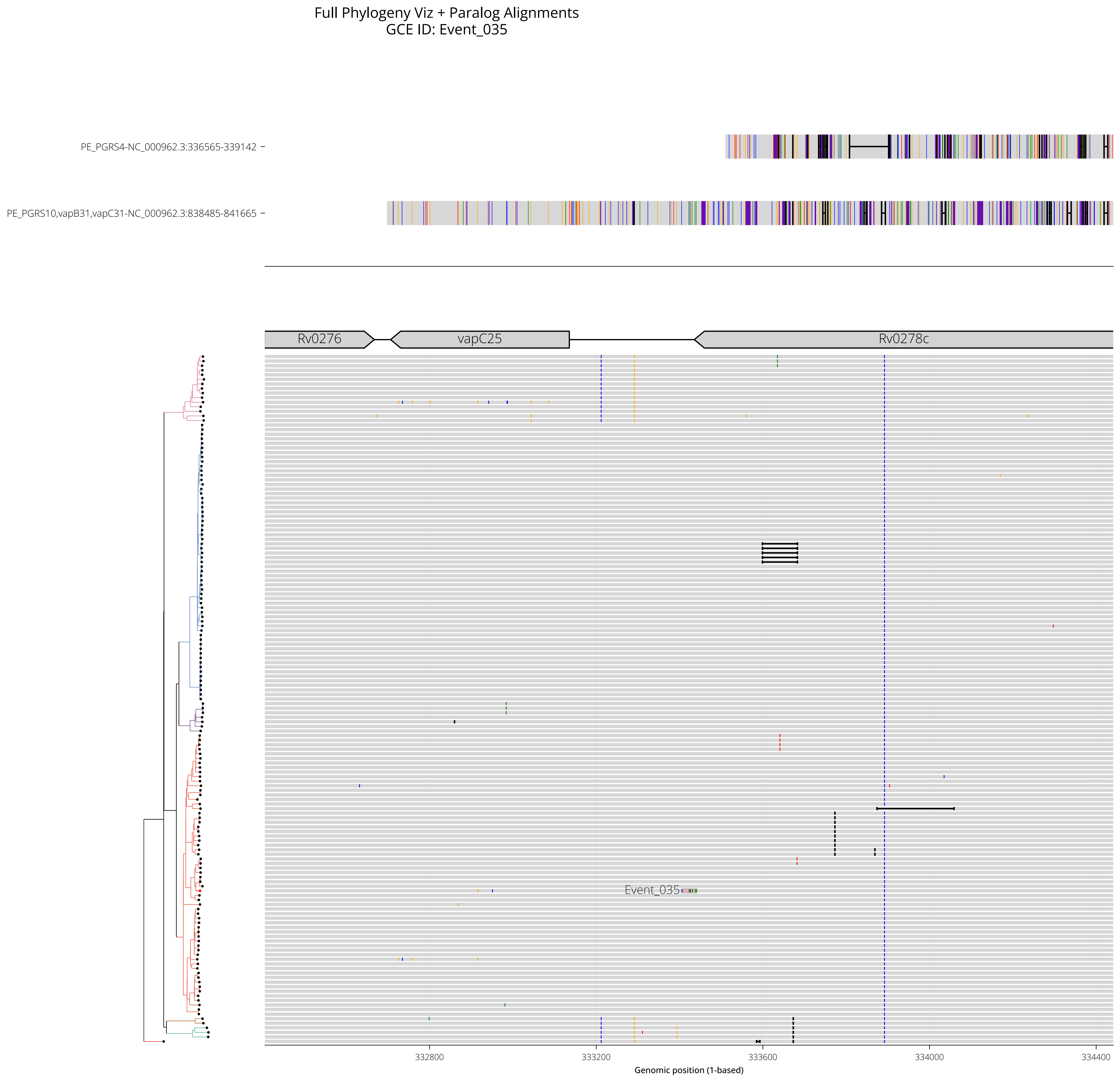
Full Phylogeny Viz + Paralog Alignments
GCE ID: Event_035
PE_PGRS4-NC_000962.3:336565-339142
PE_PGRS10,vapB31,vapC31-NC_000962.3:838485-841665
Rv0276
vapC25
Rv0278c
Event_035
332800
333200
333600
334000
334400
Genomic position (1-based)

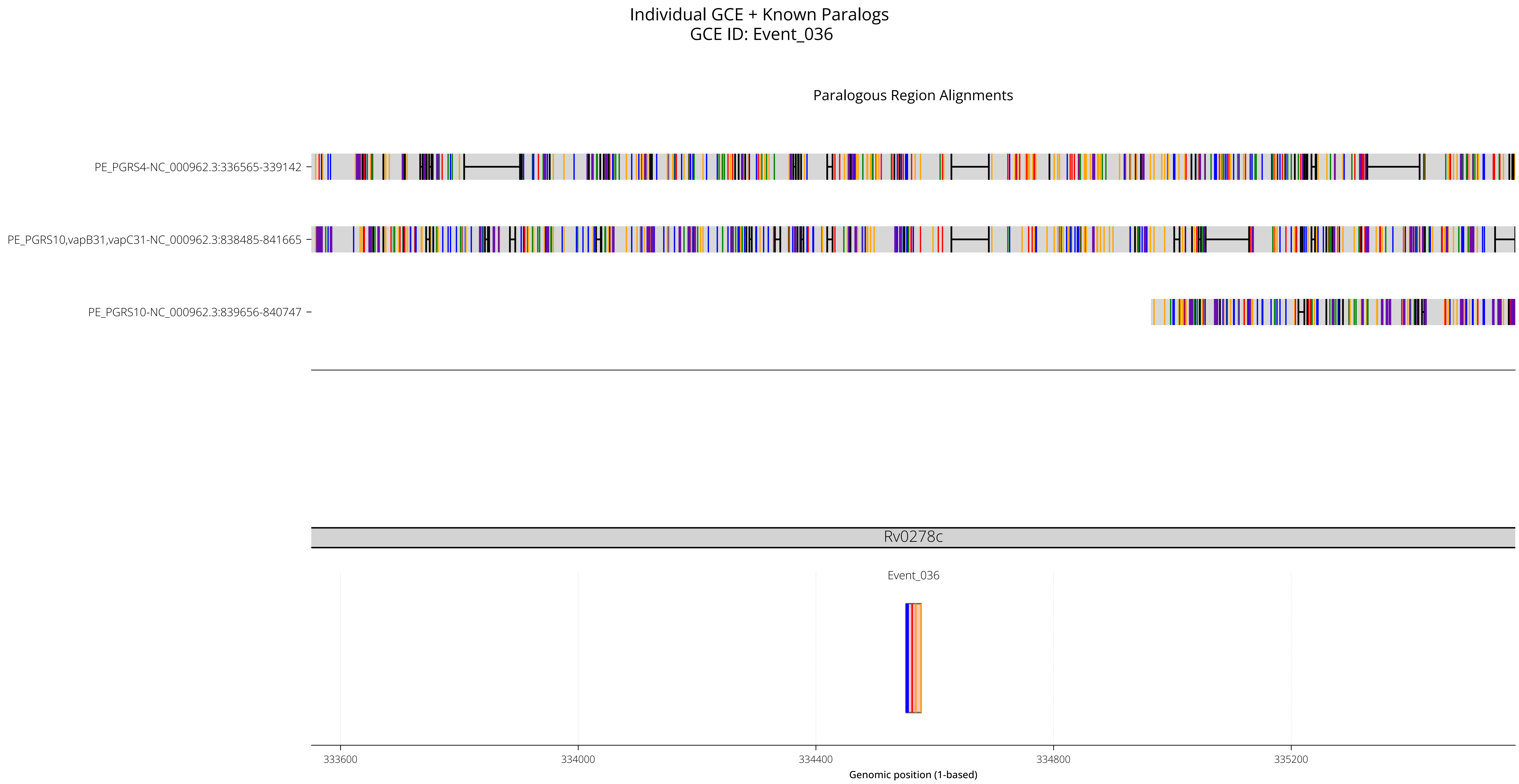
Individual GCE + Known Paralogs
GCE ID: Event_036
Paralogous Region Alignments
PE_PGRS4-NC_000962.3:336565-339142
PE_PGRS10,vapB31,vapC31-NC_000962.3:838485-841665
PE_PGRS10-NC_000962.3:839656-840747
Rv0278c
Event_036
333600
334000
334400
334800
335200
Genomic position (1-based)

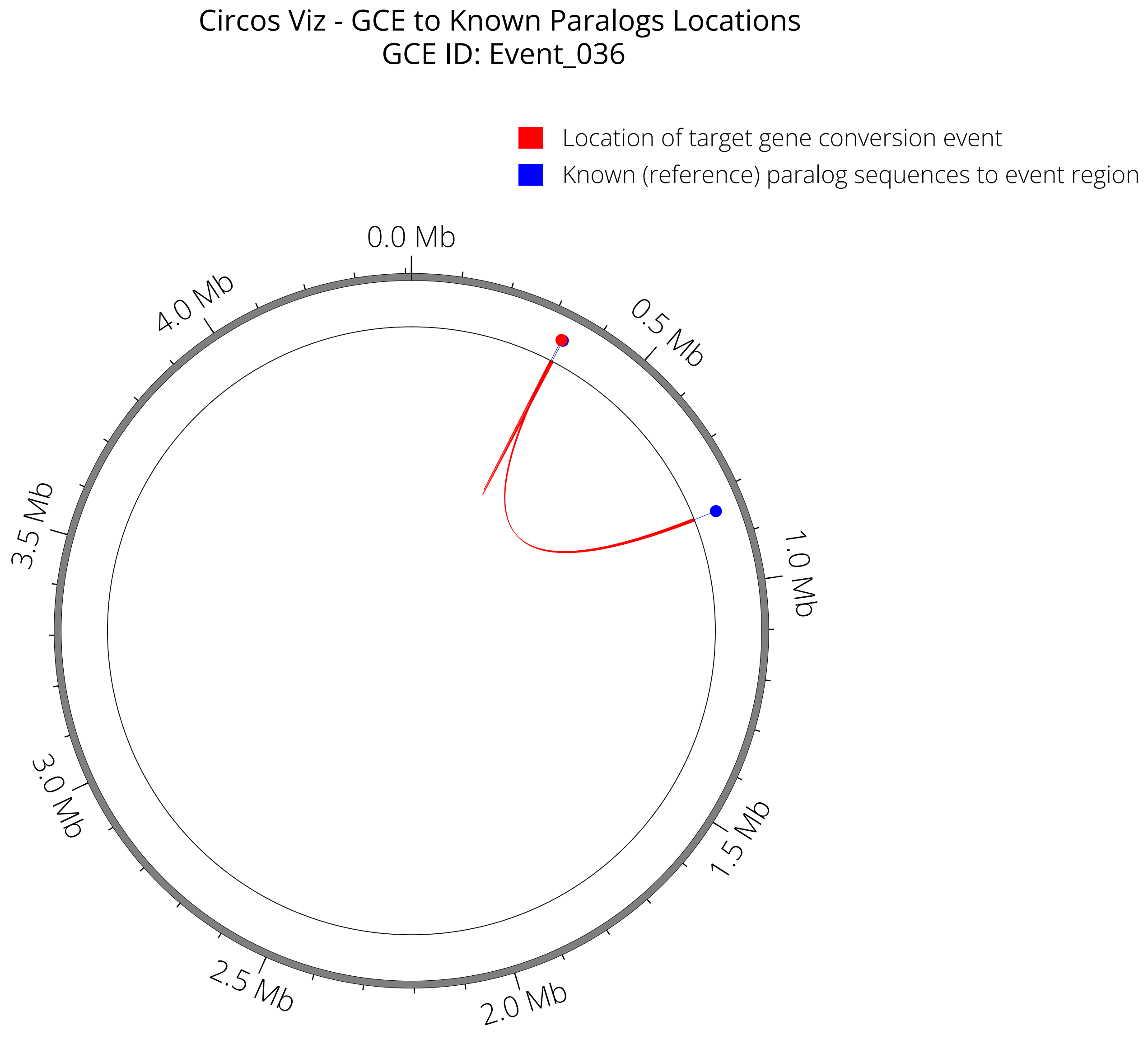

Circos Viz - GCE to Known Paralogs Locations
GCE ID: Event_036
Location of target gene conversion event
Known (reference) paralog sequences to event region
0.0 Mb
0.5 Mb
1.0 Mb
1.5 Mb
2.0 Mb
2.5 Mb
3.0 Mb
3.5 Mb
4.0 Mb

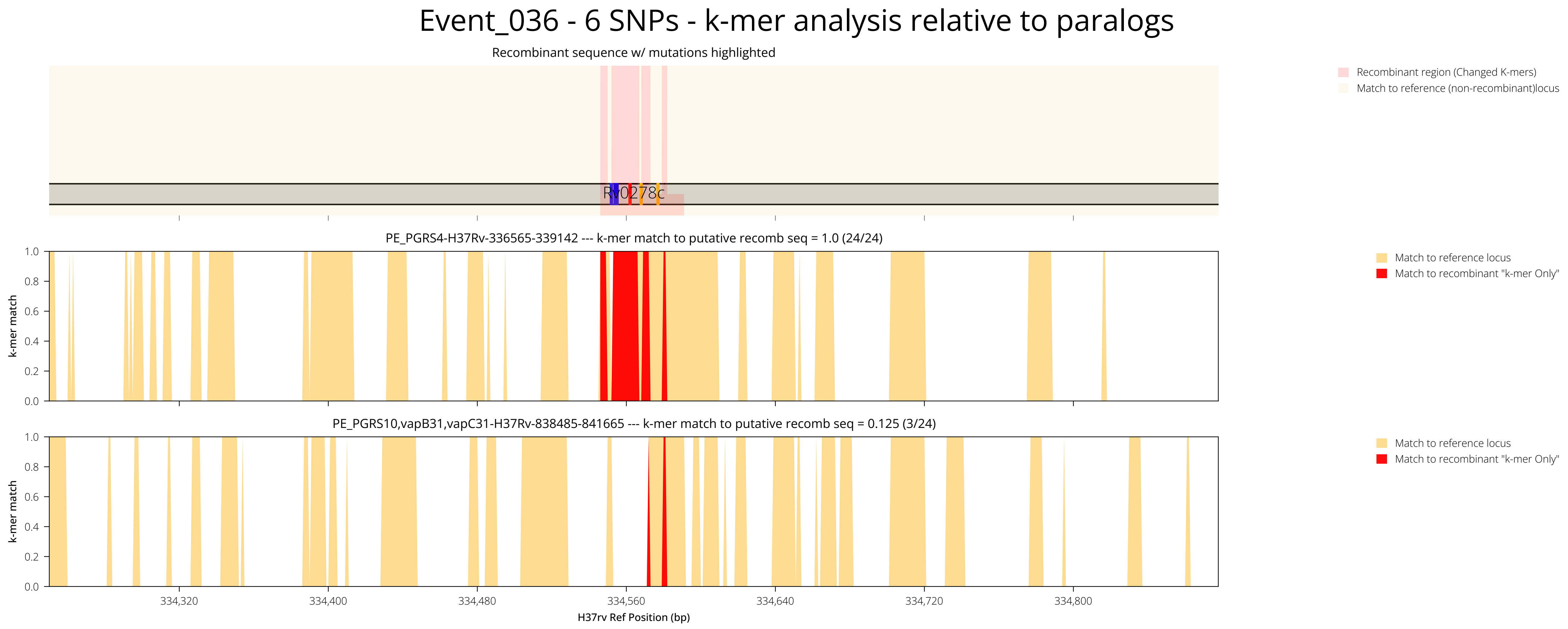
Event_036 - 6 SNPs - k-mer analysis relative to paralogs
Recombinant sequence w/ mutations highlighted
Recombinant region (Changed K-mers)
Match to reference (non-recombinant)locus
Rv0278c
PE_PGRS4-H37Rv-336565-339142 --- k-mer match to putative recomb seq = 1.0 (24/24)
Match to reference locus
Match to recombinant "k-mer Only"
k-mer match
1.0
0.8
0.6
0.4
0.2
0.0
PE_PGRS10,vapB31,vapC31-H37Rv-838485-841665 --- k-mer match to putative recomb seq = 0.125 (3/24)
Match to reference locus
Match to recombinant "k-mer Only"
k-mer match
1.0
0.8
0.6
0.4
0.2
0.0
334,320
334,400
334,480
334,560
334,640
334,720
334,800
H37rv Ref Position (bp)

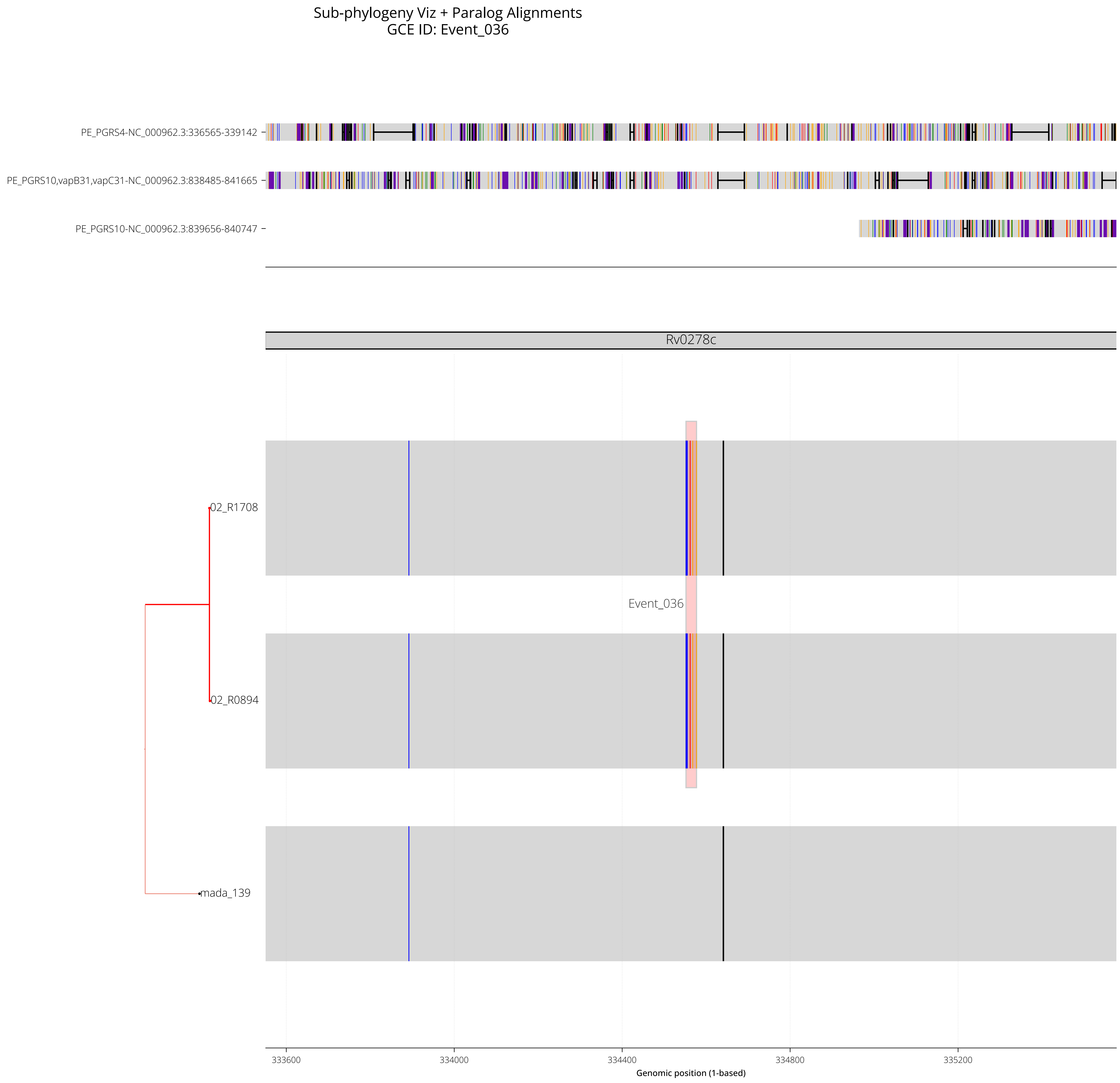

Sub-phylogeny Viz + Paralog Alignments
GCE ID: Event_036
PE_PGRS4-NC_000962.3:336565-339142
PE_PGRS10,vapB31,vapC31-NC_000962.3:838485-841665
PE_PGRS10-NC_000962.3:839656-840747
Rv0278c
02_R1708
Event_036
02_R0894
mada_139
333600
334000
334400
334800
335200
Genomic position (1-based)

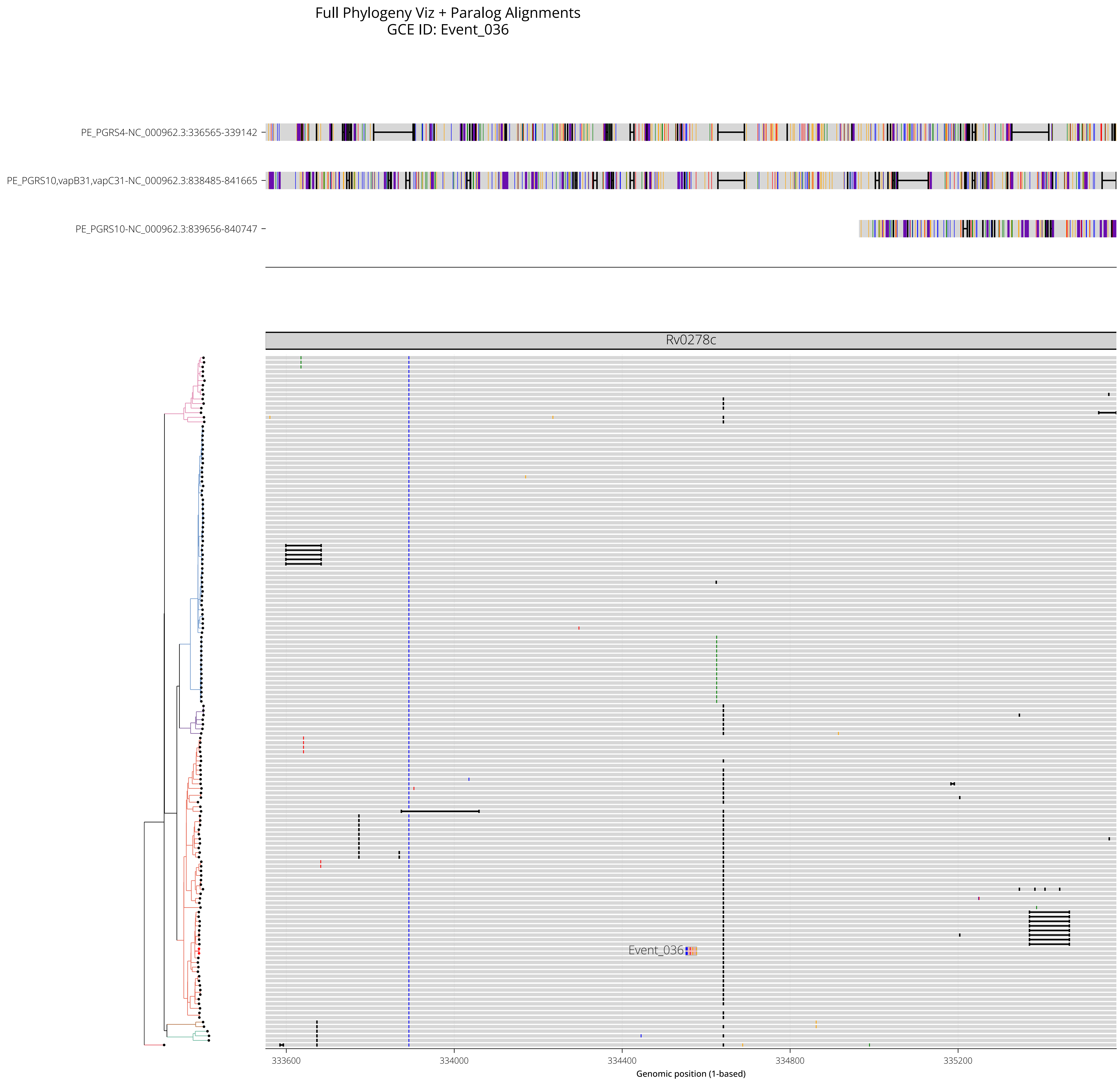

Full Phylogeny Viz + Paralog Alignments
GCE ID: Event_036
PE_PGRS4-NC_000962.3:336565-339142
PE_PGRS10,vapB31,vapC31-NC_000962.3:838485-841665
PE_PGRS10-NC_000962.3:839656-840747
Rv0278c
Event_036
333600
334000
334400
334800
335200
Genomic position (1-based)

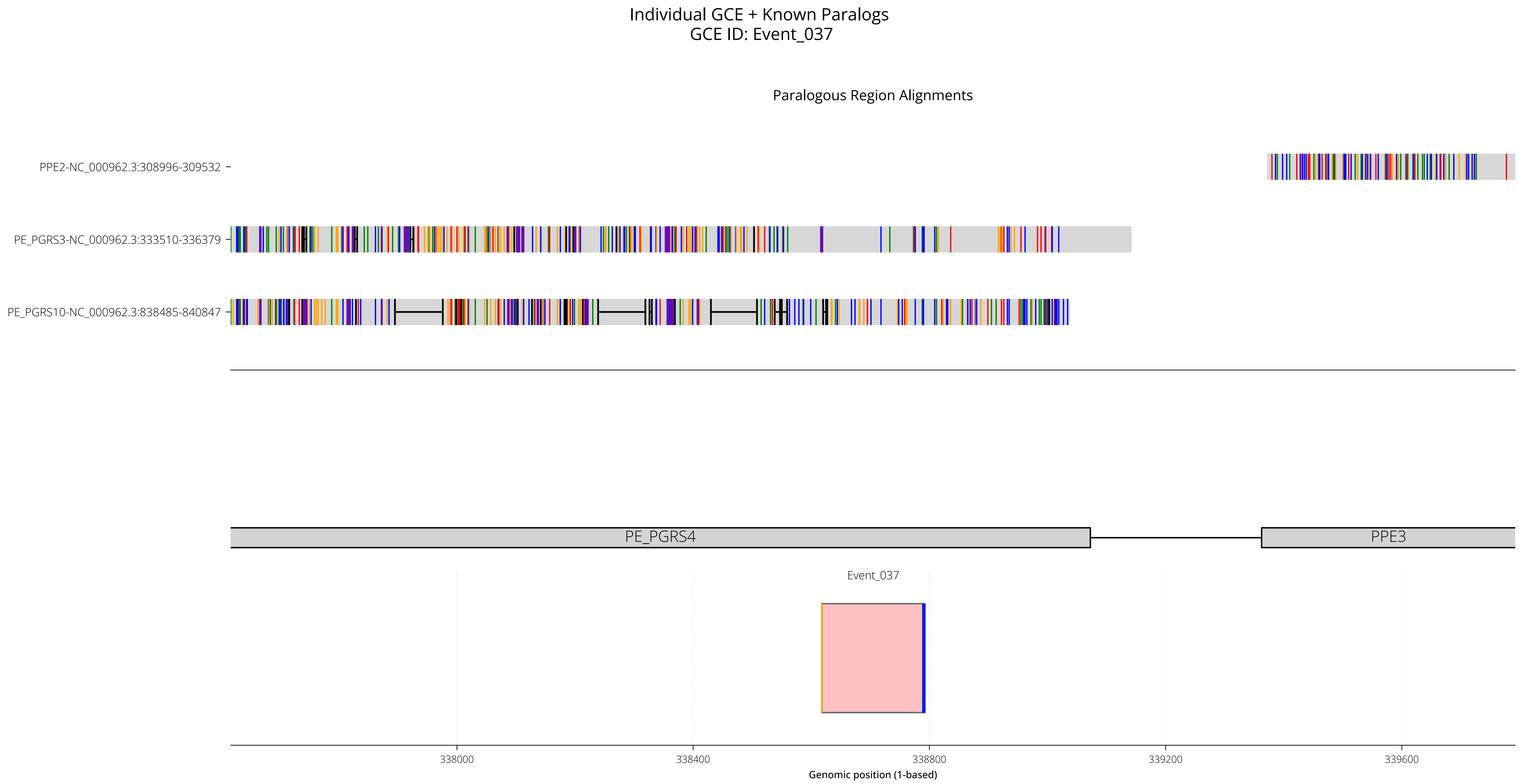
Individual GCE + Known Paralogs
GCE ID: Event_037
Paralogous Region Alignments
PPE2-NC_000962.3:308996-309532
PE_PGRS3-NC_000962.3:333510-336379
PE_PGRS10-NC_000962.3:838485-840847
PE_PGRS4
PPE3
Event_037
338000
338400
338800
339200
339600
Genomic position (1-based)

Circos Viz - GCE to Known Paralogs Locations
GCE ID: Event_037

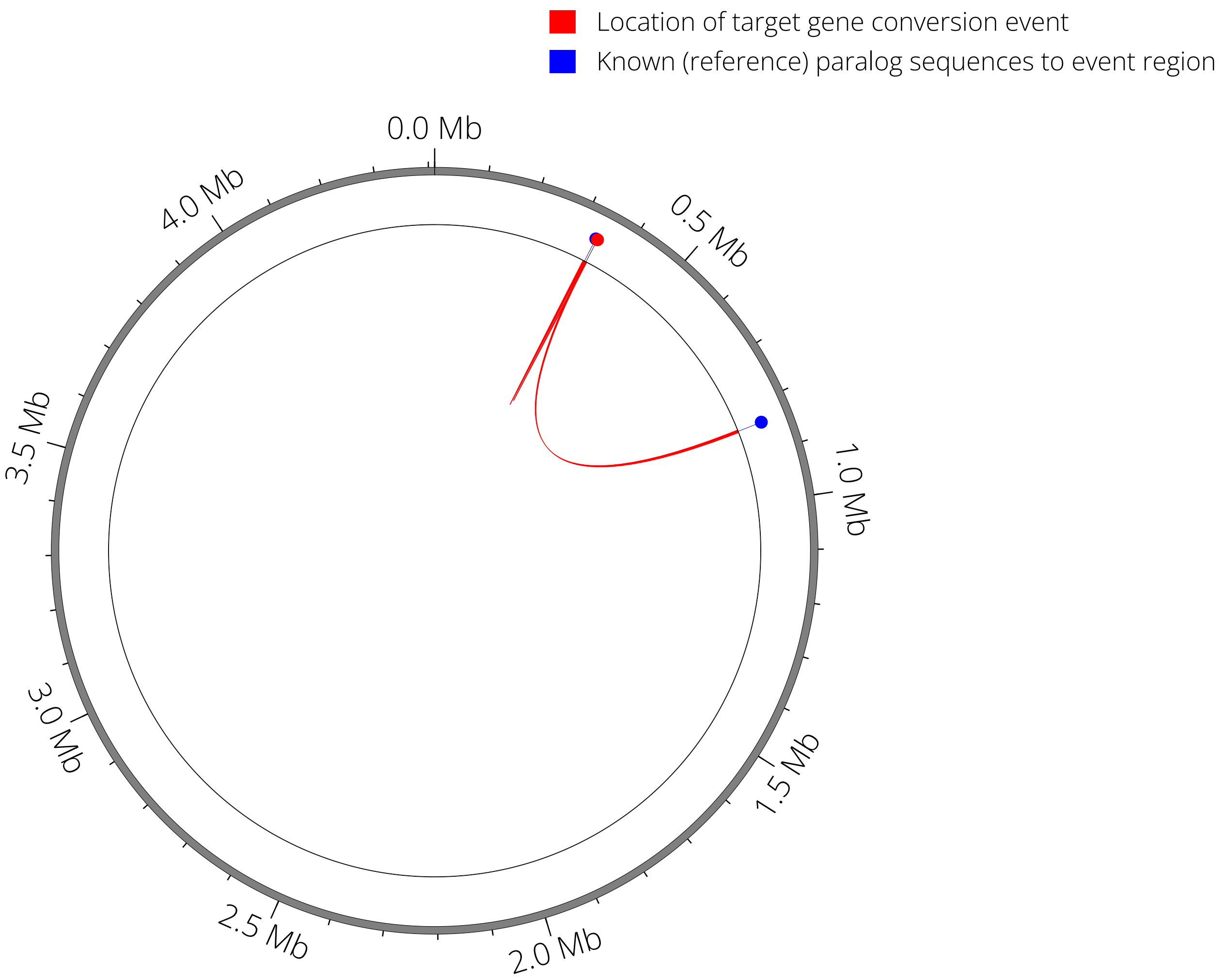

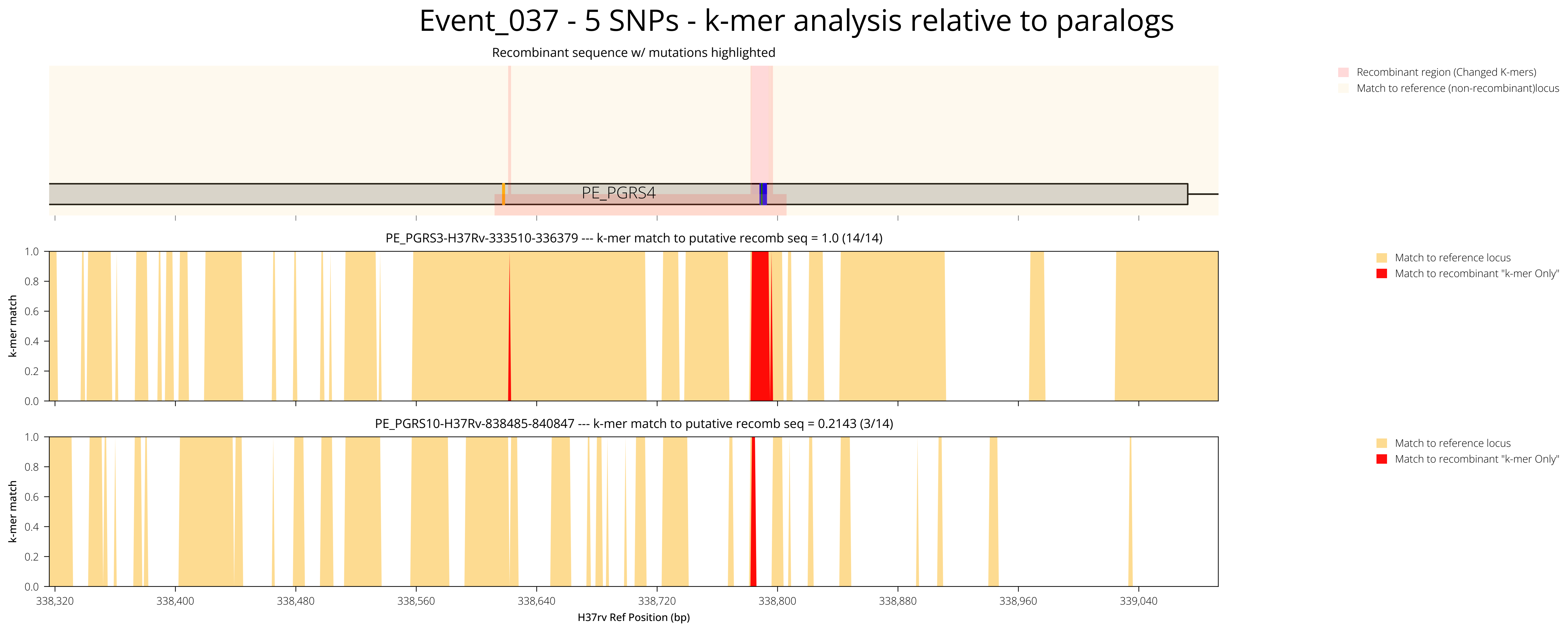

Event_037 - 5 SNPs - k-mer analysis relative to paralogs
Recombinant sequence w/ mutations highlighted
Recombinant region (Changed K-mers)
Match to reference (non-recombinant)locus
PE_PGRS4
PE_PGRS3-H37Rv-333510-336379 --- k-mer match to putative recomb seq = 1.0 (14/14)
Match to reference locus
Match to recombinant "k-mer Only"
PE_PGRS10-H37Rv-838485-840847 --- k-mer match to putative recomb seq = 0.2143 (3/14)
Match to reference locus
Match to recombinant "k-mer Only"
k-mer match
1.0
0.8
0.6
0.4
0.2
0.0
338,320
338,400
338,480
338,560
338,640
338,720
338,800
338,880
338,960
339,040
H37rv Ref Position (bp)

#### Sub-phylogeny Viz + Paralog Alignments
#### GCE ID: Event_037

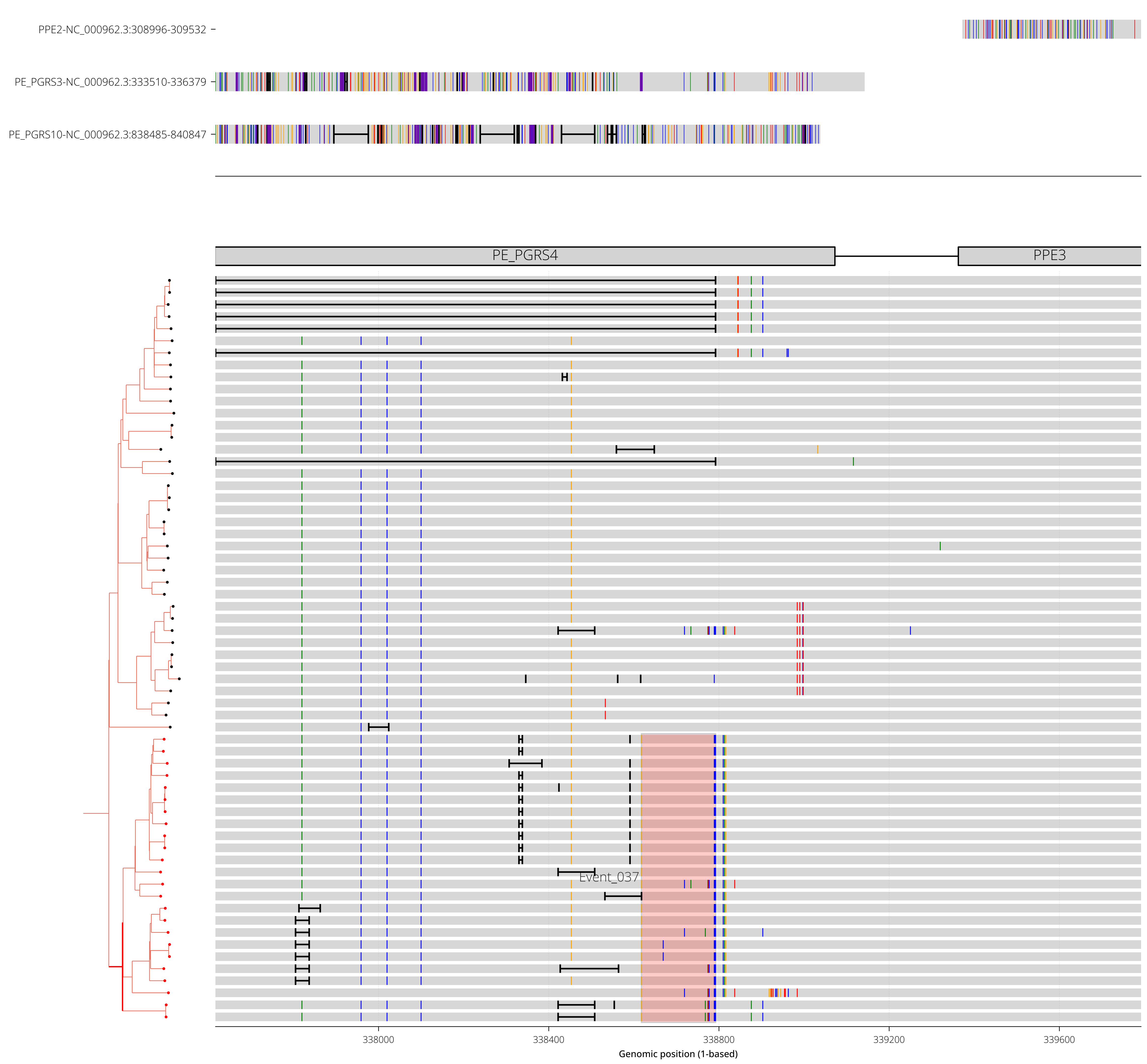

Full Phylogeny Viz + Paralog Alignments
GCE ID: Event_037

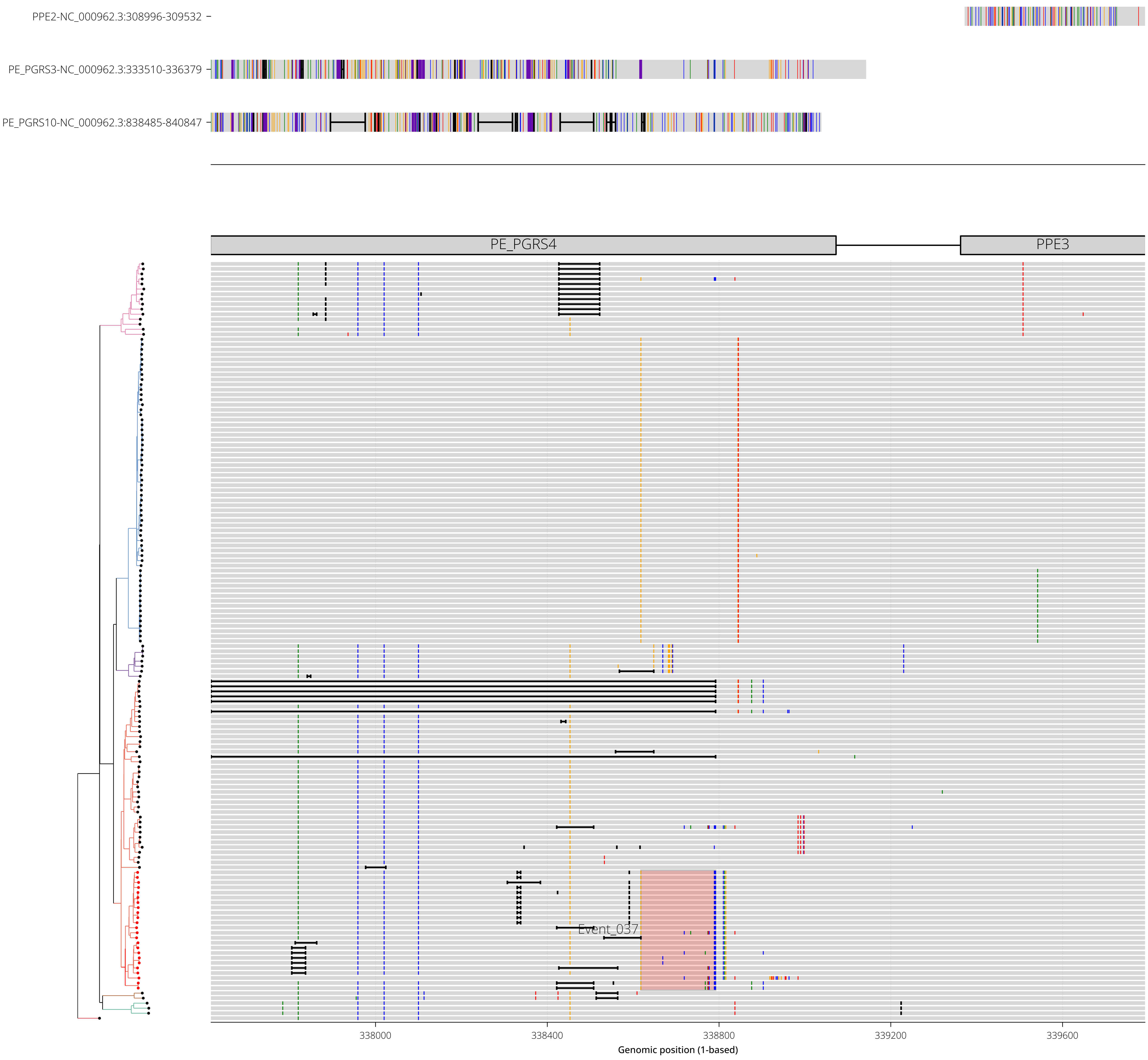

Individual GCE + Known Paralogs
GCE ID: Event_038

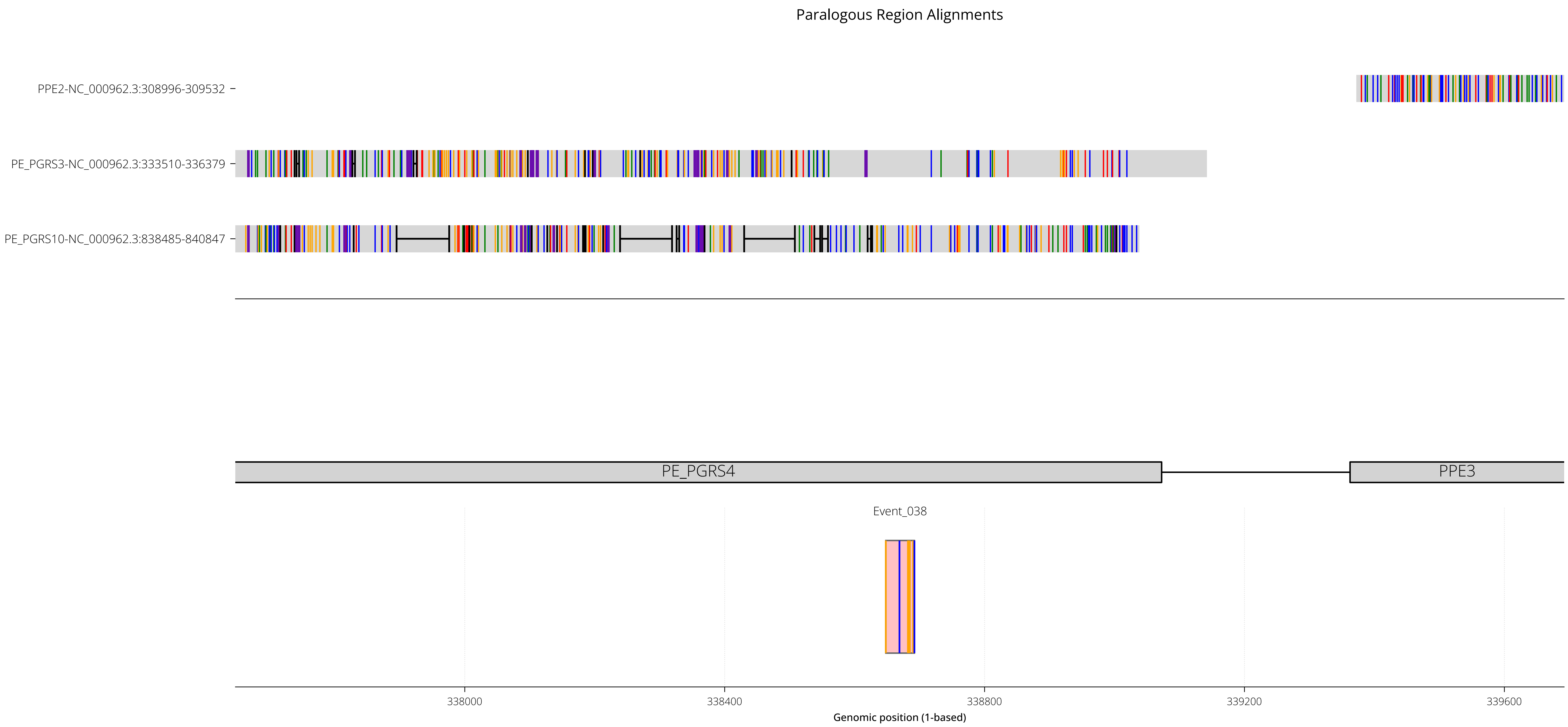

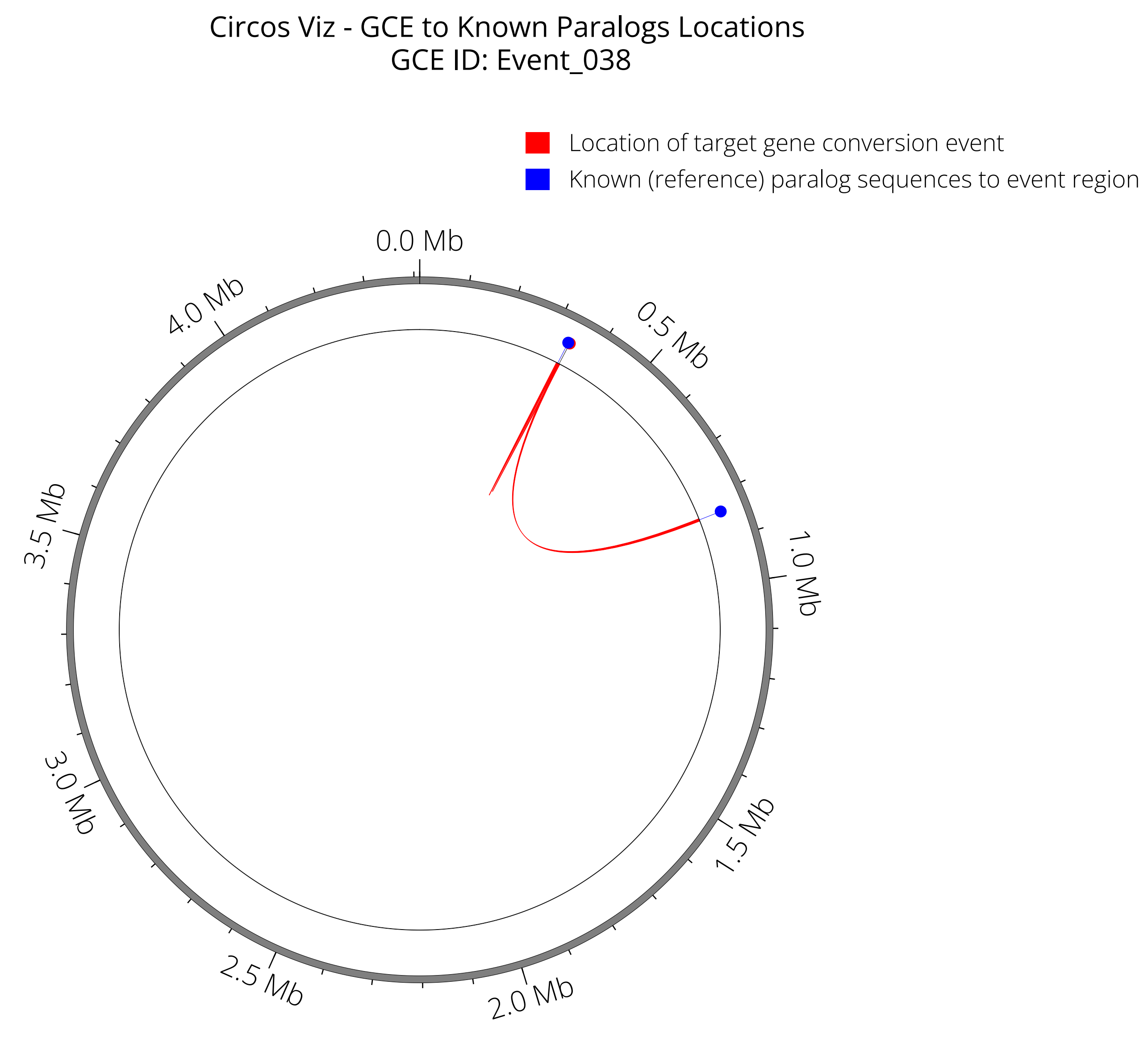
Circos Viz - GCE to Known Paralogs Locations
GCE ID: Event_038
Location of target gene conversion event
Known (reference) paralog sequences to event region
0.0 Mb
0.5 Mb
1.0 Mb
1.5 Mb
2.0 Mb
2.5 Mb
3.0 Mb
3.5 Mb
4.0 Mb

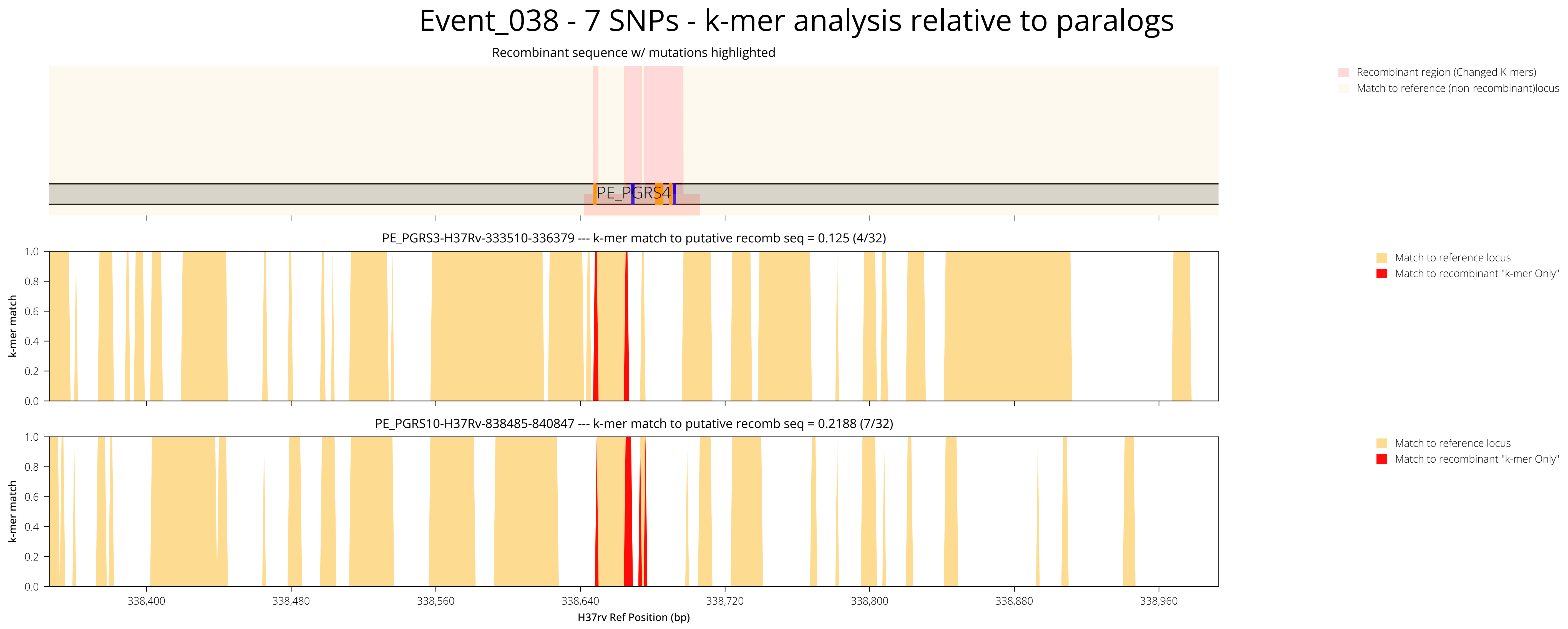
Event_038 - 7 SNPs - k-mer analysis relative to paralogs
Recombinant sequence w/ mutations highlighted
PE_PGRS4
Recombinant region (Changed K-mers)
Match to reference (non-recombinant)locus
PE_PGRS3-H37Rv-333510-336379 --- k-mer match to putative recomb seq = 0.125 (4/32)
Match to reference locus
Match to recombinant "k-mer Only"
PE_PGRS10-H37Rv-838485-840847 --- k-mer match to putative recomb seq = 0.2188 (7/32)
Match to reference locus
Match to recombinant "k-mer Only"
k-mer match
1.0
0.8
0.6
0.4
0.2
0.0
338,400
338,480
338,560
338,640
338,720
338,800
338,880
338,960
H37rv Ref Position (bp)

Sub-phylogeny Viz + Paralog Alignments
GCE ID: Event_038

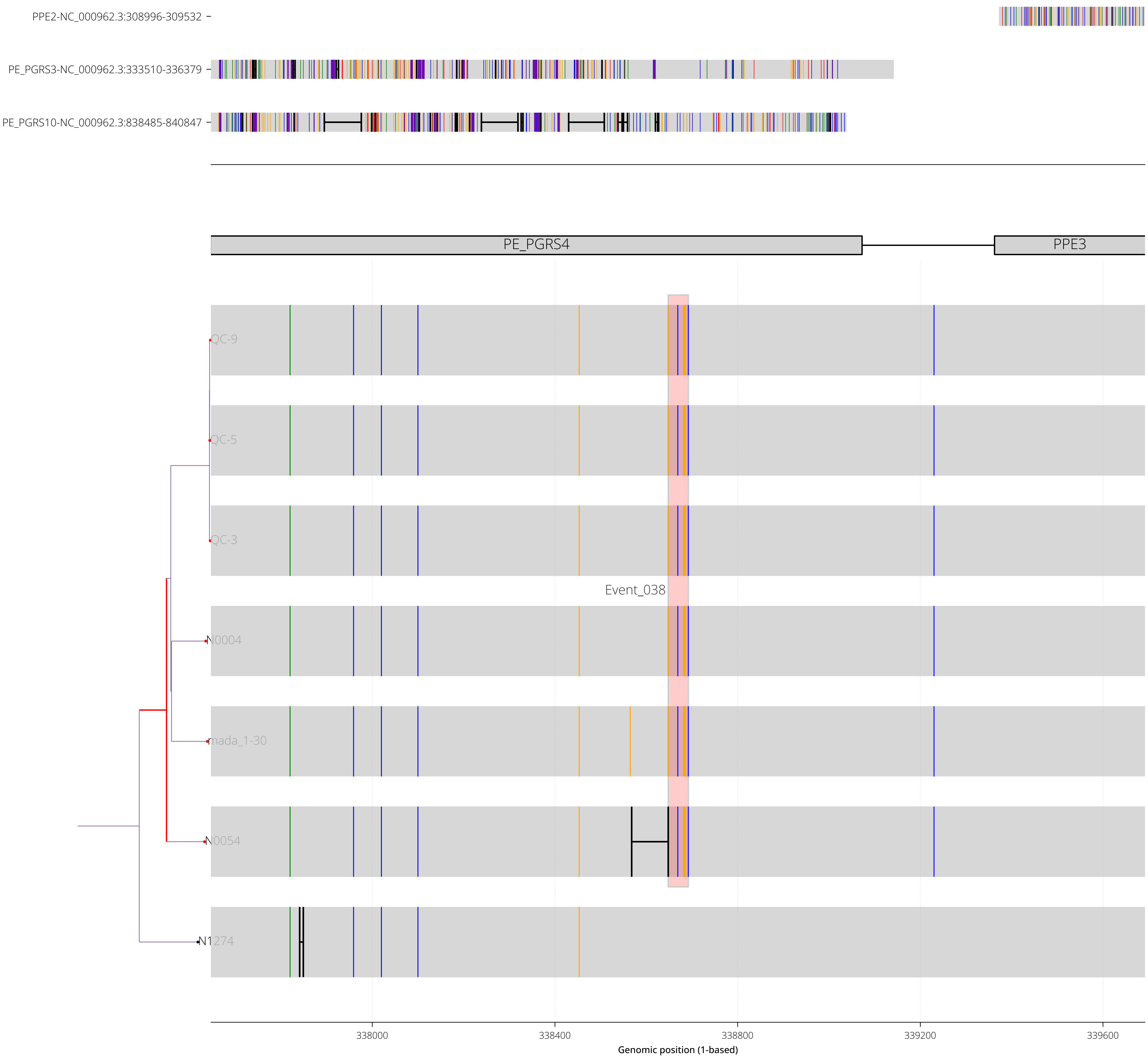

Full Phylogeny Viz + Paralog Alignments
GCE ID: Event_038

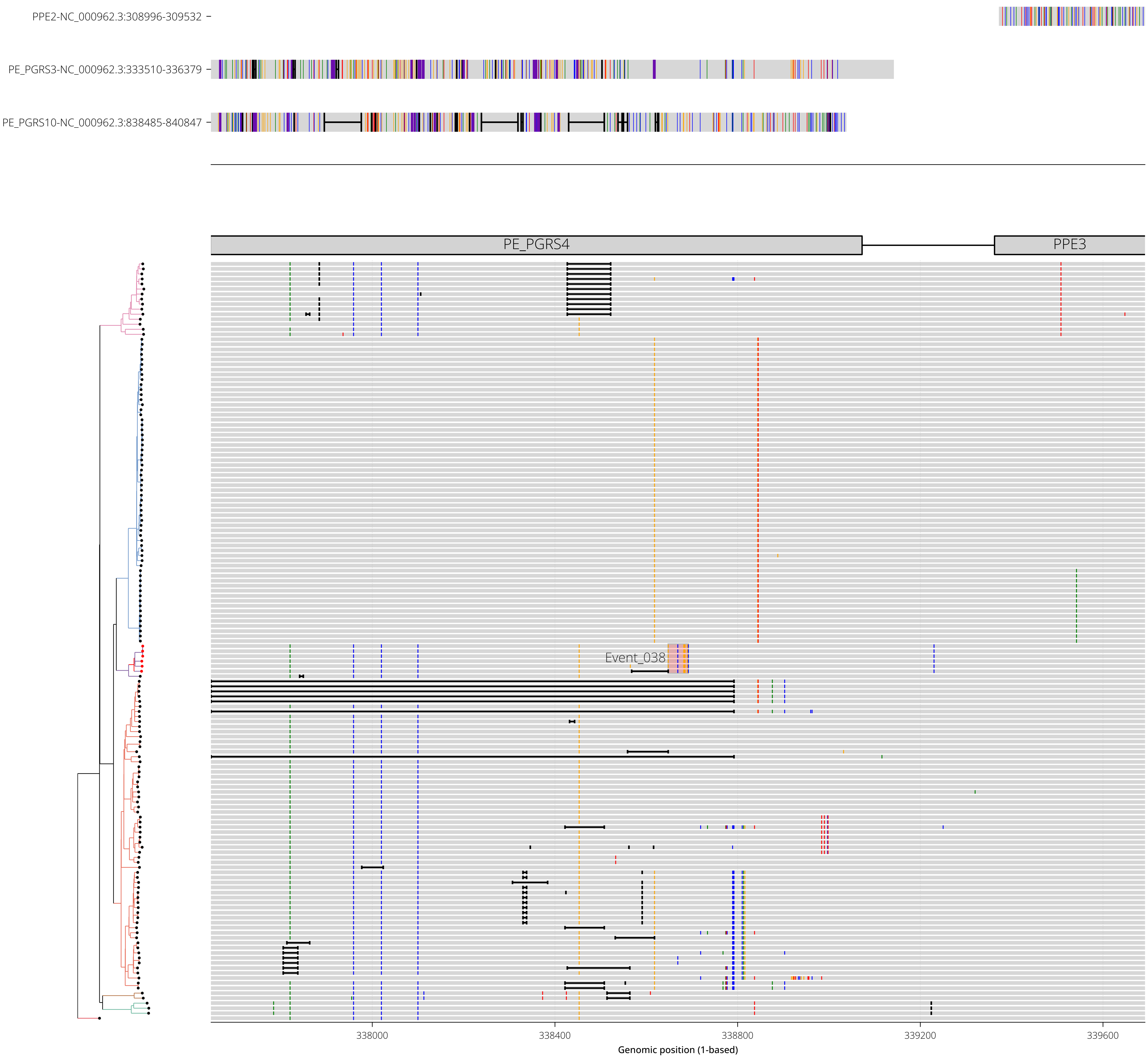

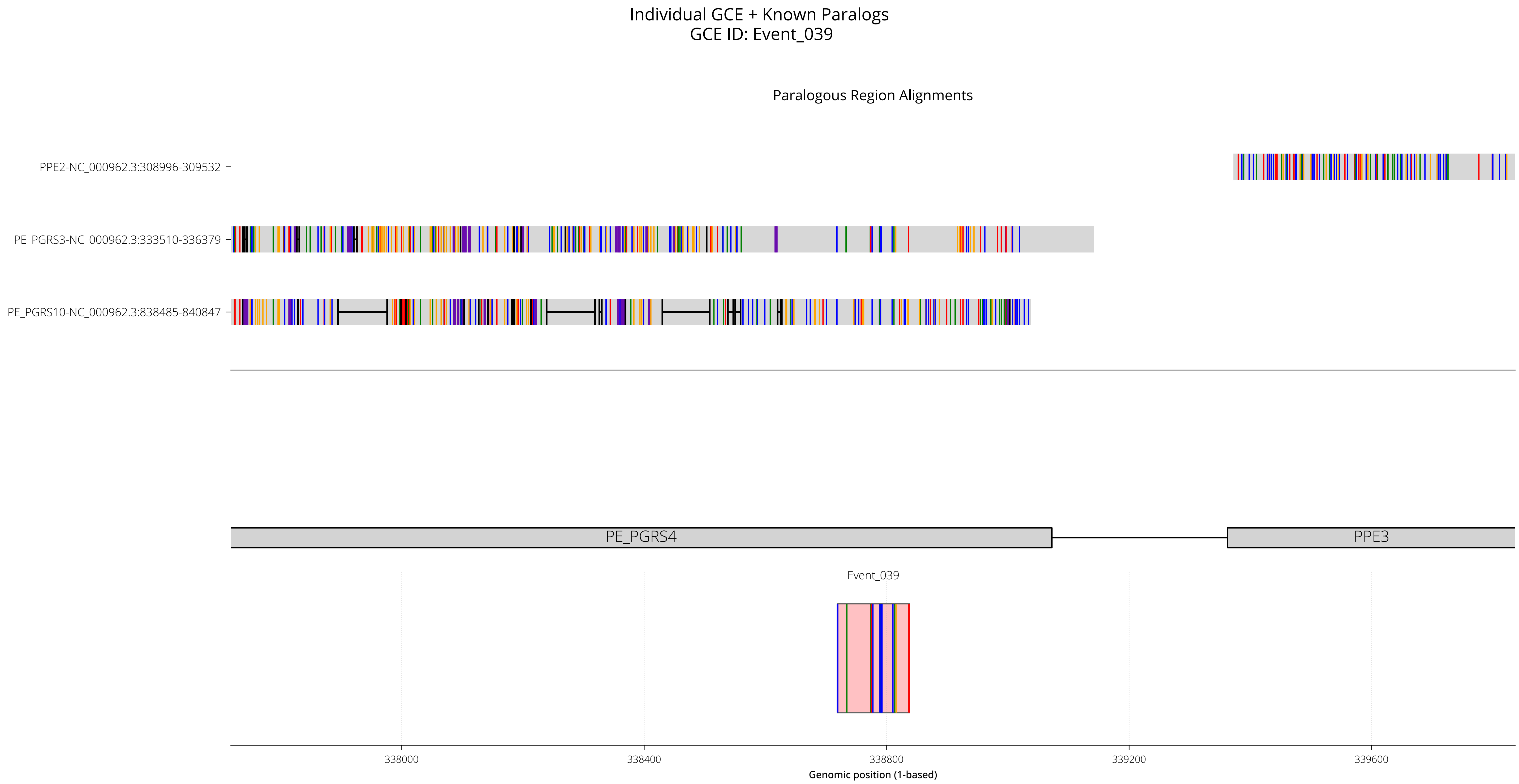

Individual GCE + Known Paralogs
GCE ID: Event_039
Paralogous Region Alignments
PPE2-NC_000962.3:308996-309532
PE_PGRS3-NC_000962.3:333510-336379
PE_PGRS10-NC_000962.3:838485-840847
PE_PGRS4
PPE3
Event_039
338000
338400
338800
339200
339600
Genomic position (1-based)

Circos Viz - GCE to Known Paralogs Locations
GCE ID: Event_039

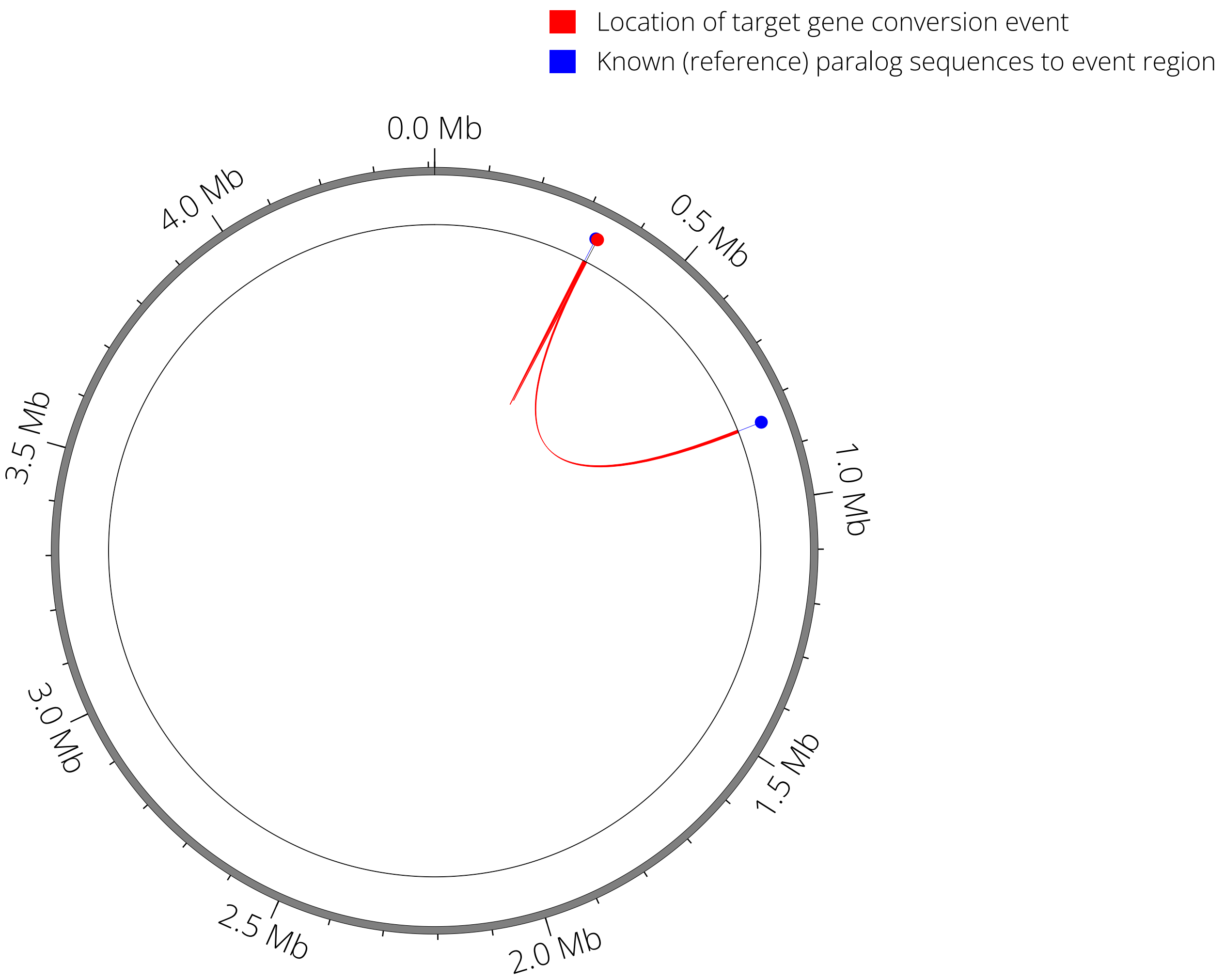

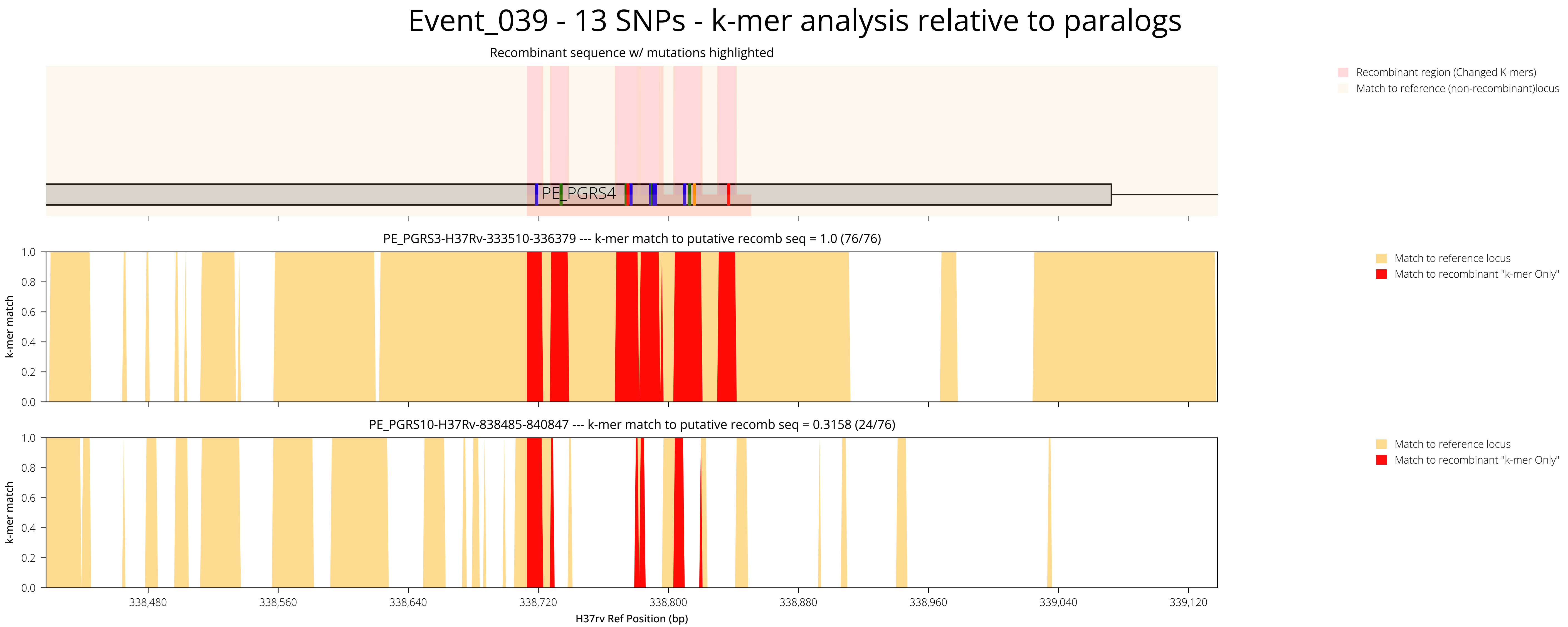

Event_039 - 13 SNPs - k-mer analysis relative to paralogs
Recombinant sequence w/ mutations highlighted
PE_PGRS4
Recombinant region (Changed K-mers)
Match to reference (non-recombinant)locus
PE_PGRS3-H37Rv-333510-336379 --- k-mer match to putative recomb seq = 1.0 (76/76)
Match to reference locus
Match to recombinant "k-mer Only"
PE_PGRS10-H37Rv-838485-840847 --- k-mer match to putative recomb seq = 0.3158 (24/76)
Match to reference locus
Match to recombinant "k-mer Only"
k-mer match
1.0
0.8
0.6
0.4
0.2
0.0
338,480
338,560
338,640
338,720
338,800
338,880
338,960
339,040
339,120
H37rv Ref Position (bp)

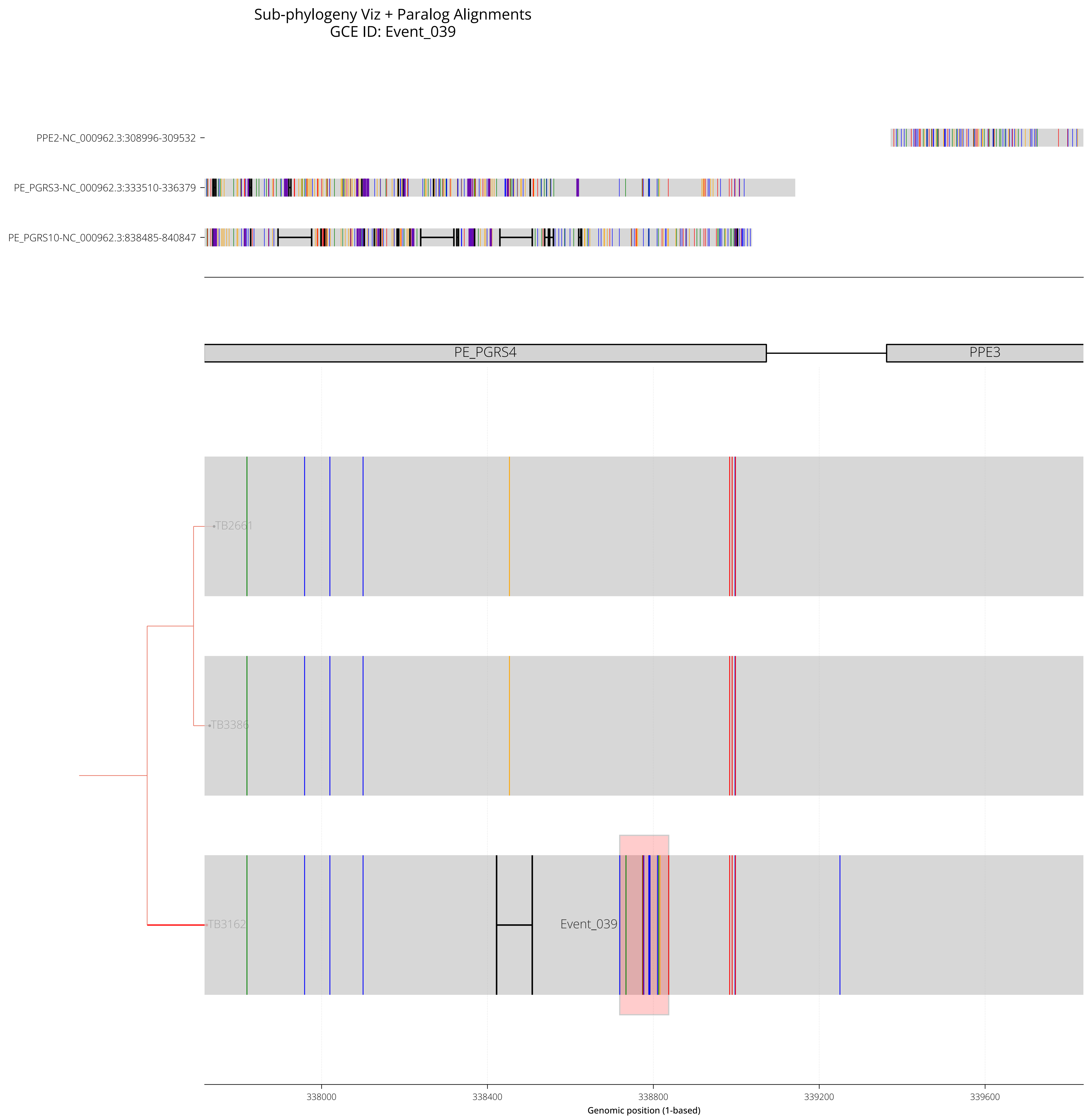

Sub-phylogeny Viz + Paralog Alignments
GCE ID: Event_039
PPE2-NC_000962.3:308996-309532
PE_PGRS3-NC_000962.3:333510-336379
PE_PGRS10-NC_000962.3:838485-840847
PE_PGRS4
PPE3
TB2661
TB3386
TB3162
Event_039
338000
338400
338800
339200
339600
Genomic position (1-based)

Full Phylogeny Viz + Paralog Alignments
GCE ID: Event_039

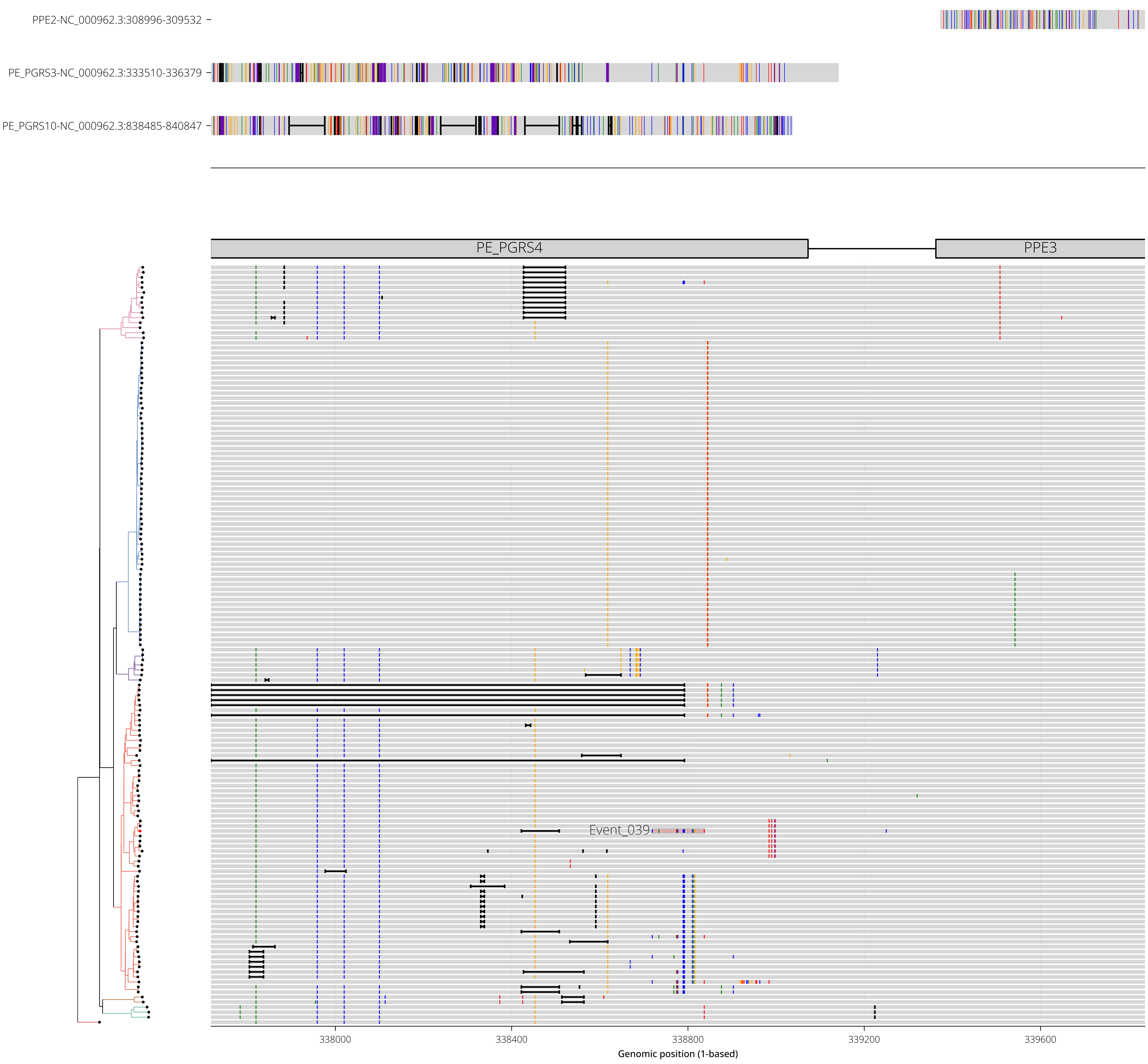

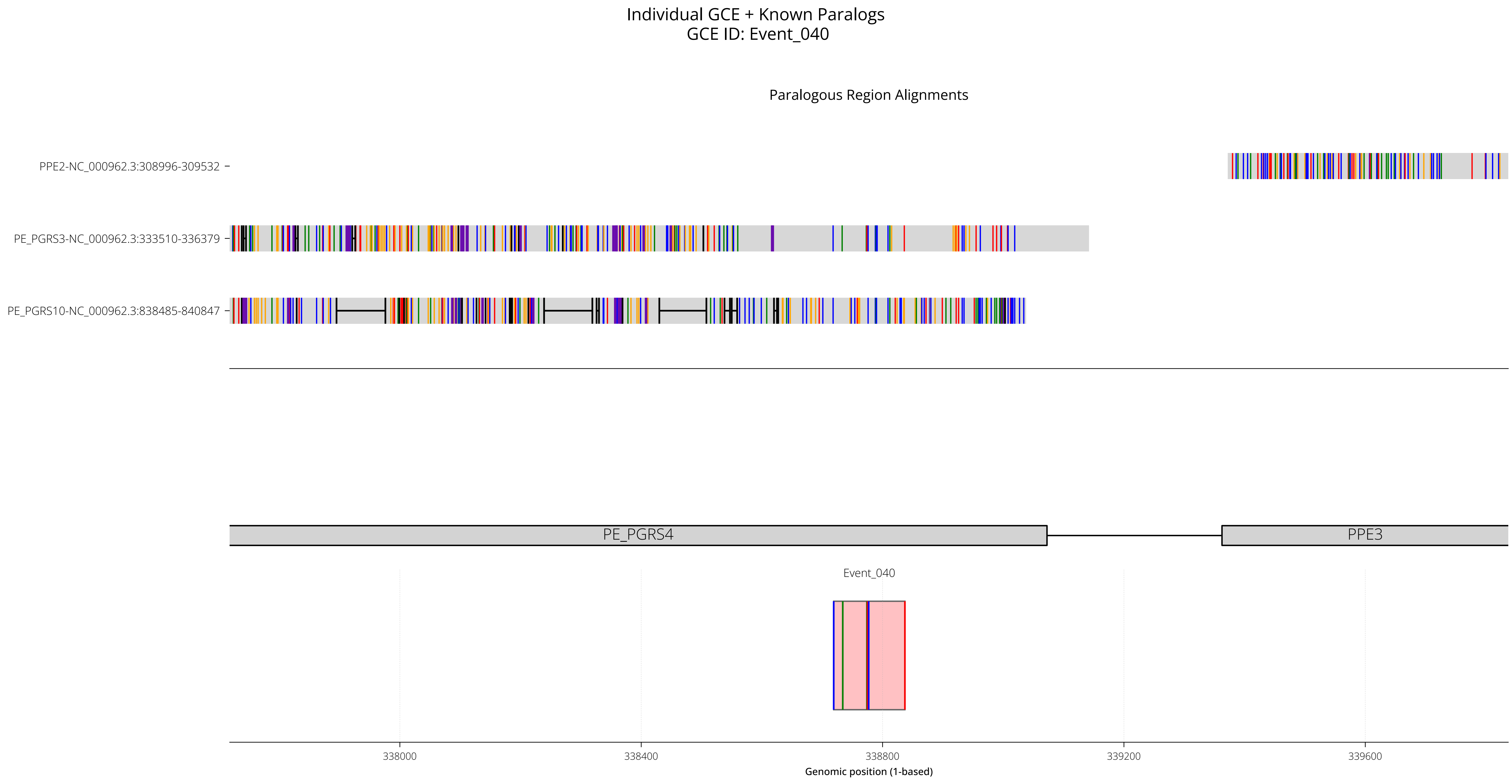
Individual GCE + Known Paralogs
GCE ID: Event_040
Paralogous Region Alignments
PPE2-NC_000962.3:308996-309532
PE_PGRS3-NC_000962.3:333510-336379
PE_PGRS10-NC_000962.3:838485-840847
PE_PGRS4
PPE3
Event_040
338000
338400
338800
339200
339600
Genomic position (1-based)

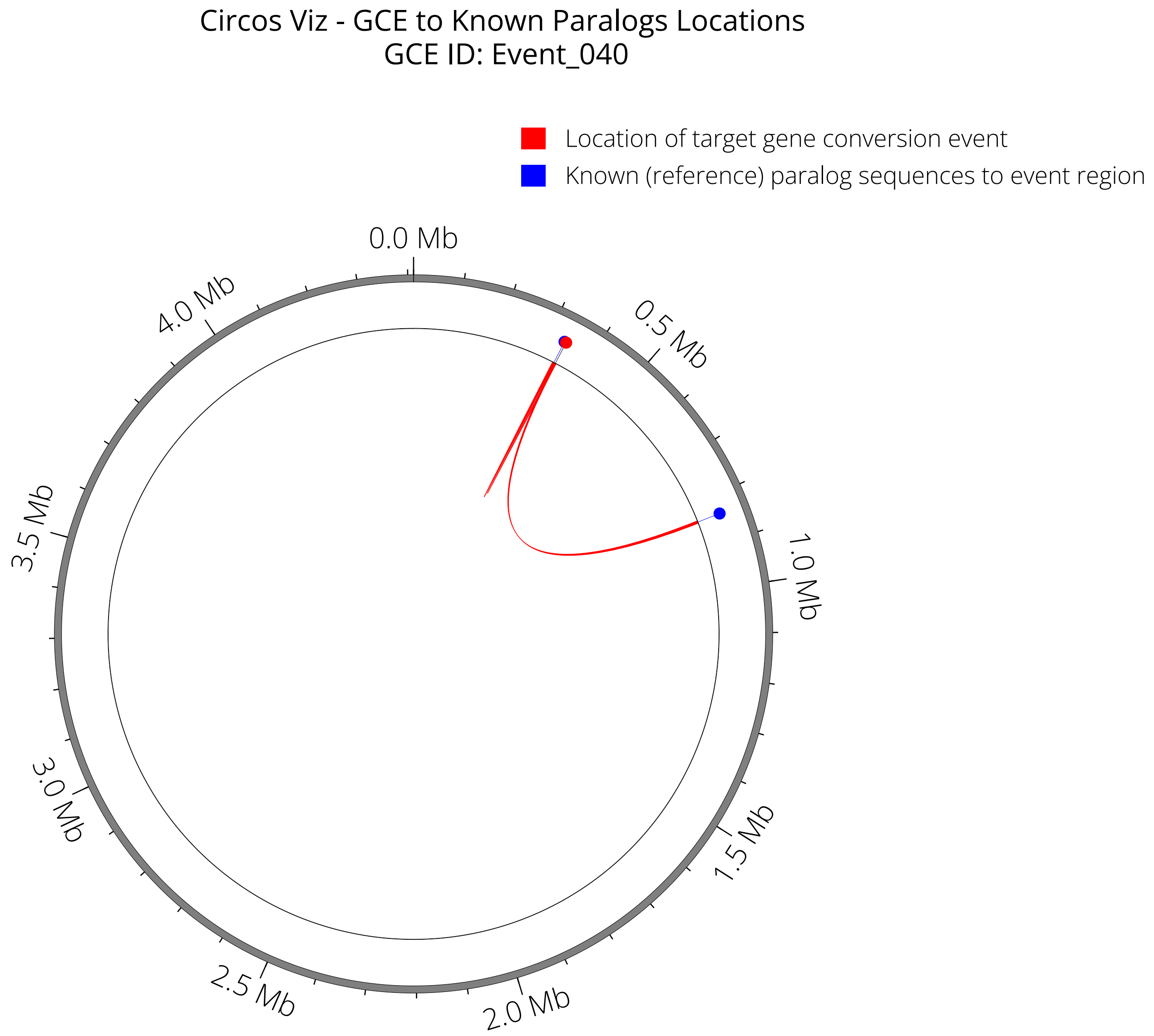
Circos Viz - GCE to Known Paralogs Locations
GCE ID: Event_040
Location of target gene conversion event
Known (reference) paralog sequences to event region
0.0 Mb
0.5 Mb
1.0 Mb
1.5 Mb
2.0 Mb
2.5 Mb
3.0 Mb
3.5 Mb
4.0 Mb

### Event_040 - 6 SNPs - k-mer analysis relative to paralogs

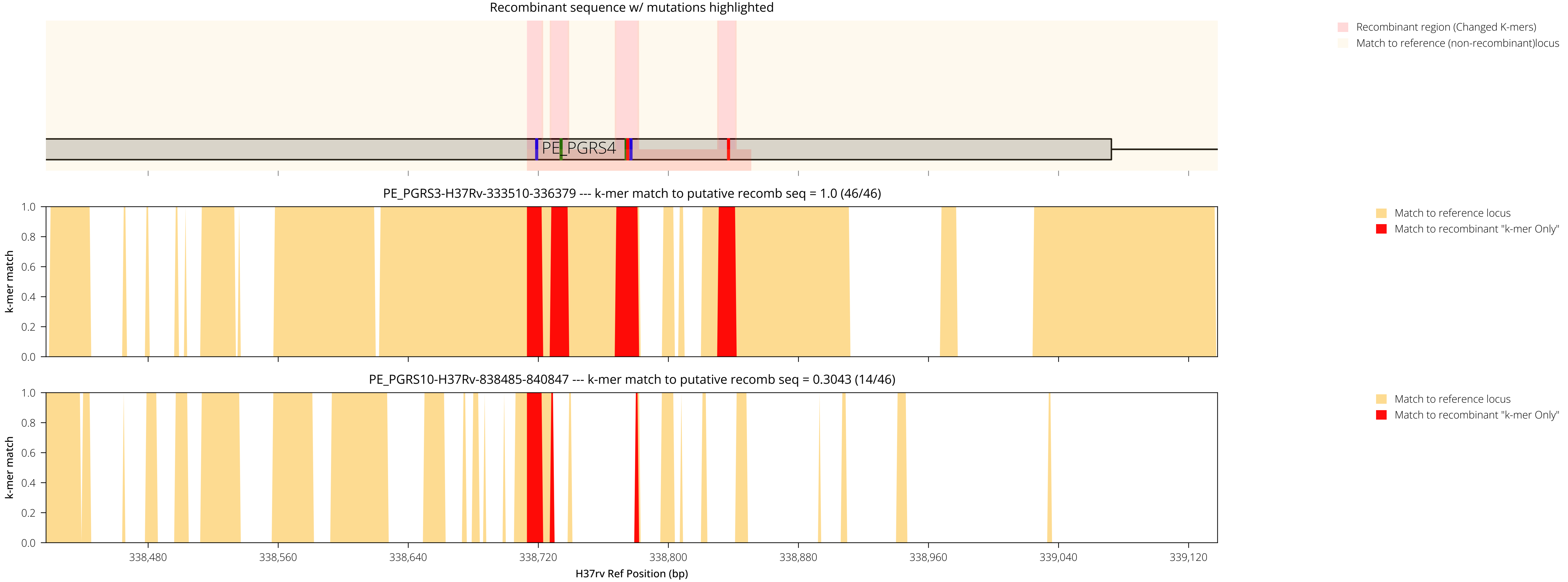

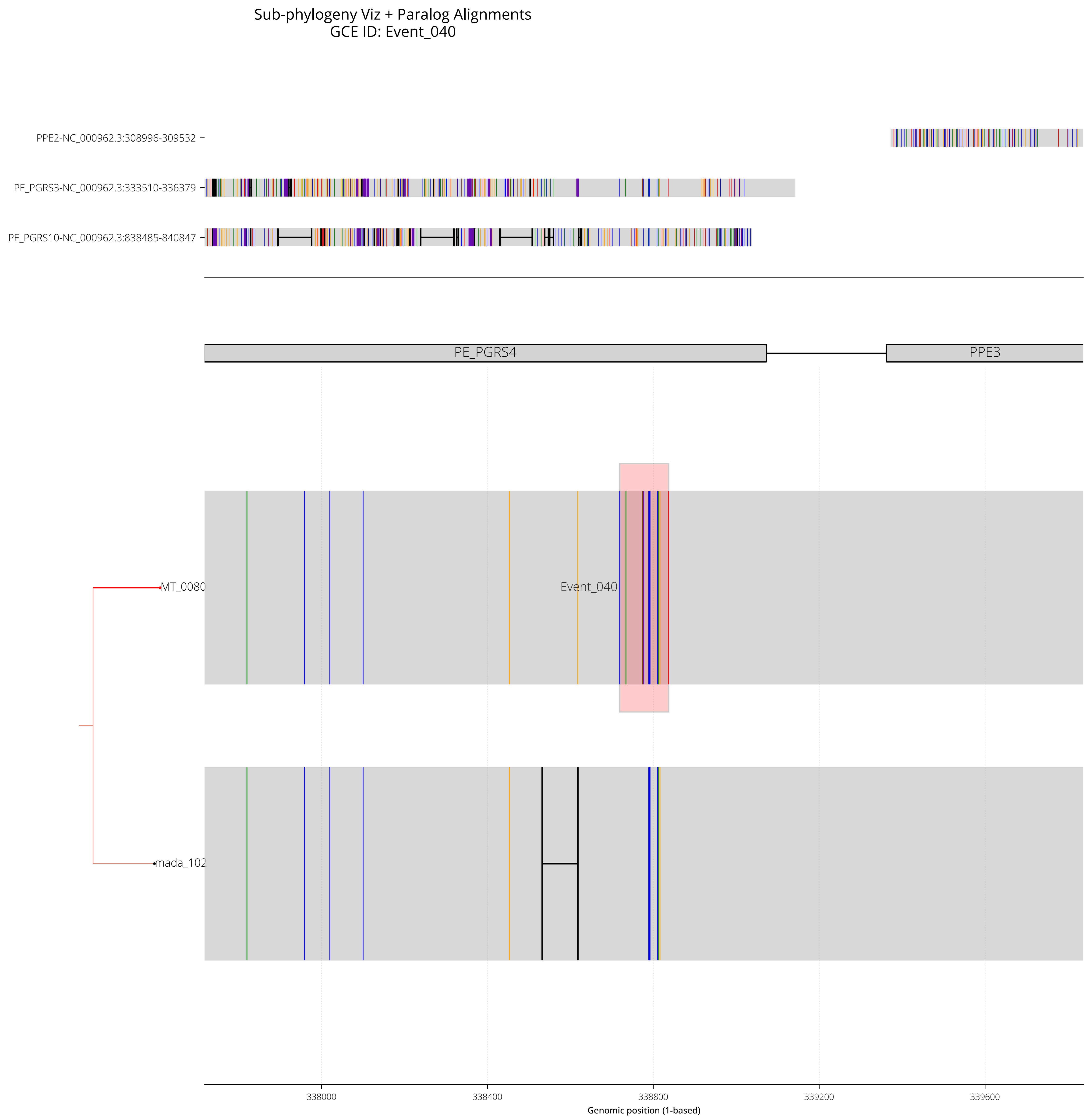
Sub-phylogeny Viz + Paralog Alignments
GCE ID: Event_040
PPE2-NC_000962.3:308996-309532
PE_PGRS3-NC_000962.3:333510-336379
PE_PGRS10-NC_000962.3:838485-840847
PE_PGRS4
PPE3
MT_0080
Event_040
mada_102
338000
338400
338800
339200
339600
Genomic position (1-based)

Full Phylogeny Viz + Paralog Alignments
GCE ID: Event_040

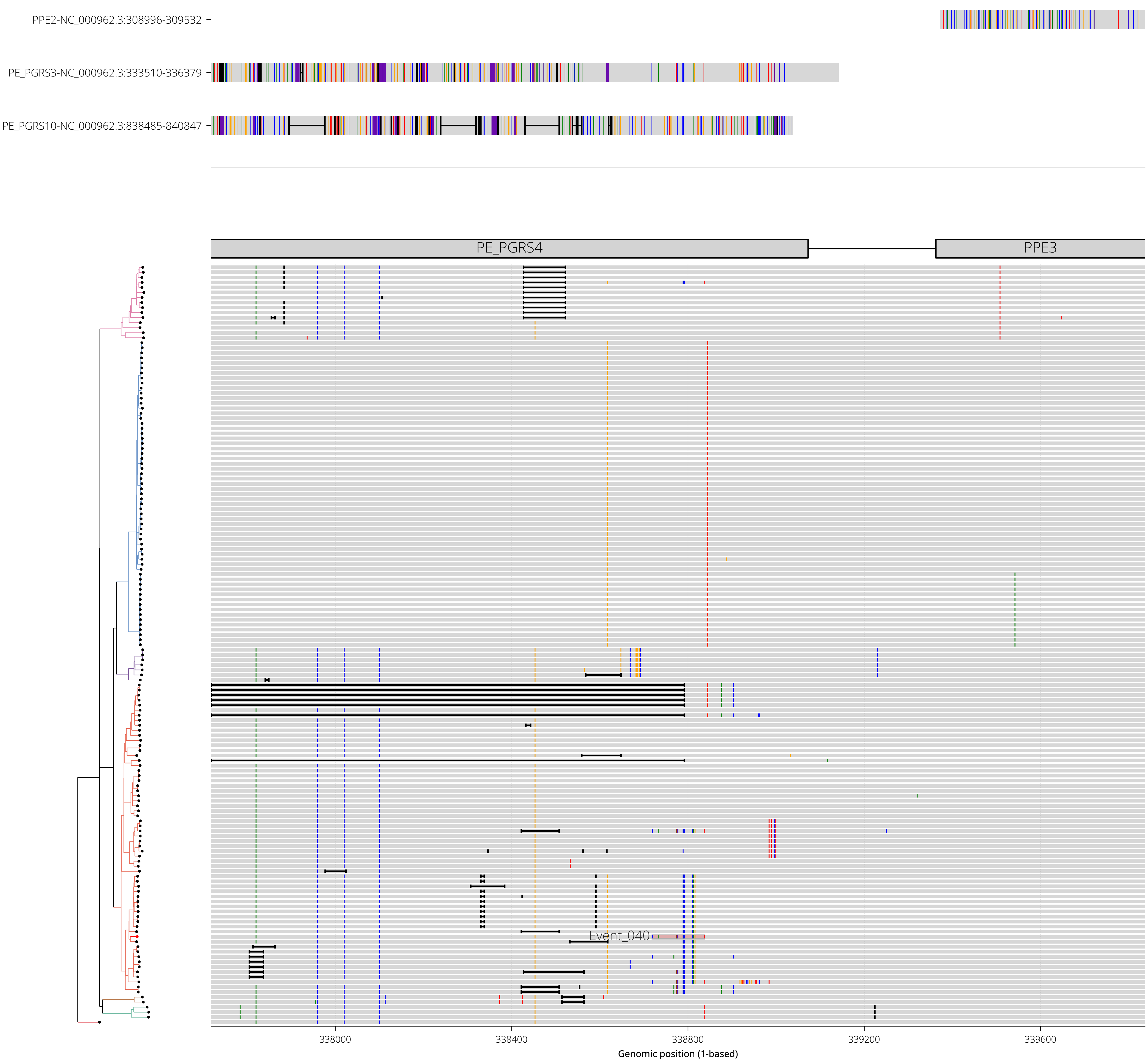

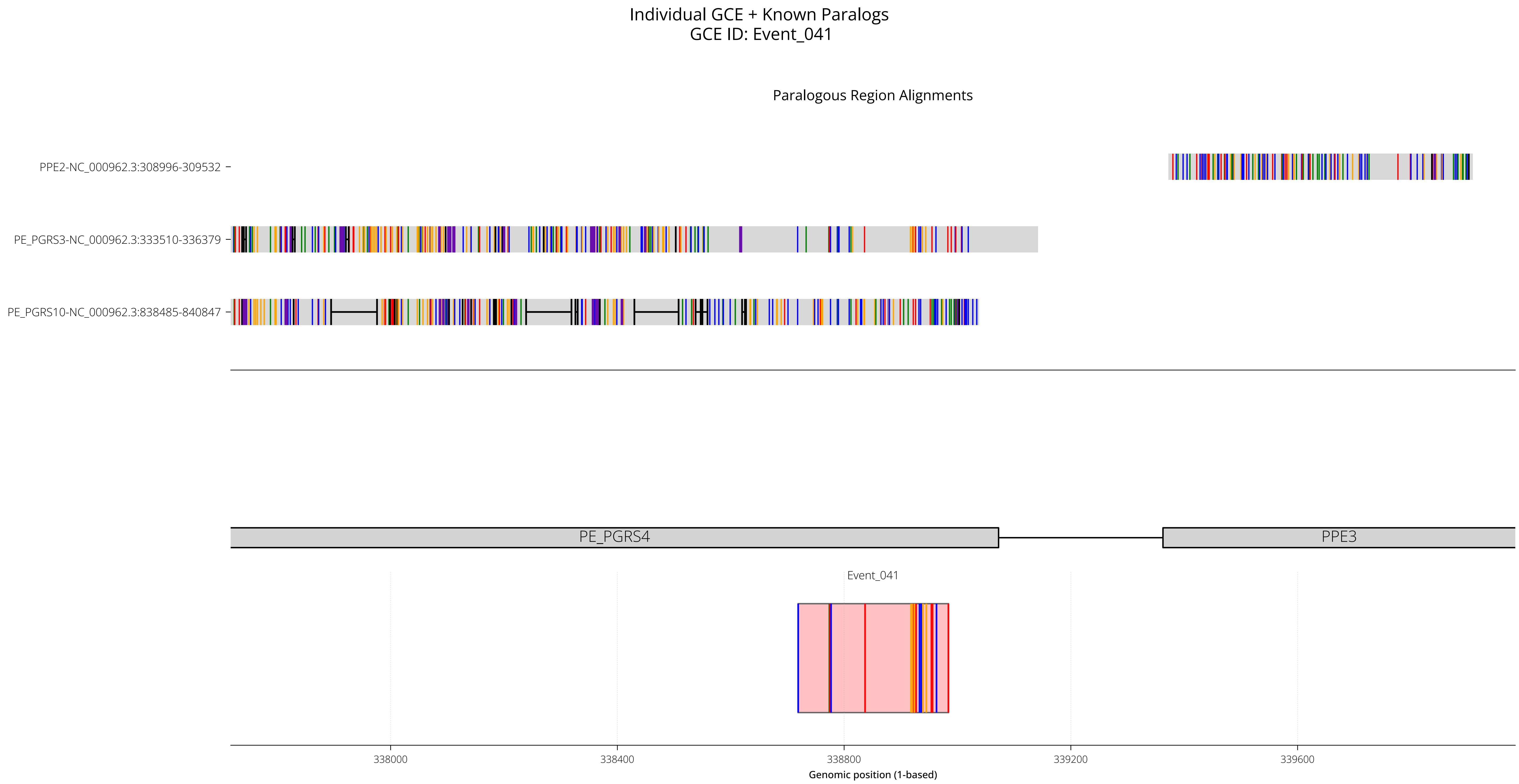

Individual GCE + Known Paralogs
GCE ID: Event_041
Paralogous Region Alignments
PPE2-NC_000962.3:308996-309532
PE_PGRS3-NC_000962.3:333510-336379
PE_PGRS10-NC_000962.3:838485-840847
PE_PGRS4
PPE3
Event_041
338000
338400
338800
339200
339600
Genomic position (1-based)

Circos Viz - GCE to Known Paralogs Locations
GCE ID: Event_041

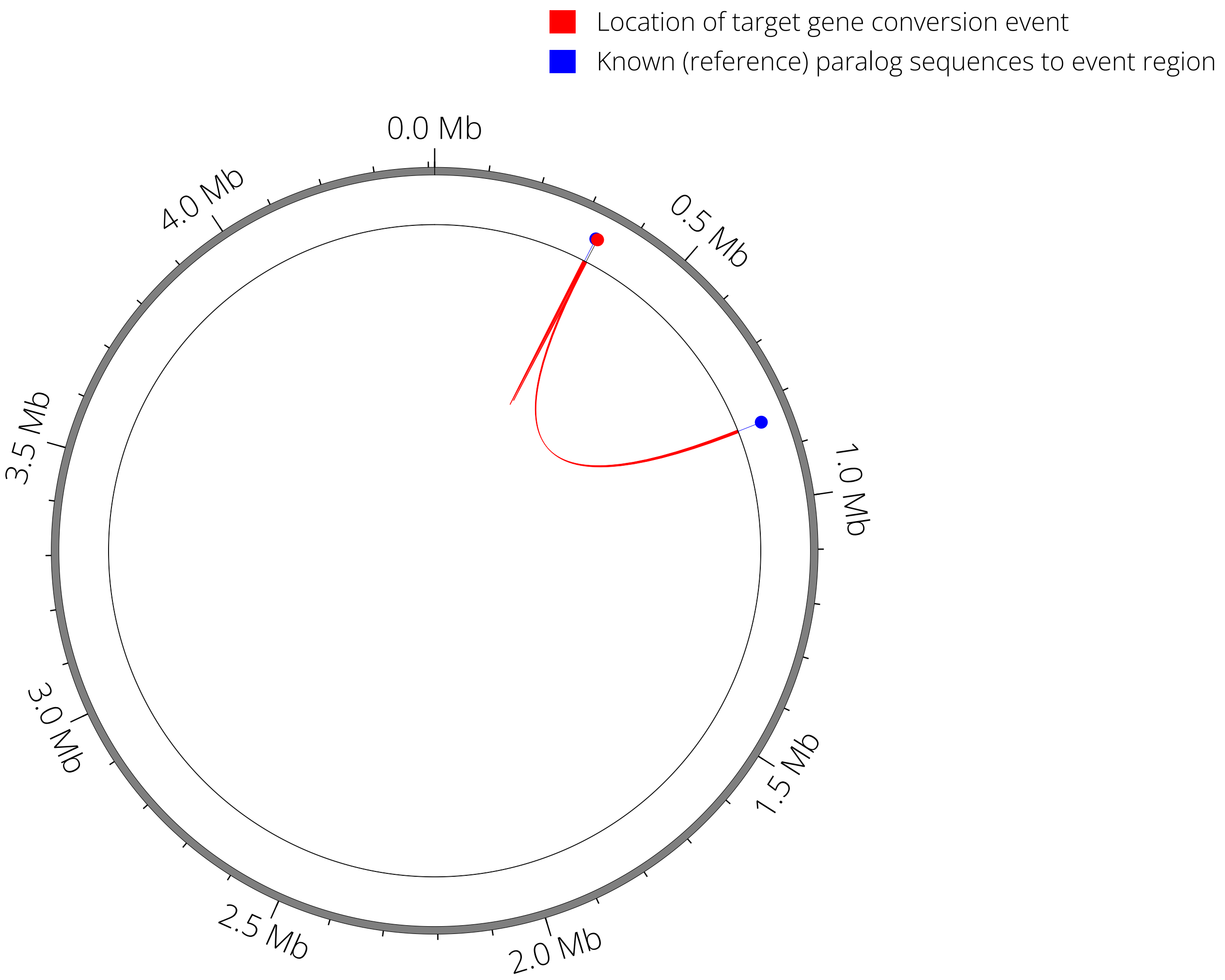

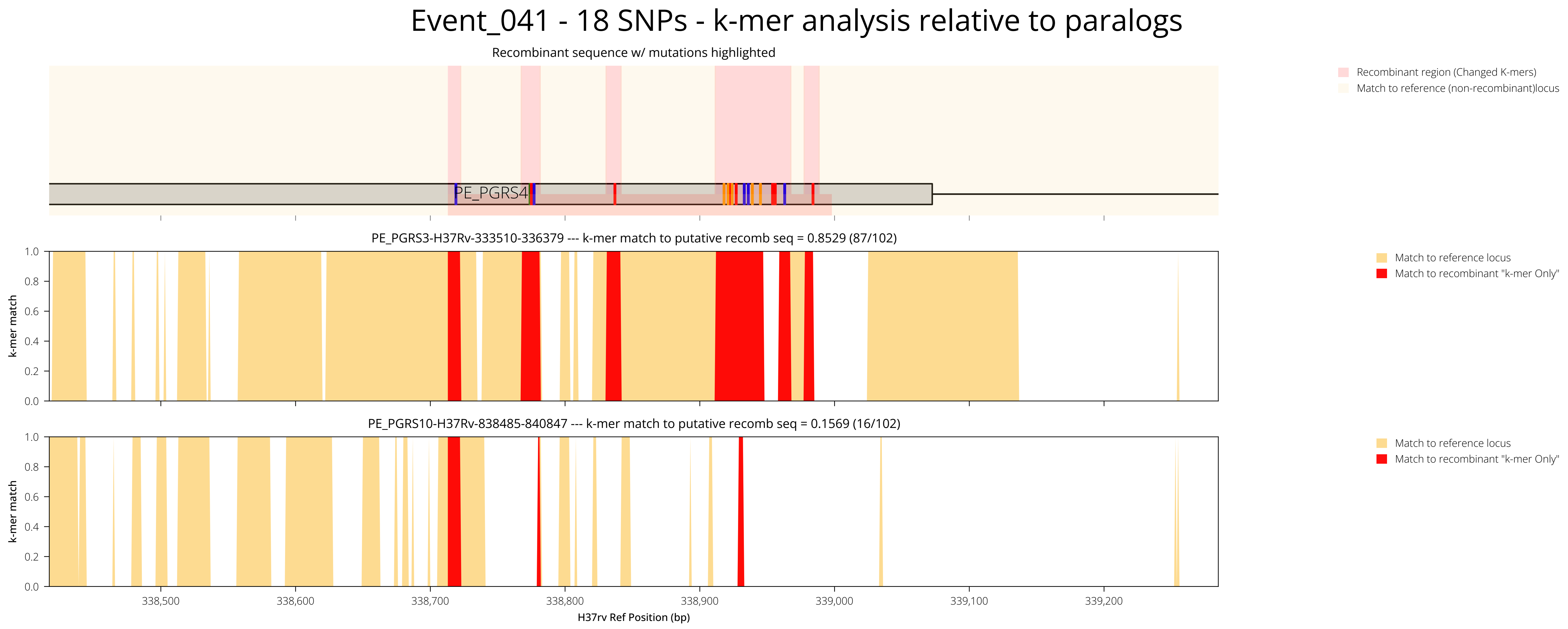

Event_041 - 18 SNPs - k-mer analysis relative to paralogs
Recombinant sequence w/ mutations highlighted
Recombinant region (Changed K-mers)
Match to reference (non-recombinant)locus
PE_PGRS4
PE_PGRS3-H37Rv-333510-336379 --- k-mer match to putative recomb seq = 0.8529 (87/102)
Match to reference locus
Match to recombinant "k-mer Only"
PE_PGRS10-H37Rv-838485-840847 --- k-mer match to putative recomb seq = 0.1569 (16/102)
Match to reference locus
Match to recombinant "k-mer Only"
k-mer match
1.0
0.8
0.6
0.4
0.2
0.0
338,500
338,600
338,700
338,800
338,900
339,000
339,100
339,200
H37rv Ref Position (bp)

Sub-phylogeny Viz + Paralog Alignments
GCE ID: Event_041

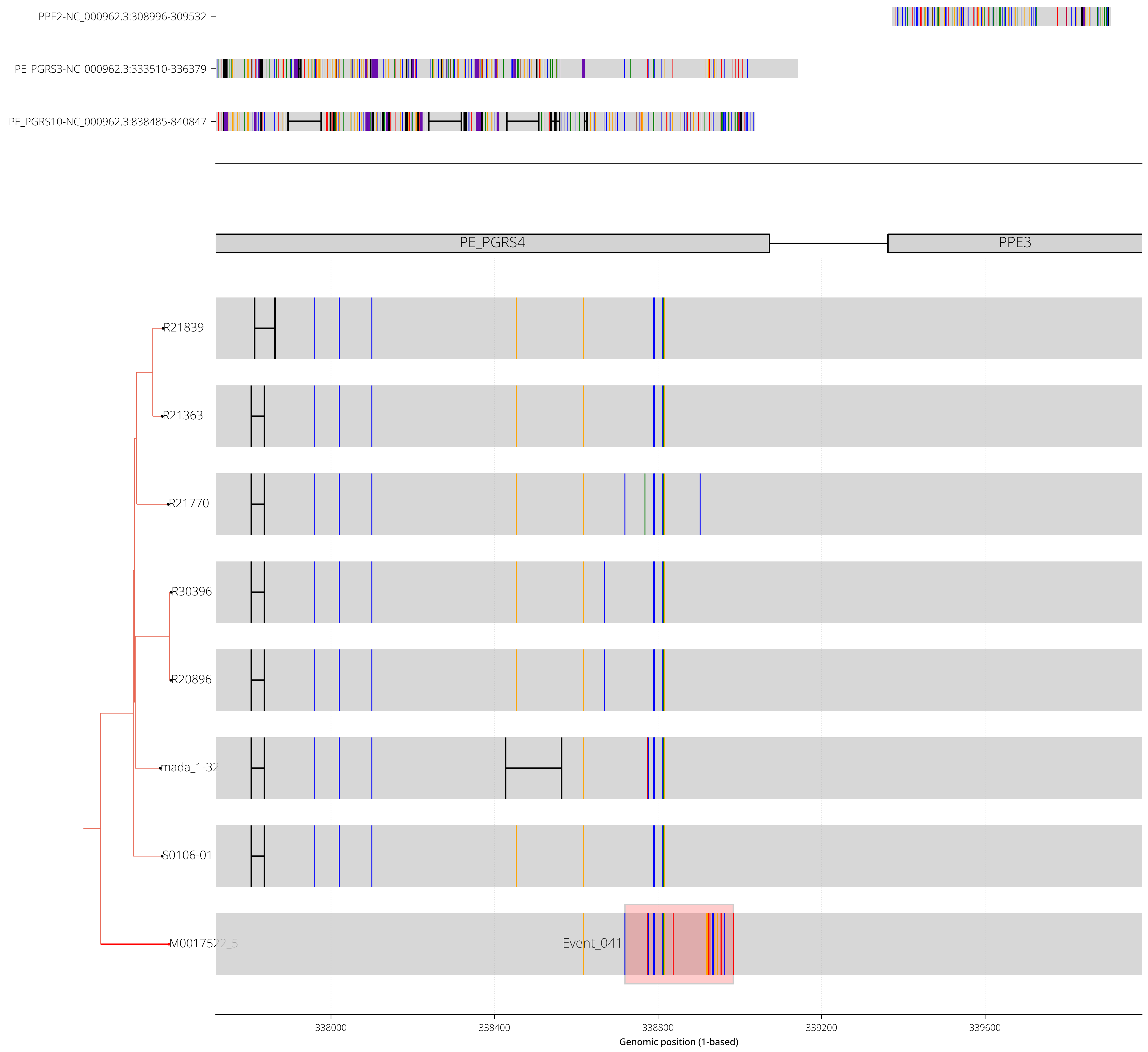

Full Phylogeny Viz + Paralog Alignments
GCE ID: Event_041

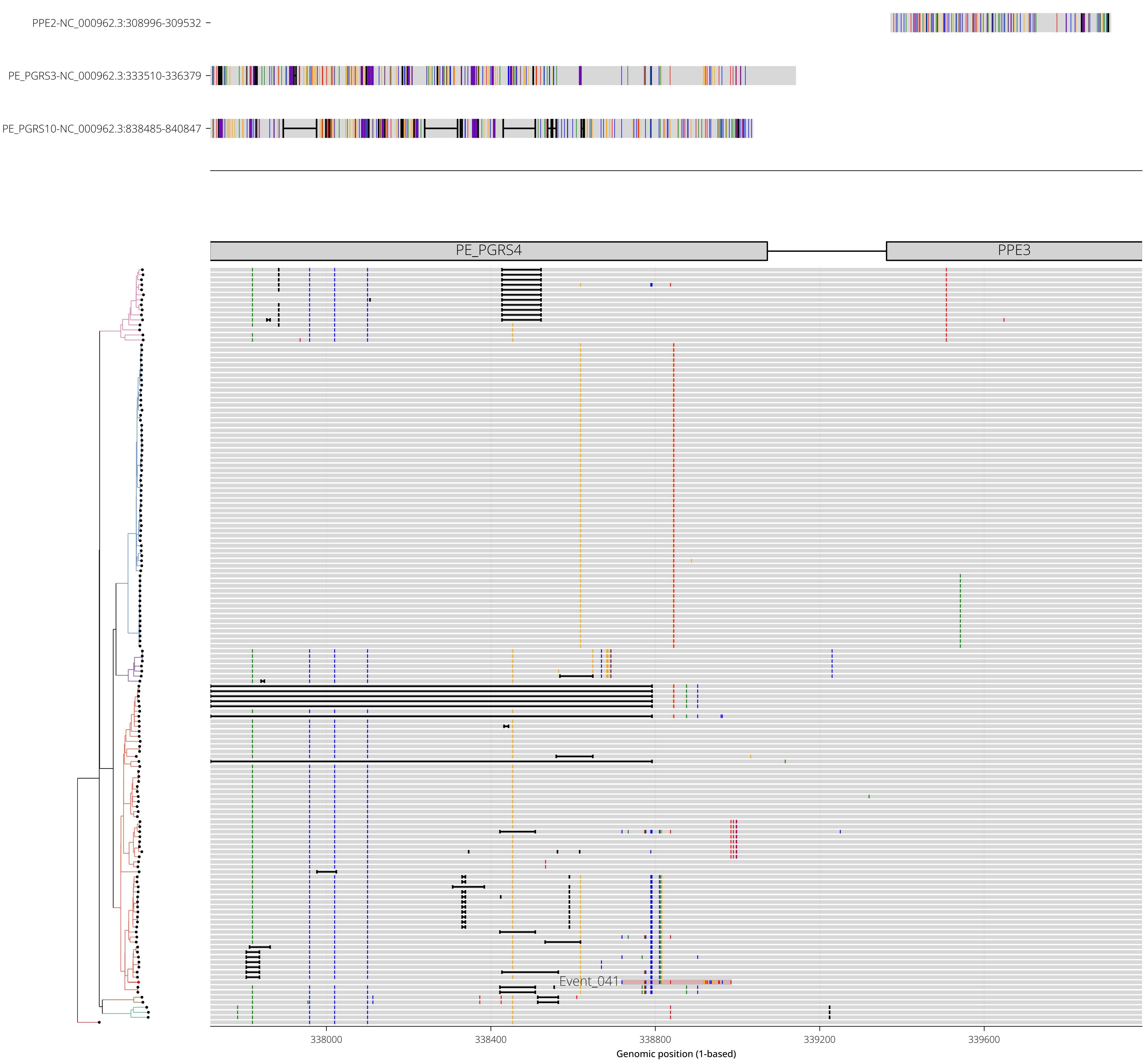

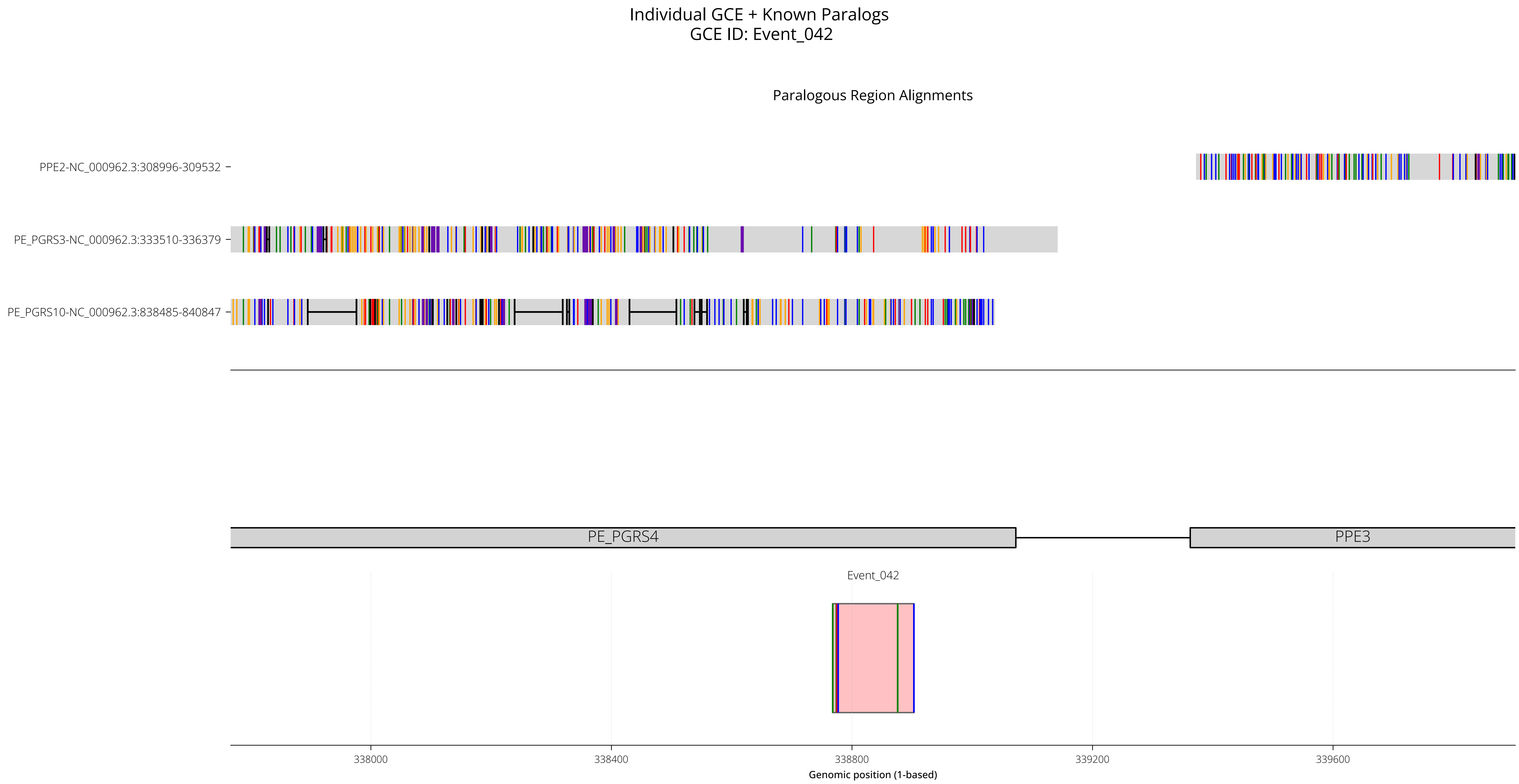

Individual GCE + Known Paralogs
GCE ID: Event_042
Paralogous Region Alignments
PPE2-NC_000962.3:308996-309532
PE_PGRS3-NC_000962.3:333510-336379
PE_PGRS10-NC_000962.3:838485-840847
PE_PGRS4
PPE3
Event_042
338000
338400
338800
339200
339600
Genomic position (1-based)

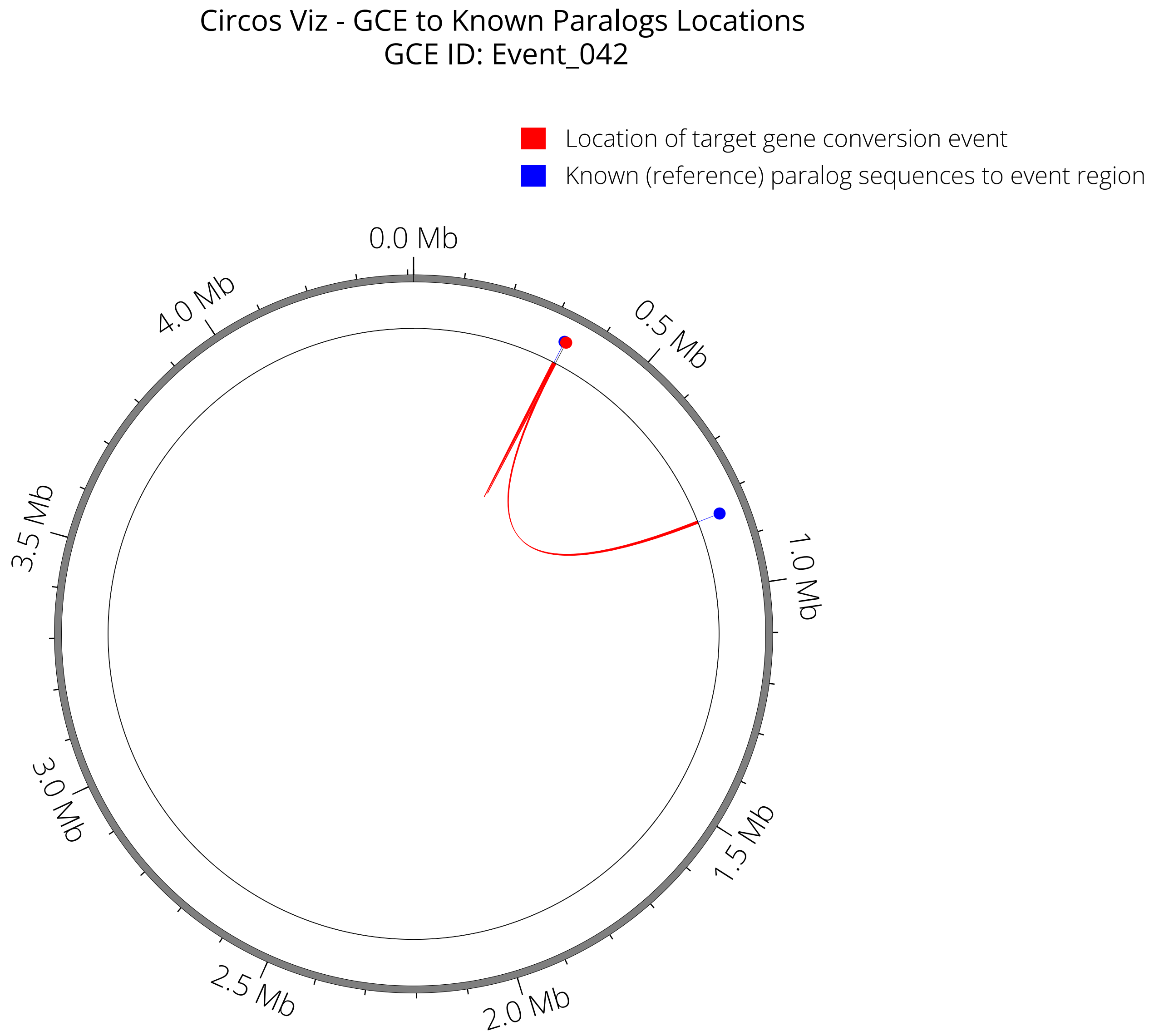
Circos Viz - GCE to Known Paralogs Locations
GCE ID: Event_042
Location of target gene conversion event
Known (reference) paralog sequences to event region
0.0 Mb
0.5 Mb
1.0 Mb
1.5 Mb
2.0 Mb
2.5 Mb
3.0 Mb
3.5 Mb
4.0 Mb

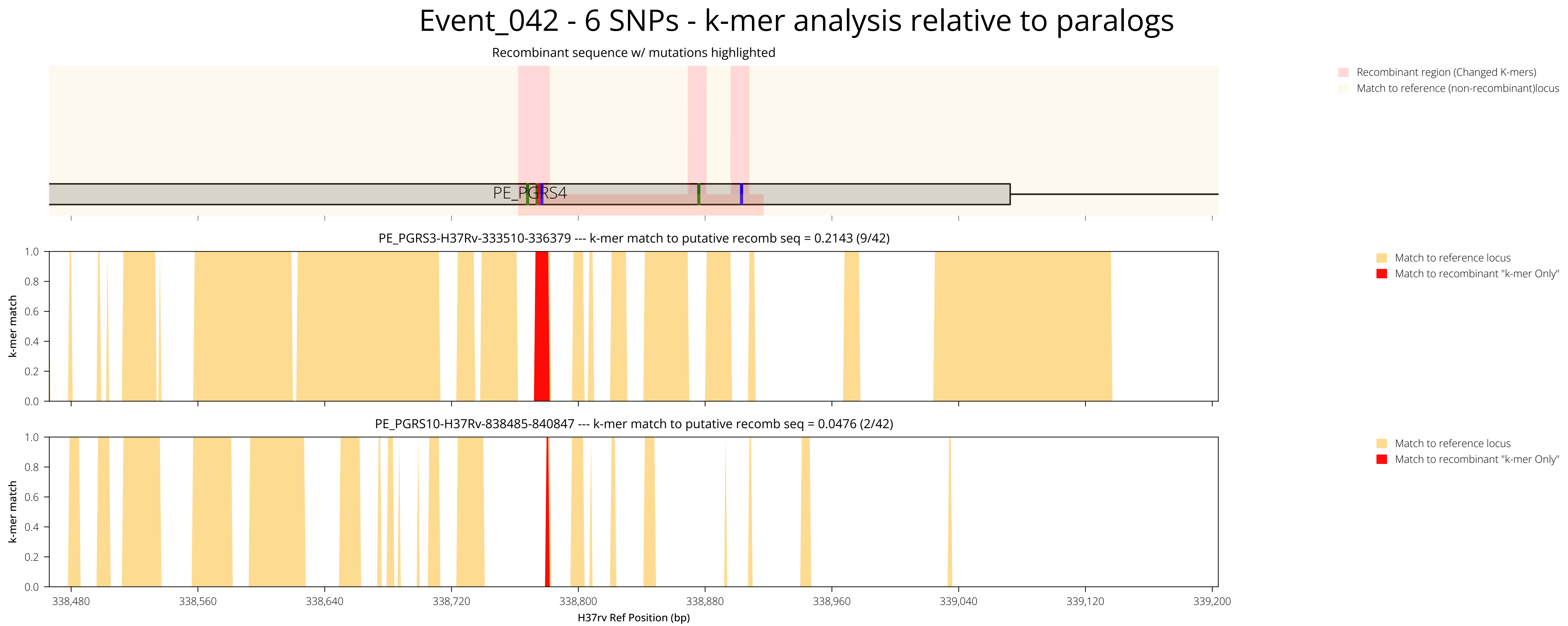

Event_042 - 6 SNPs - k-mer analysis relative to paralogs
Recombinant sequence w/ mutations highlighted
PE_PGRS4
Recombinant region (Changed K-mers)
Match to reference (non-recombinant)locus
PE_PGRS3-H37Rv-333510-336379 --- k-mer match to putative recomb seq = 0.2143 (9/42)
k-mer match
1.0
0.8
0.6
0.4
0.2
0.0
Match to reference locus
Match to recombinant "k-mer Only"
PE_PGRS10-H37Rv-838485-840847 --- k-mer match to putative recomb seq = 0.0476 (2/42)
k-mer match
1.0
0.8
0.6
0.4
0.2
0.0
Match to reference locus
Match to recombinant "k-mer Only"
338,480
338,560
338,640
338,720
338,800
338,880
338,960
339,040
339,120
339,200
H37rv Ref Position (bp)

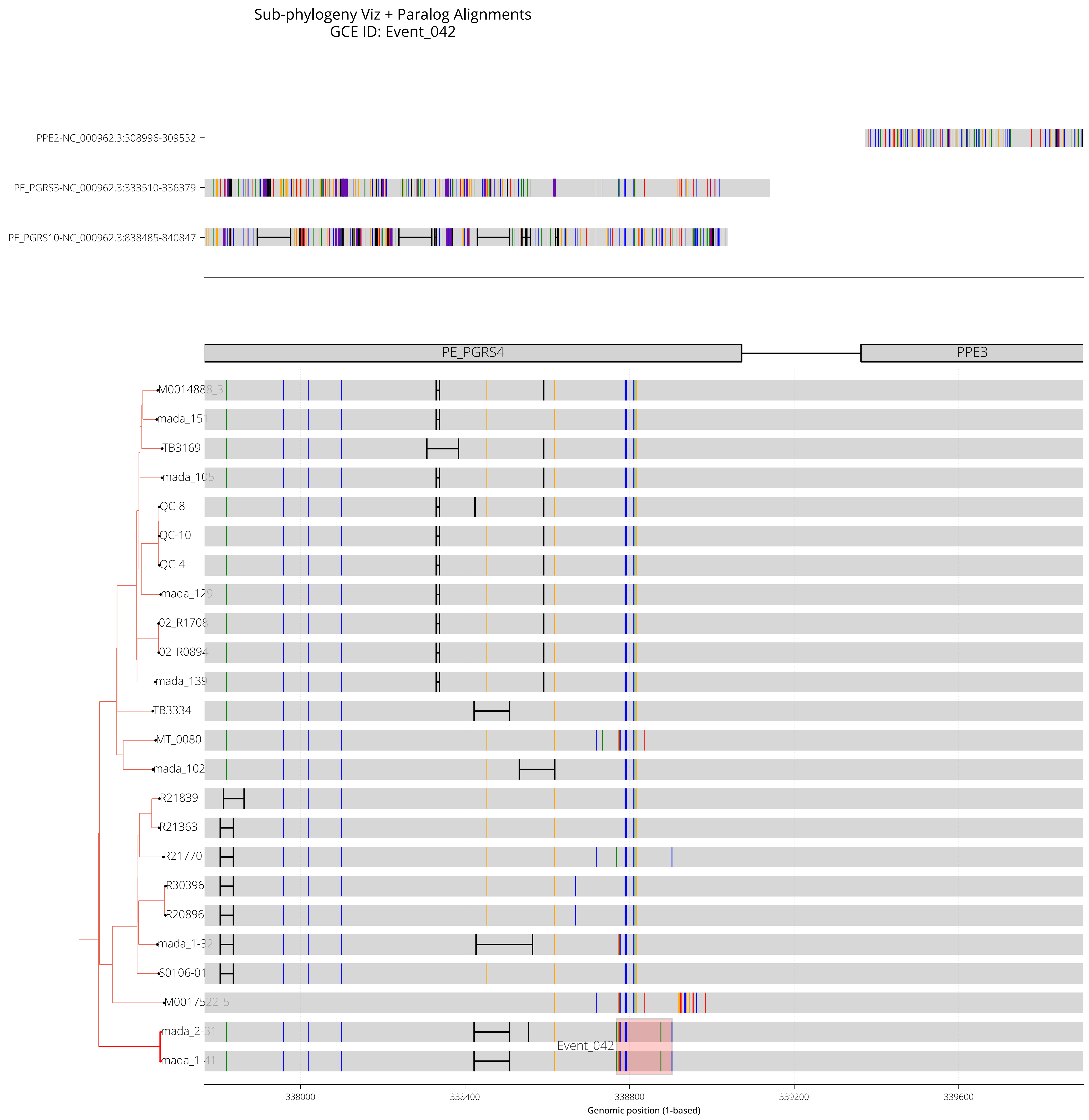

Sub-phylogeny Viz + Paralog Alignments
GCE ID: Event_042
PPE2-NC_000962.3:308996-309532
PE_PGRS3-NC_000962.3:333510-336379
PE_PGRS10-NC_000962.3:838485-840847
PE_PGRS4
PPE3
M0014888_3
mada_151
TB3169
mada_105
QC-8
QC-10
QC-4
mada_129
02_R1708
02_R0894
mada_139
TB3334
MT_0080
mada_102
R21839
R21363
R21770
R30396
R20896
mada_1-32
S0106-01
M0017522_5
mada_2-31
mada_1-41
Event_042
338000
338400
338800
339200
339600
Genomic position (1-based)

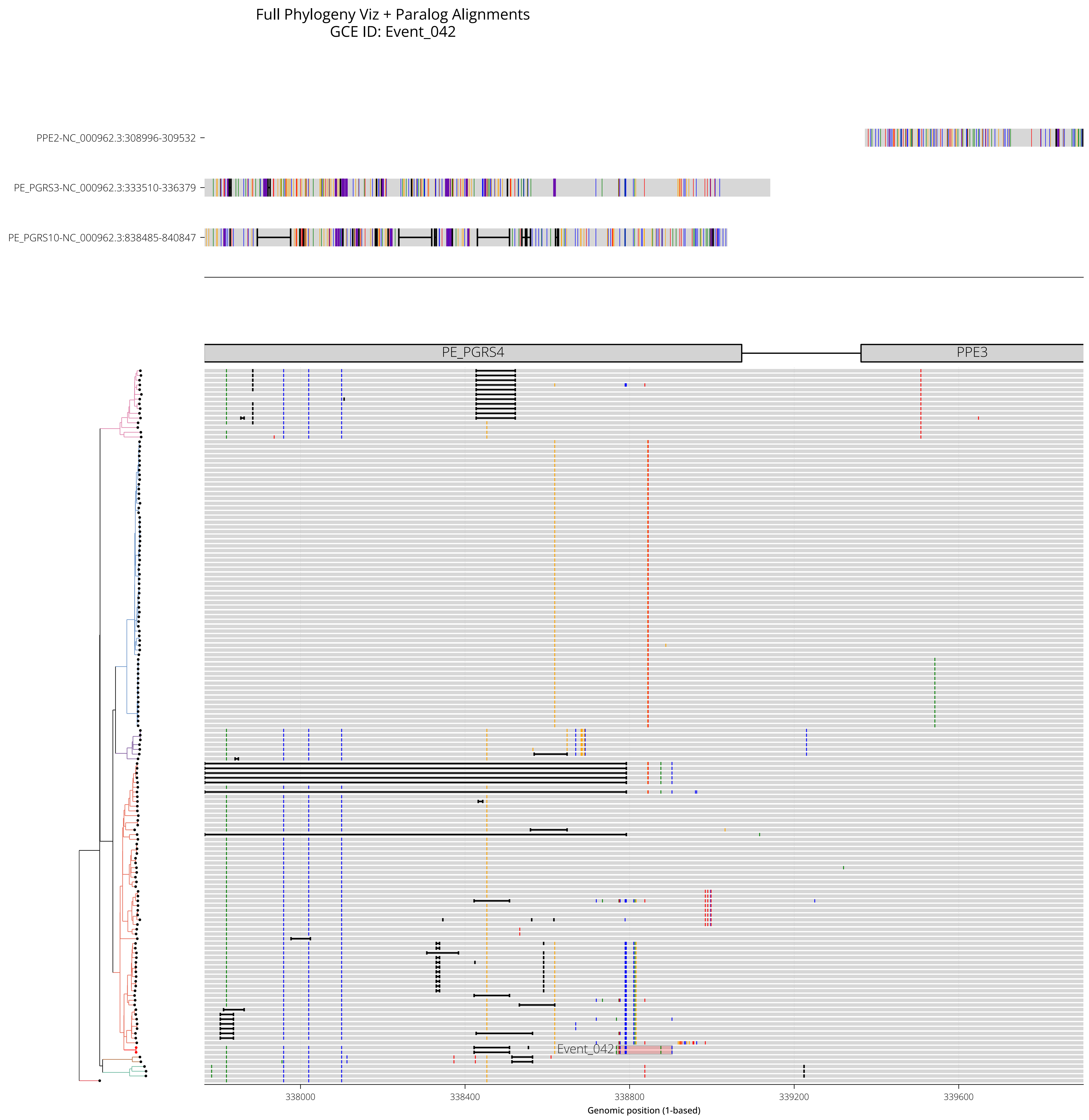

Full Phylogeny Viz + Paralog Alignments
GCE ID: Event_042
PPE2-NC_000962.3:308996-309532
PE_PGRS3-NC_000962.3:333510-336379
PE_PGRS10-NC_000962.3:838485-840847
PE_PGRS4
PPE3
Event_042
338000
338400
338800
339200
339600
Genomic position (1-based)

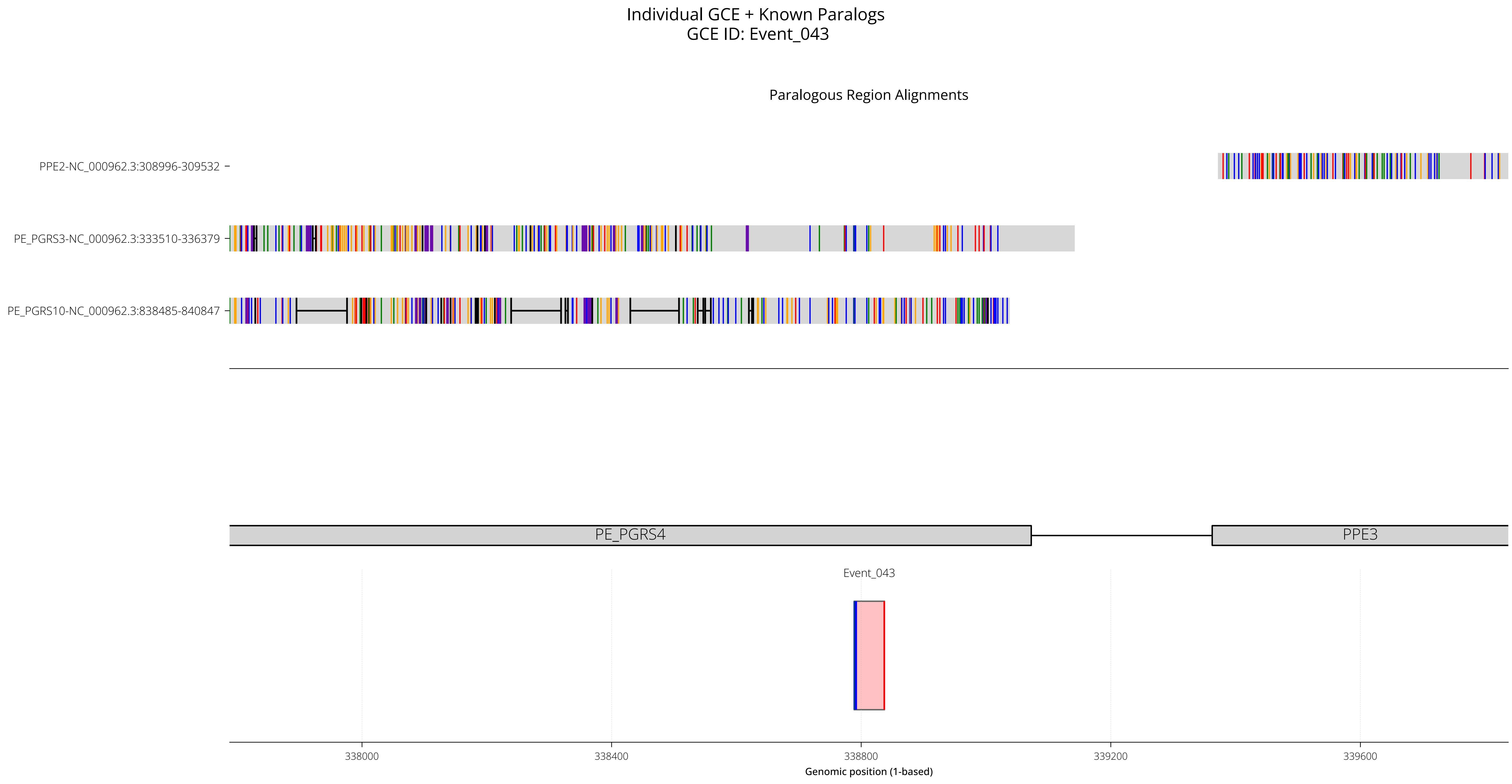
Individual GCE + Known Paralogs
GCE ID: Event_043
Paralogous Region Alignments
PPE2-NC_000962.3:308996-309532
PE_PGRS3-NC_000962.3:333510-336379
PE_PGRS10-NC_000962.3:838485-840847
PE_PGRS4
PPE3
Event_043
338000
338400
338800
339200
339600
Genomic position (1-based)

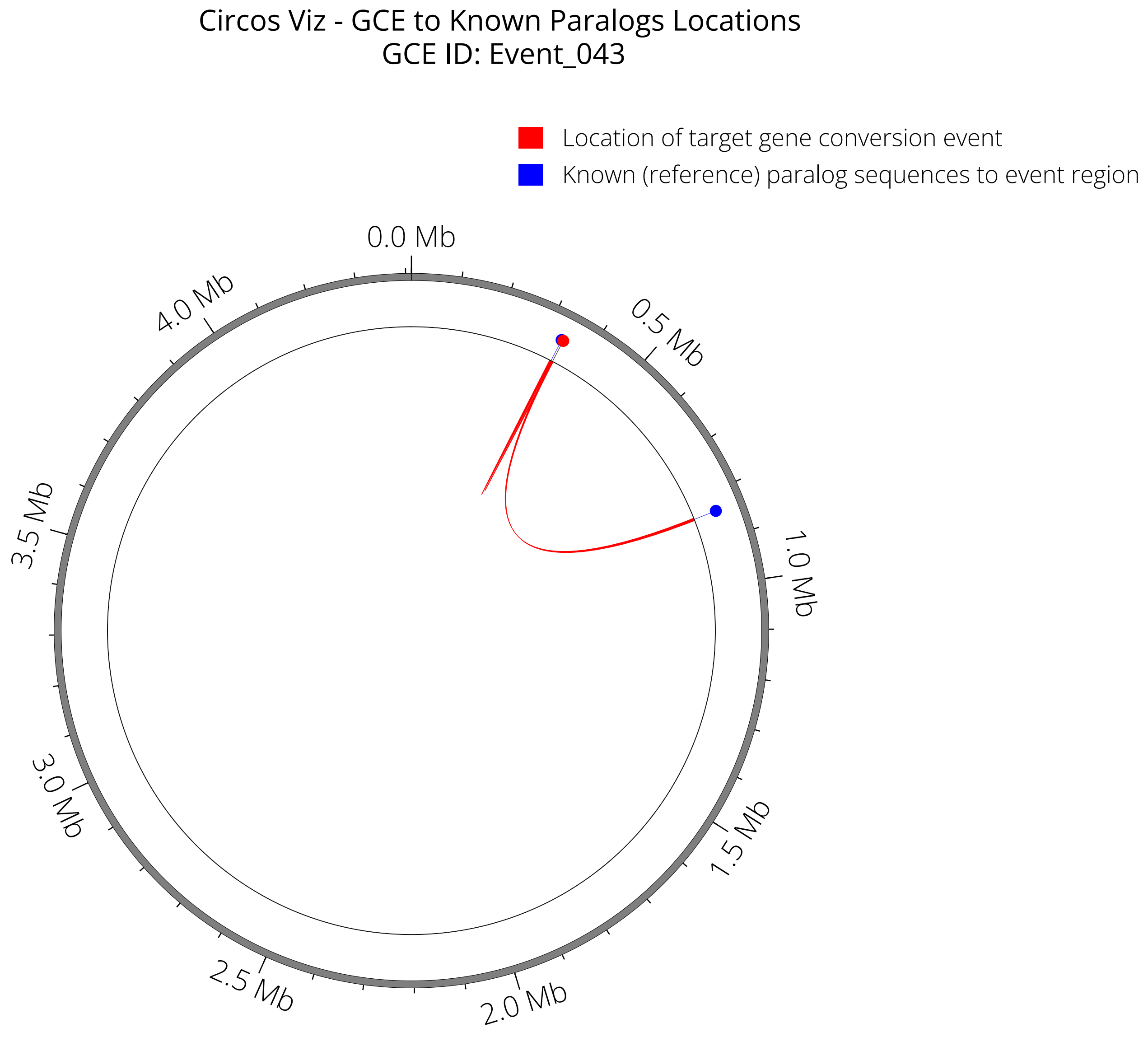

Circos Viz - GCE to Known Paralogs Locations
GCE ID: Event_043
Location of target gene conversion event
Known (reference) paralog sequences to event region
0.0 Mb
0.5 Mb
1.0 Mb
1.5 Mb
2.0 Mb
2.5 Mb
3.0 Mb
3.5 Mb
4.0 Mb

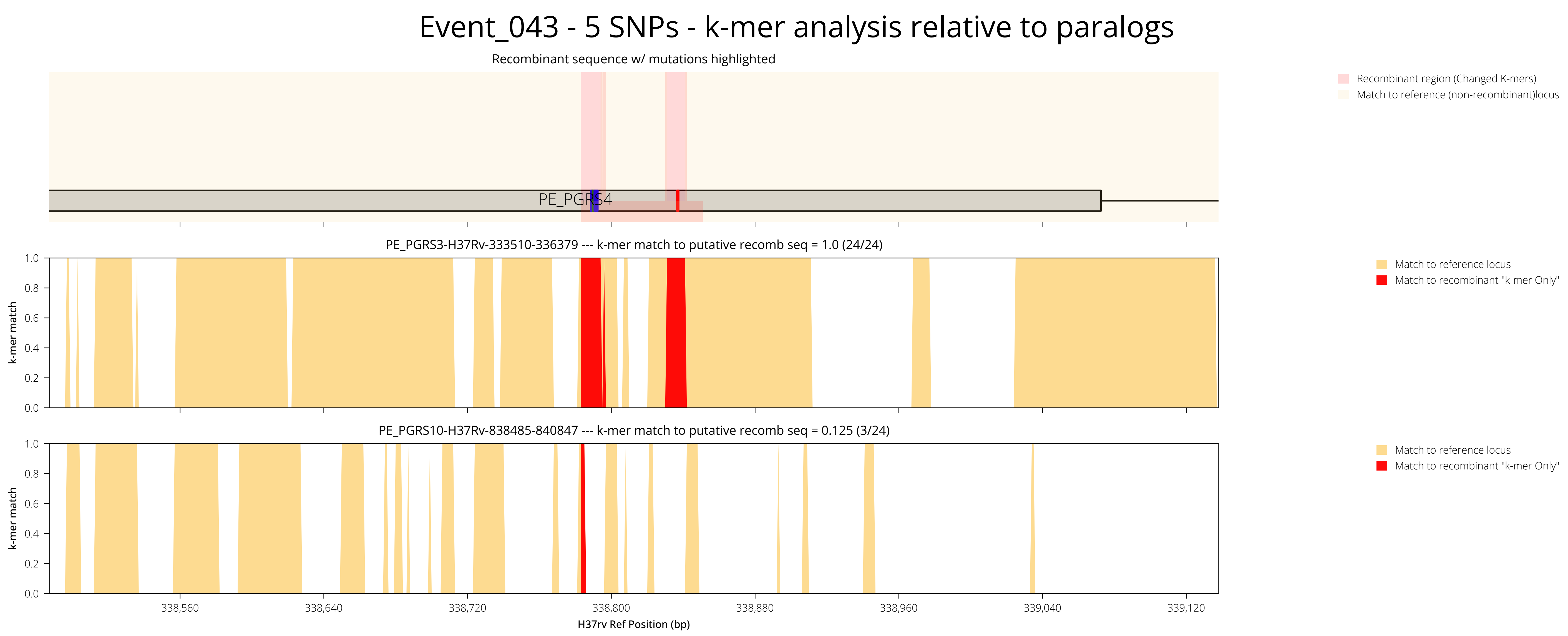
Event_043 - 5 SNPs - k-mer analysis relative to paralogs
Recombinant sequence w/ mutations highlighted
PE_PGRS4
Recombinant region (Changed K-mers)
Match to reference (non-recombinant)locus
PE_PGRS3-H37Rv-333510-336379 --- k-mer match to putative recomb seq = 1.0 (24/24)
Match to reference locus
Match to recombinant "k-mer Only"
k-mer match
1.0
0.8
0.6
0.4
0.2
0.0
PE_PGRS10-H37Rv-838485-840847 --- k-mer match to putative recomb seq = 0.125 (3/24)
Match to reference locus
Match to recombinant "k-mer Only"
k-mer match
1.0
0.8
0.6
0.4
0.2
0.0
338,560
338,640
338,720
338,800
338,880
338,960
339,040
339,120
H37rv Ref Position (bp)

Sub-phylogeny Viz + Paralog Alignments
GCE ID: Event_043

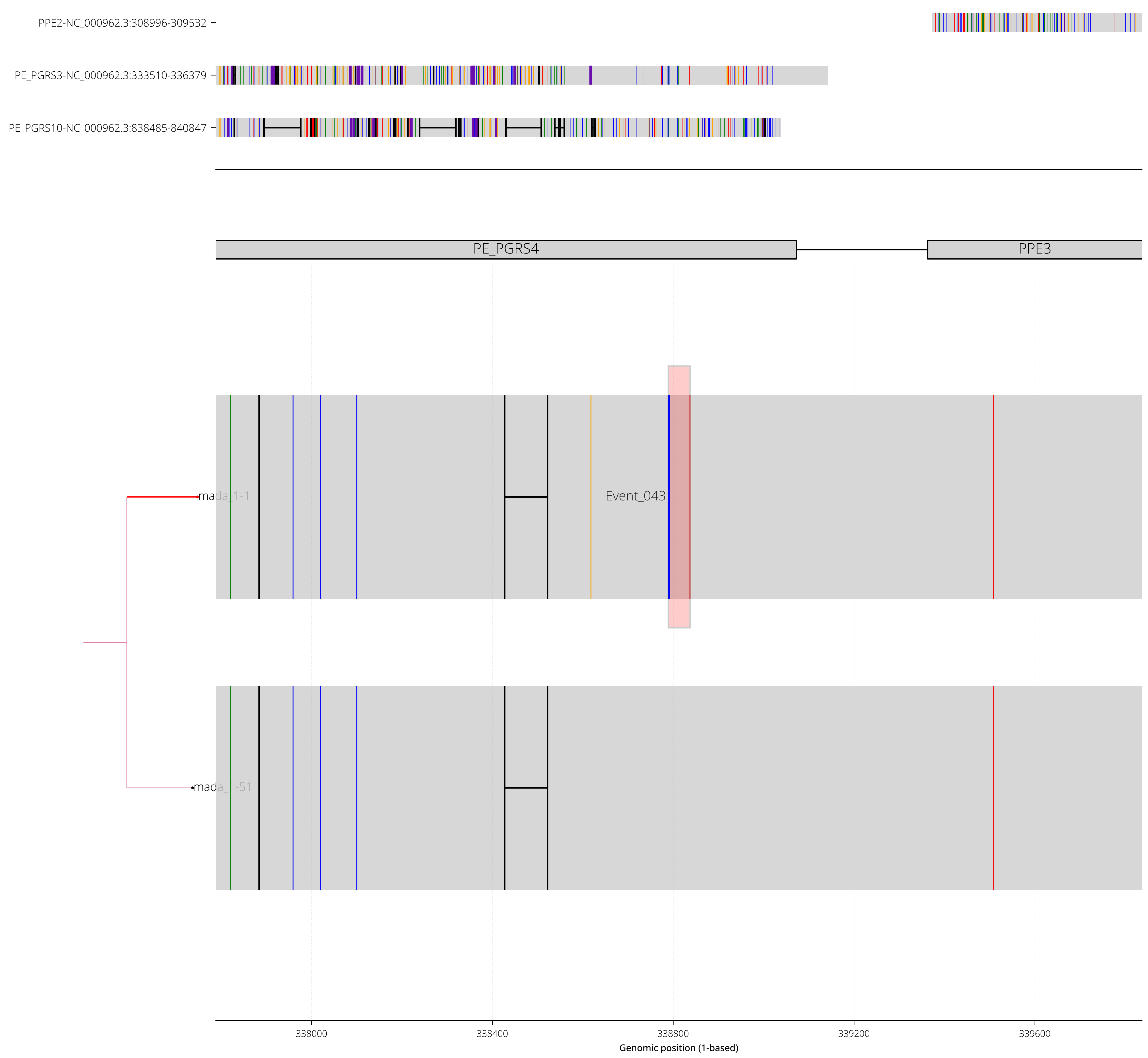

#### Full Phylogeny Viz + Paralog Alignments
#### GCE ID: Event_043

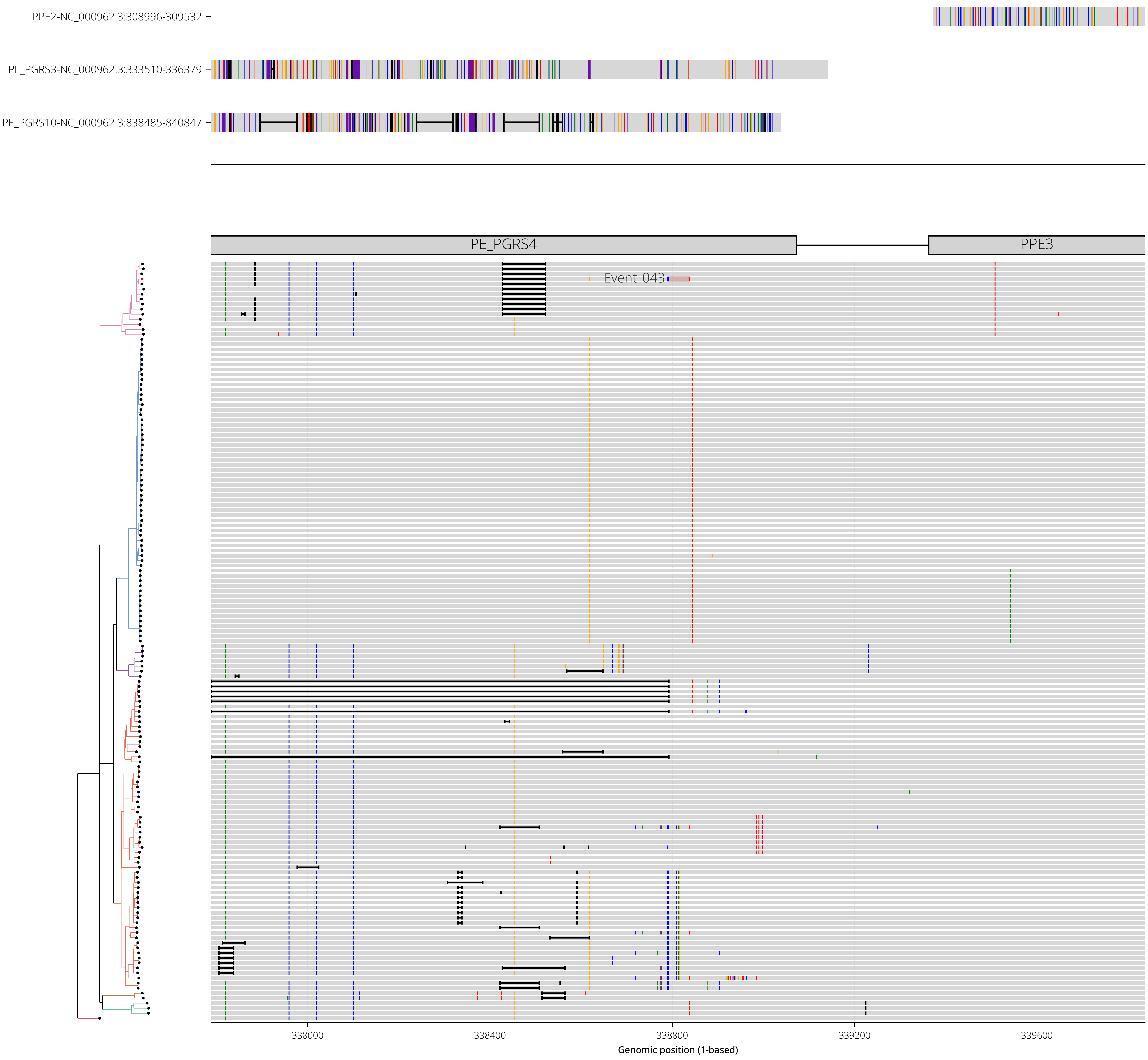

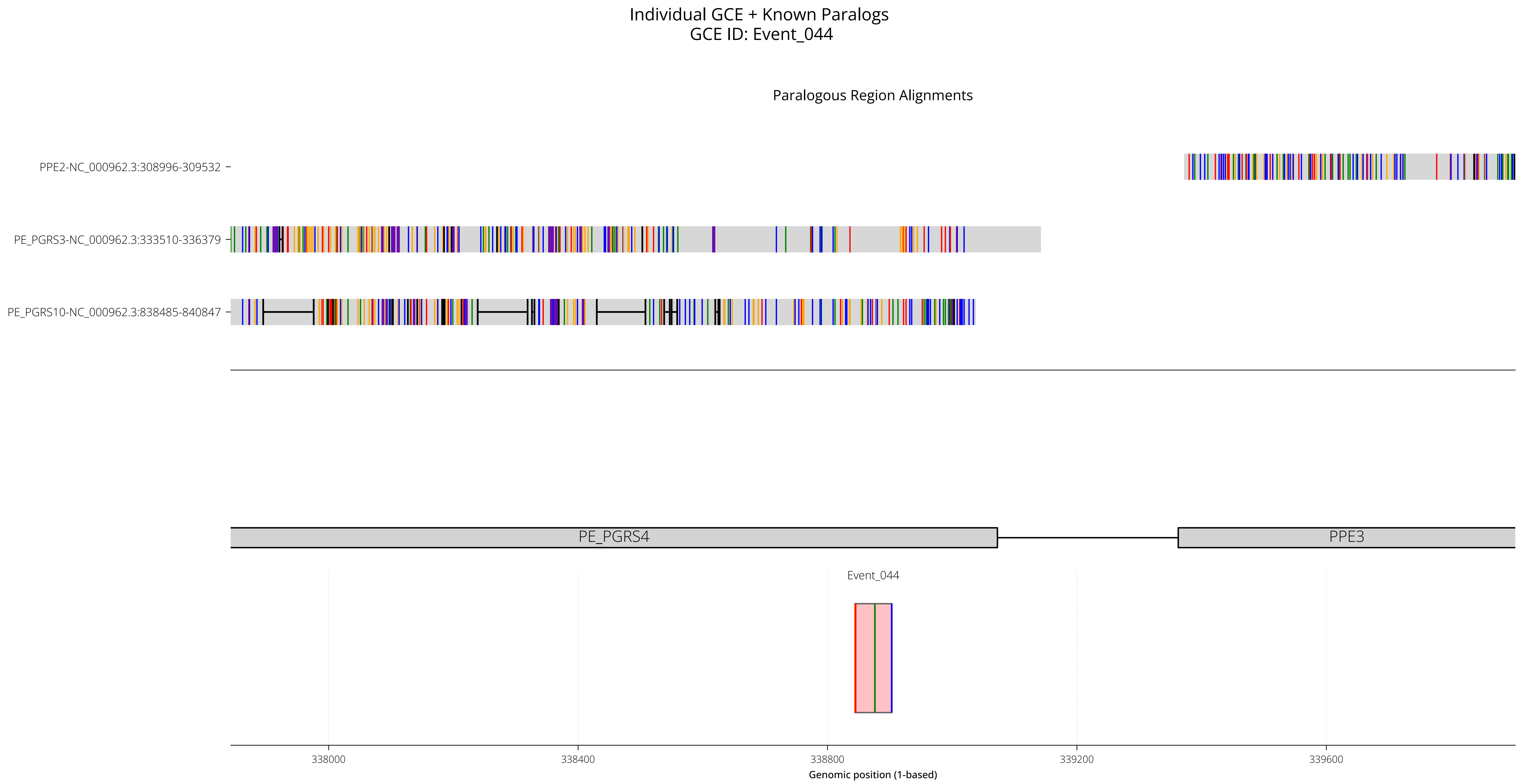

Individual GCE + Known Paralogs
GCE ID: Event_044
Paralogous Region Alignments
PPE2-NC_000962.3:308996-309532
PE_PGRS3-NC_000962.3:333510-336379
PE_PGRS10-NC_000962.3:838485-840847
PE_PGRS4
PPE3
Event_044
338000
338400
338800
339200
339600
Genomic position (1-based)

Circos Viz - GCE to Known Paralogs Locations
GCE ID: Event_044

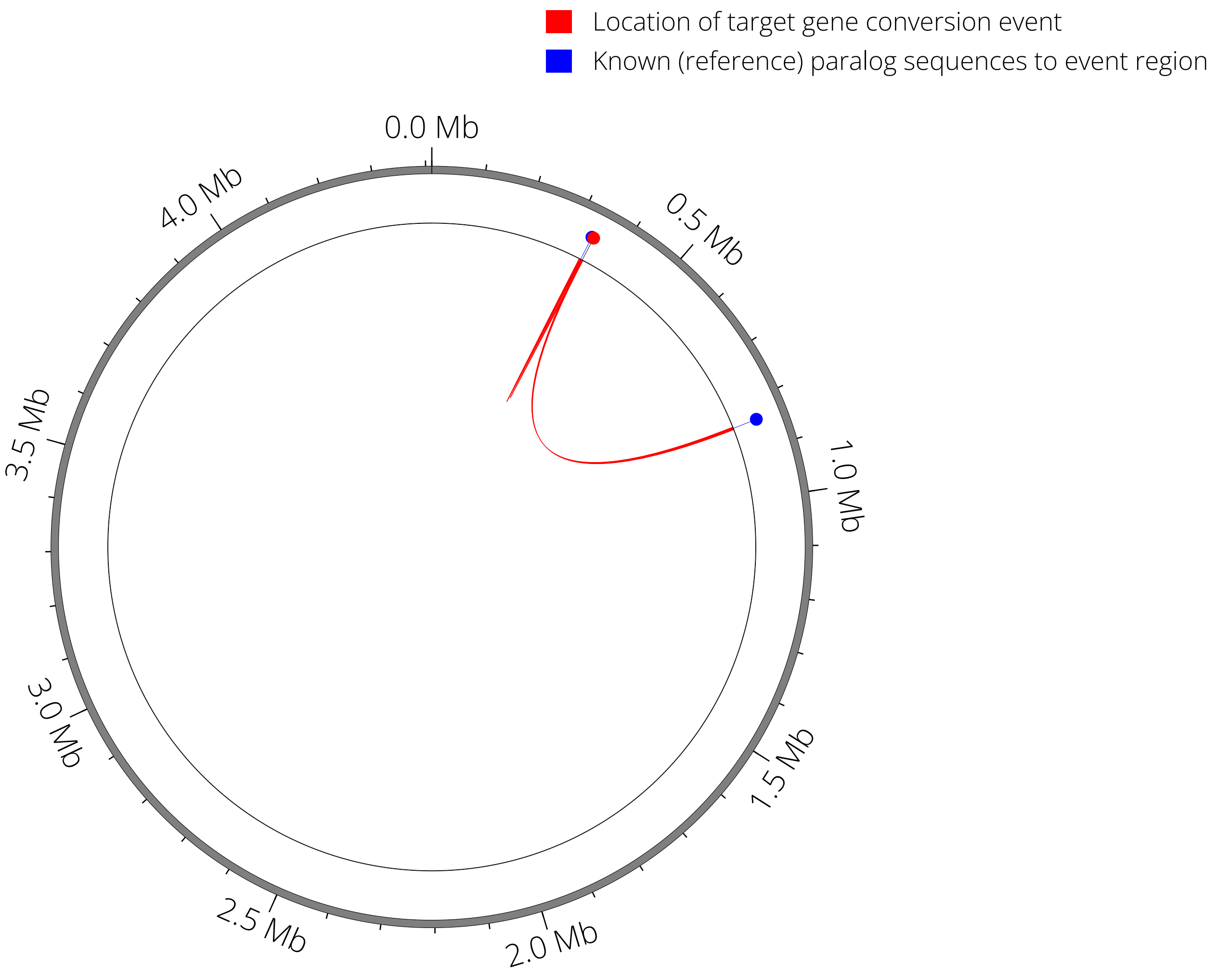

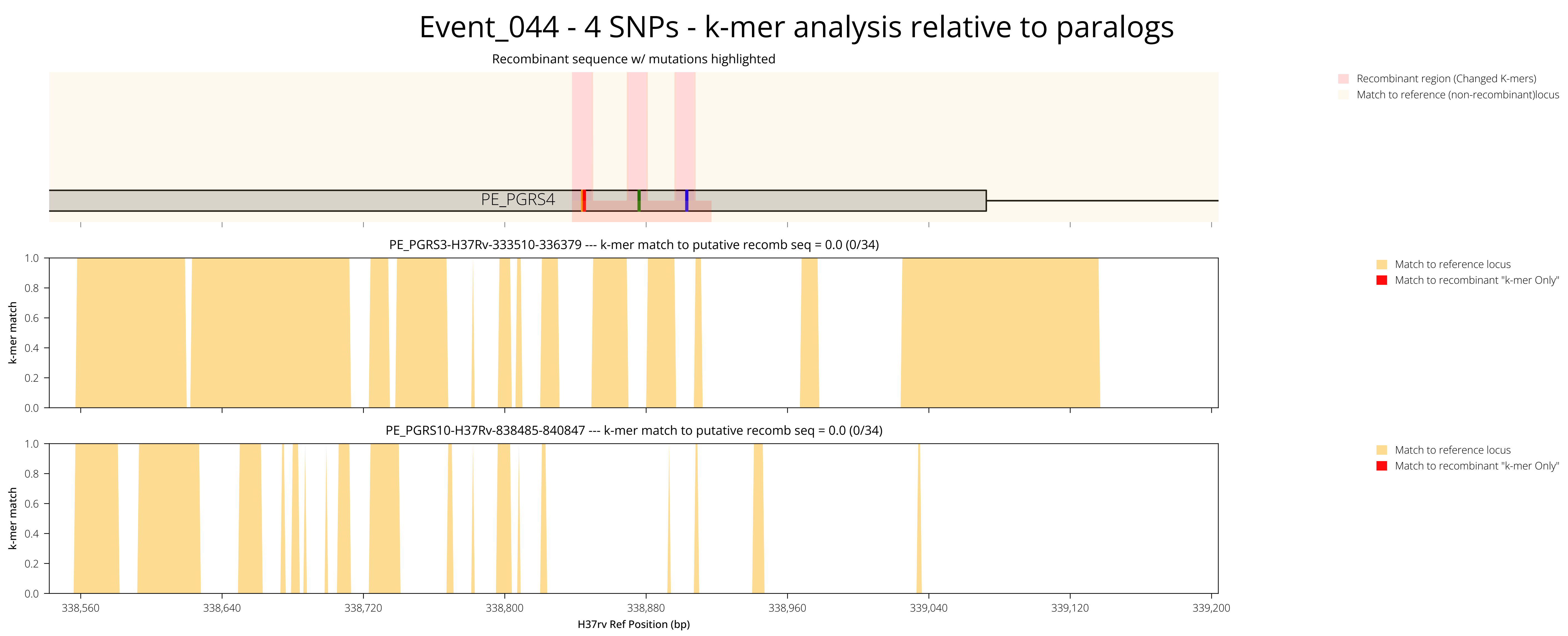
Event_044 - 4 SNPs - k-mer analysis relative to paralogs
Recombinant sequence w/ mutations highlighted
Recombinant region (Changed K-mers)
Match to reference (non-recombinant)locus
PE_PGRS4
PE_PGRS3-H37Rv-333510-336379 --- k-mer match to putative recomb seq = 0.0 (0/34)
Match to reference locus
Match to recombinant "k-mer Only"
k-mer match
1.0
0.8
0.6
0.4
0.2
0.0
PE_PGRS10-H37Rv-838485-840847 --- k-mer match to putative recomb seq = 0.0 (0/34)
Match to reference locus
Match to recombinant "k-mer Only"
k-mer match
1.0
0.8
0.6
0.4
0.2
0.0
338,560
338,640
338,720
338,800
338,880
338,960
339,040
339,120
339,200
H37rv Ref Position (bp)

Sub-phylogeny Viz + Paralog Alignments
GCE ID: Event_044

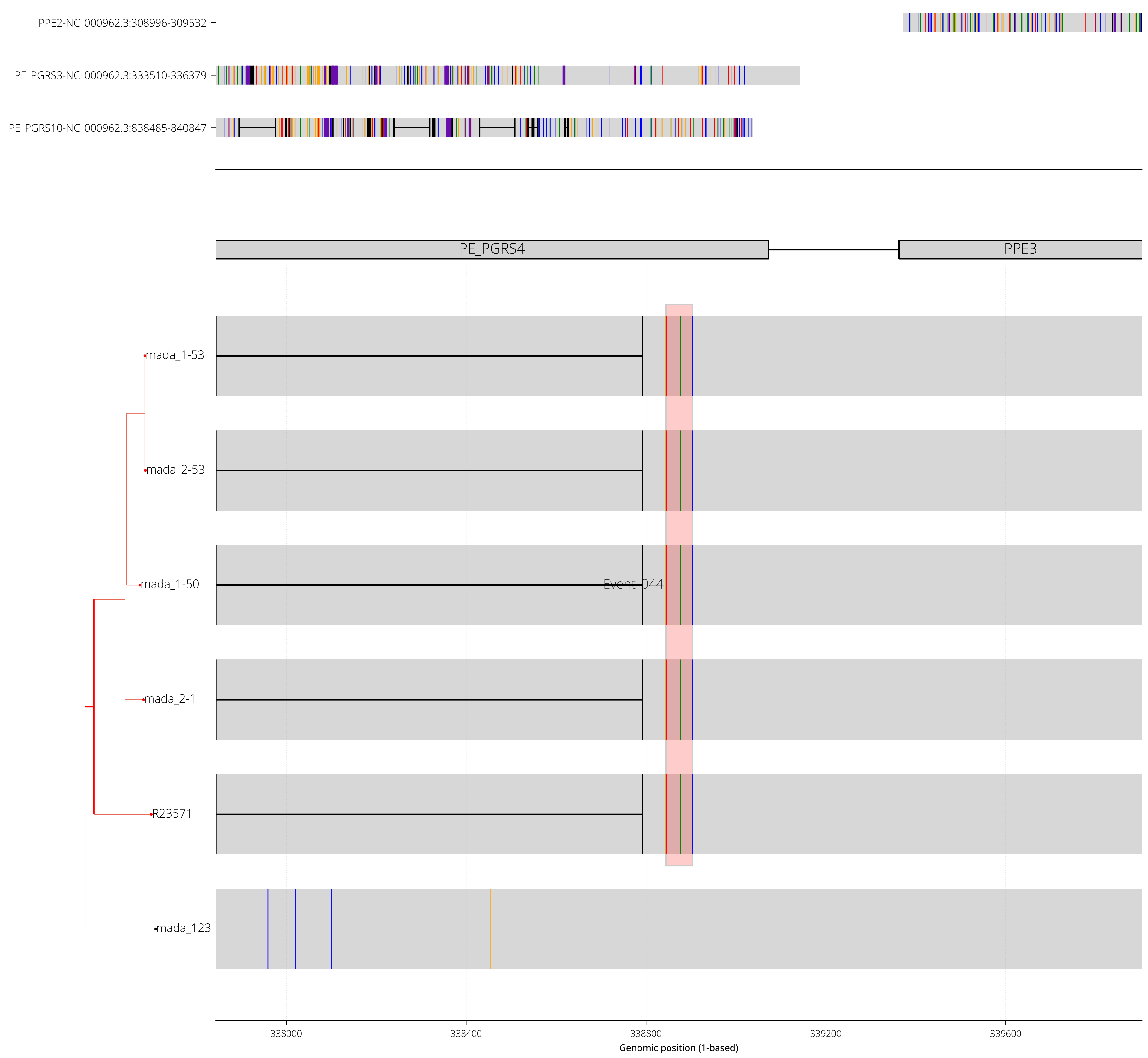

Full Phylogeny Viz + Paralog Alignments
GCE ID: Event_044

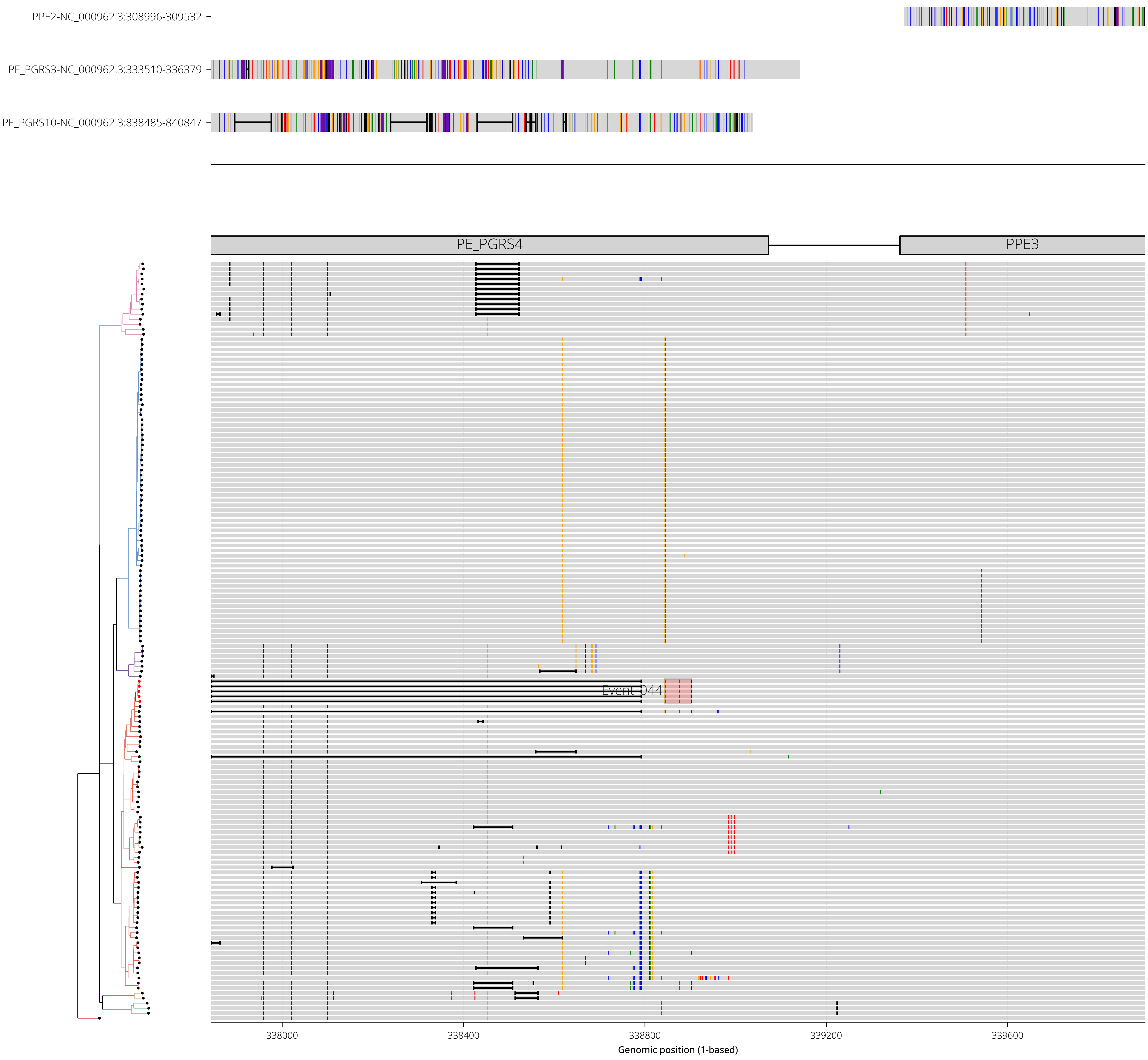

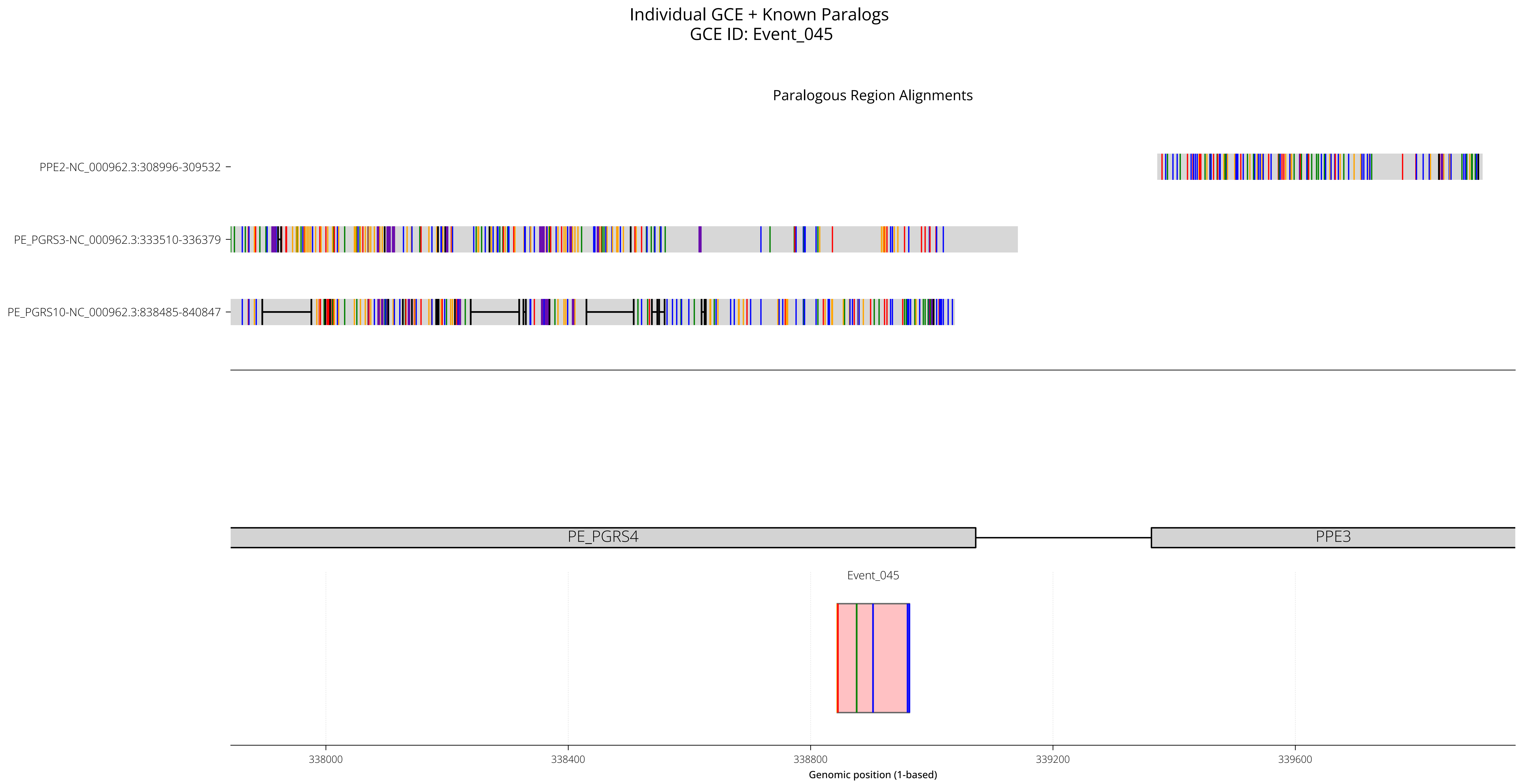

Individual GCE + Known Paralogs
GCE ID: Event_045
Paralogous Region Alignments
PPE2-NC_000962.3:308996-309532
PE_PGRS3-NC_000962.3:333510-336379
PE_PGRS10-NC_000962.3:838485-840847
PE_PGRS4
PPE3
Event_045
338000
338400
338800
339200
339600
Genomic position (1-based)

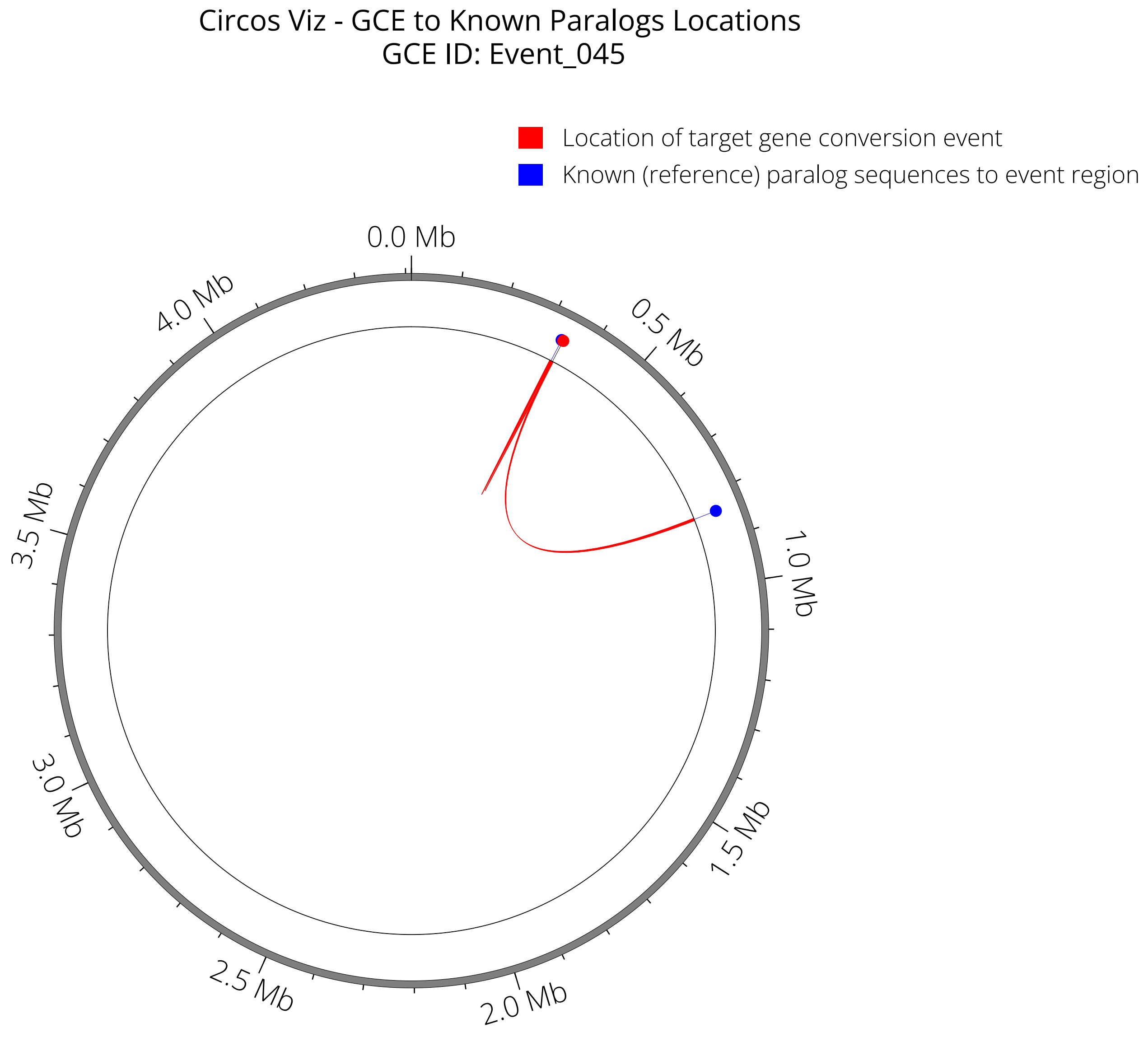

Circos Viz - GCE to Known Paralogs Locations
GCE ID: Event_045
Location of target gene conversion event
Known (reference) paralog sequences to event region
0.0 Mb
0.5 Mb
1.0 Mb
1.5 Mb
2.0 Mb
2.5 Mb
3.0 Mb
3.5 Mb
4.0 Mb

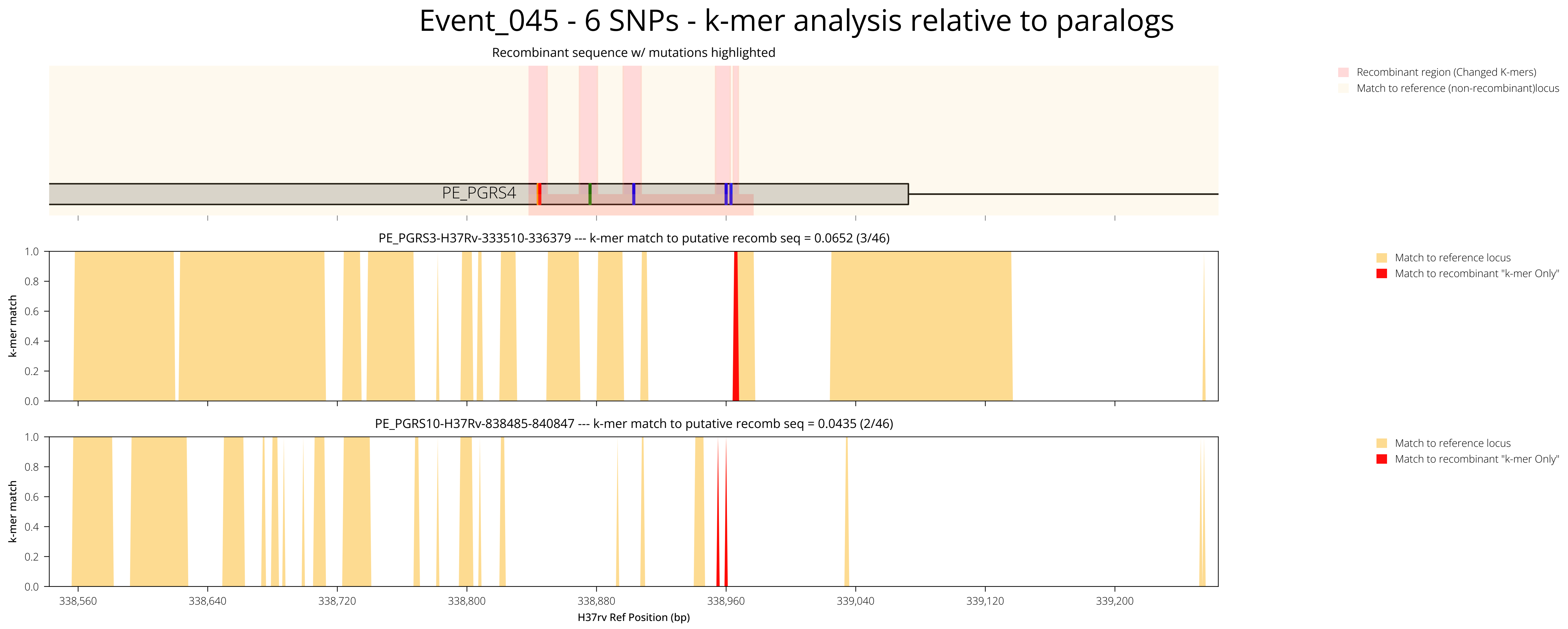

Event_045 - 6 SNPs - k-mer analysis relative to paralogs
Recombinant sequence w/ mutations highlighted
PE_PGRS4
Recombinant region (Changed K-mers)
Match to reference (non-recombinant)locus
PE_PGRS3-H37Rv-333510-336379 --- k-mer match to putative recomb seq = 0.0652 (3/46)
k-mer match
1.0
0.8
0.6
0.4
0.2
0.0
Match to reference locus
Match to recombinant "k-mer Only"
PE_PGRS10-H37Rv-838485-840847 --- k-mer match to putative recomb seq = 0.0435 (2/46)
k-mer match
Match to reference locus
Match to recombinant "k-mer Only"
338,560
338,640
338,720
338,800
338,880
338,960
339,040
339,120
339,200
H37rv Ref Position (bp)

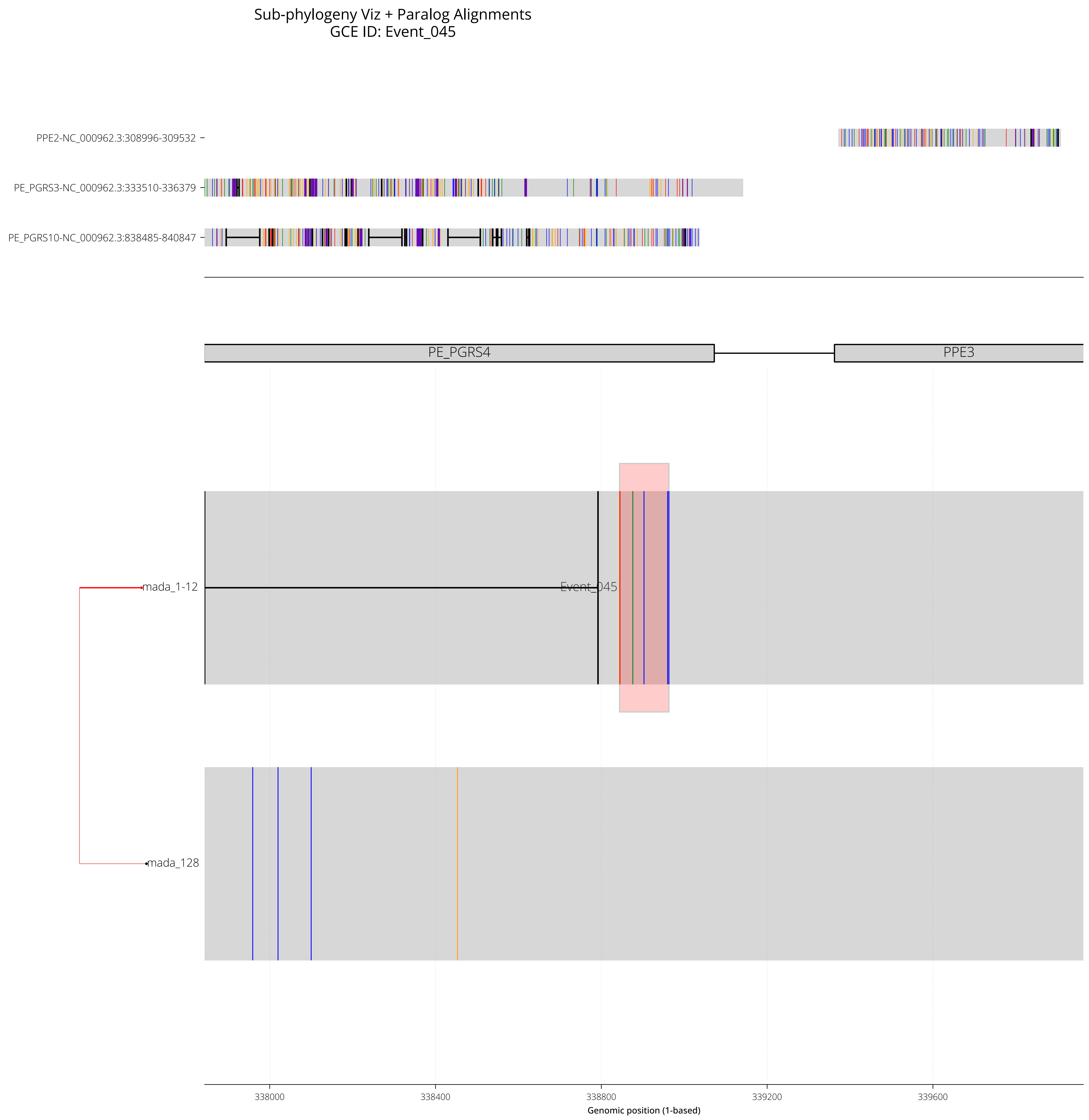
Sub-phylogeny Viz + Paralog Alignments
GCE ID: Event_045
PPE2-NC_000962.3:308996-309532
PE_PGRS3-NC_000962.3:333510-336379
PE_PGRS10-NC_000962.3:838485-840847
PE_PGRS4
PPE3
mada_1-12
Event_045
mada_128
338000
338400
338800
339200
339600
Genomic position (1-based)

Full Phylogeny Viz + Paralog Alignments
GCE ID: Event_045

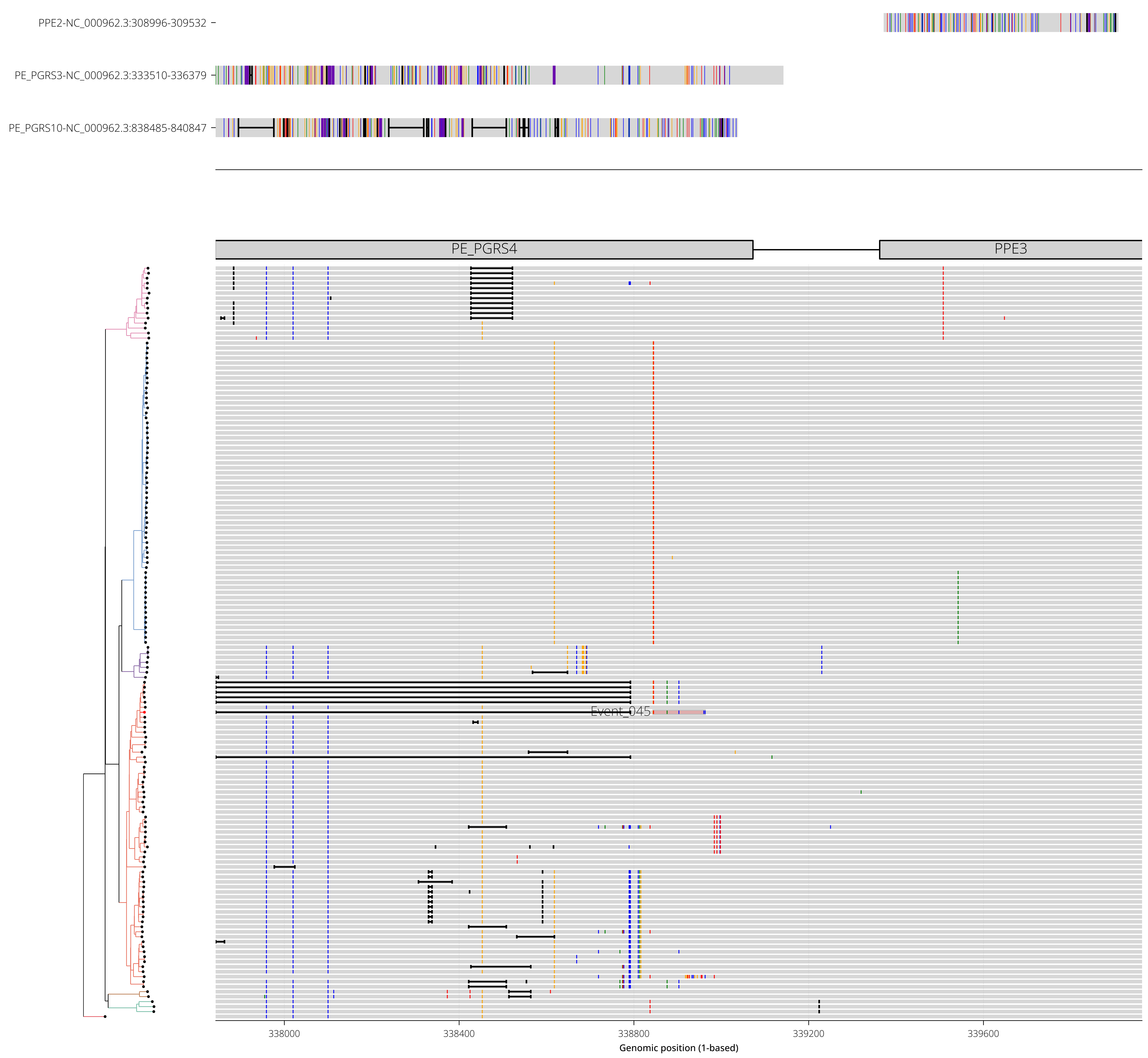

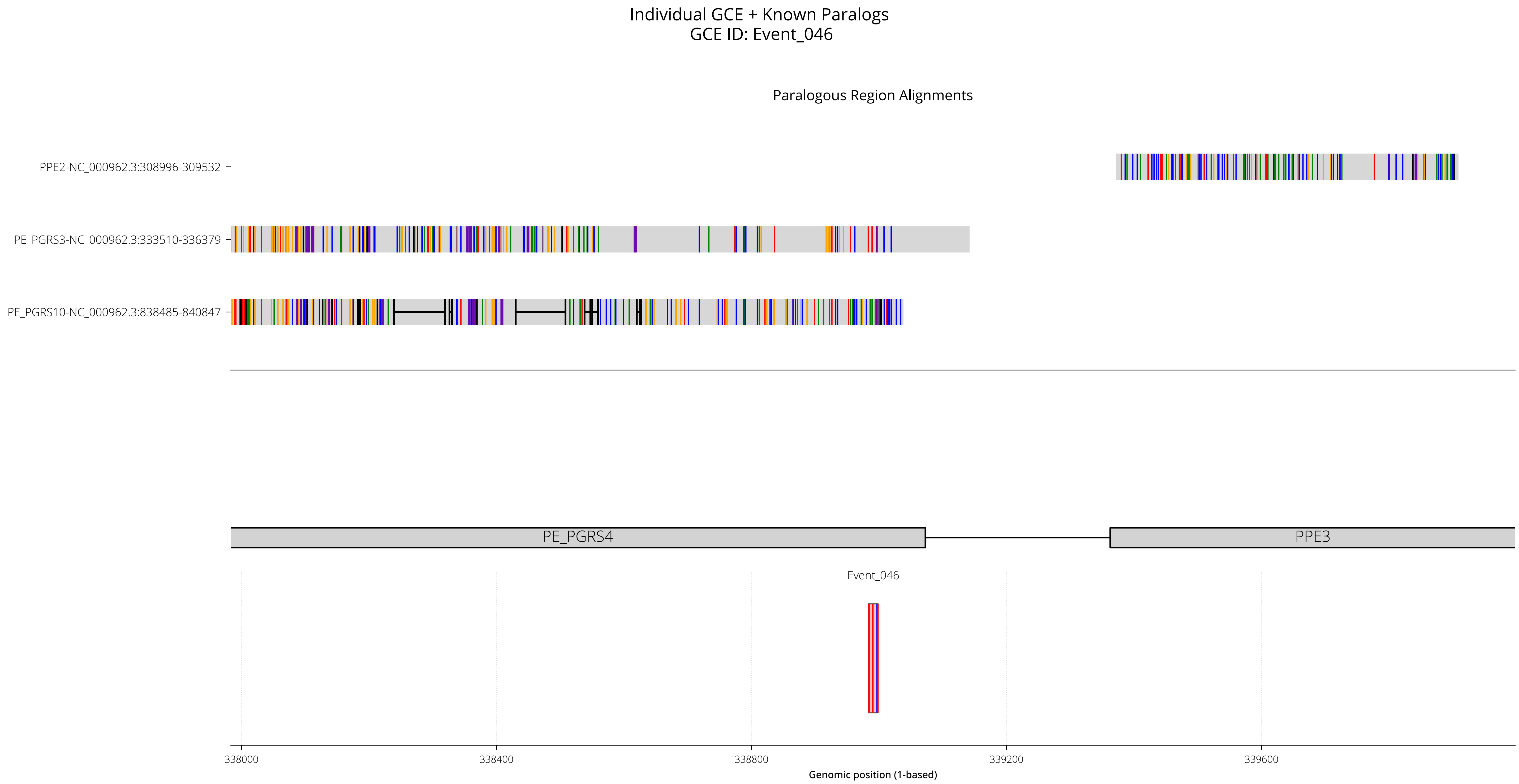
Individual GCE + Known Paralogs
GCE ID: Event_046
Paralogous Region Alignments
PPE2-NC_000962.3:308996-309532
PE_PGRS3-NC_000962.3:333510-336379
PE_PGRS10-NC_000962.3:838485-840847
PE_PGRS4
PPE3
Event_046
338000
338400
338800
339200
339600
Genomic position (1-based)

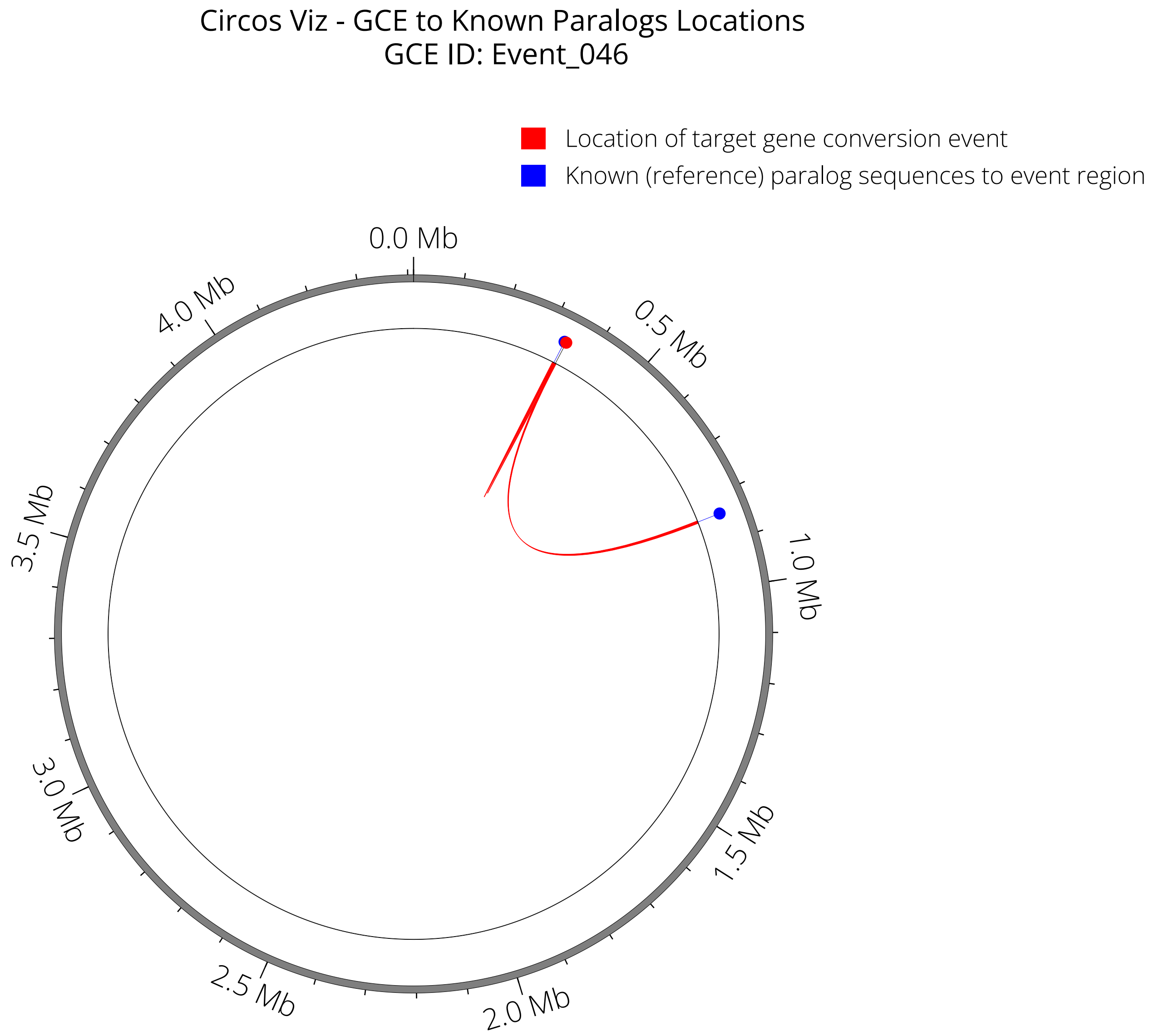
Circos Viz - GCE to Known Paralogs Locations
GCE ID: Event_046
Location of target gene conversion event
Known (reference) paralog sequences to event region
0.0 Mb
0.5 Mb
1.0 Mb
1.5 Mb
2.0 Mb
2.5 Mb
3.0 Mb
3.5 Mb
4.0 Mb

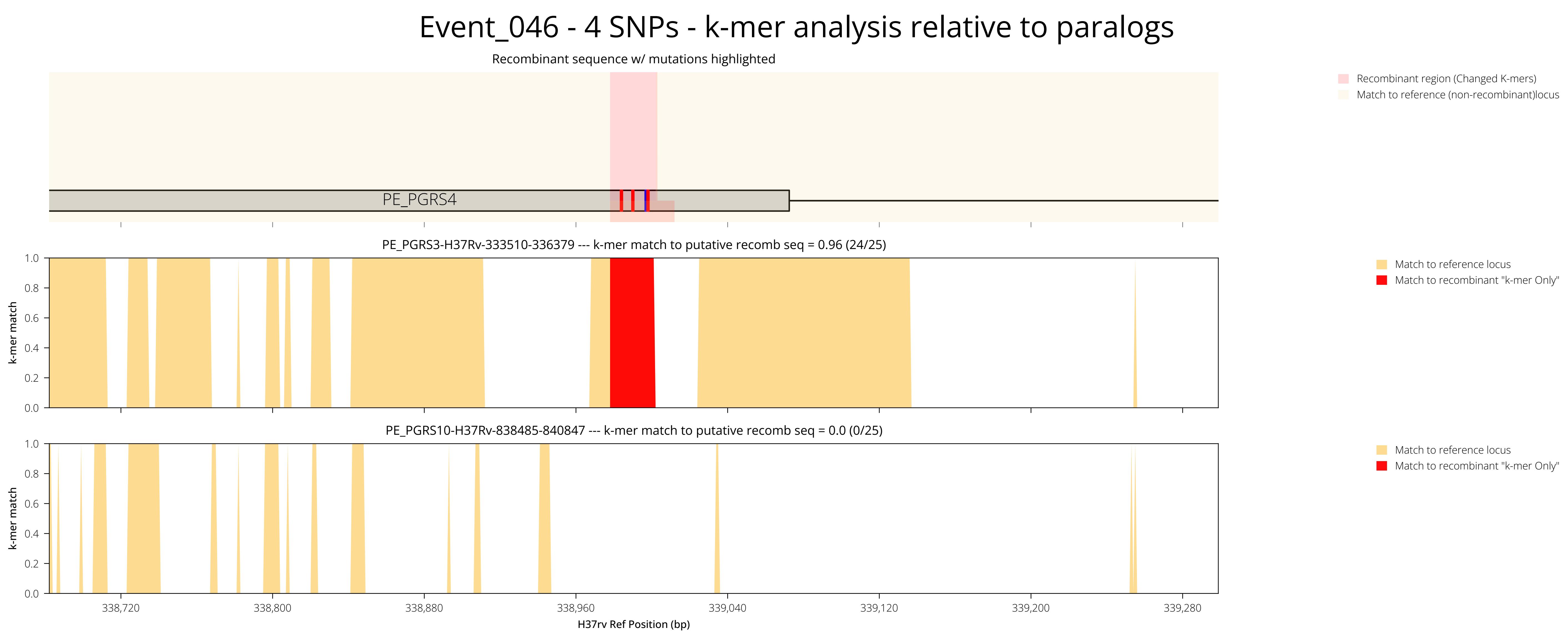

Event_046 - 4 SNPs - k-mer analysis relative to paralogs
Recombinant sequence w/ mutations highlighted
PE_PGRS4
Recombinant region (Changed K-mers)
Match to reference (non-recombinant)locus
PE_PGRS3-H37Rv-333510-336379 --- k-mer match to putative recomb seq = 0.96 (24/25)
Match to reference locus
Match to recombinant "k-mer Only"
PE_PGRS10-H37Rv-838485-840847 --- k-mer match to putative recomb seq = 0.0 (0/25)
Match to reference locus
Match to recombinant "k-mer Only"
k-mer match
1.0
0.8
0.6
0.4
0.2
0.0
338,720
338,800
338,880
338,960
339,040
339,120
339,200
339,280
H37rv Ref Position (bp)

#### Sub-phylogeny Viz + Paralog Alignments
#### GCE ID: Event_046

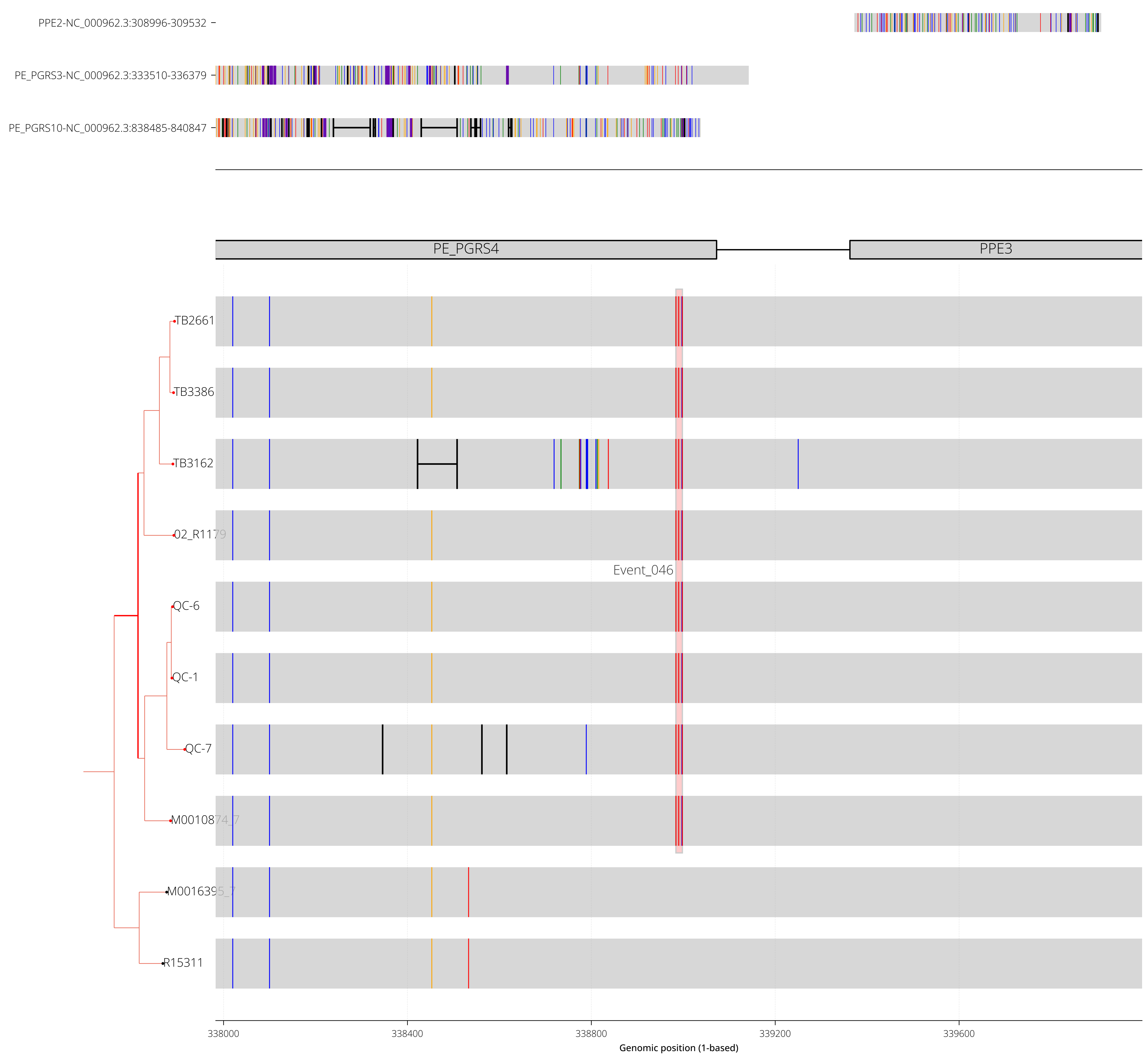

Full Phylogeny Viz + Paralog Alignments
GCE ID: Event_046

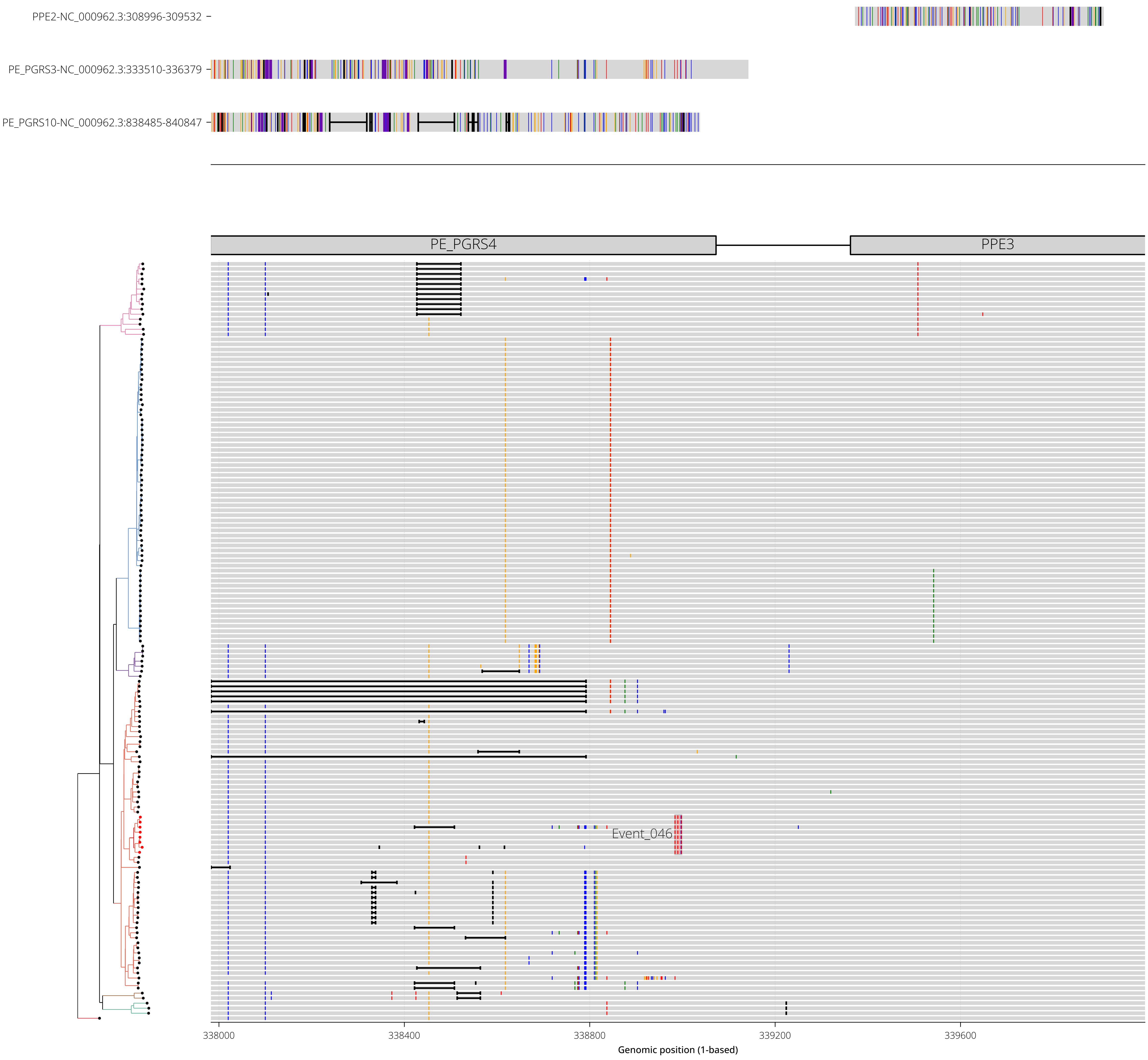

Individual GCE + Known Paralogs
GCE ID: Event_047

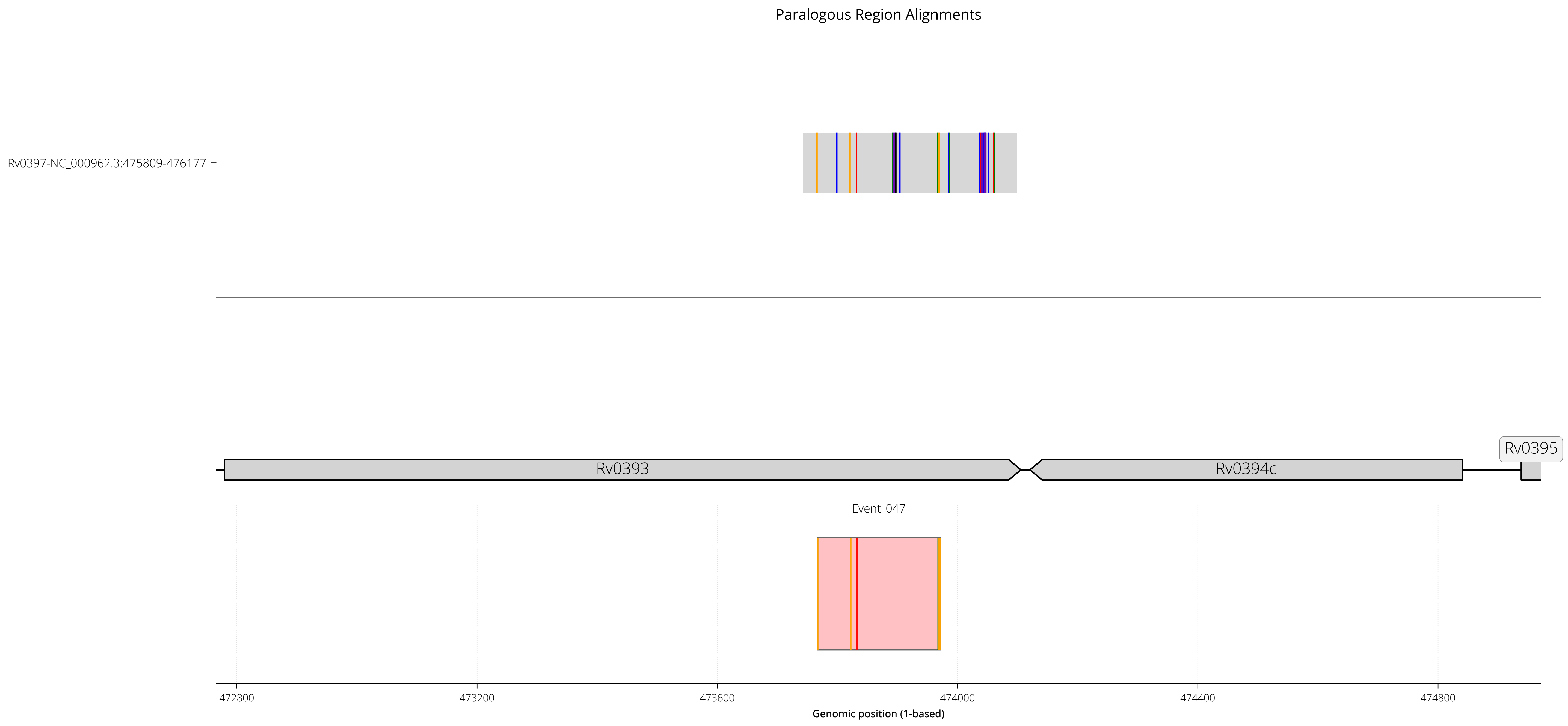

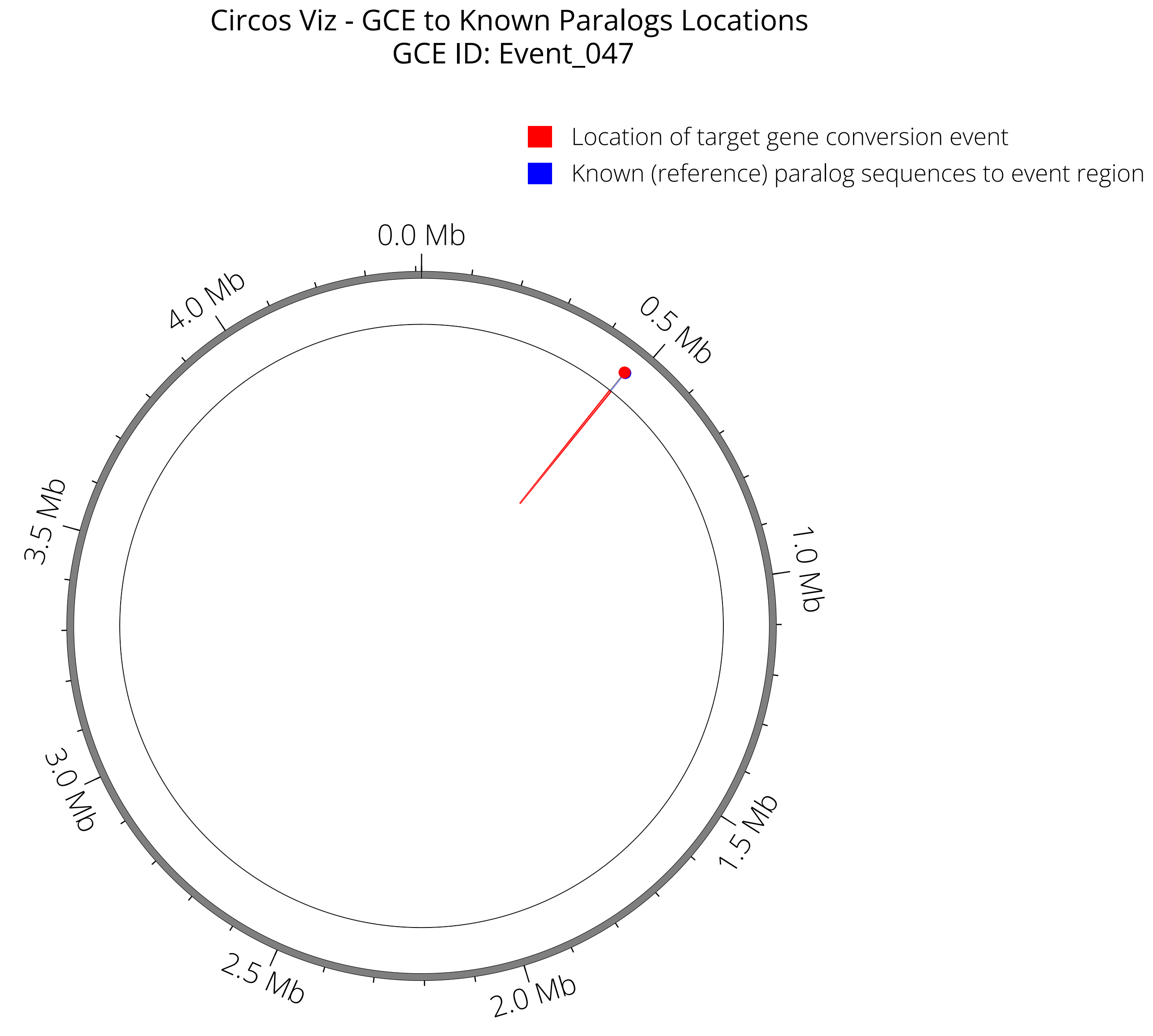
Circos Viz - GCE to Known Paralogs Locations
GCE ID: Event_047
Location of target gene conversion event
Known (reference) paralog sequences to event region
0.0 Mb
0.5 Mb
1.0 Mb
1.5 Mb
2.0 Mb
2.5 Mb
3.0 Mb
3.5 Mb
4.0 Mb

### Event_047 - 6 SNPs - k-mer analysis relative to paralogs

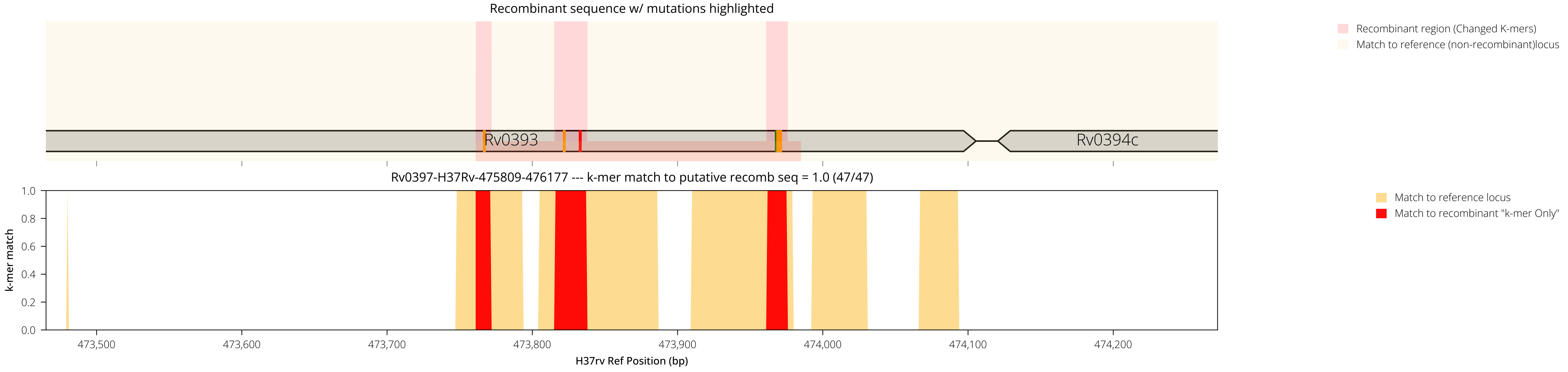

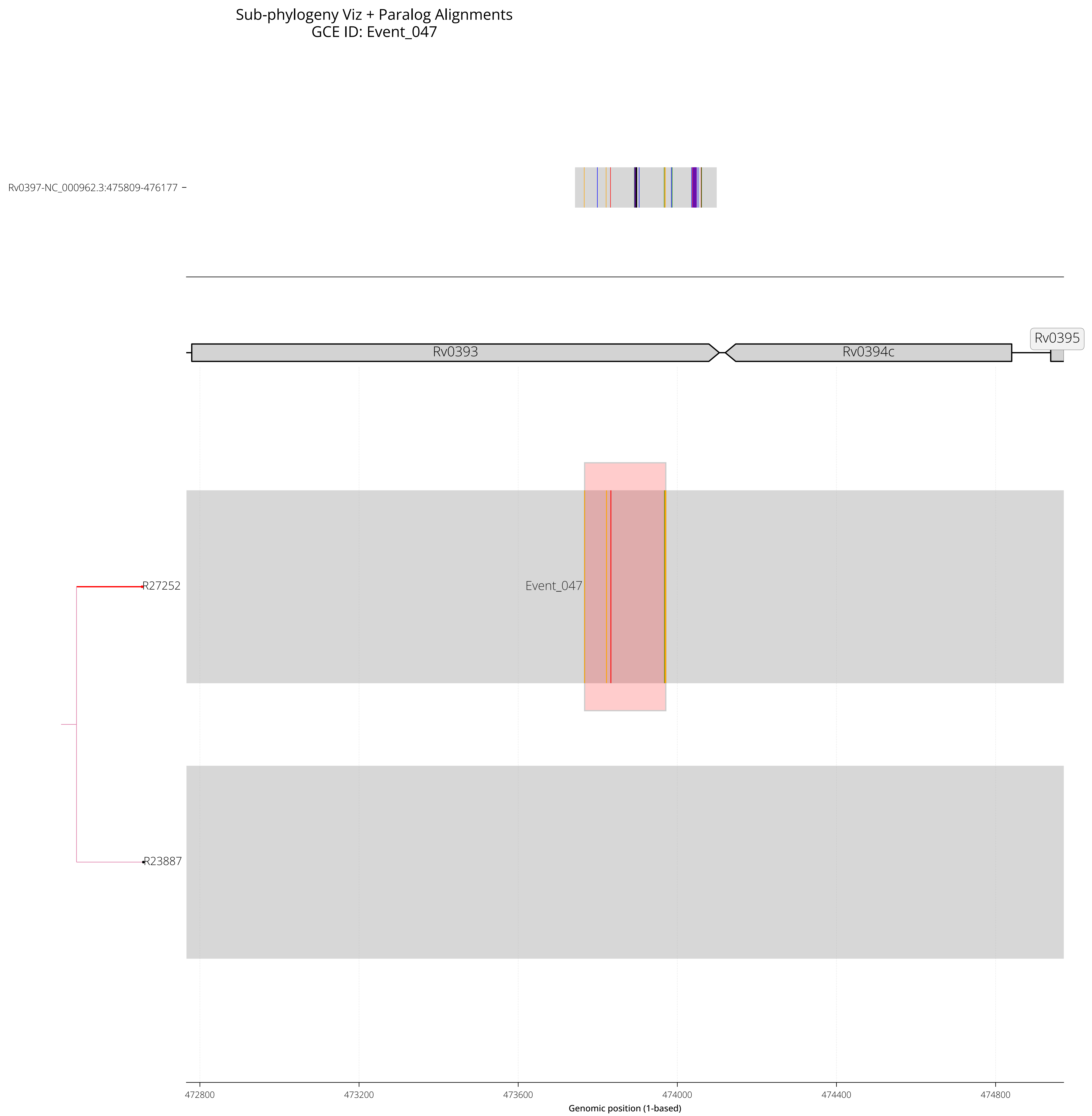

Sub-phylogeny Viz + Paralog Alignments
GCE ID: Event_047
Rv0397-NC_000962.3:475809-476177
Rv0393
Rv0394c
Rv0395
Event_047
R27252
R23887
472800
473200
473600
474000
474400
474800
Genomic position (1-based)

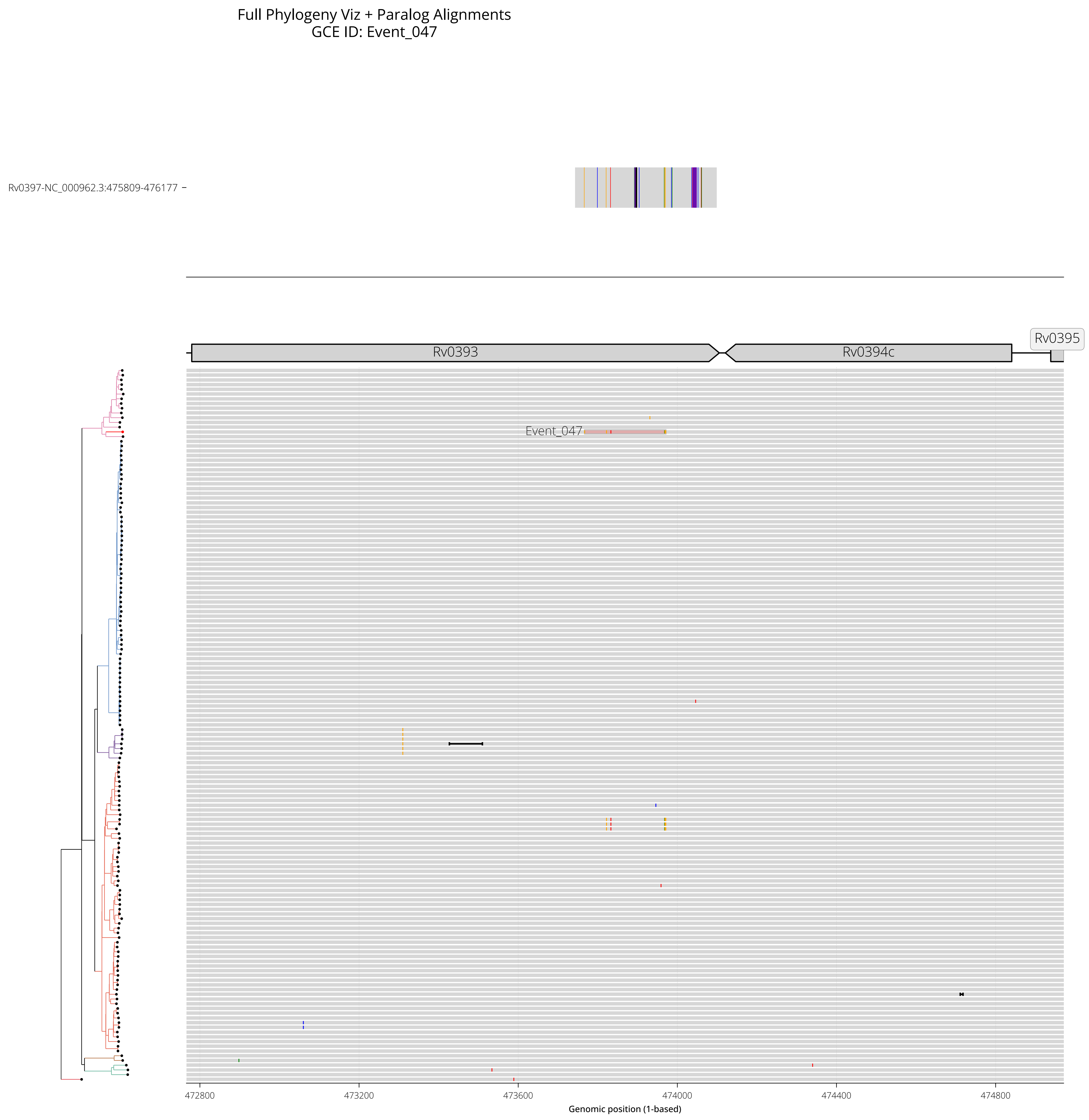

Full Phylogeny Viz + Paralog Alignments
GCE ID: Event_047
Rv0397-NC_000962.3:475809-476177
Rv0393
Rv0394c
Rv0395
Event_047
472800
473200
473600
474000
474400
474800
Genomic position (1-based)

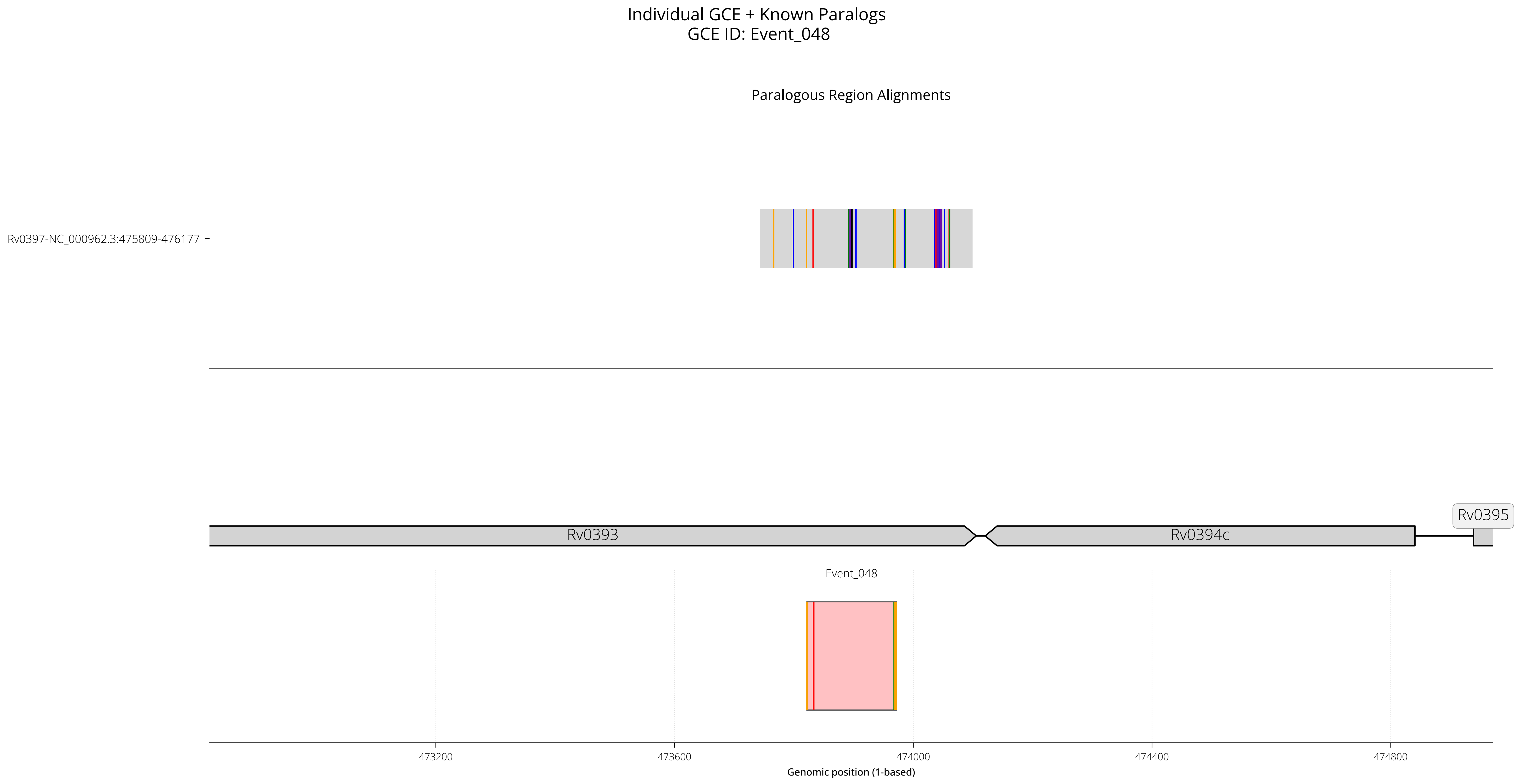

Individual GCE + Known Paralogs
GCE ID: Event_048
Paralogous Region Alignments
Rv0397-NC_000962.3:475809-476177
Rv0393
Rv0394c
Rv0395
Event_048
473200
473600
474000
474400
474800
Genomic position (1-based)

Circos Viz - GCE to Known Paralogs Locations
GCE ID: Event_048

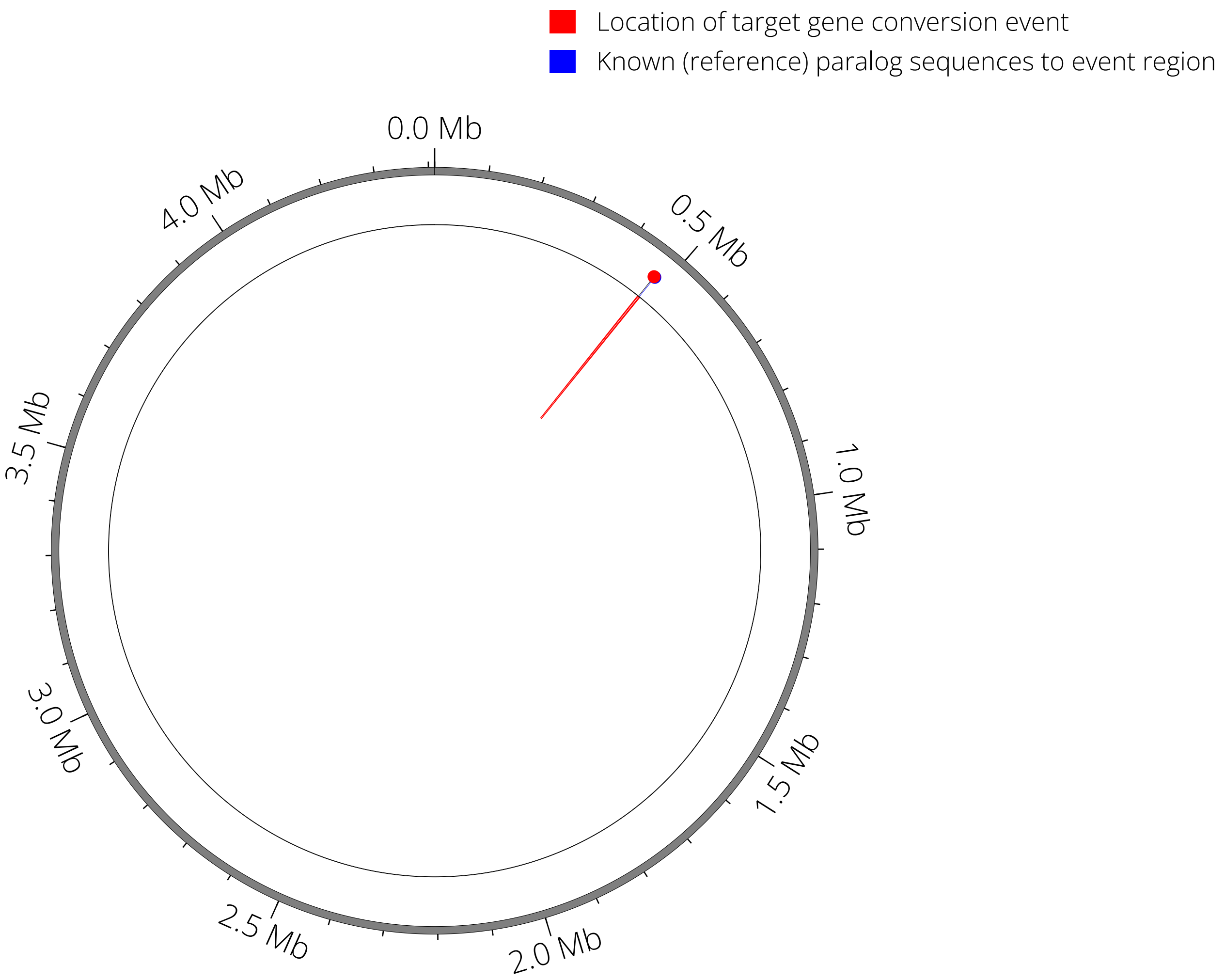

### Event_048 - 5 SNPs - k-mer analysis relative to paralogs

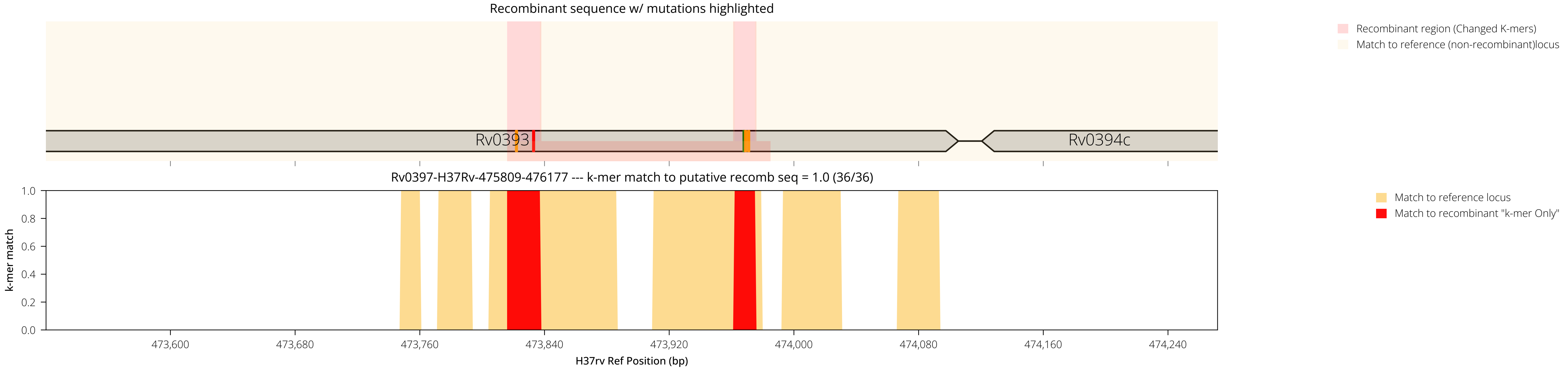

Sub-phylogeny Viz + Paralog Alignments
GCE ID: Event_048

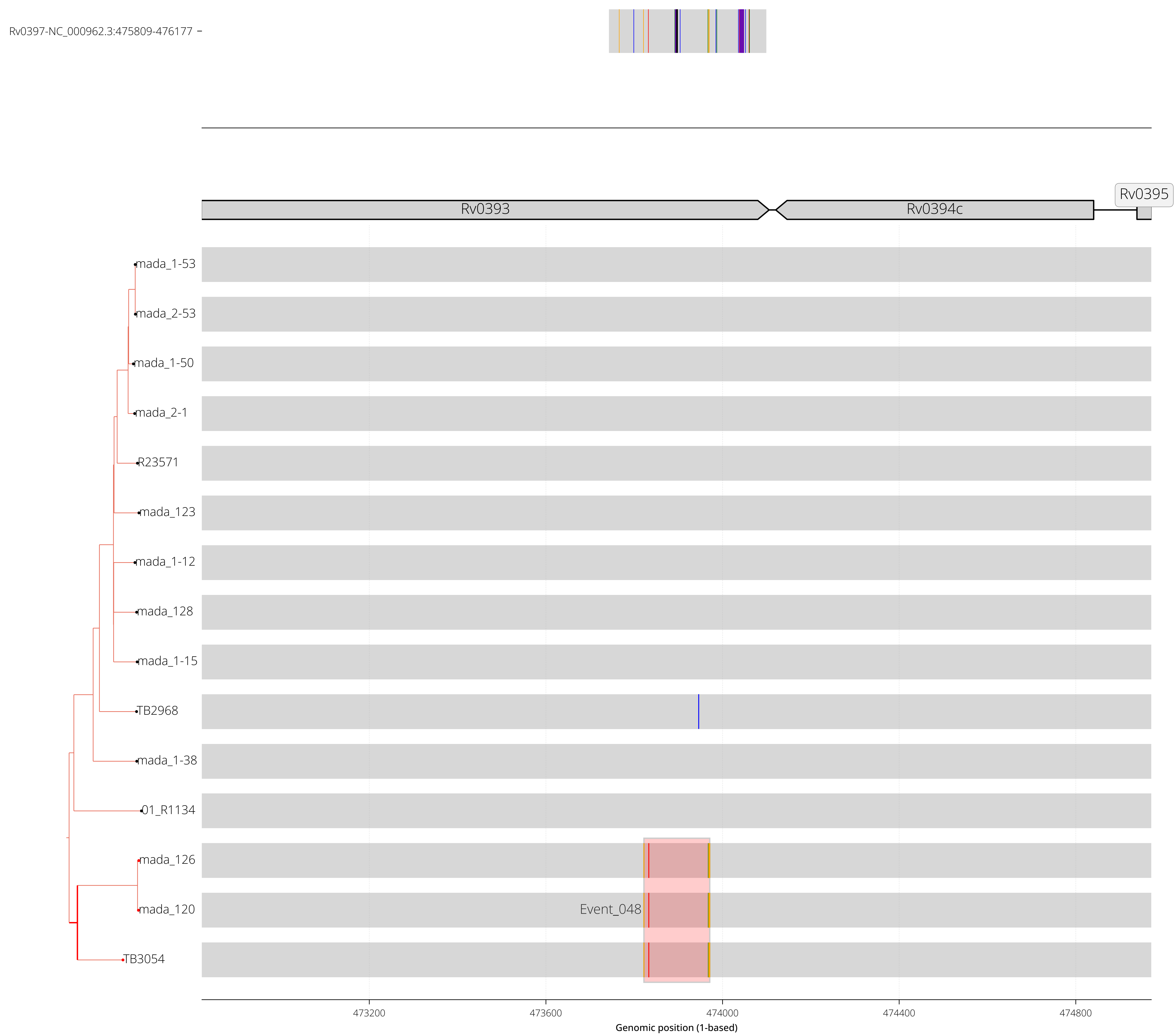

Full Phylogeny Viz + Paralog Alignments
GCE ID: Event_048

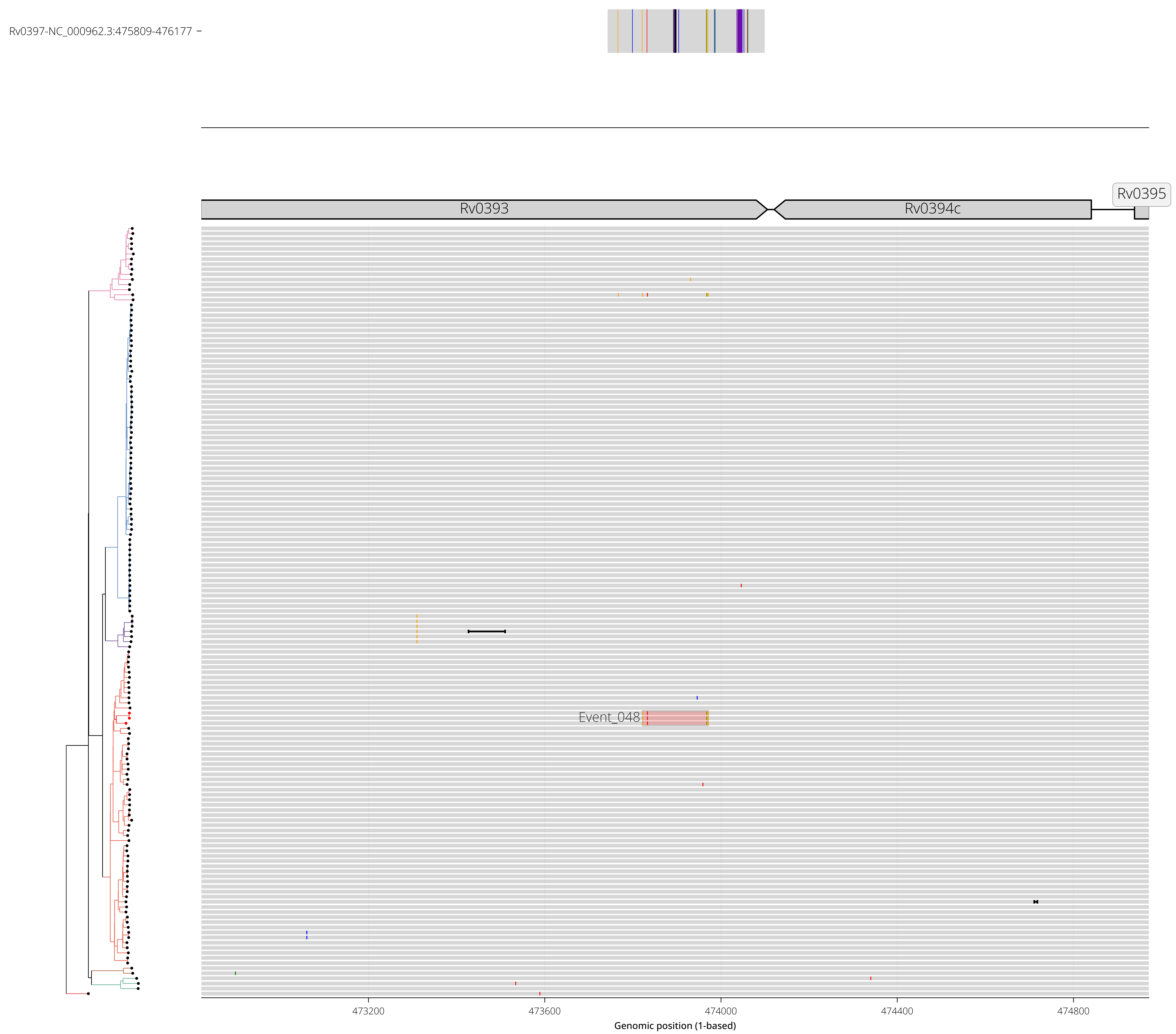

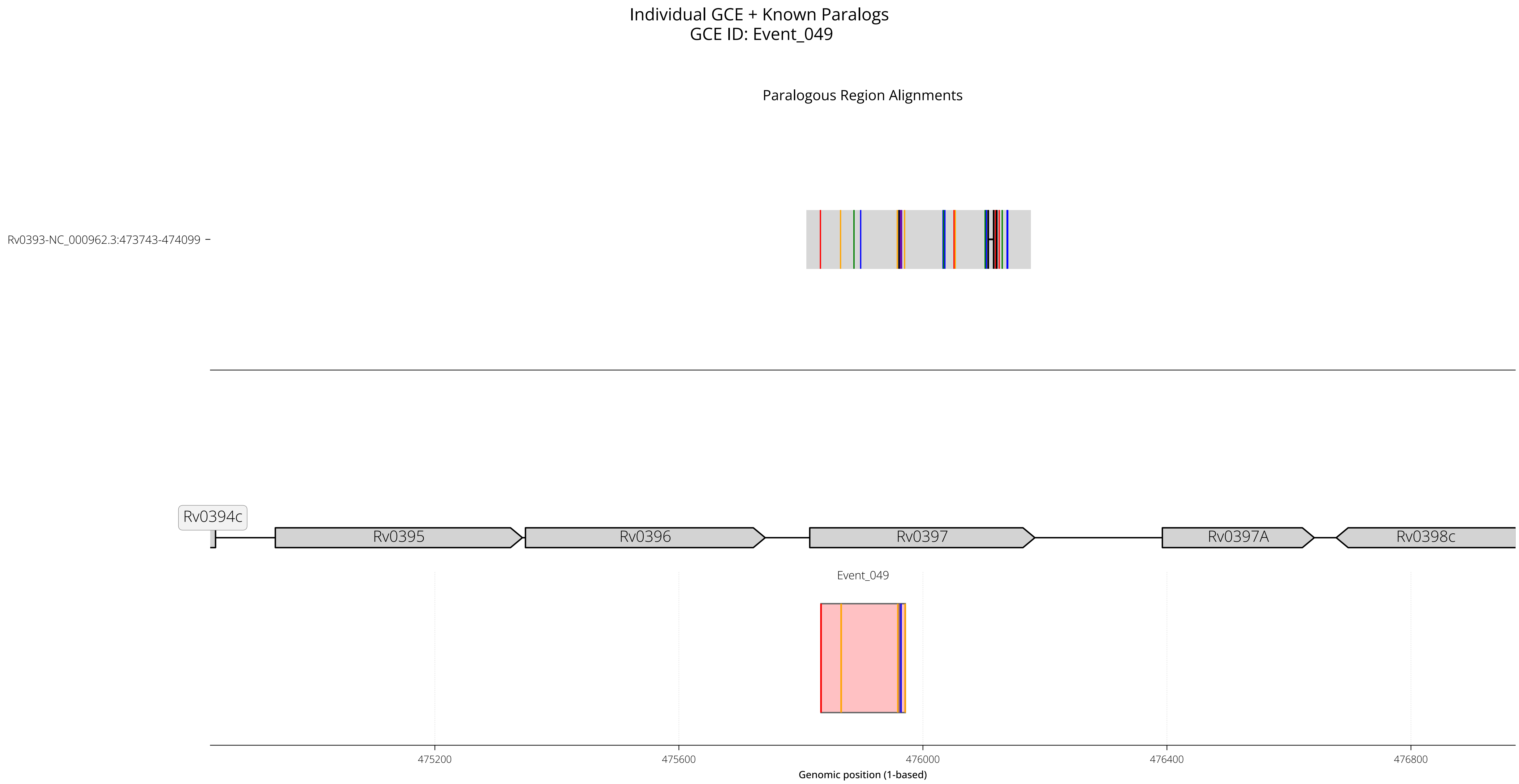

Individual GCE + Known Paralogs
GCE ID: Event_049
Paralogous Region Alignments
Rv0393-NC_000962.3:473743-474099
Rv0394c
Rv0395
Rv0396
Rv0397
Rv0397A
Rv0398c
Event_049
475200
475600
476000
476400
476800
Genomic position (1-based)

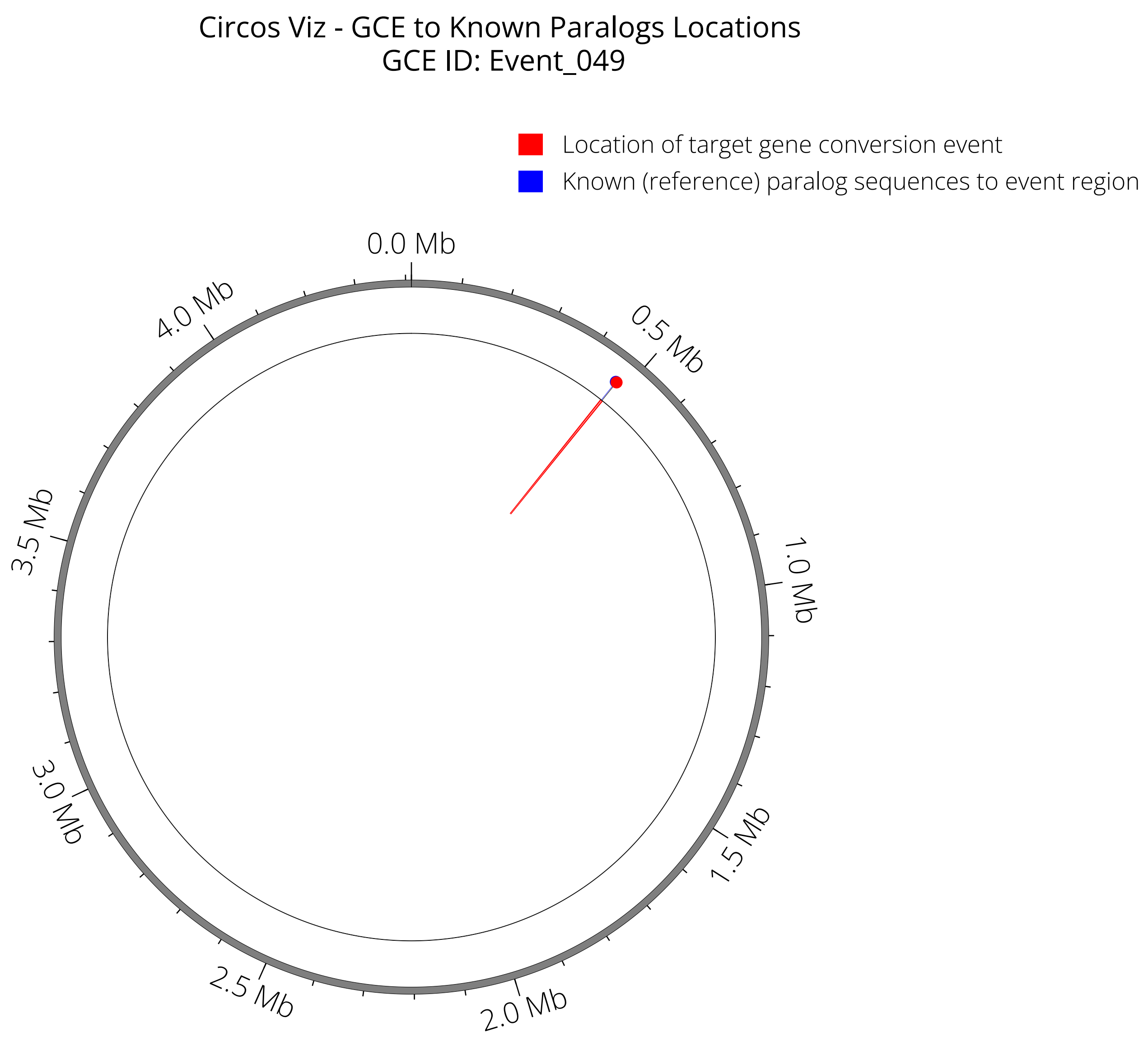
Circos Viz - GCE to Known Paralogs Locations
GCE ID: Event_049
Location of target gene conversion event
Known (reference) paralog sequences to event region
0.0 Mb
0.5 Mb
1.0 Mb
1.5 Mb
2.0 Mb
2.5 Mb
3.0 Mb
3.5 Mb
4.0 Mb

### Event_049 - 7 SNPs - k-mer analysis relative to paralogs

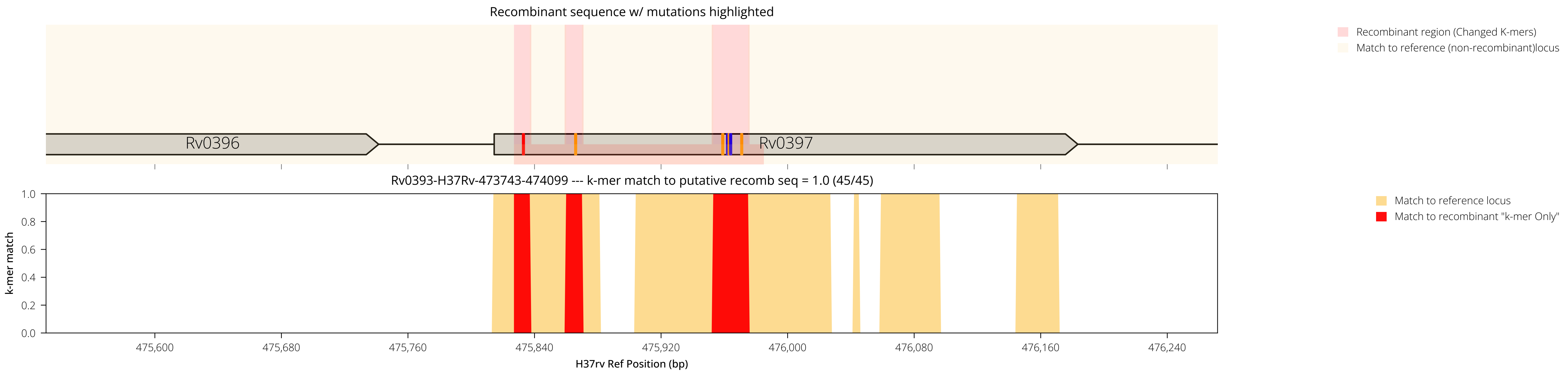

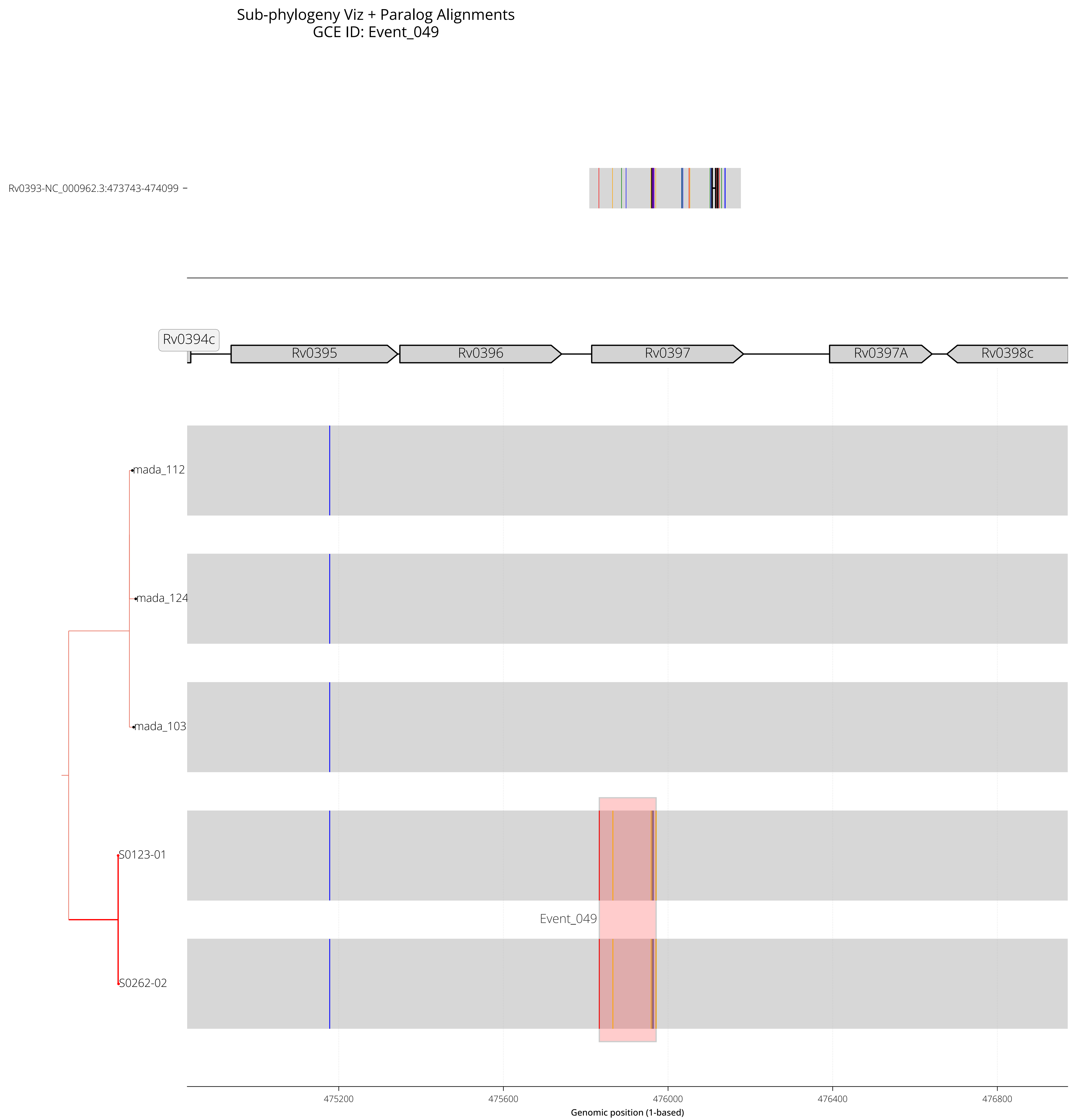

Sub-phylogeny Viz + Paralog Alignments
GCE ID: Event_049
Rv0393-NC_000962.3:473743-474099
Rv0394c
Rv0395
Rv0396
Rv0397
Rv0397A
Rv0398c
mada_112
mada_124
mada_103
S0123-01
Event_049
S0262-02
475200
475600
476000
476400
476800
Genomic position (1-based)

Full Phylogeny Viz + Paralog Alignments
GCE ID: Event_049

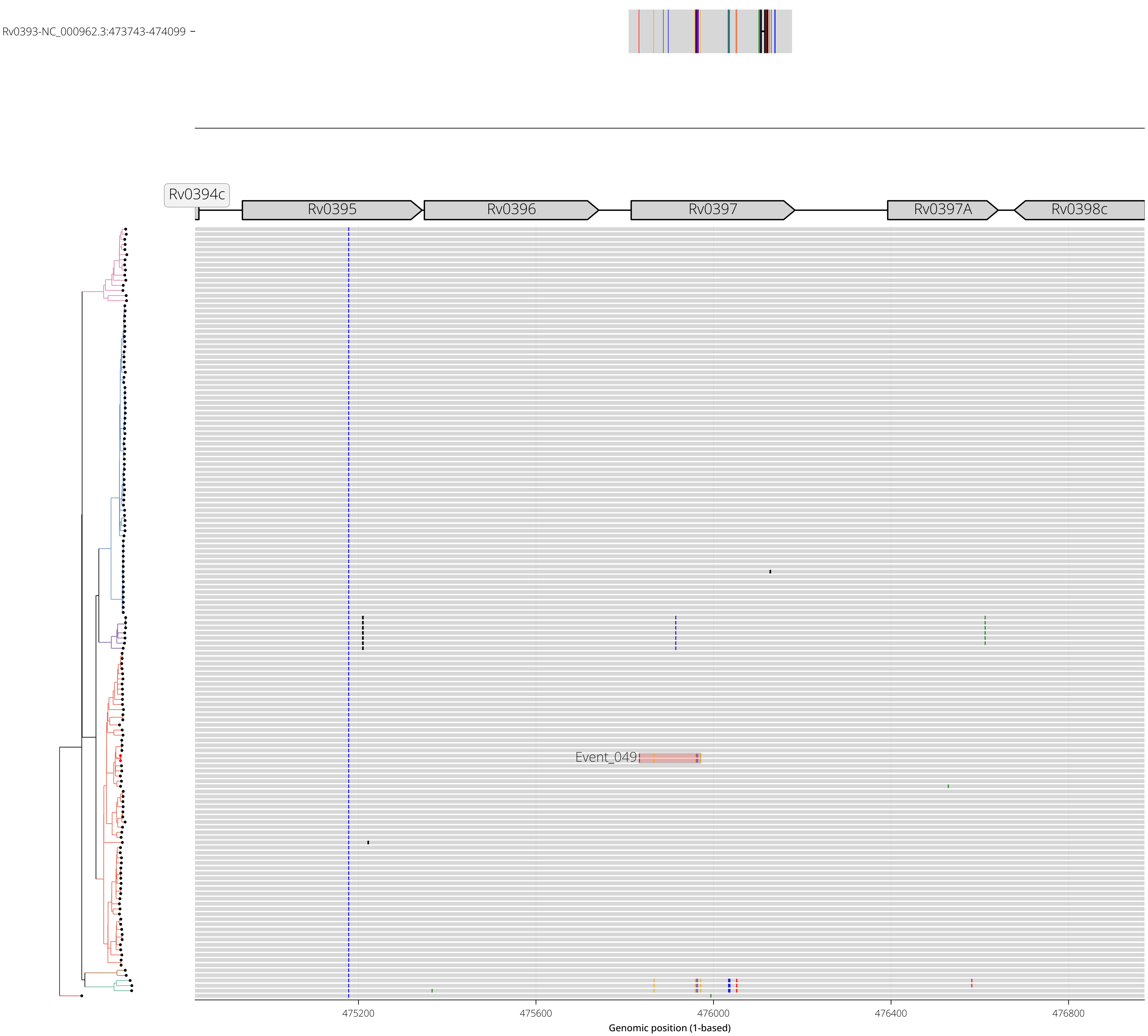

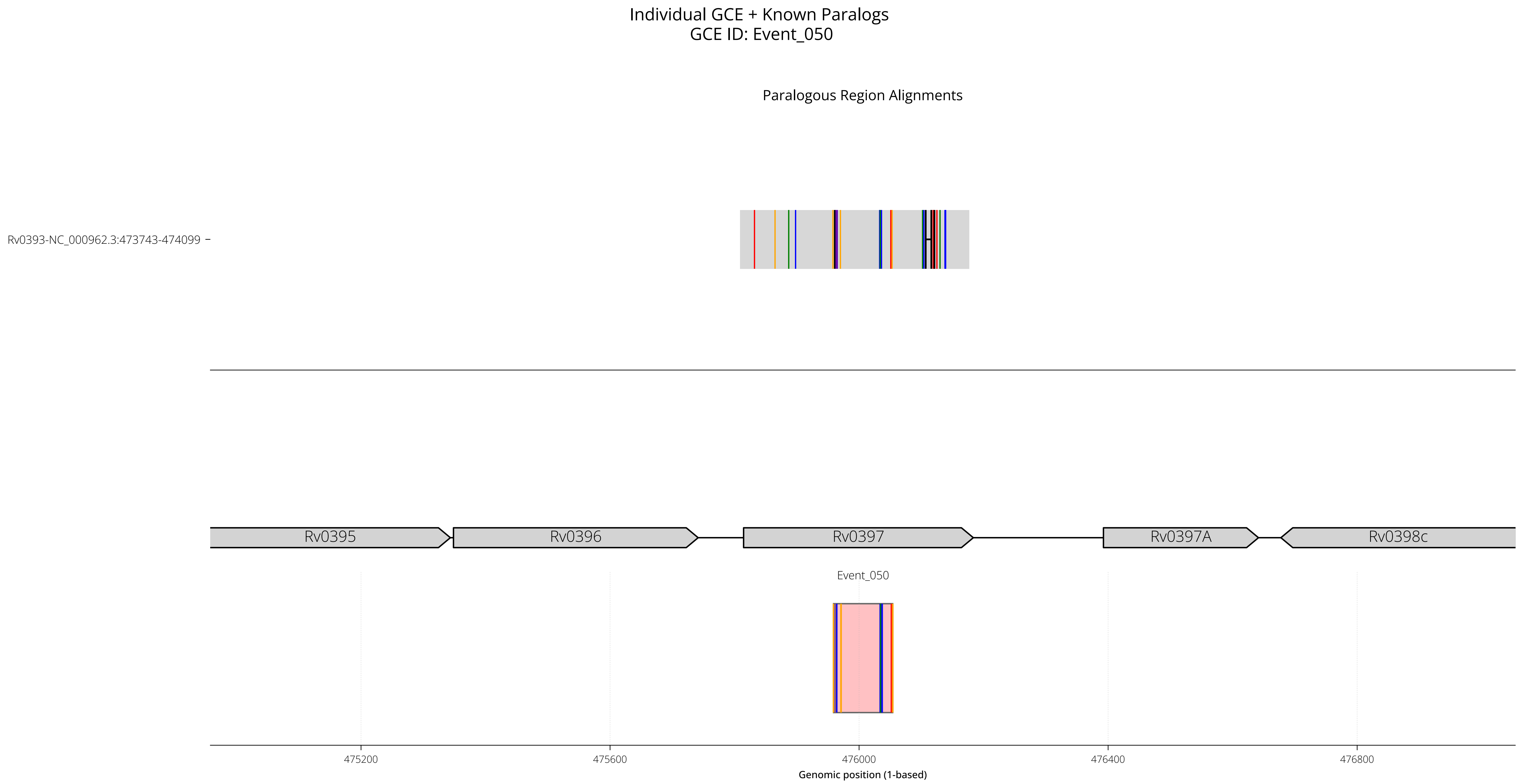

Individual GCE + Known Paralogs
GCE ID: Event_050
Paralogous Region Alignments
Rv0393-NC_000962.3:473743-474099
Rv0395
Rv0396
Rv0397
Rv0397A
Rv0398c
Event_050
475200
475600
476000
476400
476800
Genomic position (1-based)

Circos Viz - GCE to Known Paralogs Locations
GCE ID: Event_050

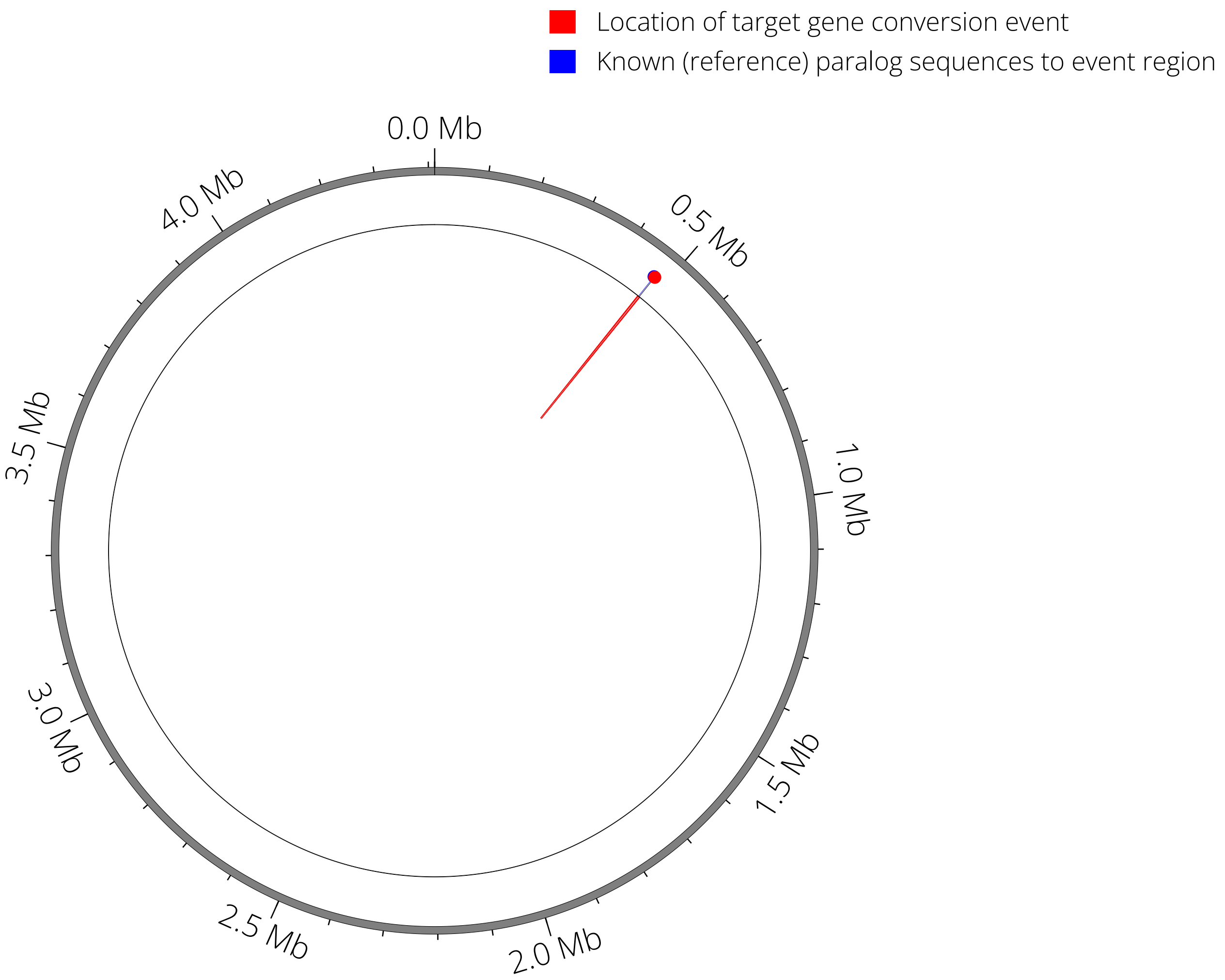

### Event_050 - 10 SNPs - k-mer analysis relative to paralogs

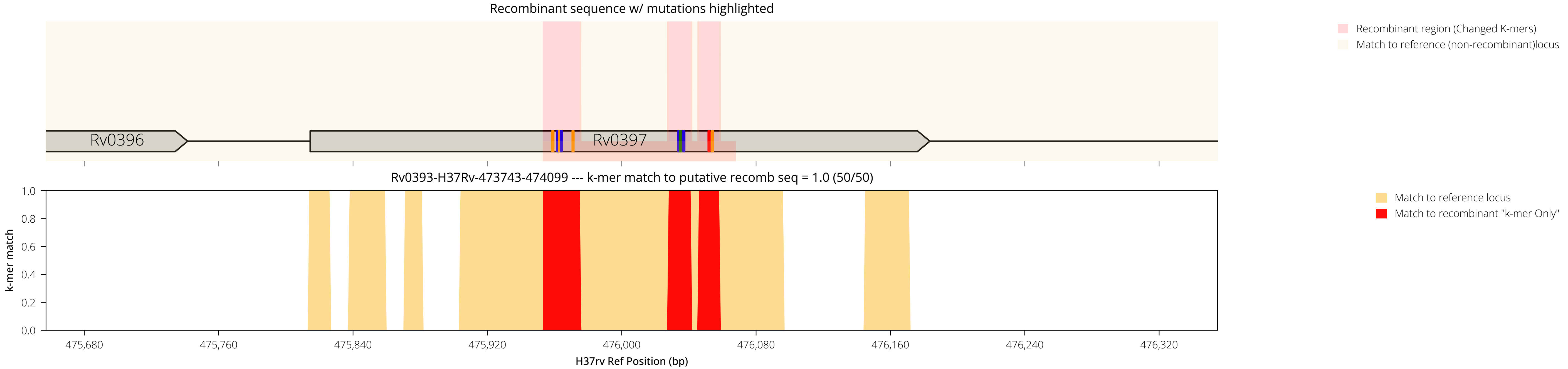

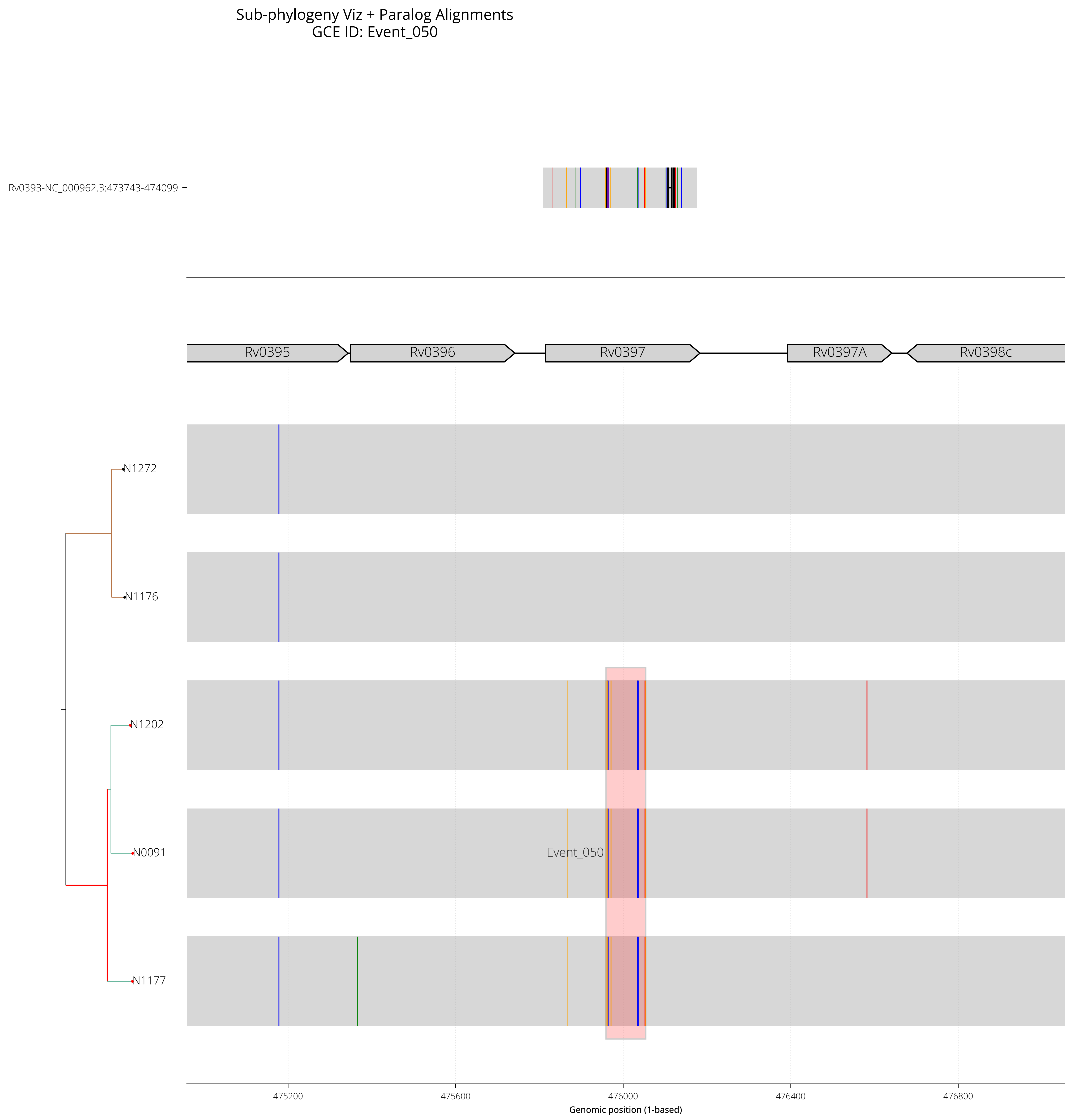

Sub-phylogeny Viz + Paralog Alignments
GCE ID: Event_050
Rv0393-NC_000962.3:473743-474099
Rv0395
Rv0396
Rv0397
Rv0397A
Rv0398c
N1272
N1176
N1202
N0091
N1177
Event_050
475200
475600
476000
476400
476800
Genomic position (1-based)

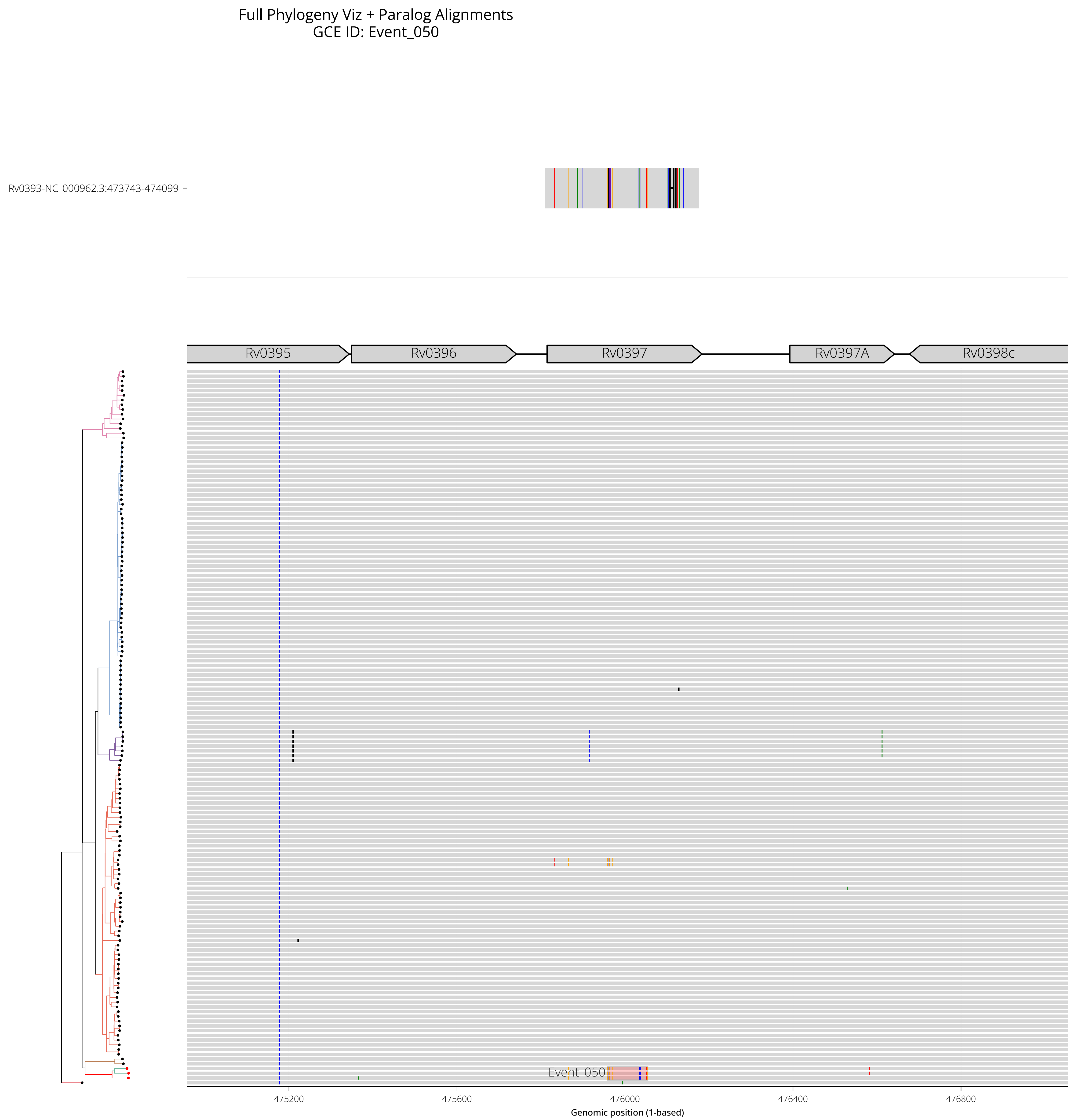

Full Phylogeny Viz + Paralog Alignments
GCE ID: Event_050
Rv0393-NC_000962.3:473743-474099
Rv0395
Rv0396
Rv0397
Rv0397A
Rv0398c
Event_050
475200
475600
476000
476400
476800
Genomic position (1-based)

Individual GCE + Known Paralogs
GCE ID: Event_051

Paralogous Region Alignments

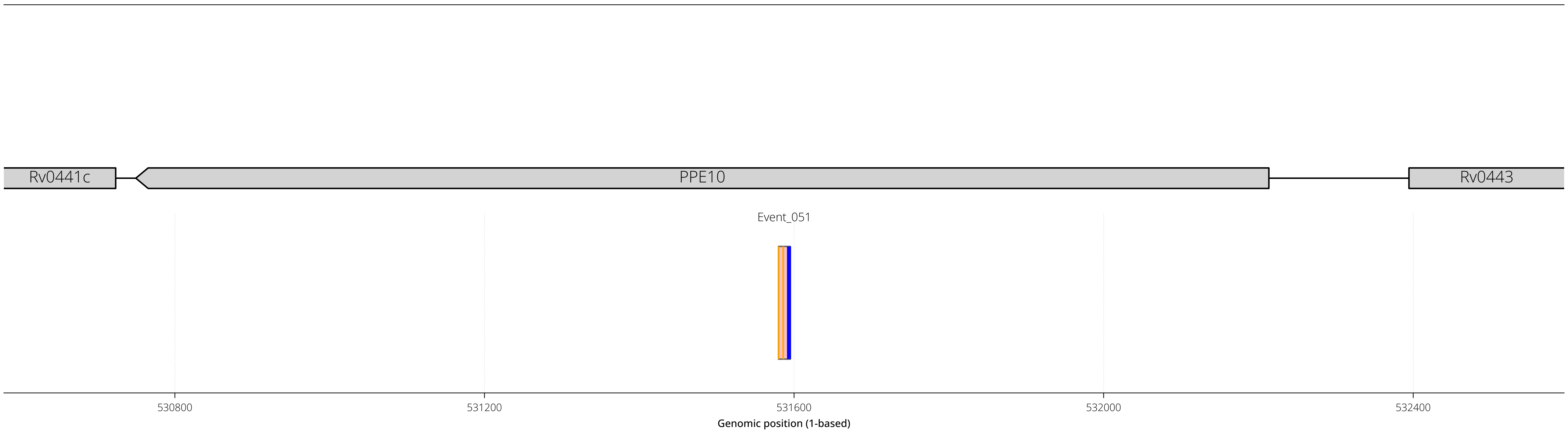

Sub-phylogeny Viz + Paralog Alignments
GCE ID: Event_051

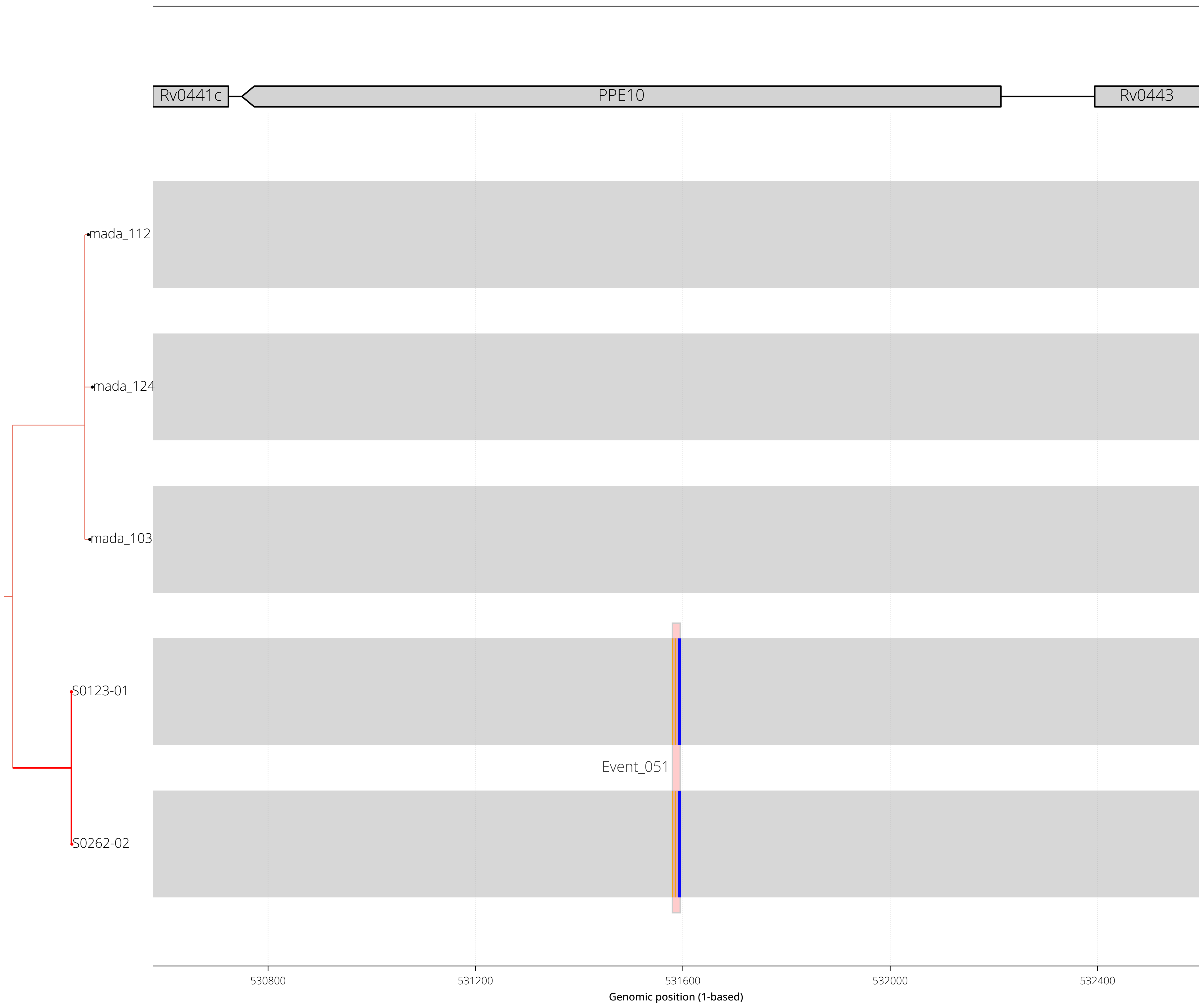

Full Phylogeny Viz + Paralog Alignments
GCE ID: Event_051

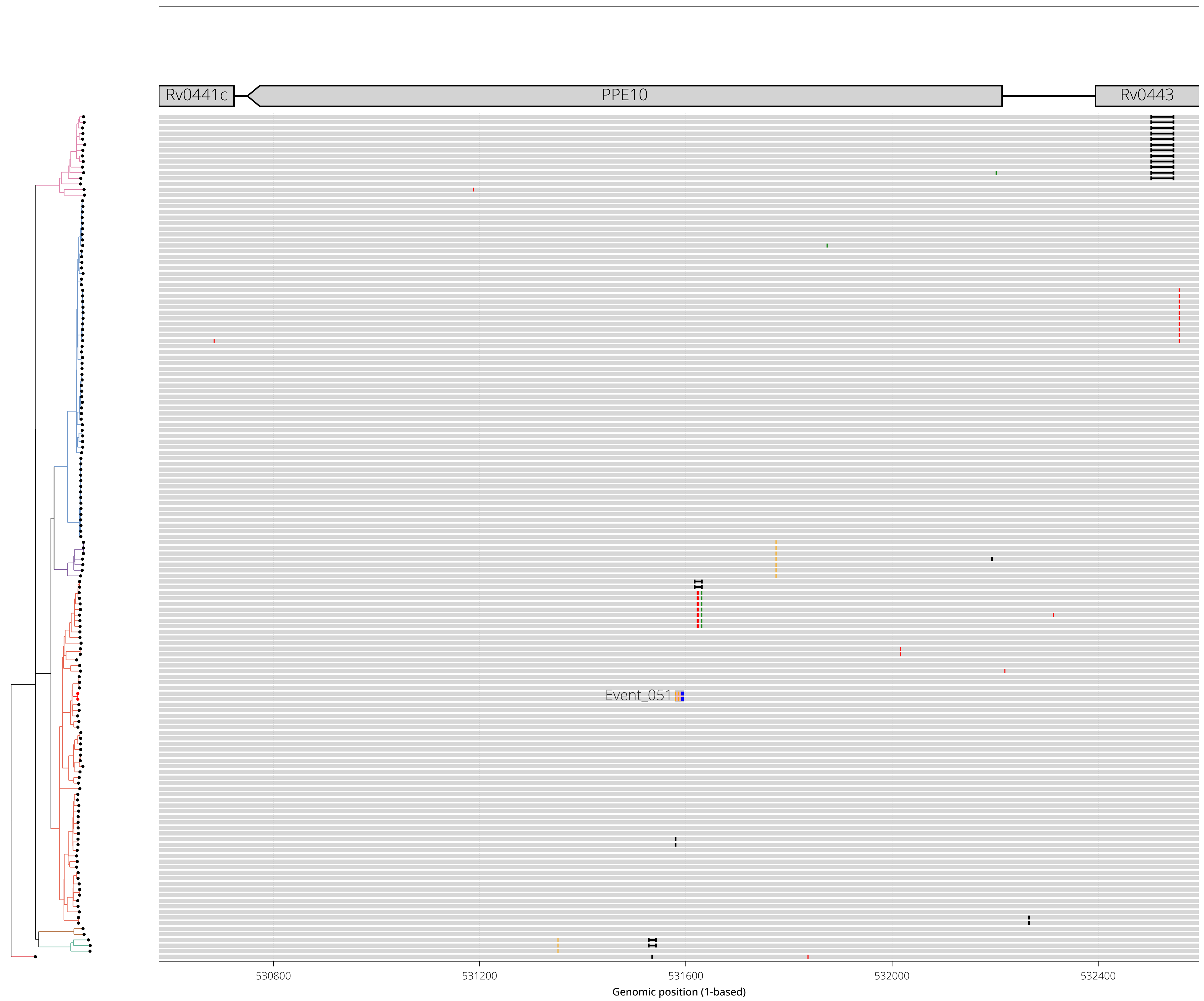

Individual GCE + Known Paralogs
GCE ID: Event_052

Paralogous Region Alignments

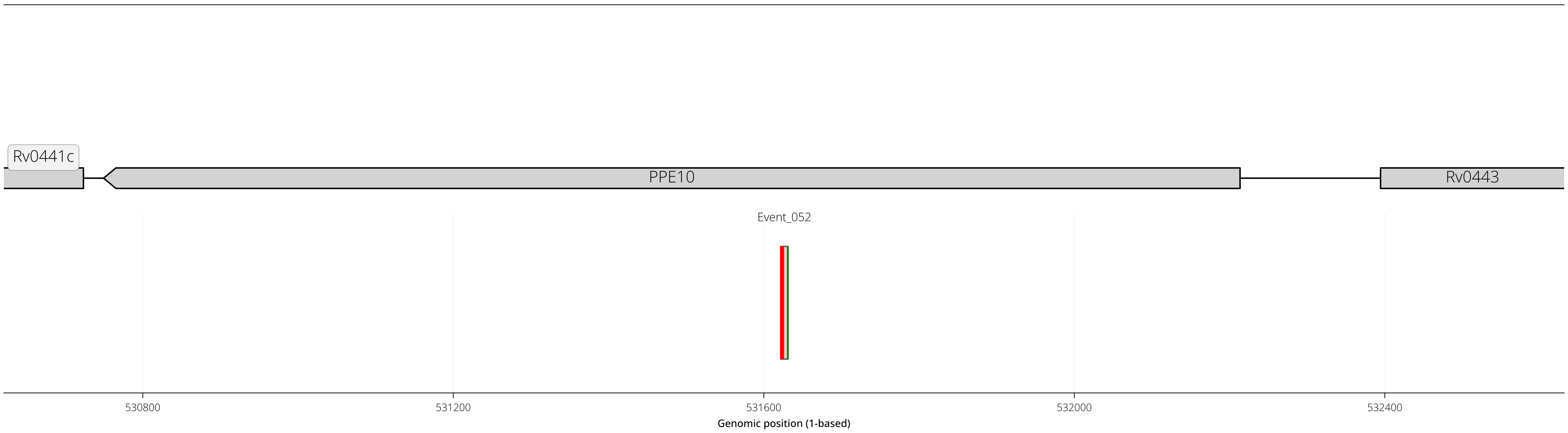

Sub-phylogeny Viz + Paralog Alignments
GCE ID: Event_052

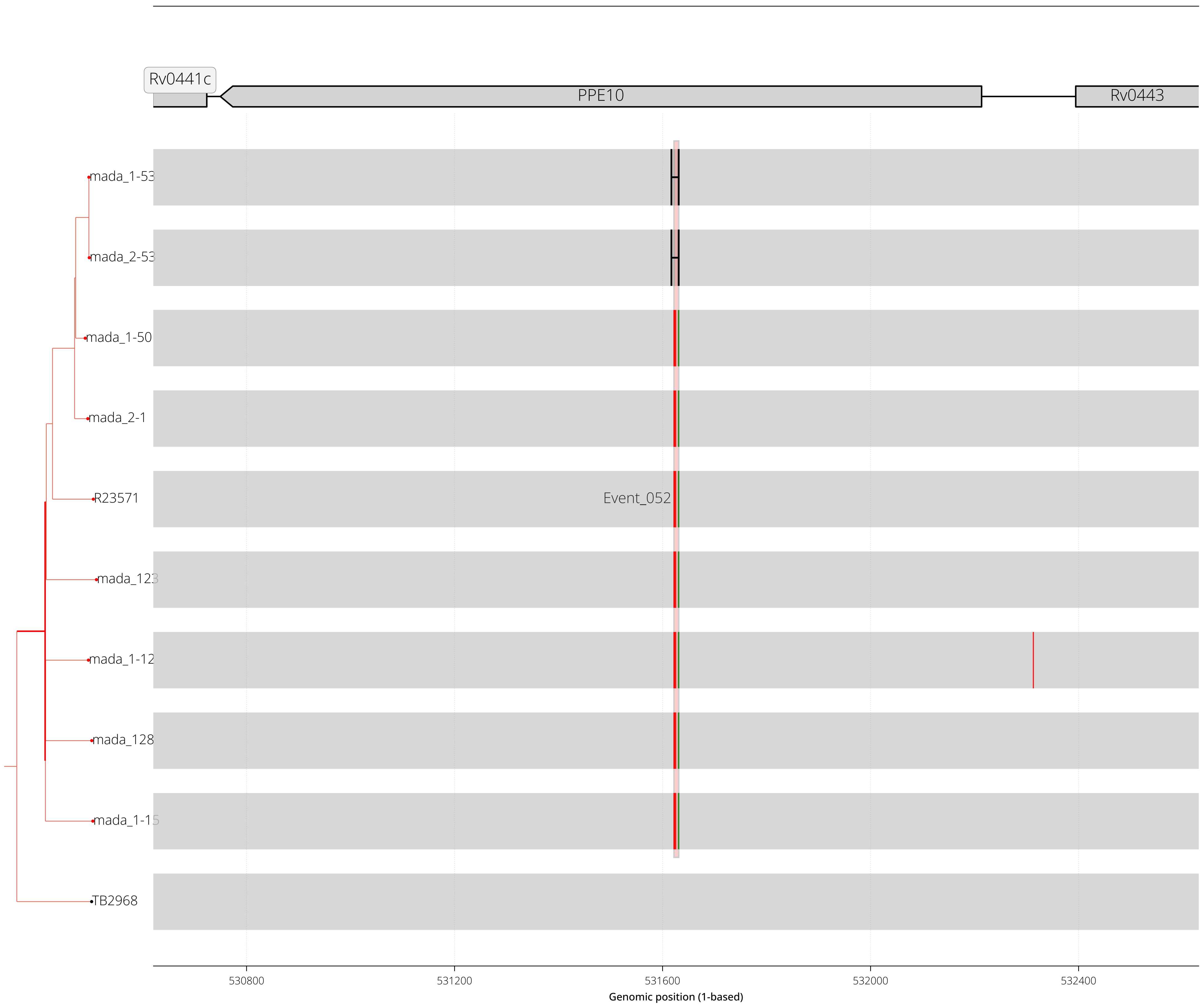

Full Phylogeny Viz + Paralog Alignments
GCE ID: Event_052

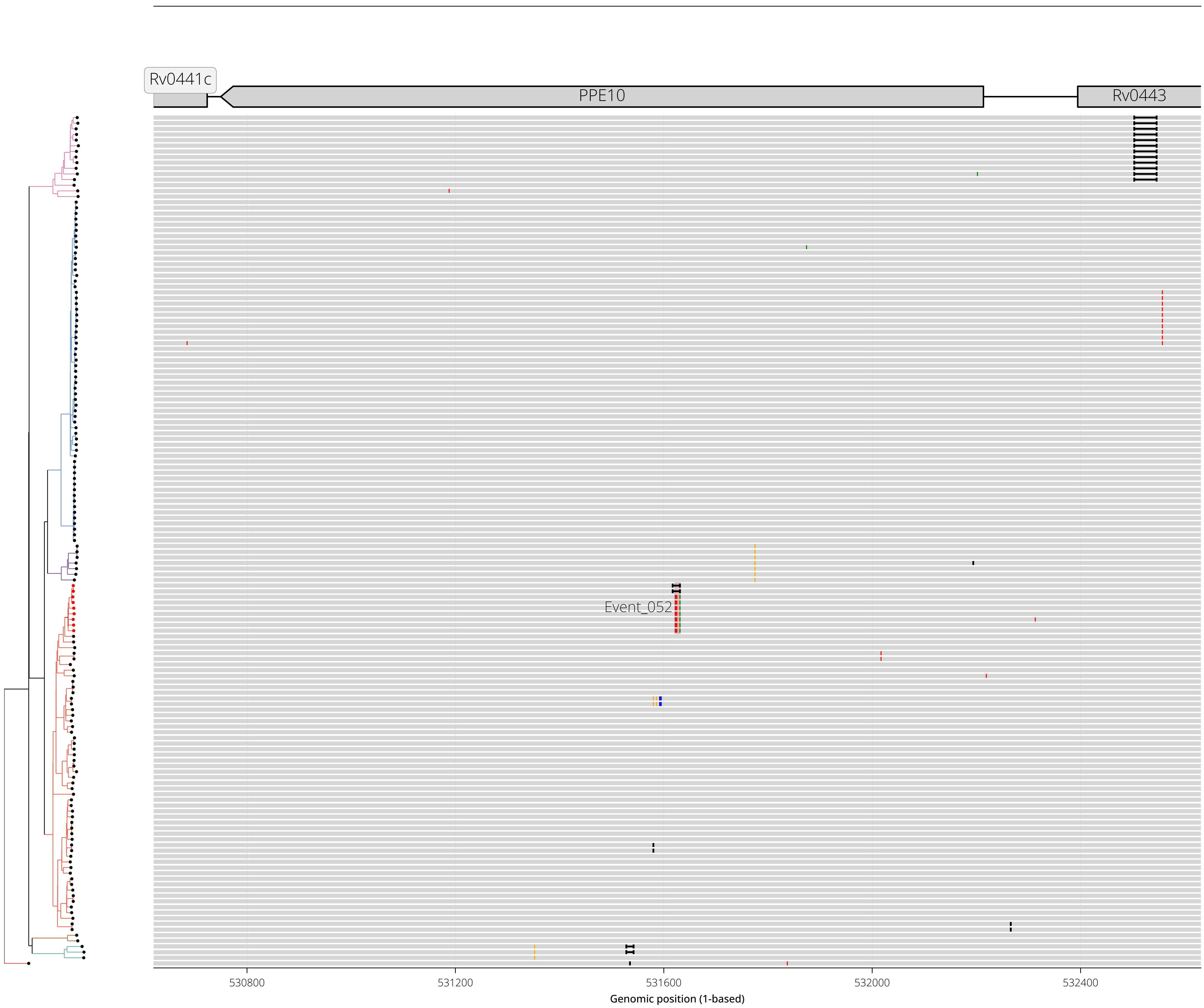

Individual GCE + Known Paralogs
GCE ID: Event_053

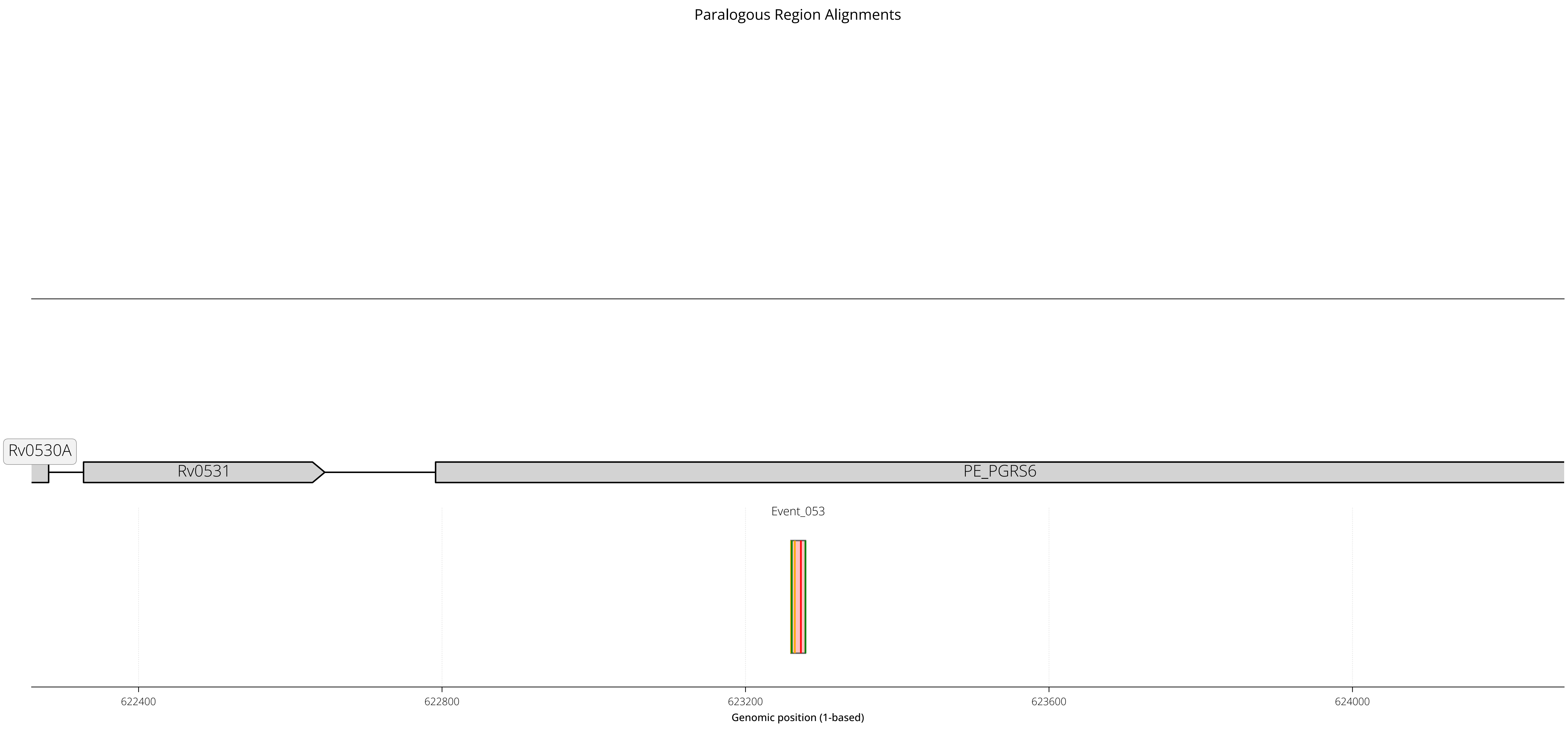

Sub-phylogeny Viz + Paralog Alignments
GCE ID: Event_053

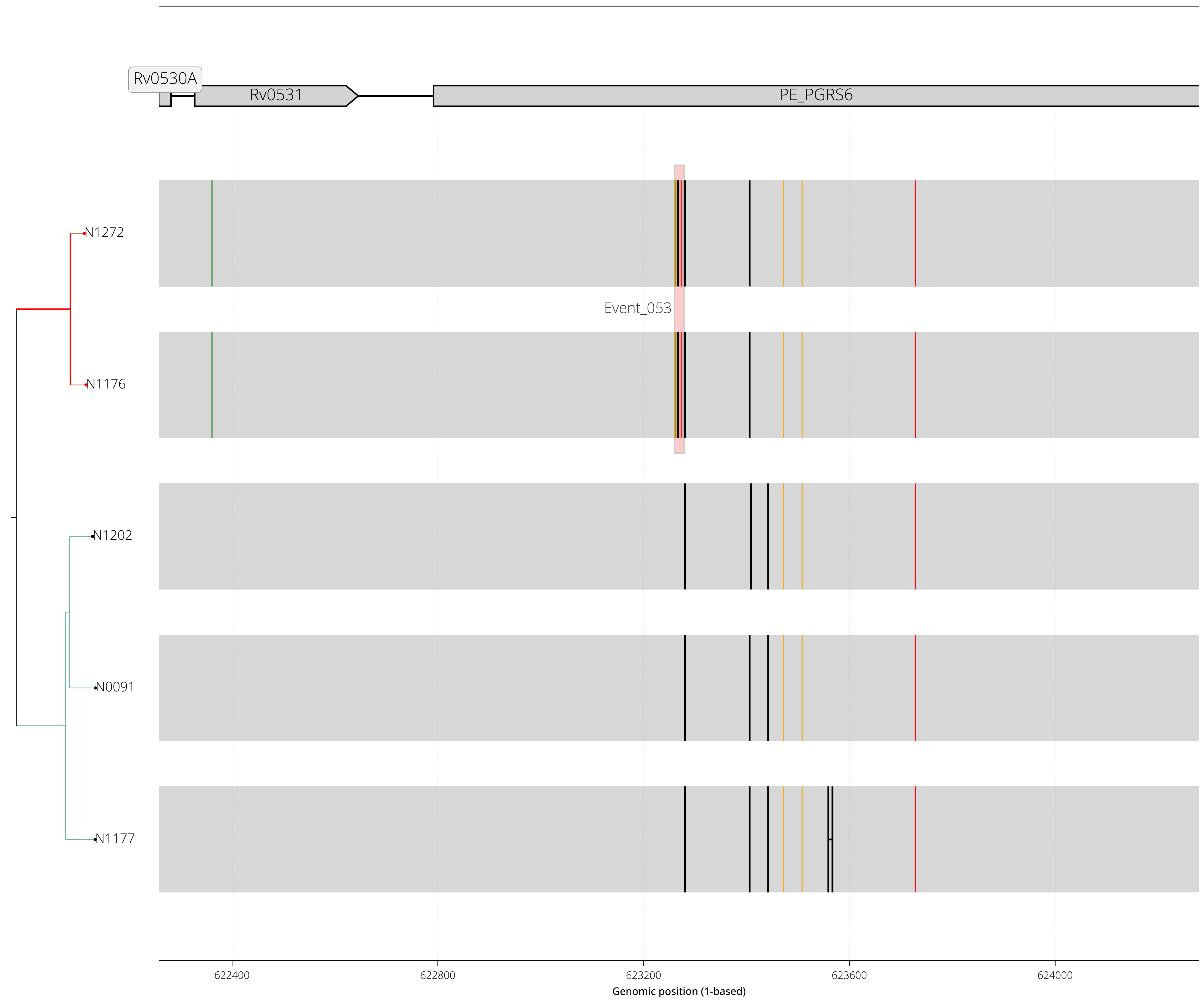

Full Phylogeny Viz + Paralog Alignments
GCE ID: Event_053

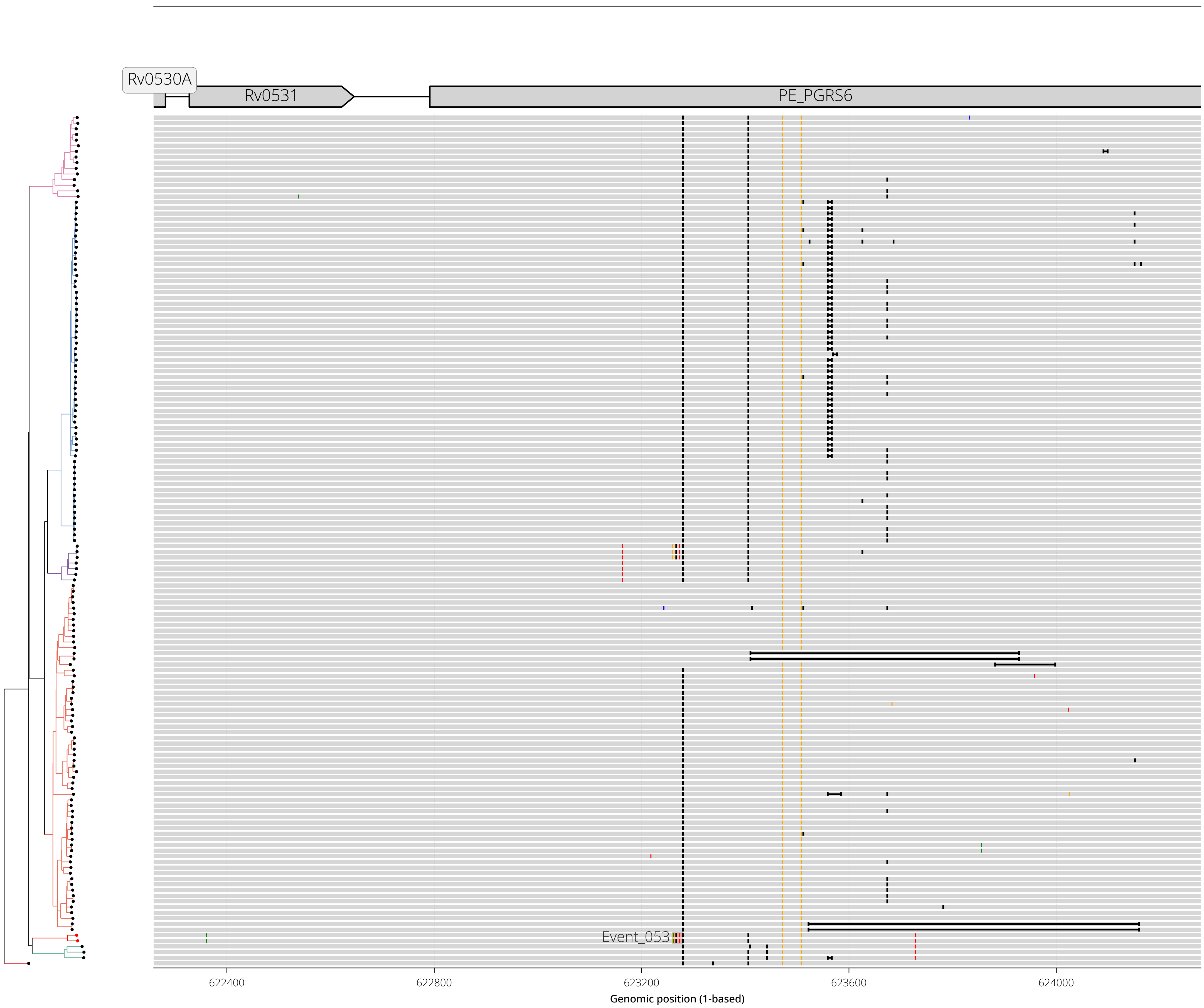

### Individual GCE + Known Paralogs
GCE ID: Event_054

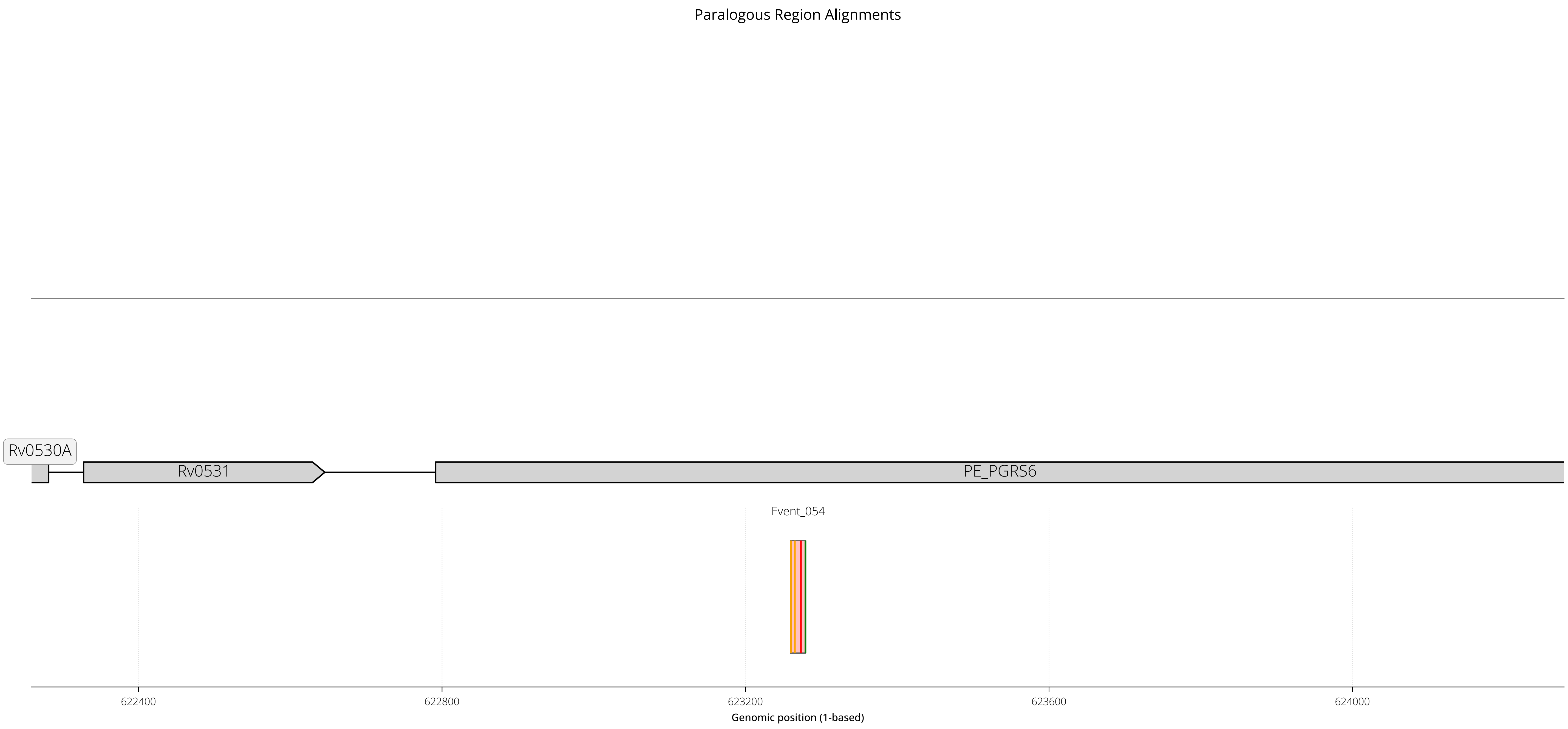

Sub-phylogeny Viz + Paralog Alignments
GCE ID: Event_054

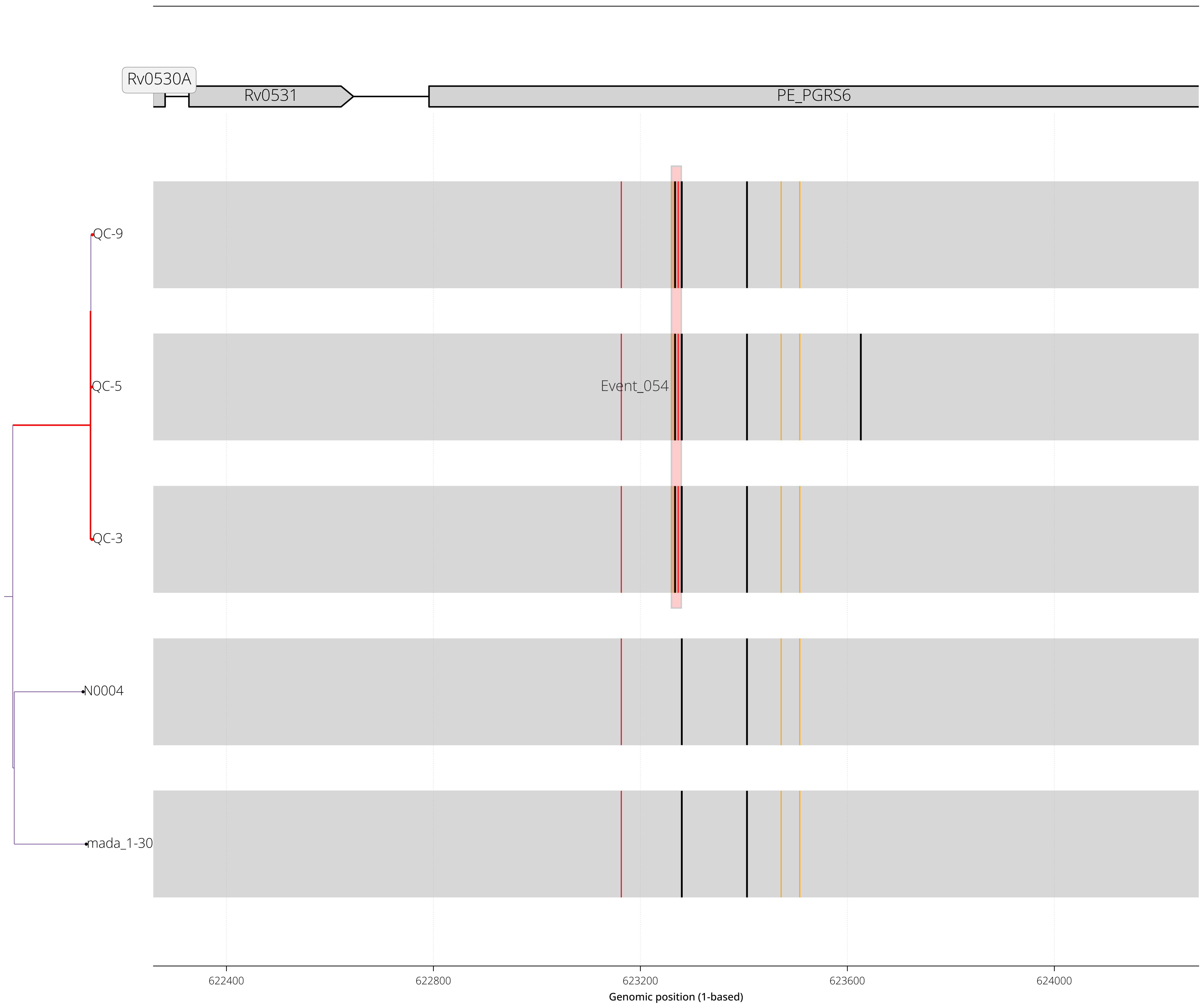

Full Phylogeny Viz + Paralog Alignments
GCE ID: Event_054

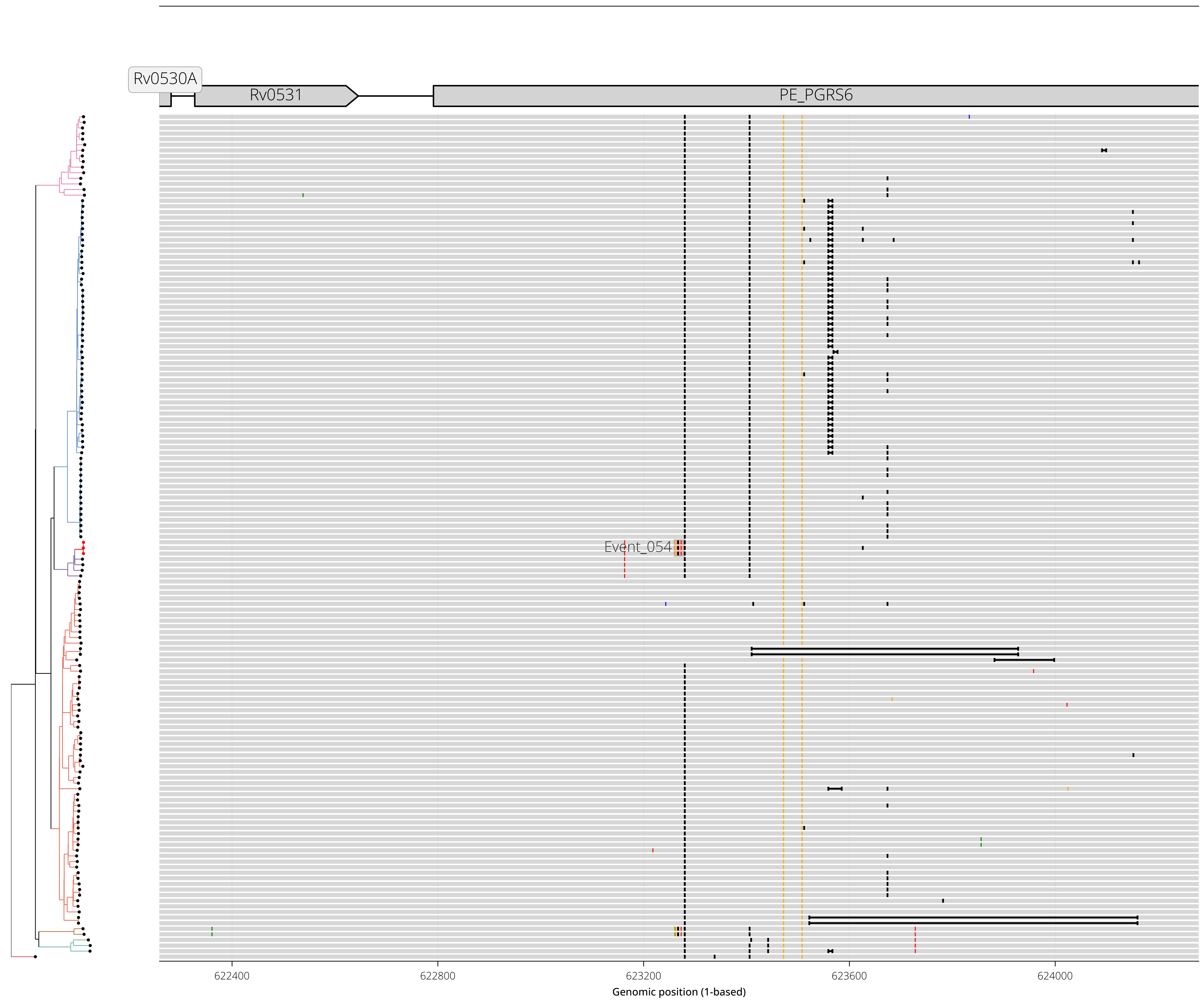

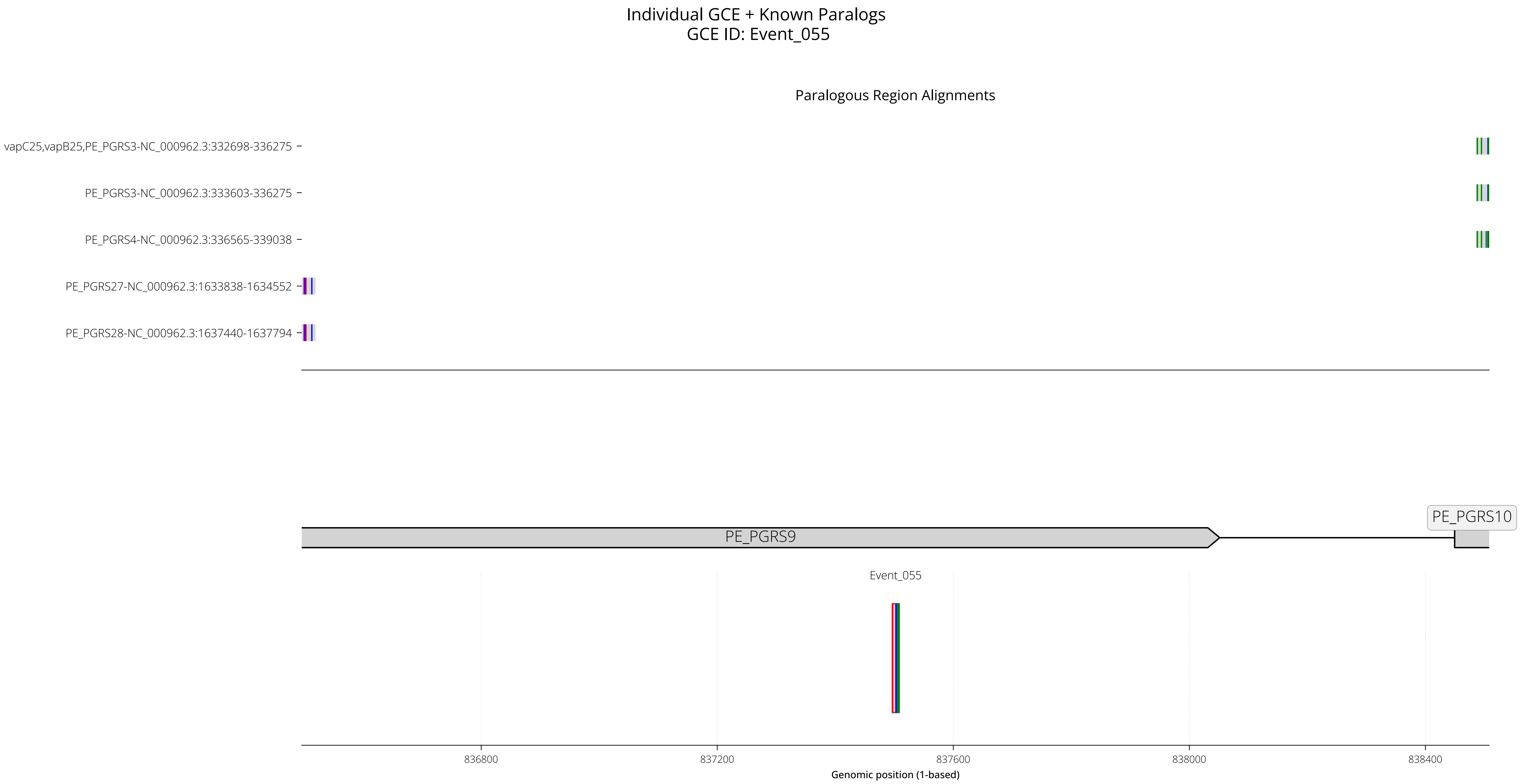
Individual GCE + Known Paralogs
GCE ID: Event_055
Paralogous Region Alignments
vapC25,vapB25,PE_PGRS3-NC_000962.3:332698-336275
PE_PGRS3-NC_000962.3:333603-336275
PE_PGRS4-NC_000962.3:336565-339038
PE_PGRS27-NC_000962.3:1633838-1634552
PE_PGRS28-NC_000962.3:1637440-1637794
PE_PGRS9
PE_PGRS10
Event_055
836800
837200
837600
838000
838400
Genomic position (1-based)

Sub-phylogeny Viz + Paralog Alignments
GCE ID: Event_055

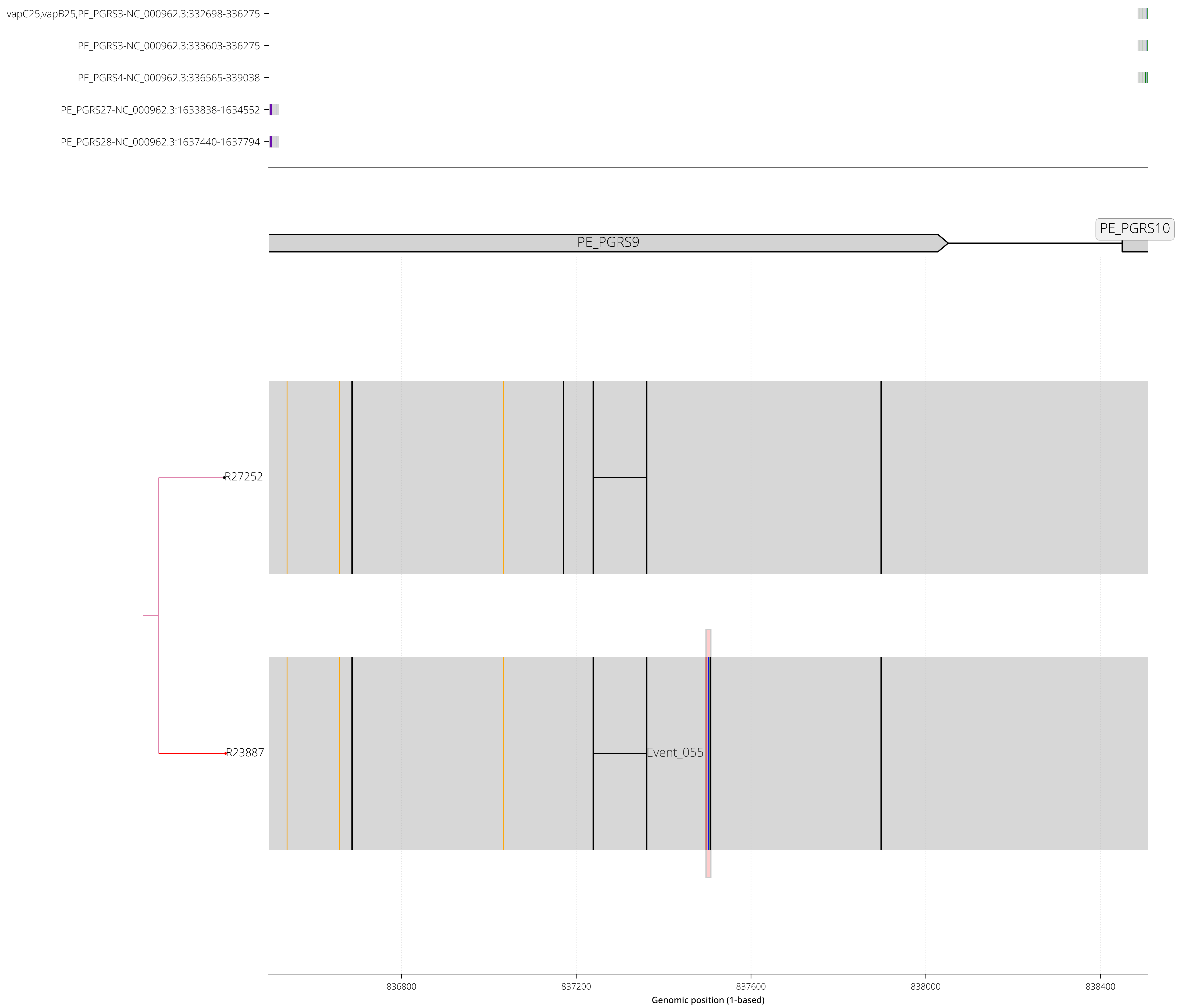

Full Phylogeny Viz + Paralog Alignments
GCE ID: Event_055

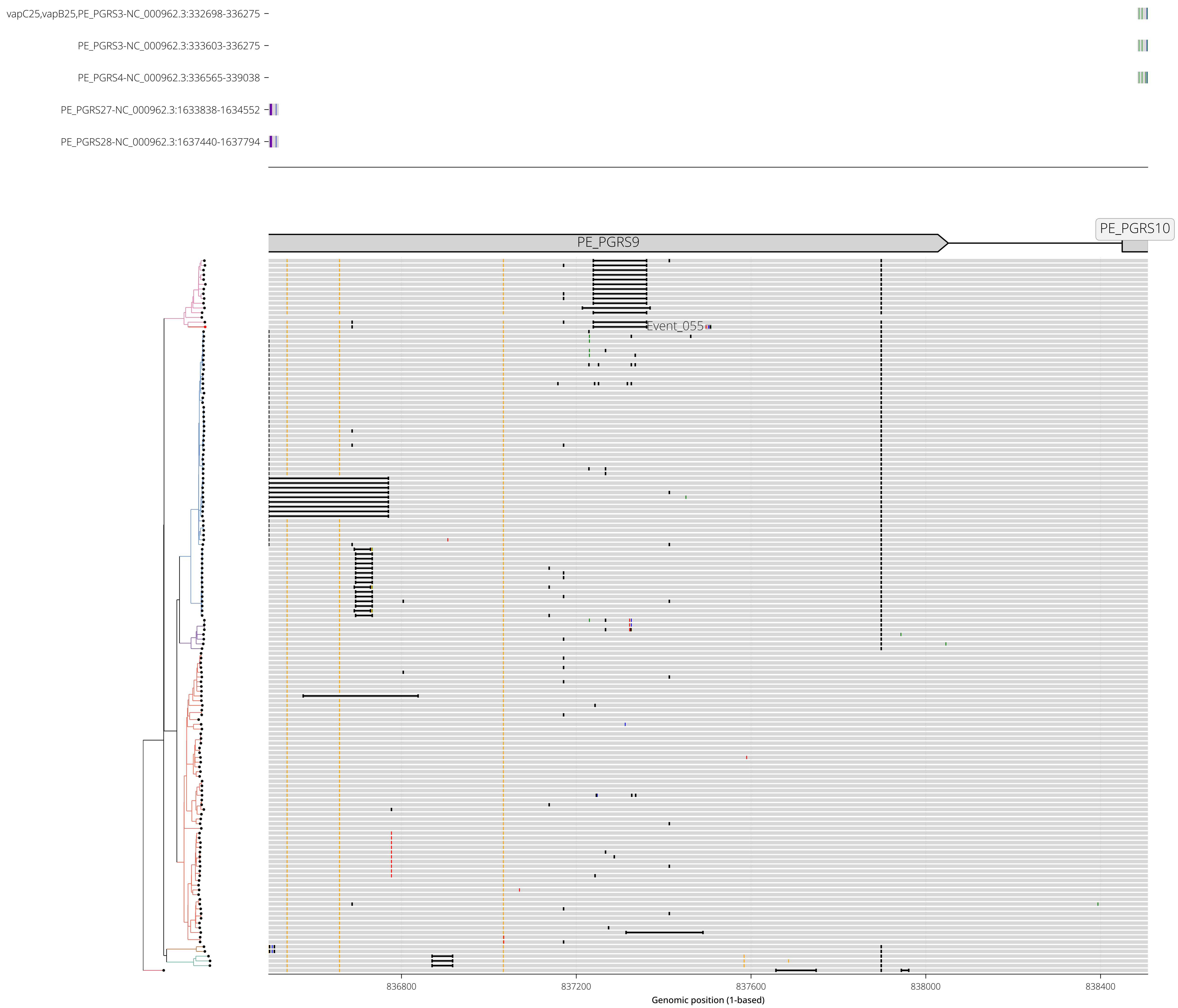

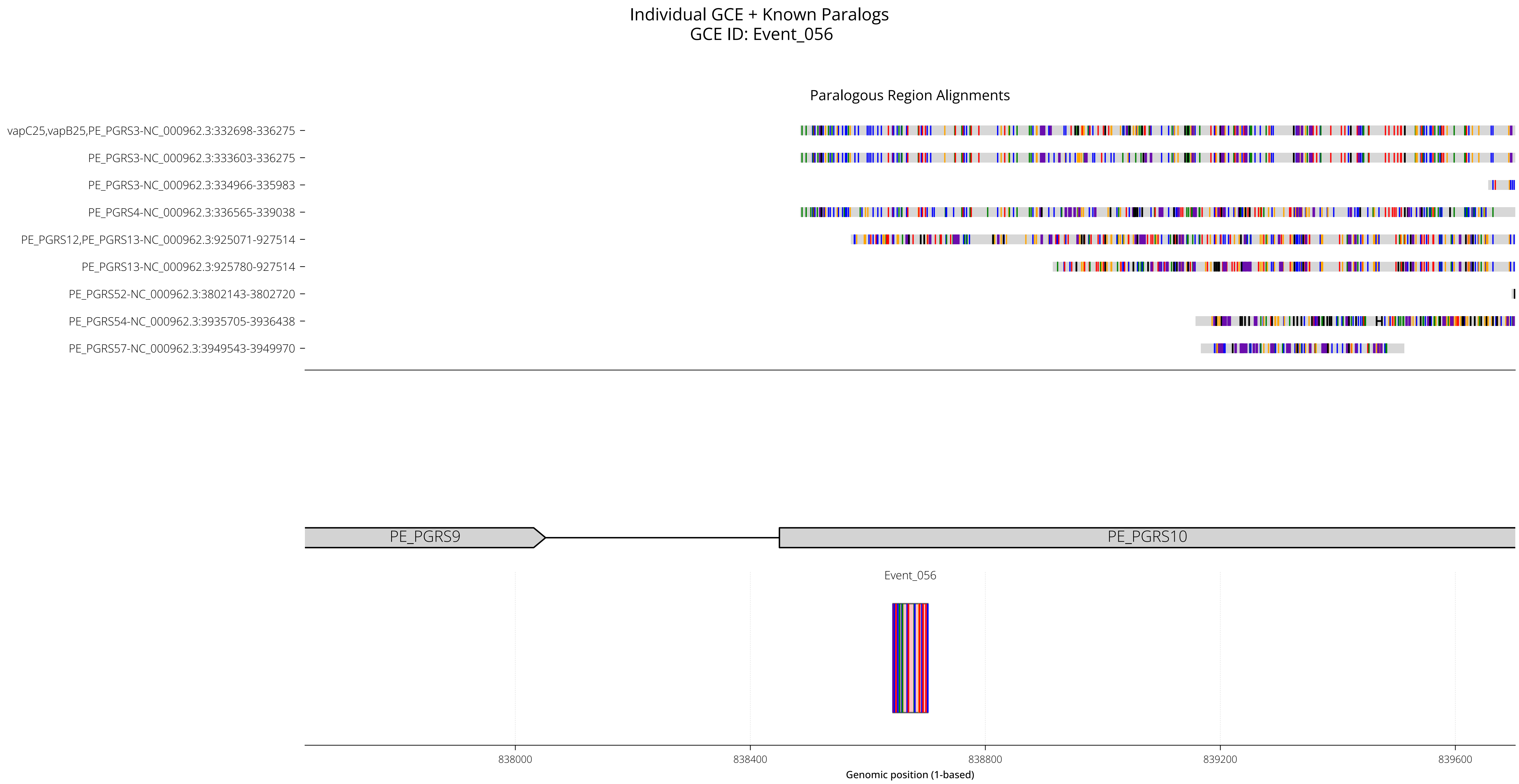
Individual GCE + Known Paralogs
GCE ID: Event_056
Paralogous Region Alignments
vapC25,vapB25,PE_PGRS3-NC_000962.3:332698-336275
PE_PGRS3-NC_000962.3:333603-336275
PE_PGRS3-NC_000962.3:334966-335983
PE_PGRS4-NC_000962.3:336565-339038
PE_PGRS12,PE_PGRS13-NC_000962.3:925071-927514
PE_PGRS13-NC_000962.3:925780-927514
PE_PGRS52-NC_000962.3:3802143-3802720
PE_PGRS54-NC_000962.3:3935705-3936438
PE_PGRS57-NC_000962.3:3949543-3949970
PE_PGRS9
PE_PGRS10
Event_056
838000
838400
838800
839200
839600
Genomic position (1-based)

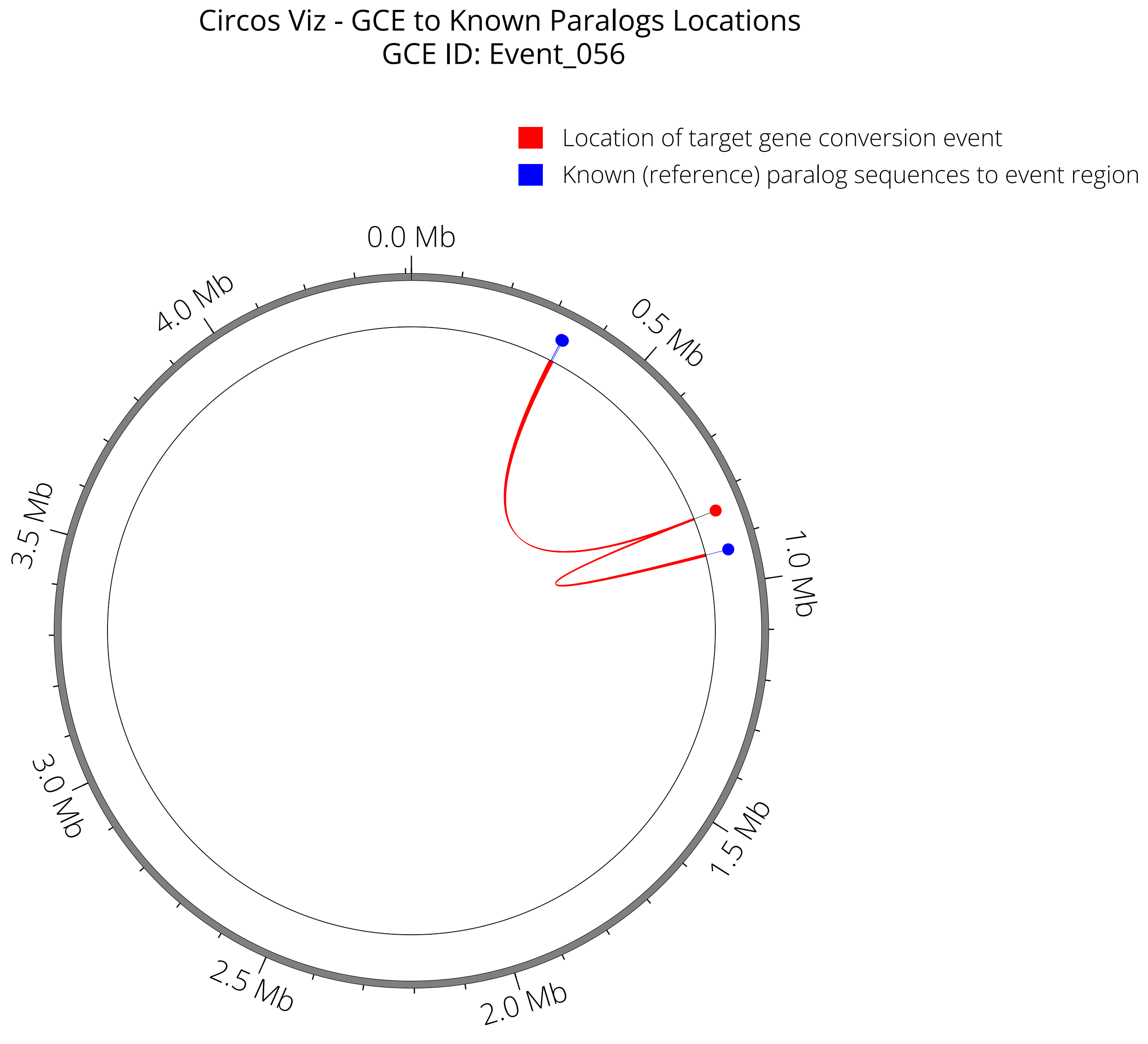

Circos Viz - GCE to Known Paralogs Locations
GCE ID: Event_056
Location of target gene conversion event
Known (reference) paralog sequences to event region
0.0 Mb
0.5 Mb
1.0 Mb
1.5 Mb
2.0 Mb
2.5 Mb
3.0 Mb
3.5 Mb
4.0 Mb

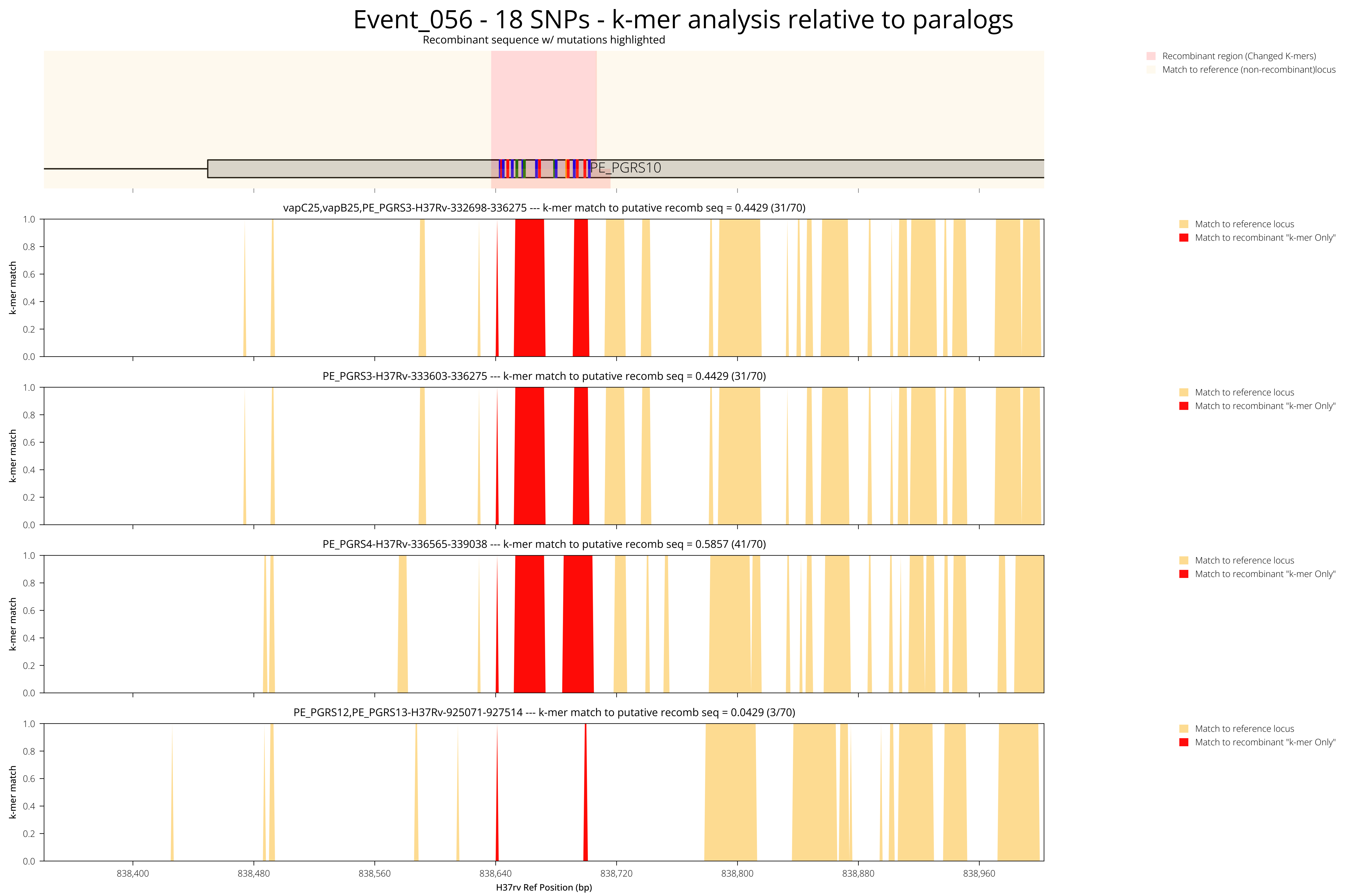

Event_056 - 18 SNPs - k-mer analysis relative to paralogs
Recombinant sequence w/ mutations highlighted
Recombinant region (Changed K-mers)
Match to reference (non-recombinant)locus
PE_PGRS10
vapC25,vapB25,PE_PGRS3-H37Rv-332698-336275 --- k-mer match to putative recomb seq = 0.4429 (31/70)
PE_PGRS3-H37Rv-333603-336275 --- k-mer match to putative recomb seq = 0.4429 (31/70)
PE_PGRS4-H37Rv-336565-339038 --- k-mer match to putative recomb seq = 0.5857 (41/70)
PE_PGRS12,PE_PGRS13-H37Rv-925071-927514 --- k-mer match to putative recomb seq = 0.0429 (3/70)
Match to reference locus
Match to recombinant "k-mer Only"
k-mer match
0.0
0.2
0.4
0.6
0.8
1.0
838,400
838,480
838,560
838,640
838,720
838,800
838,880
838,960
H37rv Ref Position (bp)

Sub-phylogeny Viz + Paralog Alignments
GCE ID: Event_056

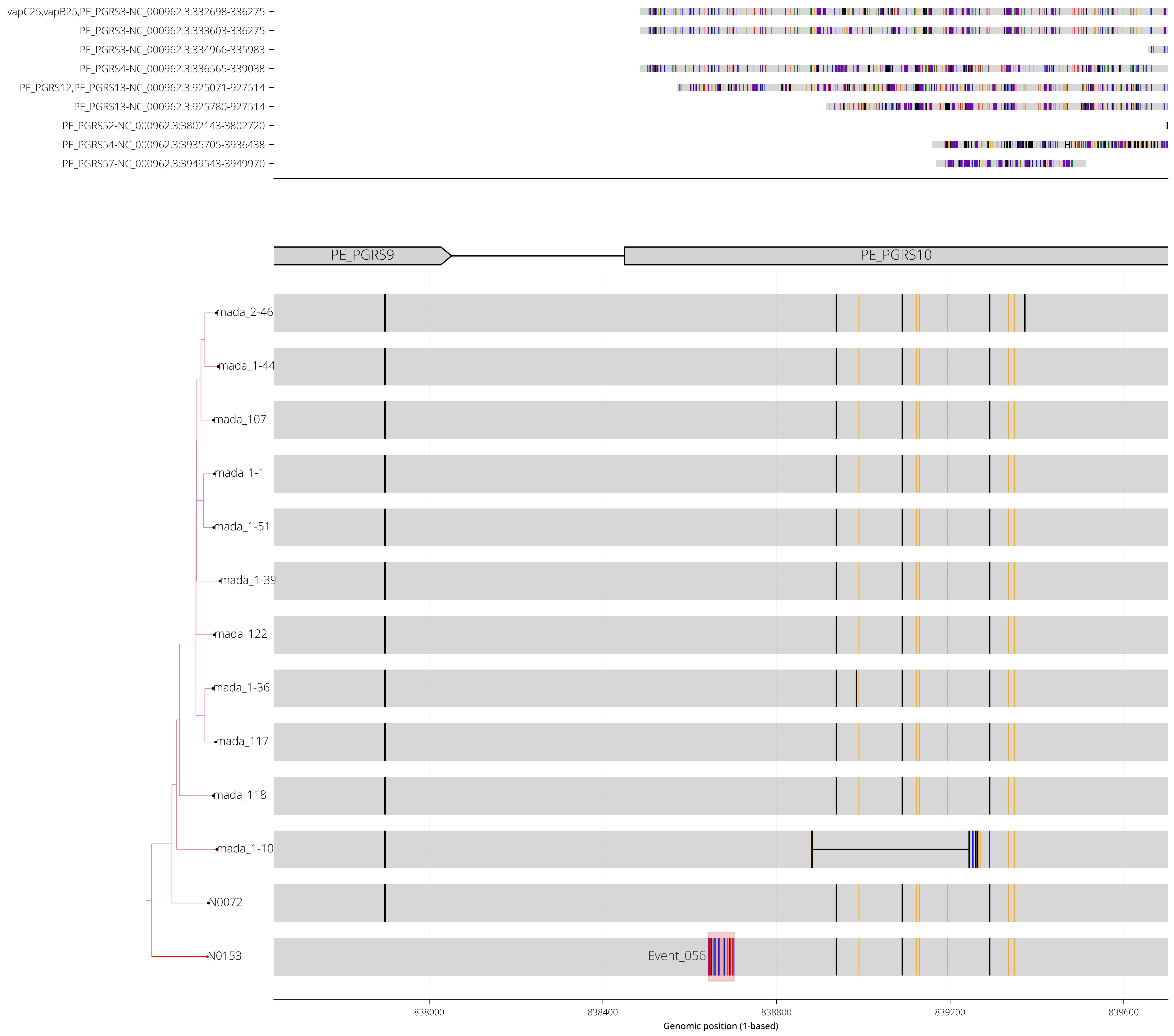

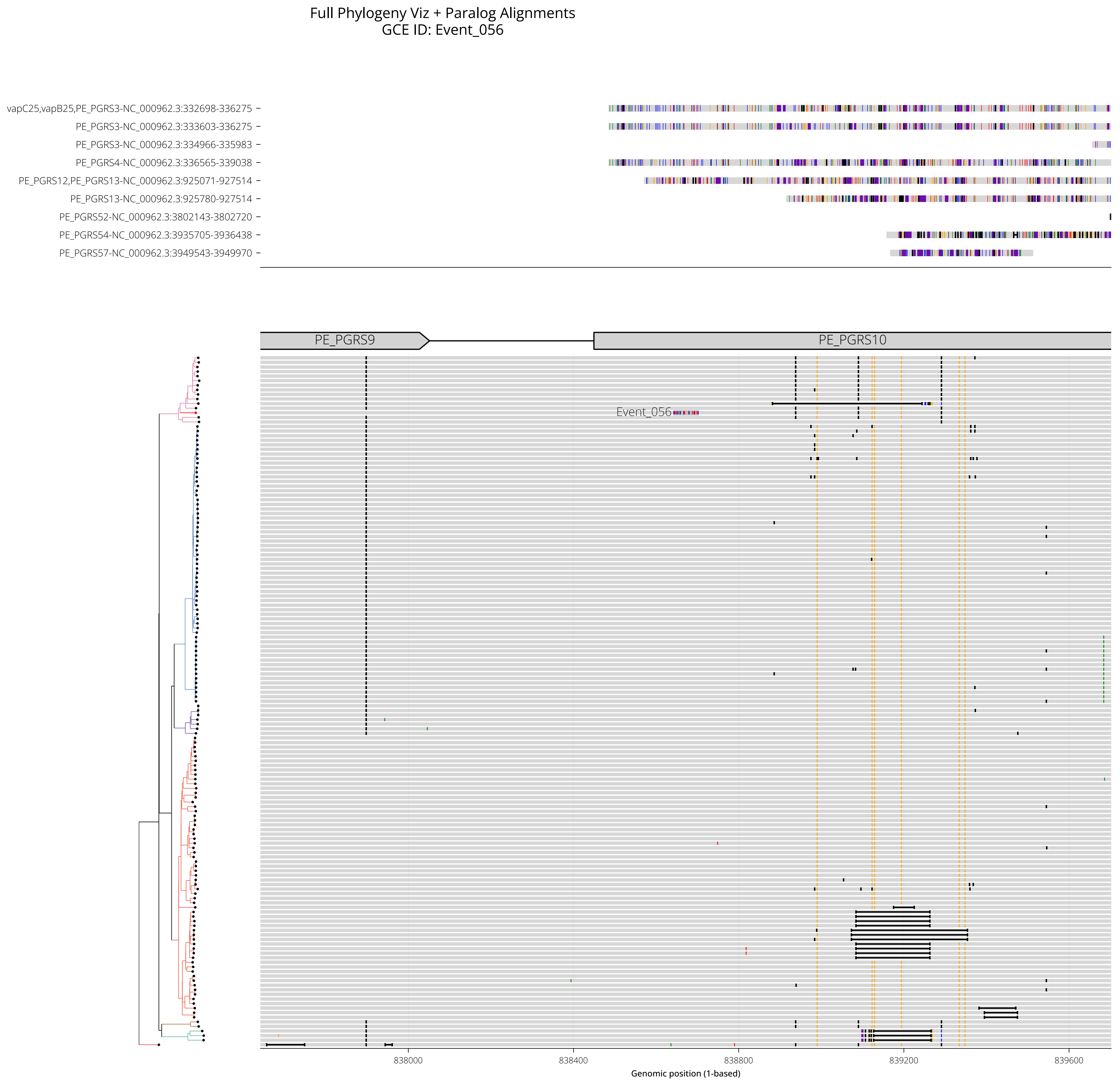
Full Phylogeny Viz + Paralog Alignments
GCE ID: Event_056
vapC25,vapB25,PE_PGRS3-NC_000962.3:332698-336275
PE_PGRS3-NC_000962.3:333603-336275
PE_PGRS3-NC_000962.3:334966-335983
PE_PGRS4-NC_000962.3:336565-339038
PE_PGRS12,PE_PGRS13-NC_000962.3:925071-927514
PE_PGRS13-NC_000962.3:925780-927514
PE_PGRS52-NC_000962.3:3802143-3802720
PE_PGRS54-NC_000962.3:3935705-3936438
PE_PGRS57-NC_000962.3:3949543-3949970
PE_PGRS9
PE_PGRS10
Event_056
838000
838400
838800
839200
839600
Genomic position (1-based)

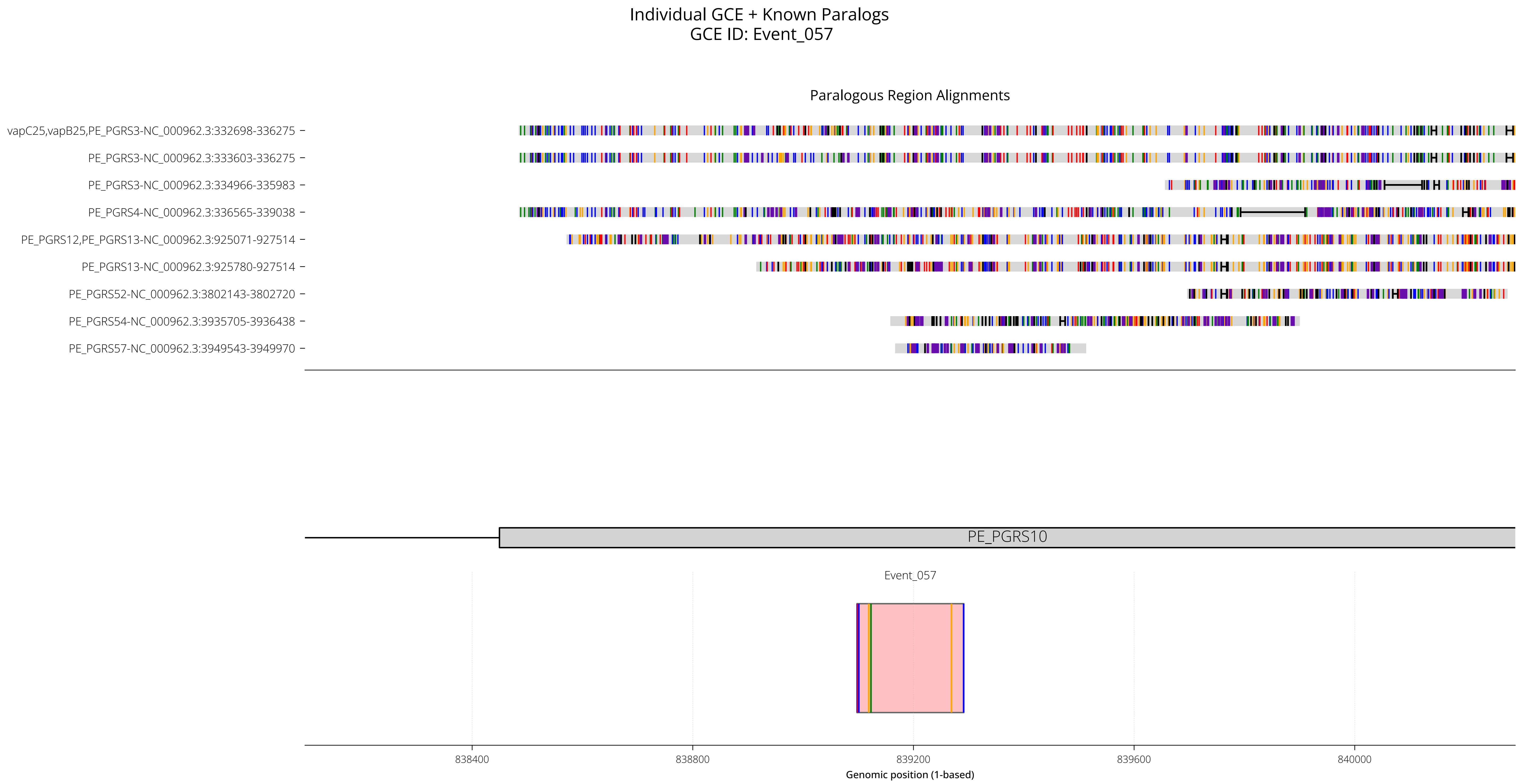

Individual GCE + Known Paralogs
GCE ID: Event_057
Paralogous Region Alignments
vapC25,vapB25,PE_PGRS3-NC_000962.3:332698-336275
PE_PGRS3-NC_000962.3:333603-336275
PE_PGRS3-NC_000962.3:334966-335983
PE_PGRS4-NC_000962.3:336565-339038
PE_PGRS12,PE_PGRS13-NC_000962.3:925071-927514
PE_PGRS13-NC_000962.3:925780-927514
PE_PGRS52-NC_000962.3:3802143-3802720
PE_PGRS54-NC_000962.3:3935705-3936438
PE_PGRS57-NC_000962.3:3949543-3949970
PE_PGRS10
Event_057
838400
838800
839200
839600
840000
Genomic position (1-based)

### Circos Viz - GCE to Known Paralogs Locations
GCE ID: Event_057

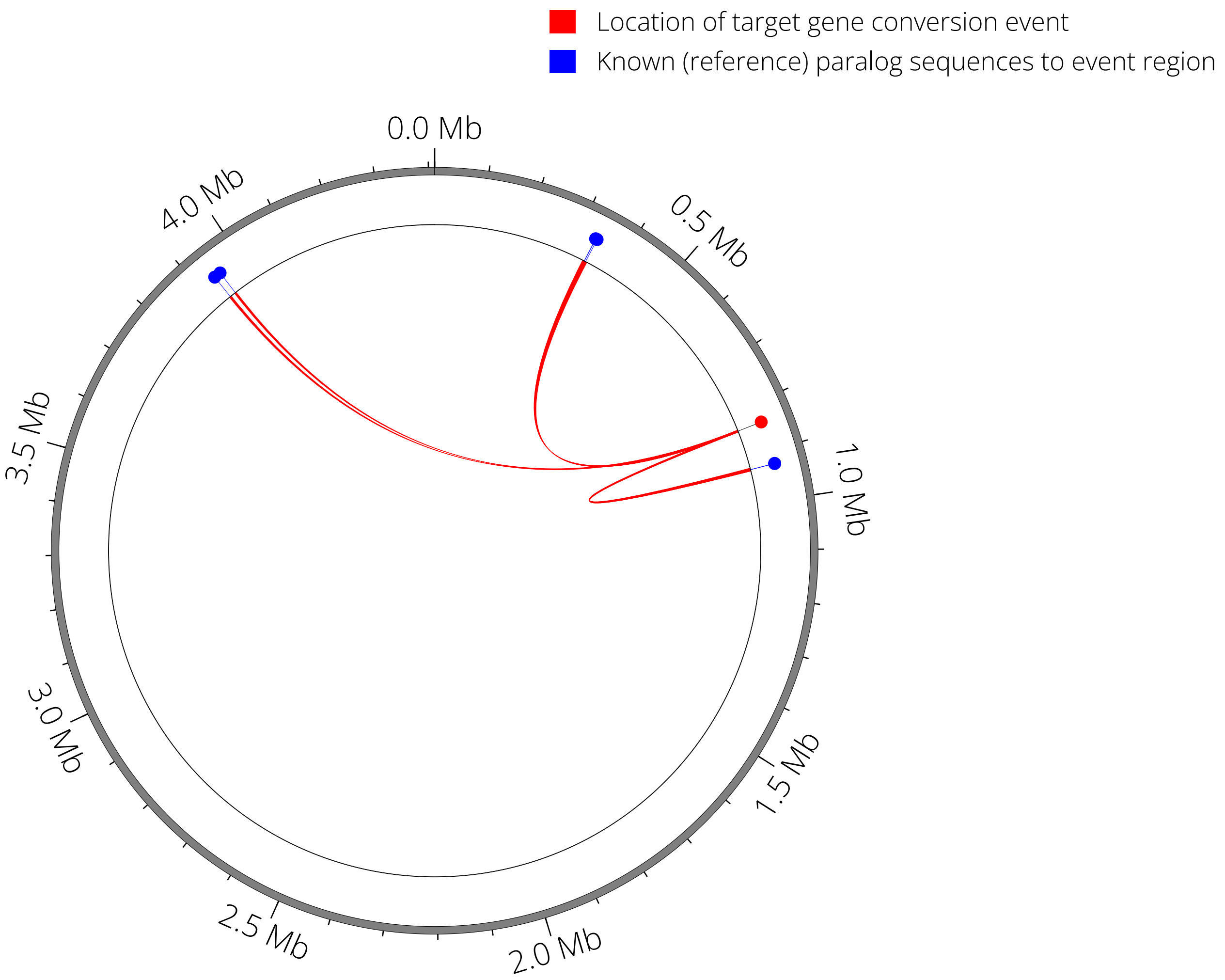

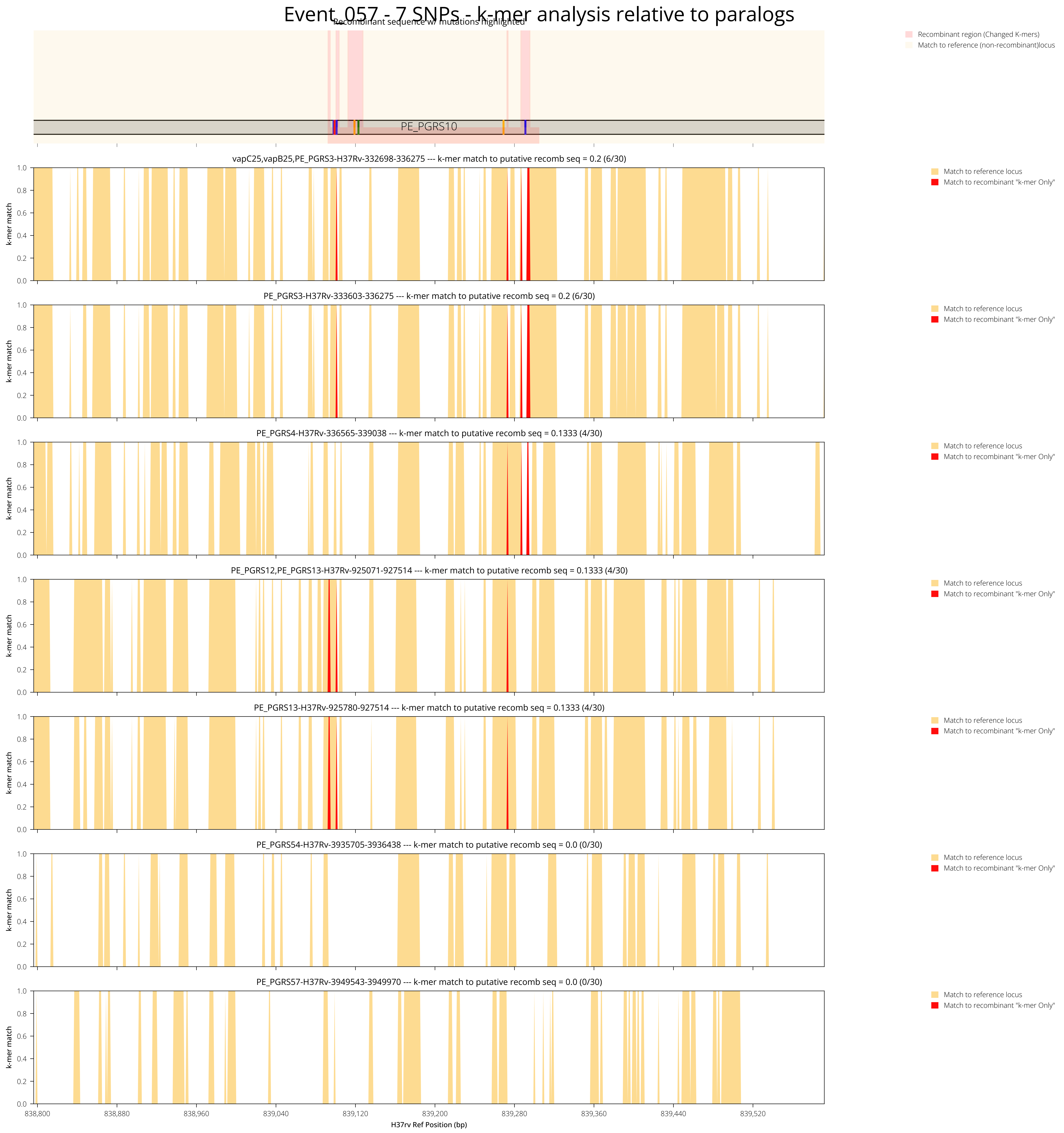

Event_057 - 7 SNPs - k-mer analysis relative to paralogs
Recombinant sequence w/ mutations highlighted
Recombinant region (Changed K-mers)
Match to reference (non-recombinant)locus
PE_PGRS10
vapC25,vapB25,PE_PGRS3-H37Rv-332698-336275 --- k-mer match to putative recomb seq = 0.2 (6/30)
PE_PGRS3-H37Rv-333603-336275 --- k-mer match to putative recomb seq = 0.2 (6/30)
PE_PGRS4-H37Rv-336565-339038 --- k-mer match to putative recomb seq = 0.1333 (4/30)
PE_PGRS12,PE_PGRS13-H37Rv-925071-927514 --- k-mer match to putative recomb seq = 0.1333 (4/30)
PE_PGRS13-H37Rv-925780-927514 --- k-mer match to putative recomb seq = 0.1333 (4/30)
PE_PGRS54-H37Rv-3935705-3936438 --- k-mer match to putative recomb seq = 0.0 (0/30)
PE_PGRS57-H37Rv-3949543-3949970 --- k-mer match to putative recomb seq = 0.0 (0/30)
Match to reference locus
Match to recombinant "k-mer Only"
k-mer match
1.0
0.8
0.6
0.4
0.2
0.0
838,800
838,880
838,960
839,040
839,120
839,200
839,280
839,360
839,440
839,520
H37rv Ref Position (bp)

Sub-phylogeny Viz + Paralog Alignments
GCE ID: Event_057

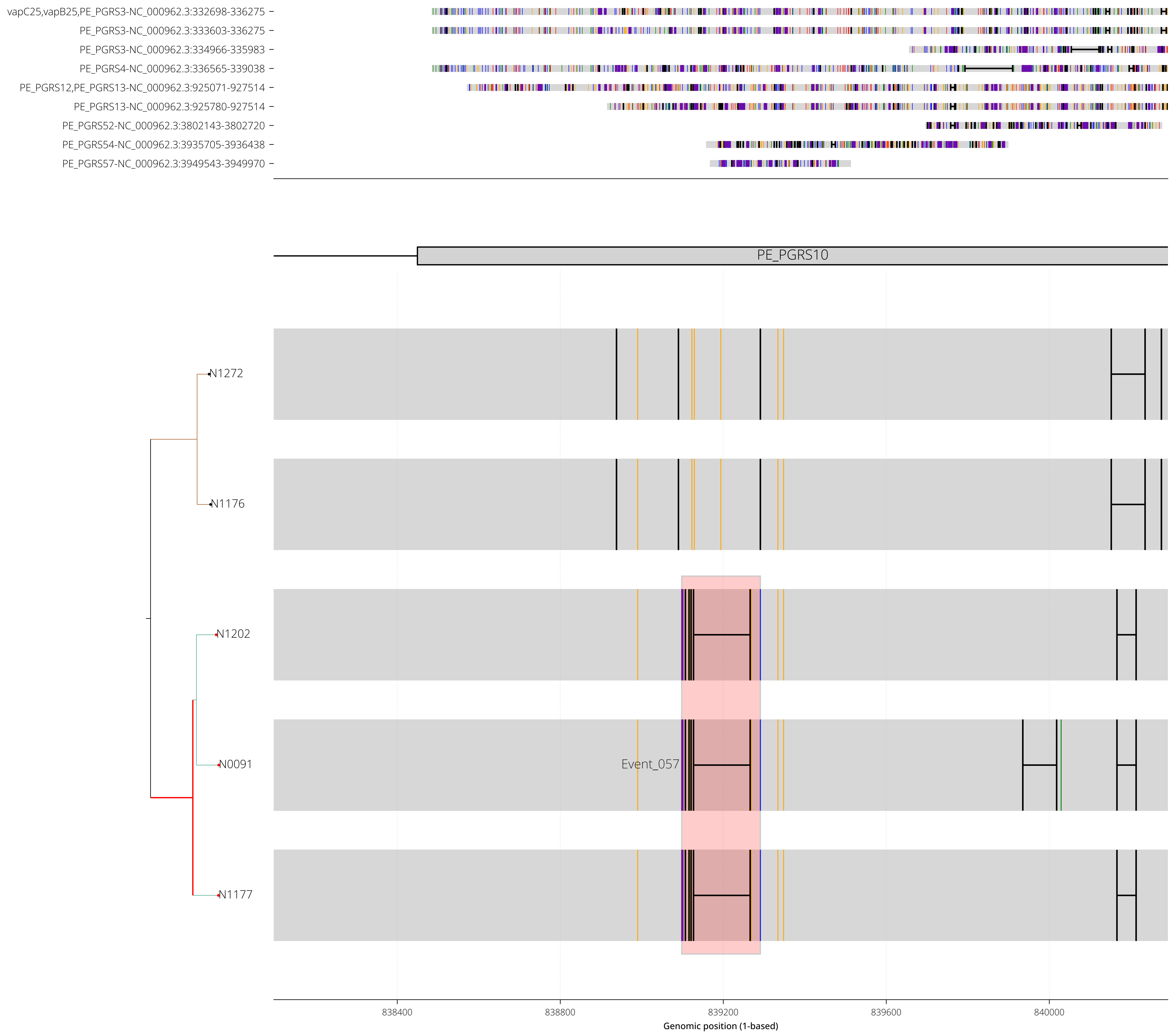

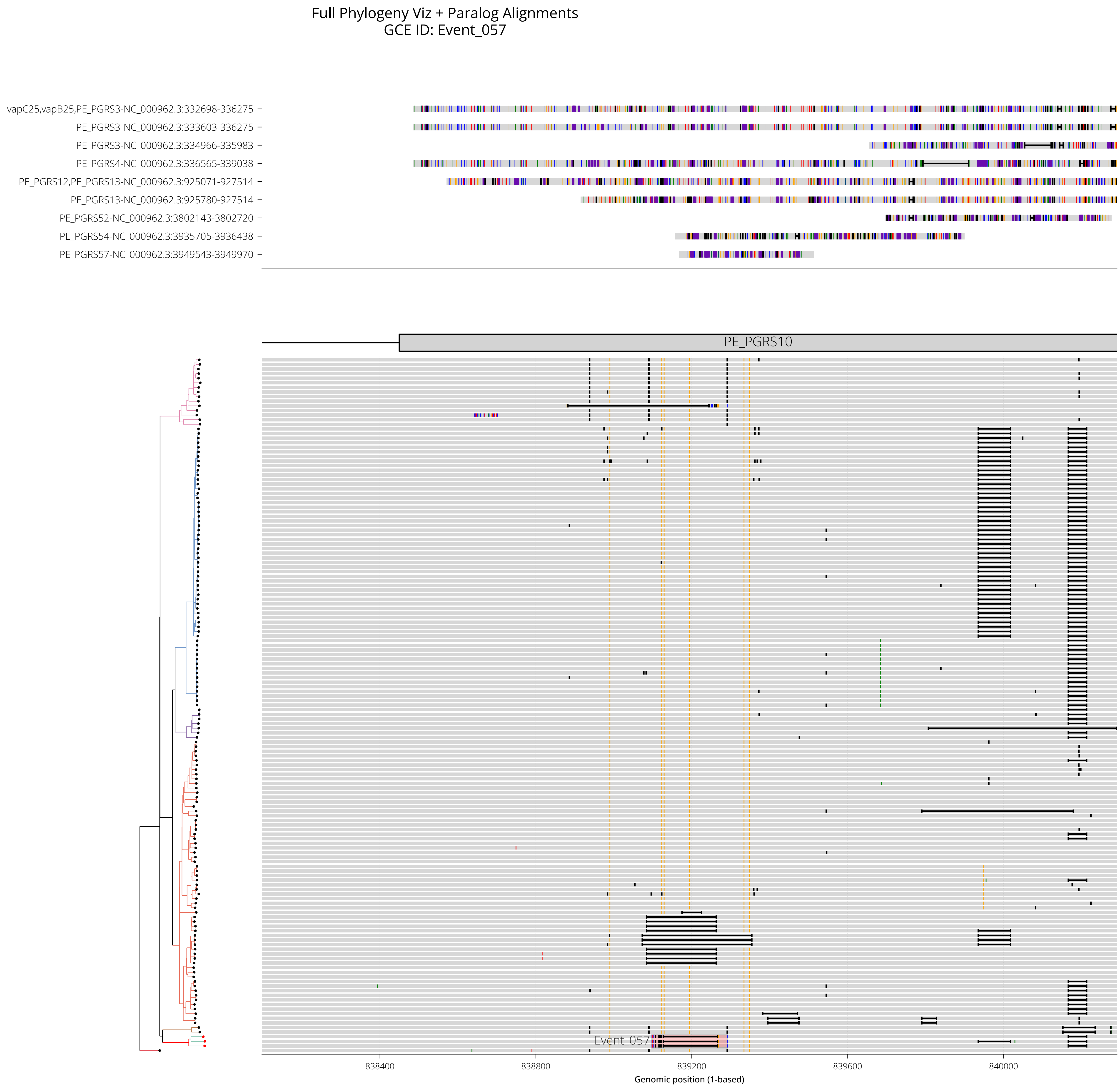

Full Phylogeny Viz + Paralog Alignments
GCE ID: Event_057
vapC25,vapB25,PE_PGRS3-NC_000962.3:332698-336275
PE_PGRS3-NC_000962.3:333603-336275
PE_PGRS3-NC_000962.3:334966-335983
PE_PGRS4-NC_000962.3:336565-339038
PE_PGRS12,PE_PGRS13-NC_000962.3:925071-927514
PE_PGRS13-NC_000962.3:925780-927514
PE_PGRS52-NC_000962.3:3802143-3802720
PE_PGRS54-NC_000962.3:3935705-3936438
PE_PGRS57-NC_000962.3:3949543-3949970
PE_PGRS10
Event_057
838400
838800
839200
839600
840000
Genomic position (1-based)

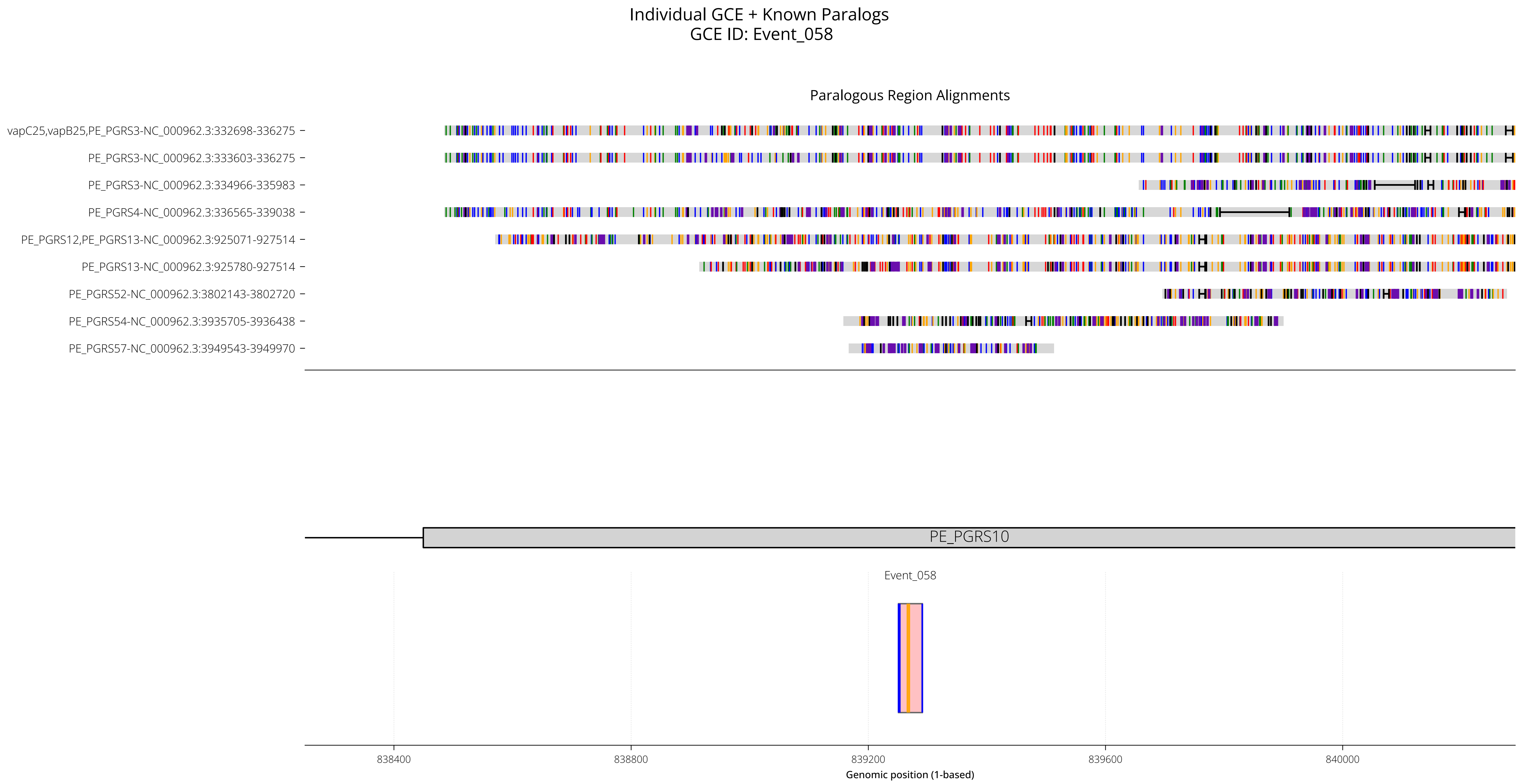

Individual GCE + Known Paralogs
GCE ID: Event_058
Paralogous Region Alignments
vapC25,vapB25,PE_PGRS3-NC_000962.3:332698-336275
PE_PGRS3-NC_000962.3:333603-336275
PE_PGRS3-NC_000962.3:334966-335983
PE_PGRS4-NC_000962.3:336565-339038
PE_PGRS12,PE_PGRS13-NC_000962.3:925071-927514
PE_PGRS13-NC_000962.3:925780-927514
PE_PGRS52-NC_000962.3:3802143-3802720
PE_PGRS54-NC_000962.3:3935705-3936438
PE_PGRS57-NC_000962.3:3949543-3949970
PE_PGRS10
Event_058
838400
838800
839200
839600
840000
Genomic position (1-based)

Circos Viz - GCE to Known Paralogs Locations
GCE ID: Event_058

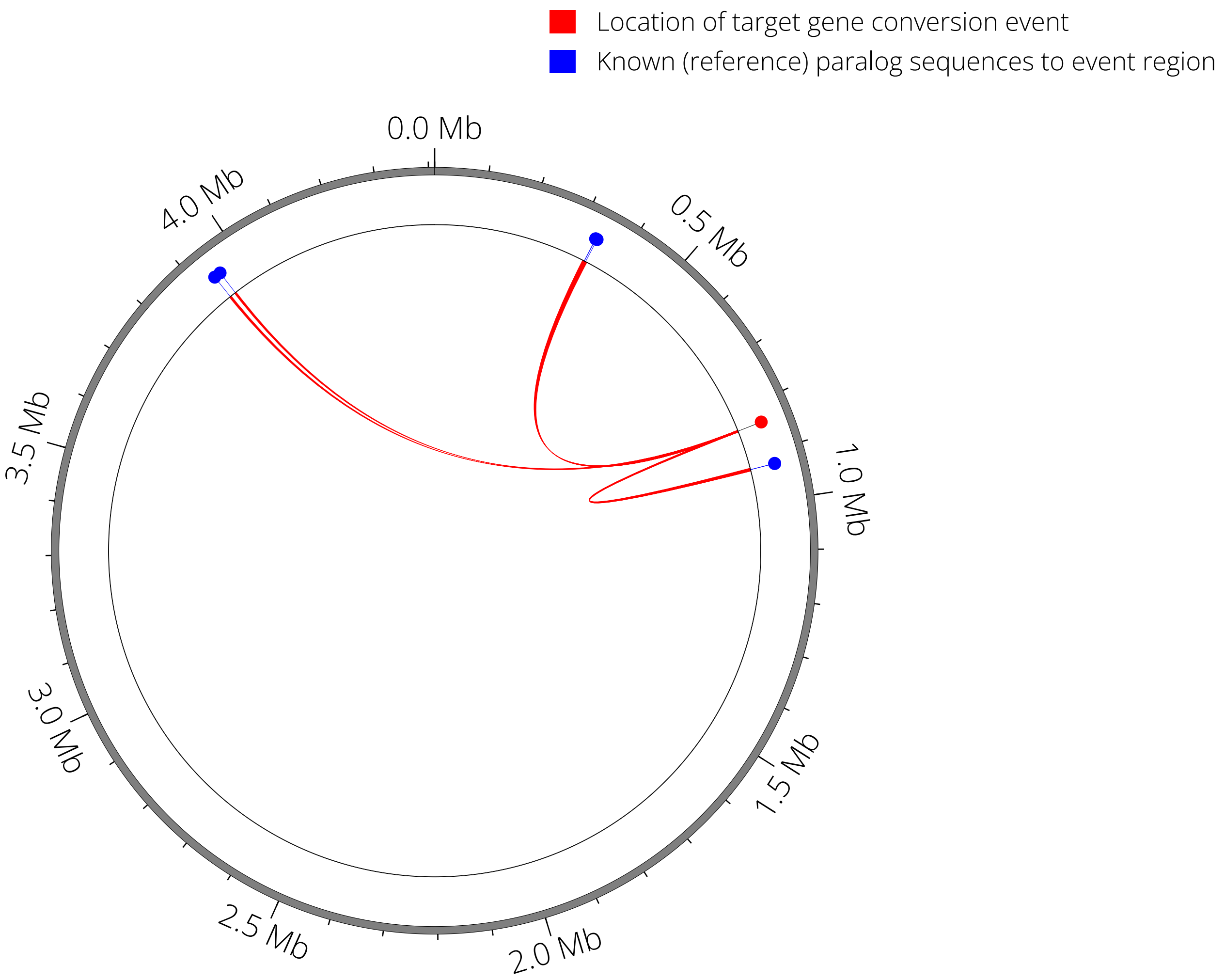

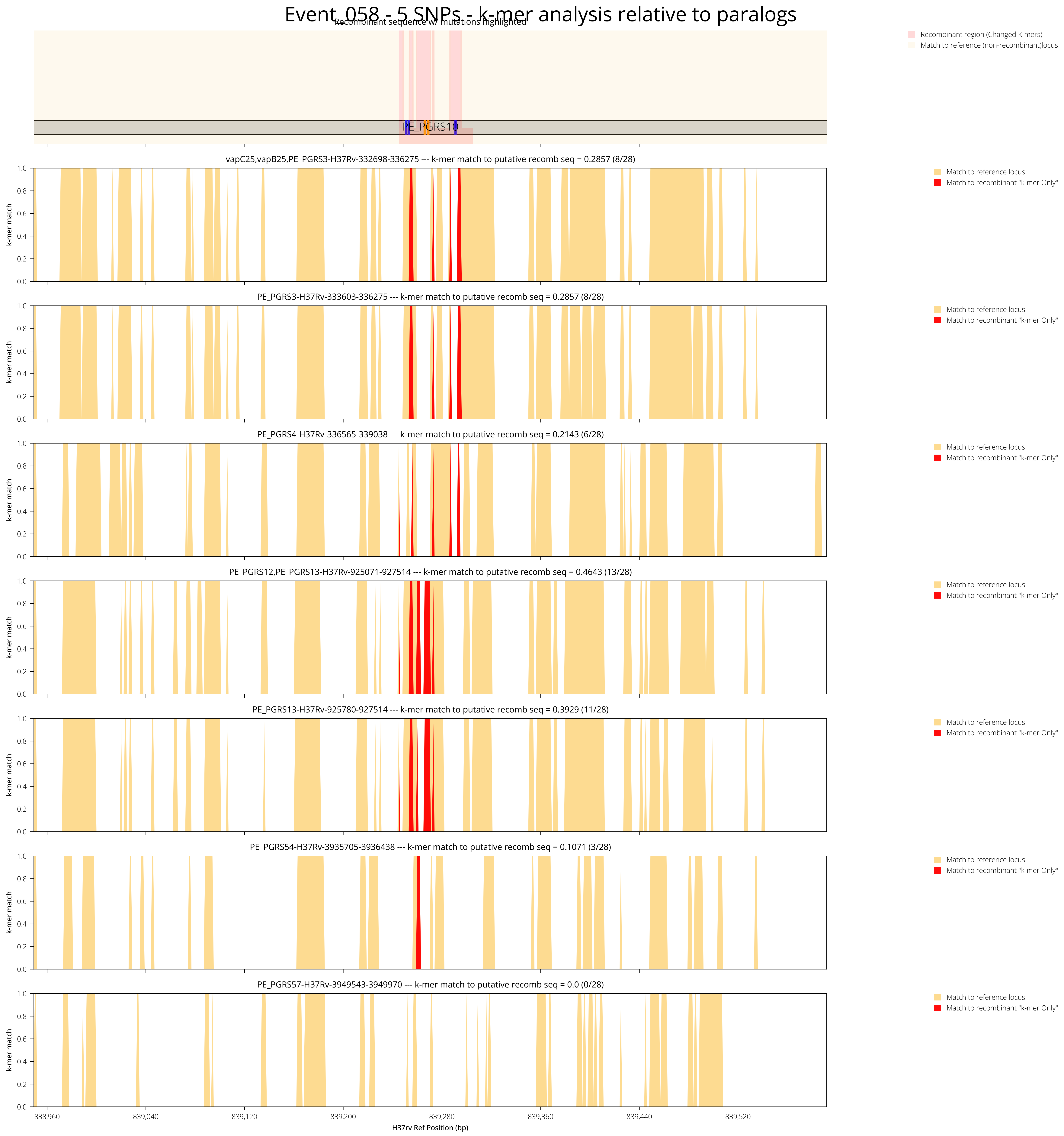
Event_058 - 5 SNPs - k-mer analysis relative to paralogs
Recombinant sequence w/ mutations highlighted
Recombinant region (Changed K-mers)
Match to reference (non-recombinant)locus
PE_PGRS10
vapC25,vapB25,PE_PGRS3-H37Rv-332698-336275 --- k-mer match to putative recomb seq = 0.2857 (8/28)
PE_PGRS3-H37Rv-333603-336275 --- k-mer match to putative recomb seq = 0.2857 (8/28)
PE_PGRS4-H37Rv-336565-339038 --- k-mer match to putative recomb seq = 0.2143 (6/28)
PE_PGRS12,PE_PGRS13-H37Rv-925071-927514 --- k-mer match to putative recomb seq = 0.4643 (13/28)
PE_PGRS13-H37Rv-925780-927514 --- k-mer match to putative recomb seq = 0.3929 (11/28)
PE_PGRS54-H37Rv-3935705-3936438 --- k-mer match to putative recomb seq = 0.1071 (3/28)
PE_PGRS57-H37Rv-3949543-3949970 --- k-mer match to putative recomb seq = 0.0 (0/28)
Match to reference locus
Match to recombinant "k-mer Only"
k-mer match
0.0
0.2
0.4
0.6
0.8
1.0
838,960
839,040
839,120
839,200
839,280
839,360
839,440
839,520
H37rv Ref Position (bp)

Sub-phylogeny Viz + Paralog Alignments
GCE ID: Event_058

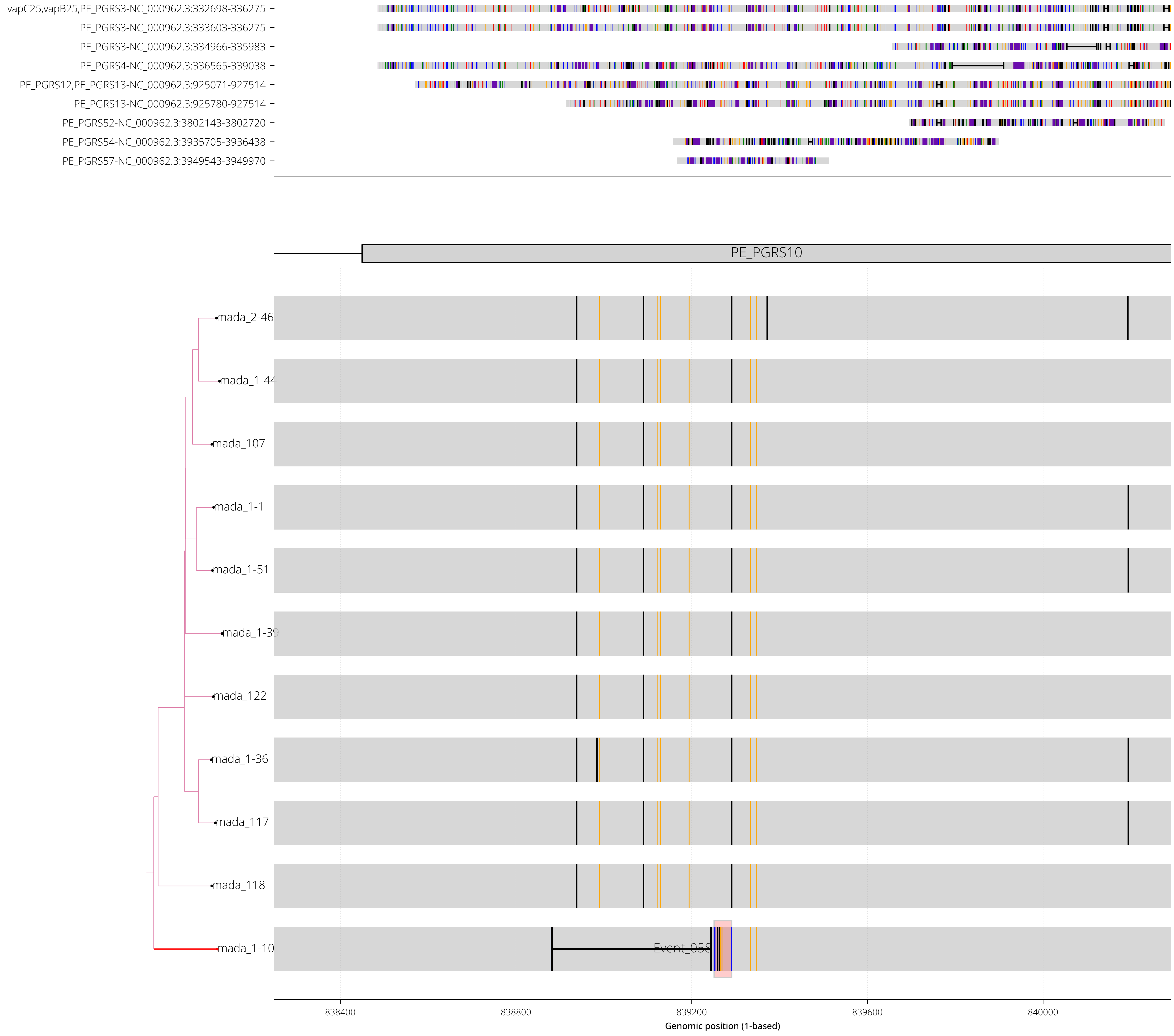

Full Phylogeny Viz + Paralog Alignments
GCE ID: Event_058

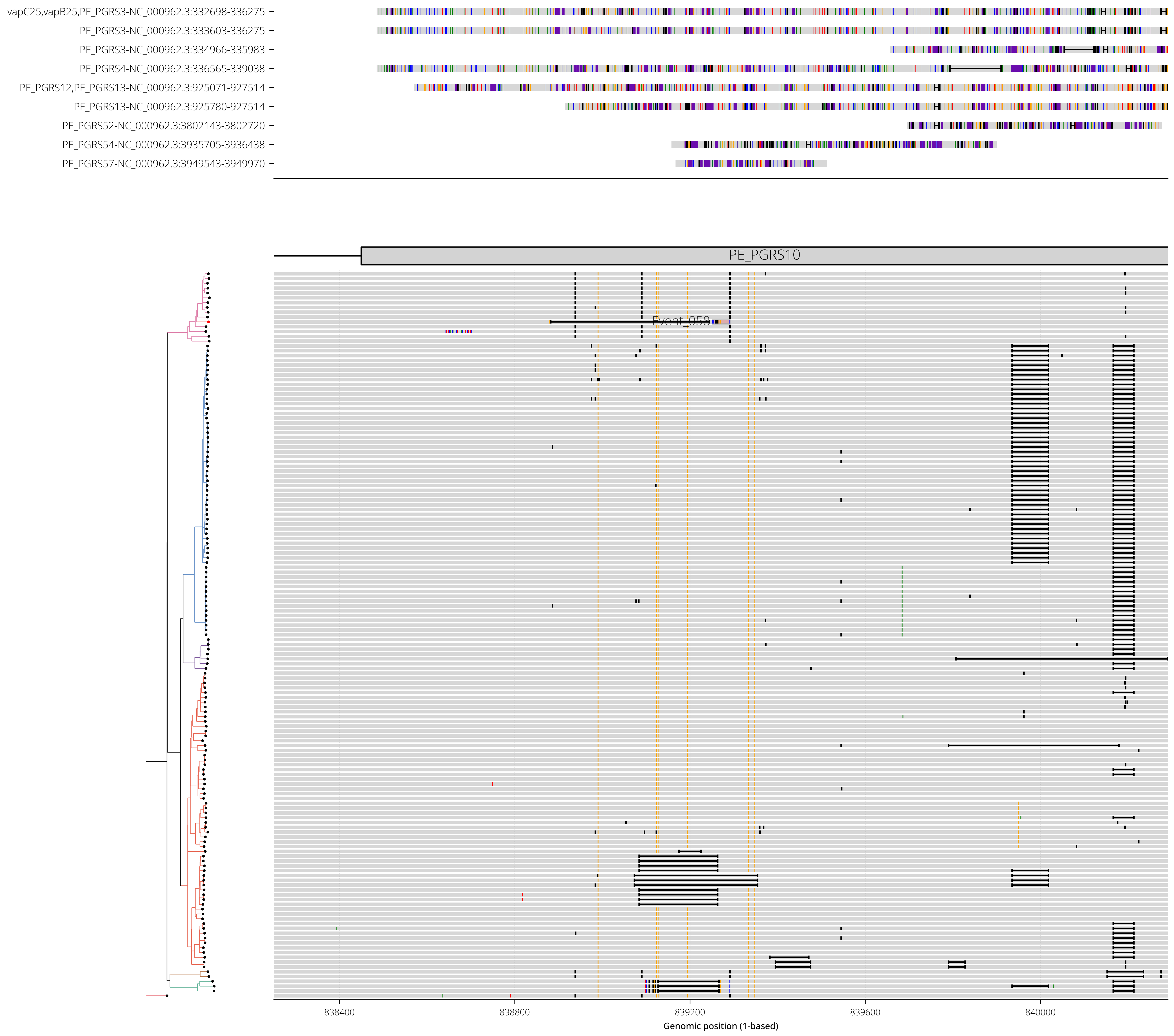

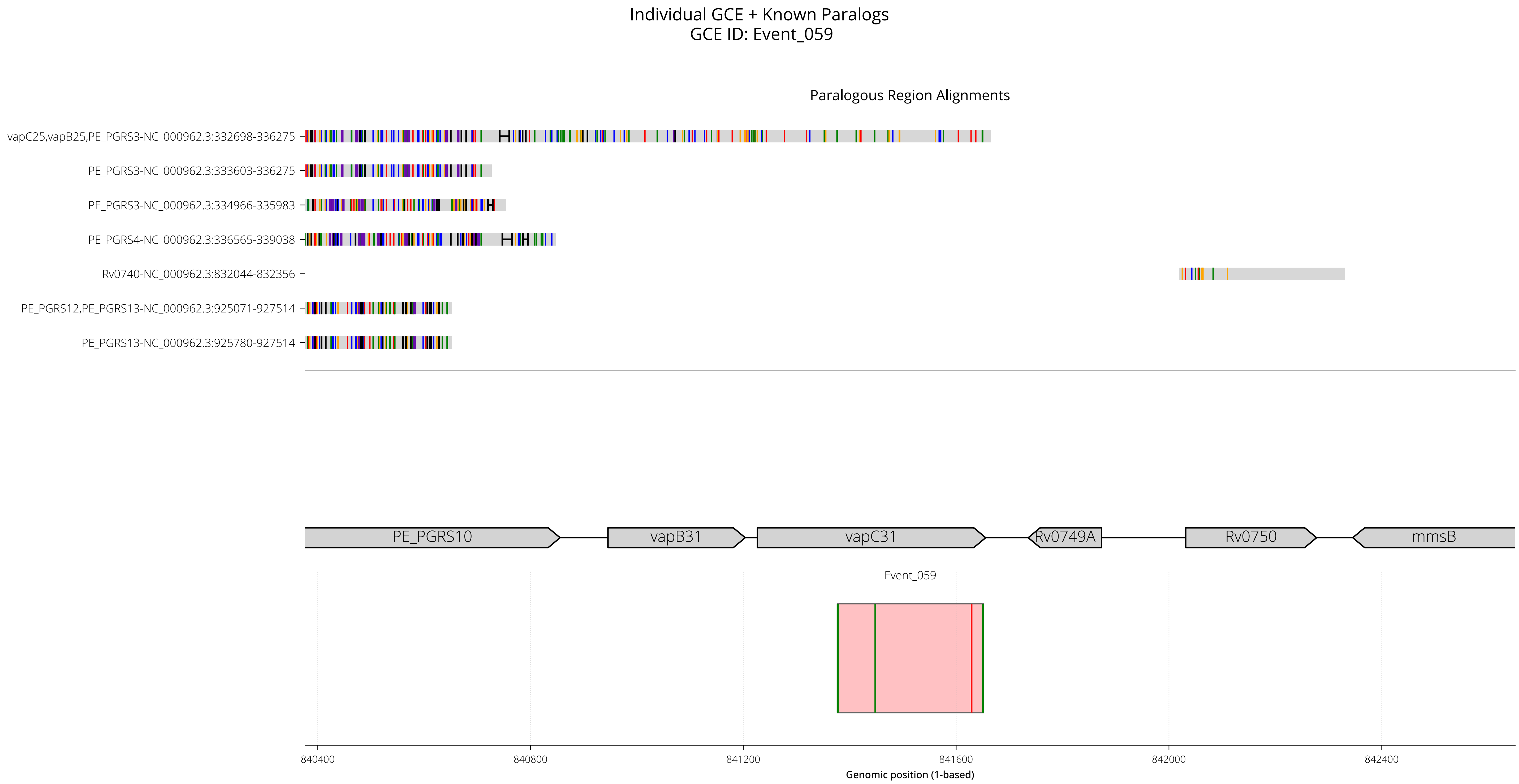
Individual GCE + Known Paralogs
GCE ID: Event_059
Paralogous Region Alignments
vapC25,vapB25,PE_PGRS3-NC_000962.3:332698-336275
PE_PGRS3-NC_000962.3:333603-336275
PE_PGRS3-NC_000962.3:334966-335983
PE_PGRS4-NC_000962.3:336565-339038
Rv0740-NC_000962.3:832044-832356
PE_PGRS12,PE_PGRS13-NC_000962.3:925071-927514
PE_PGRS13-NC_000962.3:925780-927514
PE_PGRS10
vapB31
vapC31
Rv0749A
Rv0750
mmsB
Event_059
840400
840800
841200
841600
842000
842400
Genomic position (1-based)

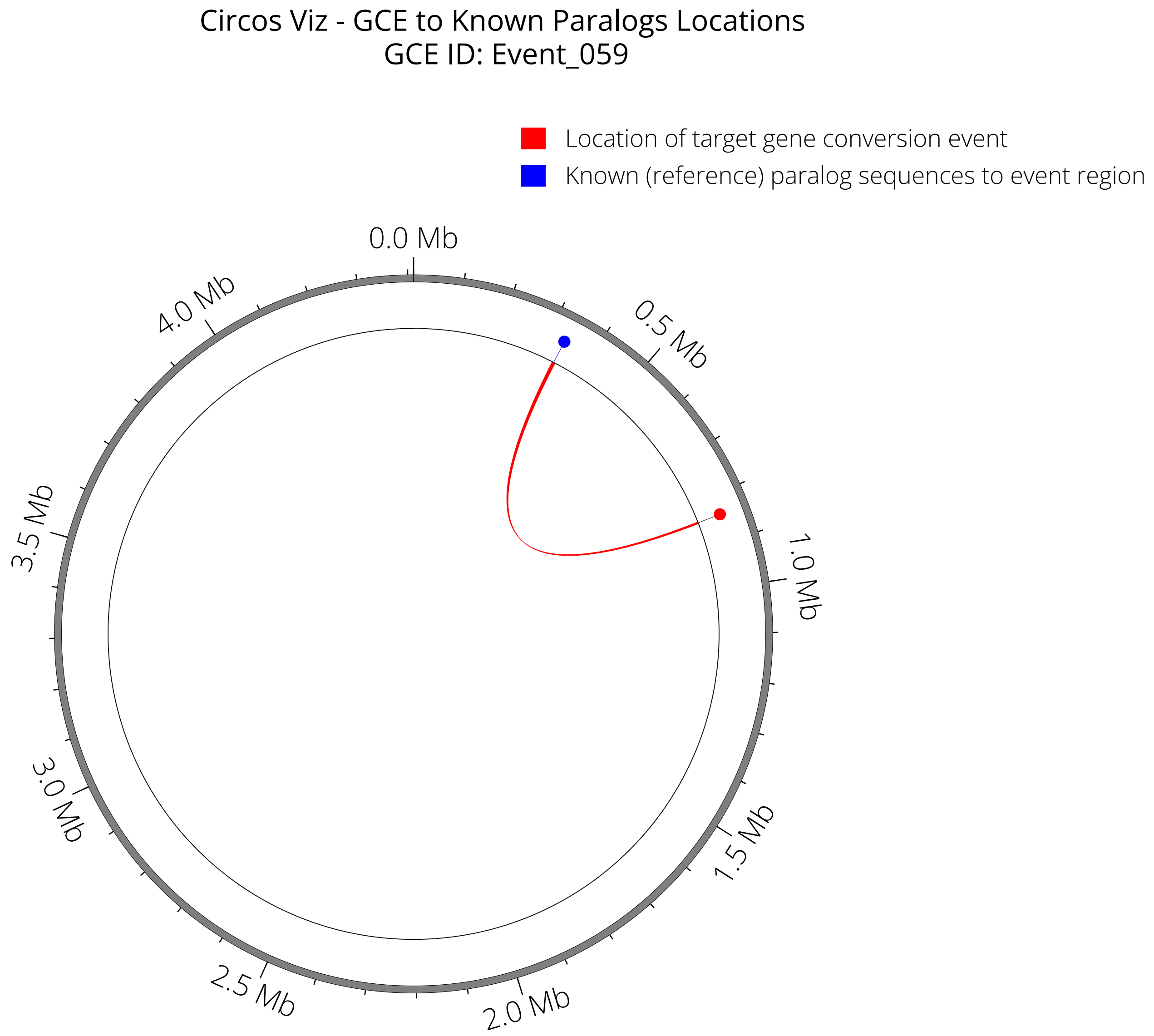
Circos Viz - GCE to Known Paralogs Locations
GCE ID: Event_059
Location of target gene conversion event
Known (reference) paralog sequences to event region
0.0 Mb
0.5 Mb
1.0 Mb
1.5 Mb
2.0 Mb
2.5 Mb
3.0 Mb
3.5 Mb
4.0 Mb

### Event_059 - 6 SNPs - k-mer analysis relative to paralogs

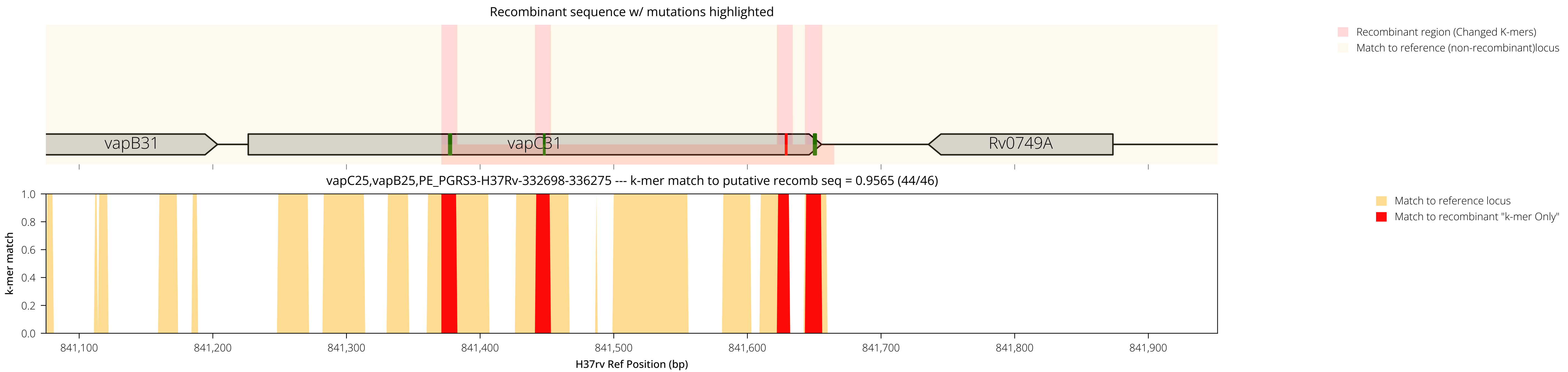

Sub-phylogeny Viz + Paralog Alignments
GCE ID: Event_059

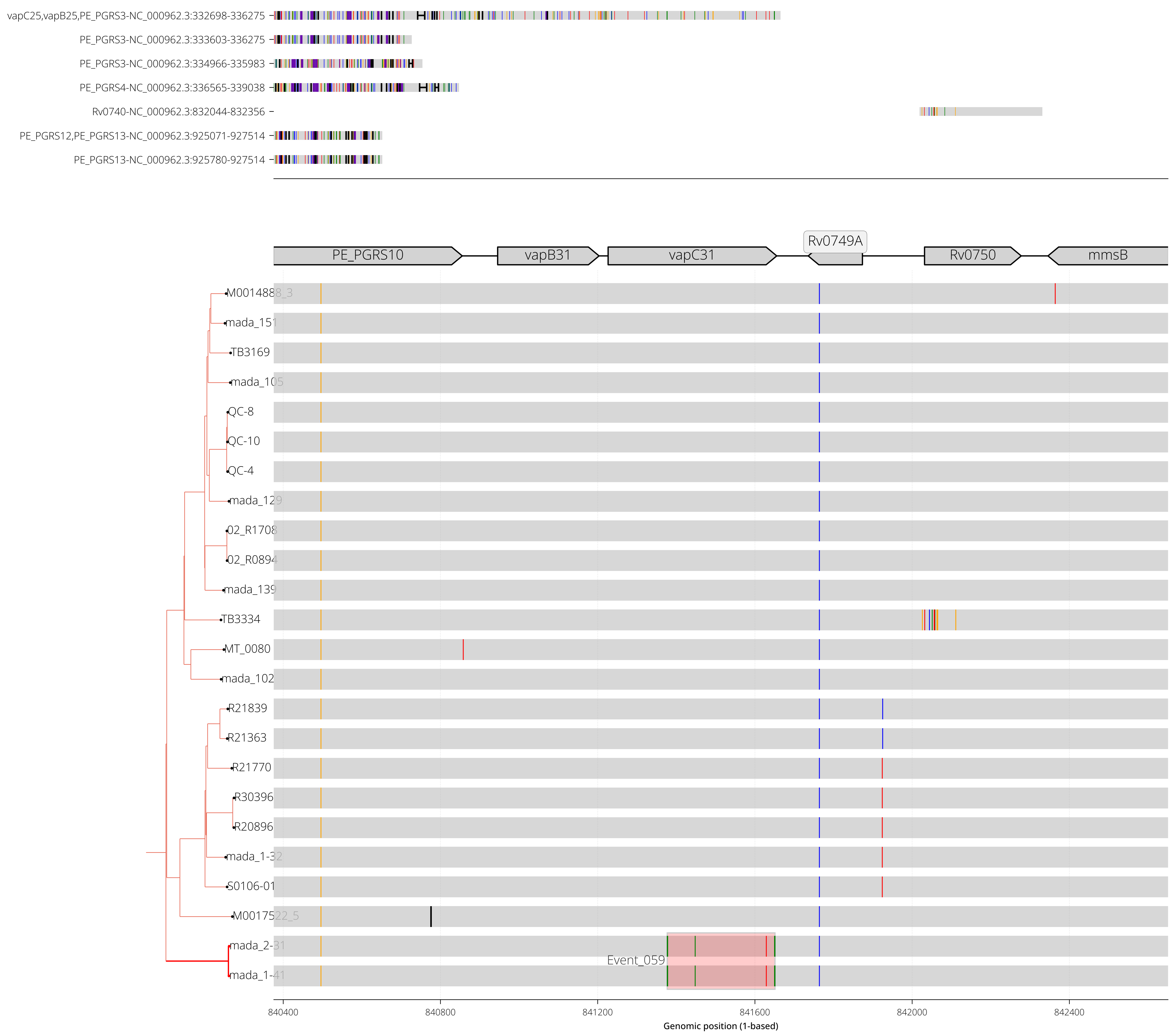

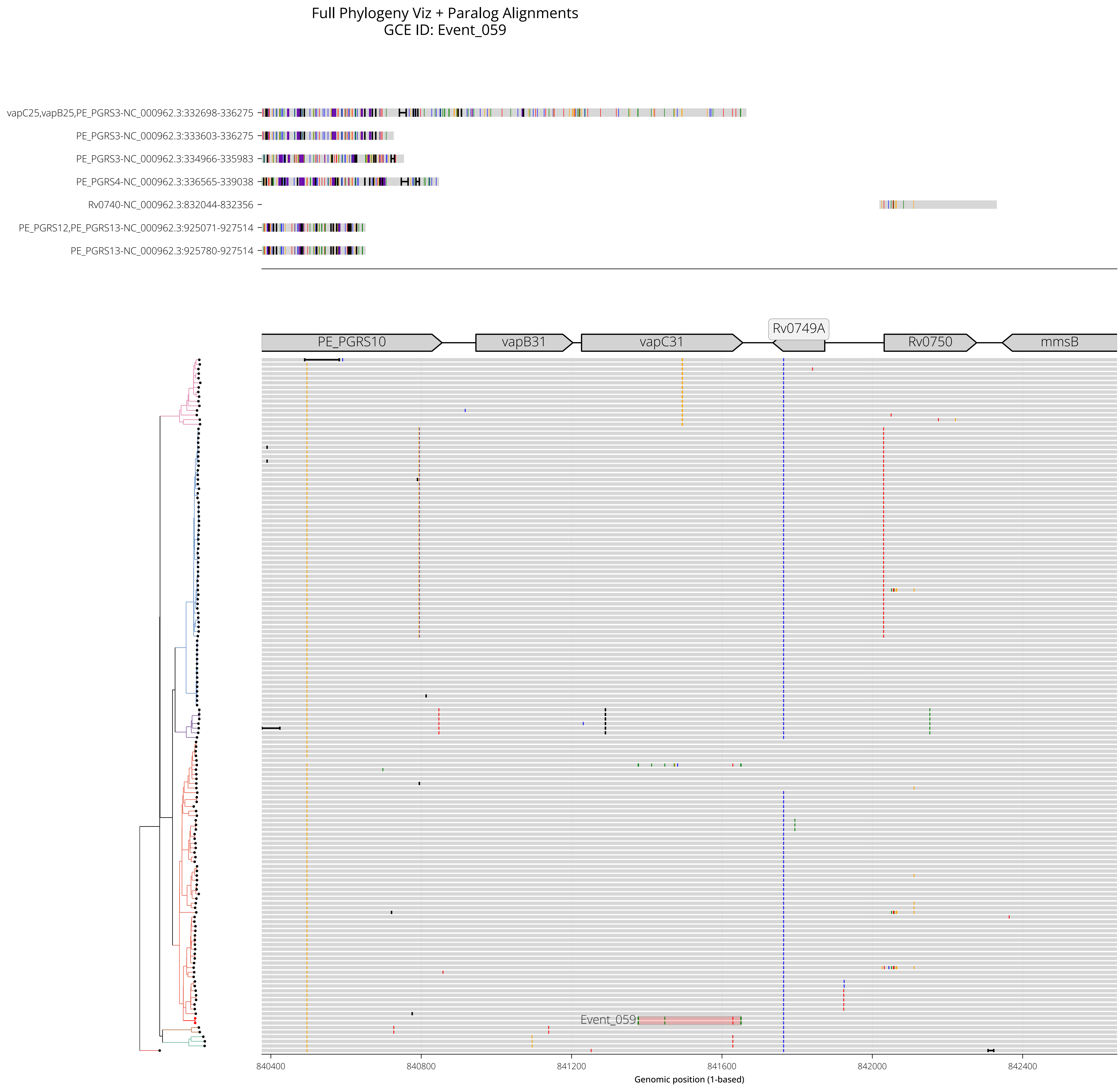

Full Phylogeny Viz + Paralog Alignments
GCE ID: Event_059
vapC25,vapB25,PE_PGRS3-NC_000962.3:332698-336275
PE_PGRS3-NC_000962.3:333603-336275
PE_PGRS3-NC_000962.3:334966-335983
PE_PGRS4-NC_000962.3:336565-339038
Rv0740-NC_000962.3:832044-832356
PE_PGRS12,PE_PGRS13-NC_000962.3:925071-927514
PE_PGRS13-NC_000962.3:925780-927514
PE_PGRS10
vapB31
vapC31
Rv0749A
Rv0750
mmsB
Event_059
840400
840800
841200
841600
842000
842400
Genomic position (1-based)

Individual GCE + Known Paralogs
GCE ID: Event_060

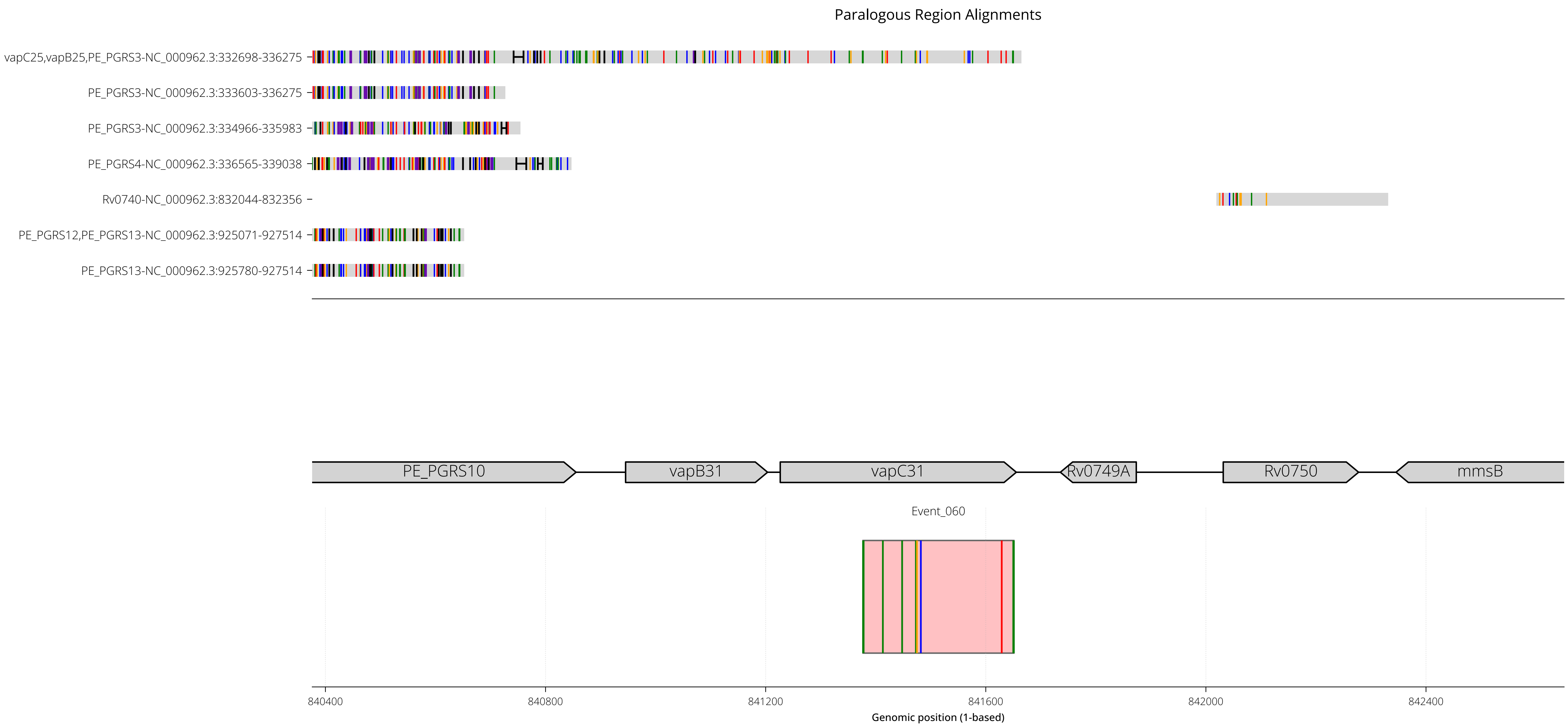

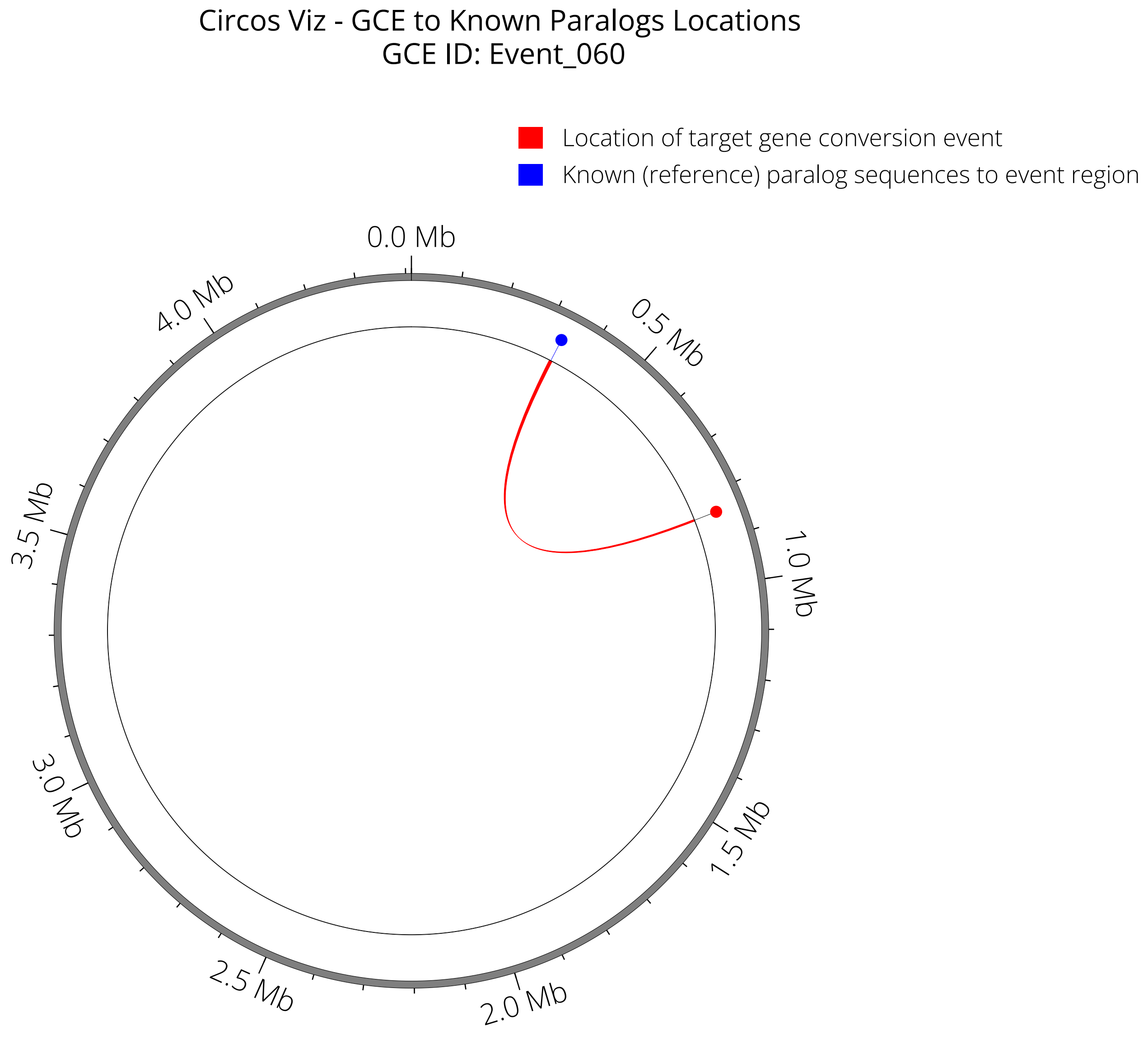

Circos Viz - GCE to Known Paralogs Locations
GCE ID: Event_060
Location of target gene conversion event
Known (reference) paralog sequences to event region
0.0 Mb
0.5 Mb
1.0 Mb
1.5 Mb
2.0 Mb
2.5 Mb
3.0 Mb
3.5 Mb
4.0 Mb

### Event_060 - 10 SNPs - k-mer analysis relative to paralogs

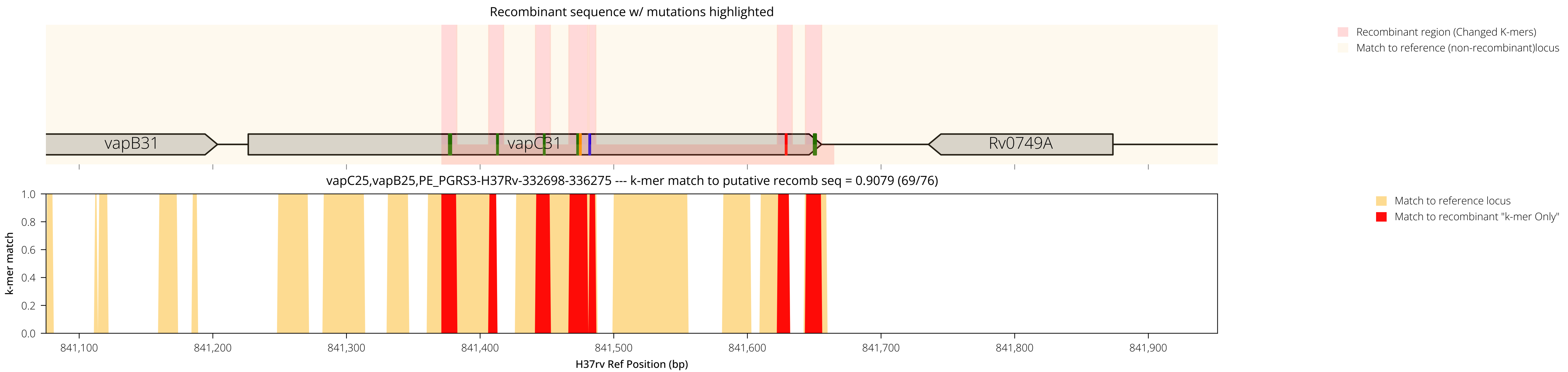

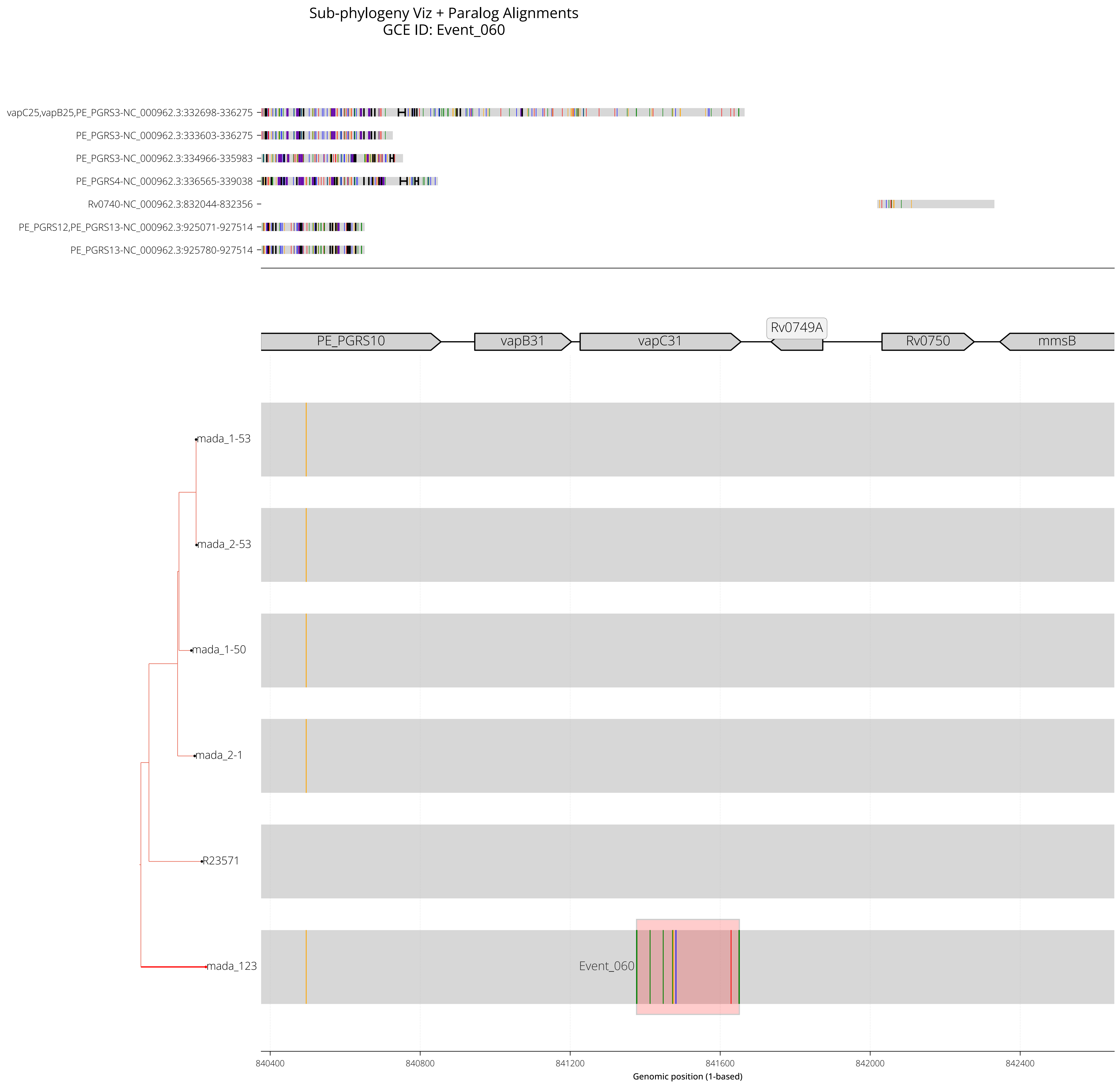

Sub-phylogeny Viz + Paralog Alignments
GCE ID: Event_060
vapC25,vapB25,PE_PGRS3-NC_000962.3:332698-336275
PE_PGRS3-NC_000962.3:333603-336275
PE_PGRS3-NC_000962.3:334966-335983
PE_PGRS4-NC_000962.3:336565-339038
Rv0740-NC_000962.3:832044-832356
PE_PGRS12,PE_PGRS13-NC_000962.3:925071-927514
PE_PGRS13-NC_000962.3:925780-927514
PE_PGRS10
vapB31
vapC31
Rv0749A
Rv0750
mmsB
mada_1-53
mada_2-53
mada_1-50
mada_2-1
R23571
mada_123
Event_060
840400
840800
841200
841600
842000
842400
Genomic position (1-based)

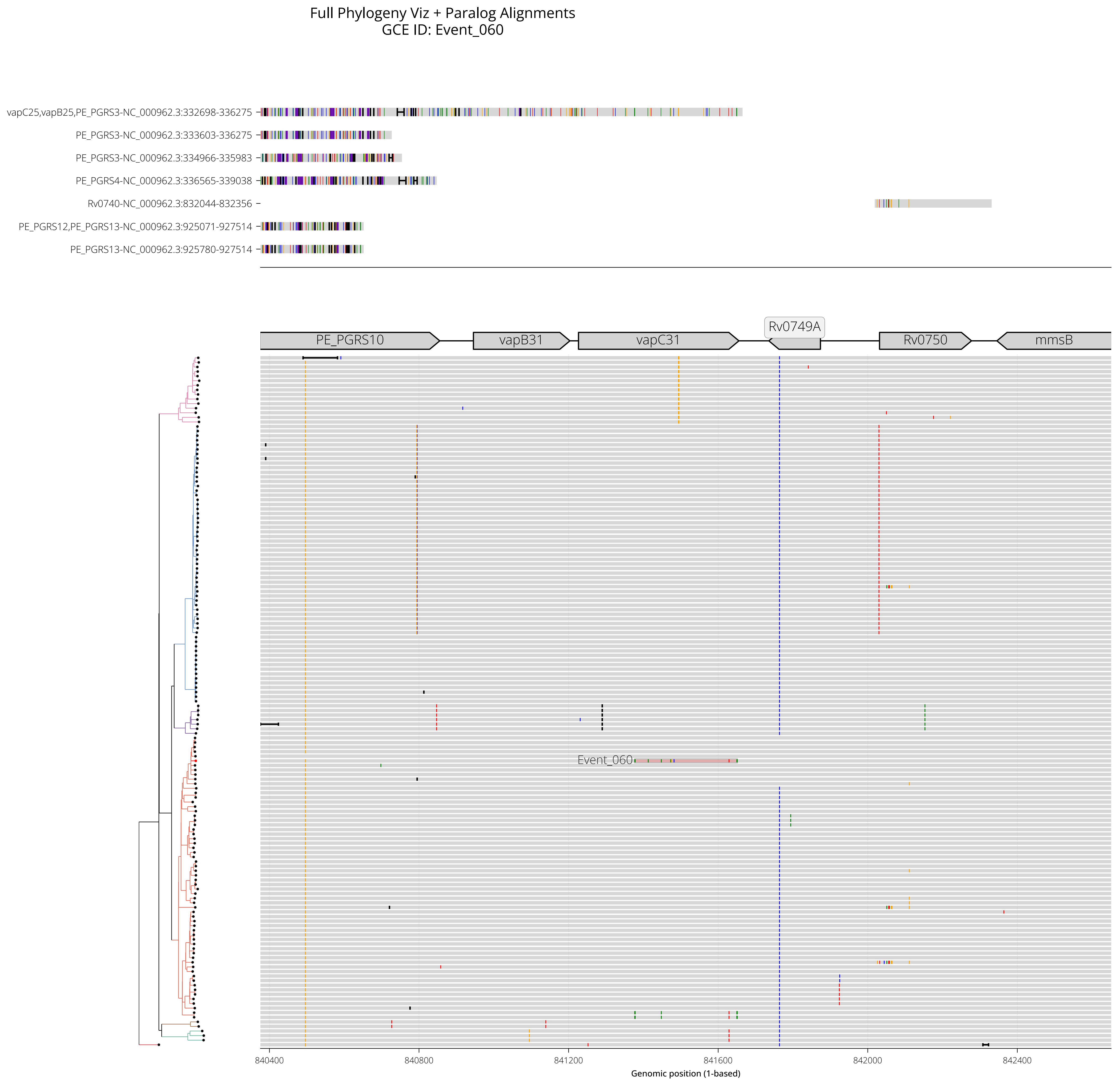

Full Phylogeny Viz + Paralog Alignments
GCE ID: Event_060
vapC25,vapB25,PE_PGRS3-NC_000962.3:332698-336275
PE_PGRS3-NC_000962.3:333603-336275
PE_PGRS3-NC_000962.3:334966-335983
PE_PGRS4-NC_000962.3:336565-339038
Rv0740-NC_000962.3:832044-832356
PE_PGRS12,PE_PGRS13-NC_000962.3:925071-927514
PE_PGRS13-NC_000962.3:925780-927514
PE_PGRS10
vapB31
vapC31
Rv0749A
Rv0750
mmsB
Event_060
840400
840800
841200
841600
842000
842400
Genomic position (1-based)

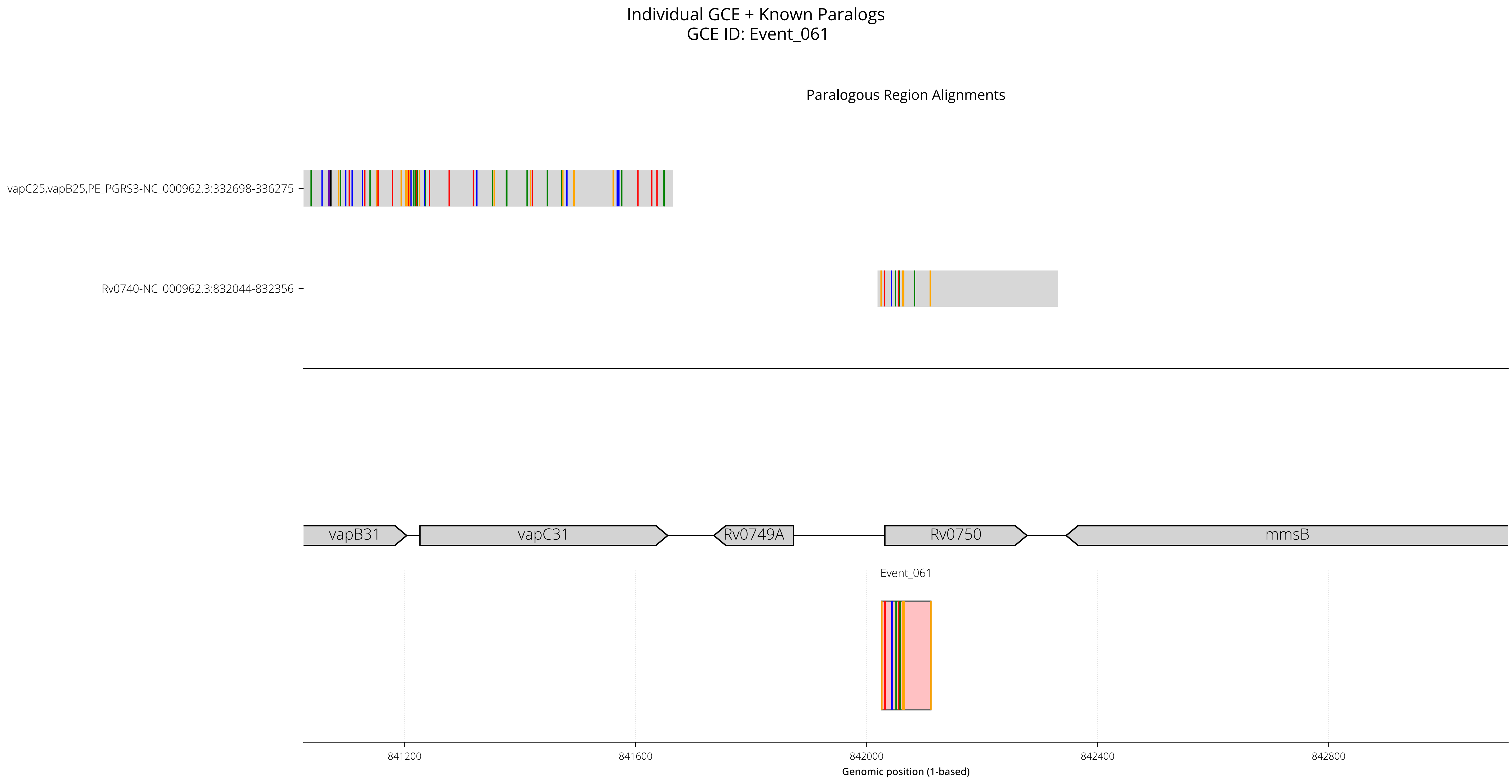
Individual GCE + Known Paralogs
GCE ID: Event_061
Paralogous Region Alignments
vapC25,vapB25,PE_PGRS3-NC_000962.3:332698-336275
Rv0740-NC_000962.3:832044-832356
vapB31
vapC31
Rv0749A
Rv0750
mmsB
Event_061
841200
841600
842000
842400
842800
Genomic position (1-based)

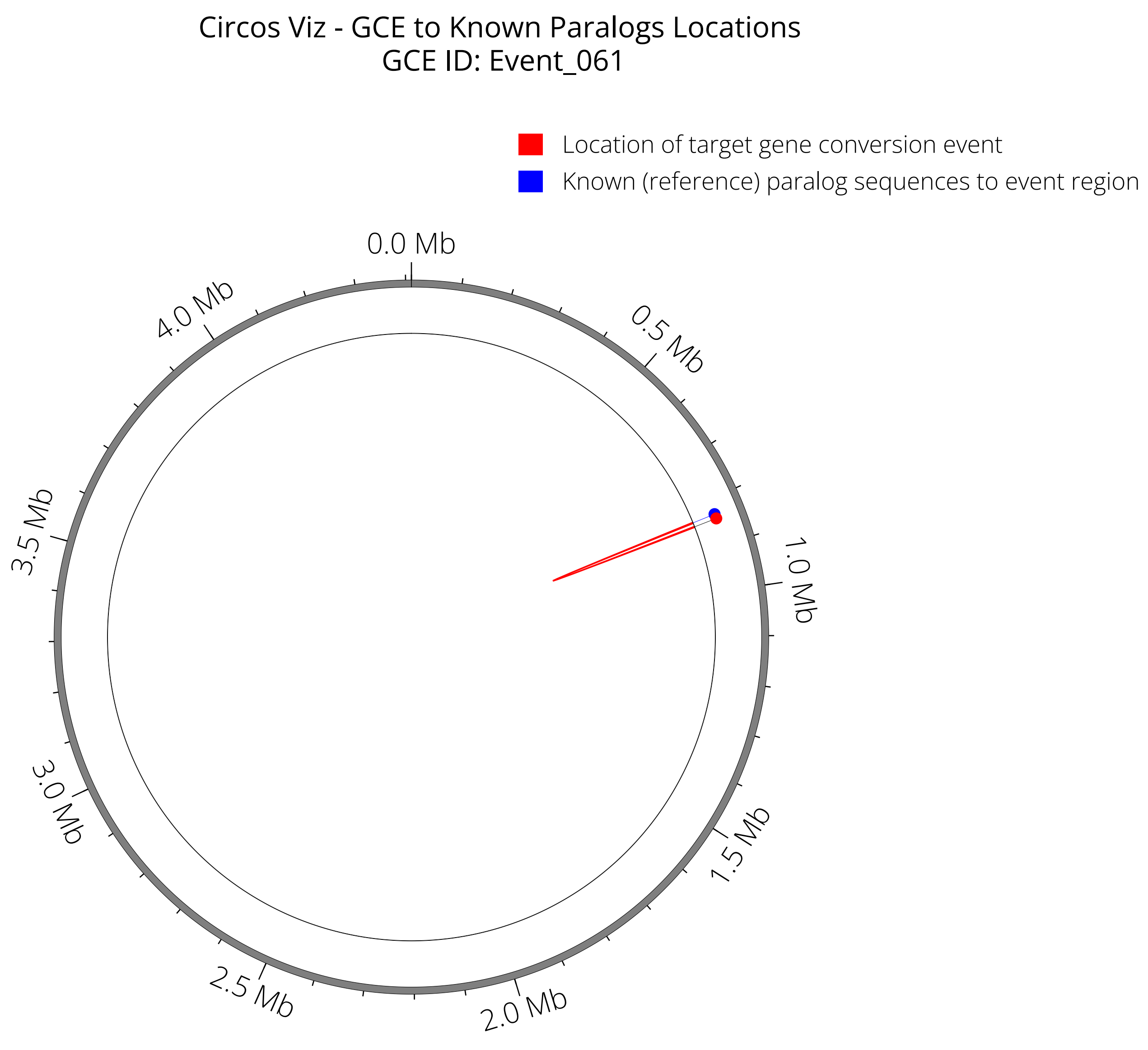
Circos Viz - GCE to Known Paralogs Locations
GCE ID: Event_061
Location of target gene conversion event
Known (reference) paralog sequences to event region
0.0 Mb
0.5 Mb
1.0 Mb
1.5 Mb
2.0 Mb
2.5 Mb
3.0 Mb
3.5 Mb
4.0 Mb

### Event_061 - 10 SNPs - k-mer analysis relative to paralogs

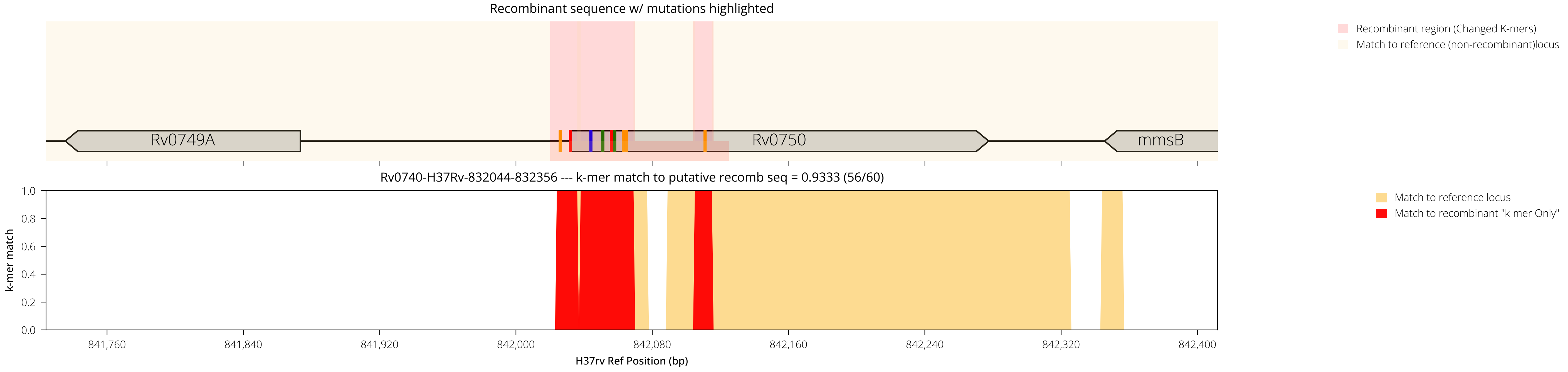

Sub-phylogeny Viz + Paralog Alignments
GCE ID: Event_061

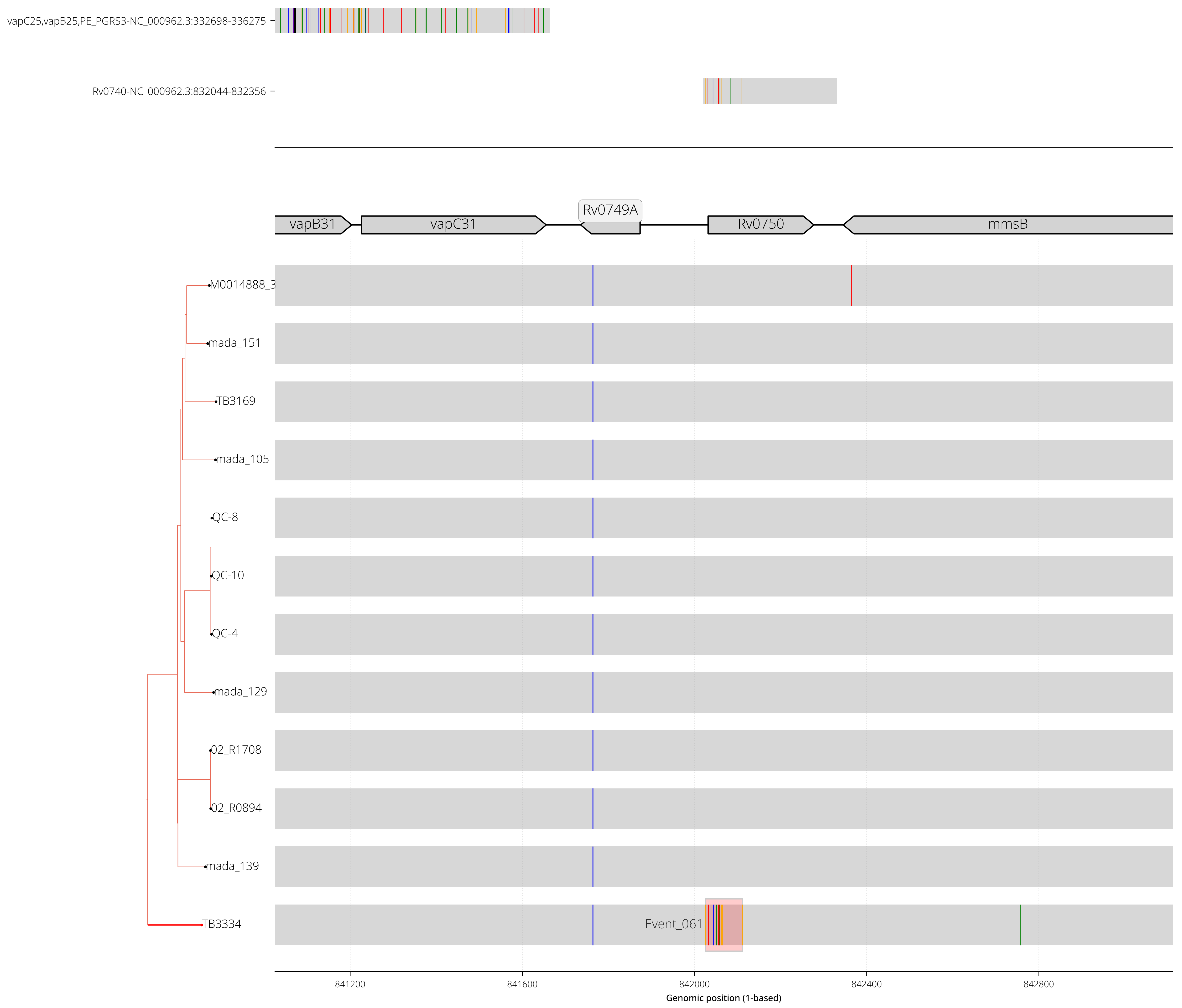

Full Phylogeny Viz + Paralog Alignments
GCE ID: Event_061

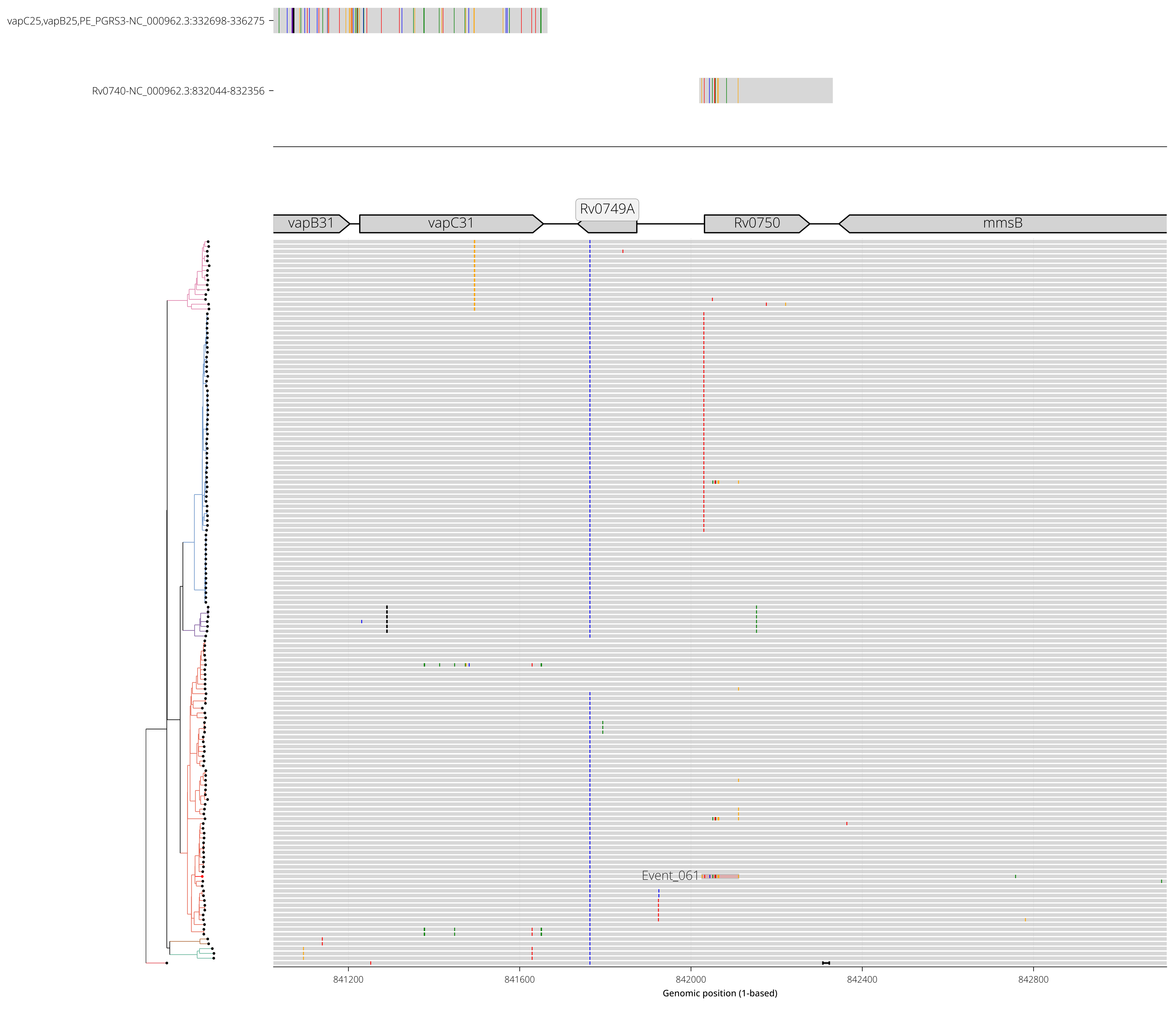

Individual GCE + Known Paralogs
GCE ID: Event_062

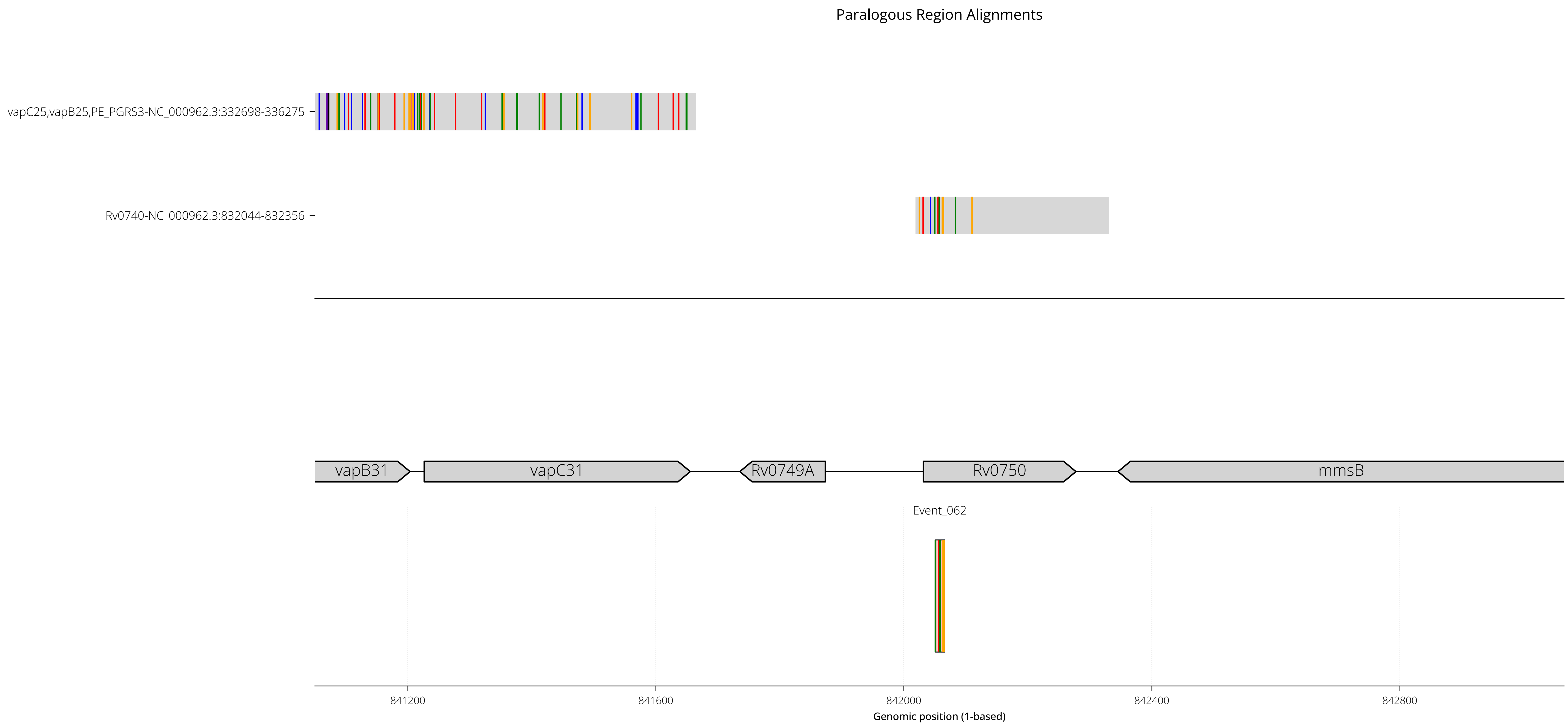

Circos Viz - GCE to Known Paralogs Locations
GCE ID: Event_062

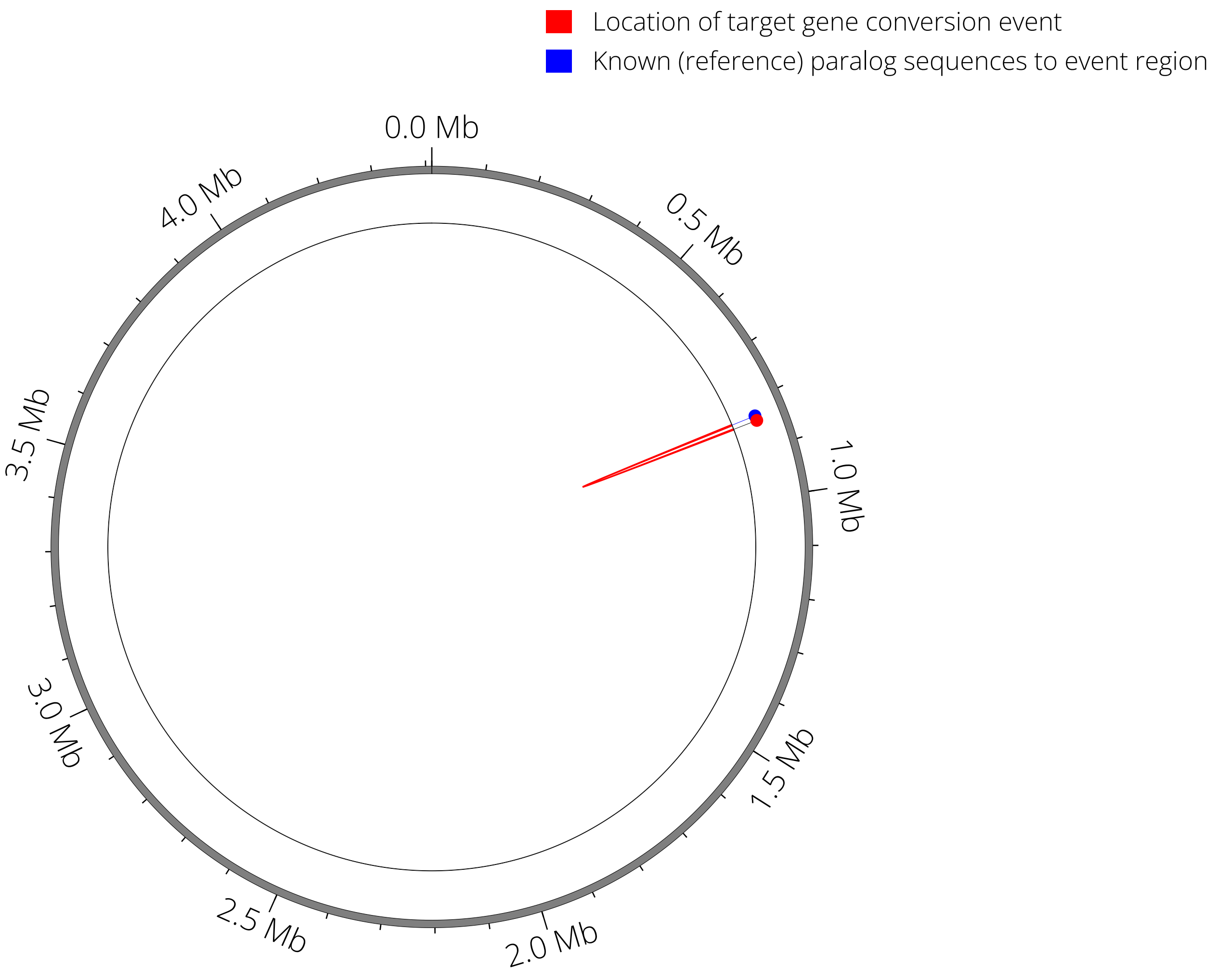

### Event_062 - 6 SNPs - k-mer analysis relative to paralogs

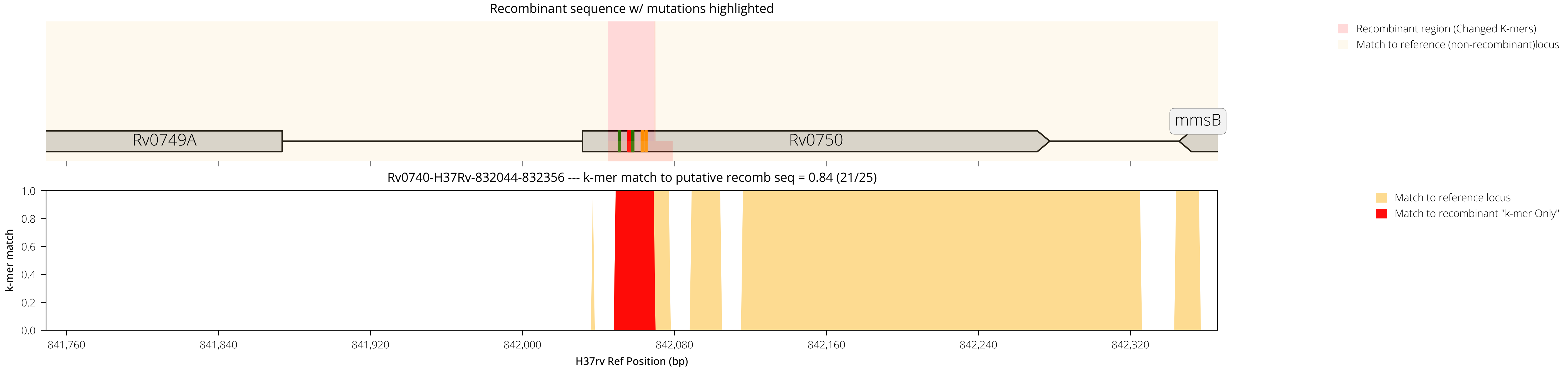

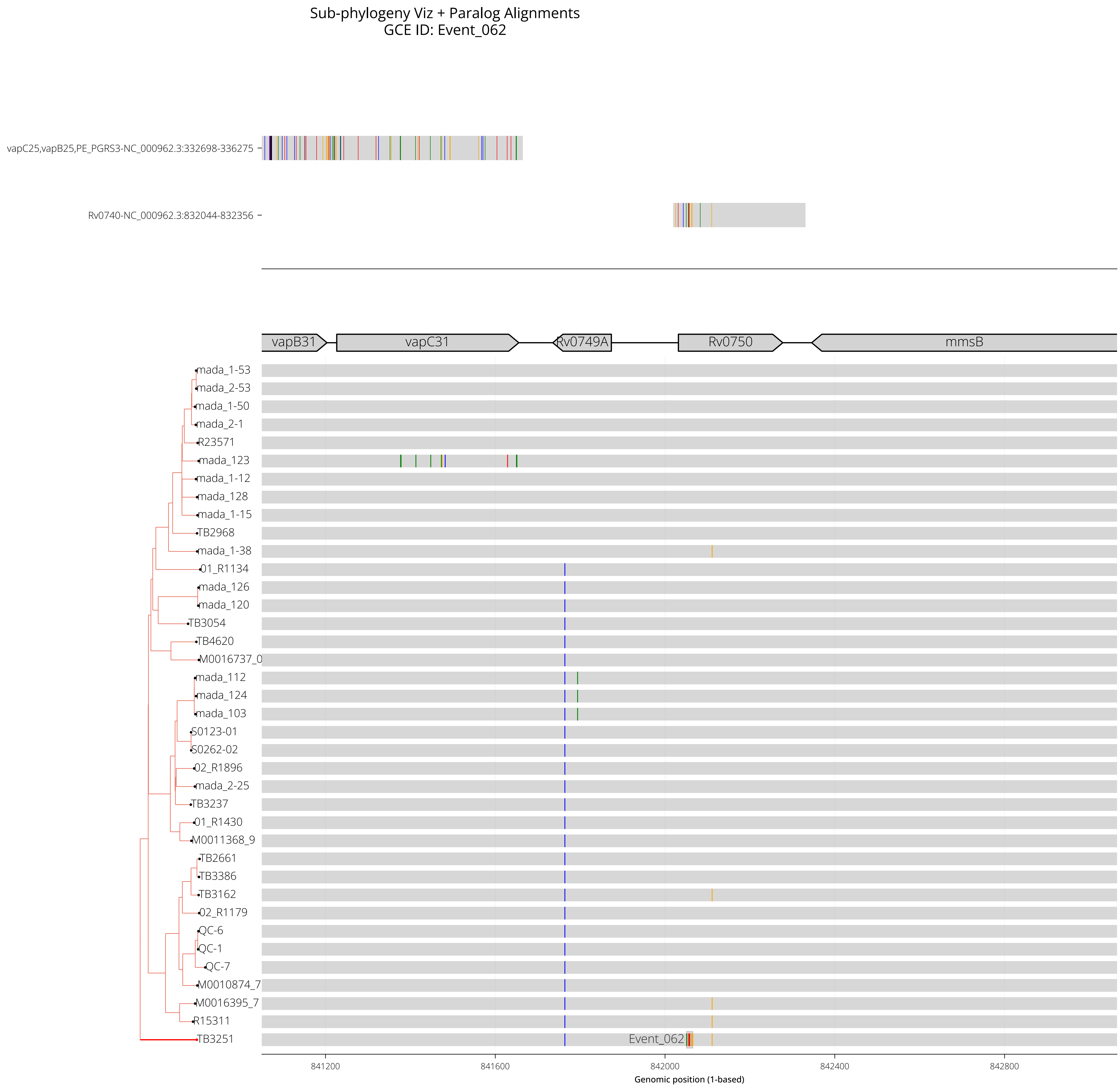
Sub-phylogeny Viz + Paralog Alignments
GCE ID: Event_062
vapC25,vapB25,PE_PGRS3-NC_000962.3:332698-336275
Rv0740-NC_000962.3:832044-832356
vapB31
vapC31
Rv0749A
Rv0750
mmsB
mada_1-53
mada_2-53
mada_1-50
mada_2-1
R23571
mada_123
mada_1-12
mada_128
mada_1-15
TB2968
mada_1-38
01_R1134
mada_126
mada_120
TB3054
TB4620
M0016737_0
mada_112
mada_124
mada_103
S0123-01
S0262-02
02_R1896
mada_2-25
TB3237
01_R1430
M0011368_9
TB2661
TB3386
TB3162
02_R1179
QC-6
QC-1
QC-7
M0010874_7
M0016395_7
R15311
TB3251
Event_062
841200
841600
842000
842400
842800
Genomic position (1-based)

Full Phylogeny Viz + Paralog Alignments
GCE ID: Event_062

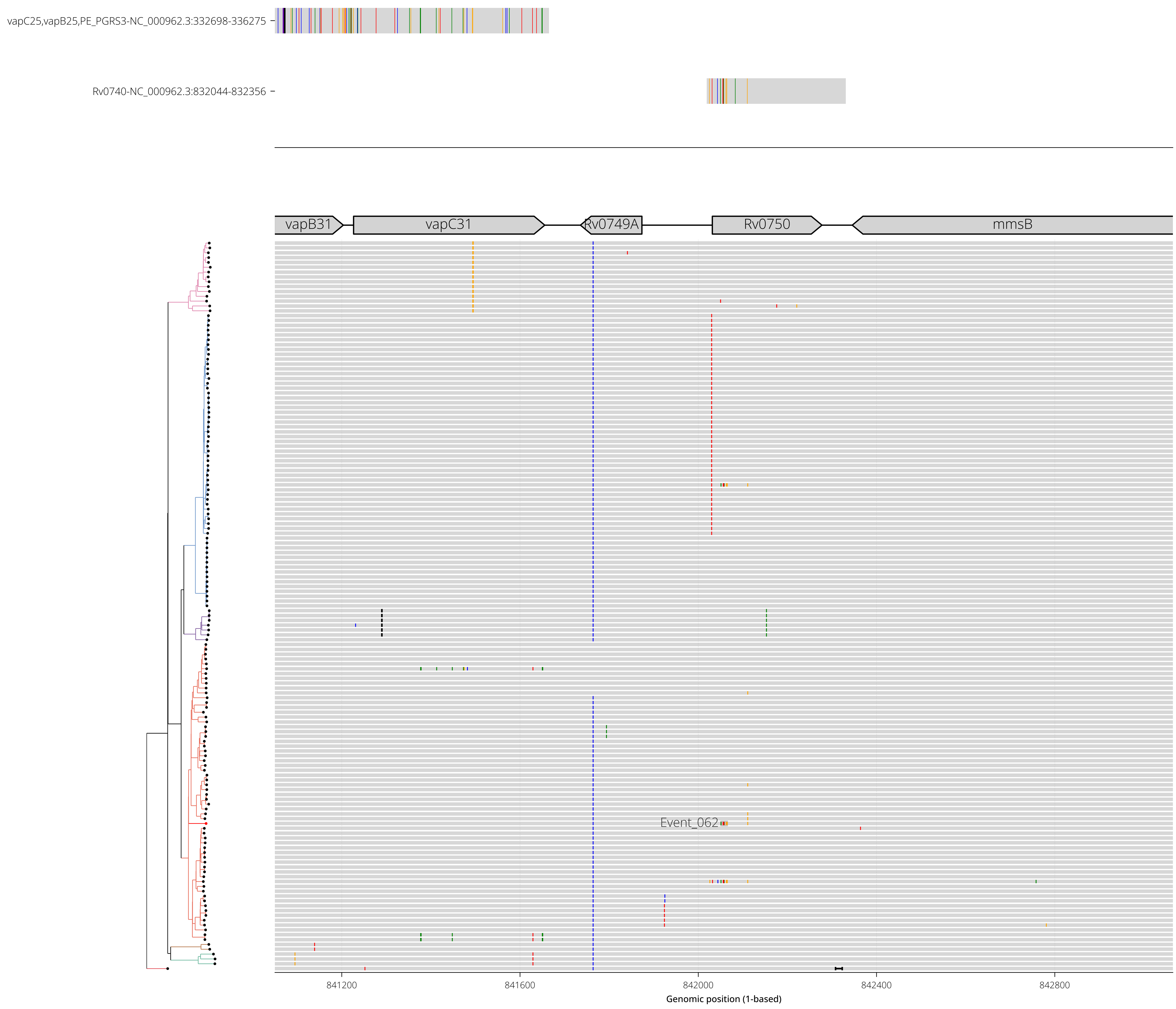

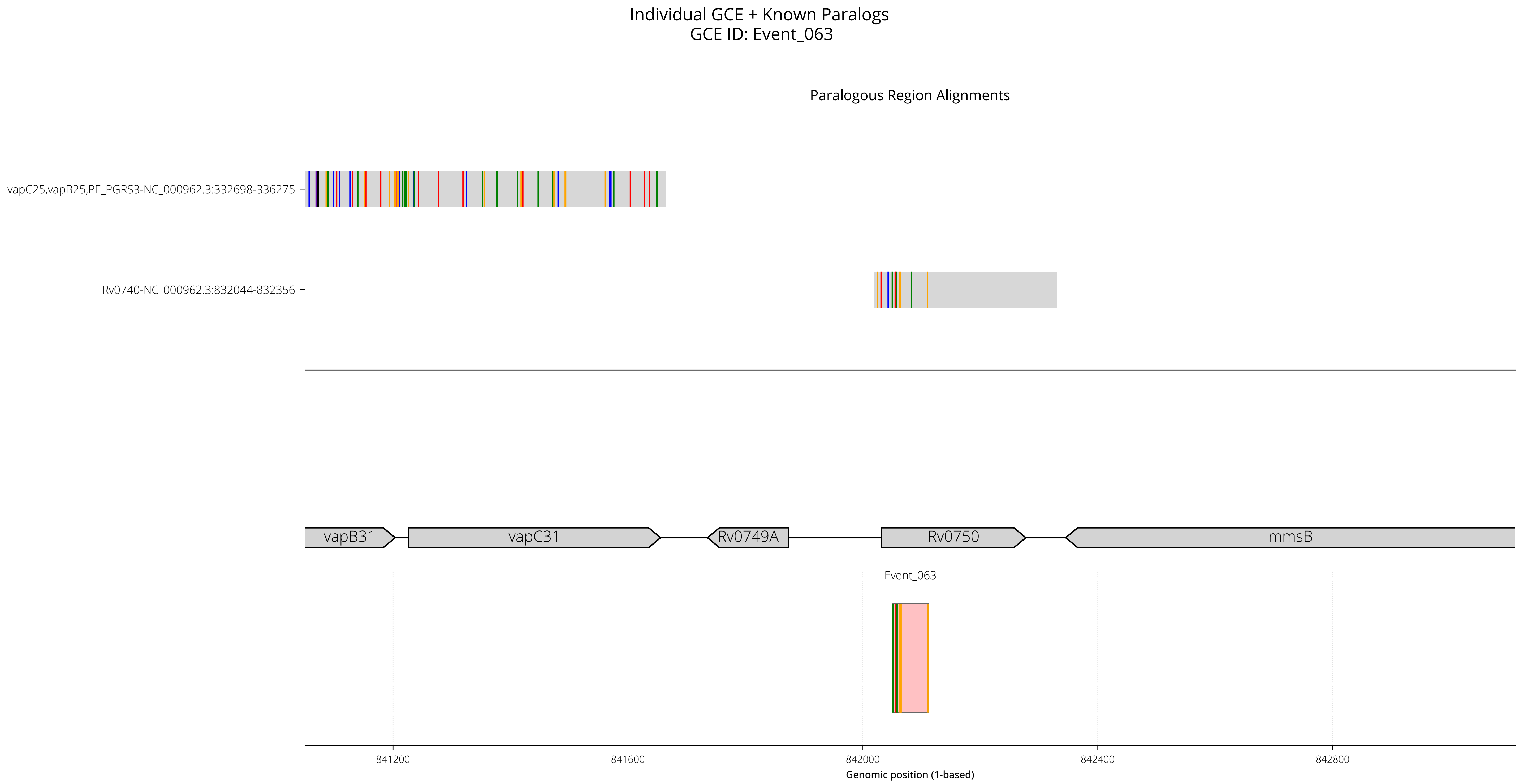
Individual GCE + Known Paralogs
GCE ID: Event_063
Paralogous Region Alignments
vapC25,vapB25,PE_PGRS3-NC_000962.3:332698-336275
Rv0740-NC_000962.3:832044-832356
vapB31
vapC31
Rv0749A
Rv0750
mmsB
Event_063
841200
841600
842000
842400
842800
Genomic position (1-based)

Circos Viz - GCE to Known Paralogs Locations
GCE ID: Event_063

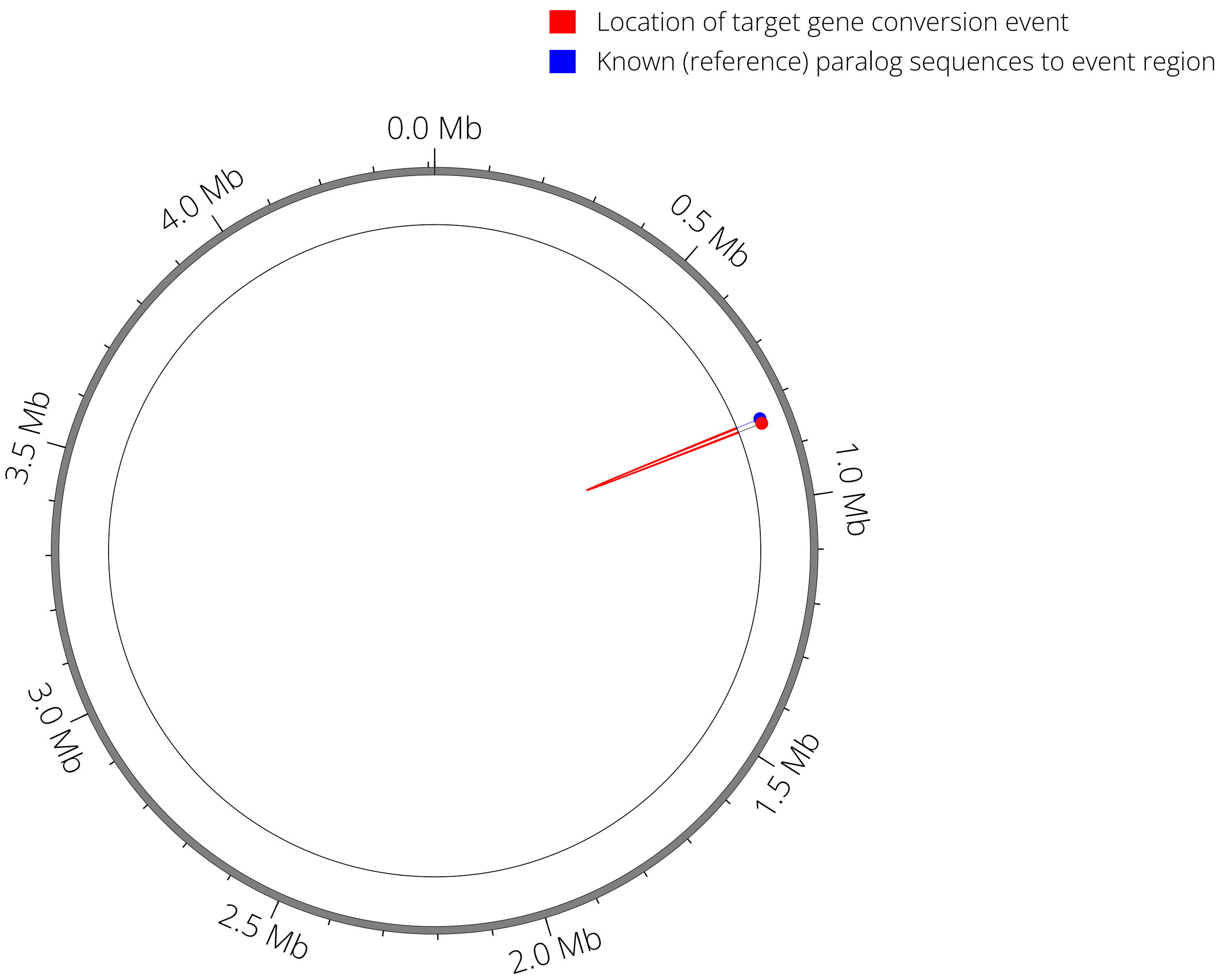

### Event_063 - 7 SNPs - k-mer analysis relative to paralogs

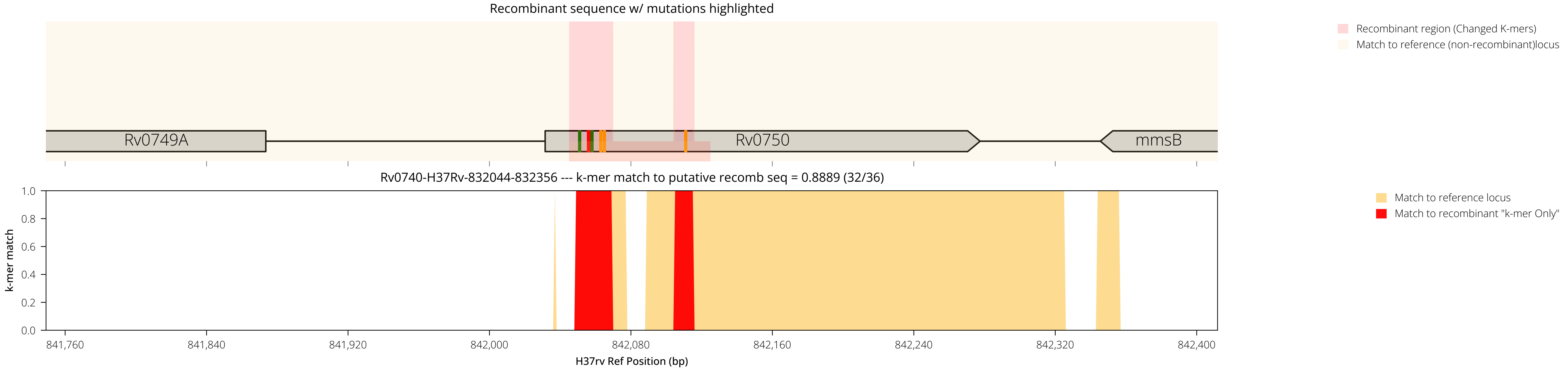

Sub-phylogeny Viz + Paralog Alignments
GCE ID: Event_063

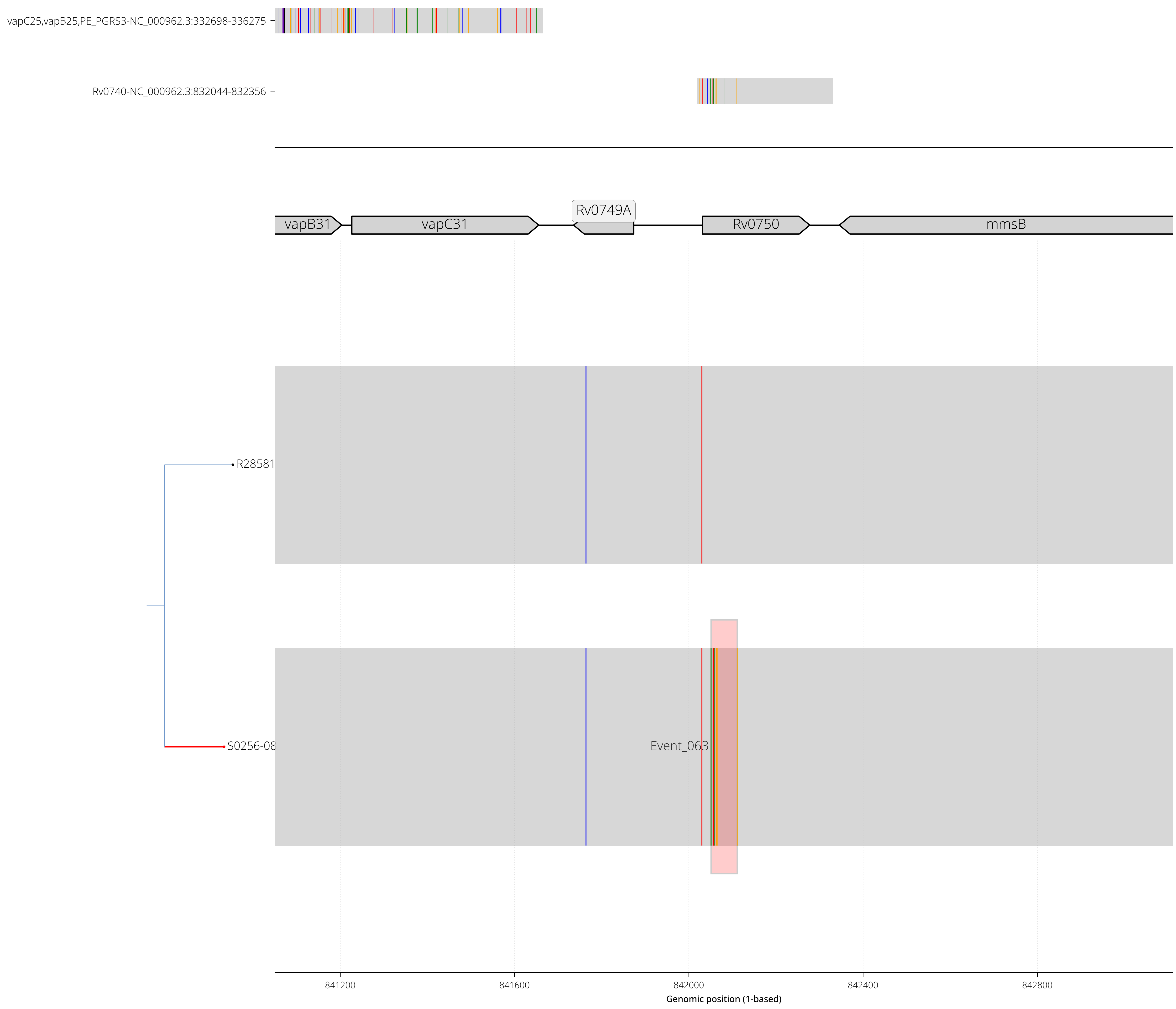

Full Phylogeny Viz + Paralog Alignments
GCE ID: Event_063

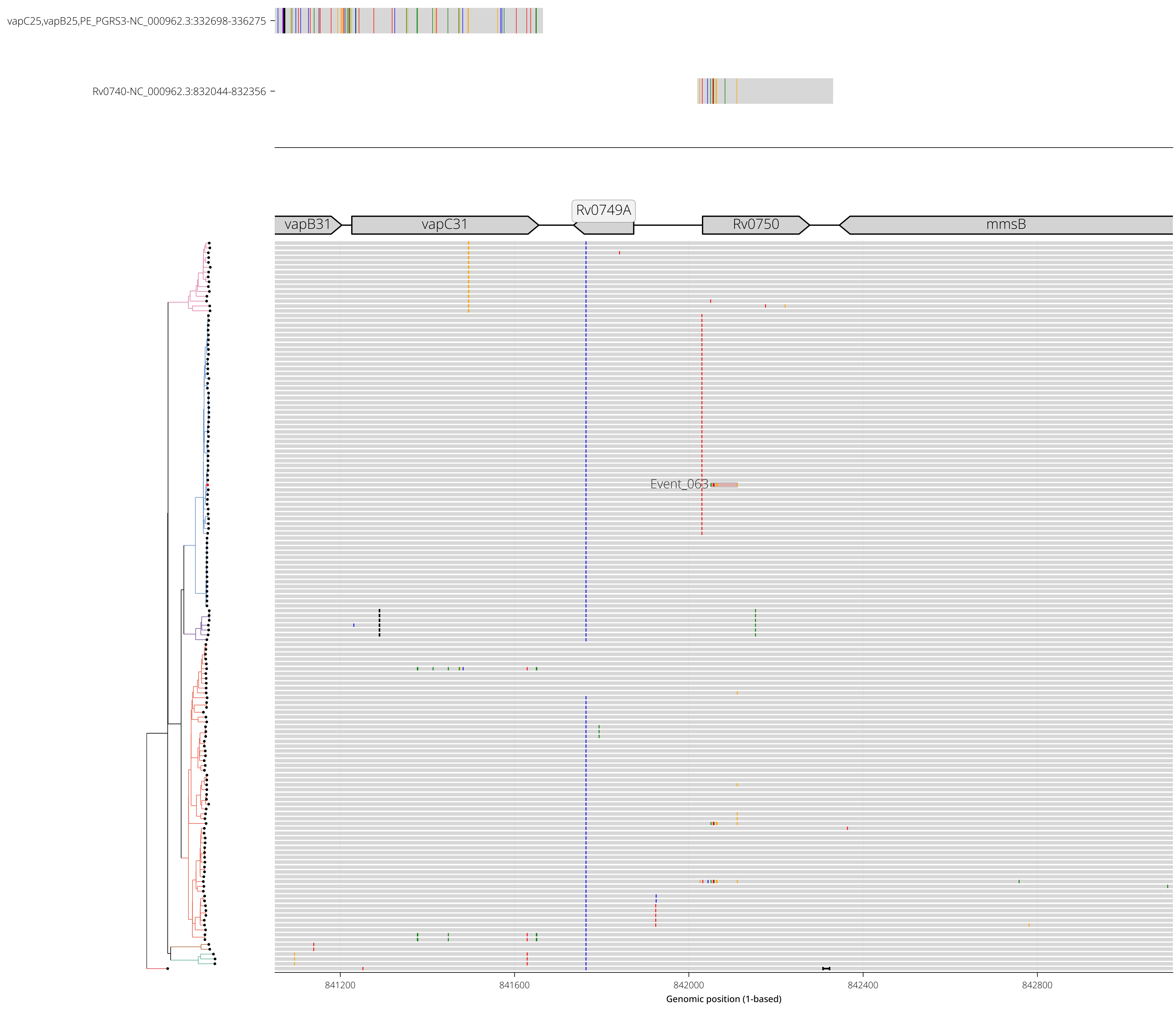

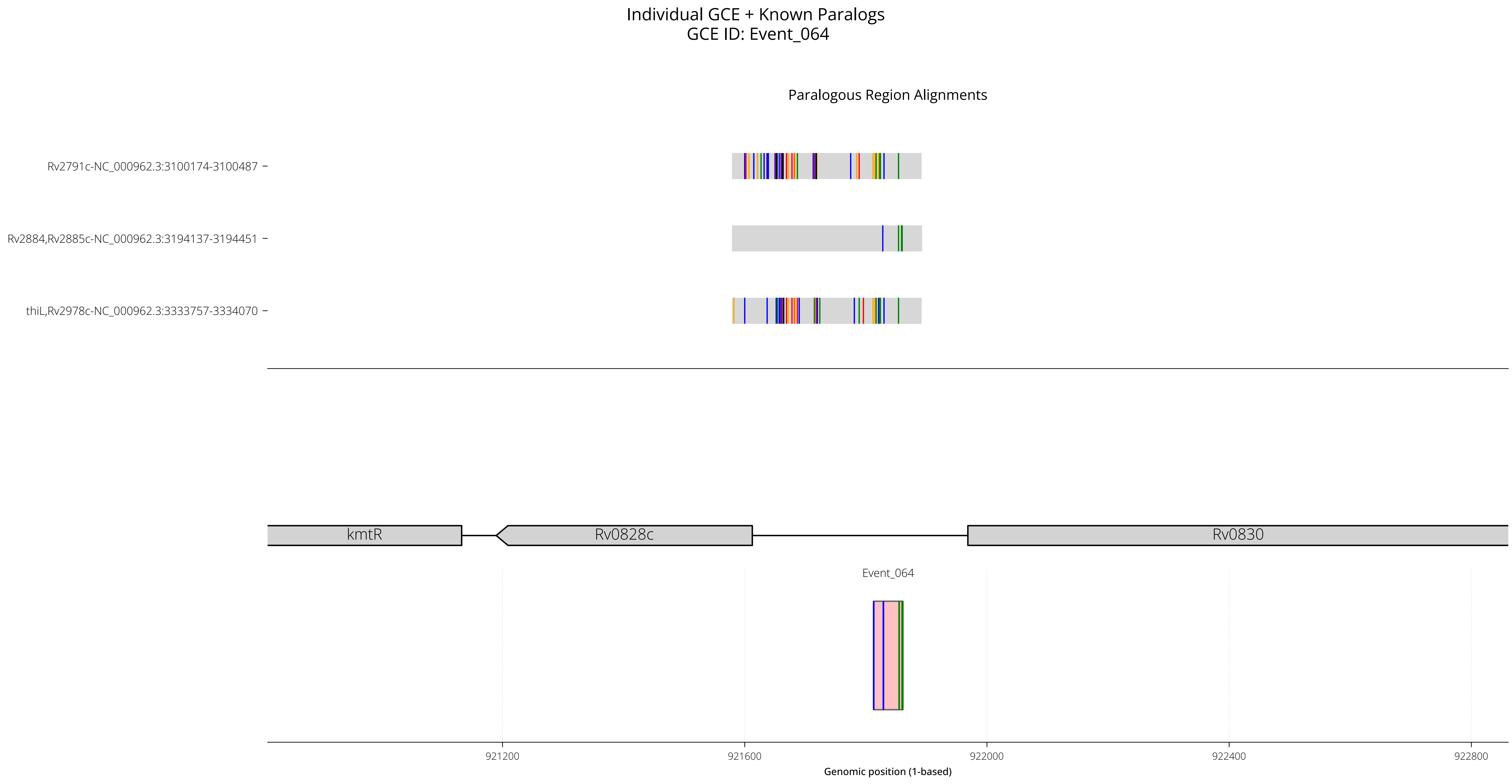
Individual GCE + Known Paralogs
GCE ID: Event_064
Paralogous Region Alignments
Rv2791c-NC_000962.3:3100174-3100487
Rv2884,Rv2885c-NC_000962.3:3194137-3194451
thiL,Rv2978c-NC_000962.3:3333757-3334070
kmtR
Rv0828c
Rv0830
Event_064
921200
921600
922000
922400
922800
Genomic position (1-based)

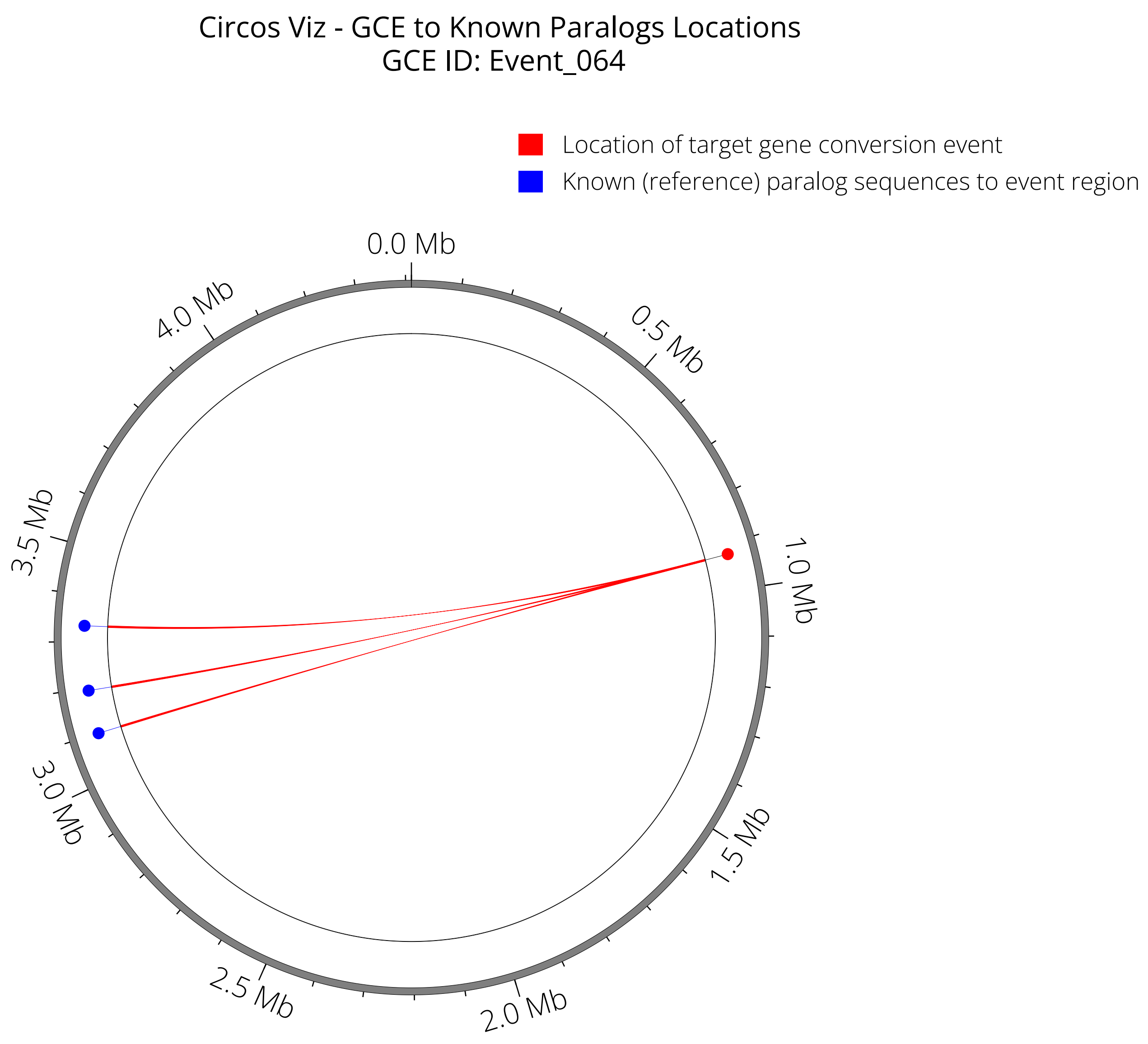
Circos Viz - GCE to Known Paralogs Locations
GCE ID: Event_064
Location of target gene conversion event
Known (reference) paralog sequences to event region
0.0 Mb
0.5 Mb
1.0 Mb
1.5 Mb
2.0 Mb
2.5 Mb
3.0 Mb
3.5 Mb
4.0 Mb

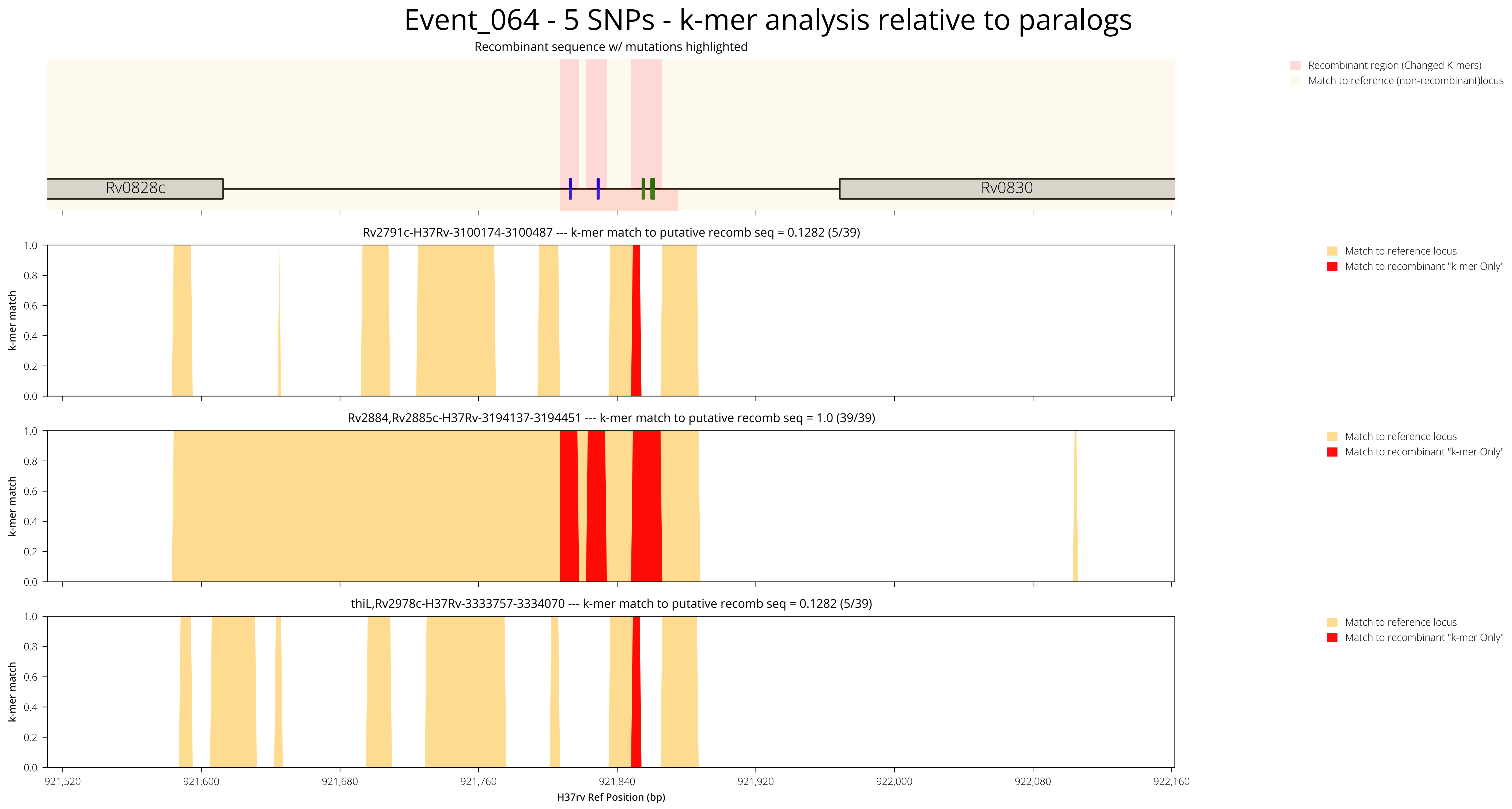

Event_064 - 5 SNPs - k-mer analysis relative to paralogs
Recombinant sequence w/ mutations highlighted
Recombinant region (Changed K-mers)
Match to reference (non-recombinant)locus
Rv0828c
Rv0830
Rv2791c-H37Rv-3100174-3100487 --- k-mer match to putative recomb seq = 0.1282 (5/39)
Rv2884,Rv2885c-H37Rv-3194137-3194451 --- k-mer match to putative recomb seq = 1.0 (39/39)
thiL,Rv2978c-H37Rv-3333757-3334070 --- k-mer match to putative recomb seq = 0.1282 (5/39)
Match to reference locus
Match to recombinant "k-mer Only"
k-mer match
1.0
0.8
0.6
0.4
0.2
0.0
921,520
921,600
921,680
921,760
921,840
921,920
922,000
922,080
922,160
H37rv Ref Position (bp)

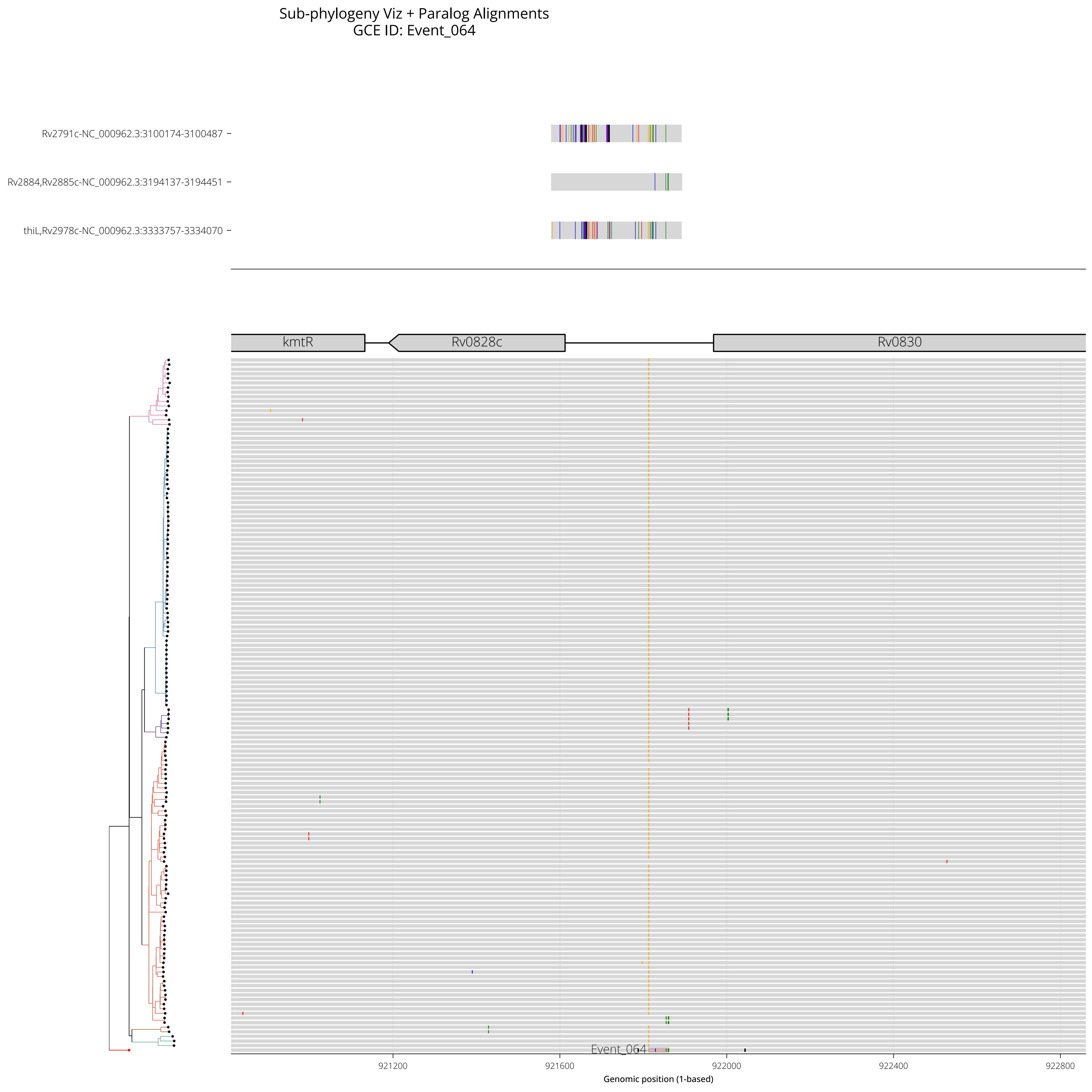

Sub-phylogeny Viz + Paralog Alignments
GCE ID: Event_064
Rv2791c-NC_000962.3:3100174-3100487
Rv2884,Rv2885c-NC_000962.3:3194137-3194451
thiL,Rv2978c-NC_000962.3:3333757-3334070
kmtR
Rv0828c
Rv0830
Event_064
921200
921600
922000
922400
922800
Genomic position (1-based)

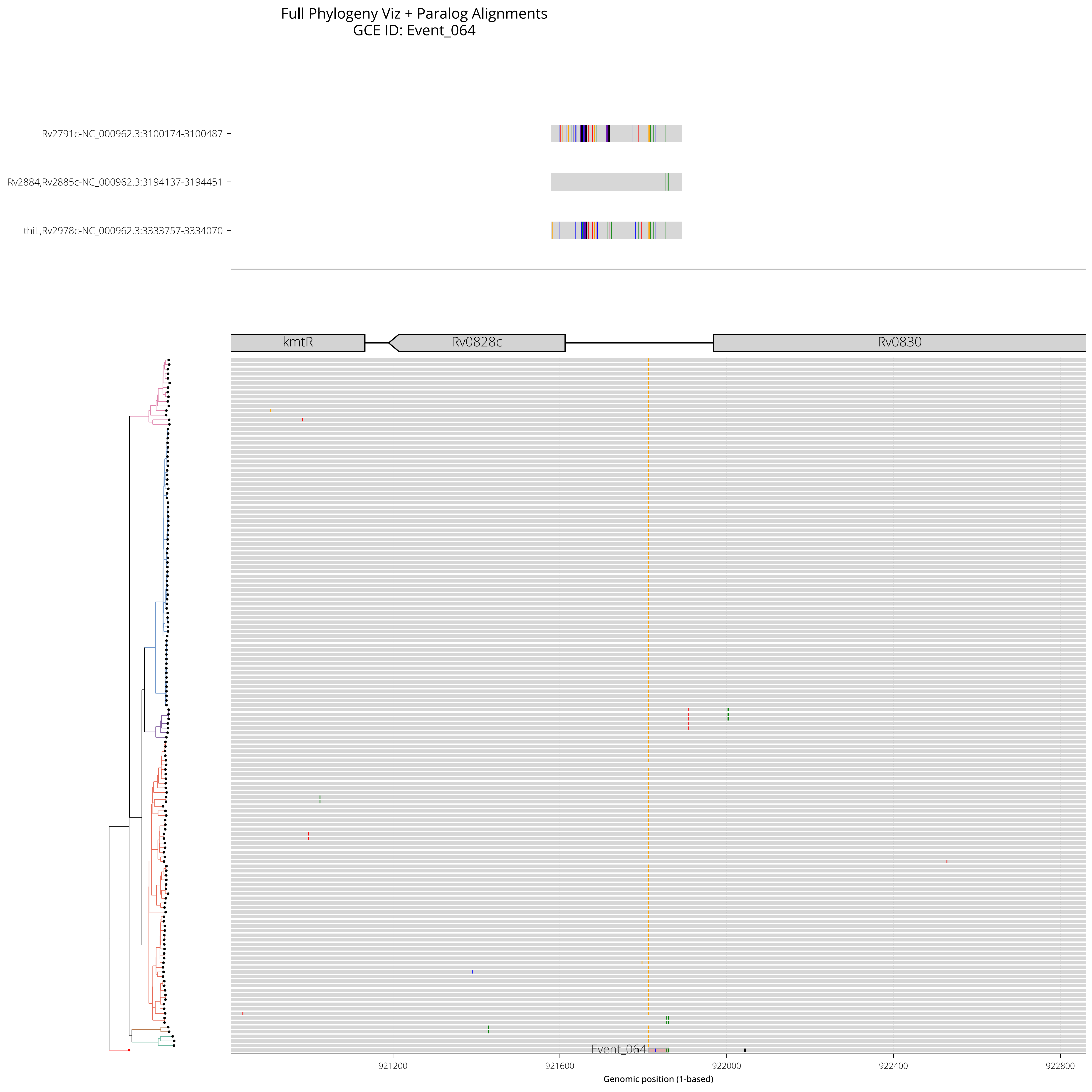

Full Phylogeny Viz + Paralog Alignments
GCE ID: Event_064
Rv2791c-NC_000962.3:3100174-3100487
Rv2884,Rv2885c-NC_000962.3:3194137-3194451
thiL,Rv2978c-NC_000962.3:3333757-3334070
kmtR
Rv0828c
Rv0830
Event_064
921200
921600
922000
922400
922800
Genomic position (1-based)

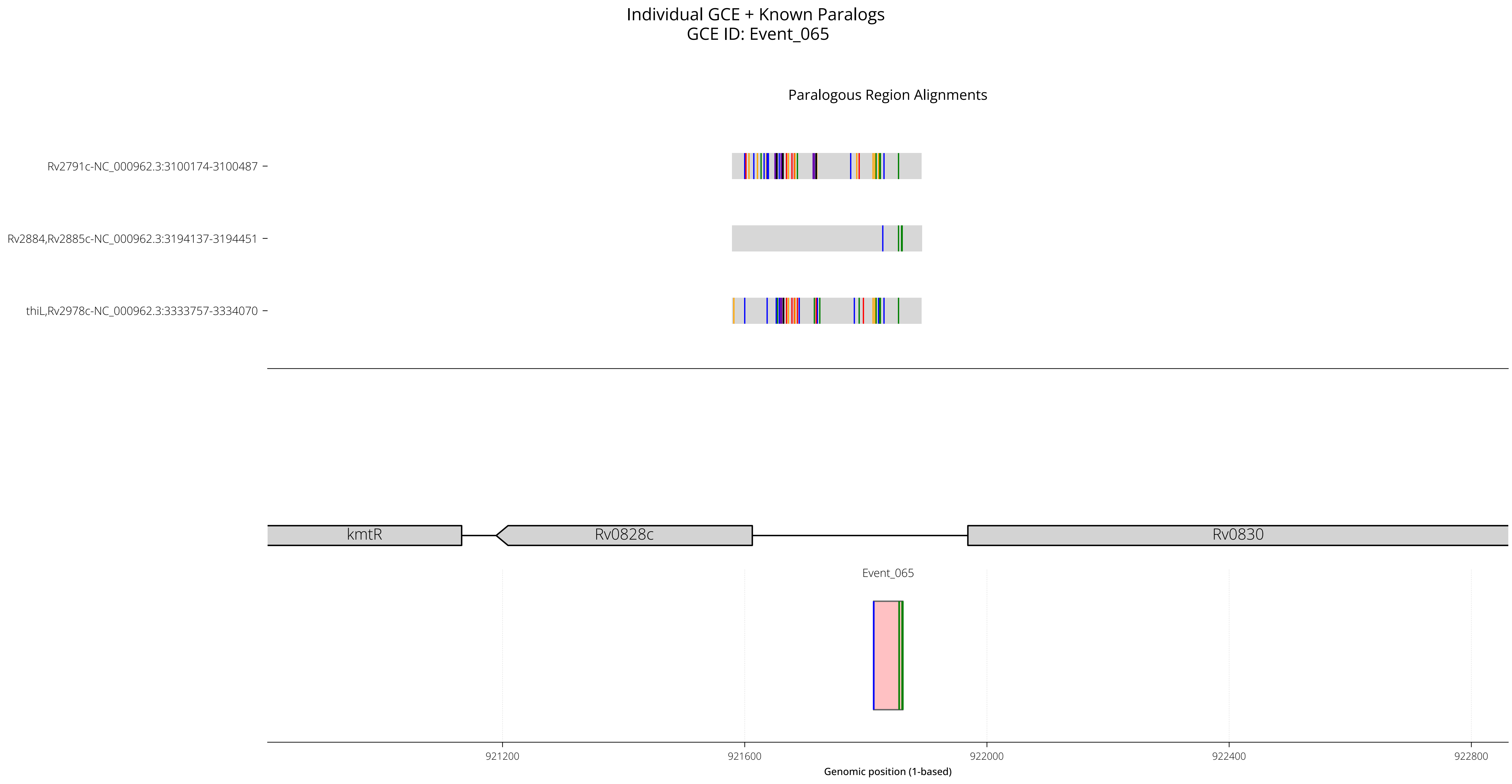

Individual GCE + Known Paralogs
GCE ID: Event_065
Paralogous Region Alignments
Rv2791c-NC_000962.3:3100174-3100487
Rv2884,Rv2885c-NC_000962.3:3194137-3194451
thiL,Rv2978c-NC_000962.3:3333757-3334070
kmtR
Rv0828c
Rv0830
Event_065
921200
921600
922000
922400
922800
Genomic position (1-based)

Circos Viz - GCE to Known Paralogs Locations
GCE ID: Event_065

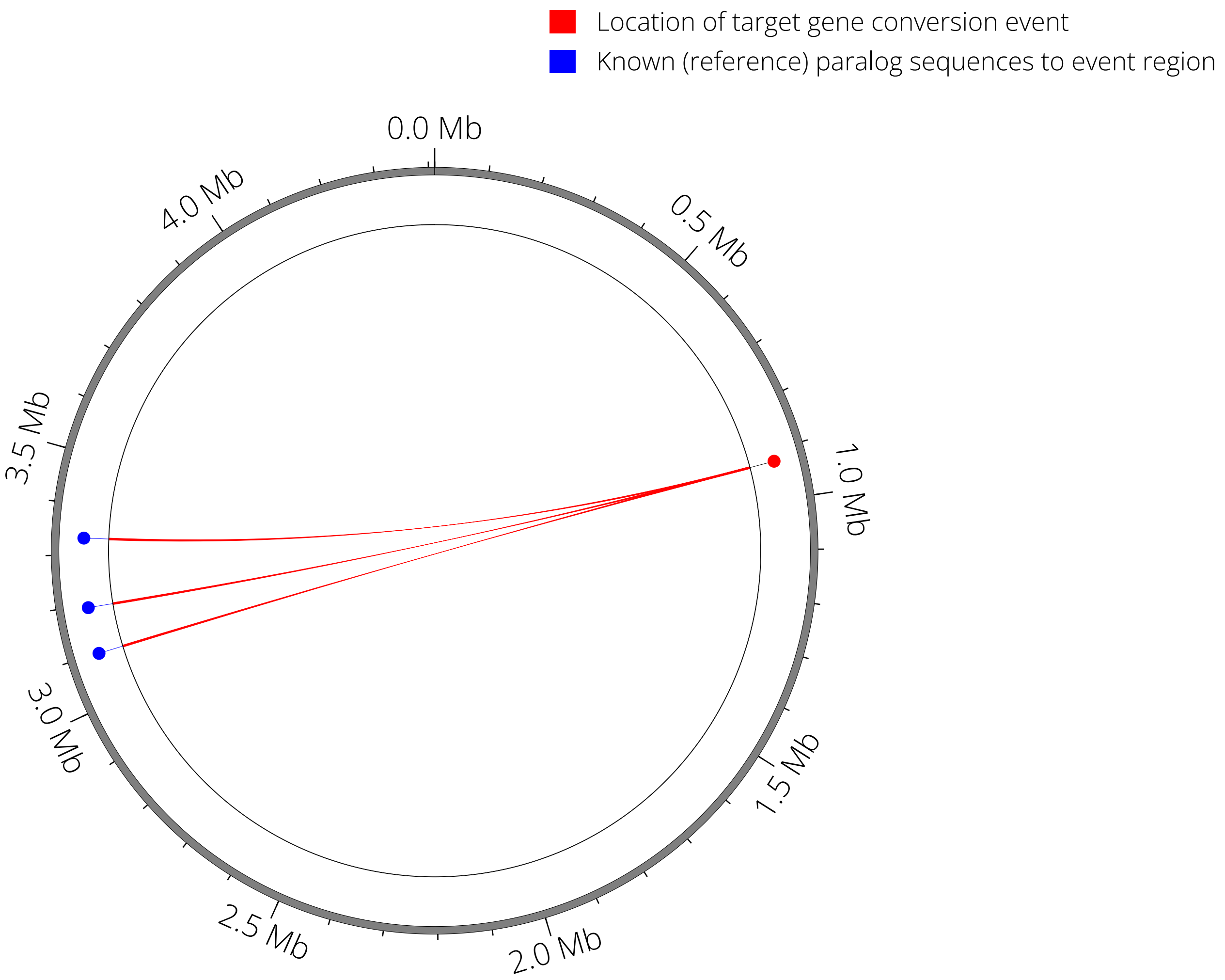

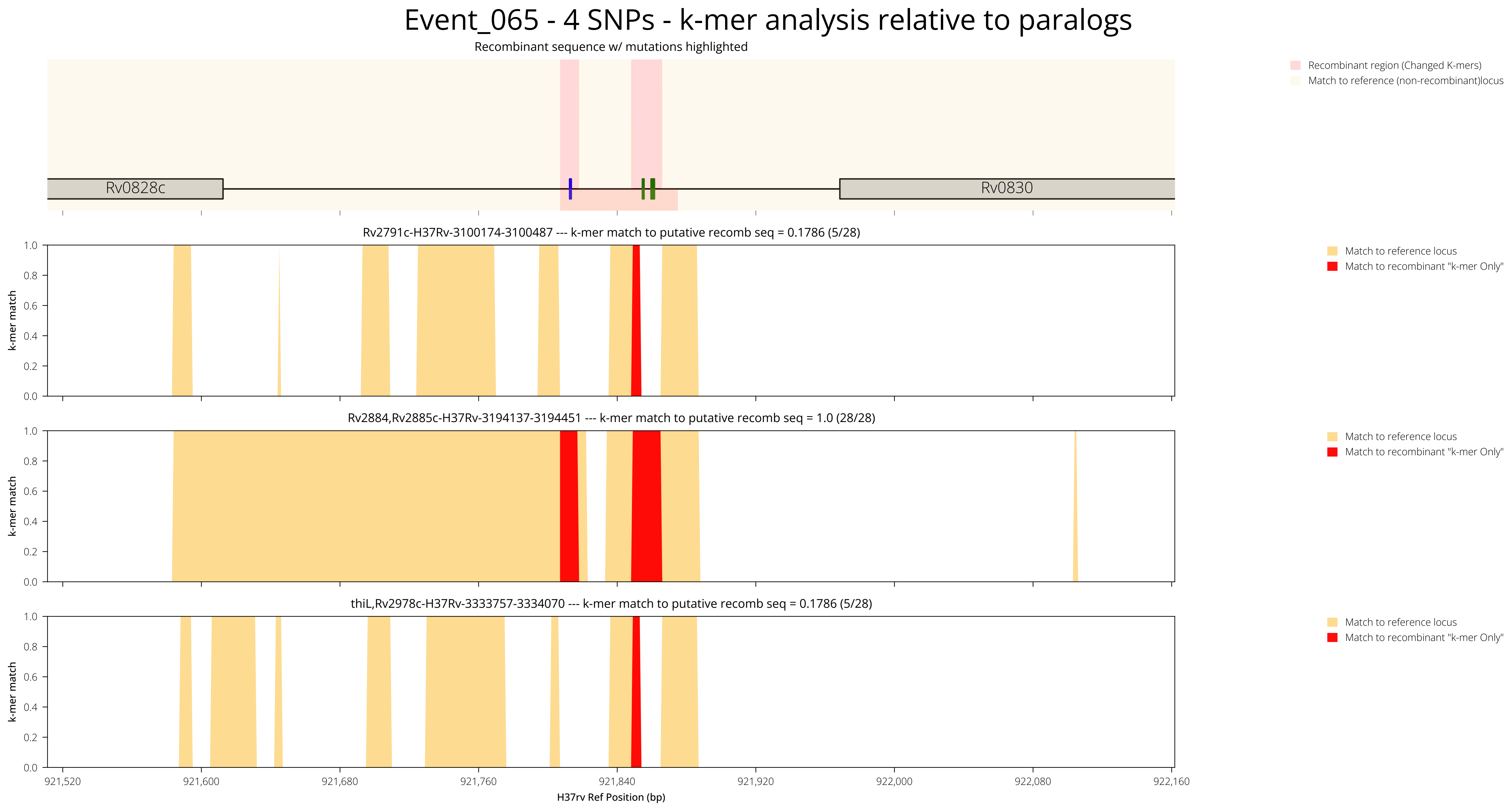
Event_065 - 4 SNPs - k-mer analysis relative to paralogs
Recombinant sequence w/ mutations highlighted
Recombinant region (Changed K-mers)
Match to reference (non-recombinant)locus
Rv0828c
Rv0830
Rv2791c-H37Rv-3100174-3100487 --- k-mer match to putative recomb seq = 0.1786 (5/28)
Match to reference locus
Match to recombinant "k-mer Only"
Rv2884,Rv2885c-H37Rv-3194137-3194451 --- k-mer match to putative recomb seq = 1.0 (28/28)
Match to reference locus
Match to recombinant "k-mer Only"
thiL,Rv2978c-H37Rv-3333757-3334070 --- k-mer match to putative recomb seq = 0.1786 (5/28)
Match to reference locus
Match to recombinant "k-mer Only"
k-mer match
1.0
0.8
0.6
0.4
0.2
0.0
921,520
921,600
921,680
921,760
921,840
921,920
922,000
922,080
922,160
H37rv Ref Position (bp)

Sub-phylogeny Viz + Paralog Alignments
GCE ID: Event_065

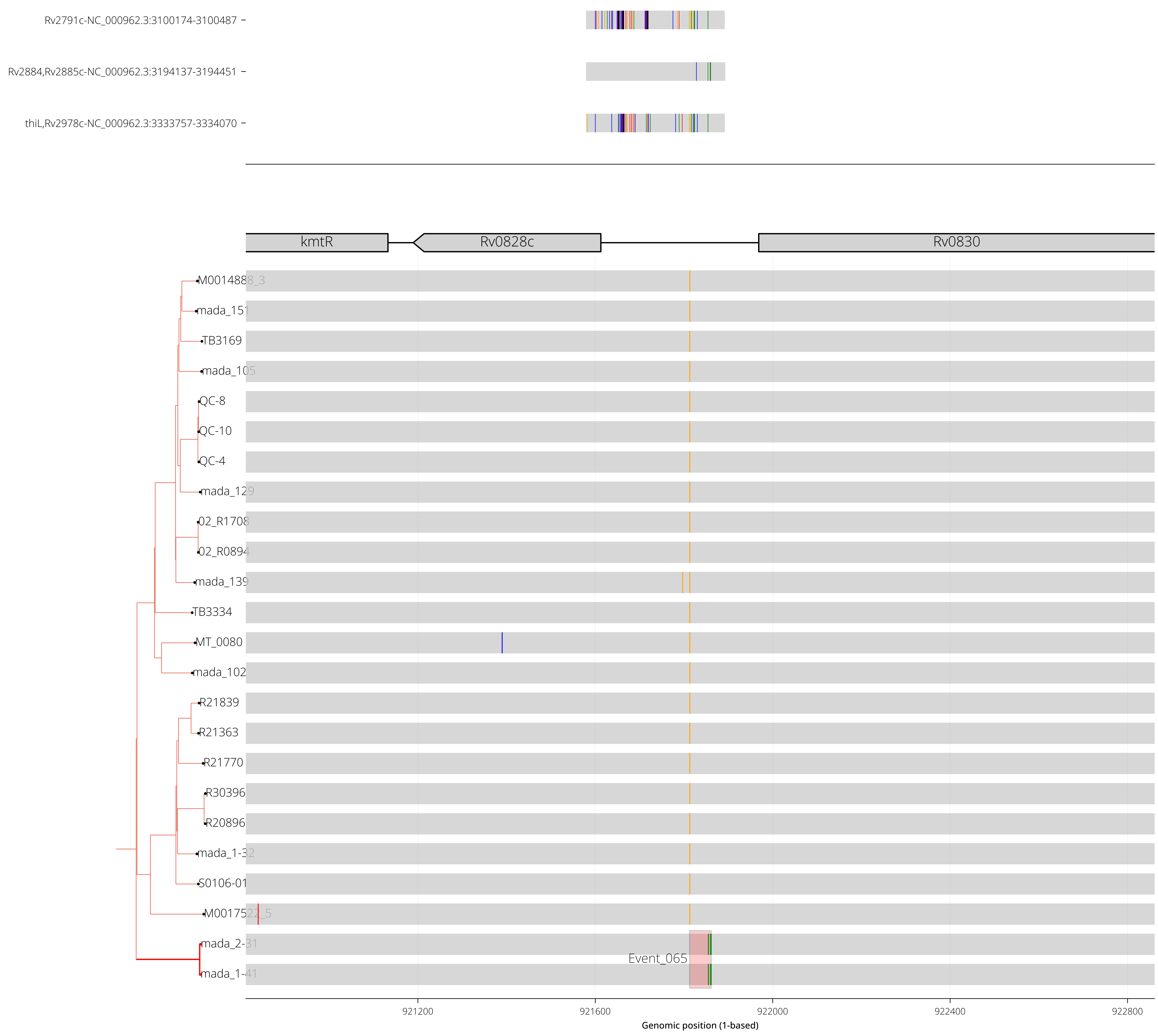

Full Phylogeny Viz + Paralog Alignments
GCE ID: Event_065

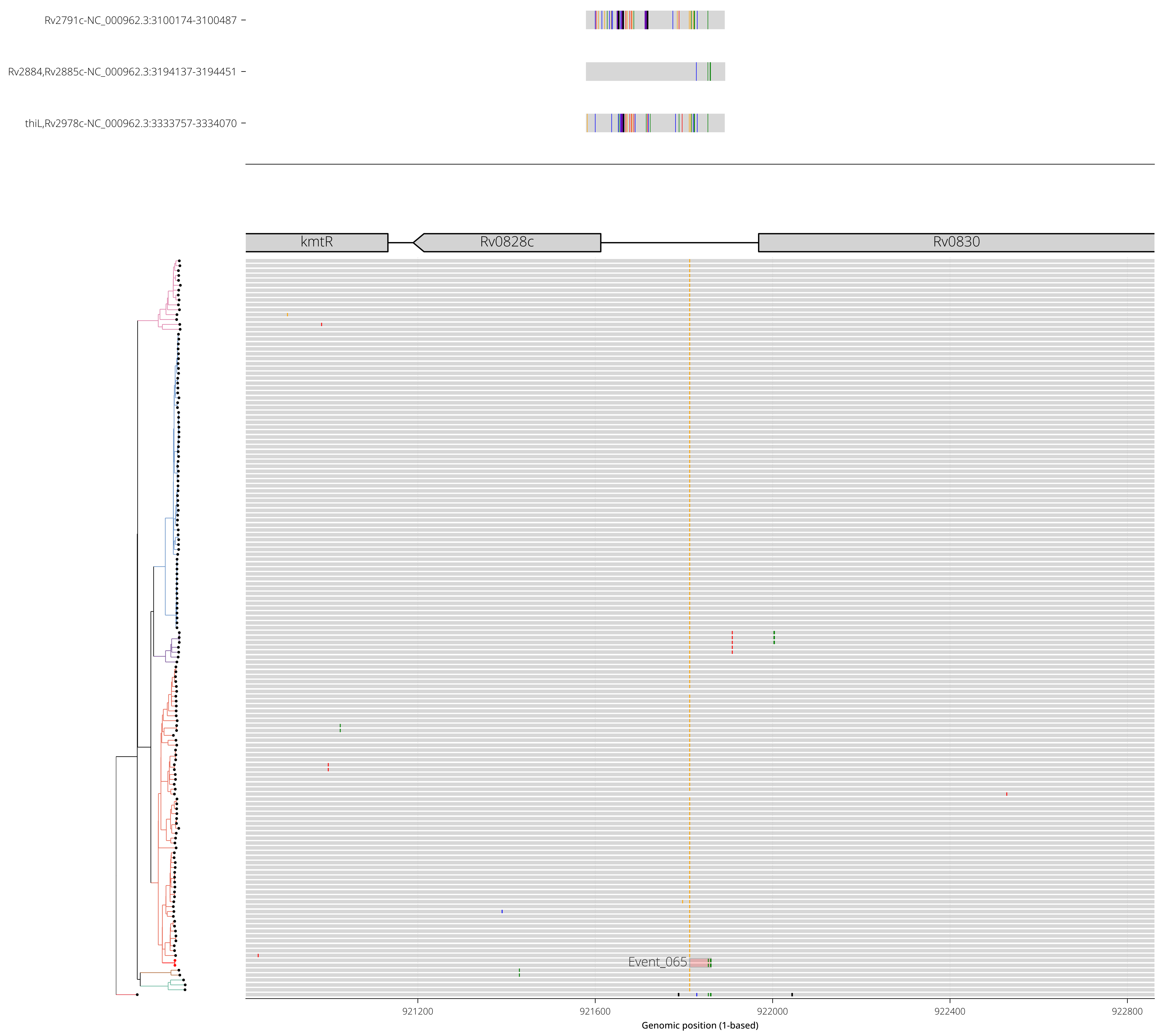

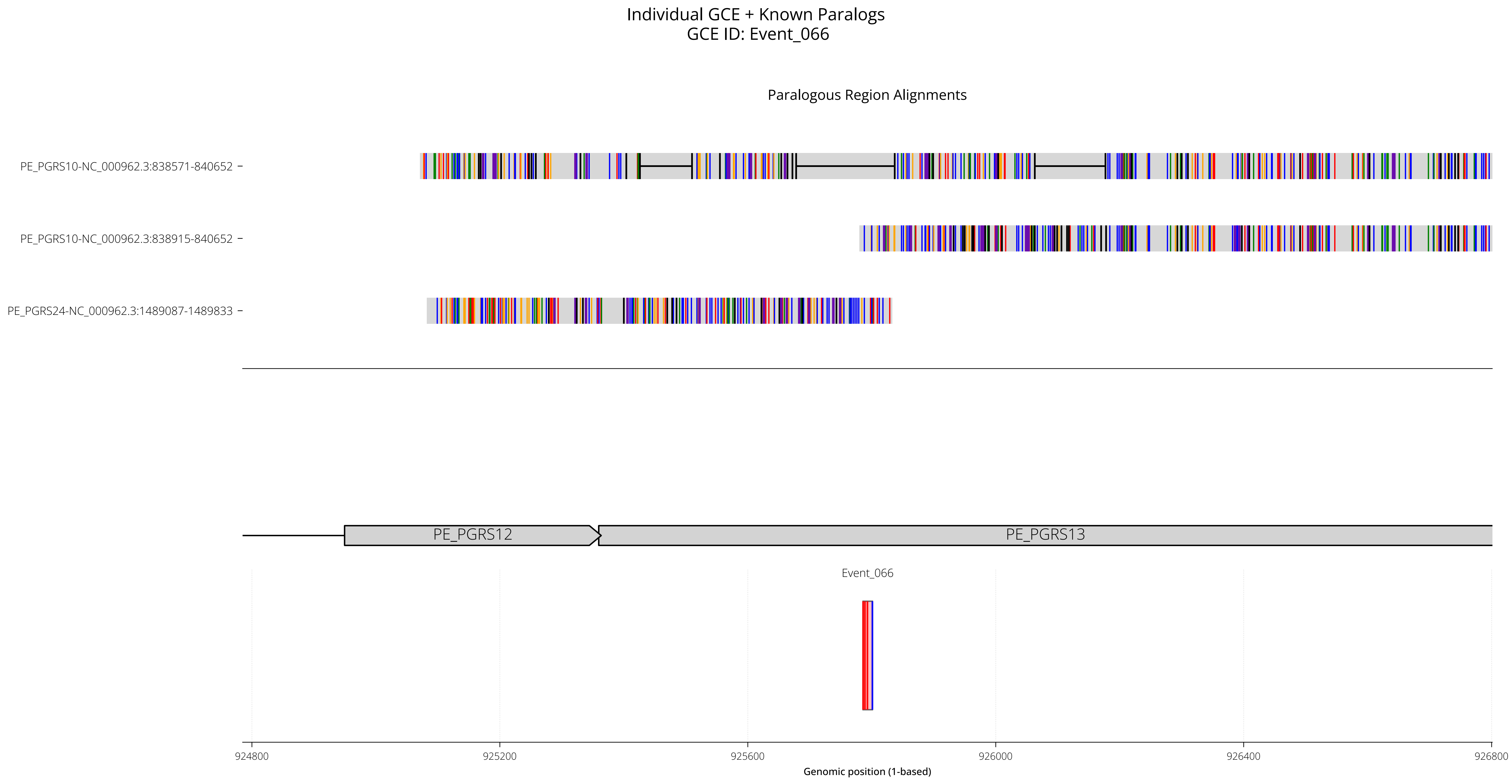
Individual GCE + Known Paralogs
GCE ID: Event_066
Paralogous Region Alignments
PE_PGRS10-NC_000962.3:838571-840652
PE_PGRS10-NC_000962.3:838915-840652
PE_PGRS24-NC_000962.3:1489087-1489833
PE_PGRS12
PE_PGRS13
Event_066
924800
925200
925600
926000
926400
926800
Genomic position (1-based)

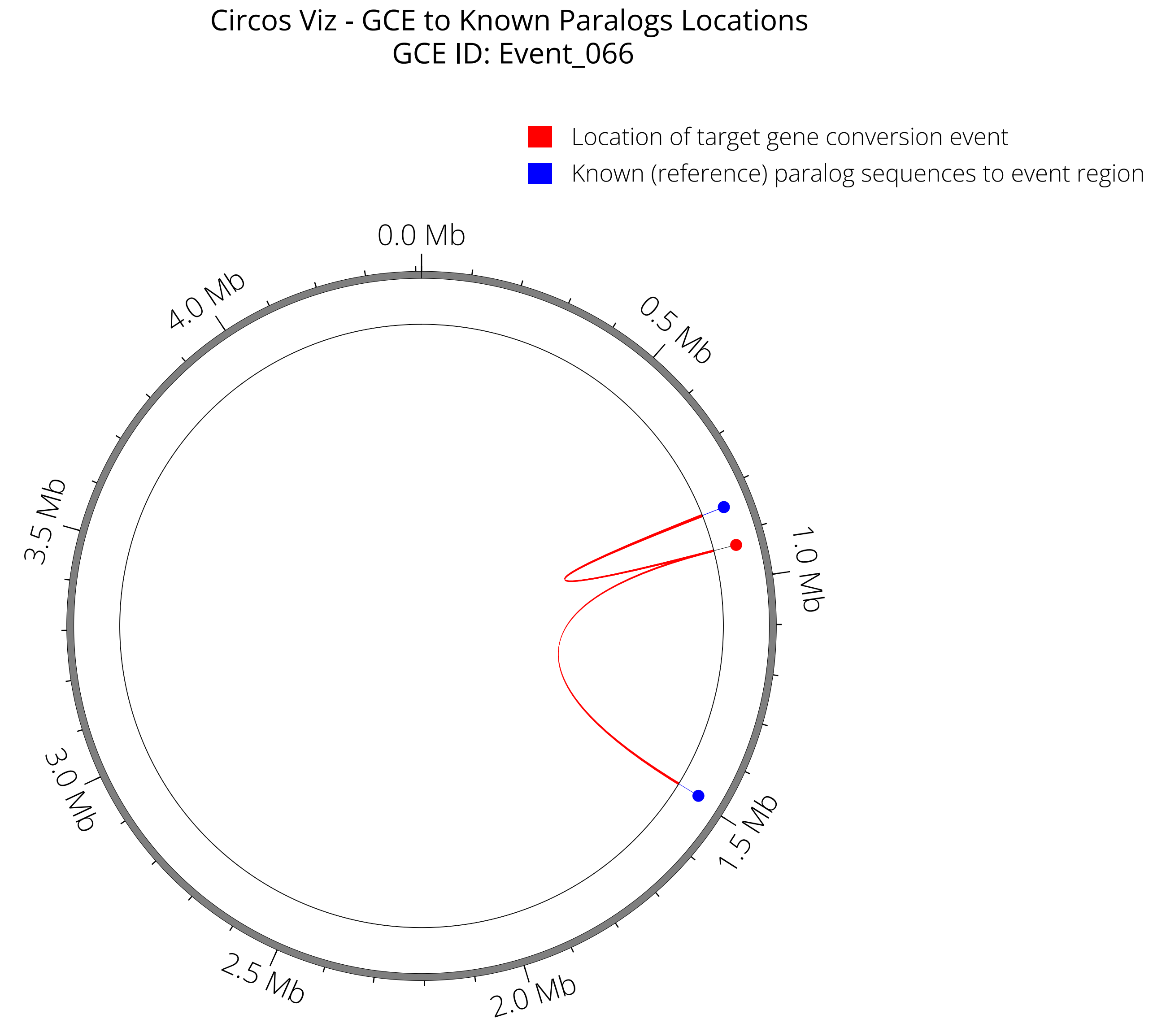
Circos Viz - GCE to Known Paralogs Locations
GCE ID: Event_066
Location of target gene conversion event
Known (reference) paralog sequences to event region
0.0 Mb
0.5 Mb
1.0 Mb
1.5 Mb
2.0 Mb
2.5 Mb
3.0 Mb
3.5 Mb
4.0 Mb

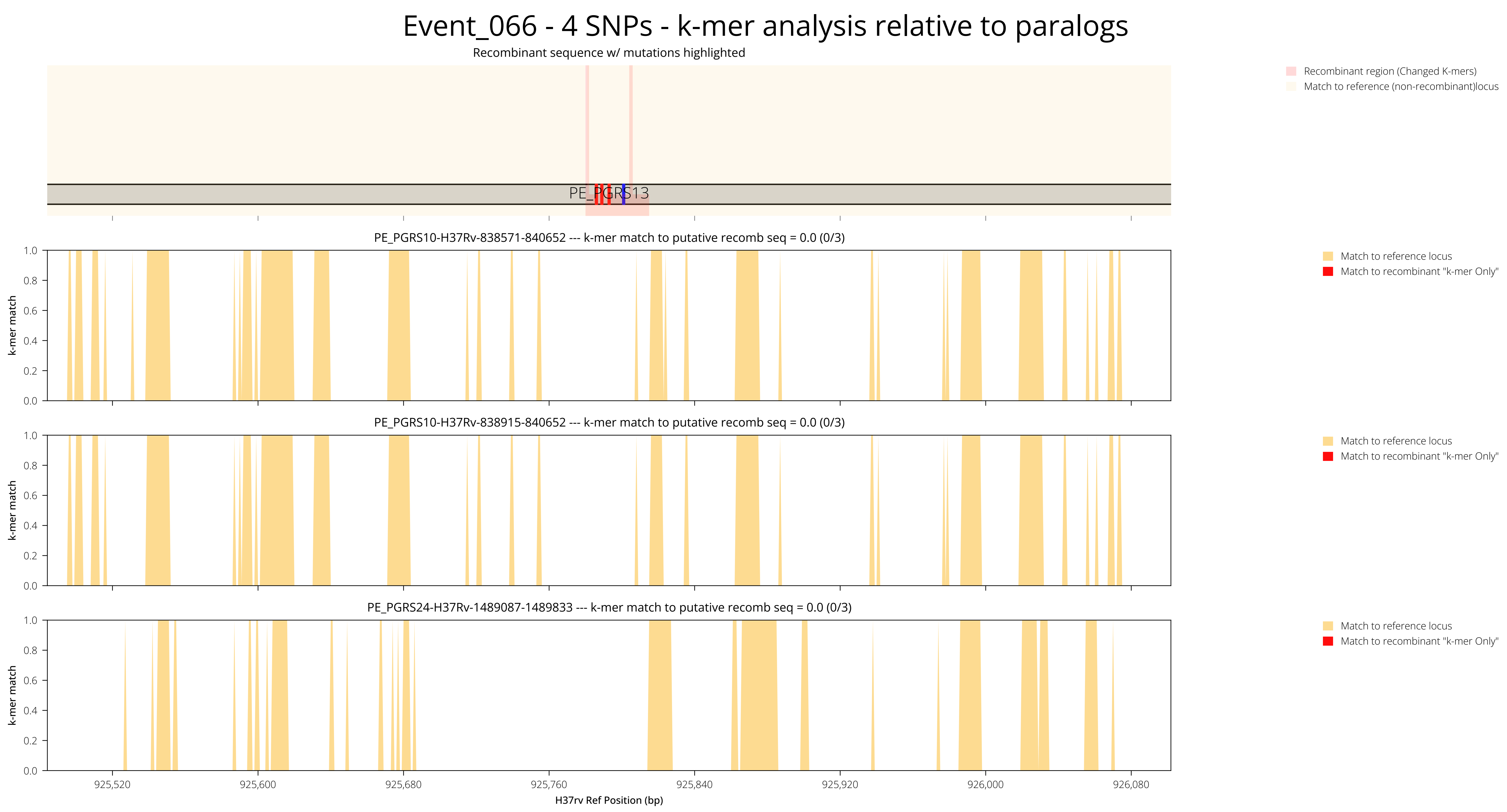
Event_066 - 4 SNPs - k-mer analysis relative to paralogs
Recombinant sequence w/ mutations highlighted
Recombinant region (Changed K-mers)
Match to reference (non-recombinant)locus
PE_PGRS13
PE_PGRS10-H37Rv-838571-840652 --- k-mer match to putative recomb seq = 0.0 (0/3)
PE_PGRS10-H37Rv-838915-840652 --- k-mer match to putative recomb seq = 0.0 (0/3)
PE_PGRS24-H37Rv-1489087-1489833 --- k-mer match to putative recomb seq = 0.0 (0/3)
Match to reference locus
Match to recombinant "k-mer Only"
k-mer match
1.0
0.8
0.6
0.4
0.2
0.0
925,520
925,600
925,680
925,760
925,840
925,920
926,000
926,080
H37rv Ref Position (bp)

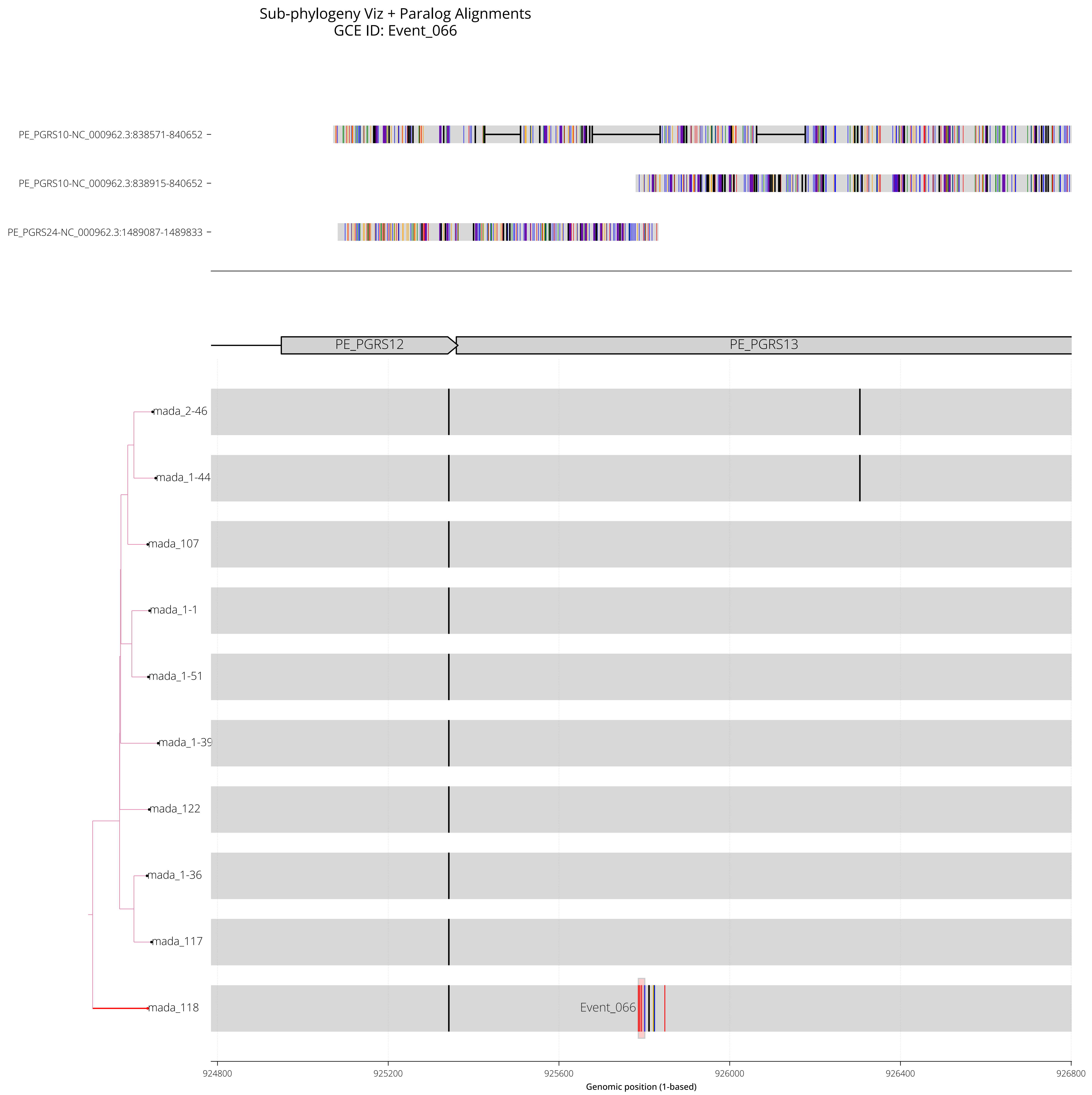
Sub-phylogeny Viz + Paralog Alignments
GCE ID: Event_066
PE_PGRS10-NC_000962.3:838571-840652
PE_PGRS10-NC_000962.3:838915-840652
PE_PGRS24-NC_000962.3:1489087-1489833
PE_PGRS12
PE_PGRS13
mada_2-46
mada_1-44
mada_107
mada_1-1
mada_1-51
mada_1-39
mada_122
mada_1-36
mada_117
mada_118
Event_066
924800
925200
925600
926000
926400
926800
Genomic position (1-based)

Full Phylogeny Viz + Paralog Alignments
GCE ID: Event_066

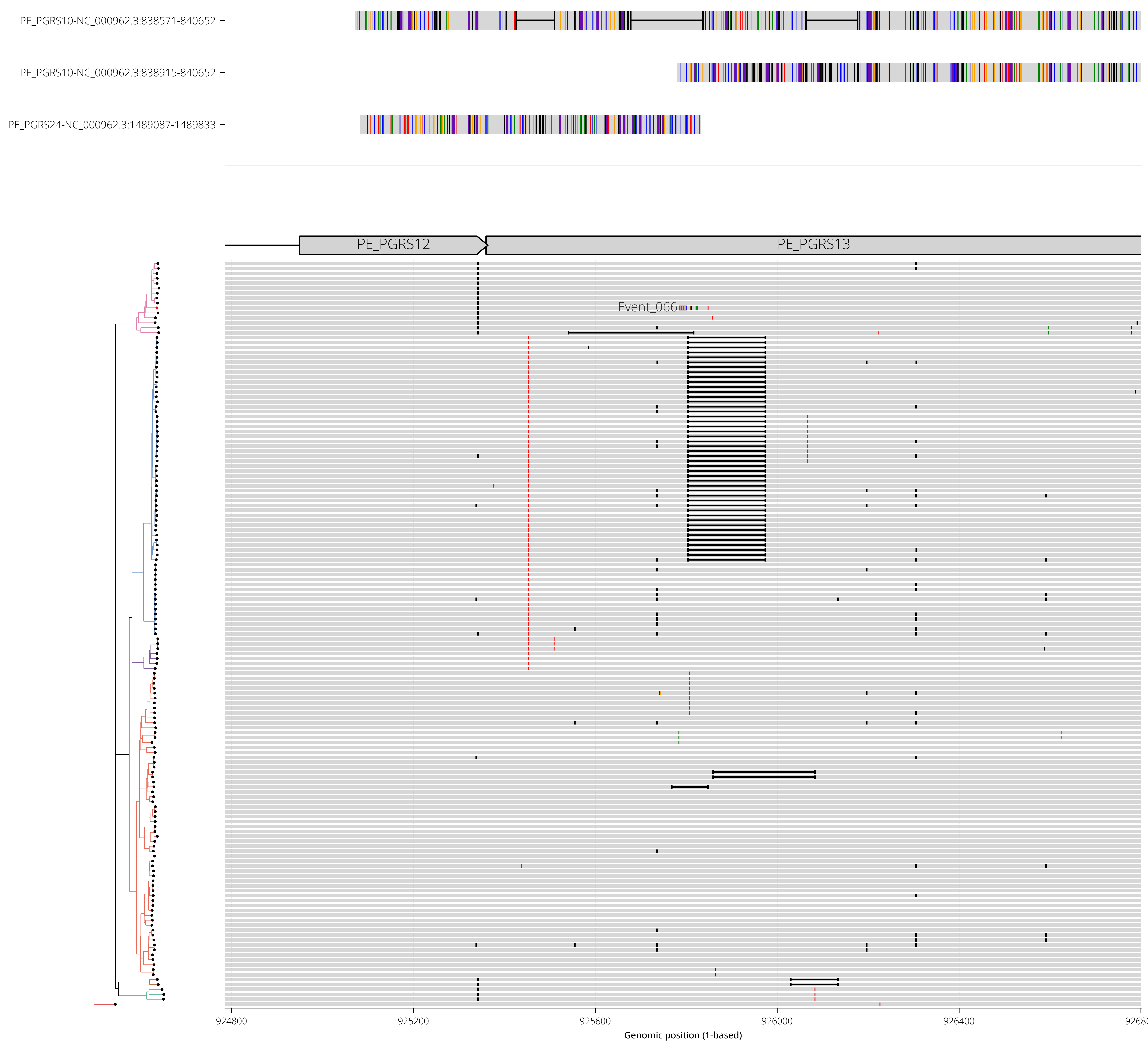

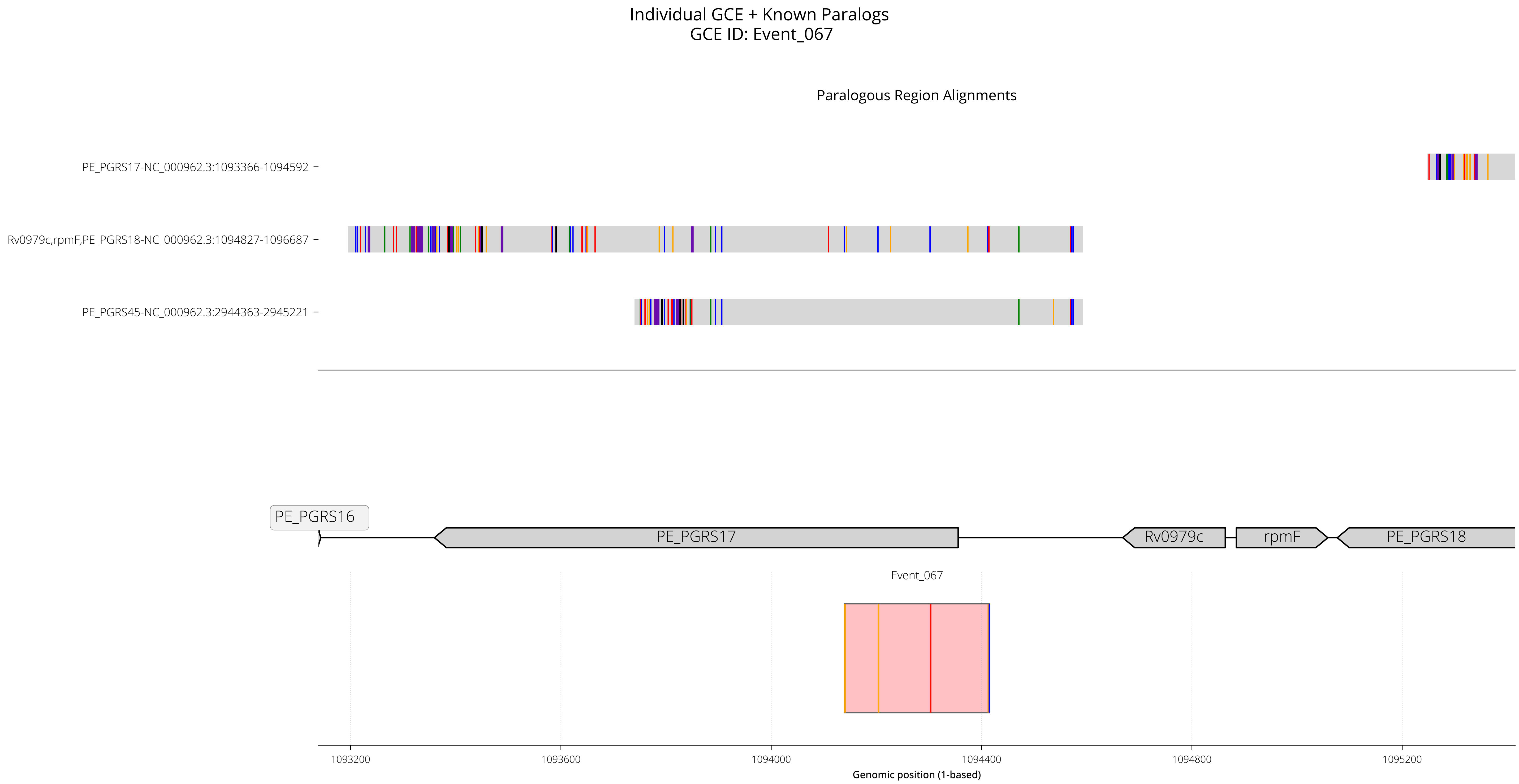
Individual GCE + Known Paralogs
GCE ID: Event_067
Paralogous Region Alignments
PE_PGRS17-NC_000962.3:1093366-1094592
Rv0979c,rpmF,PE_PGRS18-NC_000962.3:1094827-1096687
PE_PGRS45-NC_000962.3:2944363-2945221
PE_PGRS16
PE_PGRS17
Rv0979c
rpmF
PE_PGRS18
Event_067
1093200
1093600
1094000
1094400
1094800
1095200
Genomic position (1-based)

### Circos Viz - GCE to Known Paralogs Locations
GCE ID: Event_067

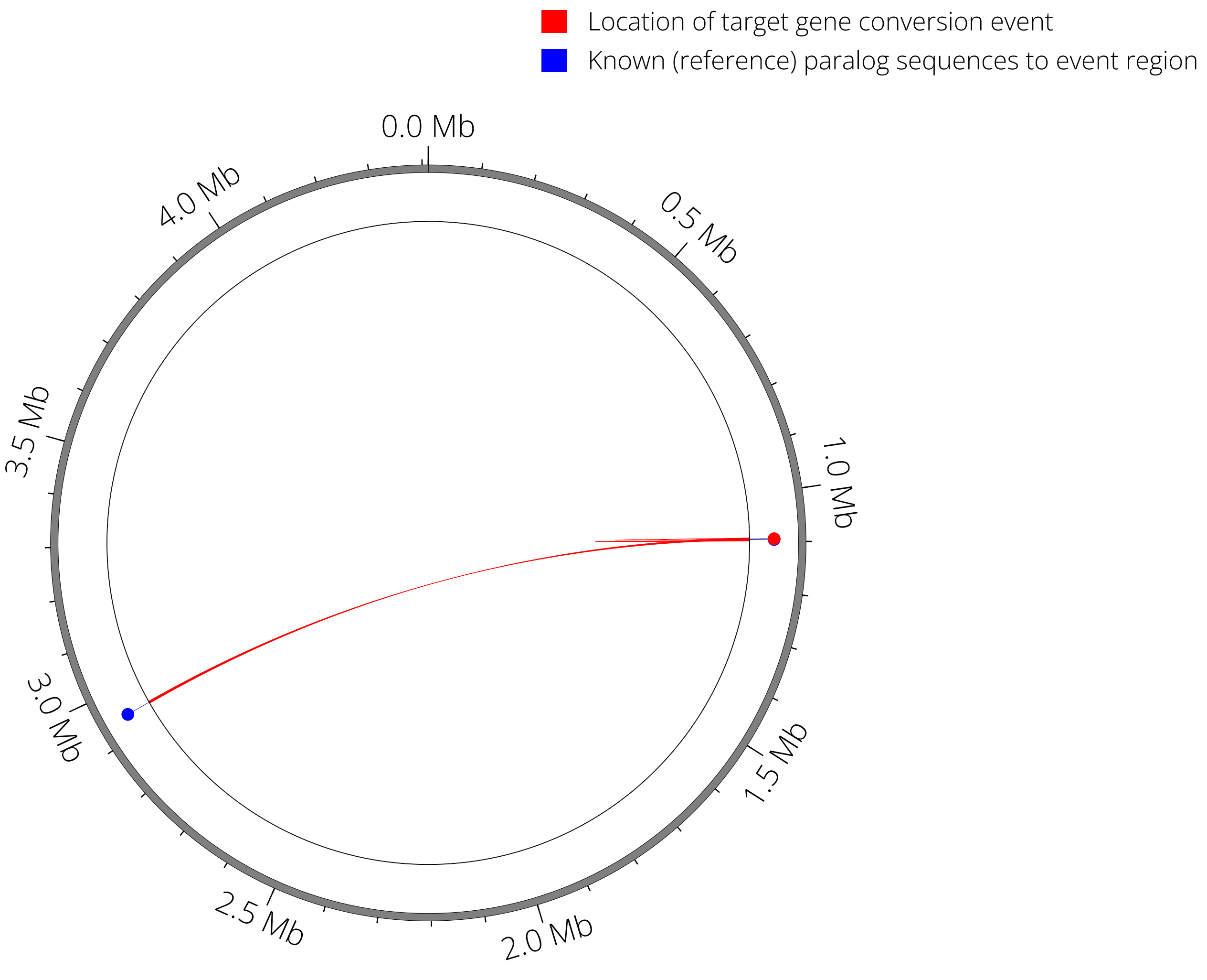

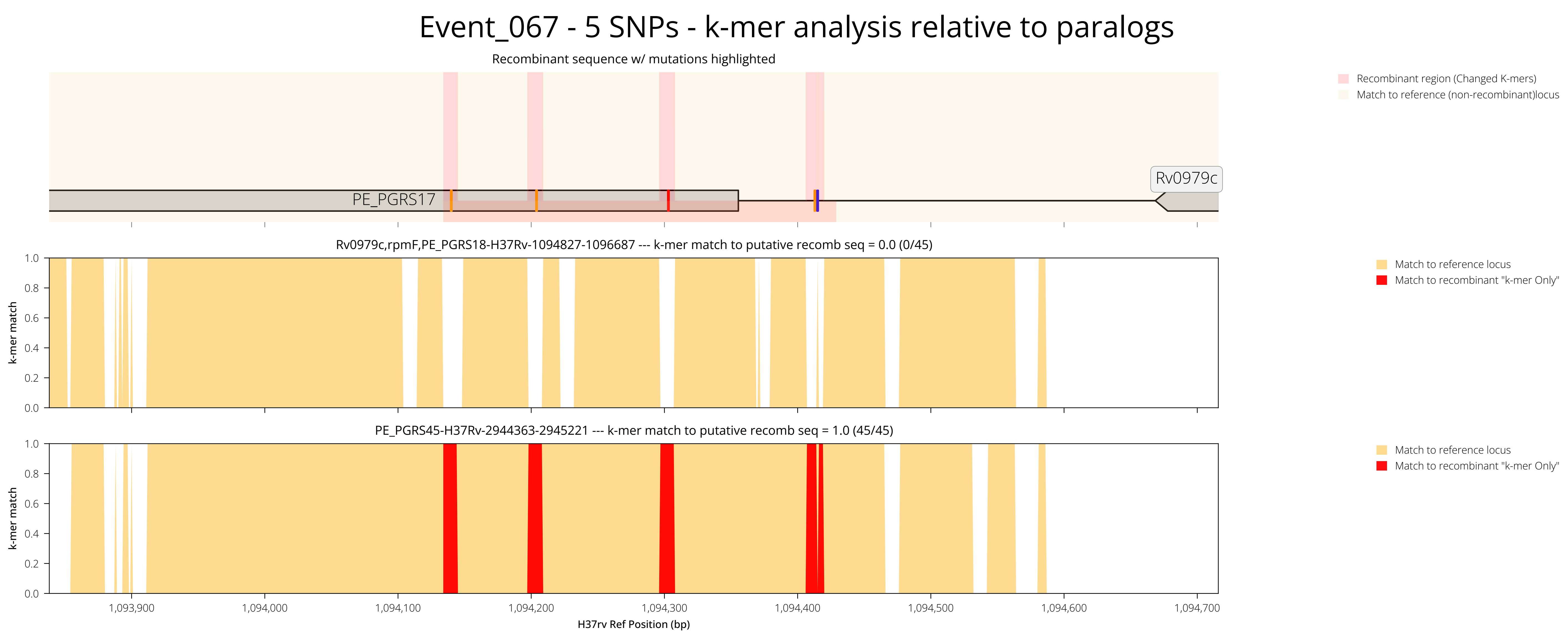

Event_067 - 5 SNPs - k-mer analysis relative to paralogs
Recombinant sequence w/ mutations highlighted
Recombinant region (Changed K-mers)
Match to reference (non-recombinant)locus
PE_PGRS17
Rv0979c
Rv0979c,rpmF,PE_PGRS18-H37Rv-1094827-1096687 --- k-mer match to putative recomb seq = 0.0 (0/45)
k-mer match
Match to reference locus
Match to recombinant "k-mer Only"
PE_PGRS45-H37Rv-2944363-2945221 --- k-mer match to putative recomb seq = 1.0 (45/45)
k-mer match
Match to reference locus
Match to recombinant "k-mer Only"
1,093,900
1,094,000
1,094,100
1,094,200
1,094,300
1,094,400
1,094,500
1,094,600
1,094,700
H37rv Ref Position (bp)

Sub-phylogeny Viz + Paralog Alignments
GCE ID: Event_067

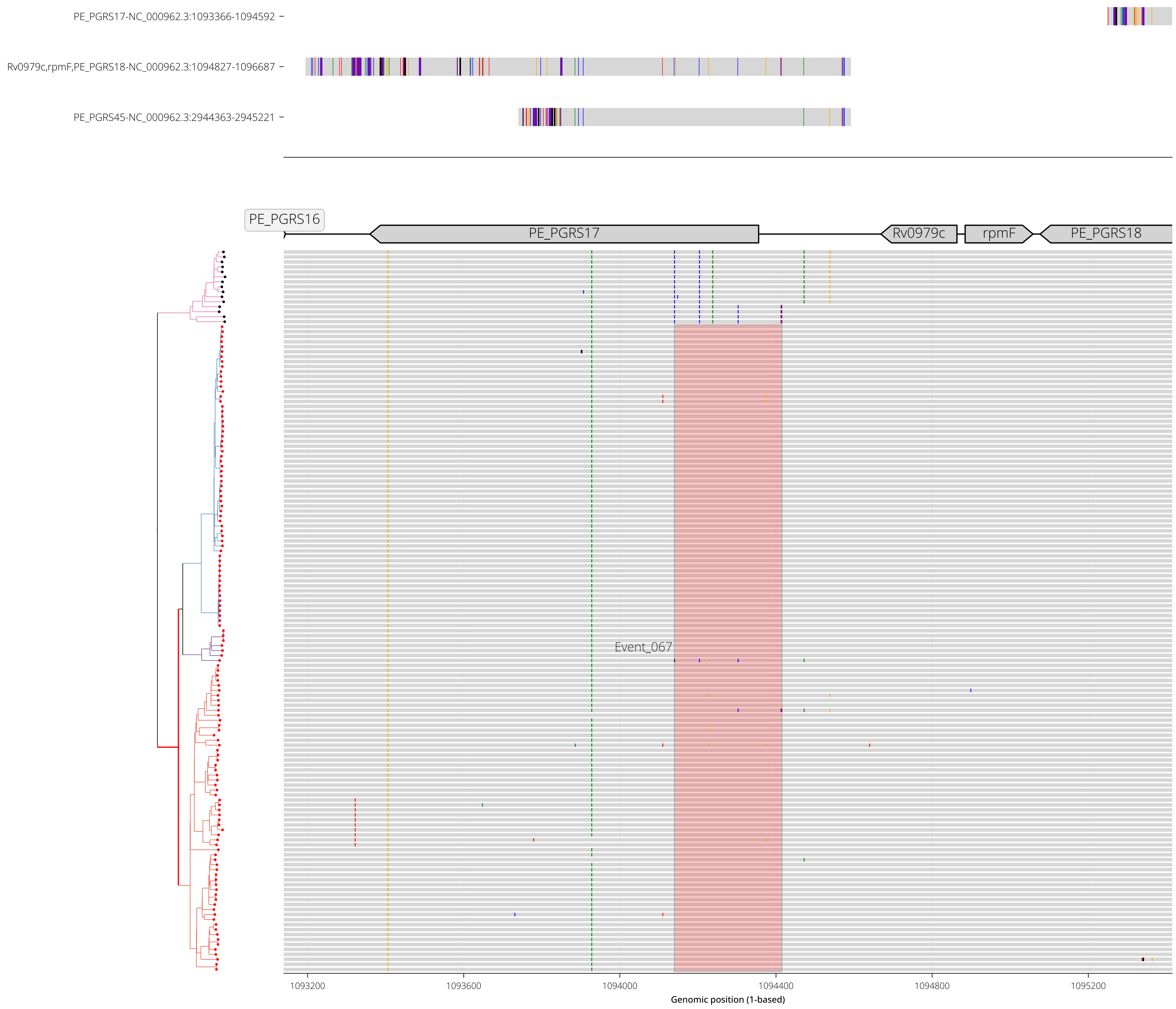

Full Phylogeny Viz + Paralog Alignments
GCE ID: Event_067

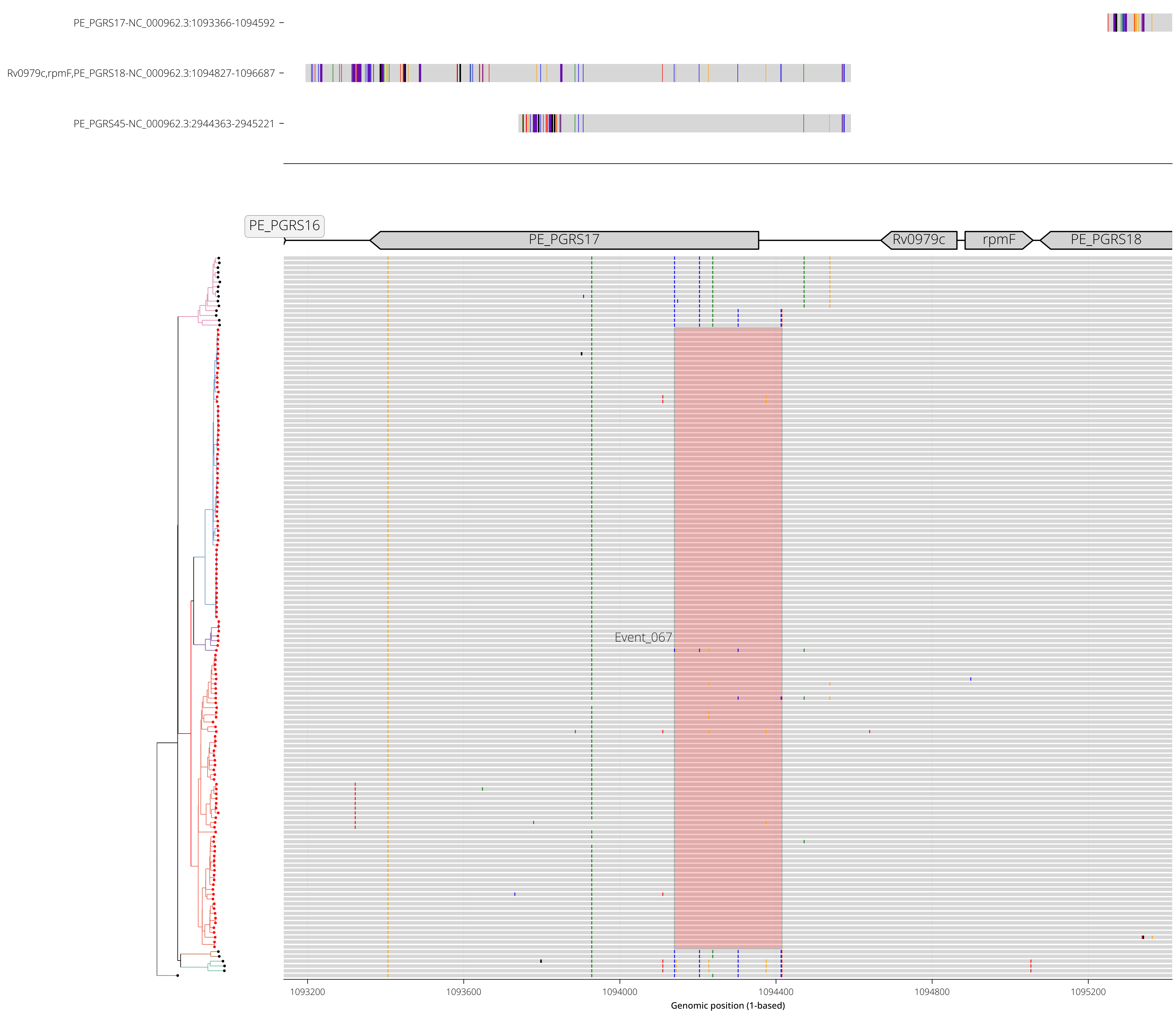

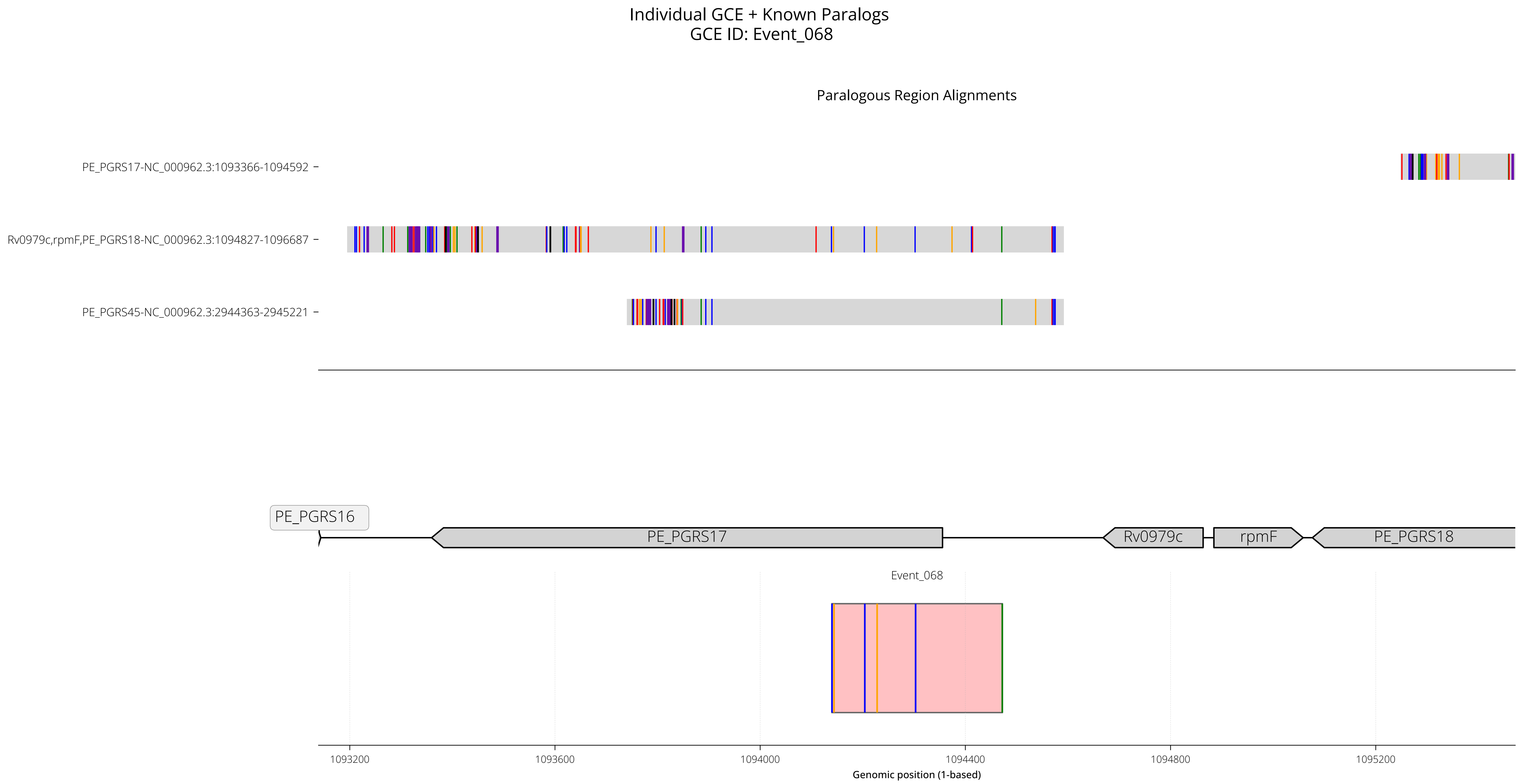

Individual GCE + Known Paralogs
GCE ID: Event_068
Paralogous Region Alignments
PE_PGRS17-NC_000962.3:1093366-1094592
Rv0979c,rpmF,PE_PGRS18-NC_000962.3:1094827-1096687
PE_PGRS45-NC_000962.3:2944363-2945221
PE_PGRS16
PE_PGRS17
Rv0979c
rpmF
PE_PGRS18
Event_068
1093200
1093600
1094000
1094400
1094800
1095200
Genomic position (1-based)

### Circos Viz - GCE to Known Paralogs Locations
GCE ID: Event_068

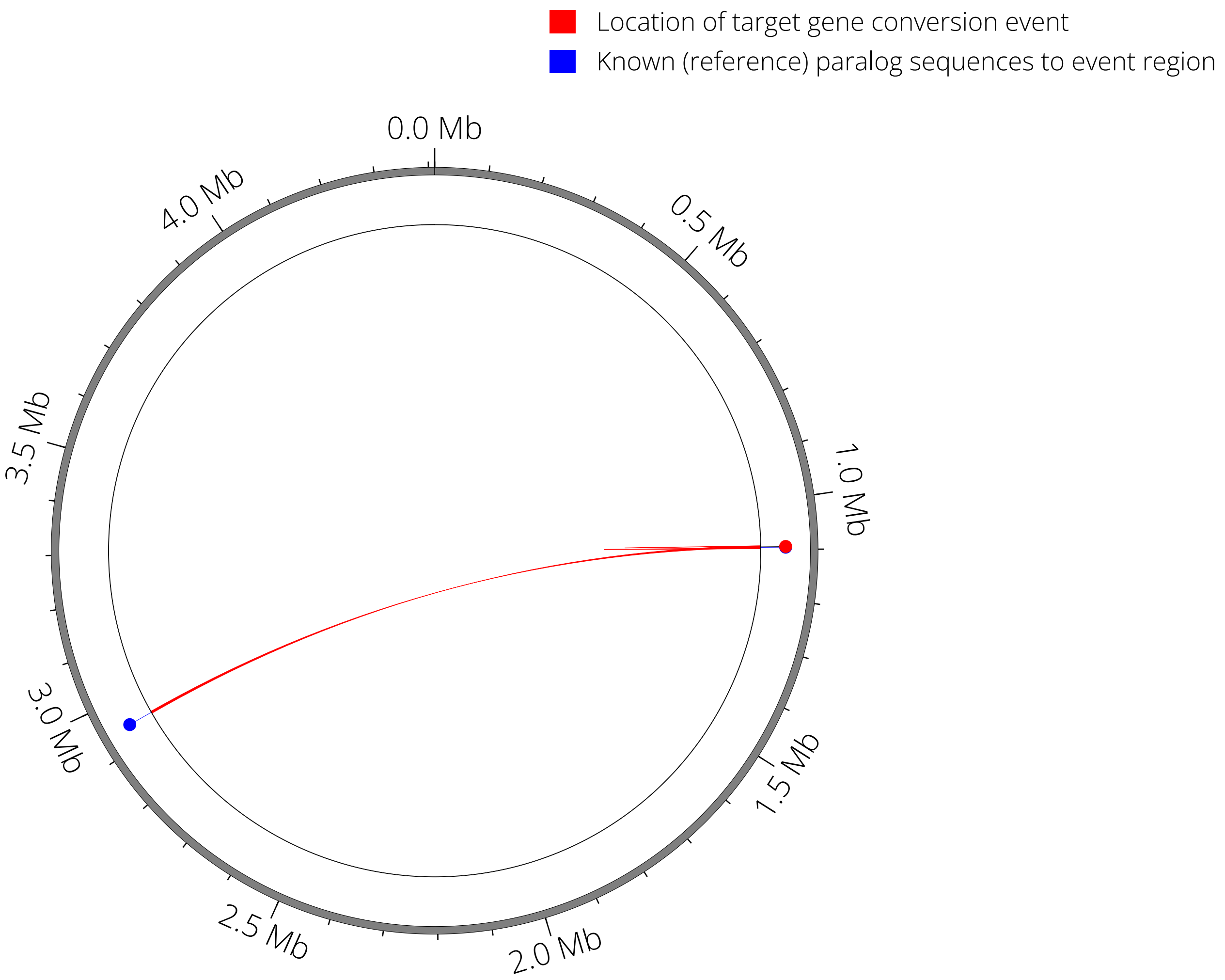

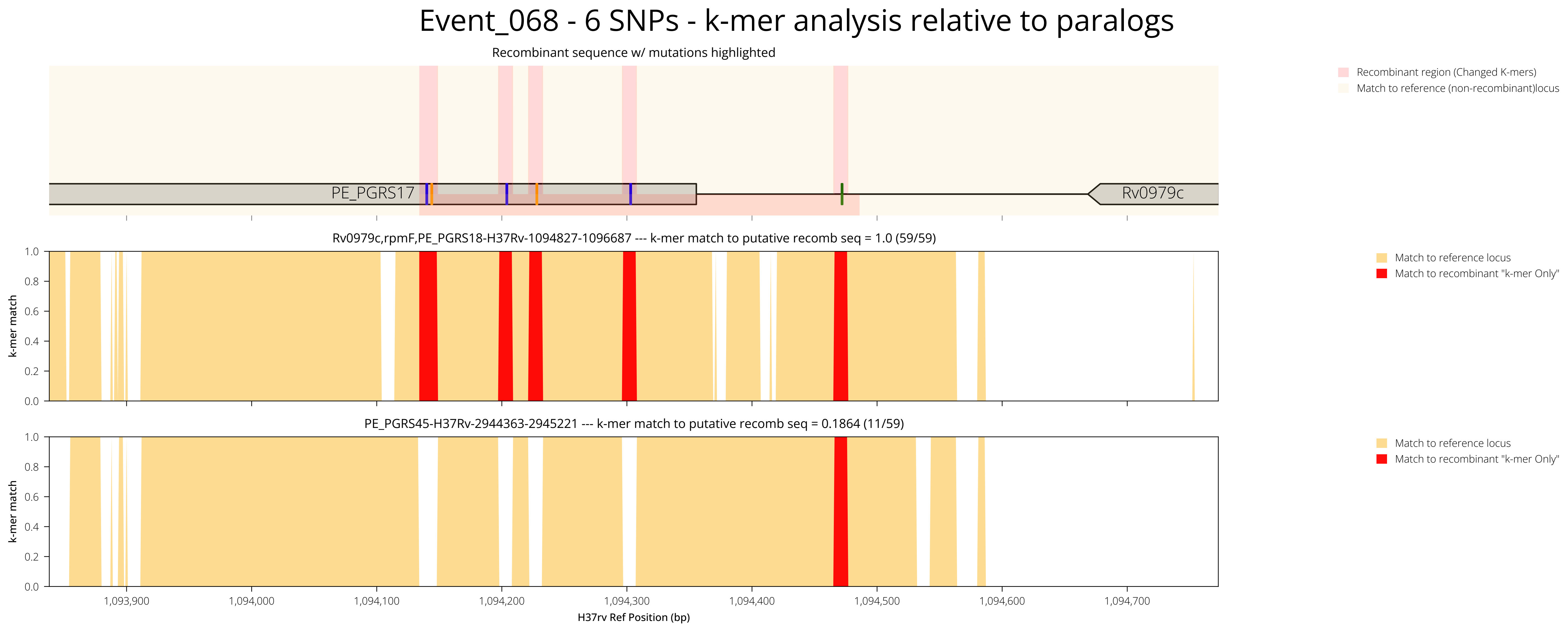

Event_068 - 6 SNPs - k-mer analysis relative to paralogs
Recombinant sequence w/ mutations highlighted
Recombinant region (Changed K-mers)
Match to reference (non-recombinant)locus
PE_PGRS17
Rv0979c
Rv0979c,rpmF,PE_PGRS18-H37Rv-1094827-1096687 --- k-mer match to putative recomb seq = 1.0 (59/59)
Match to reference locus
Match to recombinant "k-mer Only"
k-mer match
PE_PGRS45-H37Rv-2944363-2945221 --- k-mer match to putative recomb seq = 0.1864 (11/59)
Match to reference locus
Match to recombinant "k-mer Only"
k-mer match
1,093,900
1,094,000
1,094,100
1,094,200
1,094,300
1,094,400
1,094,500
1,094,600
1,094,700
H37rv Ref Position (bp)

Sub-phylogeny Viz + Paralog Alignments
GCE ID: Event_068

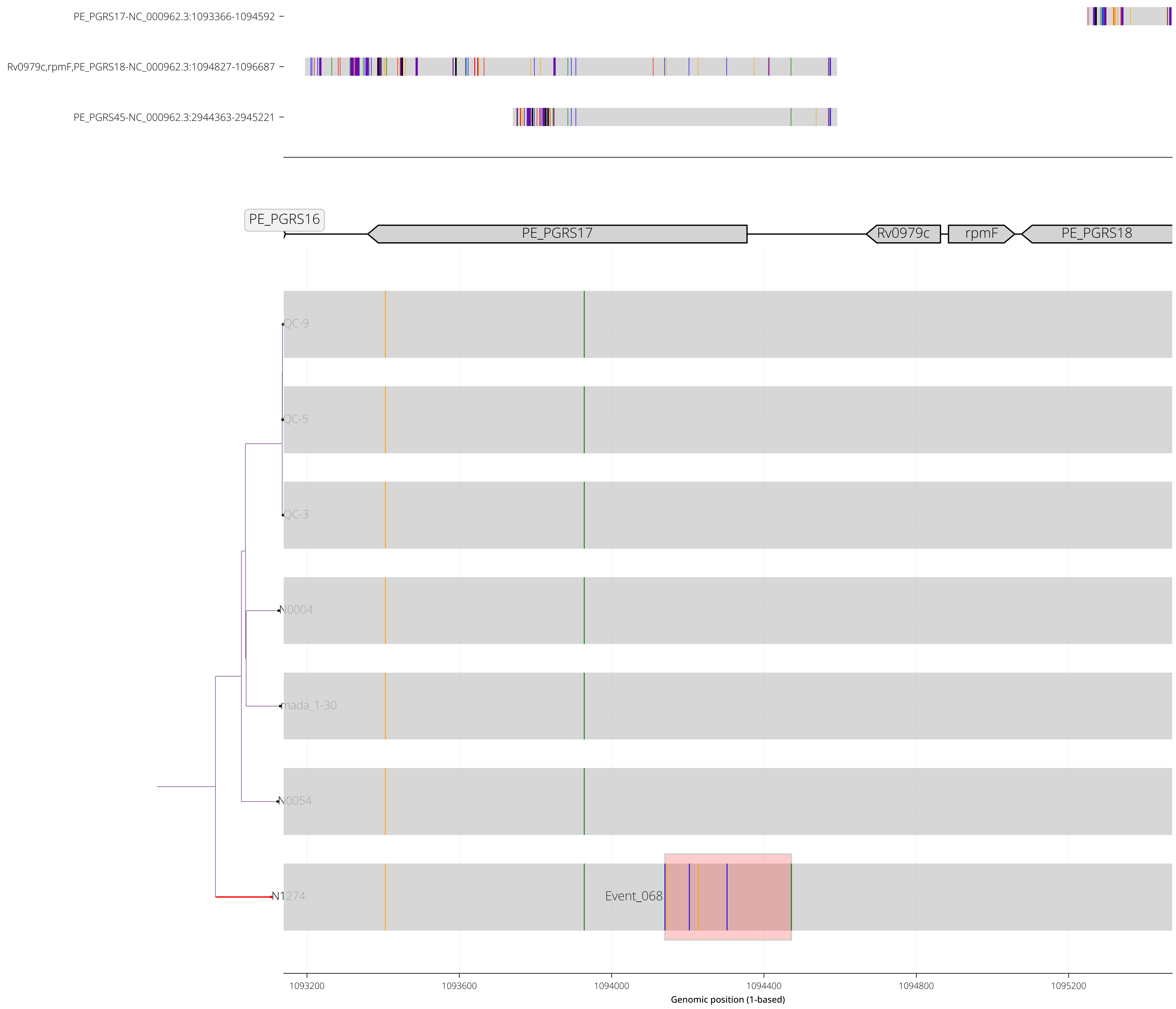

Full Phylogeny Viz + Paralog Alignments
GCE ID: Event_068

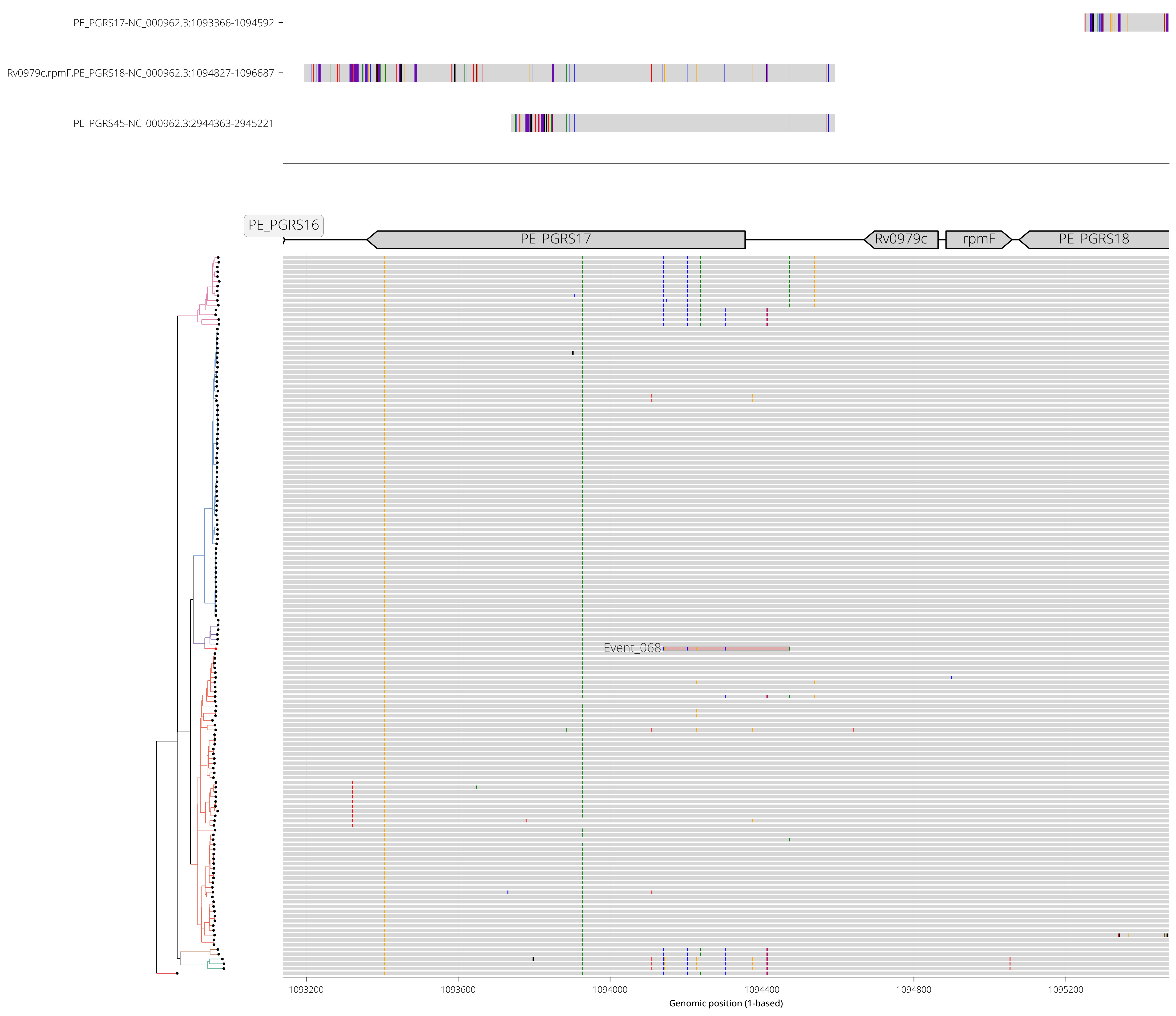

Individual GCE + Known Paralogs
GCE ID: Event_069

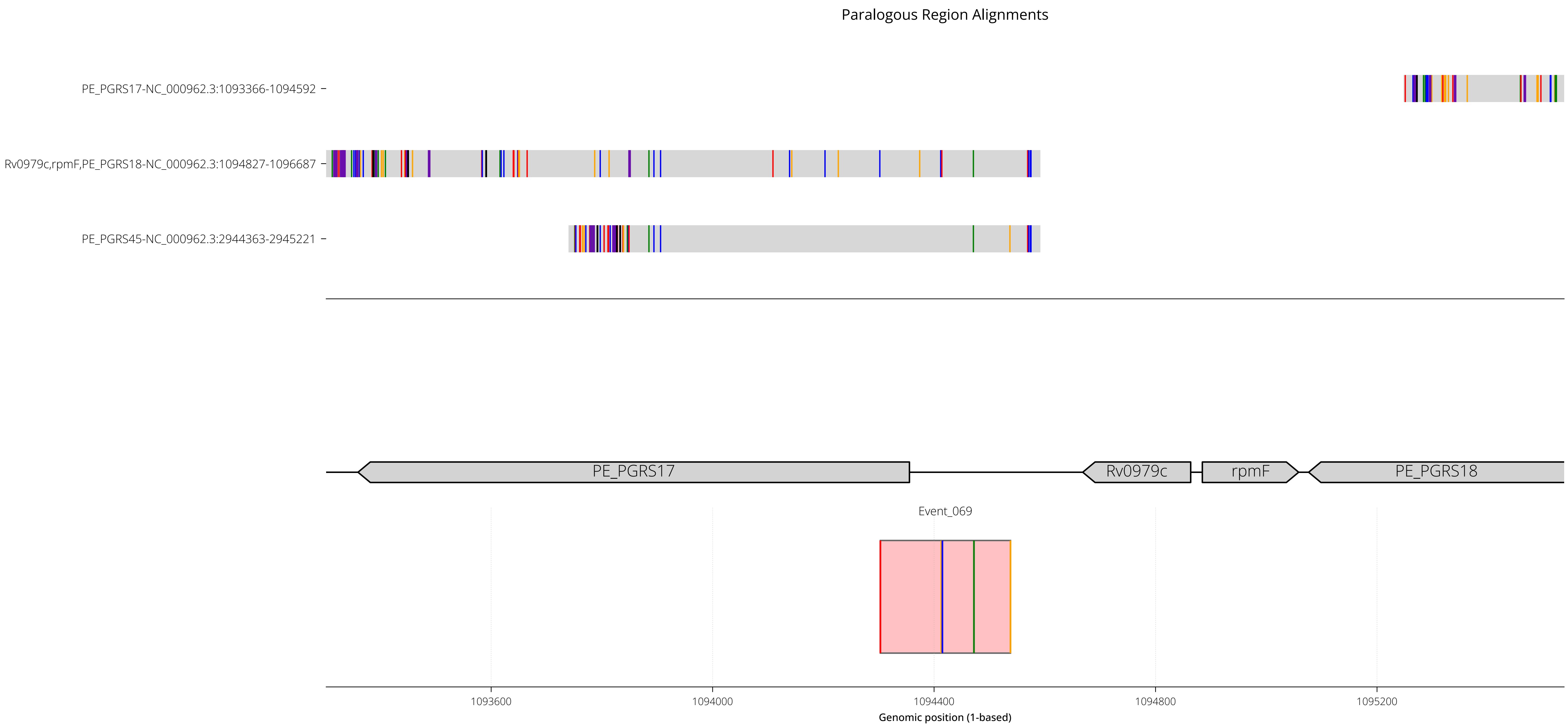

Circos Viz - GCE to Known Paralogs Locations
GCE ID: Event_069

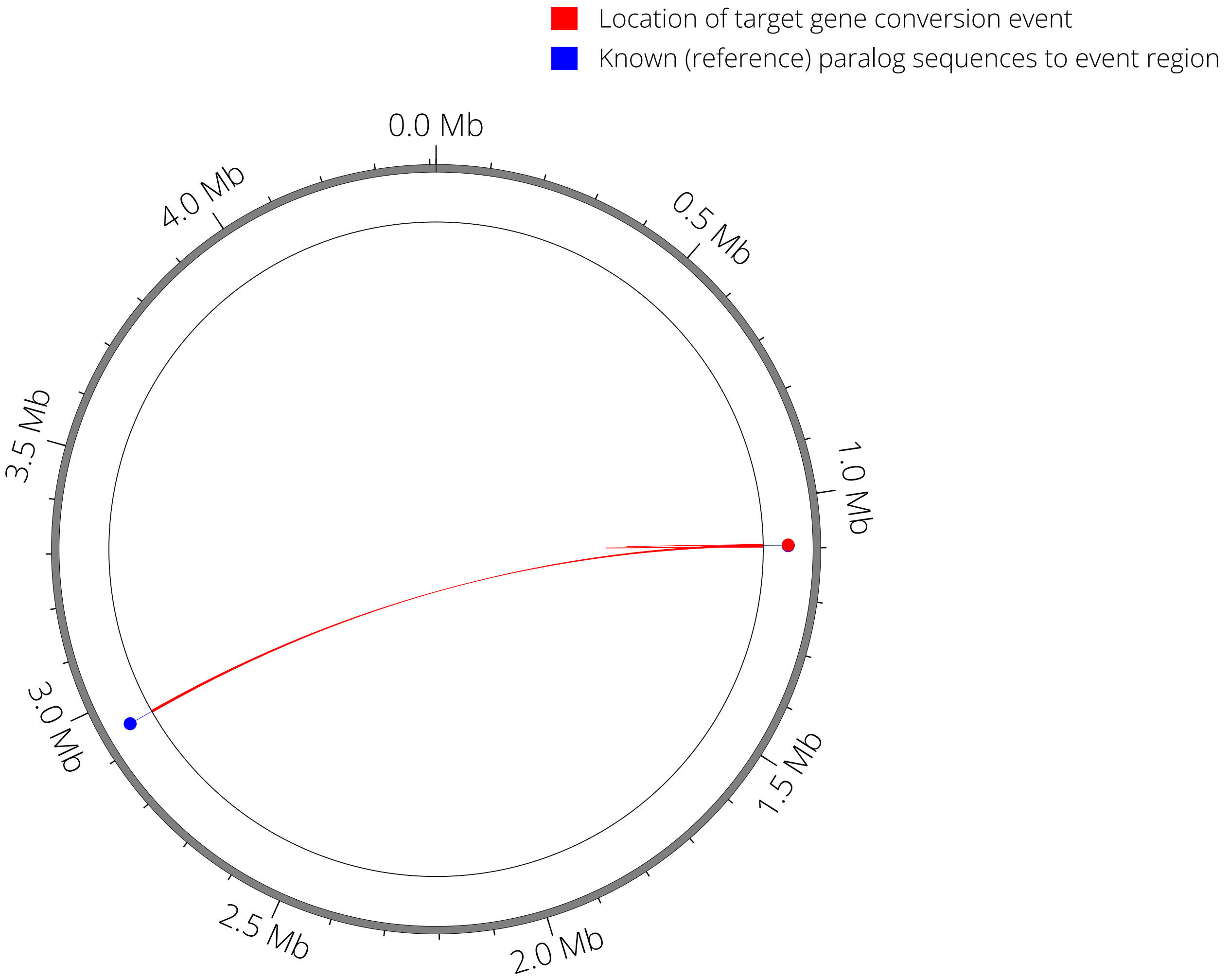

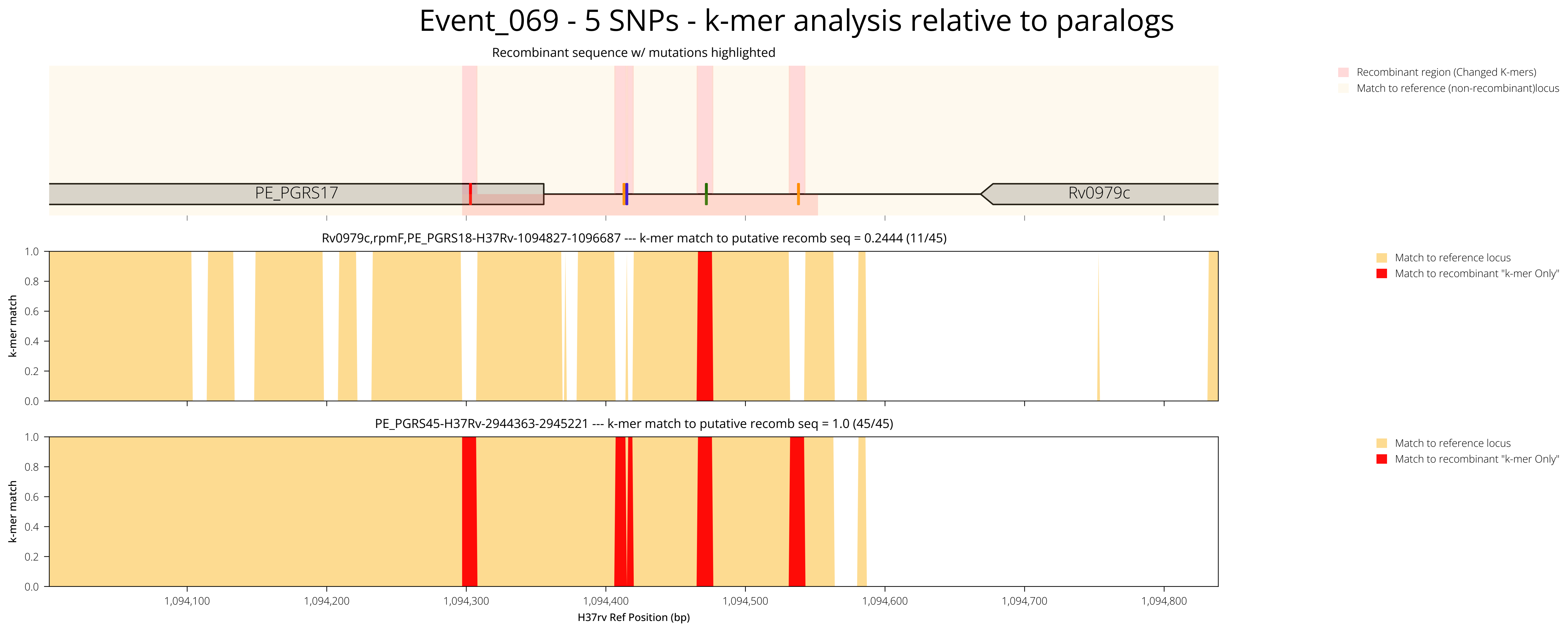

Event_069 - 5 SNPs - k-mer analysis relative to paralogs
Recombinant sequence w/ mutations highlighted
Recombinant region (Changed K-mers)
Match to reference (non-recombinant)locus
PE_PGRS17
Rv0979c
Rv0979c,rpmF,PE_PGRS18-H37Rv-1094827-1096687 --- k-mer match to putative recomb seq = 0.2444 (11/45)
k-mer match
Match to reference locus
Match to recombinant "k-mer Only"
PE_PGRS45-H37Rv-2944363-2945221 --- k-mer match to putative recomb seq = 1.0 (45/45)
k-mer match
Match to reference locus
Match to recombinant "k-mer Only"
1.0
0.8
0.6
0.4
0.2
0.0
1,094,100
1,094,200
1,094,300
1,094,400
1,094,500
1,094,600
1,094,700
1,094,800
H37rv Ref Position (bp)

Sub-phylogeny Viz + Paralog Alignments
GCE ID: Event_069

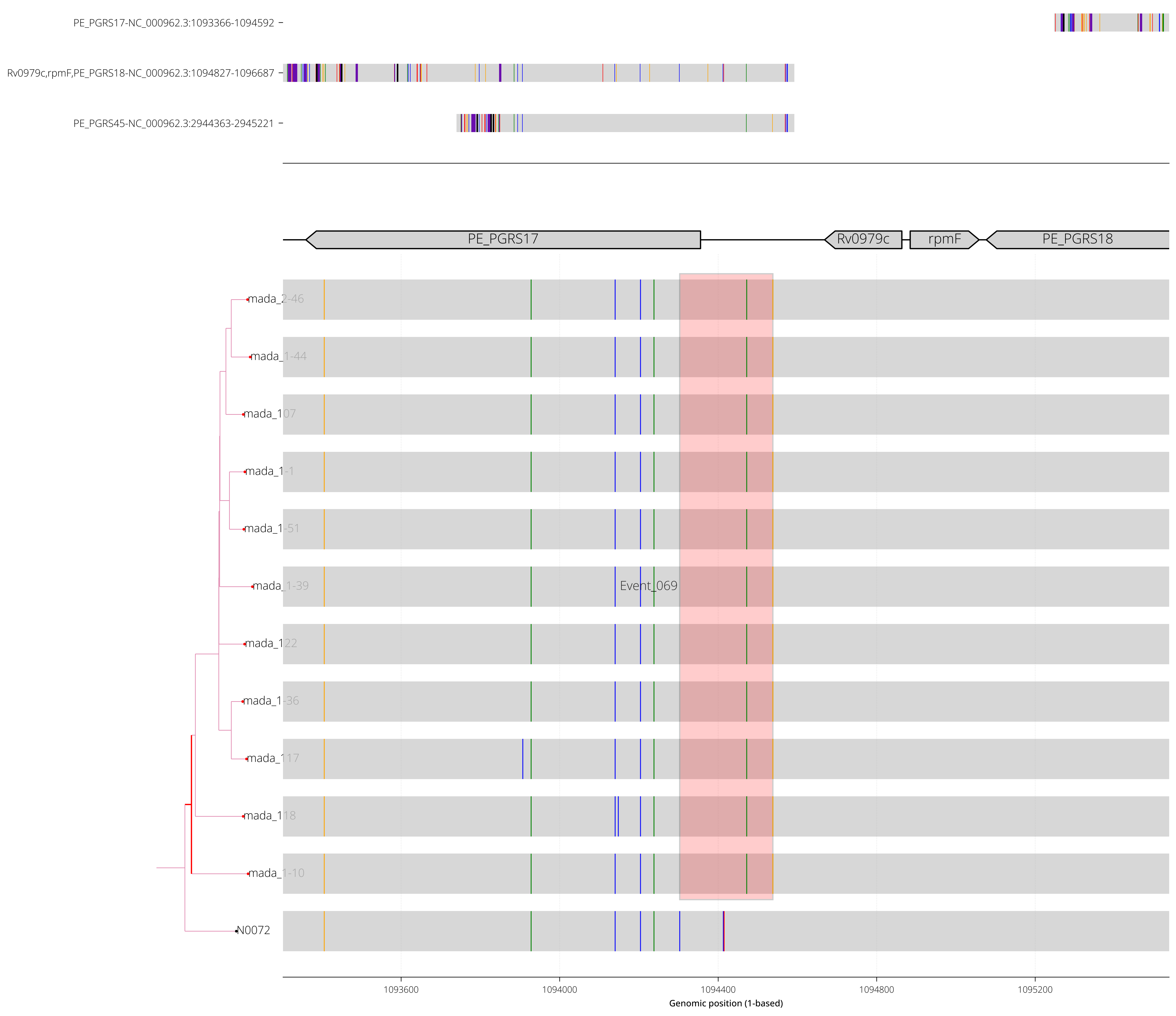

Full Phylogeny Viz + Paralog Alignments
GCE ID: Event_069

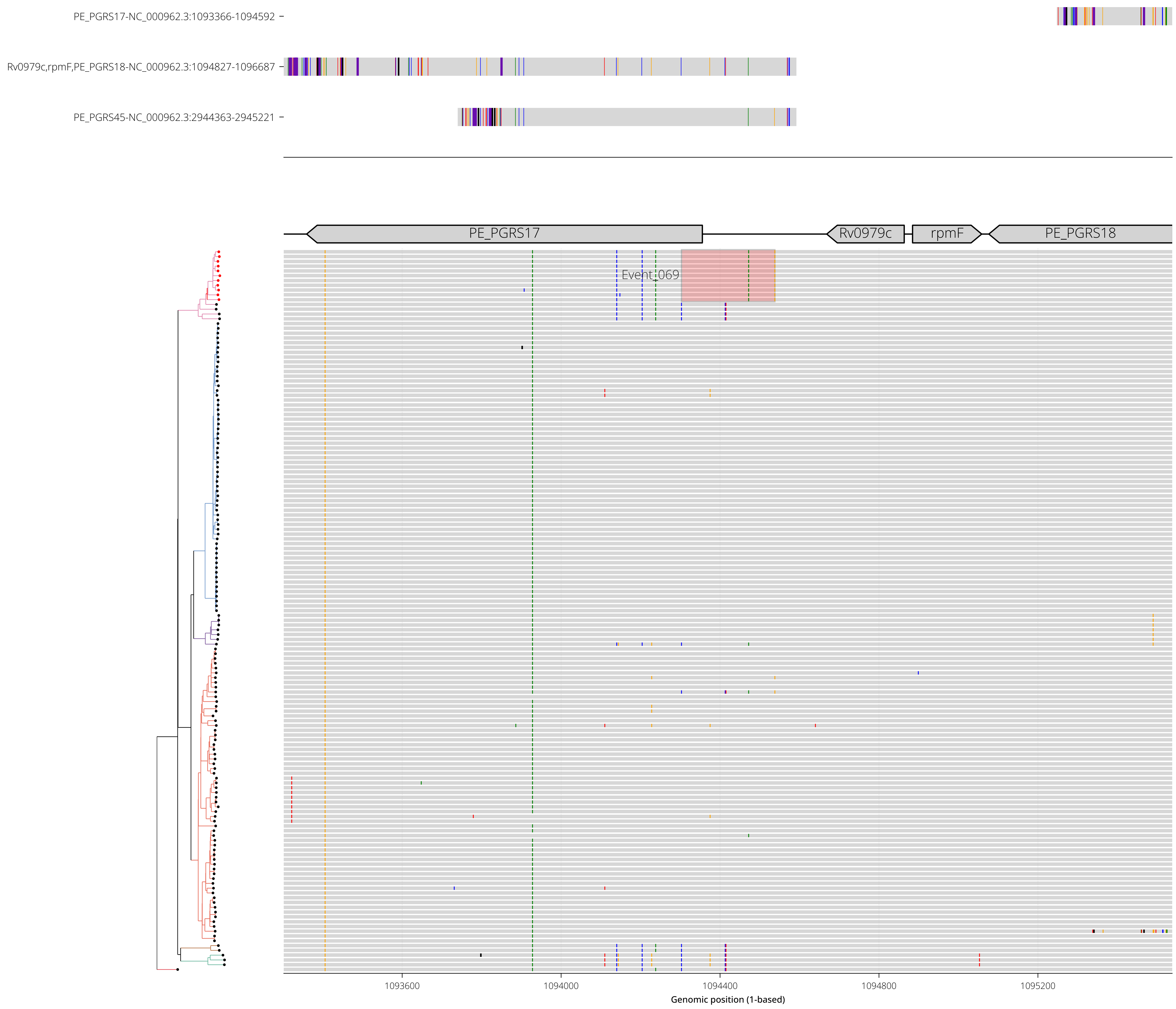

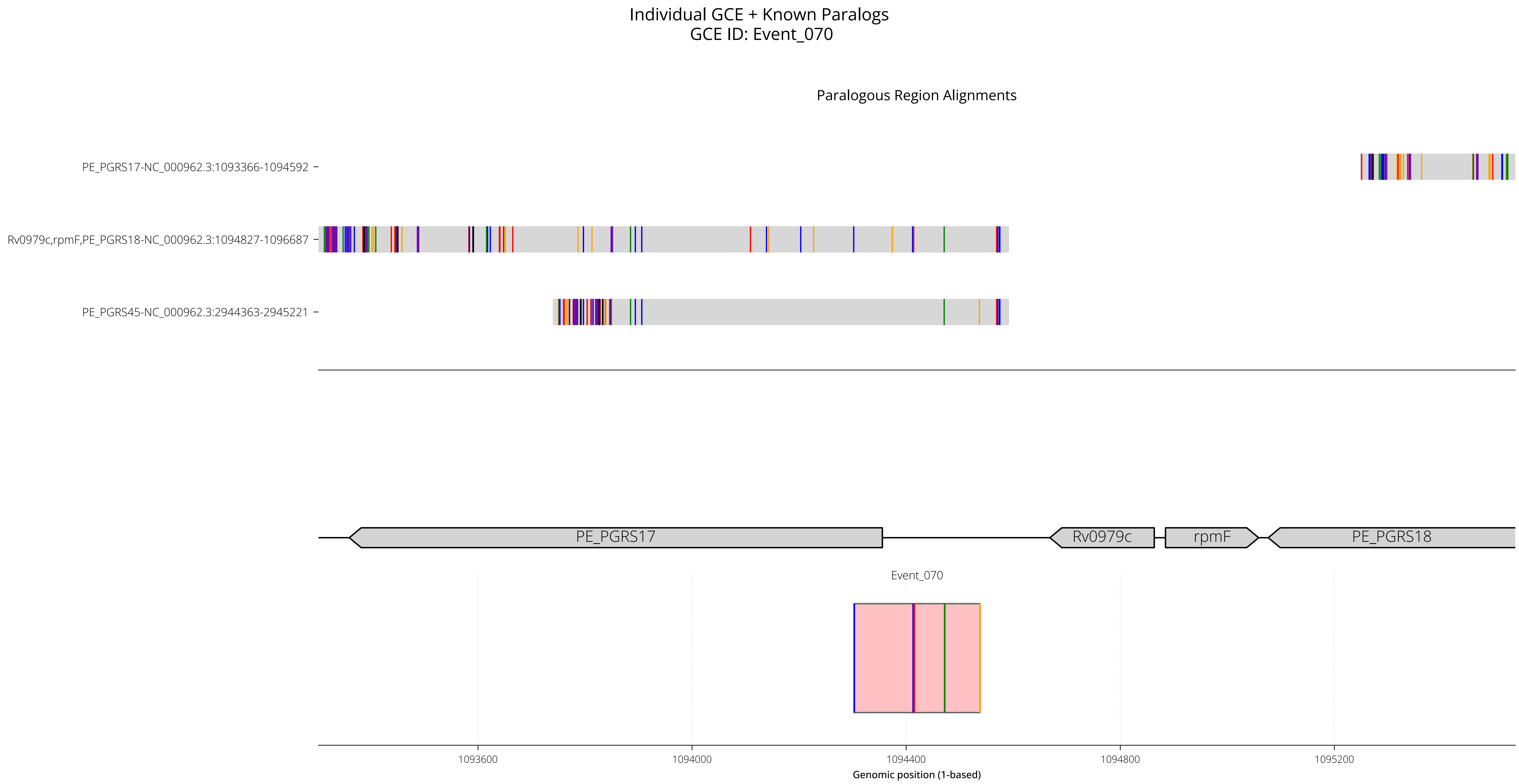

Individual GCE + Known Paralogs
GCE ID: Event_070
Paralogous Region Alignments
PE_PGRS17-NC_000962.3:1093366-1094592
Rv0979c,rpmF,PE_PGRS18-NC_000962.3:1094827-1096687
PE_PGRS45-NC_000962.3:2944363-2945221
PE_PGRS17
Rv0979c
rpmF
PE_PGRS18
Event_070
1093600
1094000
1094400
1094800
1095200
Genomic position (1-based)

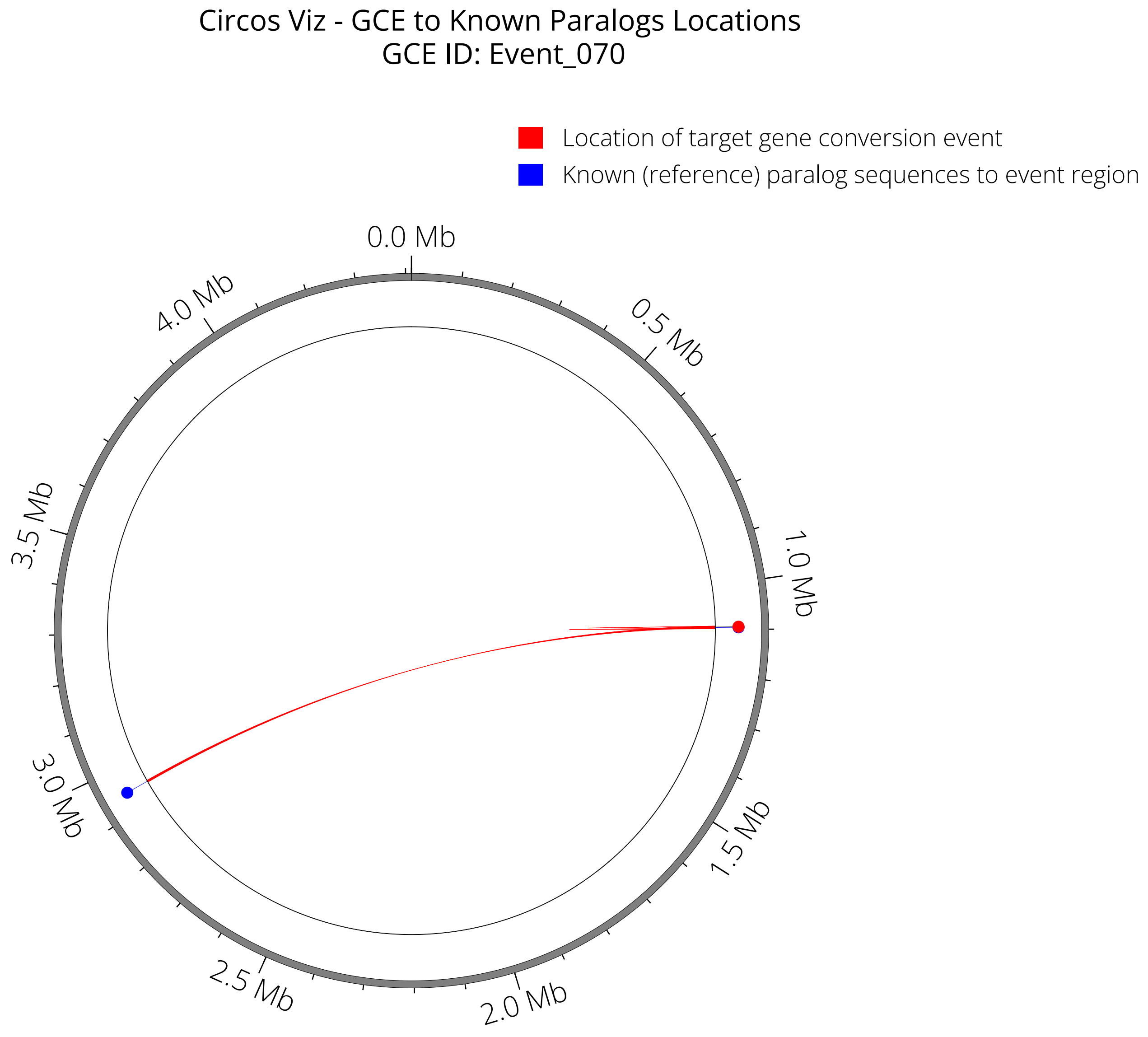

Circos Viz - GCE to Known Paralogs Locations
GCE ID: Event_070
Location of target gene conversion event
Known (reference) paralog sequences to event region
0.0 Mb
0.5 Mb
1.0 Mb
1.5 Mb
2.0 Mb
2.5 Mb
3.0 Mb
3.5 Mb
4.0 Mb

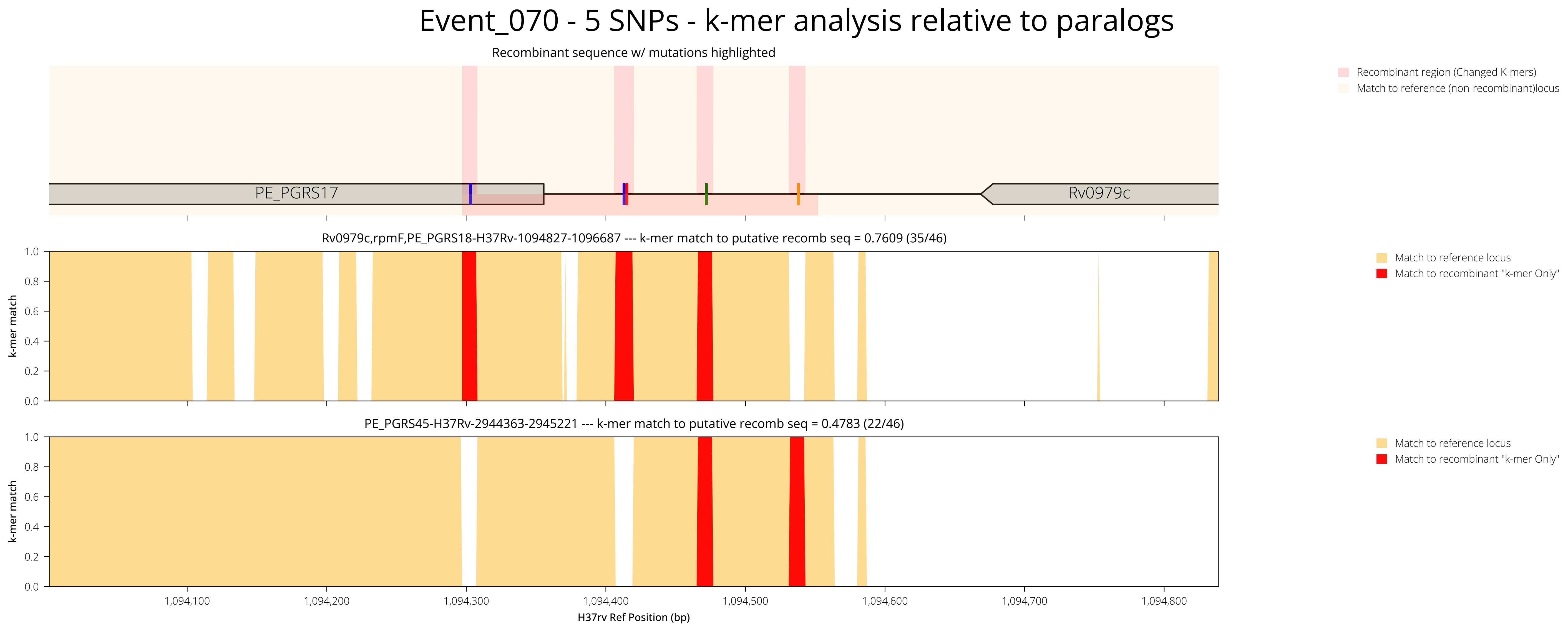

Event_070 - 5 SNPs - k-mer analysis relative to paralogs
Recombinant sequence w/ mutations highlighted
Recombinant region (Changed K-mers)
Match to reference (non-recombinant)locus
PE_PGRS17
Rv0979c
Rv0979c,rpmF,PE_PGRS18-H37Rv-1094827-1096687 --- k-mer match to putative recomb seq = 0.7609 (35/46)
Match to reference locus
Match to recombinant "k-mer Only"
k-mer match
PE_PGRS45-H37Rv-2944363-2945221 --- k-mer match to putative recomb seq = 0.4783 (22/46)
Match to reference locus
Match to recombinant "k-mer Only"
k-mer match
1.0
0.8
0.6
0.4
0.2
0.0
1,094,100
1,094,200
1,094,300
1,094,400
1,094,500
1,094,600
1,094,700
1,094,800
H37rv Ref Position (bp)

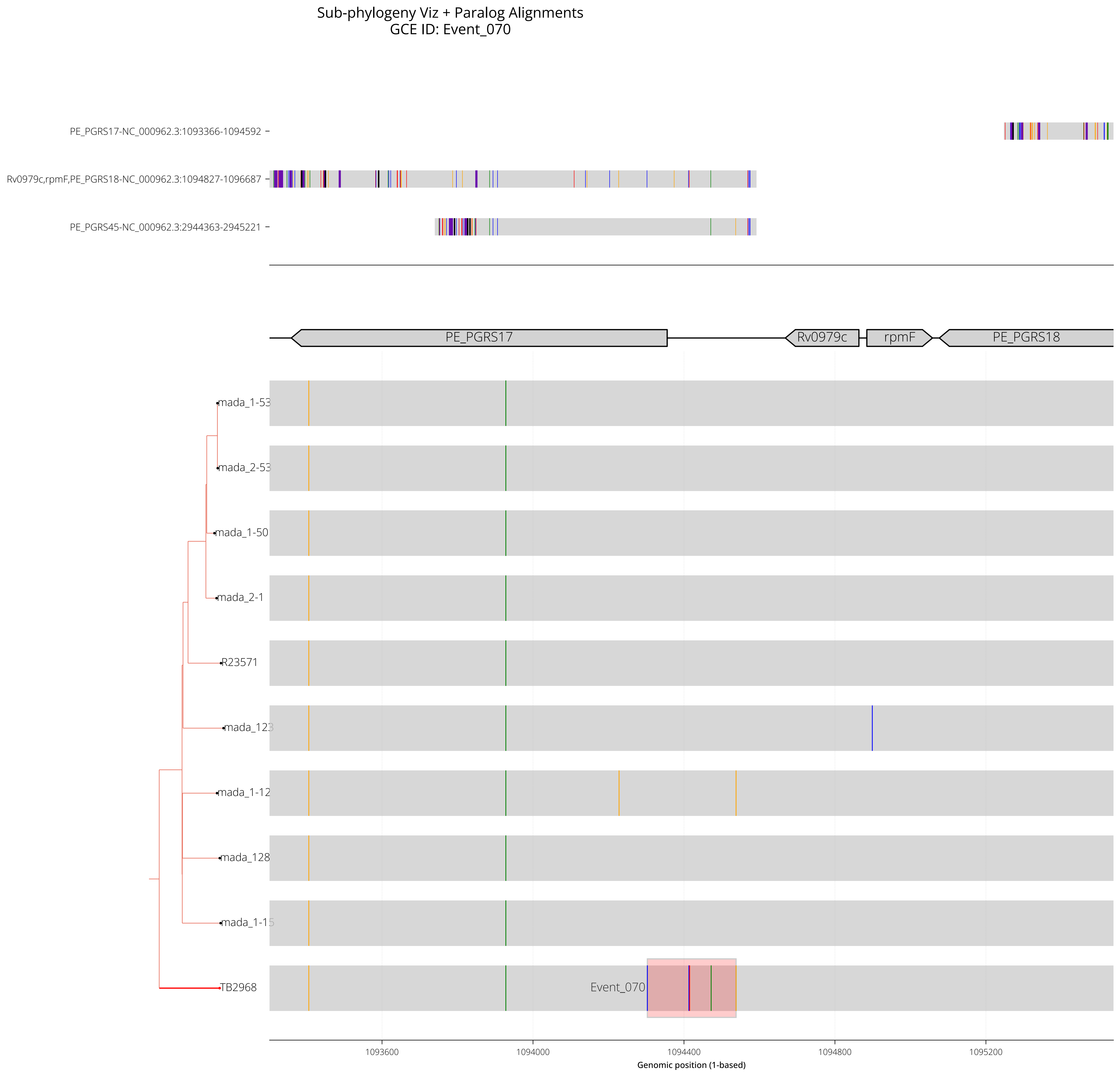
Sub-phylogeny Viz + Paralog Alignments
GCE ID: Event_070
PE_PGRS17-NC_000962.3:1093366-1094592
Rv0979c,rpmF,PE_PGRS18-NC_000962.3:1094827-1096687
PE_PGRS45-NC_000962.3:2944363-2945221
PE_PGRS17
Rv0979c
rpmF
PE_PGRS18
mada_1-53
mada_2-53
mada_1-50
mada_2-1
R23571
mada_123
mada_1-12
mada_128
mada_1-15
TB2968
Event_070
1093600
1094000
1094400
1094800
1095200
Genomic position (1-based)

Full Phylogeny Viz + Paralog Alignments
GCE ID: Event_070

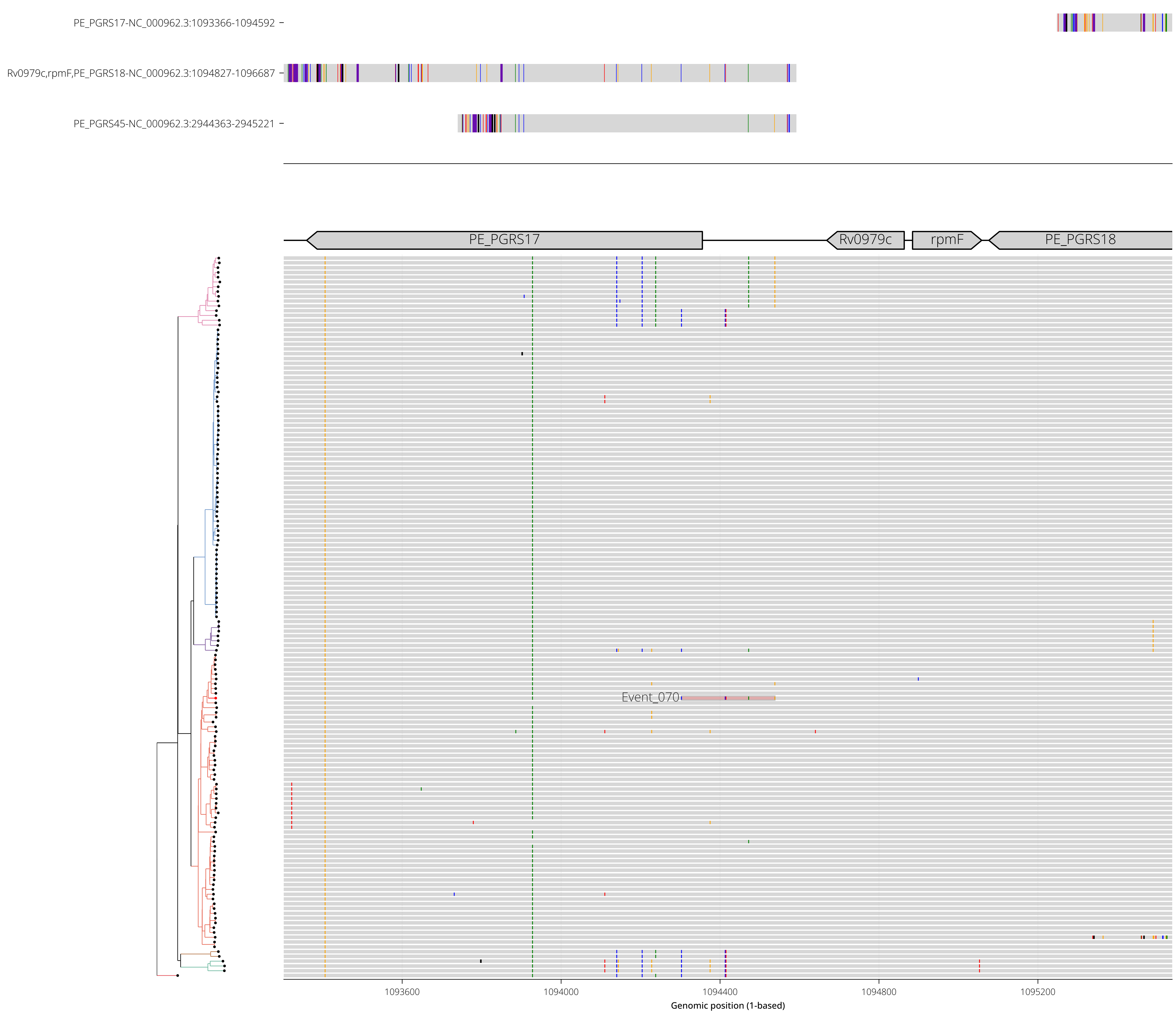

Individual GCE + Known Paralogs
GCE ID: Event_071

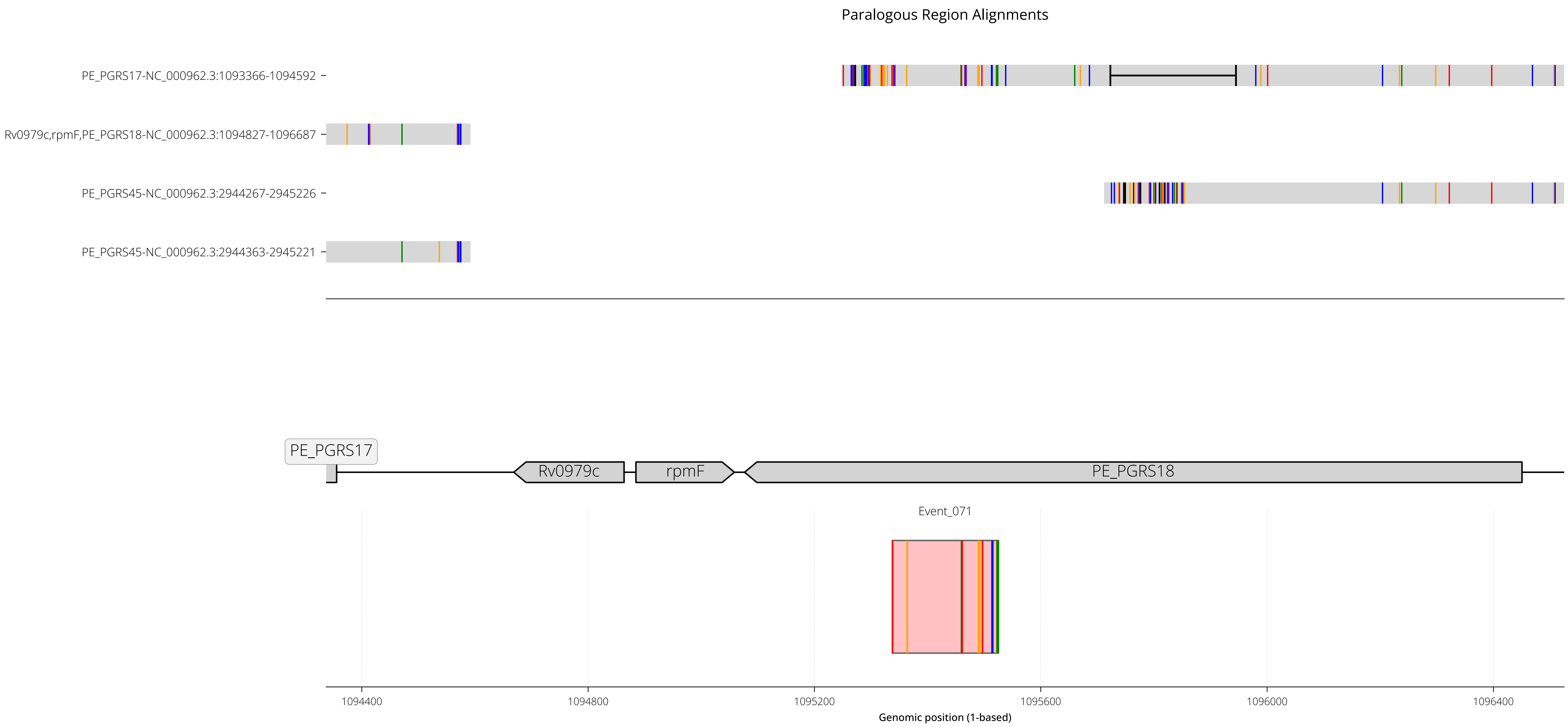

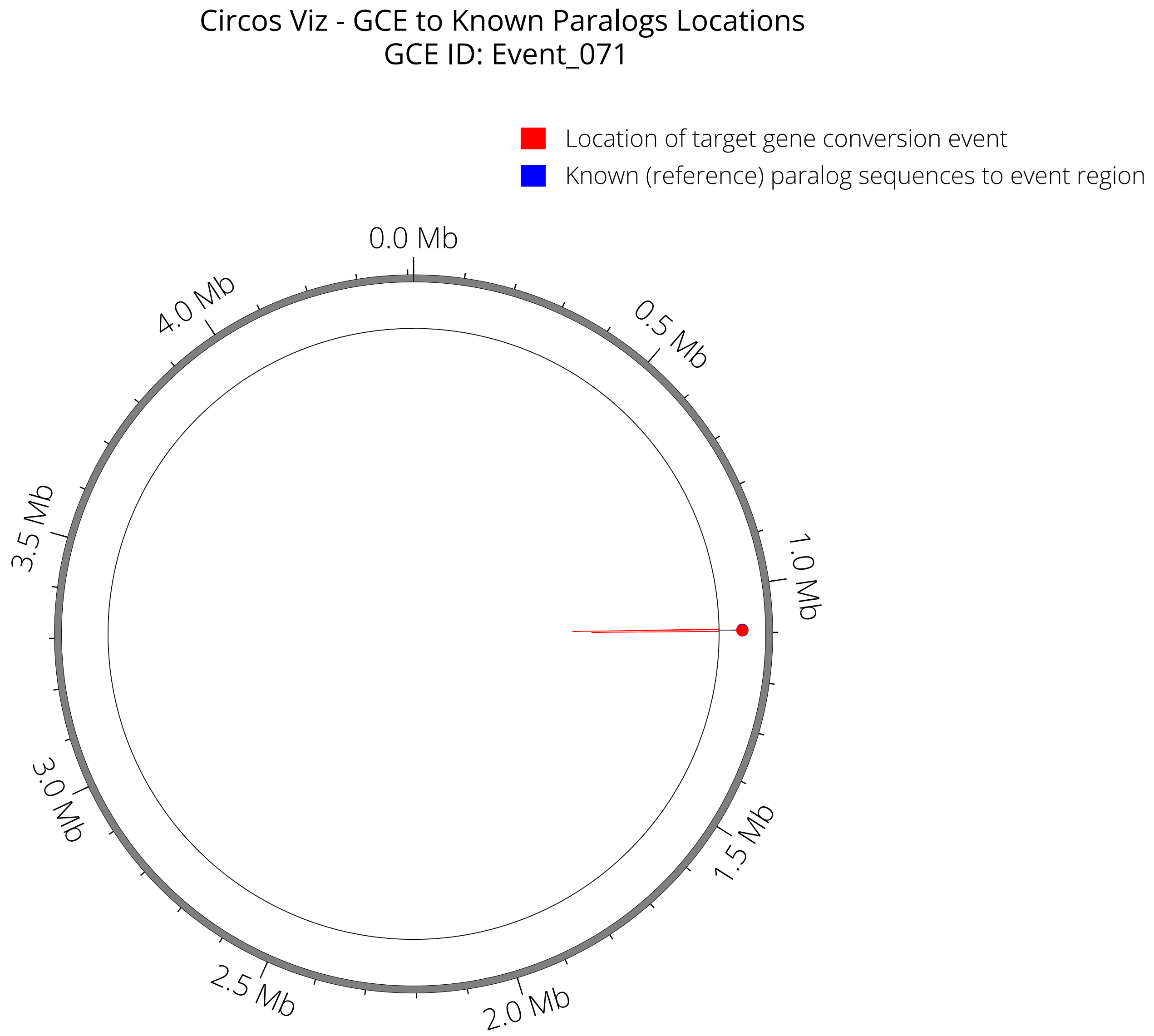

Circos Viz - GCE to Known Paralogs Locations
GCE ID: Event_071
Location of target gene conversion event
Known (reference) paralog sequences to event region
0.0 Mb
0.5 Mb
1.0 Mb
1.5 Mb
2.0 Mb
2.5 Mb
3.0 Mb
3.5 Mb
4.0 Mb

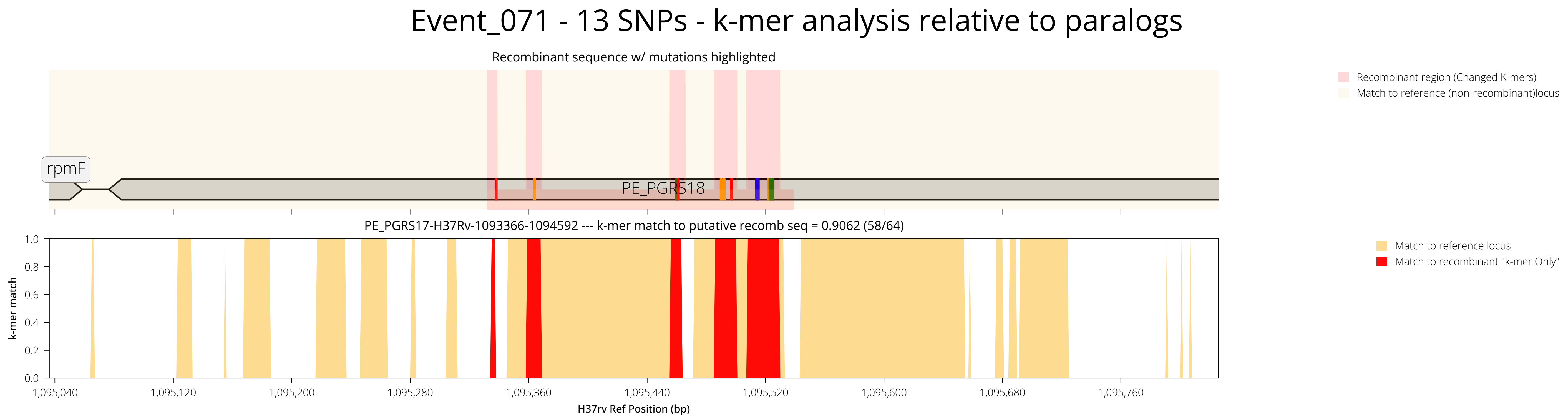

Event_071 - 13 SNPs - k-mer analysis relative to paralogs
Recombinant sequence w/ mutations highlighted
rpmF
PE_PGRS18
Recombinant region (Changed K-mers)
Match to reference (non-recombinant)locus
PE_PGRS17-H37Rv-1093366-1094592 --- k-mer match to putative recomb seq = 0.9062 (58/64)
k-mer match
1.0
0.8
0.6
0.4
0.2
0.0
1,095,040
1,095,120
1,095,200
1,095,280
1,095,360
1,095,440
1,095,520
1,095,600
1,095,680
1,095,760
H37rv Ref Position (bp)
Match to reference locus
Match to recombinant "k-mer Only"

Sub-phylogeny Viz + Paralog Alignments
GCE ID: Event_071

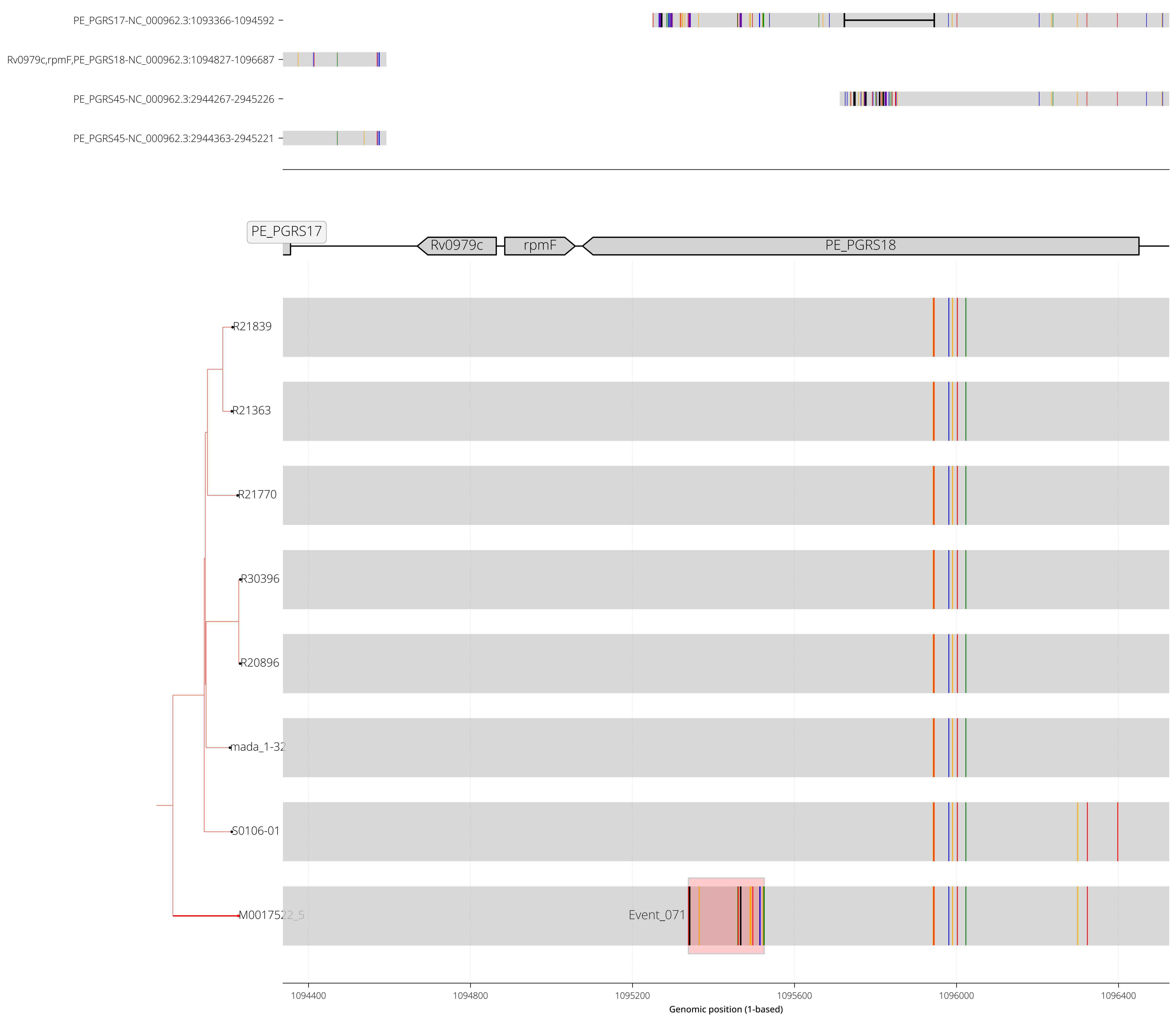

### Full Phylogeny Viz + Paralog Alignments
GCE ID: Event_071

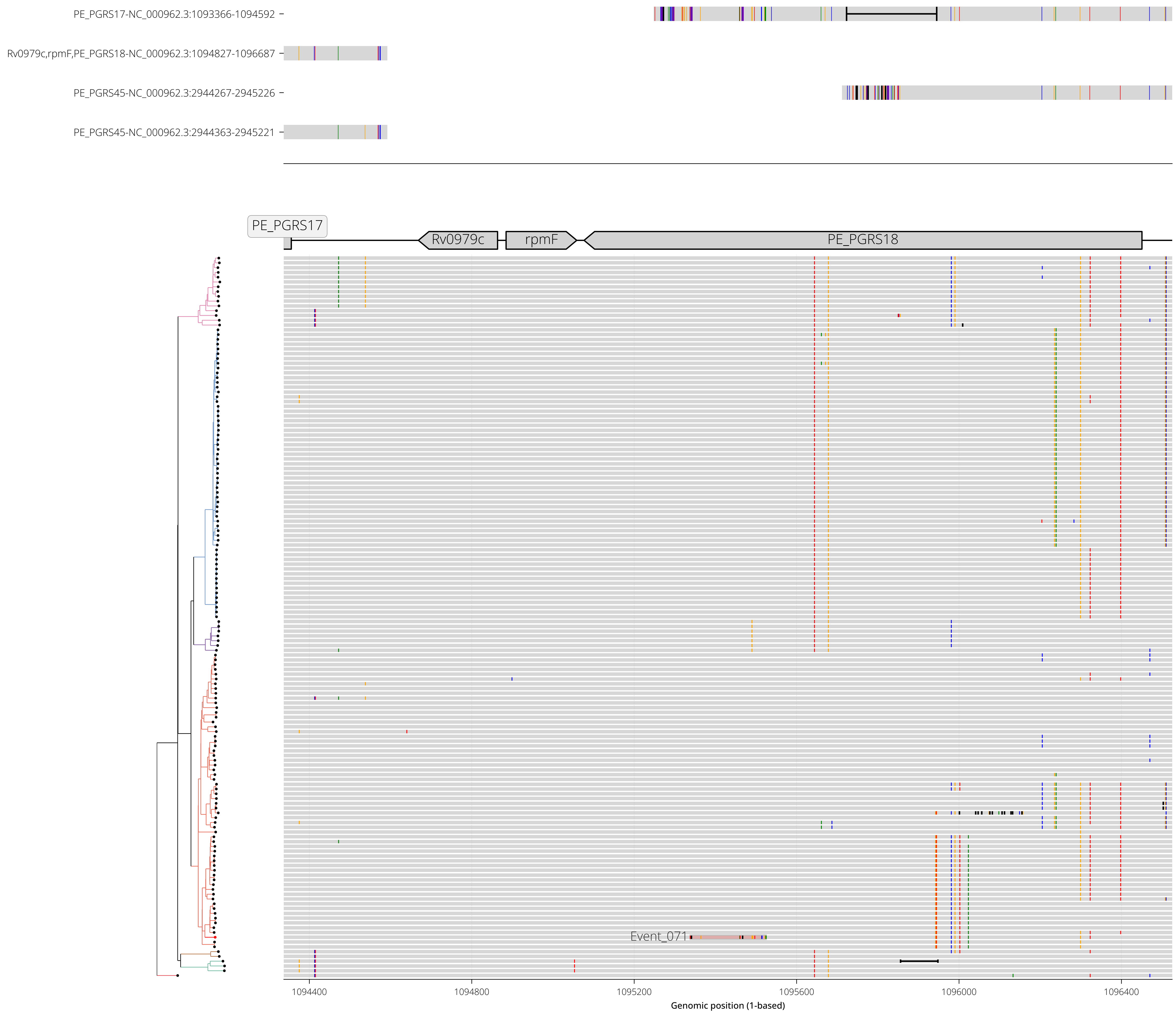

Individual GCE + Known Paralogs
GCE ID: Event_072

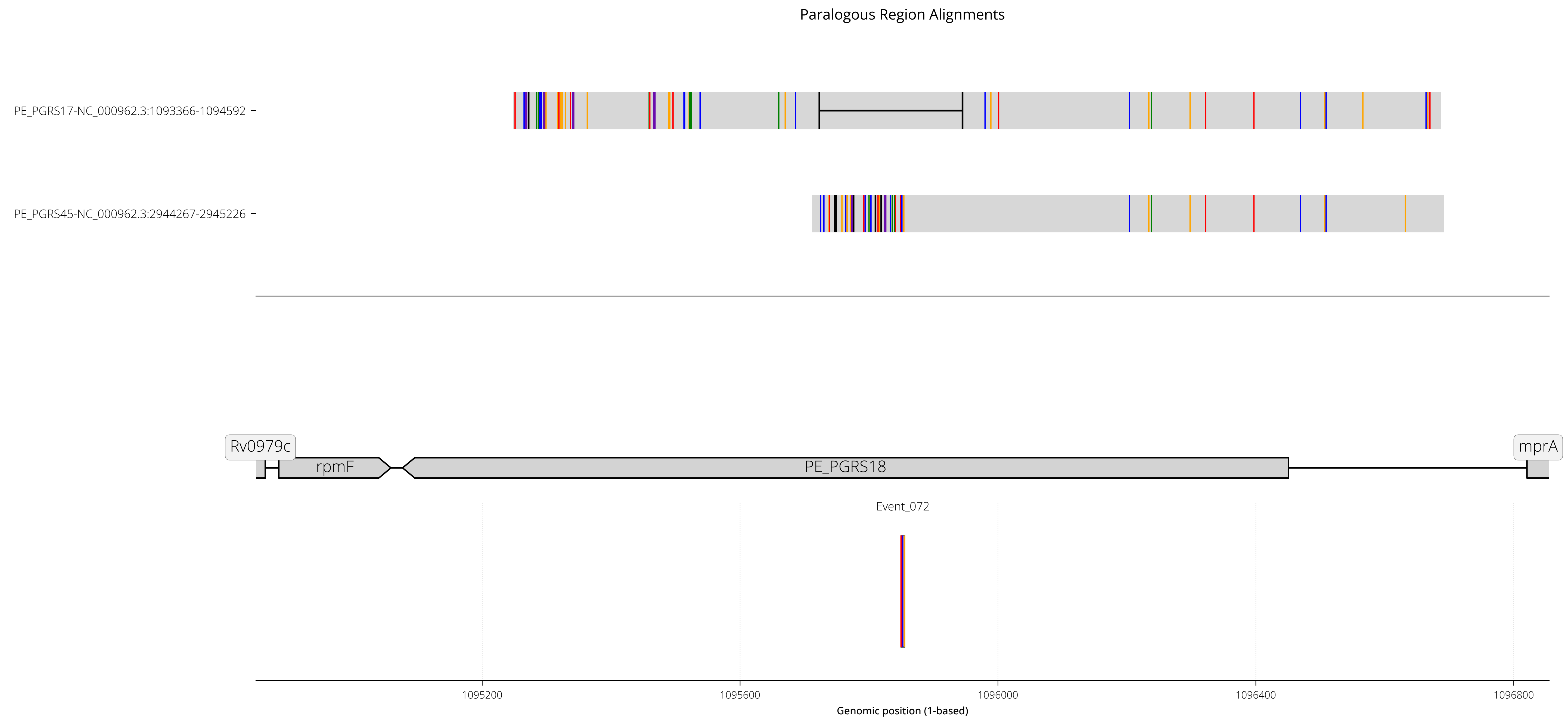

Circos Viz - GCE to Known Paralogs Locations
GCE ID: Event_072

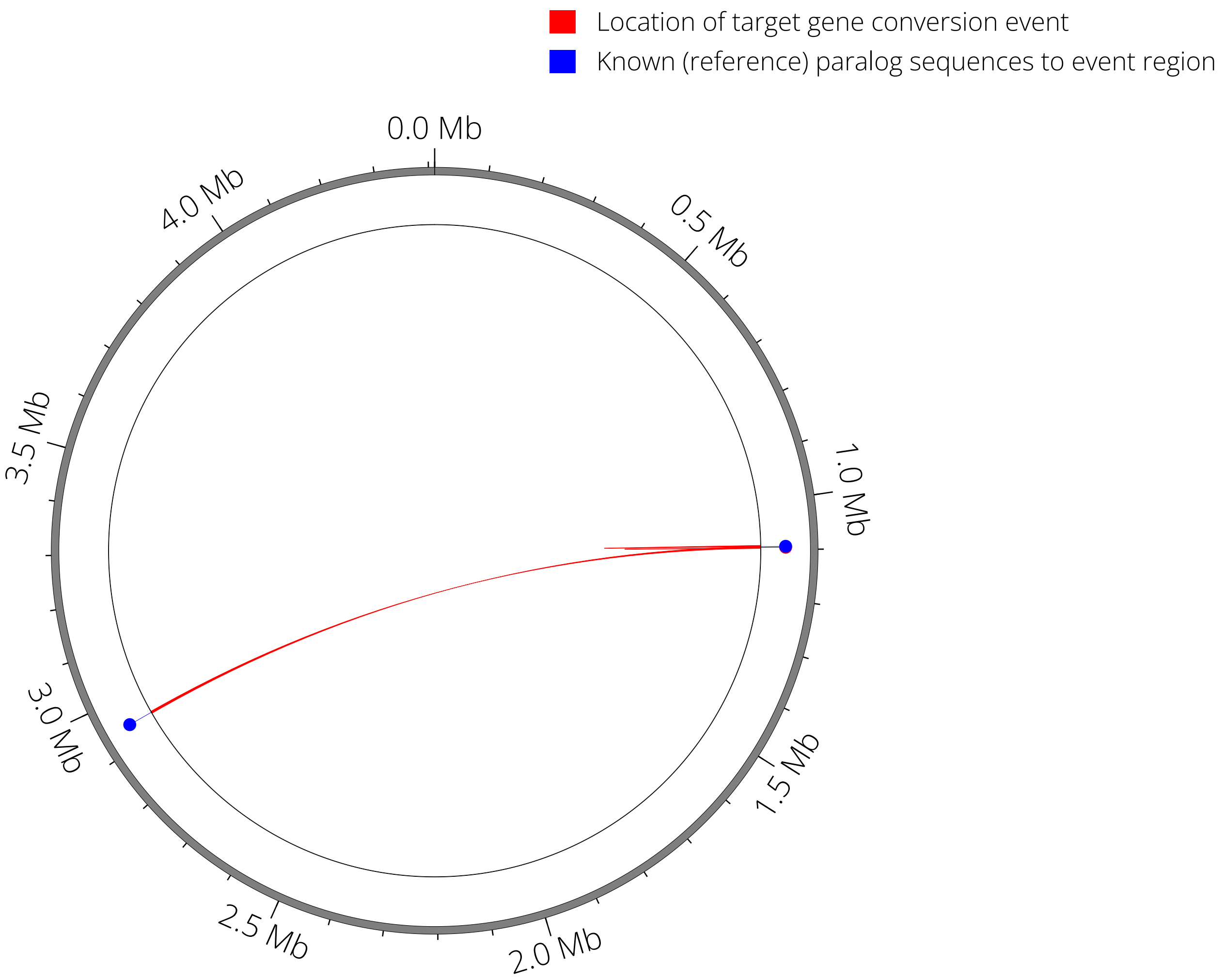

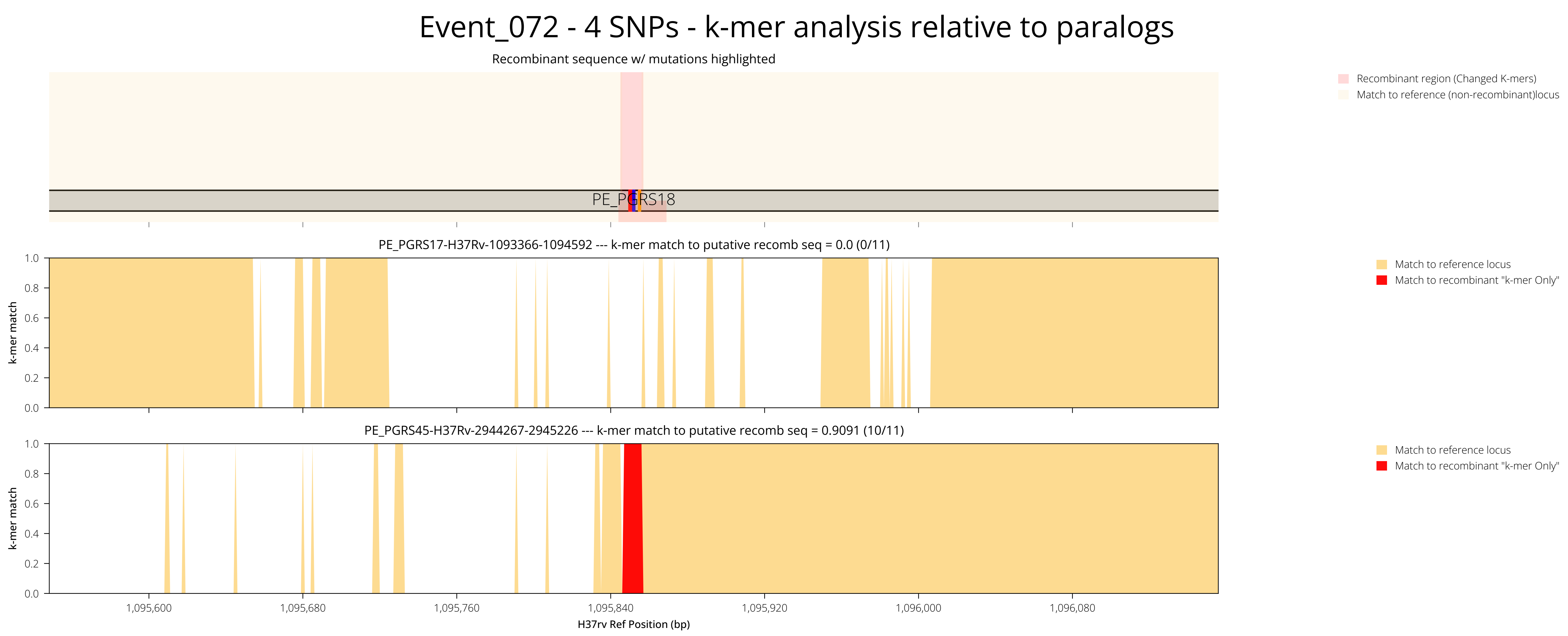

Event_072 - 4 SNPs - k-mer analysis relative to paralogs
Recombinant sequence w/ mutations highlighted
Recombinant region (Changed K-mers)
Match to reference (non-recombinant)locus
PE_PGRS18
PE_PGRS17-H37Rv-1093366-1094592 --- k-mer match to putative recomb seq = 0.0 (0/11)
Match to reference locus
Match to recombinant "k-mer Only"
k-mer match
1.0
0.8
0.6
0.4
0.2
0.0
PE_PGRS45-H37Rv-2944267-2945226 --- k-mer match to putative recomb seq = 0.9091 (10/11)
Match to reference locus
Match to recombinant "k-mer Only"
k-mer match
1,095,600
1,095,680
1,095,760
1,095,840
1,095,920
1,096,000
1,096,080
H37rv Ref Position (bp)

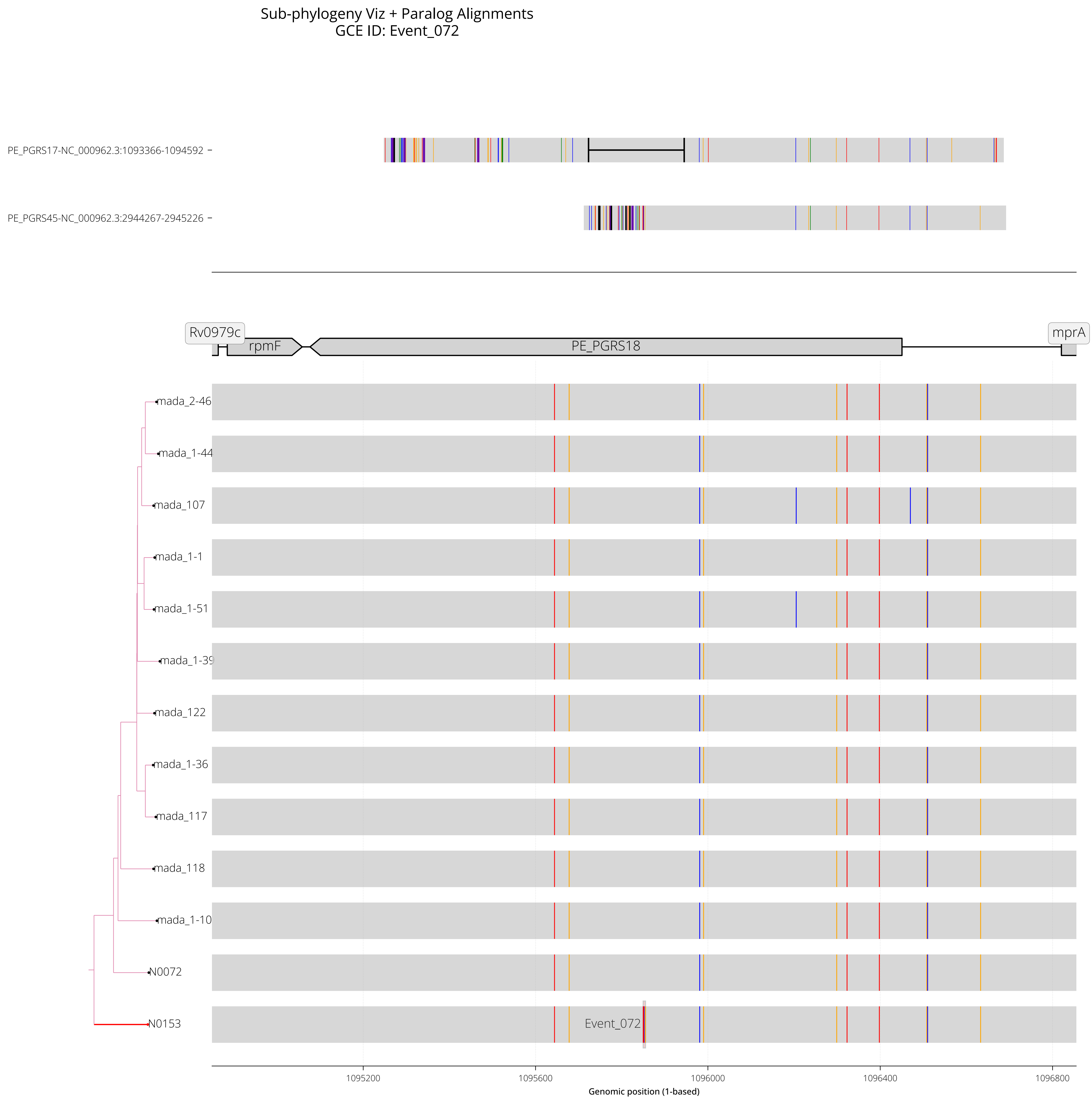

Sub-phylogeny Viz + Paralog Alignments
GCE ID: Event_072
PE_PGRS17-NC_000962.3:1093366-1094592
PE_PGRS45-NC_000962.3:2944267-2945226
Rv0979c
rpmF
PE_PGRS18
mprA
mada_2-46
mada_1-44
mada_107
mada_1-1
mada_1-51
mada_1-39
mada_122
mada_1-36
mada_117
mada_118
mada_1-10
N0072
N0153
Event_072
1095200
1095600
1096000
1096400
1096800
Genomic position (1-based)

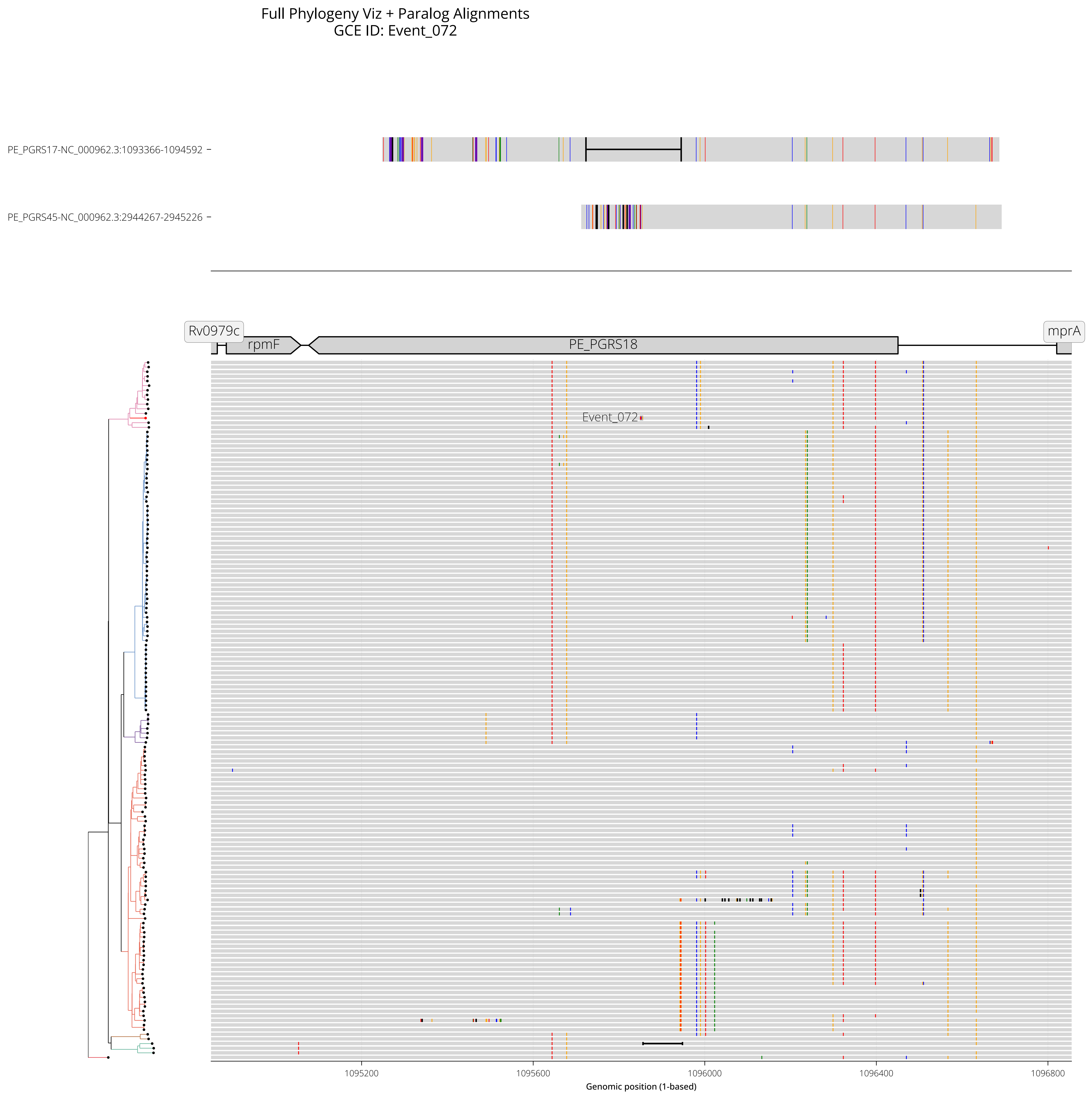

Full Phylogeny Viz + Paralog Alignments
GCE ID: Event_072
PE_PGRS17-NC_000962.3:1093366-1094592
PE_PGRS45-NC_000962.3:2944267-2945226
Rv0979c
rpmF
PE_PGRS18
mprA
Event_072
1095200
1095600
1096000
1096400
1096800
Genomic position (1-based)

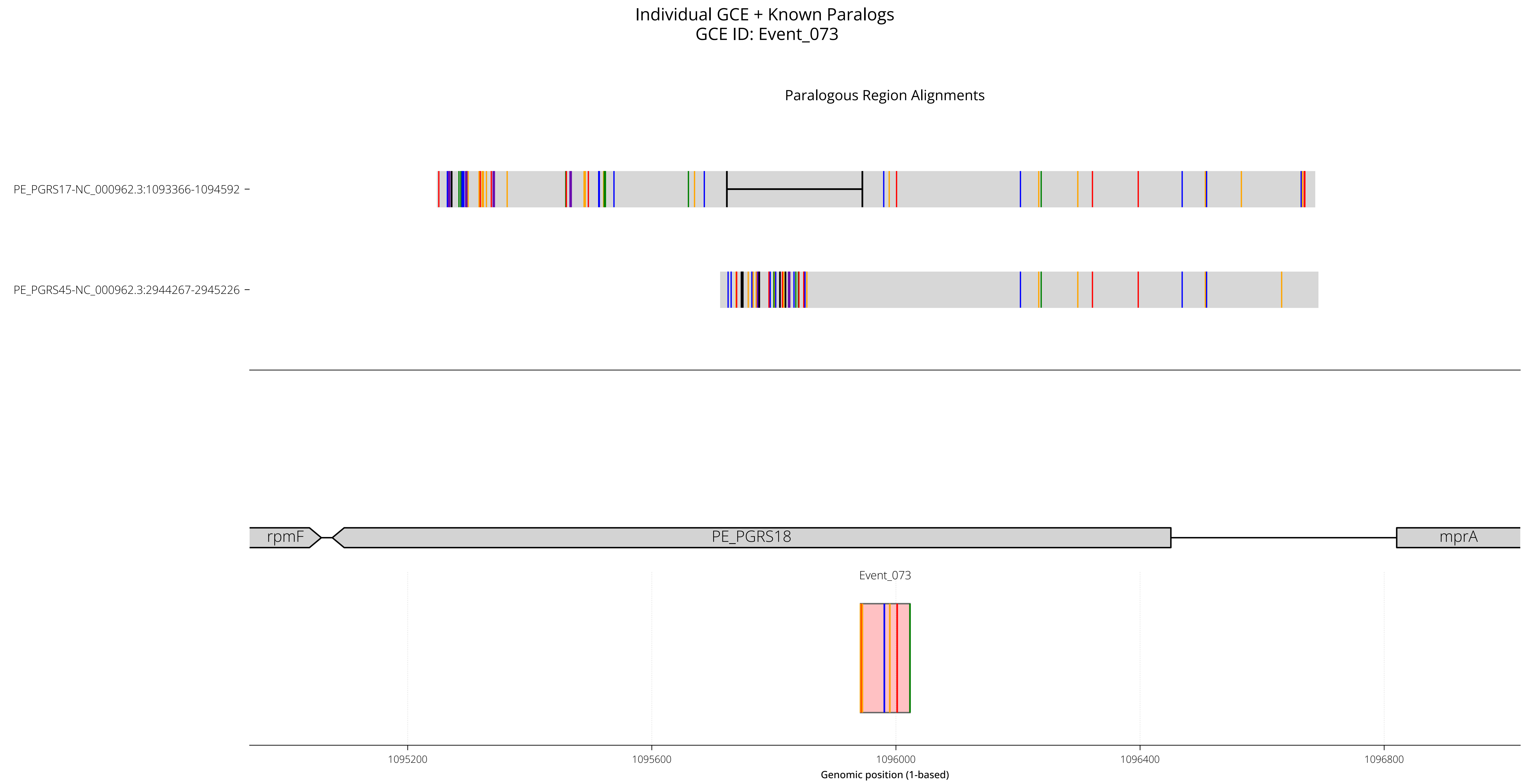

Individual GCE + Known Paralogs
GCE ID: Event_073
Paralogous Region Alignments
PE_PGRS17-NC_000962.3:1093366-1094592
PE_PGRS45-NC_000962.3:2944267-2945226
rpmF
PE_PGRS18
mprA
Event_073
1095200
1095600
1096000
1096400
1096800
Genomic position (1-based)

Circos Viz - GCE to Known Paralogs Locations
GCE ID: Event_073

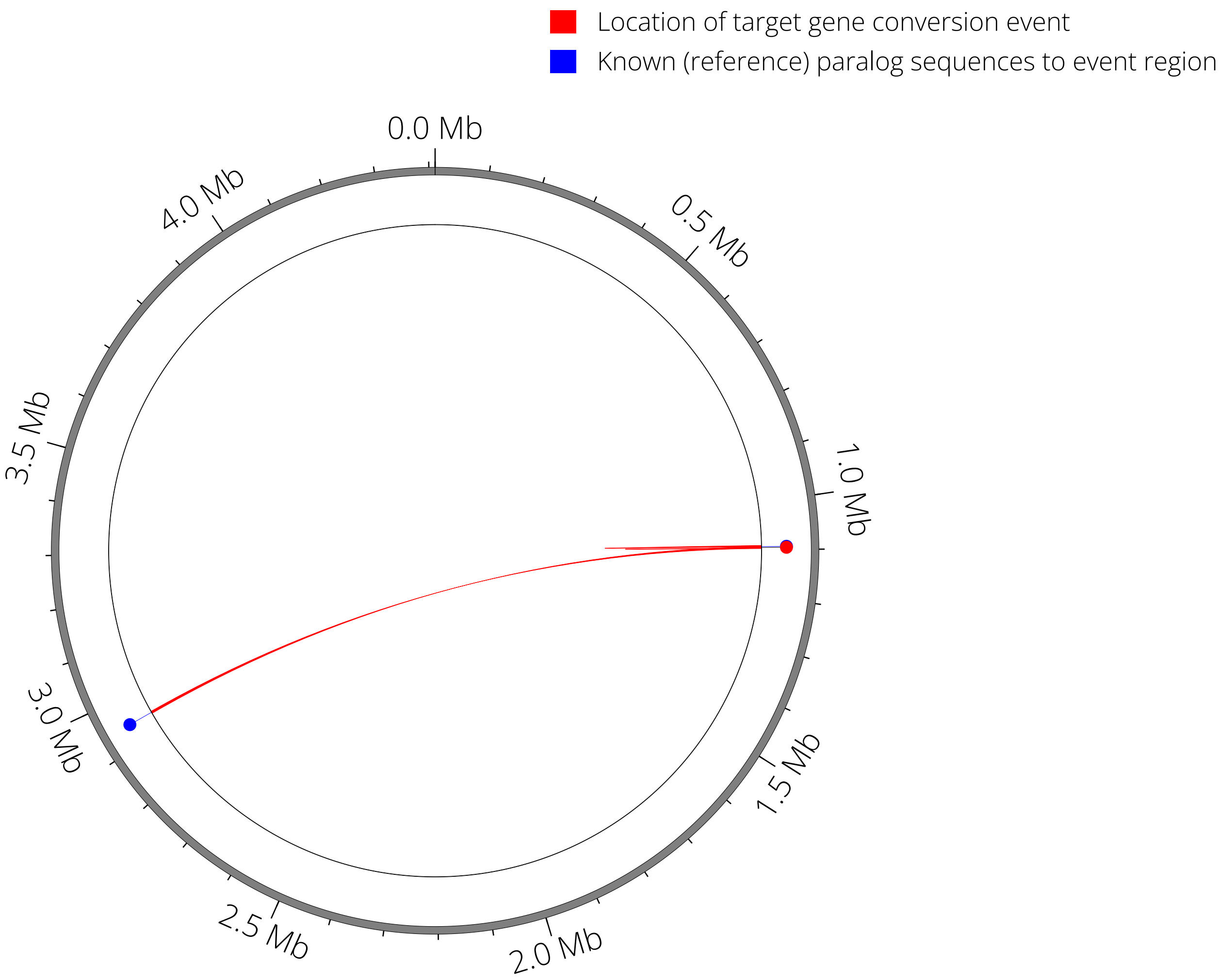

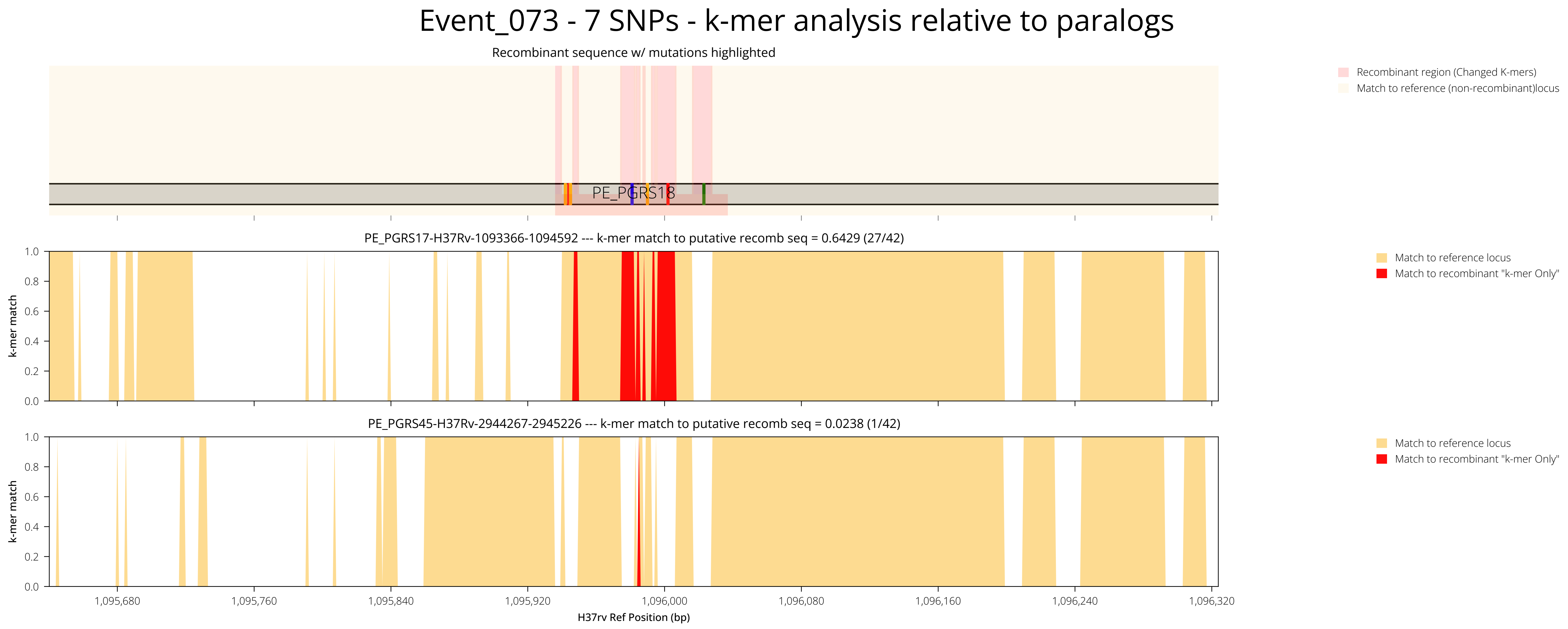

Event_073 - 7 SNPs - k-mer analysis relative to paralogs
Recombinant sequence w/ mutations highlighted
Recombinant region (Changed K-mers)
Match to reference (non-recombinant)locus
PE_PGRS18
PE_PGRS17-H37Rv-1093366-1094592 --- k-mer match to putative recomb seq = 0.6429 (27/42)
Match to reference locus
Match to recombinant "k-mer Only"
k-mer match
1.0
0.8
0.6
0.4
0.2
0.0
PE_PGRS45-H37Rv-2944267-2945226 --- k-mer match to putative recomb seq = 0.0238 (1/42)
Match to reference locus
Match to recombinant "k-mer Only"
k-mer match
1.0
0.8
0.6
0.4
0.2
0.0
1,095,680
1,095,760
1,095,840
1,095,920
1,096,000
1,096,080
1,096,160
1,096,240
1,096,320
H37rv Ref Position (bp)

Sub-phylogeny Viz + Paralog Alignments
GCE ID: Event_073

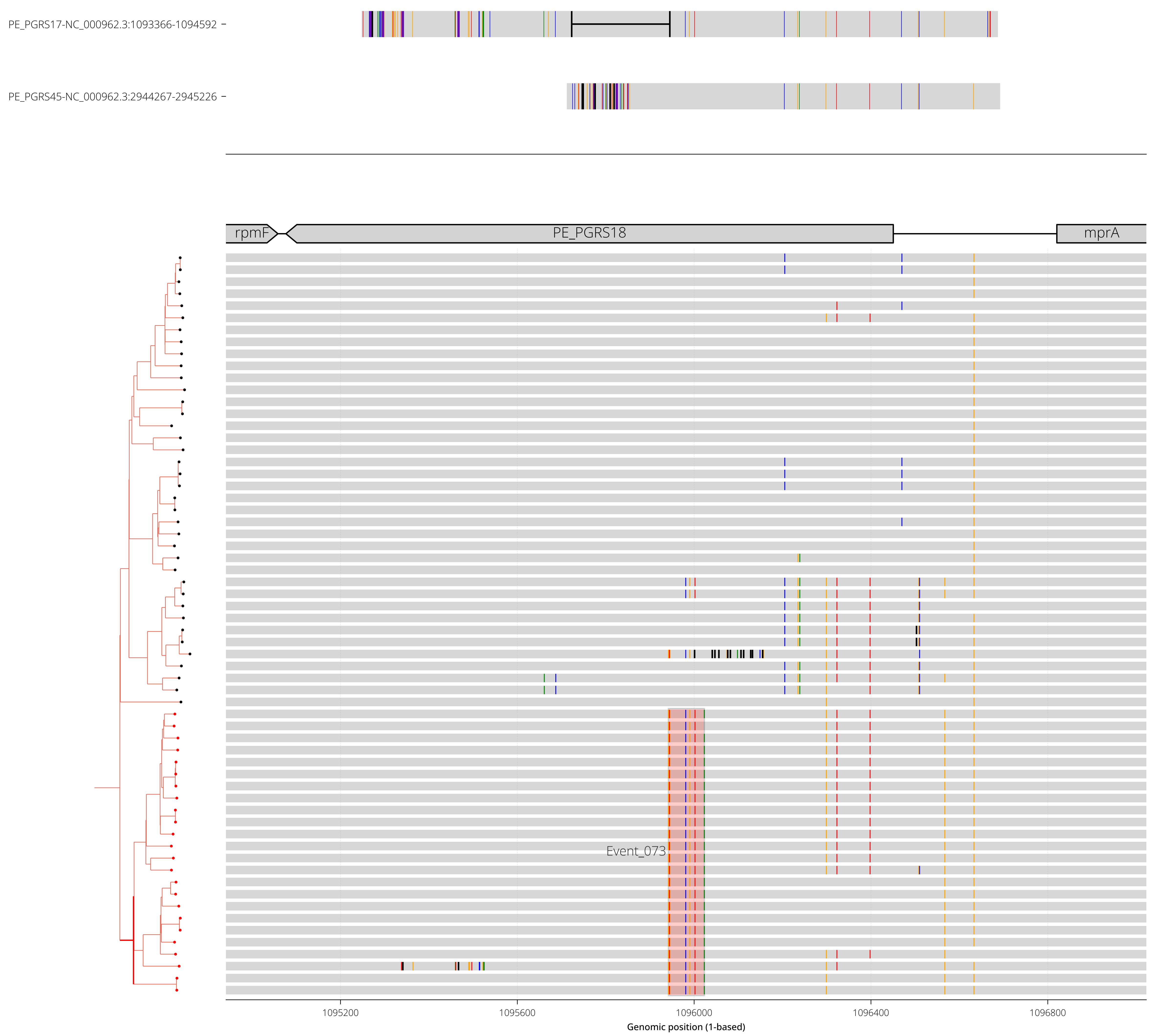

Full Phylogeny Viz + Paralog Alignments
GCE ID: Event_073

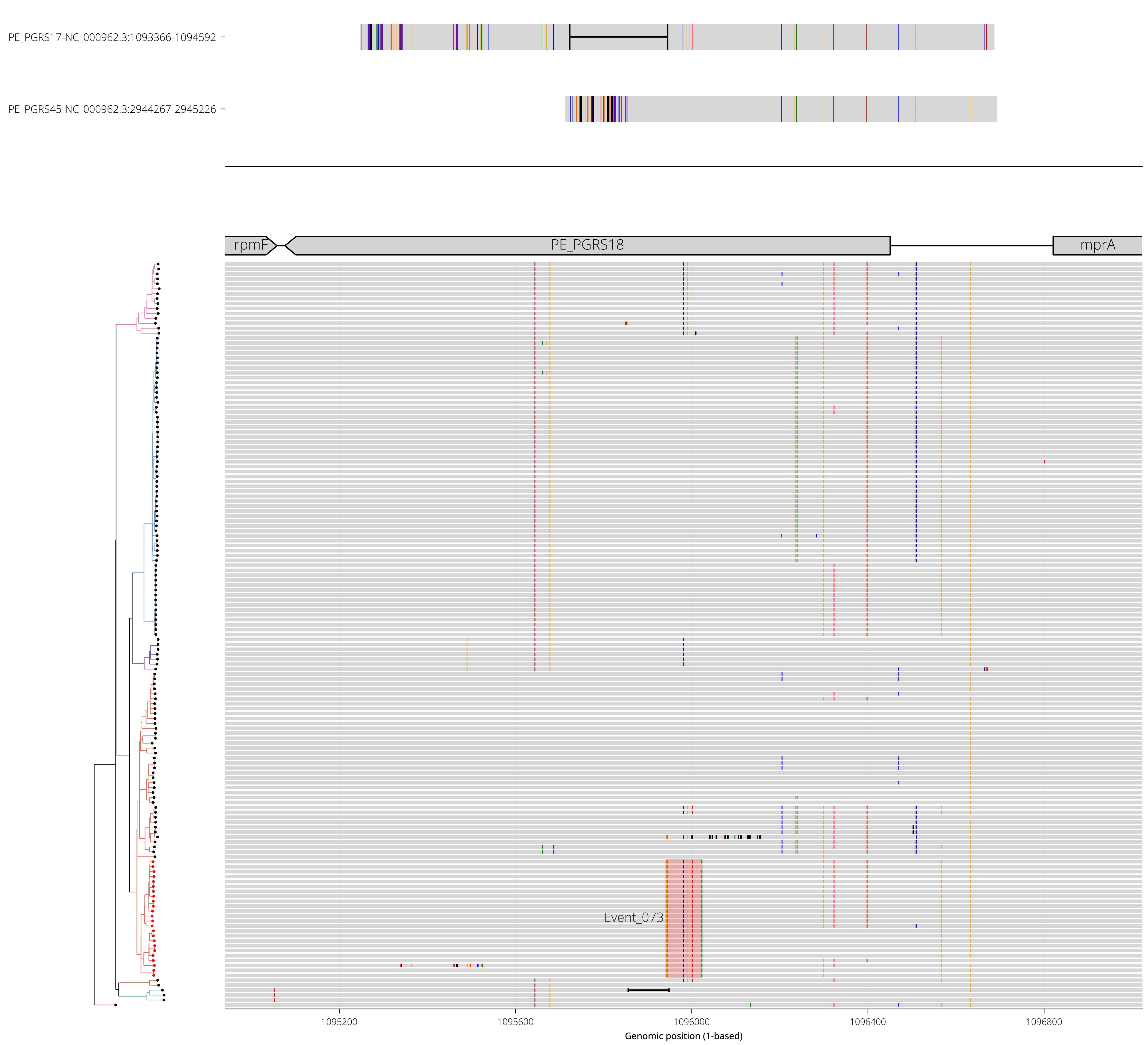

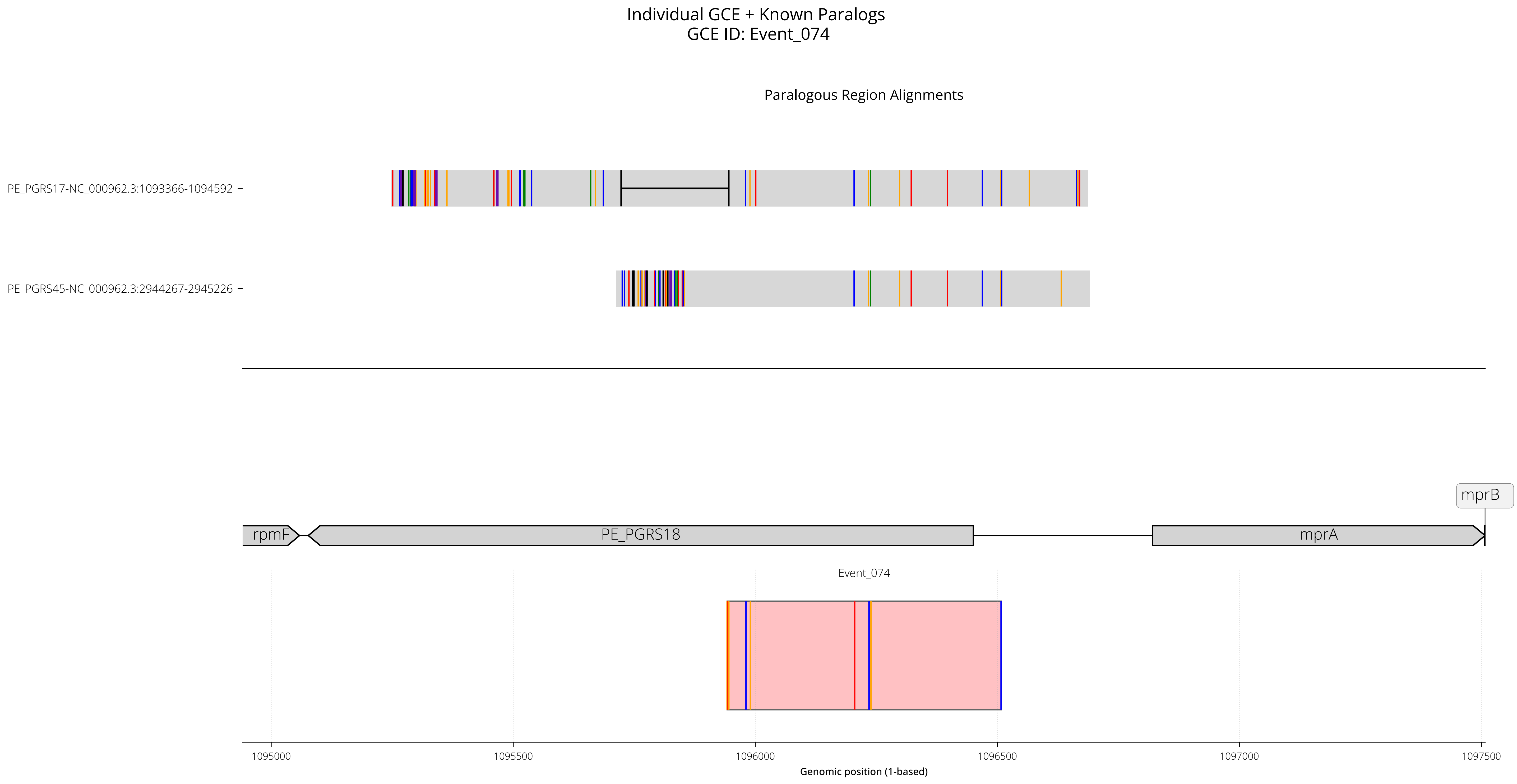
Individual GCE + Known Paralogs
GCE ID: Event_074
Paralogous Region Alignments
PE_PGRS17-NC_000962.3:1093366-1094592
PE_PGRS45-NC_000962.3:2944267-2945226
mprB
rpmF
PE_PGRS18
mprA
Event_074
1095000
1095500
1096000
1096500
1097000
1097500
Genomic position (1-based)

Circos Viz - GCE to Known Paralogs Locations
GCE ID: Event_074

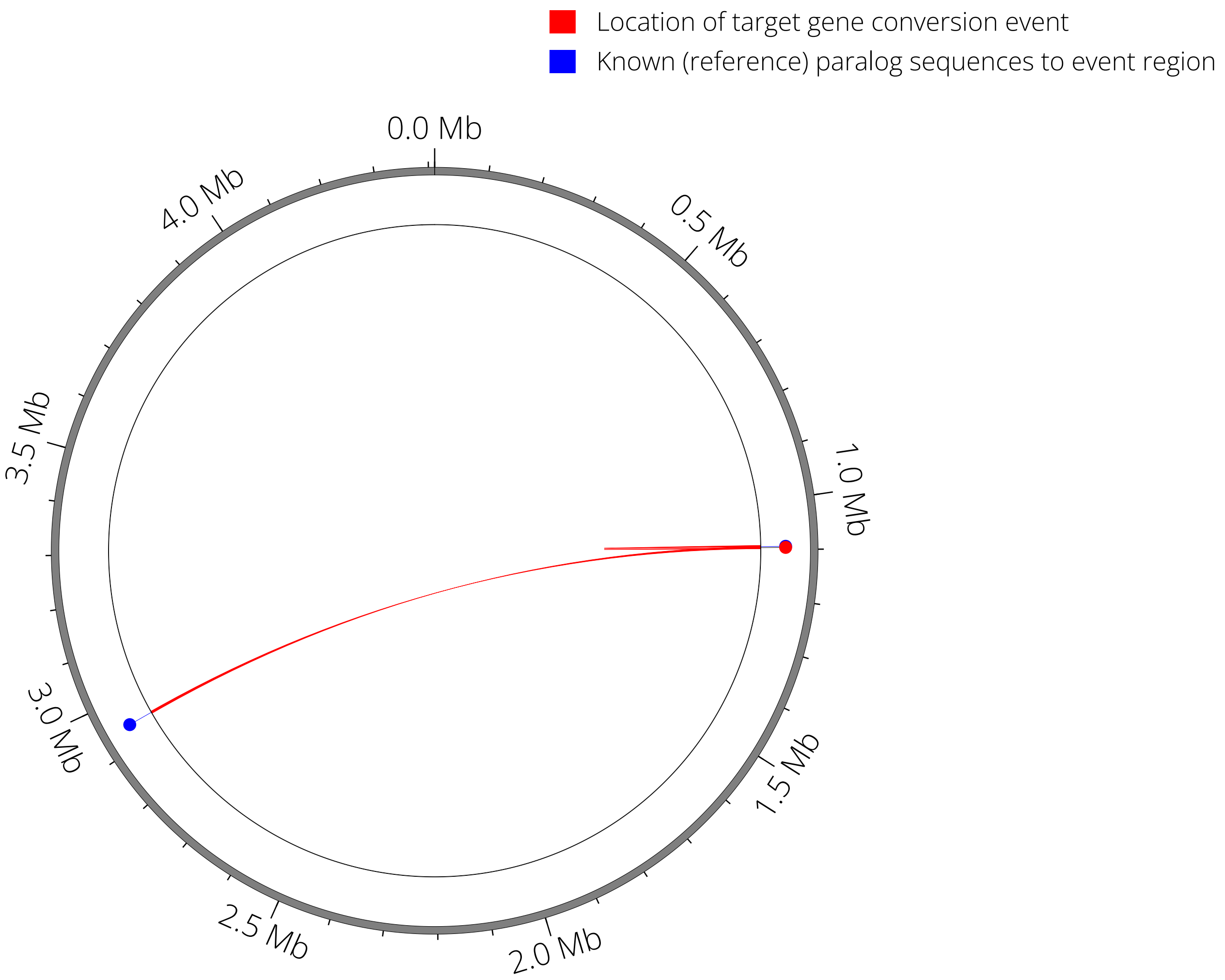

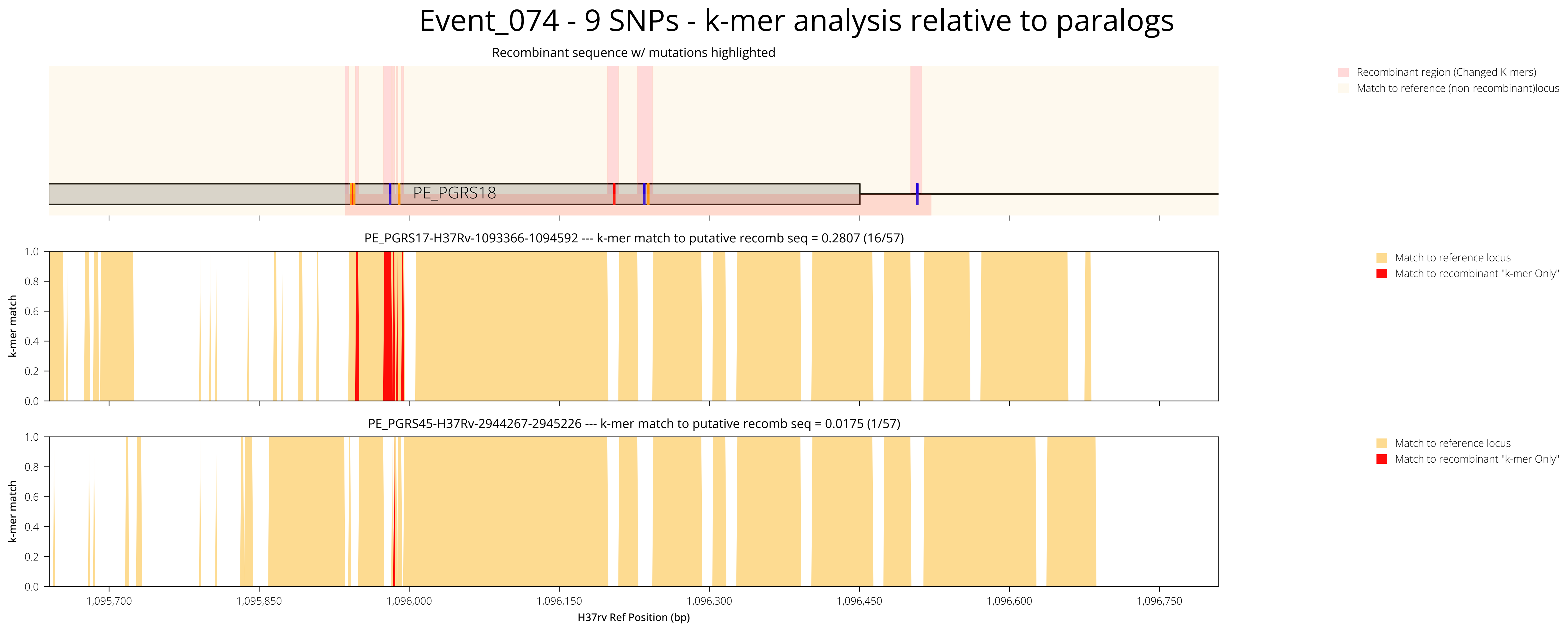

Event_074 - 9 SNPs - k-mer analysis relative to paralogs
Recombinant sequence w/ mutations highlighted
Recombinant region (Changed K-mers)
Match to reference (non-recombinant)locus
PE_PGRS18
PE_PGRS17-H37Rv-1093366-1094592 --- k-mer match to putative recomb seq = 0.2807 (16/57)
Match to reference locus
Match to recombinant "k-mer Only"
k-mer match
1.0
0.8
0.6
0.4
0.2
0.0
PE_PGRS45-H37Rv-2944267-2945226 --- k-mer match to putative recomb seq = 0.0175 (1/57)
Match to reference locus
Match to recombinant "k-mer Only"
k-mer match
1,095,700
1,095,850
1,096,000
1,096,150
1,096,300
1,096,450
1,096,600
1,096,750
H37rv Ref Position (bp)

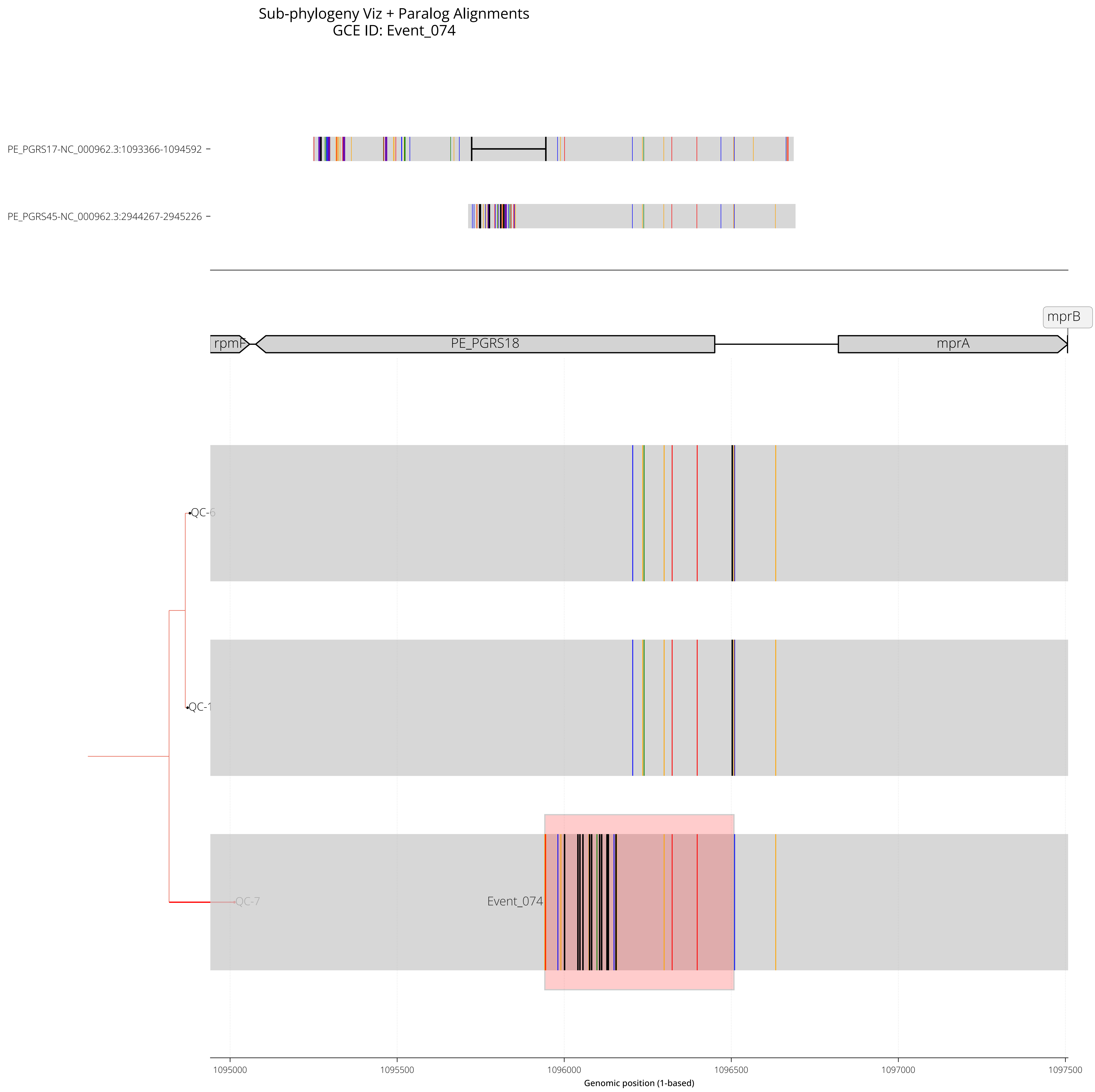

Sub-phylogeny Viz + Paralog Alignments
GCE ID: Event_074
PE_PGRS17-NC_000962.3:1093366-1094592
PE_PGRS45-NC_000962.3:2944267-2945226
mprB
rpmF
PE_PGRS18
mprA
QC-6
QC-1
QC-7
Event_074
1095000
1095500
1096000
1096500
1097000
1097500
Genomic position (1-based)

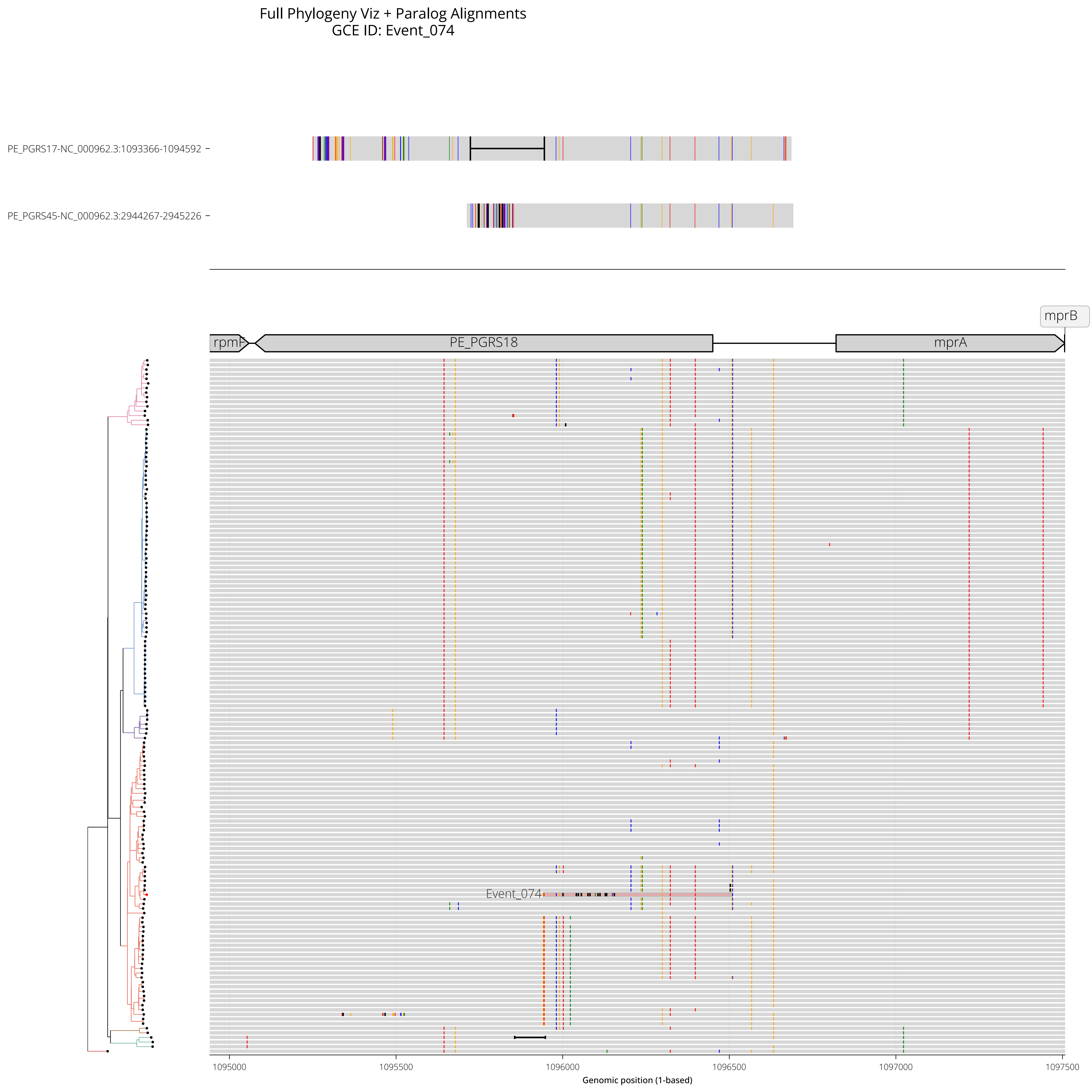
Full Phylogeny Viz + Paralog Alignments
GCE ID: Event_074
PE_PGRS17-NC_000962.3:1093366-1094592
PE_PGRS45-NC_000962.3:2944267-2945226
rpmF
PE_PGRS18
mprA
mprB
Event_074
1095000
1095500
1096000
1096500
1097000
1097500
Genomic position (1-based)

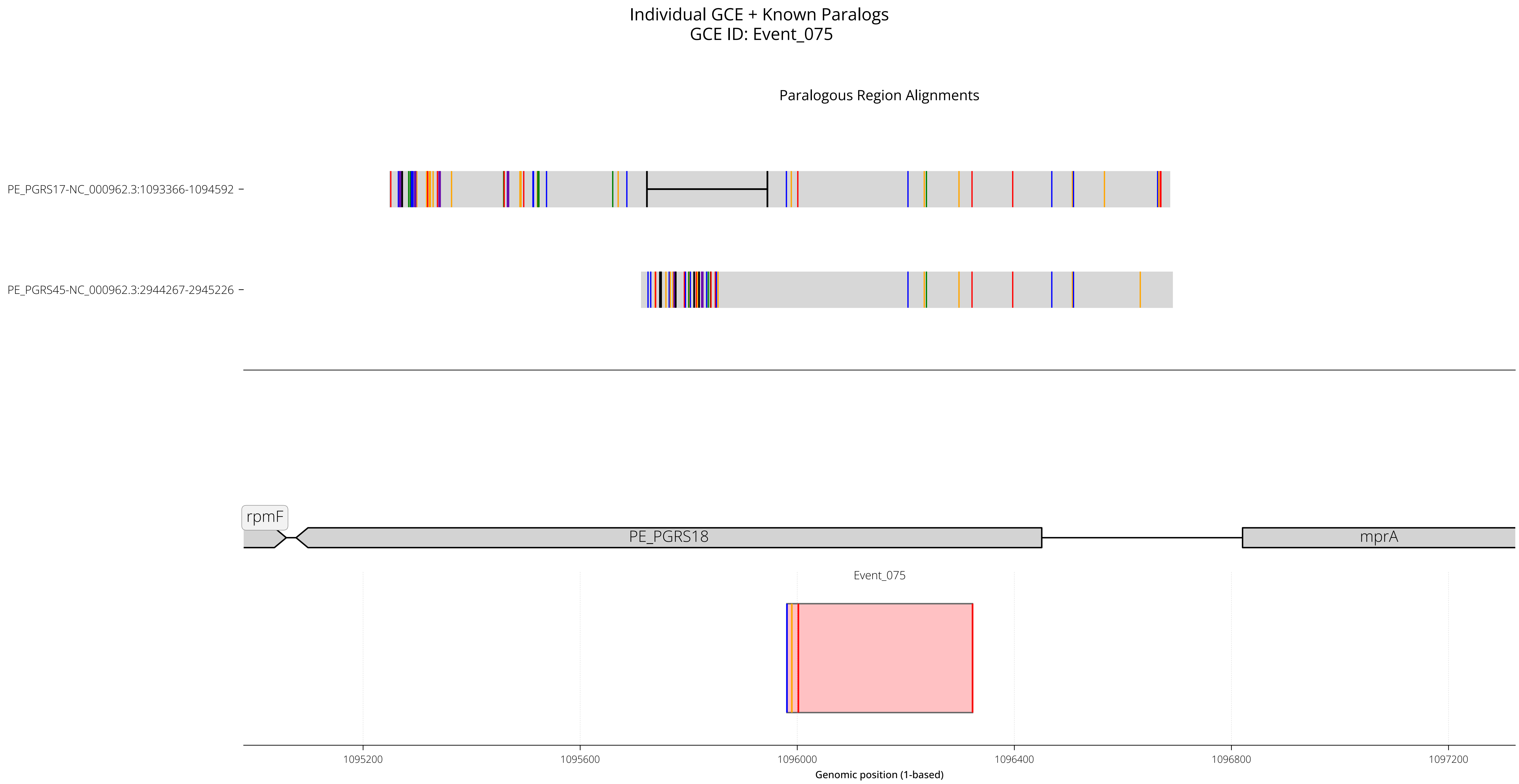

Individual GCE + Known Paralogs
GCE ID: Event_075
Paralogous Region Alignments
PE_PGRS17-NC_000962.3:1093366-1094592
PE_PGRS45-NC_000962.3:2944267-2945226
rpmF
PE_PGRS18
mprA
Event_075
1095200
1095600
1096000
1096400
1096800
1097200
Genomic position (1-based)

Circos Viz - GCE to Known Paralogs Locations
GCE ID: Event_075

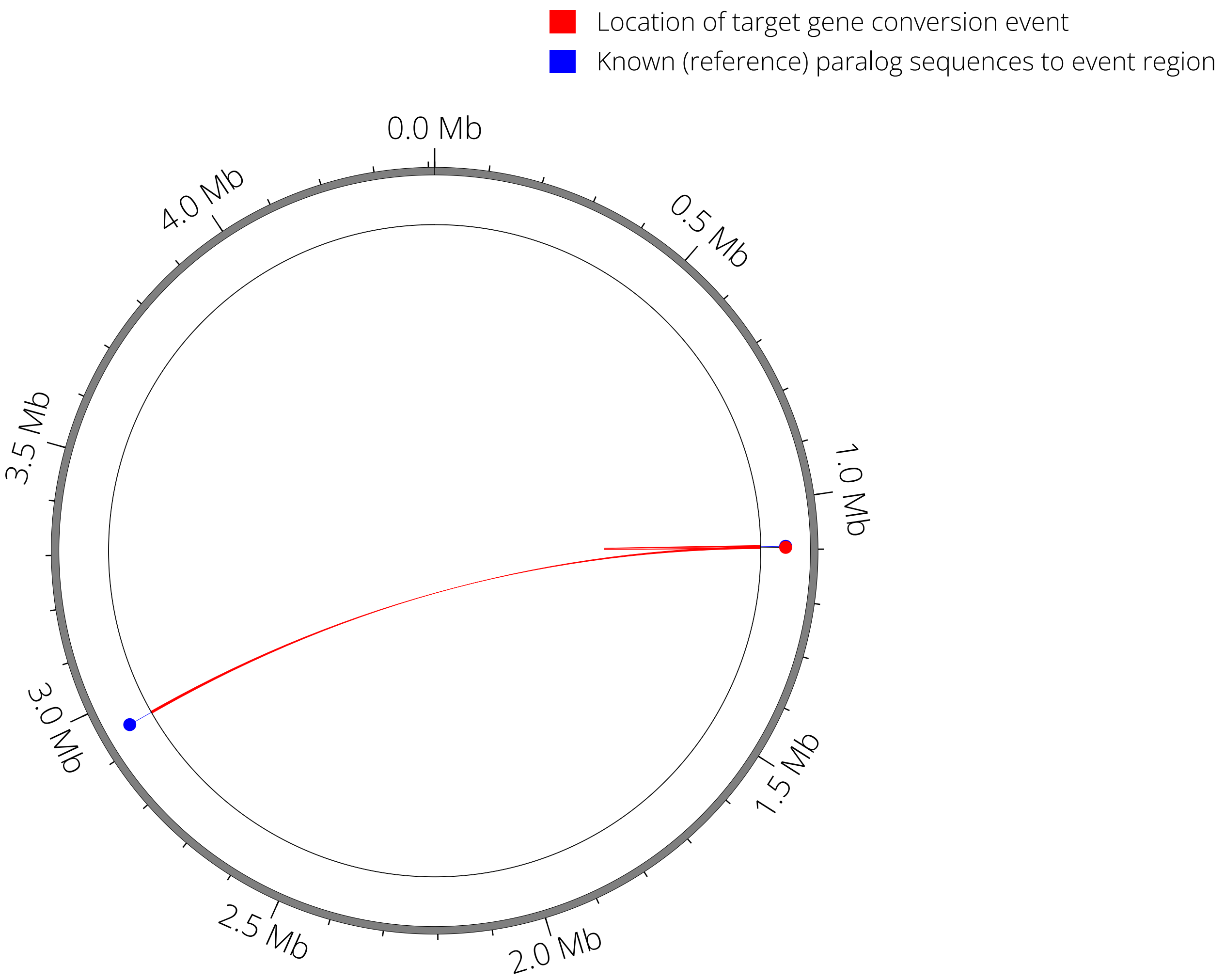

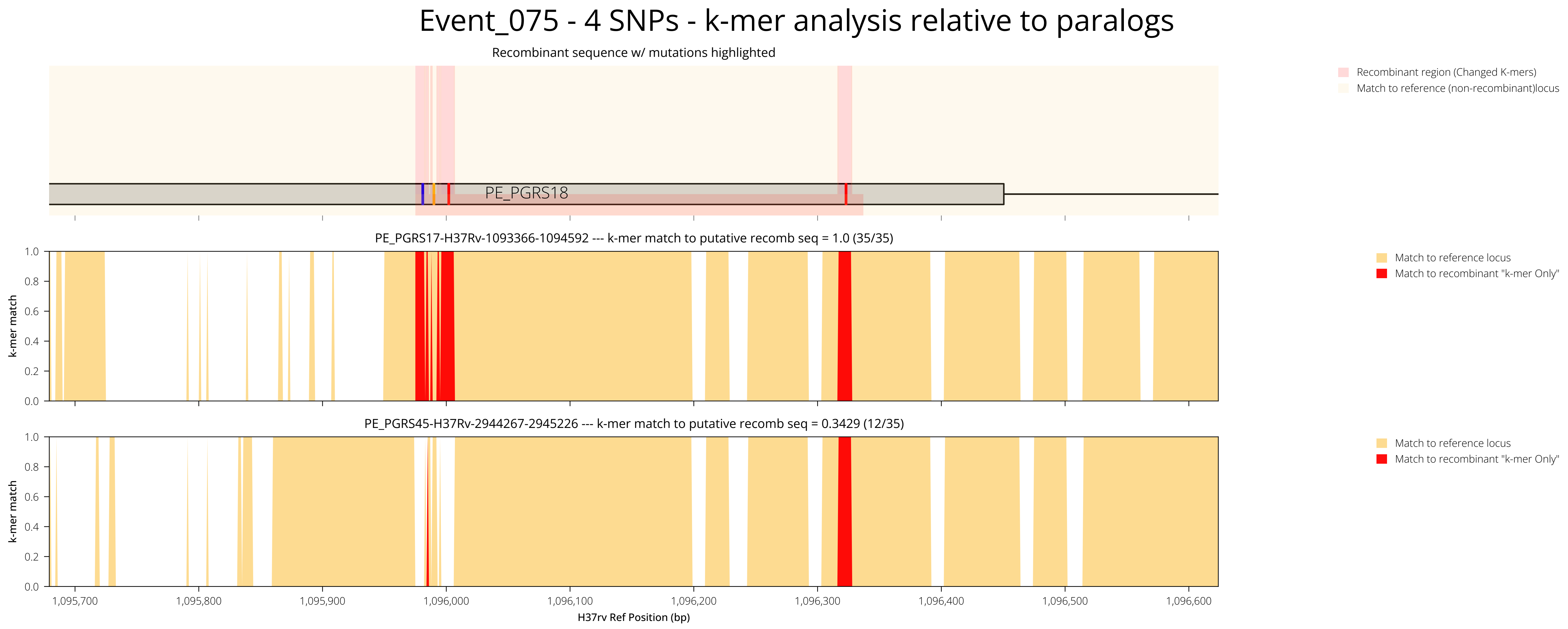

Event_075 - 4 SNPs - k-mer analysis relative to paralogs
Recombinant sequence w/ mutations highlighted
PE_PGRS18
Recombinant region (Changed K-mers)
Match to reference (non-recombinant)locus
PE_PGRS17-H37Rv-1093366-1094592 --- k-mer match to putative recomb seq = 1.0 (35/35)
Match to reference locus
Match to recombinant "k-mer Only"
PE_PGRS45-H37Rv-2944267-2945226 --- k-mer match to putative recomb seq = 0.3429 (12/35)
Match to reference locus
Match to recombinant "k-mer Only"
k-mer match
1.0
0.8
0.6
0.4
0.2
0.0
1,095,700
1,095,800
1,095,900
1,096,000
1,096,100
1,096,200
1,096,300
1,096,400
1,096,500
1,096,600
H37rv Ref Position (bp)

Sub-phylogeny Viz + Paralog Alignments
GCE ID: Event_075

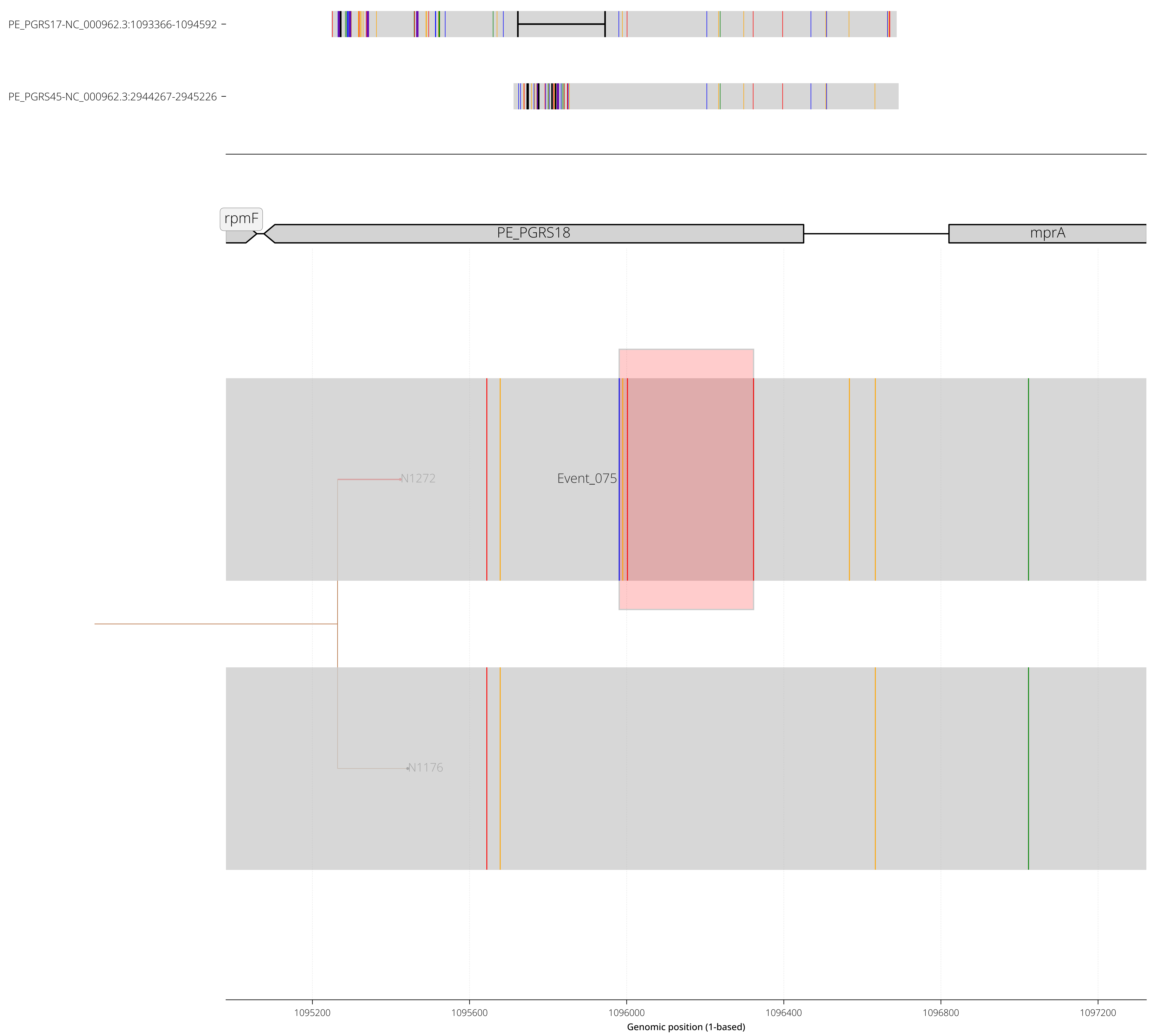

Full Phylogeny Viz + Paralog Alignments
GCE ID: Event_075

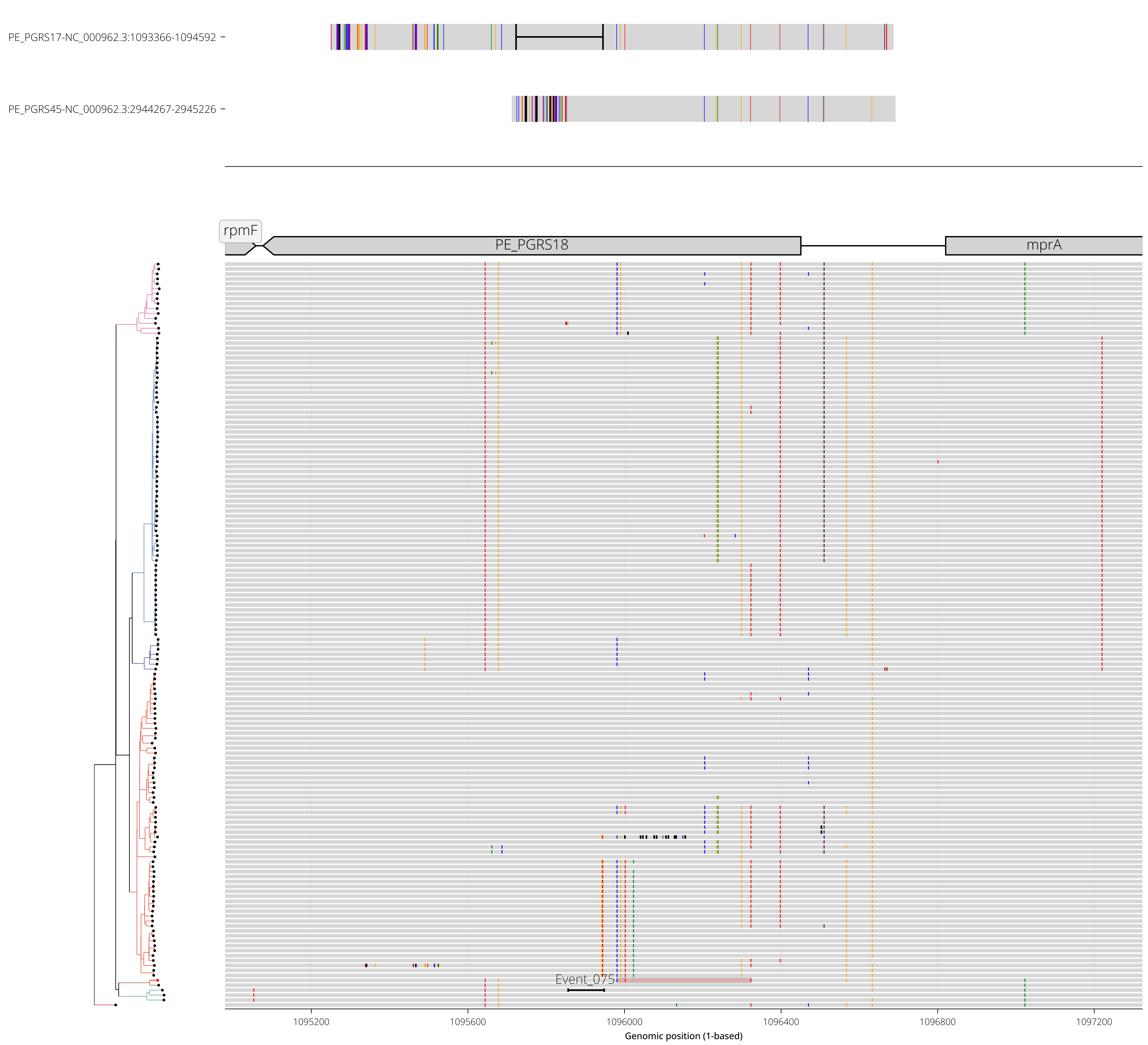

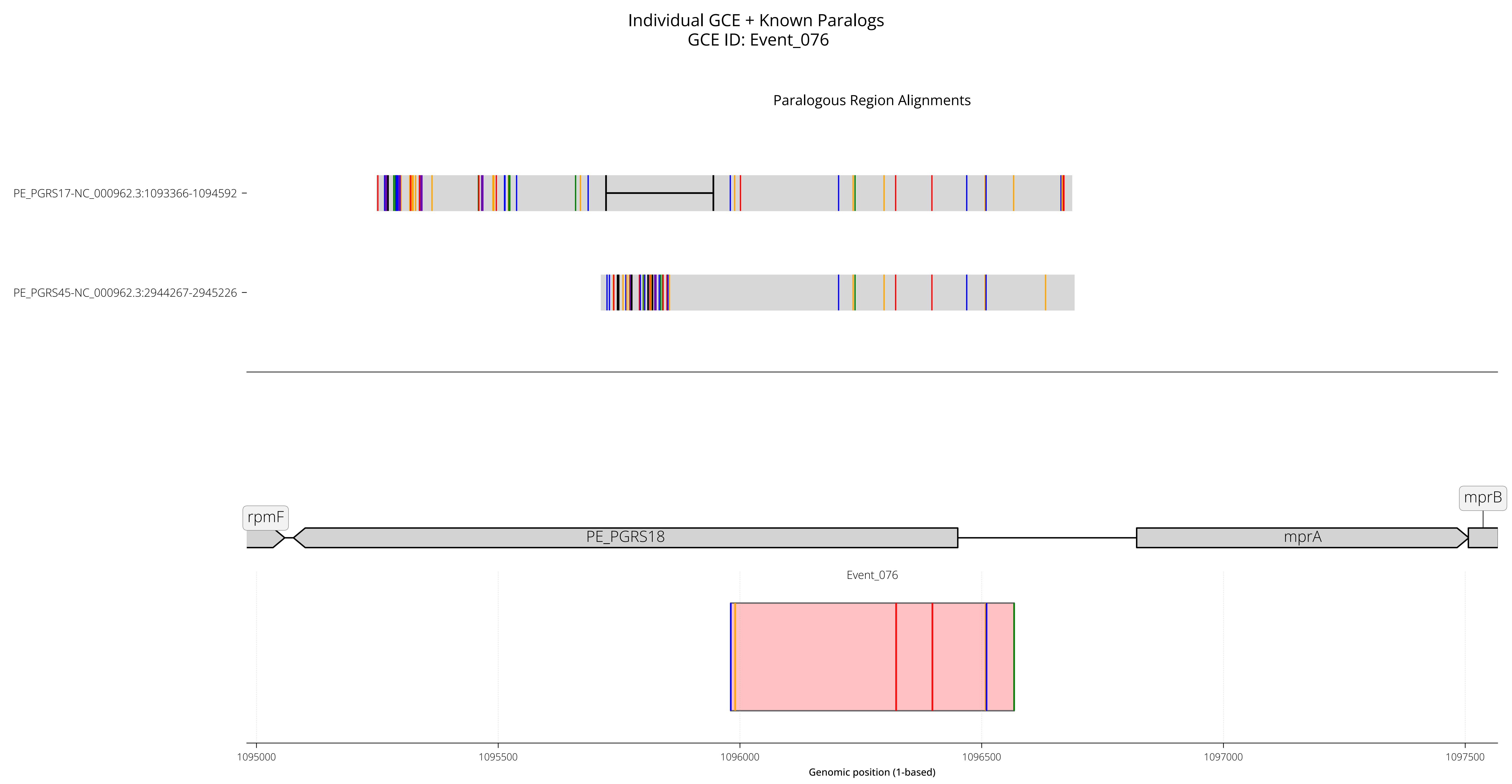

Individual GCE + Known Paralogs
GCE ID: Event_076
Paralogous Region Alignments
PE_PGRS17-NC_000962.3:1093366-1094592
PE_PGRS45-NC_000962.3:2944267-2945226
rpmF
PE_PGRS18
mprA
mprB
Event_076
1095000
1095500
1096000
1096500
1097000
1097500
Genomic position (1-based)

Circos Viz - GCE to Known Paralogs Locations
GCE ID: Event_076

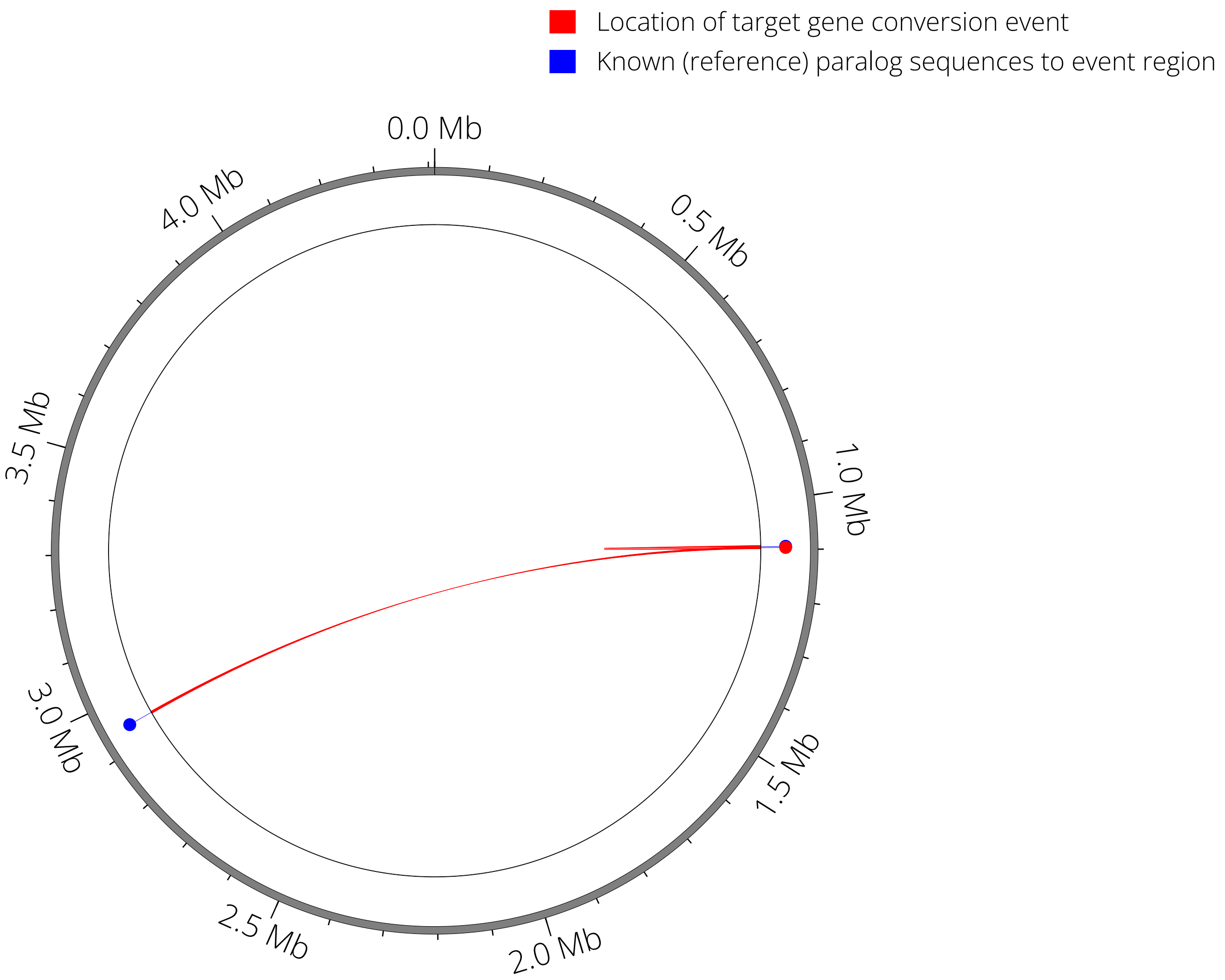

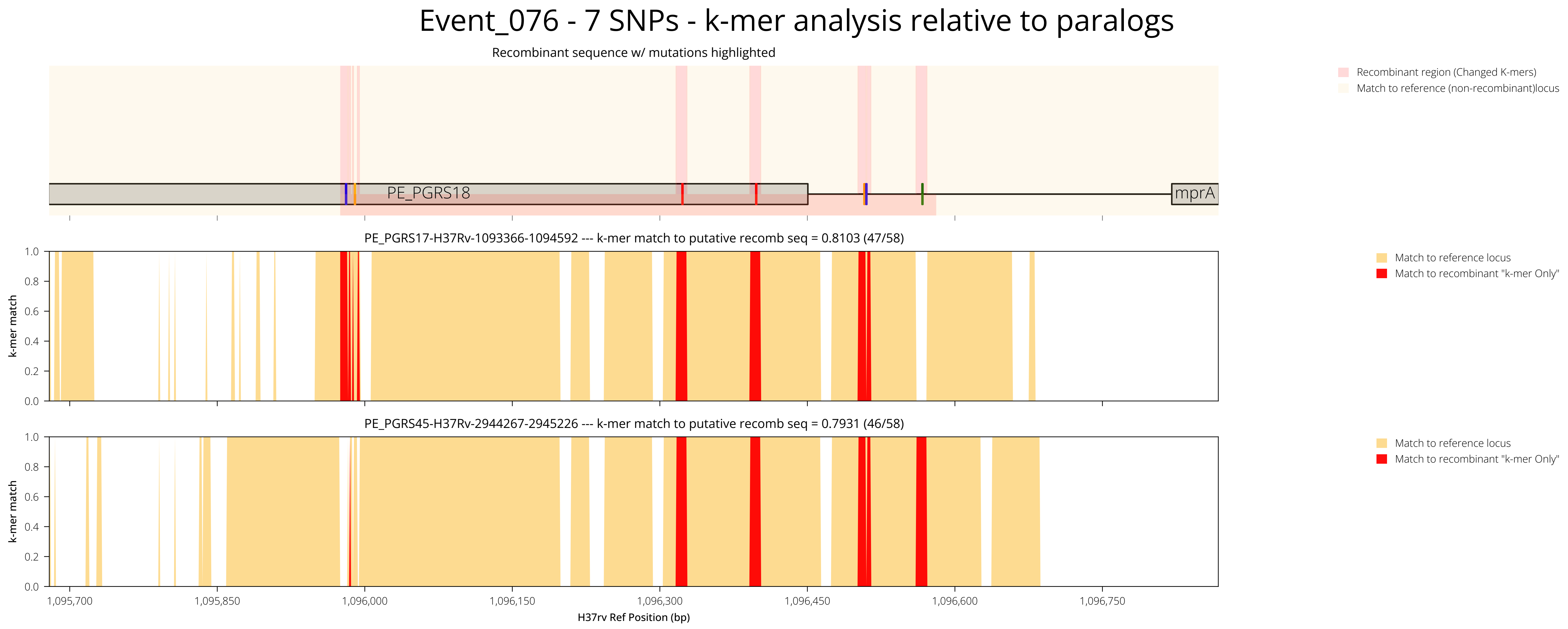

Event_076 - 7 SNPs - k-mer analysis relative to paralogs
Recombinant sequence w/ mutations highlighted
Recombinant region (Changed K-mers)
Match to reference (non-recombinant)locus
PE_PGRS18
mprA
PE_PGRS17-H37Rv-1093366-1094592 --- k-mer match to putative recomb seq = 0.8103 (47/58)
Match to reference locus
Match to recombinant "k-mer Only"
k-mer match
PE_PGRS45-H37Rv-2944267-2945226 --- k-mer match to putative recomb seq = 0.7931 (46/58)
1,095,700
1,095,850
1,096,000
1,096,150
1,096,300
1,096,450
1,096,600
1,096,750
H37rv Ref Position (bp)

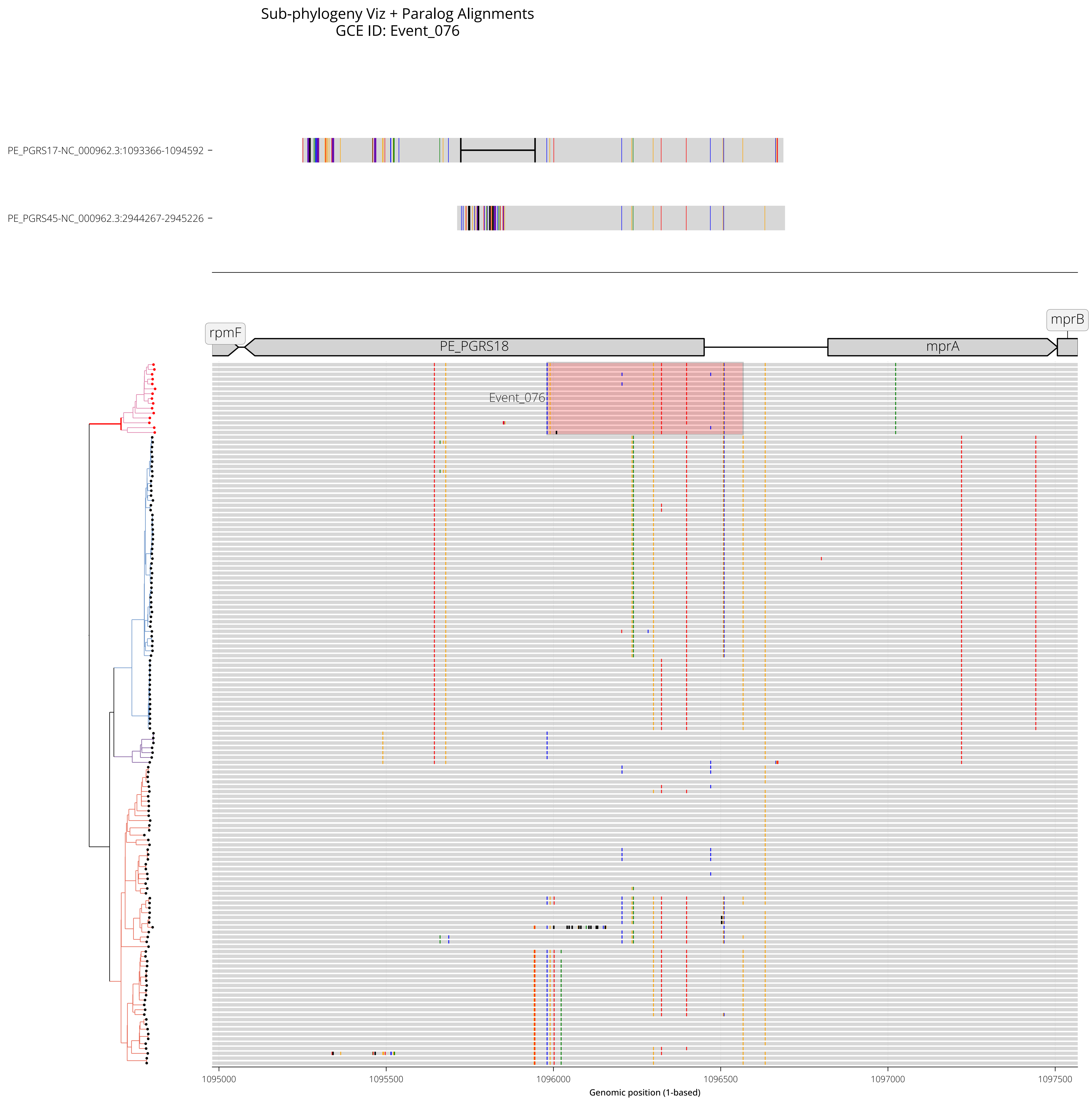

Sub-phylogeny Viz + Paralog Alignments
GCE ID: Event_076
PE_PGRS17-NC_000962.3:1093366-1094592
PE_PGRS45-NC_000962.3:2944267-2945226
rpmF
PE_PGRS18
mprA
mprB
Event_076
1095000
1095500
1096000
1096500
1097000
1097500
Genomic position (1-based)

Full Phylogeny Viz + Paralog Alignments
GCE ID: Event_076

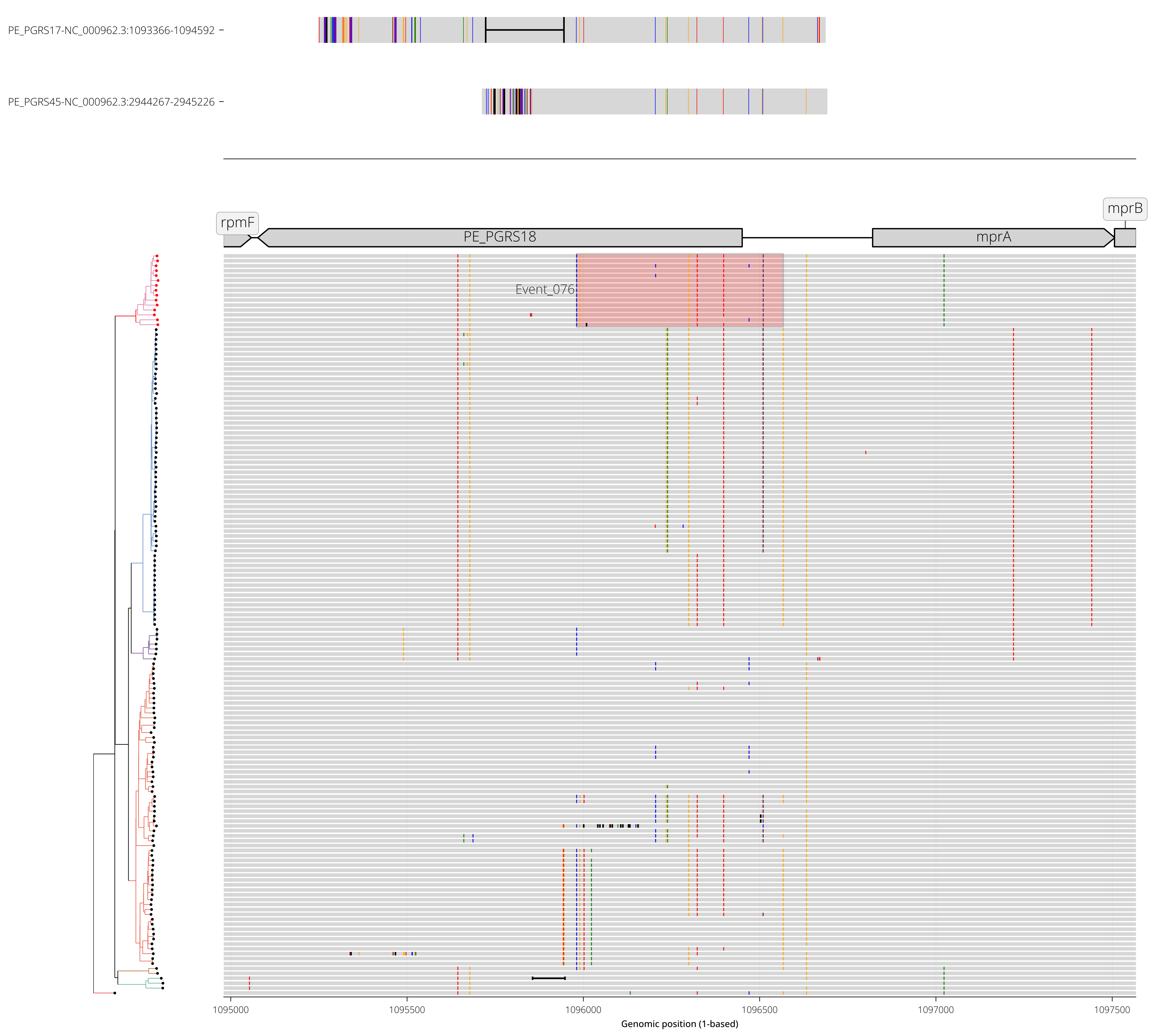

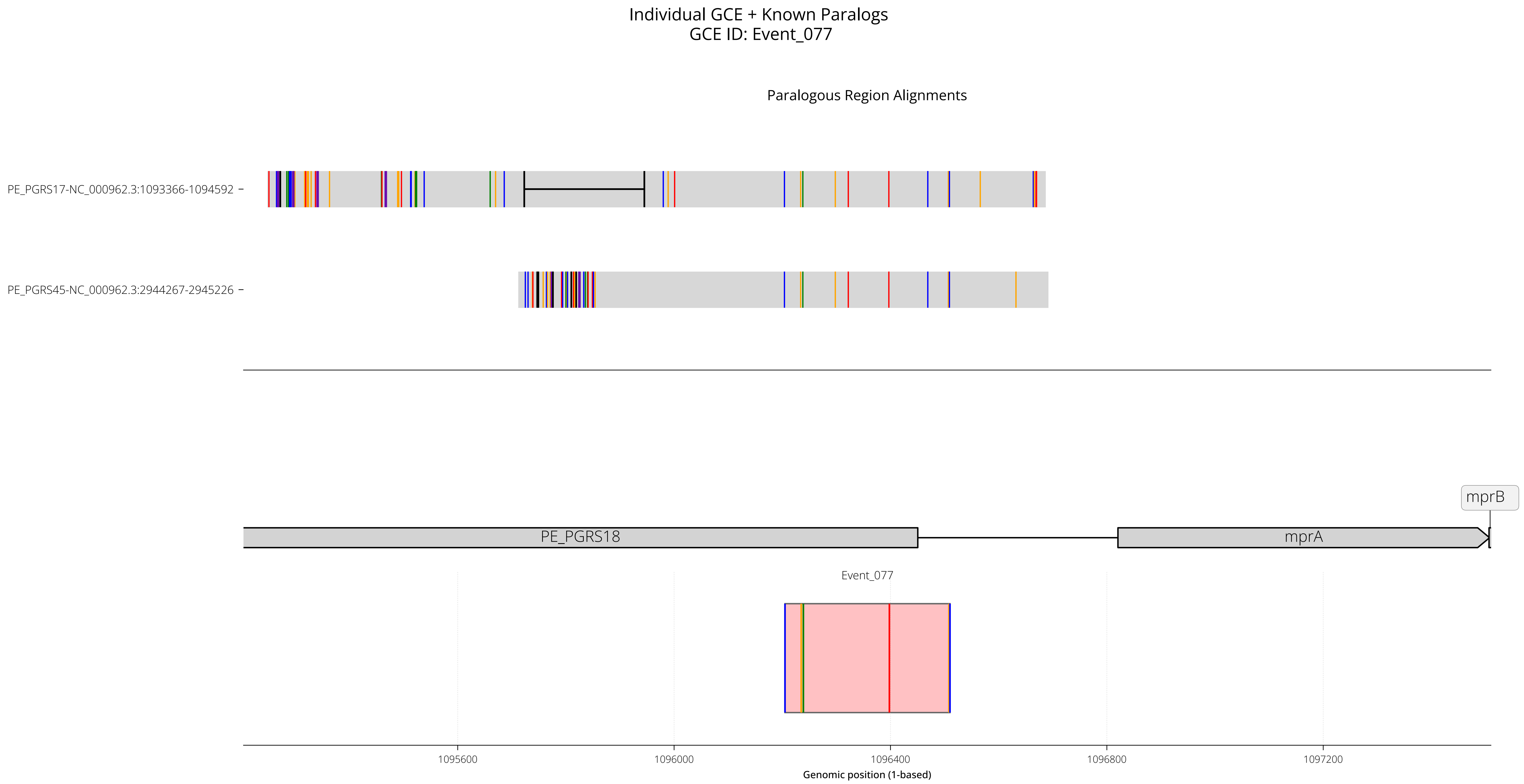

Individual GCE + Known Paralogs
GCE ID: Event_077
Paralogous Region Alignments
PE_PGRS17-NC_000962.3:1093366-1094592
PE_PGRS45-NC_000962.3:2944267-2945226
mprB
PE_PGRS18
mprA
Event_077
1095600
1096000
1096400
1096800
1097200
Genomic position (1-based)

### Circos Viz - GCE to Known Paralogs Locations
GCE ID: Event_077

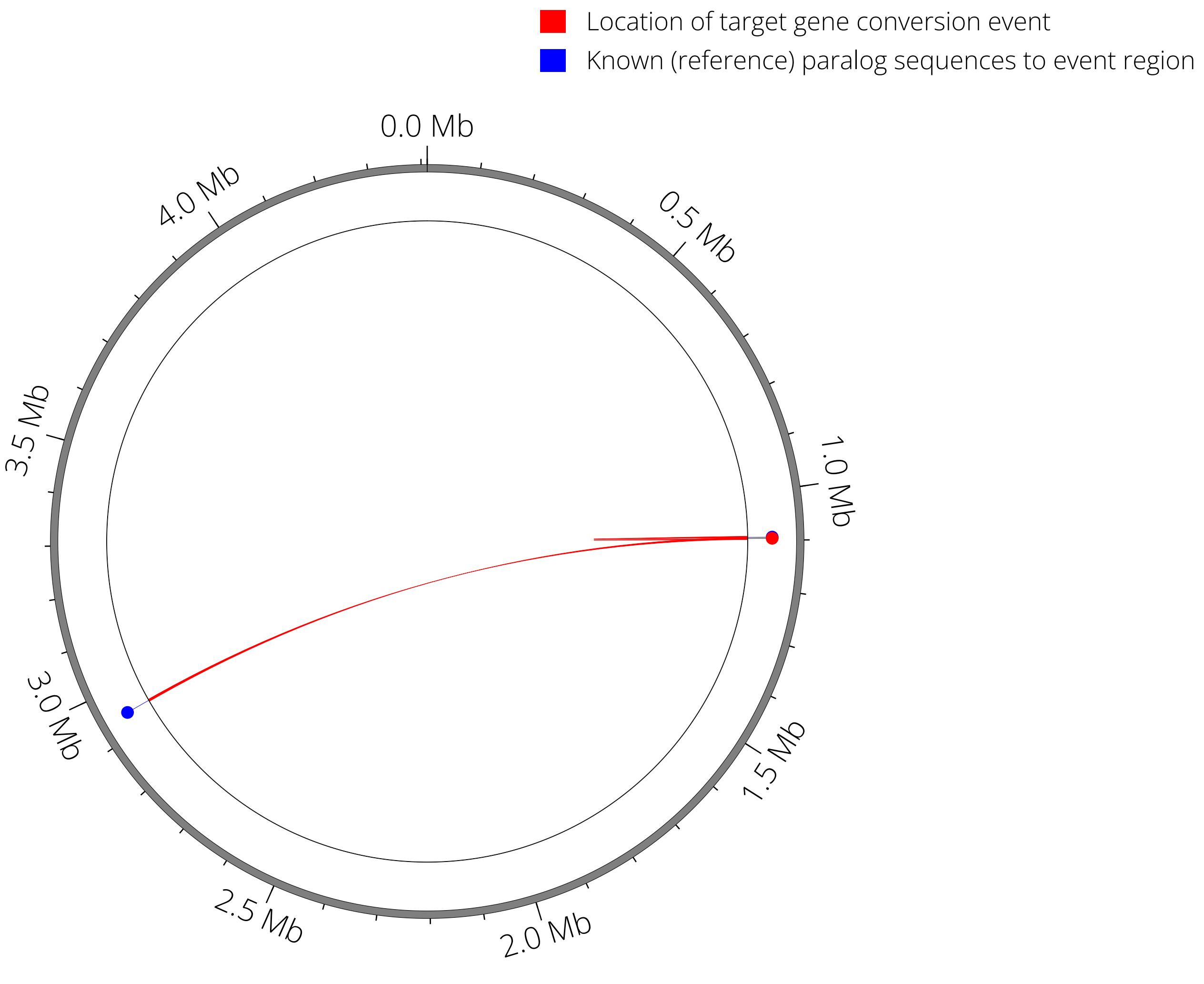

### Event_077 - 6 SNPs - k-mer analysis relative to paralogs

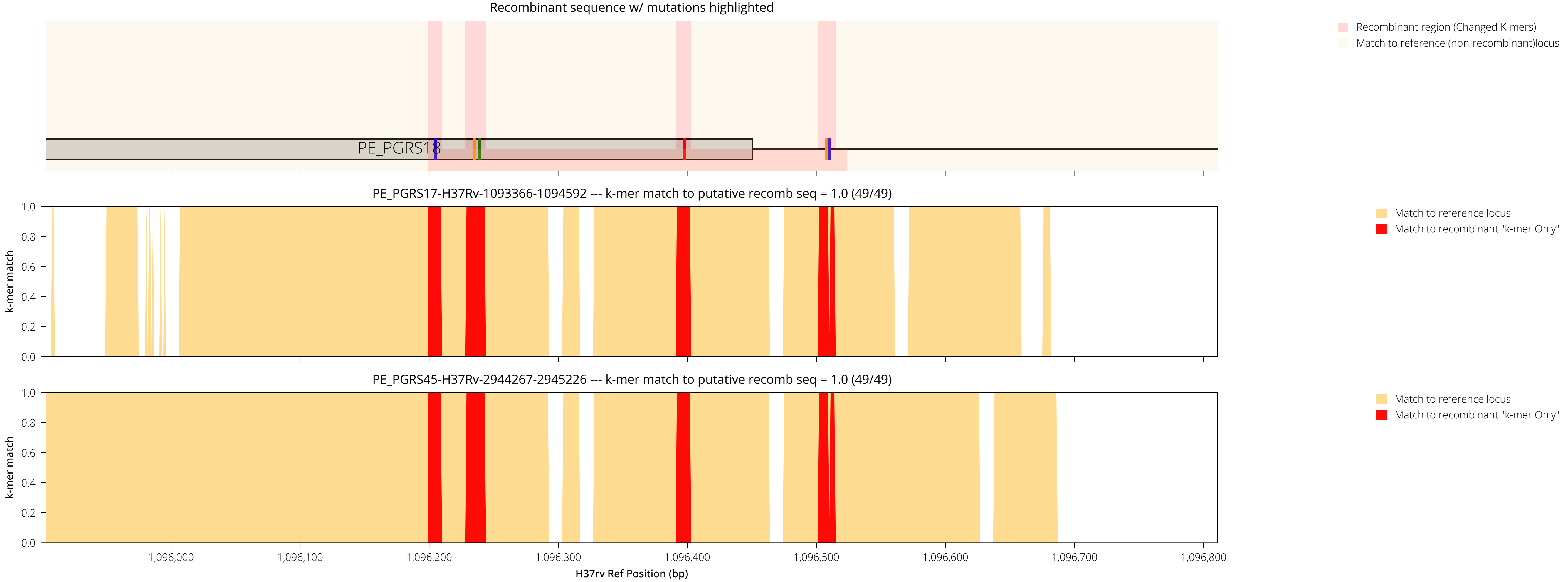

Sub-phylogeny Viz + Paralog Alignments
GCE ID: Event_077

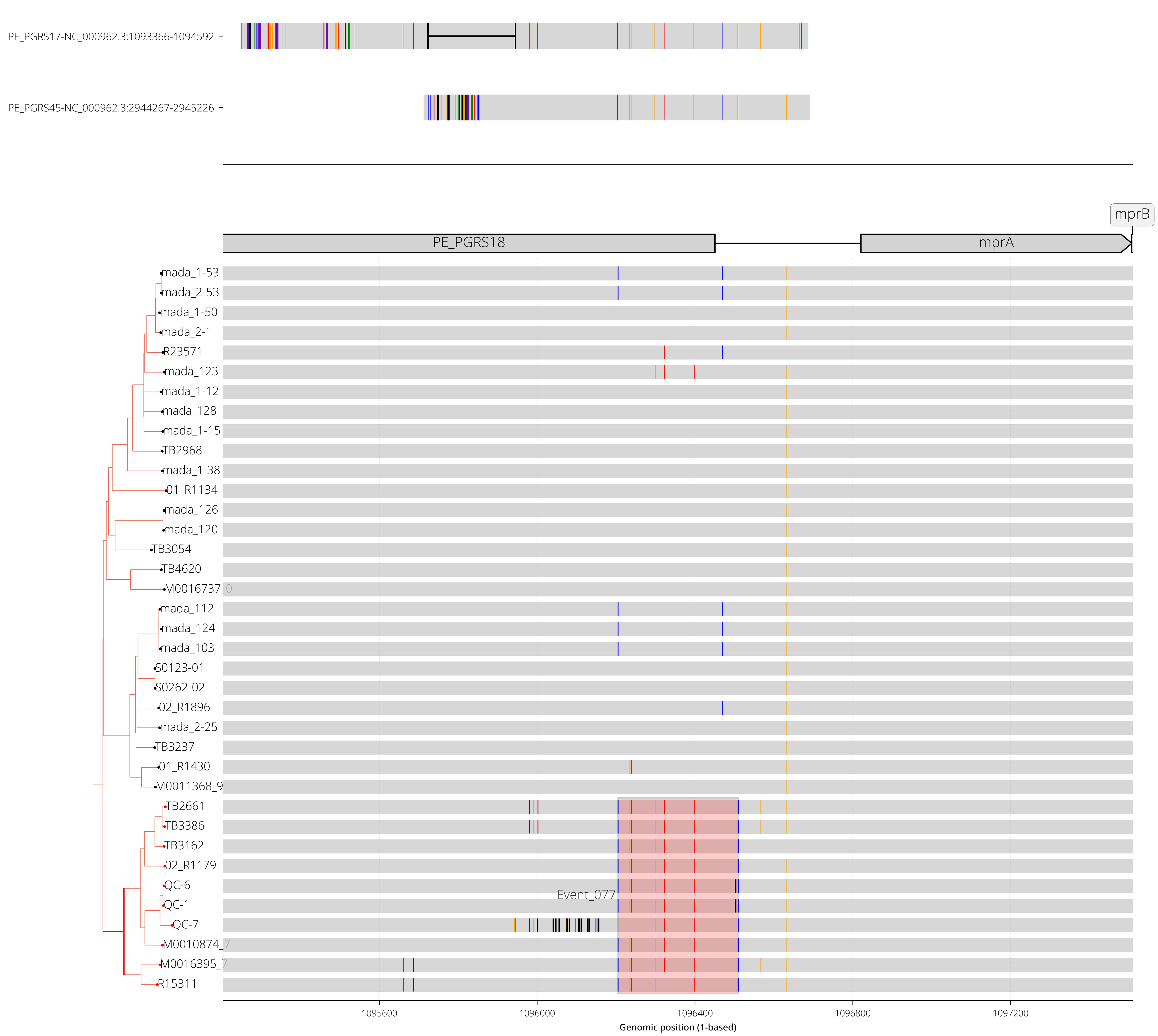

Full Phylogeny Viz + Paralog Alignments
GCE ID: Event_077

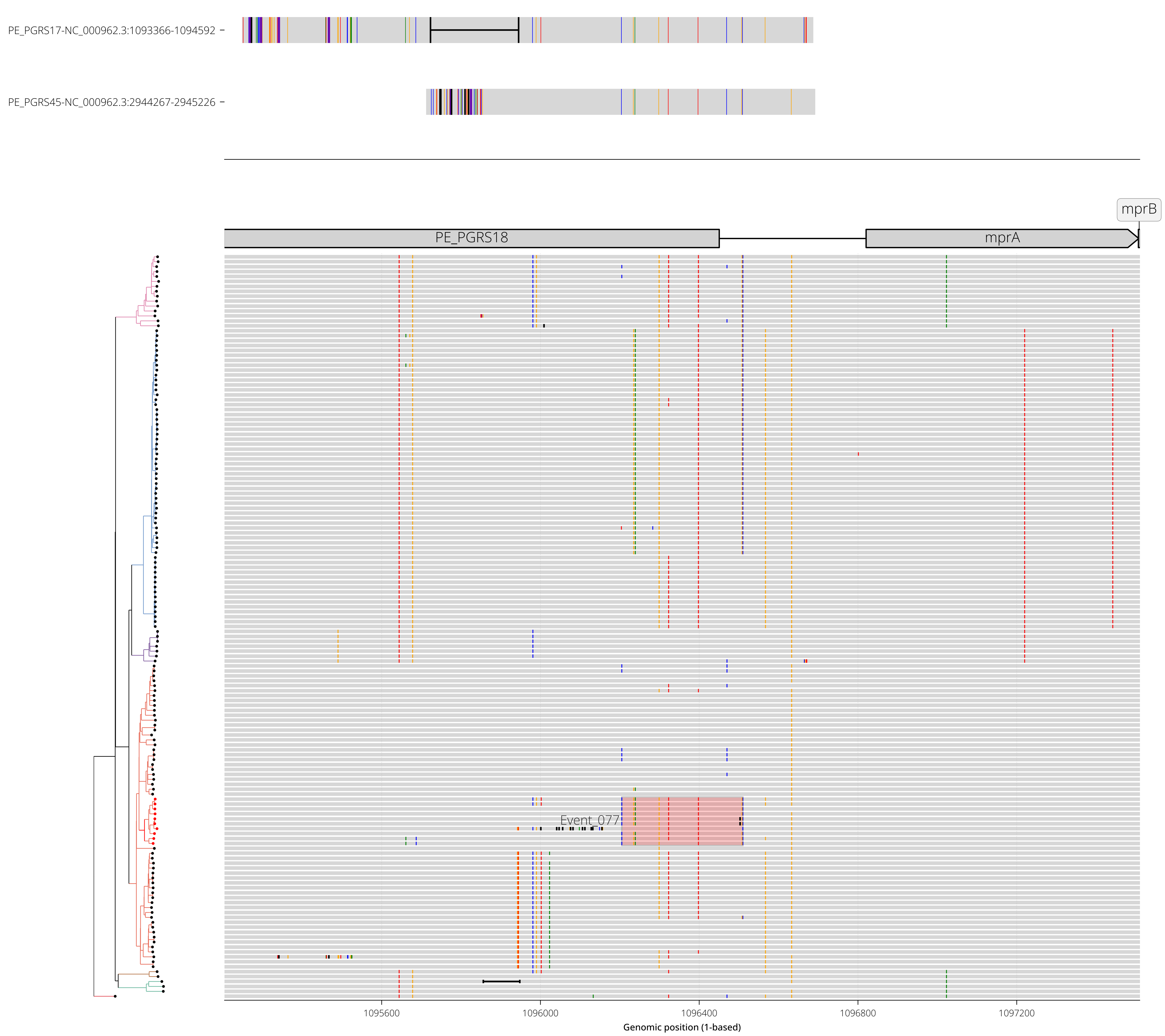

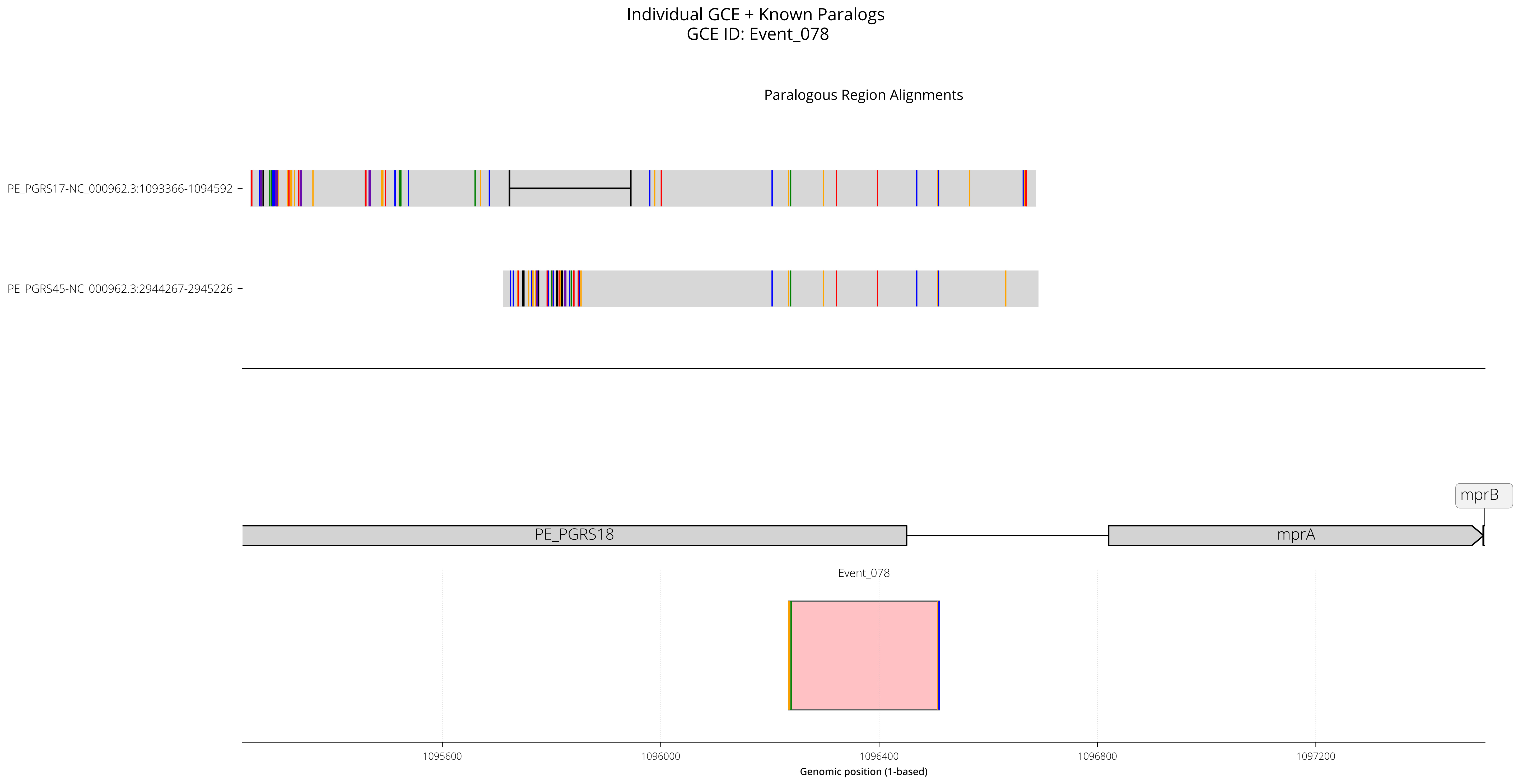

Individual GCE + Known Paralogs
GCE ID: Event_078
Paralogous Region Alignments
PE_PGRS17-NC_000962.3:1093366-1094592
PE_PGRS45-NC_000962.3:2944267-2945226
mprB
PE_PGRS18
mprA
Event_078
1095600
1096000
1096400
1096800
1097200
Genomic position (1-based)

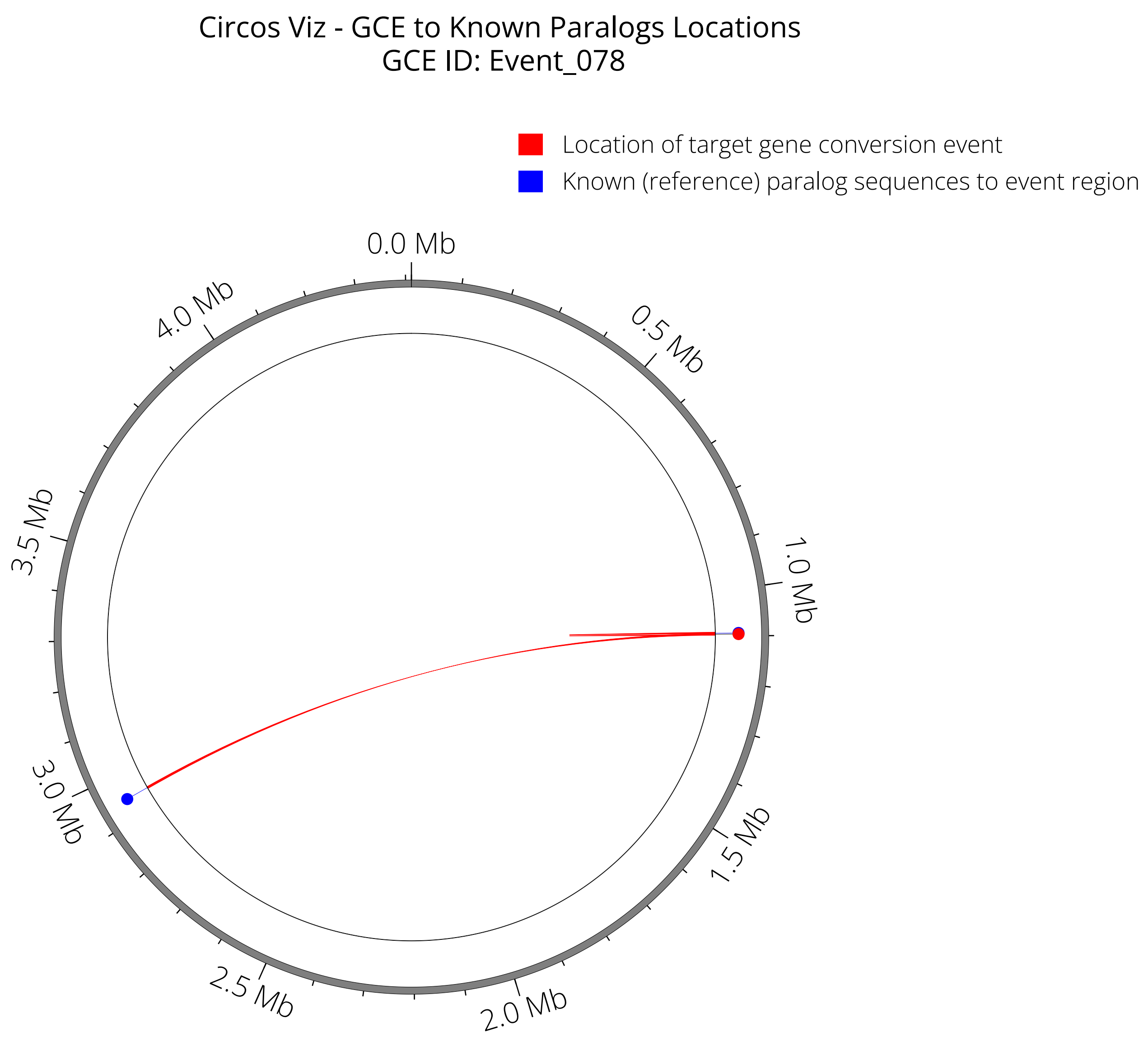
Circos Viz - GCE to Known Paralogs Locations
GCE ID: Event_078
Location of target gene conversion event
Known (reference) paralog sequences to event region
0.0 Mb
0.5 Mb
1.0 Mb
1.5 Mb
2.0 Mb
2.5 Mb
3.0 Mb
3.5 Mb
4.0 Mb

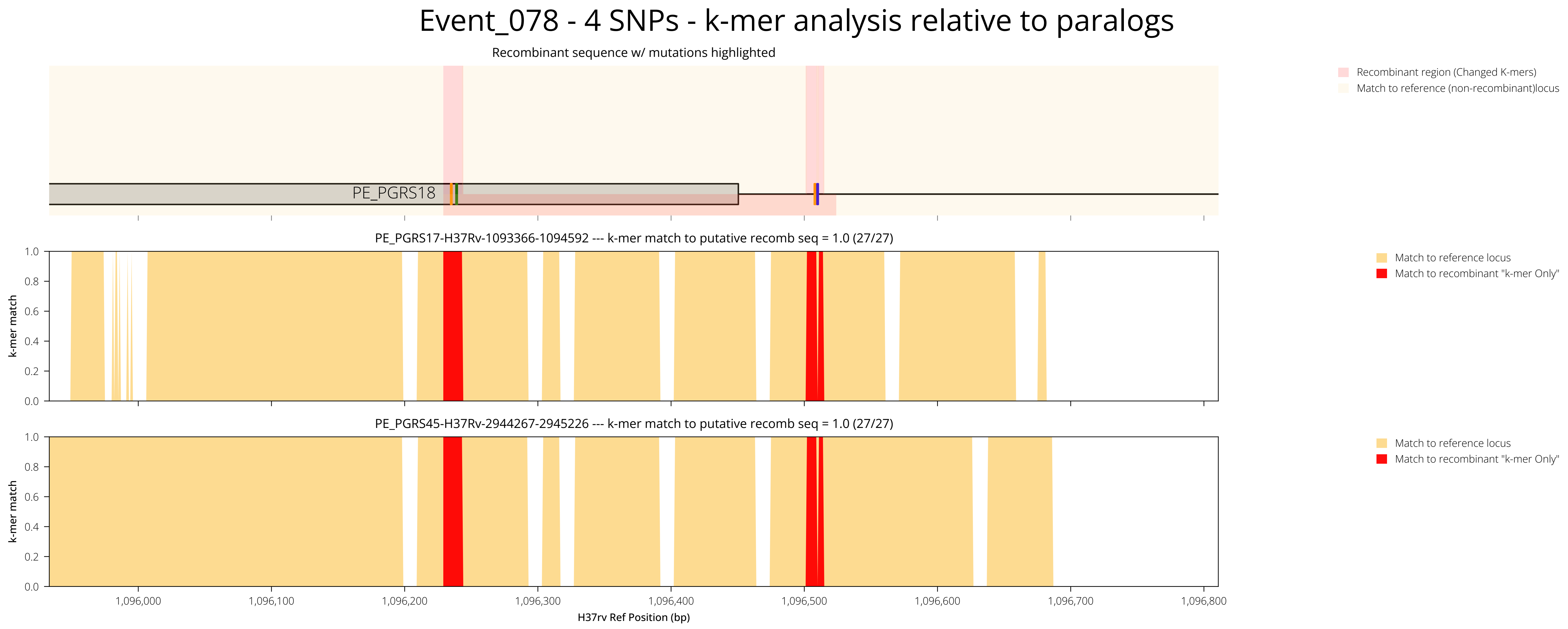

Event_078 - 4 SNPs - k-mer analysis relative to paralogs
Recombinant sequence w/ mutations highlighted
Recombinant region (Changed K-mers)
Match to reference (non-recombinant)locus
PE_PGRS18
PE_PGRS17-H37Rv-1093366-1094592 --- k-mer match to putative recomb seq = 1.0 (27/27)
Match to reference locus
Match to recombinant "k-mer Only"
k-mer match
1.0
0.8
0.6
0.4
0.2
0.0
PE_PGRS45-H37Rv-2944267-2945226 --- k-mer match to putative recomb seq = 1.0 (27/27)
Match to reference locus
Match to recombinant "k-mer Only"
k-mer match
1.0
0.8
0.6
0.4
0.2
0.0
1,096,000
1,096,100
1,096,200
1,096,300
1,096,400
1,096,500
1,096,600
1,096,700
1,096,800
H37rv Ref Position (bp)

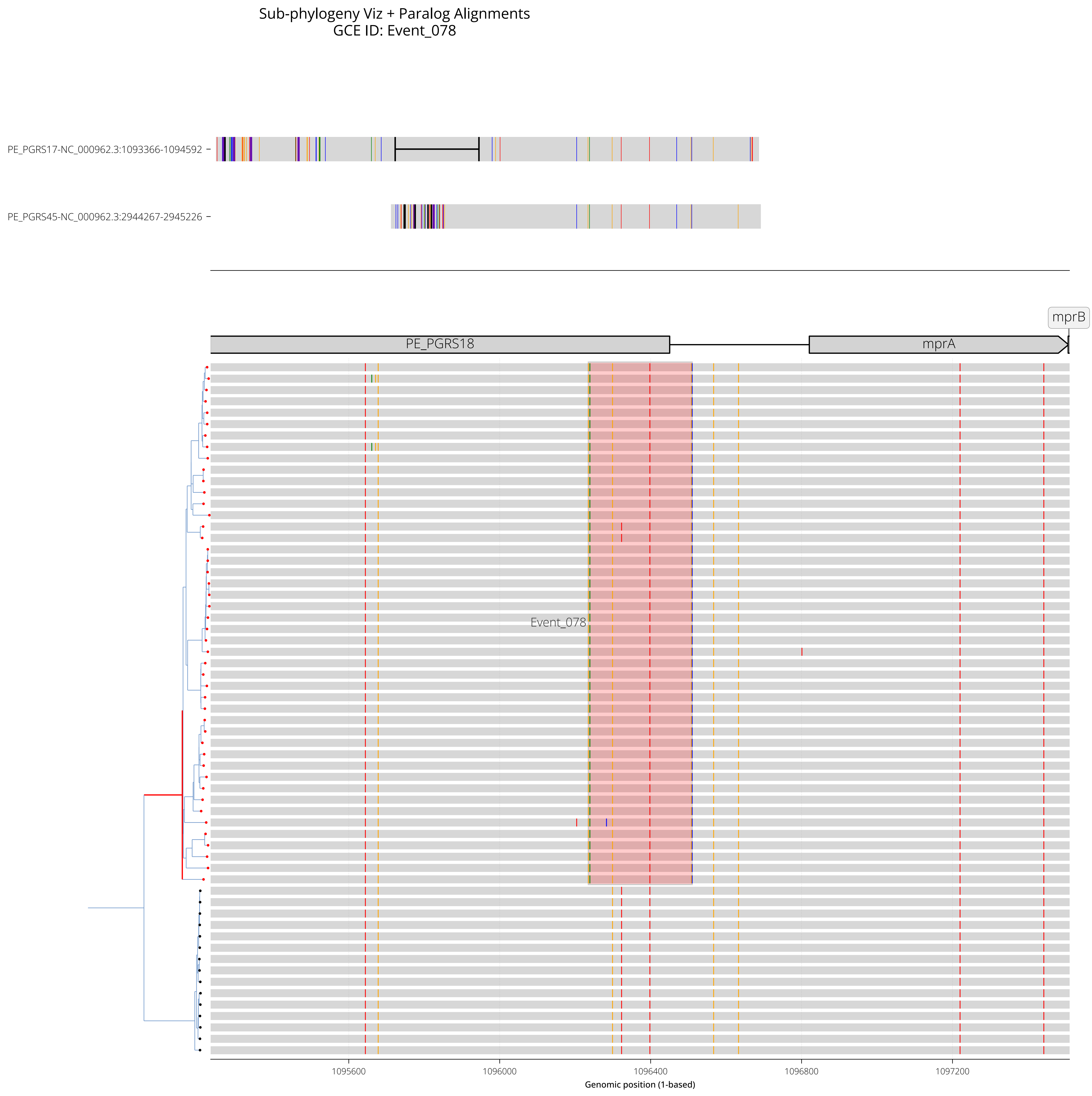

Sub-phylogeny Viz + Paralog Alignments
GCE ID: Event_078
PE_PGRS17-NC_000962.3:1093366-1094592
PE_PGRS45-NC_000962.3:2944267-2945226
mprB
PE_PGRS18
mprA
Event_078
1095600
1096000
1096400
1096800
1097200
Genomic position (1-based)

### Full Phylogeny Viz + Paralog Alignments
GCE ID: Event_078

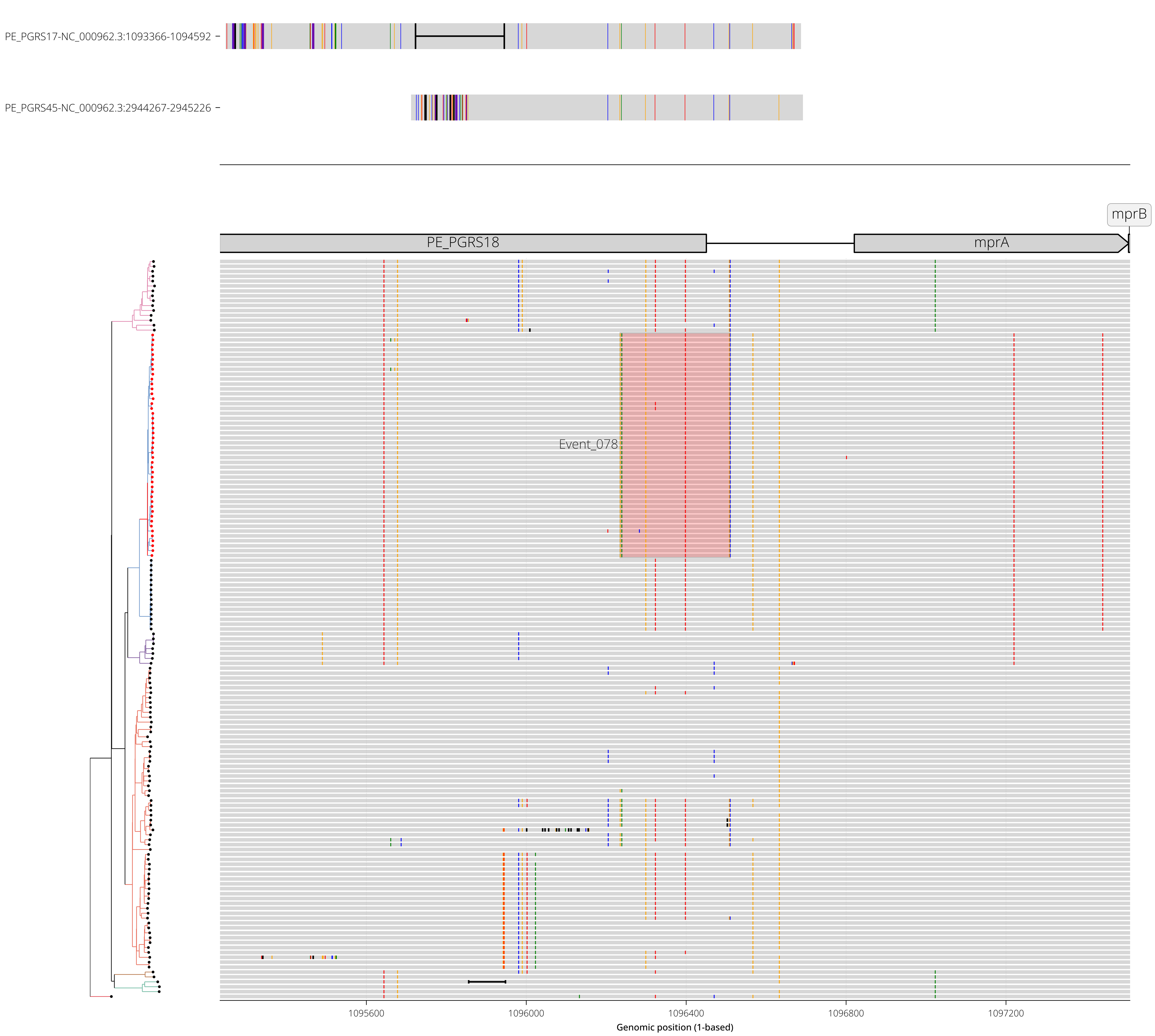

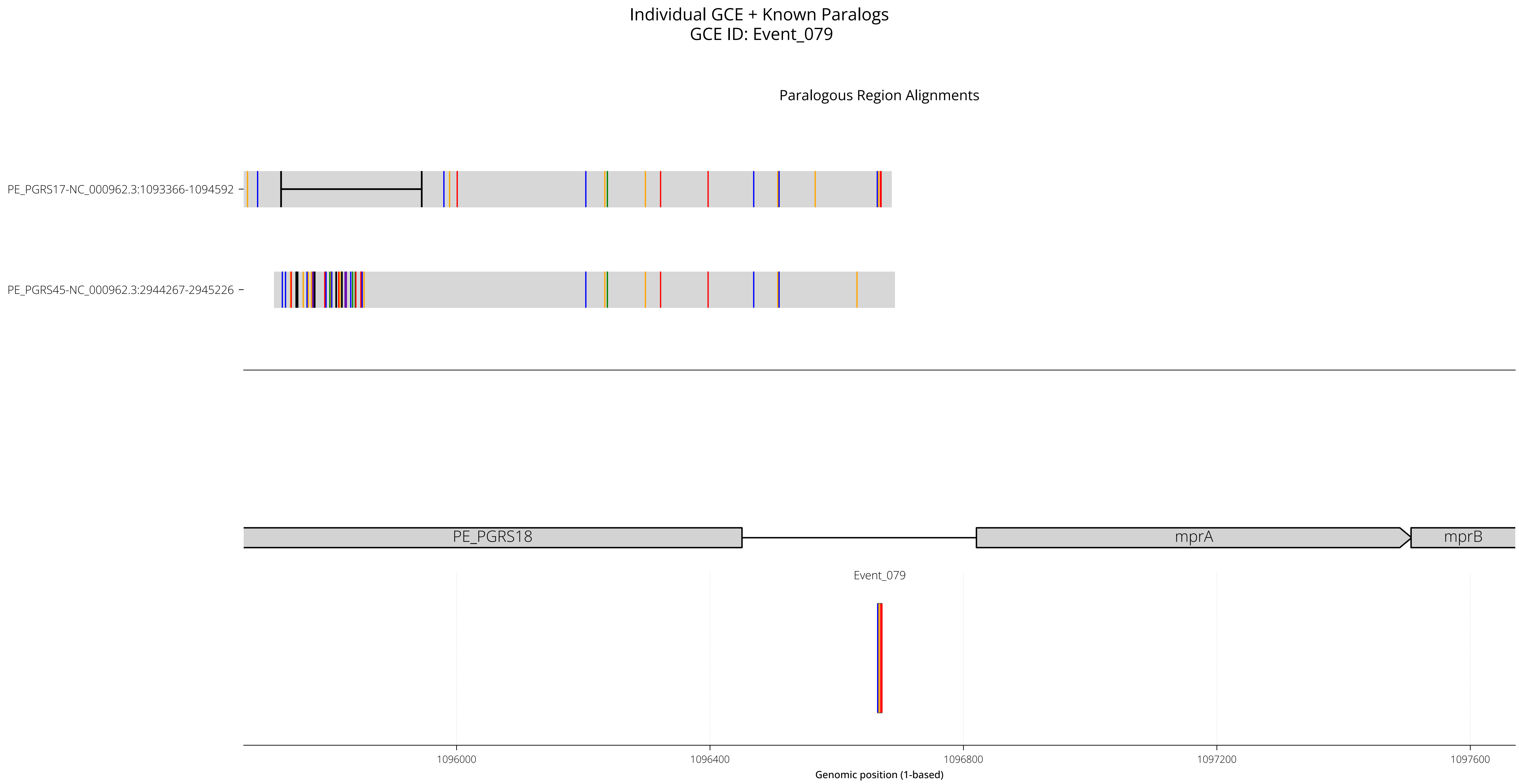
Individual GCE + Known Paralogs
GCE ID: Event_079
Paralogous Region Alignments
PE_PGRS17-NC_000962.3:1093366-1094592
PE_PGRS45-NC_000962.3:2944267-2945226
PE_PGRS18
mprA
mprB
Event_079
1096000
1096400
1096800
1097200
1097600
Genomic position (1-based)

Circos Viz - GCE to Known Paralogs Locations
GCE ID: Event_079

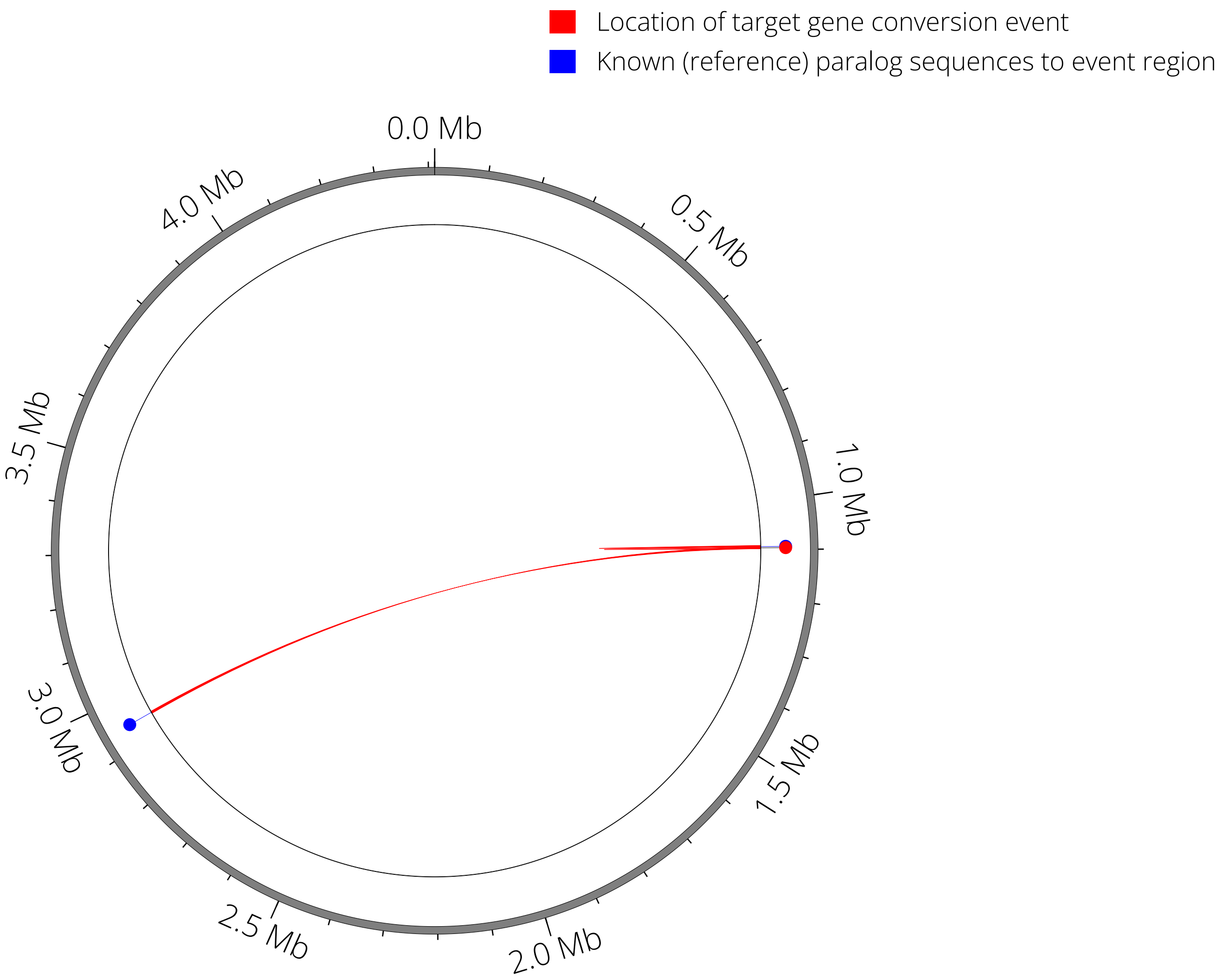

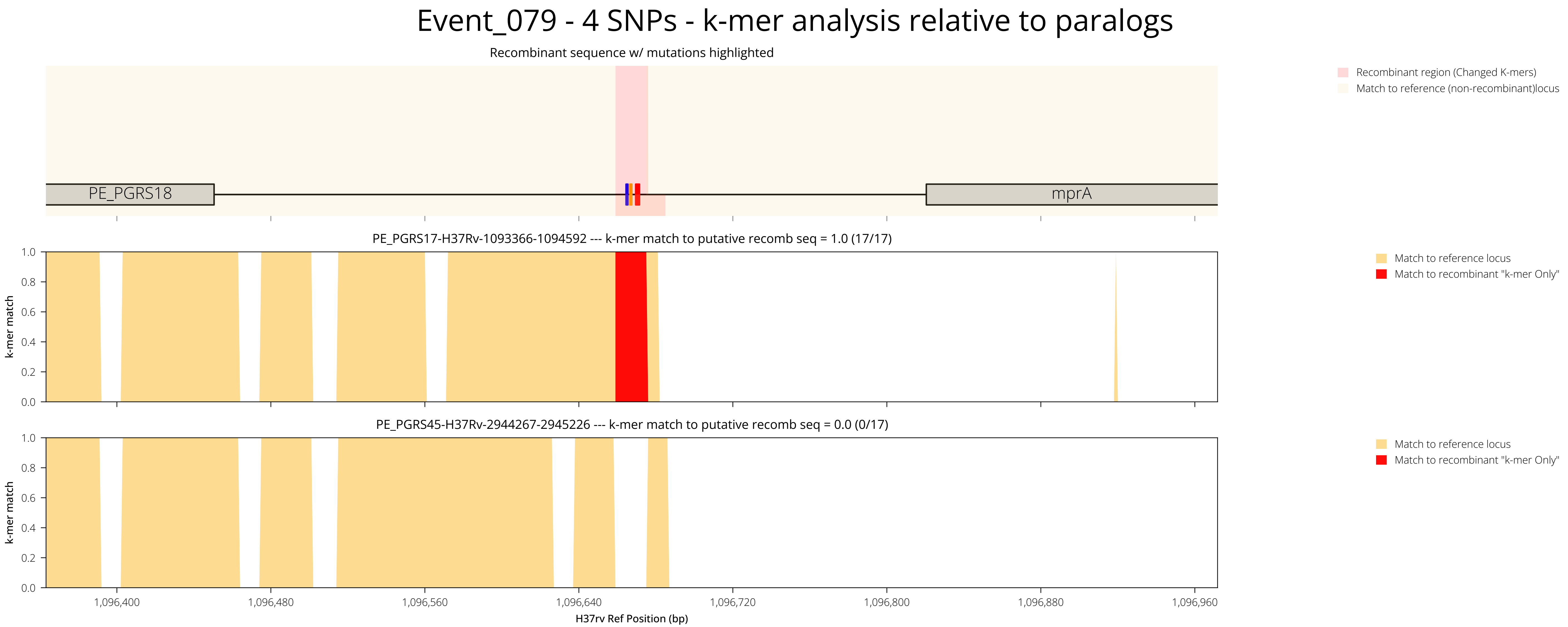

Event_079 - 4 SNPs - k-mer analysis relative to paralogs
Recombinant sequence w/ mutations highlighted
PE_PGRS18
mprA
Recombinant region (Changed K-mers)
Match to reference (non-recombinant)locus
PE_PGRS17-H37Rv-1093366-1094592 --- k-mer match to putative recomb seq = 1.0 (17/17)
Match to reference locus
Match to recombinant "k-mer Only"
PE_PGRS45-H37Rv-2944267-2945226 --- k-mer match to putative recomb seq = 0.0 (0/17)
Match to reference locus
Match to recombinant "k-mer Only"
k-mer match
1.0
0.8
0.6
0.4
0.2
0.0
1,096,400
1,096,480
1,096,560
1,096,640
1,096,720
1,096,800
1,096,880
1,096,960
H37rv Ref Position (bp)

Sub-phylogeny Viz + Paralog Alignments
GCE ID: Event_079

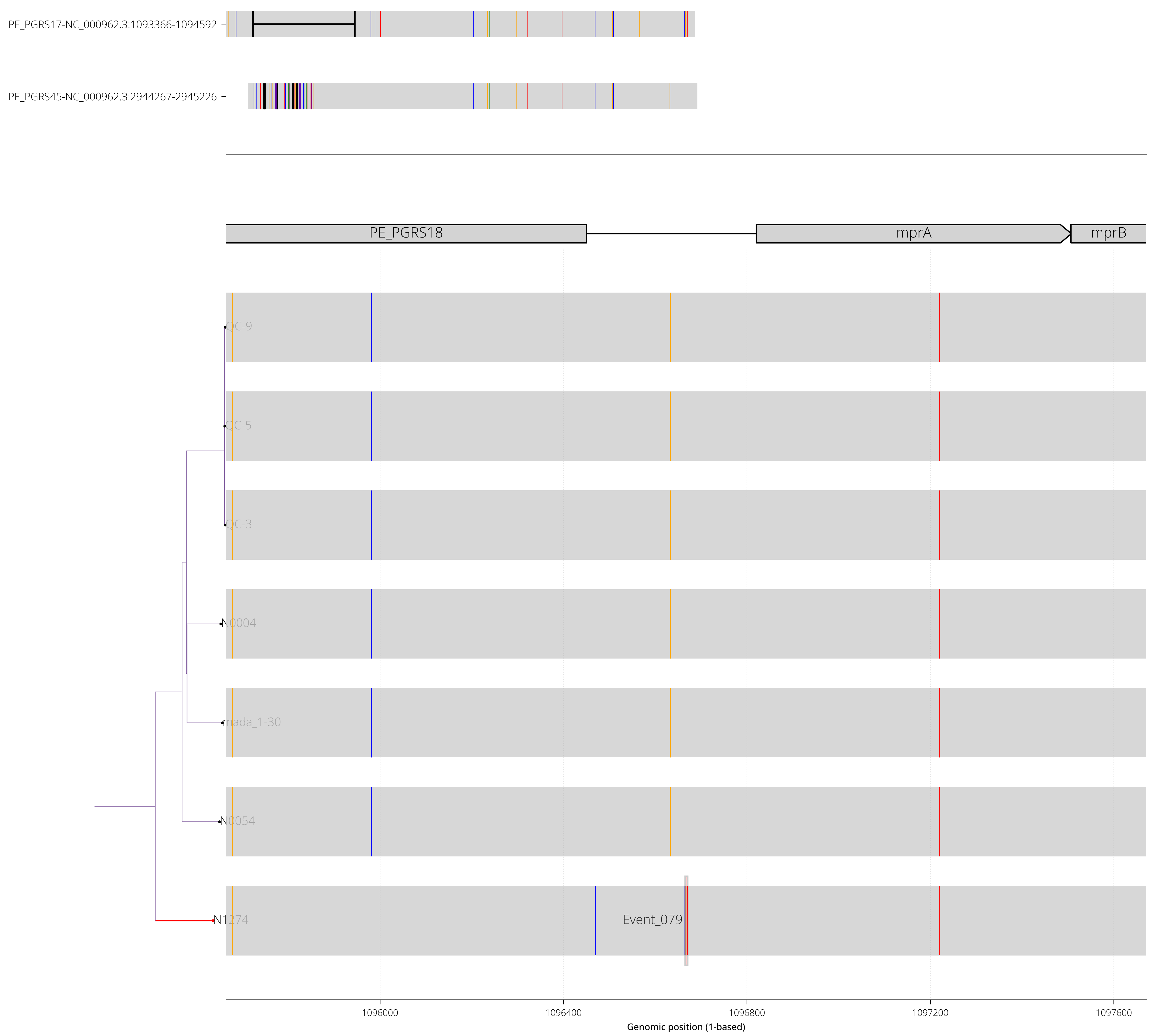

Full Phylogeny Viz + Paralog Alignments
GCE ID: Event_079

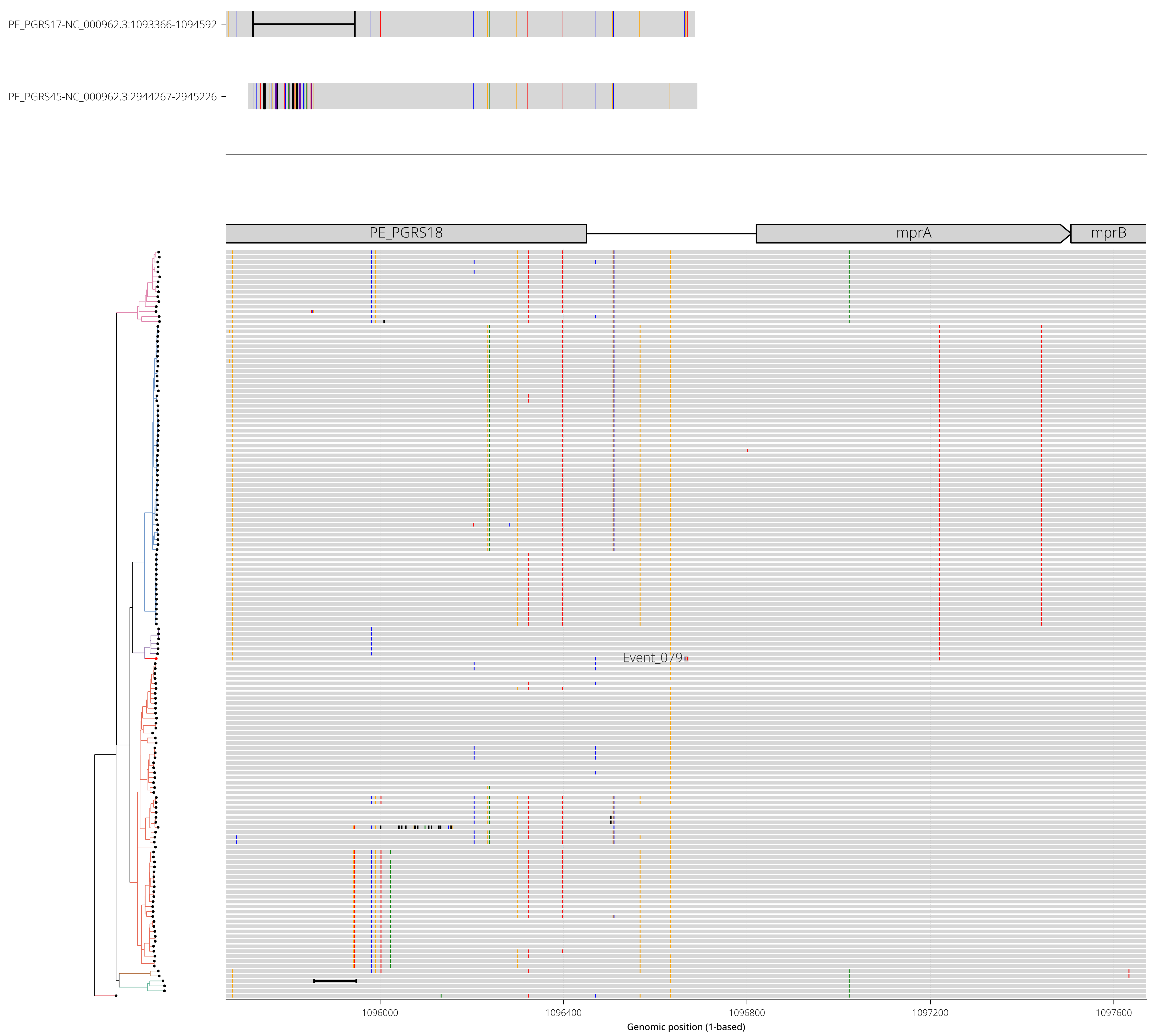

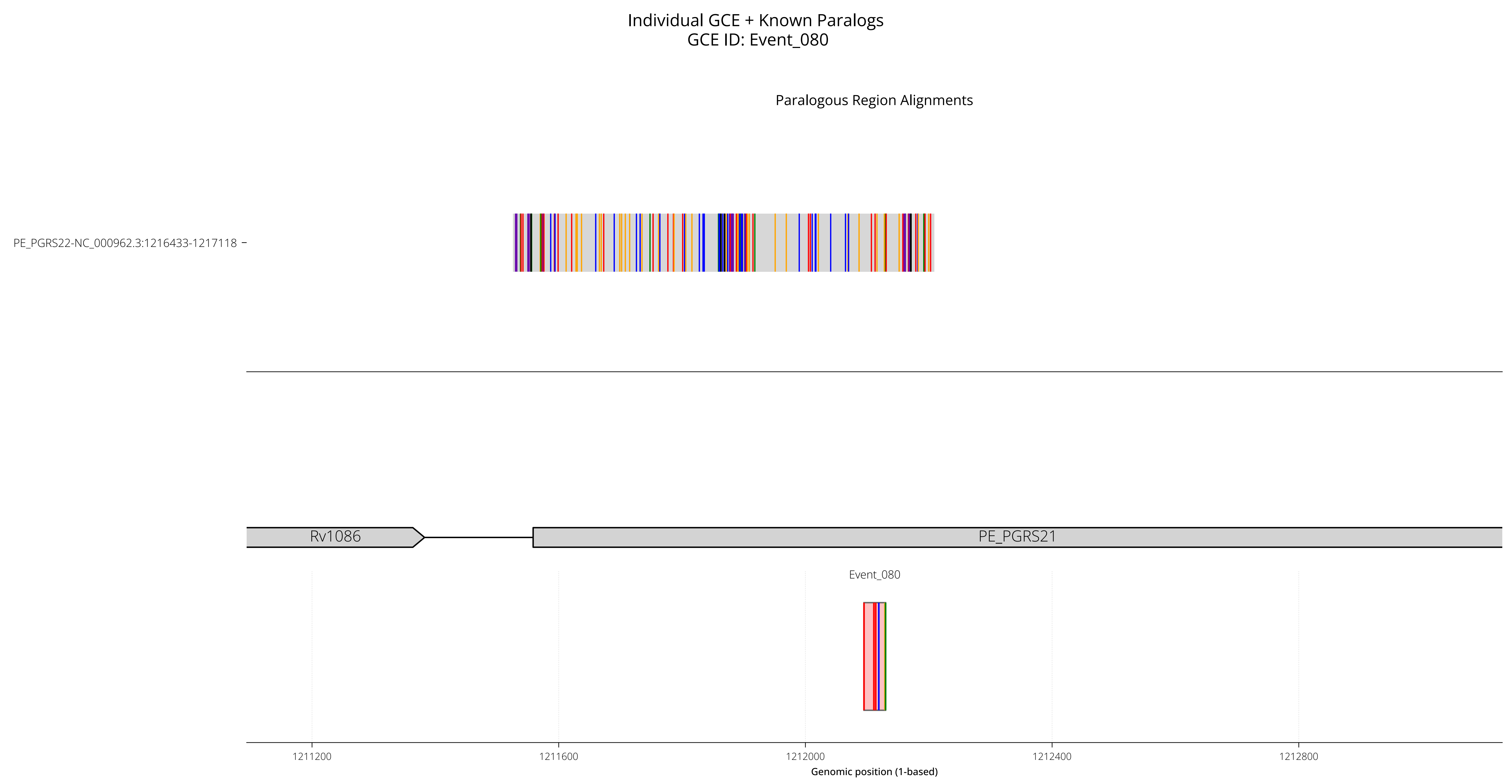

Individual GCE + Known Paralogs
GCE ID: Event_080
Paralogous Region Alignments
PE_PGRS22-NC_000962.3:1216433-1217118
Rv1086
PE_PGRS21
Event_080
1211200
1211600
1212000
1212400
1212800
Genomic position (1-based)

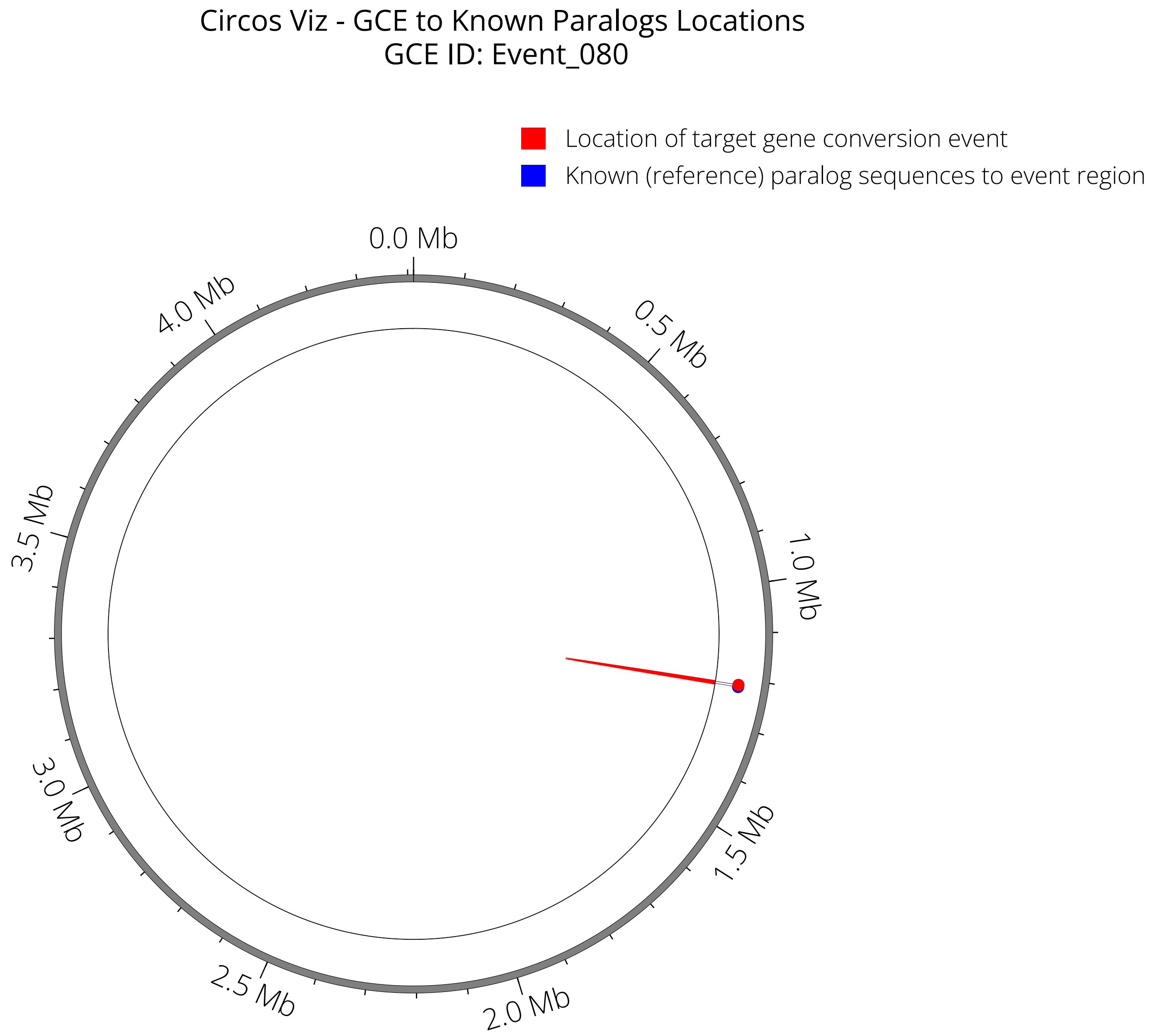
Circos Viz - GCE to Known Paralogs Locations
GCE ID: Event_080
Location of target gene conversion event
Known (reference) paralog sequences to event region
0.0 Mb
0.5 Mb
1.0 Mb
1.5 Mb
2.0 Mb
2.5 Mb
3.0 Mb
3.5 Mb
4.0 Mb

### Event_080 - 6 SNPs - k-mer analysis relative to paralogs

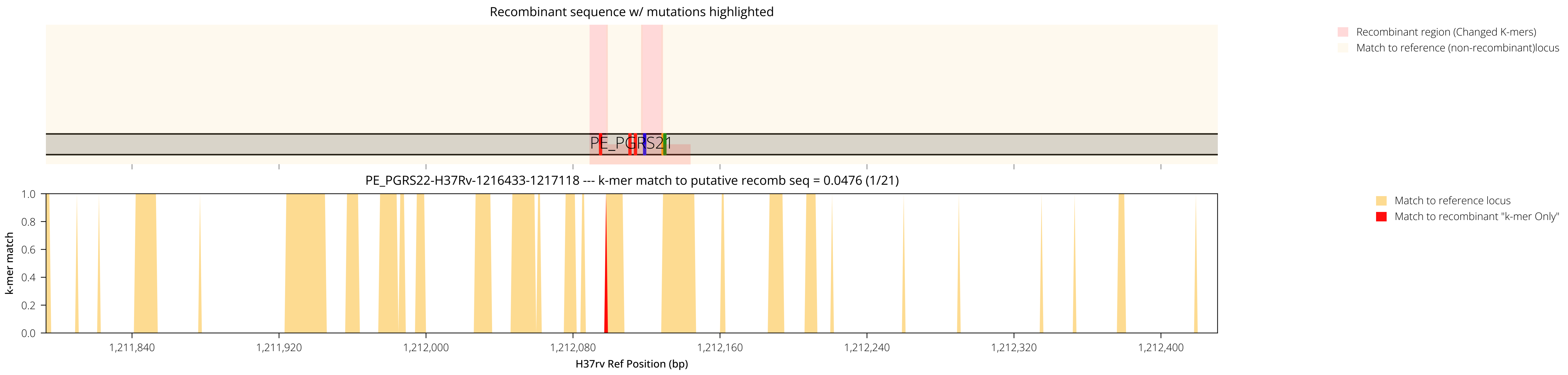

Sub-phylogeny Viz + Paralog Alignments
GCE ID: Event_080

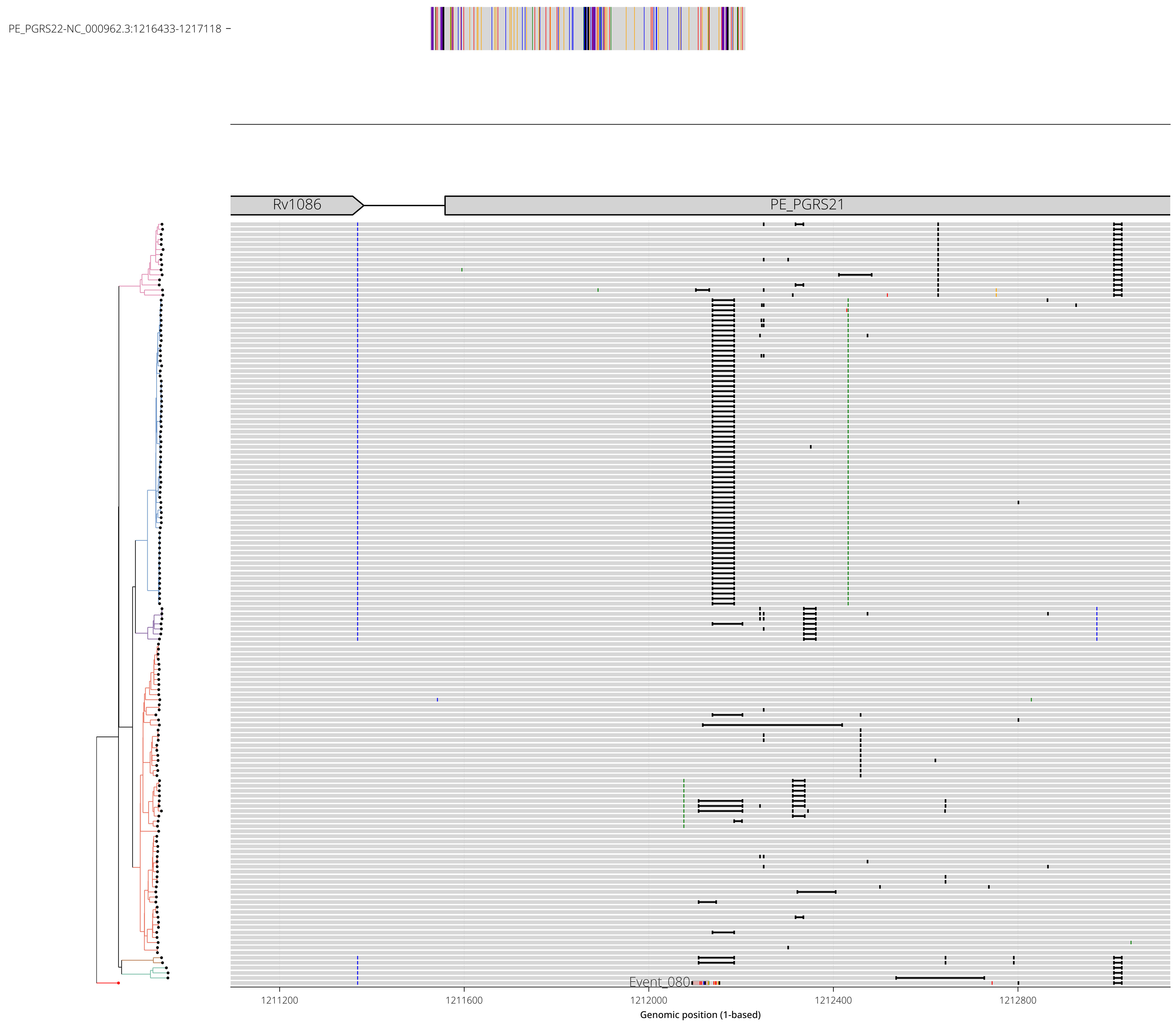

### Full Phylogeny Viz + Paralog Alignments
GCE ID: Event_080

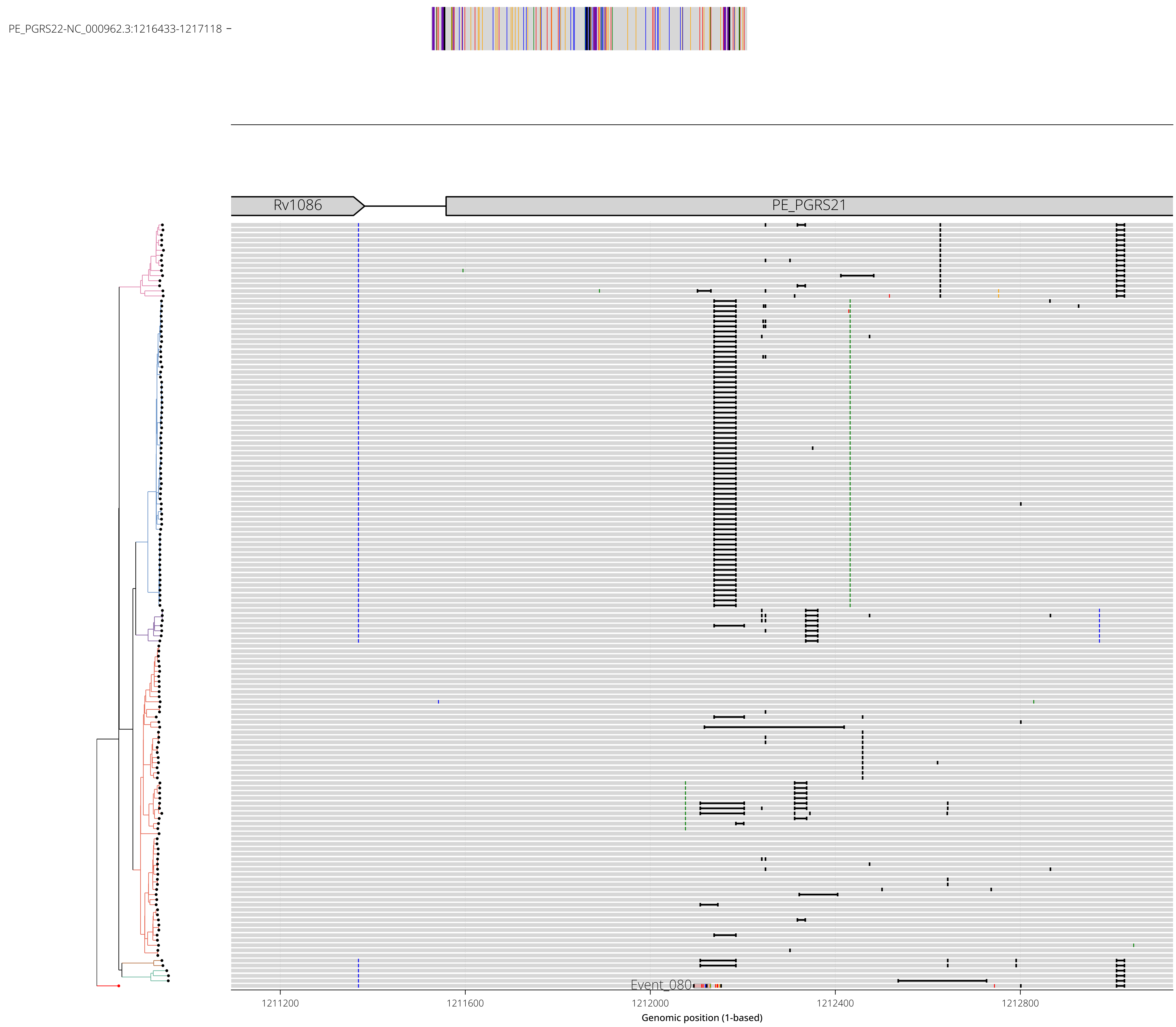

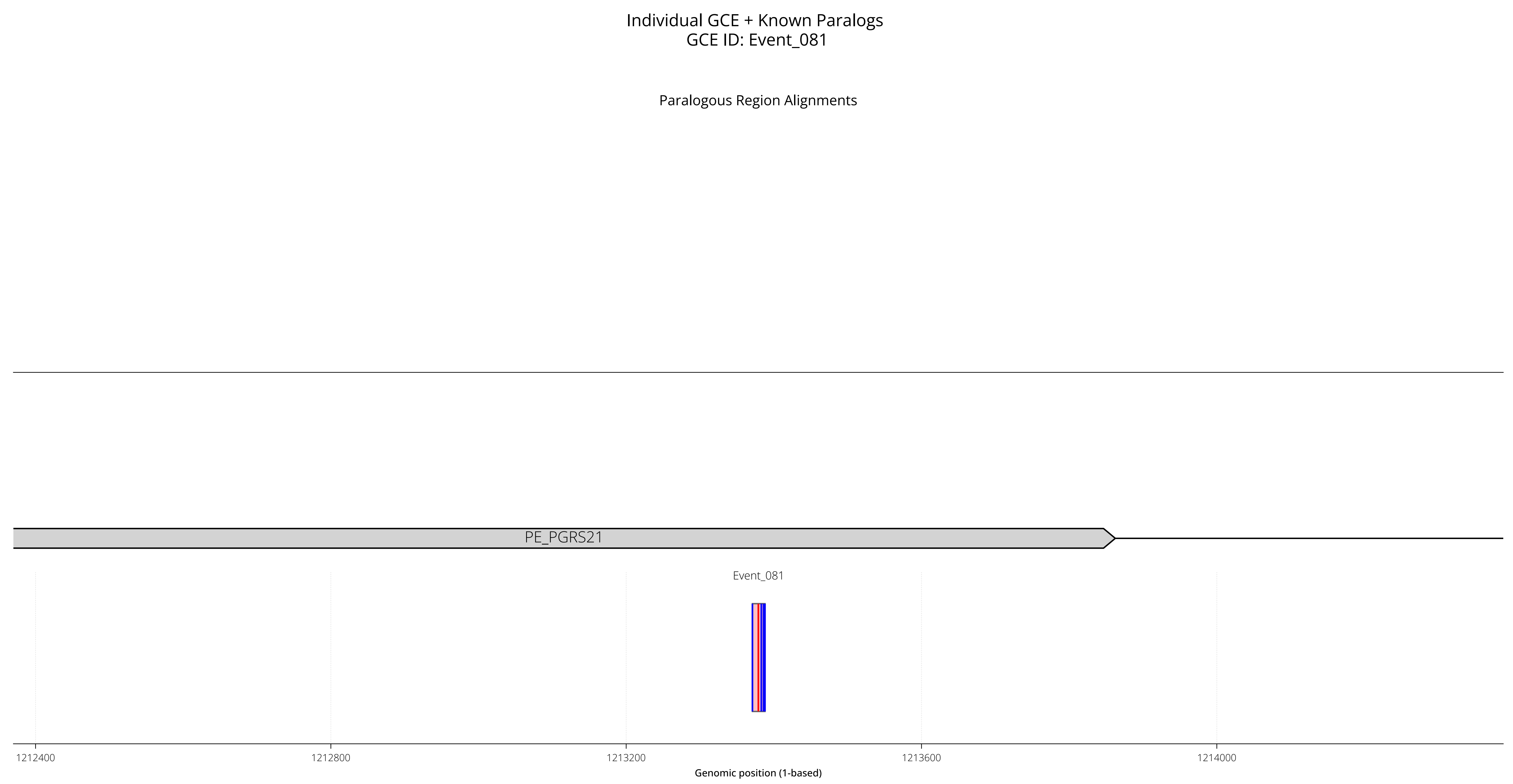
Individual GCE + Known Paralogs
GCE ID: Event_081
Paralogous Region Alignments
PE_PGRS21
Event_081
1212400
1212800
1213200
1213600
1214000
Genomic position (1-based)

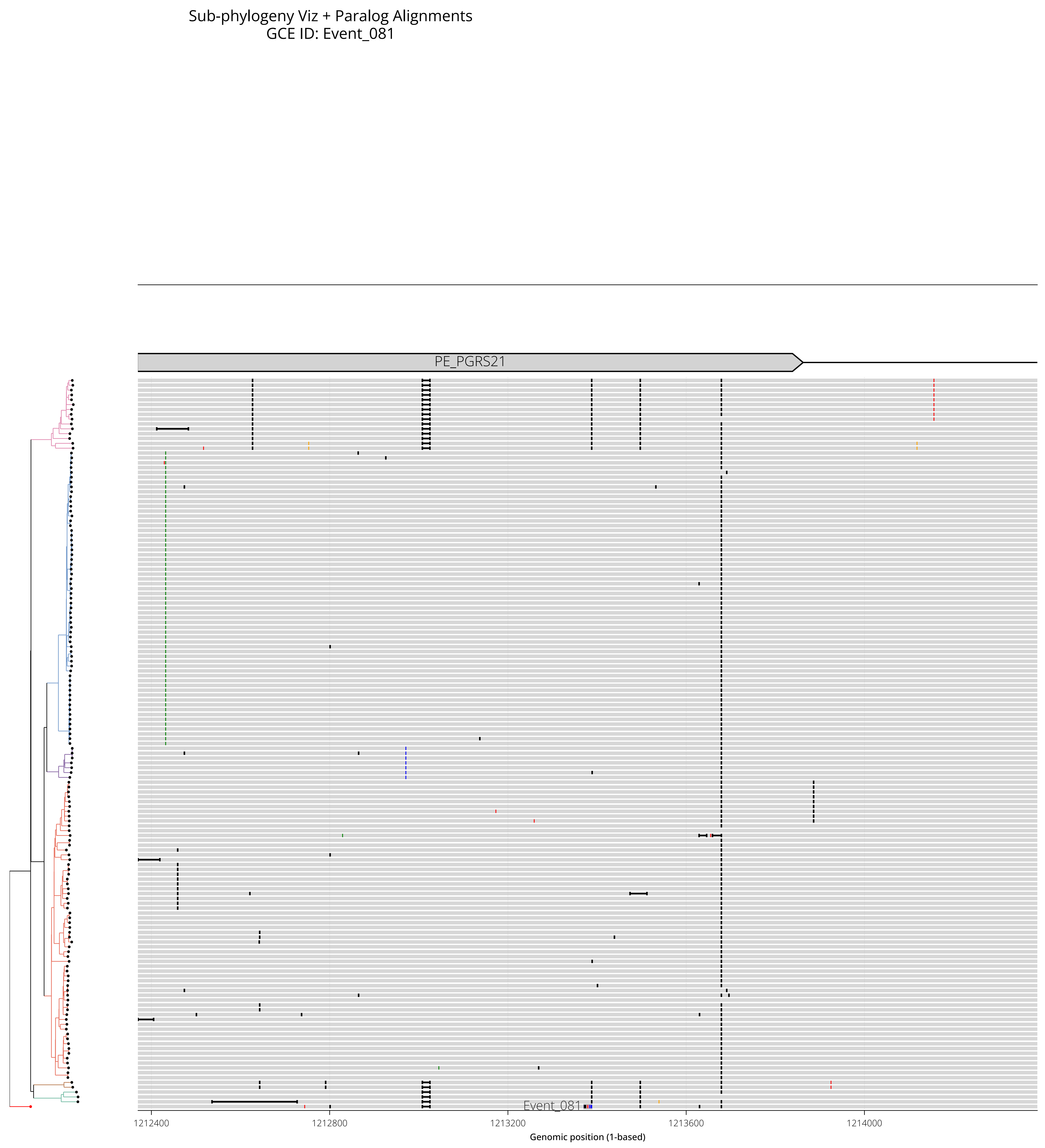

Sub-phylogeny Viz + Paralog Alignments
GCE ID: Event_081
PE_PGRS21
Event_081
1212400
1212800
1213200
1213600
1214000
Genomic position (1-based)

Full Phylogeny Viz + Paralog Alignments
GCE ID: Event_081

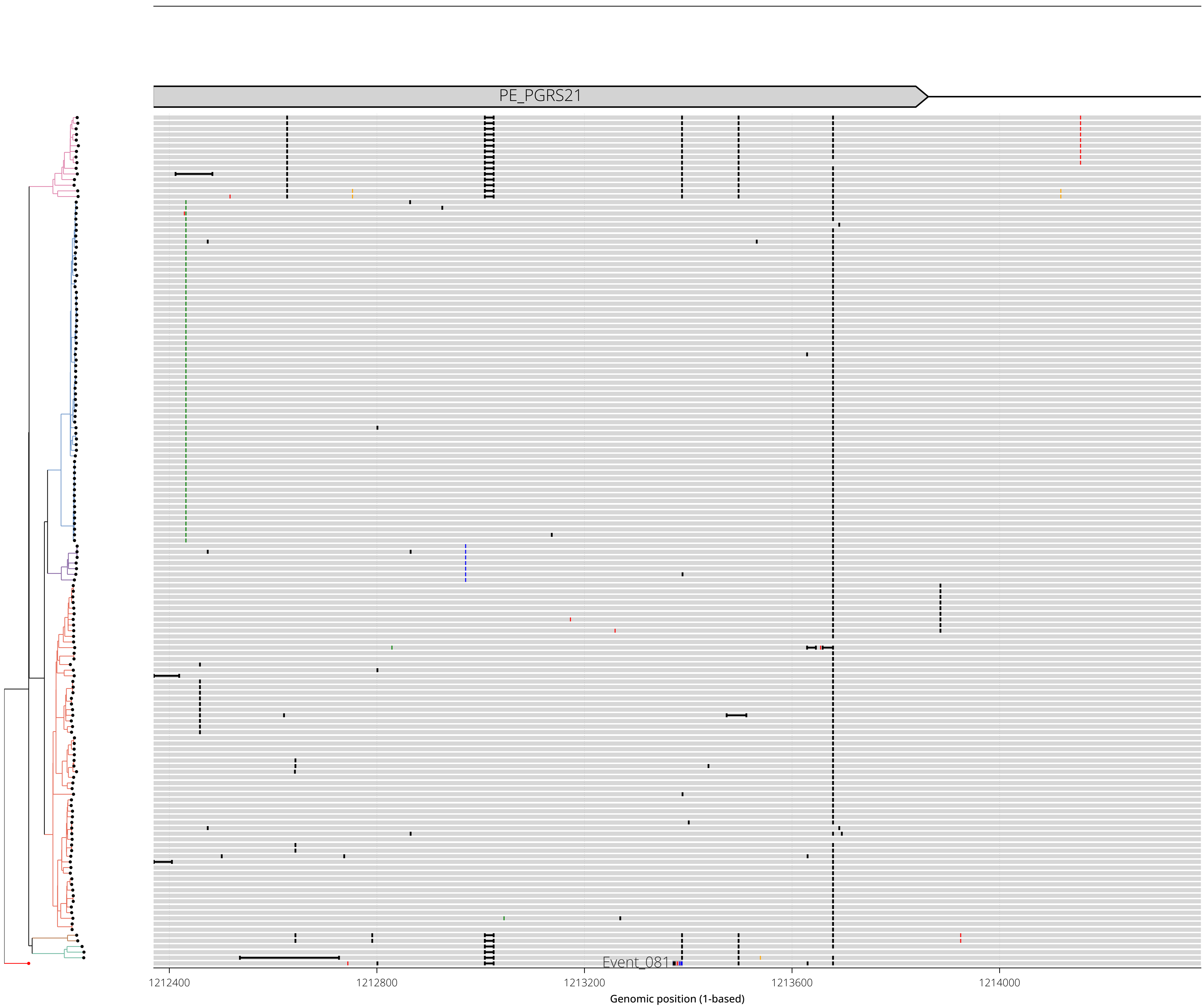

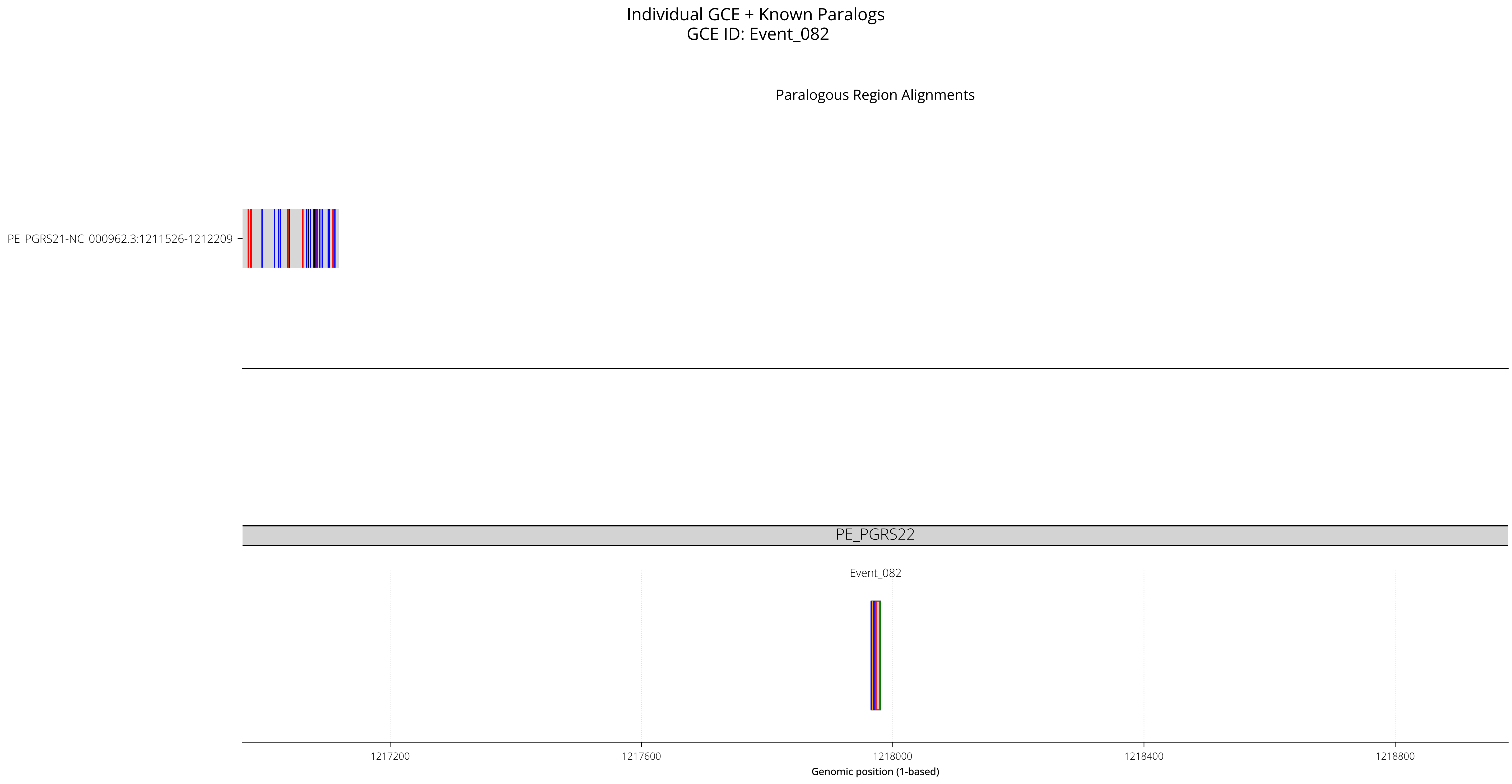

Individual GCE + Known Paralogs
GCE ID: Event_082
Paralogous Region Alignments
PE_PGRS21-NC_000962.3:1211526-1212209
PE_PGRS22
Event_082
1217200
1217600
1218000
1218400
1218800
Genomic position (1-based)

### Sub-phylogeny Viz + Paralog Alignments
GCE ID: Event_082

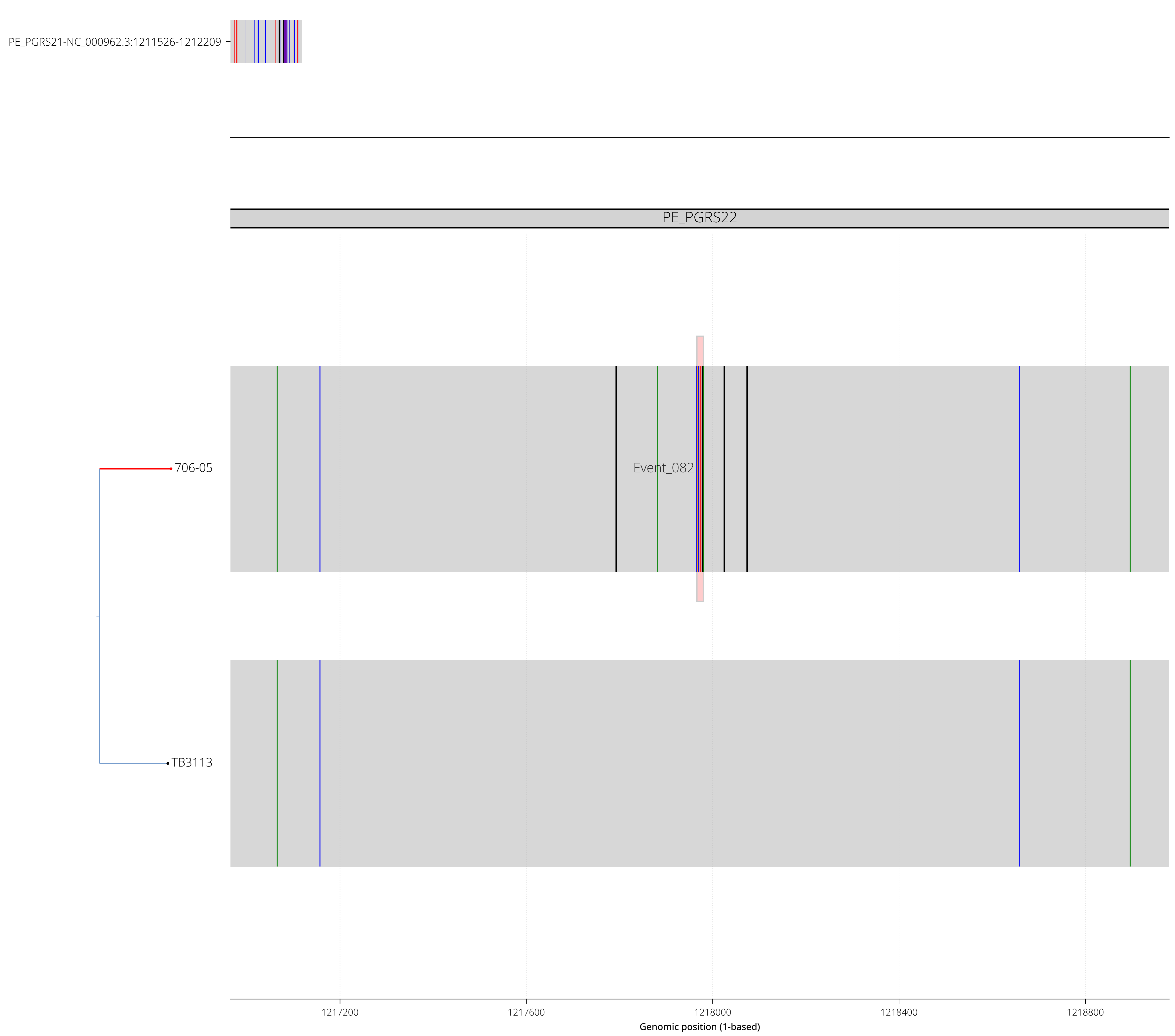

Full Phylogeny Viz + Paralog Alignments
GCE ID: Event_082

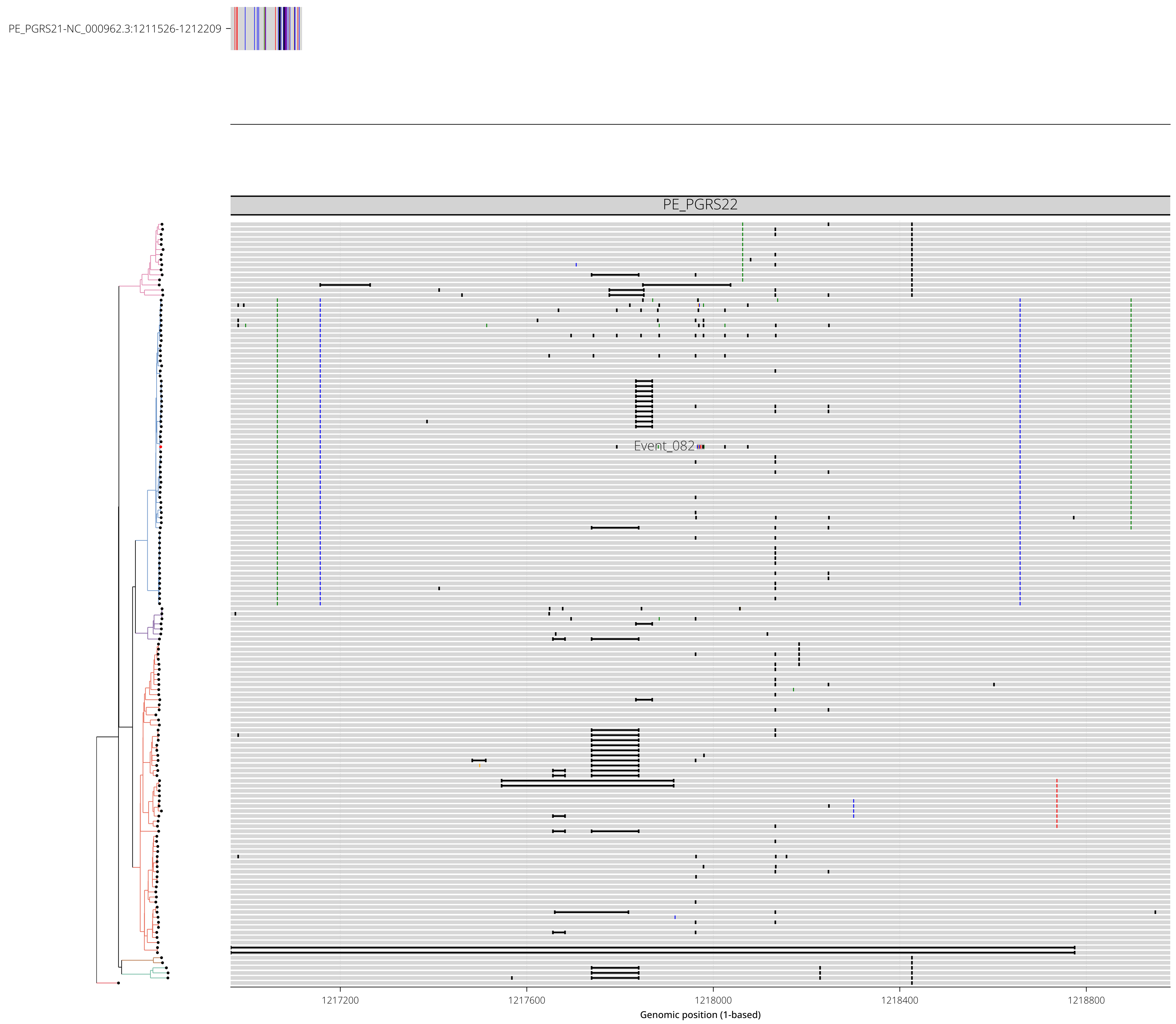

Individual GCE + Known Paralogs
GCE ID: Event_083

Paralogous Region Alignments

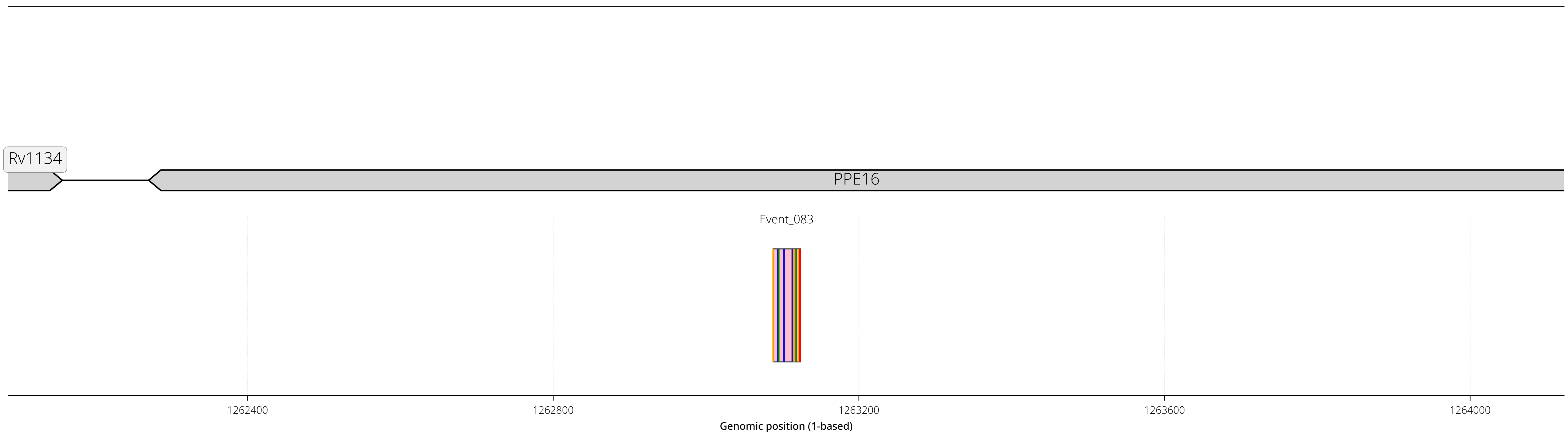

Sub-phylogeny Viz + Paralog Alignments
GCE ID: Event_083

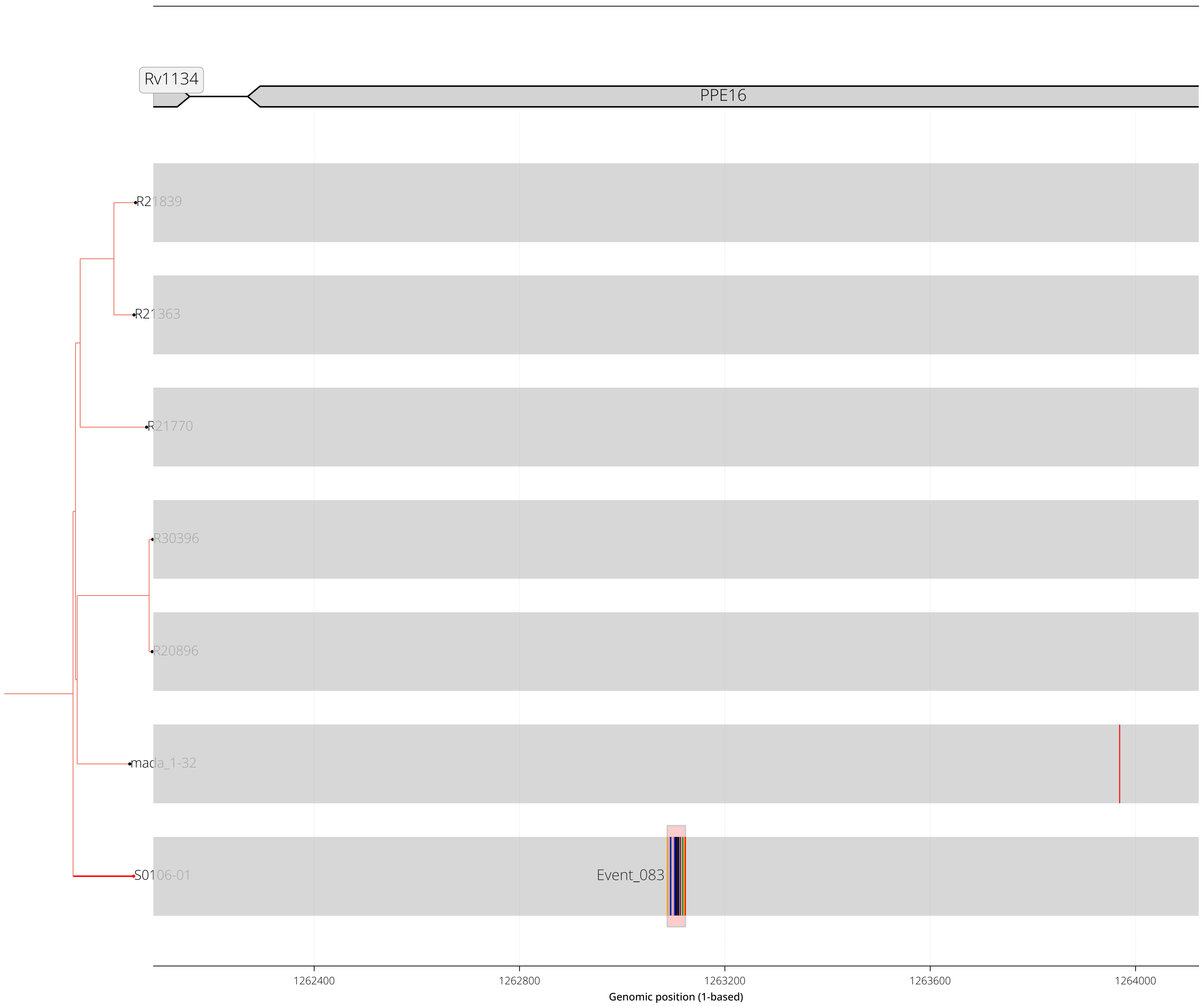

Full Phylogeny Viz + Paralog Alignments
GCE ID: Event_083

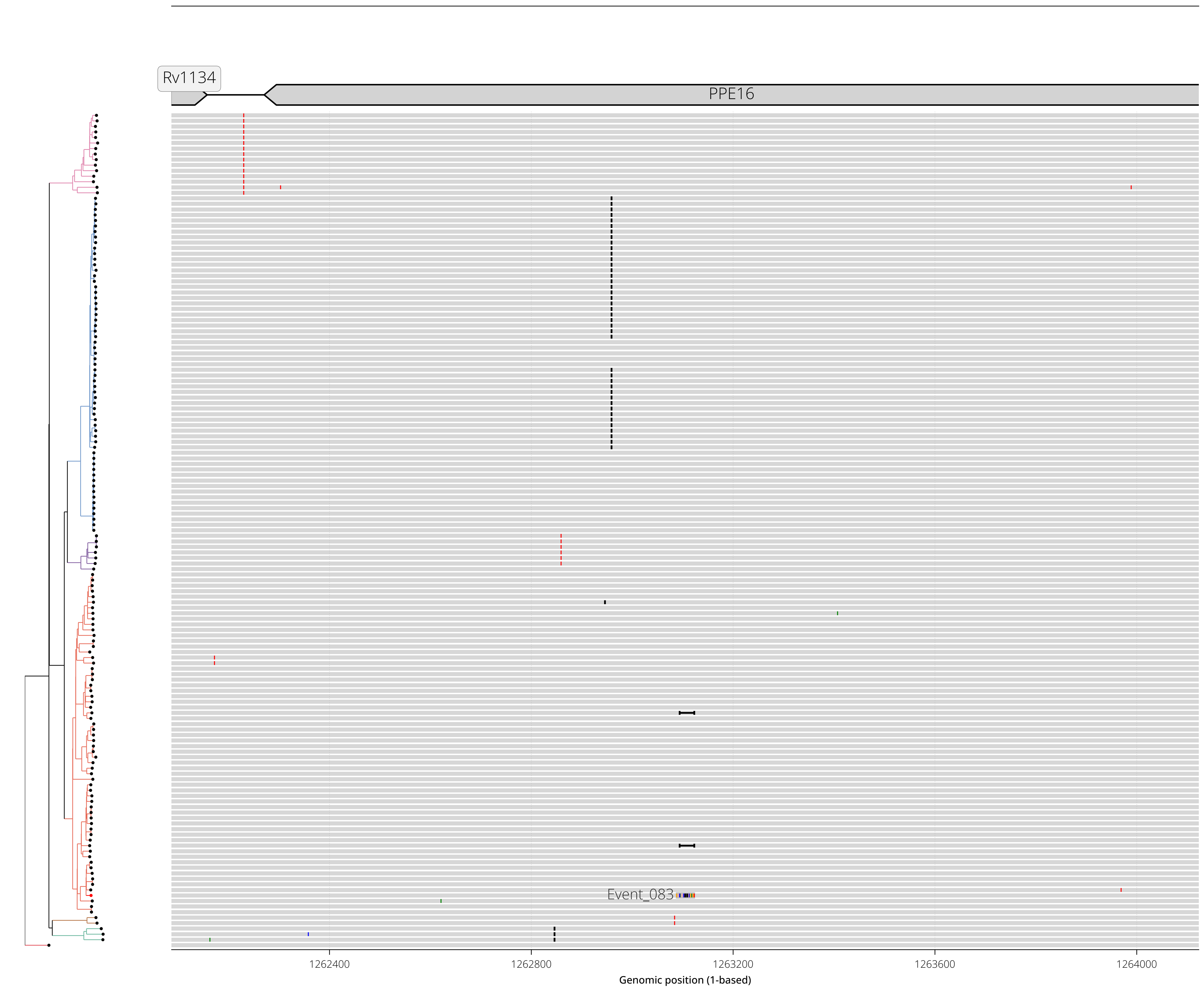

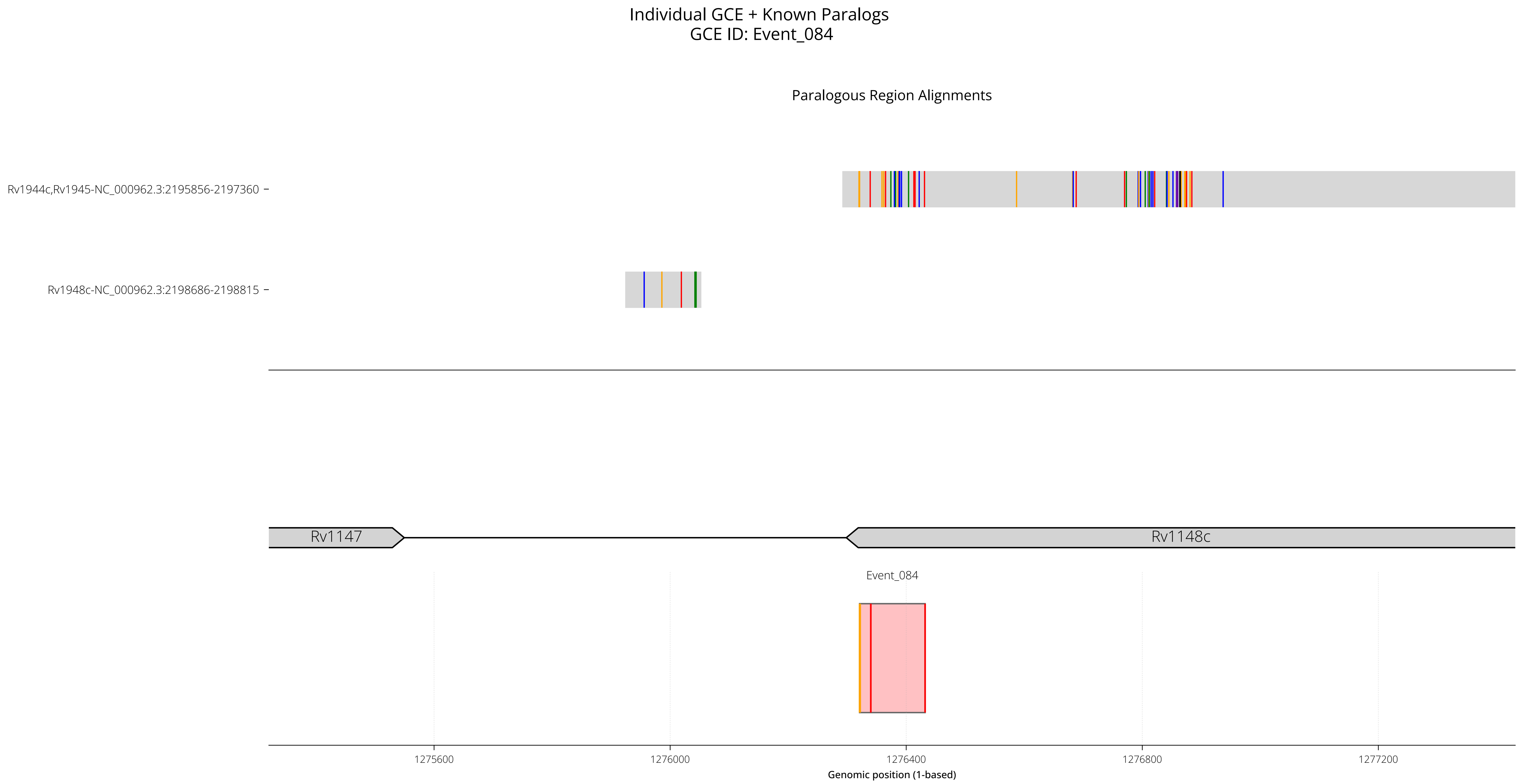
Individual GCE + Known Paralogs
GCE ID: Event_084
Paralogous Region Alignments
Rv1944c,Rv1945-NC_000962.3:2195856-2197360
Rv1948c-NC_000962.3:2198686-2198815
Rv1147
Rv1148c
Event_084
1275600
1276000
1276400
1276800
1277200
Genomic position (1-based)

Circos Viz - GCE to Known Paralogs Locations
GCE ID: Event_084

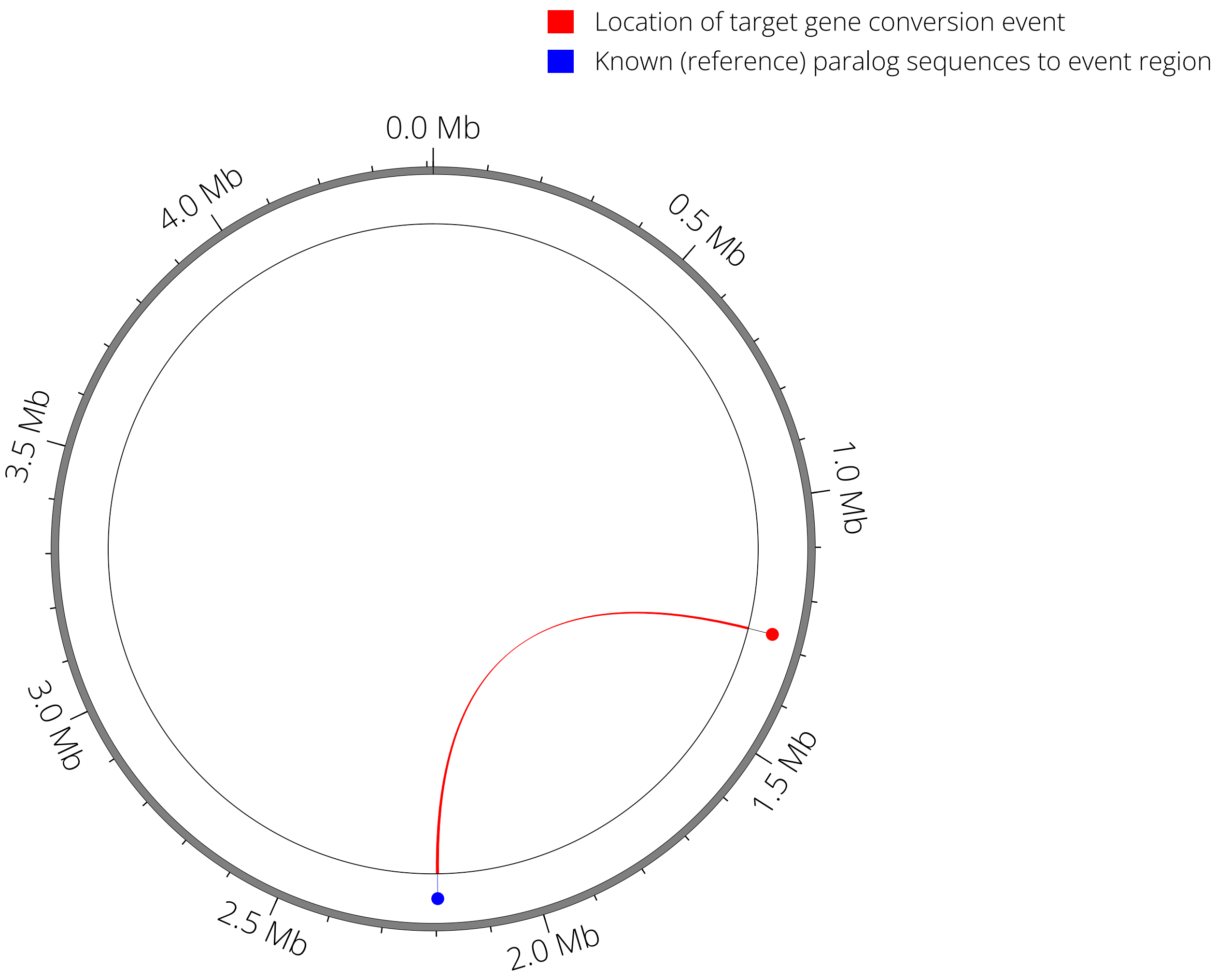

### Event_084 - 4 SNPs - k-mer analysis relative to paralogs

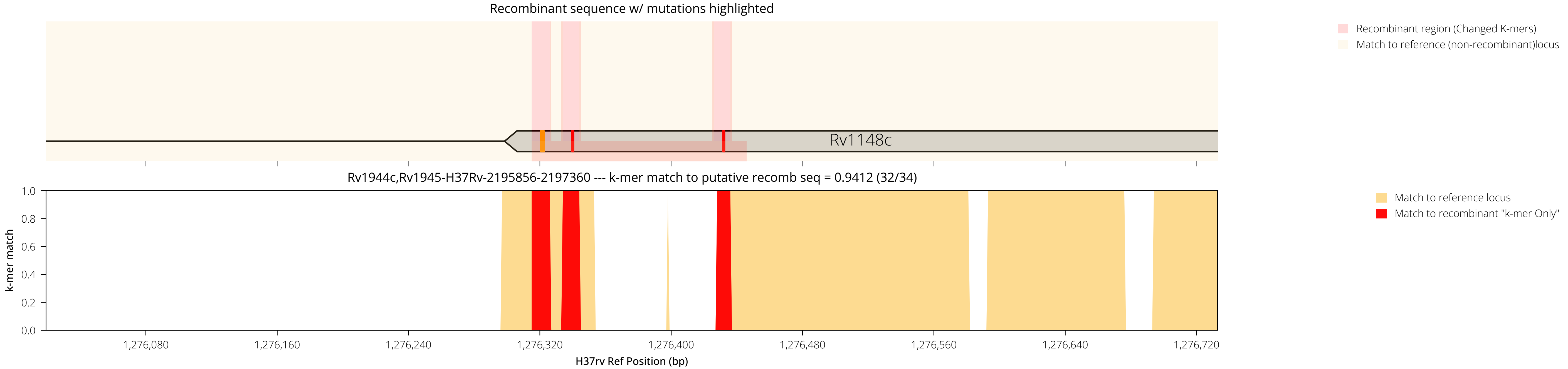

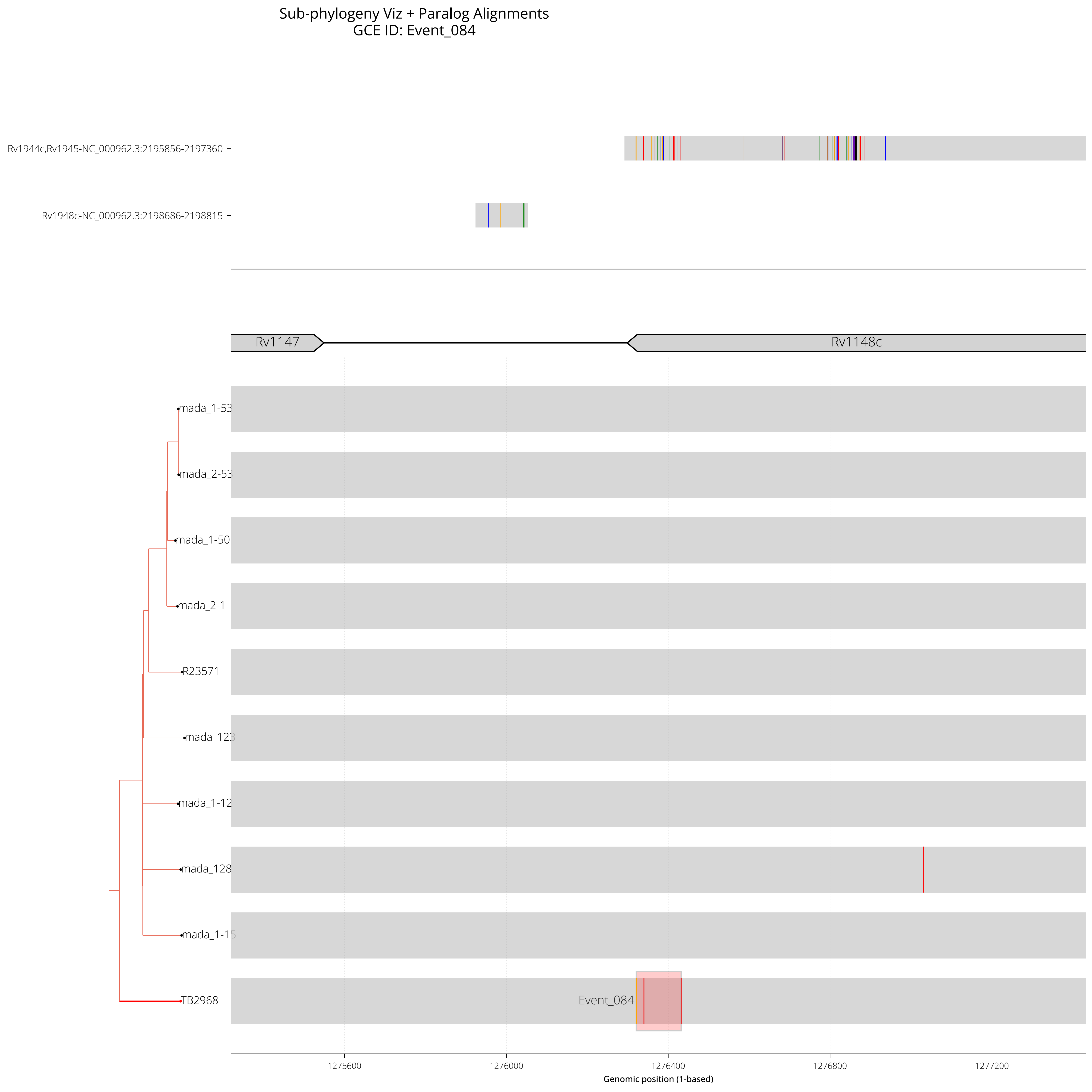
Sub-phylogeny Viz + Paralog Alignments
GCE ID: Event_084
Rv1944c,Rv1945-NC_000962.3:2195856-2197360
Rv1948c-NC_000962.3:2198686-2198815
Rv1147
Rv1148c
mada_1-53
mada_2-53
mada_1-50
mada_2-1
R23571
mada_123
mada_1-12
mada_128
mada_1-15
TB2968
Event_084
1275600
1276000
1276400
1276800
1277200
Genomic position (1-based)

Full Phylogeny Viz + Paralog Alignments
GCE ID: Event_084

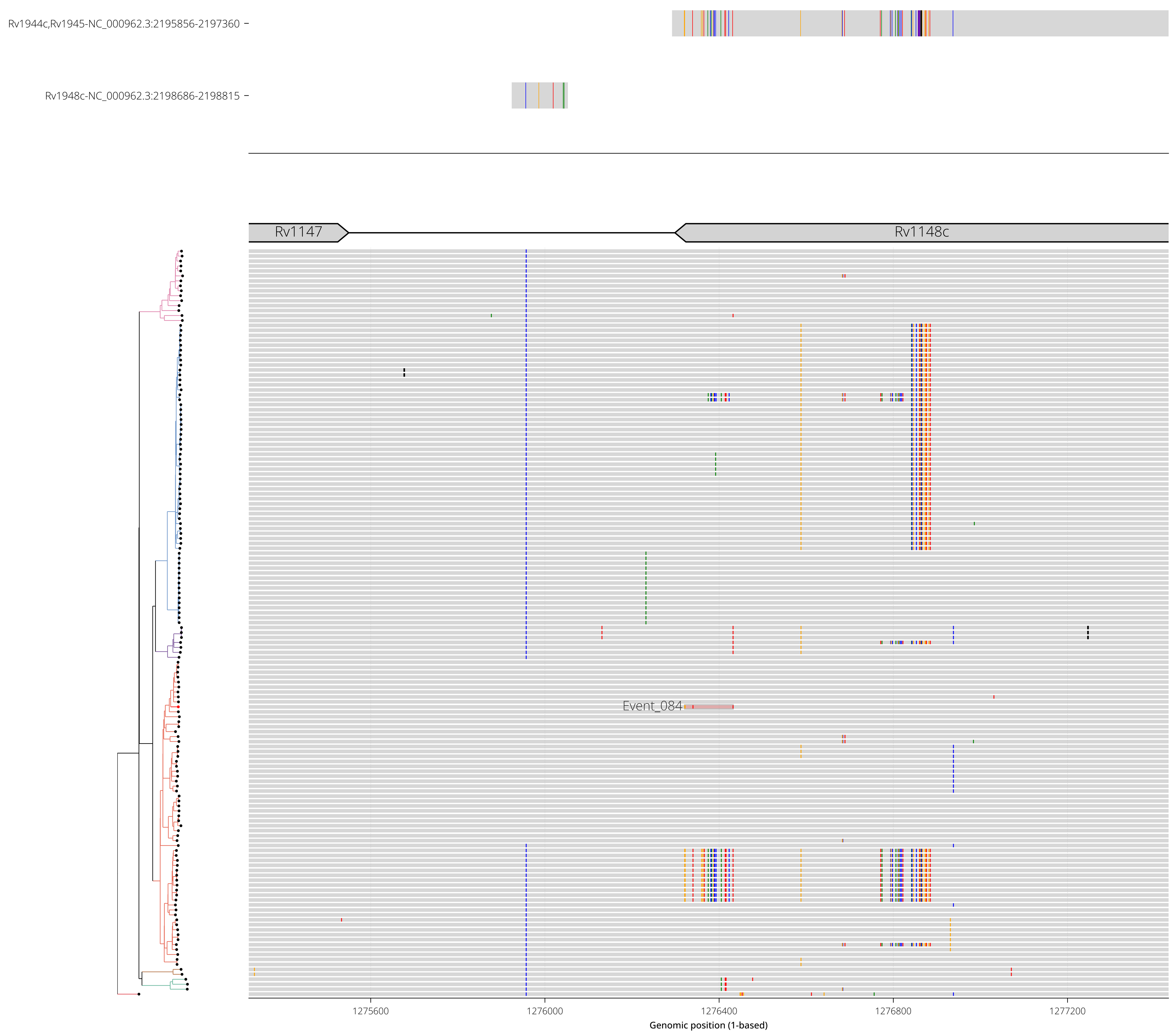

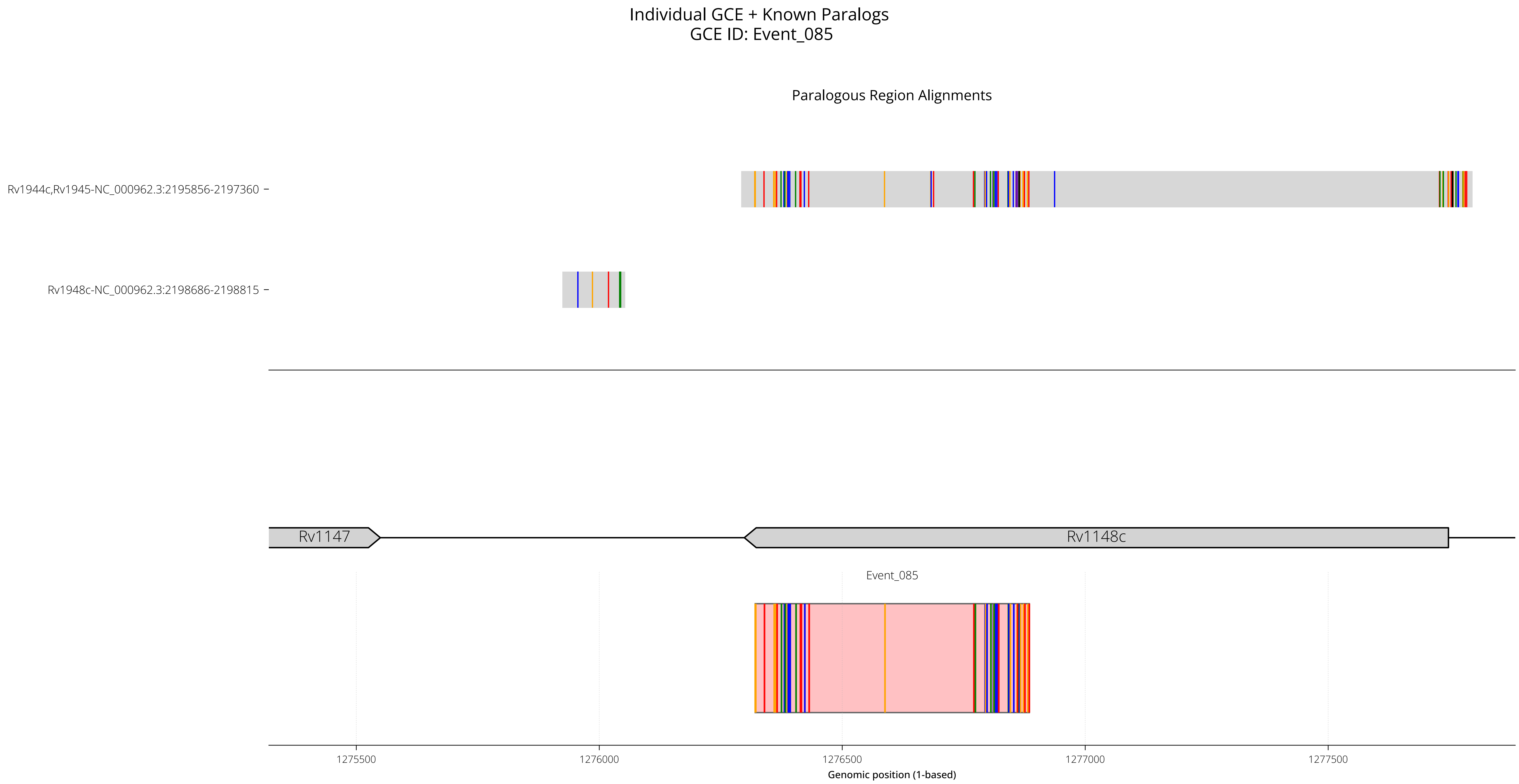

Individual GCE + Known Paralogs
GCE ID: Event_085
Paralogous Region Alignments
Rv1944c,Rv1945-NC_000962.3:2195856-2197360
Rv1948c-NC_000962.3:2198686-2198815
Rv1147
Rv1148c
Event_085
1275500
1276000
1276500
1277000
1277500
Genomic position (1-based)

Circos Viz - GCE to Known Paralogs Locations
GCE ID: Event_085

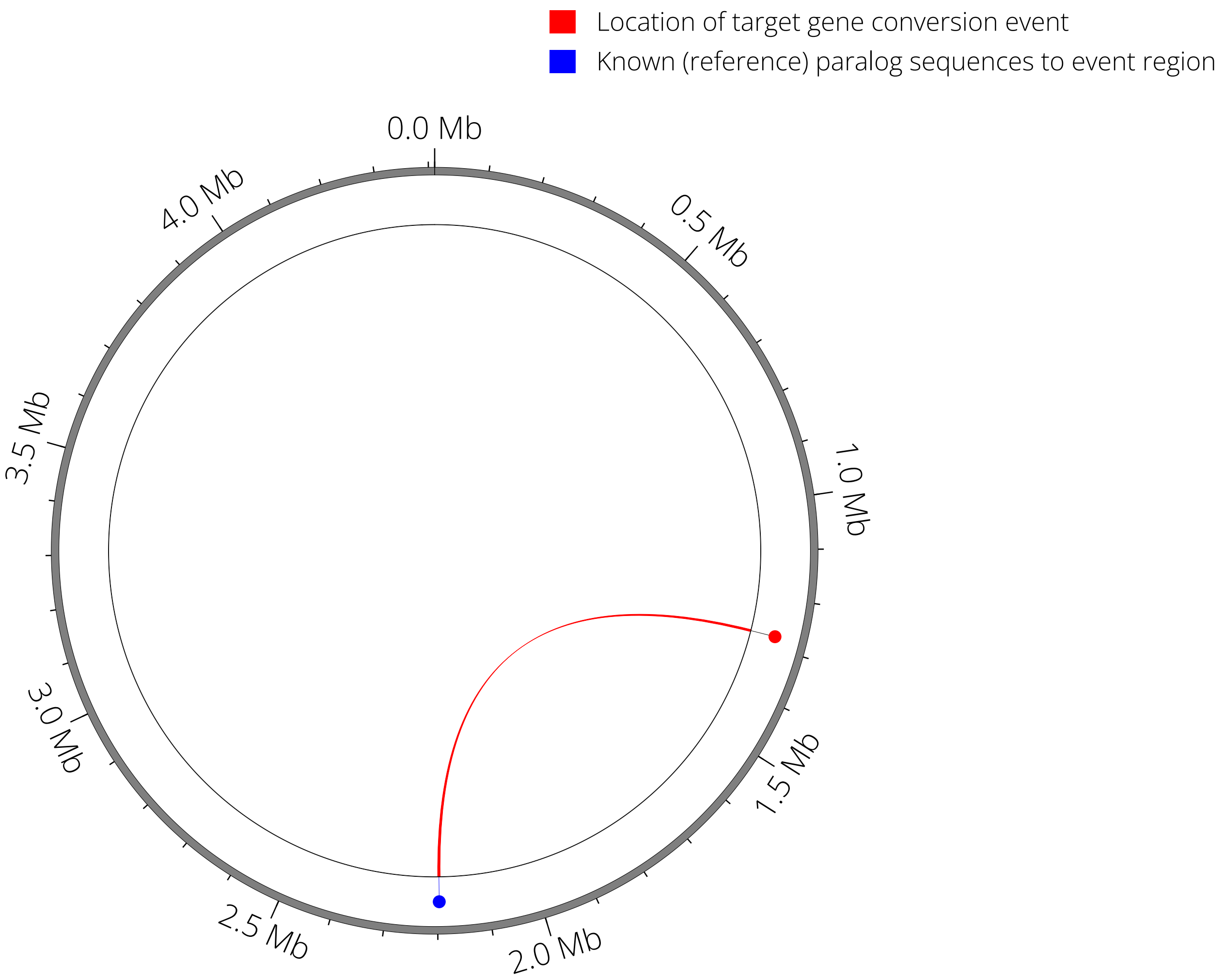

### Event_085 - 45 SNPs - k-mer analysis relative to paralogs

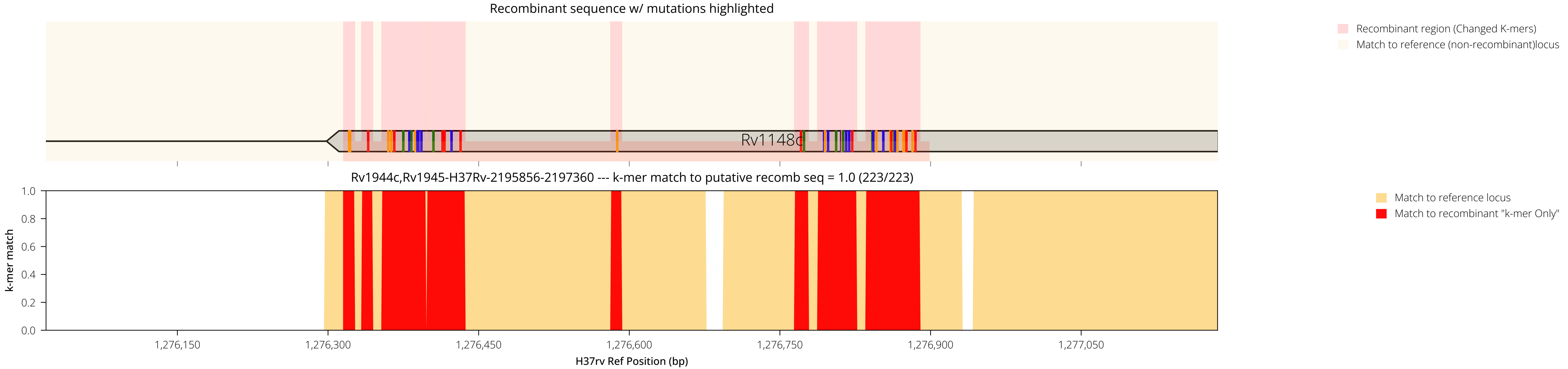

Sub-phylogeny Viz + Paralog Alignments
GCE ID: Event_085

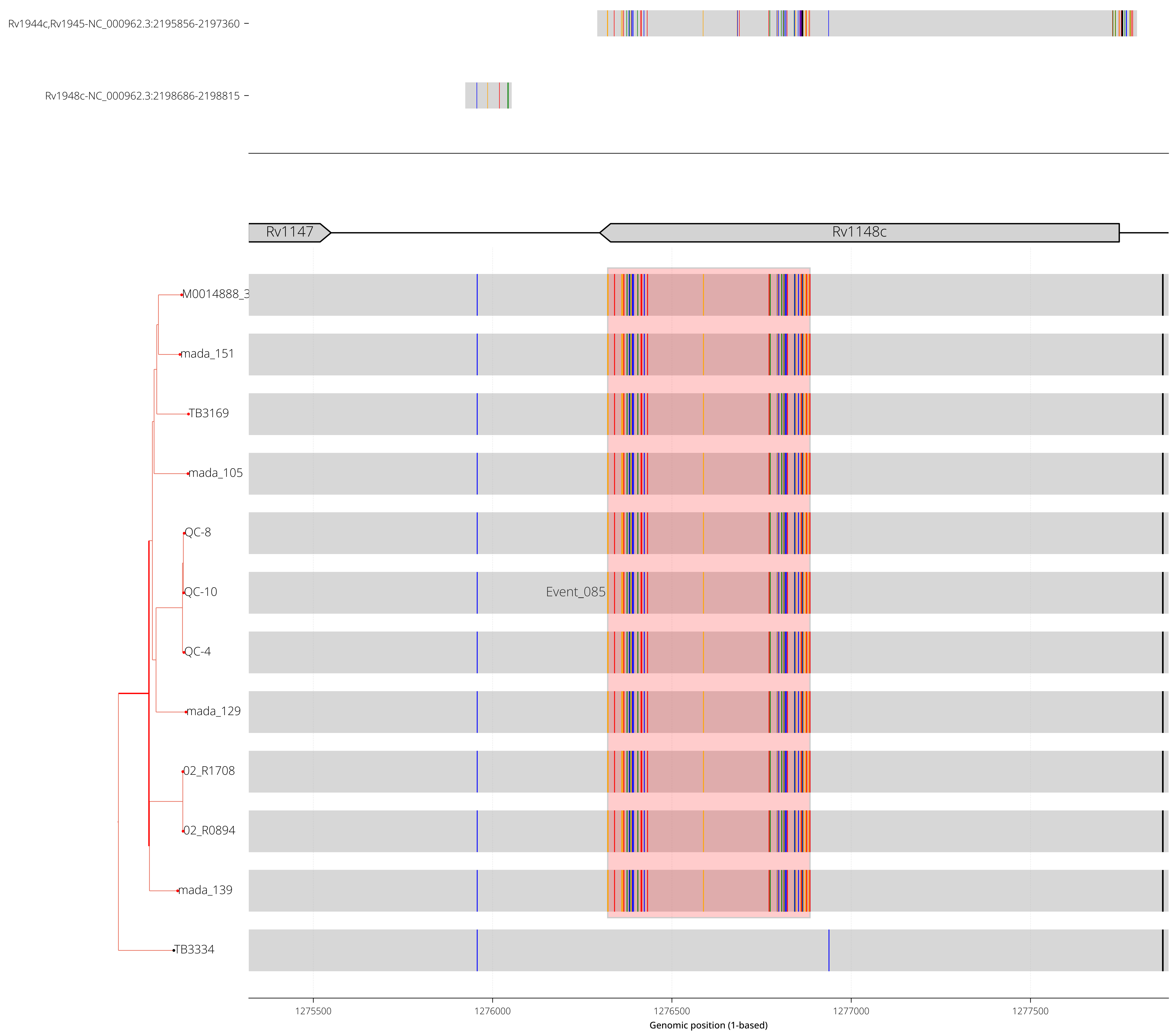

Full Phylogeny Viz + Paralog Alignments
GCE ID: Event_085

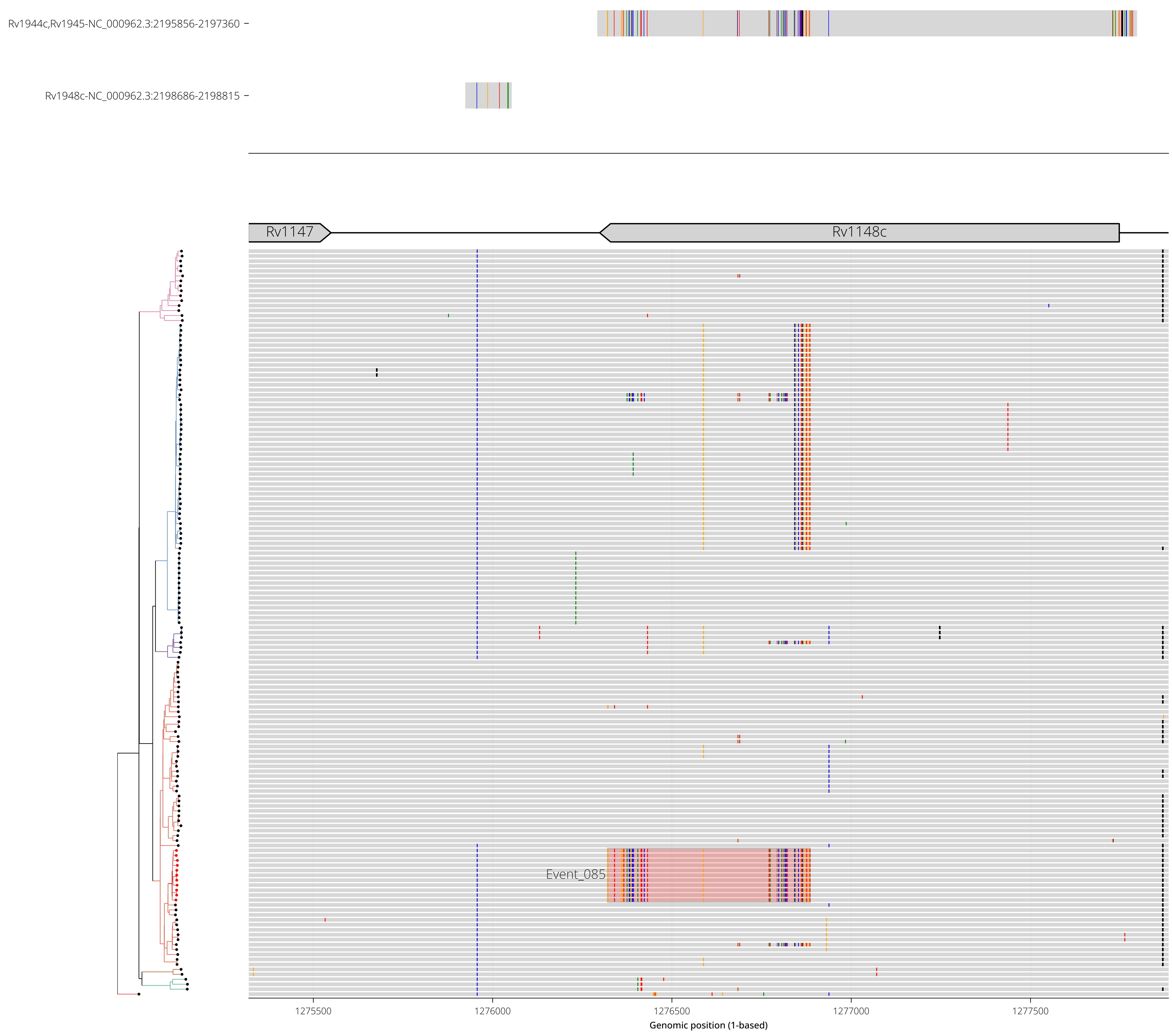

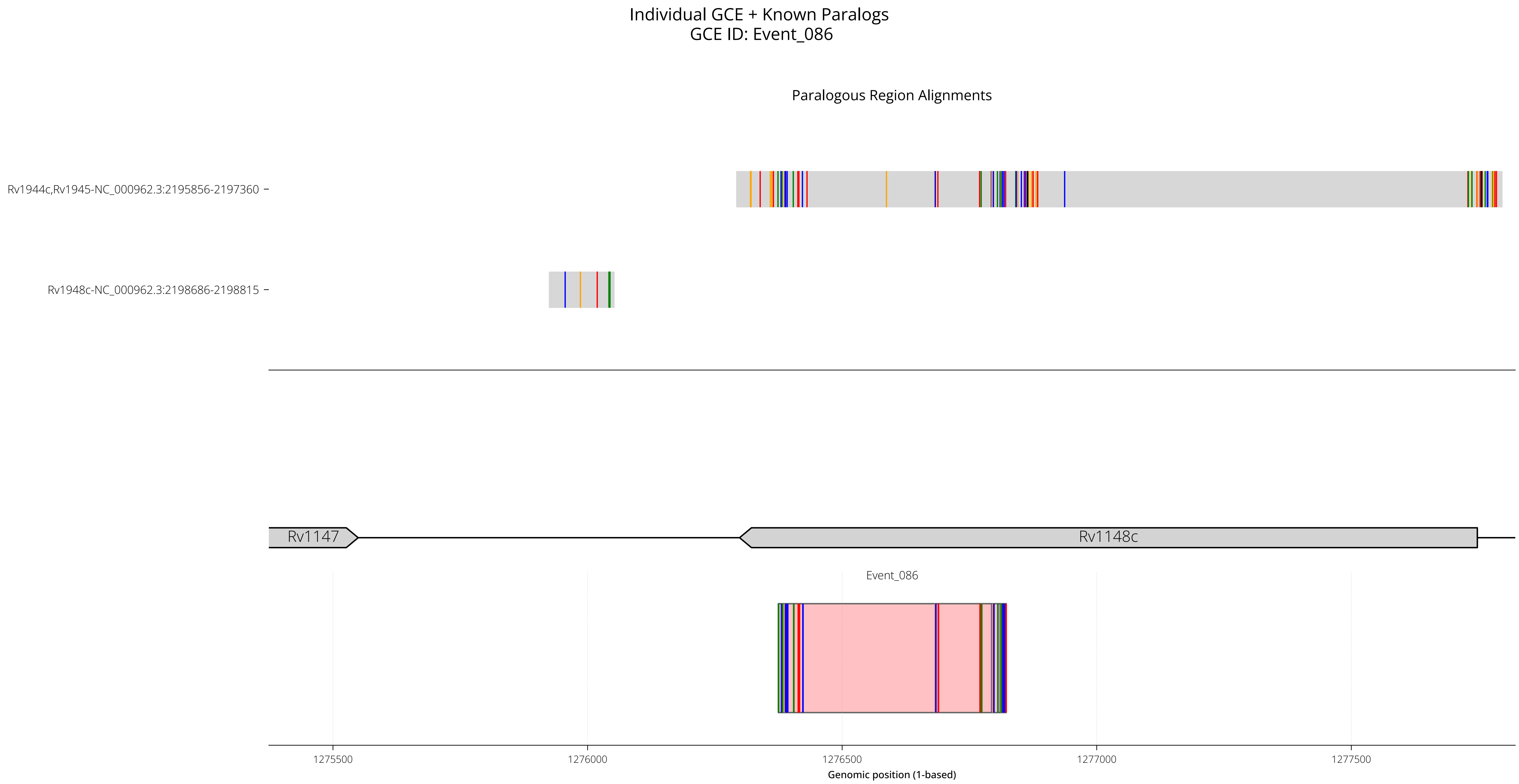
Individual GCE + Known Paralogs
GCE ID: Event_086
Paralogous Region Alignments
Rv1944c,Rv1945-NC_000962.3:2195856-2197360
Rv1948c-NC_000962.3:2198686-2198815
Rv1147
Rv1148c
Event_086
1275500
1276000
1276500
1277000
1277500
Genomic position (1-based)

Circos Viz - GCE to Known Paralogs Locations
GCE ID: Event_086

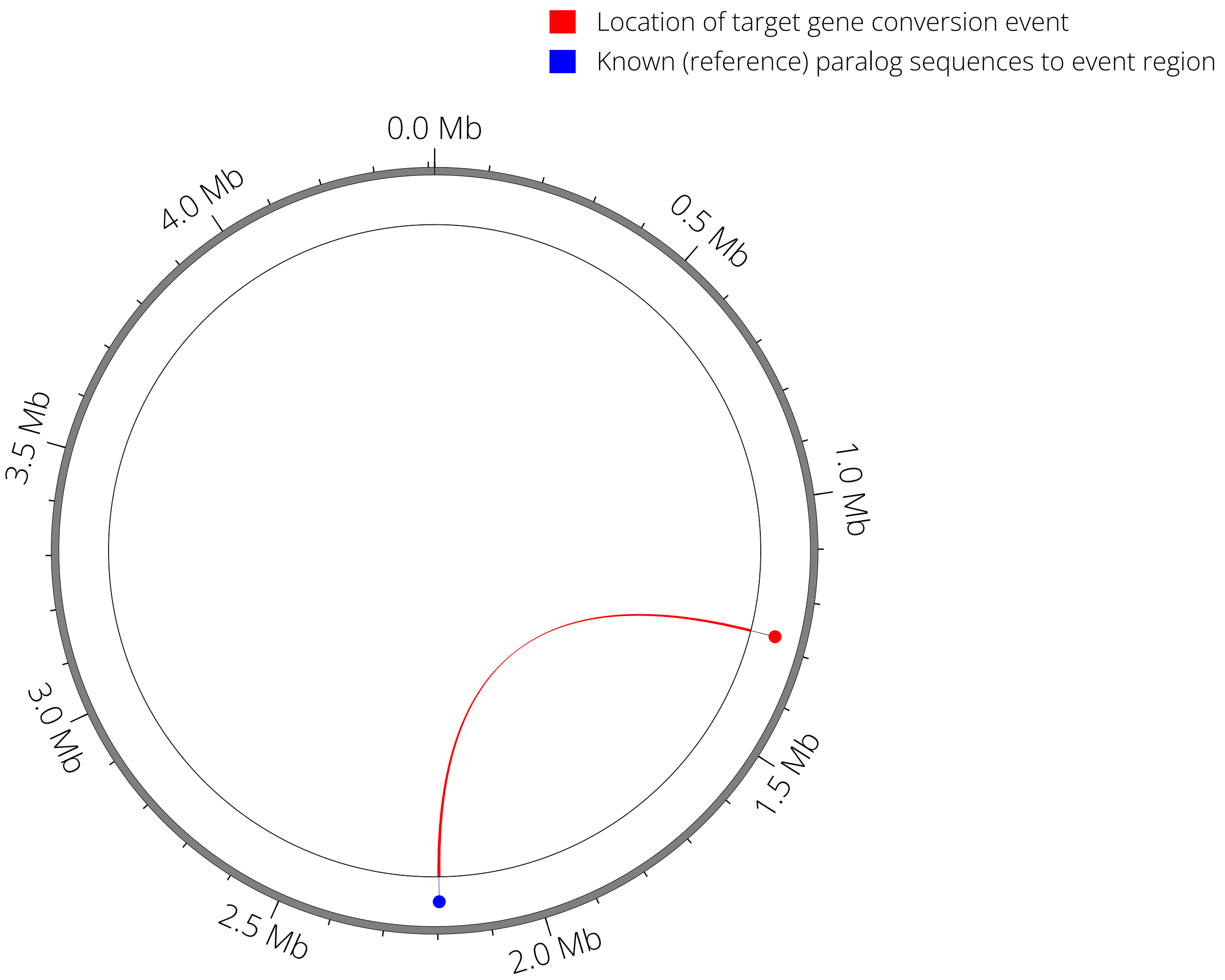

### Event_086 - 26 SNPs - k-mer analysis relative to paralogs

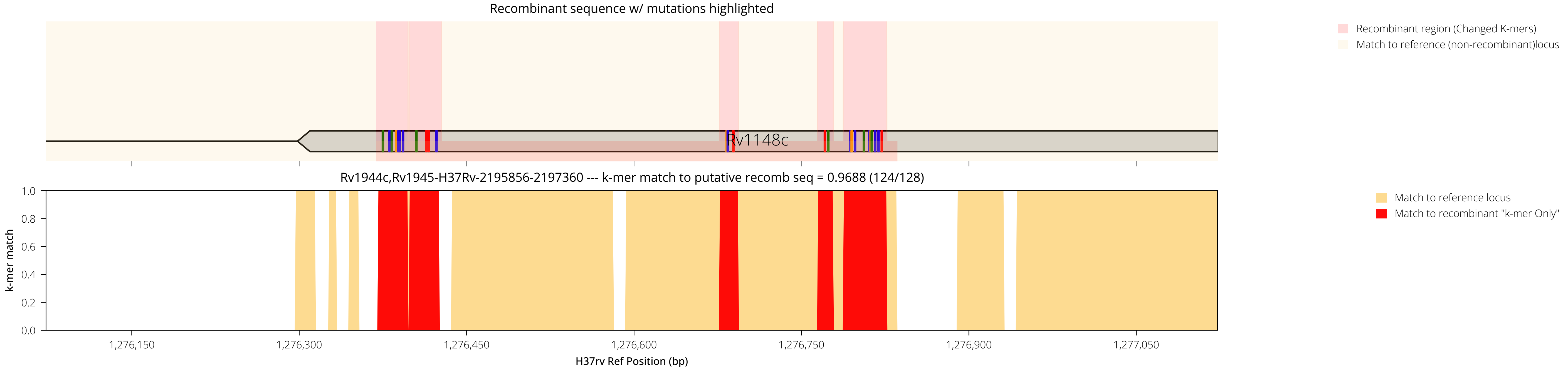

Sub-phylogeny Viz + Paralog Alignments
GCE ID: Event_086

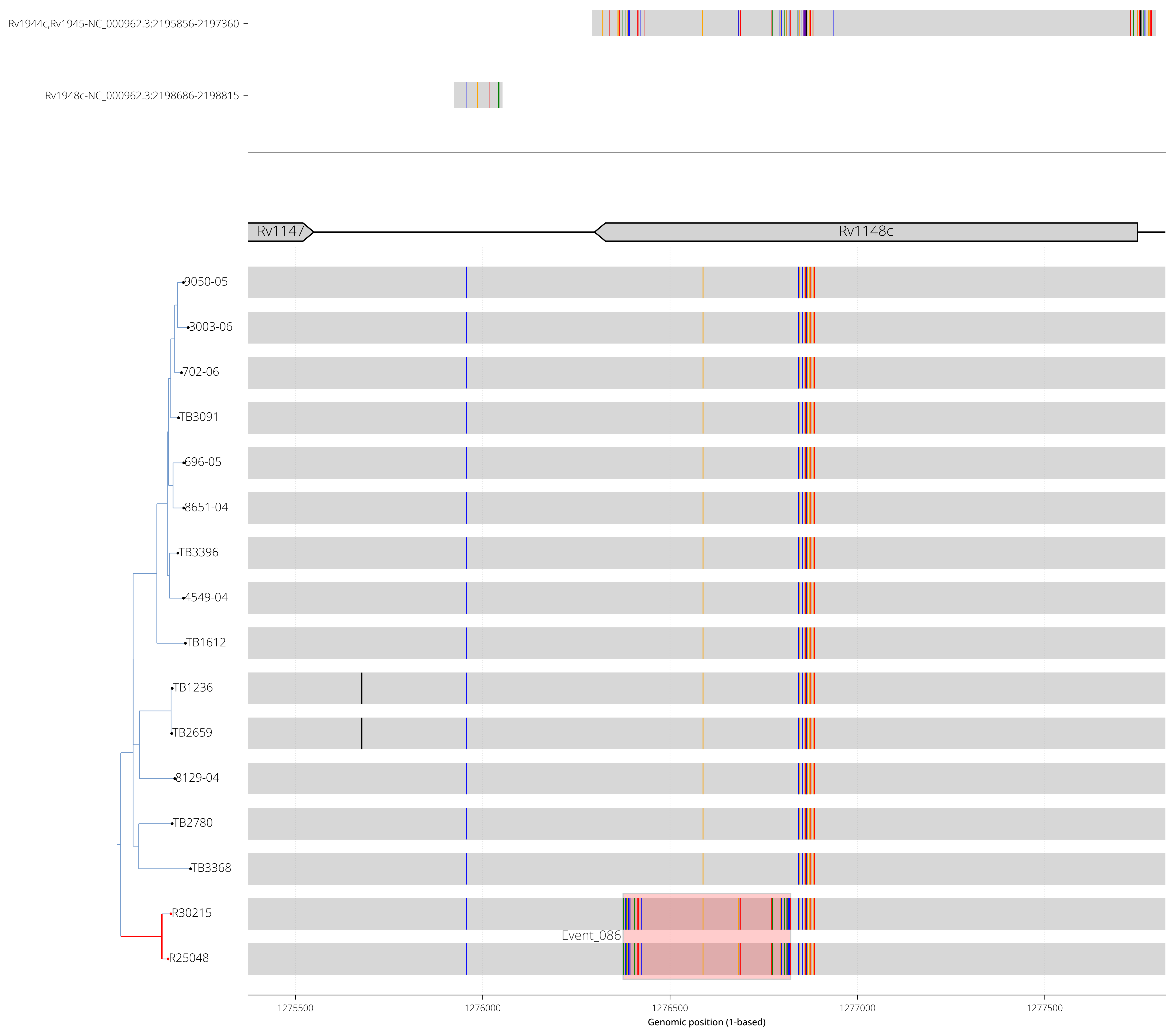

Full Phylogeny Viz + Paralog Alignments
GCE ID: Event_086

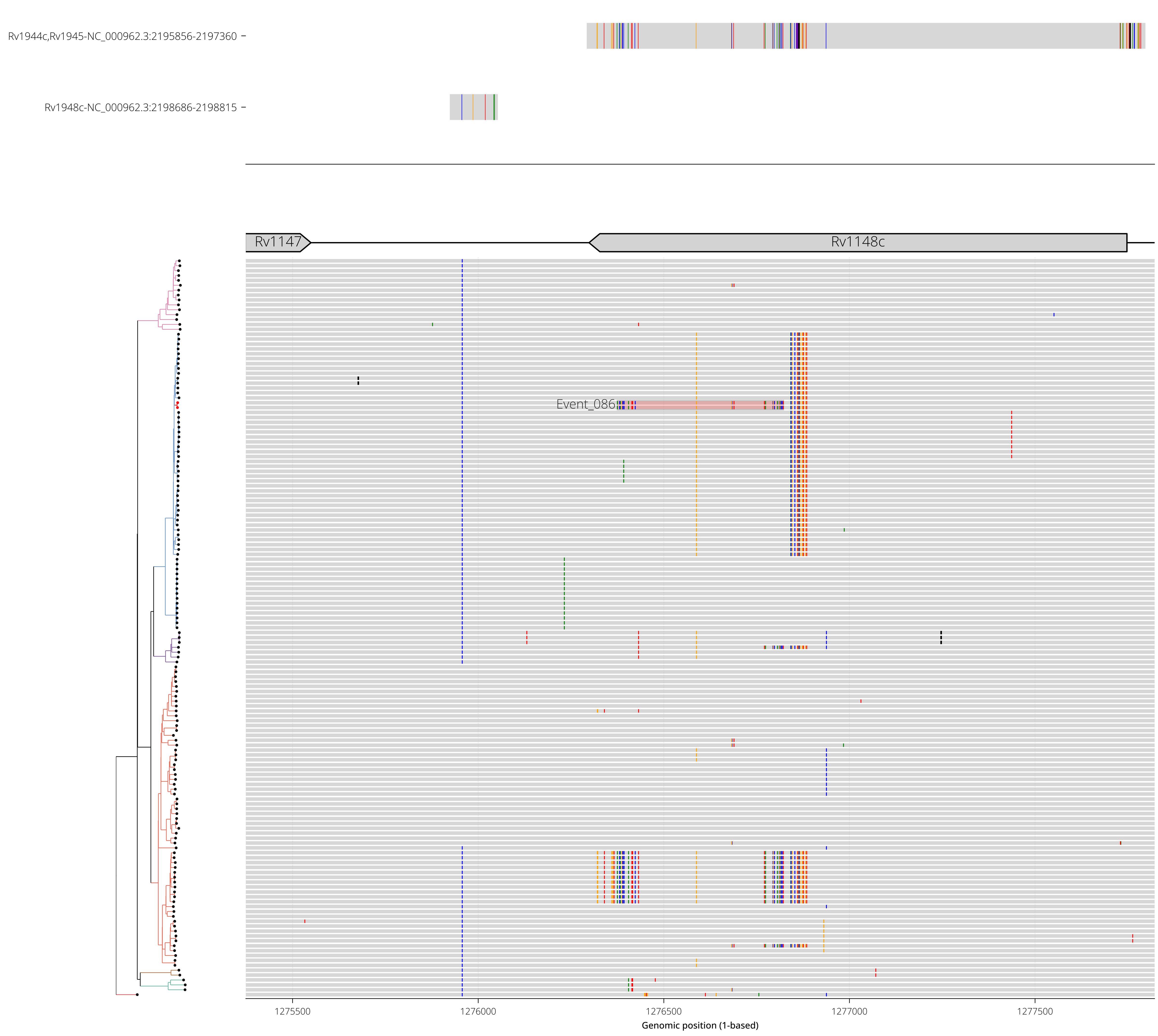

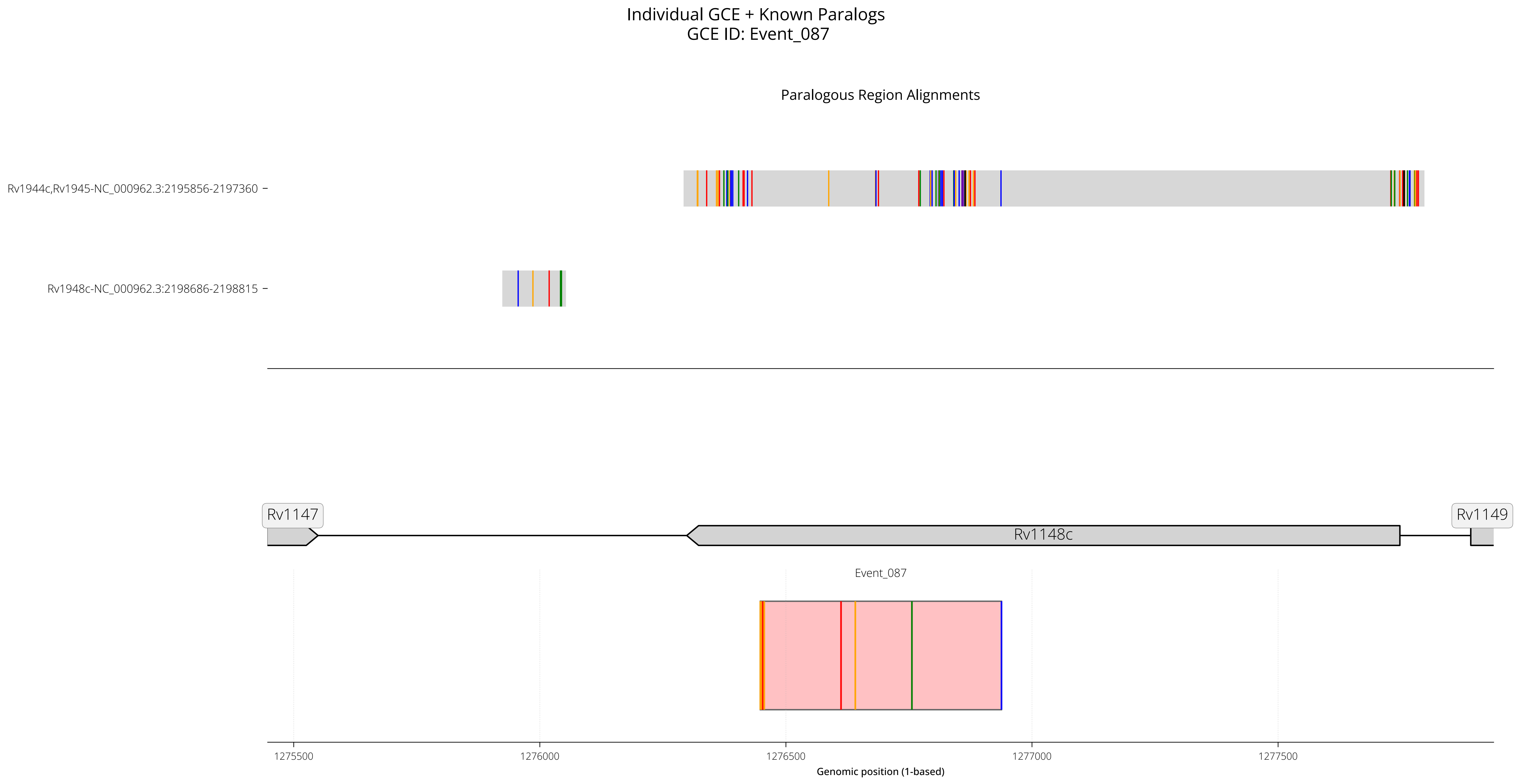
Individual GCE + Known Paralogs
GCE ID: Event_087
Paralogous Region Alignments
Rv1944c,Rv1945-NC_000962.3:2195856-2197360
Rv1948c-NC_000962.3:2198686-2198815
Rv1147
Rv1148c
Rv1149
Event_087
1275500
1276000
1276500
1277000
1277500
Genomic position (1-based)

Circos Viz - GCE to Known Paralogs Locations
GCE ID: Event_087

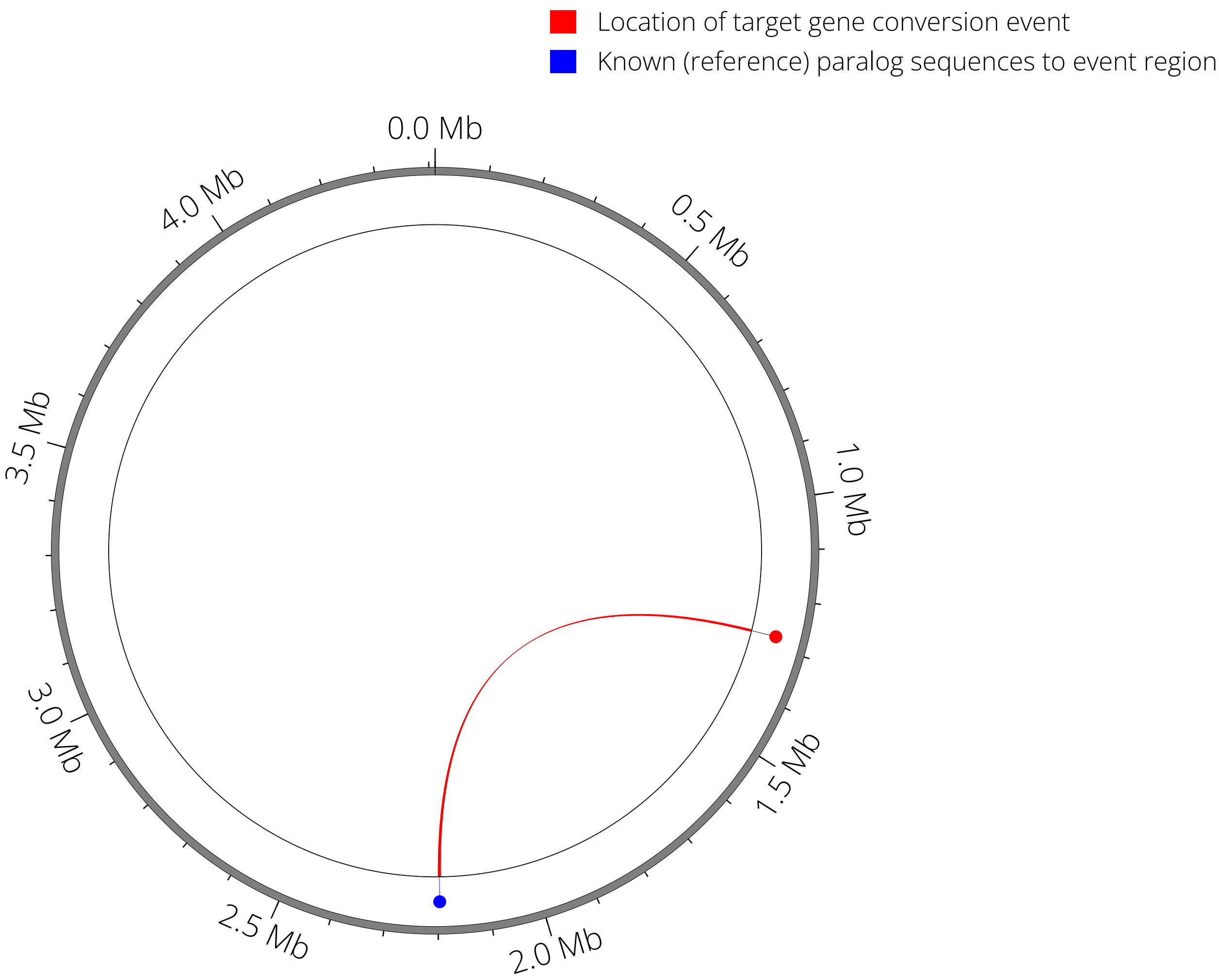

### Event_087 - 8 SNPs - k-mer analysis relative to paralogs

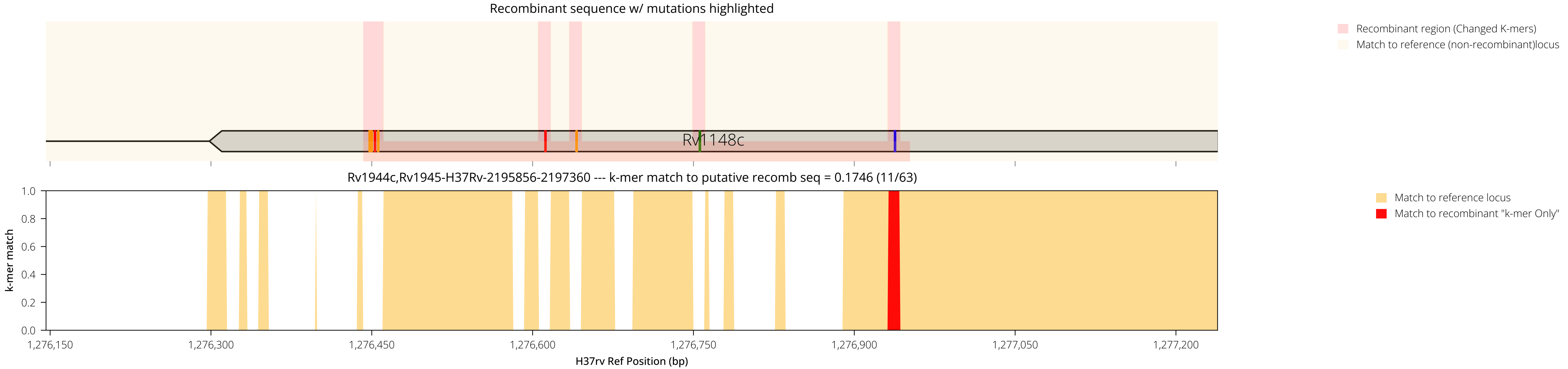

Sub-phylogeny Viz + Paralog Alignments
GCE ID: Event_087

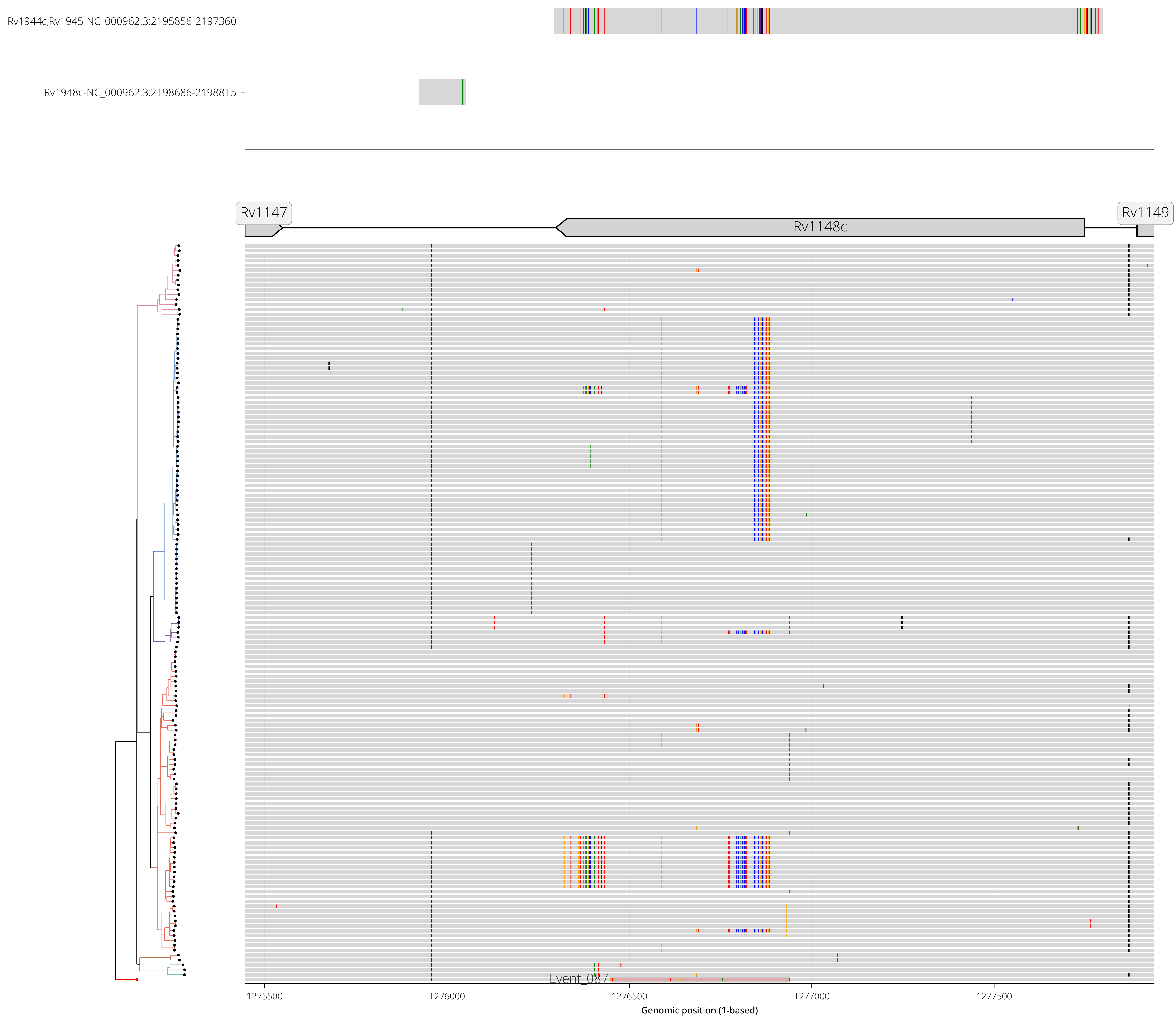

Full Phylogeny Viz + Paralog Alignments
GCE ID: Event_087

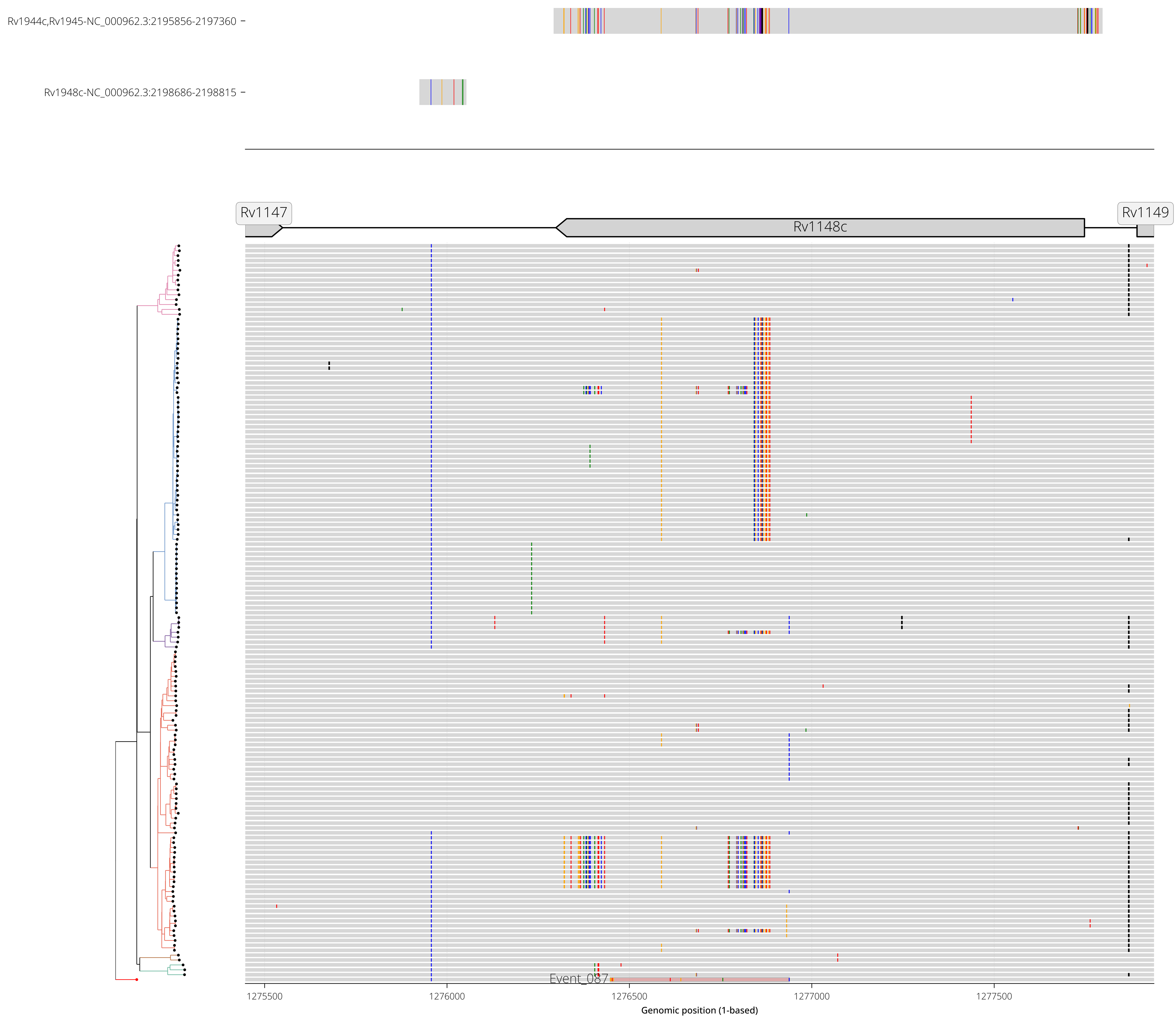

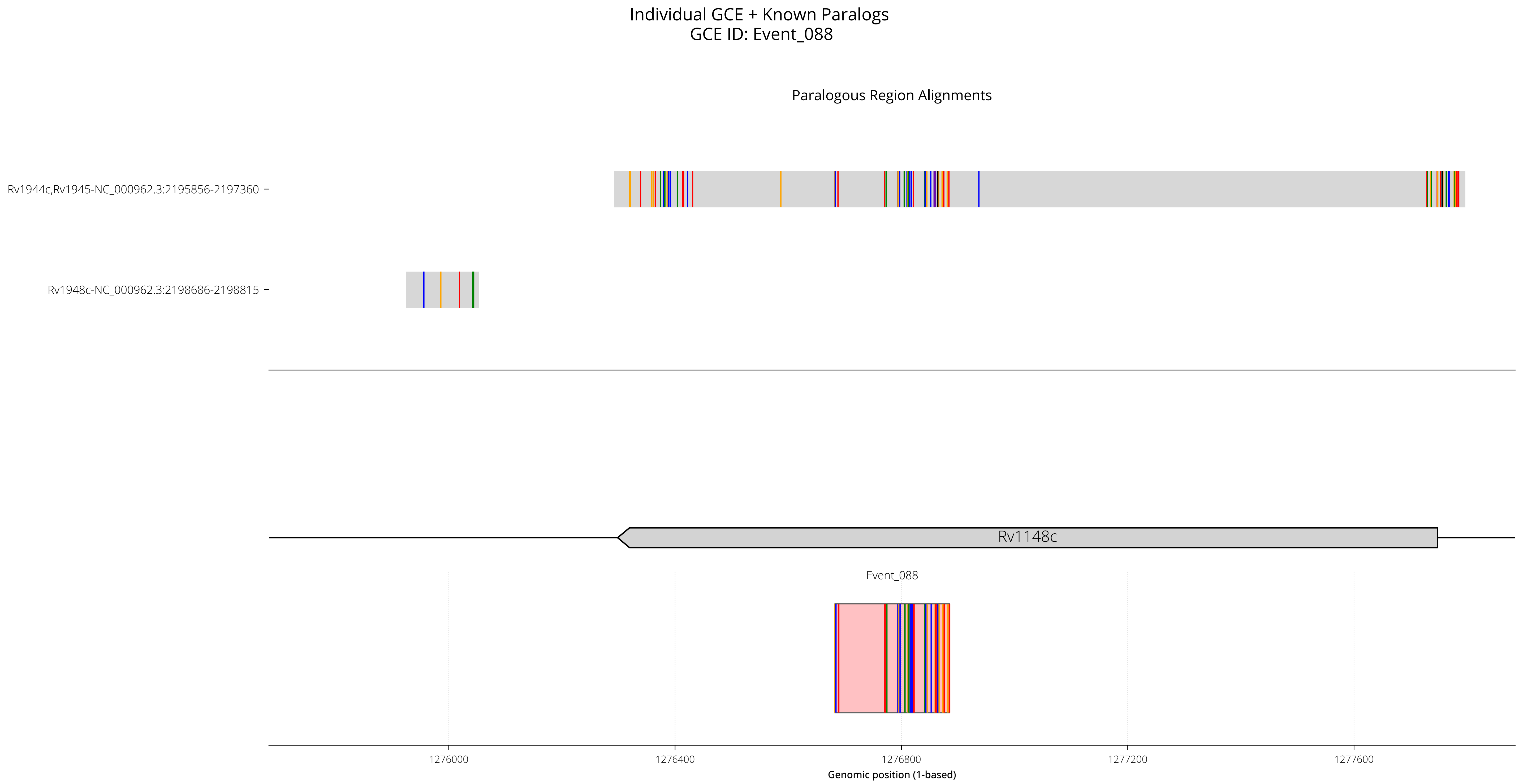

Individual GCE + Known Paralogs
GCE ID: Event_088
Paralogous Region Alignments
Rv1944c,Rv1945-NC_000962.3:2195856-2197360
Rv1948c-NC_000962.3:2198686-2198815
Rv1148c
Event_088
1276000
1276400
1276800
1277200
1277600
Genomic position (1-based)

#### Circos Viz - GCE to Known Paralogs Locations
#### GCE ID: Event_088

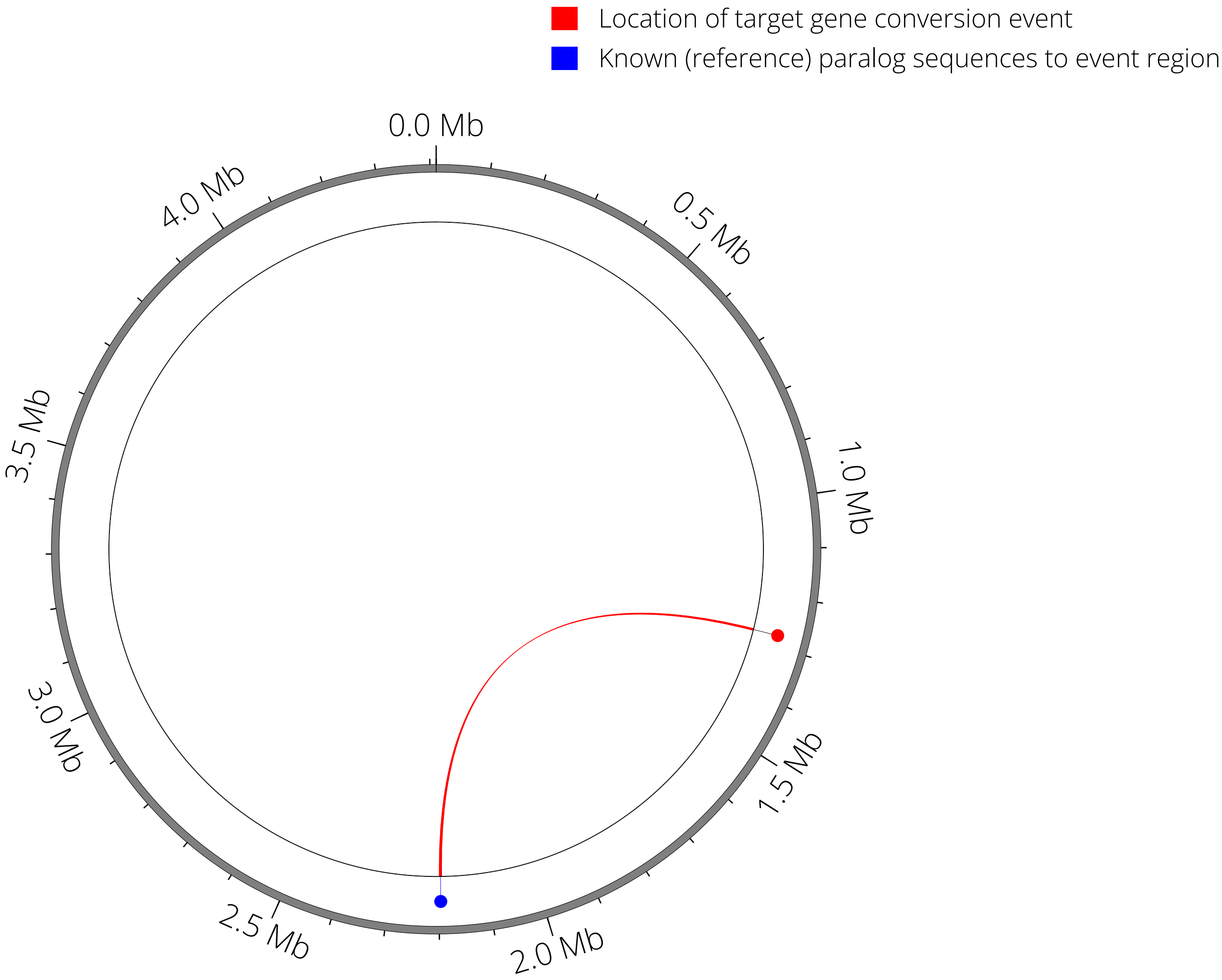

### Event_088 - 29 SNPs - k-mer analysis relative to paralogs

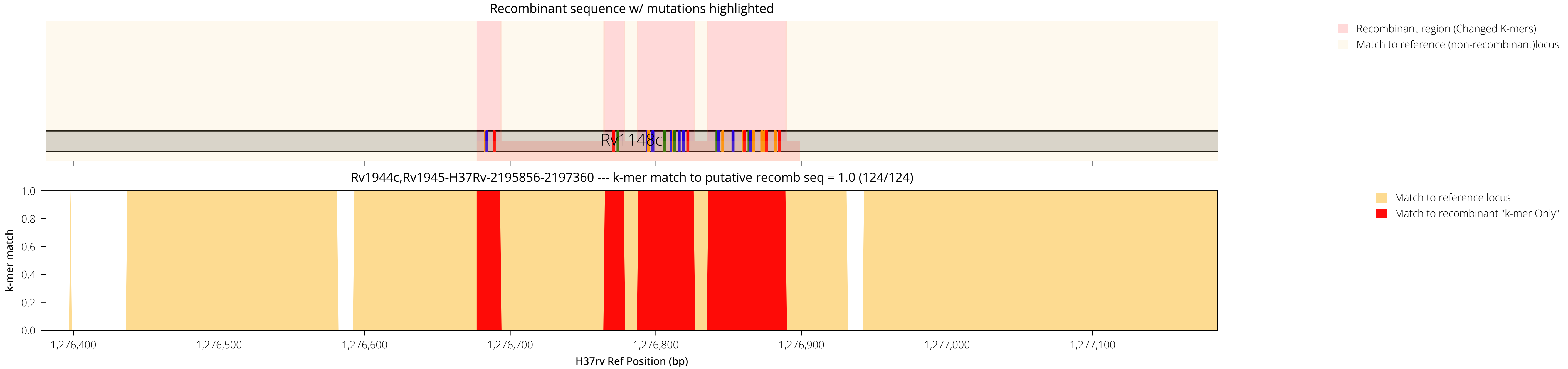

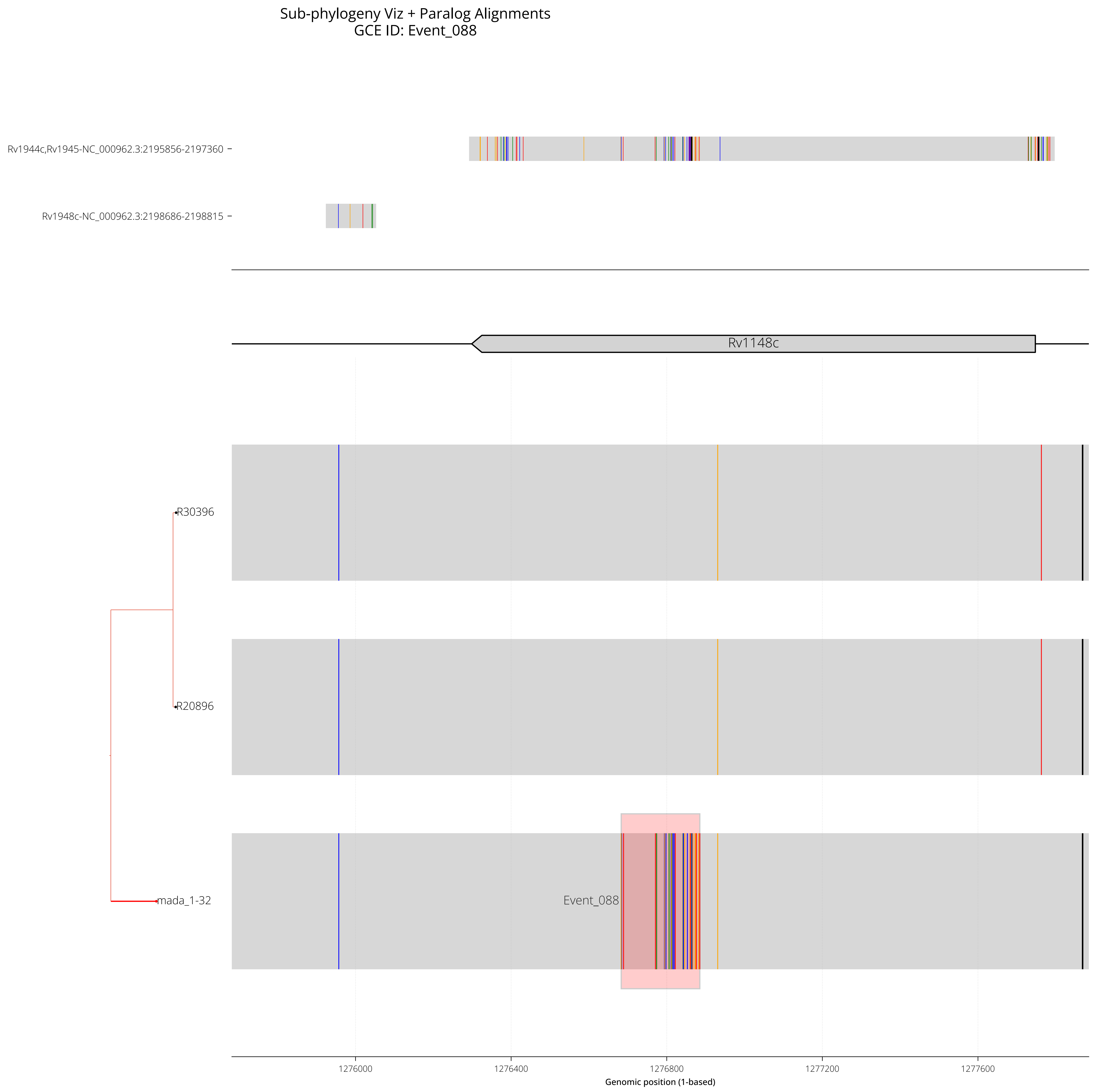

Sub-phylogeny Viz + Paralog Alignments
GCE ID: Event_088
Rv1944c,Rv1945-NC_000962.3:2195856-2197360
Rv1948c-NC_000962.3:2198686-2198815
Rv1148c
R30396
R20896
mada_1-32
Event_088
1276000
1276400
1276800
1277200
1277600
Genomic position (1-based)

Full Phylogeny Viz + Paralog Alignments
GCE ID: Event_088

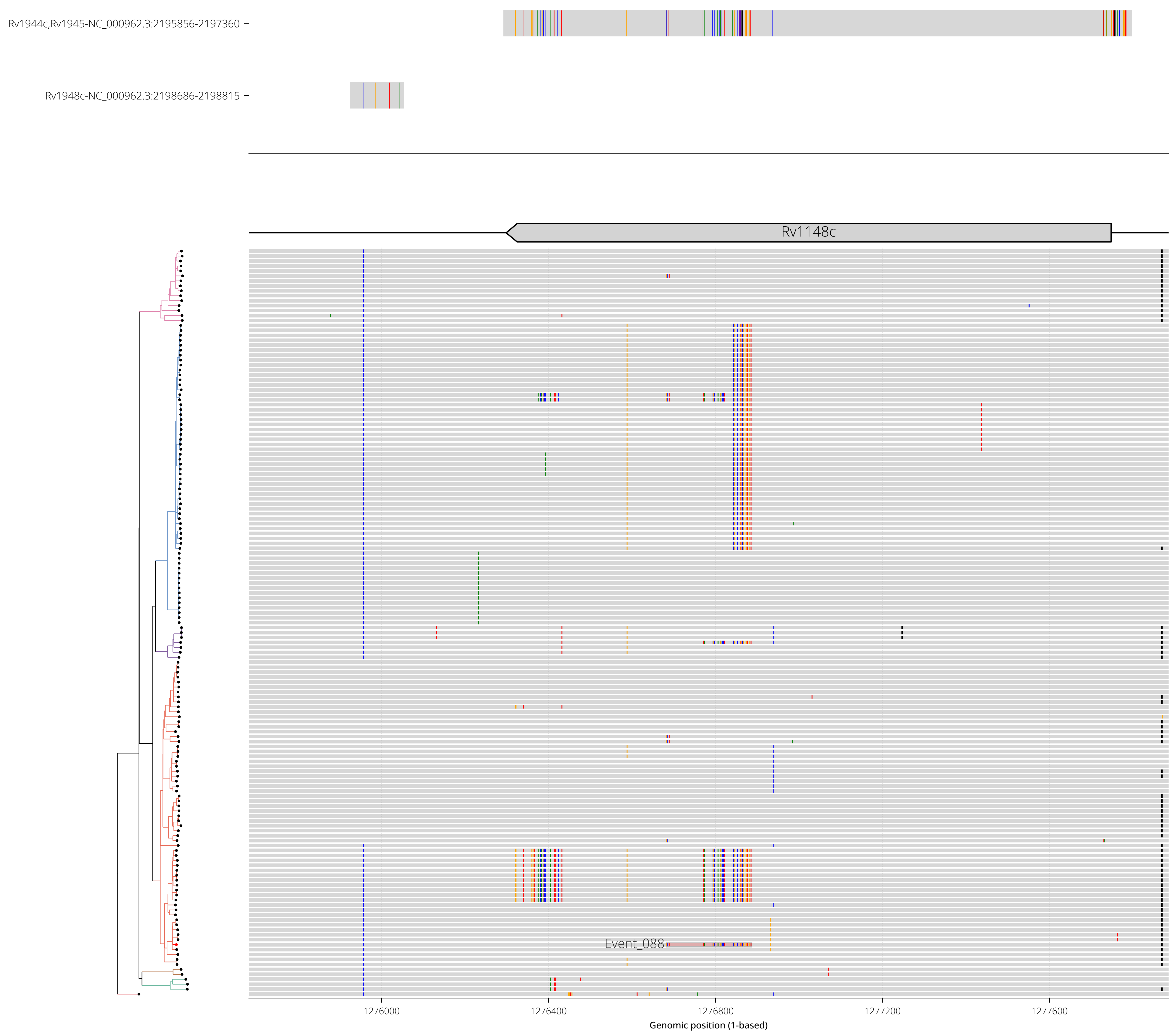

### Individual GCE + Known Paralogs
GCE ID: Event_089

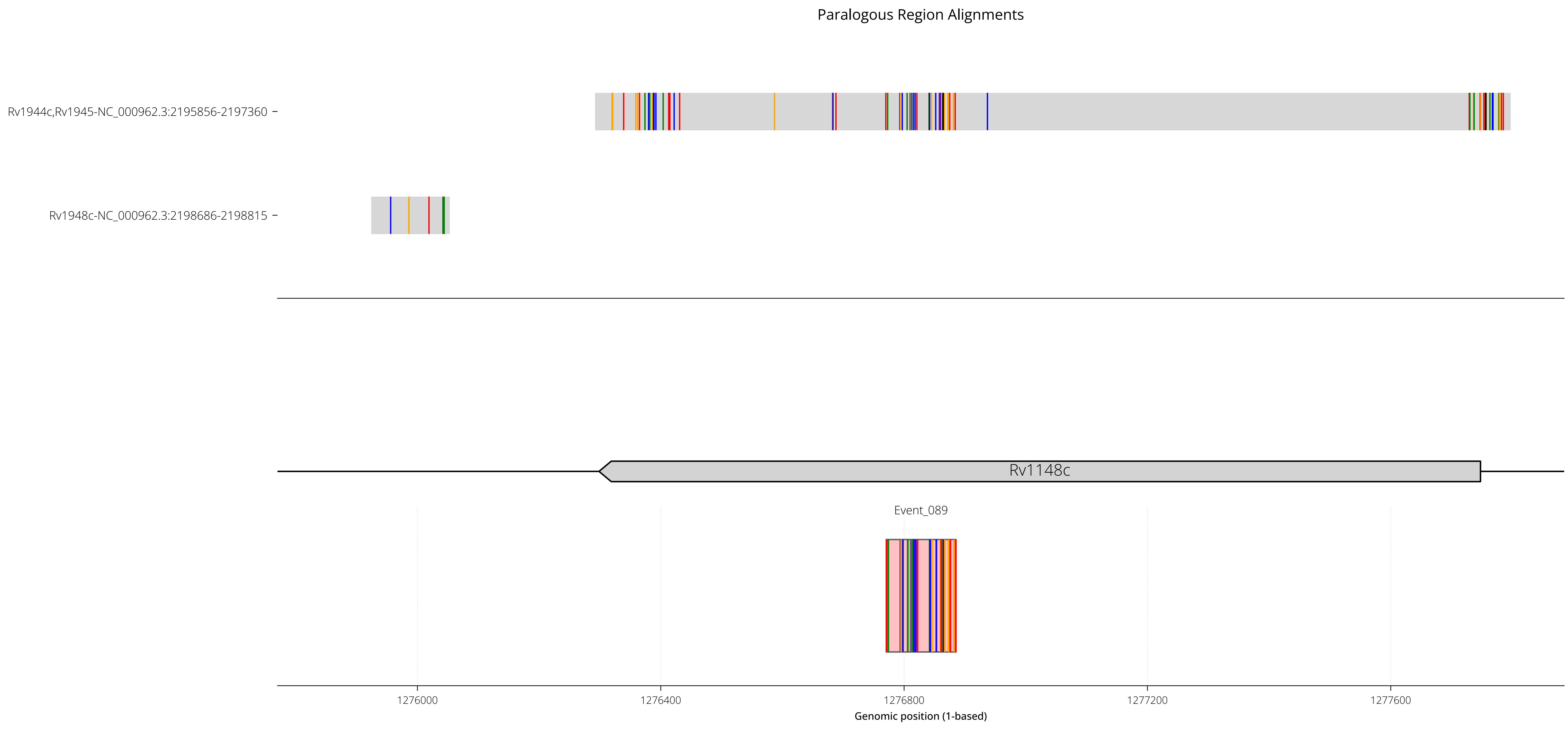

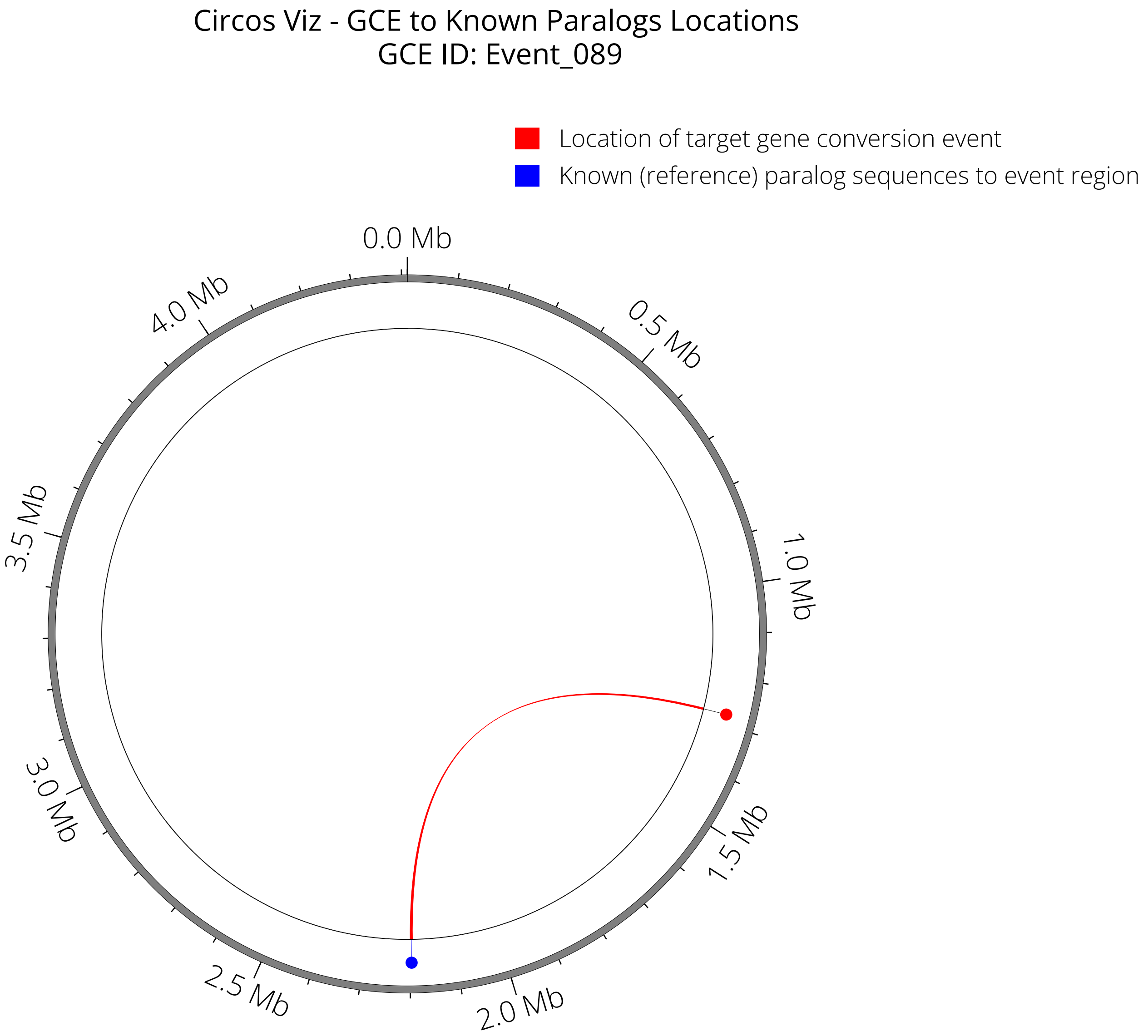
Circos Viz - GCE to Known Paralogs Locations
GCE ID: Event_089
Location of target gene conversion event
Known (reference) paralog sequences to event region
0.0 Mb
0.5 Mb
1.0 Mb
1.5 Mb
2.0 Mb
2.5 Mb
3.0 Mb
3.5 Mb
4.0 Mb

### Event_089 - 26 SNPs - k-mer analysis relative to paralogs

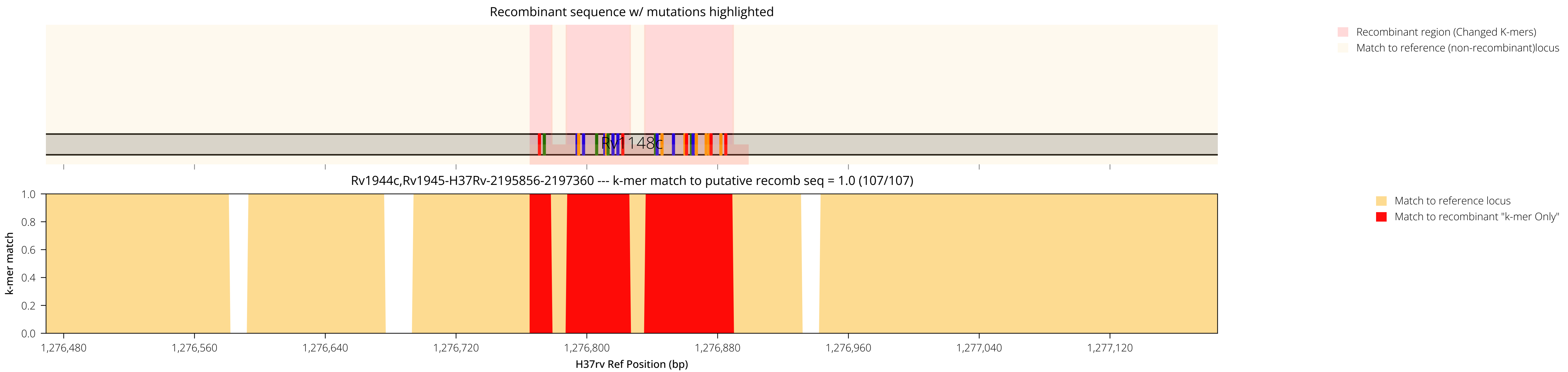

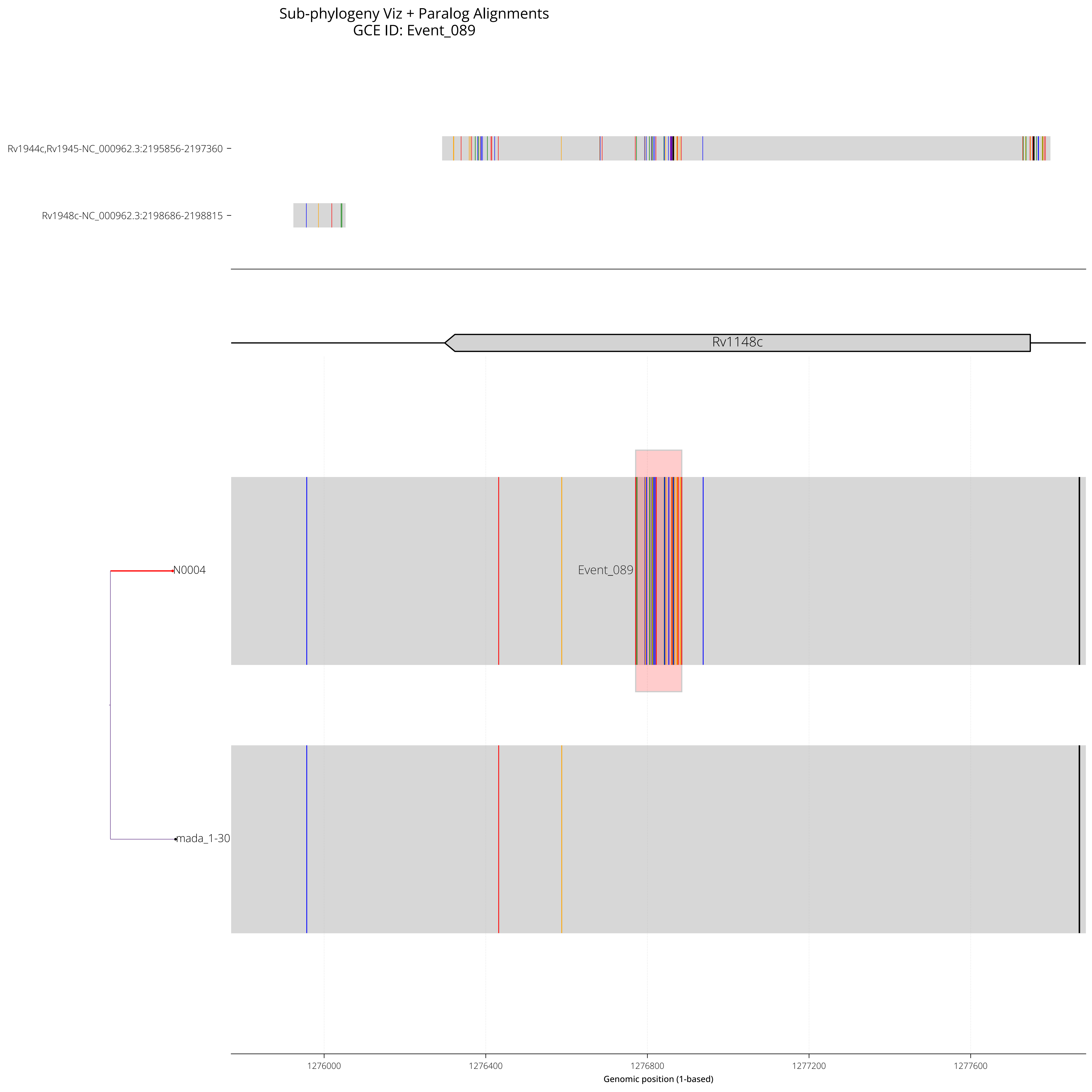
Sub-phylogeny Viz + Paralog Alignments
GCE ID: Event_089
Rv1944c,Rv1945-NC_000962.3:2195856-2197360
Rv1948c-NC_000962.3:2198686-2198815
Rv1148c
N0004
Event_089
mada_1-30
1276000
1276400
1276800
1277200
1277600
Genomic position (1-based)

Full Phylogeny Viz + Paralog Alignments
GCE ID: Event_089

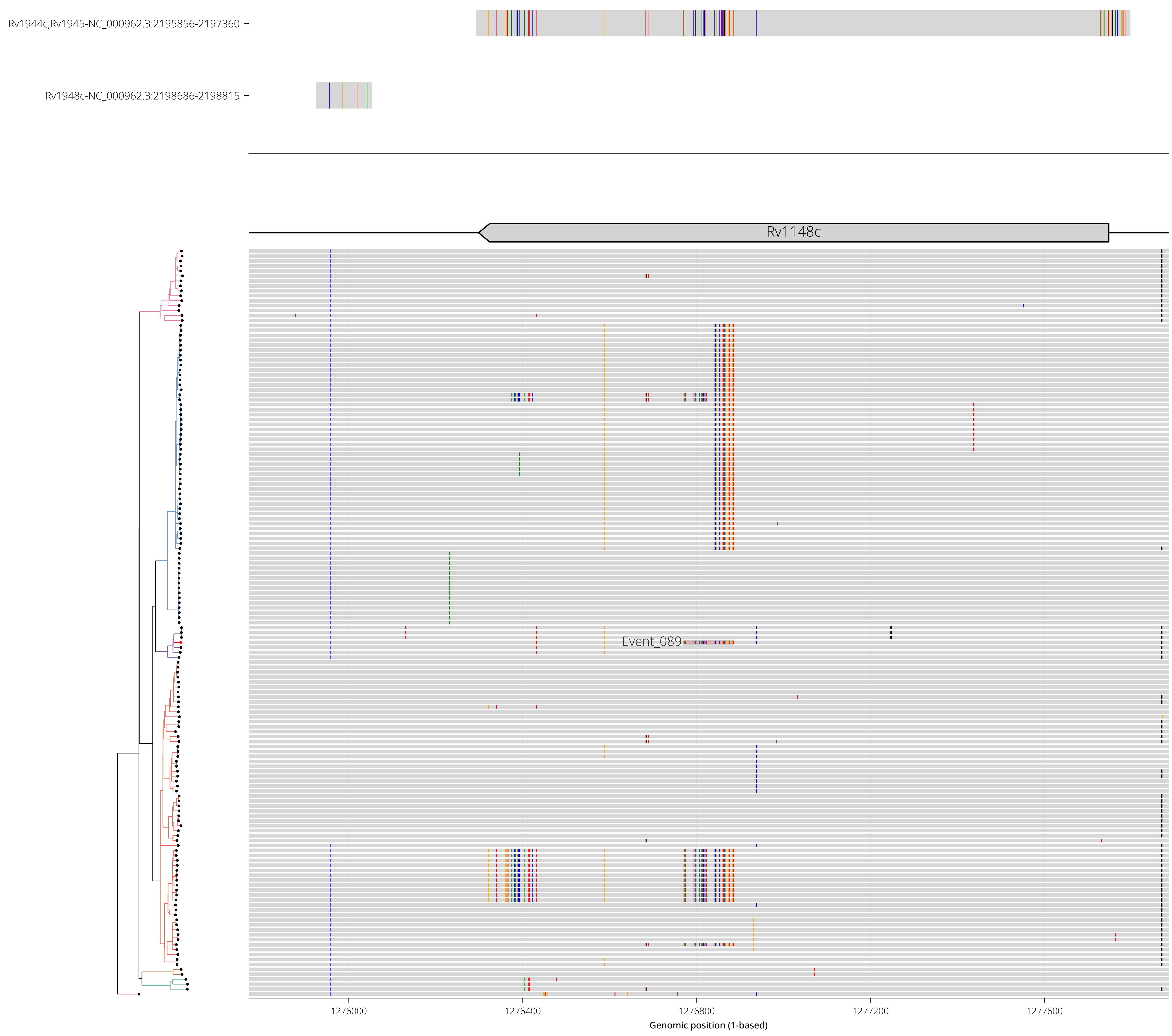

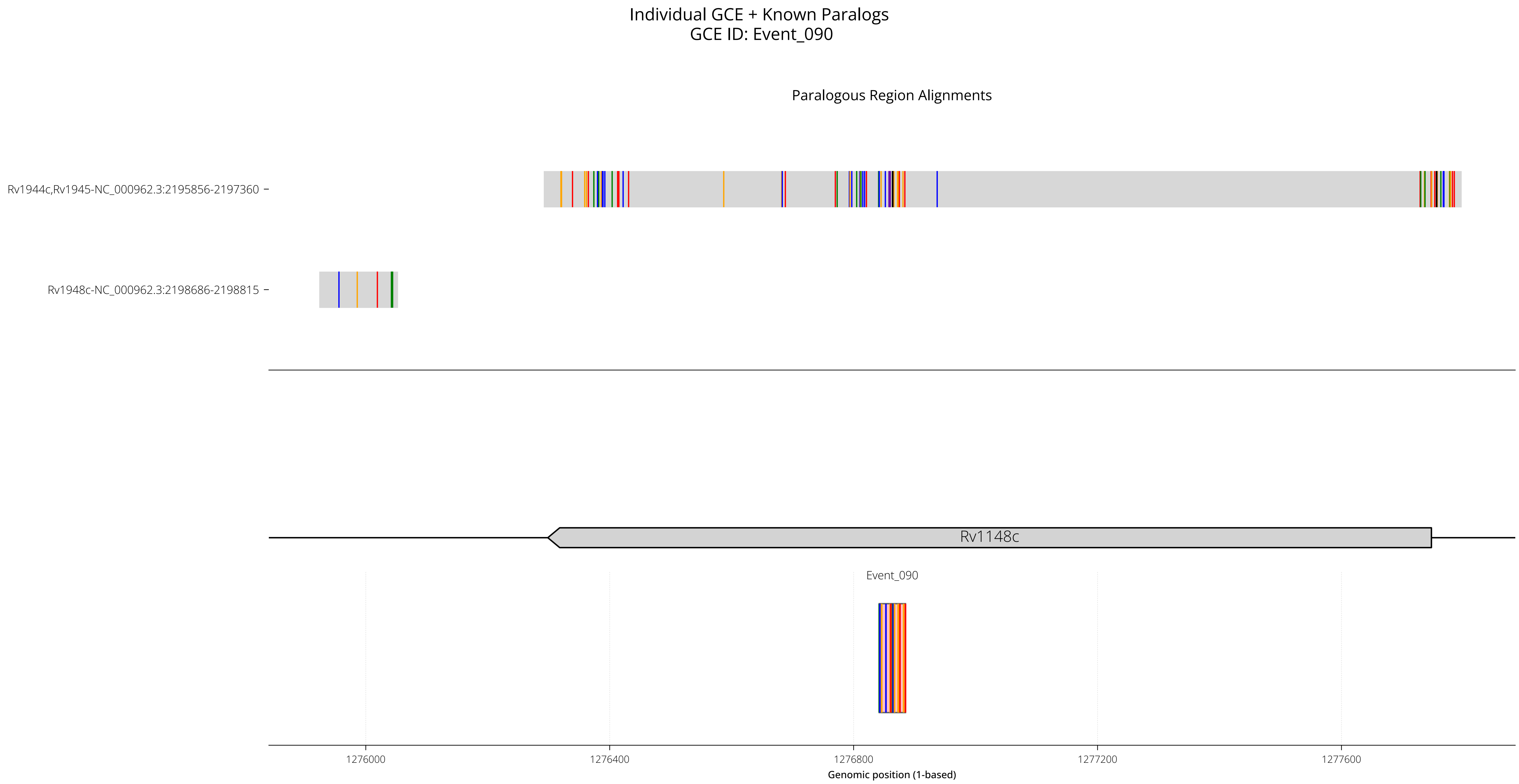
Individual GCE + Known Paralogs
GCE ID: Event_090
Paralogous Region Alignments
Rv1944c,Rv1945-NC_000962.3:2195856-2197360
Rv1948c-NC_000962.3:2198686-2198815
Rv1148c
Event_090
1276000
1276400
1276800
1277200
1277600
Genomic position (1-based)

Circos Viz - GCE to Known Paralogs Locations
GCE ID: Event_090

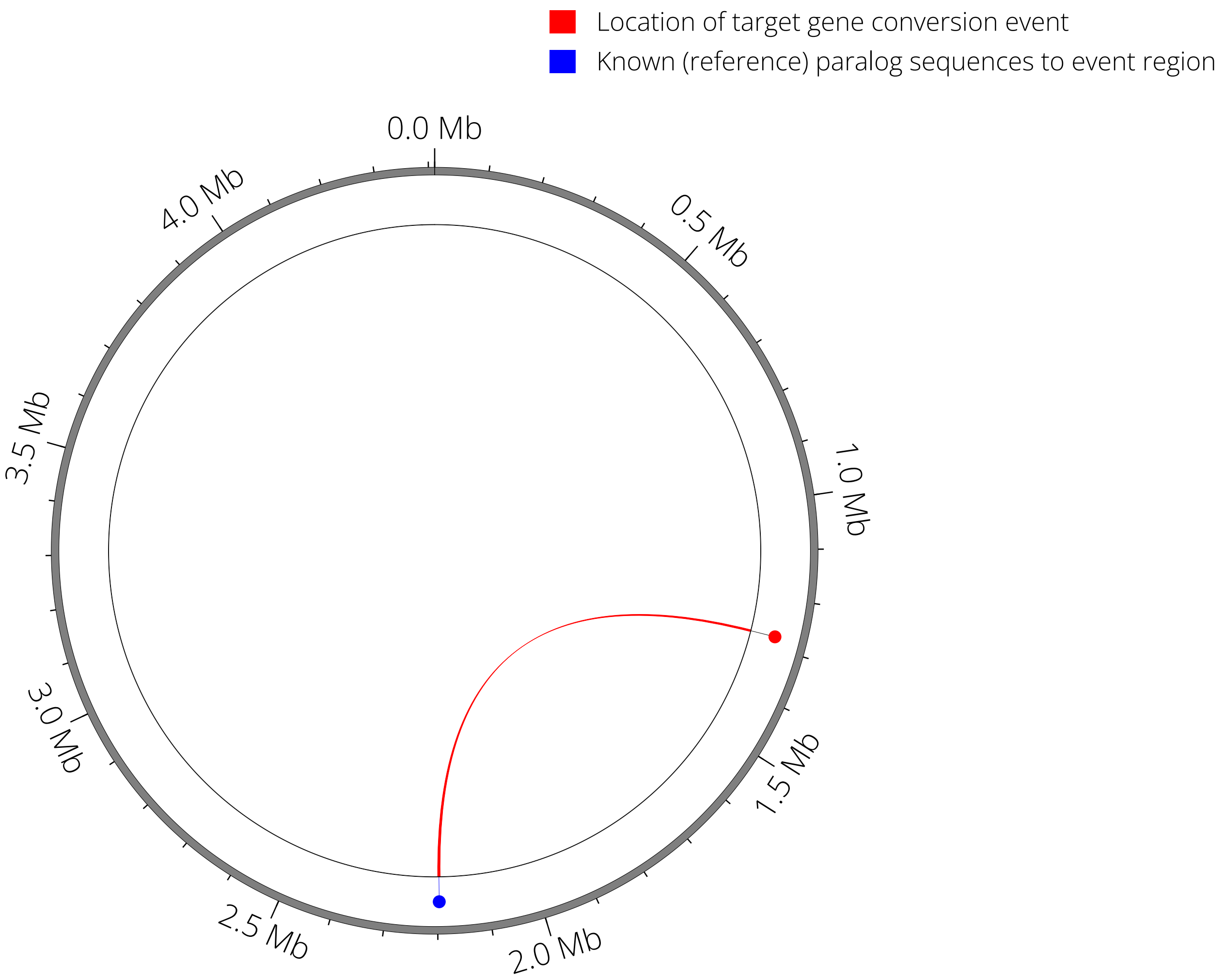

### Event_090 - 14 SNPs - k-mer analysis relative to paralogs

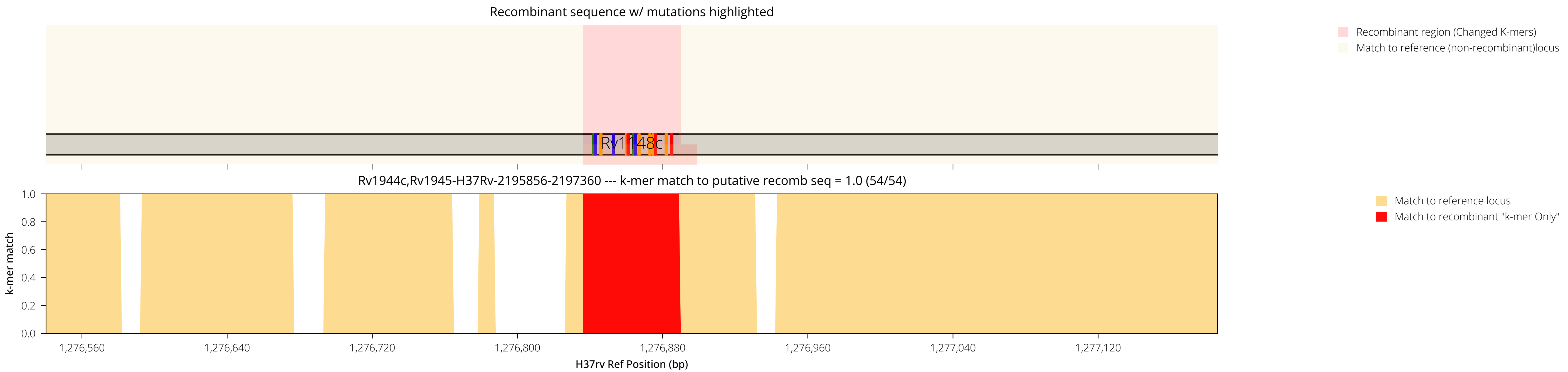

Sub-phylogeny Viz + Paralog Alignments
GCE ID: Event_090

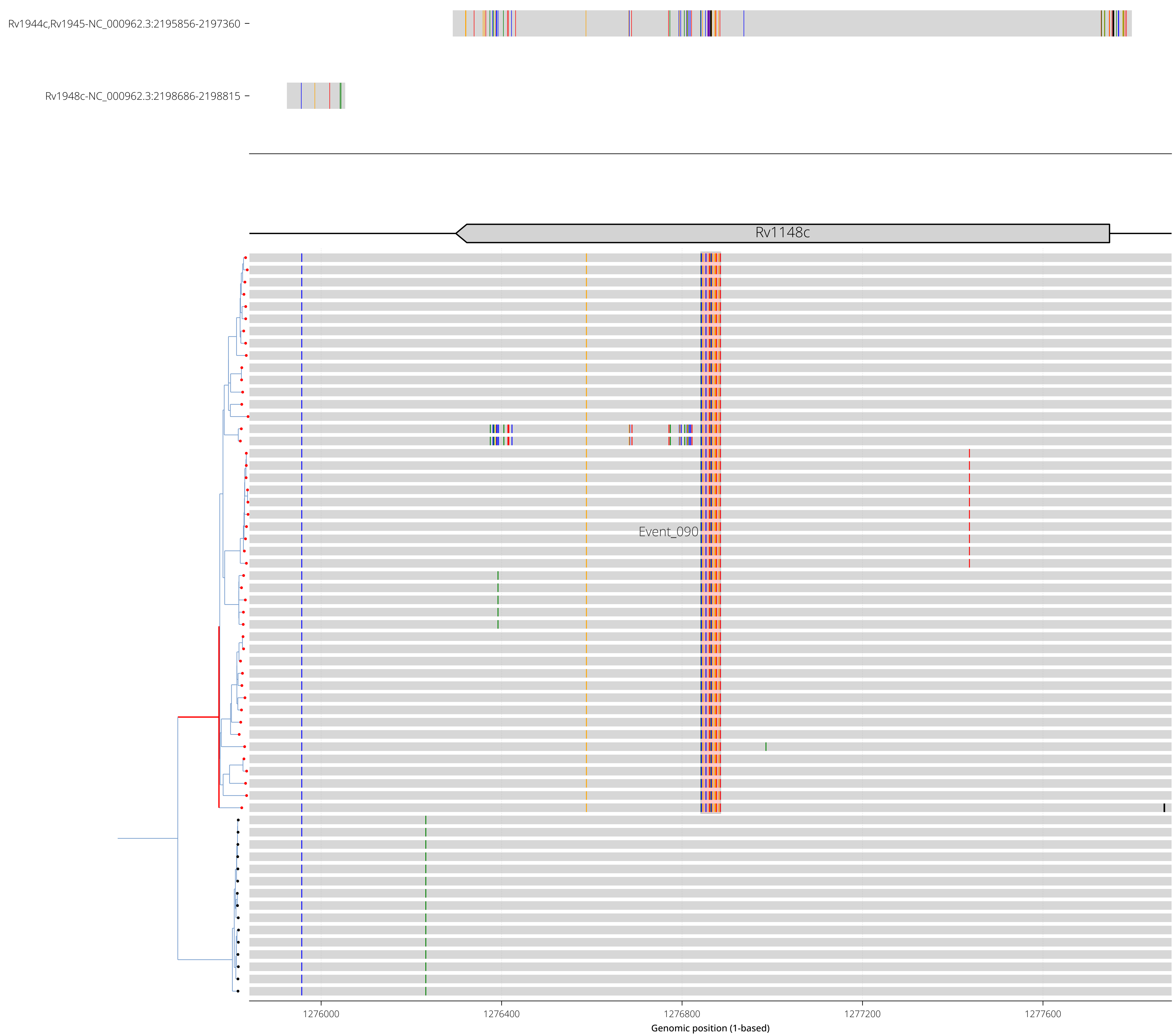

Full Phylogeny Viz + Paralog Alignments
GCE ID: Event_090

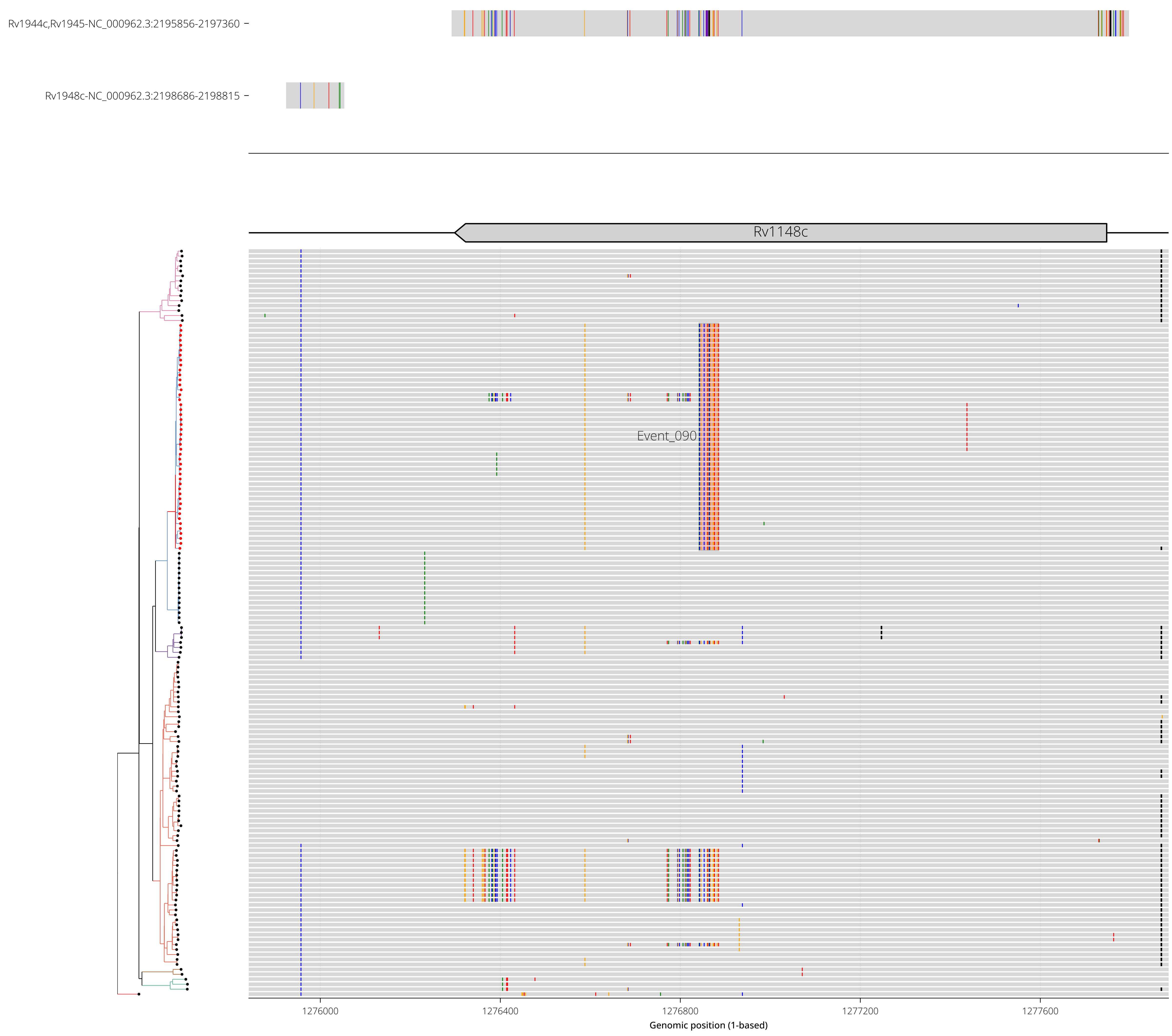

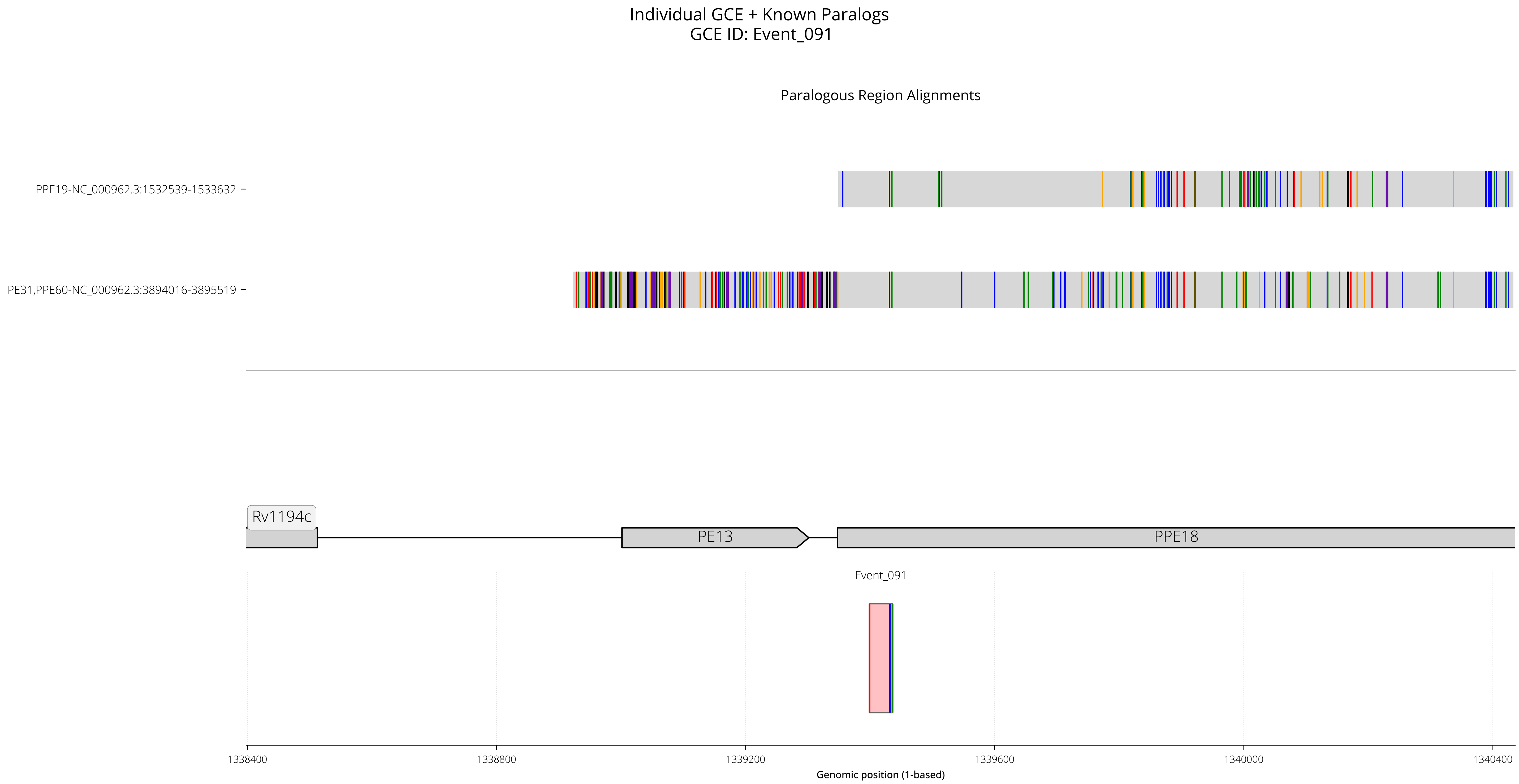

Individual GCE + Known Paralogs
GCE ID: Event_091
Paralogous Region Alignments
PPE19-NC_000962.3:1532539-1533632
PE31,PPE60-NC_000962.3:3894016-3895519
Rv1194c
PE13
PPE18
Event_091
1338400
1338800
1339200
1339600
1340000
1340400
Genomic position (1-based)

### Circos Viz - GCE to Known Paralogs Locations
GCE ID: Event_091

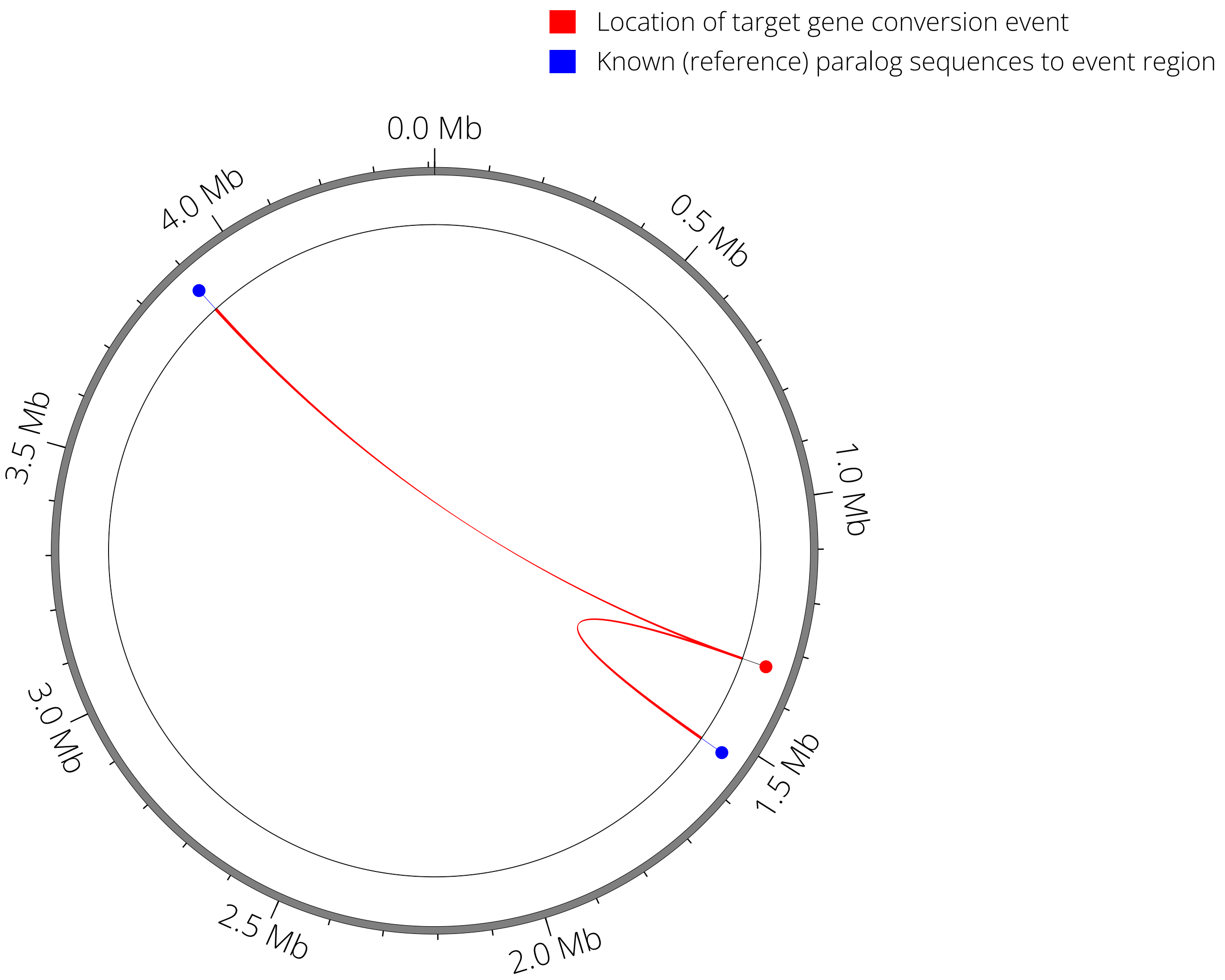

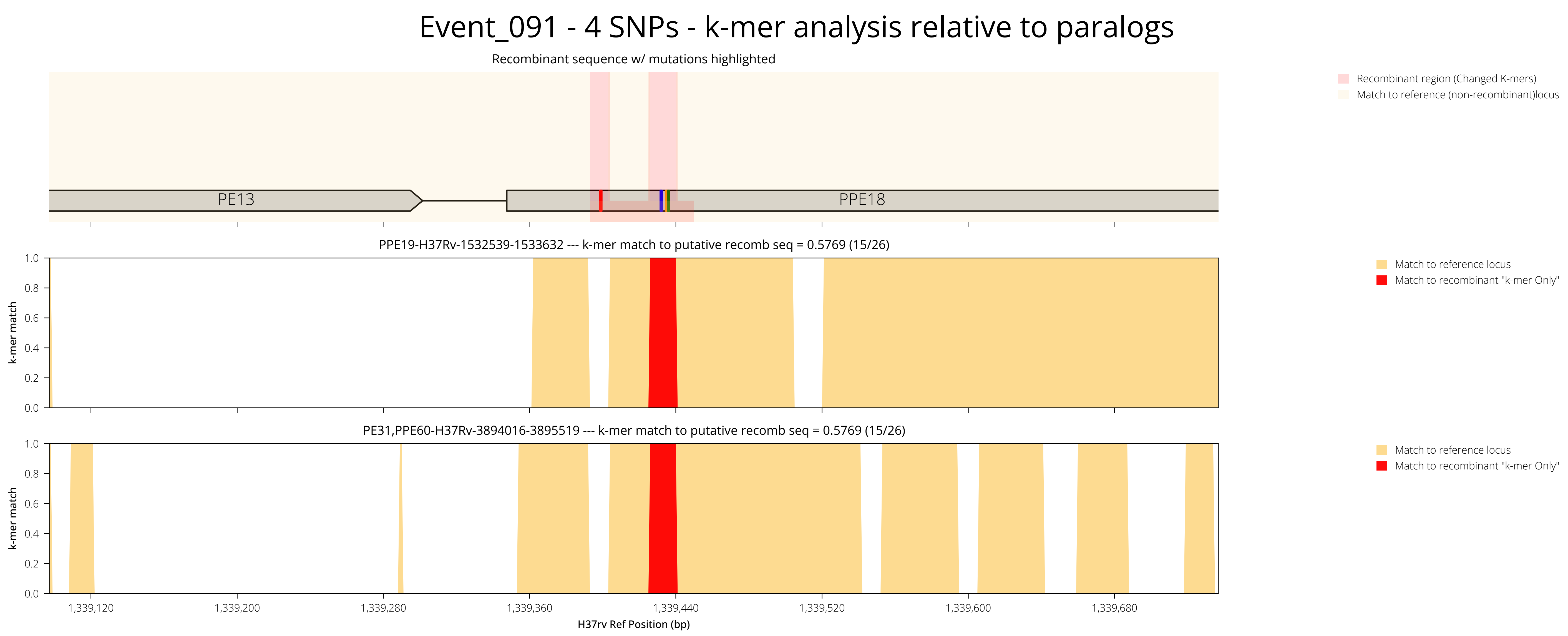

Event_091 - 4 SNPs - k-mer analysis relative to paralogs
Recombinant sequence w/ mutations highlighted
Recombinant region (Changed K-mers)
Match to reference (non-recombinant)locus
PE13
PPE18
PPE19-H37Rv-1532539-1533632 --- k-mer match to putative recomb seq = 0.5769 (15/26)
Match to reference locus
Match to recombinant "k-mer Only"
PE31,PPE60-H37Rv-3894016-3895519 --- k-mer match to putative recomb seq = 0.5769 (15/26)
Match to reference locus
Match to recombinant "k-mer Only"
k-mer match
1.0
0.8
0.6
0.4
0.2
0.0
1,339,120
1,339,200
1,339,280
1,339,360
1,339,440
1,339,520
1,339,600
1,339,680
H37rv Ref Position (bp)

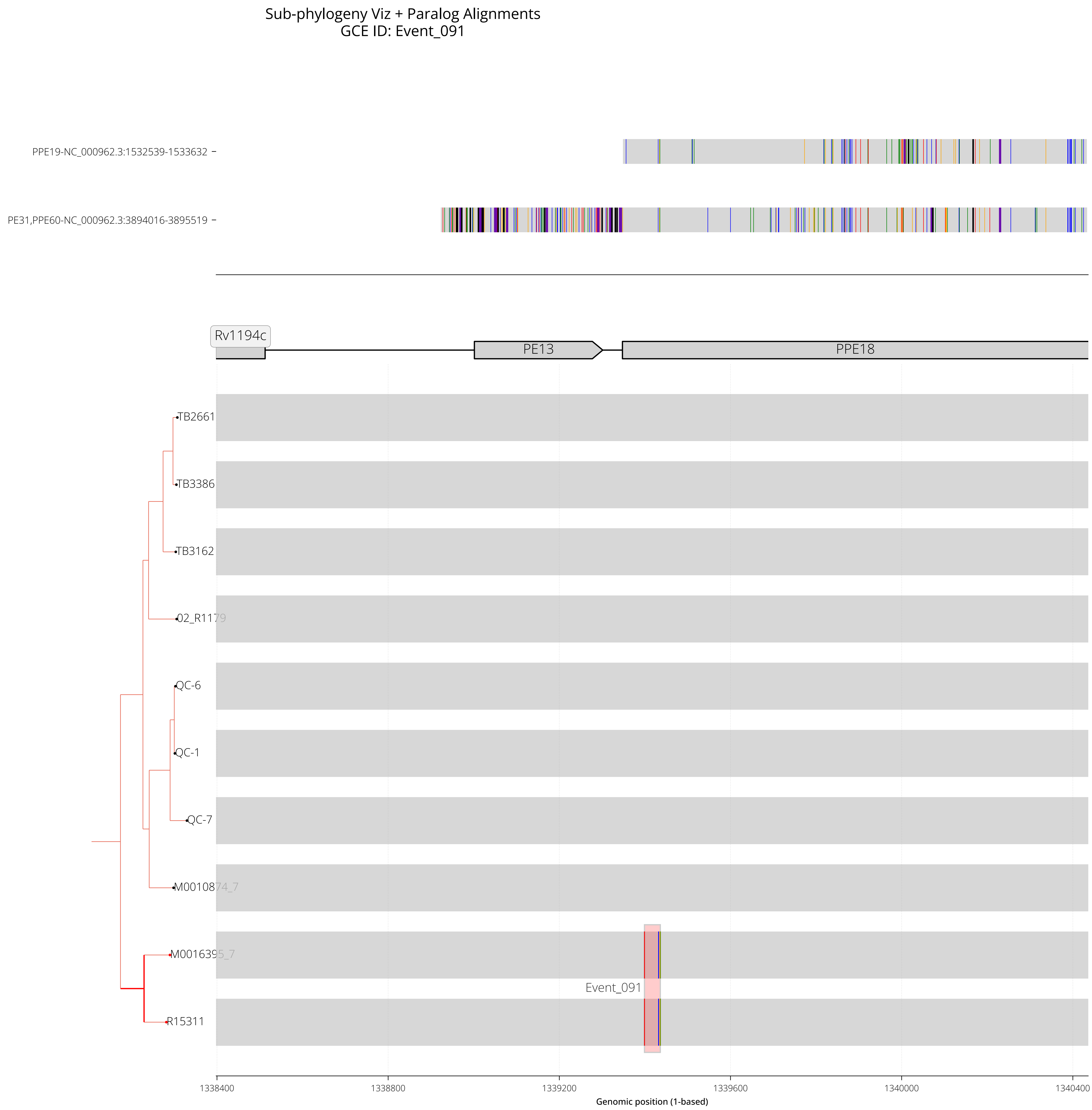
Sub-phylogeny Viz + Paralog Alignments
GCE ID: Event_091
PPE19-NC_000962.3:1532539-1533632
PE31,PPE60-NC_000962.3:3894016-3895519
Rv1194c
PE13
PPE18
TB2661
TB3386
TB3162
02_R1179
QC-6
QC-1
QC-7
M0010874_7
M0016395_7
R15311
Event_091
1338400
1338800
1339200
1339600
1340000
1340400
Genomic position (1-based)

Full Phylogeny Viz + Paralog Alignments
GCE ID: Event_091

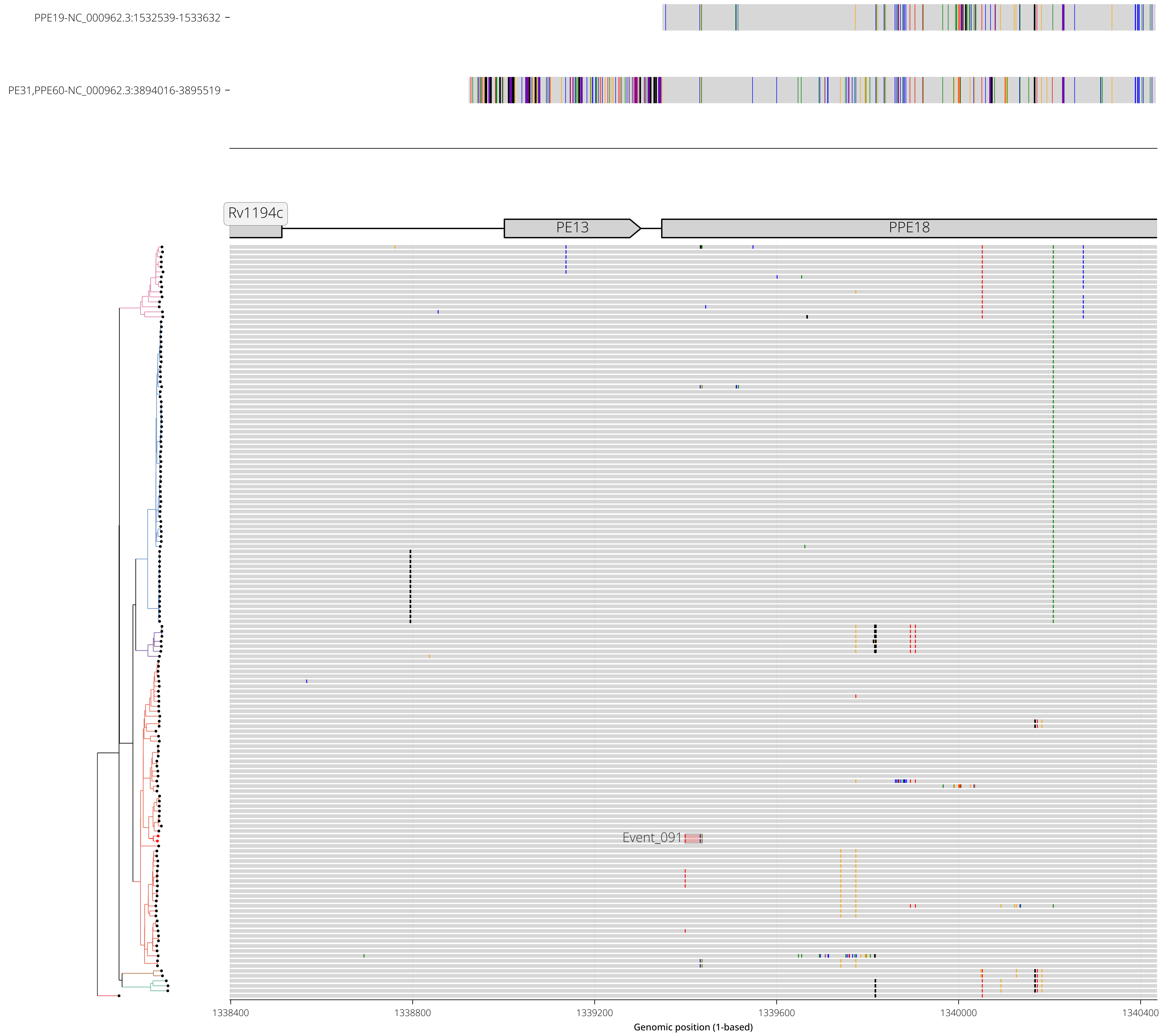

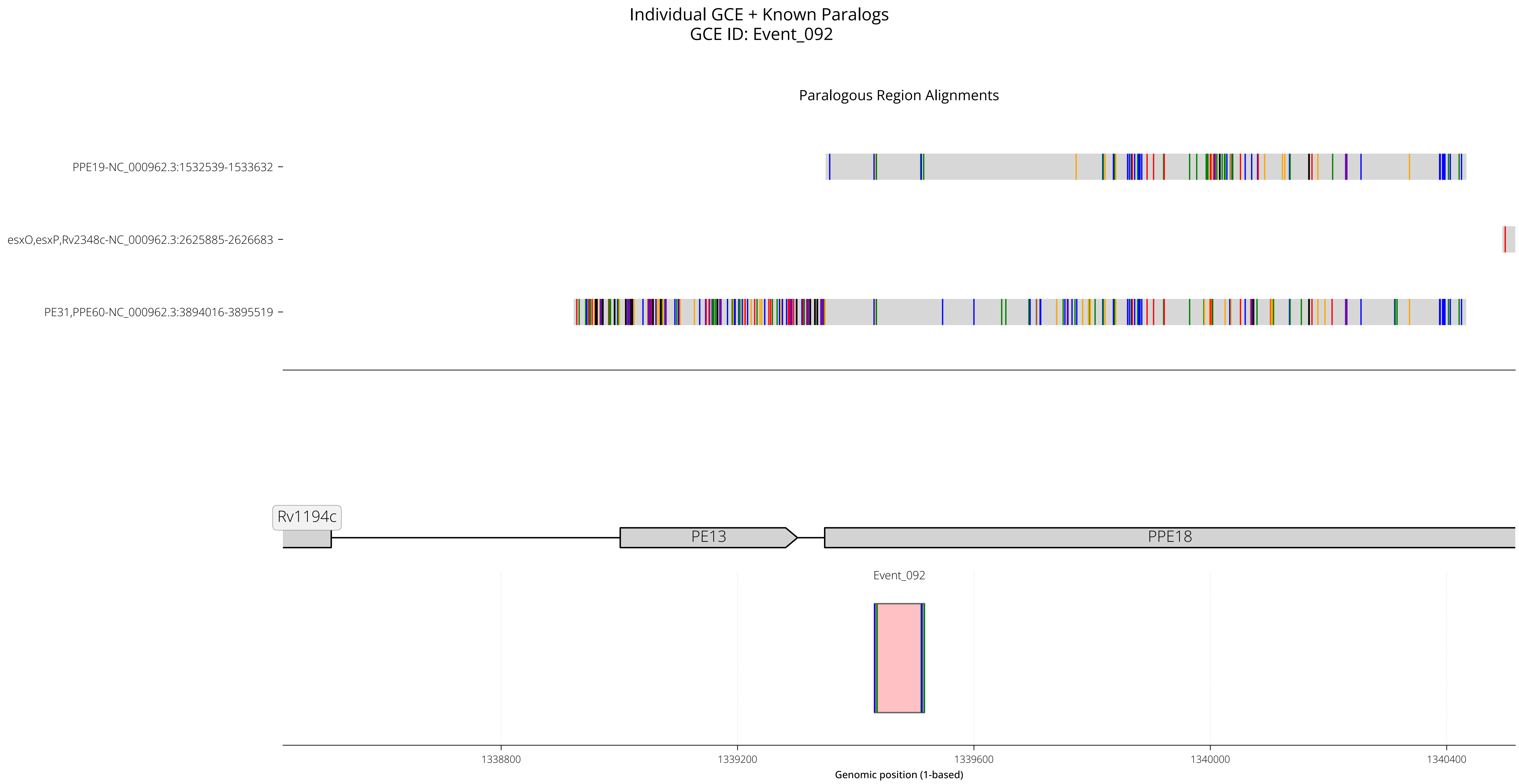
Individual GCE + Known Paralogs
GCE ID: Event_092
Paralogous Region Alignments
PPE19-NC_000962.3:1532539-1533632
esxO,esxP,Rv2348c-NC_000962.3:2625885-2626683
PE31,PPE60-NC_000962.3:3894016-3895519
Rv1194c
PE13
PPE18
Event_092
1338800
1339200
1339600
1340000
1340400
Genomic position (1-based)

Circos Viz - GCE to Known Paralogs Locations
GCE ID: Event_092

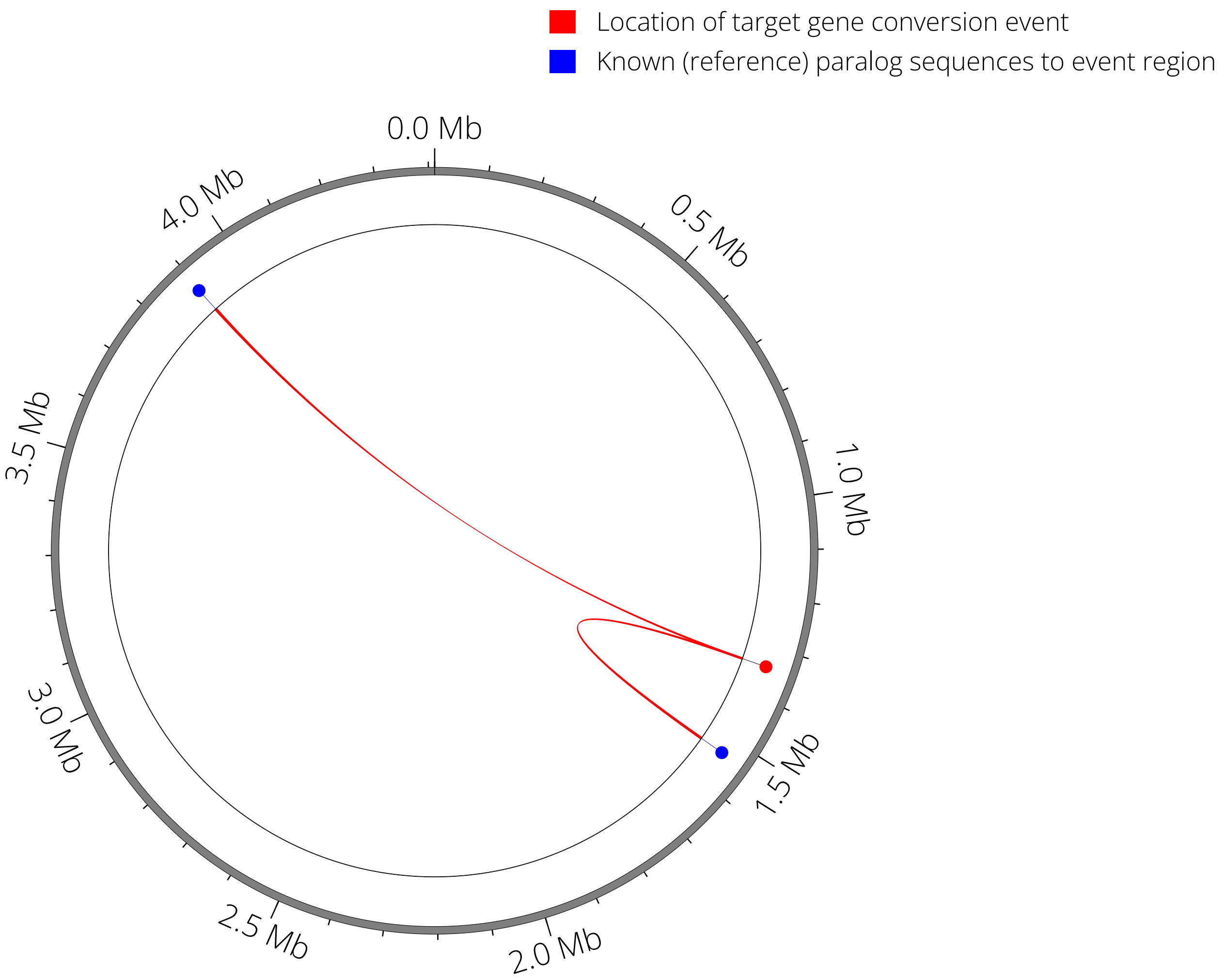

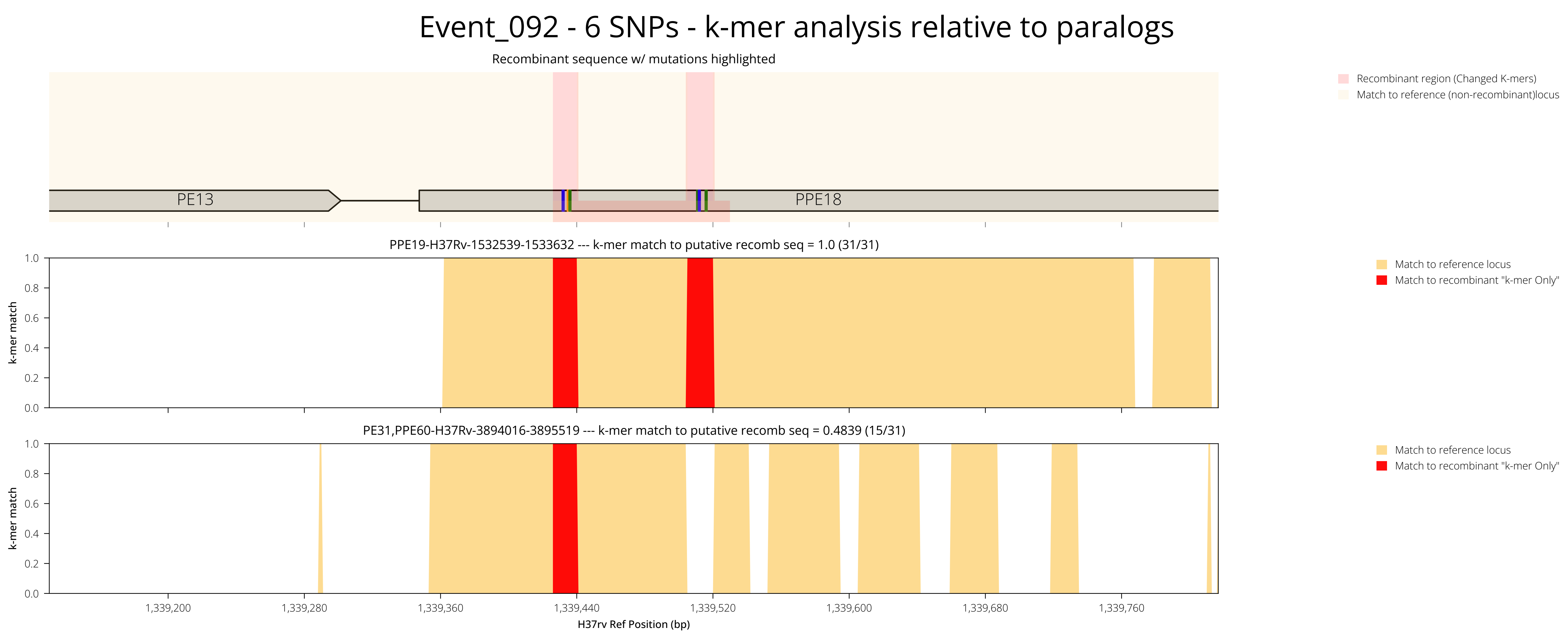
Event_092 - 6 SNPs - k-mer analysis relative to paralogs
Recombinant sequence w/ mutations highlighted
PE13
PPE18
Recombinant region (Changed K-mers)
Match to reference (non-recombinant)locus
PPE19-H37Rv-1532539-1533632 --- k-mer match to putative recomb seq = 1.0 (31/31)
Match to reference locus
Match to recombinant "k-mer Only"
PE31,PPE60-H37Rv-3894016-3895519 --- k-mer match to putative recomb seq = 0.4839 (15/31)
Match to reference locus
Match to recombinant "k-mer Only"
k-mer match
1.0
0.8
0.6
0.4
0.2
0.0
1,339,200
1,339,280
1,339,360
1,339,440
1,339,520
1,339,600
1,339,680
1,339,760
H37rv Ref Position (bp)

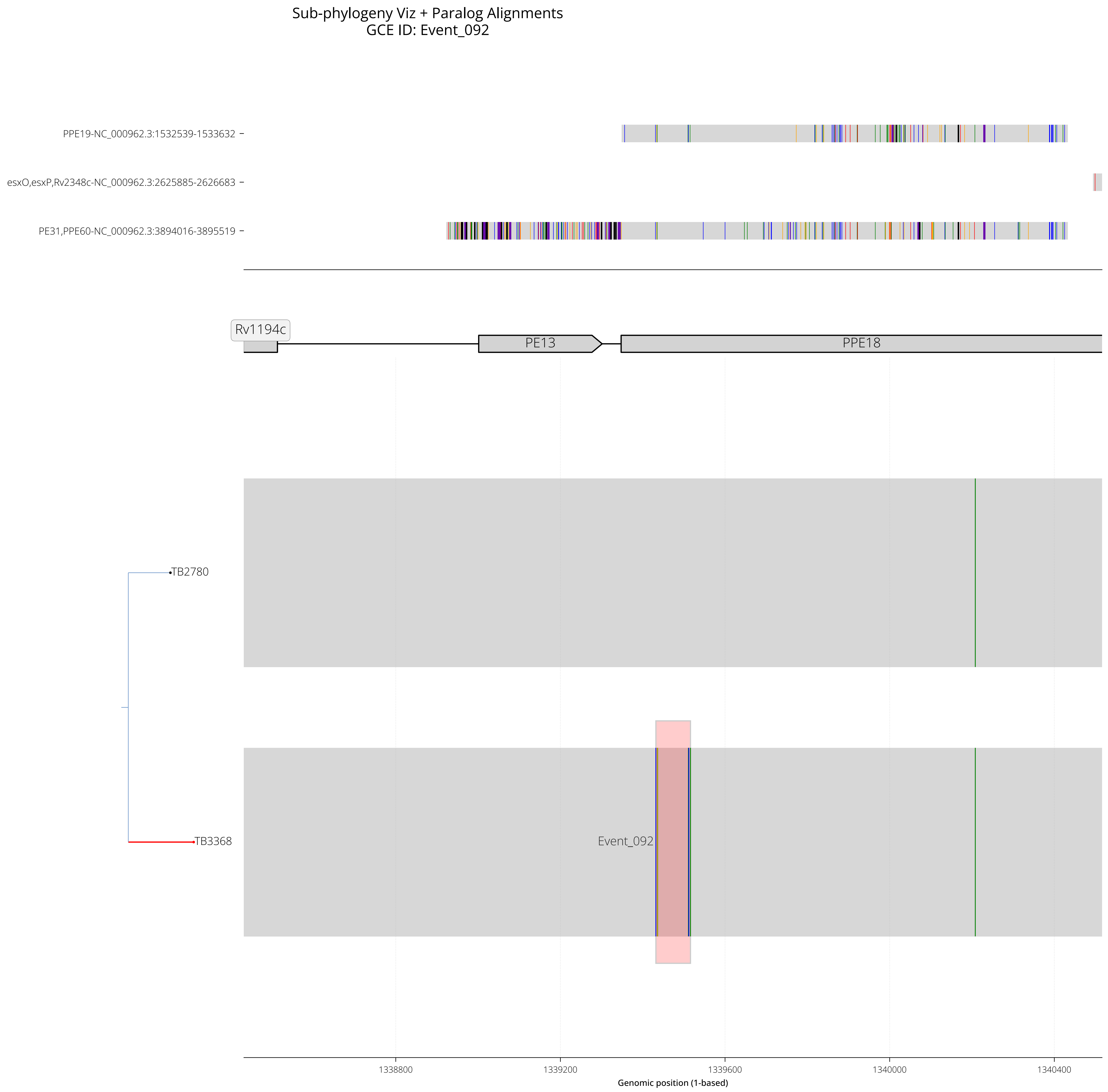

Sub-phylogeny Viz + Paralog Alignments
GCE ID: Event_092
PPE19-NC_000962.3:1532539-1533632
esxO,esxP,Rv2348c-NC_000962.3:2625885-2626683
PE31,PPE60-NC_000962.3:3894016-3895519
Rv1194c
PE13
PPE18
TB2780
TB3368
Event_092
1338800
1339200
1339600
1340000
1340400
Genomic position (1-based)

Full Phylogeny Viz + Paralog Alignments
GCE ID: Event_092

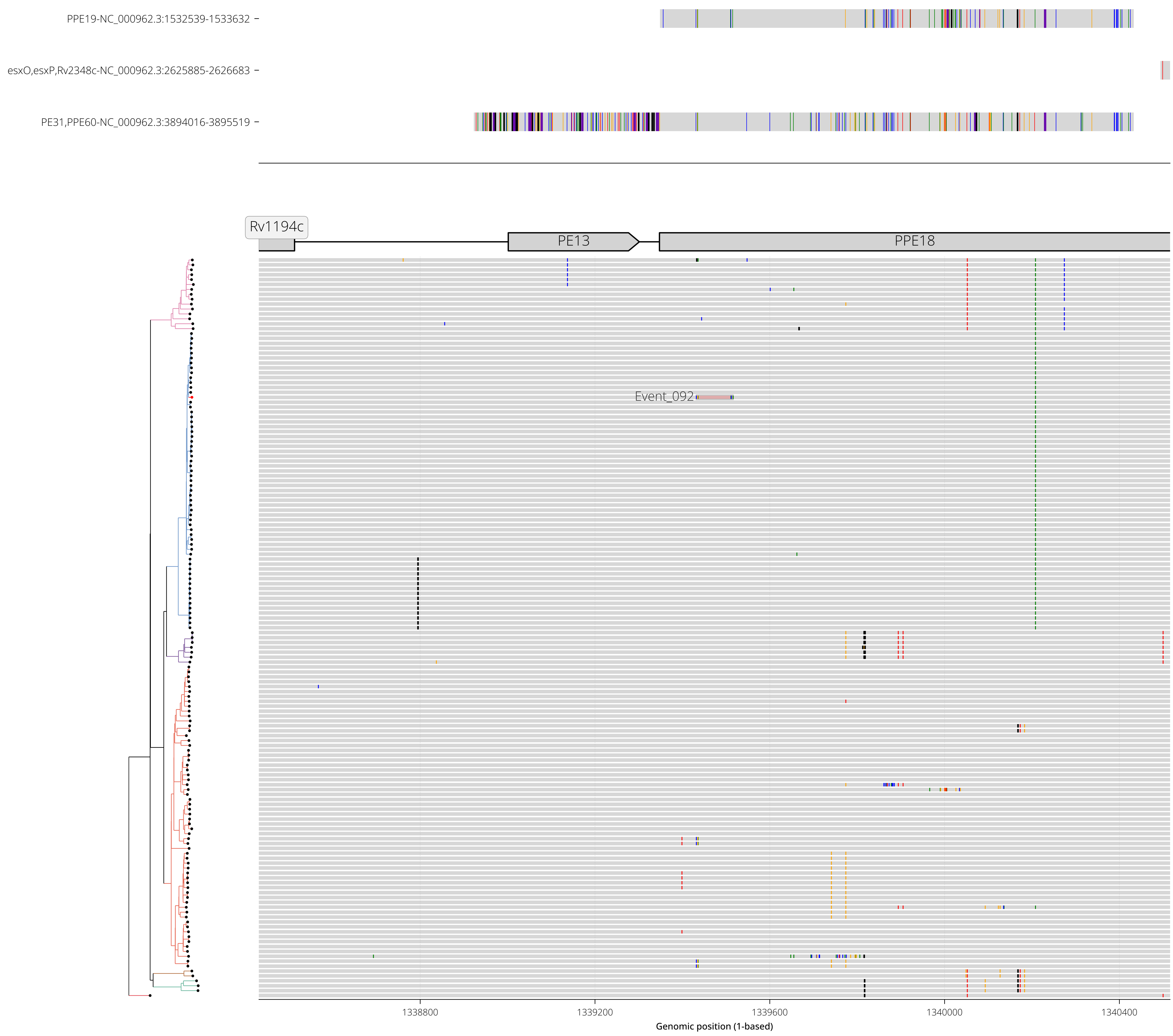

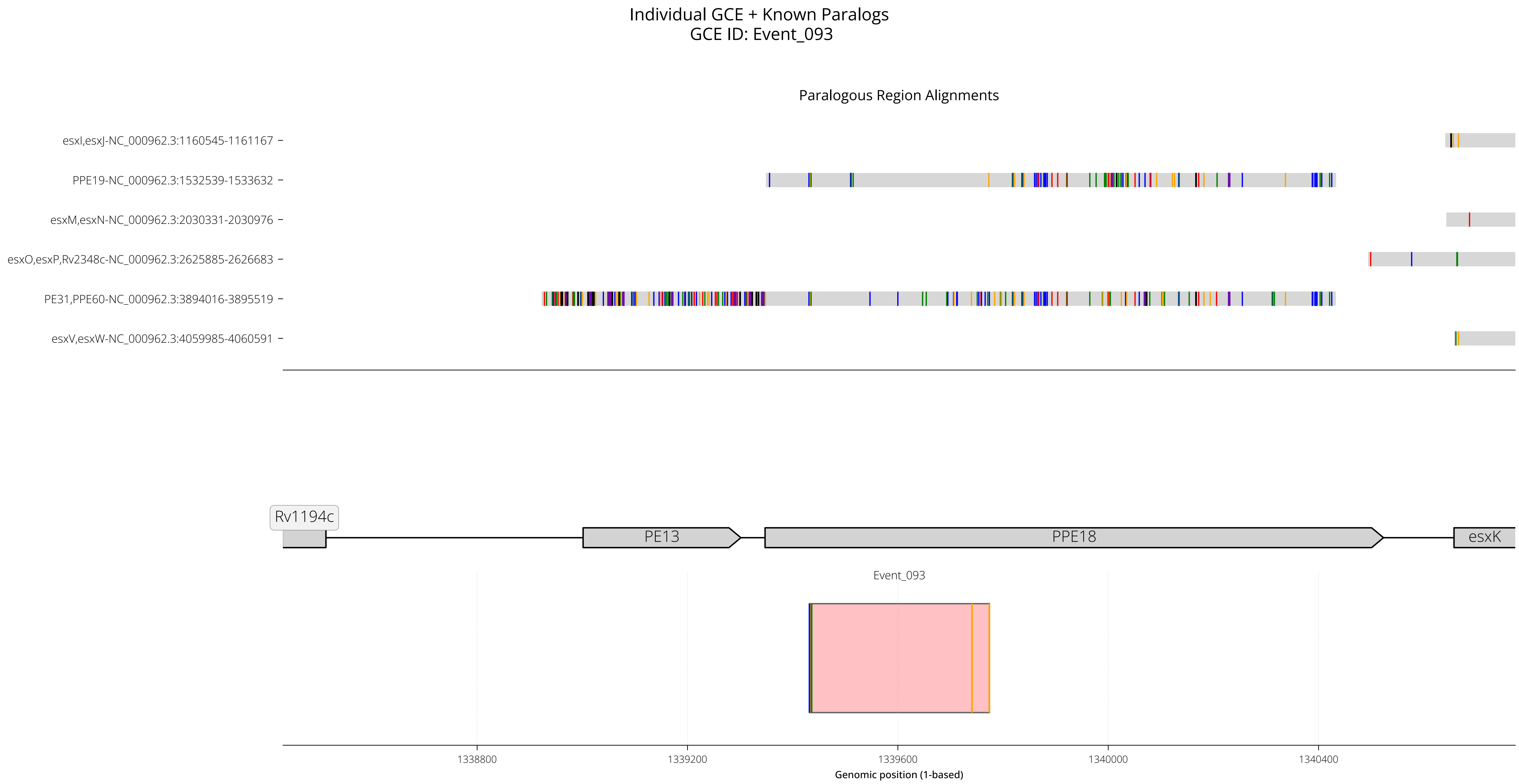

Individual GCE + Known Paralogs
GCE ID: Event_093
Paralogous Region Alignments
esxI,esxJ-NC_000962.3:1160545-1161167
PPE19-NC_000962.3:1532539-1533632
esxM,esxN-NC_000962.3:2030331-2030976
esxO,esxP,Rv2348c-NC_000962.3:2625885-2626683
PE31,PPE60-NC_000962.3:3894016-3895519
esxV,esxW-NC_000962.3:4059985-4060591
Rv1194c
PE13
PPE18
esxK
Event_093
1338800
1339200
1339600
1340000
1340400
Genomic position (1-based)

Circos Viz - GCE to Known Paralogs Locations
GCE ID: Event_093

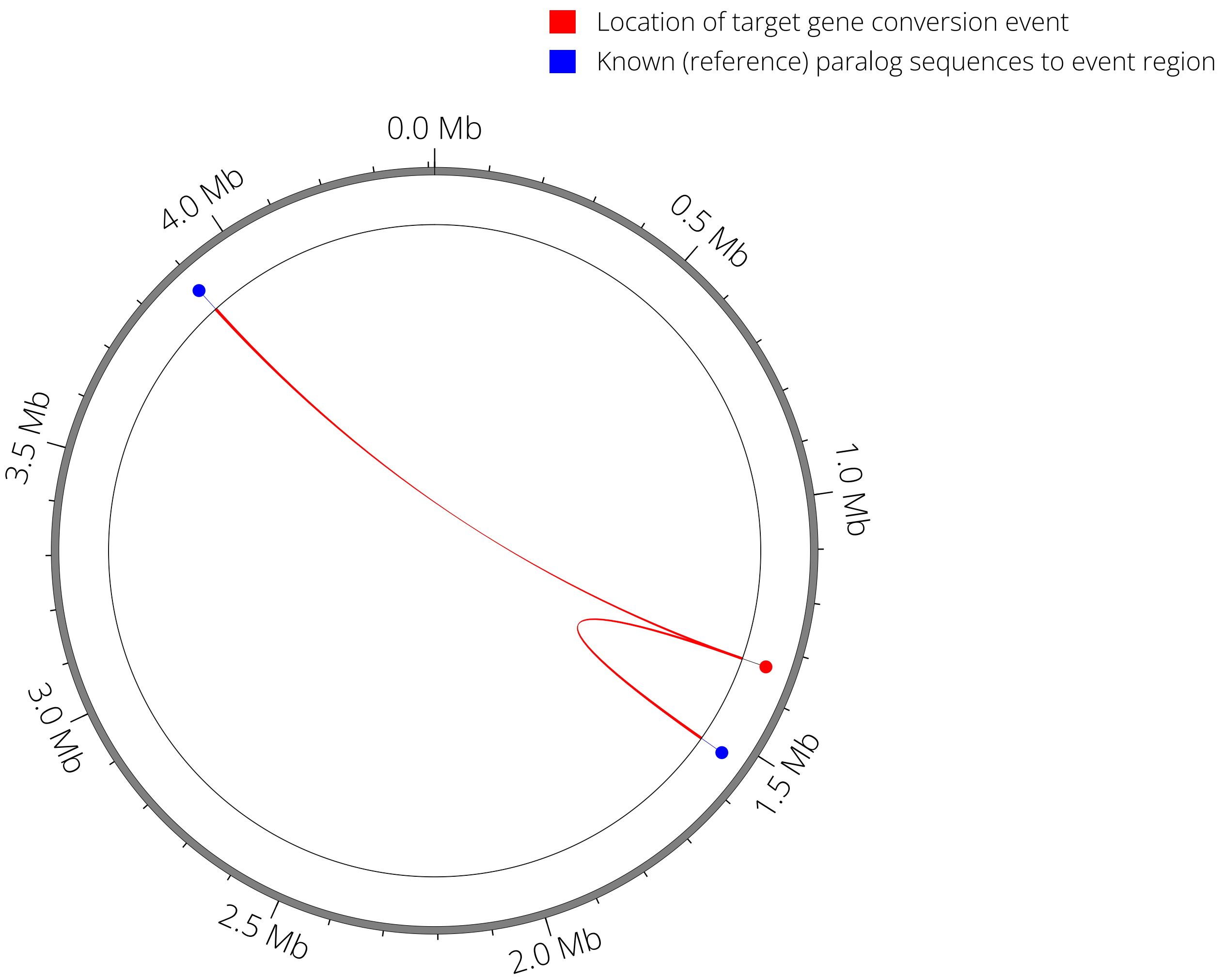

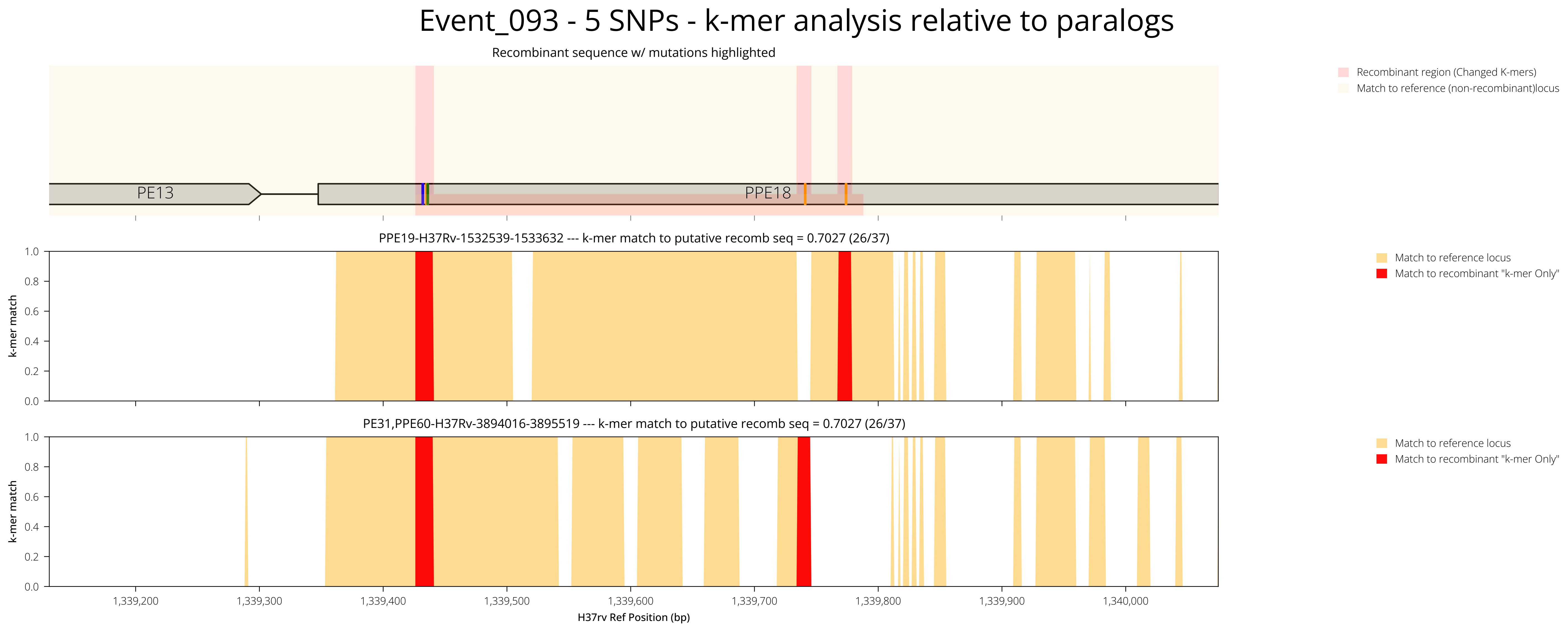

Event_093 - 5 SNPs - k-mer analysis relative to paralogs
Recombinant sequence w/ mutations highlighted
Recombinant region (Changed K-mers)
Match to reference (non-recombinant)locus
PE13
PPE18
PPE19-H37Rv-1532539-1533632 --- k-mer match to putative recomb seq = 0.7027 (26/37)
Match to reference locus
Match to recombinant "k-mer Only"
PE31,PPE60-H37Rv-3894016-3895519 --- k-mer match to putative recomb seq = 0.7027 (26/37)
Match to reference locus
Match to recombinant "k-mer Only"
k-mer match
1.0
0.8
0.6
0.4
0.2
0.0
1,339,200
1,339,300
1,339,400
1,339,500
1,339,600
1,339,700
1,339,800
1,339,900
1,340,000
H37rv Ref Position (bp)

Sub-phylogeny Viz + Paralog Alignments
GCE ID: Event_093

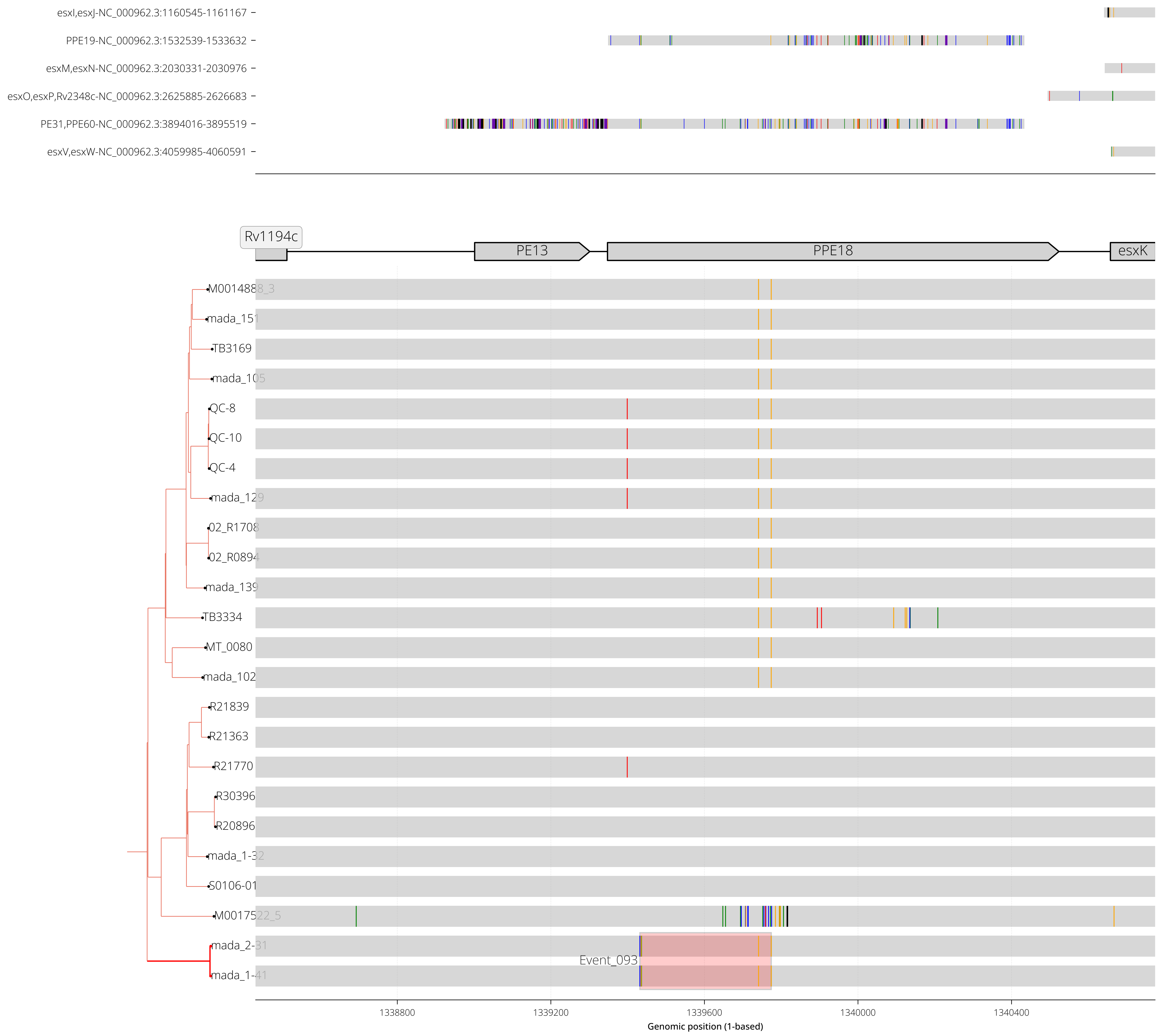

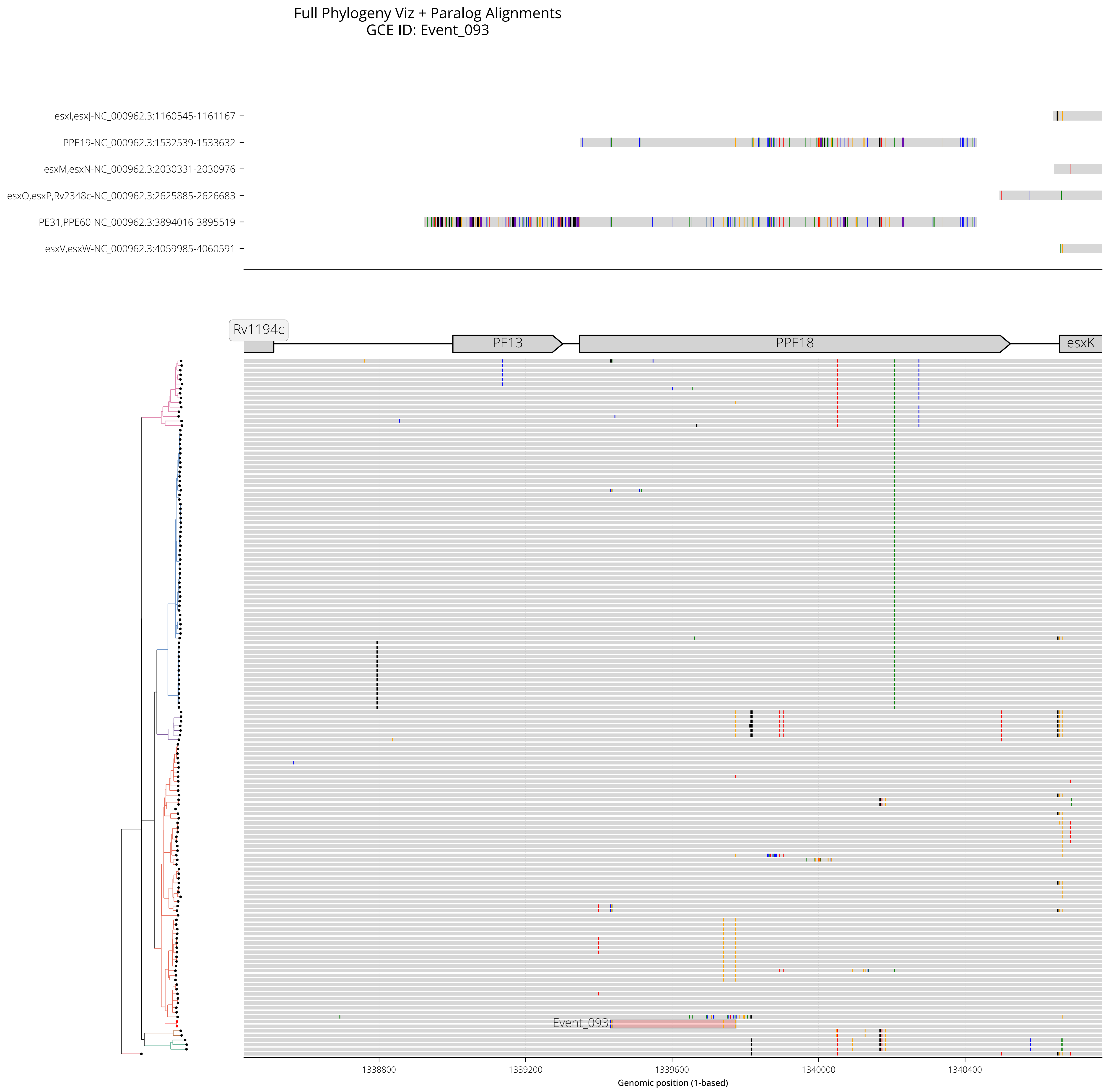

Full Phylogeny Viz + Paralog Alignments
GCE ID: Event_093
esxI,esxJ-NC_000962.3:1160545-1161167
PPE19-NC_000962.3:1532539-1533632
esxM,esxN-NC_000962.3:2030331-2030976
esxO,esxP,Rv2348c-NC_000962.3:2625885-2626683
PE31,PPE60-NC_000962.3:3894016-3895519
esxV,esxW-NC_000962.3:4059985-4060591
Rv1194c
PE13
PPE18
esxK
Event_093
1338800
1339200
1339600
1340000
1340400
Genomic position (1-based)

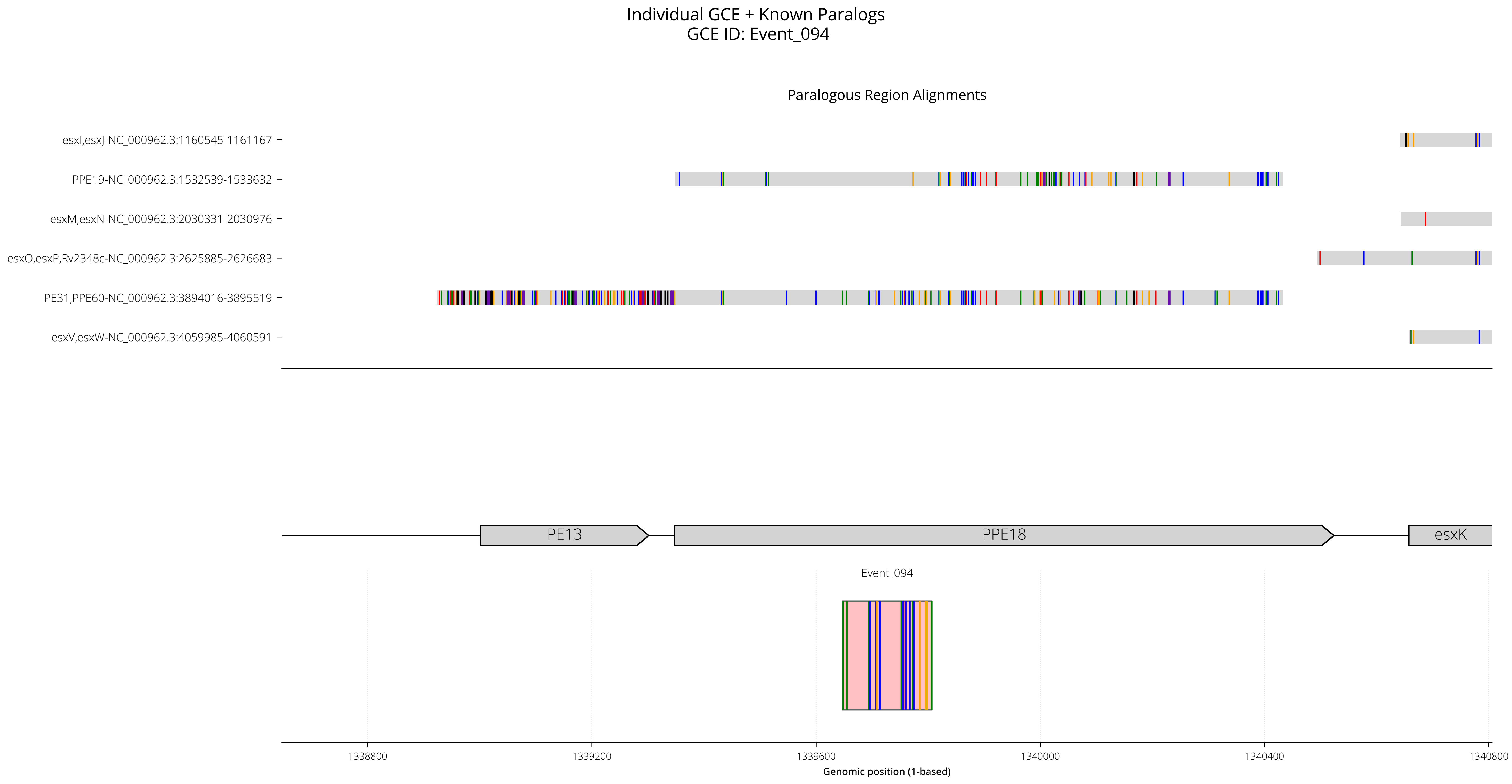

Individual GCE + Known Paralogs
GCE ID: Event_094
Paralogous Region Alignments
esxI,esxJ-NC_000962.3:1160545-1161167
PPE19-NC_000962.3:1532539-1533632
esxM,esxN-NC_000962.3:2030331-2030976
esxO,esxP,Rv2348c-NC_000962.3:2625885-2626683
PE31,PPE60-NC_000962.3:3894016-3895519
esxV,esxW-NC_000962.3:4059985-4060591
PE13
PPE18
esxK
Event_094
1338800
1339200
1339600
1340000
1340400
1340800
Genomic position (1-based)

### Circos Viz - GCE to Known Paralogs Locations
GCE ID: Event_094

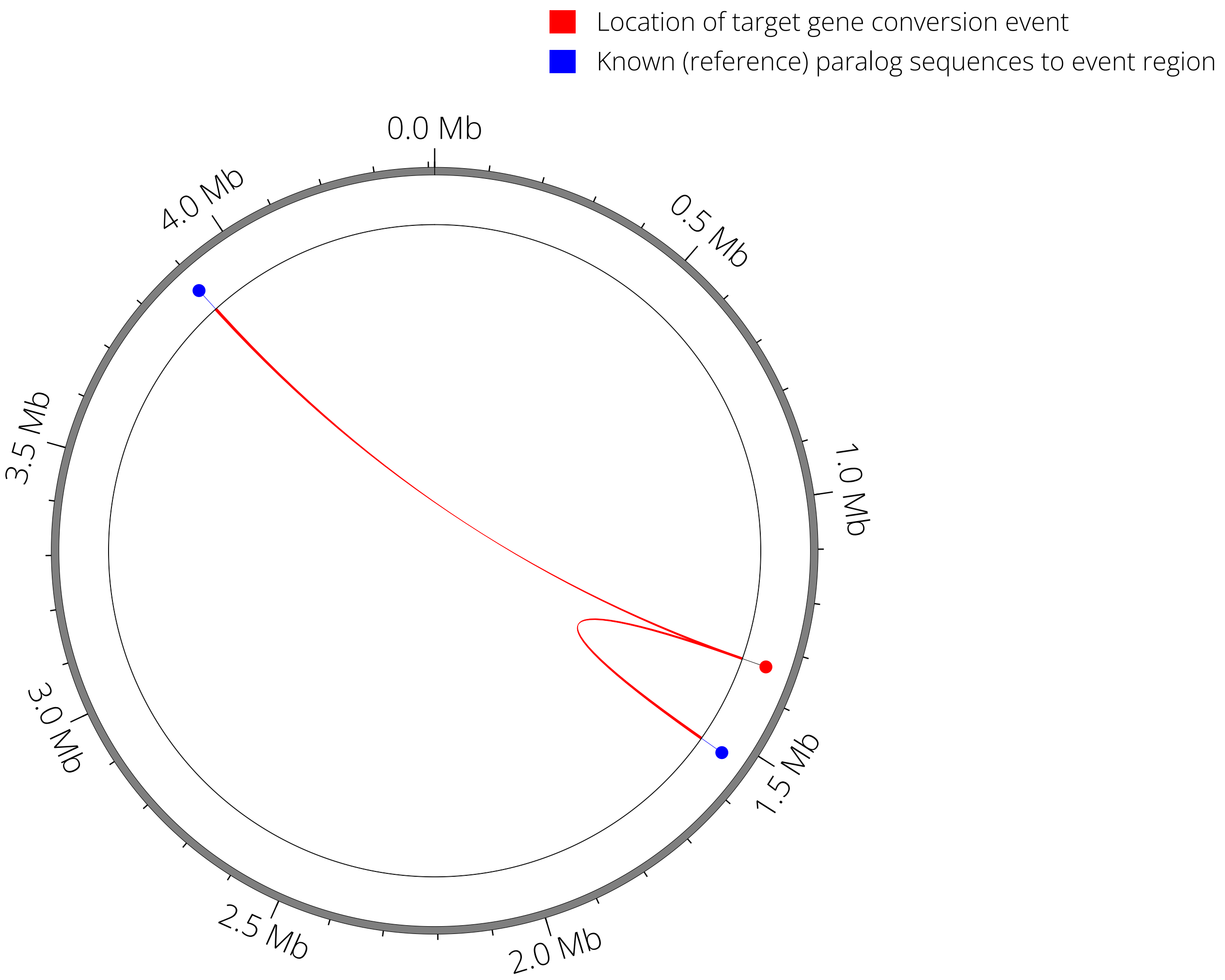

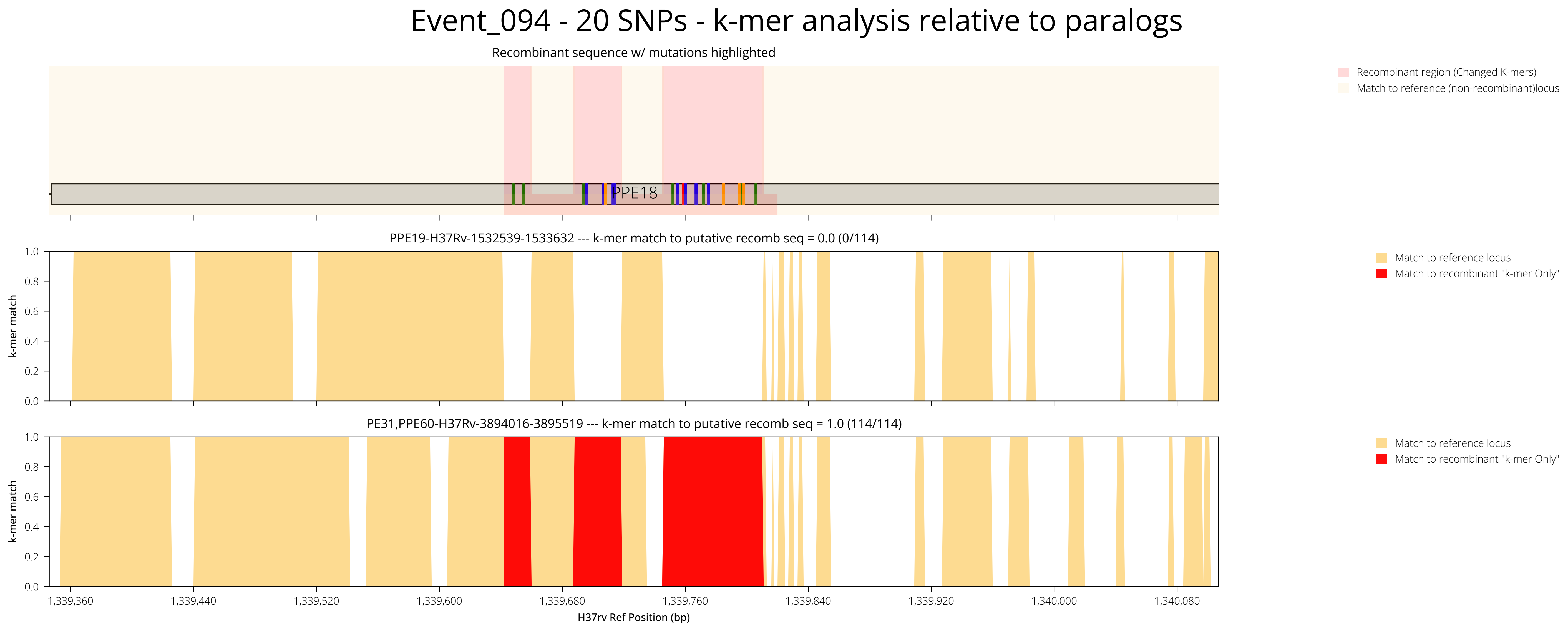
Event_094 - 20 SNPs - k-mer analysis relative to paralogs
Recombinant sequence w/ mutations highlighted
PPE18
Recombinant region (Changed K-mers)
Match to reference (non-recombinant)locus
PPE19-H37Rv-1532539-1533632 --- k-mer match to putative recomb seq = 0.0 (0/114)
Match to reference locus
Match to recombinant "k-mer Only"
PE31,PPE60-H37Rv-3894016-3895519 --- k-mer match to putative recomb seq = 1.0 (114/114)
Match to reference locus
Match to recombinant "k-mer Only"
k-mer match
1.0
0.8
0.6
0.4
0.2
0.0
1,339,360
1,339,440
1,339,520
1,339,600
1,339,680
1,339,760
1,339,840
1,339,920
1,340,000
1,340,080
H37rv Ref Position (bp)

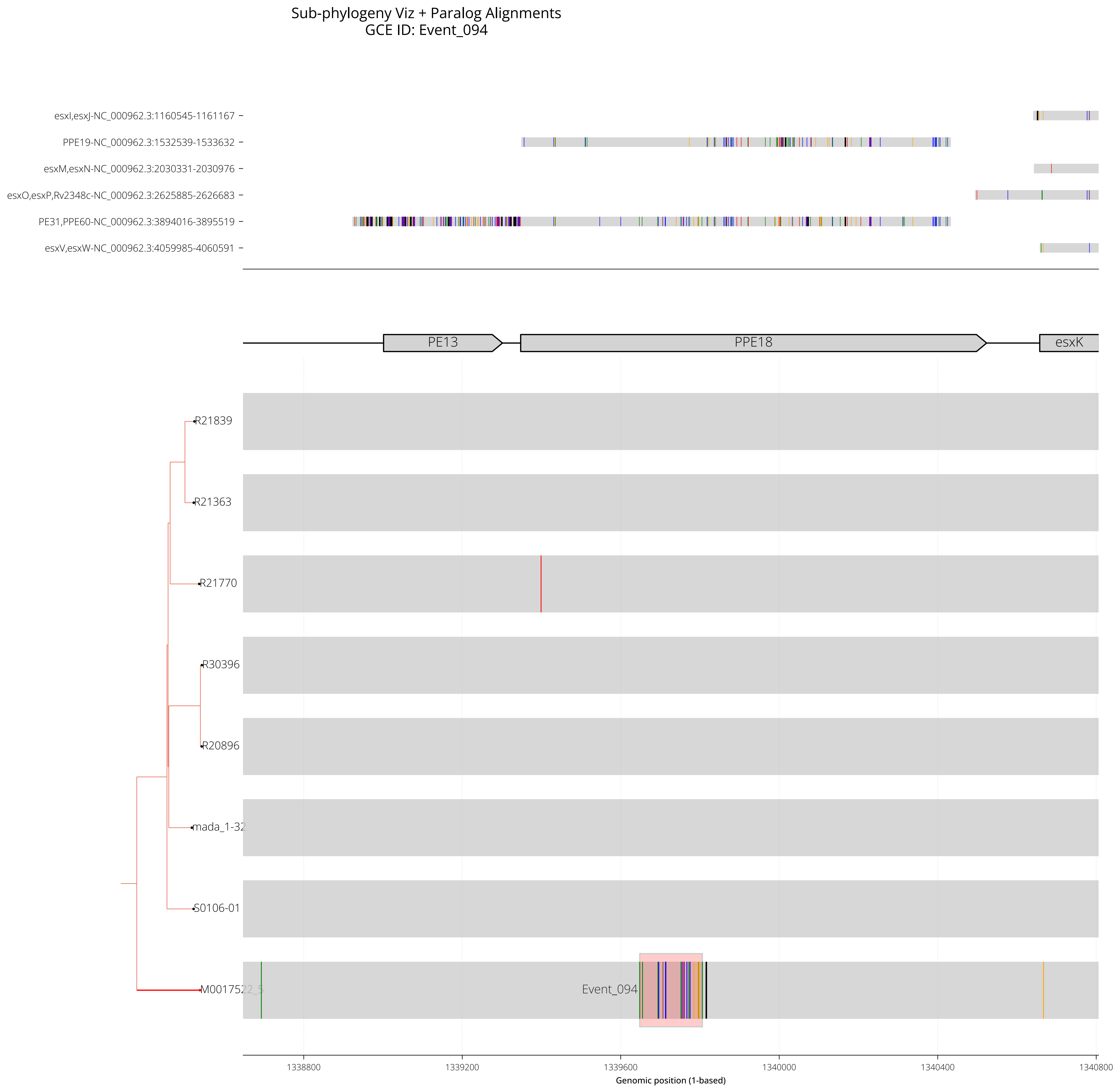

Sub-phylogeny Viz + Paralog Alignments
GCE ID: Event_094
esxI,esxJ-NC_000962.3:1160545-1161167
PPE19-NC_000962.3:1532539-1533632
esxM,esxN-NC_000962.3:2030331-2030976
esxO,esxP,Rv2348c-NC_000962.3:2625885-2626683
PE31,PPE60-NC_000962.3:3894016-3895519
esxV,esxW-NC_000962.3:4059985-4060591
PE13
PPE18
esxK
R21839
R21363
R21770
R30396
R20896
mada_1-32
S0106-01
M0017522_5
Event_094
1338800
1339200
1339600
1340000
1340400
1340800
Genomic position (1-based)

Full Phylogeny Viz + Paralog Alignments
GCE ID: Event_094

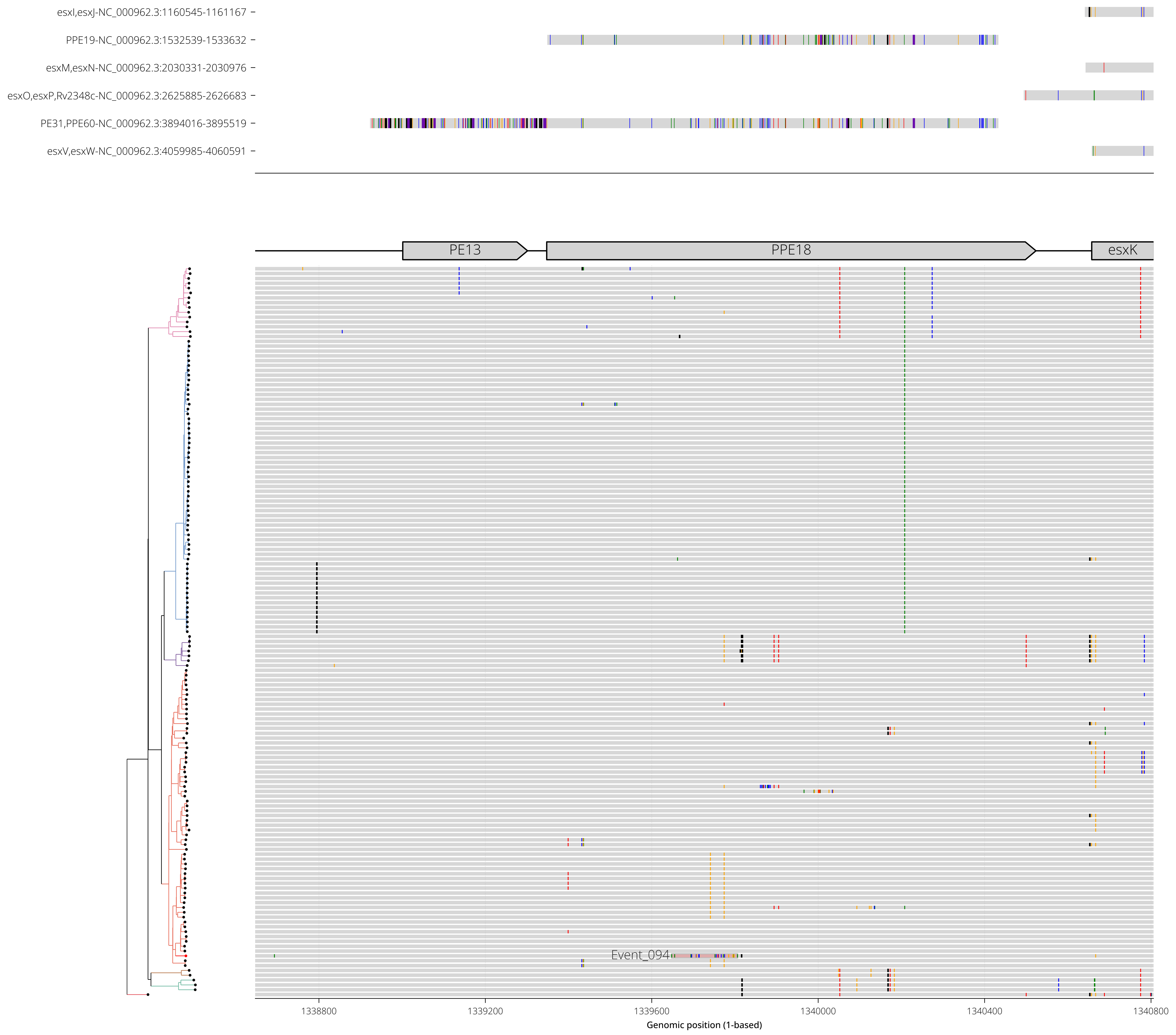

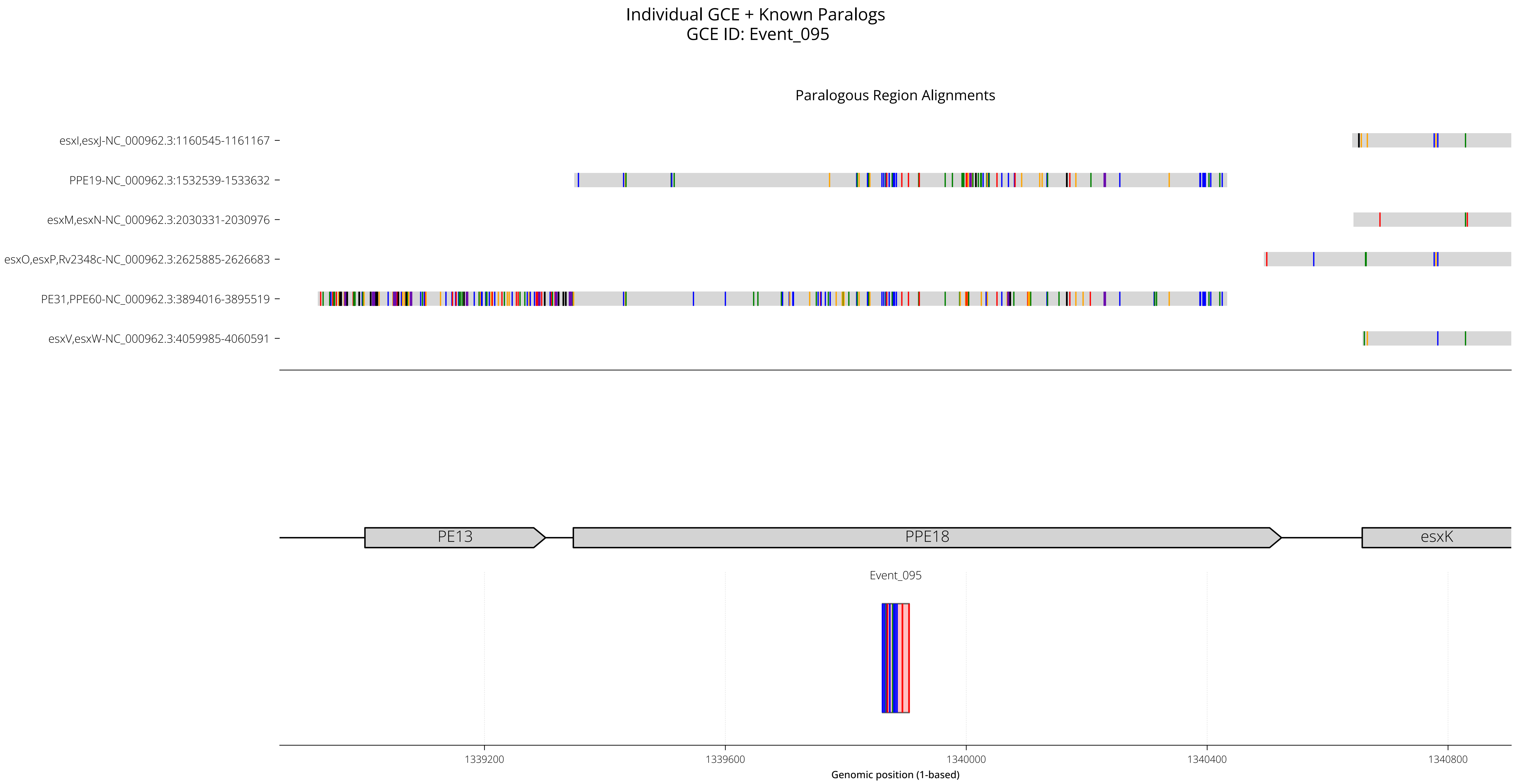

Individual GCE + Known Paralogs
GCE ID: Event_095
Paralogous Region Alignments
esxI,esxJ-NC_000962.3:1160545-1161167
PPE19-NC_000962.3:1532539-1533632
esxM,esxN-NC_000962.3:2030331-2030976
esxO,esxP,Rv2348c-NC_000962.3:2625885-2626683
PE31,PPE60-NC_000962.3:3894016-3895519
esxV,esxW-NC_000962.3:4059985-4060591
PE13
PPE18
esxK
Event_095
1339200
1339600
1340000
1340400
1340800
Genomic position (1-based)

Circos Viz - GCE to Known Paralogs Locations
GCE ID: Event_095

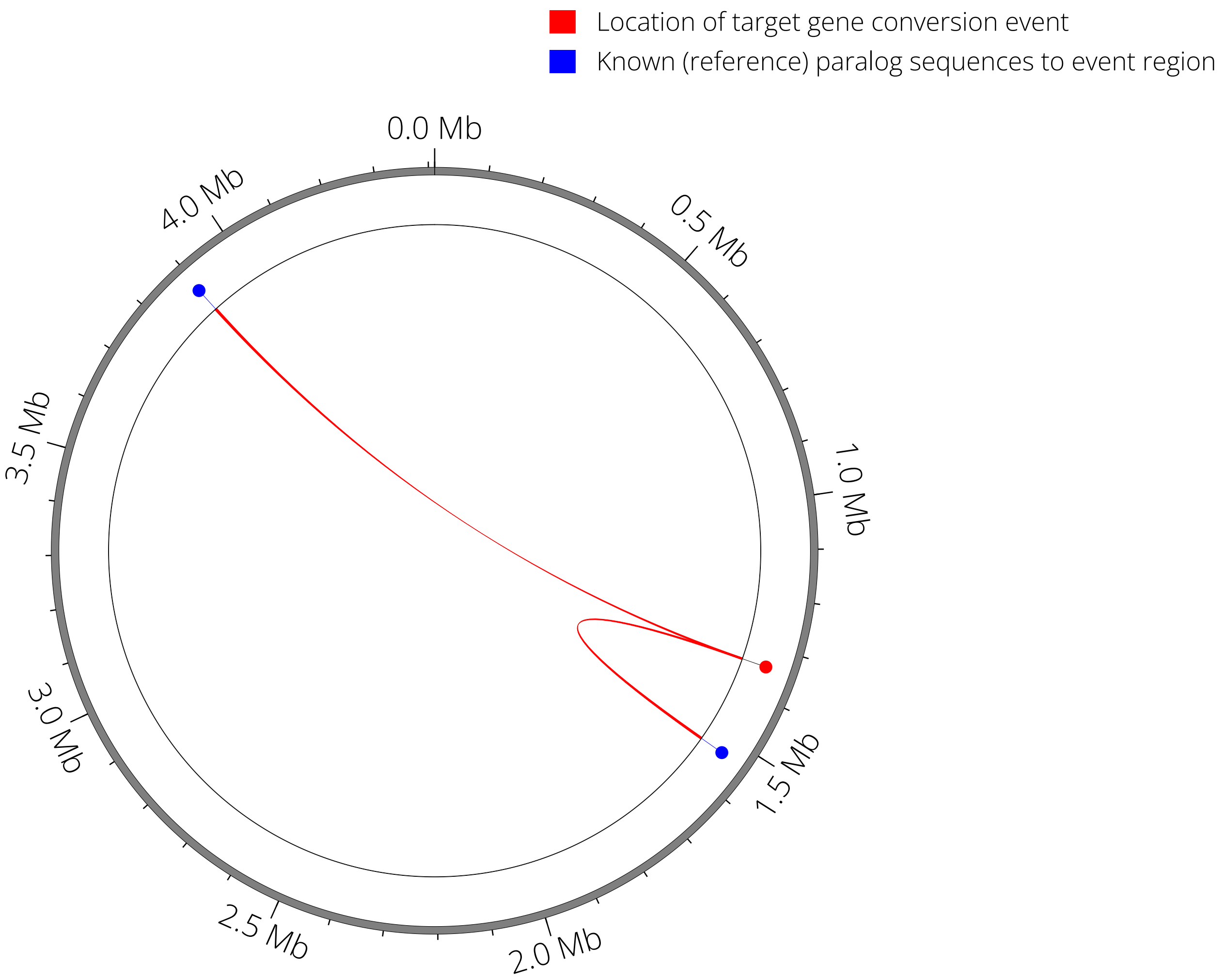

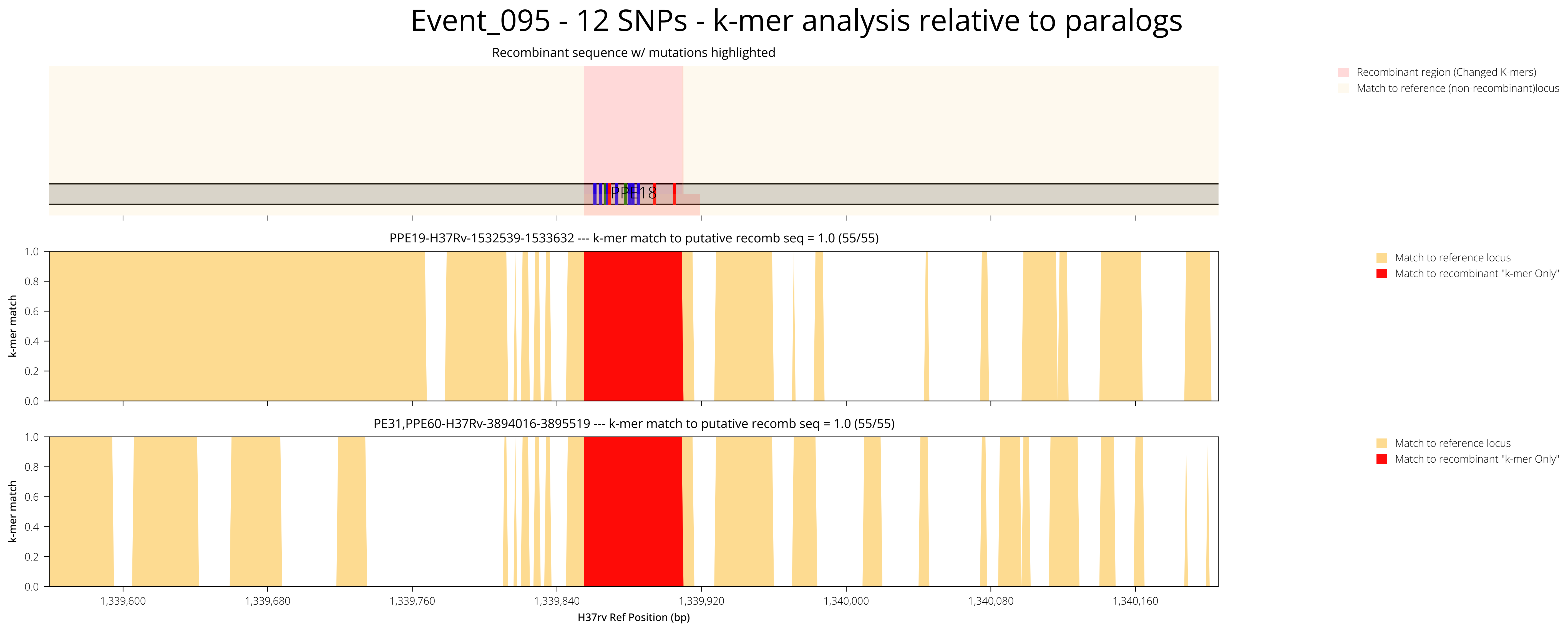

Event_095 - 12 SNPs - k-mer analysis relative to paralogs
Recombinant sequence w/ mutations highlighted
Recombinant region (Changed K-mers)
Match to reference (non-recombinant)locus
PPE18
PPE19-H37Rv-1532539-1533632 --- k-mer match to putative recomb seq = 1.0 (55/55)
Match to reference locus
Match to recombinant "k-mer Only"
PE31,PPE60-H37Rv-3894016-3895519 --- k-mer match to putative recomb seq = 1.0 (55/55)
Match to reference locus
Match to recombinant "k-mer Only"
k-mer match
1.0
0.8
0.6
0.4
0.2
0.0
1,339,600
1,339,680
1,339,760
1,339,840
1,339,920
1,340,000
1,340,080
1,340,160
H37rv Ref Position (bp)

Sub-phylogeny Viz + Paralog Alignments
GCE ID: Event_095

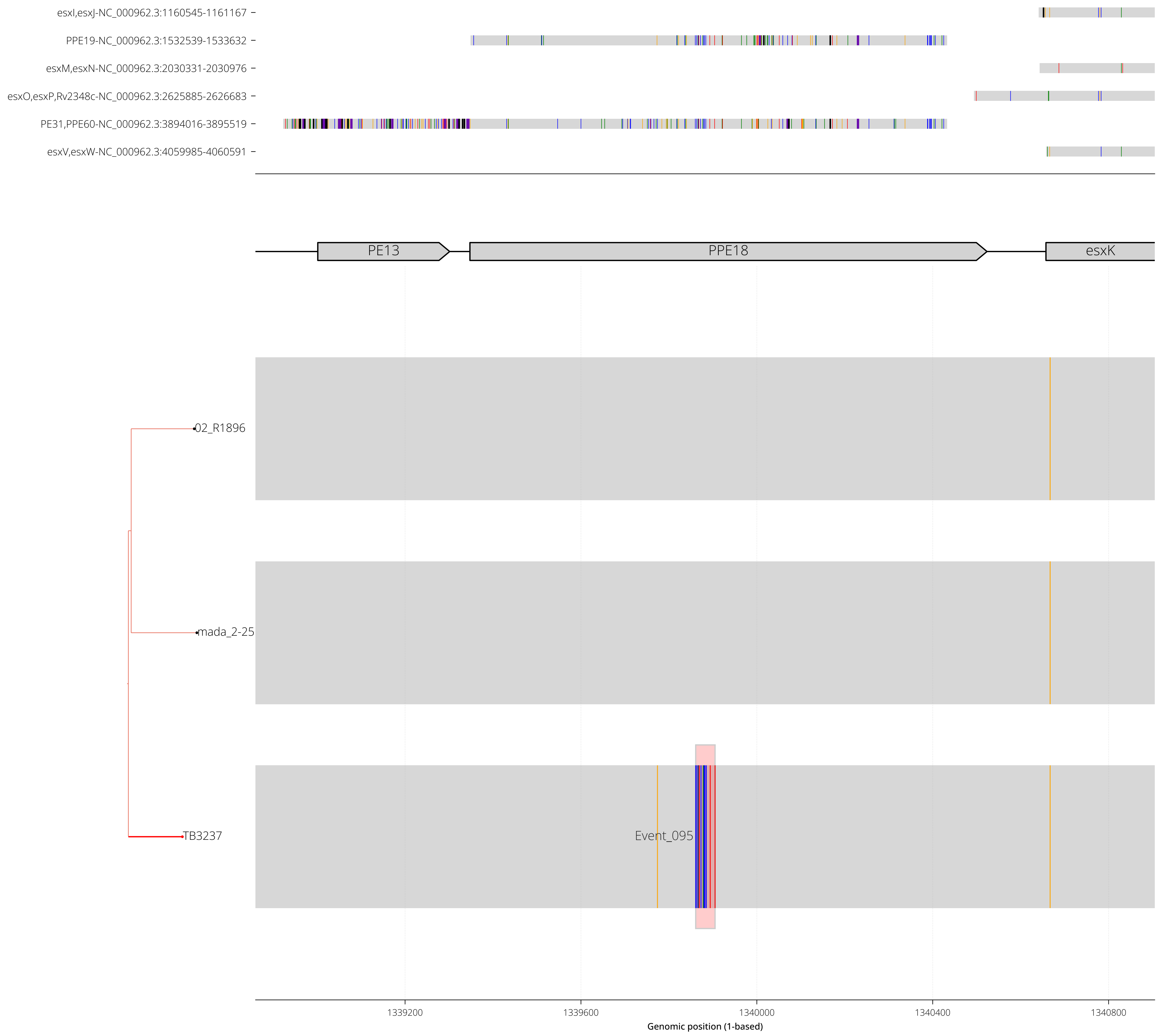

Full Phylogeny Viz + Paralog Alignments
GCE ID: Event_095

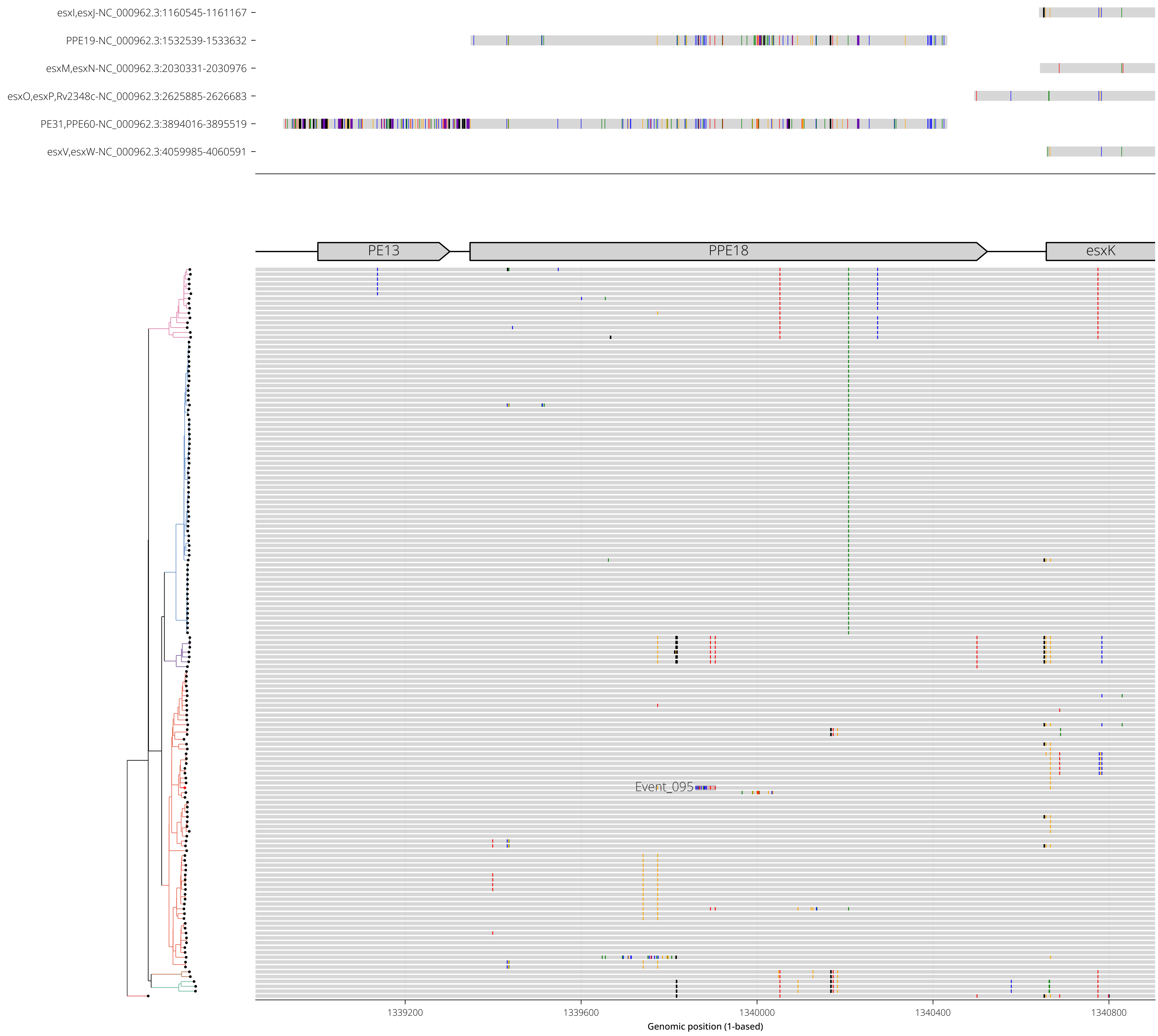

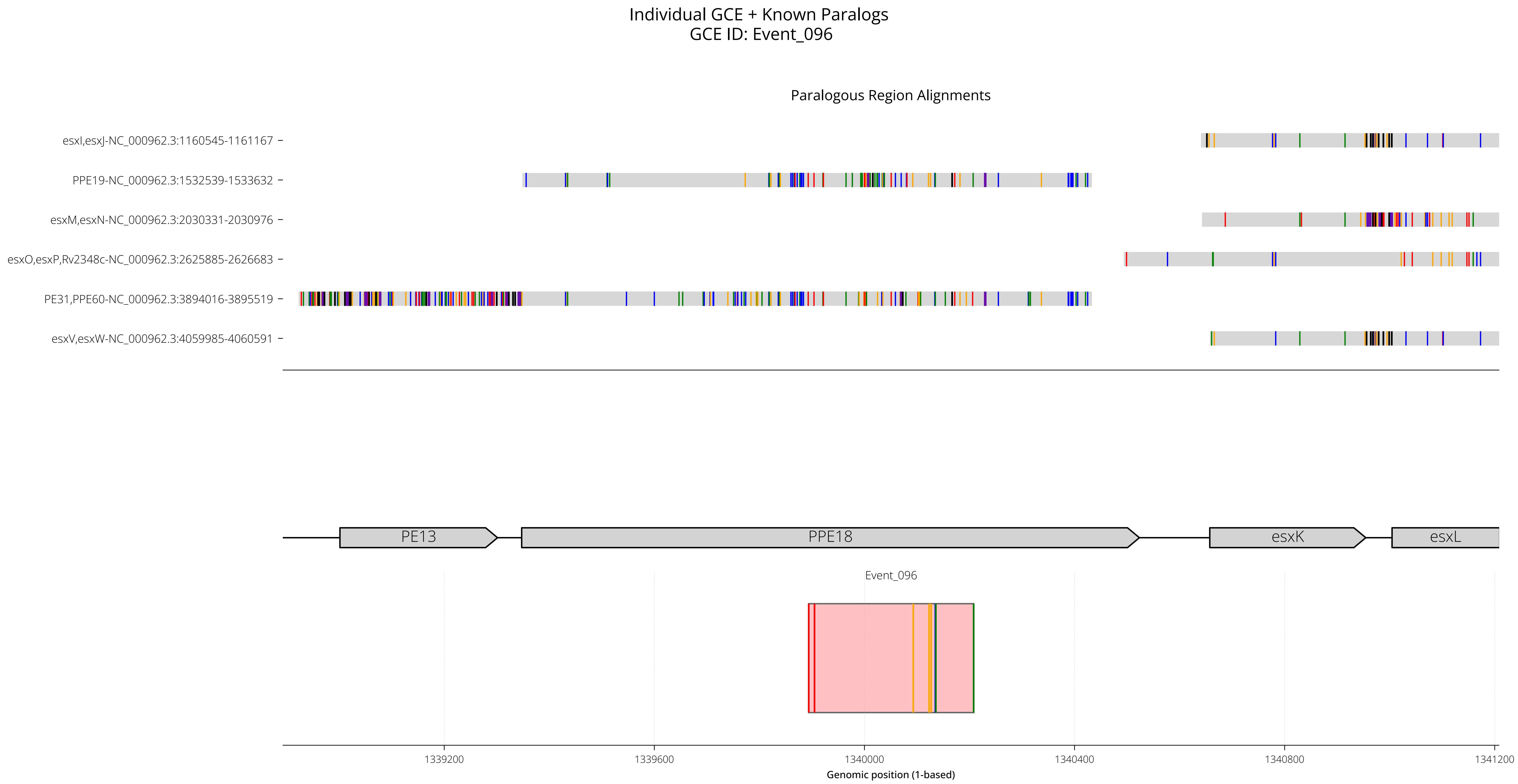

Individual GCE + Known Paralogs
GCE ID: Event_096
Paralogous Region Alignments
esxI,esxJ-NC_000962.3:1160545-1161167
PPE19-NC_000962.3:1532539-1533632
esxM,esxN-NC_000962.3:2030331-2030976
esxO,esxP,Rv2348c-NC_000962.3:2625885-2626683
PE31,PPE60-NC_000962.3:3894016-3895519
esxV,esxW-NC_000962.3:4059985-4060591
PE13
PPE18
esxK
esxL
Event_096
1339200
1339600
1340000
1340400
1340800
1341200
Genomic position (1-based)

Circos Viz - GCE to Known Paralogs Locations
GCE ID: Event_096

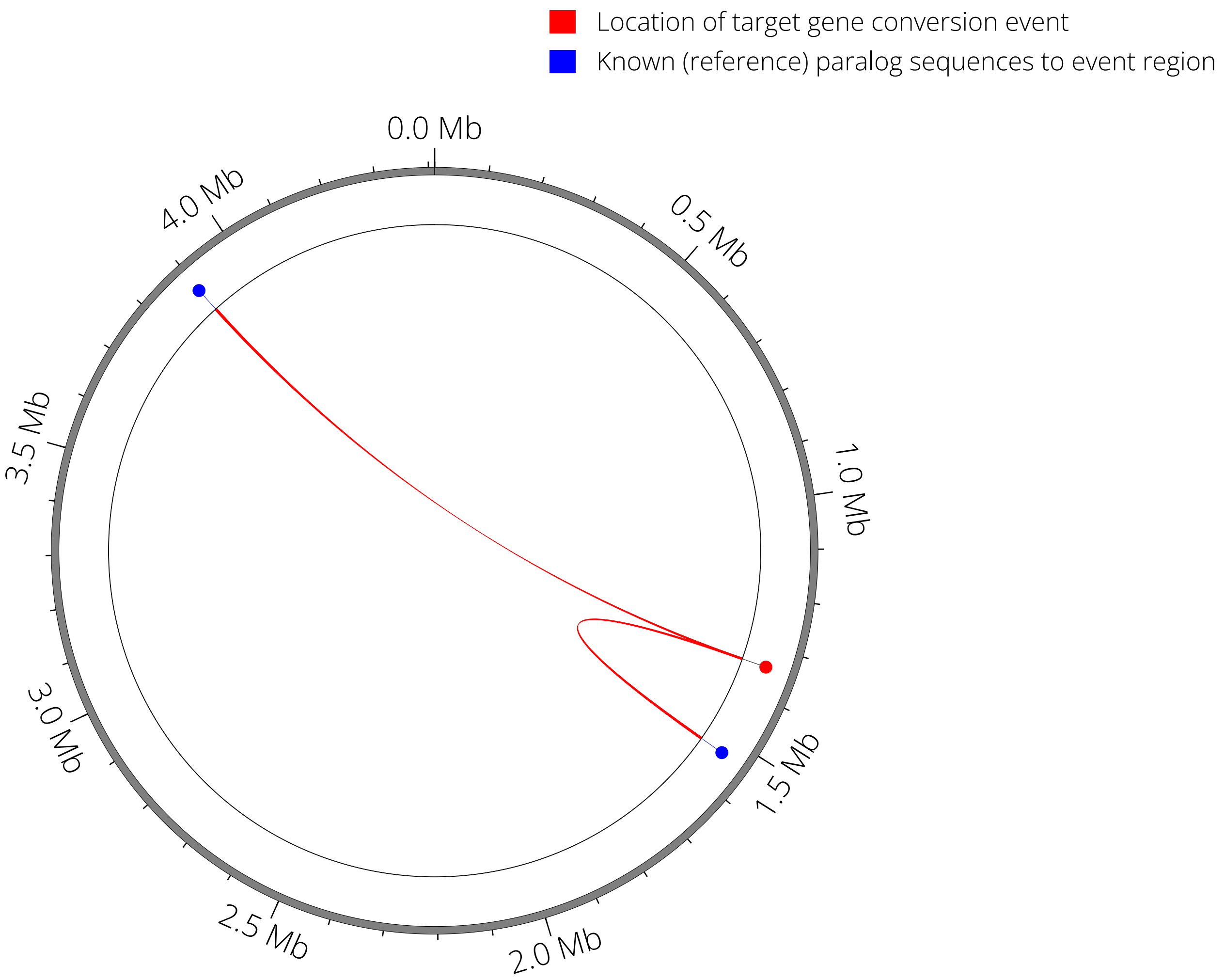

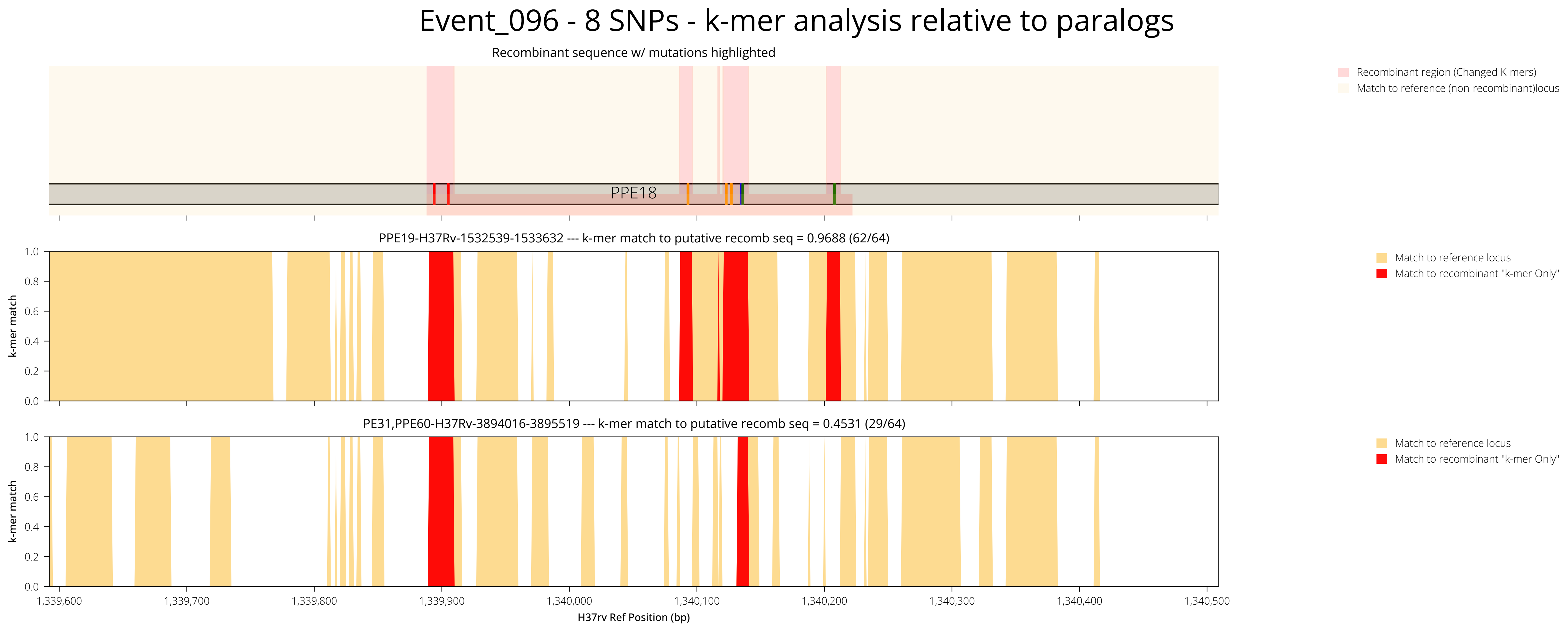

Event_096 - 8 SNPs - k-mer analysis relative to paralogs
Recombinant sequence w/ mutations highlighted
Recombinant region (Changed K-mers)
Match to reference (non-recombinant)locus
PPE18
PPE19-H37Rv-1532539-1533632 --- k-mer match to putative recomb seq = 0.9688 (62/64)
Match to reference locus
Match to recombinant "k-mer Only"
PE31,PPE60-H37Rv-3894016-3895519 --- k-mer match to putative recomb seq = 0.4531 (29/64)
Match to reference locus
Match to recombinant "k-mer Only"
k-mer match
1.0
0.8
0.6
0.4
0.2
0.0
1,339,600
1,339,700
1,339,800
1,339,900
1,340,000
1,340,100
1,340,200
1,340,300
1,340,400
1,340,500
H37rv Ref Position (bp)

Sub-phylogeny Viz + Paralog Alignments
GCE ID: Event_096

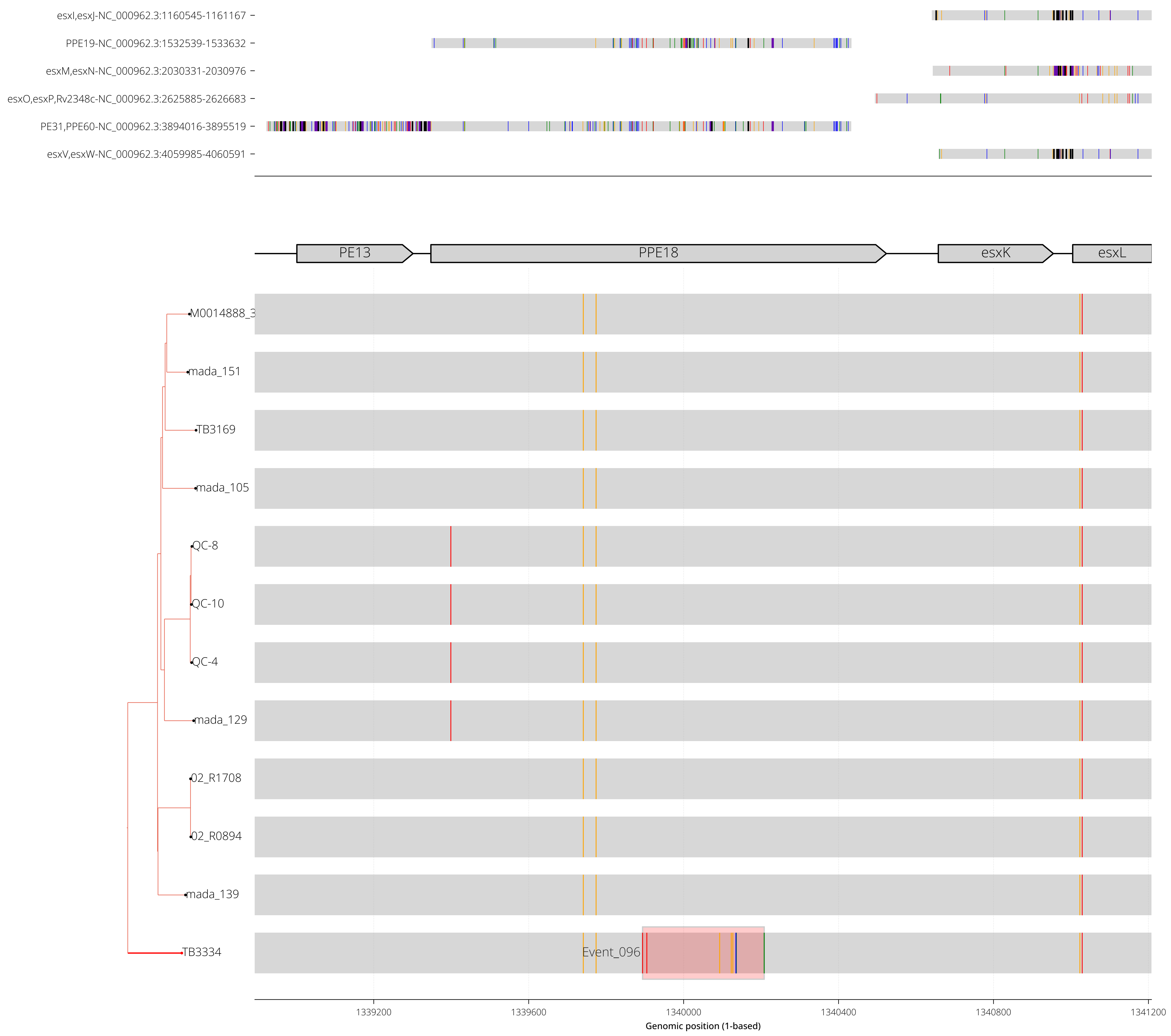

### Full Phylogeny Viz + Paralog Alignments
GCE ID: Event_096

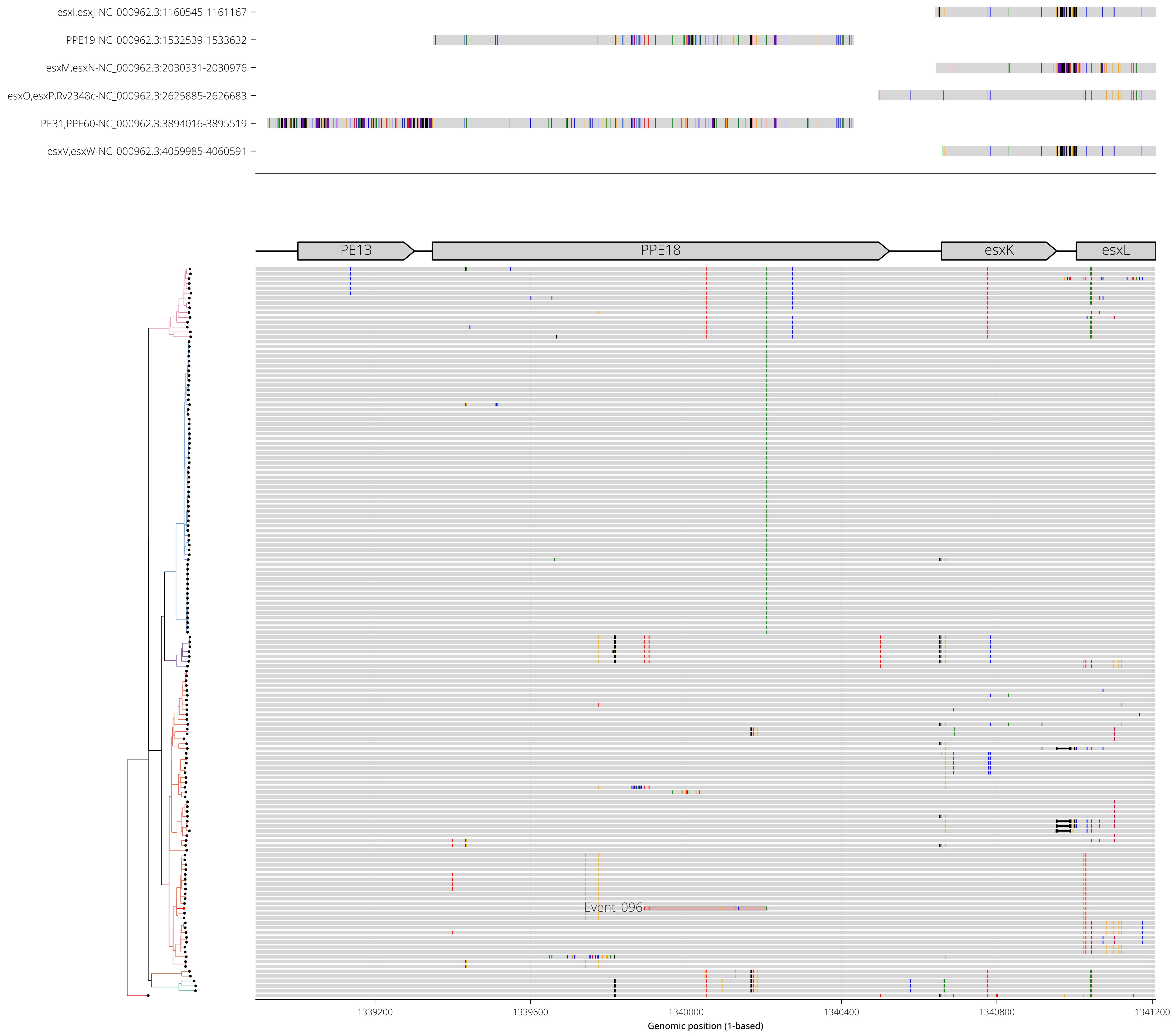

Individual GCE + Known Paralogs
GCE ID: Event_097

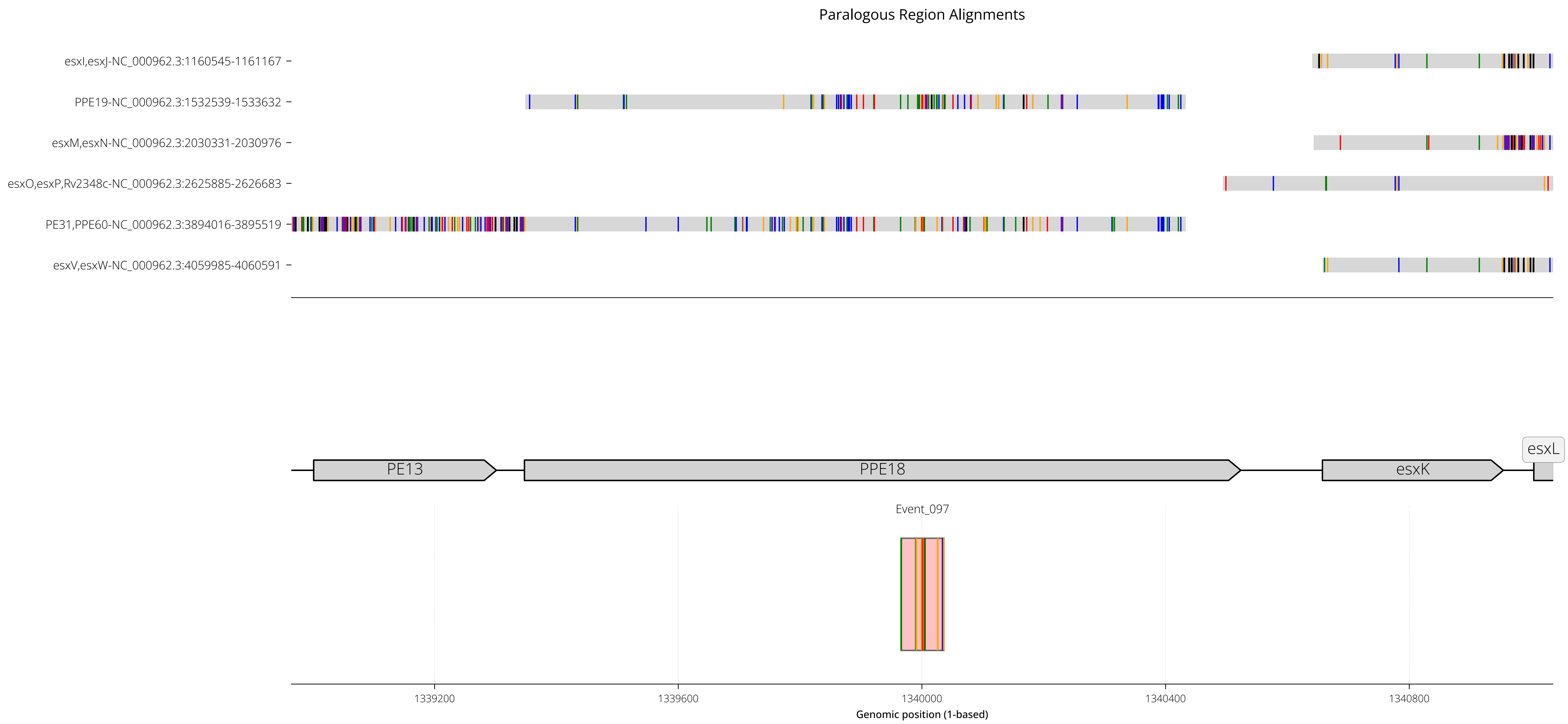

Circos Viz - GCE to Known Paralogs Locations
GCE ID: Event_097

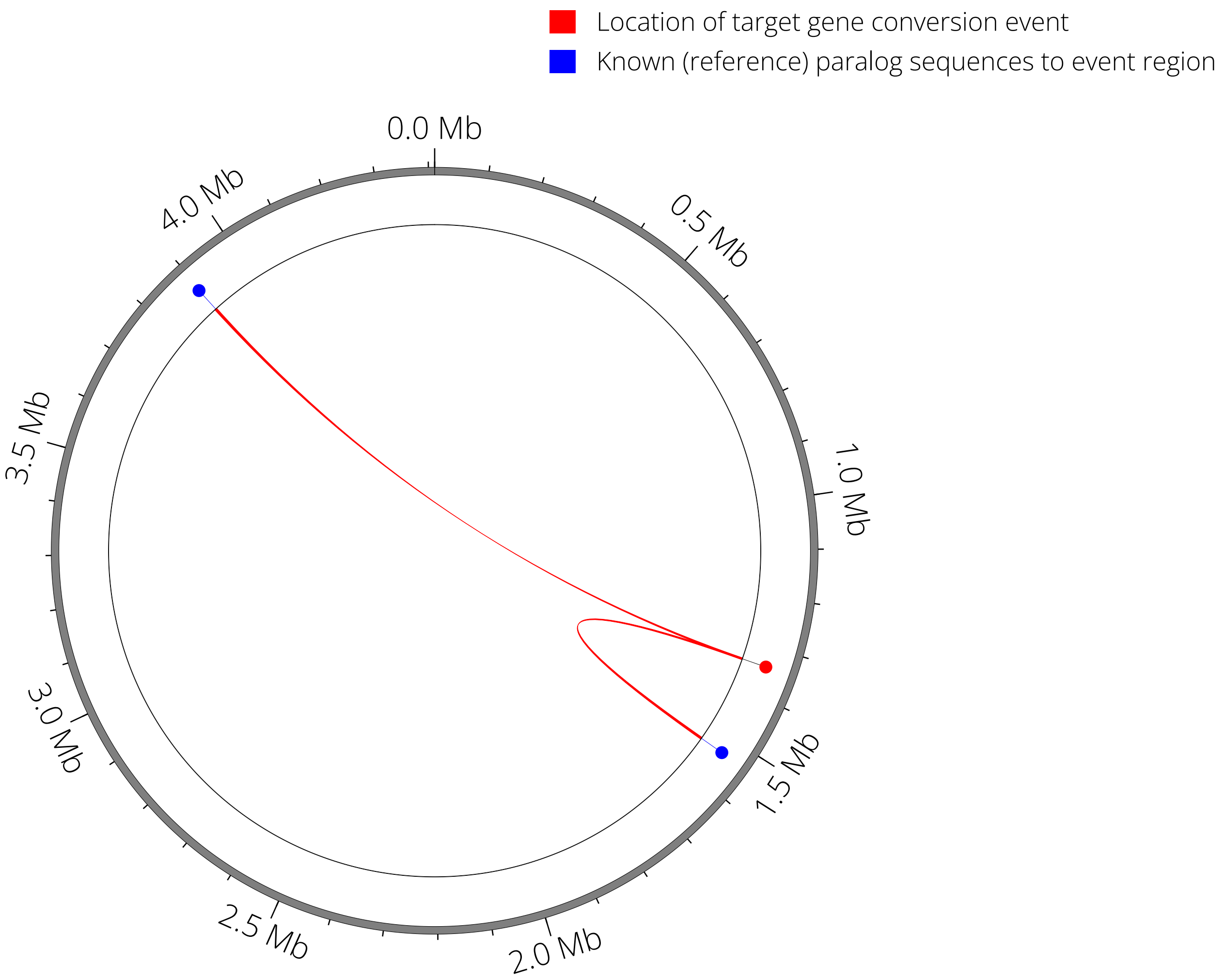

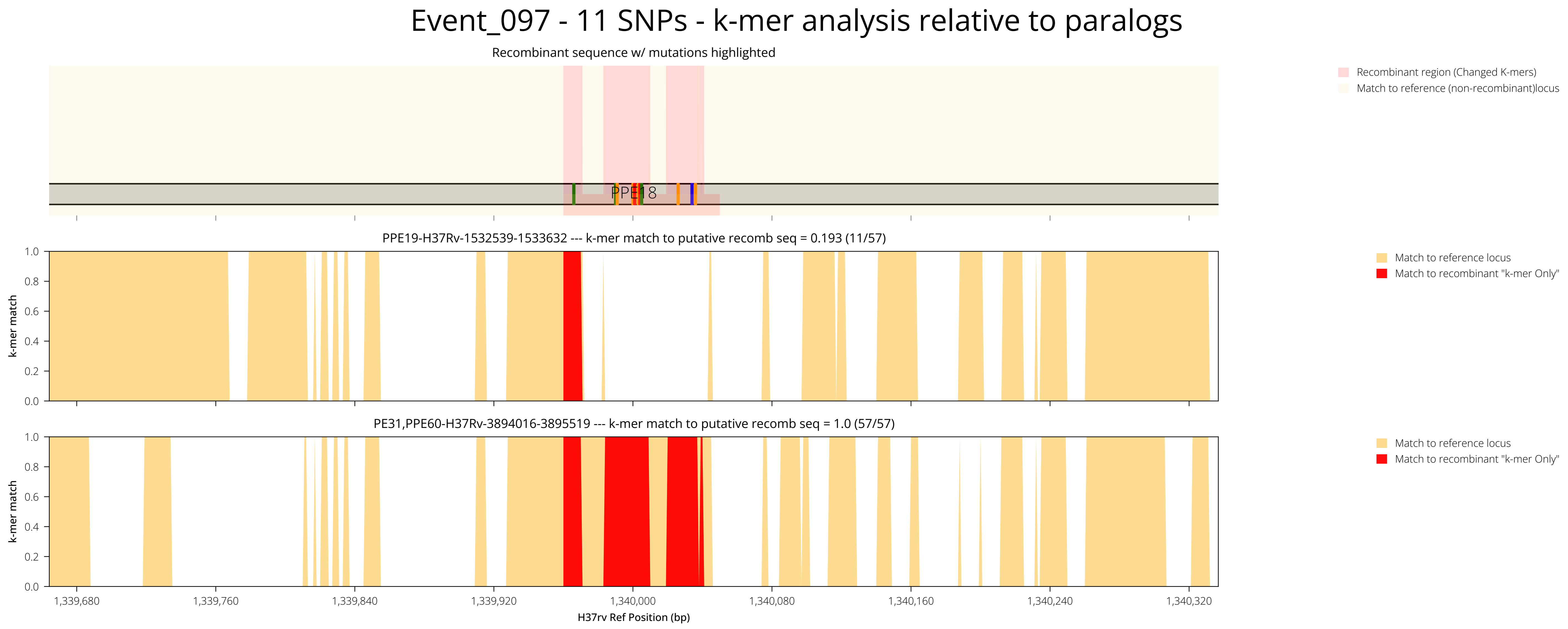

Event_097 - 11 SNPs - k-mer analysis relative to paralogs
Recombinant sequence w/ mutations highlighted
Recombinant region (Changed K-mers)
Match to reference (non-recombinant)locus
PPE18
PPE19-H37Rv-1532539-1533632 --- k-mer match to putative recomb seq = 0.193 (11/57)
Match to reference locus
Match to recombinant "k-mer Only"
k-mer match
1.0
0.8
0.6
0.4
0.2
0.0
PE31,PPE60-H37Rv-3894016-3895519 --- k-mer match to putative recomb seq = 1.0 (57/57)
Match to reference locus
Match to recombinant "k-mer Only"
k-mer match
1.0
0.8
0.6
0.4
0.2
0.0
1,339,680
1,339,760
1,339,840
1,339,920
1,340,000
1,340,080
1,340,160
1,340,240
1,340,320
H37rv Ref Position (bp)

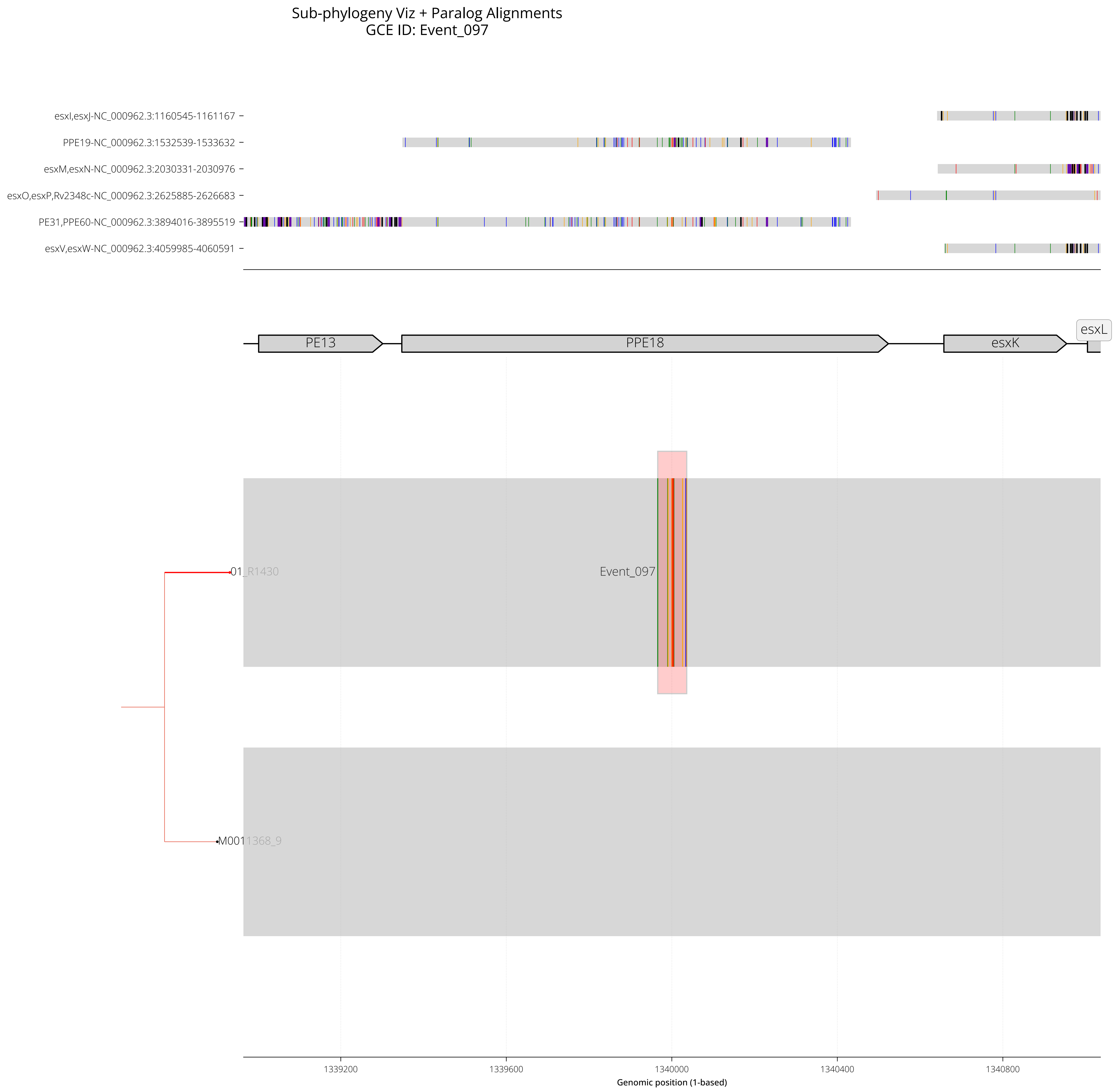

Sub-phylogeny Viz + Paralog Alignments
GCE ID: Event_097
esxI,esxJ-NC_000962.3:1160545-1161167
PPE19-NC_000962.3:1532539-1533632
esxM,esxN-NC_000962.3:2030331-2030976
esxO,esxP,Rv2348c-NC_000962.3:2625885-2626683
PE31,PPE60-NC_000962.3:3894016-3895519
esxV,esxW-NC_000962.3:4059985-4060591
PE13
PPE18
esxK
esxL
01_R1430
Event_097
M0011368_9
1339200
1339600
1340000
1340400
1340800
Genomic position (1-based)

Full Phylogeny Viz + Paralog Alignments
GCE ID: Event_097

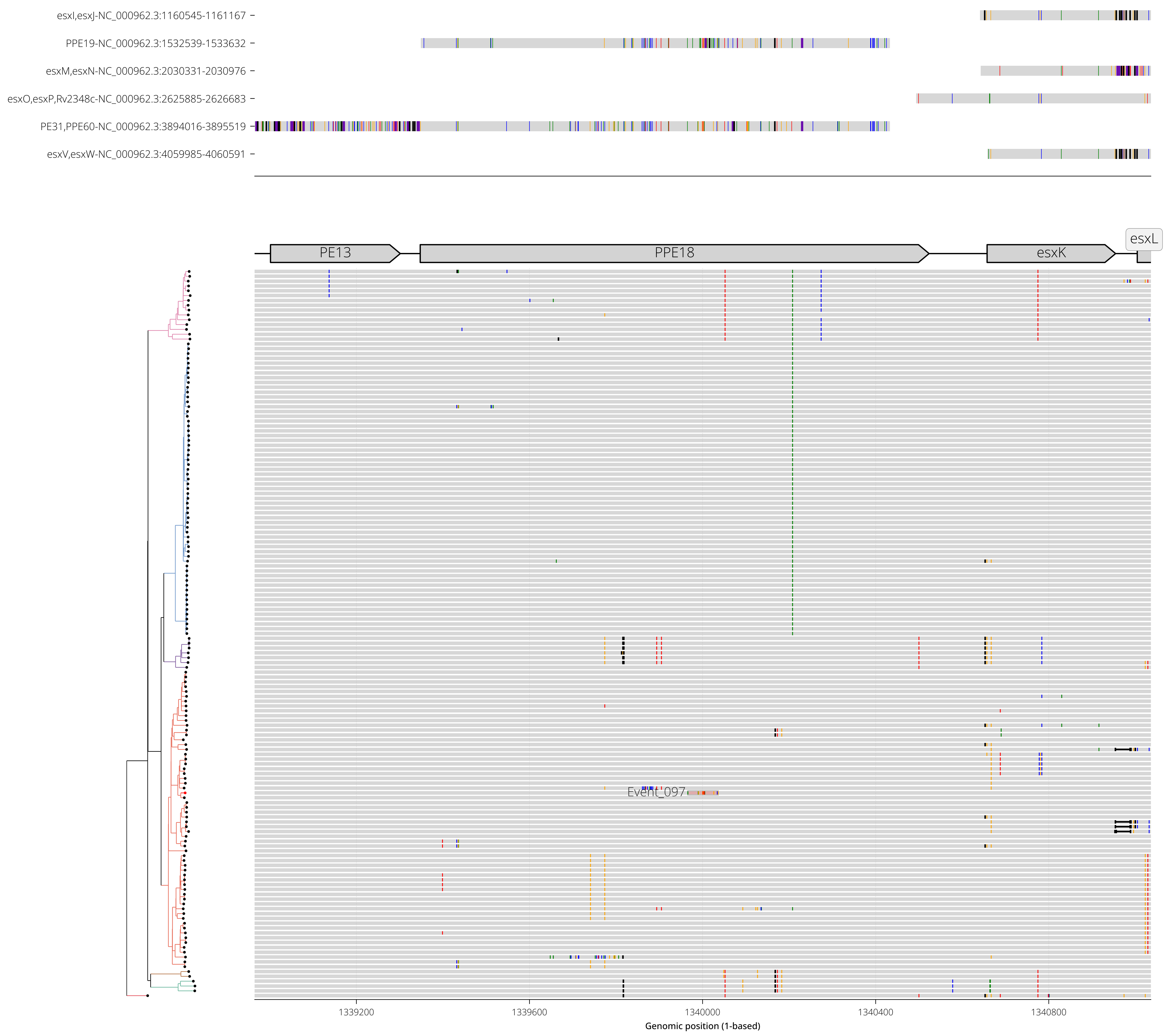

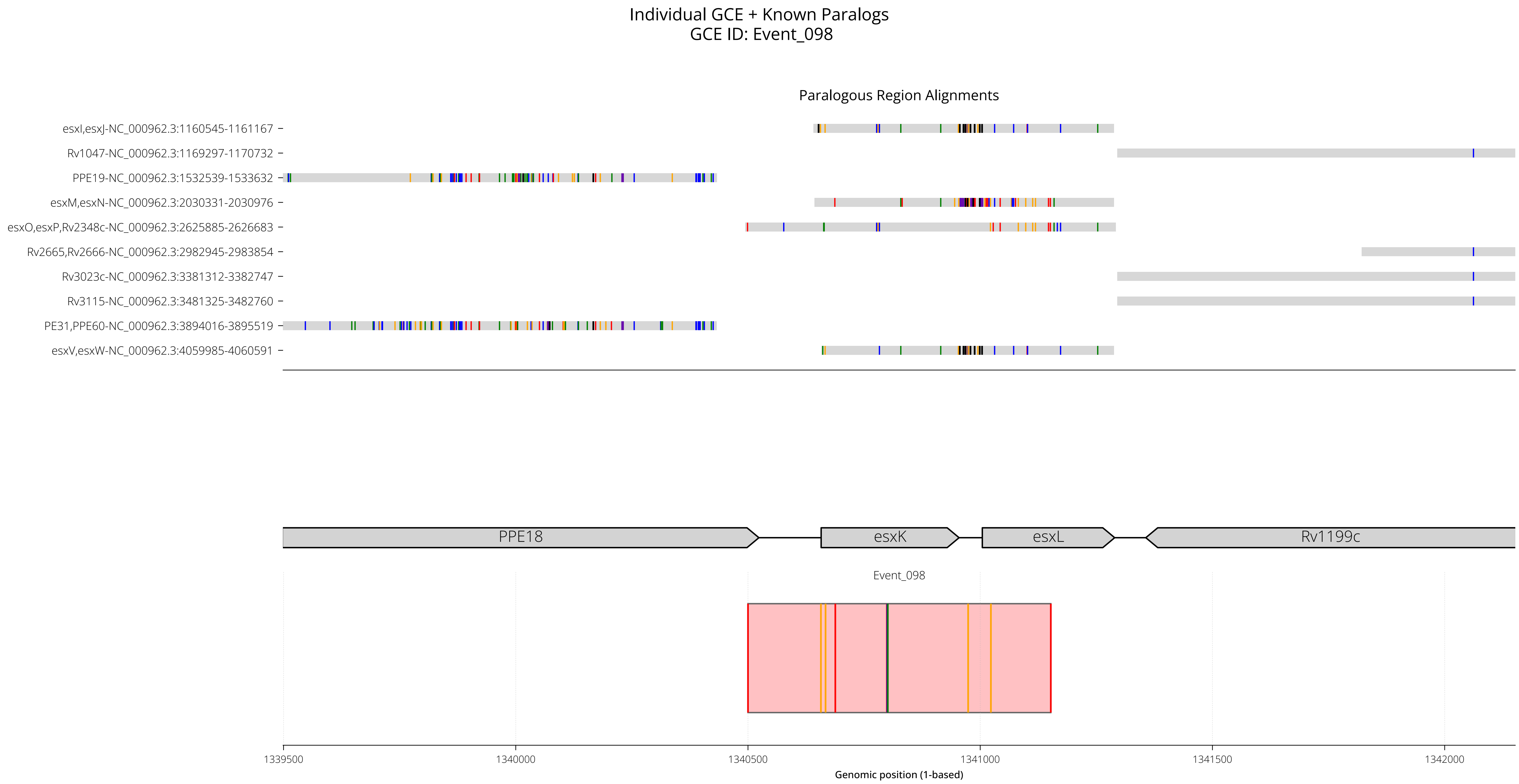

Individual GCE + Known Paralogs
GCE ID: Event_098
Paralogous Region Alignments
esxI,esxJ-NC_000962.3:1160545-1161167
Rv1047-NC_000962.3:1169297-1170732
PPE19-NC_000962.3:1532539-1533632
esxM,esxN-NC_000962.3:2030331-2030976
esxO,esxP,Rv2348c-NC_000962.3:2625885-2626683
Rv2665,Rv2666-NC_000962.3:2982945-2983854
Rv3023c-NC_000962.3:3381312-3382747
Rv3115-NC_000962.3:3481325-3482760
PE31,PPE60-NC_000962.3:3894016-3895519
esxV,esxW-NC_000962.3:4059985-4060591
PPE18
esxK
esxL
Rv1199c
Event_098
1339500
1340000
1340500
1341000
1341500
1342000
Genomic position (1-based)

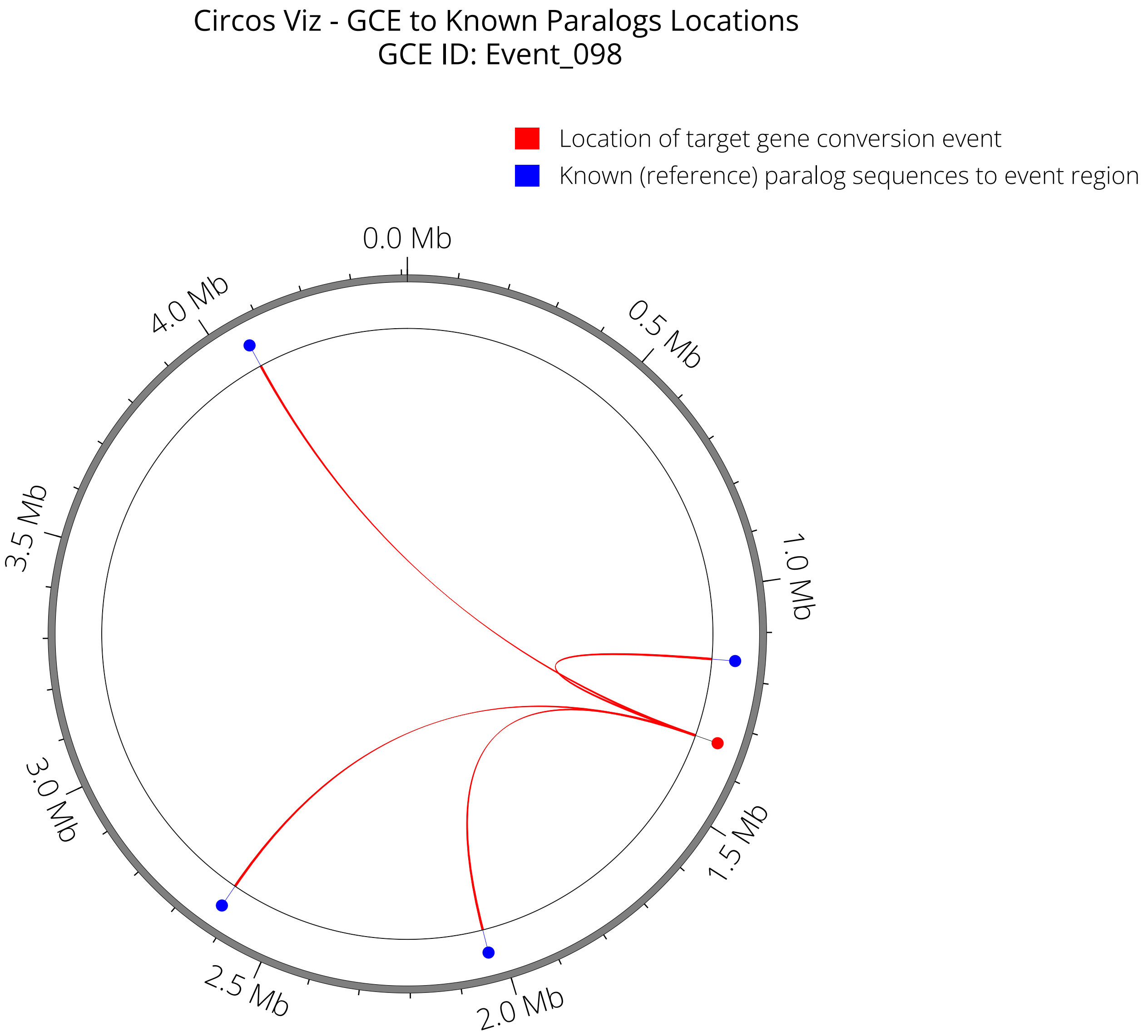

Circos Viz - GCE to Known Paralogs Locations
GCE ID: Event_098
Location of target gene conversion event
Known (reference) paralog sequences to event region
0.0 Mb
0.5 Mb
1.0 Mb
1.5 Mb
2.0 Mb
2.5 Mb
3.0 Mb
3.5 Mb
4.0 Mb

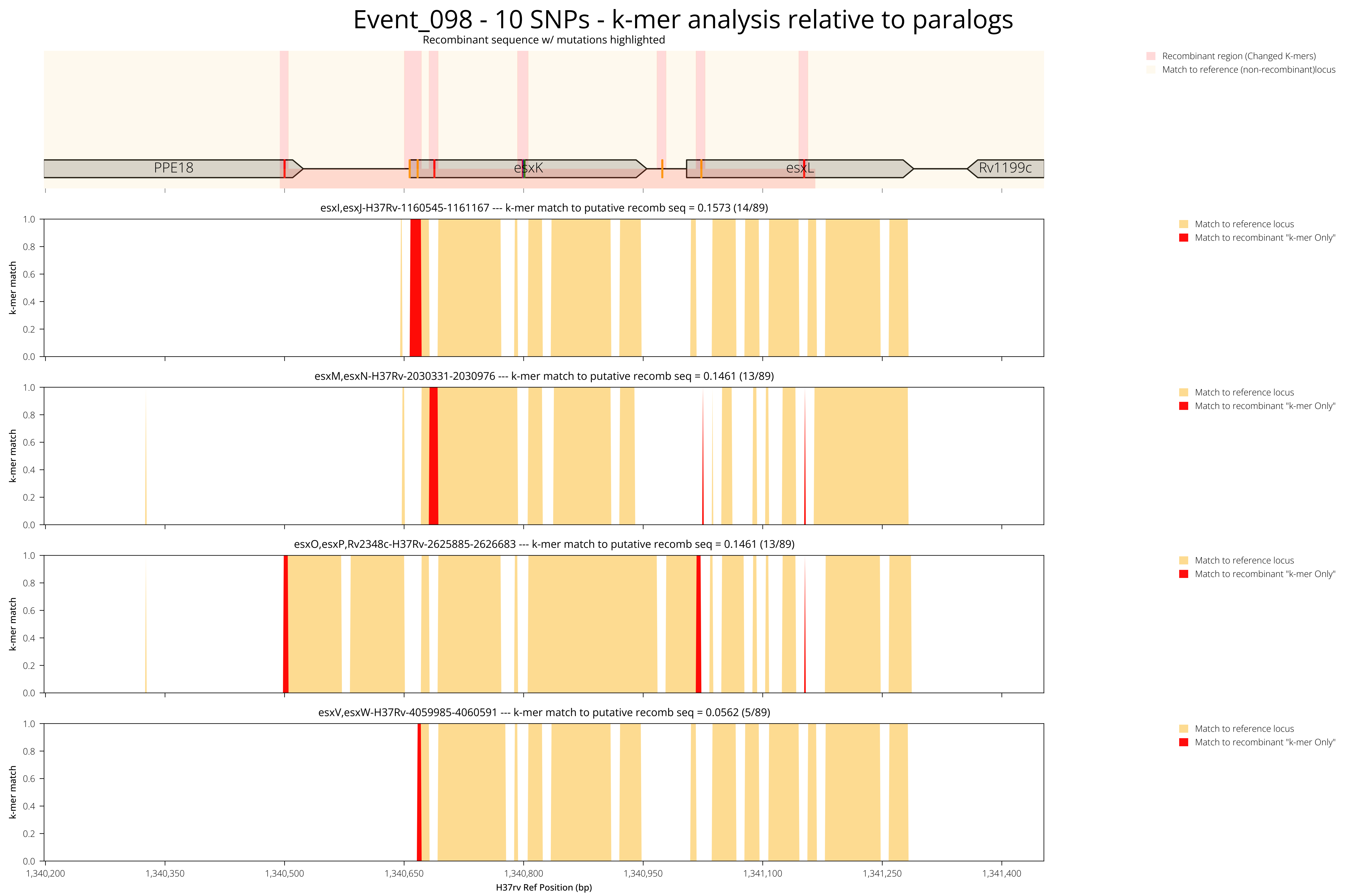
Event_098 - 10 SNPs - k-mer analysis relative to paralogs
Recombinant sequence w/ mutations highlighted
Recombinant region (Changed K-mers)
Match to reference (non-recombinant)locus
PPE18
esxK
esxL
Rv1199c
esxI,esxJ-H37Rv-1160545-1161167 --- k-mer match to putative recomb seq = 0.1573 (14/89)
esxM,esxN-H37Rv-2030331-2030976 --- k-mer match to putative recomb seq = 0.1461 (13/89)
esxO,esxP,Rv2348c-H37Rv-2625885-2626683 --- k-mer match to putative recomb seq = 0.1461 (13/89)
esxV,esxW-H37Rv-4059985-4060591 --- k-mer match to putative recomb seq = 0.0562 (5/89)
Match to reference locus
Match to recombinant "k-mer Only"
k-mer match
1.0
0.8
0.6
0.4
0.2
0.0
1,340,200
1,340,350
1,340,500
1,340,650
1,340,800
1,340,950
1,341,100
1,341,250
1,341,400
H37rv Ref Position (bp)

Sub-phylogeny Viz + Paralog Alignments
GCE ID: Event_098

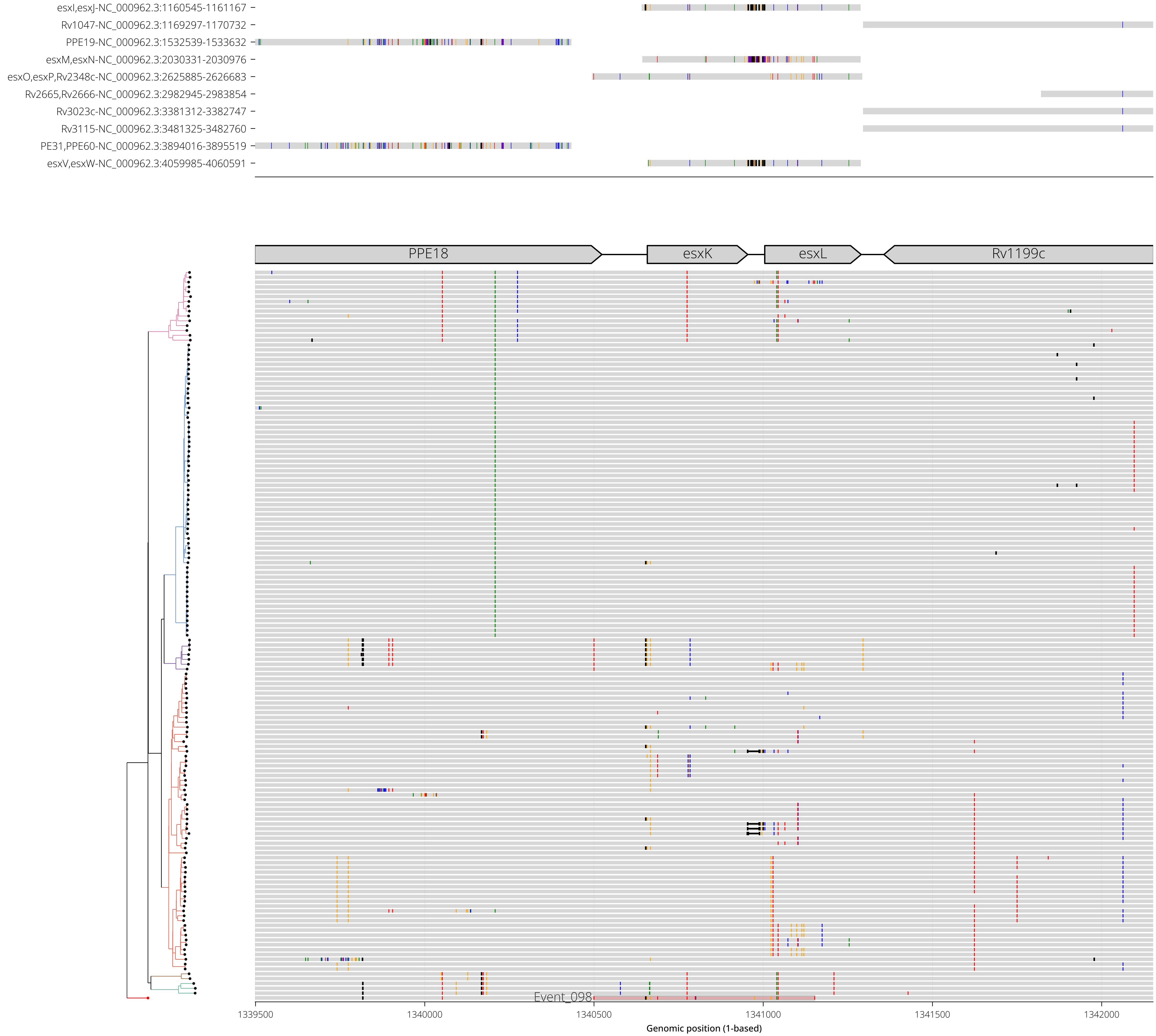

Full Phylogeny Viz + Paralog Alignments
GCE ID: Event_098

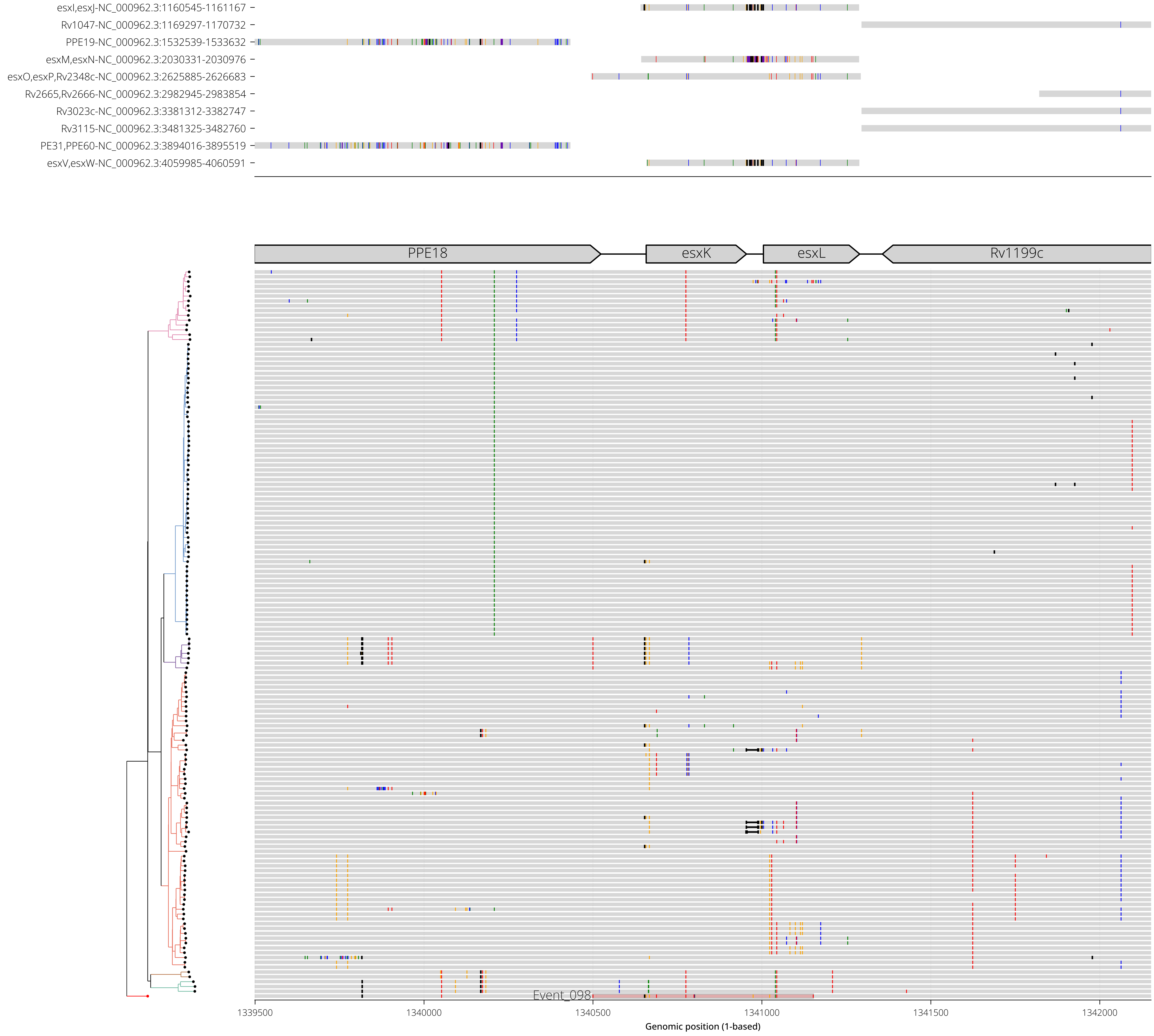

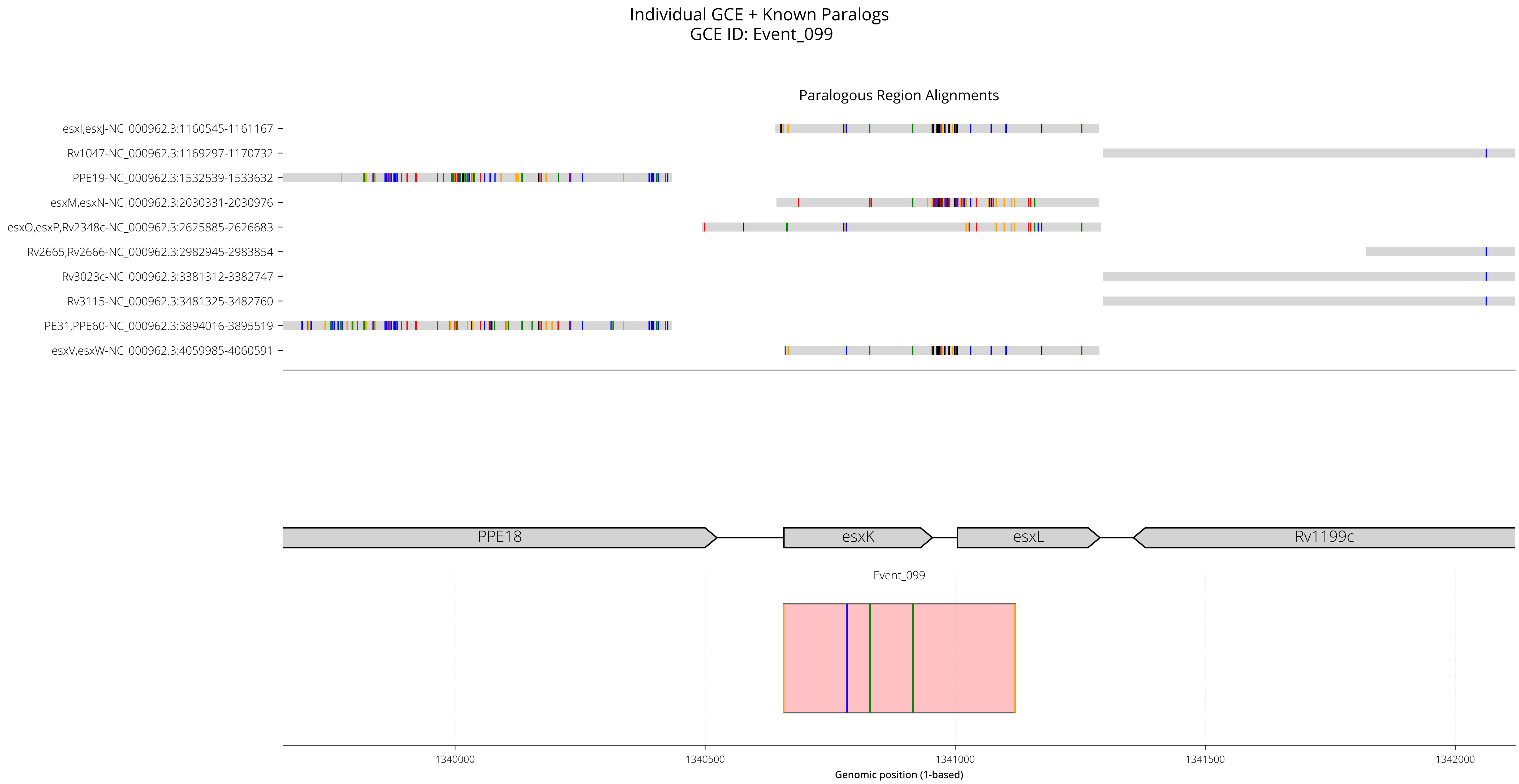

Individual GCE + Known Paralogs
GCE ID: Event_099
Paralogous Region Alignments
esxI,esxJ-NC_000962.3:1160545-1161167
Rv1047-NC_000962.3:1169297-1170732
PPE19-NC_000962.3:1532539-1533632
esxM,esxN-NC_000962.3:2030331-2030976
esxO,esxP,Rv2348c-NC_000962.3:2625885-2626683
Rv2665,Rv2666-NC_000962.3:2982945-2983854
Rv3023c-NC_000962.3:3381312-3382747
Rv3115-NC_000962.3:3481325-3482760
PE31,PPE60-NC_000962.3:3894016-3895519
esxV,esxW-NC_000962.3:4059985-4060591
PPE18
esxK
esxL
Rv1199c
Event_099
1340000
1340500
1341000
1341500
1342000
Genomic position (1-based)

### Circos Viz - GCE to Known Paralogs Locations
GCE ID: Event_099

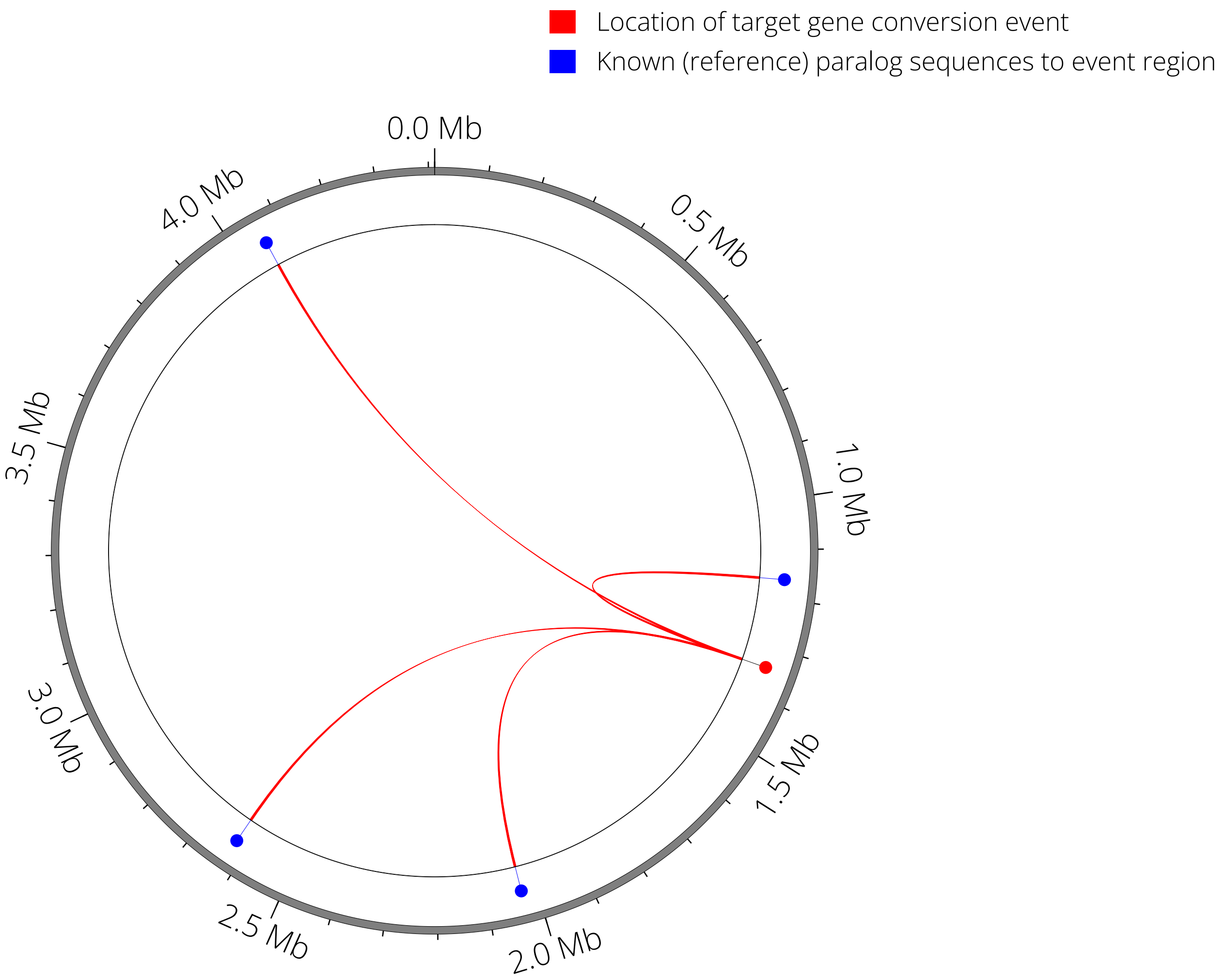

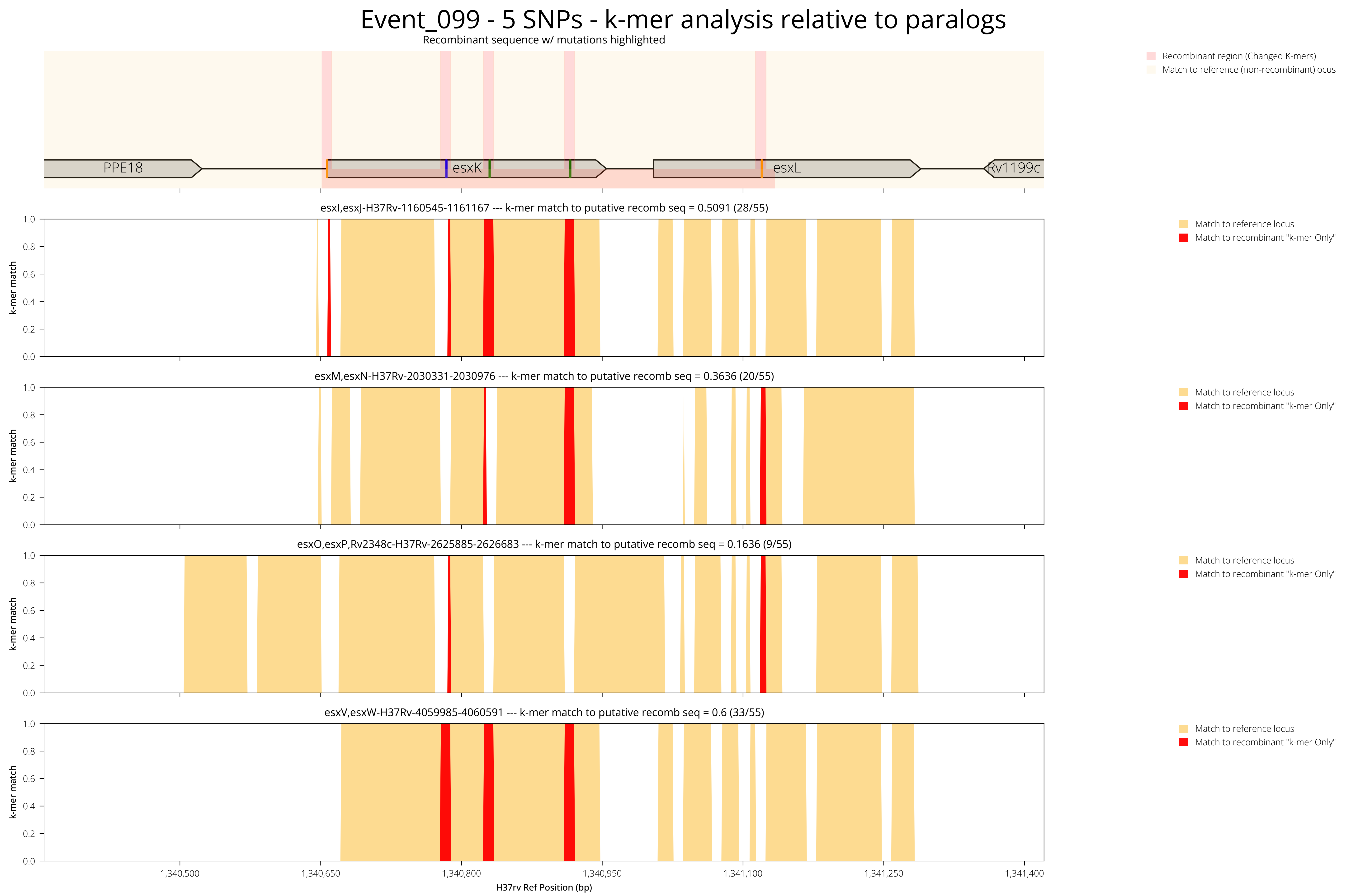
Event_099 - 5 SNPs - k-mer analysis relative to paralogs
Recombinant sequence w/ mutations highlighted
Recombinant region (Changed K-mers)
Match to reference (non-recombinant)locus
PPE18
esxK
esxL
Rv1199c
esxI,esxJ-H37Rv-1160545-1161167 --- k-mer match to putative recomb seq = 0.5091 (28/55)
esxM,esxN-H37Rv-2030331-2030976 --- k-mer match to putative recomb seq = 0.3636 (20/55)
esxO,esxP,Rv2348c-H37Rv-2625885-2626683 --- k-mer match to putative recomb seq = 0.1636 (9/55)
esxV,esxW-H37Rv-4059985-4060591 --- k-mer match to putative recomb seq = 0.6 (33/55)
Match to reference locus
Match to recombinant "k-mer Only"
k-mer match
1.0
0.8
0.6
0.4
0.2
0.0
1,340,500
1,340,650
1,340,800
1,340,950
1,341,100
1,341,250
1,341,400
H37rv Ref Position (bp)

Sub-phylogeny Viz + Paralog Alignments
GCE ID: Event_099

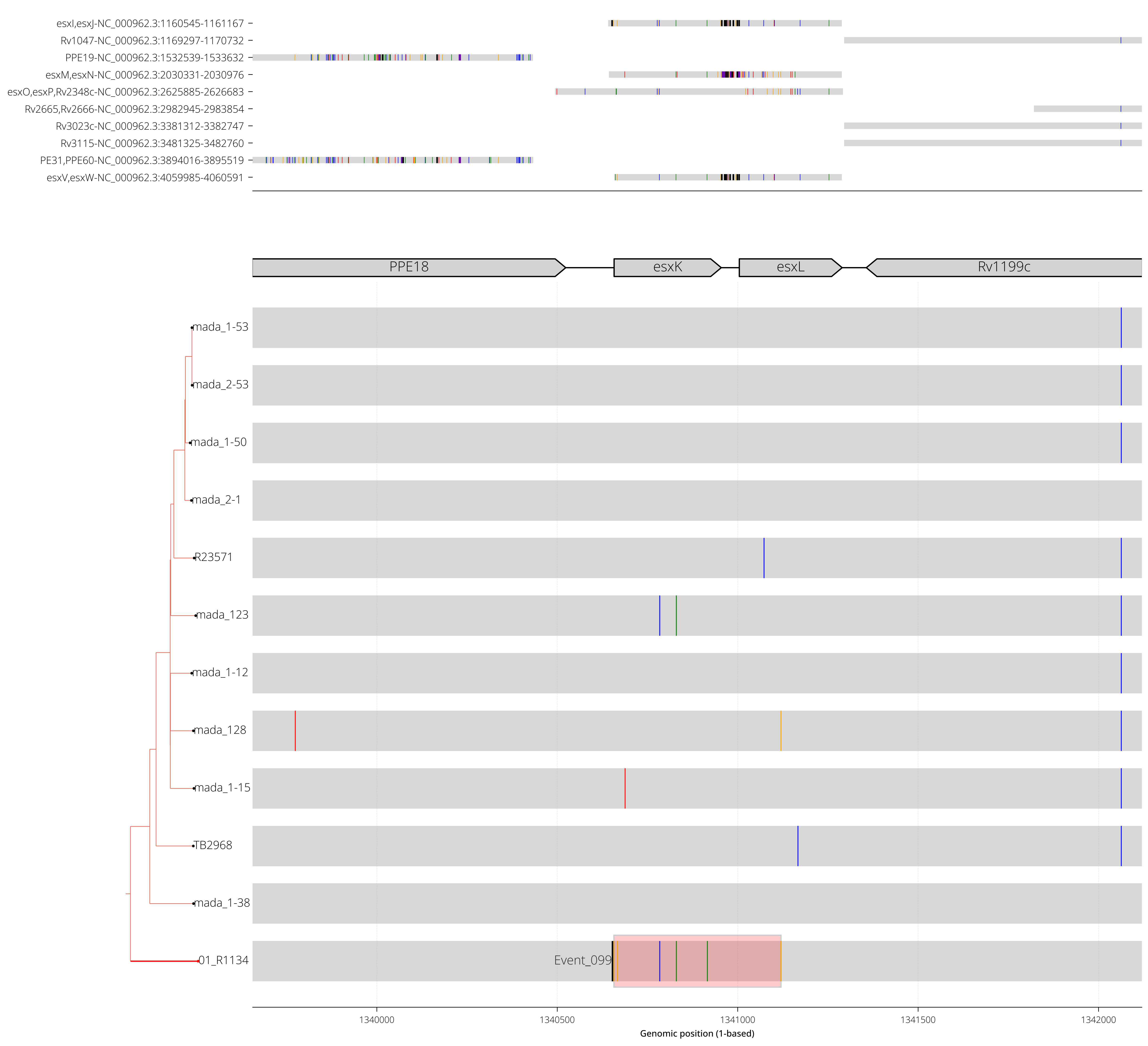

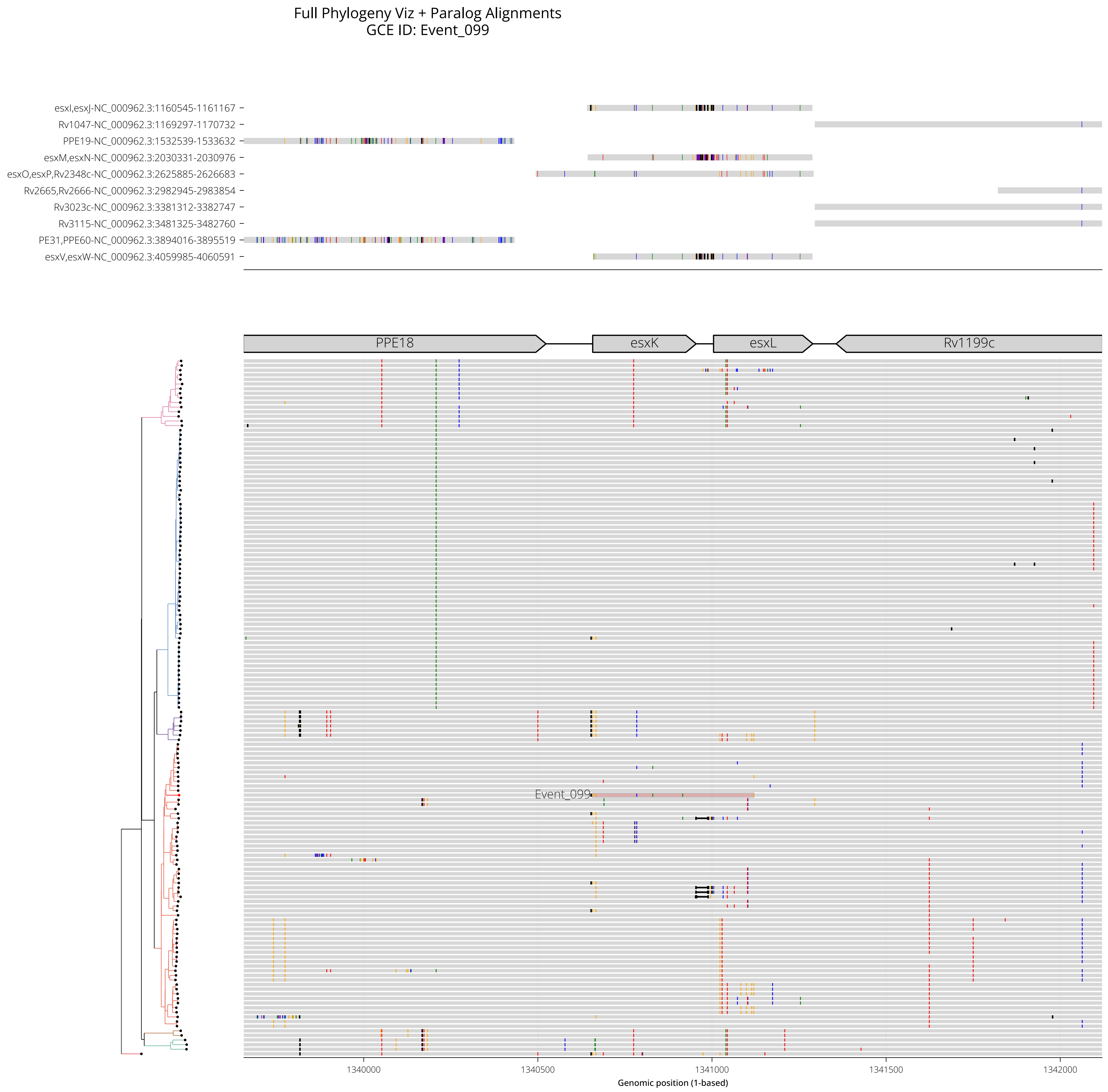

Full Phylogeny Viz + Paralog Alignments
GCE ID: Event_099
esxI,esxJ-NC_000962.3:1160545-1161167
Rv1047-NC_000962.3:1169297-1170732
PPE19-NC_000962.3:1532539-1533632
esxM,esxN-NC_000962.3:2030331-2030976
esxO,esxP,Rv2348c-NC_000962.3:2625885-2626683
Rv2665,Rv2666-NC_000962.3:2982945-2983854
Rv3023c-NC_000962.3:3381312-3382747
Rv3115-NC_000962.3:3481325-3482760
PE31,PPE60-NC_000962.3:3894016-3895519
esxV,esxW-NC_000962.3:4059985-4060591
PPE18
esxK
esxL
Rv1199c
Event_099
1340000
1340500
1341000
1341500
1342000
Genomic position (1-based)

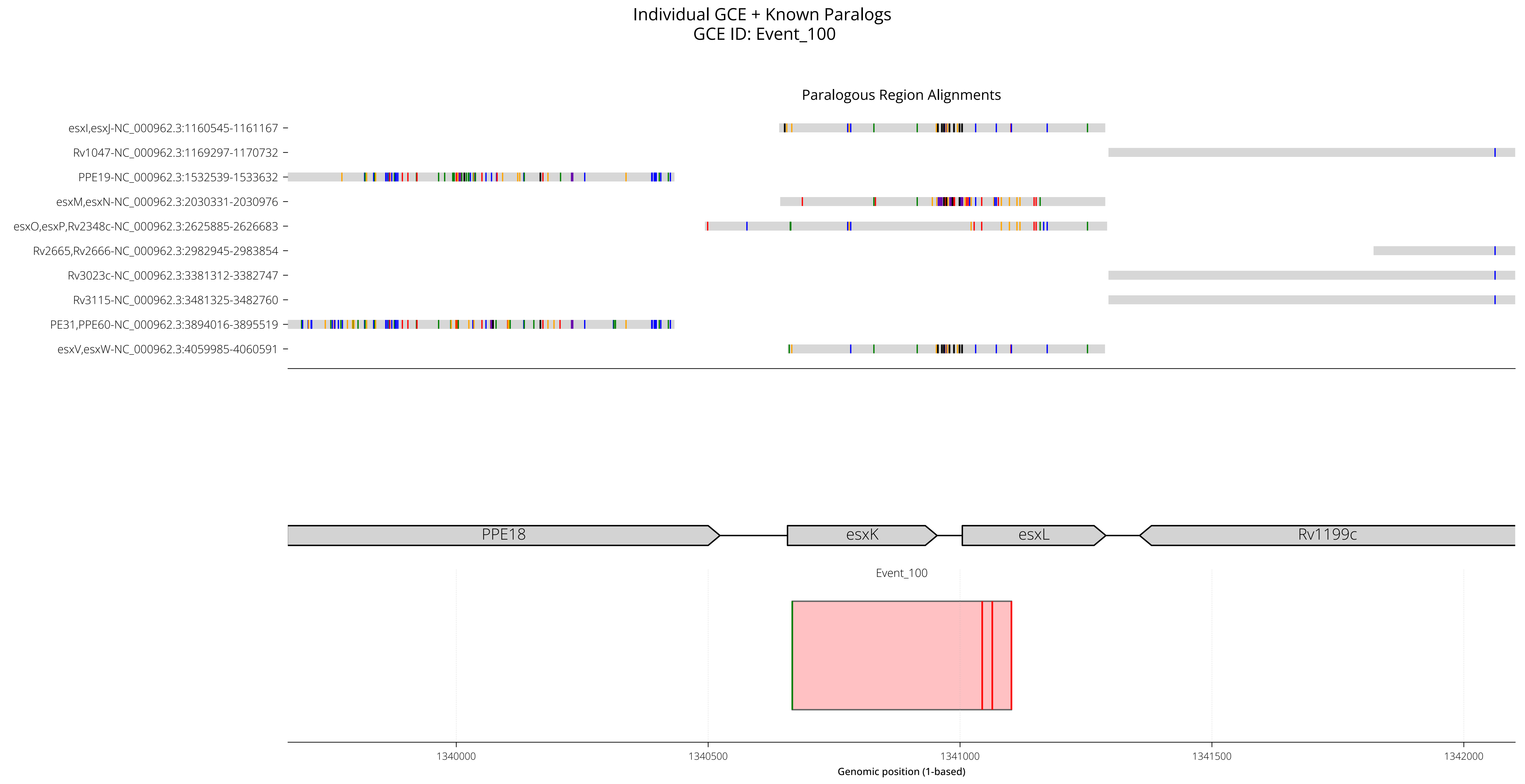

Individual GCE + Known Paralogs
GCE ID: Event_100
Paralogous Region Alignments
esxI,esxJ-NC_000962.3:1160545-1161167
Rv1047-NC_000962.3:1169297-1170732
PPE19-NC_000962.3:1532539-1533632
esxM,esxN-NC_000962.3:2030331-2030976
esxO,esxP,Rv2348c-NC_000962.3:2625885-2626683
Rv2665,Rv2666-NC_000962.3:2982945-2983854
Rv3023c-NC_000962.3:3381312-3382747
Rv3115-NC_000962.3:3481325-3482760
PE31,PPE60-NC_000962.3:3894016-3895519
esxV,esxW-NC_000962.3:4059985-4060591
PPE18
esxK
esxL
Rv1199c
Event_100
1340000
1340500
1341000
1341500
1342000
Genomic position (1-based)

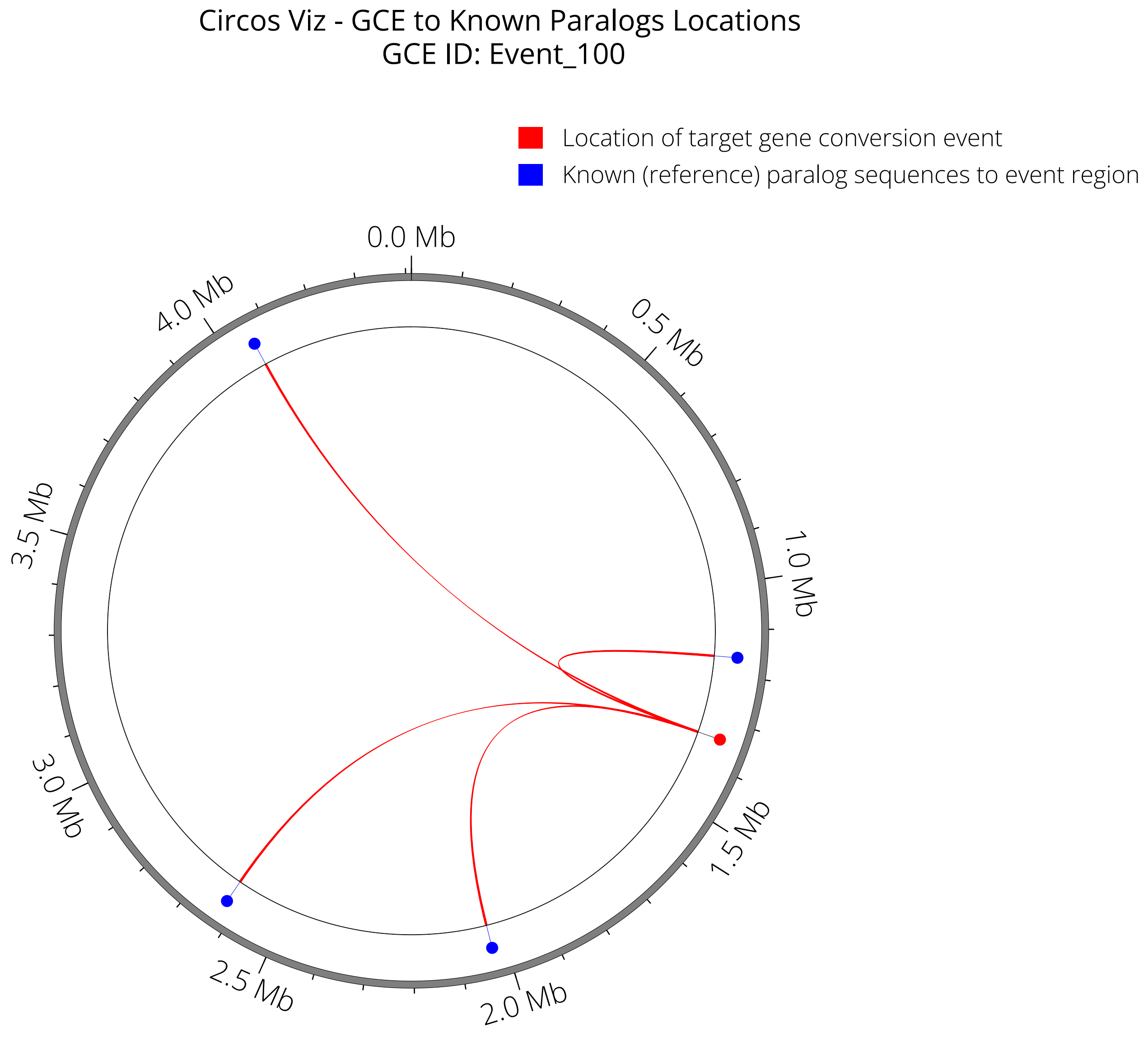

Circos Viz - GCE to Known Paralogs Locations
GCE ID: Event_100
Location of target gene conversion event
Known (reference) paralog sequences to event region
0.0 Mb
0.5 Mb
1.0 Mb
1.5 Mb
2.0 Mb
2.5 Mb
3.0 Mb
3.5 Mb
4.0 Mb

### Event_100 - 4 SNPs - k-mer analysis relative to paralogs

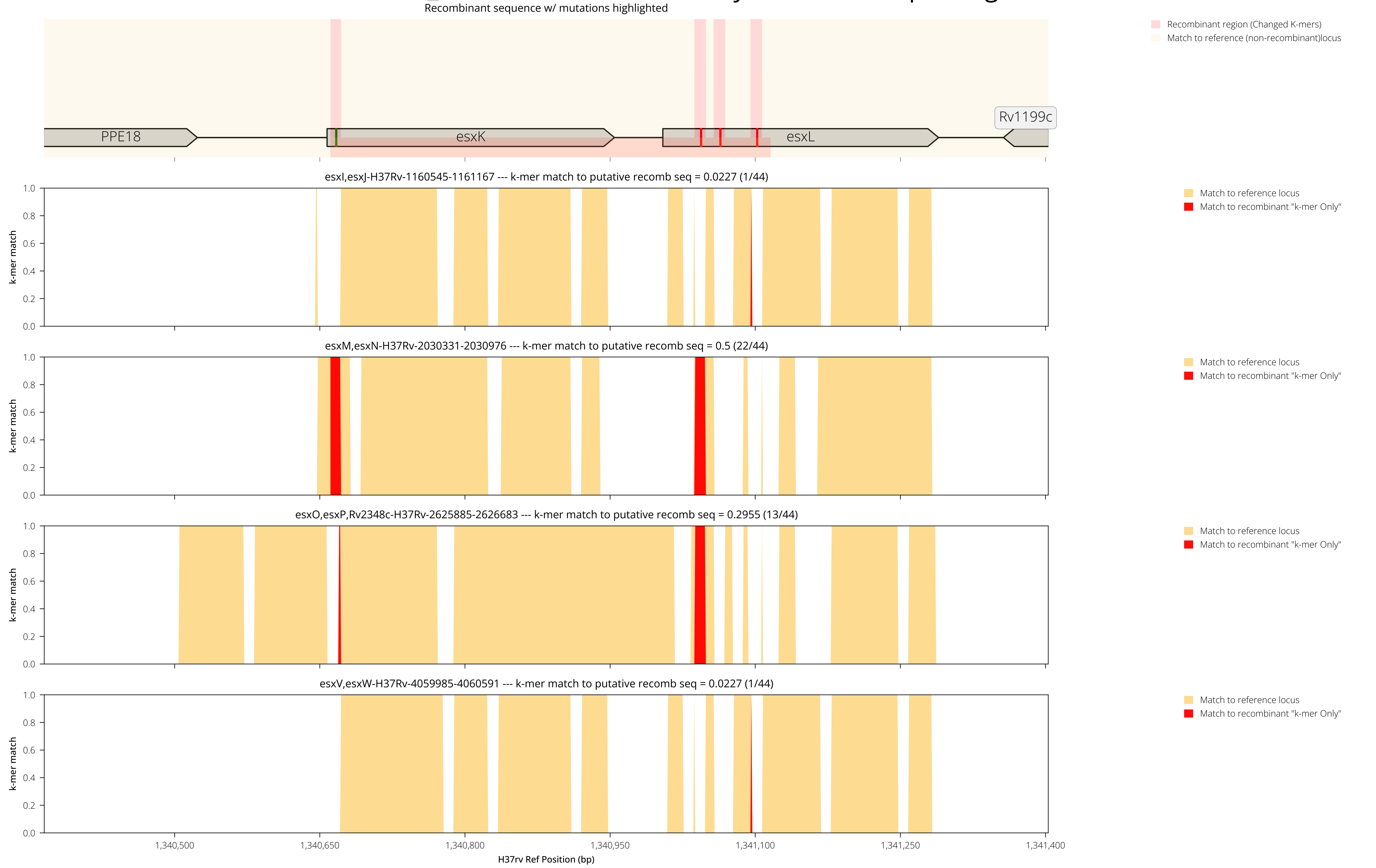

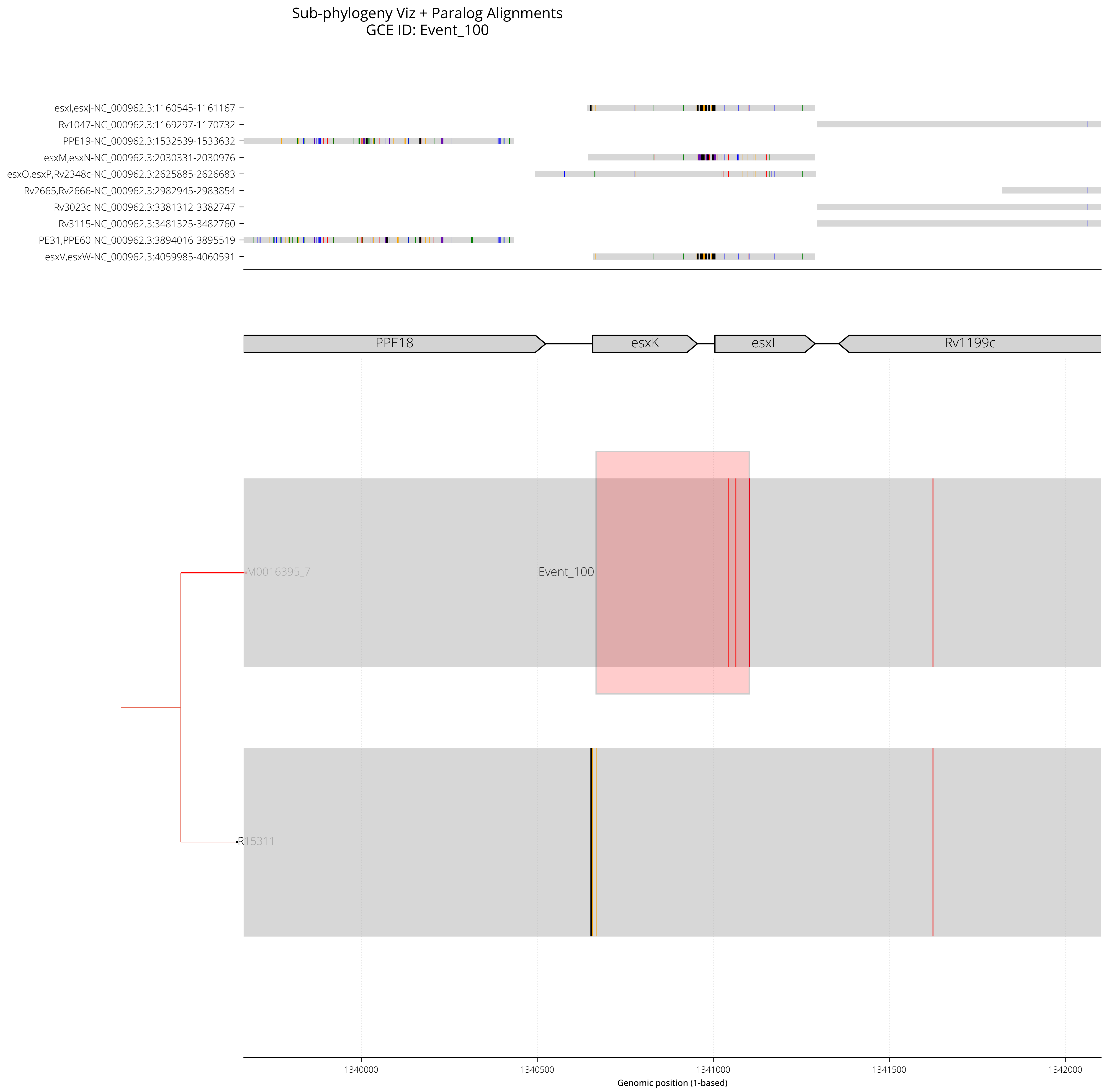

Sub-phylogeny Viz + Paralog Alignments
GCE ID: Event_100
esxI,esxJ-NC_000962.3:1160545-1161167
Rv1047-NC_000962.3:1169297-1170732
PPE19-NC_000962.3:1532539-1533632
esxM,esxN-NC_000962.3:2030331-2030976
esxO,esxP,Rv2348c-NC_000962.3:2625885-2626683
Rv2665,Rv2666-NC_000962.3:2982945-2983854
Rv3023c-NC_000962.3:3381312-3382747
Rv3115-NC_000962.3:3481325-3482760
PE31,PPE60-NC_000962.3:3894016-3895519
esxV,esxW-NC_000962.3:4059985-4060591
PPE18
esxK
esxL
Rv1199c
M0016395_7
Event_100
R15311
1340000
1340500
1341000
1341500
1342000
Genomic position (1-based)

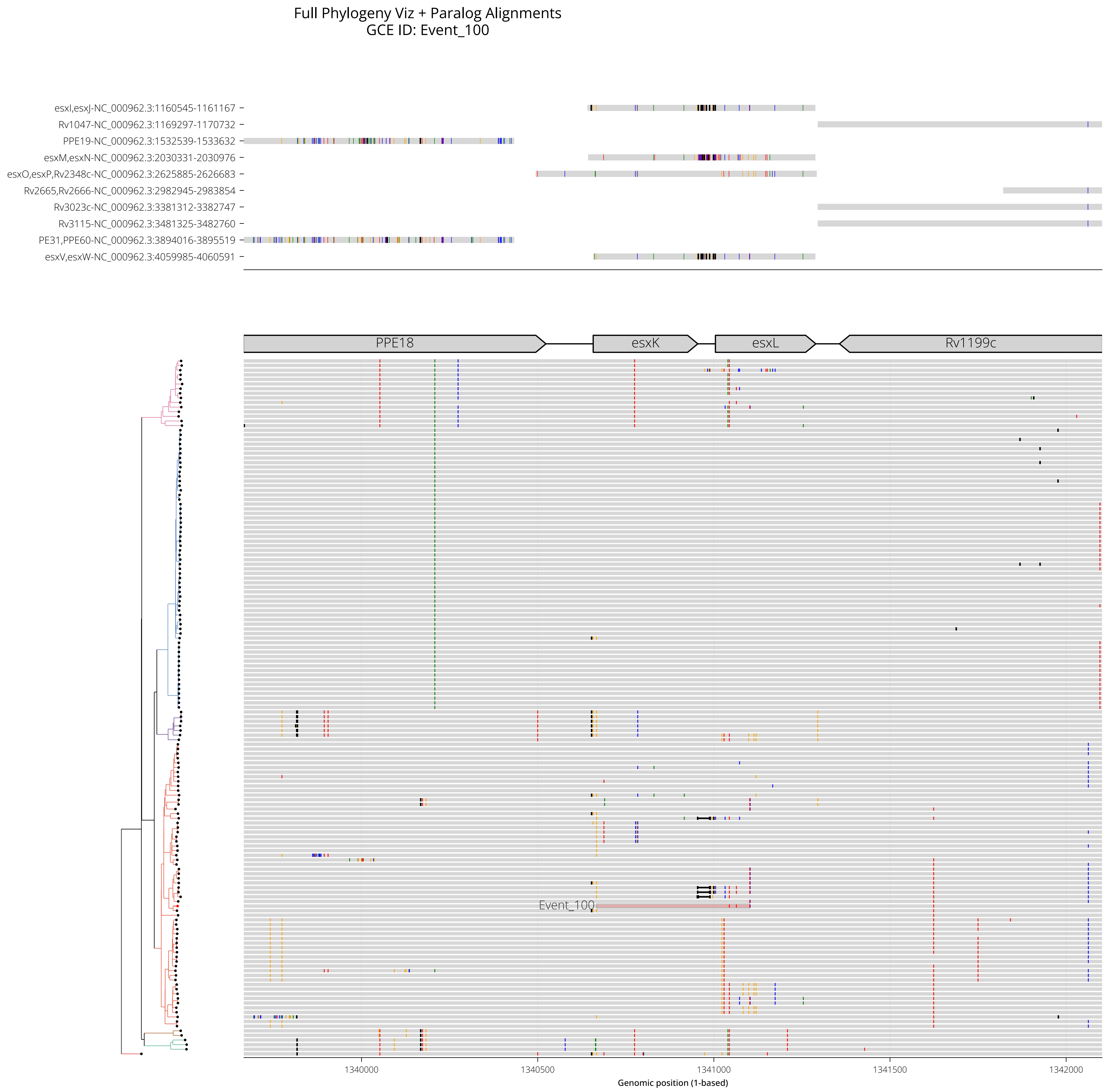

Full Phylogeny Viz + Paralog Alignments
GCE ID: Event_100
esxI,esxJ-NC_000962.3:1160545-1161167
Rv1047-NC_000962.3:1169297-1170732
PPE19-NC_000962.3:1532539-1533632
esxM,esxN-NC_000962.3:2030331-2030976
esxO,esxP,Rv2348c-NC_000962.3:2625885-2626683
Rv2665,Rv2666-NC_000962.3:2982945-2983854
Rv3023c-NC_000962.3:3381312-3382747
Rv3115-NC_000962.3:3481325-3482760
PE31,PPE60-NC_000962.3:3894016-3895519
esxV,esxW-NC_000962.3:4059985-4060591
PPE18
esxK
esxL
Rv1199c
Event_100
1340000
1340500
1341000
1341500
1342000
Genomic position (1-based)

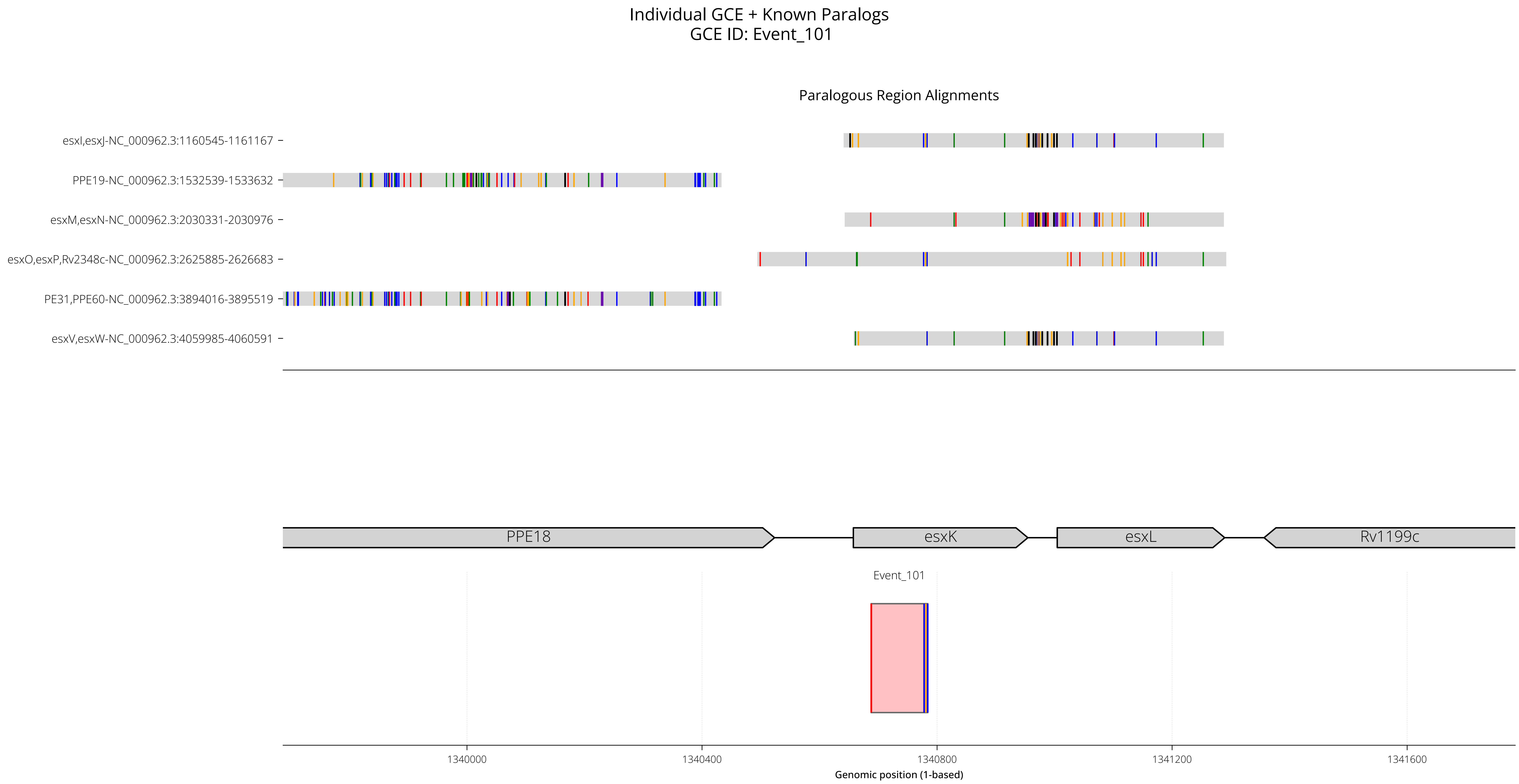

Individual GCE + Known Paralogs
GCE ID: Event_101
Paralogous Region Alignments
esxI,esxJ-NC_000962.3:1160545-1161167
PPE19-NC_000962.3:1532539-1533632
esxM,esxN-NC_000962.3:2030331-2030976
esxO,esxP,Rv2348c-NC_000962.3:2625885-2626683
PE31,PPE60-NC_000962.3:3894016-3895519
esxV,esxW-NC_000962.3:4059985-4060591
PPE18
esxK
esxL
Rv1199c
Event_101
1340000
1340400
1340800
1341200
1341600
Genomic position (1-based)

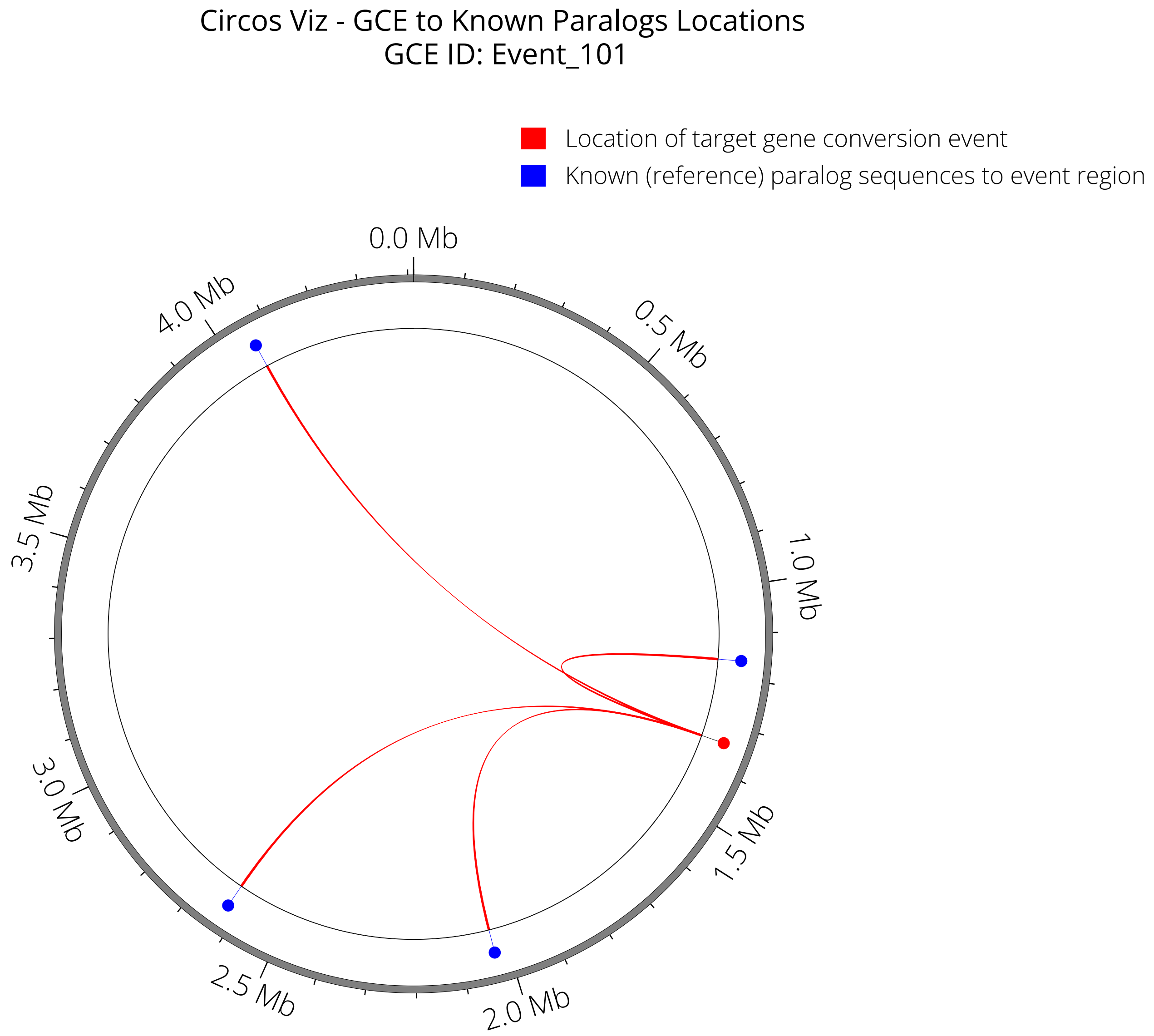
Circos Viz - GCE to Known Paralogs Locations
GCE ID: Event_101
Location of target gene conversion event
Known (reference) paralog sequences to event region
0.0 Mb
0.5 Mb
1.0 Mb
1.5 Mb
2.0 Mb
2.5 Mb
3.0 Mb
3.5 Mb
4.0 Mb

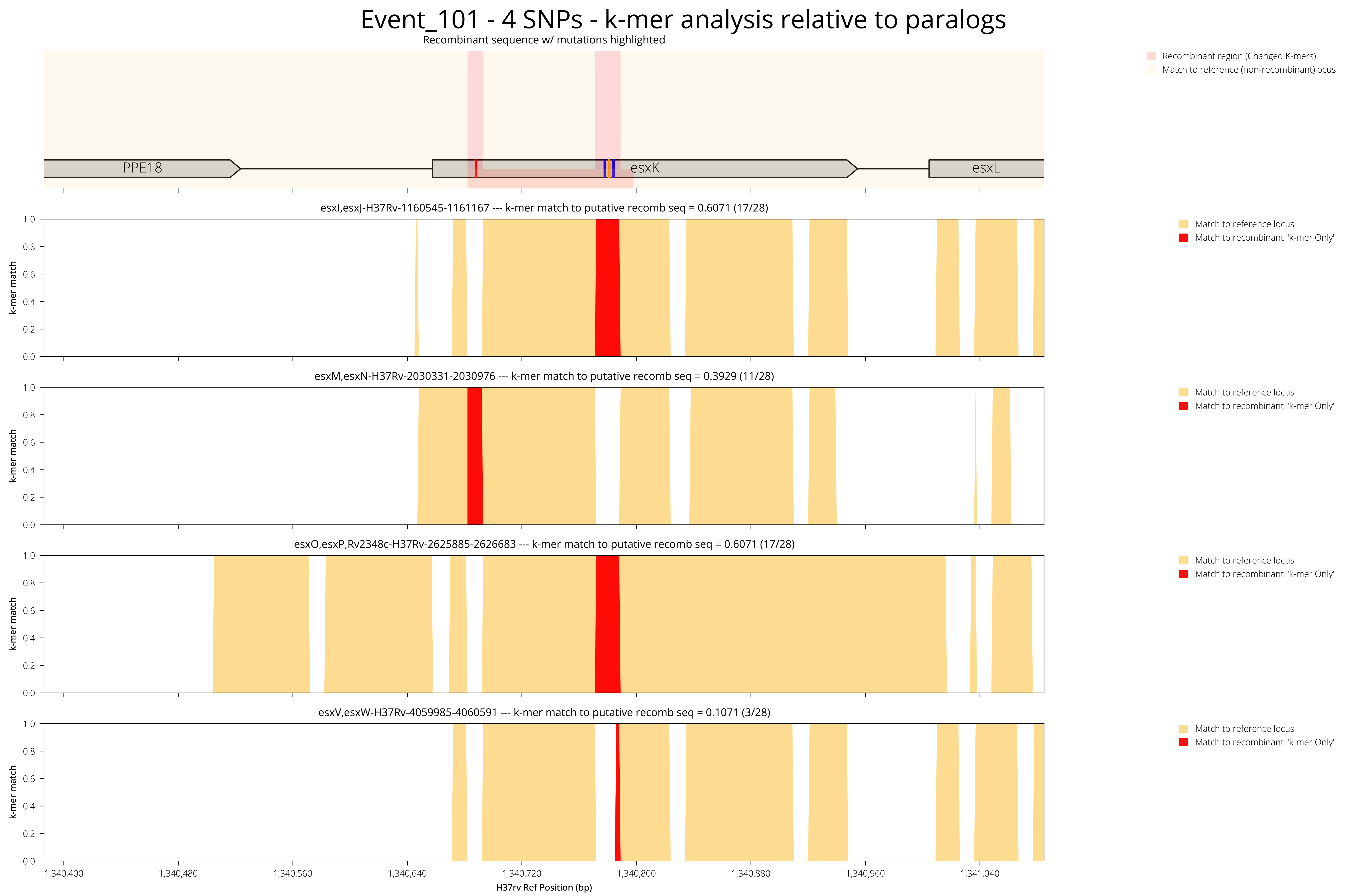

Event_101 - 4 SNPs - k-mer analysis relative to paralogs
Recombinant sequence w/ mutations highlighted
Recombinant region (Changed K-mers)
Match to reference (non-recombinant)locus
PPE18
esxK
esxL
esxI,esxJ-H37Rv-1160545-1161167 --- k-mer match to putative recomb seq = 0.6071 (17/28)
esxM,esxN-H37Rv-2030331-2030976 --- k-mer match to putative recomb seq = 0.3929 (11/28)
esxO,esxP,Rv2348c-H37Rv-2625885-2626683 --- k-mer match to putative recomb seq = 0.6071 (17/28)
esxV,esxW-H37Rv-4059985-4060591 --- k-mer match to putative recomb seq = 0.1071 (3/28)
Match to reference locus
Match to recombinant "k-mer Only"
k-mer match
1.0
0.8
0.6
0.4
0.2
0.0
1,340,400
1,340,480
1,340,560
1,340,640
1,340,720
1,340,800
1,340,880
1,340,960
1,341,040
H37rv Ref Position (bp)

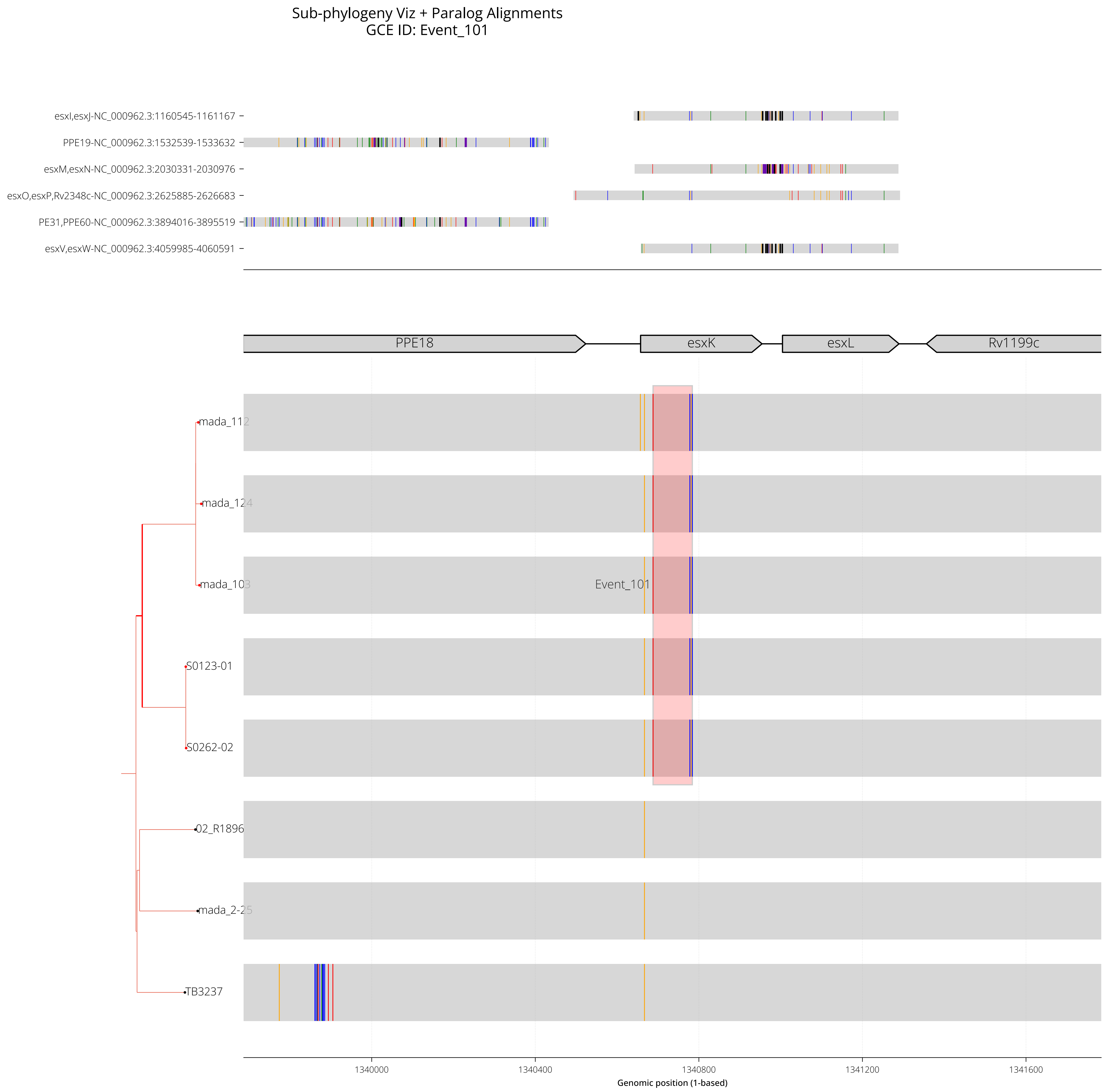

Sub-phylogeny Viz + Paralog Alignments
GCE ID: Event_101
esxI,esxJ-NC_000962.3:1160545-1161167
PPE19-NC_000962.3:1532539-1533632
esxM,esxN-NC_000962.3:2030331-2030976
esxO,esxP,Rv2348c-NC_000962.3:2625885-2626683
PE31,PPE60-NC_000962.3:3894016-3895519
esxV,esxW-NC_000962.3:4059985-4060591
PPE18
esxK
esxL
Rv1199c
mada_112
mada_124
mada_103
Event_101
S0123-01
S0262-02
02_R1896
mada_2-25
TB3237
1340000
1340400
1340800
1341200
1341600
Genomic position (1-based)

Full Phylogeny Viz + Paralog Alignments
GCE ID: Event_101

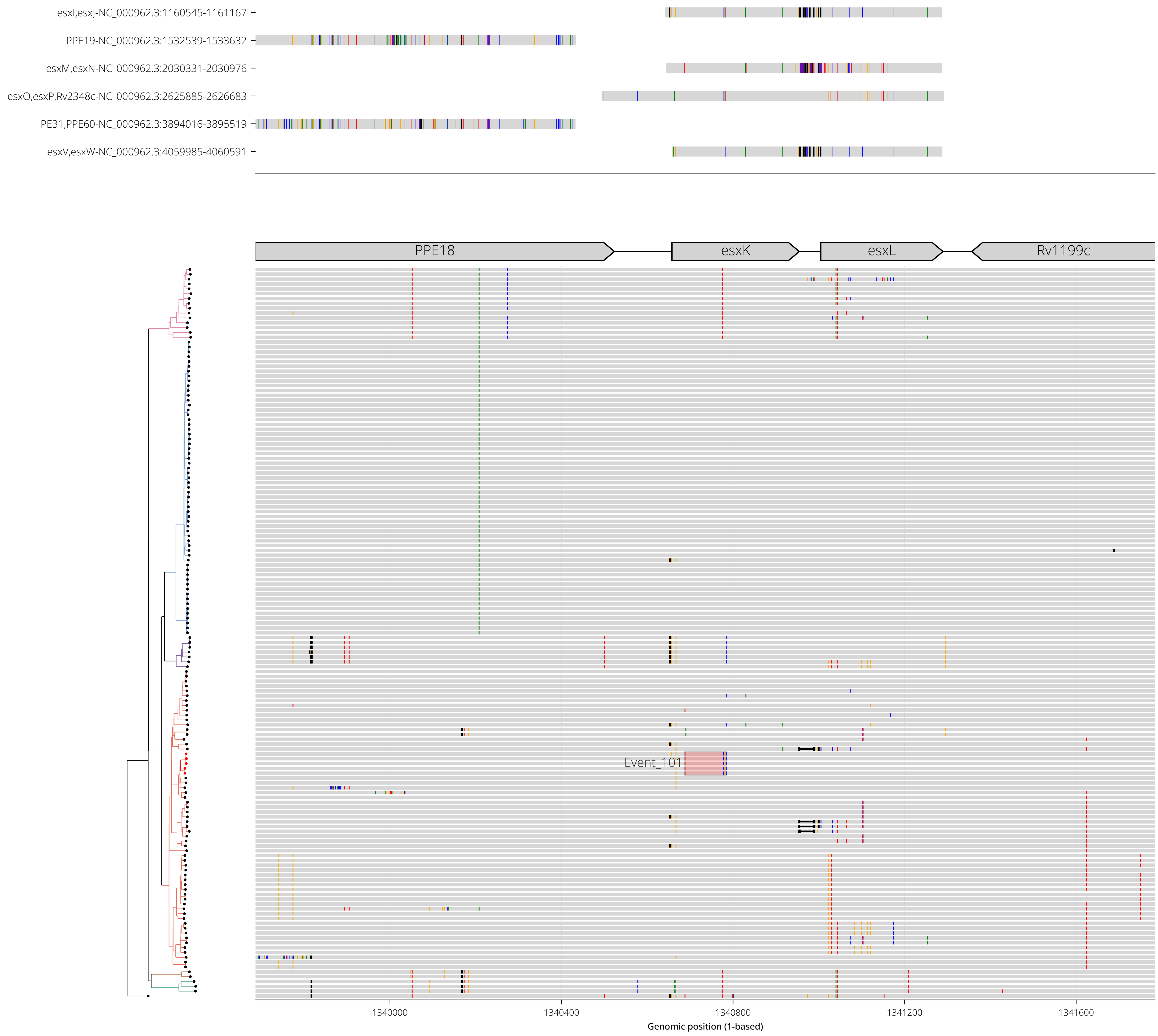

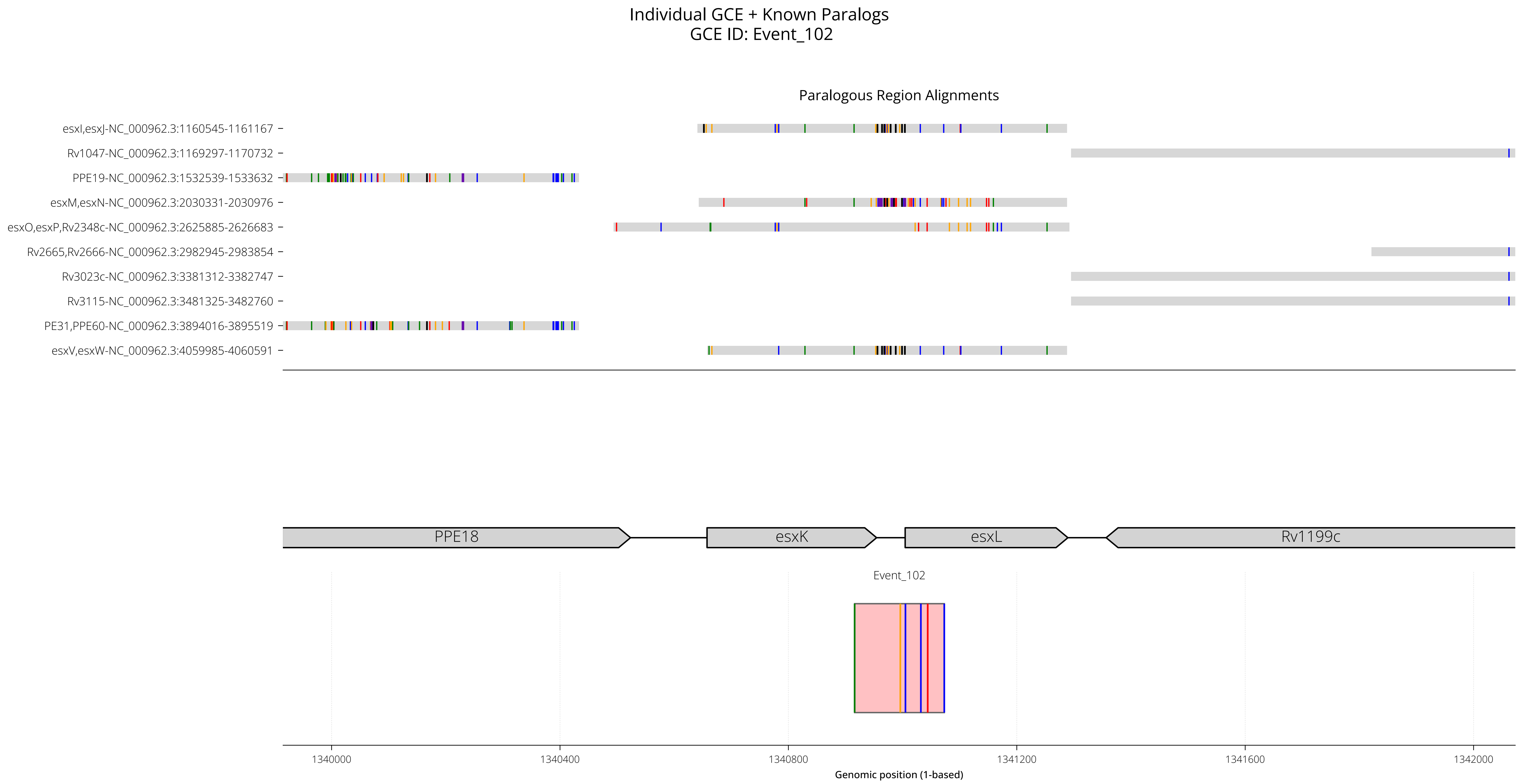
Individual GCE + Known Paralogs
GCE ID: Event_102
Paralogous Region Alignments
esxI,esxJ-NC_000962.3:1160545-1161167
Rv1047-NC_000962.3:1169297-1170732
PPE19-NC_000962.3:1532539-1533632
esxM,esxN-NC_000962.3:2030331-2030976
esxO,esxP,Rv2348c-NC_000962.3:2625885-2626683
Rv2665,Rv2666-NC_000962.3:2982945-2983854
Rv3023c-NC_000962.3:3381312-3382747
Rv3115-NC_000962.3:3481325-3482760
PE31,PPE60-NC_000962.3:3894016-3895519
esxV,esxW-NC_000962.3:4059985-4060591
PPE18
esxK
esxL
Rv1199c
Event_102
1340000
1340400
1340800
1341200
1341600
1342000
Genomic position (1-based)

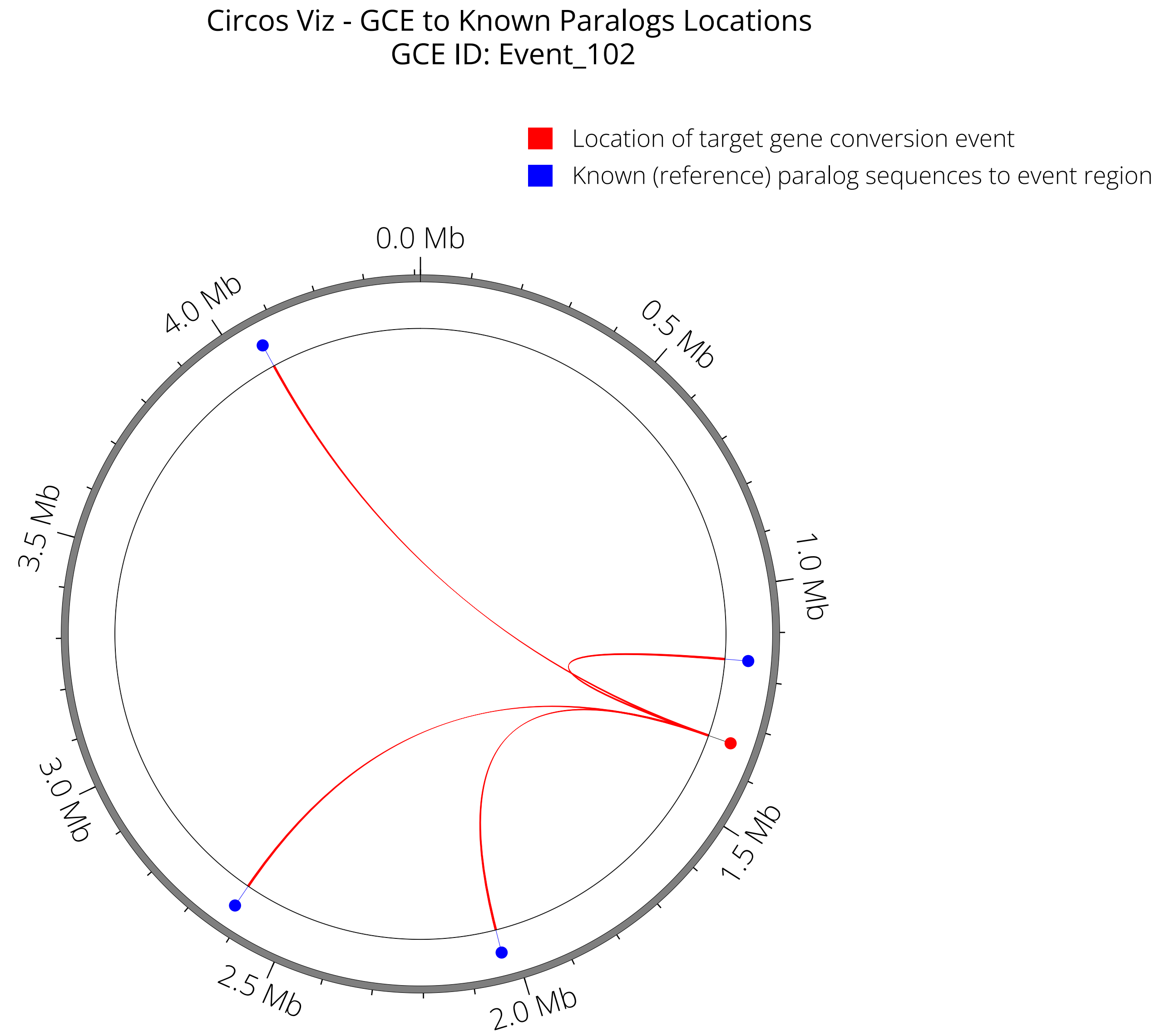
Circos Viz - GCE to Known Paralogs Locations
GCE ID: Event_102
Location of target gene conversion event
Known (reference) paralog sequences to event region
0.0 Mb
0.5 Mb
1.0 Mb
1.5 Mb
2.0 Mb
2.5 Mb
3.0 Mb
3.5 Mb
4.0 Mb

### Event_102 - 6 SNPs - k-mer analysis relative to paralogs

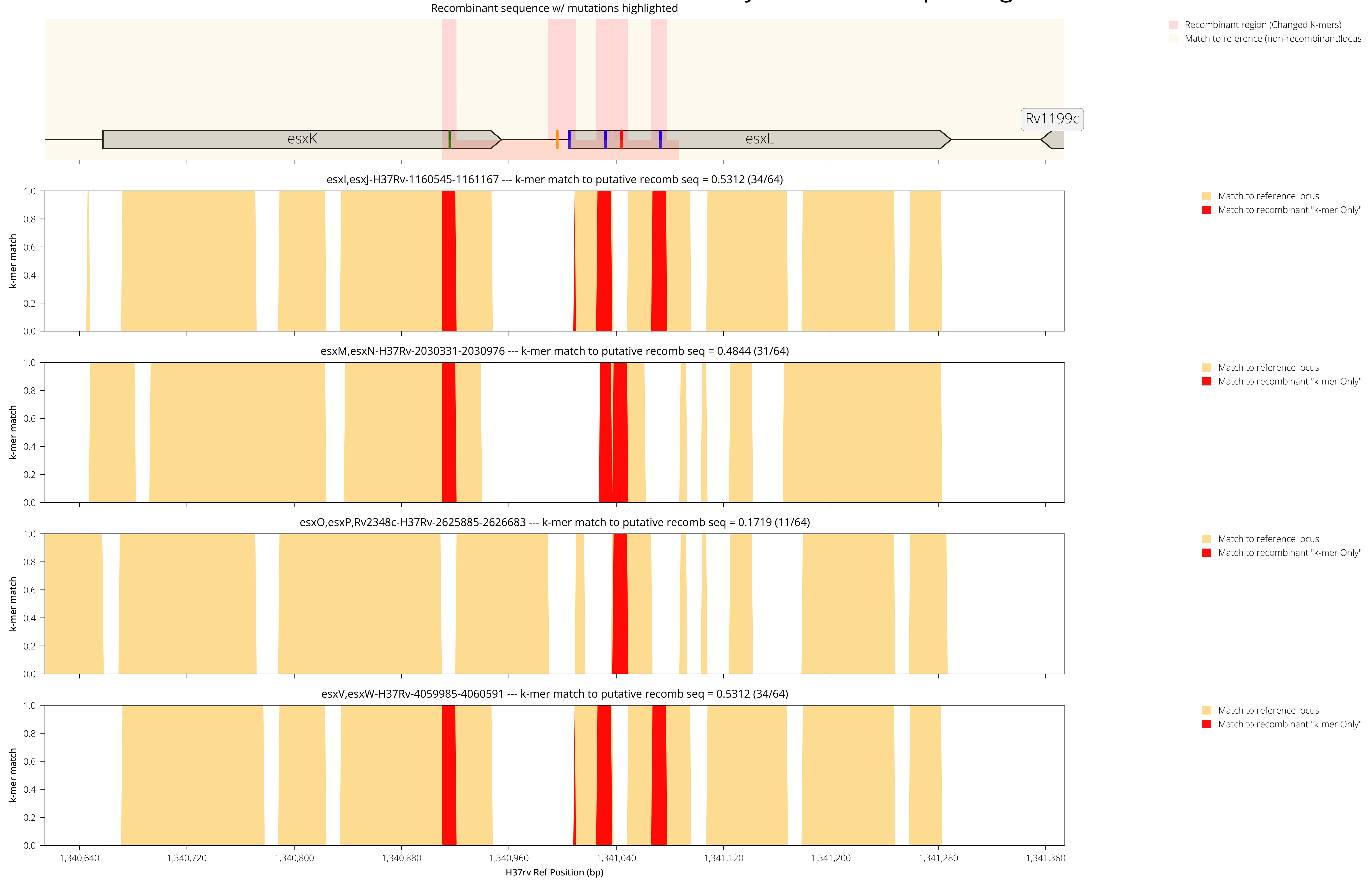

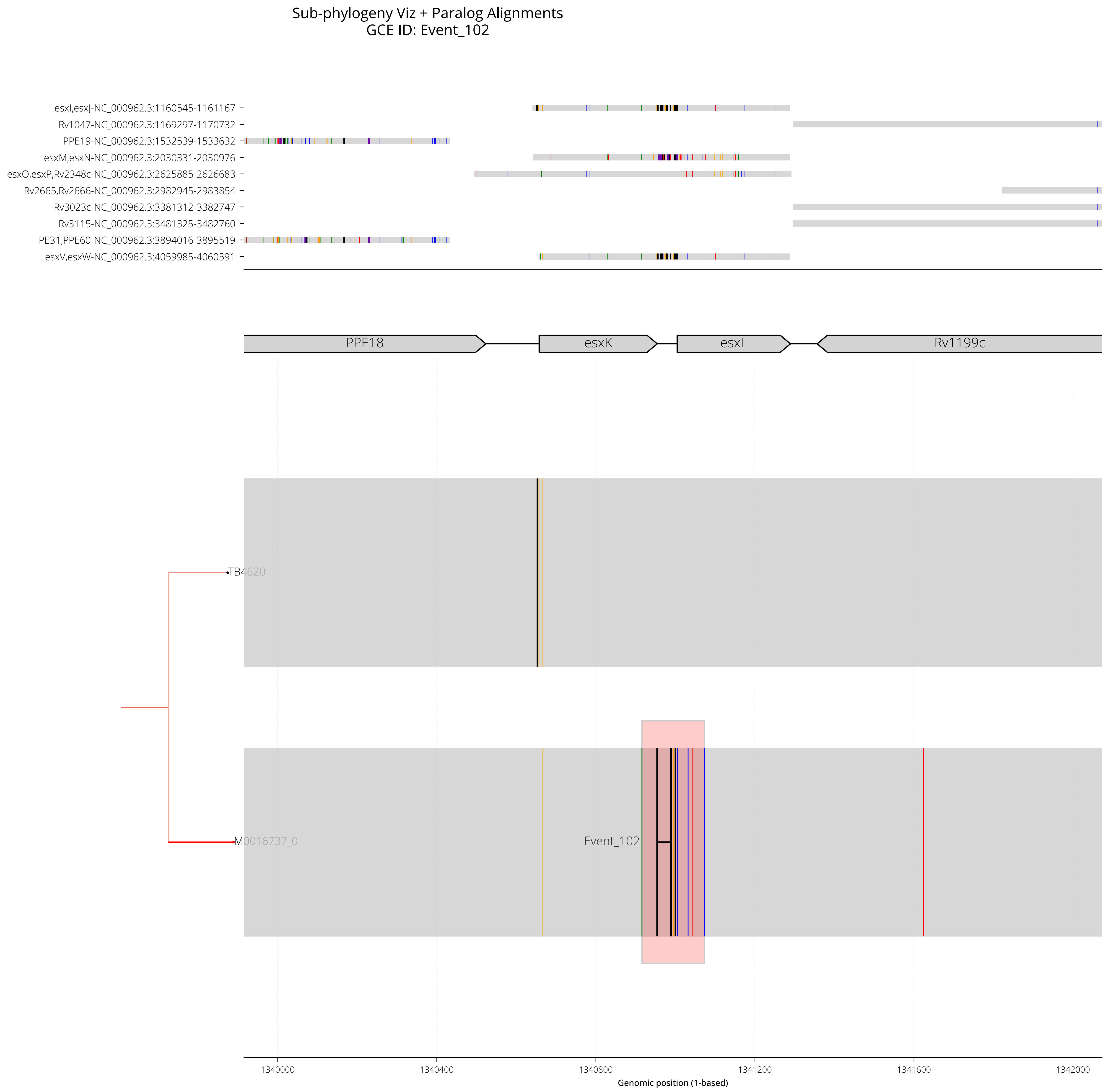

Sub-phylogeny Viz + Paralog Alignments
GCE ID: Event_102
esxI,esxJ-NC_000962.3:1160545-1161167
Rv1047-NC_000962.3:1169297-1170732
PPE19-NC_000962.3:1532539-1533632
esxM,esxN-NC_000962.3:2030331-2030976
esxO,esxP,Rv2348c-NC_000962.3:2625885-2626683
Rv2665,Rv2666-NC_000962.3:2982945-2983854
Rv3023c-NC_000962.3:3381312-3382747
Rv3115-NC_000962.3:3481325-3482760
PE31,PPE60-NC_000962.3:3894016-3895519
esxV,esxW-NC_000962.3:4059985-4060591
PPE18
esxK
esxL
Rv1199c
TB4620
M0016737_0
Event_102
1340000
1340400
1340800
1341200
1341600
1342000
Genomic position (1-based)

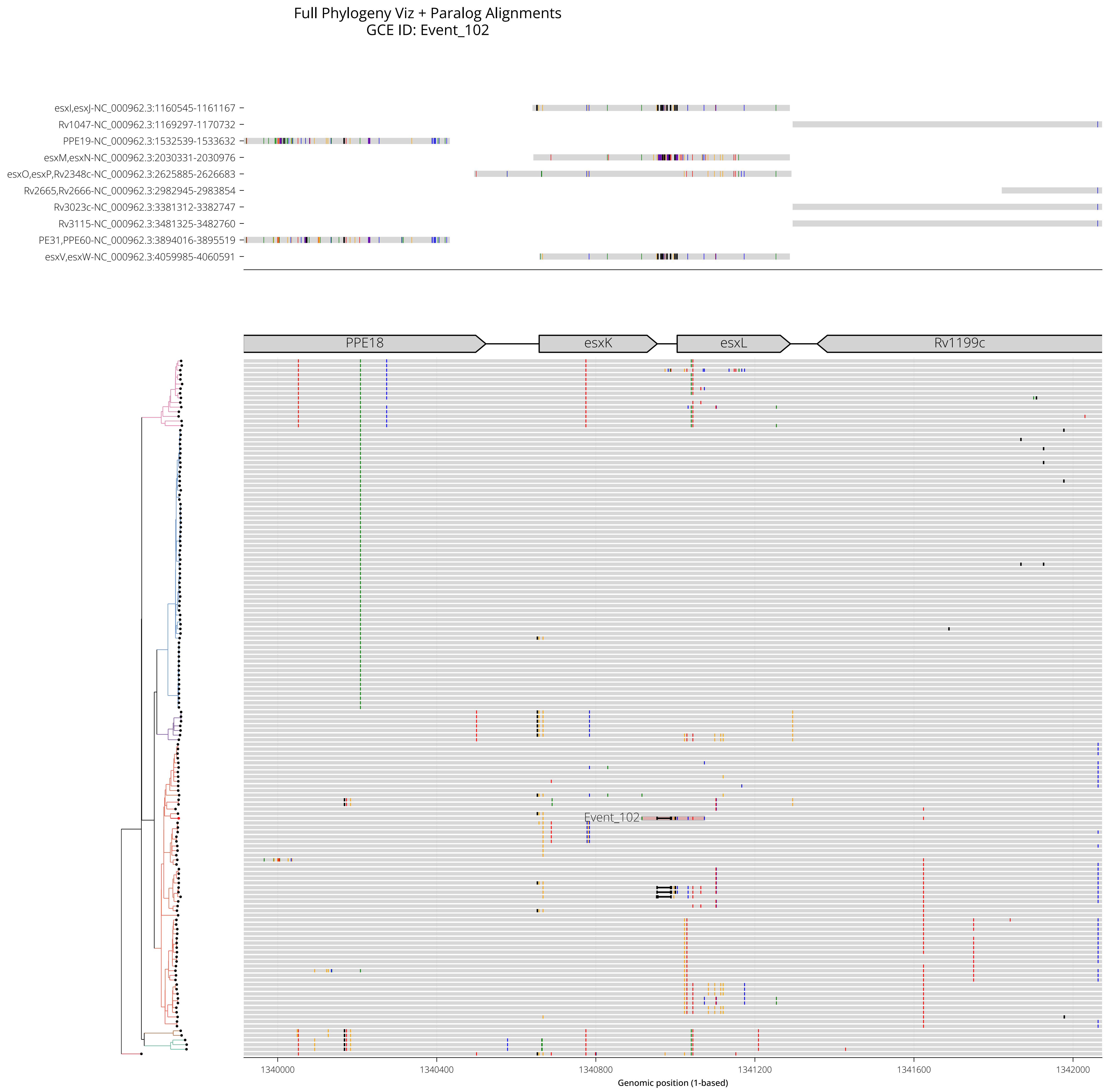

Full Phylogeny Viz + Paralog Alignments
GCE ID: Event_102
esxI,esxJ-NC_000962.3:1160545-1161167
Rv1047-NC_000962.3:1169297-1170732
PPE19-NC_000962.3:1532539-1533632
esxM,esxN-NC_000962.3:2030331-2030976
esxO,esxP,Rv2348c-NC_000962.3:2625885-2626683
Rv2665,Rv2666-NC_000962.3:2982945-2983854
Rv3023c-NC_000962.3:3381312-3382747
Rv3115-NC_000962.3:3481325-3482760
PE31,PPE60-NC_000962.3:3894016-3895519
esxV,esxW-NC_000962.3:4059985-4060591
PPE18
esxK
esxL
Rv1199c
Event_102
1340000
1340400
1340800
1341200
1341600
1342000
Genomic position (1-based)

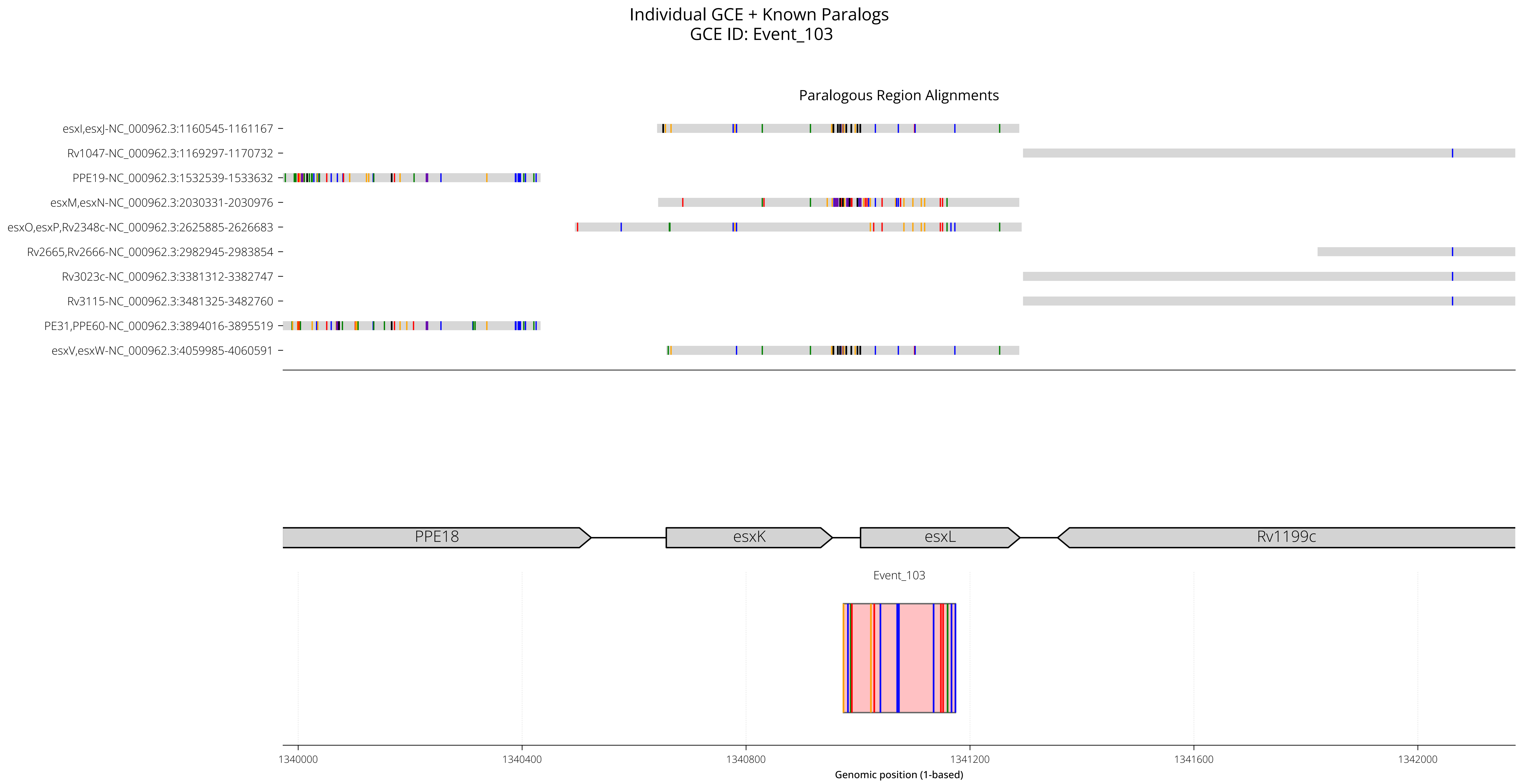
Individual GCE + Known Paralogs
GCE ID: Event_103
Paralogous Region Alignments
esxI,esxJ-NC_000962.3:1160545-1161167
Rv1047-NC_000962.3:1169297-1170732
PPE19-NC_000962.3:1532539-1533632
esxM,esxN-NC_000962.3:2030331-2030976
esxO,esxP,Rv2348c-NC_000962.3:2625885-2626683
Rv2665,Rv2666-NC_000962.3:2982945-2983854
Rv3023c-NC_000962.3:3381312-3382747
Rv3115-NC_000962.3:3481325-3482760
PE31,PPE60-NC_000962.3:3894016-3895519
esxV,esxW-NC_000962.3:4059985-4060591
PPE18
esxK
esxL
Rv1199c
Event_103
1340000
1340400
1340800
1341200
1341600
1342000
Genomic position (1-based)

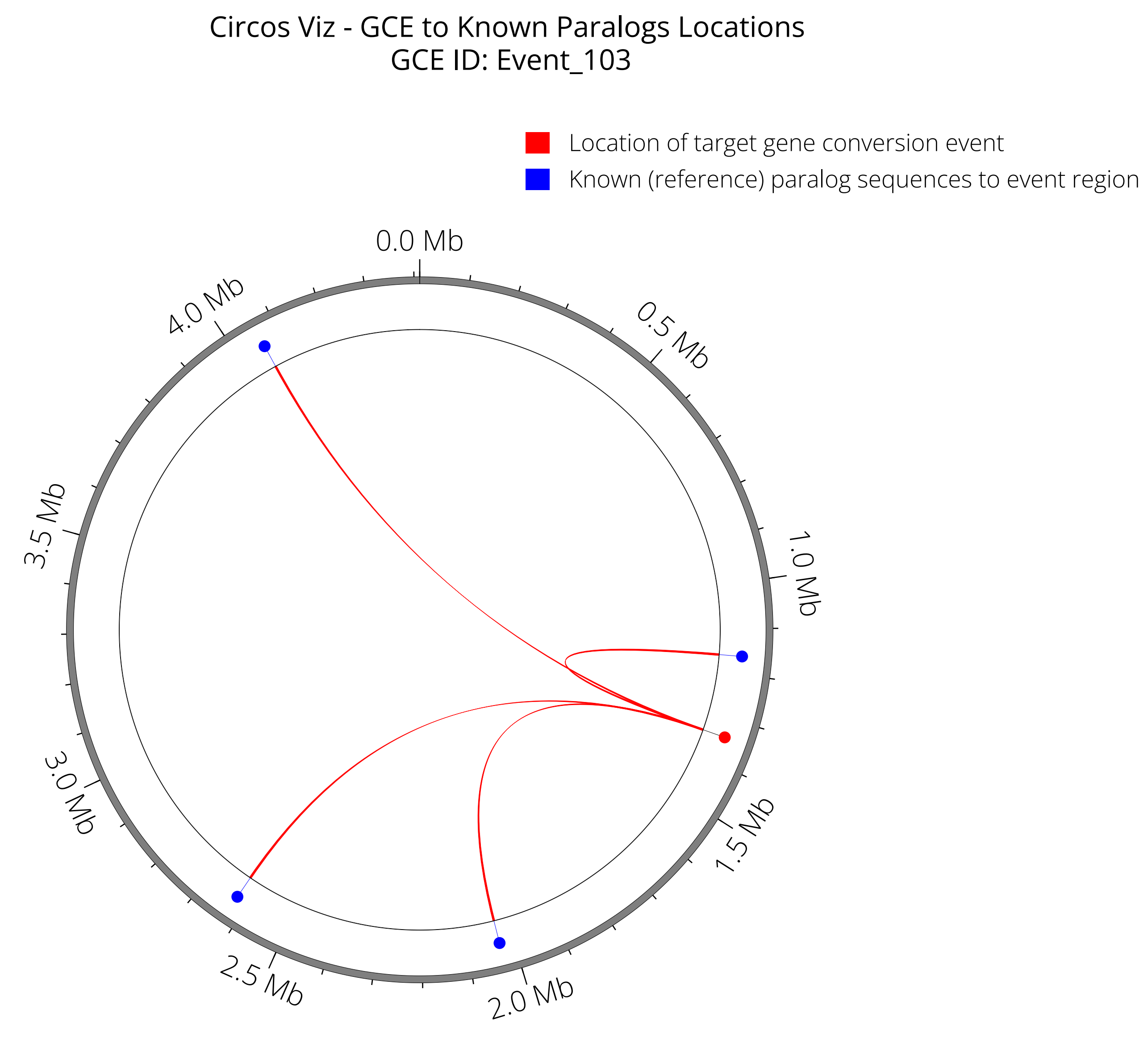
Circos Viz - GCE to Known Paralogs Locations
GCE ID: Event_103
Location of target gene conversion event
Known (reference) paralog sequences to event region
0.0 Mb
0.5 Mb
1.0 Mb
1.5 Mb
2.0 Mb
2.5 Mb
3.0 Mb
3.5 Mb
4.0 Mb

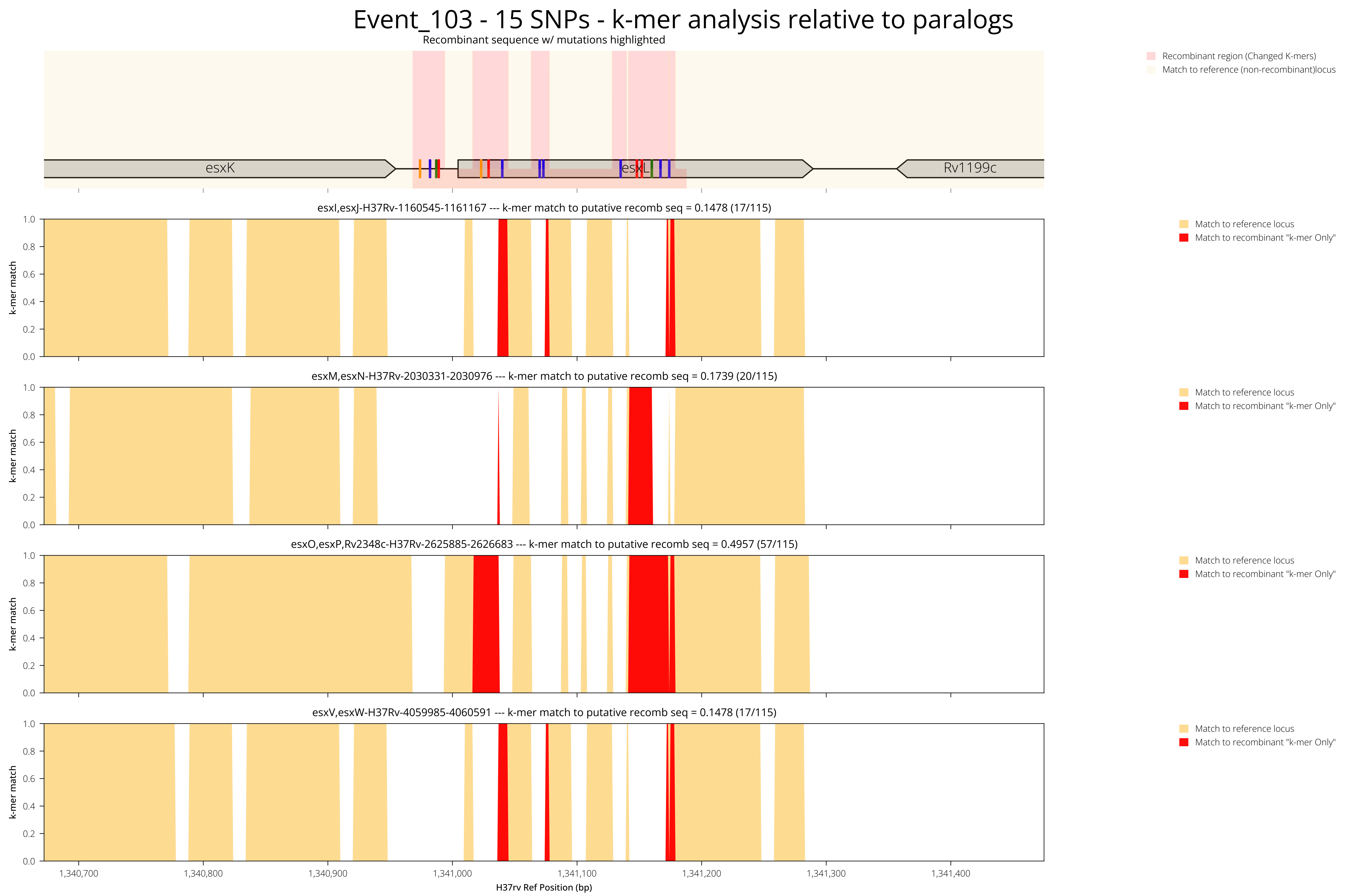
Event_103 - 15 SNPs - k-mer analysis relative to paralogs
Recombinant sequence w/ mutations highlighted
Recombinant region (Changed K-mers)
Match to reference (non-recombinant)locus
esxK
esxL
Rv1199c
esxI,esxJ-H37Rv-1160545-1161167 --- k-mer match to putative recomb seq = 0.1478 (17/115)
esxM,esxN-H37Rv-2030331-2030976 --- k-mer match to putative recomb seq = 0.1739 (20/115)
esxO,esxP,Rv2348c-H37Rv-2625885-2626683 --- k-mer match to putative recomb seq = 0.4957 (57/115)
esxV,esxW-H37Rv-4059985-4060591 --- k-mer match to putative recomb seq = 0.1478 (17/115)
Match to reference locus
Match to recombinant "k-mer Only"
k-mer match
1.0
0.8
0.6
0.4
0.2
0.0
1,340,700
1,340,800
1,340,900
1,341,000
1,341,100
1,341,200
1,341,300
1,341,400
H37rv Ref Position (bp)

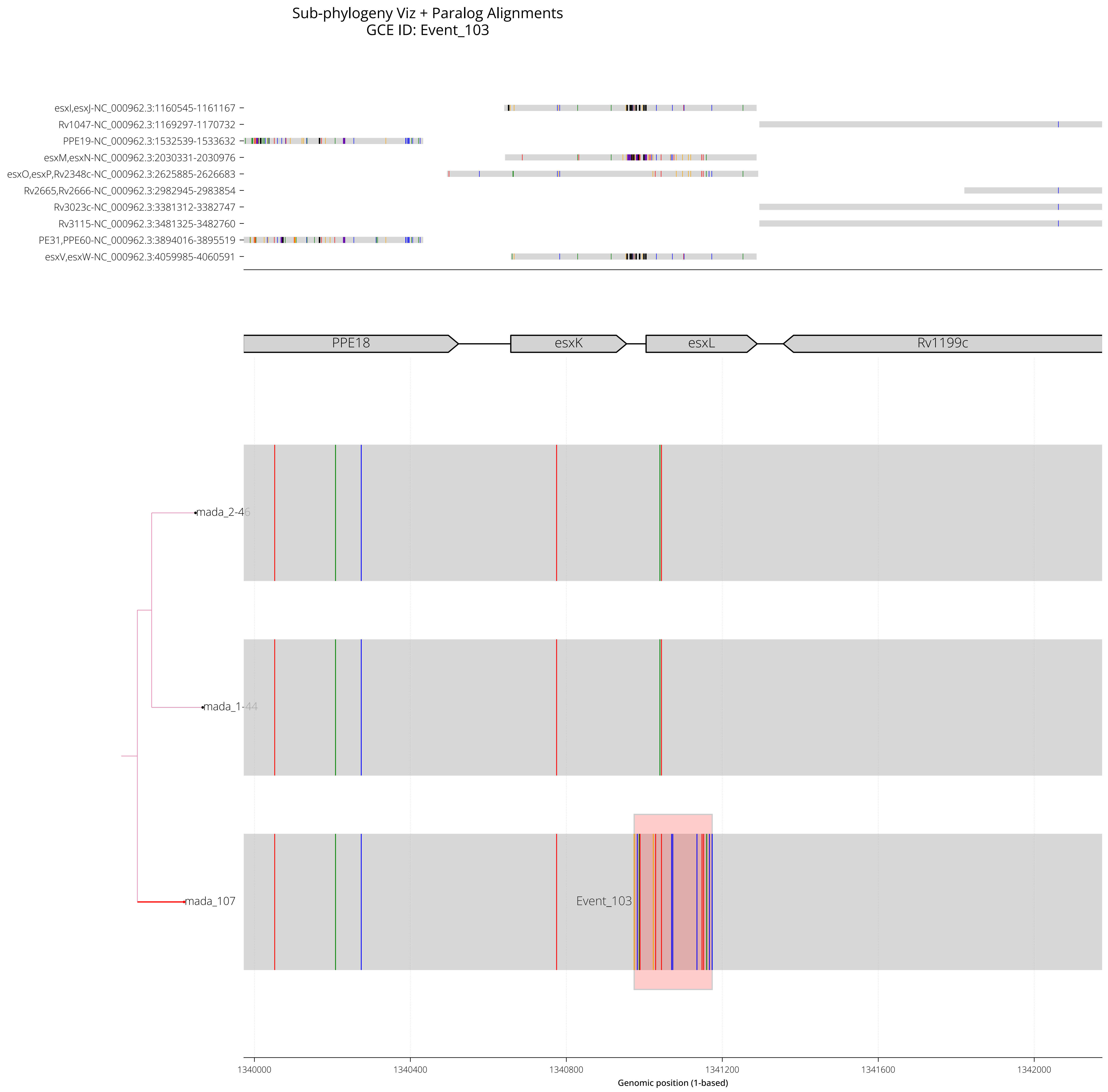

Sub-phylogeny Viz + Paralog Alignments
GCE ID: Event_103
esxI,esxJ-NC_000962.3:1160545-1161167
Rv1047-NC_000962.3:1169297-1170732
PPE19-NC_000962.3:1532539-1533632
esxM,esxN-NC_000962.3:2030331-2030976
esxO,esxP,Rv2348c-NC_000962.3:2625885-2626683
Rv2665,Rv2666-NC_000962.3:2982945-2983854
Rv3023c-NC_000962.3:3381312-3382747
Rv3115-NC_000962.3:3481325-3482760
PE31,PPE60-NC_000962.3:3894016-3895519
esxV,esxW-NC_000962.3:4059985-4060591
PPE18
esxK
esxL
Rv1199c
mada_2-46
mada_1-44
mada_107
Event_103
1340000
1340400
1340800
1341200
1341600
1342000
Genomic position (1-based)

Full Phylogeny Viz + Paralog Alignments
GCE ID: Event_103

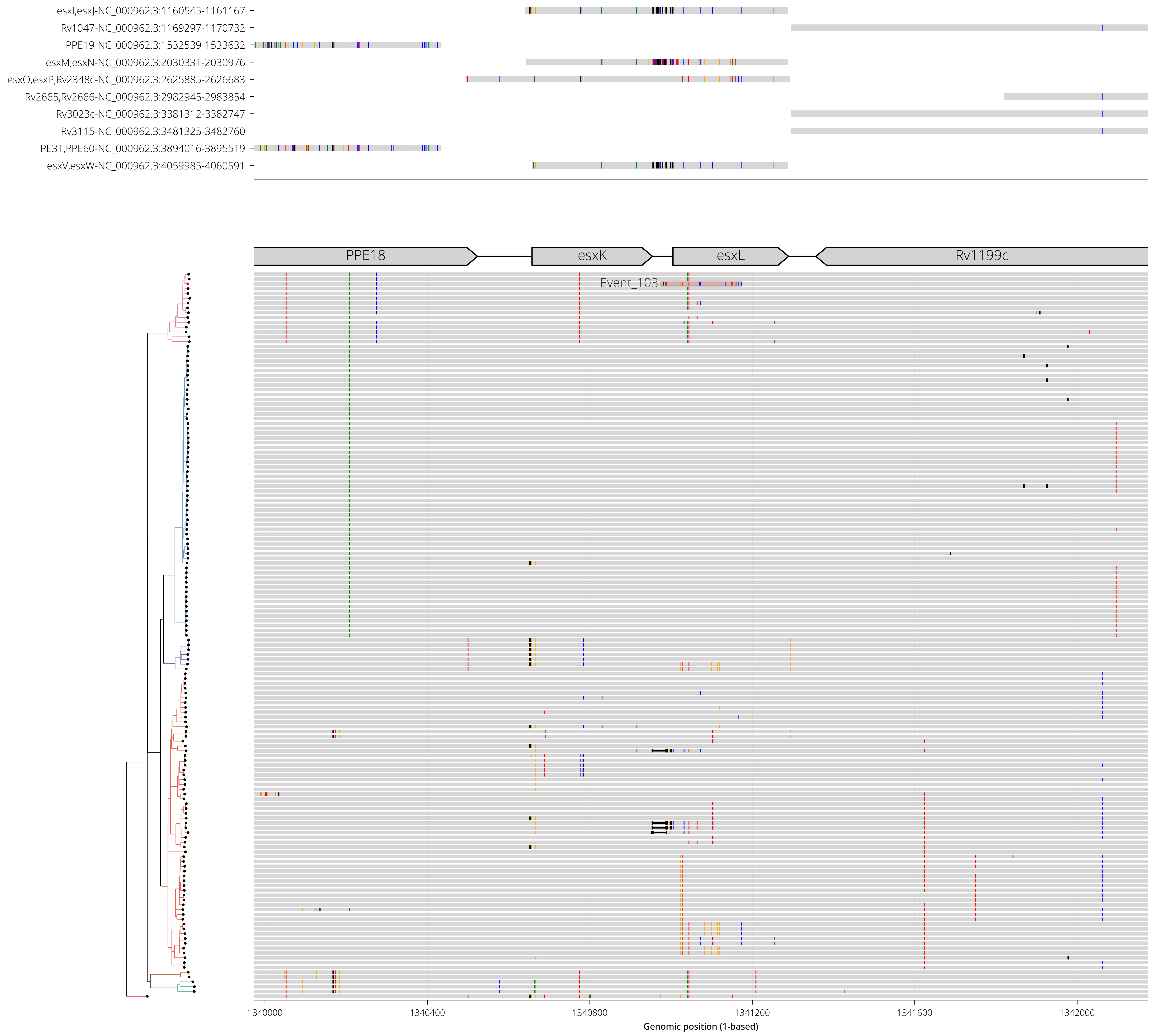

Individual GCE + Known Paralogs
GCE ID: Event_104

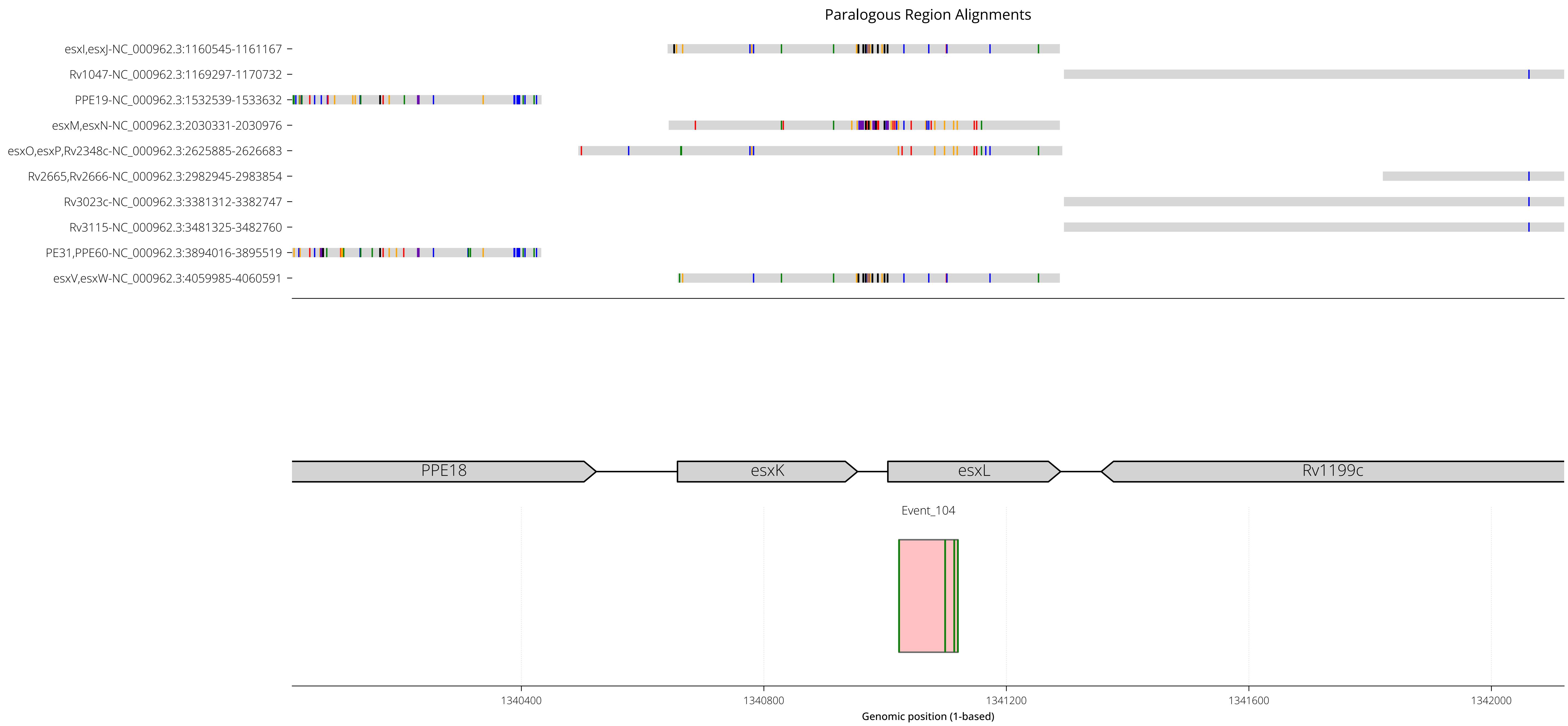

Circos Viz - GCE to Known Paralogs Locations
GCE ID: Event_104

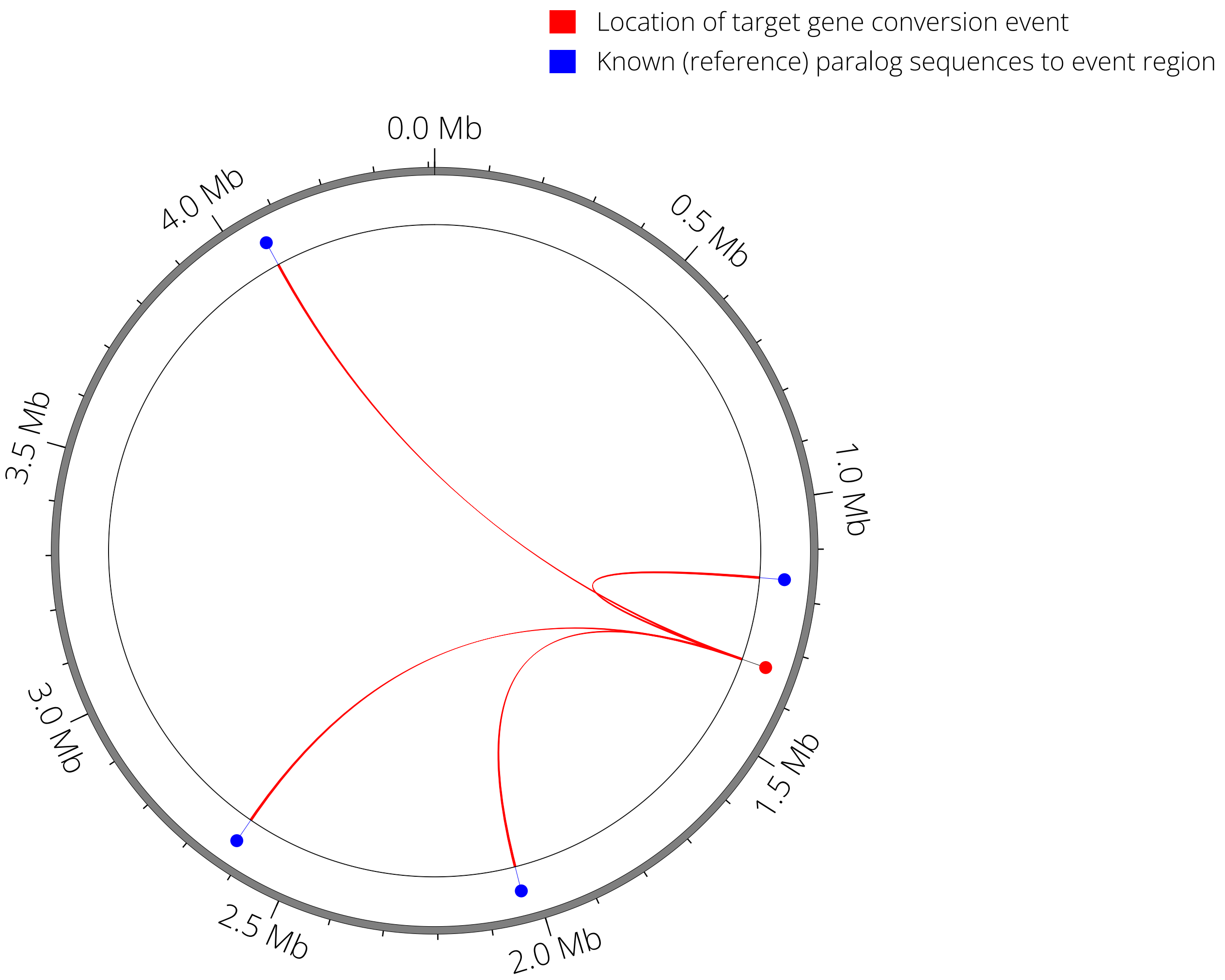

### Event_104 - 4 SNPs - k-mer analysis relative to paralogs

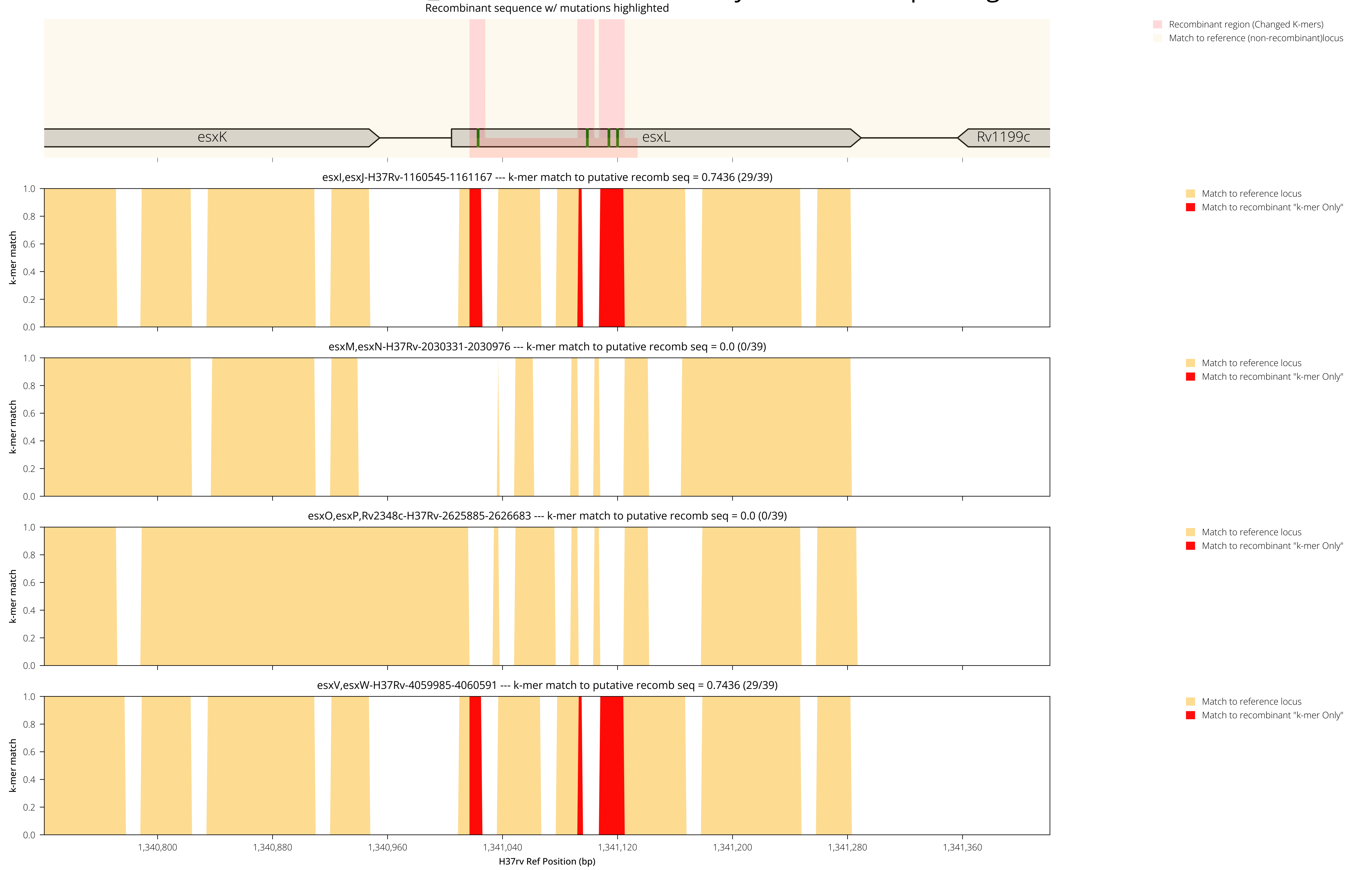

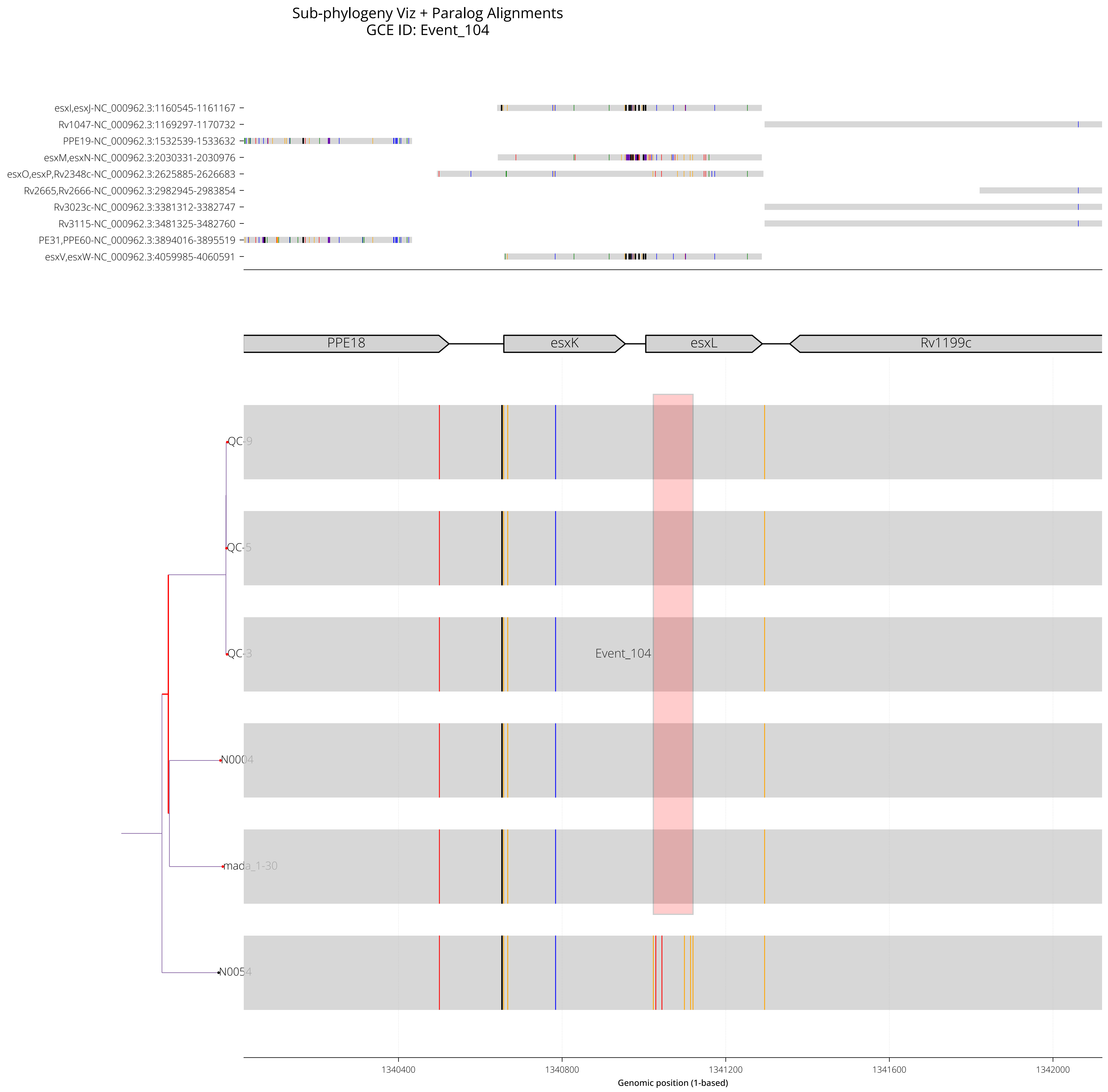

Sub-phylogeny Viz + Paralog Alignments
GCE ID: Event_104
esxI,esxJ-NC_000962.3:1160545-1161167
Rv1047-NC_000962.3:1169297-1170732
PPE19-NC_000962.3:1532539-1533632
esxM,esxN-NC_000962.3:2030331-2030976
esxO,esxP,Rv2348c-NC_000962.3:2625885-2626683
Rv2665,Rv2666-NC_000962.3:2982945-2983854
Rv3023c-NC_000962.3:3381312-3382747
Rv3115-NC_000962.3:3481325-3482760
PE31,PPE60-NC_000962.3:3894016-3895519
esxV,esxW-NC_000962.3:4059985-4060591
PPE18
esxK
esxL
Rv1199c
QC-9
QC-5
QC-3
Event_104
N0004
mada_1-30
N0054
1340400
1340800
1341200
1341600
1342000
Genomic position (1-based)

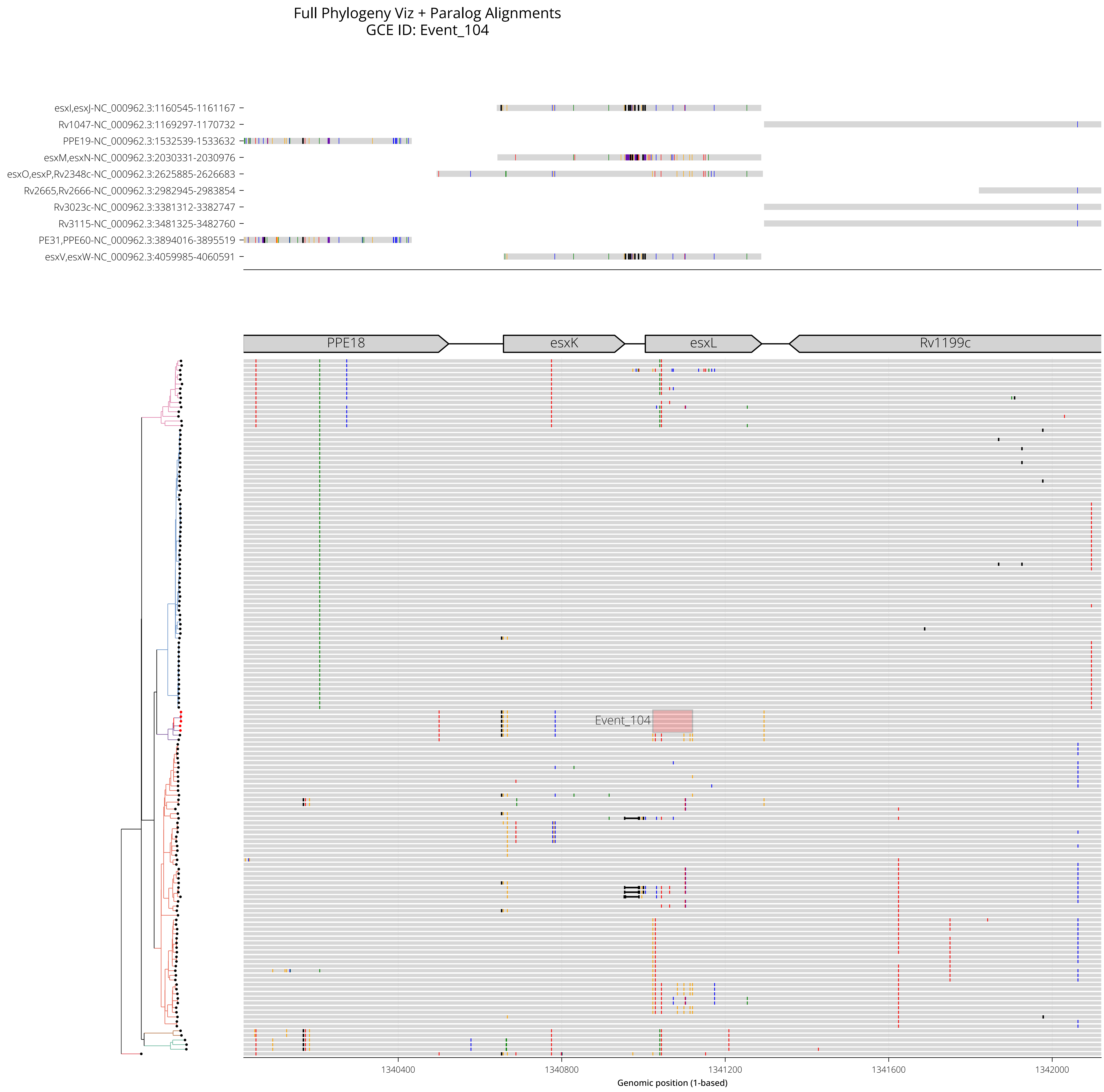

Full Phylogeny Viz + Paralog Alignments
GCE ID: Event_104
esxI,esxJ-NC_000962.3:1160545-1161167
Rv1047-NC_000962.3:1169297-1170732
PPE19-NC_000962.3:1532539-1533632
esxM,esxN-NC_000962.3:2030331-2030976
esxO,esxP,Rv2348c-NC_000962.3:2625885-2626683
Rv2665,Rv2666-NC_000962.3:2982945-2983854
Rv3023c-NC_000962.3:3381312-3382747
Rv3115-NC_000962.3:3481325-3482760
PE31,PPE60-NC_000962.3:3894016-3895519
esxV,esxW-NC_000962.3:4059985-4060591
PPE18
esxK
esxL
Rv1199c
Event_104
1340400
1340800
1341200
1341600
1342000
Genomic position (1-based)

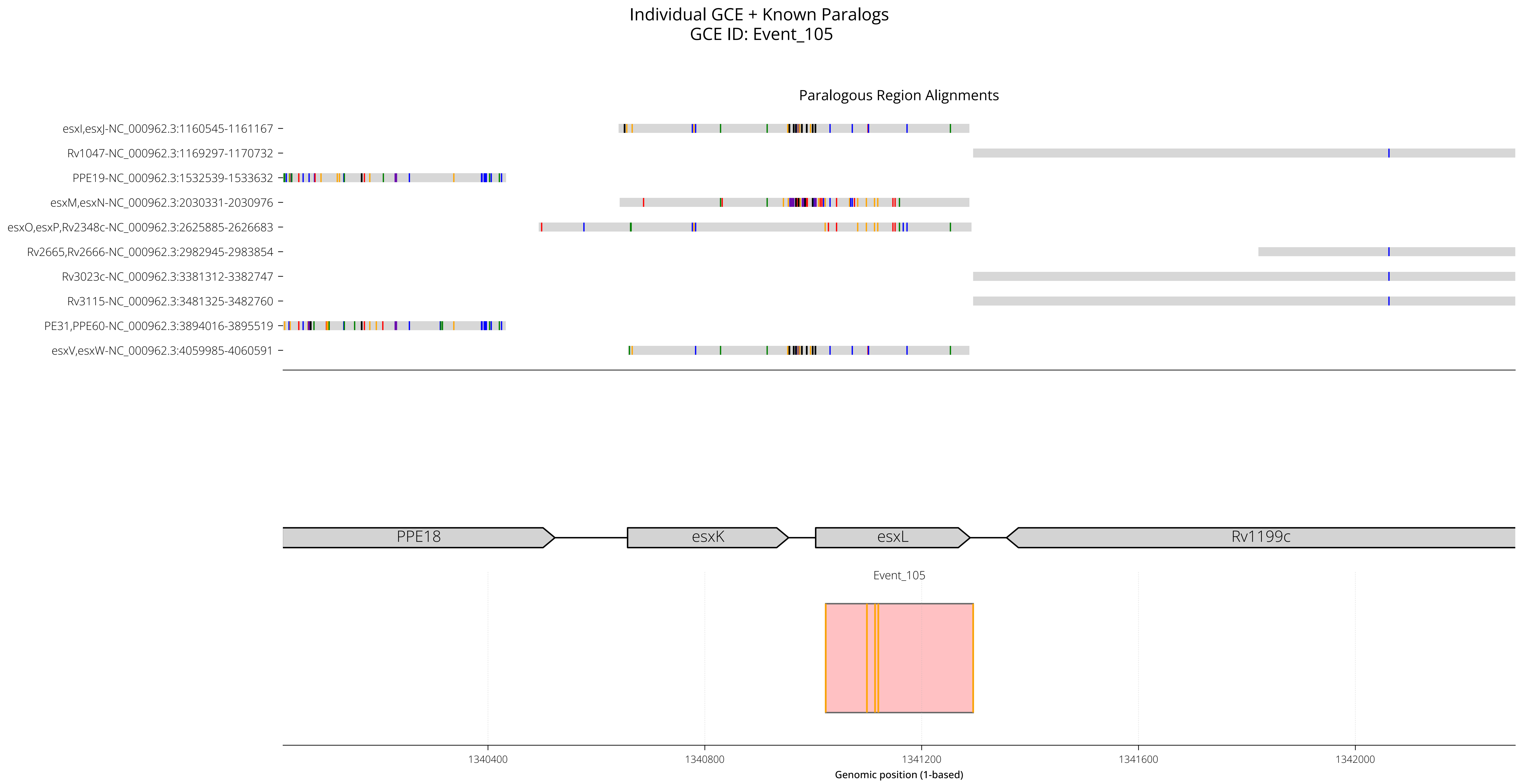

Individual GCE + Known Paralogs
GCE ID: Event_105
Paralogous Region Alignments
esxI,esxJ-NC_000962.3:1160545-1161167
Rv1047-NC_000962.3:1169297-1170732
PPE19-NC_000962.3:1532539-1533632
esxM,esxN-NC_000962.3:2030331-2030976
esxO,esxP,Rv2348c-NC_000962.3:2625885-2626683
Rv2665,Rv2666-NC_000962.3:2982945-2983854
Rv3023c-NC_000962.3:3381312-3382747
Rv3115-NC_000962.3:3481325-3482760
PE31,PPE60-NC_000962.3:3894016-3895519
esxV,esxW-NC_000962.3:4059985-4060591
PPE18
esxK
esxL
Rv1199c
Event_105
1340400
1340800
1341200
1341600
1342000
Genomic position (1-based)

Circos Viz - GCE to Known Paralogs Locations
GCE ID: Event_105

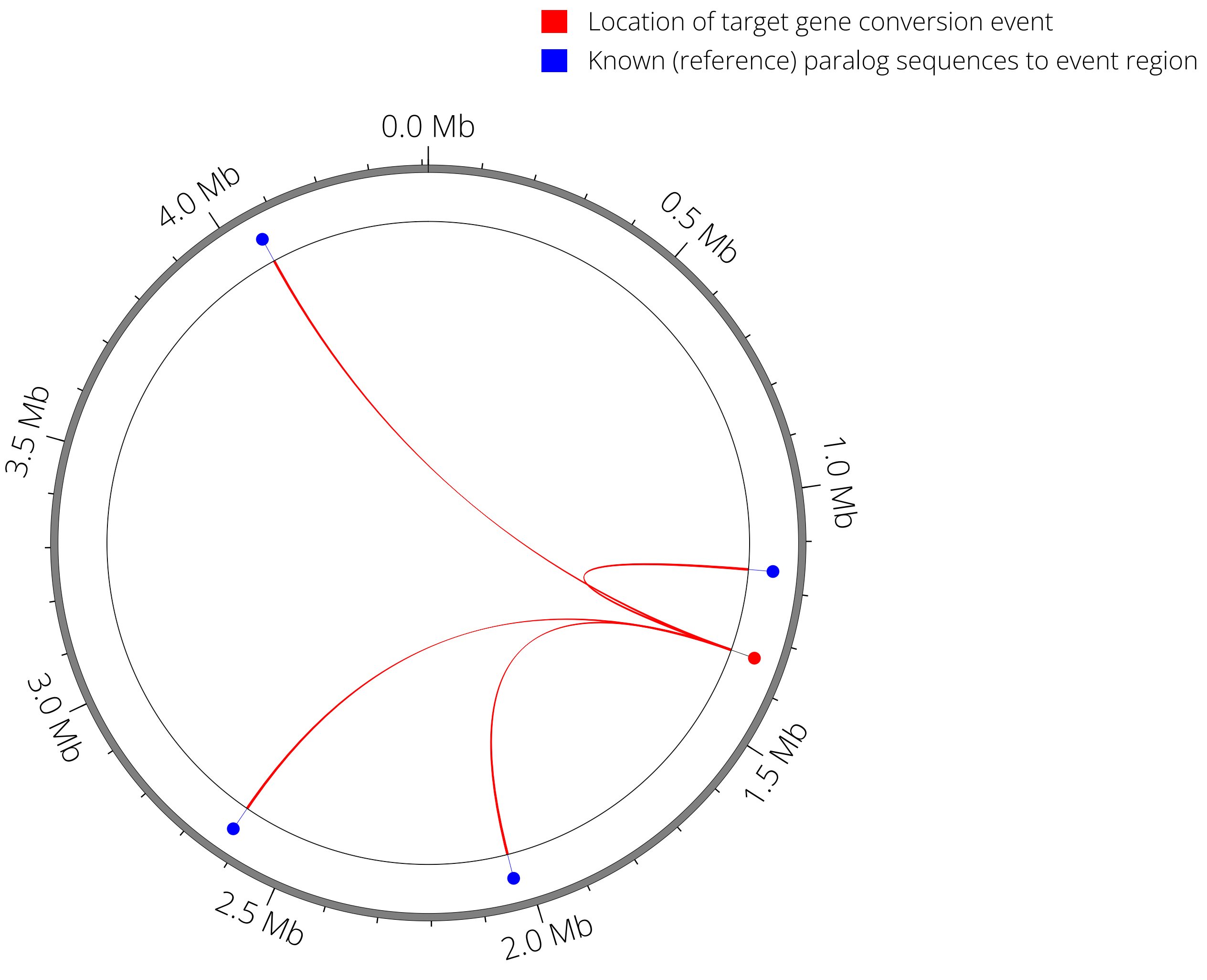

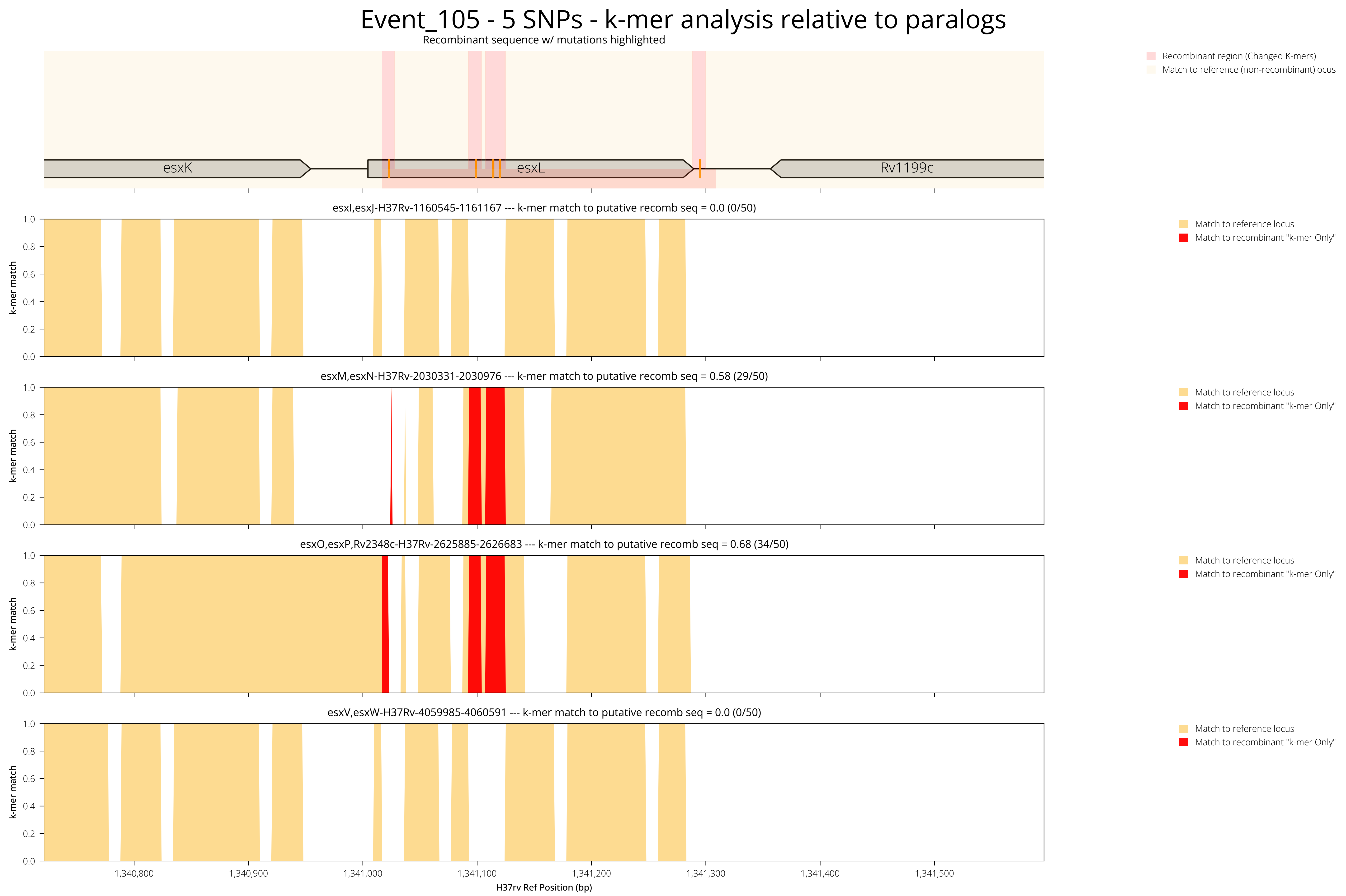

Event_105 - 5 SNPs - k-mer analysis relative to paralogs
Recombinant sequence w/ mutations highlighted
Recombinant region (Changed K-mers)
Match to reference (non-recombinant)locus
esxK
esxL
Rv1199c
esxI,esxJ-H37Rv-1160545-1161167 --- k-mer match to putative recomb seq = 0.0 (0/50)
esxM,esxN-H37Rv-2030331-2030976 --- k-mer match to putative recomb seq = 0.58 (29/50)
esxO,esxP,Rv2348c-H37Rv-2625885-2626683 --- k-mer match to putative recomb seq = 0.68 (34/50)
esxV,esxW-H37Rv-4059985-4060591 --- k-mer match to putative recomb seq = 0.0 (0/50)
Match to reference locus
Match to recombinant "k-mer Only"
k-mer match
1.0
0.8
0.6
0.4
0.2
0.0
1,340,800
1,340,900
1,341,000
1,341,100
1,341,200
1,341,300
1,341,400
1,341,500
H37rv Ref Position (bp)

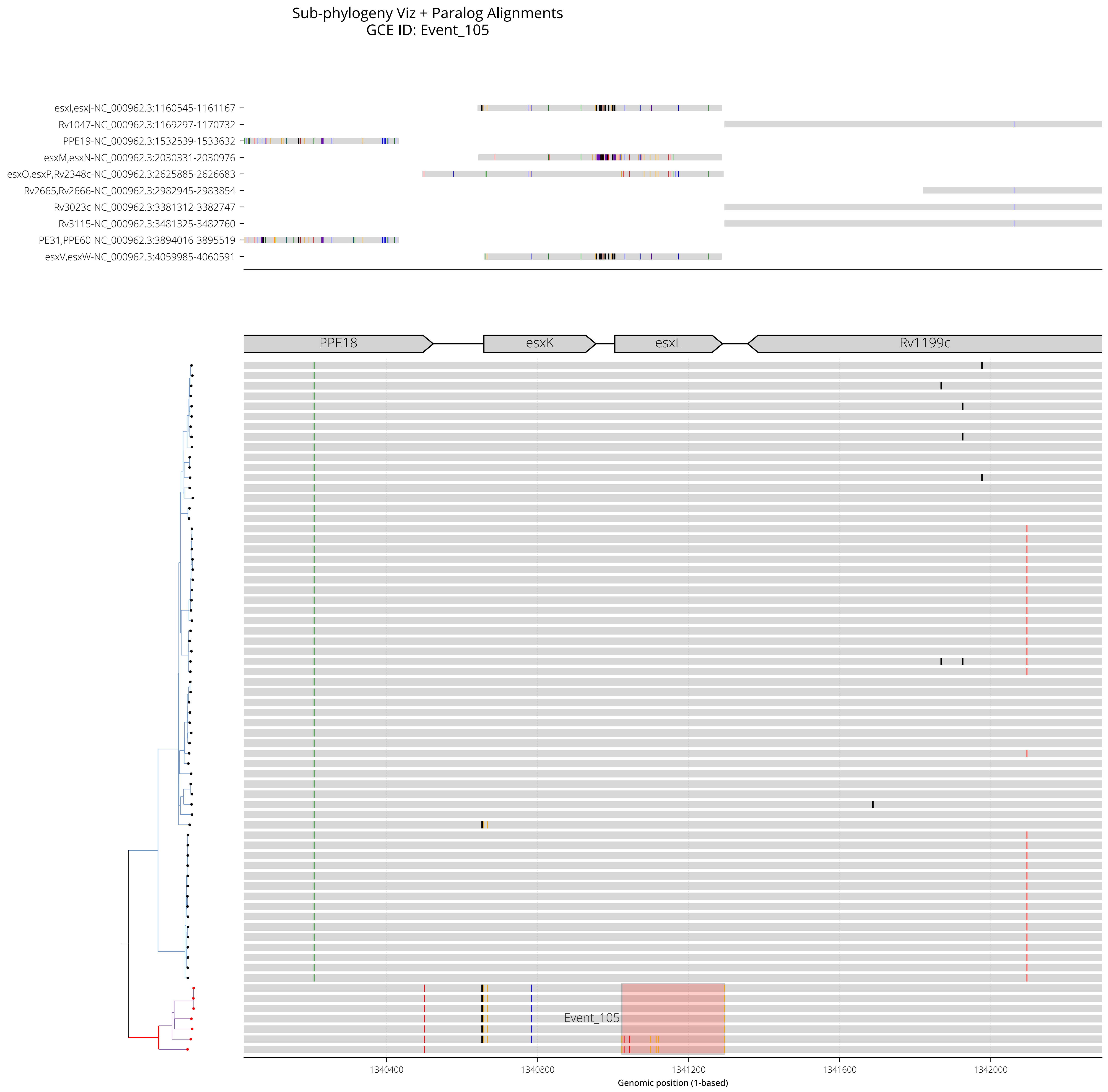

Sub-phylogeny Viz + Paralog Alignments
GCE ID: Event_105
esxI,esxJ-NC_000962.3:1160545-1161167
Rv1047-NC_000962.3:1169297-1170732
PPE19-NC_000962.3:1532539-1533632
esxM,esxN-NC_000962.3:2030331-2030976
esxO,esxP,Rv2348c-NC_000962.3:2625885-2626683
Rv2665,Rv2666-NC_000962.3:2982945-2983854
Rv3023c-NC_000962.3:3381312-3382747
Rv3115-NC_000962.3:3481325-3482760
PE31,PPE60-NC_000962.3:3894016-3895519
esxV,esxW-NC_000962.3:4059985-4060591
PPE18
esxK
esxL
Rv1199c
Event_105
1340400
1340800
1341200
1341600
1342000
Genomic position (1-based)

Full Phylogeny Viz + Paralog Alignments
GCE ID: Event_105

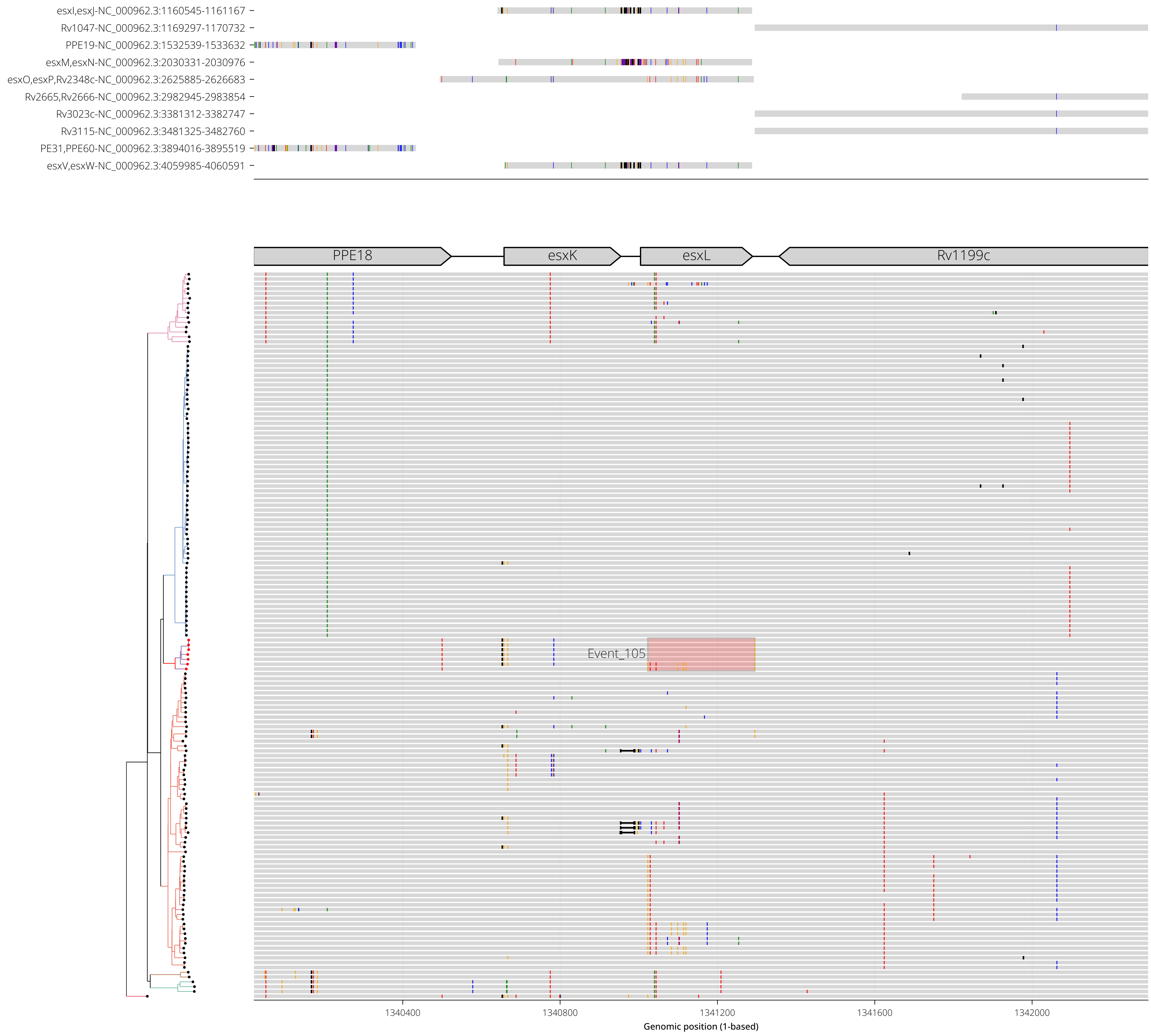

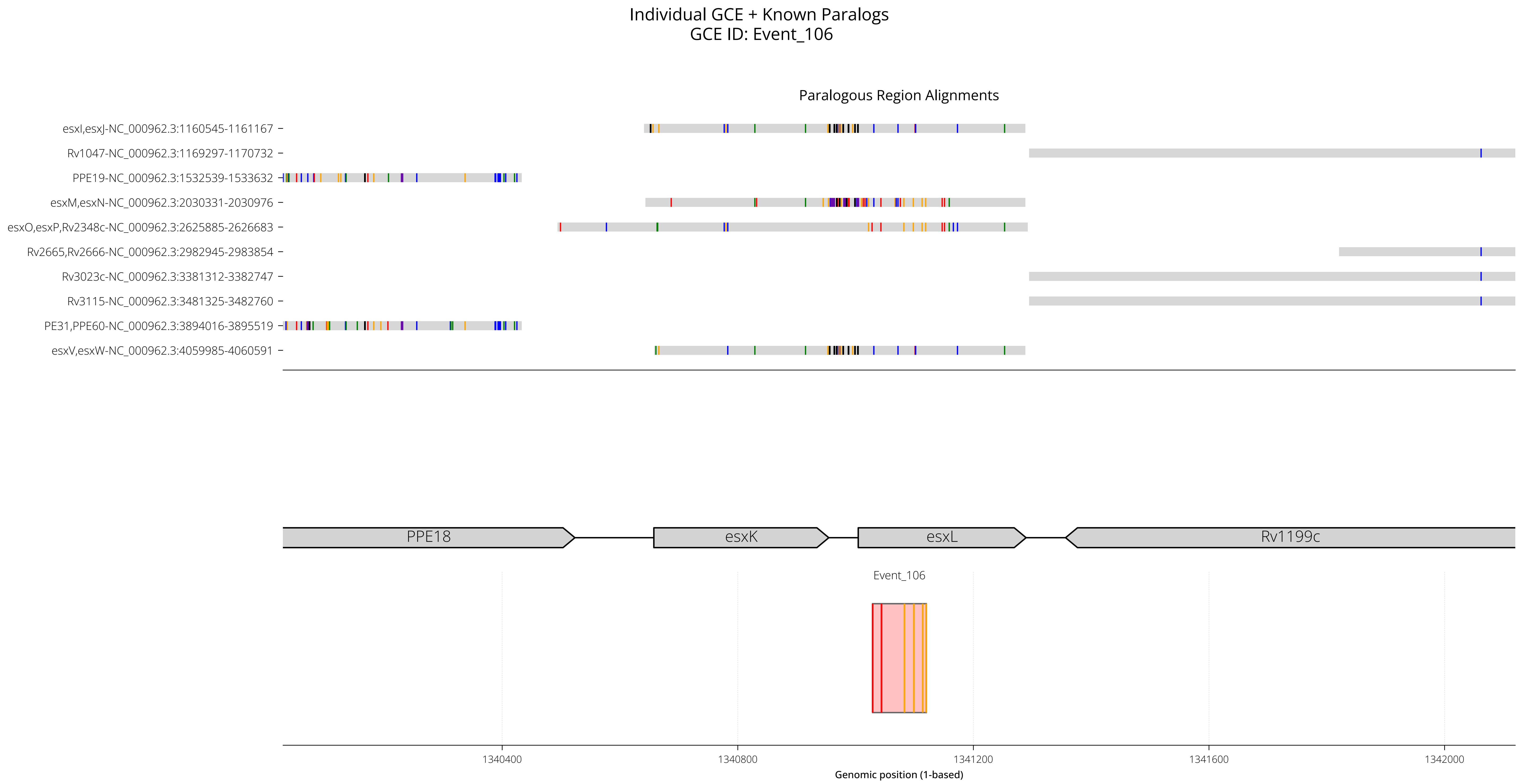
Individual GCE + Known Paralogs
GCE ID: Event_106
Paralogous Region Alignments
esxI,esxJ-NC_000962.3:1160545-1161167
Rv1047-NC_000962.3:1169297-1170732
PPE19-NC_000962.3:1532539-1533632
esxM,esxN-NC_000962.3:2030331-2030976
esxO,esxP,Rv2348c-NC_000962.3:2625885-2626683
Rv2665,Rv2666-NC_000962.3:2982945-2983854
Rv3023c-NC_000962.3:3381312-3382747
Rv3115-NC_000962.3:3481325-3482760
PE31,PPE60-NC_000962.3:3894016-3895519
esxV,esxW-NC_000962.3:4059985-4060591
PPE18
esxK
esxL
Rv1199c
Event_106
1340400
1340800
1341200
1341600
1342000
Genomic position (1-based)

Circos Viz - GCE to Known Paralogs Locations
GCE ID: Event_106

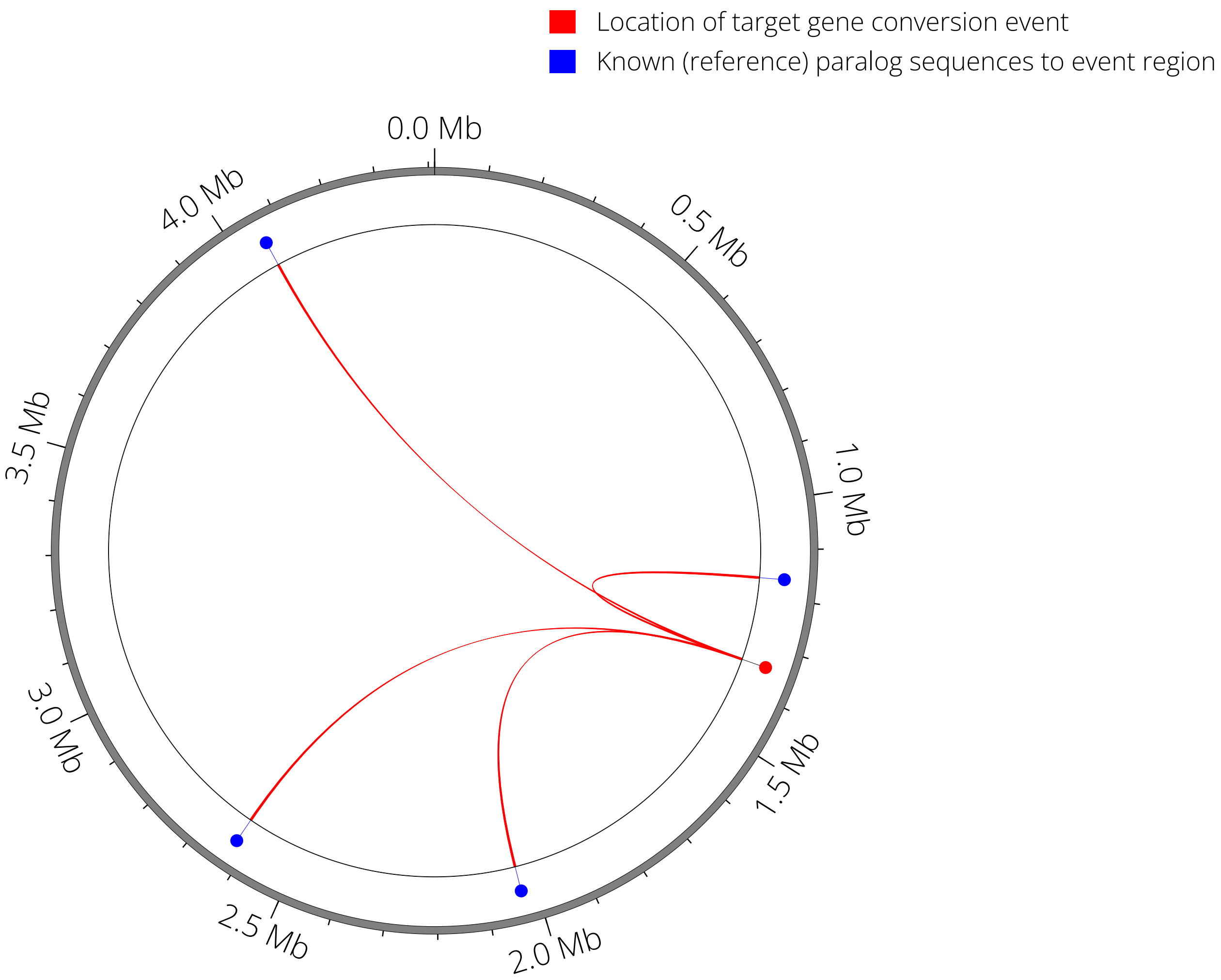

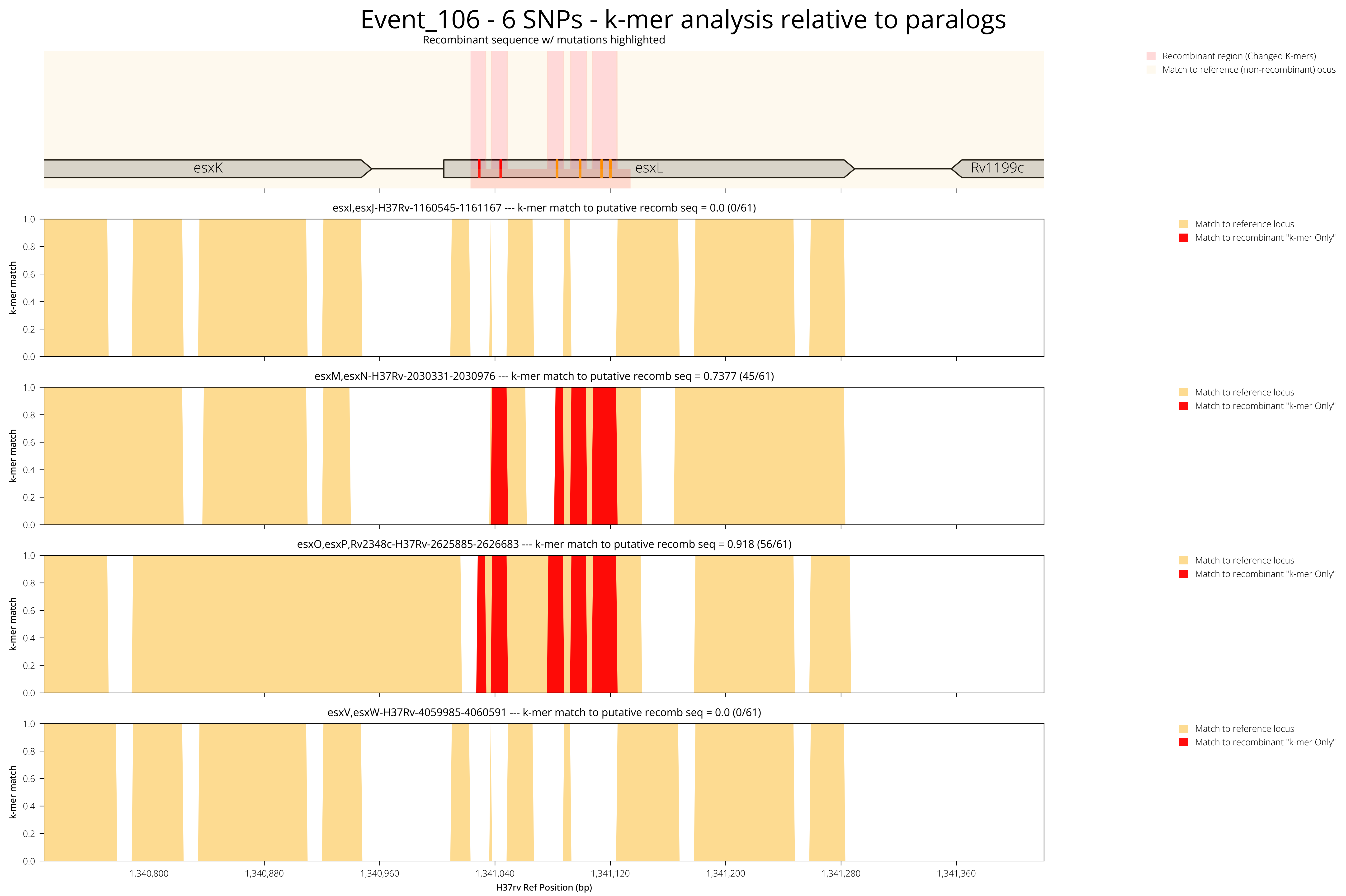

Event_106 - 6 SNPs - k-mer analysis relative to paralogs
Recombinant sequence w/ mutations highlighted
Recombinant region (Changed K-mers)
Match to reference (non-recombinant)locus
esxK
esxL
Rv1199c
esxI,esxJ-H37Rv-1160545-1161167 --- k-mer match to putative recomb seq = 0.0 (0/61)
esxM,esxN-H37Rv-2030331-2030976 --- k-mer match to putative recomb seq = 0.7377 (45/61)
esxO,esxP,Rv2348c-H37Rv-2625885-2626683 --- k-mer match to putative recomb seq = 0.918 (56/61)
esxV,esxW-H37Rv-4059985-4060591 --- k-mer match to putative recomb seq = 0.0 (0/61)
Match to reference locus
Match to recombinant "k-mer Only"
k-mer match
1.0
0.8
0.6
0.4
0.2
0.0
1,340,800
1,340,880
1,340,960
1,341,040
1,341,120
1,341,200
1,341,280
1,341,360
H37rv Ref Position (bp)

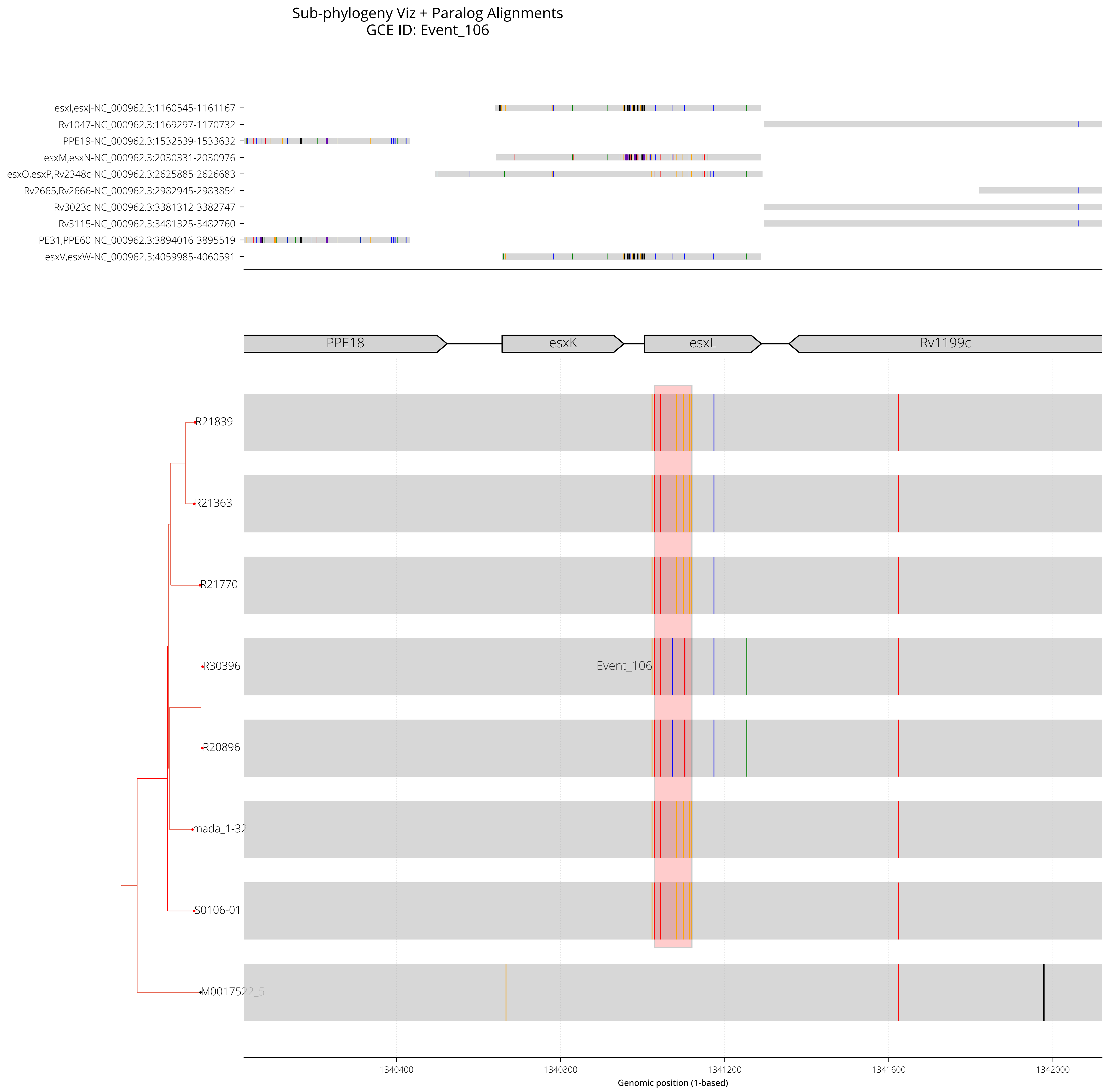

Sub-phylogeny Viz + Paralog Alignments
GCE ID: Event_106
esxI,esxJ-NC_000962.3:1160545-1161167
Rv1047-NC_000962.3:1169297-1170732
PPE19-NC_000962.3:1532539-1533632
esxM,esxN-NC_000962.3:2030331-2030976
esxO,esxP,Rv2348c-NC_000962.3:2625885-2626683
Rv2665,Rv2666-NC_000962.3:2982945-2983854
Rv3023c-NC_000962.3:3381312-3382747
Rv3115-NC_000962.3:3481325-3482760
PE31,PPE60-NC_000962.3:3894016-3895519
esxV,esxW-NC_000962.3:4059985-4060591
PPE18
esxK
esxL
Rv1199c
R21839
R21363
R21770
R30396
Event_106
R20896
mada_1-32
S0106-01
M0017522_5
1340400
1340800
1341200
1341600
1342000
Genomic position (1-based)

Full Phylogeny Viz + Paralog Alignments
GCE ID: Event_106

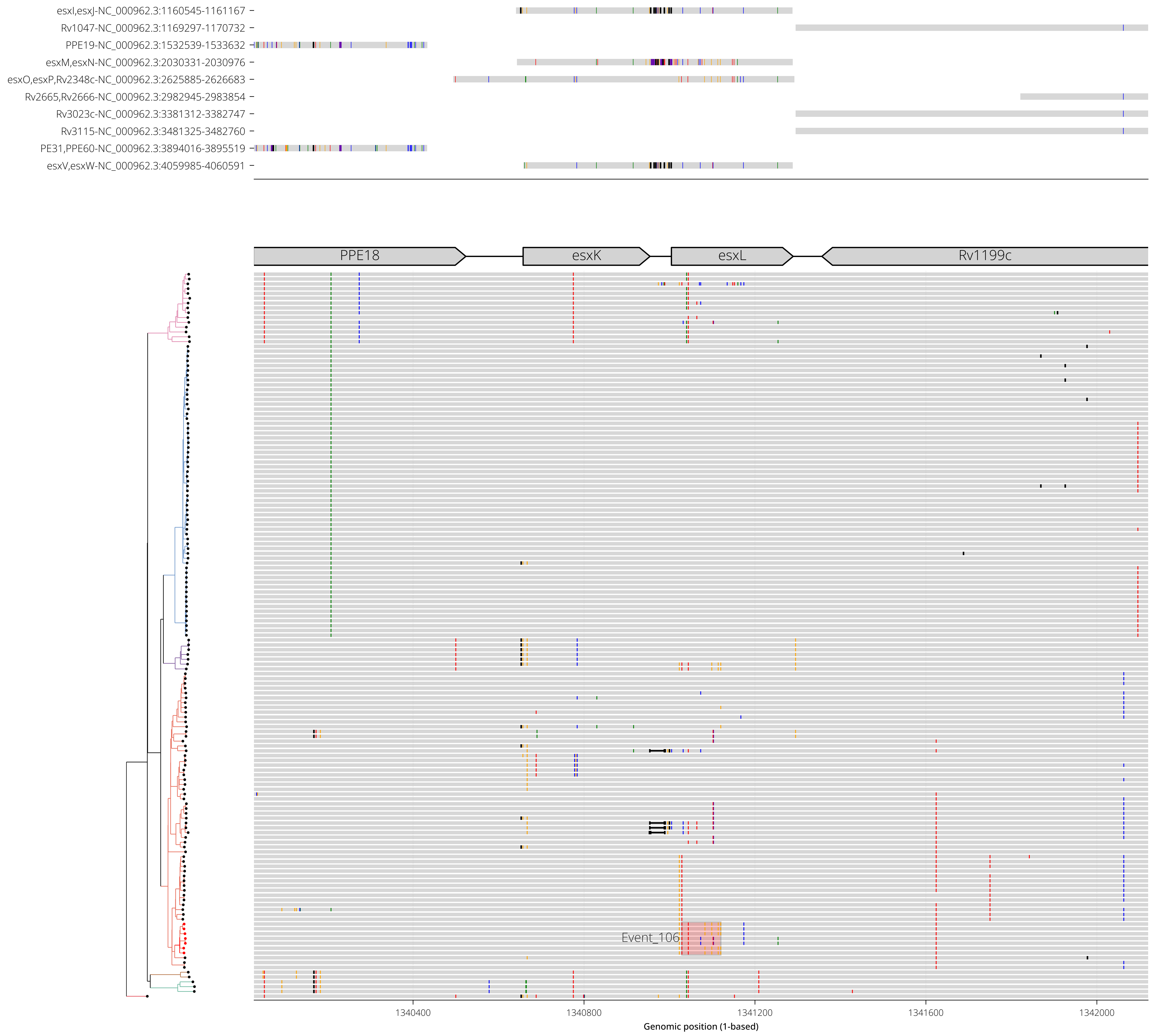

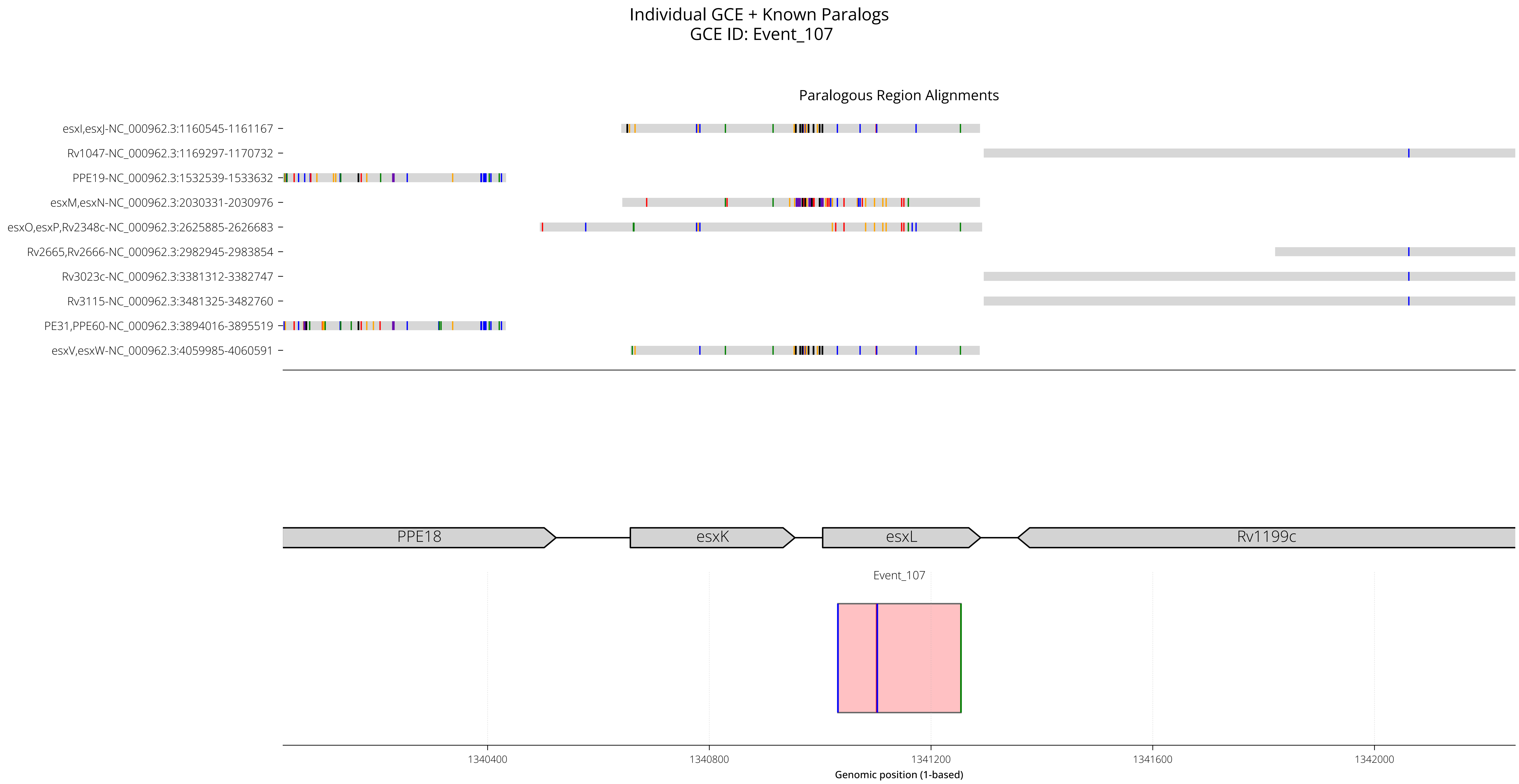
Individual GCE + Known Paralogs
GCE ID: Event_107
Paralogous Region Alignments
esxI,esxJ-NC_000962.3:1160545-1161167
Rv1047-NC_000962.3:1169297-1170732
PPE19-NC_000962.3:1532539-1533632
esxM,esxN-NC_000962.3:2030331-2030976
esxO,esxP,Rv2348c-NC_000962.3:2625885-2626683
Rv2665,Rv2666-NC_000962.3:2982945-2983854
Rv3023c-NC_000962.3:3381312-3382747
Rv3115-NC_000962.3:3481325-3482760
PE31,PPE60-NC_000962.3:3894016-3895519
esxV,esxW-NC_000962.3:4059985-4060591
PPE18
esxK
esxL
Rv1199c
Event_107
1340400
1340800
1341200
1341600
1342000
Genomic position (1-based)

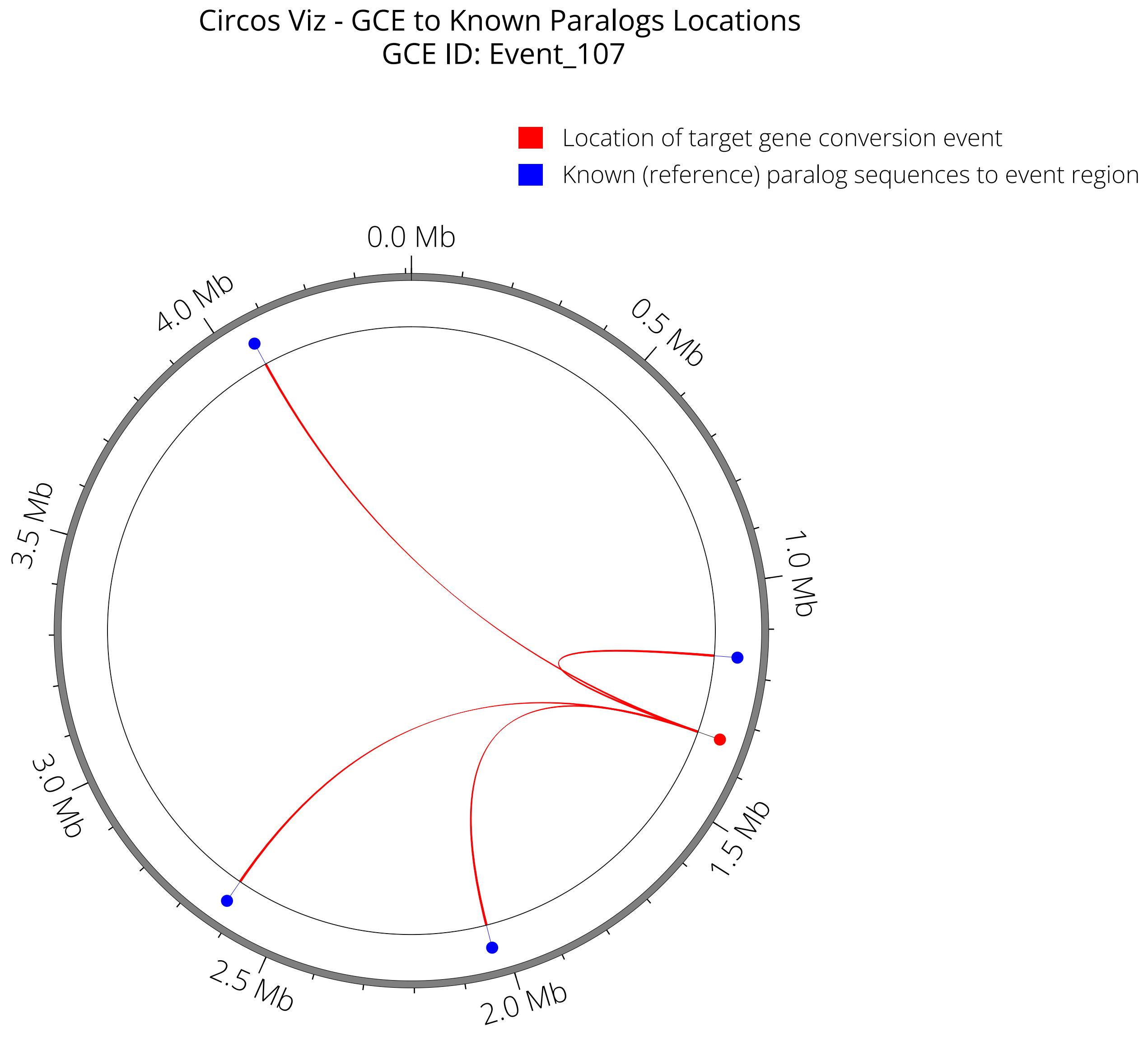
Circos Viz - GCE to Known Paralogs Locations
GCE ID: Event_107
Location of target gene conversion event
Known (reference) paralog sequences to event region
0.0 Mb
0.5 Mb
1.0 Mb
1.5 Mb
2.0 Mb
2.5 Mb
3.0 Mb
3.5 Mb
4.0 Mb

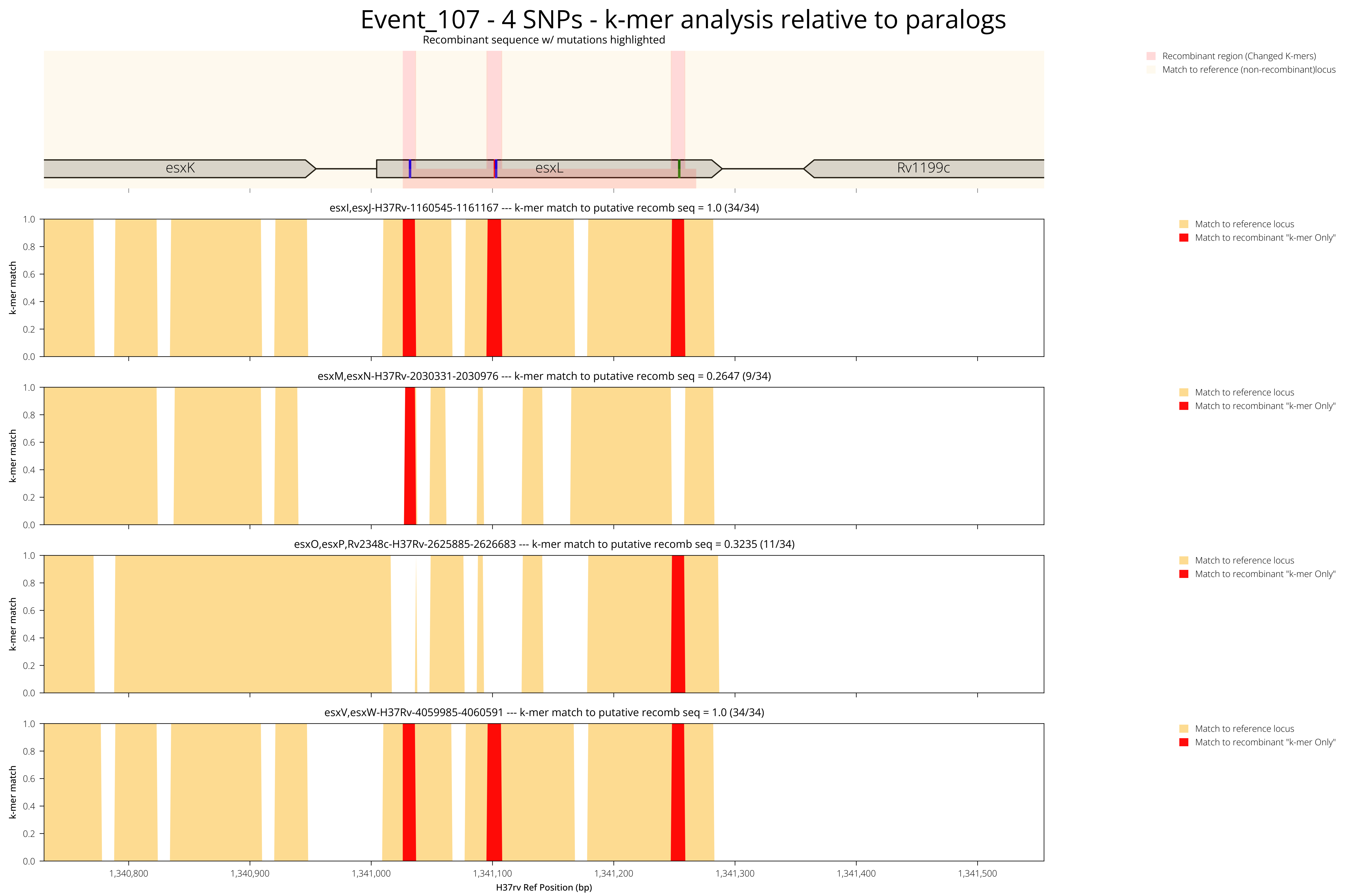
Event_107 - 4 SNPs - k-mer analysis relative to paralogs
Recombinant sequence w/ mutations highlighted
Recombinant region (Changed K-mers)
Match to reference (non-recombinant)locus
esxK
esxL
Rv1199c
esxI,esxJ-H37Rv-1160545-1161167 --- k-mer match to putative recomb seq = 1.0 (34/34)
esxM,esxN-H37Rv-2030331-2030976 --- k-mer match to putative recomb seq = 0.2647 (9/34)
esxO,esxP,Rv2348c-H37Rv-2625885-2626683 --- k-mer match to putative recomb seq = 0.3235 (11/34)
esxV,esxW-H37Rv-4059985-4060591 --- k-mer match to putative recomb seq = 1.0 (34/34)
Match to reference locus
Match to recombinant "k-mer Only"
k-mer match
0.0
0.2
0.4
0.6
0.8
1.0
1,340,800
1,340,900
1,341,000
1,341,100
1,341,200
1,341,300
1,341,400
1,341,500
H37rv Ref Position (bp)

Sub-phylogeny Viz + Paralog Alignments
GCE ID: Event_107

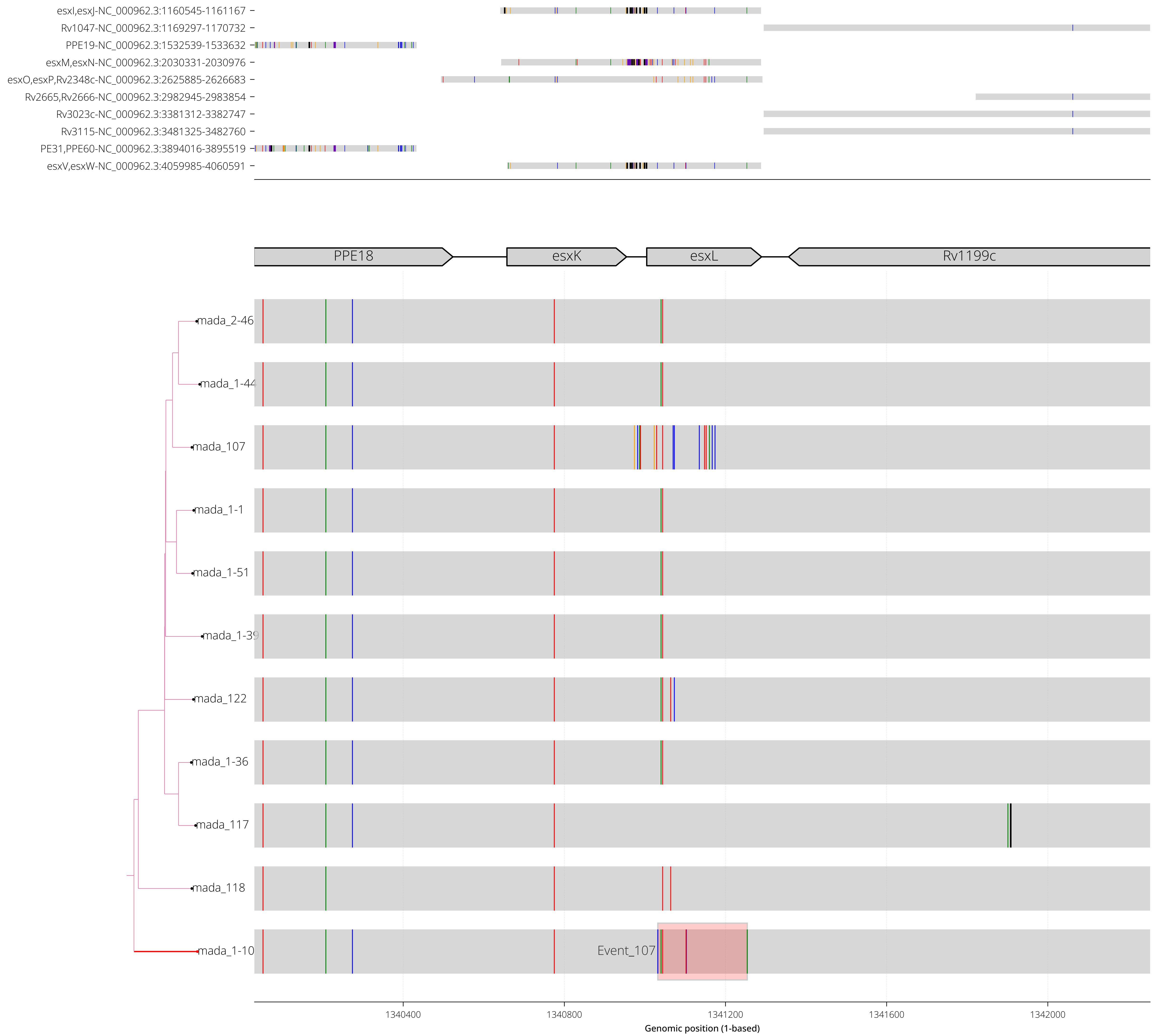

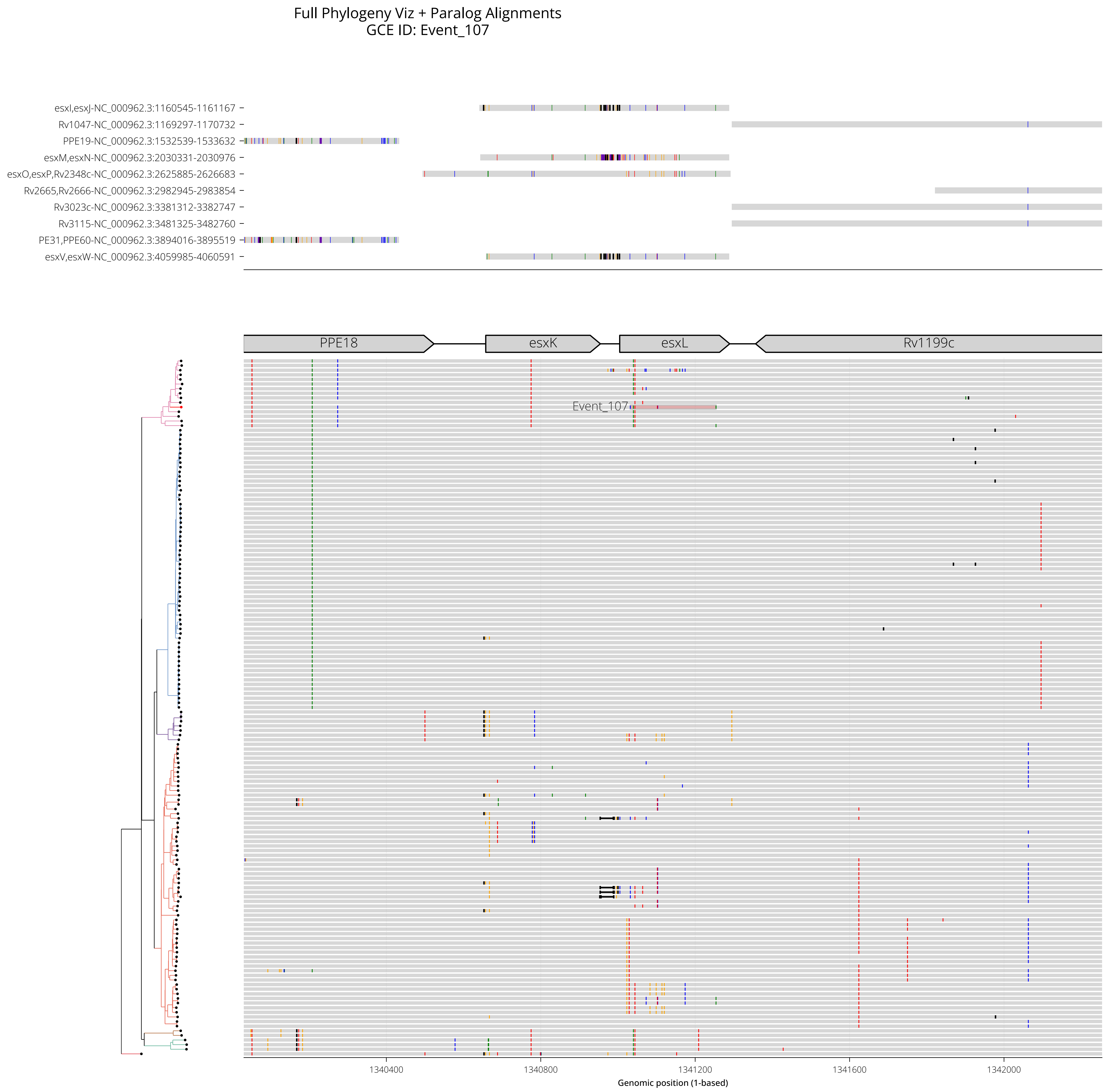

Full Phylogeny Viz + Paralog Alignments
GCE ID: Event_107
esxI,esxJ-NC_000962.3:1160545-1161167
Rv1047-NC_000962.3:1169297-1170732
PPE19-NC_000962.3:1532539-1533632
esxM,esxN-NC_000962.3:2030331-2030976
esxO,esxP,Rv2348c-NC_000962.3:2625885-2626683
Rv2665,Rv2666-NC_000962.3:2982945-2983854
Rv3023c-NC_000962.3:3381312-3382747
Rv3115-NC_000962.3:3481325-3482760
PE31,PPE60-NC_000962.3:3894016-3895519
esxV,esxW-NC_000962.3:4059985-4060591
PPE18
esxK
esxL
Rv1199c
Event_107
1340400
1340800
1341200
1341600
1342000
Genomic position (1-based)

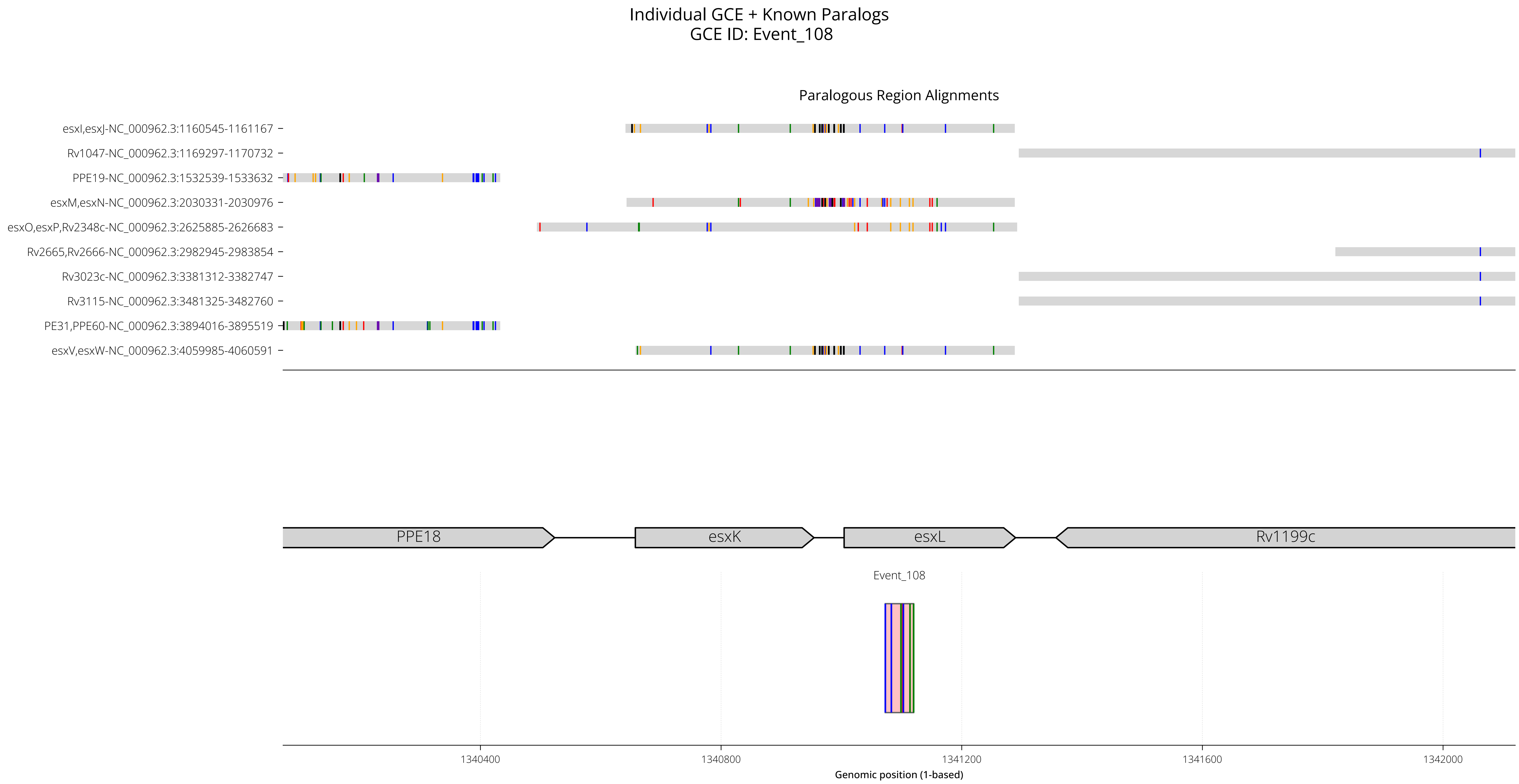

Individual GCE + Known Paralogs
GCE ID: Event_108
Paralogous Region Alignments
esxI,esxJ-NC_000962.3:1160545-1161167
Rv1047-NC_000962.3:1169297-1170732
PPE19-NC_000962.3:1532539-1533632
esxM,esxN-NC_000962.3:2030331-2030976
esxO,esxP,Rv2348c-NC_000962.3:2625885-2626683
Rv2665,Rv2666-NC_000962.3:2982945-2983854
Rv3023c-NC_000962.3:3381312-3382747
Rv3115-NC_000962.3:3481325-3482760
PE31,PPE60-NC_000962.3:3894016-3895519
esxV,esxW-NC_000962.3:4059985-4060591
PPE18
esxK
esxL
Rv1199c
Event_108
1340400
1340800
1341200
1341600
1342000
Genomic position (1-based)

Circos Viz - GCE to Known Paralogs Locations
GCE ID: Event_108

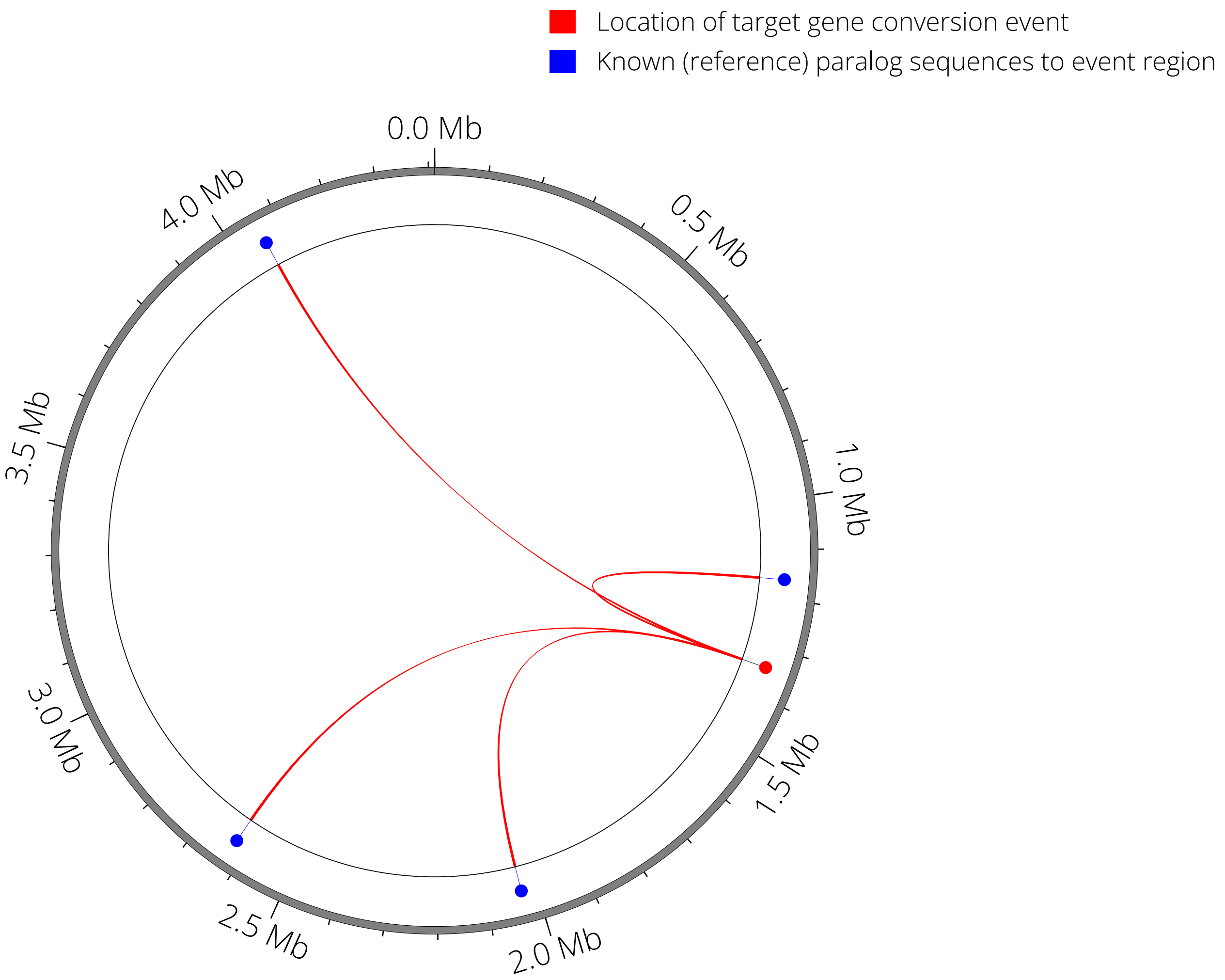

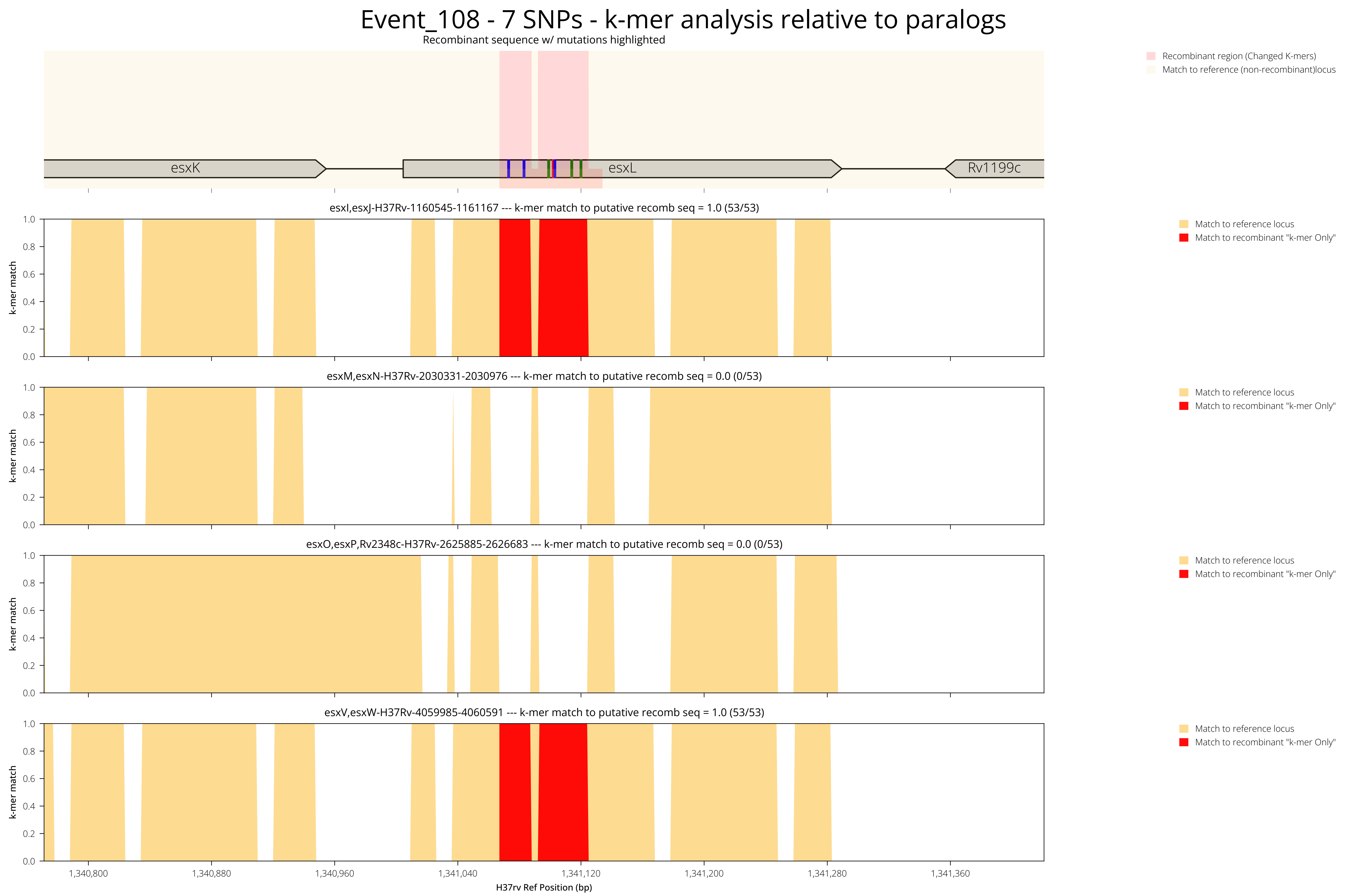

Event_108 - 7 SNPs - k-mer analysis relative to paralogs
Recombinant sequence w/ mutations highlighted
Recombinant region (Changed K-mers)
Match to reference (non-recombinant)locus
esxK
esxL
Rv1199c
esxI,esxJ-H37Rv-1160545-1161167 --- k-mer match to putative recomb seq = 1.0 (53/53)
esxM,esxN-H37Rv-2030331-2030976 --- k-mer match to putative recomb seq = 0.0 (0/53)
esxO,esxP,Rv2348c-H37Rv-2625885-2626683 --- k-mer match to putative recomb seq = 0.0 (0/53)
esxV,esxW-H37Rv-4059985-4060591 --- k-mer match to putative recomb seq = 1.0 (53/53)
Match to reference locus
Match to recombinant "k-mer Only"
k-mer match
0.0
0.2
0.4
0.6
0.8
1.0
1,340,800
1,340,880
1,340,960
1,341,040
1,341,120
1,341,200
1,341,280
1,341,360
H37rv Ref Position (bp)

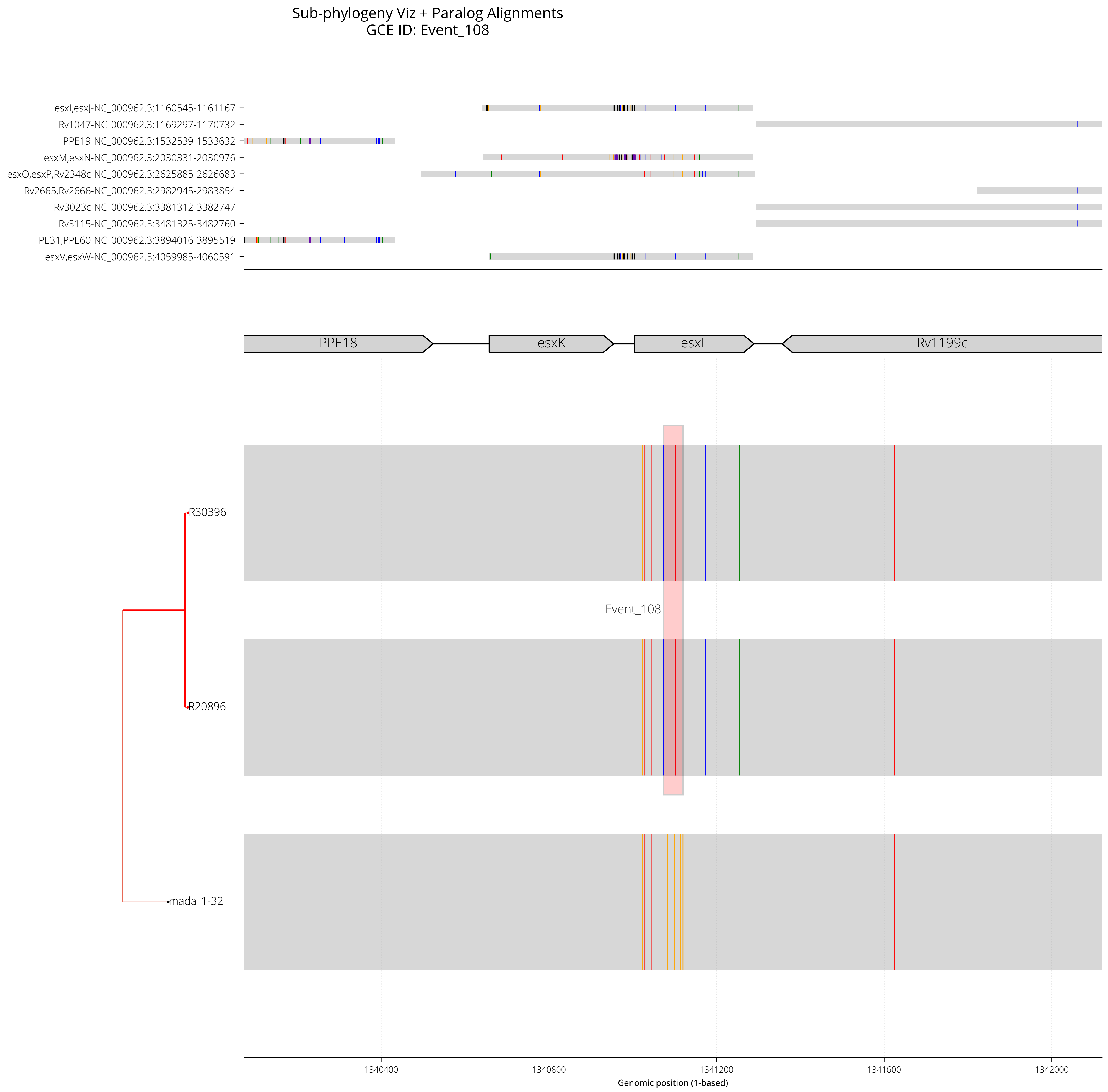
Sub-phylogeny Viz + Paralog Alignments
GCE ID: Event_108
esxI,esxJ-NC_000962.3:1160545-1161167
Rv1047-NC_000962.3:1169297-1170732
PPE19-NC_000962.3:1532539-1533632
esxM,esxN-NC_000962.3:2030331-2030976
esxO,esxP,Rv2348c-NC_000962.3:2625885-2626683
Rv2665,Rv2666-NC_000962.3:2982945-2983854
Rv3023c-NC_000962.3:3381312-3382747
Rv3115-NC_000962.3:3481325-3482760
PE31,PPE60-NC_000962.3:3894016-3895519
esxV,esxW-NC_000962.3:4059985-4060591
PPE18
esxK
esxL
Rv1199c
R30396
Event_108
R20896
mada_1-32
1340400
1340800
1341200
1341600
1342000
Genomic position (1-based)

Full Phylogeny Viz + Paralog Alignments
GCE ID: Event_108

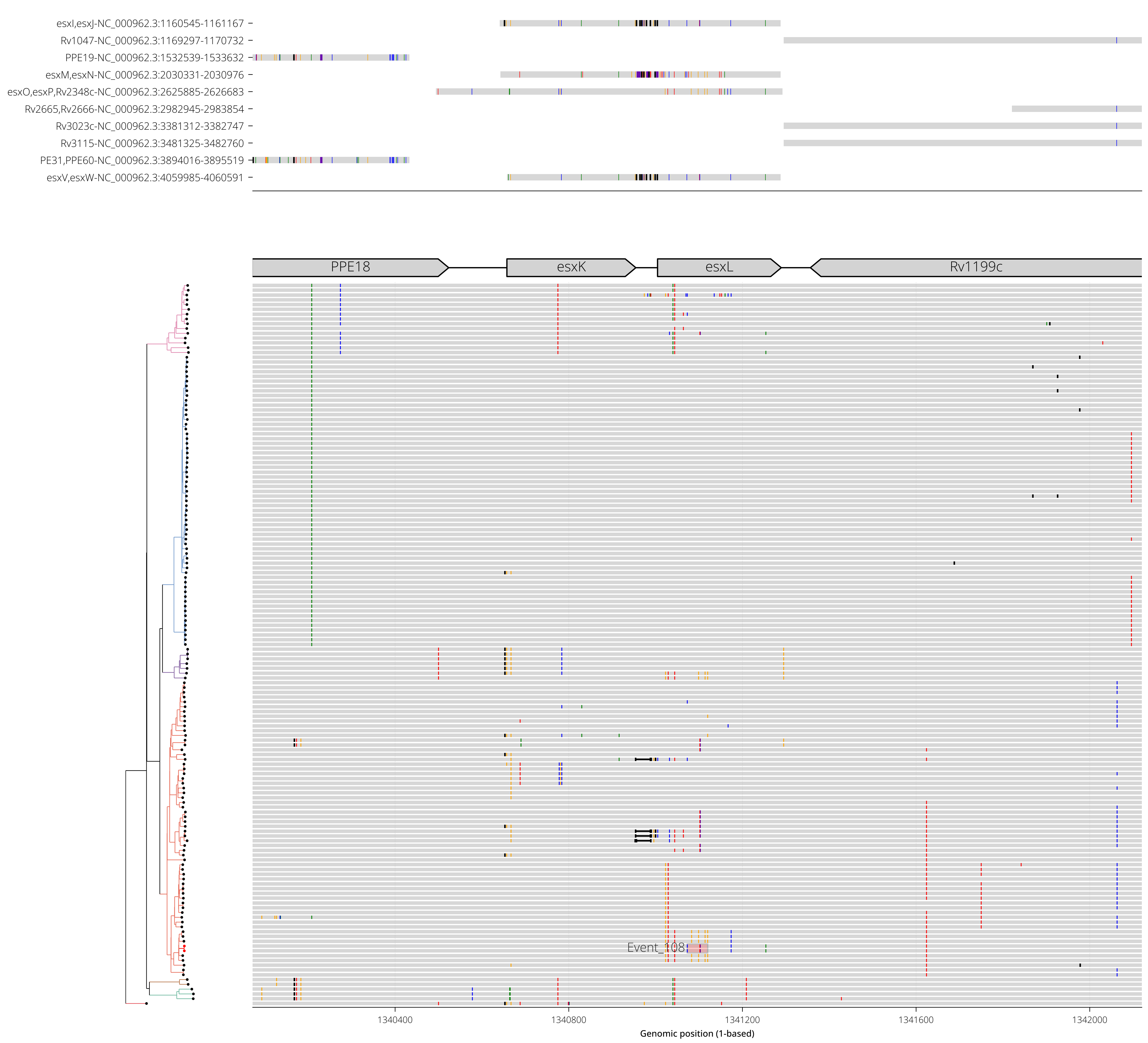

Individual GCE + Known Paralogs
GCE ID: Event_109

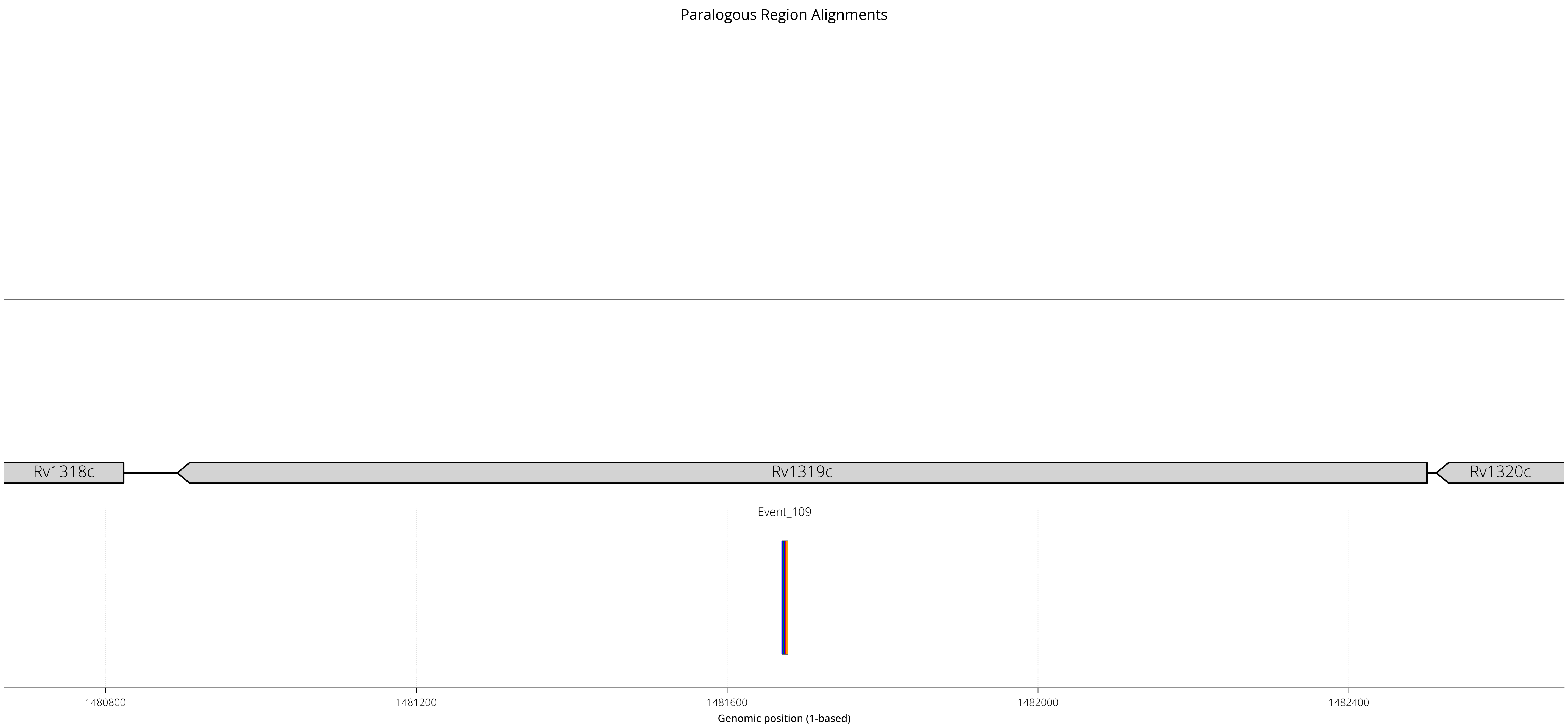

Sub-phylogeny Viz + Paralog Alignments
GCE ID: Event_109

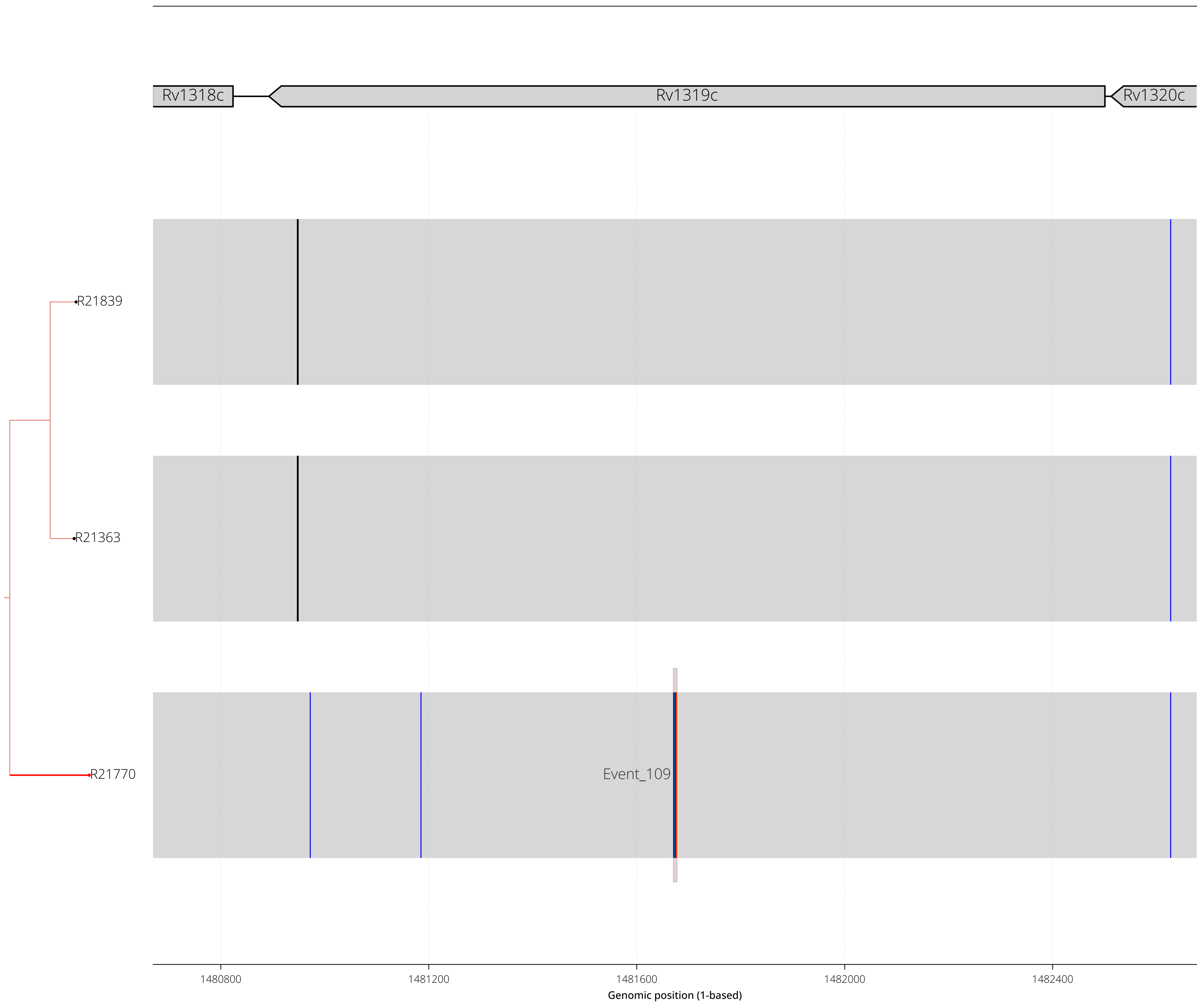

Full Phylogeny Viz + Paralog Alignments
GCE ID: Event_109

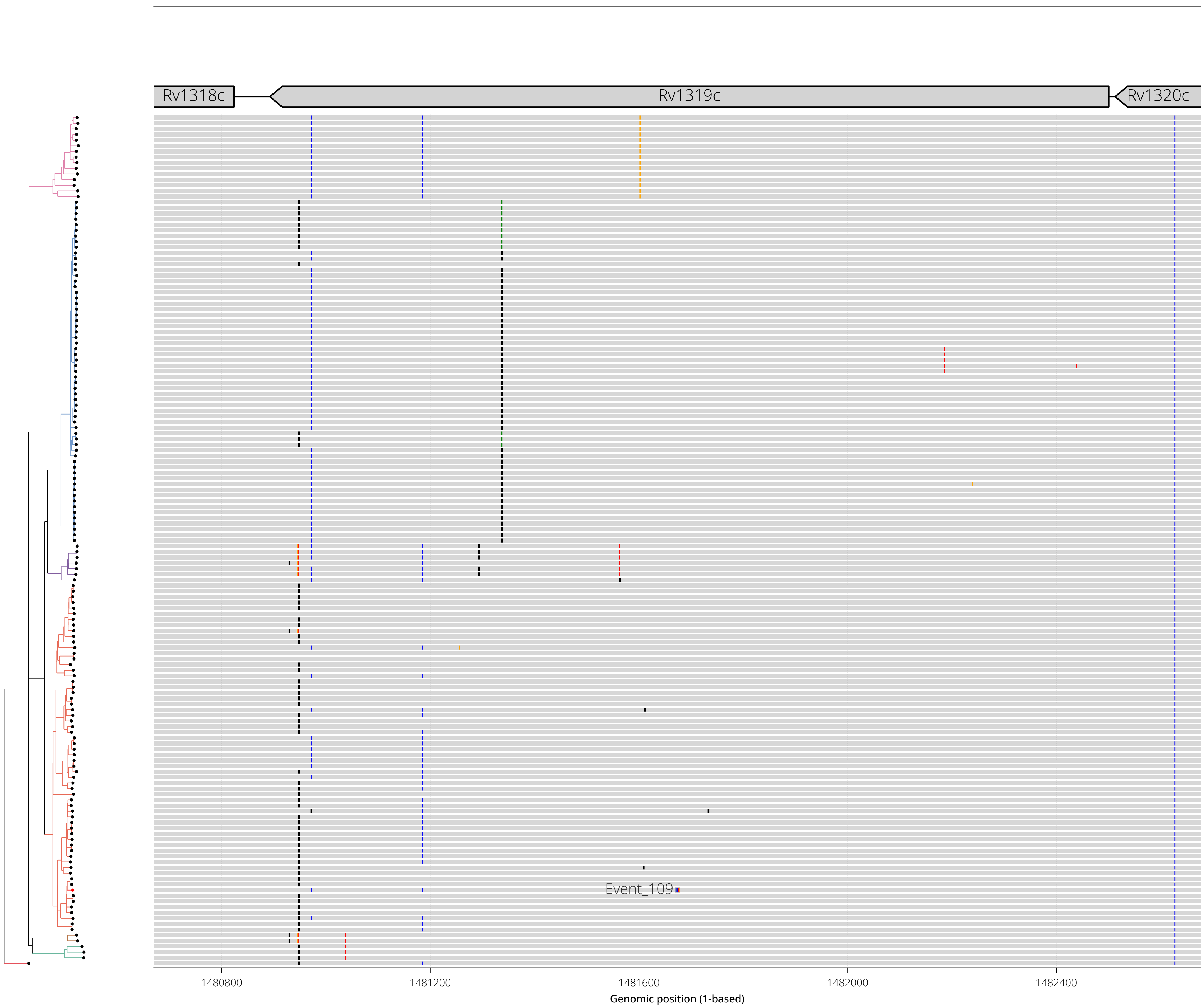

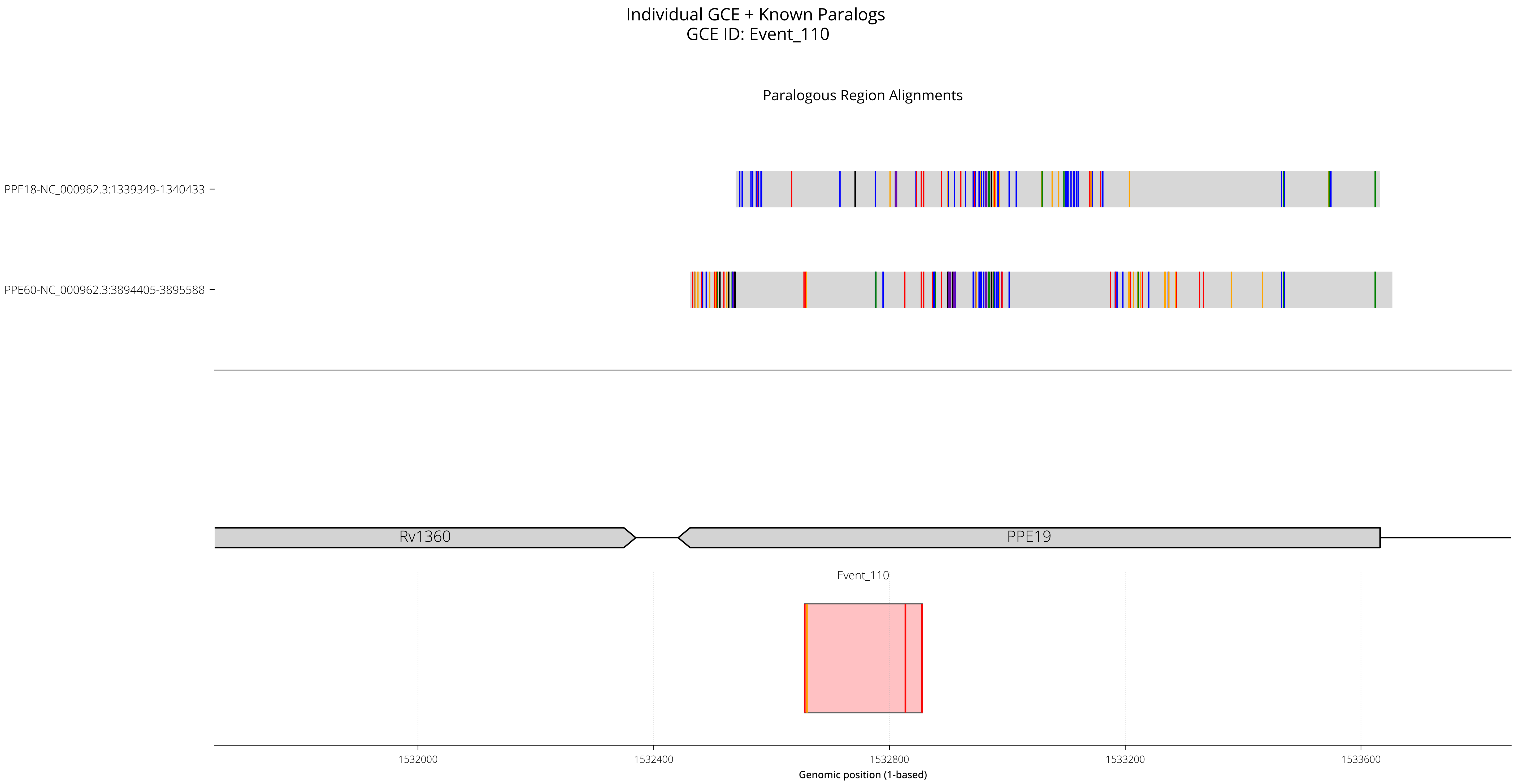
Individual GCE + Known Paralogs
GCE ID: Event_110
Paralogous Region Alignments
PPE18-NC_000962.3:1339349-1340433
PPE60-NC_000962.3:3894405-3895588
Rv1360
PPE19
Event_110
1532000
1532400
1532800
1533200
1533600
Genomic position (1-based)

Circos Viz - GCE to Known Paralogs Locations
GCE ID: Event_110

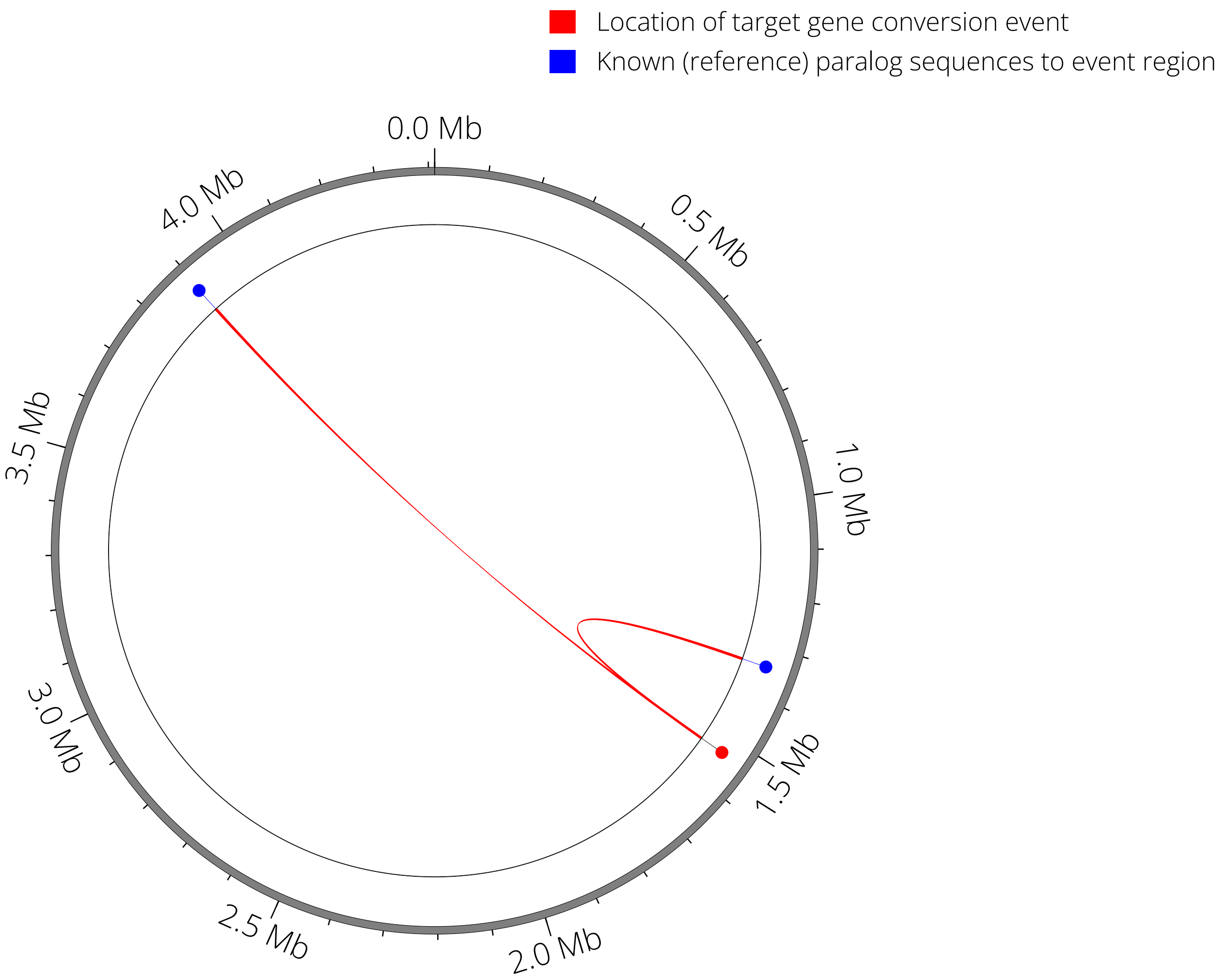

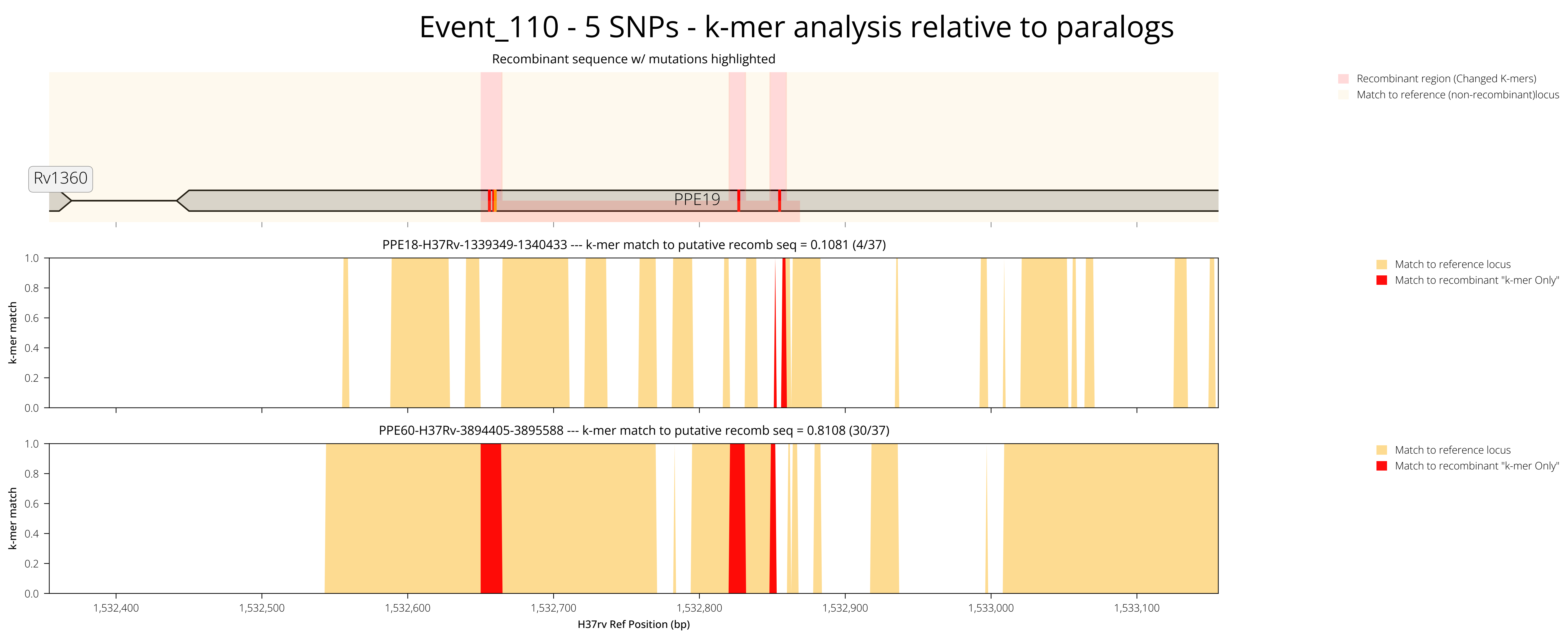
Event_110 - 5 SNPs - k-mer analysis relative to paralogs
Recombinant sequence w/ mutations highlighted
Recombinant region (Changed K-mers)
Match to reference (non-recombinant)locus
Rv1360
PPE19
PPE18-H37Rv-1339349-1340433 --- k-mer match to putative recomb seq = 0.1081 (4/37)
Match to reference locus
Match to recombinant "k-mer Only"
k-mer match
PPE60-H37Rv-3894405-3895588 --- k-mer match to putative recomb seq = 0.8108 (30/37)
1,532,400
1,532,500
1,532,600
1,532,700
1,532,800
1,532,900
1,533,000
1,533,100
H37rv Ref Position (bp)

Sub-phylogeny Viz + Paralog Alignments
GCE ID: Event_110

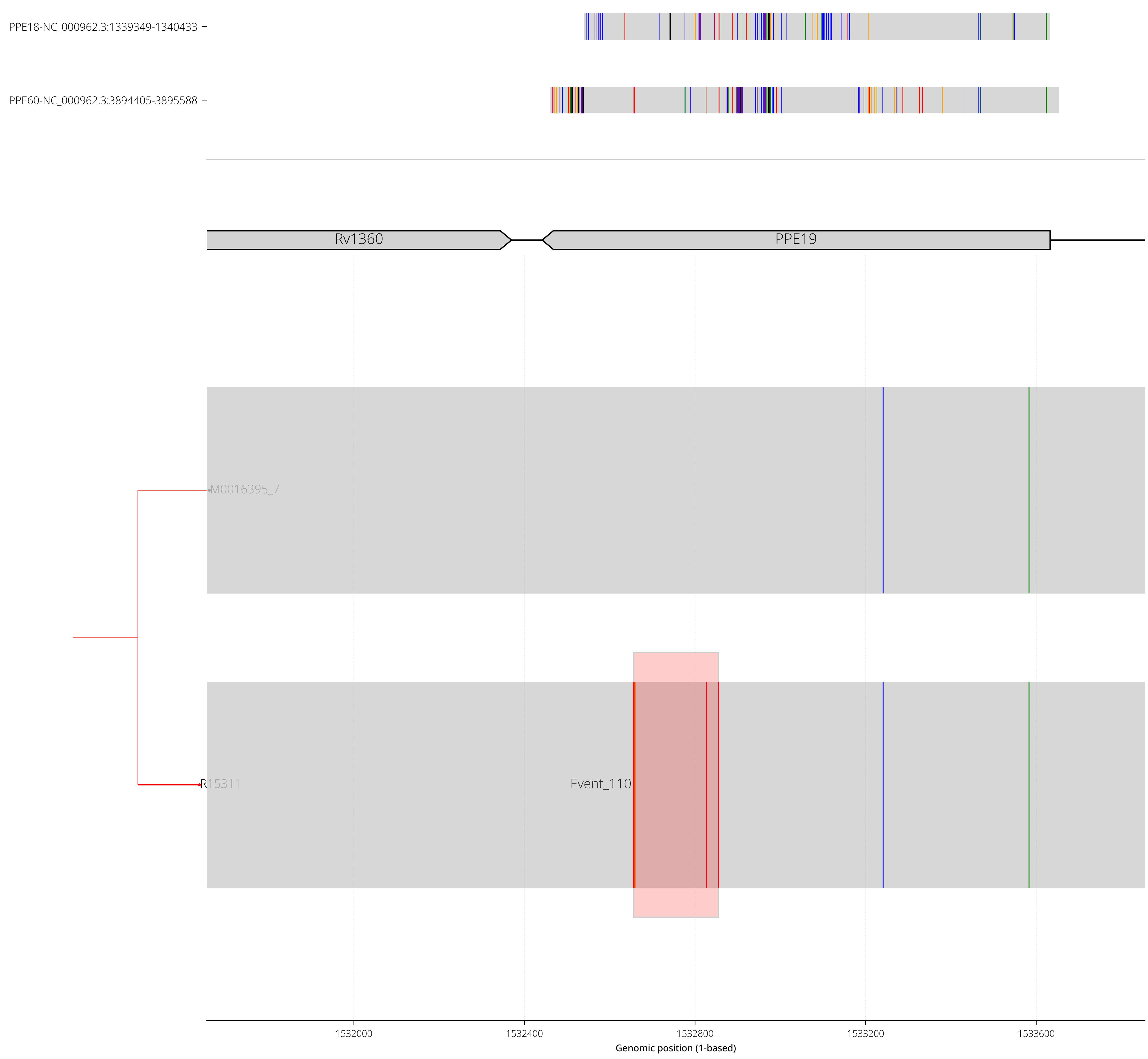

Full Phylogeny Viz + Paralog Alignments
GCE ID: Event_110

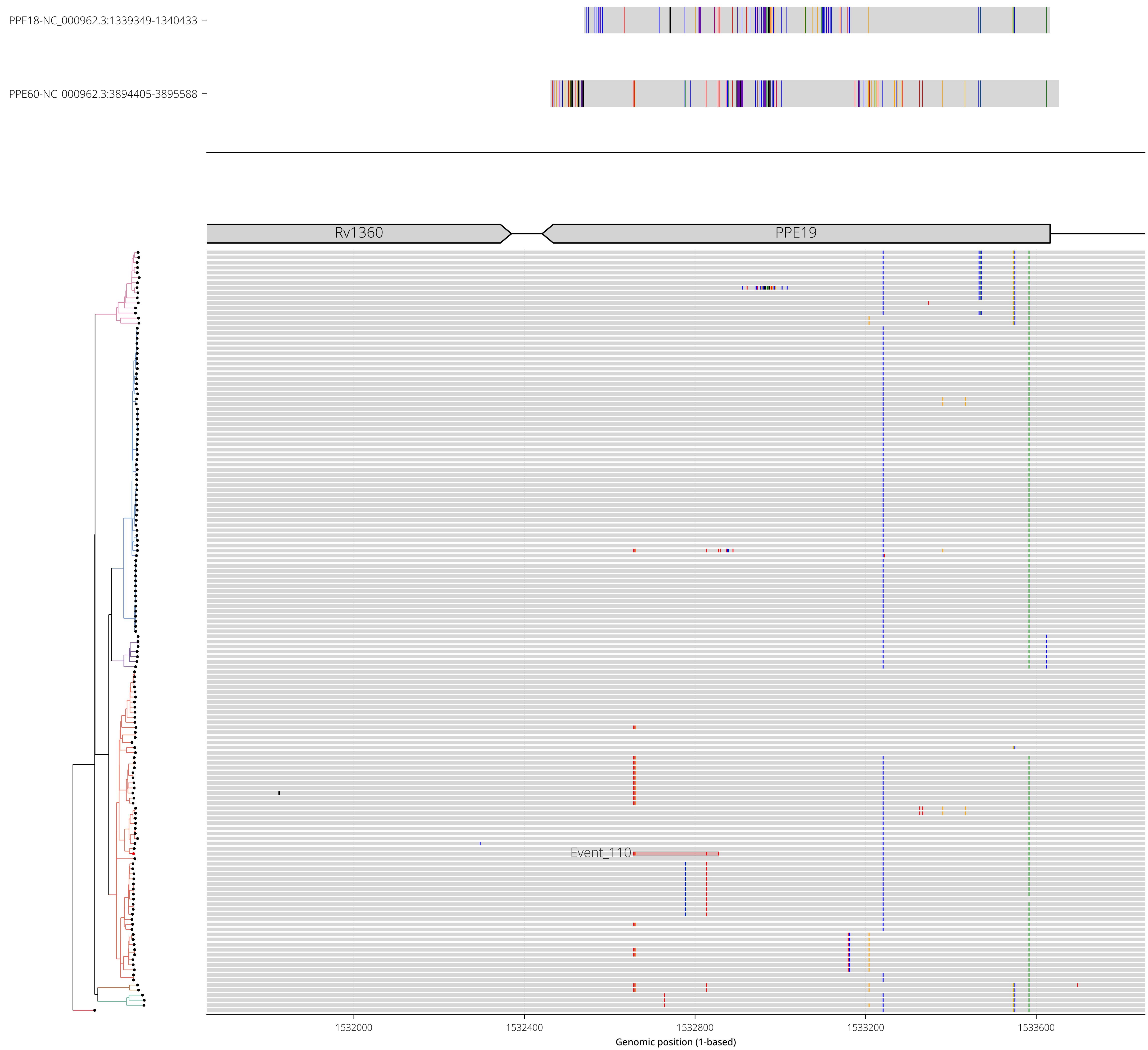

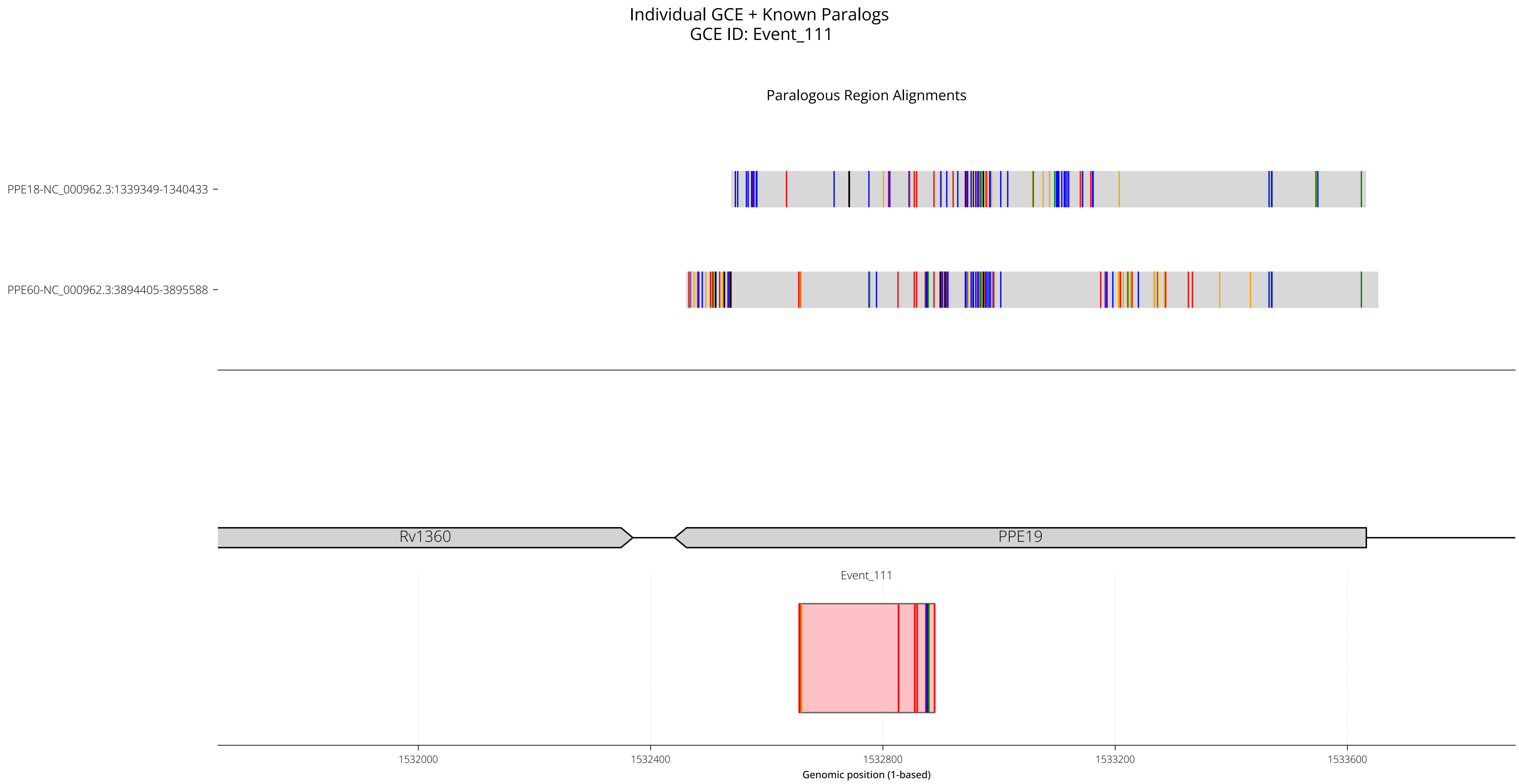
Individual GCE + Known Paralogs
GCE ID: Event_111
Paralogous Region Alignments
PPE18-NC_000962.3:1339349-1340433
PPE60-NC_000962.3:3894405-3895588
Rv1360
PPE19
Event_111
1532000
1532400
1532800
1533200
1533600
Genomic position (1-based)

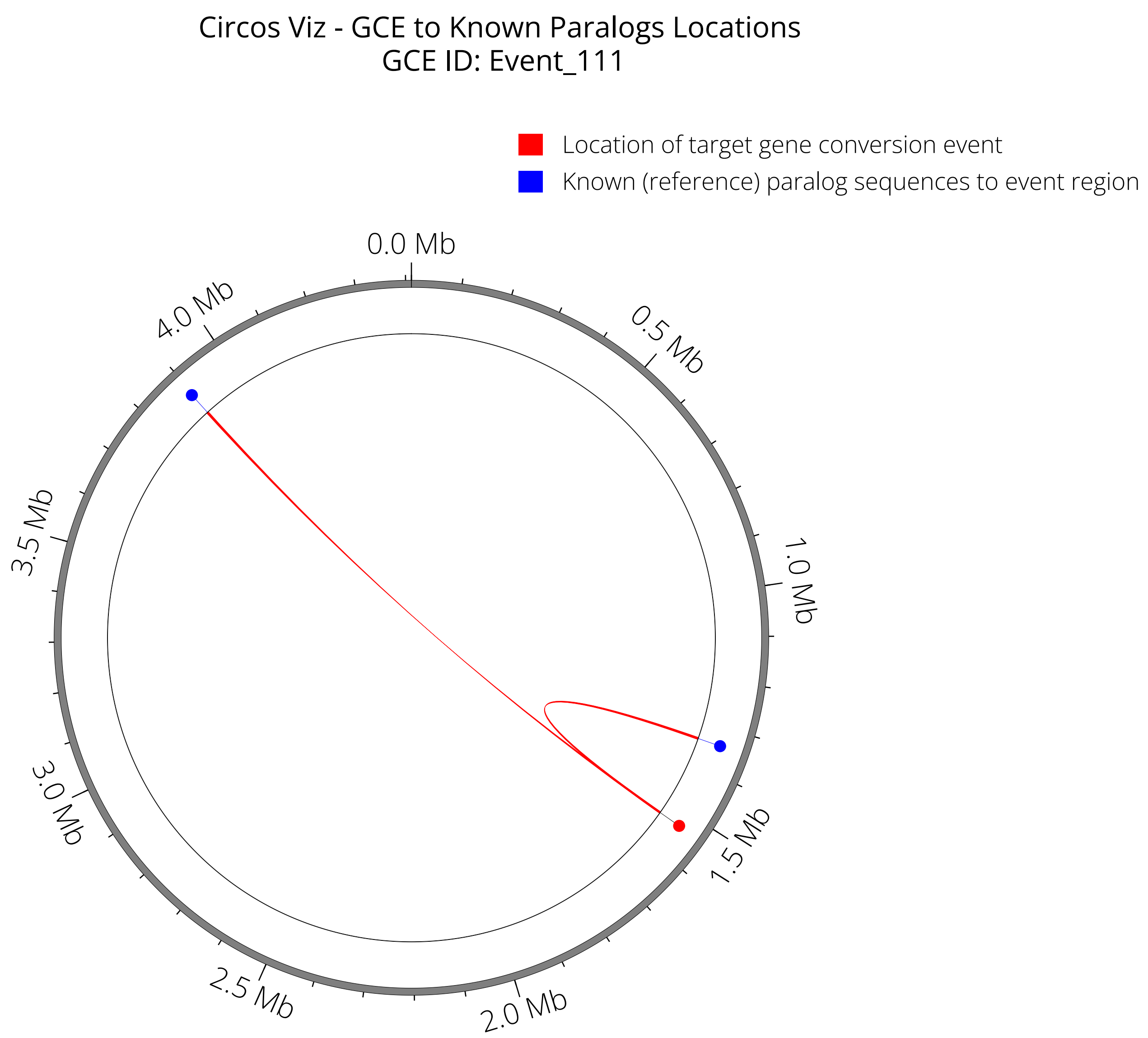

Circos Viz - GCE to Known Paralogs Locations
GCE ID: Event_111
Location of target gene conversion event
Known (reference) paralog sequences to event region
0.0 Mb
0.5 Mb
1.0 Mb
1.5 Mb
2.0 Mb
2.5 Mb
3.0 Mb
3.5 Mb
4.0 Mb

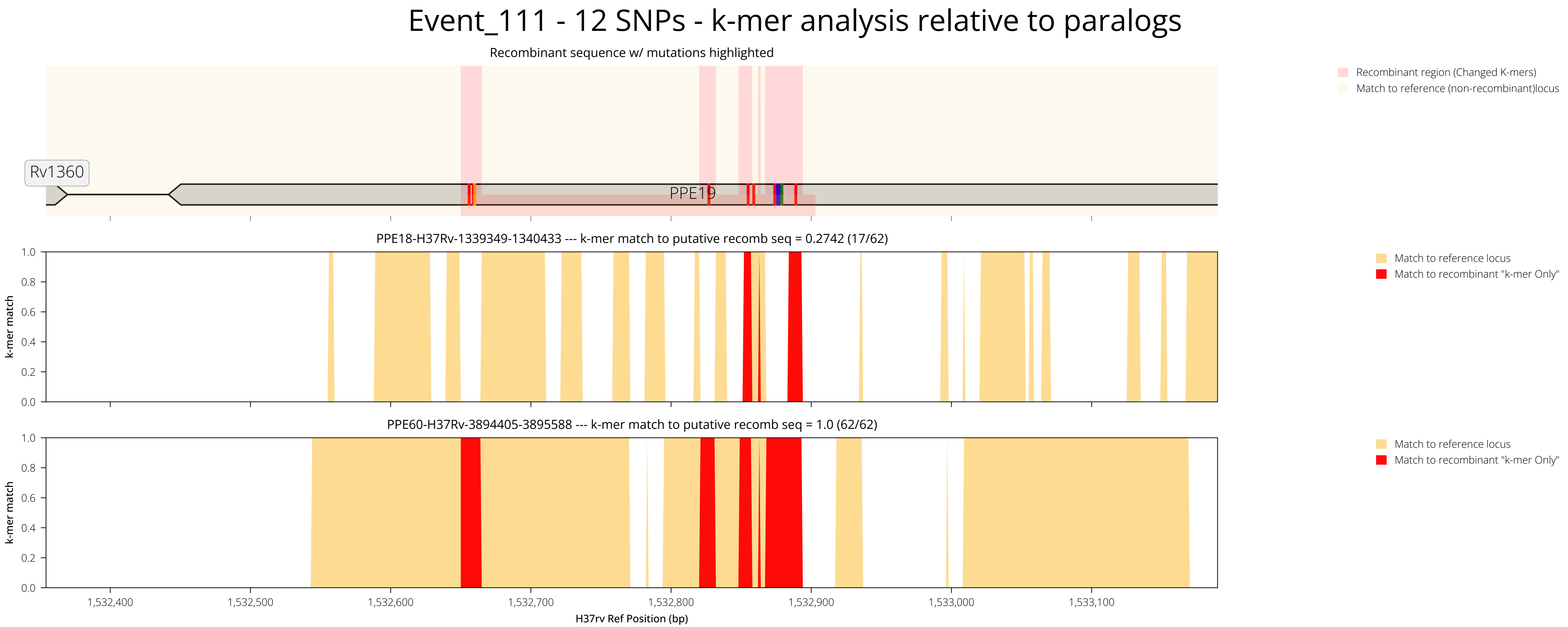

Event_111 - 12 SNPs - k-mer analysis relative to paralogs
Recombinant sequence w/ mutations highlighted
Recombinant region (Changed K-mers)
Match to reference (non-recombinant)locus
Rv1360
PPE19
PPE18-H37Rv-1339349-1340433 --- k-mer match to putative recomb seq = 0.2742 (17/62)
Match to reference locus
Match to recombinant "k-mer Only"
k-mer match
1.0
0.8
0.6
0.4
0.2
0.0
PPE60-H37Rv-3894405-3895588 --- k-mer match to putative recomb seq = 1.0 (62/62)
Match to reference locus
Match to recombinant "k-mer Only"
k-mer match
1,532,400
1,532,500
1,532,600
1,532,700
1,532,800
1,532,900
1,533,000
1,533,100
H37rv Ref Position (bp)

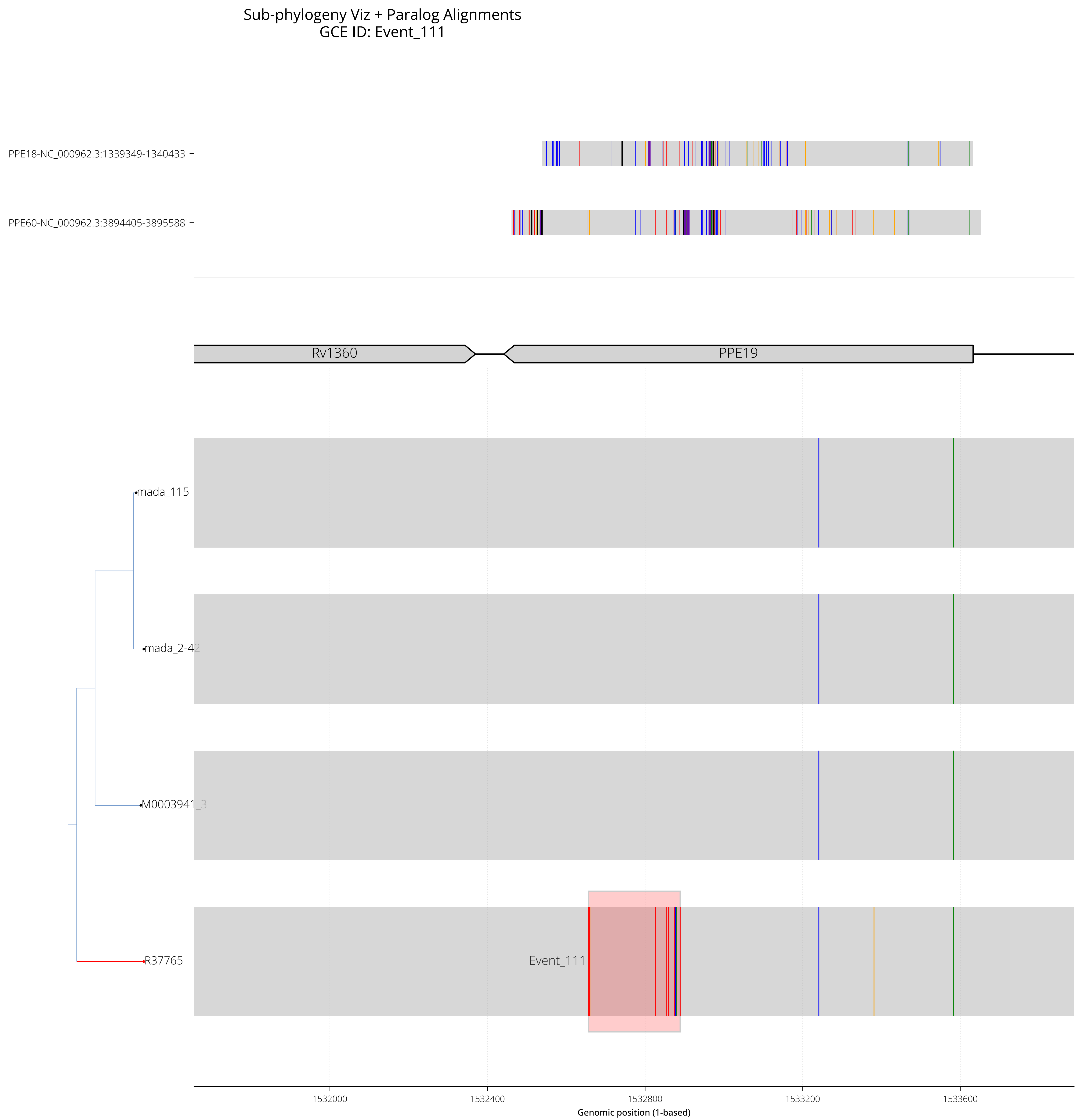

Sub-phylogeny Viz + Paralog Alignments
GCE ID: Event_111
PPE18-NC_000962.3:1339349-1340433
PPE60-NC_000962.3:3894405-3895588
Rv1360
PPE19
mada_115
mada_2-42
M0003941_3
R37765
Event_111
1532000
1532400
1532800
1533200
1533600
Genomic position (1-based)

Full Phylogeny Viz + Paralog Alignments
GCE ID: Event_111

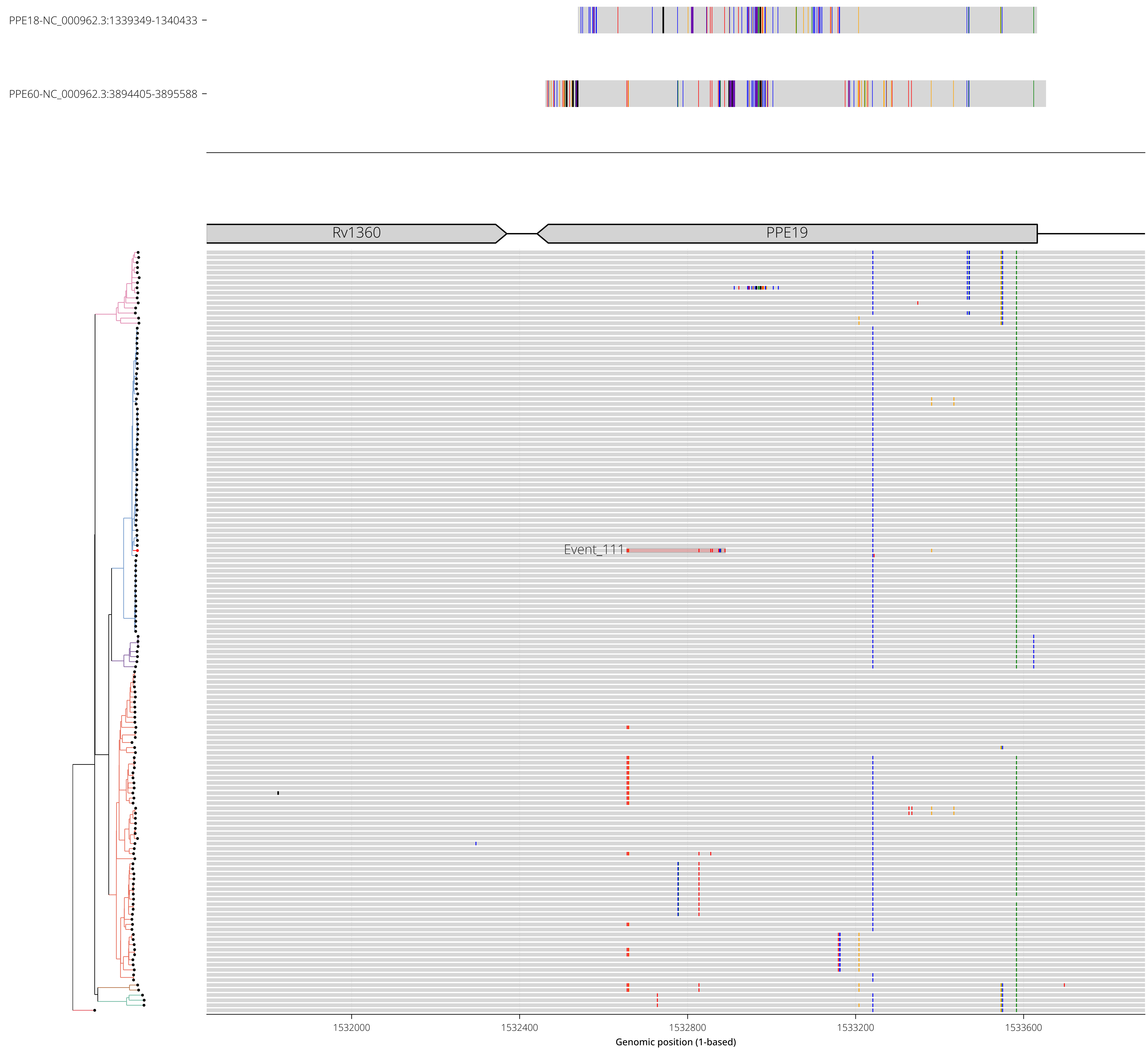

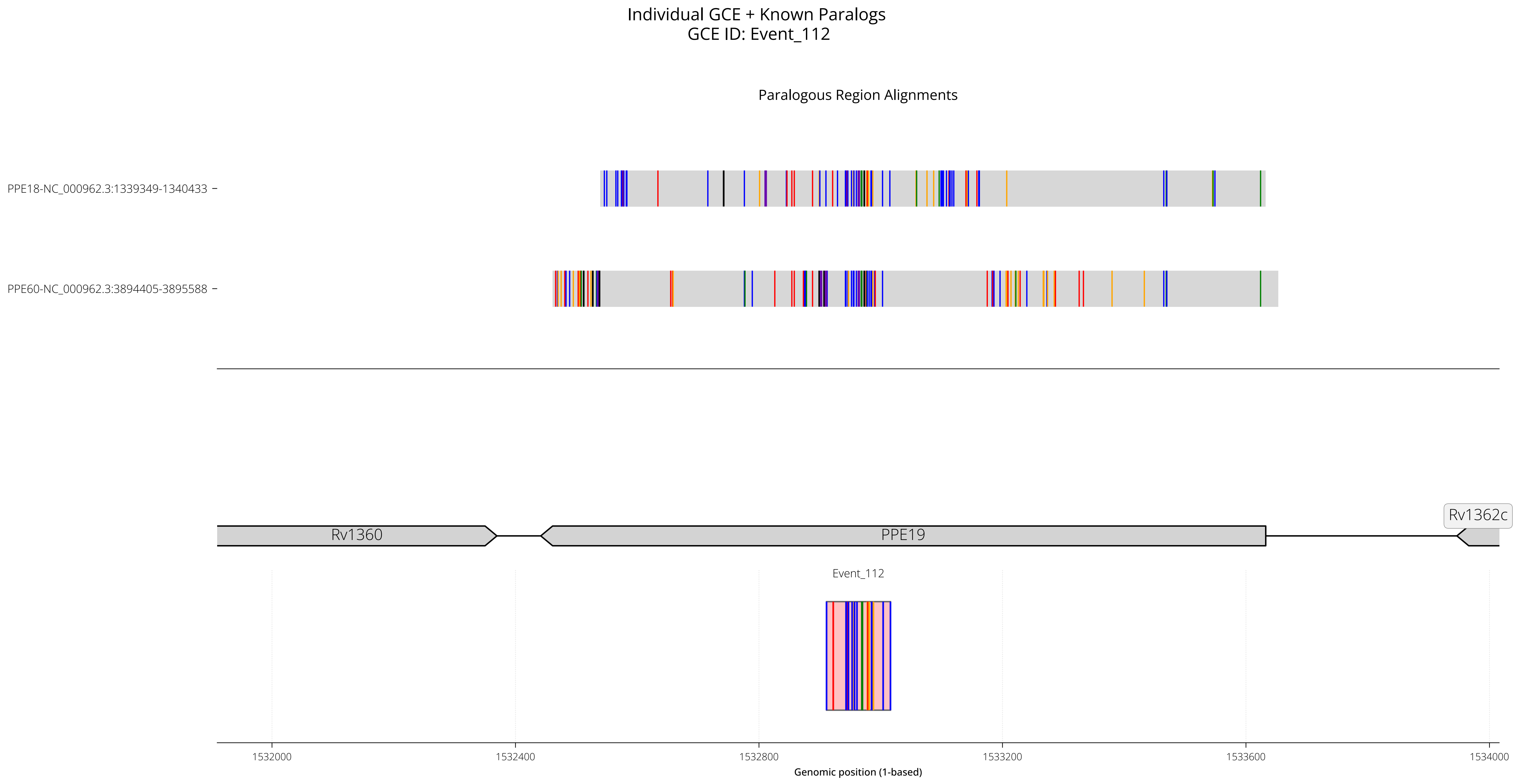

Individual GCE + Known Paralogs
GCE ID: Event_112
Paralogous Region Alignments
PPE18-NC_000962.3:1339349-1340433
PPE60-NC_000962.3:3894405-3895588
Rv1360
PPE19
Rv1362c
Event_112
1532000
1532400
1532800
1533200
1533600
1534000
Genomic position (1-based)

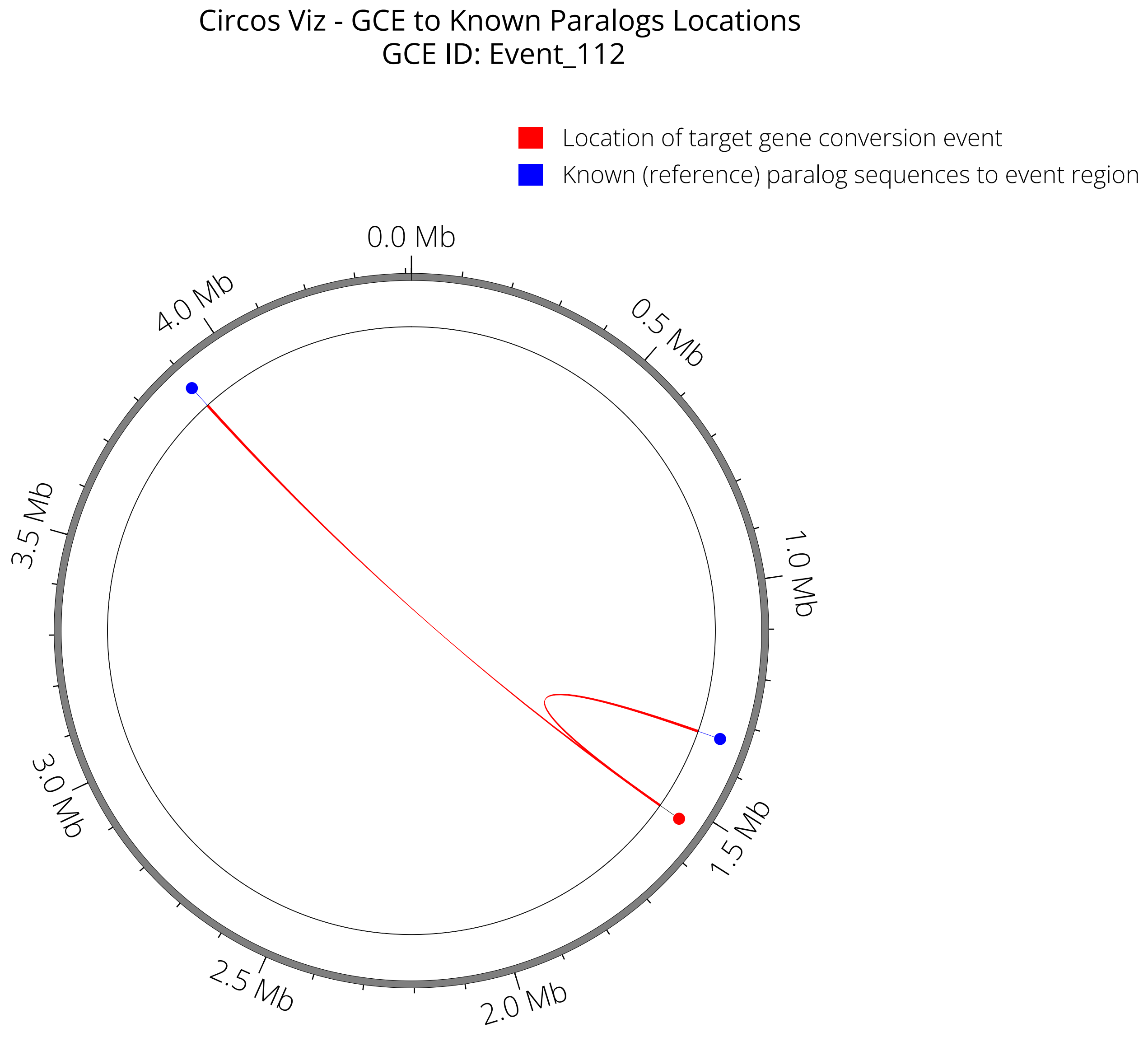

Circos Viz - GCE to Known Paralogs Locations
GCE ID: Event_112
Location of target gene conversion event
Known (reference) paralog sequences to event region
0.0 Mb
0.5 Mb
1.0 Mb
1.5 Mb
2.0 Mb
2.5 Mb
3.0 Mb
3.5 Mb
4.0 Mb

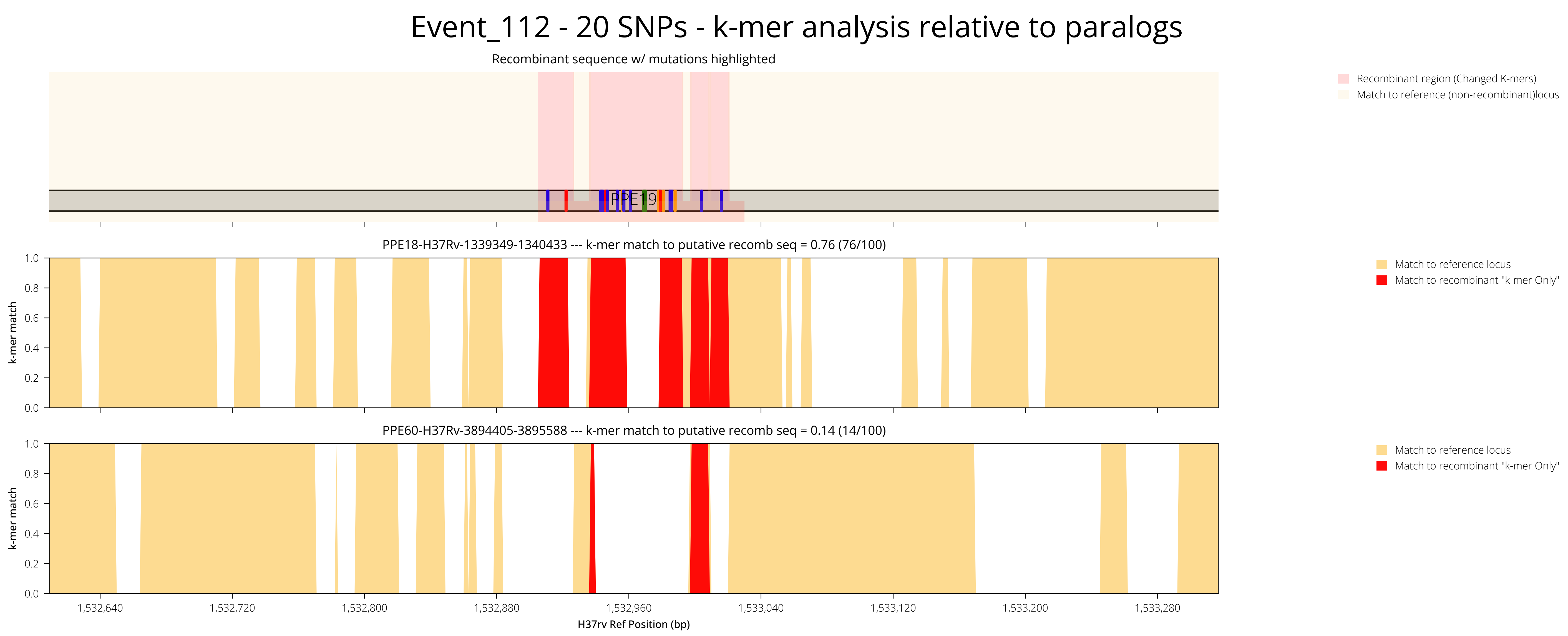
Event_112 - 20 SNPs - k-mer analysis relative to paralogs
Recombinant sequence w/ mutations highlighted
Recombinant region (Changed K-mers)
Match to reference (non-recombinant)locus
PPE19
PPE18-H37Rv-1339349-1340433 --- k-mer match to putative recomb seq = 0.76 (76/100)
Match to reference locus
Match to recombinant "k-mer Only"
k-mer match
1.0
0.8
0.6
0.4
0.2
0.0
PPE60-H37Rv-3894405-3895588 --- k-mer match to putative recomb seq = 0.14 (14/100)
Match to reference locus
Match to recombinant "k-mer Only"
k-mer match
1.0
0.8
0.6
0.4
0.2
0.0
1,532,640
1,532,720
1,532,800
1,532,880
1,532,960
1,533,040
1,533,120
1,533,200
1,533,280
H37rv Ref Position (bp)

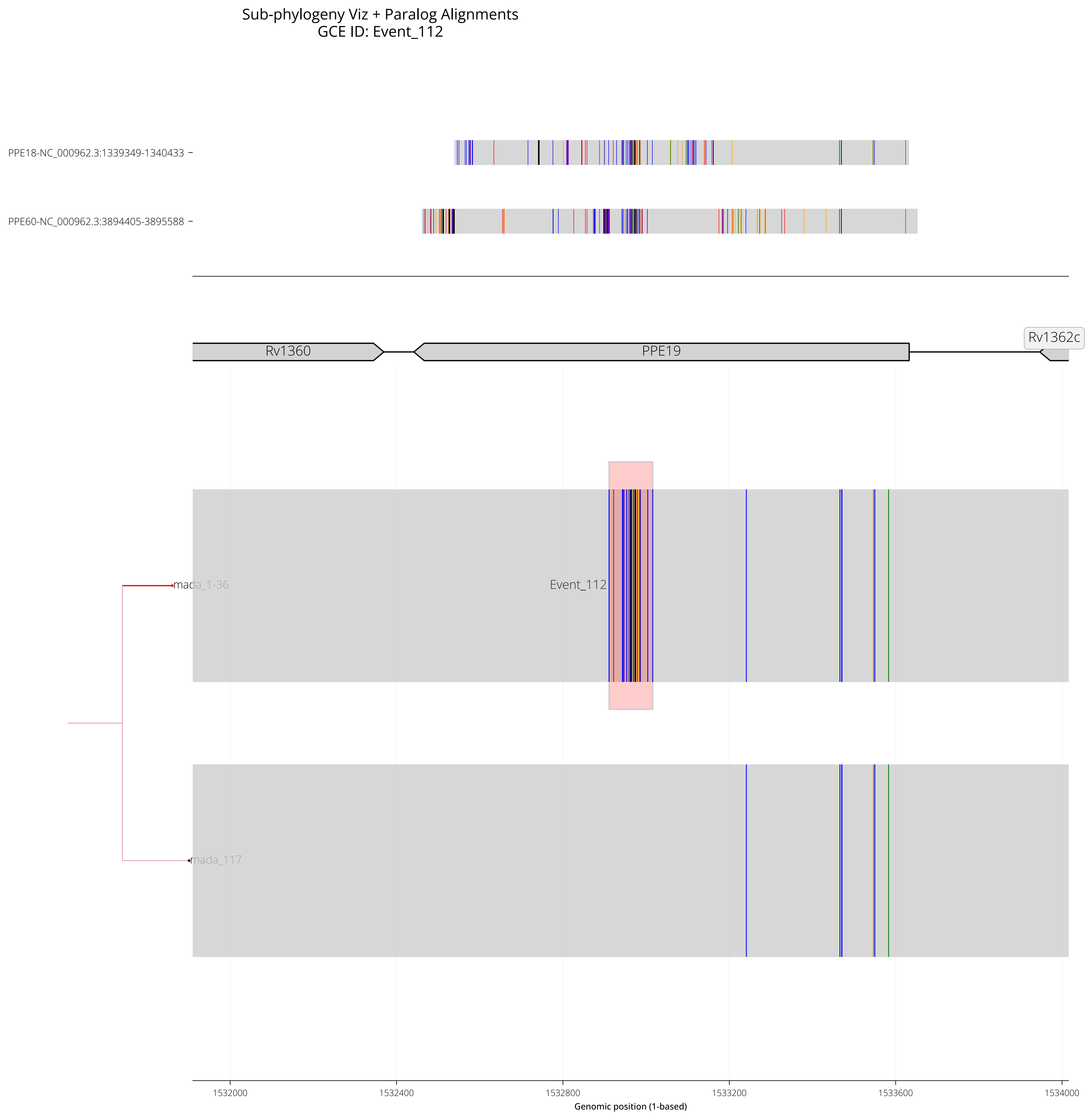

Sub-phylogeny Viz + Paralog Alignments
GCE ID: Event_112
PPE18-NC_000962.3:1339349-1340433
PPE60-NC_000962.3:3894405-3895588
Rv1360
PPE19
Rv1362c
Event_112
mada_1-36
mada_117
1532000
1532400
1532800
1533200
1533600
1534000
Genomic position (1-based)

Full Phylogeny Viz + Paralog Alignments
GCE ID: Event_112

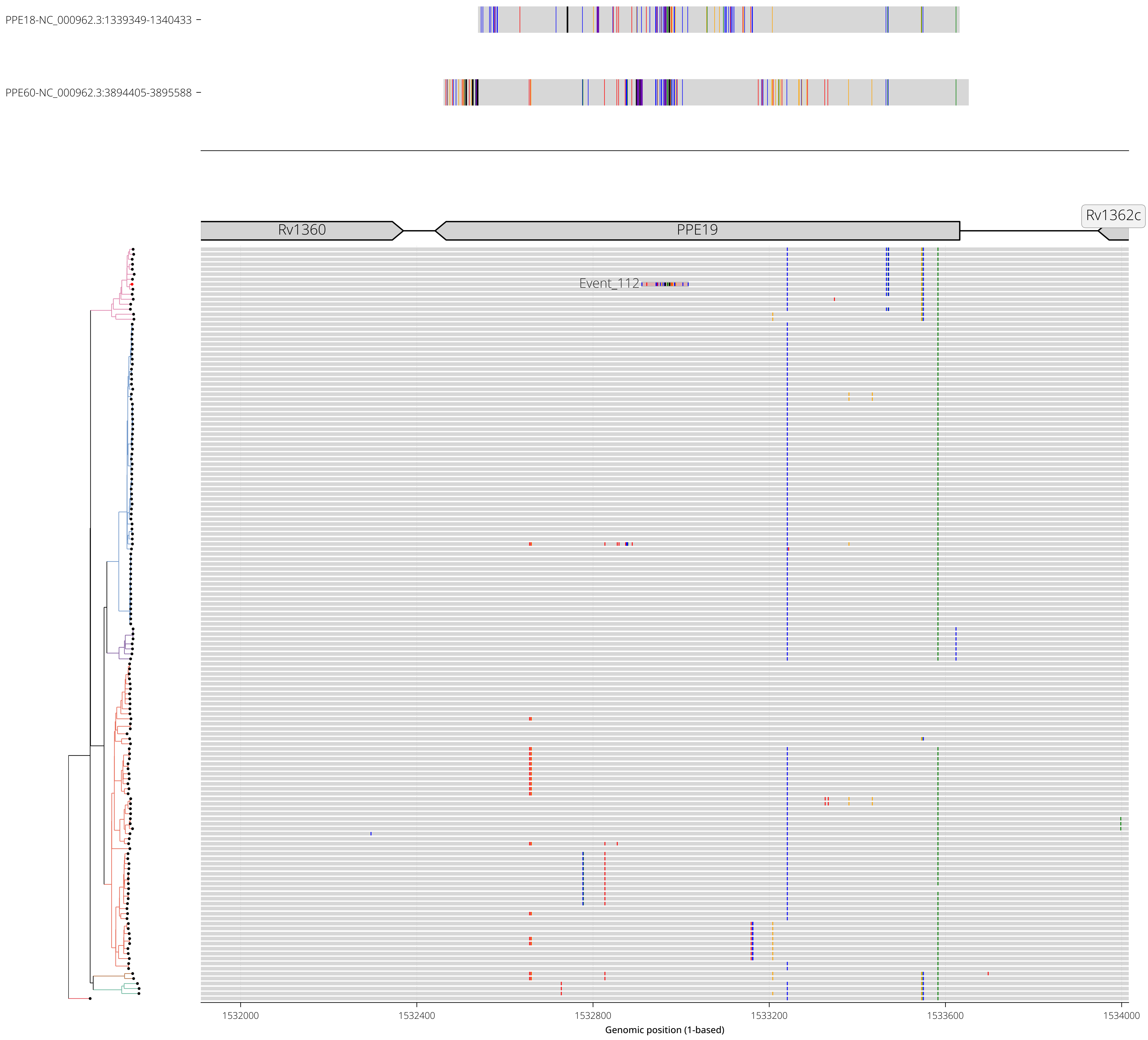

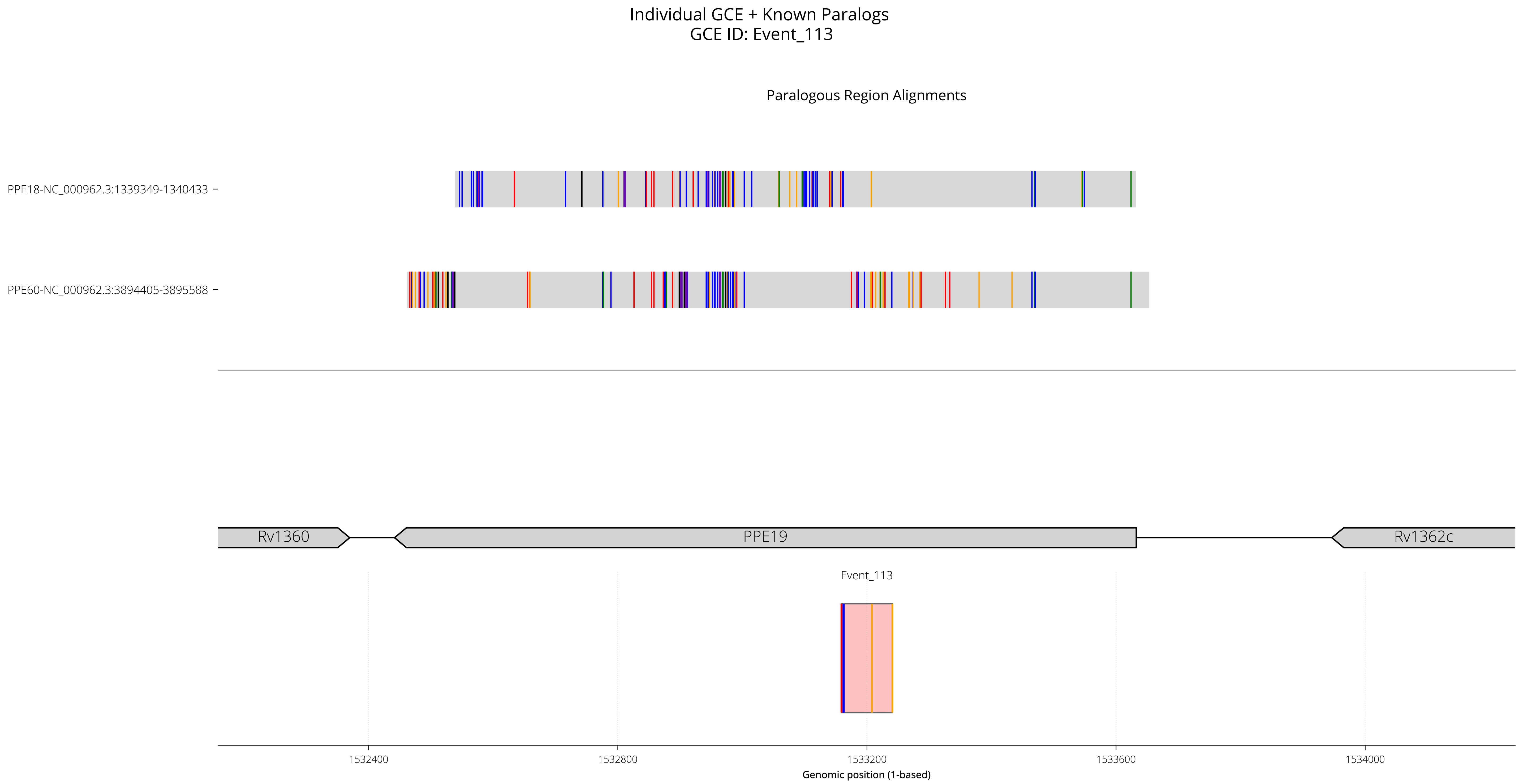

Individual GCE + Known Paralogs
GCE ID: Event_113
Paralogous Region Alignments
PPE18-NC_000962.3:1339349-1340433
PPE60-NC_000962.3:3894405-3895588
Rv1360
PPE19
Rv1362c
Event_113
1532400
1532800
1533200
1533600
1534000
Genomic position (1-based)

Circos Viz - GCE to Known Paralogs Locations
GCE ID: Event_113

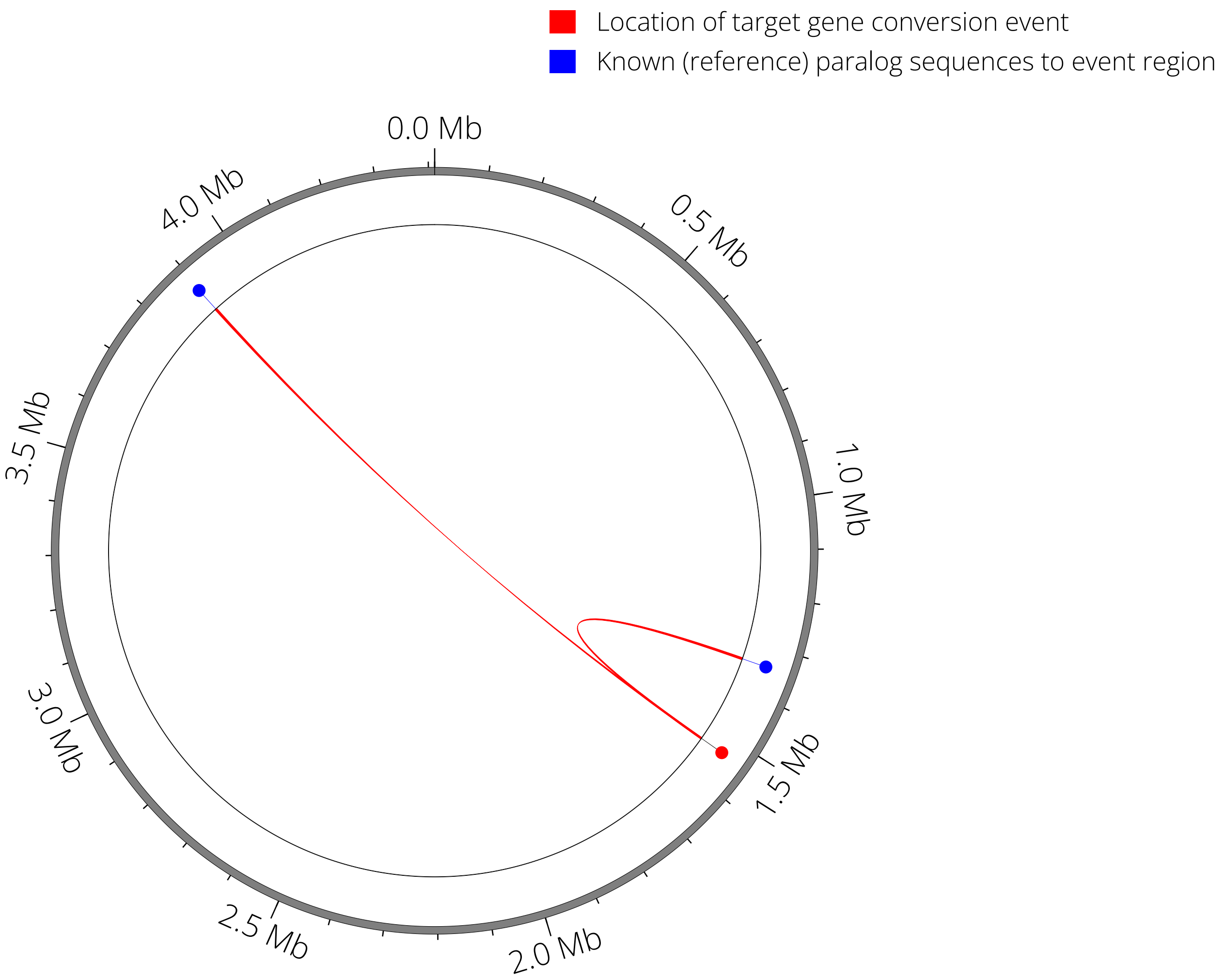

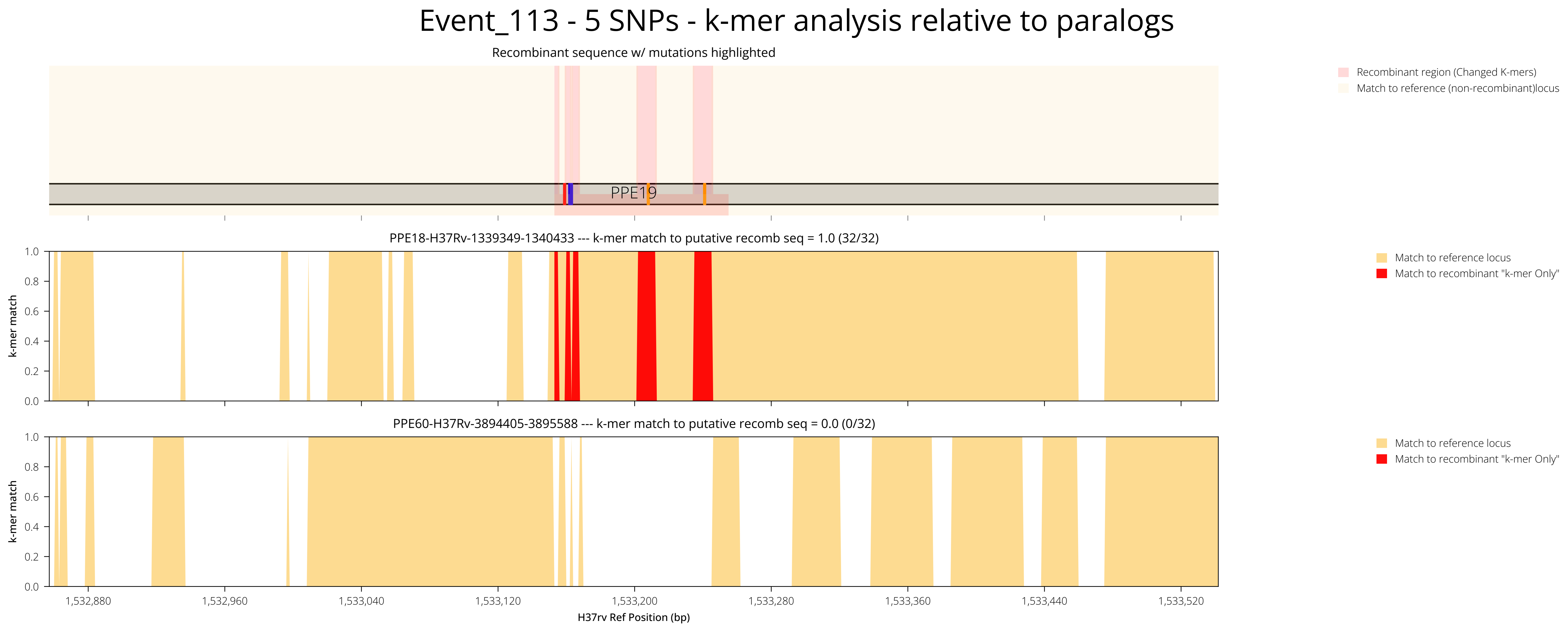
Event_113 - 5 SNPs - k-mer analysis relative to paralogs
Recombinant sequence w/ mutations highlighted
PPE19
Recombinant region (Changed K-mers)
Match to reference (non-recombinant)locus
PPE18-H37Rv-1339349-1340433 --- k-mer match to putative recomb seq = 1.0 (32/32)
Match to reference locus
Match to recombinant "k-mer Only"
PPE60-H37Rv-3894405-3895588 --- k-mer match to putative recomb seq = 0.0 (0/32)
Match to reference locus
Match to recombinant "k-mer Only"
k-mer match
k-mer match
1.0
0.8
0.6
0.4
0.2
0.0
1,532,880
1,532,960
1,533,040
1,533,120
1,533,200
1,533,280
1,533,360
1,533,440
1,533,520
H37rv Ref Position (bp)

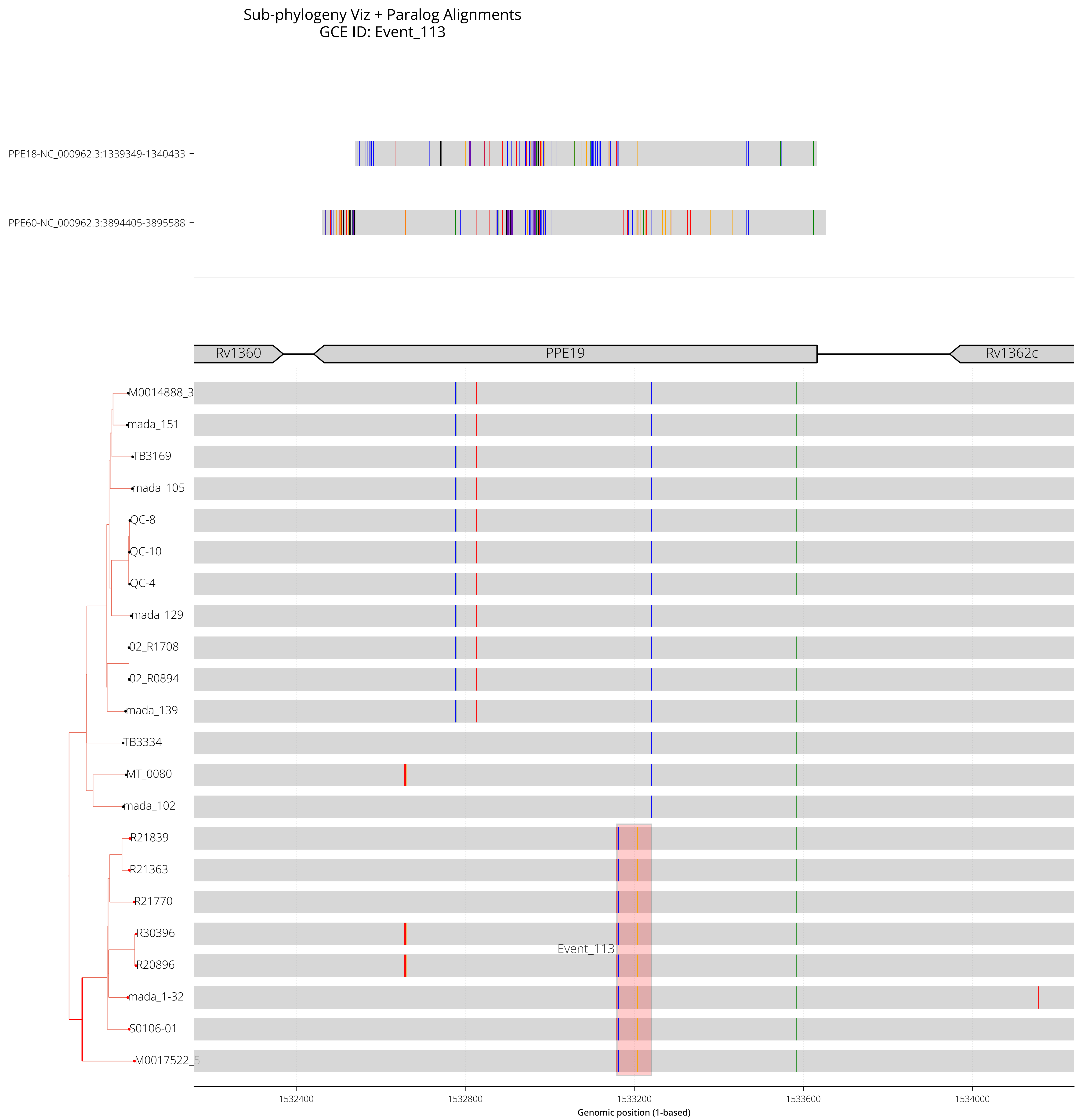

Sub-phylogeny Viz + Paralog Alignments
GCE ID: Event_113
PPE18-NC_000962.3:1339349-1340433
PPE60-NC_000962.3:3894405-3895588
Rv1360
PPE19
Rv1362c
M0014888_3
mada_151
TB3169
mada_105
QC-8
QC-10
QC-4
mada_129
02_R1708
02_R0894
mada_139
TB3334
MT_0080
mada_102
R21839
R21363
R21770
R30396
R20896
mada_1-32
S0106-01
M0017522_5
Event_113
1532400
1532800
1533200
1533600
1534000
Genomic position (1-based)

Full Phylogeny Viz + Paralog Alignments
GCE ID: Event_113

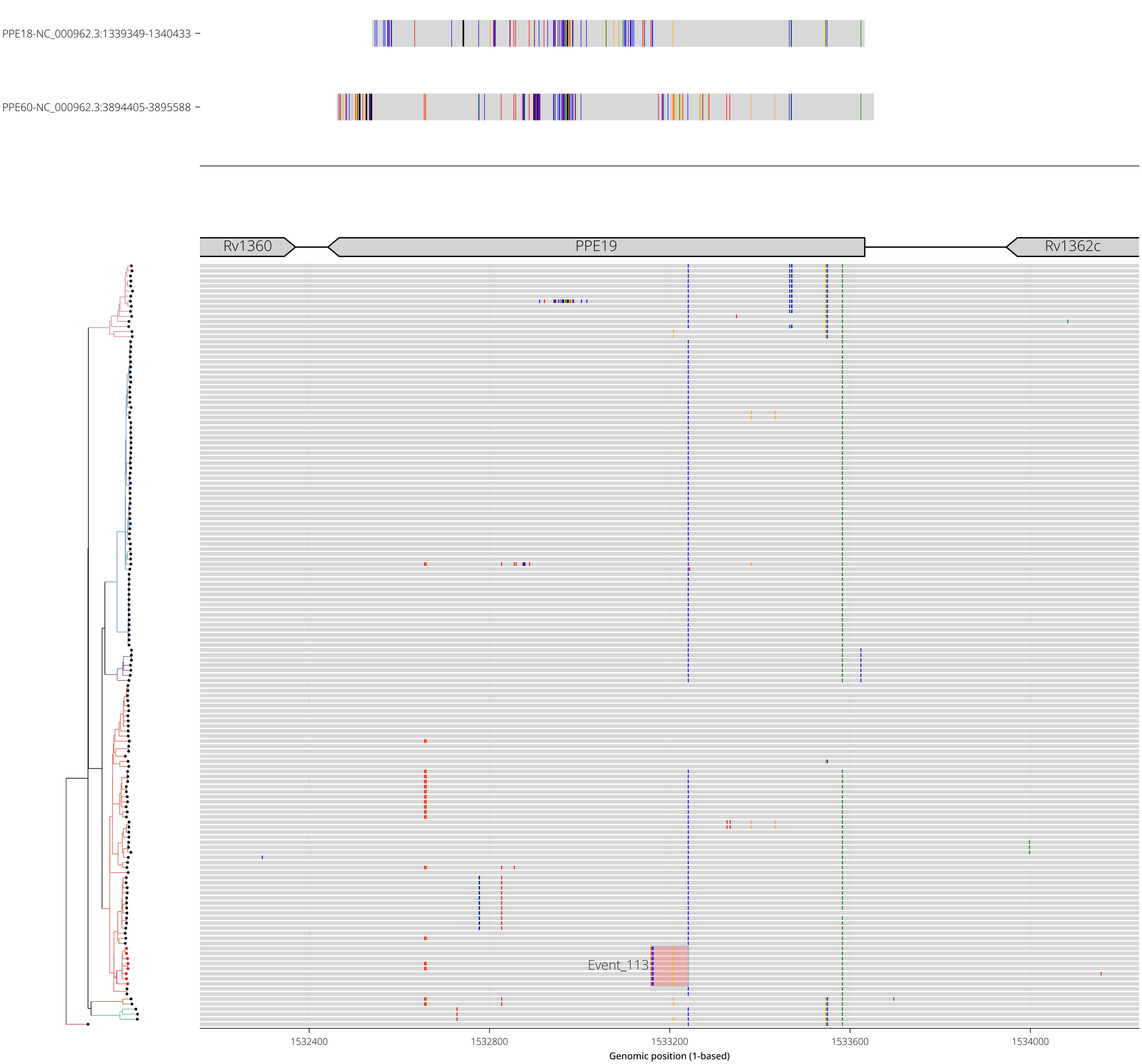

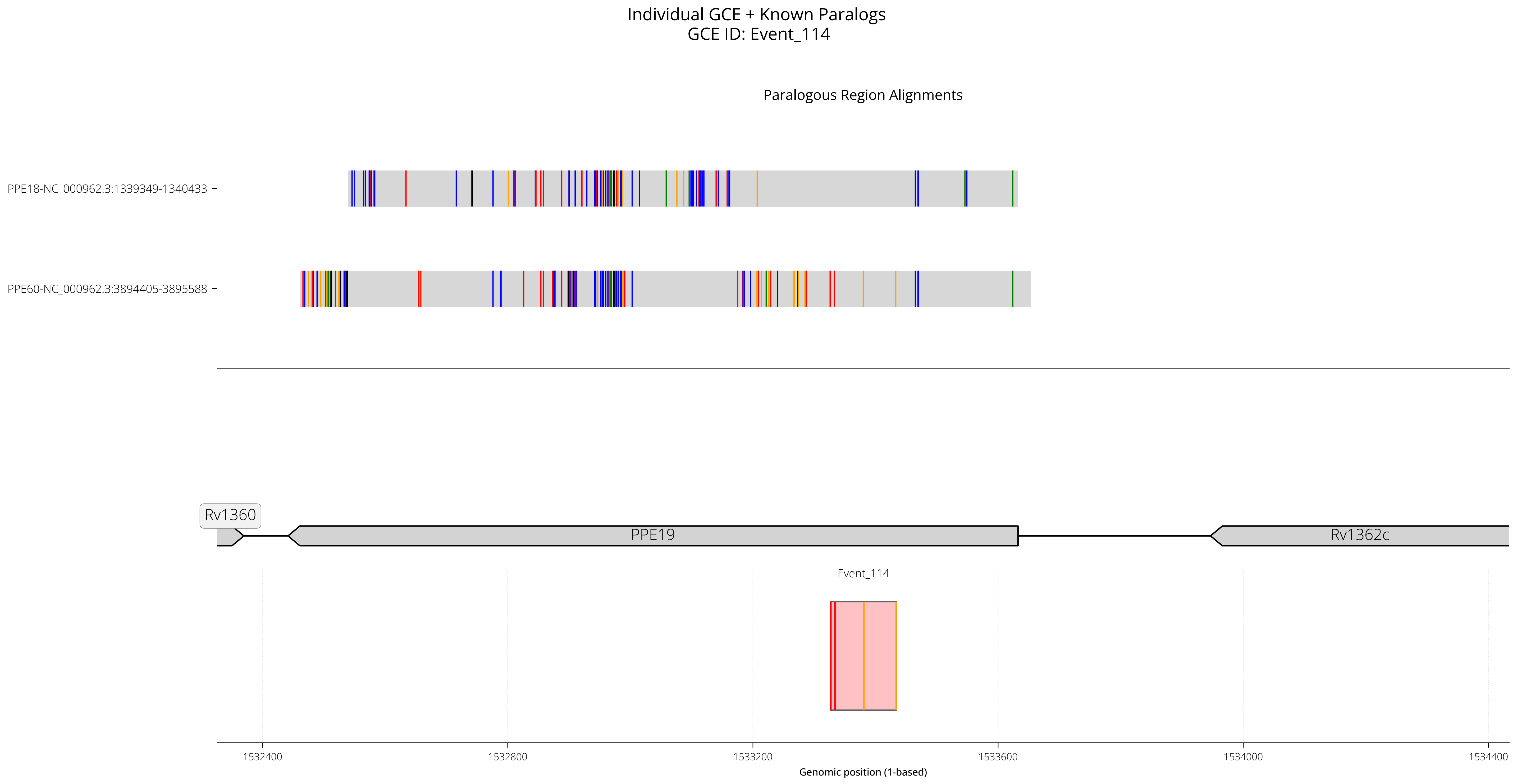

Individual GCE + Known Paralogs
GCE ID: Event_114
Paralogous Region Alignments
PPE18-NC_000962.3:1339349-1340433
PPE60-NC_000962.3:3894405-3895588
Rv1360
PPE19
Rv1362c
Event_114
1532400
1532800
1533200
1533600
1534000
1534400
Genomic position (1-based)

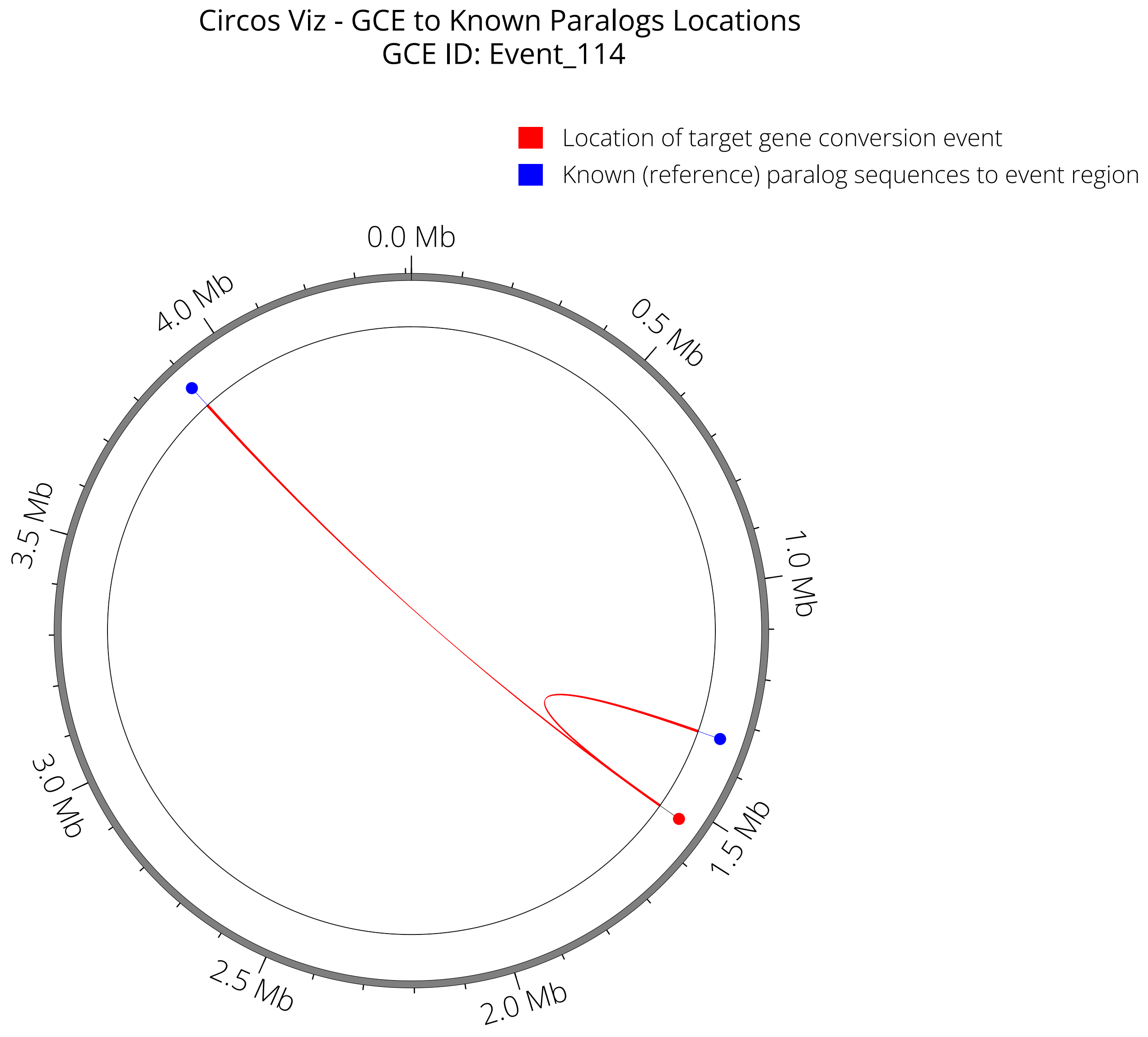
Circos Viz - GCE to Known Paralogs Locations
GCE ID: Event_114
Location of target gene conversion event
Known (reference) paralog sequences to event region
0.0 Mb
0.5 Mb
1.0 Mb
1.5 Mb
2.0 Mb
2.5 Mb
3.0 Mb
3.5 Mb
4.0 Mb

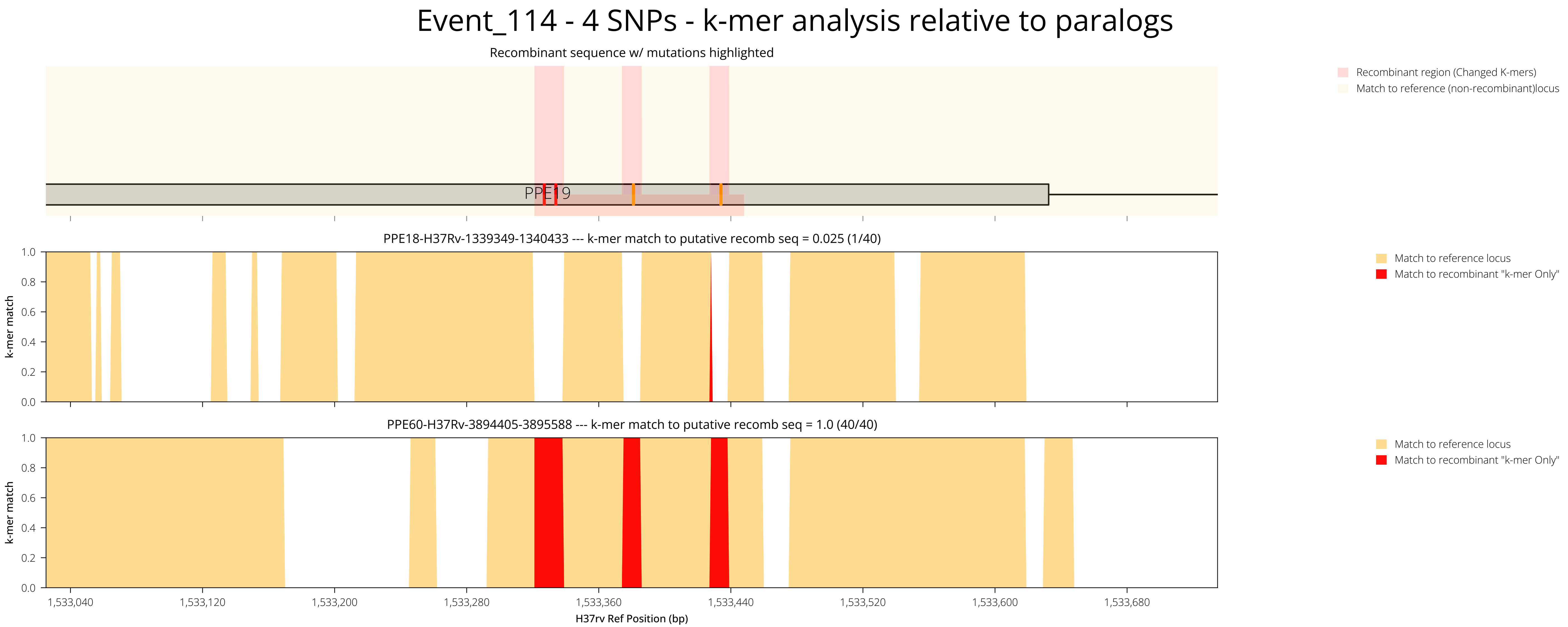
Event_114 - 4 SNPs - k-mer analysis relative to paralogs
Recombinant sequence w/ mutations highlighted
PPE19
Recombinant region (Changed K-mers)
Match to reference (non-recombinant)locus
PPE18-H37Rv-1339349-1340433 --- k-mer match to putative recomb seq = 0.025 (1/40)
Match to reference locus
Match to recombinant "k-mer Only"
k-mer match
1.0
0.8
0.6
0.4
0.2
0.0
PPE60-H37Rv-3894405-3895588 --- k-mer match to putative recomb seq = 1.0 (40/40)
Match to reference locus
Match to recombinant "k-mer Only"
k-mer match
1,533,040
1,533,120
1,533,200
1,533,280
1,533,360
1,533,440
1,533,520
1,533,600
1,533,680
H37rv Ref Position (bp)

Sub-phylogeny Viz + Paralog Alignments
GCE ID: Event_114

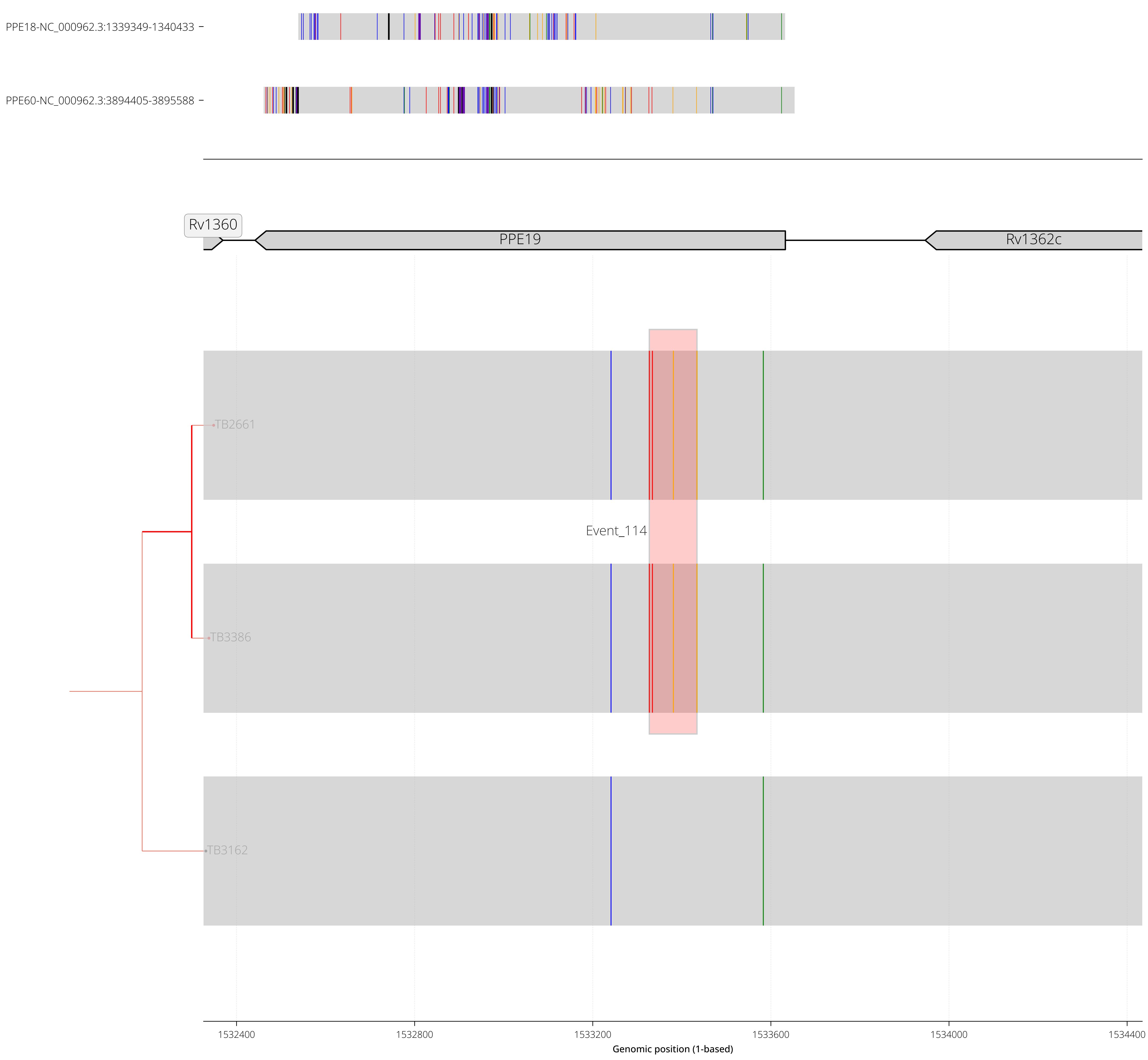

Full Phylogeny Viz + Paralog Alignments
GCE ID: Event_114

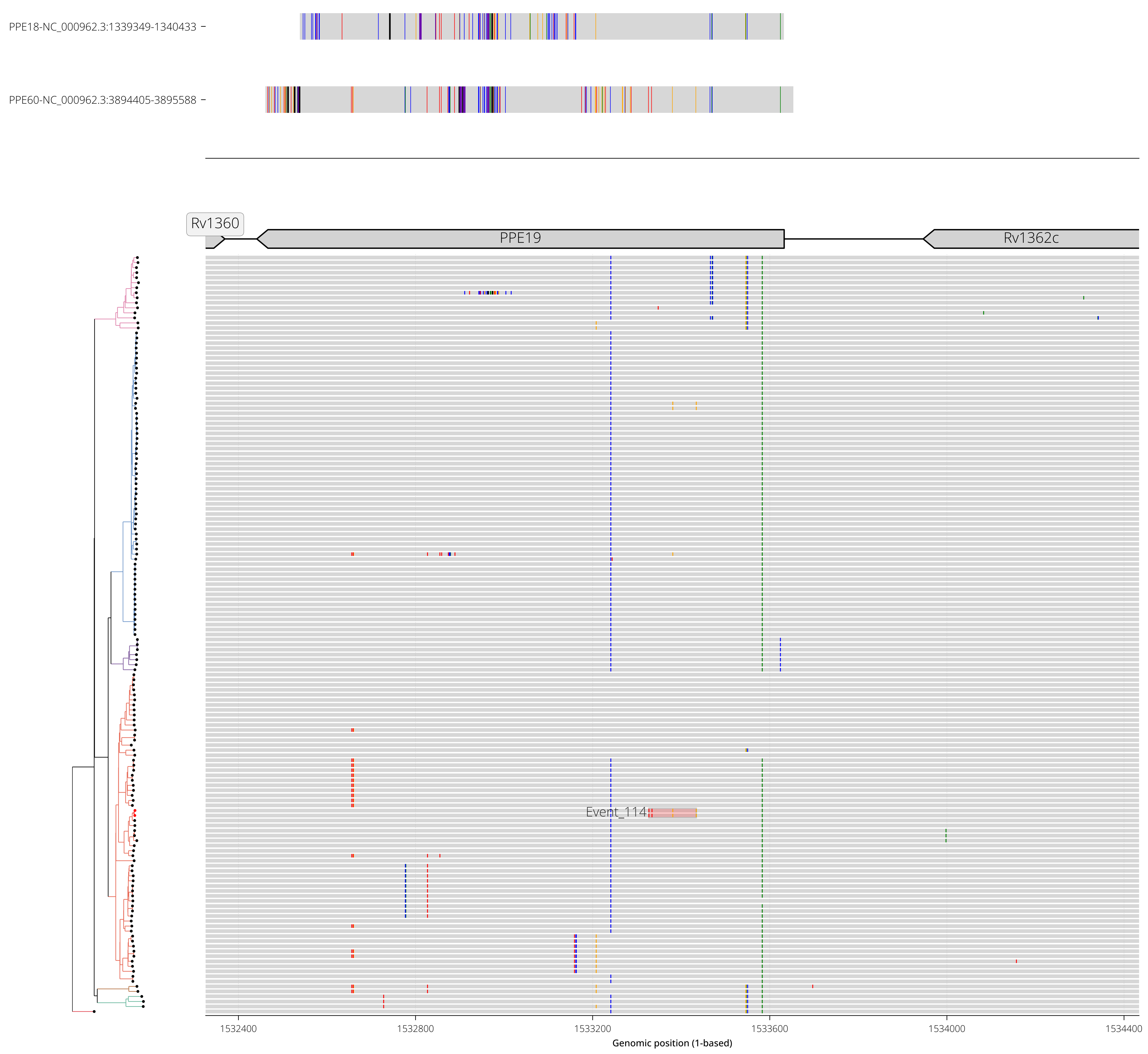

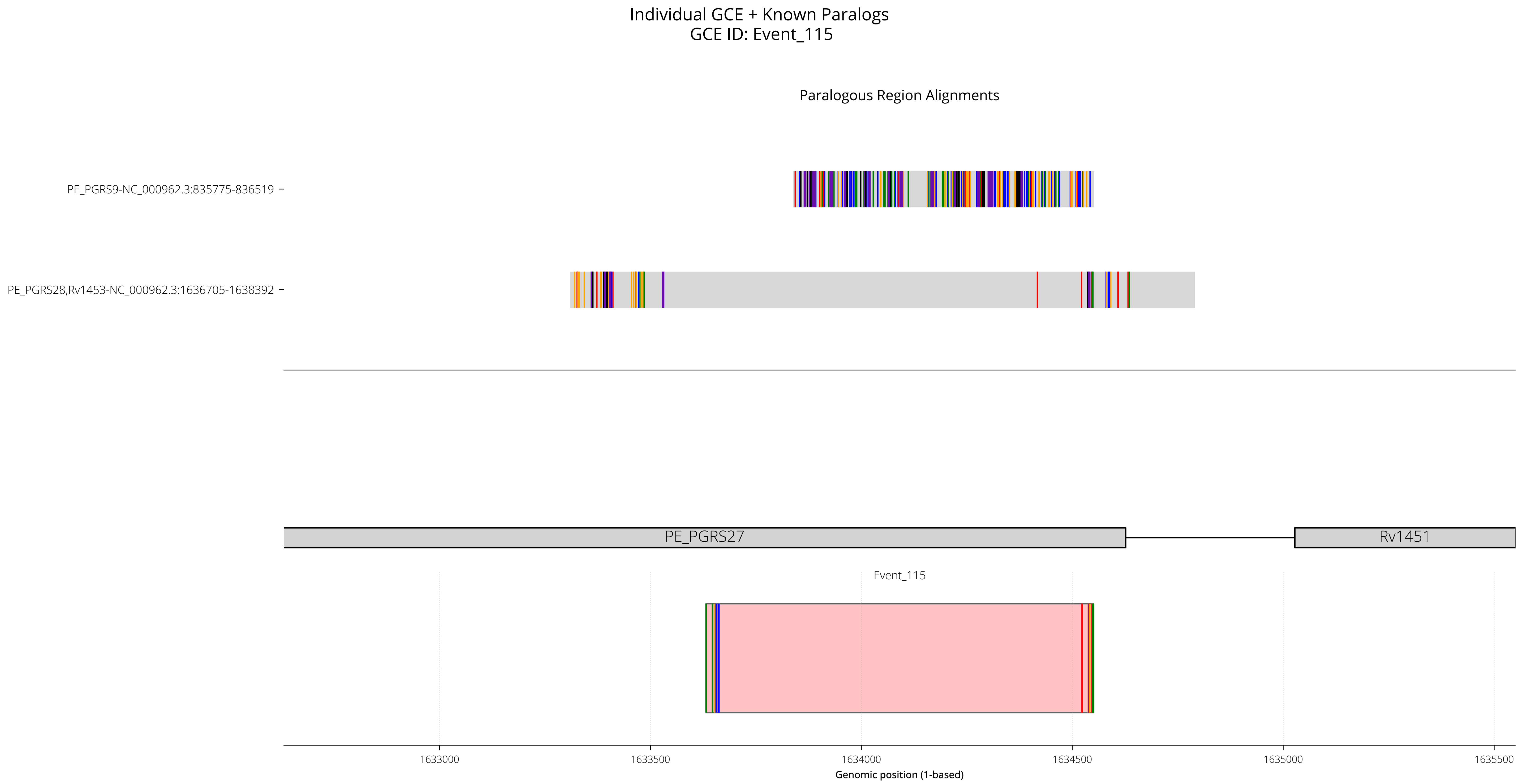
Individual GCE + Known Paralogs
GCE ID: Event_115
Paralogous Region Alignments
PE_PGRS9-NC_000962.3:835775-836519
PE_PGRS28,Rv1453-NC_000962.3:1636705-1638392
PE_PGRS27
Rv1451
Event_115
1633000
1633500
1634000
1634500
1635000
1635500
Genomic position (1-based)

Circos Viz - GCE to Known Paralogs Locations
GCE ID: Event_115

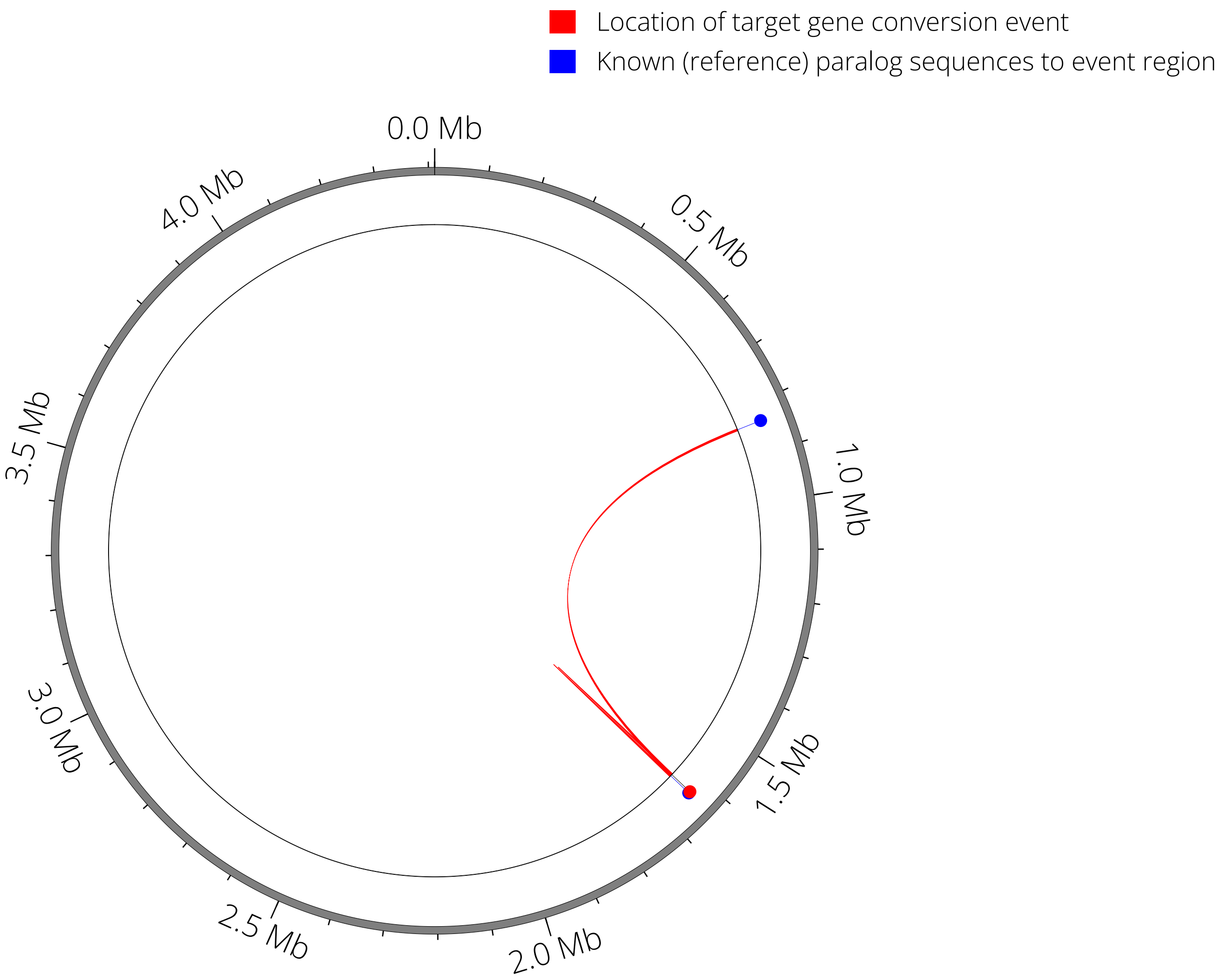

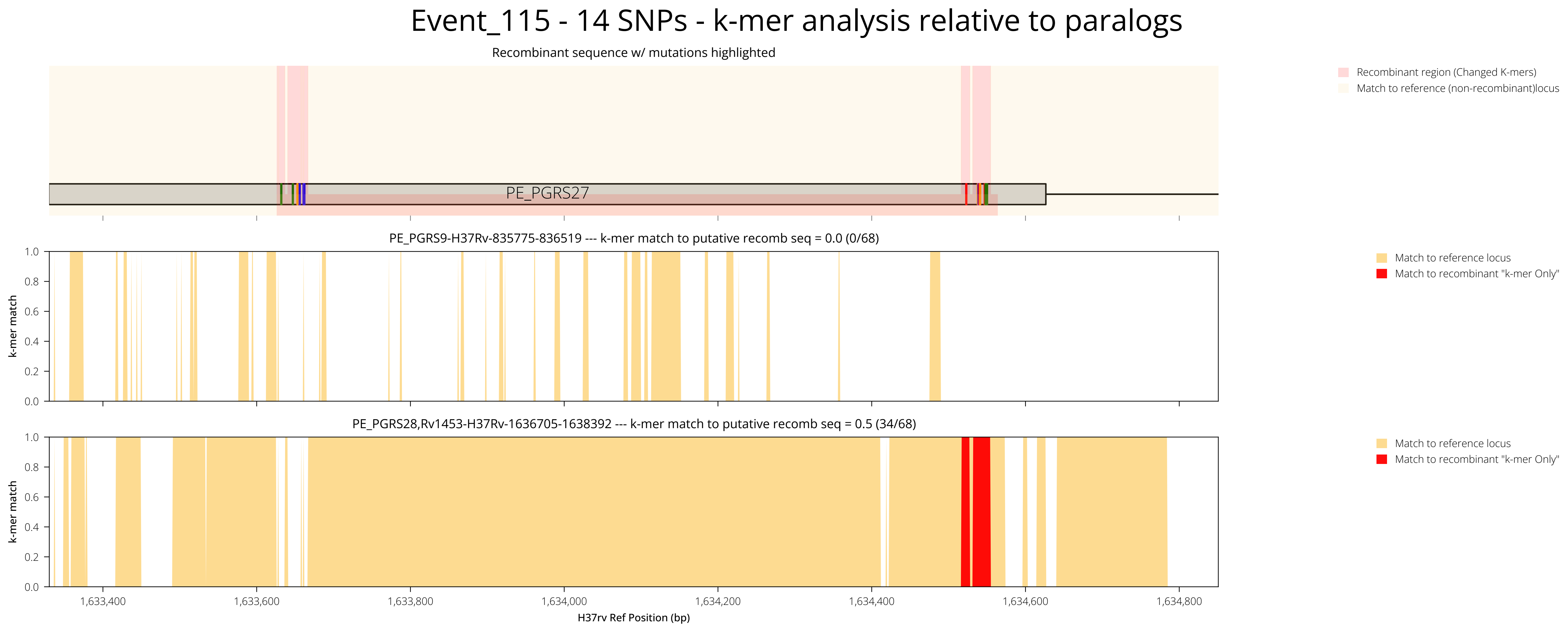

Event_115 - 14 SNPs - k-mer analysis relative to paralogs
Recombinant sequence w/ mutations highlighted
Recombinant region (Changed K-mers)
Match to reference (non-recombinant)locus
PE_PGRS27
PE_PGRS9-H37Rv-835775-836519 --- k-mer match to putative recomb seq = 0.0 (0/68)
Match to reference locus
Match to recombinant "k-mer Only"
PE_PGRS28,Rv1453-H37Rv-1636705-1638392 --- k-mer match to putative recomb seq = 0.5 (34/68)
Match to reference locus
Match to recombinant "k-mer Only"
k-mer match
1.0
0.8
0.6
0.4
0.2
0.0
1,633,400
1,633,600
1,633,800
1,634,000
1,634,200
1,634,400
1,634,600
1,634,800
H37rv Ref Position (bp)

Sub-phylogeny Viz + Paralog Alignments
GCE ID: Event_115

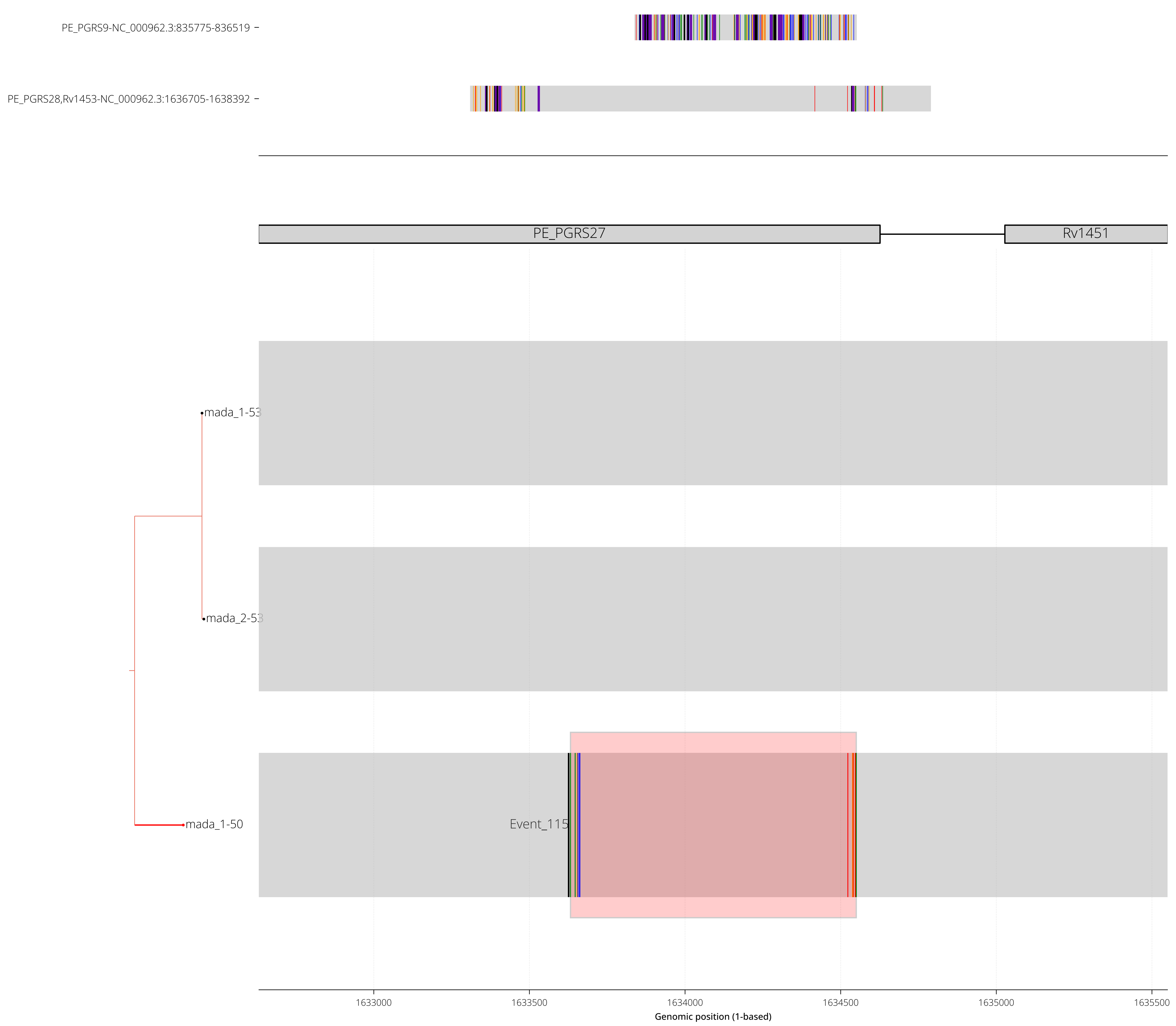

Full Phylogeny Viz + Paralog Alignments
GCE ID: Event_115

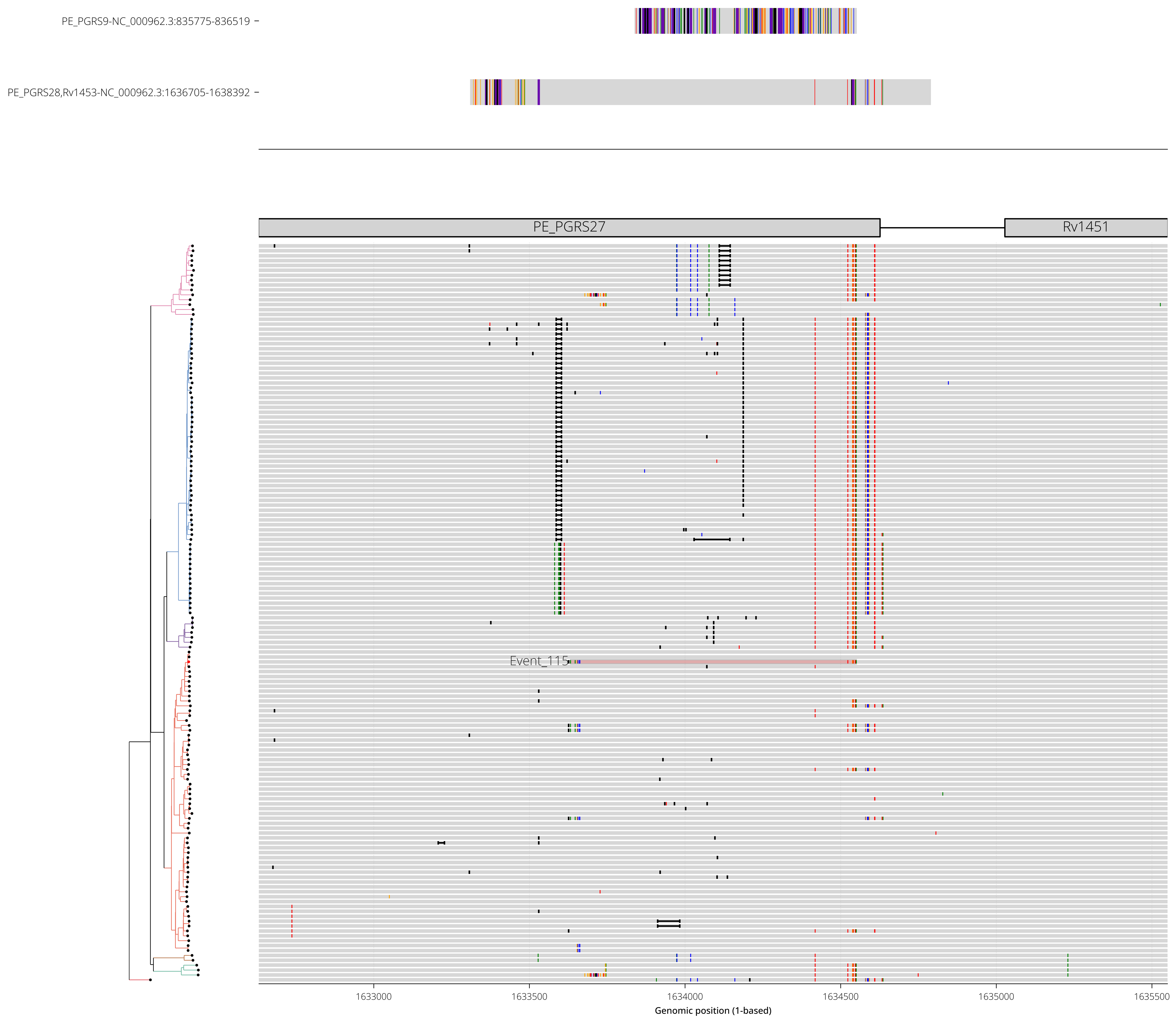

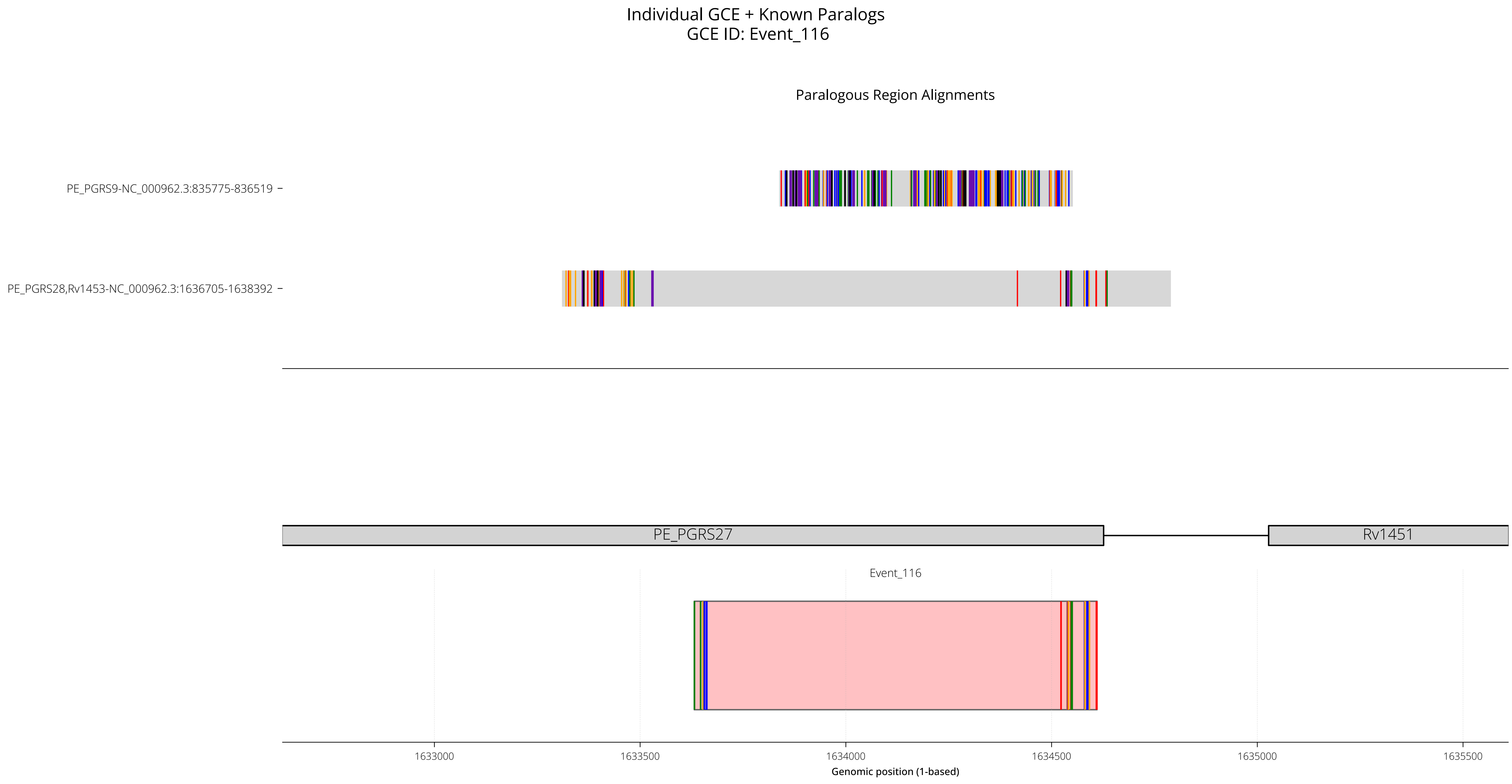
Individual GCE + Known Paralogs
GCE ID: Event_116
Paralogous Region Alignments
PE_PGRS9-NC_000962.3:835775-836519
PE_PGRS28,Rv1453-NC_000962.3:1636705-1638392
PE_PGRS27
Rv1451
Event_116
1633000
1633500
1634000
1634500
1635000
1635500
Genomic position (1-based)

### Circos Viz - GCE to Known Paralogs Locations
#### GCE ID: Event_116

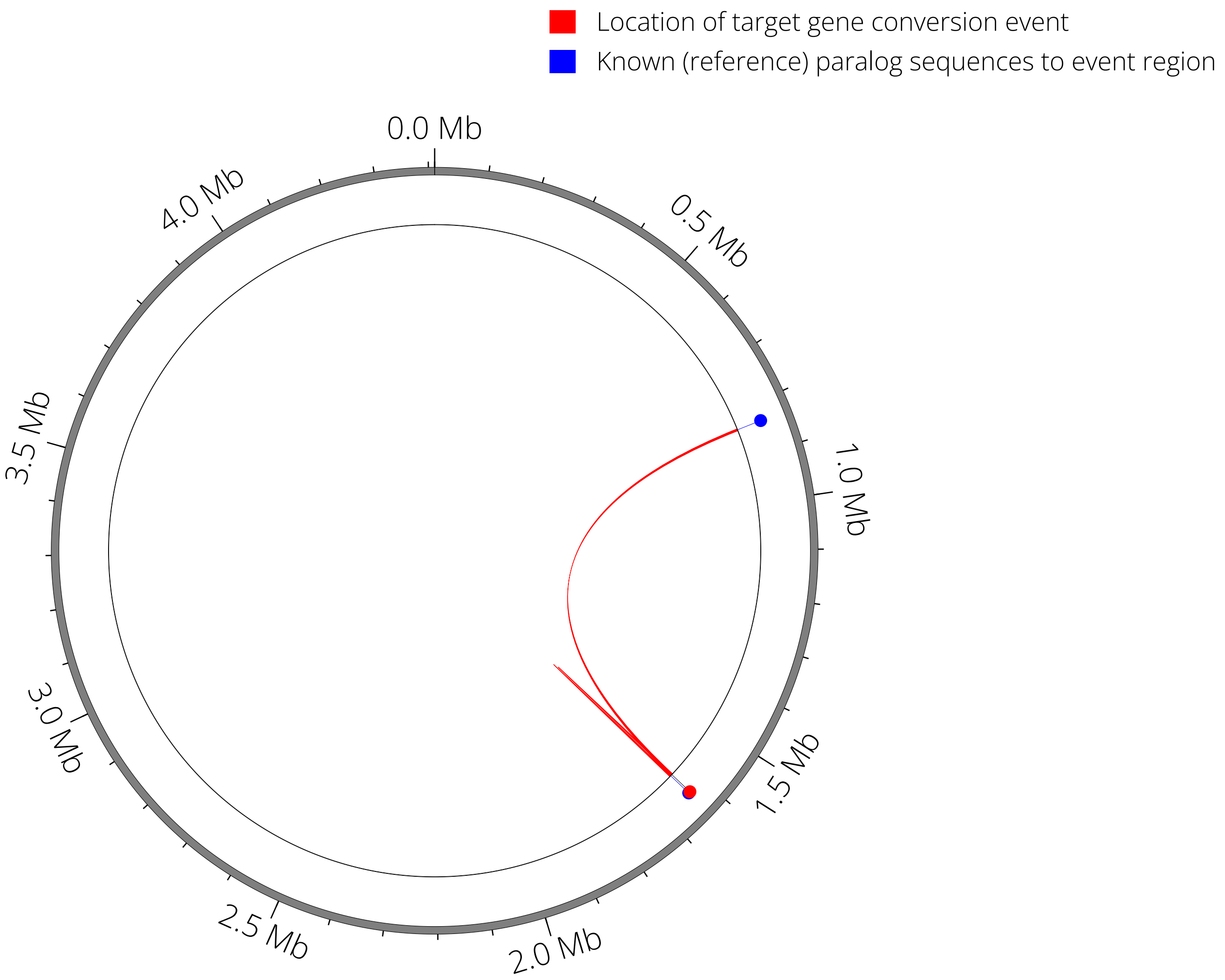

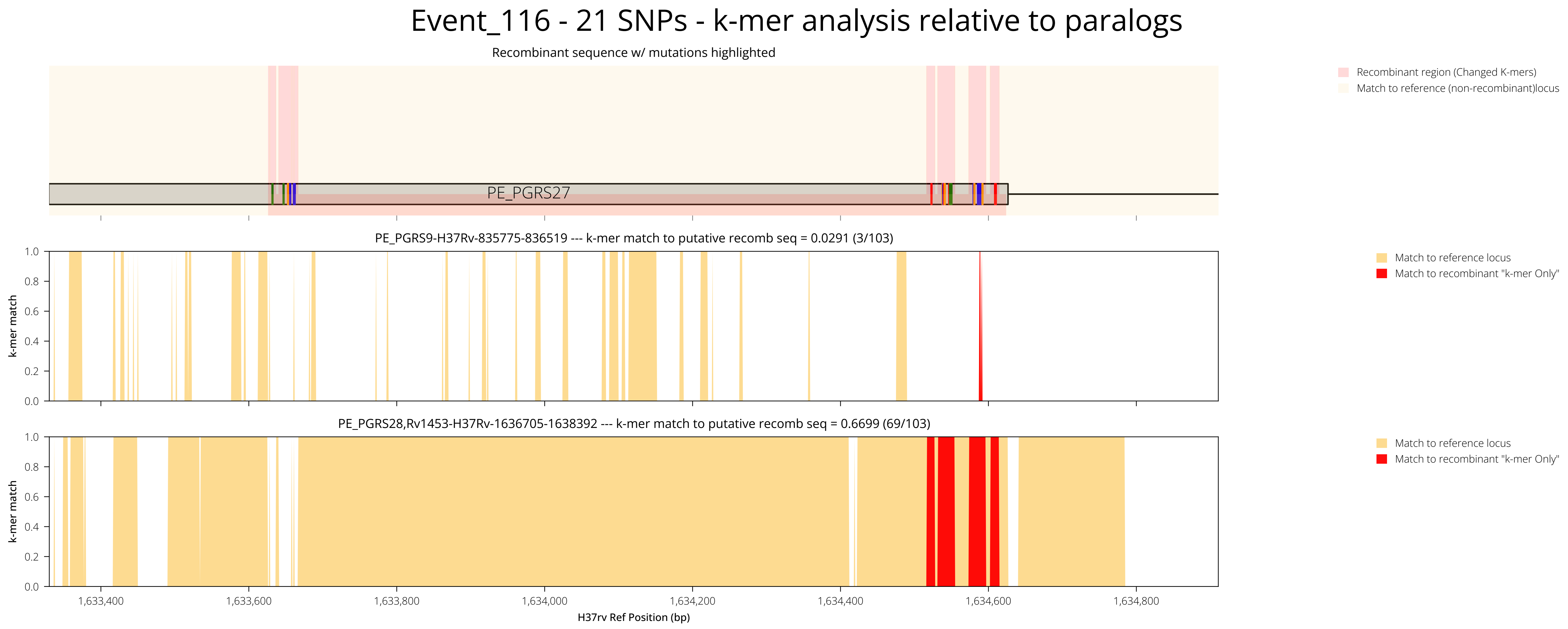

Event_116 - 21 SNPs - k-mer analysis relative to paralogs
Recombinant sequence w/ mutations highlighted
PE_PGRS27
Recombinant region (Changed K-mers)
Match to reference (non-recombinant)locus
PE_PGRS9-H37Rv-835775-836519 --- k-mer match to putative recomb seq = 0.0291 (3/103)
Match to reference locus
Match to recombinant "k-mer Only"
PE_PGRS28,Rv1453-H37Rv-1636705-1638392 --- k-mer match to putative recomb seq = 0.6699 (69/103)
Match to reference locus
Match to recombinant "k-mer Only"
k-mer match
1.0
0.8
0.6
0.4
0.2
0.0
1,633,400
1,633,600
1,633,800
1,634,000
1,634,200
1,634,400
1,634,600
1,634,800
H37rv Ref Position (bp)

Sub-phylogeny Viz + Paralog Alignments
GCE ID: Event_116

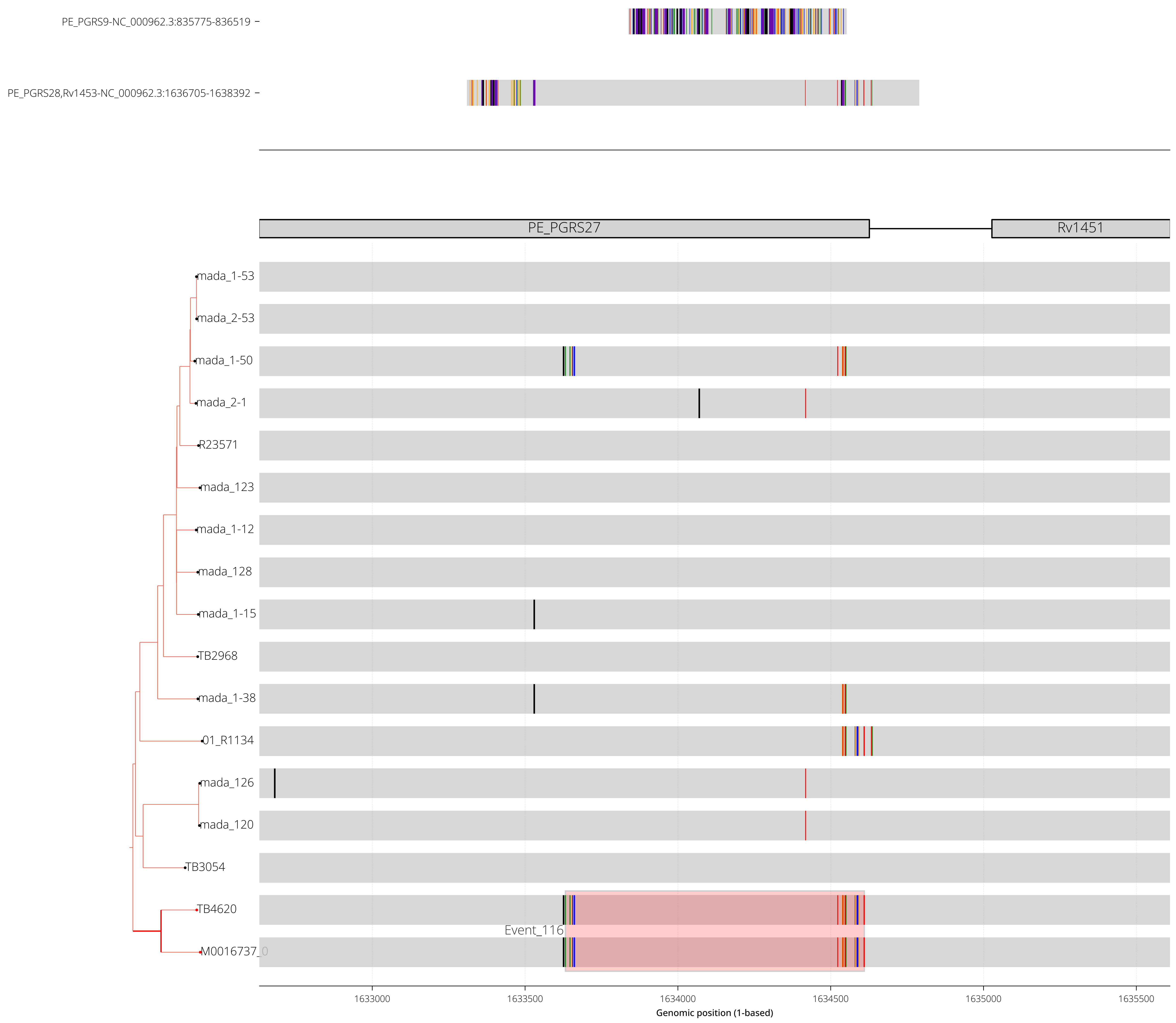

Full Phylogeny Viz + Paralog Alignments
GCE ID: Event_116

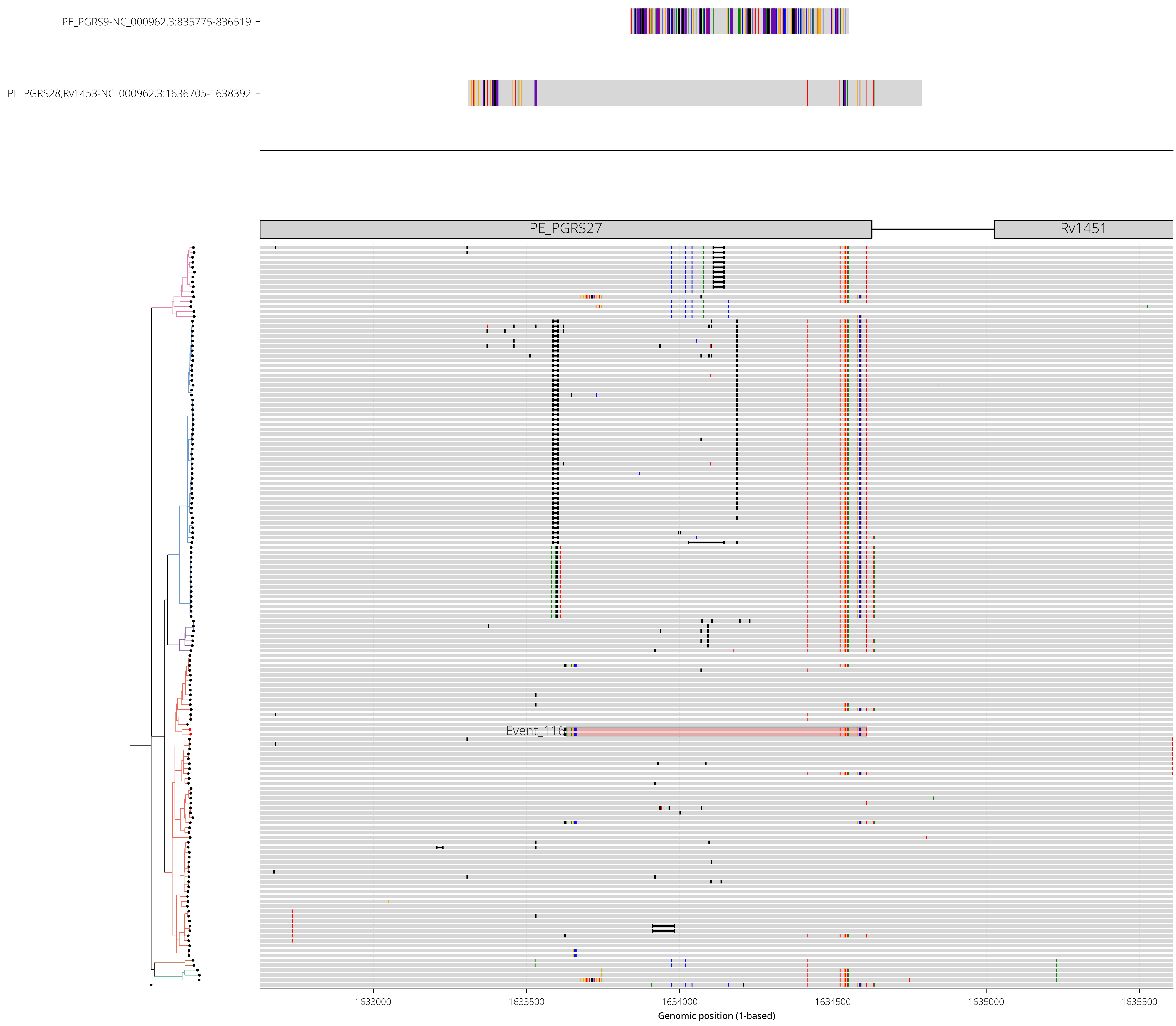

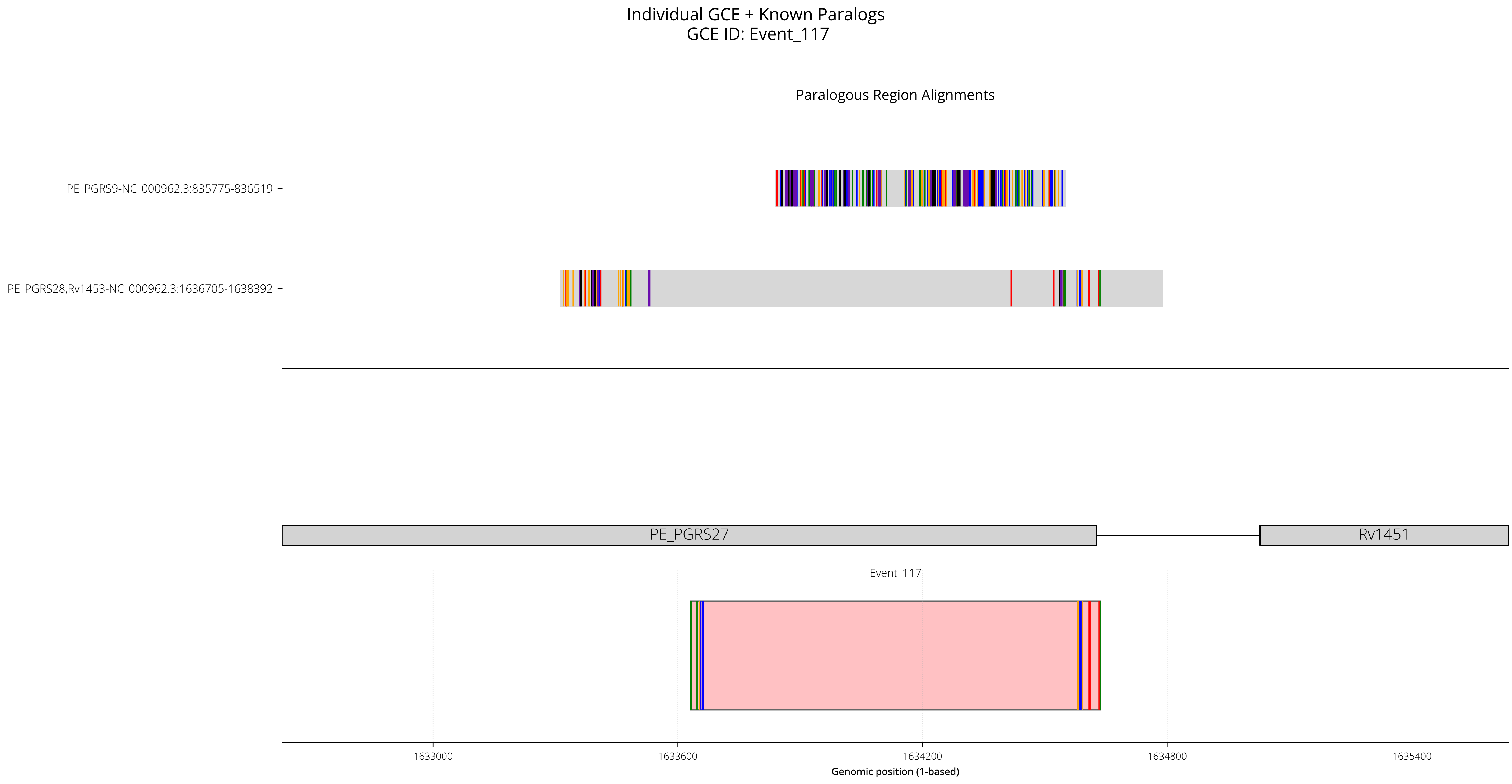

Individual GCE + Known Paralogs
GCE ID: Event_117
Paralogous Region Alignments
PE_PGRS9-NC_000962.3:835775-836519
PE_PGRS28,Rv1453-NC_000962.3:1636705-1638392
PE_PGRS27
Rv1451
Event_117
1633000
1633600
1634200
1634800
1635400
Genomic position (1-based)

Circos Viz - GCE to Known Paralogs Locations
GCE ID: Event_117

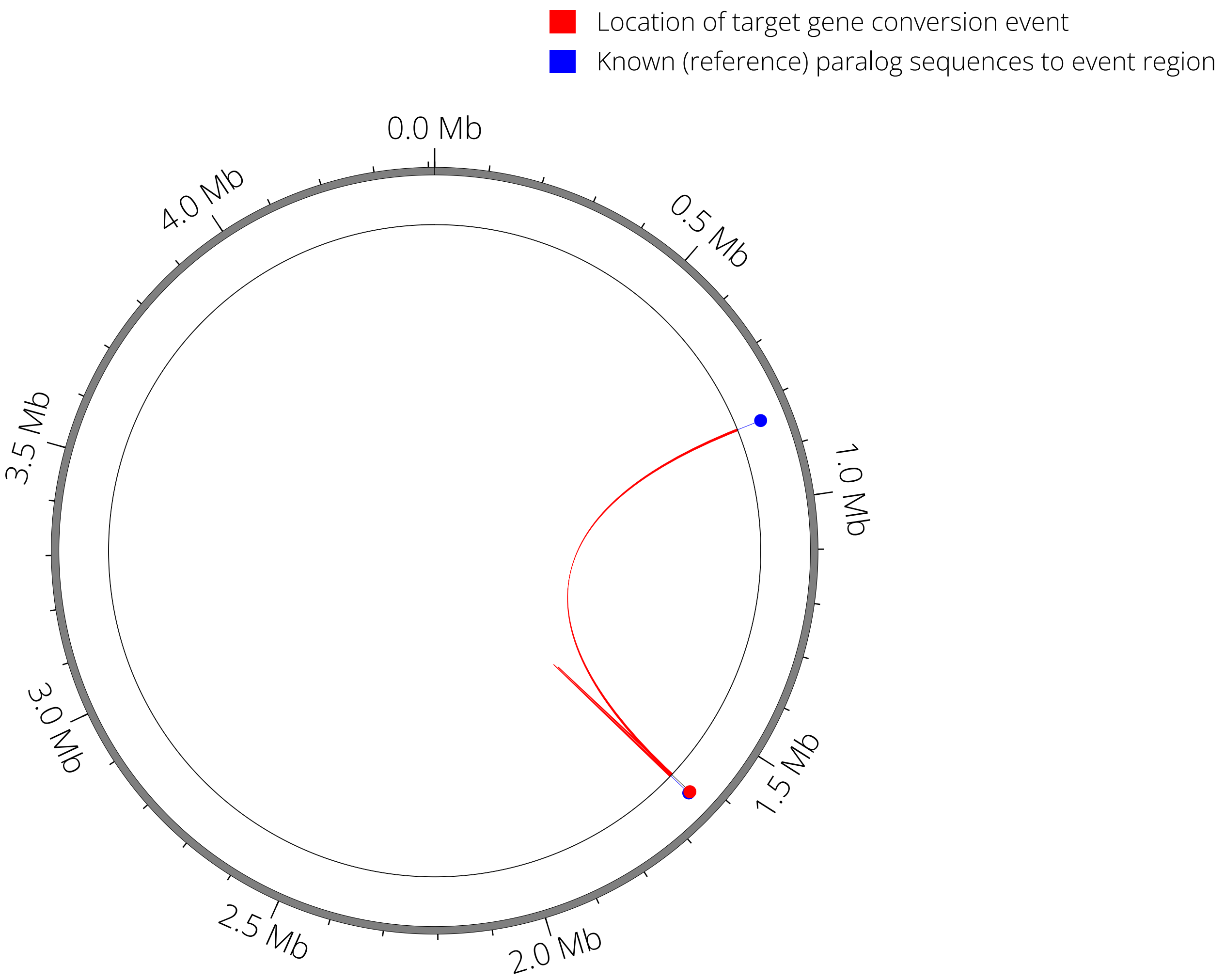

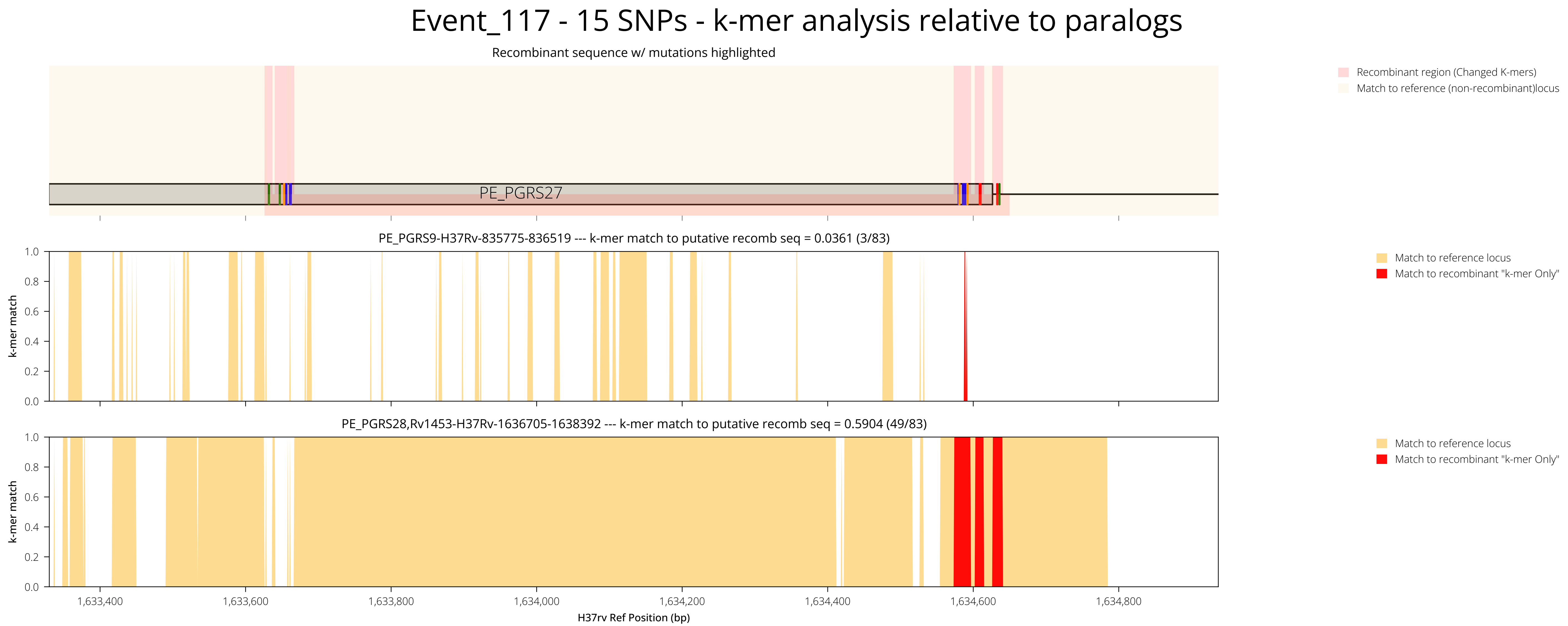
Event_117 - 15 SNPs - k-mer analysis relative to paralogs
Recombinant sequence w/ mutations highlighted
Recombinant region (Changed K-mers)
Match to reference (non-recombinant)locus
PE_PGRS27
PE_PGRS9-H37Rv-835775-836519 --- k-mer match to putative recomb seq = 0.0361 (3/83)
Match to reference locus
Match to recombinant "k-mer Only"
k-mer match
1.0
0.8
0.6
0.4
0.2
0.0
PE_PGRS28,Rv1453-H37Rv-1636705-1638392 --- k-mer match to putative recomb seq = 0.5904 (49/83)
Match to reference locus
Match to recombinant "k-mer Only"
k-mer match
1,633,400
1,633,600
1,633,800
1,634,000
1,634,200
1,634,400
1,634,600
1,634,800
H37rv Ref Position (bp)

Sub-phylogeny Viz + Paralog Alignments
GCE ID: Event_117

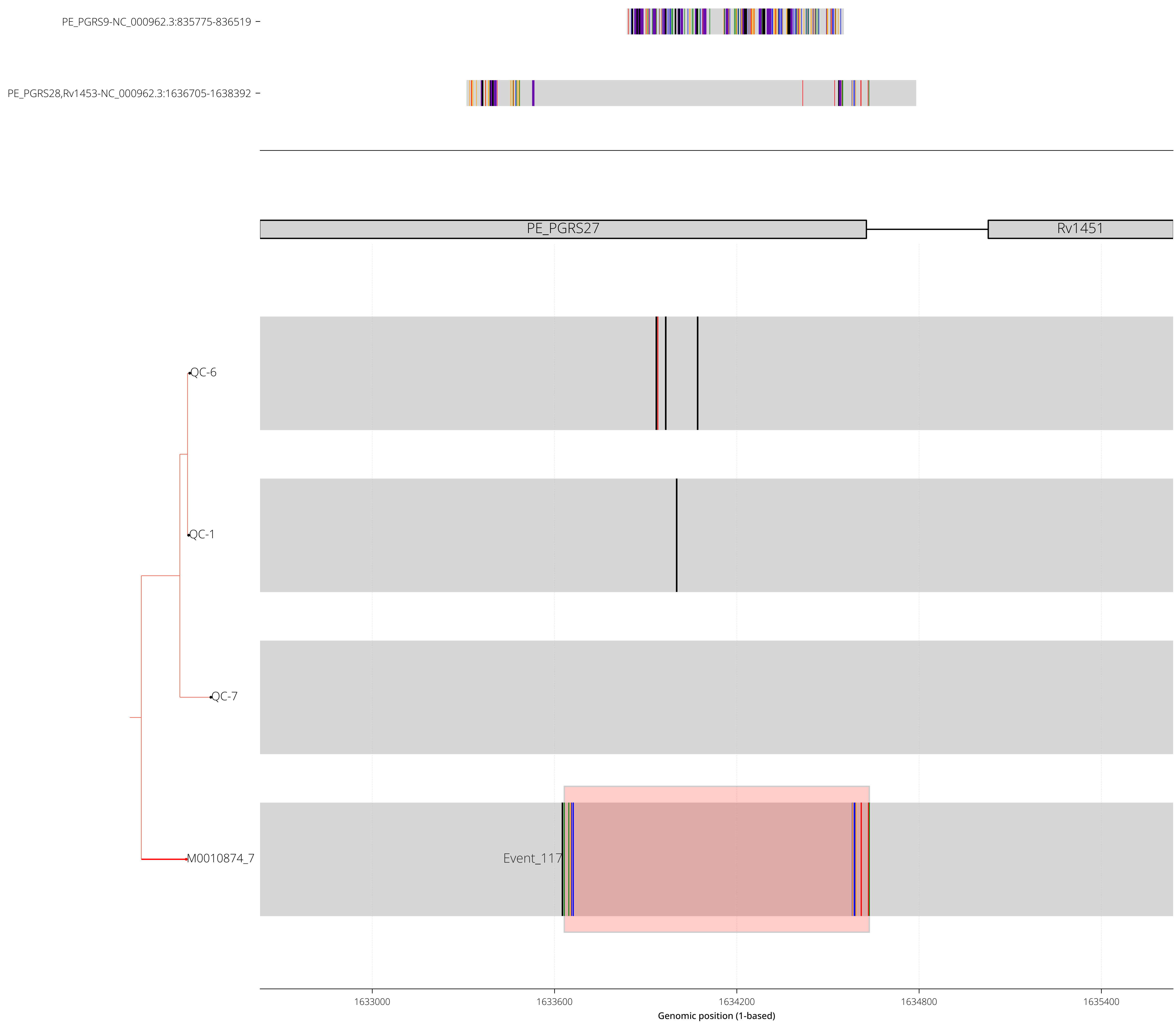

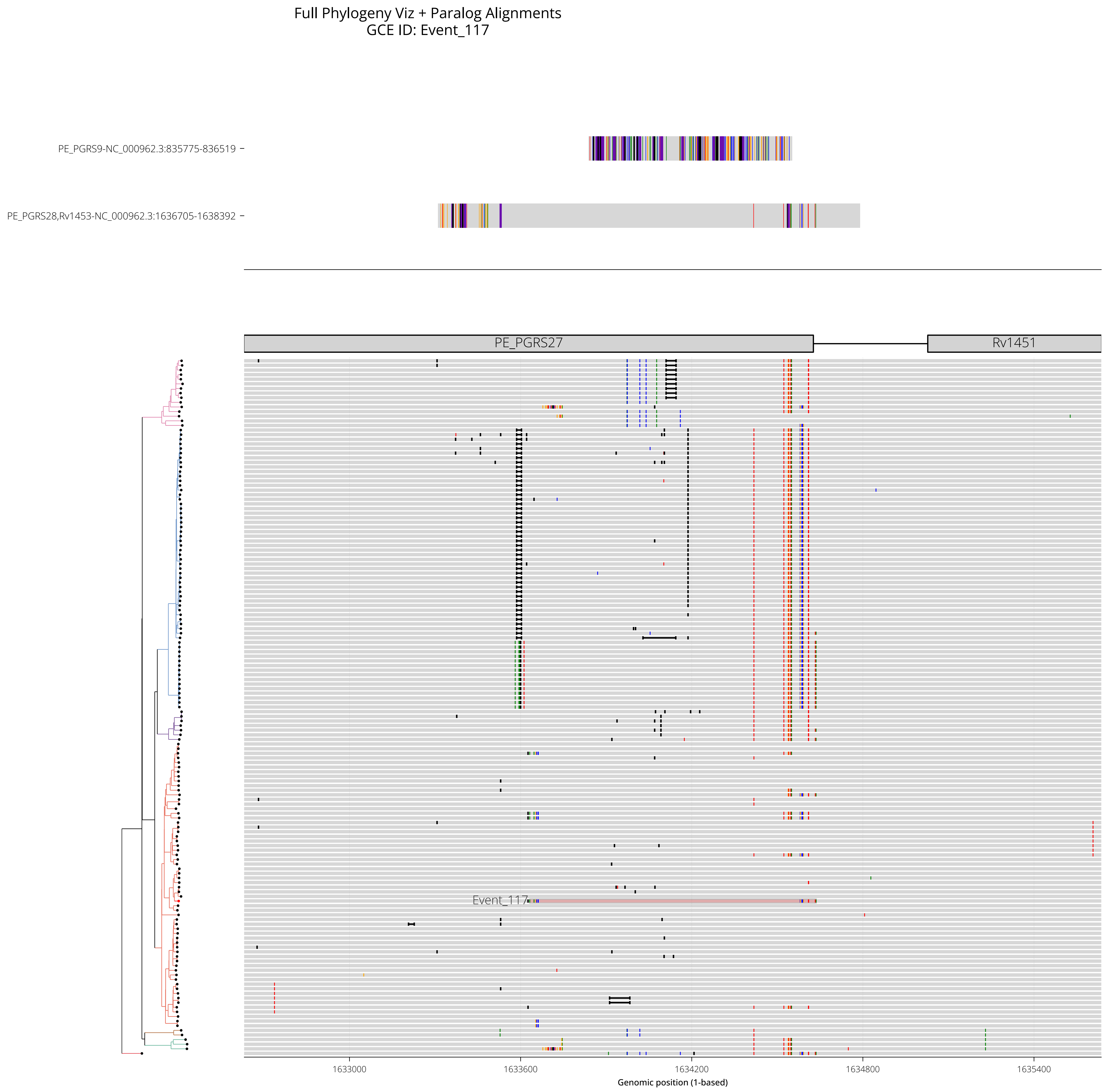
Full Phylogeny Viz + Paralog Alignments
GCE ID: Event_117
PE_PGRS9-NC_000962.3:835775-836519
PE_PGRS28,Rv1453-NC_000962.3:1636705-1638392
PE_PGRS27
Rv1451
Event_117
1633000
1633600
1634200
1634800
1635400
Genomic position (1-based)

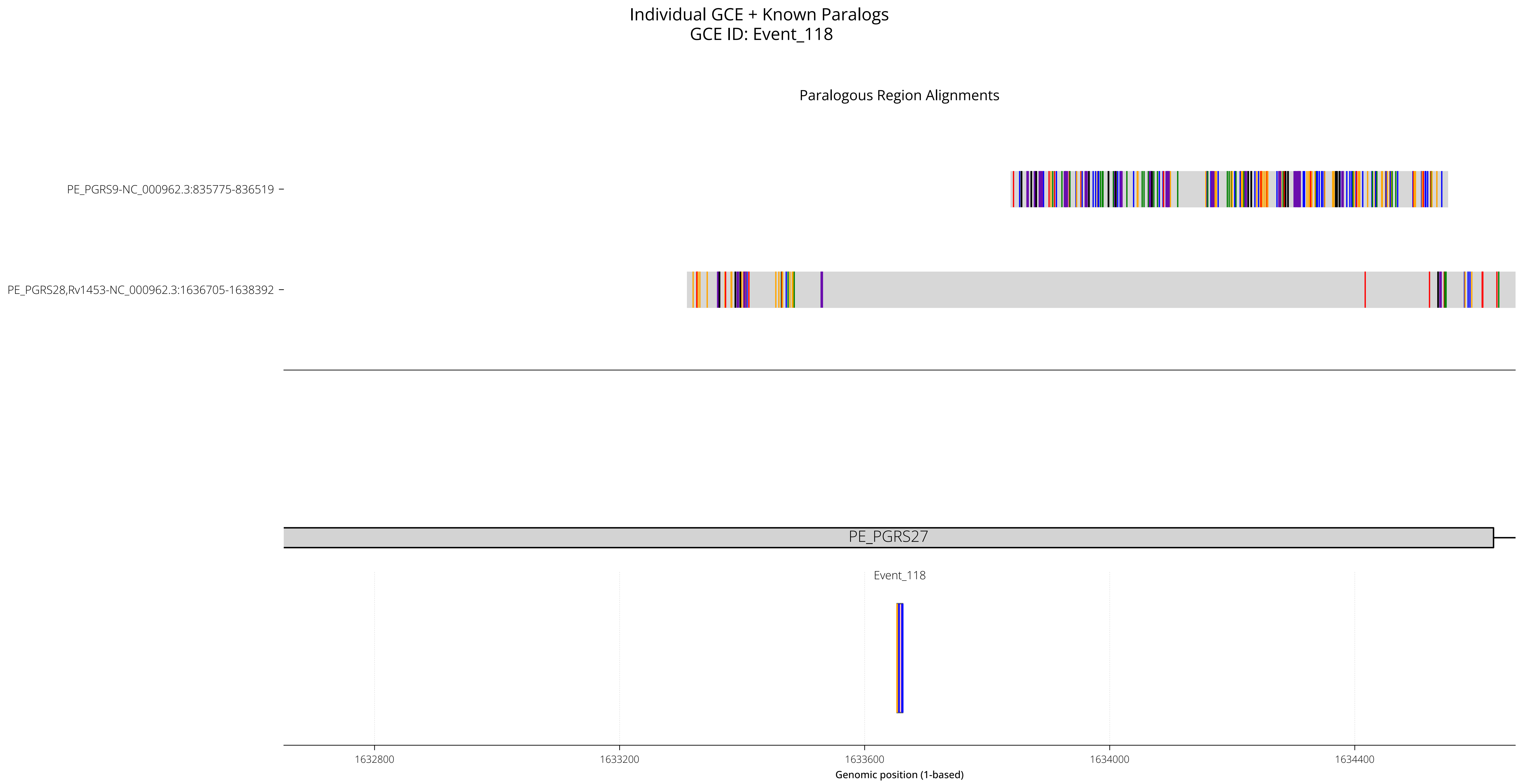

Individual GCE + Known Paralogs
GCE ID: Event_118
Paralogous Region Alignments
PE_PGRS9-NC_000962.3:835775-836519
PE_PGRS28,Rv1453-NC_000962.3:1636705-1638392
PE_PGRS27
Event_118
1632800
1633200
1633600
1634000
1634400
Genomic position (1-based)

Circos Viz - GCE to Known Paralogs Locations
GCE ID: Event_118

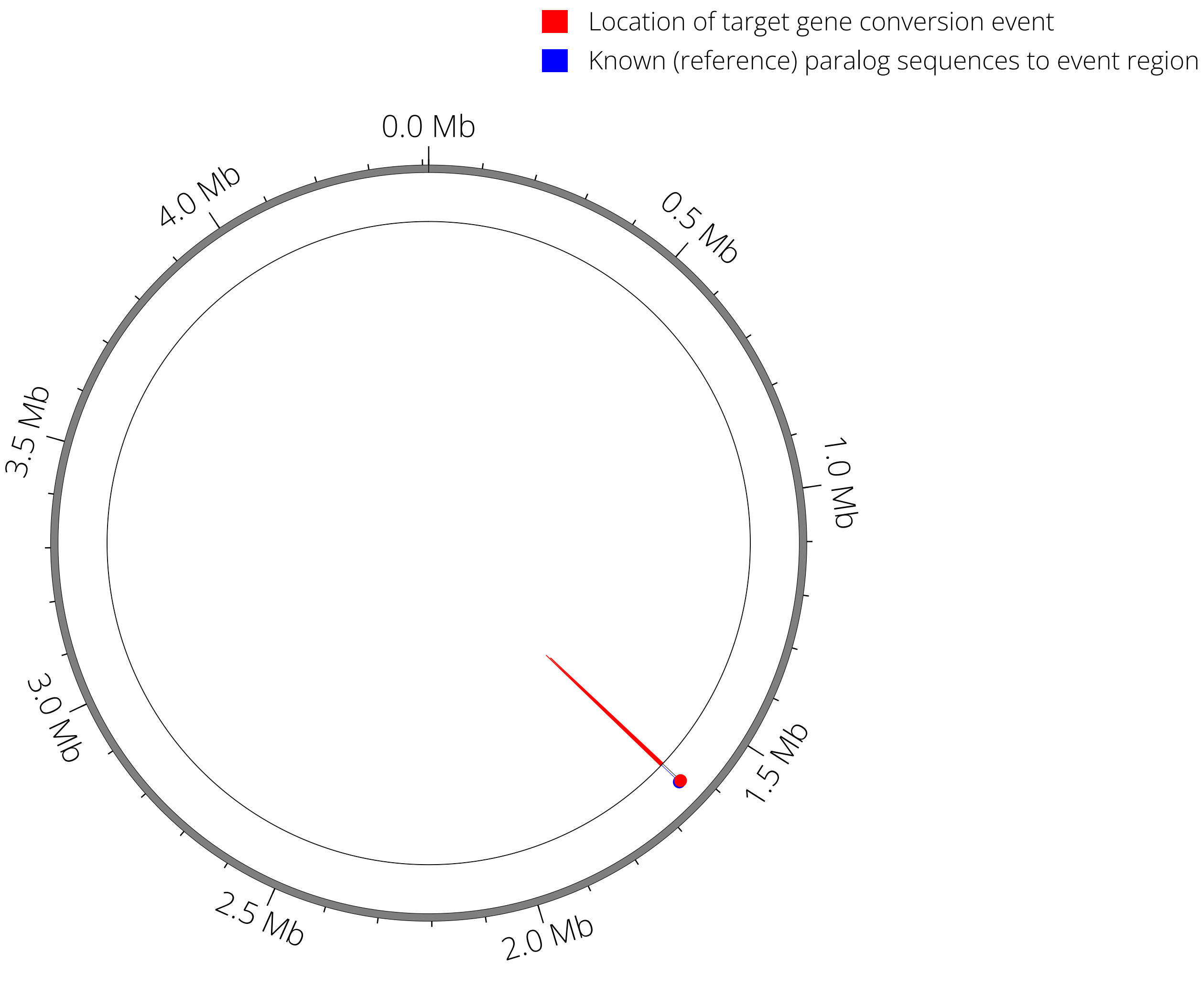

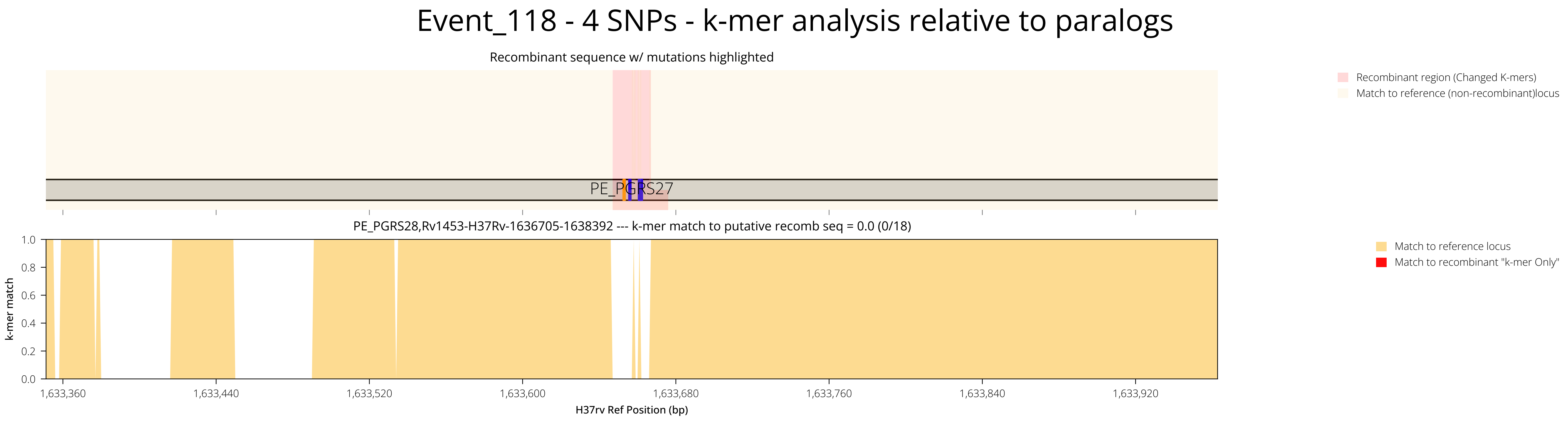
Event_118 - 4 SNPs - k-mer analysis relative to paralogs
Recombinant sequence w/ mutations highlighted
Recombinant region (Changed K-mers)
Match to reference (non-recombinant)locus
PE_PGRS27
PE_PGRS28,Rv1453-H37Rv-1636705-1638392 --- k-mer match to putative recomb seq = 0.0 (0/18)
Match to reference locus
Match to recombinant "k-mer Only"
k-mer match
1.0
0.8
0.6
0.4
0.2
0.0
1,633,360
1,633,440
1,633,520
1,633,600
1,633,680
1,633,760
1,633,840
1,633,920
H37rv Ref Position (bp)

Sub-phylogeny Viz + Paralog Alignments
GCE ID: Event_118

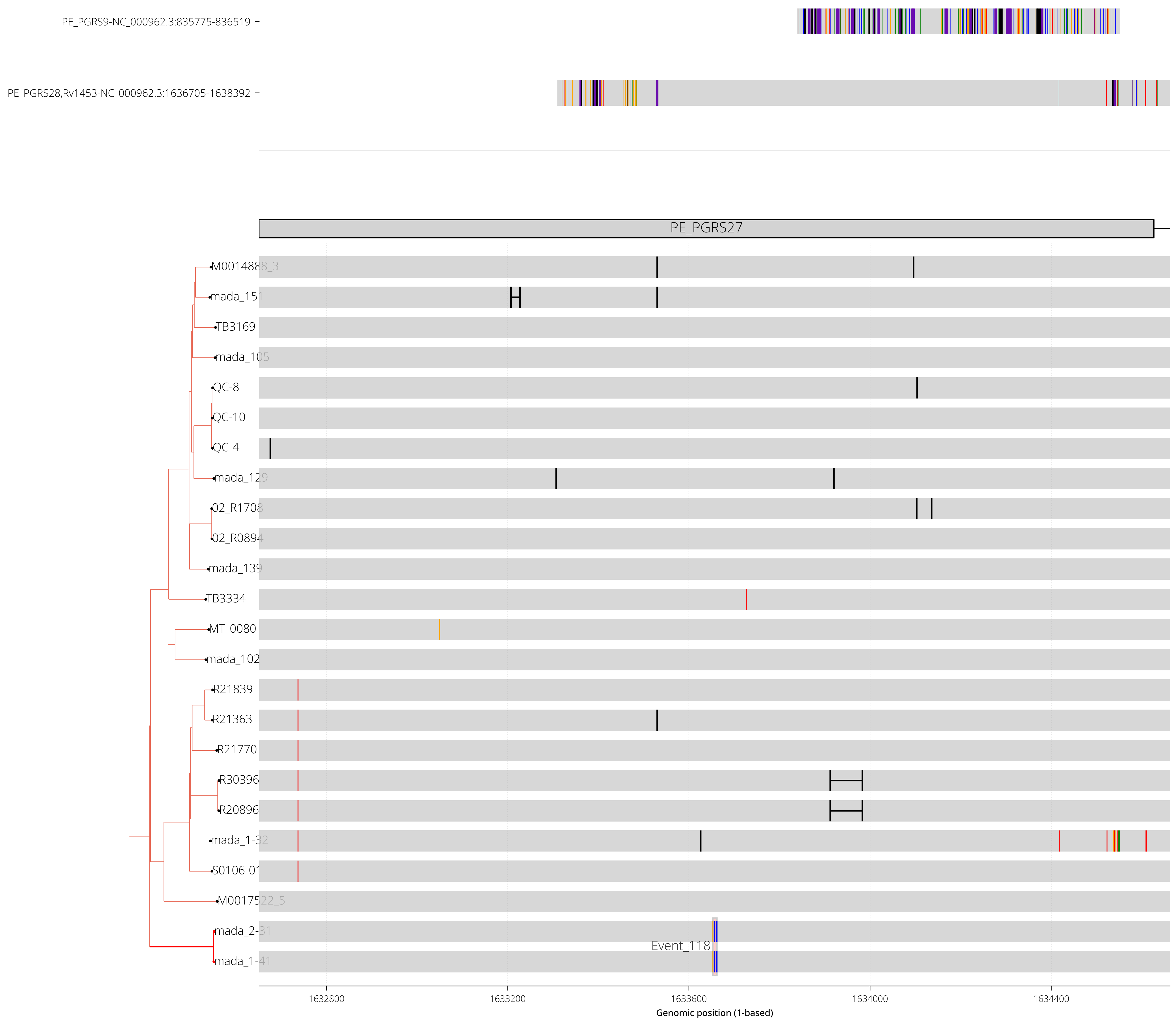

Full Phylogeny Viz + Paralog Alignments
GCE ID: Event_118

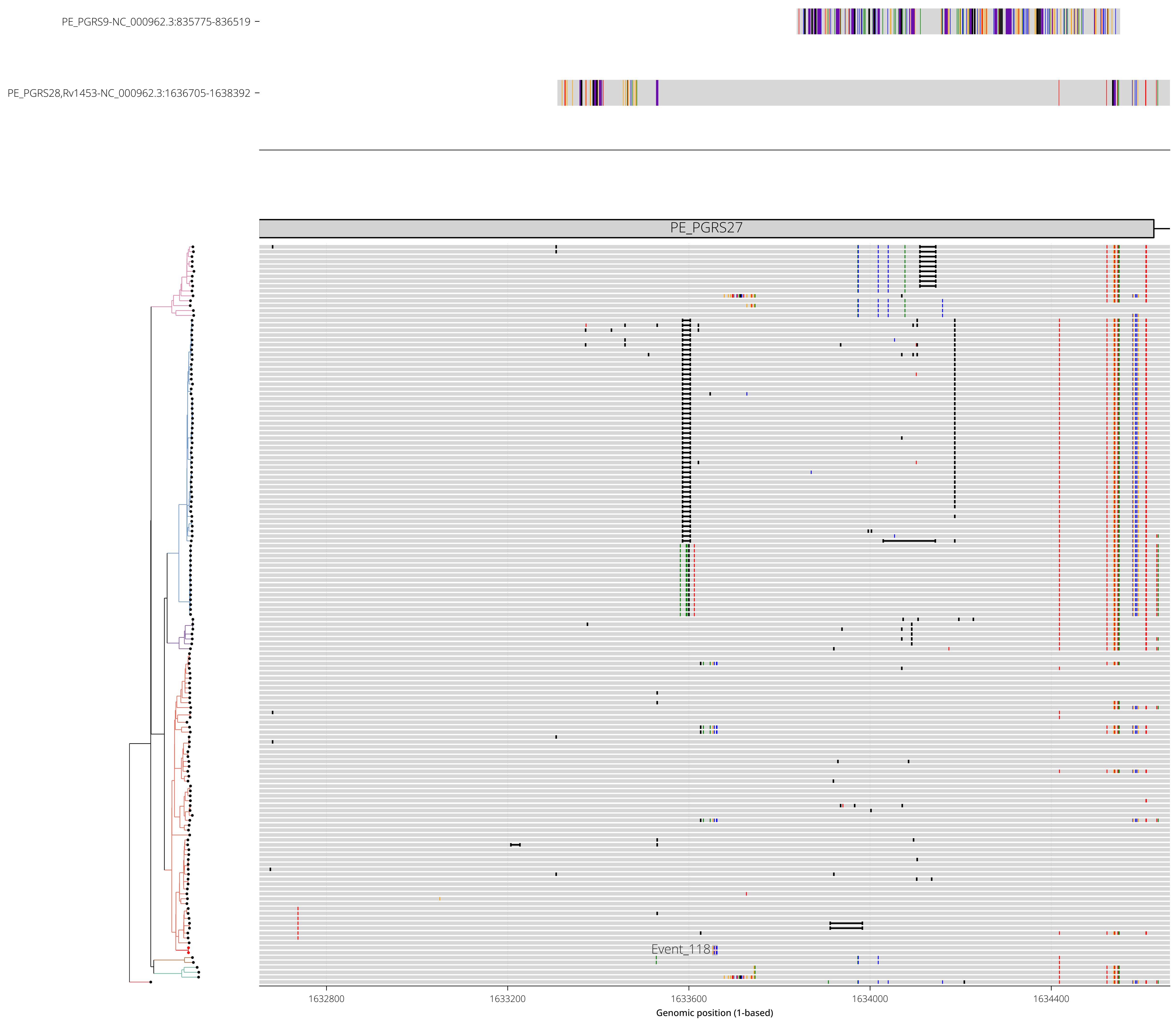

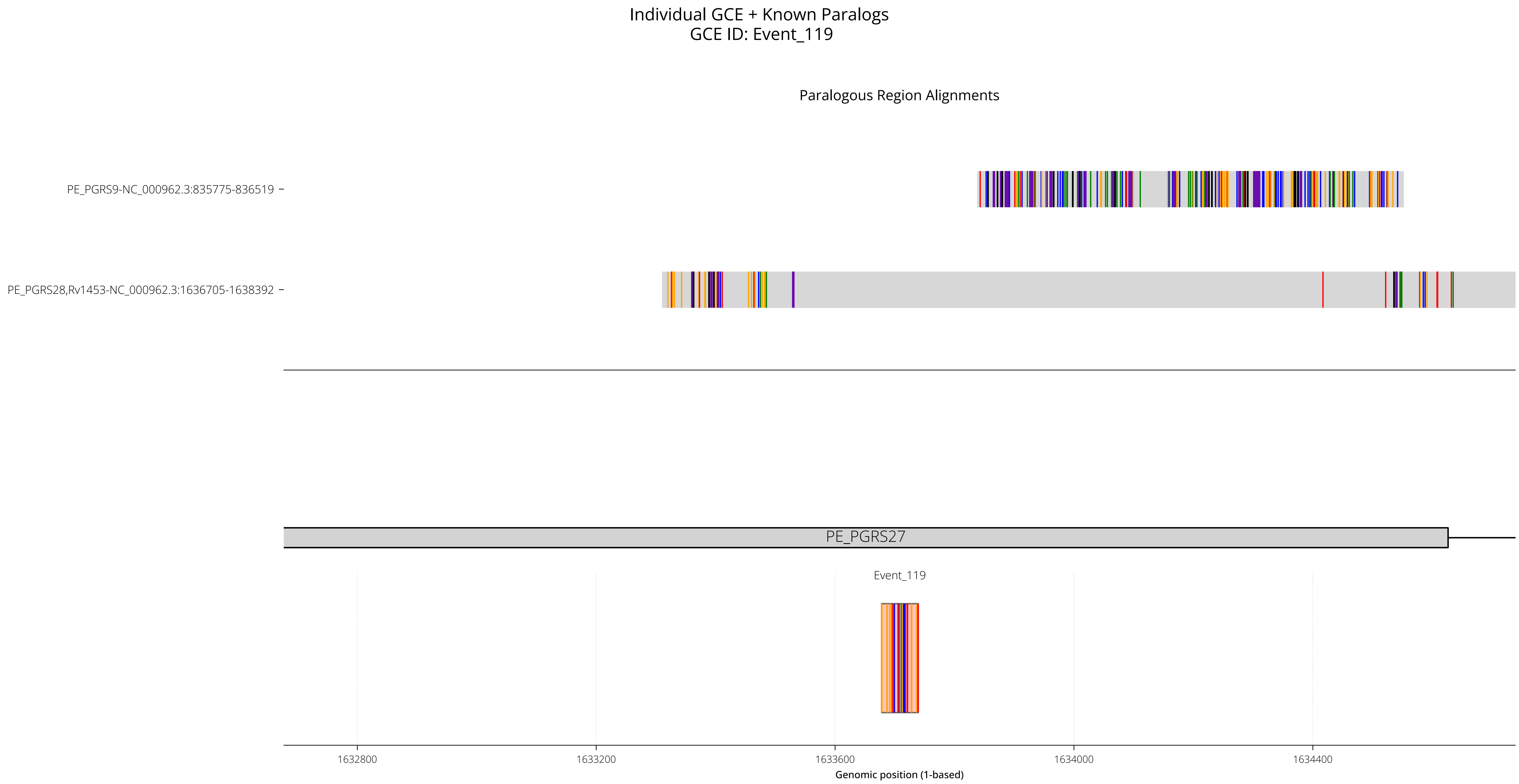

Individual GCE + Known Paralogs
GCE ID: Event_119
Paralogous Region Alignments
PE_PGRS9-NC_000962.3:835775-836519
PE_PGRS28,Rv1453-NC_000962.3:1636705-1638392
PE_PGRS27
Event_119
1632800
1633200
1633600
1634000
1634400
Genomic position (1-based)

Circos Viz - GCE to Known Paralogs Locations
GCE ID: Event_119

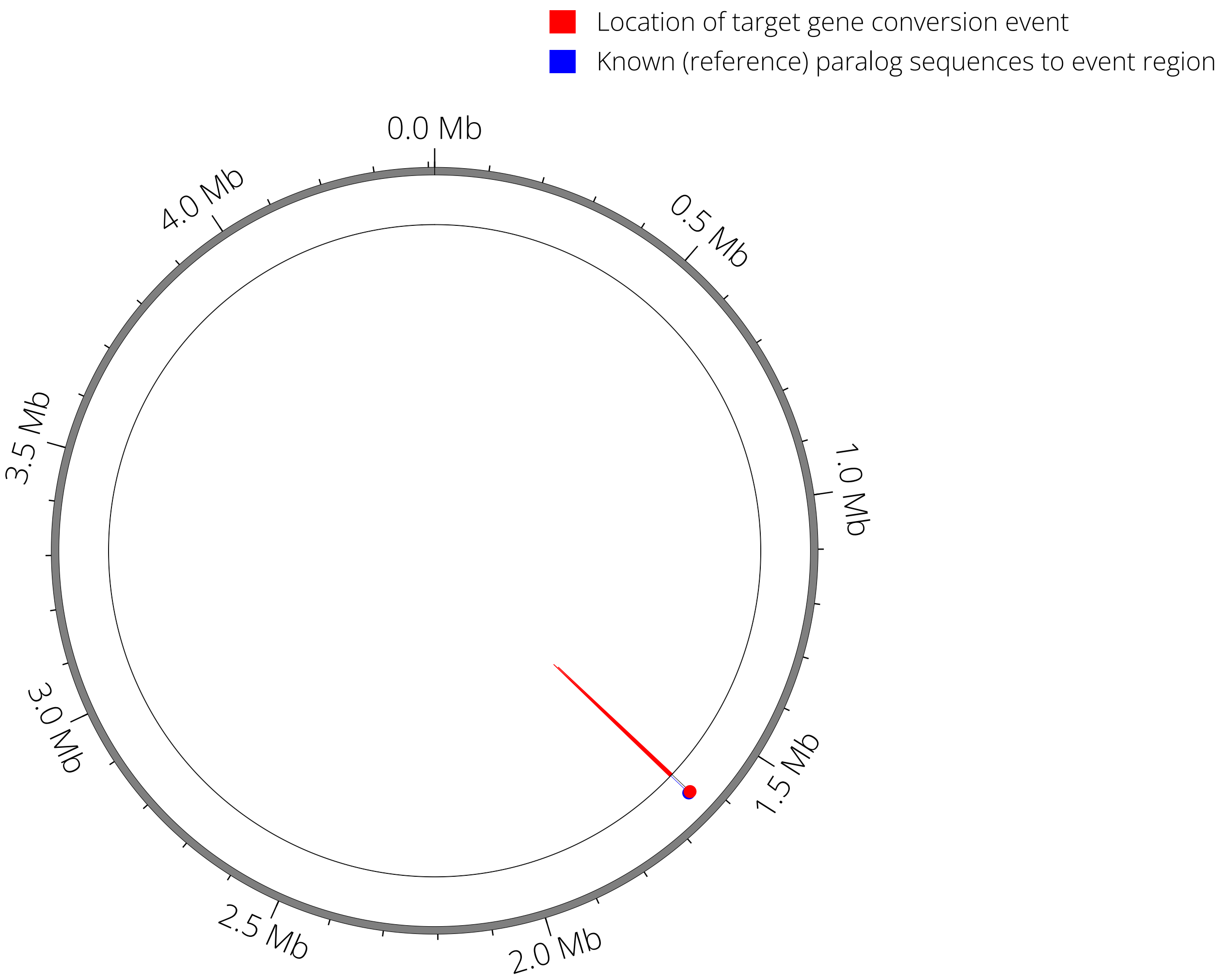

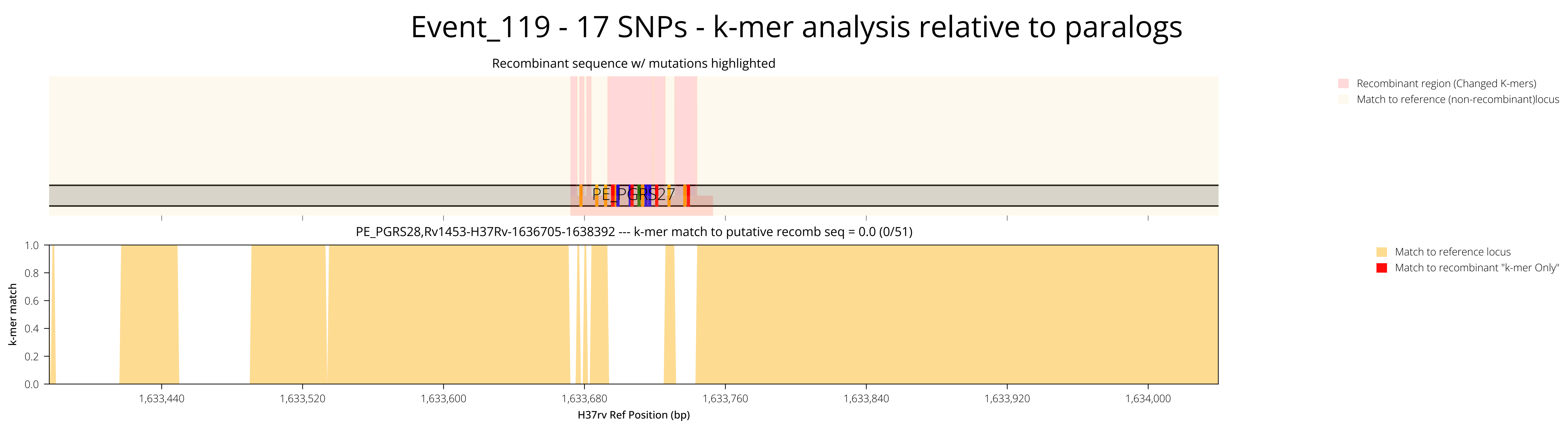
Event_119 - 17 SNPs - k-mer analysis relative to paralogs
Recombinant sequence w/ mutations highlighted
PE_PGRS27
Recombinant region (Changed K-mers)
Match to reference (non-recombinant)locus
PE_PGRS28,Rv1453-H37Rv-1636705-1638392 --- k-mer match to putative recomb seq = 0.0 (0/51)
Match to reference locus
Match to recombinant "k-mer Only"
k-mer match
1.0
0.8
0.6
0.4
0.2
0.0
1,633,440
1,633,520
1,633,600
1,633,680
1,633,760
1,633,840
1,633,920
1,634,000
H37rv Ref Position (bp)

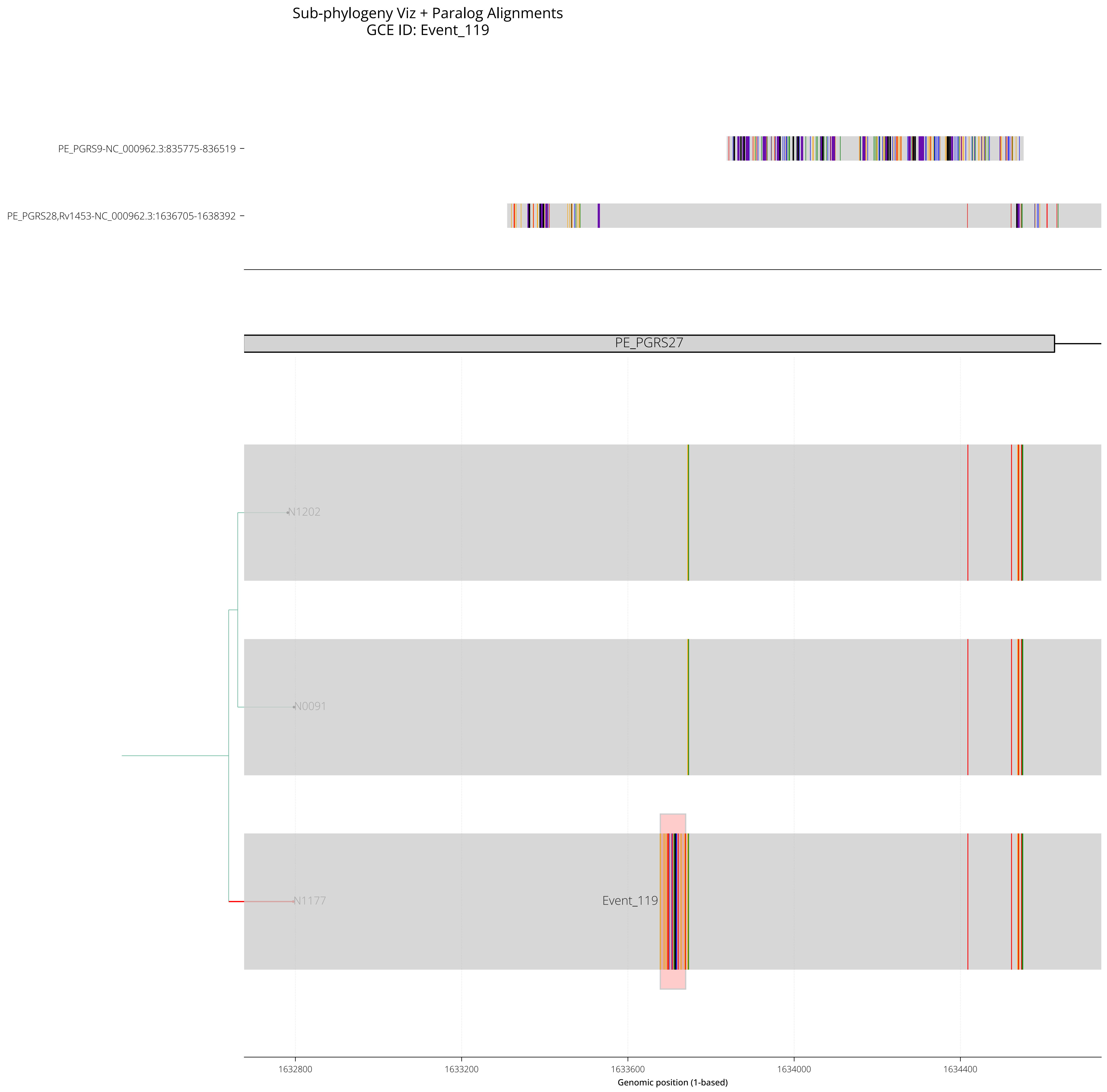

Sub-phylogeny Viz + Paralog Alignments
GCE ID: Event_119
PE_PGRS9-NC_000962.3:835775-836519
PE_PGRS28,Rv1453-NC_000962.3:1636705-1638392
PE_PGRS27
N1202
N0091
N1177
Event_119
1632800
1633200
1633600
1634000
1634400
Genomic position (1-based)

Full Phylogeny Viz + Paralog Alignments
GCE ID: Event_119

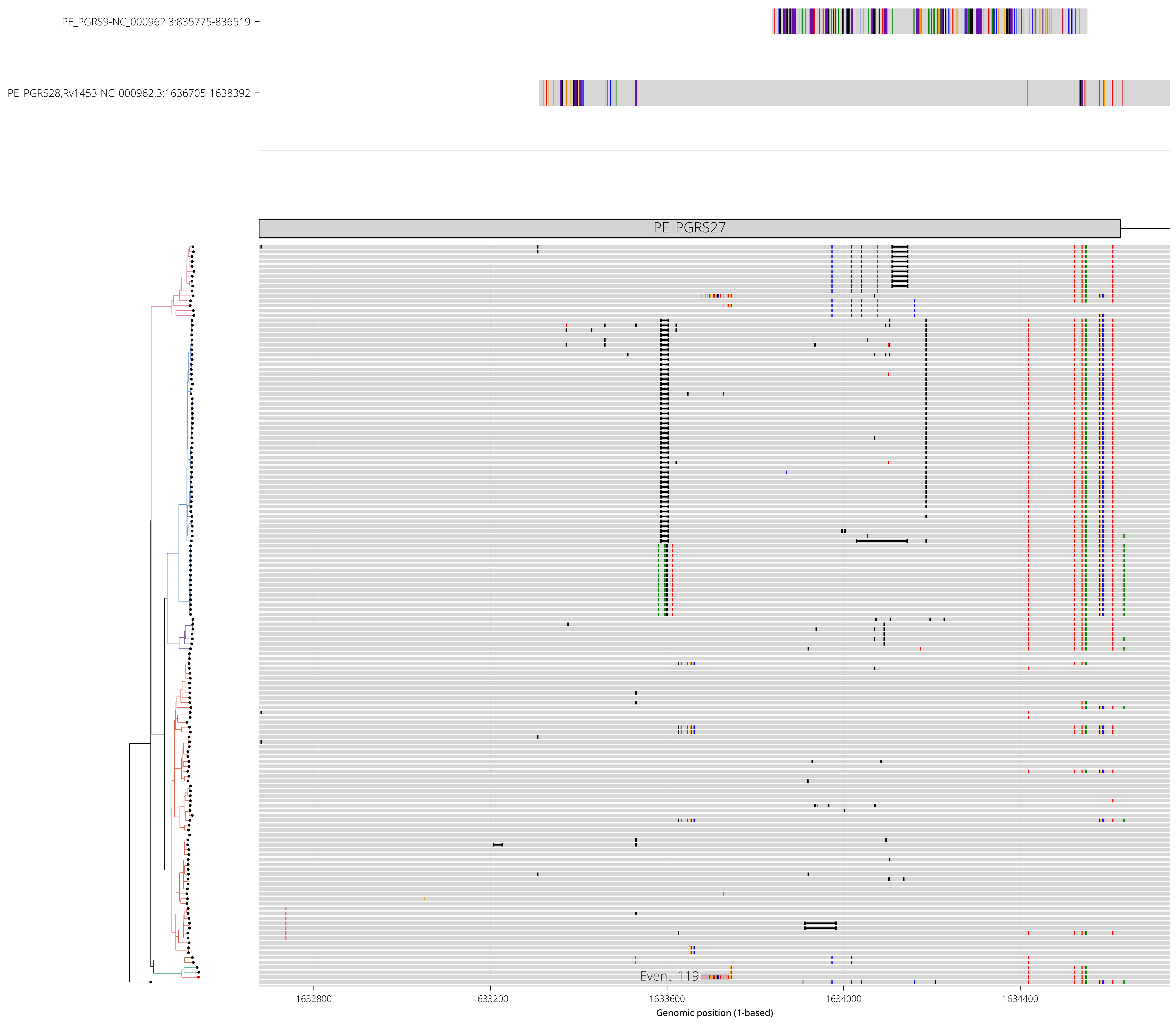

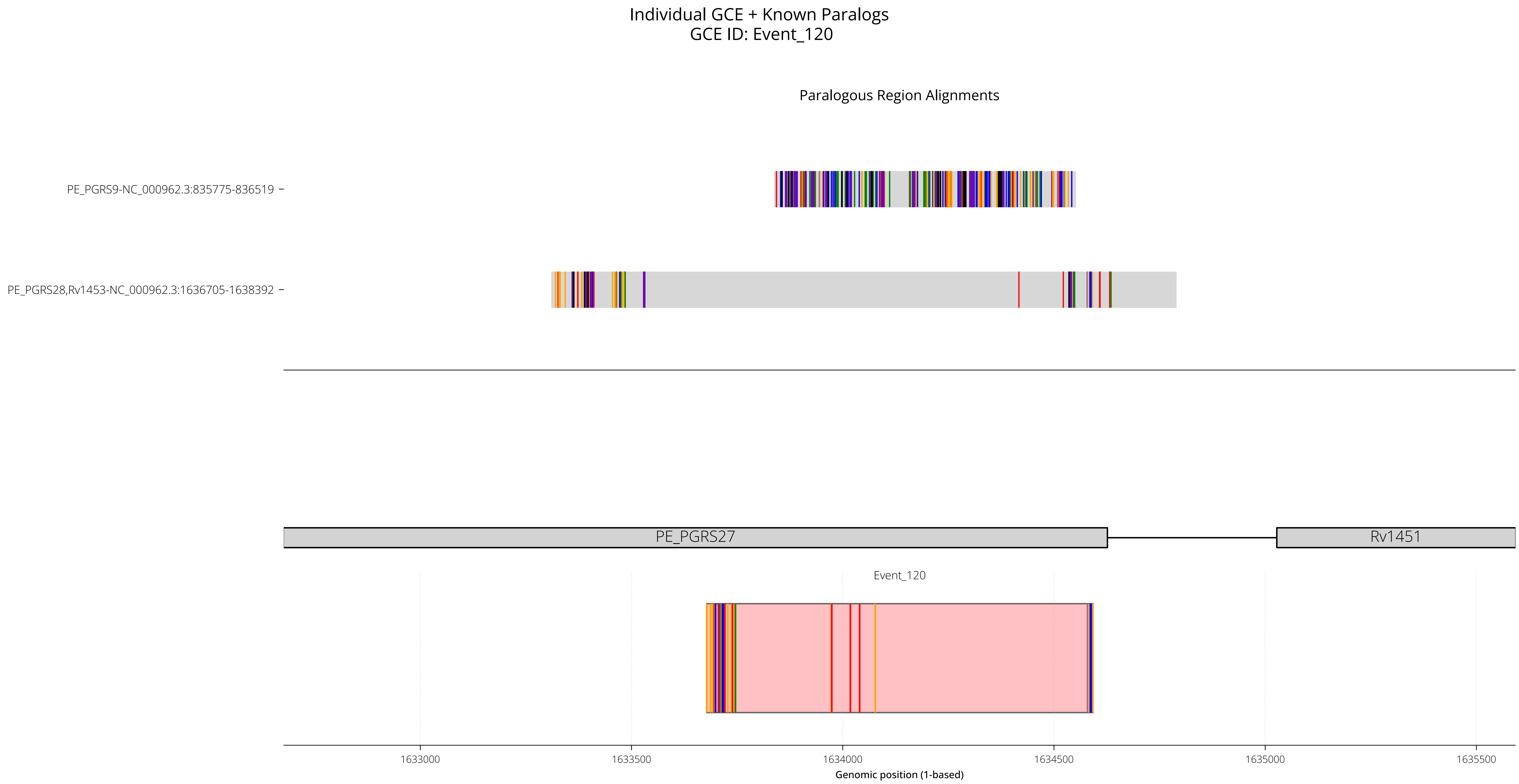

Individual GCE + Known Paralogs
GCE ID: Event_120
Paralogous Region Alignments
PE_PGRS9-NC_000962.3:835775-836519
PE_PGRS28,Rv1453-NC_000962.3:1636705-1638392
PE_PGRS27
Rv1451
Event_120
1633000
1633500
1634000
1634500
1635000
1635500
Genomic position (1-based)

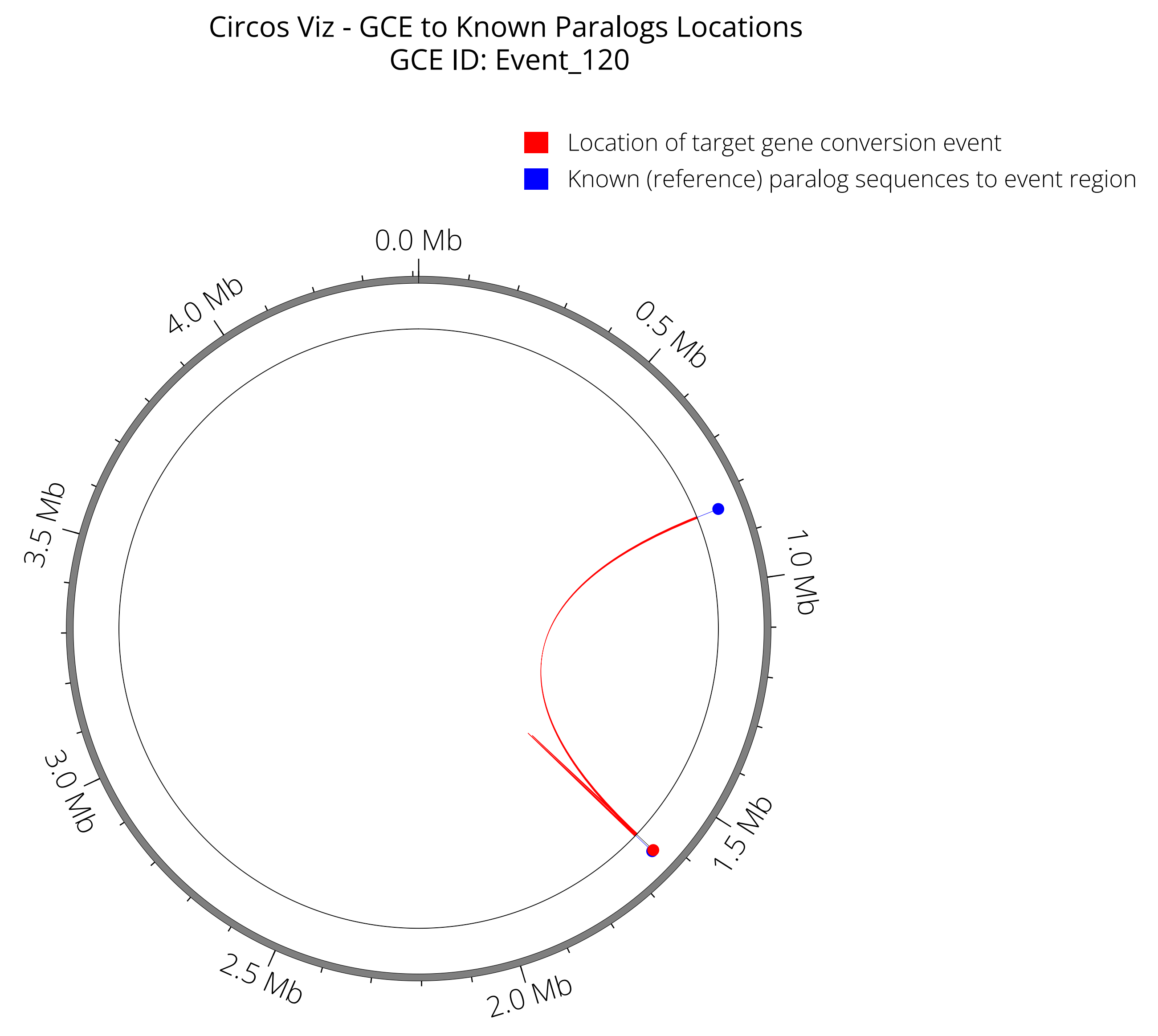
Circos Viz - GCE to Known Paralogs Locations
GCE ID: Event_120
Location of target gene conversion event
Known (reference) paralog sequences to event region
0.0 Mb
0.5 Mb
1.0 Mb
1.5 Mb
2.0 Mb
2.5 Mb
3.0 Mb
3.5 Mb
4.0 Mb

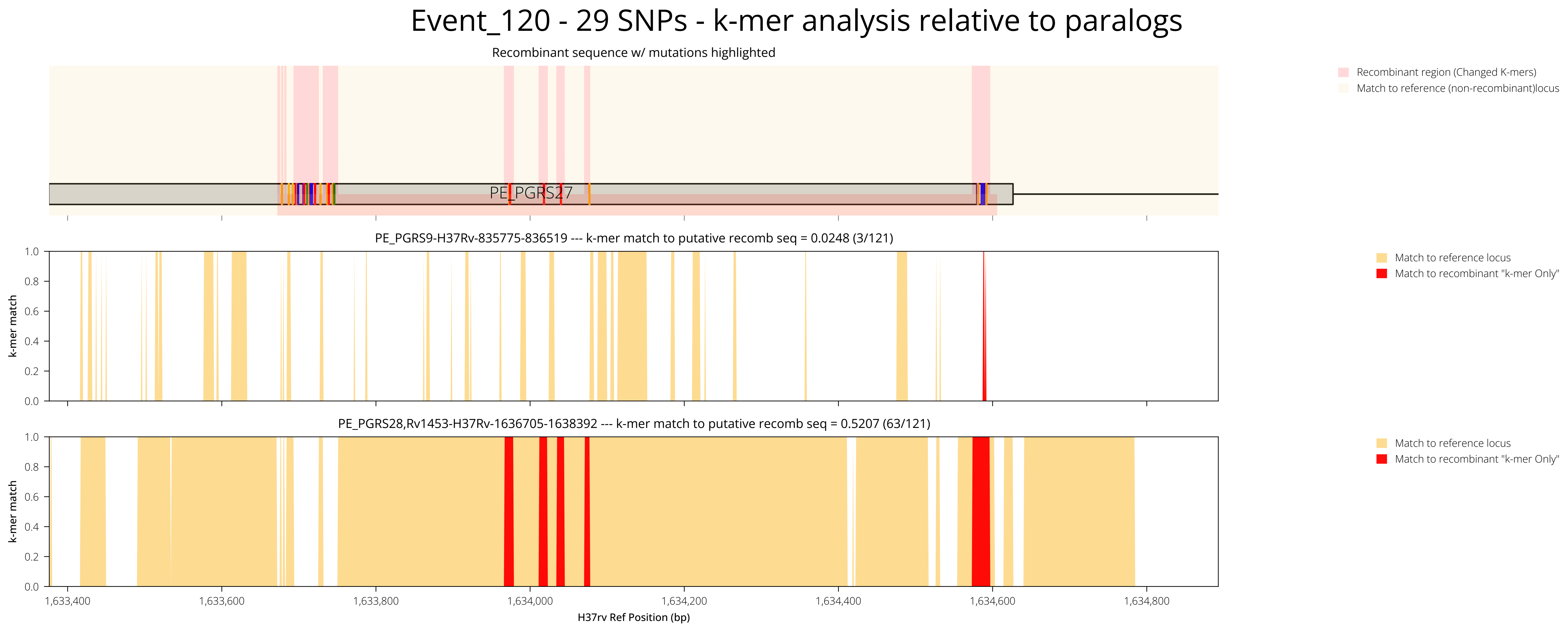

Event_120 - 29 SNPs - k-mer analysis relative to paralogs
Recombinant sequence w/ mutations highlighted
PE_PGRS27
Recombinant region (Changed K-mers)
Match to reference (non-recombinant)locus
PE_PGRS9-H37Rv-835775-836519 --- k-mer match to putative recomb seq = 0.0248 (3/121)
Match to reference locus
Match to recombinant "k-mer Only"
PE_PGRS28,Rv1453-H37Rv-1636705-1638392 --- k-mer match to putative recomb seq = 0.5207 (63/121)
Match to reference locus
Match to recombinant "k-mer Only"
k-mer match
0.0
0.2
0.4
0.6
0.8
1.0
1,633,400
1,633,600
1,633,800
1,634,000
1,634,200
1,634,400
1,634,600
1,634,800
H37rv Ref Position (bp)

Sub-phylogeny Viz + Paralog Alignments
GCE ID: Event_120

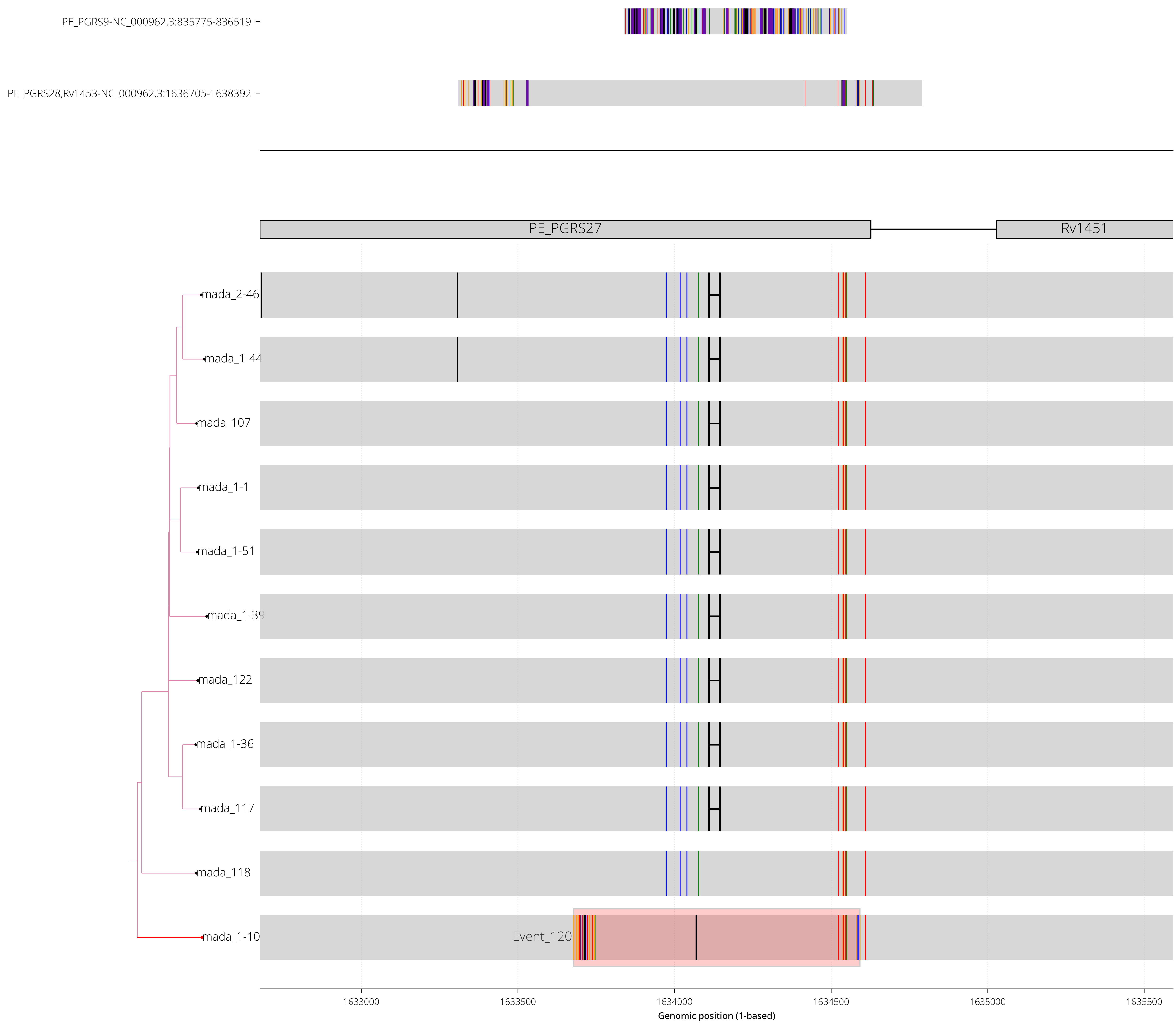

Full Phylogeny Viz + Paralog Alignments
GCE ID: Event_120

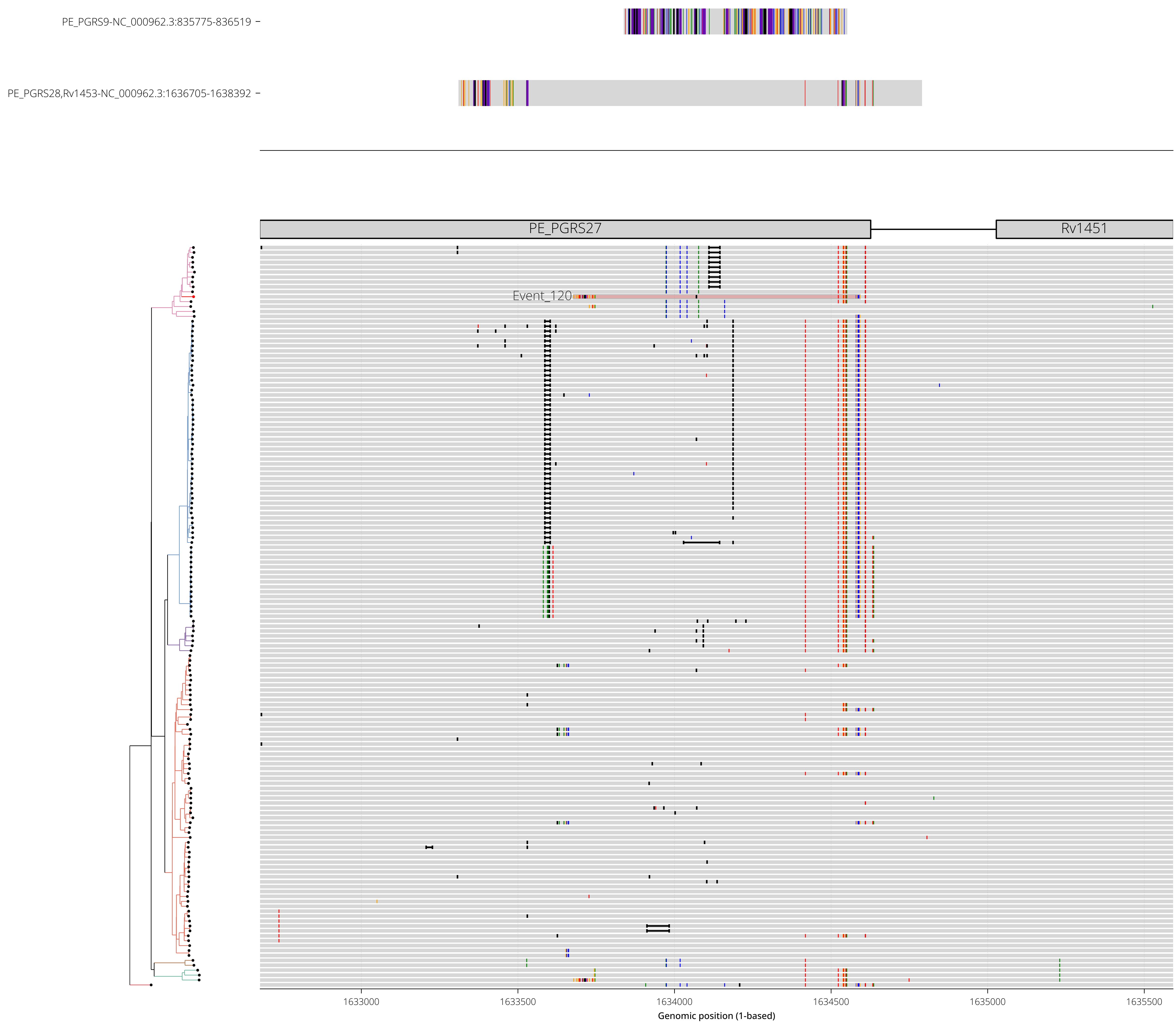

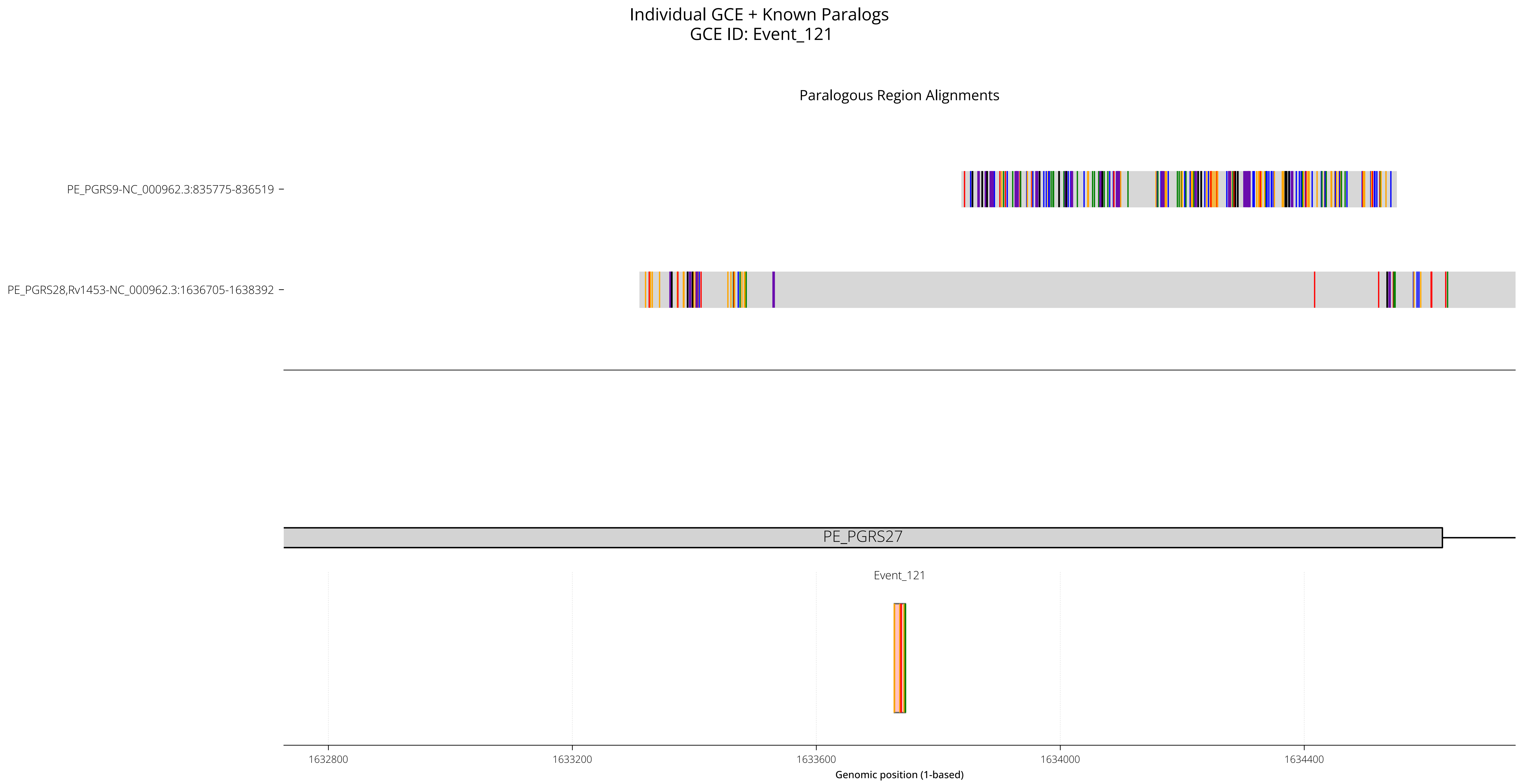
Individual GCE + Known Paralogs
GCE ID: Event_121
Paralogous Region Alignments
PE_PGRS9-NC_000962.3:835775-836519
PE_PGRS28,Rv1453-NC_000962.3:1636705-1638392
PE_PGRS27
Event_121
1632800
1633200
1633600
1634000
1634400
Genomic position (1-based)

Circos Viz - GCE to Known Paralogs Locations
GCE ID: Event_121

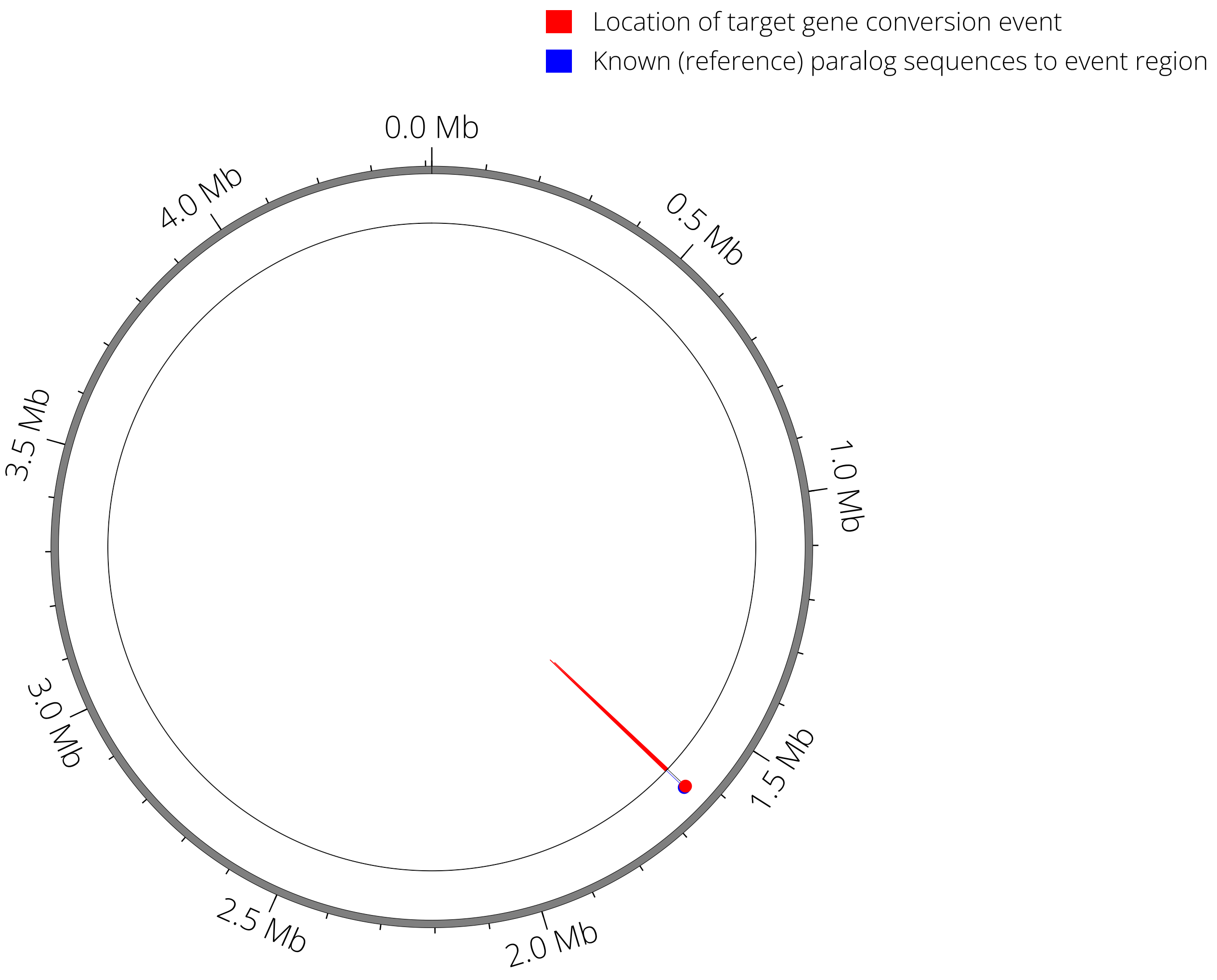

### Event_121 - 5 SNPs - k-mer analysis relative to paralogs

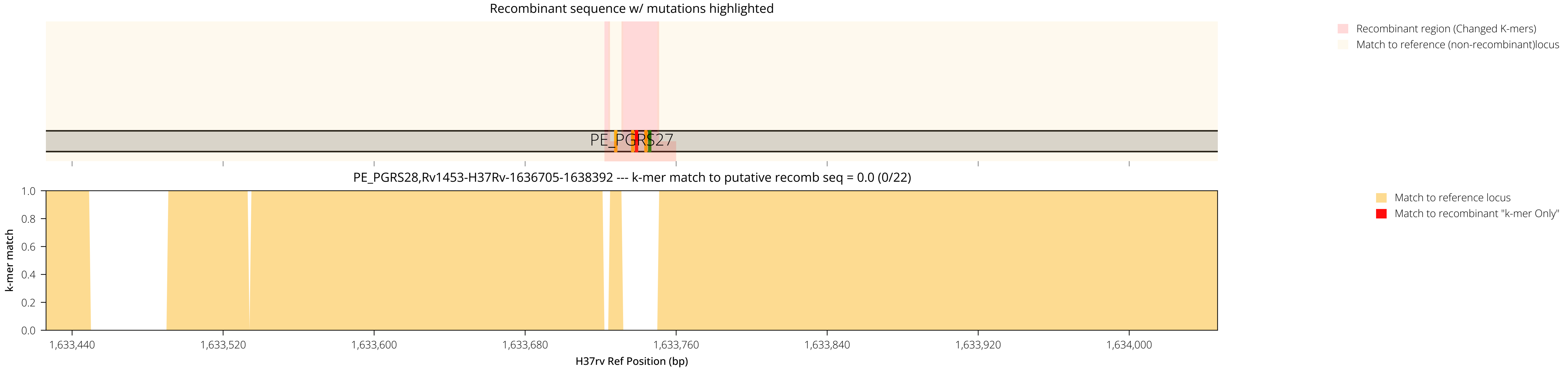

Sub-phylogeny Viz + Paralog Alignments
GCE ID: Event_121

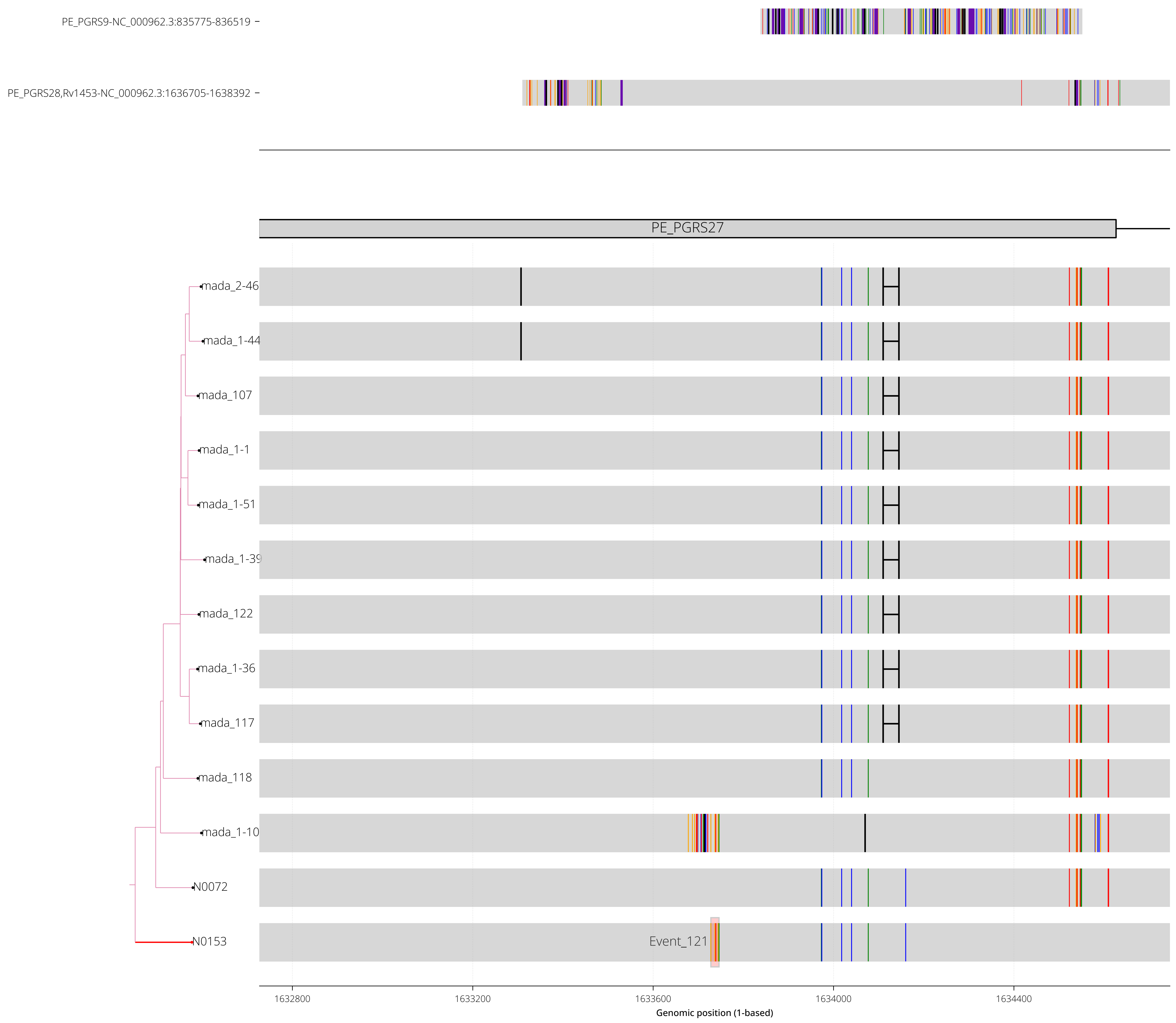

Full Phylogeny Viz + Paralog Alignments
GCE ID: Event_121

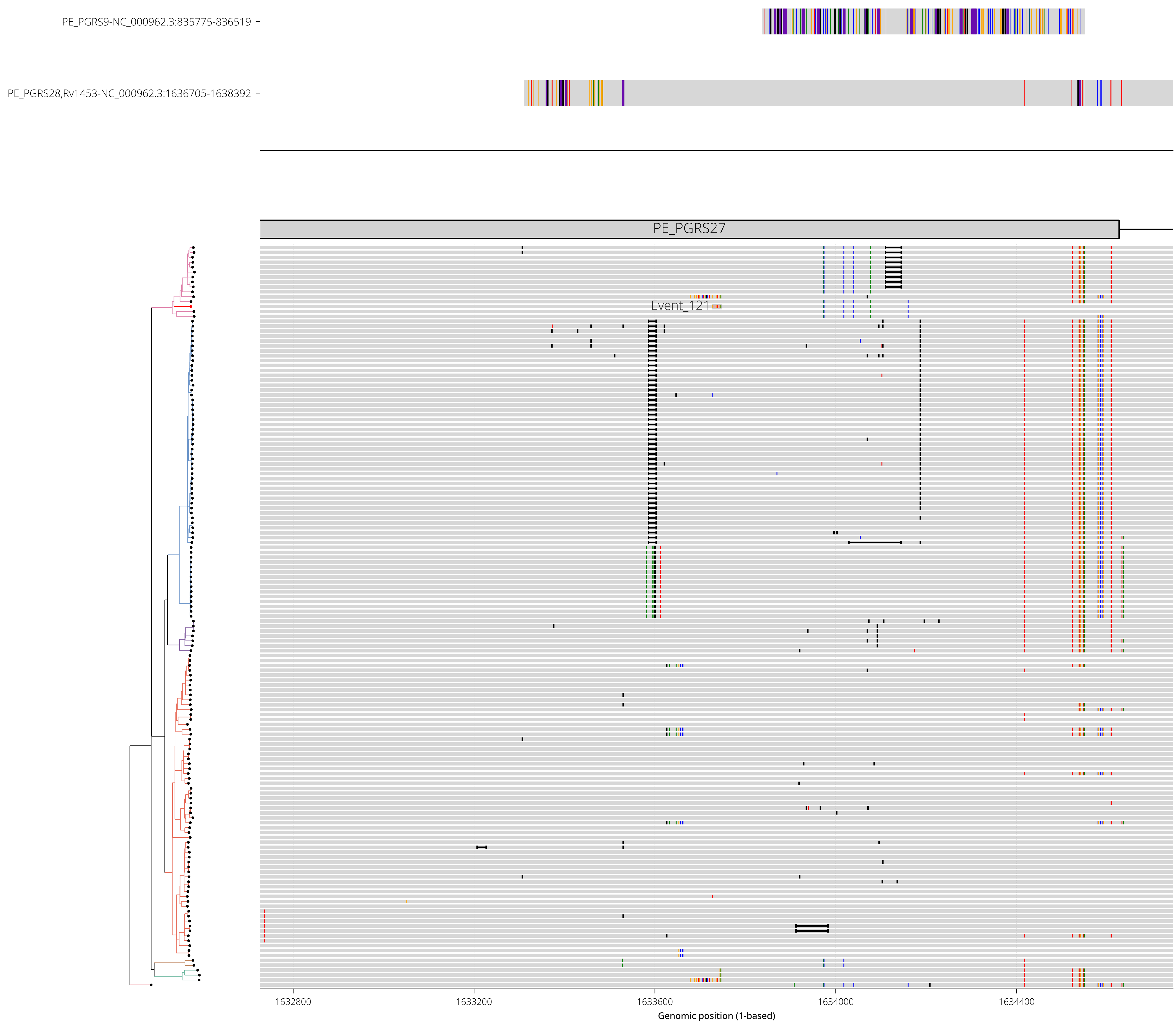

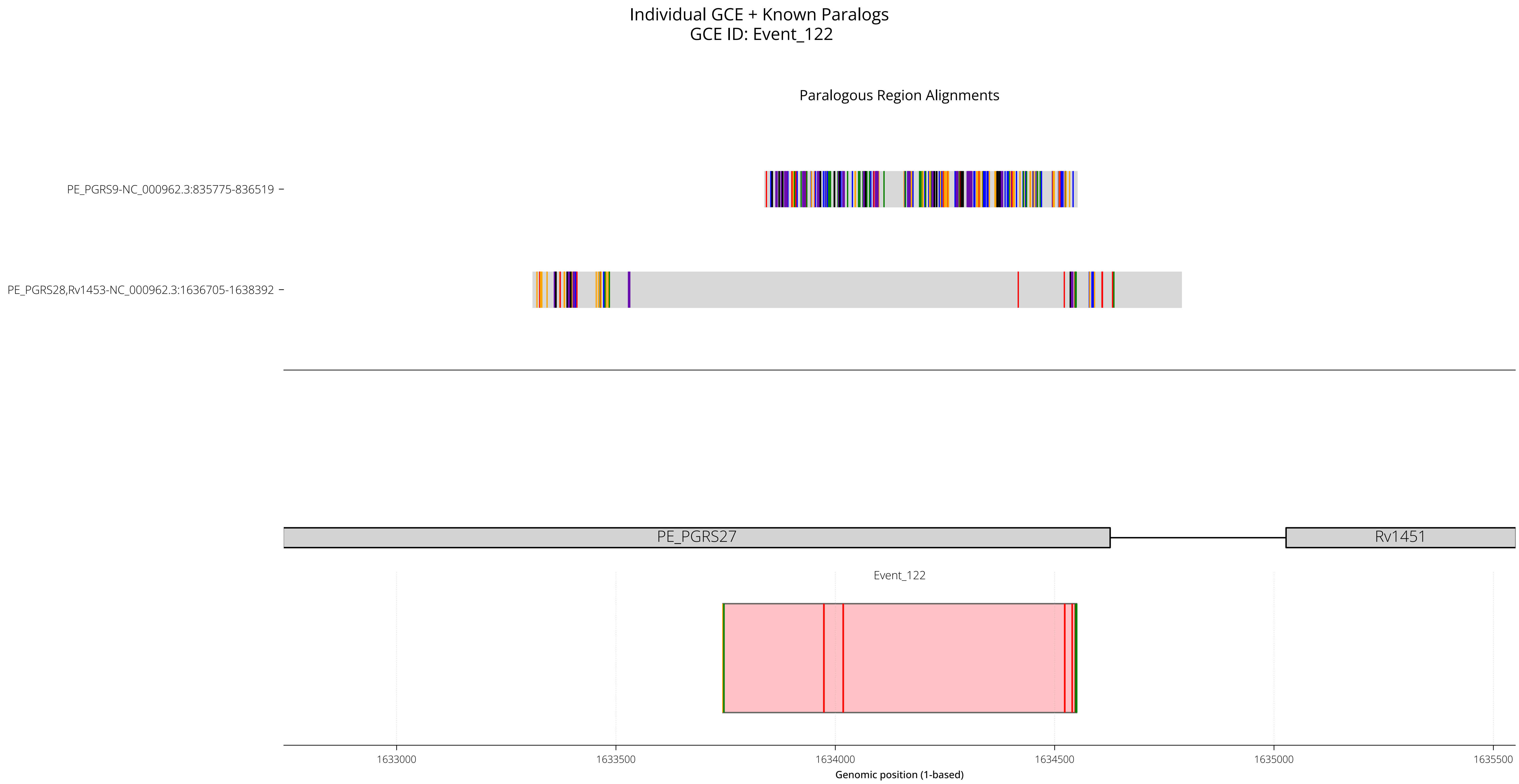

Individual GCE + Known Paralogs
GCE ID: Event_122
Paralogous Region Alignments
PE_PGRS9-NC_000962.3:835775-836519
PE_PGRS28,Rv1453-NC_000962.3:1636705-1638392
PE_PGRS27
Rv1451
Event_122
1633000
1633500
1634000
1634500
1635000
1635500
Genomic position (1-based)

### Circos Viz - GCE to Known Paralogs Locations
GCE ID: Event_122

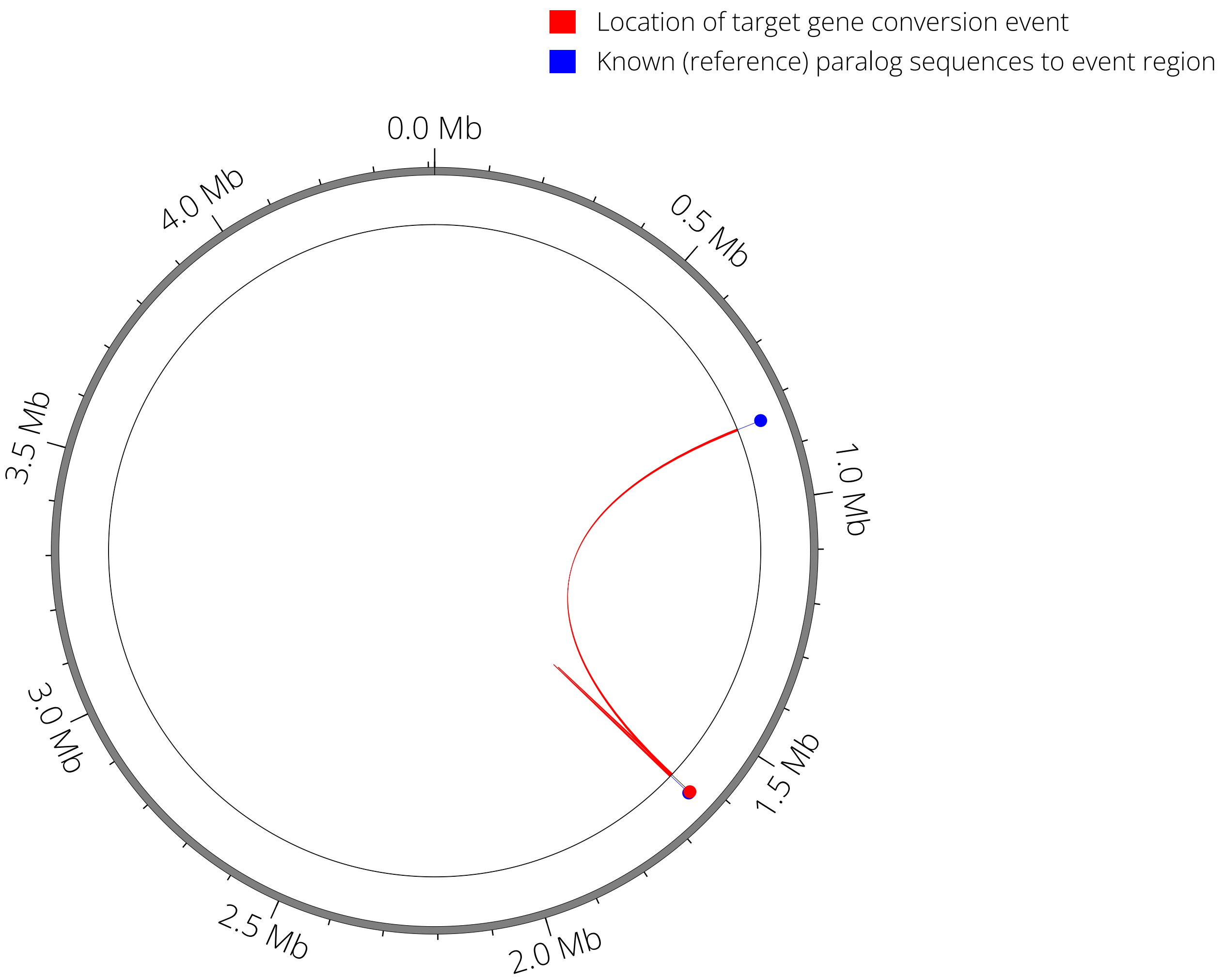

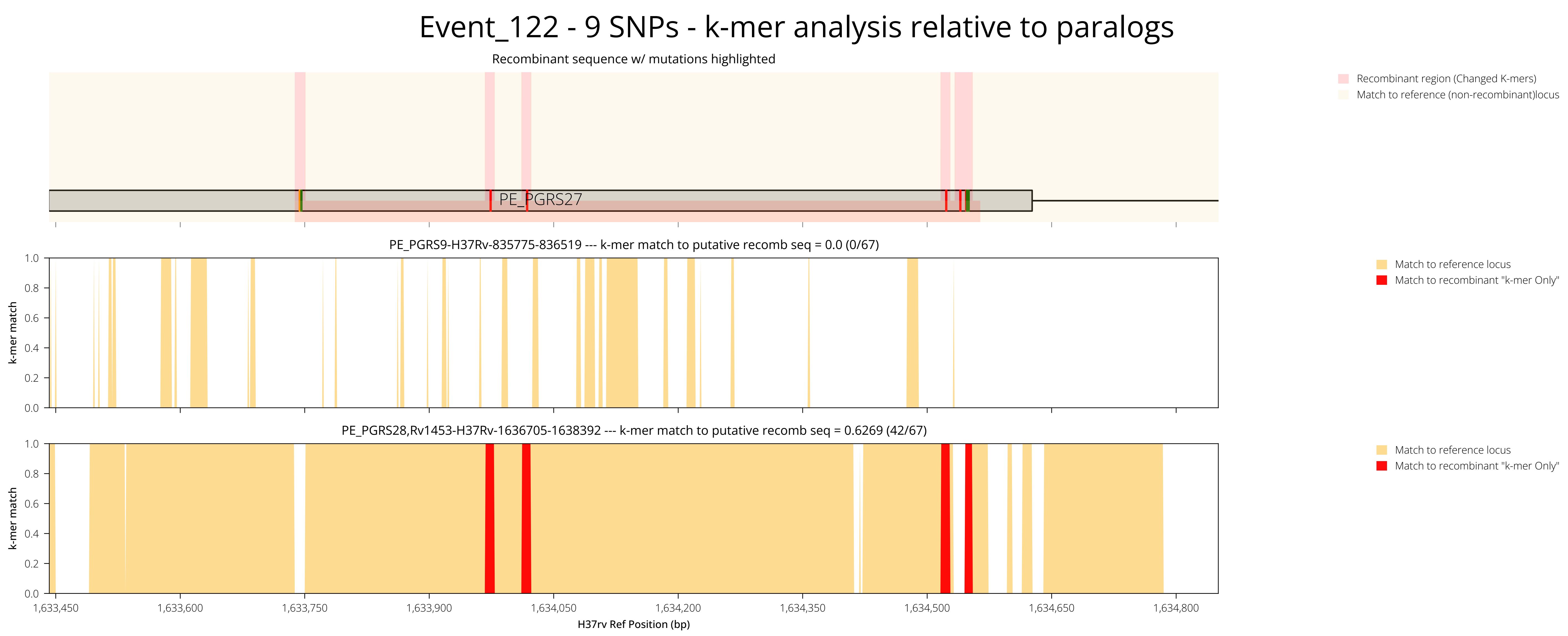

Event_122 - 9 SNPs - k-mer analysis relative to paralogs
Recombinant sequence w/ mutations highlighted
PE_PGRS27
Recombinant region (Changed K-mers)
Match to reference (non-recombinant)locus
PE_PGRS9-H37Rv-835775-836519 --- k-mer match to putative recomb seq = 0.0 (0/67)
Match to reference locus
Match to recombinant "k-mer Only"
k-mer match
1.0
0.8
0.6
0.4
0.2
0.0
PE_PGRS28,Rv1453-H37Rv-1636705-1638392 --- k-mer match to putative recomb seq = 0.6269 (42/67)
Match to reference locus
Match to recombinant "k-mer Only"
k-mer match
1,633,450
1,633,600
1,633,750
1,633,900
1,634,050
1,634,200
1,634,350
1,634,500
1,634,650
1,634,800
H37rv Ref Position (bp)

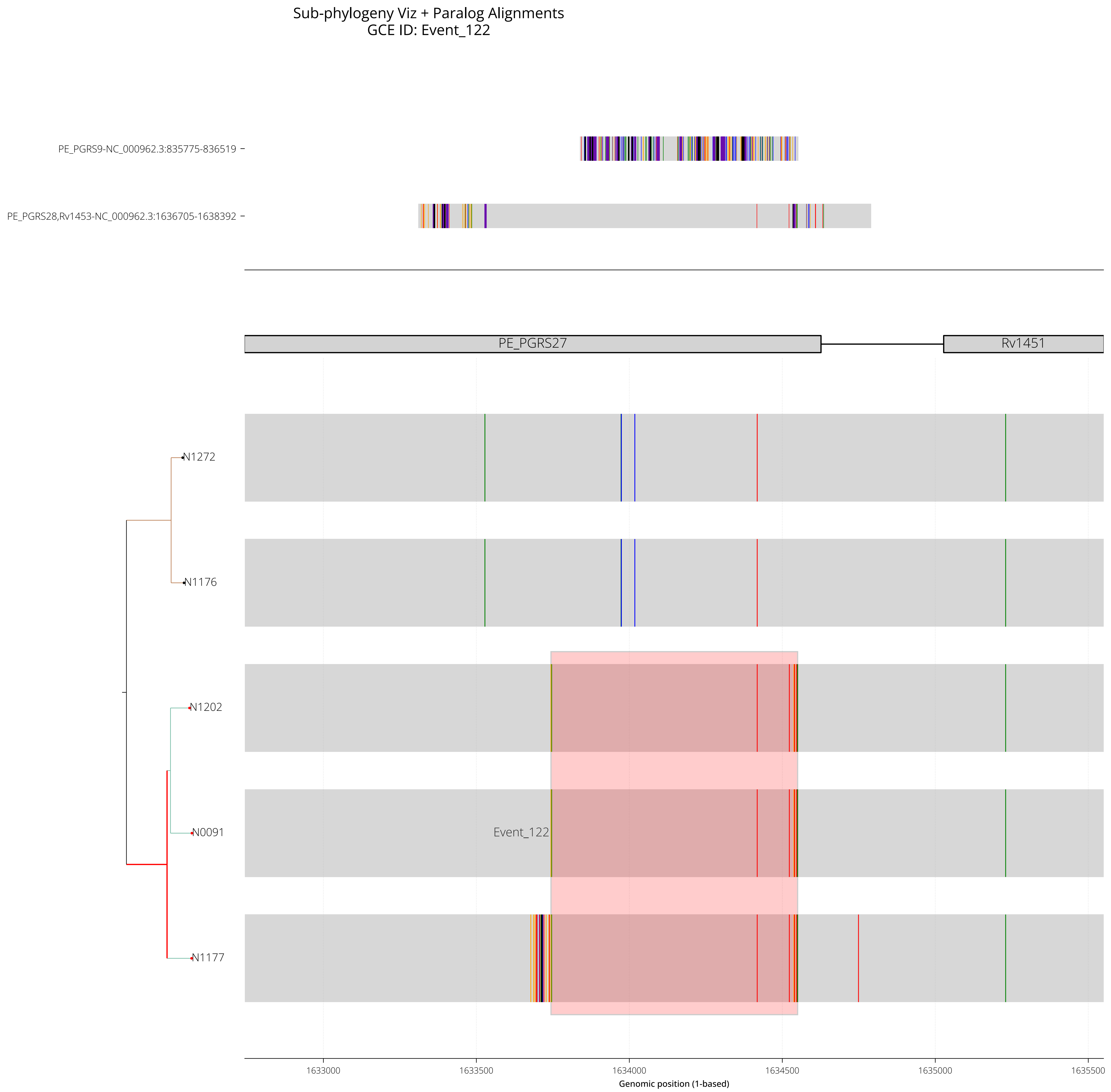

Sub-phylogeny Viz + Paralog Alignments
GCE ID: Event_122
PE_PGRS9-NC_000962.3:835775-836519
PE_PGRS28,Rv1453-NC_000962.3:1636705-1638392
PE_PGRS27
Rv1451
N1272
N1176
N1202
N0091
N1177
Event_122
1633000
1633500
1634000
1634500
1635000
1635500
Genomic position (1-based)

Full Phylogeny Viz + Paralog Alignments
GCE ID: Event_122

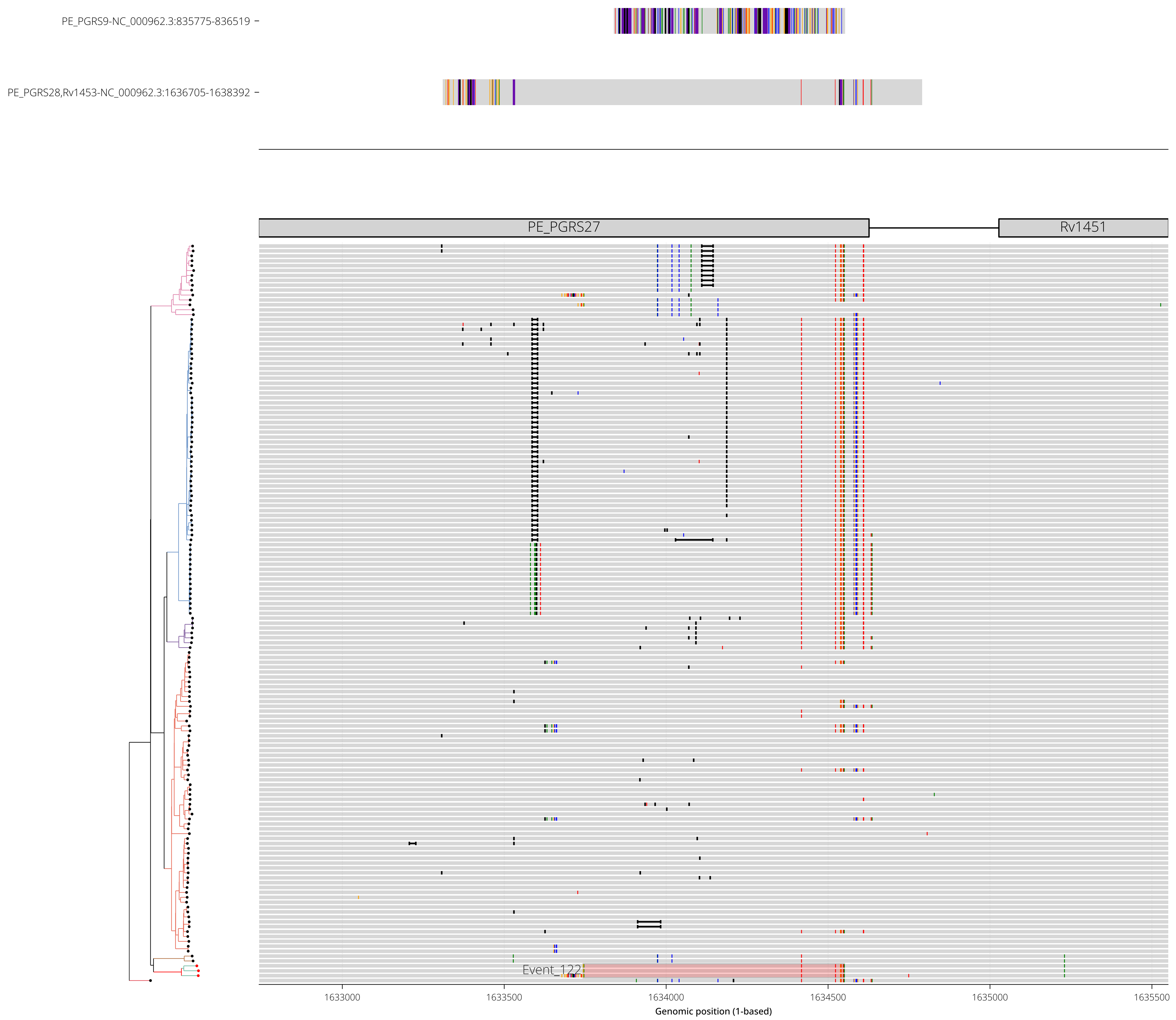

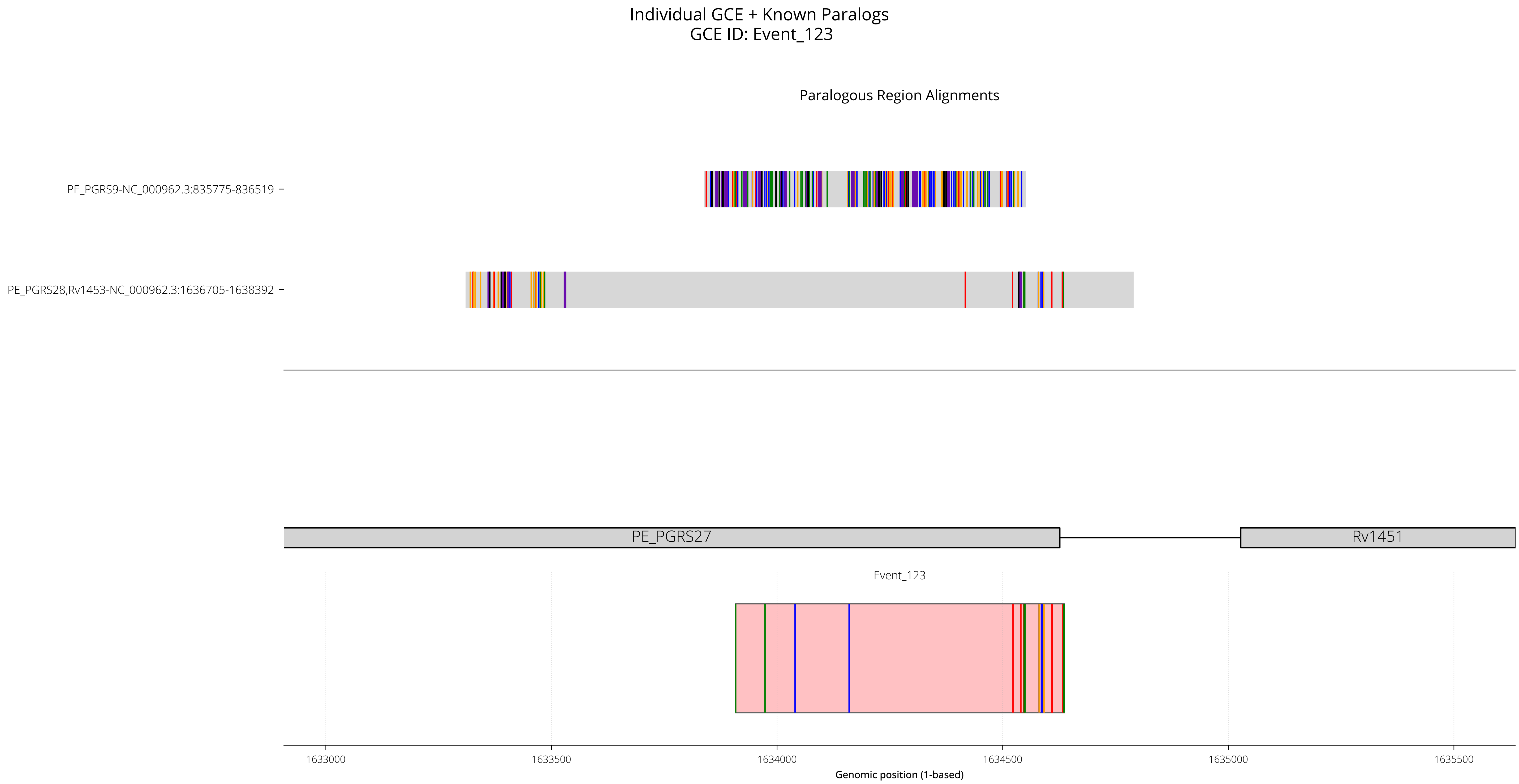
Individual GCE + Known Paralogs
GCE ID: Event_123
Paralogous Region Alignments
PE_PGRS9-NC_000962.3:835775-836519
PE_PGRS28,Rv1453-NC_000962.3:1636705-1638392
PE_PGRS27
Rv1451
Event_123
1633000
1633500
1634000
1634500
1635000
1635500
Genomic position (1-based)

Circos Viz - GCE to Known Paralogs Locations
GCE ID: Event_123

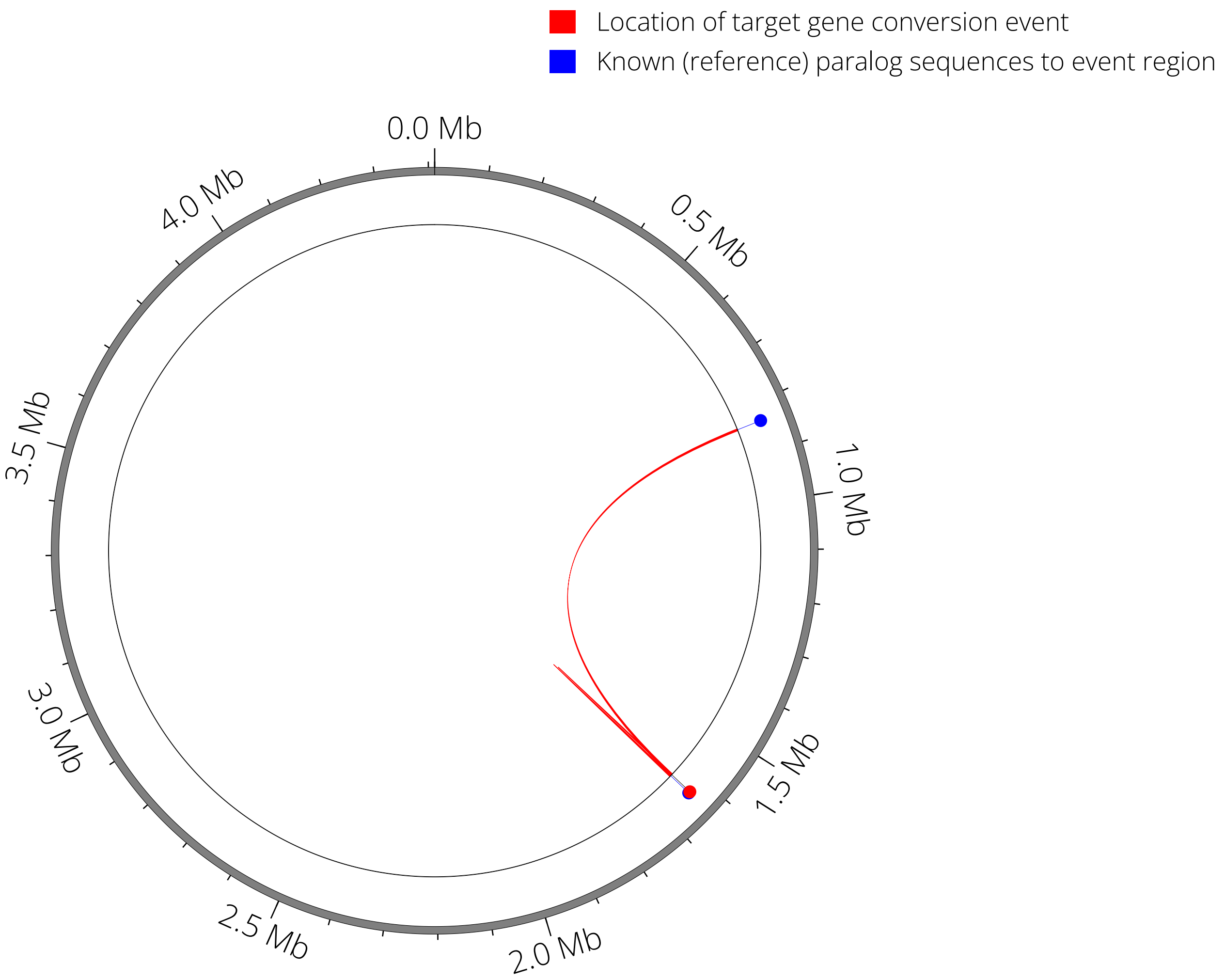

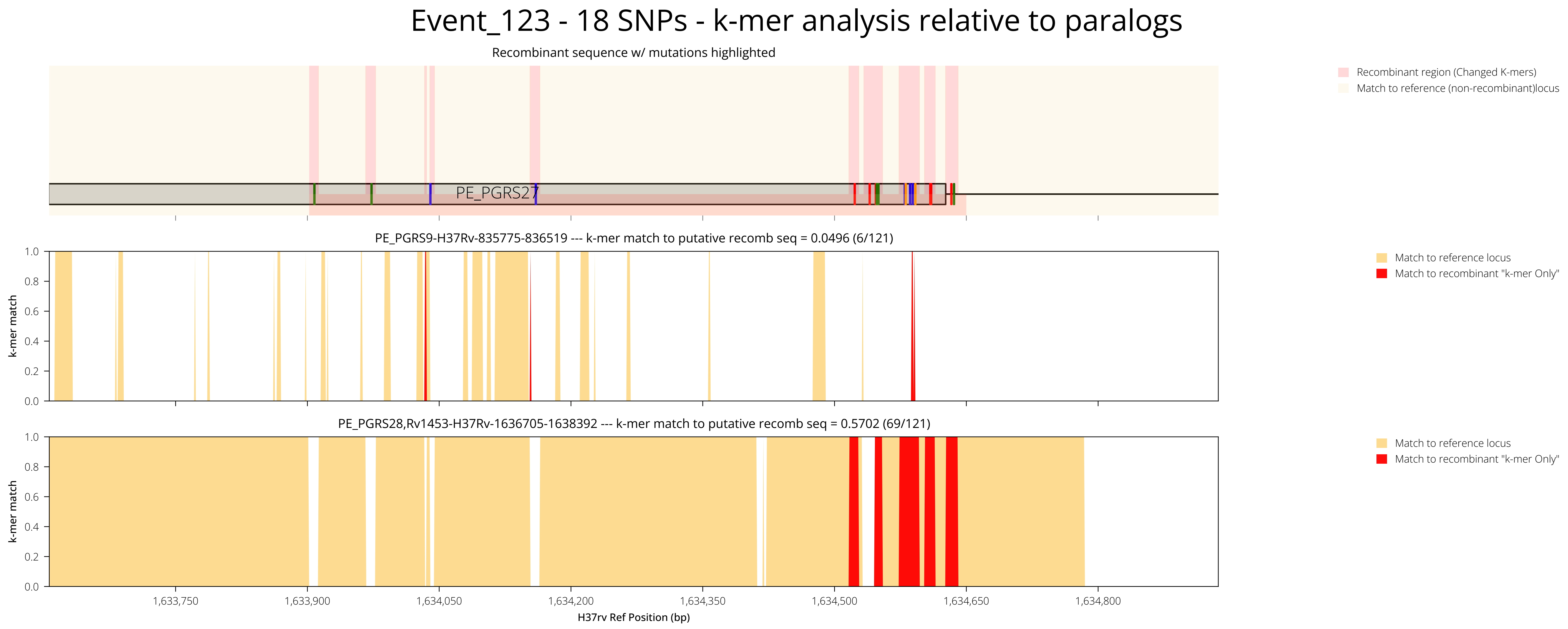

Event_123 - 18 SNPs - k-mer analysis relative to paralogs
Recombinant sequence w/ mutations highlighted
Recombinant region (Changed K-mers)
Match to reference (non-recombinant)locus
PE_PGRS27
PE_PGRS9-H37Rv-835775-836519 --- k-mer match to putative recomb seq = 0.0496 (6/121)
Match to reference locus
Match to recombinant "k-mer Only"
k-mer match
1.0
0.8
0.6
0.4
0.2
0.0
PE_PGRS28,Rv1453-H37Rv-1636705-1638392 --- k-mer match to putative recomb seq = 0.5702 (69/121)
Match to reference locus
Match to recombinant "k-mer Only"
k-mer match
1,633,750
1,633,900
1,634,050
1,634,200
1,634,350
1,634,500
1,634,650
1,634,800
H37rv Ref Position (bp)

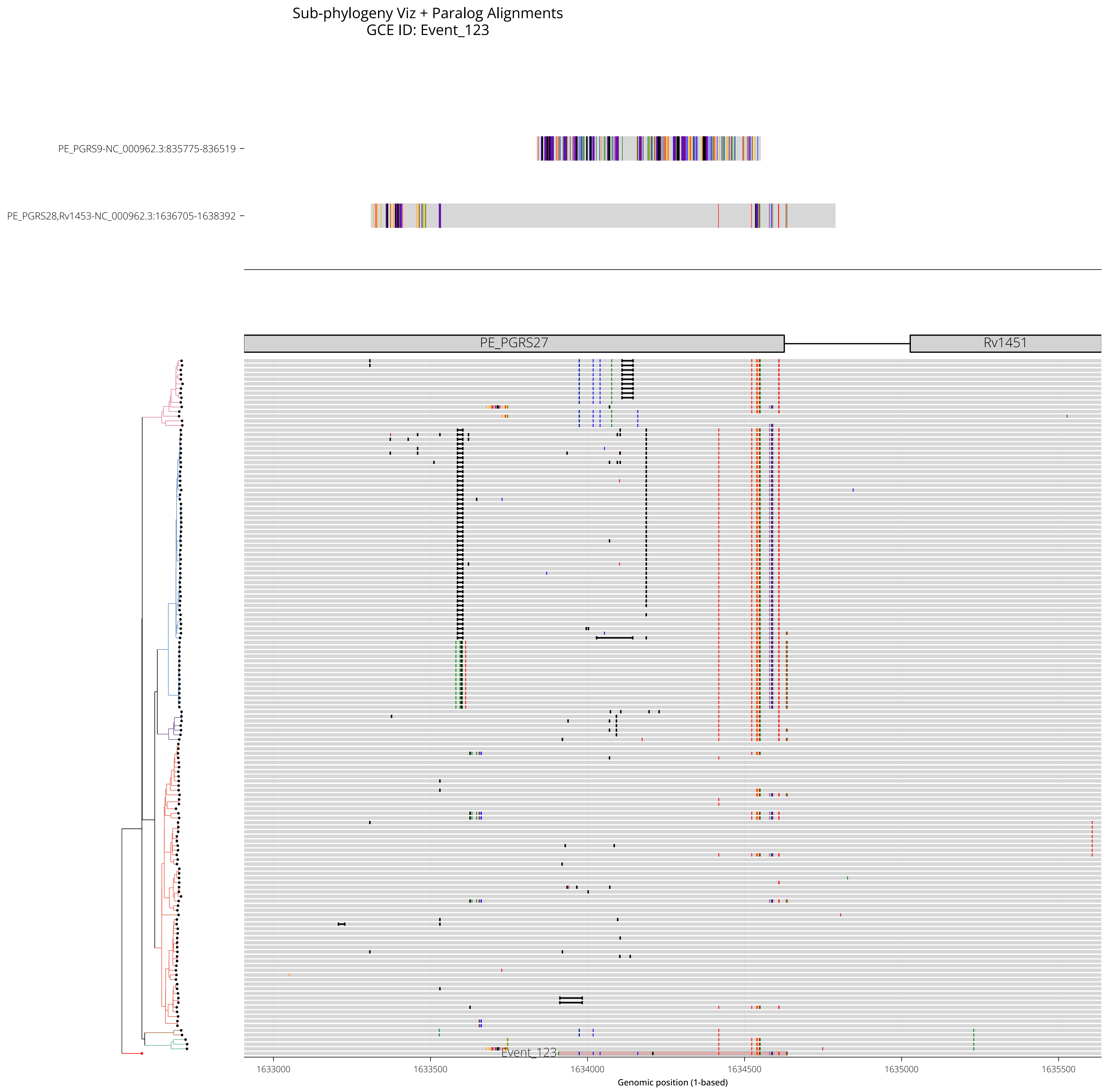

Sub-phylogeny Viz + Paralog Alignments
GCE ID: Event_123
PE_PGRS9-NC_000962.3:835775-836519
PE_PGRS28,Rv1453-NC_000962.3:1636705-1638392
PE_PGRS27
Rv1451
Event_123
1633000
1633500
1634000
1634500
1635000
1635500
Genomic position (1-based)

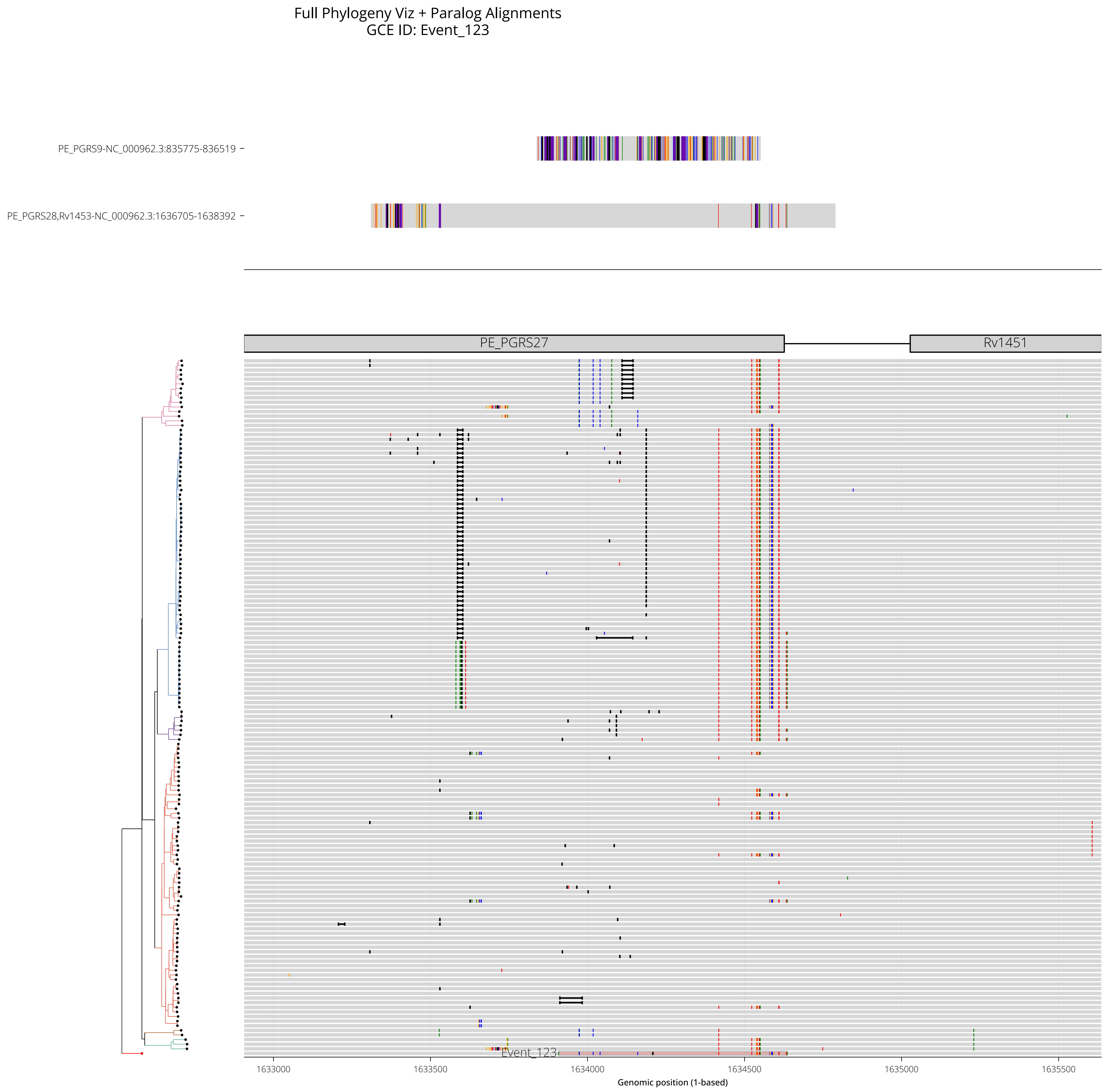

Full Phylogeny Viz + Paralog Alignments
GCE ID: Event_123
PE_PGRS9-NC_000962.3:835775-836519
PE_PGRS28,Rv1453-NC_000962.3:1636705-1638392
PE_PGRS27
Rv1451
Event_123
1633000
1633500
1634000
1634500
1635000
1635500
Genomic position (1-based)

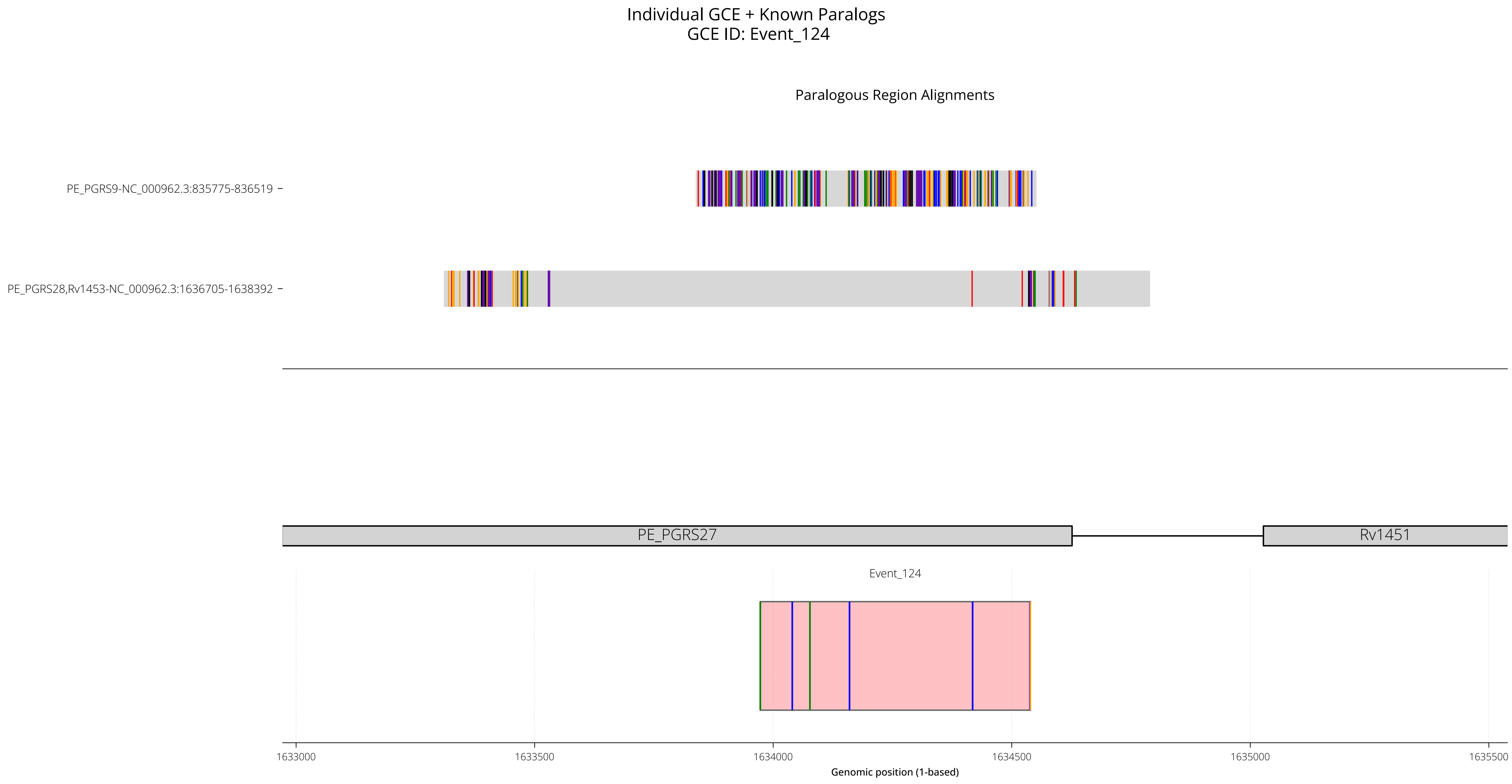
Individual GCE + Known Paralogs
GCE ID: Event_124
Paralogous Region Alignments
PE_PGRS9-NC_000962.3:835775-836519
PE_PGRS28,Rv1453-NC_000962.3:1636705-1638392
PE_PGRS27
Rv1451
Event_124
1633000
1633500
1634000
1634500
1635000
1635500
Genomic position (1-based)

Circos Viz - GCE to Known Paralogs Locations
GCE ID: Event_124

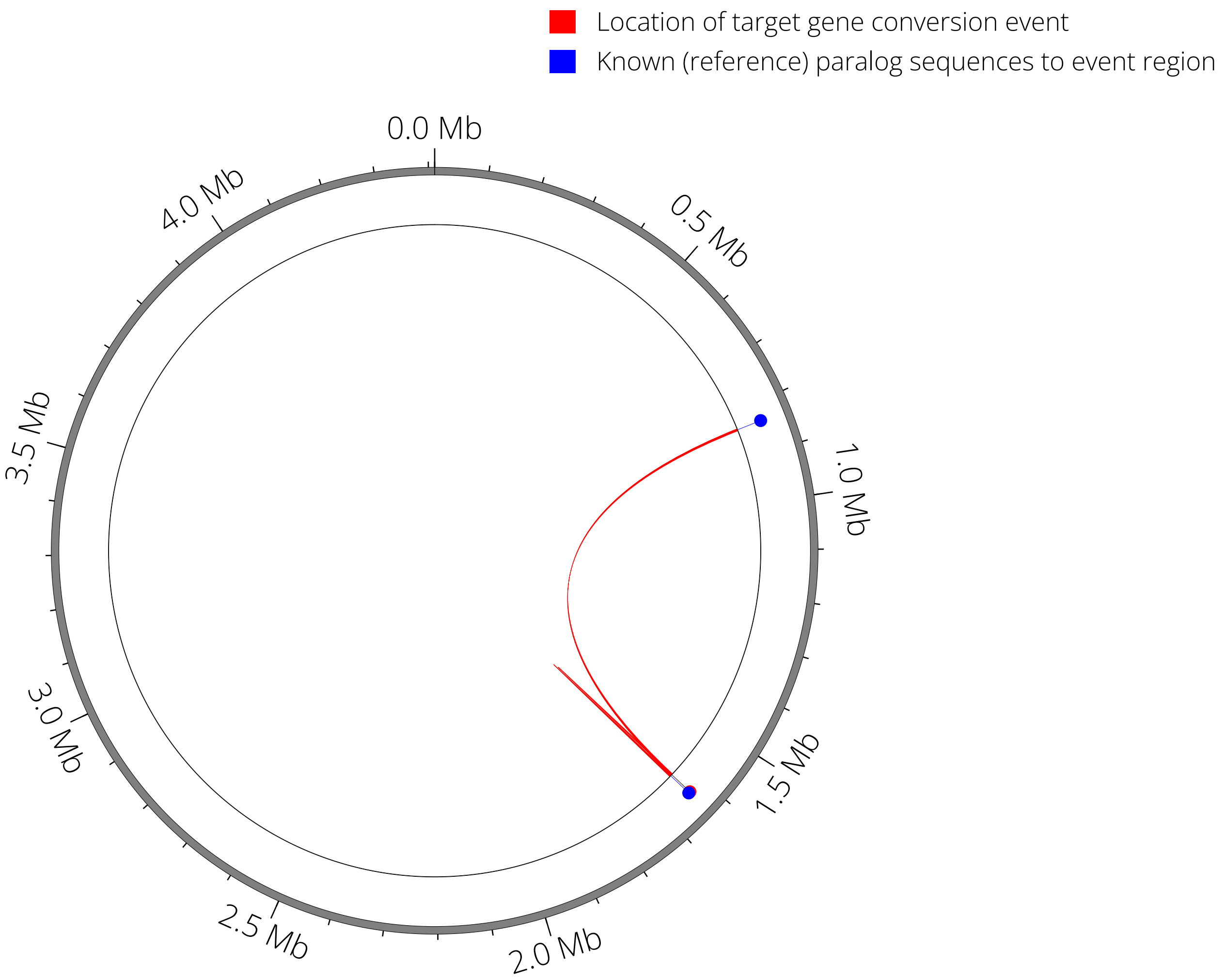

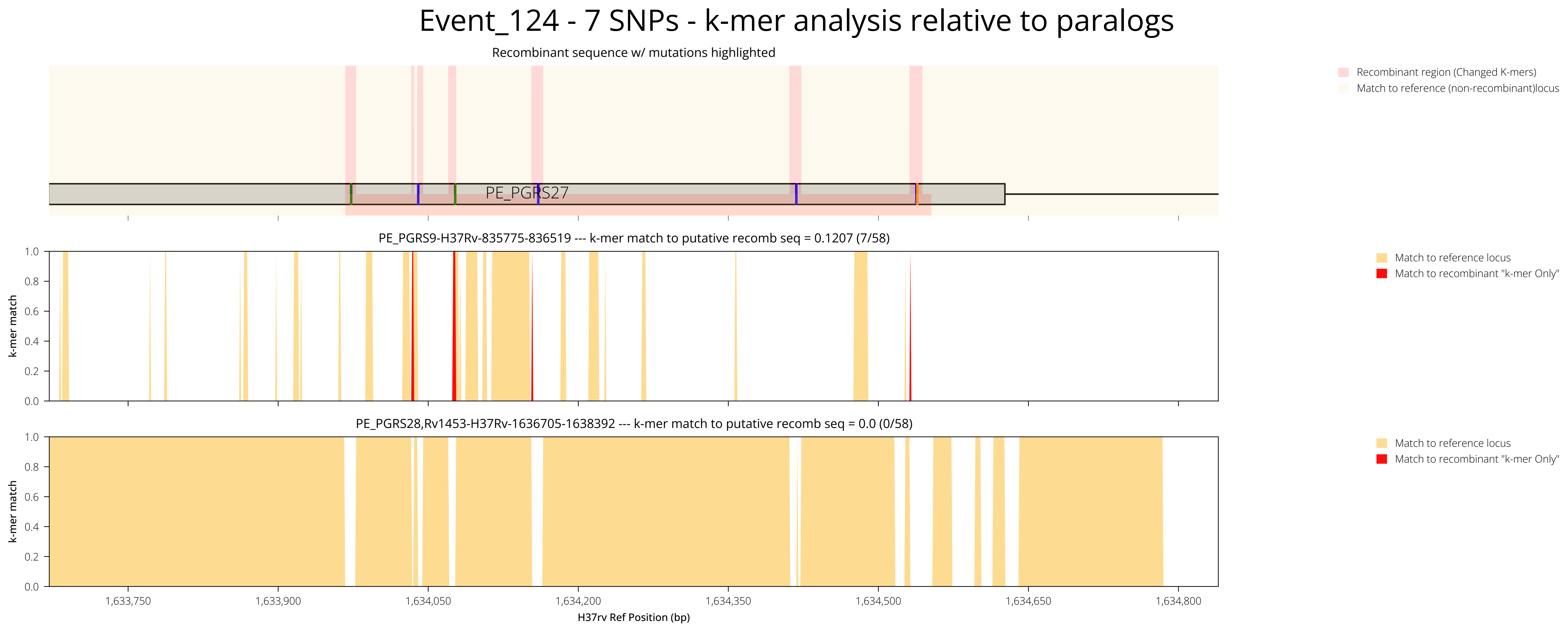

Event_124 - 7 SNPs - k-mer analysis relative to paralogs
Recombinant sequence w/ mutations highlighted
Recombinant region (Changed K-mers)
Match to reference (non-recombinant)locus
PE_PGRS27
PE_PGRS9-H37Rv-835775-836519 --- k-mer match to putative recomb seq = 0.1207 (7/58)
Match to reference locus
Match to recombinant "k-mer Only"
k-mer match
1.0
0.8
0.6
0.4
0.2
0.0
PE_PGRS28,Rv1453-H37Rv-1636705-1638392 --- k-mer match to putative recomb seq = 0.0 (0/58)
Match to reference locus
Match to recombinant "k-mer Only"
k-mer match
1.0
0.8
0.6
0.4
0.2
0.0
1,633,750
1,633,900
1,634,050
1,634,200
1,634,350
1,634,500
1,634,650
1,634,800
H37rv Ref Position (bp)

Sub-phylogeny Viz + Paralog Alignments
GCE ID: Event_124

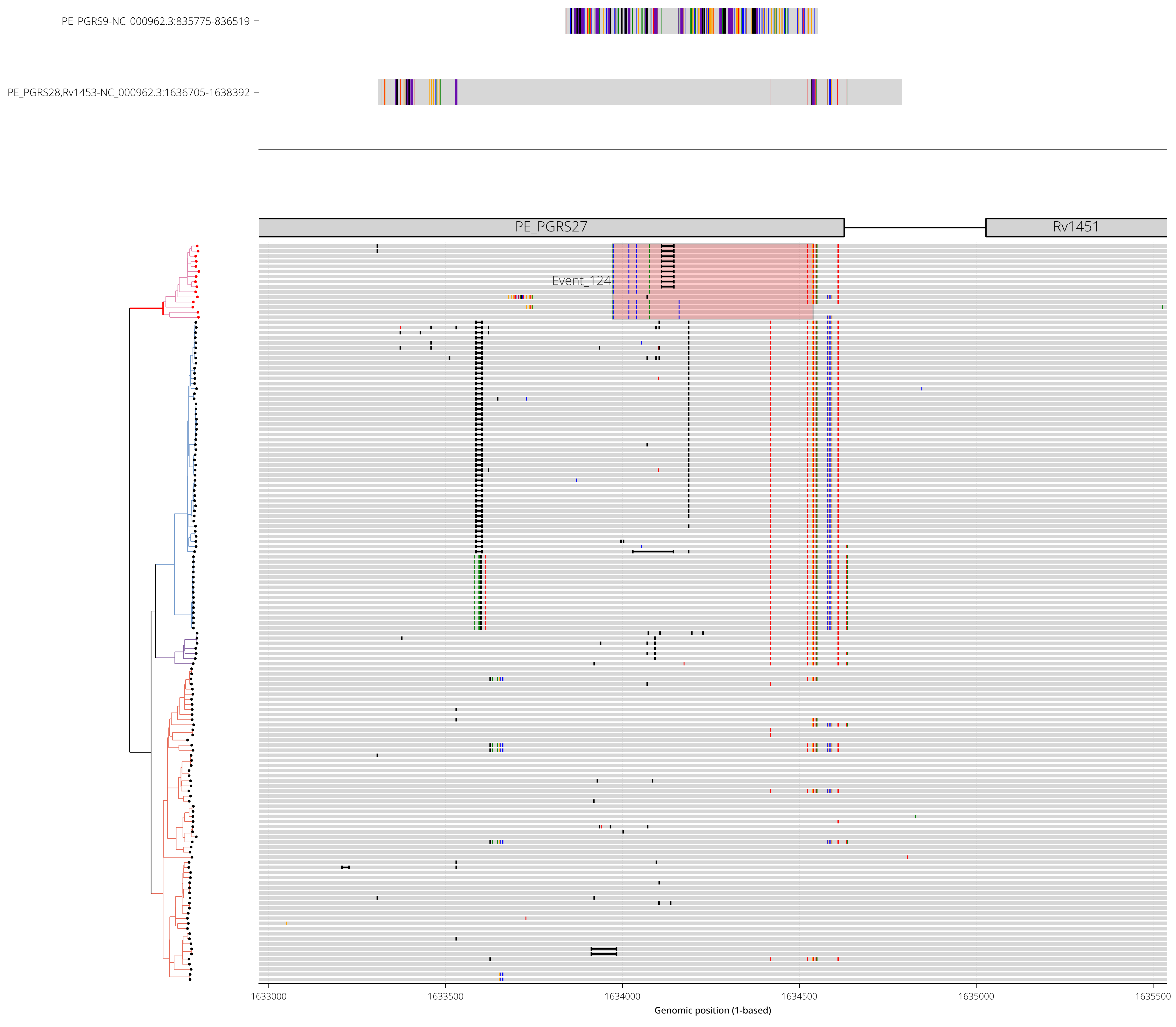

Full Phylogeny Viz + Paralog Alignments
GCE ID: Event_124

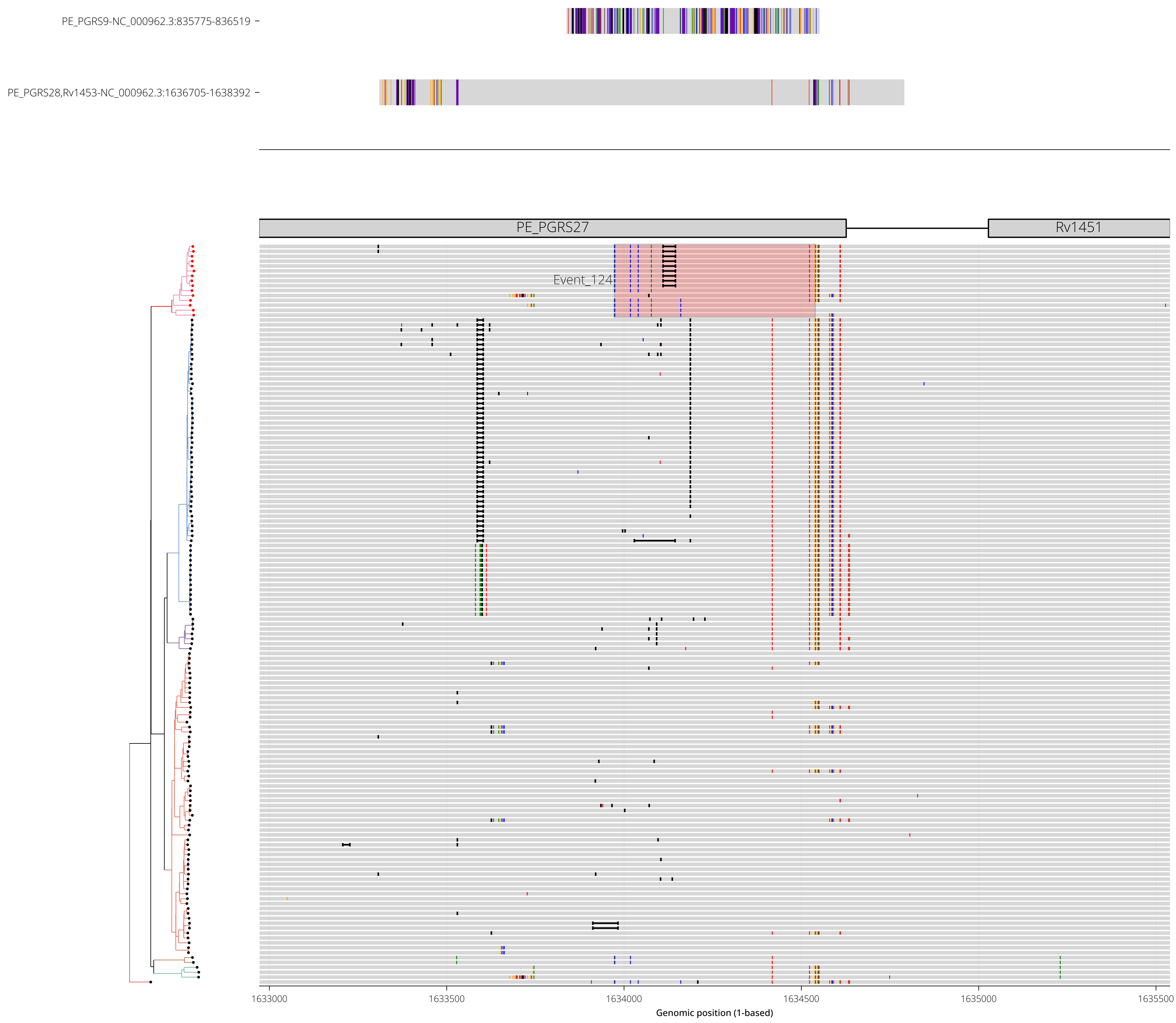

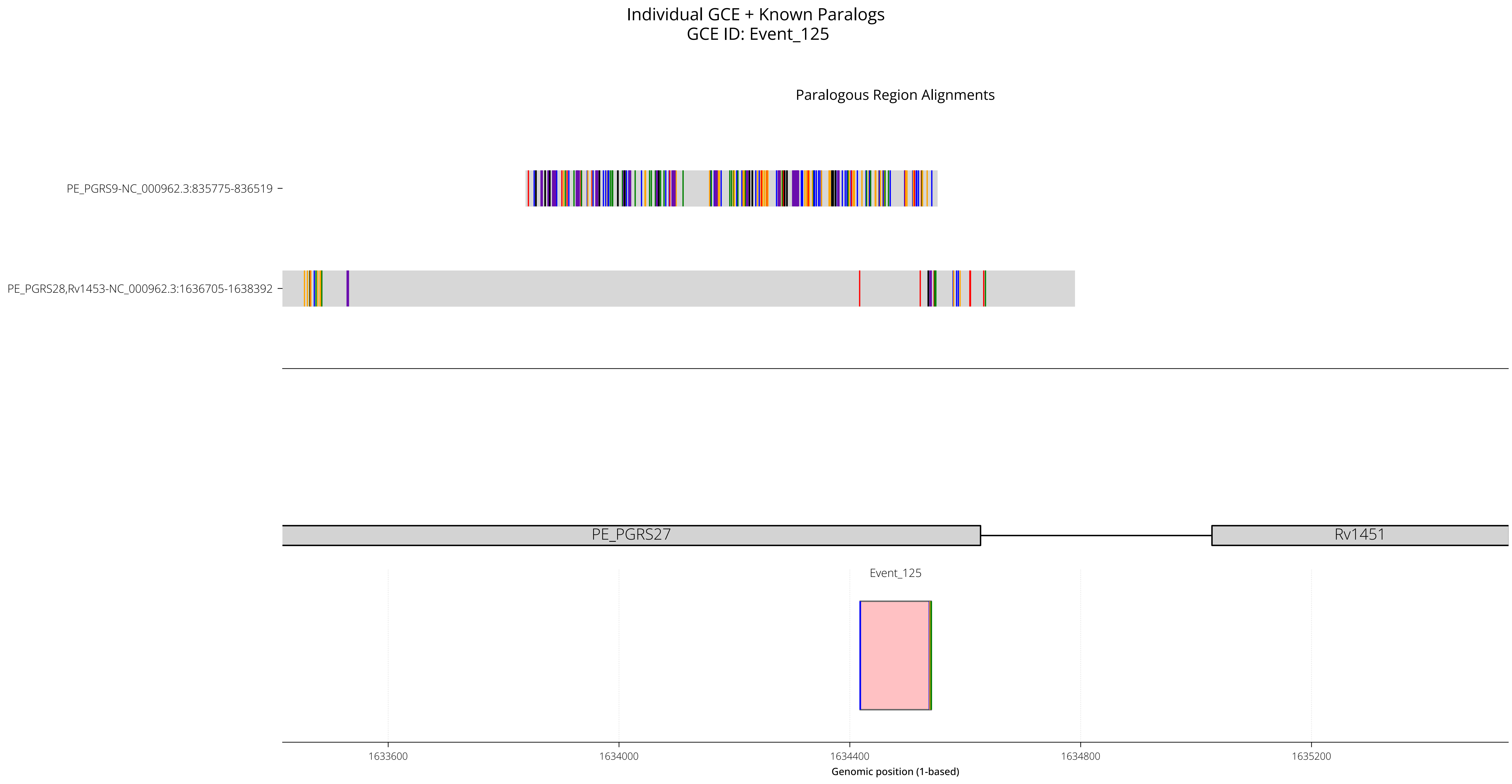
Individual GCE + Known Paralogs
GCE ID: Event_125
Paralogous Region Alignments
PE_PGRS9-NC_000962.3:835775-836519
PE_PGRS28,Rv1453-NC_000962.3:1636705-1638392
PE_PGRS27
Rv1451
Event_125
1633600
1634000
1634400
1634800
1635200
Genomic position (1-based)

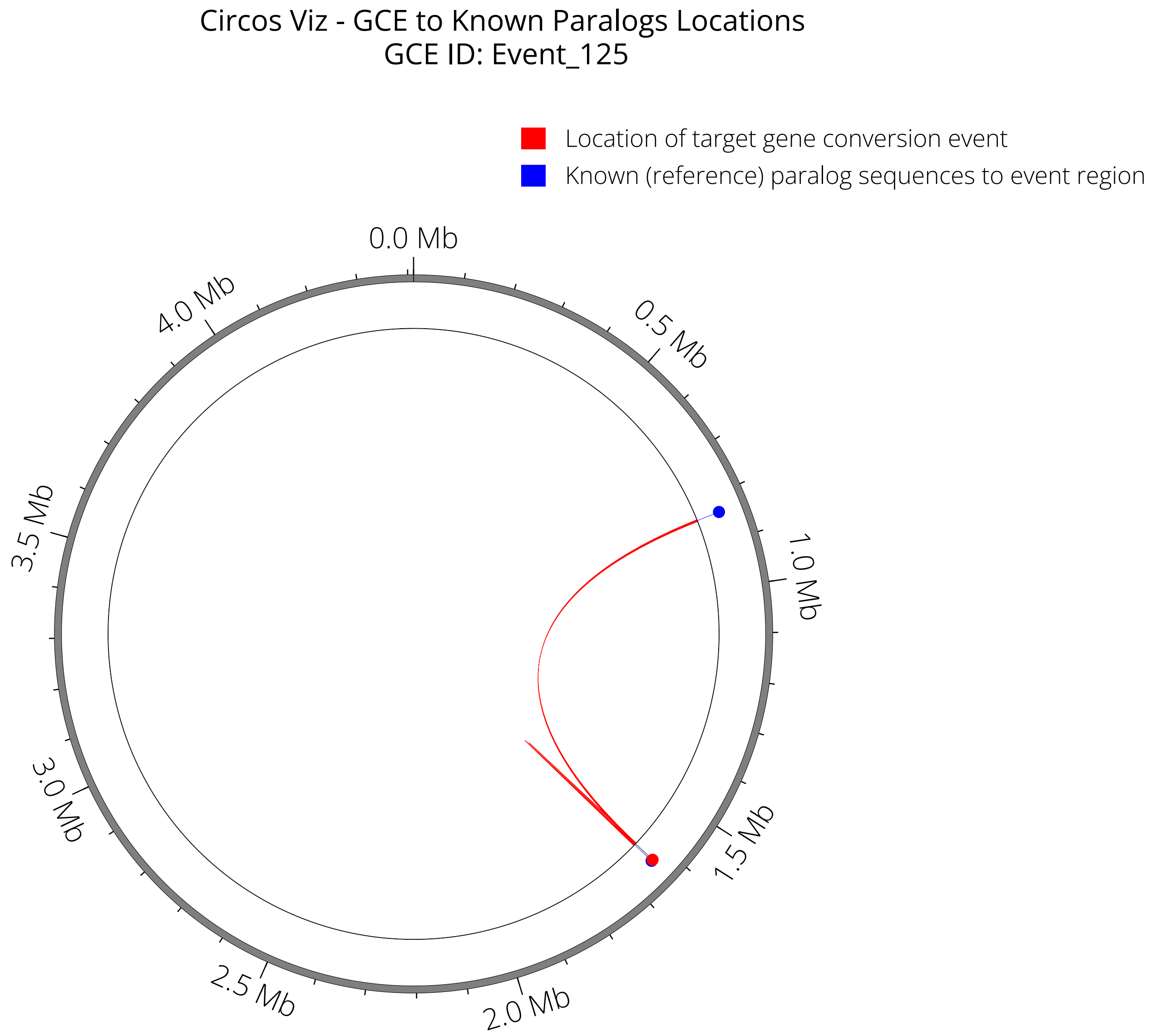
Circos Viz - GCE to Known Paralogs Locations
GCE ID: Event_125
Location of target gene conversion event
Known (reference) paralog sequences to event region
0.0 Mb
0.5 Mb
1.0 Mb
1.5 Mb
2.0 Mb
2.5 Mb
3.0 Mb
3.5 Mb
4.0 Mb

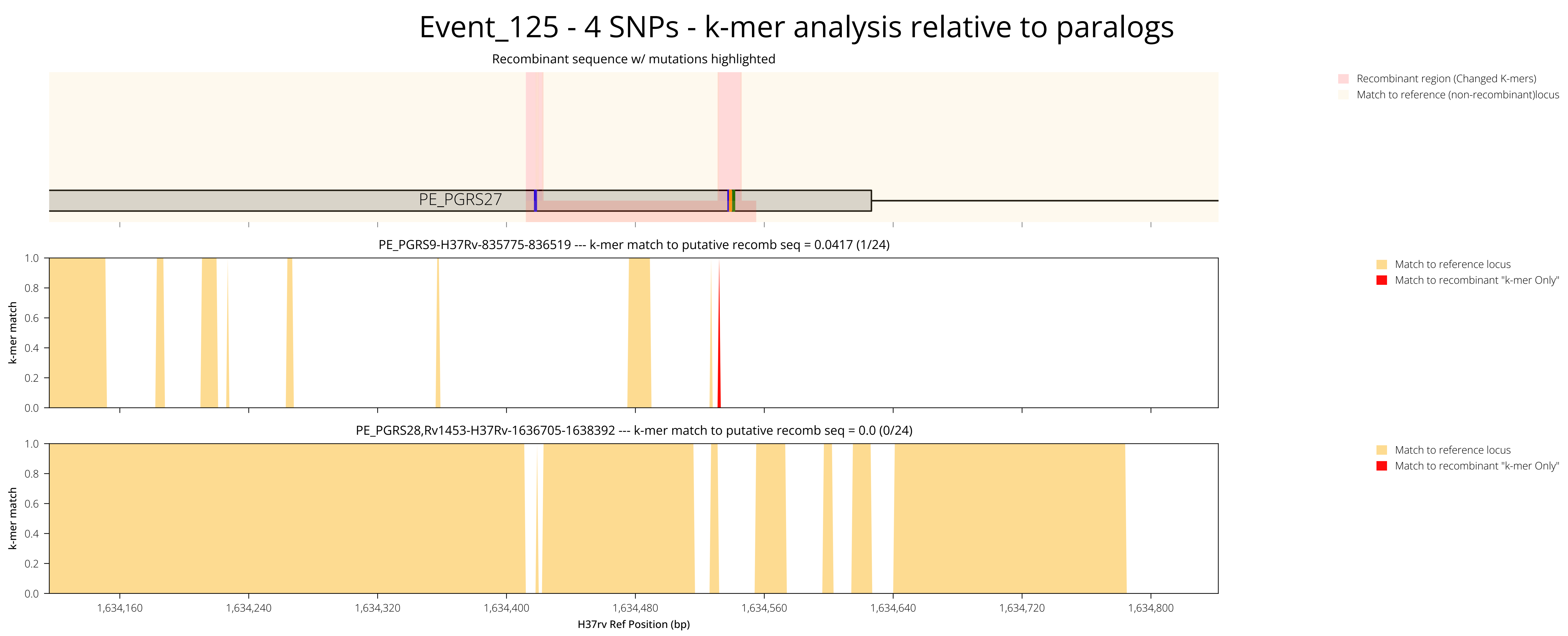
Event_125 - 4 SNPs - k-mer analysis relative to paralogs
Recombinant sequence w/ mutations highlighted
Recombinant region (Changed K-mers)
Match to reference (non-recombinant)locus
PE_PGRS27
PE_PGRS9-H37Rv-835775-836519 --- k-mer match to putative recomb seq = 0.0417 (1/24)
Match to reference locus
Match to recombinant "k-mer Only"
k-mer match
1.0
0.8
0.6
0.4
0.2
0.0
PE_PGRS28,Rv1453-H37Rv-1636705-1638392 --- k-mer match to putative recomb seq = 0.0 (0/24)
Match to reference locus
Match to recombinant "k-mer Only"
k-mer match
1,634,160
1,634,240
1,634,320
1,634,400
1,634,480
1,634,560
1,634,640
1,634,720
1,634,800
H37rv Ref Position (bp)

Sub-phylogeny Viz + Paralog Alignments
GCE ID: Event_125

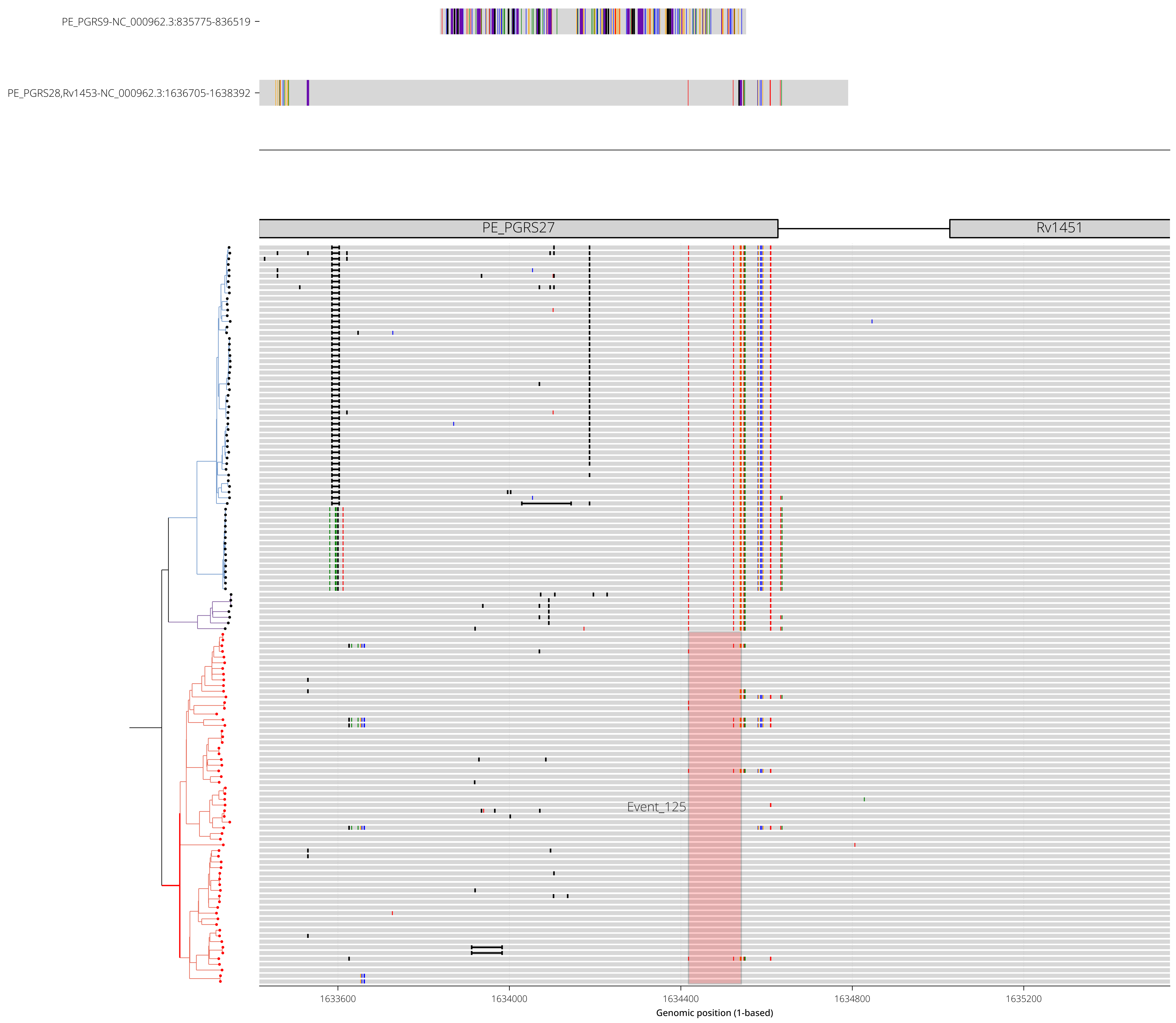

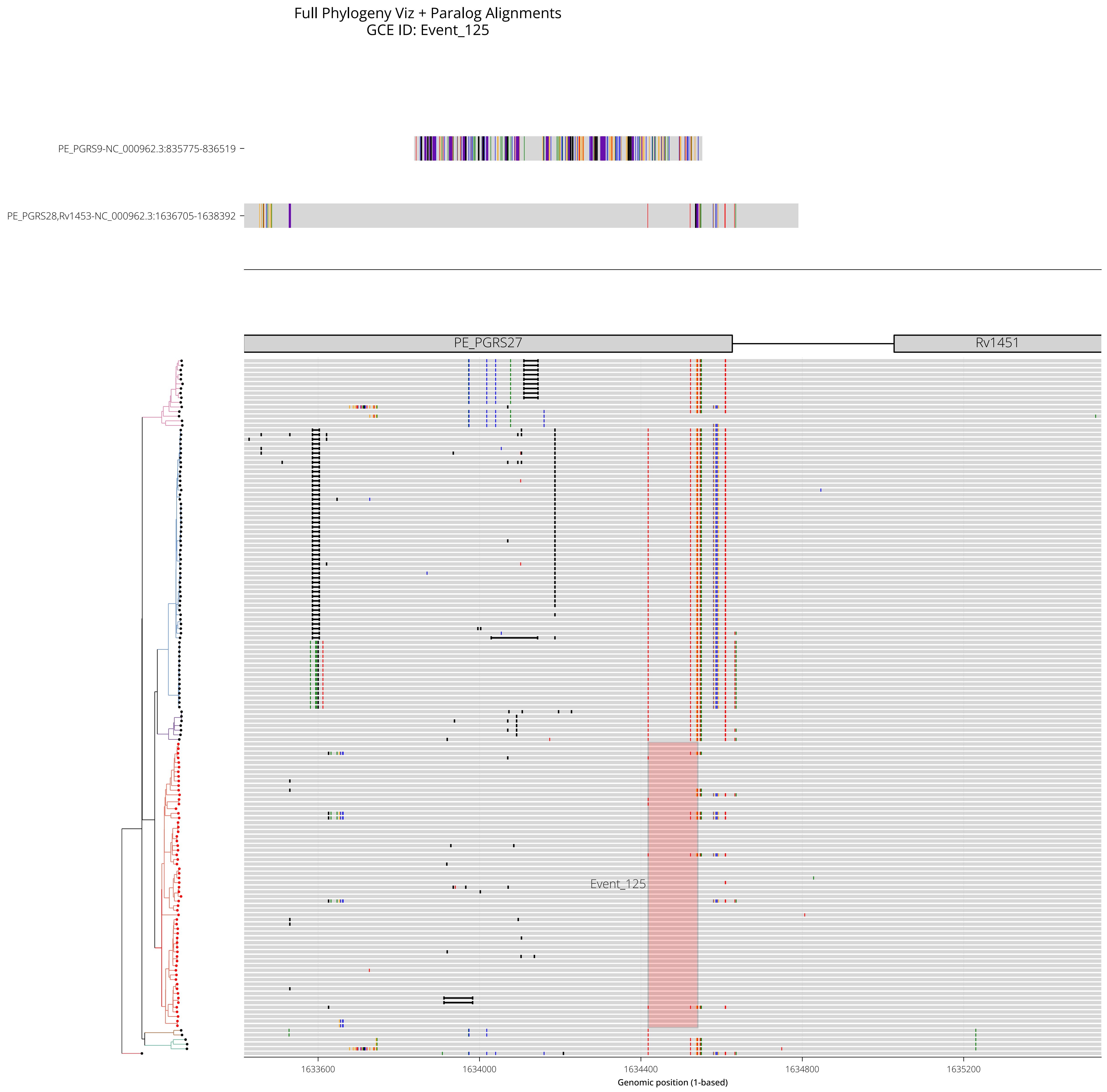
Full Phylogeny Viz + Paralog Alignments
GCE ID: Event_125
PE_PGRS9-NC_000962.3:835775-836519
PE_PGRS28,Rv1453-NC_000962.3:1636705-1638392
PE_PGRS27
Rv1451
Event_125
1633600
1634000
1634400
1634800
1635200
Genomic position (1-based)

Individual GCE + Known Paralogs
GCE ID: Event_126

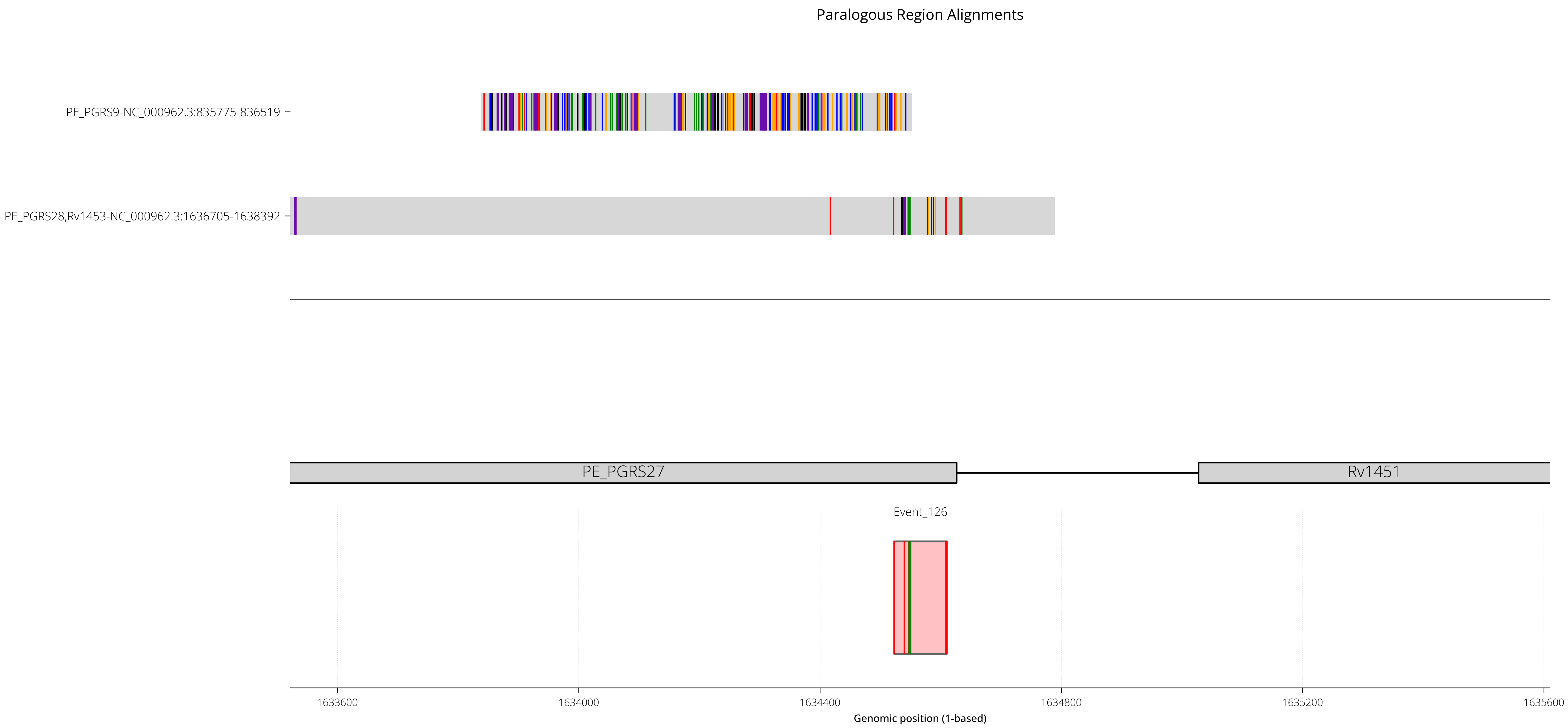

Circos Viz - GCE to Known Paralogs Locations
GCE ID: Event_126

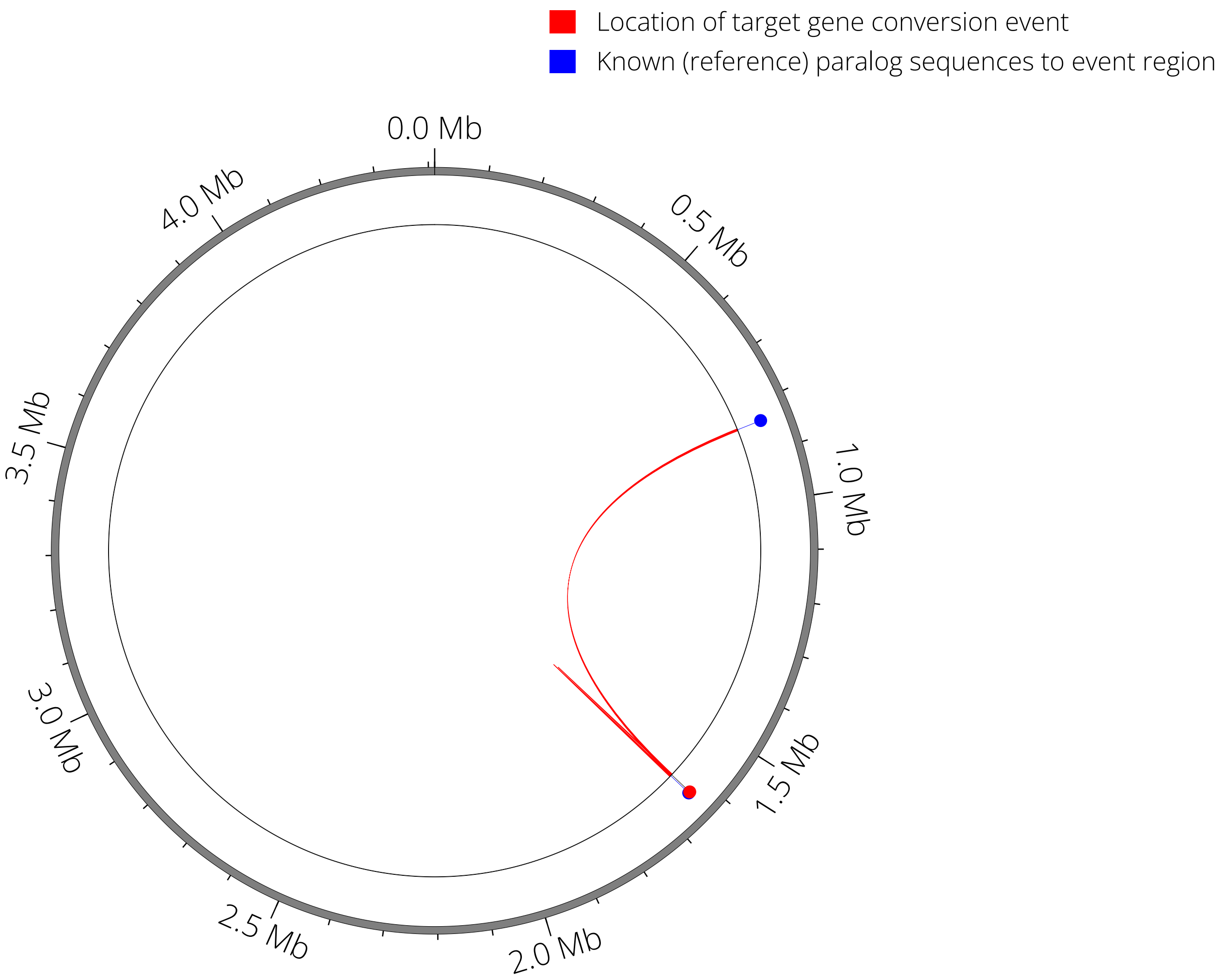

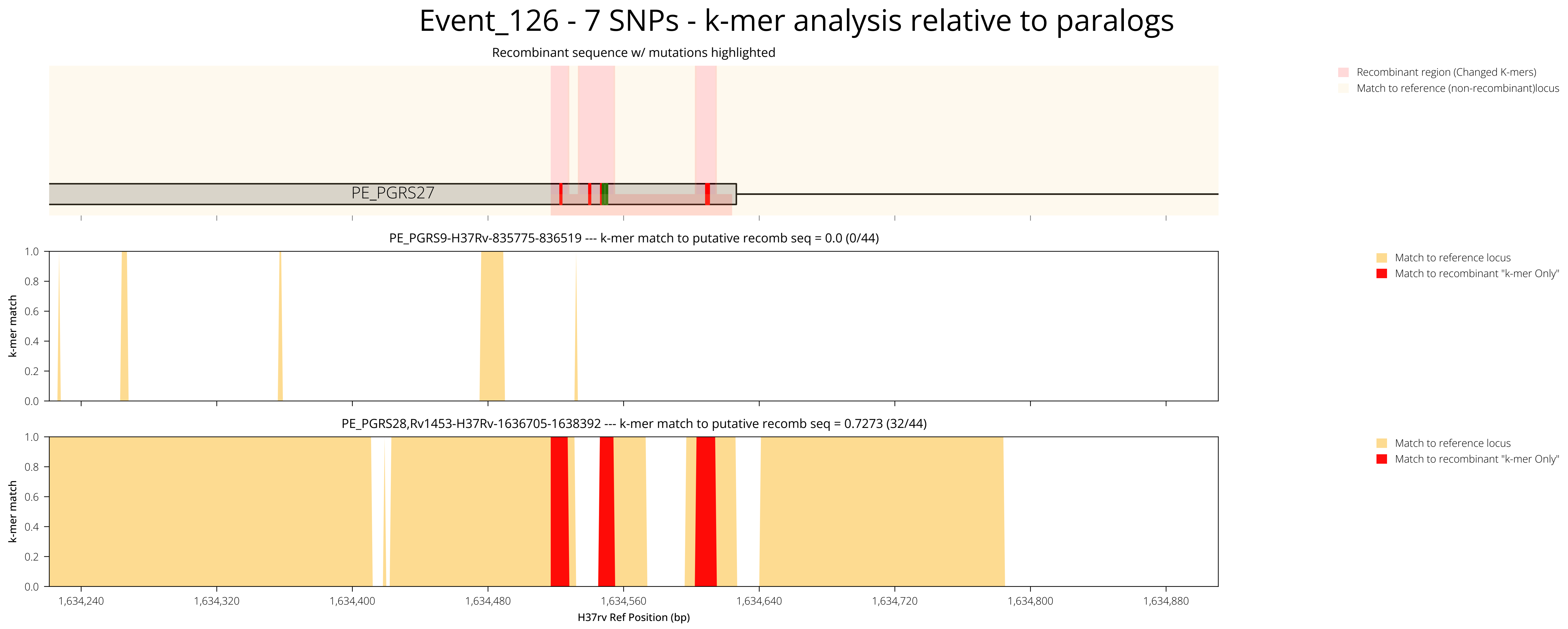

Event_126 - 7 SNPs - k-mer analysis relative to paralogs
Recombinant sequence w/ mutations highlighted
PE_PGRS27
Recombinant region (Changed K-mers)
Match to reference (non-recombinant)locus
PE_PGRS9-H37Rv-835775-836519 --- k-mer match to putative recomb seq = 0.0 (0/44)
k-mer match
1.0
0.8
0.6
0.4
0.2
0.0
Match to reference locus
Match to recombinant "k-mer Only"
PE_PGRS28,Rv1453-H37Rv-1636705-1638392 --- k-mer match to putative recomb seq = 0.7273 (32/44)
k-mer match
1.0
0.8
0.6
0.4
0.2
0.0
Match to reference locus
Match to recombinant "k-mer Only"
1,634,240
1,634,320
1,634,400
1,634,480
1,634,560
1,634,640
1,634,720
1,634,800
1,634,880
H37rv Ref Position (bp)

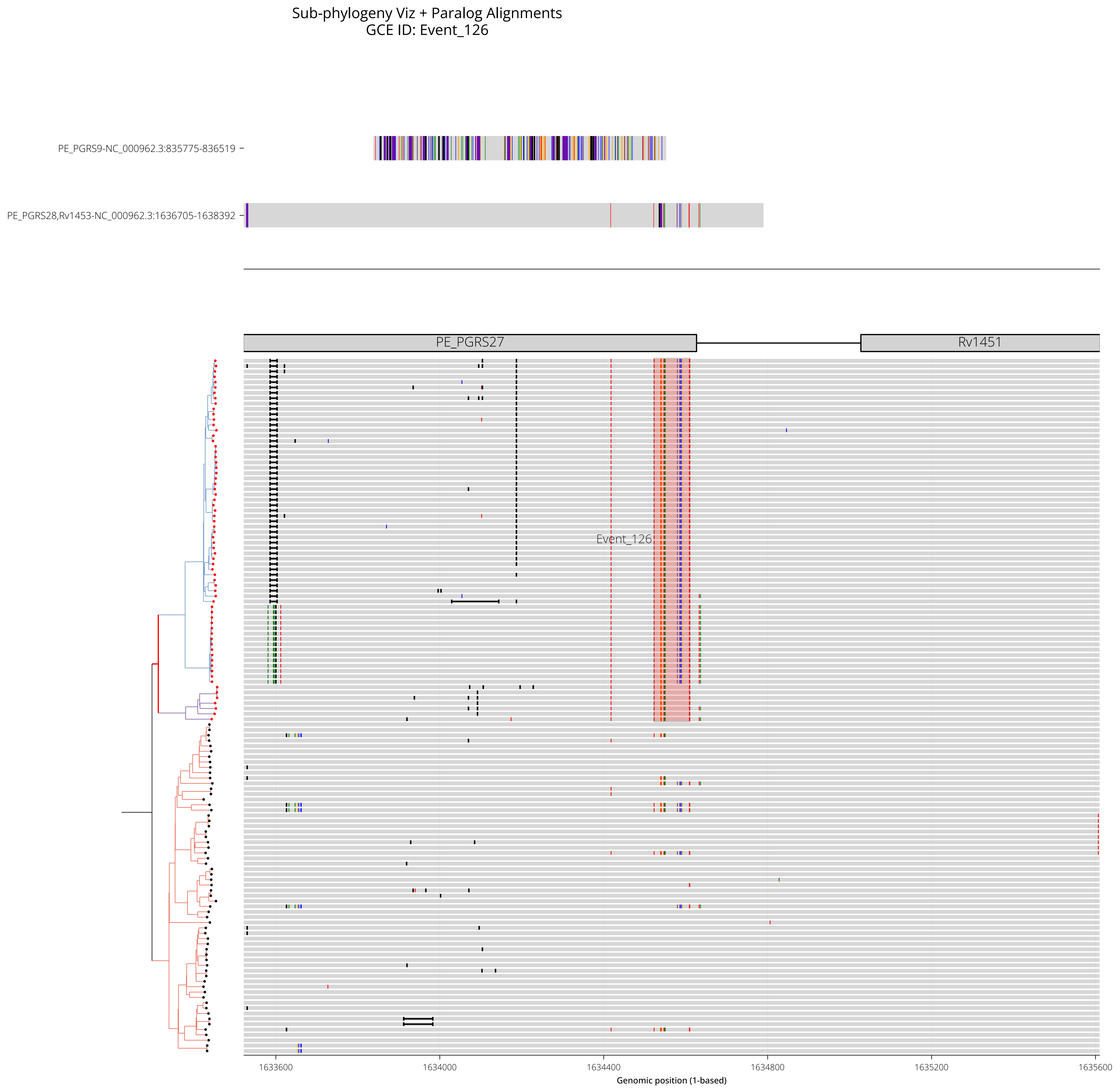

Sub-phylogeny Viz + Paralog Alignments
GCE ID: Event_126
PE_PGRS9-NC_000962.3:835775-836519
PE_PGRS28,Rv1453-NC_000962.3:1636705-1638392
PE_PGRS27
Rv1451
Event_126
1633600
1634000
1634400
1634800
1635200
1635600
Genomic position (1-based)

Full Phylogeny Viz + Paralog Alignments
GCE ID: Event_126

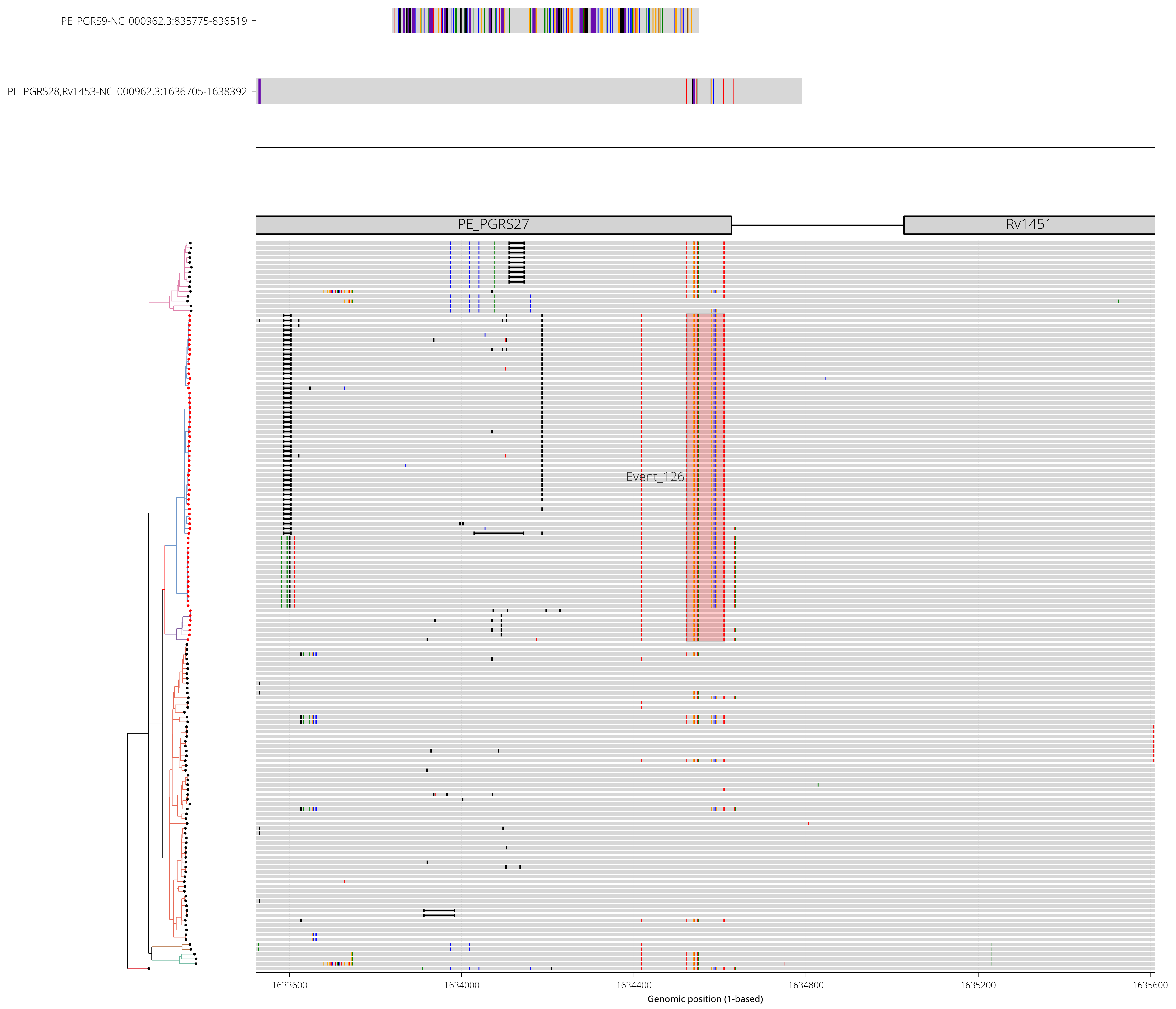

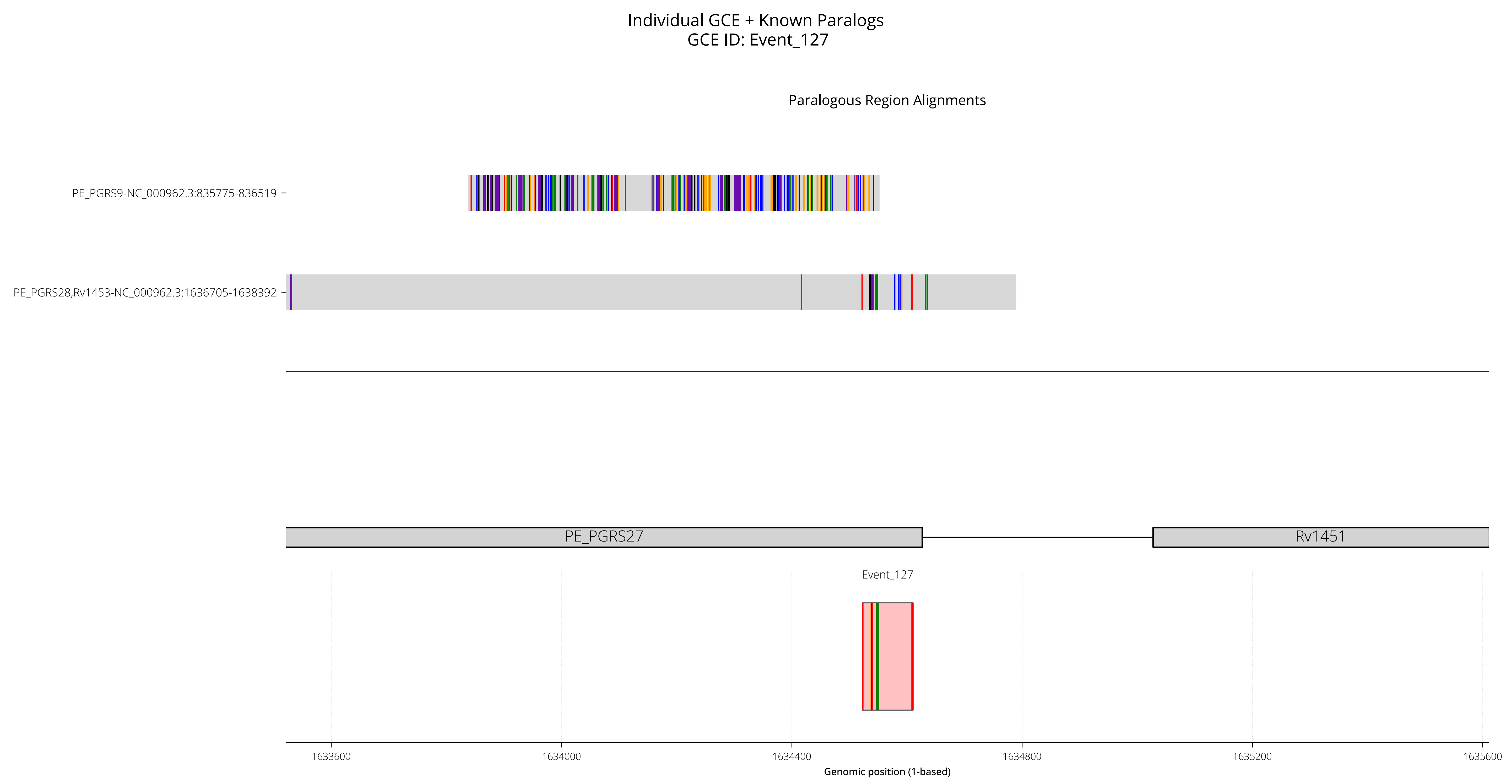

Individual GCE + Known Paralogs
GCE ID: Event_127
Paralogous Region Alignments
PE_PGRS9-NC_000962.3:835775-836519
PE_PGRS28,Rv1453-NC_000962.3:1636705-1638392
PE_PGRS27
Rv1451
Event_127
1633600
1634000
1634400
1634800
1635200
1635600
Genomic position (1-based)

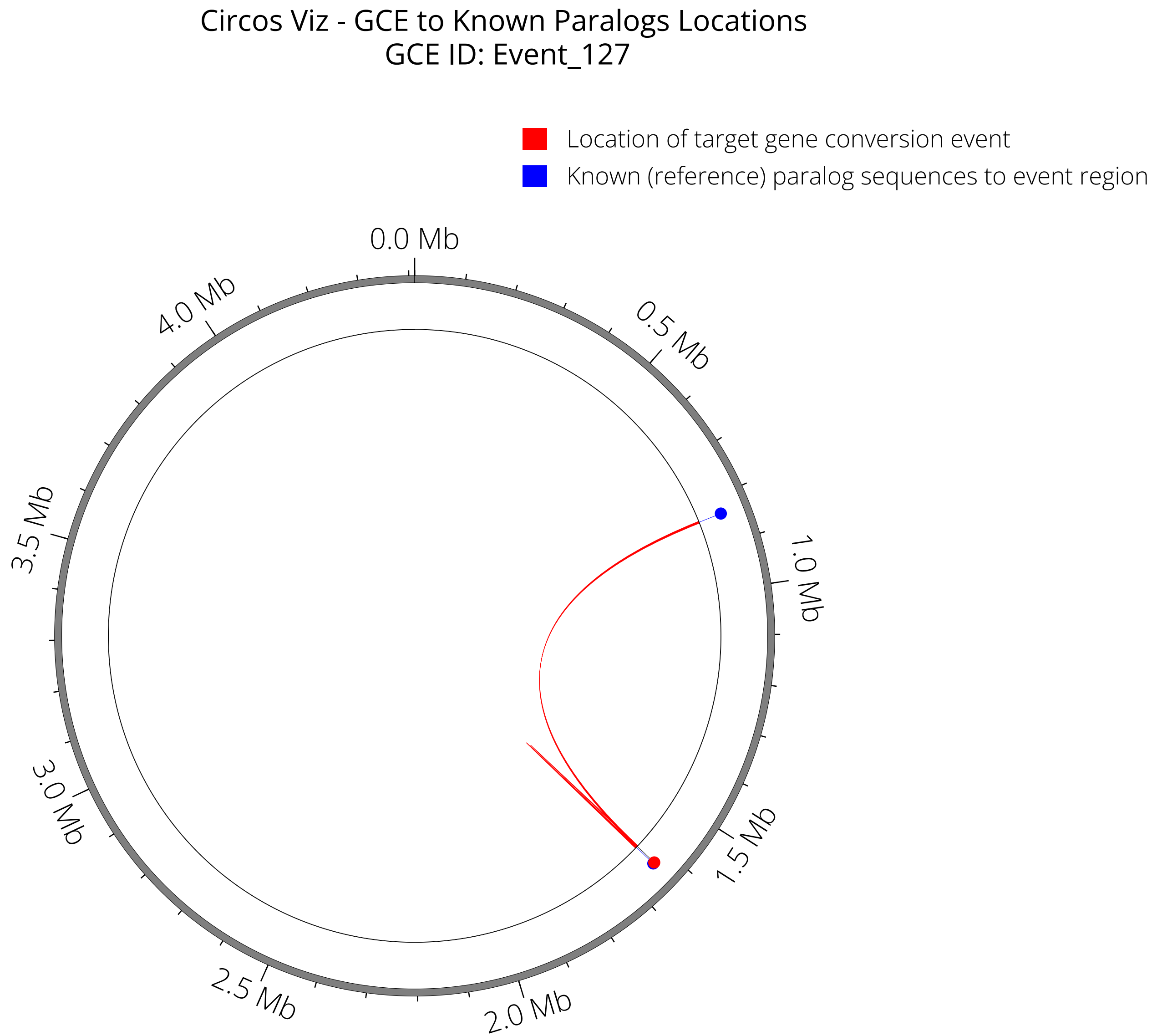
Circos Viz - GCE to Known Paralogs Locations
GCE ID: Event_127
Location of target gene conversion event
Known (reference) paralog sequences to event region
0.0 Mb
0.5 Mb
1.0 Mb
1.5 Mb
2.0 Mb
2.5 Mb
3.0 Mb
3.5 Mb
4.0 Mb

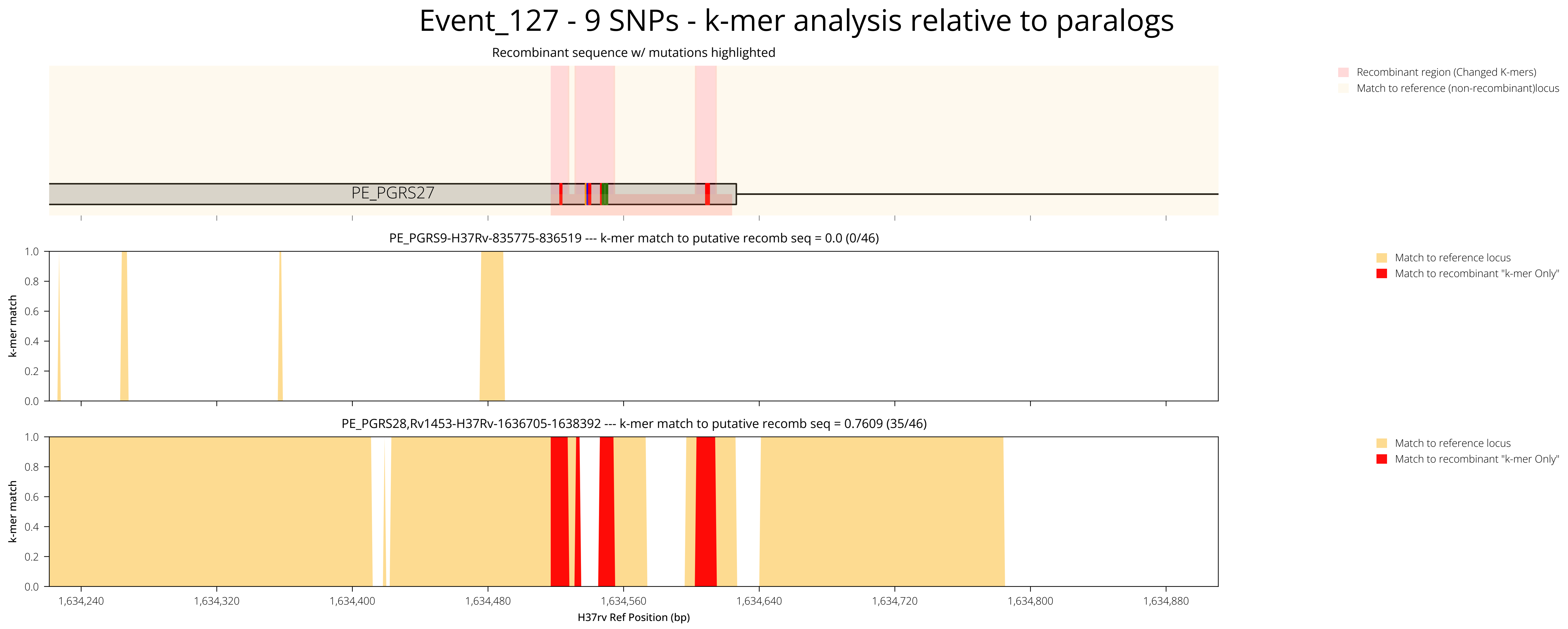

Event_127 - 9 SNPs - k-mer analysis relative to paralogs
Recombinant sequence w/ mutations highlighted
PE_PGRS27
Recombinant region (Changed K-mers)
Match to reference (non-recombinant)locus
PE_PGRS9-H37Rv-835775-836519 --- k-mer match to putative recomb seq = 0.0 (0/46)
k-mer match
1.0
0.8
0.6
0.4
0.2
0.0
Match to reference locus
Match to recombinant "k-mer Only"
PE_PGRS28,Rv1453-H37Rv-1636705-1638392 --- k-mer match to putative recomb seq = 0.7609 (35/46)
1,634,240
1,634,320
1,634,400
1,634,480
1,634,560
1,634,640
1,634,720
1,634,800
1,634,880
H37rv Ref Position (bp)

Sub-phylogeny Viz + Paralog Alignments
GCE ID: Event_127

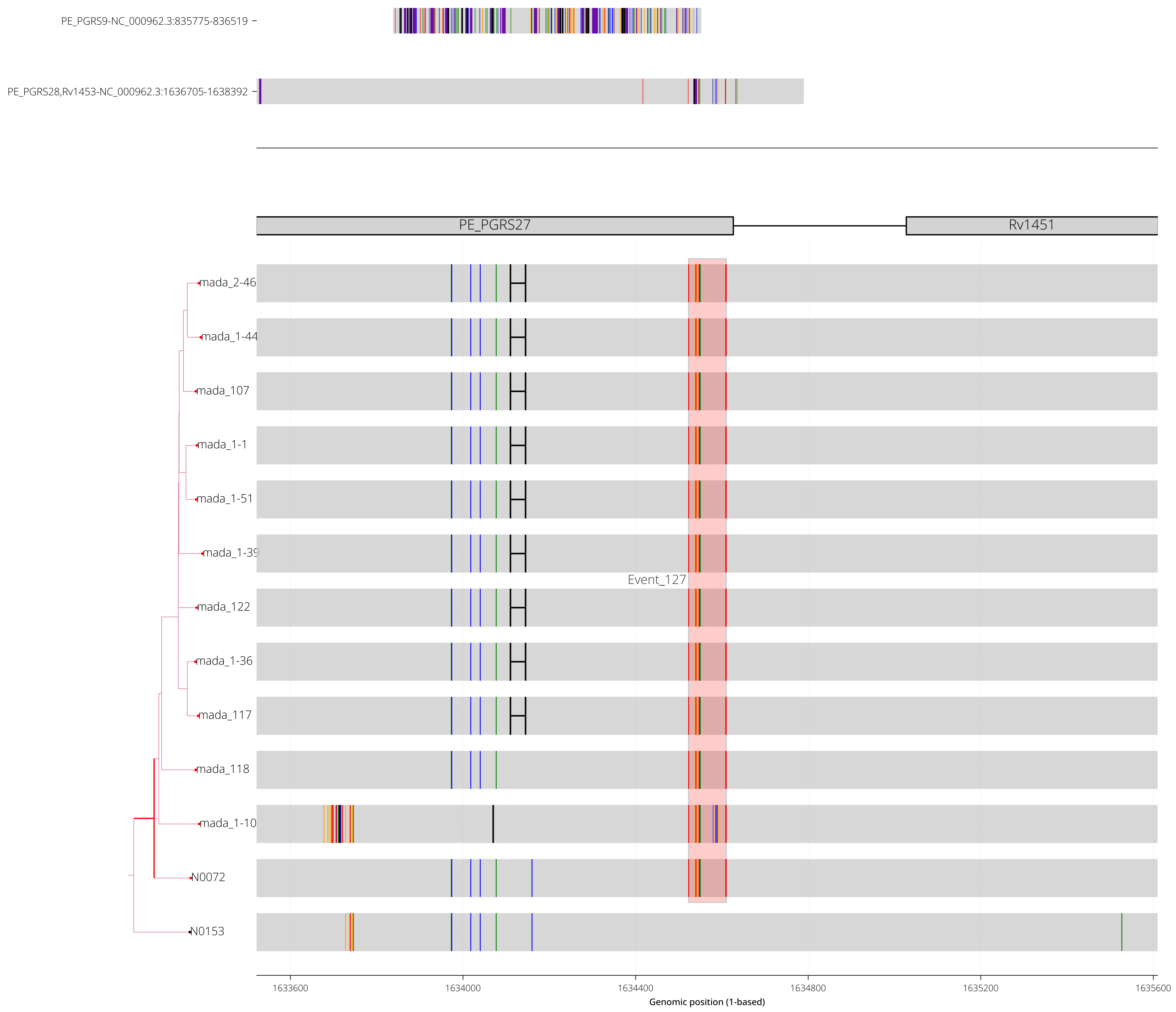

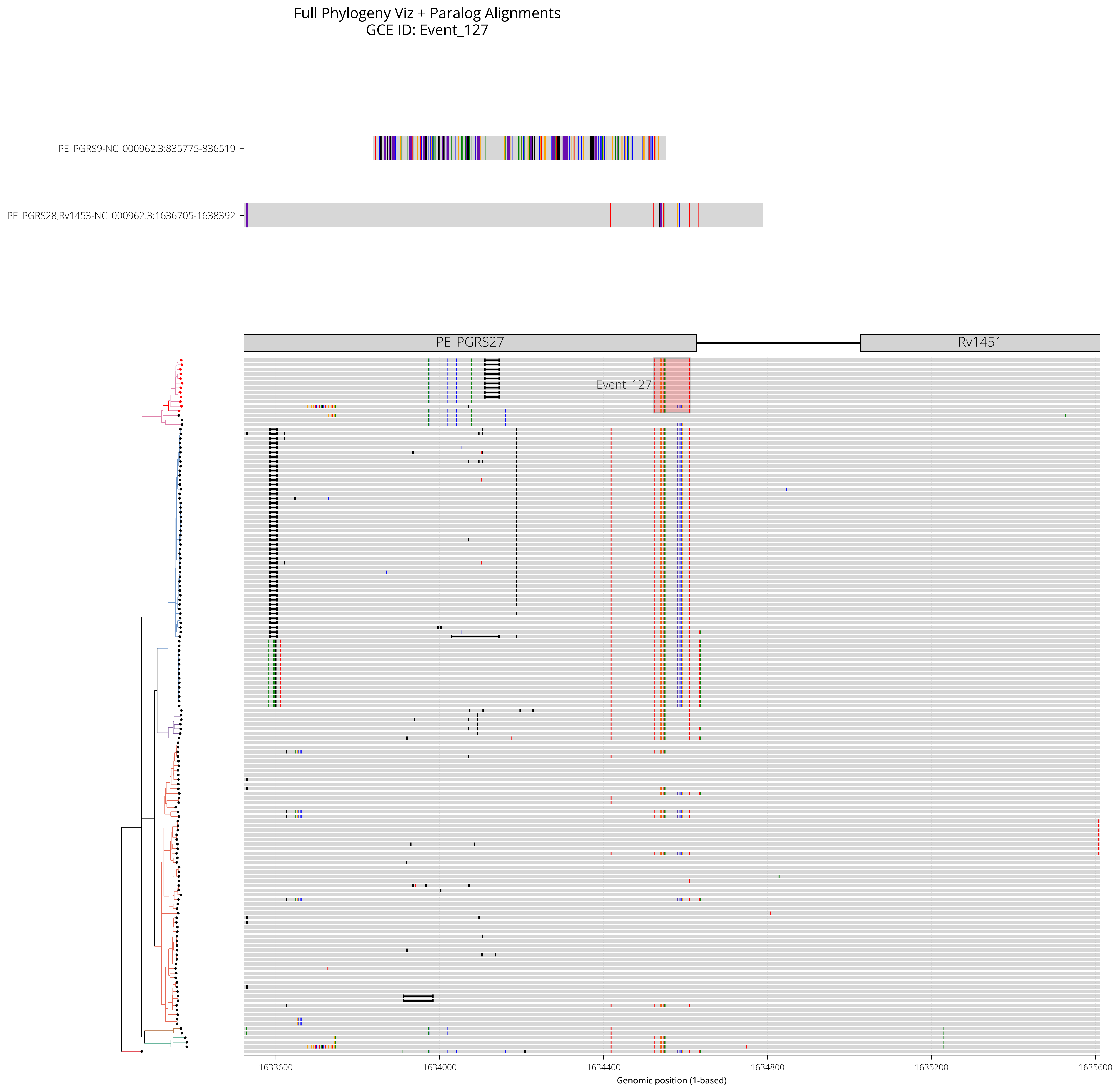
Full Phylogeny Viz + Paralog Alignments
GCE ID: Event_127
PE_PGRS9-NC_000962.3:835775-836519
PE_PGRS28,Rv1453-NC_000962.3:1636705-1638392
PE_PGRS27
Rv1451
Event_127
1633600
1634000
1634400
1634800
1635200
1635600
Genomic position (1-based)

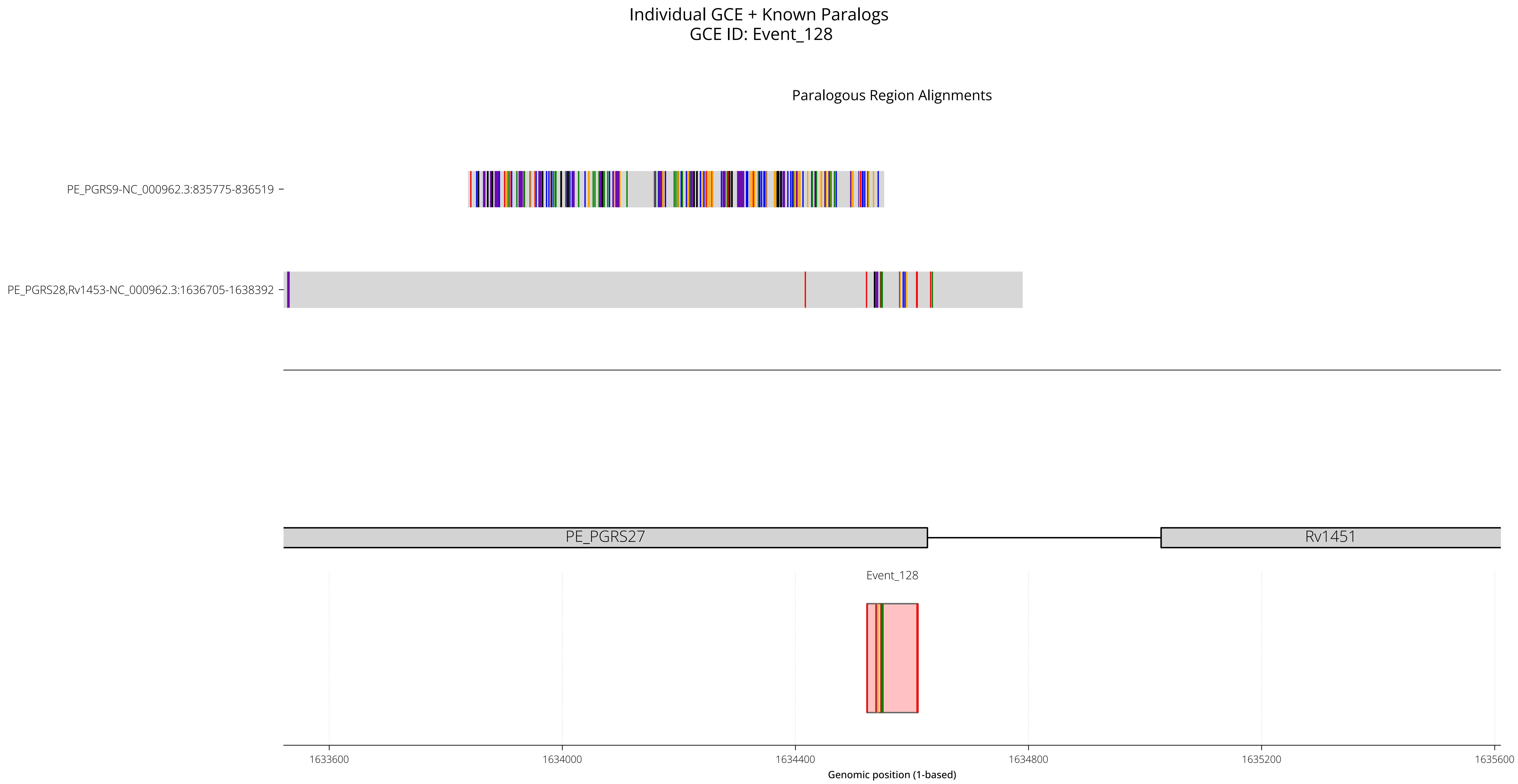

Individual GCE + Known Paralogs
GCE ID: Event_128
Paralogous Region Alignments
PE_PGRS9-NC_000962.3:835775-836519
PE_PGRS28,Rv1453-NC_000962.3:1636705-1638392
PE_PGRS27
Rv1451
Event_128
1633600
1634000
1634400
1634800
1635200
1635600
Genomic position (1-based)

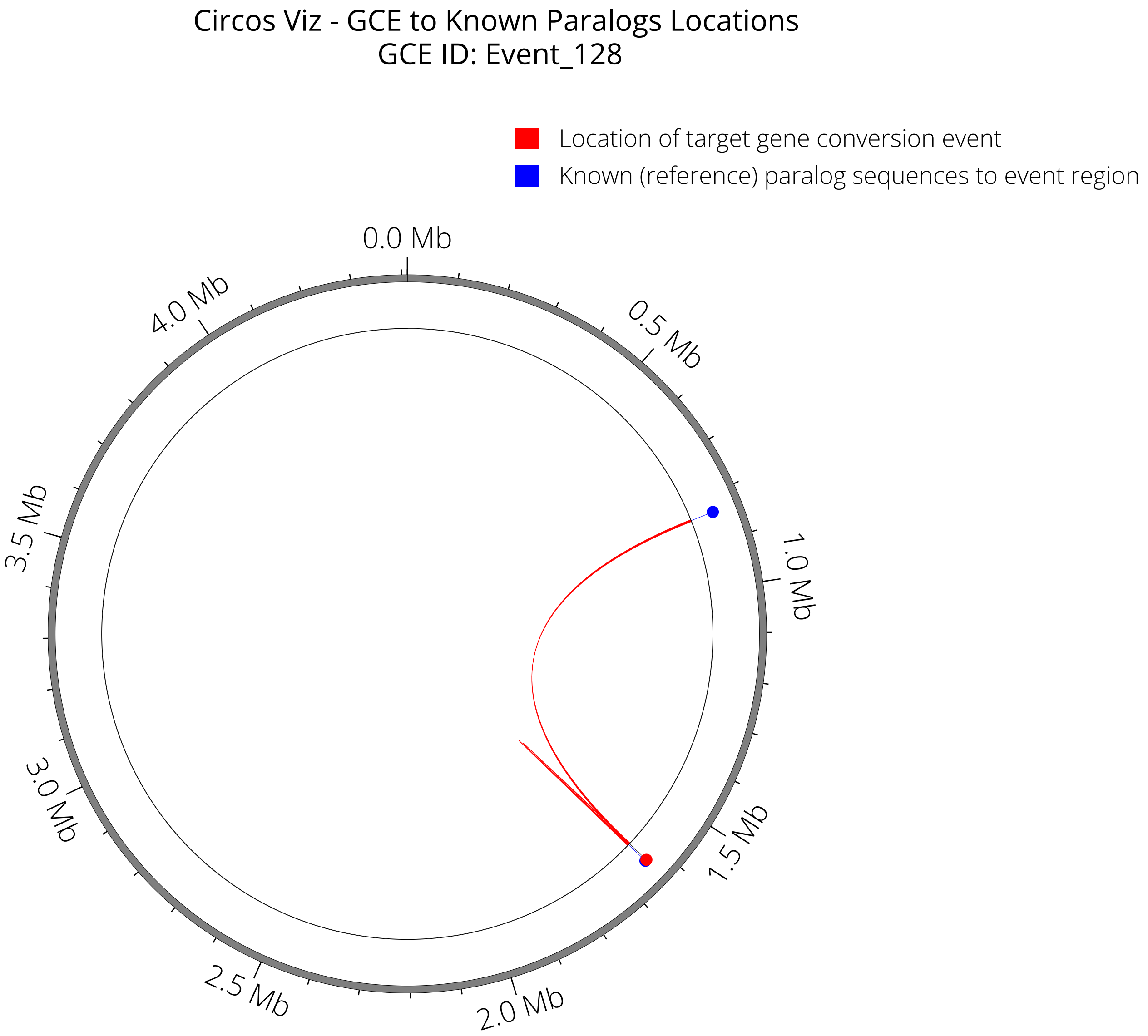
Circos Viz - GCE to Known Paralogs Locations
GCE ID: Event_128
Location of target gene conversion event
Known (reference) paralog sequences to event region
0.0 Mb
0.5 Mb
1.0 Mb
1.5 Mb
2.0 Mb
2.5 Mb
3.0 Mb
3.5 Mb
4.0 Mb

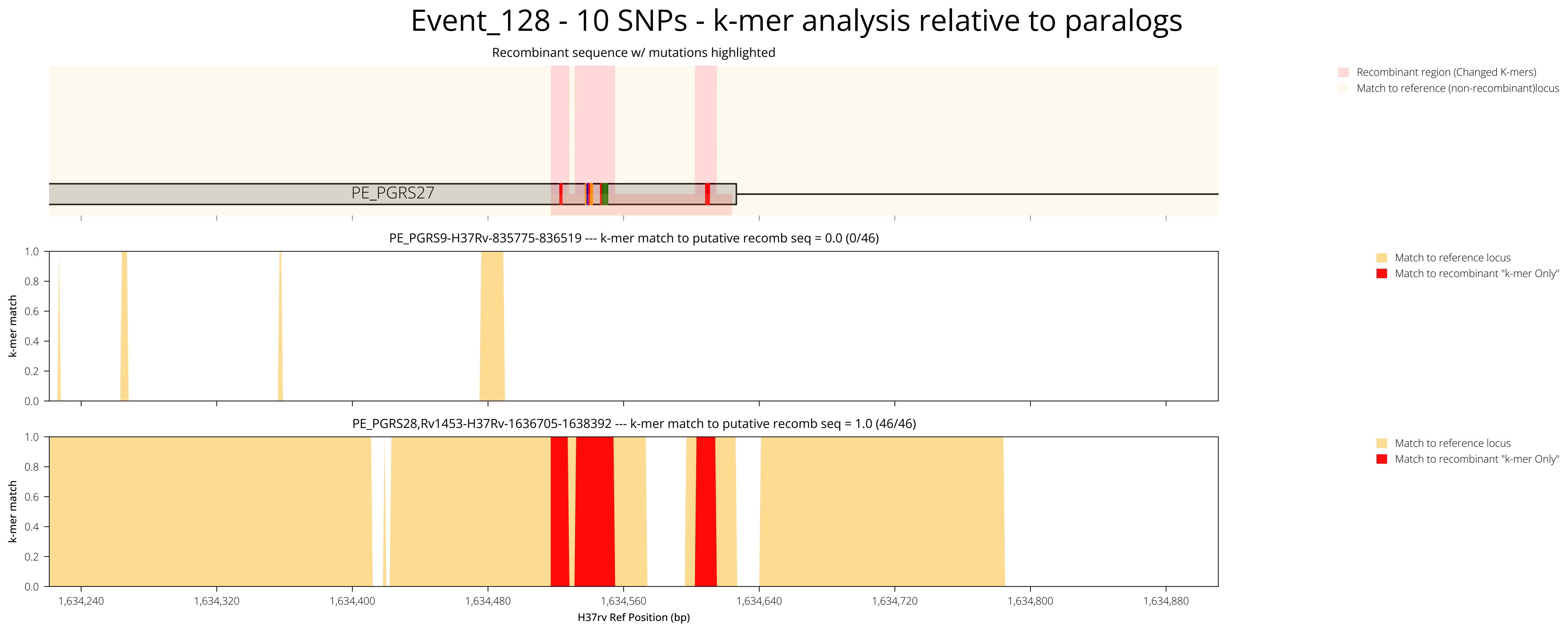

Event_128 - 10 SNPs - k-mer analysis relative to paralogs
Recombinant sequence w/ mutations highlighted
PE_PGRS27
Recombinant region (Changed K-mers)
Match to reference (non-recombinant)locus
PE_PGRS9-H37Rv-835775-836519 --- k-mer match to putative recomb seq = 0.0 (0/46)
Match to reference locus
Match to recombinant "k-mer Only"
k-mer match
1.0
0.8
0.6
0.4
0.2
0.0
PE_PGRS28,Rv1453-H37Rv-1636705-1638392 --- k-mer match to putative recomb seq = 1.0 (46/46)
Match to reference locus
Match to recombinant "k-mer Only"
k-mer match
1.0
0.8
0.6
0.4
0.2
0.0
1,634,240
1,634,320
1,634,400
1,634,480
1,634,560
1,634,640
1,634,720
1,634,800
1,634,880
H37rv Ref Position (bp)

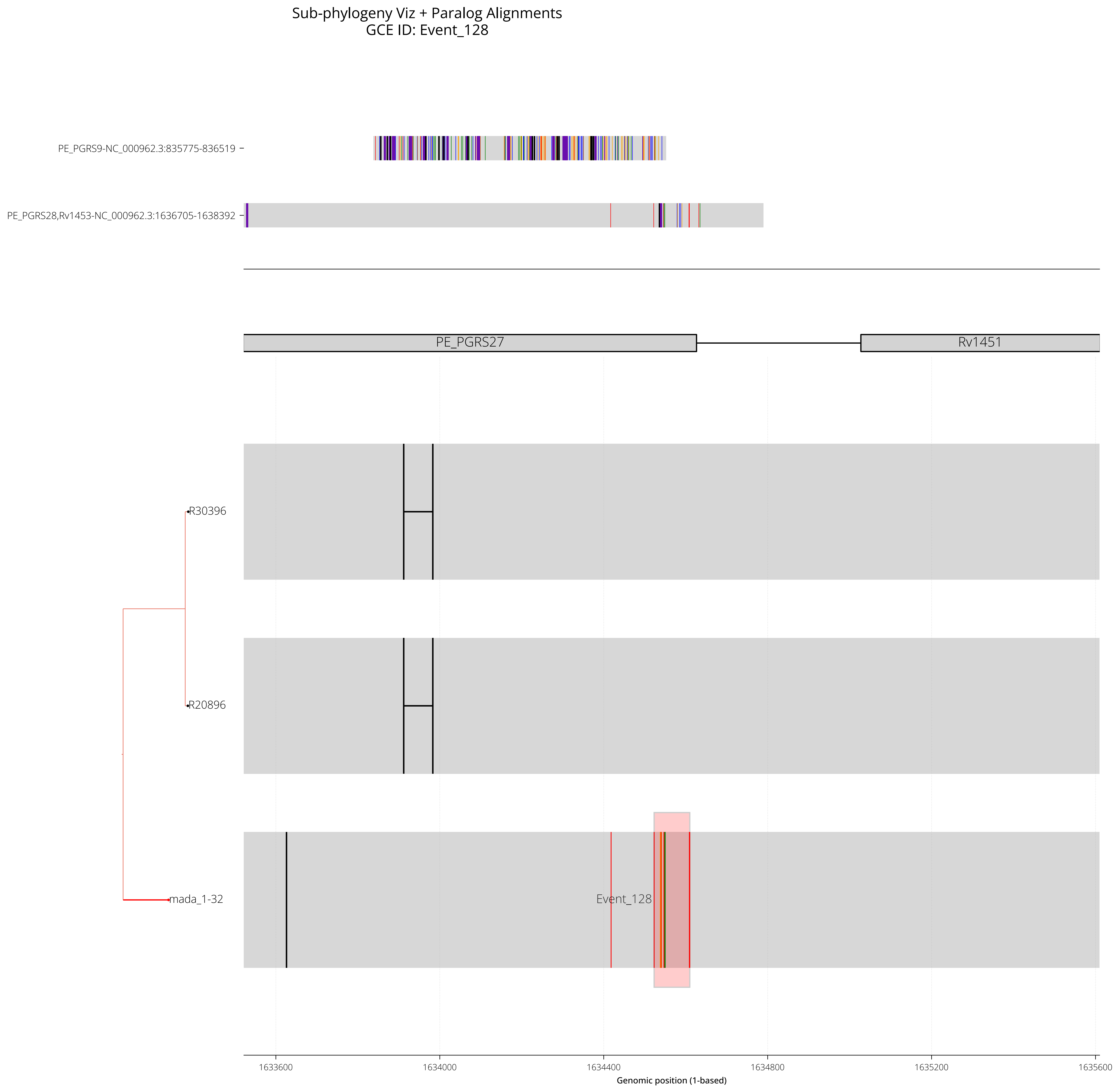

Sub-phylogeny Viz + Paralog Alignments
GCE ID: Event_128
PE_PGRS9-NC_000962.3:835775-836519
PE_PGRS28,Rv1453-NC_000962.3:1636705-1638392
PE_PGRS27
Rv1451
R30396
R20896
mada_1-32
Event_128
1633600
1634000
1634400
1634800
1635200
1635600
Genomic position (1-based)

Full Phylogeny Viz + Paralog Alignments
GCE ID: Event_128

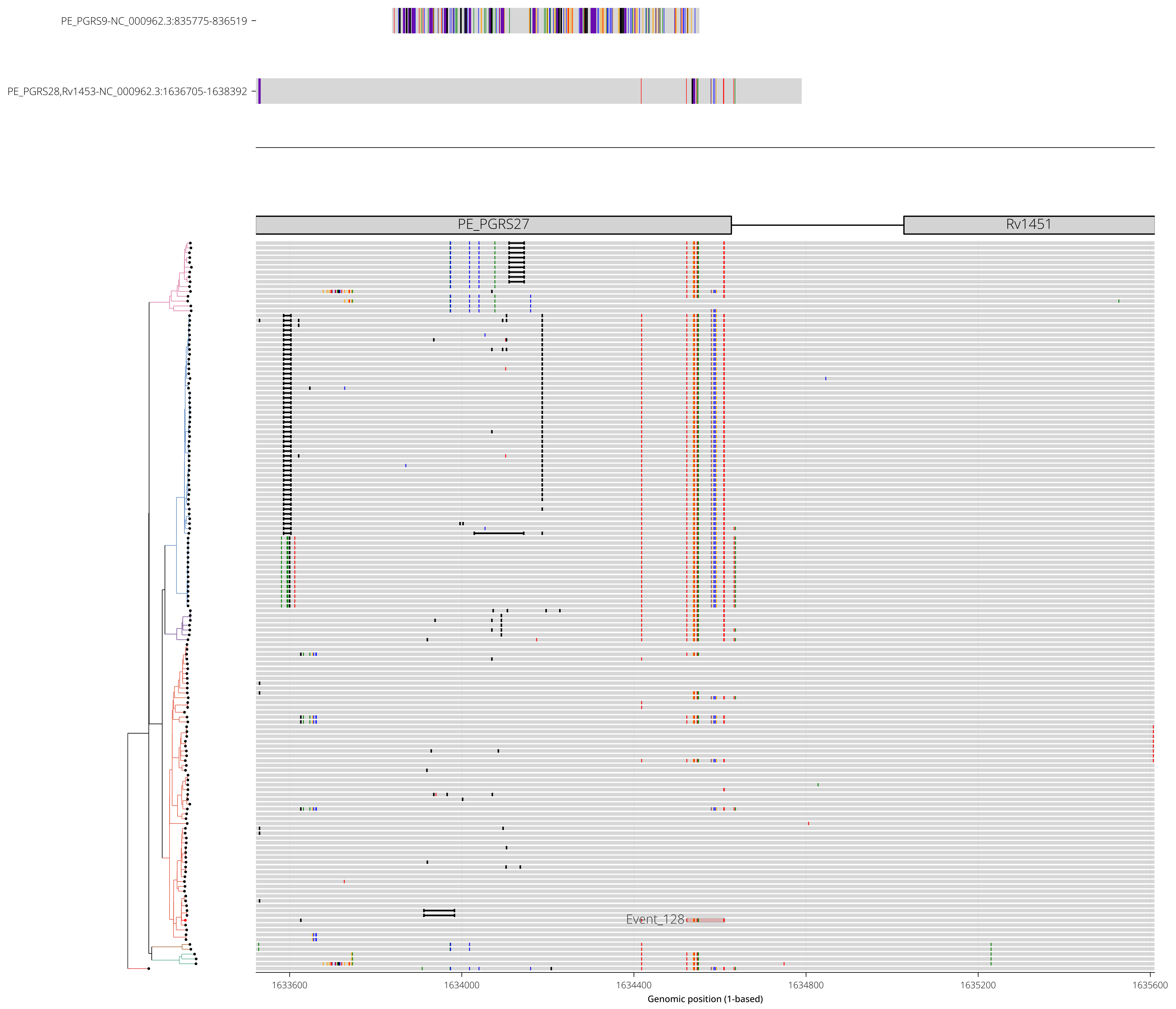

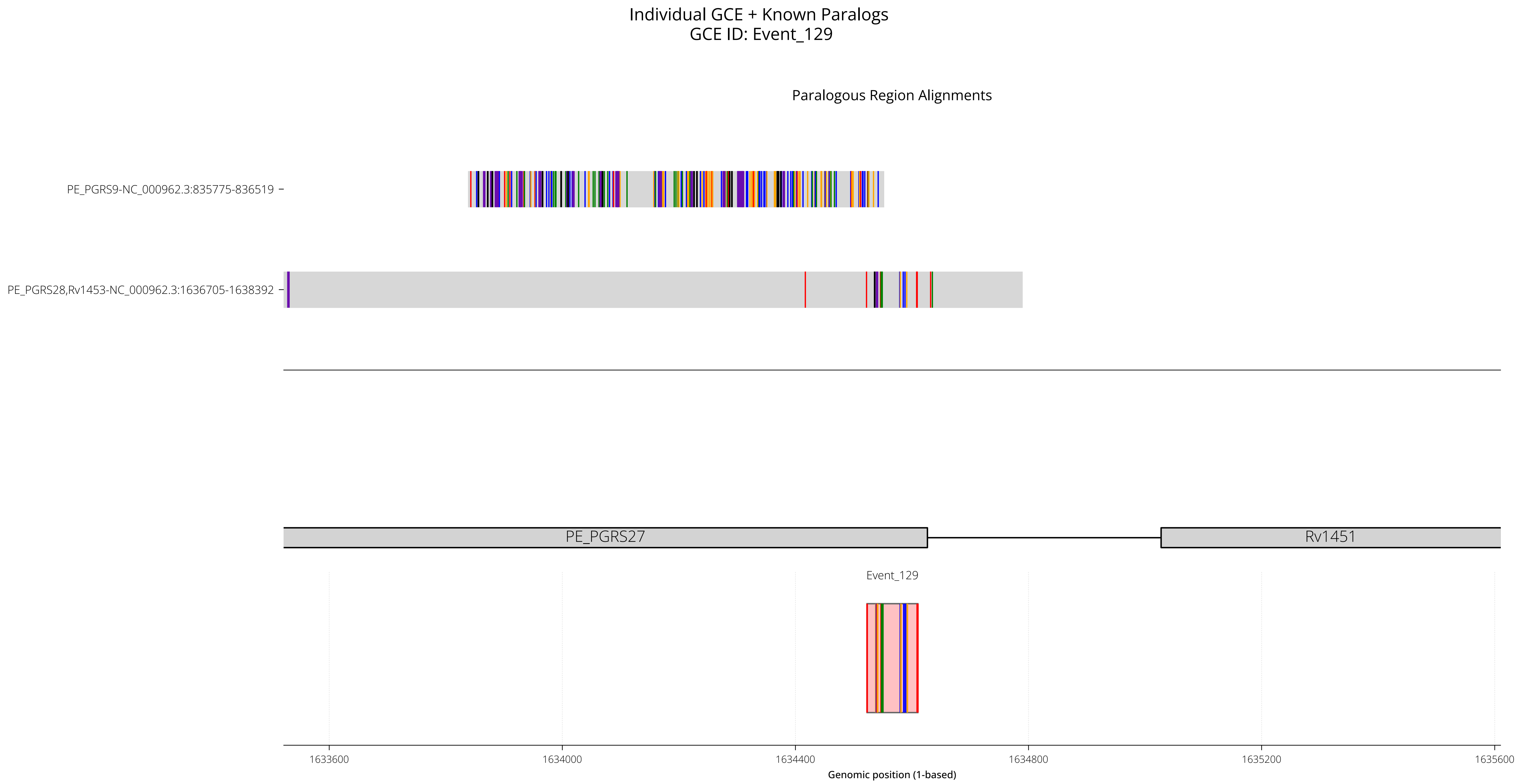
Individual GCE + Known Paralogs
GCE ID: Event_129
Paralogous Region Alignments
PE_PGRS9-NC_000962.3:835775-836519
PE_PGRS28,Rv1453-NC_000962.3:1636705-1638392
PE_PGRS27
Rv1451
Event_129
1633600
1634000
1634400
1634800
1635200
1635600
Genomic position (1-based)

Circos Viz - GCE to Known Paralogs Locations
GCE ID: Event_129

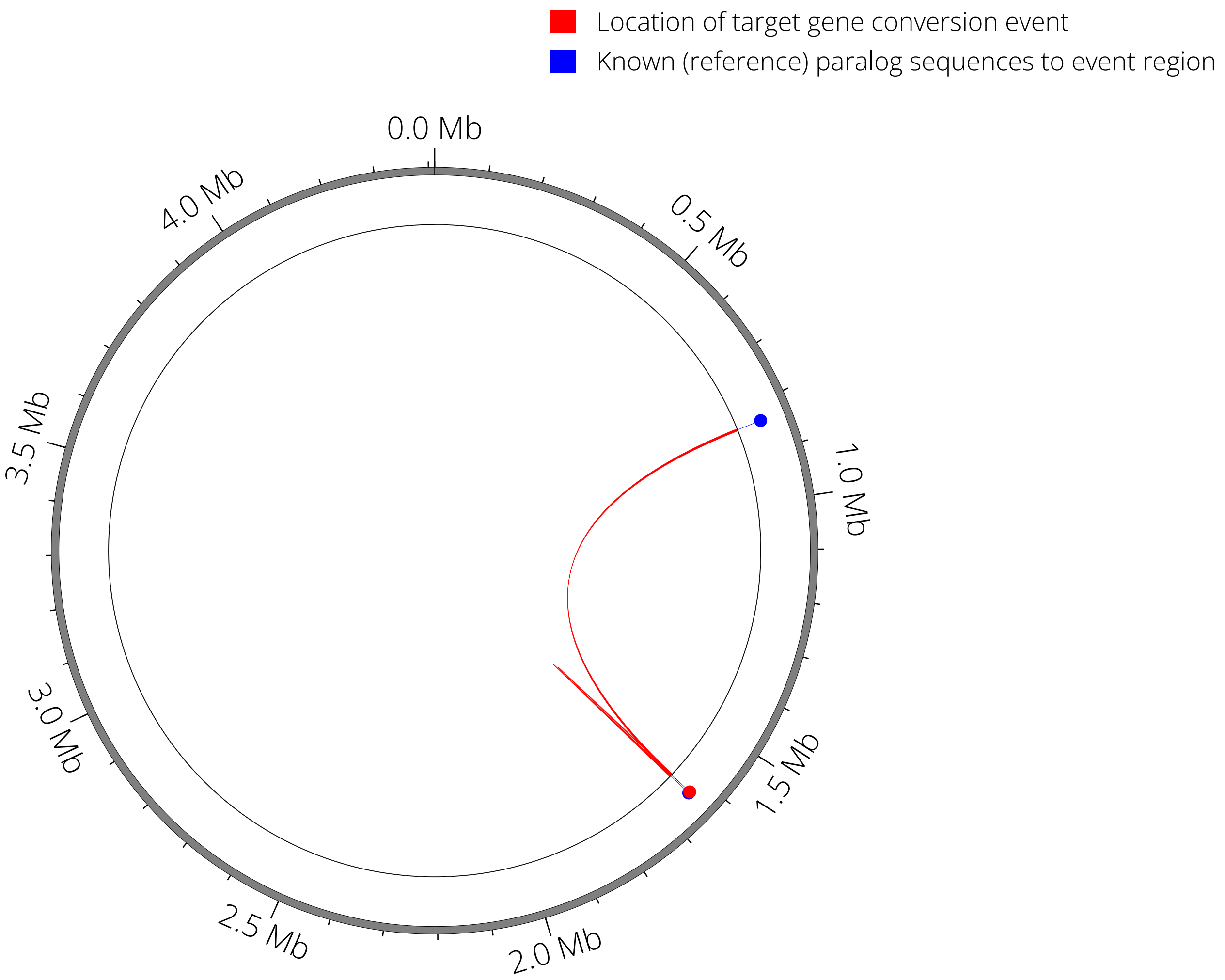

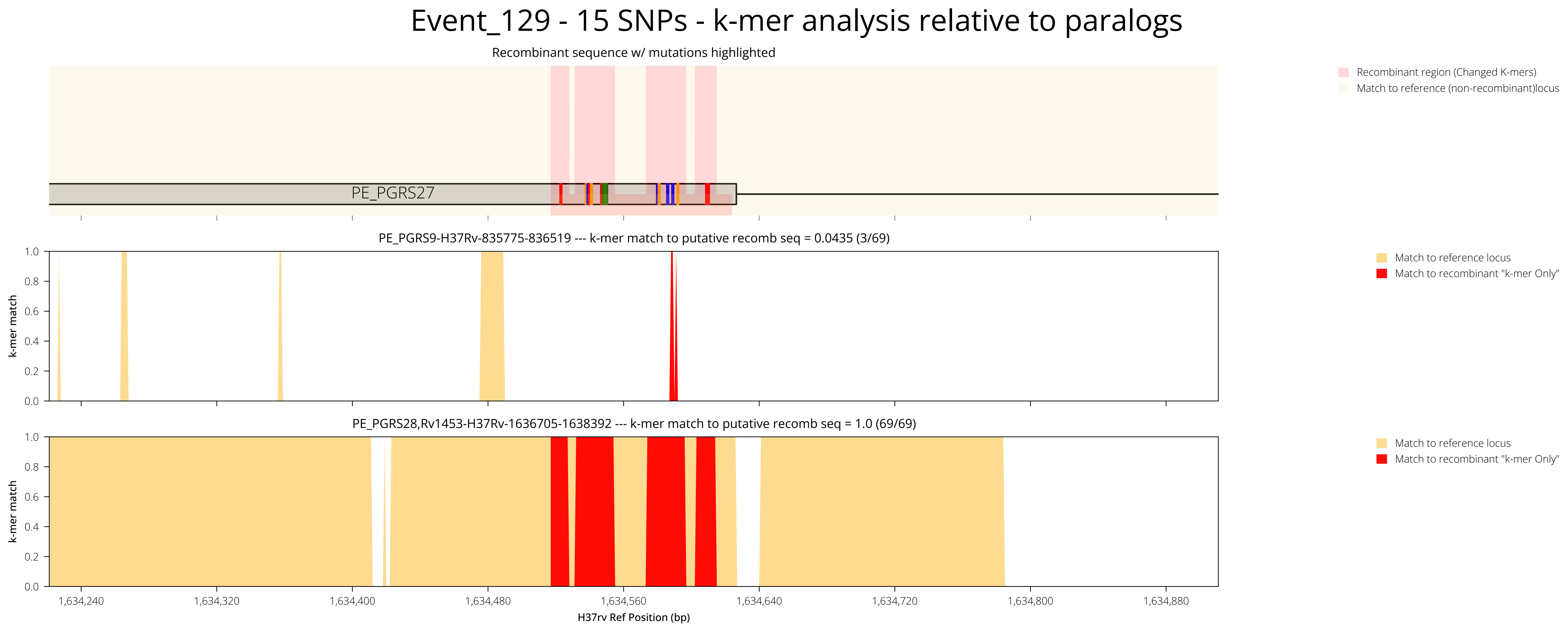

Event_129 - 15 SNPs - k-mer analysis relative to paralogs
Recombinant sequence w/ mutations highlighted
PE_PGRS27
Recombinant region (Changed K-mers)
Match to reference (non-recombinant)locus
PE_PGRS9-H37Rv-835775-836519 --- k-mer match to putative recomb seq = 0.0435 (3/69)
Match to reference locus
Match to recombinant "k-mer Only"
k-mer match
1.0
0.8
0.6
0.4
0.2
0.0
PE_PGRS28,Rv1453-H37Rv-1636705-1638392 --- k-mer match to putative recomb seq = 1.0 (69/69)
Match to reference locus
Match to recombinant "k-mer Only"
k-mer match
1.0
0.8
0.6
0.4
0.2
0.0
1,634,240
1,634,320
1,634,400
1,634,480
1,634,560
1,634,640
1,634,720
1,634,800
1,634,880
H37rv Ref Position (bp)

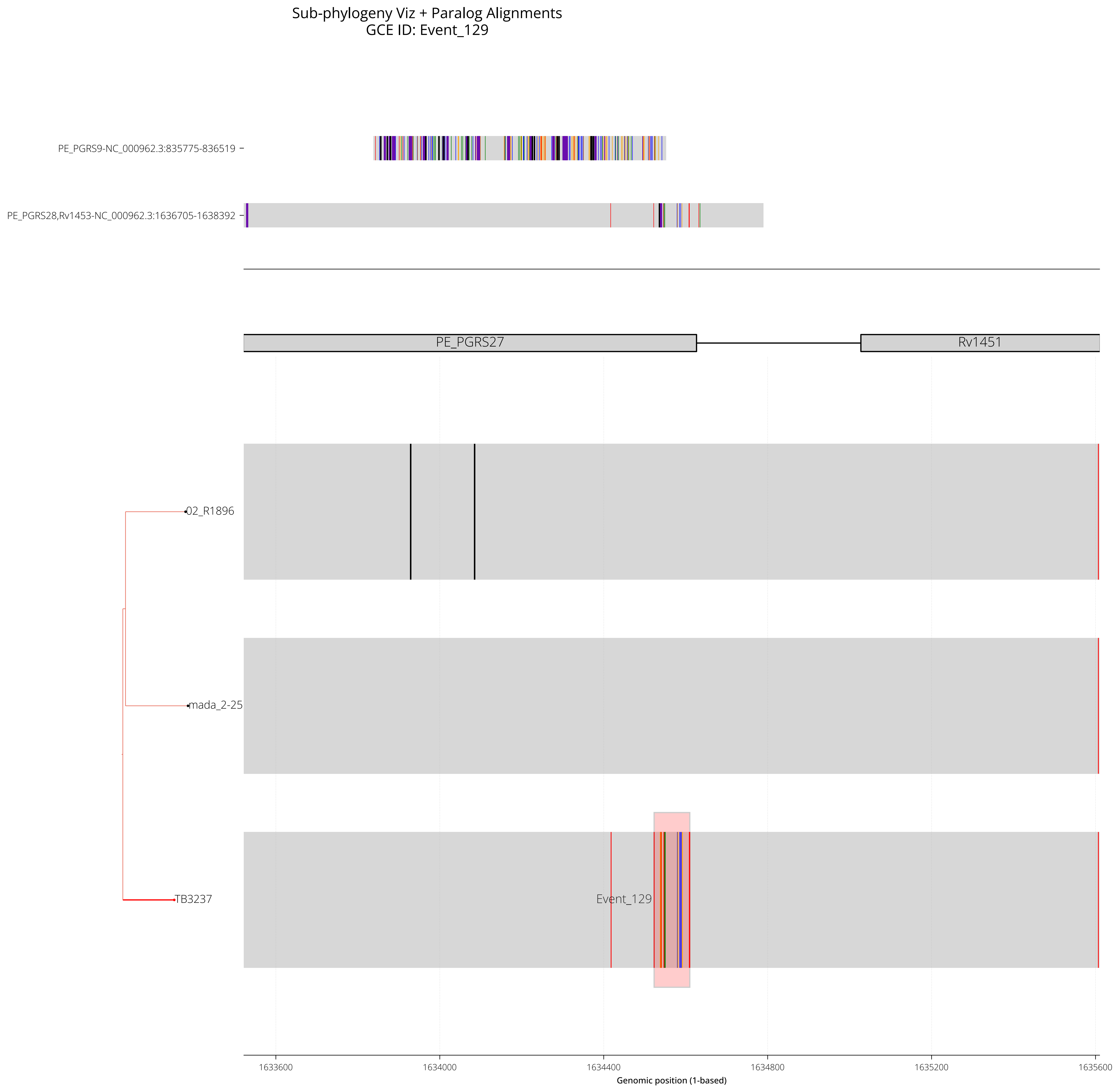

Sub-phylogeny Viz + Paralog Alignments
GCE ID: Event_129
PE_PGRS9-NC_000962.3:835775-836519
PE_PGRS28,Rv1453-NC_000962.3:1636705-1638392
PE_PGRS27
Rv1451
02_R1896
mada_2-25
TB3237
Event_129
1633600
1634000
1634400
1634800
1635200
1635600
Genomic position (1-based)

Full Phylogeny Viz + Paralog Alignments
GCE ID: Event_129

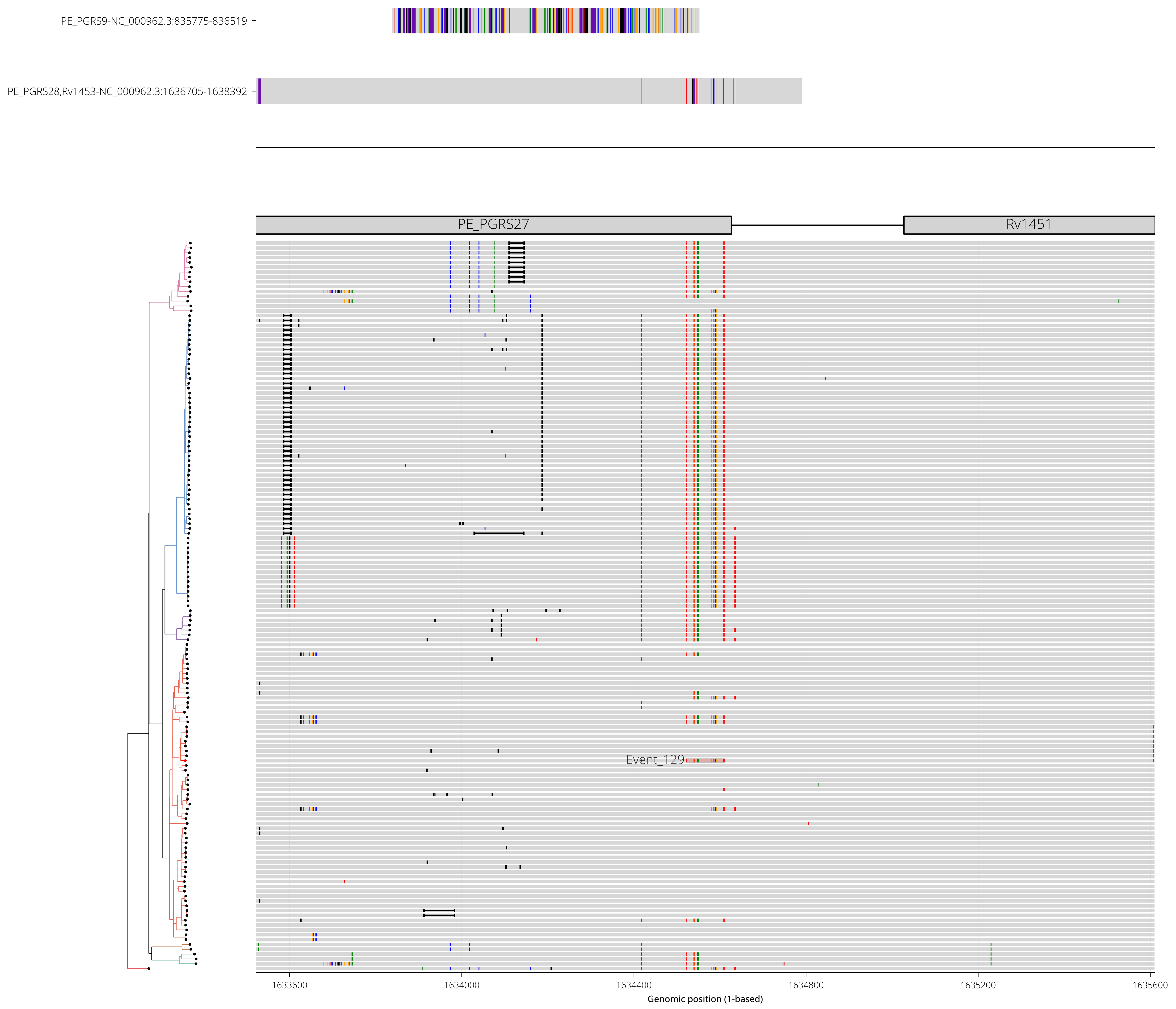

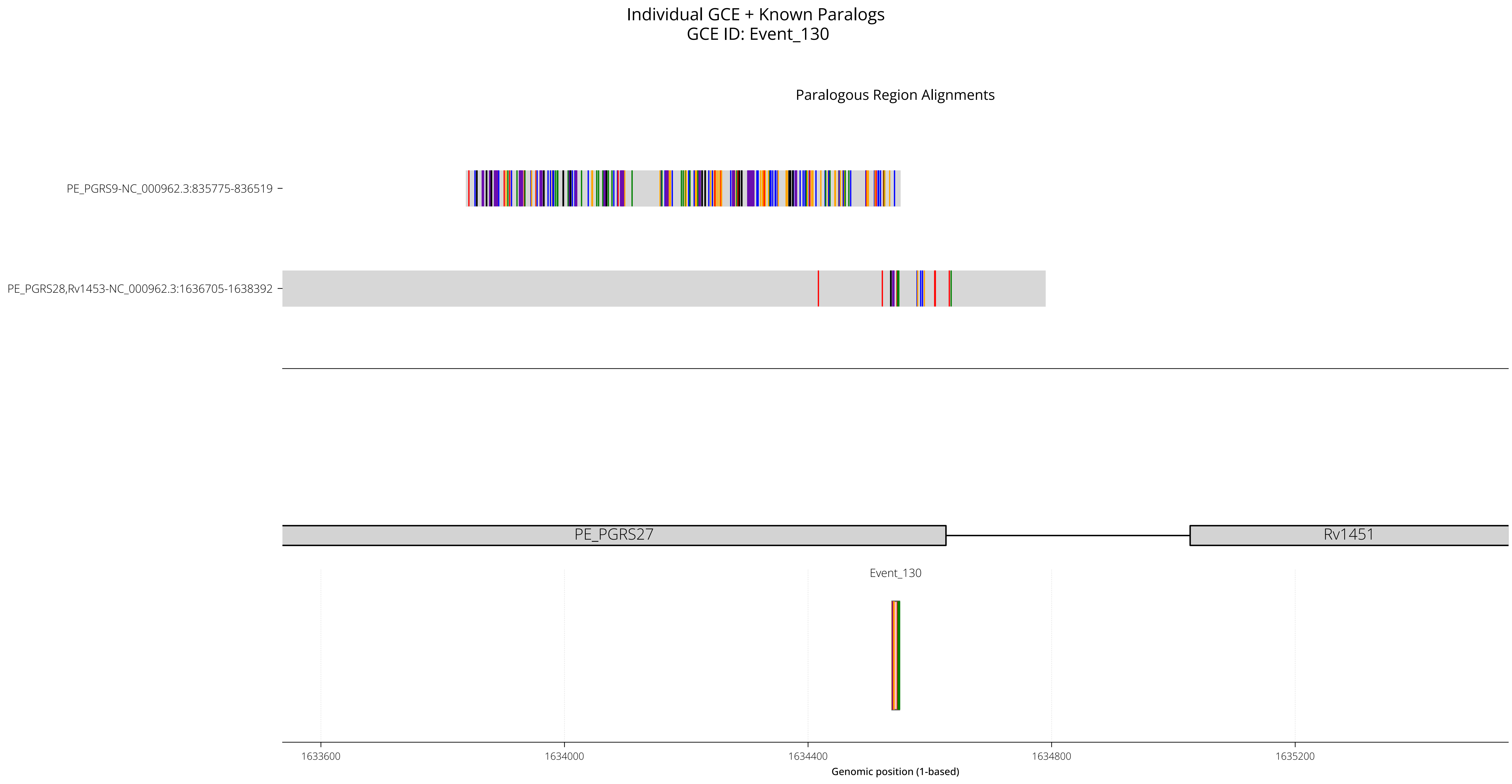
Individual GCE + Known Paralogs
GCE ID: Event_130
Paralogous Region Alignments
PE_PGRS9-NC_000962.3:835775-836519
PE_PGRS28,Rv1453-NC_000962.3:1636705-1638392
PE_PGRS27
Rv1451
Event_130
1633600
1634000
1634400
1634800
1635200
Genomic position (1-based)

Circos Viz - GCE to Known Paralogs Locations
GCE ID: Event_130

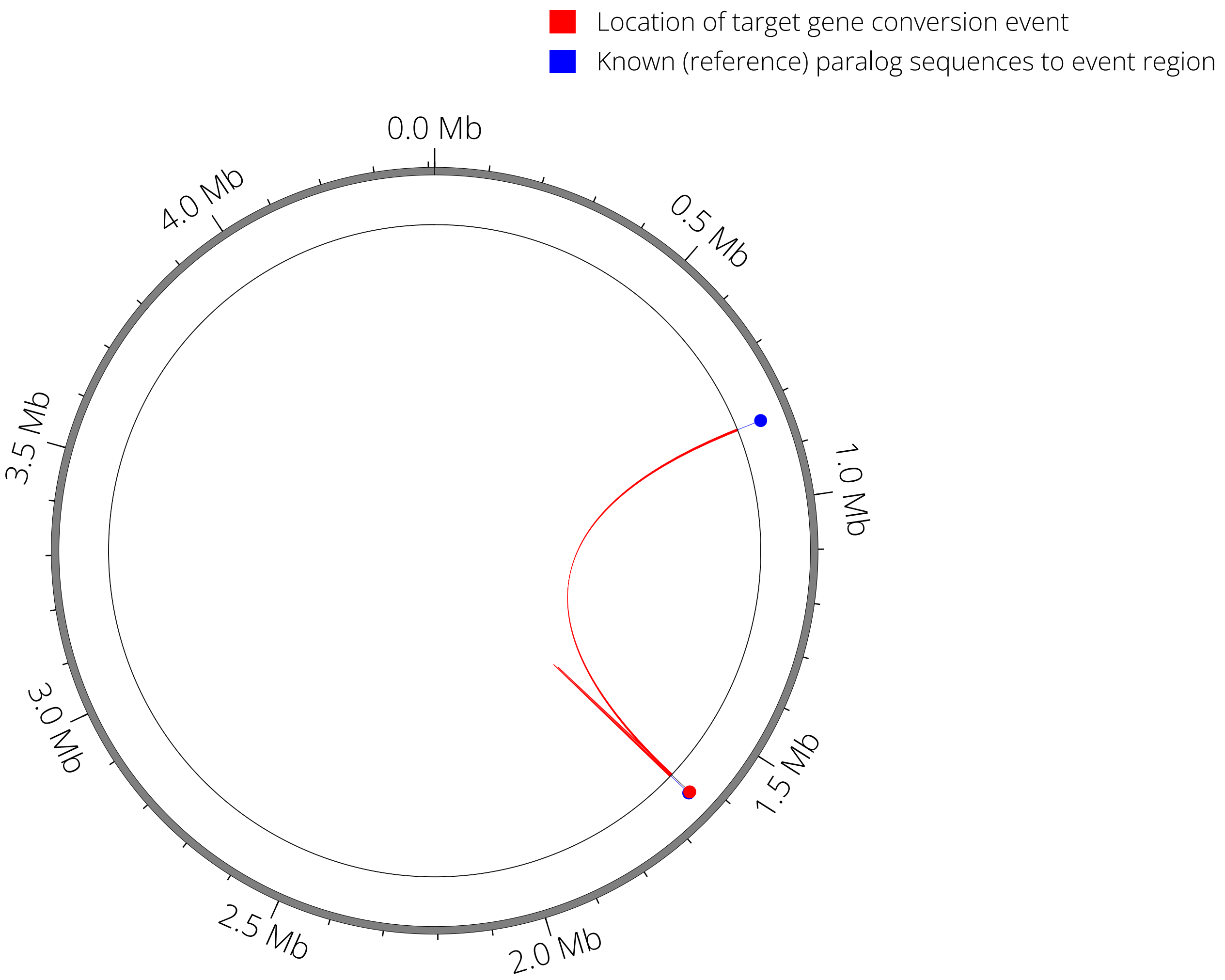

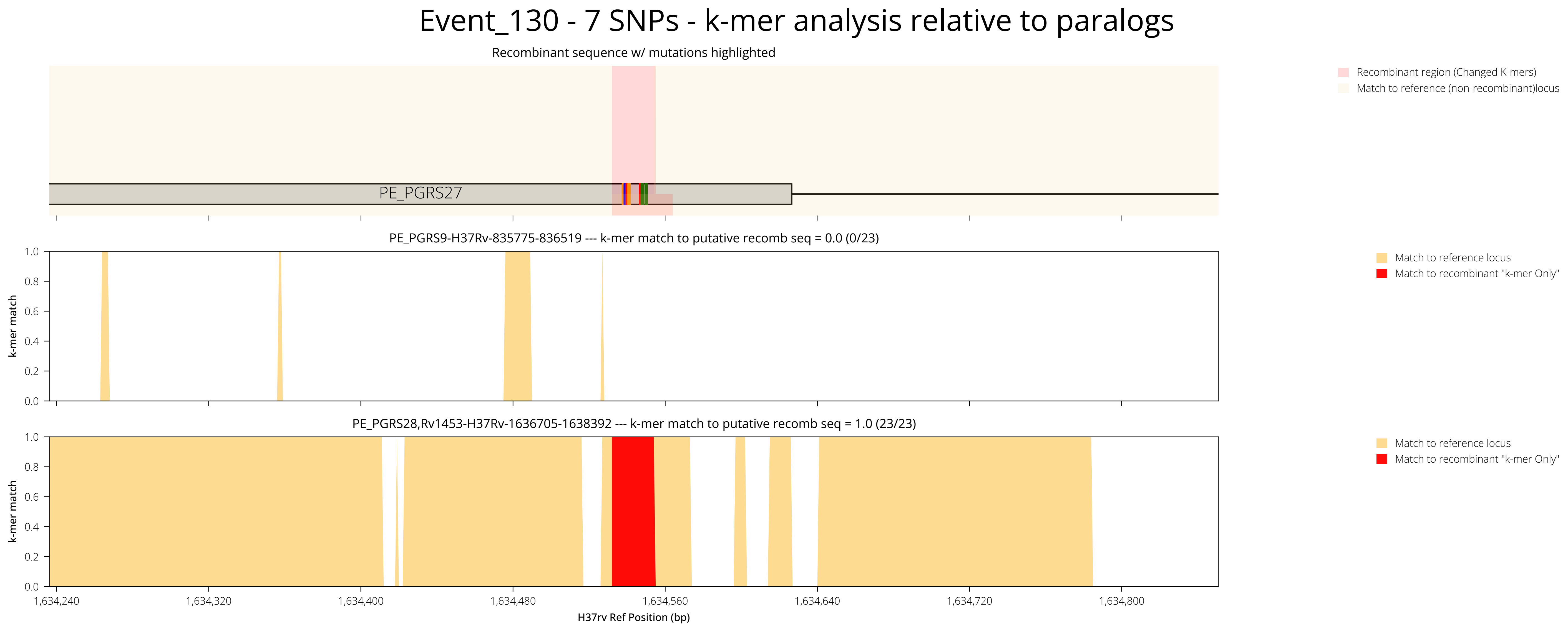
Event_130 - 7 SNPs - k-mer analysis relative to paralogs
Recombinant sequence w/ mutations highlighted
Recombinant region (Changed K-mers)
Match to reference (non-recombinant)locus
PE_PGRS27
PE_PGRS9-H37Rv-835775-836519 --- k-mer match to putative recomb seq = 0.0 (0/23)
Match to reference locus
Match to recombinant "k-mer Only"
PE_PGRS28,Rv1453-H37Rv-1636705-1638392 --- k-mer match to putative recomb seq = 1.0 (23/23)
Match to reference locus
Match to recombinant "k-mer Only"
k-mer match
1.0
0.8
0.6
0.4
0.2
0.0
1,634,240
1,634,320
1,634,400
1,634,480
1,634,560
1,634,640
1,634,720
1,634,800
H37rv Ref Position (bp)

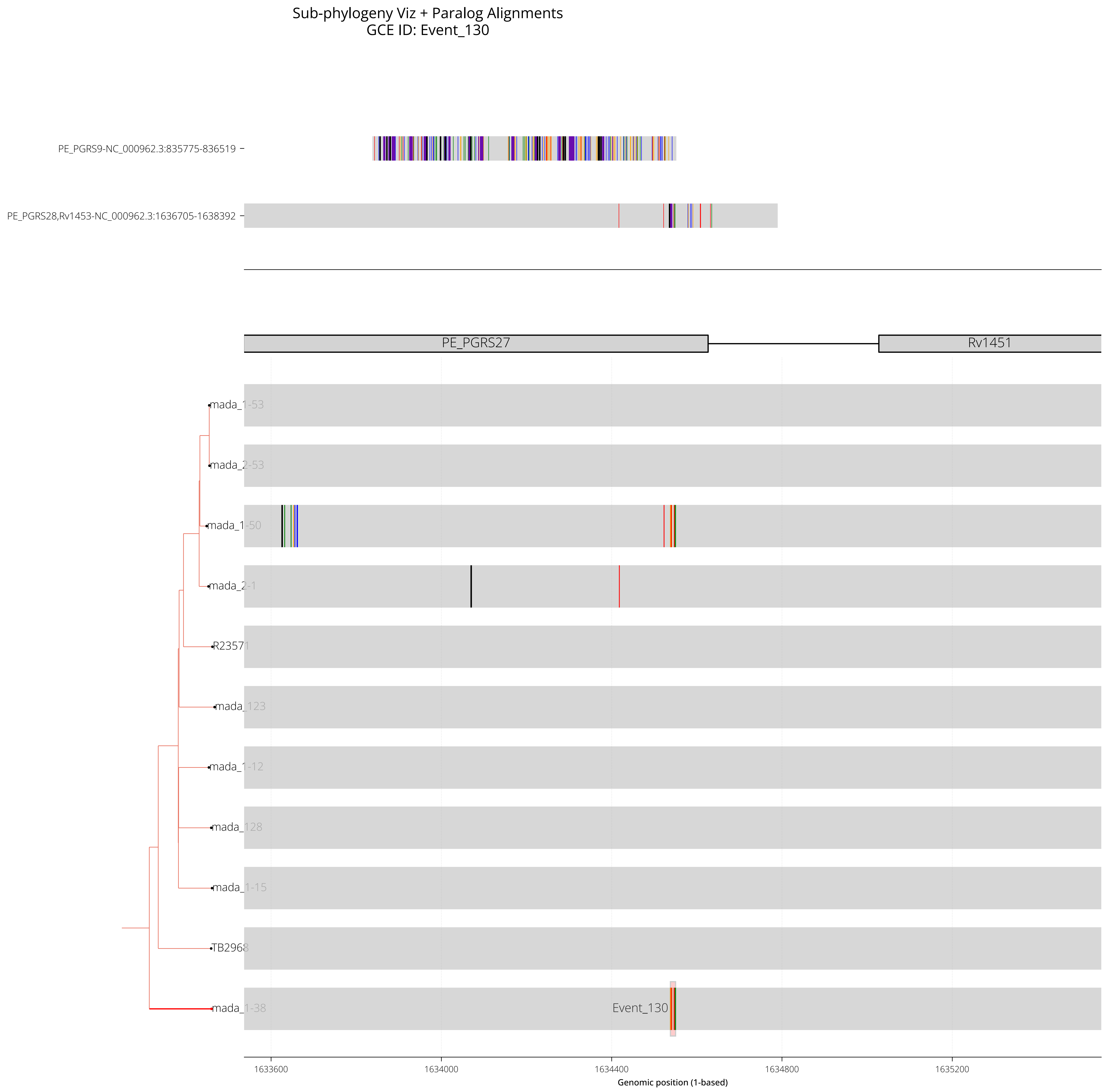

Sub-phylogeny Viz + Paralog Alignments
GCE ID: Event_130
PE_PGRS9-NC_000962.3:835775-836519
PE_PGRS28,Rv1453-NC_000962.3:1636705-1638392
PE_PGRS27
Rv1451
mada_1-53
mada_2-53
mada_1-50
mada_2-1
R23571
mada_123
mada_1-12
mada_128
mada_1-15
TB2968
mada_1-38
Event_130
1633600
1634000
1634400
1634800
1635200
Genomic position (1-based)

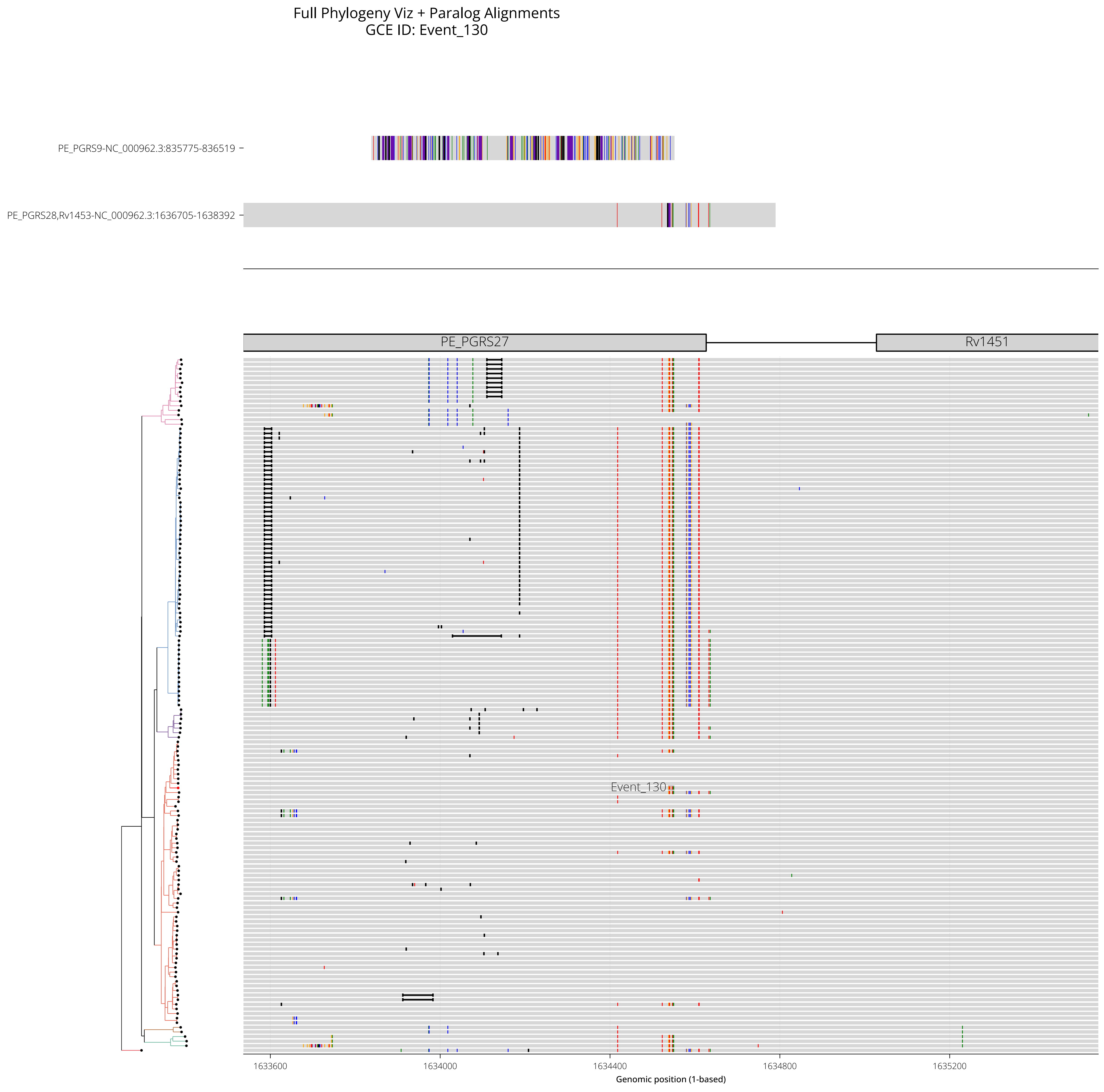
Full Phylogeny Viz + Paralog Alignments
GCE ID: Event_130
PE_PGRS9-NC_000962.3:835775-836519
PE_PGRS28,Rv1453-NC_000962.3:1636705-1638392
PE_PGRS27
Rv1451
Event_130
1633600
1634000
1634400
1634800
1635200
Genomic position (1-based)

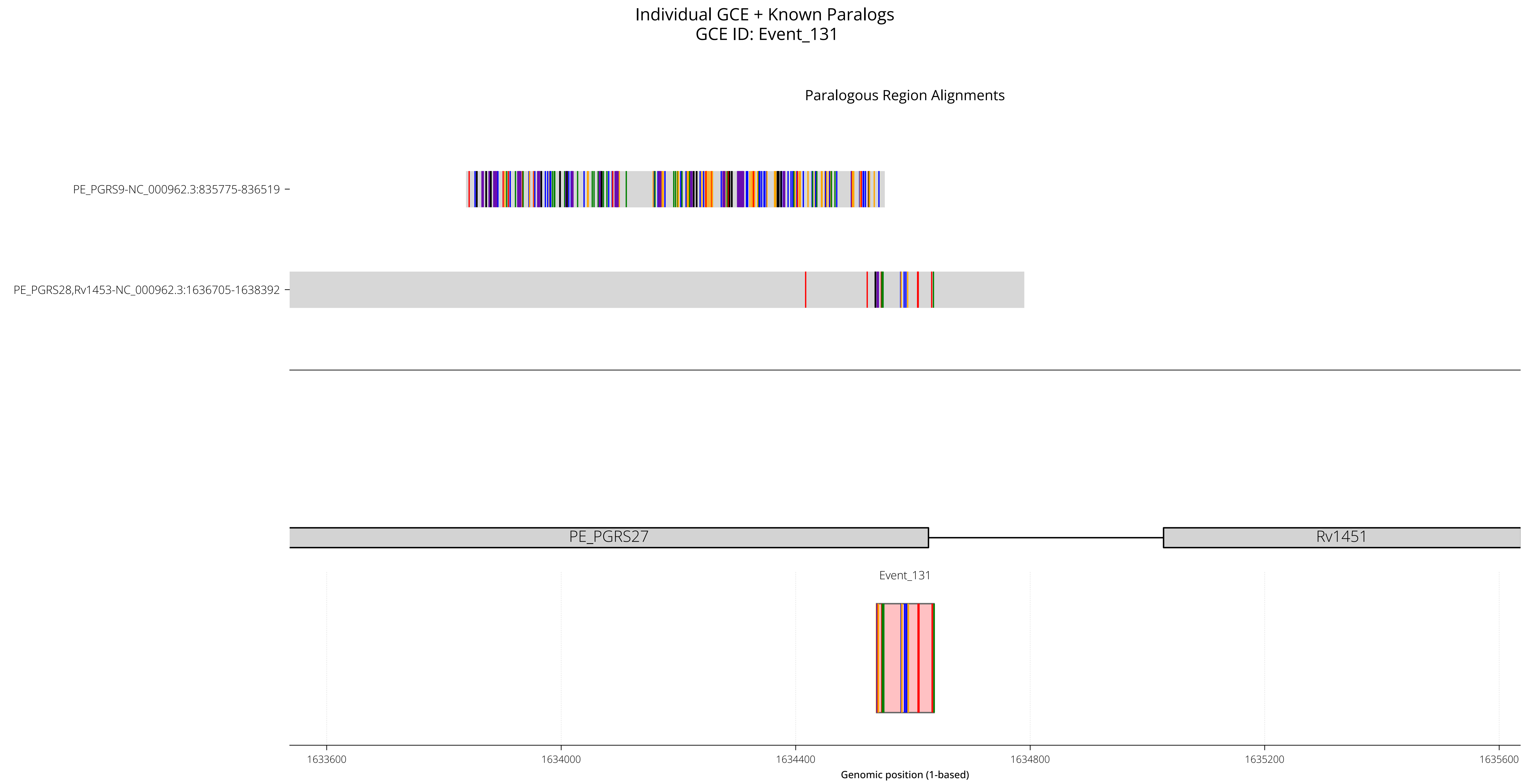

Individual GCE + Known Paralogs
GCE ID: Event_131
Paralogous Region Alignments
PE_PGRS9-NC_000962.3:835775-836519
PE_PGRS28,Rv1453-NC_000962.3:1636705-1638392
PE_PGRS27
Rv1451
Event_131
1633600
1634000
1634400
1634800
1635200
1635600
Genomic position (1-based)

Circos Viz - GCE to Known Paralogs Locations
GCE ID: Event_131

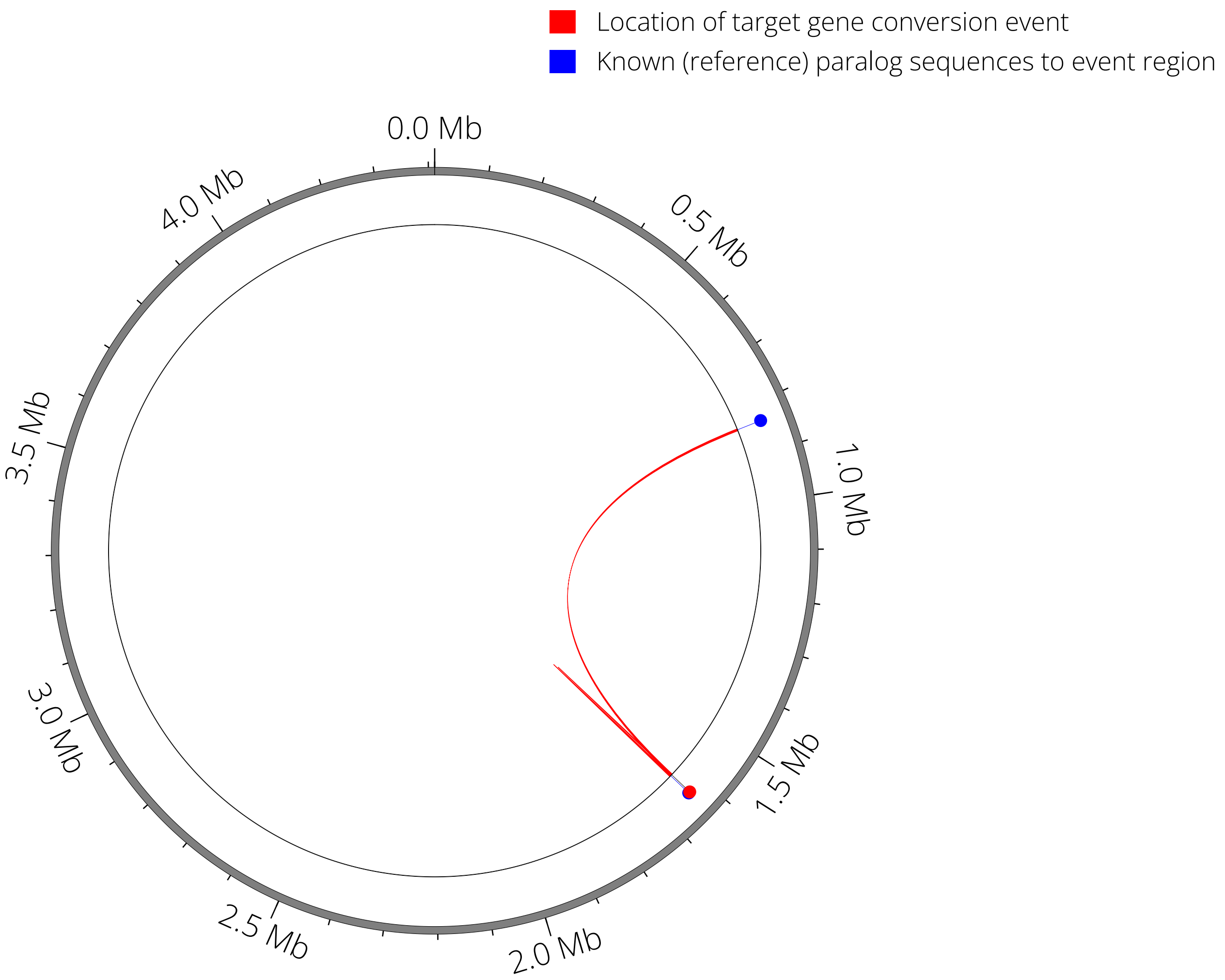

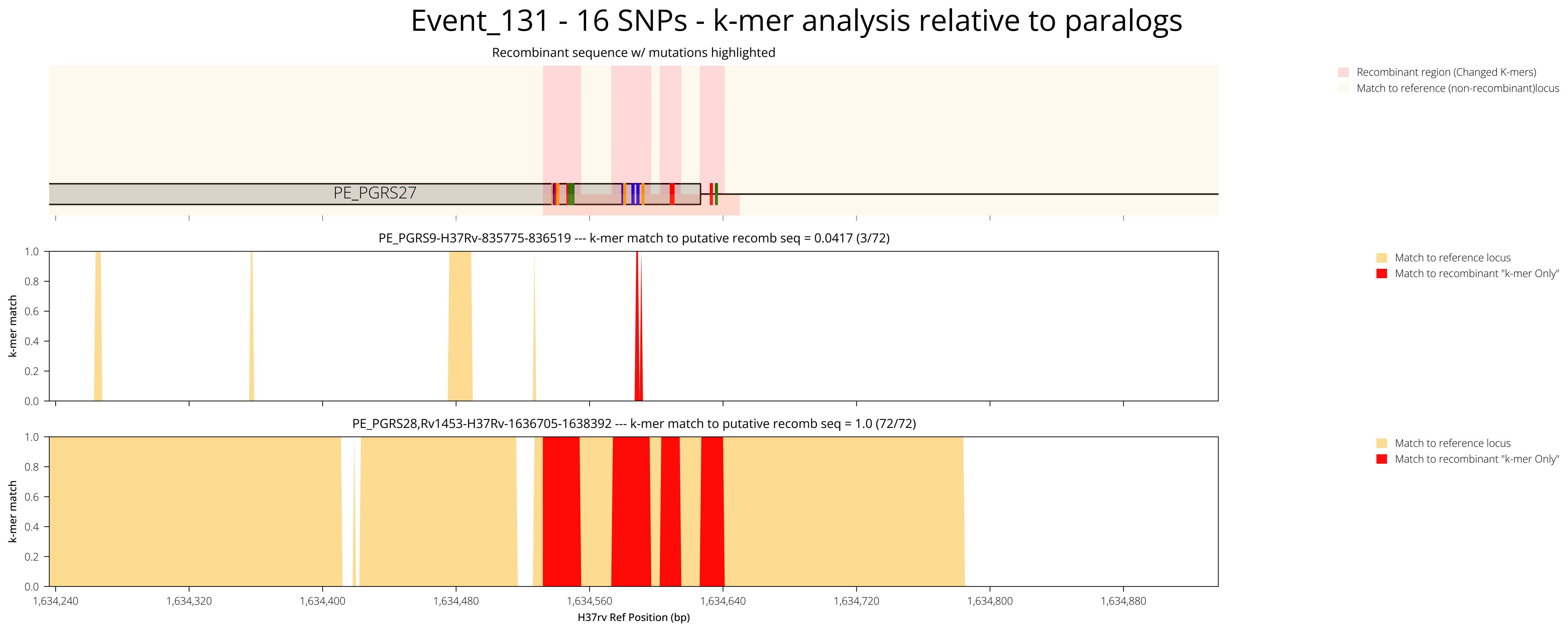
Event_131 - 16 SNPs - k-mer analysis relative to paralogs
Recombinant sequence w/ mutations highlighted
PE_PGRS27
Recombinant region (Changed K-mers)
Match to reference (non-recombinant)locus
PE_PGRS9-H37Rv-835775-836519 --- k-mer match to putative recomb seq = 0.0417 (3/72)
Match to reference locus
Match to recombinant "k-mer Only"
k-mer match
PE_PGRS28,Rv1453-H37Rv-1636705-1638392 --- k-mer match to putative recomb seq = 1.0 (72/72)
1.0
0.8
0.6
0.4
0.2
0.0
1,634,240
1,634,320
1,634,400
1,634,480
1,634,560
1,634,640
1,634,720
1,634,800
1,634,880
H37rv Ref Position (bp)

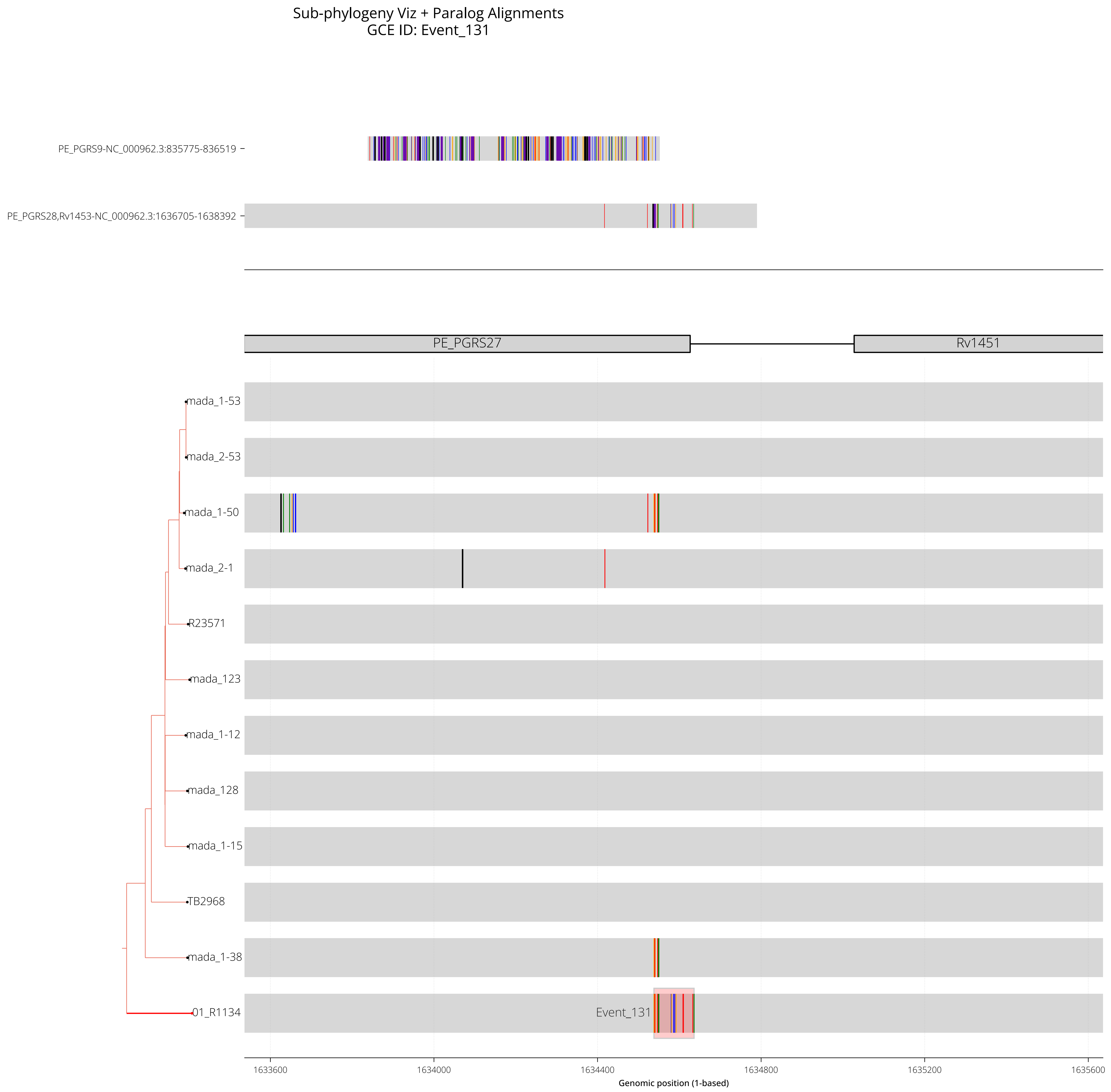

Sub-phylogeny Viz + Paralog Alignments
GCE ID: Event_131
PE_PGRS9-NC_000962.3:835775-836519
PE_PGRS28,Rv1453-NC_000962.3:1636705-1638392
PE_PGRS27
Rv1451
mada_1-53
mada_2-53
mada_1-50
mada_2-1
R23571
mada_123
mada_1-12
mada_128
mada_1-15
TB2968
mada_1-38
01_R1134
Event_131
1633600
1634000
1634400
1634800
1635200
1635600
Genomic position (1-based)

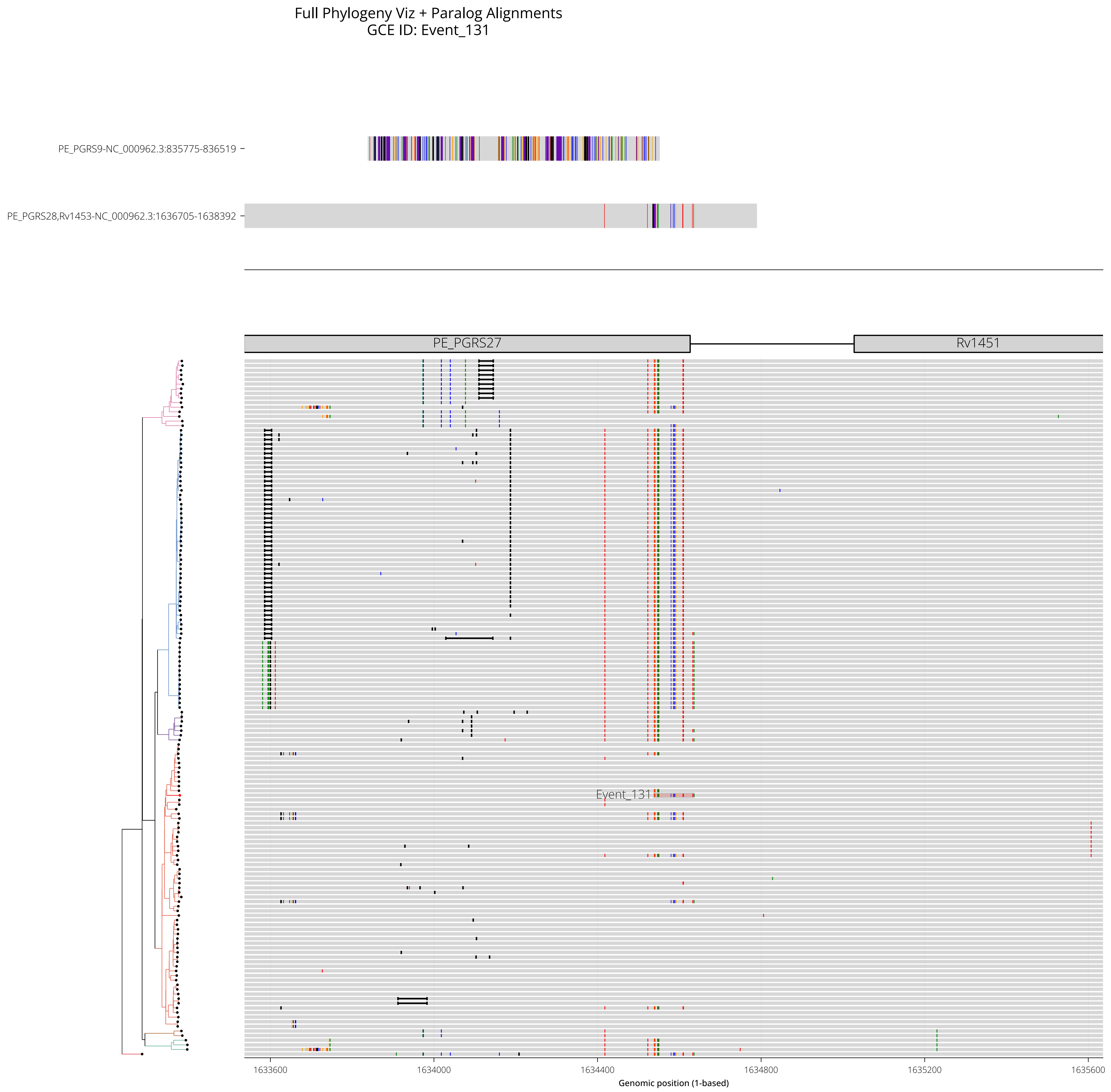

Full Phylogeny Viz + Paralog Alignments
GCE ID: Event_131
PE_PGRS9-NC_000962.3:835775-836519
PE_PGRS28,Rv1453-NC_000962.3:1636705-1638392
PE_PGRS27
Rv1451
Event_131
1633600
1634000
1634400
1634800
1635200
1635600
Genomic position (1-based)

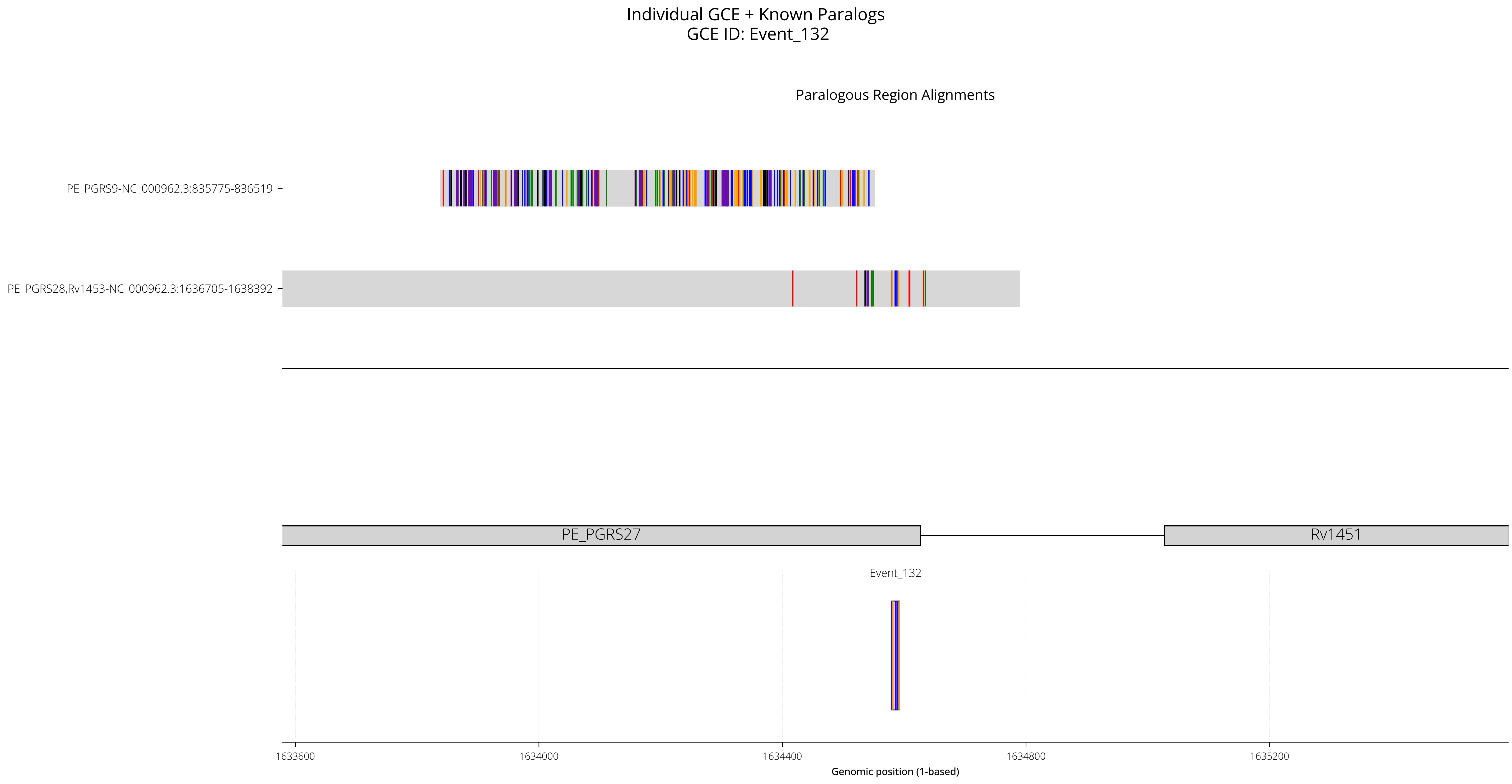
Individual GCE + Known Paralogs
GCE ID: Event_132
Paralogous Region Alignments
PE_PGRS9-NC_000962.3:835775-836519
PE_PGRS28,Rv1453-NC_000962.3:1636705-1638392
PE_PGRS27
Rv1451
Event_132
1633600
1634000
1634400
1634800
1635200
Genomic position (1-based)

Circos Viz - GCE to Known Paralogs Locations
GCE ID: Event_132

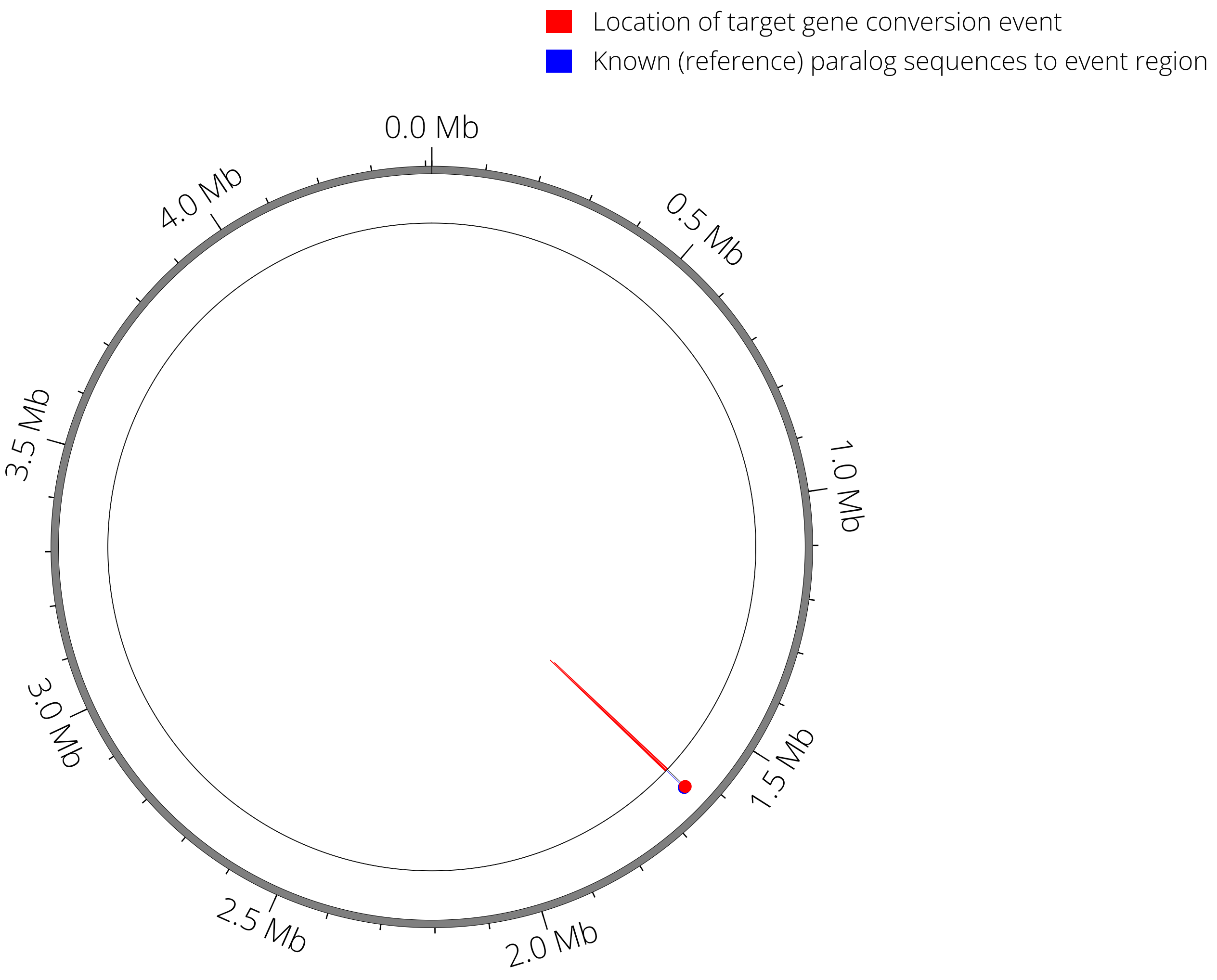

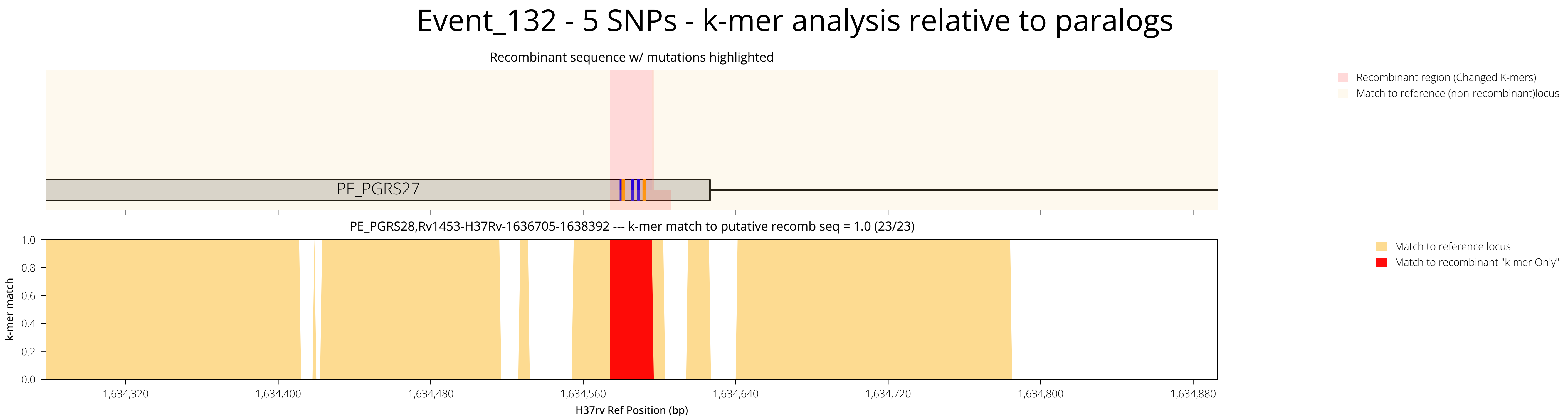

Event_132 - 5 SNPs - k-mer analysis relative to paralogs
Recombinant sequence w/ mutations highlighted
PE_PGRS27
Recombinant region (Changed K-mers)
Match to reference (non-recombinant)locus
PE_PGRS28,Rv1453-H37Rv-1636705-1638392 --- k-mer match to putative recomb seq = 1.0 (23/23)
k-mer match
1.0
0.8
0.6
0.4
0.2
0.0
1,634,320
1,634,400
1,634,480
1,634,560
1,634,640
1,634,720
1,634,800
1,634,880
H37rv Ref Position (bp)
Match to reference locus
Match to recombinant "k-mer Only"

Sub-phylogeny Viz + Paralog Alignments
GCE ID: Event_132

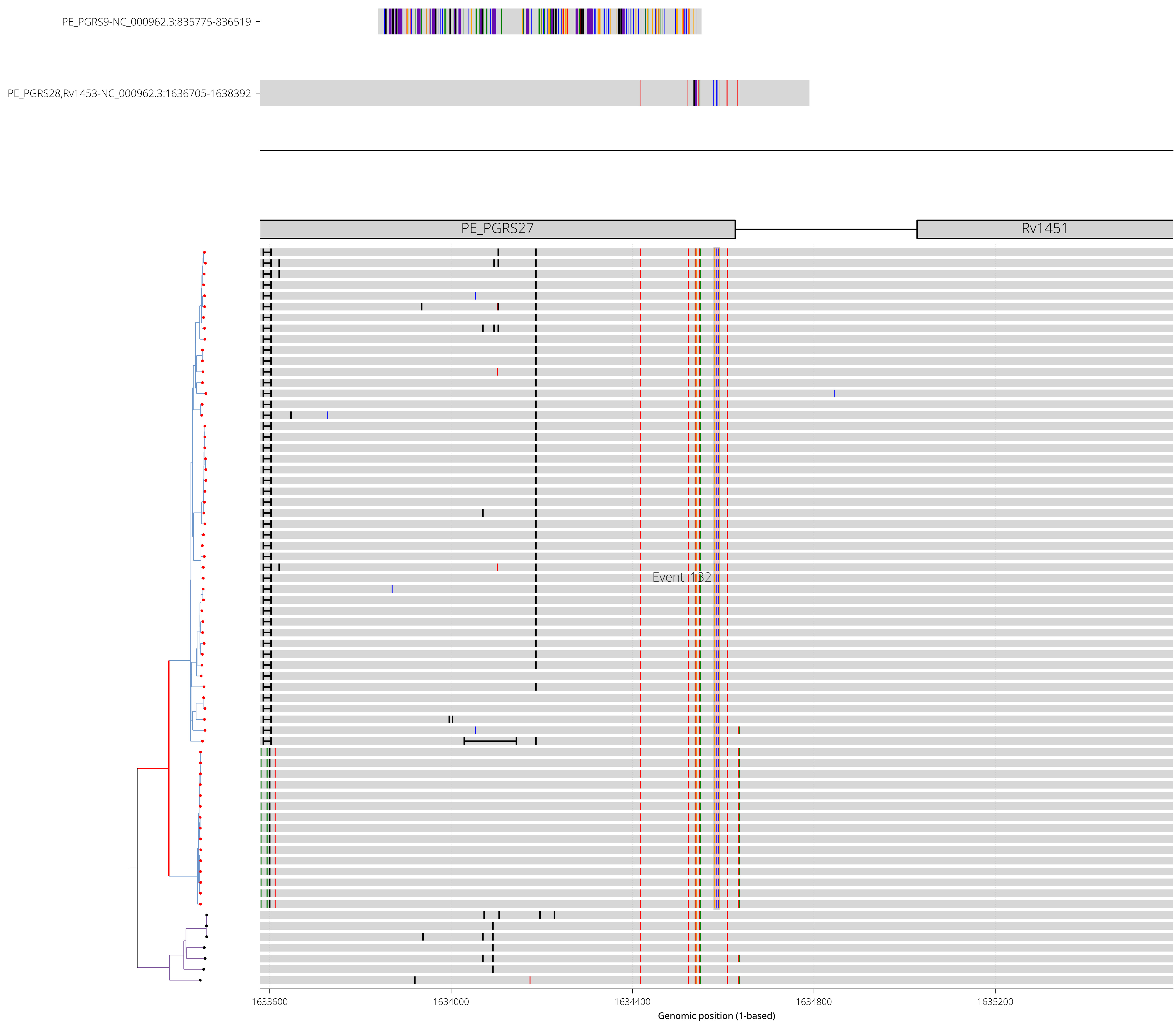

Full Phylogeny Viz + Paralog Alignments
GCE ID: Event_132

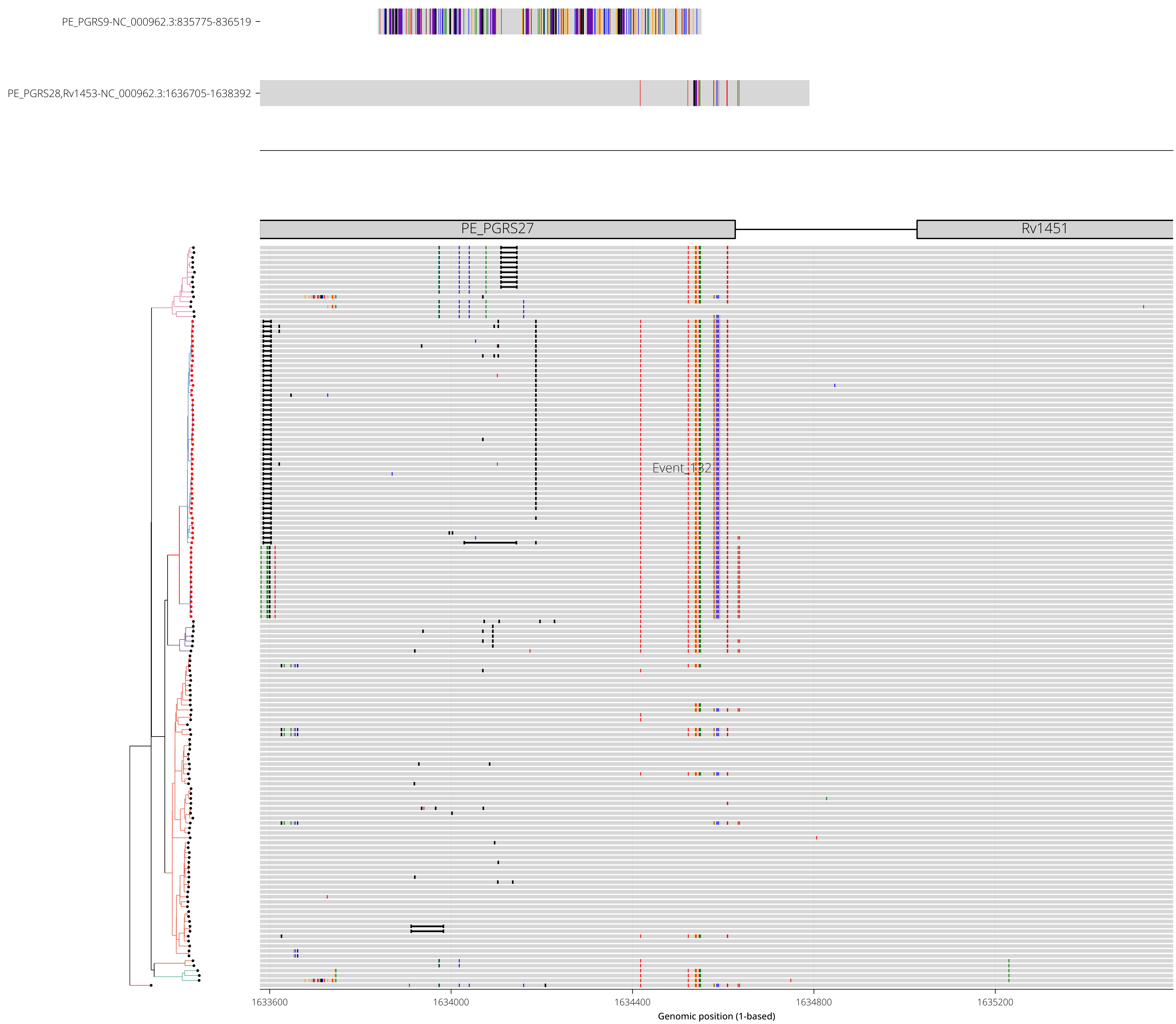

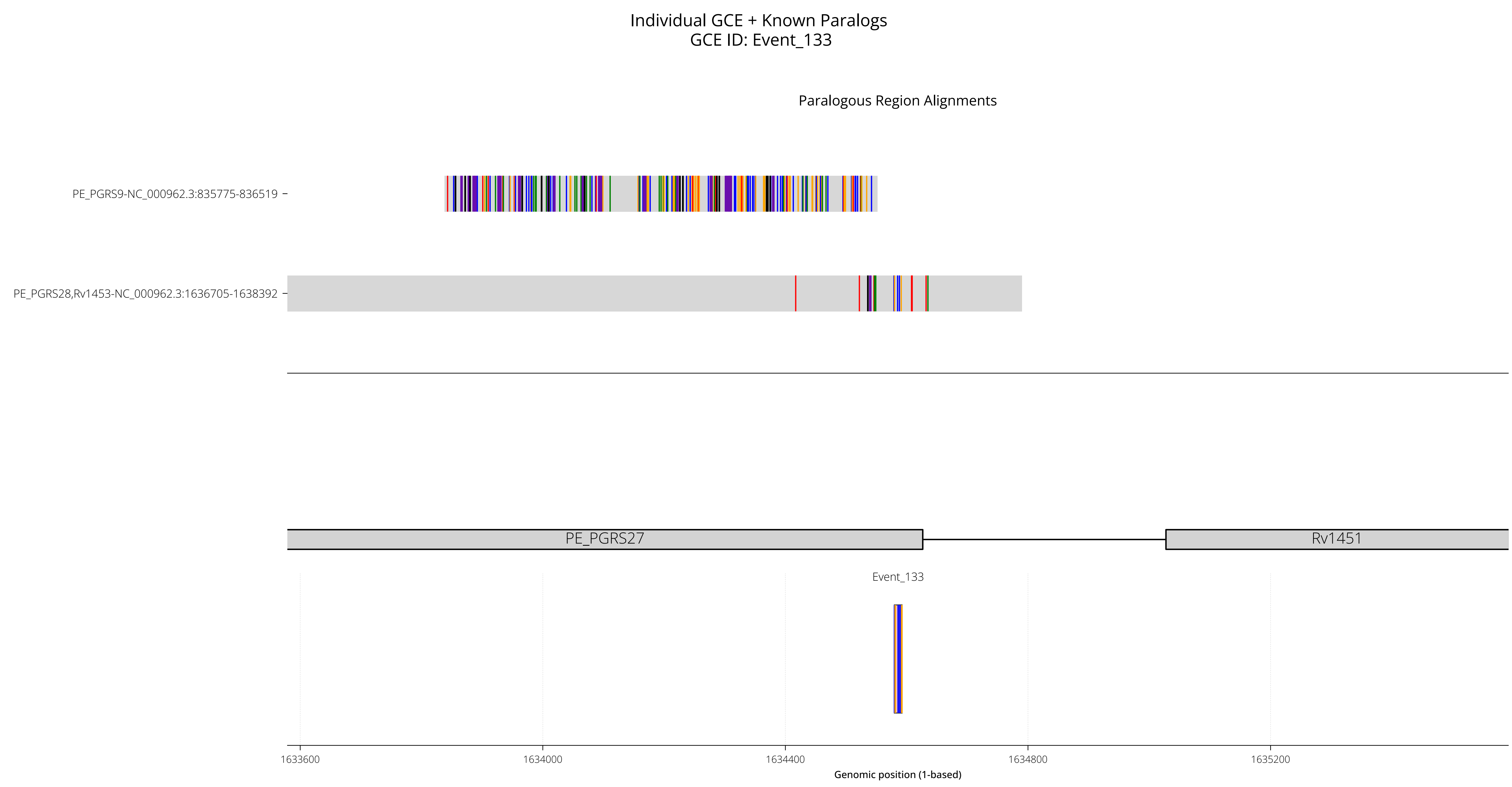
Individual GCE + Known Paralogs
GCE ID: Event_133
Paralogous Region Alignments
PE_PGRS9-NC_000962.3:835775-836519
PE_PGRS28,Rv1453-NC_000962.3:1636705-1638392
PE_PGRS27
Rv1451
Event_133
1633600
1634000
1634400
1634800
1635200
Genomic position (1-based)

### Circos Viz - GCE to Known Paralogs Locations
GCE ID: Event_133

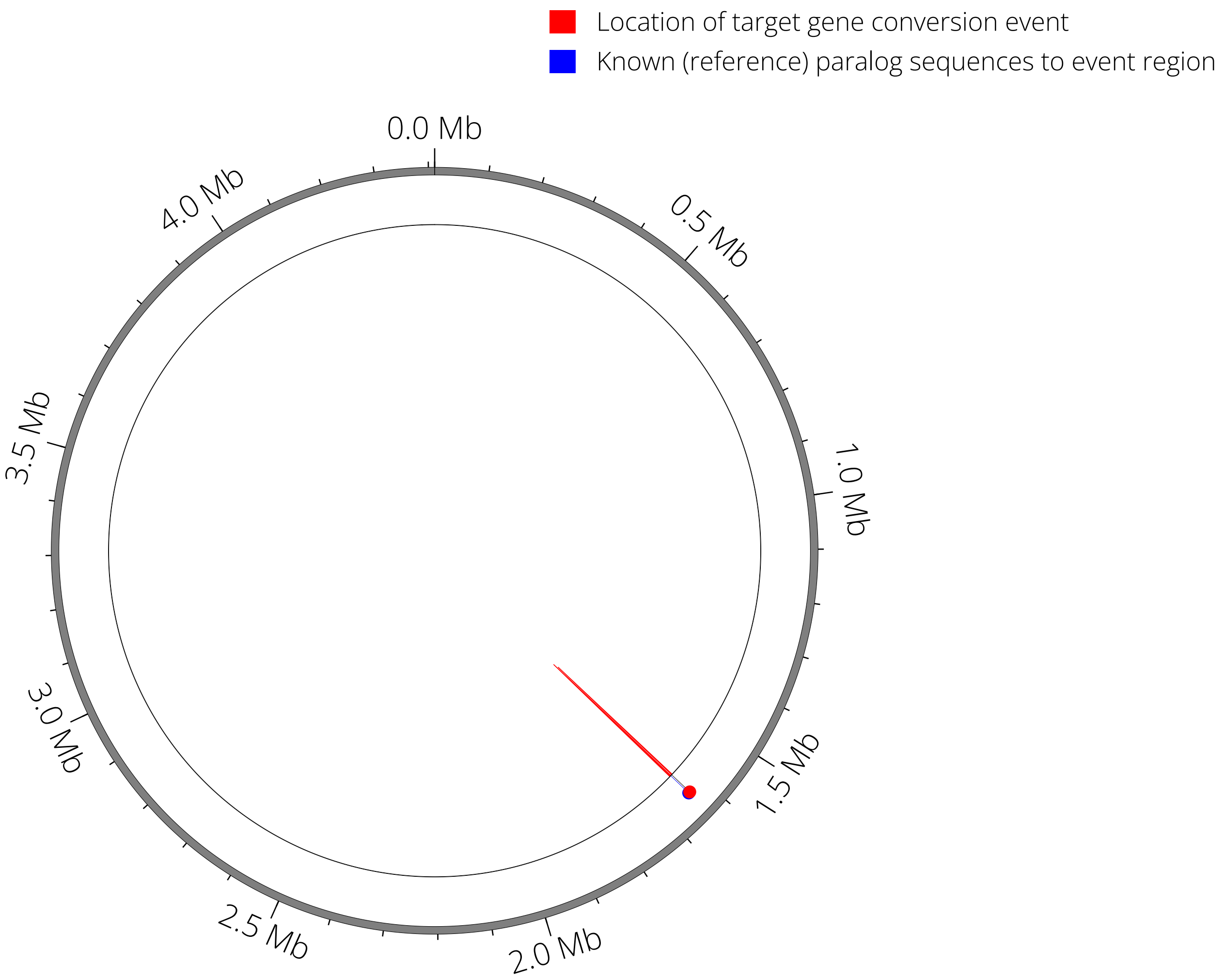

### Event_133 - 5 SNPs - k-mer analysis relative to paralogs

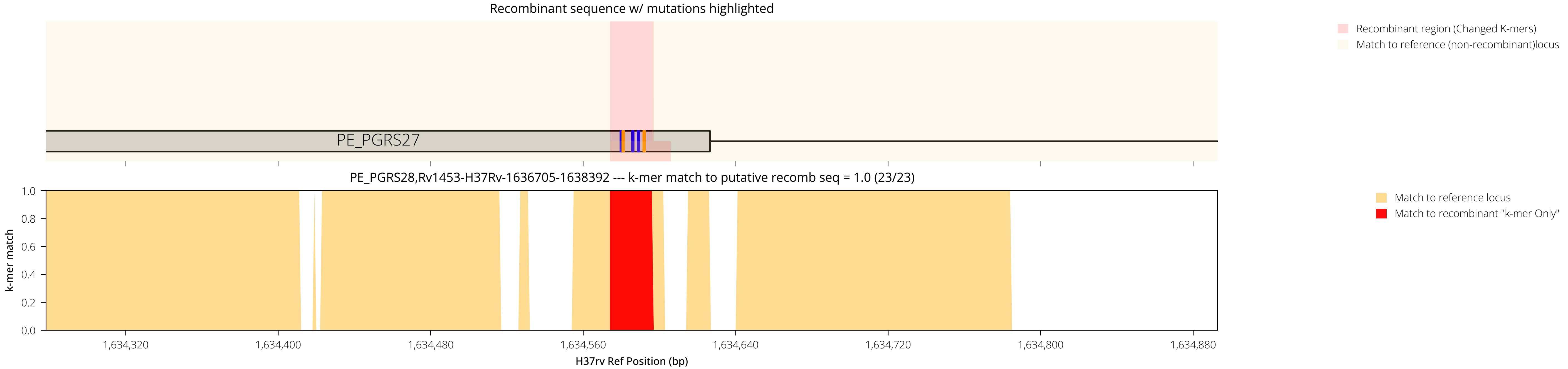

Sub-phylogeny Viz + Paralog Alignments
GCE ID: Event_133

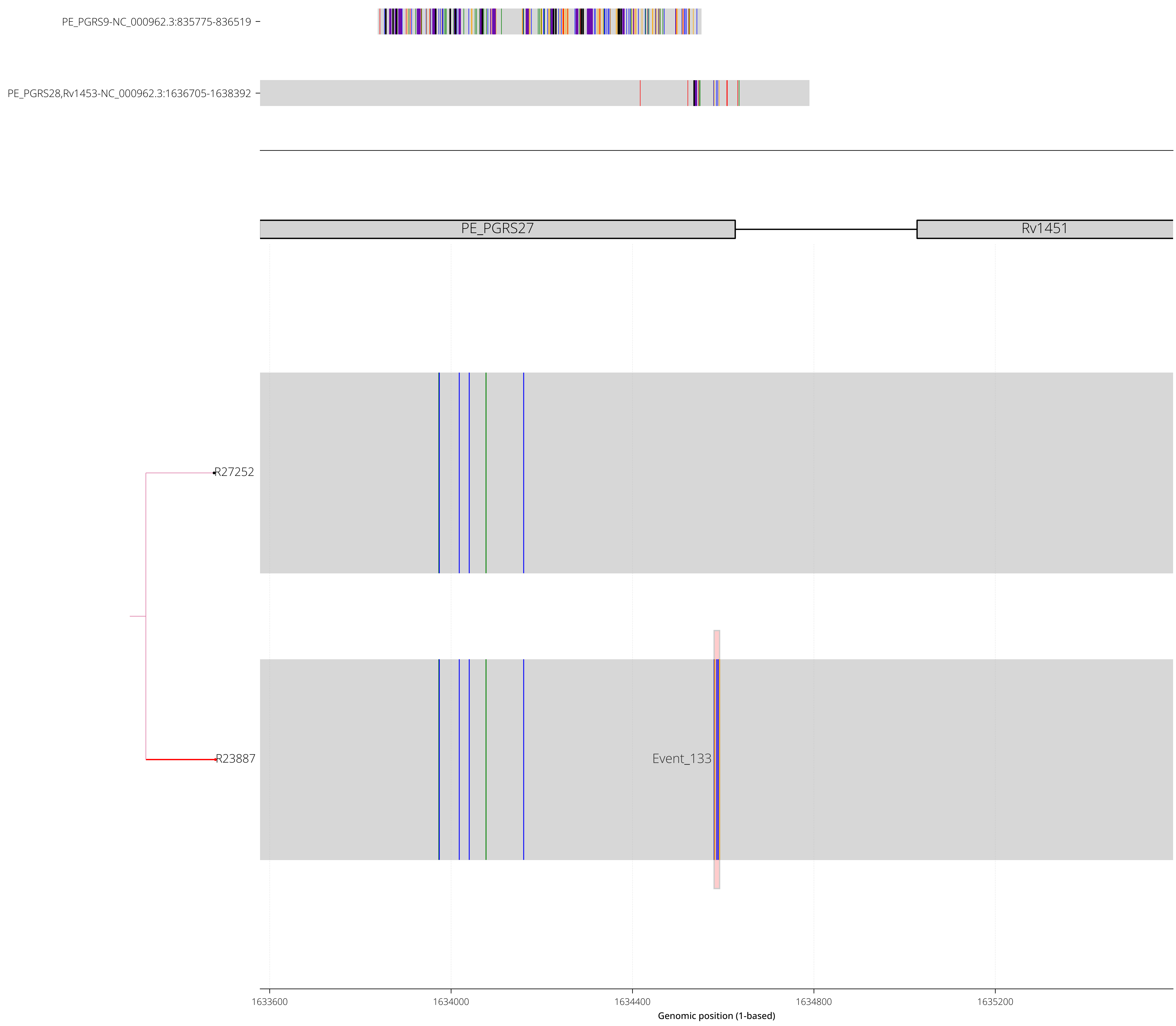

Full Phylogeny Viz + Paralog Alignments
GCE ID: Event_133

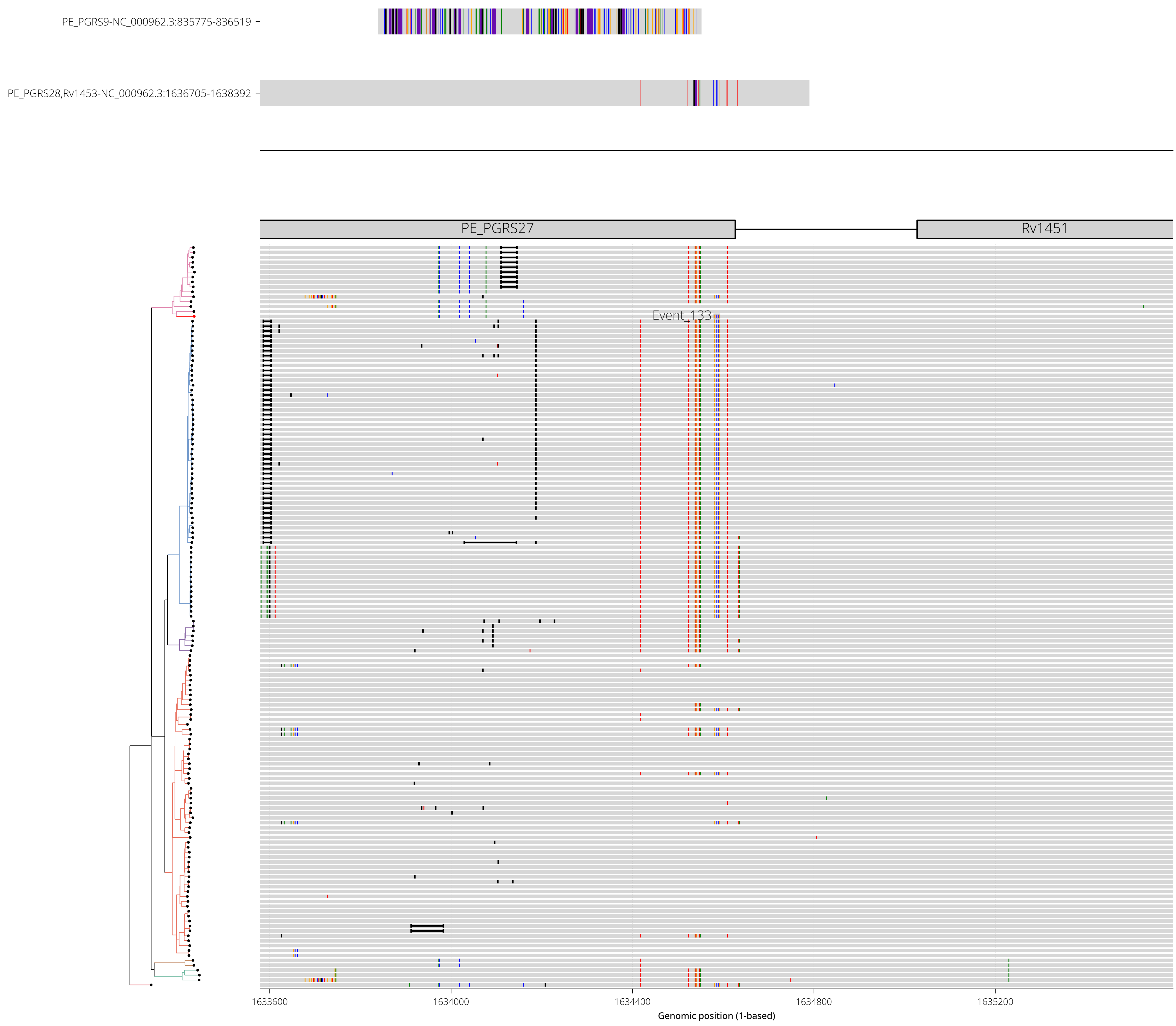

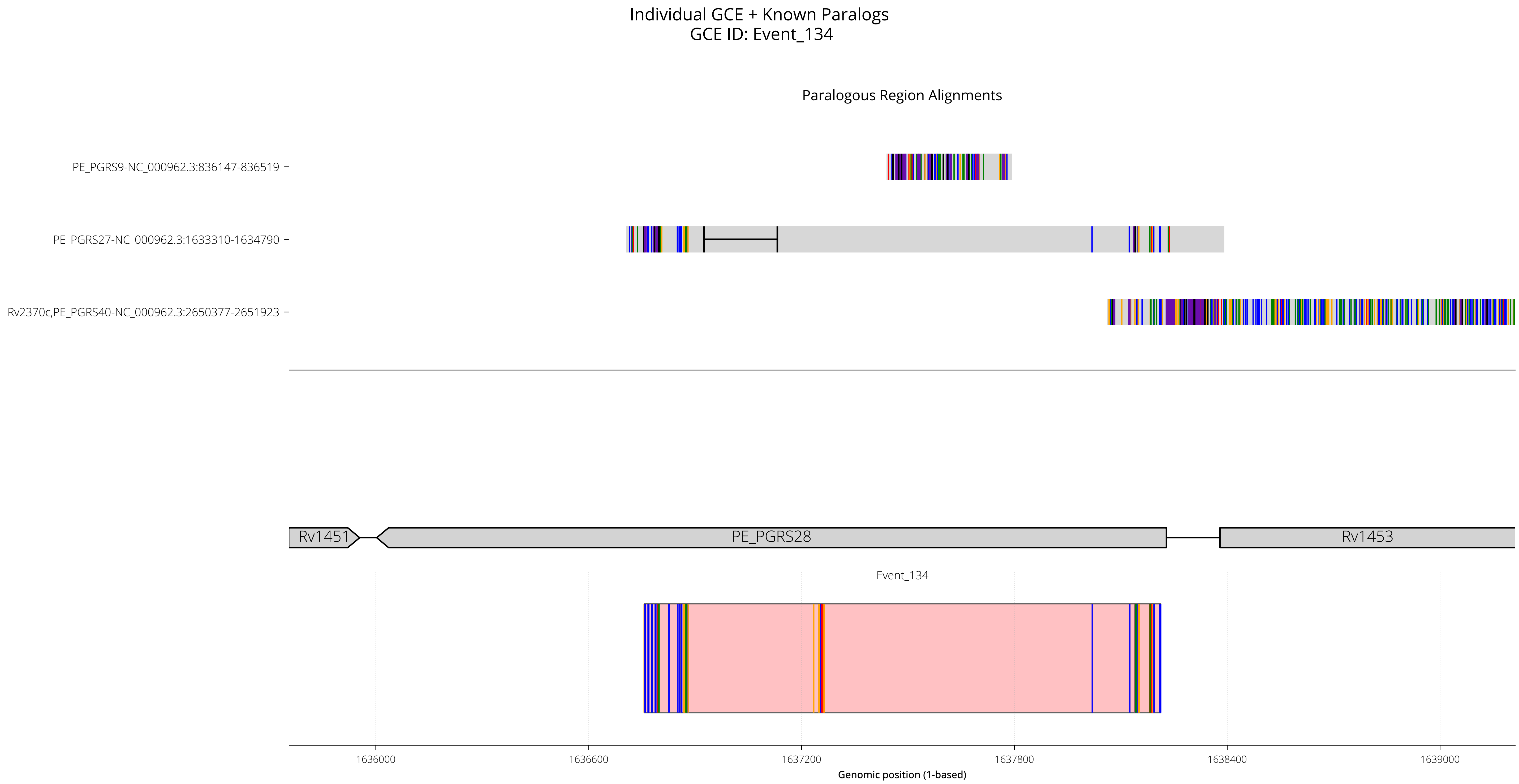

Individual GCE + Known Paralogs
GCE ID: Event_134
Paralogous Region Alignments
PE_PGRS9-NC_000962.3:836147-836519
PE_PGRS27-NC_000962.3:1633310-1634790
Rv2370c,PE_PGRS40-NC_000962.3:2650377-2651923
Rv1451
PE_PGRS28
Rv1453
Event_134
1636000
1636600
1637200
1637800
1638400
1639000
Genomic position (1-based)

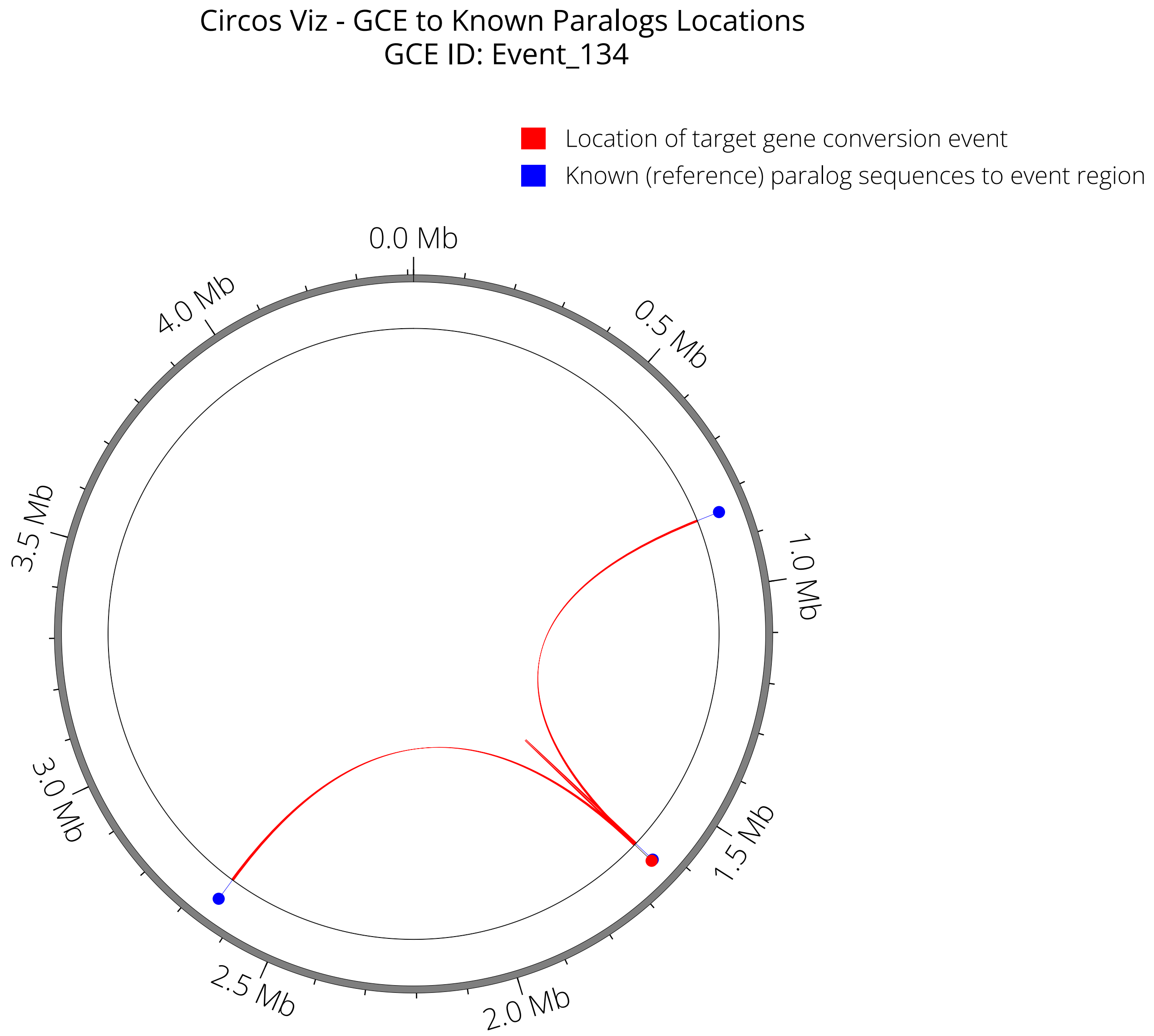

Circos Viz - GCE to Known Paralogs Locations
GCE ID: Event_134
Location of target gene conversion event
Known (reference) paralog sequences to event region
0.0 Mb
0.5 Mb
1.0 Mb
1.5 Mb
2.0 Mb
2.5 Mb
3.0 Mb
3.5 Mb
4.0 Mb

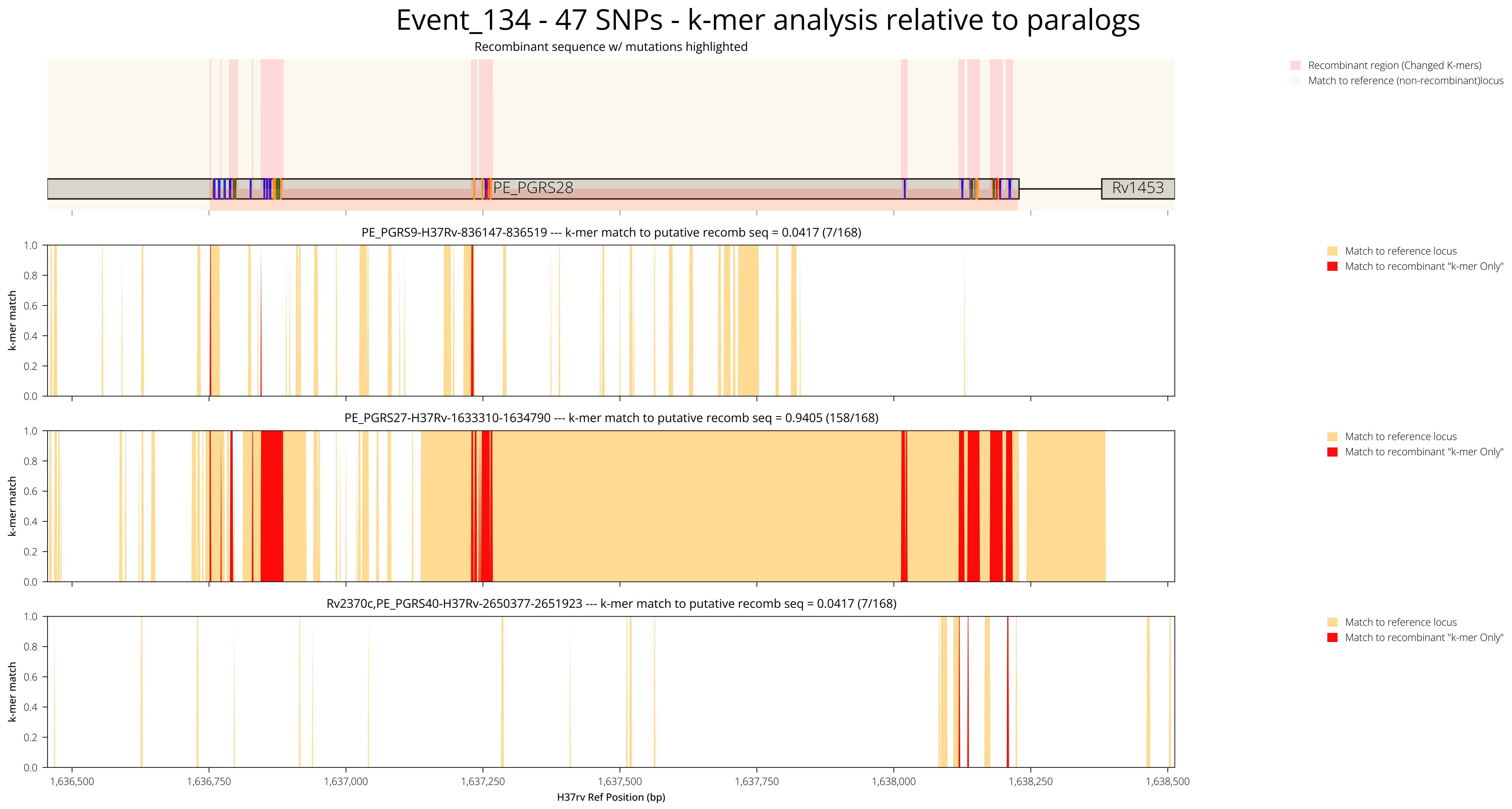

Event_134 - 47 SNPs - k-mer analysis relative to paralogs
Recombinant sequence w/ mutations highlighted
Recombinant region (Changed K-mers)
Match to reference (non-recombinant)locus
PE_PGRS28
Rv1453
PE_PGRS9-H37Rv-836147-836519 --- k-mer match to putative recomb seq = 0.0417 (7/168)
PE_PGRS27-H37Rv-1633310-1634790 --- k-mer match to putative recomb seq = 0.9405 (158/168)
Rv2370c,PE_PGRS40-H37Rv-2650377-2651923 --- k-mer match to putative recomb seq = 0.0417 (7/168)
Match to reference locus
Match to recombinant "k-mer Only"
k-mer match
1.0
0.8
0.6
0.4
0.2
0.0
1,636,500
1,636,750
1,637,000
1,637,250
1,637,500
1,637,750
1,638,000
1,638,250
1,638,500
H37rv Ref Position (bp)

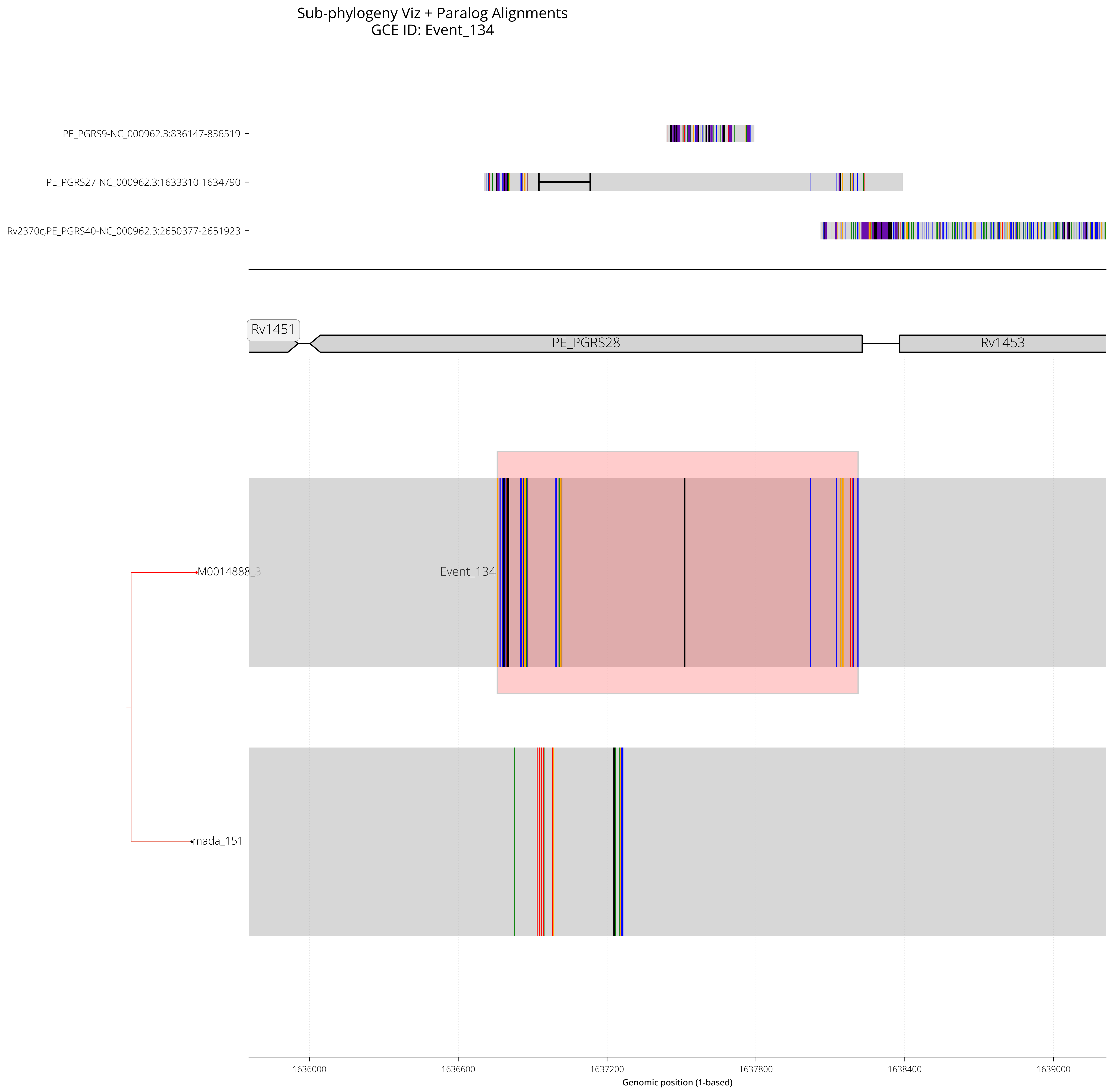
Sub-phylogeny Viz + Paralog Alignments
GCE ID: Event_134
PE_PGRS9-NC_000962.3:836147-836519
PE_PGRS27-NC_000962.3:1633310-1634790
Rv2370c,PE_PGRS40-NC_000962.3:2650377-2651923
Rv1451
PE_PGRS28
Rv1453
M0014888_3
Event_134
mada_151
1636000
1636600
1637200
1637800
1638400
1639000
Genomic position (1-based)

Full Phylogeny Viz + Paralog Alignments
GCE ID: Event_134

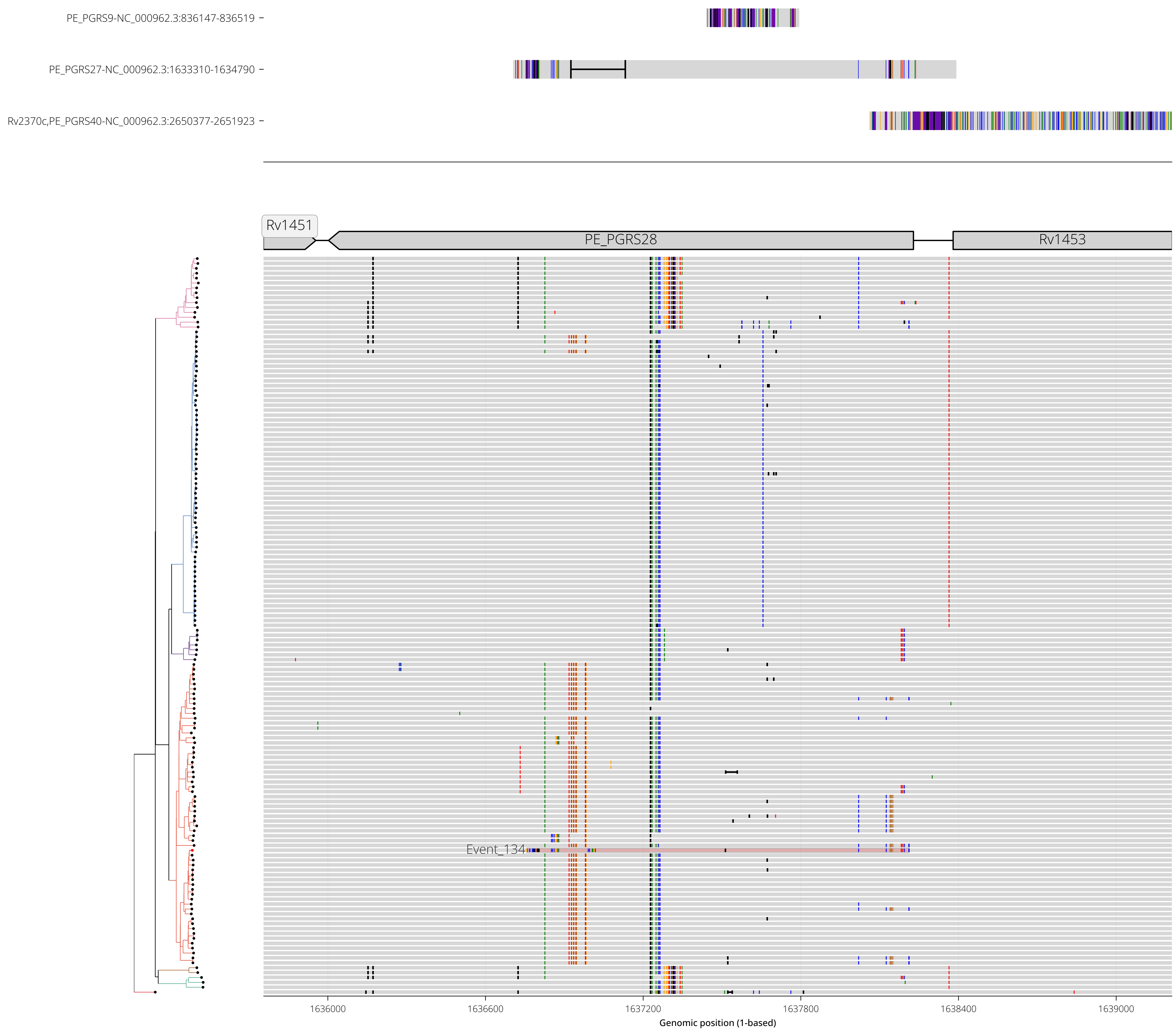

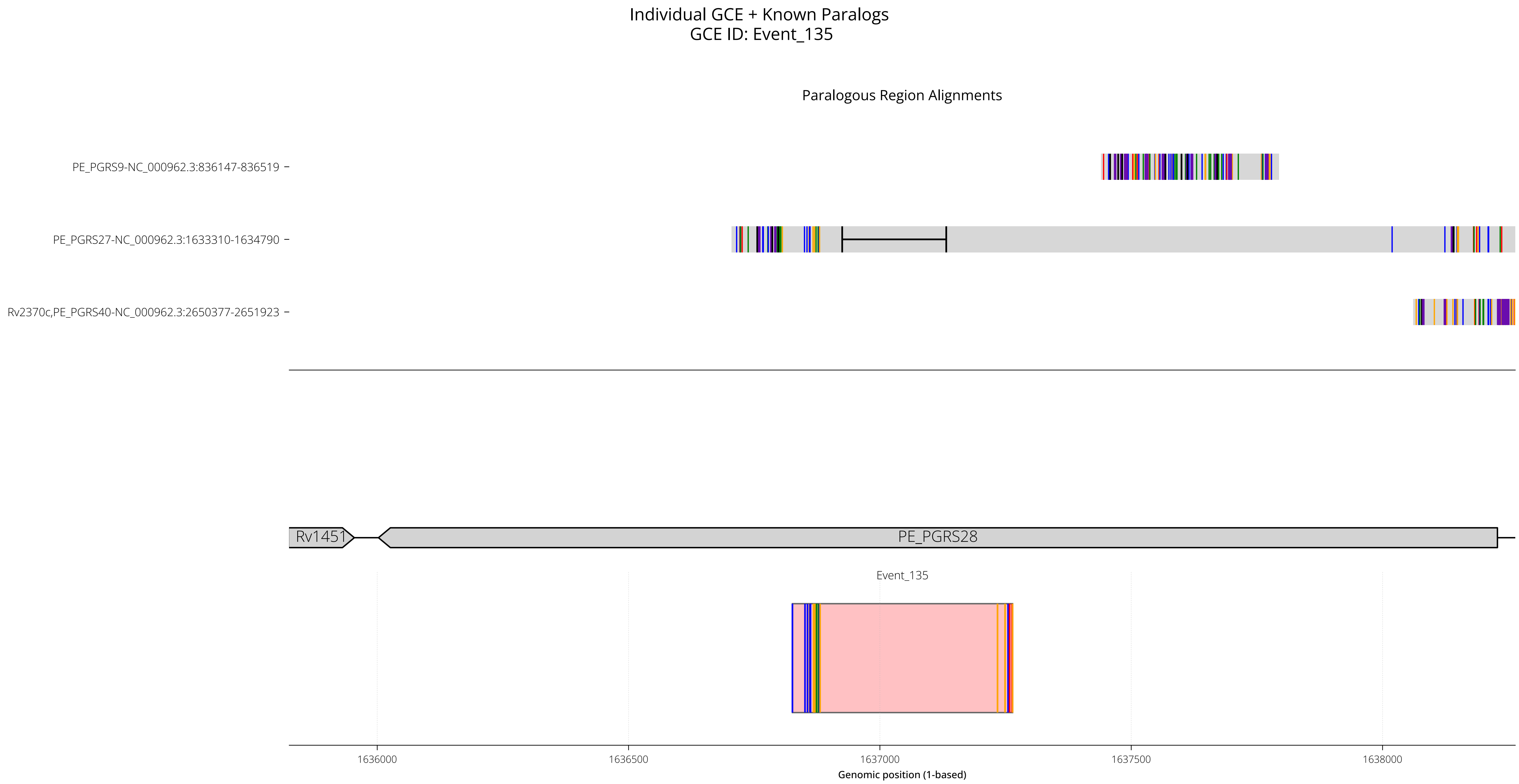
Individual GCE + Known Paralogs
GCE ID: Event_135
Paralogous Region Alignments
PE_PGRS9-NC_000962.3:836147-836519
PE_PGRS27-NC_000962.3:1633310-1634790
Rv2370c,PE_PGRS40-NC_000962.3:2650377-2651923
Rv1451
PE_PGRS28
Event_135
1636000
1636500
1637000
1637500
1638000
Genomic position (1-based)

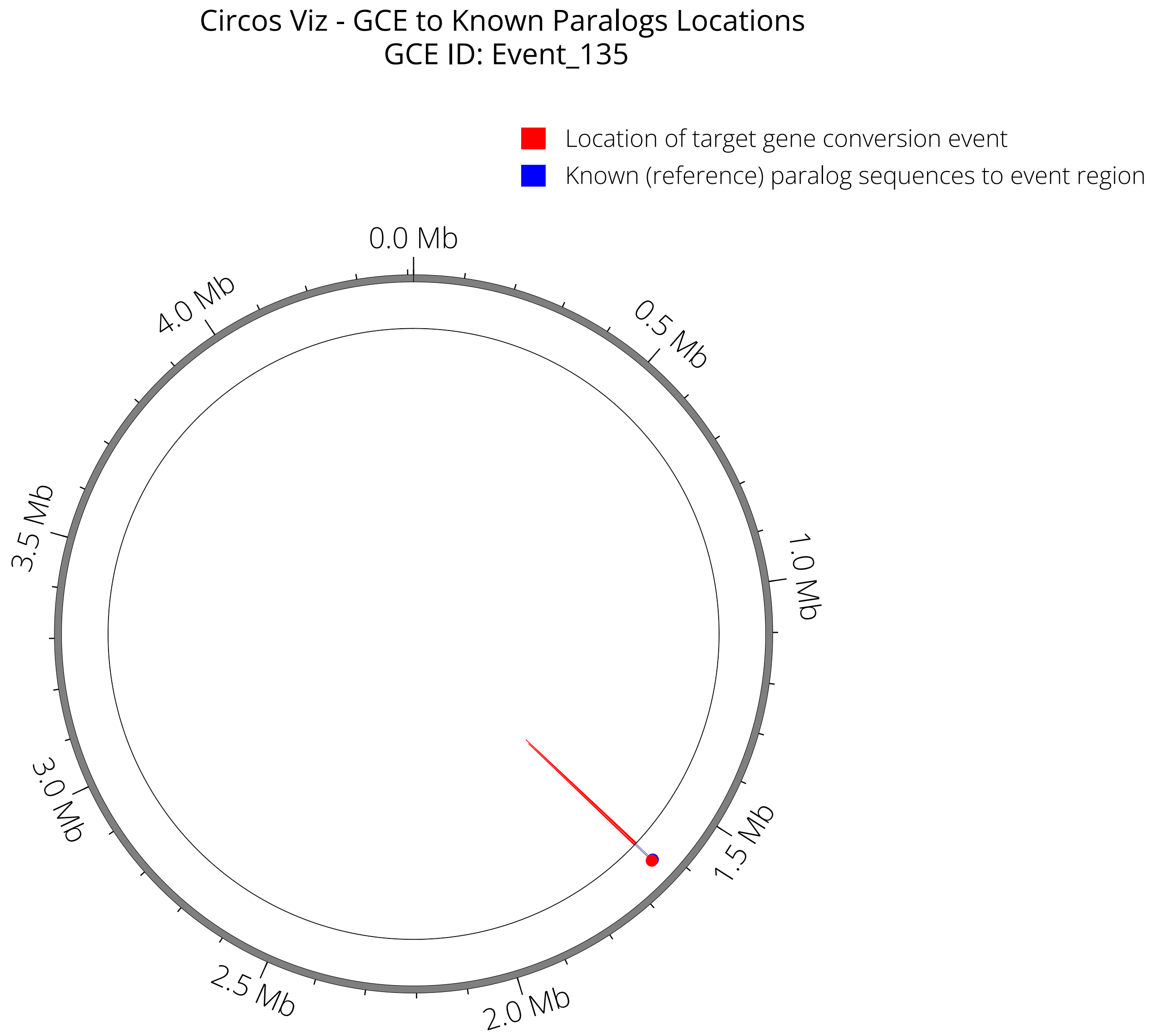
Circos Viz - GCE to Known Paralogs Locations
GCE ID: Event_135
Location of target gene conversion event
Known (reference) paralog sequences to event region
0.0 Mb
0.5 Mb
1.0 Mb
1.5 Mb
2.0 Mb
2.5 Mb
3.0 Mb
3.5 Mb
4.0 Mb

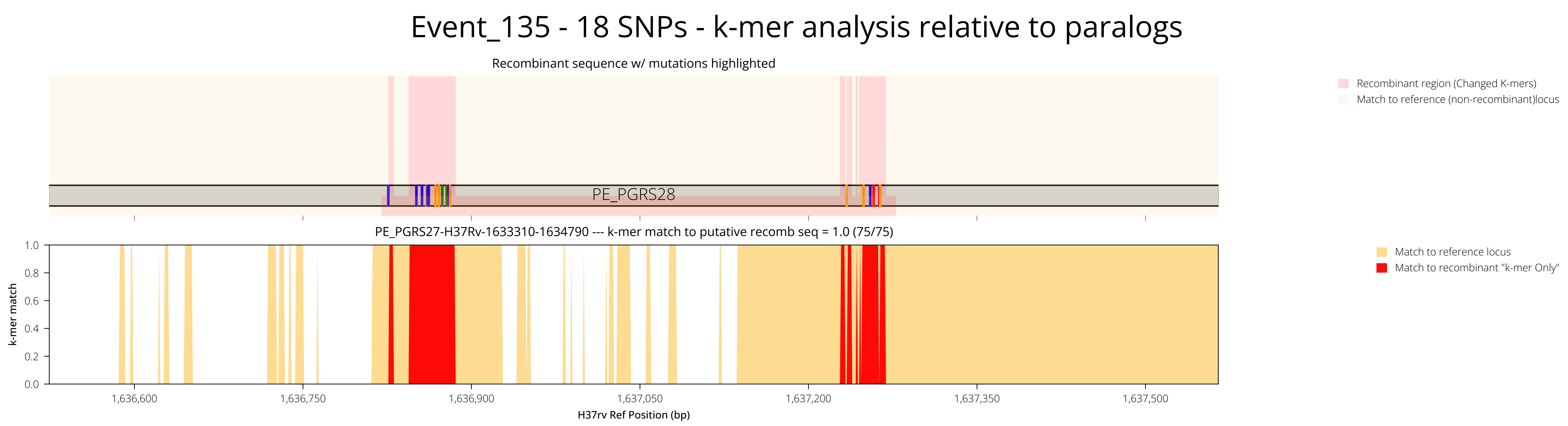

Event_135 - 18 SNPs - k-mer analysis relative to paralogs
Recombinant sequence w/ mutations highlighted
PE_PGRS28
Recombinant region (Changed K-mers)
Match to reference (non-recombinant)locus
PE_PGRS27-H37Rv-1633310-1634790 --- k-mer match to putative recomb seq = 1.0 (75/75)
Match to reference locus
Match to recombinant "k-mer Only"
k-mer match
1.0
0.8
0.6
0.4
0.2
0.0
1,636,600
1,636,750
1,636,900
1,637,050
1,637,200
1,637,350
1,637,500
H37rv Ref Position (bp)

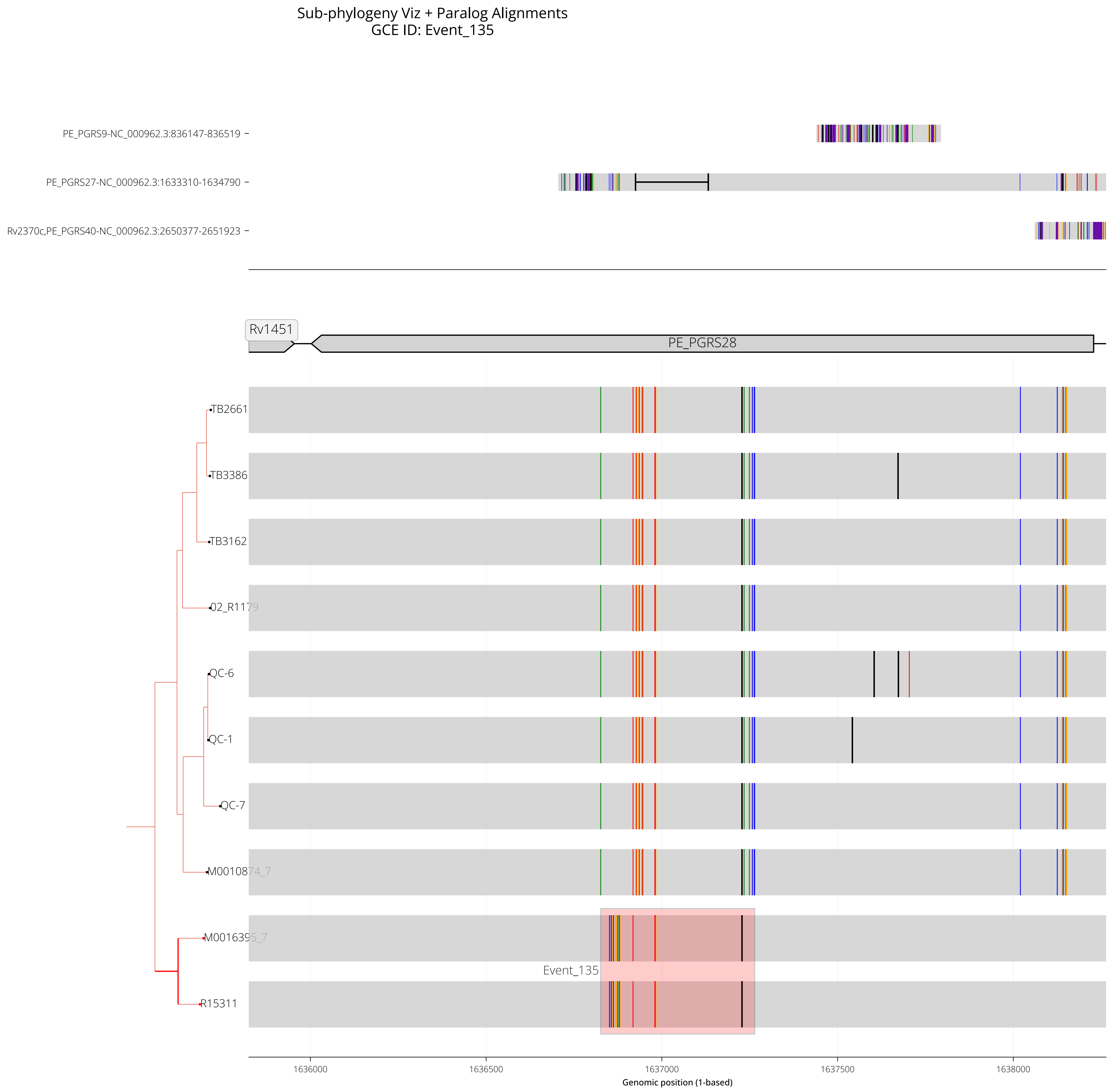

Sub-phylogeny Viz + Paralog Alignments
GCE ID: Event_135
PE_PGRS9-NC_000962.3:836147-836519
PE_PGRS27-NC_000962.3:1633310-1634790
Rv2370c,PE_PGRS40-NC_000962.3:2650377-2651923
Rv1451
PE_PGRS28
TB2661
TB3386
TB3162
02_R1179
QC-6
QC-1
QC-7
M0010874_7
M0016395_7
R15311
Event_135
1636000
1636500
1637000
1637500
1638000
Genomic position (1-based)

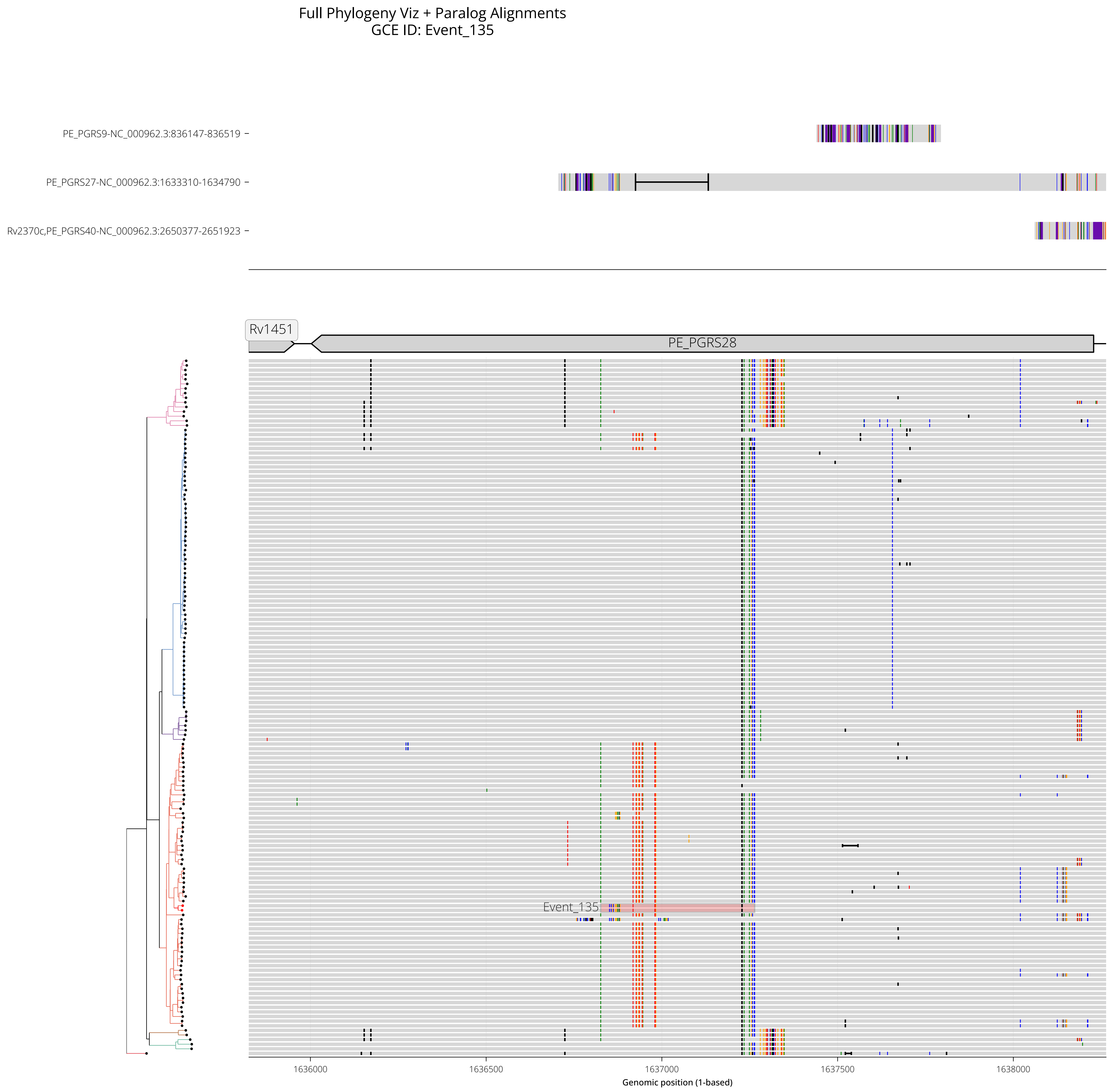
Full Phylogeny Viz + Paralog Alignments
GCE ID: Event_135
PE_PGRS9-NC_000962.3:836147-836519
PE_PGRS27-NC_000962.3:1633310-1634790
Rv2370c,PE_PGRS40-NC_000962.3:2650377-2651923
Rv1451
PE_PGRS28
Event_135
1636000
1636500
1637000
1637500
1638000
Genomic position (1-based)

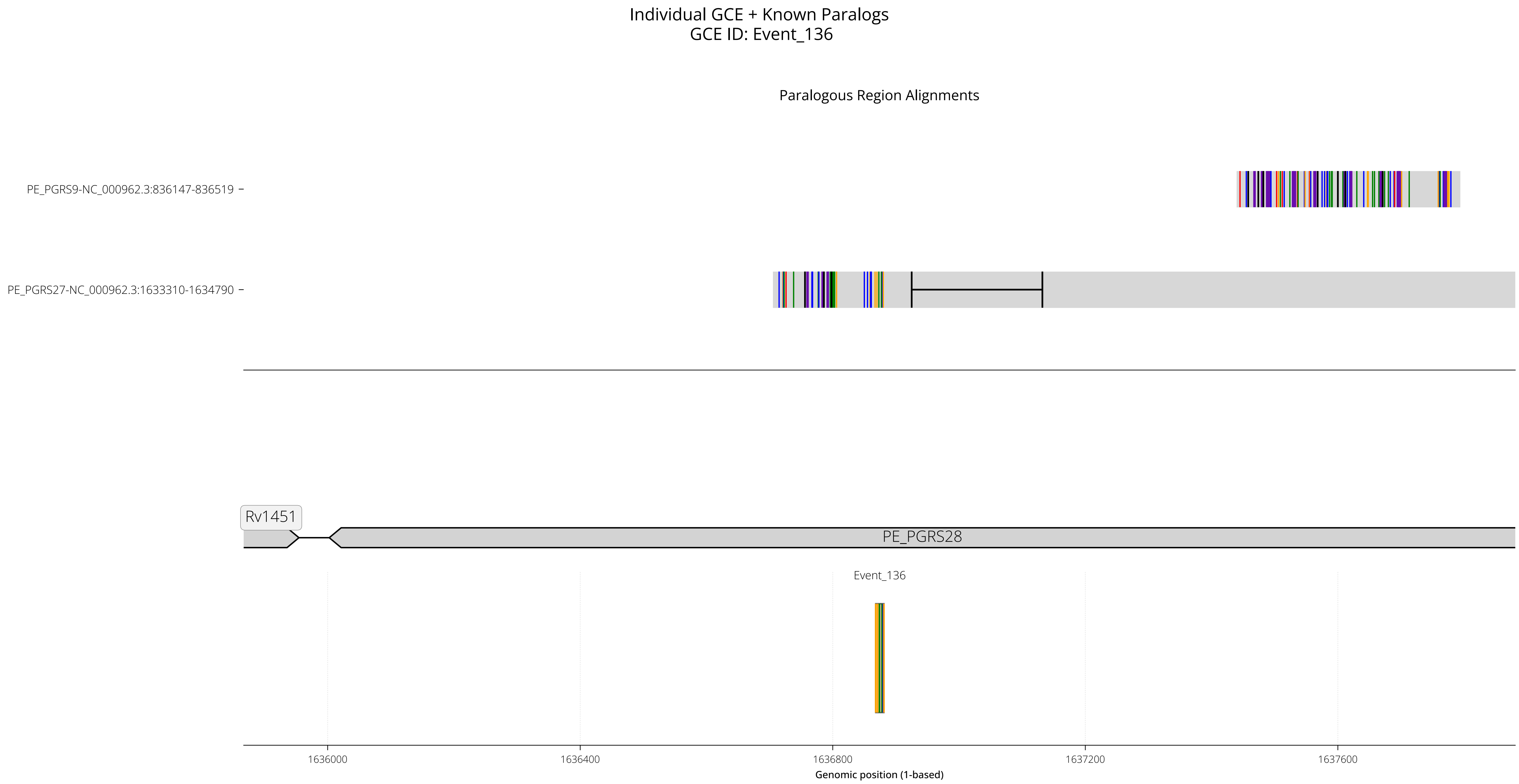
Individual GCE + Known Paralogs
GCE ID: Event_136
Paralogous Region Alignments
PE_PGRS9-NC_000962.3:836147-836519
PE_PGRS27-NC_000962.3:1633310-1634790
Rv1451
PE_PGRS28
Event_136
1636000
1636400
1636800
1637200
1637600
Genomic position (1-based)

Circos Viz - GCE to Known Paralogs Locations
GCE ID: Event_136

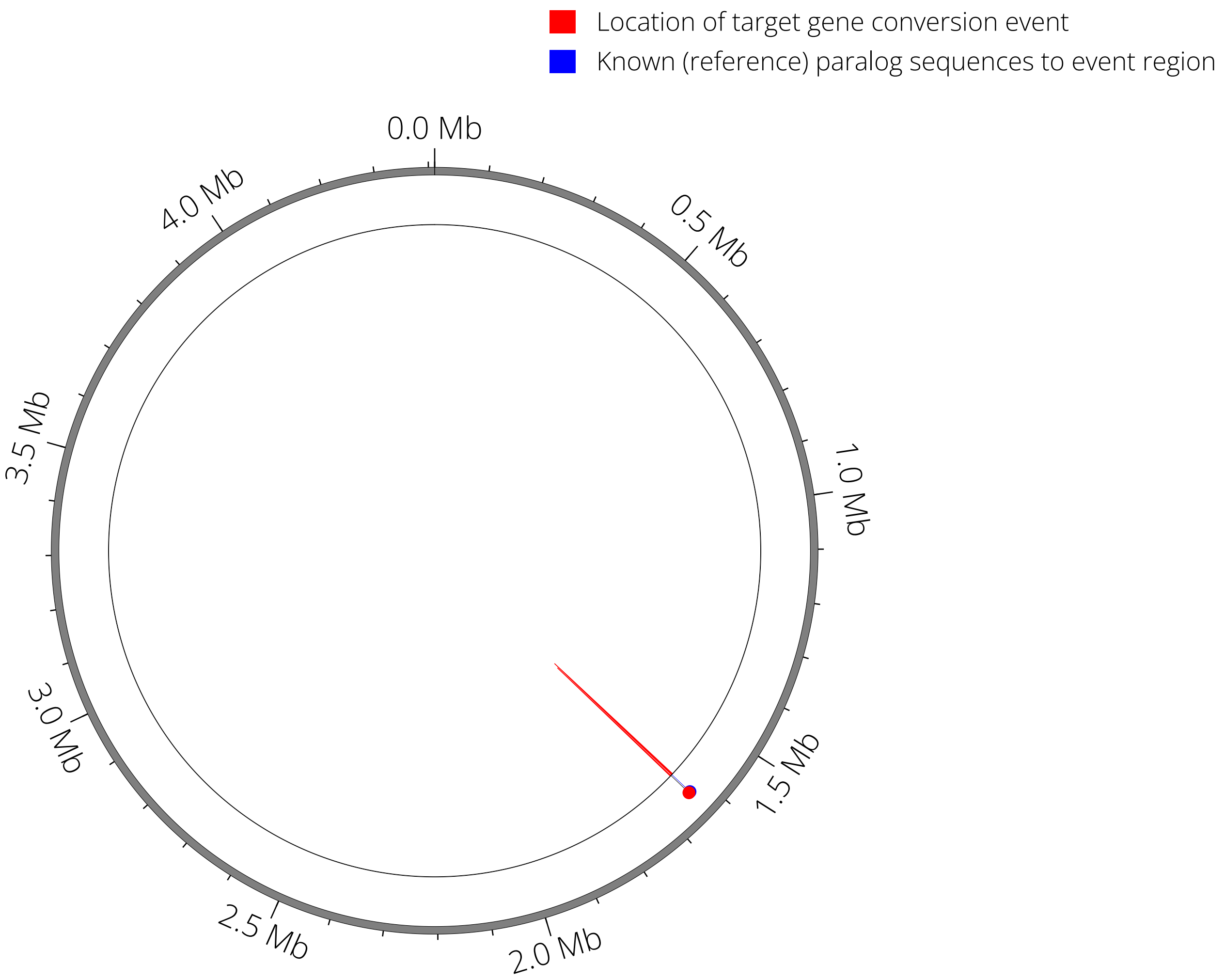

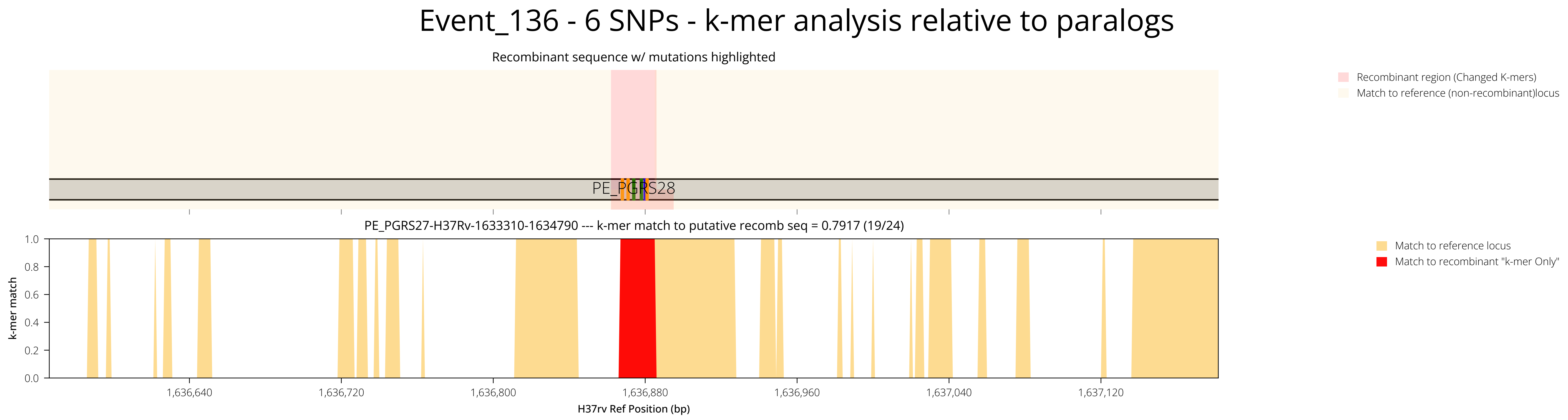
Event_136 - 6 SNPs - k-mer analysis relative to paralogs
Recombinant sequence w/ mutations highlighted
Recombinant region (Changed K-mers)
Match to reference (non-recombinant)locus
PE_PGRS28
PE_PGRS27-H37Rv-1633310-1634790 --- k-mer match to putative recomb seq = 0.7917 (19/24)
Match to reference locus
Match to recombinant "k-mer Only"
k-mer match
1.0
0.8
0.6
0.4
0.2
0.0
1,636,640
1,636,720
1,636,800
1,636,880
1,636,960
1,637,040
1,637,120
H37rv Ref Position (bp)

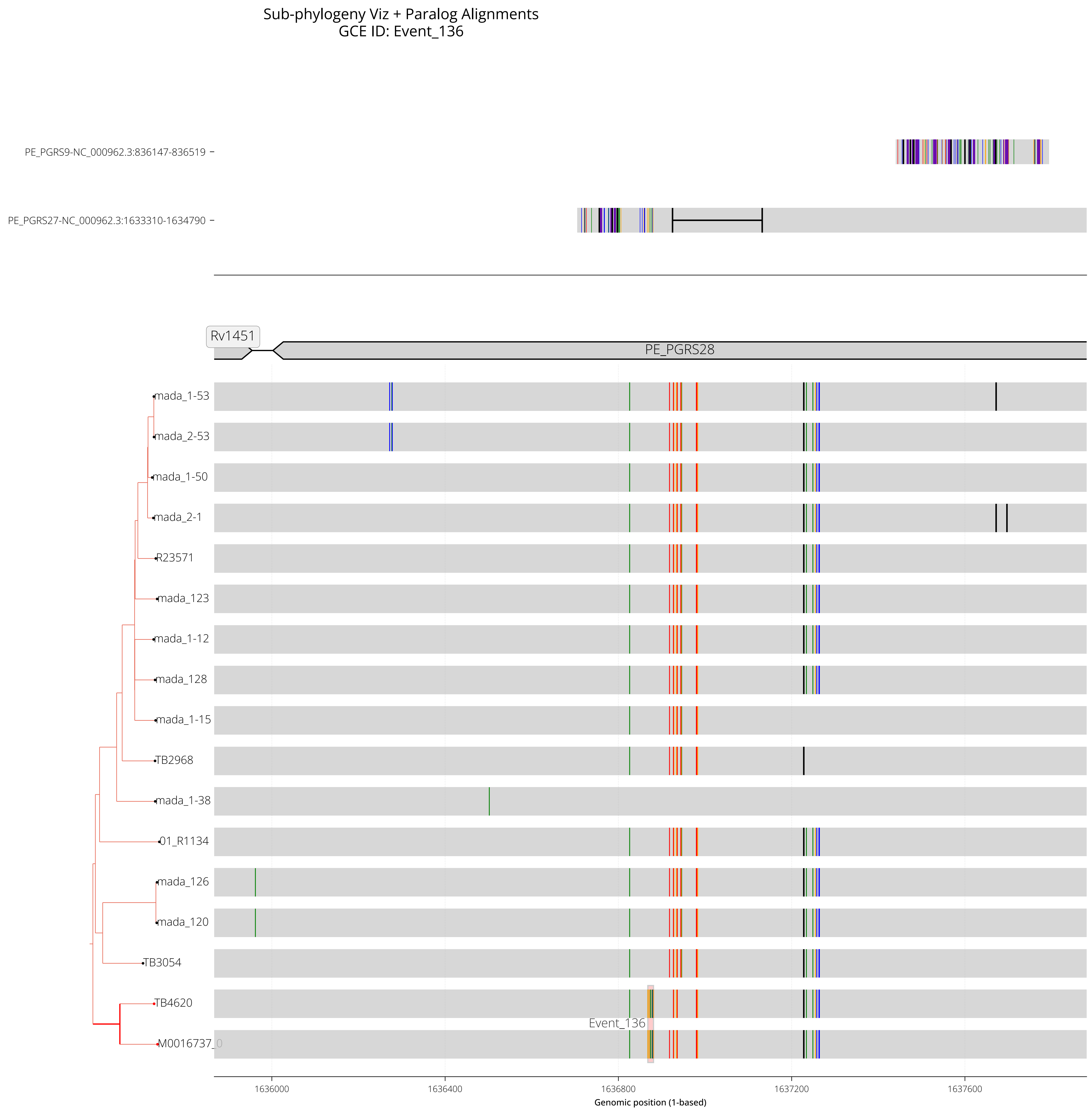

Sub-phylogeny Viz + Paralog Alignments
GCE ID: Event_136
PE_PGRS9-NC_000962.3:836147-836519
PE_PGRS27-NC_000962.3:1633310-1634790
Rv1451
PE_PGRS28
mada_1-53
mada_2-53
mada_1-50
mada_2-1
R23571
mada_123
mada_1-12
mada_128
mada_1-15
TB2968
mada_1-38
01_R1134
mada_126
mada_120
TB3054
TB4620
Event_136
M0016737_0
1636000
1636400
1636800
1637200
1637600
Genomic position (1-based)

Full Phylogeny Viz + Paralog Alignments
GCE ID: Event_136

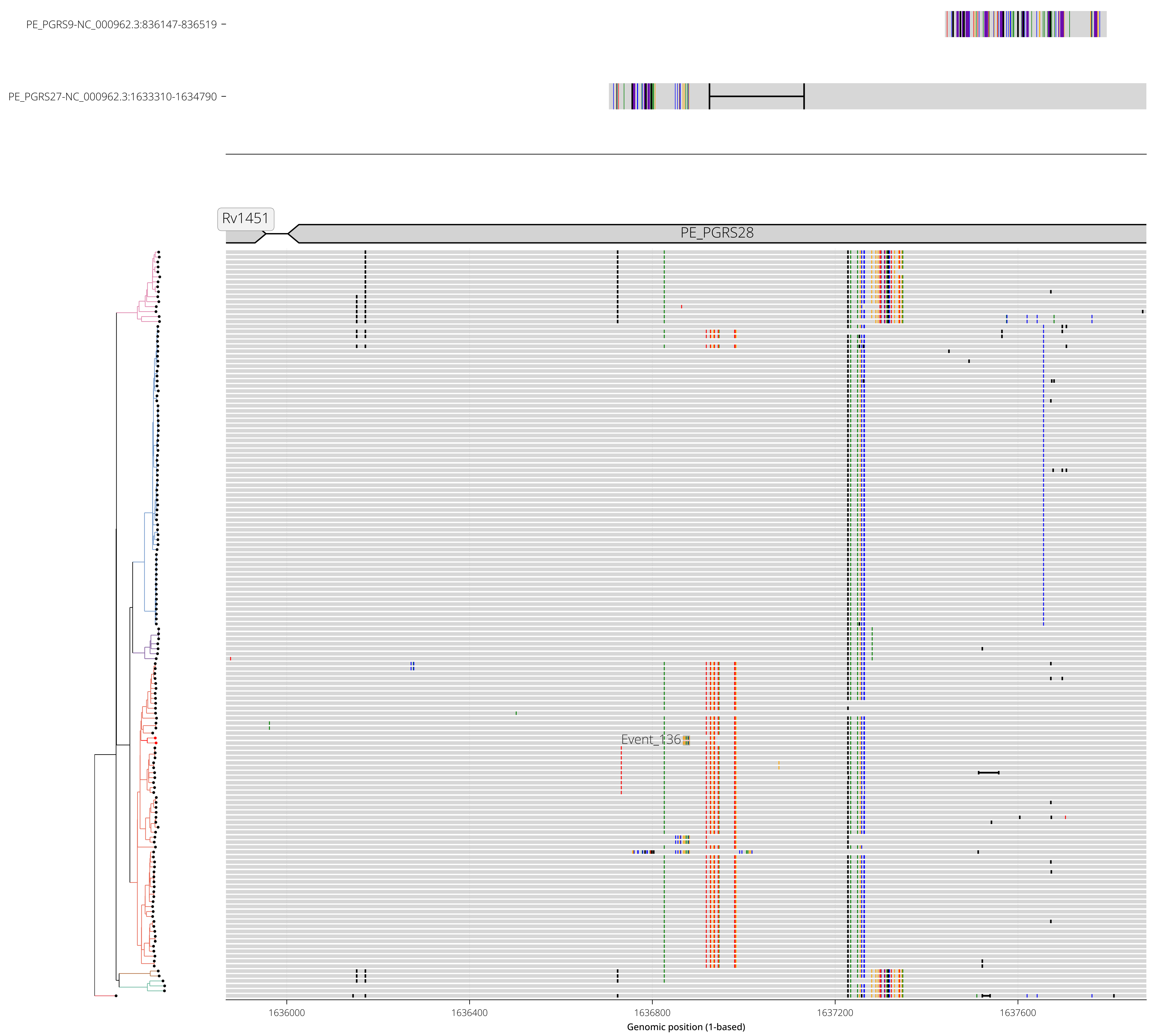

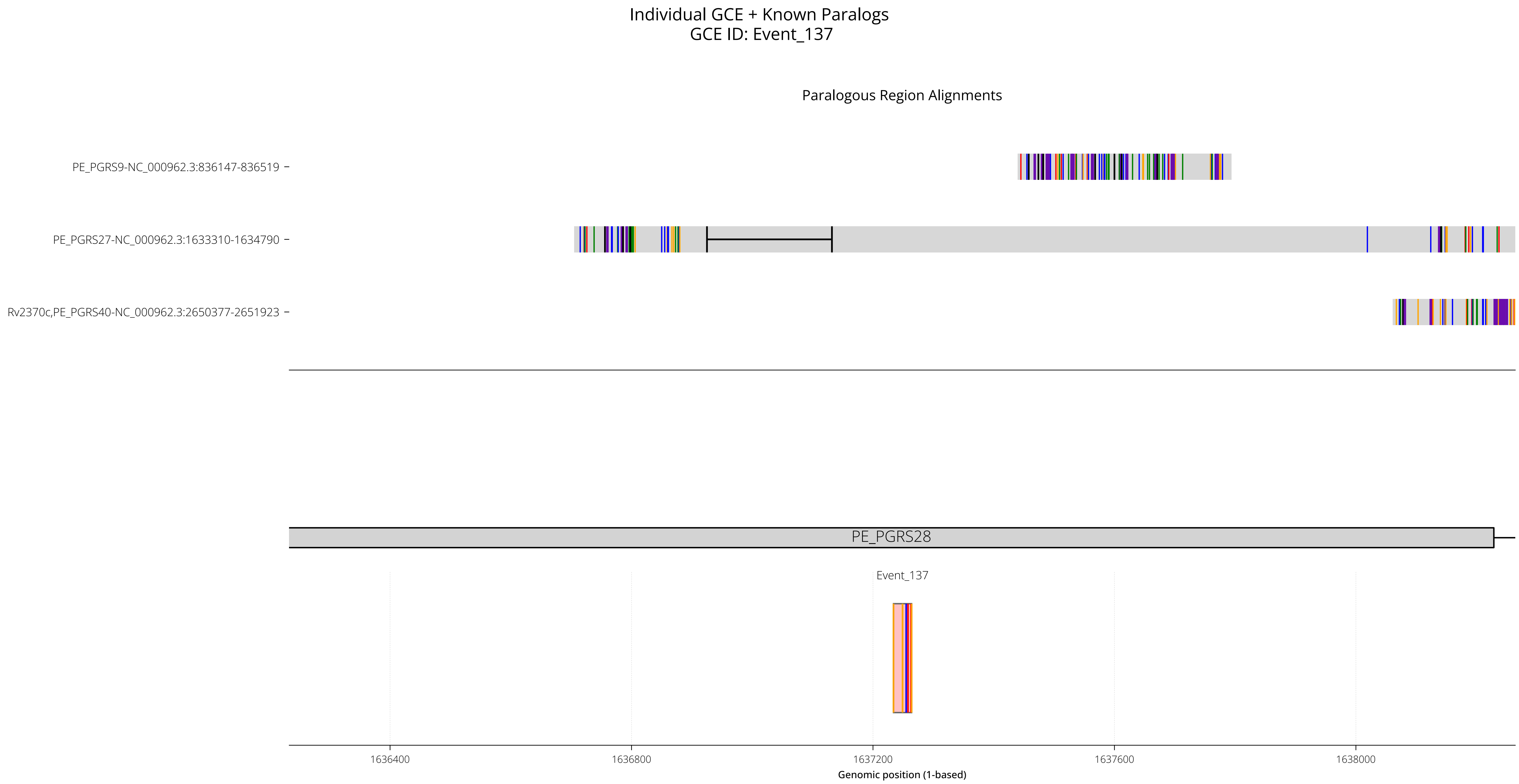
Individual GCE + Known Paralogs
GCE ID: Event_137
Paralogous Region Alignments
PE_PGRS9-NC_000962.3:836147-836519
PE_PGRS27-NC_000962.3:1633310-1634790
Rv2370c,PE_PGRS40-NC_000962.3:2650377-2651923
PE_PGRS28
Event_137
1636400
1636800
1637200
1637600
1638000
Genomic position (1-based)

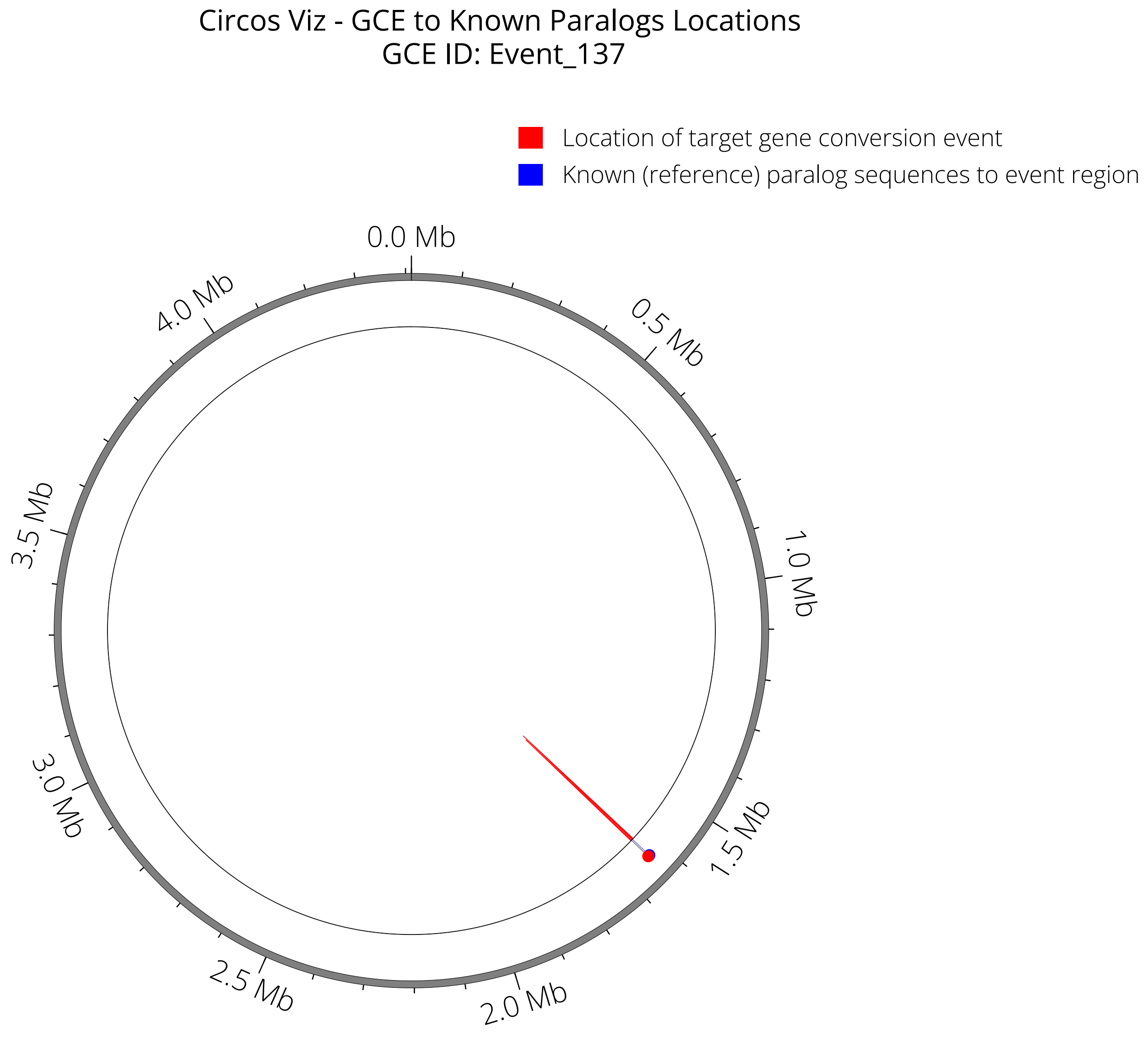
Circos Viz - GCE to Known Paralogs Locations
GCE ID: Event_137
Location of target gene conversion event
Known (reference) paralog sequences to event region
0.0 Mb
0.5 Mb
1.0 Mb
1.5 Mb
2.0 Mb
2.5 Mb
3.0 Mb
3.5 Mb
4.0 Mb

### Event_137 - 6 SNPs - k-mer analysis relative to paralogs

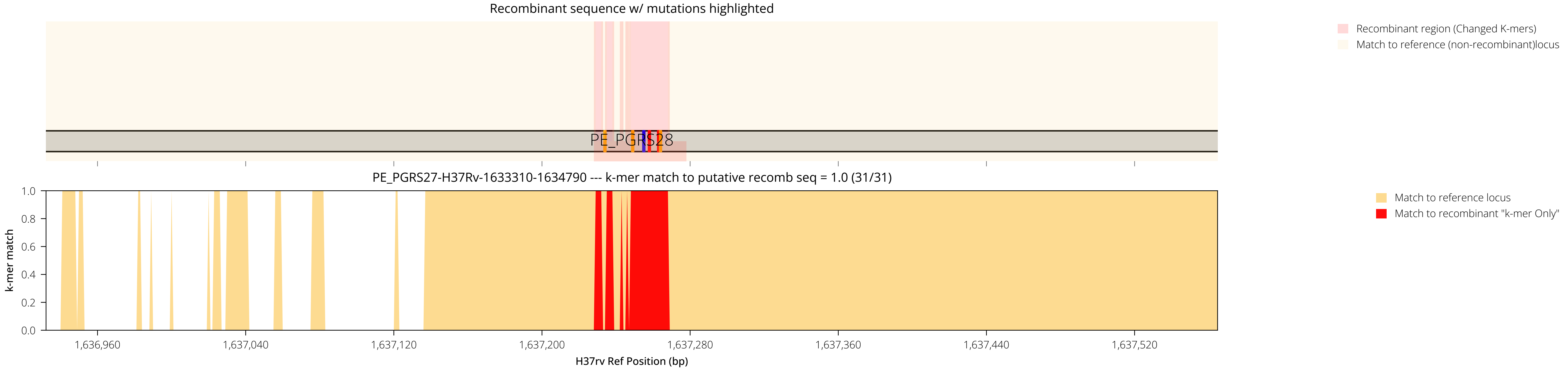

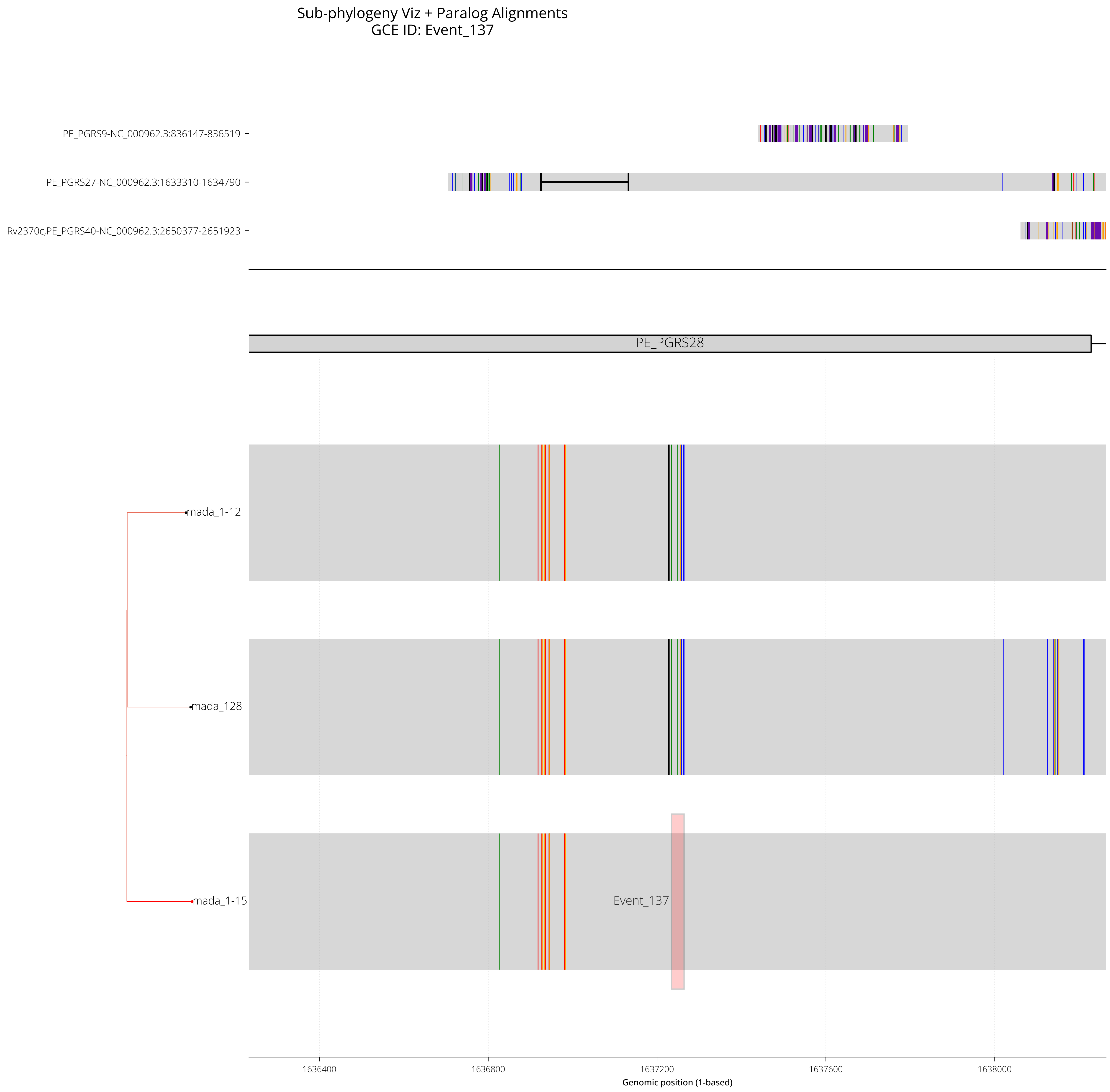
Sub-phylogeny Viz + Paralog Alignments
GCE ID: Event_137
PE_PGRS9-NC_000962.3:836147-836519
PE_PGRS27-NC_000962.3:1633310-1634790
Rv2370c,PE_PGRS40-NC_000962.3:2650377-2651923
PE_PGRS28
mada_1-12
mada_128
mada_1-15
Event_137
1636400
1636800
1637200
1637600
1638000
Genomic position (1-based)

Full Phylogeny Viz + Paralog Alignments
GCE ID: Event_137

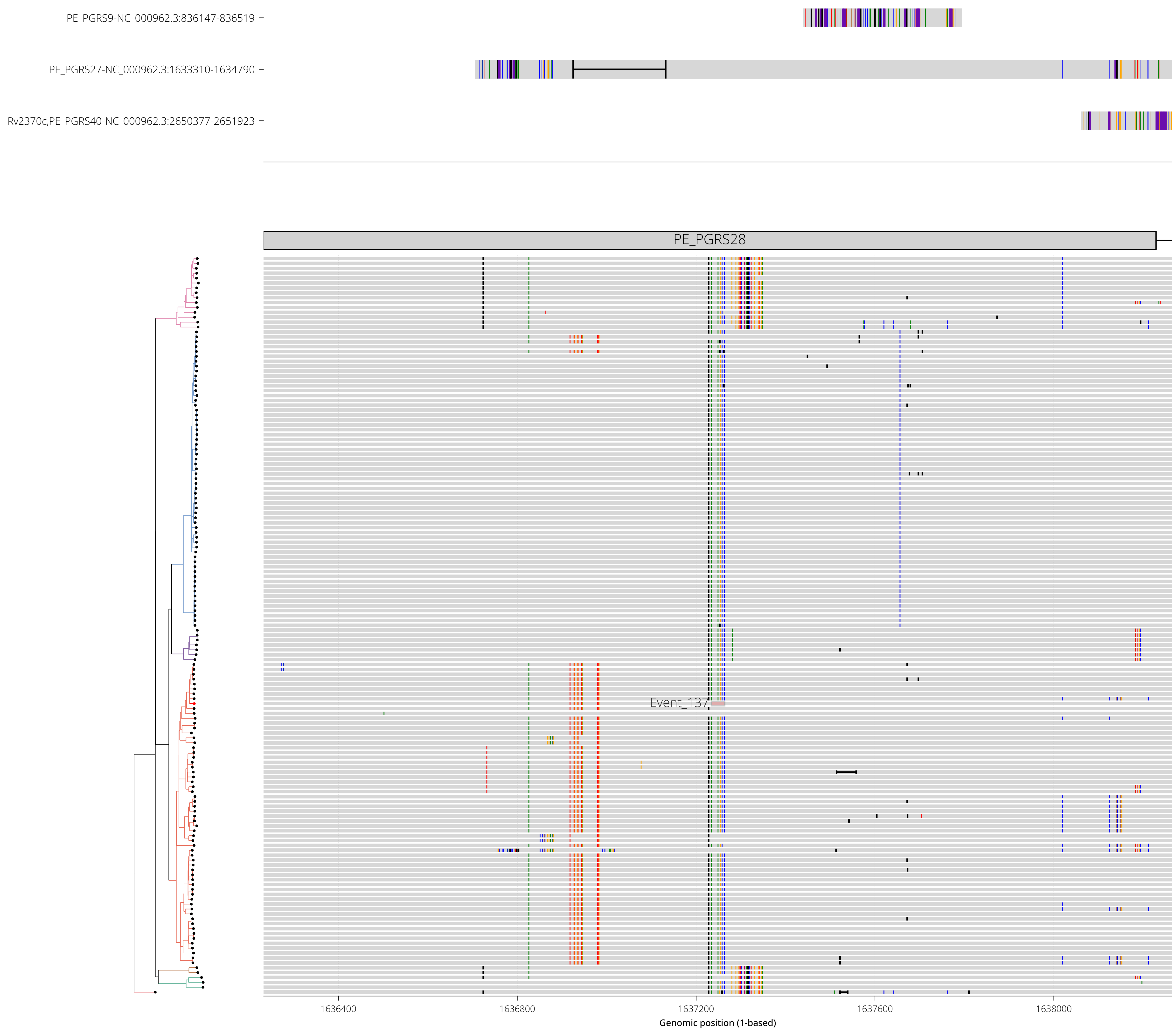

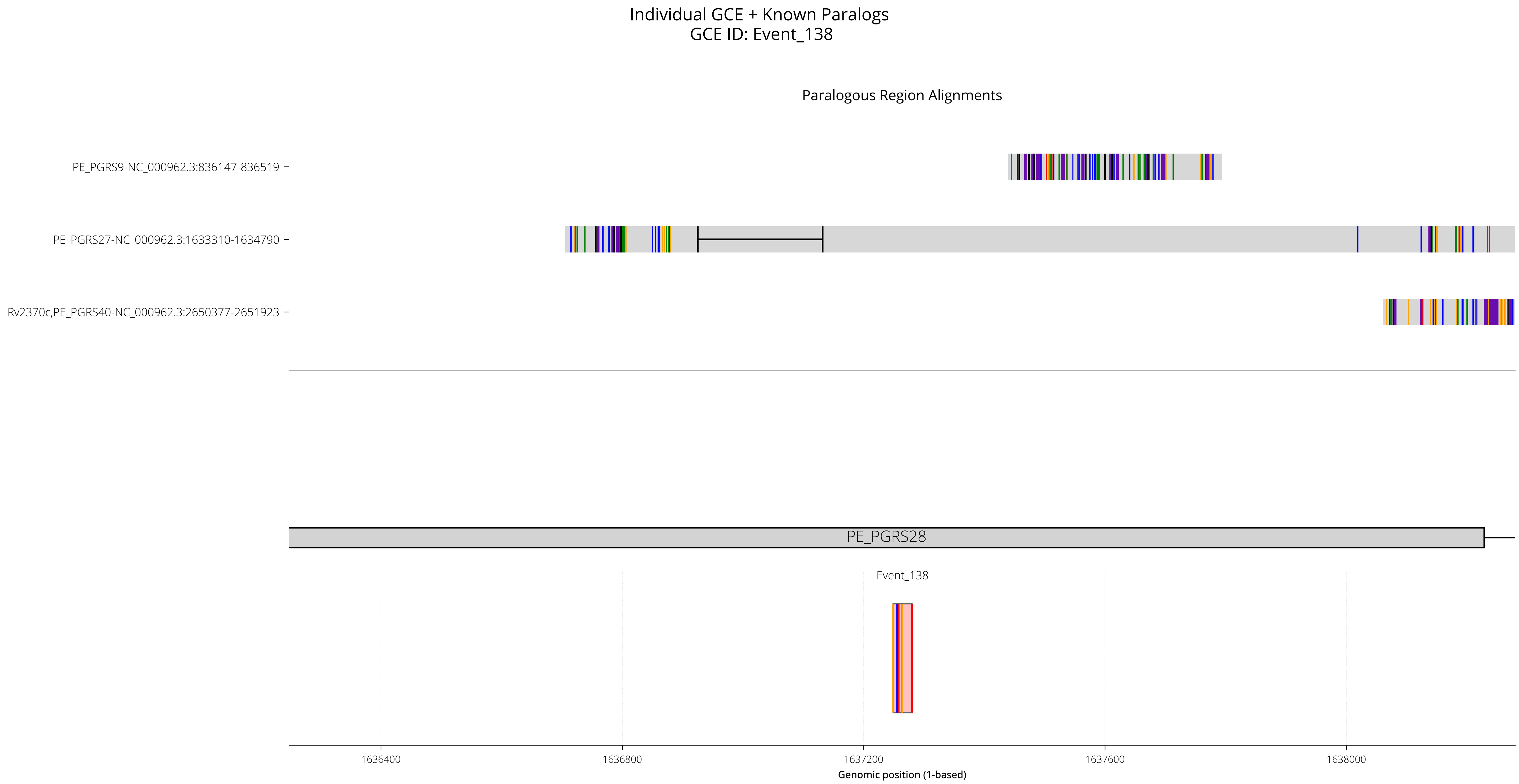
Individual GCE + Known Paralogs
GCE ID: Event_138
Paralogous Region Alignments
PE_PGRS9-NC_000962.3:836147-836519
PE_PGRS27-NC_000962.3:1633310-1634790
Rv2370c,PE_PGRS40-NC_000962.3:2650377-2651923
PE_PGRS28
Event_138
1636400
1636800
1637200
1637600
1638000
Genomic position (1-based)

Circos Viz - GCE to Known Paralogs Locations
GCE ID: Event_138

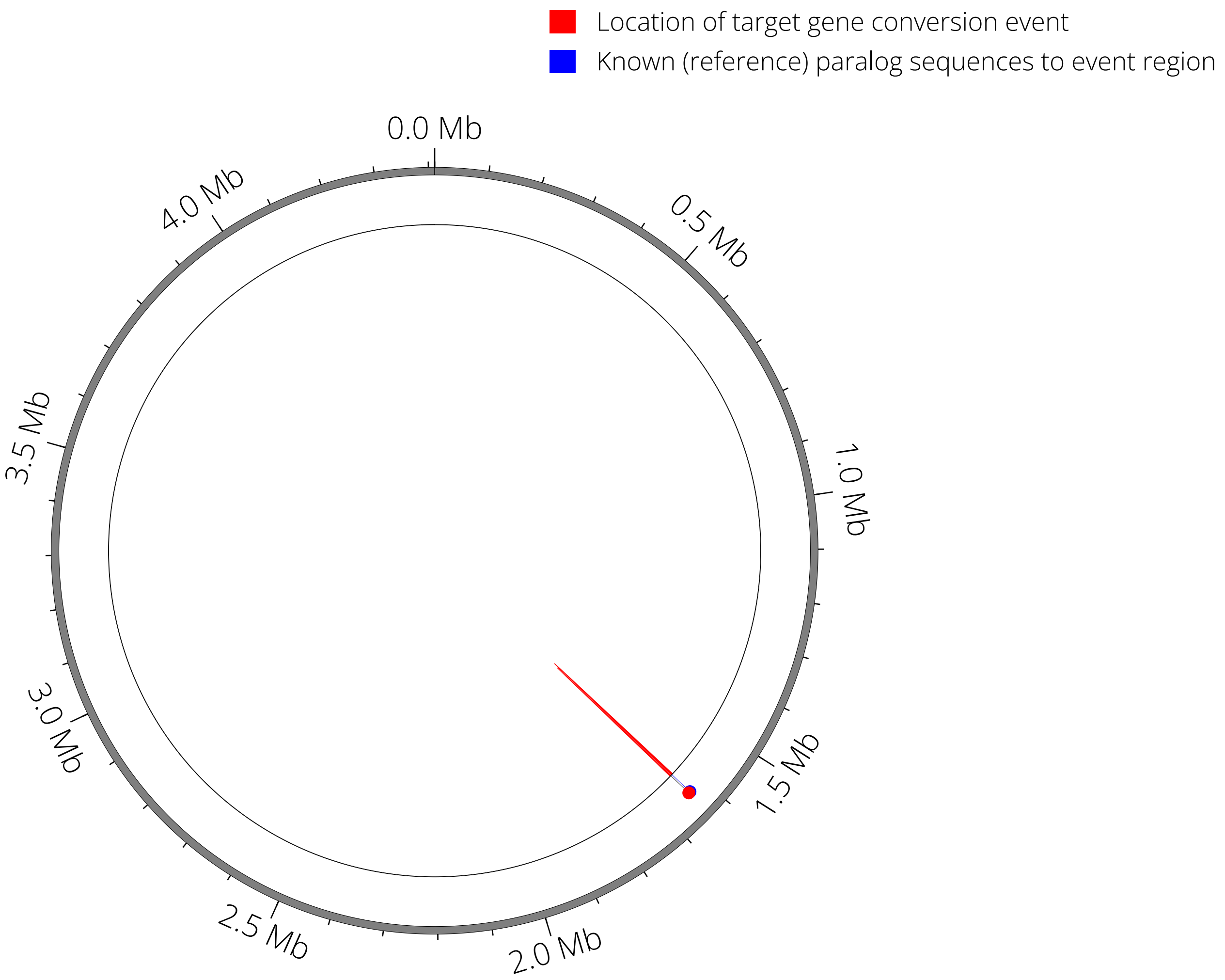

### Event_138 - 6 SNPs - k-mer analysis relative to paralogs

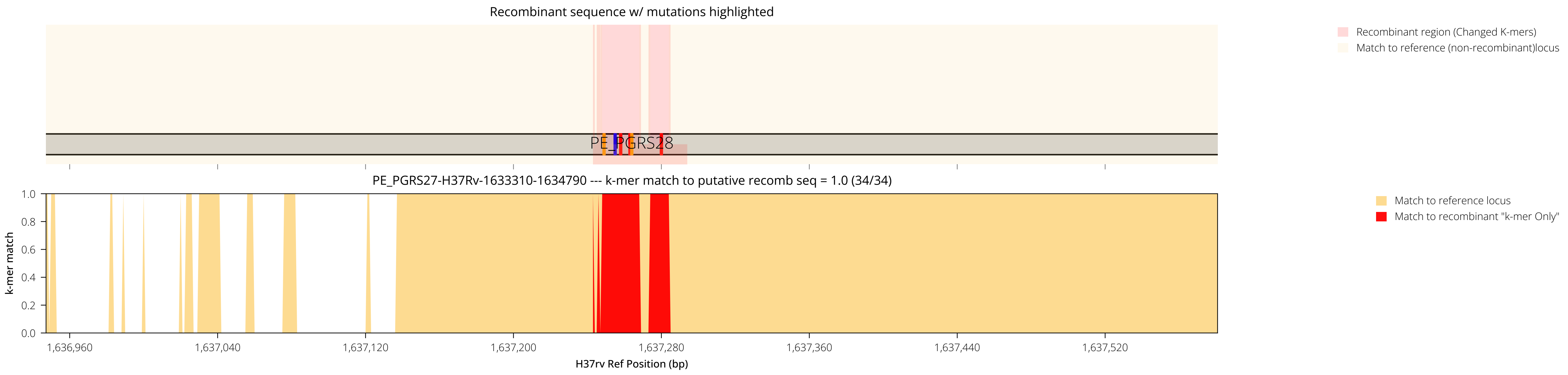

Sub-phylogeny Viz + Paralog Alignments
GCE ID: Event_138

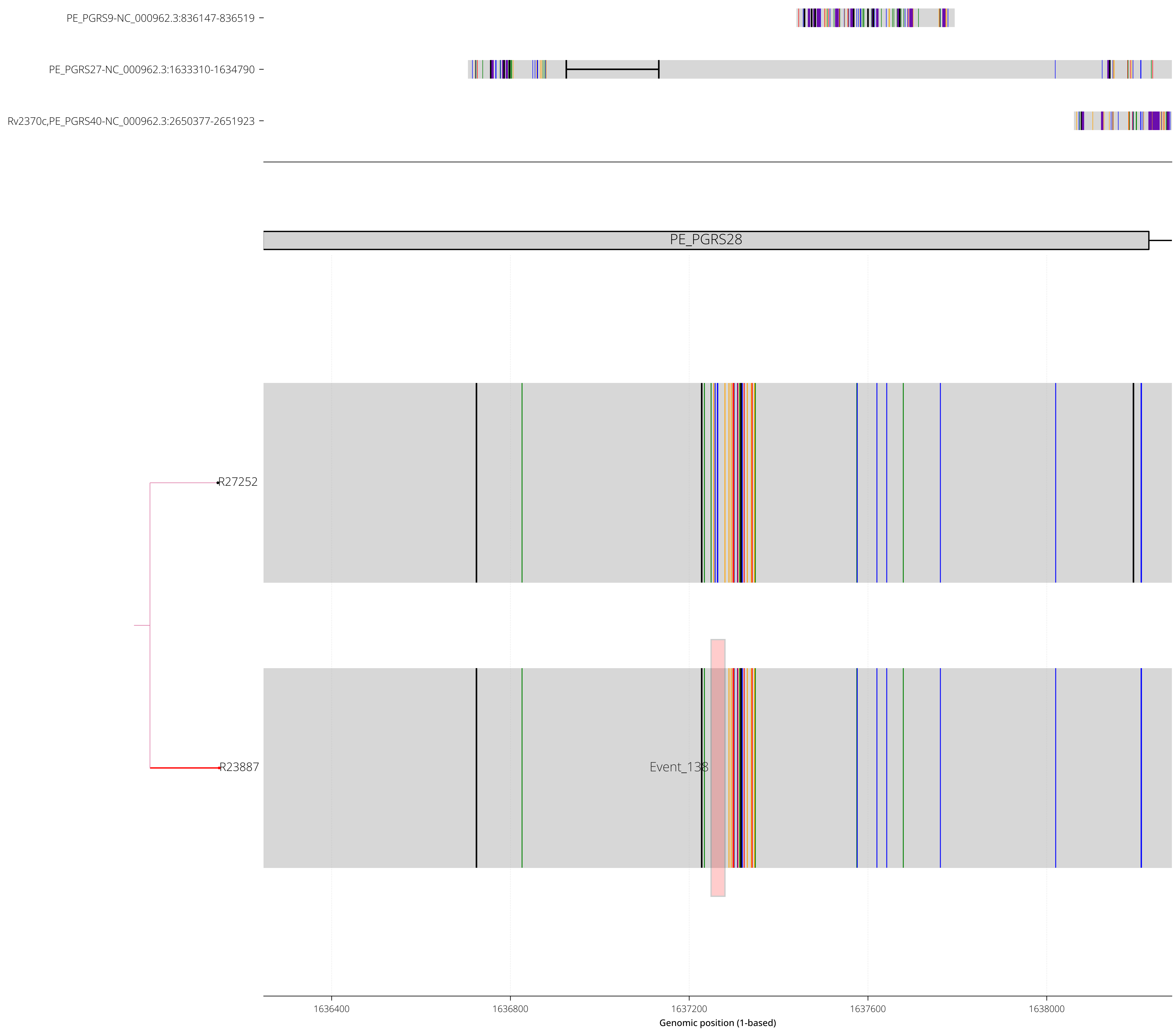

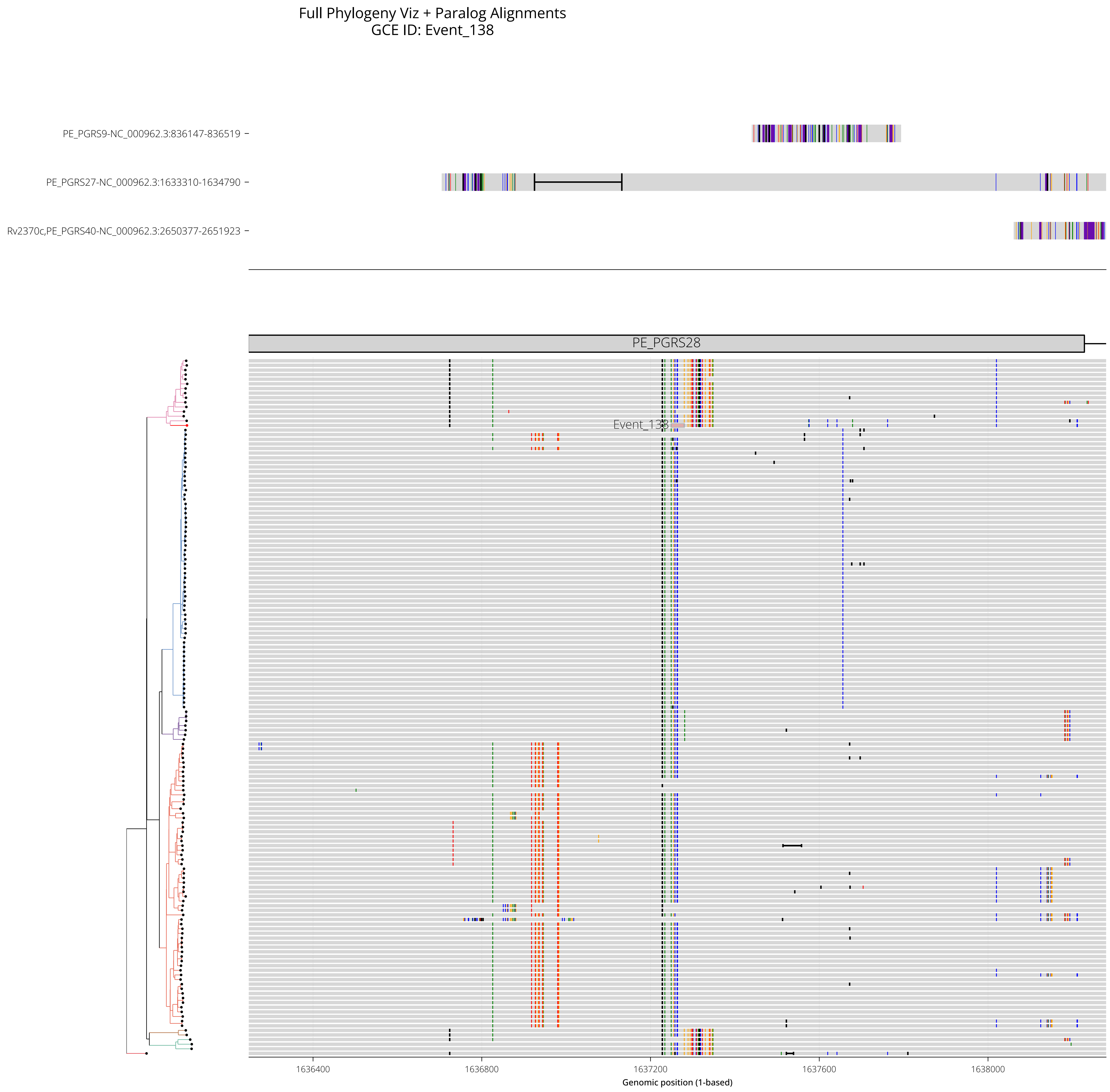

Full Phylogeny Viz + Paralog Alignments
GCE ID: Event_138
PE_PGRS9-NC_000962.3:836147-836519
PE_PGRS27-NC_000962.3:1633310-1634790
Rv2370c,PE_PGRS40-NC_000962.3:2650377-2651923
PE_PGRS28
Event_138
1636400
1636800
1637200
1637600
1638000
Genomic position (1-based)

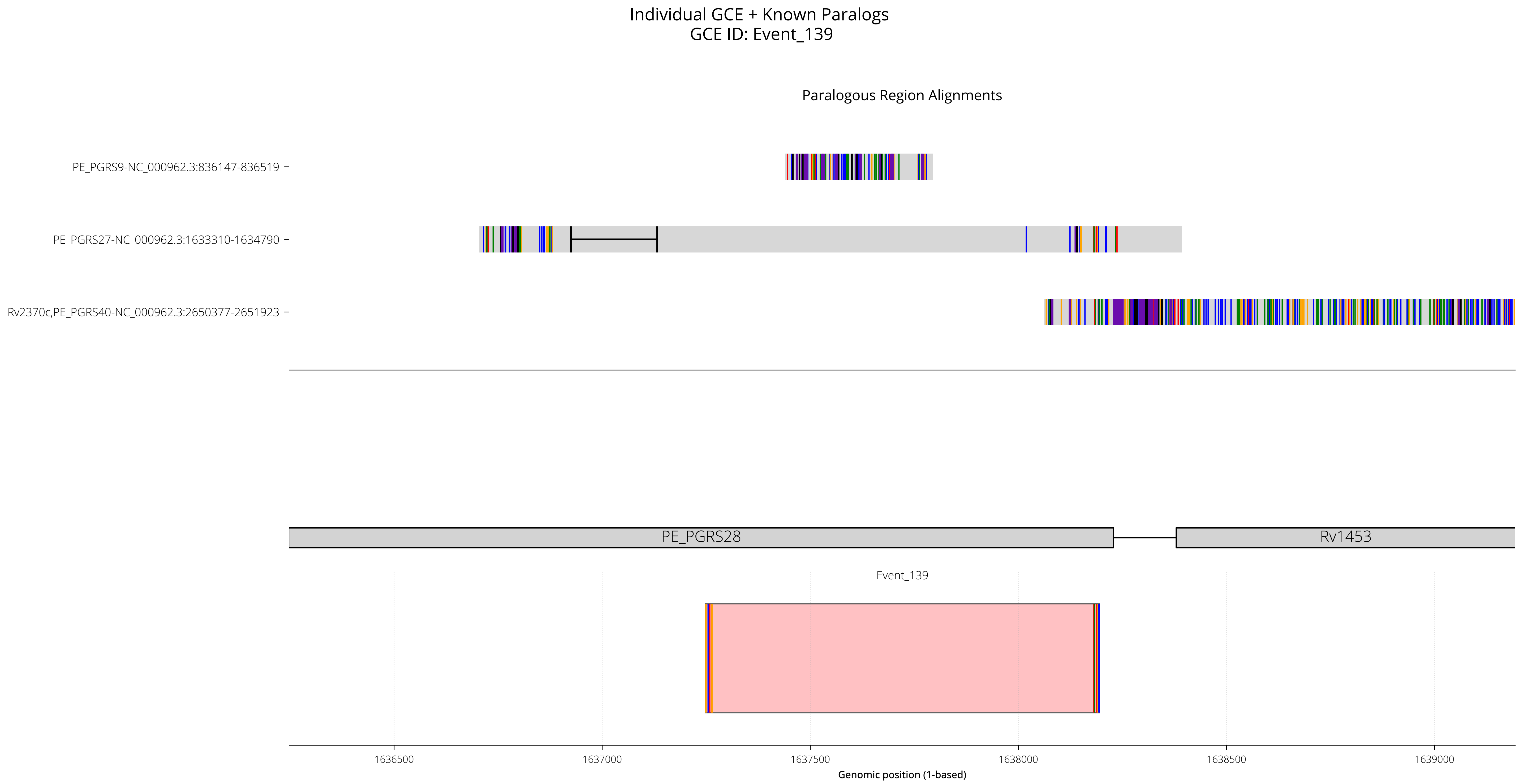
Individual GCE + Known Paralogs
GCE ID: Event_139
Paralogous Region Alignments
PE_PGRS9-NC_000962.3:836147-836519
PE_PGRS27-NC_000962.3:1633310-1634790
Rv2370c,PE_PGRS40-NC_000962.3:2650377-2651923
PE_PGRS28
Rv1453
Event_139
1636500
1637000
1637500
1638000
1638500
1639000
Genomic position (1-based)

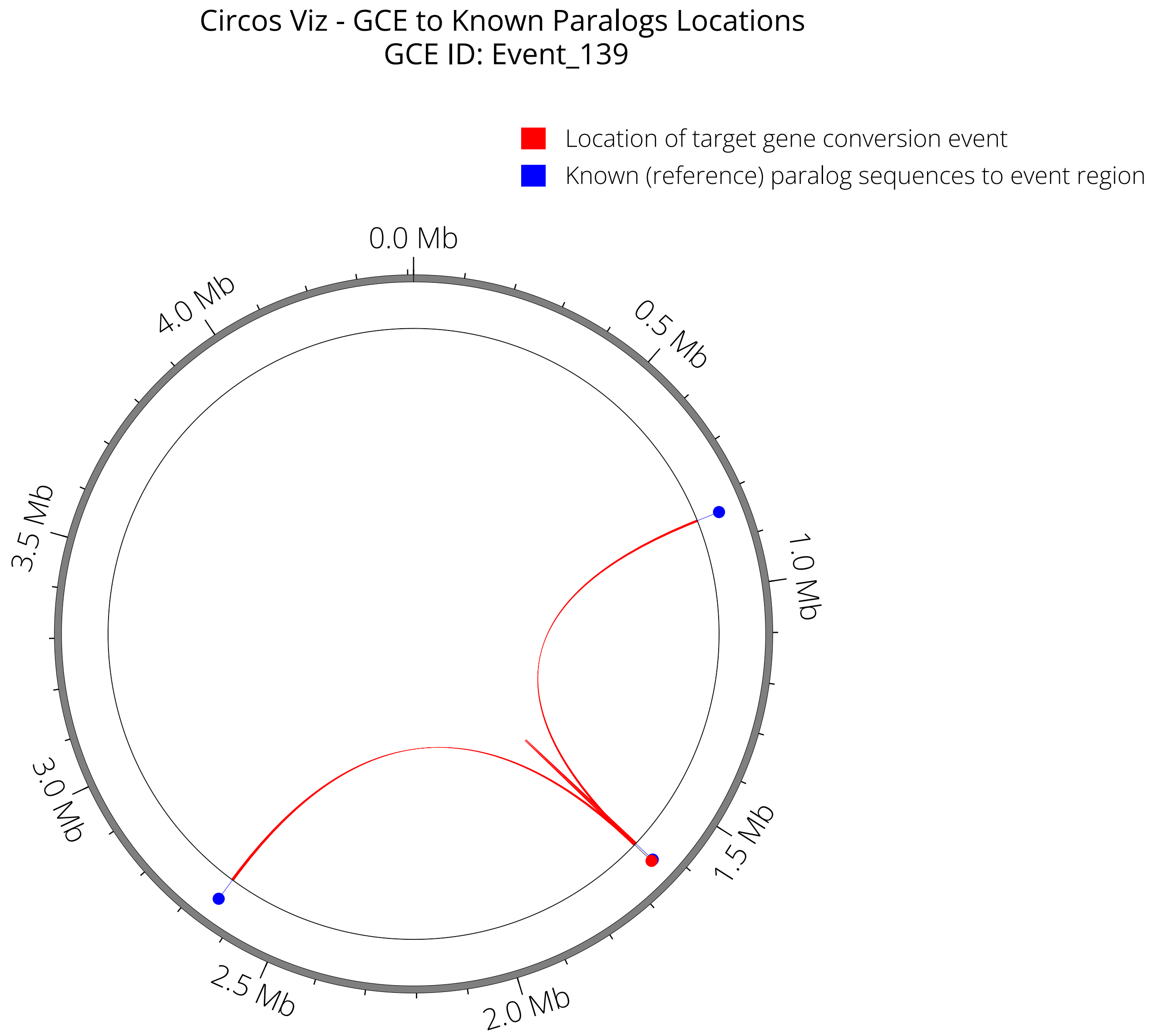
Circos Viz - GCE to Known Paralogs Locations
GCE ID: Event_139
Location of target gene conversion event
Known (reference) paralog sequences to event region
0.0 Mb
0.5 Mb
1.0 Mb
1.5 Mb
2.0 Mb
2.5 Mb
3.0 Mb
3.5 Mb
4.0 Mb

### Event_139 - 10 SNPs - k-mer analysis relative to paralogs

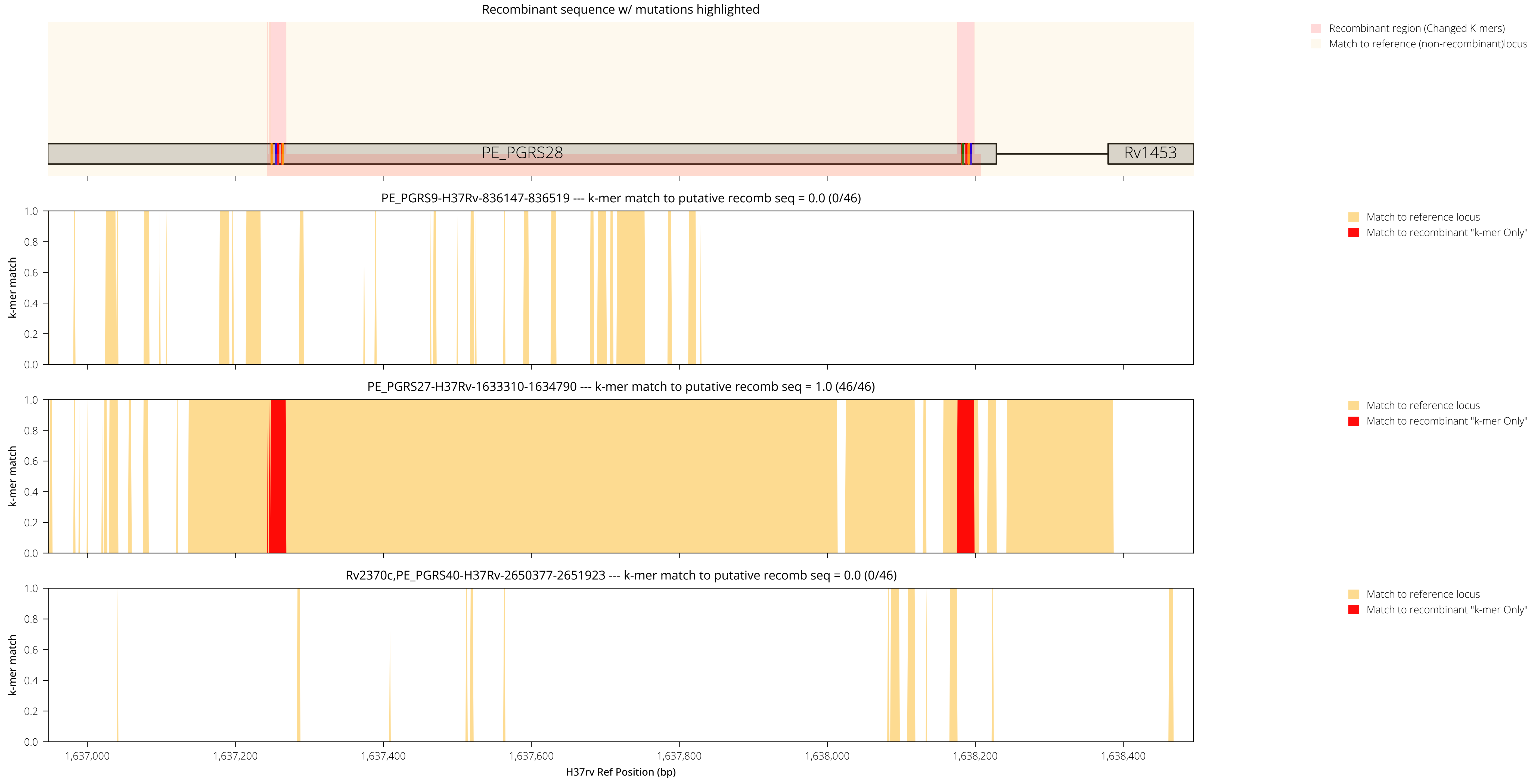

Sub-phylogeny Viz + Paralog Alignments
GCE ID: Event_139

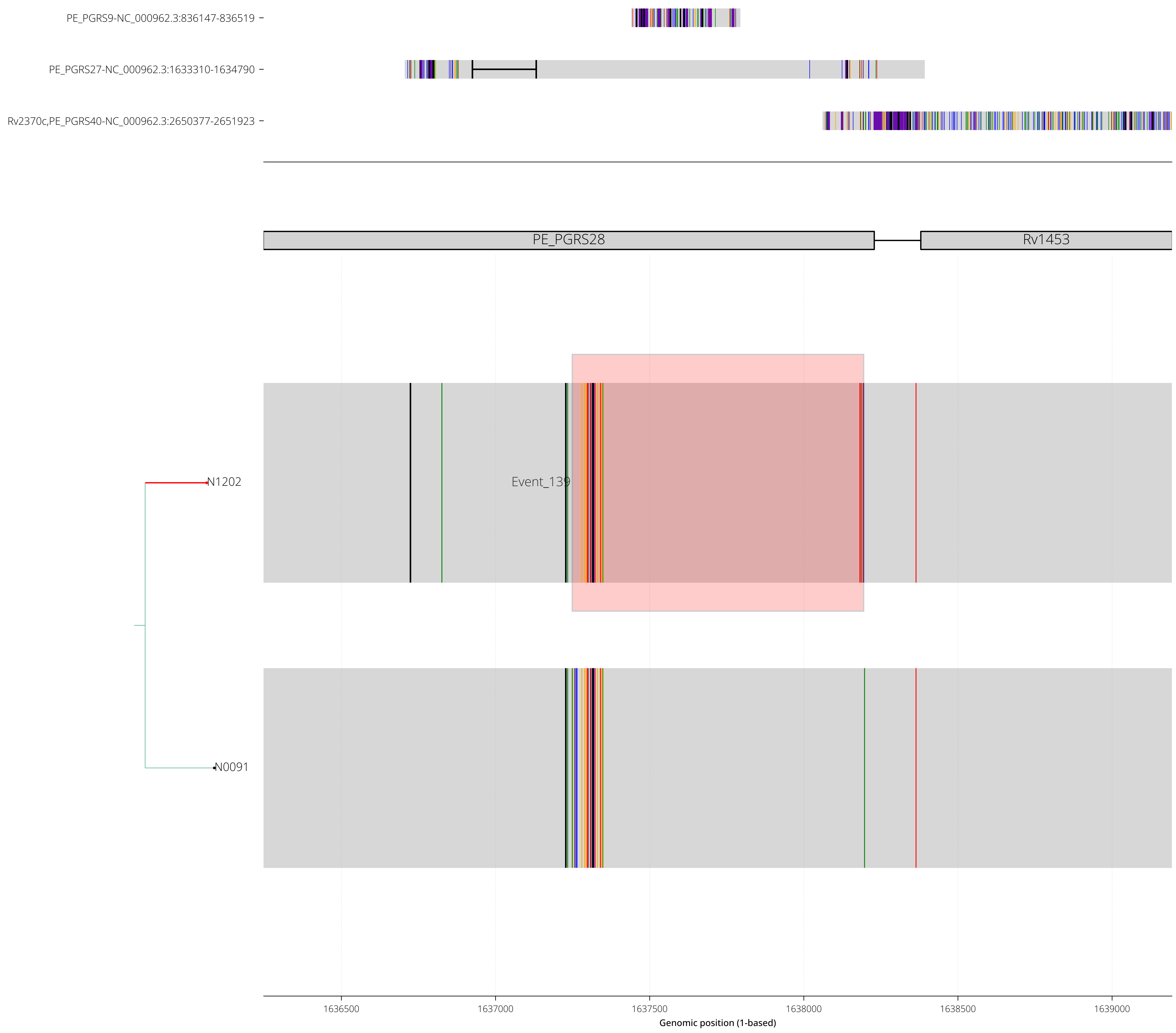

Full Phylogeny Viz + Paralog Alignments
GCE ID: Event_139

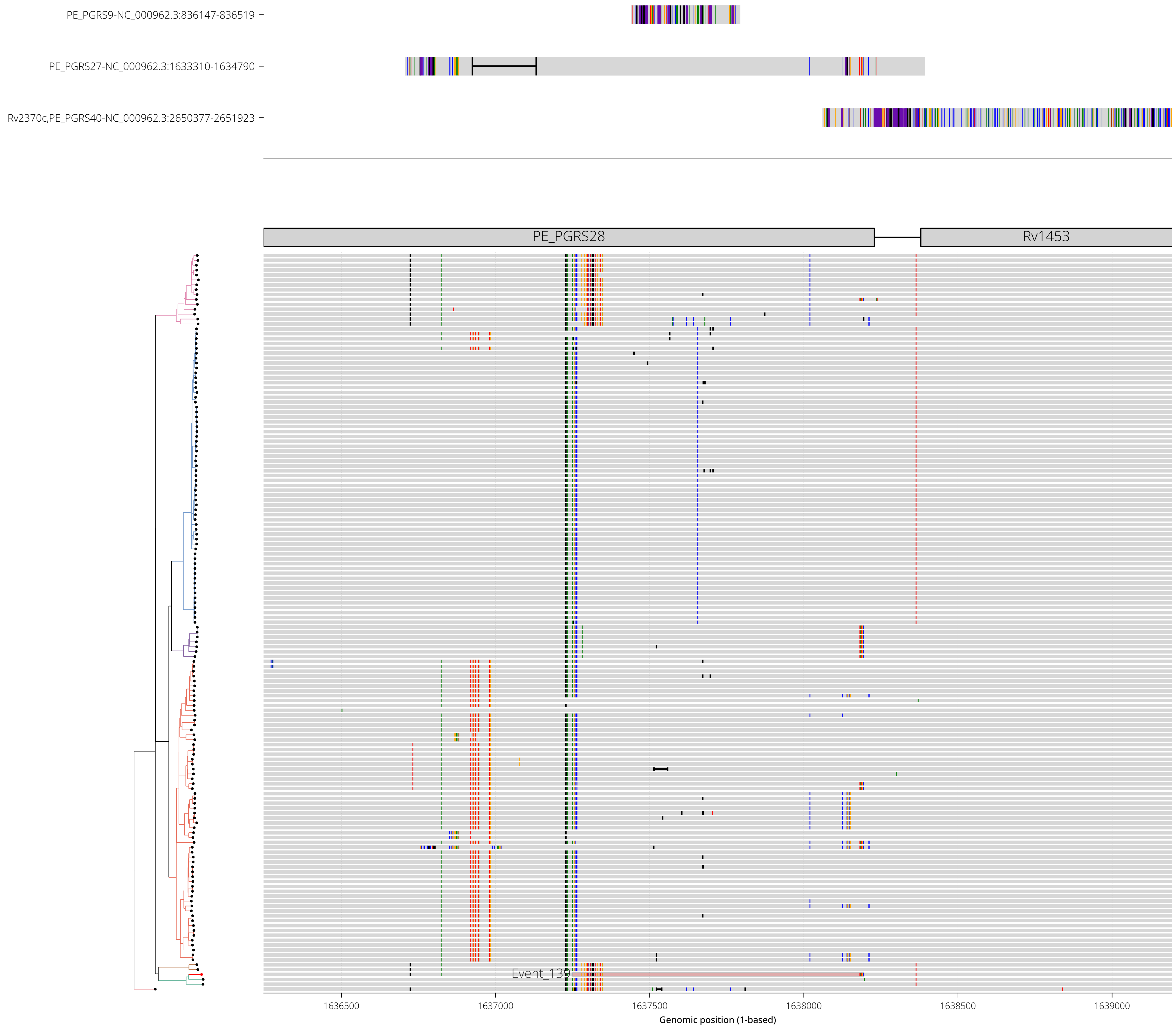

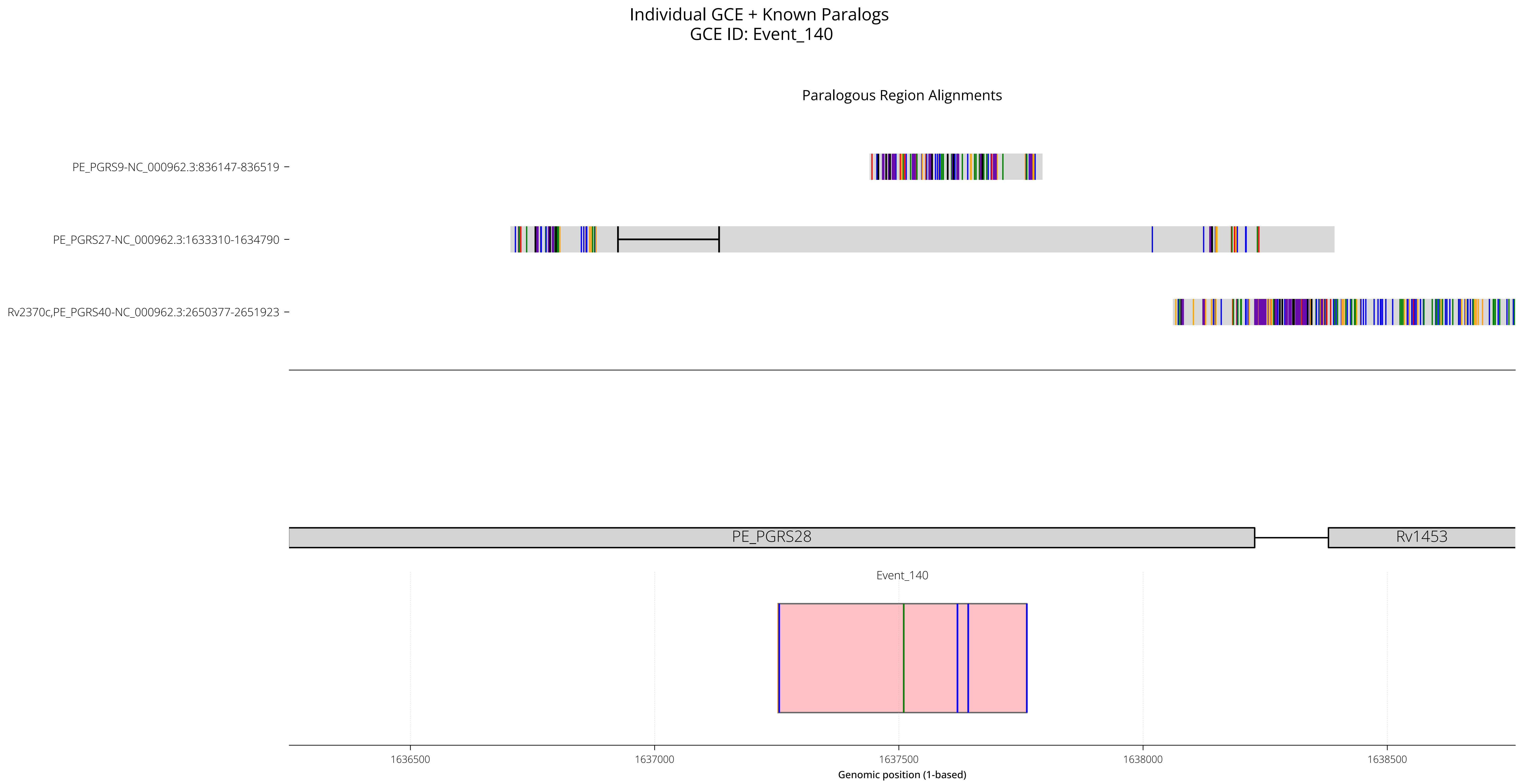

Individual GCE + Known Paralogs
GCE ID: Event_140
Paralogous Region Alignments
PE_PGRS9-NC_000962.3:836147-836519
PE_PGRS27-NC_000962.3:1633310-1634790
Rv2370c,PE_PGRS40-NC_000962.3:2650377-2651923
PE_PGRS28
Rv1453
Event_140
1636500
1637000
1637500
1638000
1638500
Genomic position (1-based)

Circos Viz - GCE to Known Paralogs Locations
GCE ID: Event_140

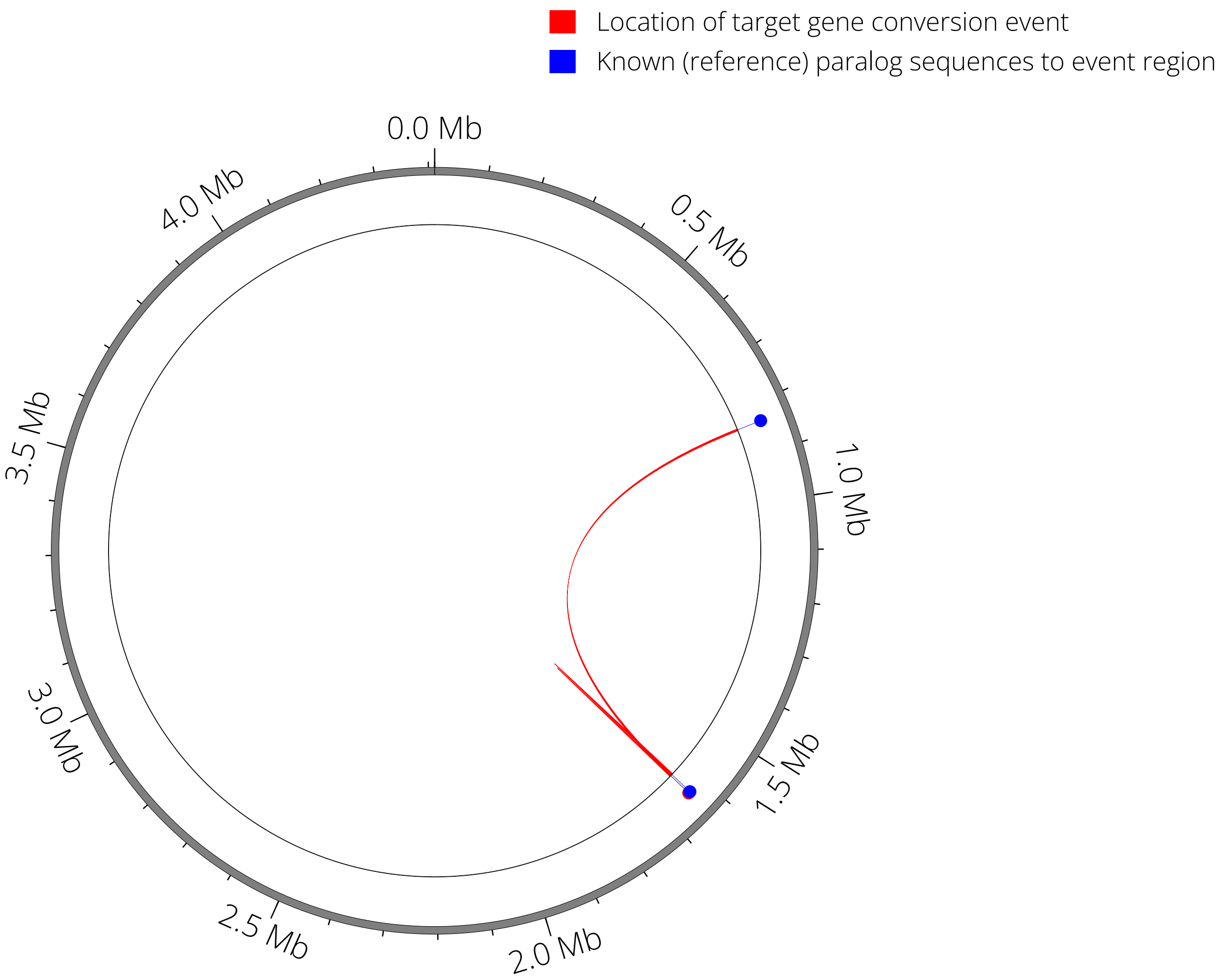

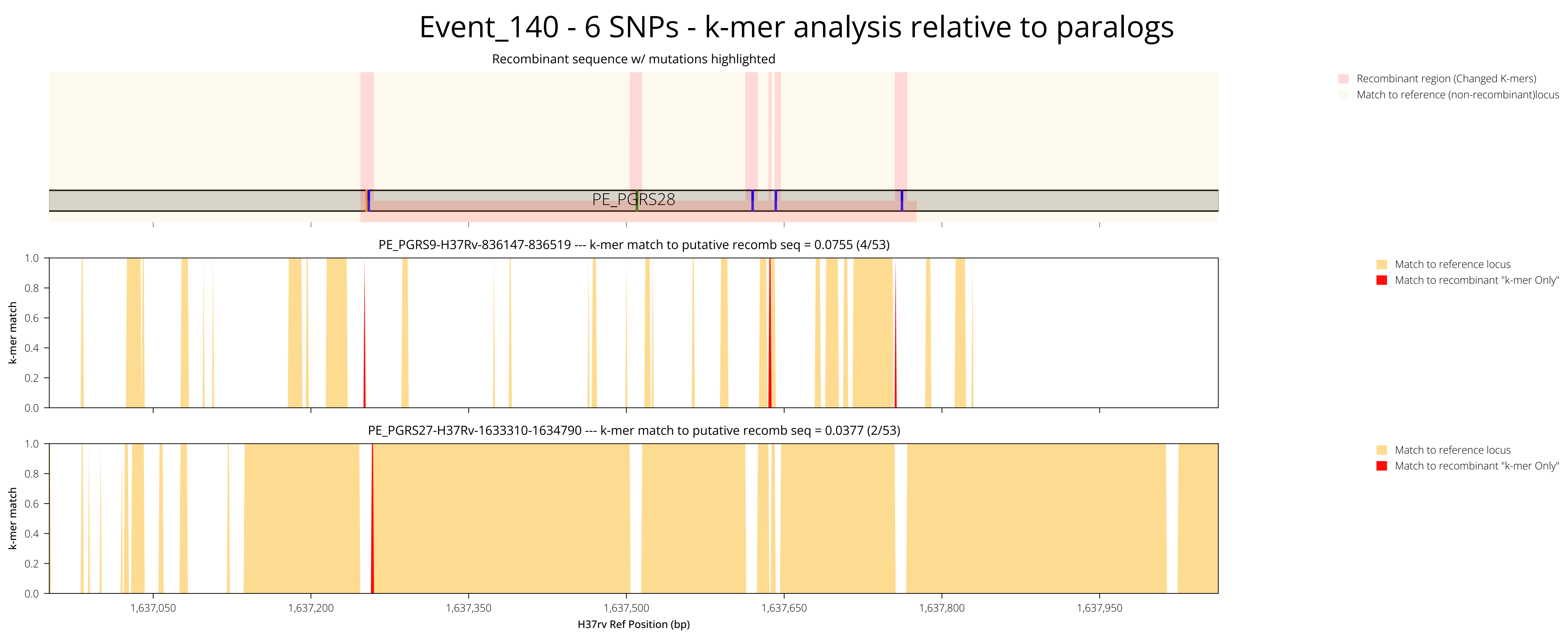

Event_140 - 6 SNPs - k-mer analysis relative to paralogs
Recombinant sequence w/ mutations highlighted
Recombinant region (Changed K-mers)
Match to reference (non-recombinant)locus
PE_PGRS28
PE_PGRS9-H37Rv-836147-836519 --- k-mer match to putative recomb seq = 0.0755 (4/53)
Match to reference locus
Match to recombinant "k-mer Only"
PE_PGRS27-H37Rv-1633310-1634790 --- k-mer match to putative recomb seq = 0.0377 (2/53)
Match to reference locus
Match to recombinant "k-mer Only"
k-mer match
1.0
0.8
0.6
0.4
0.2
0.0
1,637,050
1,637,200
1,637,350
1,637,500
1,637,650
1,637,800
1,637,950
H37rv Ref Position (bp)

Sub-phylogeny Viz + Paralog Alignments
GCE ID: Event_140

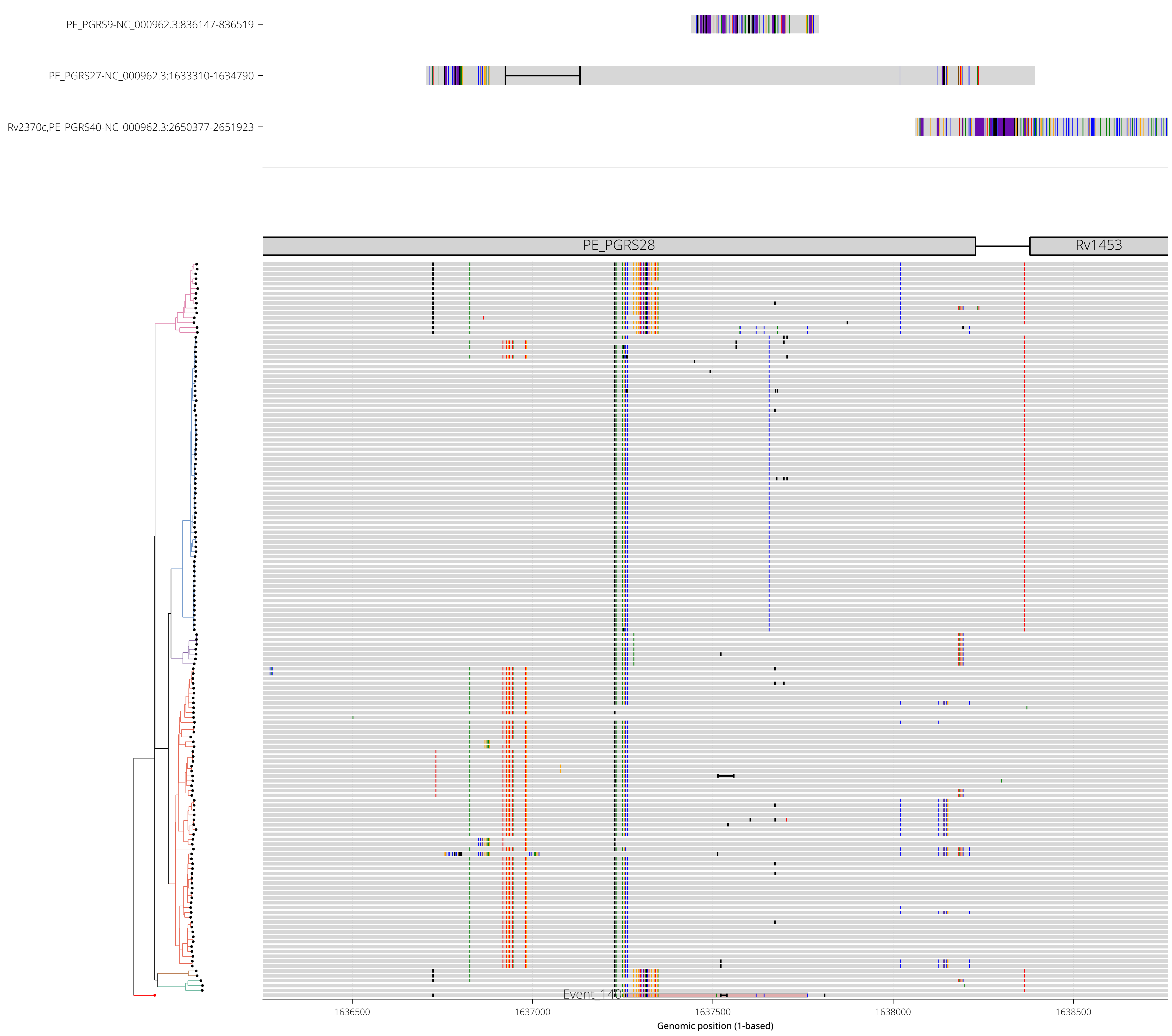

Full Phylogeny Viz + Paralog Alignments
GCE ID: Event_140

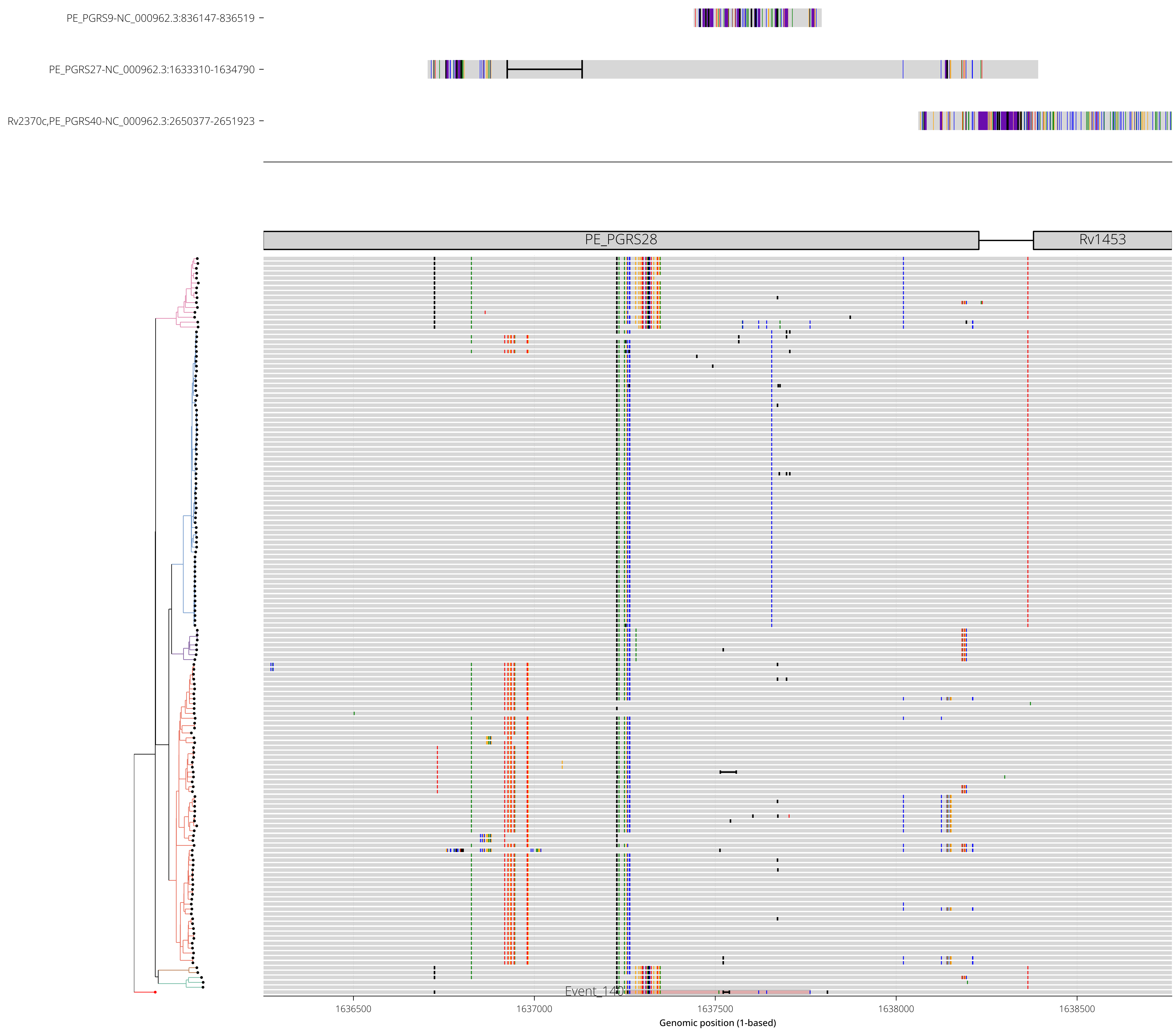

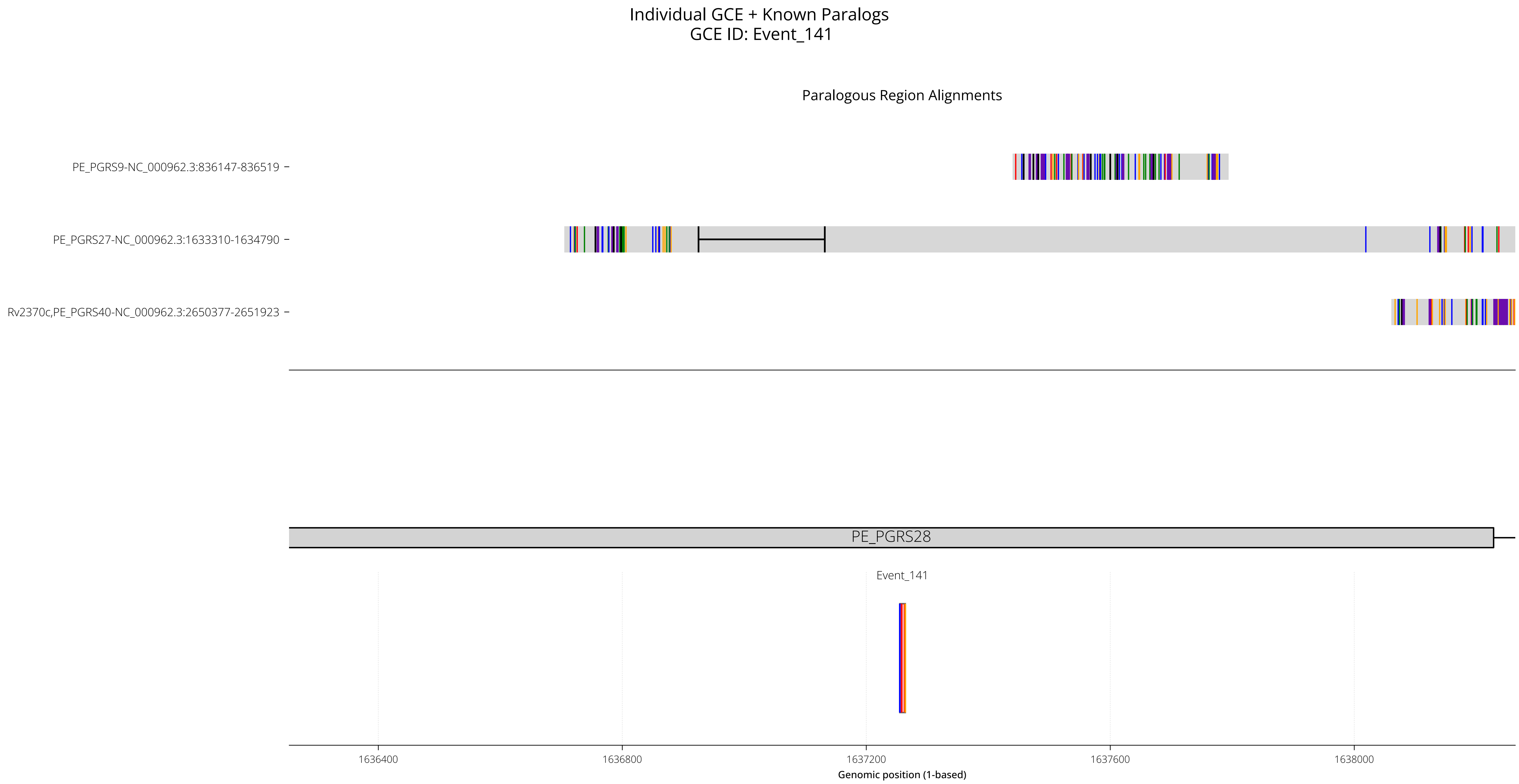

Individual GCE + Known Paralogs
GCE ID: Event_141
Paralogous Region Alignments
PE_PGRS9-NC_000962.3:836147-836519
PE_PGRS27-NC_000962.3:1633310-1634790
Rv2370c,PE_PGRS40-NC_000962.3:2650377-2651923
PE_PGRS28
Event_141
1636400
1636800
1637200
1637600
1638000
Genomic position (1-based)

### Circos Viz - GCE to Known Paralogs Locations
GCE ID: Event_141

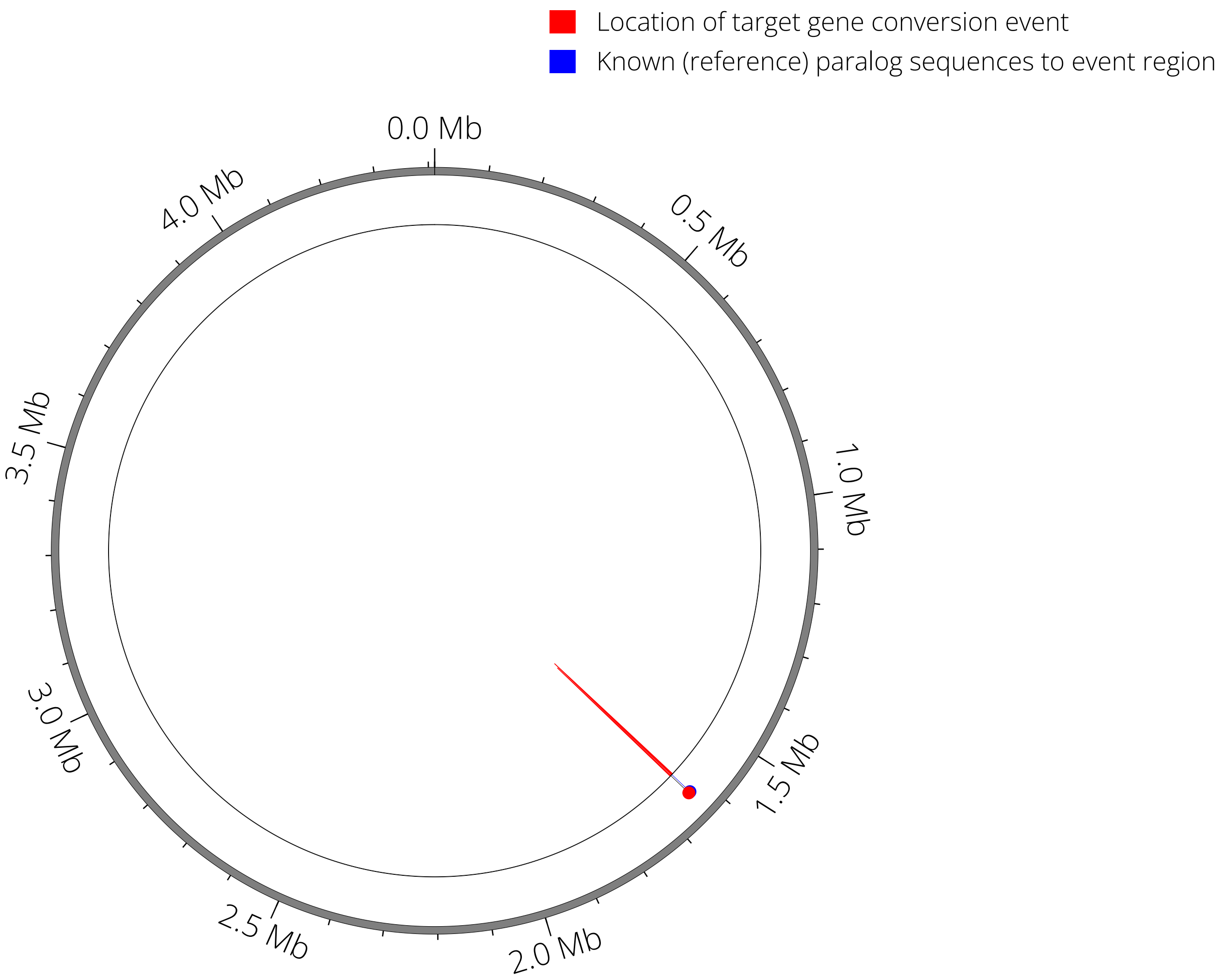

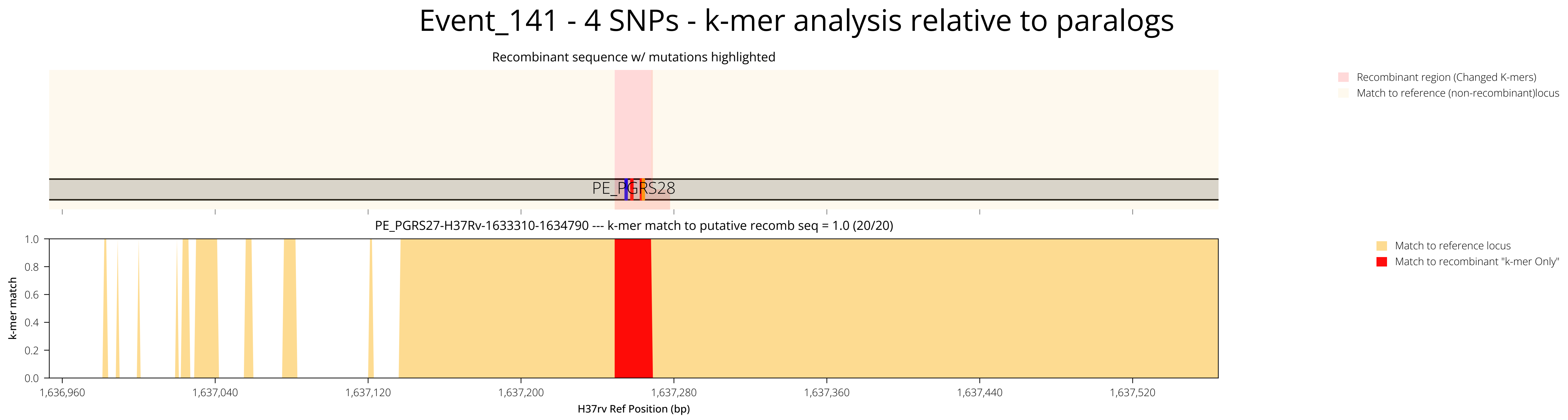

Event_141 - 4 SNPs - k-mer analysis relative to paralogs
Recombinant sequence w/ mutations highlighted
Recombinant region (Changed K-mers)
Match to reference (non-recombinant)locus
PE_PGRS28
PE_PGRS27-H37Rv-1633310-1634790 --- k-mer match to putative recomb seq = 1.0 (20/20)
Match to reference locus
Match to recombinant "k-mer Only"
k-mer match
1.0
0.8
0.6
0.4
0.2
0.0
1,636,960
1,637,040
1,637,120
1,637,200
1,637,280
1,637,360
1,637,440
1,637,520
H37rv Ref Position (bp)

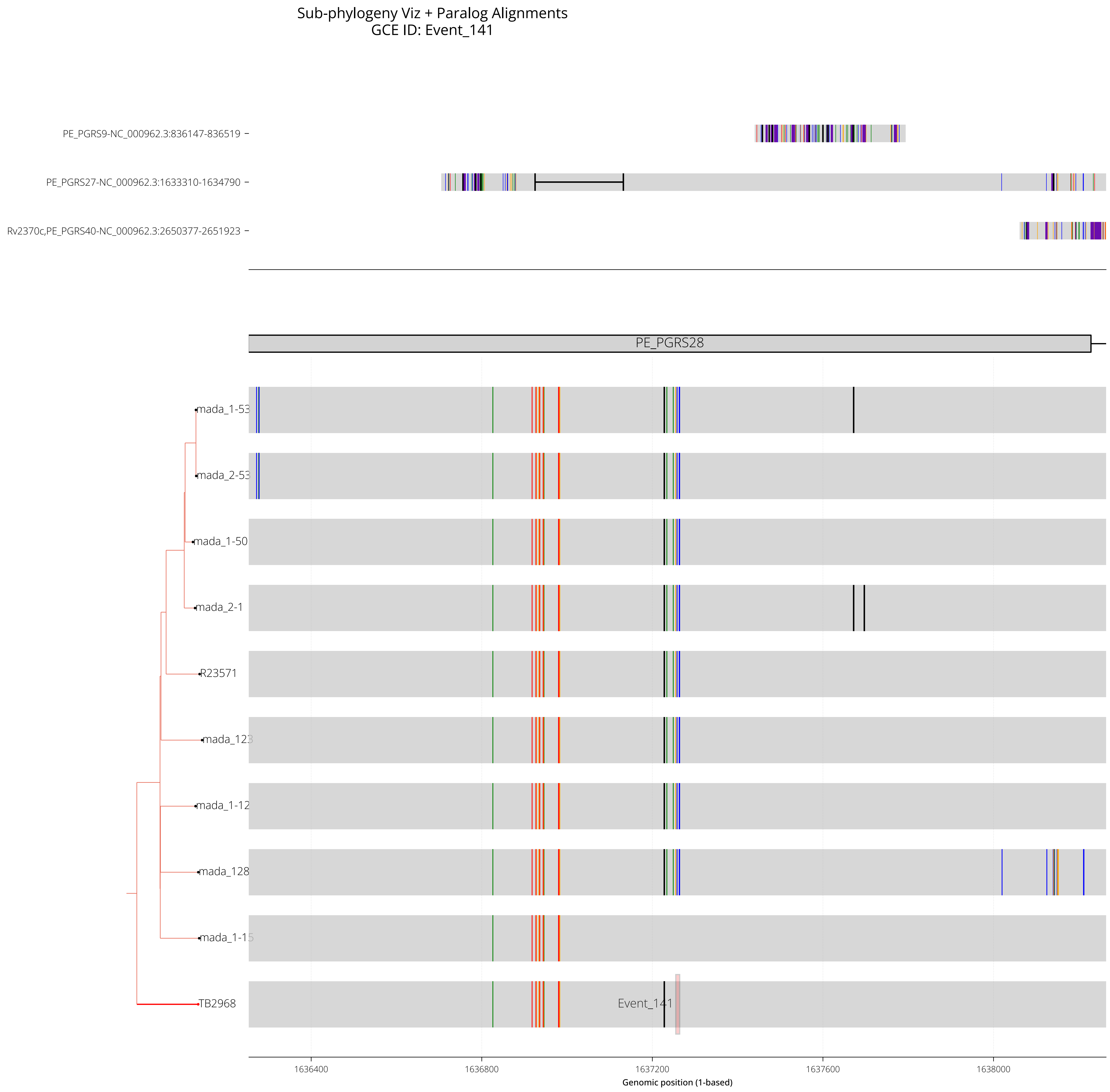
Sub-phylogeny Viz + Paralog Alignments
GCE ID: Event_141
PE_PGRS9-NC_000962.3:836147-836519
PE_PGRS27-NC_000962.3:1633310-1634790
Rv2370c,PE_PGRS40-NC_000962.3:2650377-2651923
PE_PGRS28
mada_1-53
mada_2-53
mada_1-50
mada_2-1
R23571
mada_1-12
mada_128
TB2968
Event_141
1636400
1636800
1637200
1637600
1638000
Genomic position (1-based)

### Full Phylogeny Viz + Paralog Alignments
GCE ID: Event_141

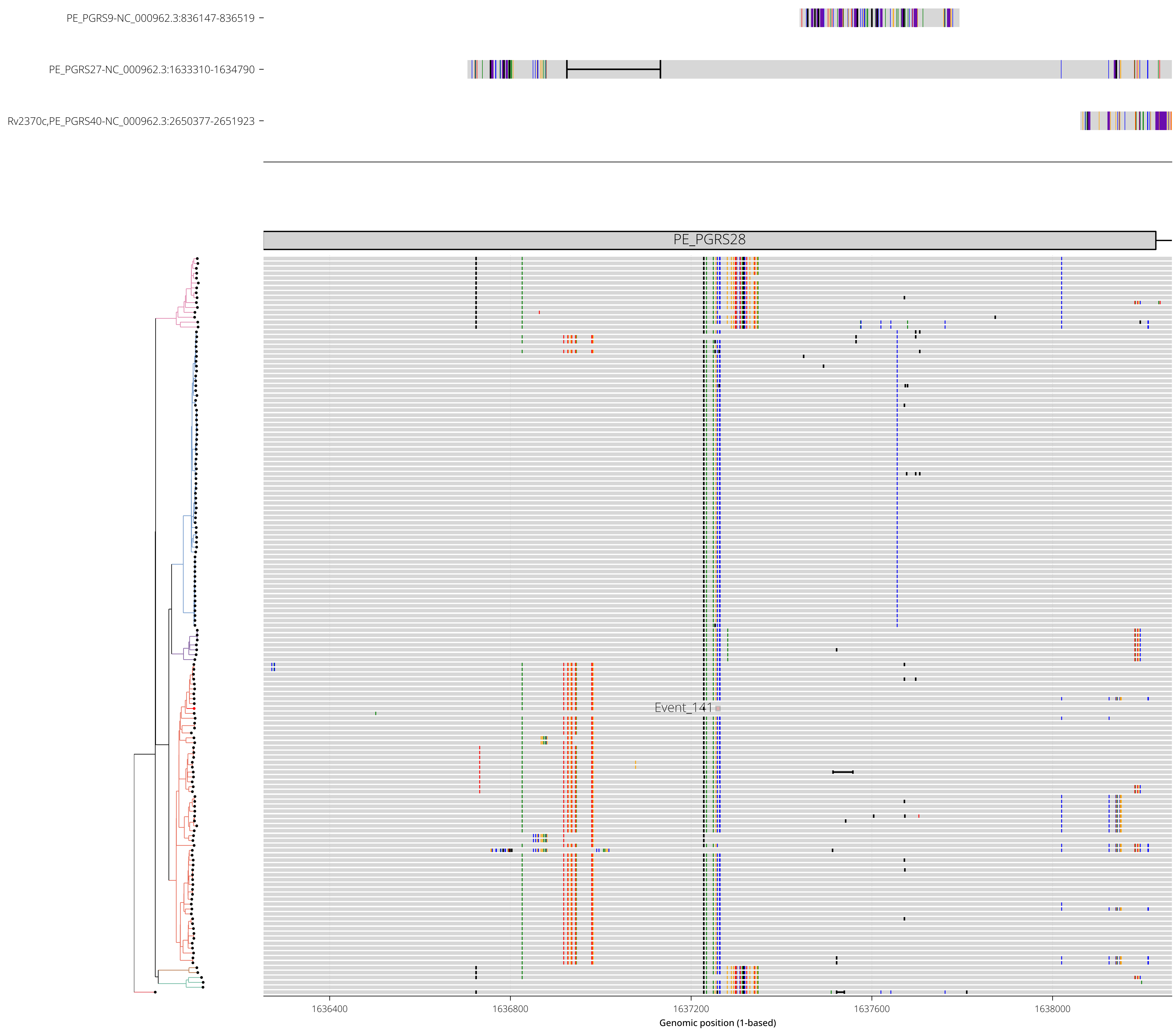

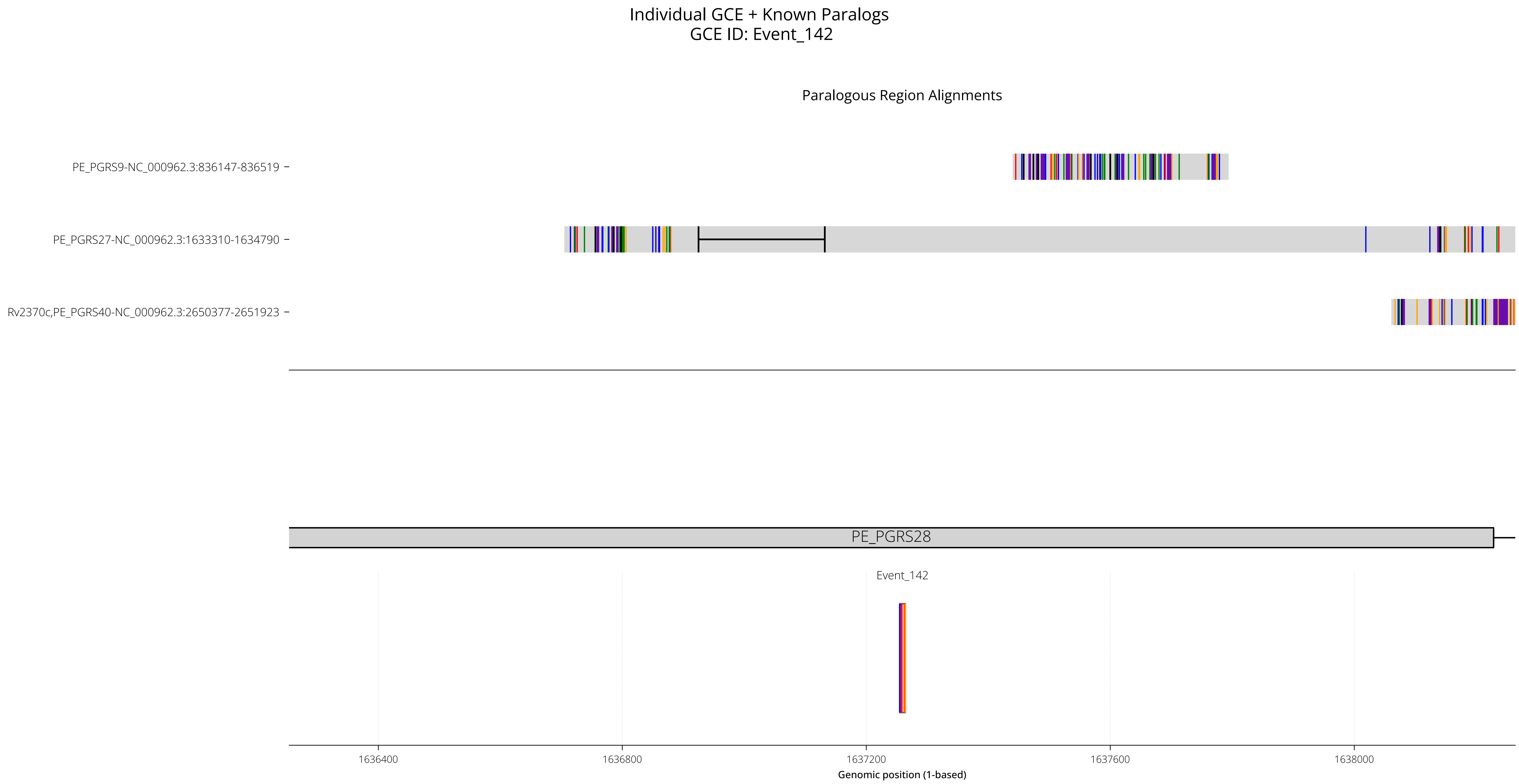

Individual GCE + Known Paralogs
GCE ID: Event_142
Paralogous Region Alignments
PE_PGRS9-NC_000962.3:836147-836519
PE_PGRS27-NC_000962.3:1633310-1634790
Rv2370c,PE_PGRS40-NC_000962.3:2650377-2651923
PE_PGRS28
Event_142
1636400
1636800
1637200
1637600
1638000
Genomic position (1-based)

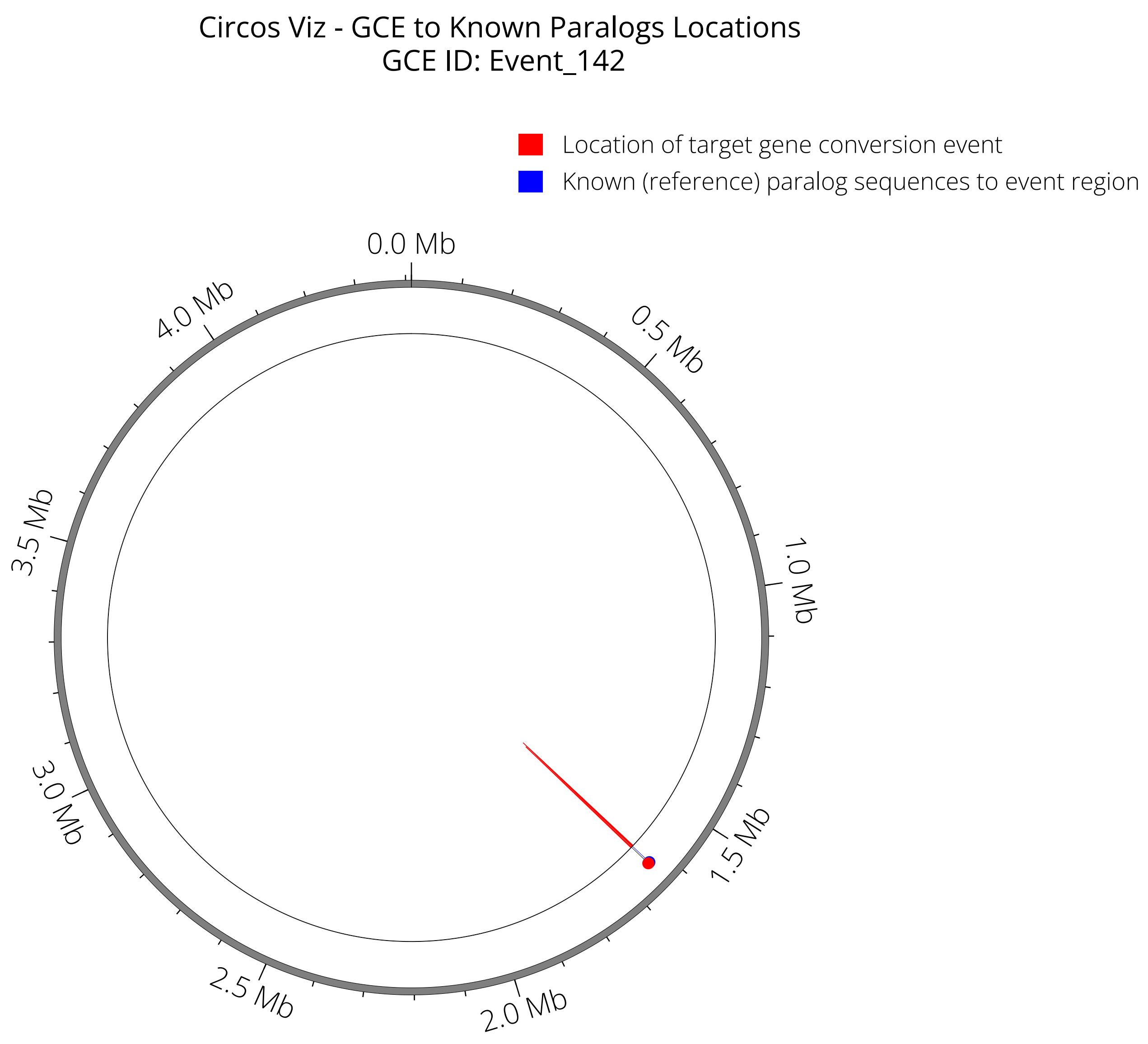

Circos Viz - GCE to Known Paralogs Locations
GCE ID: Event_142
Location of target gene conversion event
Known (reference) paralog sequences to event region
0.0 Mb
0.5 Mb
1.0 Mb
1.5 Mb
2.0 Mb
2.5 Mb
3.0 Mb
3.5 Mb
4.0 Mb

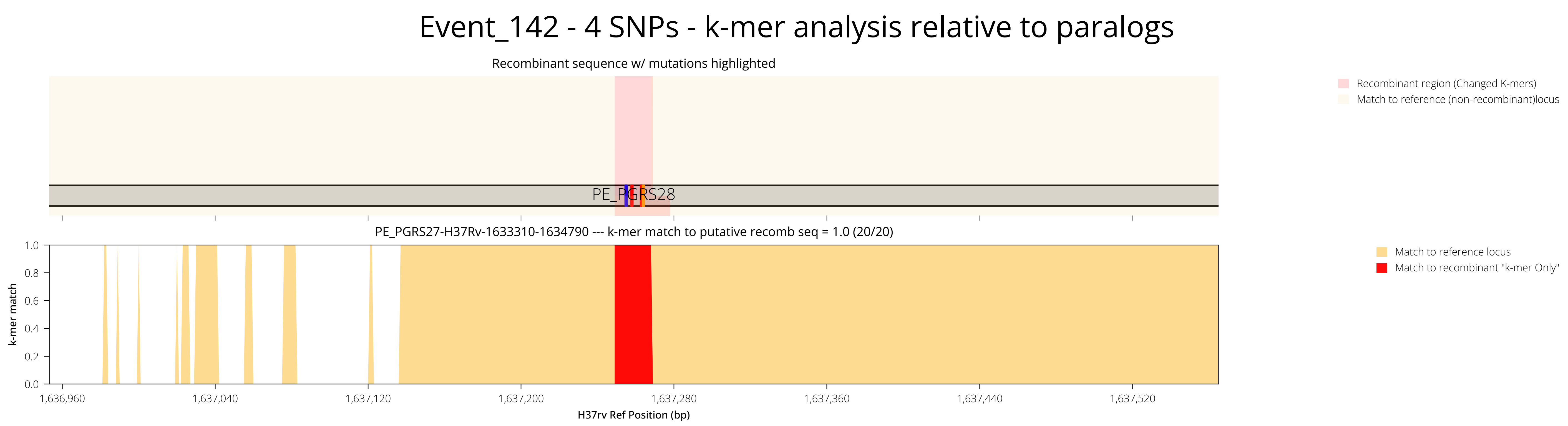

Event_142 - 4 SNPs - k-mer analysis relative to paralogs
Recombinant sequence w/ mutations highlighted
Recombinant region (Changed K-mers)
Match to reference (non-recombinant)locus
PE_PGRS28
PE_PGRS27-H37Rv-1633310-1634790 --- k-mer match to putative recomb seq = 1.0 (20/20)
Match to reference locus
Match to recombinant "k-mer Only"
k-mer match
1.0
0.8
0.6
0.4
0.2
0.0
1,636,960
1,637,040
1,637,120
1,637,200
1,637,280
1,637,360
1,637,440
1,637,520
H37rv Ref Position (bp)

Sub-phylogeny Viz + Paralog Alignments
GCE ID: Event_142

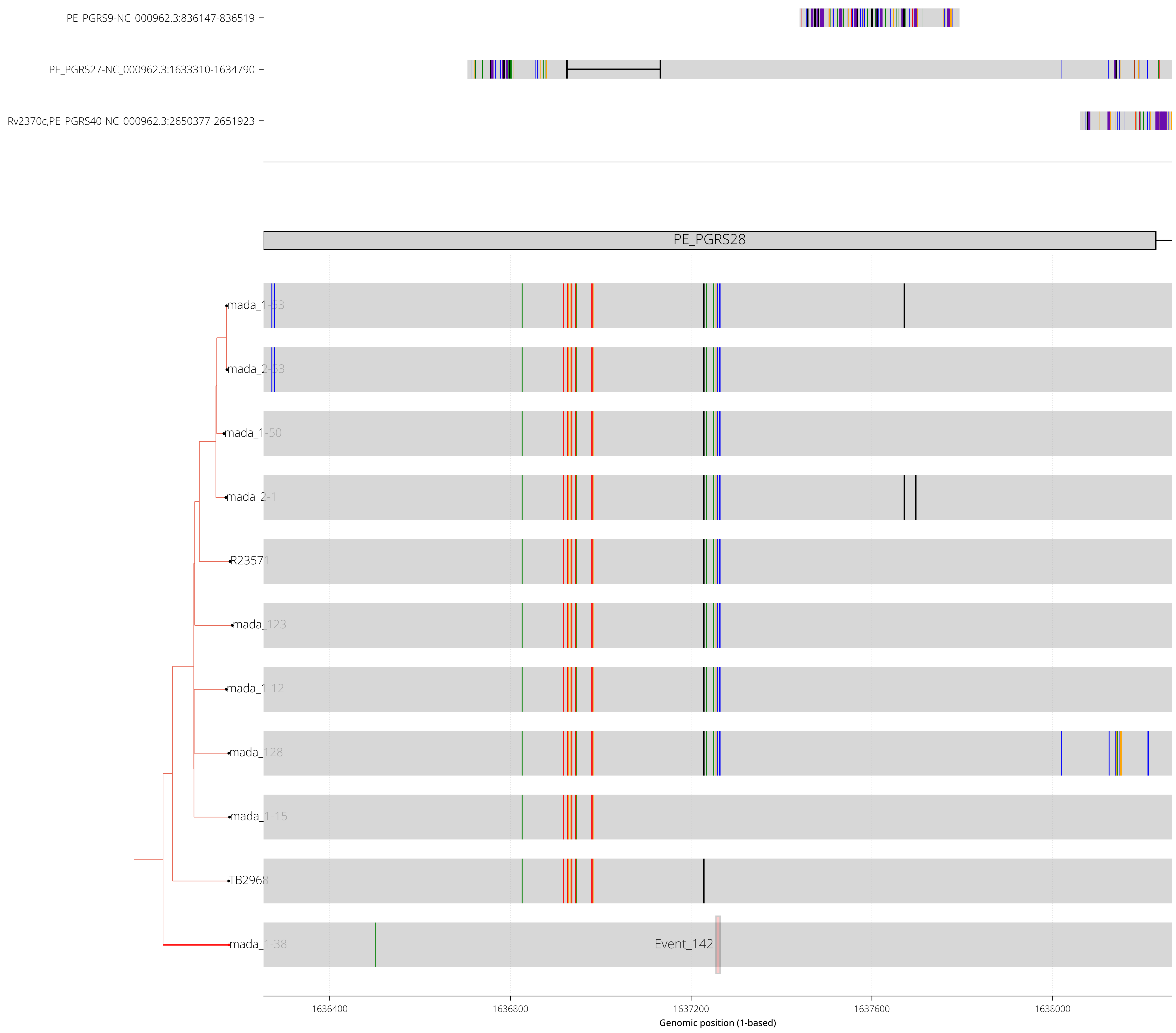

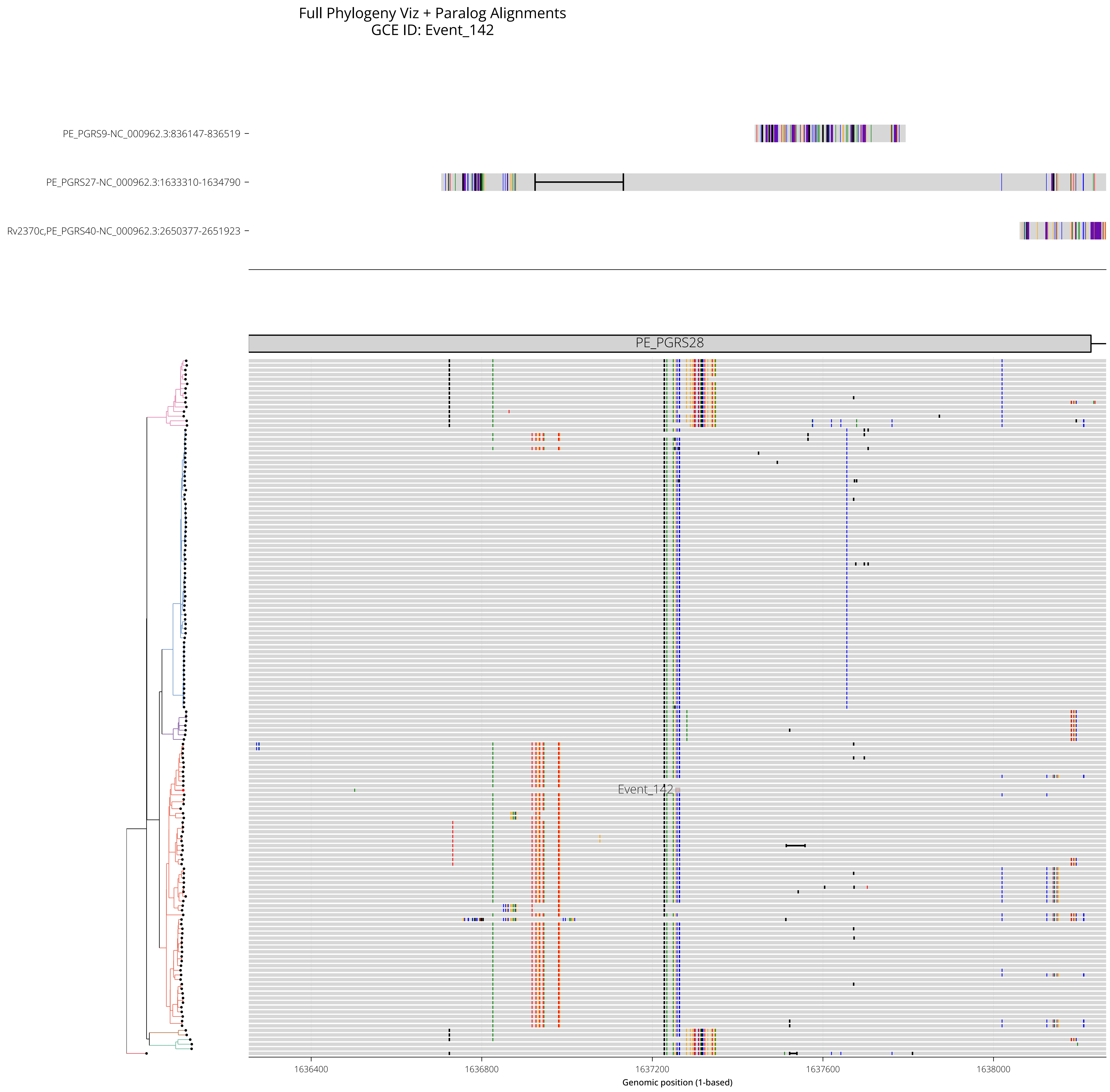

Full Phylogeny Viz + Paralog Alignments
GCE ID: Event_142
PE_PGRS9-NC_000962.3:836147-836519
PE_PGRS27-NC_000962.3:1633310-1634790
Rv2370c,PE_PGRS40-NC_000962.3:2650377-2651923
PE_PGRS28
Event_142
1636400
1636800
1637200
1637600
1638000
Genomic position (1-based)

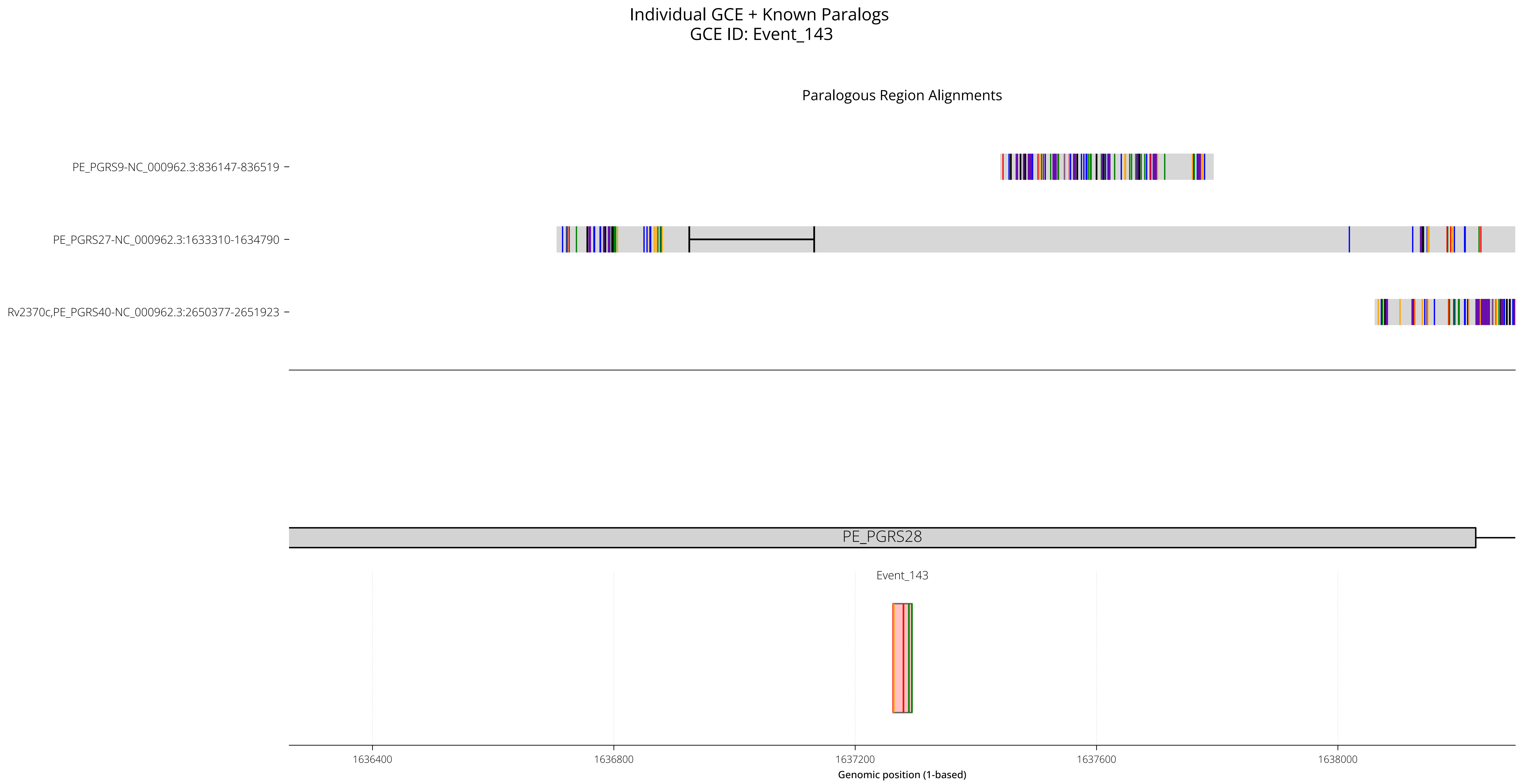

Individual GCE + Known Paralogs
GCE ID: Event_143
Paralogous Region Alignments
PE_PGRS9-NC_000962.3:836147-836519
PE_PGRS27-NC_000962.3:1633310-1634790
Rv2370c,PE_PGRS40-NC_000962.3:2650377-2651923
PE_PGRS28
Event_143
1636400
1636800
1637200
1637600
1638000
Genomic position (1-based)

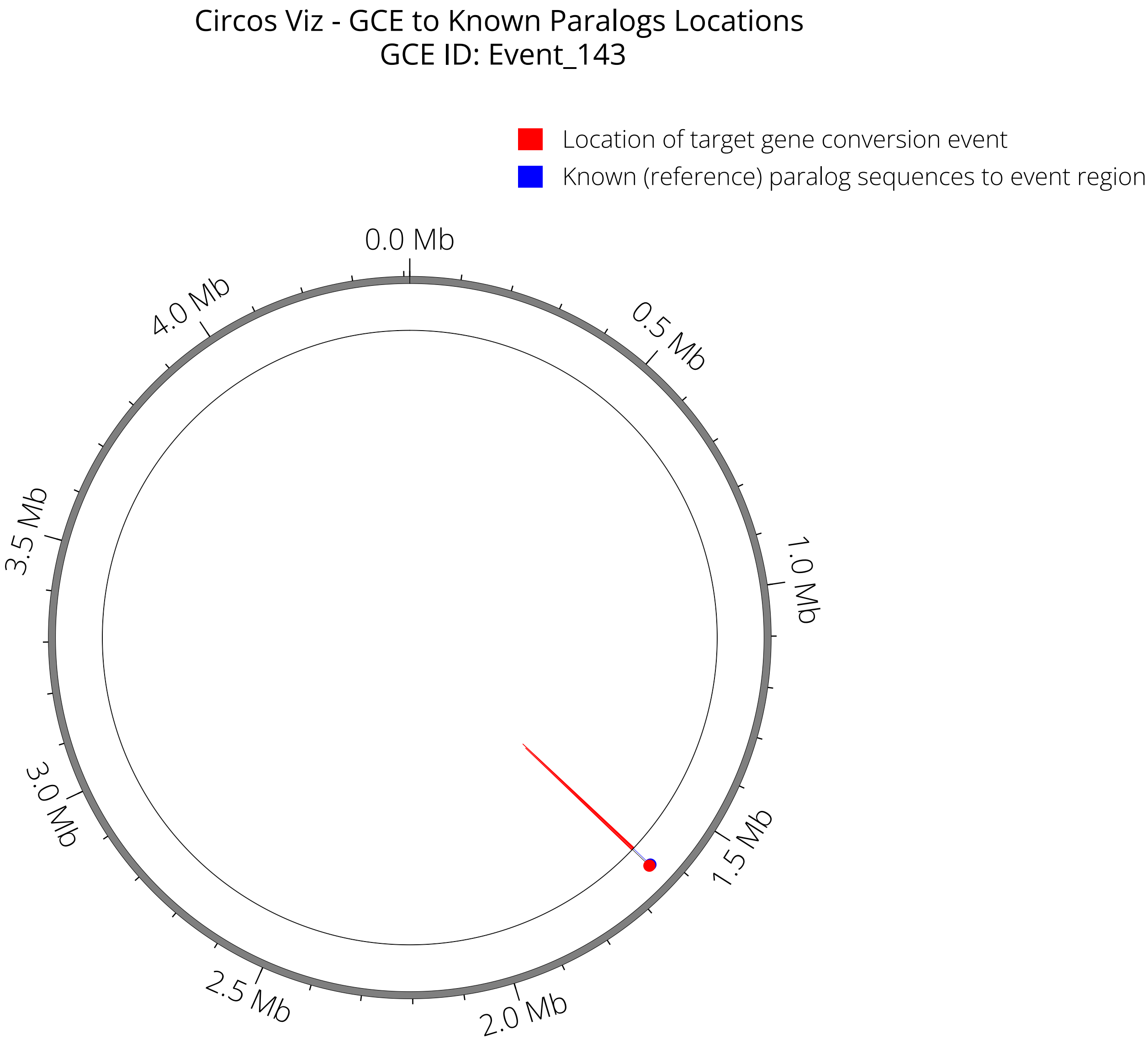
Circos Viz - GCE to Known Paralogs Locations
GCE ID: Event_143
Location of target gene conversion event
Known (reference) paralog sequences to event region
0.0 Mb
0.5 Mb
1.0 Mb
1.5 Mb
2.0 Mb
2.5 Mb
3.0 Mb
3.5 Mb
4.0 Mb

### Event_143 - 5 SNPs - k-mer analysis relative to paralogs

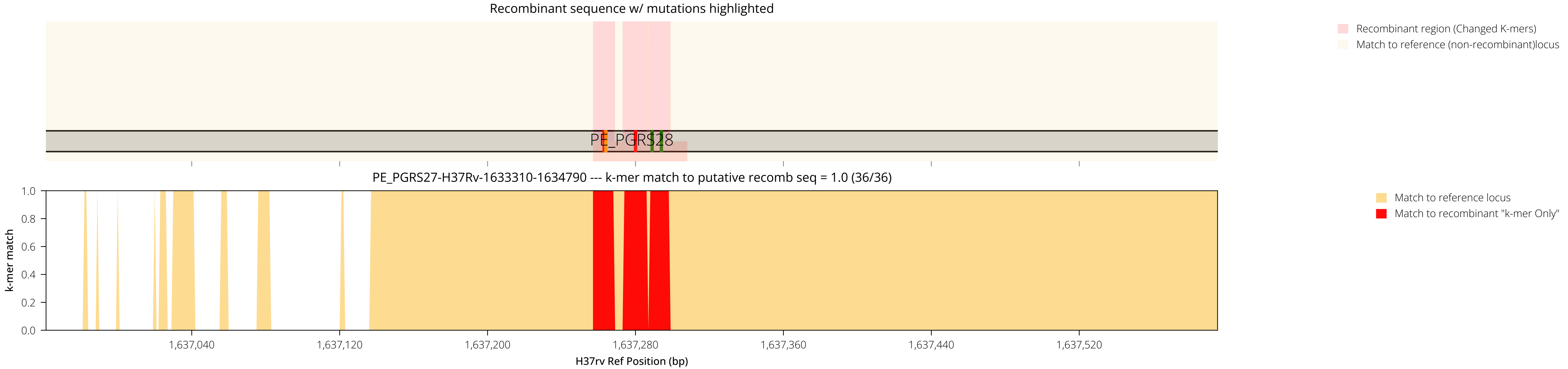

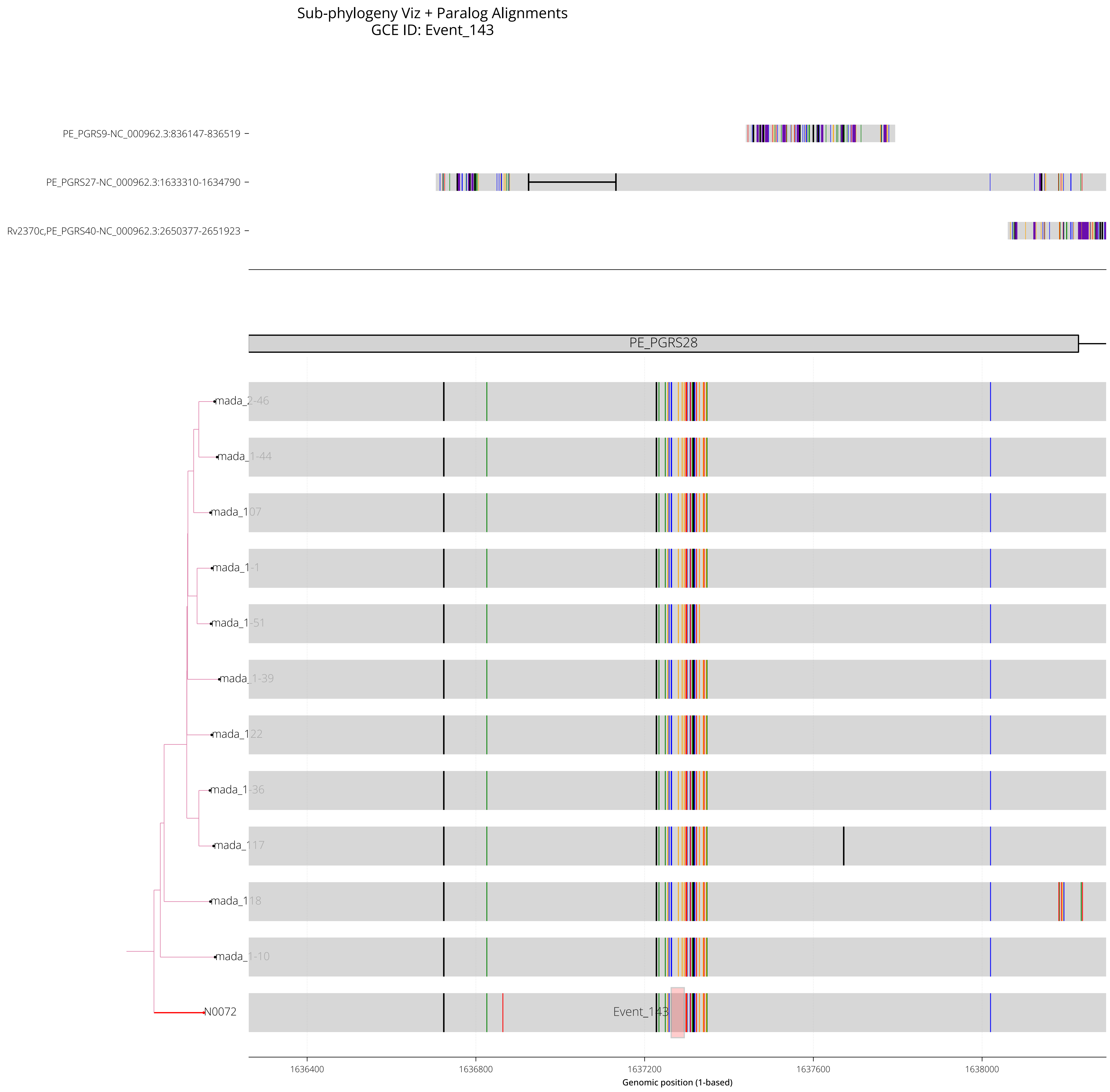
Sub-phylogeny Viz + Paralog Alignments
GCE ID: Event_143
PE_PGRS9-NC_000962.3:836147-836519
PE_PGRS27-NC_000962.3:1633310-1634790
Rv2370c,PE_PGRS40-NC_000962.3:2650377-2651923
PE_PGRS28
mada_2-46
mada_1-44
mada_107
mada_1-1
mada_1-51
mada_1-39
mada_122
mada_1-36
mada_117
mada_118
mada_1-10
N0072
Event_143
1636400
1636800
1637200
1637600
1638000
Genomic position (1-based)

Full Phylogeny Viz + Paralog Alignments
GCE ID: Event_143

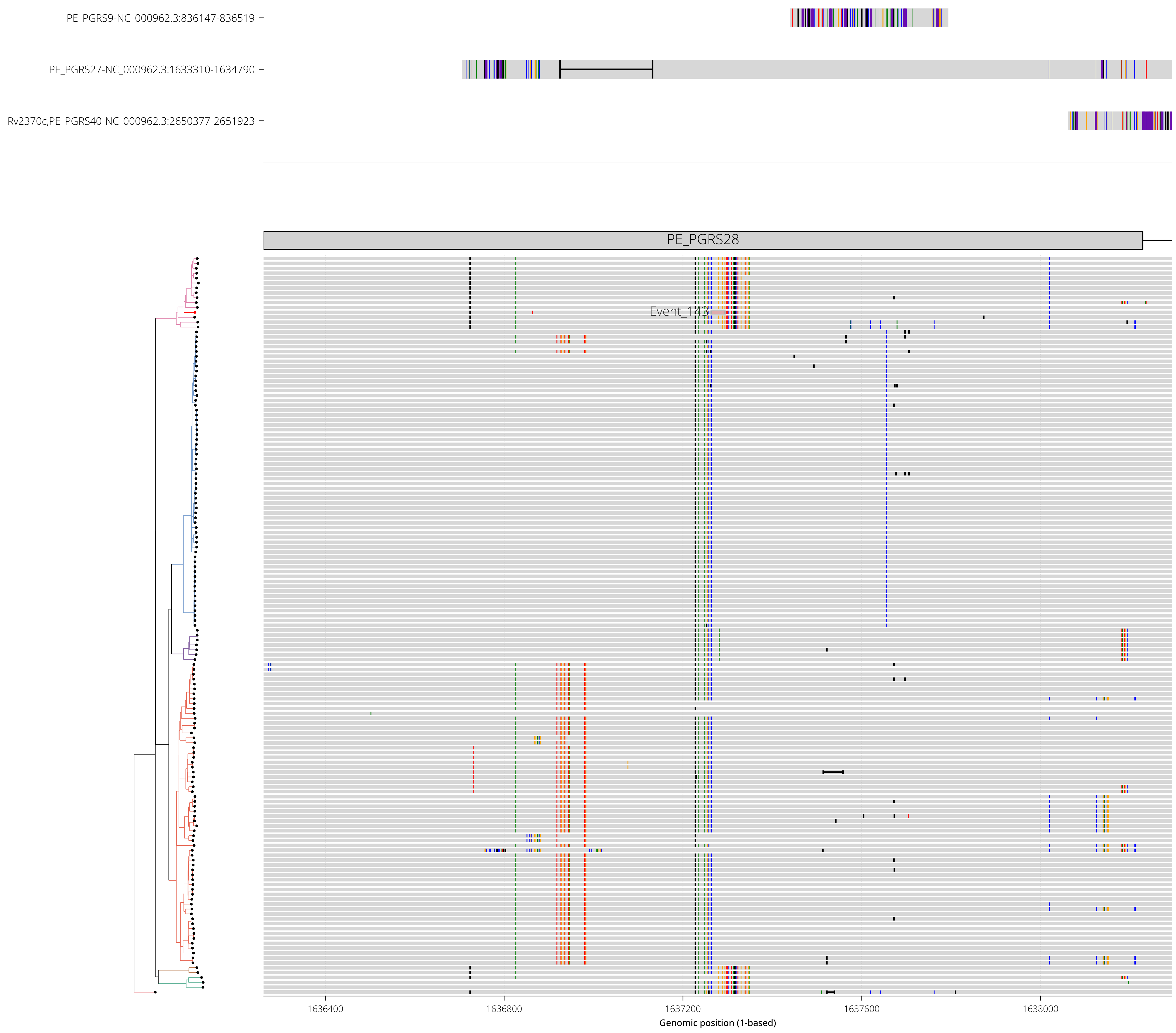

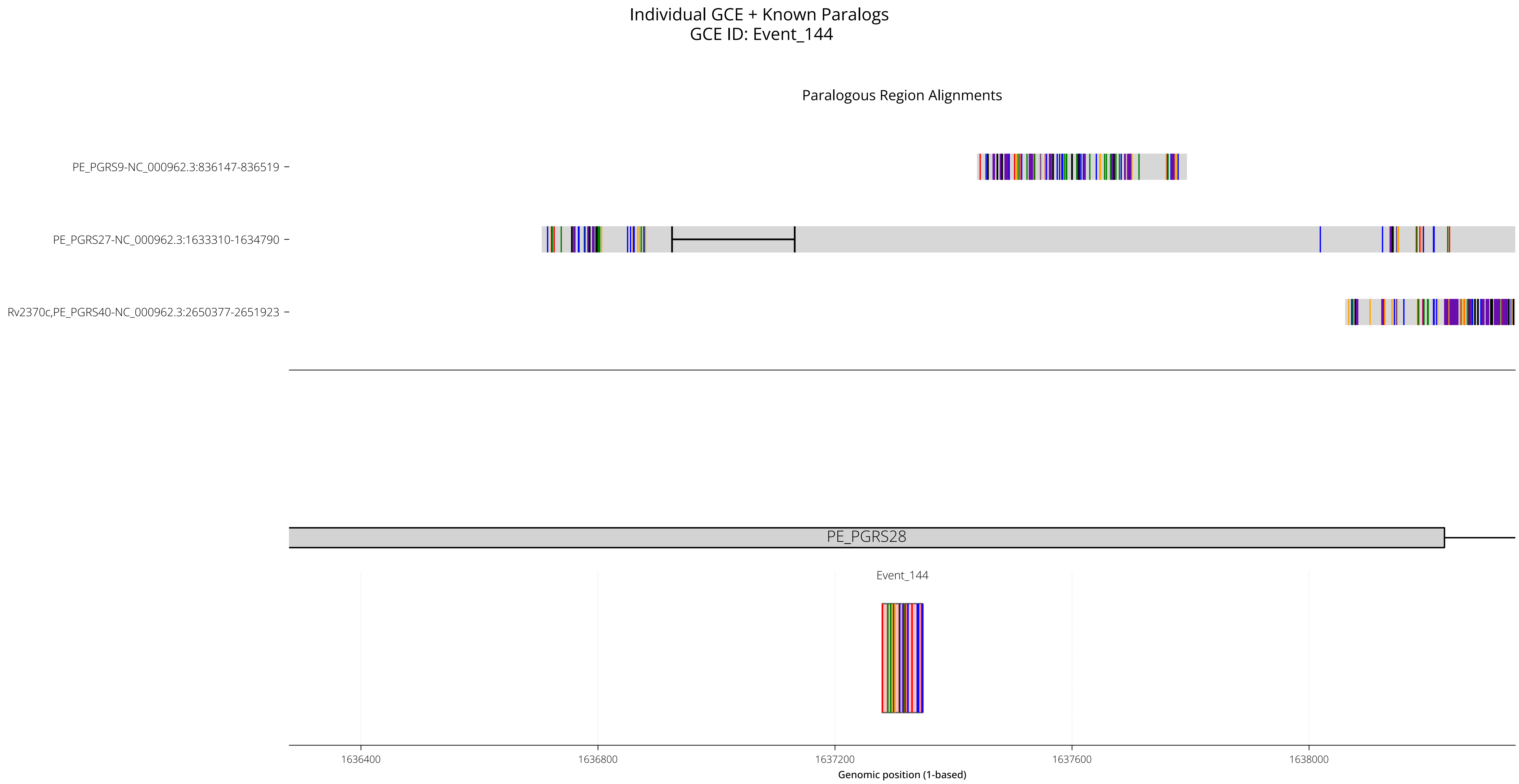

Individual GCE + Known Paralogs
GCE ID: Event_144
Paralogous Region Alignments
PE_PGRS9-NC_000962.3:836147-836519
PE_PGRS27-NC_000962.3:1633310-1634790
Rv2370c,PE_PGRS40-NC_000962.3:2650377-2651923
PE_PGRS28
Event_144
1636400
1636800
1637200
1637600
1638000
Genomic position (1-based)

Circos Viz - GCE to Known Paralogs Locations
GCE ID: Event_144

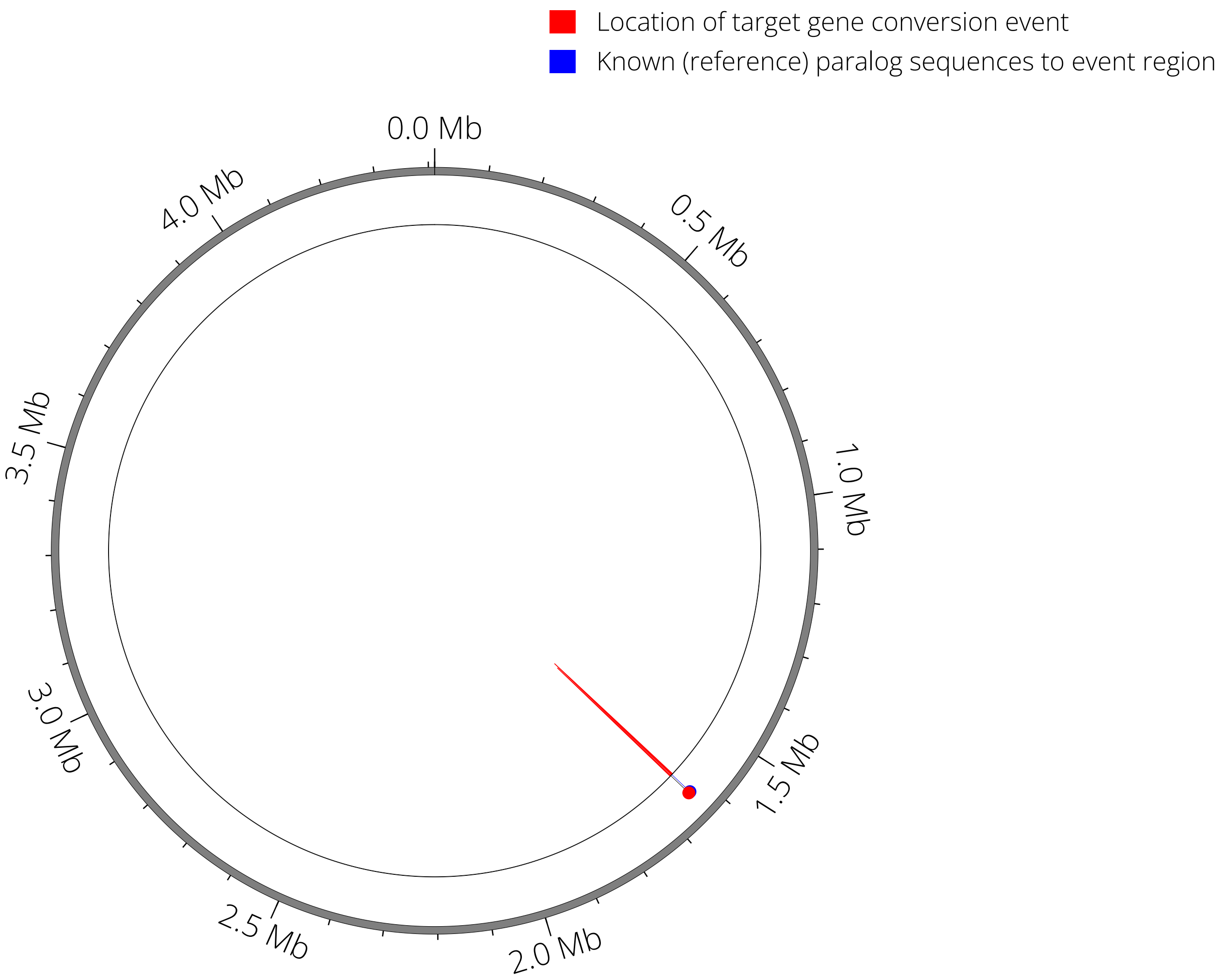

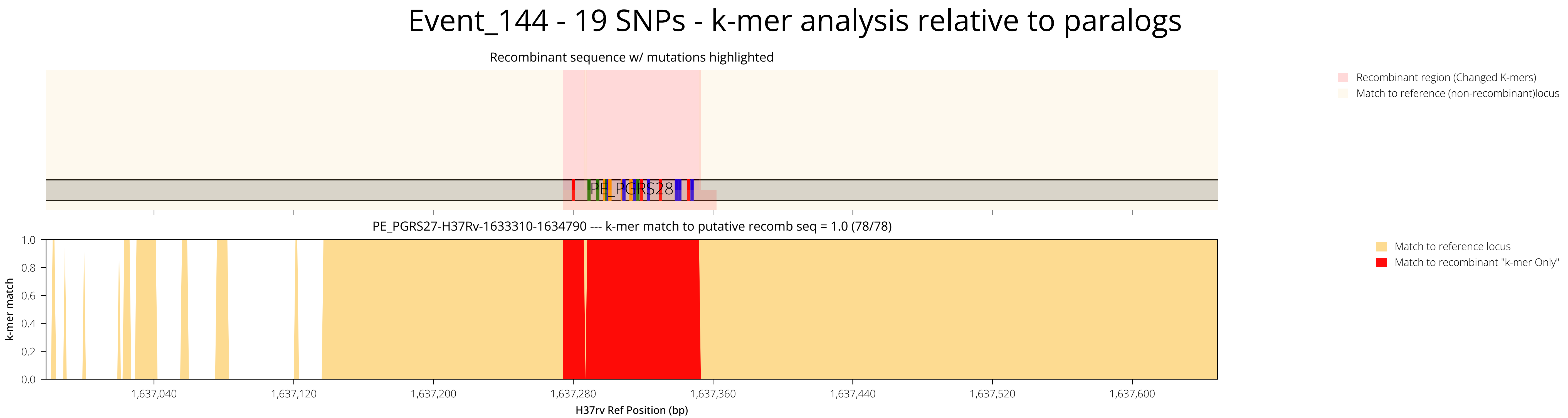
Event_144 - 19 SNPs - k-mer analysis relative to paralogs
Recombinant sequence w/ mutations highlighted
Recombinant region (Changed K-mers)
Match to reference (non-recombinant)locus
PE_PGRS28
PE_PGRS27-H37Rv-1633310-1634790 --- k-mer match to putative recomb seq = 1.0 (78/78)
Match to reference locus
Match to recombinant "k-mer Only"
k-mer match
1.0
0.8
0.6
0.4
0.2
0.0
1,637,040
1,637,120
1,637,200
1,637,280
1,637,360
1,637,440
1,637,520
1,637,600
H37rv Ref Position (bp)

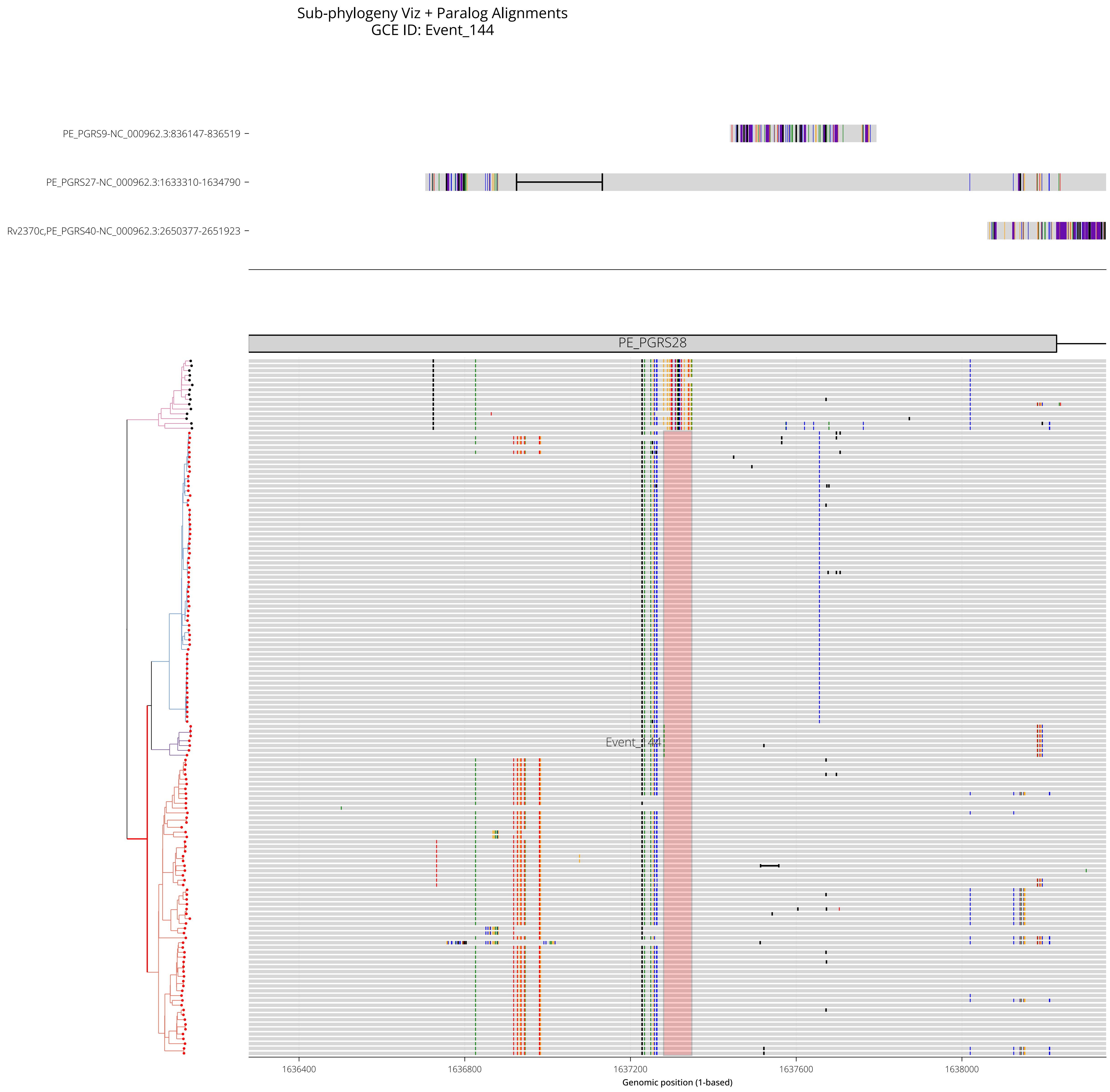

Sub-phylogeny Viz + Paralog Alignments
GCE ID: Event_144
PE_PGRS9-NC_000962.3:836147-836519
PE_PGRS27-NC_000962.3:1633310-1634790
Rv2370c,PE_PGRS40-NC_000962.3:2650377-2651923
PE_PGRS28
Event_144
1636400
1636800
1637200
1637600
1638000
Genomic position (1-based)

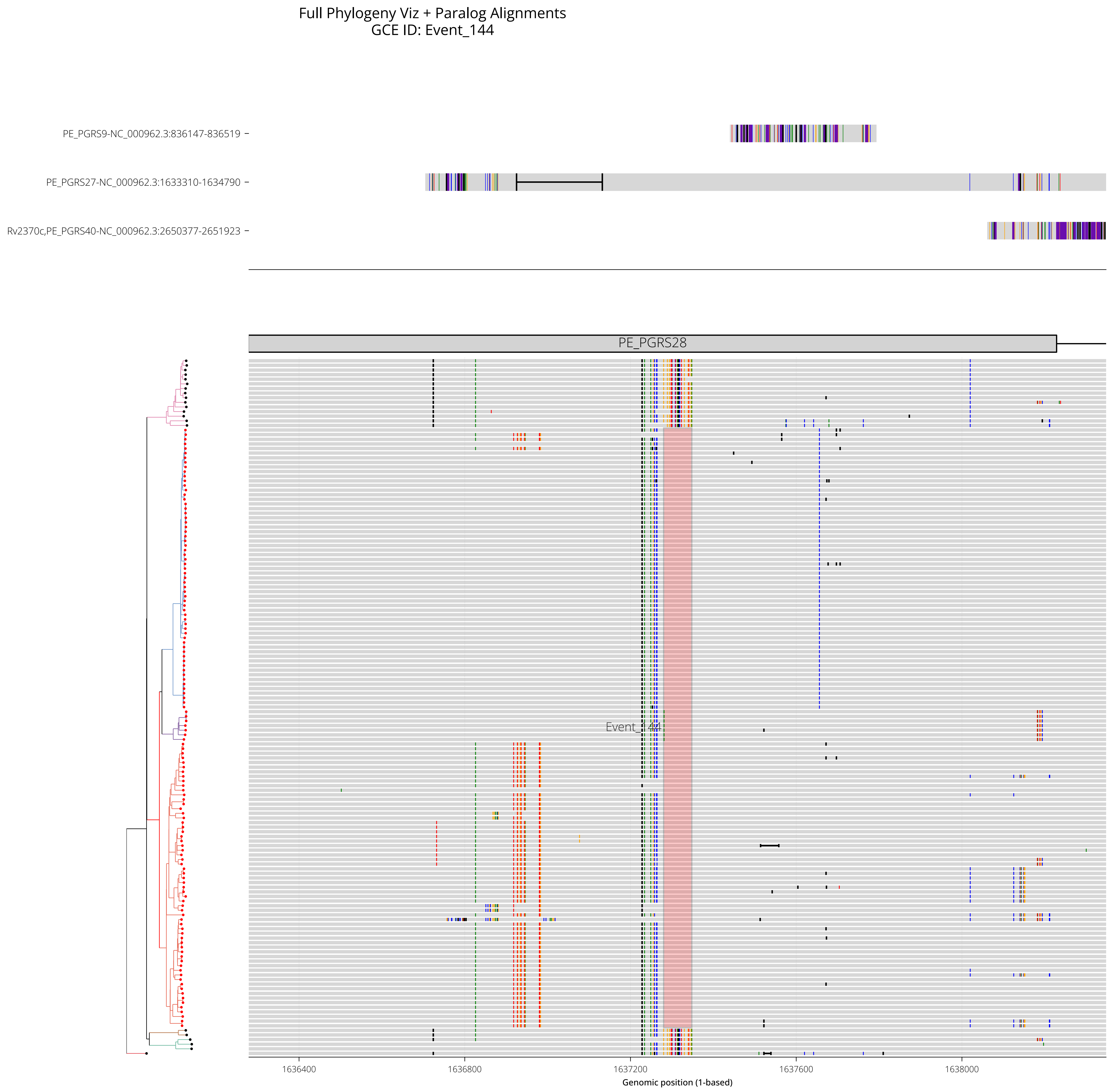
Full Phylogeny Viz + Paralog Alignments
GCE ID: Event_144
PE_PGRS9-NC_000962.3:836147-836519
PE_PGRS27-NC_000962.3:1633310-1634790
Rv2370c,PE_PGRS40-NC_000962.3:2650377-2651923
PE_PGRS28
Event_144
1636400
1636800
1637200
1637600
1638000
Genomic position (1-based)

Individual GCE + Known Paralogs
GCE ID: Event_145

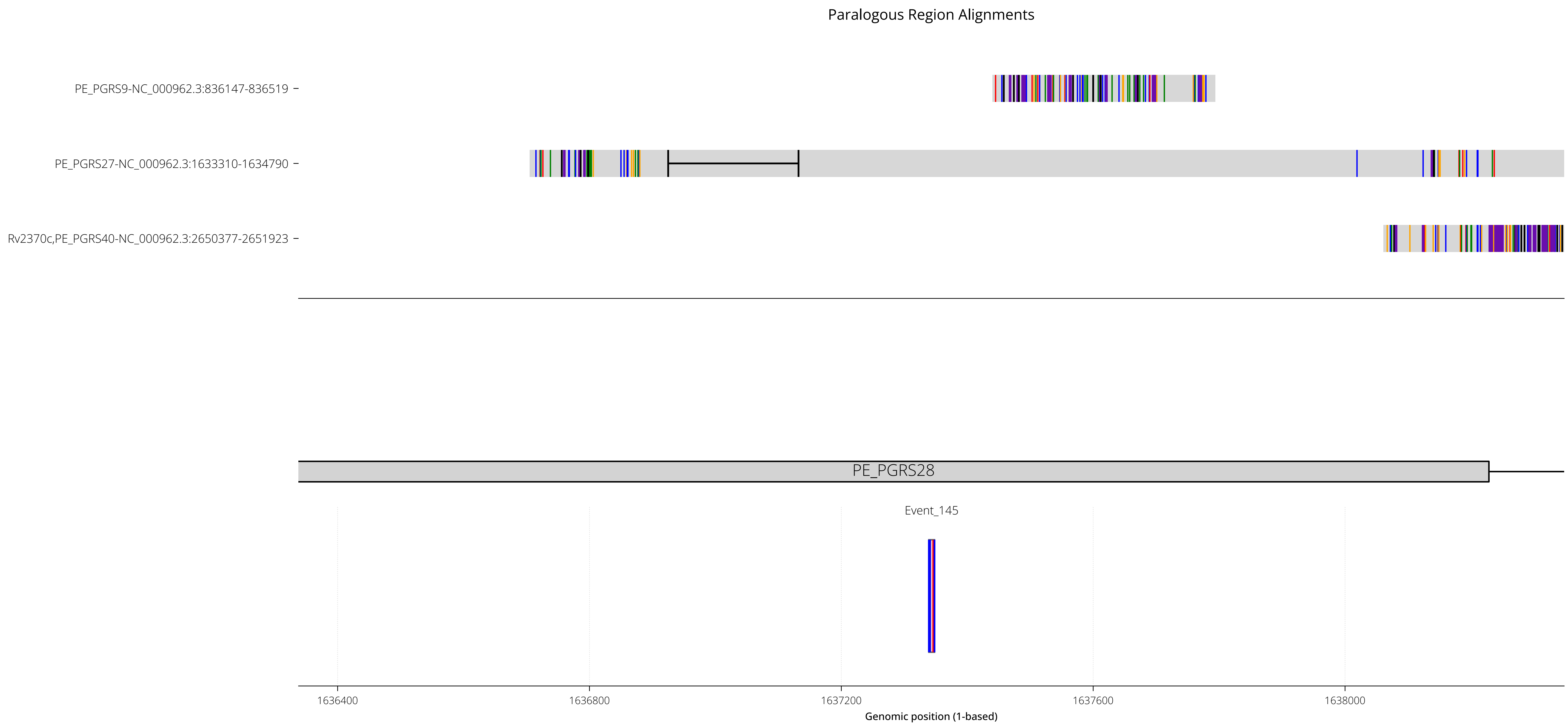

Circos Viz - GCE to Known Paralogs Locations
GCE ID: Event_145

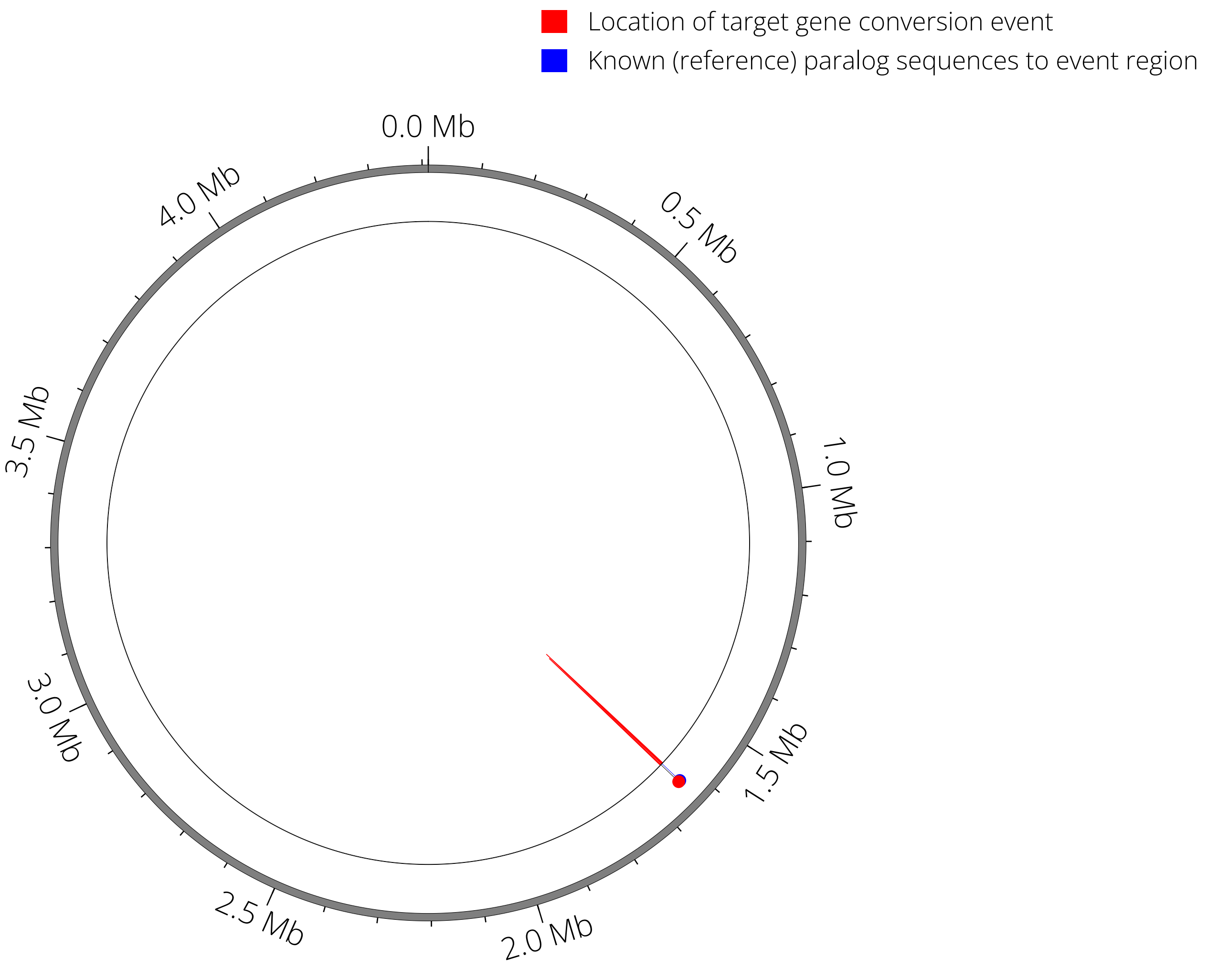

### Event_145 - 4 SNPs - k-mer analysis relative to paralogs

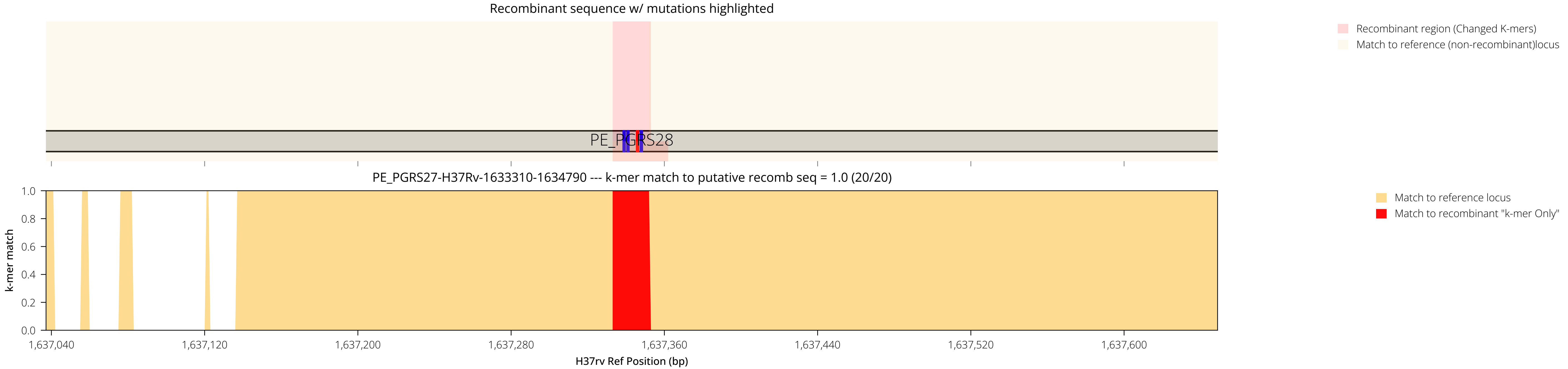

Sub-phylogeny Viz + Paralog Alignments
GCE ID: Event_145

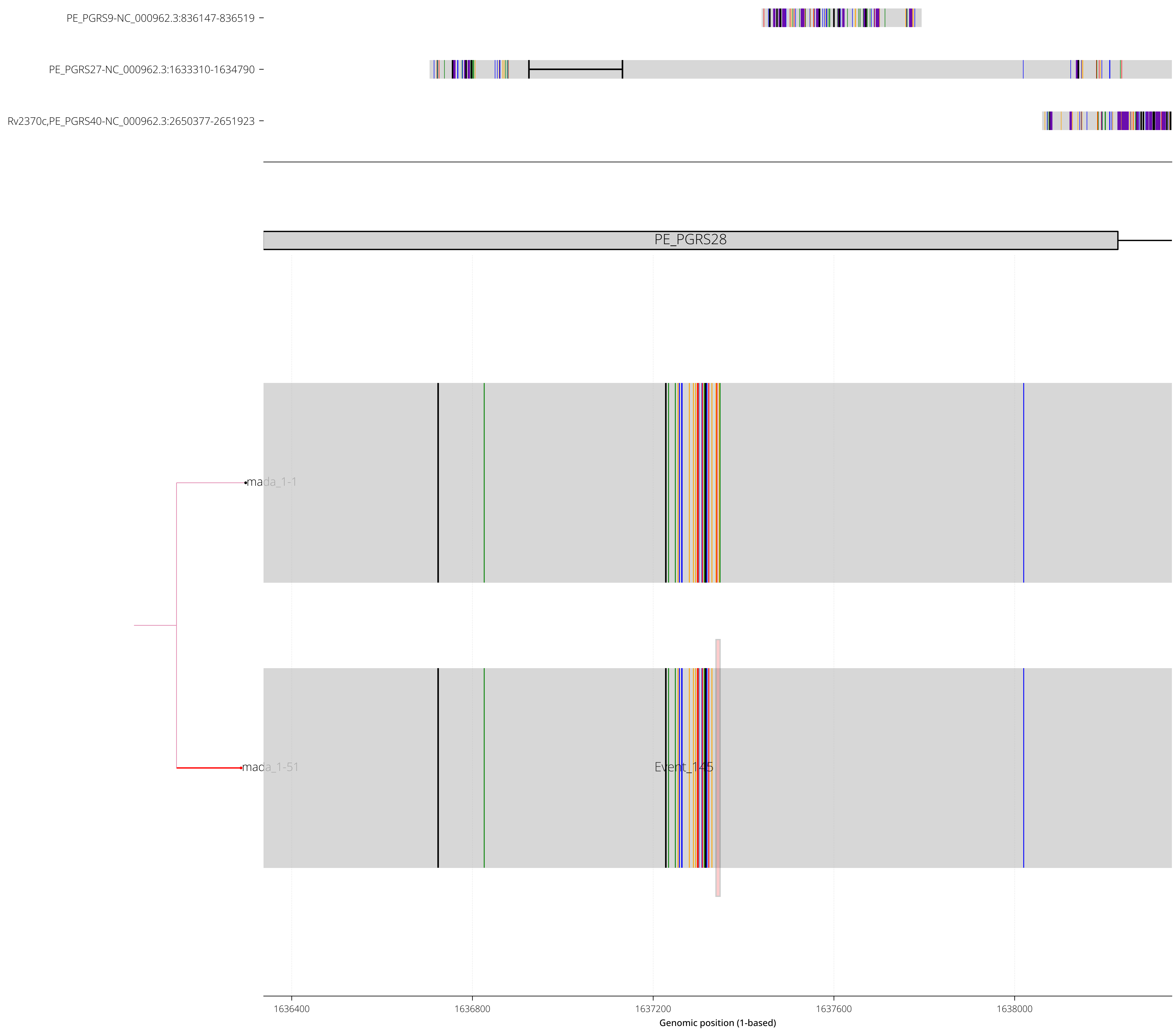

Full Phylogeny Viz + Paralog Alignments
GCE ID: Event_145

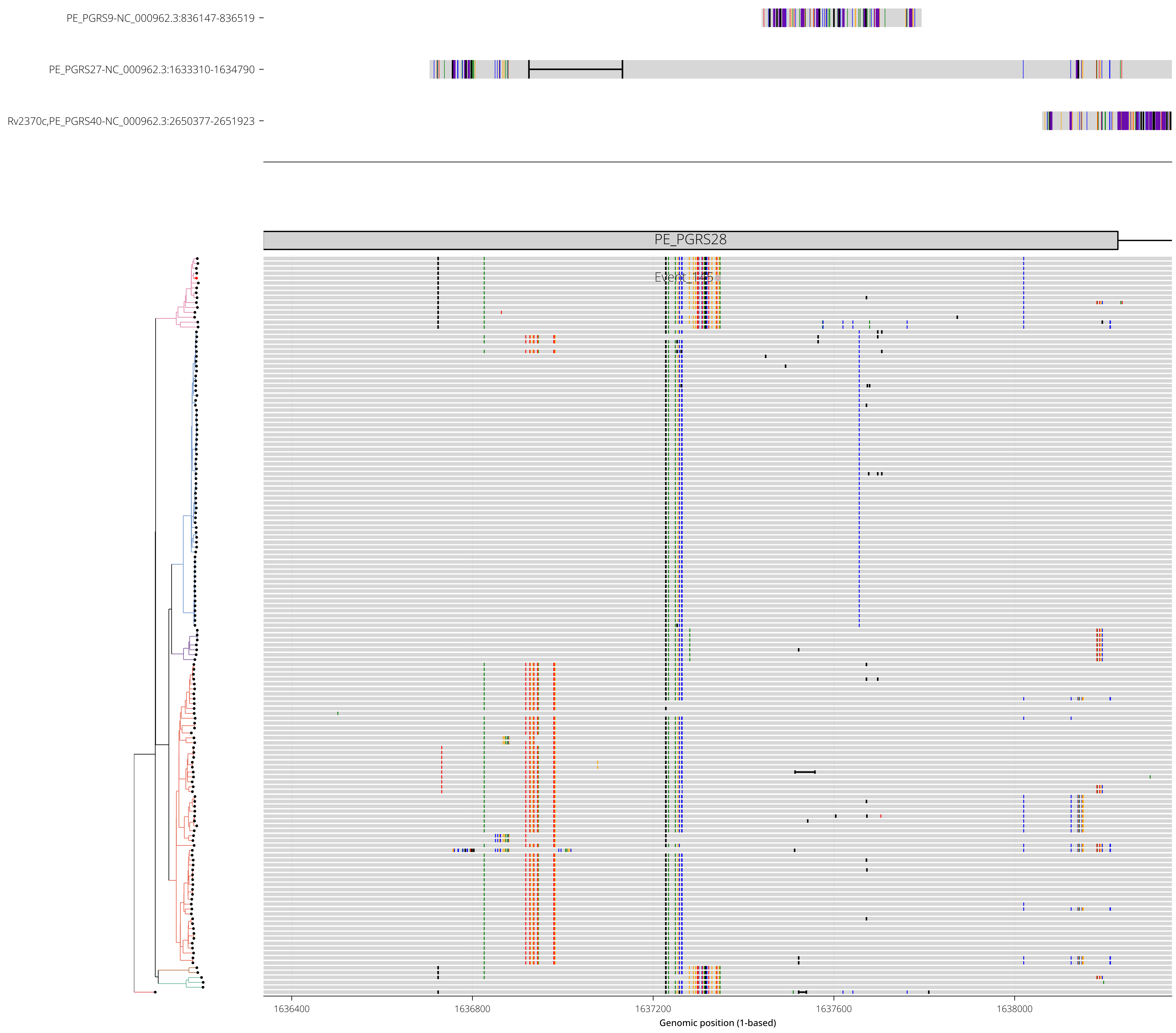

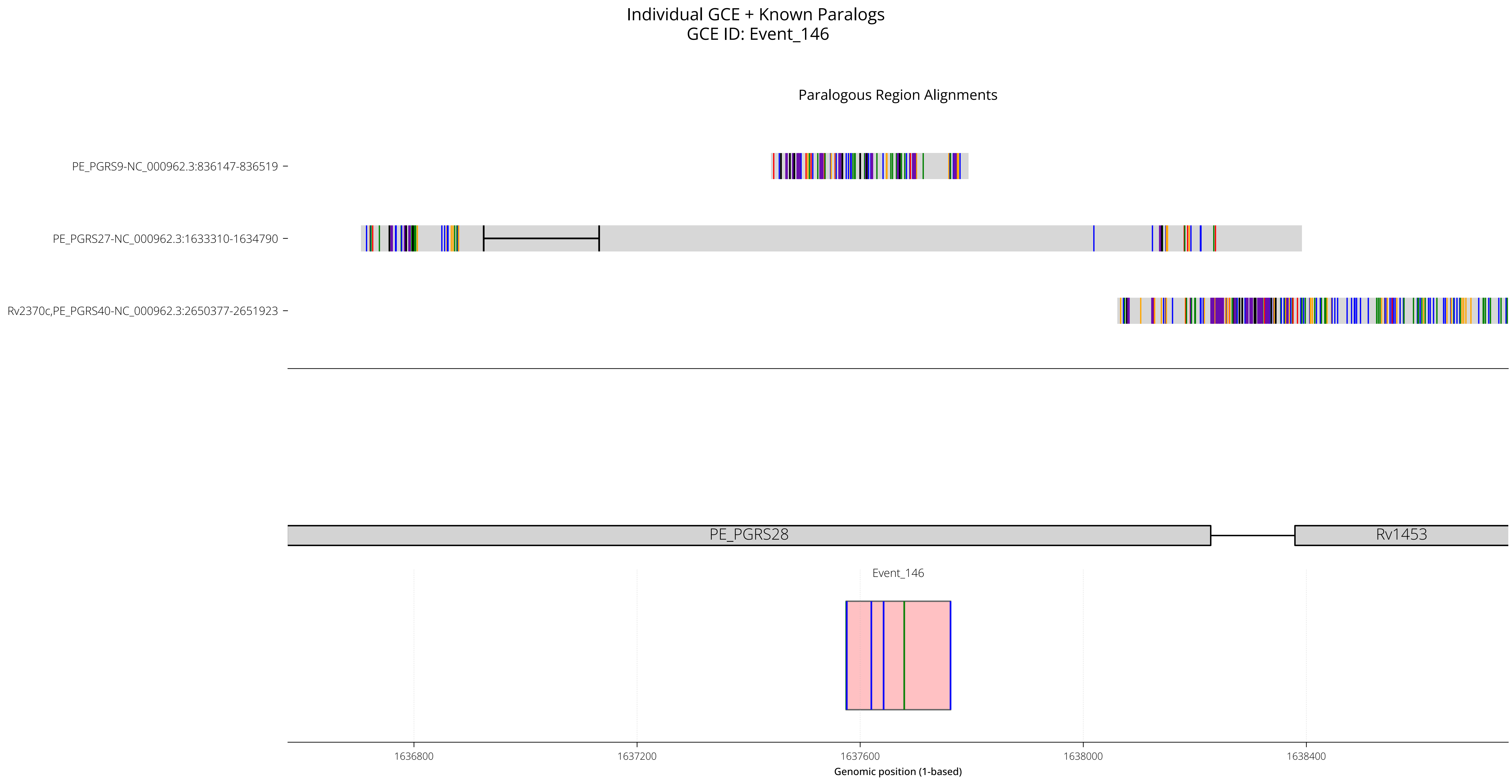

Individual GCE + Known Paralogs
GCE ID: Event_146
Paralogous Region Alignments
PE_PGRS9-NC_000962.3:836147-836519
PE_PGRS27-NC_000962.3:1633310-1634790
Rv2370c,PE_PGRS40-NC_000962.3:2650377-2651923
PE_PGRS28
Rv1453
Event_146
1636800
1637200
1637600
1638000
1638400
Genomic position (1-based)

Circos Viz - GCE to Known Paralogs Locations
GCE ID: Event_146

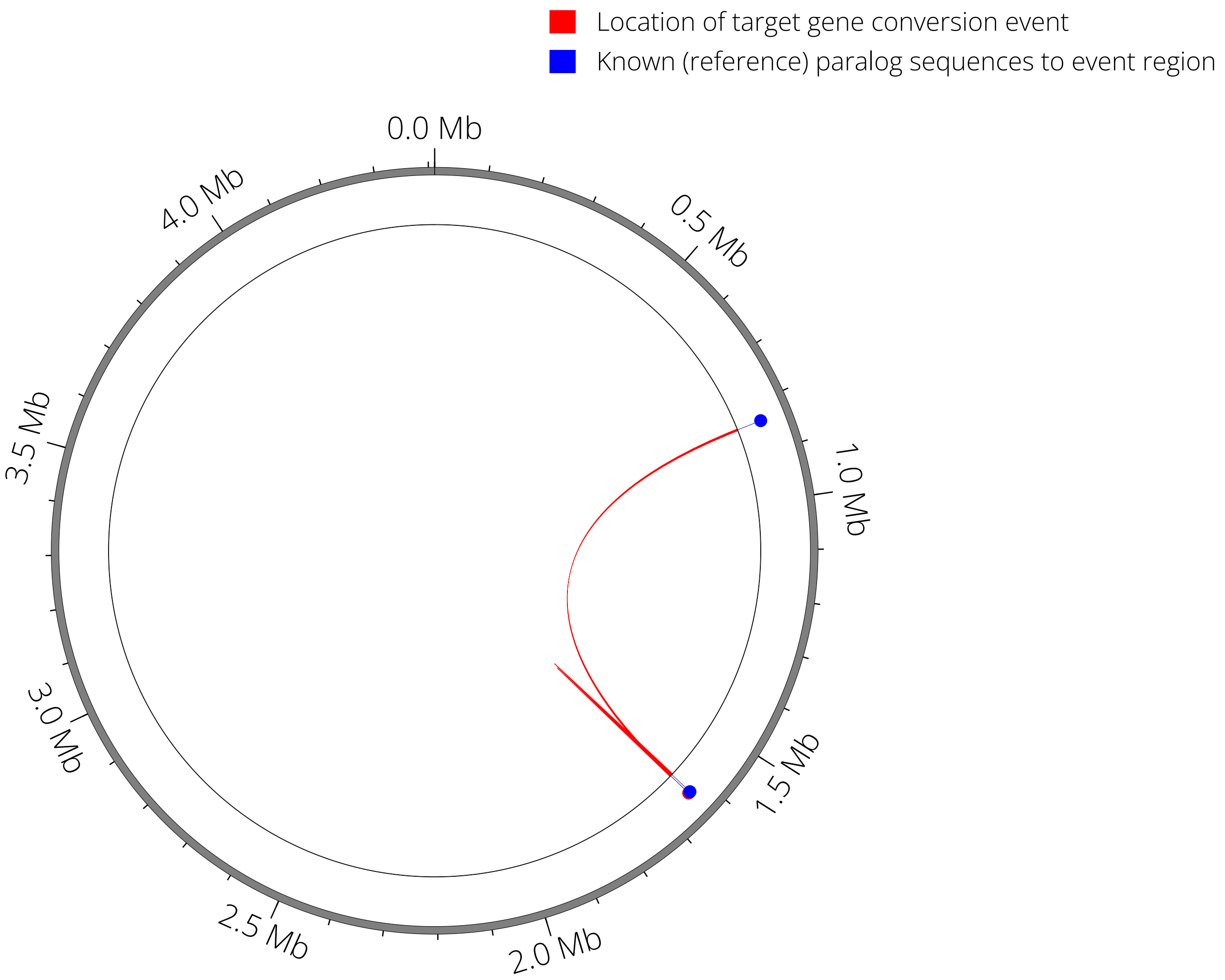

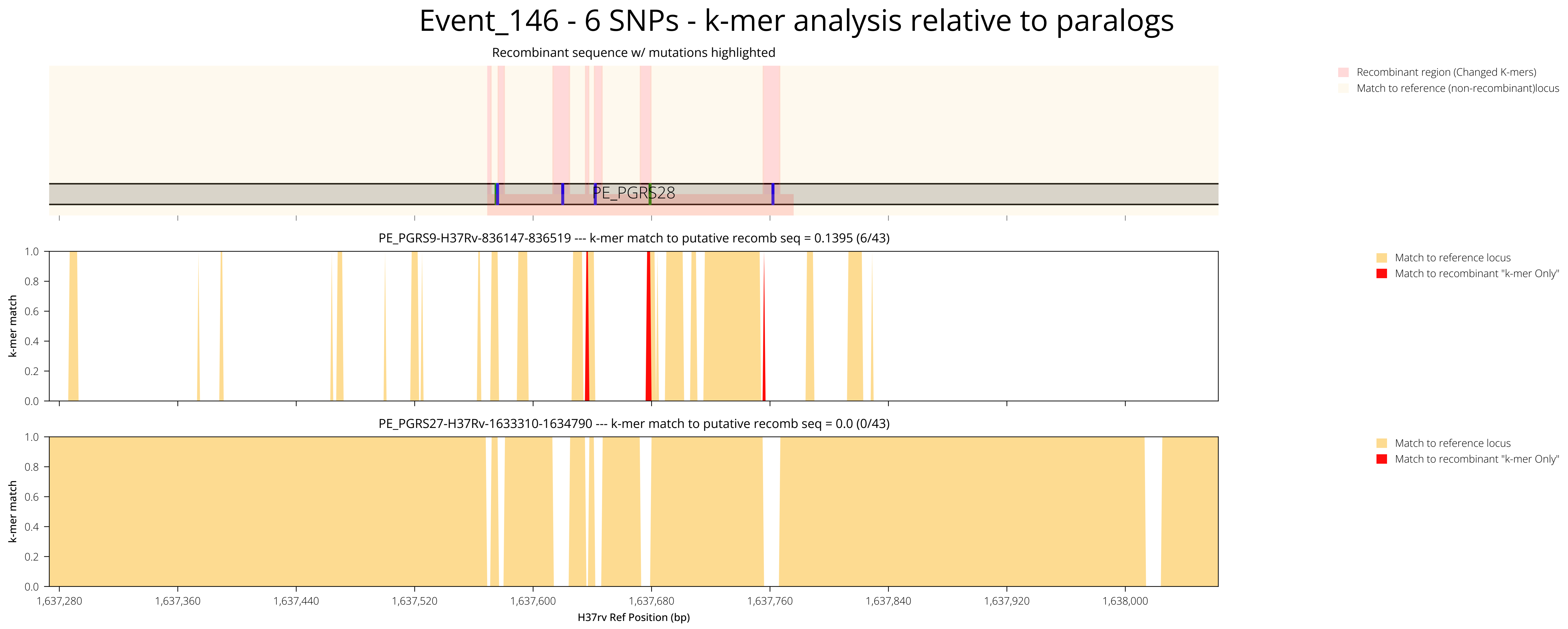

Event_146 - 6 SNPs - k-mer analysis relative to paralogs
Recombinant sequence w/ mutations highlighted
PE_PGRS28
Recombinant region (Changed K-mers)
Match to reference (non-recombinant)locus
PE_PGRS9-H37Rv-836147-836519 --- k-mer match to putative recomb seq = 0.1395 (6/43)
k-mer match
1.0
0.8
0.6
0.4
0.2
0.0
Match to reference locus
Match to recombinant "k-mer Only"
PE_PGRS27-H37Rv-1633310-1634790 --- k-mer match to putative recomb seq = 0.0 (0/43)
k-mer match
Match to reference locus
Match to recombinant "k-mer Only"
1,637,280
1,637,360
1,637,440
1,637,520
1,637,600
1,637,680
1,637,760
1,637,840
1,637,920
1,638,000
H37rv Ref Position (bp)

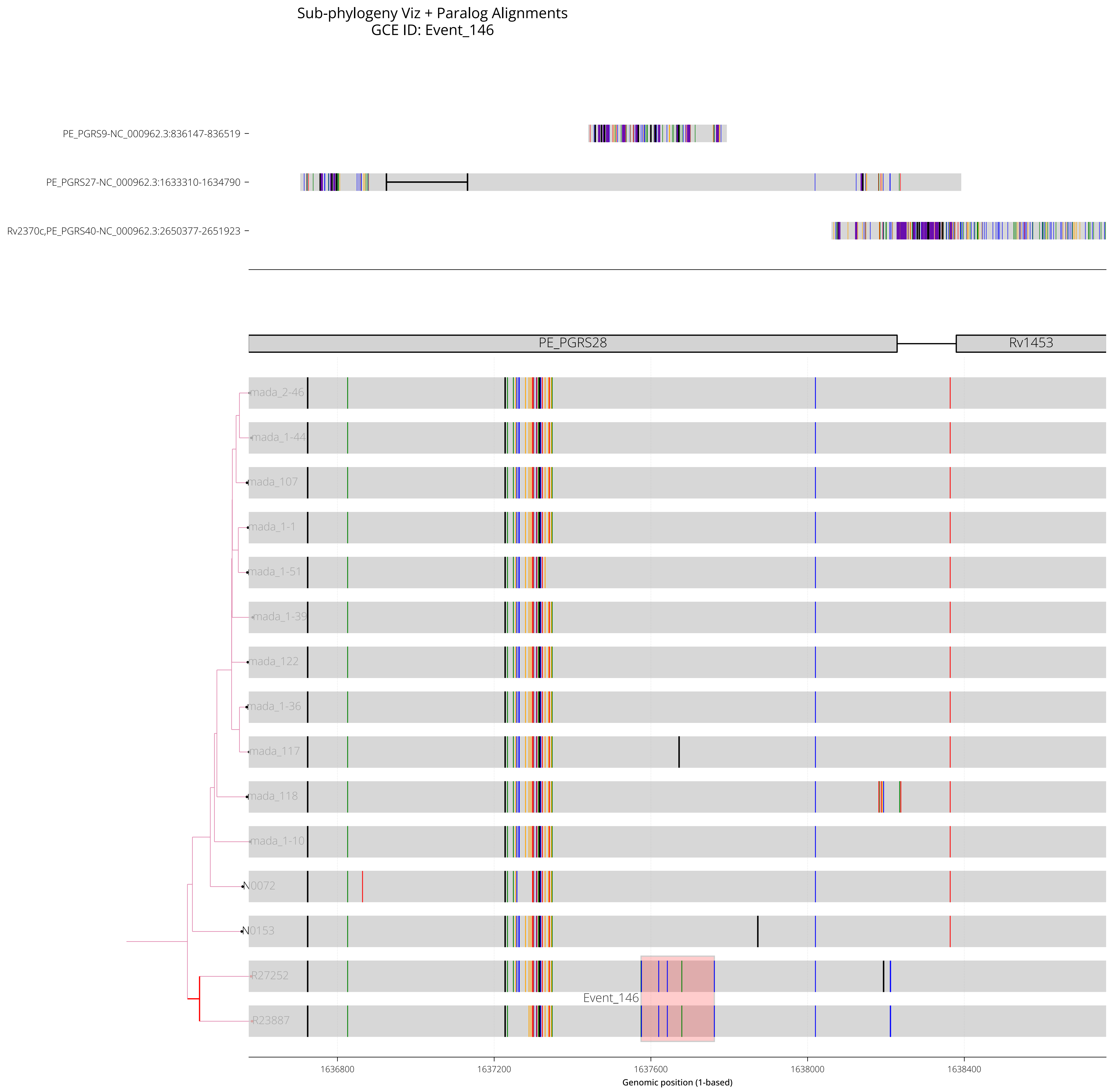

Sub-phylogeny Viz + Paralog Alignments
GCE ID: Event_146
PE_PGRS9-NC_000962.3:836147-836519
PE_PGRS27-NC_000962.3:1633310-1634790
Rv2370c,PE_PGRS40-NC_000962.3:2650377-2651923
PE_PGRS28
Rv1453
mada_2-46
mada_1-44
mada_107
mada_1-1
mada_1-51
mada_1-39
mada_122
mada_1-36
mada_117
mada_118
mada_1-10
N0072
N0153
R27252
R23887
Event_146
1636800
1637200
1637600
1638000
1638400
Genomic position (1-based)

Full Phylogeny Viz + Paralog Alignments
GCE ID: Event_146

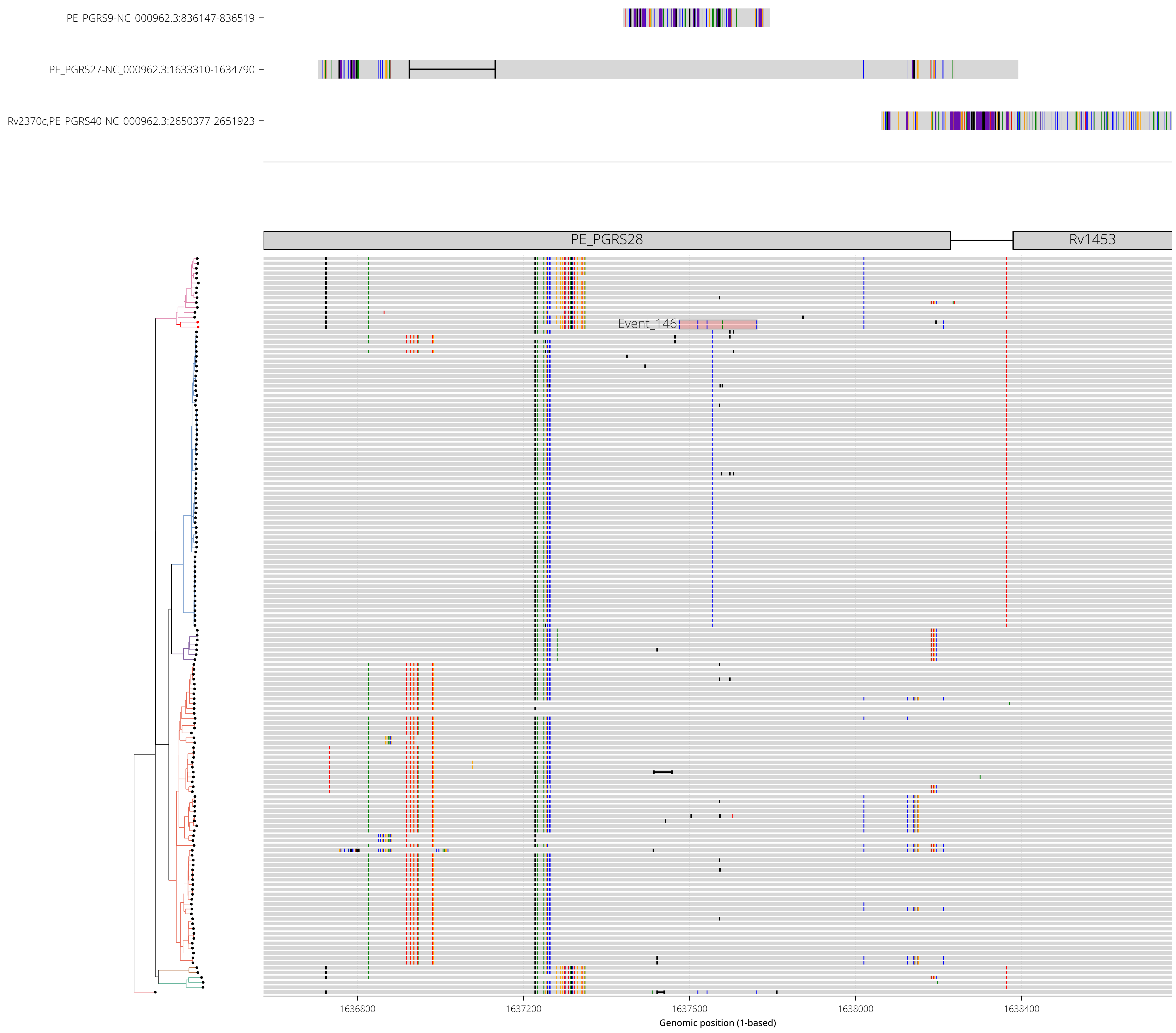

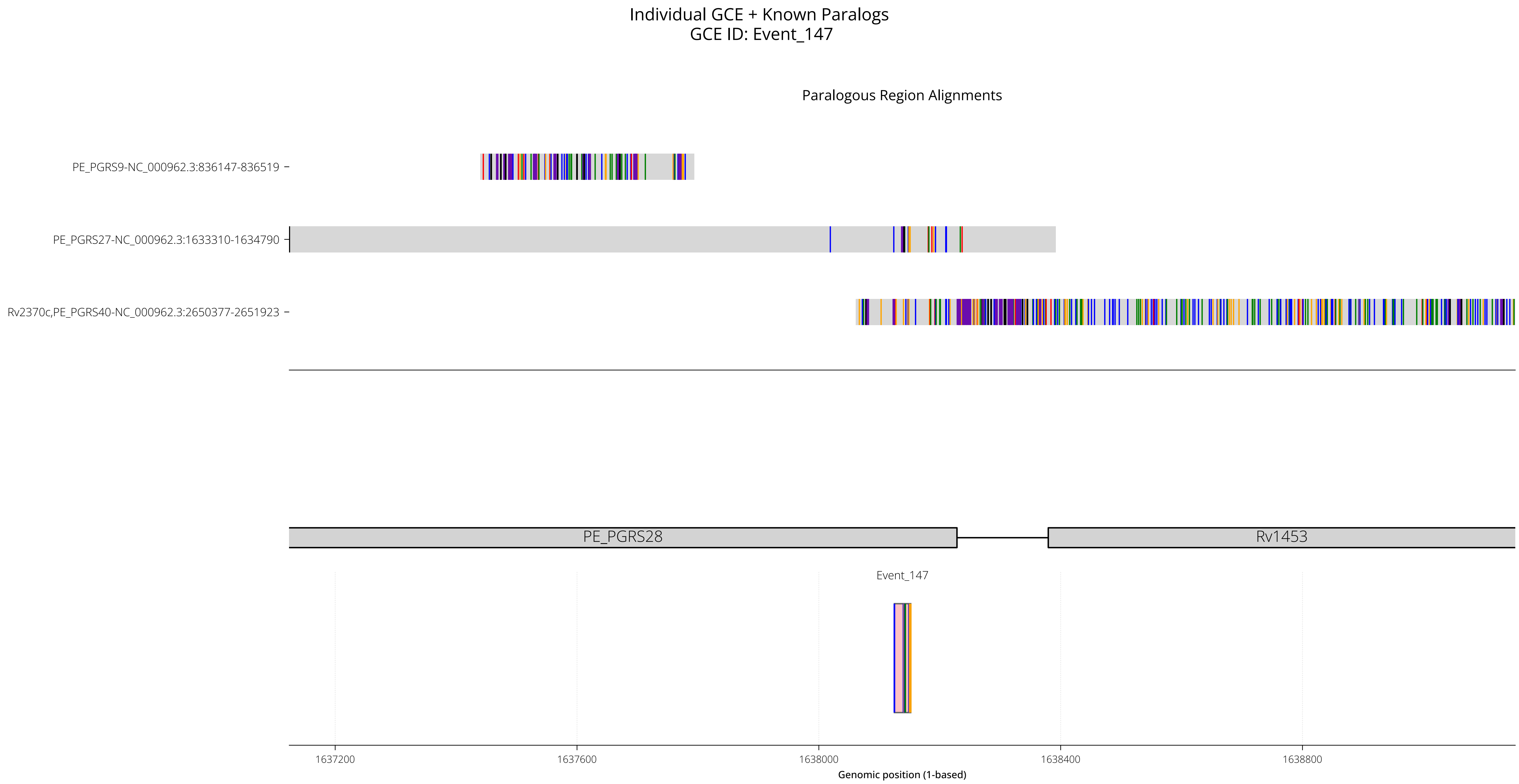
Individual GCE + Known Paralogs
GCE ID: Event_147
Paralogous Region Alignments
PE_PGRS9-NC_000962.3:836147-836519
PE_PGRS27-NC_000962.3:1633310-1634790
Rv2370c,PE_PGRS40-NC_000962.3:2650377-2651923
PE_PGRS28
Rv1453
Event_147
1637200
1637600
1638000
1638400
1638800
Genomic position (1-based)

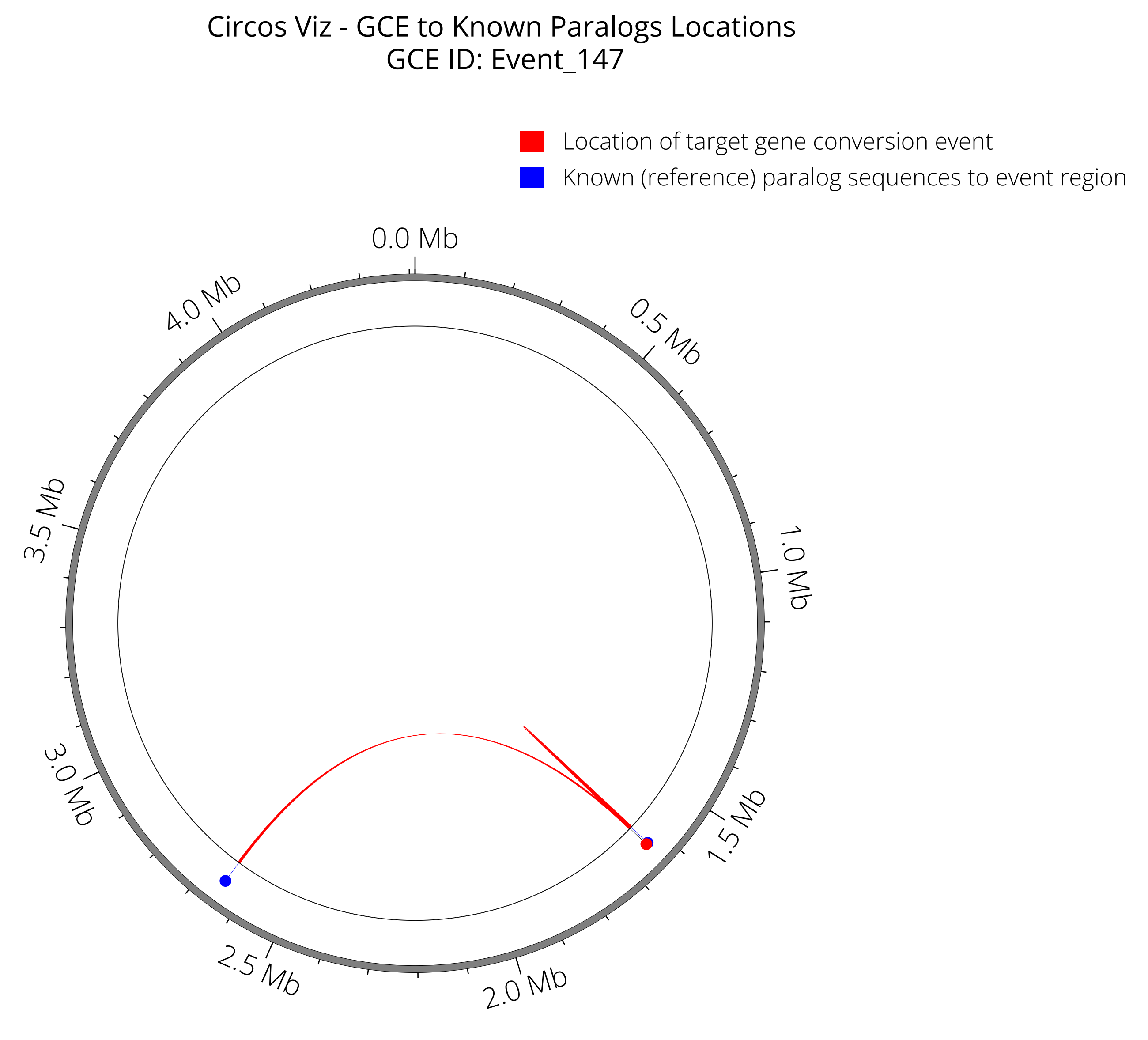

Circos Viz - GCE to Known Paralogs Locations
GCE ID: Event_147
Location of target gene conversion event
Known (reference) paralog sequences to event region
0.0 Mb
0.5 Mb
1.0 Mb
1.5 Mb
2.0 Mb
2.5 Mb
3.0 Mb
3.5 Mb
4.0 Mb

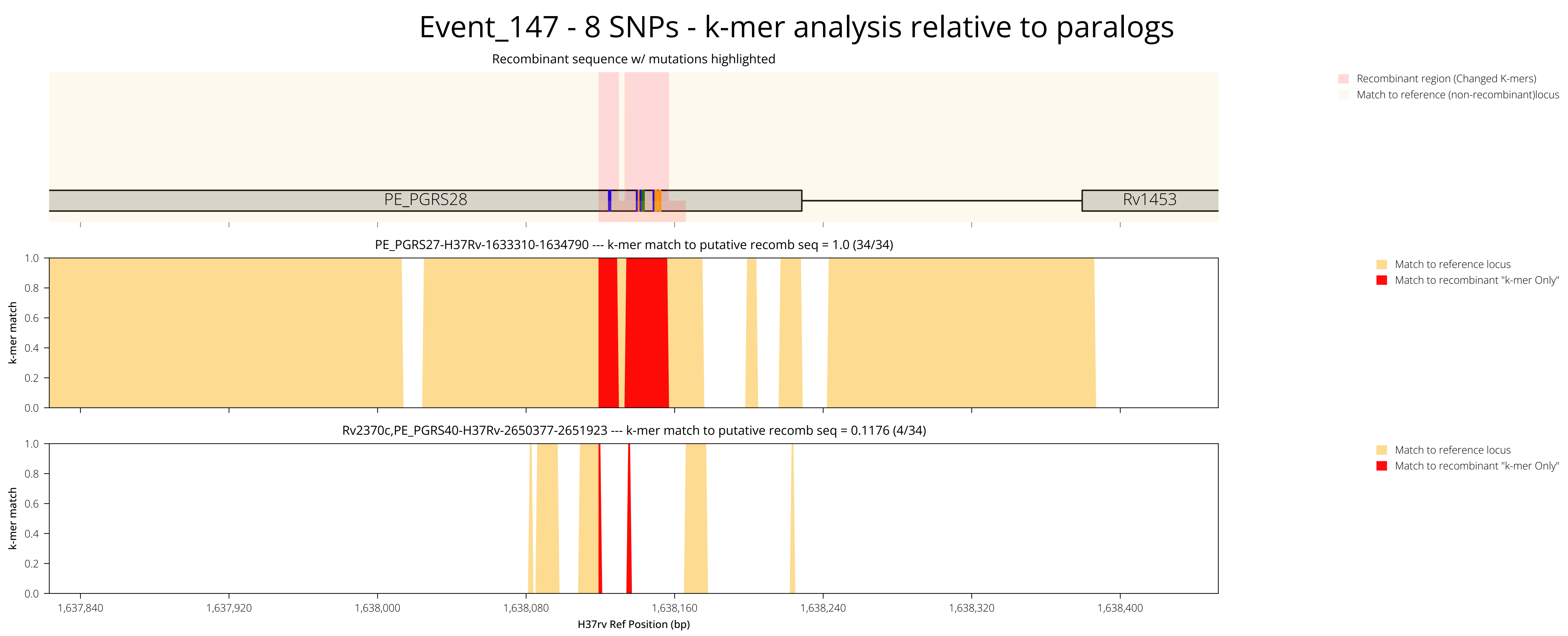

Event_147 - 8 SNPs - k-mer analysis relative to paralogs
Recombinant sequence w/ mutations highlighted
Recombinant region (Changed K-mers)
Match to reference (non-recombinant)locus
PE_PGRS28
Rv1453
PE_PGRS27-H37Rv-1633310-1634790 --- k-mer match to putative recomb seq = 1.0 (34/34)
Match to reference locus
Match to recombinant "k-mer Only"
Rv2370c,PE_PGRS40-H37Rv-2650377-2651923 --- k-mer match to putative recomb seq = 0.1176 (4/34)
Match to reference locus
Match to recombinant "k-mer Only"
k-mer match
0.0
0.2
0.4
0.6
0.8
1.0
1,637,840
1,637,920
1,638,000
1,638,080
1,638,160
1,638,240
1,638,320
1,638,400
H37rv Ref Position (bp)

Sub-phylogeny Viz + Paralog Alignments
GCE ID: Event_147

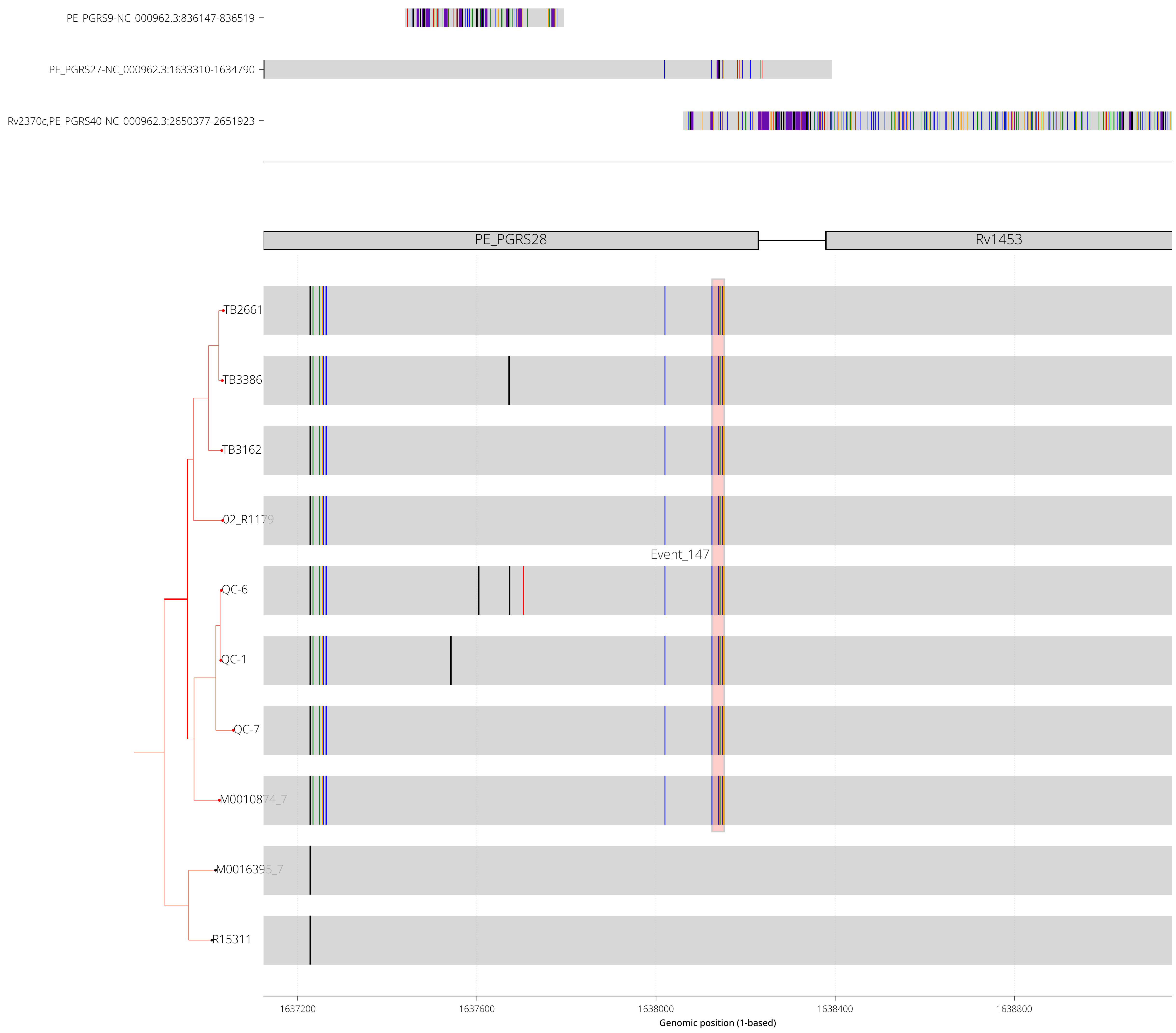

Full Phylogeny Viz + Paralog Alignments
GCE ID: Event_147

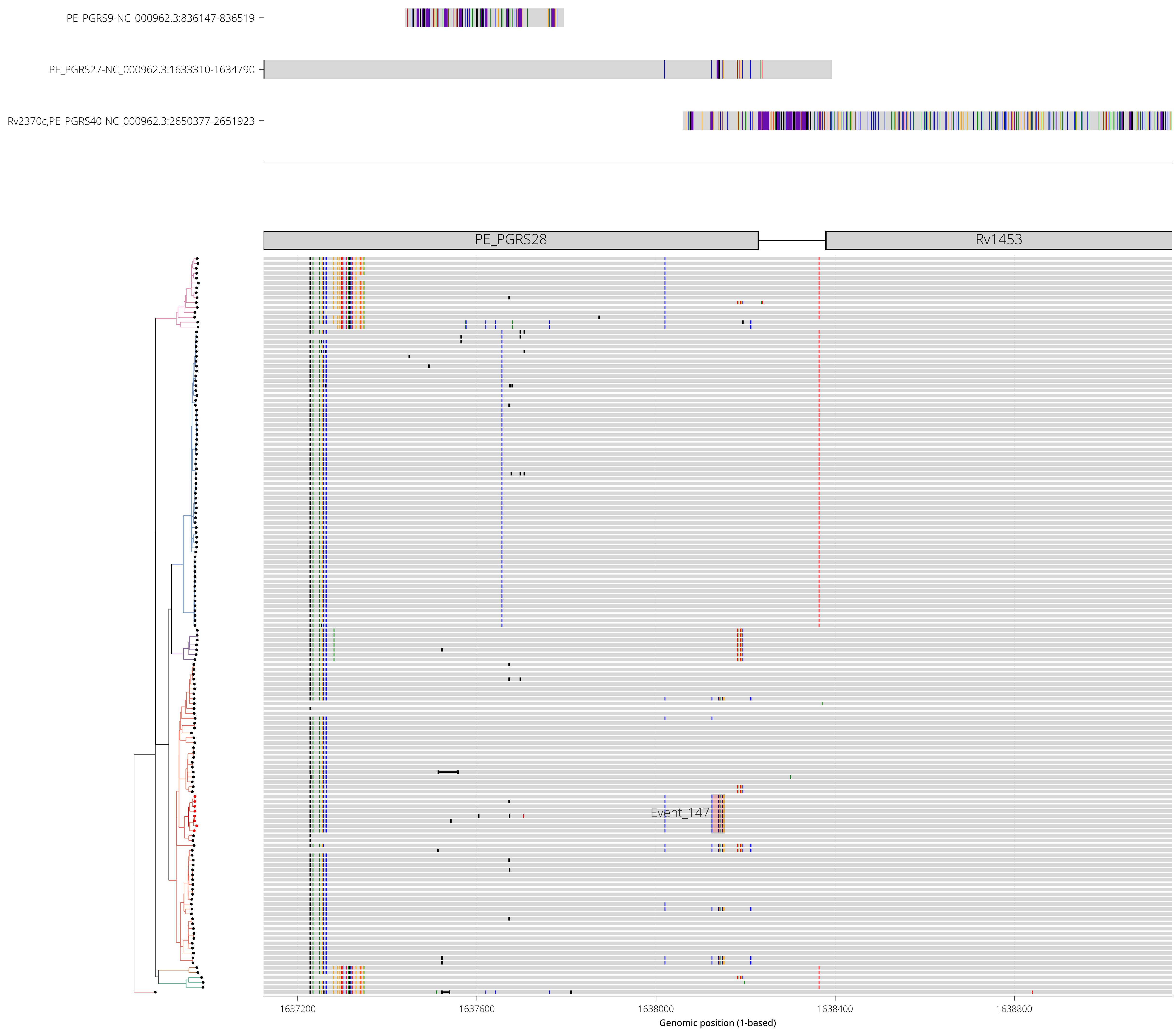

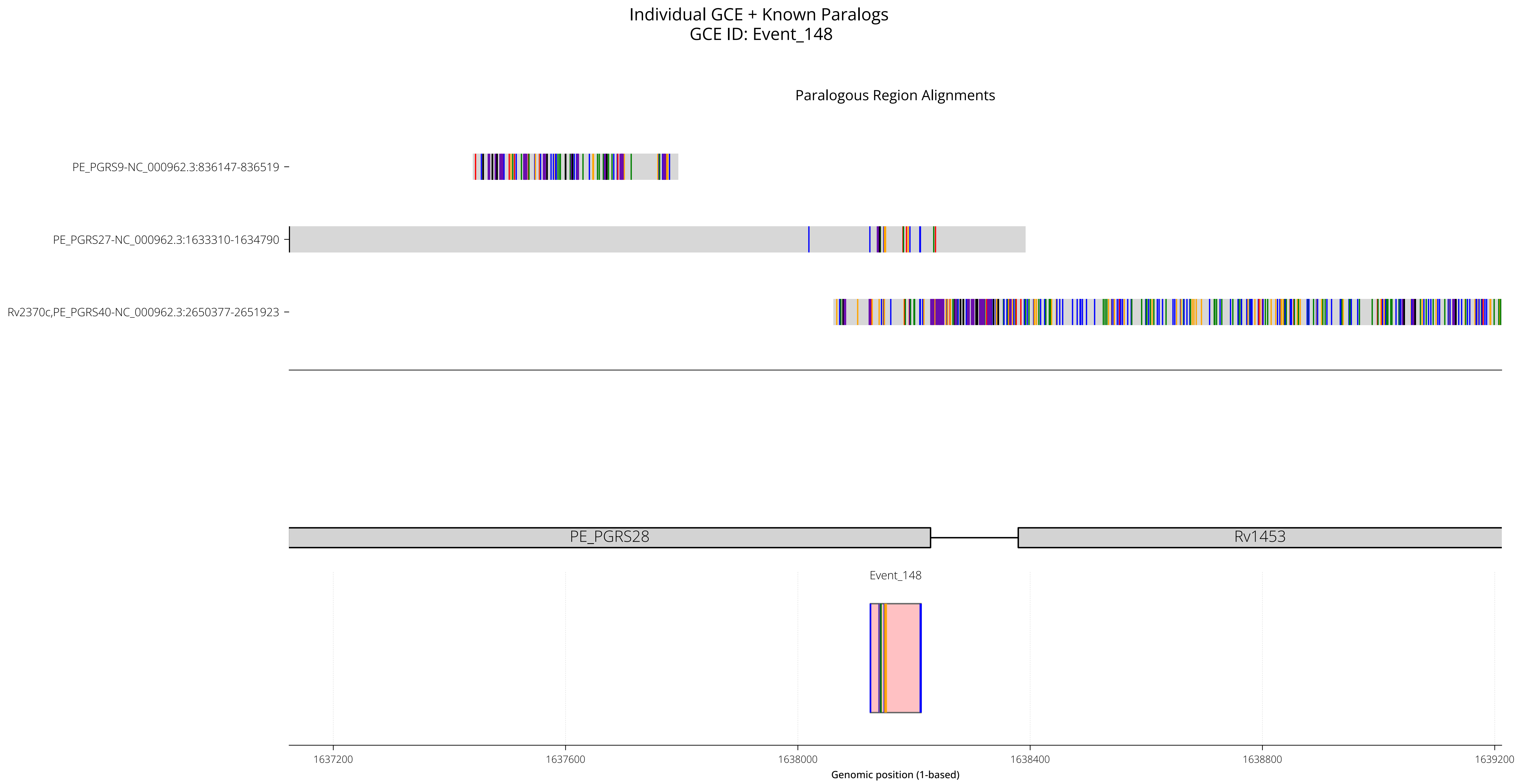
Individual GCE + Known Paralogs
GCE ID: Event_148
Paralogous Region Alignments
PE_PGRS9-NC_000962.3:836147-836519
PE_PGRS27-NC_000962.3:1633310-1634790
Rv2370c,PE_PGRS40-NC_000962.3:2650377-2651923
PE_PGRS28
Rv1453
Event_148
1637200
1637600
1638000
1638400
1638800
1639200
Genomic position (1-based)

Circos Viz - GCE to Known Paralogs Locations
GCE ID: Event_148

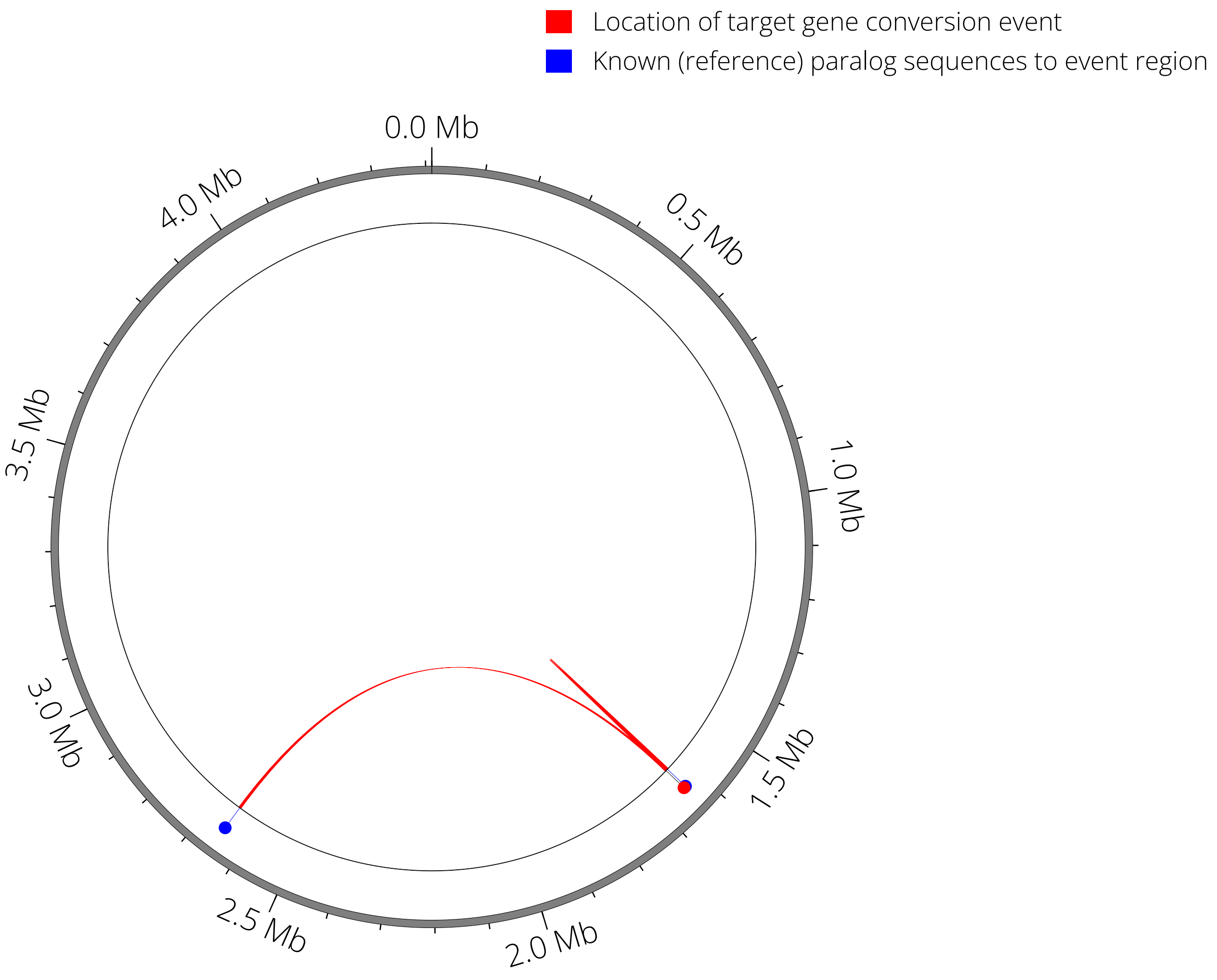

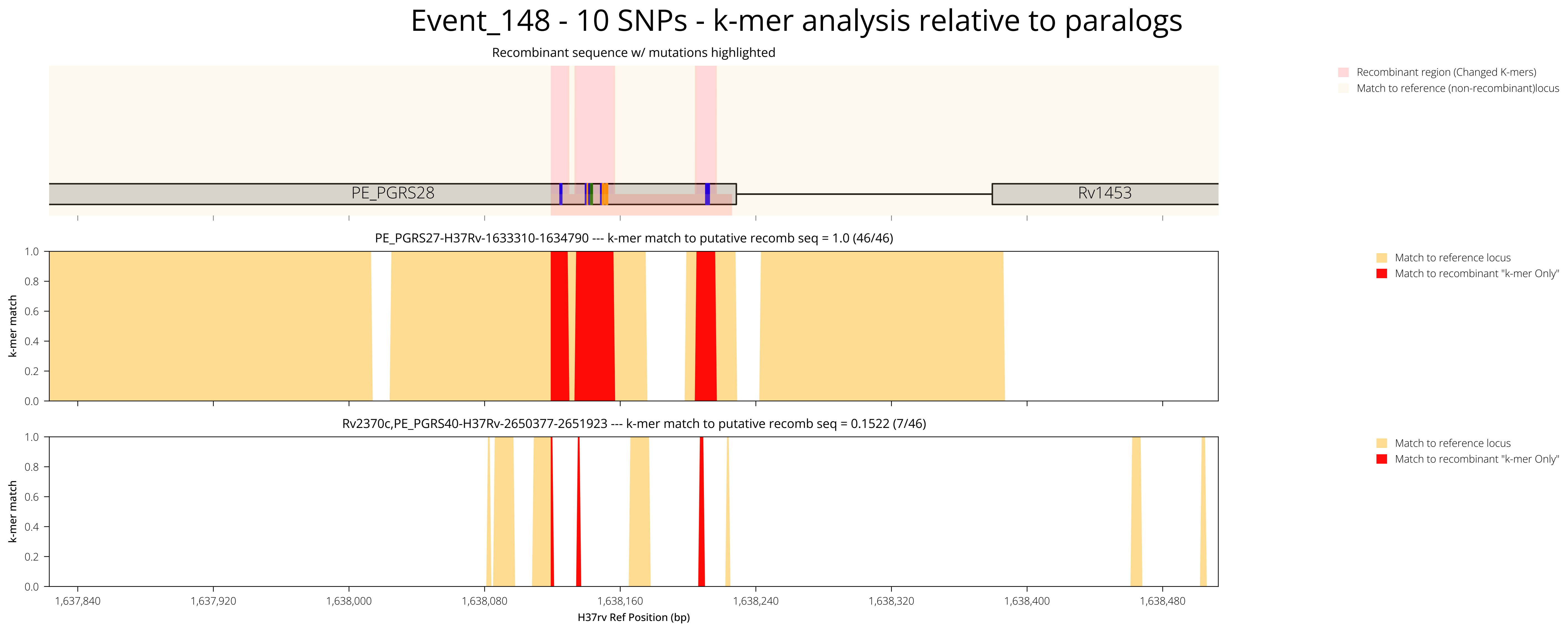
Event_148 - 10 SNPs - k-mer analysis relative to paralogs
Recombinant sequence w/ mutations highlighted
PE_PGRS28
Rv1453
Recombinant region (Changed K-mers)
Match to reference (non-recombinant)locus
PE_PGRS27-H37Rv-1633310-1634790 --- k-mer match to putative recomb seq = 1.0 (46/46)
Match to reference locus
Match to recombinant "k-mer Only"
Rv2370c,PE_PGRS40-H37Rv-2650377-2651923 --- k-mer match to putative recomb seq = 0.1522 (7/46)
Match to reference locus
Match to recombinant "k-mer Only"
k-mer match
1.0
0.8
0.6
0.4
0.2
0.0
1,637,840
1,637,920
1,638,000
1,638,080
1,638,160
1,638,240
1,638,320
1,638,400
1,638,480
H37rv Ref Position (bp)

Sub-phylogeny Viz + Paralog Alignments
GCE ID: Event_148

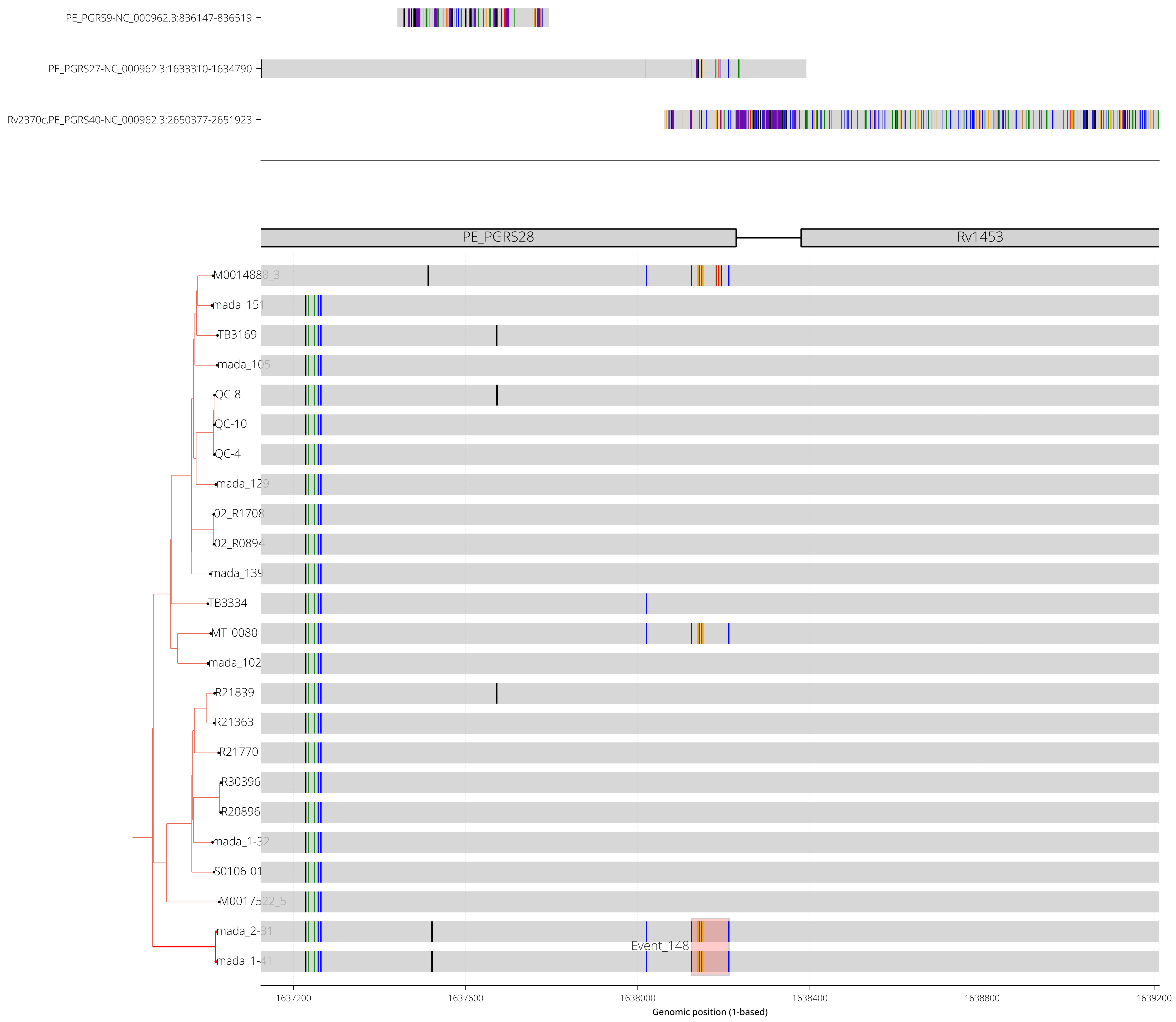

Full Phylogeny Viz + Paralog Alignments
GCE ID: Event_148

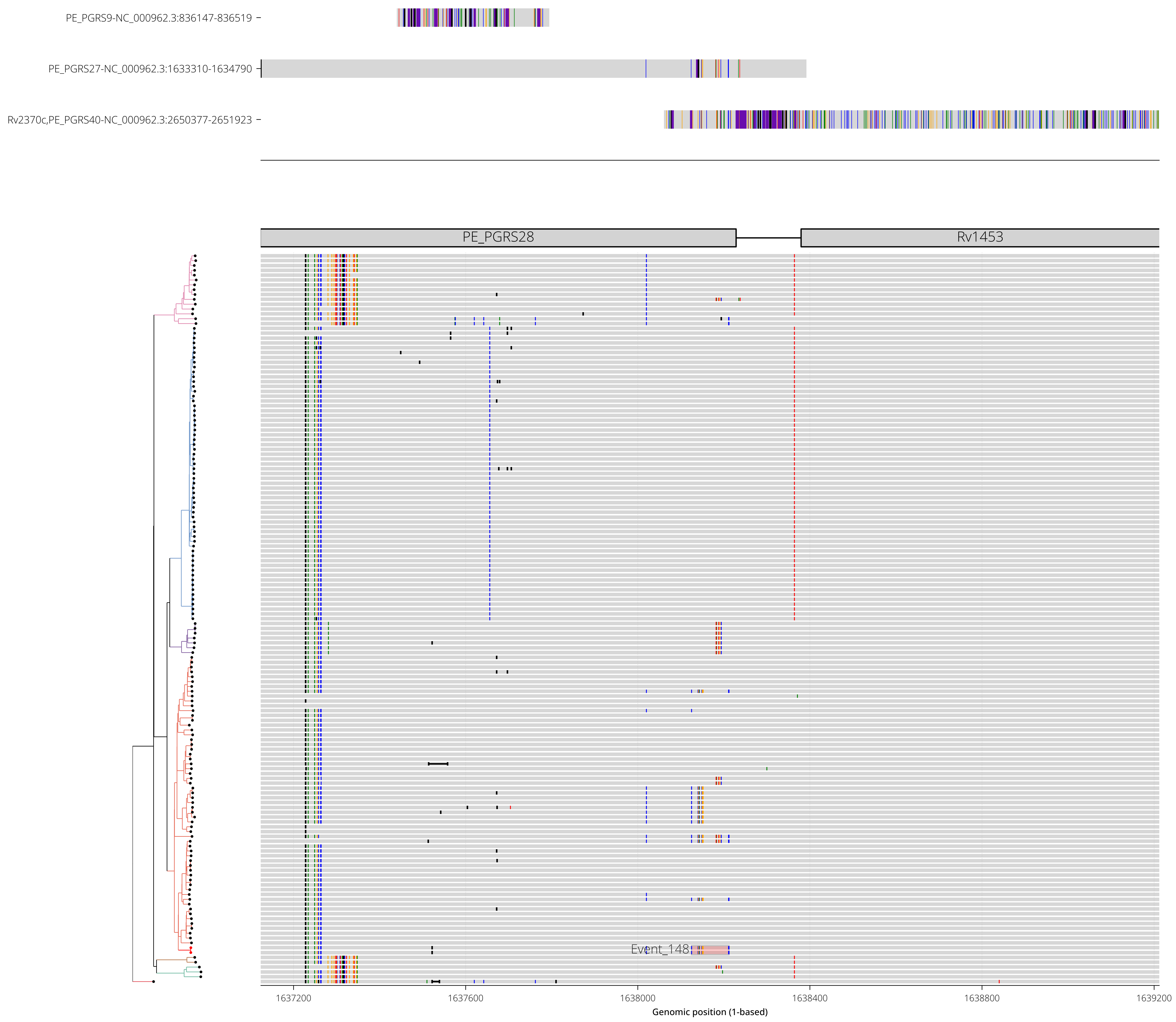

#### Individual GCE + Known Paralogs
#### GCE ID: Event_149

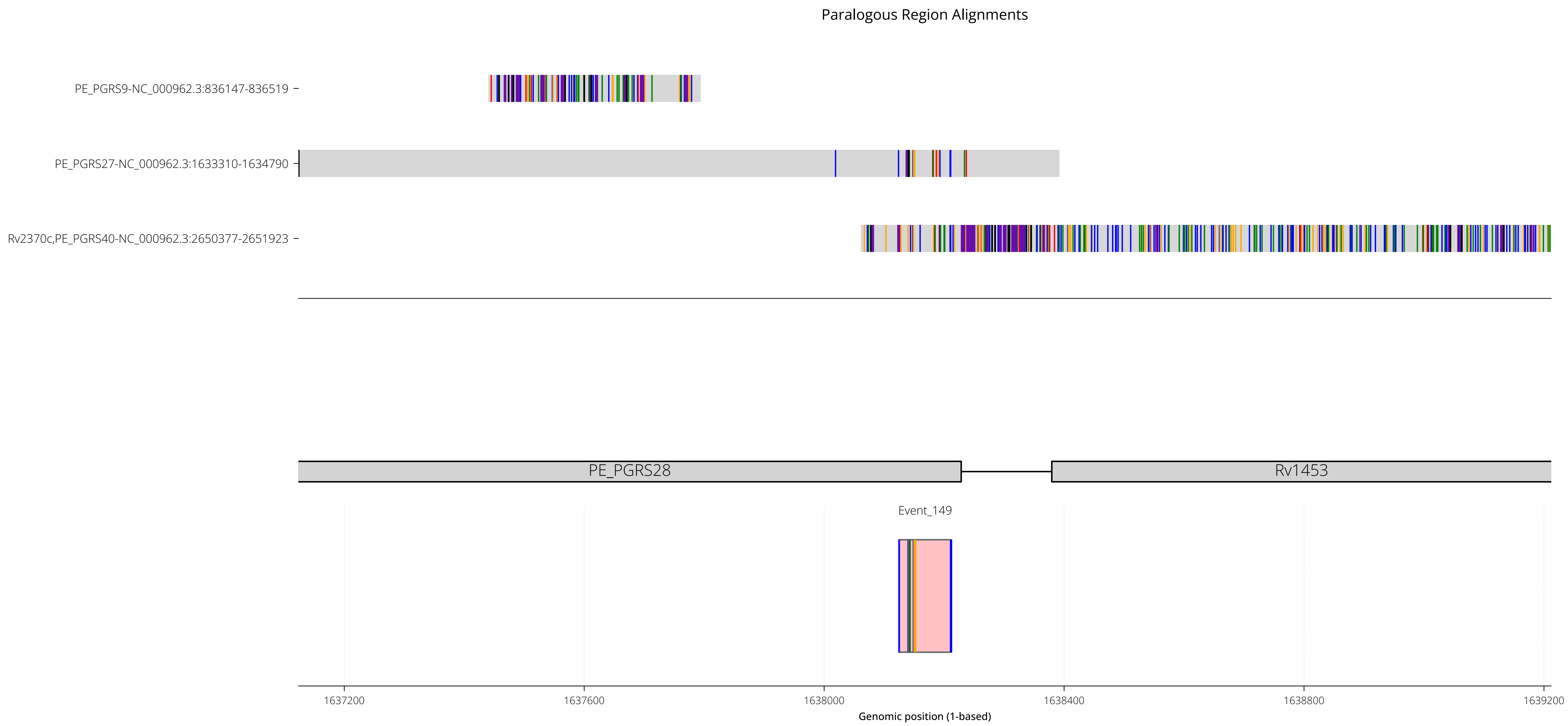

Circos Viz - GCE to Known Paralogs Locations
GCE ID: Event_149

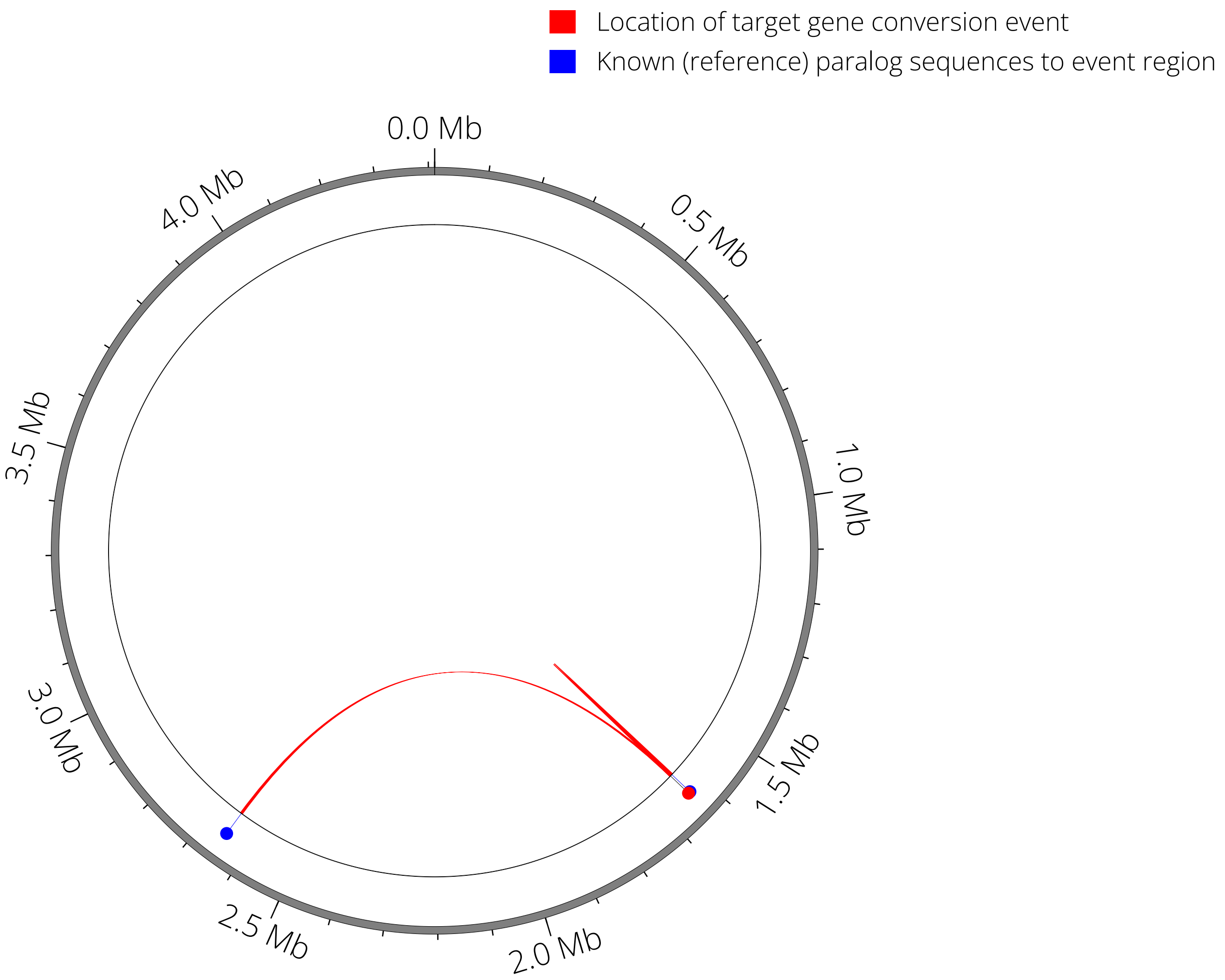

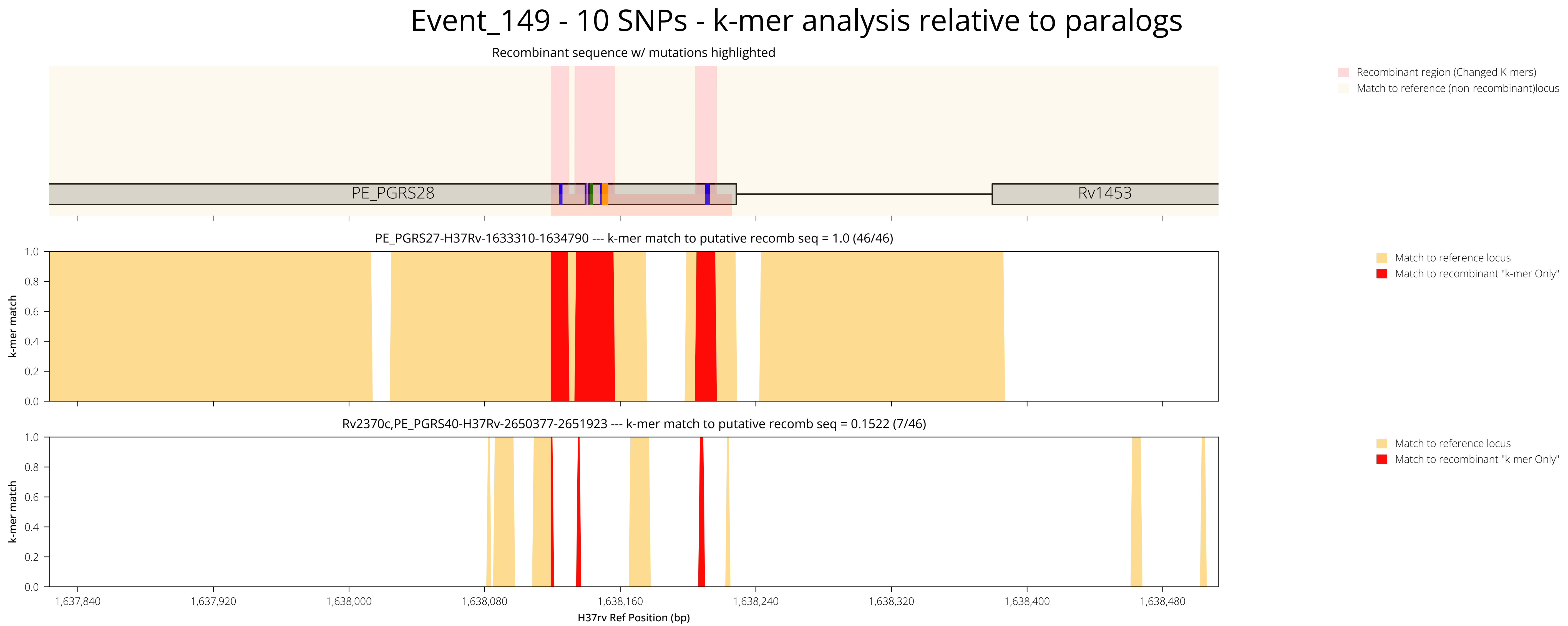
Event_149 - 10 SNPs - k-mer analysis relative to paralogs
Recombinant sequence w/ mutations highlighted
Recombinant region (Changed K-mers)
Match to reference (non-recombinant)locus
PE_PGRS28
Rv1453
PE_PGRS27-H37Rv-1633310-1634790 --- k-mer match to putative recomb seq = 1.0 (46/46)
Match to reference locus
Match to recombinant "k-mer Only"
k-mer match
1.0
0.8
0.6
0.4
0.2
0.0
Rv2370c,PE_PGRS40-H37Rv-2650377-2651923 --- k-mer match to putative recomb seq = 0.1522 (7/46)
Match to reference locus
Match to recombinant "k-mer Only"
k-mer match
1,637,840
1,637,920
1,638,000
1,638,080
1,638,160
1,638,240
1,638,320
1,638,400
1,638,480
H37rv Ref Position (bp)

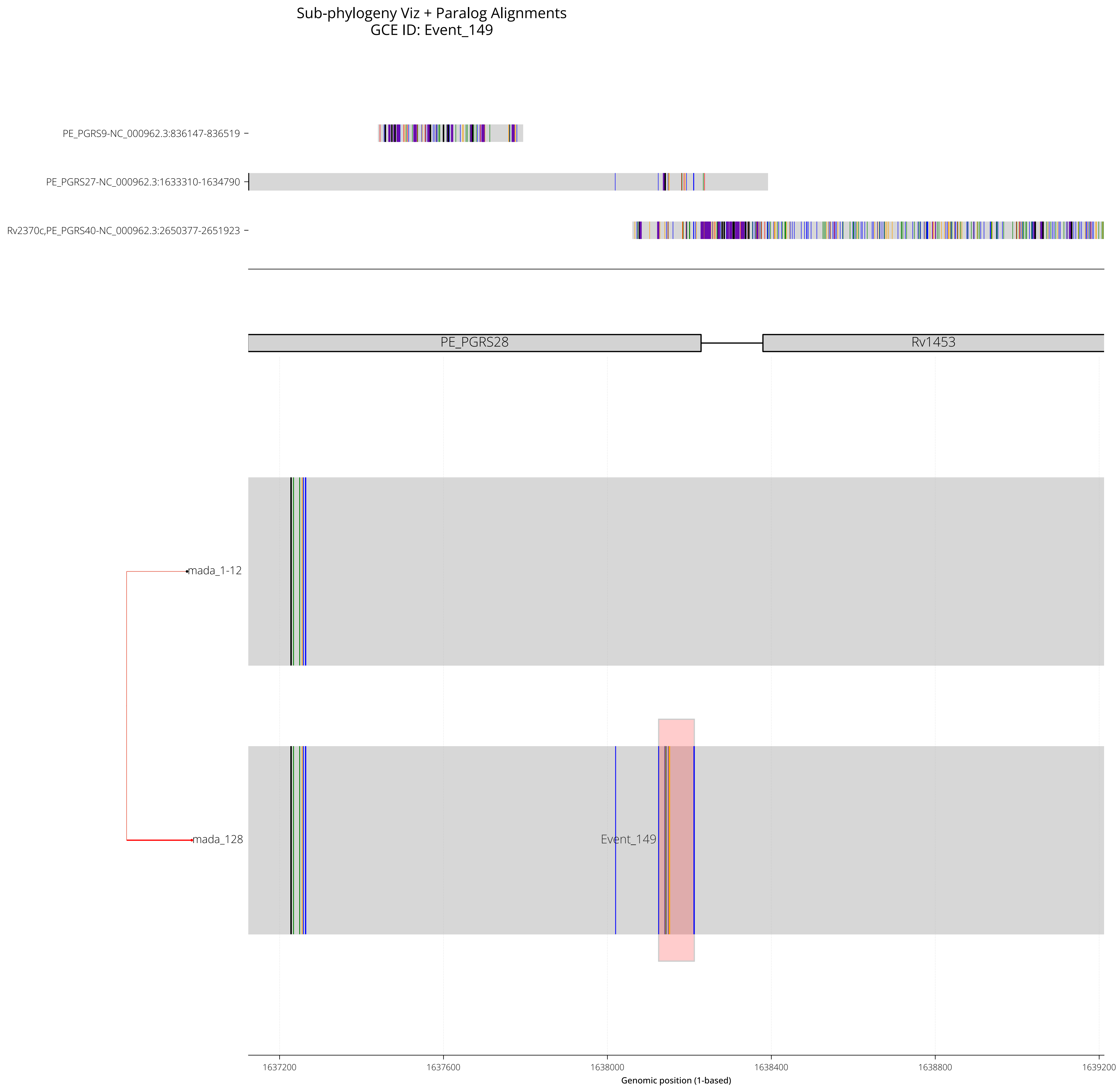

Sub-phylogeny Viz + Paralog Alignments
GCE ID: Event_149
PE_PGRS9-NC_000962.3:836147-836519
PE_PGRS27-NC_000962.3:1633310-1634790
Rv2370c,PE_PGRS40-NC_000962.3:2650377-2651923
PE_PGRS28
Rv1453
mada_1-12
mada_128
Event_149
1637200
1637600
1638000
1638400
1638800
1639200
Genomic position (1-based)

Full Phylogeny Viz + Paralog Alignments
GCE ID: Event_149

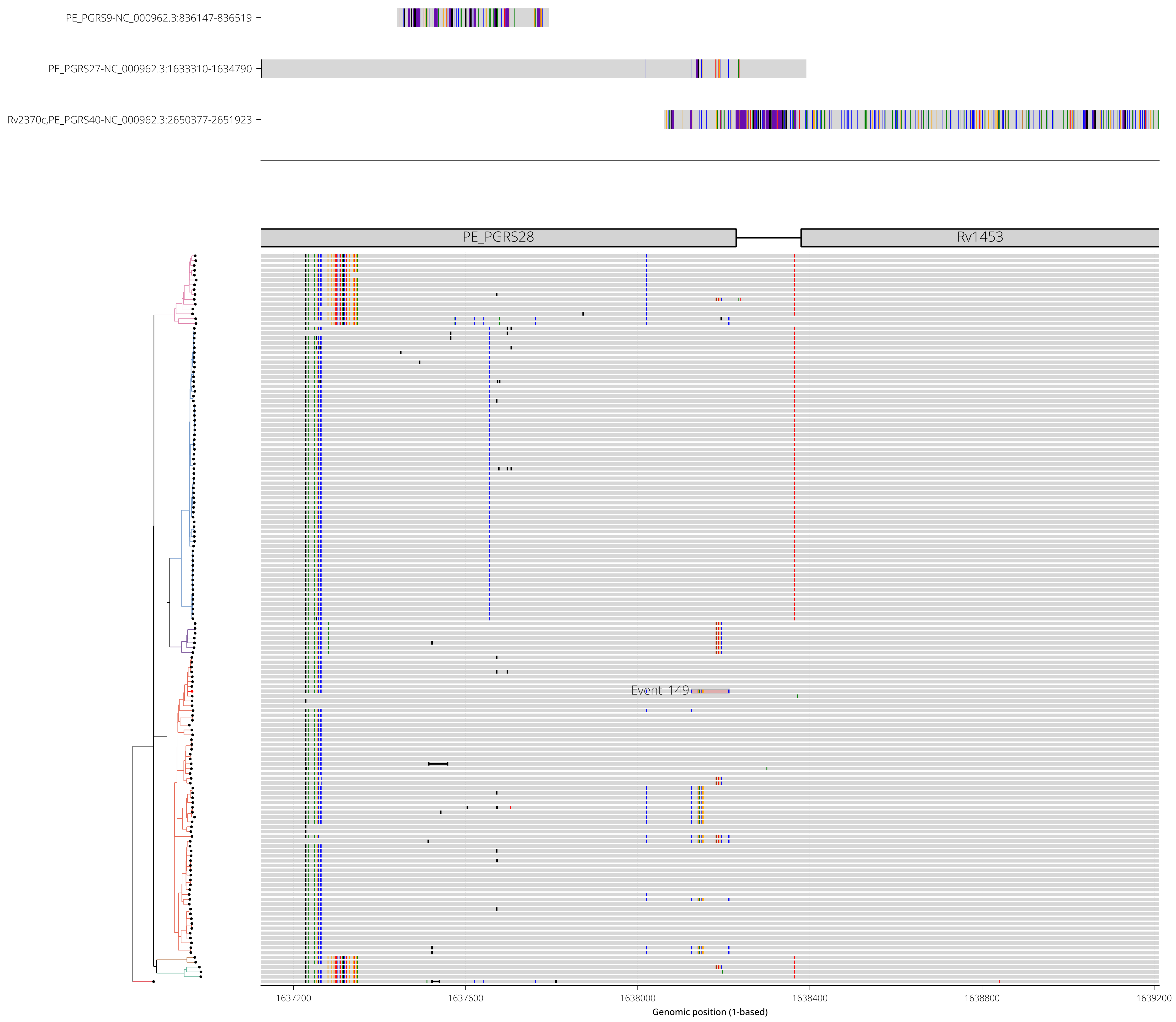

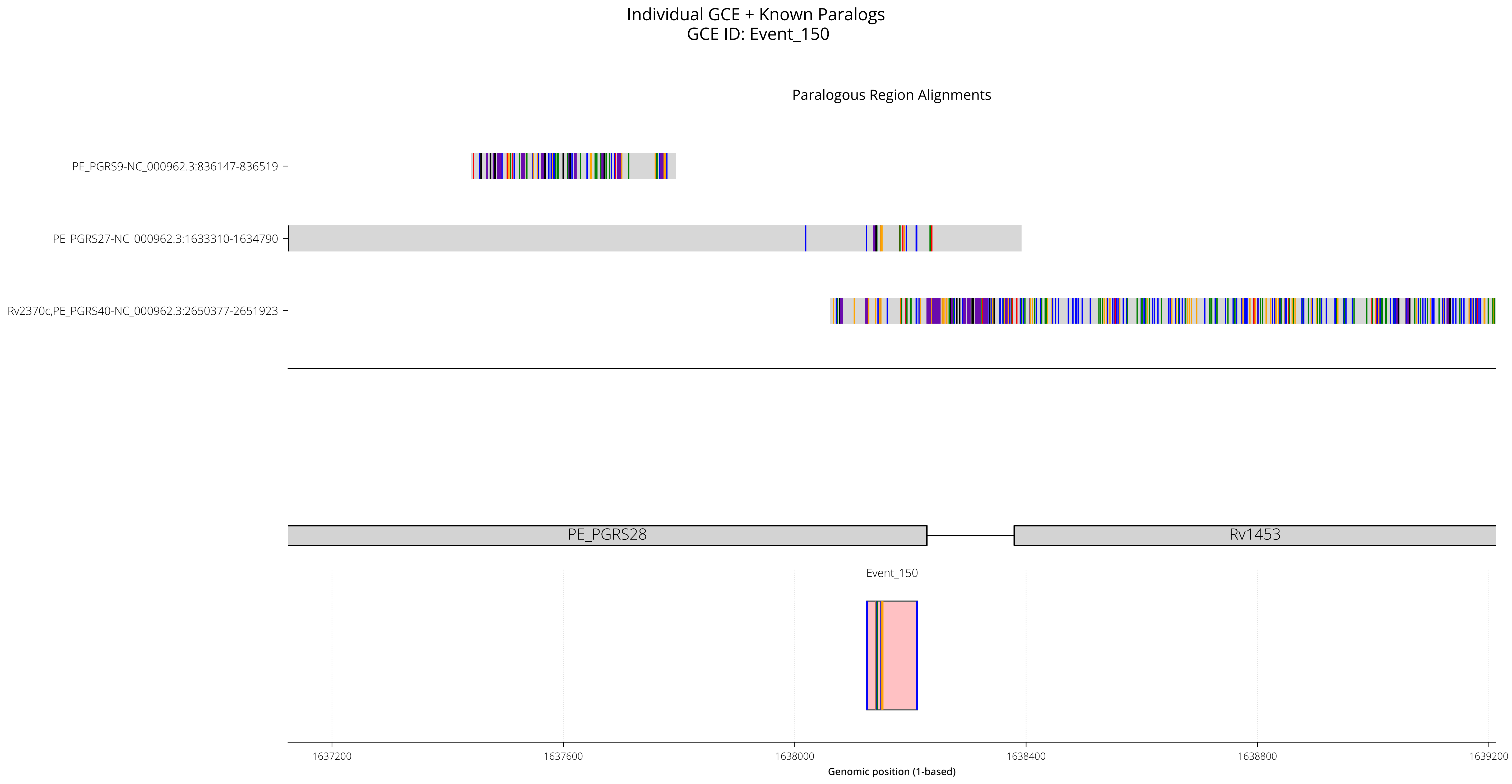
Individual GCE + Known Paralogs
GCE ID: Event_150
Paralogous Region Alignments
PE_PGRS9-NC_000962.3:836147-836519
PE_PGRS27-NC_000962.3:1633310-1634790
Rv2370c,PE_PGRS40-NC_000962.3:2650377-2651923
PE_PGRS28
Rv1453
Event_150
1637200
1637600
1638000
1638400
1638800
1639200
Genomic position (1-based)

Circos Viz - GCE to Known Paralogs Locations
GCE ID: Event_150

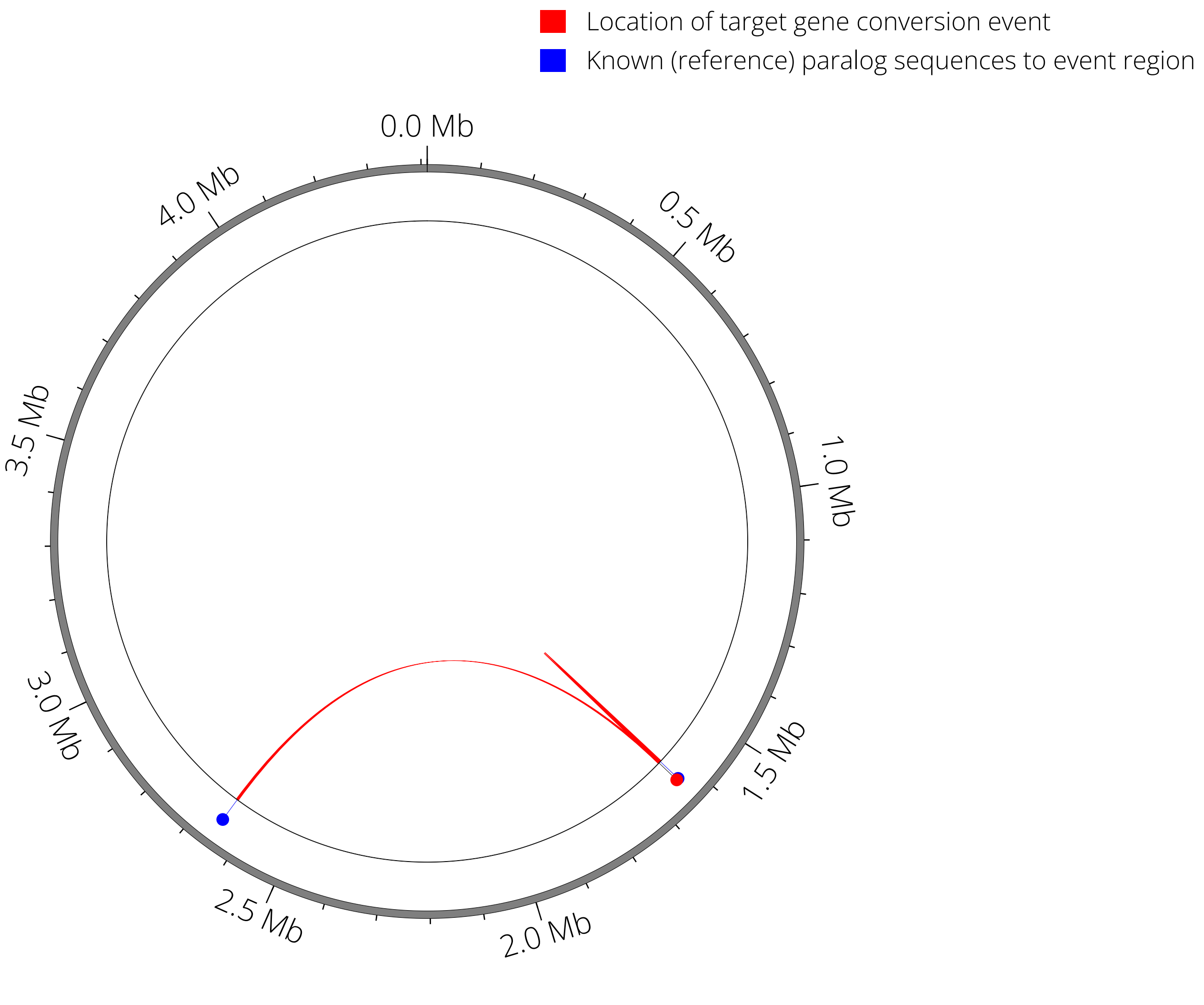

### Event_150 - 10 SNPs - k-mer analysis relative to paralogs

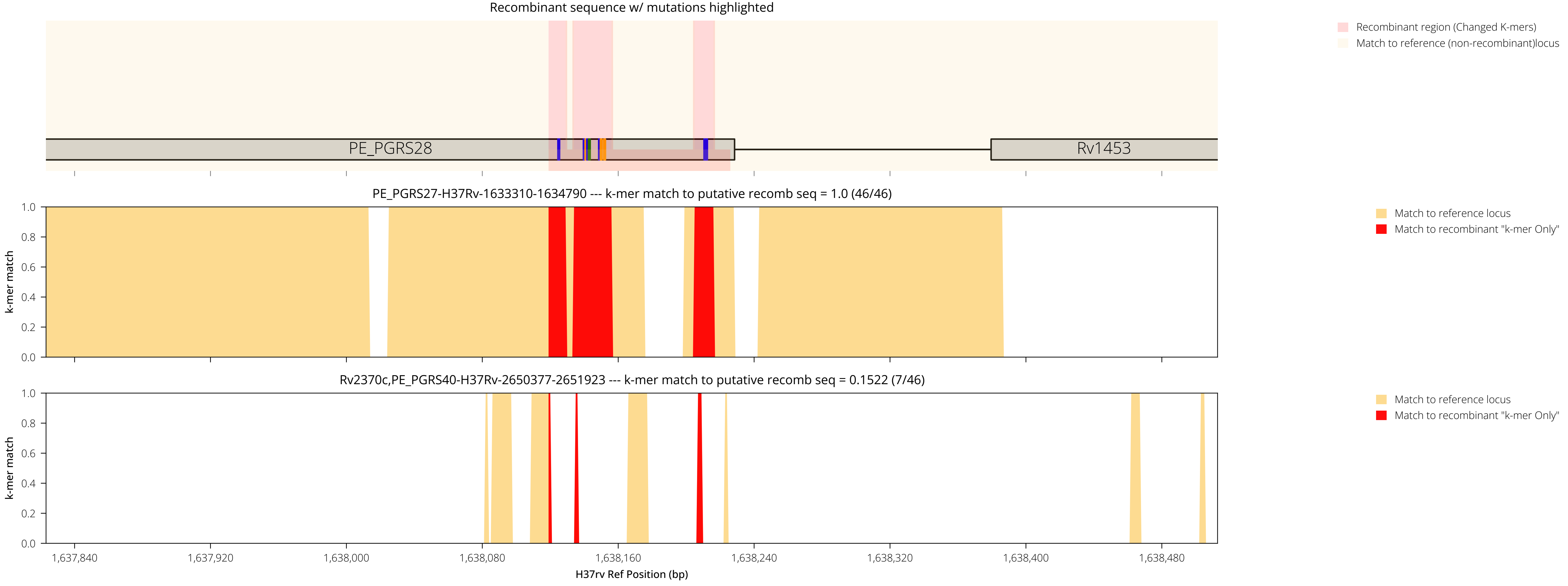

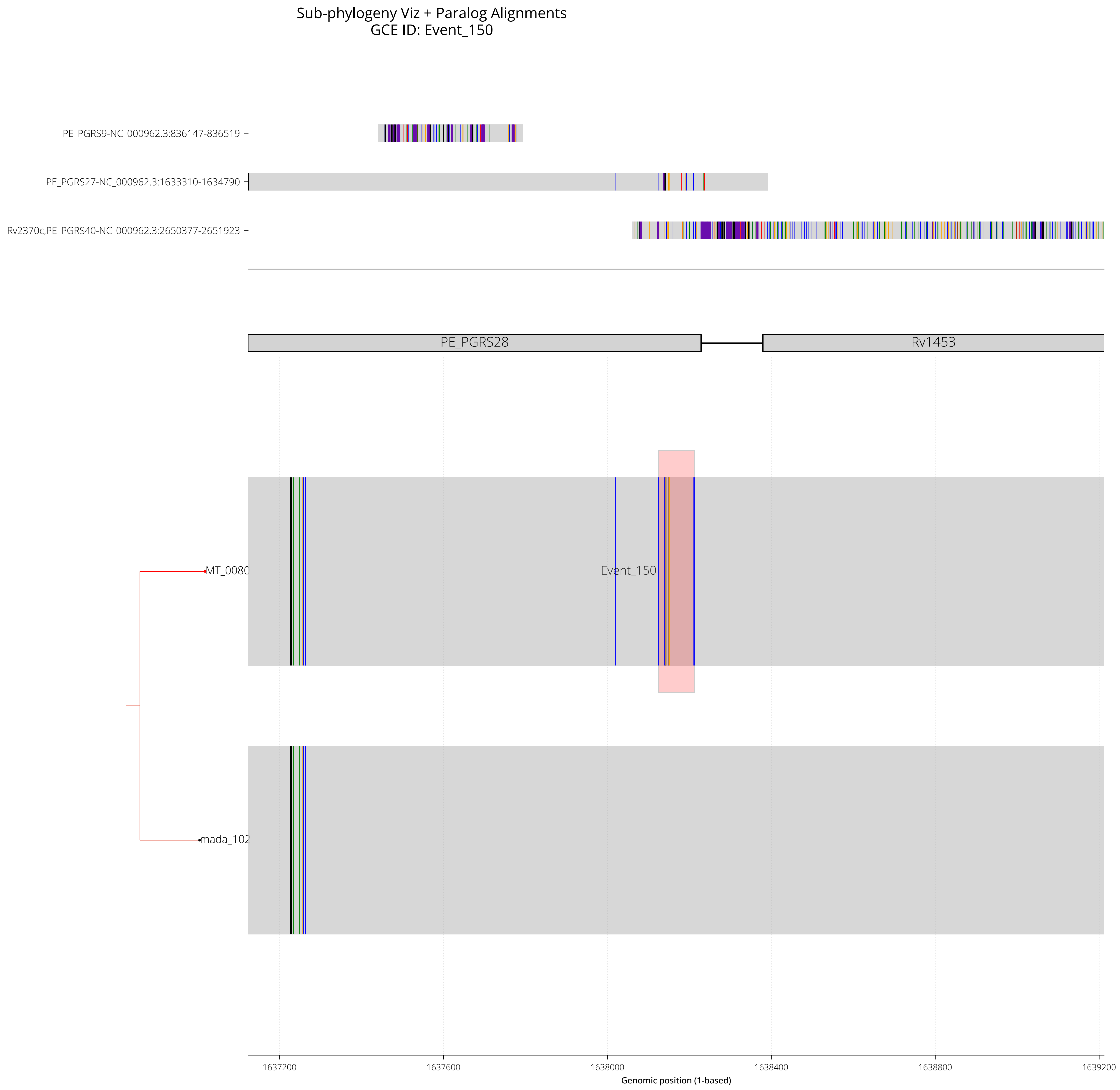

Sub-phylogeny Viz + Paralog Alignments
GCE ID: Event_150
PE_PGRS9-NC_000962.3:836147-836519
PE_PGRS27-NC_000962.3:1633310-1634790
Rv2370c,PE_PGRS40-NC_000962.3:2650377-2651923
PE_PGRS28
Rv1453
MT_0080
Event_150
mada_102
1637200
1637600
1638000
1638400
1638800
1639200
Genomic position (1-based)

Full Phylogeny Viz + Paralog Alignments
GCE ID: Event_150

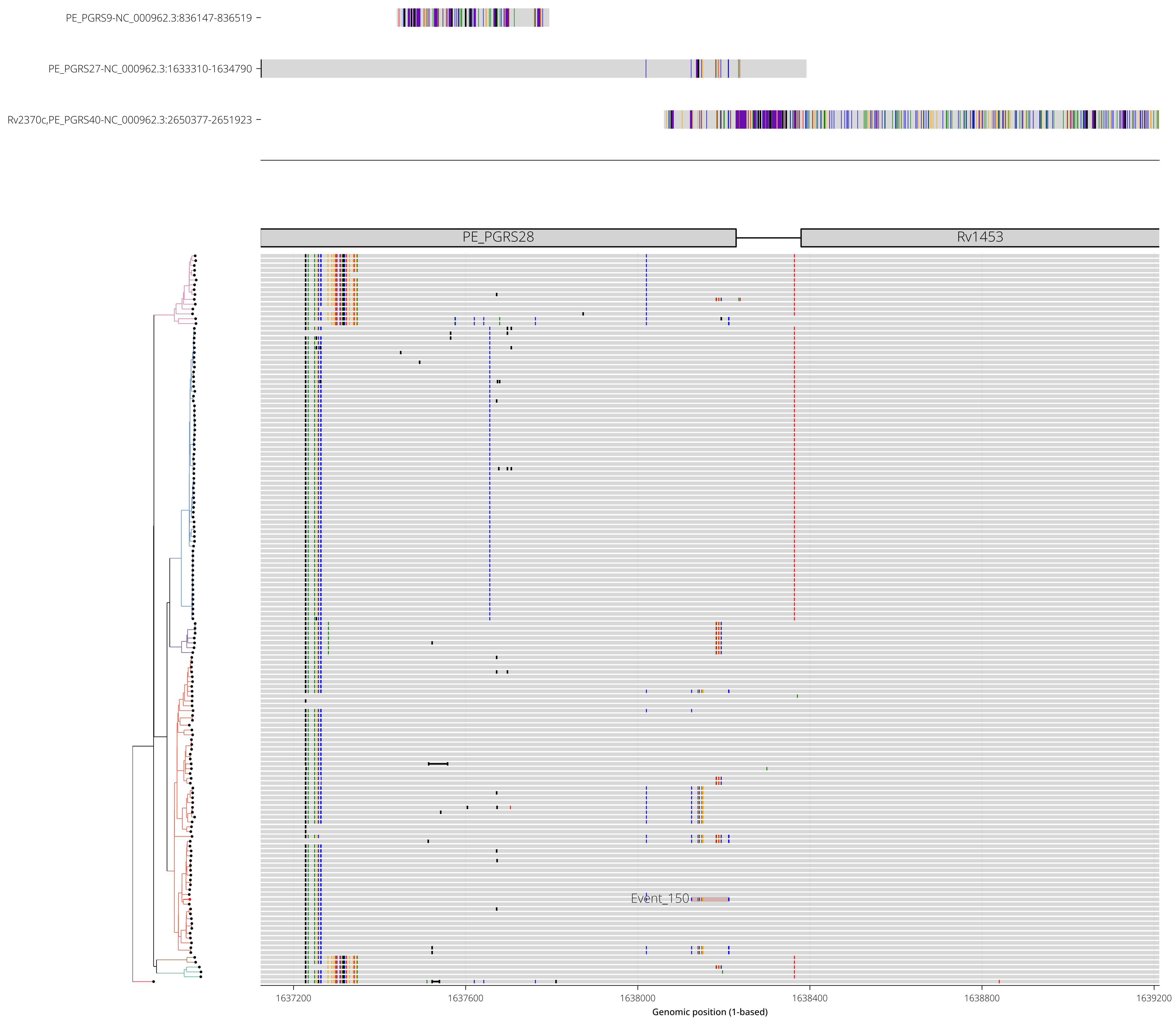

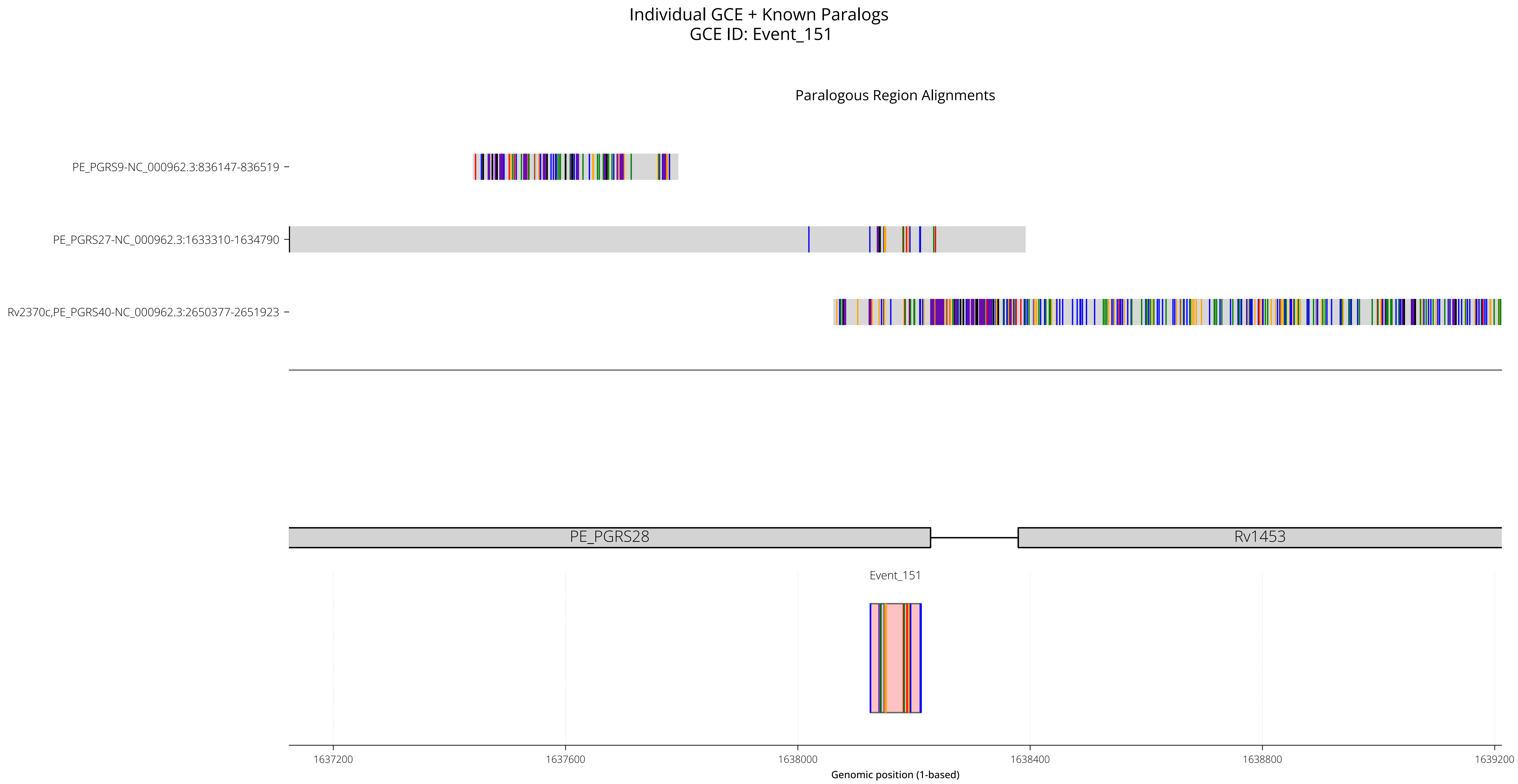

Individual GCE + Known Paralogs
GCE ID: Event_151
Paralogous Region Alignments
PE_PGRS9-NC_000962.3:836147-836519
PE_PGRS27-NC_000962.3:1633310-1634790
Rv2370c,PE_PGRS40-NC_000962.3:2650377-2651923
PE_PGRS28
Rv1453
Event_151
1637200
1637600
1638000
1638400
1638800
1639200
Genomic position (1-based)

Circos Viz - GCE to Known Paralogs Locations
GCE ID: Event_151

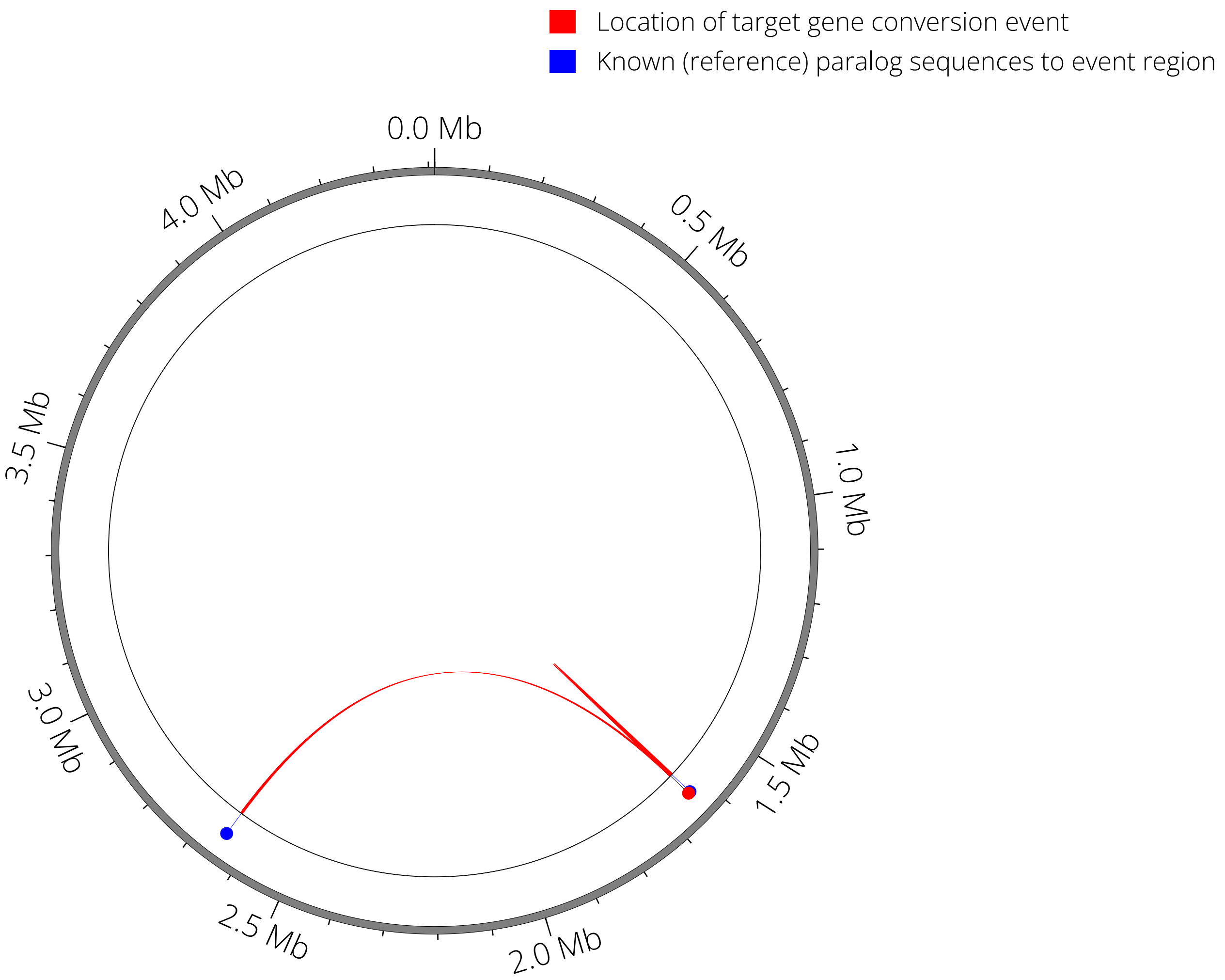

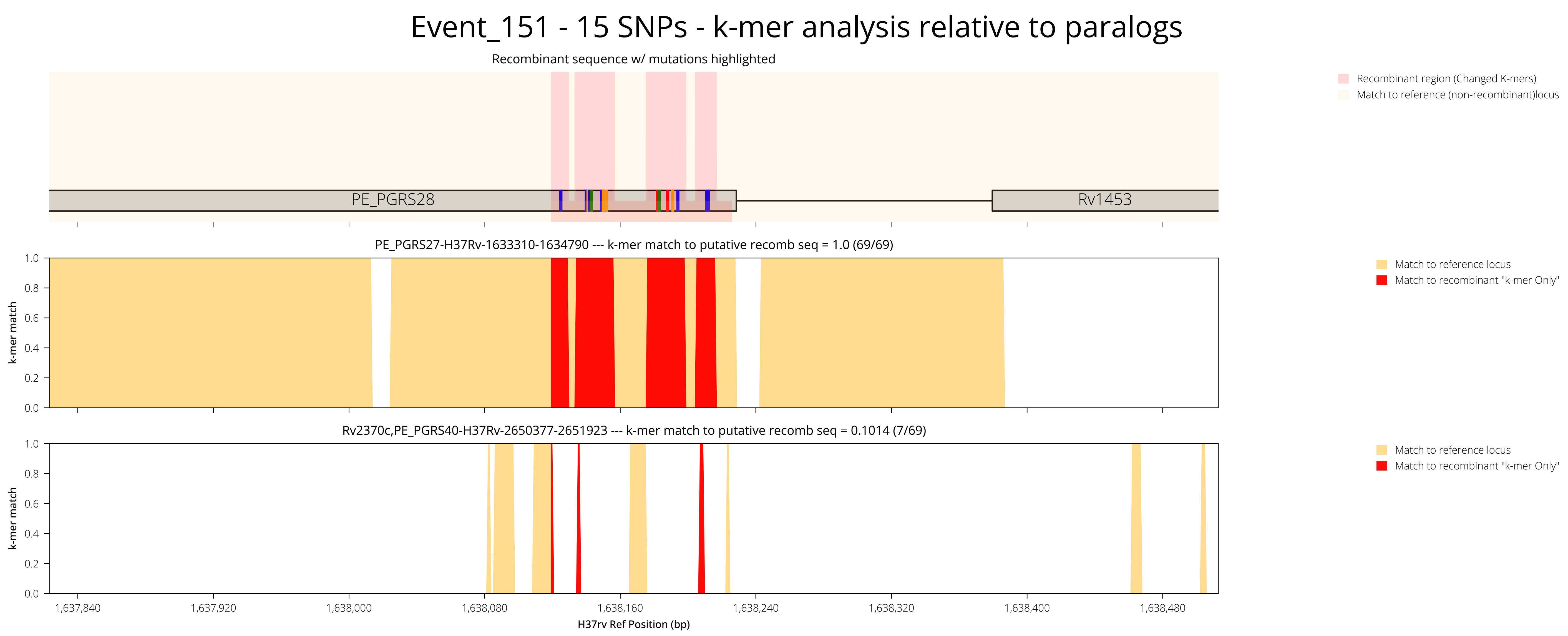

Event_151 - 15 SNPs - k-mer analysis relative to paralogs
Recombinant sequence w/ mutations highlighted
Recombinant region (Changed K-mers)
Match to reference (non-recombinant)locus
PE_PGRS28
Rv1453
PE_PGRS27-H37Rv-1633310-1634790 --- k-mer match to putative recomb seq = 1.0 (69/69)
Match to reference locus
Match to recombinant "k-mer Only"
k-mer match
1.0
0.8
0.6
0.4
0.2
0.0
Rv2370c,PE_PGRS40-H37Rv-2650377-2651923 --- k-mer match to putative recomb seq = 0.1014 (7/69)
Match to reference locus
Match to recombinant "k-mer Only"
k-mer match
1,637,840
1,637,920
1,638,000
1,638,080
1,638,160
1,638,240
1,638,320
1,638,400
1,638,480
H37rv Ref Position (bp)

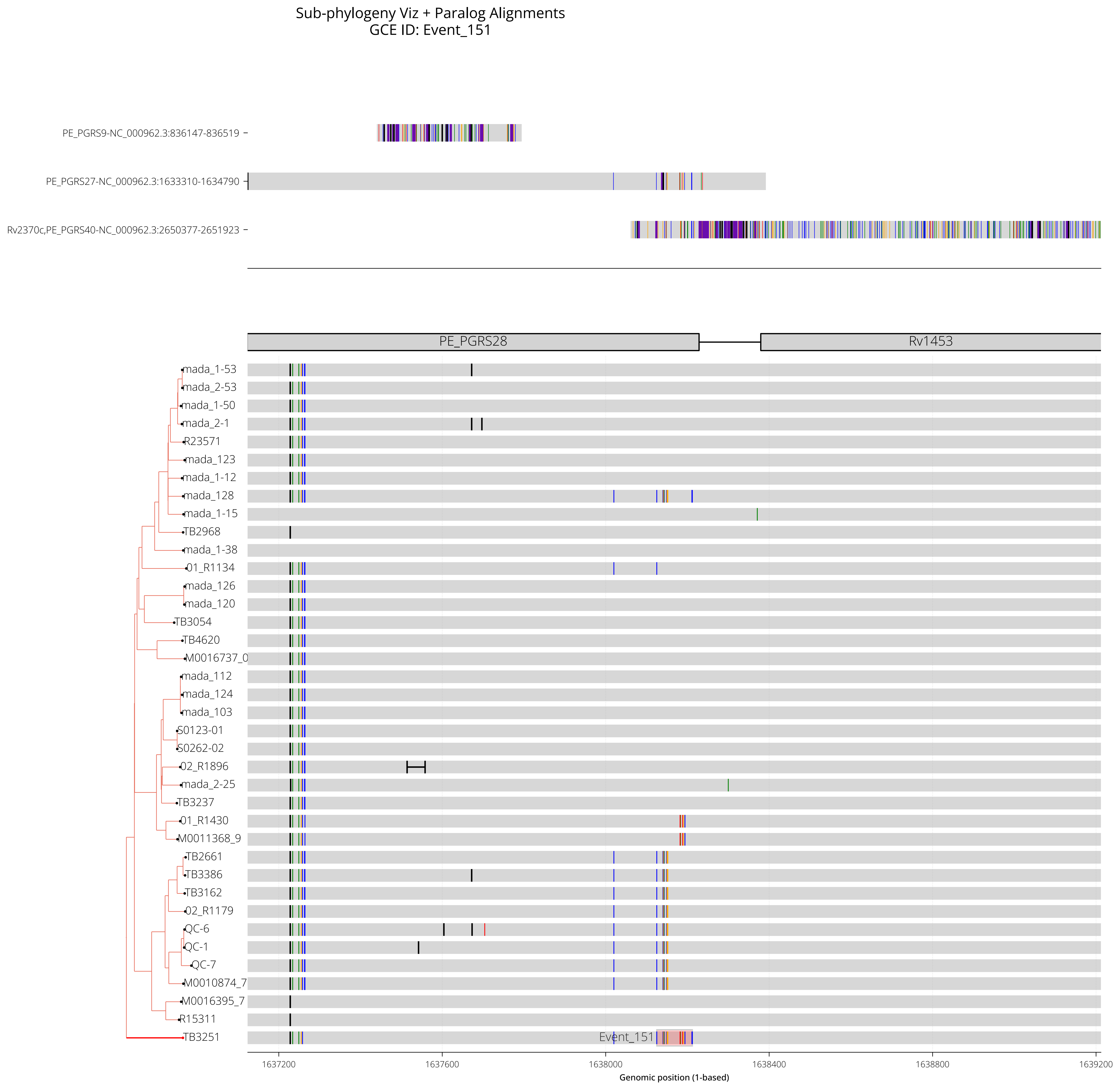

Sub-phylogeny Viz + Paralog Alignments
GCE ID: Event_151
PE_PGRS9-NC_000962.3:836147-836519
PE_PGRS27-NC_000962.3:1633310-1634790
Rv2370c,PE_PGRS40-NC_000962.3:2650377-2651923
PE_PGRS28
Rv1453
mada_1-53
mada_2-53
mada_1-50
mada_2-1
R23571
mada_123
mada_1-12
mada_128
mada_1-15
TB2968
mada_1-38
01_R1134
mada_126
mada_120
TB3054
TB4620
M0016737_0
mada_112
mada_124
mada_103
S0123-01
S0262-02
02_R1896
mada_2-25
TB3237
01_R1430
M0011368_9
TB2661
TB3386
TB3162
02_R1179
QC-6
QC-1
QC-7
M0010874_7
M0016395_7
R15311
TB3251
Event_151
1637200
1637600
1638000
1638400
1638800
1639200
Genomic position (1-based)

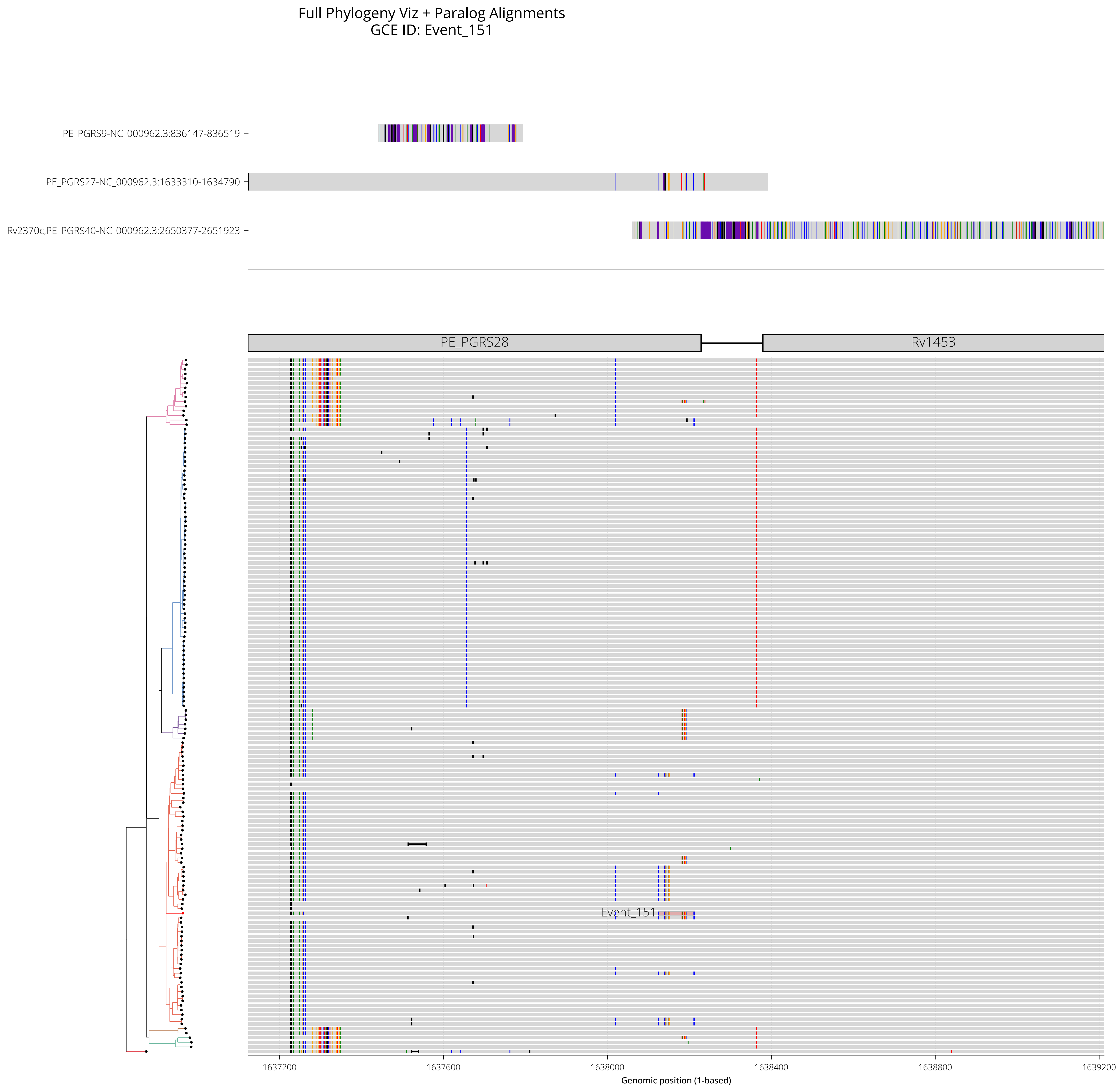

Full Phylogeny Viz + Paralog Alignments
GCE ID: Event_151
PE_PGRS9-NC_000962.3:836147-836519
PE_PGRS27-NC_000962.3:1633310-1634790
Rv2370c,PE_PGRS40-NC_000962.3:2650377-2651923
PE_PGRS28
Rv1453
Event_151
1637200
1637600
1638000
1638400
1638800
1639200
Genomic position (1-based)

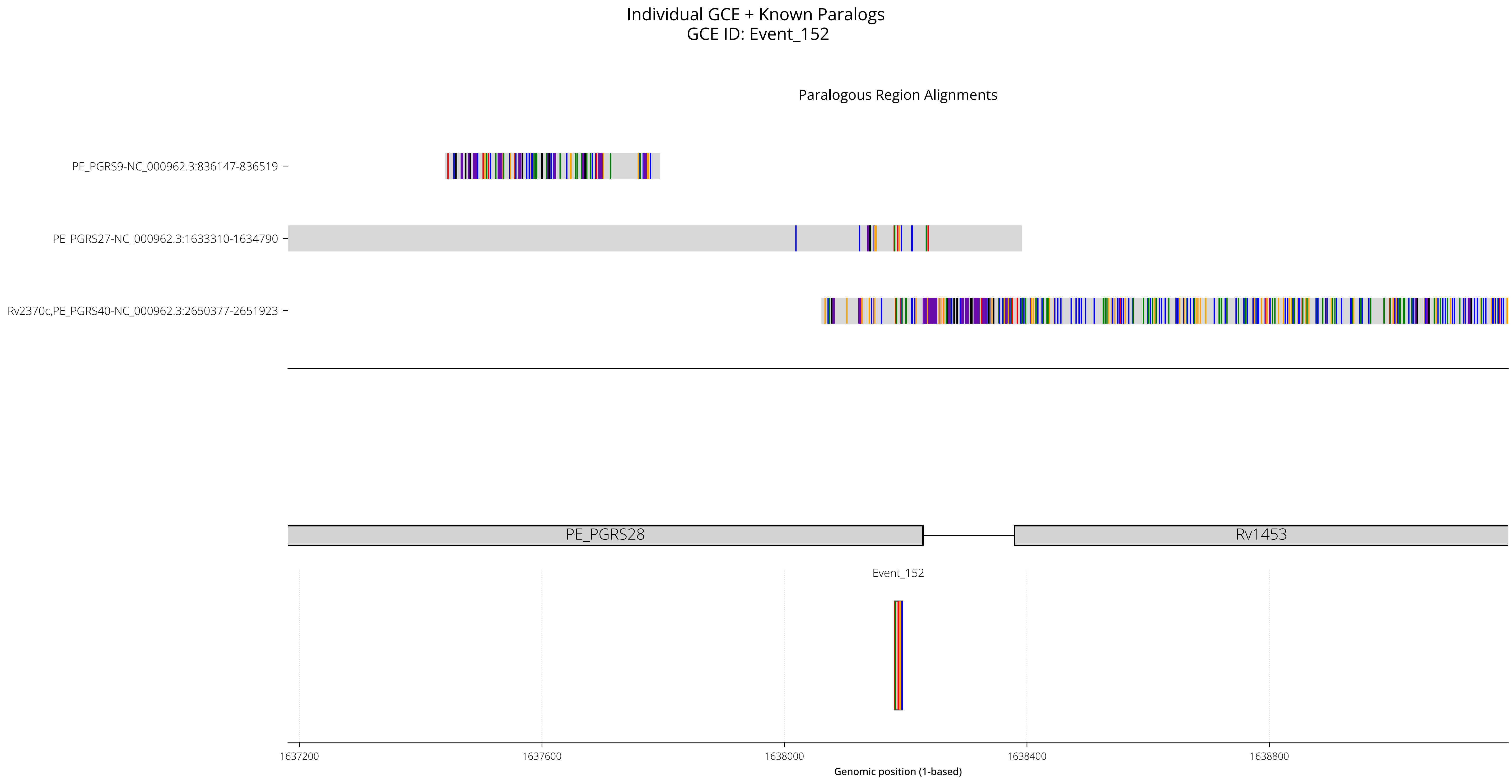
Individual GCE + Known Paralogs
GCE ID: Event_152
Paralogous Region Alignments
PE_PGRS9-NC_000962.3:836147-836519
PE_PGRS27-NC_000962.3:1633310-1634790
Rv2370c,PE_PGRS40-NC_000962.3:2650377-2651923
PE_PGRS28
Rv1453
Event_152
1637200
1637600
1638000
1638400
1638800
Genomic position (1-based)

Circos Viz - GCE to Known Paralogs Locations
GCE ID: Event_152

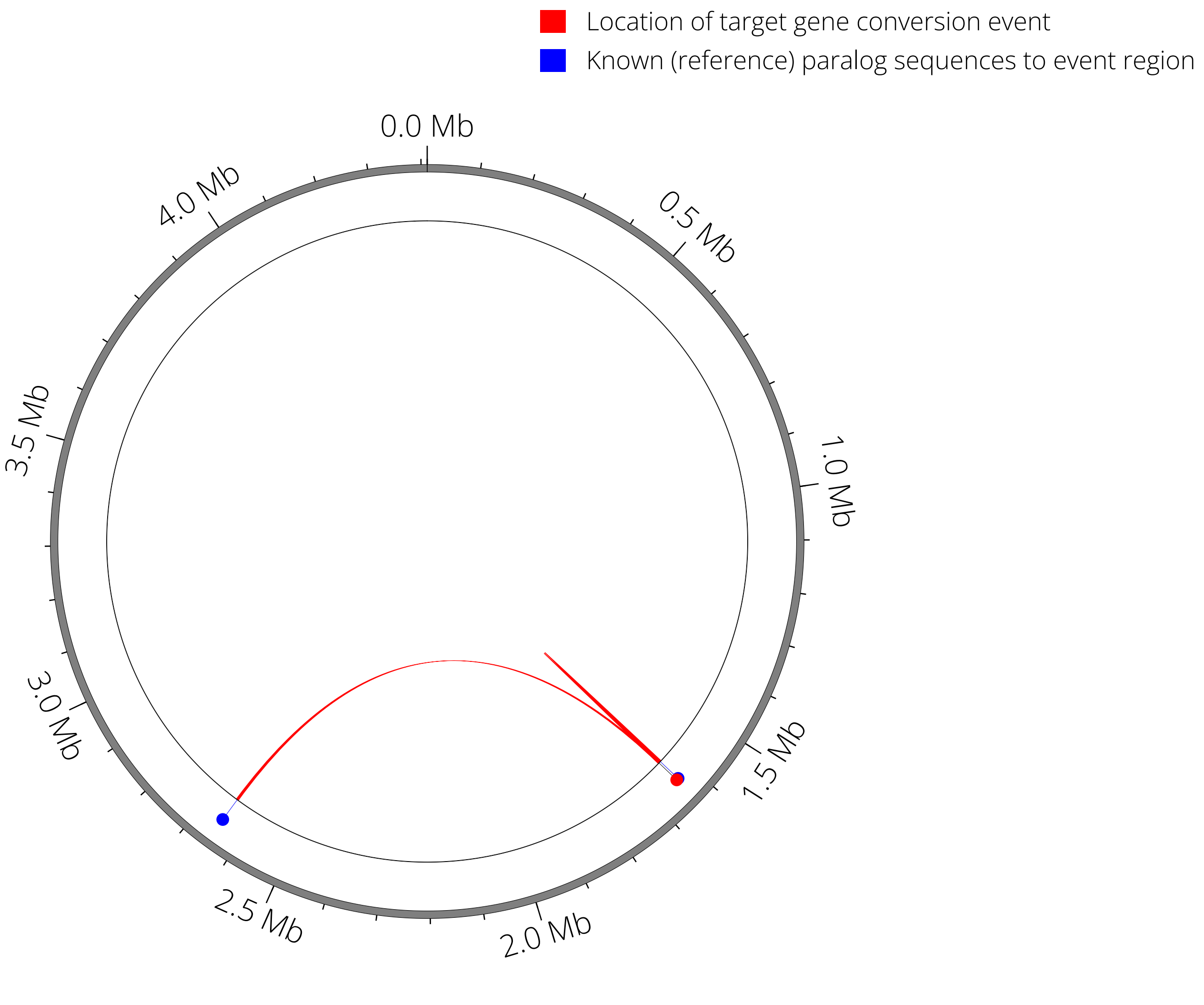

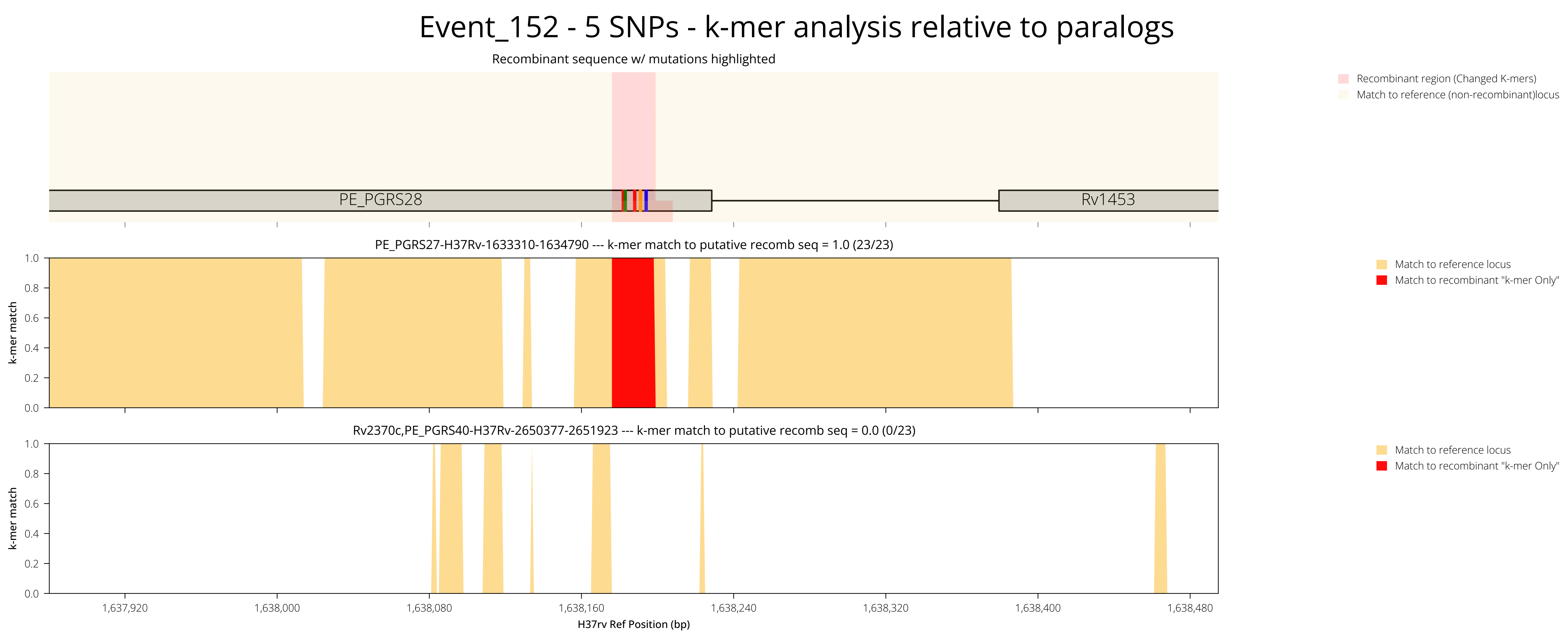

Event_152 - 5 SNPs - k-mer analysis relative to paralogs
Recombinant sequence w/ mutations highlighted
Recombinant region (Changed K-mers)
Match to reference (non-recombinant)locus
PE_PGRS28
Rv1453
PE_PGRS27-H37Rv-1633310-1634790 --- k-mer match to putative recomb seq = 1.0 (23/23)
Match to reference locus
Match to recombinant "k-mer Only"
Rv2370c,PE_PGRS40-H37Rv-2650377-2651923 --- k-mer match to putative recomb seq = 0.0 (0/23)
Match to reference locus
Match to recombinant "k-mer Only"
k-mer match
1.0
0.8
0.6
0.4
0.2
0.0
1,637,920
1,638,000
1,638,080
1,638,160
1,638,240
1,638,320
1,638,400
1,638,480
H37rv Ref Position (bp)

Sub-phylogeny Viz + Paralog Alignments
GCE ID: Event_152

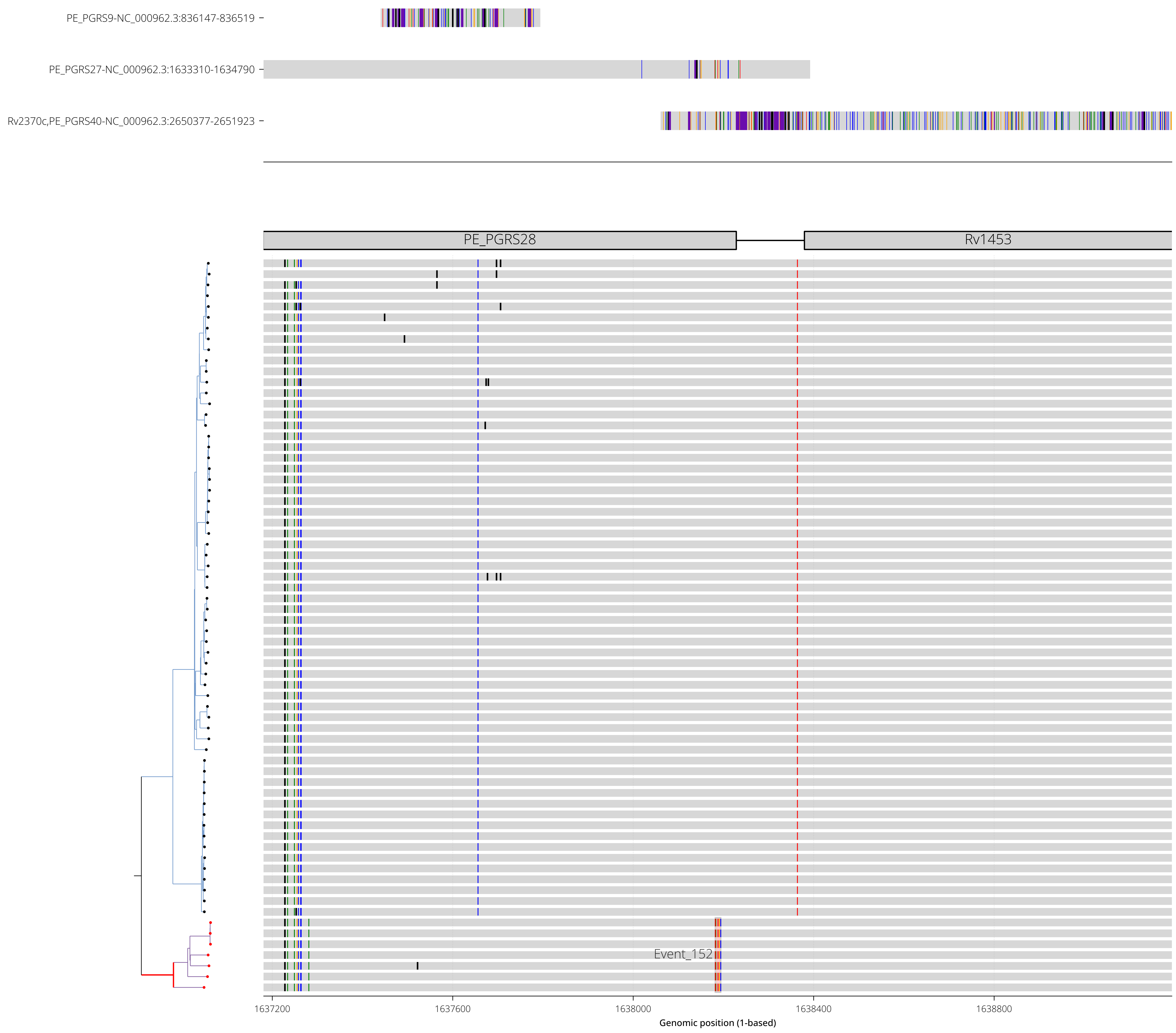

#### Full Phylogeny Viz + Paralog Alignments
#### GCE ID: Event_152

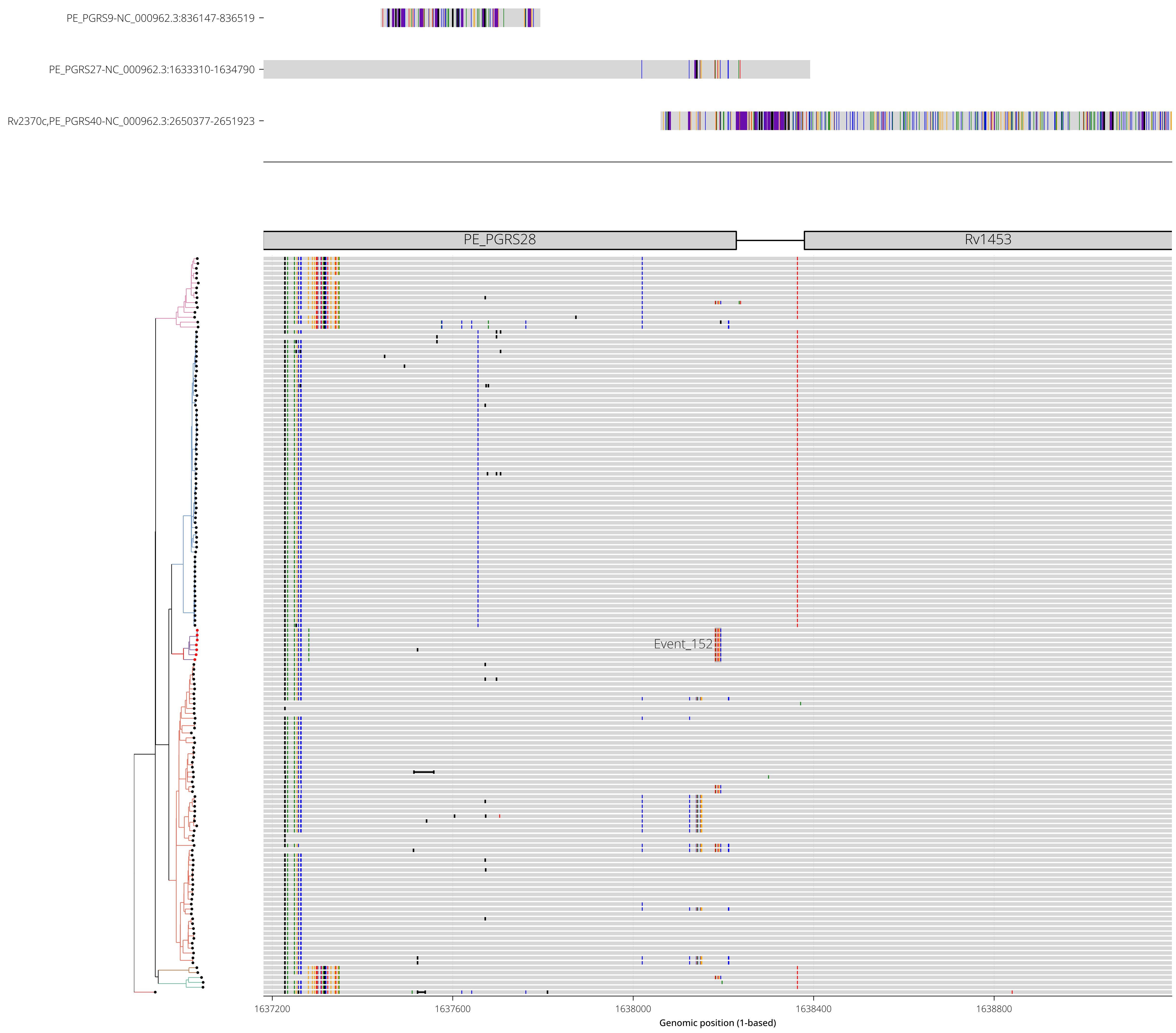

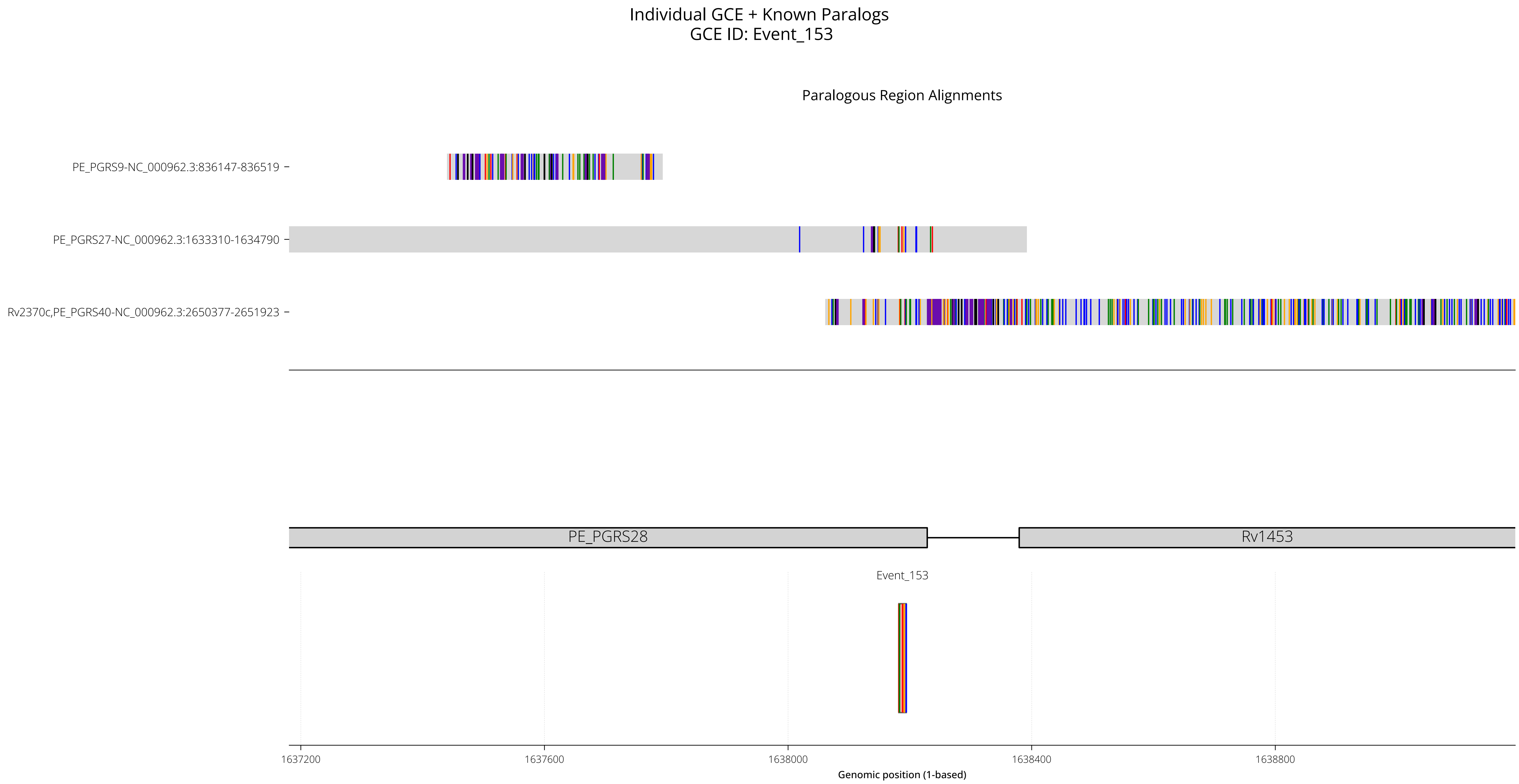

Individual GCE + Known Paralogs
GCE ID: Event_153
Paralogous Region Alignments
PE_PGRS9-NC_000962.3:836147-836519
PE_PGRS27-NC_000962.3:1633310-1634790
Rv2370c,PE_PGRS40-NC_000962.3:2650377-2651923
PE_PGRS28
Rv1453
Event_153
1637200
1637600
1638000
1638400
1638800
Genomic position (1-based)

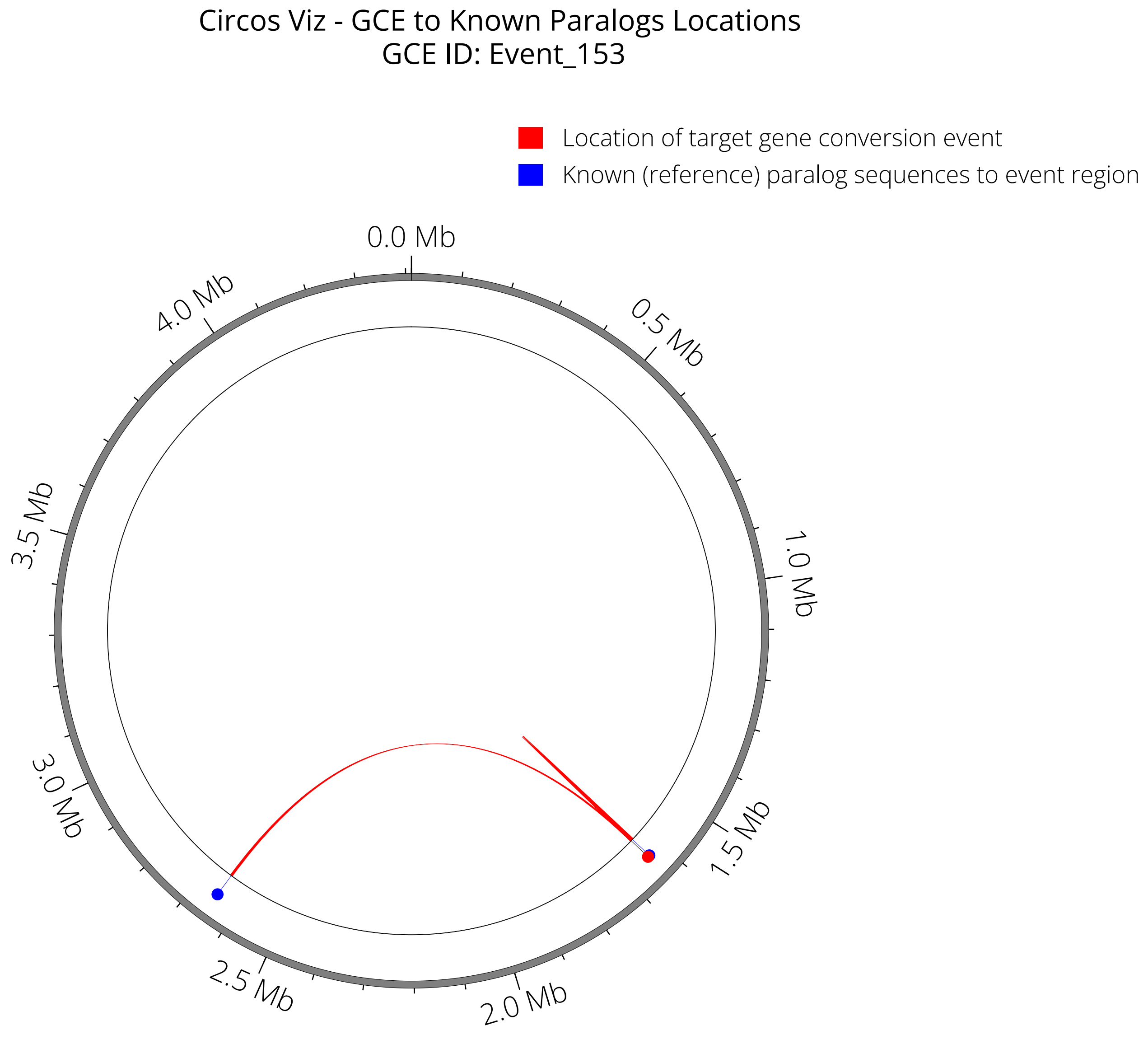
Circos Viz - GCE to Known Paralogs Locations
GCE ID: Event_153
Location of target gene conversion event
Known (reference) paralog sequences to event region
0.0 Mb
0.5 Mb
1.0 Mb
1.5 Mb
2.0 Mb
2.5 Mb
3.0 Mb
3.5 Mb
4.0 Mb

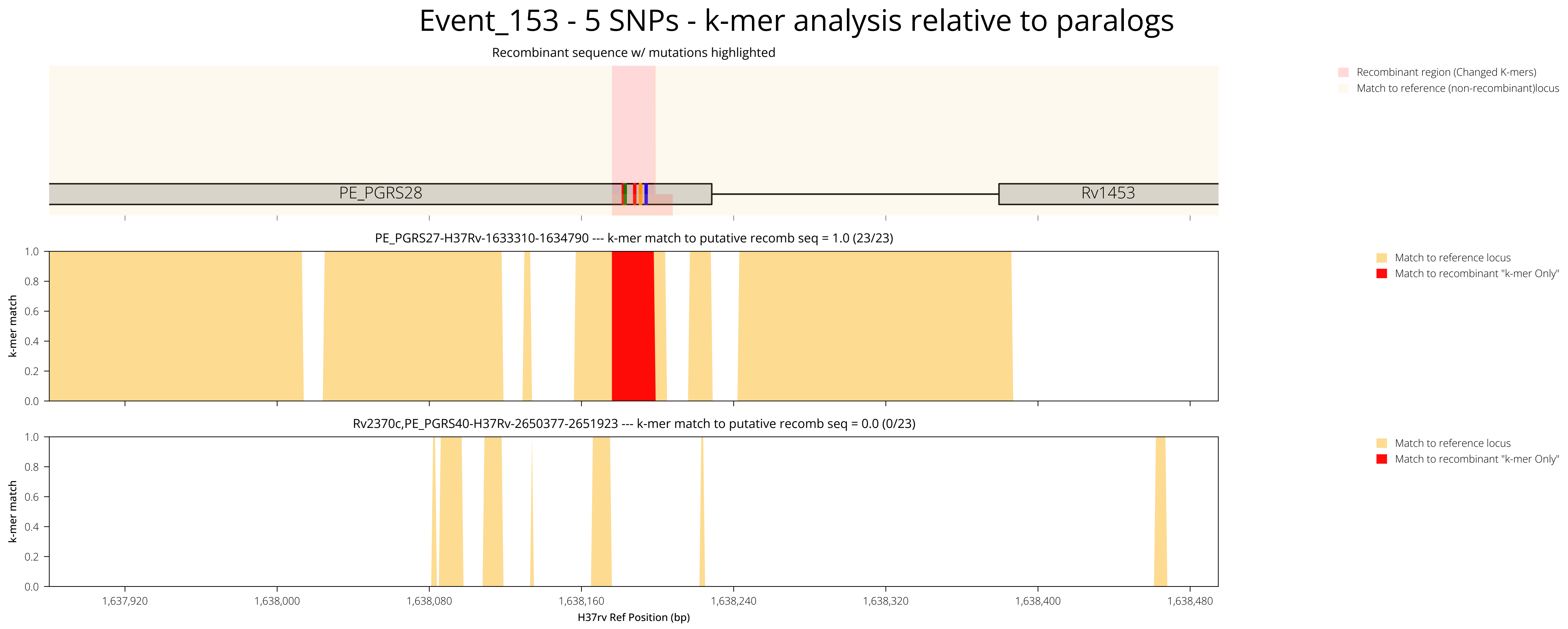

Event_153 - 5 SNPs - k-mer analysis relative to paralogs
Recombinant sequence w/ mutations highlighted
Recombinant region (Changed K-mers)
Match to reference (non-recombinant)locus
PE_PGRS28
Rv1453
PE_PGRS27-H37Rv-1633310-1634790 --- k-mer match to putative recomb seq = 1.0 (23/23)
Match to reference locus
Match to recombinant "k-mer Only"
k-mer match
1.0
0.8
0.6
0.4
0.2
0.0
Rv2370c,PE_PGRS40-H37Rv-2650377-2651923 --- k-mer match to putative recomb seq = 0.0 (0/23)
Match to reference locus
Match to recombinant "k-mer Only"
k-mer match
1,637,920
1,638,000
1,638,080
1,638,160
1,638,240
1,638,320
1,638,400
1,638,480
H37rv Ref Position (bp)

Sub-phylogeny Viz + Paralog Alignments
GCE ID: Event_153

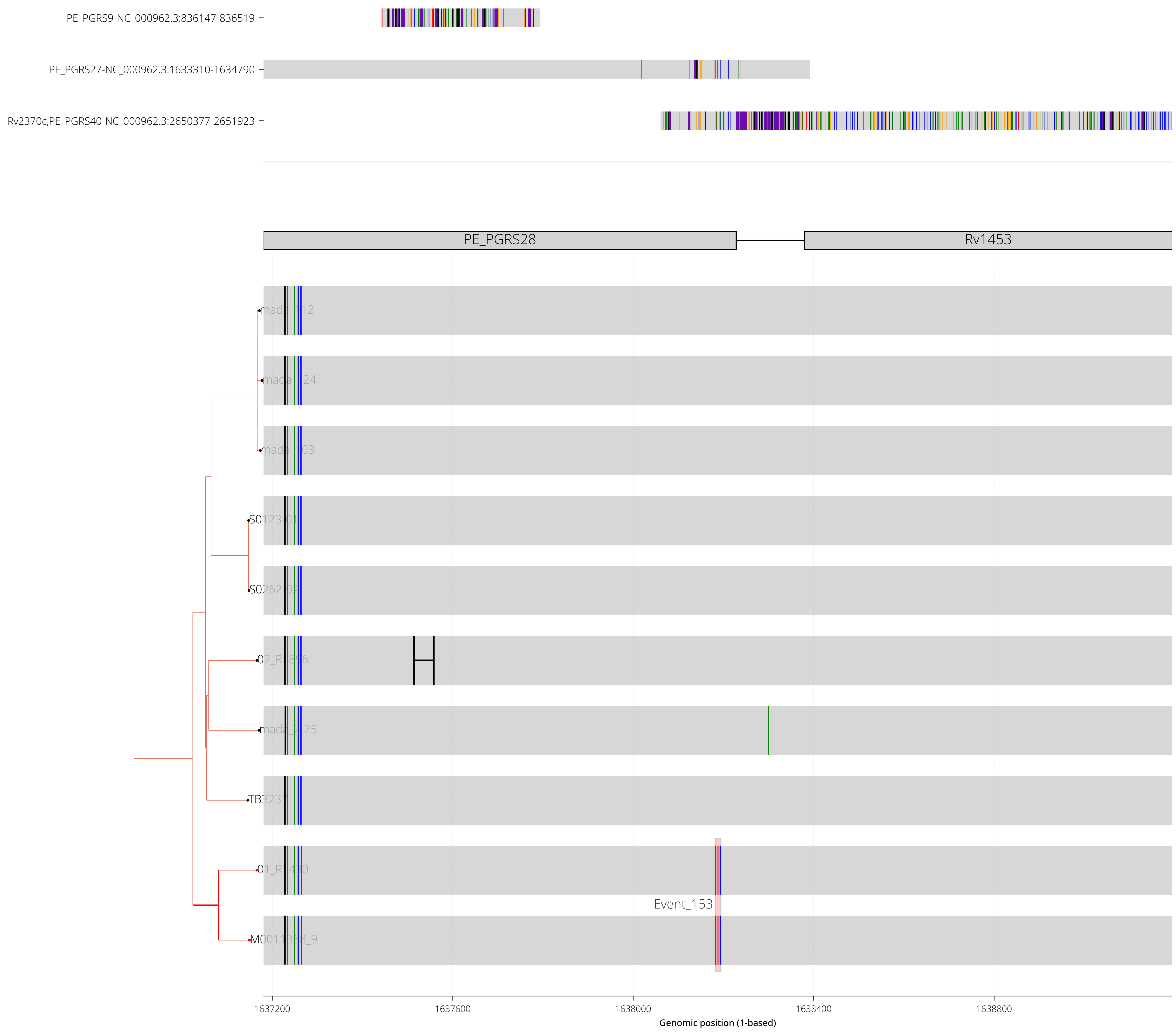

Full Phylogeny Viz + Paralog Alignments
GCE ID: Event_153

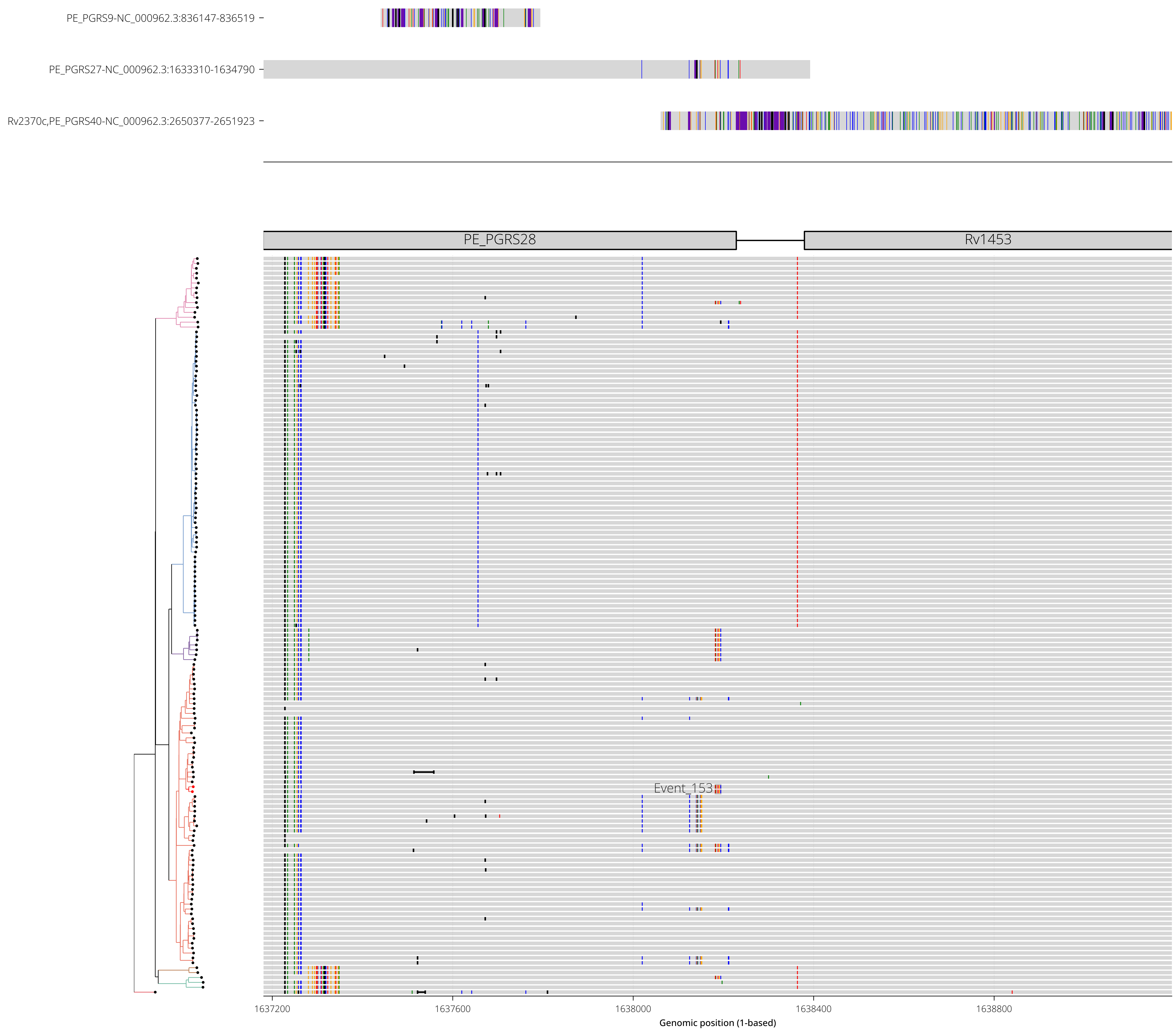

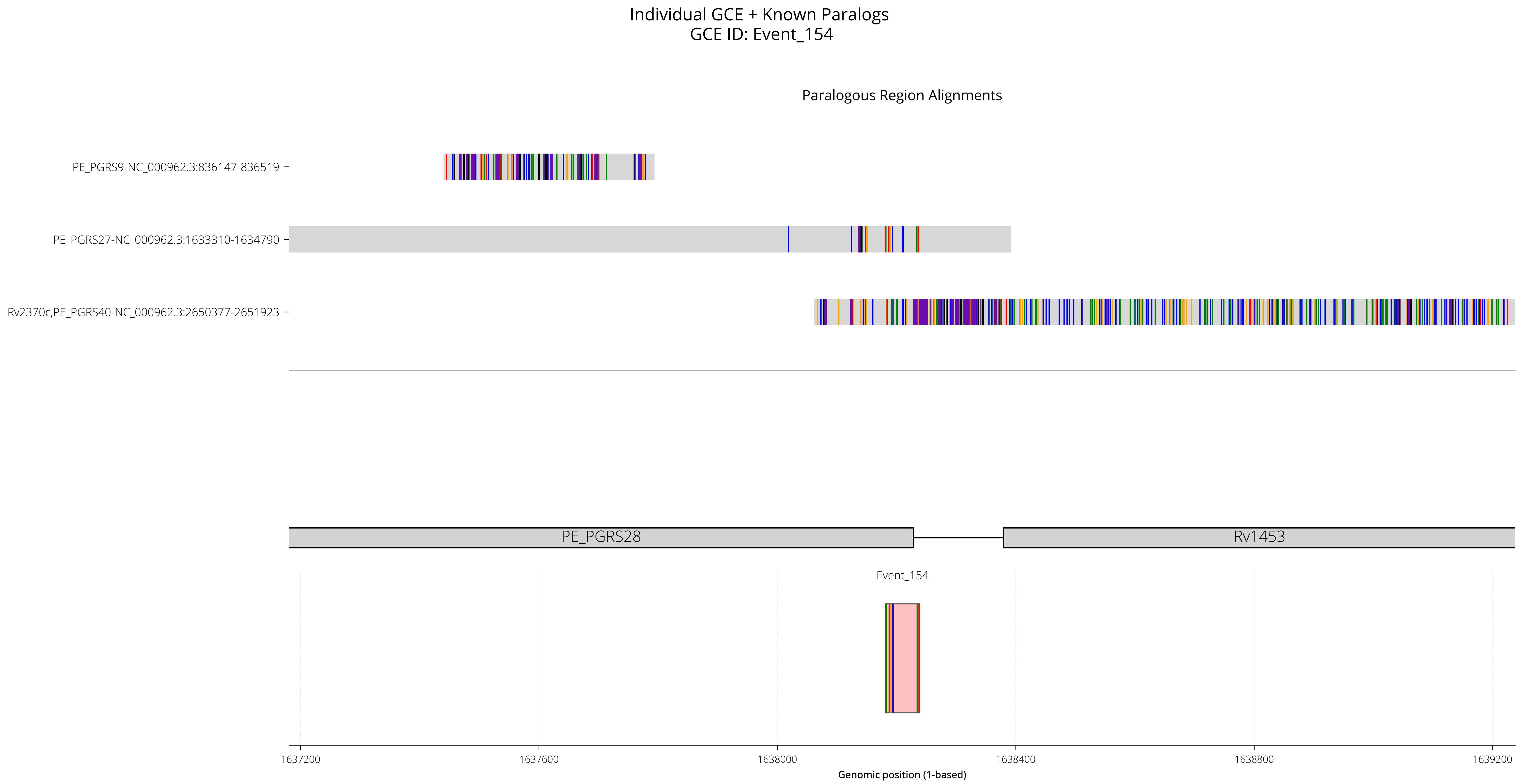
Individual GCE + Known Paralogs
GCE ID: Event_154
Paralogous Region Alignments
PE_PGRS9-NC_000962.3:836147-836519
PE_PGRS27-NC_000962.3:1633310-1634790
Rv2370c,PE_PGRS40-NC_000962.3:2650377-2651923
PE_PGRS28
Rv1453
Event_154
1637200
1637600
1638000
1638400
1638800
1639200
Genomic position (1-based)

Circos Viz - GCE to Known Paralogs Locations
GCE ID: Event_154

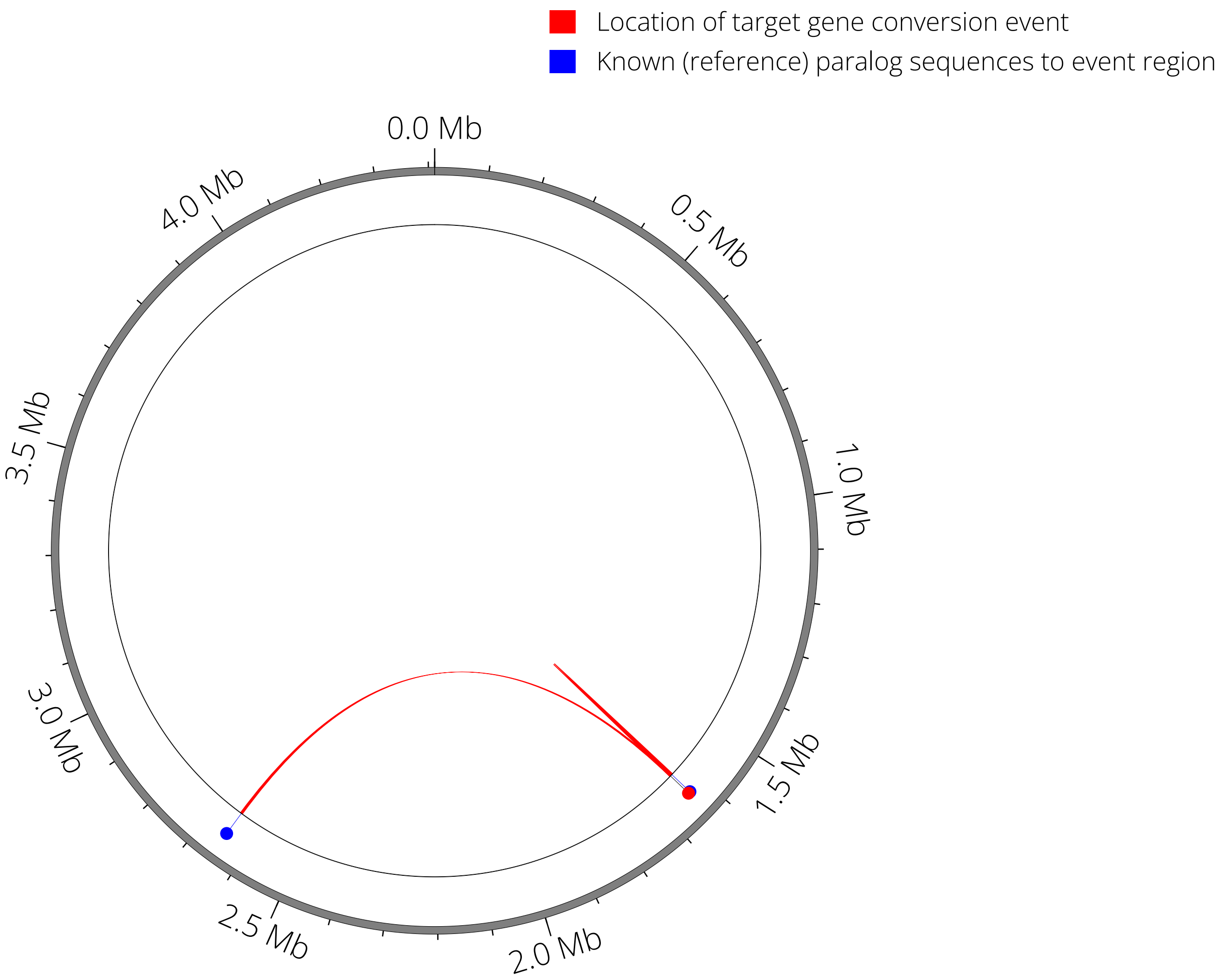

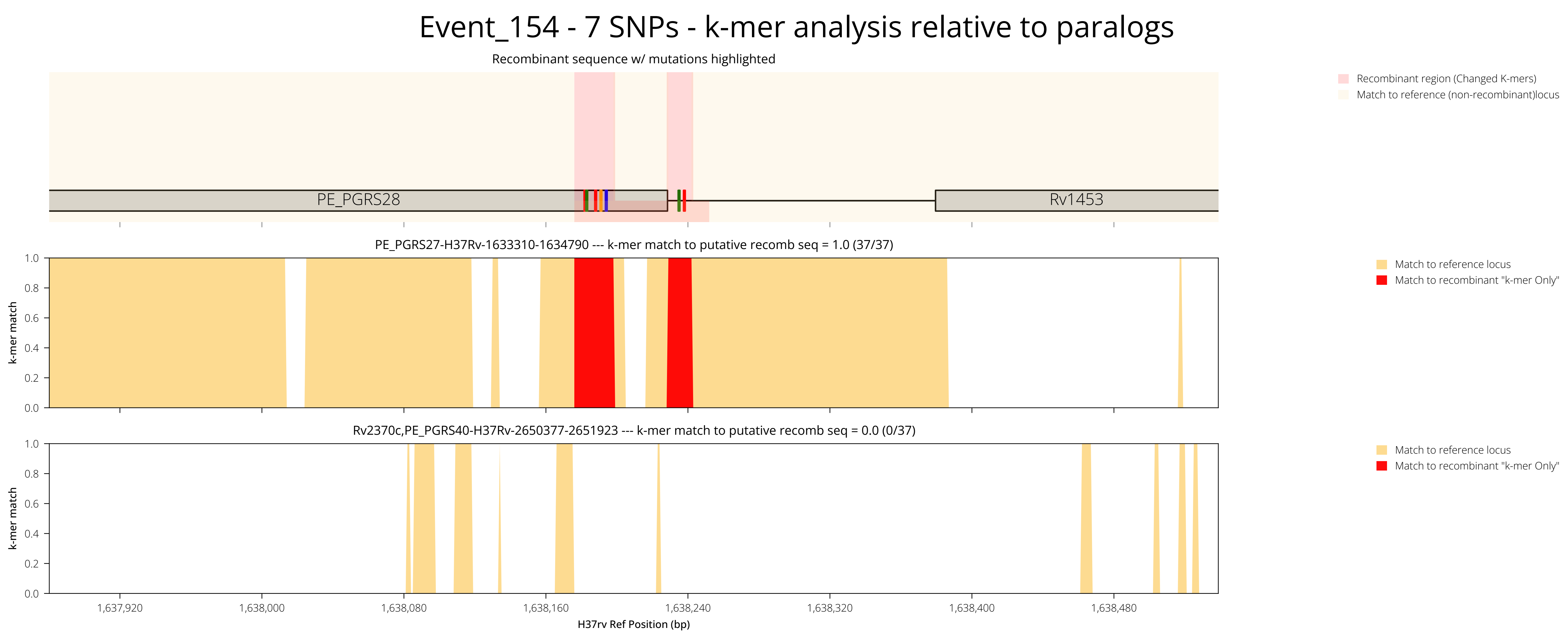

Event_154 - 7 SNPs - k-mer analysis relative to paralogs
Recombinant sequence w/ mutations highlighted
Recombinant region (Changed K-mers)
Match to reference (non-recombinant)locus
PE_PGRS28
Rv1453
PE_PGRS27-H37Rv-1633310-1634790 --- k-mer match to putative recomb seq = 1.0 (37/37)
Match to reference locus
Match to recombinant "k-mer Only"
k-mer match
1.0
0.8
0.6
0.4
0.2
0.0
Rv2370c,PE_PGRS40-H37Rv-2650377-2651923 --- k-mer match to putative recomb seq = 0.0 (0/37)
Match to reference locus
Match to recombinant "k-mer Only"
k-mer match
1,637,920
1,638,000
1,638,080
1,638,160
1,638,240
1,638,320
1,638,400
1,638,480
H37rv Ref Position (bp)

Sub-phylogeny Viz + Paralog Alignments
GCE ID: Event_154

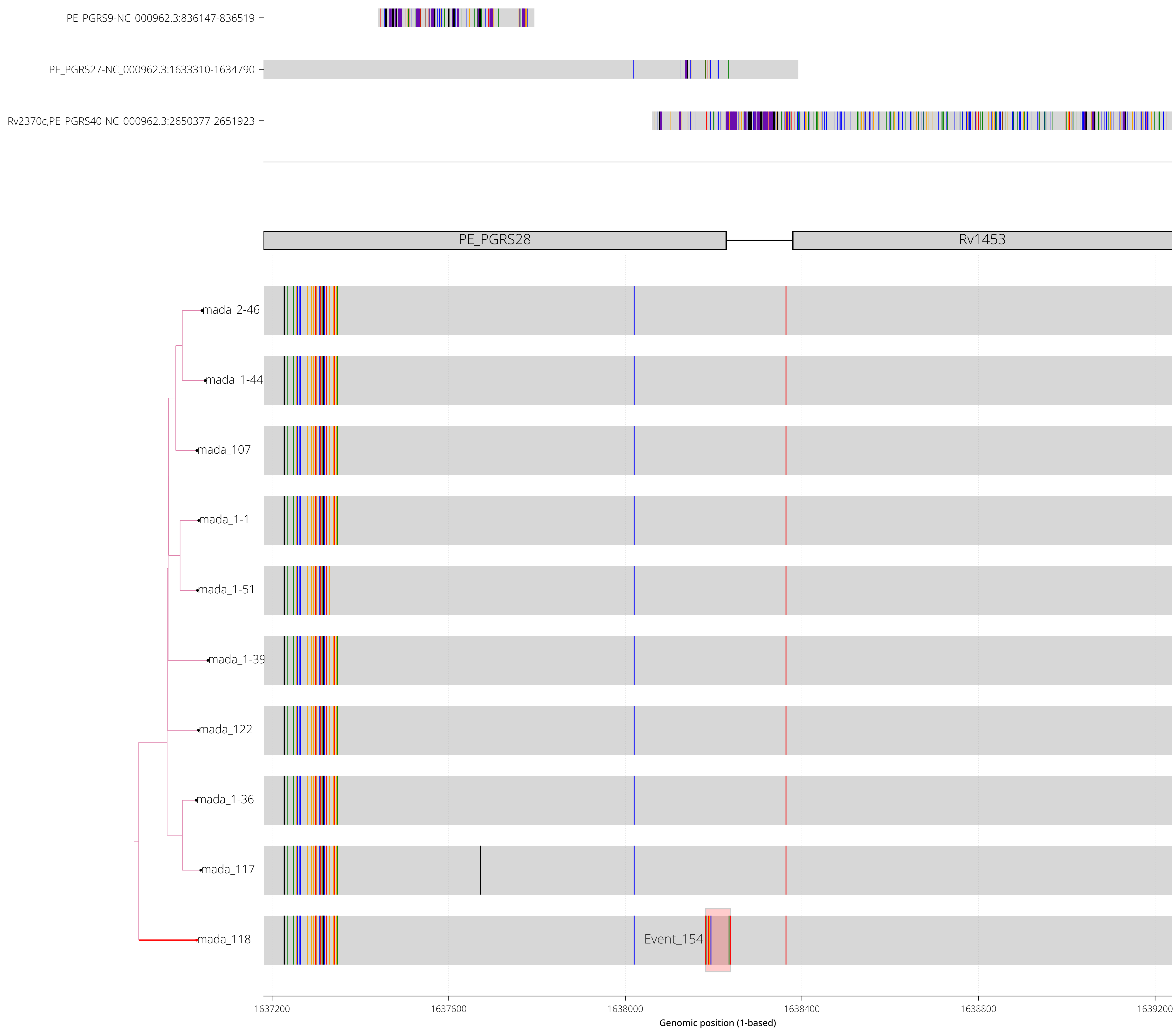

### Full Phylogeny Viz + Paralog Alignments
GCE ID: Event_154

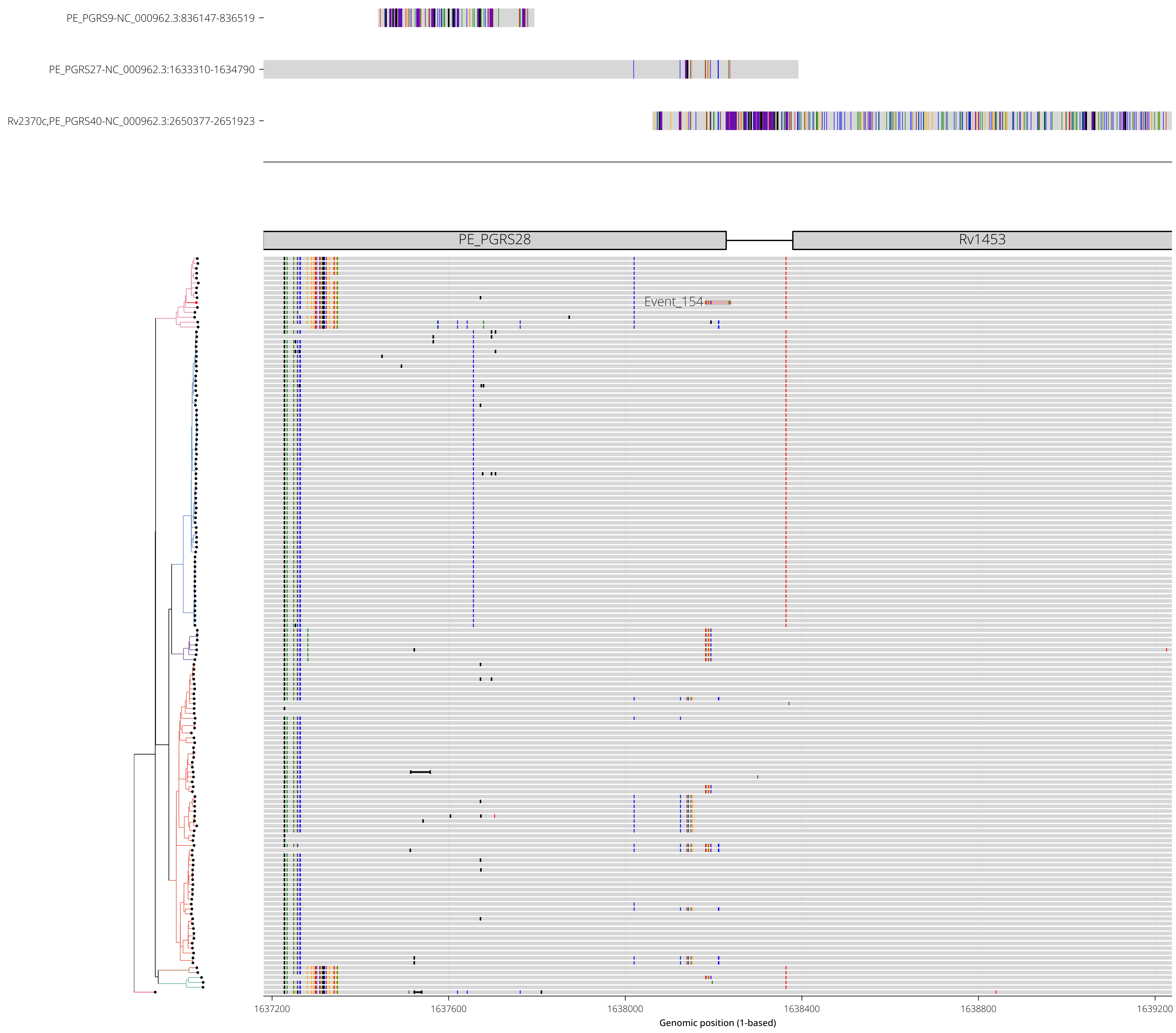

Individual GCE + Known Paralogs
GCE ID: Event_155

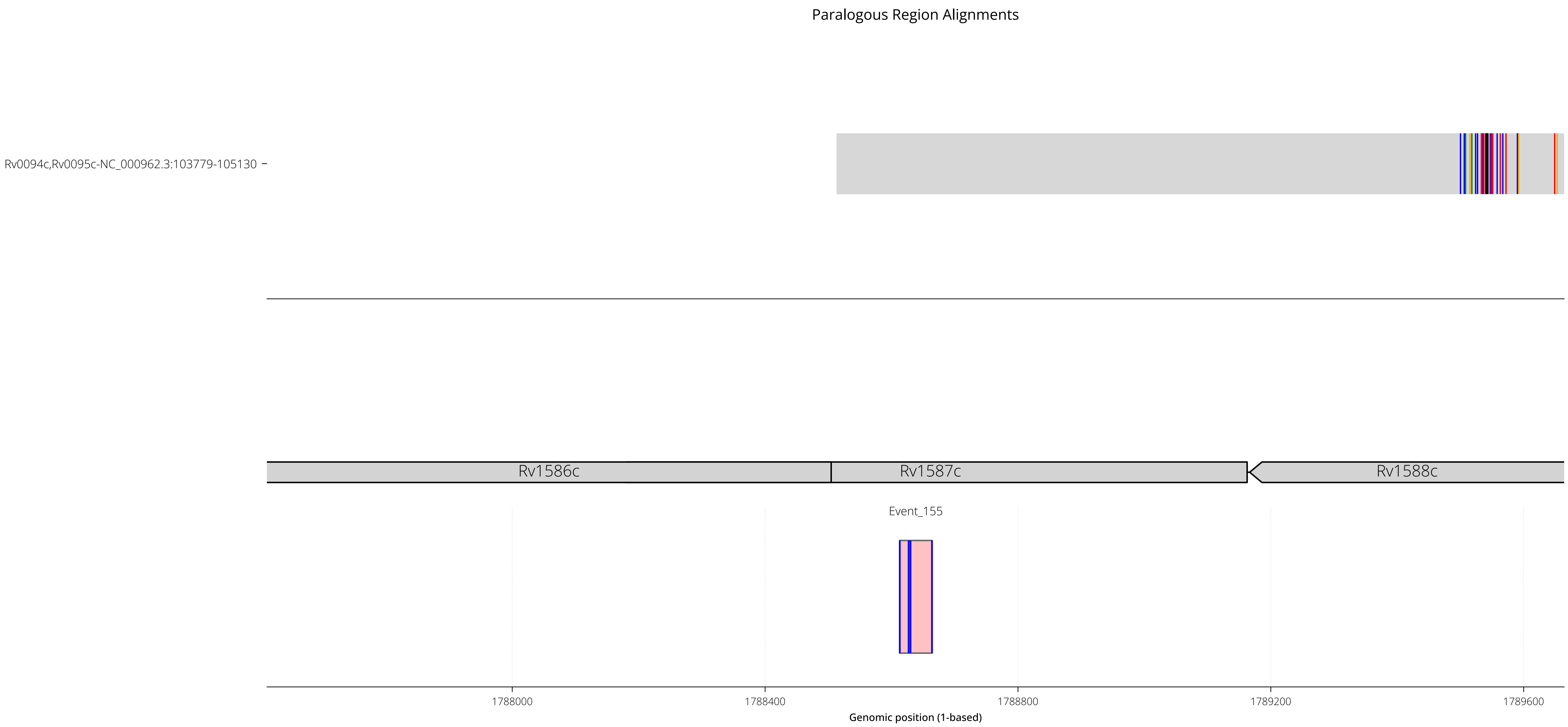

### Circos Viz - GCE to Known Paralogs Locations
GCE ID: Event_155

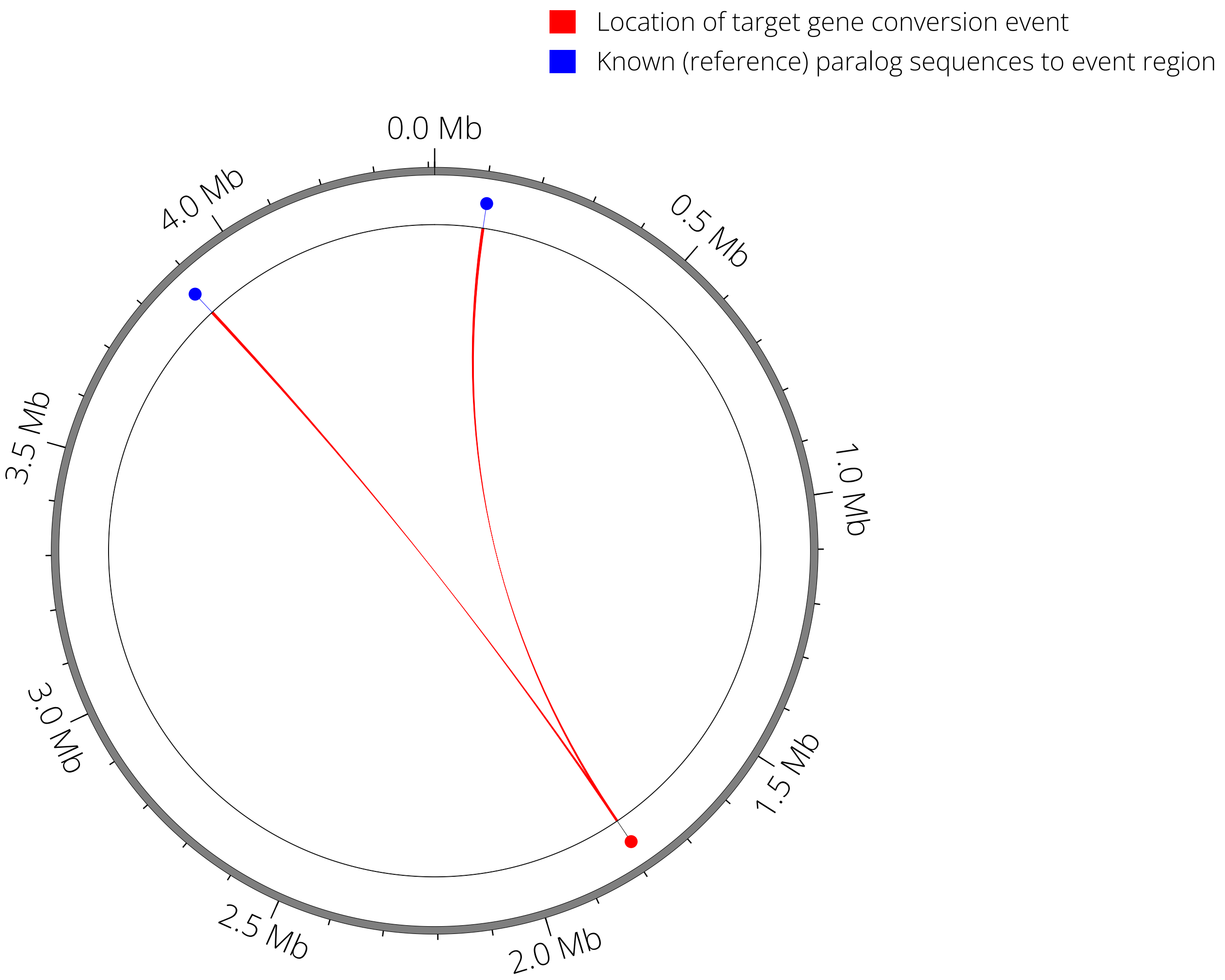

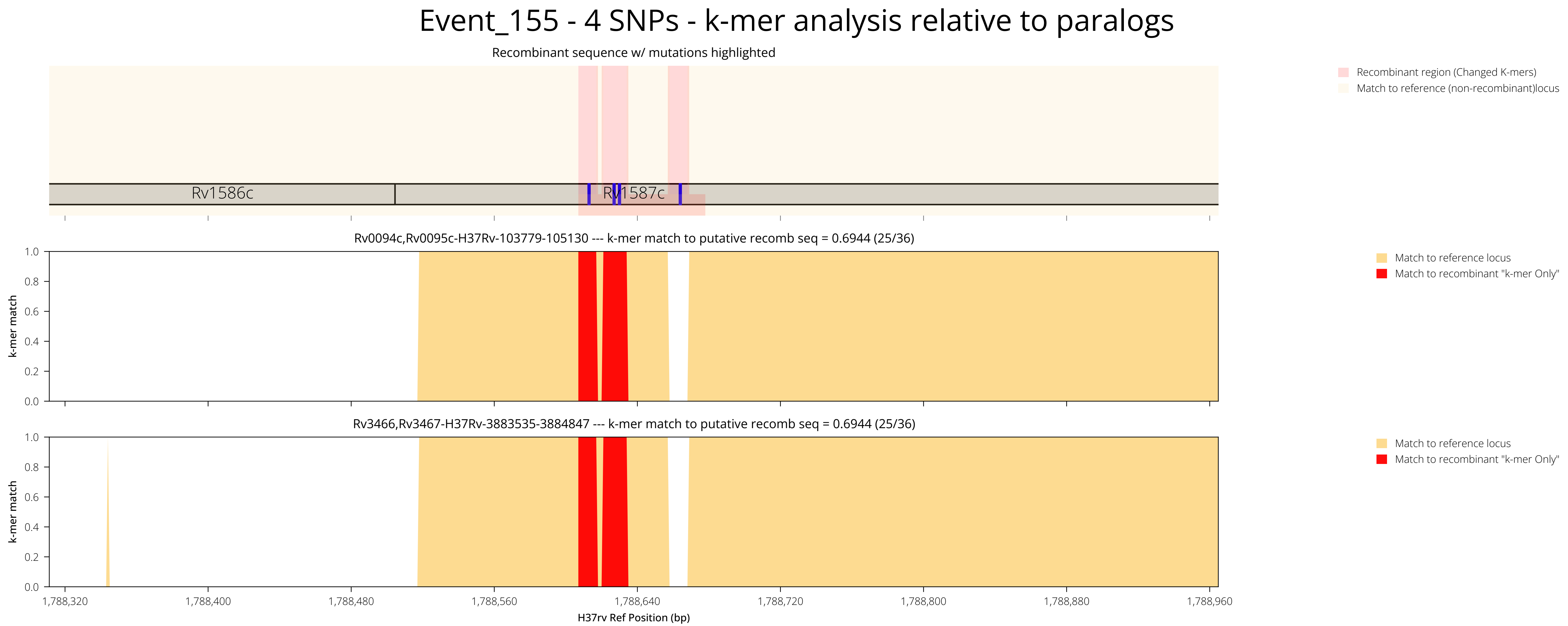
Event_155 - 4 SNPs - k-mer analysis relative to paralogs
Recombinant sequence w/ mutations highlighted
Rv1586c
Rv1587c
Recombinant region (Changed K-mers)
Match to reference (non-recombinant)locus
Rv0094c,Rv0095c-H37Rv-103779-105130 --- k-mer match to putative recomb seq = 0.6944 (25/36)
Match to reference locus
Match to recombinant "k-mer Only"
Rv3466,Rv3467-H37Rv-3883535-3884847 --- k-mer match to putative recomb seq = 0.6944 (25/36)
Match to reference locus
Match to recombinant "k-mer Only"
k-mer match
1.0
0.8
0.6
0.4
0.2
0.0
1,788,320
1,788,400
1,788,480
1,788,560
1,788,640
1,788,720
1,788,800
1,788,880
1,788,960
H37rv Ref Position (bp)

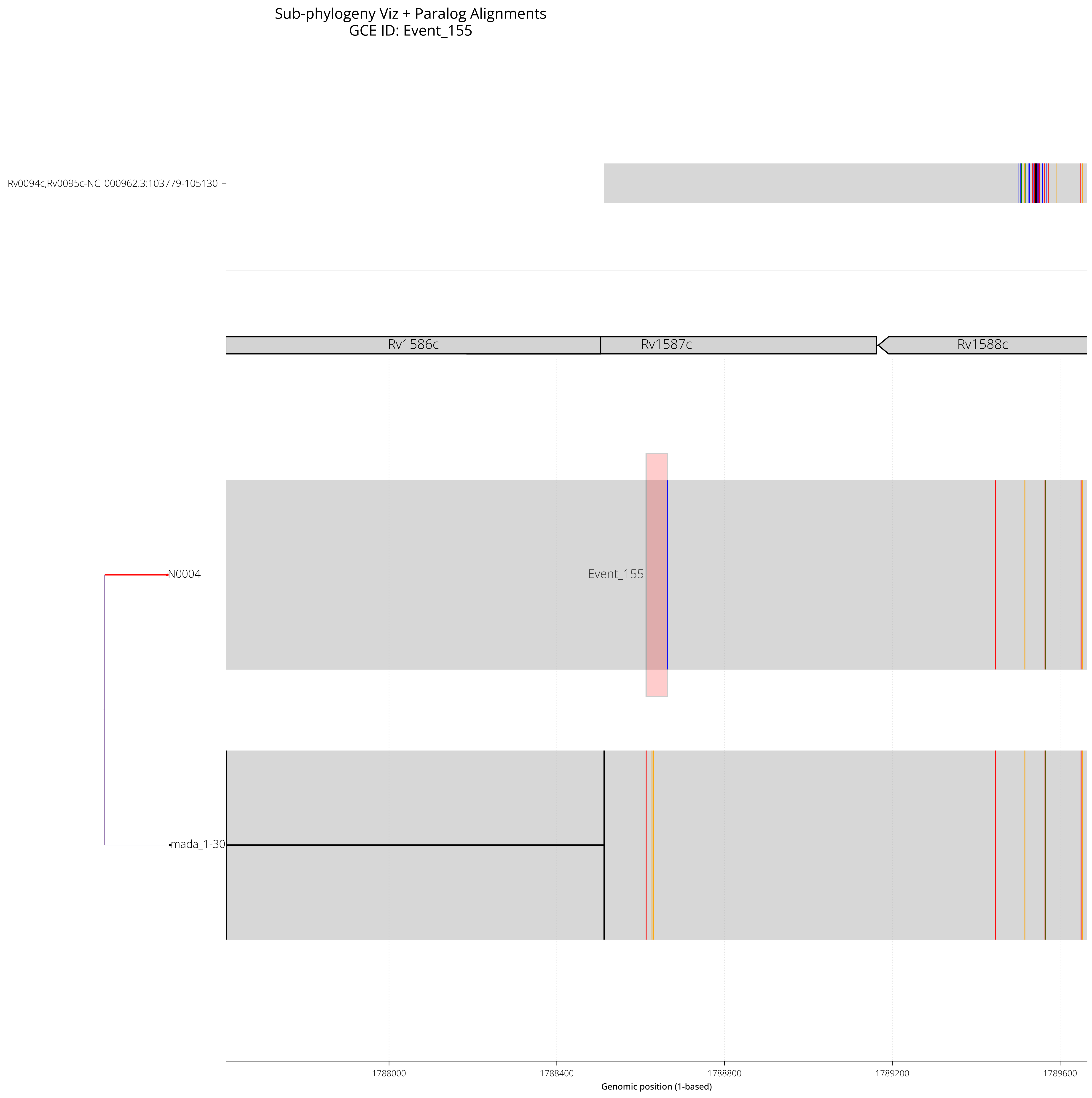

Sub-phylogeny Viz + Paralog Alignments
GCE ID: Event_155
Rv0094c,Rv0095c-NC_000962.3:103779-105130
Rv1586c
Rv1587c
Rv1588c
N0004
Event_155
mada_1-30
1788000
1788400
1788800
1789200
1789600
Genomic position (1-based)

Full Phylogeny Viz + Paralog Alignments
GCE ID: Event_155

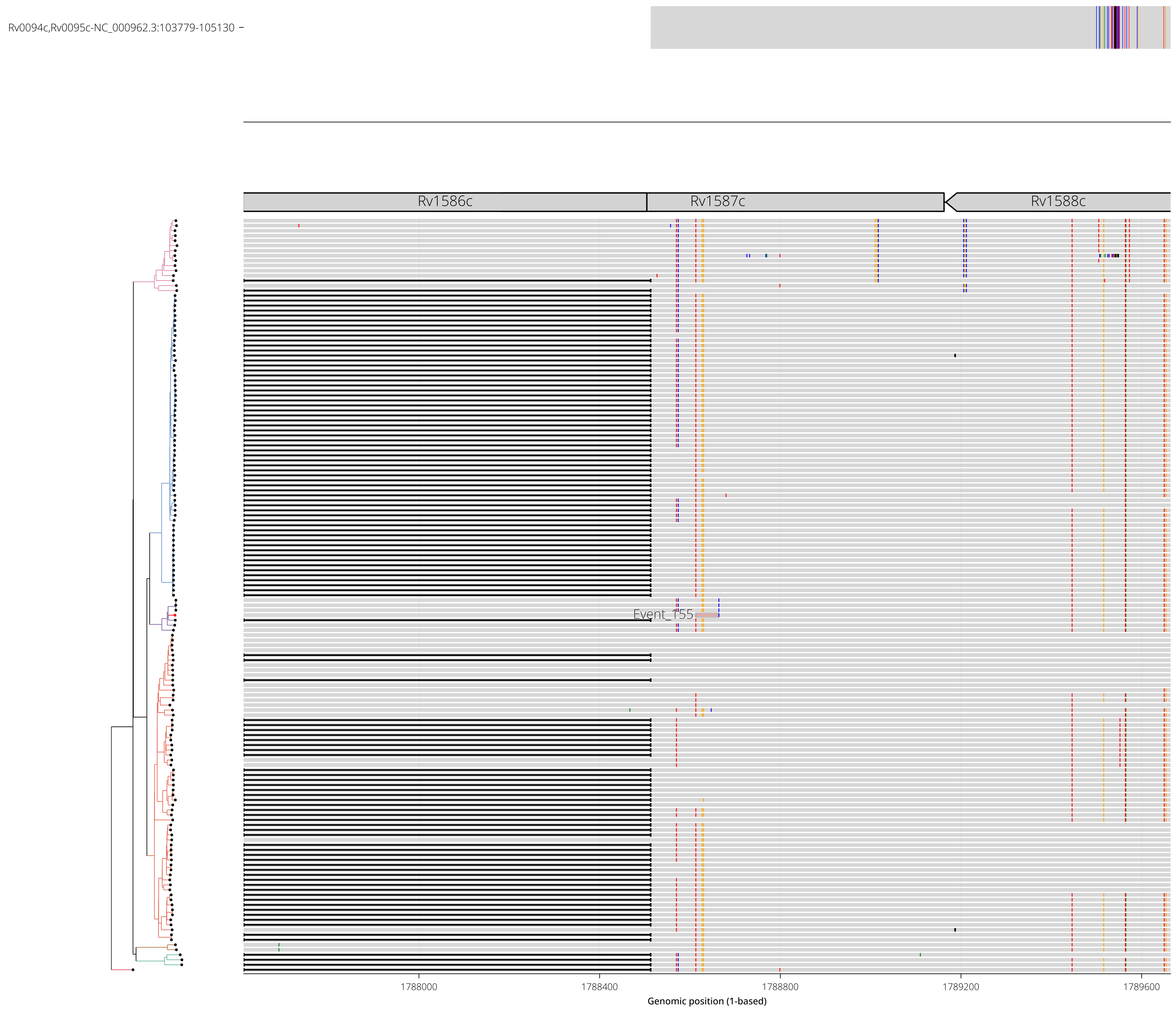

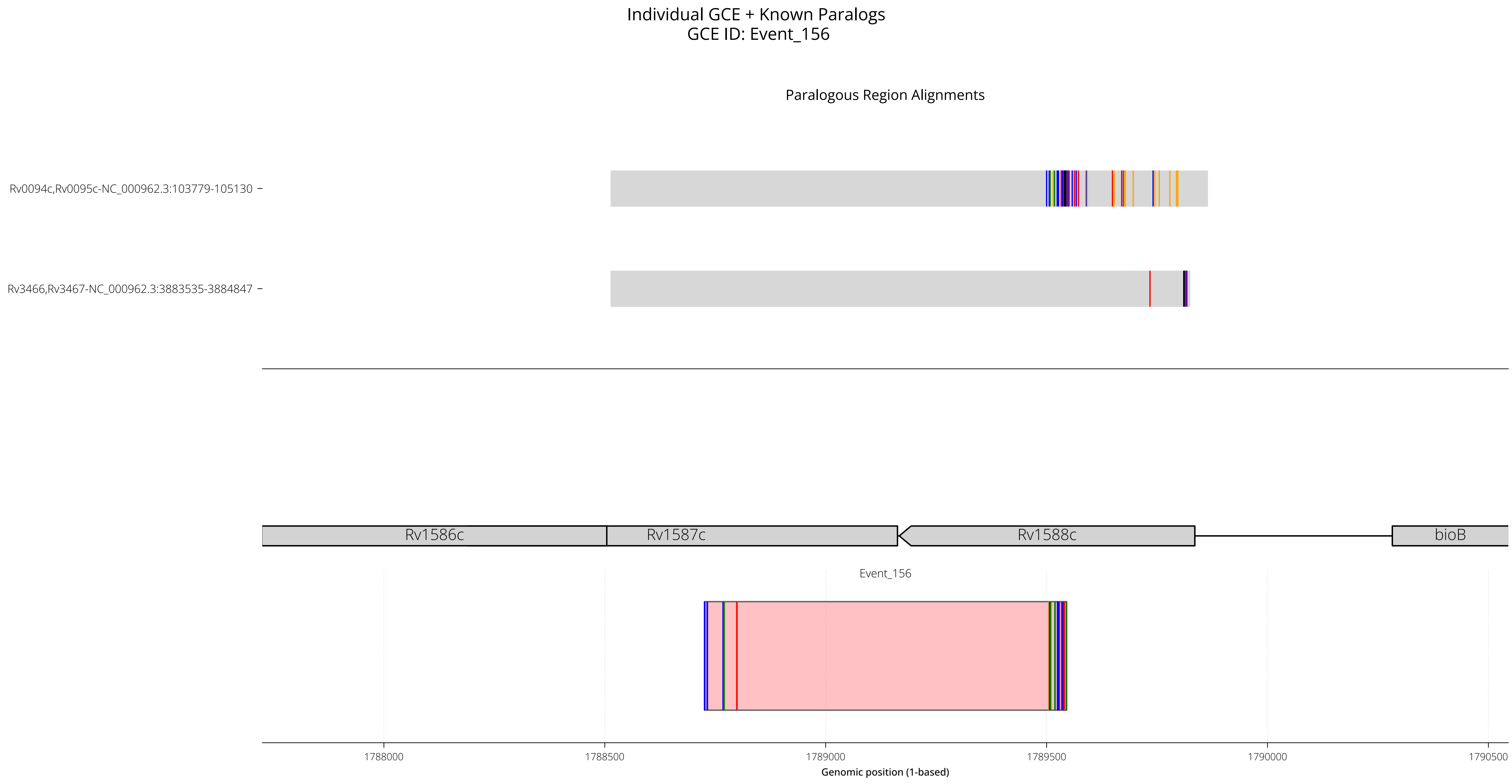

Individual GCE + Known Paralogs
GCE ID: Event_156
Paralogous Region Alignments
Rv0094c,Rv0095c-NC_000962.3:103779-105130
Rv3466,Rv3467-NC_000962.3:3883535-3884847
Rv1586c
Rv1587c
Rv1588c
bioB
Event_156
1788000
1788500
1789000
1789500
1790000
1790500
Genomic position (1-based)

Circos Viz - GCE to Known Paralogs Locations
GCE ID: Event_156

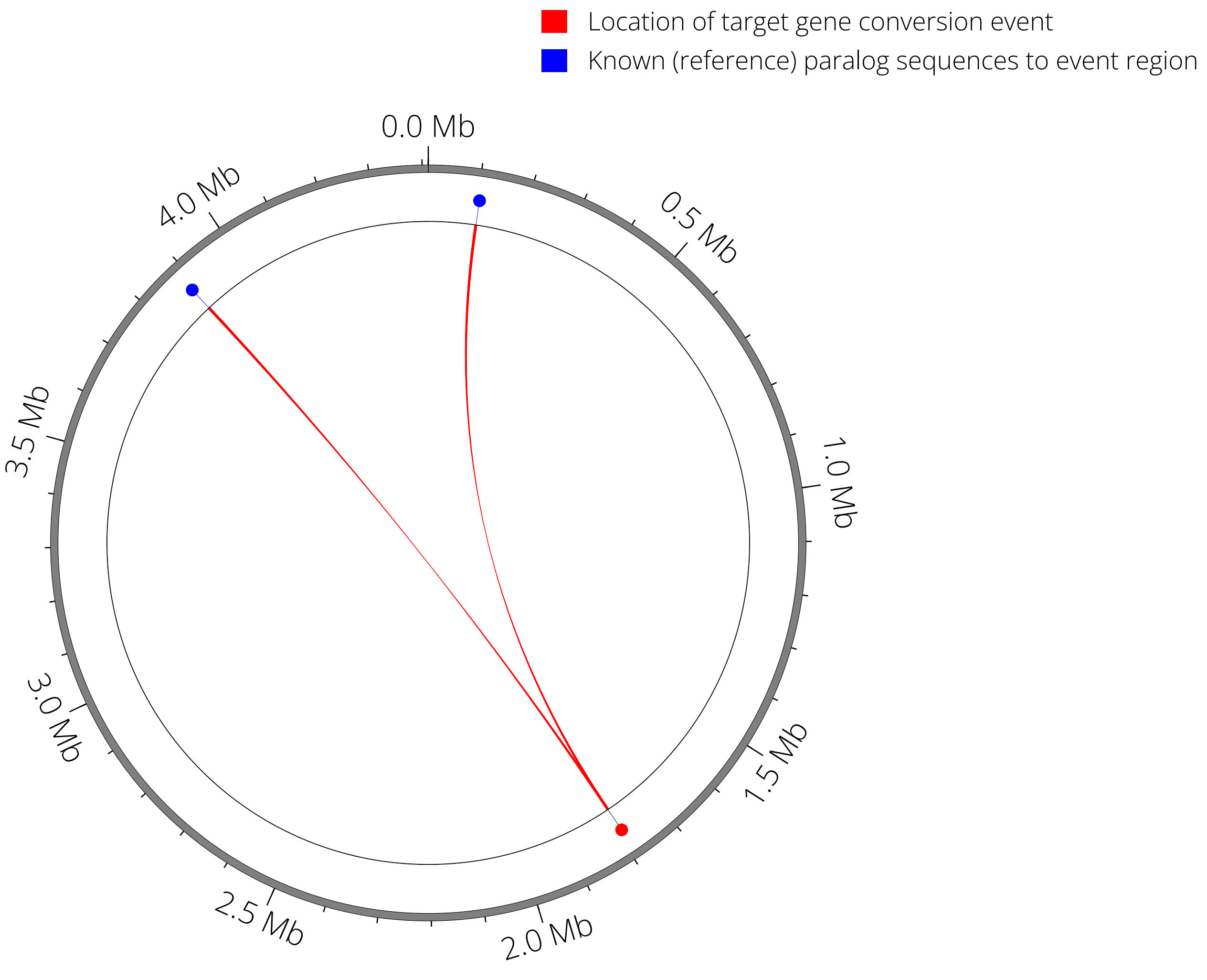

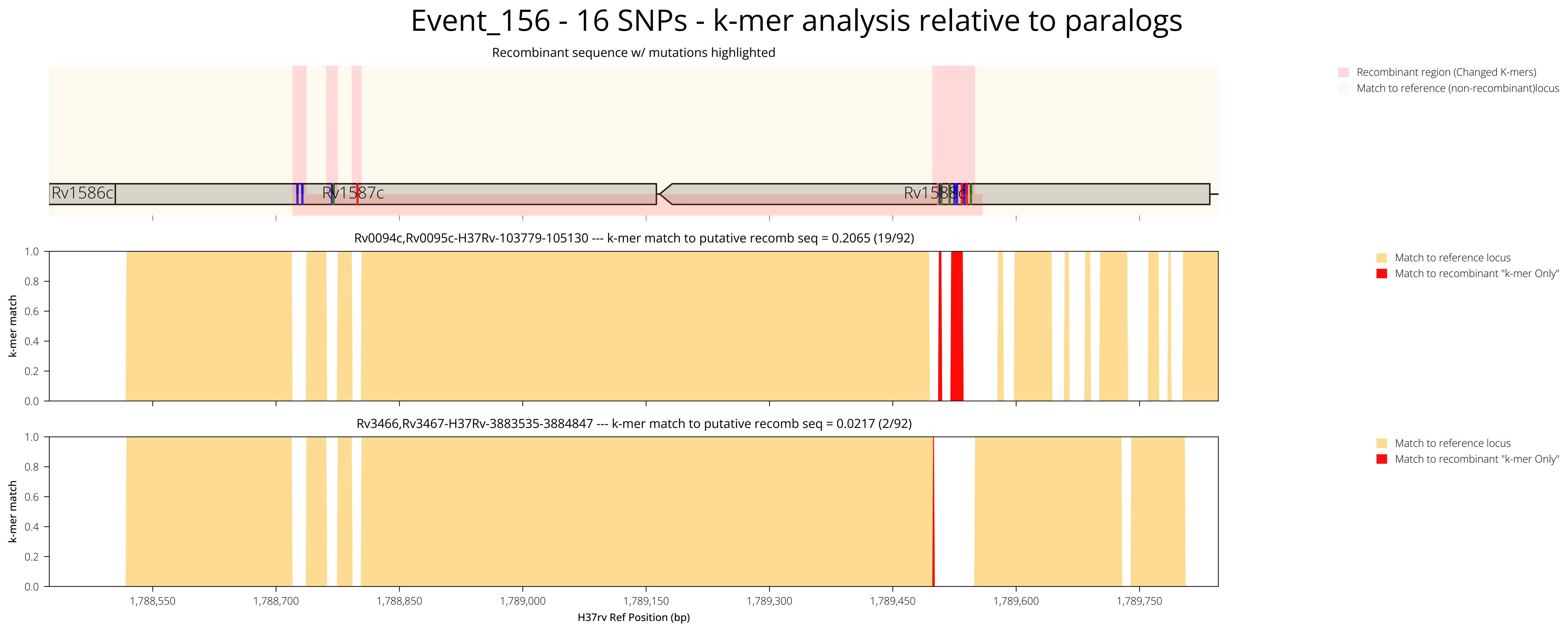
Event_156 - 16 SNPs - k-mer analysis relative to paralogs
Recombinant sequence w/ mutations highlighted
Recombinant region (Changed K-mers)
Match to reference (non-recombinant)locus
Rv1586c
Rv1587c
Rv1588c
Rv0094c,Rv0095c-H37Rv-103779-105130 --- k-mer match to putative recomb seq = 0.2065 (19/92)
Match to reference locus
Match to recombinant "k-mer Only"
k-mer match
Rv3466,Rv3467-H37Rv-3883535-3884847 --- k-mer match to putative recomb seq = 0.0217 (2/92)
Match to reference locus
Match to recombinant "k-mer Only"
k-mer match
1,788,550
1,788,700
1,788,850
1,789,000
1,789,150
1,789,300
1,789,450
1,789,600
1,789,750
H37rv Ref Position (bp)

Sub-phylogeny Viz + Paralog Alignments
GCE ID: Event_156

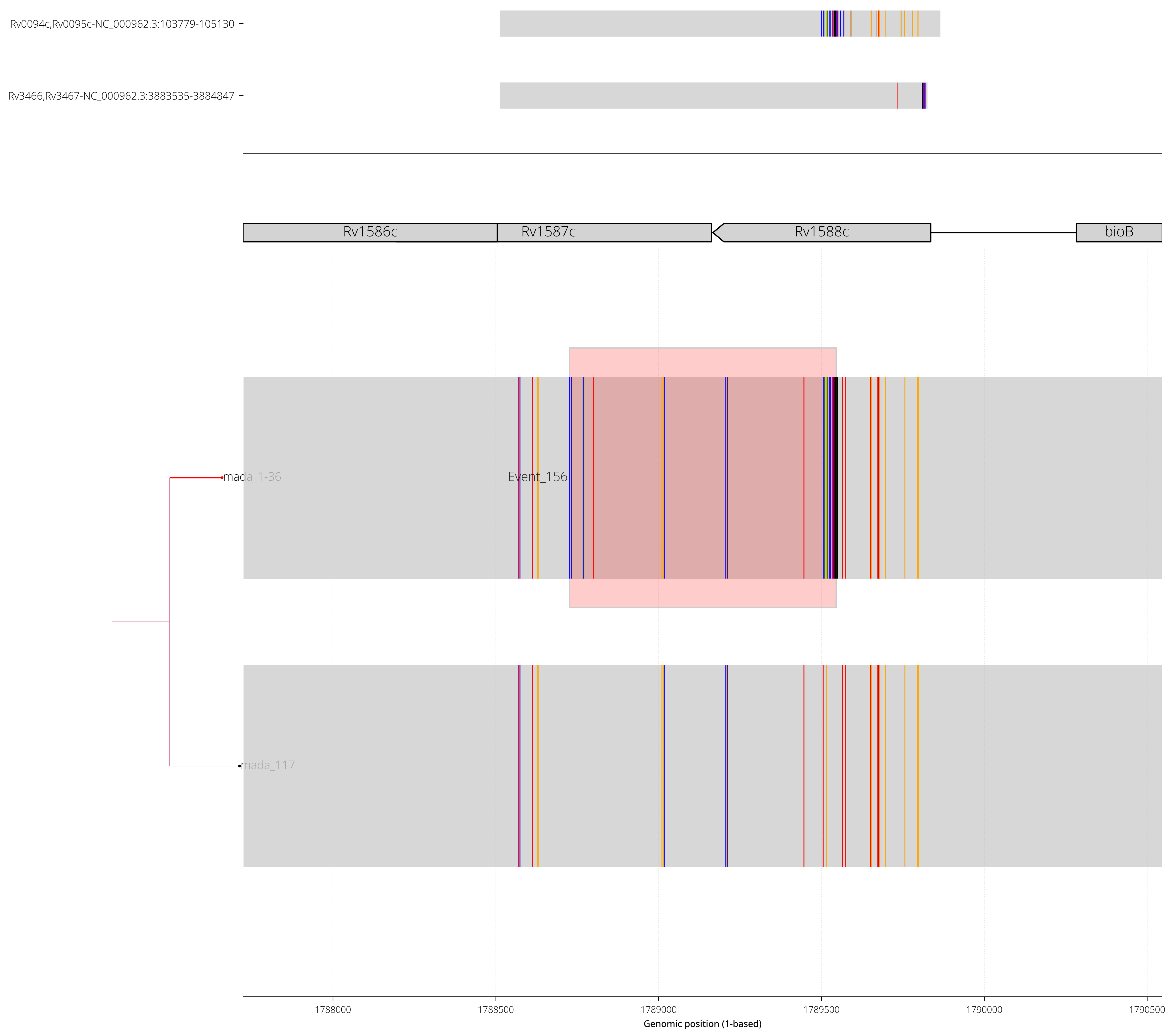

Full Phylogeny Viz + Paralog Alignments
GCE ID: Event_156

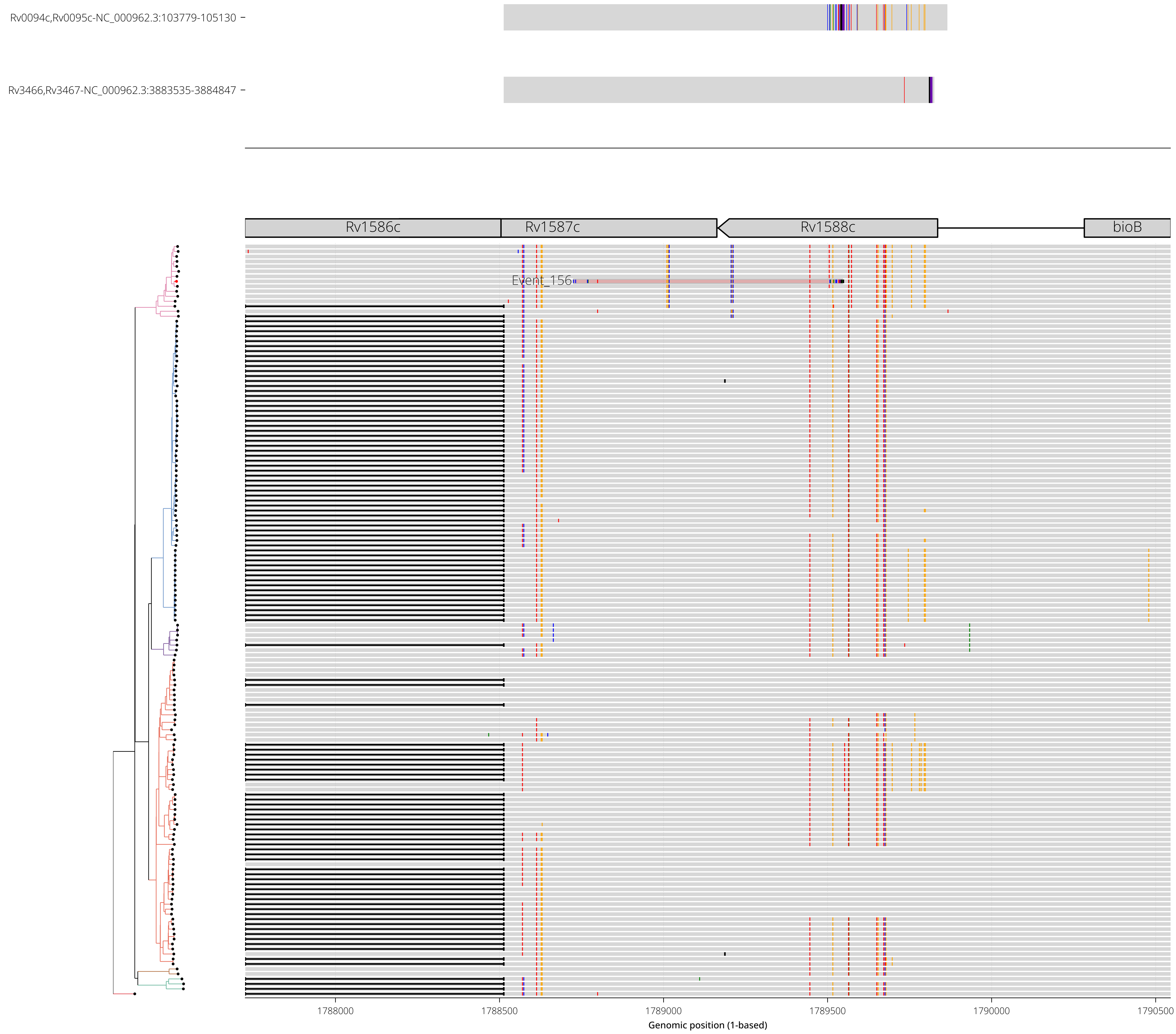

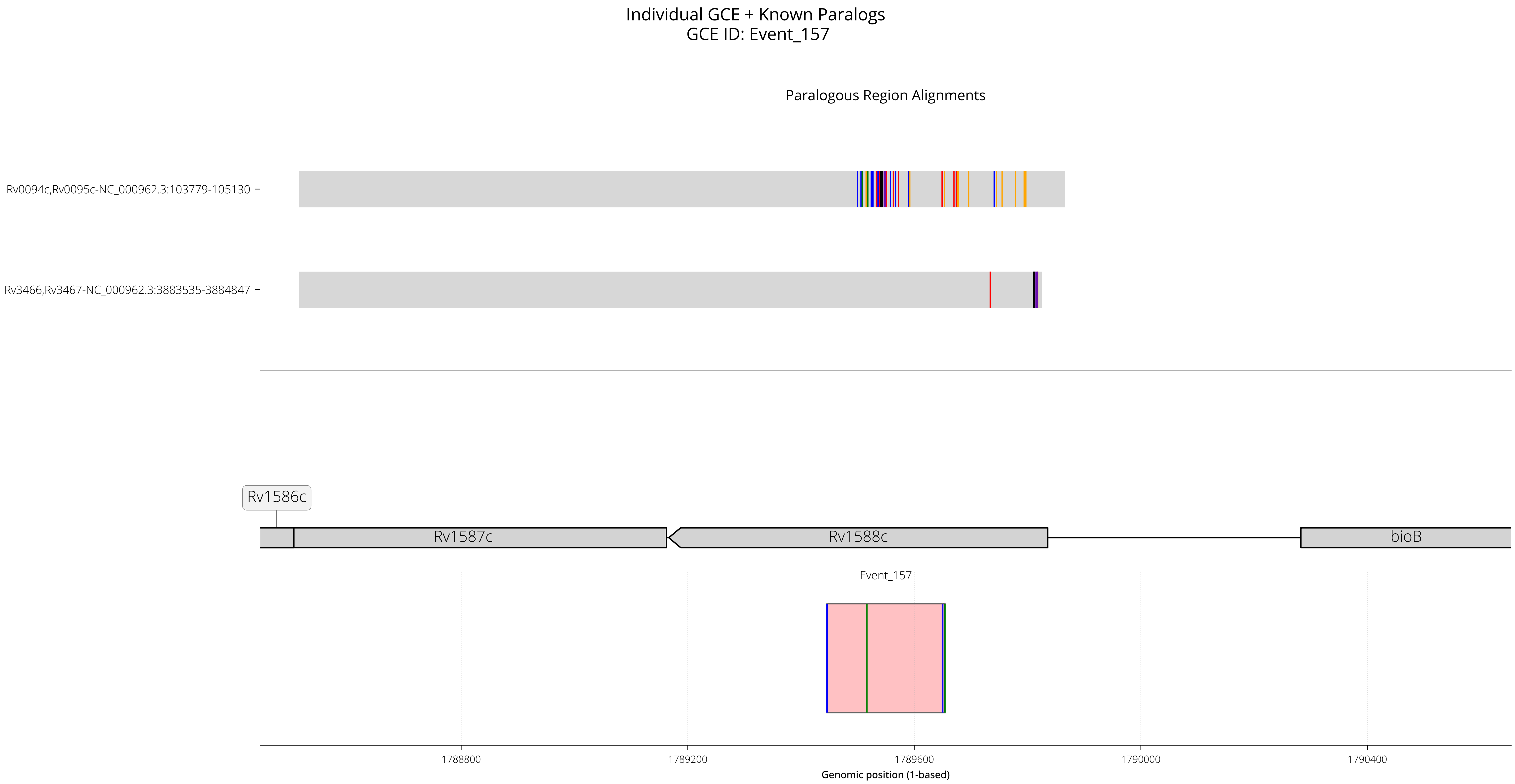
Individual GCE + Known Paralogs
GCE ID: Event_157
Paralogous Region Alignments
Rv0094c,Rv0095c-NC_000962.3:103779-105130
Rv3466,Rv3467-NC_000962.3:3883535-3884847
Rv1586c
Rv1587c
Rv1588c
bioB
Event_157
1788800
1789200
1789600
1790000
1790400
Genomic position (1-based)

Circos Viz - GCE to Known Paralogs Locations
GCE ID: Event_157

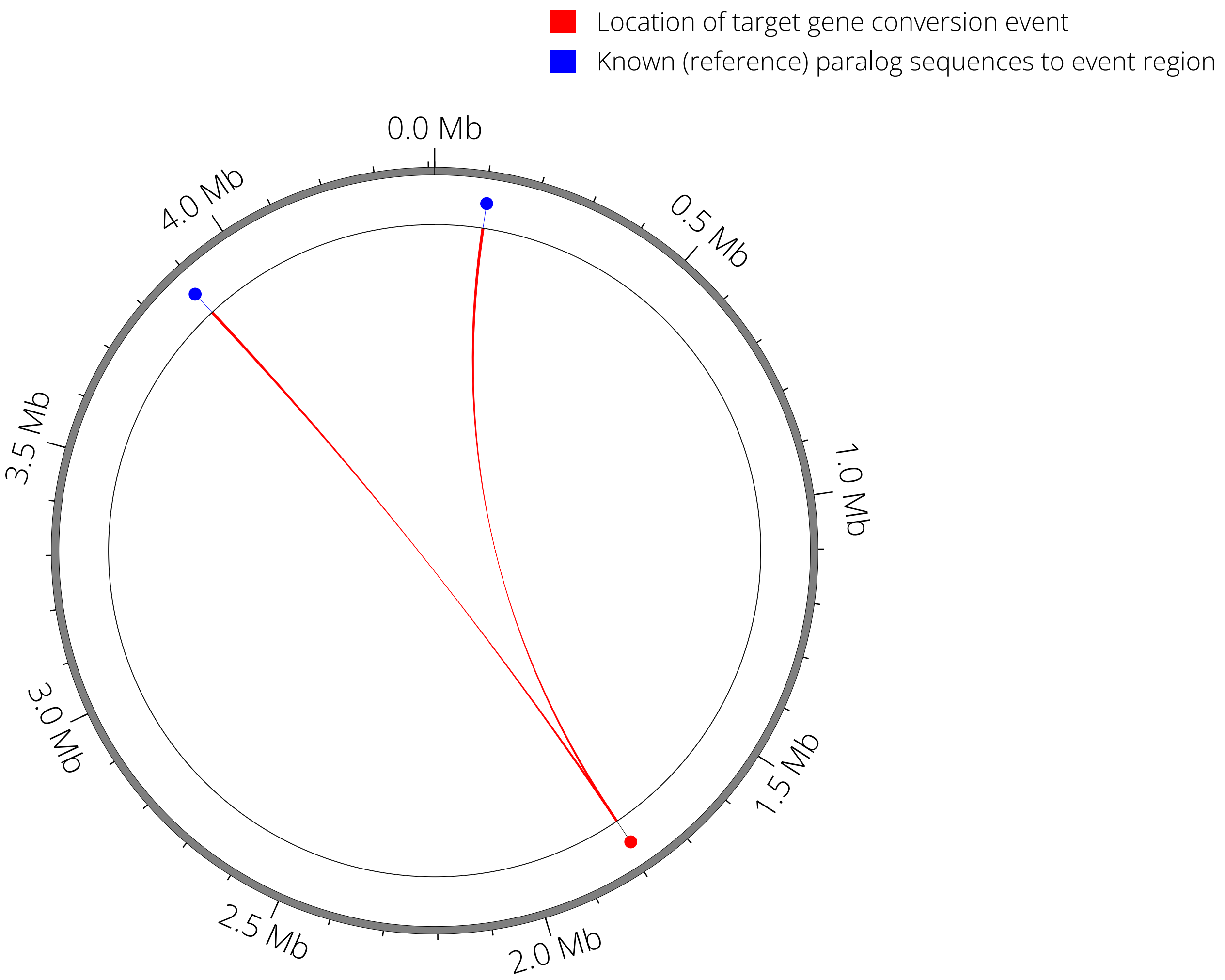

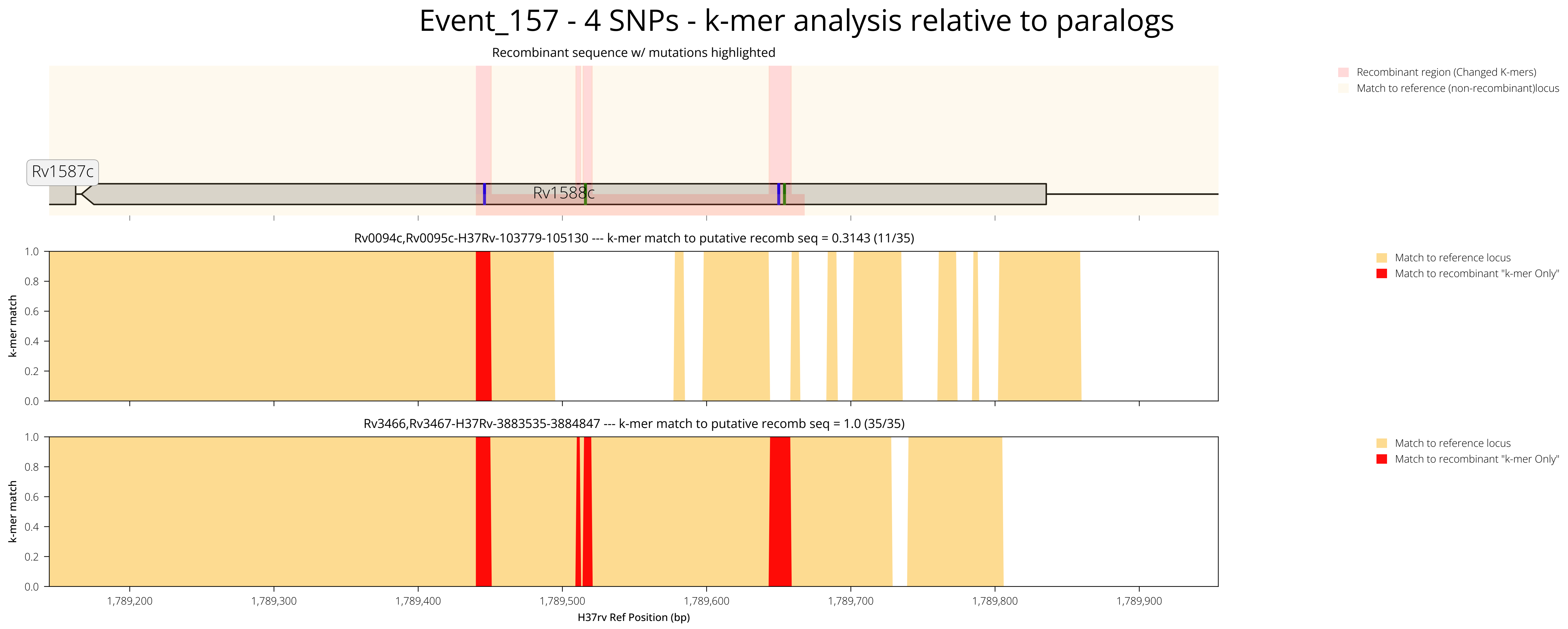

Event_157 - 4 SNPs - k-mer analysis relative to paralogs
Recombinant sequence w/ mutations highlighted
Recombinant region (Changed K-mers)
Match to reference (non-recombinant)locus
Rv1587c
Rv1588c
Rv0094c,Rv0095c-H37Rv-103779-105130 --- k-mer match to putative recomb seq = 0.3143 (11/35)
Match to reference locus
Match to recombinant "k-mer Only"
k-mer match
1.0
0.8
0.6
0.4
0.2
0.0
Rv3466,Rv3467-H37Rv-3883535-3884847 --- k-mer match to putative recomb seq = 1.0 (35/35)
Match to reference locus
Match to recombinant "k-mer Only"
k-mer match
1.0
0.8
0.6
0.4
0.2
0.0
1,789,200
1,789,300
1,789,400
1,789,500
1,789,600
1,789,700
1,789,800
1,789,900
H37rv Ref Position (bp)

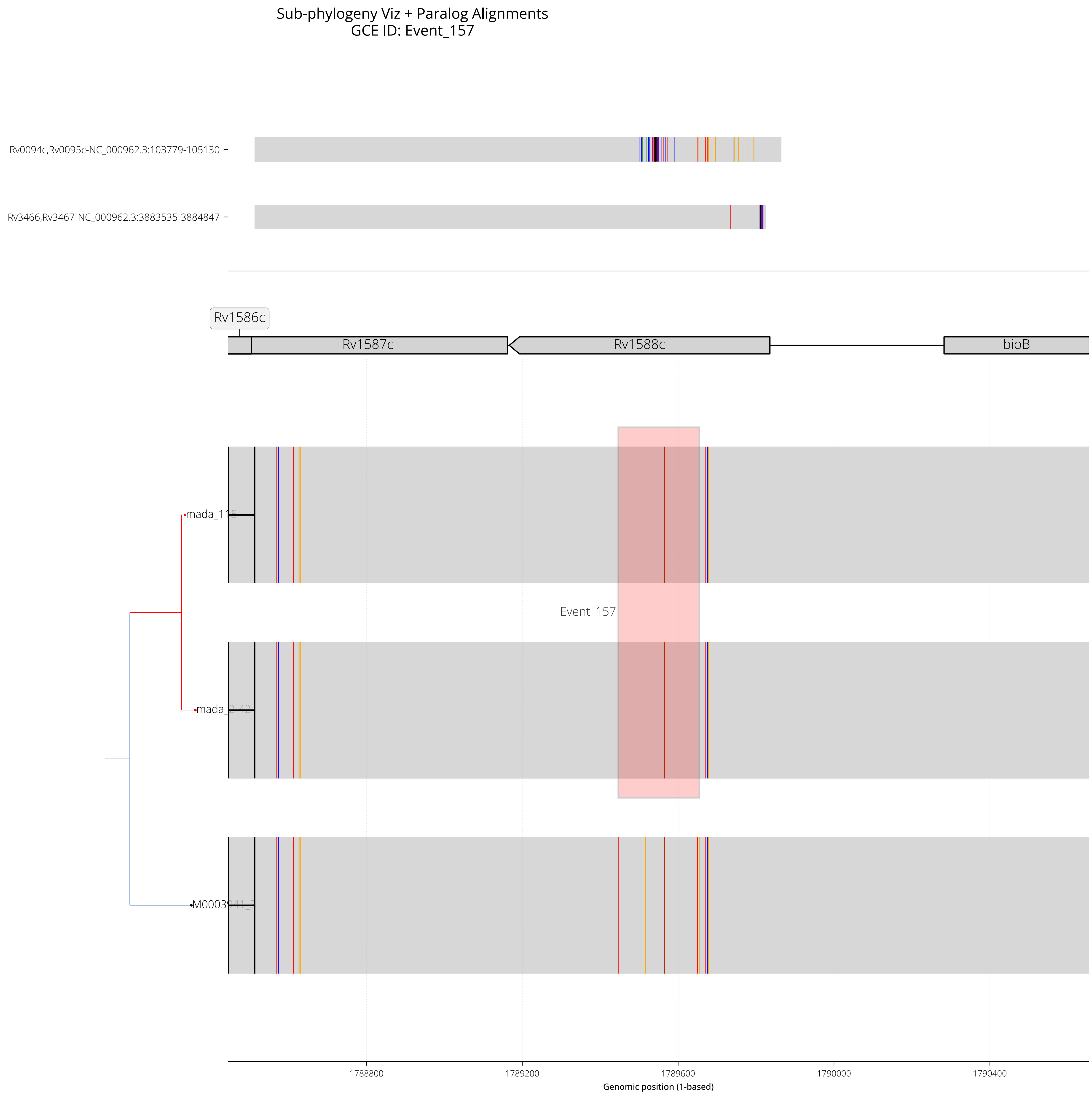

Sub-phylogeny Viz + Paralog Alignments
GCE ID: Event_157
Rv0094c,Rv0095c-NC_000962.3:103779-105130
Rv3466,Rv3467-NC_000962.3:3883535-3884847
Rv1586c
Rv1587c
Rv1588c
bioB
mada_115
Event_157
mada_2-42
M0003941_3
1788800
1789200
1789600
1790000
1790400
Genomic position (1-based)

Full Phylogeny Viz + Paralog Alignments
GCE ID: Event_157

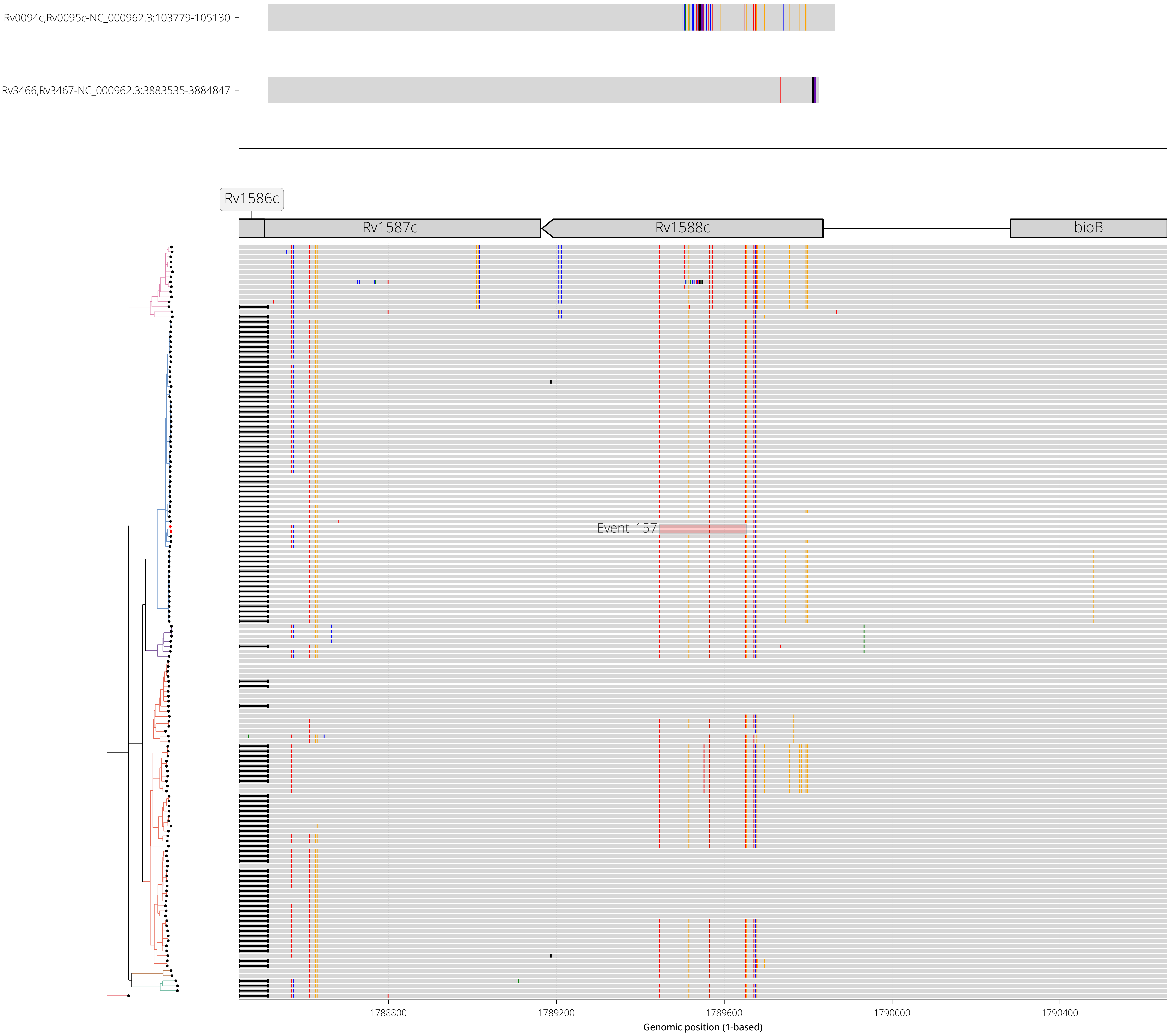

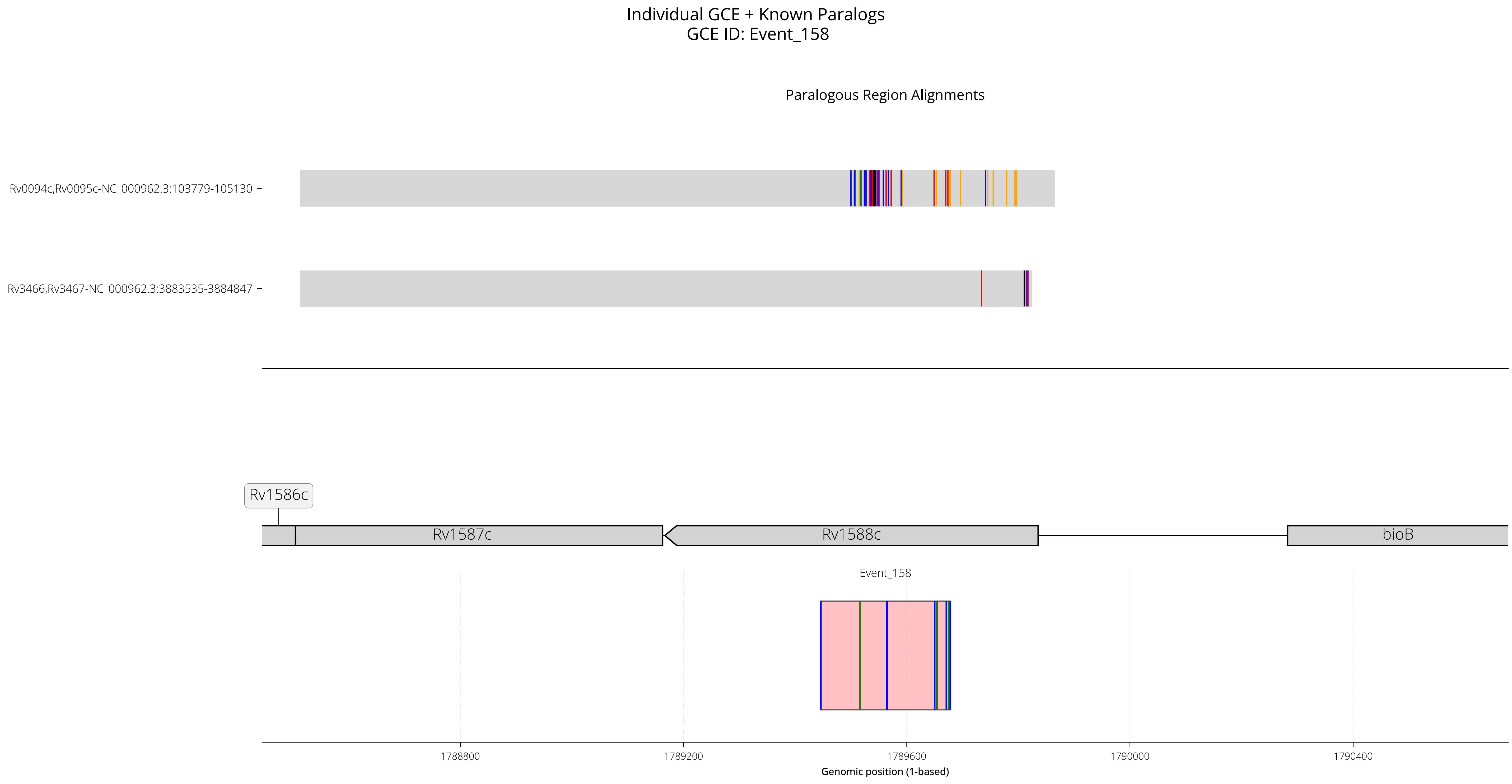

Individual GCE + Known Paralogs
GCE ID: Event_158
Paralogous Region Alignments
Rv0094c,Rv0095c-NC_000962.3:103779-105130
Rv3466,Rv3467-NC_000962.3:3883535-3884847
Rv1586c
Rv1587c
Rv1588c
bioB
Event_158
1788800
1789200
1789600
1790000
1790400
Genomic position (1-based)

Circos Viz - GCE to Known Paralogs Locations
GCE ID: Event_158

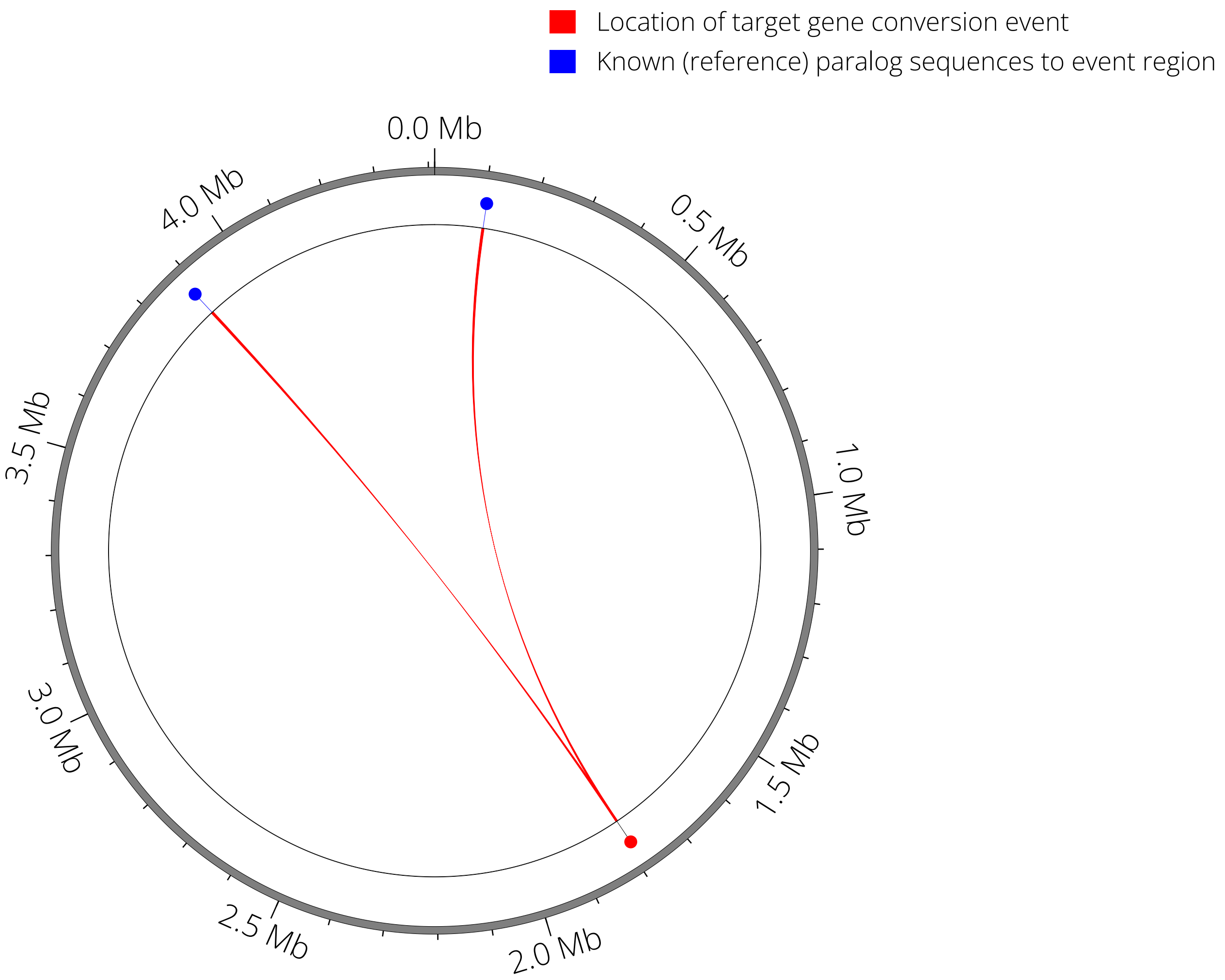

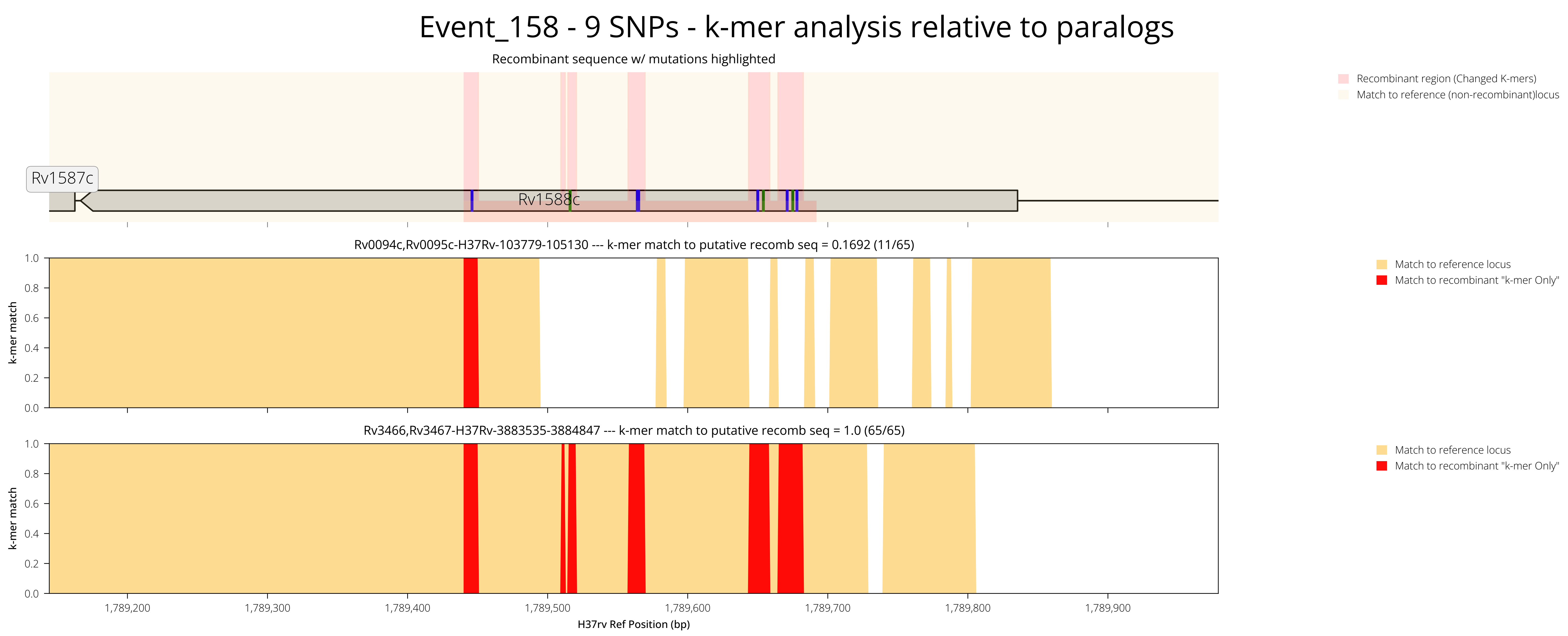

Event_158 - 9 SNPs - k-mer analysis relative to paralogs
Recombinant sequence w/ mutations highlighted
Rv1587c
Rv1588c
Recombinant region (Changed K-mers)
Match to reference (non-recombinant)locus
Rv0094c,Rv0095c-H37Rv-103779-105130 --- k-mer match to putative recomb seq = 0.1692 (11/65)
Match to reference locus
Match to recombinant "k-mer Only"
Rv3466,Rv3467-H37Rv-3883535-3884847 --- k-mer match to putative recomb seq = 1.0 (65/65)
Match to reference locus
Match to recombinant "k-mer Only"
k-mer match
1.0
0.8
0.6
0.4
0.2
0.0
1,789,200
1,789,300
1,789,400
1,789,500
1,789,600
1,789,700
1,789,800
1,789,900
H37rv Ref Position (bp)

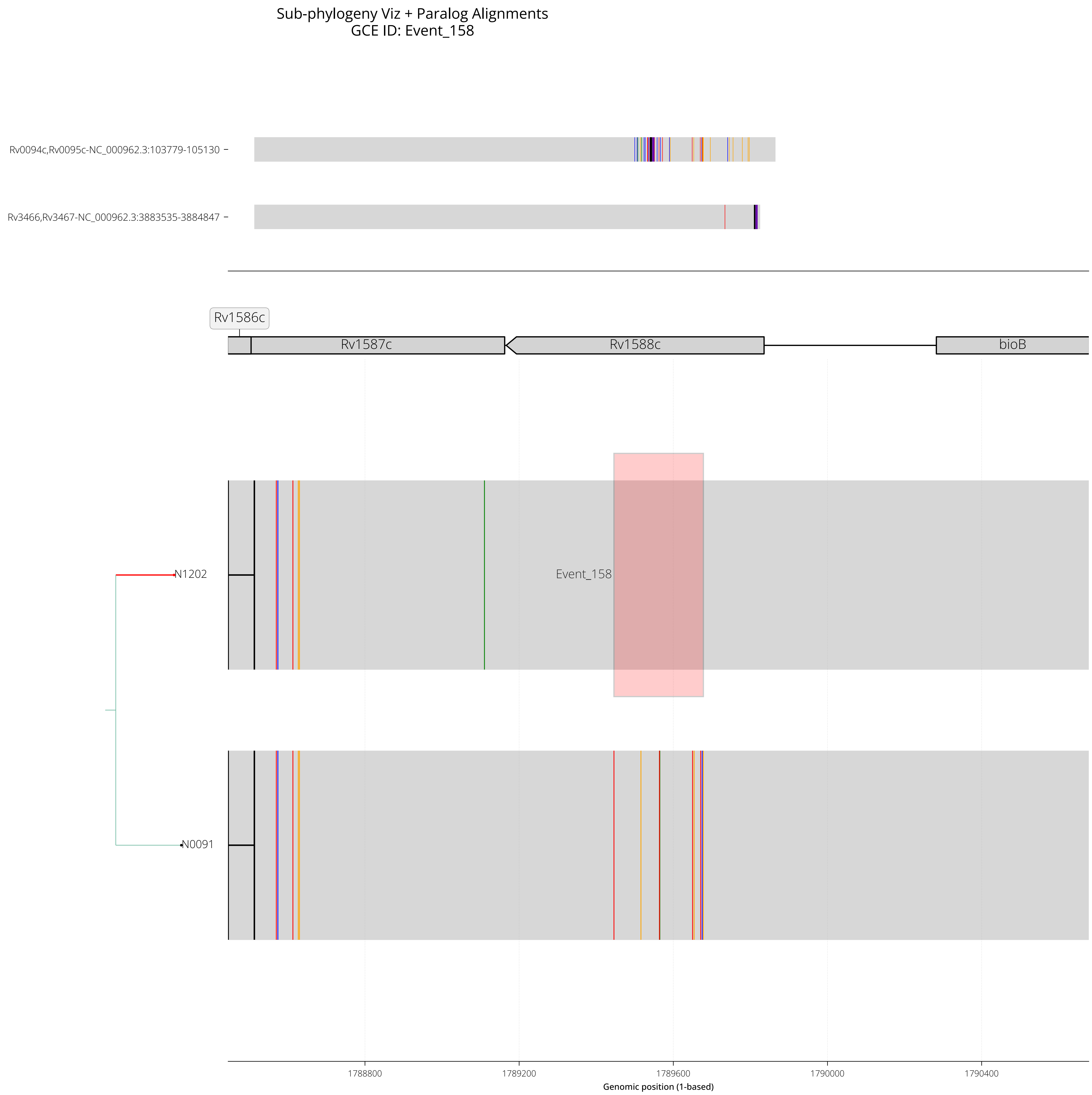
Sub-phylogeny Viz + Paralog Alignments
GCE ID: Event_158
Rv0094c,Rv0095c-NC_000962.3:103779-105130
Rv3466,Rv3467-NC_000962.3:3883535-3884847
Rv1586c
Rv1587c
Rv1588c
bioB
N1202
Event_158
N0091
1788800
1789200
1789600
1790000
1790400
Genomic position (1-based)

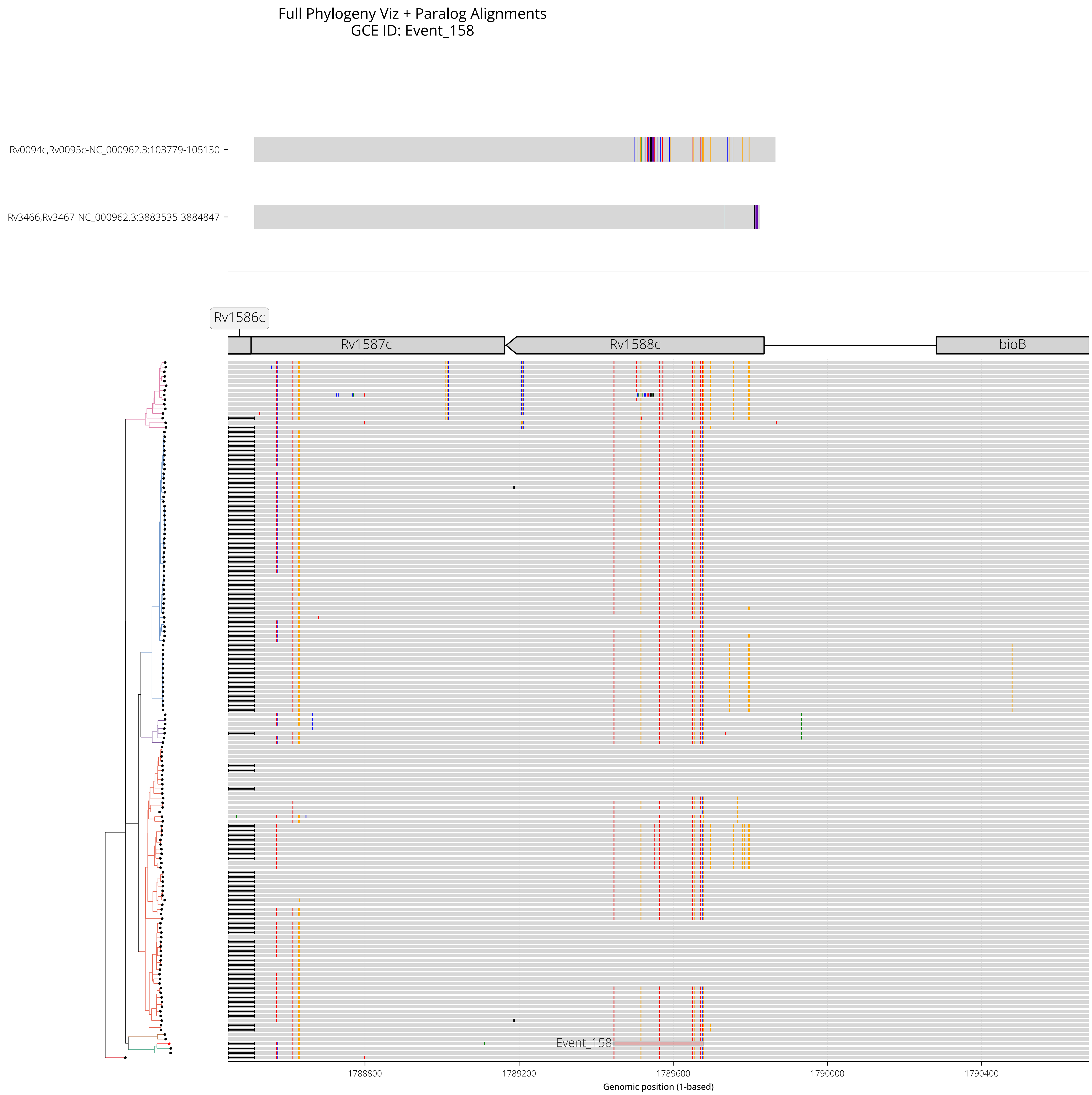

Full Phylogeny Viz + Paralog Alignments
GCE ID: Event_158
Rv0094c,Rv0095c-NC_000962.3:103779-105130
Rv3466,Rv3467-NC_000962.3:3883535-3884847
Rv1586c
Rv1587c
Rv1588c
bioB
Event_158
1788800
1789200
1789600
1790000
1790400
Genomic position (1-based)

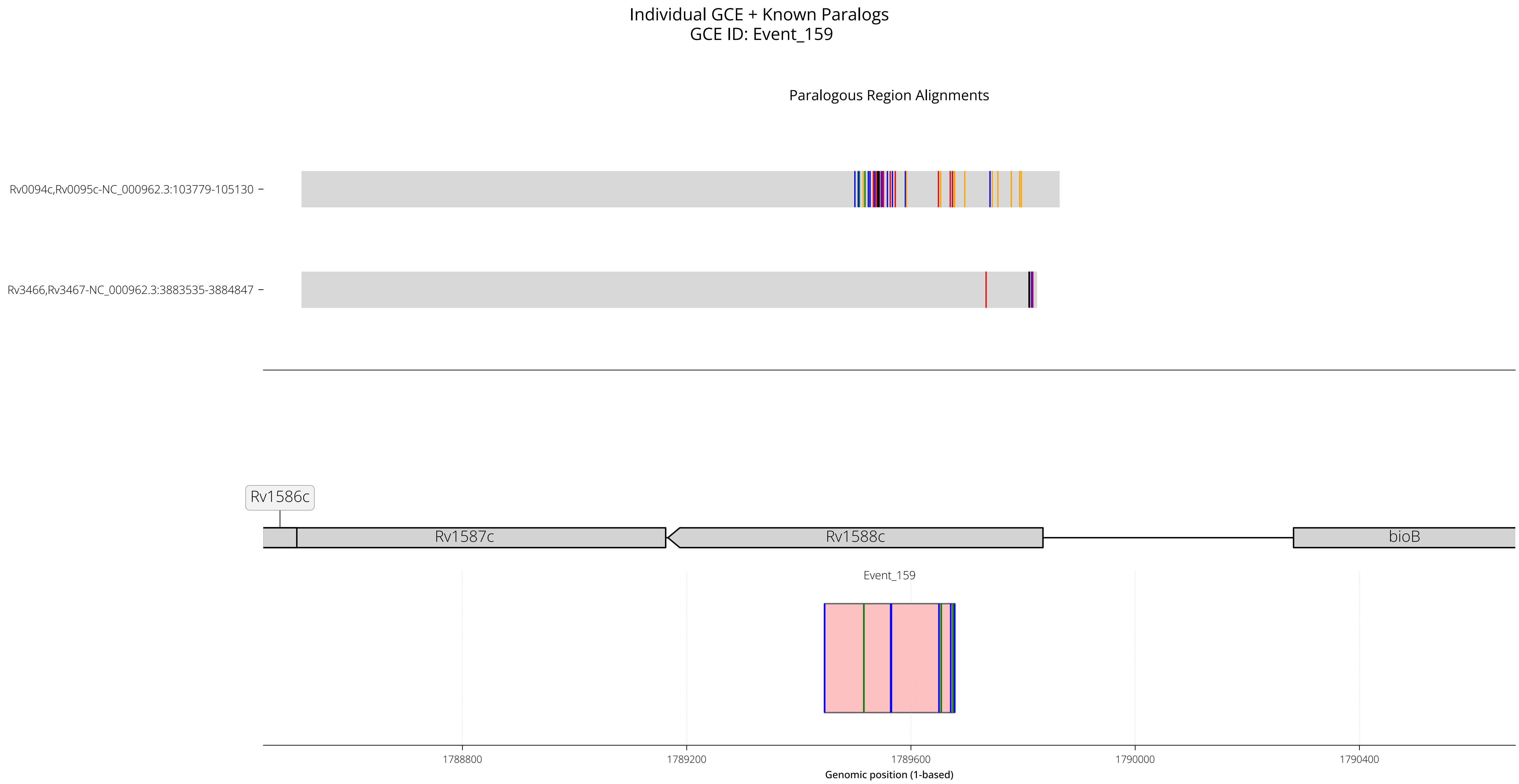

Individual GCE + Known Paralogs
GCE ID: Event_159
Paralogous Region Alignments
Rv0094c,Rv0095c-NC_000962.3:103779-105130
Rv3466,Rv3467-NC_000962.3:3883535-3884847
Rv1586c
Rv1587c
Rv1588c
bioB
Event_159
1788800
1789200
1789600
1790000
1790400
Genomic position (1-based)

Circos Viz - GCE to Known Paralogs Locations
GCE ID: Event_159

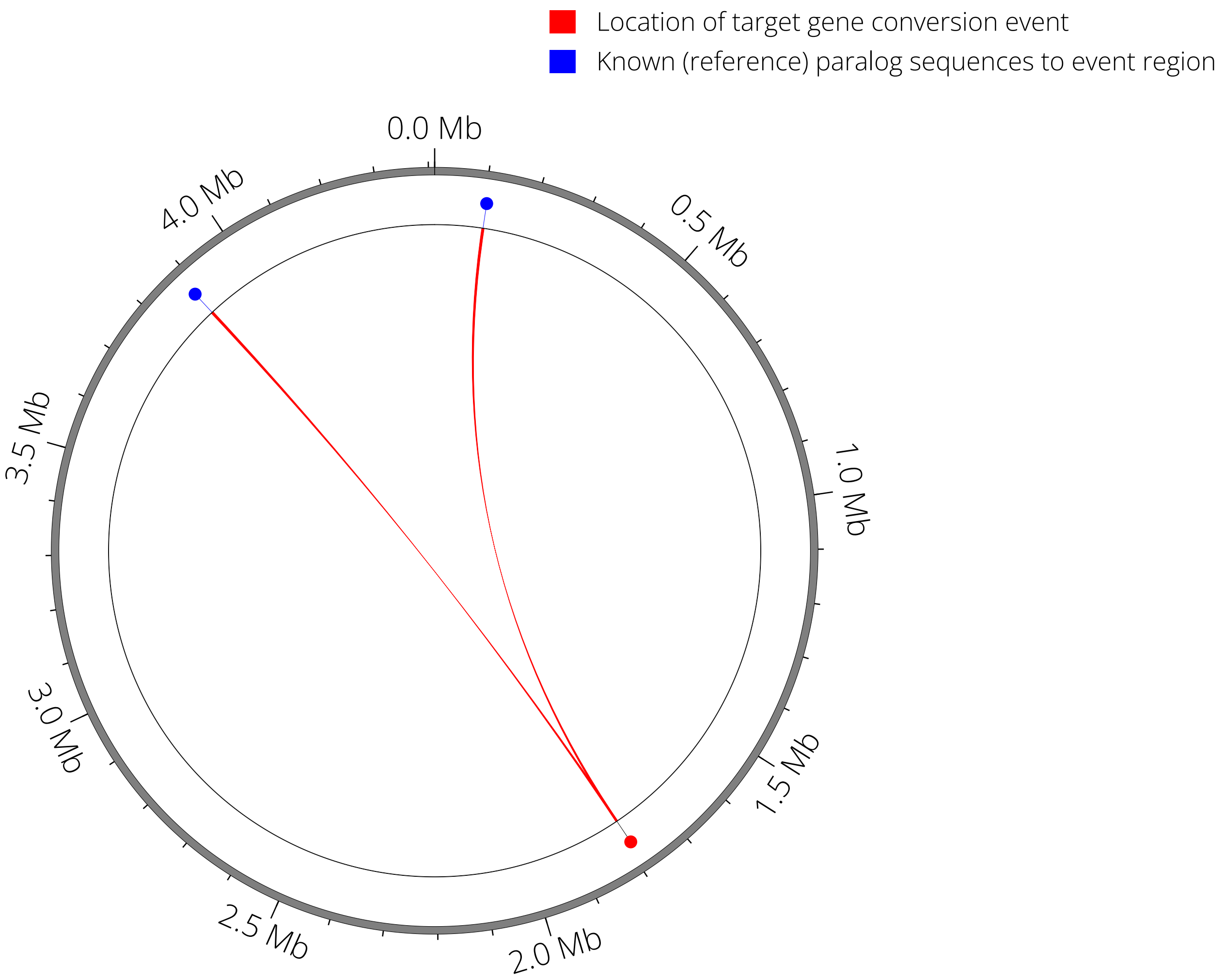

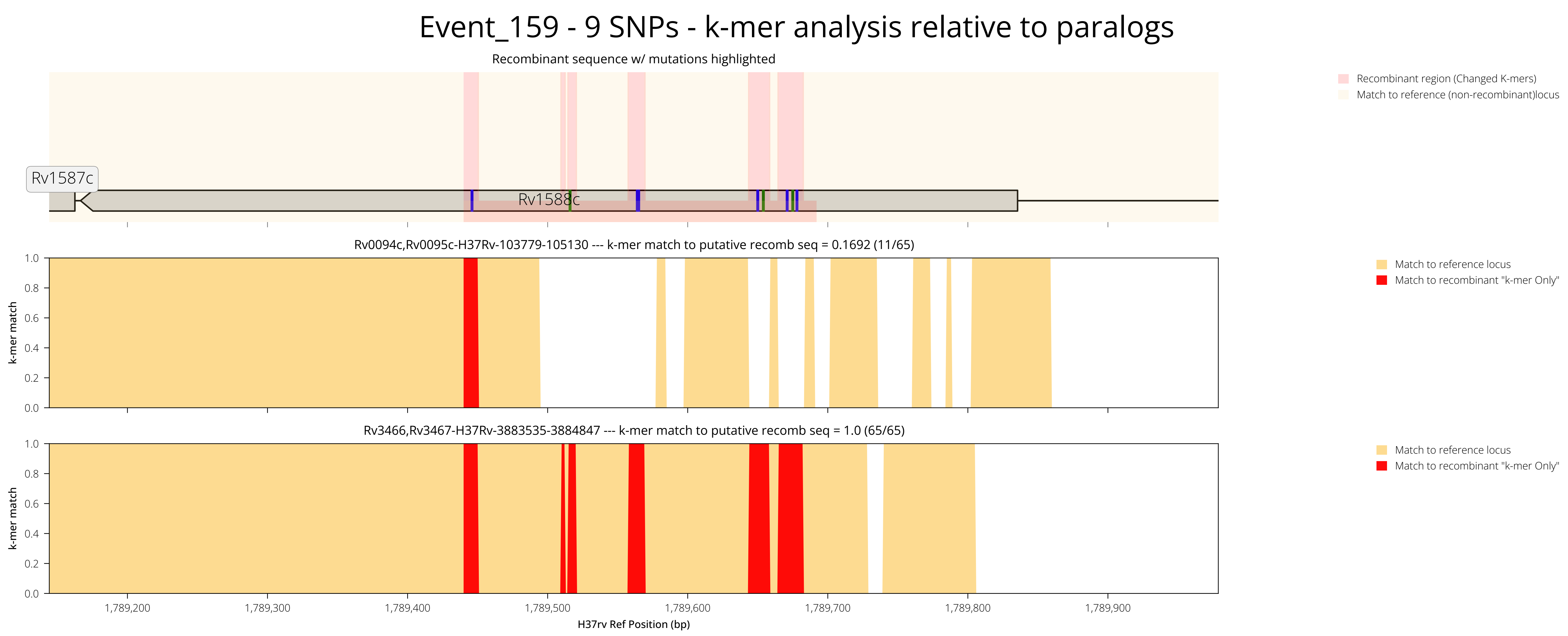
Event_159 - 9 SNPs - k-mer analysis relative to paralogs
Recombinant sequence w/ mutations highlighted
Rv1587c
Rv1588c
Recombinant region (Changed K-mers)
Match to reference (non-recombinant)locus
Rv0094c,Rv0095c-H37Rv-103779-105130 --- k-mer match to putative recomb seq = 0.1692 (11/65)
Match to reference locus
Match to recombinant "k-mer Only"
k-mer match
Rv3466,Rv3467-H37Rv-3883535-3884847 --- k-mer match to putative recomb seq = 1.0 (65/65)
1,789,200
1,789,300
1,789,400
1,789,500
1,789,600
1,789,700
1,789,800
1,789,900
H37rv Ref Position (bp)

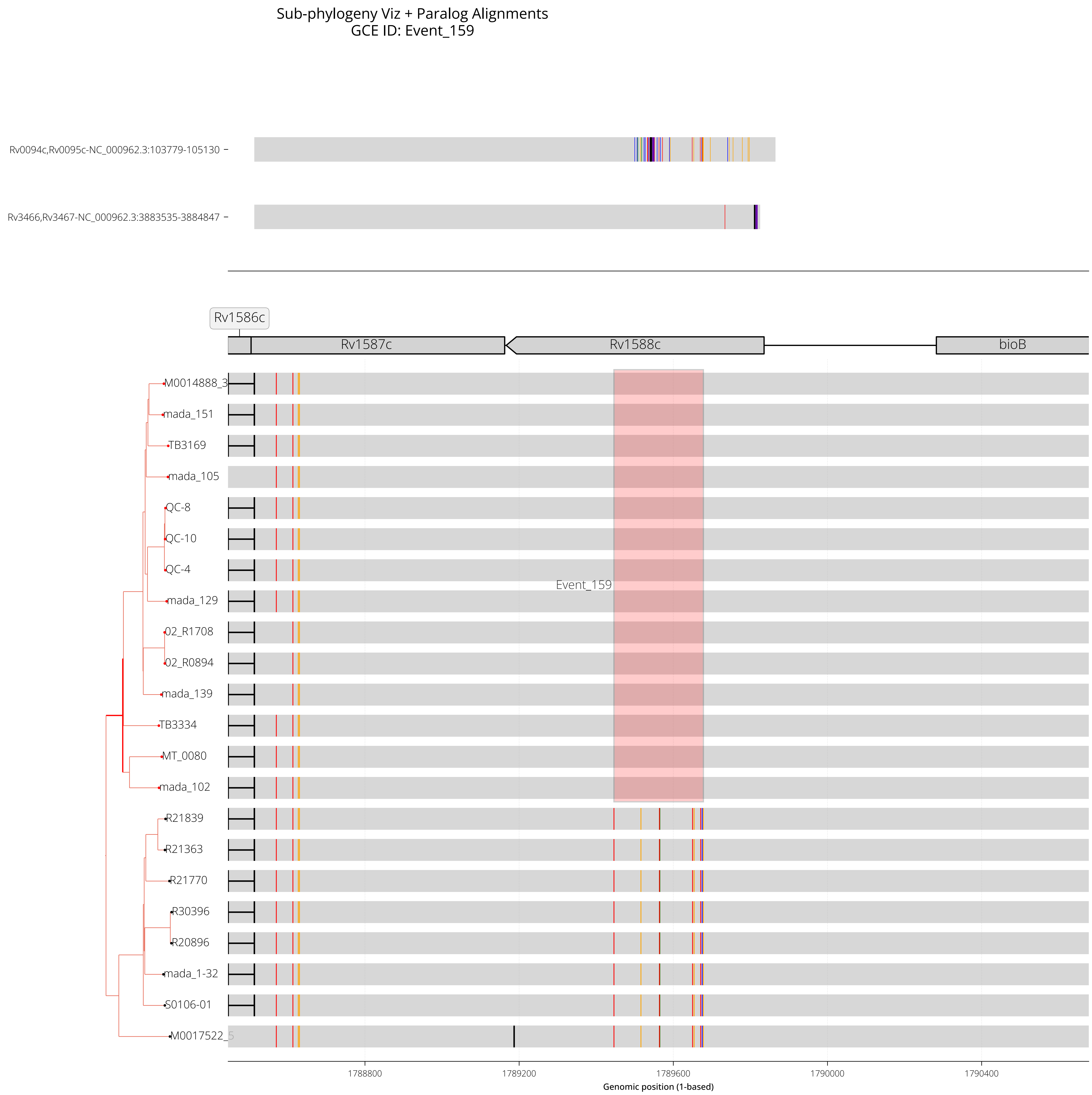

Sub-phylogeny Viz + Paralog Alignments
GCE ID: Event_159
Rv0094c,Rv0095c-NC_000962.3:103779-105130
Rv3466,Rv3467-NC_000962.3:3883535-3884847
Rv1586c
Rv1587c
Rv1588c
bioB
M0014888_3
mada_151
TB3169
mada_105
QC-8
QC-10
QC-4
mada_129
02_R1708
02_R0894
mada_139
TB3334
MT_0080
mada_102
R21839
R21363
R21770
R30396
R20896
mada_1-32
S0106-01
M0017522_5
Event_159
1788800
1789200
1789600
1790000
1790400
Genomic position (1-based)

Full Phylogeny Viz + Paralog Alignments
GCE ID: Event_159

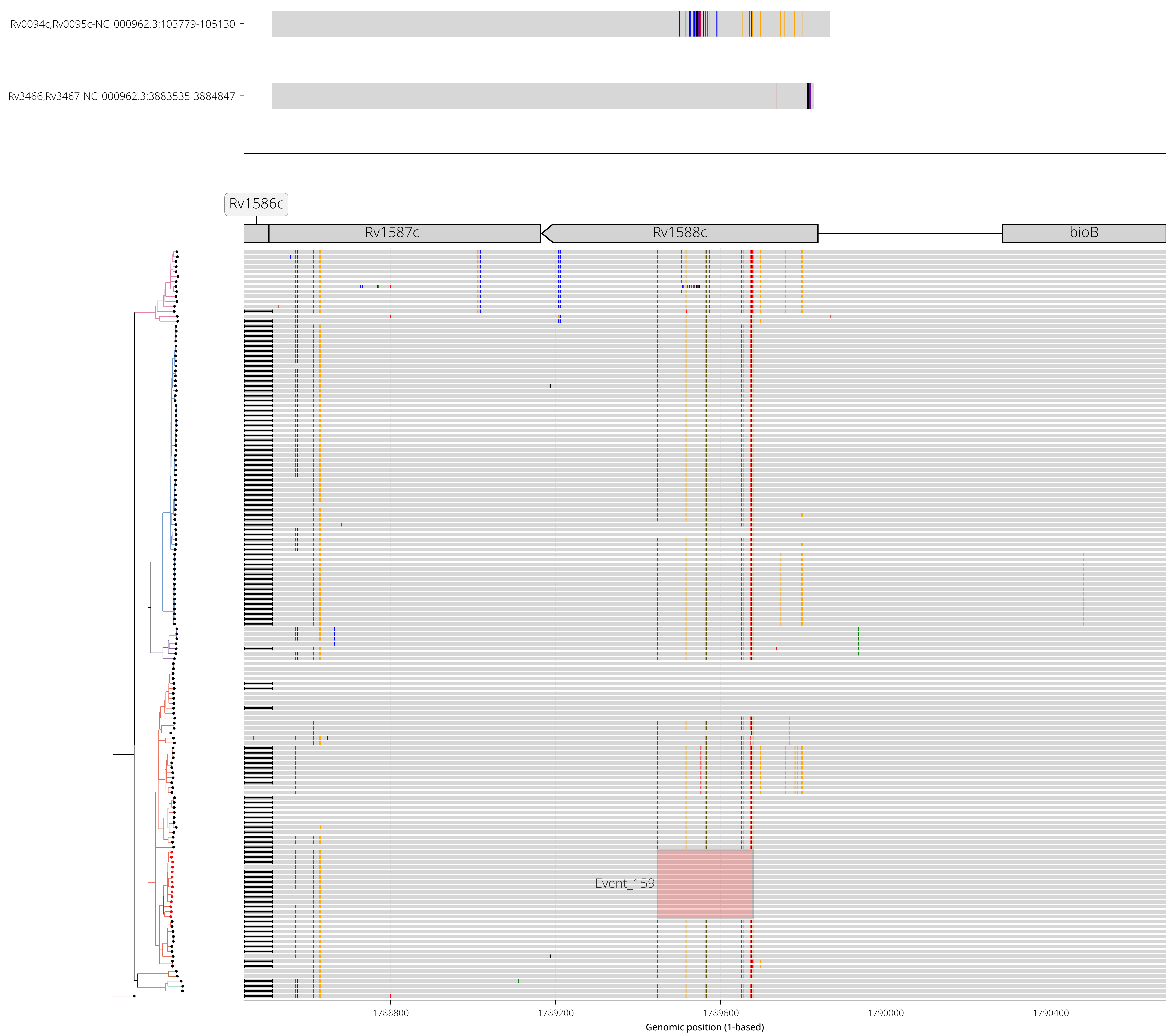

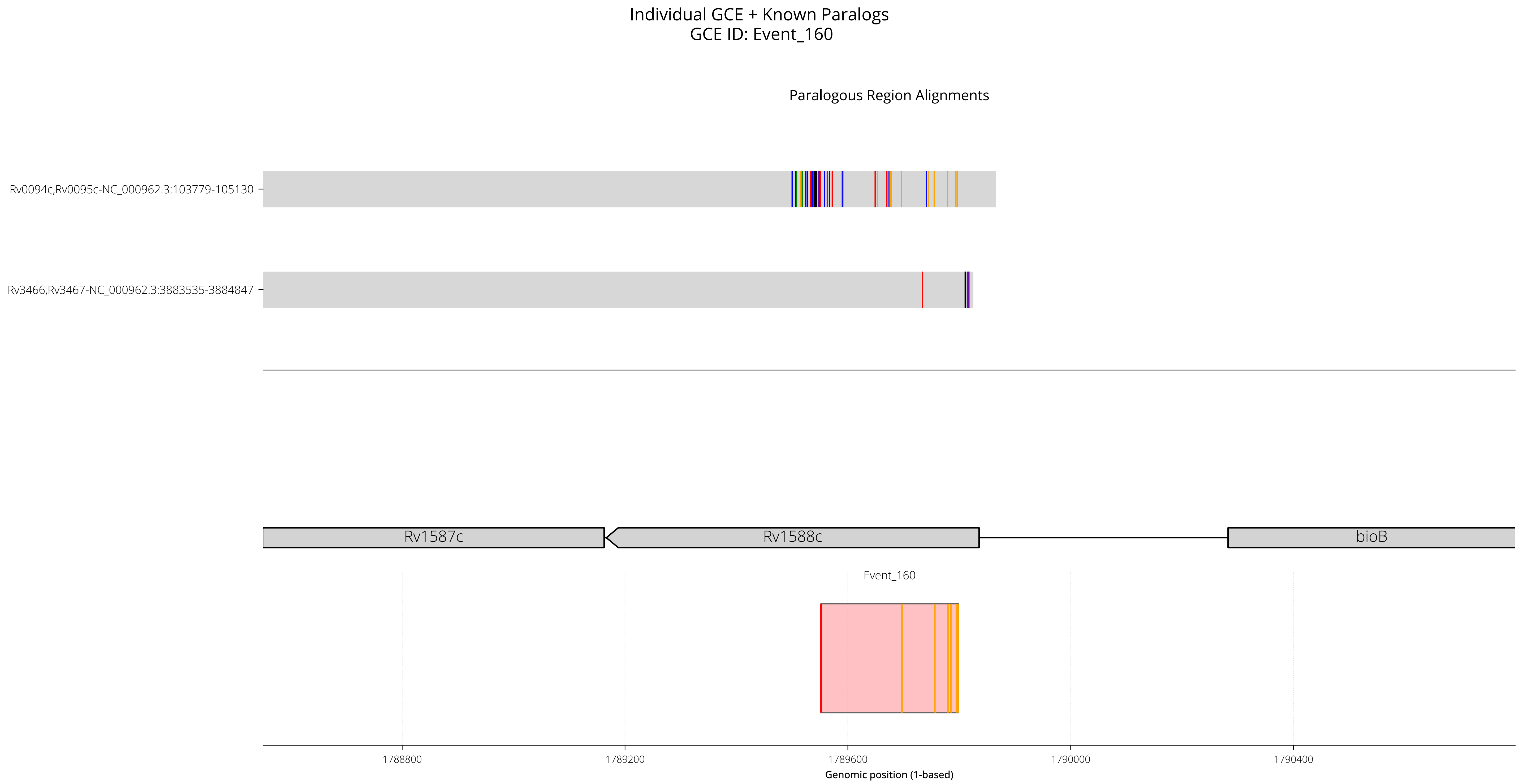

Individual GCE + Known Paralogs
GCE ID: Event_160
Paralogous Region Alignments
Rv0094c,Rv0095c-NC_000962.3:103779-105130
Rv3466,Rv3467-NC_000962.3:3883535-3884847
Rv1587c
Rv1588c
bioB
Event_160
1788800
1789200
1789600
1790000
1790400
Genomic position (1-based)

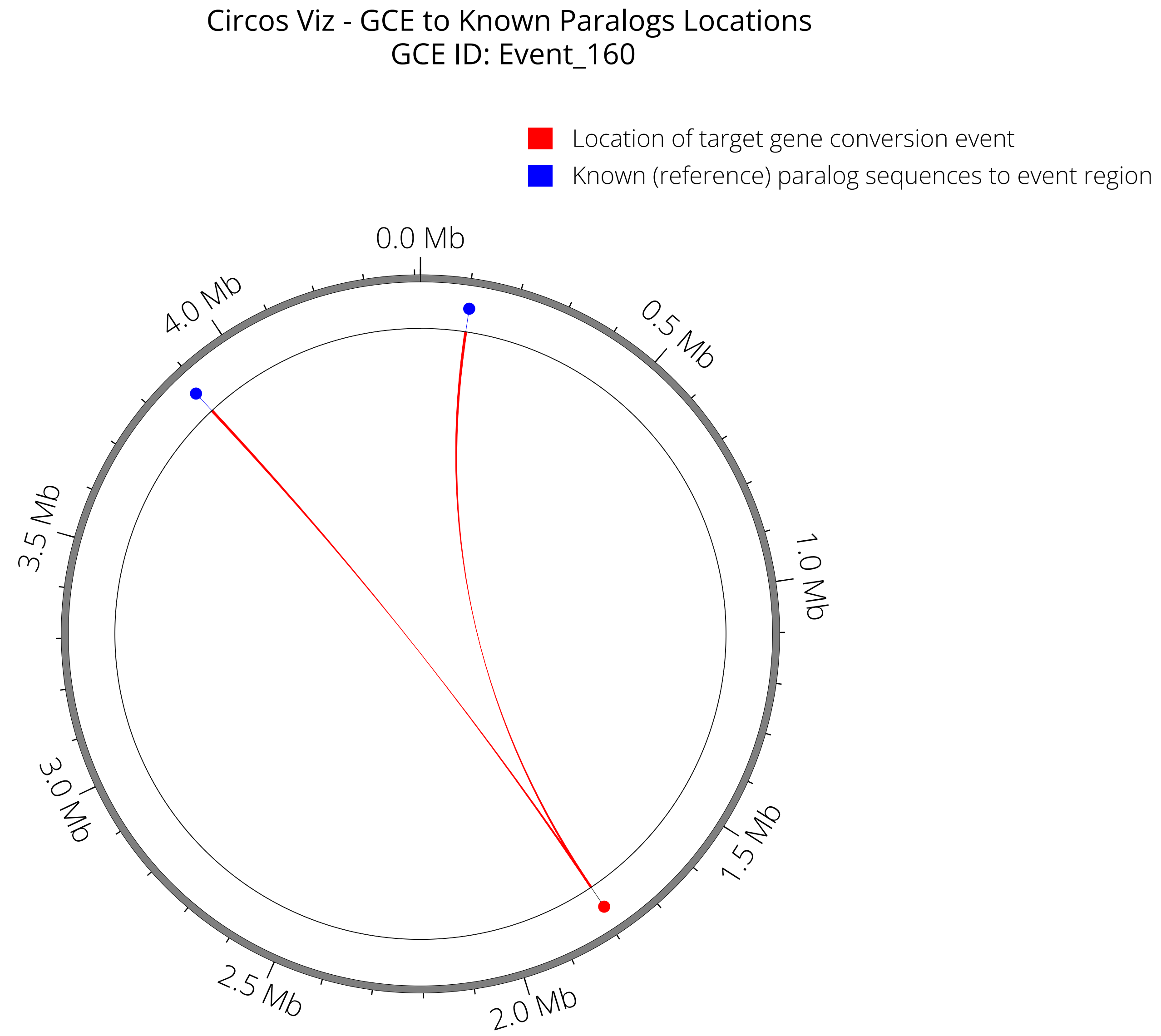
Circos Viz - GCE to Known Paralogs Locations
GCE ID: Event_160
Location of target gene conversion event
Known (reference) paralog sequences to event region
0.0 Mb
0.5 Mb
1.0 Mb
1.5 Mb
2.0 Mb
2.5 Mb
3.0 Mb
3.5 Mb
4.0 Mb

### Event_160 - 7 SNPs - k-mer analysis relative to paralogs

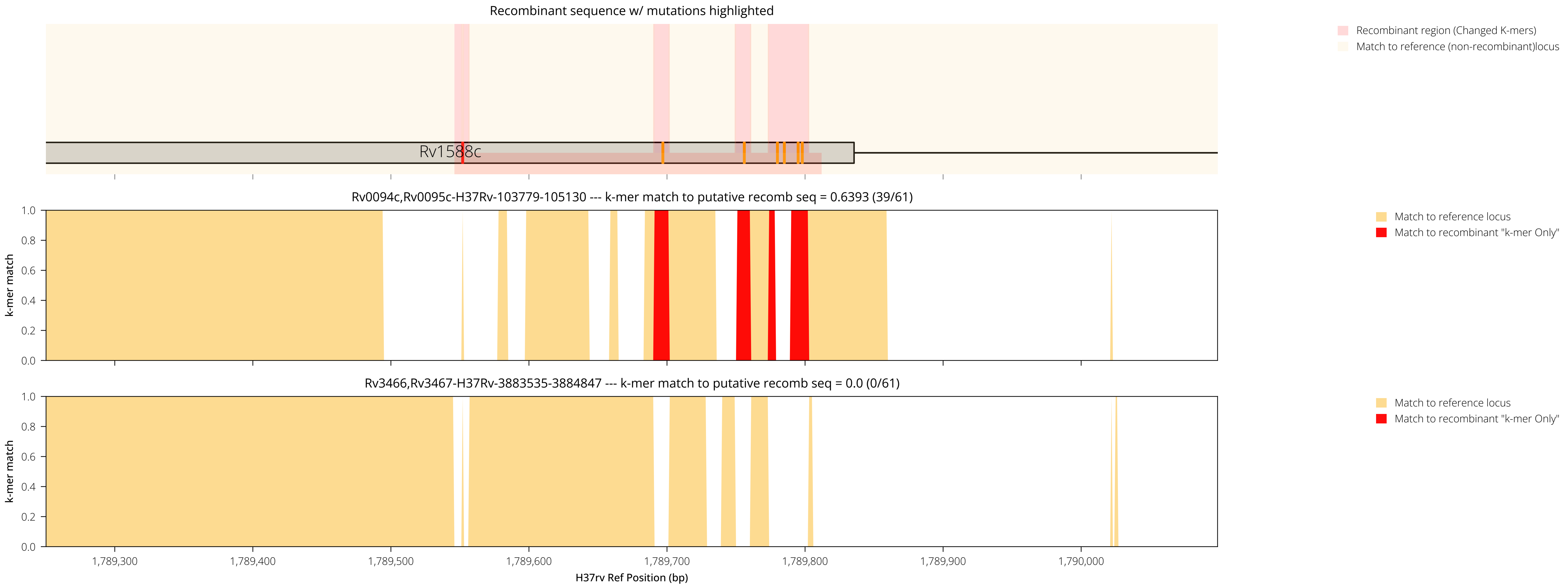

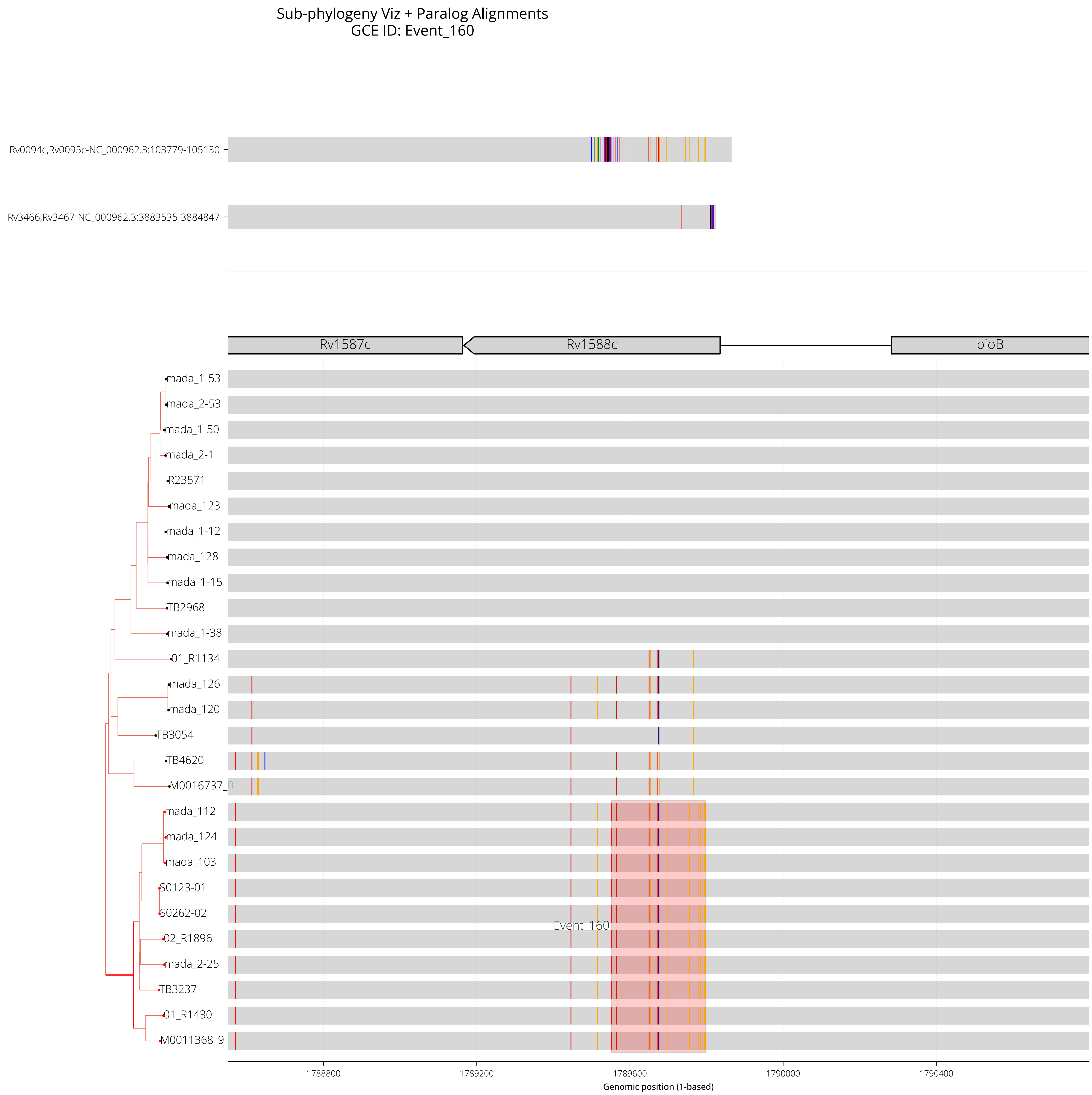

Sub-phylogeny Viz + Paralog Alignments
GCE ID: Event_160
Rv0094c,Rv0095c-NC_000962.3:103779-105130
Rv3466,Rv3467-NC_000962.3:3883535-3884847
Rv1587c
Rv1588c
bioB
mada_1-53
mada_2-53
mada_1-50
mada_2-1
R23571
mada_123
mada_1-12
mada_128
mada_1-15
TB2968
mada_1-38
01_R1134
mada_126
mada_120
TB3054
TB4620
M0016737_0
mada_112
mada_124
mada_103
S0123-01
S0262-02
02_R1896
mada_2-25
TB3237
01_R1430
M0011368_9
Event_160
1788800
1789200
1789600
1790000
1790400
Genomic position (1-based)

Full Phylogeny Viz + Paralog Alignments
GCE ID: Event_160

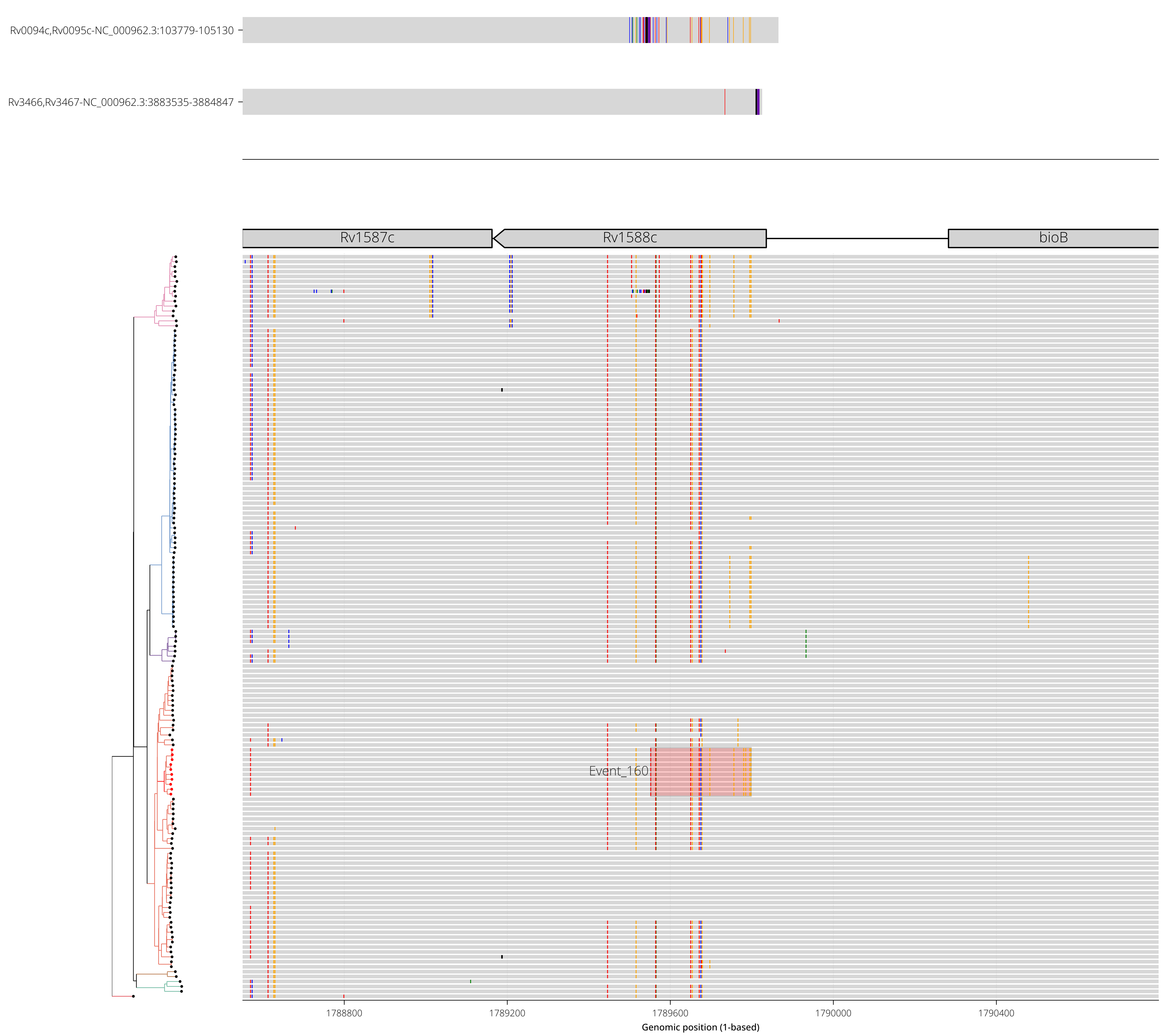

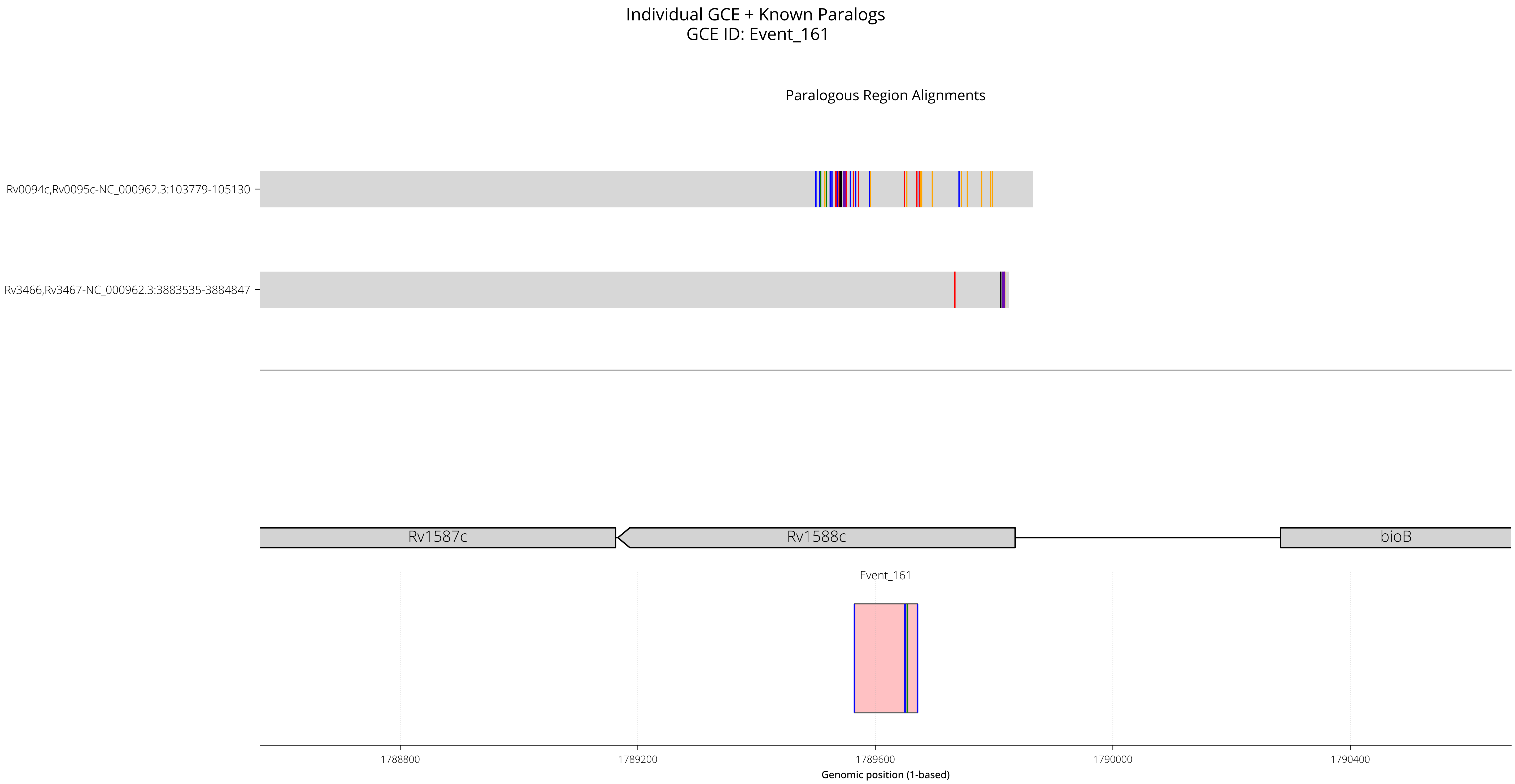

Individual GCE + Known Paralogs
GCE ID: Event_161
Paralogous Region Alignments
Rv0094c,Rv0095c-NC_000962.3:103779-105130
Rv3466,Rv3467-NC_000962.3:3883535-3884847
Rv1587c
Rv1588c
bioB
Event_161
1788800
1789200
1789600
1790000
1790400
Genomic position (1-based)

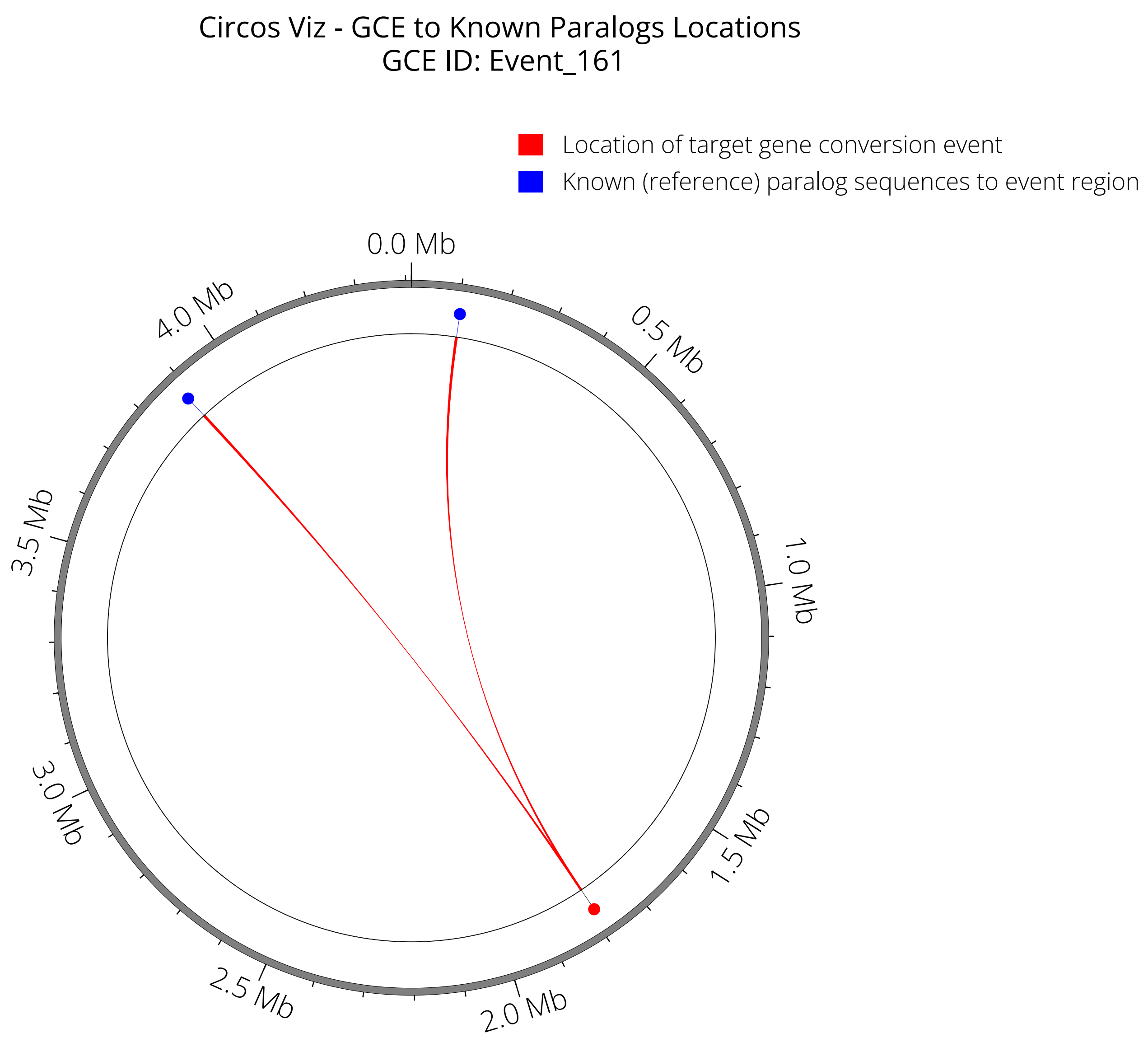
Circos Viz - GCE to Known Paralogs Locations
GCE ID: Event_161
Location of target gene conversion event
Known (reference) paralog sequences to event region
0.0 Mb
0.5 Mb
1.0 Mb
1.5 Mb
2.0 Mb
2.5 Mb
3.0 Mb
3.5 Mb
4.0 Mb

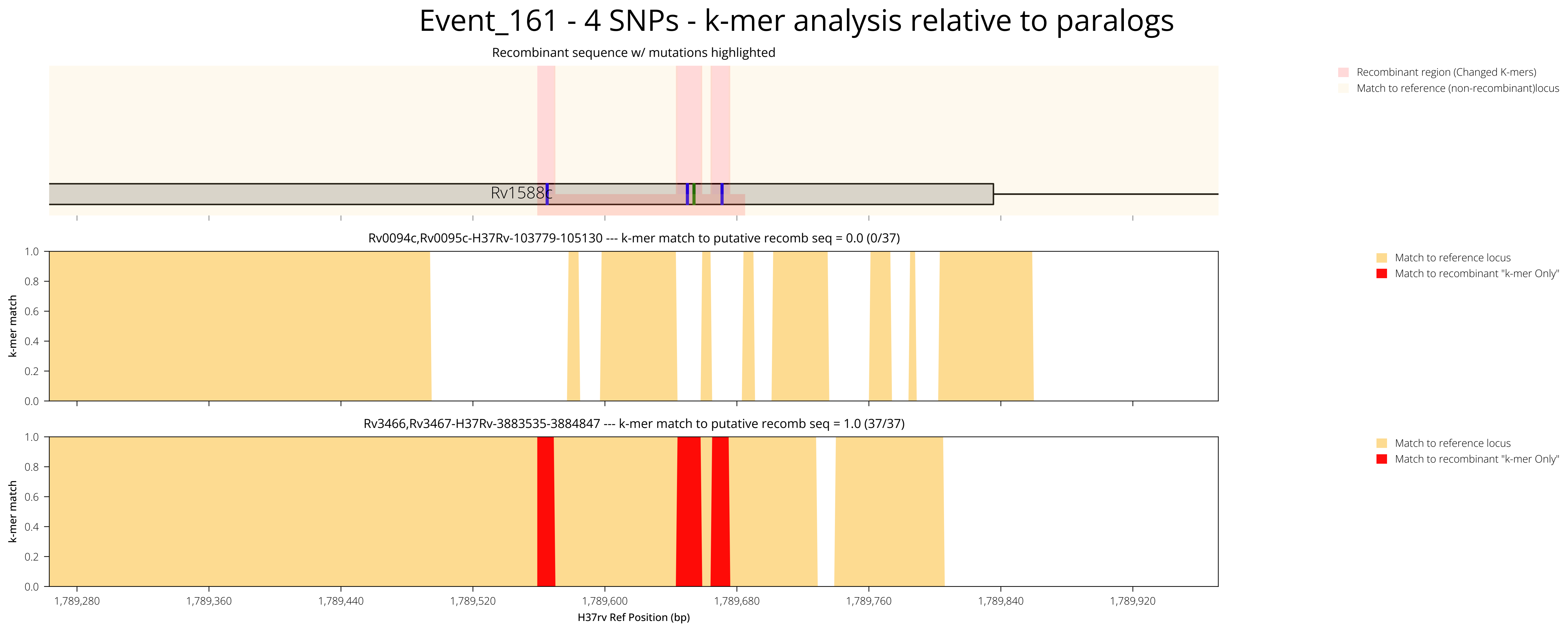

Event_161 - 4 SNPs - k-mer analysis relative to paralogs
Recombinant sequence w/ mutations highlighted
Recombinant region (Changed K-mers)
Match to reference (non-recombinant)locus
Rv1588c
Rv0094c,Rv0095c-H37Rv-103779-105130 --- k-mer match to putative recomb seq = 0.0 (0/37)
Match to reference locus
Match to recombinant "k-mer Only"
Rv3466,Rv3467-H37Rv-3883535-3884847 --- k-mer match to putative recomb seq = 1.0 (37/37)
Match to reference locus
Match to recombinant "k-mer Only"
k-mer match
1.0
0.8
0.6
0.4
0.2
0.0
1,789,280
1,789,360
1,789,440
1,789,520
1,789,600
1,789,680
1,789,760
1,789,840
1,789,920
H37rv Ref Position (bp)

### Sub-phylogeny Viz + Paralog Alignments
GCE ID: Event_161

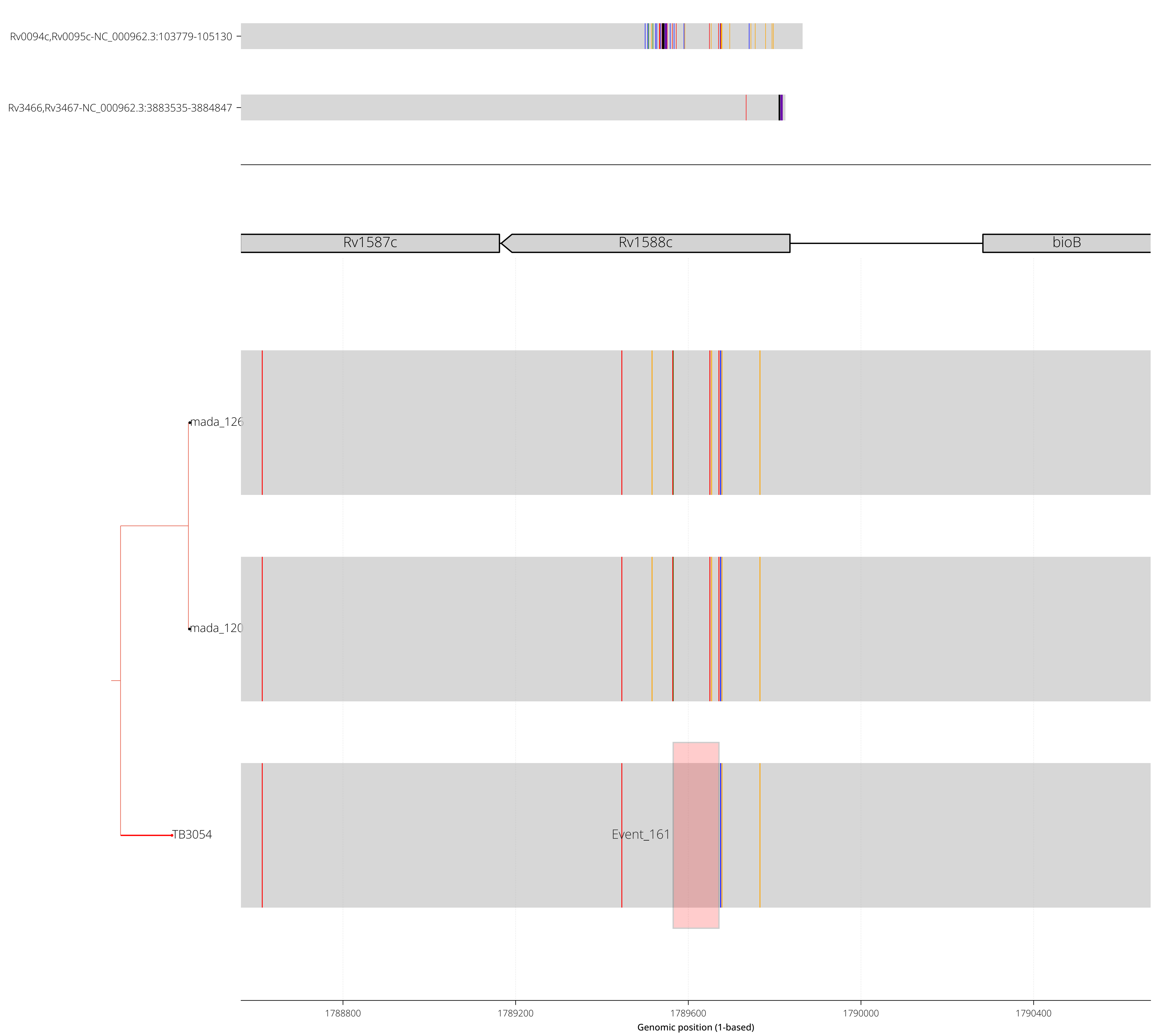

Full Phylogeny Viz + Paralog Alignments
GCE ID: Event_161

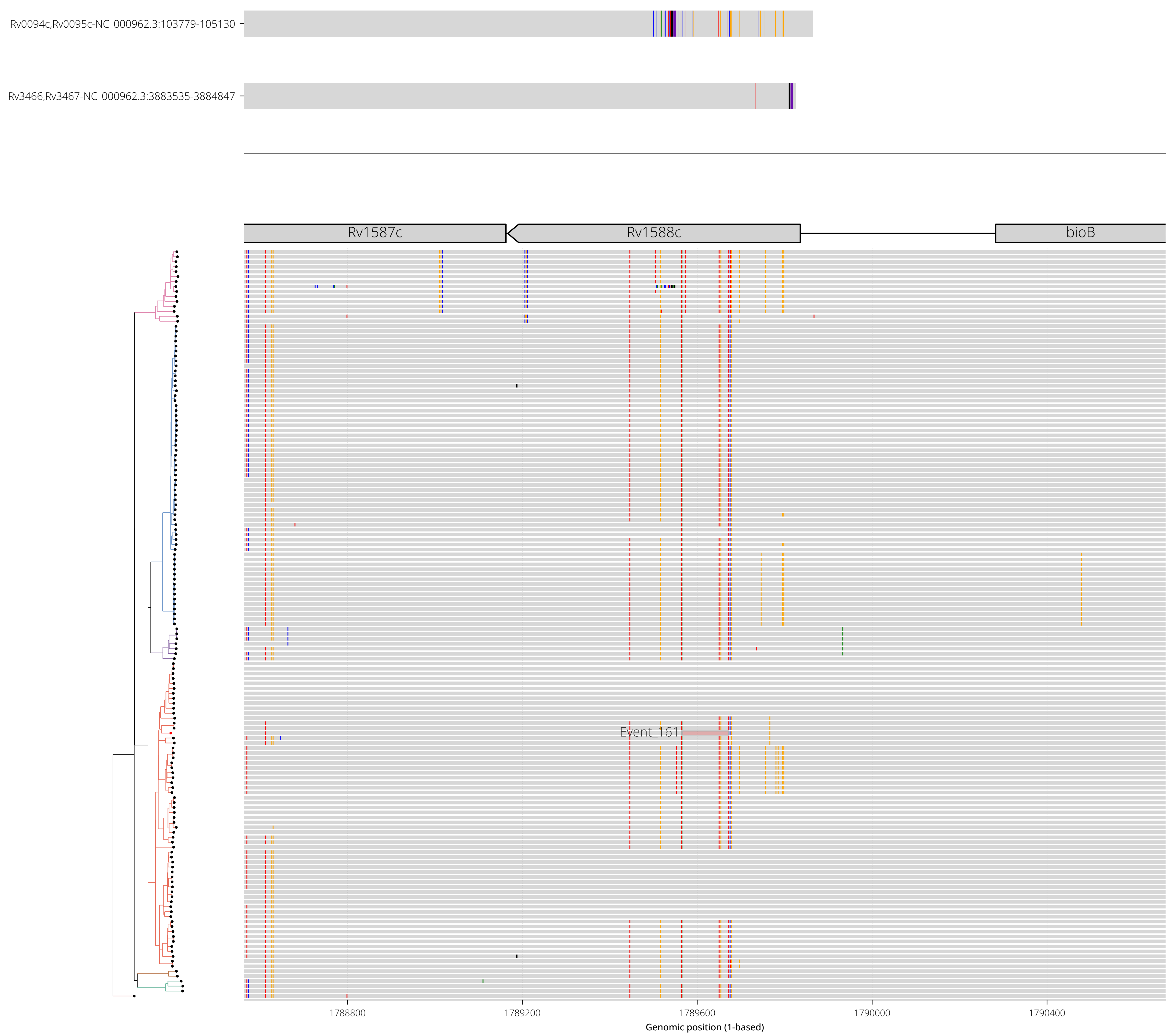

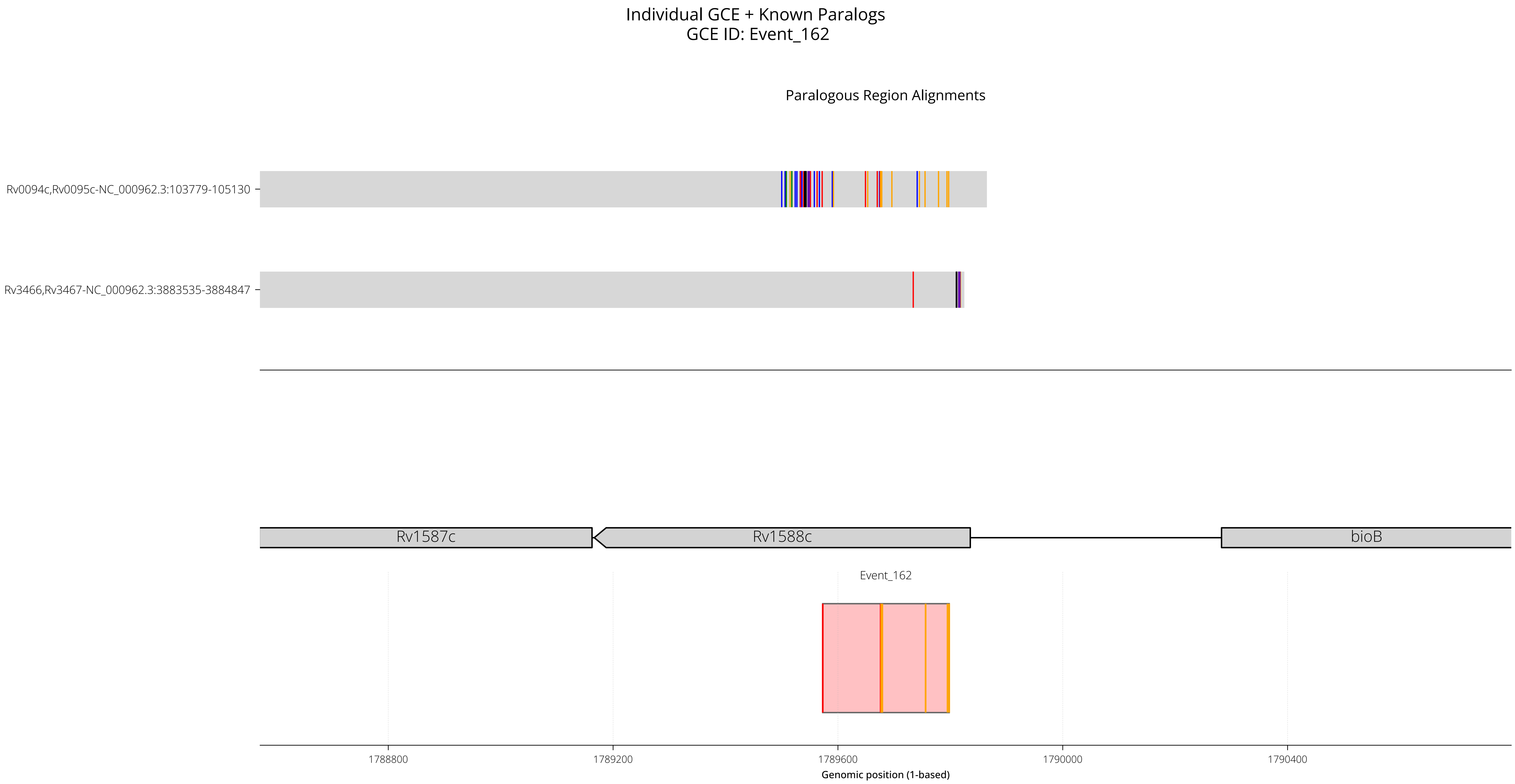

Individual GCE + Known Paralogs
GCE ID: Event_162
Paralogous Region Alignments
Rv0094c,Rv0095c-NC_000962.3:103779-105130
Rv3466,Rv3467-NC_000962.3:3883535-3884847
Rv1587c
Rv1588c
bioB
Event_162
1788800
1789200
1789600
1790000
1790400
Genomic position (1-based)

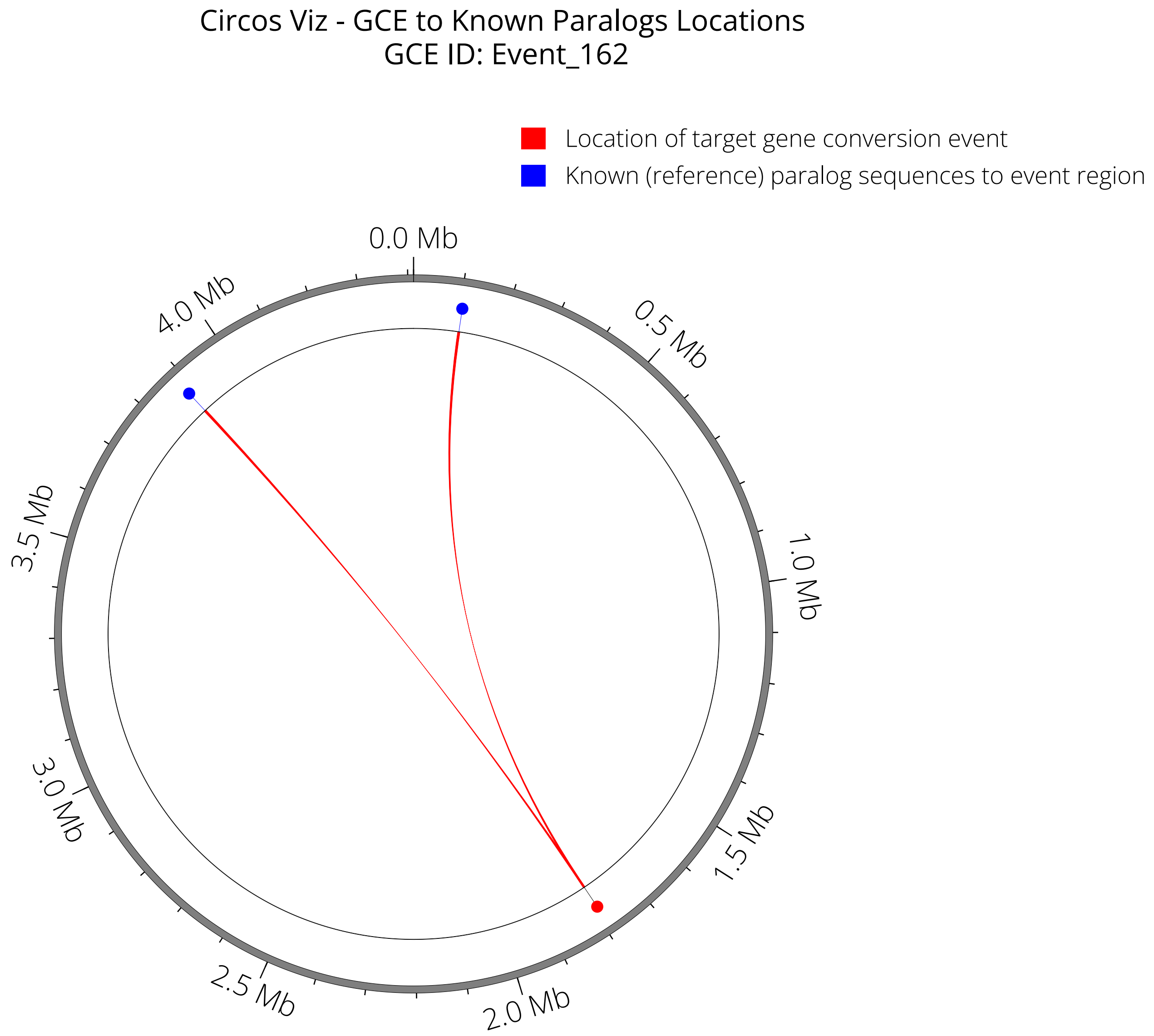
Circos Viz - GCE to Known Paralogs Locations
GCE ID: Event_162
Location of target gene conversion event
Known (reference) paralog sequences to event region
0.0 Mb
0.5 Mb
1.0 Mb
1.5 Mb
2.0 Mb
2.5 Mb
3.0 Mb
3.5 Mb
4.0 Mb

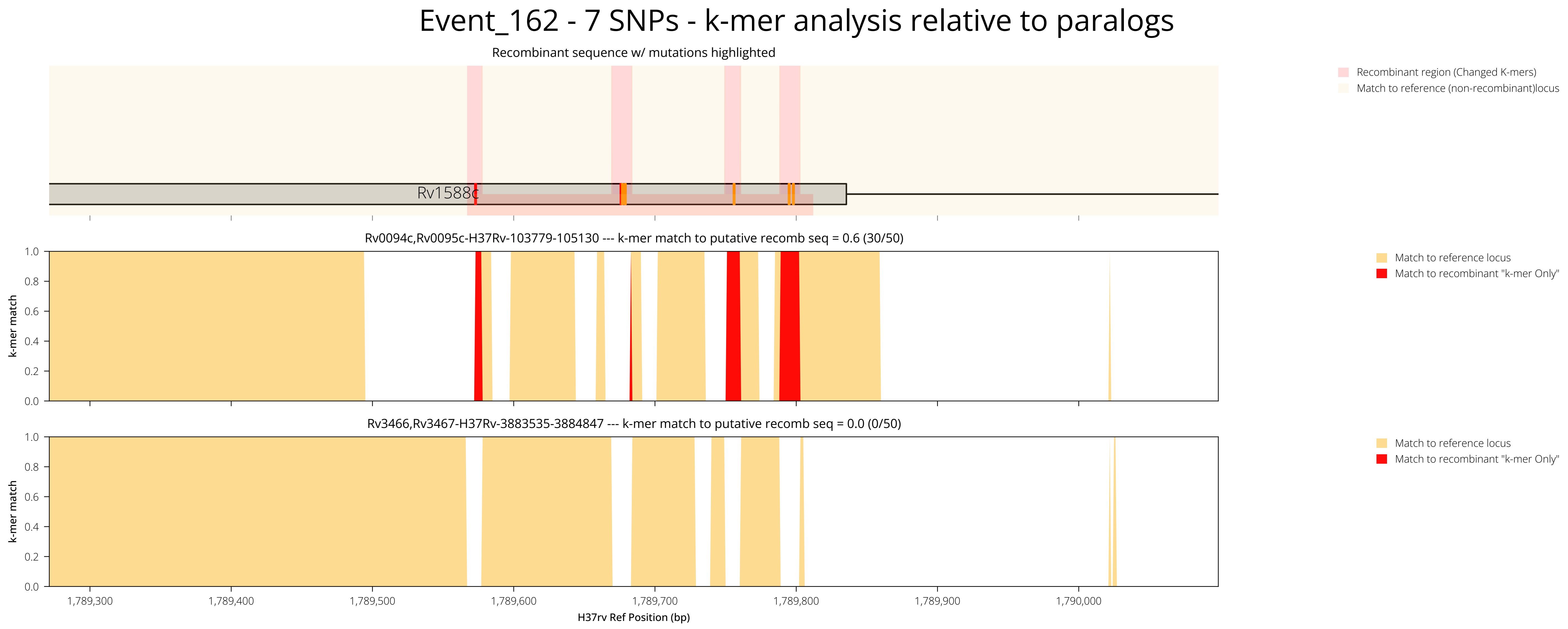
Event_162 - 7 SNPs - k-mer analysis relative to paralogs
Recombinant sequence w/ mutations highlighted
Rv1588c
Recombinant region (Changed K-mers)
Match to reference (non-recombinant)locus
Rv0094c,Rv0095c-H37Rv-103779-105130 --- k-mer match to putative recomb seq = 0.6 (30/50)
Match to reference locus
Match to recombinant "k-mer Only"
k-mer match
1.0
0.8
0.6
0.4
0.2
0.0
Rv3466,Rv3467-H37Rv-3883535-3884847 --- k-mer match to putative recomb seq = 0.0 (0/50)
Match to reference locus
Match to recombinant "k-mer Only"
k-mer match
1,789,300
1,789,400
1,789,500
1,789,600
1,789,700
1,789,800
1,789,900
1,790,000
H37rv Ref Position (bp)

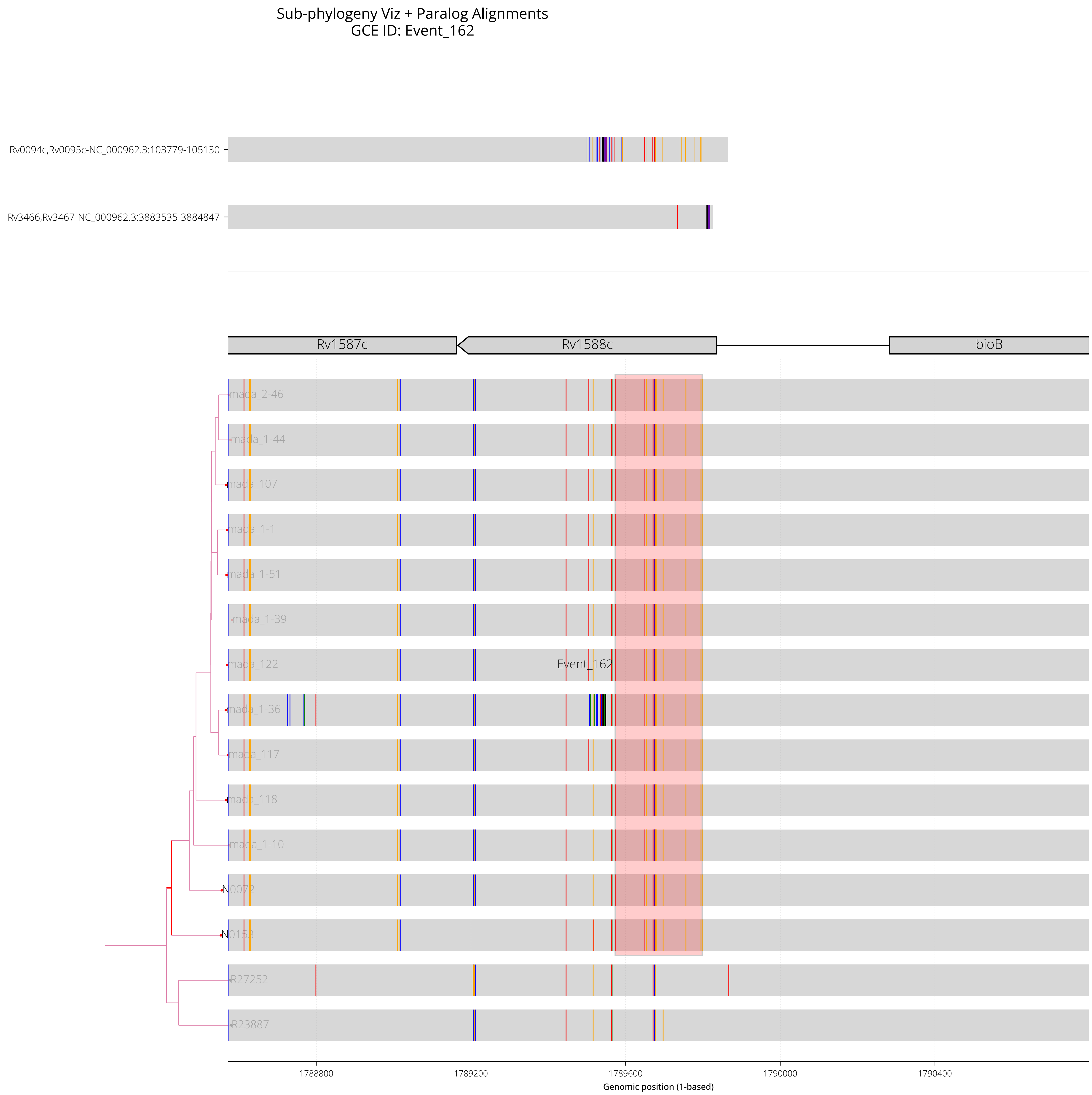
Sub-phylogeny Viz + Paralog Alignments
GCE ID: Event_162
Rv0094c,Rv0095c-NC_000962.3:103779-105130
Rv3466,Rv3467-NC_000962.3:3883535-3884847
Rv1587c
Rv1588c
bioB
mada_2-46
mada_1-44
mada_107
mada_1-1
mada_1-51
mada_1-39
mada_122
Event_162
mada_1-36
mada_117
mada_118
mada_1-10
N0072
N0153
R27252
R23887
1788800
1789200
1789600
1790000
1790400
Genomic position (1-based)

Full Phylogeny Viz + Paralog Alignments
GCE ID: Event_162

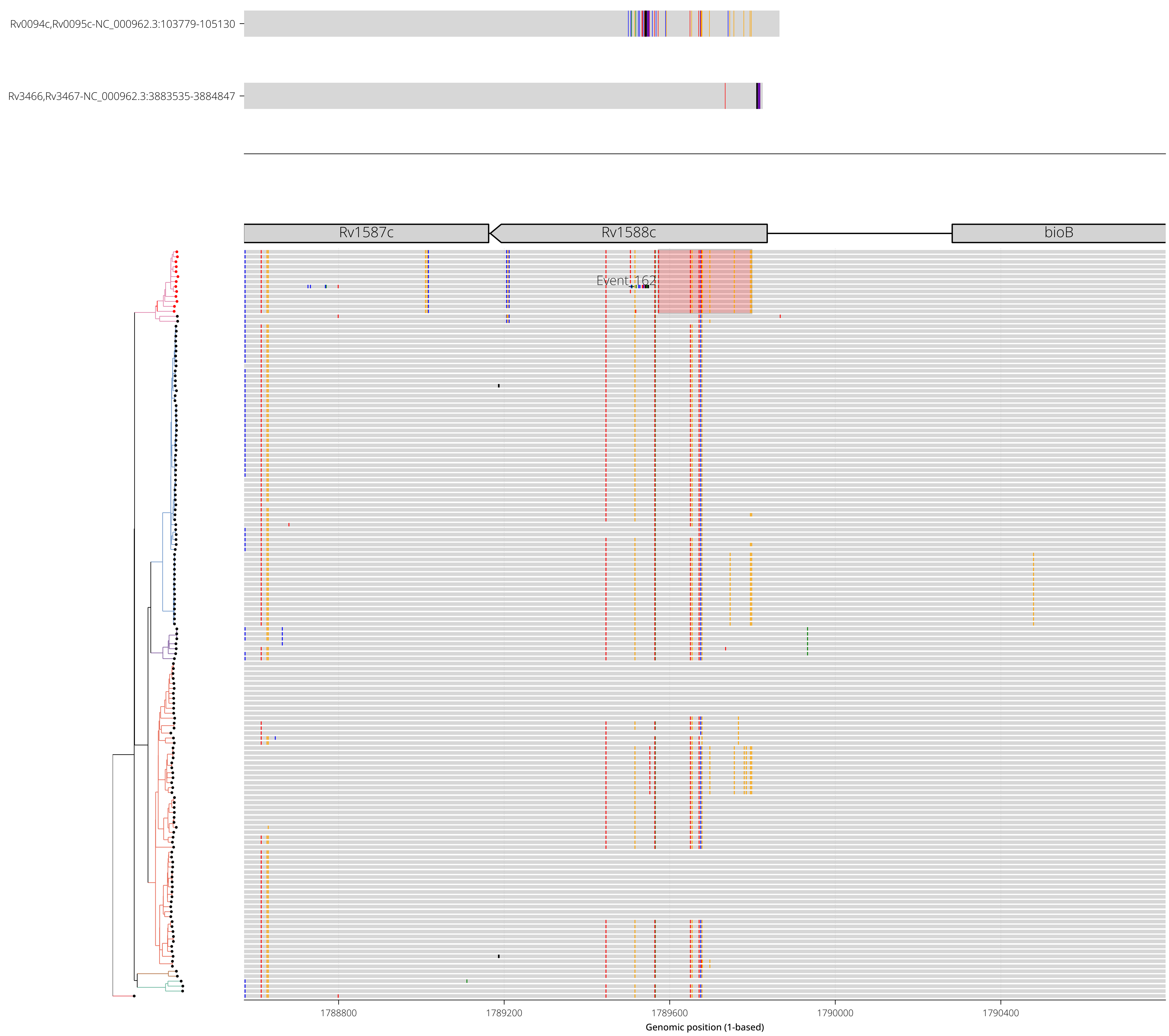

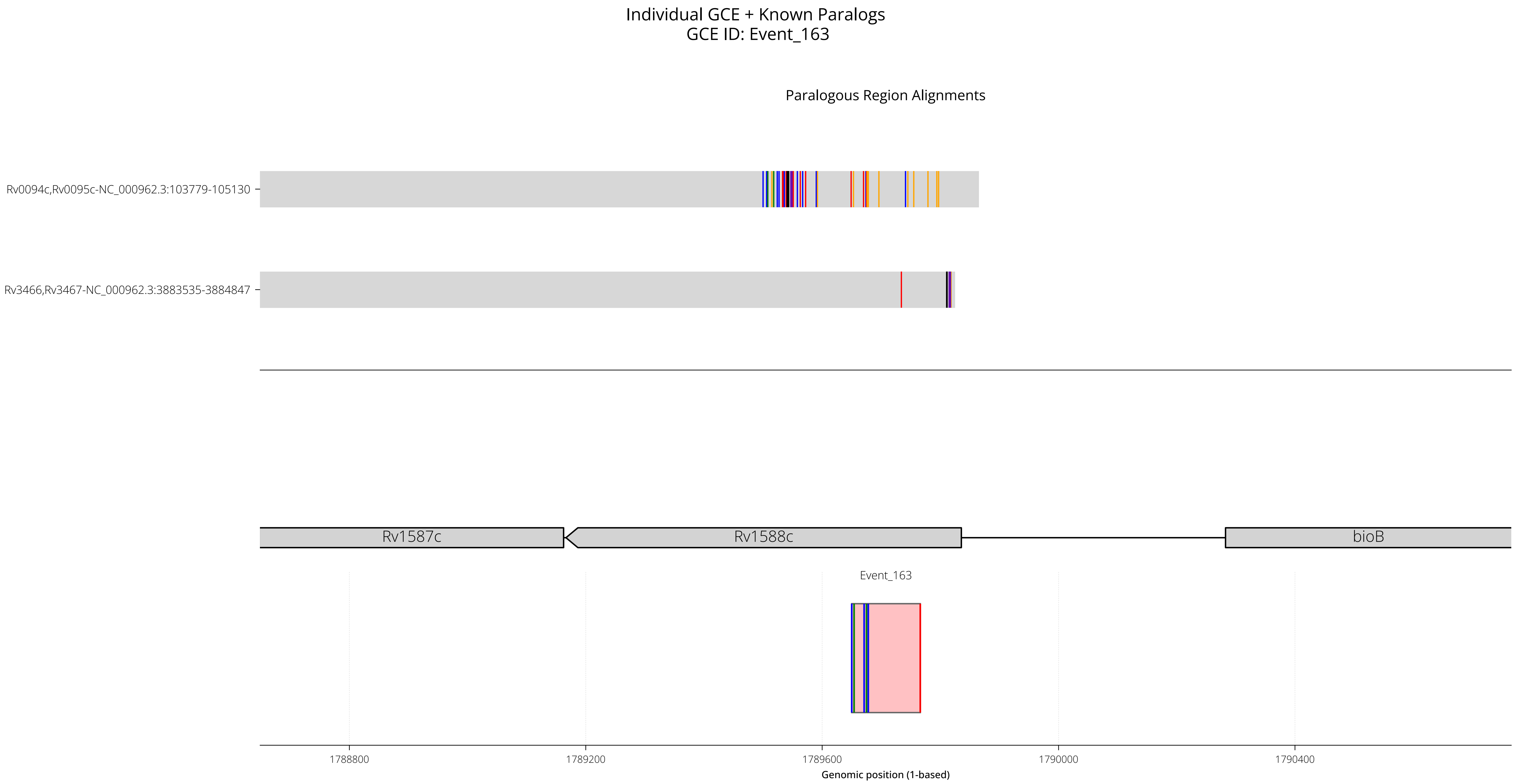
Individual GCE + Known Paralogs
GCE ID: Event_163
Paralogous Region Alignments
Rv0094c,Rv0095c-NC_000962.3:103779-105130
Rv3466,Rv3467-NC_000962.3:3883535-3884847
Rv1587c
Rv1588c
bioB
Event_163
1788800
1789200
1789600
1790000
1790400
Genomic position (1-based)

Circos Viz - GCE to Known Paralogs Locations
GCE ID: Event_163

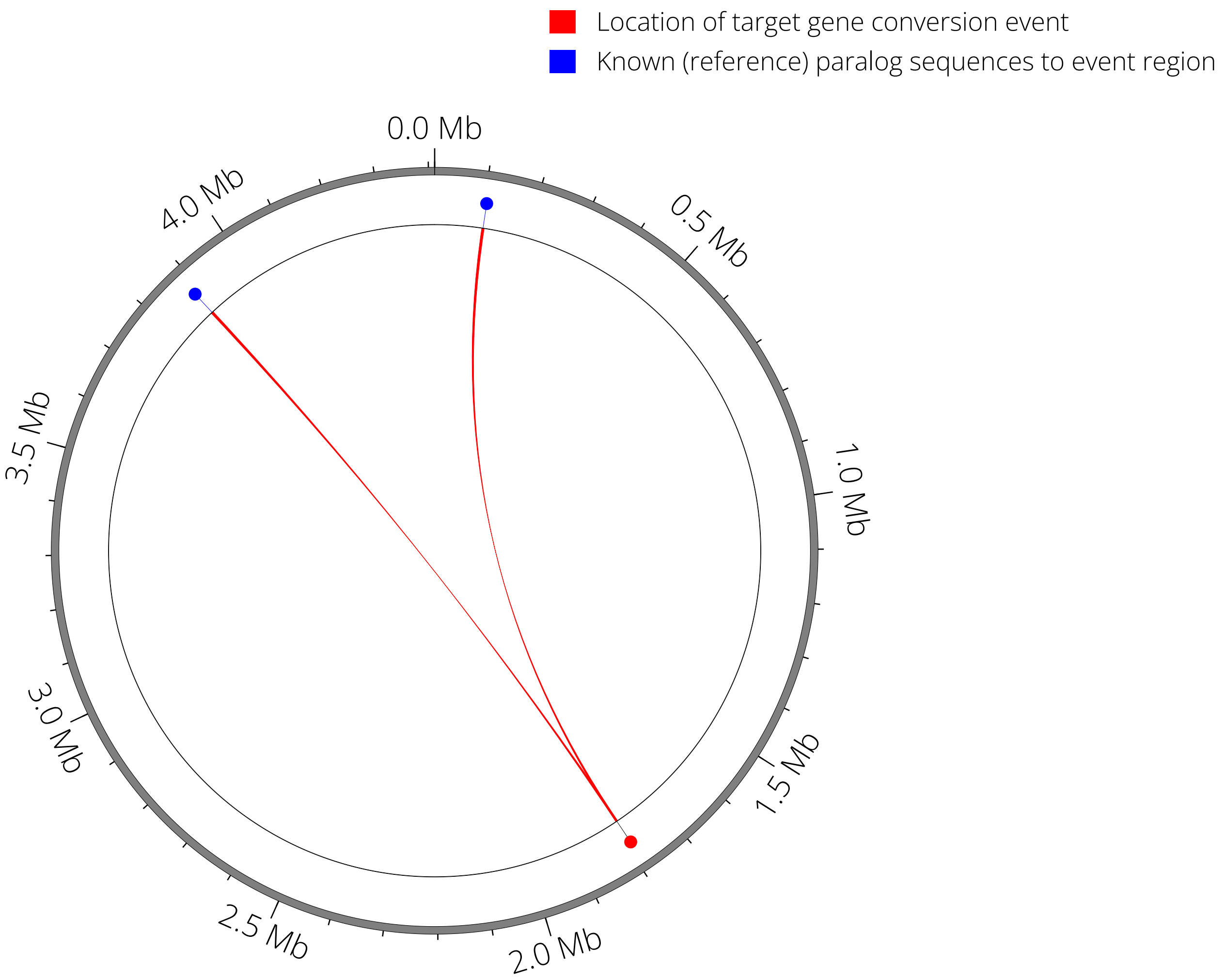

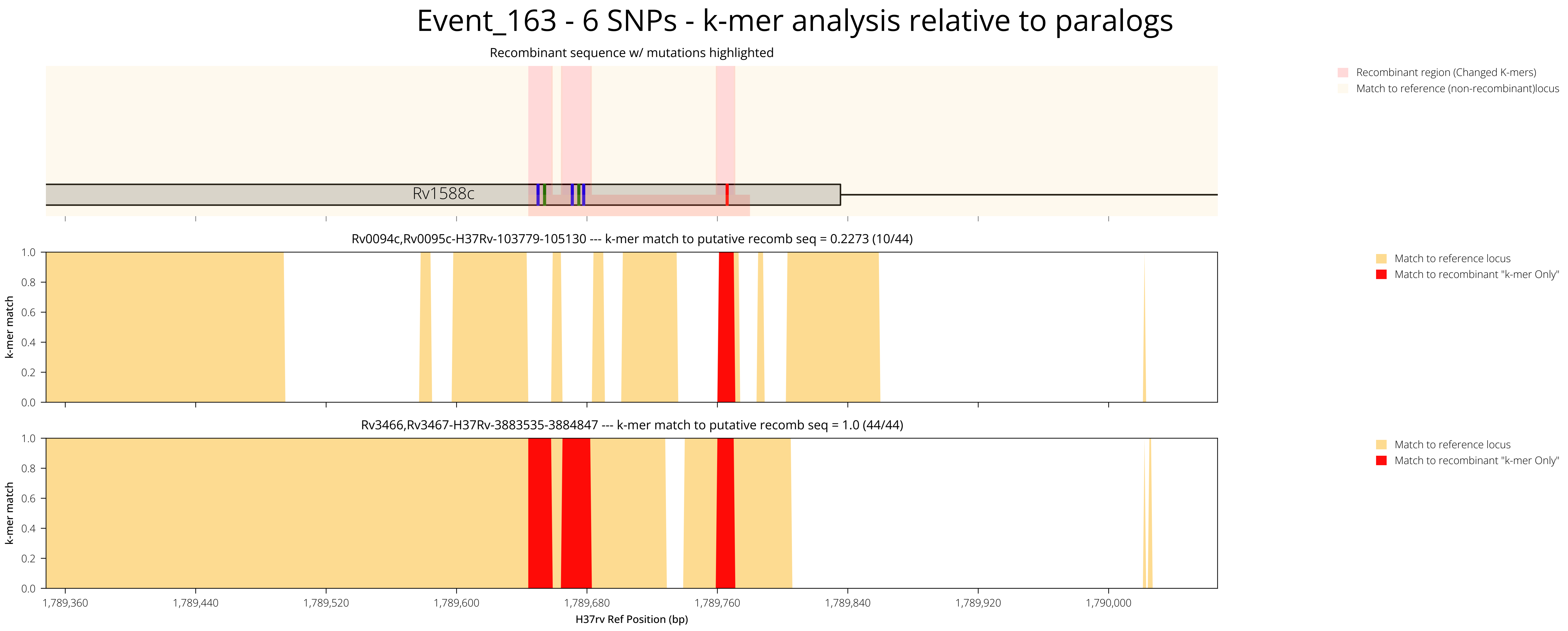

Event_163 - 6 SNPs - k-mer analysis relative to paralogs
Recombinant sequence w/ mutations highlighted
Rv1588c
Recombinant region (Changed K-mers)
Match to reference (non-recombinant)locus
Rv0094c,Rv0095c-H37Rv-103779-105130 --- k-mer match to putative recomb seq = 0.2273 (10/44)
Match to reference locus
Match to recombinant "k-mer Only"
Rv3466,Rv3467-H37Rv-3883535-3884847 --- k-mer match to putative recomb seq = 1.0 (44/44)
Match to reference locus
Match to recombinant "k-mer Only"
k-mer match
1.0
0.8
0.6
0.4
0.2
0.0
1,789,360
1,789,440
1,789,520
1,789,600
1,789,680
1,789,760
1,789,840
1,789,920
1,790,000
H37rv Ref Position (bp)

Sub-phylogeny Viz + Paralog Alignments
GCE ID: Event_163

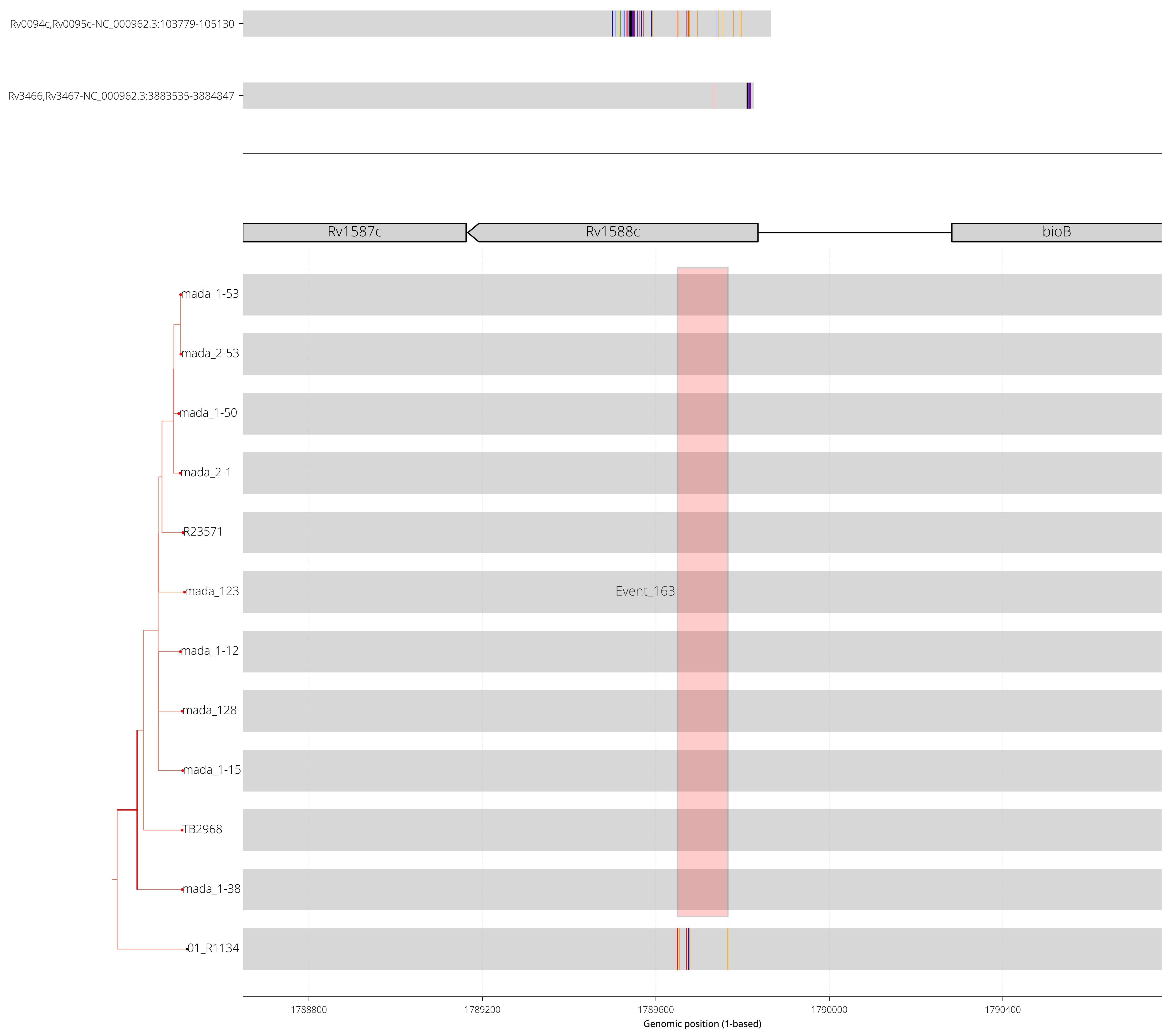

Full Phylogeny Viz + Paralog Alignments
GCE ID: Event_163

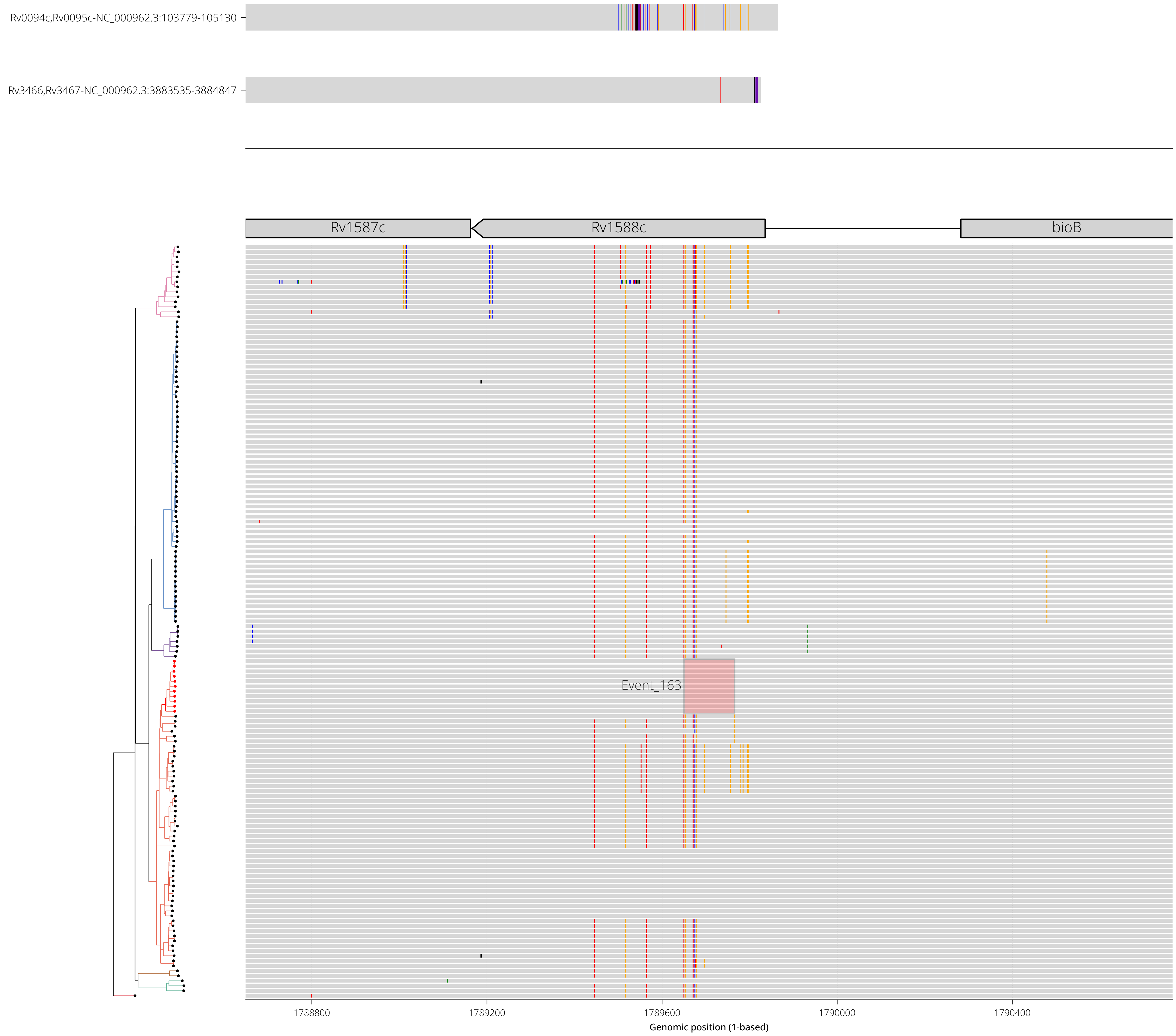

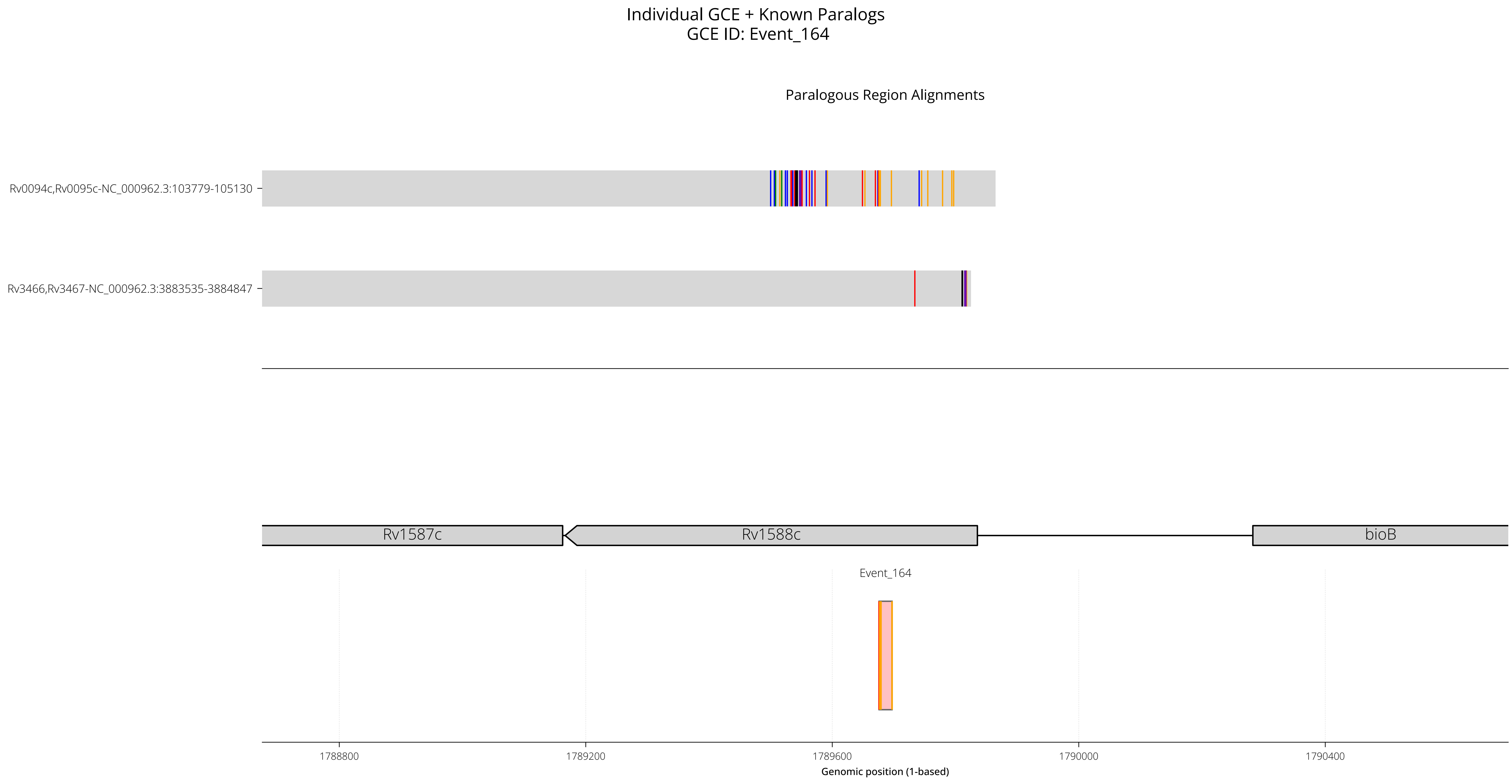

Individual GCE + Known Paralogs
GCE ID: Event_164
Paralogous Region Alignments
Rv0094c,Rv0095c-NC_000962.3:103779-105130
Rv3466,Rv3467-NC_000962.3:3883535-3884847
Rv1587c
Rv1588c
bioB
Event_164
1788800
1789200
1789600
1790000
1790400
Genomic position (1-based)

Circos Viz - GCE to Known Paralogs Locations
GCE ID: Event_164

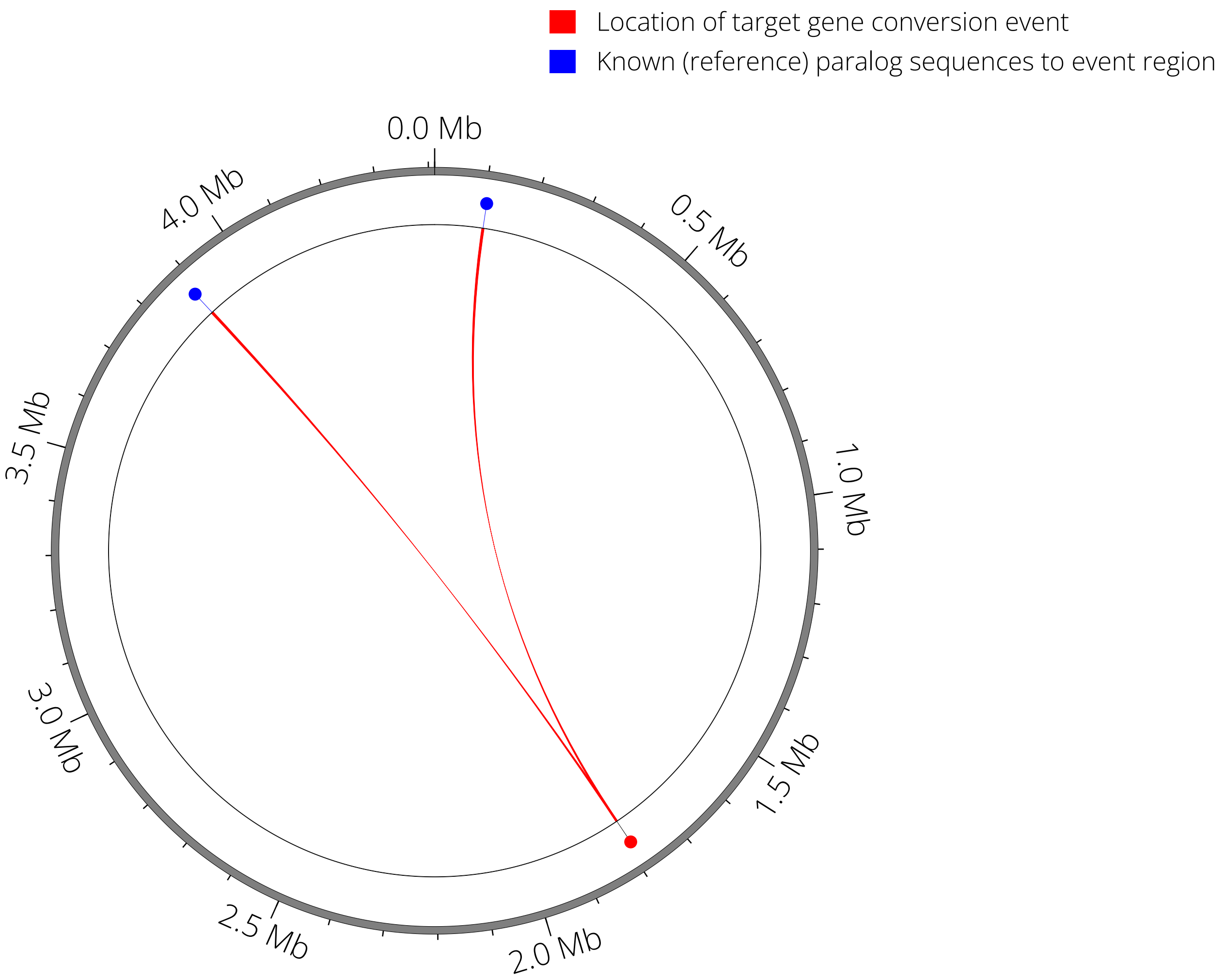

### Event_164 - 4 SNPs - k-mer analysis relative to paralogs

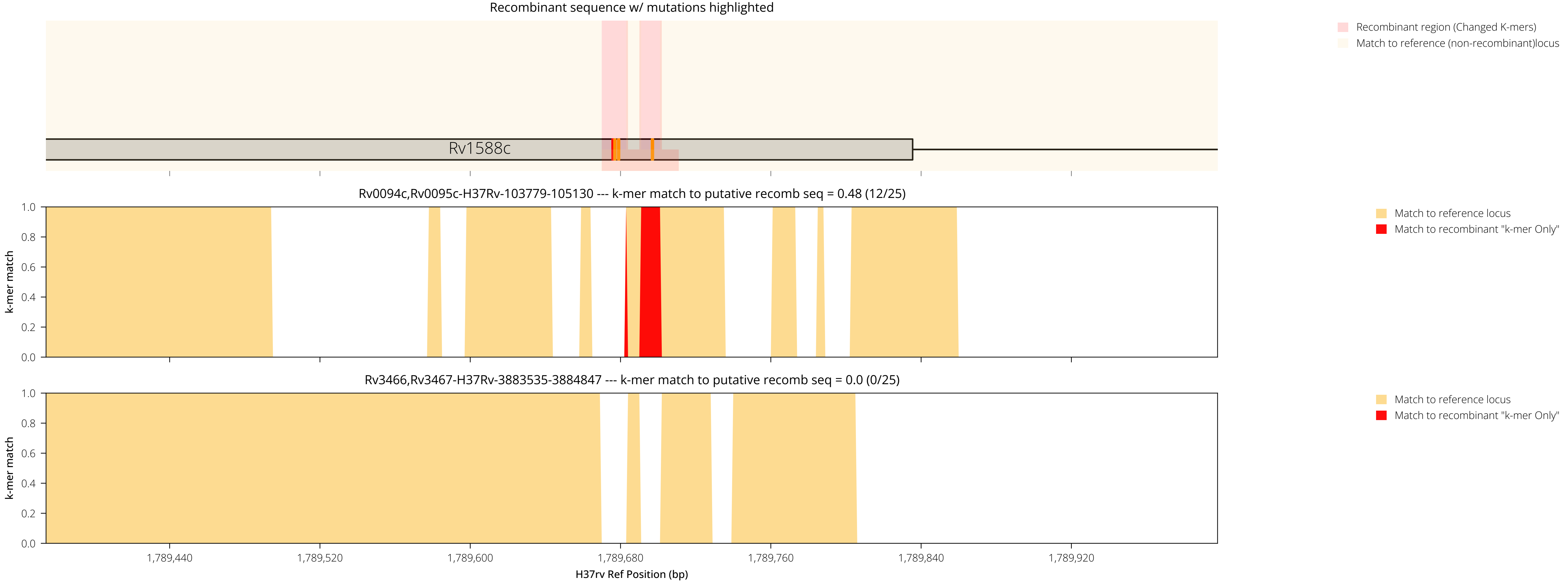

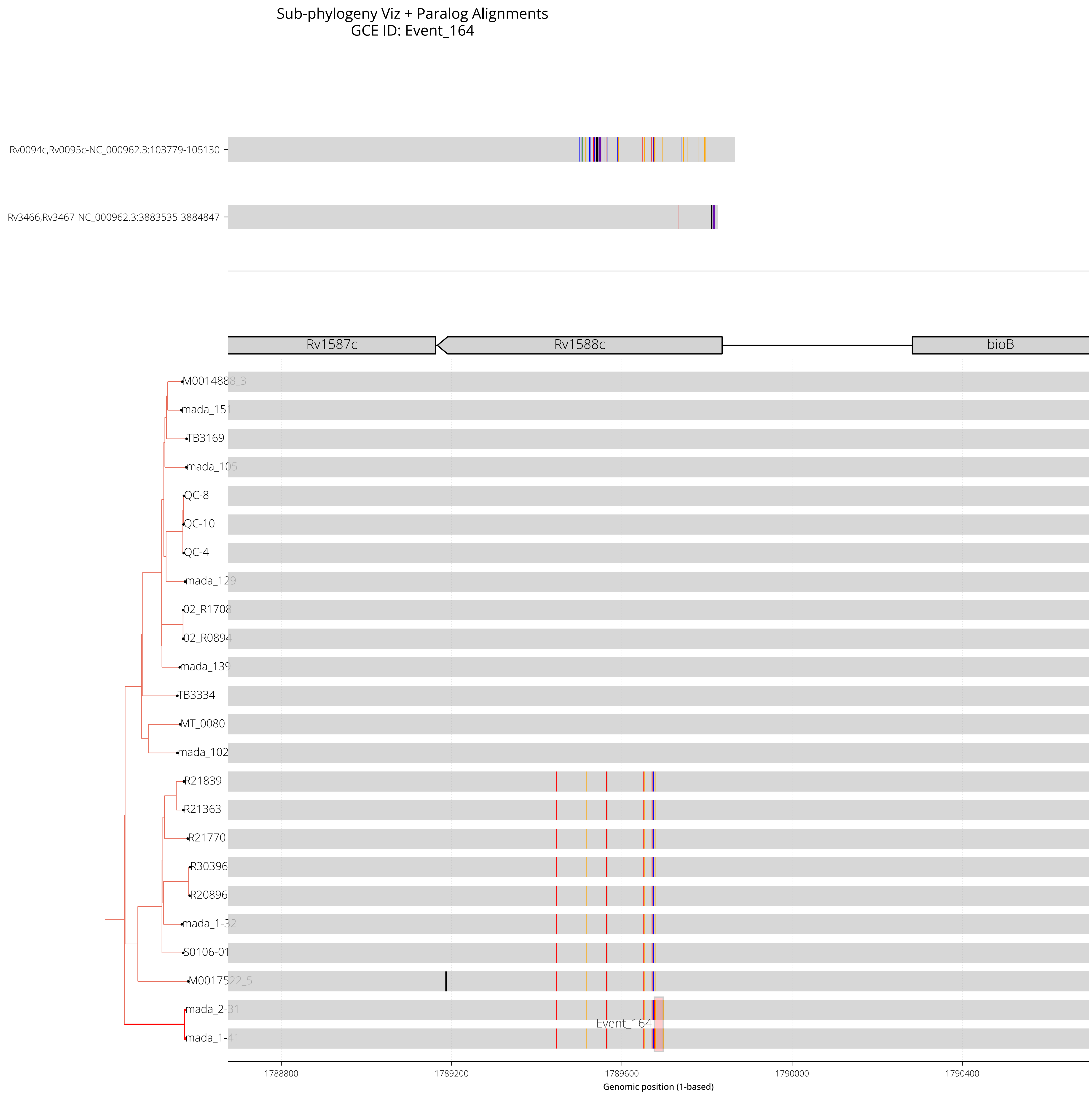

Sub-phylogeny Viz + Paralog Alignments
GCE ID: Event_164
Rv0094c,Rv0095c-NC_000962.3:103779-105130
Rv3466,Rv3467-NC_000962.3:3883535-3884847
Rv1587c
Rv1588c
bioB
M0014888_3
mada_151
TB3169
mada_105
QC-8
QC-10
QC-4
mada_129
02_R1708
02_R0894
mada_139
TB3334
MT_0080
mada_102
R21839
R21363
R21770
R30396
R20896
mada_1-32
S0106-01
M0017522_5
mada_2-31
Event_164
mada_1-41
1788800
1789200
1789600
1790000
1790400
Genomic position (1-based)

Full Phylogeny Viz + Paralog Alignments
GCE ID: Event_164

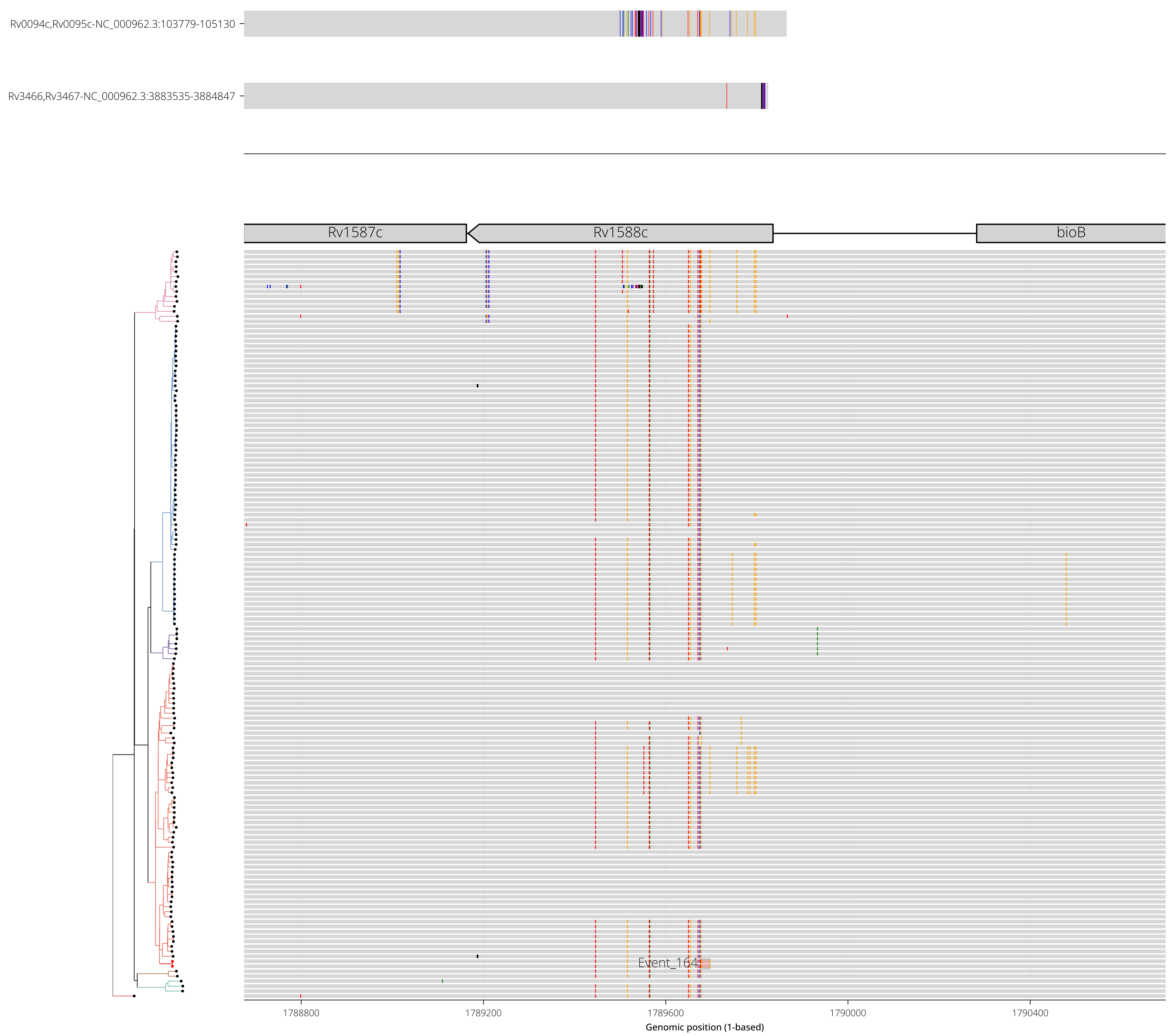

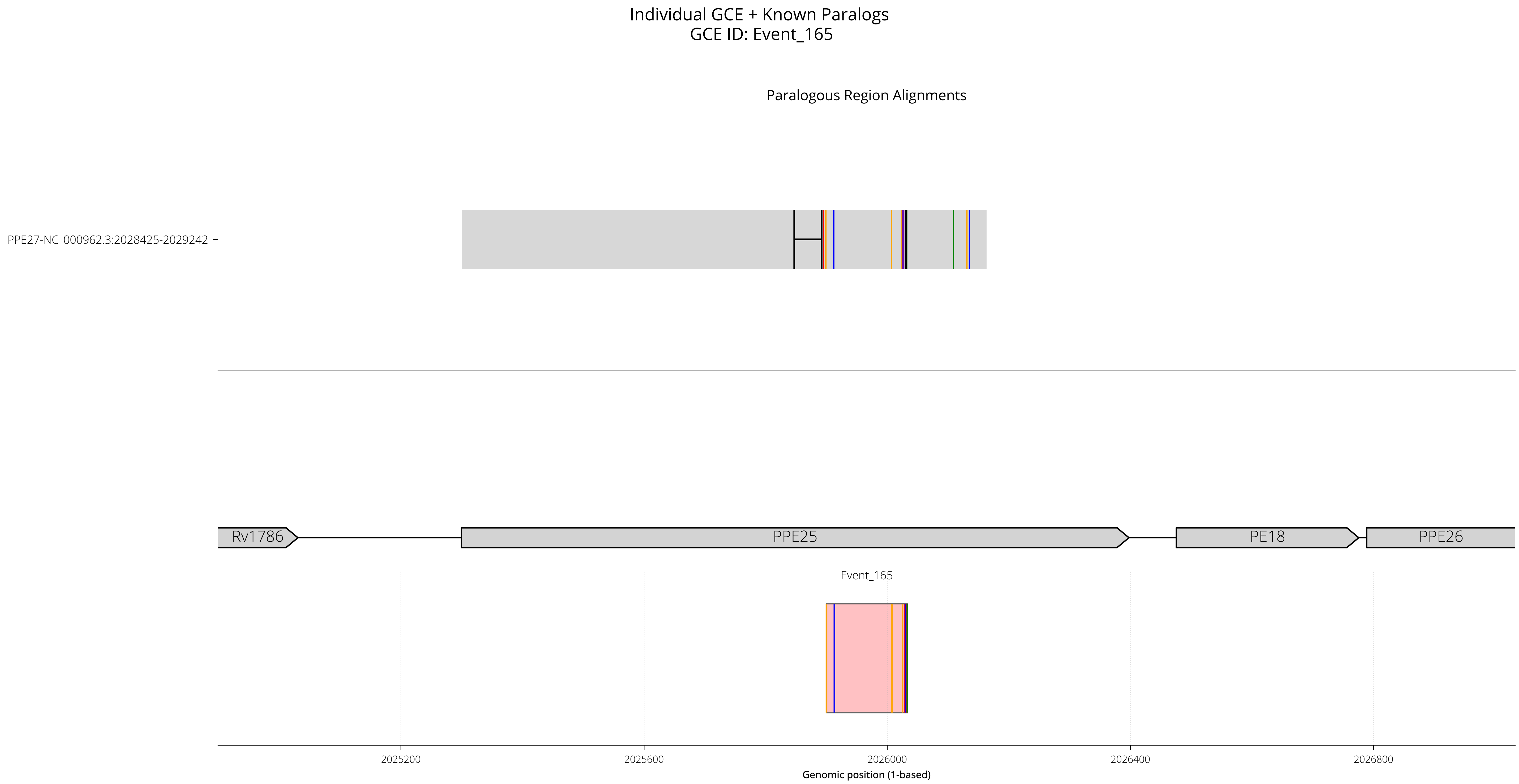
Individual GCE + Known Paralogs
GCE ID: Event_165
Paralogous Region Alignments
PPE27-NC_000962.3:2028425-2029242
Rv1786
PPE25
PE18
PPE26
Event_165
2025200
2025600
2026000
2026400
2026800
Genomic position (1-based)

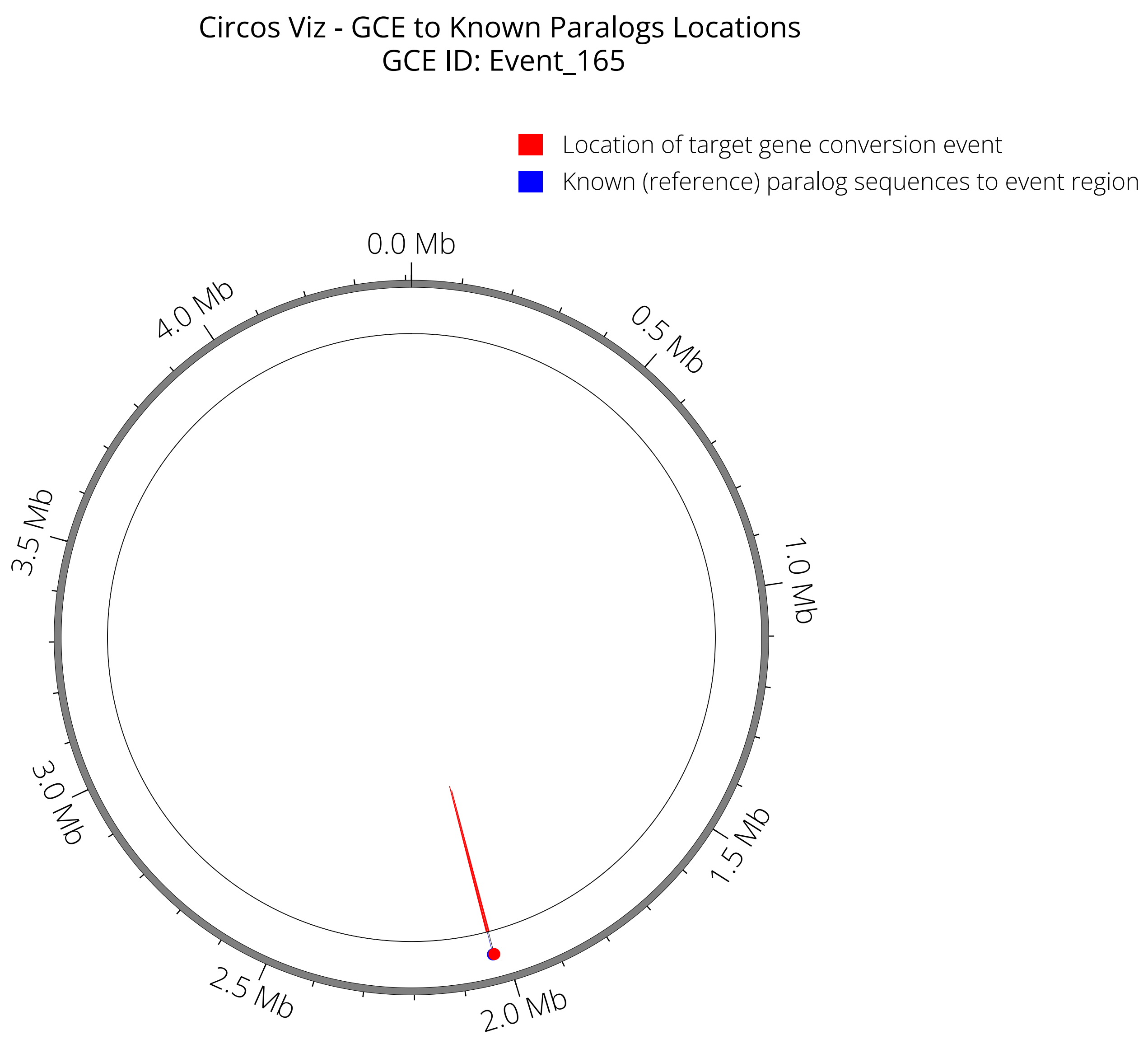
Circos Viz - GCE to Known Paralogs Locations
GCE ID: Event_165
Location of target gene conversion event
Known (reference) paralog sequences to event region
0.0 Mb
0.5 Mb
1.0 Mb
1.5 Mb
2.0 Mb
2.5 Mb
3.0 Mb
3.5 Mb
4.0 Mb

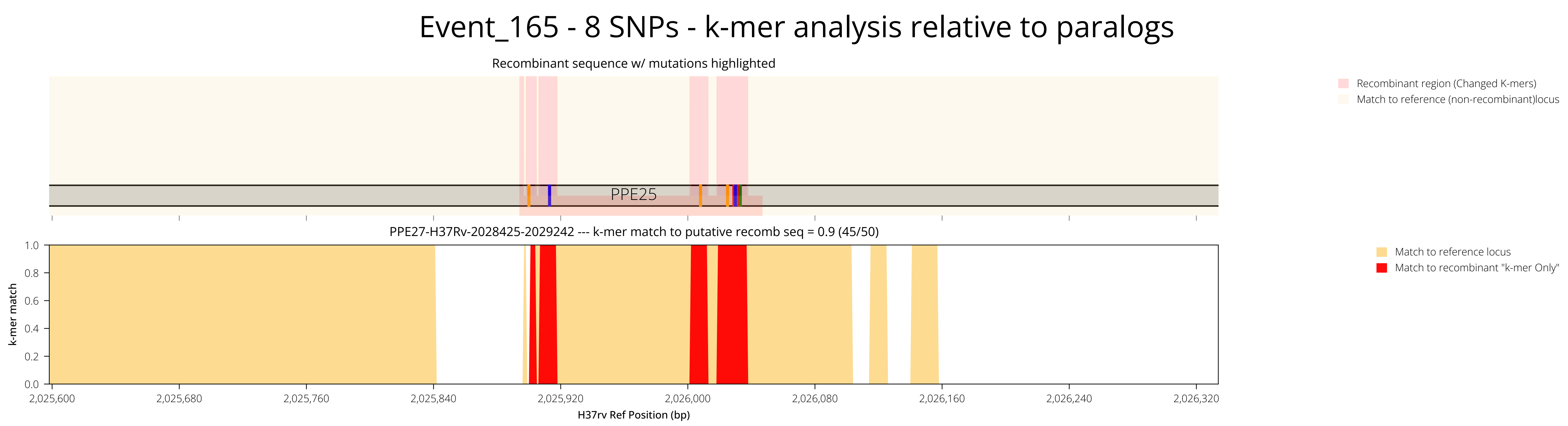
Event_165 - 8 SNPs - k-mer analysis relative to paralogs
Recombinant sequence w/ mutations highlighted
PPE25
Recombinant region (Changed K-mers)
Match to reference (non-recombinant)locus
PPE27-H37Rv-2028425-2029242 --- k-mer match to putative recomb seq = 0.9 (45/50)
Match to reference locus
Match to recombinant "k-mer Only"
k-mer match
1.0
0.8
0.6
0.4
0.2
0.0
2,025,600
2,025,680
2,025,760
2,025,840
2,025,920
2,026,000
2,026,080
2,026,160
2,026,240
2,026,320
H37rv Ref Position (bp)

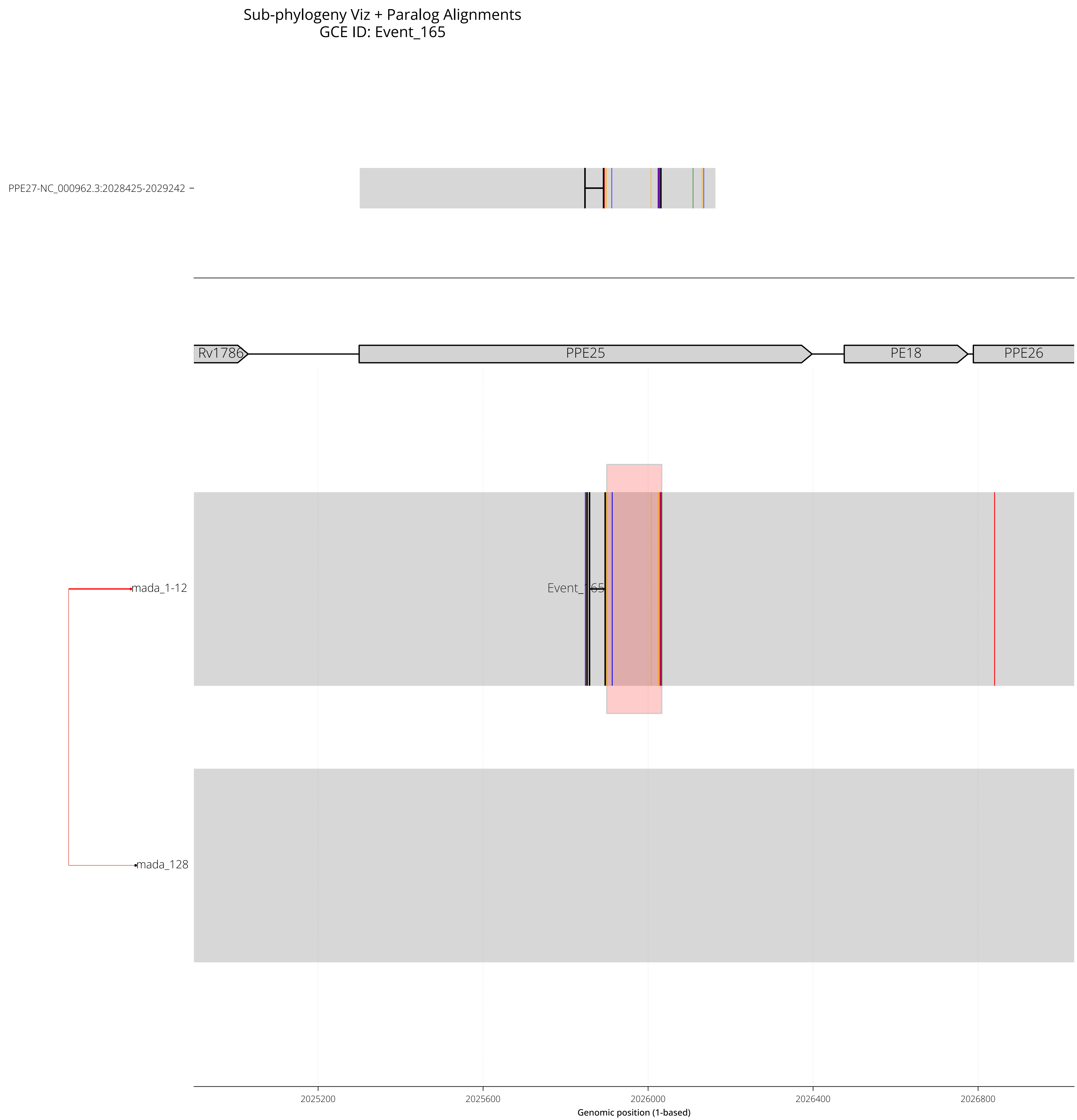

Sub-phylogeny Viz + Paralog Alignments
GCE ID: Event_165
PPE27-NC_000962.3:2028425-2029242
Rv1786
PPE25
PE18
PPE26
mada_1-12
Event_165
mada_128
2025200
2025600
2026000
2026400
2026800
Genomic position (1-based)

Full Phylogeny Viz + Paralog Alignments
GCE ID: Event_165

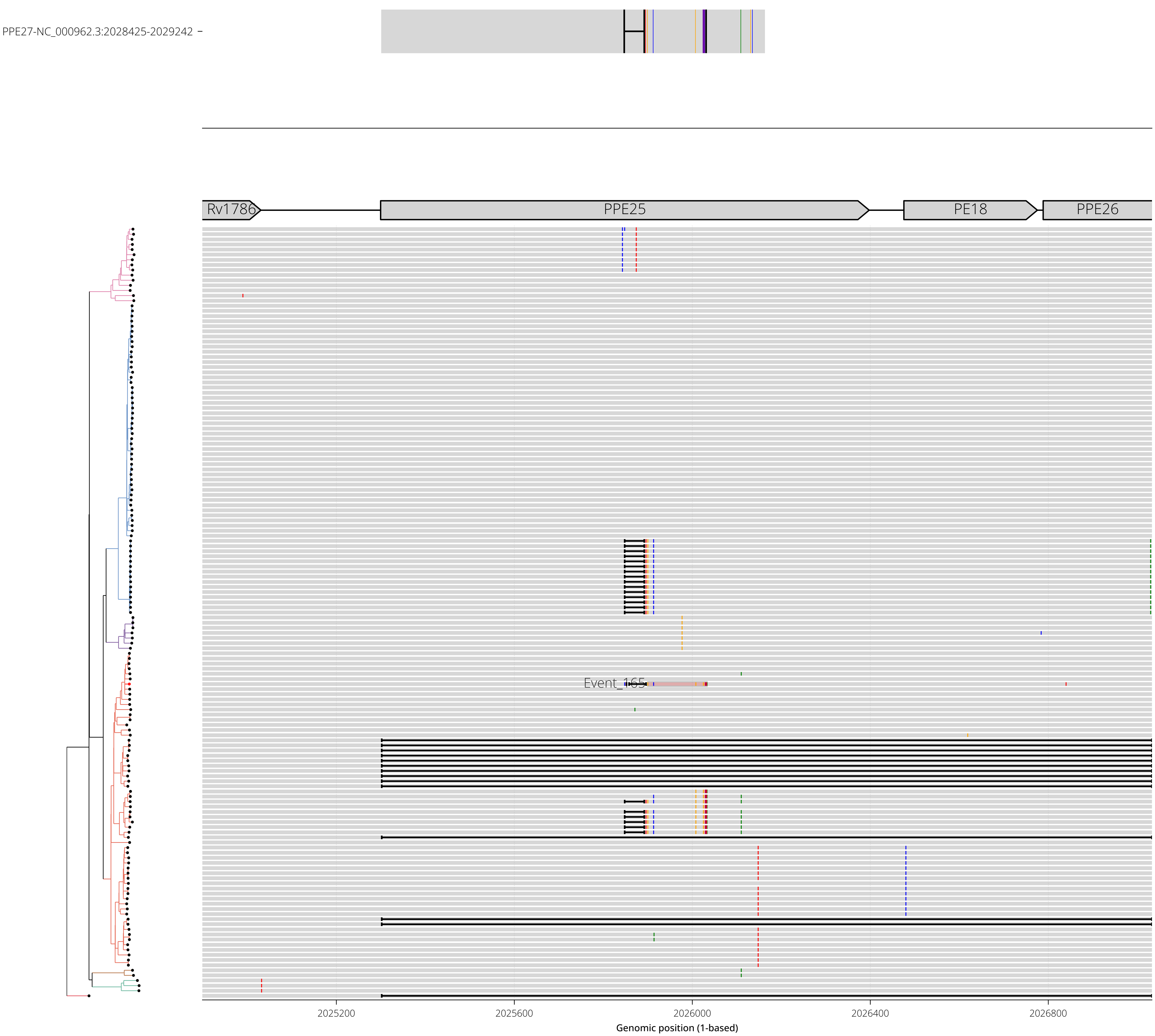

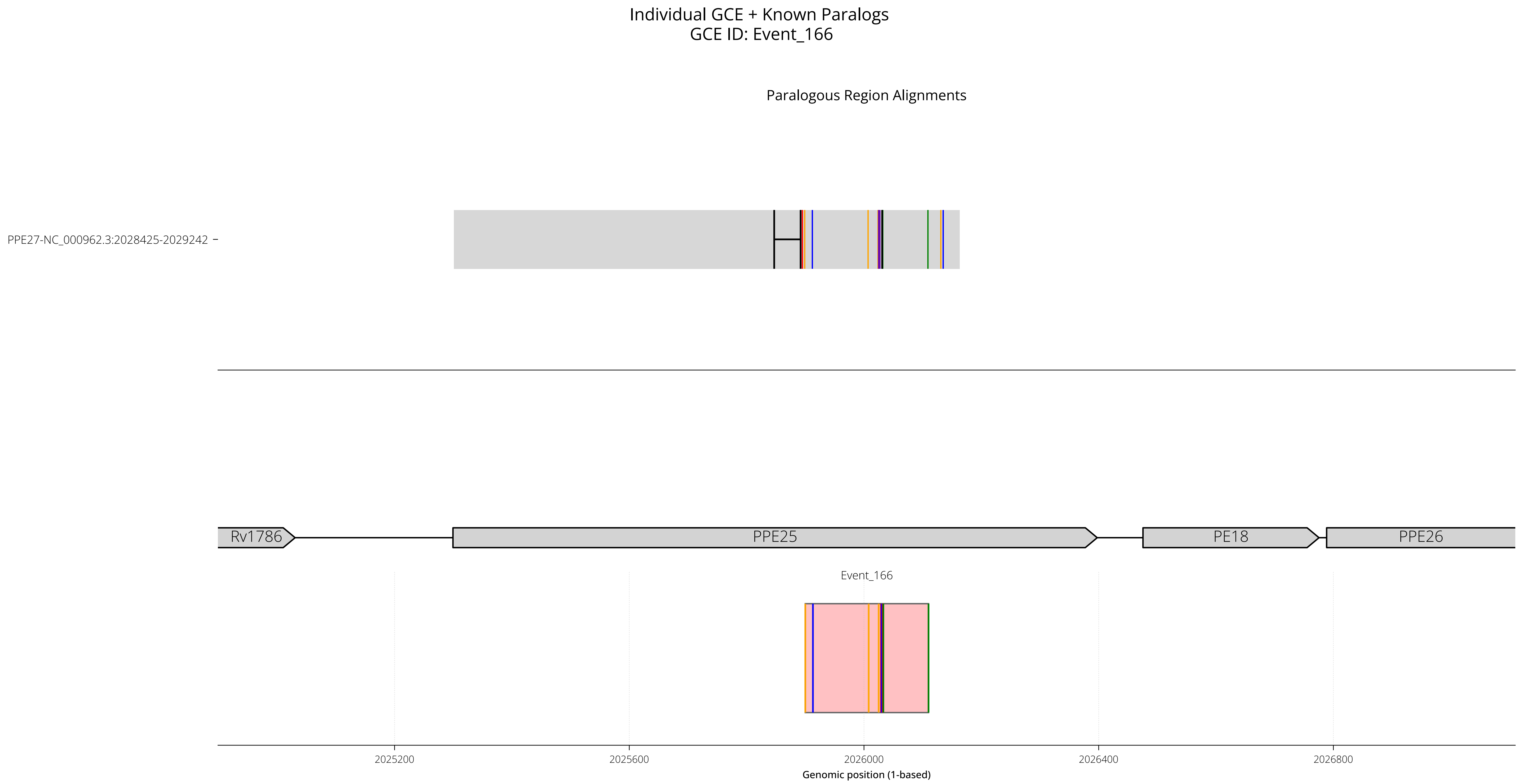

Individual GCE + Known Paralogs
GCE ID: Event_166
Paralogous Region Alignments
PPE27-NC_000962.3:2028425-2029242
Rv1786
PPE25
PE18
PPE26
Event_166
2025200
2025600
2026000
2026400
2026800
Genomic position (1-based)

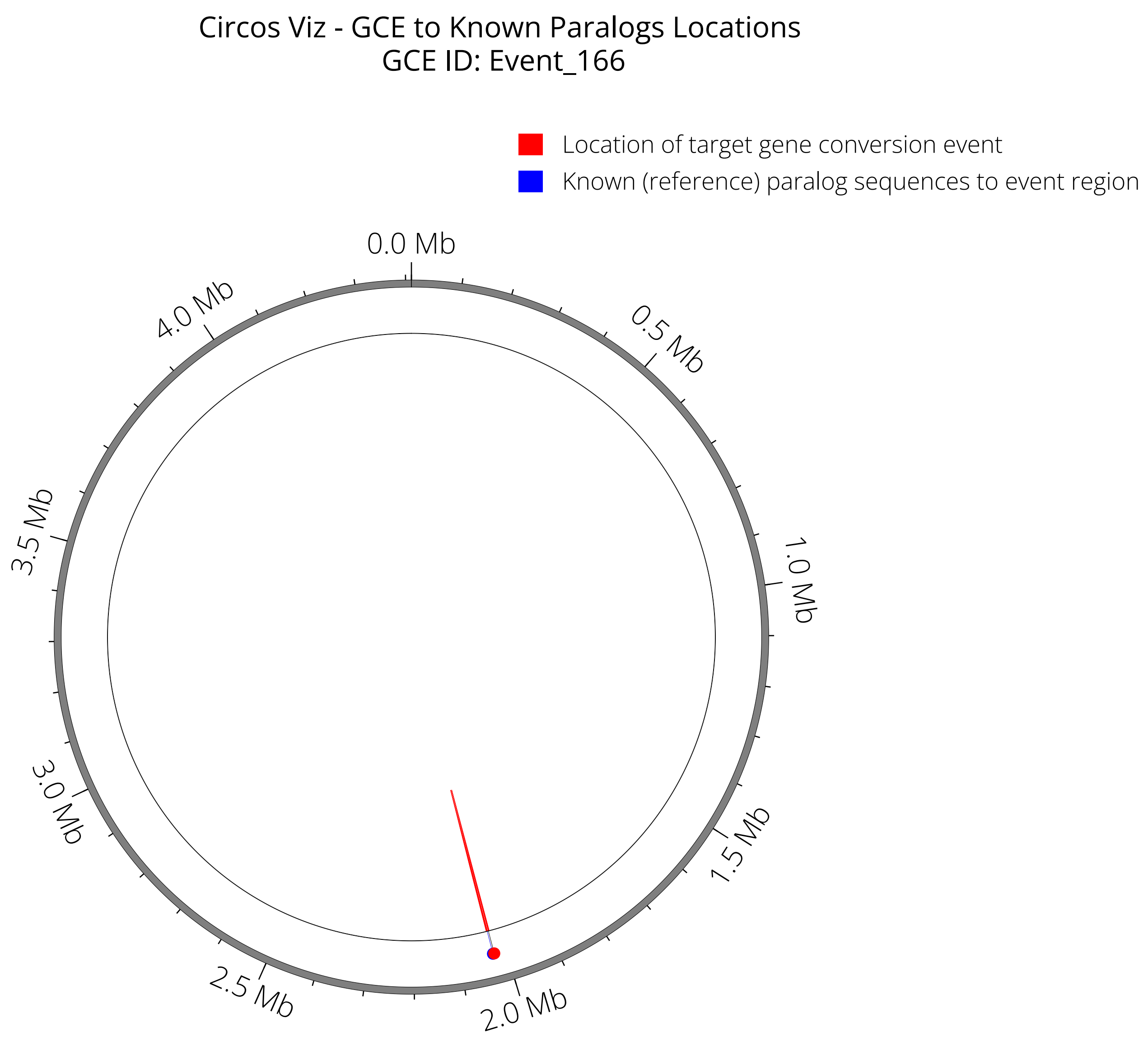

Circos Viz - GCE to Known Paralogs Locations
GCE ID: Event_166
Location of target gene conversion event
Known (reference) paralog sequences to event region
0.0 Mb
0.5 Mb
1.0 Mb
1.5 Mb
2.0 Mb
2.5 Mb
3.0 Mb
3.5 Mb
4.0 Mb

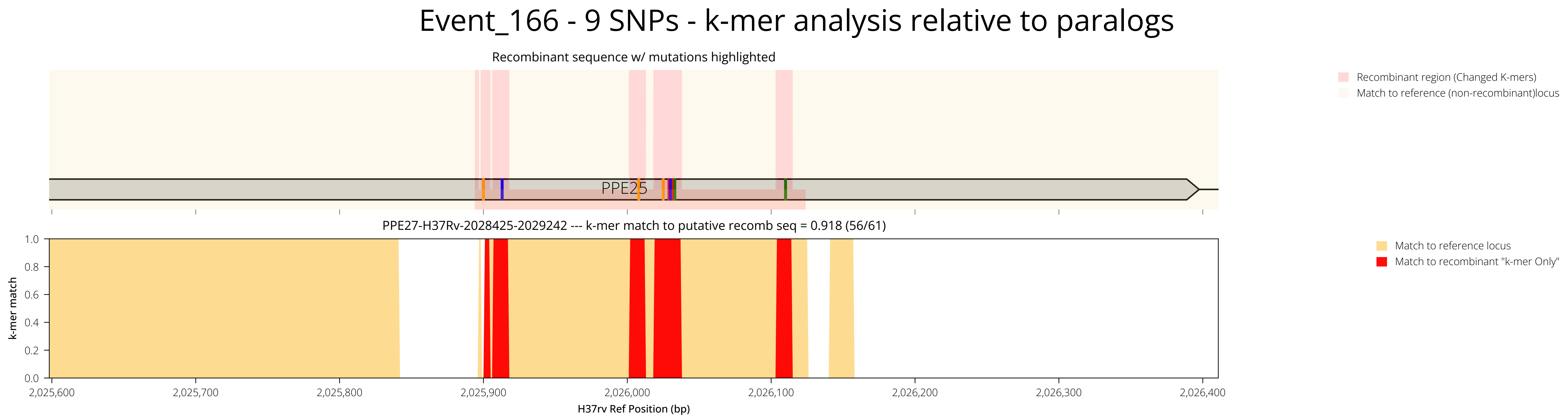

Event_166 - 9 SNPs - k-mer analysis relative to paralogs
Recombinant sequence w/ mutations highlighted
PPE25
Recombinant region (Changed K-mers)
Match to reference (non-recombinant)locus
PPE27-H37Rv-2028425-2029242 --- k-mer match to putative recomb seq = 0.918 (56/61)
Match to reference locus
Match to recombinant "k-mer Only"
k-mer match
1.0
0.8
0.6
0.4
0.2
0.0
2,025,600
2,025,700
2,025,800
2,025,900
2,026,000
2,026,100
2,026,200
2,026,300
2,026,400
H37rv Ref Position (bp)

Sub-phylogeny Viz + Paralog Alignments
GCE ID: Event_166

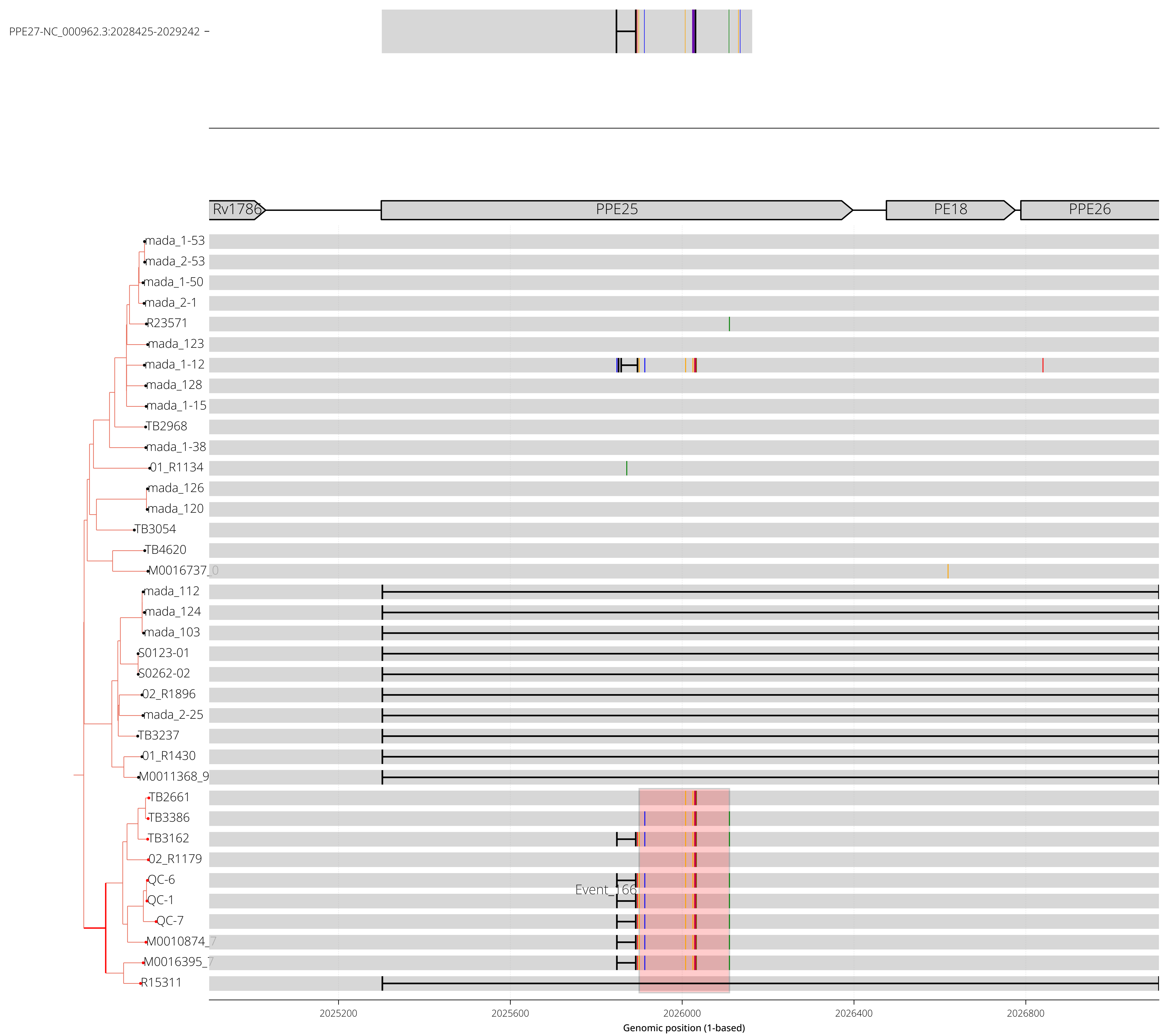

Full Phylogeny Viz + Paralog Alignments
GCE ID: Event_166

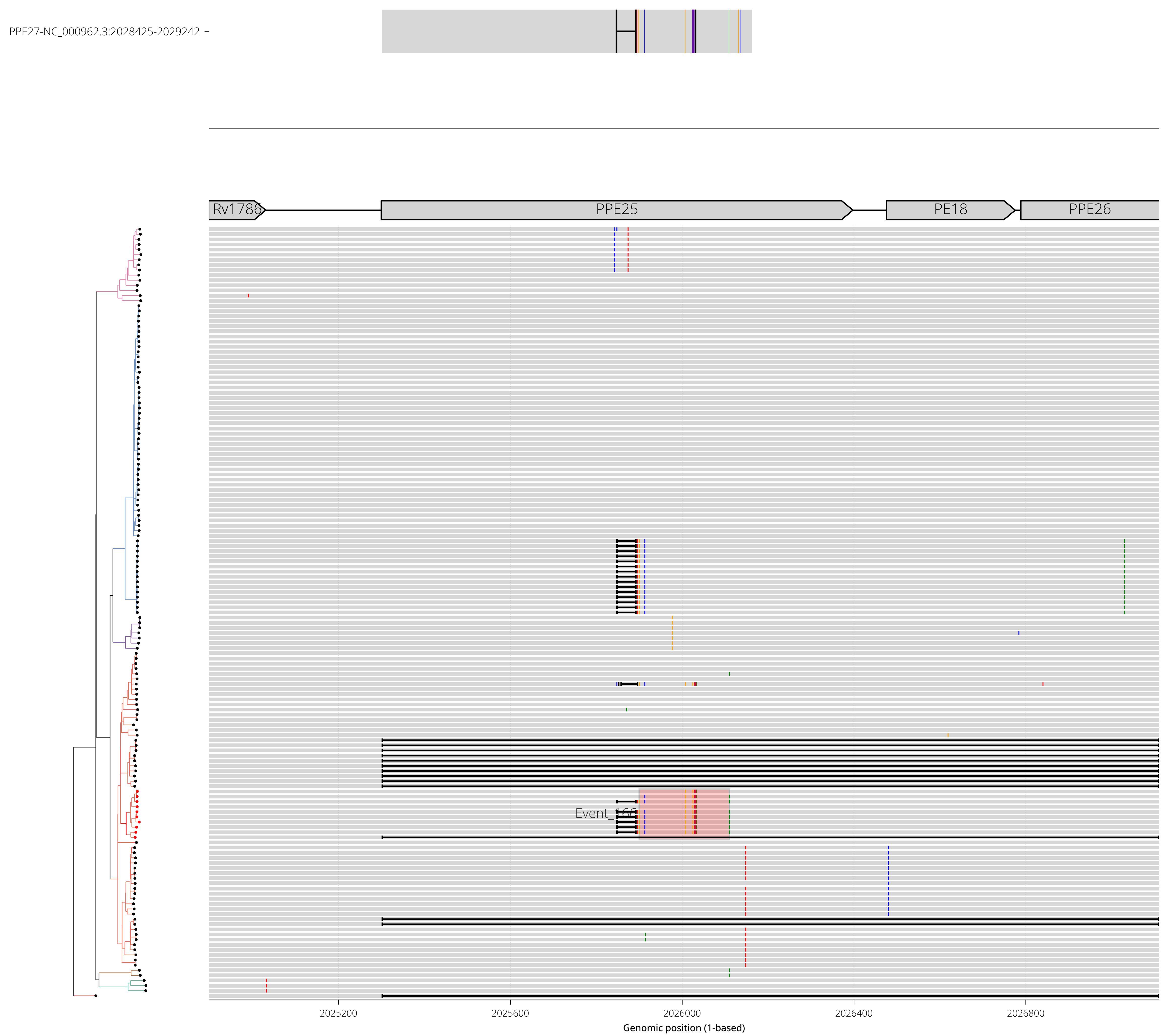

### Individual GCE + Known Paralogs
#### GCE ID: Event_167

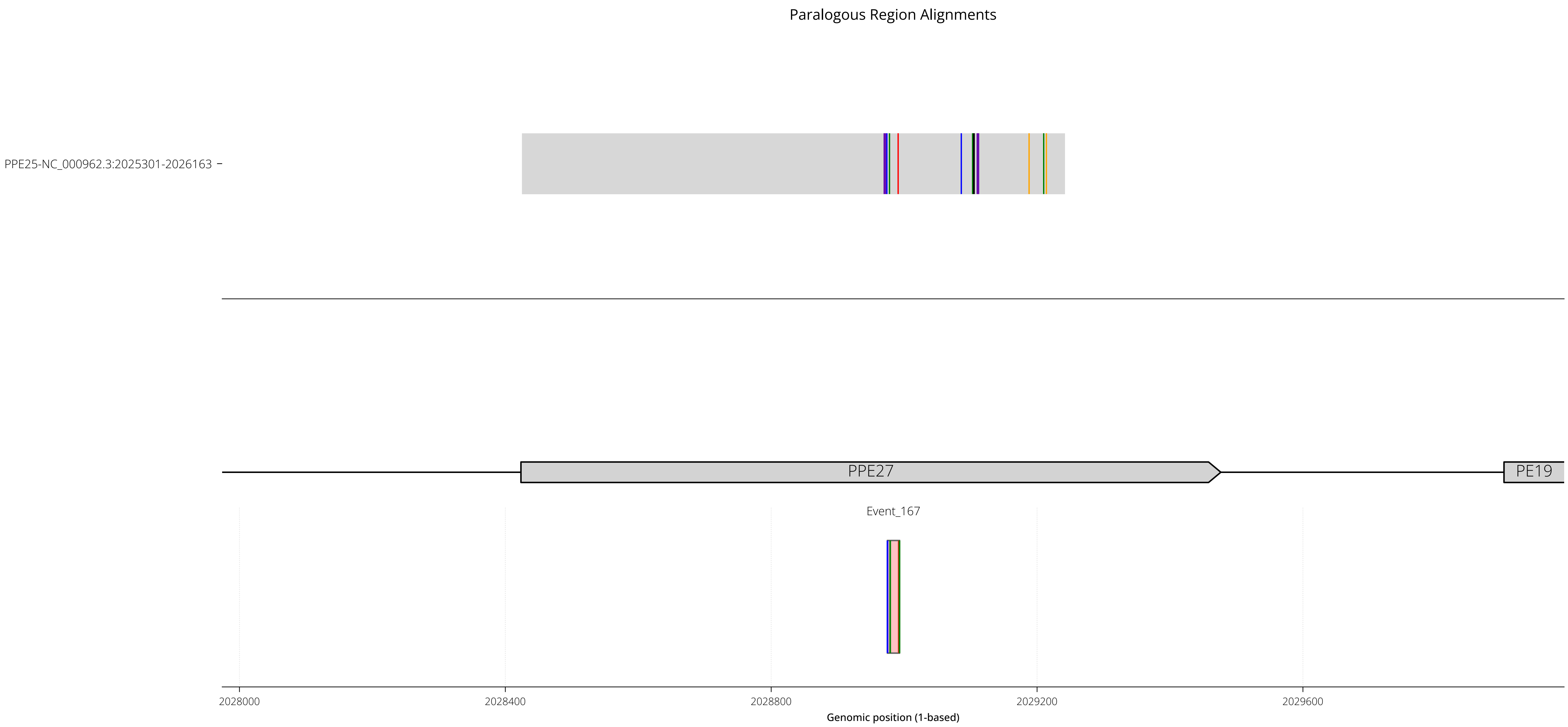

Circos Viz - GCE to Known Paralogs Locations
GCE ID: Event_167

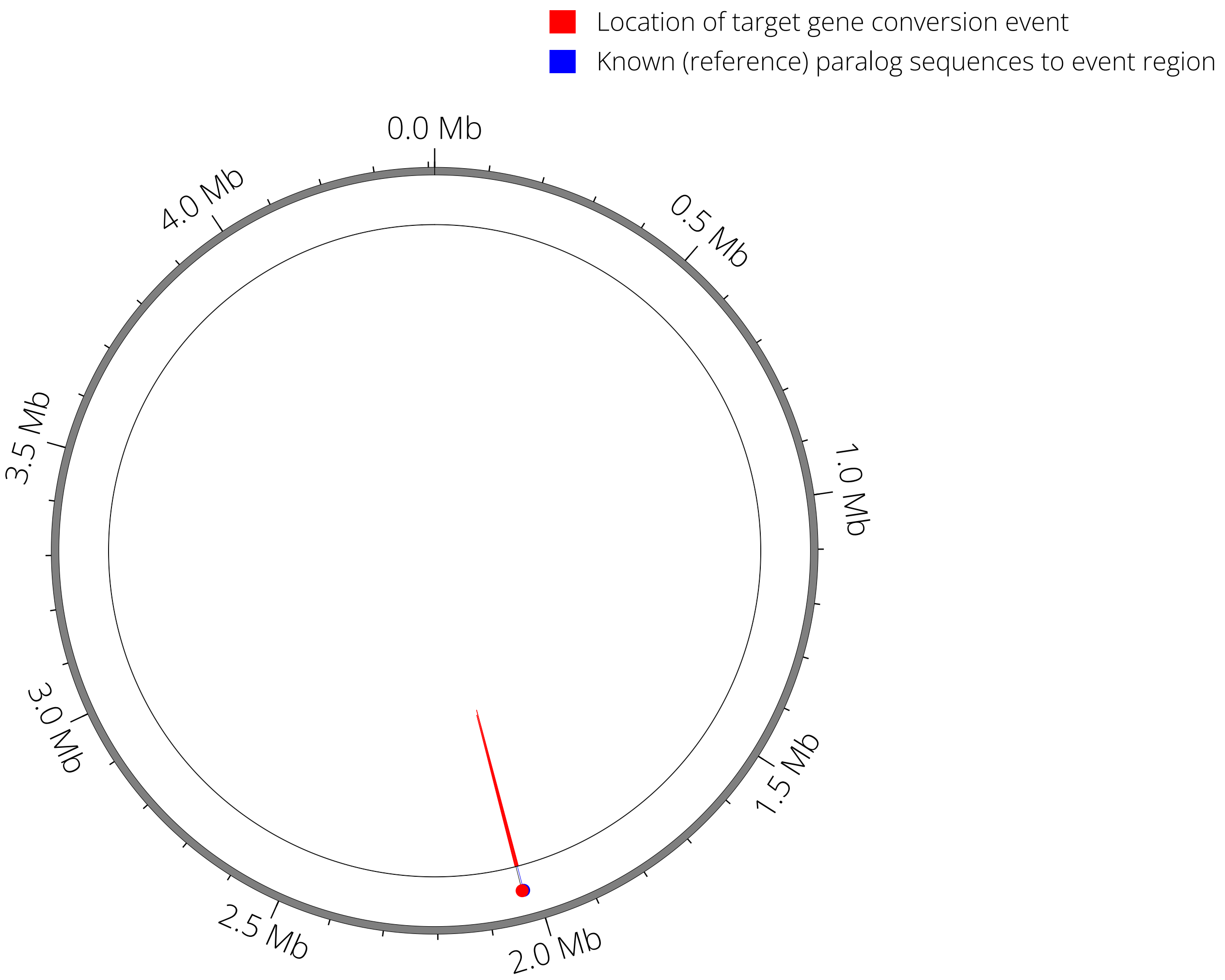

### Event_167 - 4 SNPs - k-mer analysis relative to paralogs

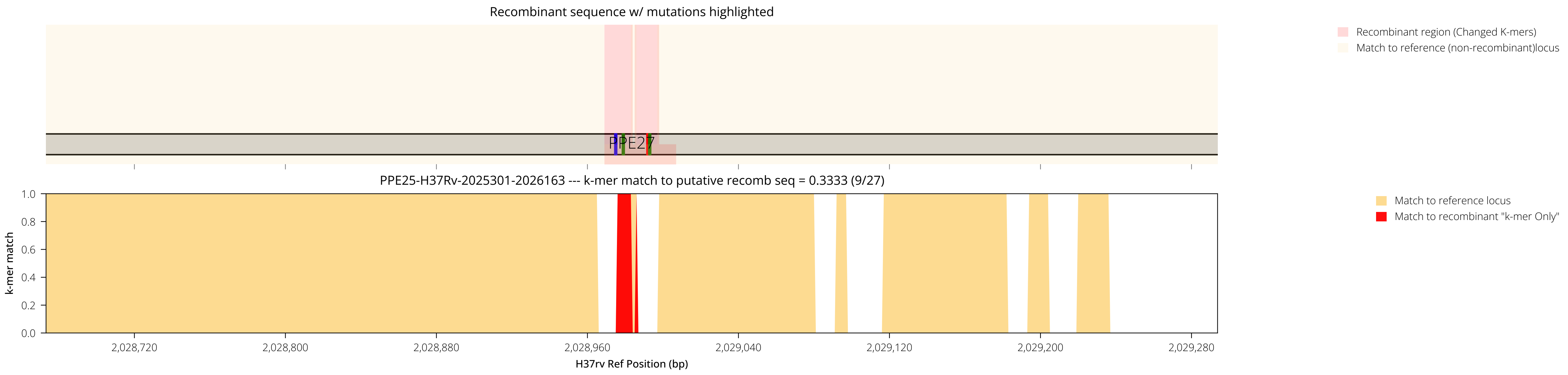

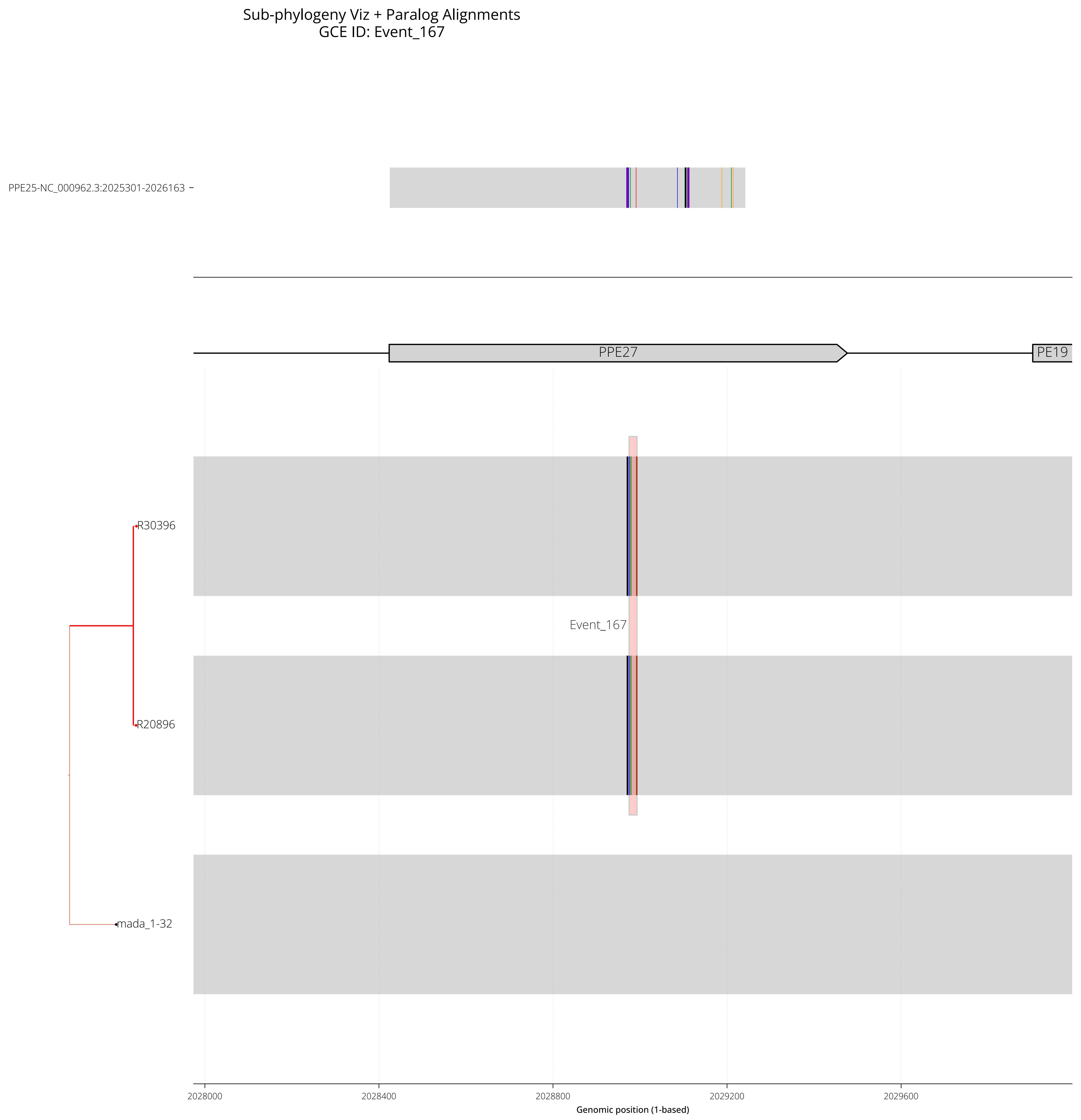

Sub-phylogeny Viz + Paralog Alignments
GCE ID: Event_167
PPE25-NC_000962.3:2025301-2026163
PPE27
PE19
R30396
Event_167
R20896
mada_1-32
2028000
2028400
2028800
2029200
2029600
Genomic position (1-based)

### Full Phylogeny Viz + Paralog Alignments
GCE ID: Event_167

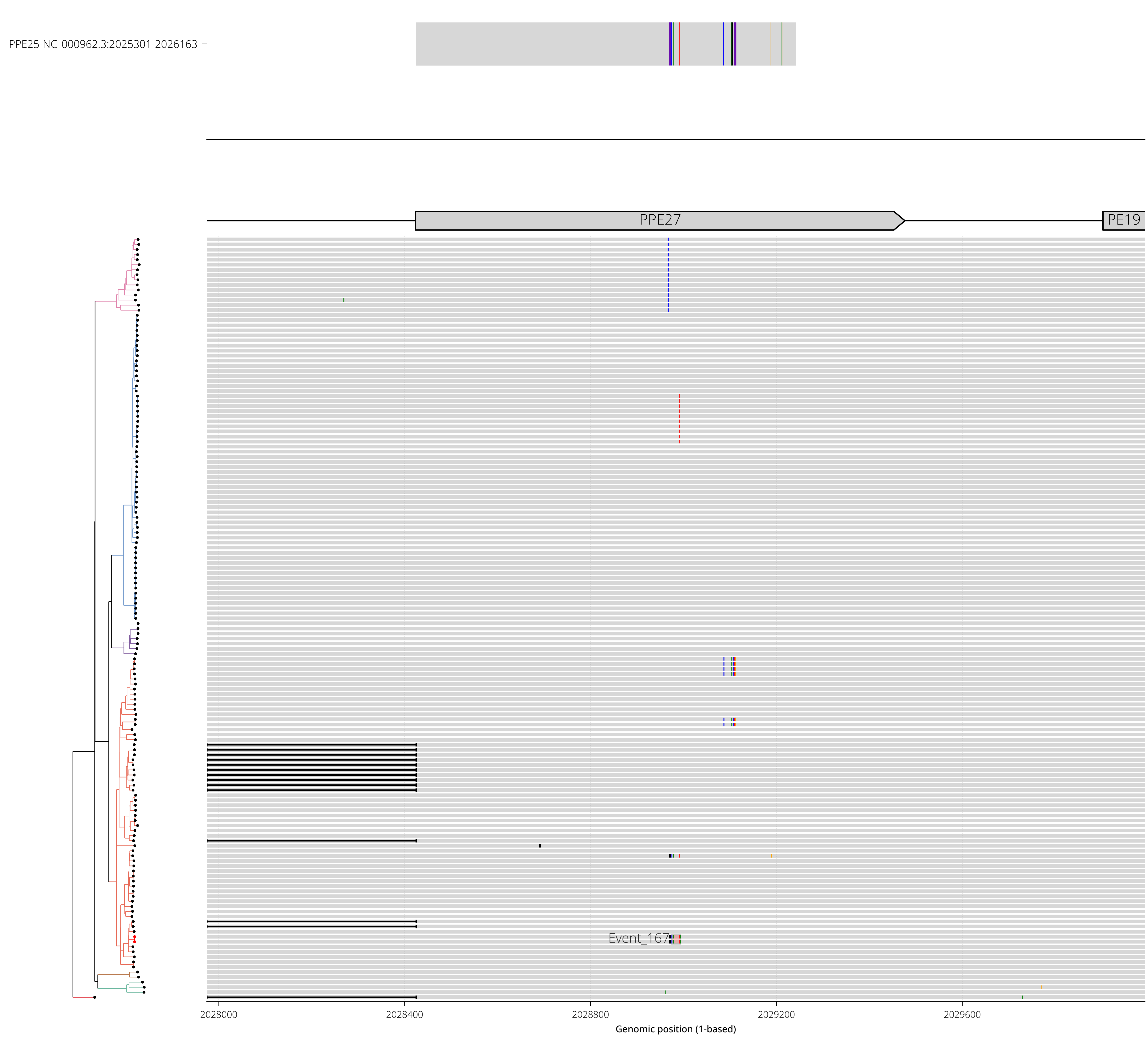

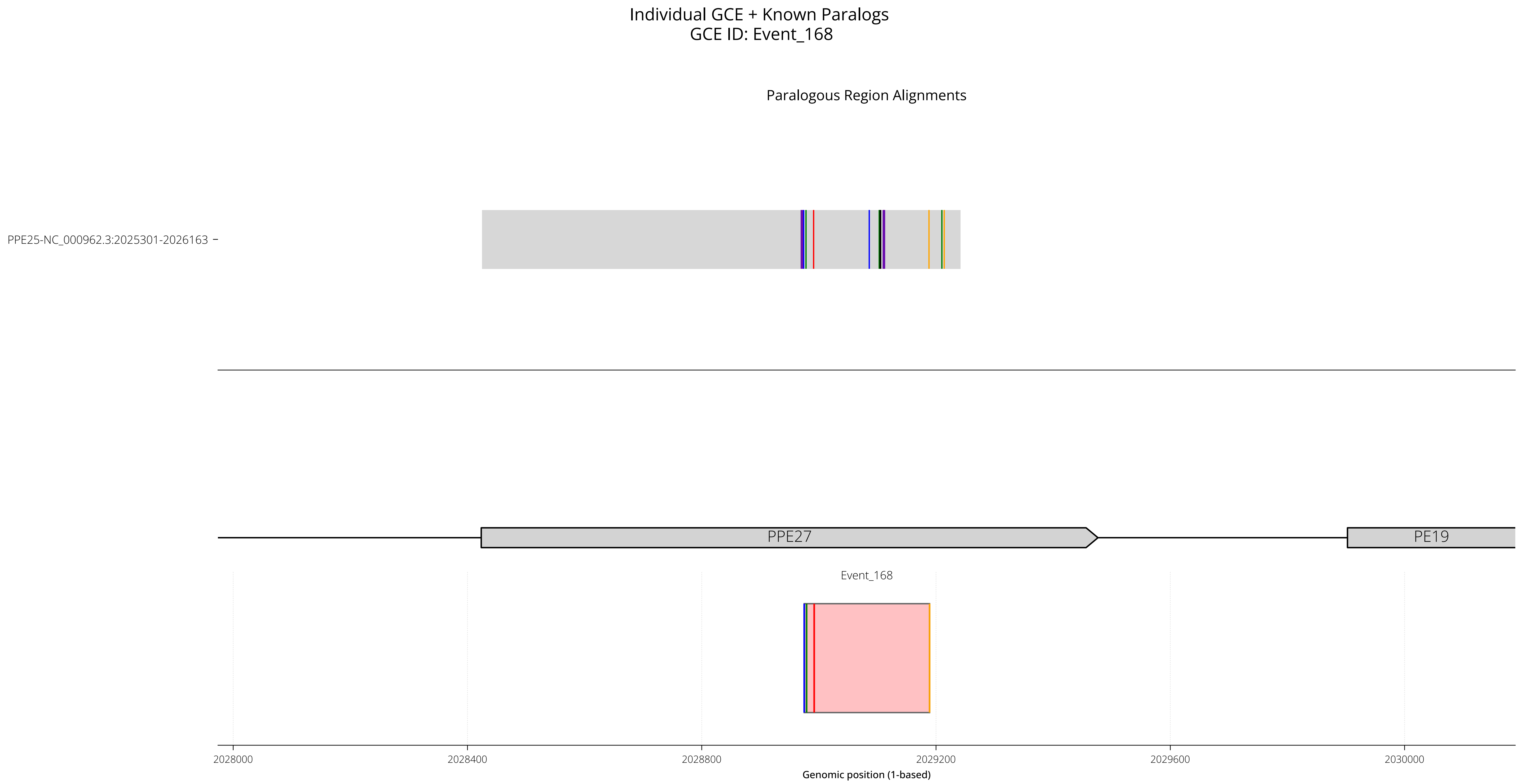
Individual GCE + Known Paralogs
GCE ID: Event_168
Paralogous Region Alignments
PPE25-NC_000962.3:2025301-2026163
PPE27
PE19
Event_168
2028000
2028400
2028800
2029200
2029600
2030000
Genomic position (1-based)

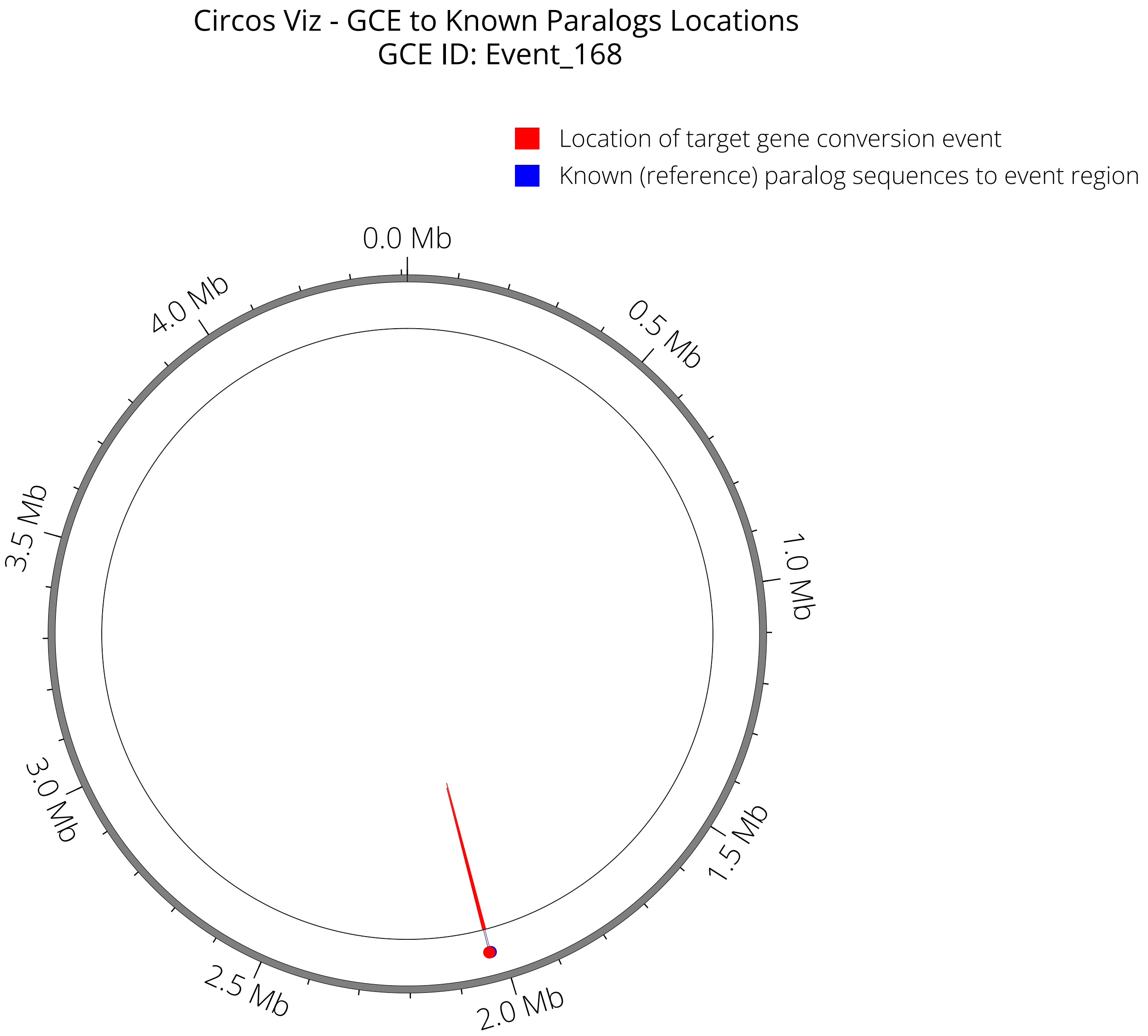

Circos Viz - GCE to Known Paralogs Locations
GCE ID: Event_168
Location of target gene conversion event
Known (reference) paralog sequences to event region
0.0 Mb
0.5 Mb
1.0 Mb
1.5 Mb
2.0 Mb
2.5 Mb
3.0 Mb
3.5 Mb
4.0 Mb

### Event_168 - 4 SNPs - k-mer analysis relative to paralogs

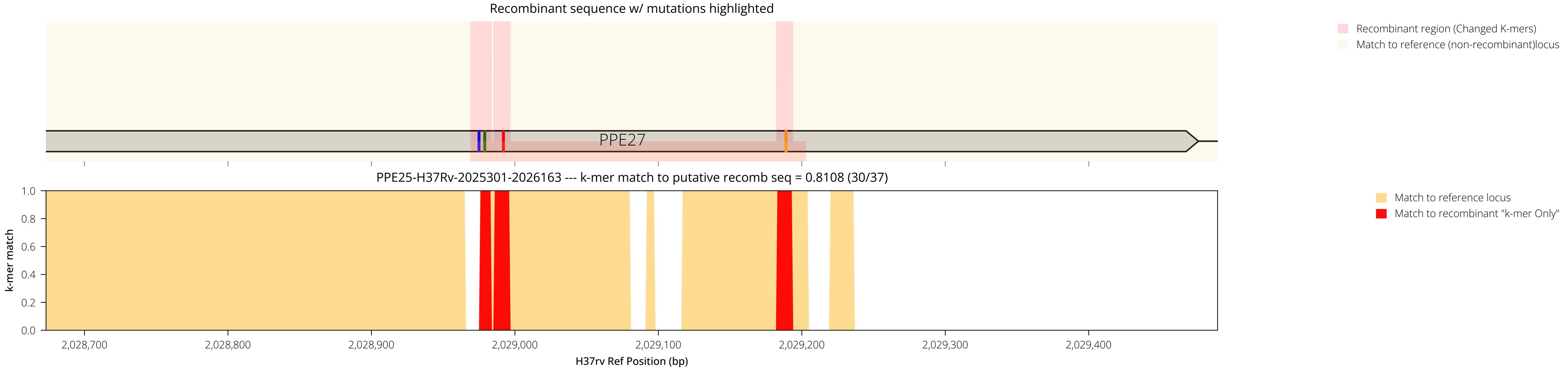

Sub-phylogeny Viz + Paralog Alignments
GCE ID: Event_168

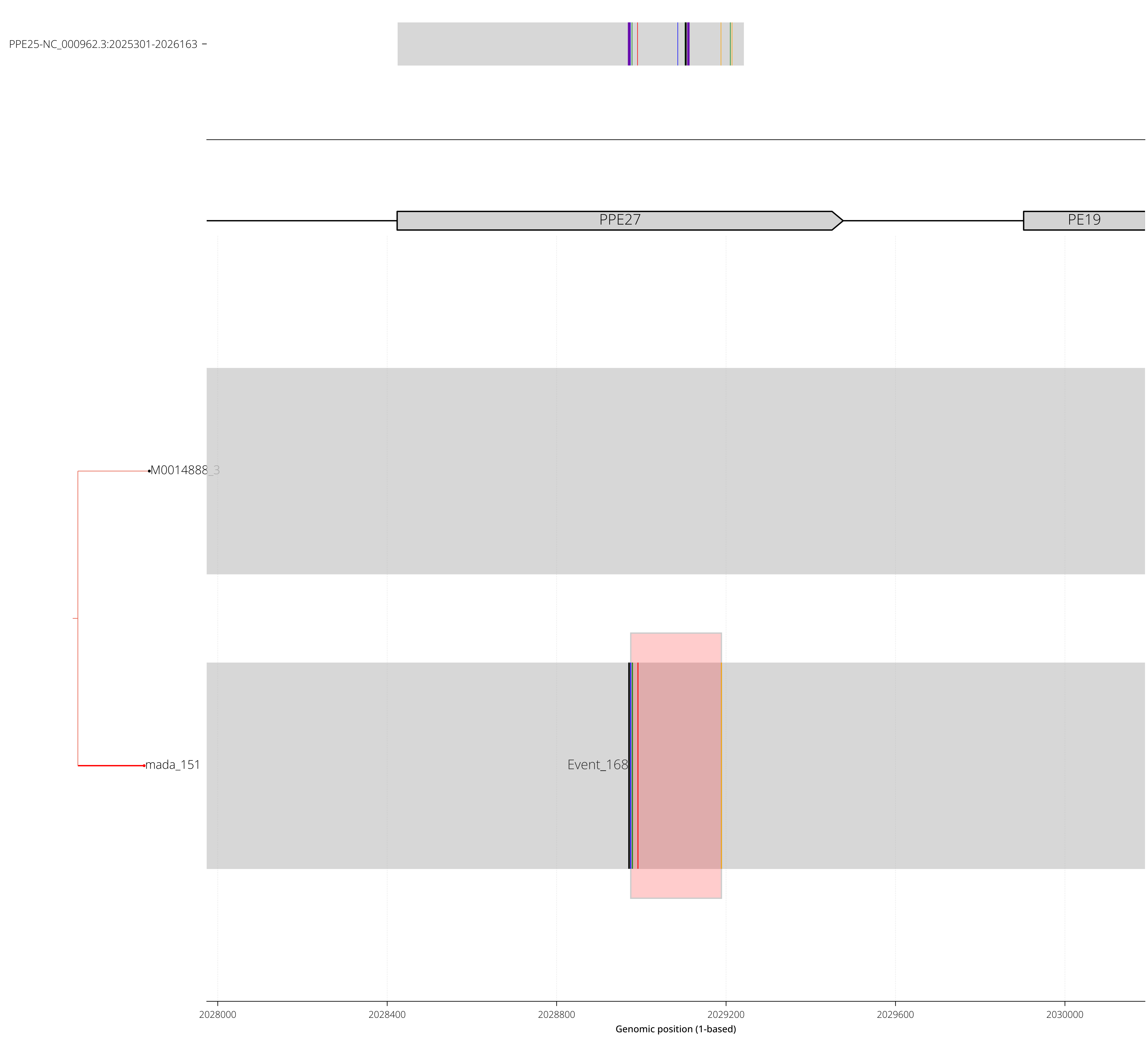

Full Phylogeny Viz + Paralog Alignments
GCE ID: Event_168

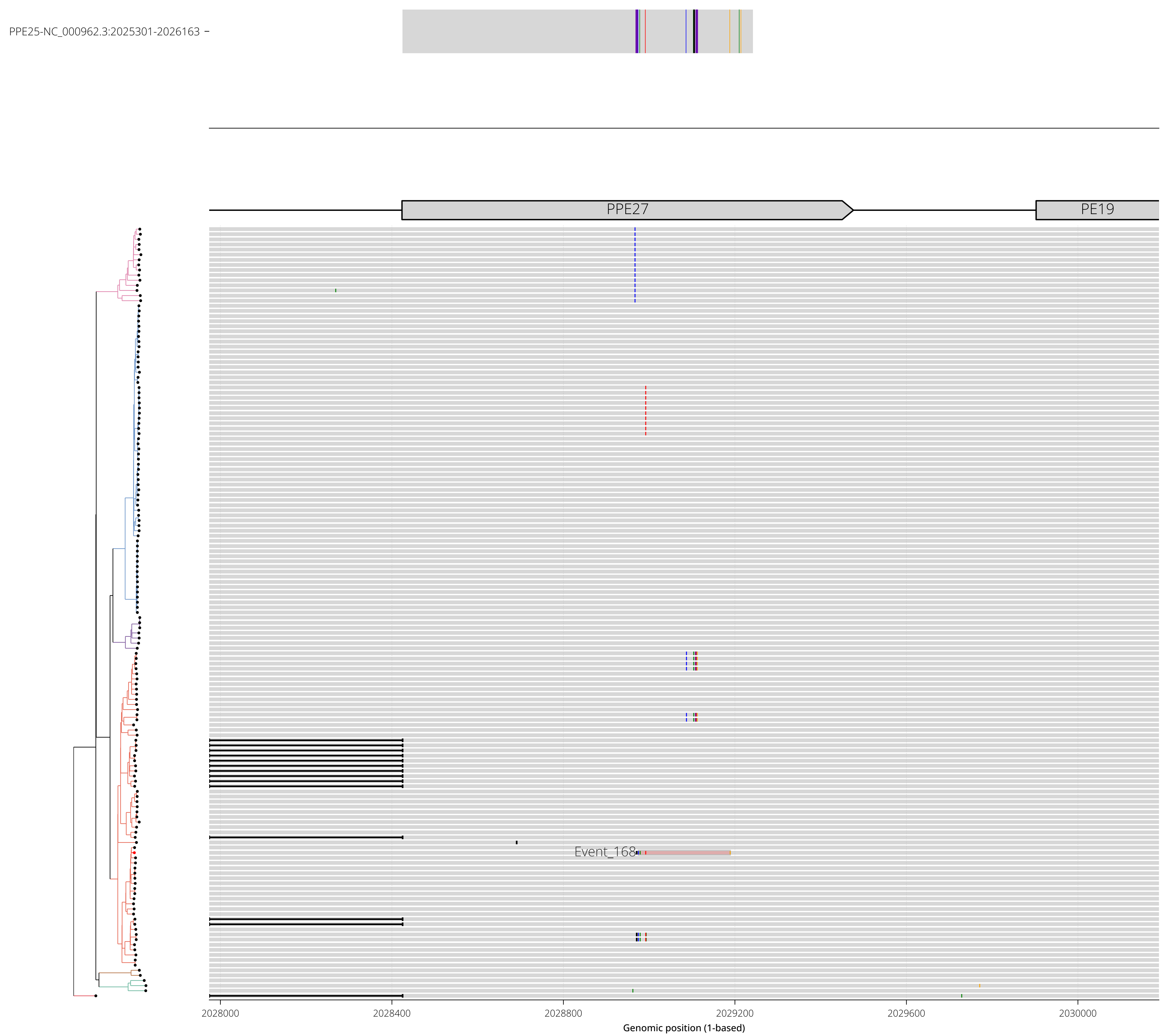

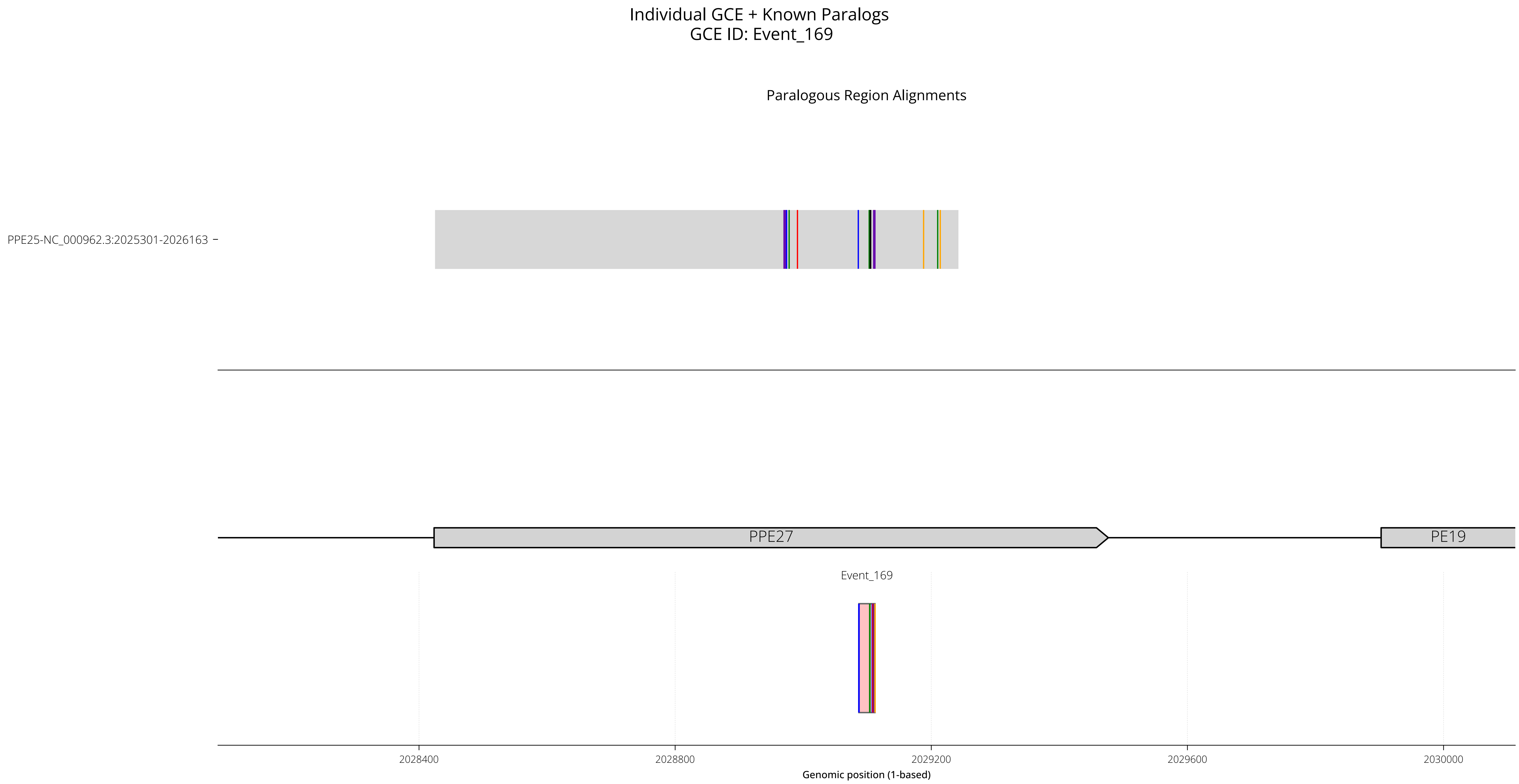

Individual GCE + Known Paralogs
GCE ID: Event_169
Paralogous Region Alignments
PPE25-NC_000962.3:2025301-2026163
PPE27
PE19
Event_169
2028400
2028800
2029200
2029600
2030000
Genomic position (1-based)

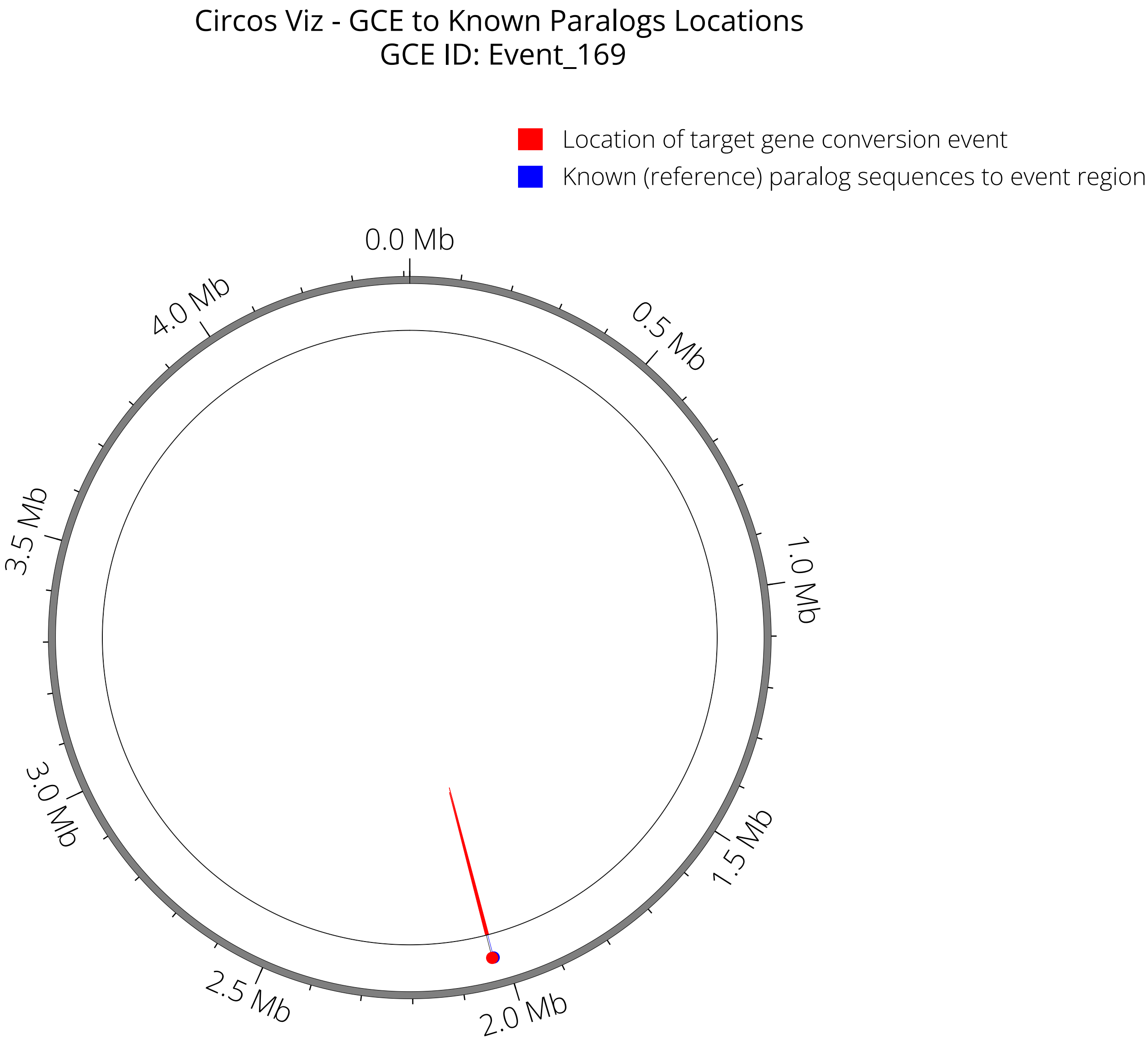
Circos Viz - GCE to Known Paralogs Locations
GCE ID: Event_169
Location of target gene conversion event
Known (reference) paralog sequences to event region
0.0 Mb
0.5 Mb
1.0 Mb
1.5 Mb
2.0 Mb
2.5 Mb
3.0 Mb
3.5 Mb
4.0 Mb

### Event_169 - 6 SNPs - k-mer analysis relative to paralogs

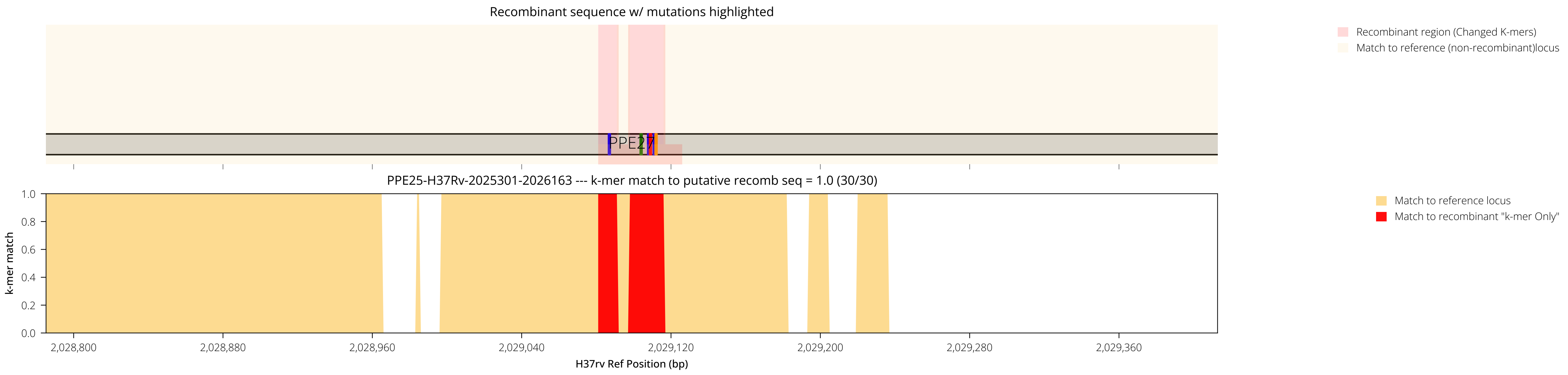

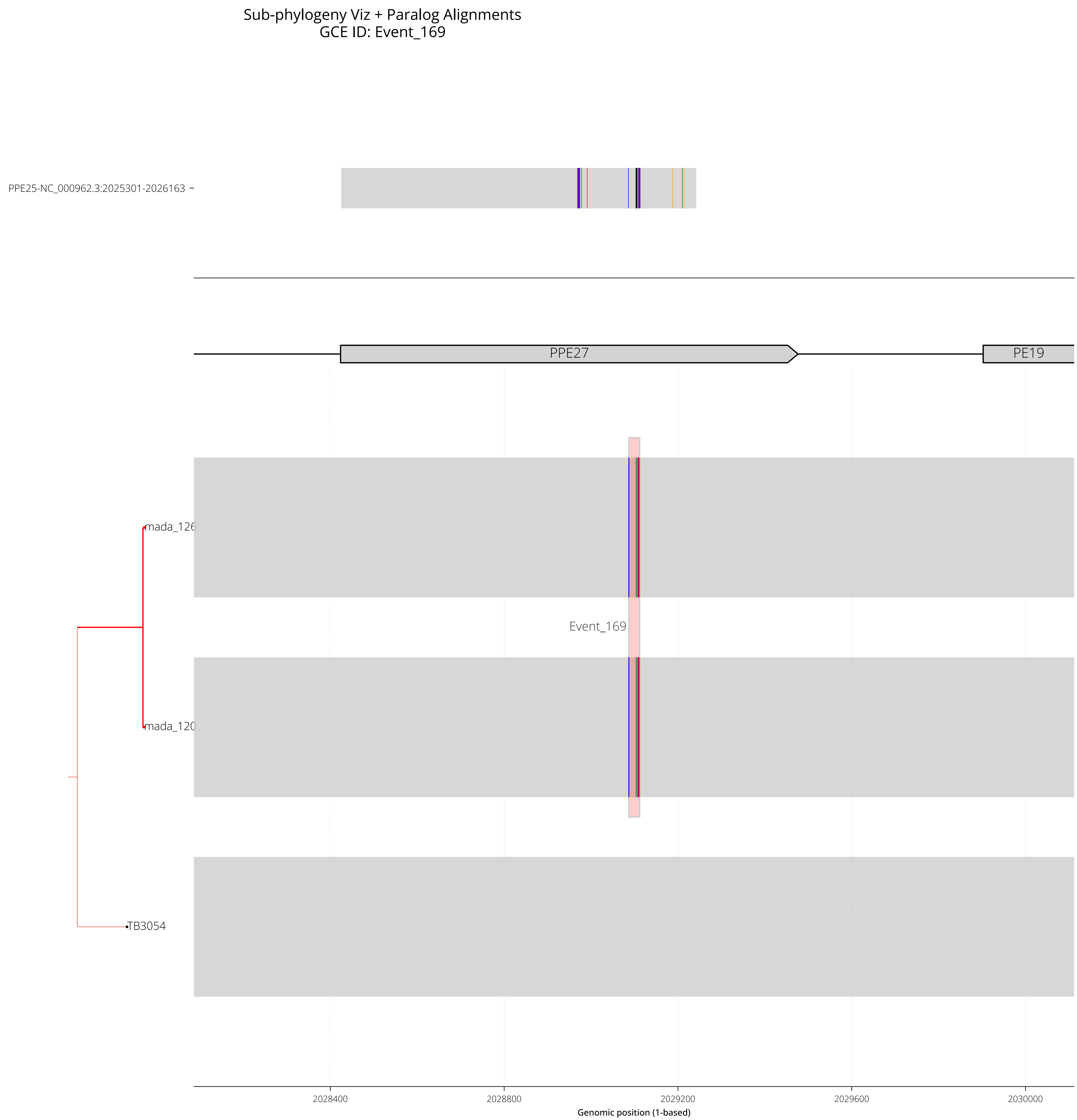

Sub-phylogeny Viz + Paralog Alignments
GCE ID: Event_169
PPE25-NC_000962.3:2025301-2026163
PPE27
PE19
mada_126
Event_169
mada_120
TB3054
2028400
2028800
2029200
2029600
2030000
Genomic position (1-based)

Full Phylogeny Viz + Paralog Alignments
GCE ID: Event_169

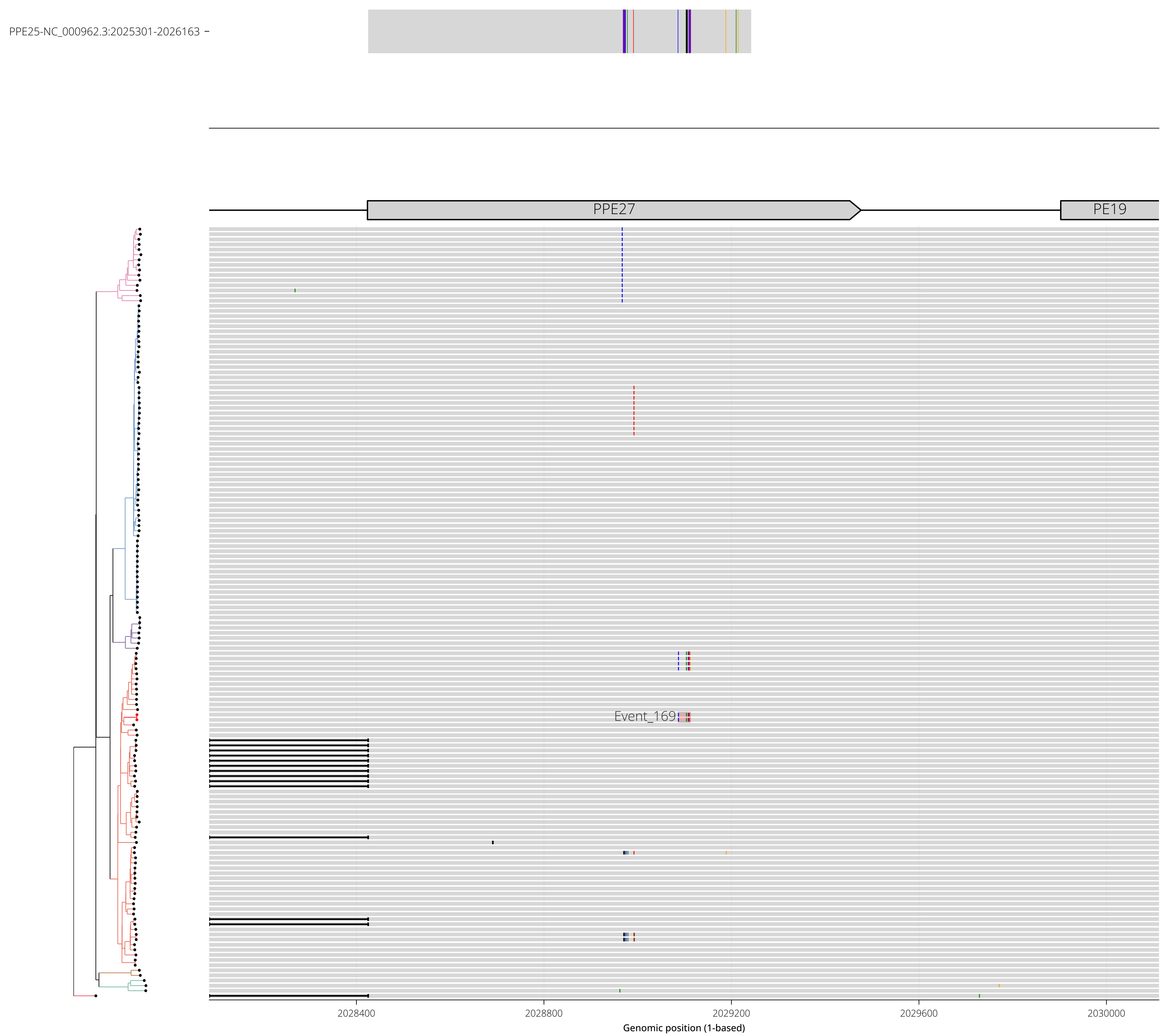

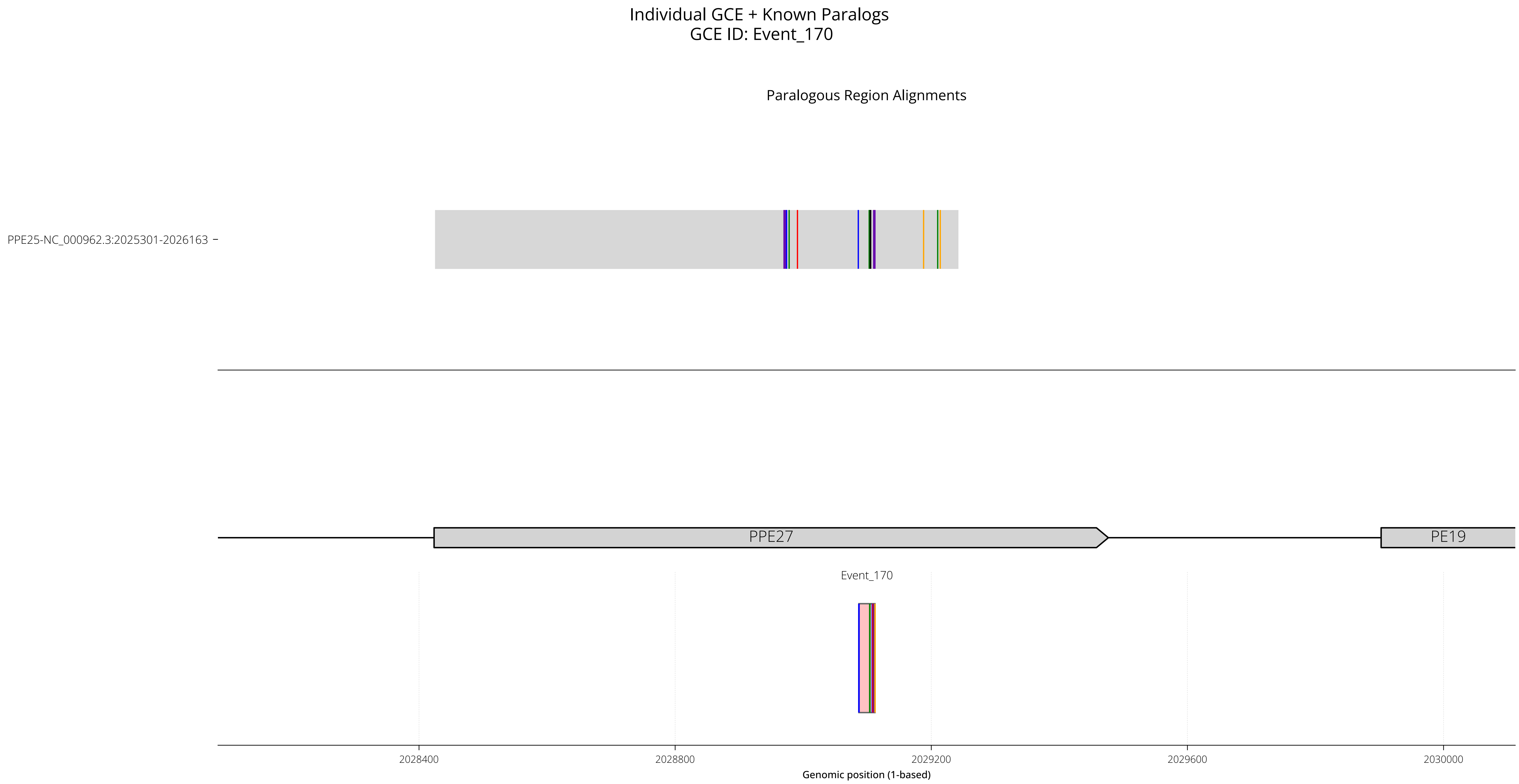
Individual GCE + Known Paralogs
GCE ID: Event_170
Paralogous Region Alignments
PPE25-NC_000962.3:2025301-2026163
PPE27
PE19
Event_170
2028400
2028800
2029200
2029600
2030000
Genomic position (1-based)

Circos Viz - GCE to Known Paralogs Locations
GCE ID: Event_170

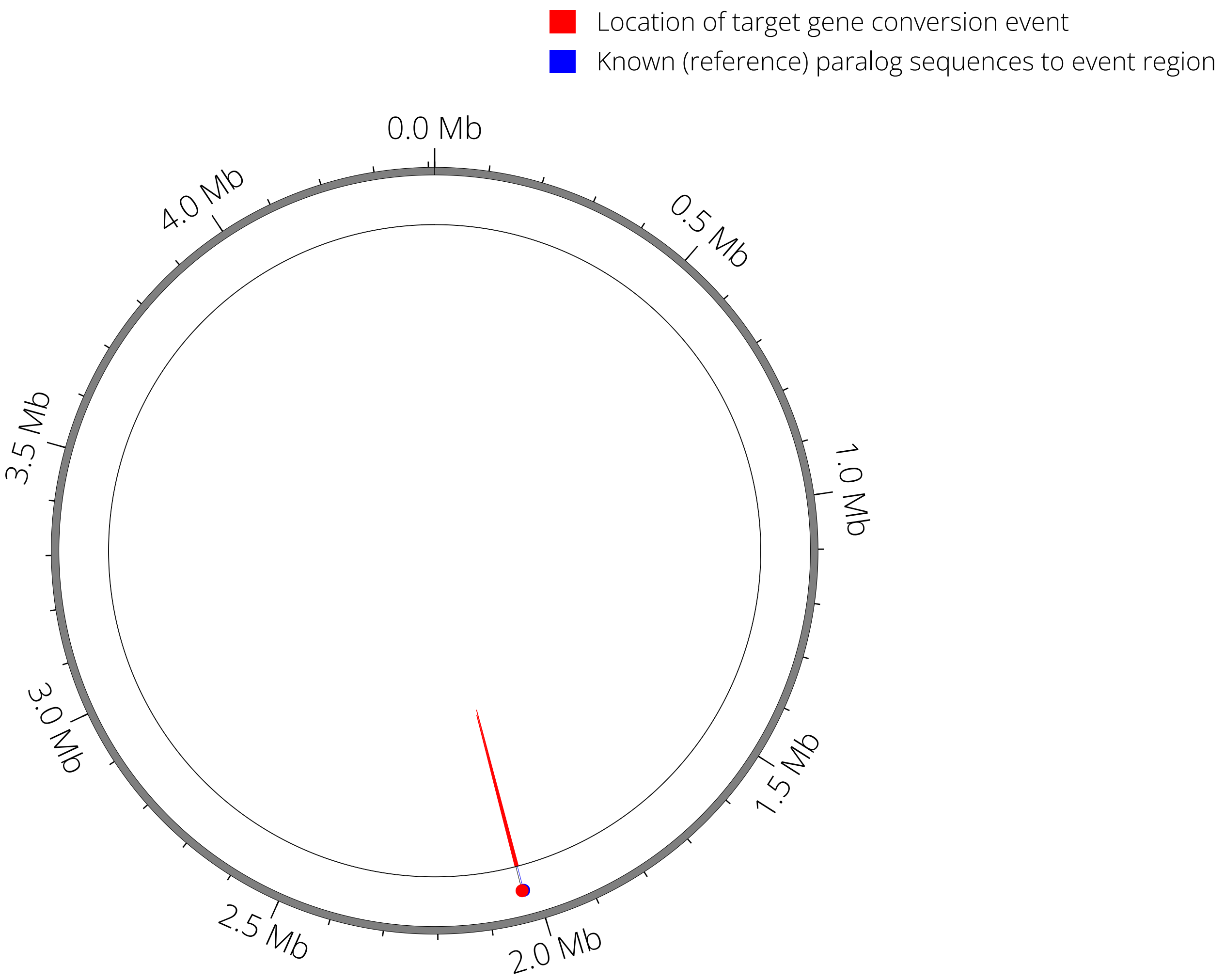

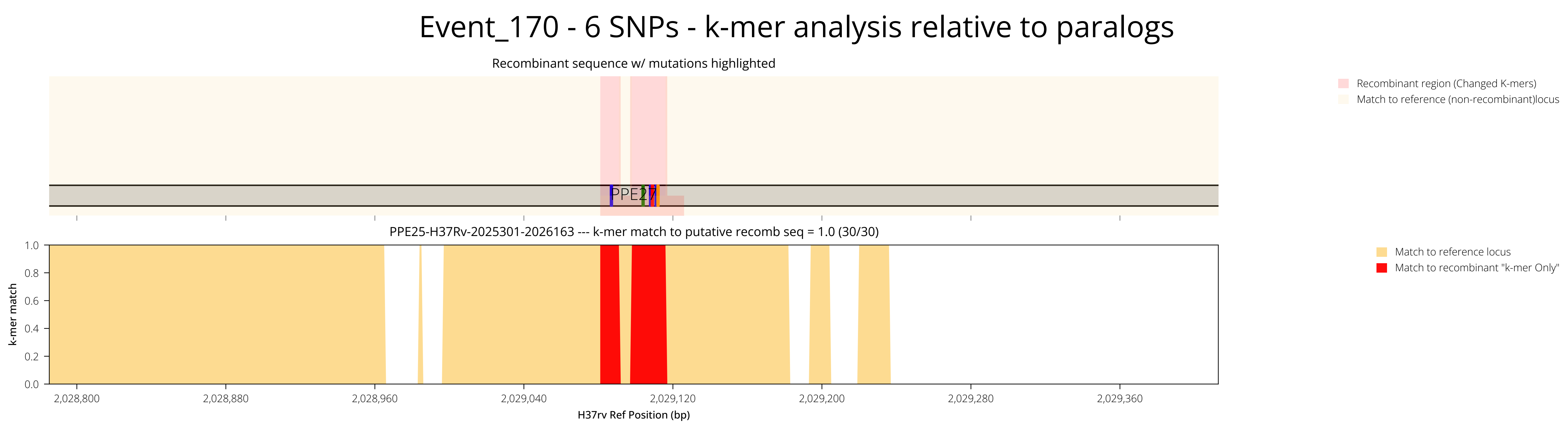
Event_170 - 6 SNPs - k-mer analysis relative to paralogs
Recombinant sequence w/ mutations highlighted
Recombinant region (Changed K-mers)
Match to reference (non-recombinant)locus
PPE27
PPE25-H37Rv-2025301-2026163 --- k-mer match to putative recomb seq = 1.0 (30/30)
Match to reference locus
Match to recombinant "k-mer Only"
k-mer match
1.0
0.8
0.6
0.4
0.2
0.0
2,028,800
2,028,880
2,028,960
2,029,040
2,029,120
2,029,200
2,029,280
2,029,360
H37rv Ref Position (bp)

Sub-phylogeny Viz + Paralog Alignments
GCE ID: Event_170

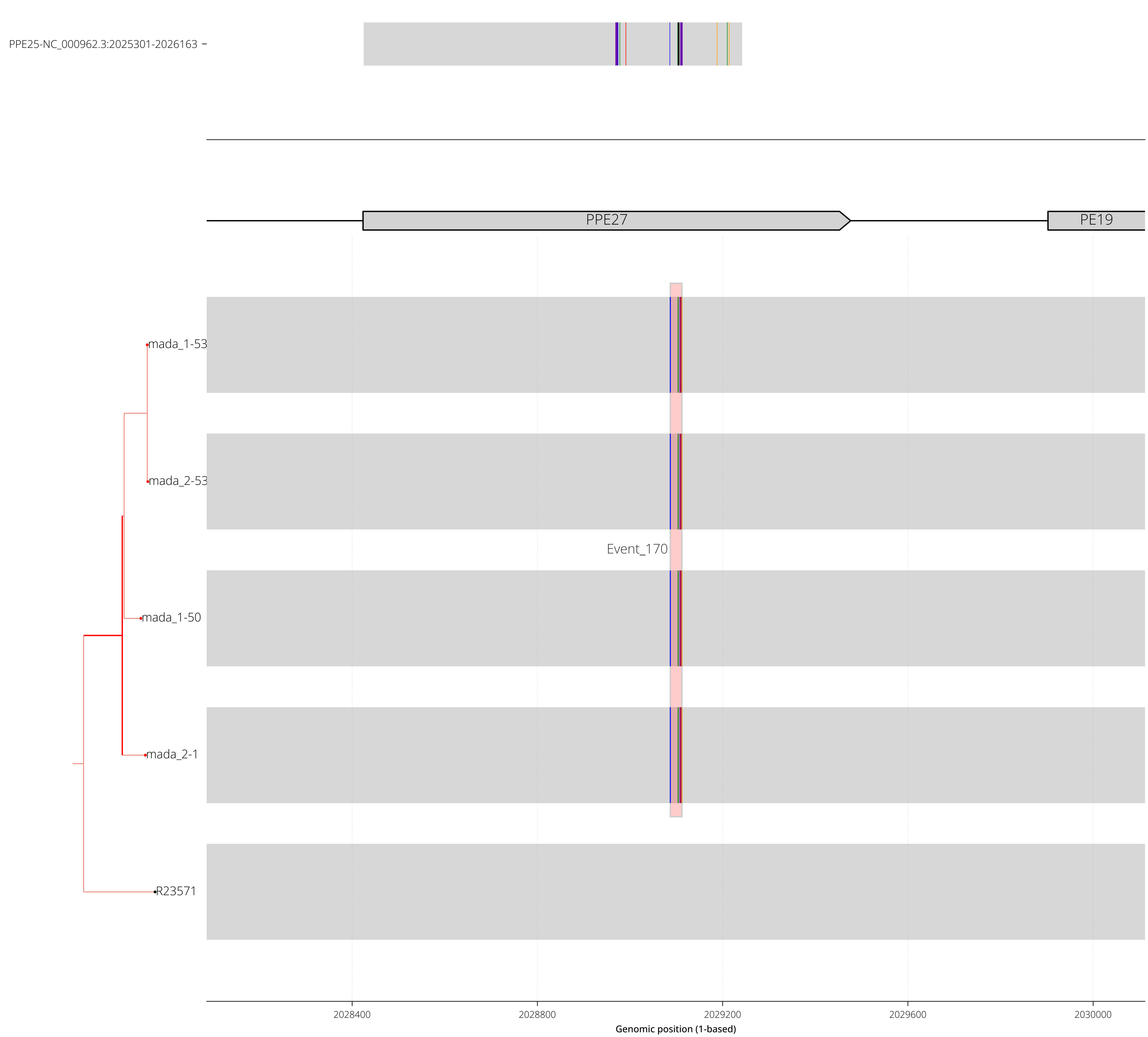

Full Phylogeny Viz + Paralog Alignments
GCE ID: Event_170

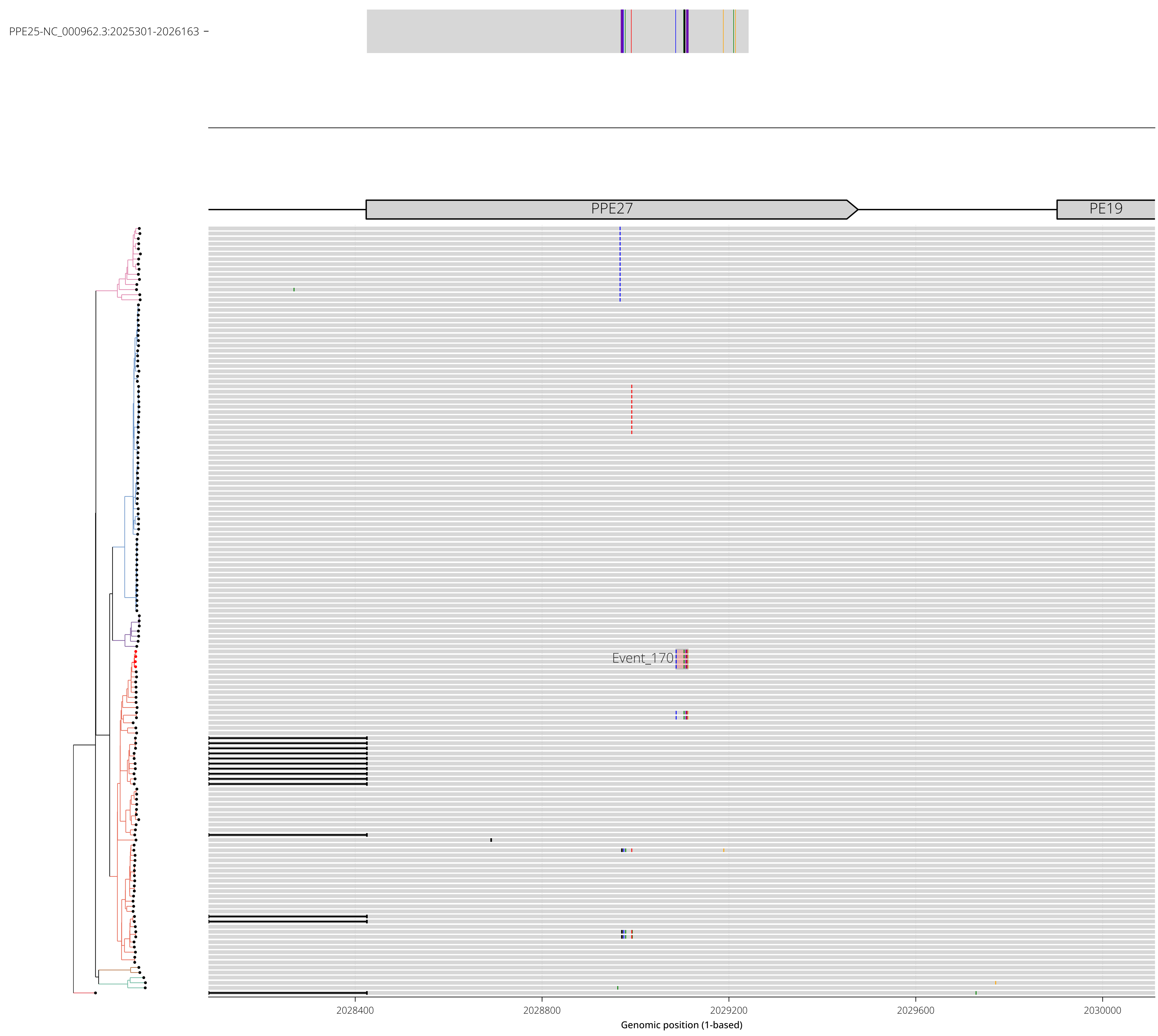

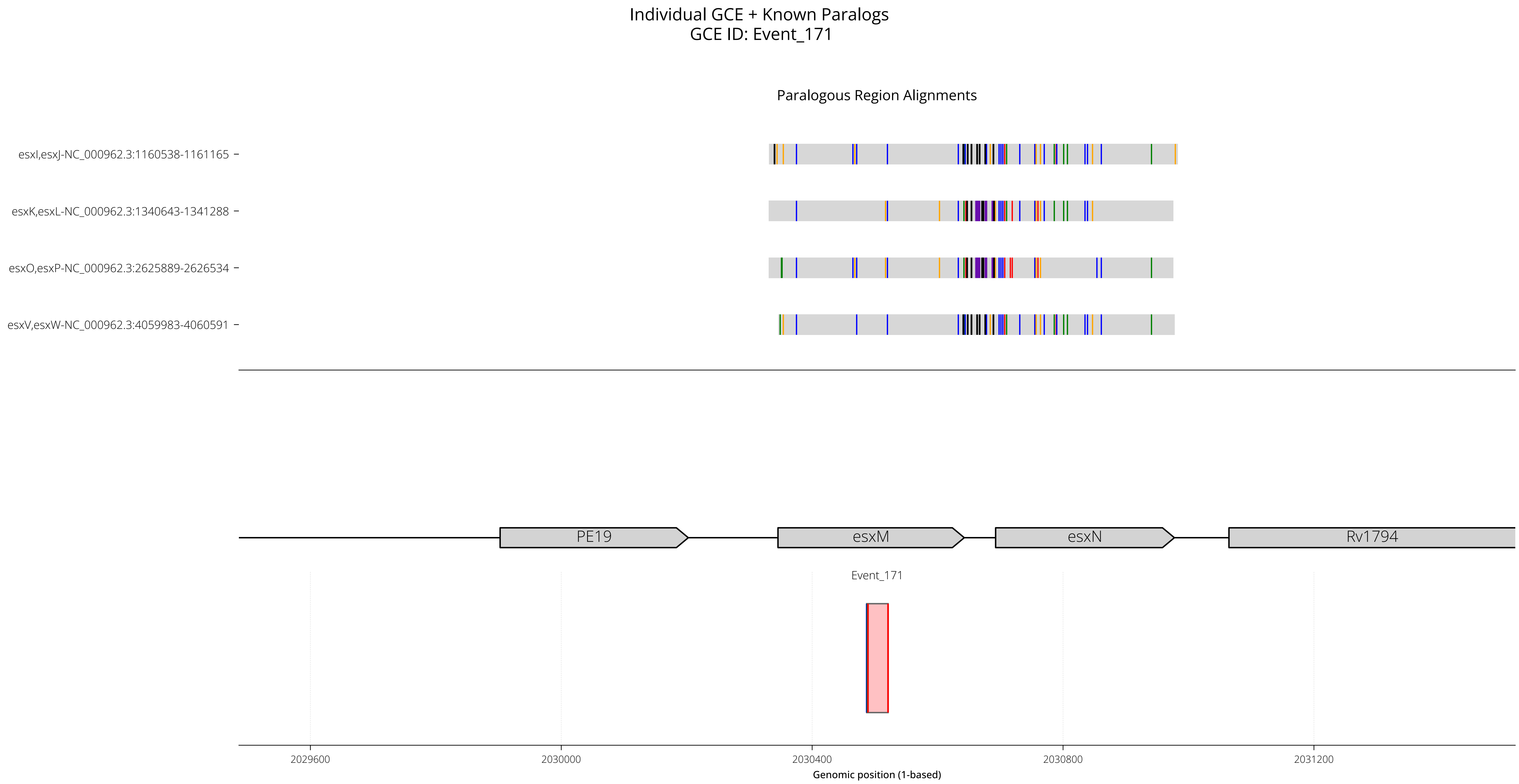

Individual GCE + Known Paralogs
GCE ID: Event_171
Paralogous Region Alignments
esxI,esxJ-NC_000962.3:1160538-1161165
esxK,esxL-NC_000962.3:1340643-1341288
esxO,esxP-NC_000962.3:2625889-2626534
esxV,esxW-NC_000962.3:4059983-4060591
PE19
esxM
esxN
Rv1794
Event_171
2029600
2030000
2030400
2030800
2031200
Genomic position (1-based)

Circos Viz - GCE to Known Paralogs Locations
GCE ID: Event_171

Location of target gene conversion event
Known (reference) paralog sequences to event region

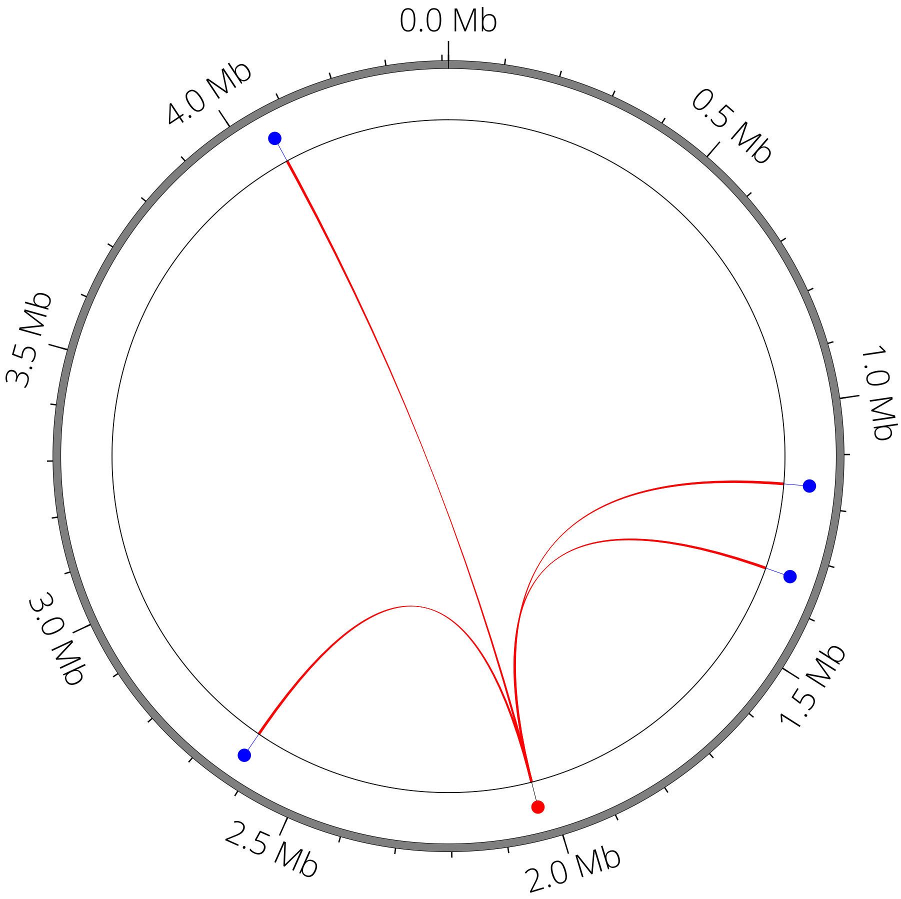

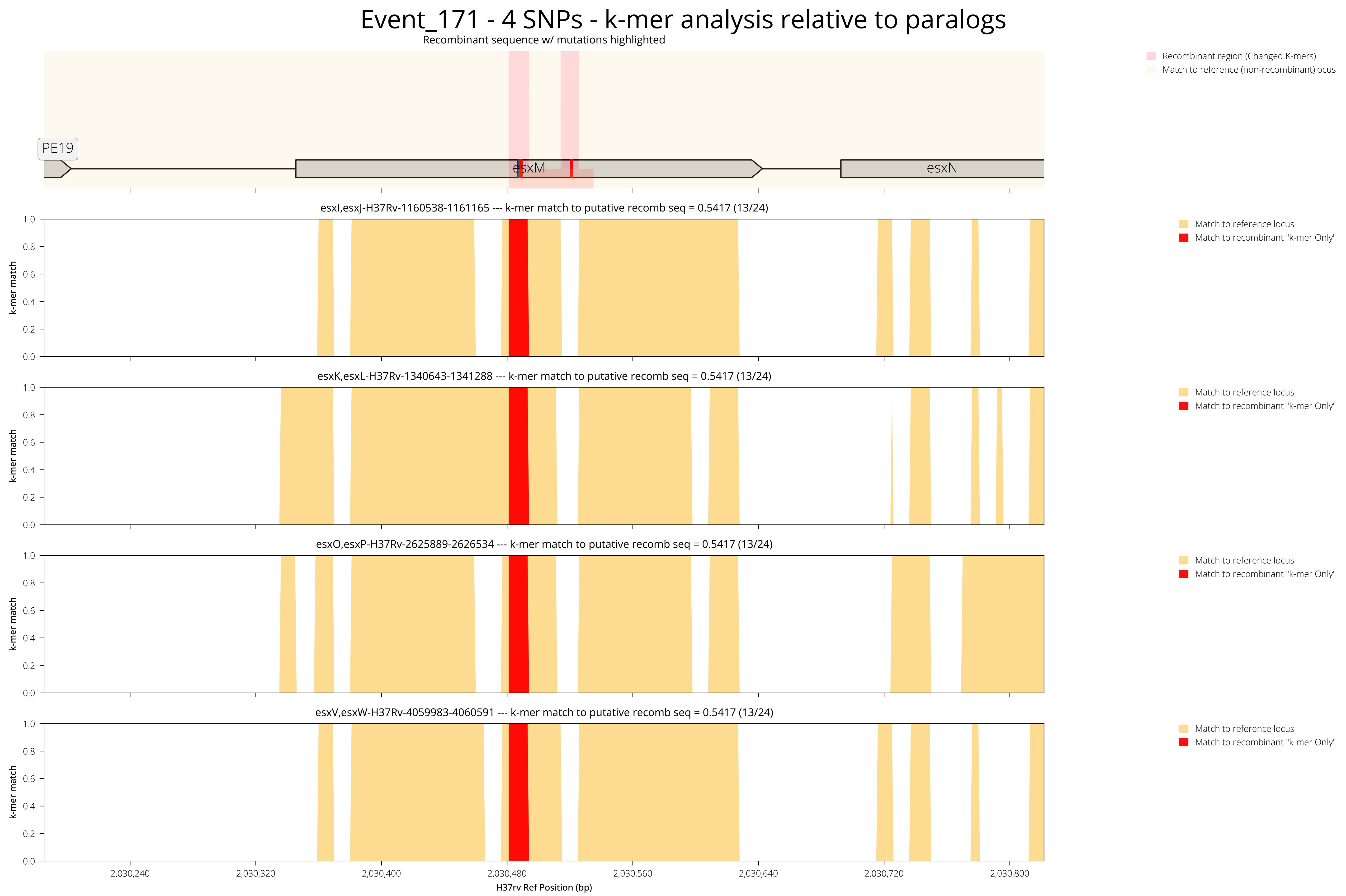
Event_171 - 4 SNPs - k-mer analysis relative to paralogs
Recombinant sequence w/ mutations highlighted
Recombinant region (Changed K-mers)
Match to reference (non-recombinant)locus
PE19
esxM
esxN
esxI,esxJ-H37Rv-1160538-1161165 --- k-mer match to putative recomb seq = 0.5417 (13/24)
esxK,esxL-H37Rv-1340643-1341288 --- k-mer match to putative recomb seq = 0.5417 (13/24)
esxO,esxP-H37Rv-2625889-2626534 --- k-mer match to putative recomb seq = 0.5417 (13/24)
esxV,esxW-H37Rv-4059983-4060591 --- k-mer match to putative recomb seq = 0.5417 (13/24)
Match to reference locus
Match to recombinant "k-mer Only"
k-mer match
1.0
0.8
0.6
0.4
0.2
0.0
2,030,240
2,030,320
2,030,400
2,030,480
2,030,560
2,030,640
2,030,720
2,030,800
H37rv Ref Position (bp)

Sub-phylogeny Viz + Paralog Alignments
GCE ID: Event_171

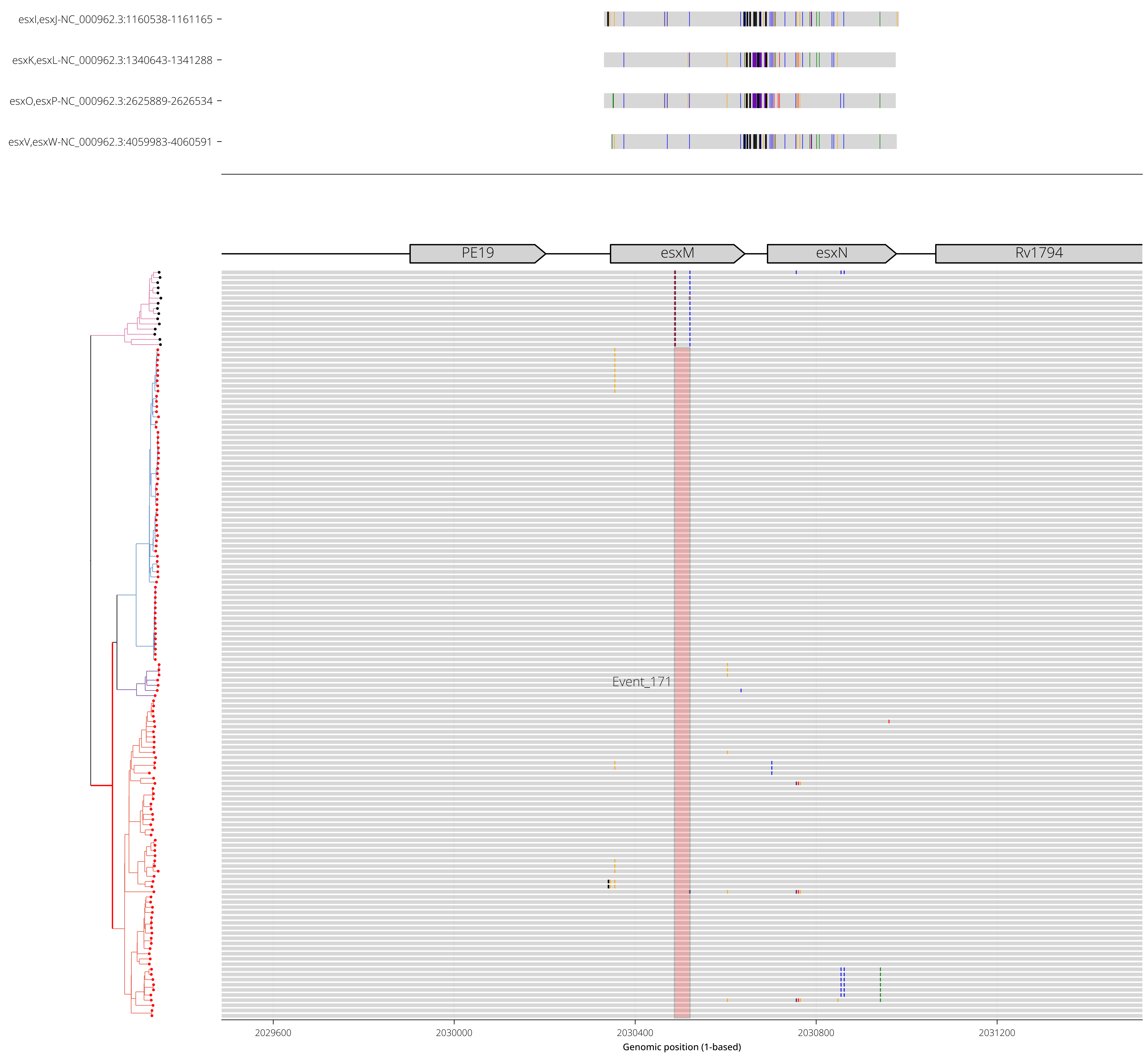

Full Phylogeny Viz + Paralog Alignments
GCE ID: Event_171

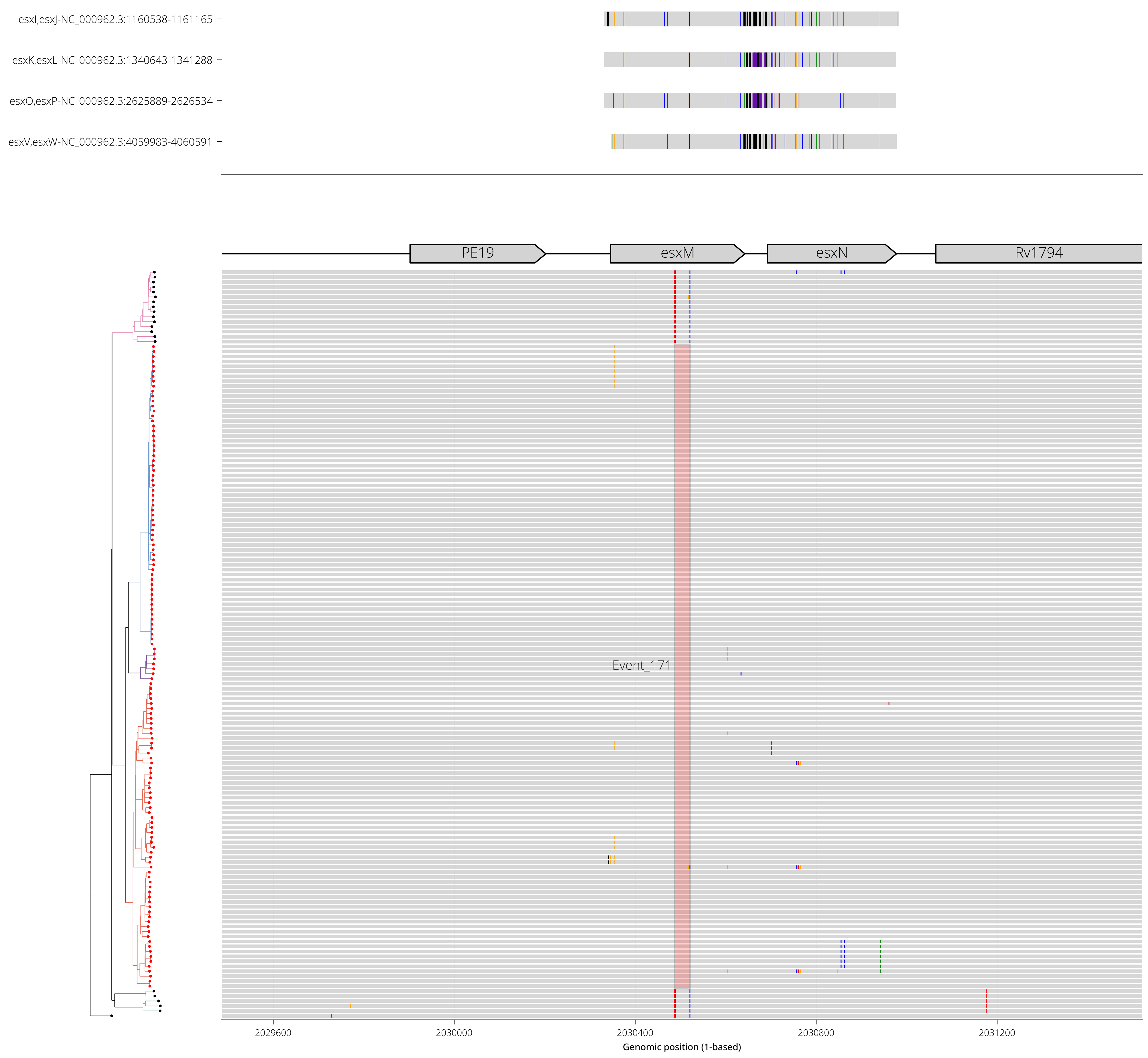

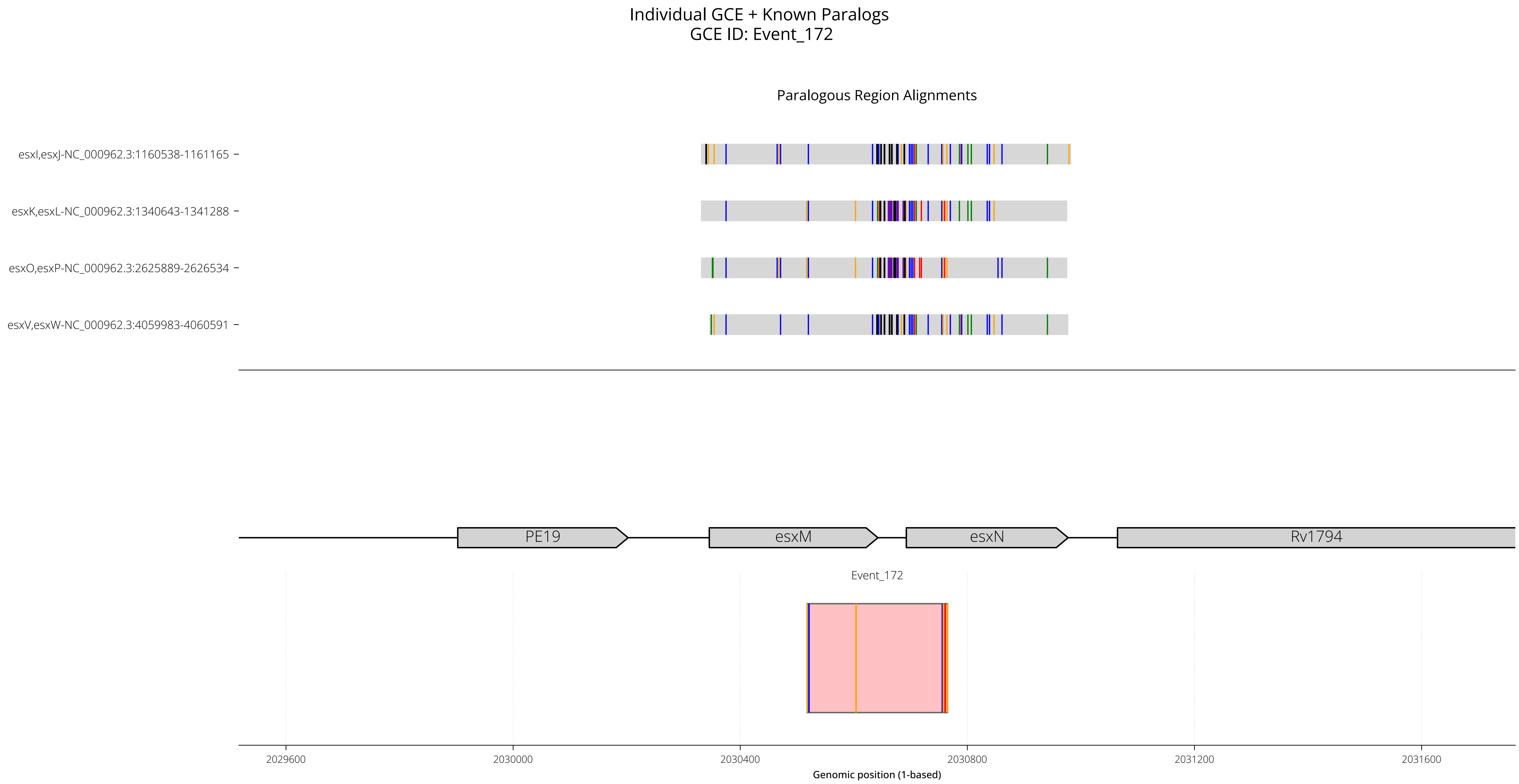
Individual GCE + Known Paralogs
GCE ID: Event_172
Paralogous Region Alignments
esxI,esxJ-NC_000962.3:1160538-1161165
esxK,esxL-NC_000962.3:1340643-1341288
esxO,esxP-NC_000962.3:2625889-2626534
esxV,esxW-NC_000962.3:4059983-4060591
PE19
esxM
esxN
Rv1794
Event_172
2029600
2030000
2030400
2030800
2031200
2031600
Genomic position (1-based)

Circos Viz - GCE to Known Paralogs Locations
GCE ID: Event_172

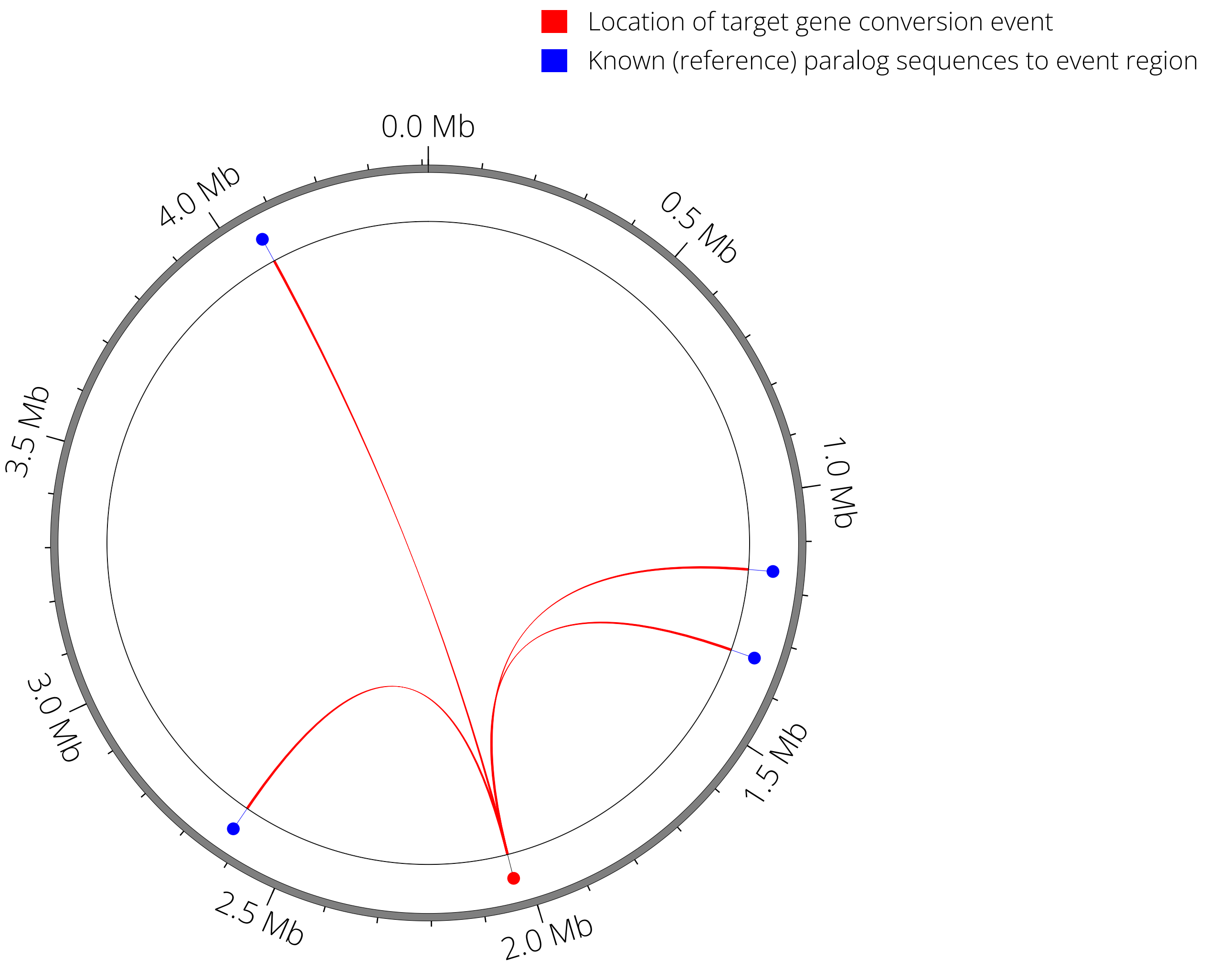

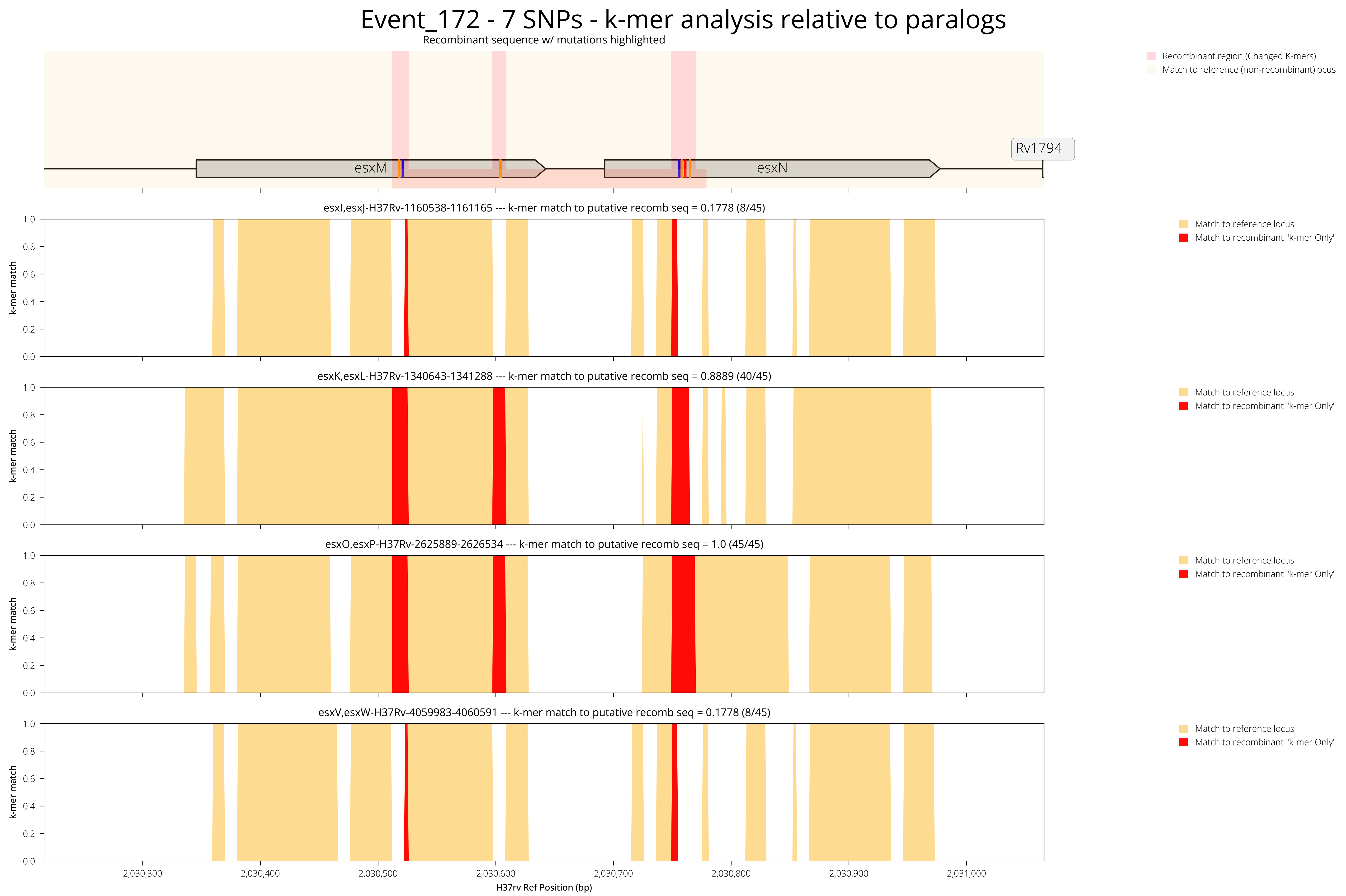
Event_172 - 7 SNPs - k-mer analysis relative to paralogs
Recombinant sequence w/ mutations highlighted
Recombinant region (Changed K-mers)
Match to reference (non-recombinant)locus
esxM
esxN
Rv1794
esxI,esxJ-H37Rv-1160538-1161165 --- k-mer match to putative recomb seq = 0.1778 (8/45)
esxK,esxL-H37Rv-1340643-1341288 --- k-mer match to putative recomb seq = 0.8889 (40/45)
esxO,esxP-H37Rv-2625889-2626534 --- k-mer match to putative recomb seq = 1.0 (45/45)
esxV,esxW-H37Rv-4059983-4060591 --- k-mer match to putative recomb seq = 0.1778 (8/45)
k-mer match
Match to reference locus
Match to recombinant "k-mer Only"
2,030,300
2,030,400
2,030,500
2,030,600
2,030,700
2,030,800
2,030,900
2,031,000
H37rv Ref Position (bp)

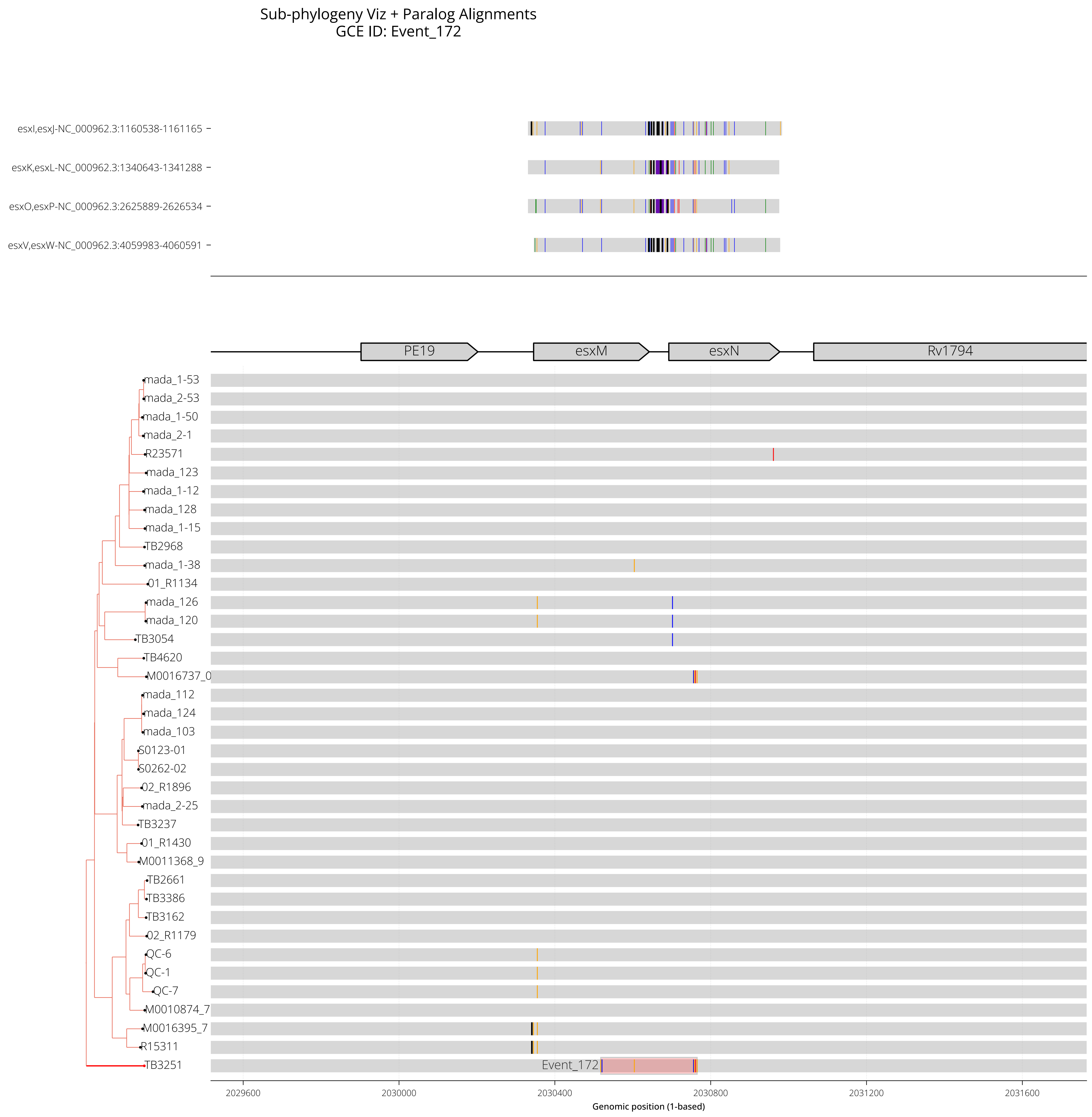
Sub-phylogeny Viz + Paralog Alignments
GCE ID: Event_172
esxI,esxJ-NC_000962.3:1160538-1161165
esxK,esxL-NC_000962.3:1340643-1341288
esxO,esxP-NC_000962.3:2625889-2626534
esxV,esxW-NC_000962.3:4059983-4060591
PE19
esxM
esxN
Rv1794
mada_1-53
mada_2-53
mada_1-50
mada_2-1
R23571
mada_123
mada_1-12
mada_128
mada_1-15
TB2968
mada_1-38
01_R1134
mada_126
mada_120
TB3054
TB4620
M0016737_0
mada_112
mada_124
mada_103
S0123-01
S0262-02
02_R1896
mada_2-25
TB3237
01_R1430
M0011368_9
TB2661
TB3386
TB3162
02_R1179
QC-6
QC-1
QC-7
M0010874_7
M0016395_7
R15311
TB3251
Event_172
2029600
2030000
2030400
2030800
2031200
2031600
Genomic position (1-based)

Full Phylogeny Viz + Paralog Alignments
GCE ID: Event_172

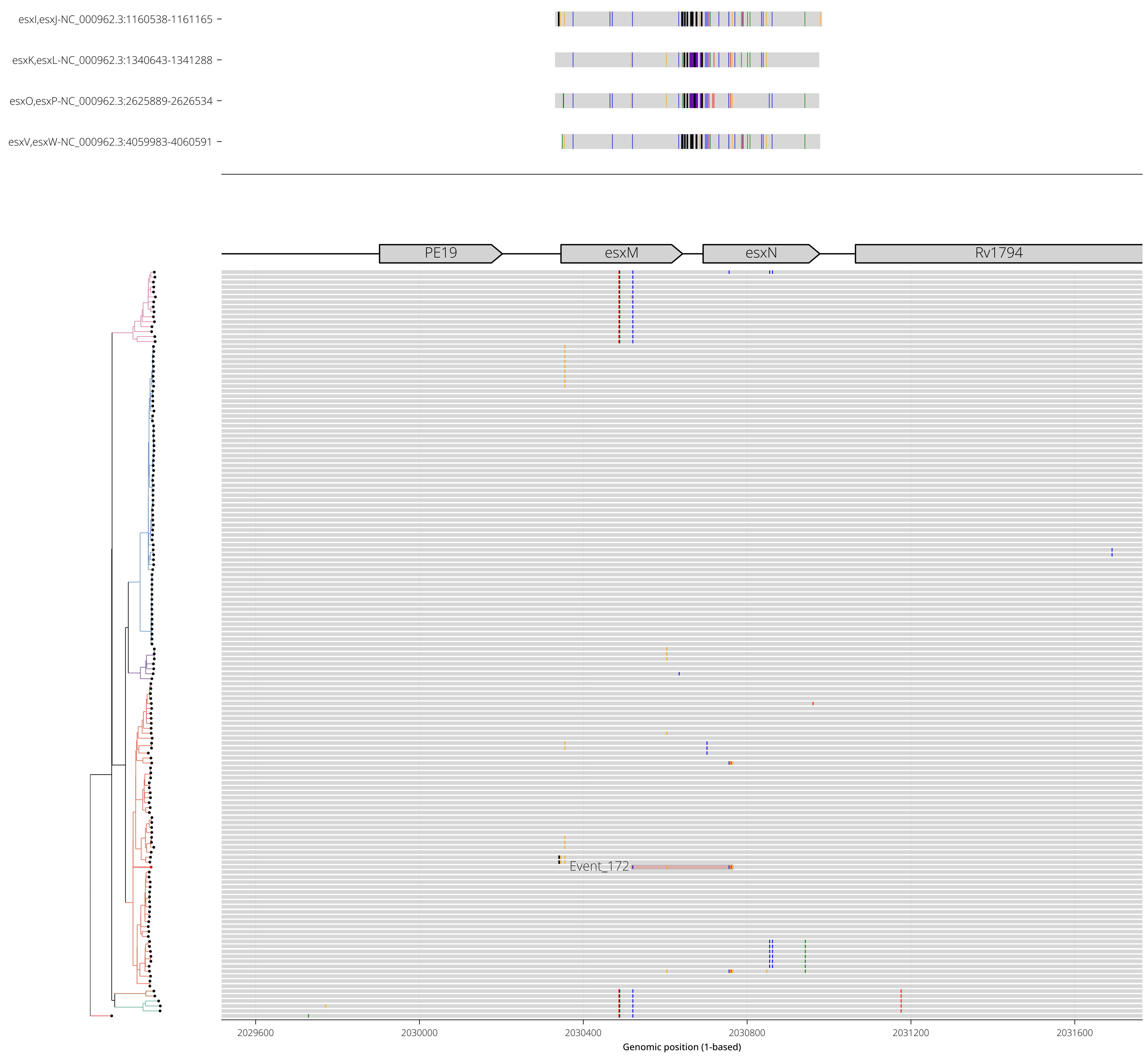

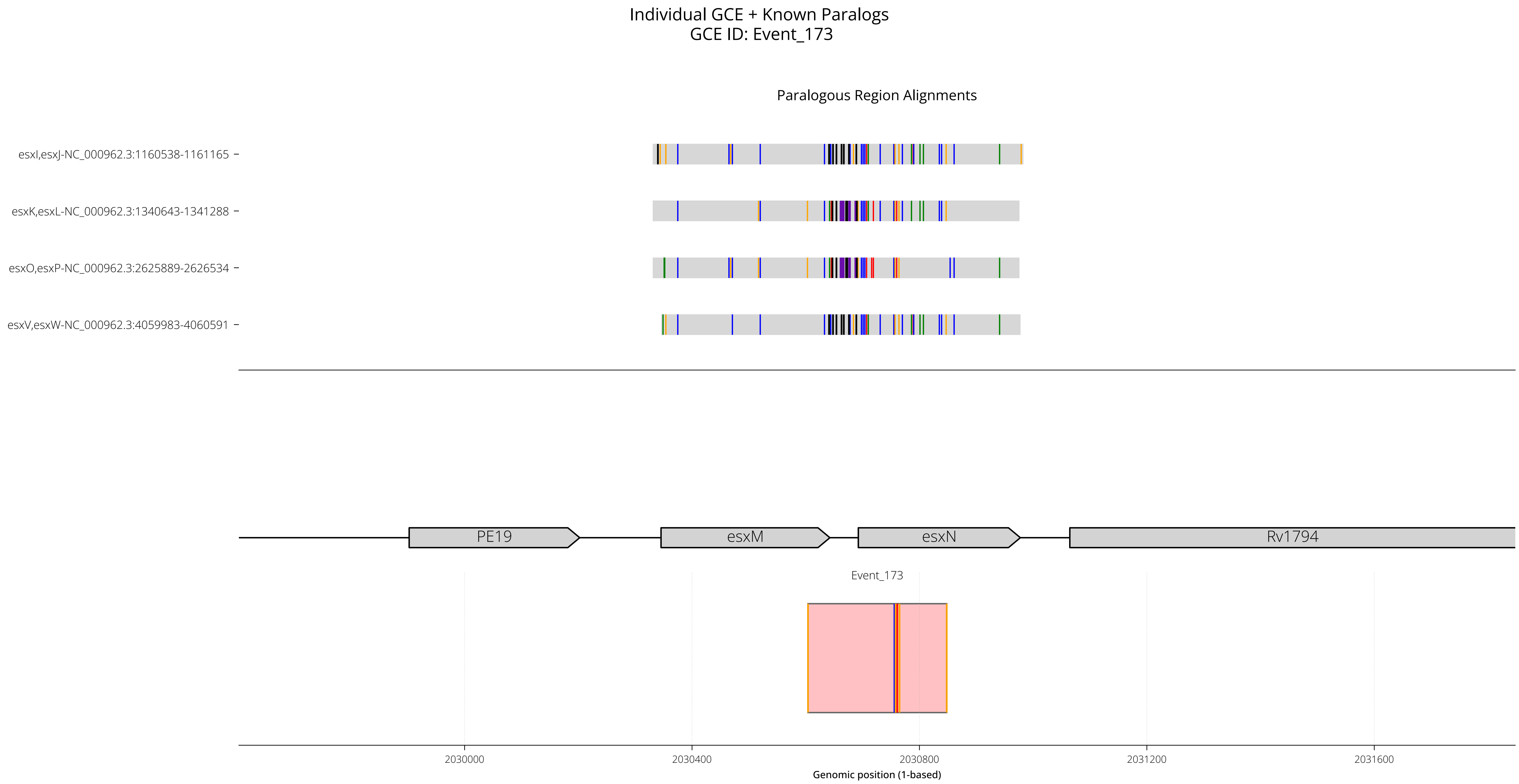

Individual GCE + Known Paralogs
GCE ID: Event_173
Paralogous Region Alignments
esxI,esxJ-NC_000962.3:1160538-1161165
esxK,esxL-NC_000962.3:1340643-1341288
esxO,esxP-NC_000962.3:2625889-2626534
esxV,esxW-NC_000962.3:4059983-4060591
PE19
esxM
esxN
Rv1794
Event_173
2030000
2030400
2030800
2031200
2031600
Genomic position (1-based)

Circos Viz - GCE to Known Paralogs Locations
GCE ID: Event_173

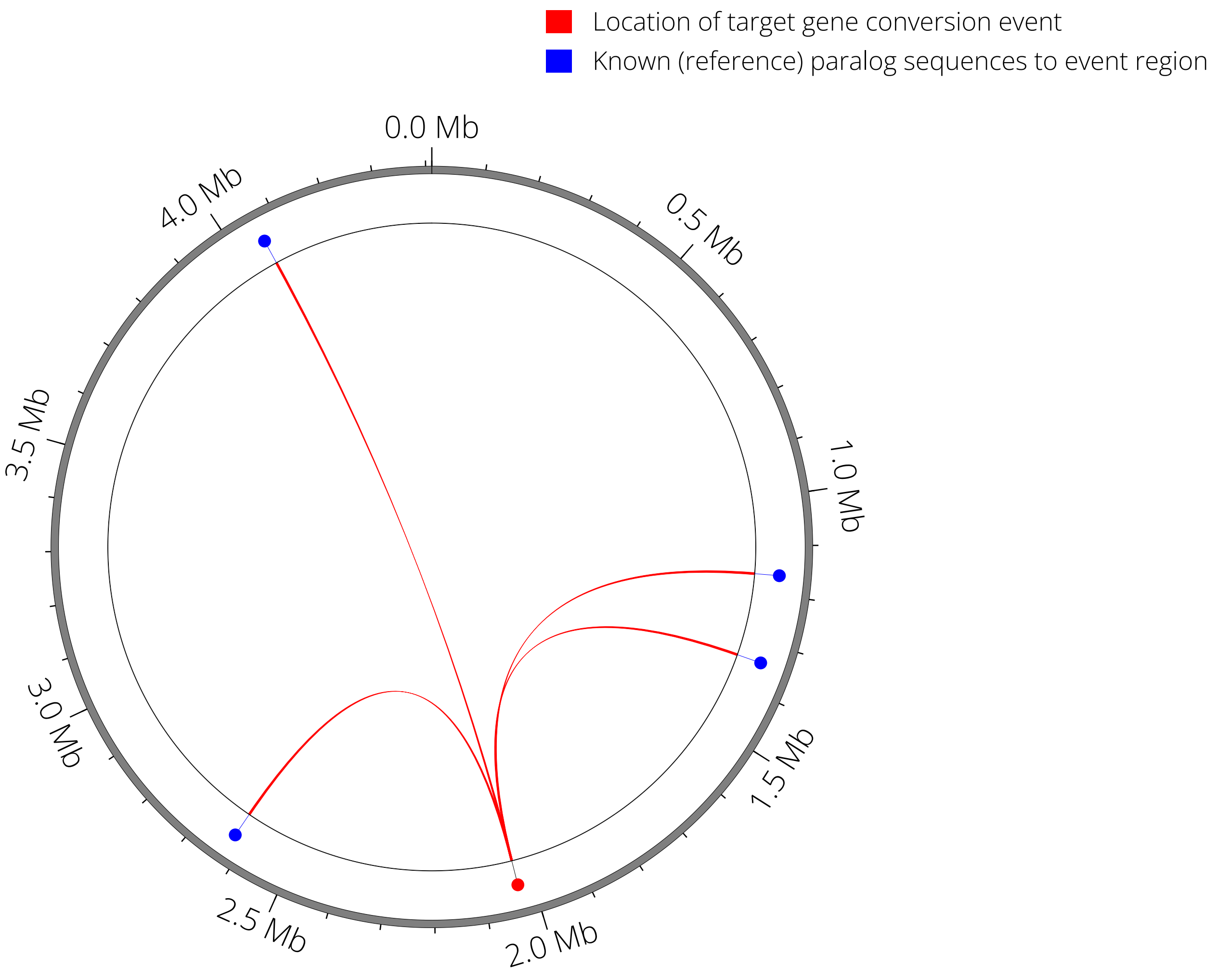

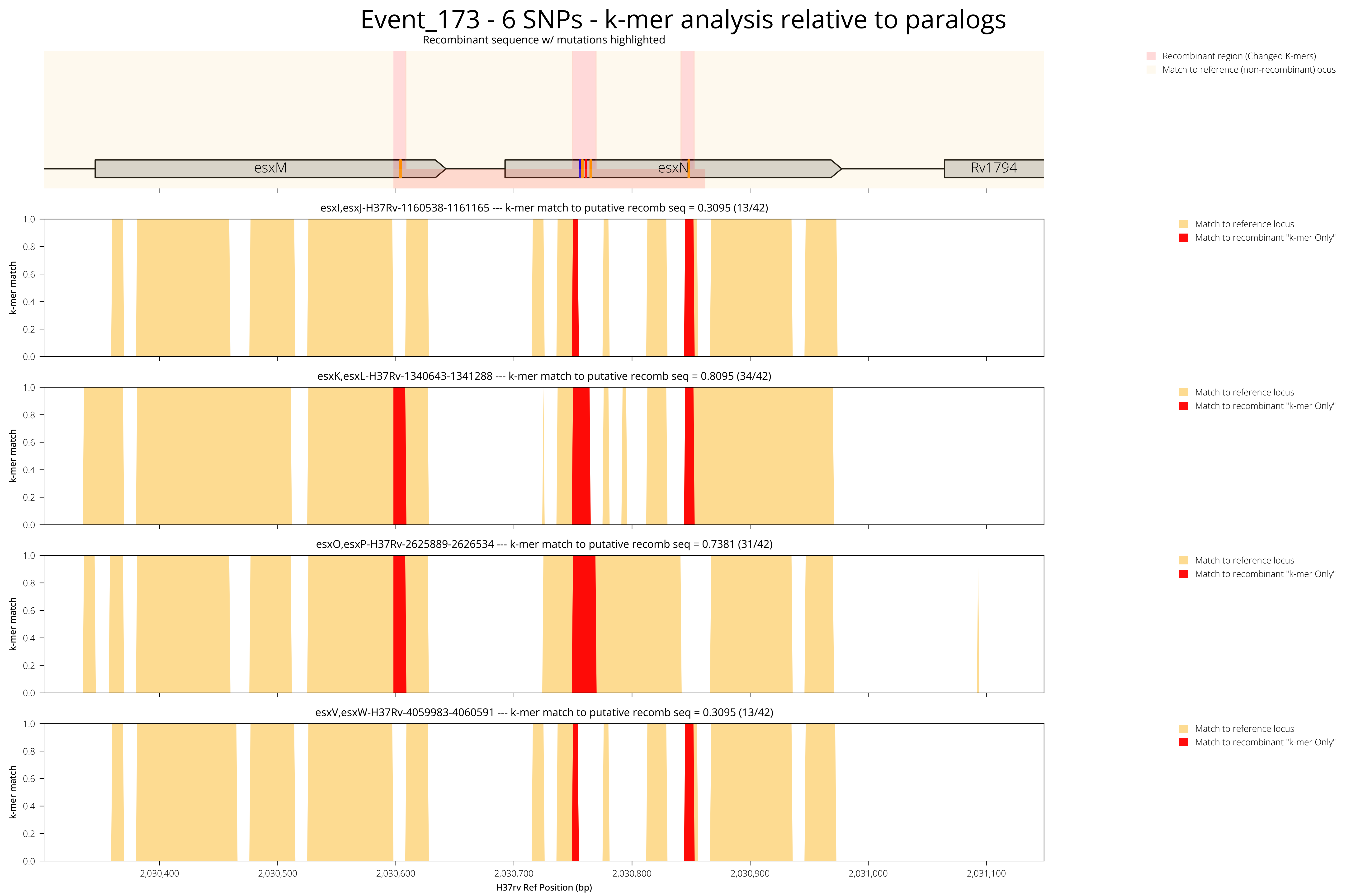

Event_173 - 6 SNPs - k-mer analysis relative to paralogs
Recombinant sequence w/ mutations highlighted
Recombinant region (Changed K-mers)
Match to reference (non-recombinant)locus
esxM
esxN
Rv1794
esxI,esxJ-H37Rv-1160538-1161165 --- k-mer match to putative recomb seq = 0.3095 (13/42)
esxK,esxL-H37Rv-1340643-1341288 --- k-mer match to putative recomb seq = 0.8095 (34/42)
esxO,esxP-H37Rv-2625889-2626534 --- k-mer match to putative recomb seq = 0.7381 (31/42)
esxV,esxW-H37Rv-4059983-4060591 --- k-mer match to putative recomb seq = 0.3095 (13/42)
Match to reference locus
Match to recombinant "k-mer Only"
k-mer match
1.0
0.8
0.6
0.4
0.2
0.0
2,030,400
2,030,500
2,030,600
2,030,700
2,030,800
2,030,900
2,031,000
2,031,100
H37rv Ref Position (bp)

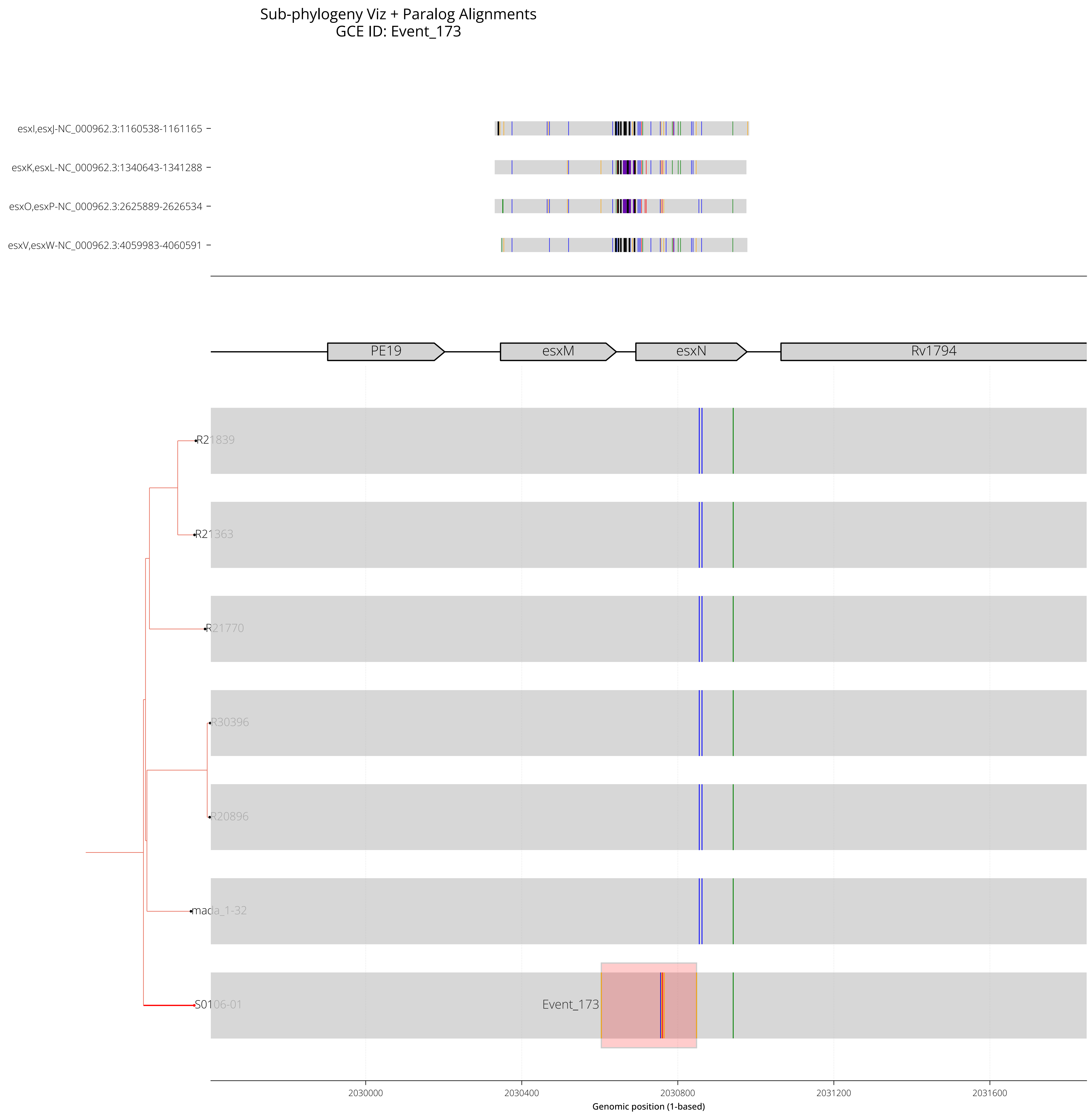
Sub-phylogeny Viz + Paralog Alignments
GCE ID: Event_173
esxI,esxJ-NC_000962.3:1160538-1161165
esxK,esxL-NC_000962.3:1340643-1341288
esxO,esxP-NC_000962.3:2625889-2626534
esxV,esxW-NC_000962.3:4059983-4060591
PE19
esxM
esxN
Rv1794
R21839
R21363
R21770
R30396
R20896
mada_1-32
S0106-01
Event_173
2030000
2030400
2030800
2031200
2031600
Genomic position (1-based)

Full Phylogeny Viz + Paralog Alignments
GCE ID: Event_173

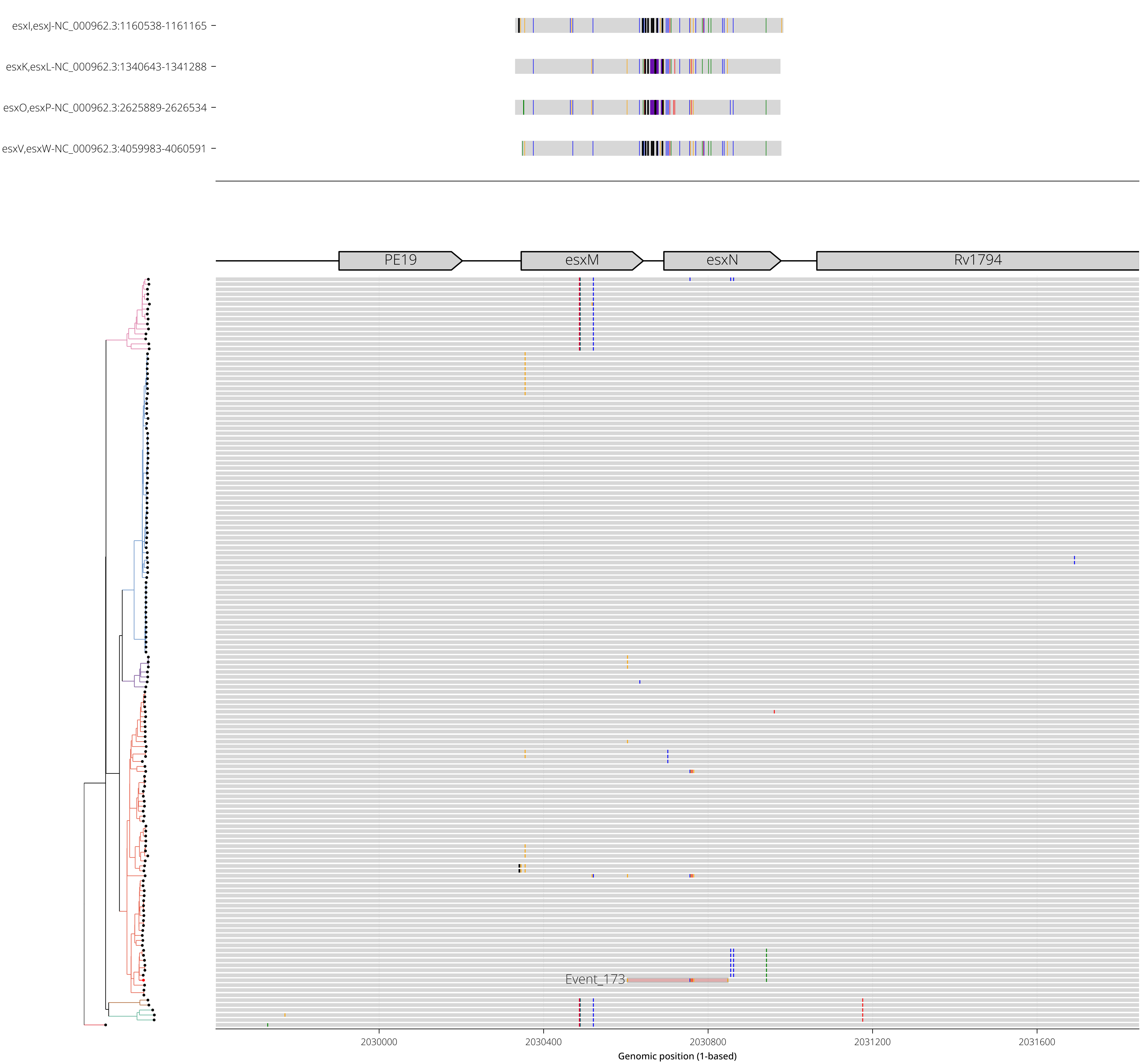

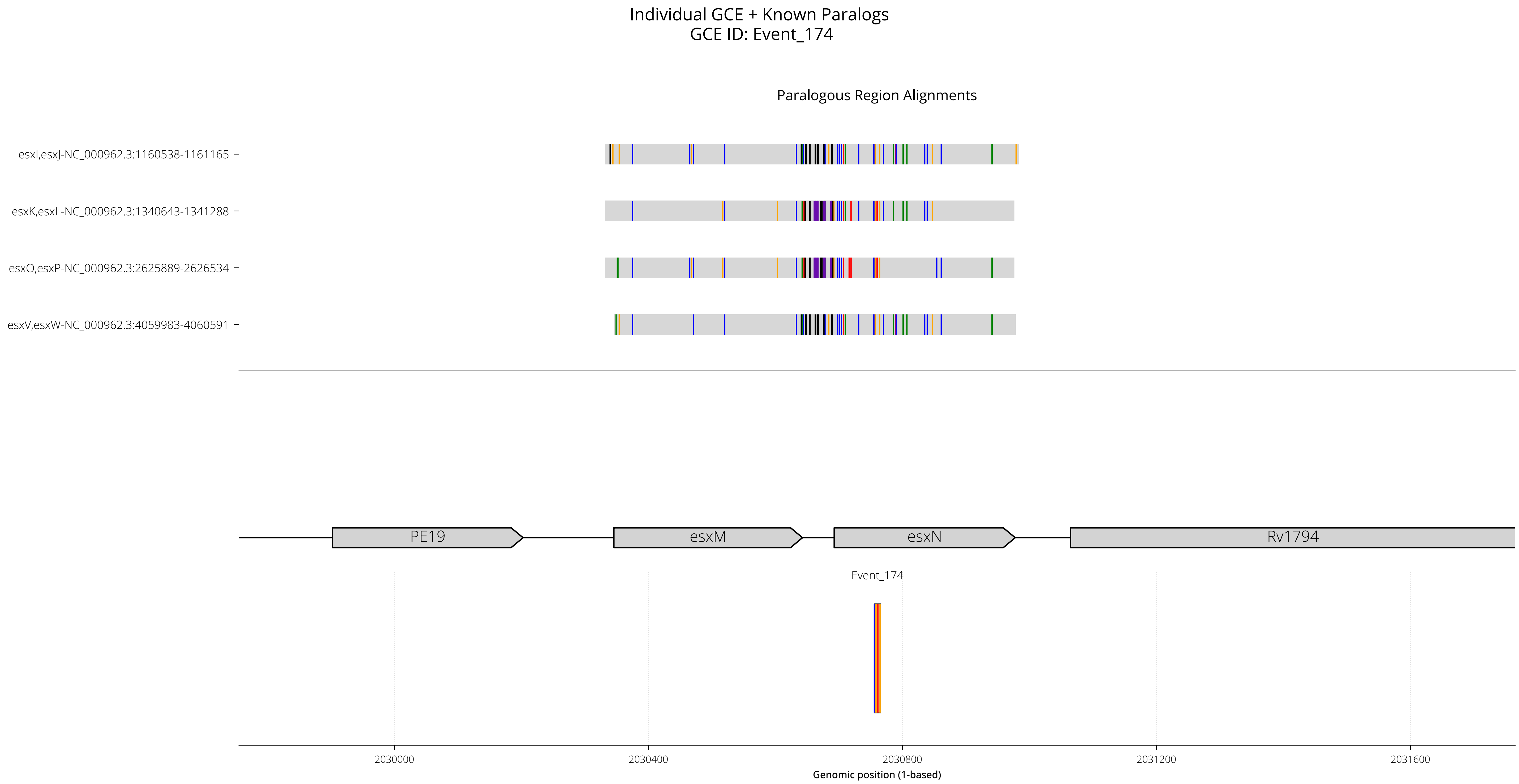
Individual GCE + Known Paralogs
GCE ID: Event_174
Paralogous Region Alignments
esxI,esxJ-NC_000962.3:1160538-1161165
esxK,esxL-NC_000962.3:1340643-1341288
esxO,esxP-NC_000962.3:2625889-2626534
esxV,esxW-NC_000962.3:4059983-4060591
PE19
esxM
esxN
Rv1794
Event_174
2030000
2030400
2030800
2031200
2031600
Genomic position (1-based)

Circos Viz - GCE to Known Paralogs Locations
GCE ID: Event_174

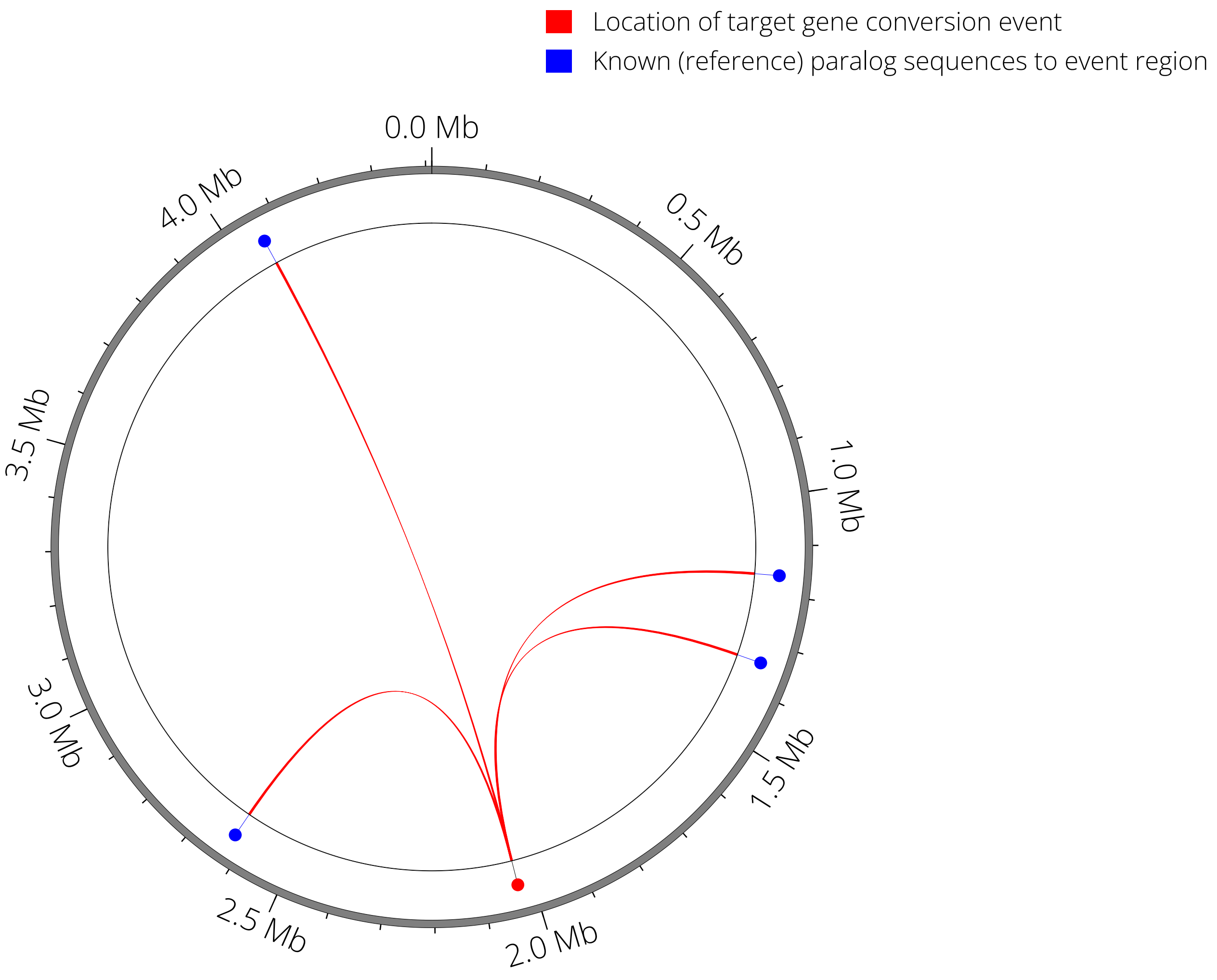

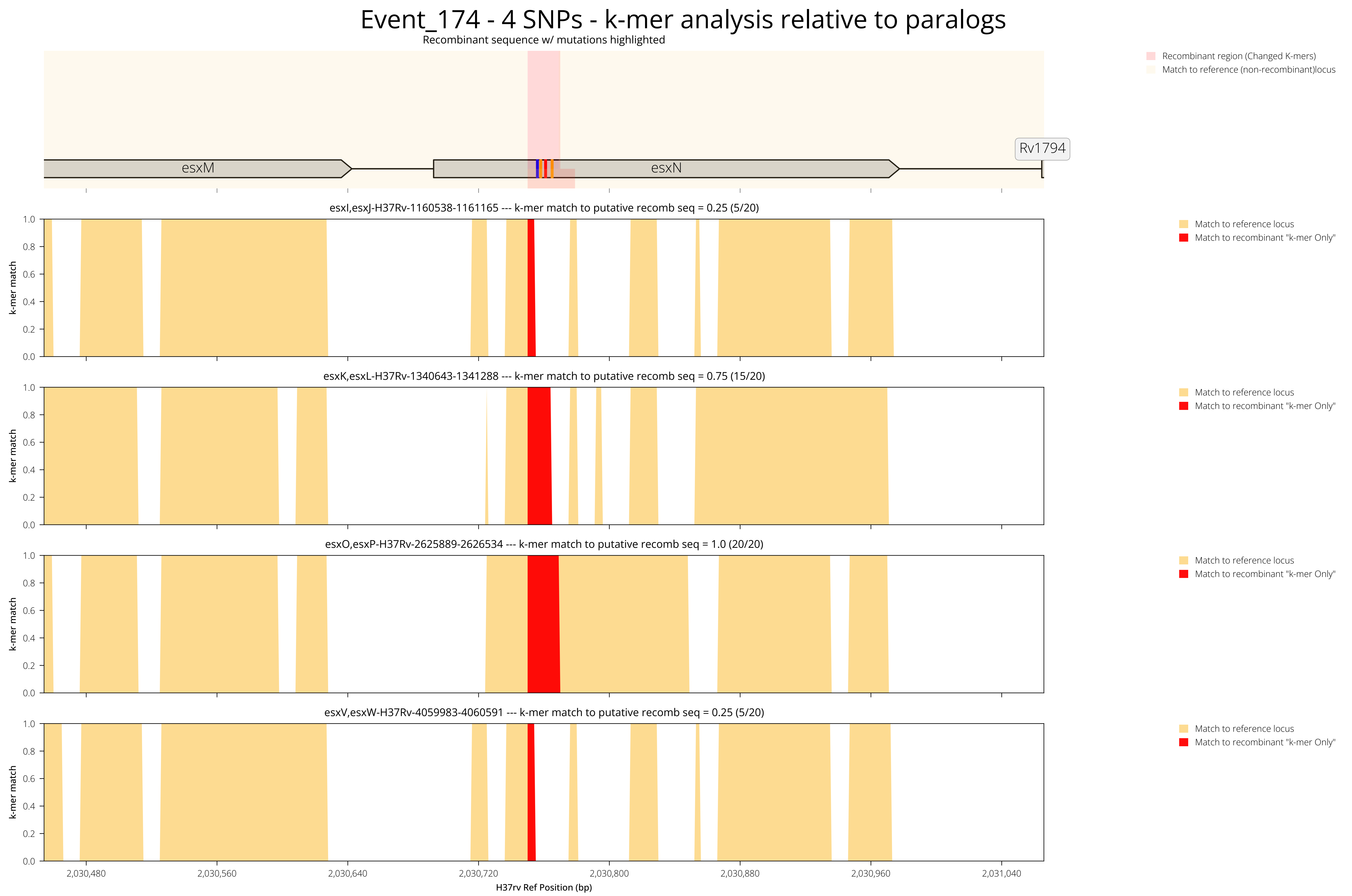

Event_174 - 4 SNPs - k-mer analysis relative to paralogs
Recombinant sequence w/ mutations highlighted
Recombinant region (Changed K-mers)
Match to reference (non-recombinant)locus
esxM
esxN
Rv1794
esxI,esxJ-H37Rv-1160538-1161165 --- k-mer match to putative recomb seq = 0.25 (5/20)
esxK,esxL-H37Rv-1340643-1341288 --- k-mer match to putative recomb seq = 0.75 (15/20)
esxO,esxP-H37Rv-2625889-2626534 --- k-mer match to putative recomb seq = 1.0 (20/20)
esxV,esxW-H37Rv-4059983-4060591 --- k-mer match to putative recomb seq = 0.25 (5/20)
k-mer match
Match to reference locus
Match to recombinant "k-mer Only"
2,030,480
2,030,560
2,030,640
2,030,720
2,030,800
2,030,880
2,030,960
2,031,040
H37rv Ref Position (bp)

Sub-phylogeny Viz + Paralog Alignments
GCE ID: Event_174

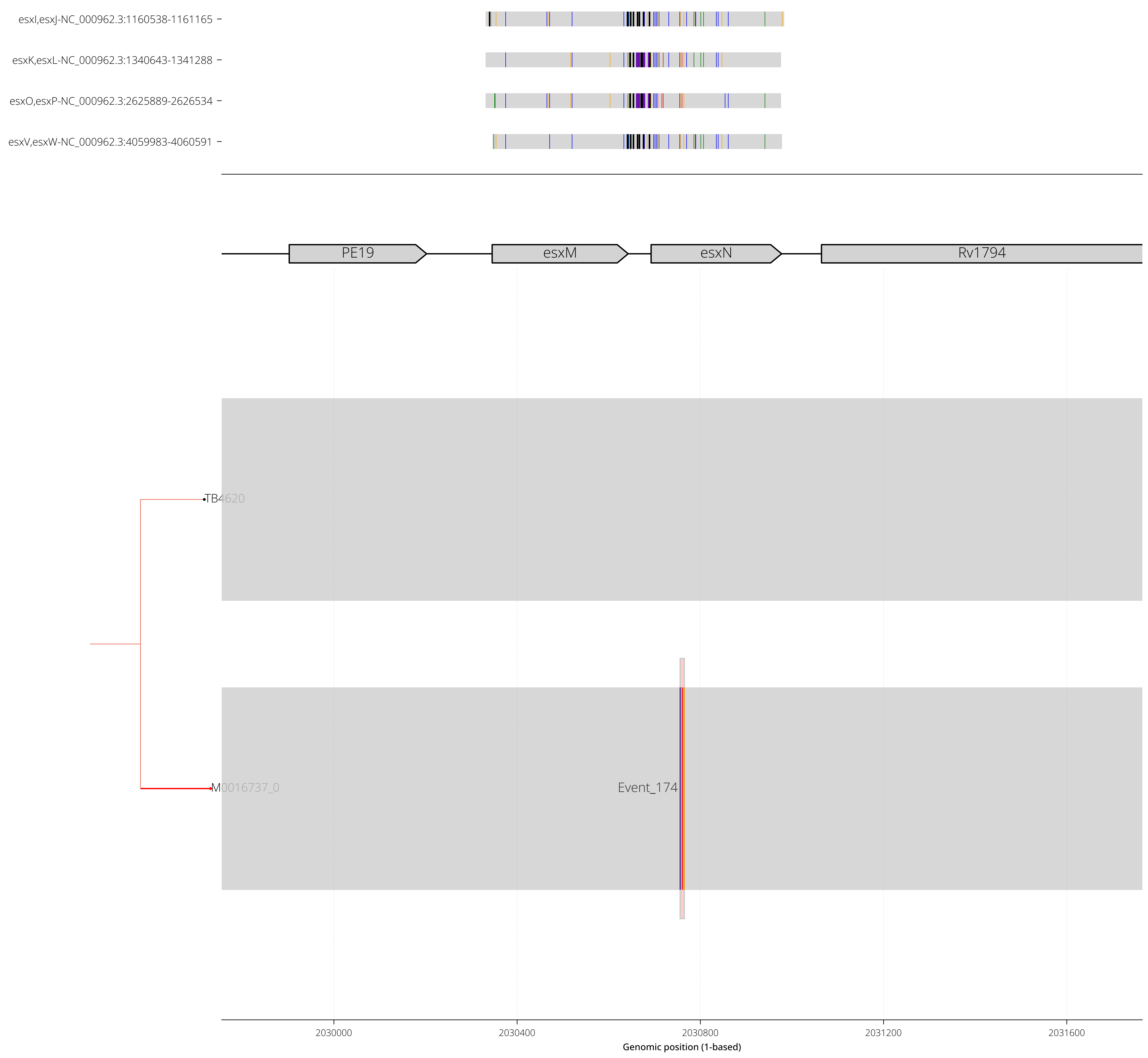

Full Phylogeny Viz + Paralog Alignments
GCE ID: Event_174

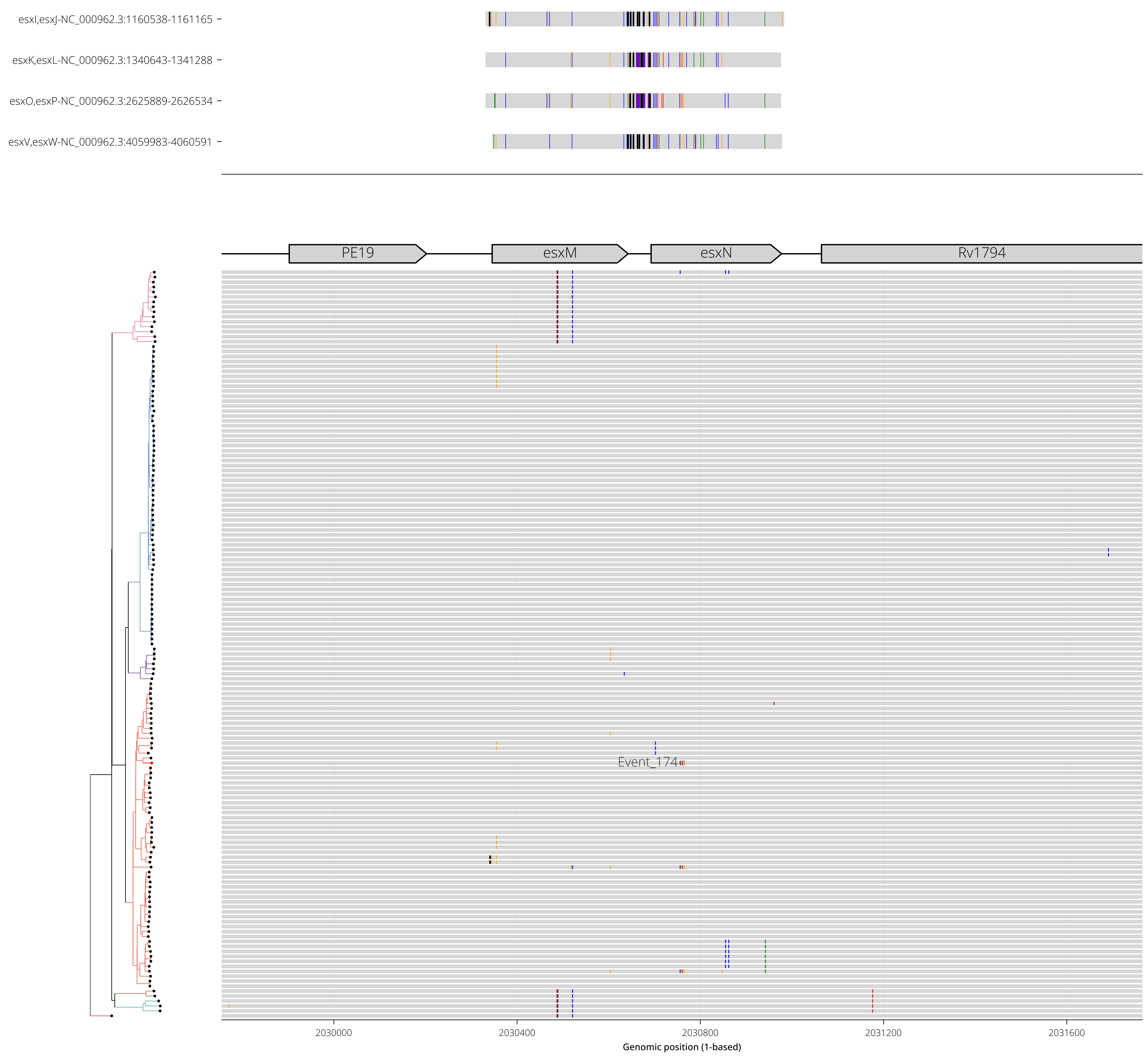

Individual GCE + Known Paralogs
GCE ID: Event_175

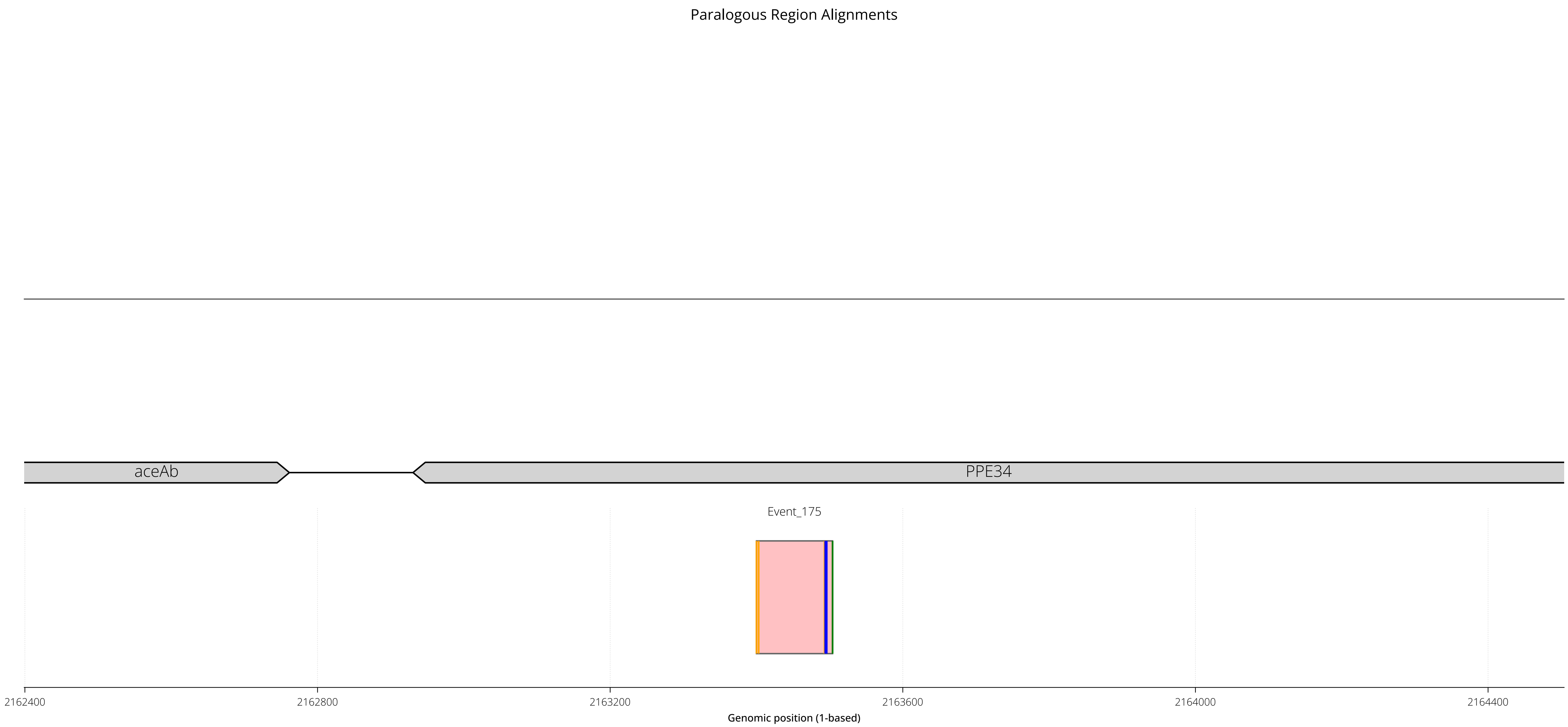

Sub-phylogeny Viz + Paralog Alignments
GCE ID: Event_175

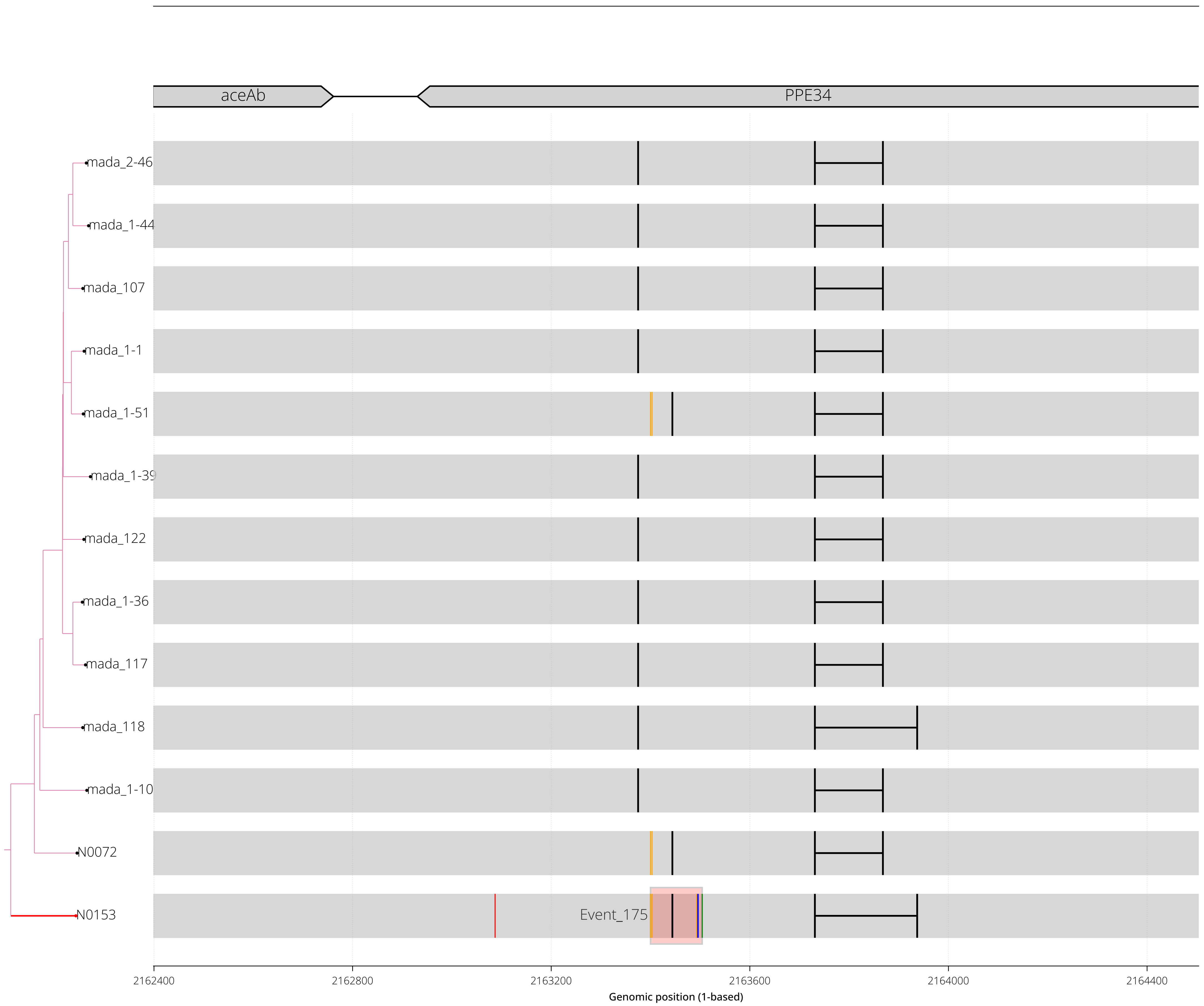

Full Phylogeny Viz + Paralog Alignments
GCE ID: Event_175

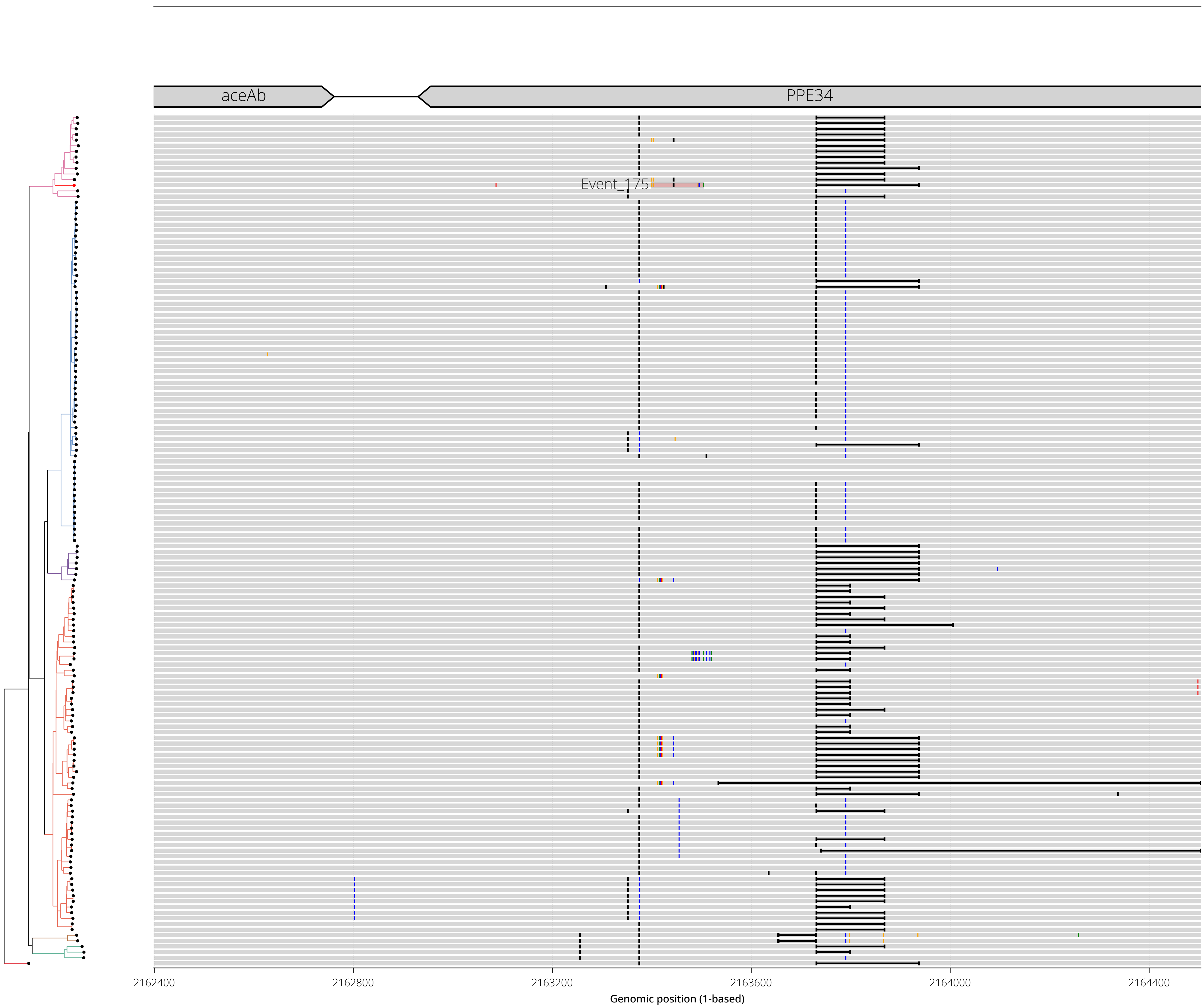

Individual GCE + Known Paralogs
GCE ID: Event_176

Paralogous Region Alignments

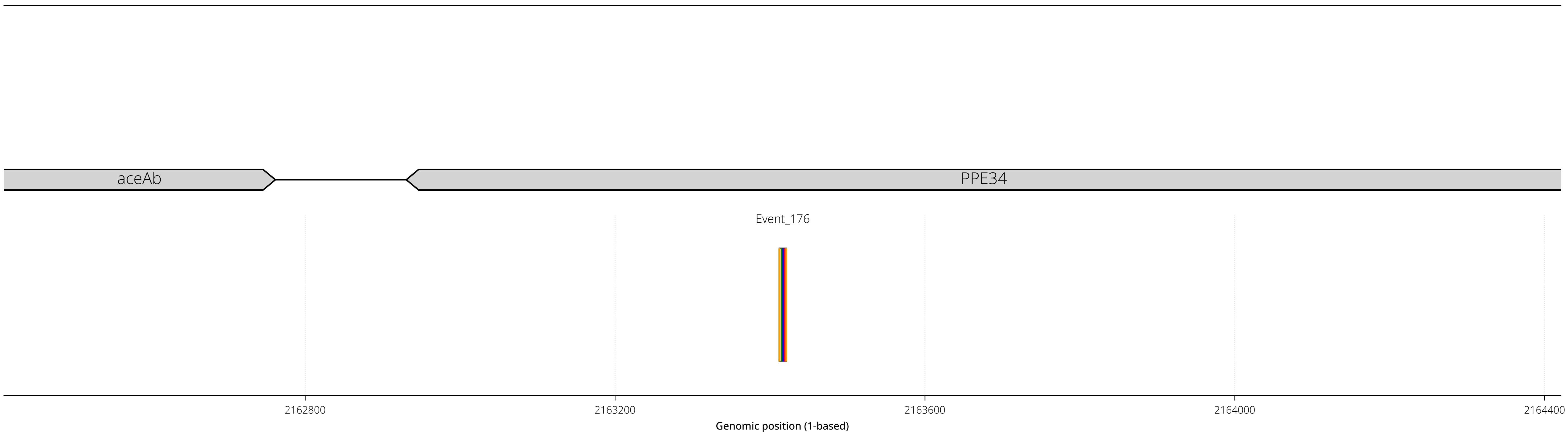

Sub-phylogeny Viz + Paralog Alignments
GCE ID: Event_176

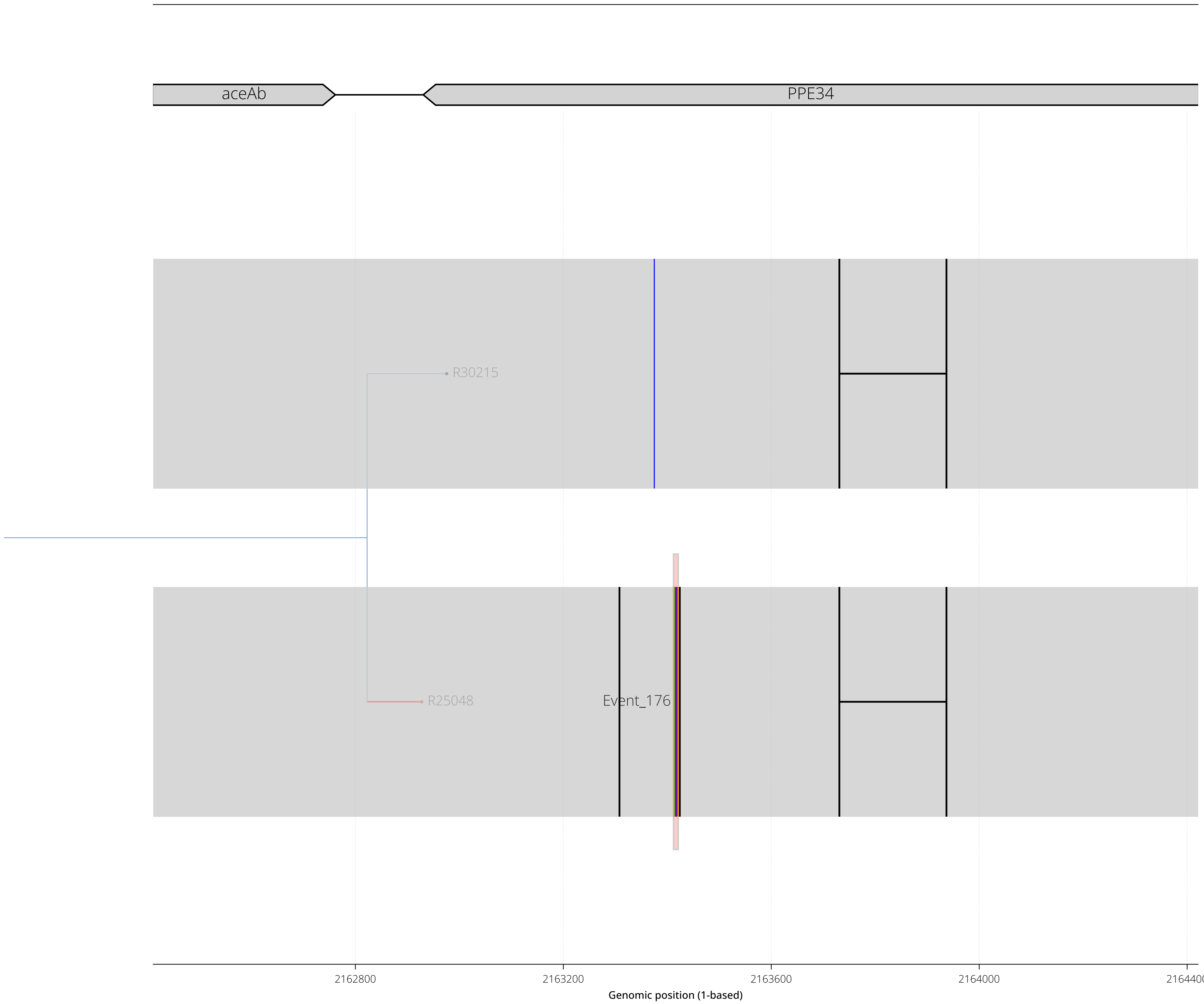

Full Phylogeny Viz + Paralog Alignments
GCE ID: Event_176

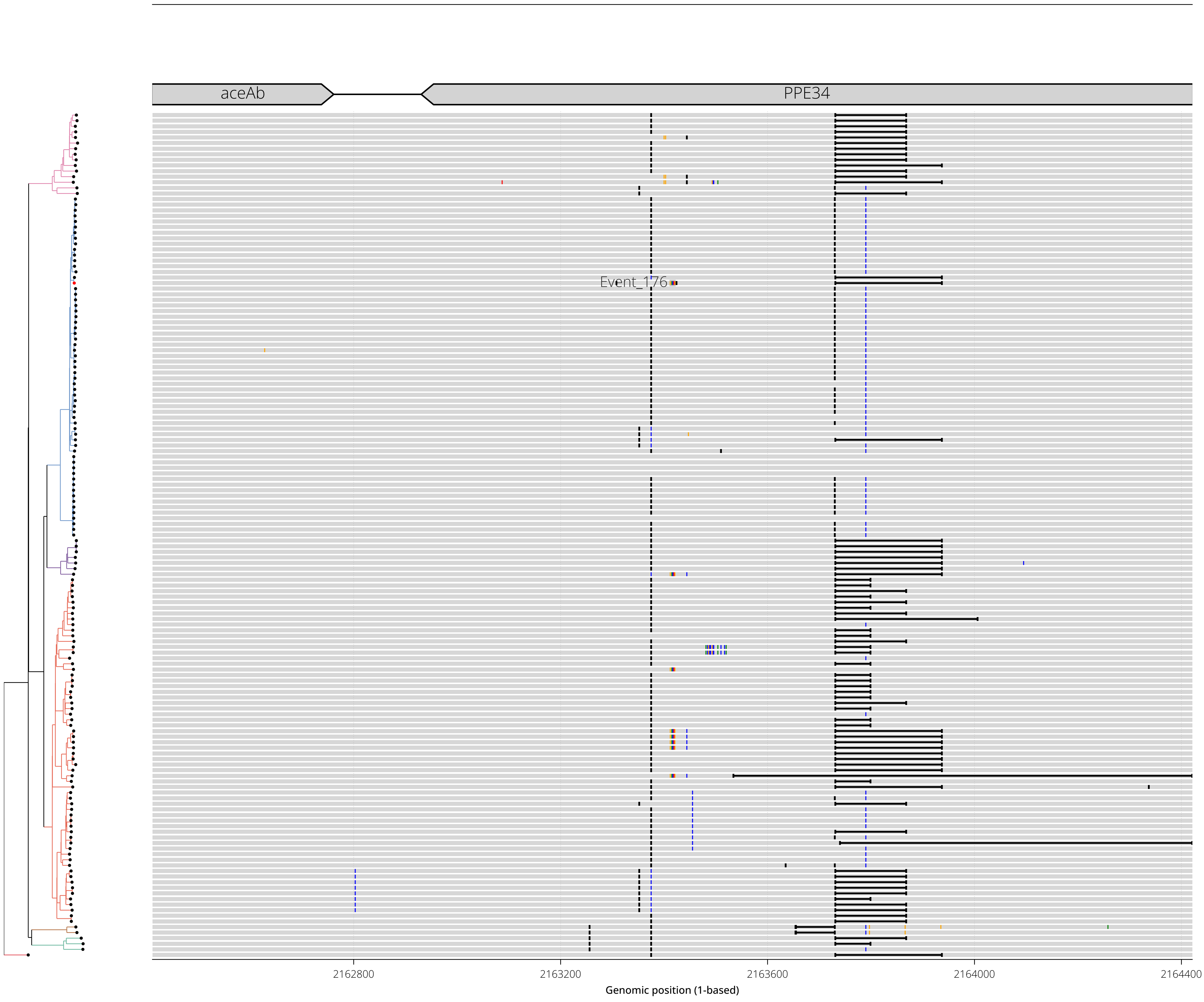

### Individual GCE + Known Paralogs
GCE ID: Event_177

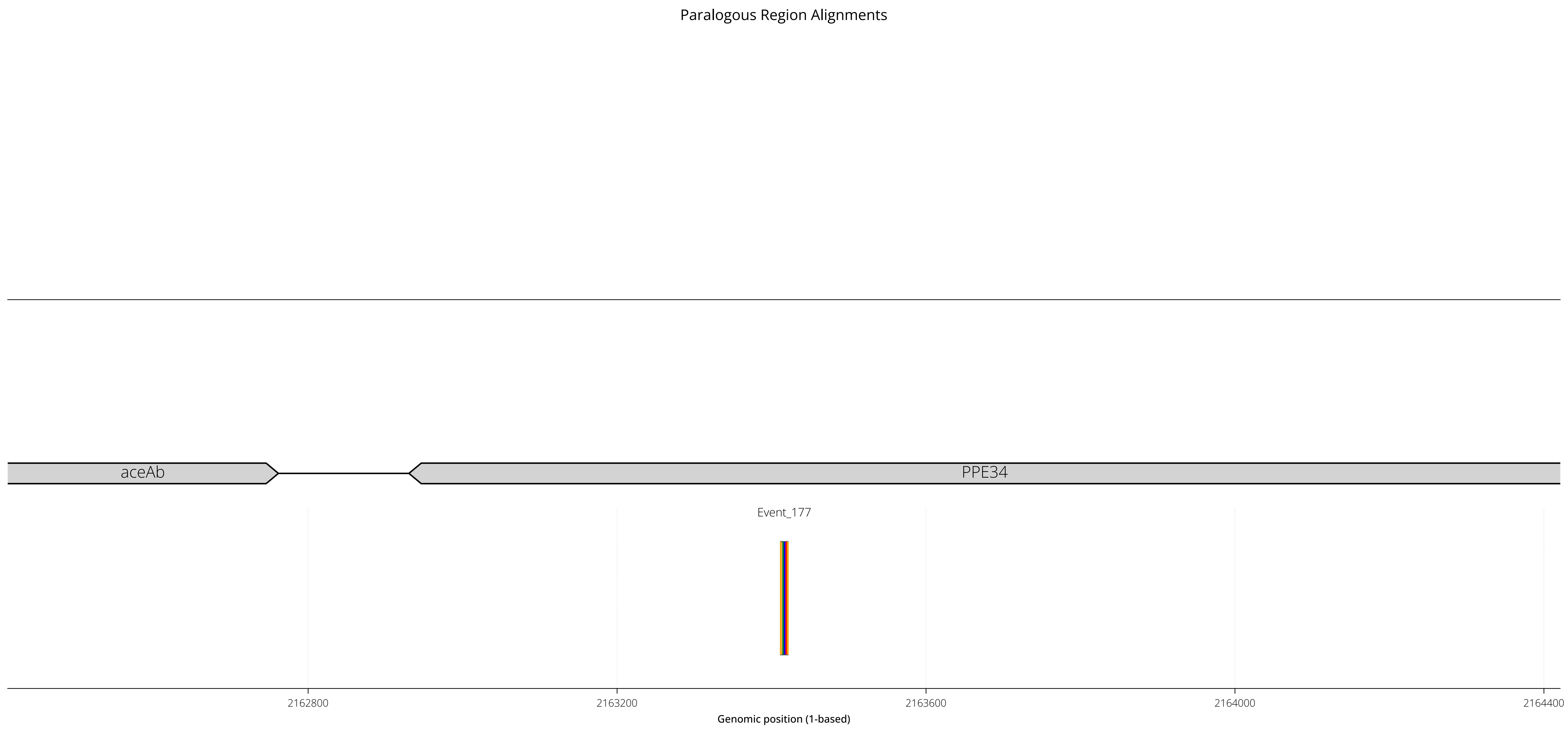

Sub-phylogeny Viz + Paralog Alignments
GCE ID: Event_177

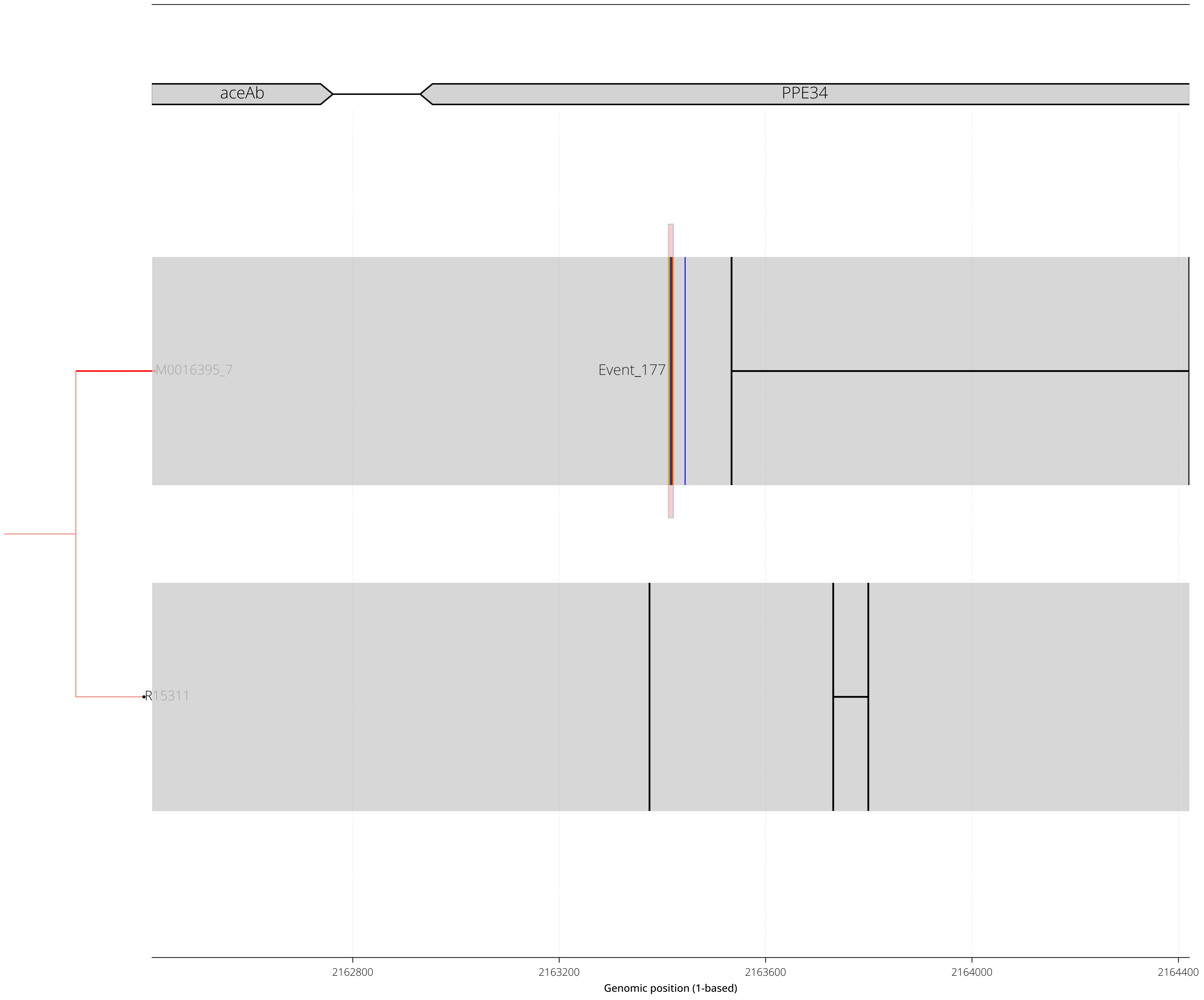

Full Phylogeny Viz + Paralog Alignments
GCE ID: Event_177

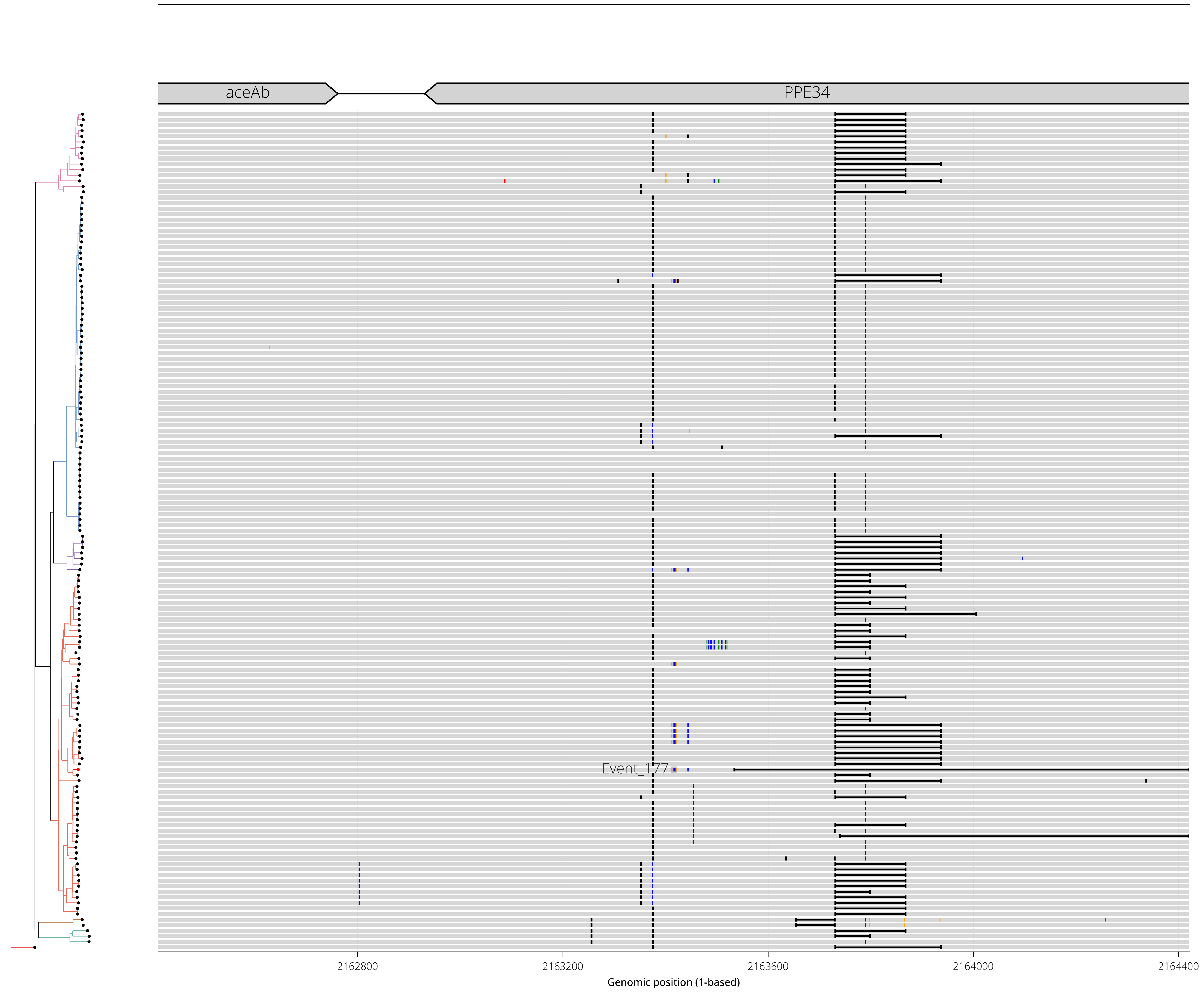

Individual GCE + Known Paralogs
GCE ID: Event_178

Paralogous Region Alignments

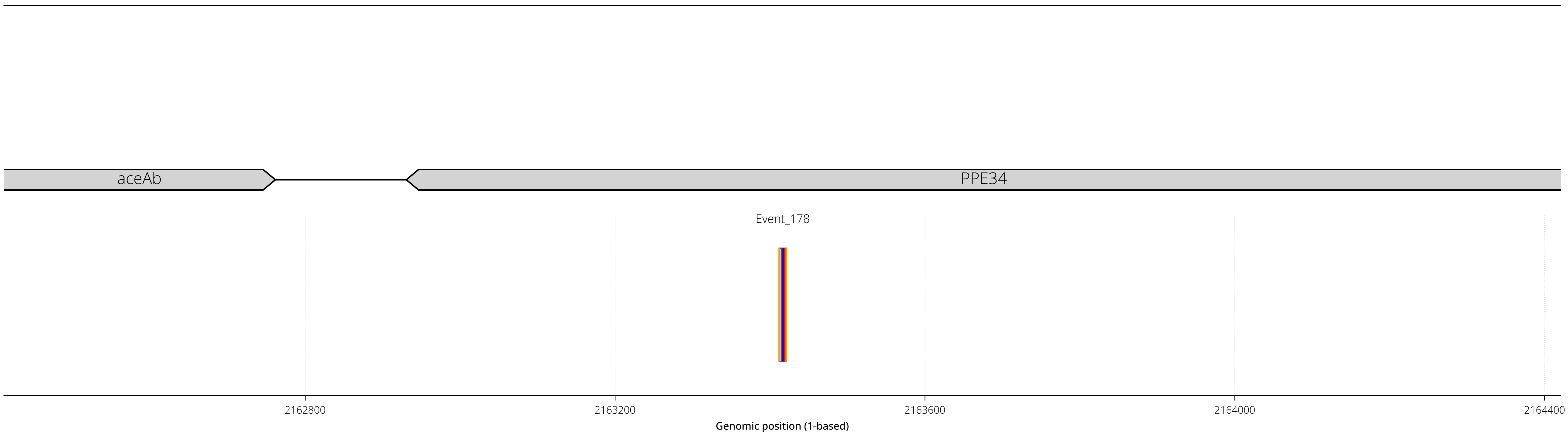

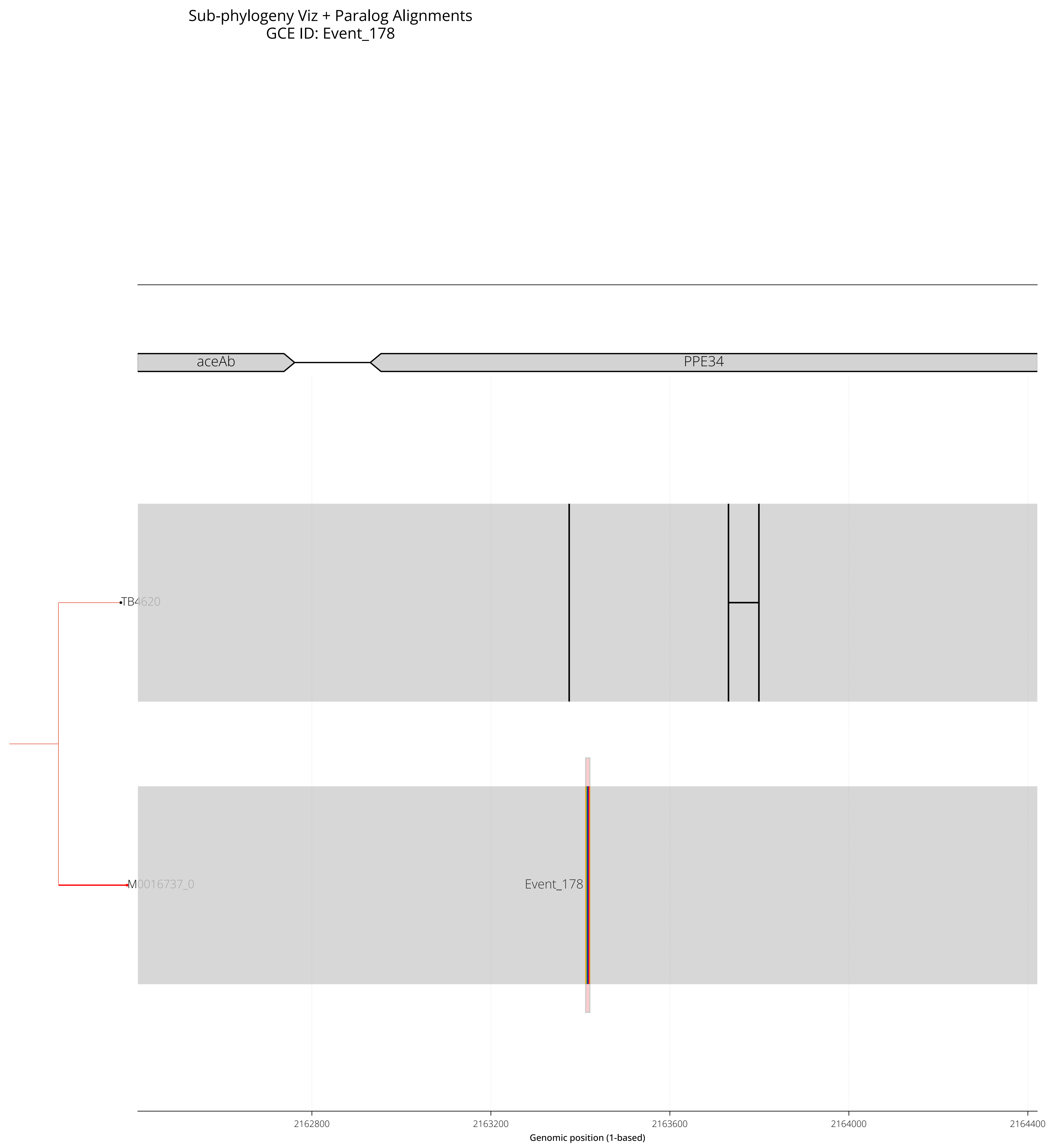
Sub-phylogeny Viz + Paralog Alignments
GCE ID: Event_178
aceAb
PPE34
TB4620
M0016737_0
Event_178
2162800
2163200
2163600
2164000
2164400
Genomic position (1-based)

Full Phylogeny Viz + Paralog Alignments
GCE ID: Event_178

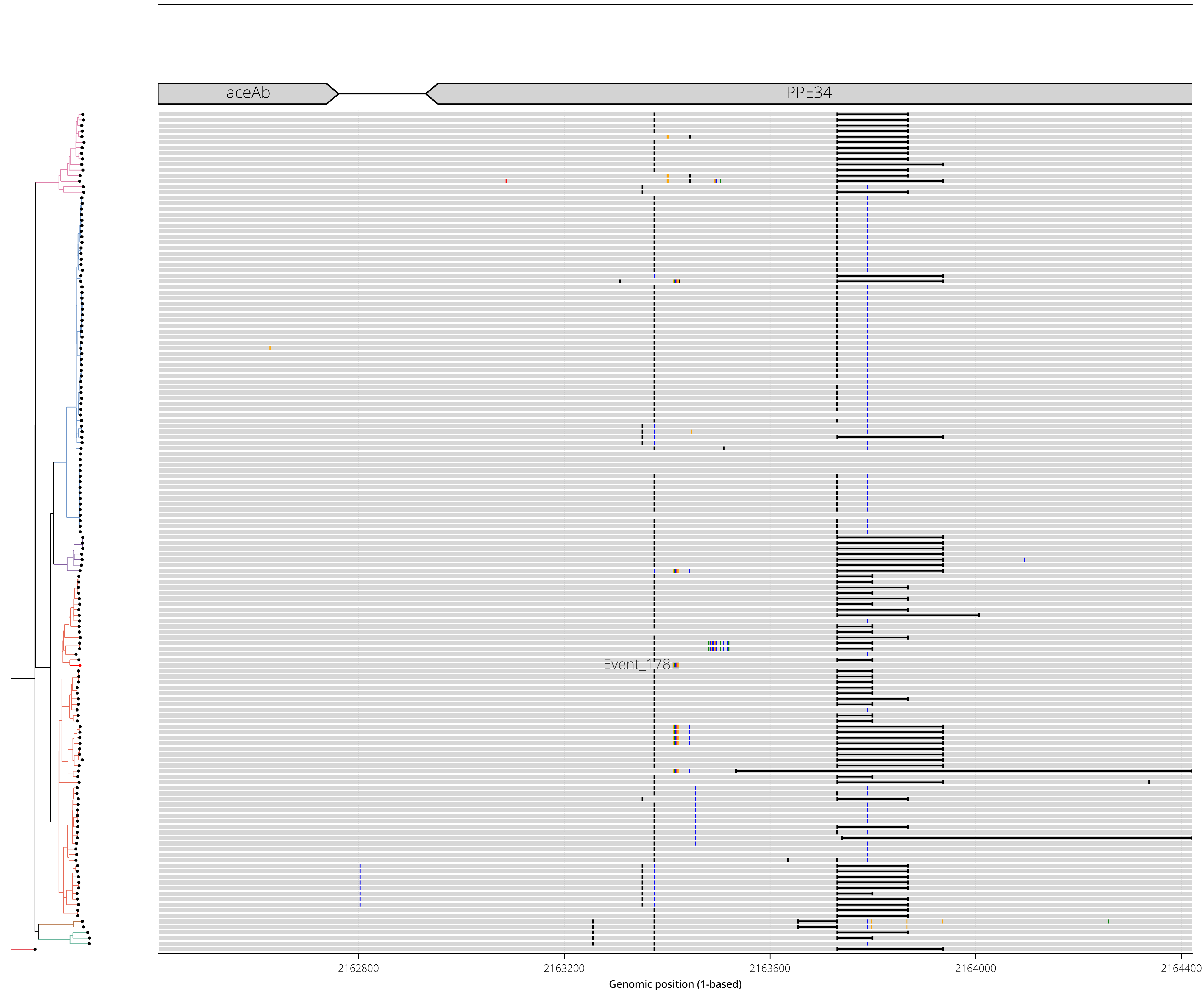

Individual GCE + Known Paralogs
GCE ID: Event_179

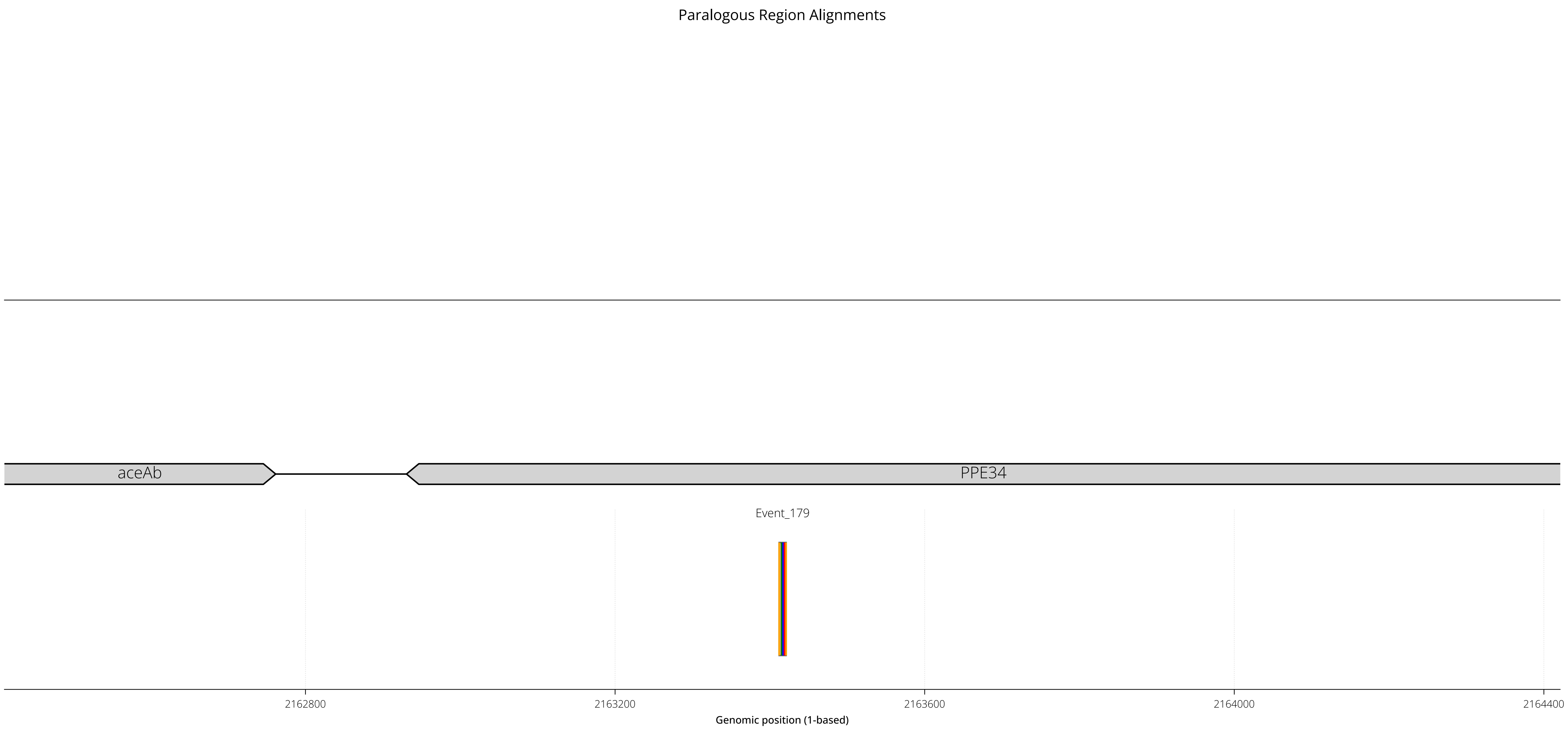

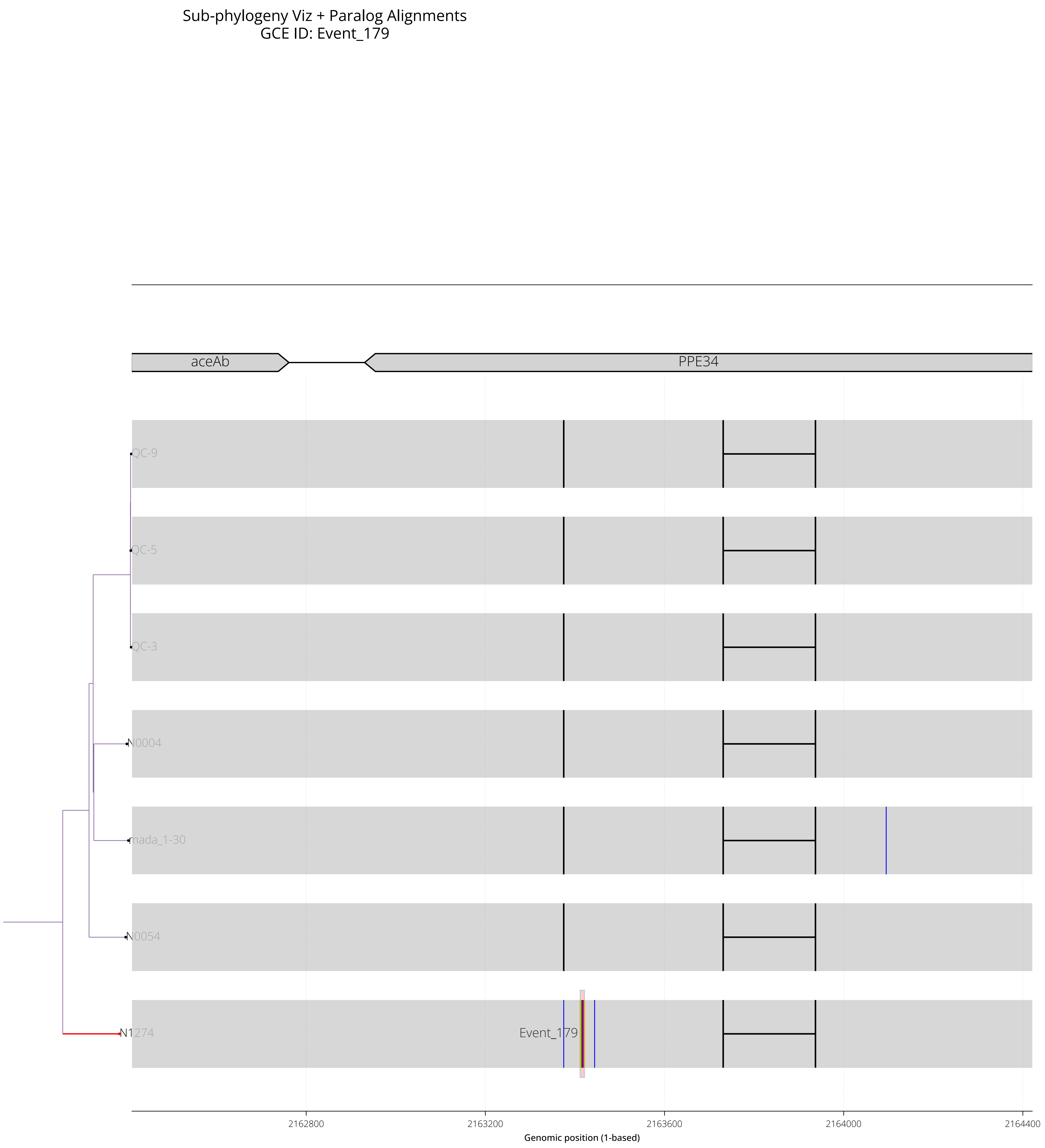

Sub-phylogeny Viz + Paralog Alignments
GCE ID: Event_179
aceAb
PPE34
QC-9
QC-5
QC-3
N0004
mada_1-30
N0054
N1274
Event_179
2162800
2163200
2163600
2164000
2164400
Genomic position (1-based)

Full Phylogeny Viz + Paralog Alignments
GCE ID: Event_179

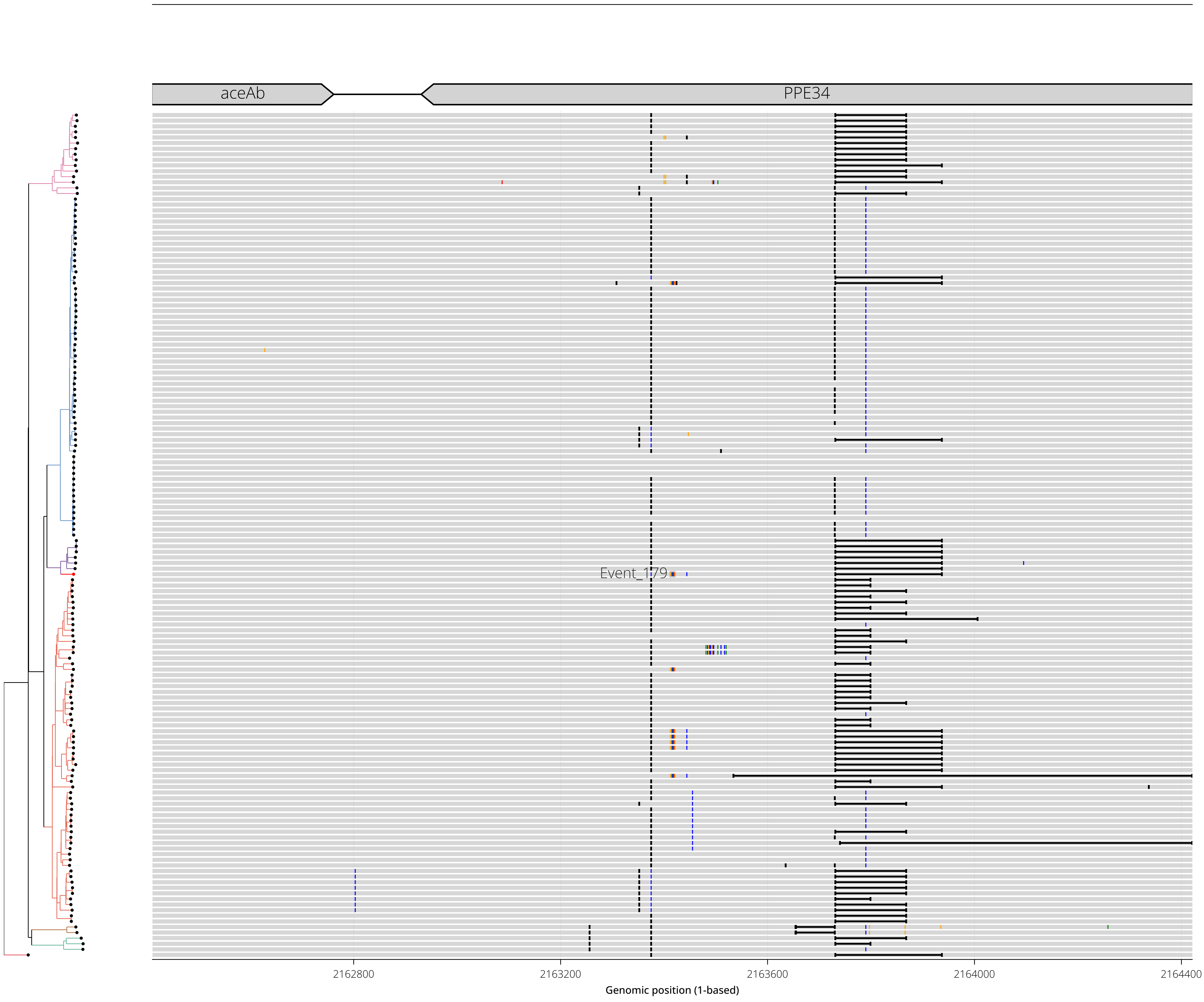

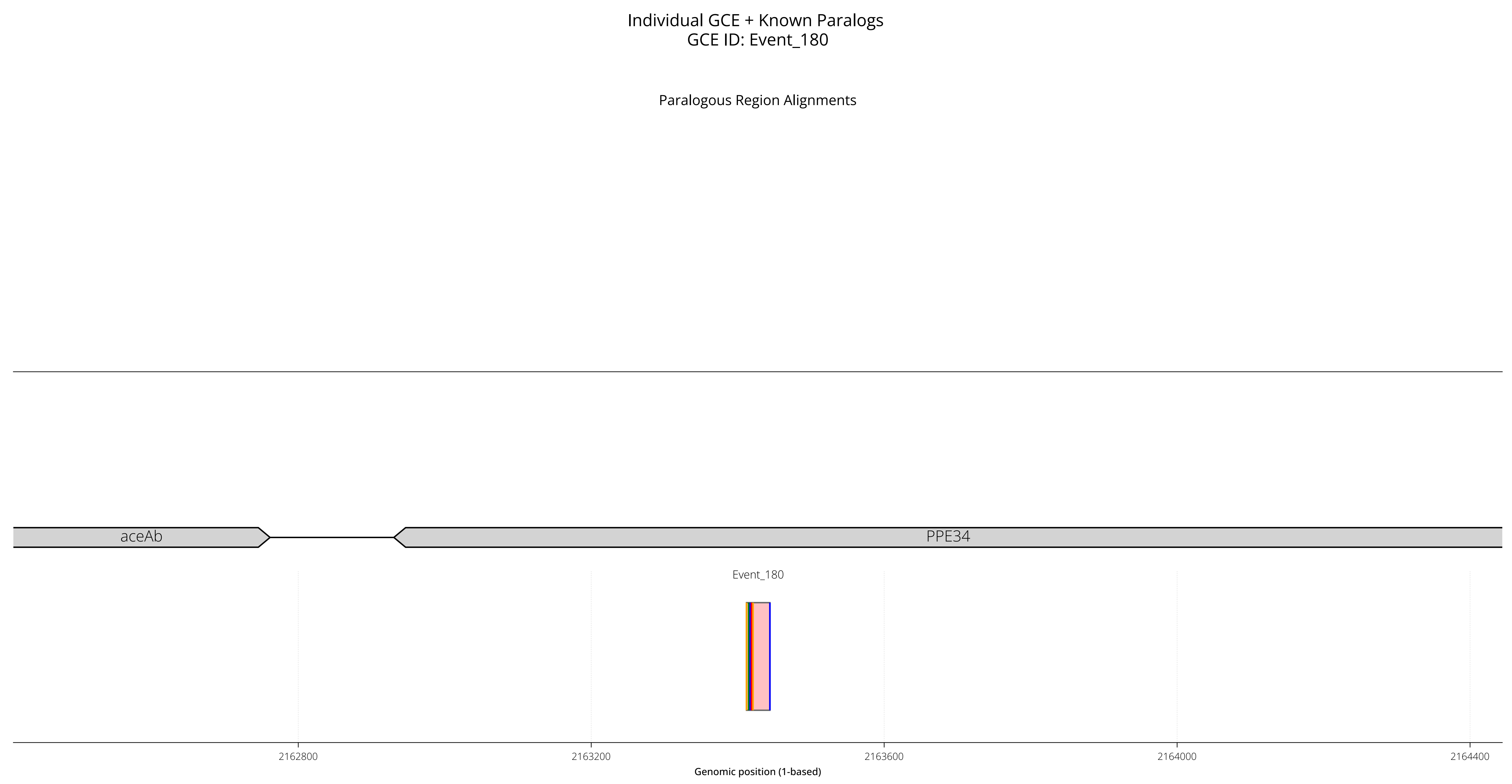
Individual GCE + Known Paralogs
GCE ID: Event_180
Paralogous Region Alignments
aceAb
PPE34
Event_180
2162800
2163200
2163600
2164000
2164400
Genomic position (1-based)

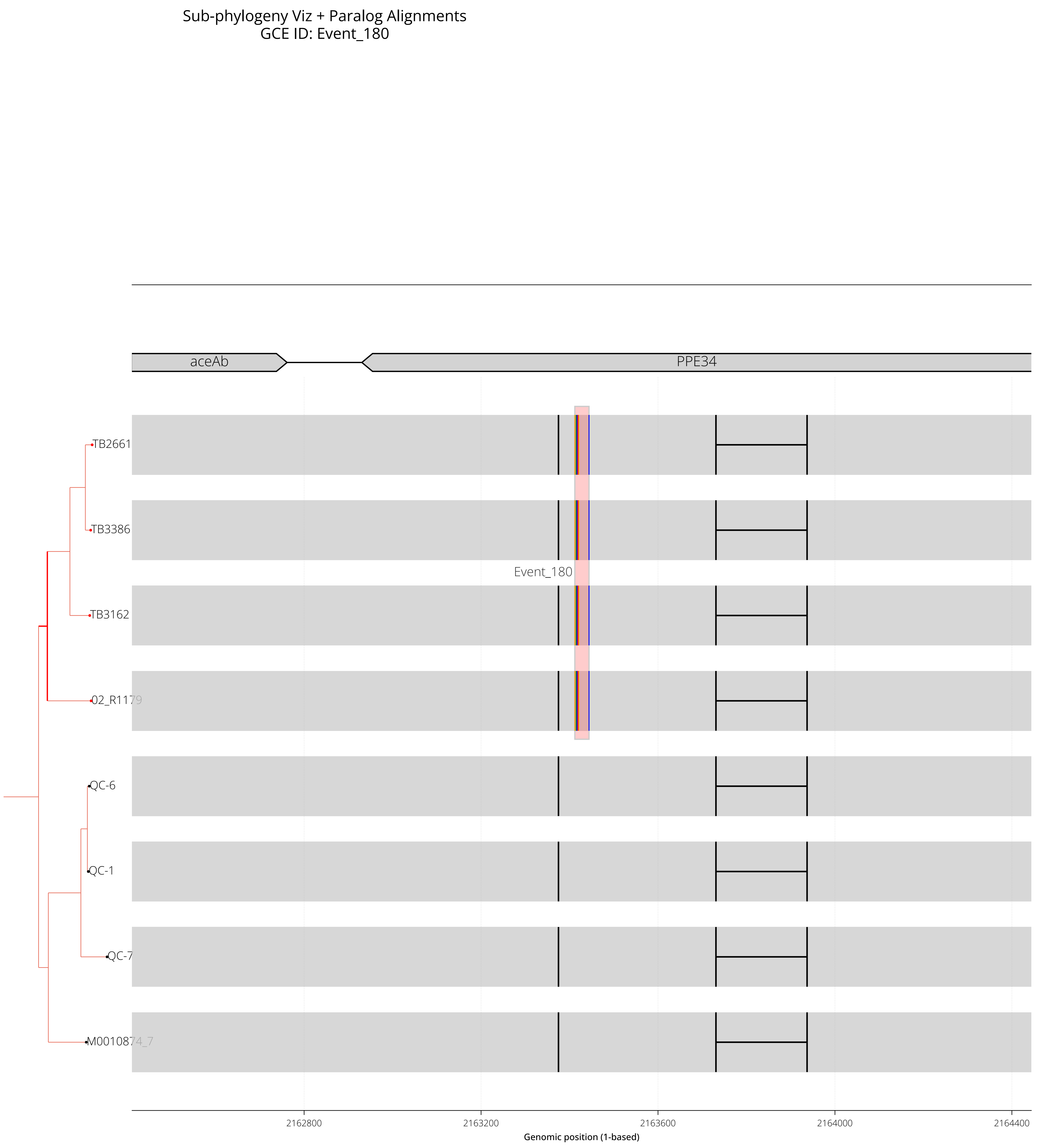

Sub-phylogeny Viz + Paralog Alignments
GCE ID: Event_180
aceAb
PPE34
TB2661
TB3386
Event_180
TB3162
02_R1179
QC-6
QC-1
QC-7
M0010874_7
2162800
2163200
2163600
2164000
2164400
Genomic position (1-based)

Full Phylogeny Viz + Paralog Alignments
GCE ID: Event_180

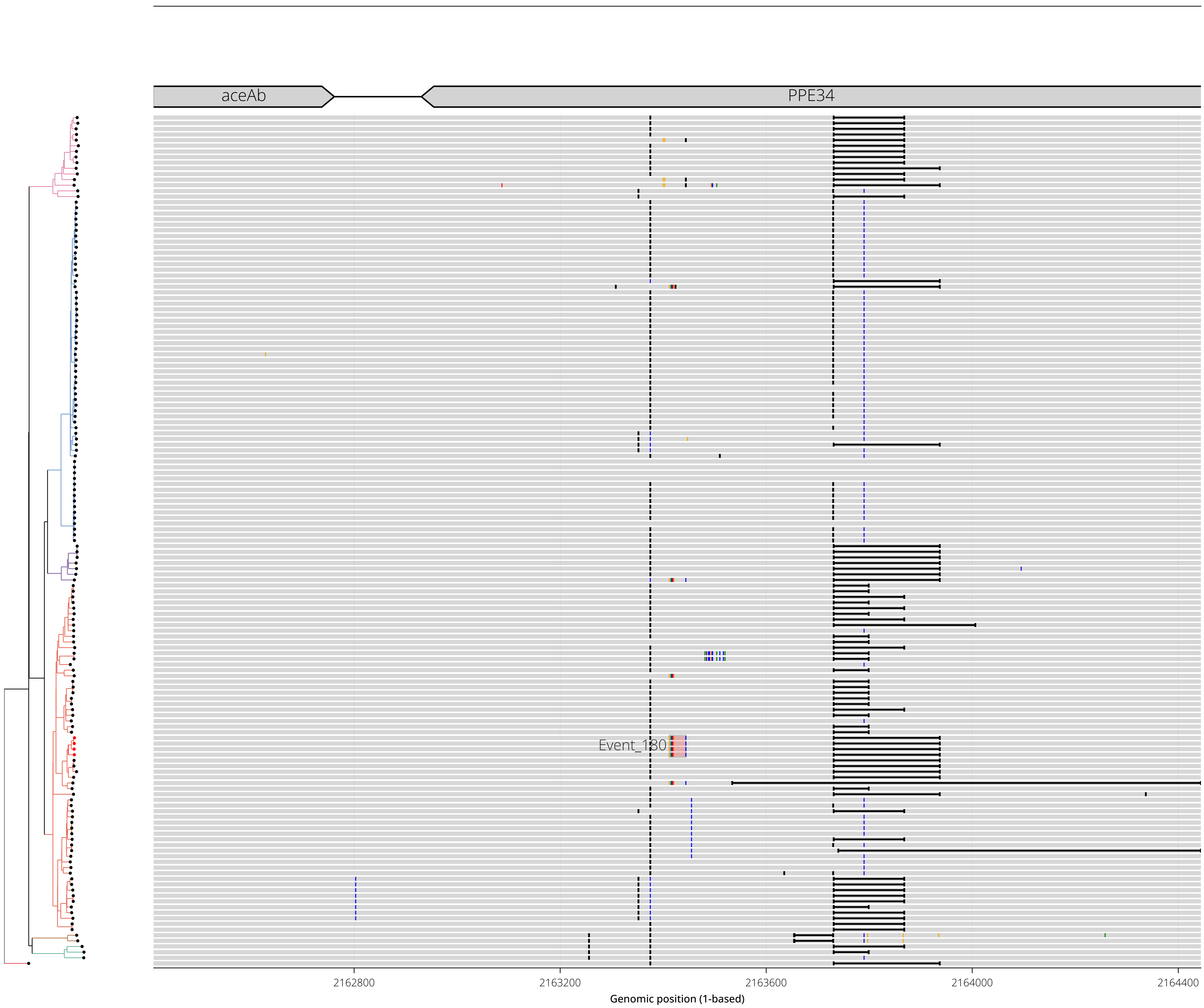

Individual GCE + Known Paralogs
GCE ID: Event_181

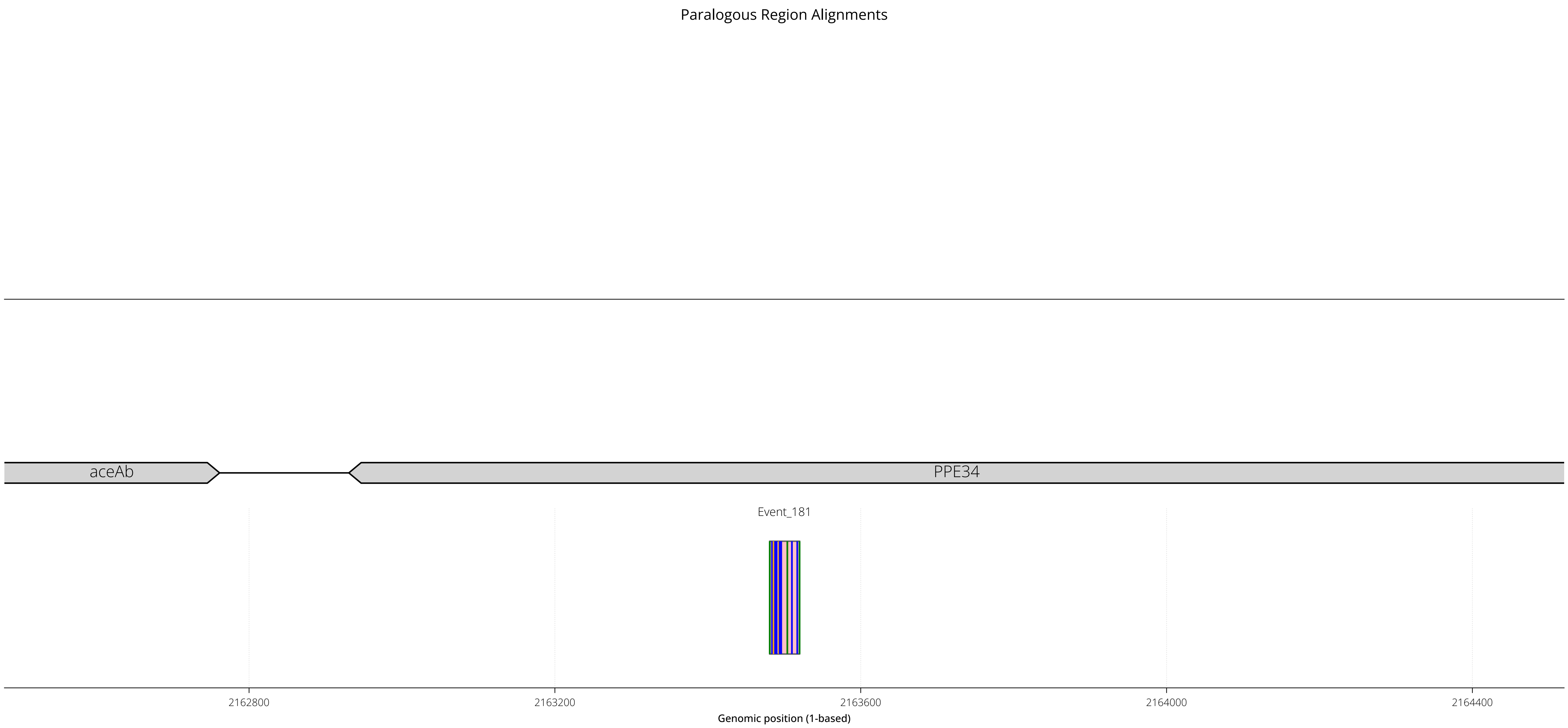

Sub-phylogeny Viz + Paralog Alignments
GCE ID: Event_181

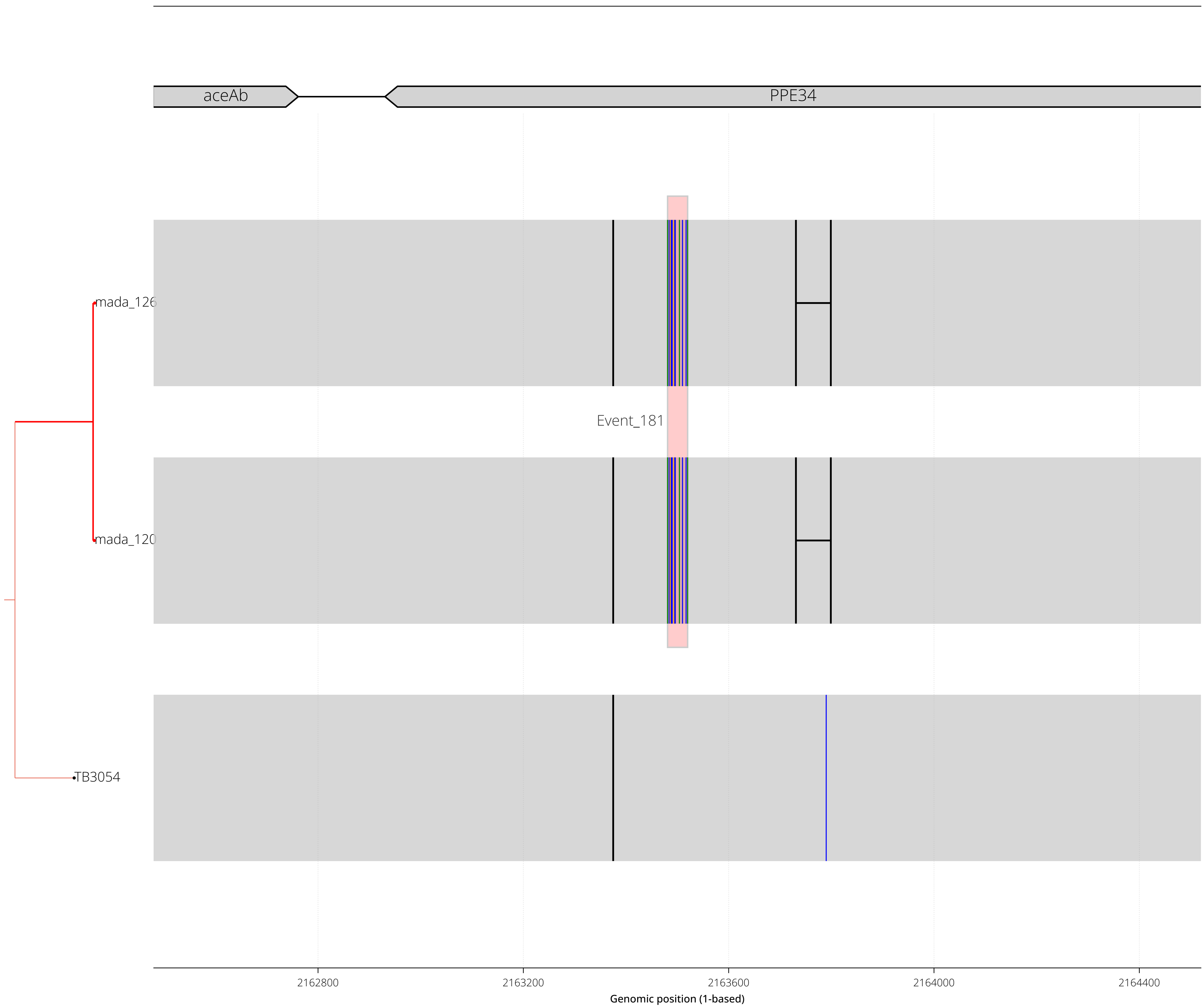

Full Phylogeny Viz + Paralog Alignments
GCE ID: Event_181

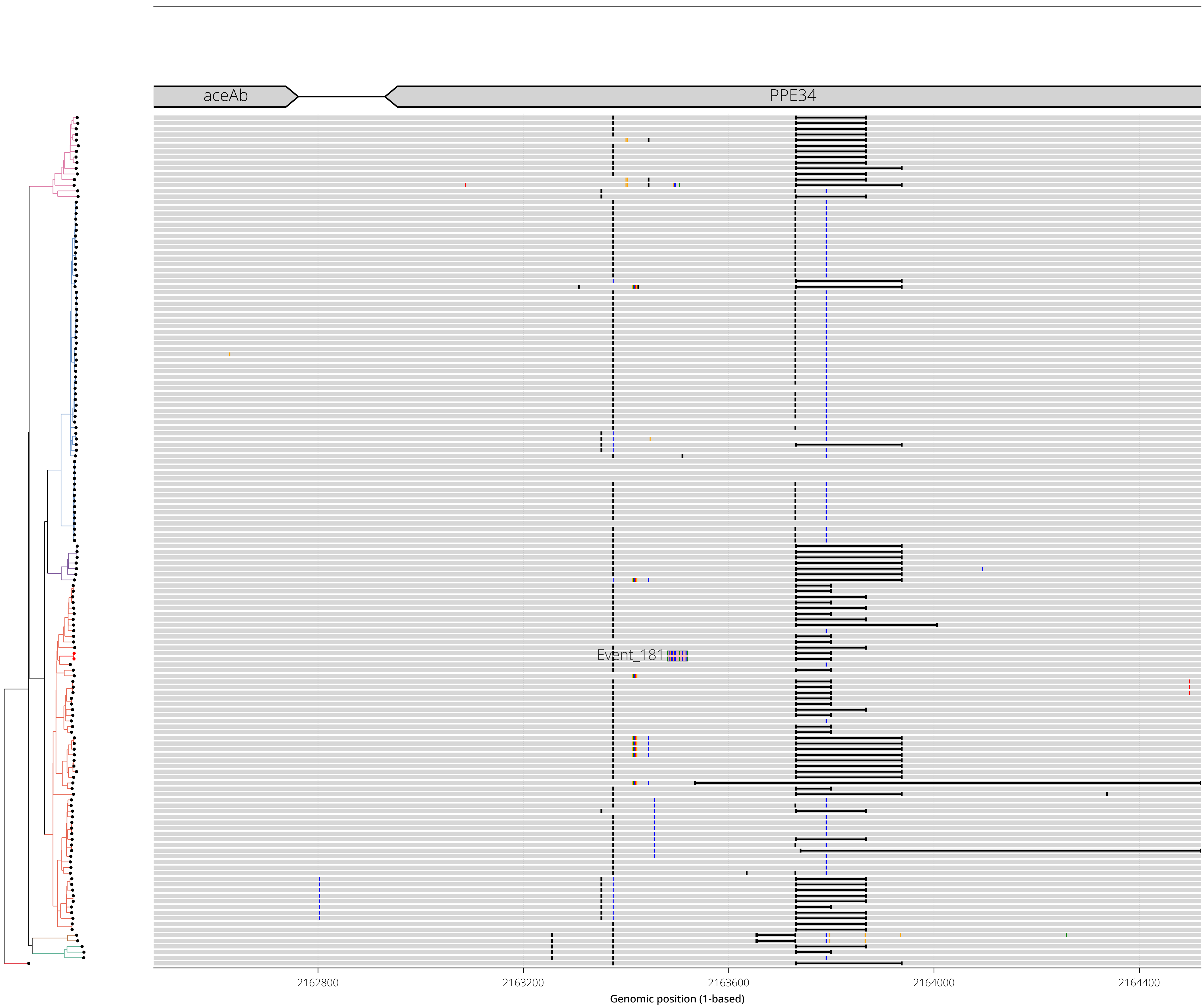

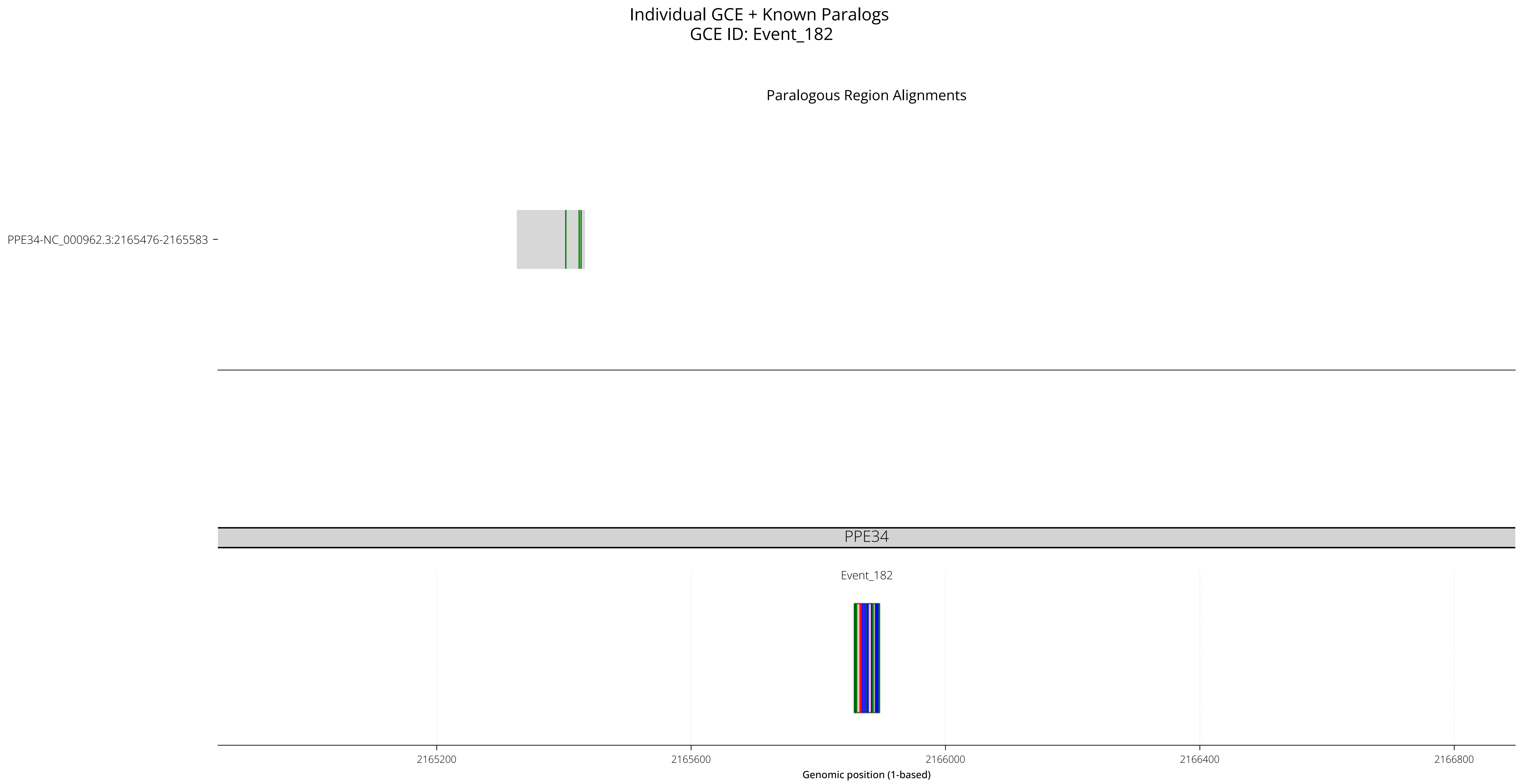

Individual GCE + Known Paralogs
GCE ID: Event_182
Paralogous Region Alignments
PPE34-NC_000962.3:2165476-2165583
PPE34
Event_182
2165200
2165600
2166000
2166400
2166800
Genomic position (1-based)

Sub-phylogeny Viz + Paralog Alignments
GCE ID: Event_182

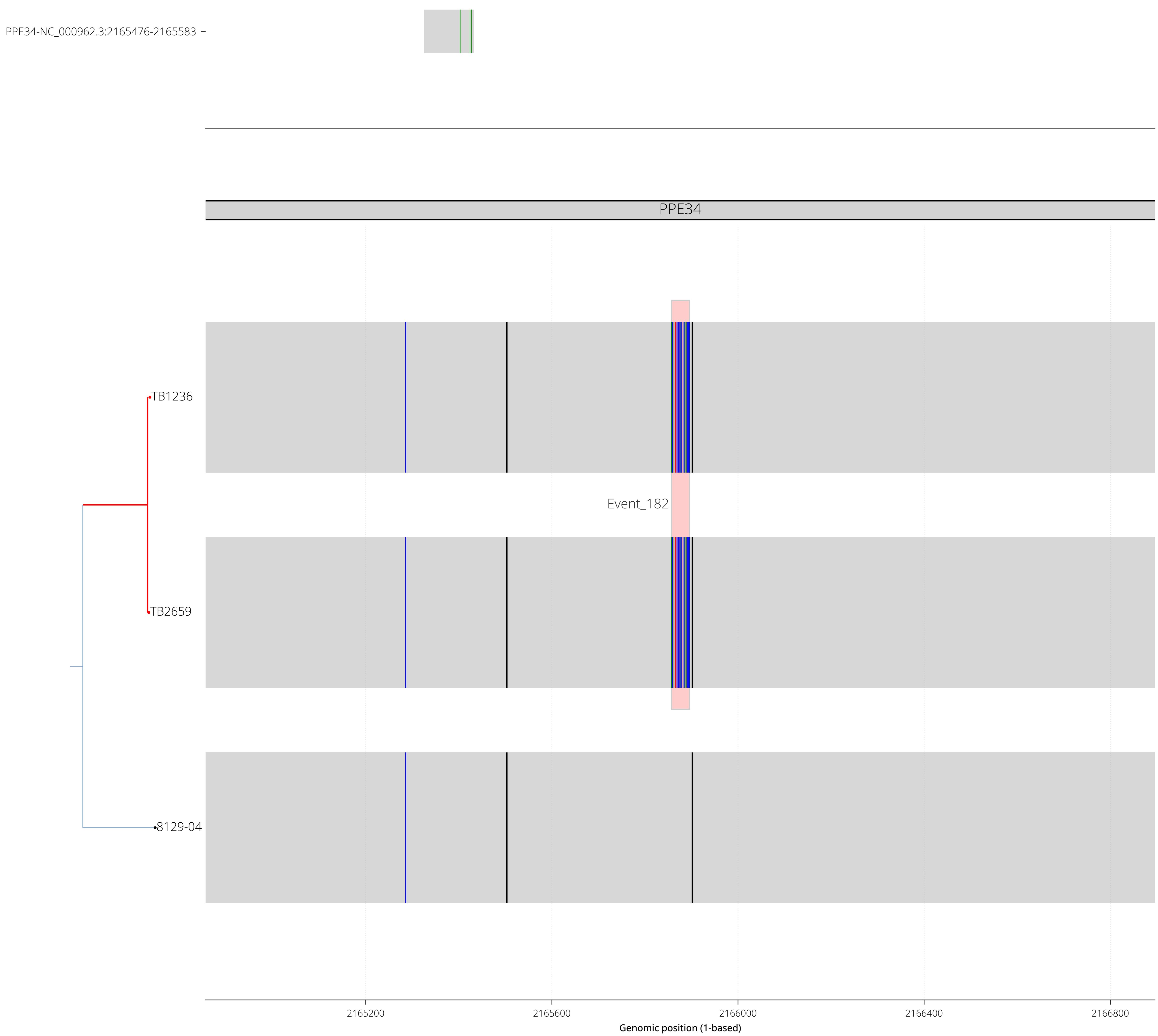

### Full Phylogeny Viz + Paralog Alignments
GCE ID: Event_182

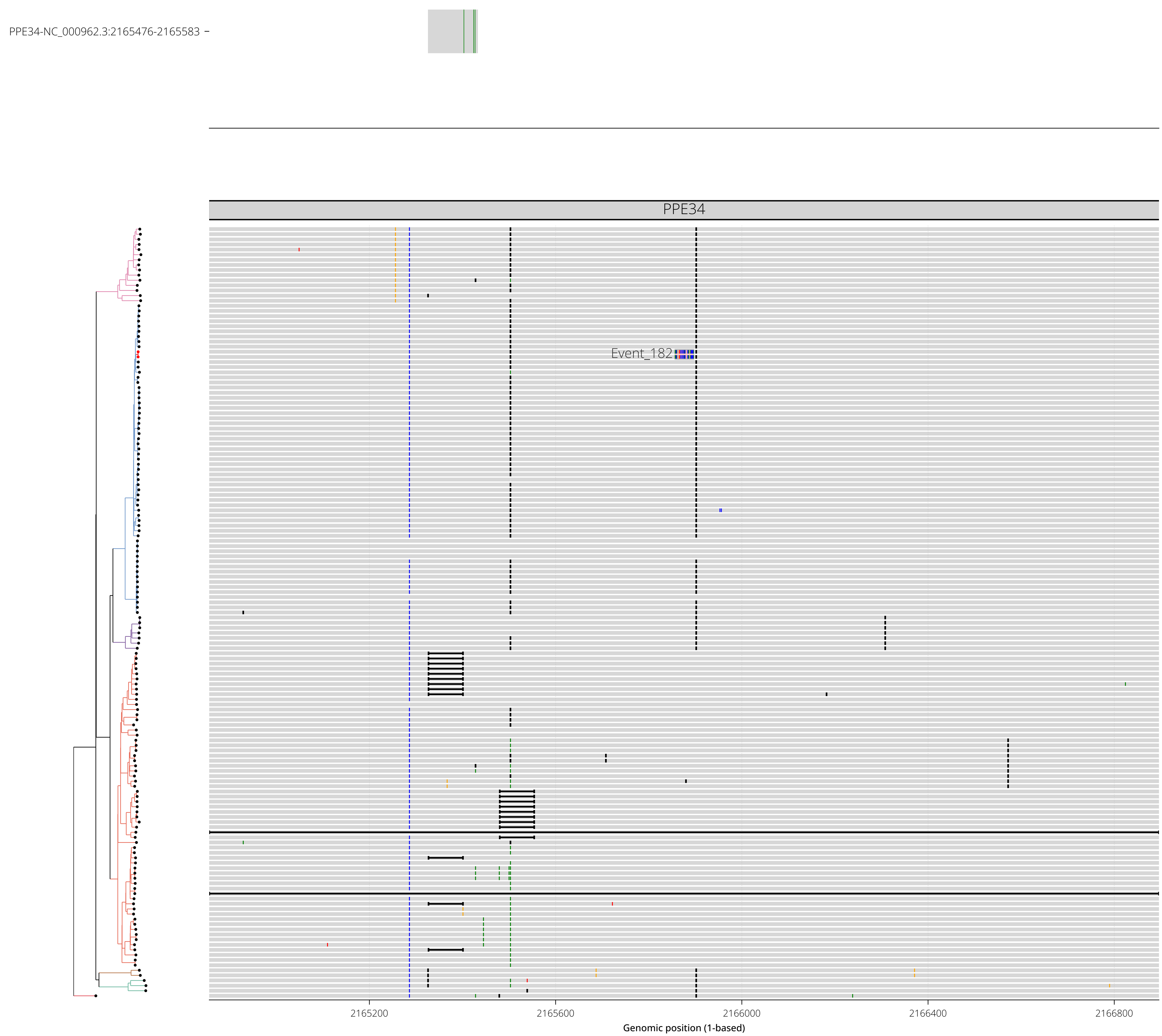

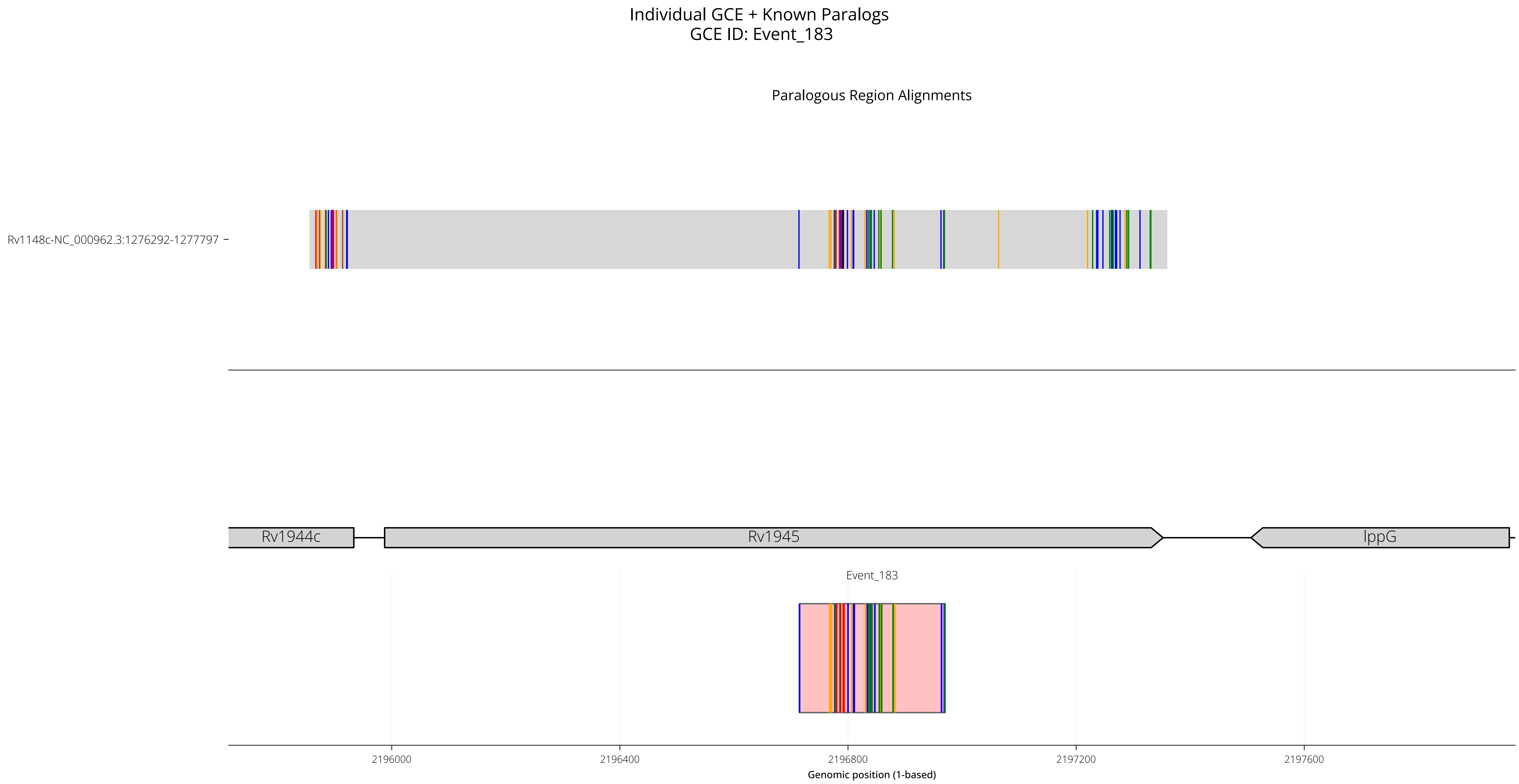
Individual GCE + Known Paralogs
GCE ID: Event_183
Paralogous Region Alignments
Rv1148c-NC_000962.3:1276292-1277797
Rv1944c
Rv1945
lppG
Event_183
2196000
2196400
2196800
2197200
2197600
Genomic position (1-based)

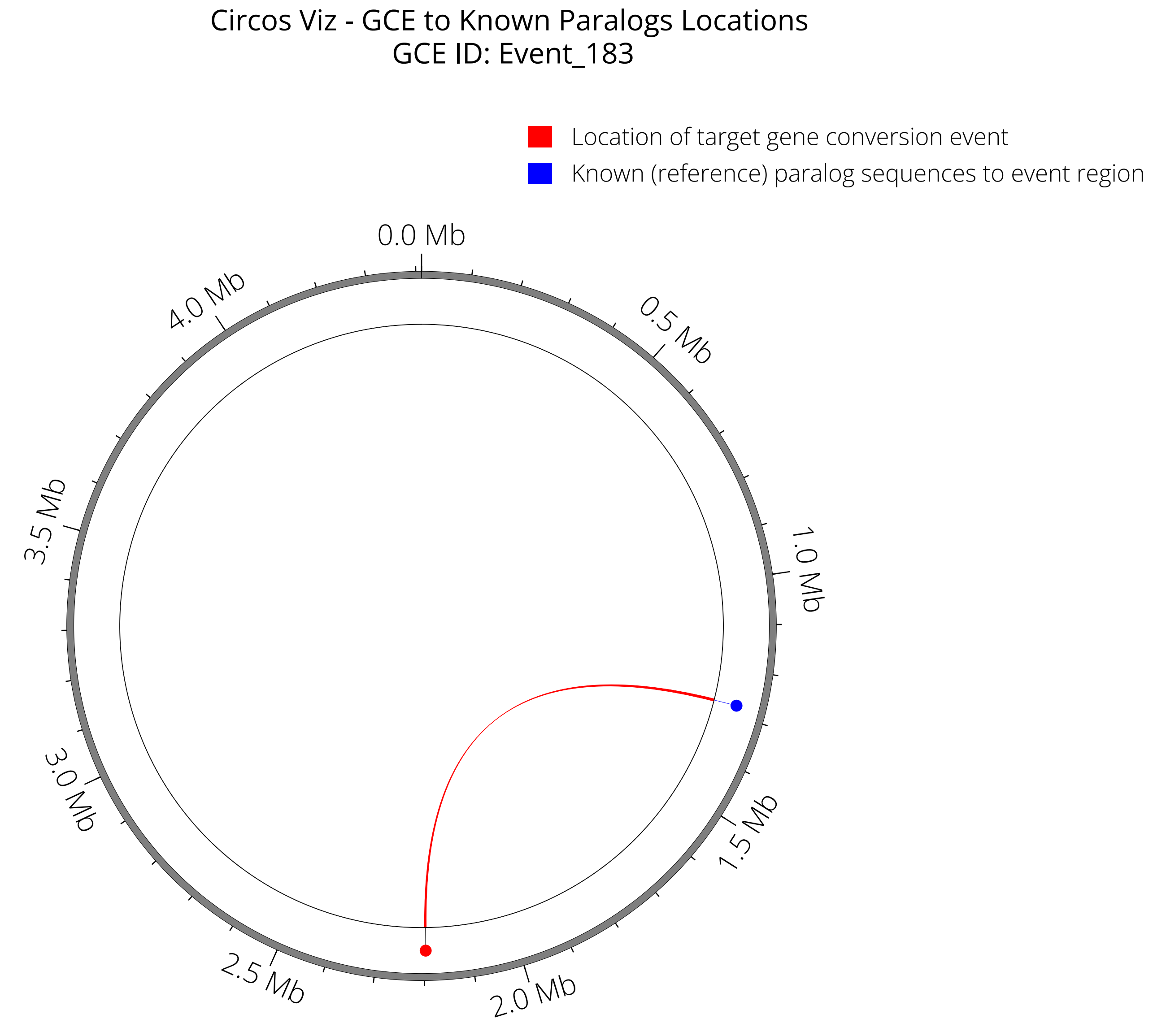
Circos Viz - GCE to Known Paralogs Locations
GCE ID: Event_183
Location of target gene conversion event
Known (reference) paralog sequences to event region
0.0 Mb
0.5 Mb
1.0 Mb
1.5 Mb
2.0 Mb
2.5 Mb
3.0 Mb
3.5 Mb
4.0 Mb

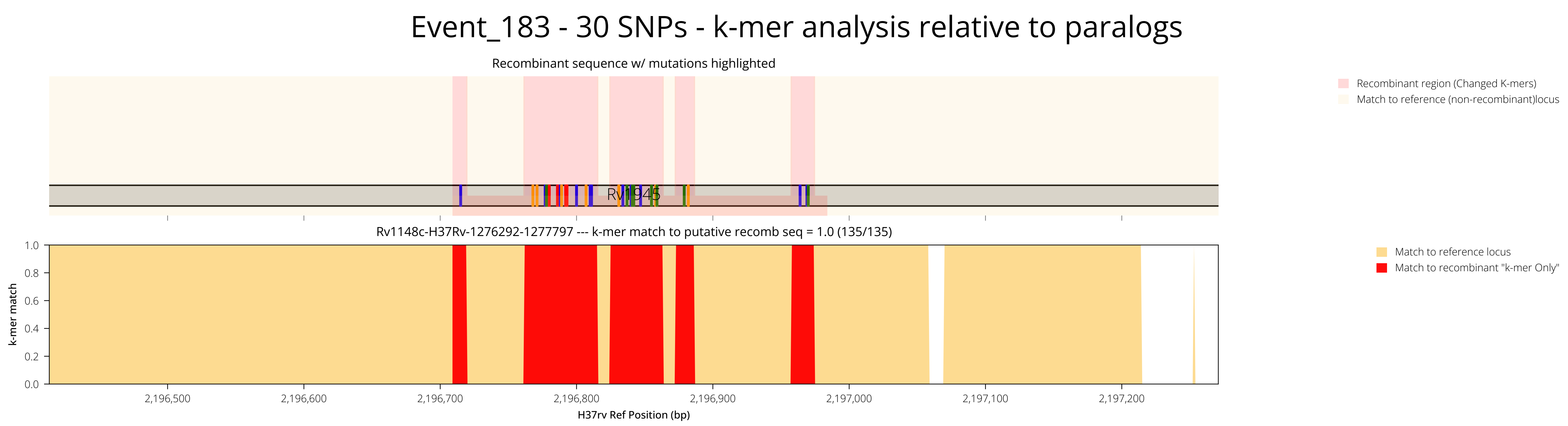

Event_183 - 30 SNPs - k-mer analysis relative to paralogs
Recombinant sequence w/ mutations highlighted
Rv1945
Recombinant region (Changed K-mers)
Match to reference (non-recombinant)locus
Rv1148c-H37Rv-1276292-1277797 --- k-mer match to putative recomb seq = 1.0 (135/135)
Match to reference locus
Match to recombinant "k-mer Only"
k-mer match
1.0
0.8
0.6
0.4
0.2
0.0
2,196,500
2,196,600
2,196,700
2,196,800
2,196,900
2,197,000
2,197,100
2,197,200
H37rv Ref Position (bp)

Sub-phylogeny Viz + Paralog Alignments
GCE ID: Event_183

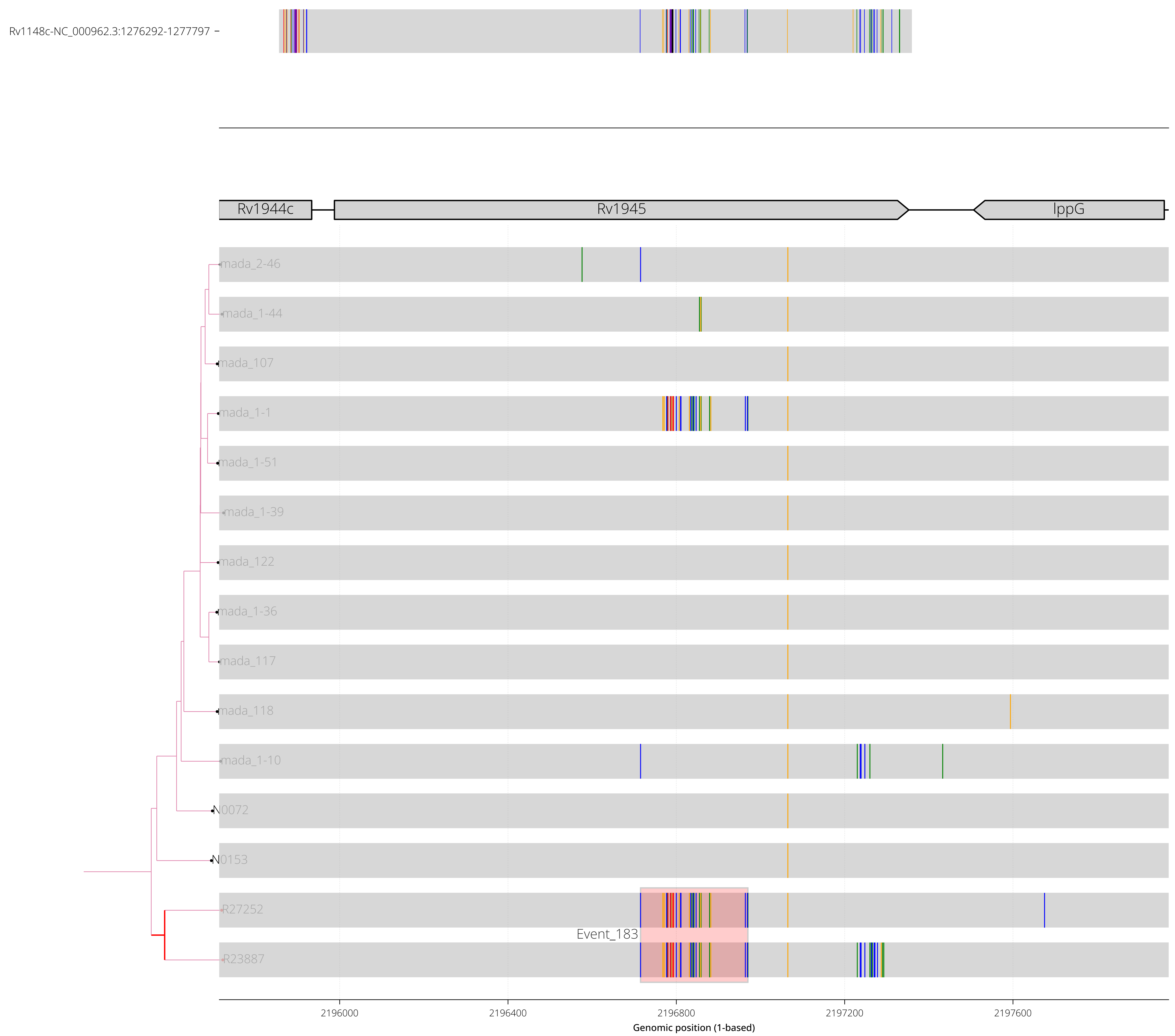

Full Phylogeny Viz + Paralog Alignments
GCE ID: Event_183

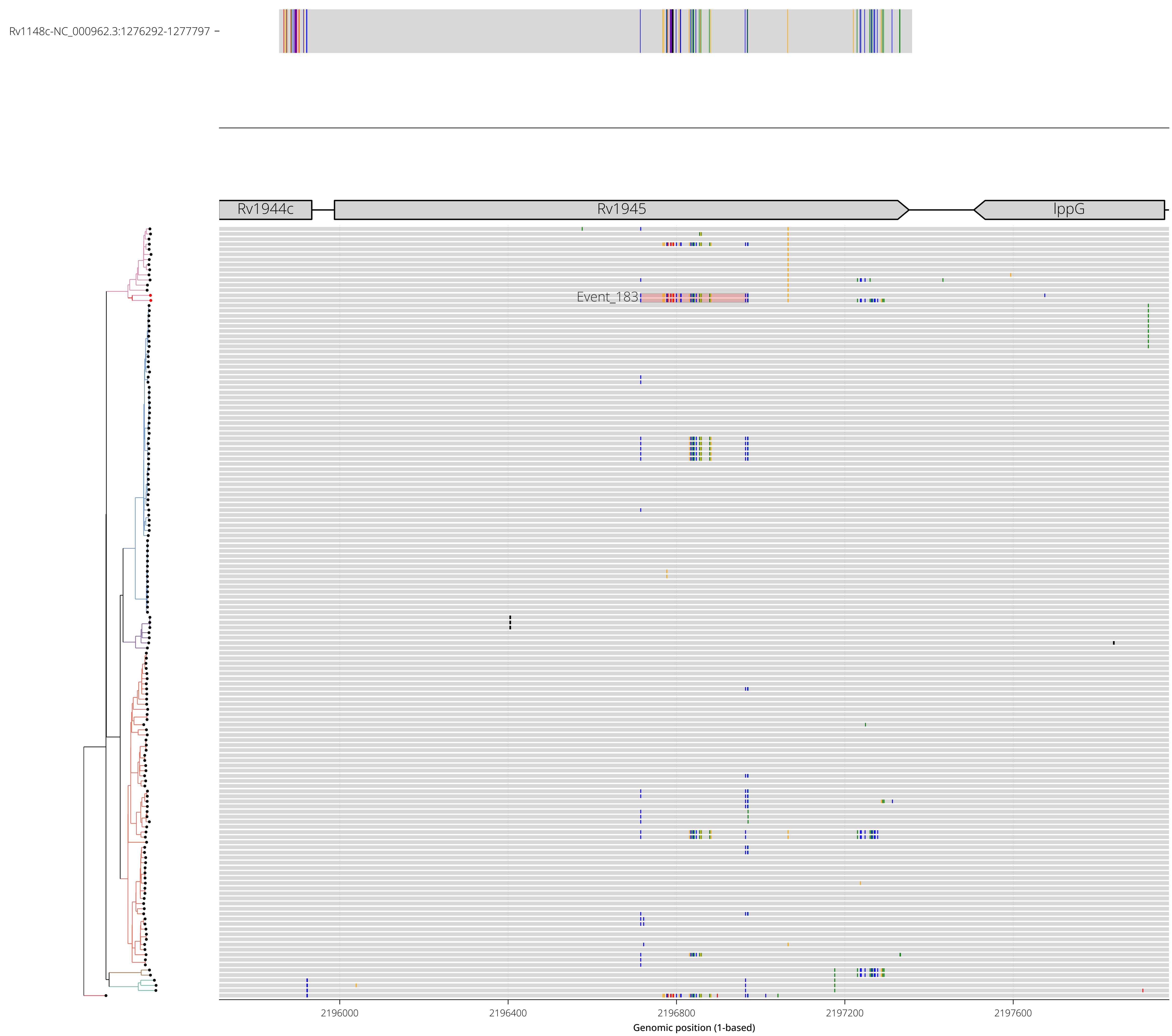

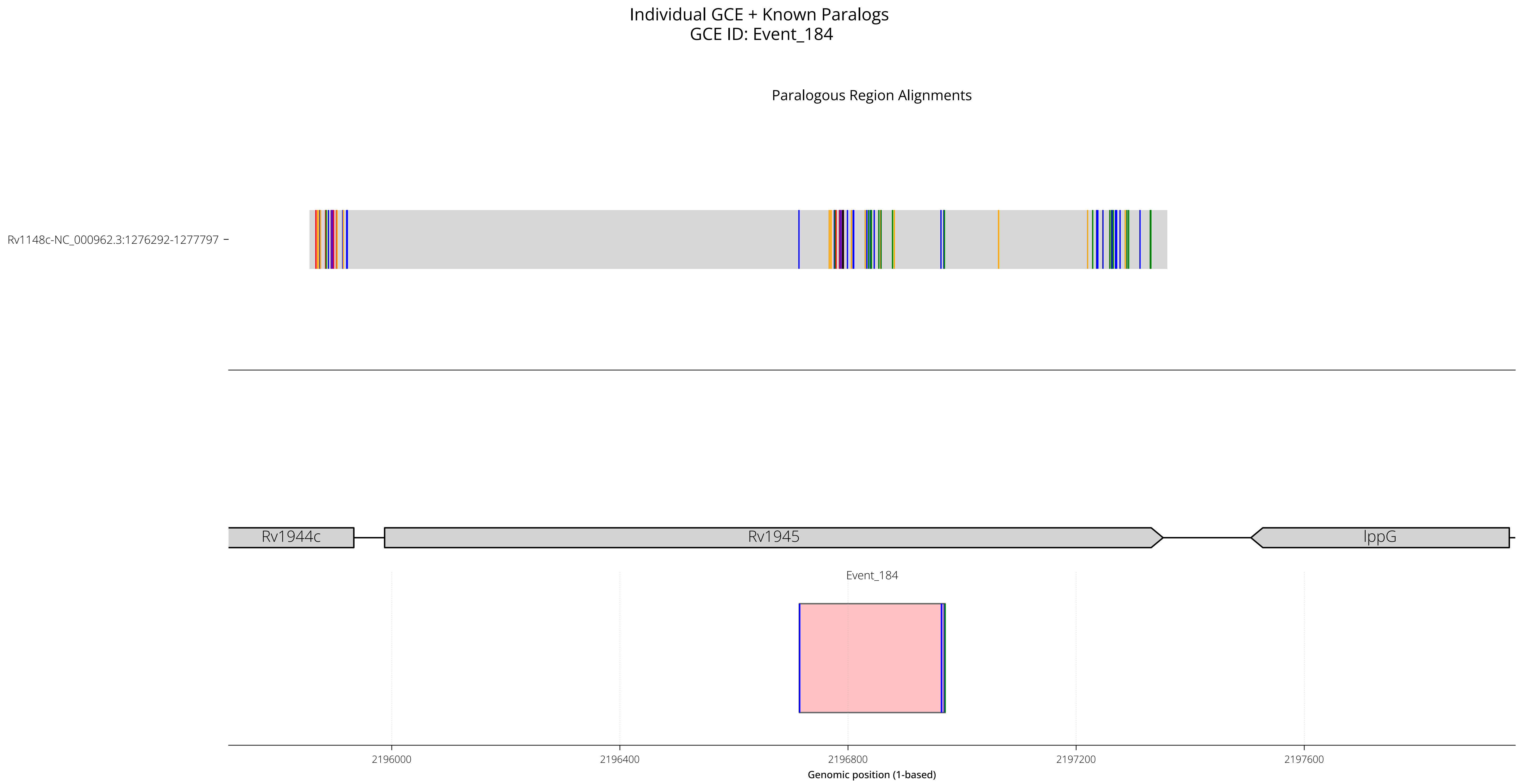
Individual GCE + Known Paralogs
GCE ID: Event_184
Paralogous Region Alignments
Rv1148c-NC_000962.3:1276292-1277797
Rv1944c
Rv1945
lppG
Event_184
2196000
2196400
2196800
2197200
2197600
Genomic position (1-based)

Circos Viz - GCE to Known Paralogs Locations
GCE ID: Event_184

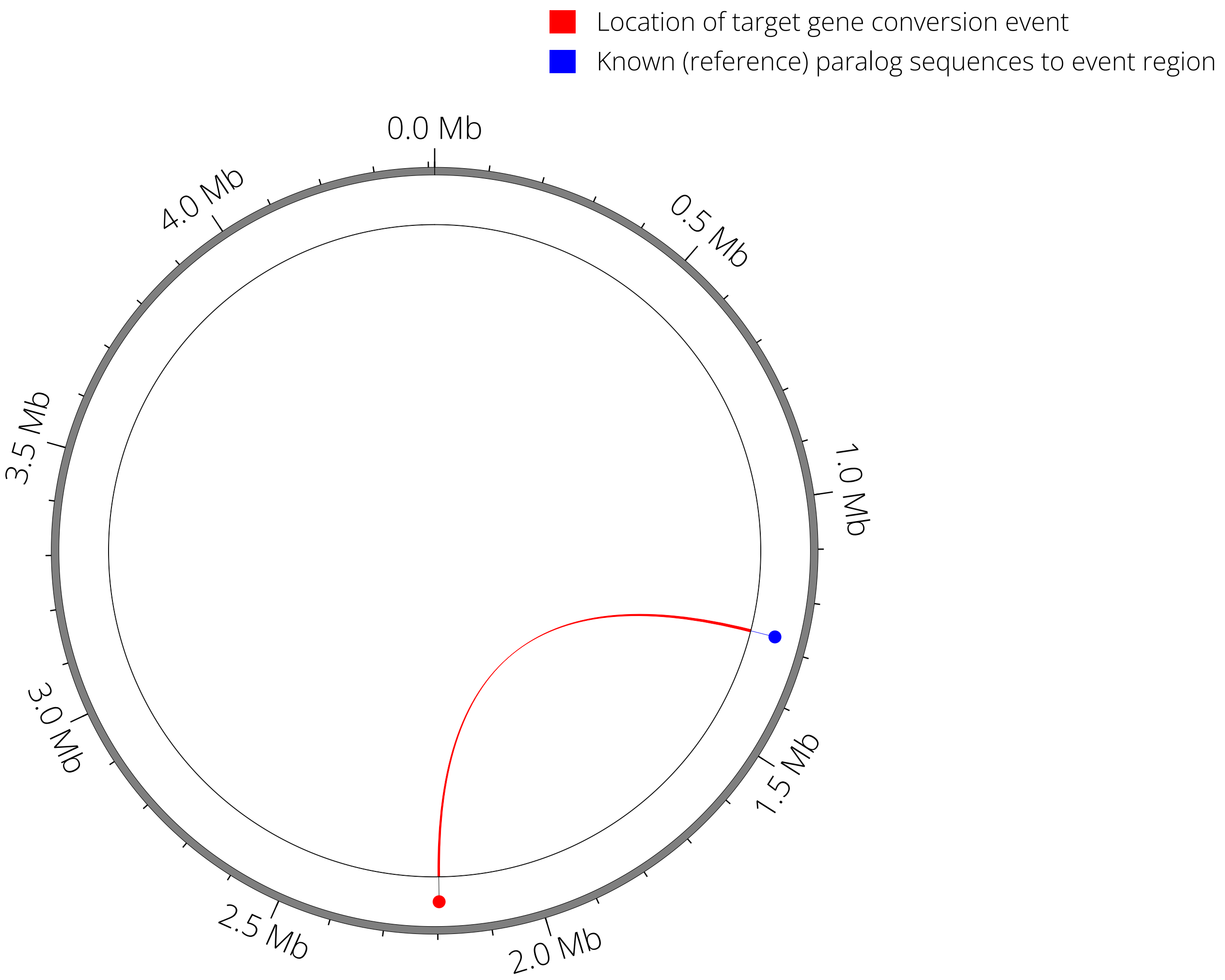

### Event_184 - 4 SNPs - k-mer analysis relative to paralogs

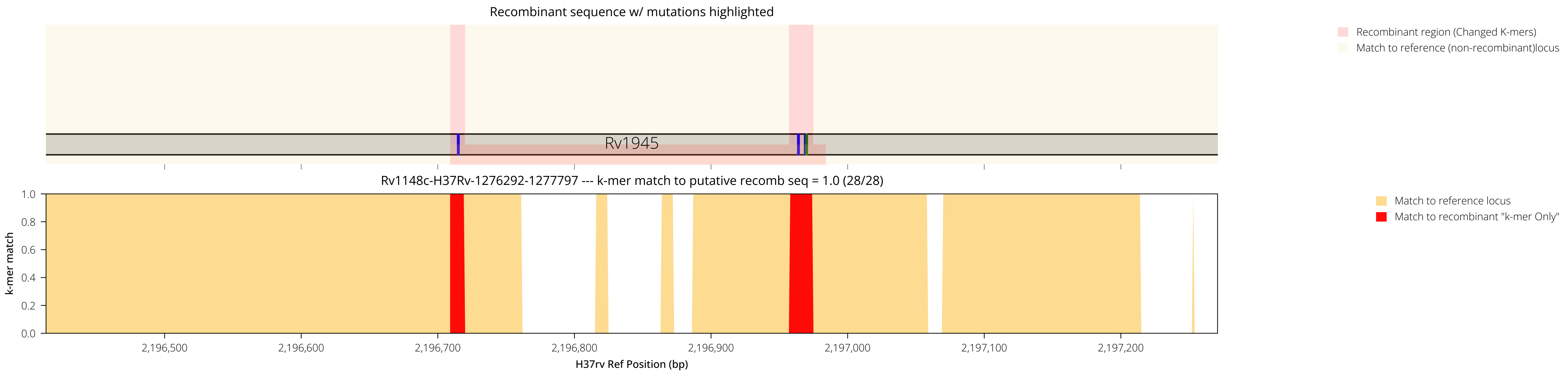

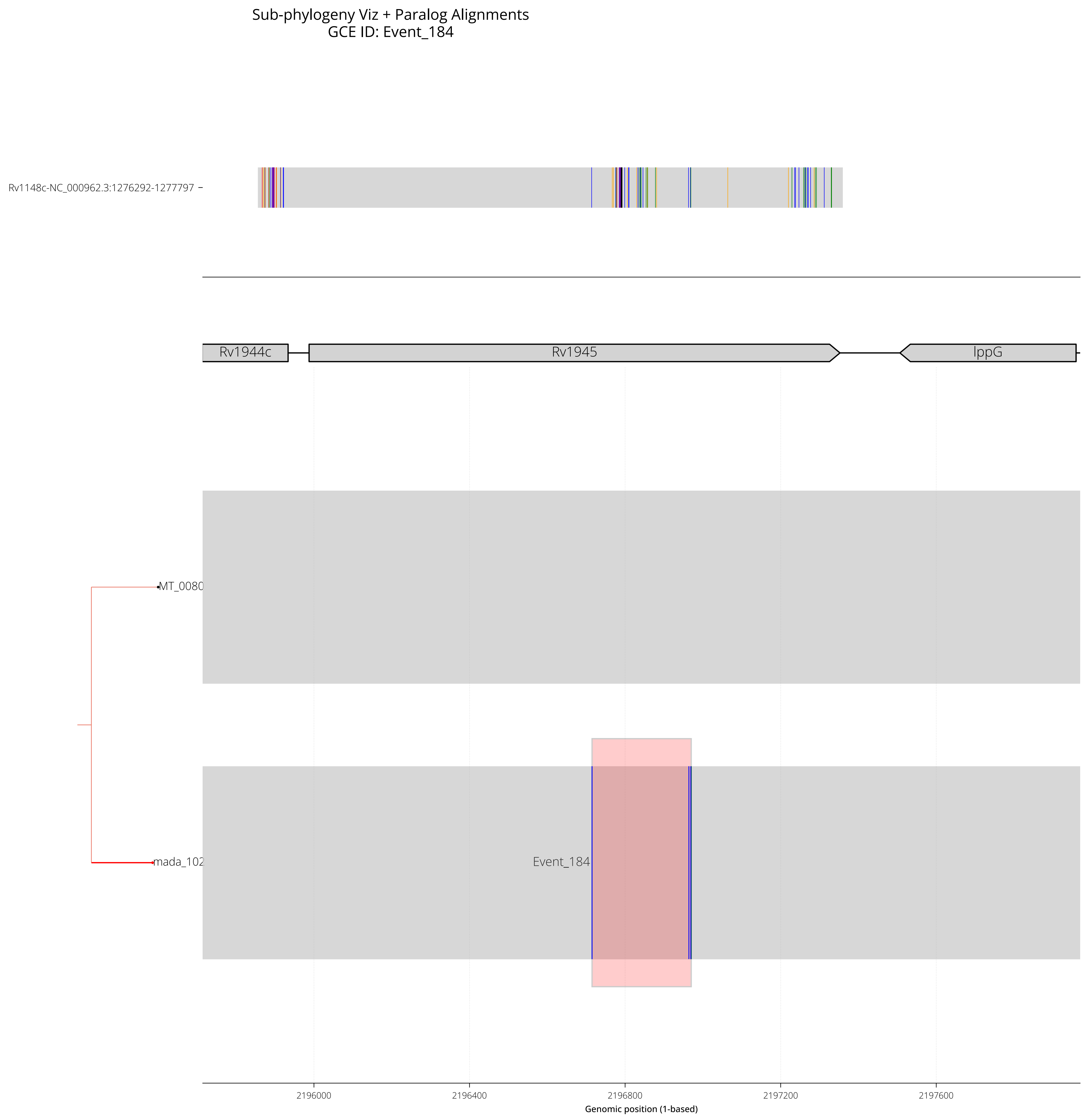
Sub-phylogeny Viz + Paralog Alignments
GCE ID: Event_184
Rv1148c-NC_000962.3:1276292-1277797
Rv1944c
Rv1945
lppG
MT_0080
mada_102
Event_184
2196000
2196400
2196800
2197200
2197600
Genomic position (1-based)

Full Phylogeny Viz + Paralog Alignments
GCE ID: Event_184

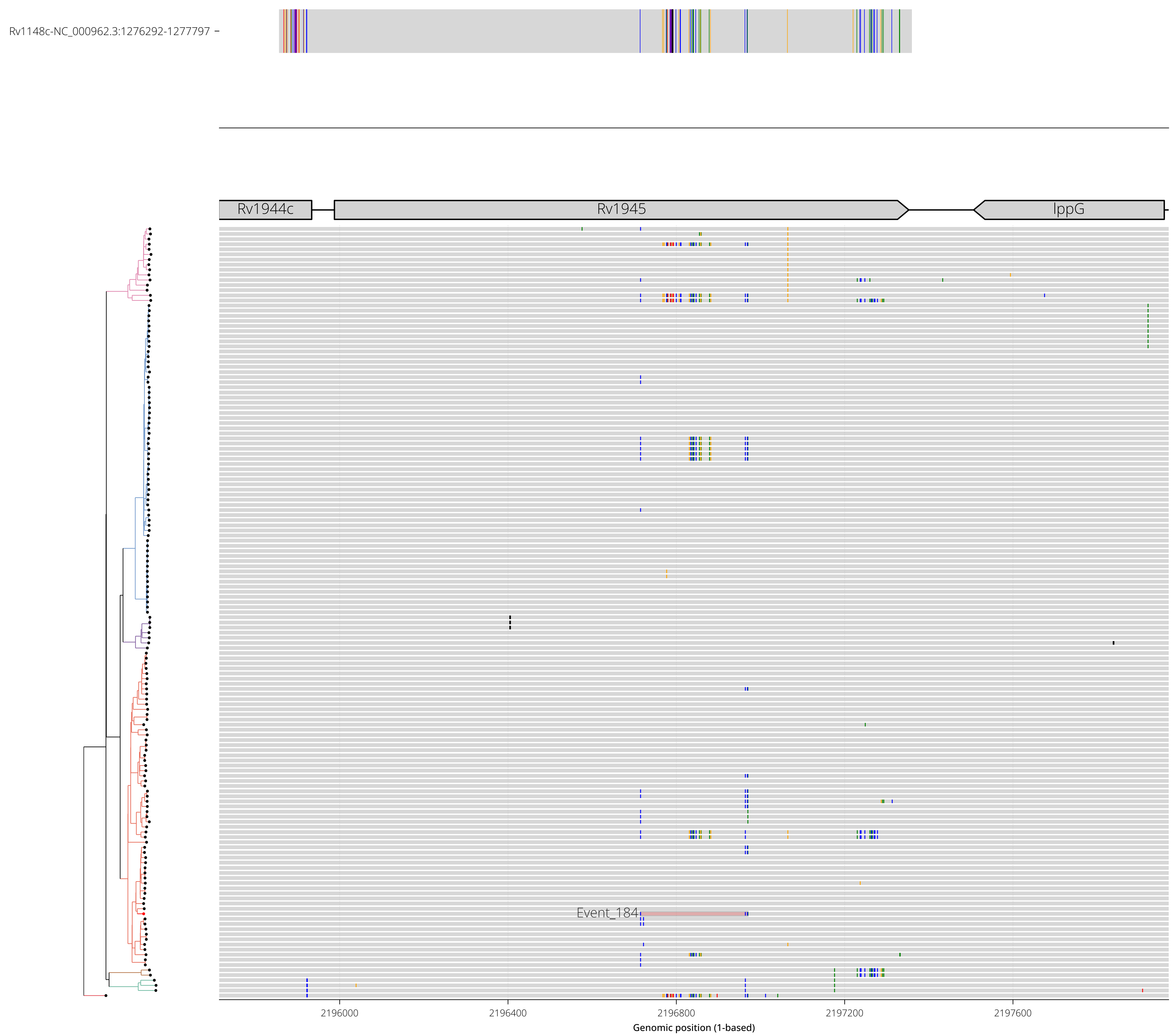

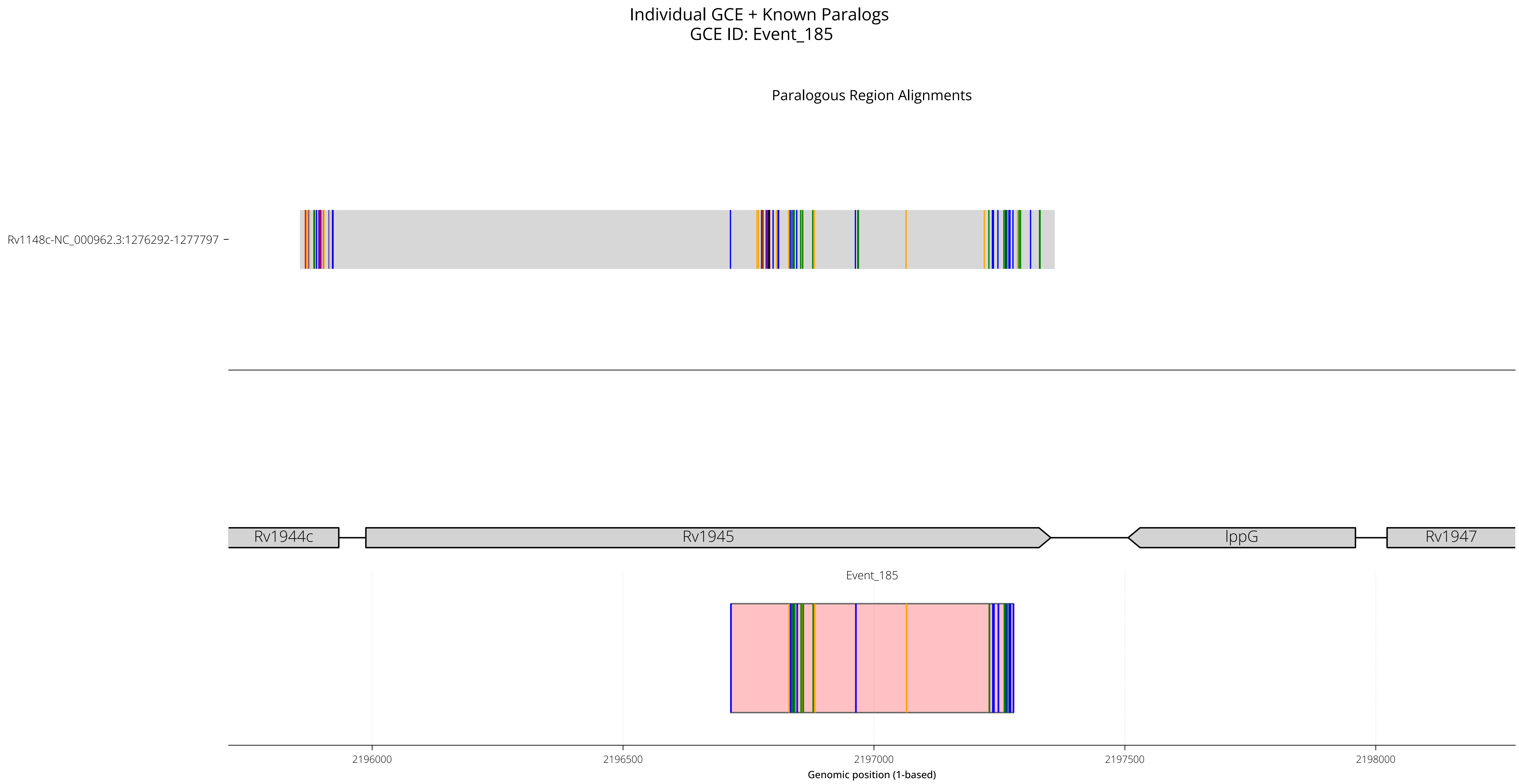

Individual GCE + Known Paralogs
GCE ID: Event_185
Paralogous Region Alignments
Rv1148c-NC_000962.3:1276292-1277797
Rv1944c
Rv1945
lppG
Rv1947
Event_185
2196000
2196500
2197000
2197500
2198000
Genomic position (1-based)

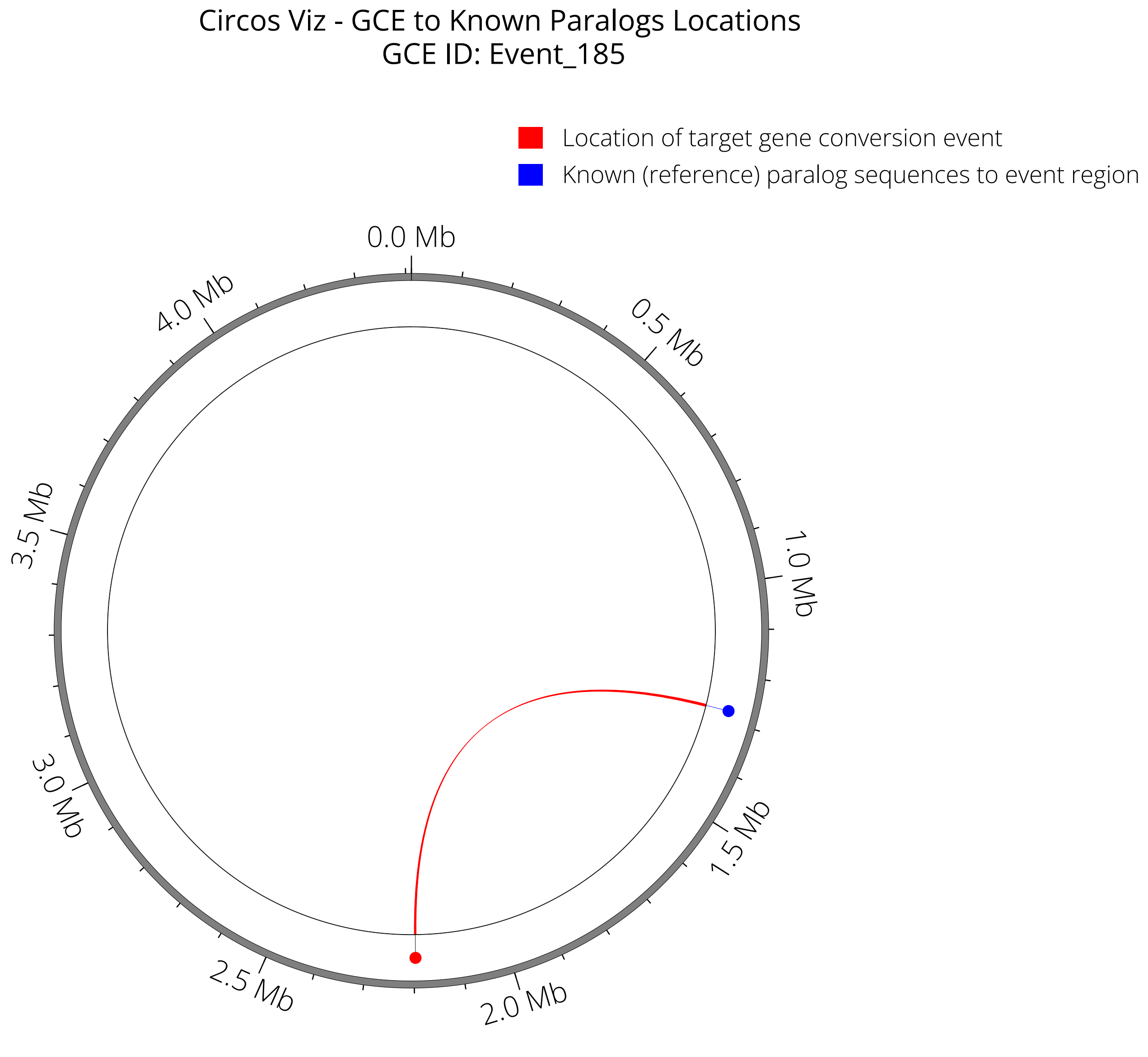
Circos Viz - GCE to Known Paralogs Locations
GCE ID: Event_185
Location of target gene conversion event
Known (reference) paralog sequences to event region
0.0 Mb
0.5 Mb
1.0 Mb
1.5 Mb
2.0 Mb
2.5 Mb
3.0 Mb
3.5 Mb
4.0 Mb

### Event_185 - 26 SNPs - k-mer analysis relative to paralogs

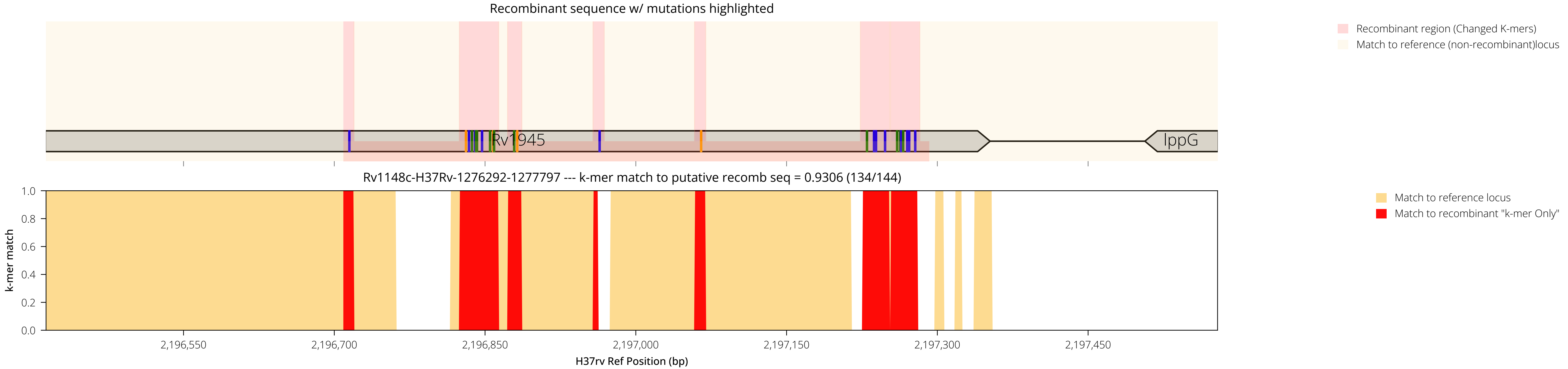

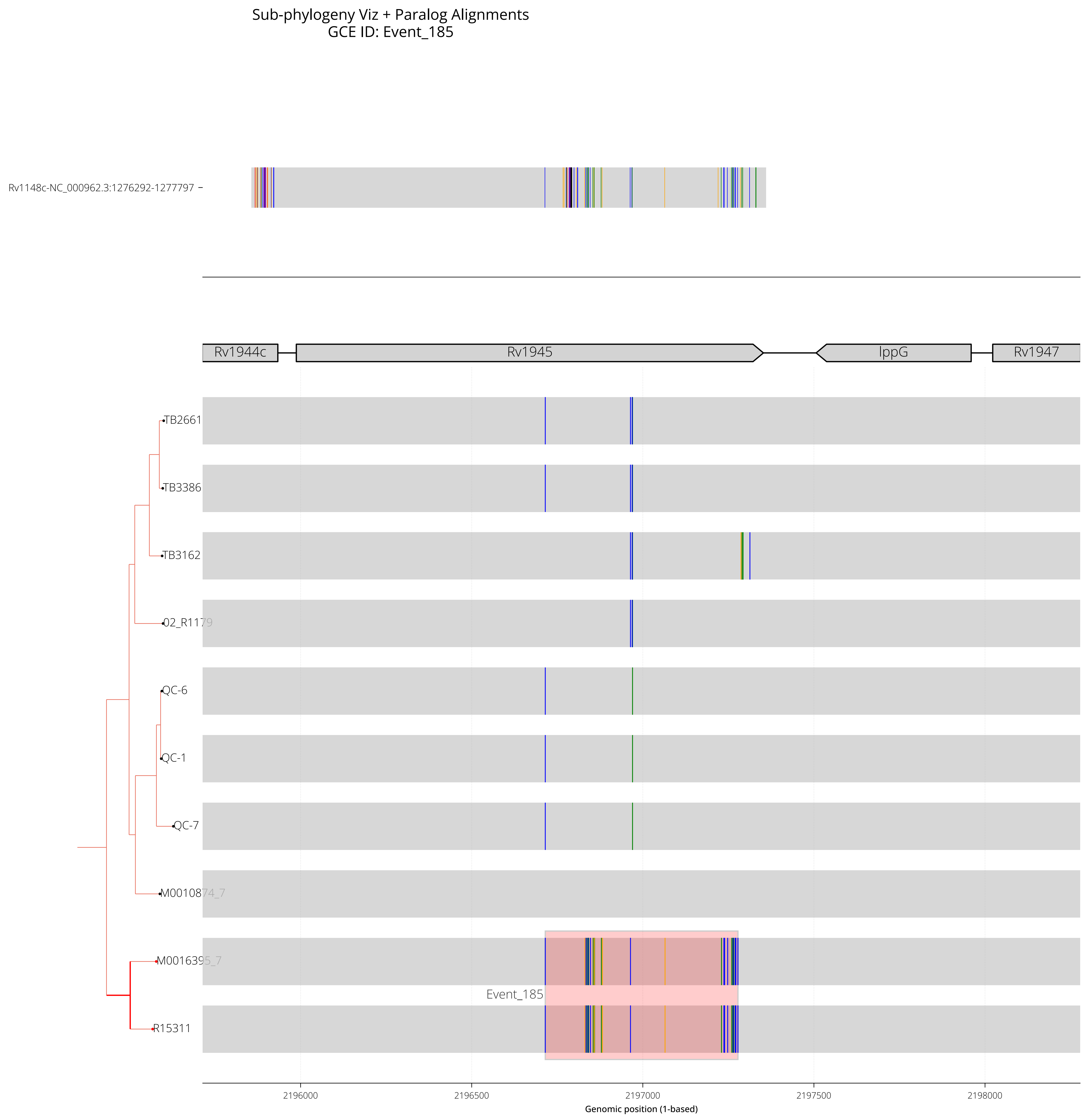

Sub-phylogeny Viz + Paralog Alignments
GCE ID: Event_185
Rv1148c-NC_000962.3:1276292-1277797
Rv1944c
Rv1945
lppG
Rv1947
TB2661
TB3386
TB3162
02_R1179
QC-6
QC-1
QC-7
M0010874_7
M0016395_7
Event_185
R15311
2196000
2196500
2197000
2197500
2198000
Genomic position (1-based)

Full Phylogeny Viz + Paralog Alignments
GCE ID: Event_185

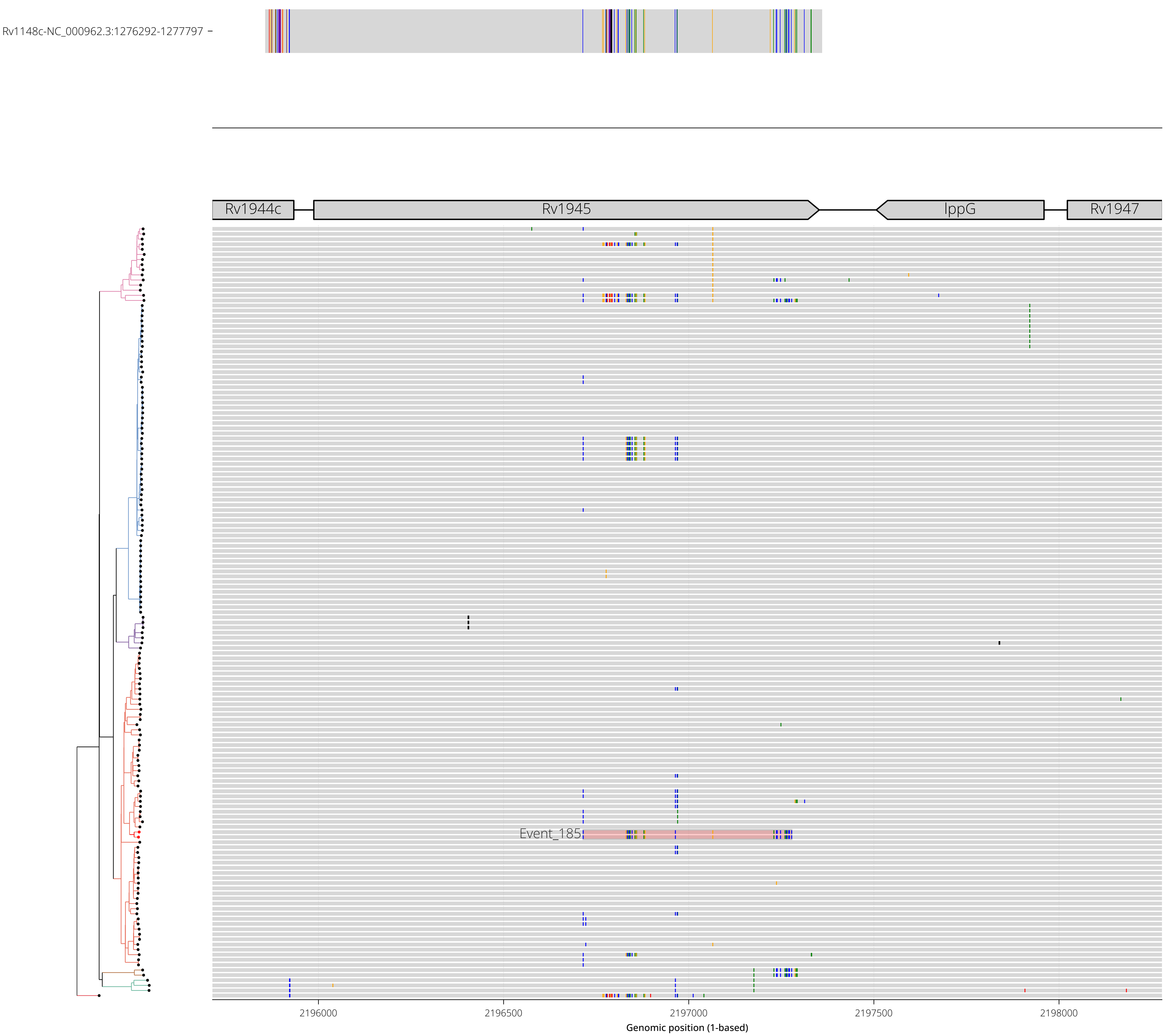

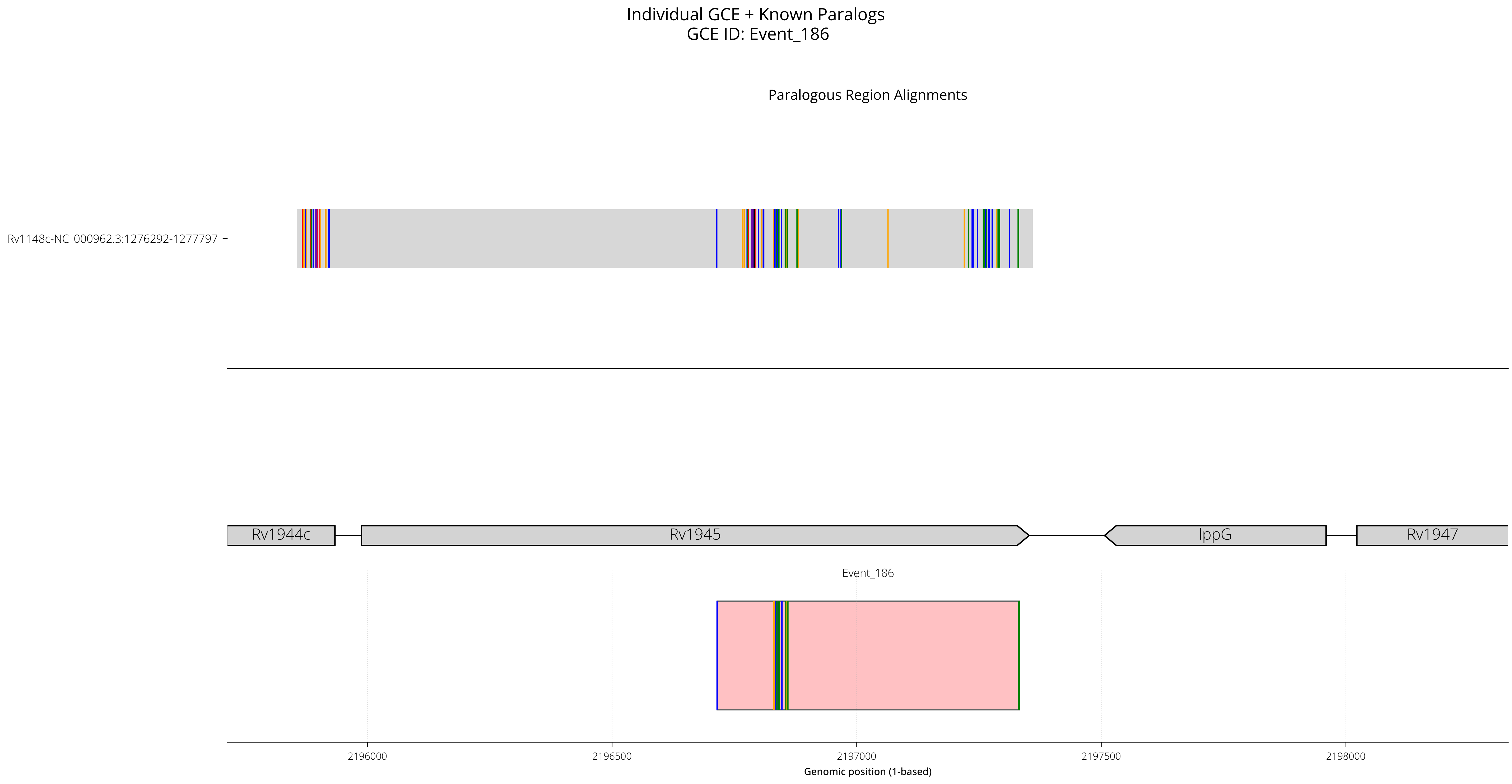

Individual GCE + Known Paralogs
GCE ID: Event_186
Paralogous Region Alignments
Rv1148c-NC_000962.3:1276292-1277797
Rv1944c
Rv1945
lppG
Rv1947
Event_186
2196000
2196500
2197000
2197500
2198000
Genomic position (1-based)

Circos Viz - GCE to Known Paralogs Locations
GCE ID: Event_186

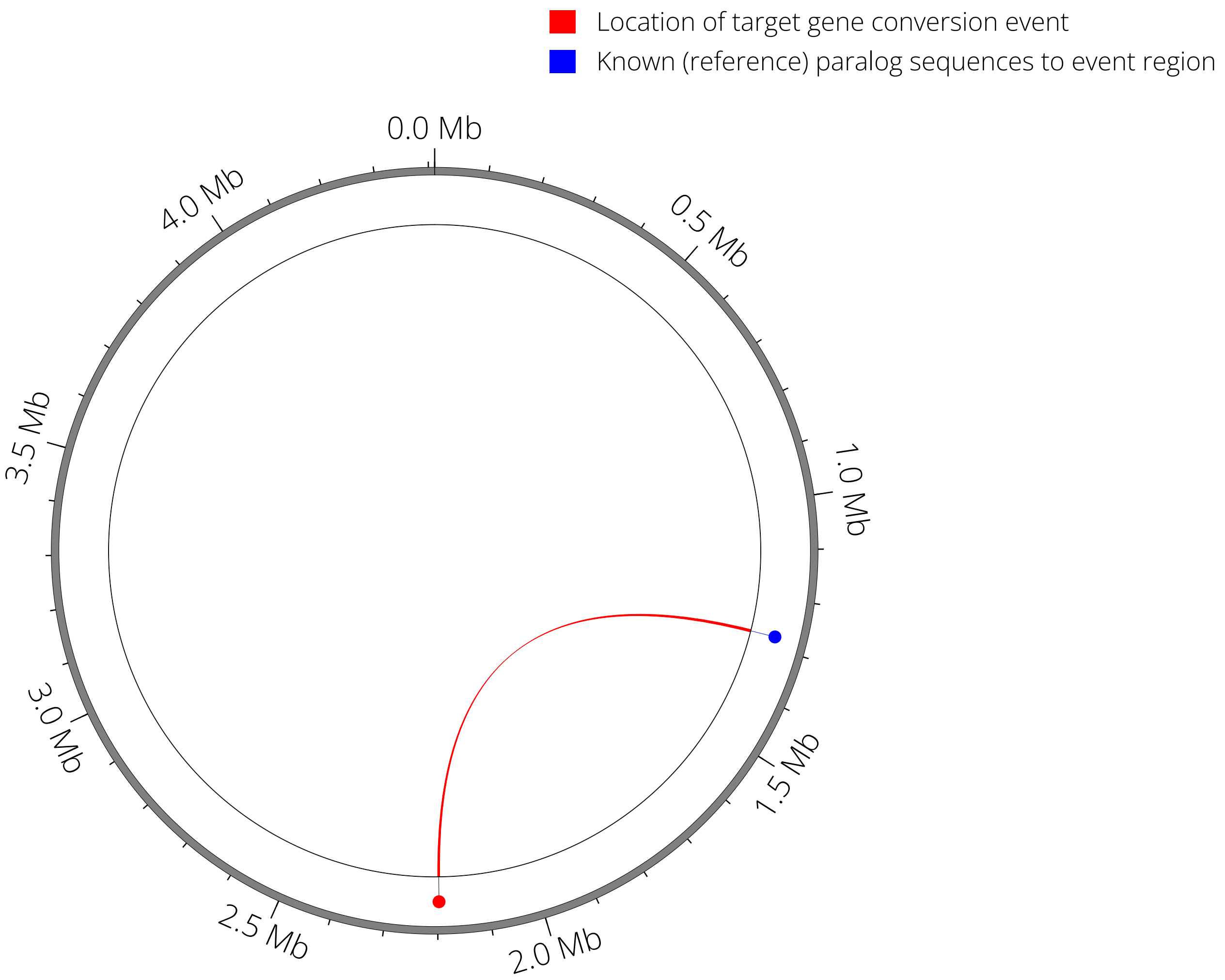

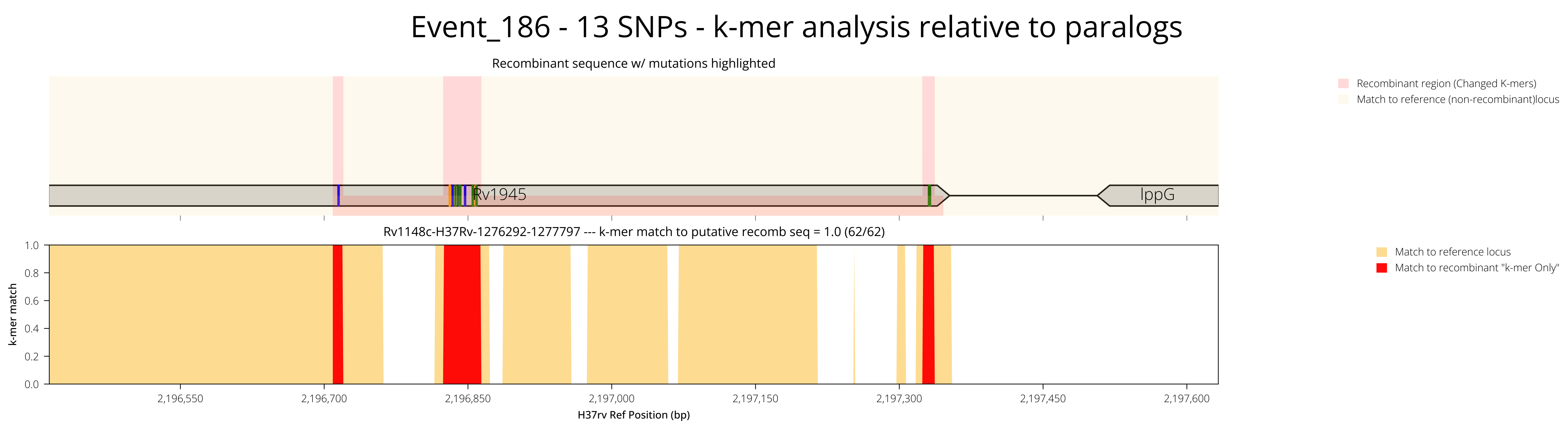
Event_186 - 13 SNPs - k-mer analysis relative to paralogs
Recombinant sequence w/ mutations highlighted
Recombinant region (Changed K-mers)
Match to reference (non-recombinant)locus
Rv1945
lppG
Rv1148c-H37Rv-1276292-1277797 --- k-mer match to putative recomb seq = 1.0 (62/62)
Match to reference locus
Match to recombinant "k-mer Only"
k-mer match
1.0
0.8
0.6
0.4
0.2
0.0
2,196,550
2,196,700
2,196,850
2,197,000
2,197,150
2,197,300
2,197,450
2,197,600
H37rv Ref Position (bp)

Sub-phylogeny Viz + Paralog Alignments
GCE ID: Event_186

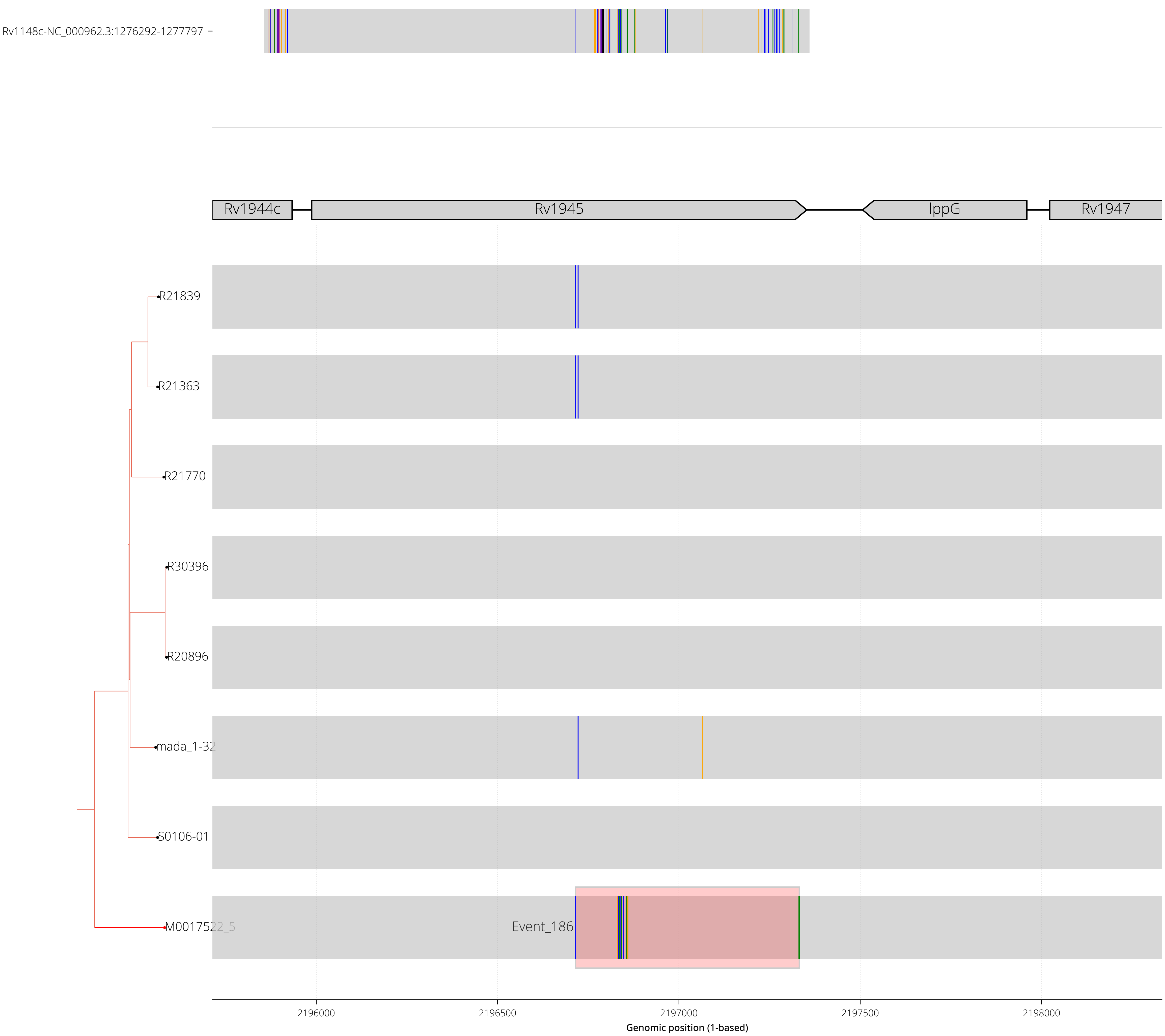

Full Phylogeny Viz + Paralog Alignments
GCE ID: Event_186

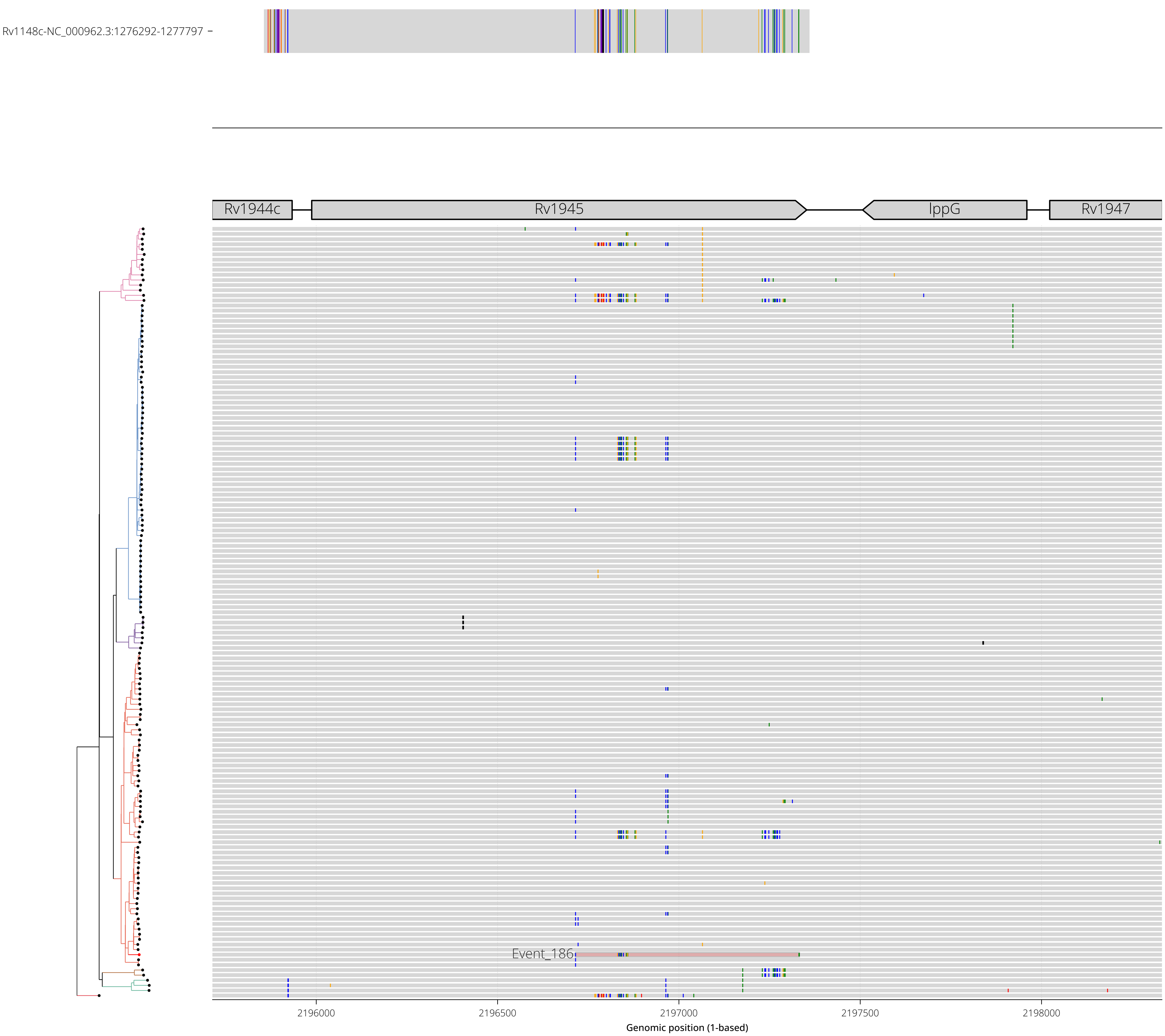

#### Individual GCE + Known Paralogs
#### GCE ID: Event_187

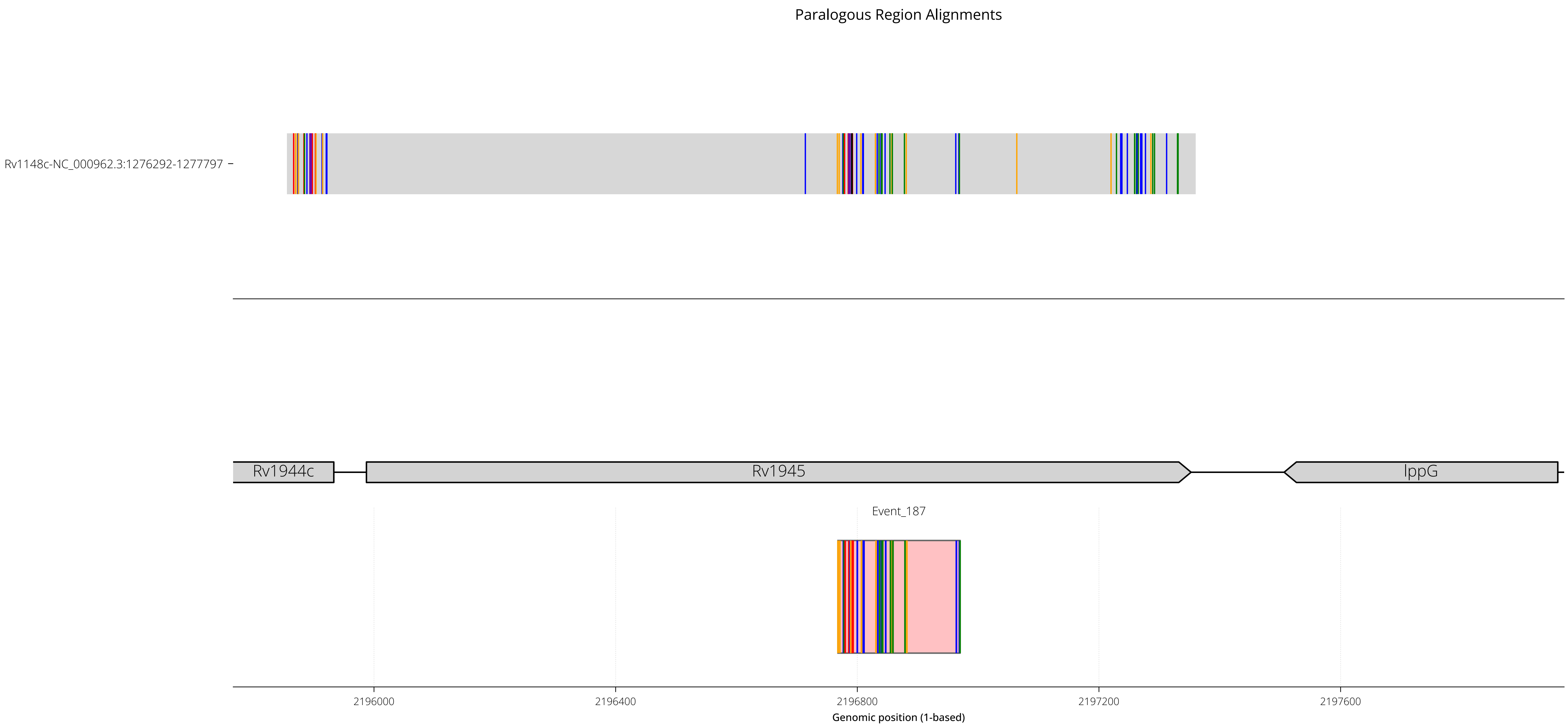

Circos Viz - GCE to Known Paralogs Locations
GCE ID: Event_187

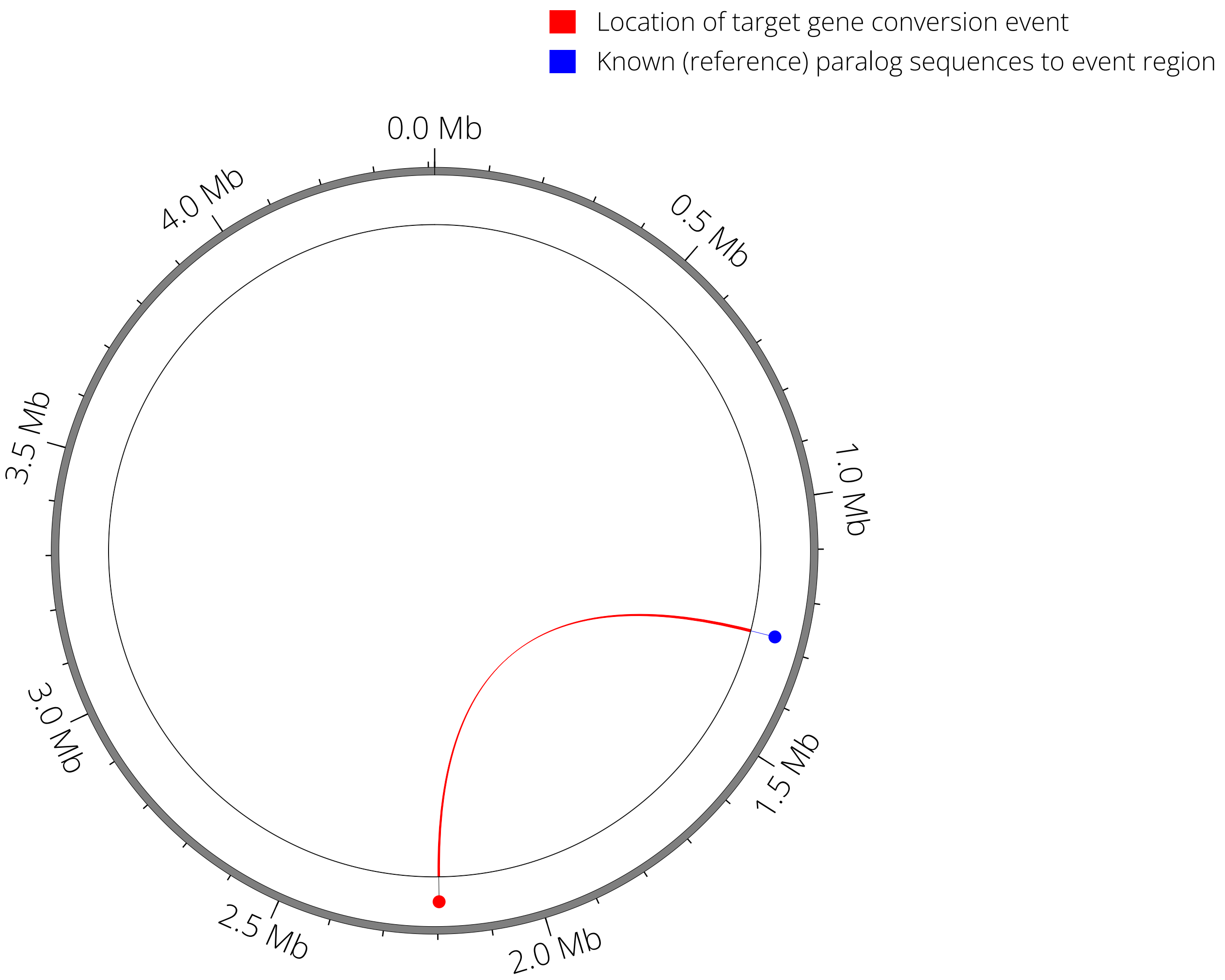

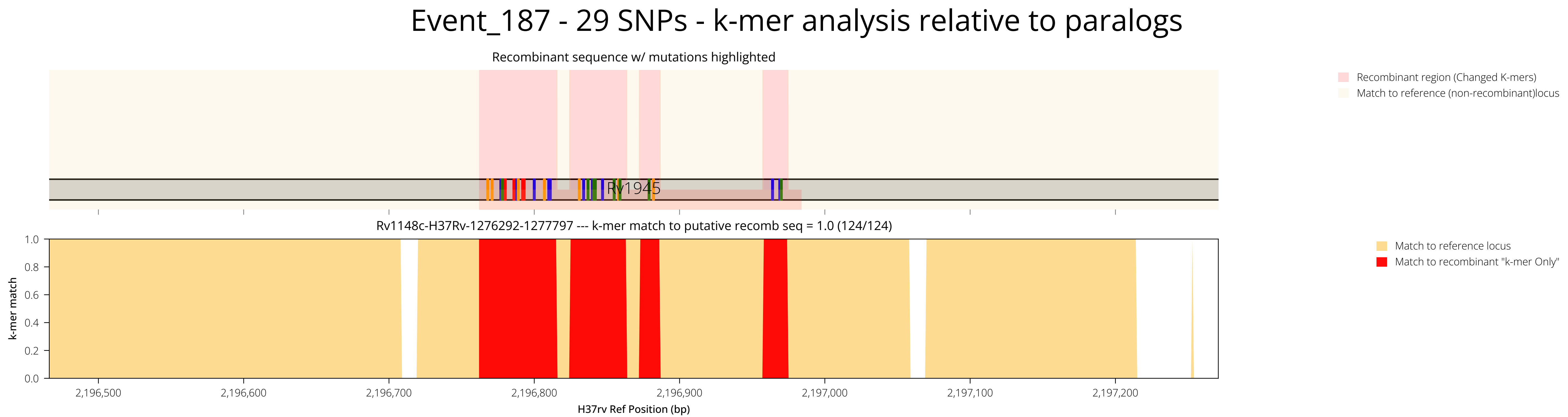
Event_187 - 29 SNPs - k-mer analysis relative to paralogs
Recombinant sequence w/ mutations highlighted
Recombinant region (Changed K-mers)
Match to reference (non-recombinant)locus
Rv1945
Rv1148c-H37Rv-1276292-1277797 --- k-mer match to putative recomb seq = 1.0 (124/124)
Match to reference locus
Match to recombinant "k-mer Only"
k-mer match
1.0
0.8
0.6
0.4
0.2
0.0
2,196,500
2,196,600
2,196,700
2,196,800
2,196,900
2,197,000
2,197,100
2,197,200
H37rv Ref Position (bp)

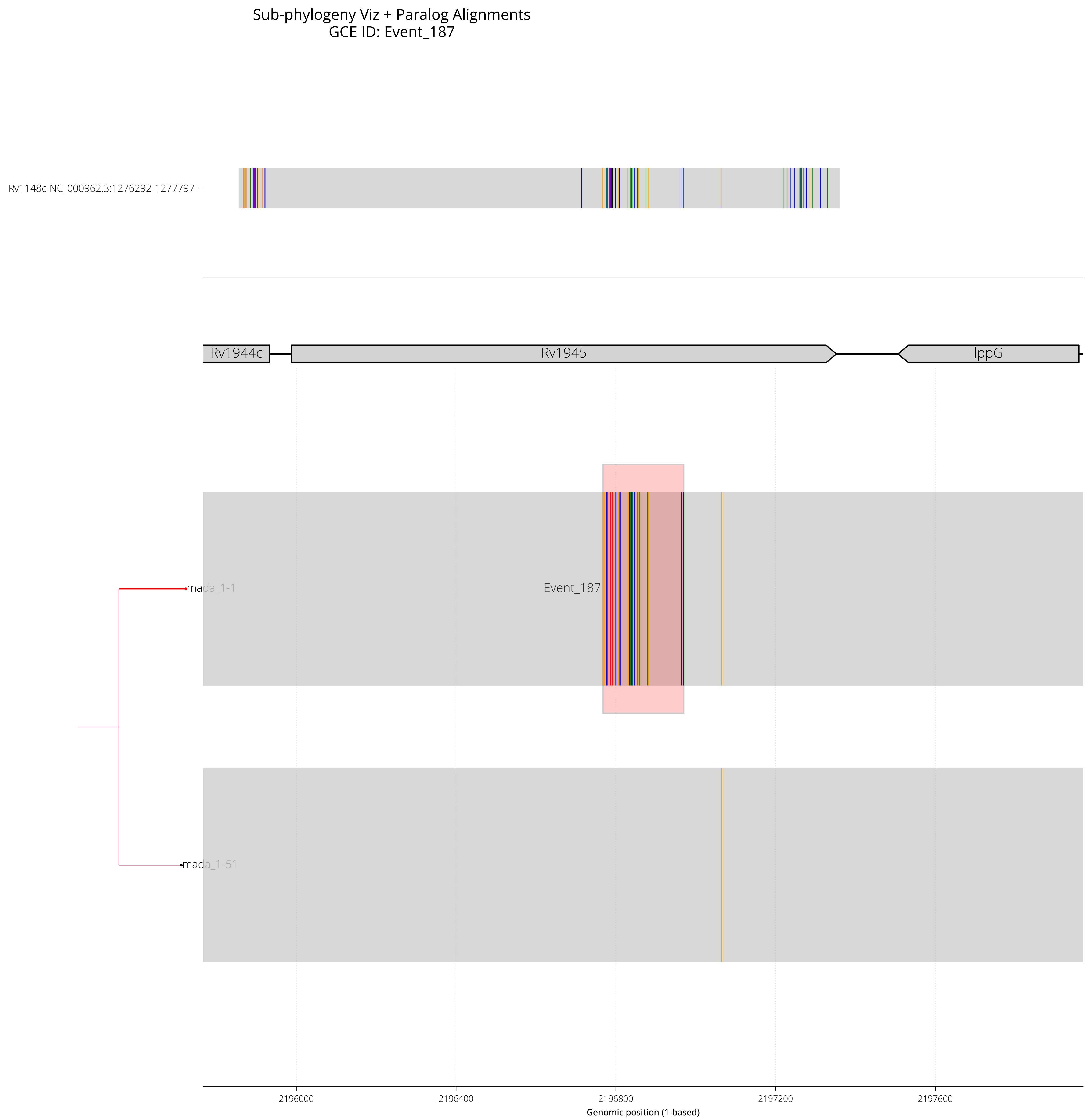
Sub-phylogeny Viz + Paralog Alignments
GCE ID: Event_187
Rv1148c-NC_000962.3:1276292-1277797
Rv1944c
Rv1945
lppG
mada_1-1
Event_187
mada_1-51
2196000
2196400
2196800
2197200
2197600
Genomic position (1-based)

Full Phylogeny Viz + Paralog Alignments
GCE ID: Event_187

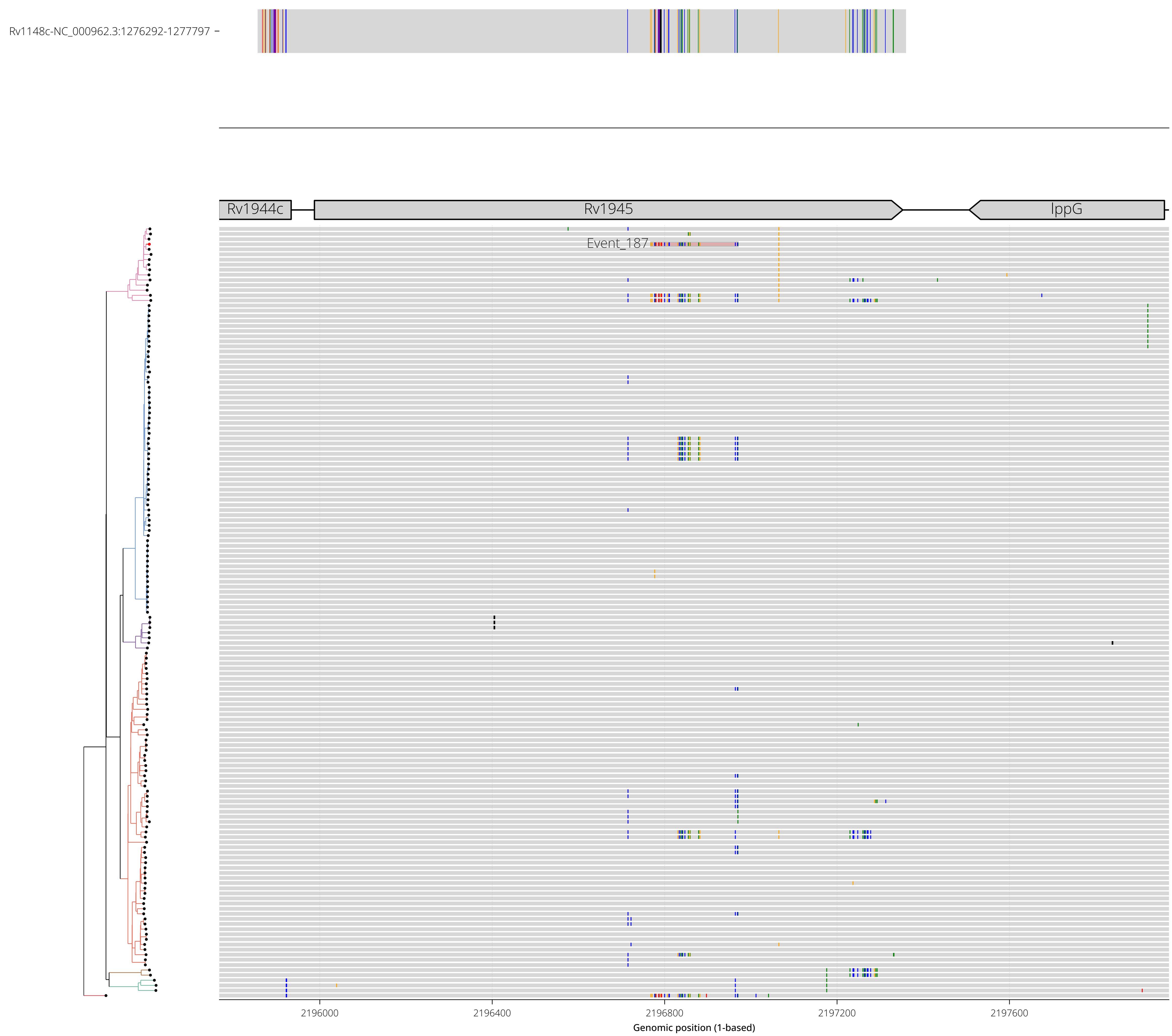

Individual GCE + Known Paralogs
GCE ID: Event_188

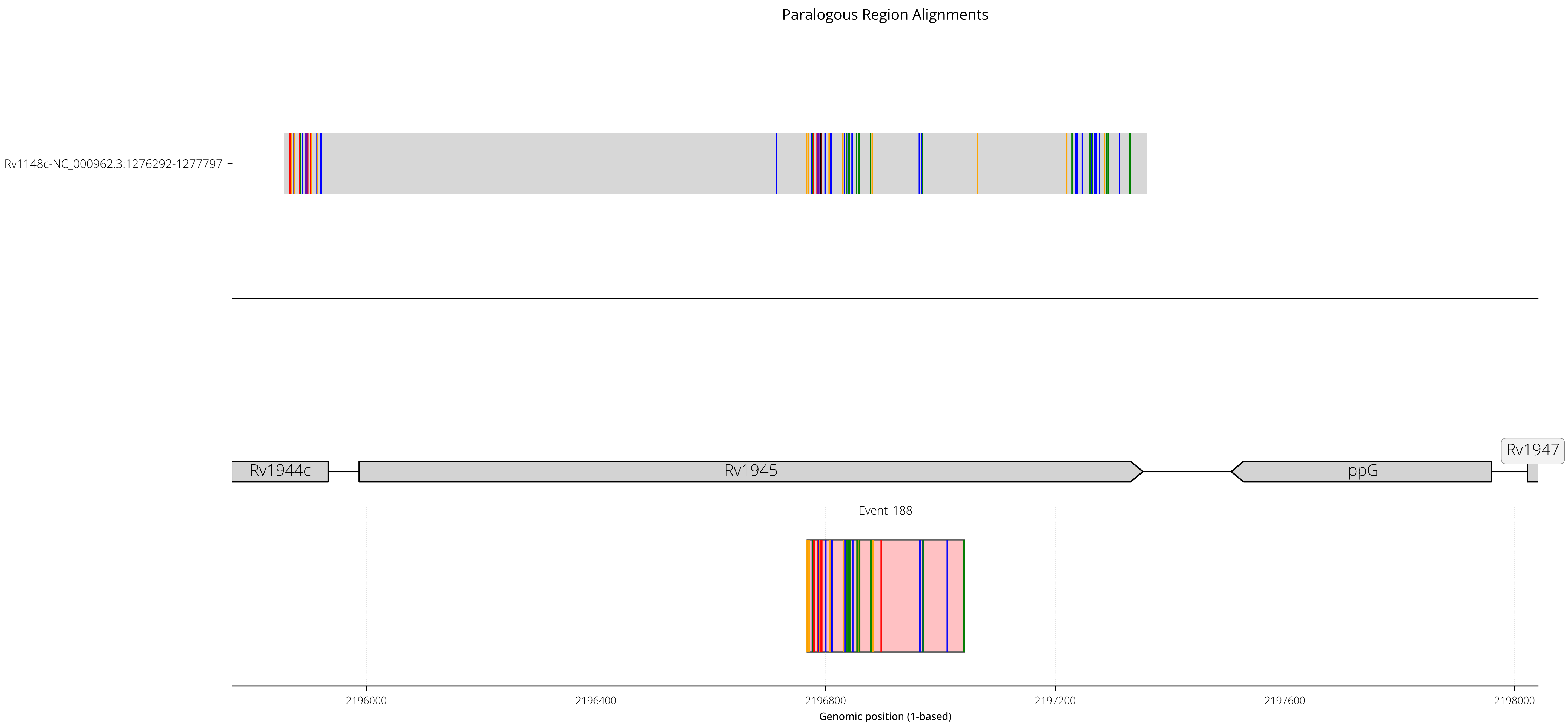

Circos Viz - GCE to Known Paralogs Locations
GCE ID: Event_188

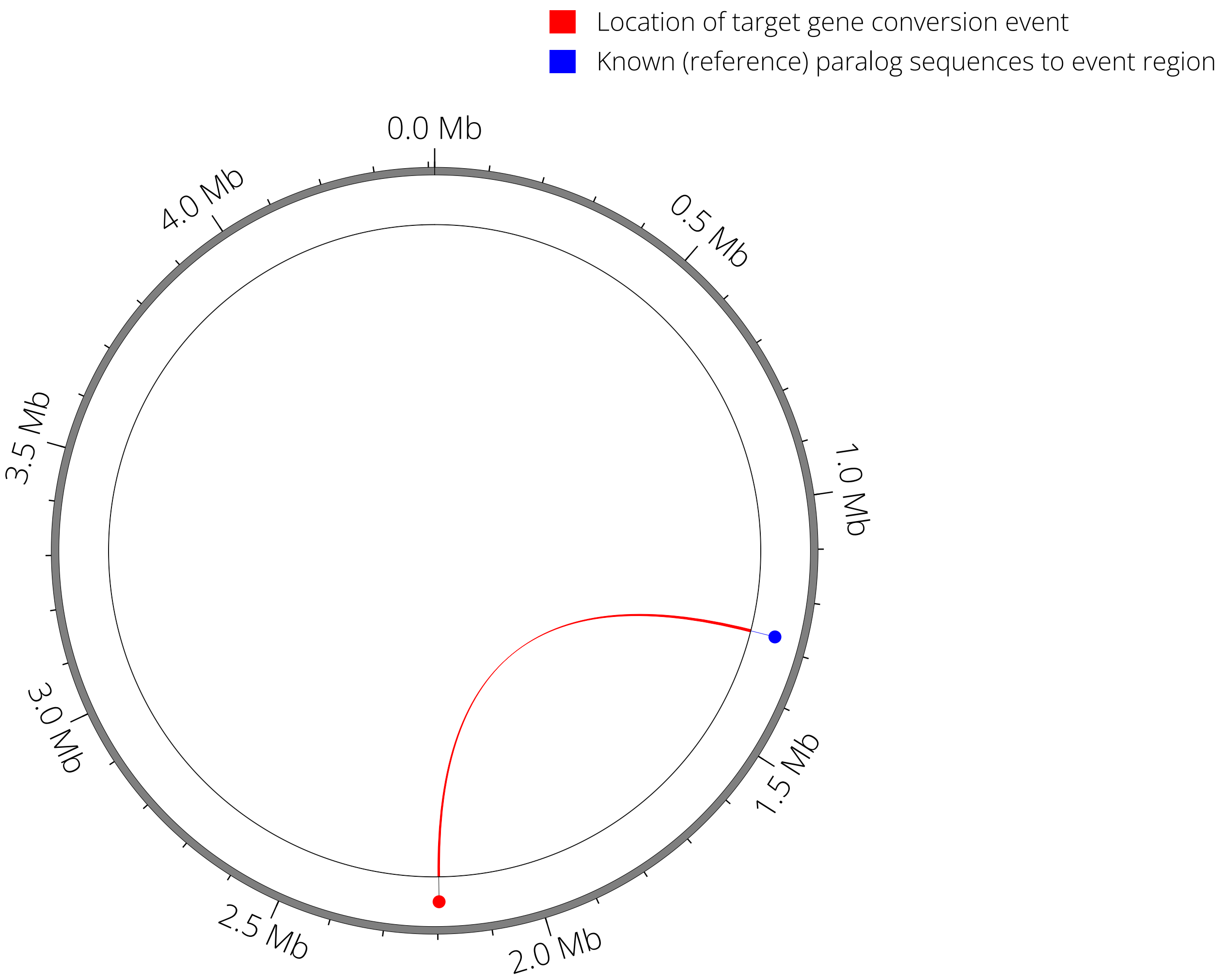

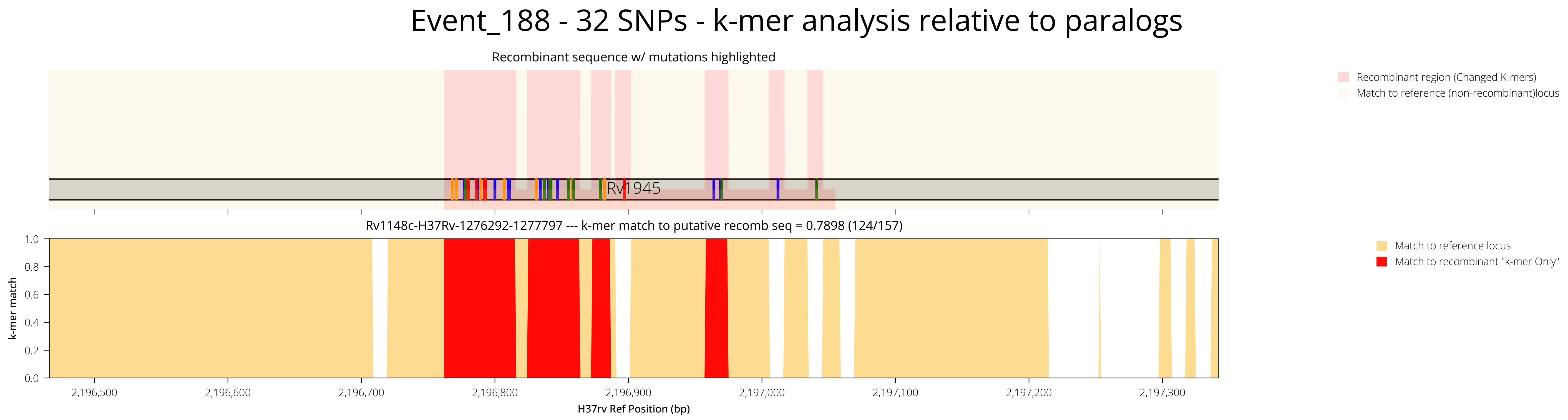

Event_188 - 32 SNPs - k-mer analysis relative to paralogs
Recombinant sequence w/ mutations highlighted
Recombinant region (Changed K-mers)
Match to reference (non-recombinant)locus
Rv1945
Rv1148c-H37Rv-1276292-1277797 --- k-mer match to putative recomb seq = 0.7898 (124/157)
Match to reference locus
Match to recombinant "k-mer Only"
k-mer match
1.0
0.8
0.6
0.4
0.2
0.0
2,196,500
2,196,600
2,196,700
2,196,800
2,196,900
2,197,000
2,197,100
2,197,200
2,197,300
H37rv Ref Position (bp)

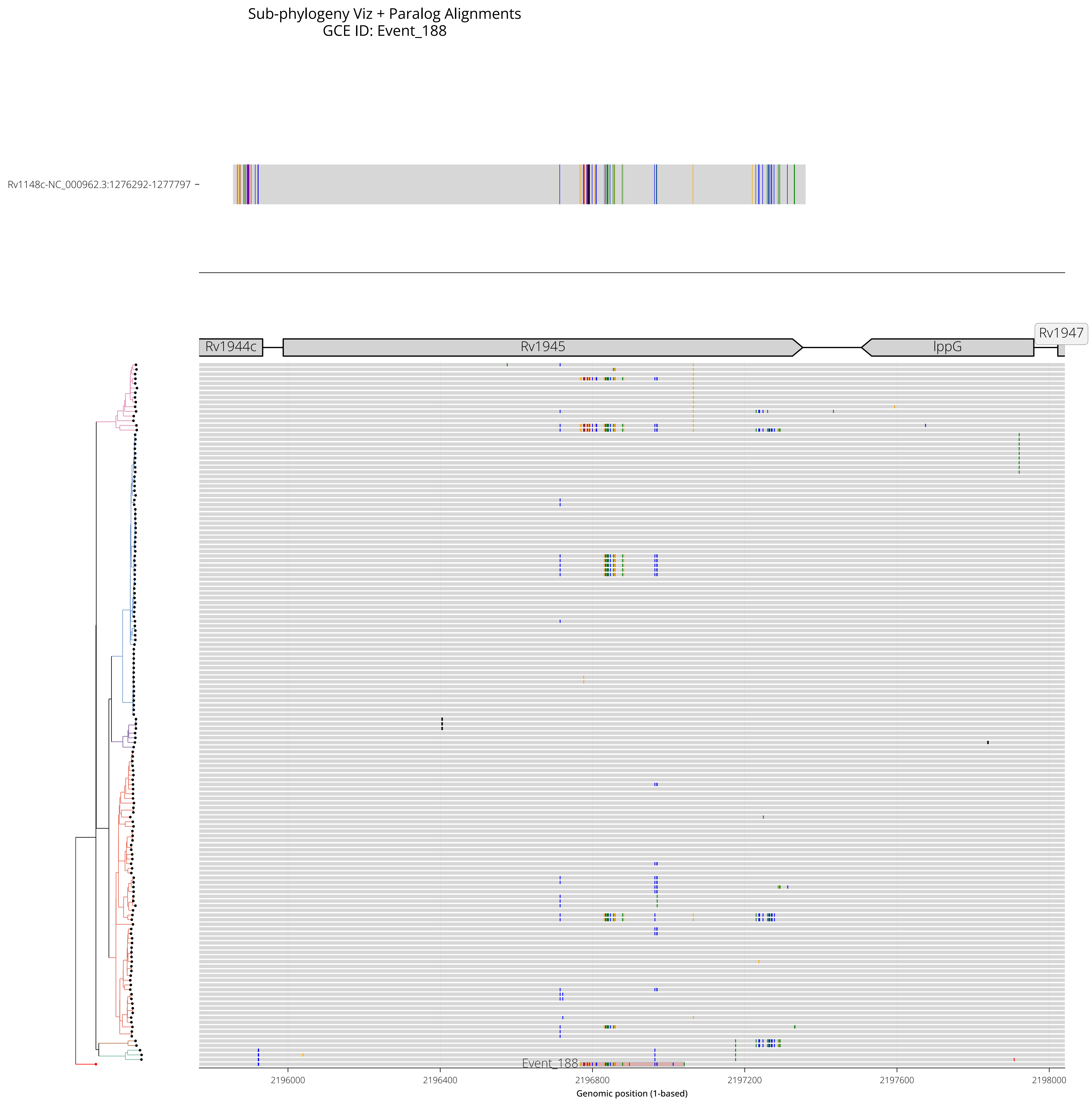
Sub-phylogeny Viz + Paralog Alignments
GCE ID: Event_188
Rv1148c-NC_000962.3:1276292-1277797
Rv1944c
Rv1945
lppG
Rv1947
Event_188
2196000
2196400
2196800
2197200
2197600
2198000
Genomic position (1-based)

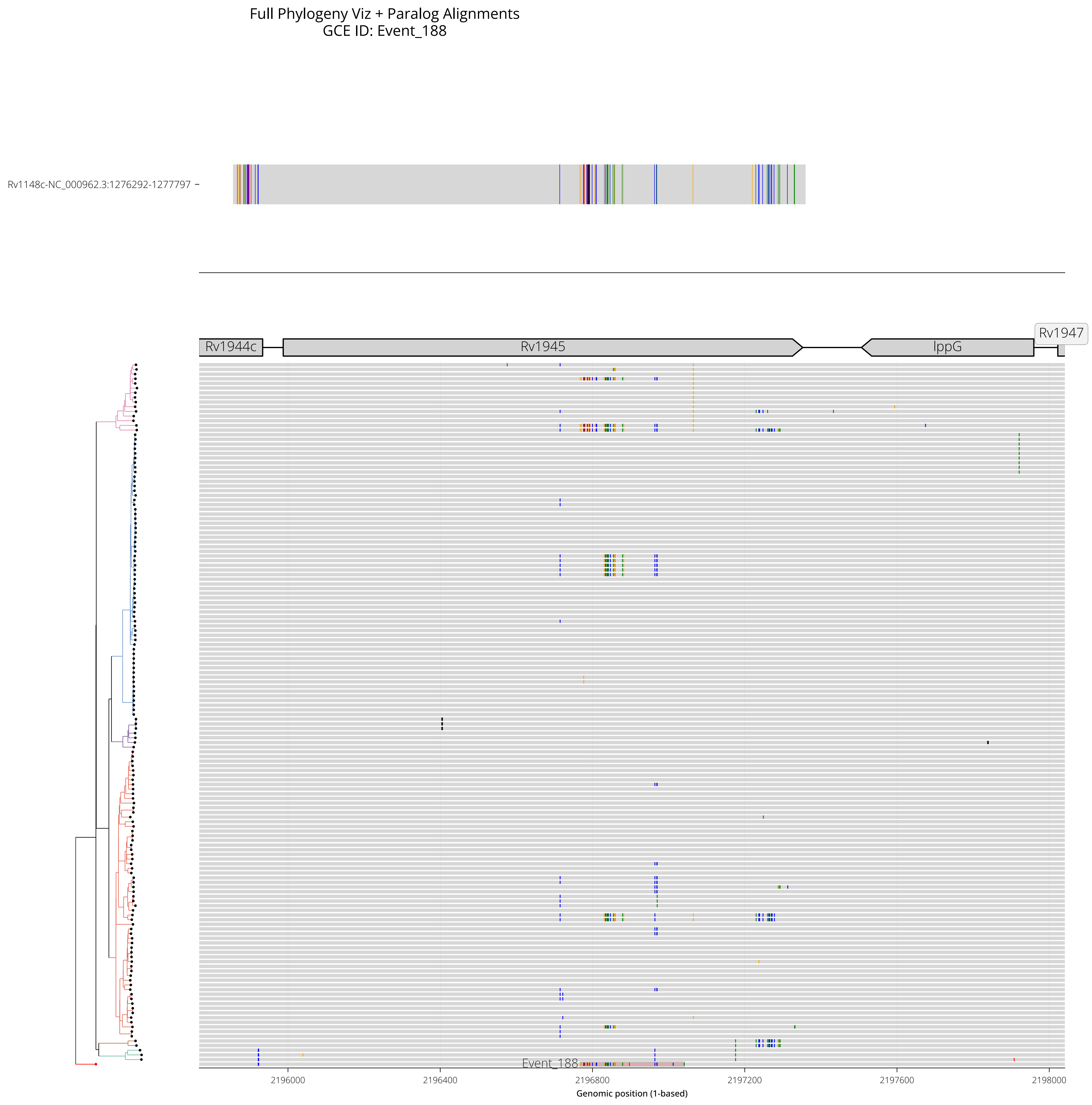

Full Phylogeny Viz + Paralog Alignments
GCE ID: Event_188
Rv1148c-NC_000962.3:1276292-1277797
Rv1944c
Rv1945
lppG
Rv1947
Event_188
2196000
2196400
2196800
2197200
2197600
2198000
Genomic position (1-based)

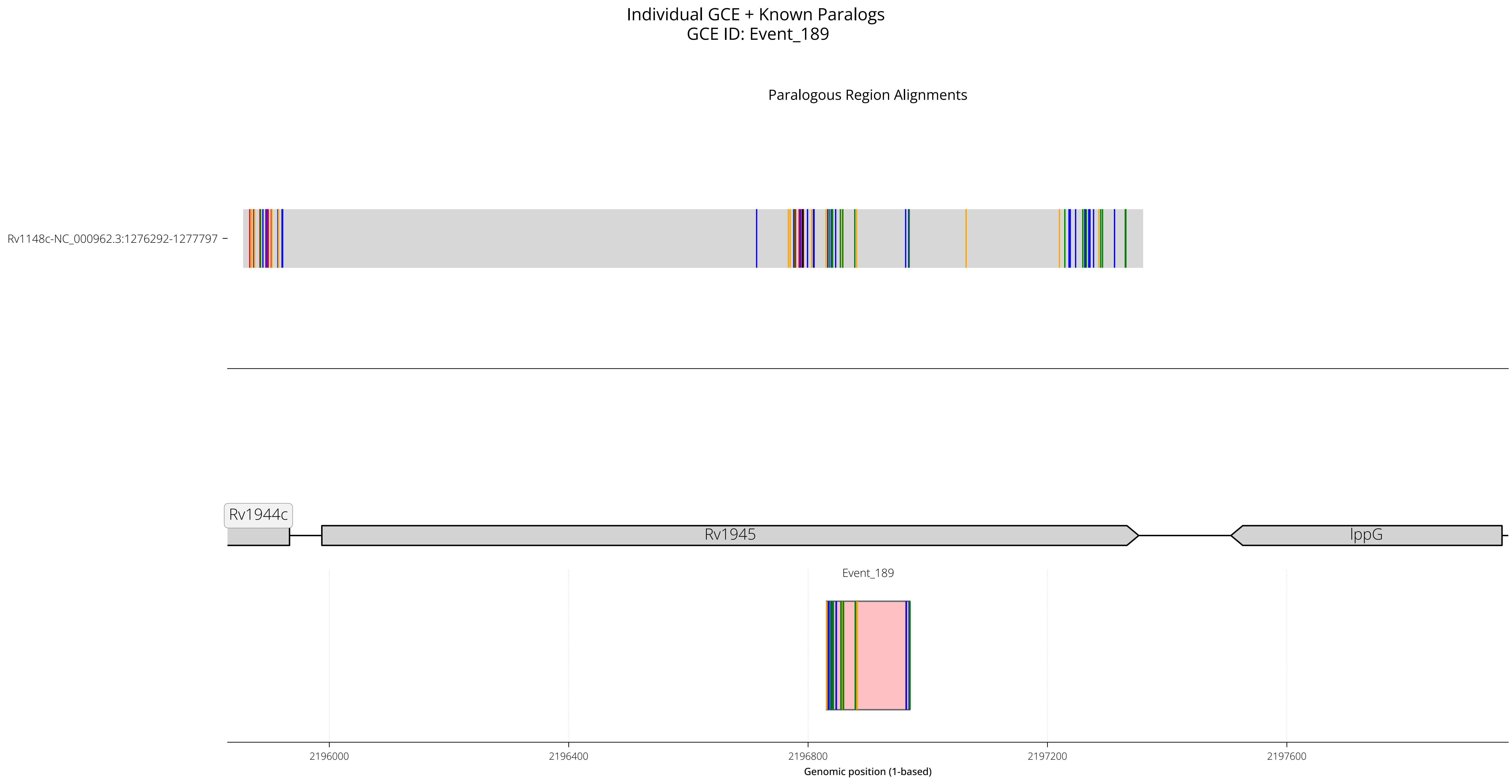
Individual GCE + Known Paralogs
GCE ID: Event_189
Paralogous Region Alignments
Rv1148c-NC_000962.3:1276292-1277797
Rv1944c
Rv1945
lppG
Event_189
2196000
2196400
2196800
2197200
2197600
Genomic position (1-based)

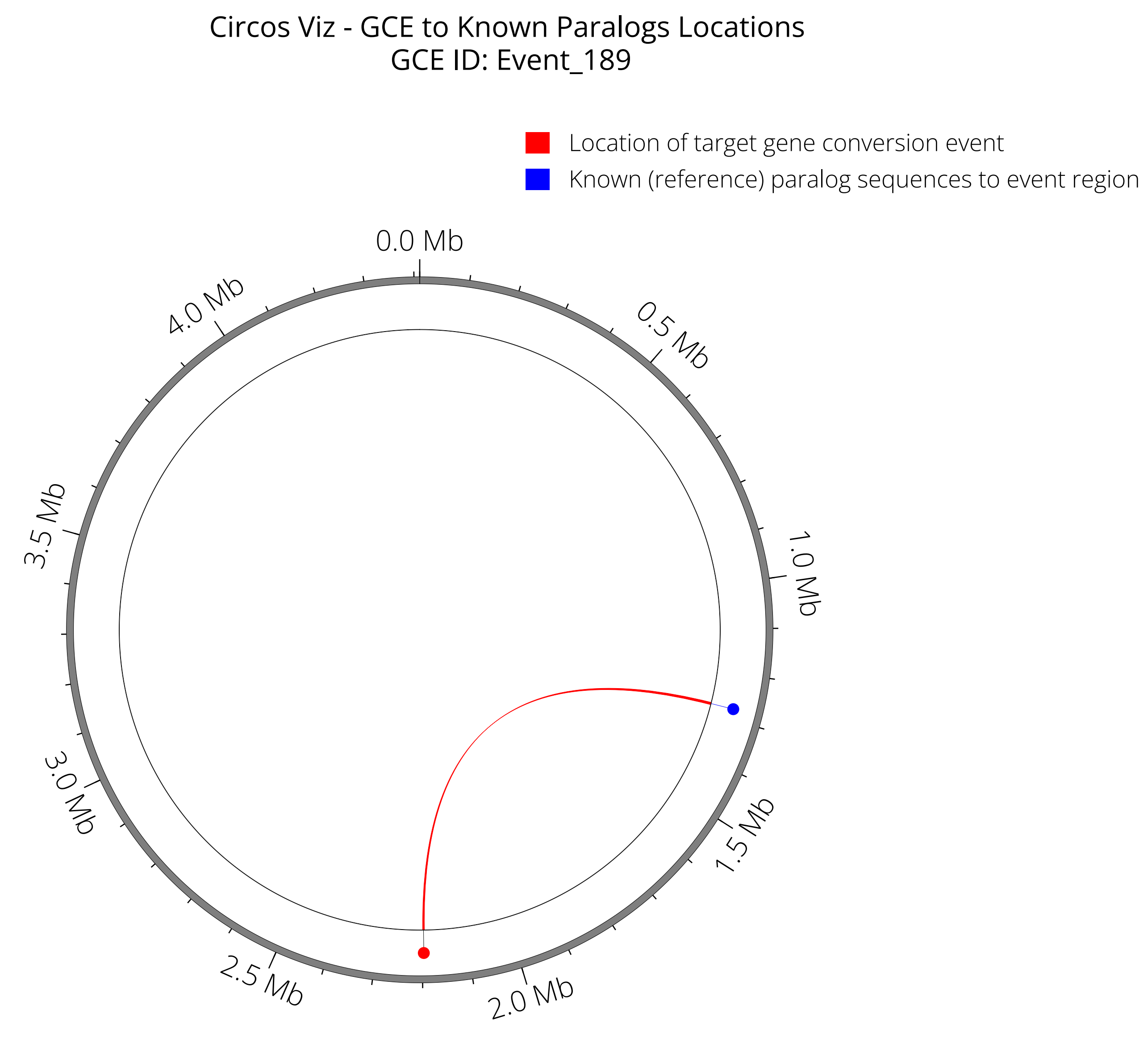
Circos Viz - GCE to Known Paralogs Locations
GCE ID: Event_189
Location of target gene conversion event
Known (reference) paralog sequences to event region
0.0 Mb
0.5 Mb
1.0 Mb
1.5 Mb
2.0 Mb
2.5 Mb
3.0 Mb
3.5 Mb
4.0 Mb

### Event_189 - 15 SNPs - k-mer analysis relative to paralogs

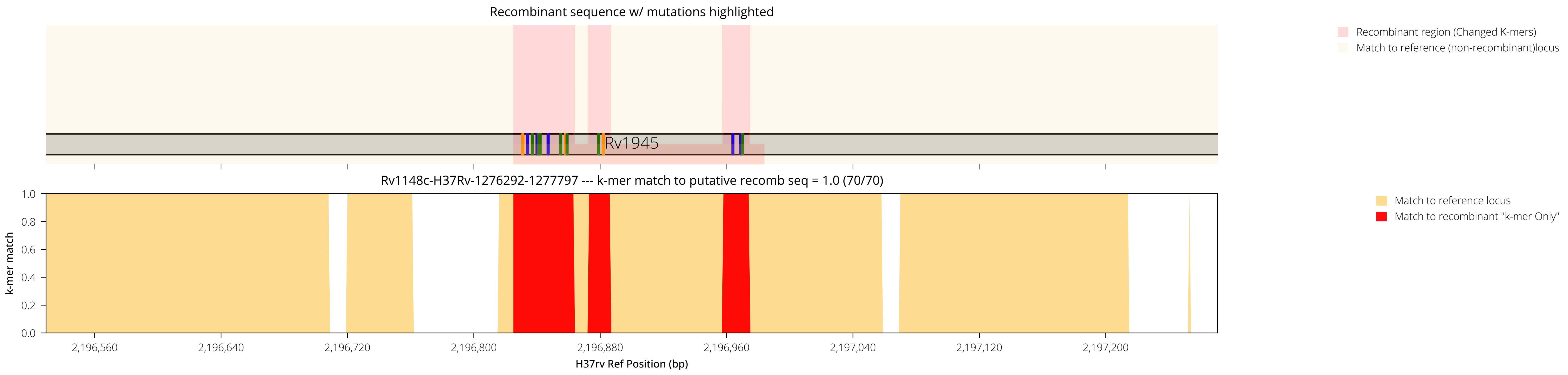

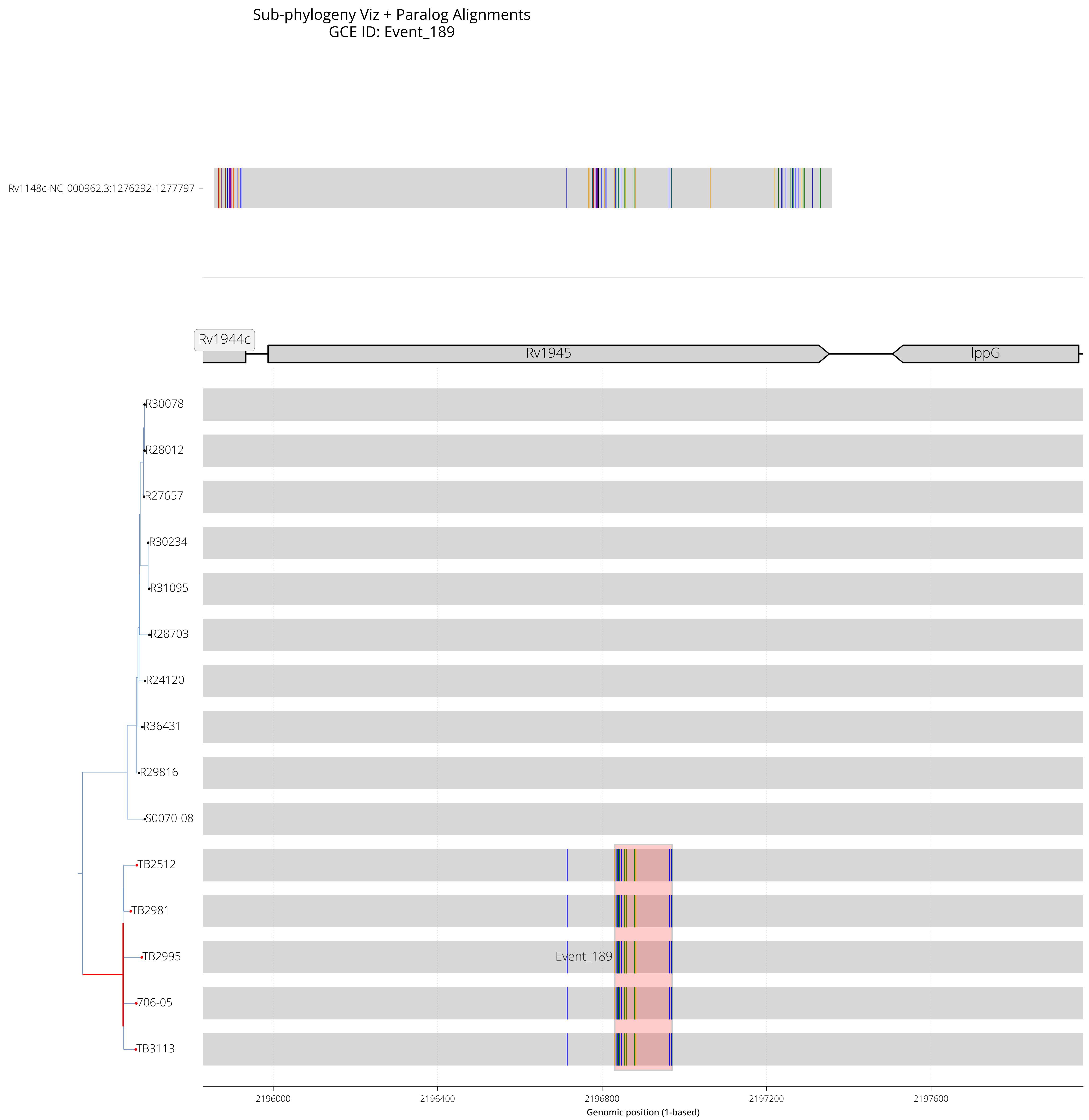

Sub-phylogeny Viz + Paralog Alignments
GCE ID: Event_189
Rv1148c-NC_000962.3:1276292-1277797
Rv1944c
Rv1945
lppG
R30078
R28012
R27657
R30234
R31095
R28703
R24120
R36431
R29816
S0070-08
TB2512
TB2981
TB2995
706-05
TB3113
Event_189
2196000
2196400
2196800
2197200
2197600
Genomic position (1-based)

#### Full Phylogeny Viz + Paralog Alignments
#### GCE ID: Event_189

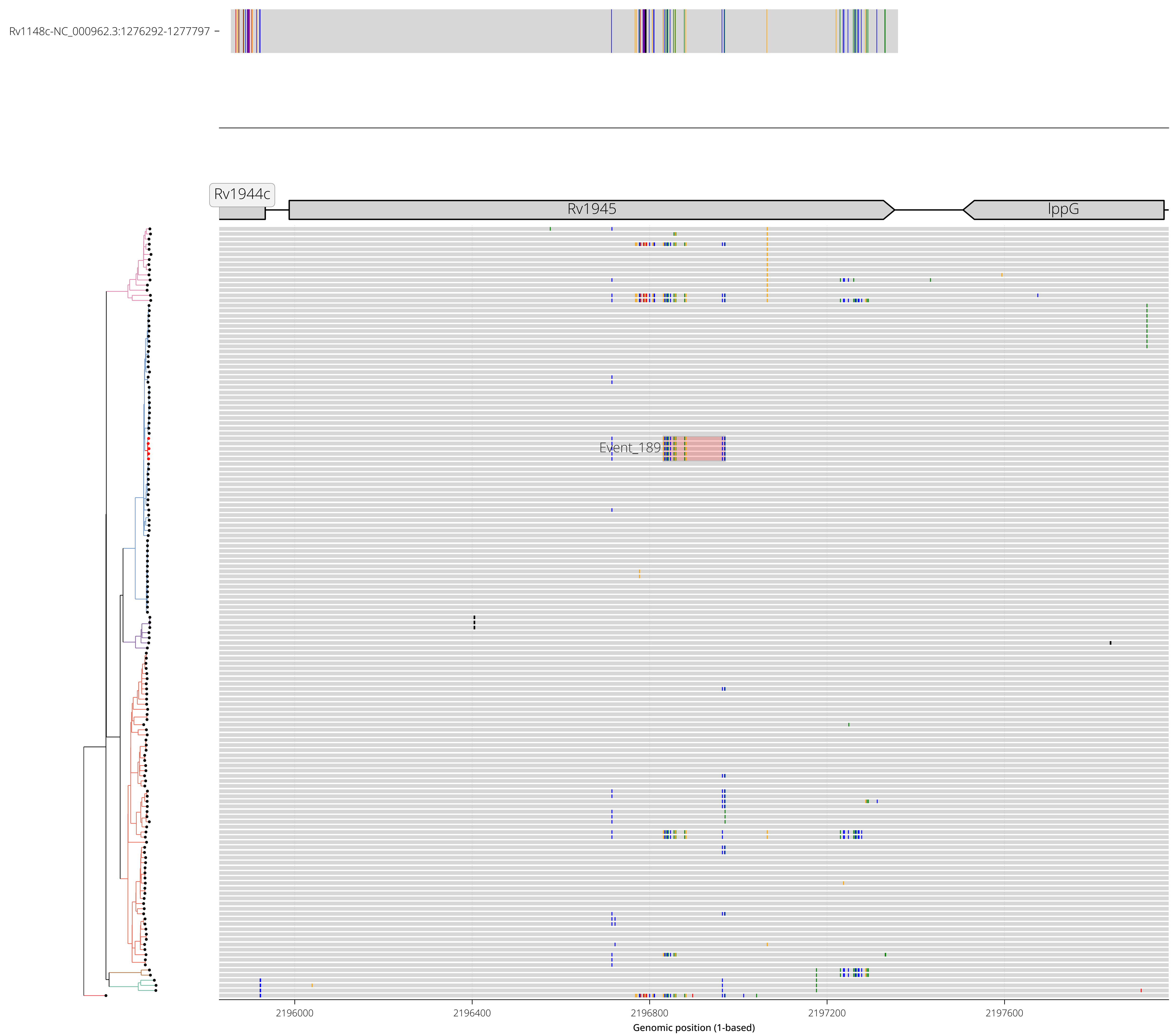

Individual GCE + Known Paralogs
GCE ID: Event_190

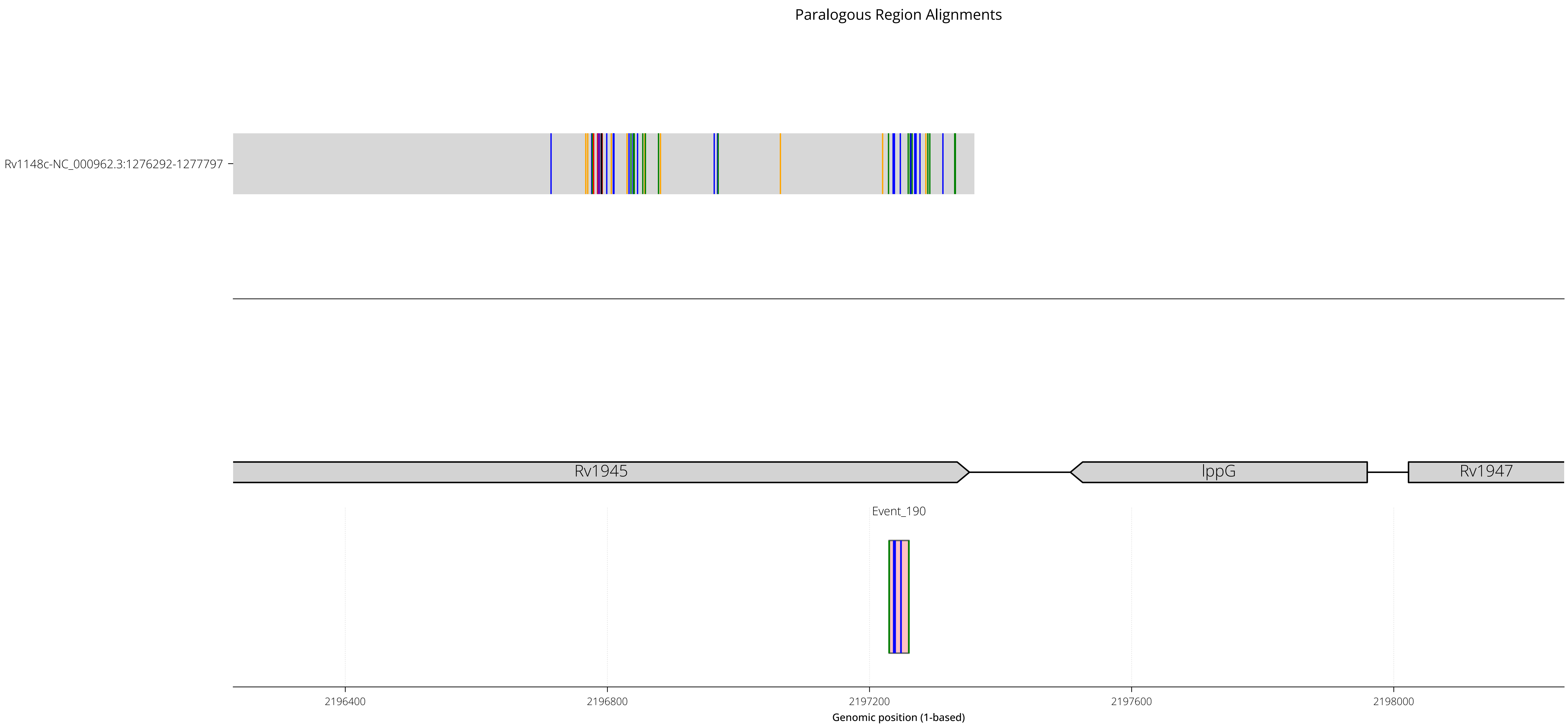

Circos Viz - GCE to Known Paralogs Locations
GCE ID: Event_190

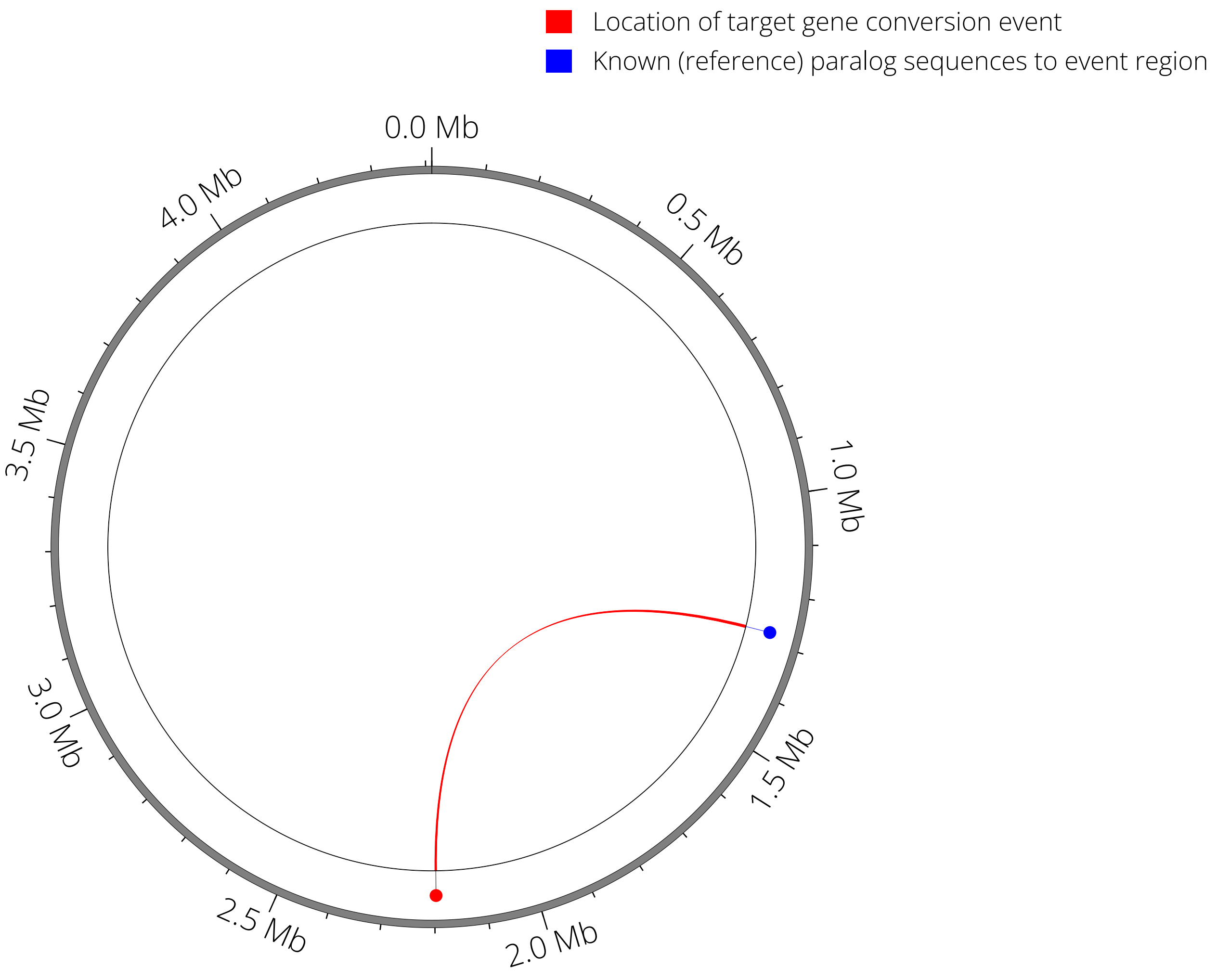

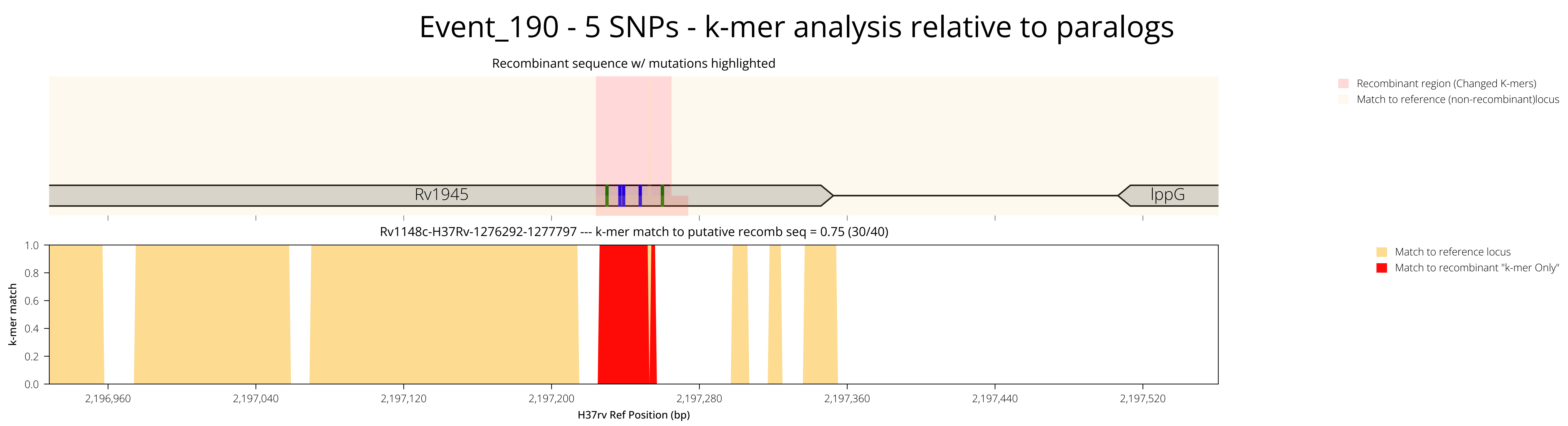

Event_190 - 5 SNPs - k-mer analysis relative to paralogs
Recombinant sequence w/ mutations highlighted
Rv1945
lppG
Recombinant region (Changed K-mers)
Match to reference (non-recombinant)locus
Rv1148c-H37Rv-1276292-1277797 --- k-mer match to putative recomb seq = 0.75 (30/40)
k-mer match
1.0
0.8
0.6
0.4
0.2
0.0
2,196,960
2,197,040
2,197,120
2,197,200
2,197,280
2,197,360
2,197,440
2,197,520
H37rv Ref Position (bp)
Match to reference locus
Match to recombinant "k-mer Only"

Sub-phylogeny Viz + Paralog Alignments
GCE ID: Event_190

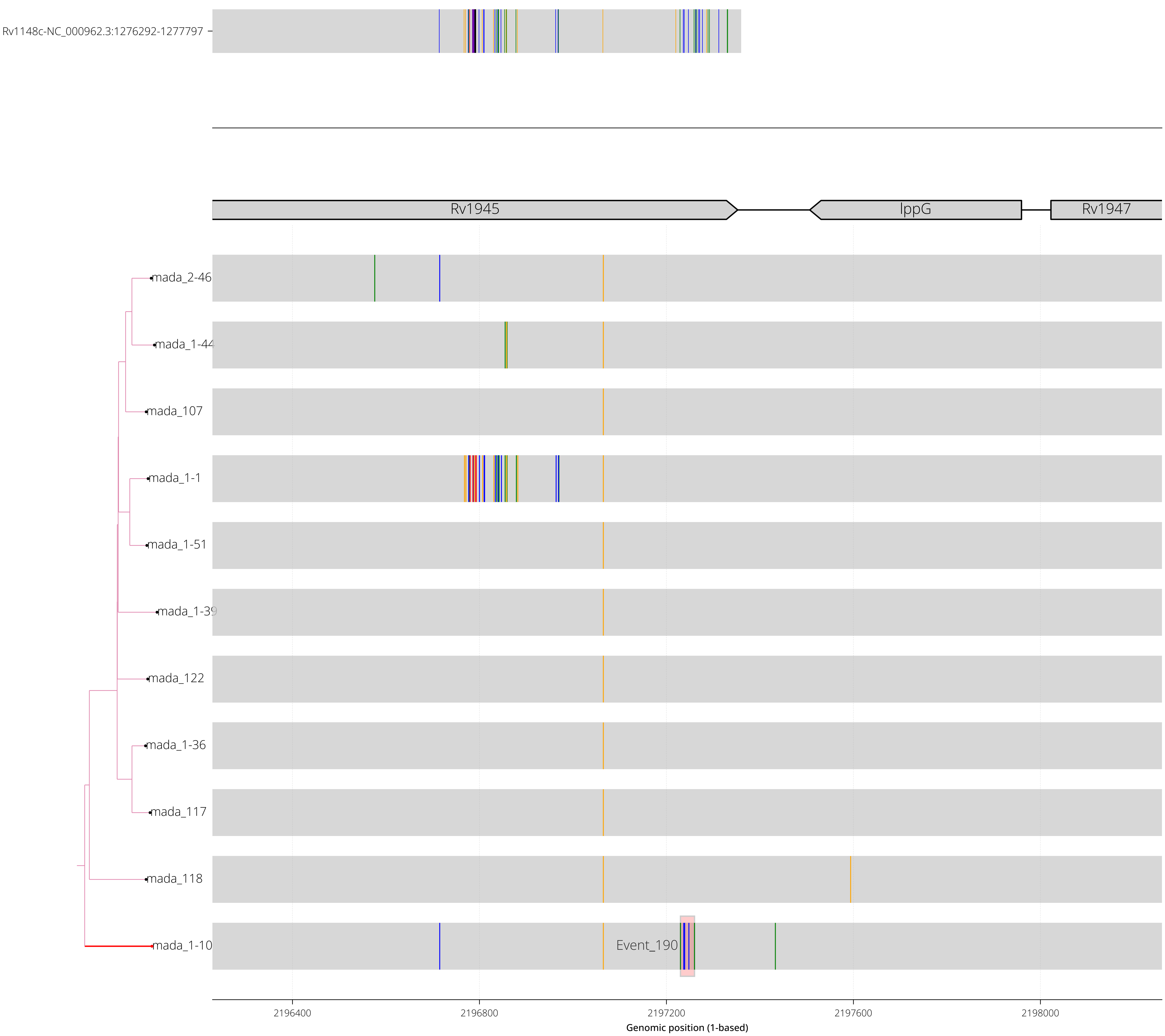

Full Phylogeny Viz + Paralog Alignments
GCE ID: Event_190

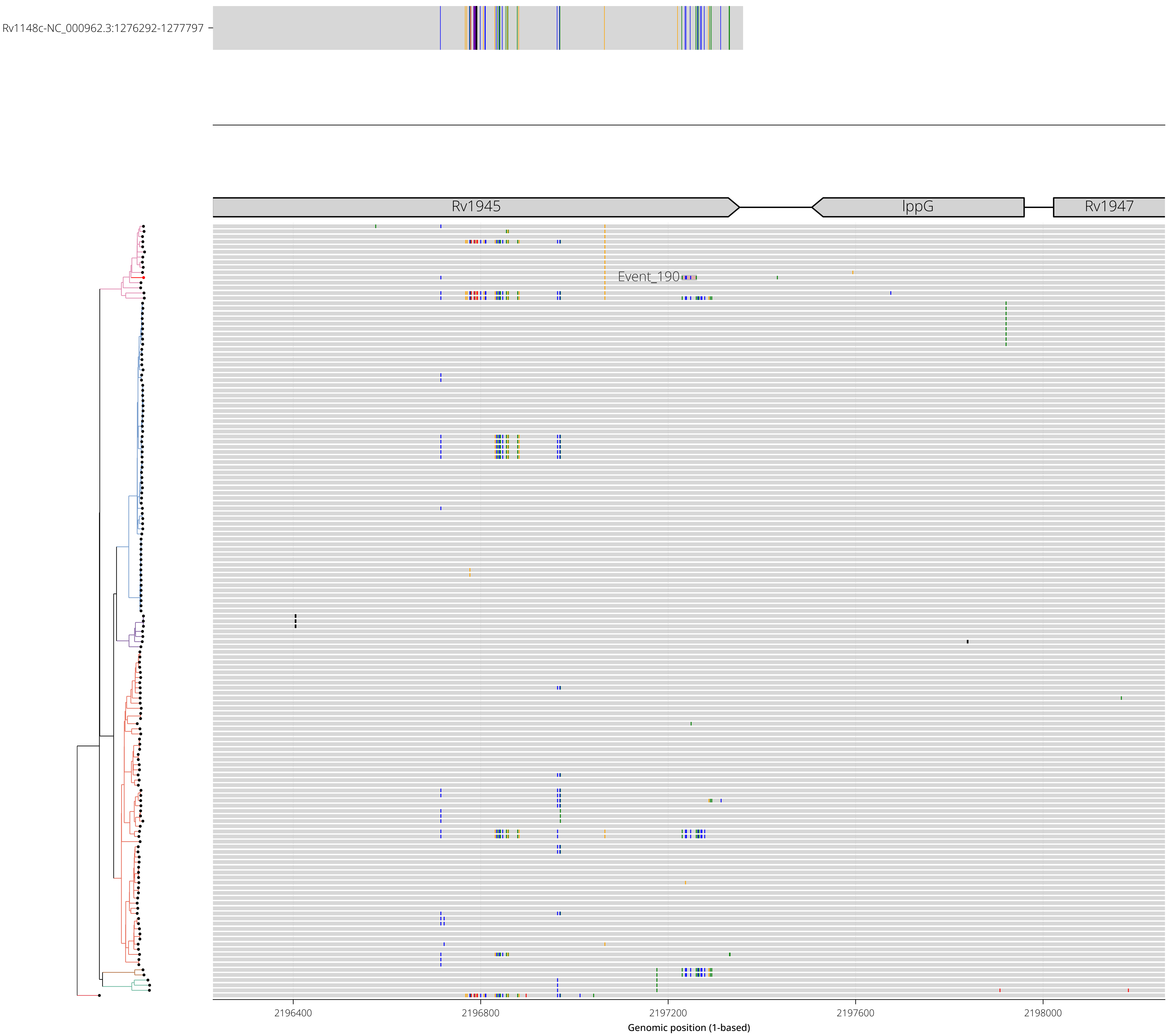

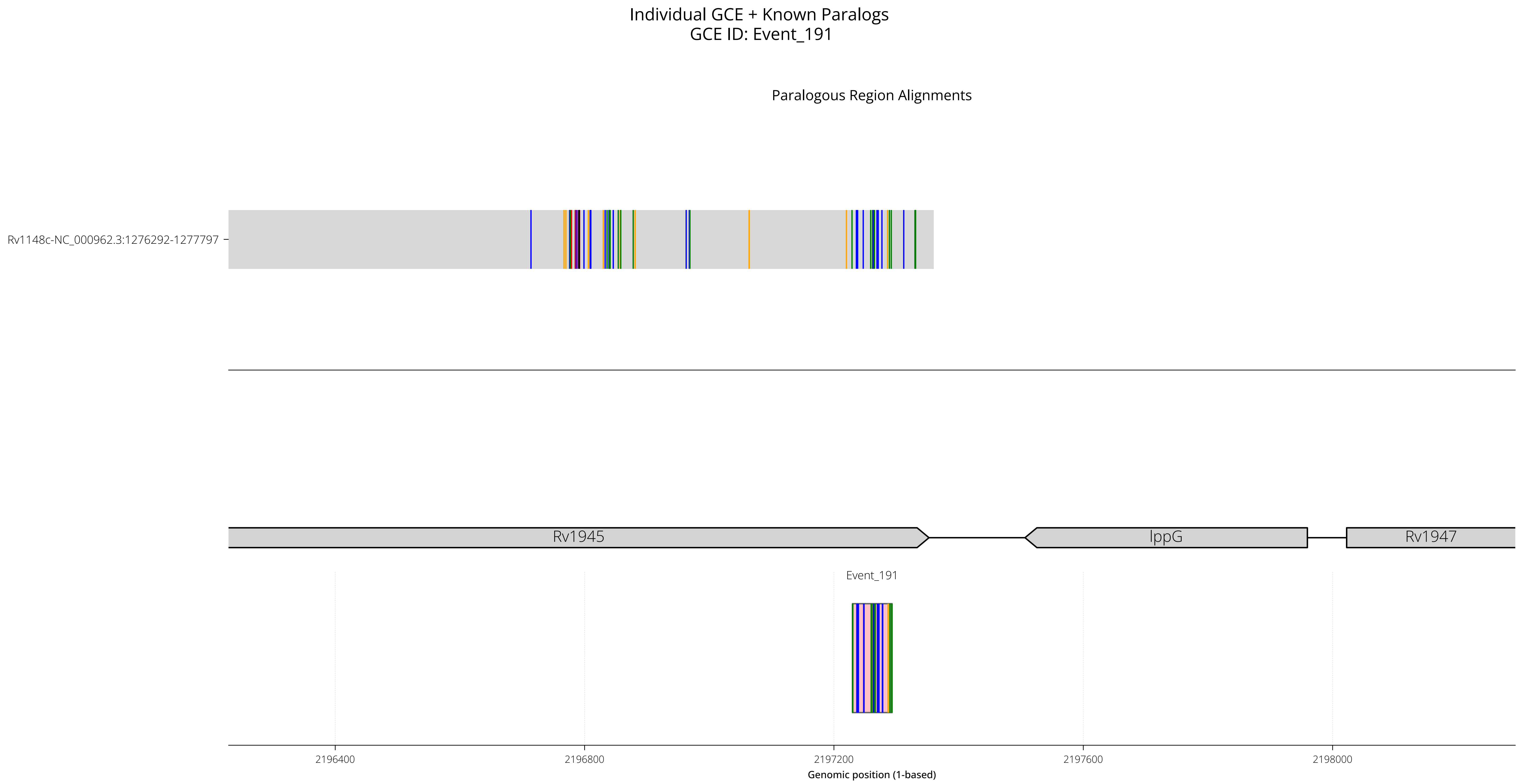

Individual GCE + Known Paralogs
GCE ID: Event_191
Paralogous Region Alignments
Rv1148c-NC_000962.3:1276292-1277797
Rv1945
lppG
Rv1947
Event_191
2196400
2196800
2197200
2197600
2198000
Genomic position (1-based)

Circos Viz - GCE to Known Paralogs Locations
GCE ID: Event_191

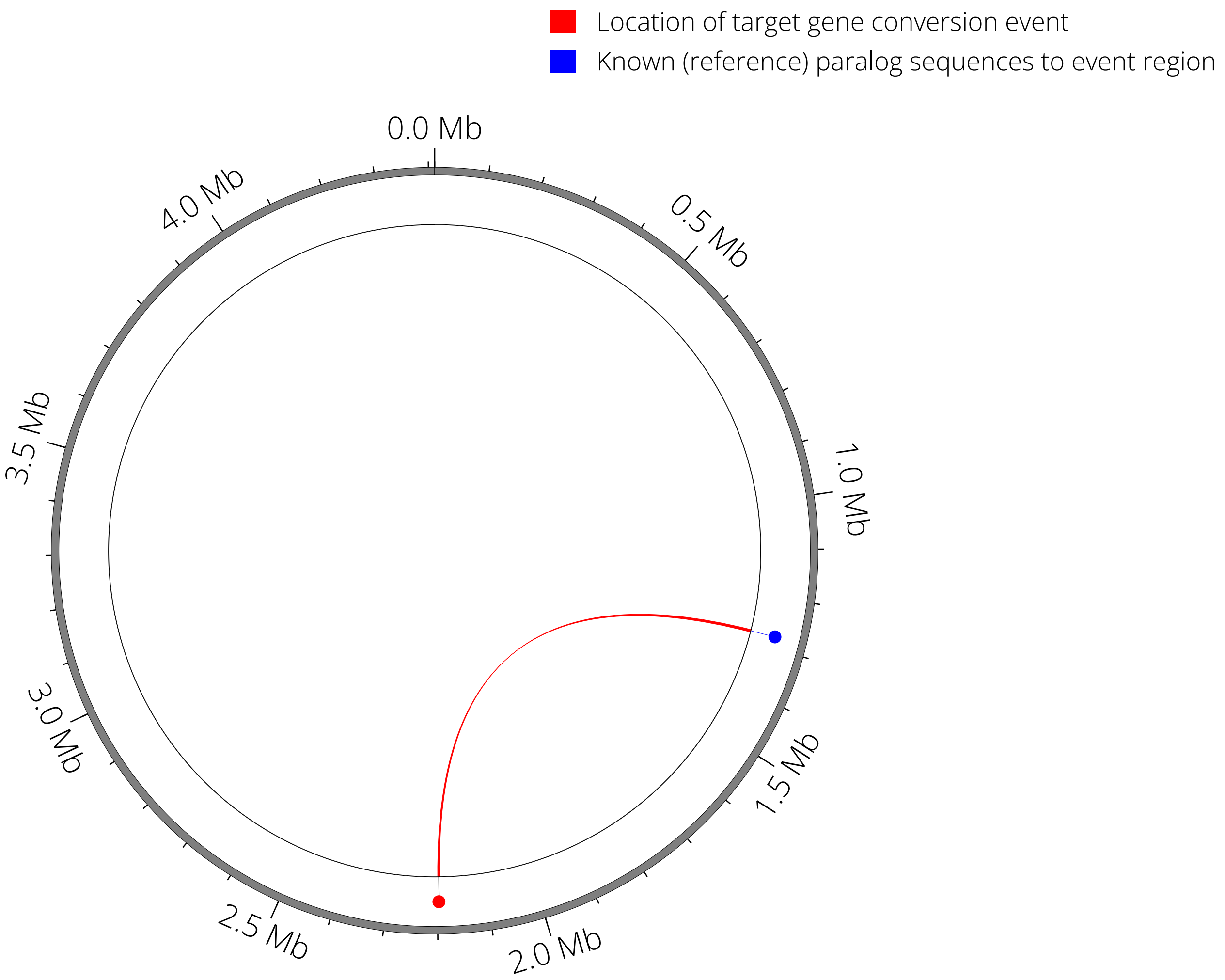

### Event_191 - 14 SNPs - k-mer analysis relative to paralogs

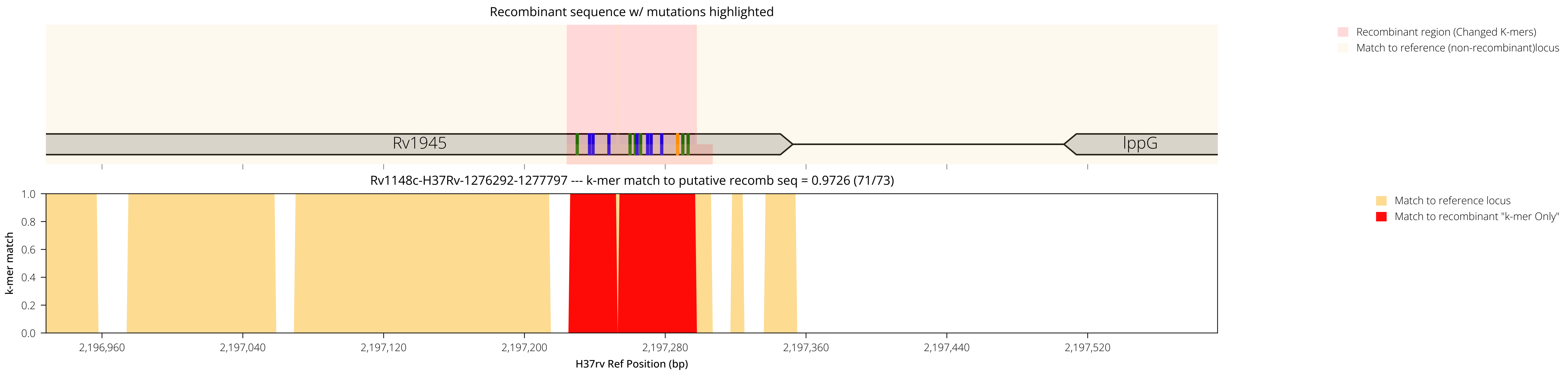

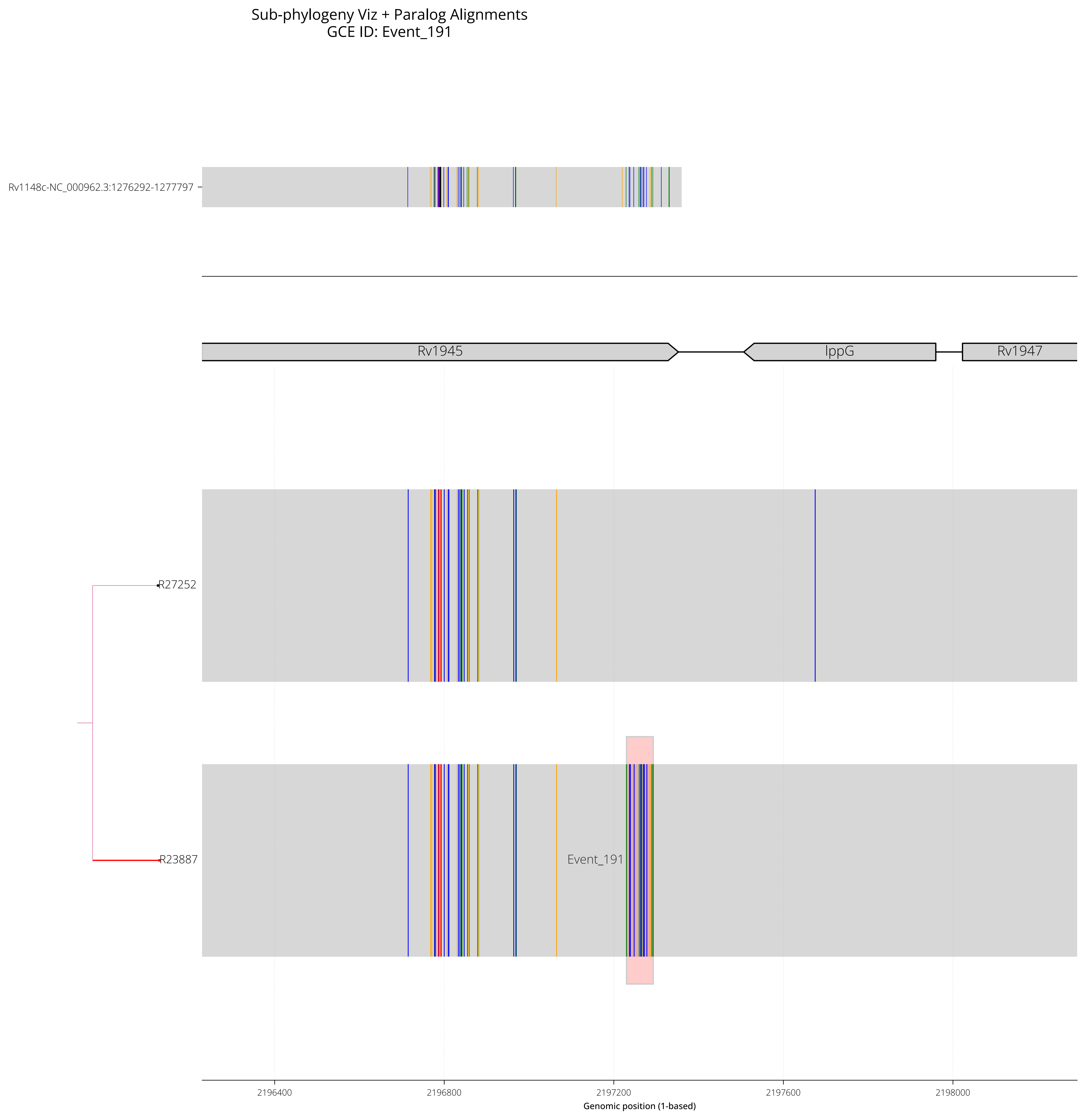

Sub-phylogeny Viz + Paralog Alignments
GCE ID: Event_191
Rv1148c-NC_000962.3:1276292-1277797
Rv1945
lppG
Rv1947
R27252
R23887
Event_191
2196400
2196800
2197200
2197600
2198000
Genomic position (1-based)

Full Phylogeny Viz + Paralog Alignments
GCE ID: Event_191

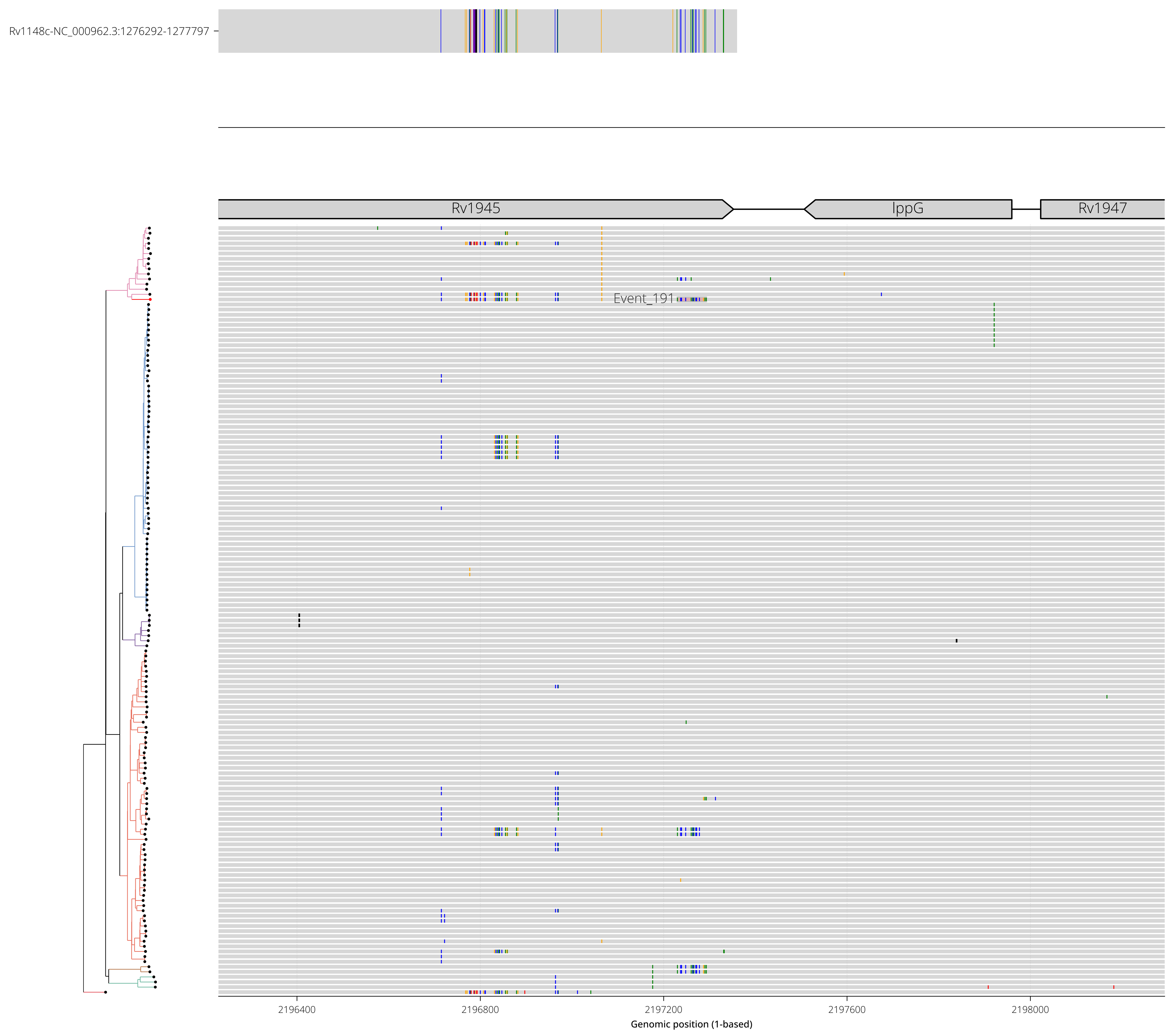

### Individual GCE + Known Paralogs
GCE ID: Event_192

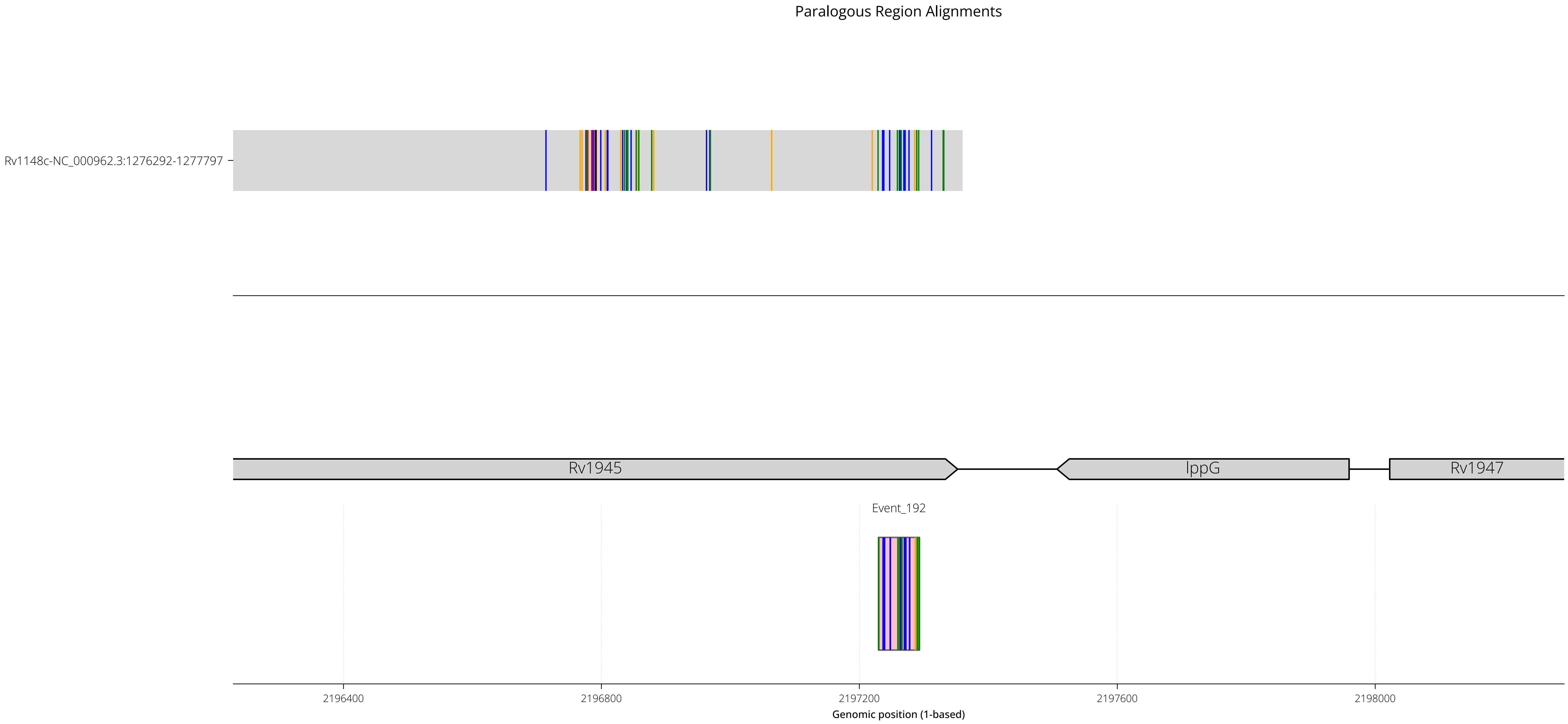

Circos Viz - GCE to Known Paralogs Locations
GCE ID: Event_192

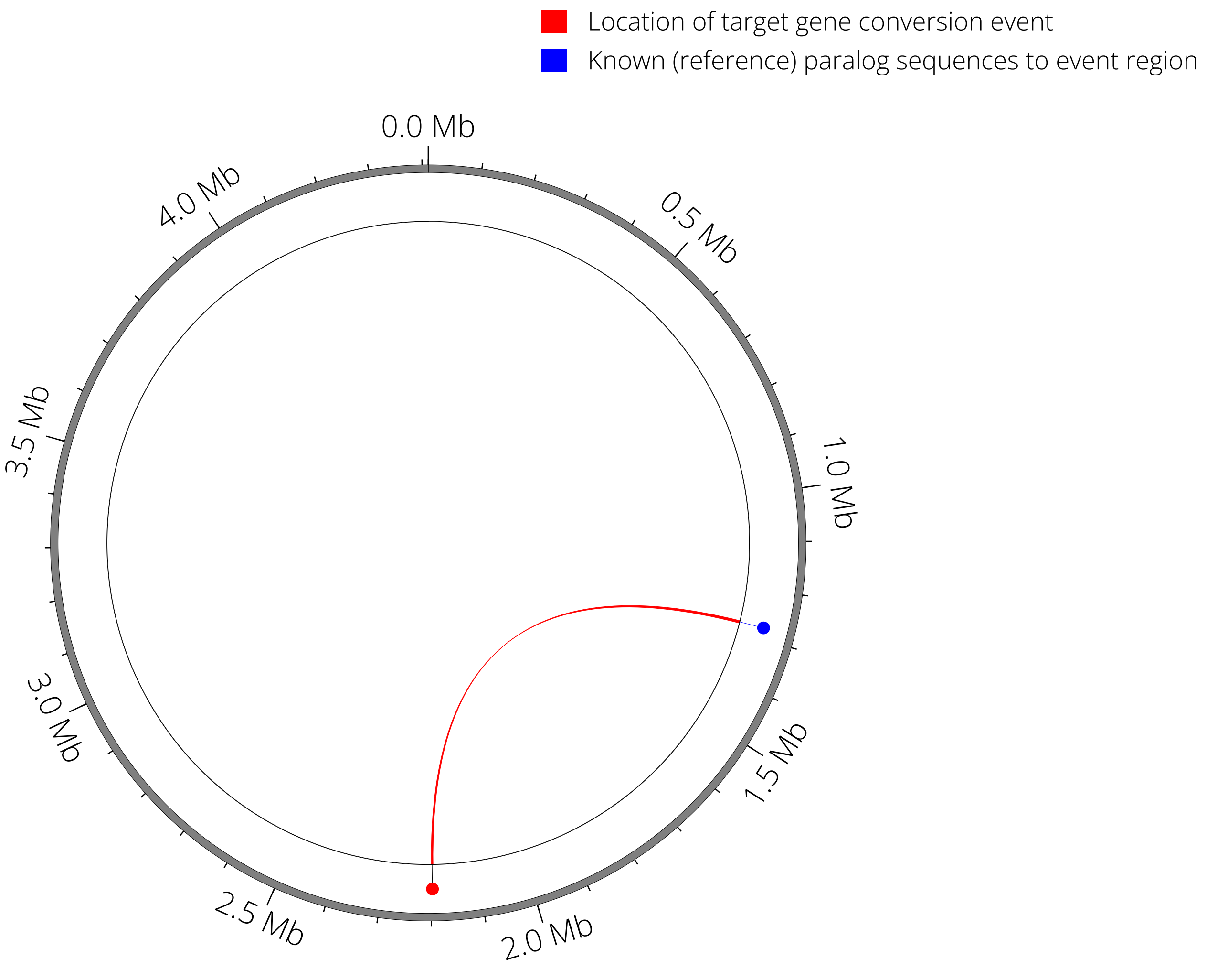

### Event_192 - 14 SNPs - k-mer analysis relative to paralogs

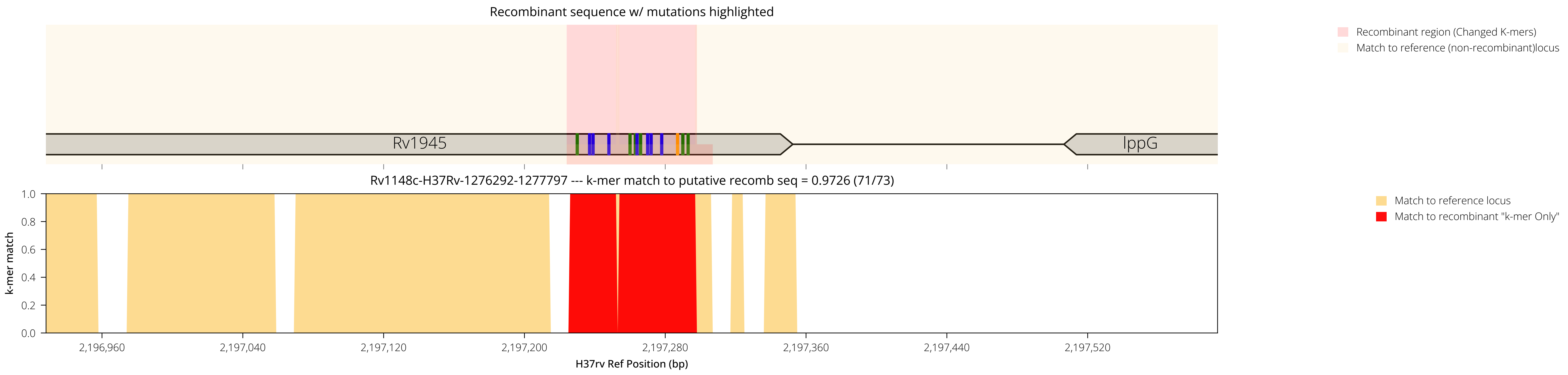

Sub-phylogeny Viz + Paralog Alignments
GCE ID: Event_192

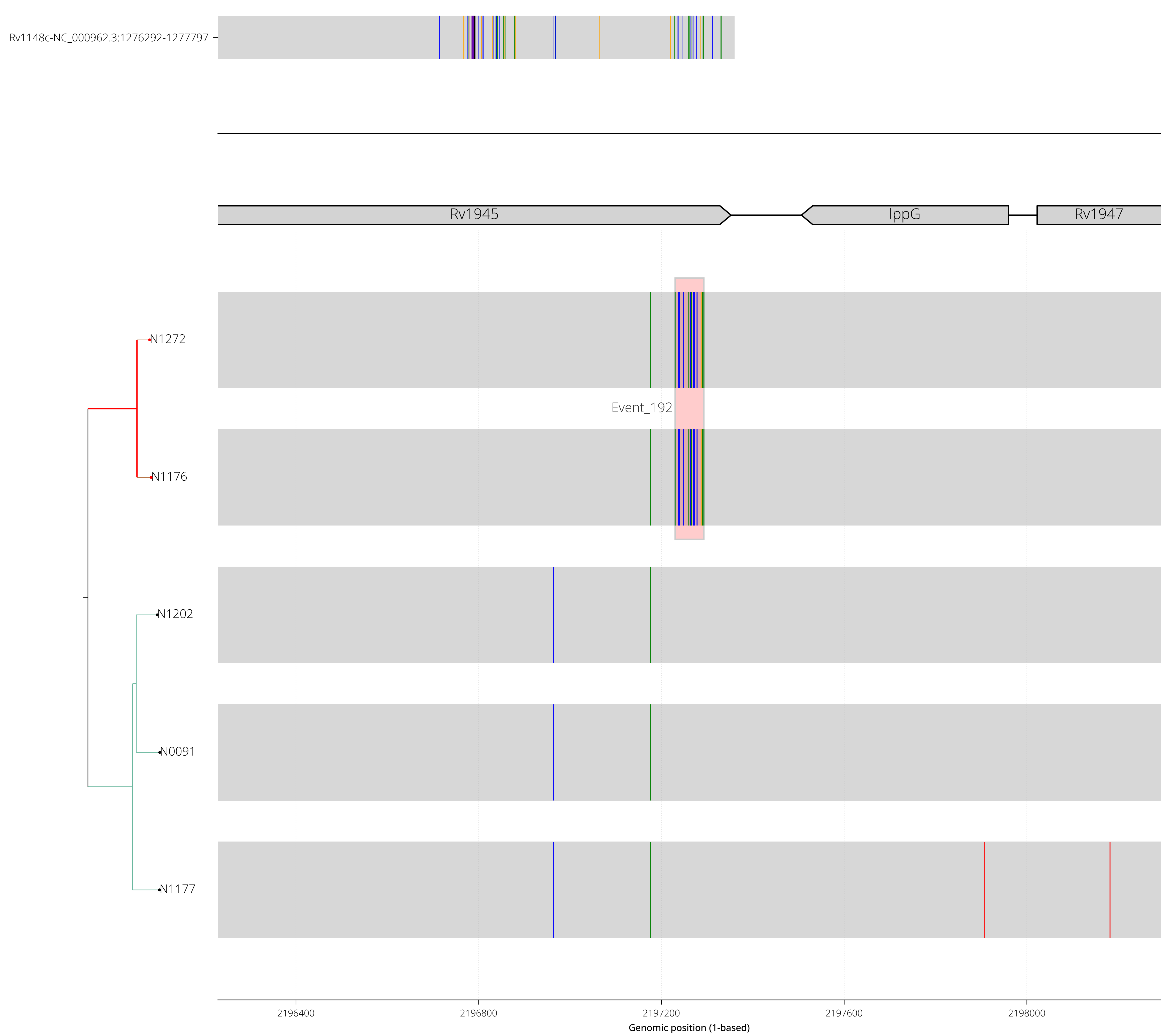

Full Phylogeny Viz + Paralog Alignments
GCE ID: Event_192

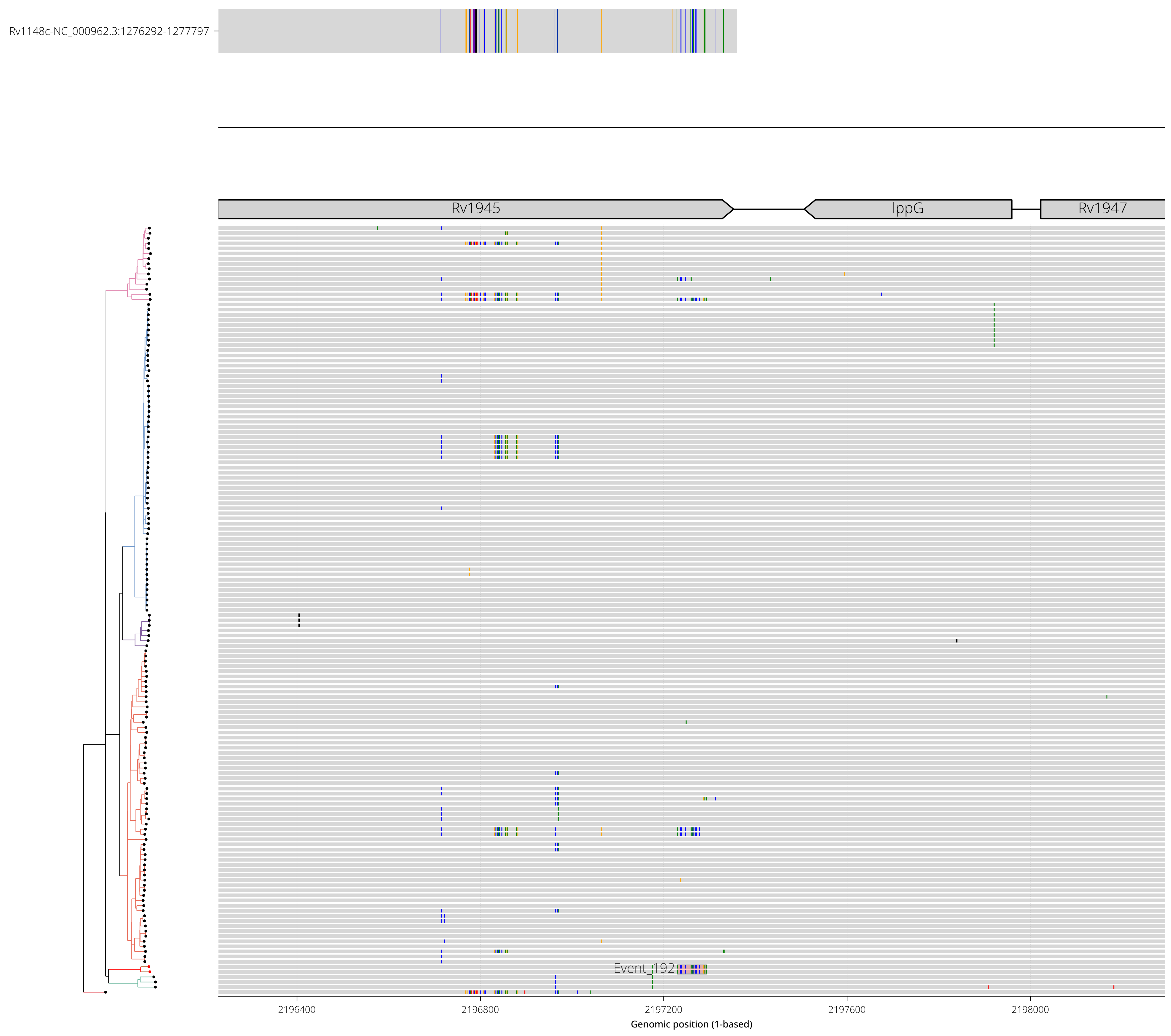

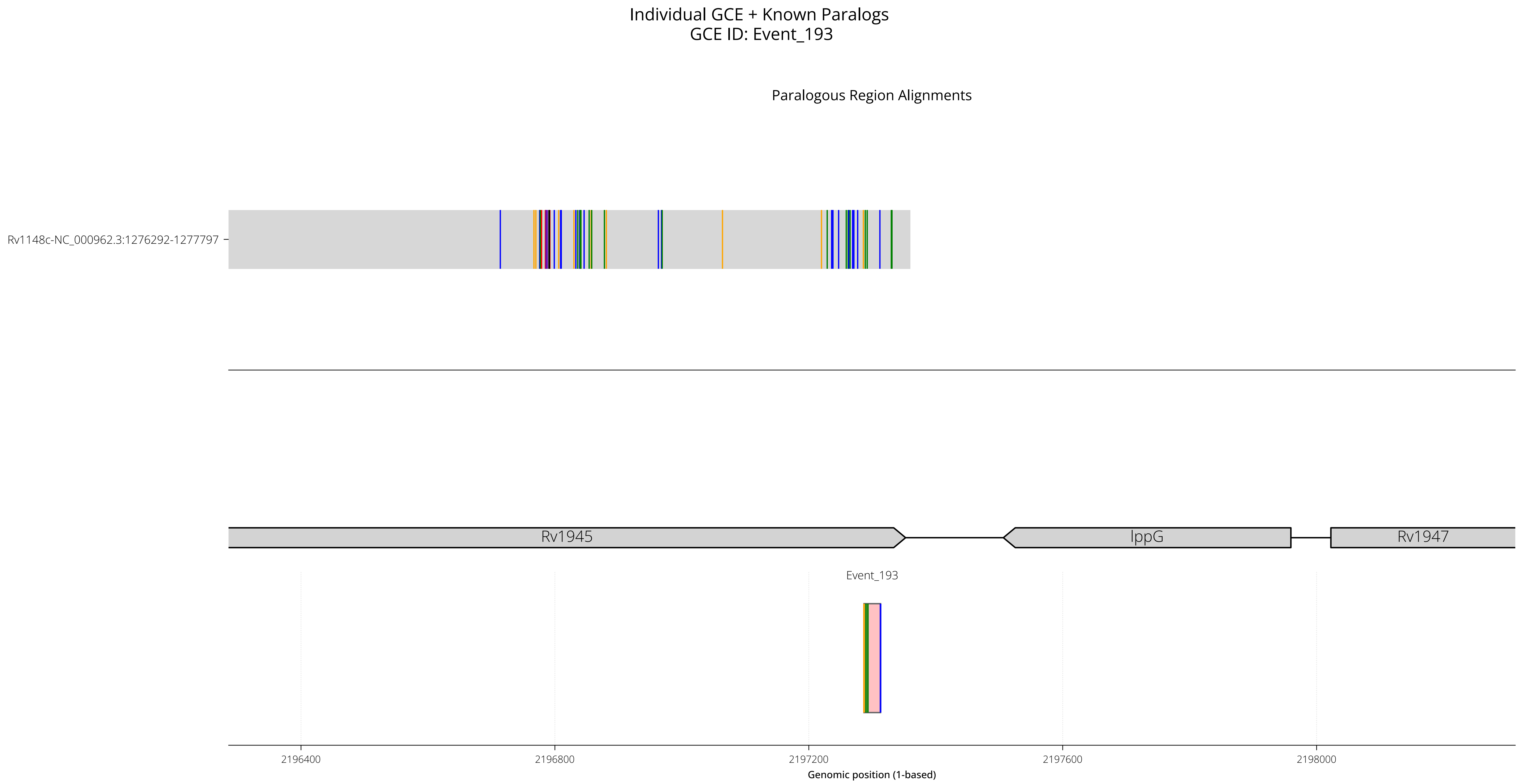

Individual GCE + Known Paralogs
GCE ID: Event_193
Paralogous Region Alignments
Rv1148c-NC_000962.3:1276292-1277797
Rv1945
lppG
Rv1947
Event_193
2196400
2196800
2197200
2197600
2198000
Genomic position (1-based)

Circos Viz - GCE to Known Paralogs Locations
GCE ID: Event_193

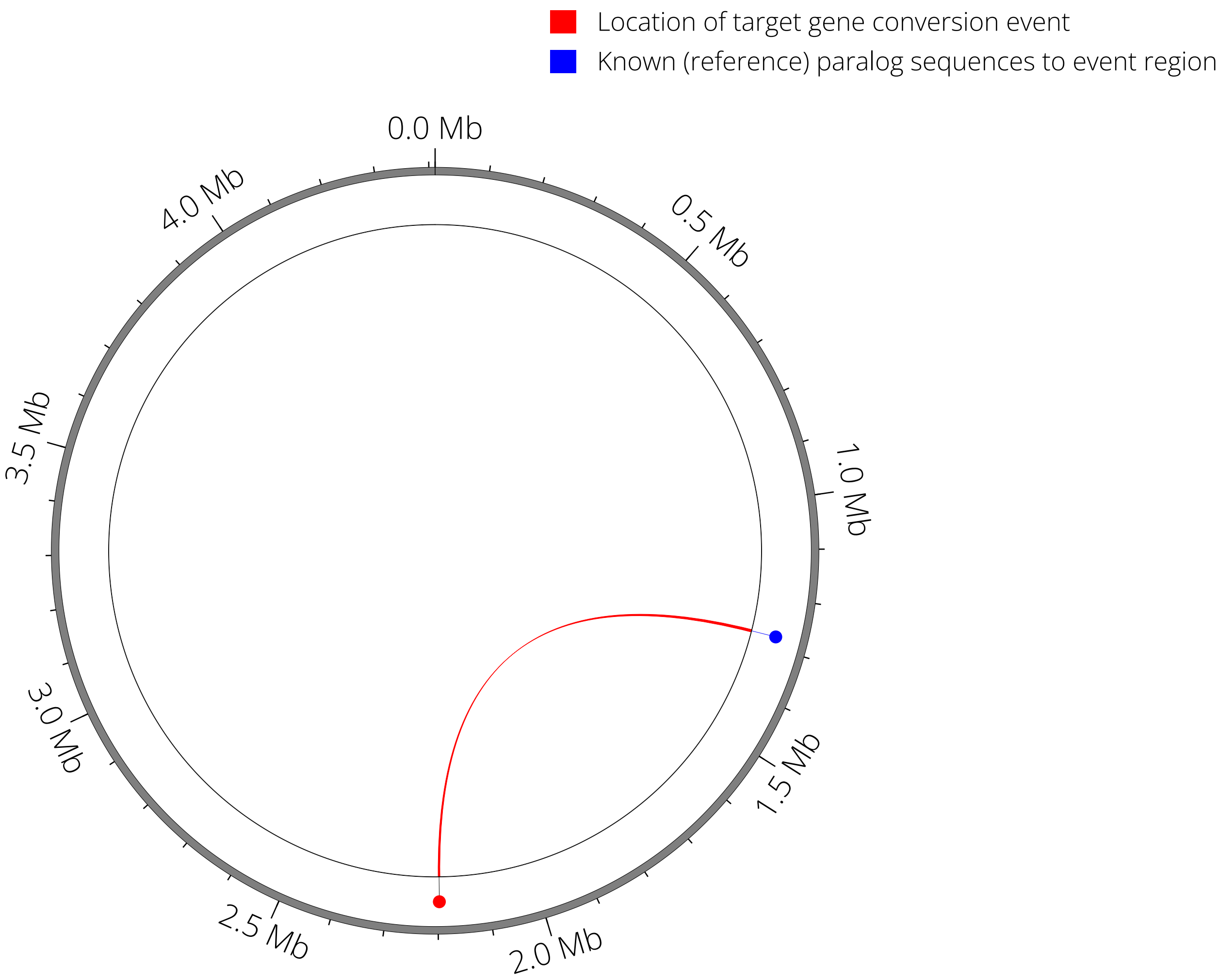

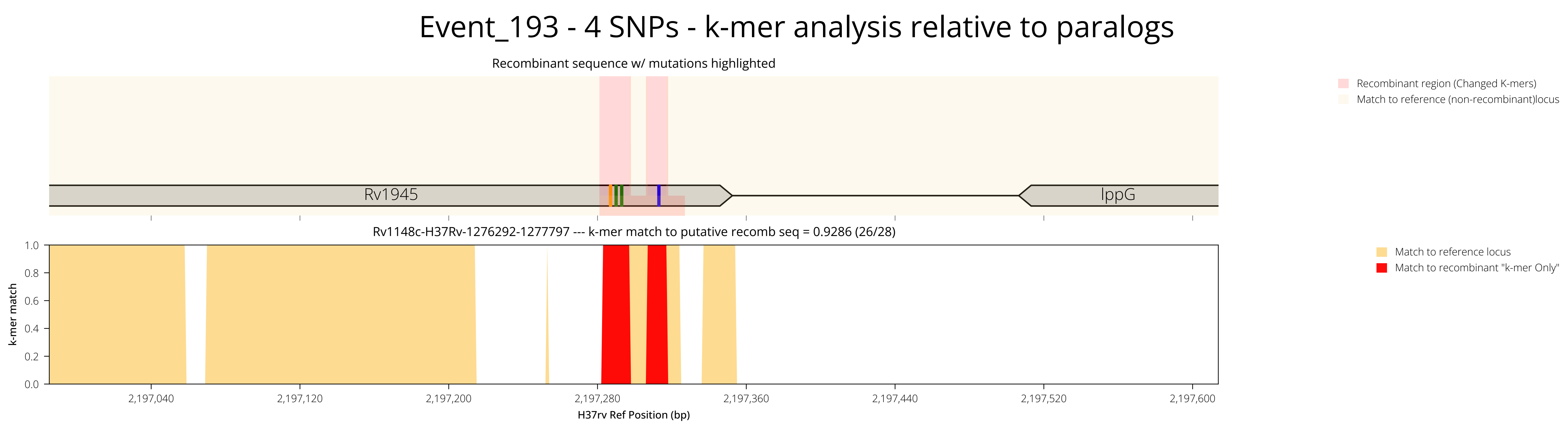

Event_193 - 4 SNPs - k-mer analysis relative to paralogs
Recombinant sequence w/ mutations highlighted
Recombinant region (Changed K-mers)
Match to reference (non-recombinant)locus
Rv1945
lppG
Rv1148c-H37Rv-1276292-1277797 --- k-mer match to putative recomb seq = 0.9286 (26/28)
Match to reference locus
Match to recombinant "k-mer Only"
k-mer match
1.0
0.8
0.6
0.4
0.2
0.0
2,197,040
2,197,120
2,197,200
2,197,280
2,197,360
2,197,440
2,197,520
2,197,600
H37rv Ref Position (bp)

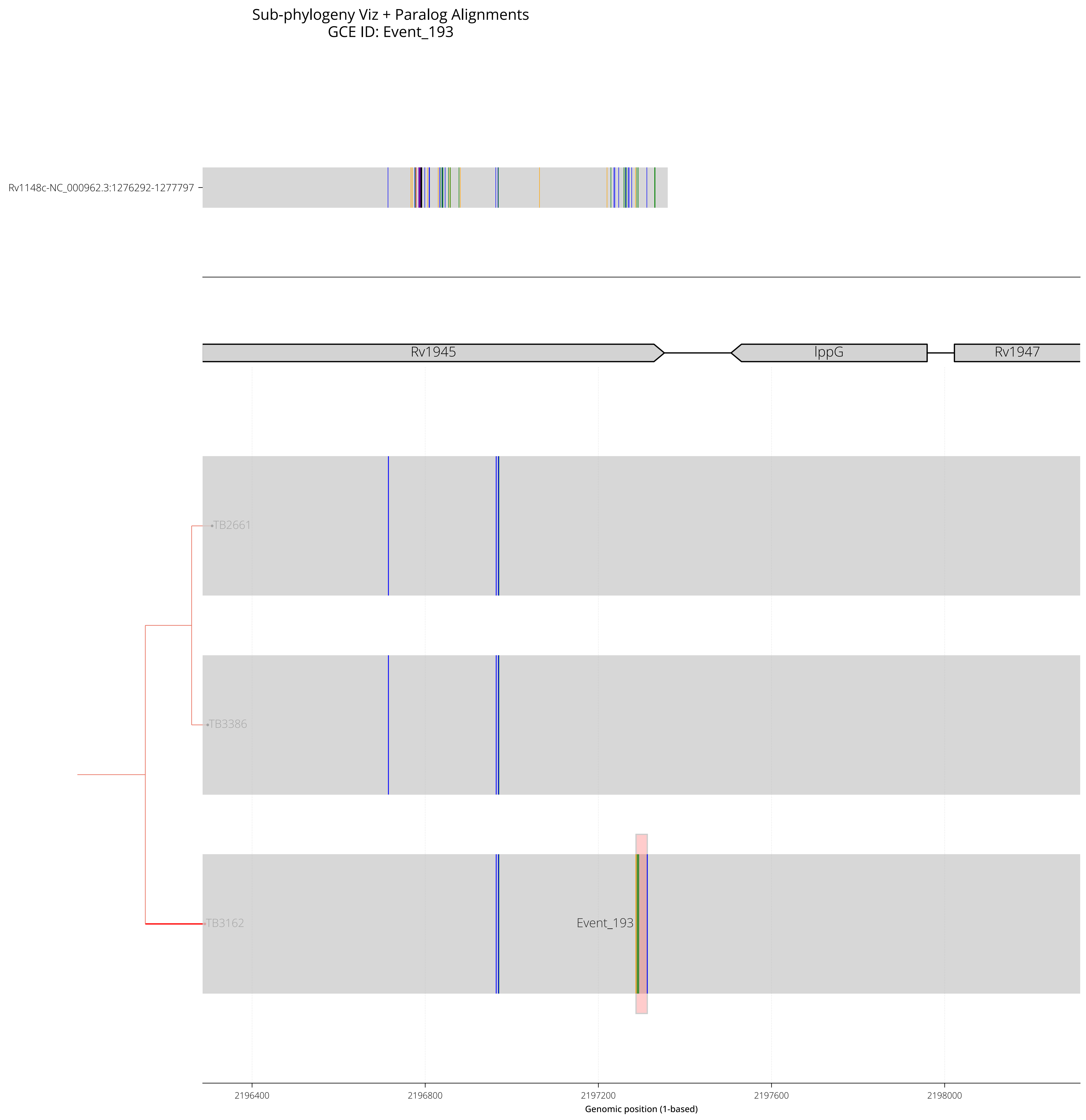

Sub-phylogeny Viz + Paralog Alignments
GCE ID: Event_193
Rv1148c-NC_000962.3:1276292-1277797
Rv1945
lppG
Rv1947
TB2661
TB3386
TB3162
Event_193
2196400
2196800
2197200
2197600
2198000
Genomic position (1-based)

Full Phylogeny Viz + Paralog Alignments
GCE ID: Event_193

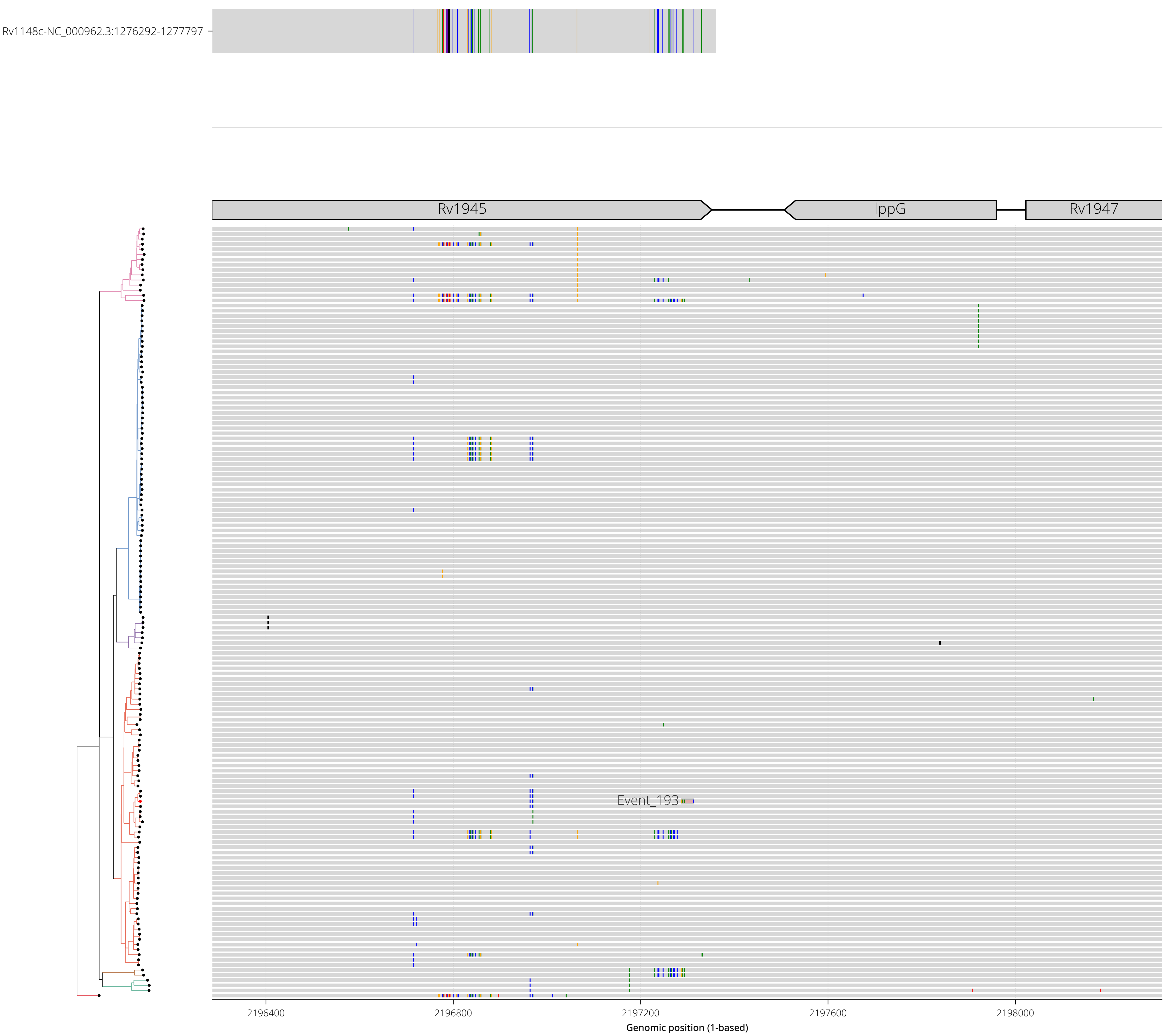

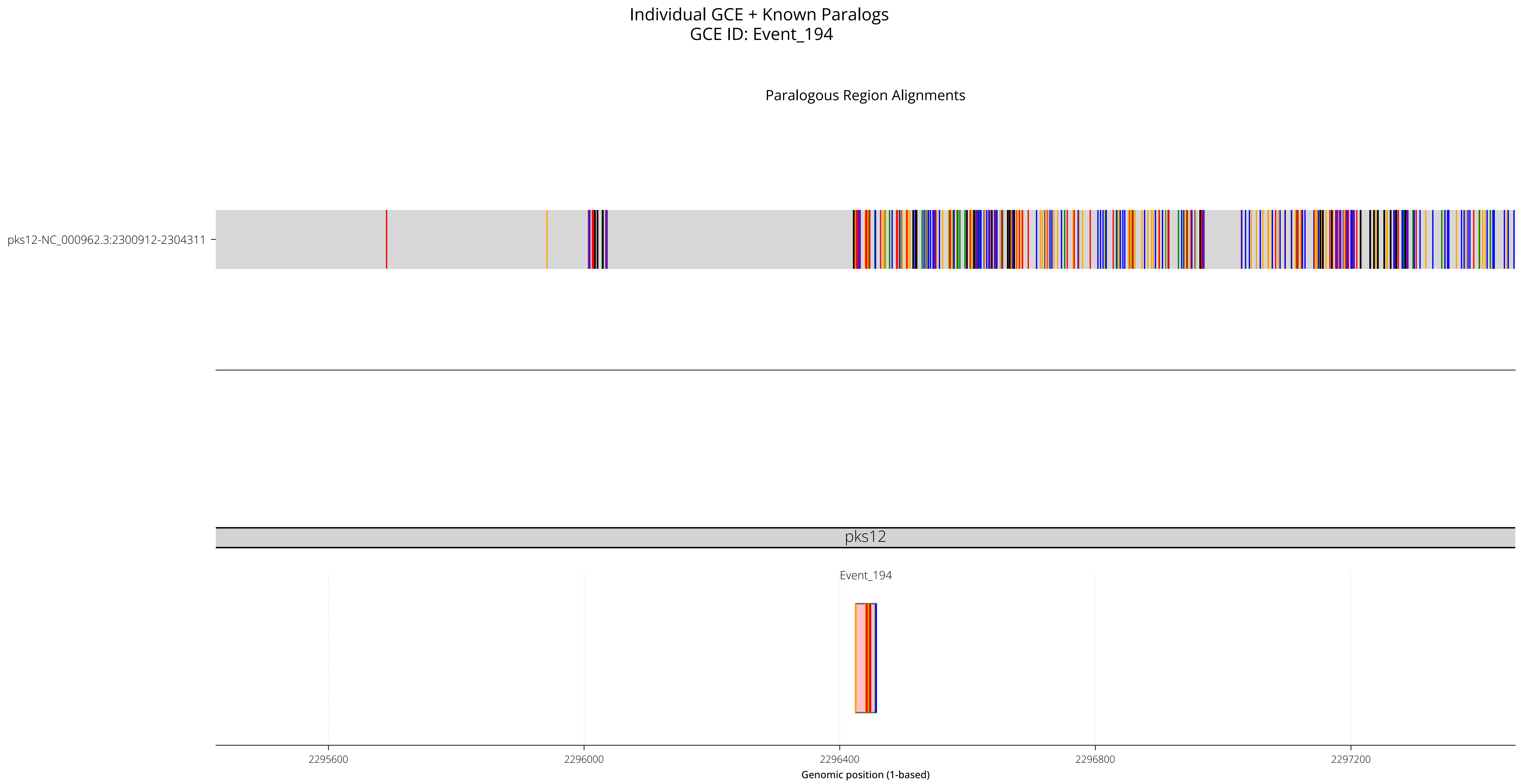

Individual GCE + Known Paralogs
GCE ID: Event_194
Paralogous Region Alignments
pks12-NC_000962.3:2300912-2304311
pks12
Event_194
2295600
2296000
2296400
2296800
2297200
Genomic position (1-based)

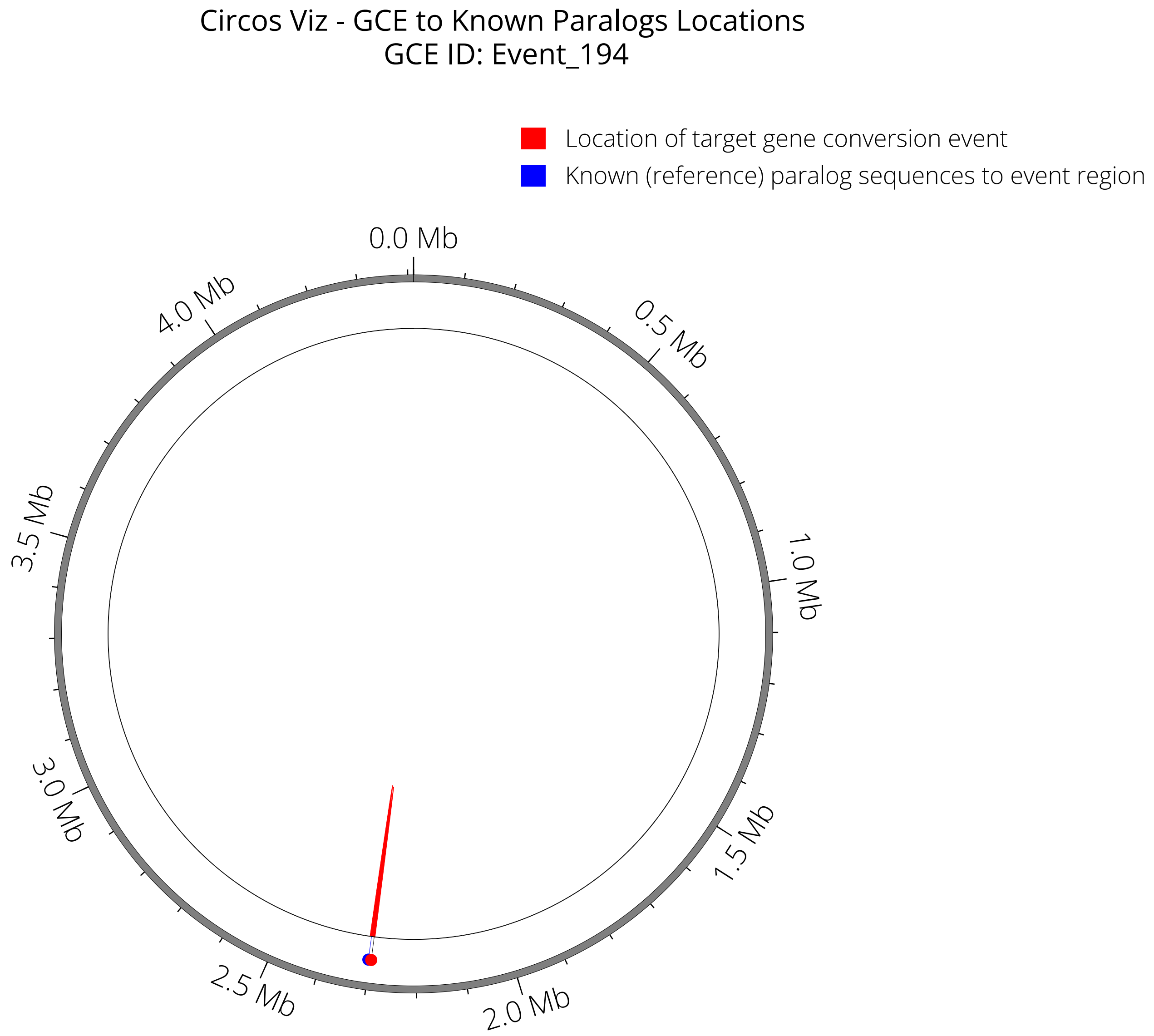
Circos Viz - GCE to Known Paralogs Locations
GCE ID: Event_194
Location of target gene conversion event
Known (reference) paralog sequences to event region
0.0 Mb
0.5 Mb
1.0 Mb
1.5 Mb
2.0 Mb
2.5 Mb
3.0 Mb
3.5 Mb
4.0 Mb

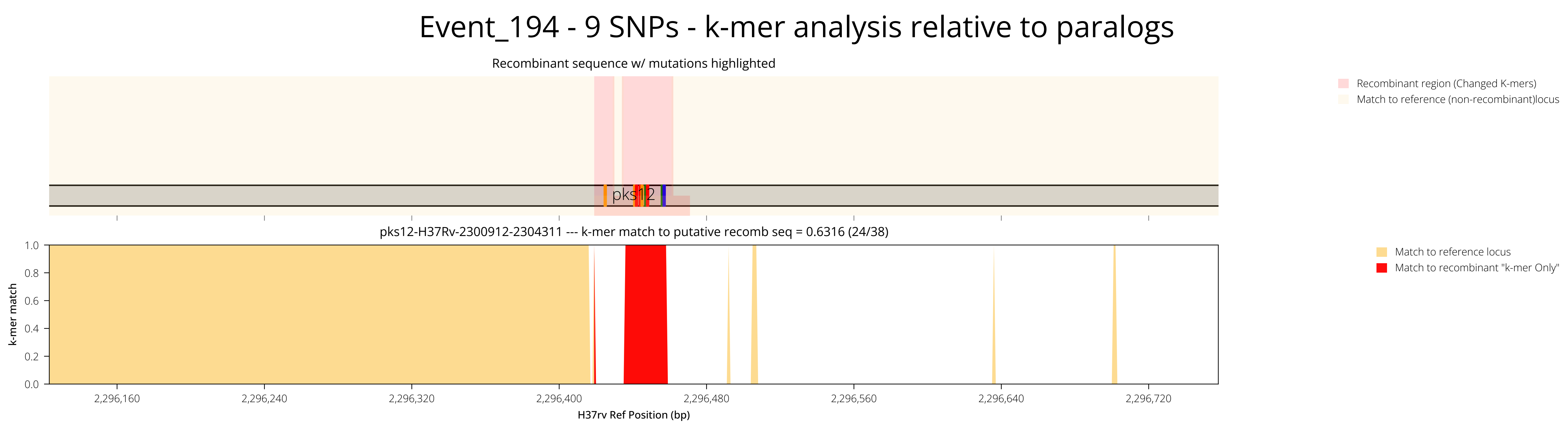
Event_194 - 9 SNPs - k-mer analysis relative to paralogs
Recombinant sequence w/ mutations highlighted
Recombinant region (Changed K-mers)
Match to reference (non-recombinant)locus
pks12
pks12-H37Rv-2300912-2304311 --- k-mer match to putative recomb seq = 0.6316 (24/38)
Match to reference locus
Match to recombinant "k-mer Only"
k-mer match
1.0
0.8
0.6
0.4
0.2
0.0
2,296,160
2,296,240
2,296,320
2,296,400
2,296,480
2,296,560
2,296,640
2,296,720
H37rv Ref Position (bp)

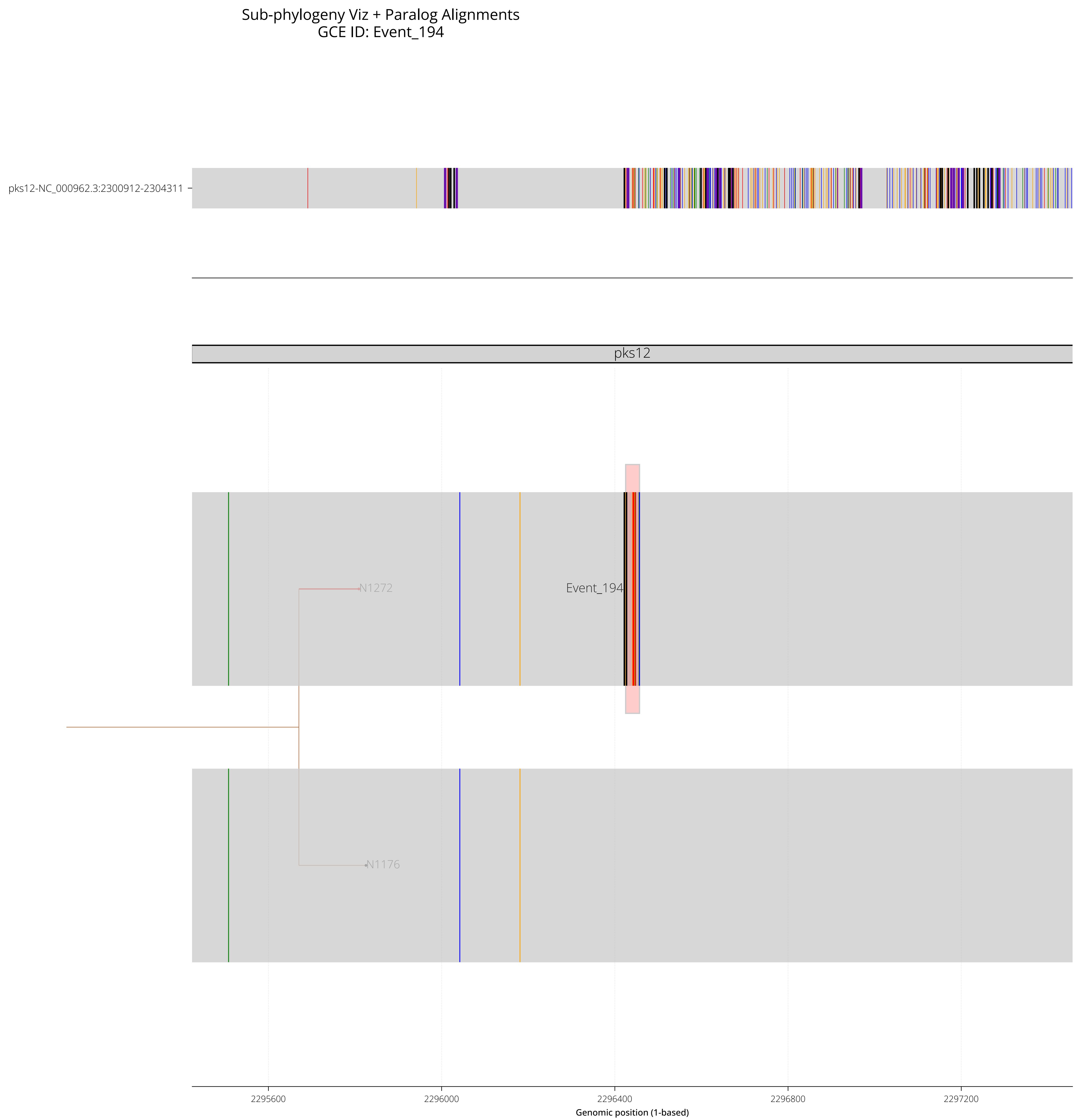

Sub-phylogeny Viz + Paralog Alignments
GCE ID: Event_194
pks12-NC_000962.3:2300912-2304311
pks12
N1272
Event_194
N1176
2295600
2296000
2296400
2296800
2297200
Genomic position (1-based)

Full Phylogeny Viz + Paralog Alignments
GCE ID: Event_194

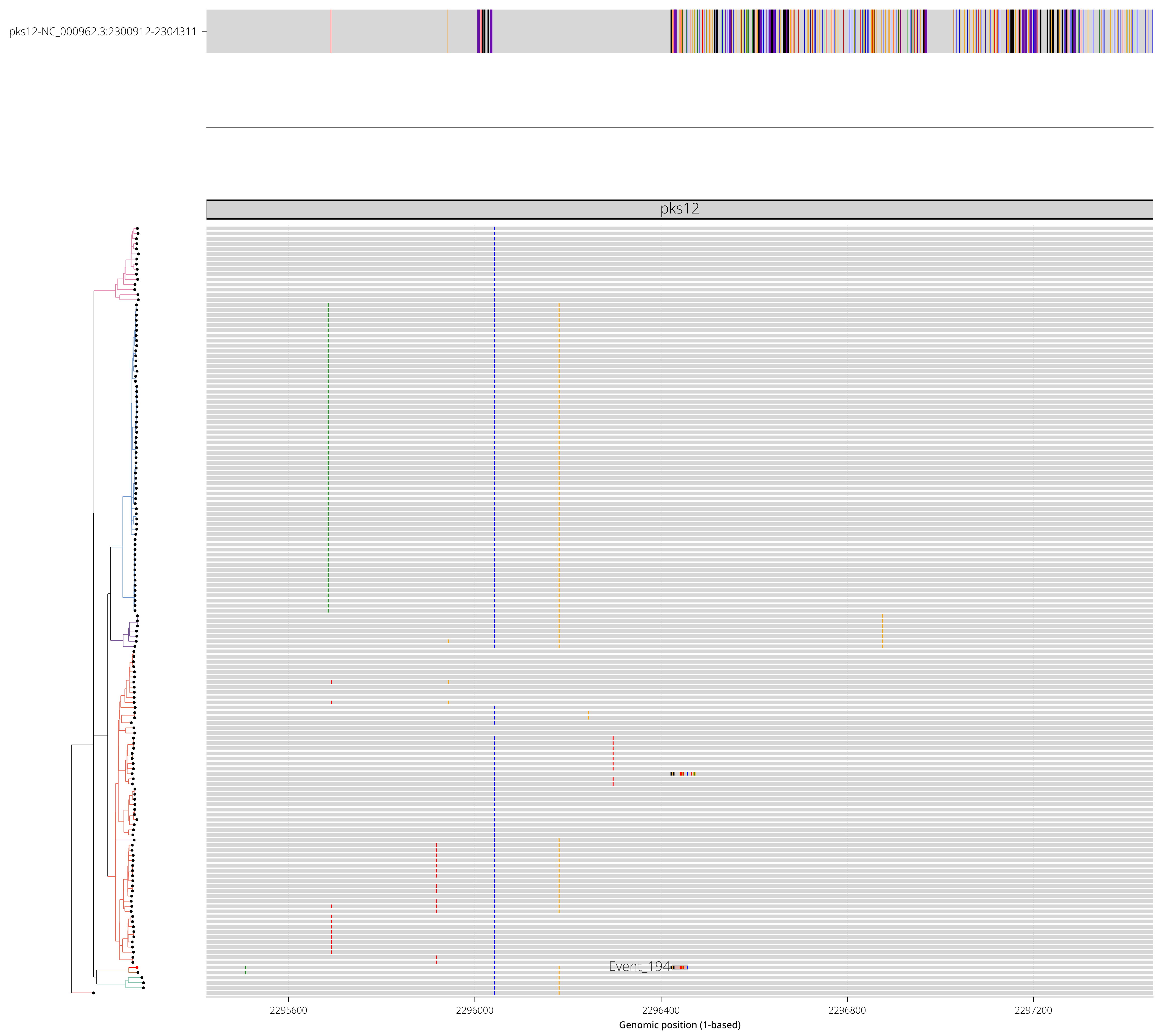

Individual GCE + Known Paralogs
GCE ID: Event_195

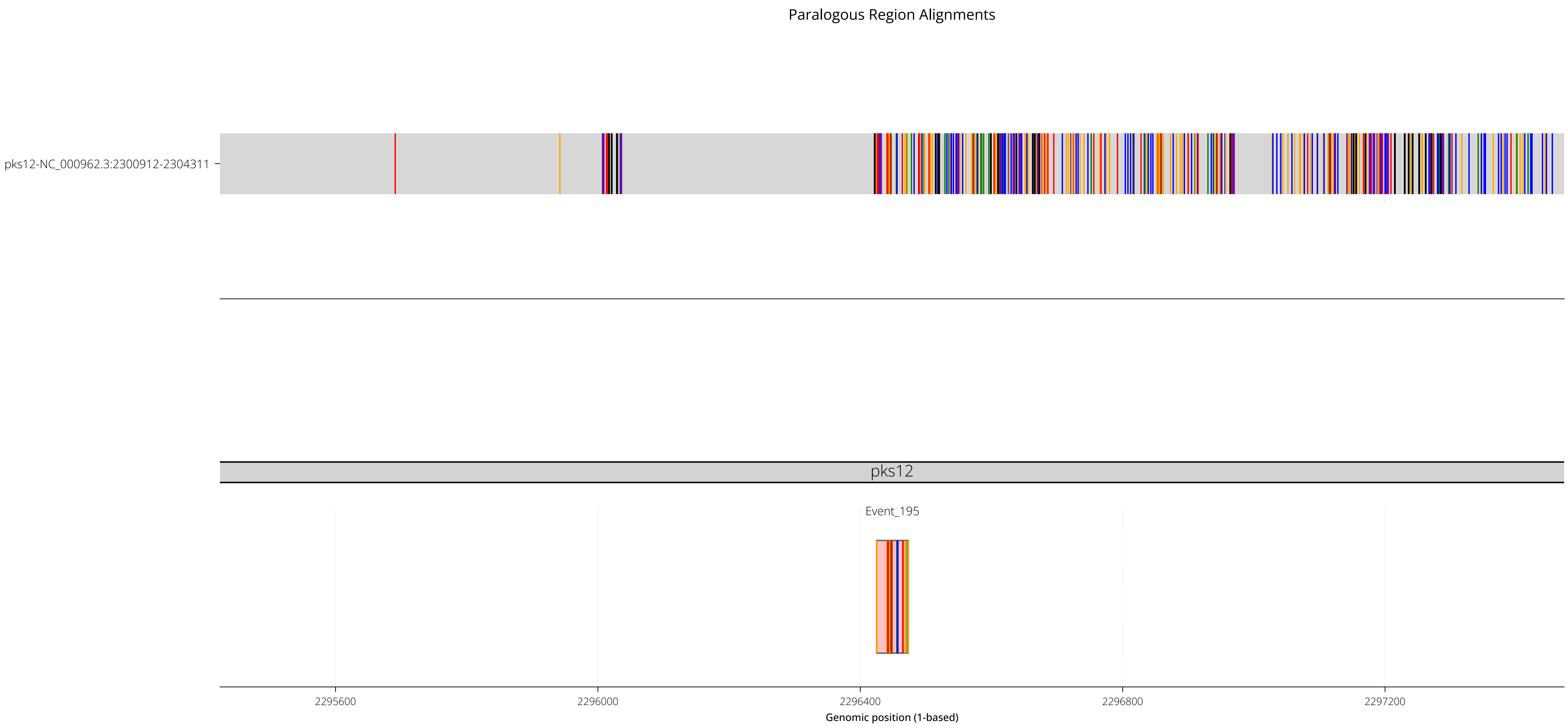

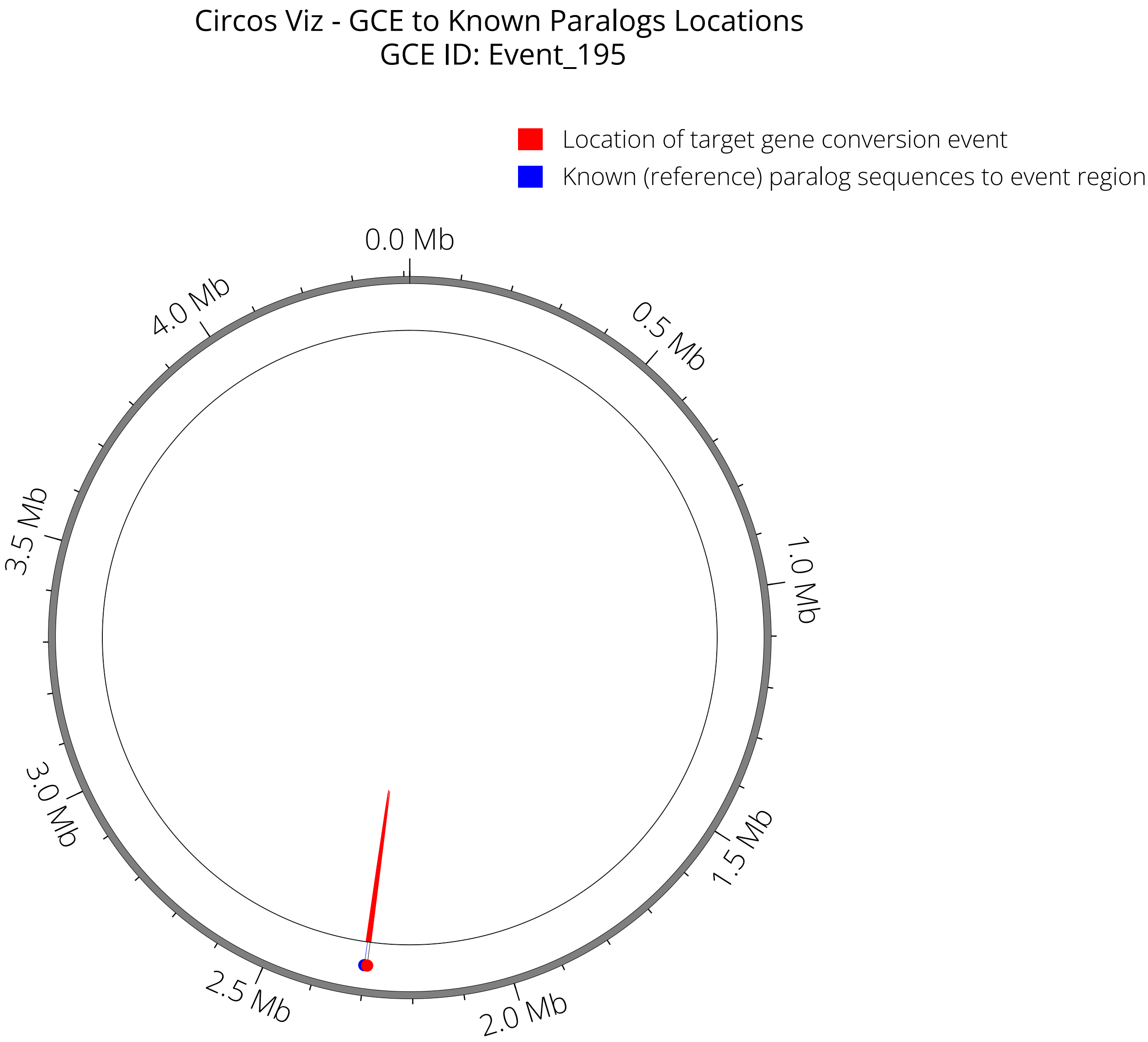

Circos Viz - GCE to Known Paralogs Locations
GCE ID: Event_195
Location of target gene conversion event
Known (reference) paralog sequences to event region
0.0 Mb
0.5 Mb
1.0 Mb
1.5 Mb
2.0 Mb
2.5 Mb
3.0 Mb
3.5 Mb
4.0 Mb

### Event_195 - 13 SNPs - k-mer analysis relative to paralogs

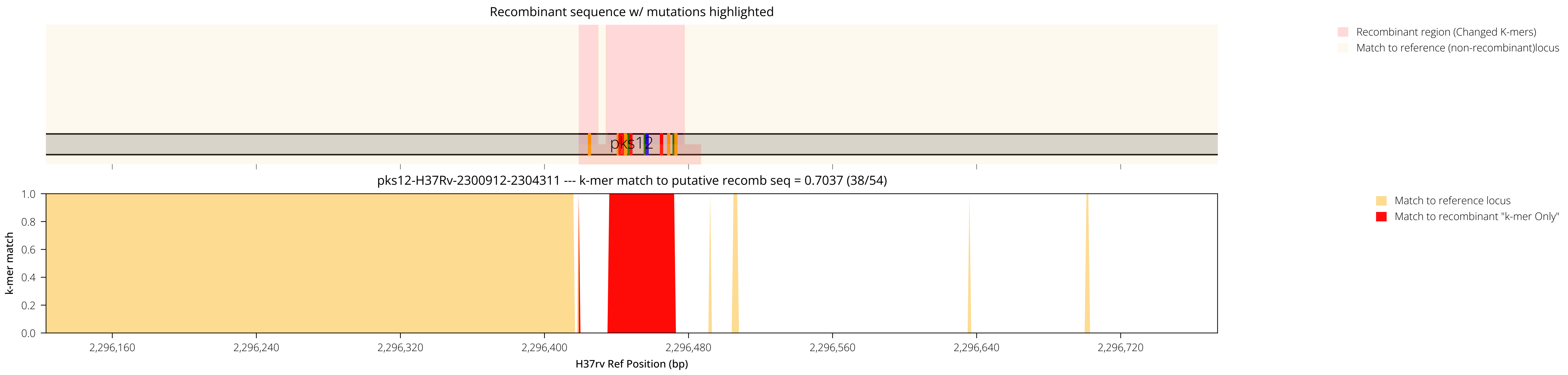

### Sub-phylogeny Viz + Paralog Alignments
GCE ID: Event_195

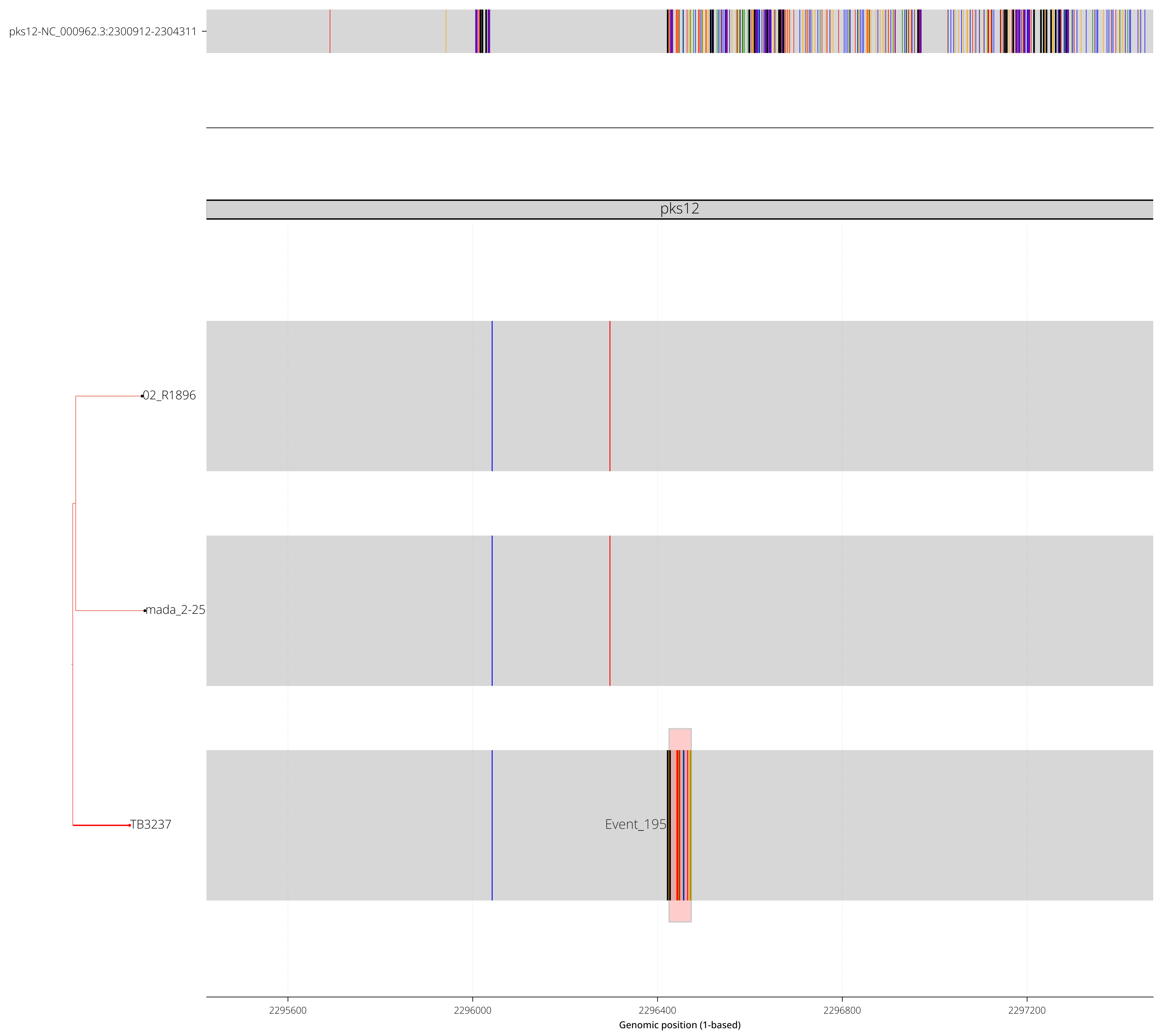

Full Phylogeny Viz + Paralog Alignments
GCE ID: Event_195

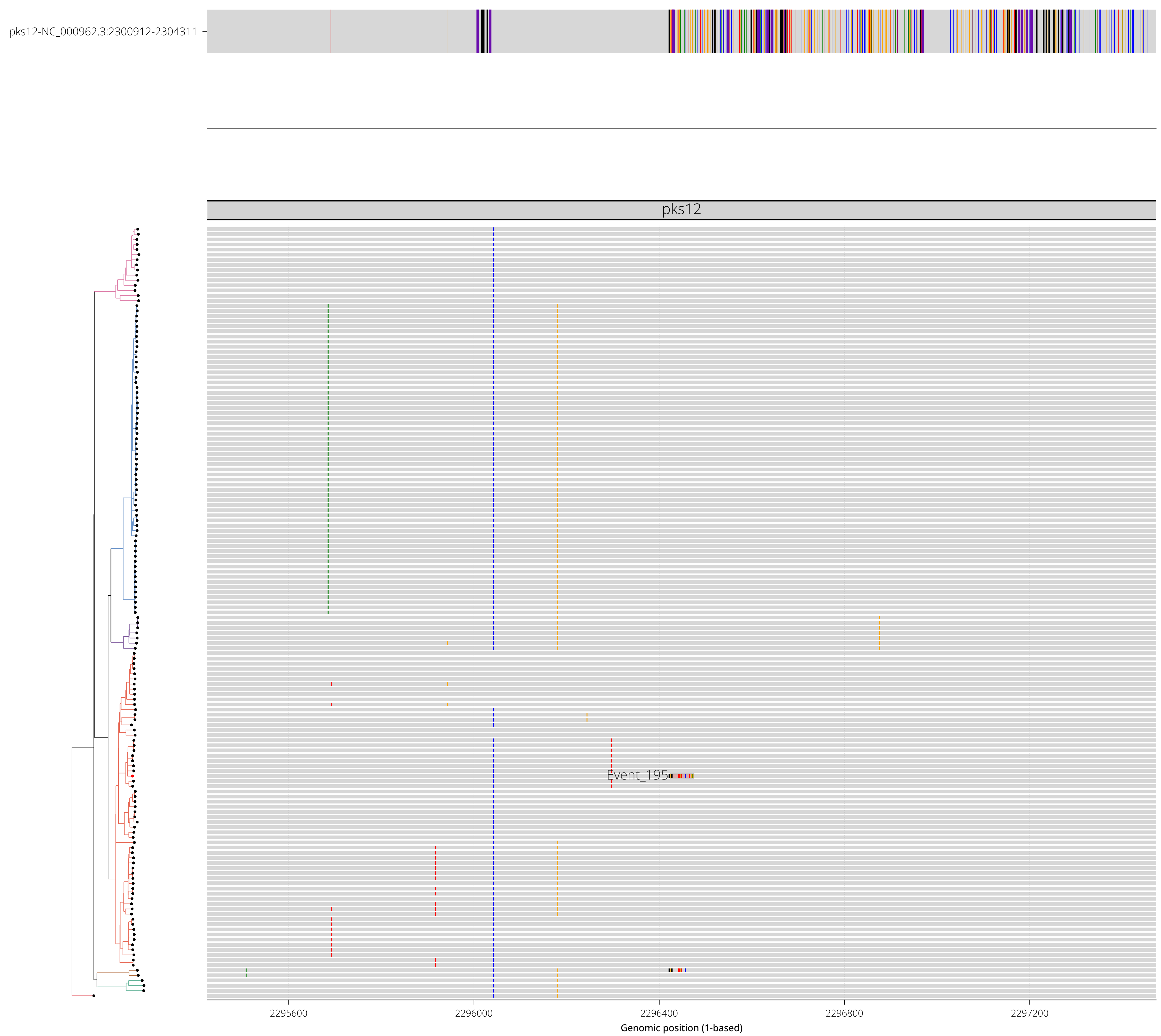

Individual GCE + Known Paralogs
GCE ID: Event_196

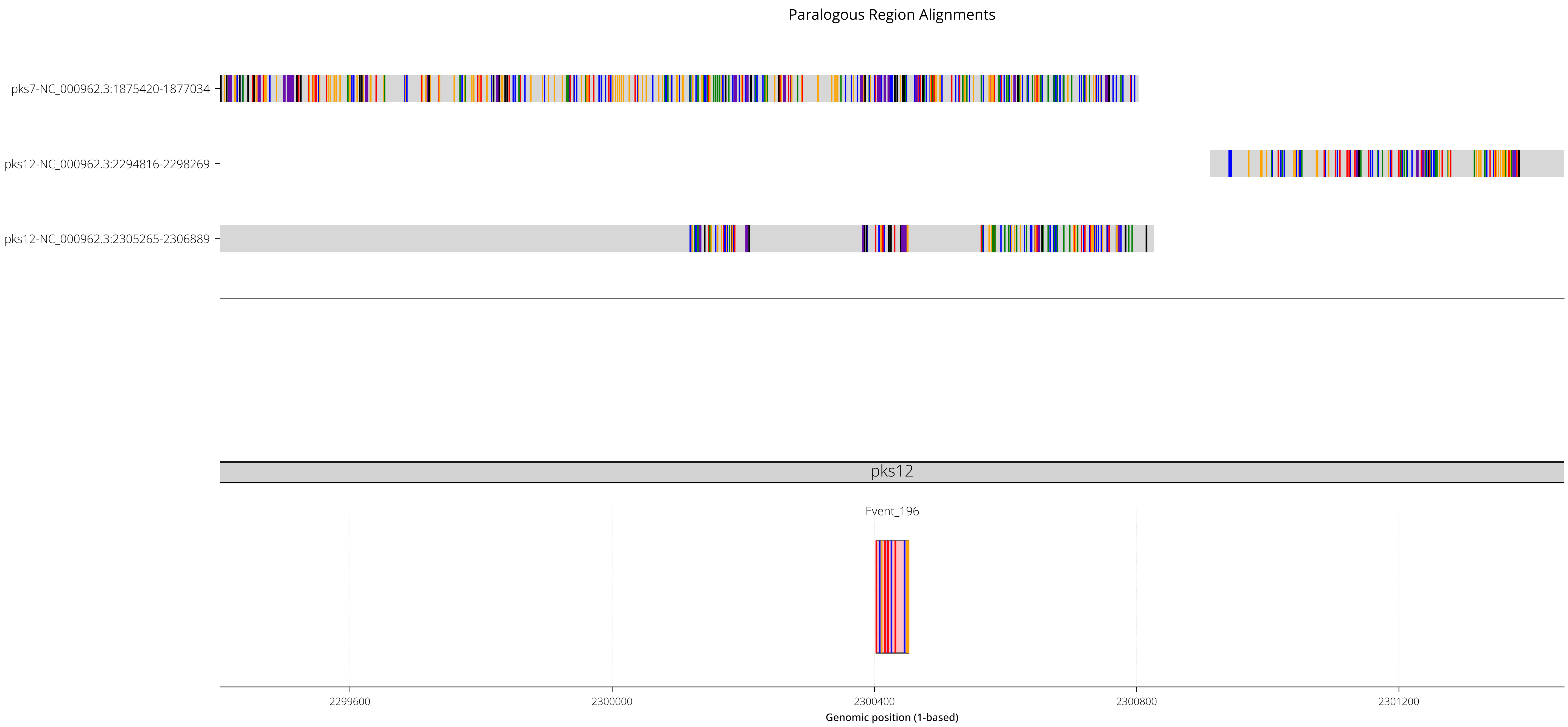

Circos Viz - GCE to Known Paralogs Locations
GCE ID: Event_196

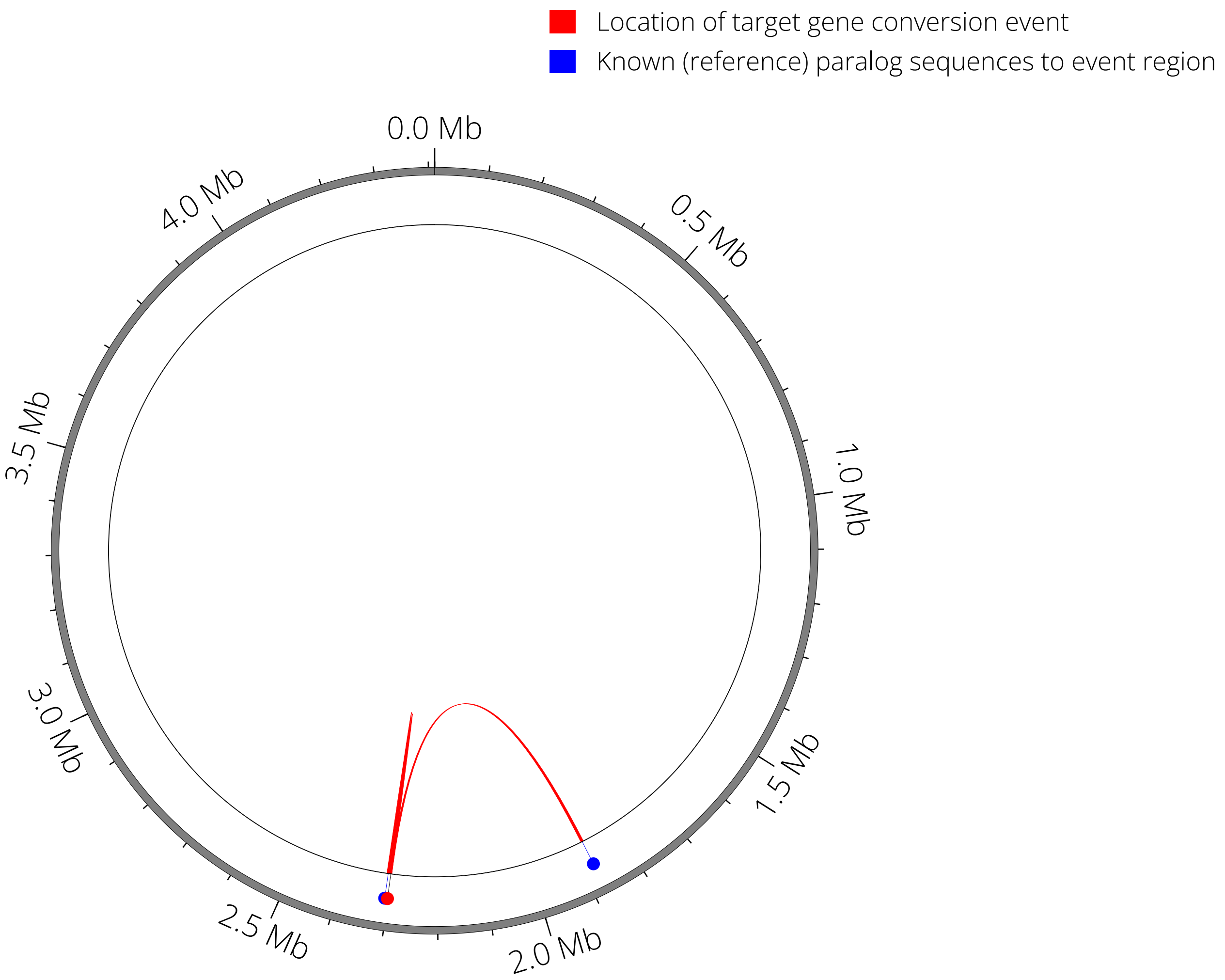

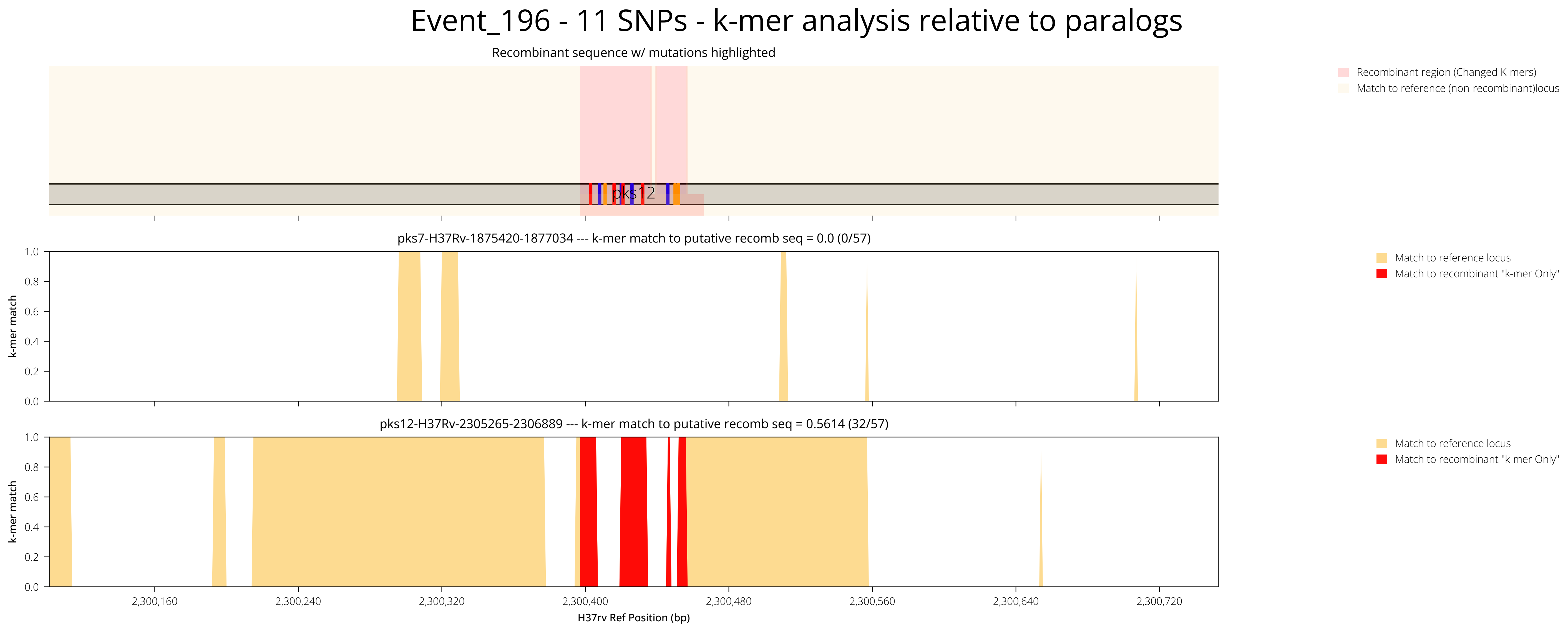

Event_196 - 11 SNPs - k-mer analysis relative to paralogs
Recombinant sequence w/ mutations highlighted
pks12
Recombinant region (Changed K-mers)
Match to reference (non-recombinant)locus
pks7-H37Rv-1875420-1877034 --- k-mer match to putative recomb seq = 0.0 (0/57)
Match to reference locus
Match to recombinant "k-mer Only"
pks12-H37Rv-2305265-2306889 --- k-mer match to putative recomb seq = 0.5614 (32/57)
k-mer match
1.0
0.8
0.6
0.4
0.2
0.0
2,300,160
2,300,240
2,300,320
2,300,400
2,300,480
2,300,560
2,300,640
2,300,720
H37rv Ref Position (bp)

Sub-phylogeny Viz + Paralog Alignments
GCE ID: Event_196

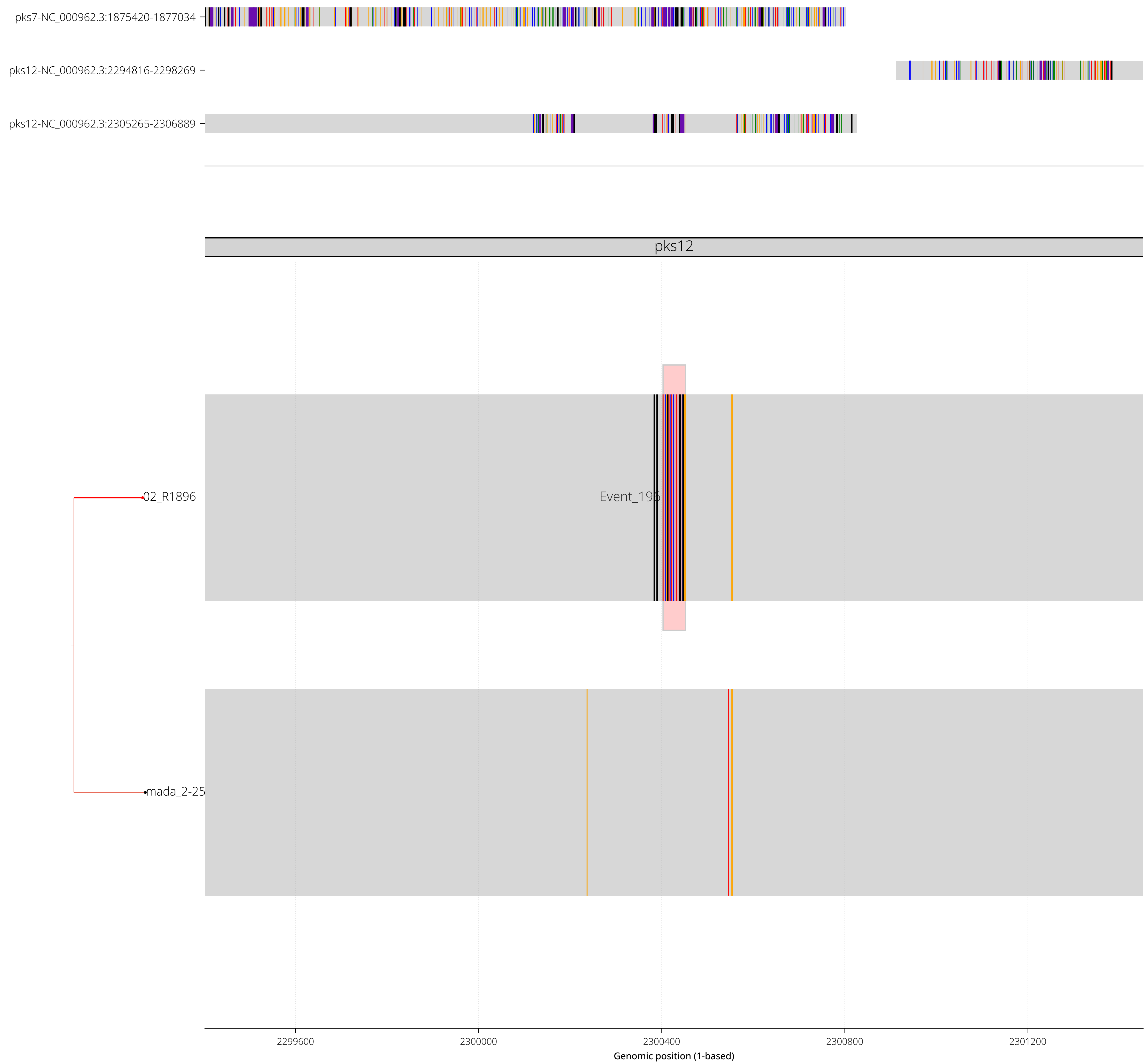

Full Phylogeny Viz + Paralog Alignments
GCE ID: Event_196

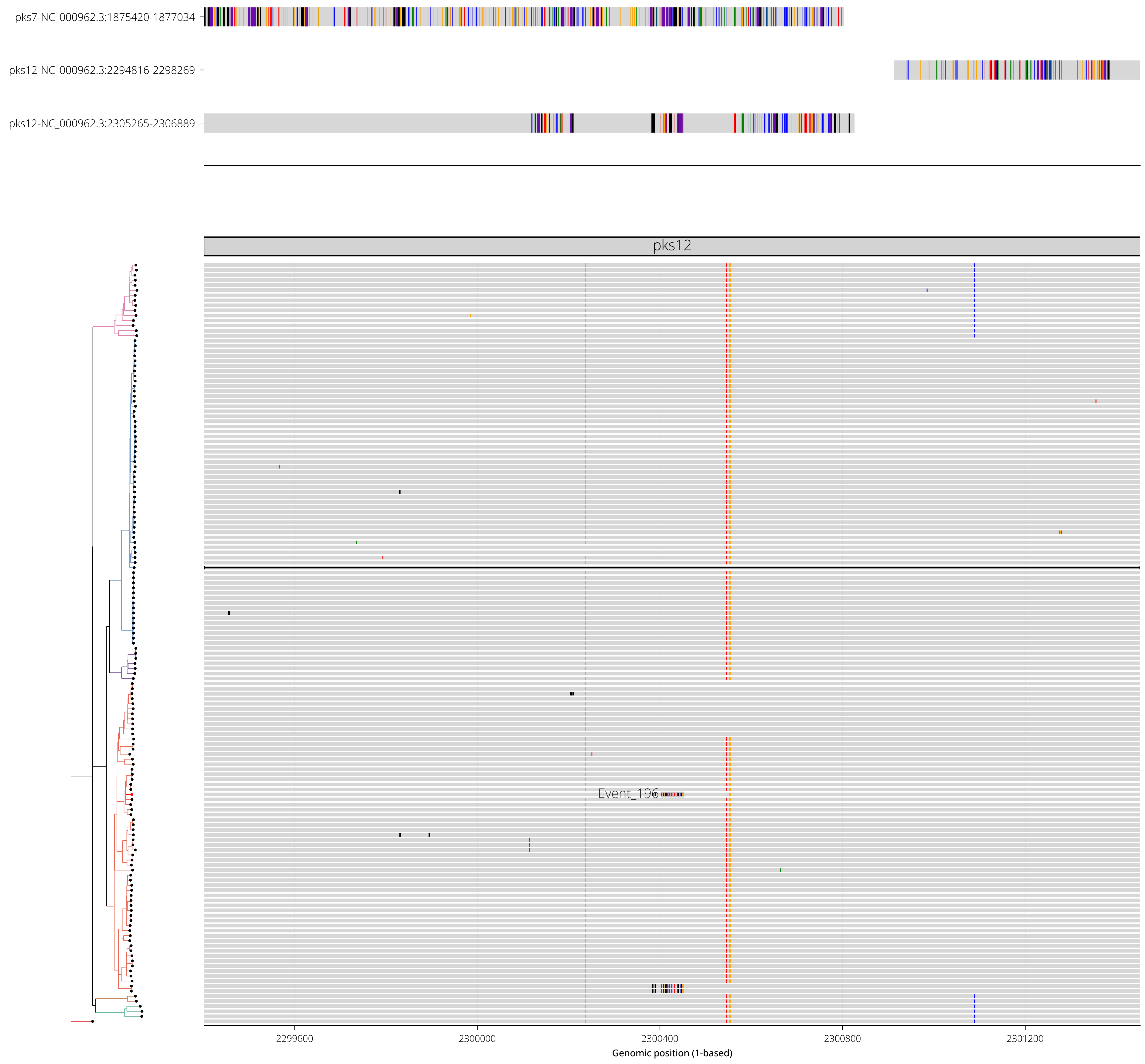

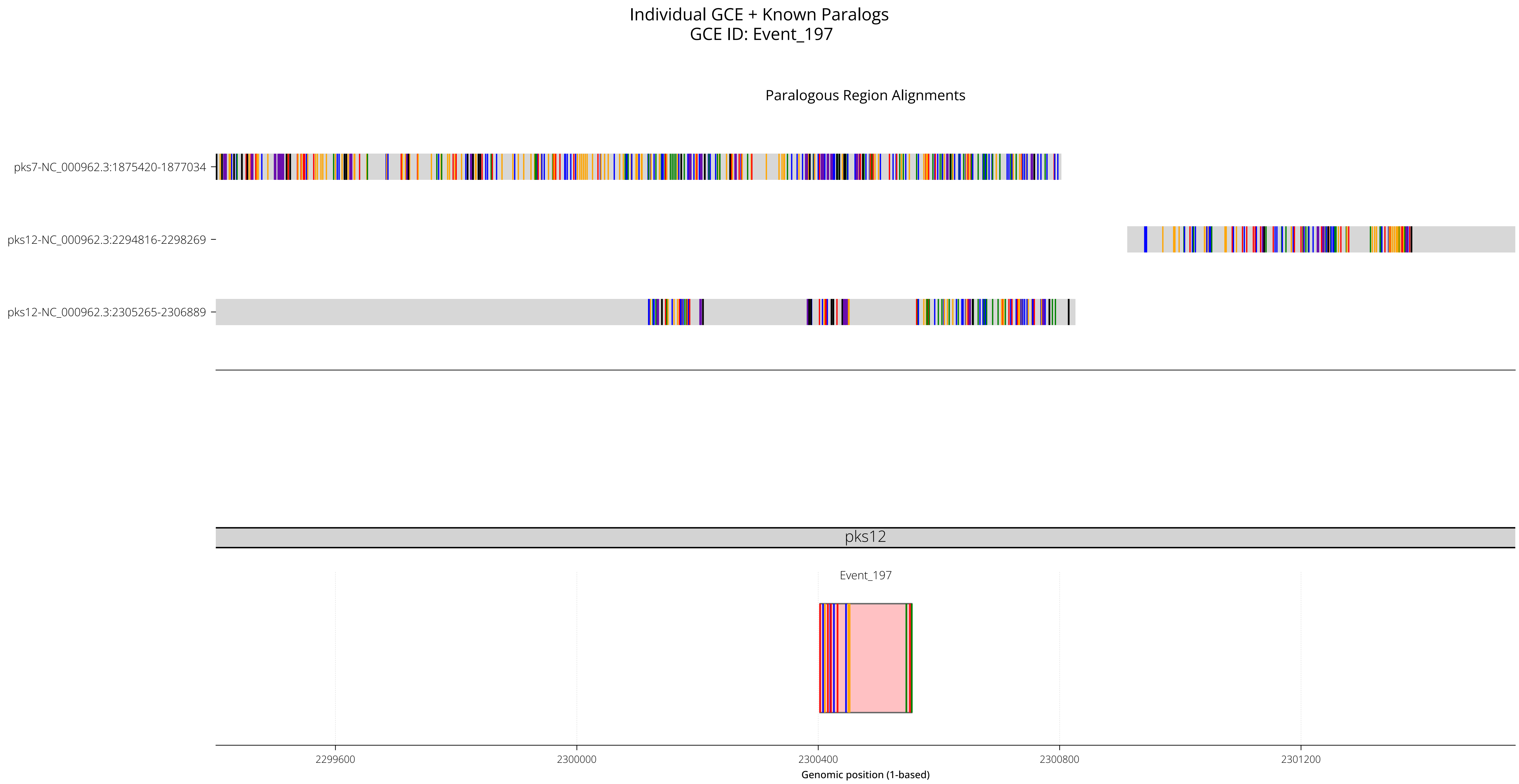

Individual GCE + Known Paralogs
GCE ID: Event_197
Paralogous Region Alignments
pks7-NC_000962.3:1875420-1877034
pks12-NC_000962.3:2294816-2298269
pks12-NC_000962.3:2305265-2306889
pks12
Event_197
2299600
2300000
2300400
2300800
2301200
Genomic position (1-based)

Circos Viz - GCE to Known Paralogs Locations
GCE ID: Event_197

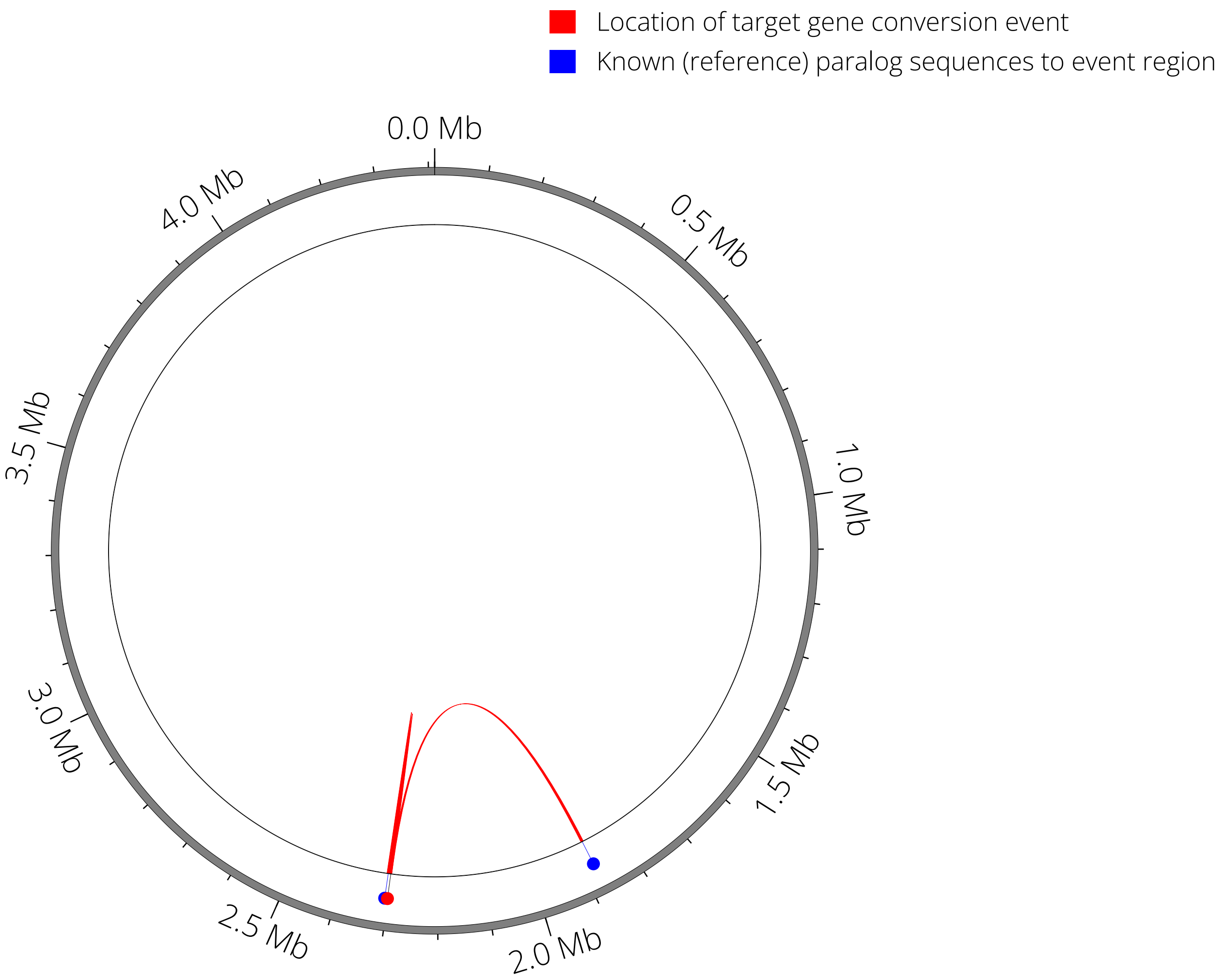

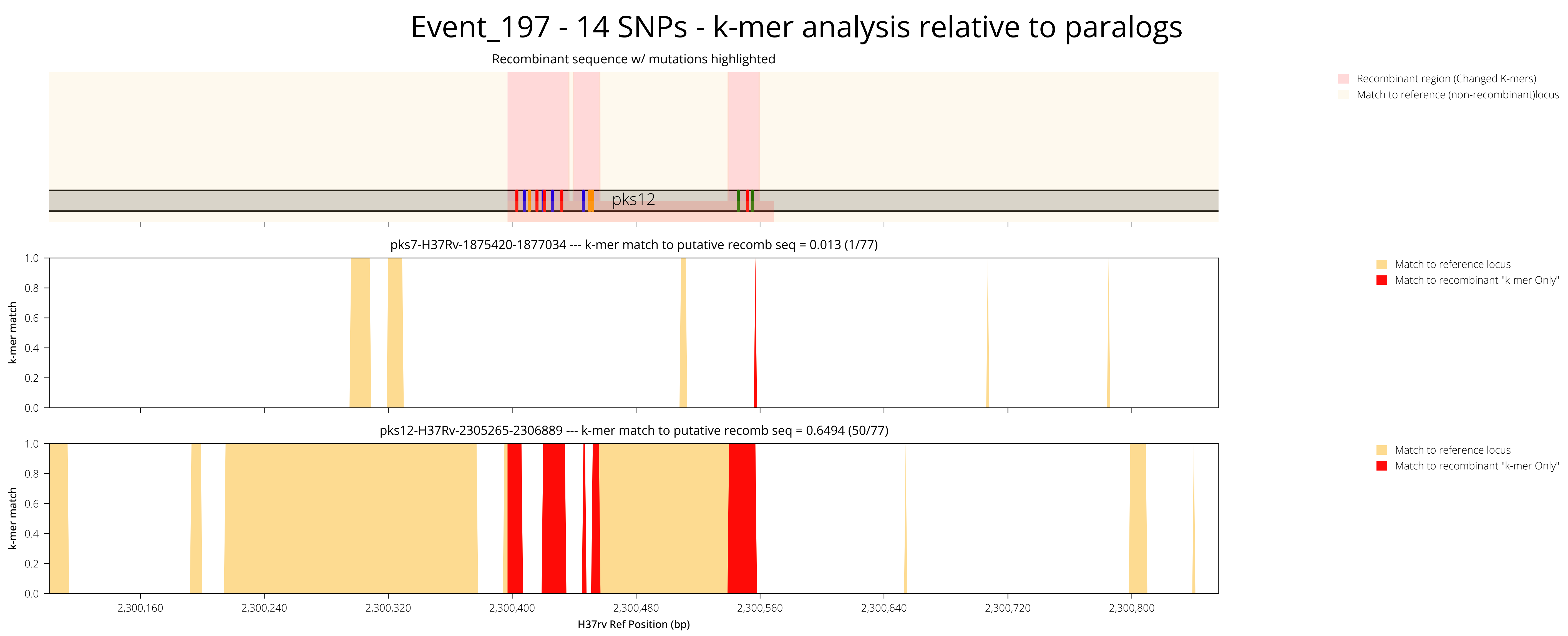

Event_197 - 14 SNPs - k-mer analysis relative to paralogs
Recombinant sequence w/ mutations highlighted
Recombinant region (Changed K-mers)
Match to reference (non-recombinant)locus
pks12
pks7-H37Rv-1875420-1877034 --- k-mer match to putative recomb seq = 0.013 (1/77)
Match to reference locus
Match to recombinant "k-mer Only"
k-mer match
1.0
0.8
0.6
0.4
0.2
0.0
pks12-H37Rv-2305265-2306889 --- k-mer match to putative recomb seq = 0.6494 (50/77)
Match to reference locus
Match to recombinant "k-mer Only"
k-mer match
2,300,160
2,300,240
2,300,320
2,300,400
2,300,480
2,300,560
2,300,640
2,300,720
2,300,800
H37rv Ref Position (bp)

Sub-phylogeny Viz + Paralog Alignments
GCE ID: Event_197

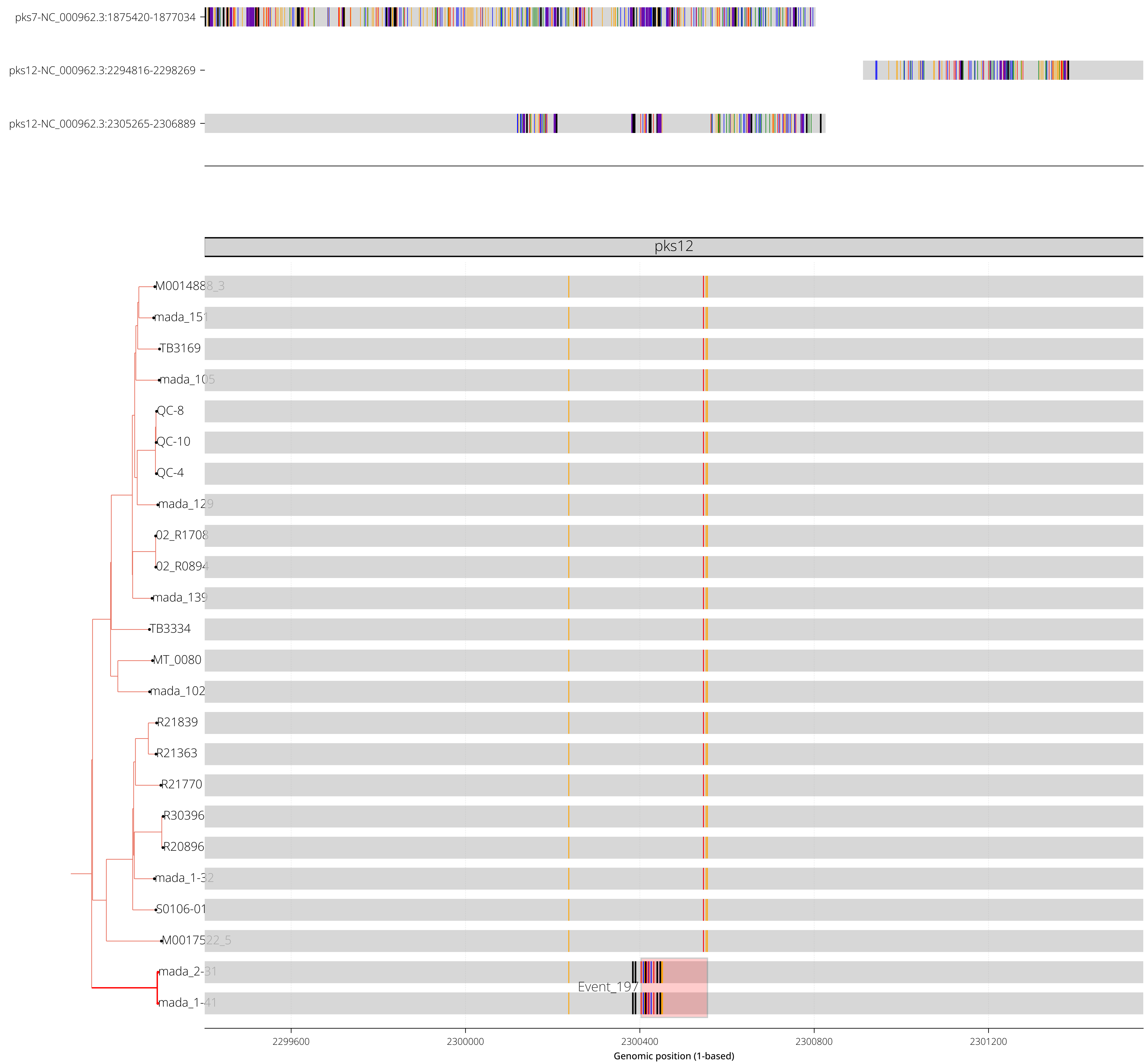

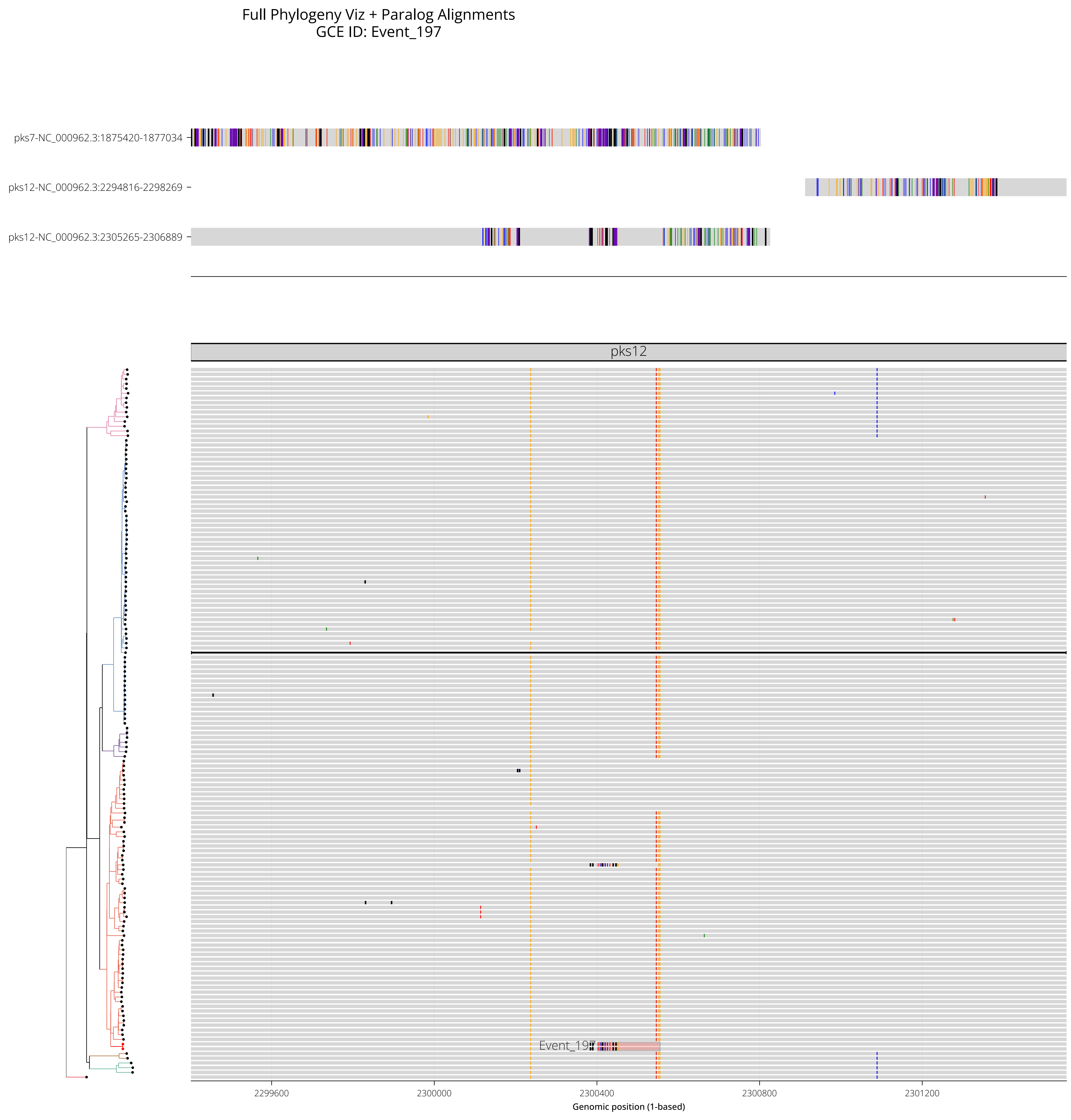

Full Phylogeny Viz + Paralog Alignments
GCE ID: Event_197
pks7-NC_000962.3:1875420-1877034
pks12-NC_000962.3:2294816-2298269
pks12-NC_000962.3:2305265-2306889
pks12
Event_197
2299600
2300000
2300400
2300800
2301200
Genomic position (1-based)

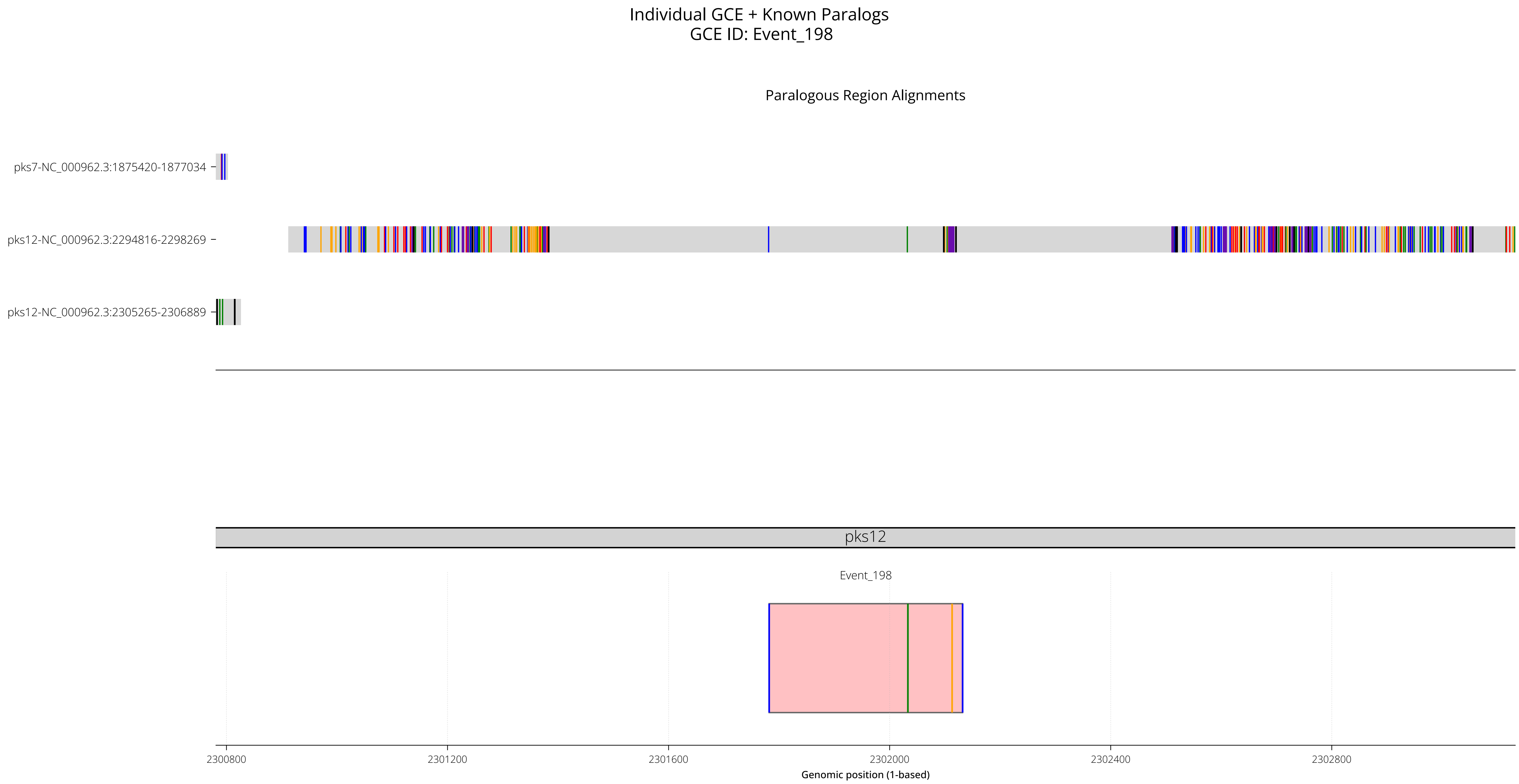
Individual GCE + Known Paralogs
GCE ID: Event_198
Paralogous Region Alignments
pks7-NC_000962.3:1875420-1877034
pks12-NC_000962.3:2294816-2298269
pks12-NC_000962.3:2305265-2306889
pks12
Event_198
2300800
2301200
2301600
2302000
2302400
2302800
Genomic position (1-based)

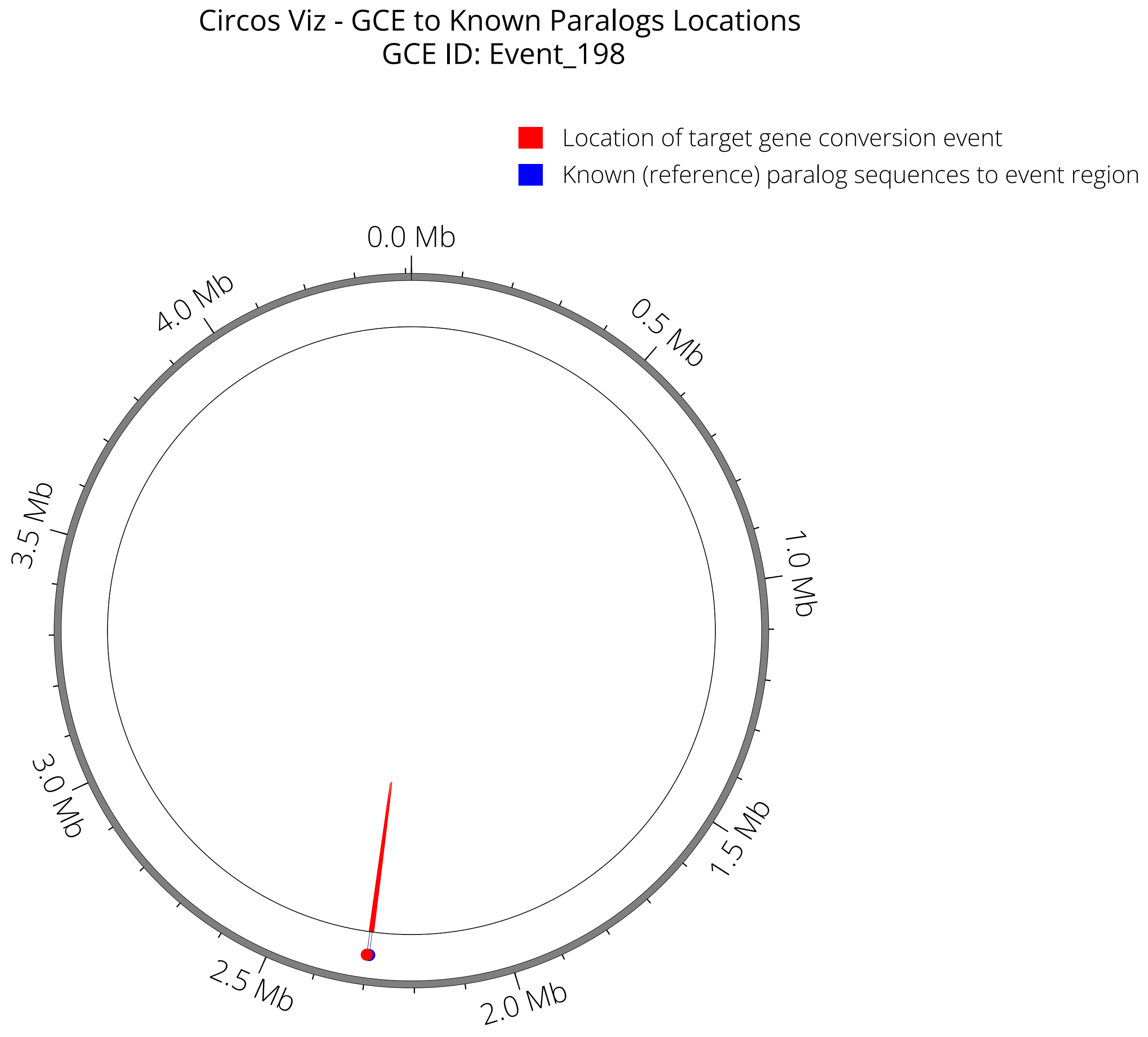
Circos Viz - GCE to Known Paralogs Locations
GCE ID: Event_198
Location of target gene conversion event
Known (reference) paralog sequences to event region
0.0 Mb
0.5 Mb
1.0 Mb
1.5 Mb
2.0 Mb
2.5 Mb
3.0 Mb
3.5 Mb
4.0 Mb

### Event_198 - 4 SNPs - k-mer analysis relative to paralogs

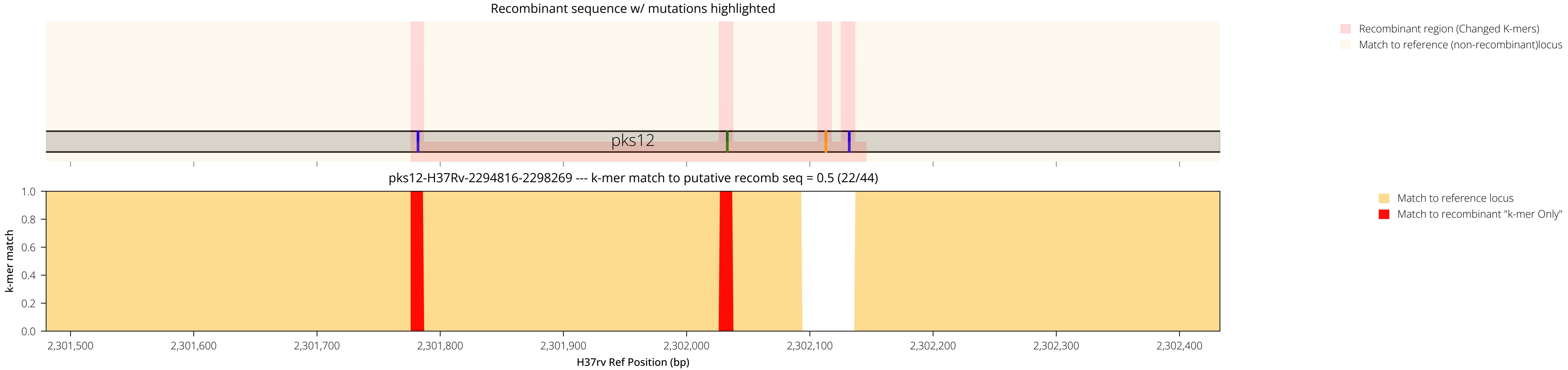

Sub-phylogeny Viz + Paralog Alignments
GCE ID: Event_198

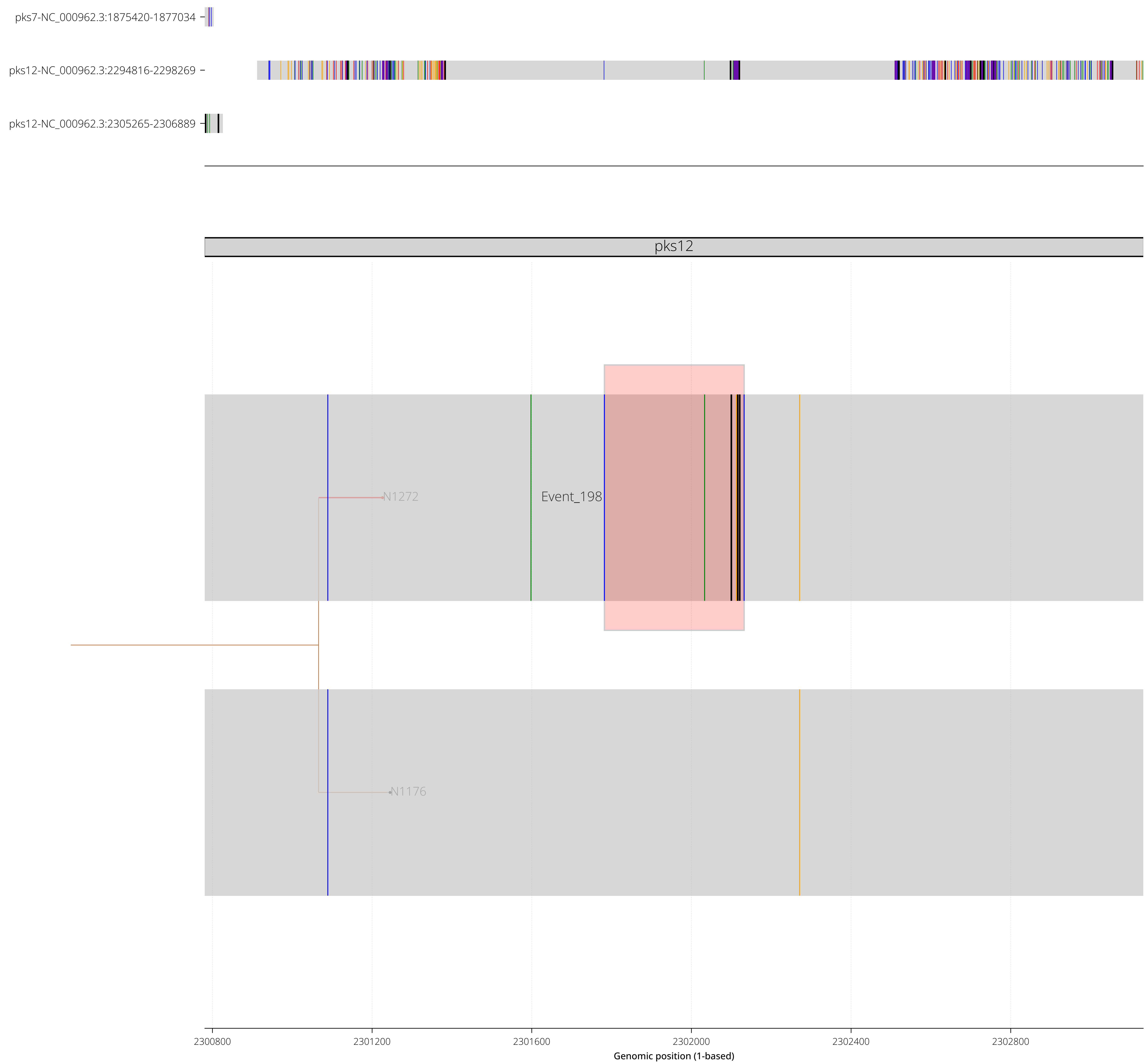

Full Phylogeny Viz + Paralog Alignments
GCE ID: Event_198

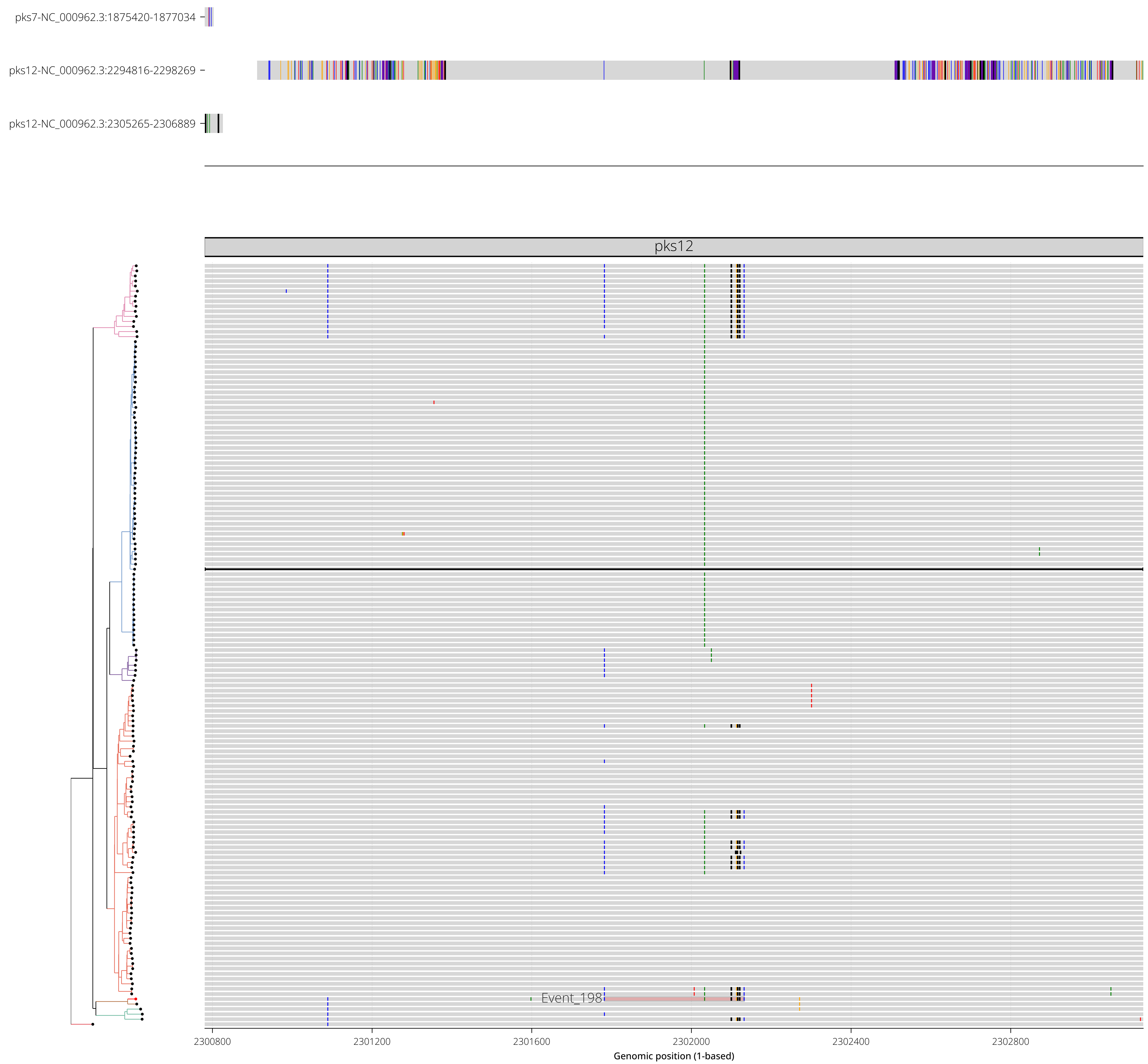

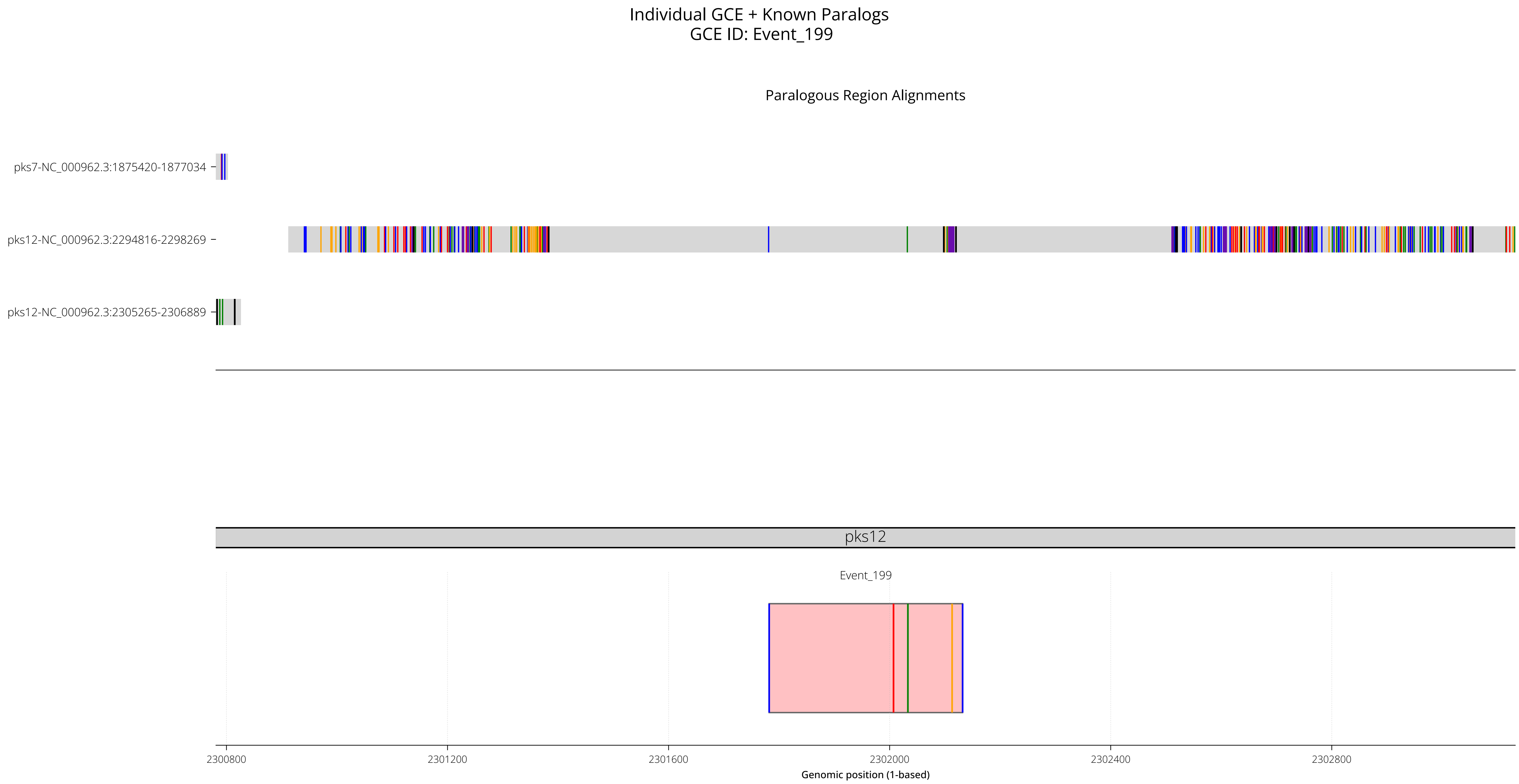

Individual GCE + Known Paralogs
GCE ID: Event_199
Paralogous Region Alignments
pks7-NC_000962.3:1875420-1877034
pks12-NC_000962.3:2294816-2298269
pks12-NC_000962.3:2305265-2306889
pks12
Event_199
2300800
2301200
2301600
2302000
2302400
2302800
Genomic position (1-based)

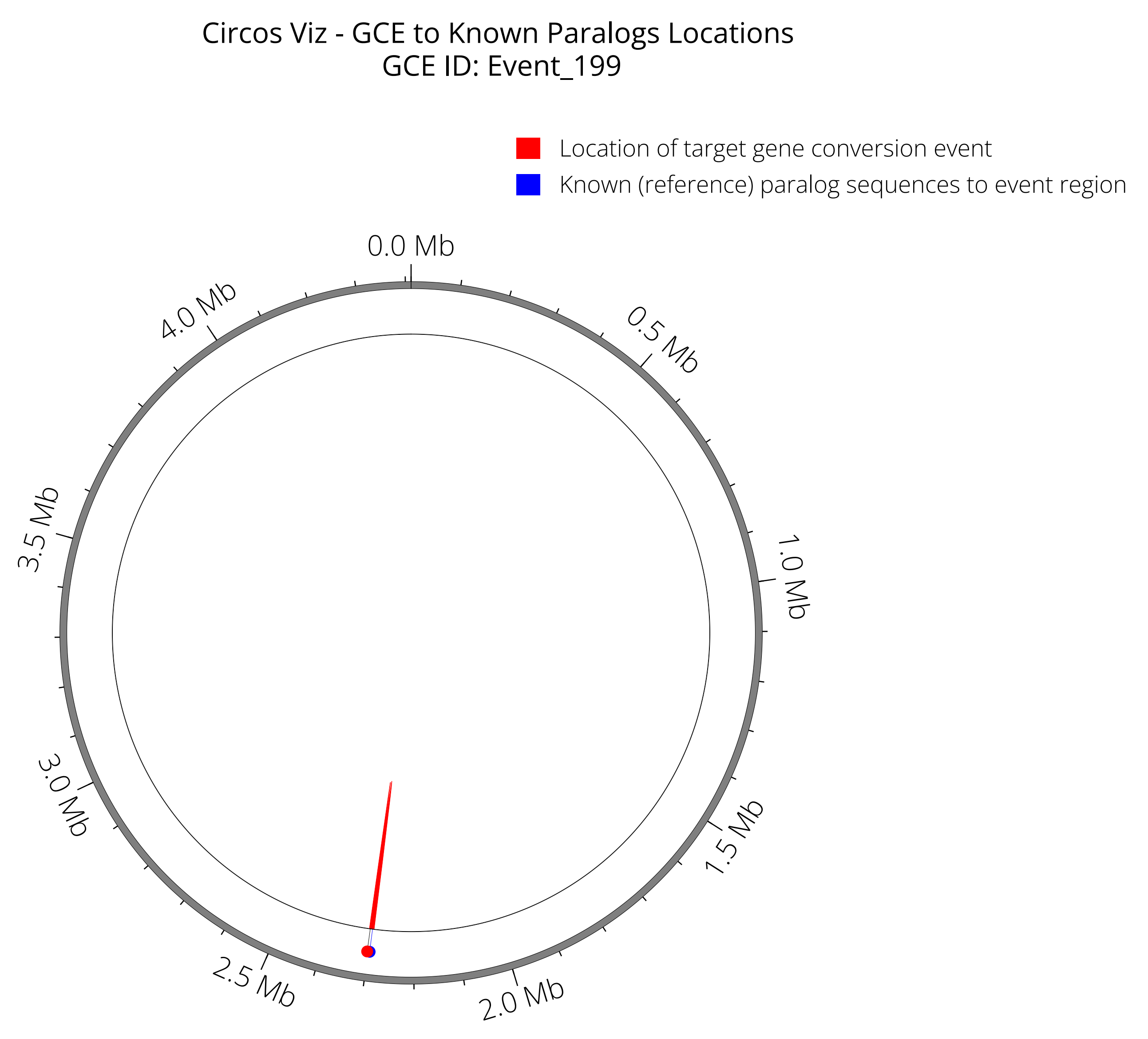
Circos Viz - GCE to Known Paralogs Locations
GCE ID: Event_199
Location of target gene conversion event
Known (reference) paralog sequences to event region
0.0 Mb
0.5 Mb
1.0 Mb
1.5 Mb
2.0 Mb
2.5 Mb
3.0 Mb
3.5 Mb
4.0 Mb

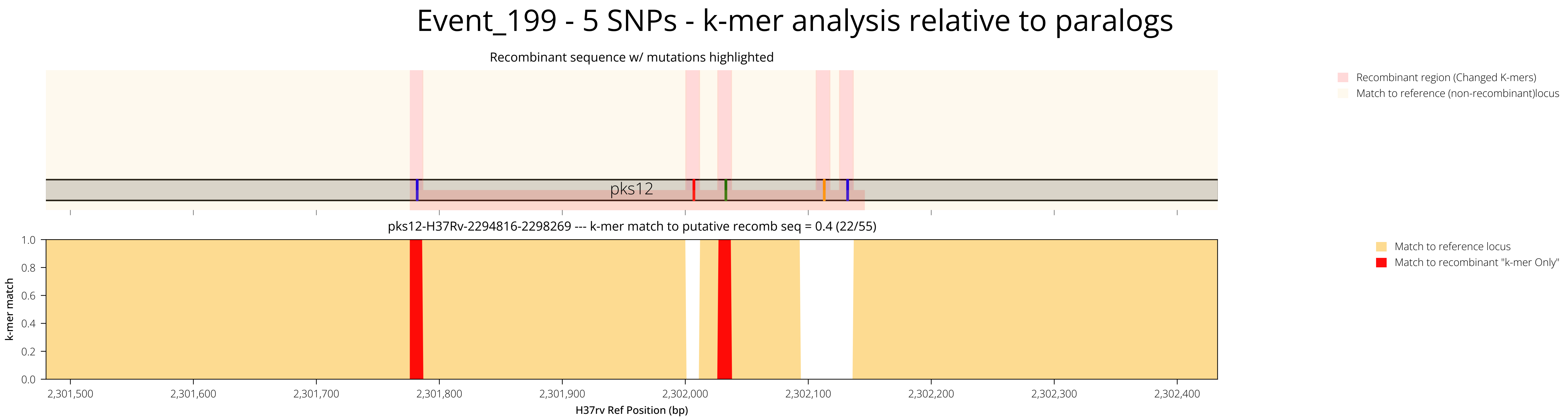
Event_199 - 5 SNPs - k-mer analysis relative to paralogs
Recombinant sequence w/ mutations highlighted
pks12
Recombinant region (Changed K-mers)
Match to reference (non-recombinant)locus
pks12-H37Rv-2294816-2298269 --- k-mer match to putative recomb seq = 0.4 (22/55)
Match to reference locus
Match to recombinant "k-mer Only"
k-mer match
1.0
0.8
0.6
0.4
0.2
0.0
2,301,500
2,301,600
2,301,700
2,301,800
2,301,900
2,302,000
2,302,100
2,302,200
2,302,300
2,302,400
H37rv Ref Position (bp)

Sub-phylogeny Viz + Paralog Alignments
GCE ID: Event_199

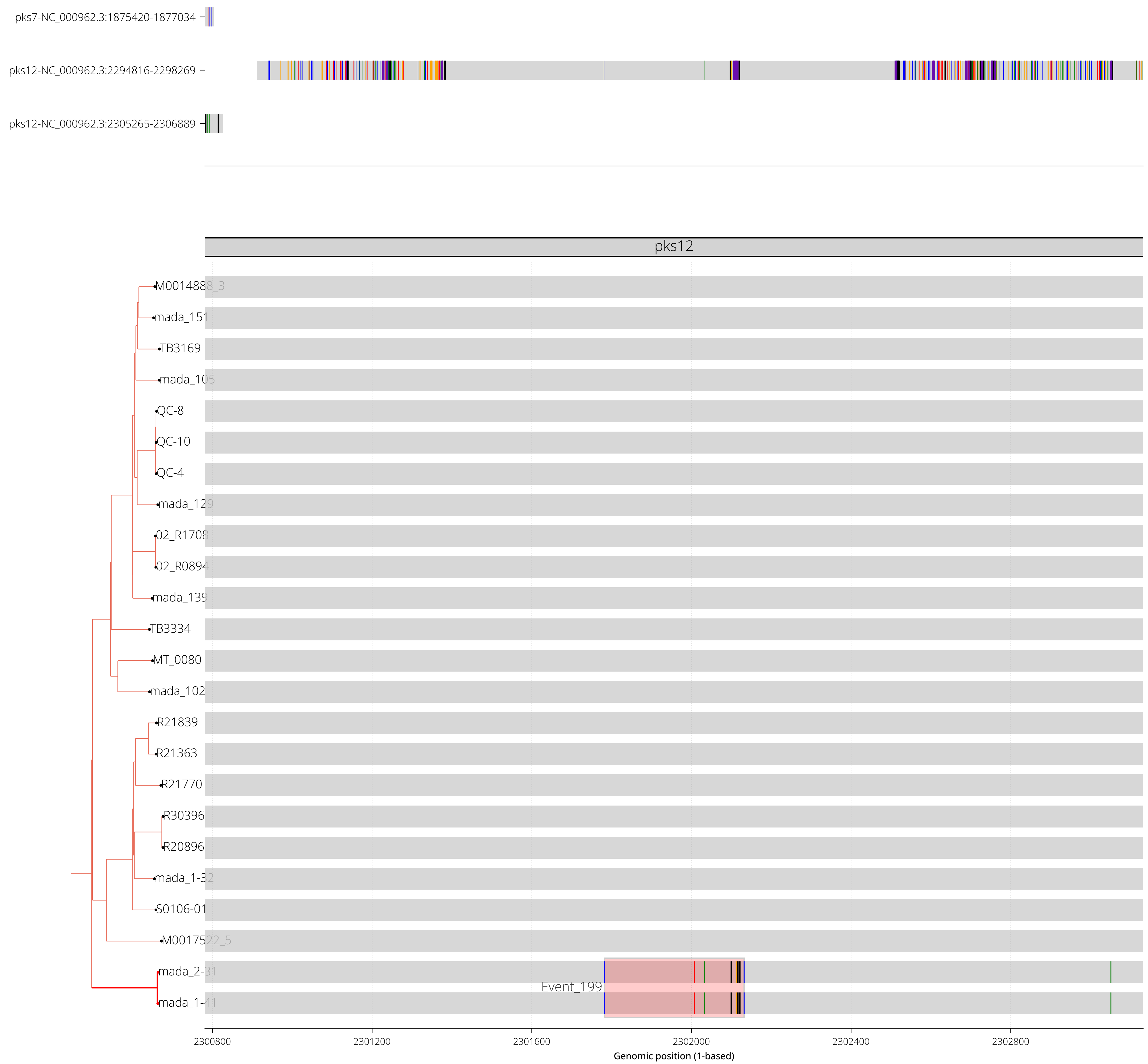

Full Phylogeny Viz + Paralog Alignments
GCE ID: Event_199

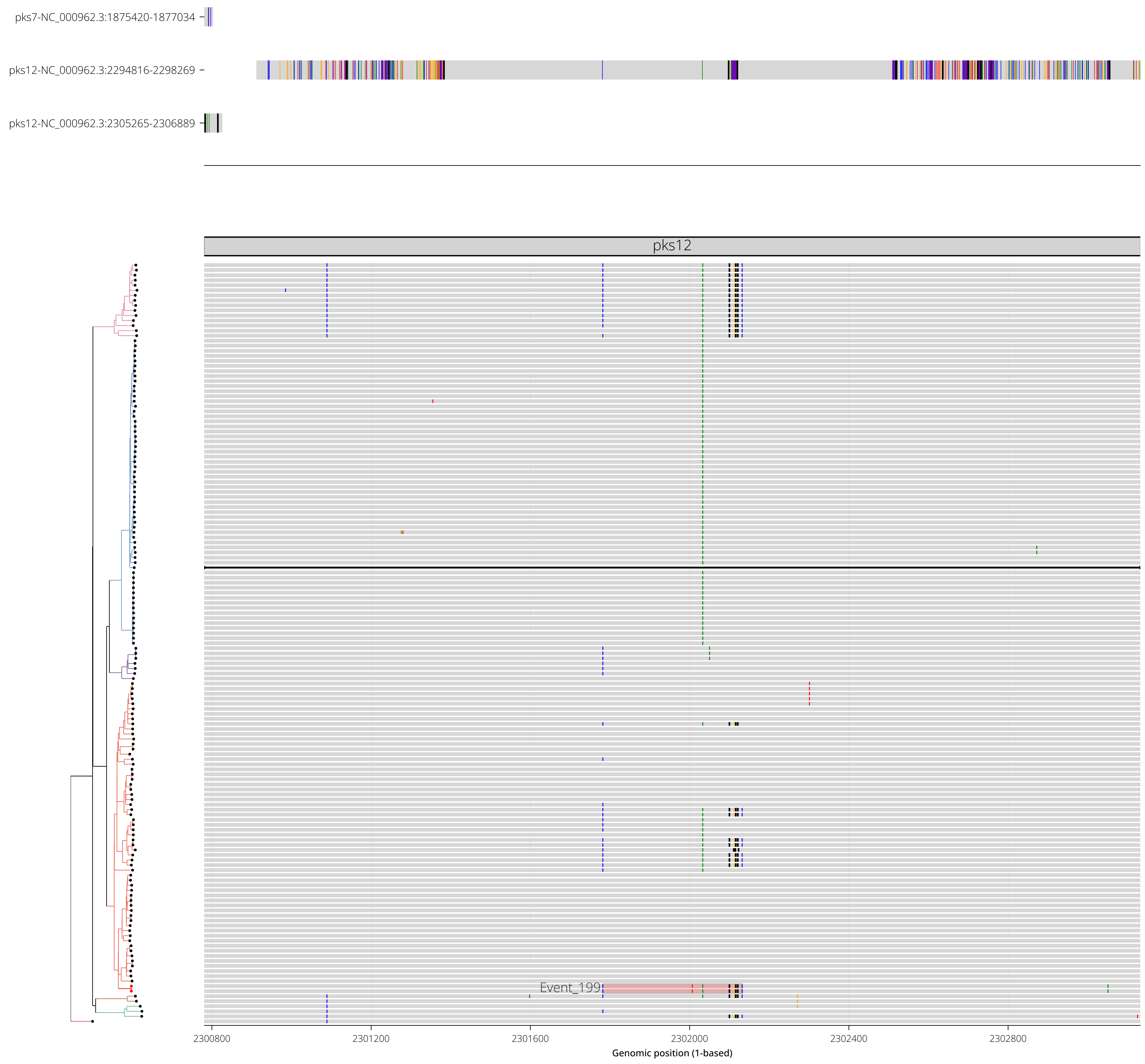

#### Individual GCE + Known Paralogs
#### GCE ID: Event_200

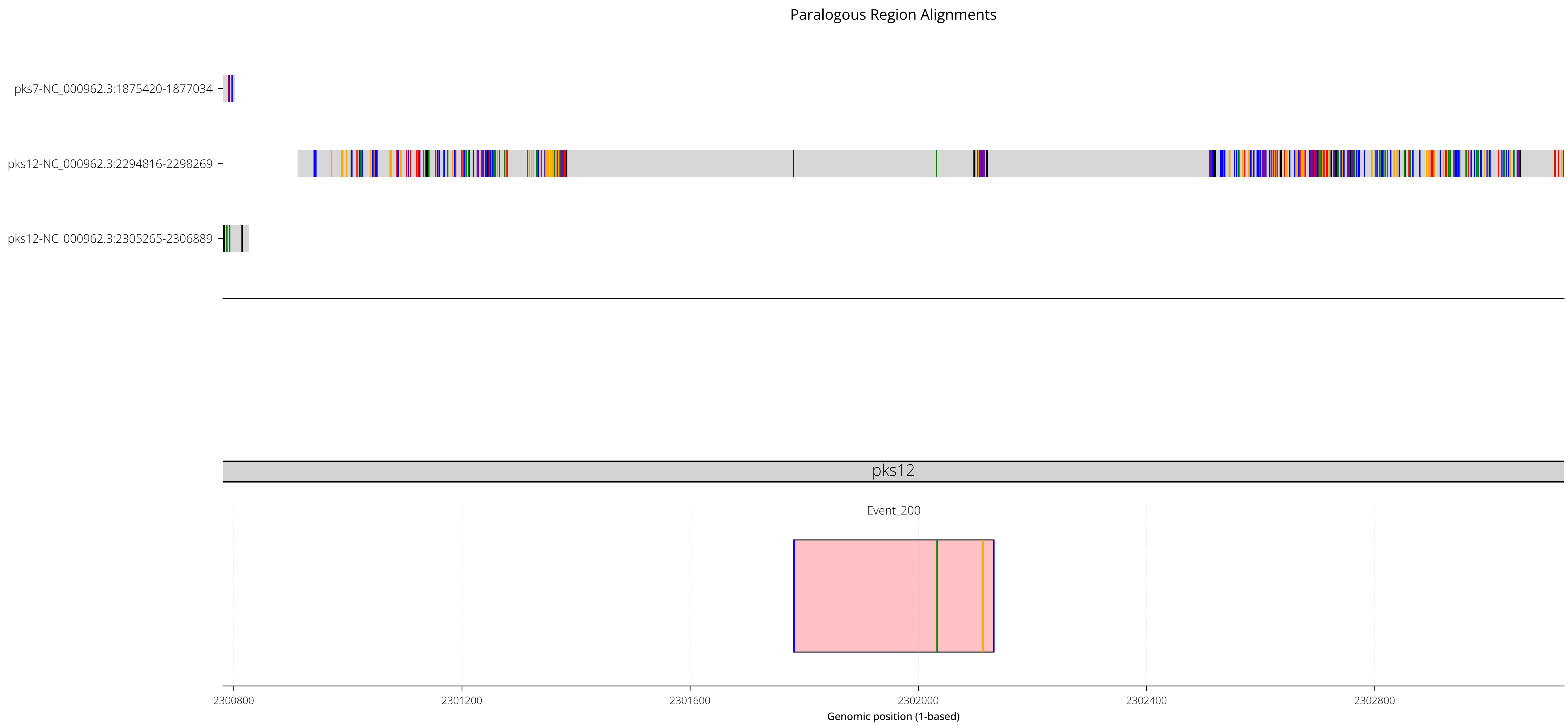

### Circos Viz - GCE to Known Paralogs Locations
GCE ID: Event_200

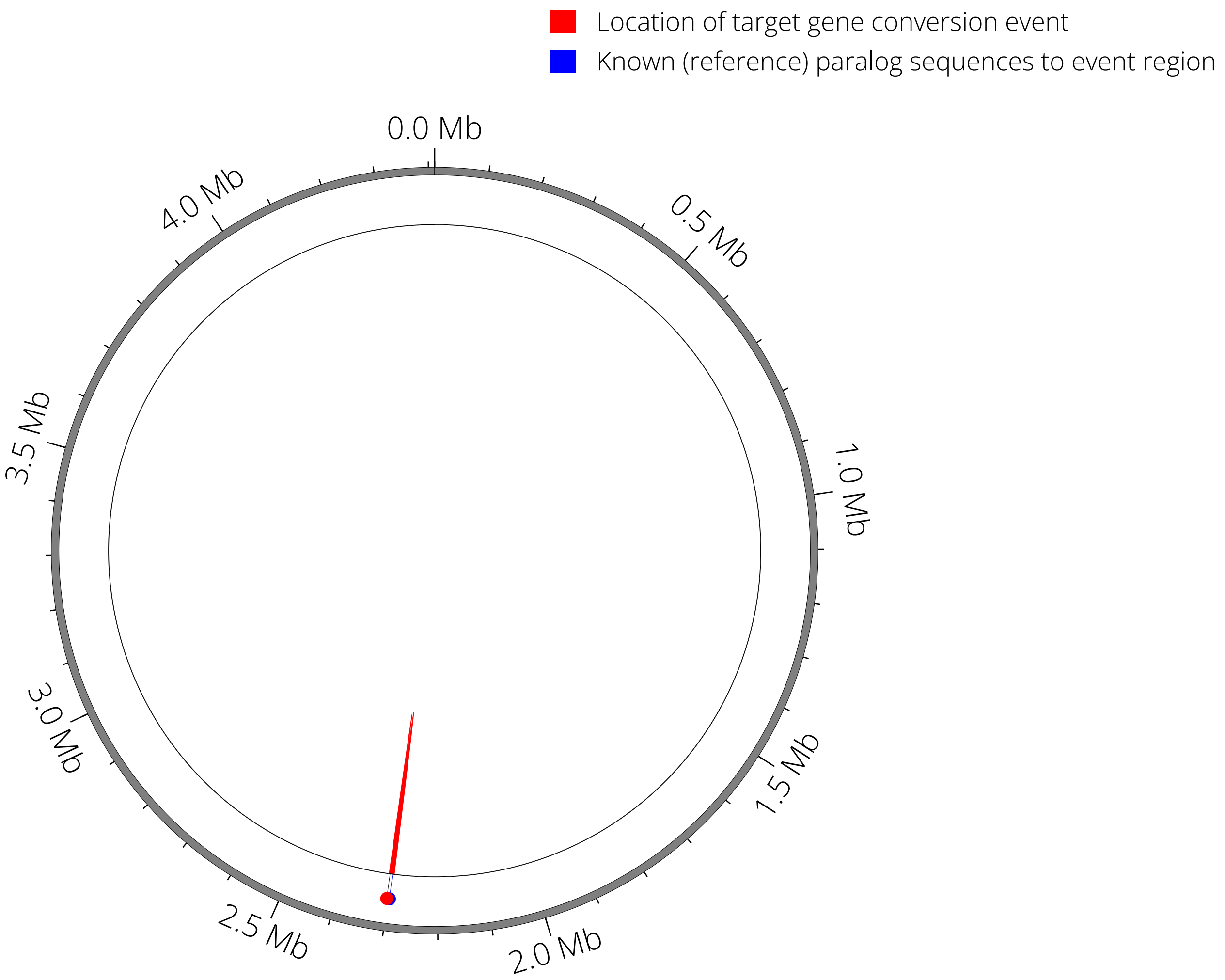

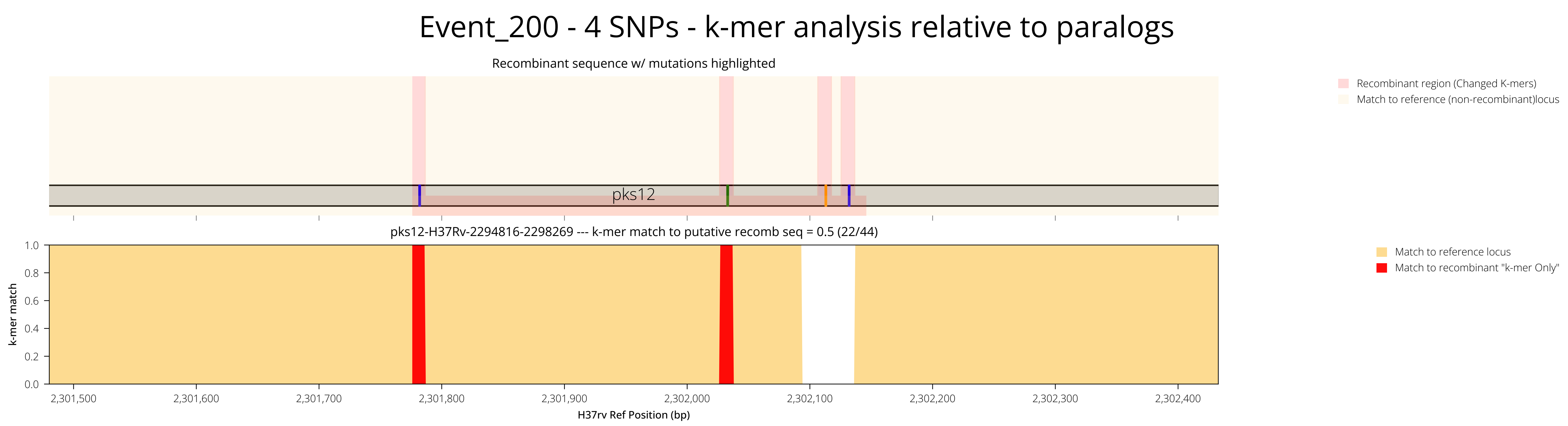

Event_200 - 4 SNPs - k-mer analysis relative to paralogs
Recombinant sequence w/ mutations highlighted
Recombinant region (Changed K-mers)
Match to reference (non-recombinant)locus
pks12
pks12-H37Rv-2294816-2298269 --- k-mer match to putative recomb seq = 0.5 (22/44)
Match to reference locus
Match to recombinant "k-mer Only"
k-mer match
1.0
0.8
0.6
0.4
0.2
0.0
2,301,500
2,301,600
2,301,700
2,301,800
2,301,900
2,302,000
2,302,100
2,302,200
2,302,300
2,302,400
H37rv Ref Position (bp)

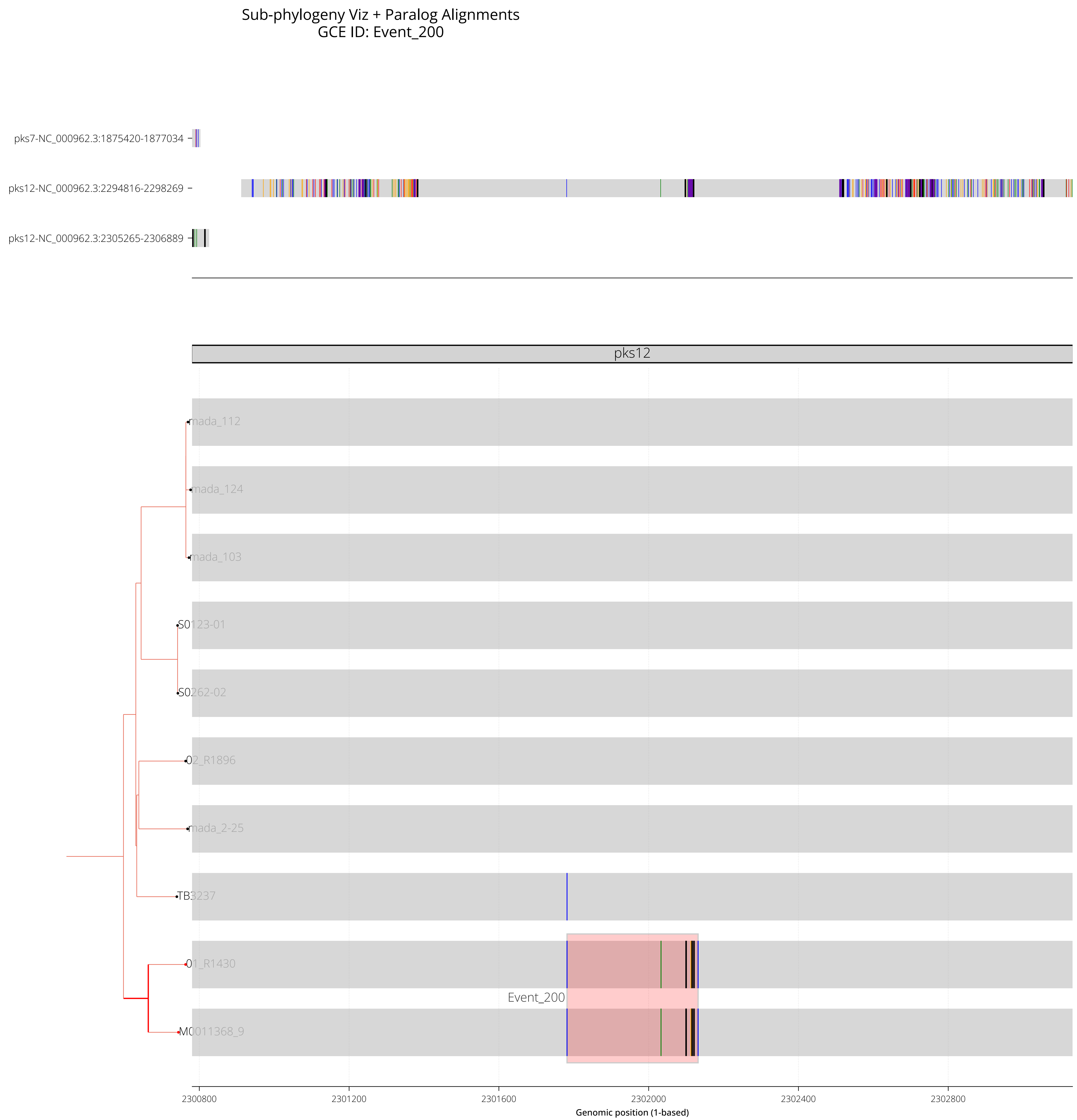

Sub-phylogeny Viz + Paralog Alignments
GCE ID: Event_200
pks7-NC_000962.3:1875420-1877034
pks12-NC_000962.3:2294816-2298269
pks12-NC_000962.3:2305265-2306889
pks12
mada_112
mada_124
mada_103
S0123-01
S0262-02
02_R1896
mada_2-25
TB3237
01_R1430
Event_200
M0011368_9
2300800
2301200
2301600
2302000
2302400
2302800
Genomic position (1-based)

Full Phylogeny Viz + Paralog Alignments
GCE ID: Event_200

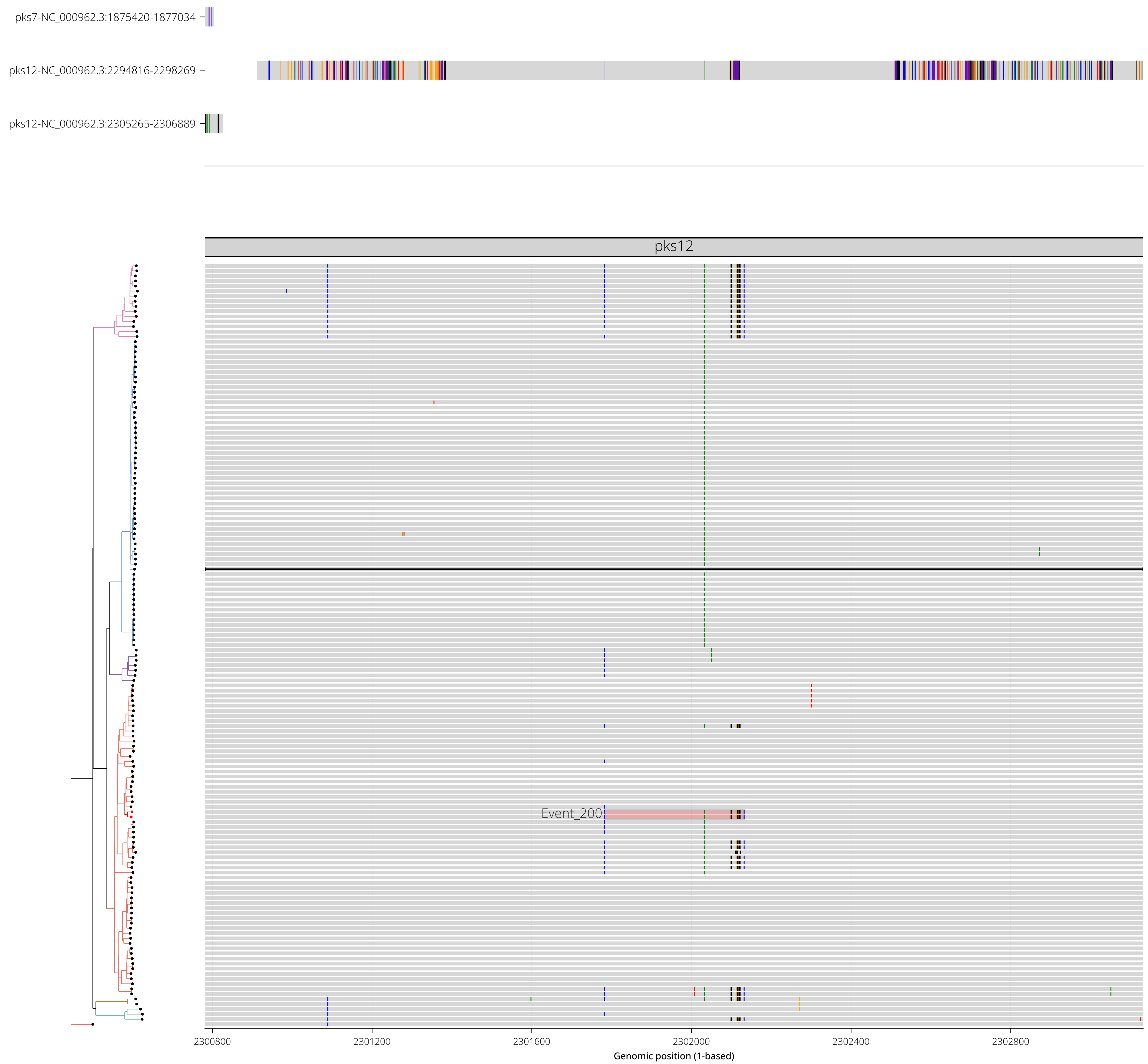

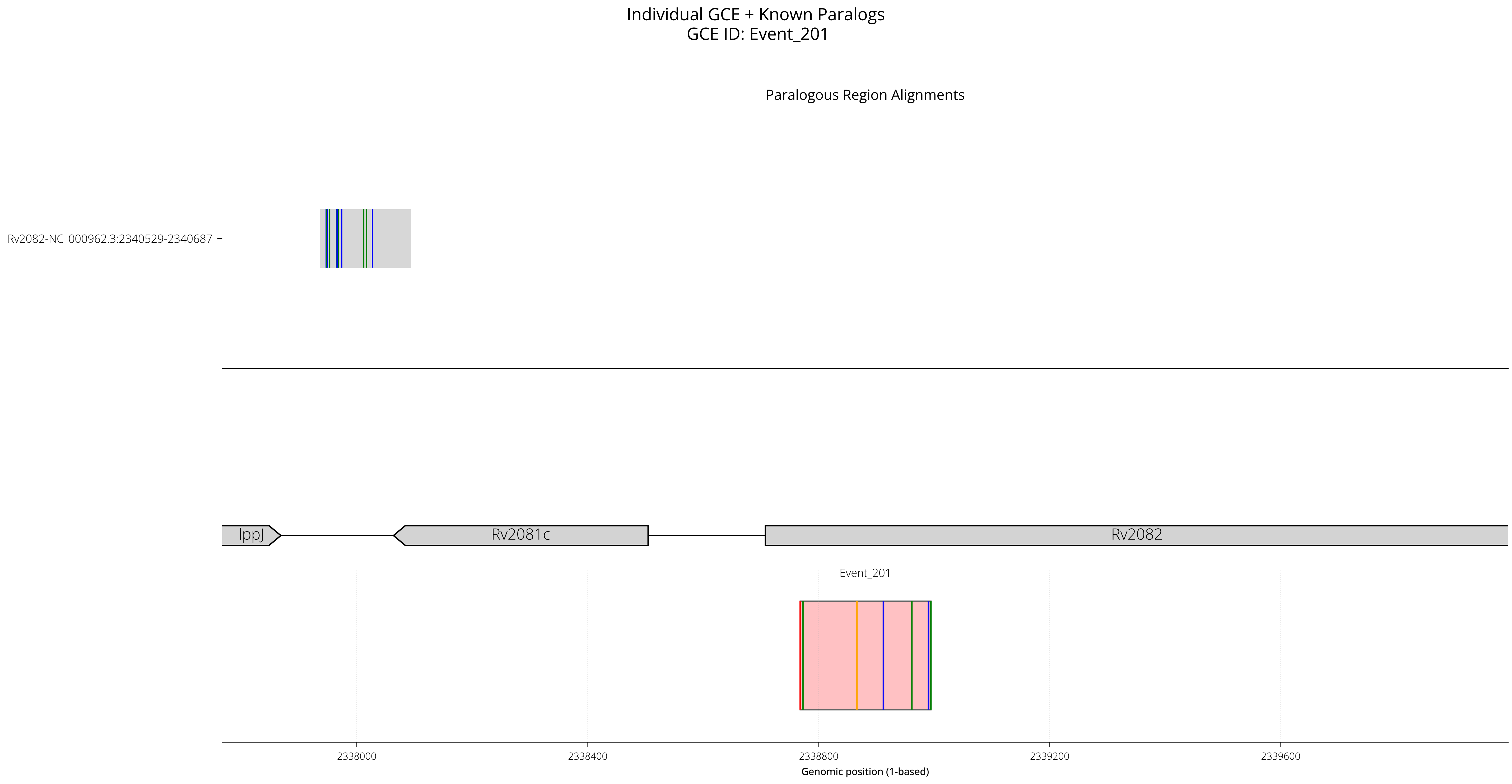

Individual GCE + Known Paralogs
GCE ID: Event_201
Paralogous Region Alignments
Rv2082-NC_000962.3:2340529-2340687
lppJ
Rv2081c
Rv2082
Event_201
2338000
2338400
2338800
2339200
2339600
Genomic position (1-based)

Sub-phylogeny Viz + Paralog Alignments
GCE ID: Event_201

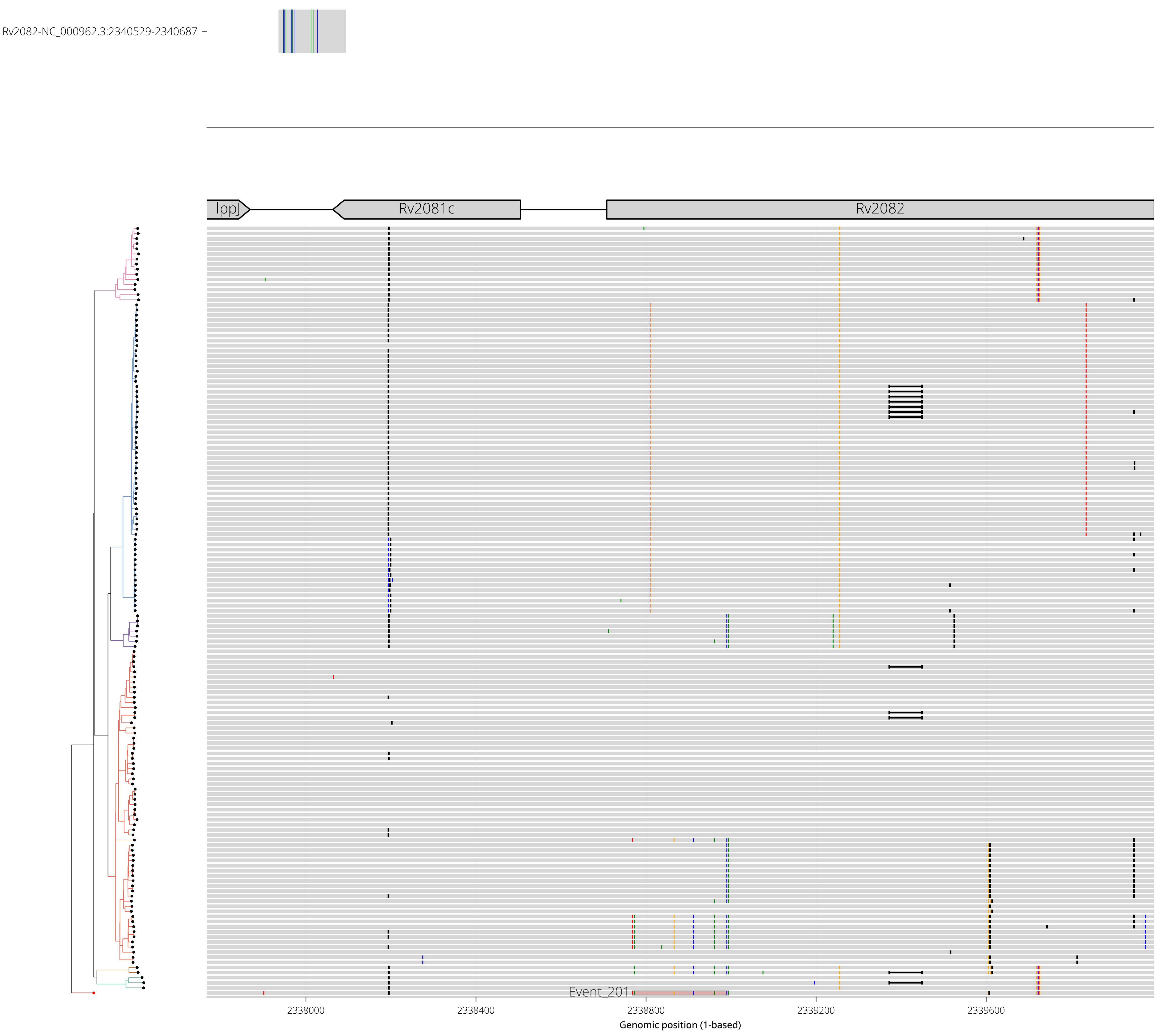

Full Phylogeny Viz + Paralog Alignments
GCE ID: Event_201

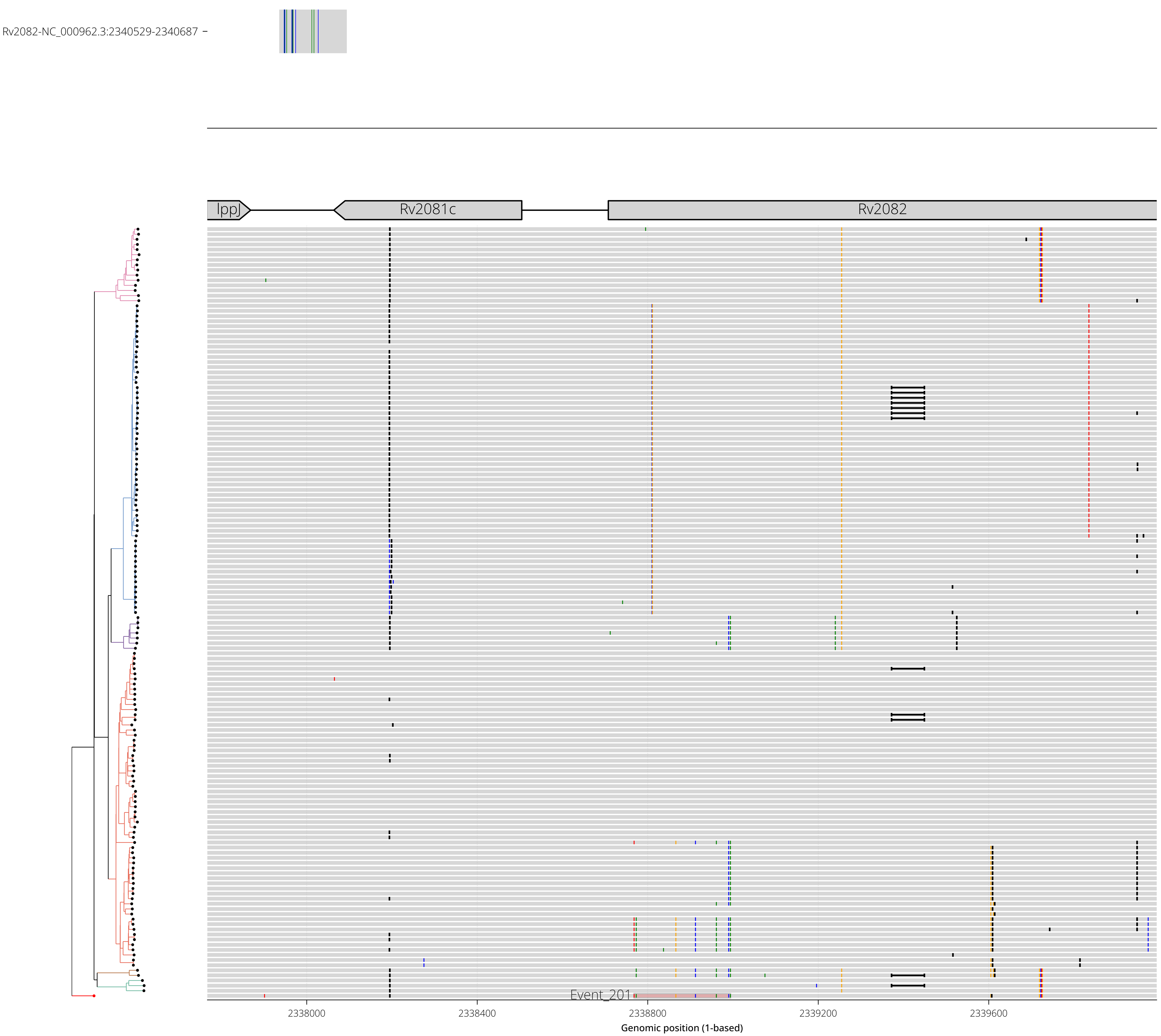

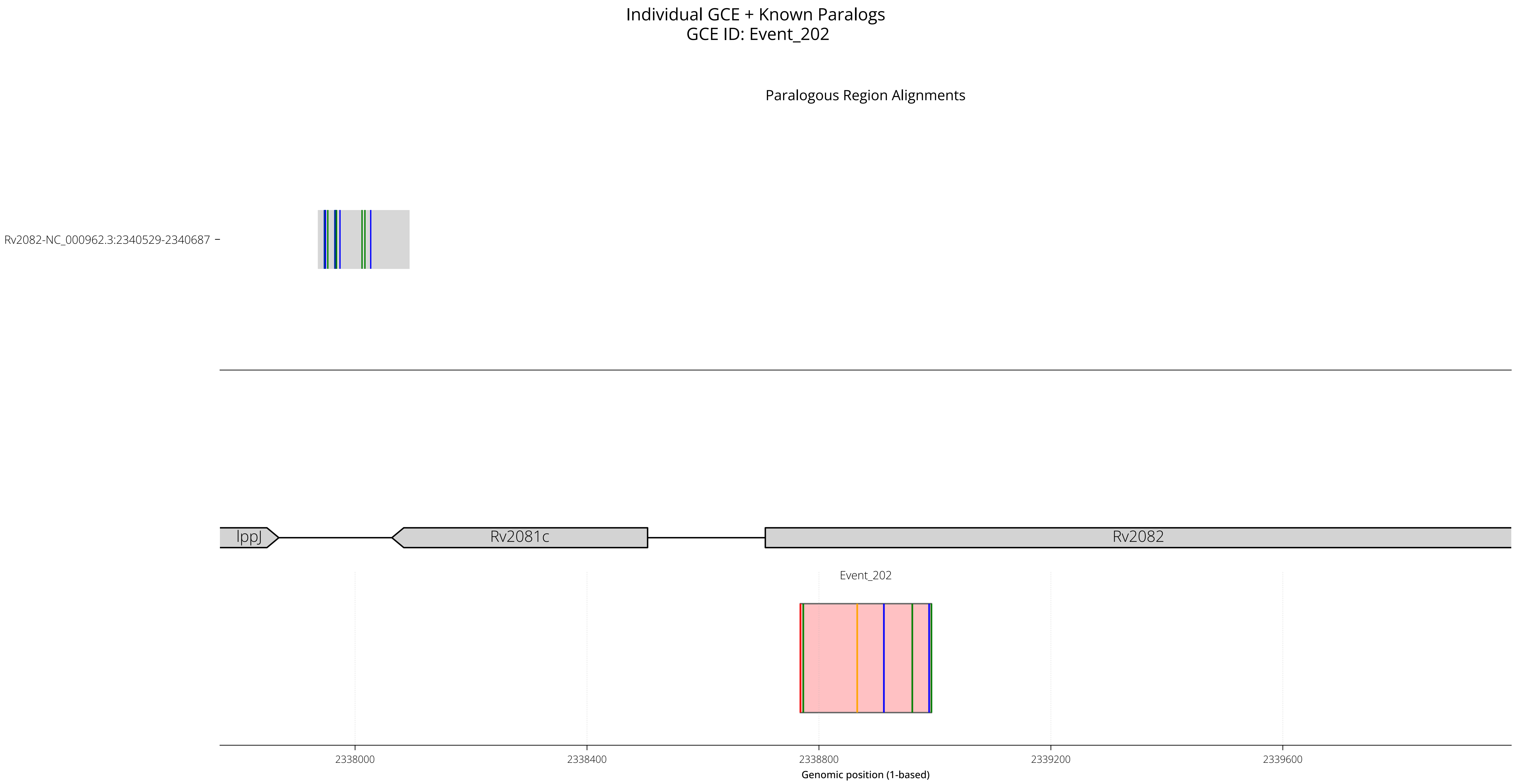

Individual GCE + Known Paralogs
GCE ID: Event_202
Paralogous Region Alignments
Rv2082-NC_000962.3:2340529-2340687
lppJ
Rv2081c
Rv2082
Event_202
2338000
2338400
2338800
2339200
2339600
Genomic position (1-based)

### Sub-phylogeny Viz + Paralog Alignments
GCE ID: Event_202

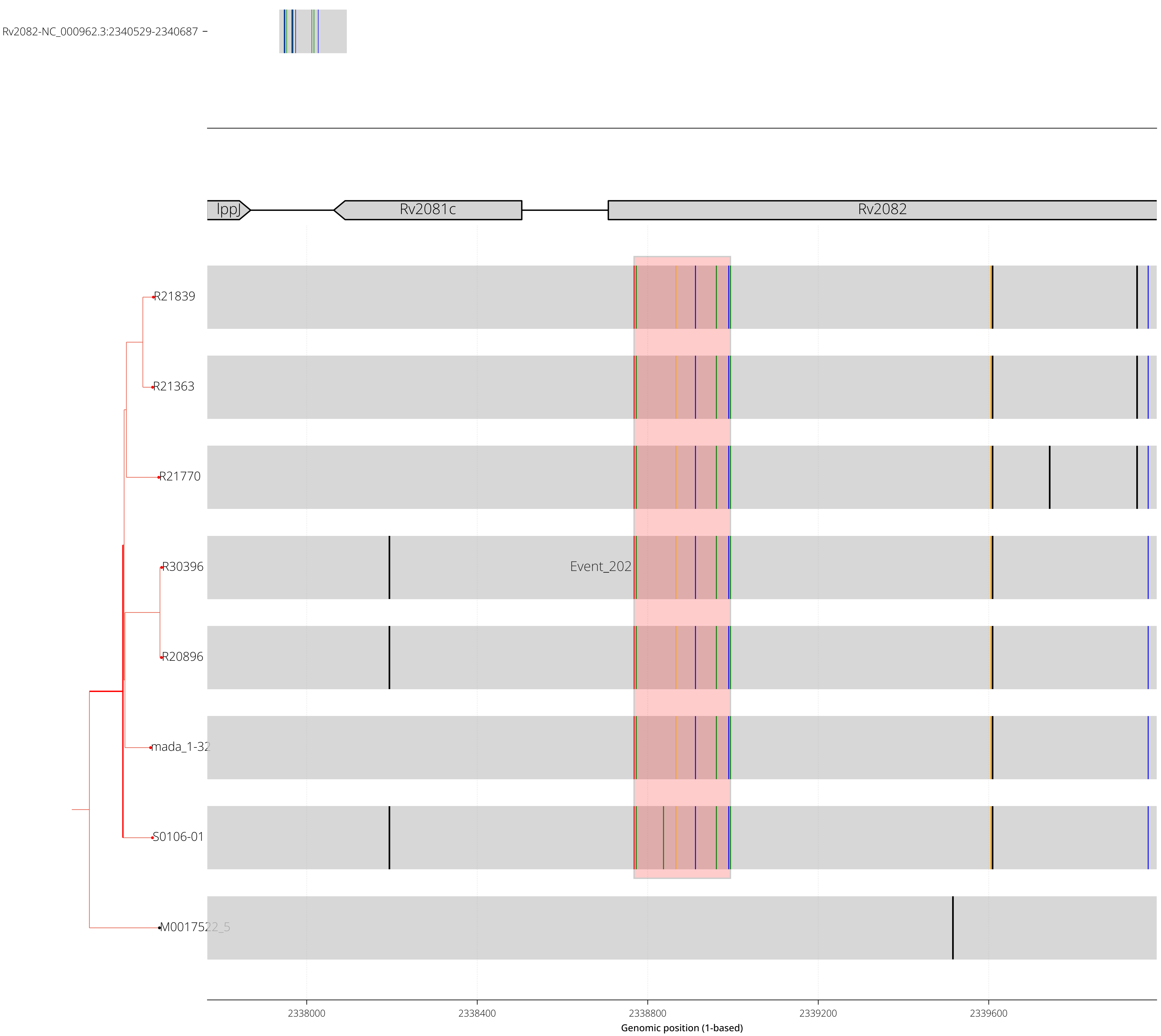

Full Phylogeny Viz + Paralog Alignments
GCE ID: Event_202

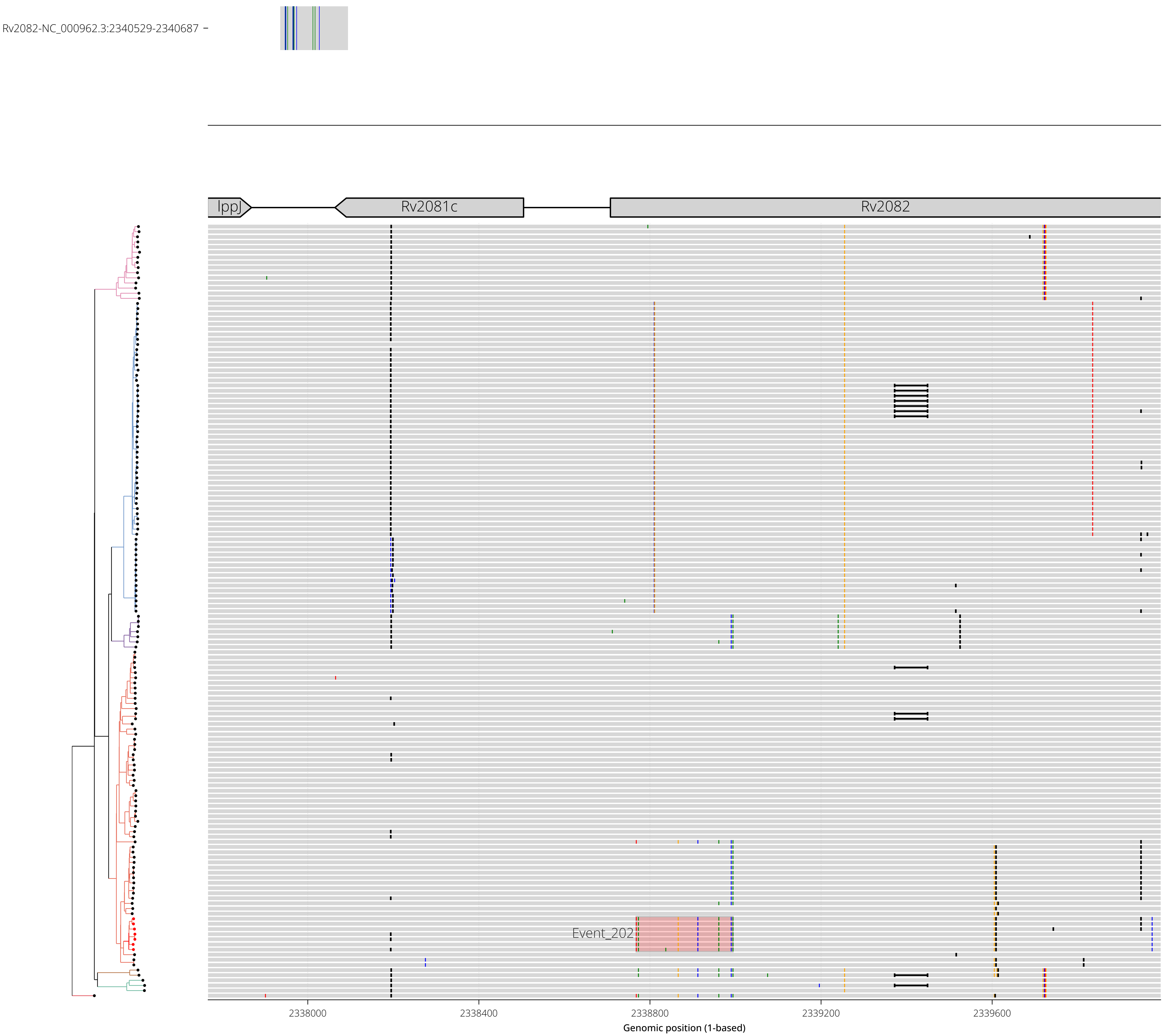

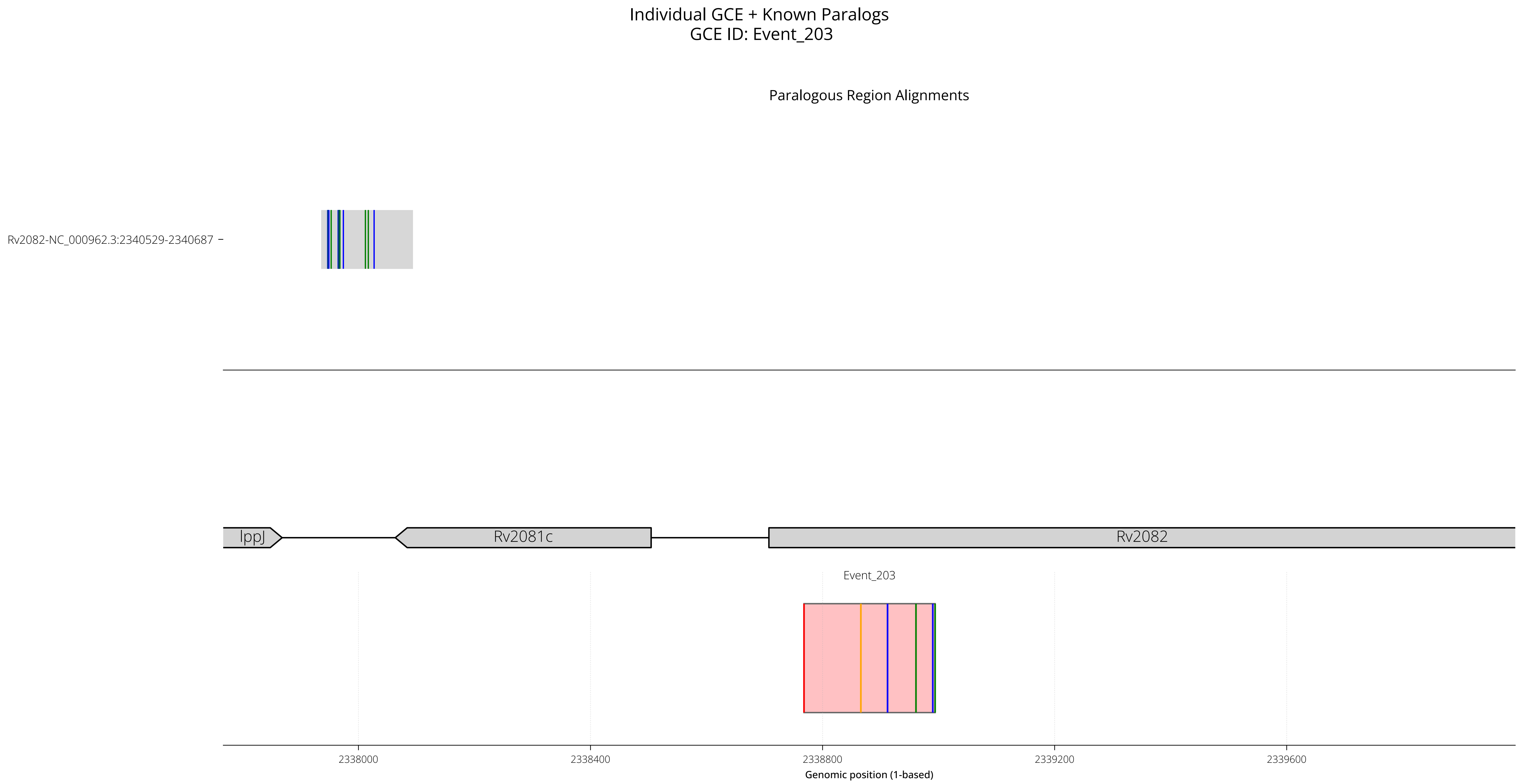
Individual GCE + Known Paralogs
GCE ID: Event_203
Paralogous Region Alignments
Rv2082-NC_000962.3:2340529-2340687
lppJ
Rv2081c
Rv2082
Event_203
2338000
2338400
2338800
2339200
2339600
Genomic position (1-based)

Sub-phylogeny Viz + Paralog Alignments
GCE ID: Event_203

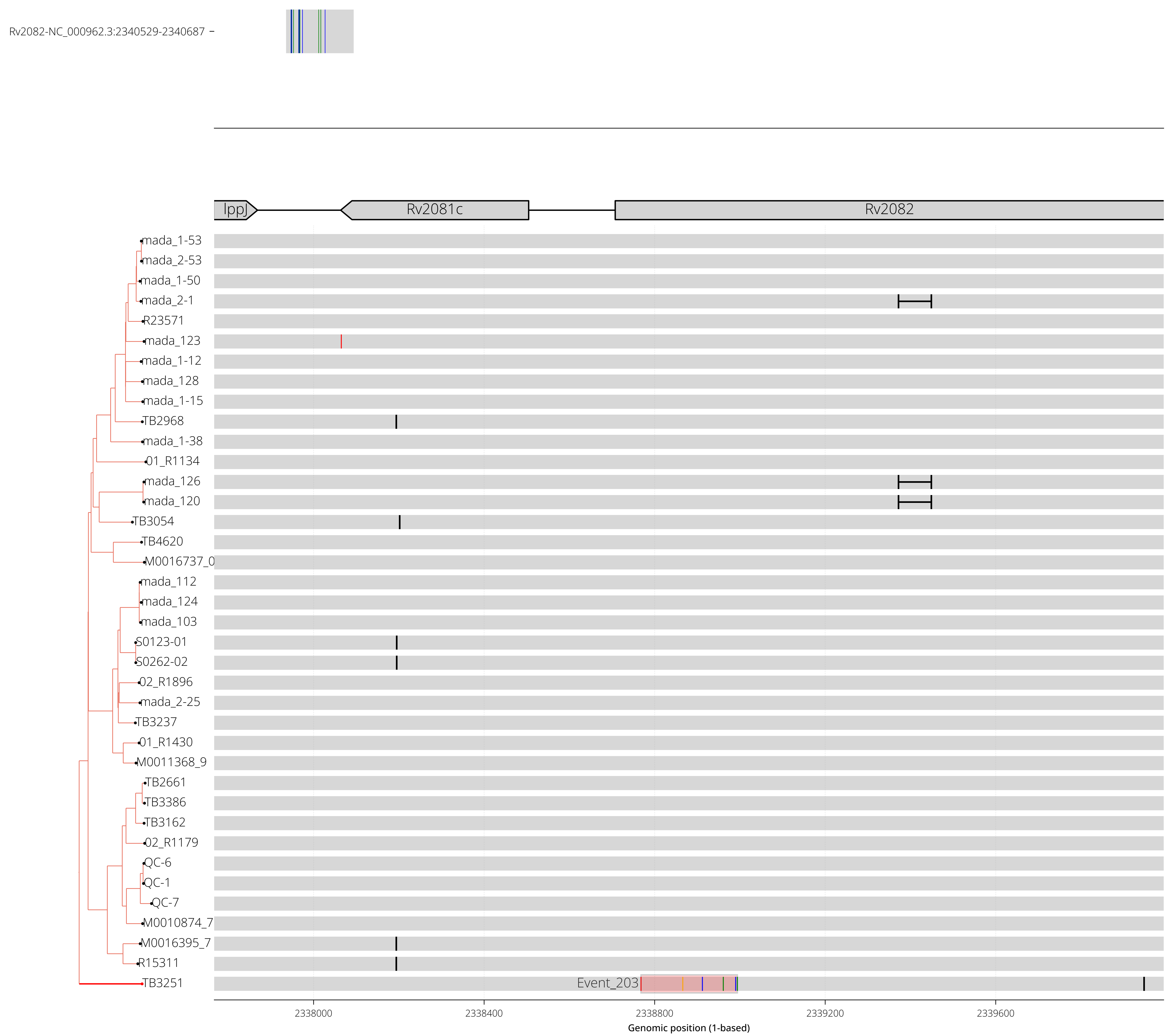

Full Phylogeny Viz + Paralog Alignments
GCE ID: Event_203

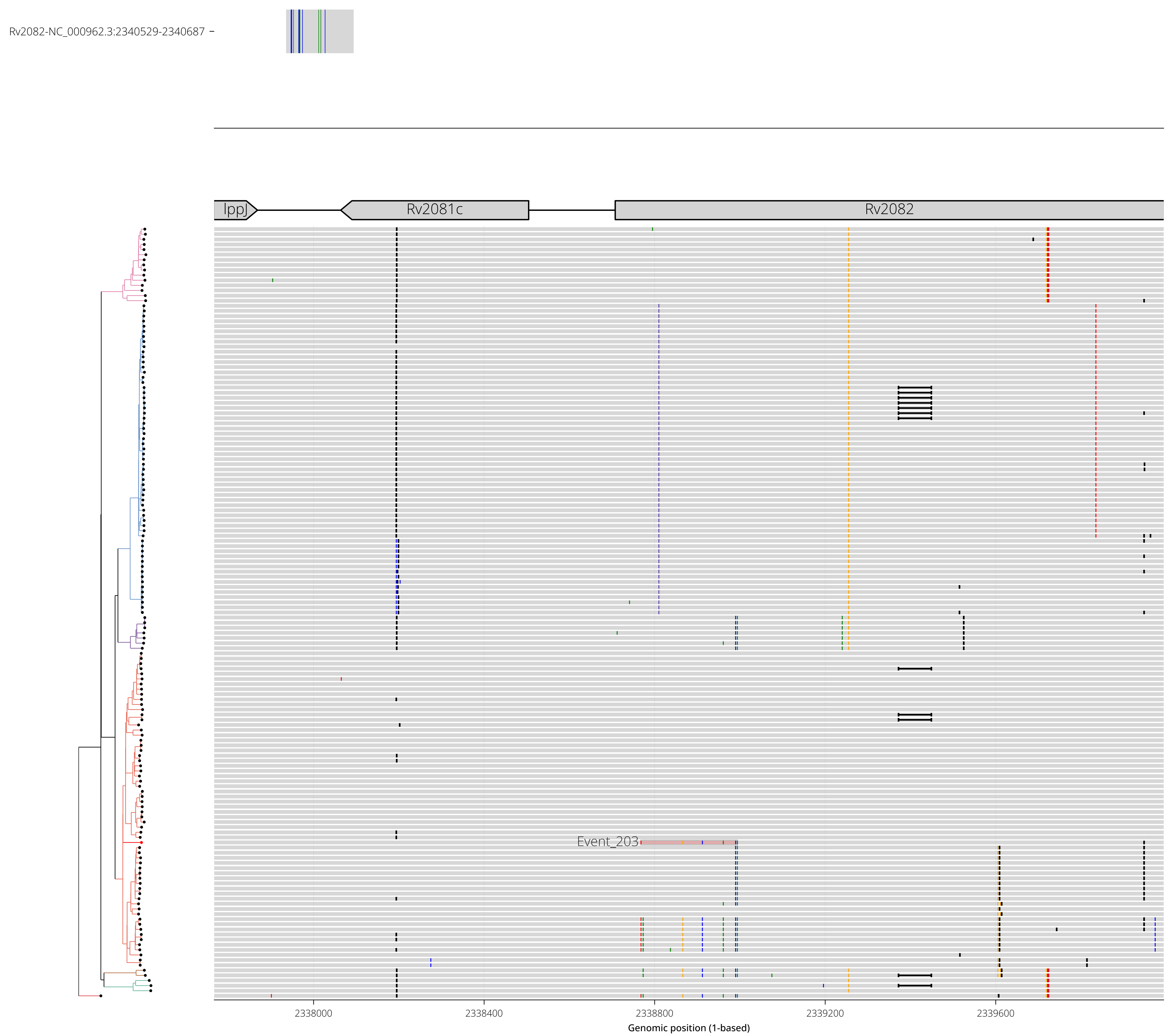

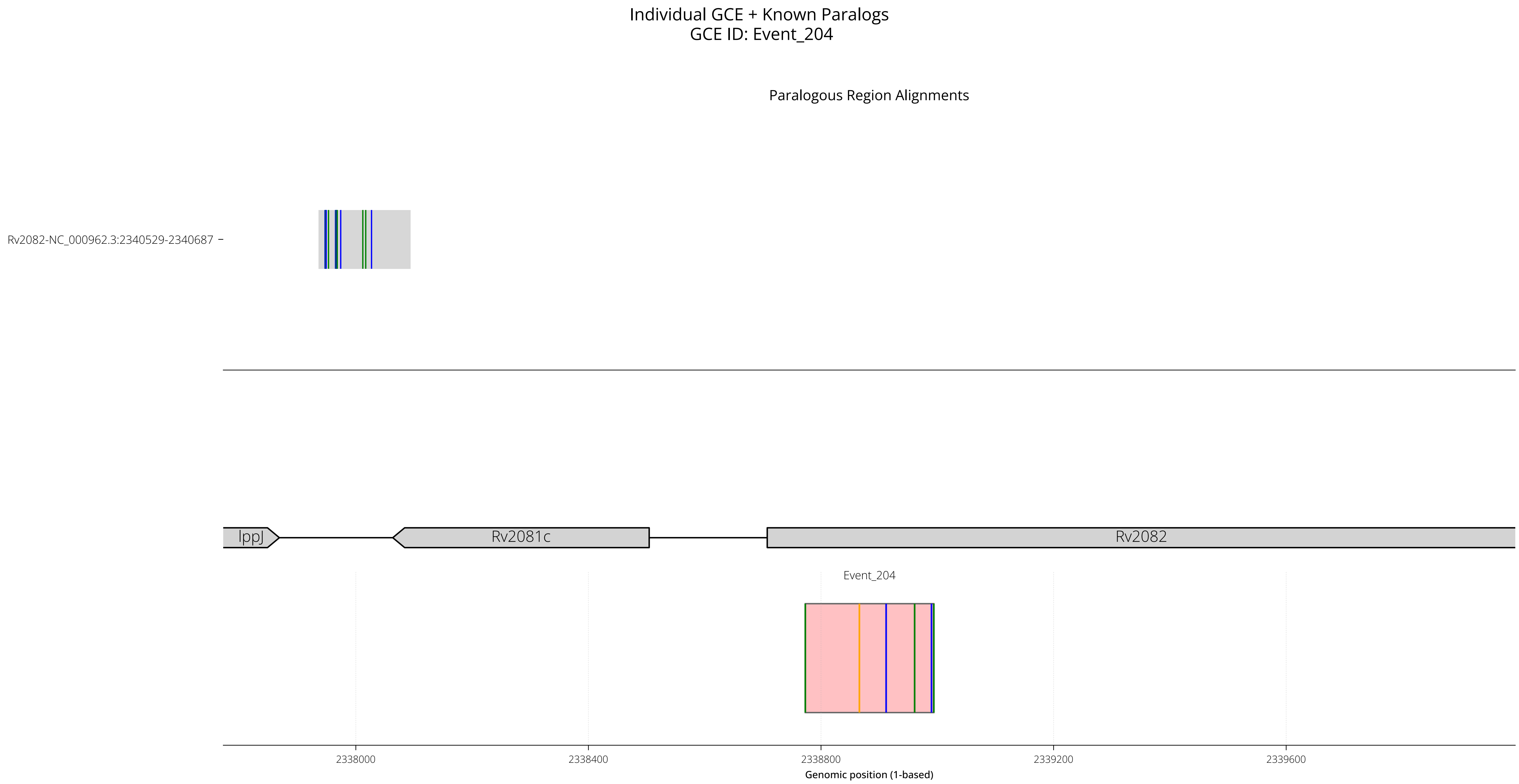

Individual GCE + Known Paralogs
GCE ID: Event_204
Paralogous Region Alignments
Rv2082-NC_000962.3:2340529-2340687
lppJ
Rv2081c
Rv2082
Event_204
2338000
2338400
2338800
2339200
2339600
Genomic position (1-based)

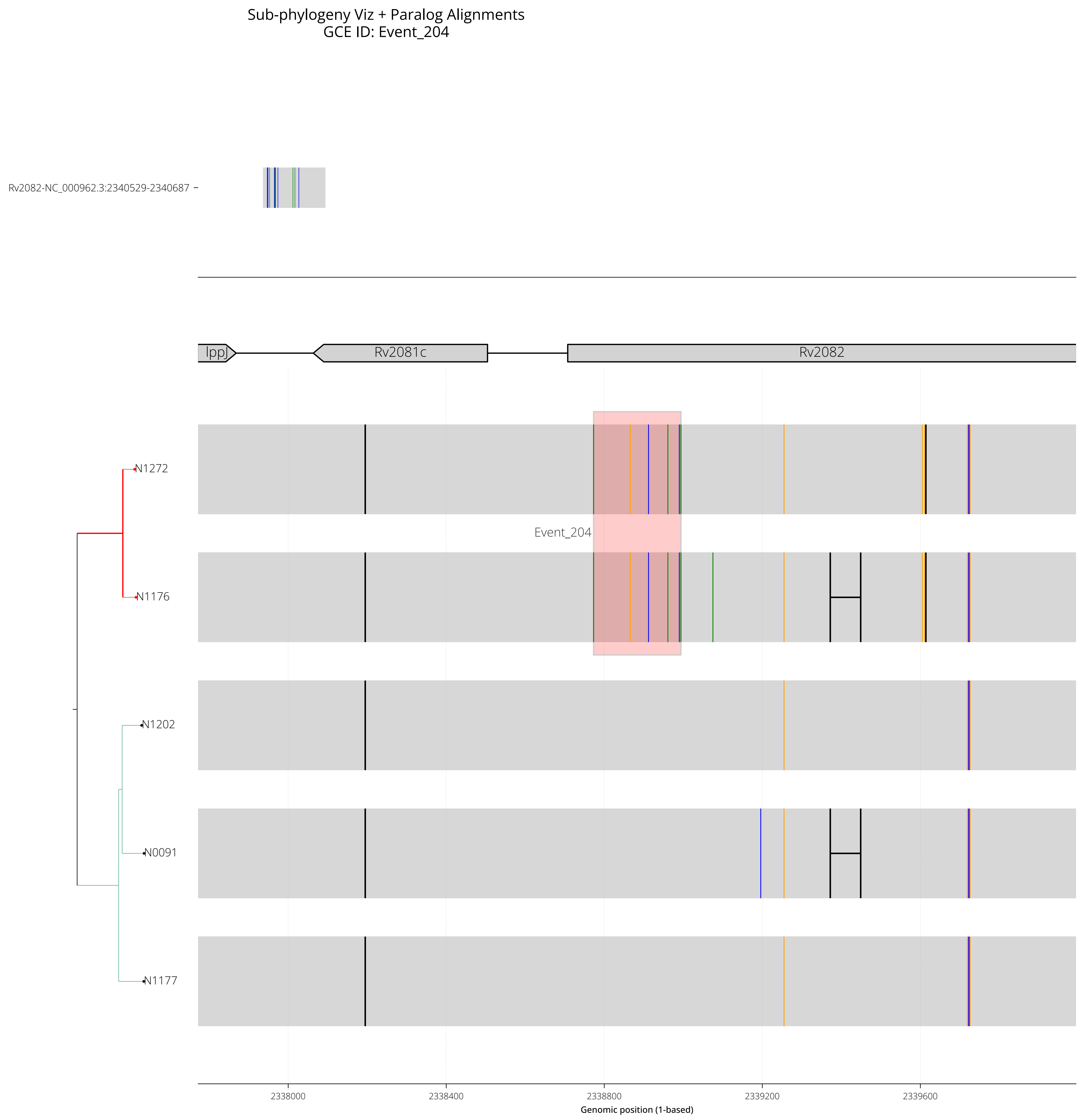
Sub-phylogeny Viz + Paralog Alignments
GCE ID: Event_204
Rv2082-NC_000962.3:2340529-2340687
lppJ
Rv2081c
Rv2082
Event_204
N1272
N1176
N1202
N0091
N1177
2338000
2338400
2338800
2339200
2339600
Genomic position (1-based)

Full Phylogeny Viz + Paralog Alignments
GCE ID: Event_204

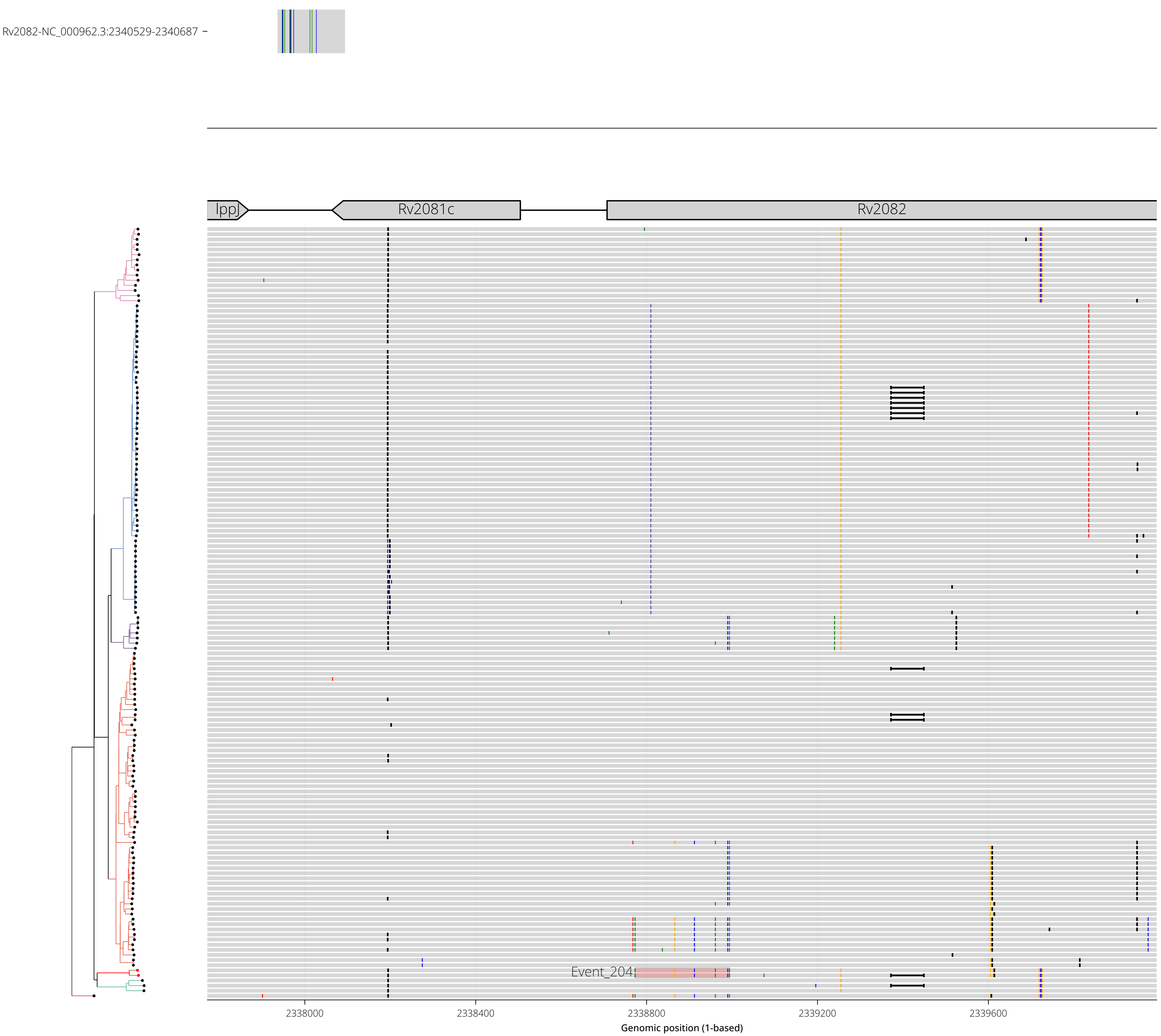

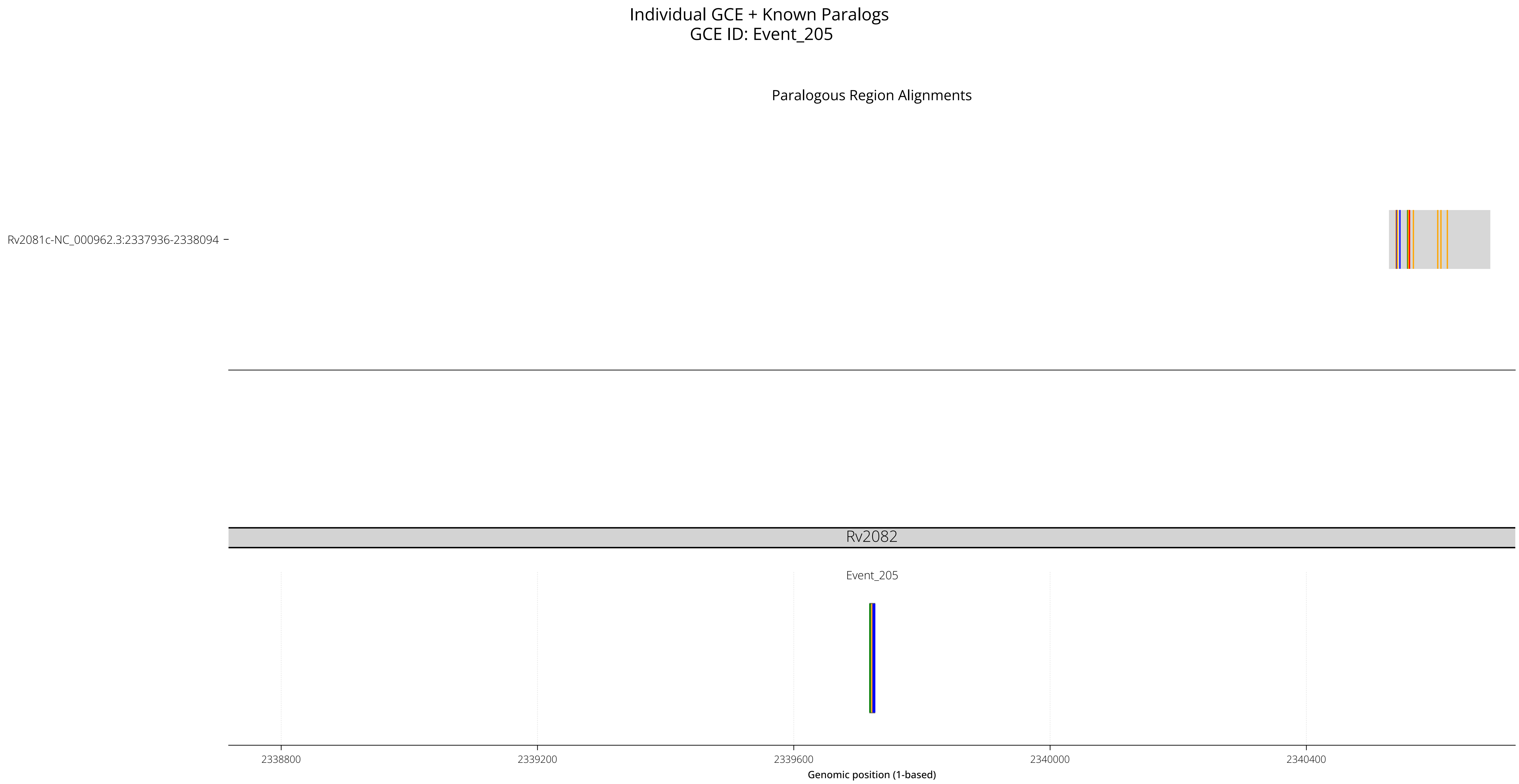

Individual GCE + Known Paralogs
GCE ID: Event_205
Paralogous Region Alignments
Rv2081c-NC_000962.3:2337936-2338094
Rv2082
Event_205
2338800
2339200
2339600
2340000
2340400
Genomic position (1-based)

Sub-phylogeny Viz + Paralog Alignments
GCE ID: Event_205

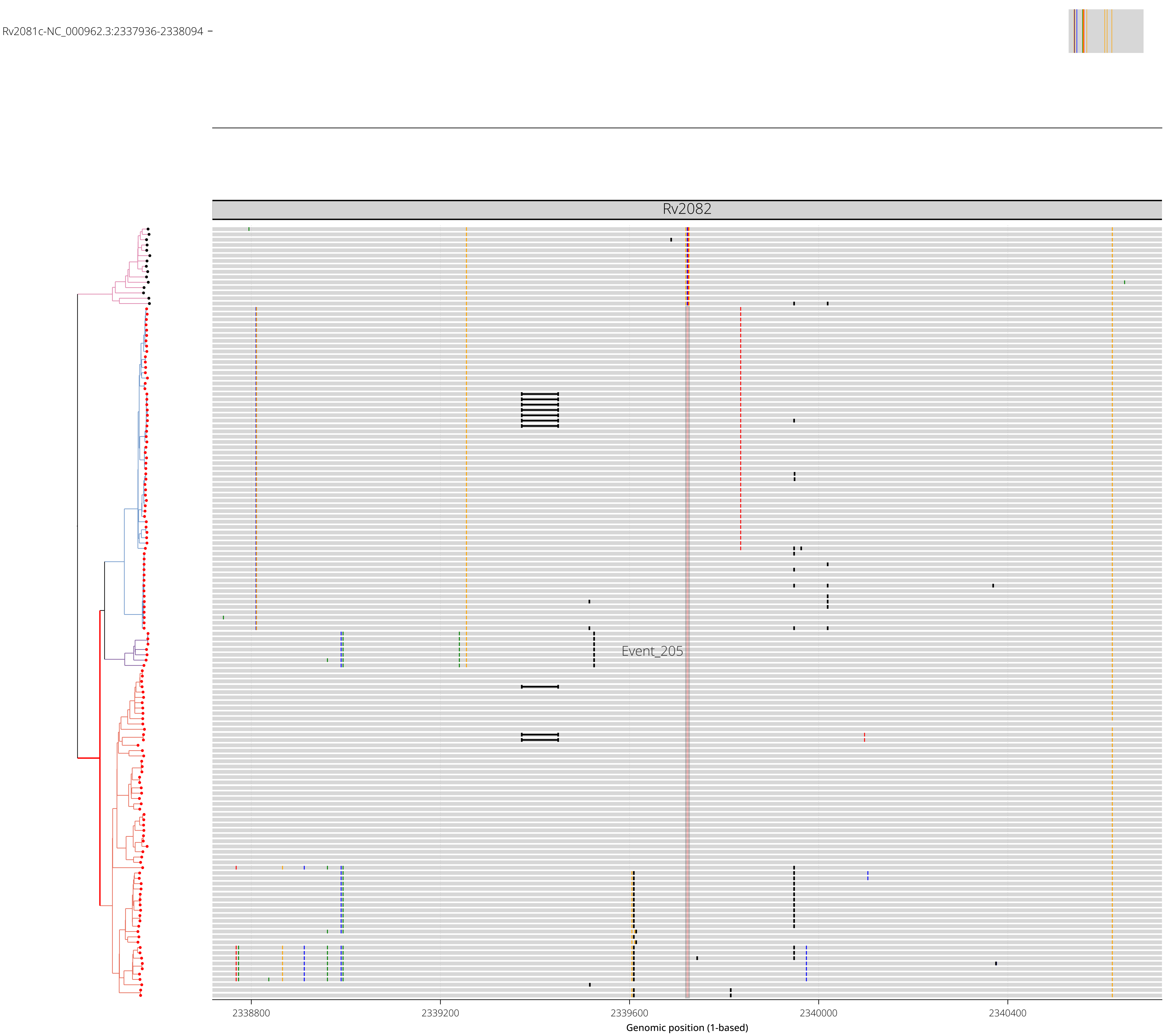

Full Phylogeny Viz + Paralog Alignments
GCE ID: Event_205

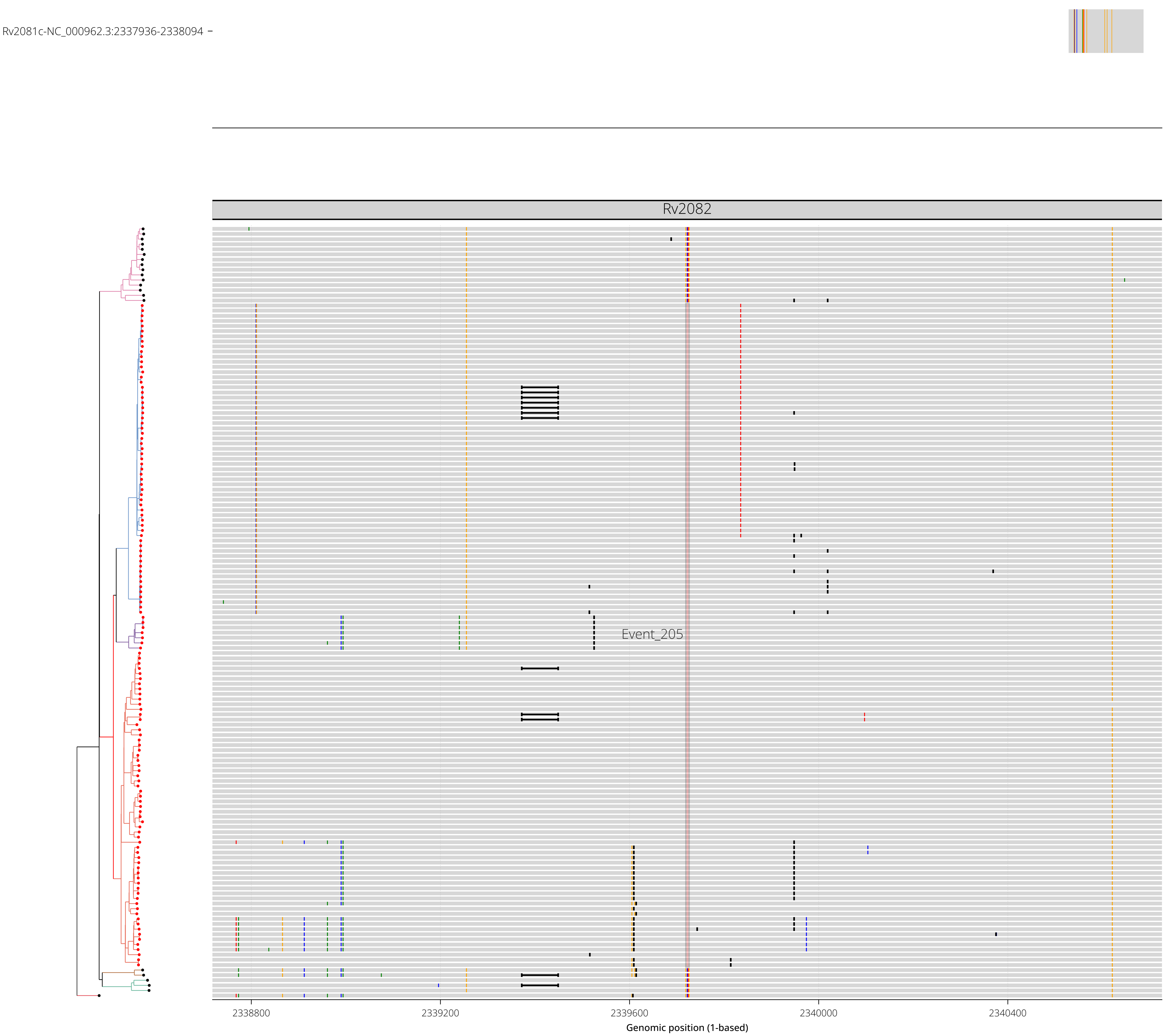

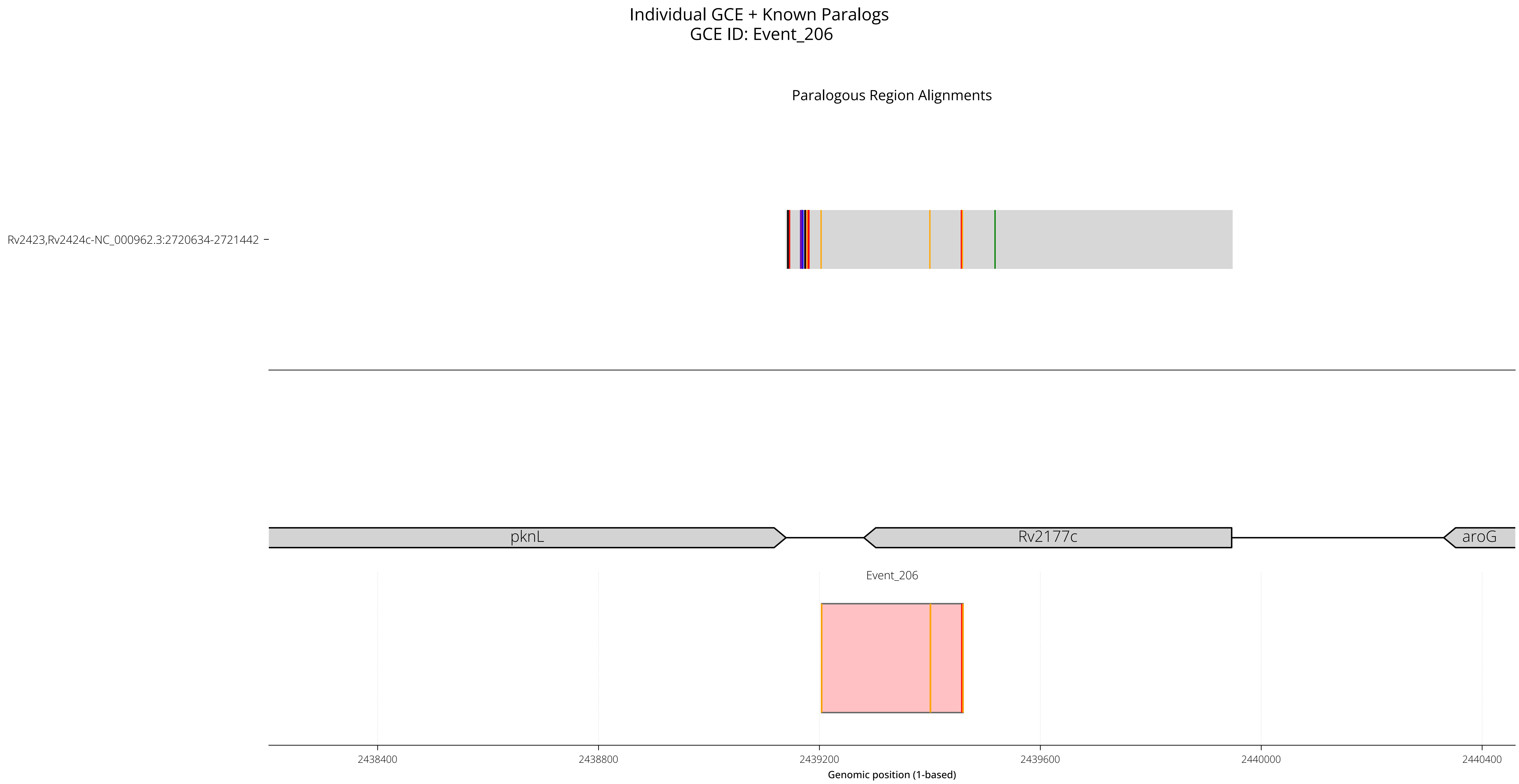

Individual GCE + Known Paralogs
GCE ID: Event_206
Paralogous Region Alignments
Rv2423,Rv2424c-NC_000962.3:2720634-2721442
pknL
Rv2177c
aroG
Event_206
2438400
2438800
2439200
2439600
2440000
2440400
Genomic position (1-based)

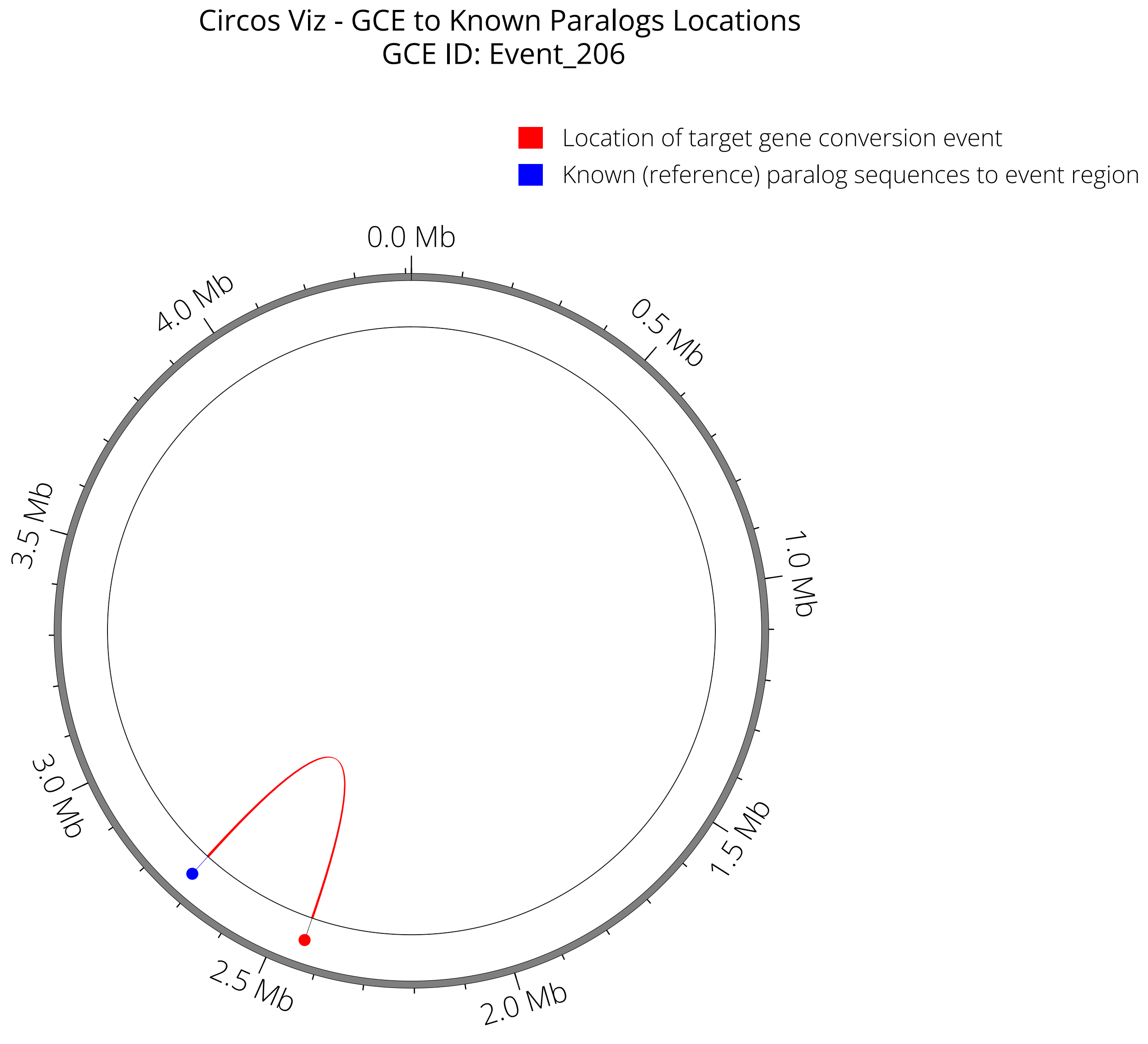

Circos Viz - GCE to Known Paralogs Locations
GCE ID: Event_206
Location of target gene conversion event
Known (reference) paralog sequences to event region
0.0 Mb
0.5 Mb
1.0 Mb
1.5 Mb
2.0 Mb
2.5 Mb
3.0 Mb
3.5 Mb
4.0 Mb

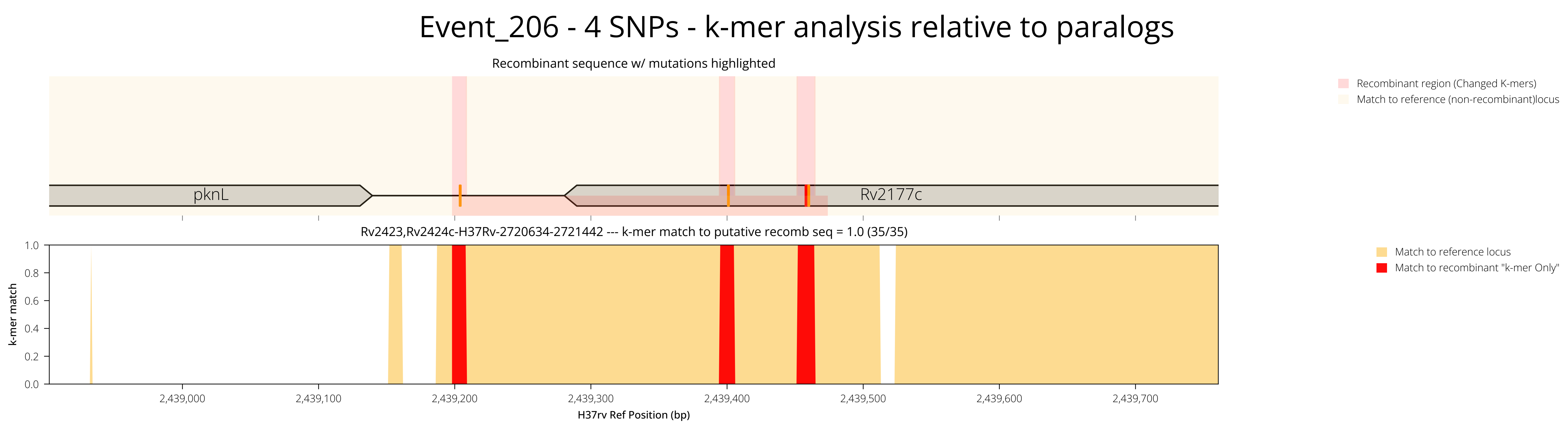
Event_206 - 4 SNPs - k-mer analysis relative to paralogs
Recombinant sequence w/ mutations highlighted
pknL
Rv2177c
Recombinant region (Changed K-mers)
Match to reference (non-recombinant)locus
Rv2423,Rv2424c-H37Rv-2720634-2721442 --- k-mer match to putative recomb seq = 1.0 (35/35)
k-mer match
1.0
0.8
0.6
0.4
0.2
0.0
2,439,000
2,439,100
2,439,200
2,439,300
2,439,400
2,439,500
2,439,600
2,439,700
H37rv Ref Position (bp)
Match to reference locus
Match to recombinant "k-mer Only"

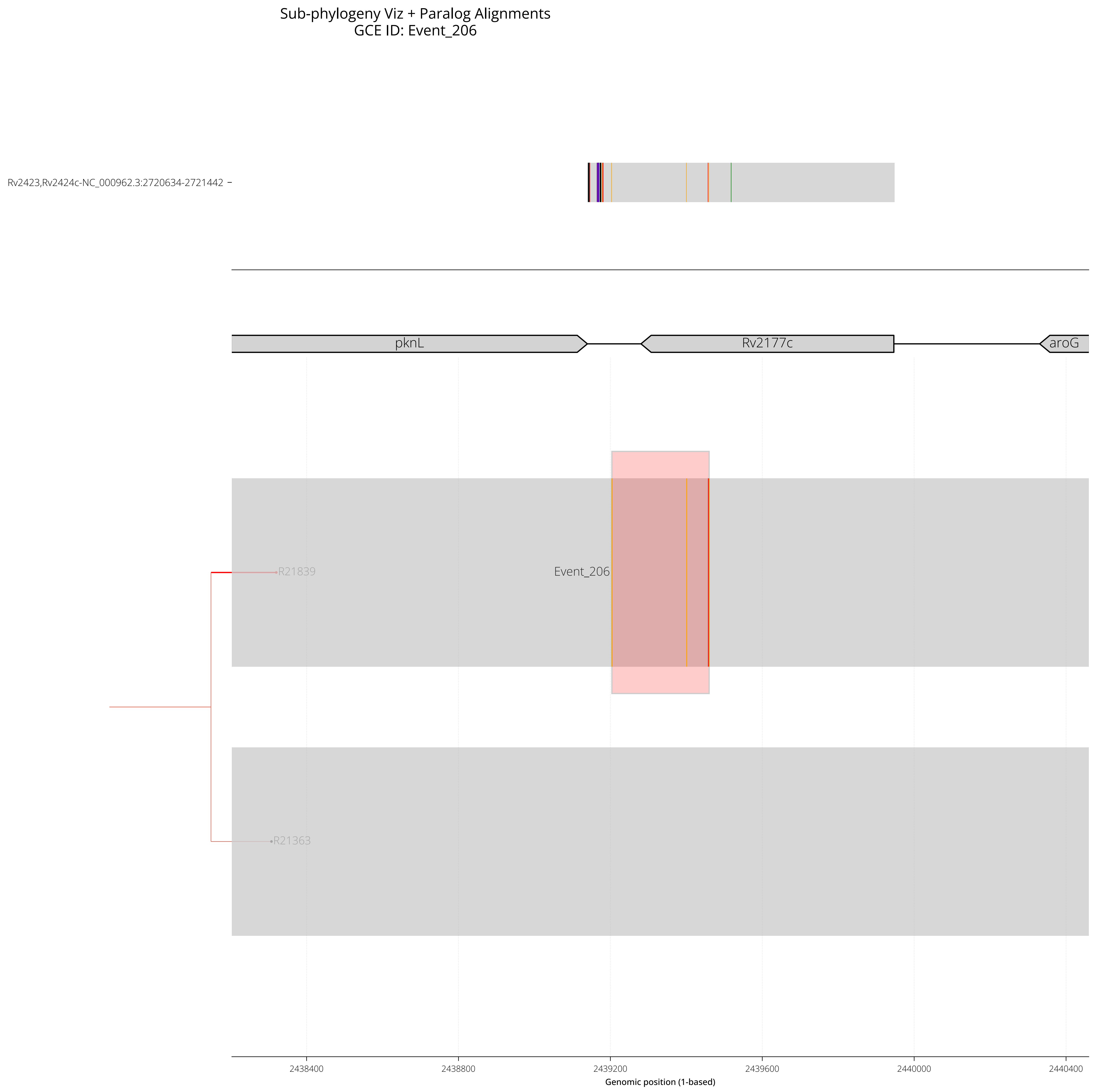
Sub-phylogeny Viz + Paralog Alignments
GCE ID: Event_206
Rv2423,Rv2424c-NC_000962.3:2720634-2721442
pknL
Rv2177c
aroG
R21839
Event_206
R21363
2438400
2438800
2439200
2439600
2440000
2440400
Genomic position (1-based)

Full Phylogeny Viz + Paralog Alignments
GCE ID: Event_206

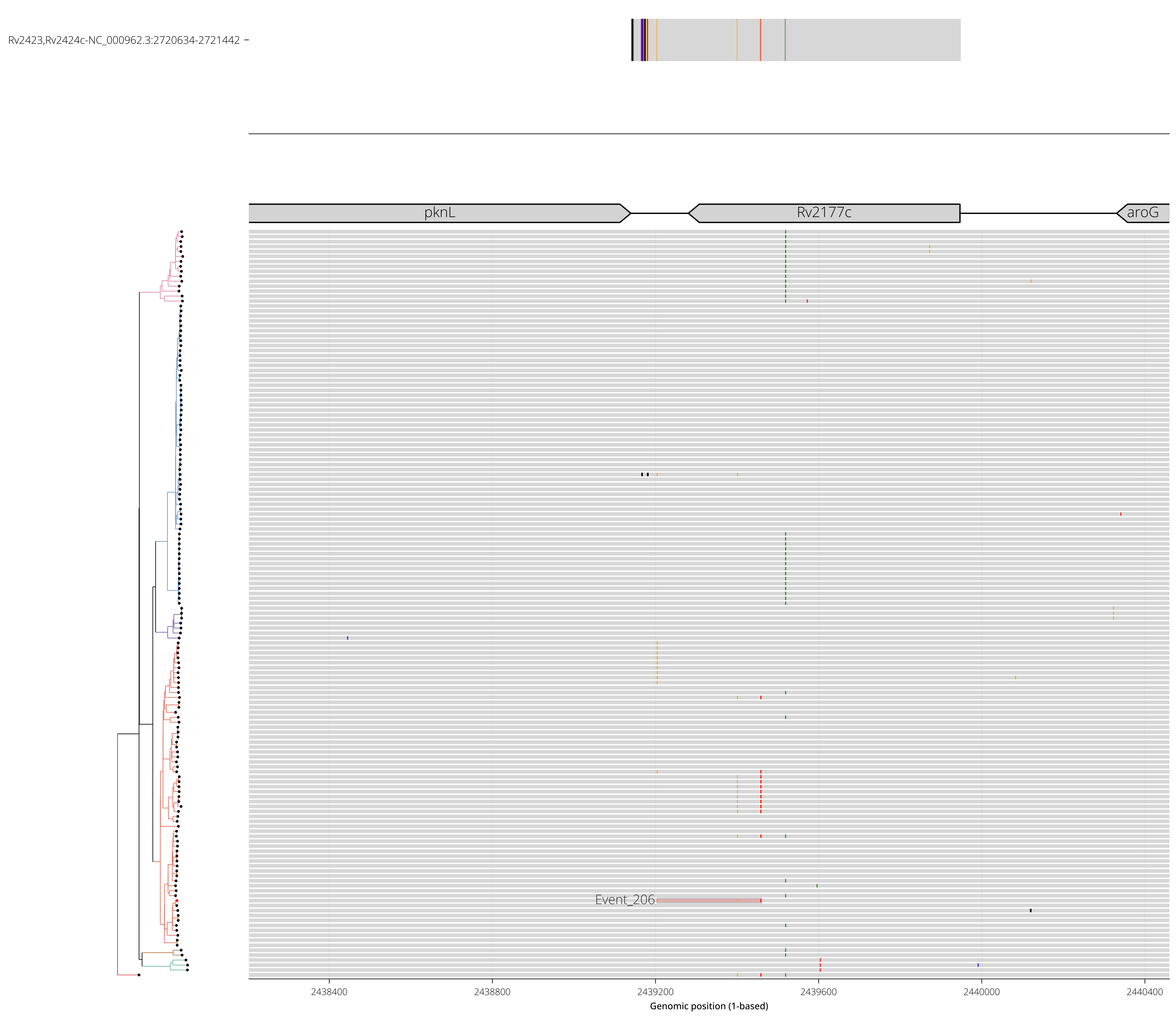

Individual GCE + Known Paralogs
GCE ID: Event_207

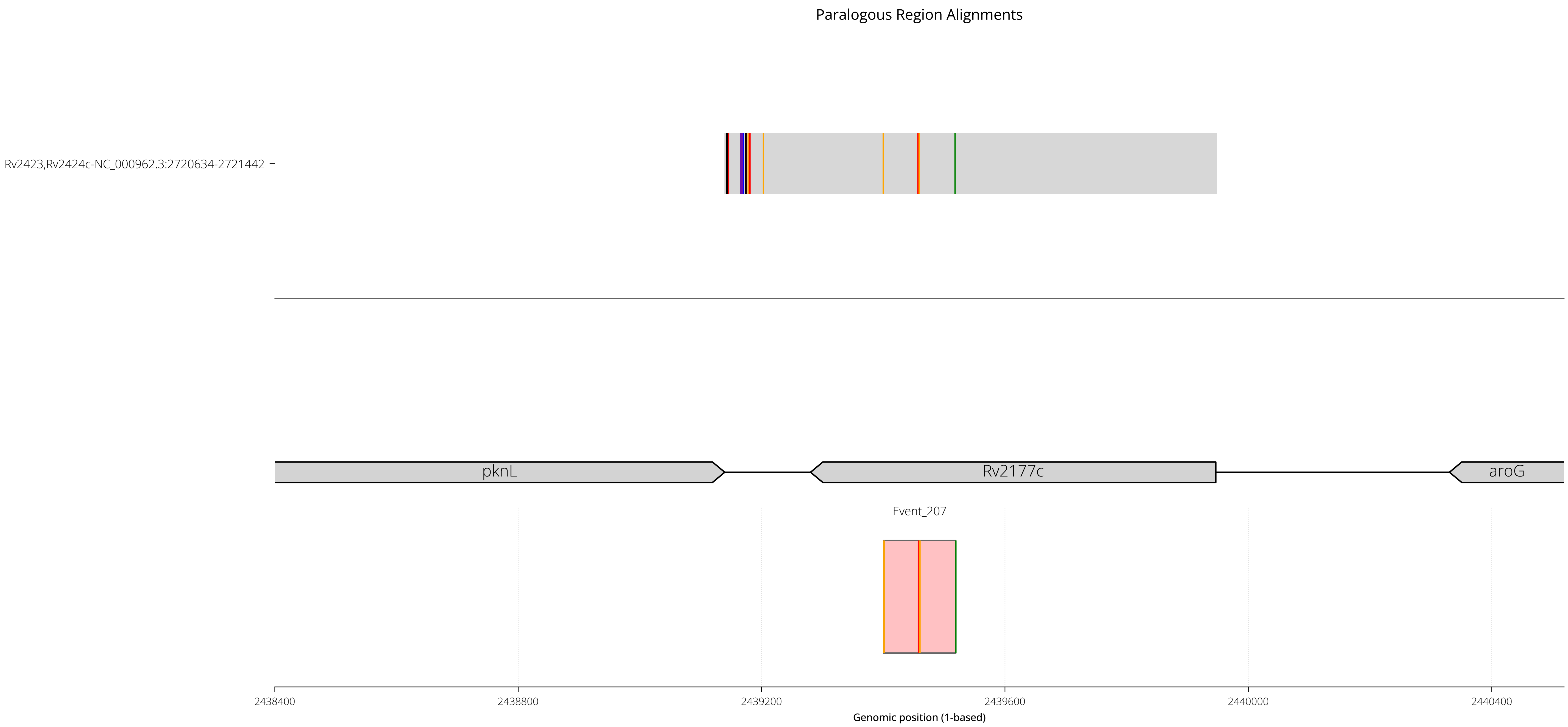

Circos Viz - GCE to Known Paralogs Locations
GCE ID: Event_207

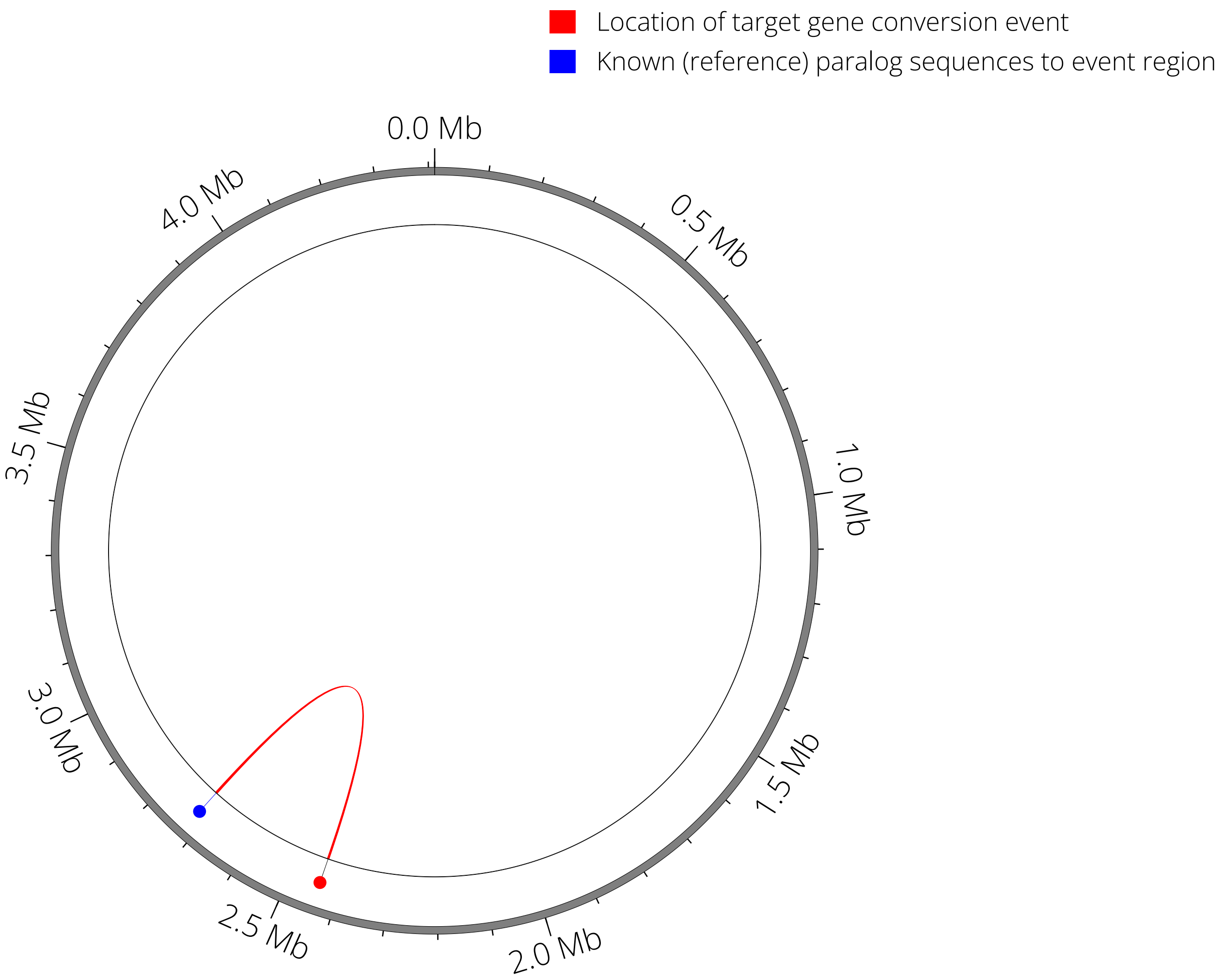

### Event_207 - 4 SNPs - k-mer analysis relative to paralogs

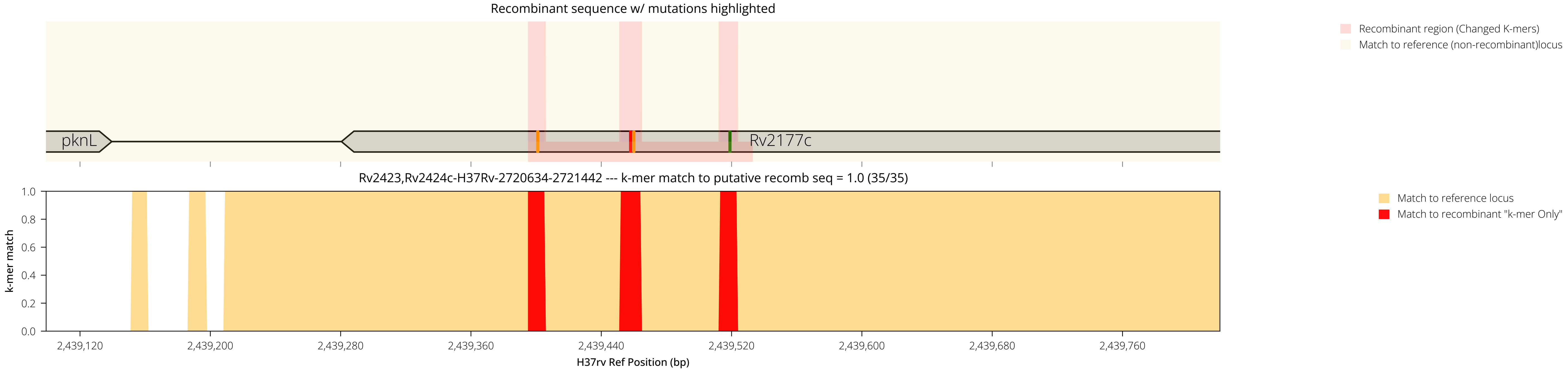

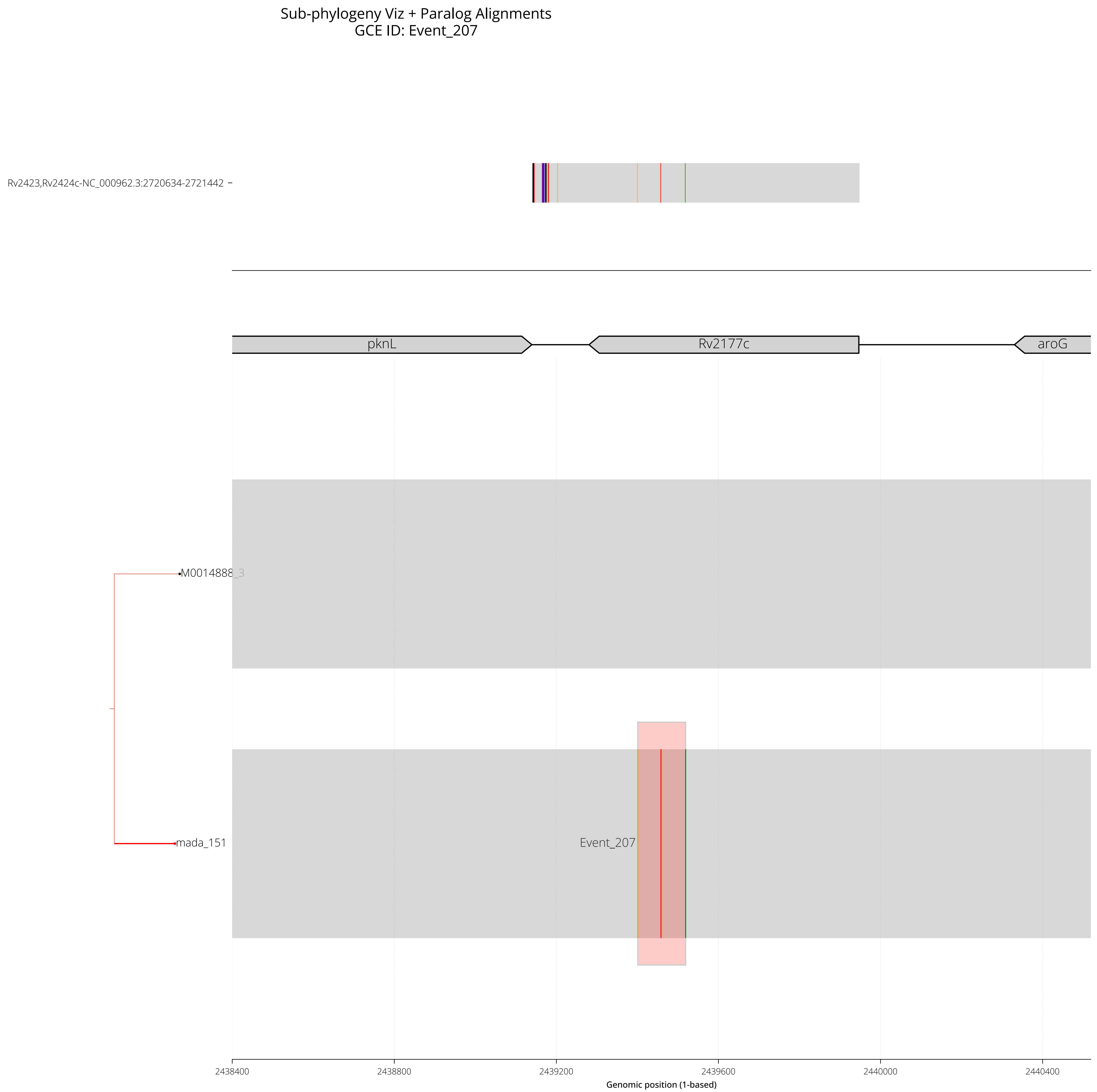
Sub-phylogeny Viz + Paralog Alignments
GCE ID: Event_207
Rv2423,Rv2424c-NC_000962.3:2720634-2721442
pknL
Rv2177c
aroG
M0014888_3
mada_151
Event_207
2438400
2438800
2439200
2439600
2440000
2440400
Genomic position (1-based)

Full Phylogeny Viz + Paralog Alignments
GCE ID: Event_207

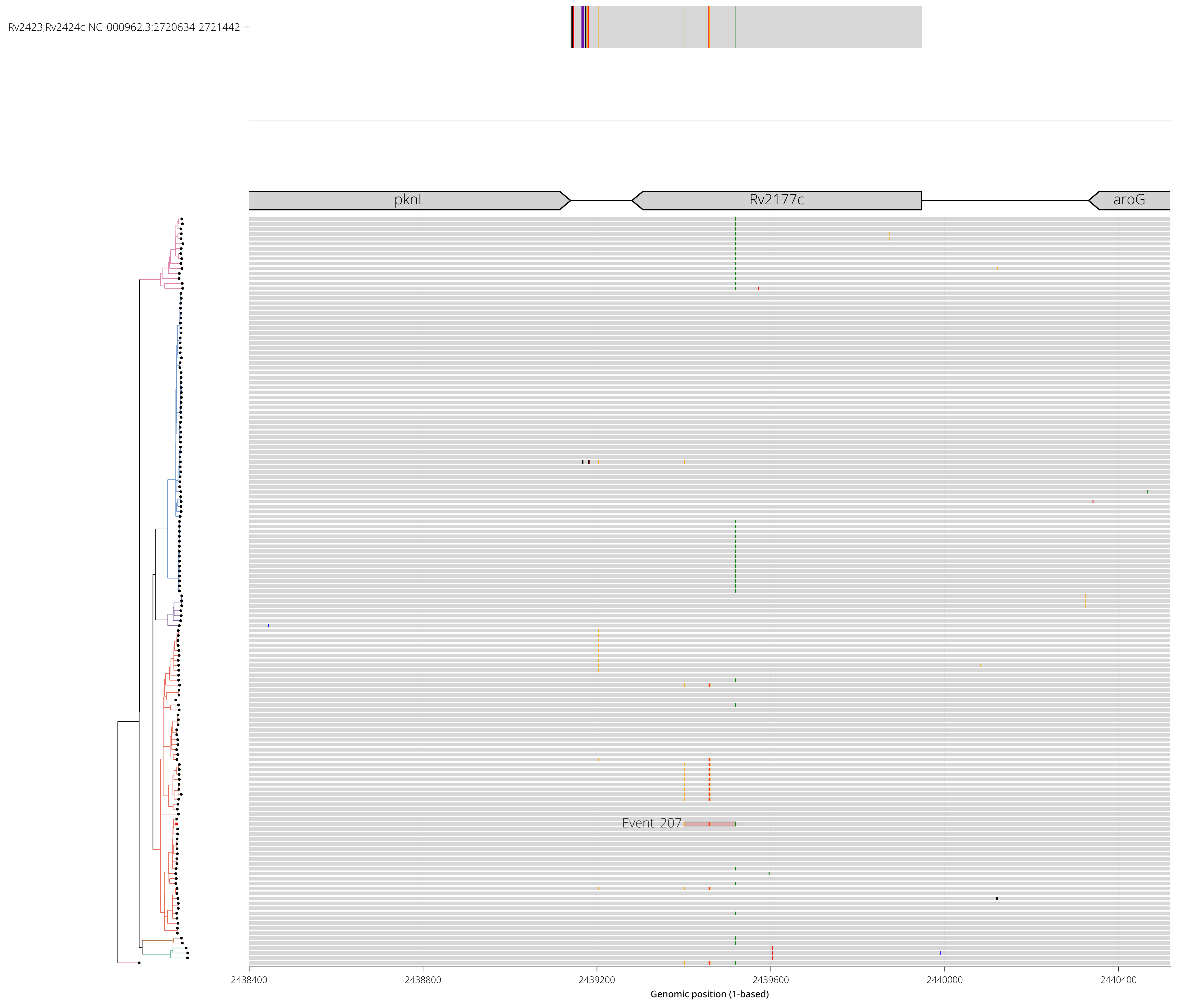

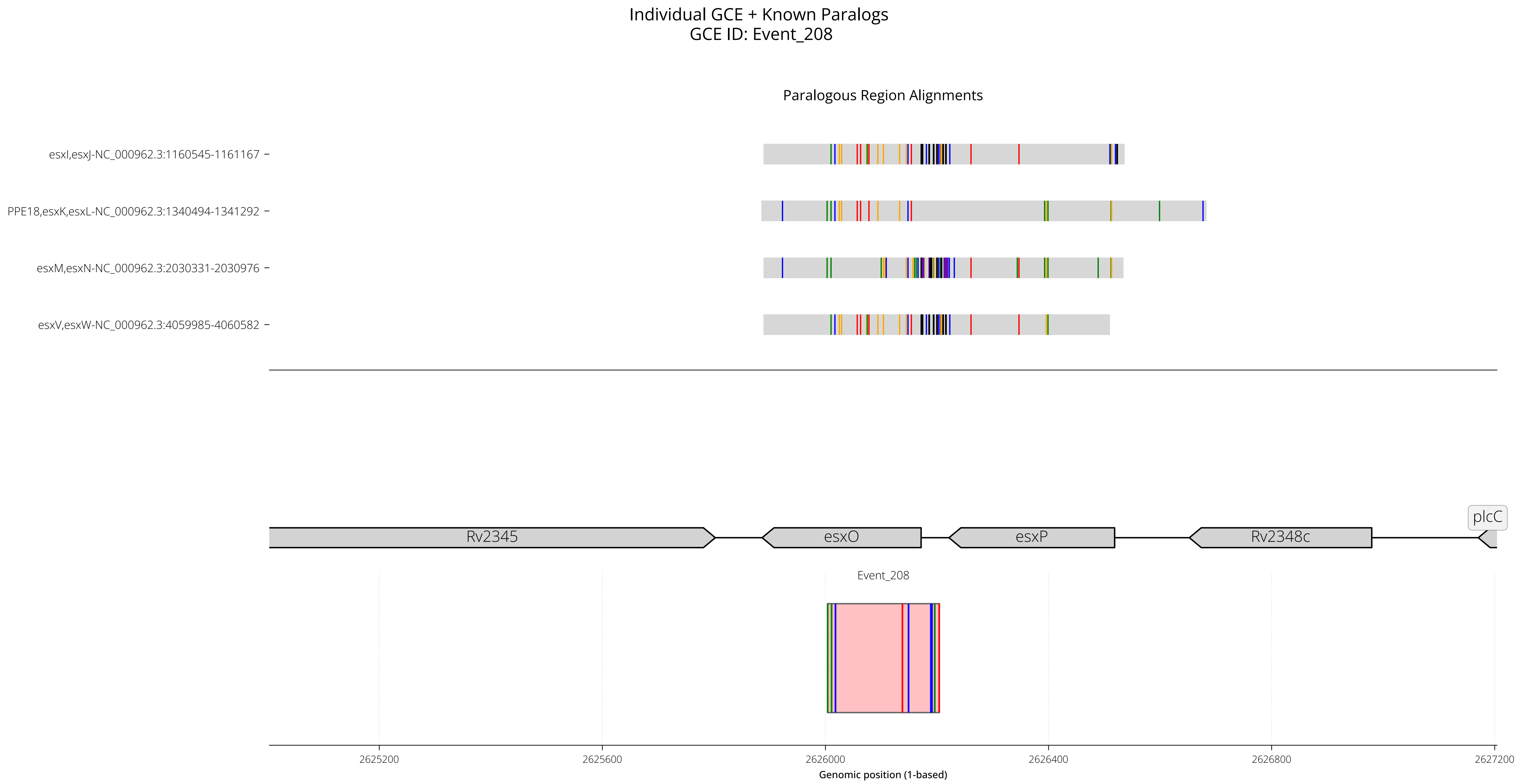
Individual GCE + Known Paralogs
GCE ID: Event_208
Paralogous Region Alignments
esxI,esxJ-NC_000962.3:1160545-1161167
PPE18,esxK,esxL-NC_000962.3:1340494-1341292
esxM,esxN-NC_000962.3:2030331-2030976
esxV,esxW-NC_000962.3:4059985-4060582
Rv2345
esxO
esxP
Rv2348c
plcC
Event_208
2625200
2625600
2626000
2626400
2626800
2627200
Genomic position (1-based)

Circos Viz - GCE to Known Paralogs Locations
GCE ID: Event_208

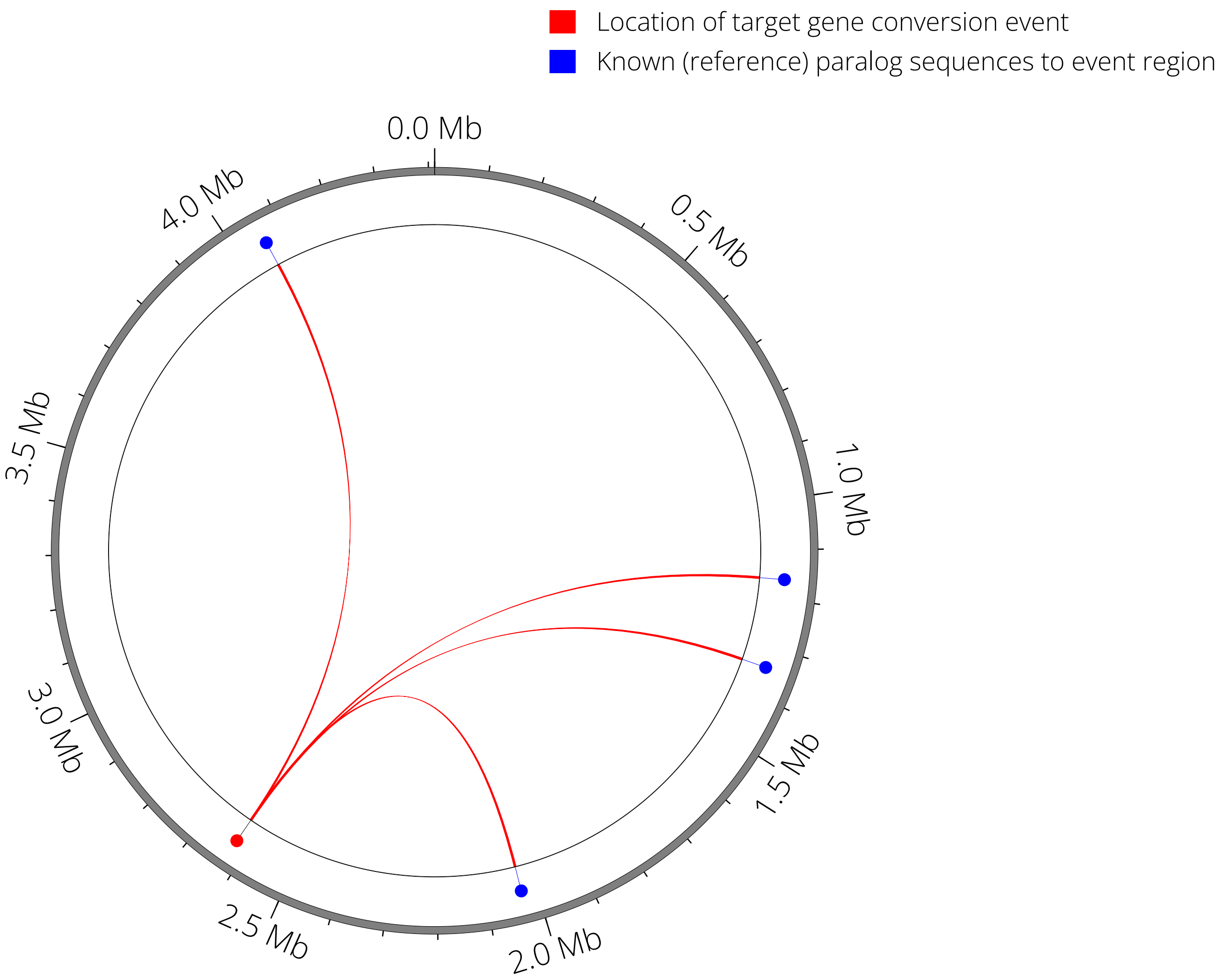

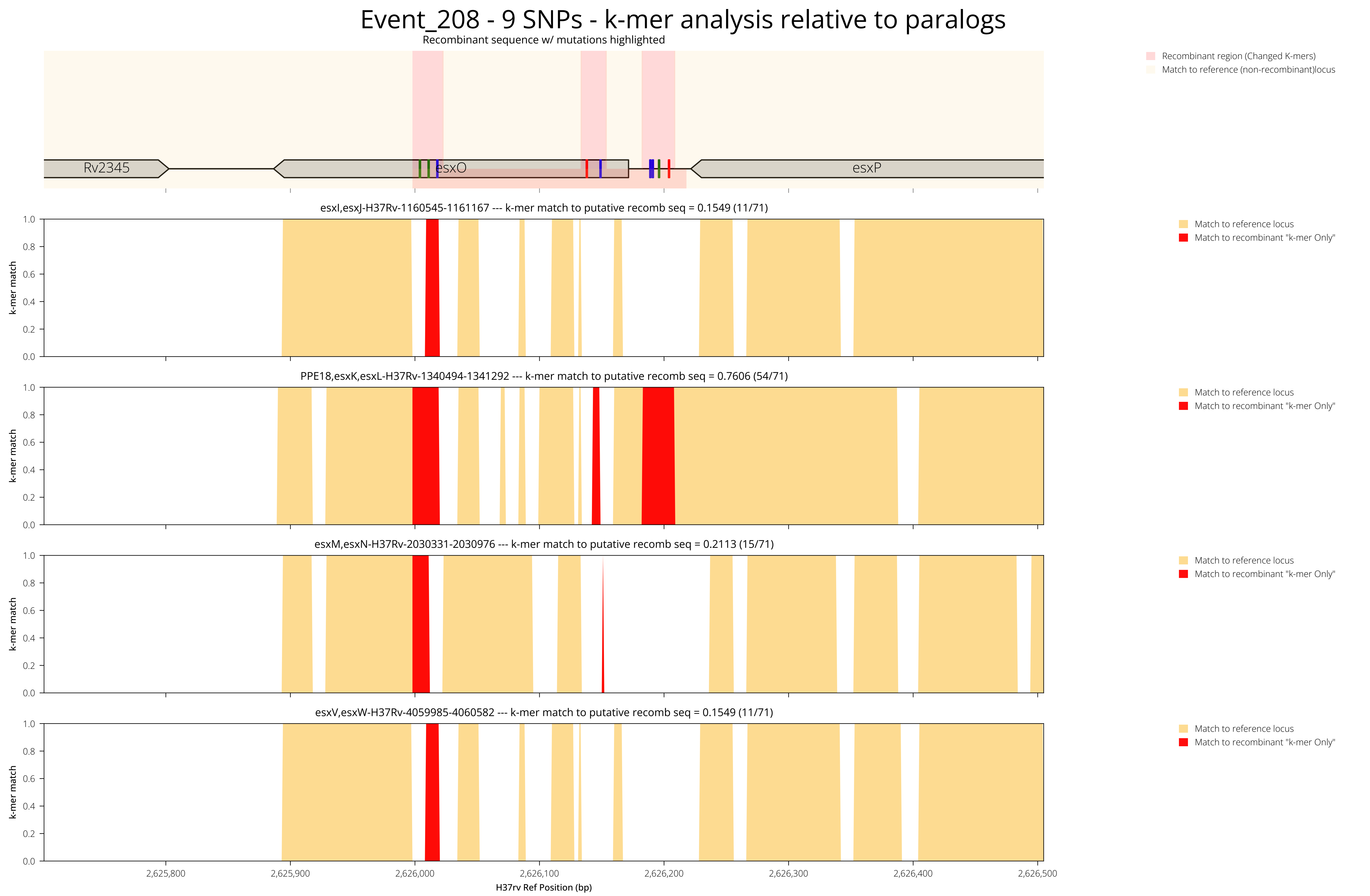

Event_208 - 9 SNPs - k-mer analysis relative to paralogs
Recombinant sequence w/ mutations highlighted
Rv2345
esxO
esxP
Recombinant region (Changed K-mers)
Match to reference (non-recombinant)locus
esxI,esxJ-H37Rv-1160545-1161167 --- k-mer match to putative recomb seq = 0.1549 (11/71)
PPE18,esxK,esxL-H37Rv-1340494-1341292 --- k-mer match to putative recomb seq = 0.7606 (54/71)
esxM,esxN-H37Rv-2030331-2030976 --- k-mer match to putative recomb seq = 0.2113 (15/71)
esxV,esxW-H37Rv-4059985-4060582 --- k-mer match to putative recomb seq = 0.1549 (11/71)
k-mer match
0.0
0.2
0.4
0.6
0.8
1.0
Match to reference locus
Match to recombinant "k-mer Only"
2,625,800
2,625,900
2,626,000
2,626,100
2,626,200
2,626,300
2,626,400
2,626,500
H37rv Ref Position (bp)

Sub-phylogeny Viz + Paralog Alignments
GCE ID: Event_208

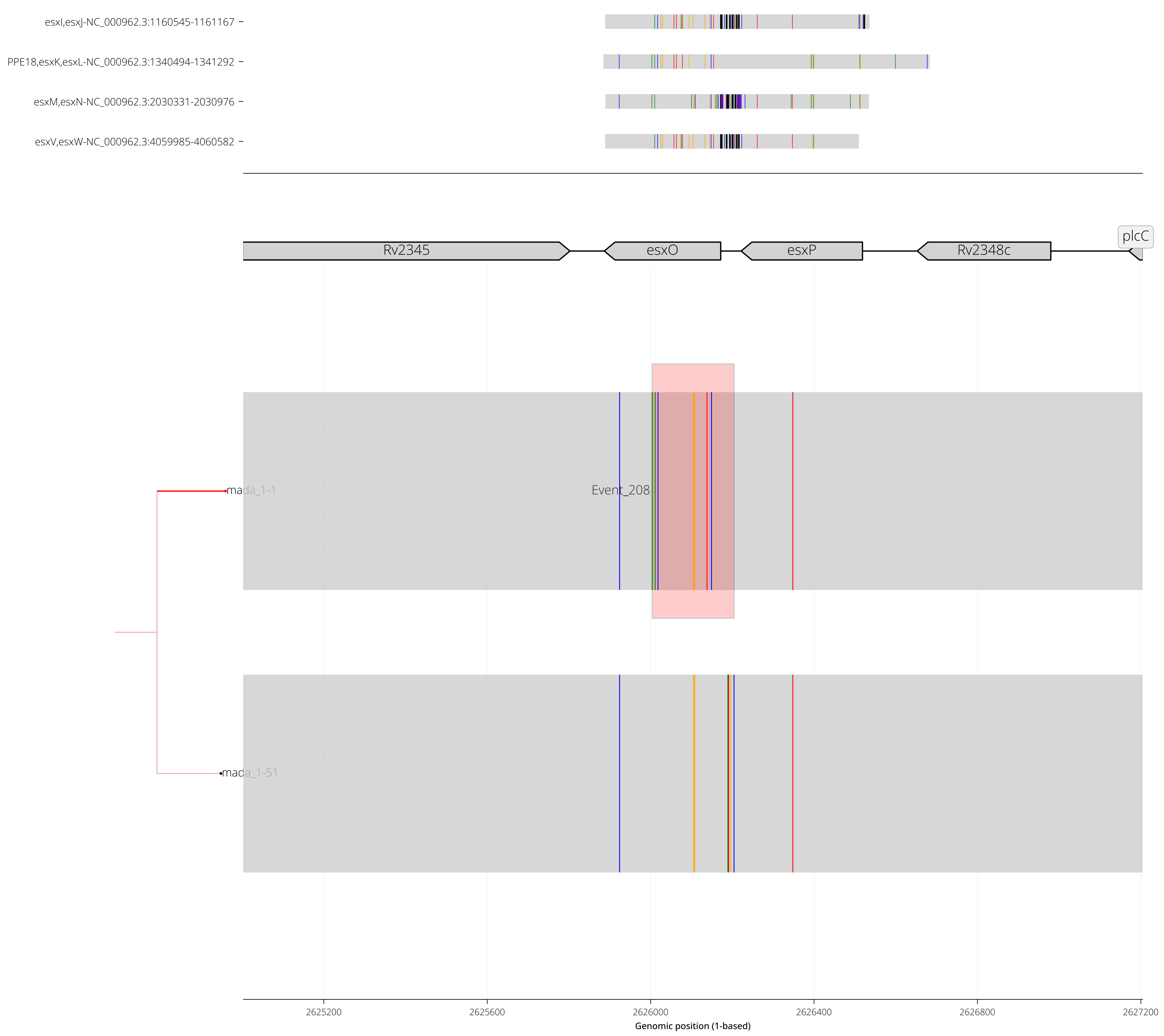

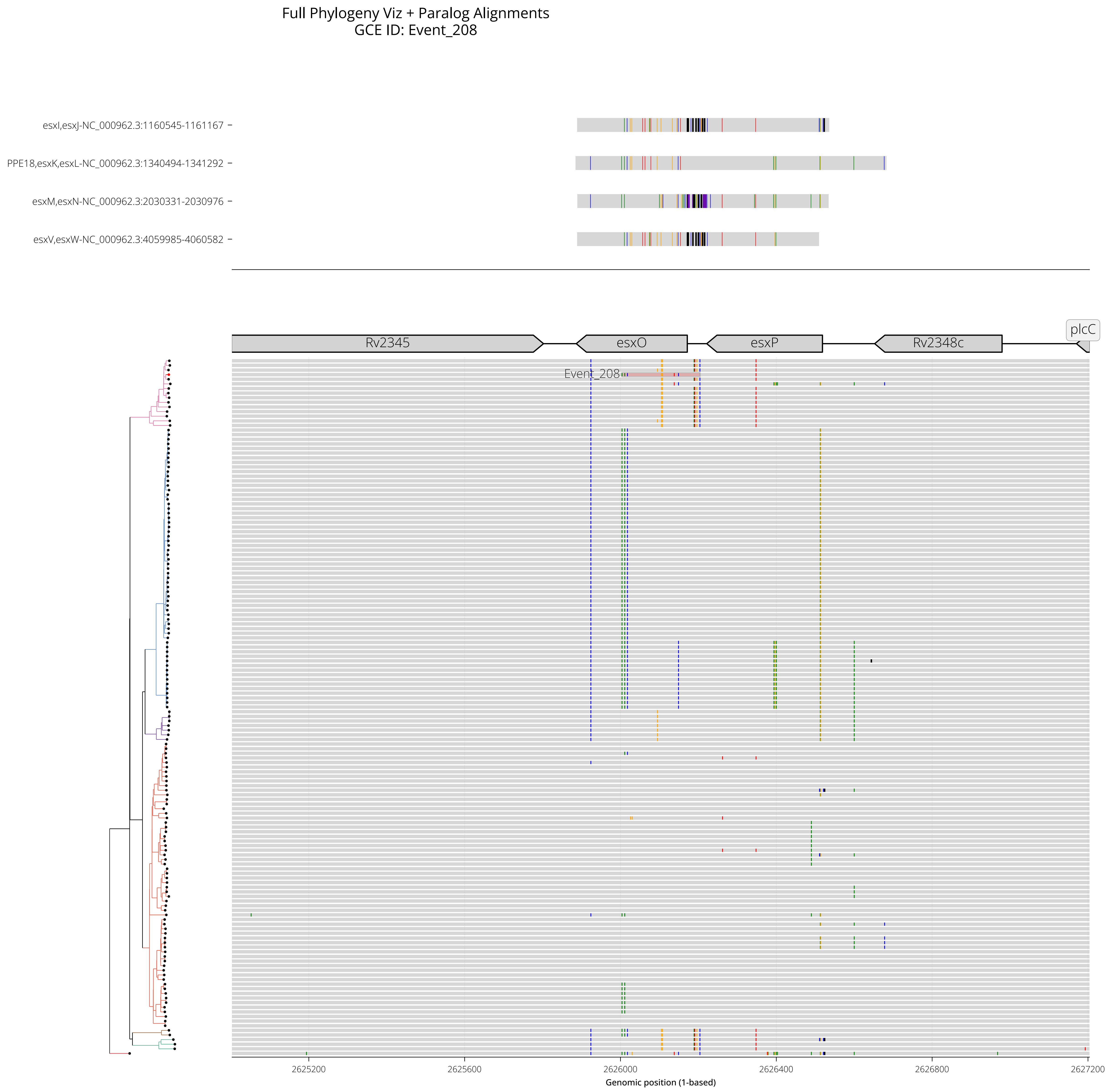

Full Phylogeny Viz + Paralog Alignments
GCE ID: Event_208
esxI,esxJ-NC_000962.3:1160545-1161167
PPE18,esxK,esxL-NC_000962.3:1340494-1341292
esxM,esxN-NC_000962.3:2030331-2030976
esxV,esxW-NC_000962.3:4059985-4060582
Rv2345
esxO
esxP
Rv2348c
plcC
Event_208
2625200
2625600
2626000
2626400
2626800
2627200
Genomic position (1-based)

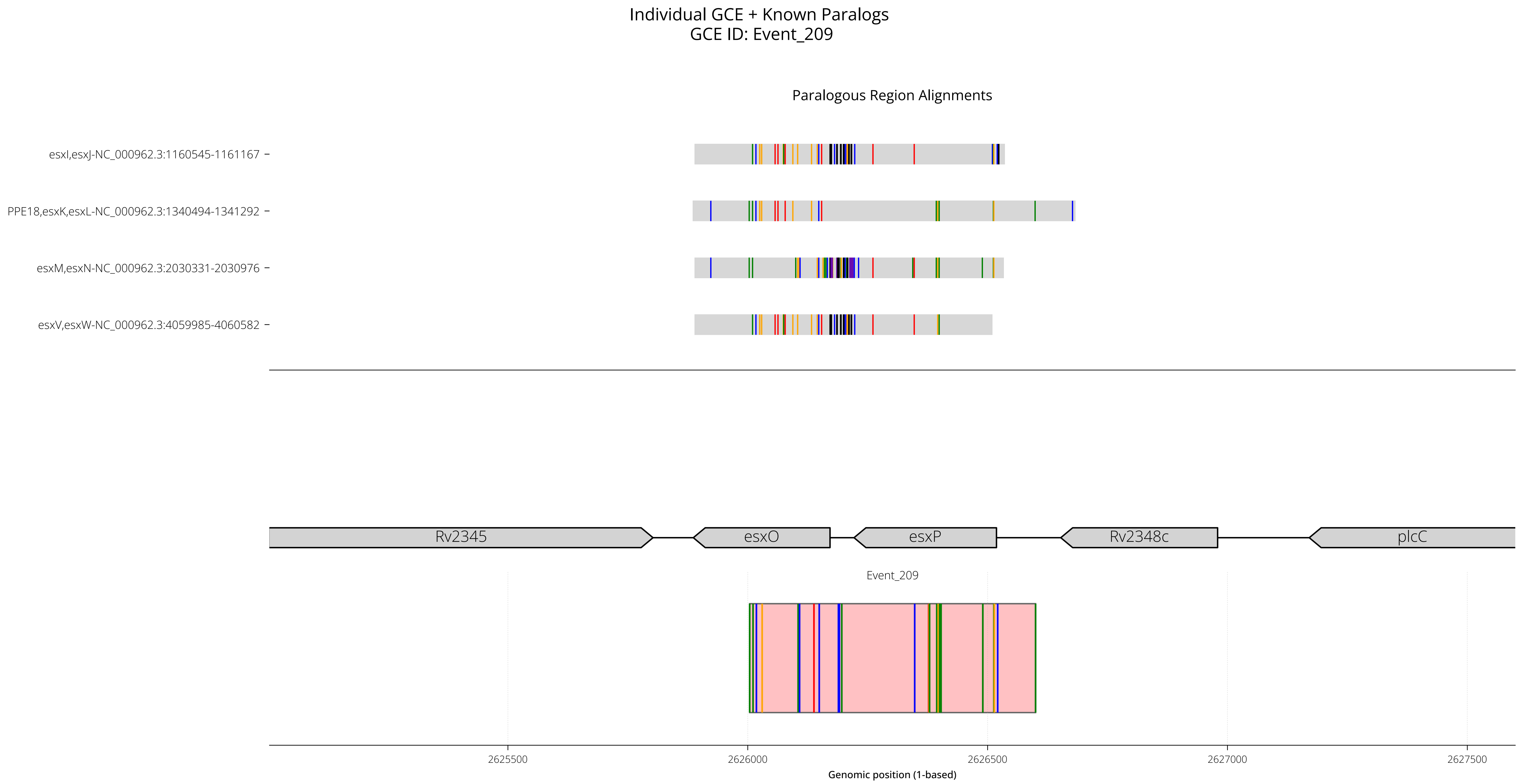
Individual GCE + Known Paralogs
GCE ID: Event_209
Paralogous Region Alignments
esxI,esxJ-NC_000962.3:1160545-1161167
PPE18,esxK,esxL-NC_000962.3:1340494-1341292
esxM,esxN-NC_000962.3:2030331-2030976
esxV,esxW-NC_000962.3:4059985-4060582
Rv2345
esxO
esxP
Rv2348c
plcC
Event_209
2625500
2626000
2626500
2627000
2627500
Genomic position (1-based)

Circos Viz - GCE to Known Paralogs Locations
GCE ID: Event_209

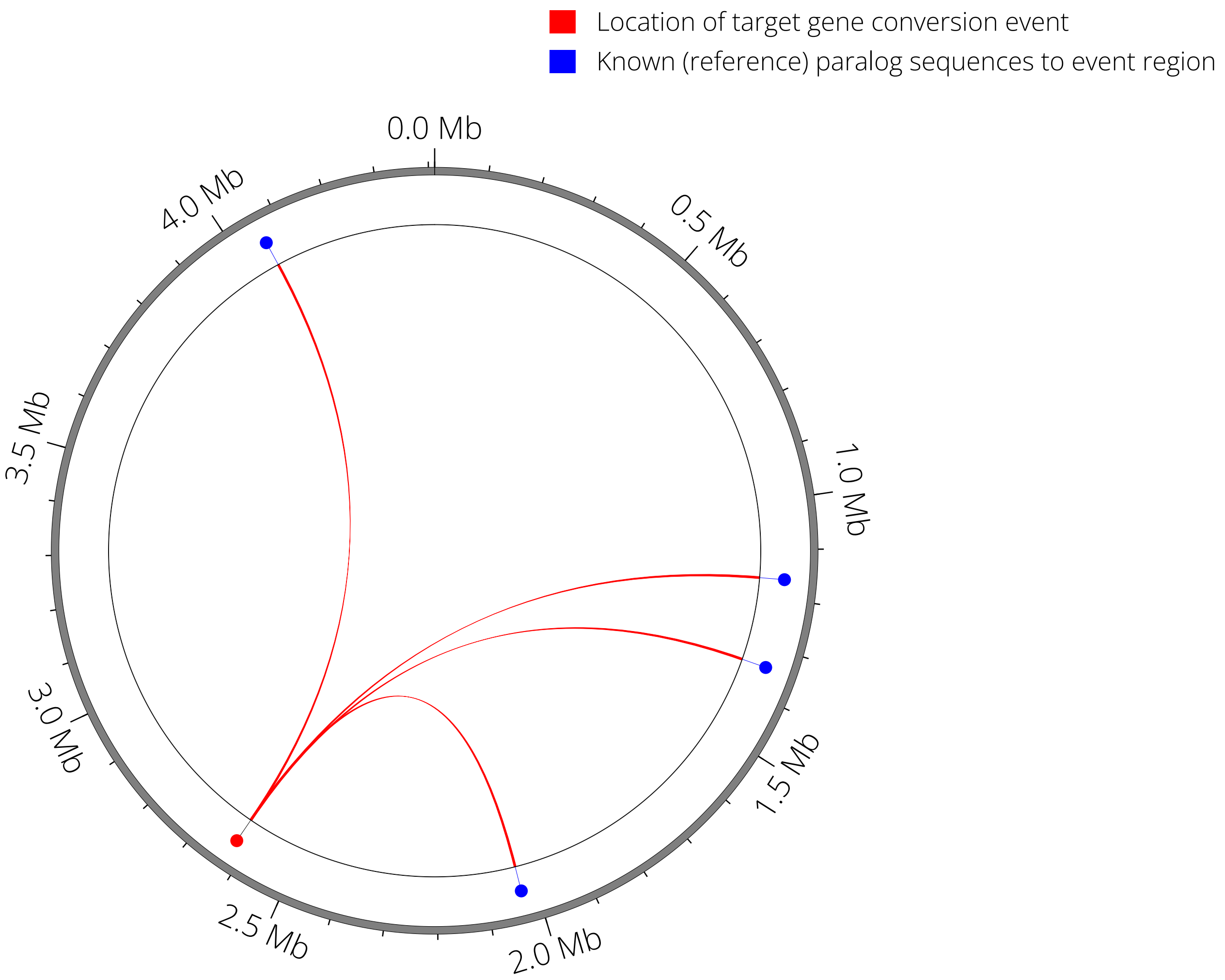

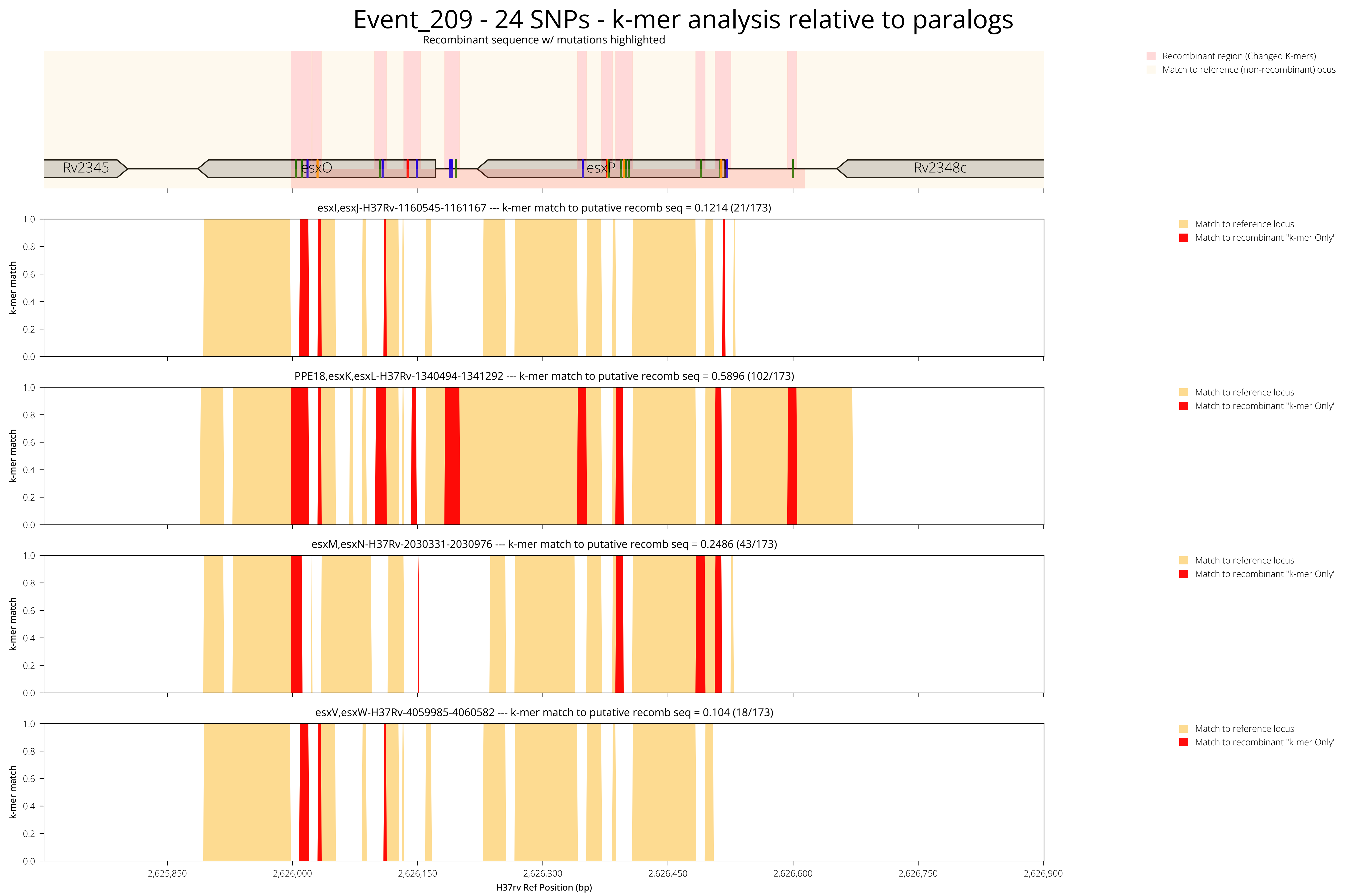

Event_209 - 24 SNPs - k-mer analysis relative to paralogs
Recombinant sequence w/ mutations highlighted
Recombinant region (Changed K-mers)
Match to reference (non-recombinant)locus
Rv2345
esxO
esxP
Rv2348c
esxI,esxJ-H37Rv-1160545-1161167 --- k-mer match to putative recomb seq = 0.1214 (21/173)
PPE18,esxK,esxL-H37Rv-1340494-1341292 --- k-mer match to putative recomb seq = 0.5896 (102/173)
esxM,esxN-H37Rv-2030331-2030976 --- k-mer match to putative recomb seq = 0.2486 (43/173)
esxV,esxW-H37Rv-4059985-4060582 --- k-mer match to putative recomb seq = 0.104 (18/173)
Match to reference locus
Match to recombinant "k-mer Only"
k-mer match
1.0
0.8
0.6
0.4
0.2
0.0
2,625,850
2,626,000
2,626,150
2,626,300
2,626,450
2,626,600
2,626,750
2,626,900
H37rv Ref Position (bp)

Sub-phylogeny Viz + Paralog Alignments
GCE ID: Event_209

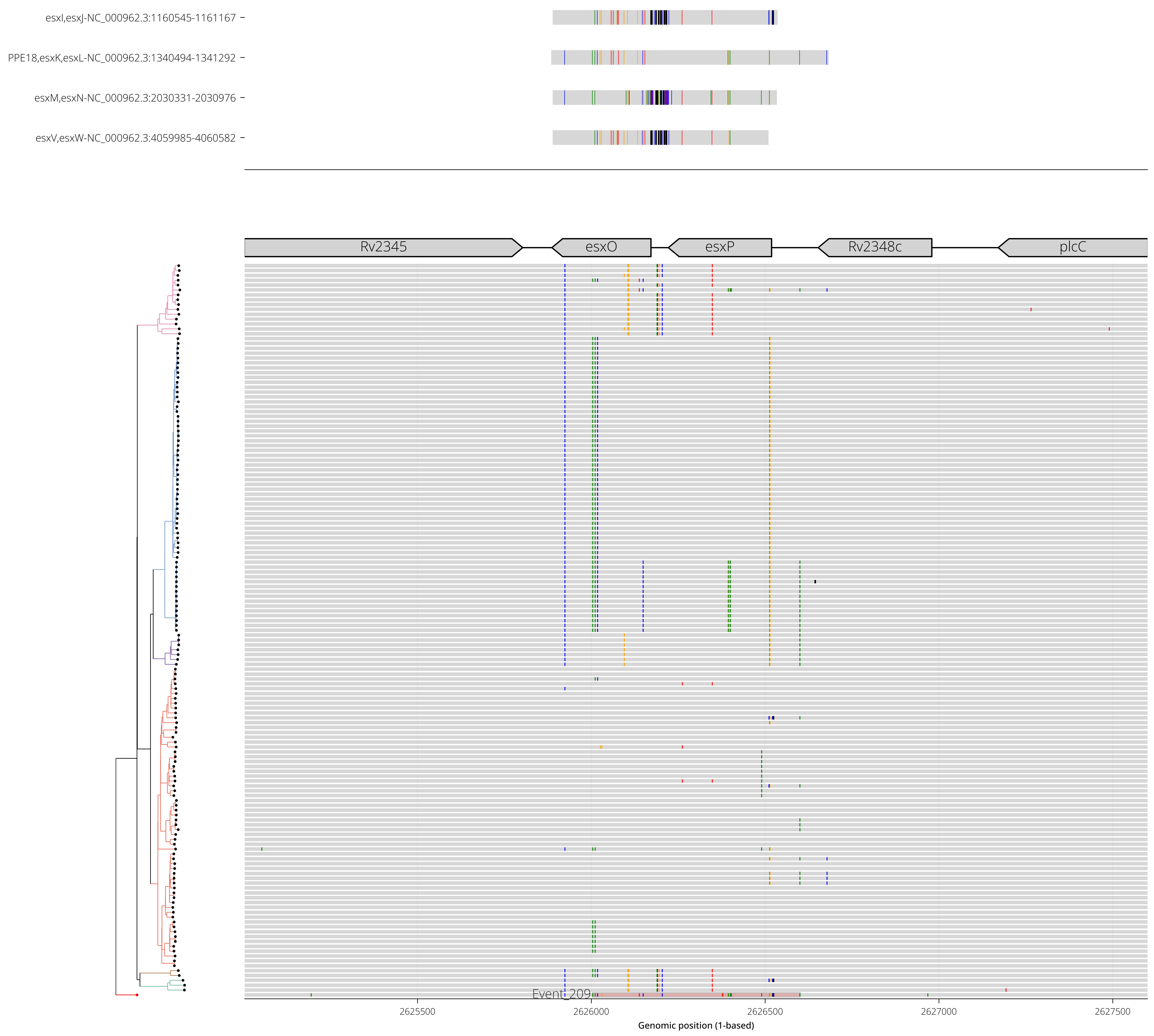

Full Phylogeny Viz + Paralog Alignments
GCE ID: Event_209

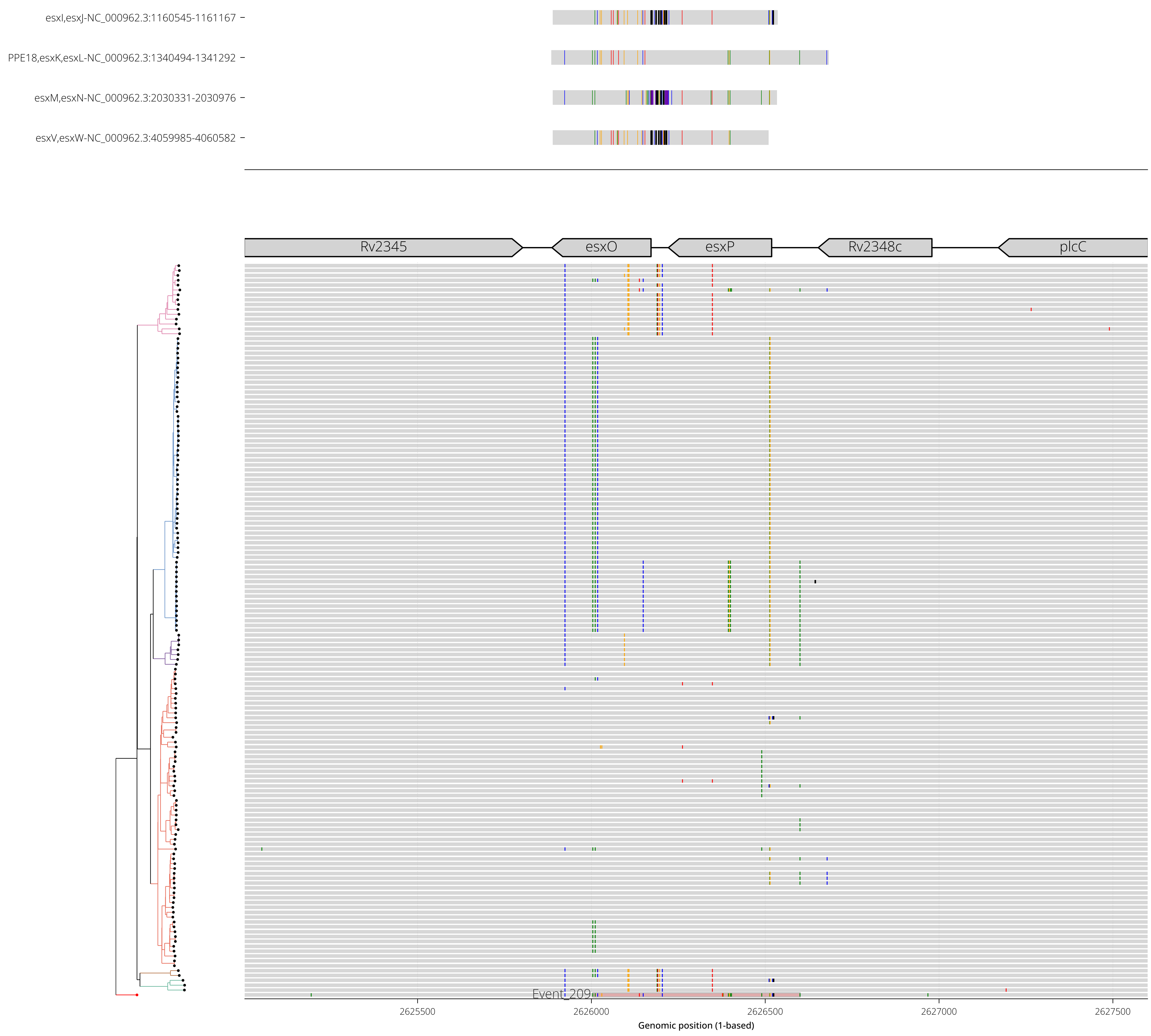

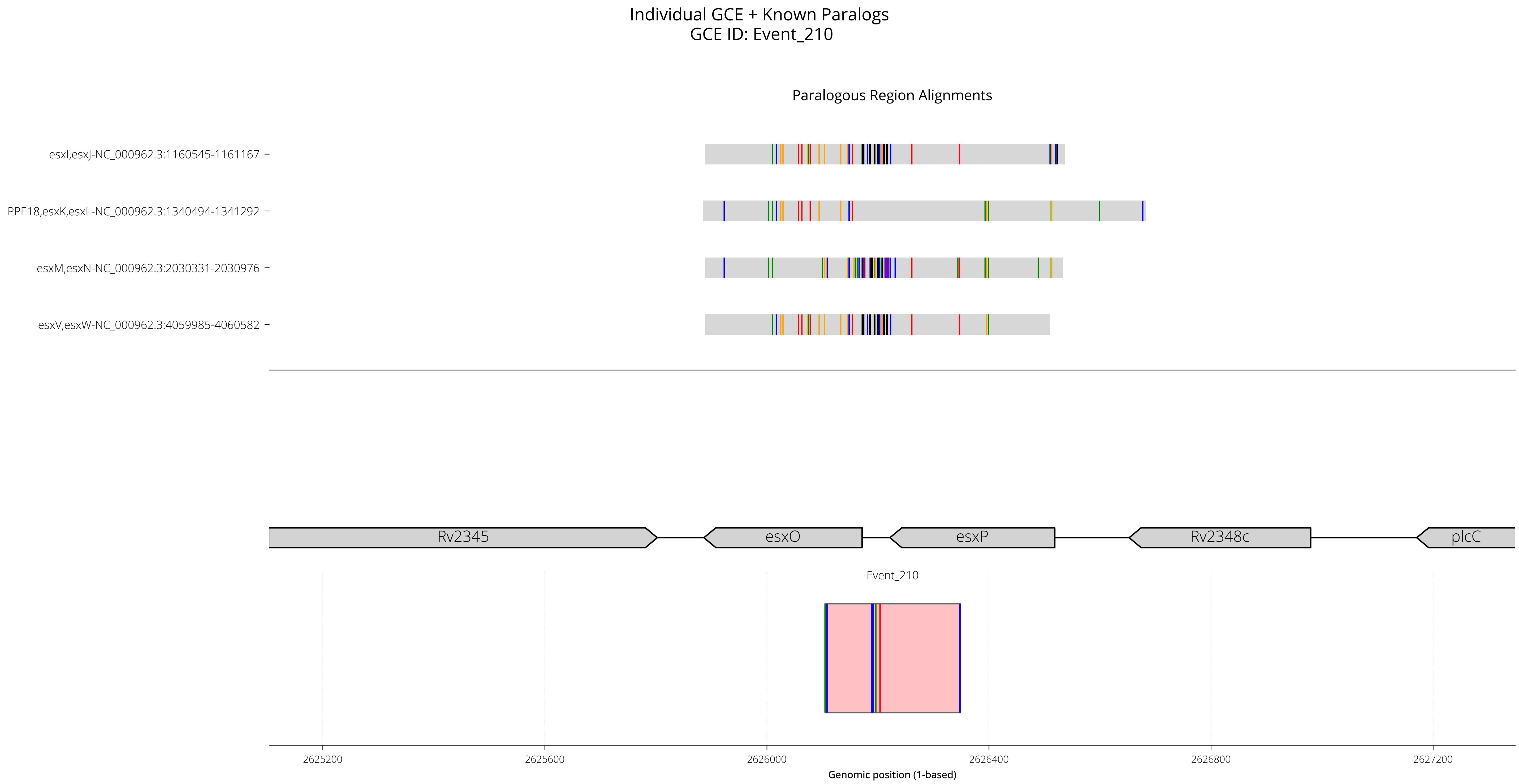
Individual GCE + Known Paralogs
GCE ID: Event_210
Paralogous Region Alignments
esxI,esxJ-NC_000962.3:1160545-1161167
PPE18,esxK,esxL-NC_000962.3:1340494-1341292
esxM,esxN-NC_000962.3:2030331-2030976
esxV,esxW-NC_000962.3:4059985-4060582
Rv2345
esxO
esxP
Rv2348c
plcC
Event_210
2625200
2625600
2626000
2626400
2626800
2627200
Genomic position (1-based)

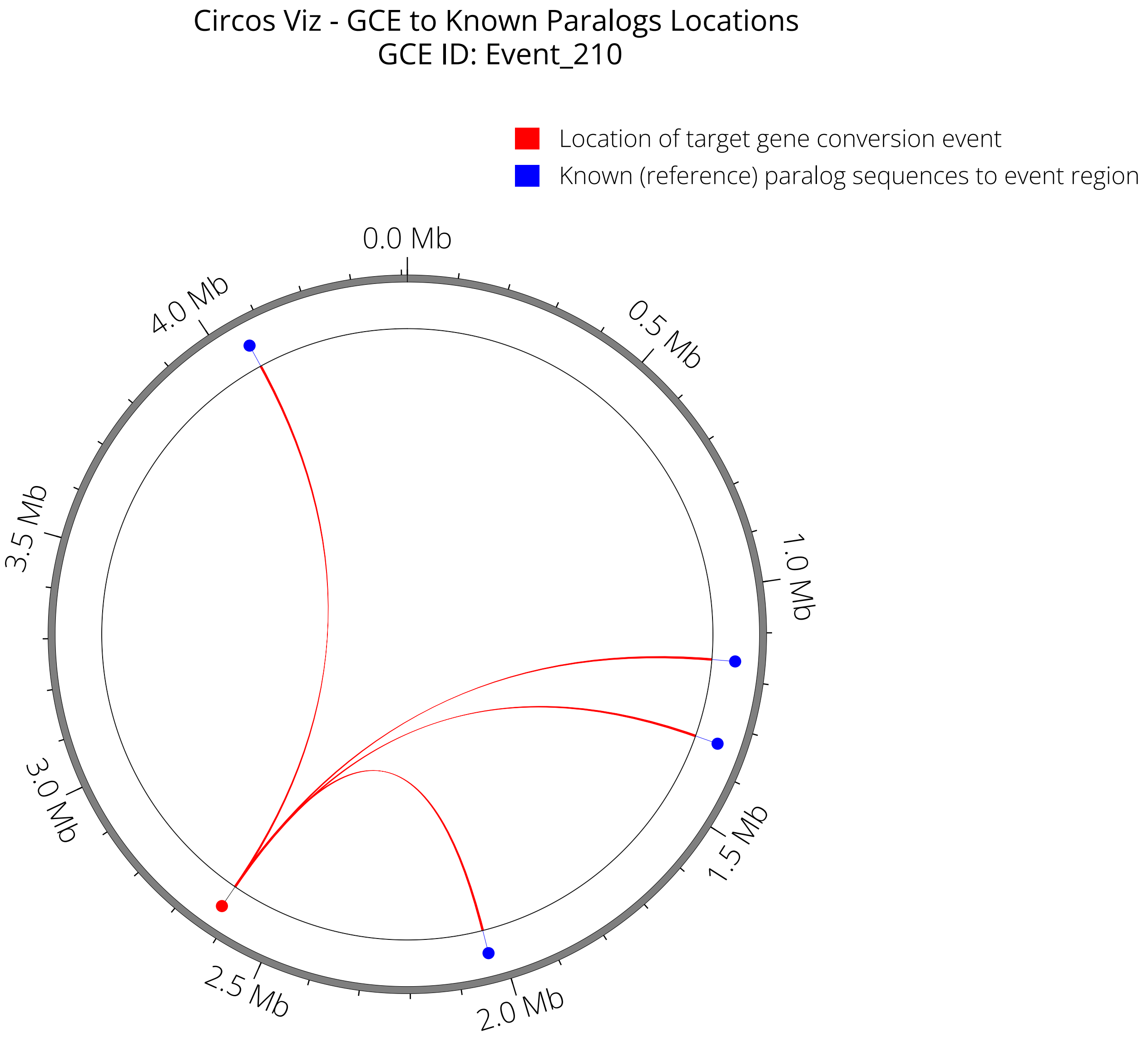

Circos Viz - GCE to Known Paralogs Locations
GCE ID: Event_210
Location of target gene conversion event
Known (reference) paralog sequences to event region
0.0 Mb
0.5 Mb
1.0 Mb
1.5 Mb
2.0 Mb
2.5 Mb
3.0 Mb
3.5 Mb
4.0 Mb

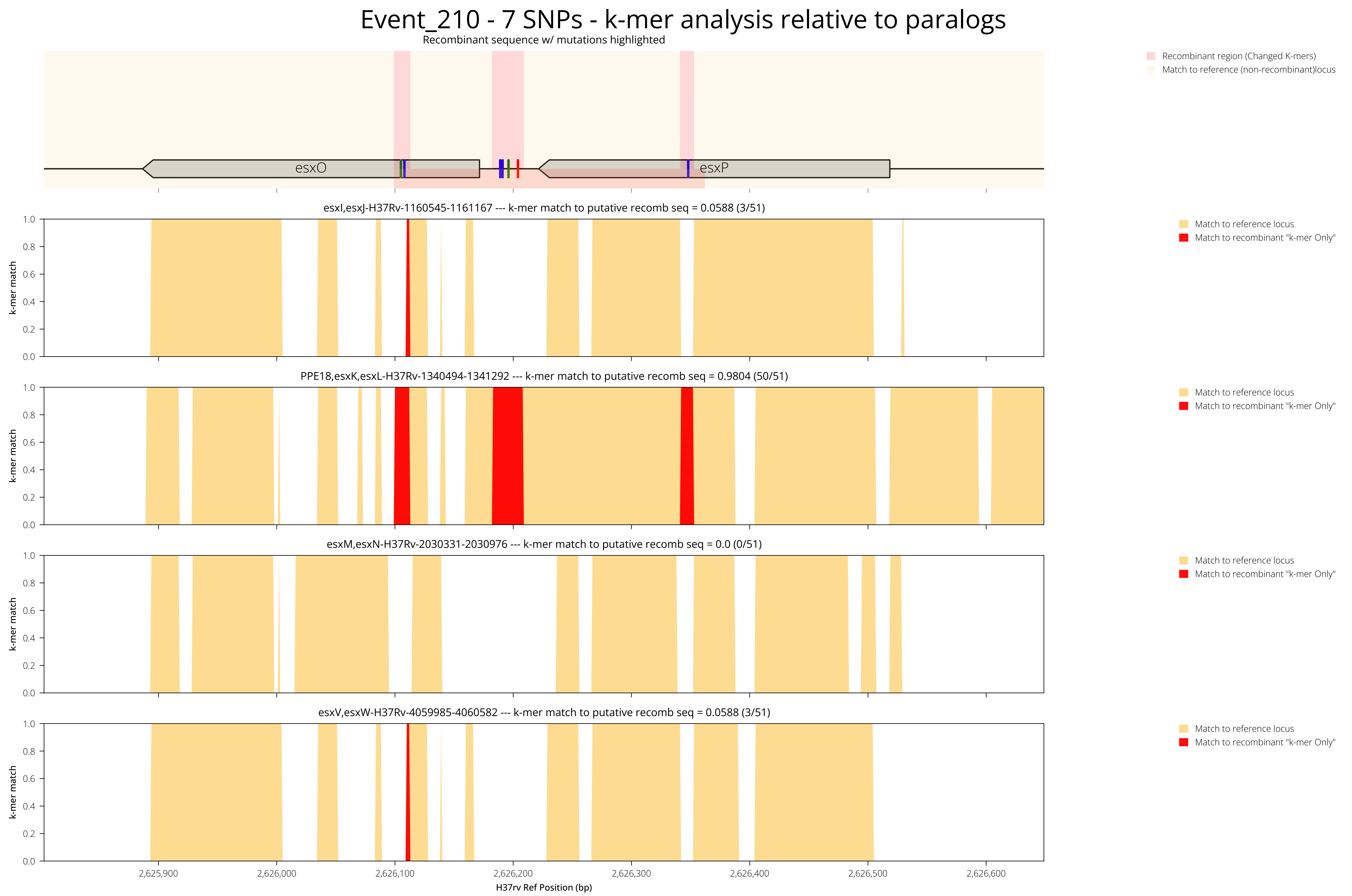

Event_210 - 7 SNPs - k-mer analysis relative to paralogs
Recombinant sequence w/ mutations highlighted
Recombinant region (Changed K-mers)
Match to reference (non-recombinant)locus
esxO
esxP
esxI,esxJ-H37Rv-1160545-1161167 --- k-mer match to putative recomb seq = 0.0588 (3/51)
PPE18,esxK,esxL-H37Rv-1340494-1341292 --- k-mer match to putative recomb seq = 0.9804 (50/51)
esxM,esxN-H37Rv-2030331-2030976 --- k-mer match to putative recomb seq = 0.0 (0/51)
esxV,esxW-H37Rv-4059985-4060582 --- k-mer match to putative recomb seq = 0.0588 (3/51)
Match to reference locus
Match to recombinant "k-mer Only"
k-mer match
1.0
0.8
0.6
0.4
0.2
0.0
2,625,900
2,626,000
2,626,100
2,626,200
2,626,300
2,626,400
2,626,500
2,626,600
H37rv Ref Position (bp)

Sub-phylogeny Viz + Paralog Alignments
GCE ID: Event_210

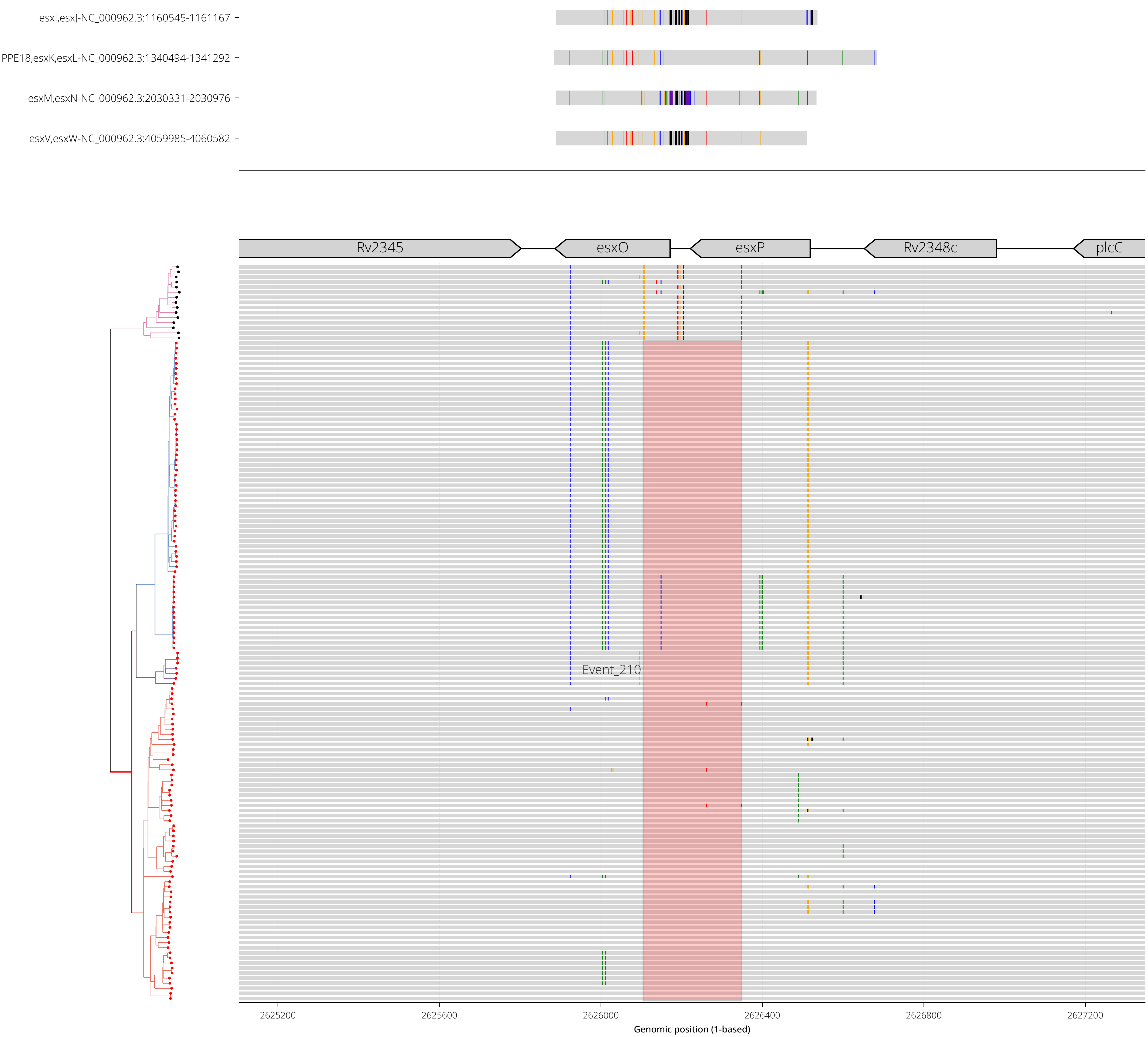

Full Phylogeny Viz + Paralog Alignments
GCE ID: Event_210

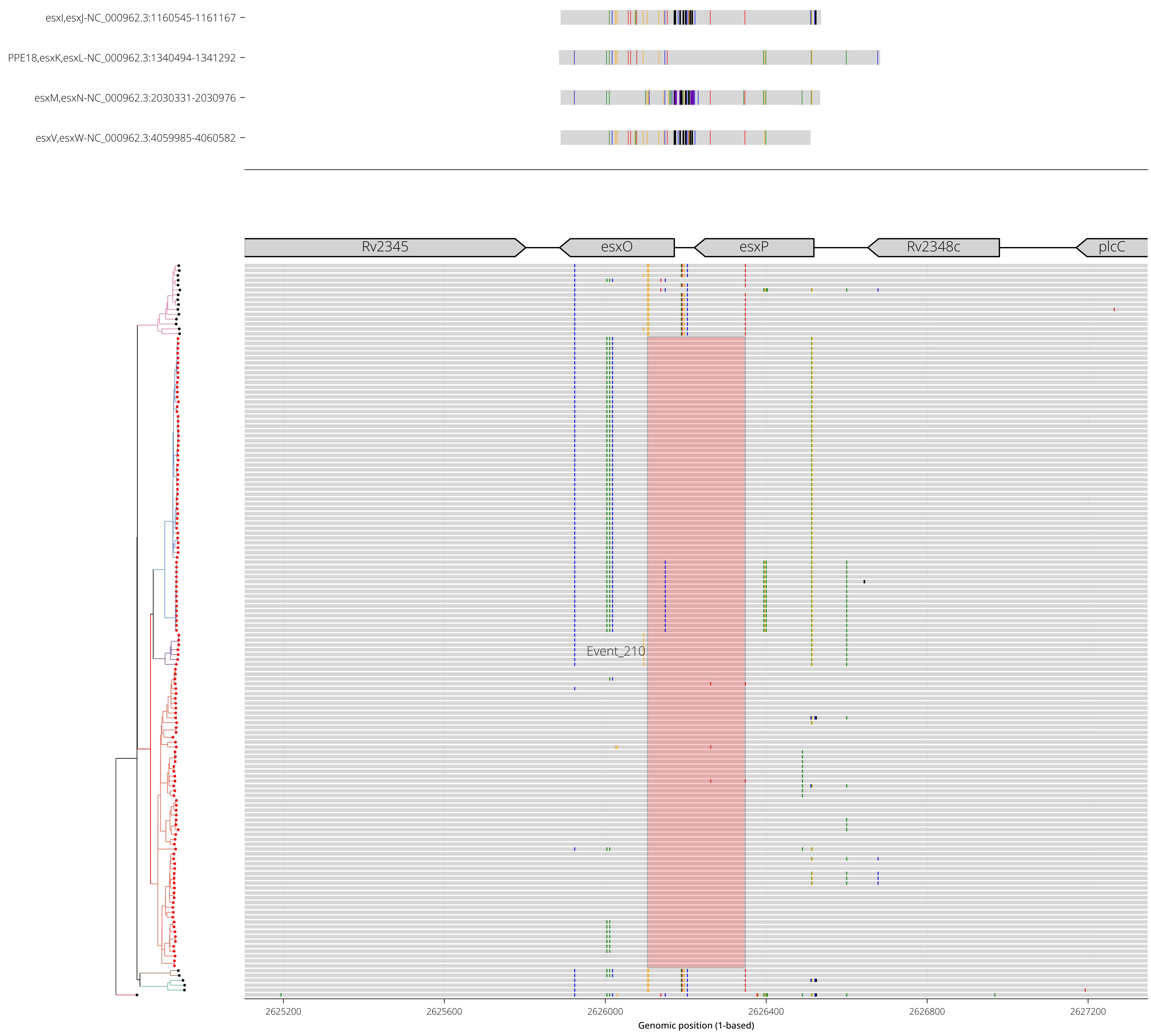

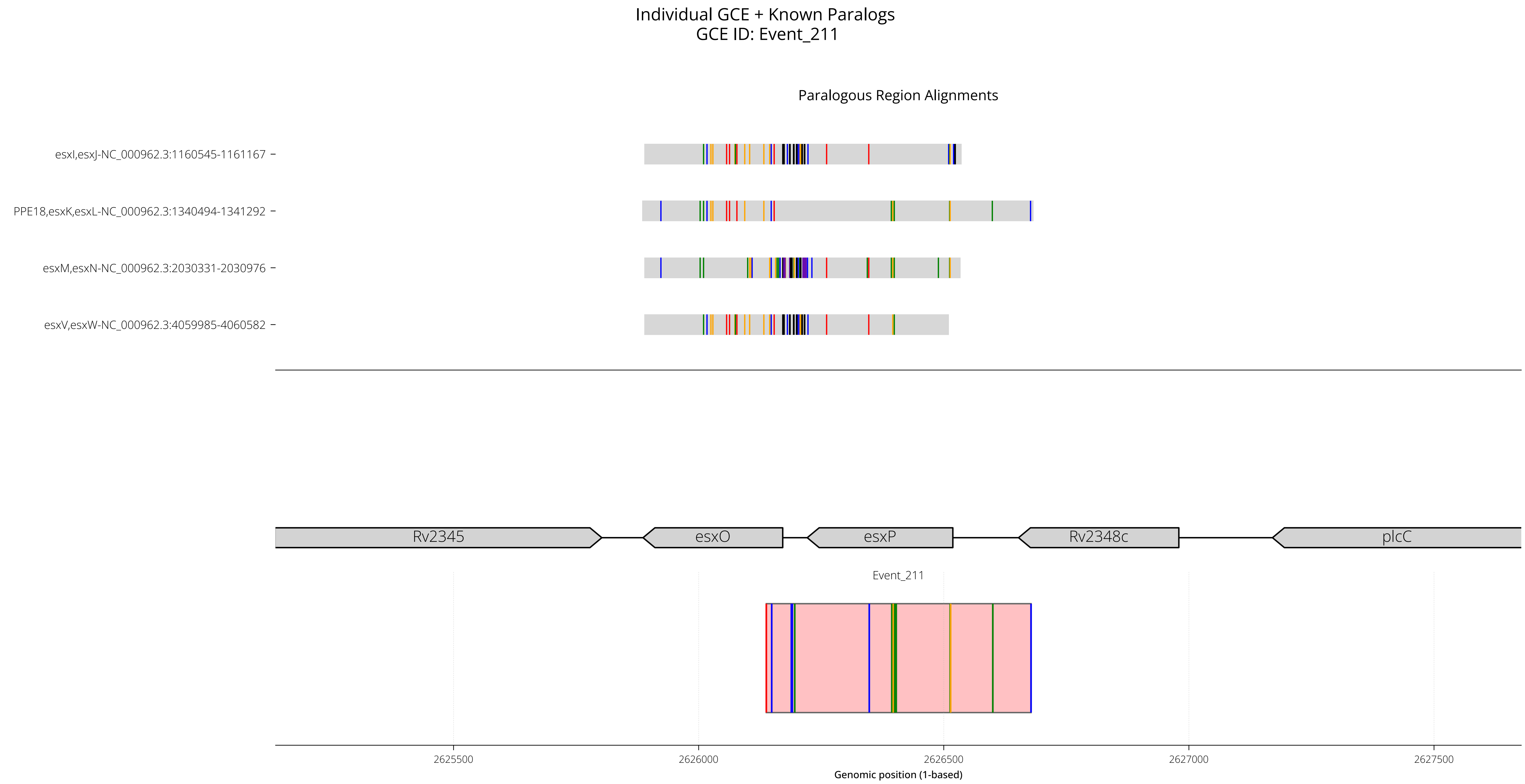
Individual GCE + Known Paralogs
GCE ID: Event_211
Paralogous Region Alignments
esxI,esxJ-NC_000962.3:1160545-1161167
PPE18,esxK,esxL-NC_000962.3:1340494-1341292
esxM,esxN-NC_000962.3:2030331-2030976
esxV,esxW-NC_000962.3:4059985-4060582
Rv2345
esxO
esxP
Rv2348c
plcC
Event_211
2625500
2626000
2626500
2627000
2627500
Genomic position (1-based)

Circos Viz - GCE to Known Paralogs Locations
GCE ID: Event_211

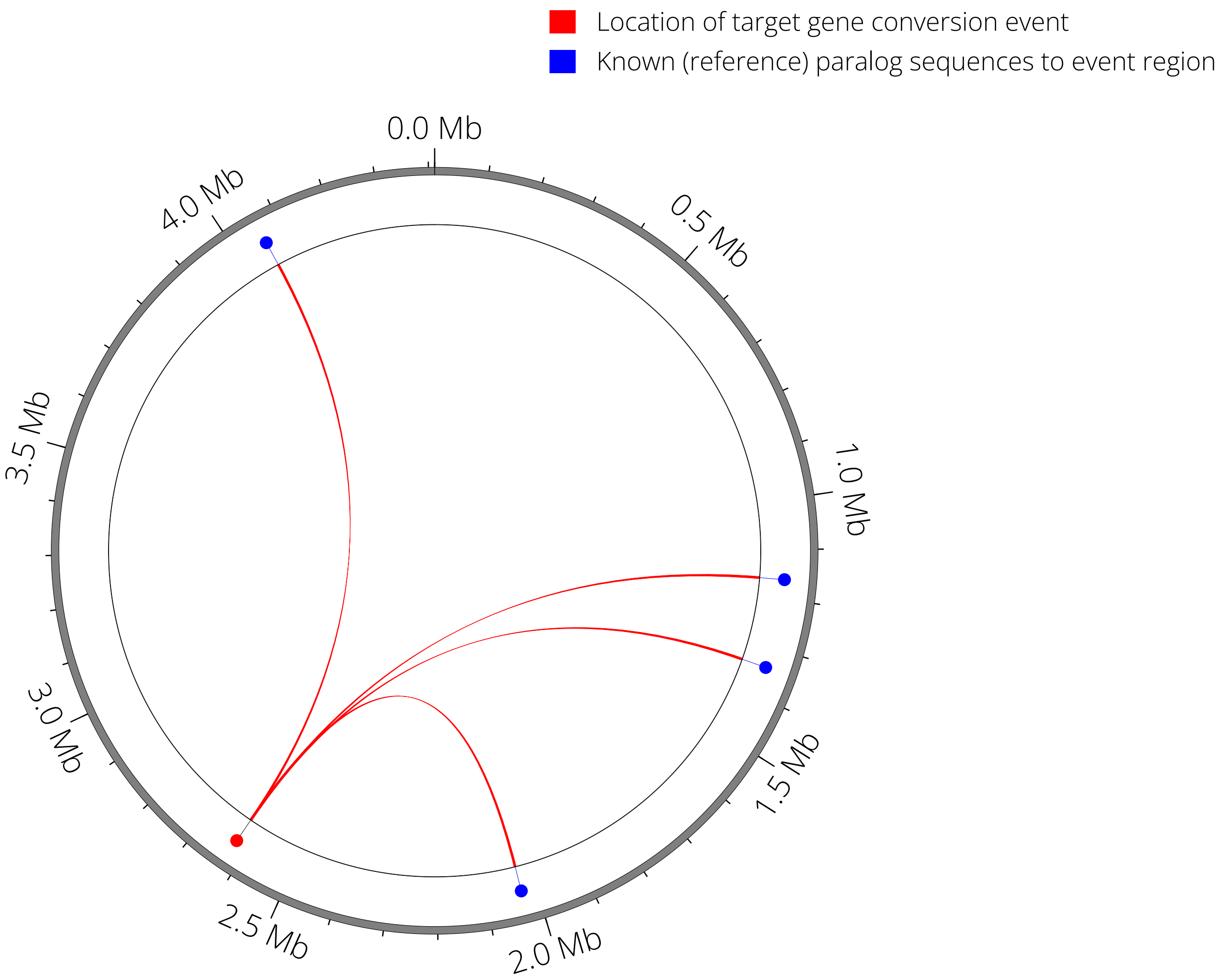

### Event_211 - 14 SNPs - k-mer analysis relative to paralogs

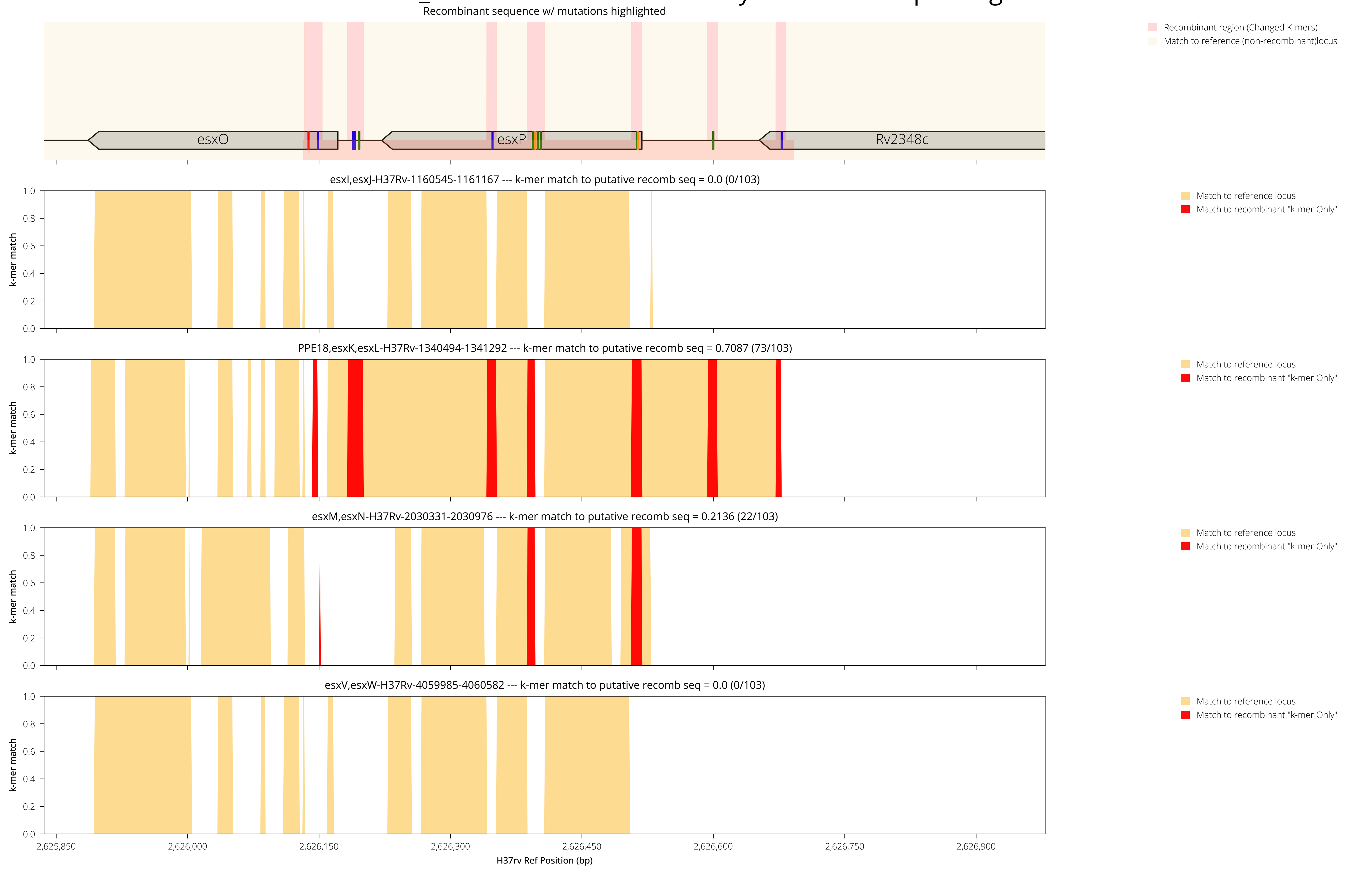

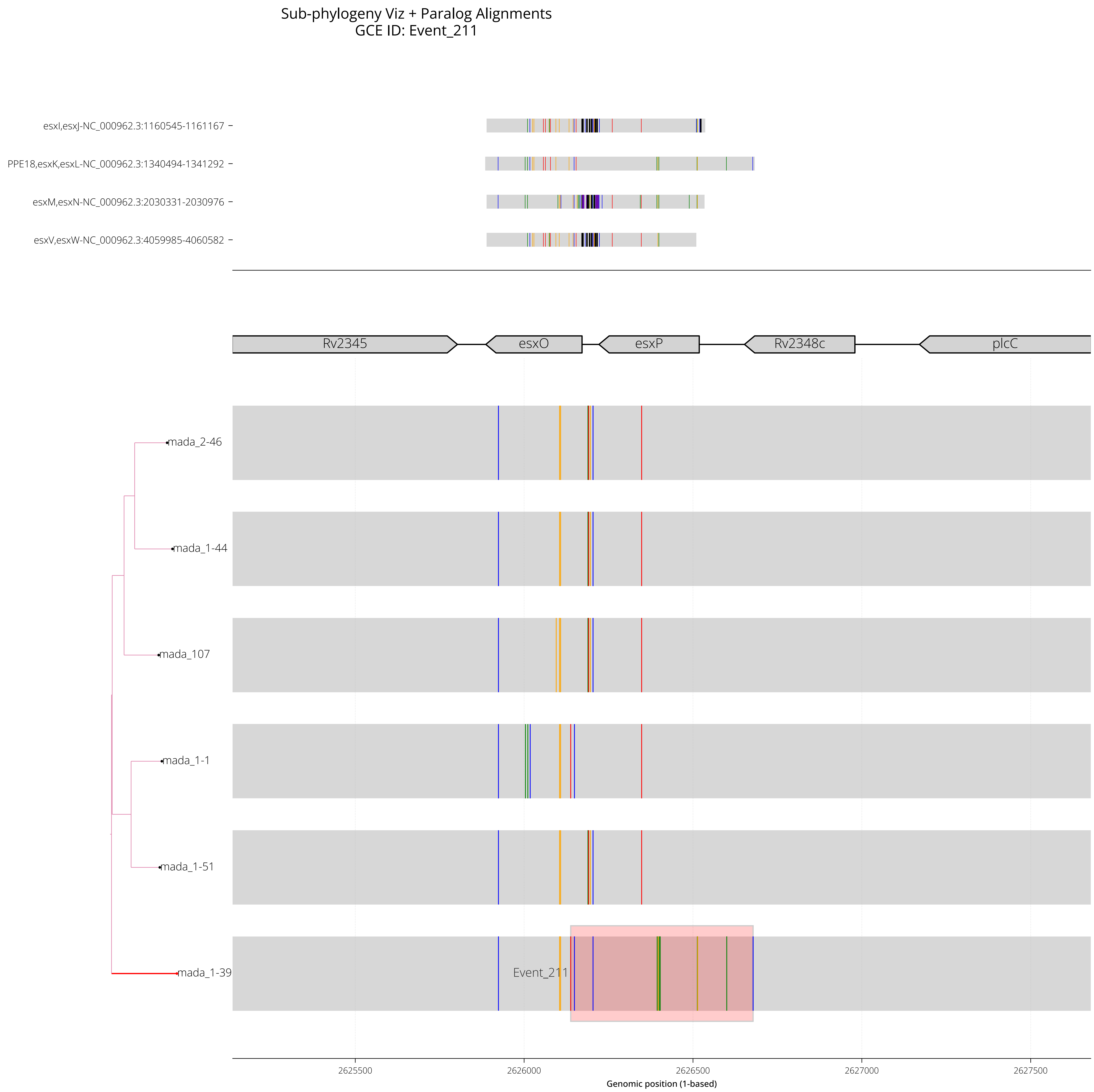

Sub-phylogeny Viz + Paralog Alignments
GCE ID: Event_211
esxI,esxJ-NC_000962.3:1160545-1161167
PPE18,esxK,esxL-NC_000962.3:1340494-1341292
esxM,esxN-NC_000962.3:2030331-2030976
esxV,esxW-NC_000962.3:4059985-4060582
Rv2345
esxO
esxP
Rv2348c
plcC
mada_2-46
mada_1-44
mada_107
mada_1-1
mada_1-51
mada_1-39
Event_211
2625500
2626000
2626500
2627000
2627500
Genomic position (1-based)

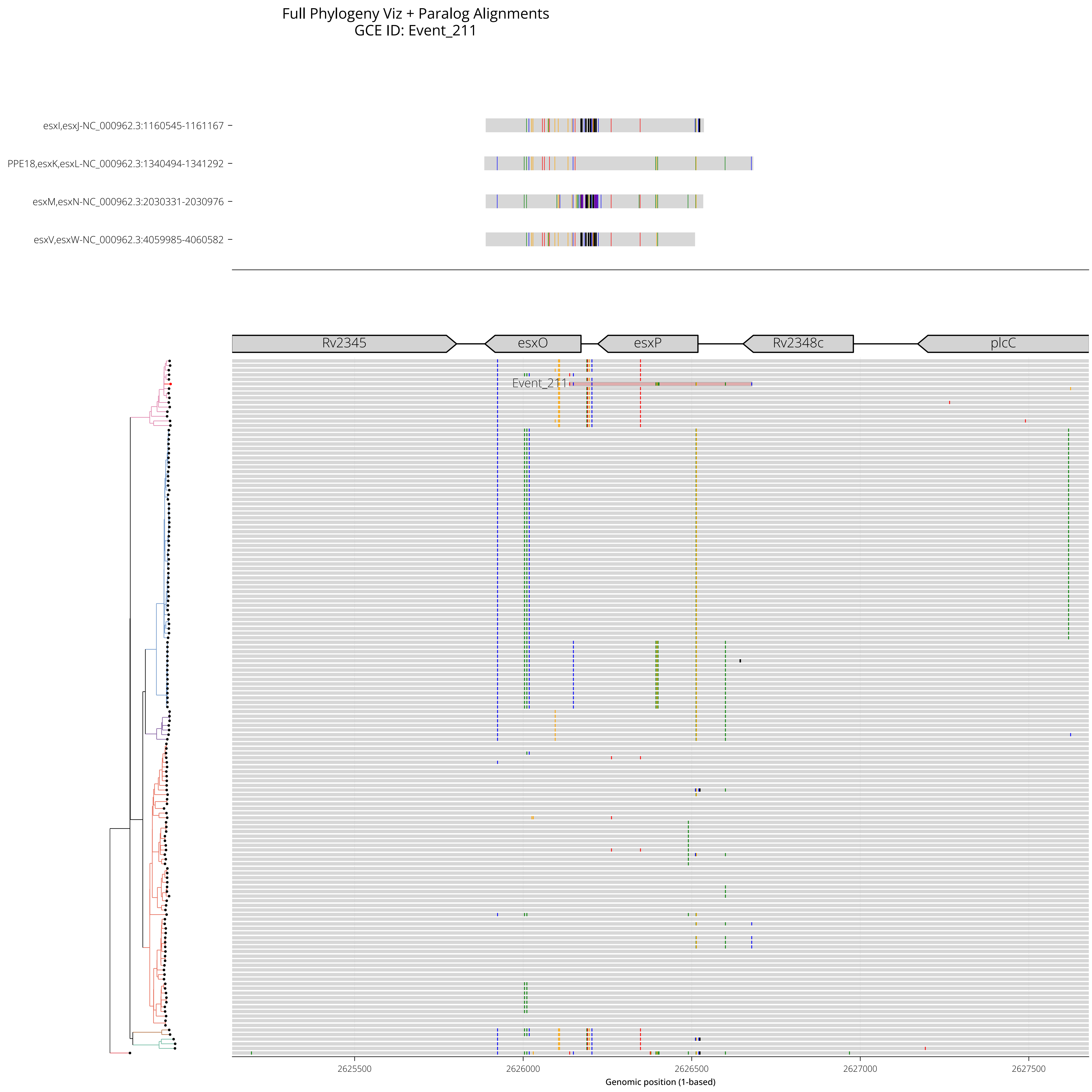
Full Phylogeny Viz + Paralog Alignments
GCE ID: Event_211
esxI,esxJ-NC_000962.3:1160545-1161167
PPE18,esxK,esxL-NC_000962.3:1340494-1341292
esxM,esxN-NC_000962.3:2030331-2030976
esxV,esxW-NC_000962.3:4059985-4060582
Rv2345
esxO
esxP
Rv2348c
plcC
Event_211
2625500
2626000
2626500
2627000
2627500
Genomic position (1-based)

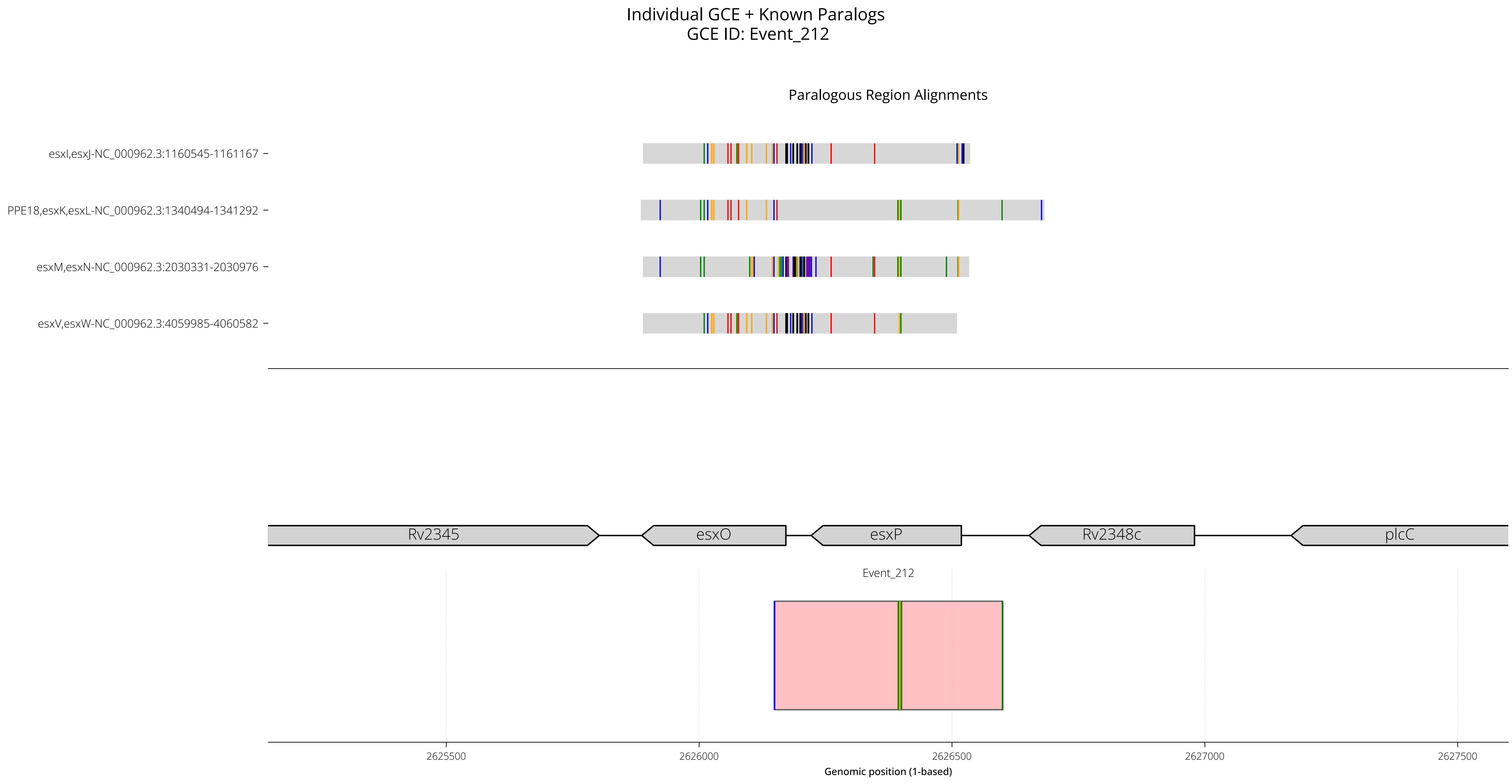

Individual GCE + Known Paralogs
GCE ID: Event_212
Paralogous Region Alignments
esxI,esxJ-NC_000962.3:1160545-1161167
PPE18,esxK,esxL-NC_000962.3:1340494-1341292
esxM,esxN-NC_000962.3:2030331-2030976
esxV,esxW-NC_000962.3:4059985-4060582
Rv2345
esxO
esxP
Rv2348c
plcC
Event_212
2625500
2626000
2626500
2627000
2627500
Genomic position (1-based)

Circos Viz - GCE to Known Paralogs Locations
GCE ID: Event_212

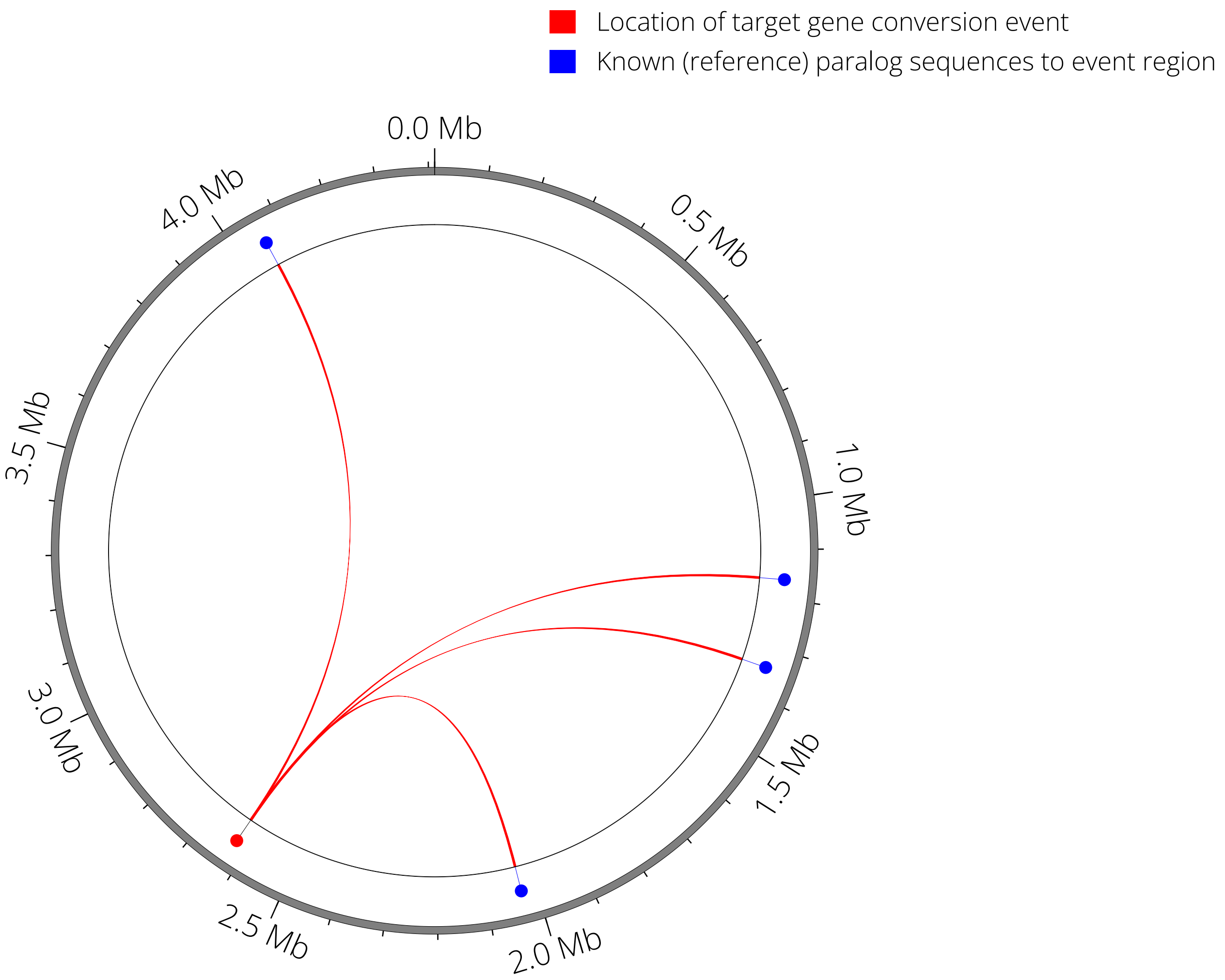

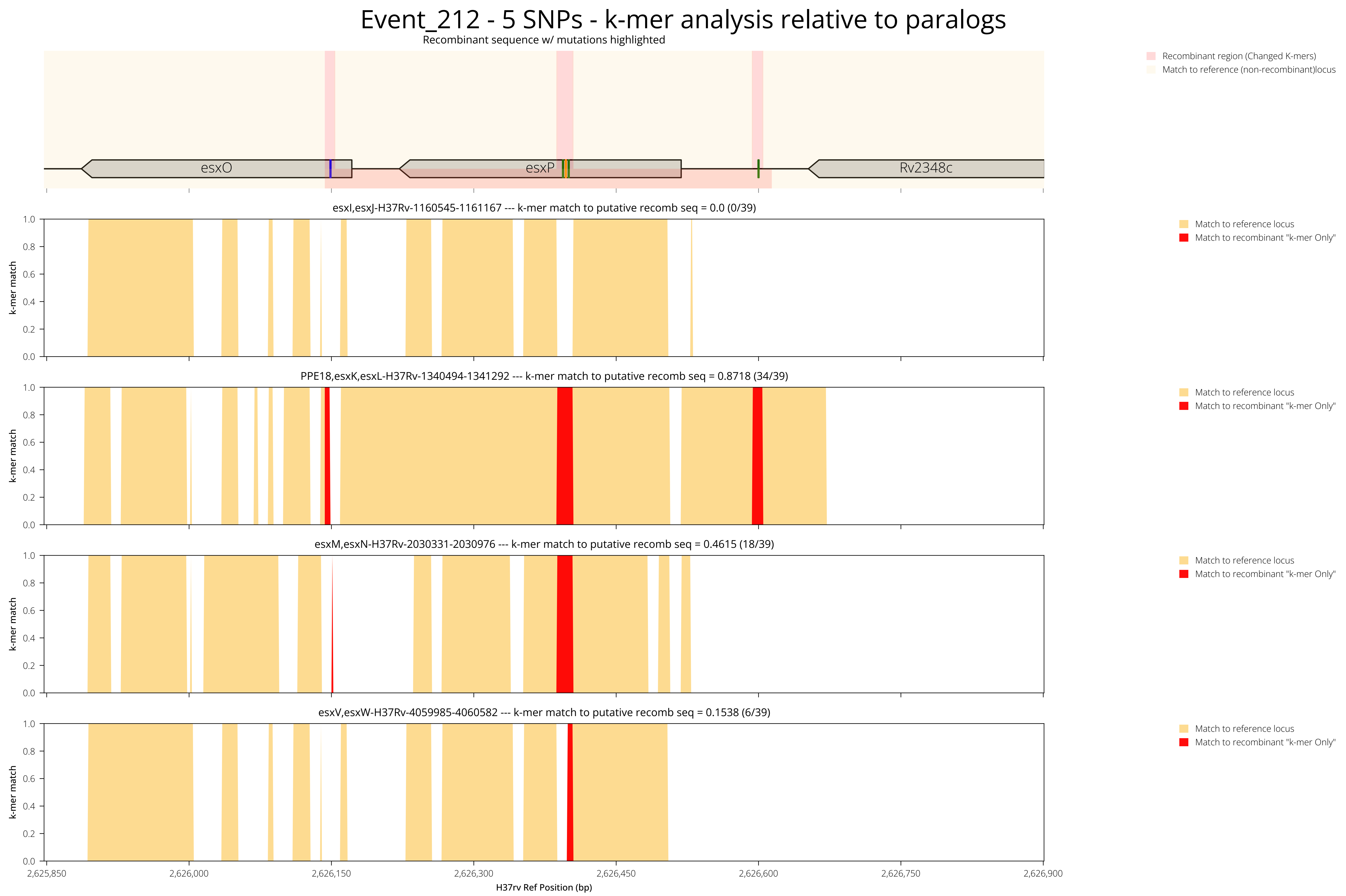
Event_212 - 5 SNPs - k-mer analysis relative to paralogs
Recombinant sequence w/ mutations highlighted
Recombinant region (Changed K-mers)
Match to reference (non-recombinant)locus
esxO
esxP
Rv2348c
esxI,esxJ-H37Rv-1160545-1161167 --- k-mer match to putative recomb seq = 0.0 (0/39)
PPE18,esxK,esxL-H37Rv-1340494-1341292 --- k-mer match to putative recomb seq = 0.8718 (34/39)
esxM,esxN-H37Rv-2030331-2030976 --- k-mer match to putative recomb seq = 0.4615 (18/39)
esxV,esxW-H37Rv-4059985-4060582 --- k-mer match to putative recomb seq = 0.1538 (6/39)
Match to reference locus
Match to recombinant "k-mer Only"
k-mer match
1.0
0.8
0.6
0.4
0.2
0.0
2,625,850
2,626,000
2,626,150
2,626,300
2,626,450
2,626,600
2,626,750
2,626,900
H37rv Ref Position (bp)

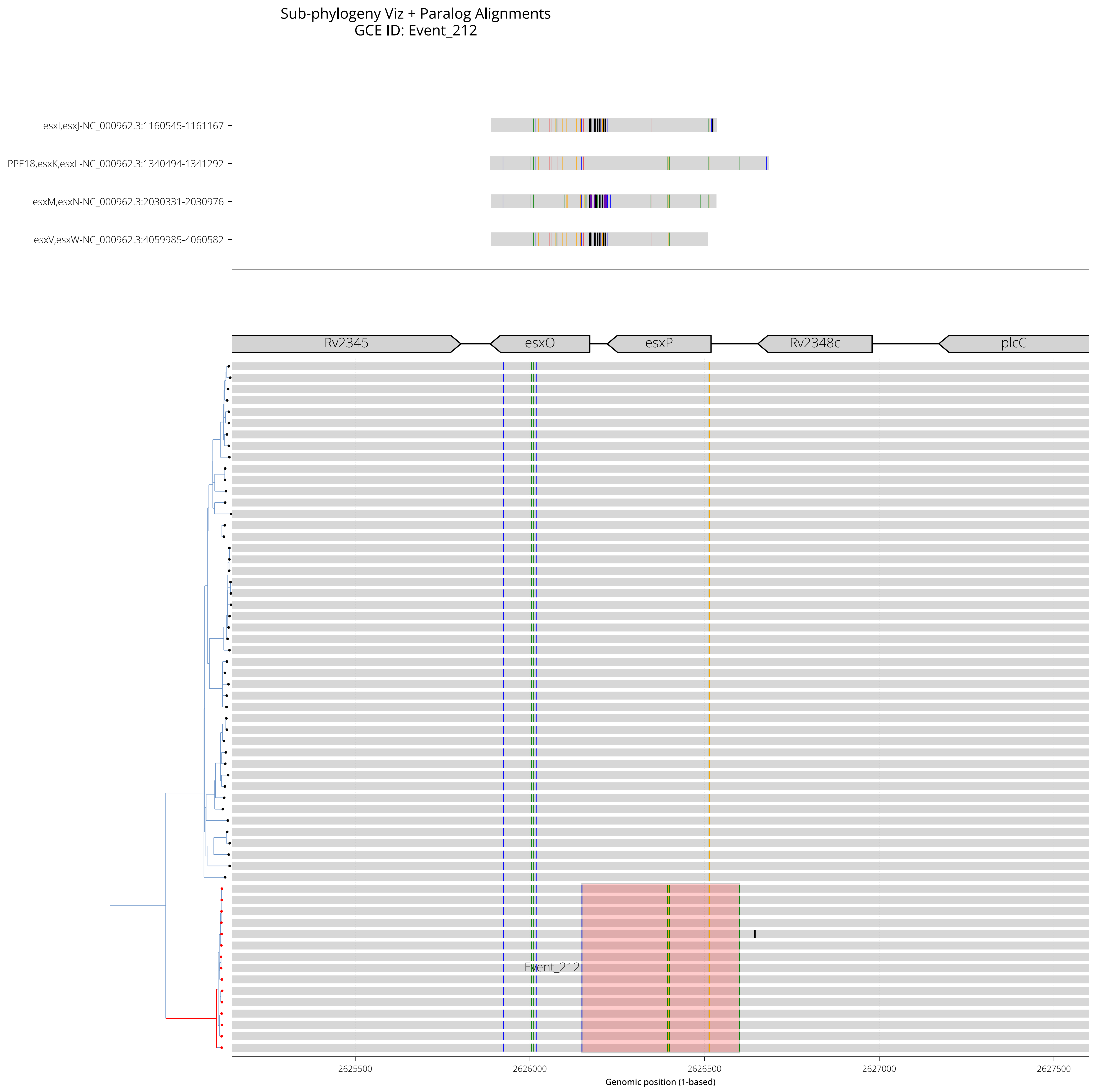

Sub-phylogeny Viz + Paralog Alignments
GCE ID: Event_212
esxI,esxJ-NC_000962.3:1160545-1161167
PPE18,esxK,esxL-NC_000962.3:1340494-1341292
esxM,esxN-NC_000962.3:2030331-2030976
esxV,esxW-NC_000962.3:4059985-4060582
Rv2345
esxO
esxP
Rv2348c
plcC
Event_212
2625500
2626000
2626500
2627000
2627500
Genomic position (1-based)

Full Phylogeny Viz + Paralog Alignments
GCE ID: Event_212

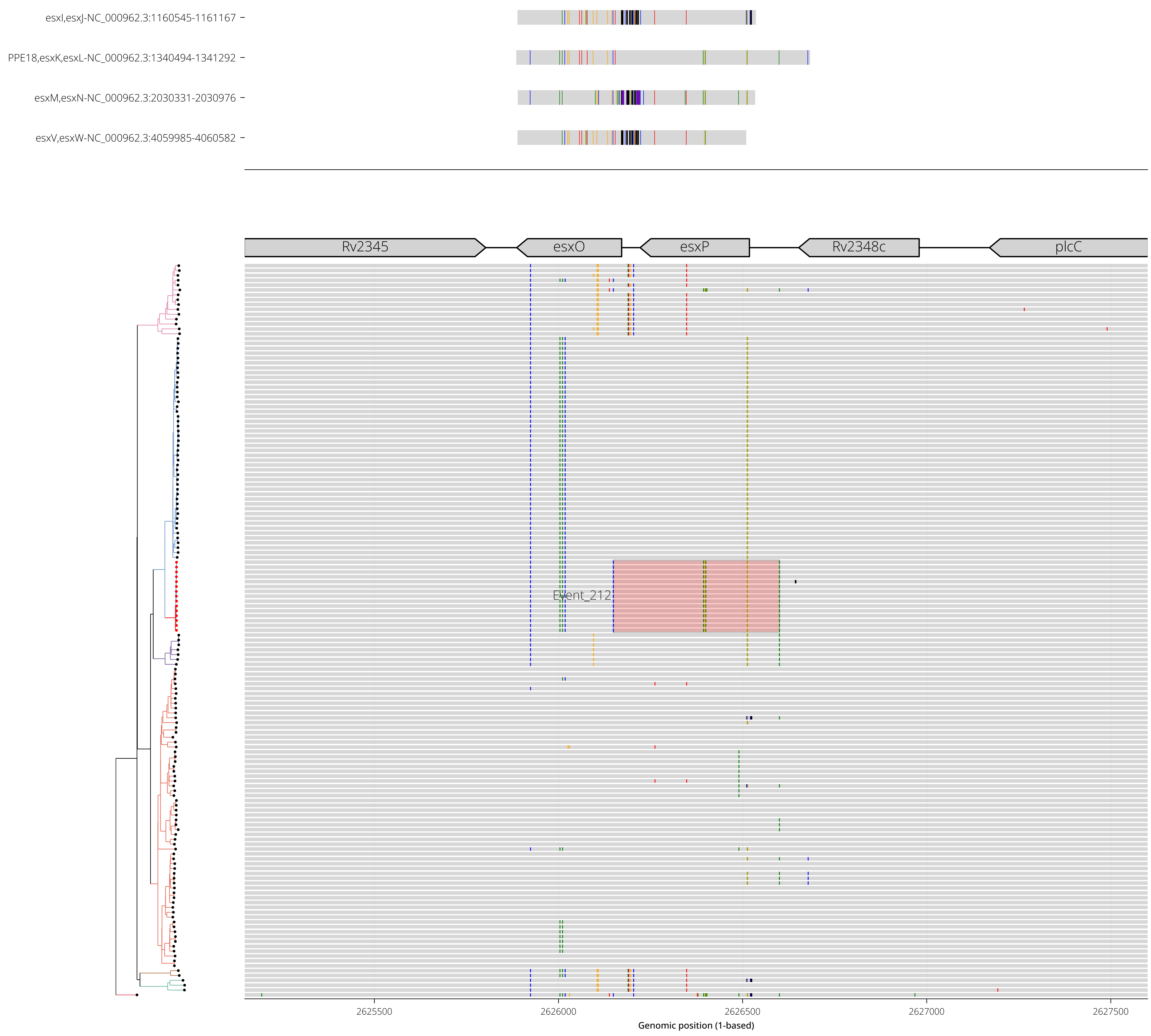

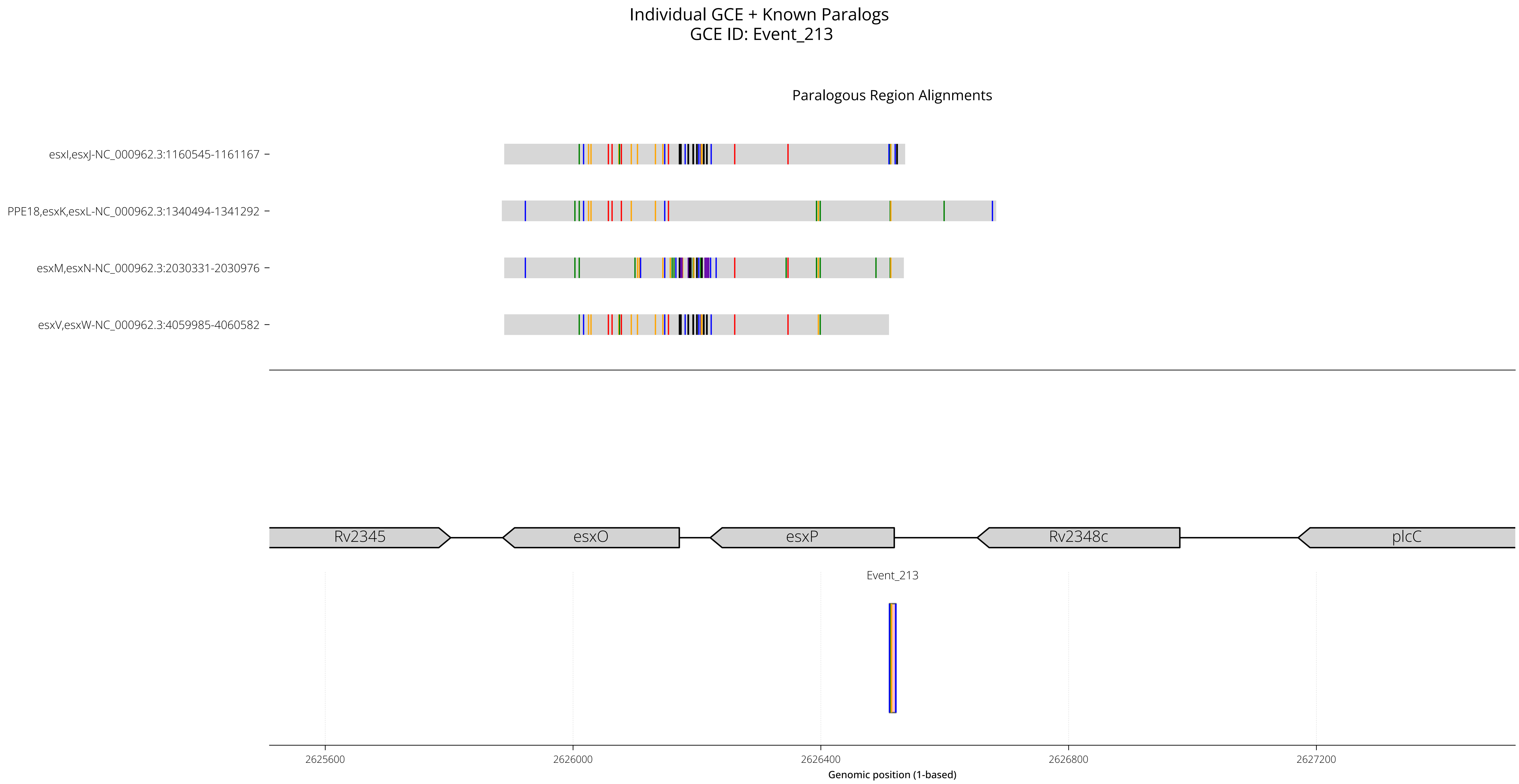

Individual GCE + Known Paralogs
GCE ID: Event_213
Paralogous Region Alignments
esxI,esxJ-NC_000962.3:1160545-1161167
PPE18,esxK,esxL-NC_000962.3:1340494-1341292
esxM,esxN-NC_000962.3:2030331-2030976
esxV,esxW-NC_000962.3:4059985-4060582
Rv2345
esxO
esxP
Rv2348c
plcC
Event_213
2625600
2626000
2626400
2626800
2627200
Genomic position (1-based)

Circos Viz - GCE to Known Paralogs Locations
GCE ID: Event_213

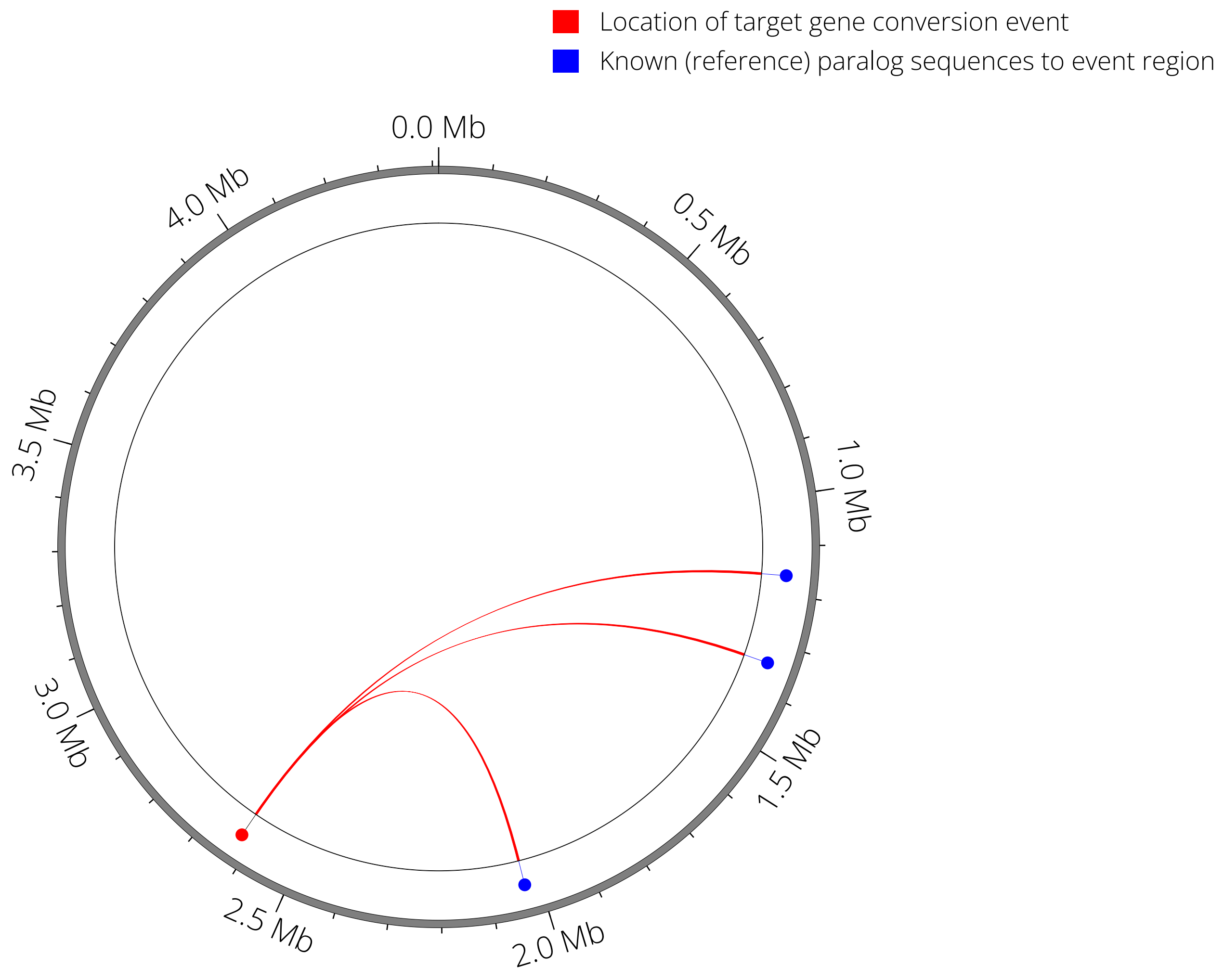

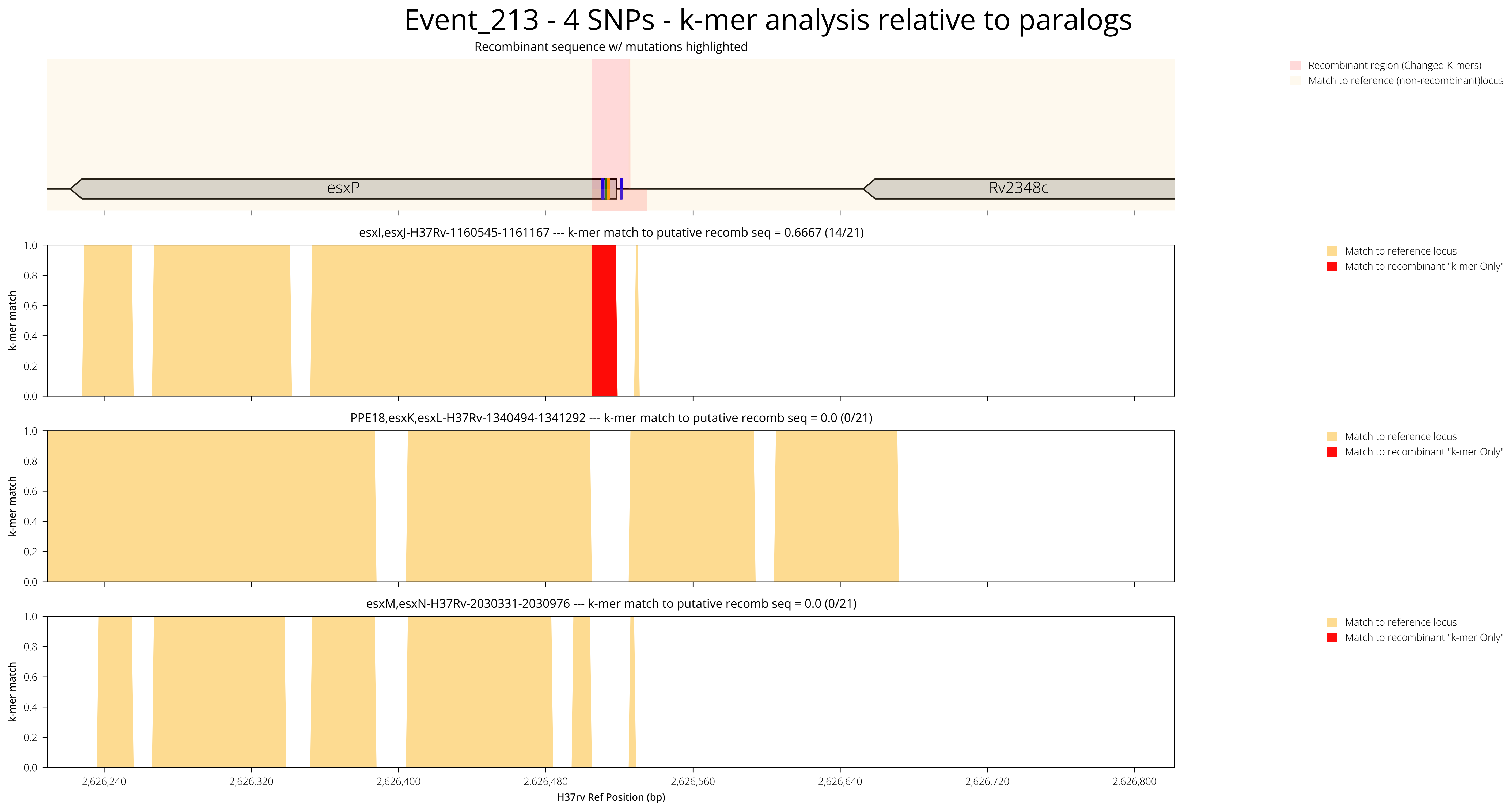

Event_213 - 4 SNPs - k-mer analysis relative to paralogs
Recombinant sequence w/ mutations highlighted
esxP
Rv2348c
Recombinant region (Changed K-mers)
Match to reference (non-recombinant)locus
esxI,esxJ-H37Rv-1160545-1161167 --- k-mer match to putative recomb seq = 0.6667 (14/21)
PPE18,esxK,esxL-H37Rv-1340494-1341292 --- k-mer match to putative recomb seq = 0.0 (0/21)
esxM,esxN-H37Rv-2030331-2030976 --- k-mer match to putative recomb seq = 0.0 (0/21)
Match to reference locus
Match to recombinant "k-mer Only"
k-mer match
1.0
0.8
0.6
0.4
0.2
0.0
2,626,240
2,626,320
2,626,400
2,626,480
2,626,560
2,626,640
2,626,720
2,626,800
H37rv Ref Position (bp)

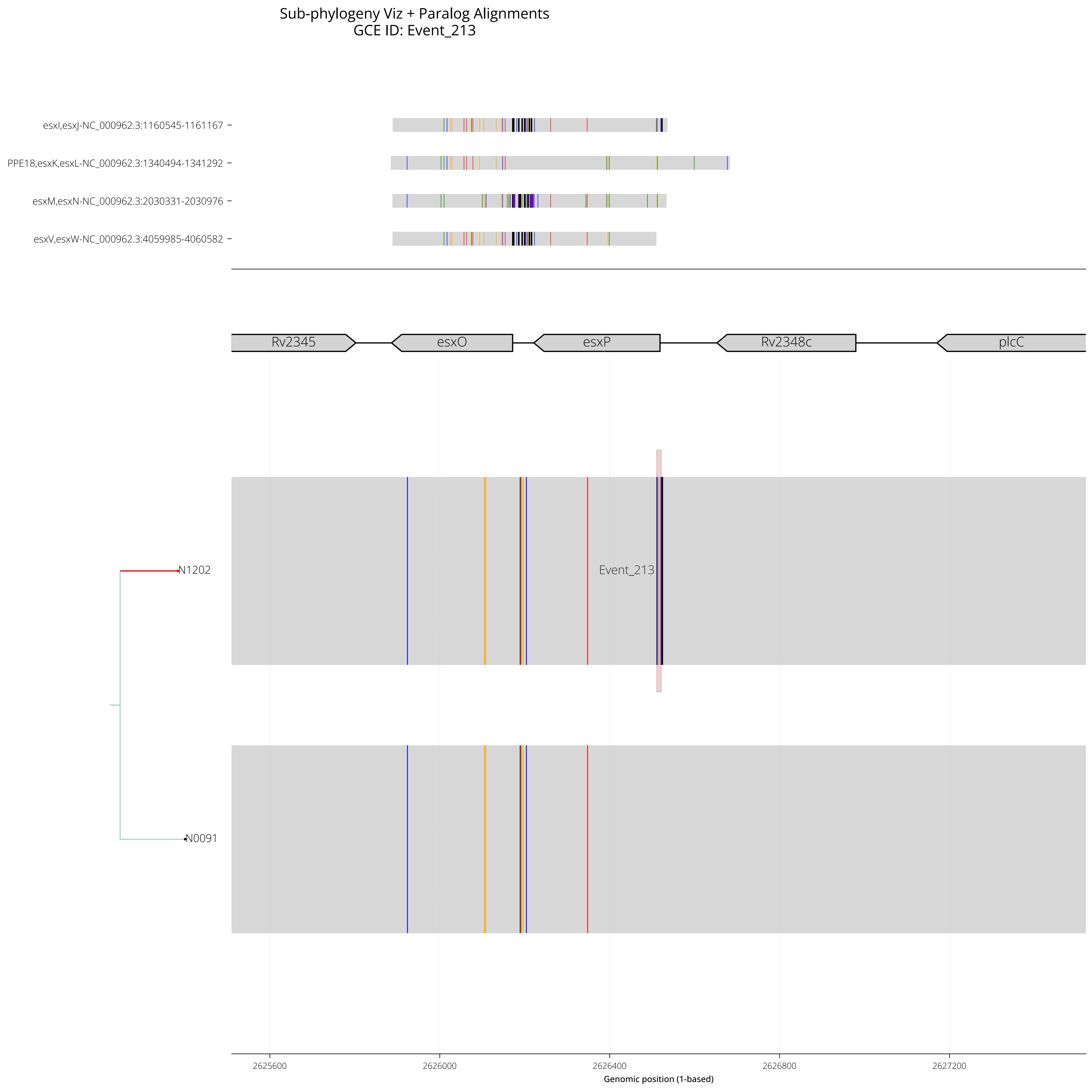

Sub-phylogeny Viz + Paralog Alignments
GCE ID: Event_213
esxI,esxJ-NC_000962.3:1160545-1161167
PPE18,esxK,esxL-NC_000962.3:1340494-1341292
esxM,esxN-NC_000962.3:2030331-2030976
esxV,esxW-NC_000962.3:4059985-4060582
Rv2345
esxO
esxP
Rv2348c
plcC
N1202
Event_213
N0091
2625600
2626000
2626400
2626800
2627200
Genomic position (1-based)

Full Phylogeny Viz + Paralog Alignments
GCE ID: Event_213

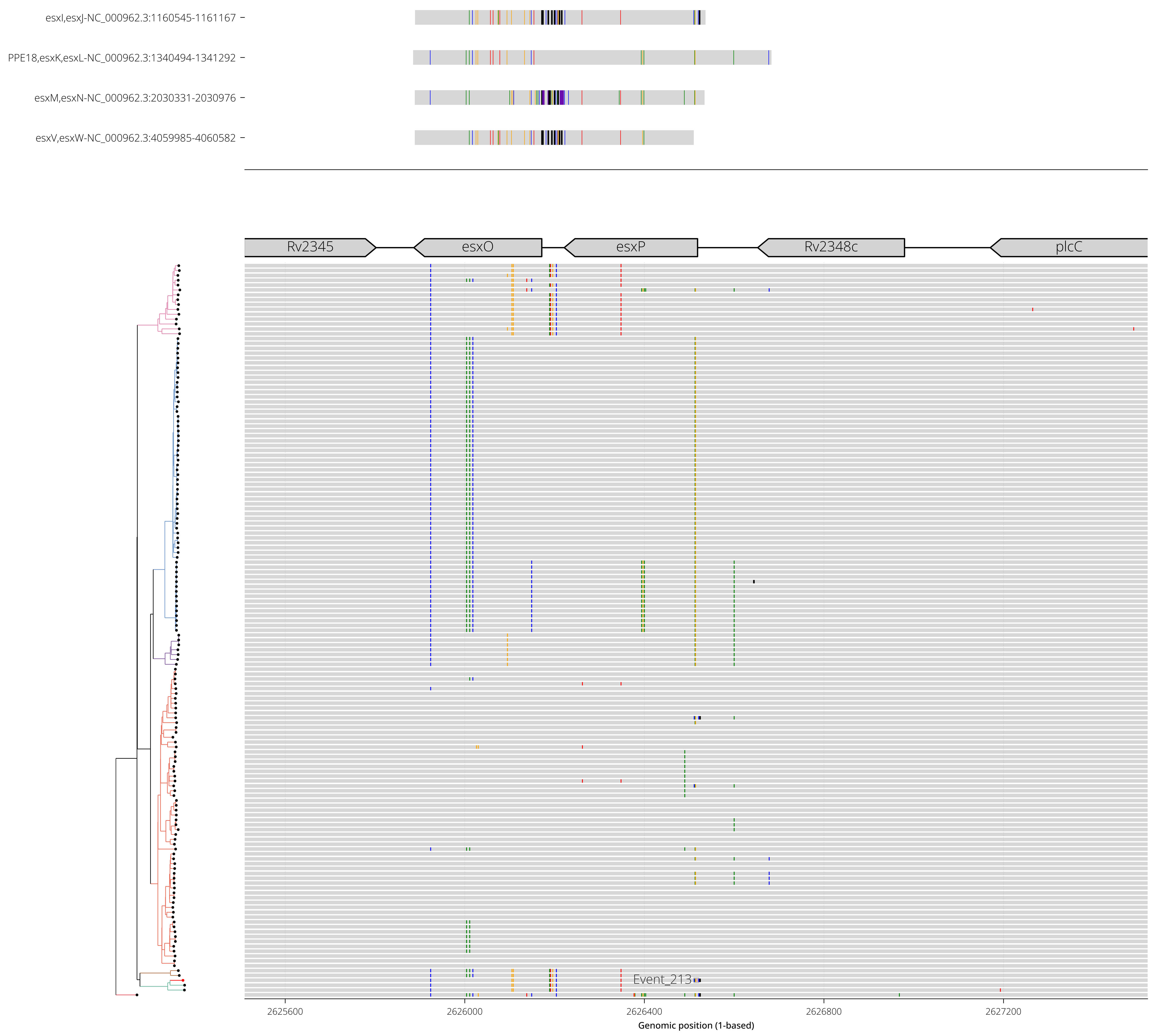

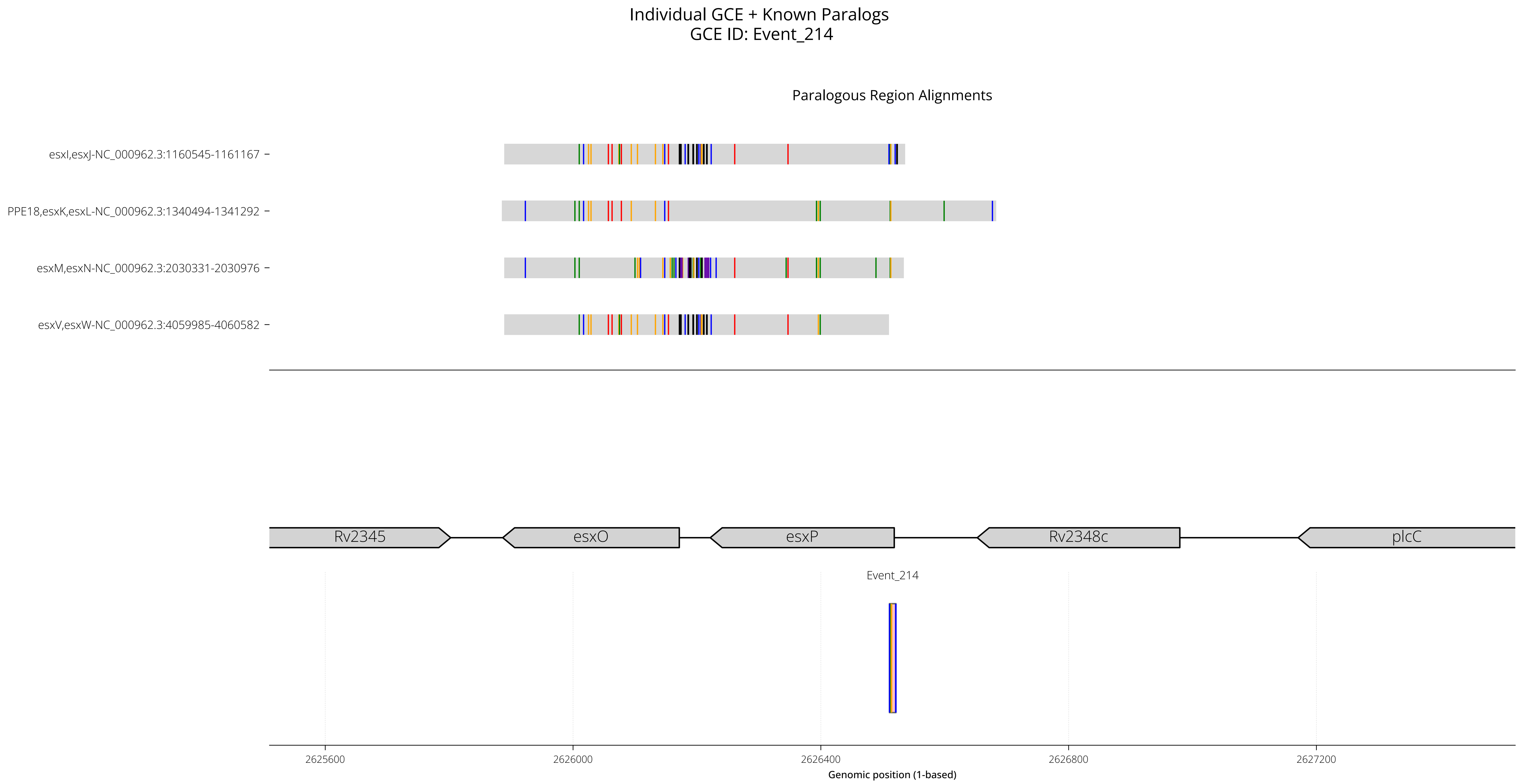

Individual GCE + Known Paralogs
GCE ID: Event_214
Paralogous Region Alignments
esxI,esxJ-NC_000962.3:1160545-1161167
PPE18,esxK,esxL-NC_000962.3:1340494-1341292
esxM,esxN-NC_000962.3:2030331-2030976
esxV,esxW-NC_000962.3:4059985-4060582
Rv2345
esxO
esxP
Rv2348c
plcC
Event_214
2625600
2626000
2626400
2626800
2627200
Genomic position (1-based)

### Circos Viz - GCE to Known Paralogs Locations
GCE ID: Event_214

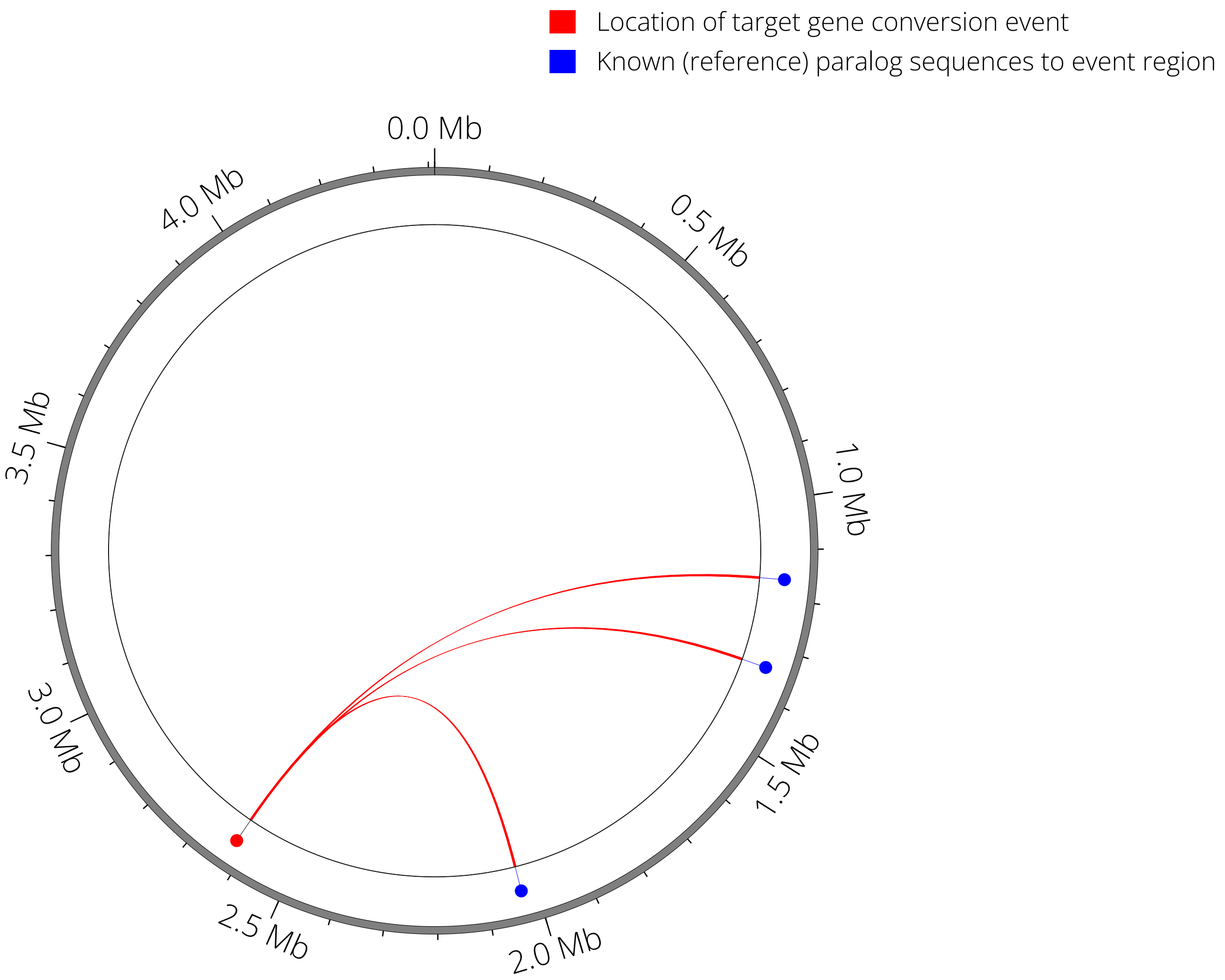

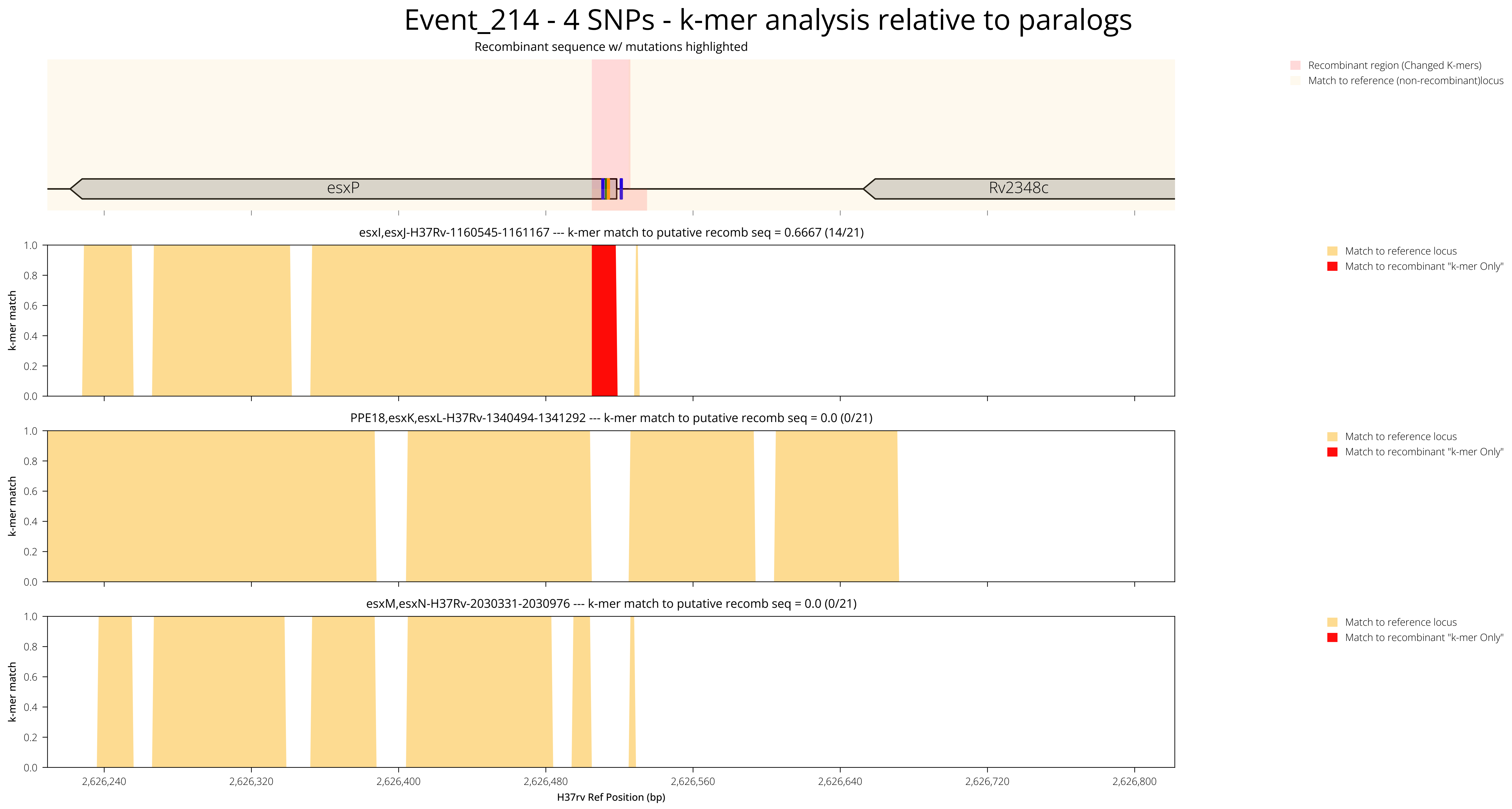
Event_214 - 4 SNPs - k-mer analysis relative to paralogs
Recombinant sequence w/ mutations highlighted
Recombinant region (Changed K-mers)
Match to reference (non-recombinant)locus
esxP
Rv2348c
esxI,esxJ-H37Rv-1160545-1161167 --- k-mer match to putative recomb seq = 0.6667 (14/21)
PPE18,esxK,esxL-H37Rv-1340494-1341292 --- k-mer match to putative recomb seq = 0.0 (0/21)
esxM,esxN-H37Rv-2030331-2030976 --- k-mer match to putative recomb seq = 0.0 (0/21)
Match to reference locus
Match to recombinant "k-mer Only"
k-mer match
1.0
0.8
0.6
0.4
0.2
0.0
2,626,240
2,626,320
2,626,400
2,626,480
2,626,560
2,626,640
2,626,720
2,626,800
H37rv Ref Position (bp)

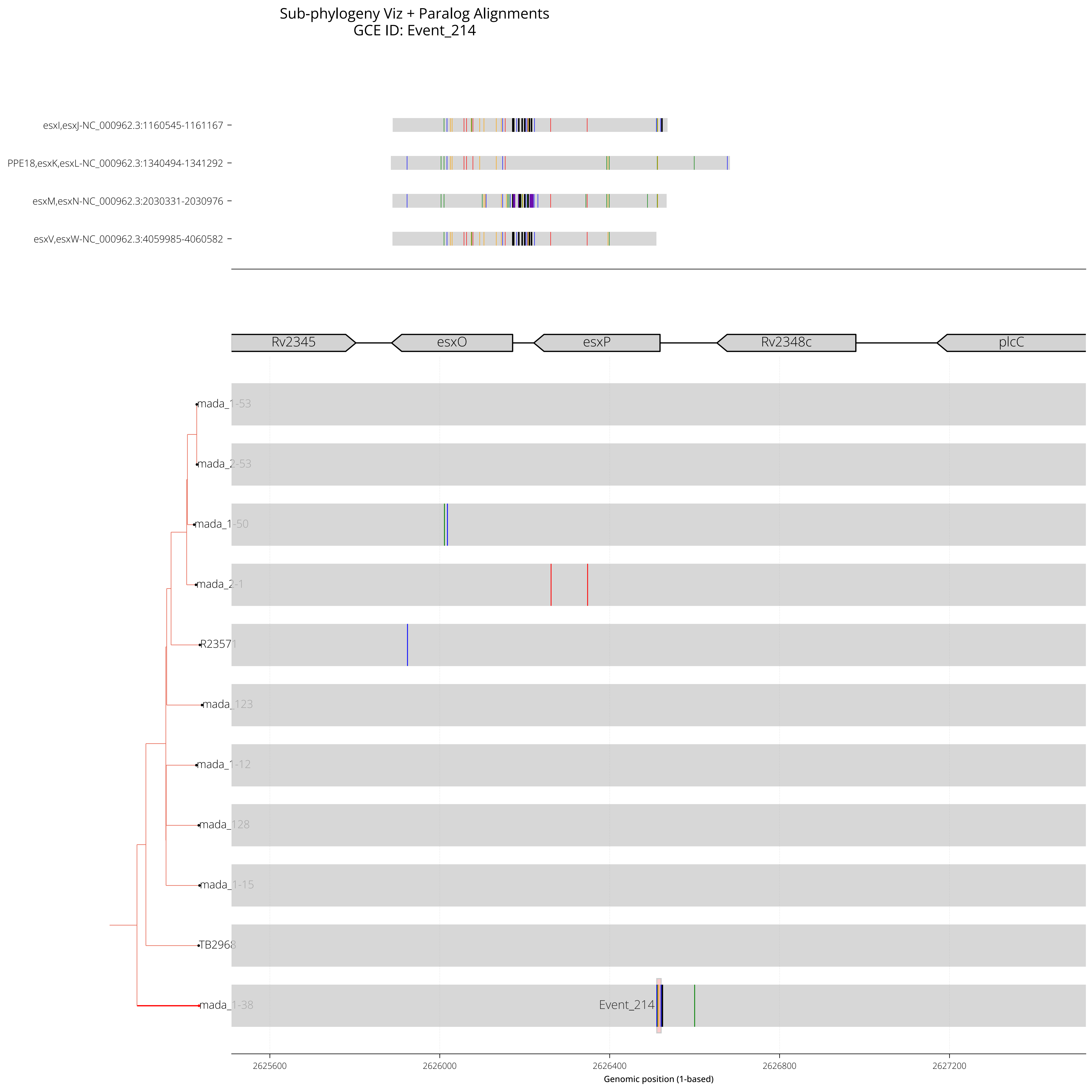
Sub-phylogeny Viz + Paralog Alignments
GCE ID: Event_214
esxI,esxJ-NC_000962.3:1160545-1161167
PPE18,esxK,esxL-NC_000962.3:1340494-1341292
esxM,esxN-NC_000962.3:2030331-2030976
esxV,esxW-NC_000962.3:4059985-4060582
Rv2345
esxO
esxP
Rv2348c
plcC
mada_1-53
mada_2-53
mada_1-50
mada_2-1
R23571
mada_123
mada_1-12
mada_128
mada_1-15
TB2968
mada_1-38
Event_214
2625600
2626000
2626400
2626800
2627200
Genomic position (1-based)

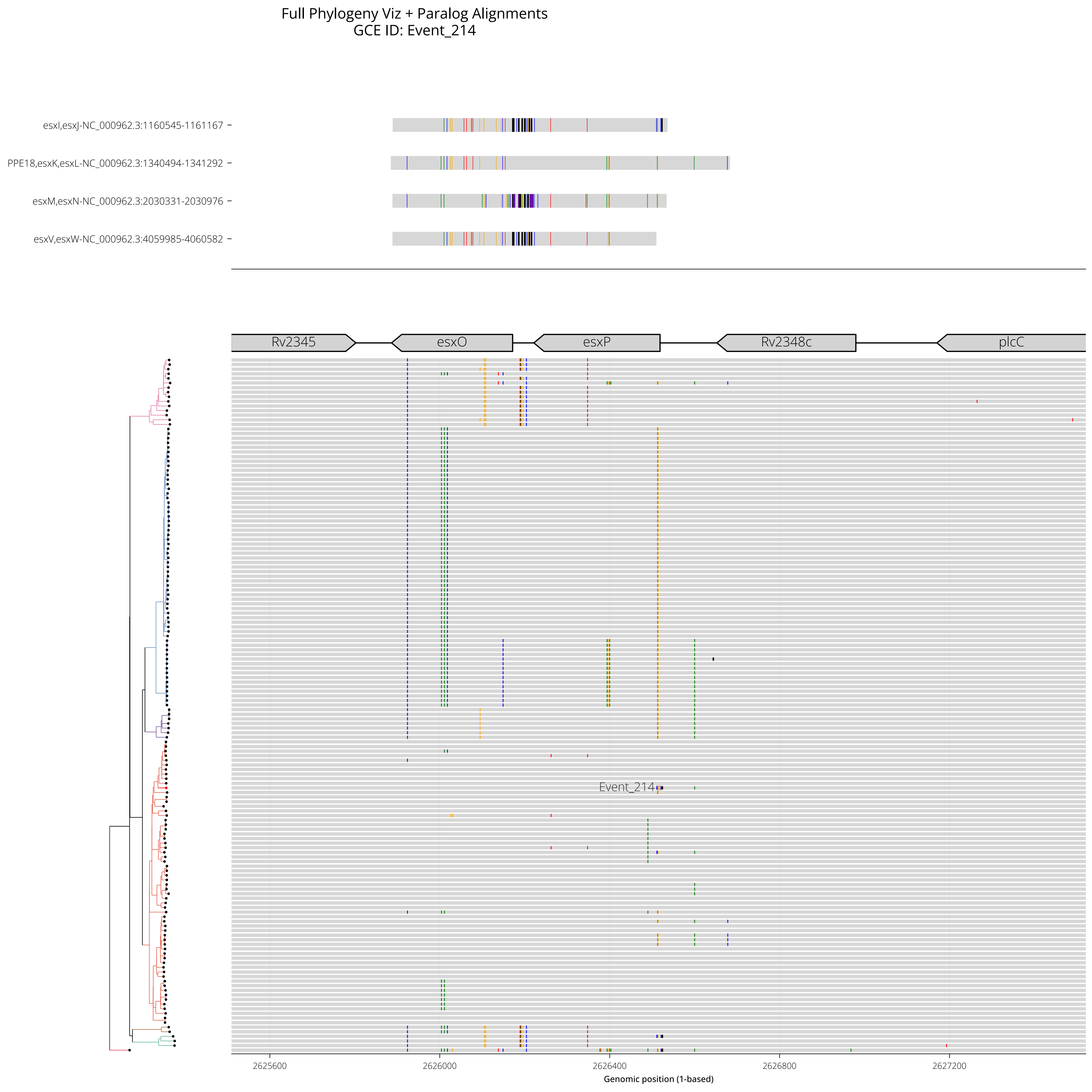

Full Phylogeny Viz + Paralog Alignments
GCE ID: Event_214
esxI,esxJ-NC_000962.3:1160545-1161167
PPE18,esxK,esxL-NC_000962.3:1340494-1341292
esxM,esxN-NC_000962.3:2030331-2030976
esxV,esxW-NC_000962.3:4059985-4060582
Rv2345
esxO
esxP
Rv2348c
plcC
Event_214
2625600
2626000
2626400
2626800
2627200
Genomic position (1-based)

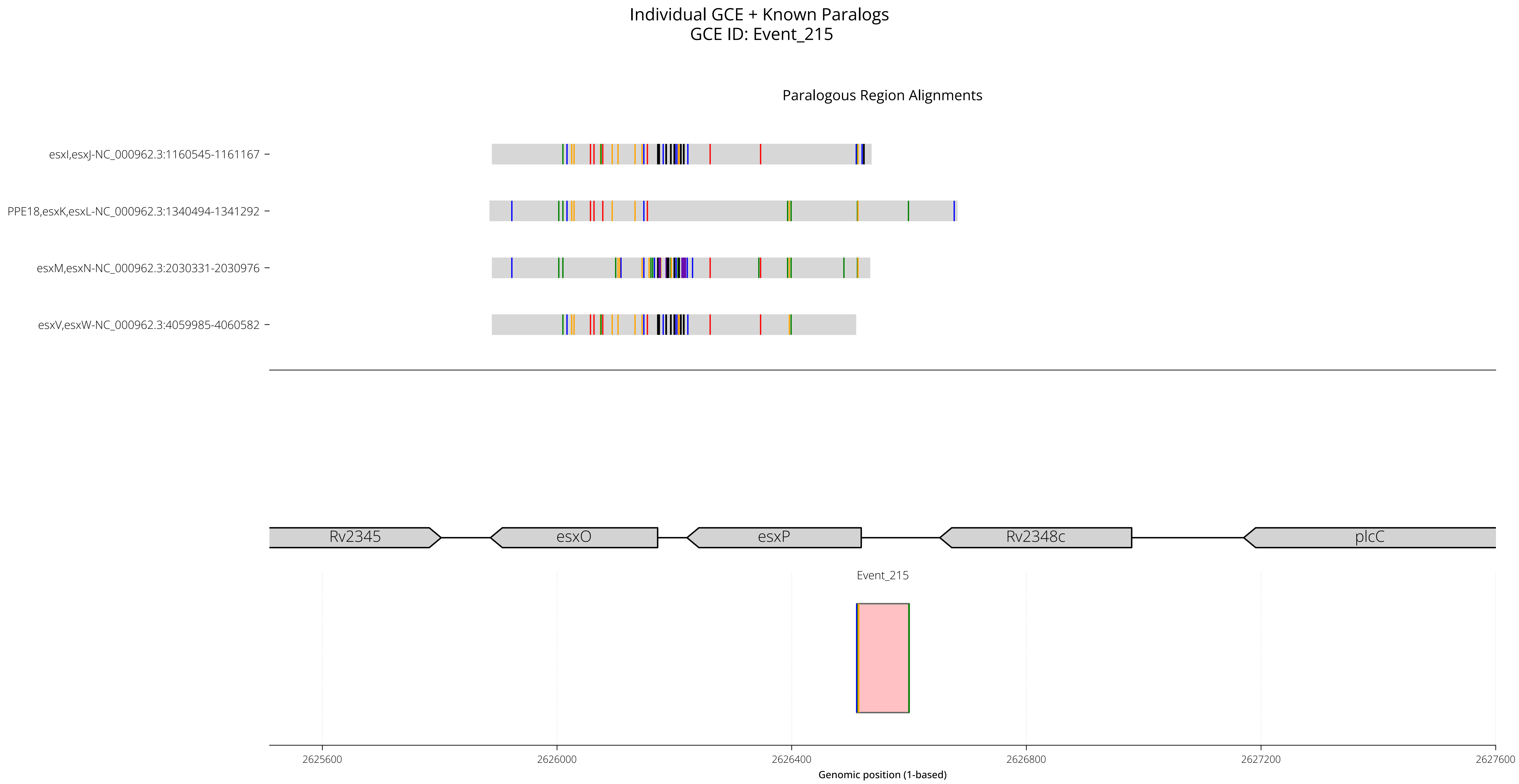

Individual GCE + Known Paralogs
GCE ID: Event_215
Paralogous Region Alignments
esxI,esxJ-NC_000962.3:1160545-1161167
PPE18,esxK,esxL-NC_000962.3:1340494-1341292
esxM,esxN-NC_000962.3:2030331-2030976
esxV,esxW-NC_000962.3:4059985-4060582
Rv2345
esxO
esxP
Rv2348c
plcC
Event_215
2625600
2626000
2626400
2626800
2627200
2627600
Genomic position (1-based)

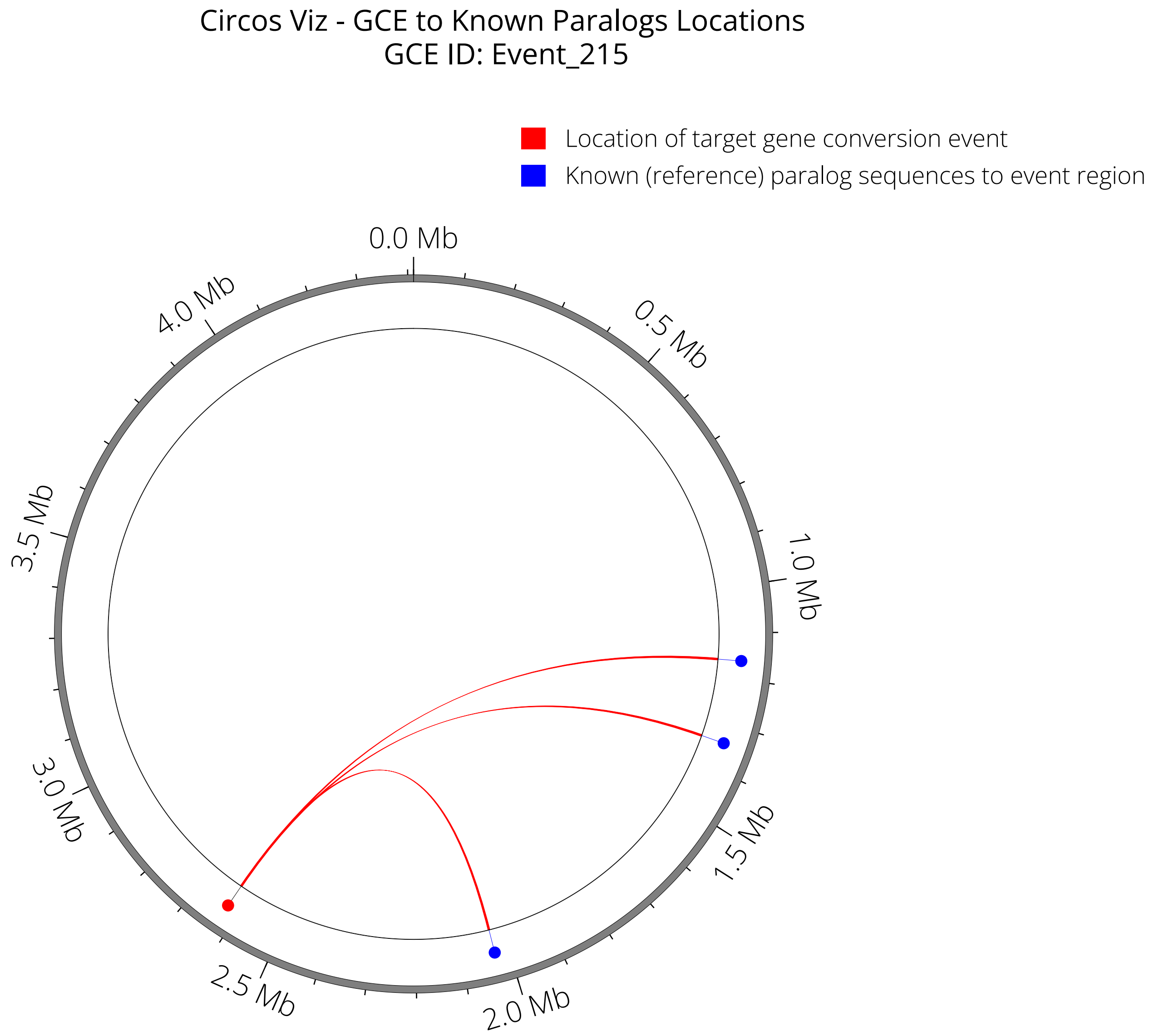
Circos Viz - GCE to Known Paralogs Locations
GCE ID: Event_215
Location of target gene conversion event
Known (reference) paralog sequences to event region
0.0 Mb
0.5 Mb
1.0 Mb
1.5 Mb
2.0 Mb
2.5 Mb
3.0 Mb
3.5 Mb
4.0 Mb

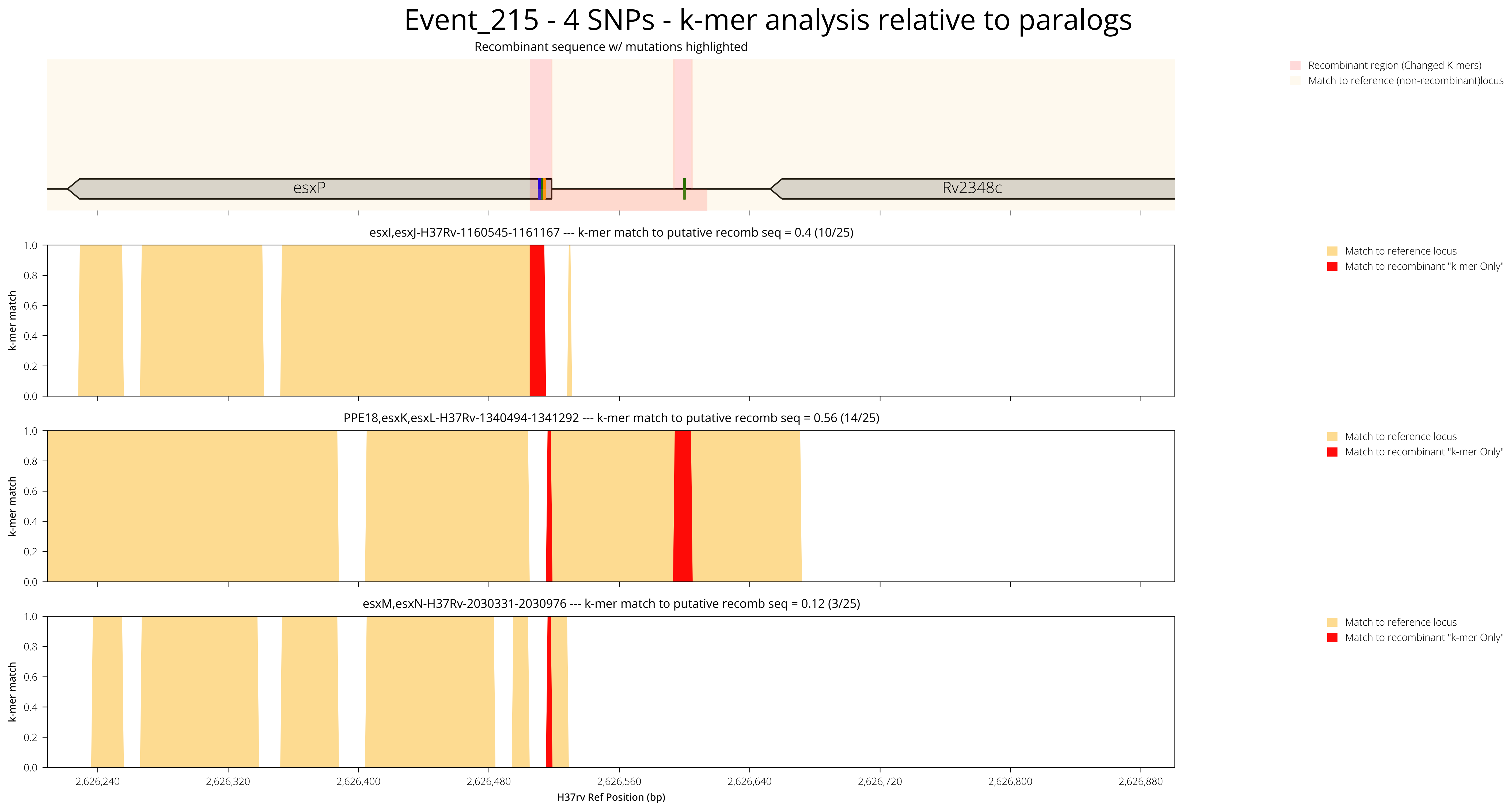

Event_215 - 4 SNPs - k-mer analysis relative to paralogs
Recombinant sequence w/ mutations highlighted
Recombinant region (Changed K-mers)
Match to reference (non-recombinant)locus
esxP
Rv2348c
esxI,esxJ-H37Rv-1160545-1161167 --- k-mer match to putative recomb seq = 0.4 (10/25)
PPE18,esxK,esxL-H37Rv-1340494-1341292 --- k-mer match to putative recomb seq = 0.56 (14/25)
esxM,esxN-H37Rv-2030331-2030976 --- k-mer match to putative recomb seq = 0.12 (3/25)
Match to reference locus
Match to recombinant "k-mer Only"
k-mer match
1.0
0.8
0.6
0.4
0.2
0.0
2,626,240
2,626,320
2,626,400
2,626,480
2,626,560
2,626,640
2,626,720
2,626,800
2,626,880
H37rv Ref Position (bp)

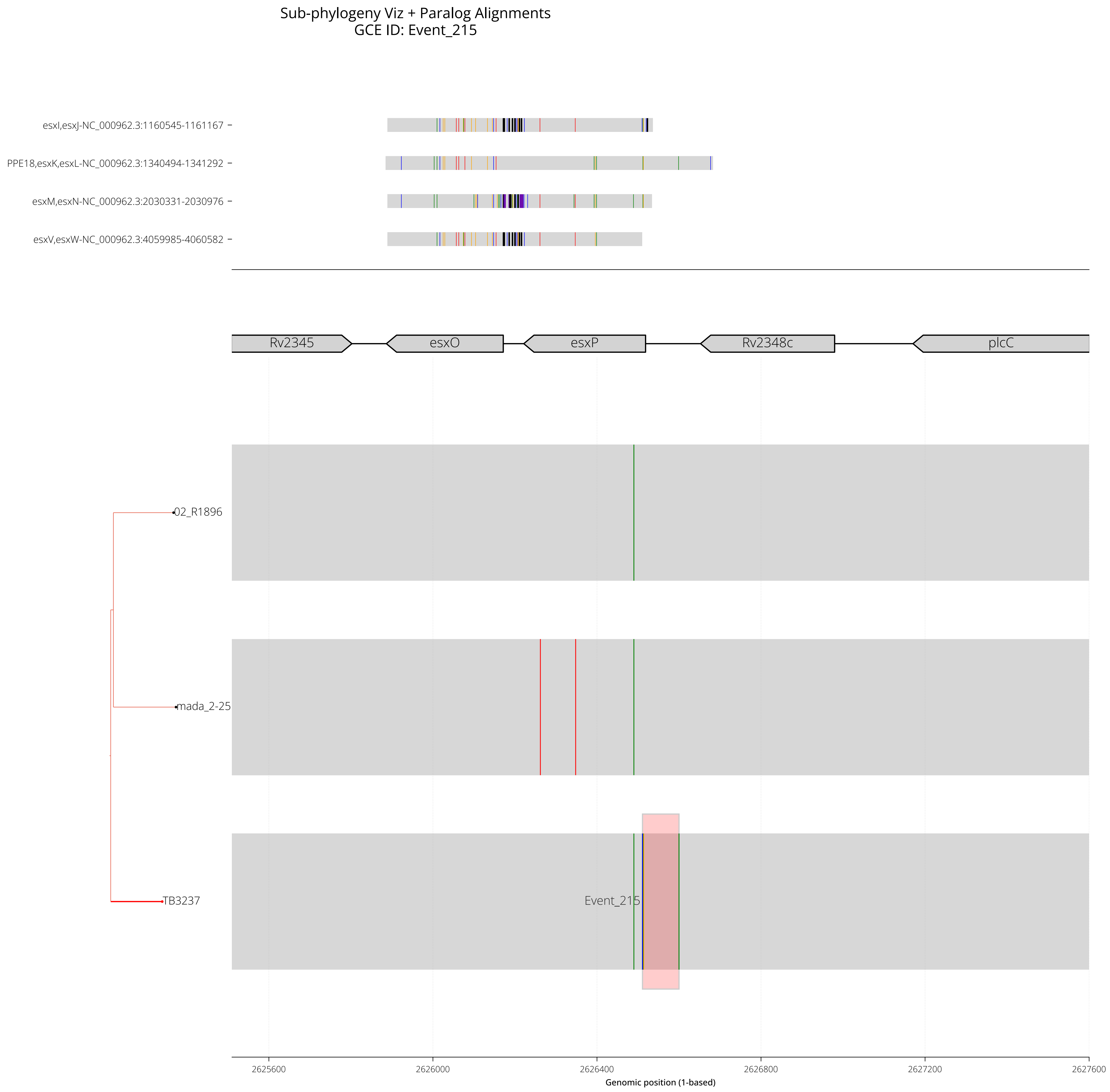

Sub-phylogeny Viz + Paralog Alignments
GCE ID: Event_215
esxI,esxJ-NC_000962.3:1160545-1161167
PPE18,esxK,esxL-NC_000962.3:1340494-1341292
esxM,esxN-NC_000962.3:2030331-2030976
esxV,esxW-NC_000962.3:4059985-4060582
Rv2345
esxO
esxP
Rv2348c
plcC
02_R1896
mada_2-25
TB3237
Event_215
2625600
2626000
2626400
2626800
2627200
2627600
Genomic position (1-based)

Full Phylogeny Viz + Paralog Alignments
GCE ID: Event_215

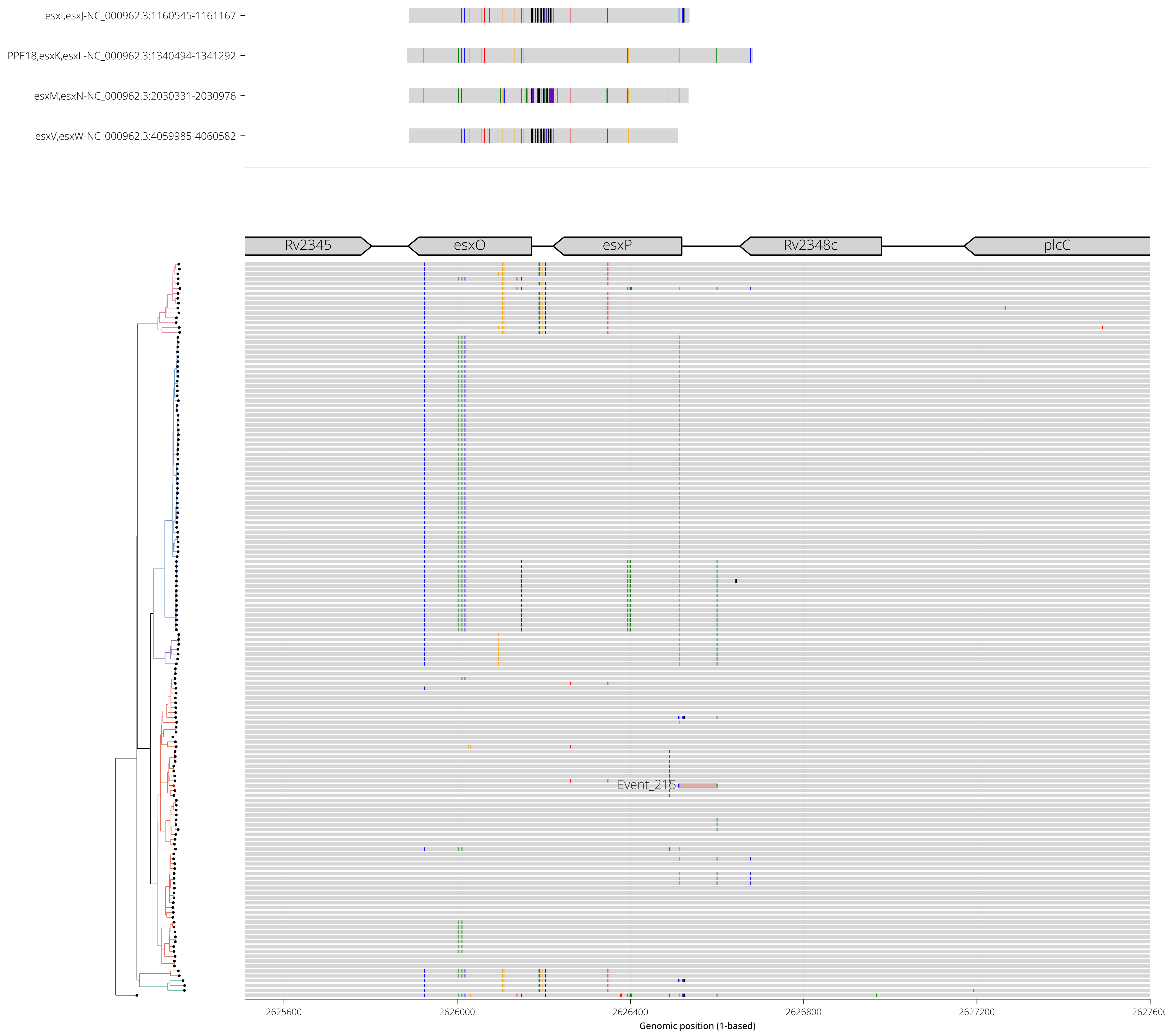

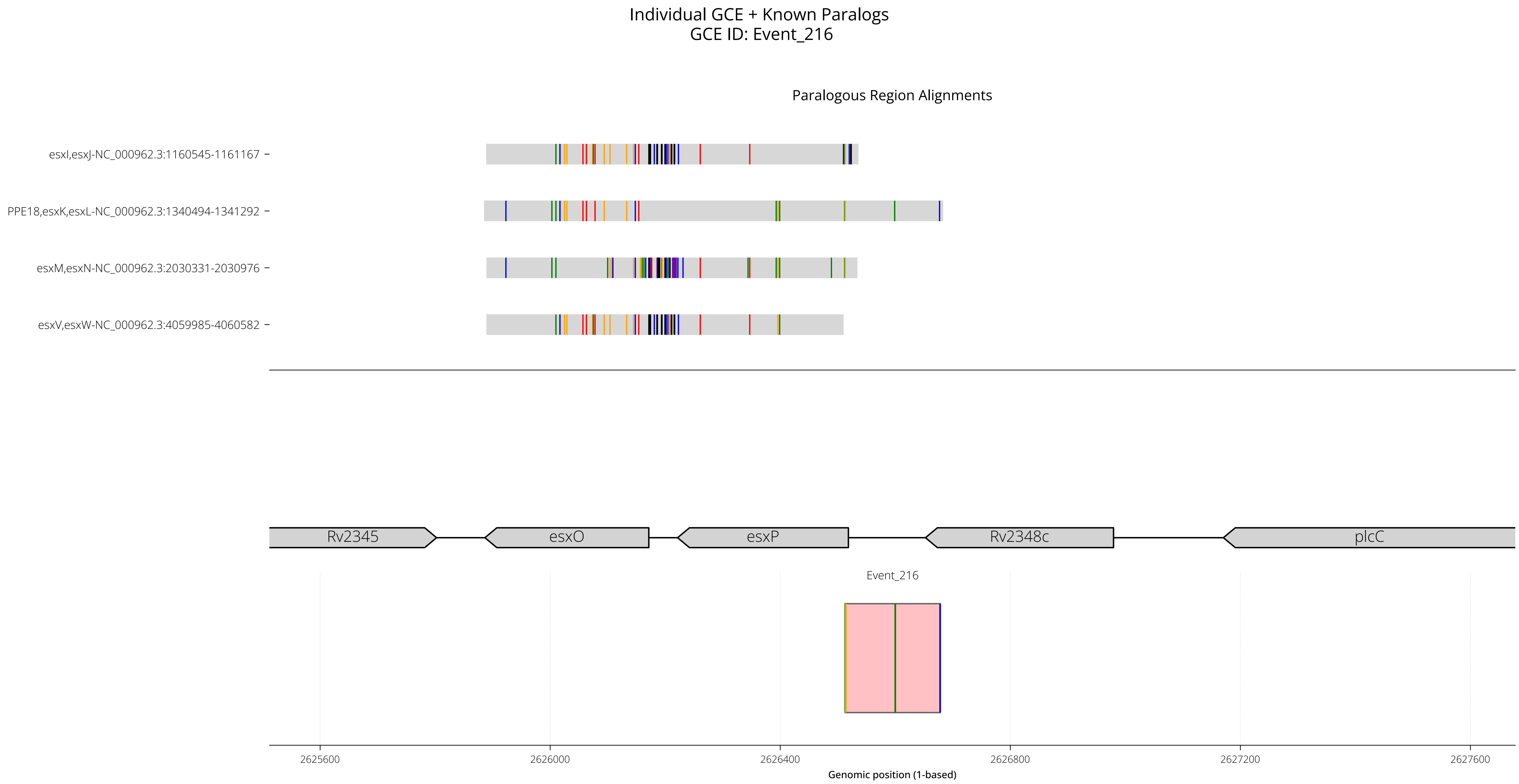
Individual GCE + Known Paralogs
GCE ID: Event_216
Paralogous Region Alignments
esxI,esxJ-NC_000962.3:1160545-1161167
PPE18,esxK,esxL-NC_000962.3:1340494-1341292
esxM,esxN-NC_000962.3:2030331-2030976
esxV,esxW-NC_000962.3:4059985-4060582
Rv2345
esxO
esxP
Rv2348c
plcC
Event_216
2625600
2626000
2626400
2626800
2627200
2627600
Genomic position (1-based)

Circos Viz - GCE to Known Paralogs Locations
GCE ID: Event_216

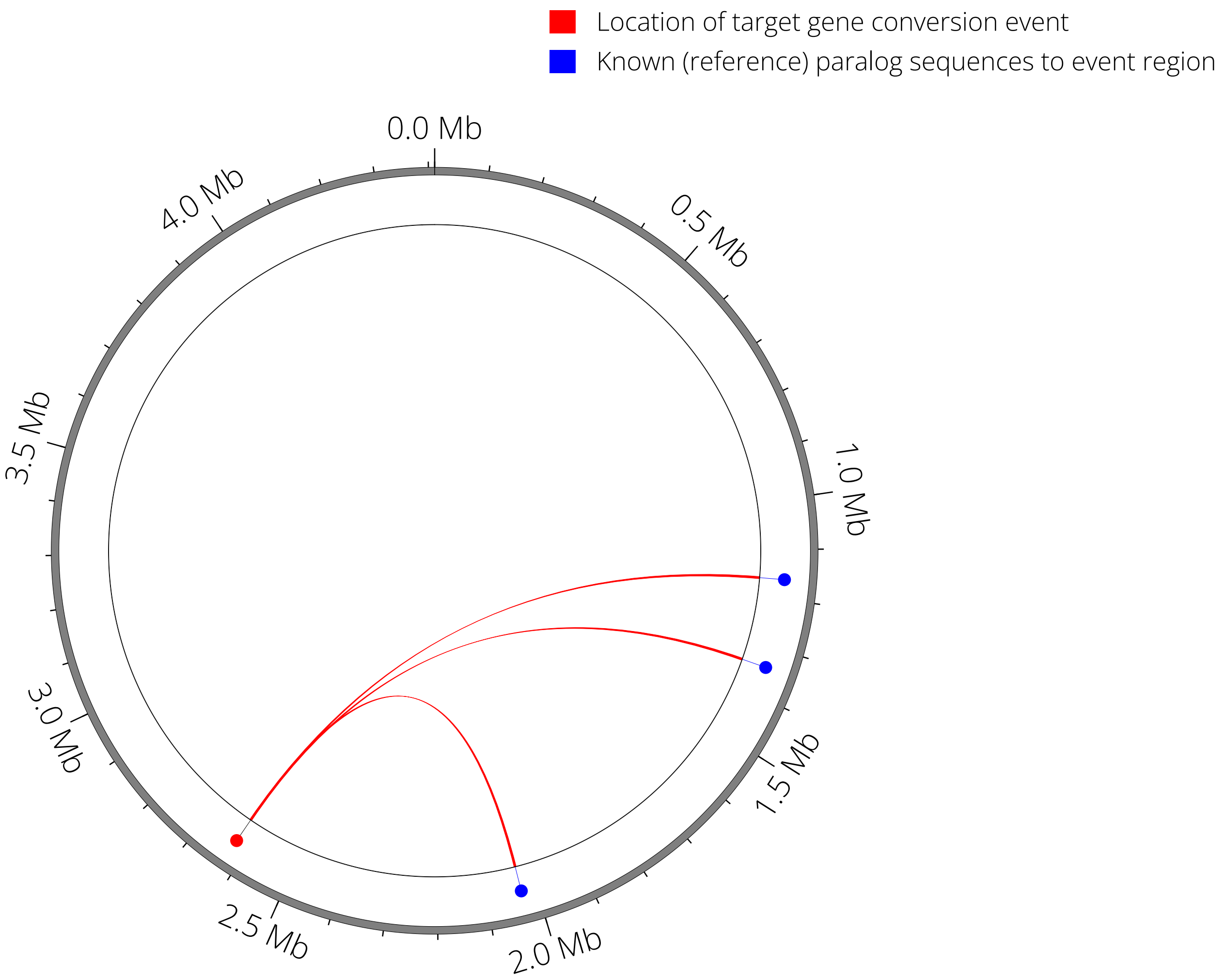

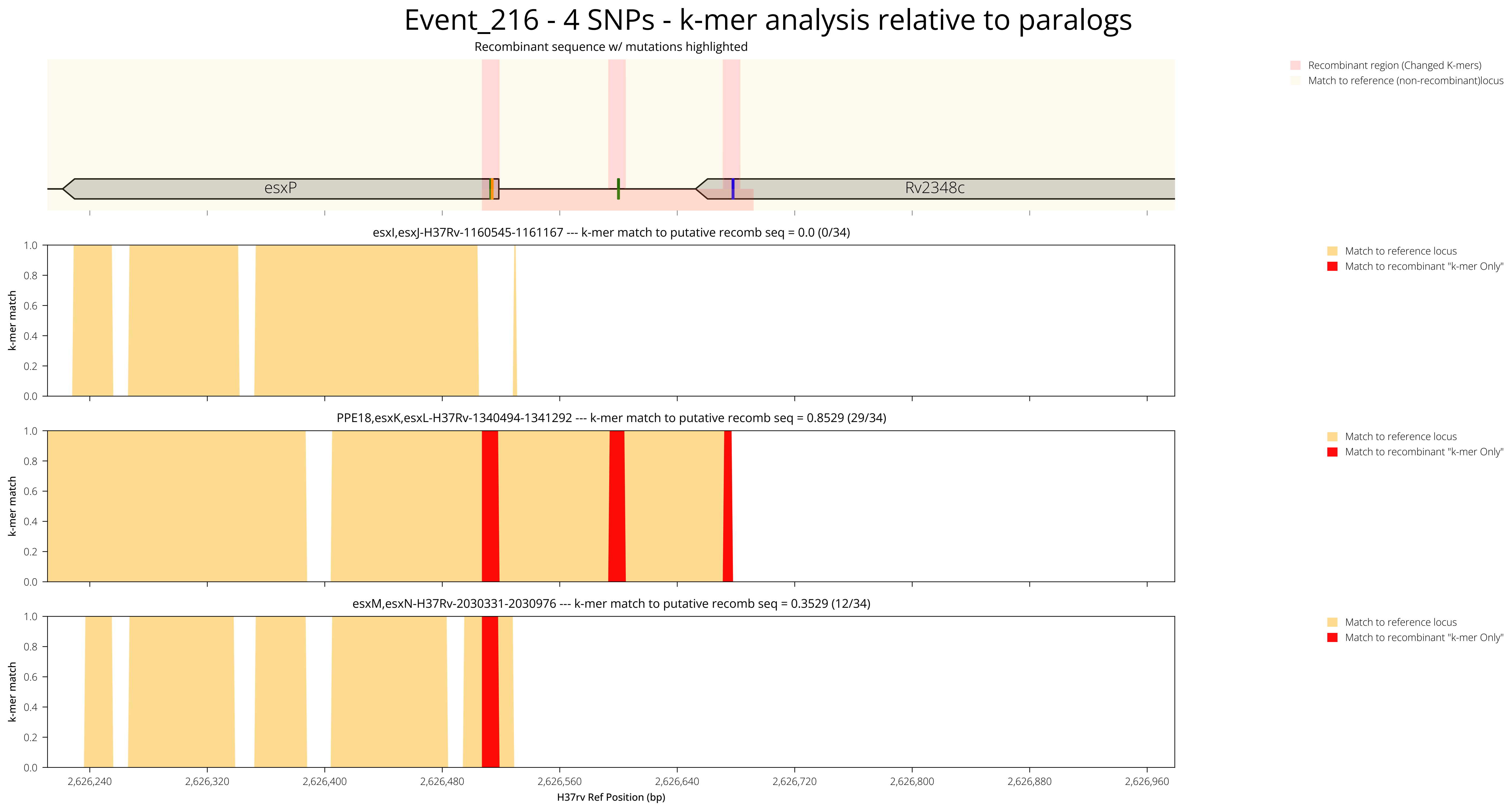

Event_216 - 4 SNPs - k-mer analysis relative to paralogs
Recombinant sequence w/ mutations highlighted
Recombinant region (Changed K-mers)
Match to reference (non-recombinant)locus
esxP
Rv2348c
esxI,esxJ-H37Rv-1160545-1161167 --- k-mer match to putative recomb seq = 0.0 (0/34)
PPE18,esxK,esxL-H37Rv-1340494-1341292 --- k-mer match to putative recomb seq = 0.8529 (29/34)
esxM,esxN-H37Rv-2030331-2030976 --- k-mer match to putative recomb seq = 0.3529 (12/34)
k-mer match
Match to reference locus
Match to recombinant "k-mer Only"
1.0
0.8
0.6
0.4
0.2
0.0
2,626,240
2,626,320
2,626,400
2,626,480
2,626,560
2,626,640
2,626,720
2,626,800
2,626,880
2,626,960
H37rv Ref Position (bp)

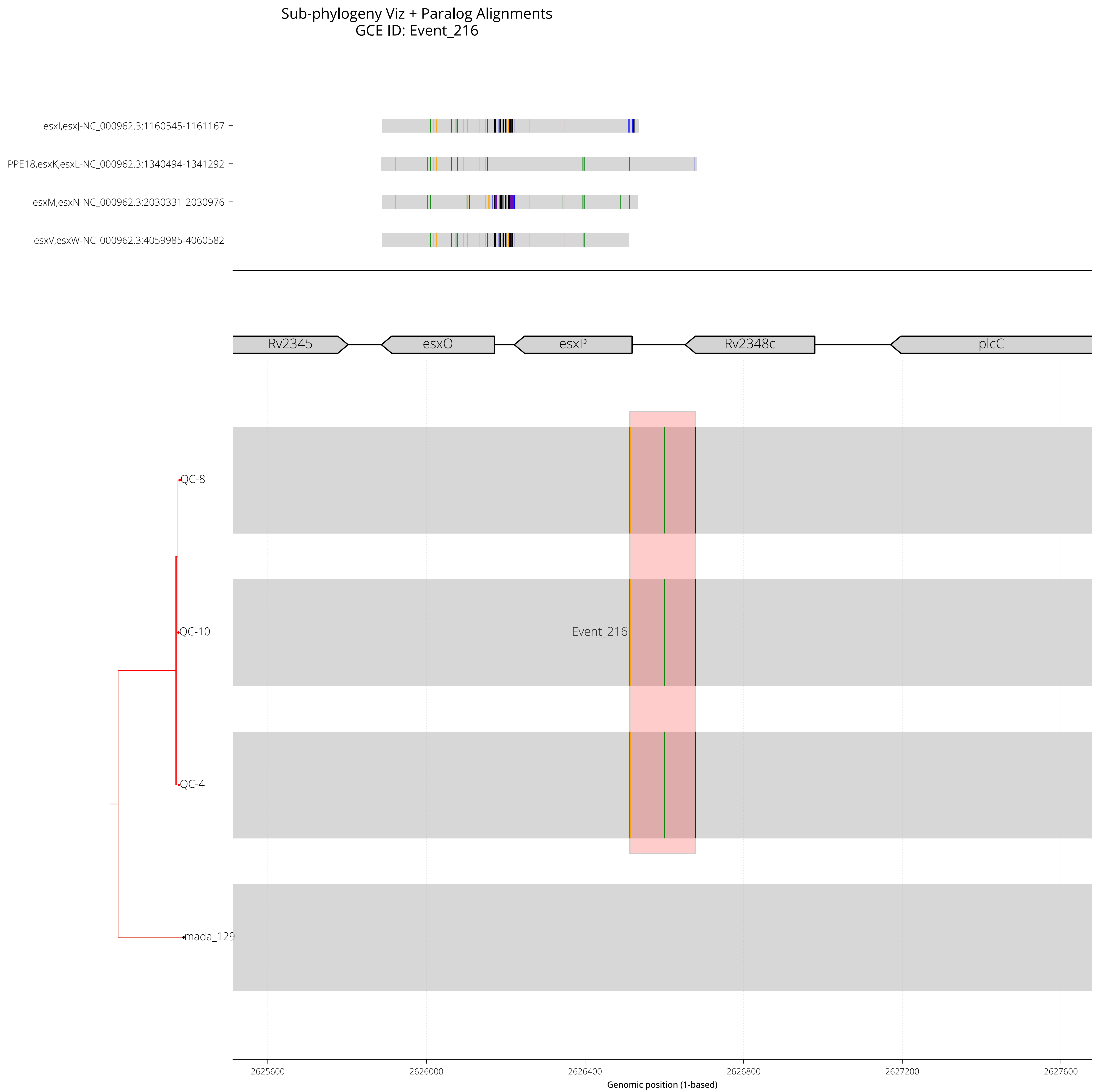
Sub-phylogeny Viz + Paralog Alignments
GCE ID: Event_216
esxI,esxJ-NC_000962.3:1160545-1161167
PPE18,esxK,esxL-NC_000962.3:1340494-1341292
esxM,esxN-NC_000962.3:2030331-2030976
esxV,esxW-NC_000962.3:4059985-4060582
Rv2345
esxO
esxP
Rv2348c
plcC
QC-8
QC-10
Event_216
QC-4
mada_129
2625600
2626000
2626400
2626800
2627200
2627600
Genomic position (1-based)

Full Phylogeny Viz + Paralog Alignments
GCE ID: Event_216

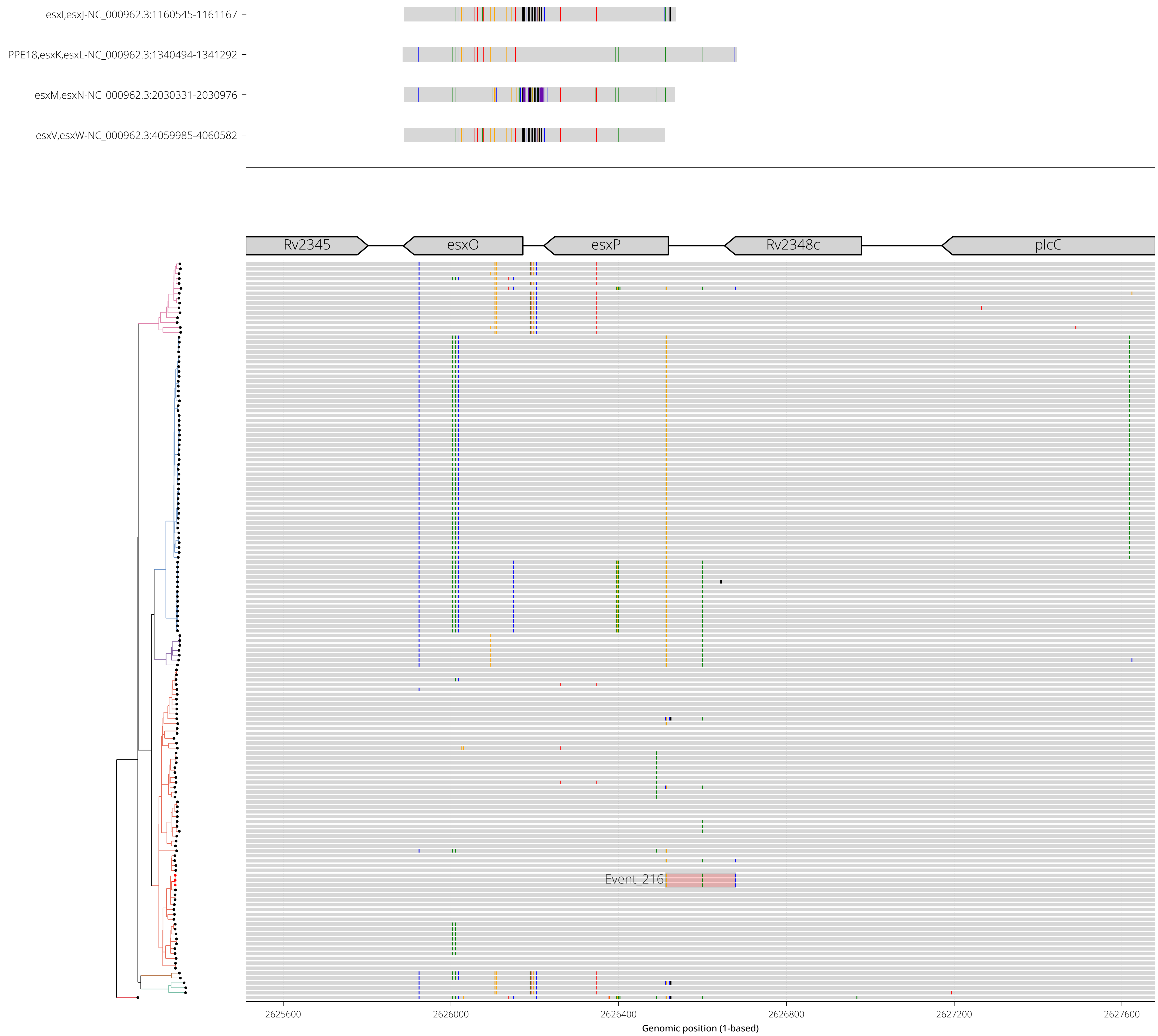

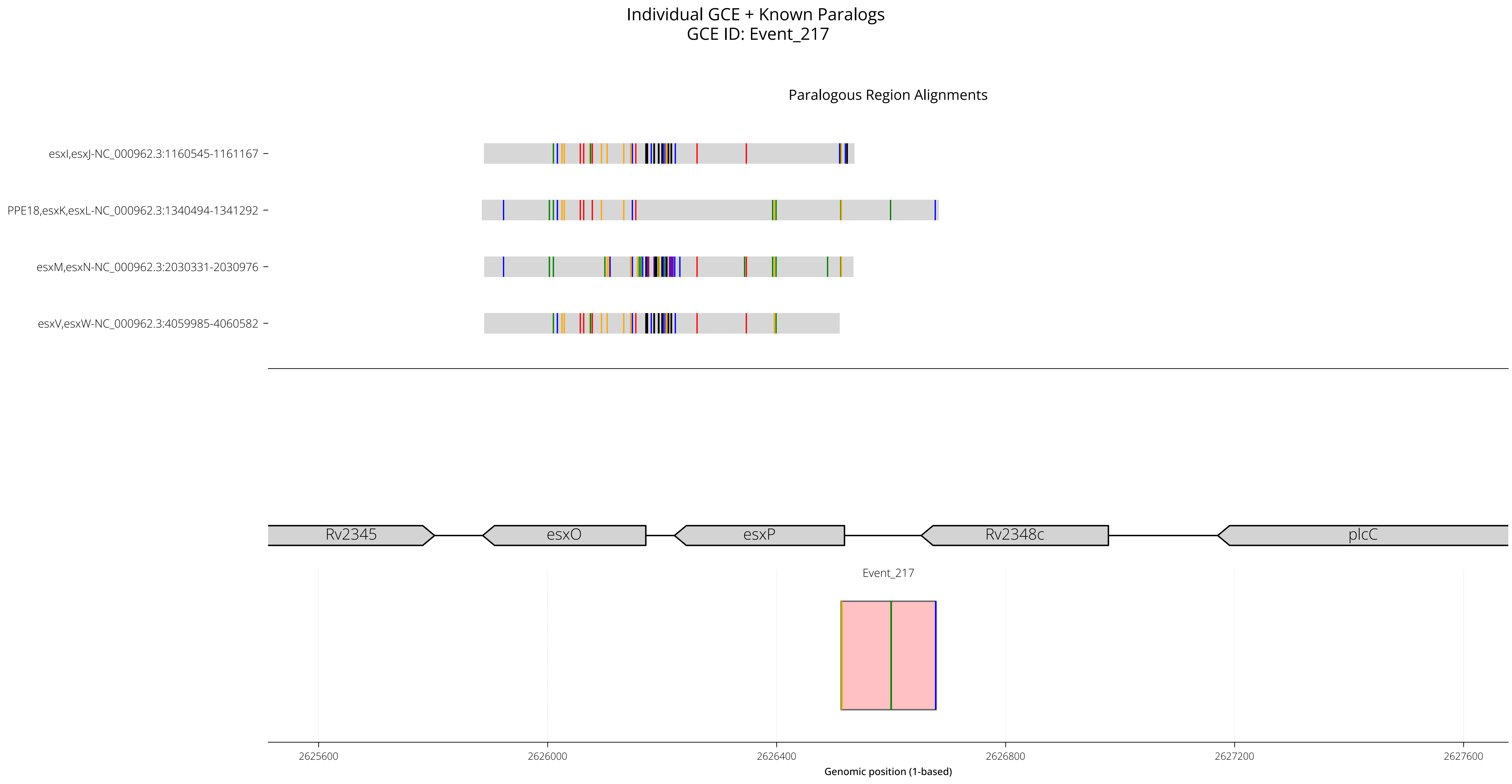
Individual GCE + Known Paralogs
GCE ID: Event_217
Paralogous Region Alignments
esxI,esxJ-NC_000962.3:1160545-1161167
PPE18,esxK,esxL-NC_000962.3:1340494-1341292
esxM,esxN-NC_000962.3:2030331-2030976
esxV,esxW-NC_000962.3:4059985-4060582
Rv2345
esxO
esxP
Rv2348c
plcC
Event_217
2625600
2626000
2626400
2626800
2627200
2627600
Genomic position (1-based)

Circos Viz - GCE to Known Paralogs Locations
GCE ID: Event_217

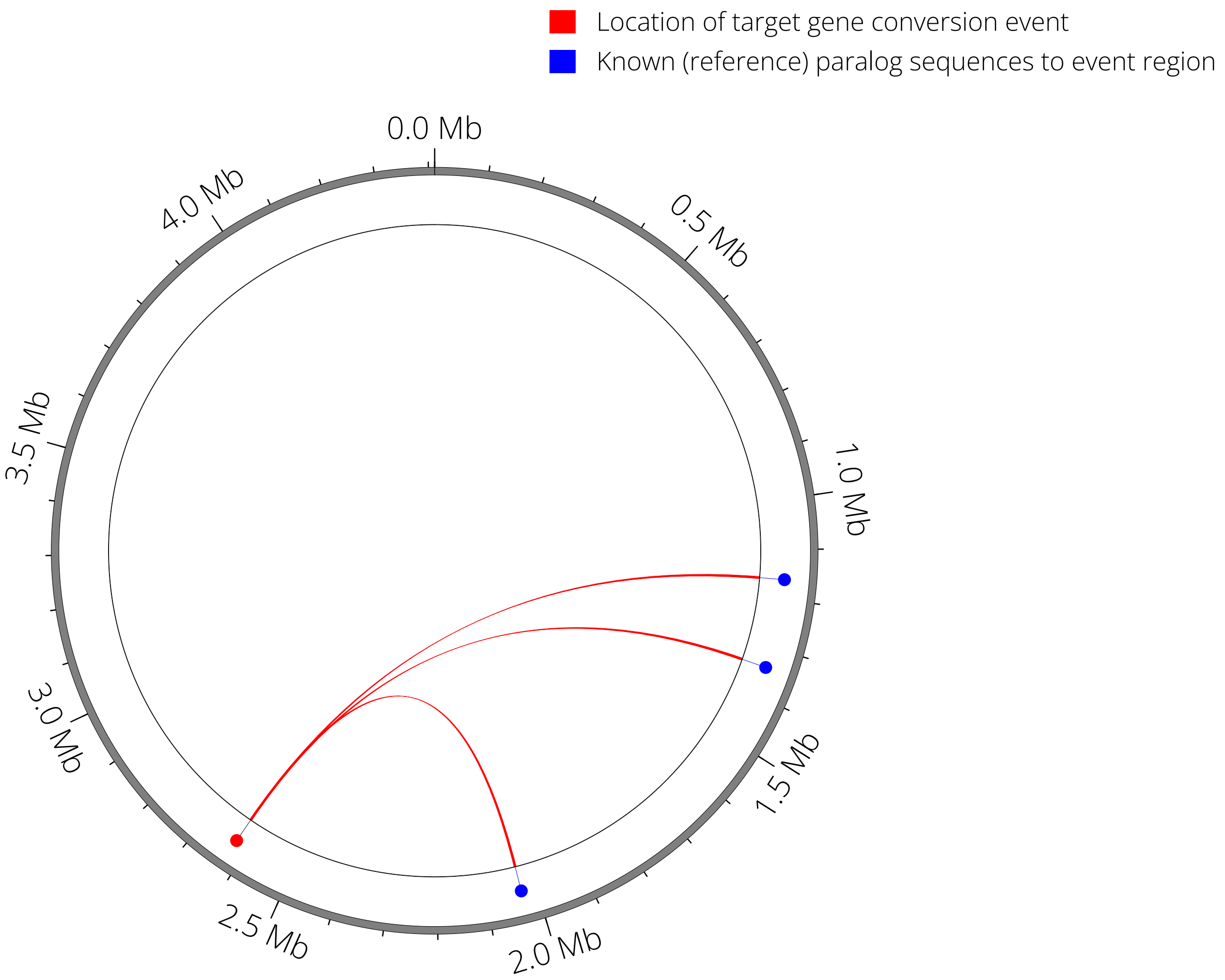

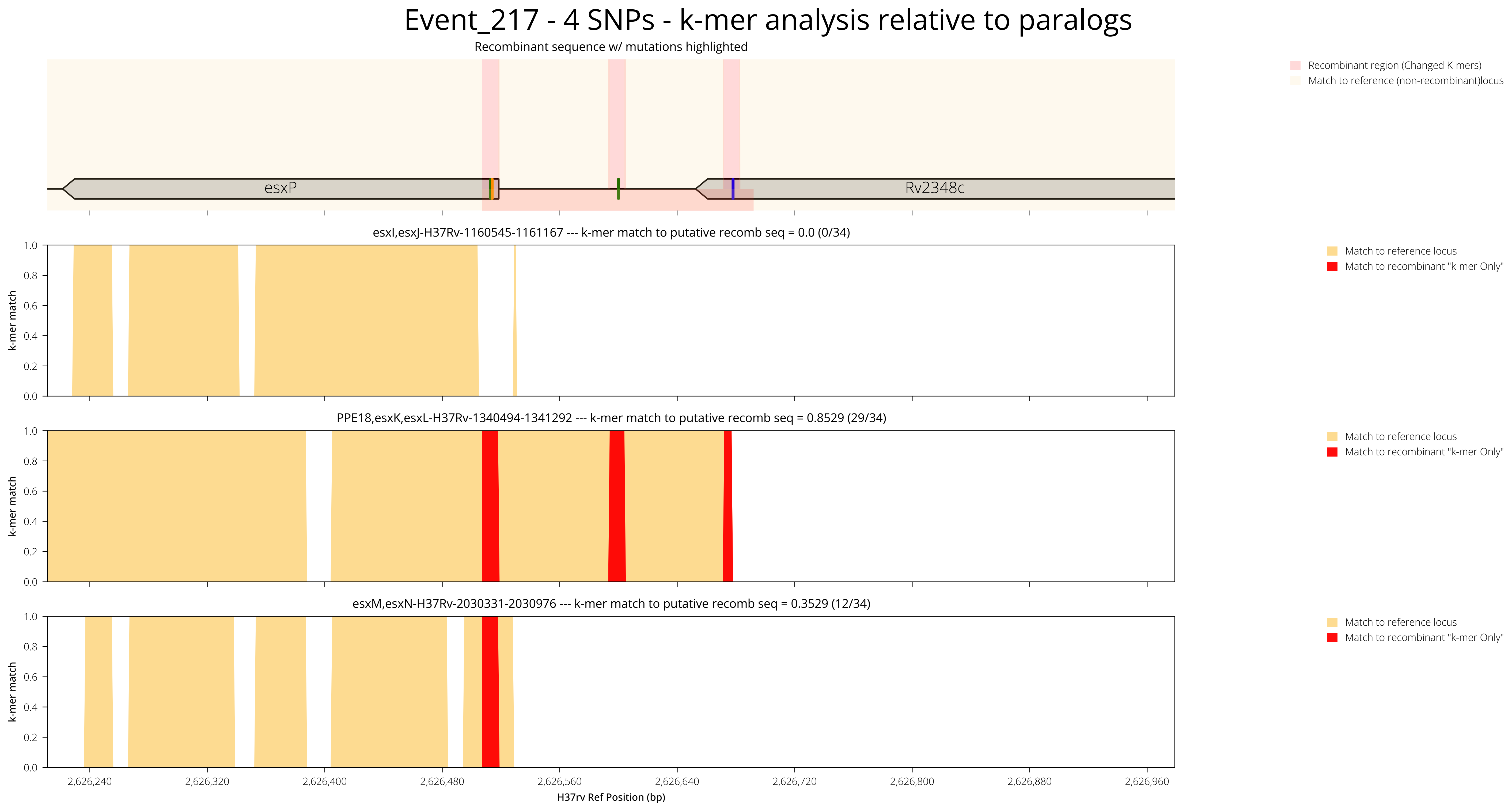

Event_217 - 4 SNPs - k-mer analysis relative to paralogs
Recombinant sequence w/ mutations highlighted
Recombinant region (Changed K-mers)
Match to reference (non-recombinant)locus
esxP
Rv2348c
esxI,esxJ-H37Rv-1160545-1161167 --- k-mer match to putative recomb seq = 0.0 (0/34)
PPE18,esxK,esxL-H37Rv-1340494-1341292 --- k-mer match to putative recomb seq = 0.8529 (29/34)
esxM,esxN-H37Rv-2030331-2030976 --- k-mer match to putative recomb seq = 0.3529 (12/34)
Match to reference locus
Match to recombinant "k-mer Only"
k-mer match
1.0
0.8
0.6
0.4
0.2
0.0
2,626,240
2,626,320
2,626,400
2,626,480
2,626,560
2,626,640
2,626,720
2,626,800
2,626,880
2,626,960
H37rv Ref Position (bp)

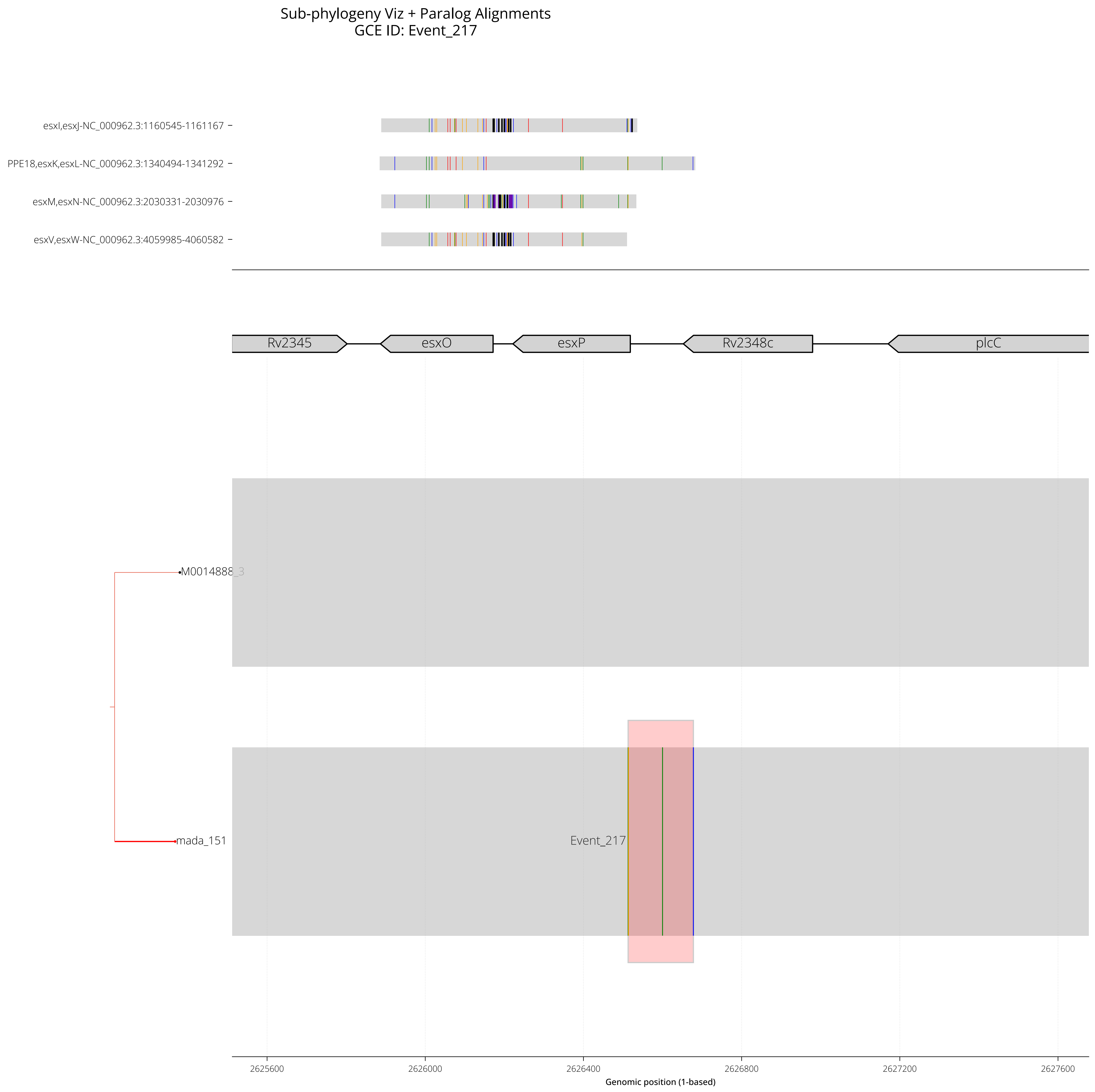
Sub-phylogeny Viz + Paralog Alignments
GCE ID: Event_217
esxI,esxJ-NC_000962.3:1160545-1161167
PPE18,esxK,esxL-NC_000962.3:1340494-1341292
esxM,esxN-NC_000962.3:2030331-2030976
esxV,esxW-NC_000962.3:4059985-4060582
Rv2345
esxO
esxP
Rv2348c
plcC
M0014888_3
mada_151
Event_217
2625600
2626000
2626400
2626800
2627200
2627600
Genomic position (1-based)

Full Phylogeny Viz + Paralog Alignments
GCE ID: Event_217

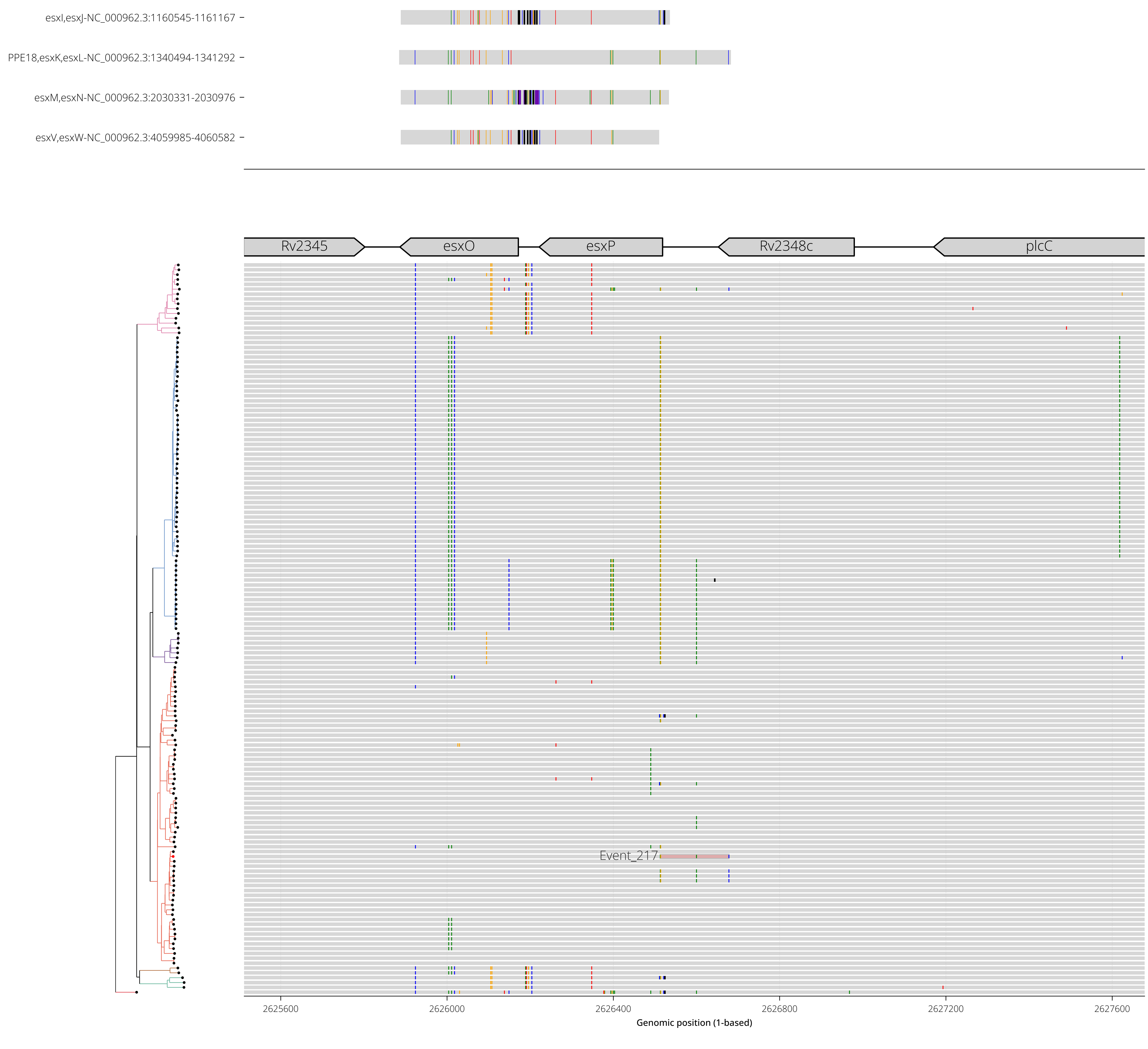

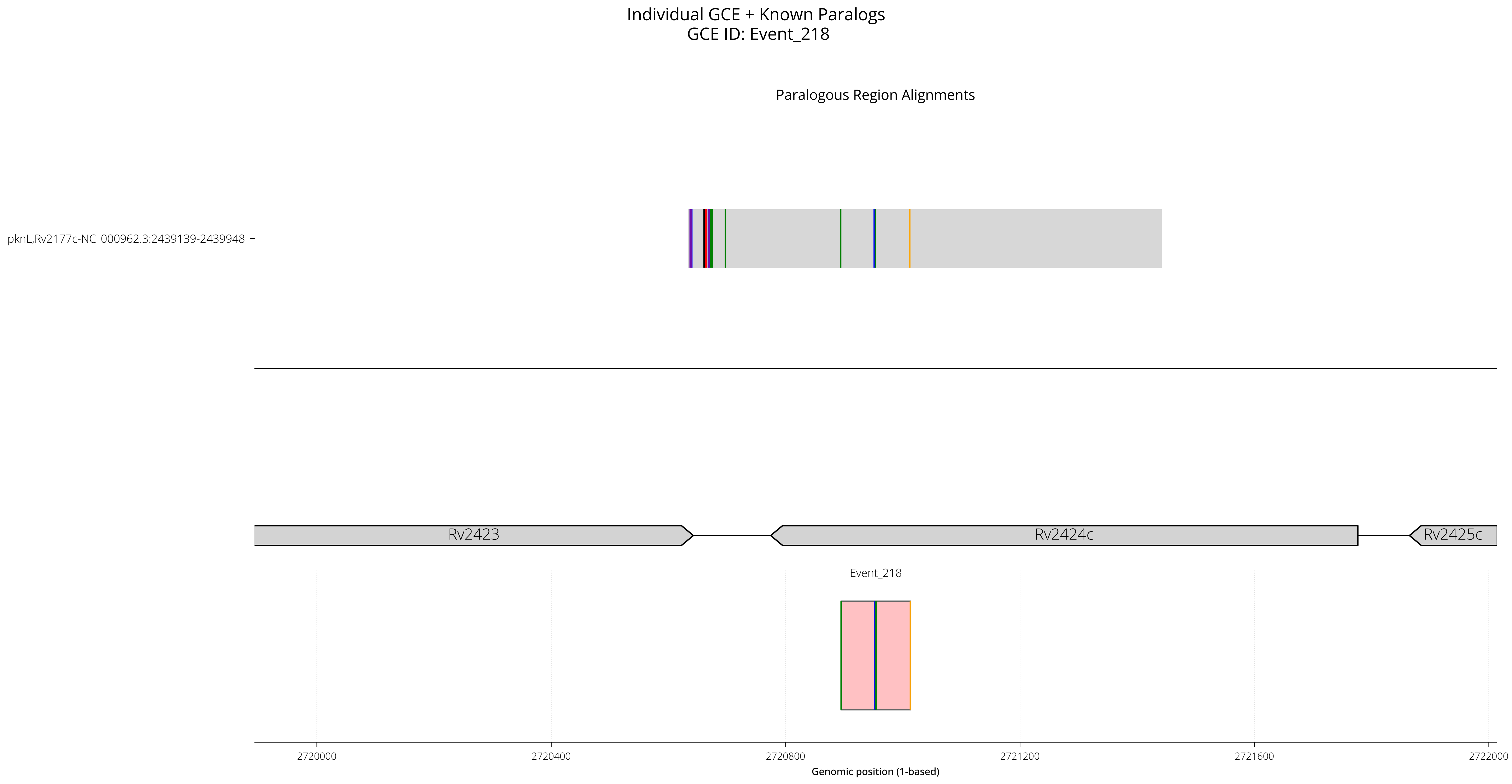

Individual GCE + Known Paralogs
GCE ID: Event_218
Paralogous Region Alignments
pknL,Rv2177c-NC_000962.3:2439139-2439948
Rv2423
Rv2424c
Rv2425c
Event_218
2720000
2720400
2720800
2721200
2721600
2722000
Genomic position (1-based)

Circos Viz - GCE to Known Paralogs Locations
GCE ID: Event_218

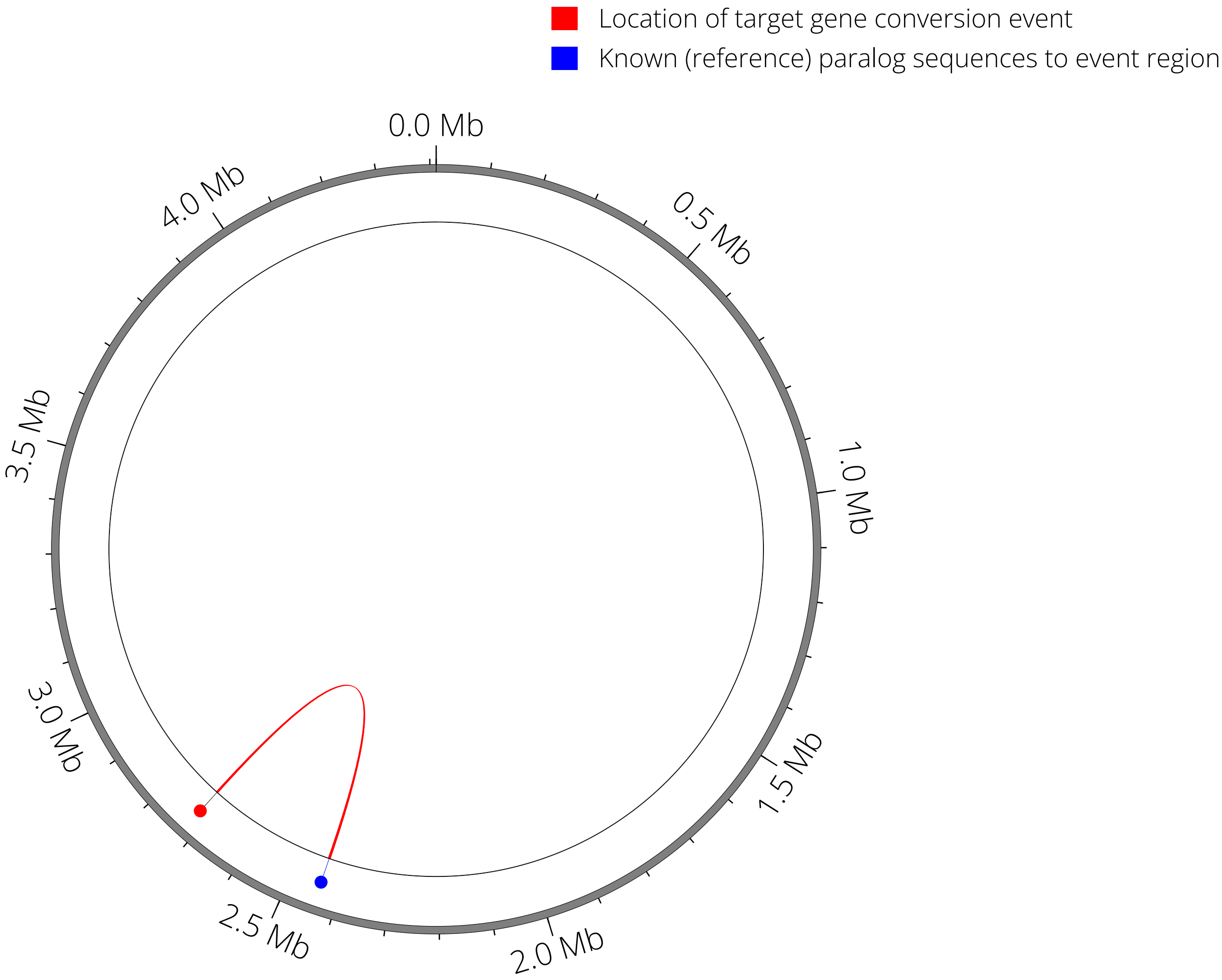

### Event_218 - 4 SNPs - k-mer analysis relative to paralogs

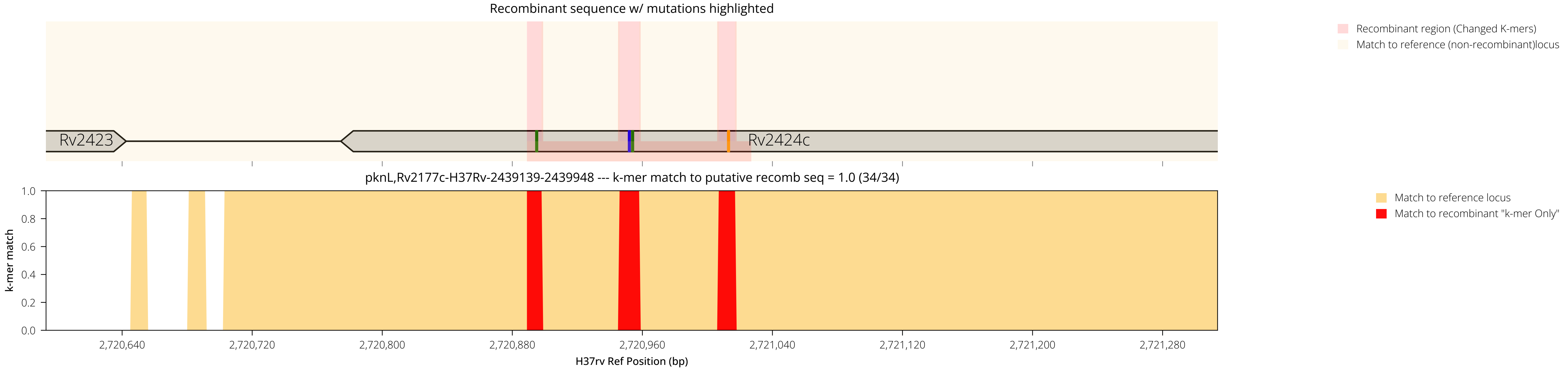

Sub-phylogeny Viz + Paralog Alignments
GCE ID: Event_218

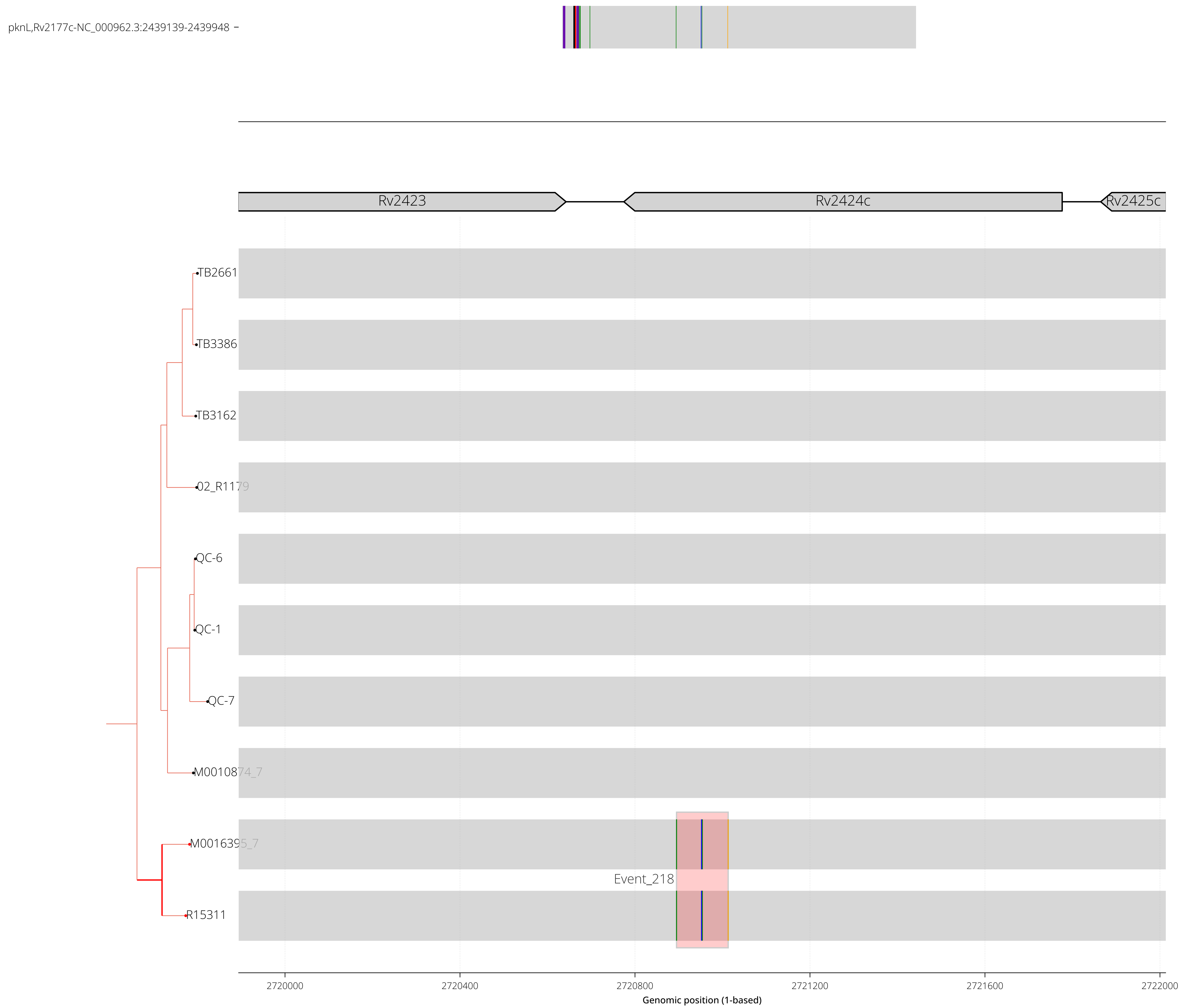

Full Phylogeny Viz + Paralog Alignments
GCE ID: Event_218

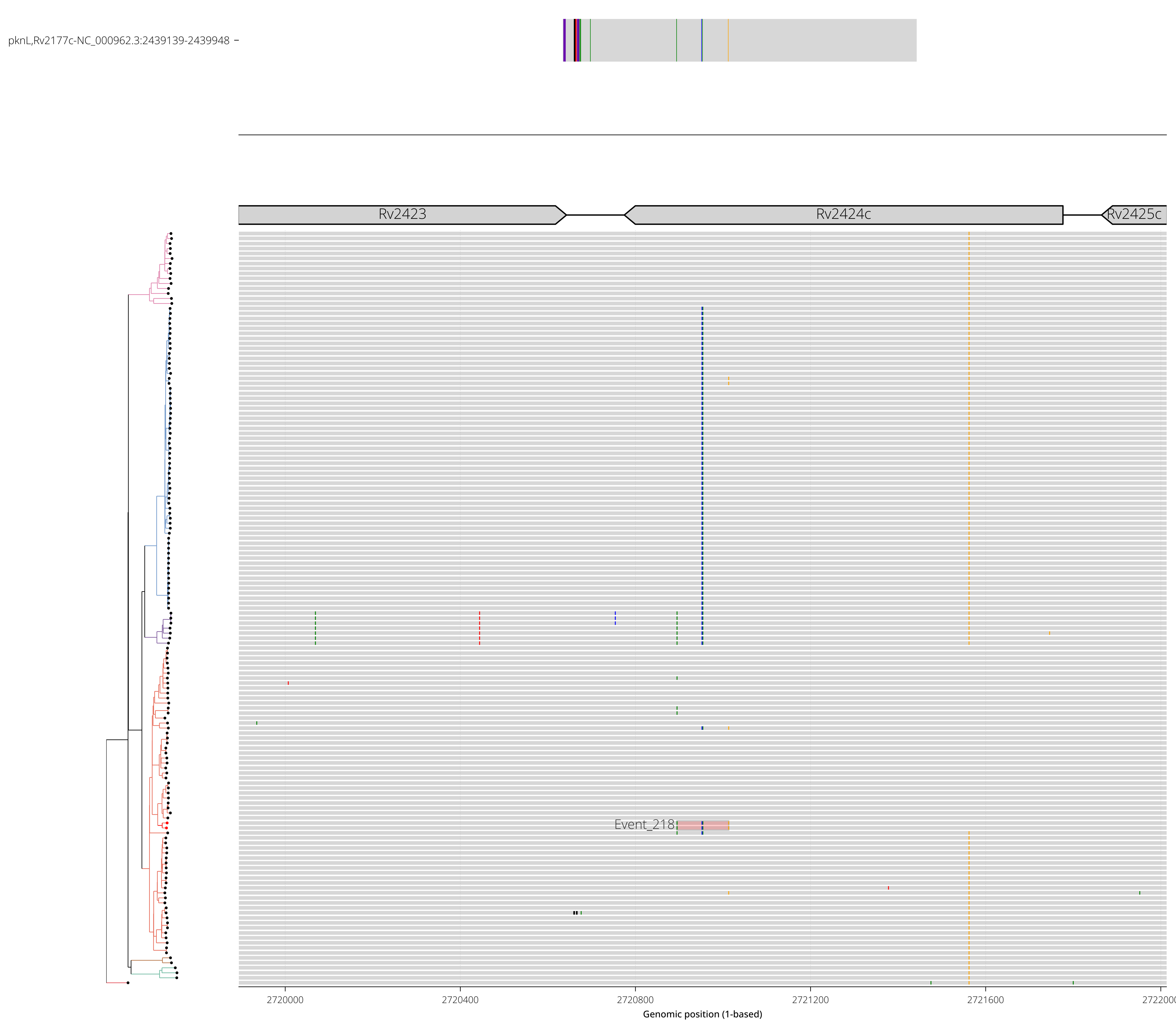

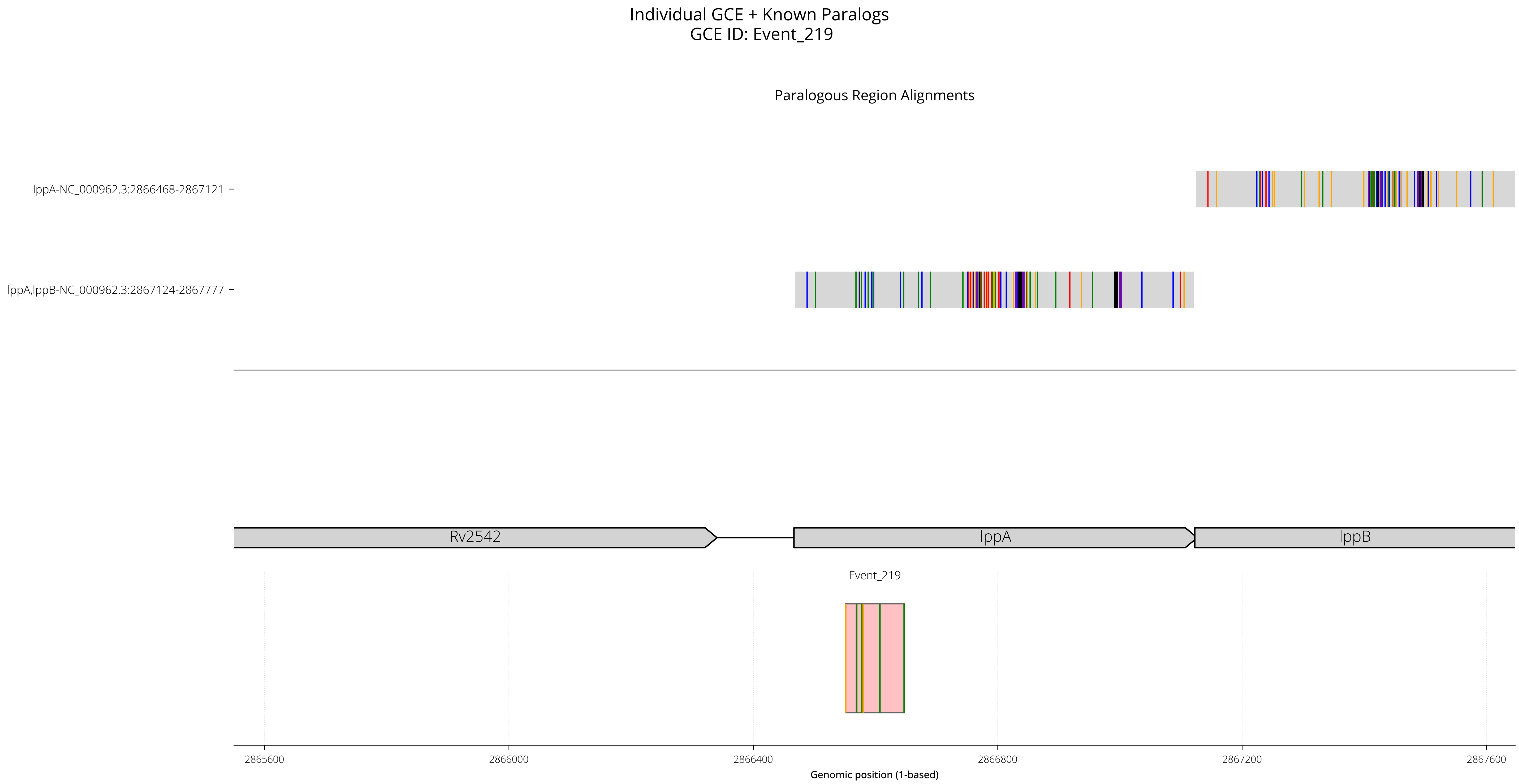

Individual GCE + Known Paralogs
GCE ID: Event_219
Paralogous Region Alignments
lppA-NC_000962.3:2866468-2867121
lppA,lppB-NC_000962.3:2867124-2867777
Rv2542
lppA
lppB
Event_219
2865600
2866000
2866400
2866800
2867200
2867600
Genomic position (1-based)

Circos Viz - GCE to Known Paralogs Locations
GCE ID: Event_219

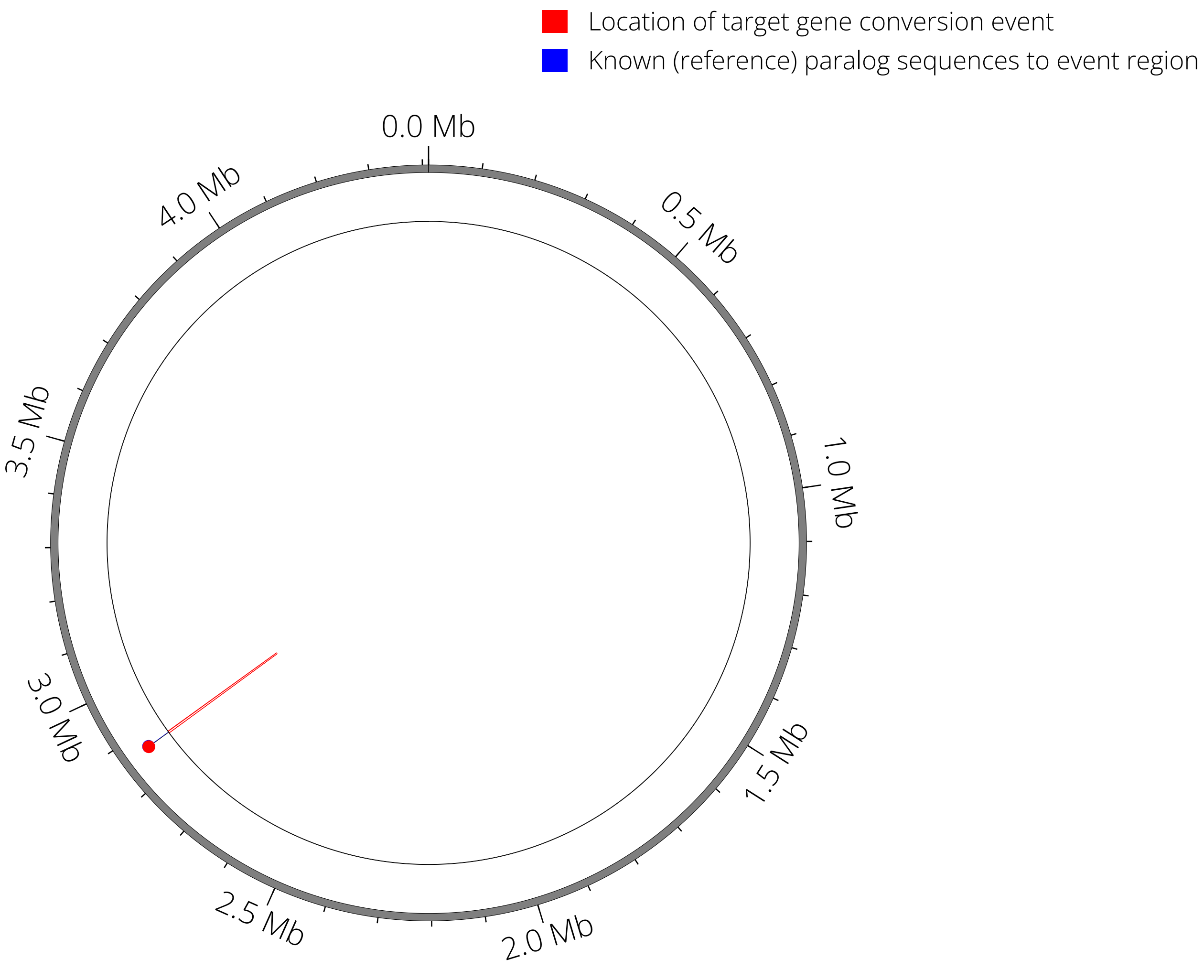

### Event_219 - 6 SNPs - k-mer analysis relative to paralogs

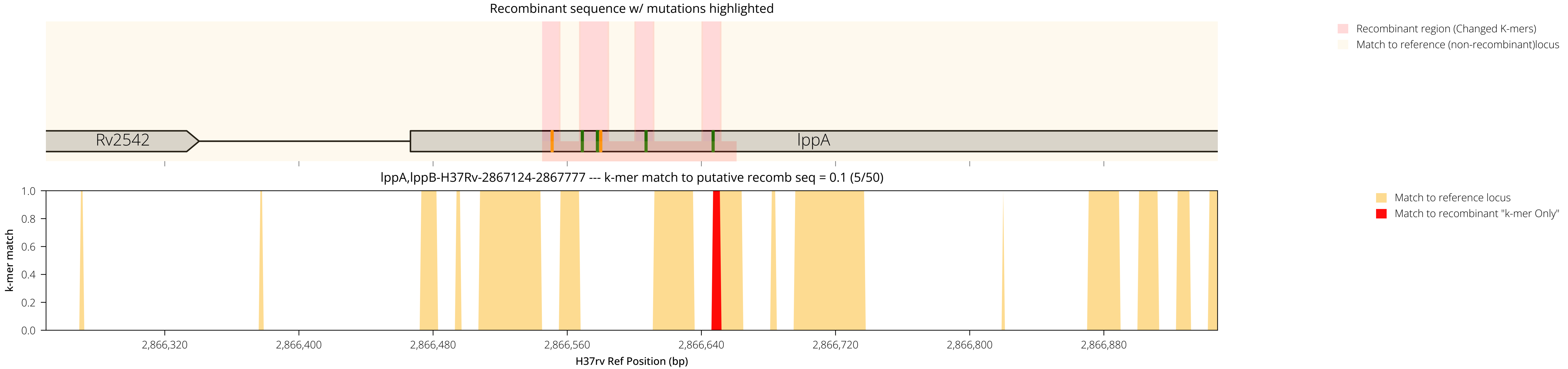

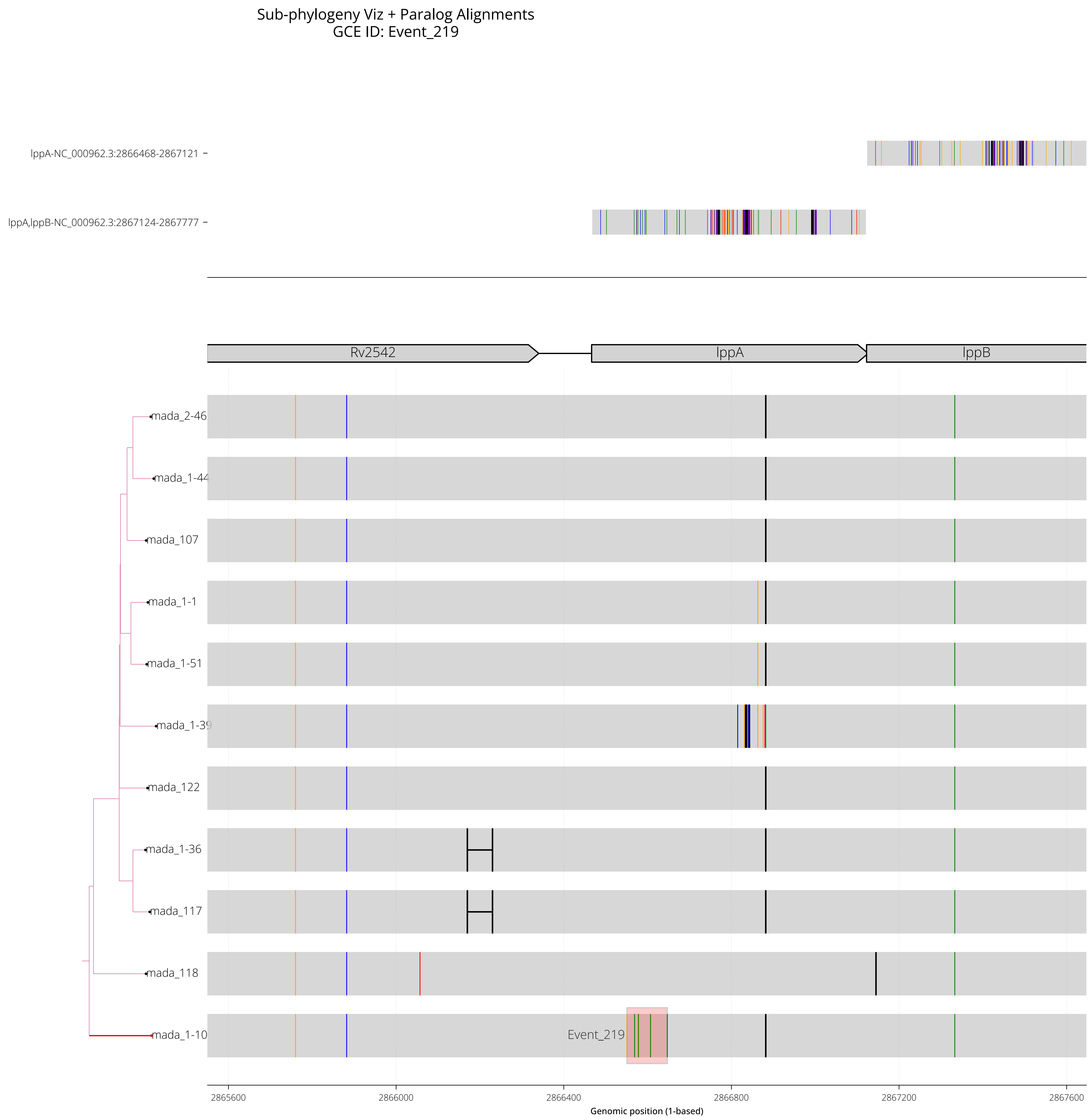

Sub-phylogeny Viz + Paralog Alignments
GCE ID: Event_219
lppA-NC_000962.3:2866468-2867121
lppA,lppB-NC_000962.3:2867124-2867777
Rv2542
lppA
lppB
mada_2-46
mada_1-44
mada_107
mada_1-1
mada_1-51
mada_1-39
mada_122
mada_1-36
mada_117
mada_118
mada_1-10
Event_219
2865600
2866000
2866400
2866800
2867200
2867600
Genomic position (1-based)

Full Phylogeny Viz + Paralog Alignments
GCE ID: Event_219

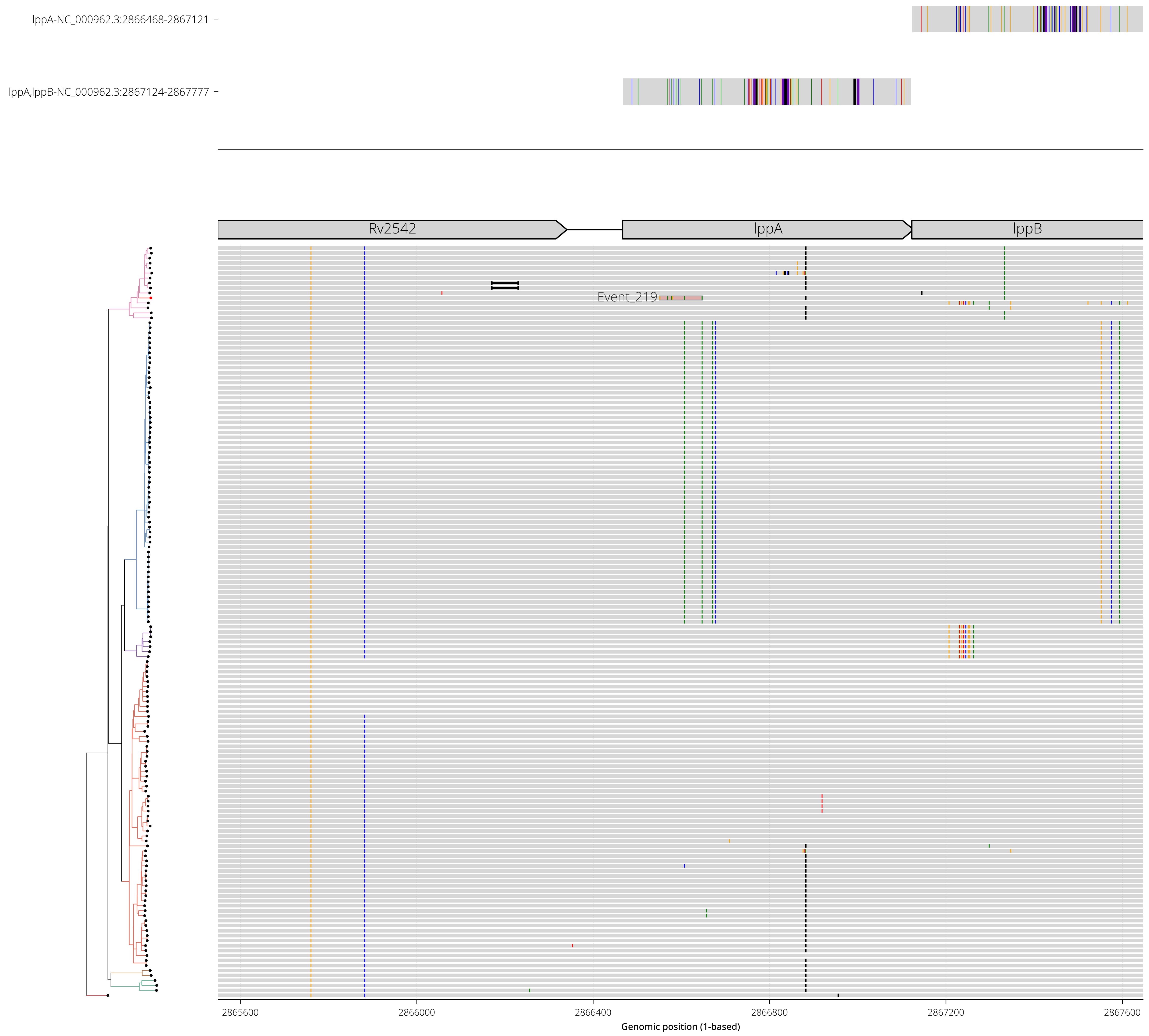

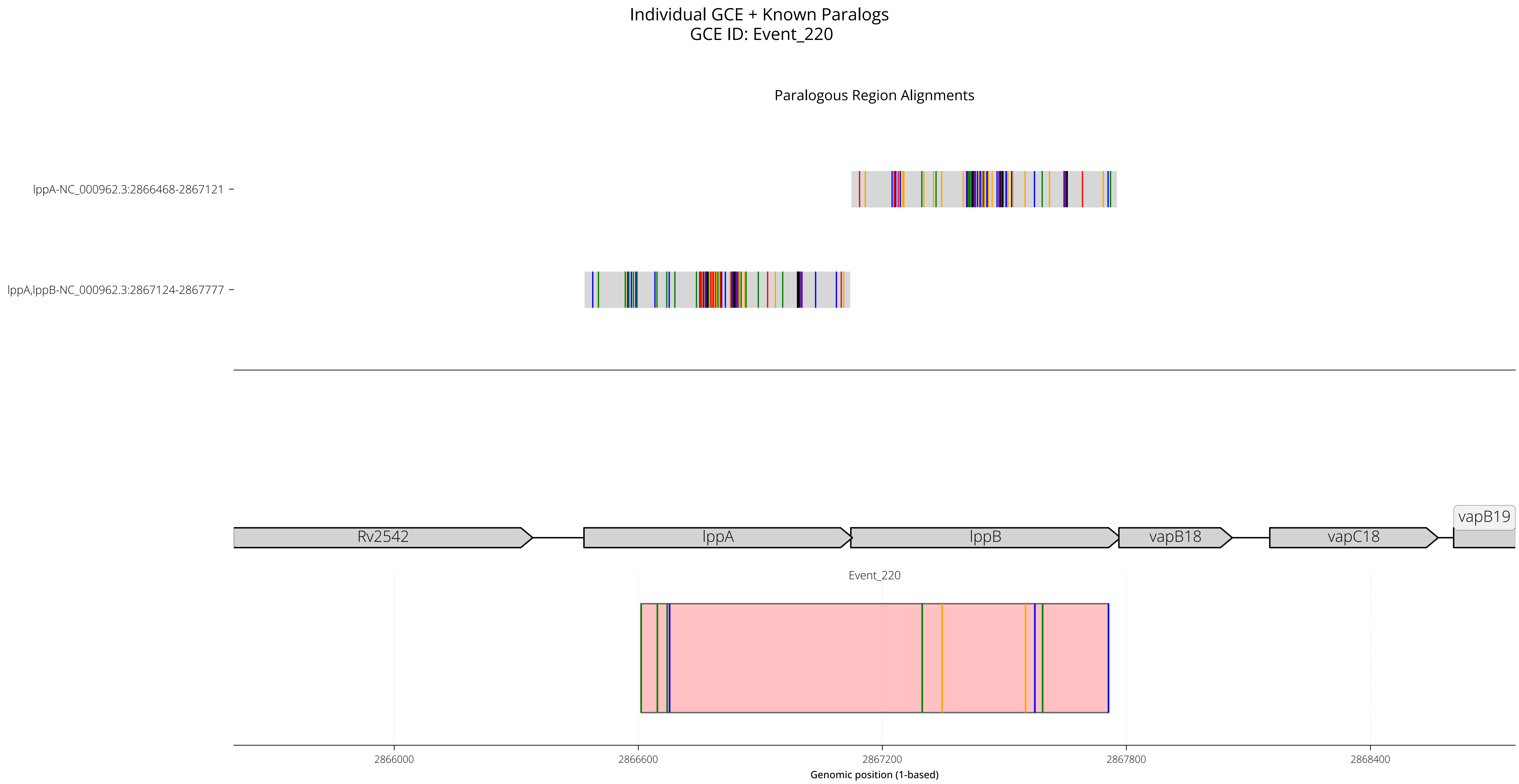
Individual GCE + Known Paralogs
GCE ID: Event_220
Paralogous Region Alignments
lppA-NC_000962.3:2866468-2867121
lppA,lppB-NC_000962.3:2867124-2867777
Rv2542
lppA
lppB
vapB18
vapC18
vapB19
Event_220
2866000
2866600
2867200
2867800
2868400
Genomic position (1-based)

Circos Viz - GCE to Known Paralogs Locations
GCE ID: Event_220

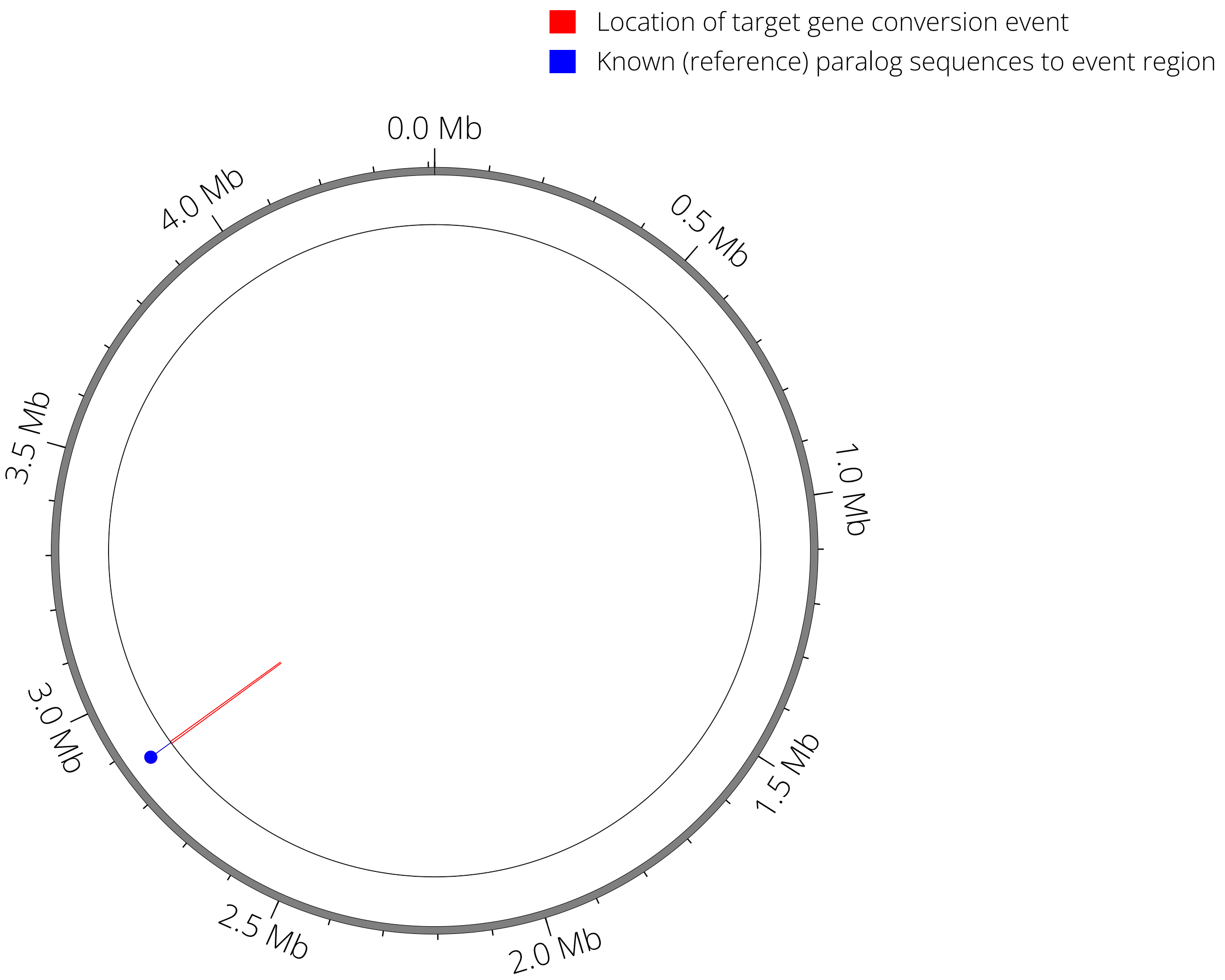

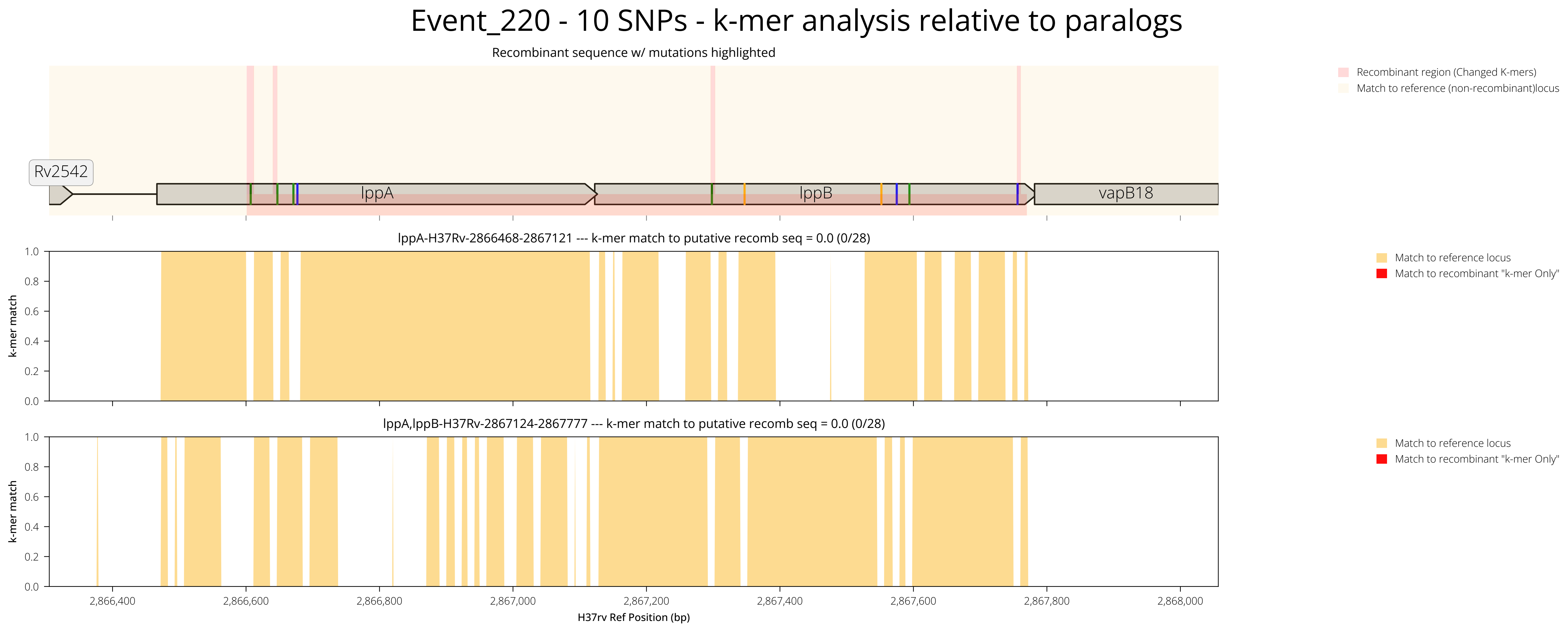
Event_220 - 10 SNPs - k-mer analysis relative to paralogs
Recombinant sequence w/ mutations highlighted
Recombinant region (Changed K-mers)
Match to reference (non-recombinant)locus
Rv2542
lppA
lppB
vapB18
lppA-H37Rv-2866468-2867121 --- k-mer match to putative recomb seq = 0.0 (0/28)
Match to reference locus
Match to recombinant "k-mer Only"
k-mer match
1.0
0.8
0.6
0.4
0.2
0.0
lppA,lppB-H37Rv-2867124-2867777 --- k-mer match to putative recomb seq = 0.0 (0/28)
Match to reference locus
Match to recombinant "k-mer Only"
k-mer match
2,866,400
2,866,600
2,866,800
2,867,000
2,867,200
2,867,400
2,867,600
2,867,800
2,868,000
H37rv Ref Position (bp)

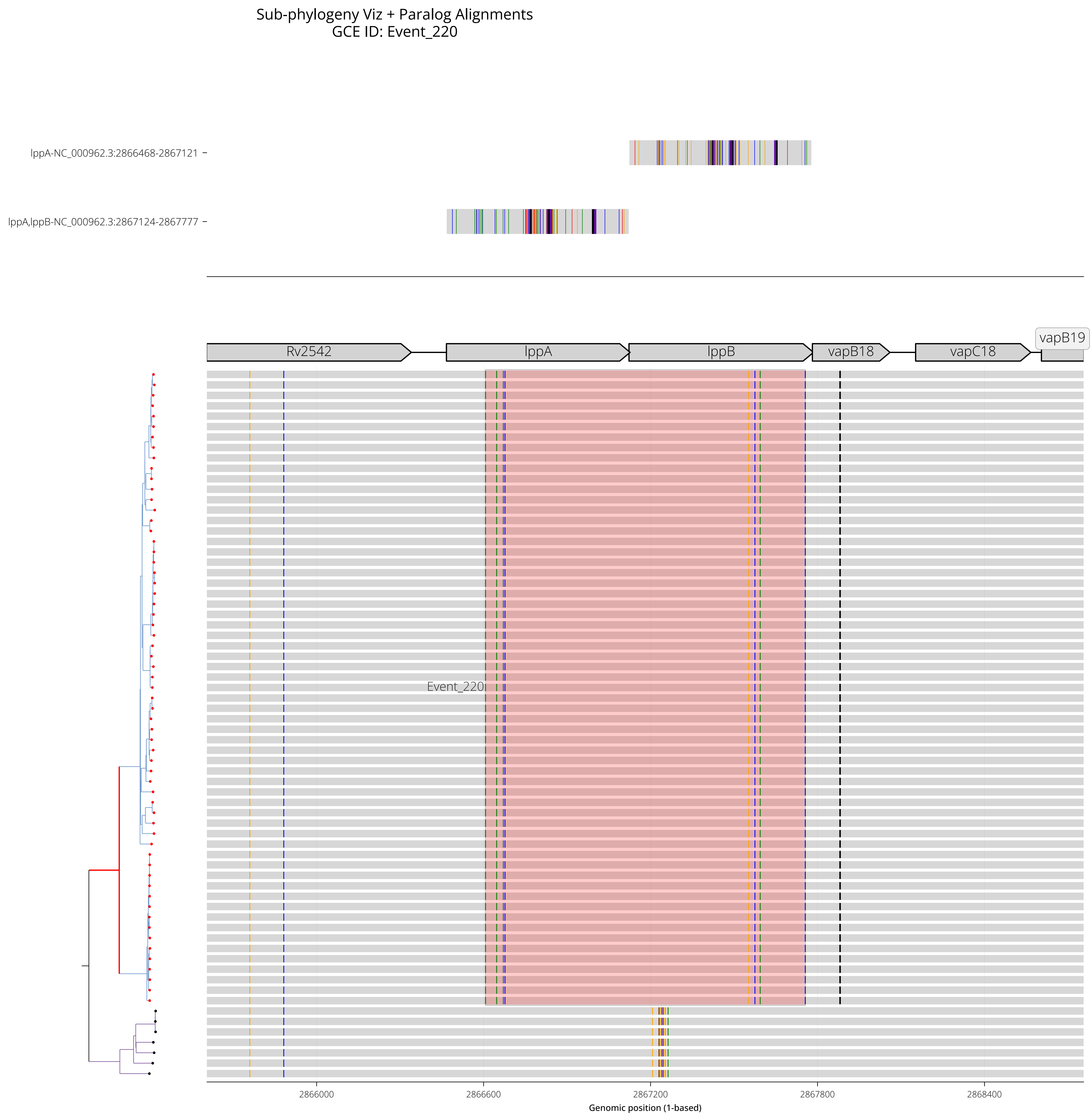
Sub-phylogeny Viz + Paralog Alignments
GCE ID: Event_220
lppA-NC_000962.3:2866468-2867121
lppA,lppB-NC_000962.3:2867124-2867777
Rv2542
lppA
lppB
vapB18
vapC18
vapB19
Event_220
2866000
2866600
2867200
2867800
2868400
Genomic position (1-based)

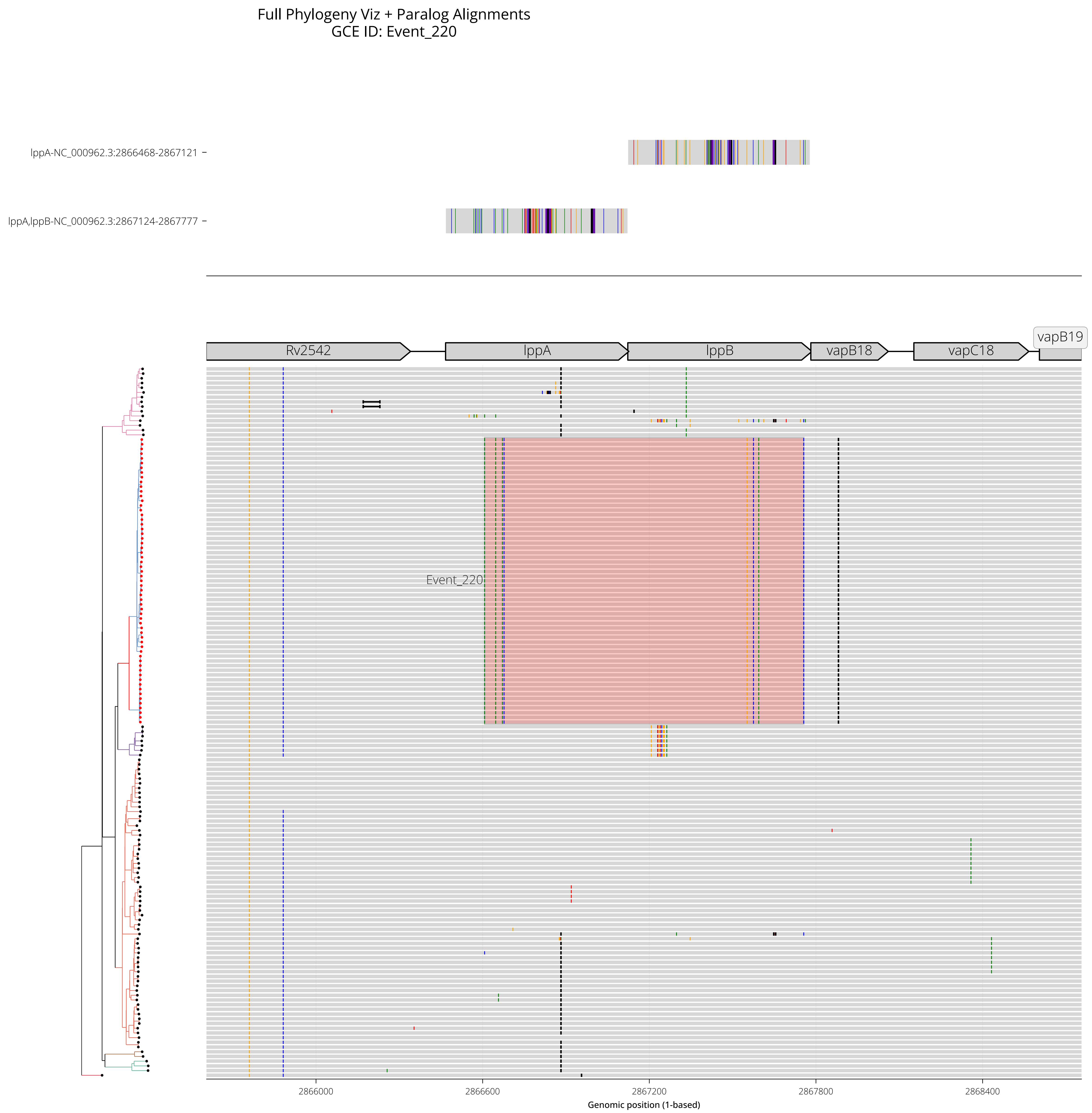

Full Phylogeny Viz + Paralog Alignments
GCE ID: Event_220
lppA-NC_000962.3:2866468-2867121
lppA,lppB-NC_000962.3:2867124-2867777
Rv2542
lppA
lppB
vapB18
vapC18
vapB19
Event_220
2866000
2866600
2867200
2867800
2868400
Genomic position (1-based)

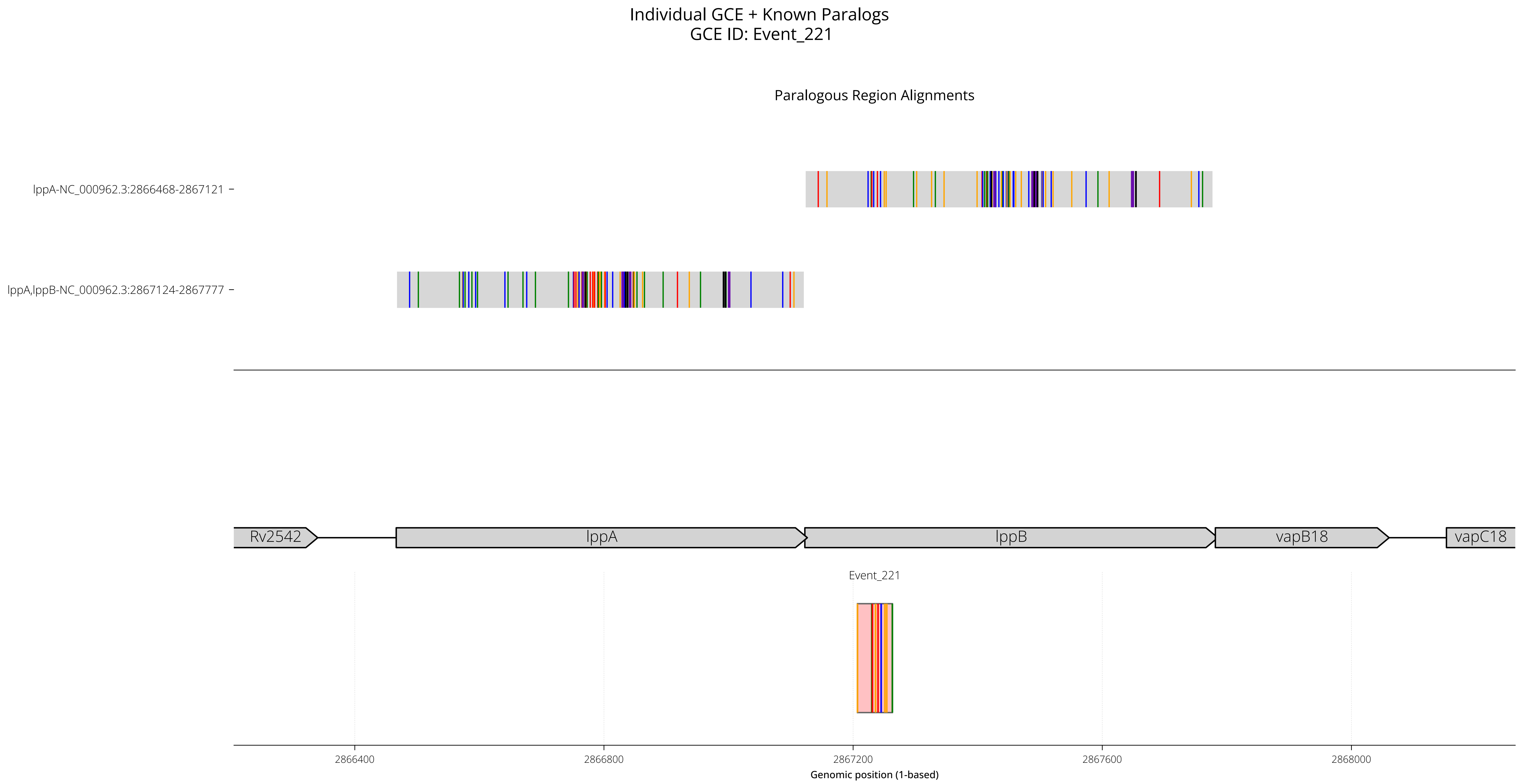

Individual GCE + Known Paralogs
GCE ID: Event_221
Paralogous Region Alignments
lppA-NC_000962.3:2866468-2867121
lppA,lppB-NC_000962.3:2867124-2867777
Rv2542
lppA
lppB
vapB18
vapC18
Event_221
2866400
2866800
2867200
2867600
2868000
Genomic position (1-based)

Circos Viz - GCE to Known Paralogs Locations
GCE ID: Event_221

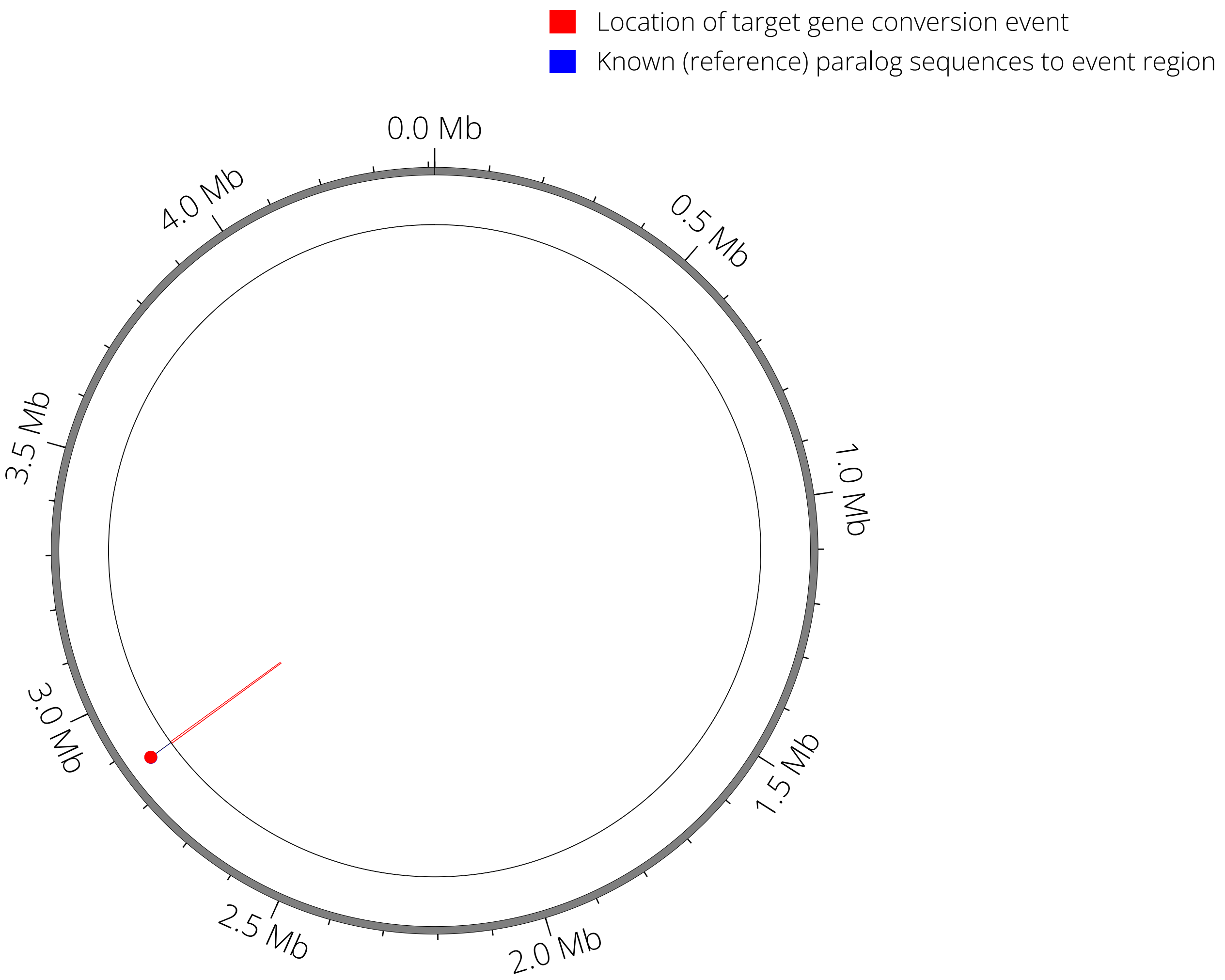

### Event_221 - 9 SNPs - k-mer analysis relative to paralogs

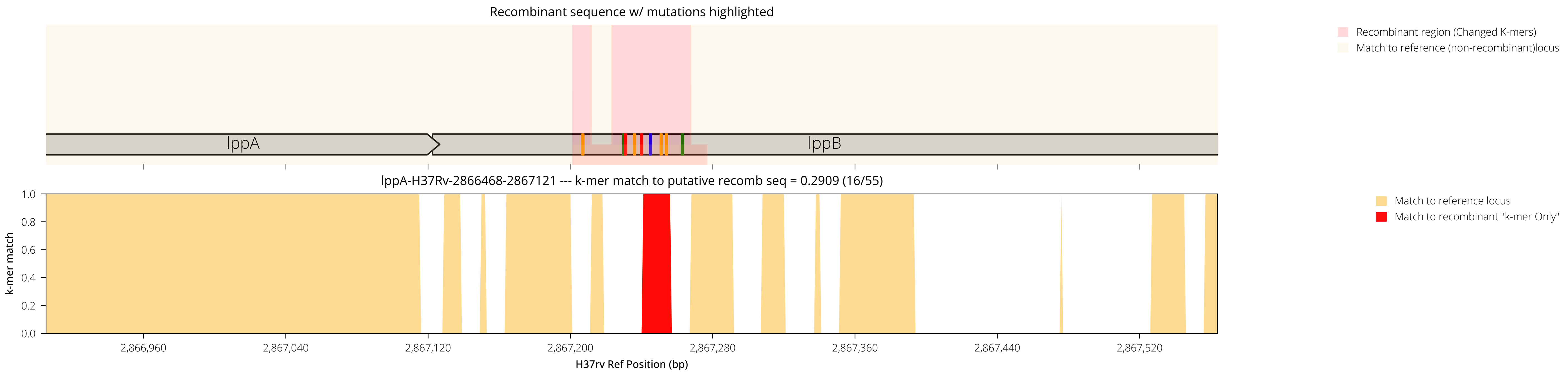

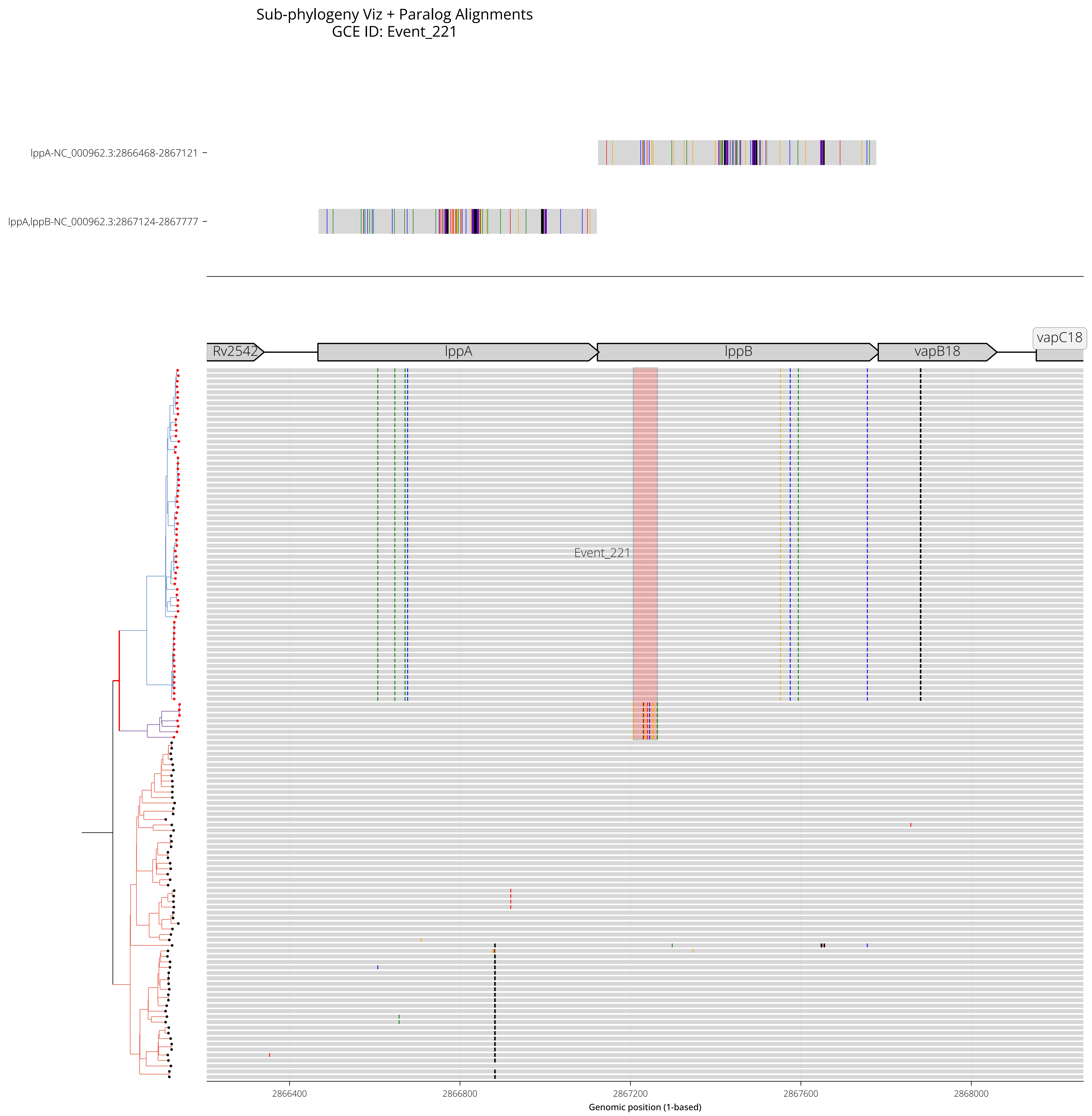
Sub-phylogeny Viz + Paralog Alignments
GCE ID: Event_221
lppA-NC_000962.3:2866468-2867121
lppA,lppB-NC_000962.3:2867124-2867777
Rv2542
lppA
lppB
vapB18
vapC18
Event_221
2866400
2866800
2867200
2867600
2868000
Genomic position (1-based)

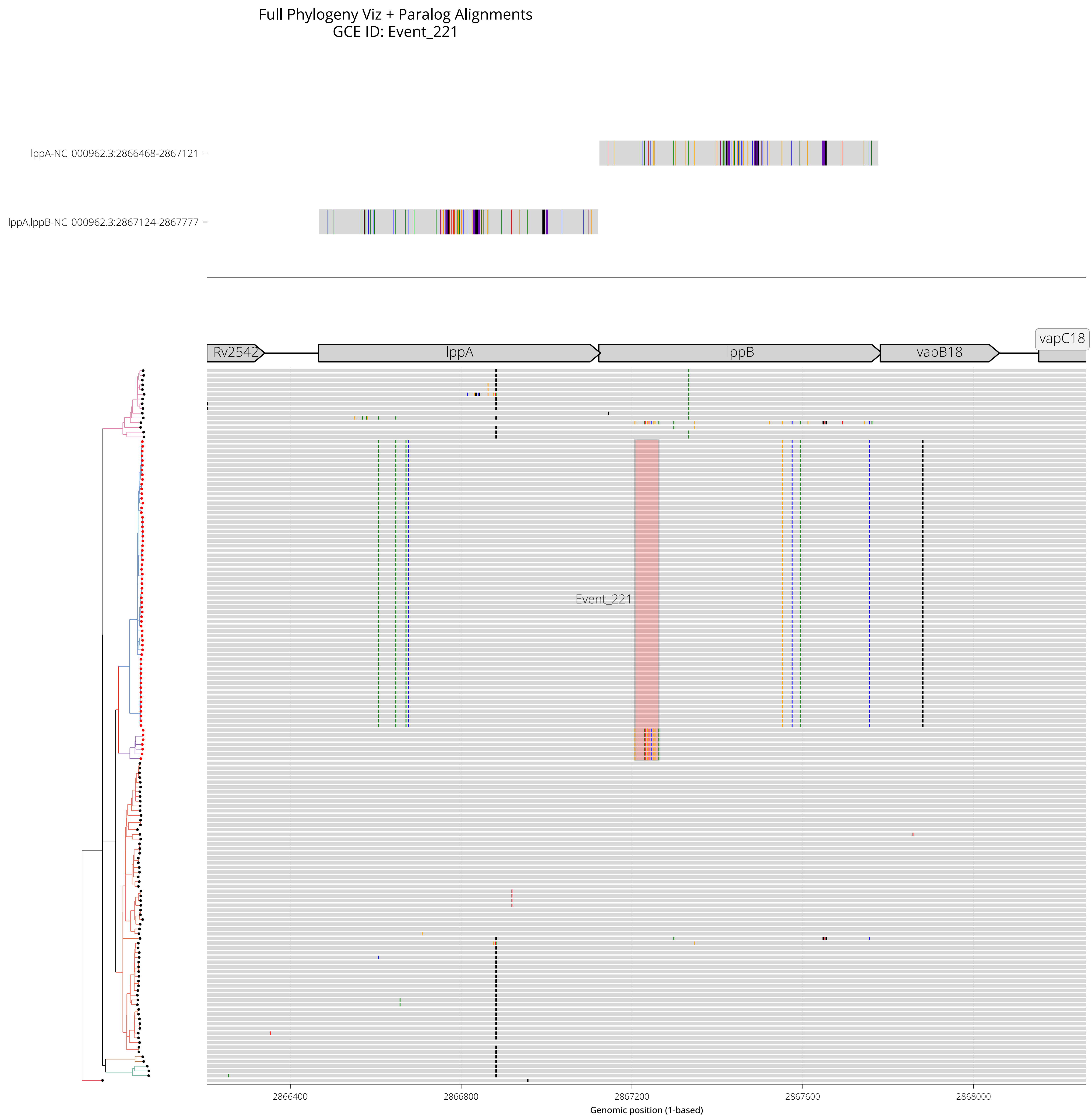
Full Phylogeny Viz + Paralog Alignments
GCE ID: Event_221
lppA-NC_000962.3:2866468-2867121
lppA,lppB-NC_000962.3:2867124-2867777
Rv2542
lppA
lppB
vapB18
vapC18
Event_221
2866400
2866800
2867200
2867600
2868000
Genomic position (1-based)

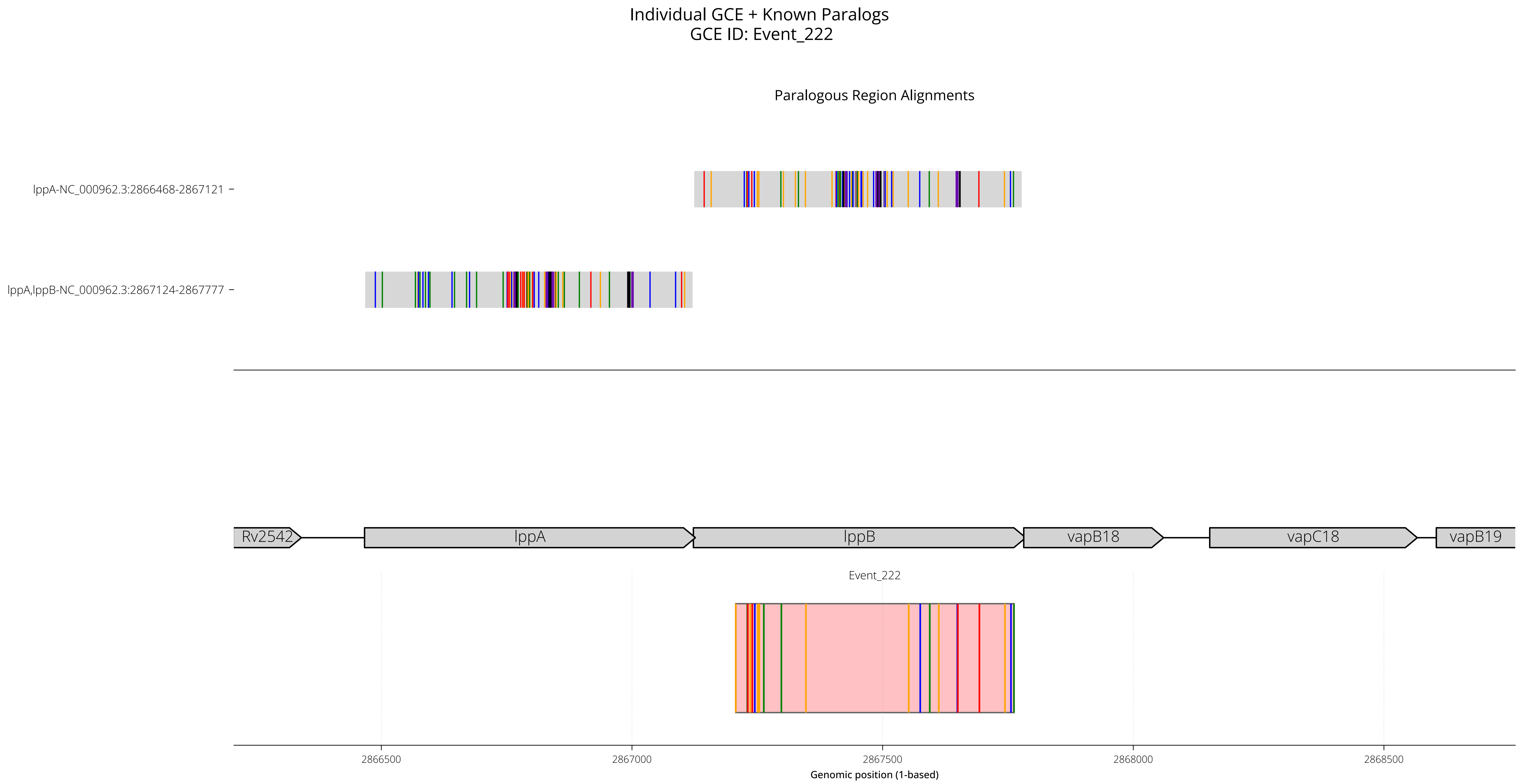

Individual GCE + Known Paralogs
GCE ID: Event_222
Paralogous Region Alignments
lppA-NC_000962.3:2866468-2867121
lppA,lppB-NC_000962.3:2867124-2867777
Rv2542
lppA
lppB
vapB18
vapC18
vapB19
Event_222
2866500
2867000
2867500
2868000
2868500
Genomic position (1-based)

### Circos Viz - GCE to Known Paralogs Locations
GCE ID: Event_222

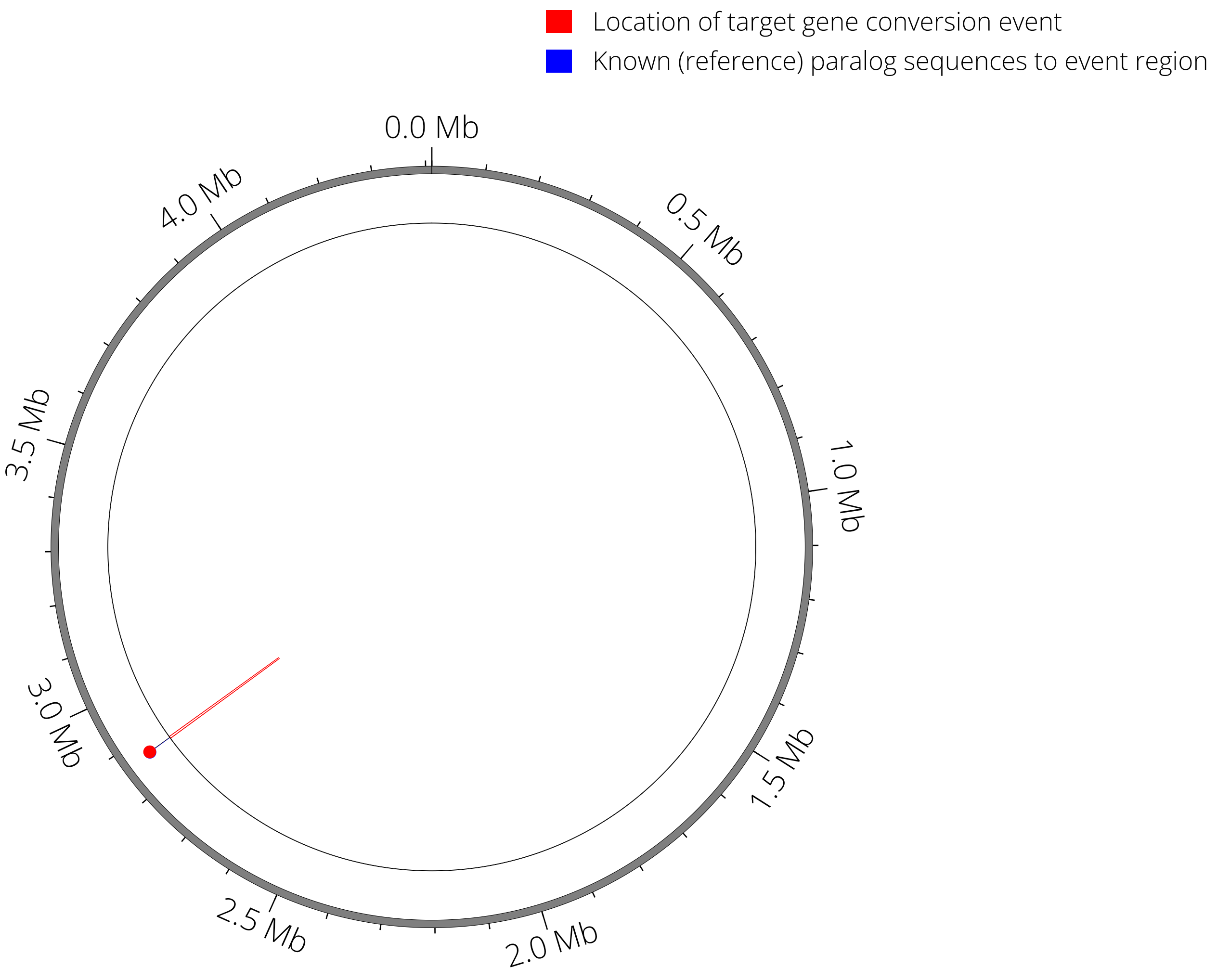

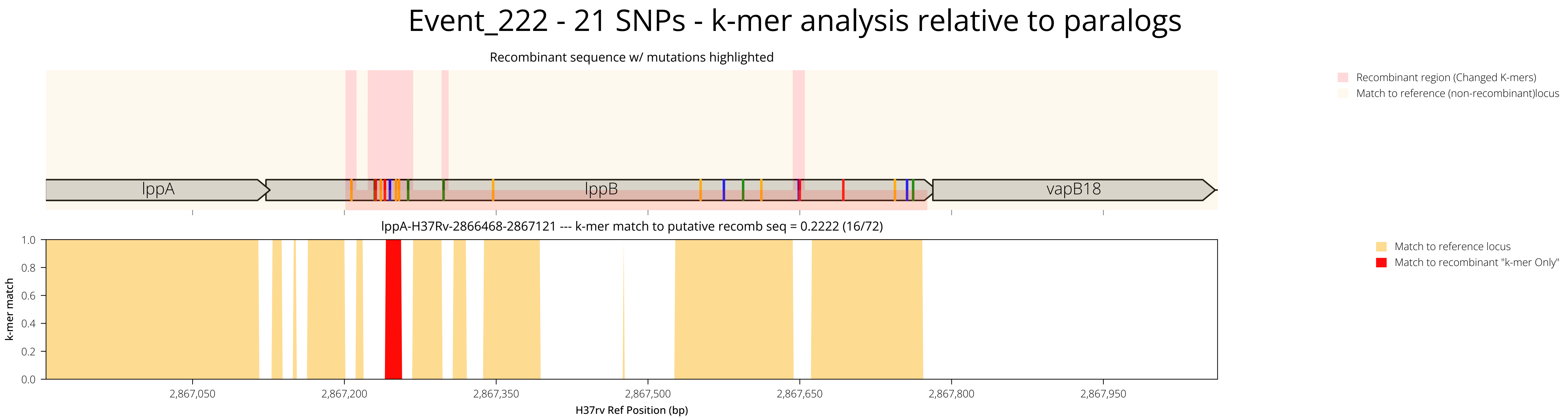
Event_222 - 21 SNPs - k-mer analysis relative to paralogs
Recombinant sequence w/ mutations highlighted
Recombinant region (Changed K-mers)
Match to reference (non-recombinant)locus
lppA
lppB
vapB18
lppA-H37Rv-2866468-2867121 --- k-mer match to putative recomb seq = 0.2222 (16/72)
Match to reference locus
Match to recombinant "k-mer Only"
k-mer match
1.0
0.8
0.6
0.4
0.2
0.0
2,867,050
2,867,200
2,867,350
2,867,500
2,867,650
2,867,800
2,867,950
H37rv Ref Position (bp)

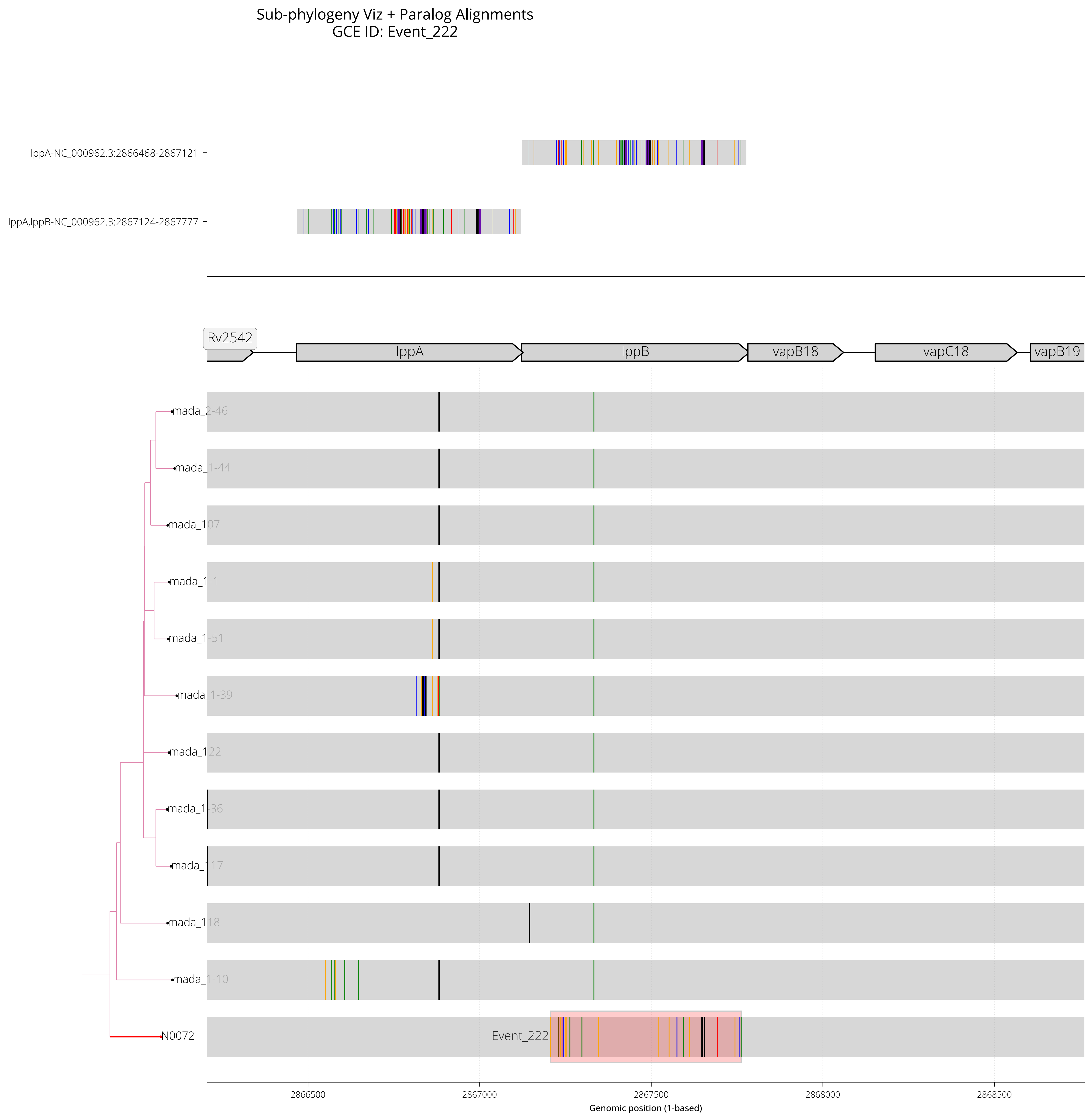
Sub-phylogeny Viz + Paralog Alignments
GCE ID: Event_222
lppA-NC_000962.3:2866468-2867121
lppA,lppB-NC_000962.3:2867124-2867777
Rv2542
lppA
lppB
vapB18
vapC18
vapB19
mada_2-46
mada_1-44
mada_107
mada_1-1
mada_1-51
mada_1-39
mada_122
mada_1-36
mada_117
mada_118
mada_1-10
N0072
Event_222
2866500
2867000
2867500
2868000
2868500
Genomic position (1-based)

Full Phylogeny Viz + Paralog Alignments
GCE ID: Event_222

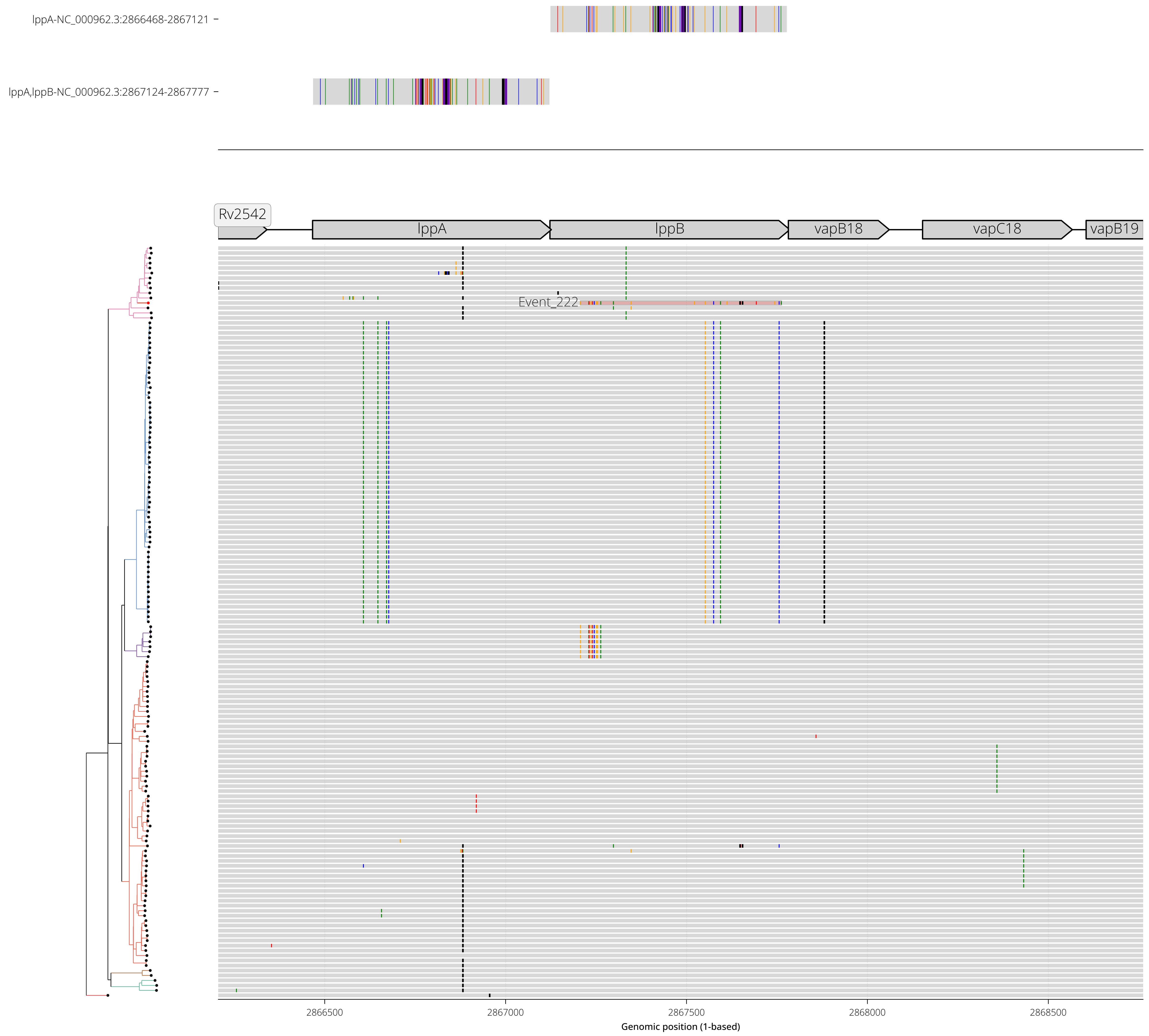

Individual GCE + Known Paralogs
GCE ID: Event_223

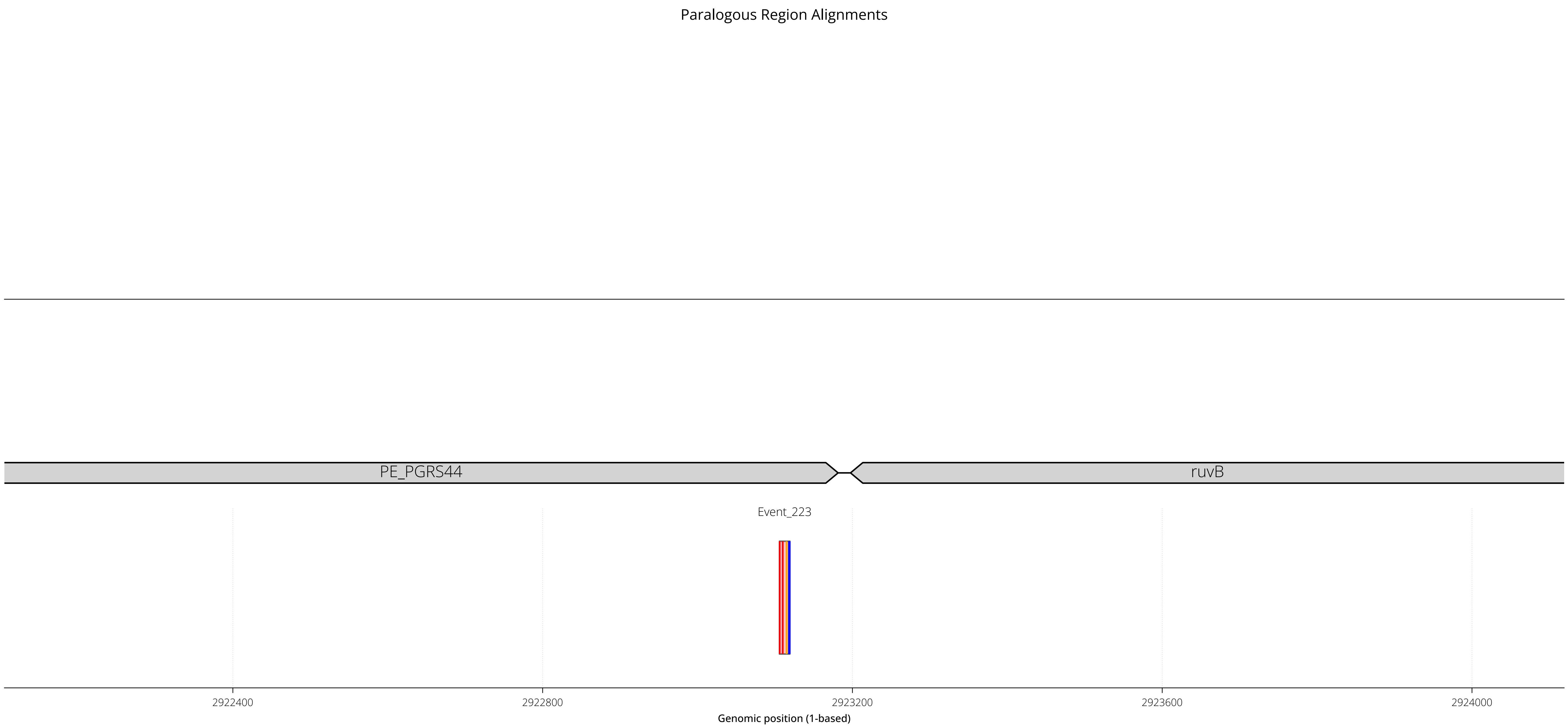

Sub-phylogeny Viz + Paralog Alignments
GCE ID: Event_223

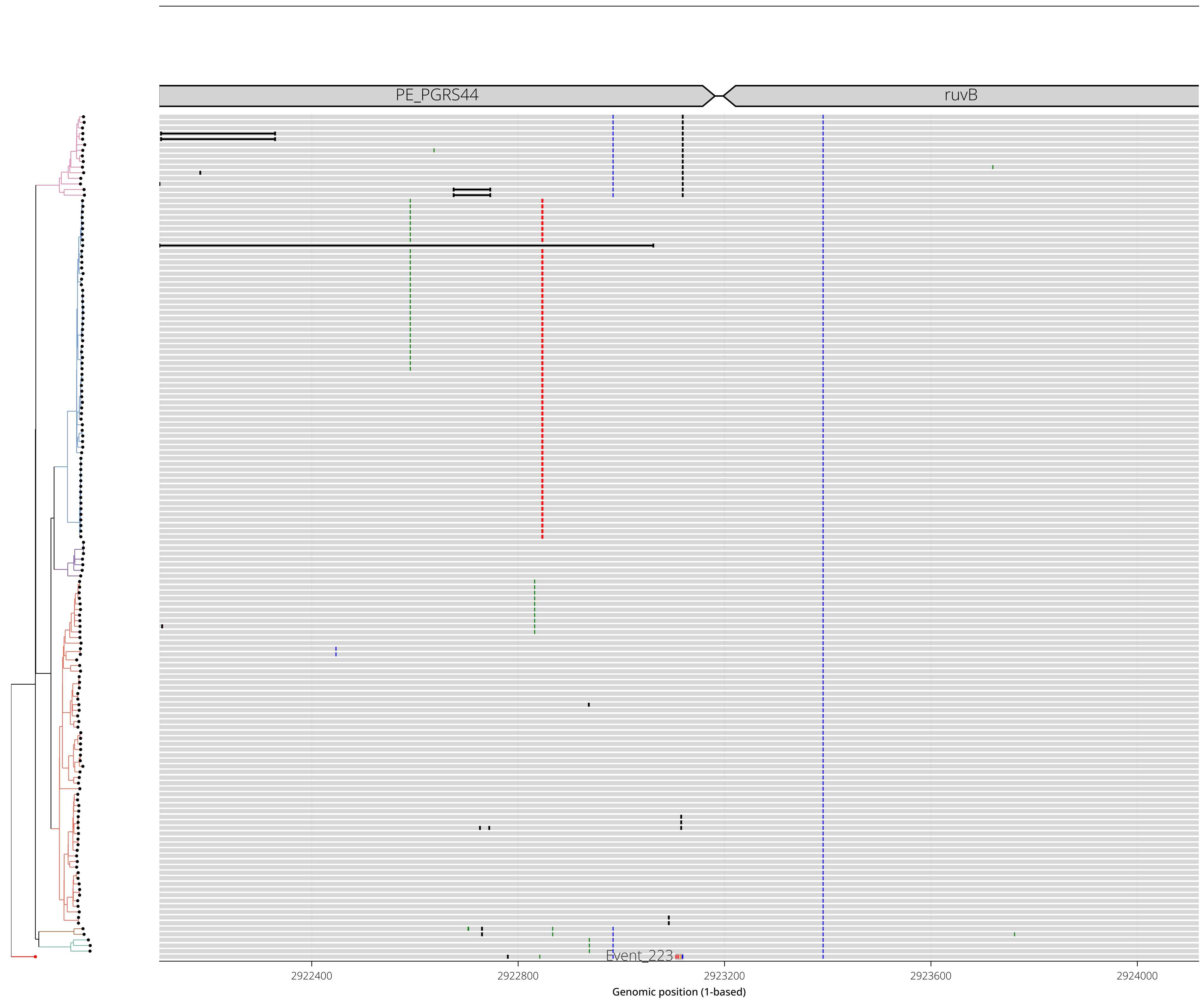

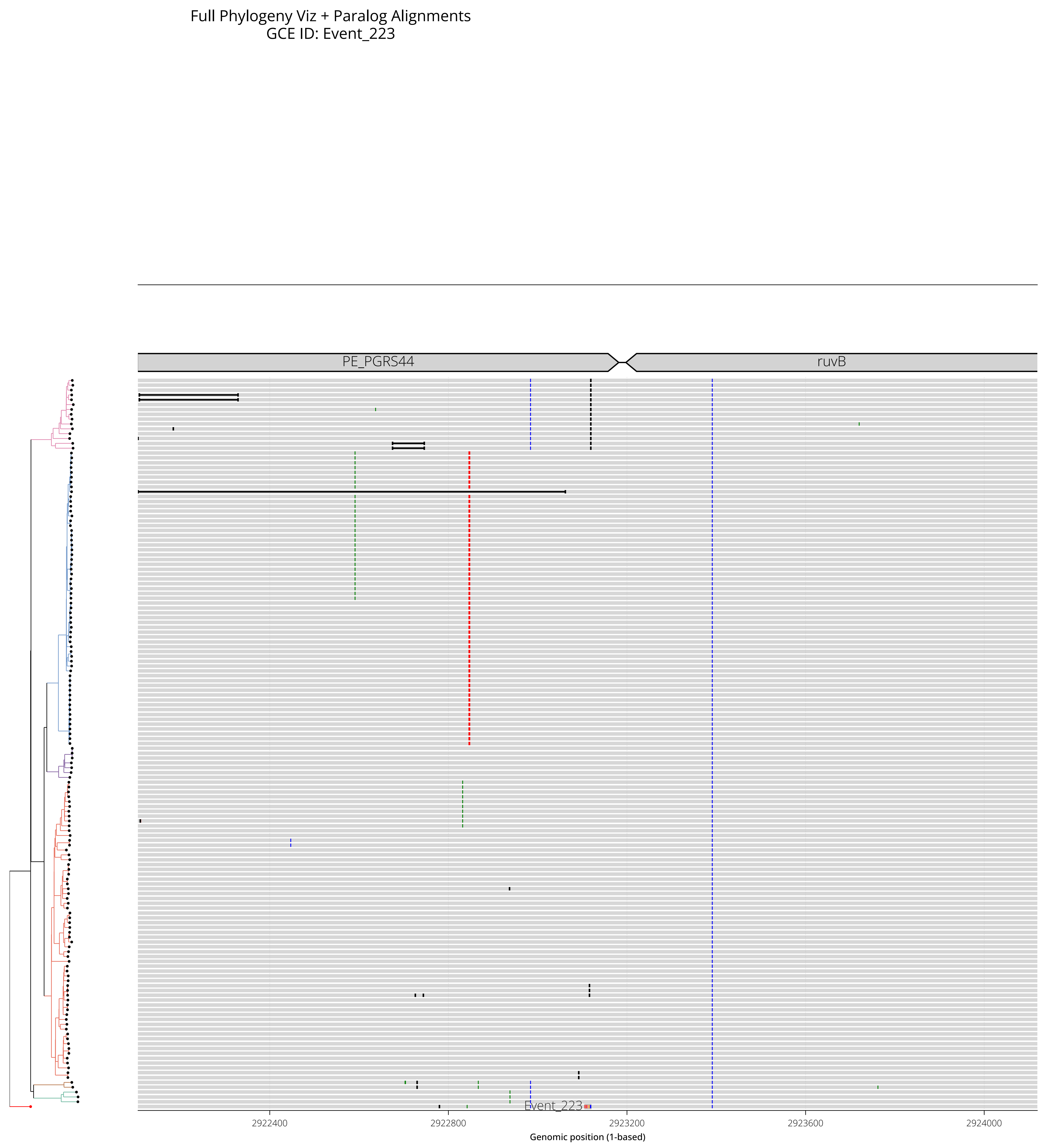
Full Phylogeny Viz + Paralog Alignments
GCE ID: Event_223
PE_PGRS44
ruvB
Event_223
2922400
2922800
2923200
2923600
2924000
Genomic position (1-based)

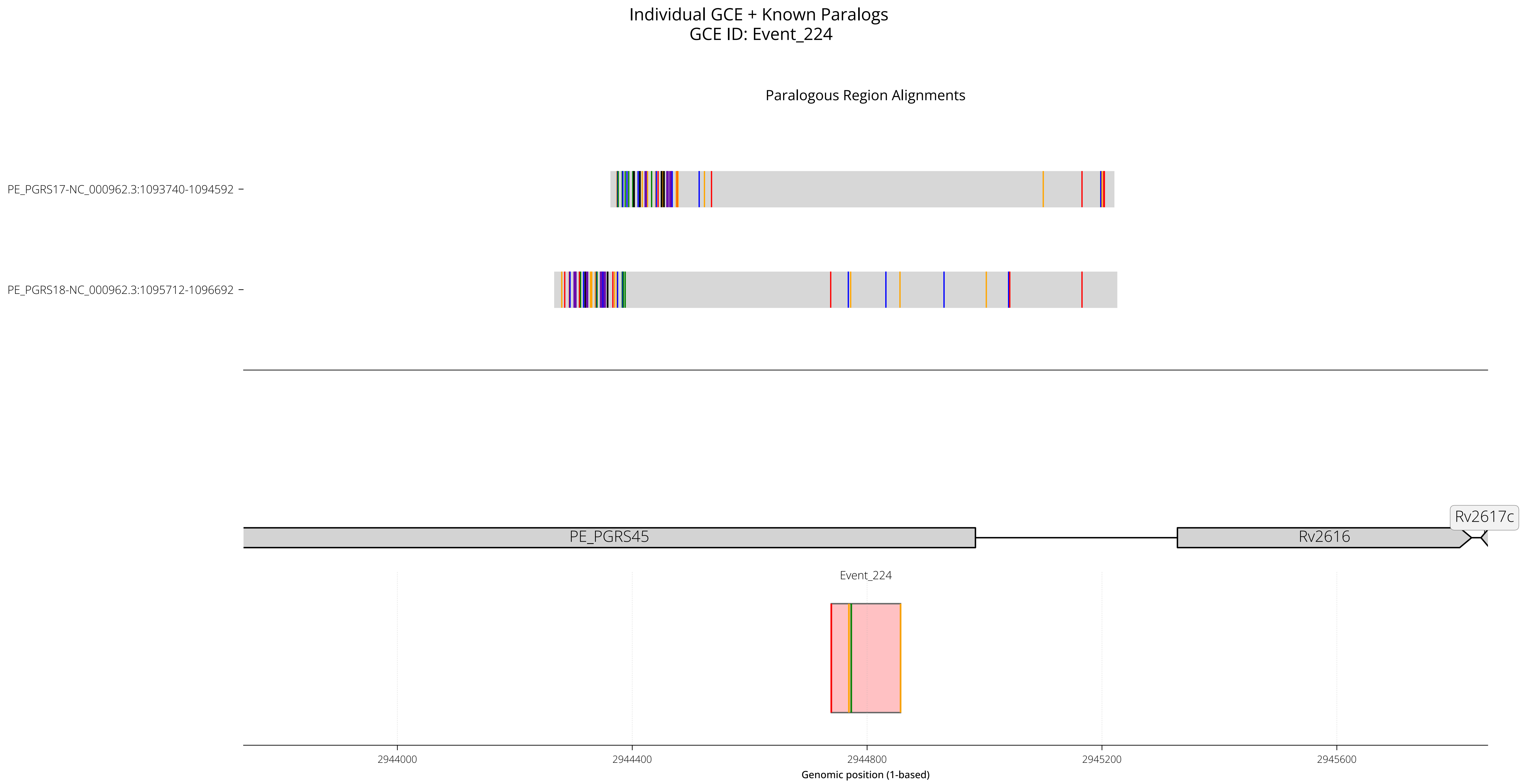

Individual GCE + Known Paralogs
GCE ID: Event_224
Paralogous Region Alignments
PE_PGRS17-NC_000962.3:1093740-1094592
PE_PGRS18-NC_000962.3:1095712-1096692
PE_PGRS45
Rv2616
Rv2617c
Event_224
2944000
2944400
2944800
2945200
2945600
Genomic position (1-based)

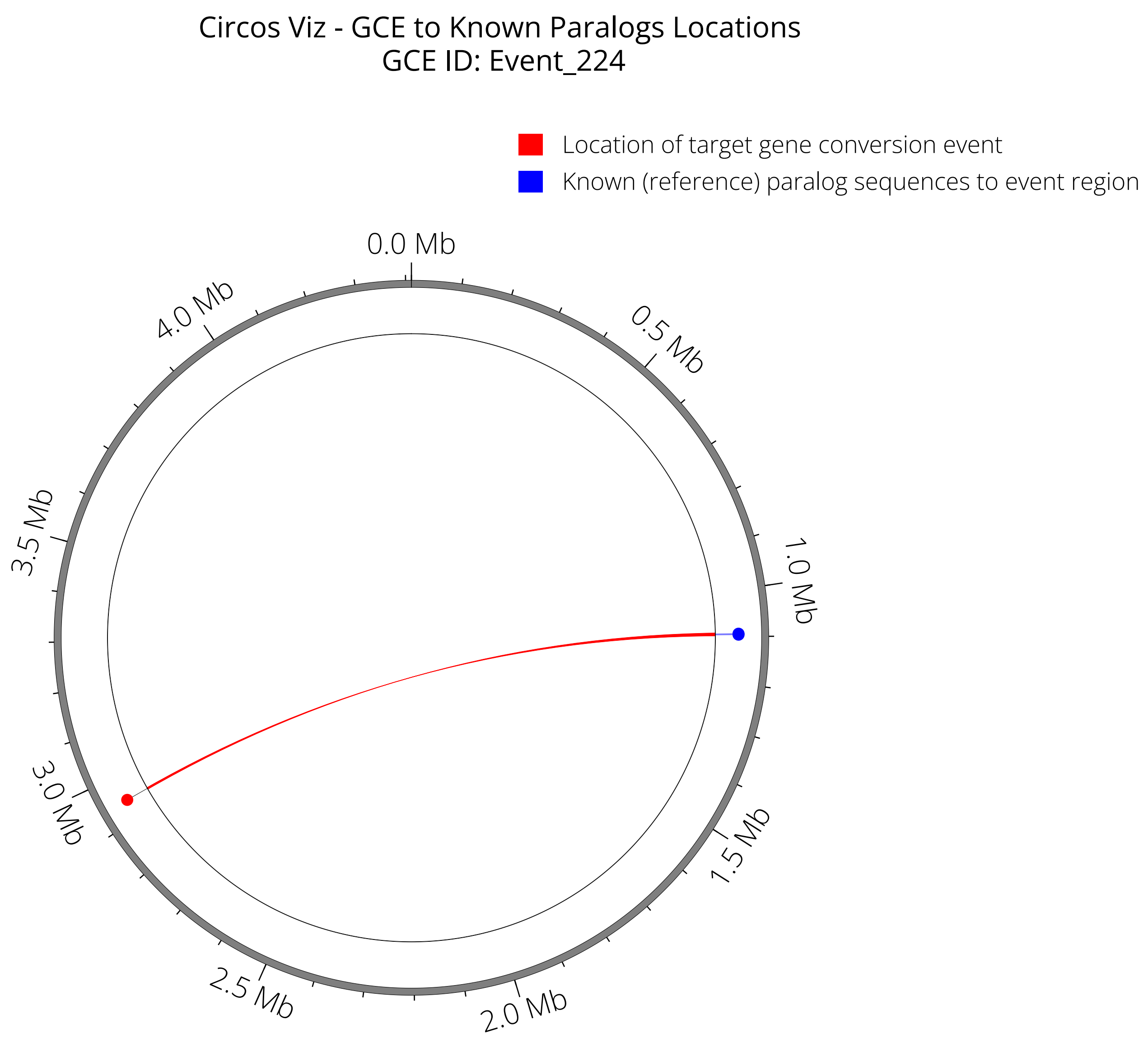
Circos Viz - GCE to Known Paralogs Locations
GCE ID: Event_224
Location of target gene conversion event
Known (reference) paralog sequences to event region
0.0 Mb
0.5 Mb
1.0 Mb
1.5 Mb
2.0 Mb
2.5 Mb
3.0 Mb
3.5 Mb
4.0 Mb

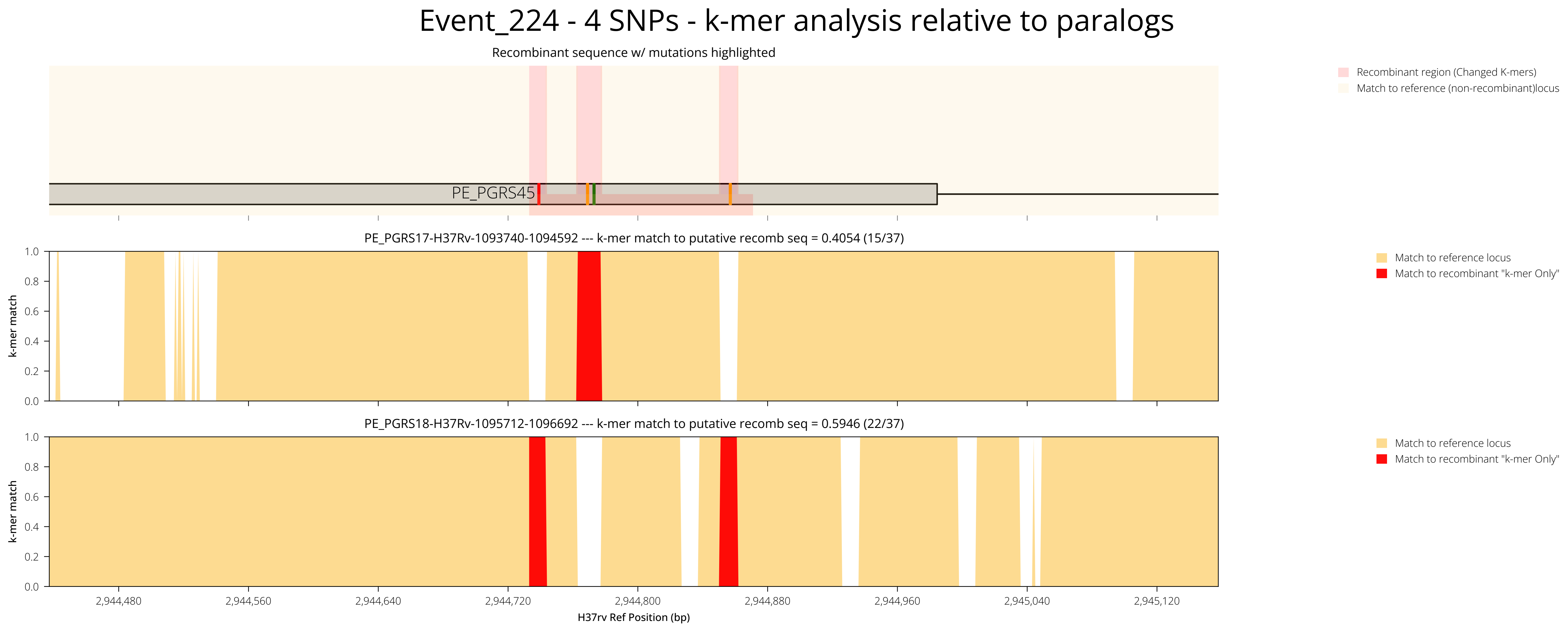
Event_224 - 4 SNPs - k-mer analysis relative to paralogs
Recombinant sequence w/ mutations highlighted
PE_PGRS45
Recombinant region (Changed K-mers)
Match to reference (non-recombinant)locus
PE_PGRS17-H37Rv-1093740-1094592 --- k-mer match to putative recomb seq = 0.4054 (15/37)
Match to reference locus
Match to recombinant "k-mer Only"
PE_PGRS18-H37Rv-1095712-1096692 --- k-mer match to putative recomb seq = 0.5946 (22/37)
Match to reference locus
Match to recombinant "k-mer Only"
k-mer match
1.0
0.8
0.6
0.4
0.2
0.0
2,944,480
2,944,560
2,944,640
2,944,720
2,944,800
2,944,880
2,944,960
2,945,040
2,945,120
H37rv Ref Position (bp)

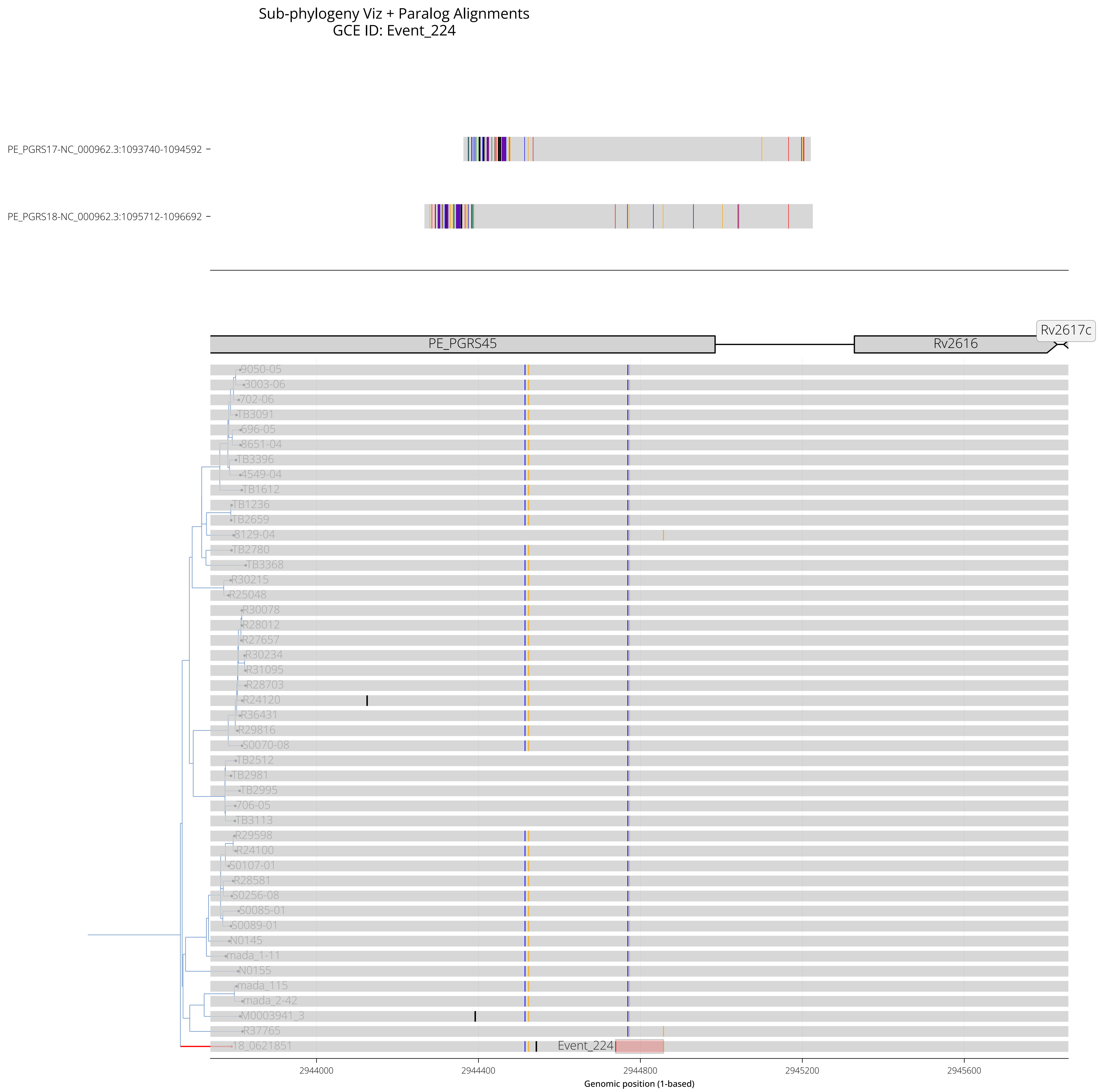

Sub-phylogeny Viz + Paralog Alignments
GCE ID: Event_224
PE_PGRS17-NC_000962.3:1093740-1094592
PE_PGRS18-NC_000962.3:1095712-1096692
PE_PGRS45
Rv2616
Rv2617c
9050-05
3003-06
702-06
TB3091
696-05
8651-04
TB3396
4549-04
TB1612
TB1236
TB2659
8129-04
TB2780
TB3368
R30215
R25048
R30078
R28012
R27657
R30234
R31095
R28703
R24120
R36431
R29816
S0070-08
TB2512
TB2981
TB2995
706-05
TB3113
R29598
R24100
S0107-01
R28581
S0256-08
S0085-01
S0089-01
N0145
mada_1-11
N0155
mada_115
mada_2-42
M0003941_3
R37765
18_0621851
Event_224
2944000
2944400
2944800
2945200
2945600
Genomic position (1-based)

Full Phylogeny Viz + Paralog Alignments
GCE ID: Event_224

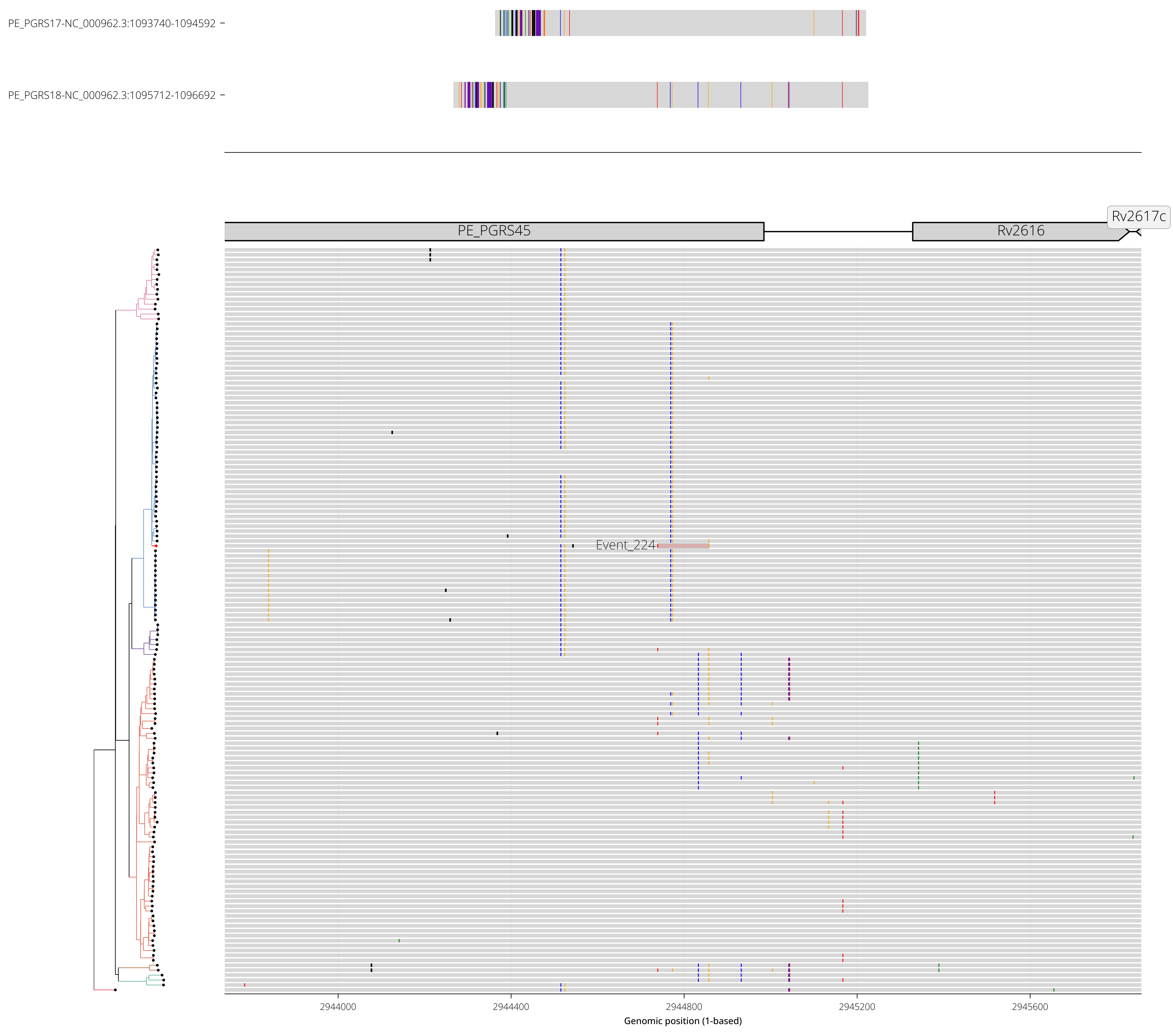

### Individual GCE + Known Paralogs
GCE ID: Event_225

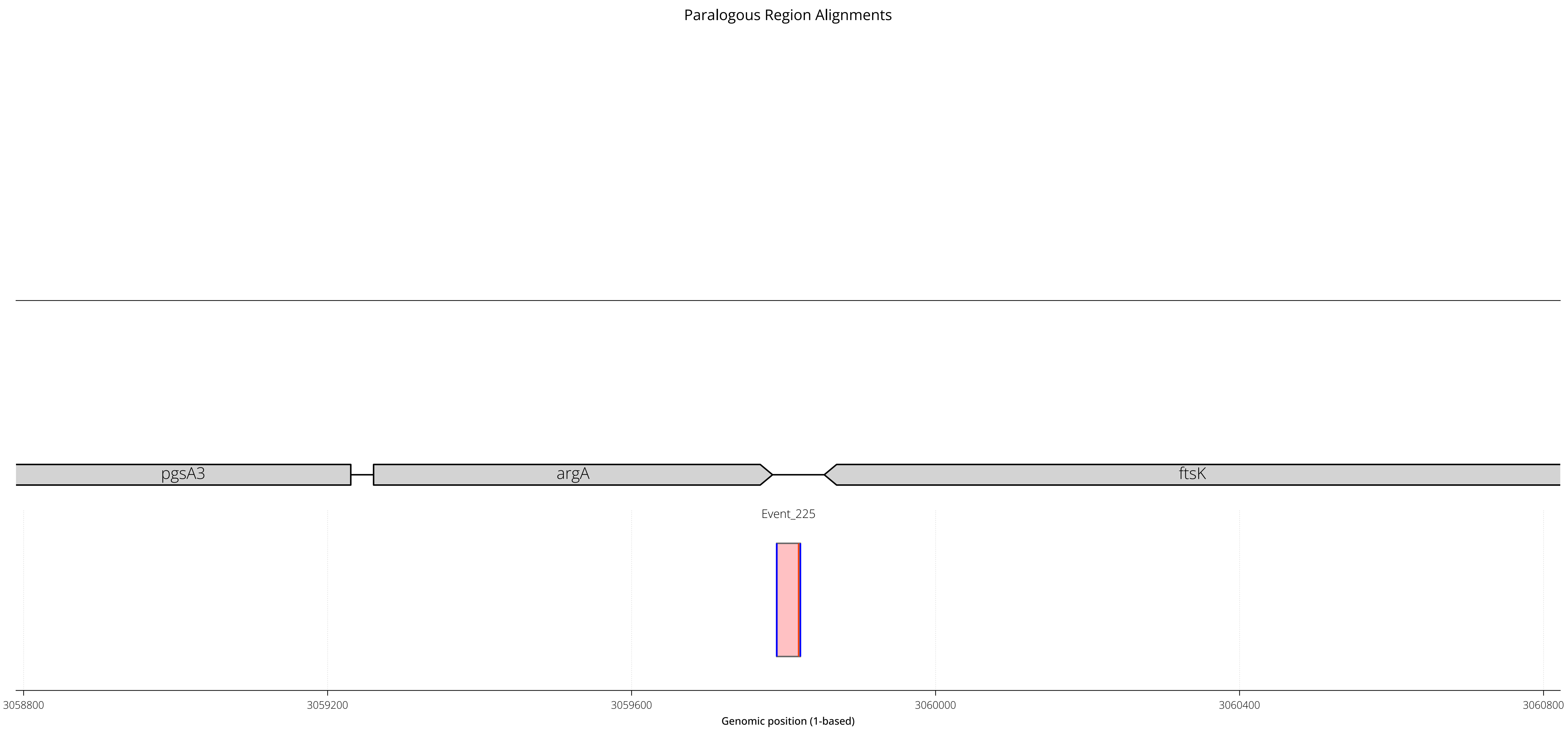

Sub-phylogeny Viz + Paralog Alignments
GCE ID: Event_225

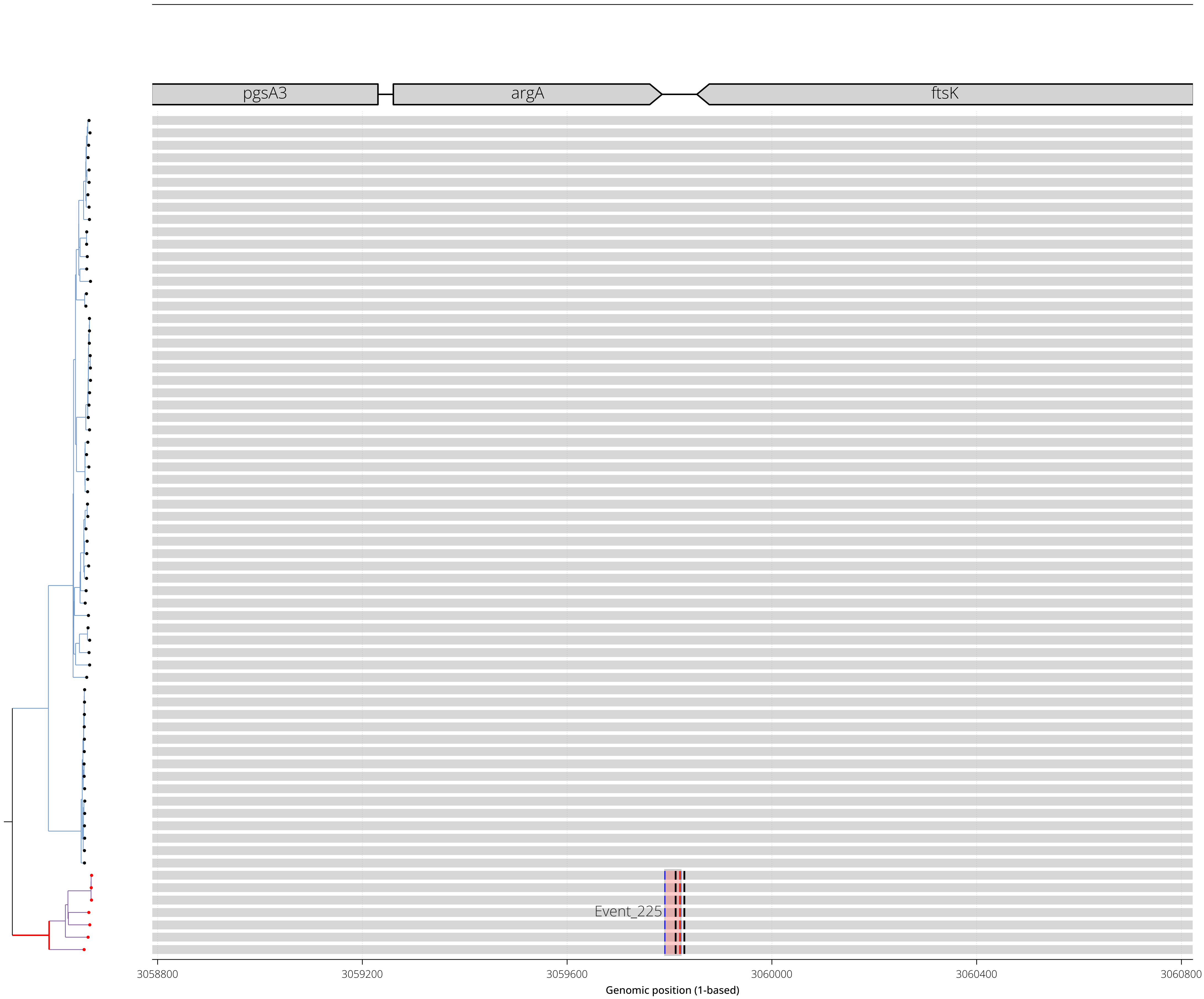

Full Phylogeny Viz + Paralog Alignments
GCE ID: Event_225

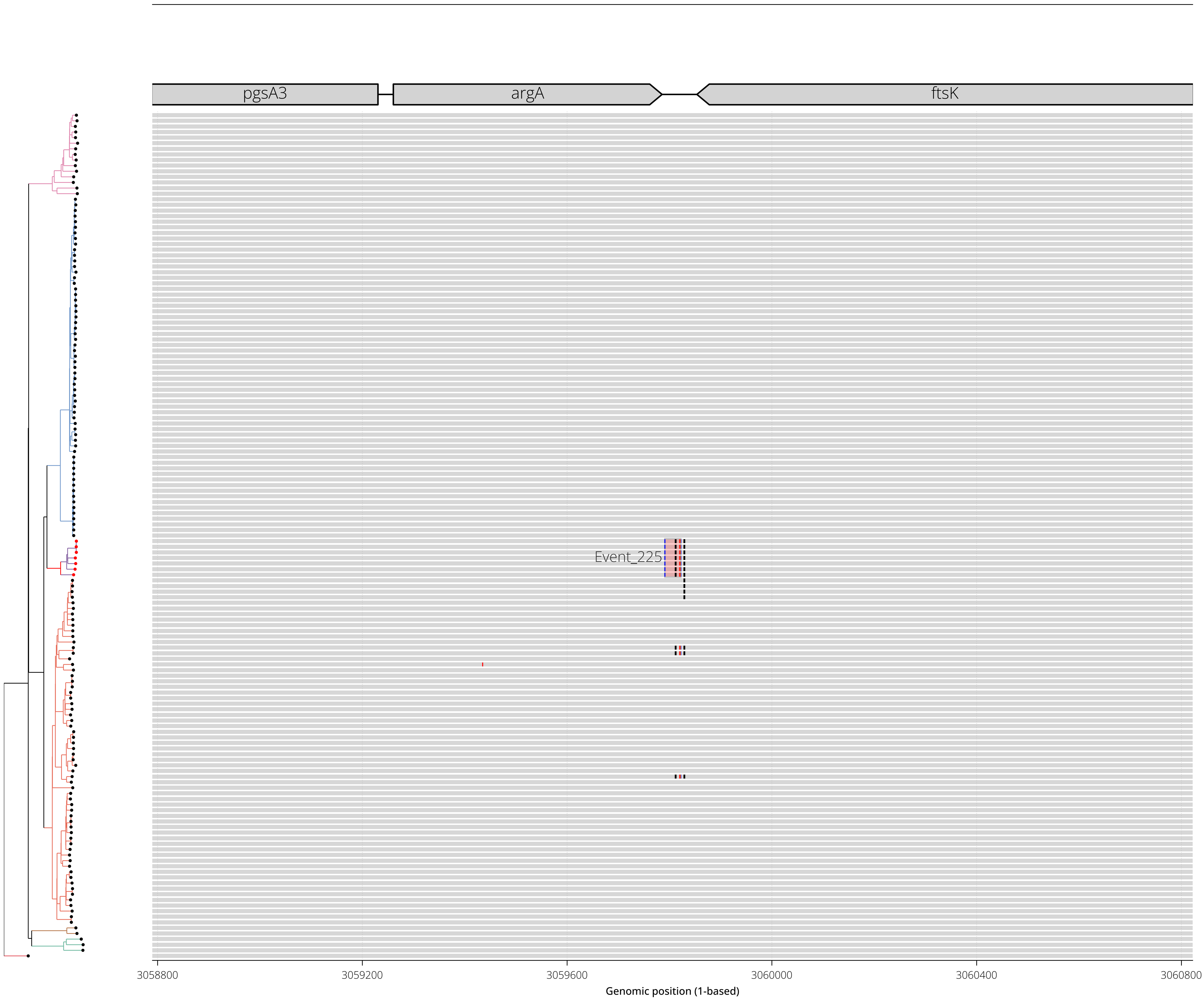

Individual GCE + Known Paralogs
GCE ID: Event_226

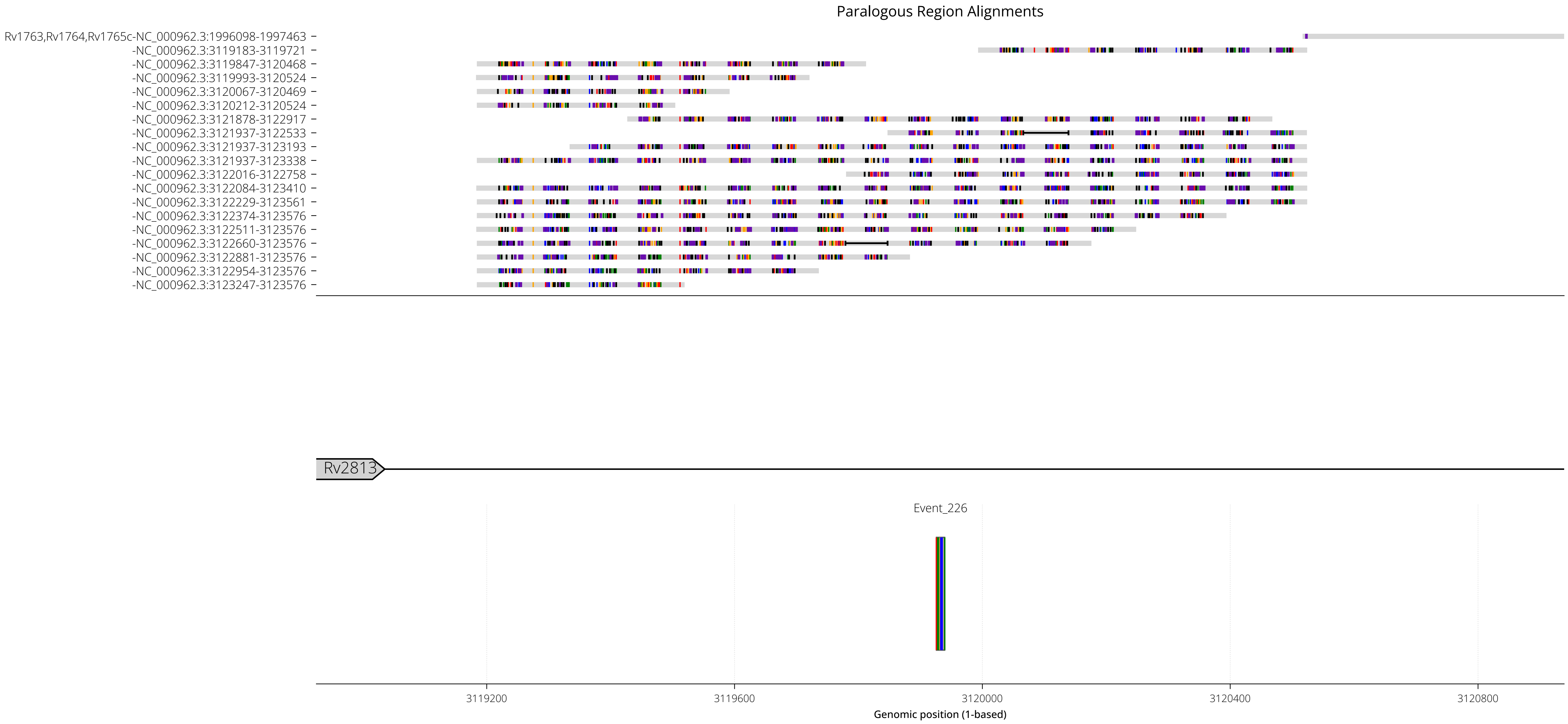

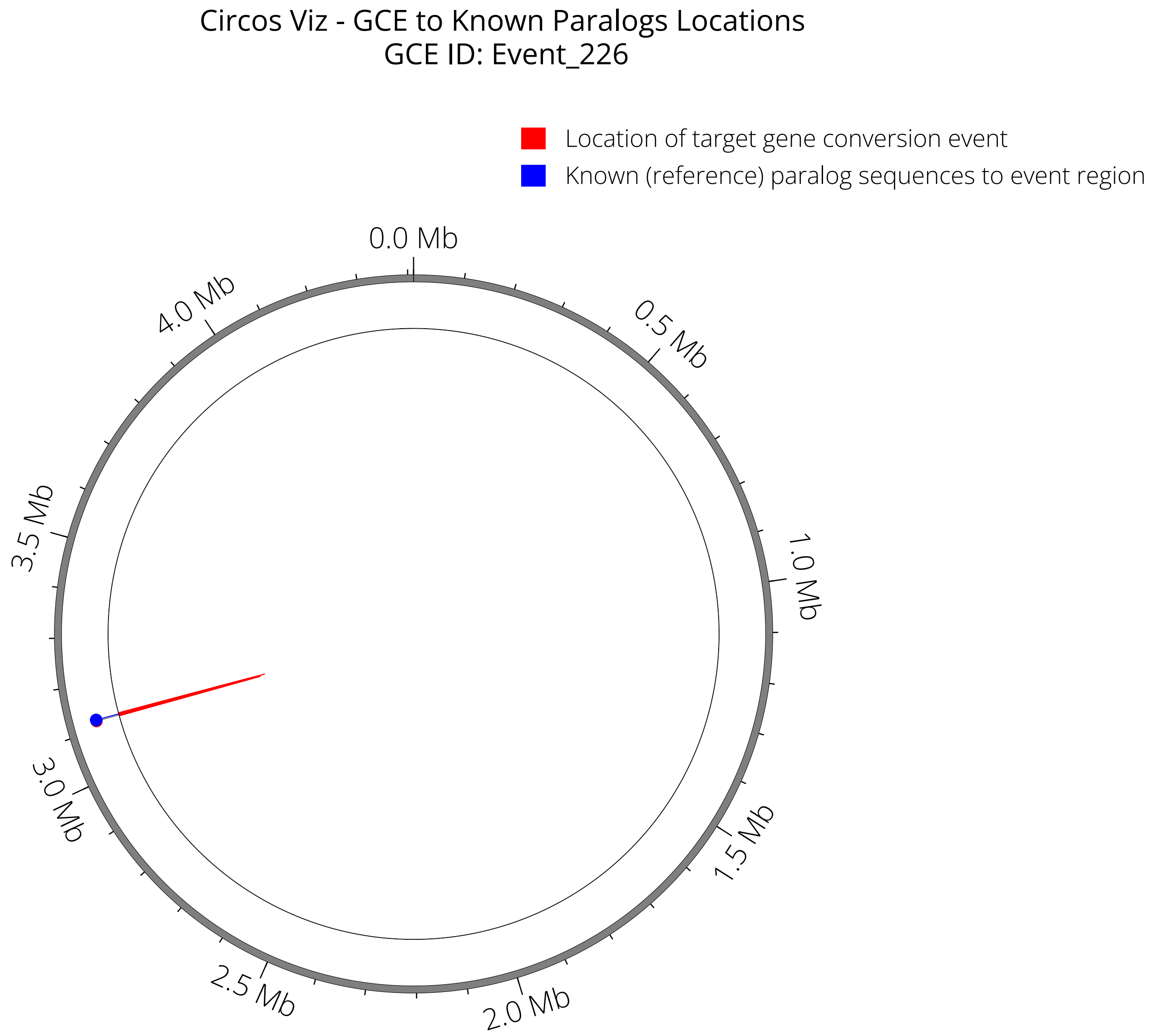
Circos Viz - GCE to Known Paralogs Locations
GCE ID: Event_226
Location of target gene conversion event
Known (reference) paralog sequences to event region
0.0 Mb
0.5 Mb
1.0 Mb
1.5 Mb
2.0 Mb
2.5 Mb
3.0 Mb
3.5 Mb
4.0 Mb

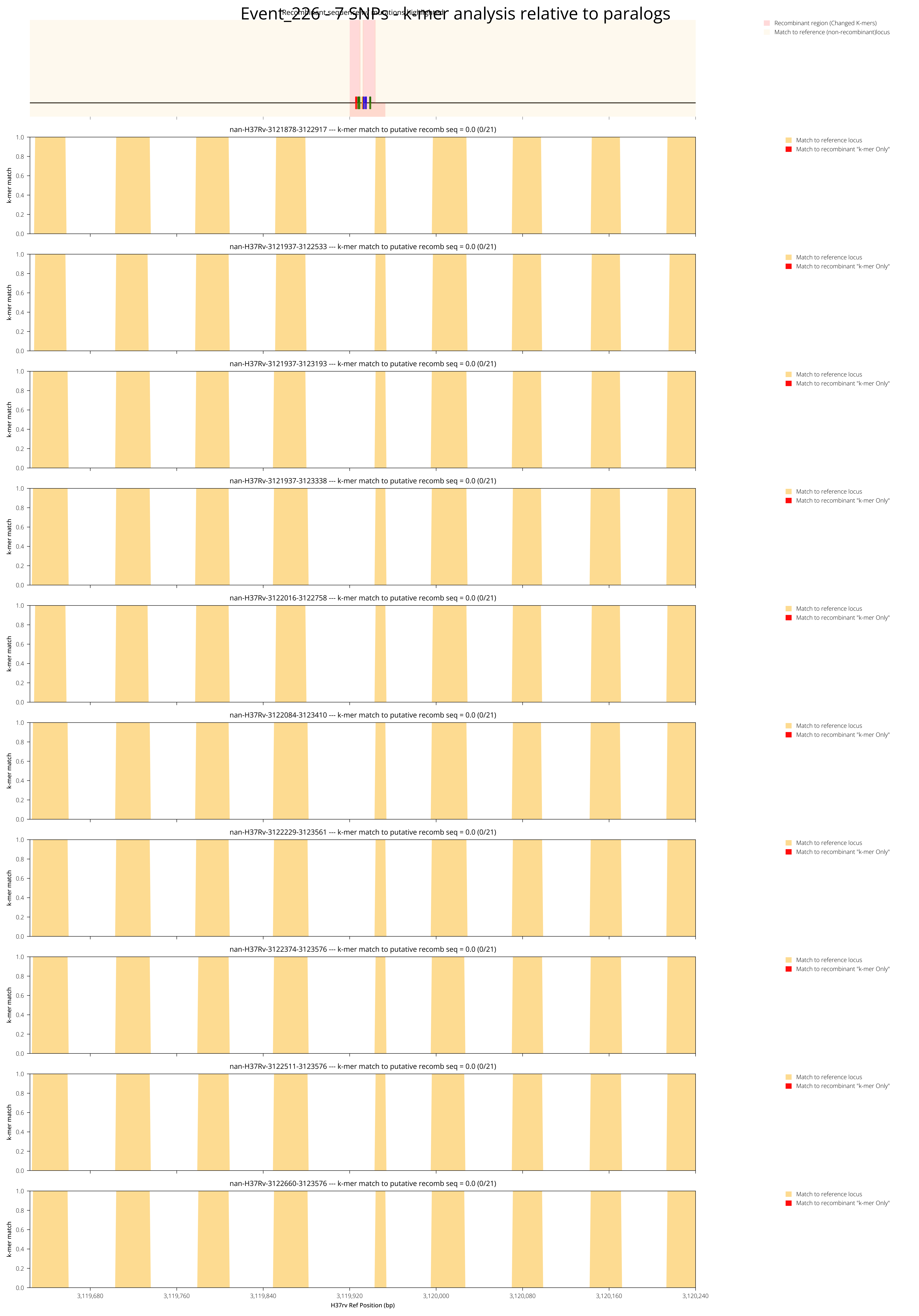

Event_226 - 7 SNPs - k-mer analysis relative to paralogs
Recombinant sequence w/ mutations highlighted
Recombinant region (Changed K-mers)
Match to reference (non-recombinant)locus
nan-H37Rv-3121878-3122917 --- k-mer match to putative recomb seq = 0.0 (0/21)
nan-H37Rv-3121937-3122533 --- k-mer match to putative recomb seq = 0.0 (0/21)
nan-H37Rv-3121937-3123193 --- k-mer match to putative recomb seq = 0.0 (0/21)
nan-H37Rv-3121937-3123338 --- k-mer match to putative recomb seq = 0.0 (0/21)
nan-H37Rv-3122016-3122758 --- k-mer match to putative recomb seq = 0.0 (0/21)
nan-H37Rv-3122084-3123410 --- k-mer match to putative recomb seq = 0.0 (0/21)
nan-H37Rv-3122229-3123561 --- k-mer match to putative recomb seq = 0.0 (0/21)
nan-H37Rv-3122374-3123576 --- k-mer match to putative recomb seq = 0.0 (0/21)
nan-H37Rv-3122511-3123576 --- k-mer match to putative recomb seq = 0.0 (0/21)
nan-H37Rv-3122660-3123576 --- k-mer match to putative recomb seq = 0.0 (0/21)
Match to reference locus
Match to recombinant "k-mer Only"
k-mer match
1.0
0.8
0.6
0.4
0.2
0.0
3,119,680
3,119,760
3,119,840
3,119,920
3,120,000
3,120,080
3,120,160
3,120,240
H37rv Ref Position (bp)

Sub-phylogeny Viz + Paralog Alignments
GCE ID: Event_226

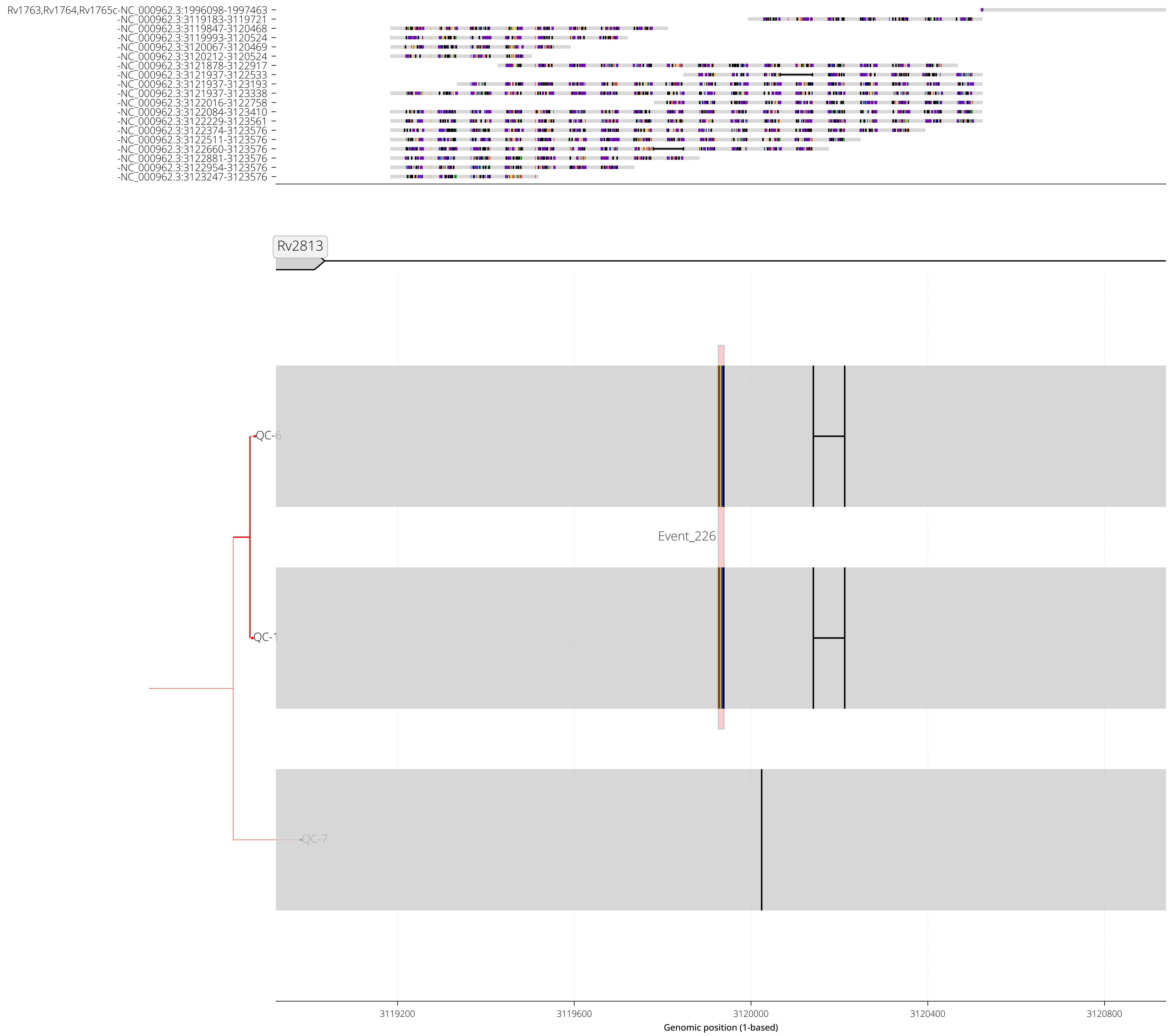

Full Phylogeny Viz + Paralog Alignments
GCE ID: Event_226

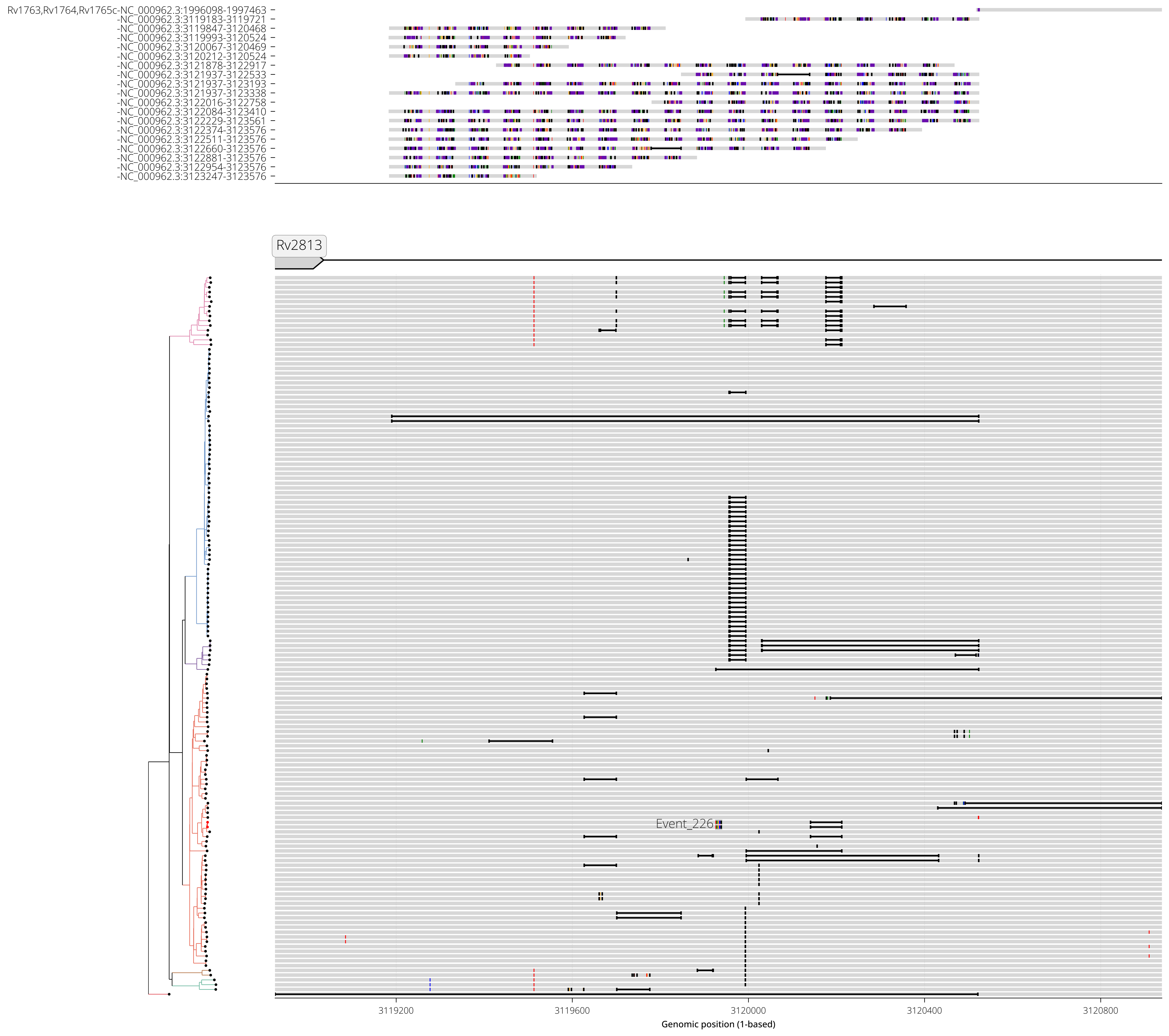

Individual GCE + Known Paralogs
GCE ID: Event_227

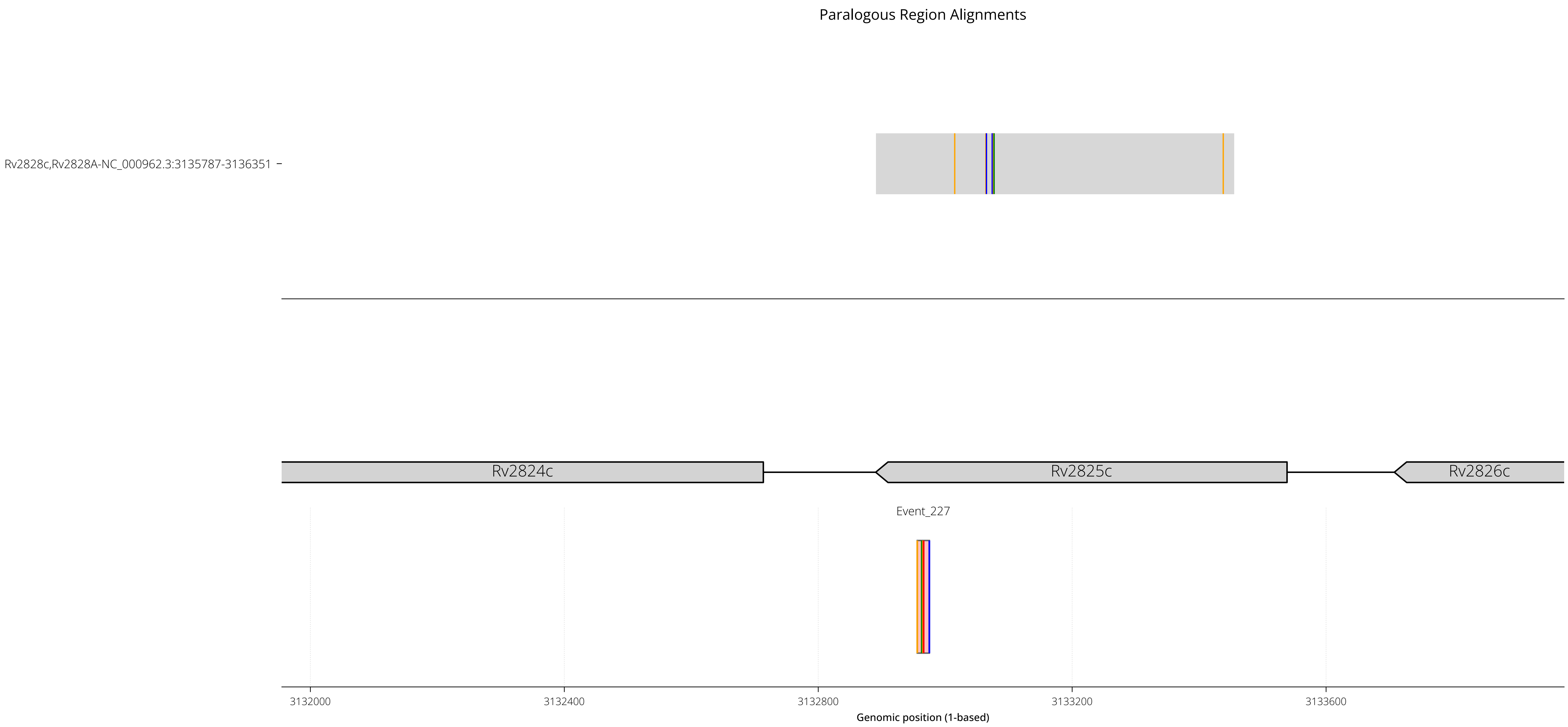

### Circos Viz - GCE to Known Paralogs Locations
GCE ID: Event_227

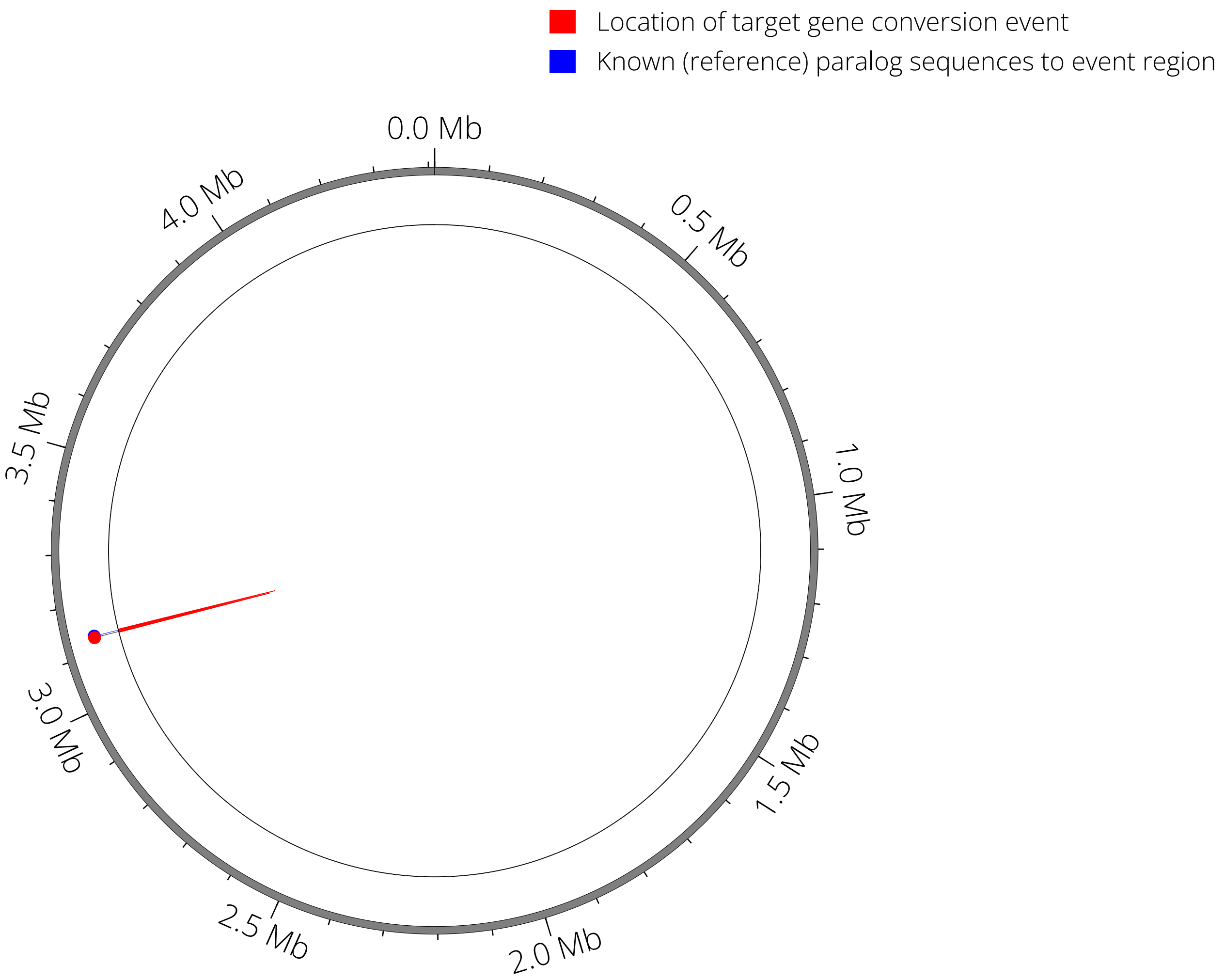

### Event_227 - 5 SNPs - k-mer analysis relative to paralogs

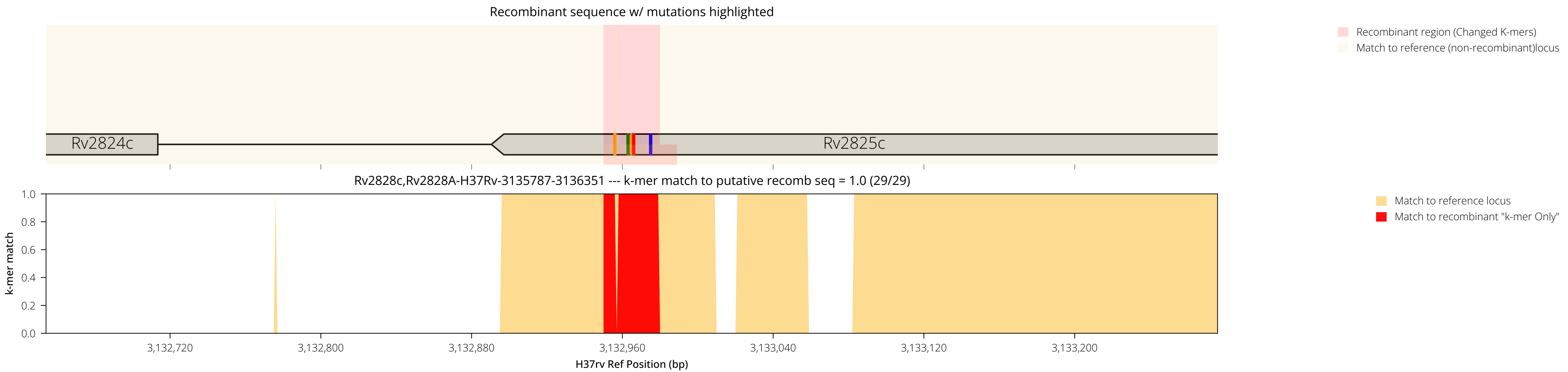

Sub-phylogeny Viz + Paralog Alignments
GCE ID: Event_227

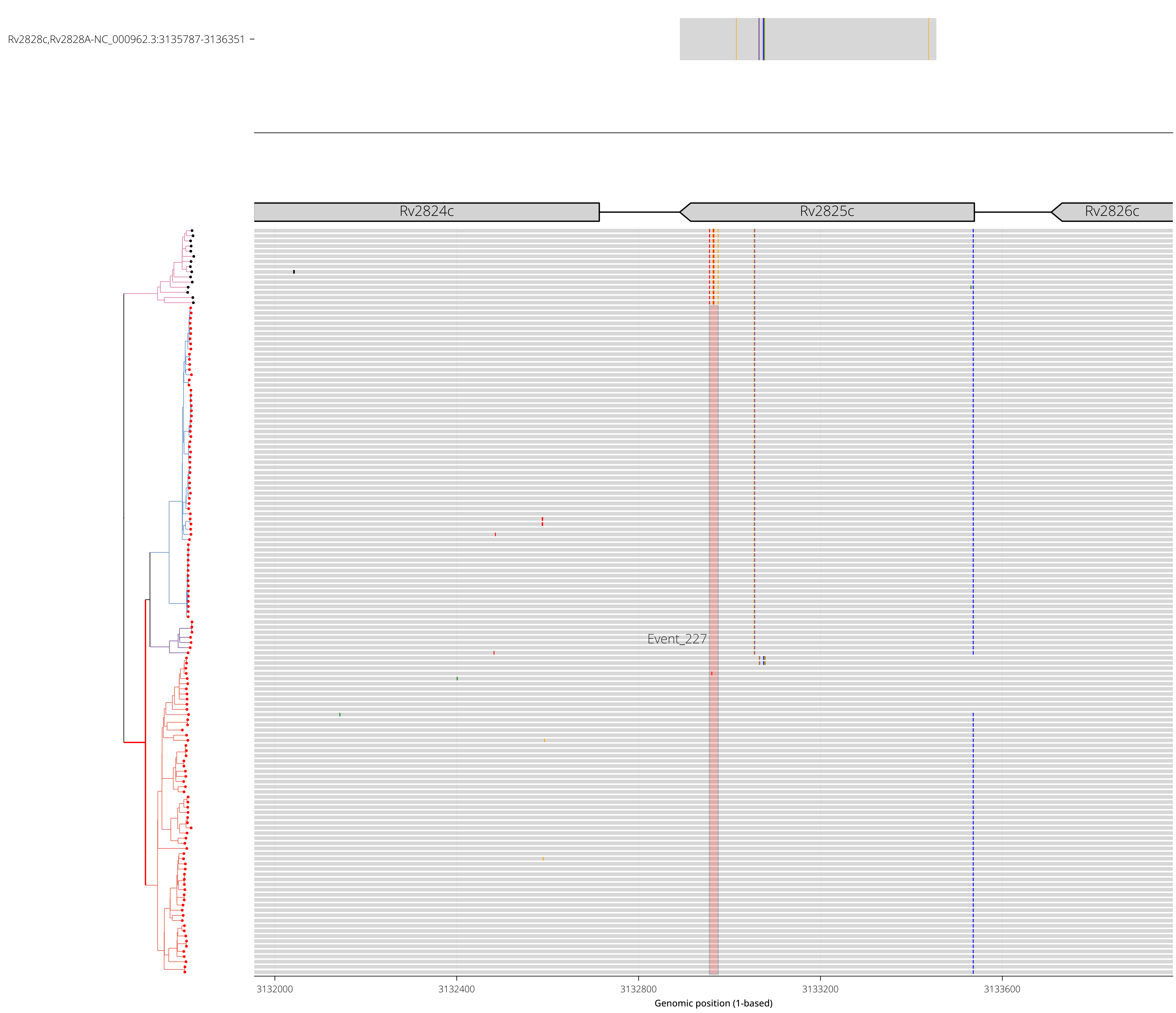

Full Phylogeny Viz + Paralog Alignments
GCE ID: Event_227

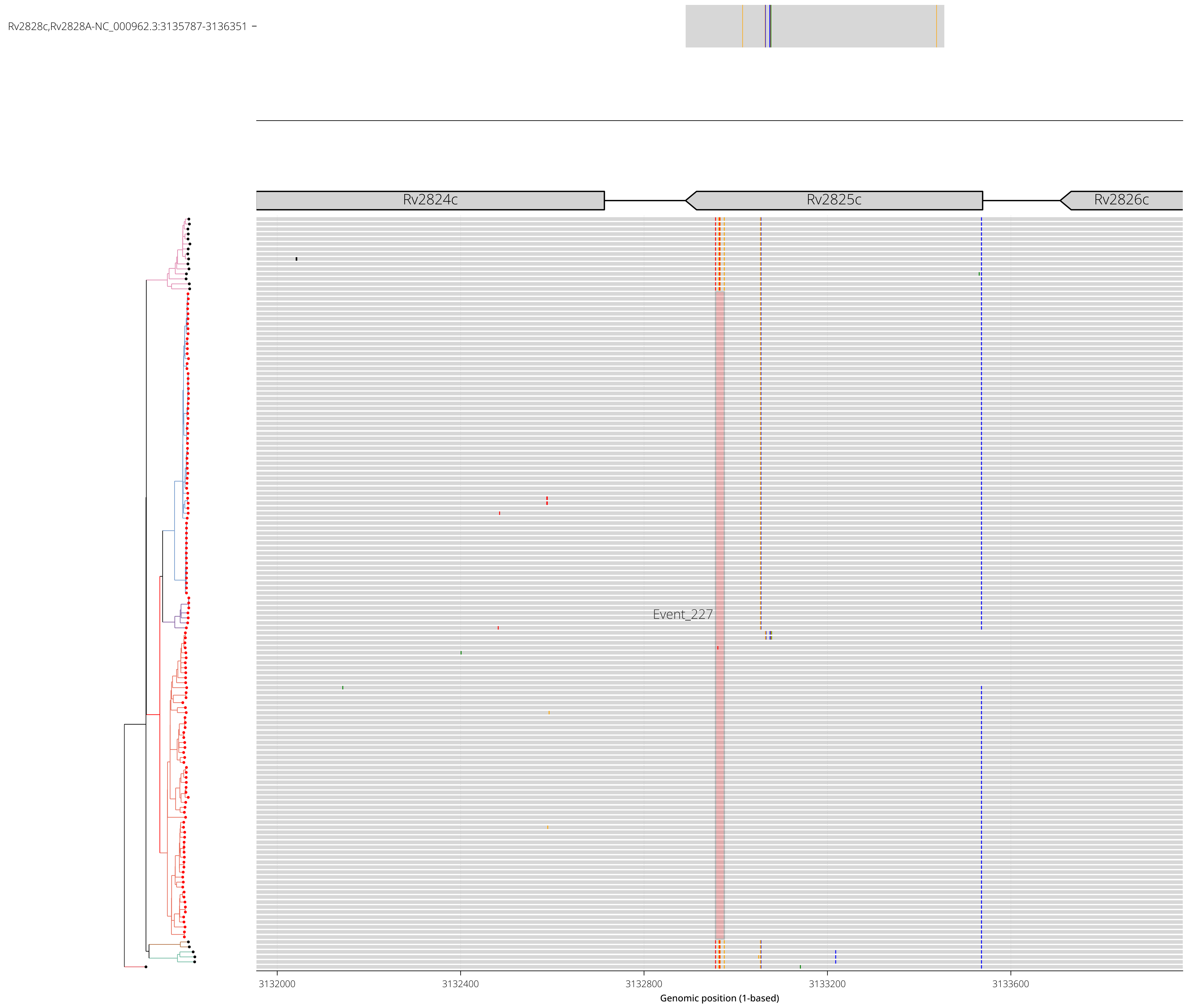

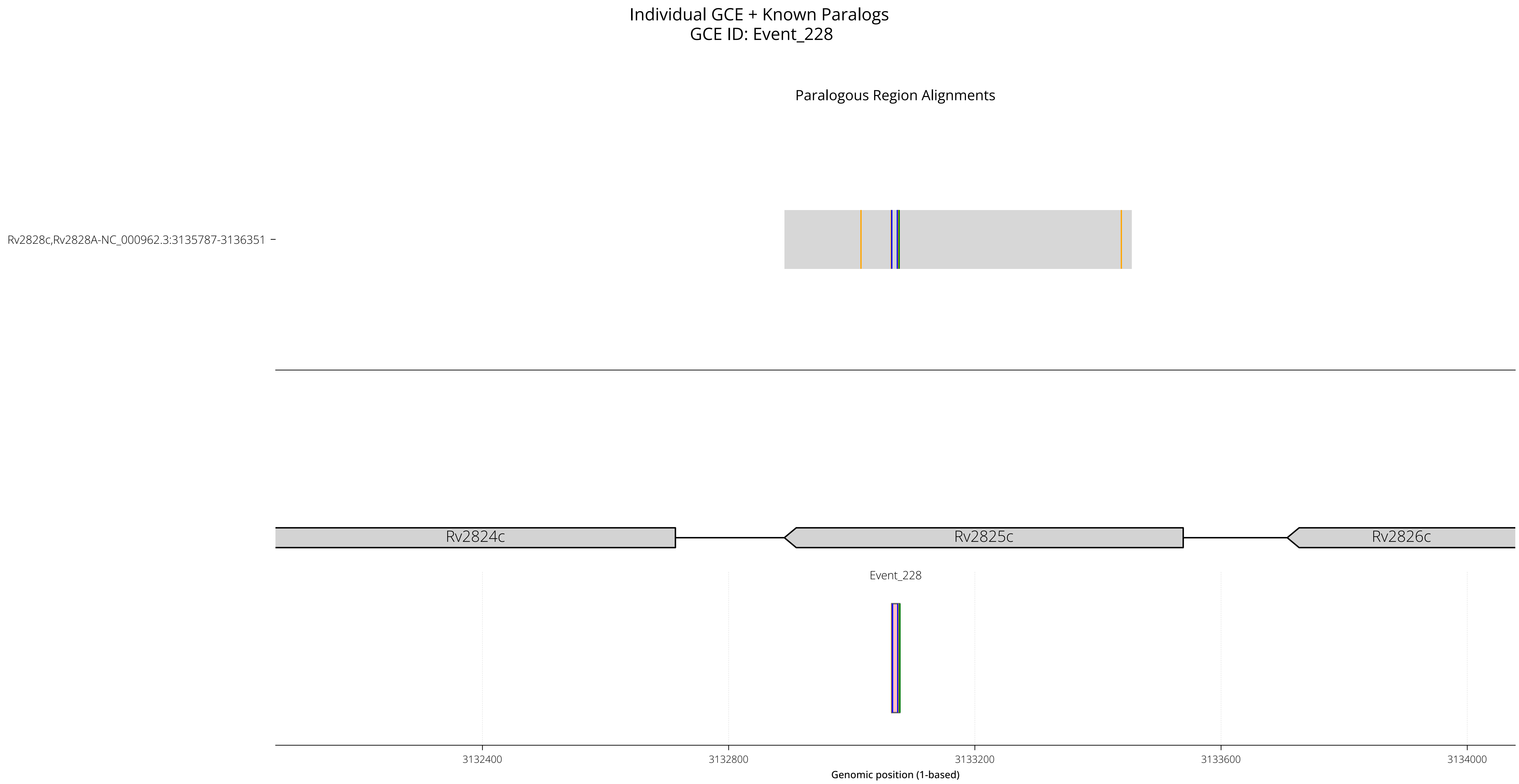
Individual GCE + Known Paralogs
GCE ID: Event_228
Paralogous Region Alignments
Rv2828c,Rv2828A-NC_000962.3:3135787-3136351
Rv2824c
Rv2825c
Rv2826c
Event_228
3132400
3132800
3133200
3133600
3134000
Genomic position (1-based)

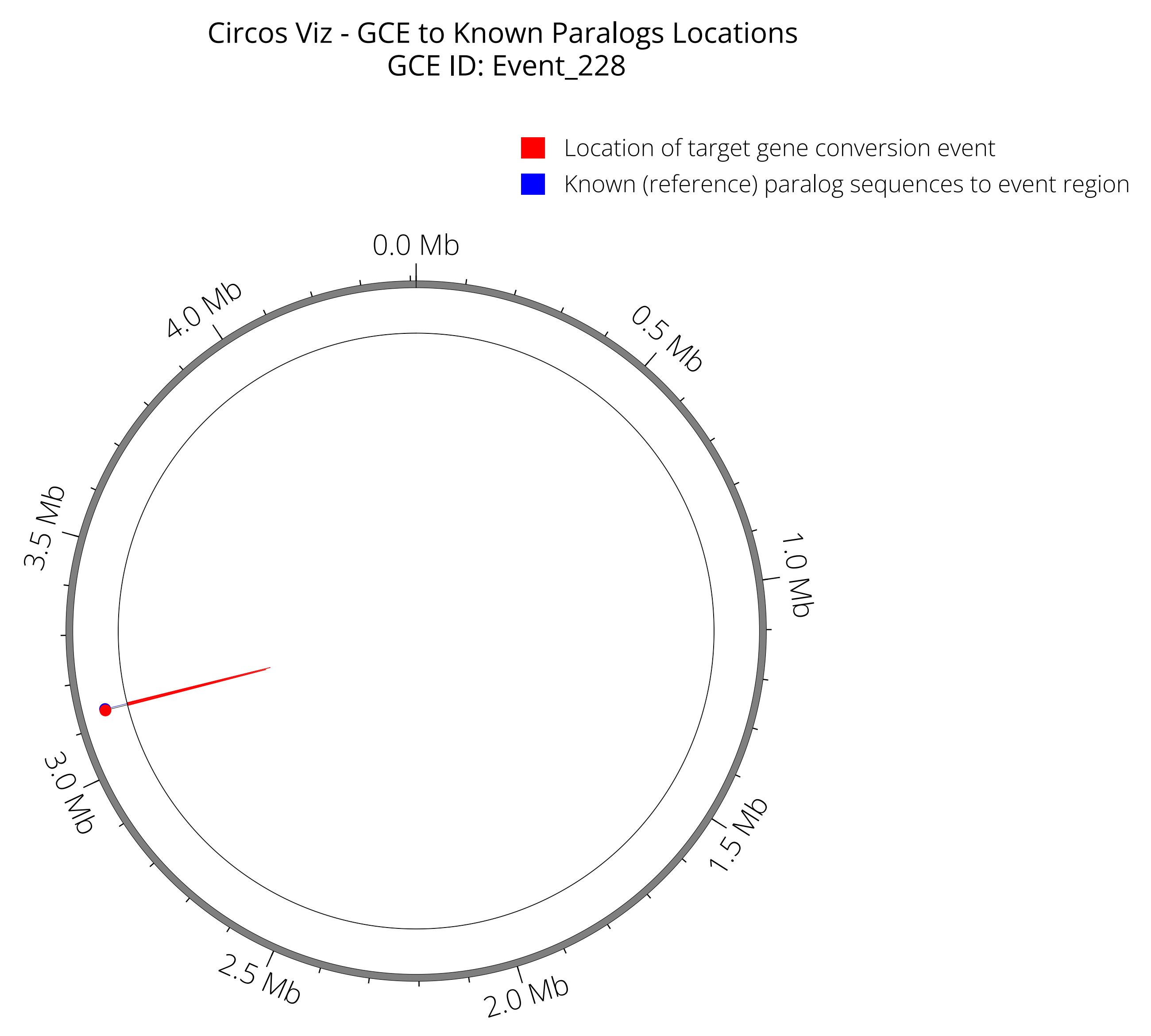

Circos Viz - GCE to Known Paralogs Locations
GCE ID: Event_228
Location of target gene conversion event
Known (reference) paralog sequences to event region
0.0 Mb
0.5 Mb
1.0 Mb
1.5 Mb
2.0 Mb
2.5 Mb
3.0 Mb
3.5 Mb
4.0 Mb

### Event_228 - 6 SNPs - k-mer analysis relative to paralogs

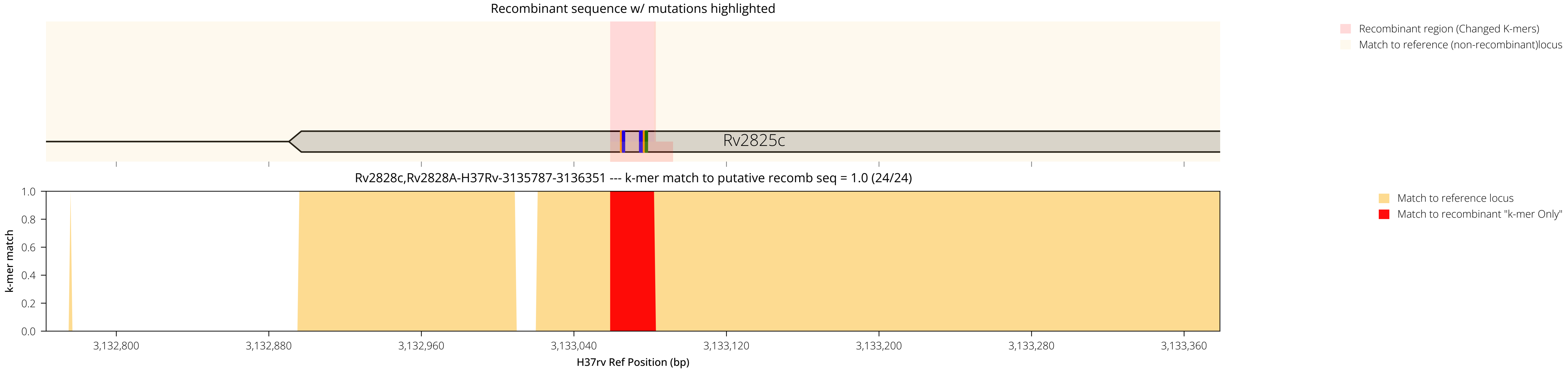

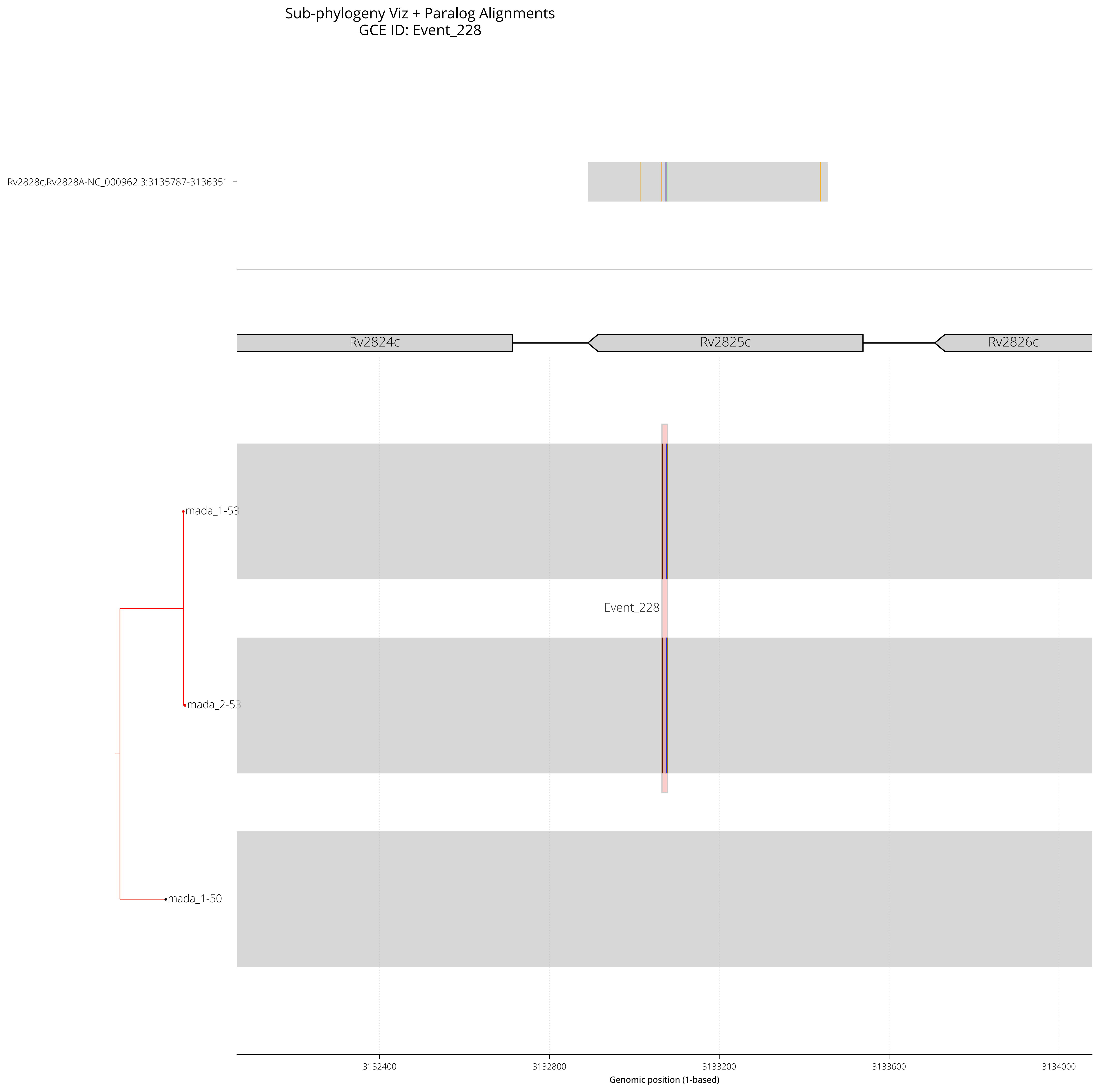

Sub-phylogeny Viz + Paralog Alignments
GCE ID: Event_228
Rv2828c,Rv2828A-NC_000962.3:3135787-3136351
Rv2824c
Rv2825c
Rv2826c
mada_1-53
Event_228
mada_2-53
mada_1-50
3132400
3132800
3133200
3133600
3134000
Genomic position (1-based)

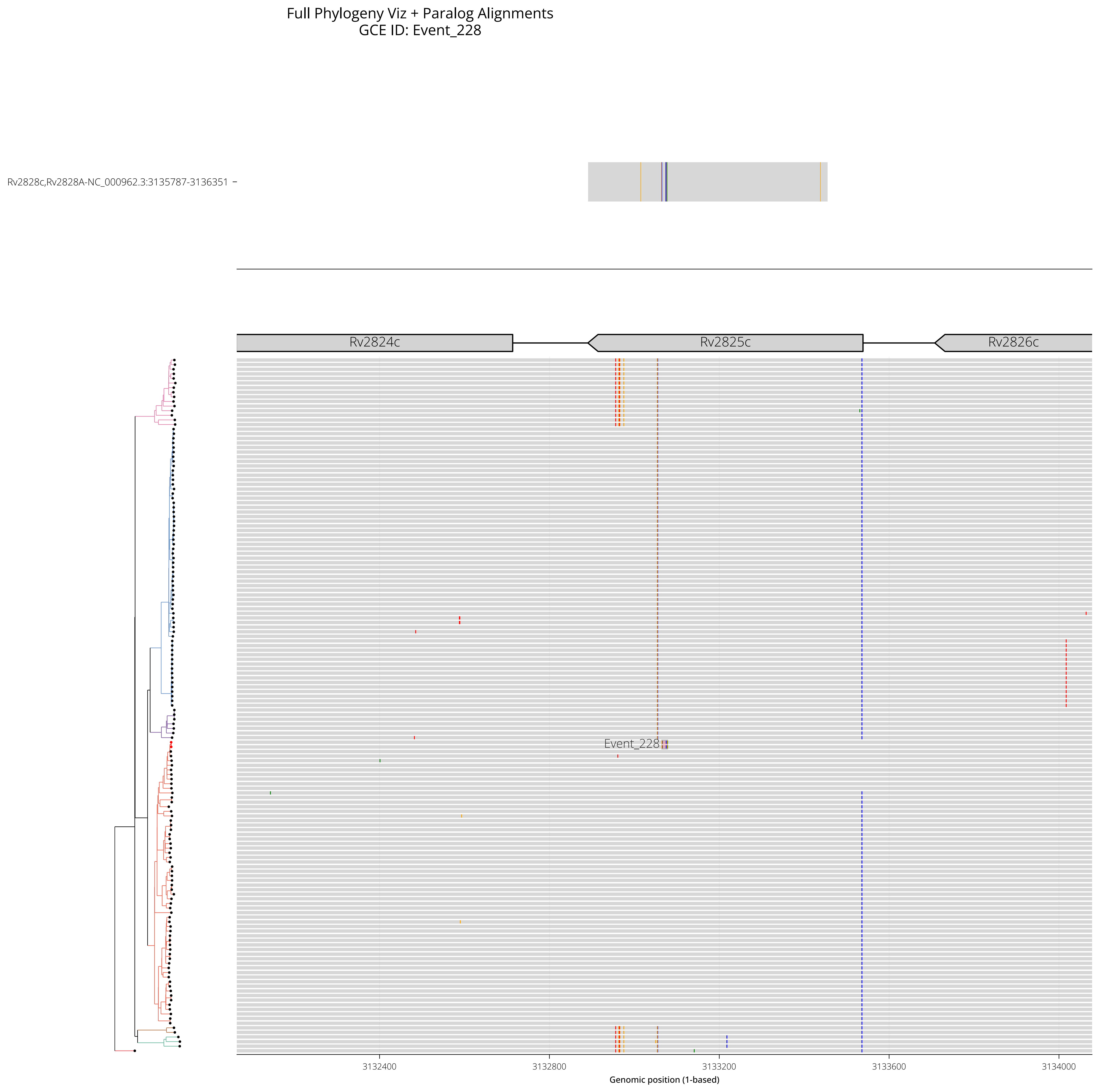

Full Phylogeny Viz + Paralog Alignments
GCE ID: Event_228
Rv2828c,Rv2828A-NC_000962.3:3135787-3136351
Rv2824c
Rv2825c
Rv2826c
Event_228
3132400
3132800
3133200
3133600
3134000
Genomic position (1-based)

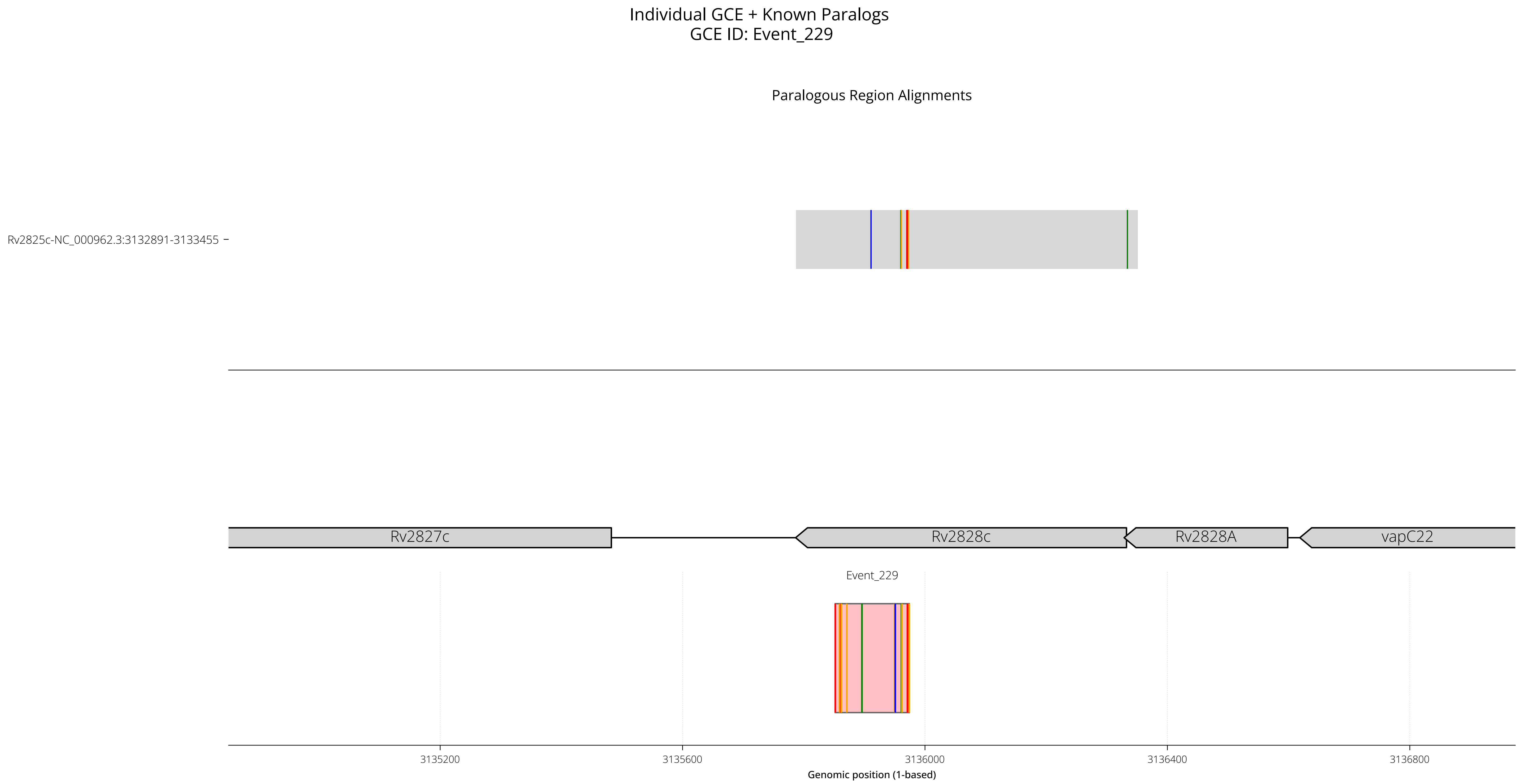

Individual GCE + Known Paralogs
GCE ID: Event_229
Paralogous Region Alignments
Rv2825c-NC_000962.3:3132891-3133455
Rv2827c
Rv2828c
Rv2828A
vapC22
Event_229
3135200
3135600
3136000
3136400
3136800
Genomic position (1-based)

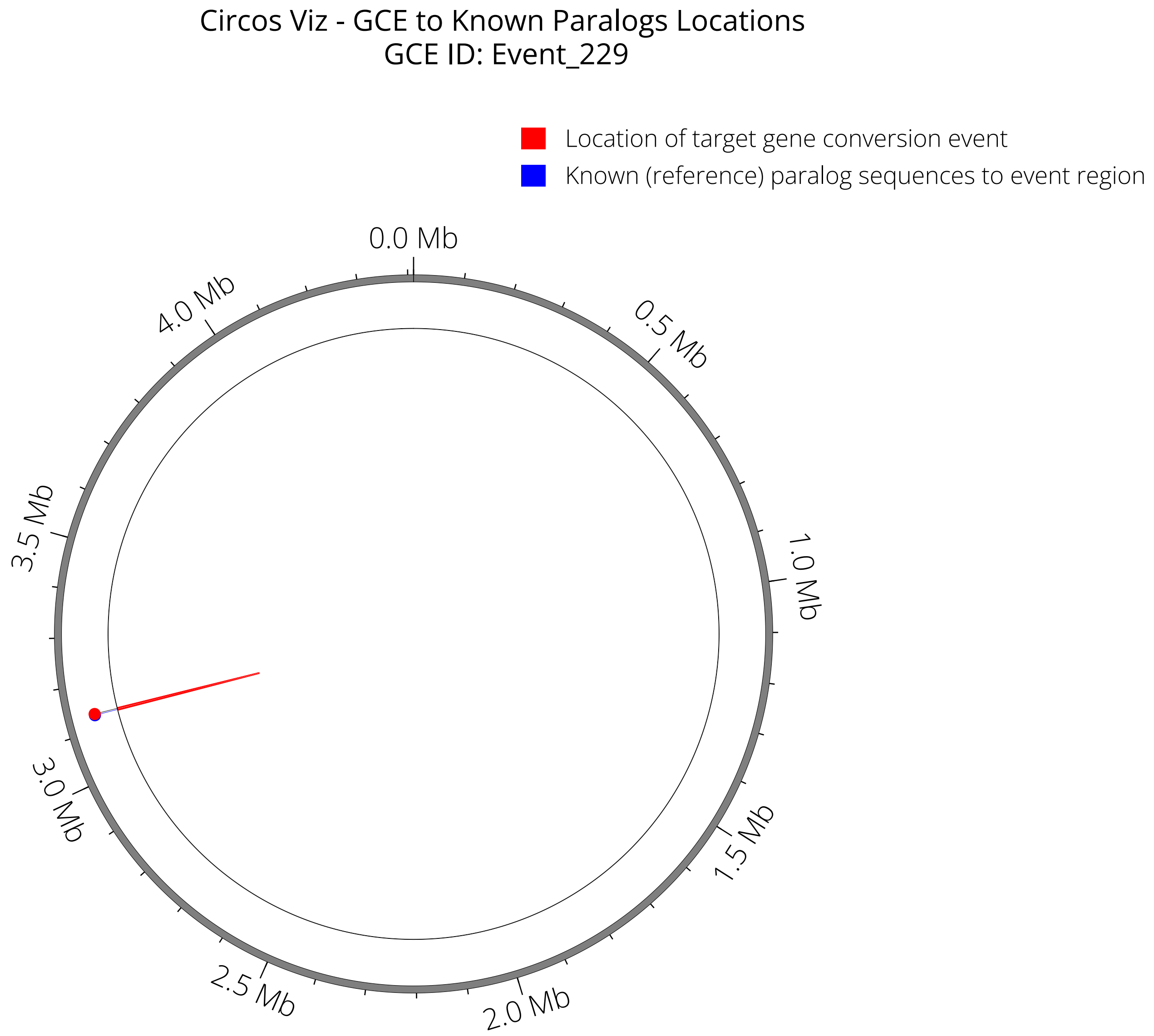

Circos Viz - GCE to Known Paralogs Locations
GCE ID: Event_229
Location of target gene conversion event
Known (reference) paralog sequences to event region
0.0 Mb
0.5 Mb
1.0 Mb
1.5 Mb
2.0 Mb
2.5 Mb
3.0 Mb
3.5 Mb
4.0 Mb

### Event_229 - 14 SNPs - k-mer analysis relative to paralogs

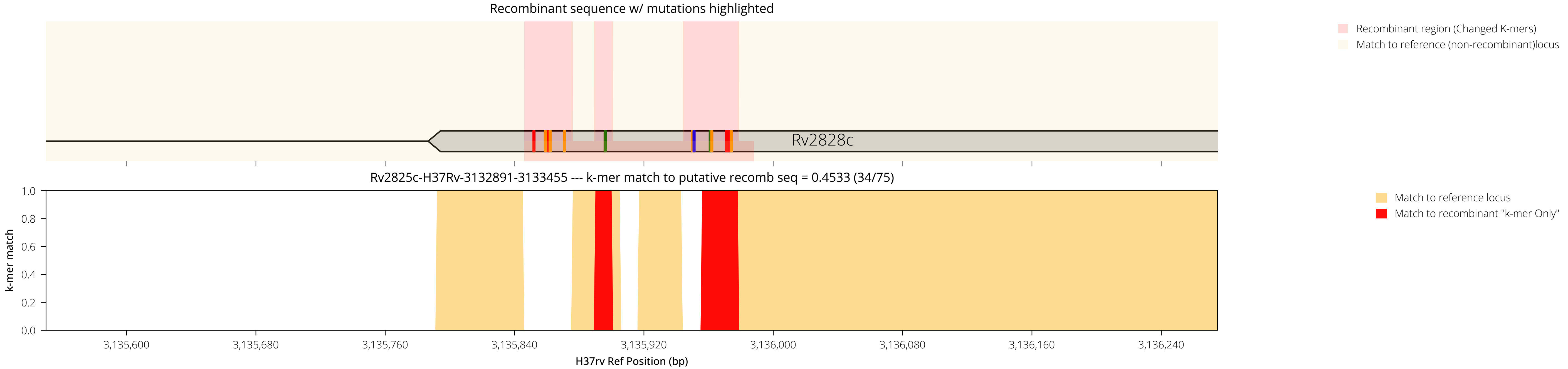

Sub-phylogeny Viz + Paralog Alignments
GCE ID: Event_229

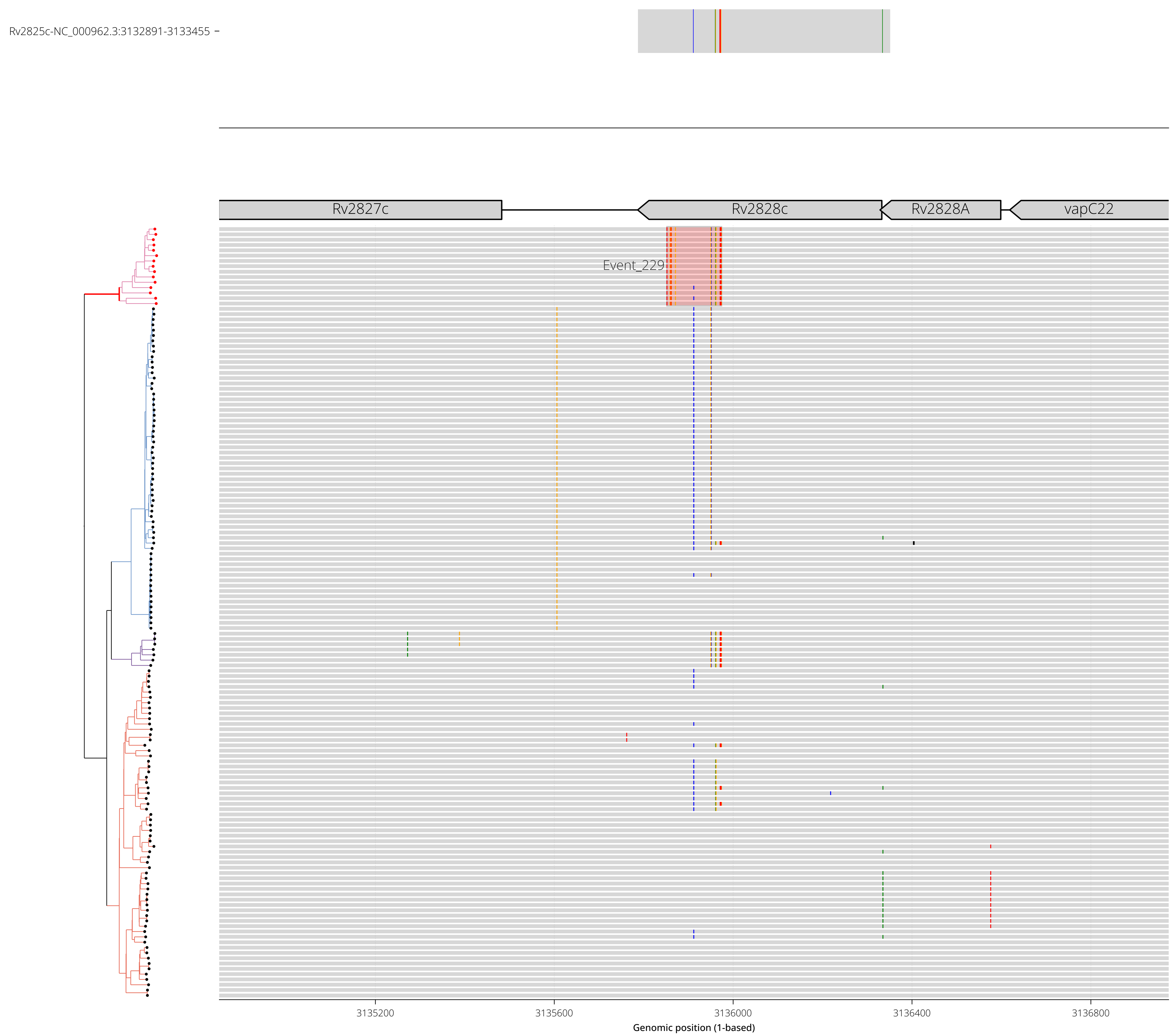

Full Phylogeny Viz + Paralog Alignments
GCE ID: Event_229

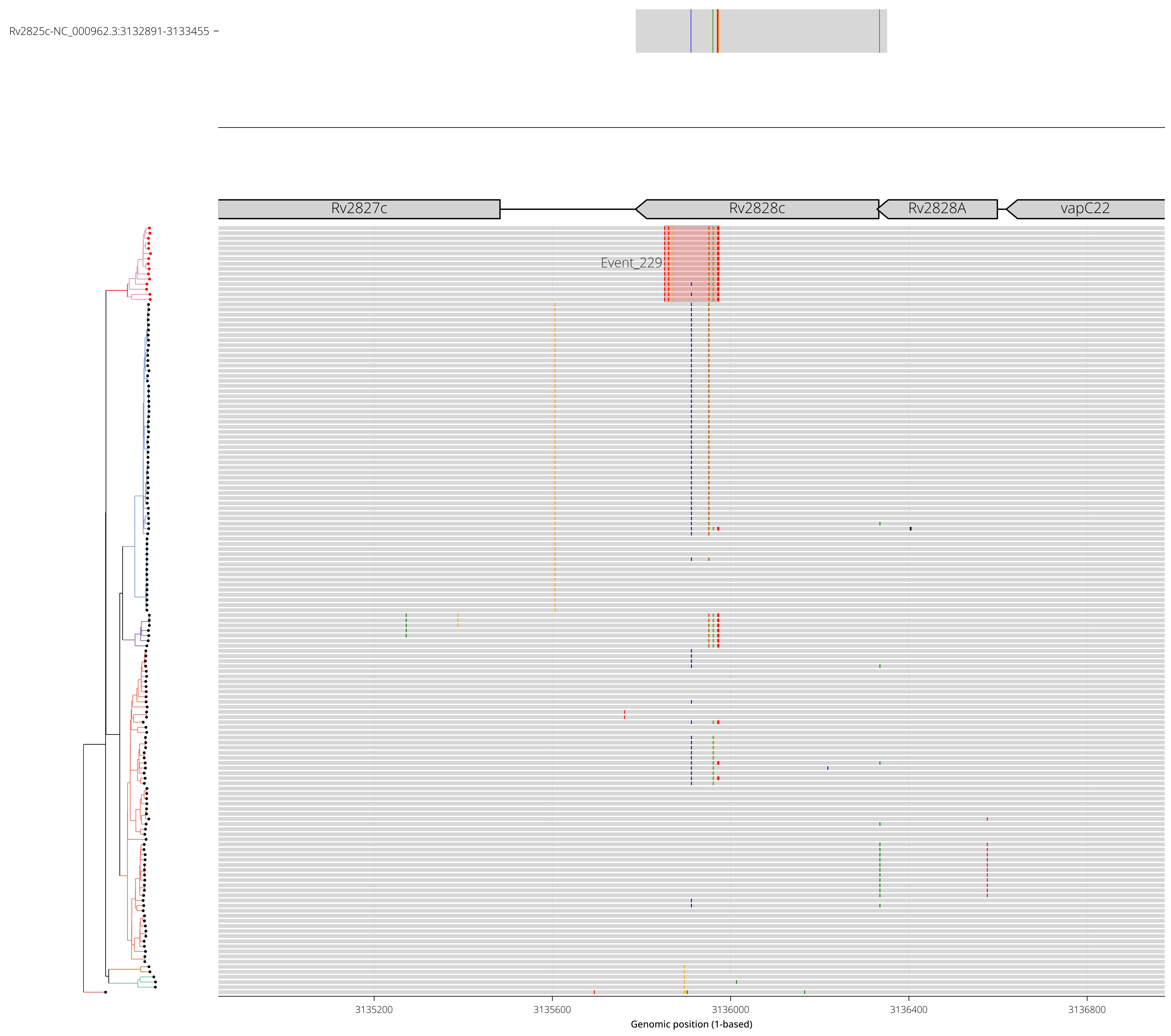

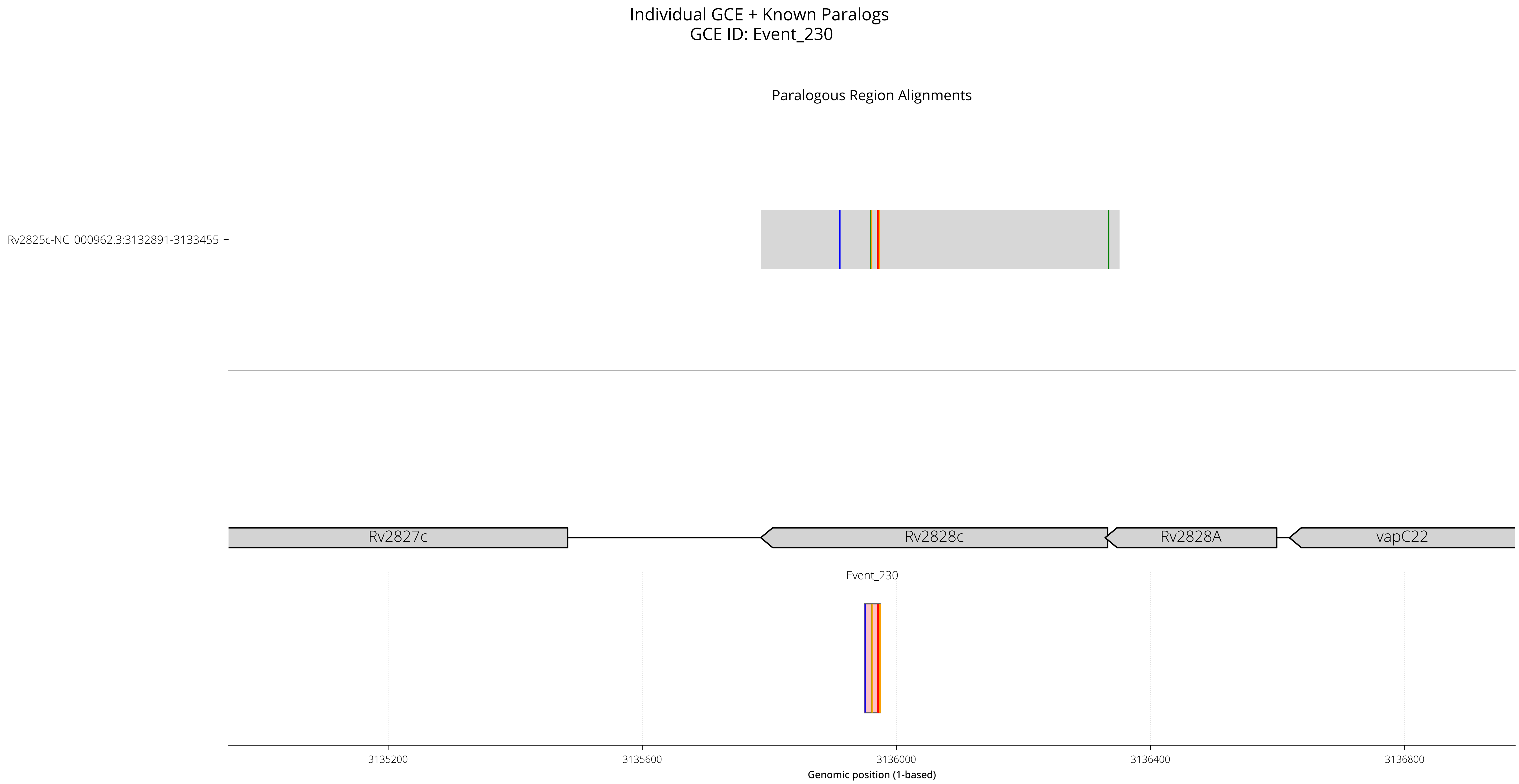
Individual GCE + Known Paralogs
GCE ID: Event_230
Paralogous Region Alignments
Rv2825c-NC_000962.3:3132891-3133455
Rv2827c
Rv2828c
Rv2828A
vapC22
Event_230
3135200
3135600
3136000
3136400
3136800
Genomic position (1-based)

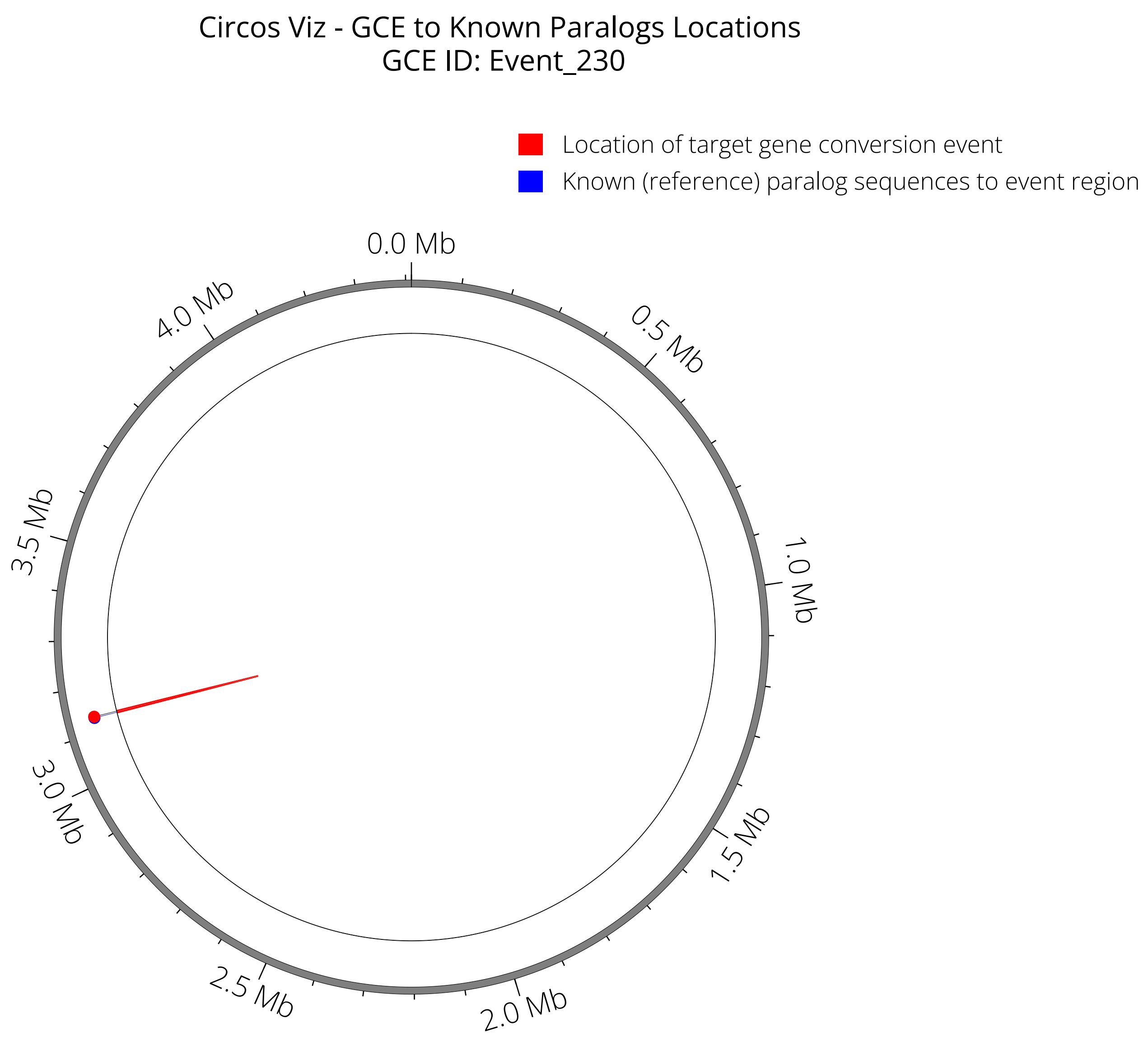
Circos Viz - GCE to Known Paralogs Locations
GCE ID: Event_230
Location of target gene conversion event
Known (reference) paralog sequences to event region
0.0 Mb
0.5 Mb
1.0 Mb
1.5 Mb
2.0 Mb
2.5 Mb
3.0 Mb
3.5 Mb
4.0 Mb

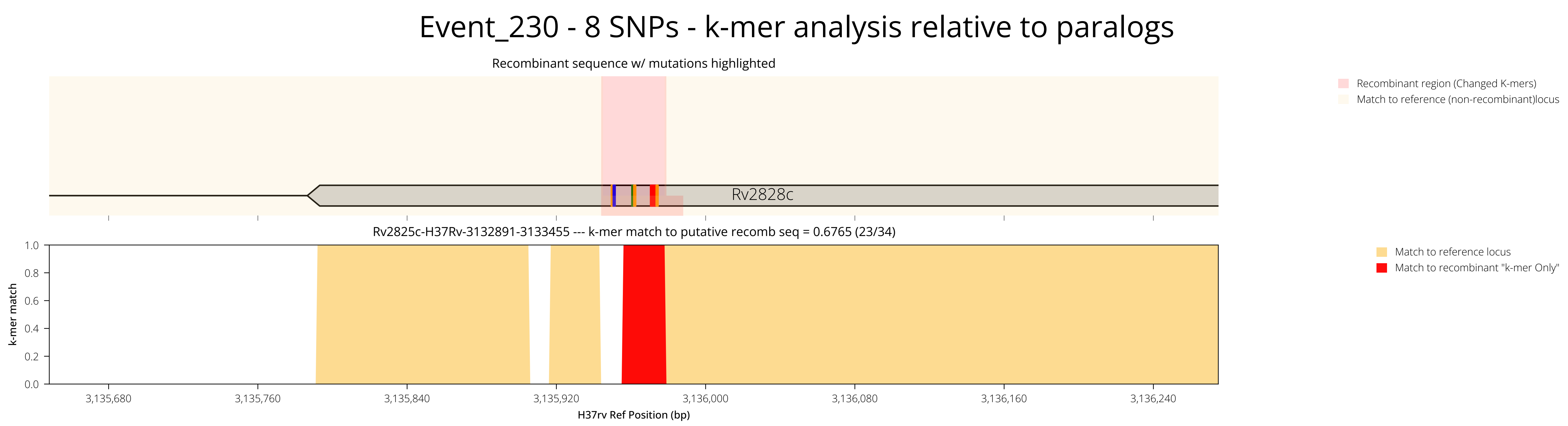
Event_230 - 8 SNPs - k-mer analysis relative to paralogs
Recombinant sequence w/ mutations highlighted
Rv2828c
Recombinant region (Changed K-mers)
Match to reference (non-recombinant)locus
Rv2825c-H37Rv-3132891-3133455 --- k-mer match to putative recomb seq = 0.6765 (23/34)
k-mer match
1.0
0.8
0.6
0.4
0.2
0.0
3,135,680
3,135,760
3,135,840
3,135,920
3,136,000
3,136,080
3,136,160
3,136,240
H37rv Ref Position (bp)
Match to reference locus
Match to recombinant "k-mer Only"

Sub-phylogeny Viz + Paralog Alignments
GCE ID: Event_230

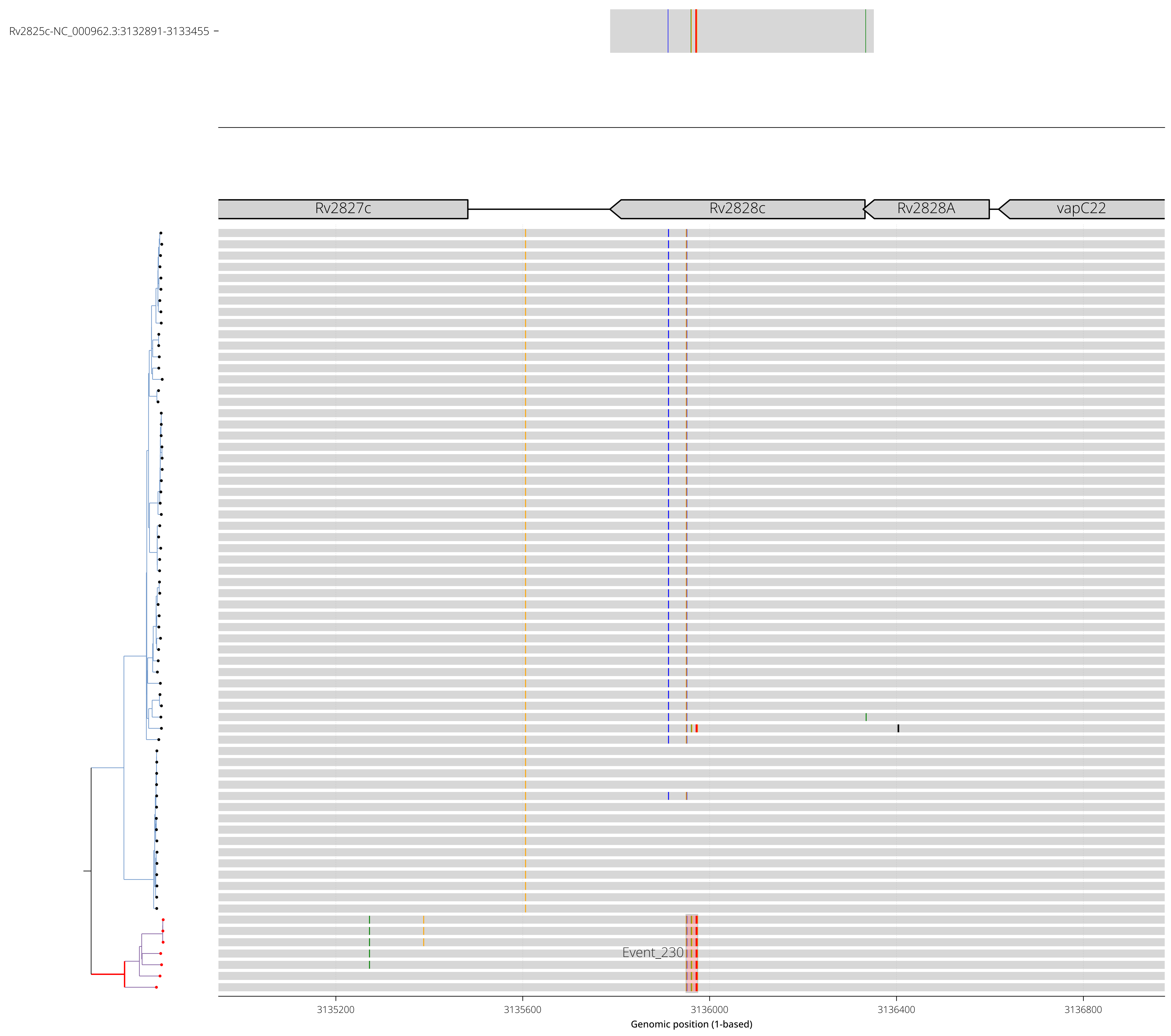

Full Phylogeny Viz + Paralog Alignments
GCE ID: Event_230

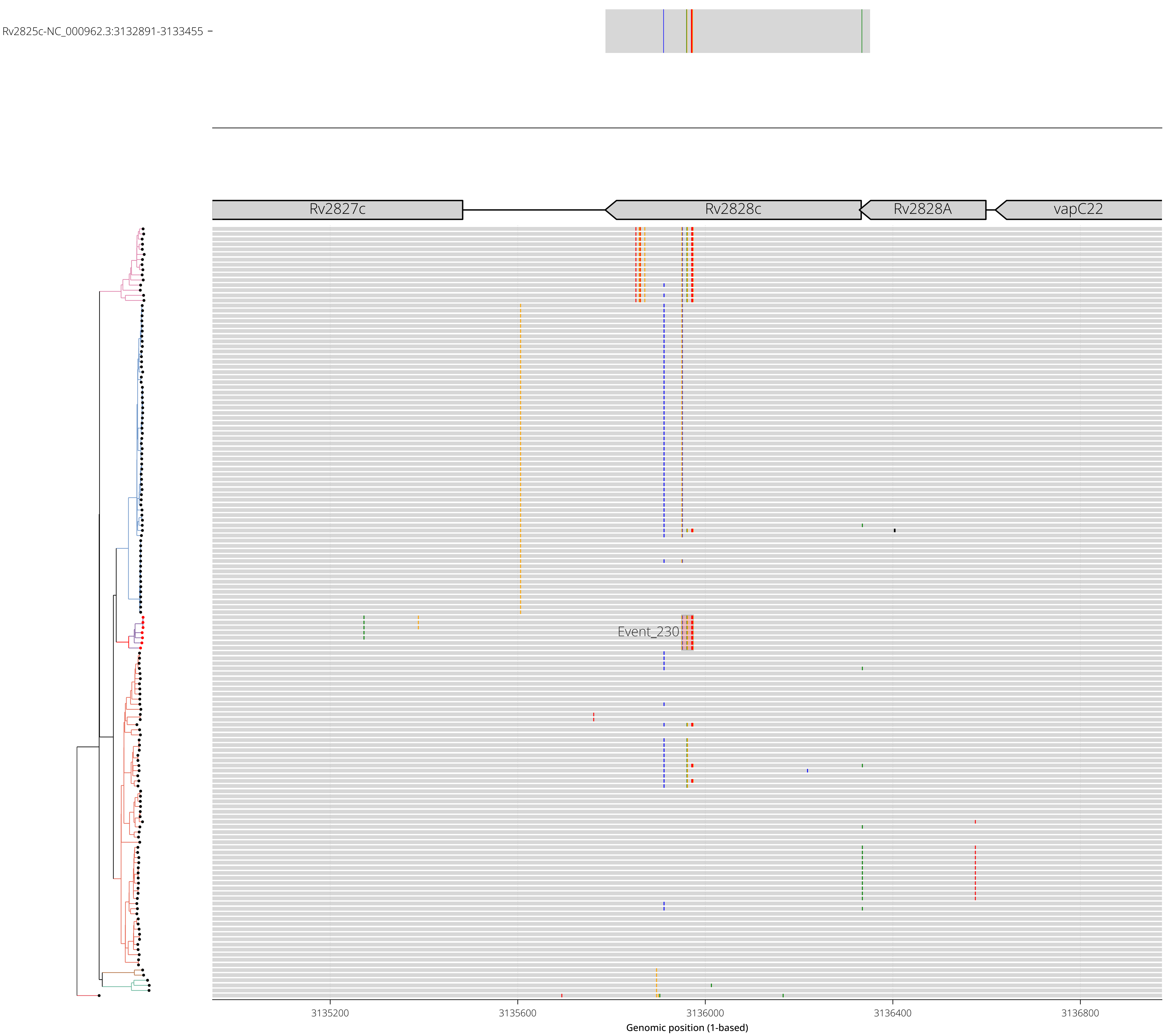

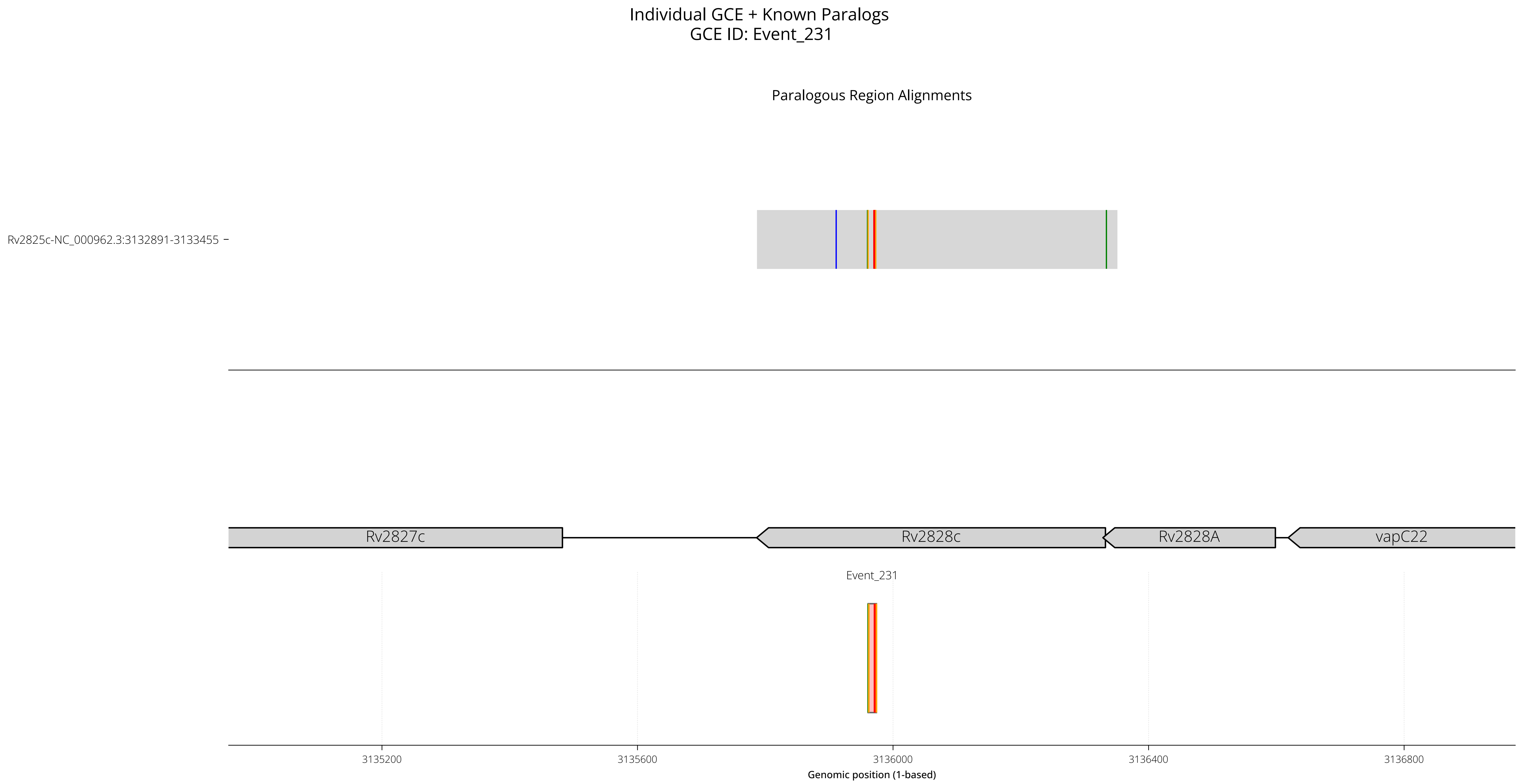
Individual GCE + Known Paralogs
GCE ID: Event_231
Paralogous Region Alignments
Rv2825c-NC_000962.3:3132891-3133455
Rv2827c
Rv2828c
Rv2828A
vapC22
Event_231
3135200
3135600
3136000
3136400
3136800
Genomic position (1-based)

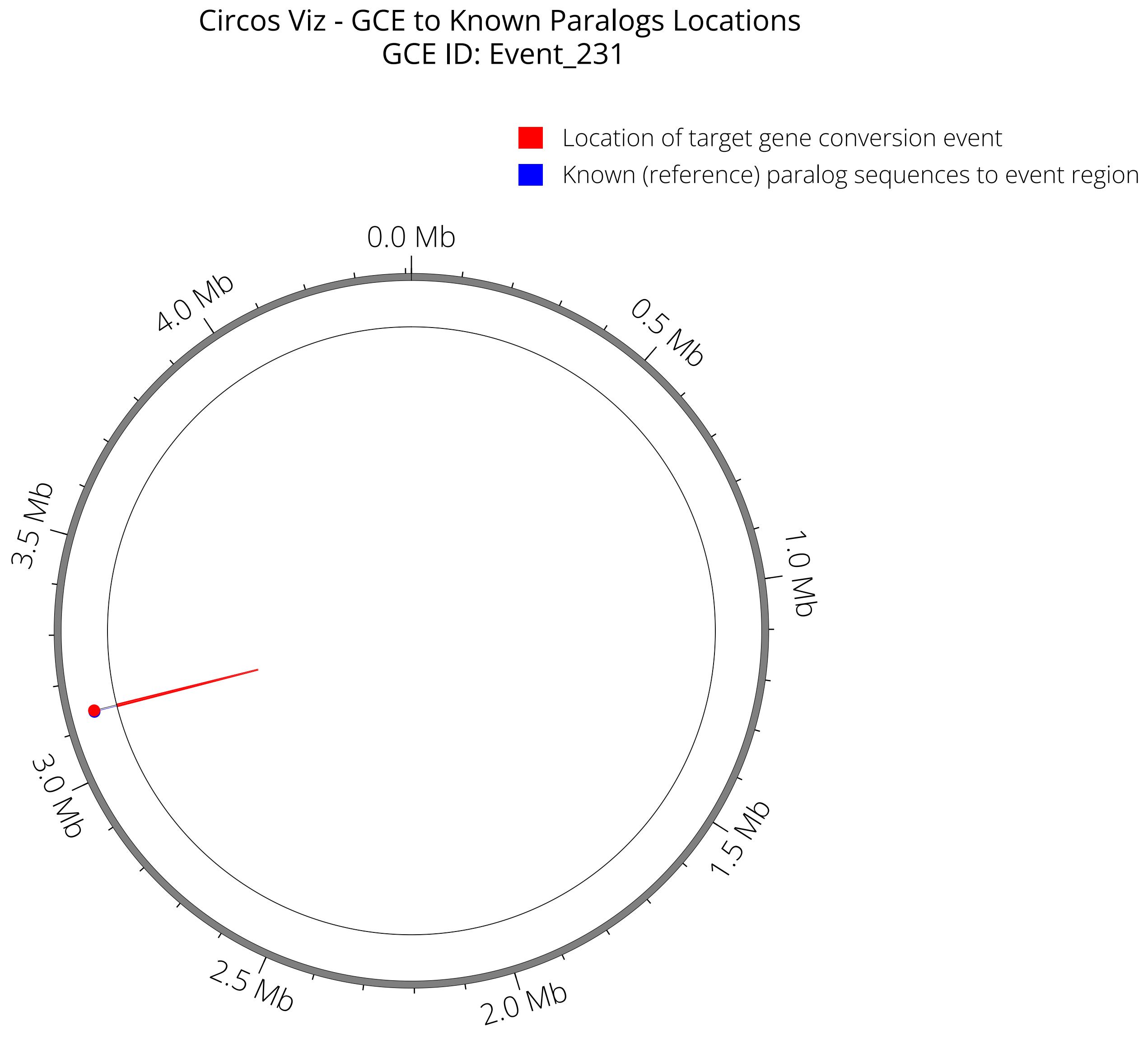
Circos Viz - GCE to Known Paralogs Locations
GCE ID: Event_231
Location of target gene conversion event
Known (reference) paralog sequences to event region
0.0 Mb
0.5 Mb
1.0 Mb
1.5 Mb
2.0 Mb
2.5 Mb
3.0 Mb
3.5 Mb
4.0 Mb

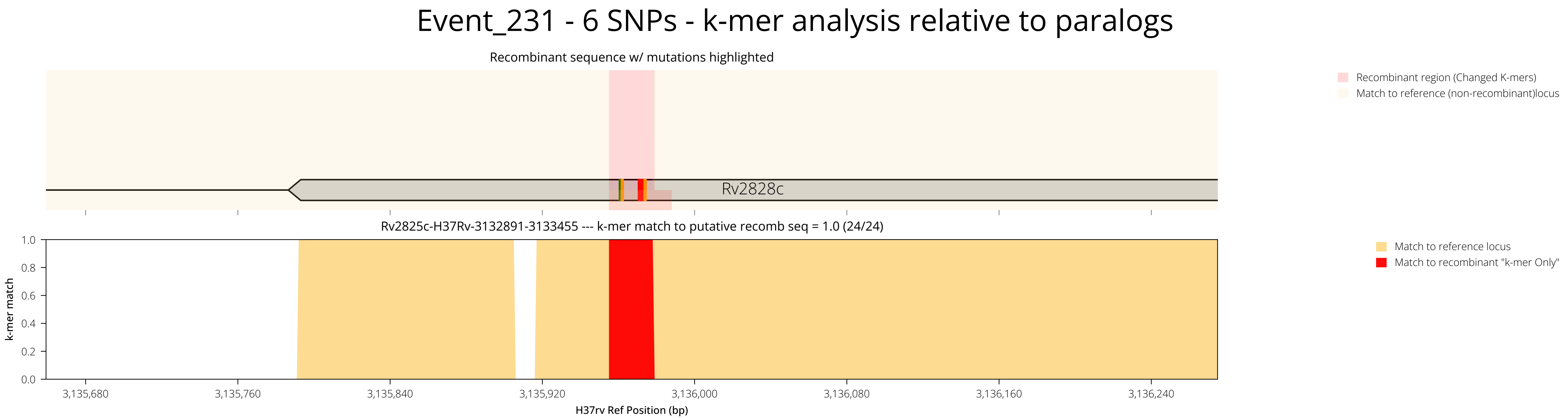
Event_231 - 6 SNPs - k-mer analysis relative to paralogs
Recombinant sequence w/ mutations highlighted
Rv2828c
Recombinant region (Changed K-mers)
Match to reference (non-recombinant)locus
Rv2825c-H37Rv-3132891-3133455 --- k-mer match to putative recomb seq = 1.0 (24/24)
Match to reference locus
Match to recombinant "k-mer Only"
k-mer match
1.0
0.8
0.6
0.4
0.2
0.0
3,135,680
3,135,760
3,135,840
3,135,920
3,136,000
3,136,080
3,136,160
3,136,240
H37rv Ref Position (bp)

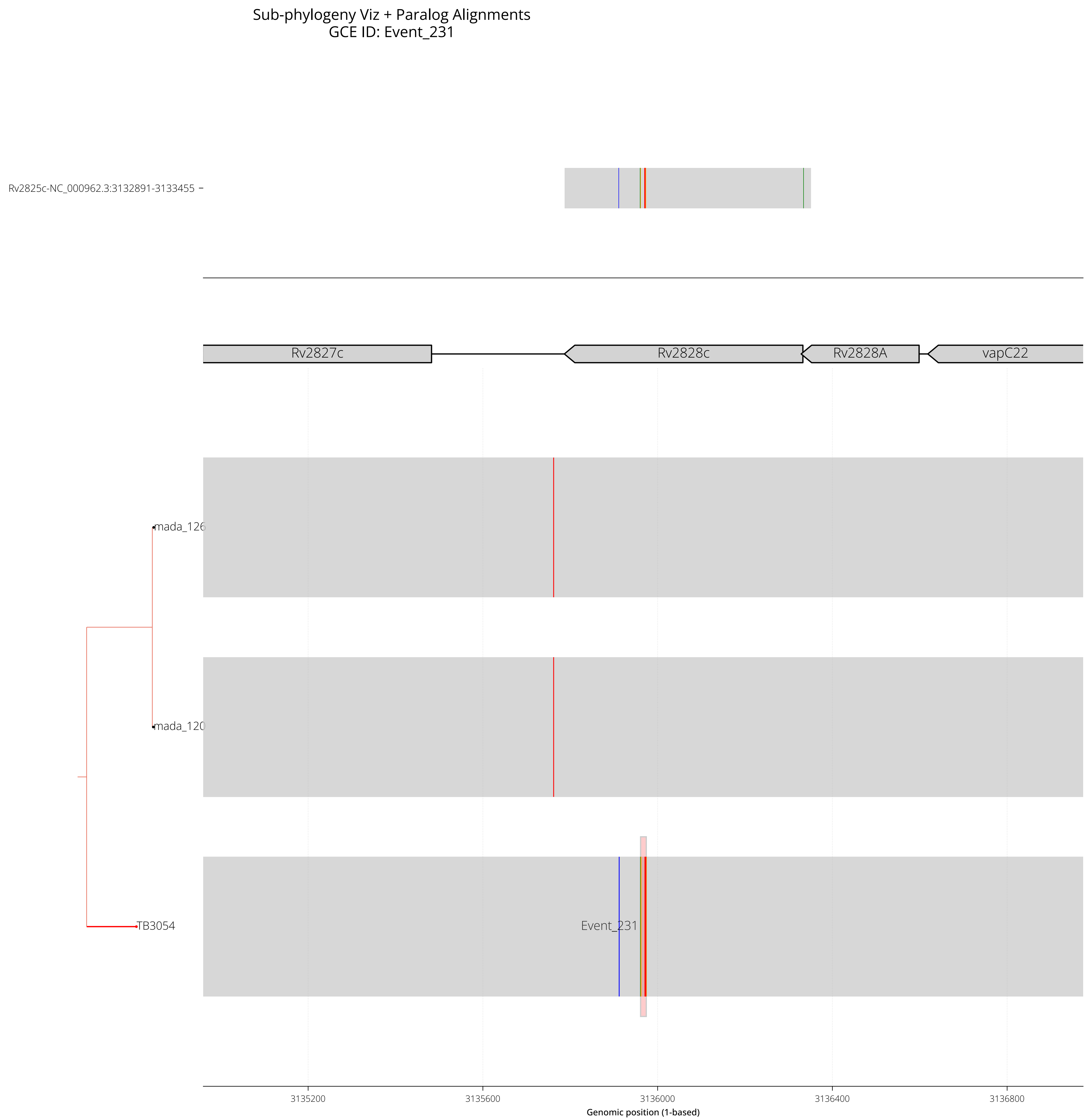

Sub-phylogeny Viz + Paralog Alignments
GCE ID: Event_231
Rv2825c-NC_000962.3:3132891-3133455
Rv2827c
Rv2828c
Rv2828A
vapC22
mada_126
mada_120
TB3054
Event_231
3135200
3135600
3136000
3136400
3136800
Genomic position (1-based)

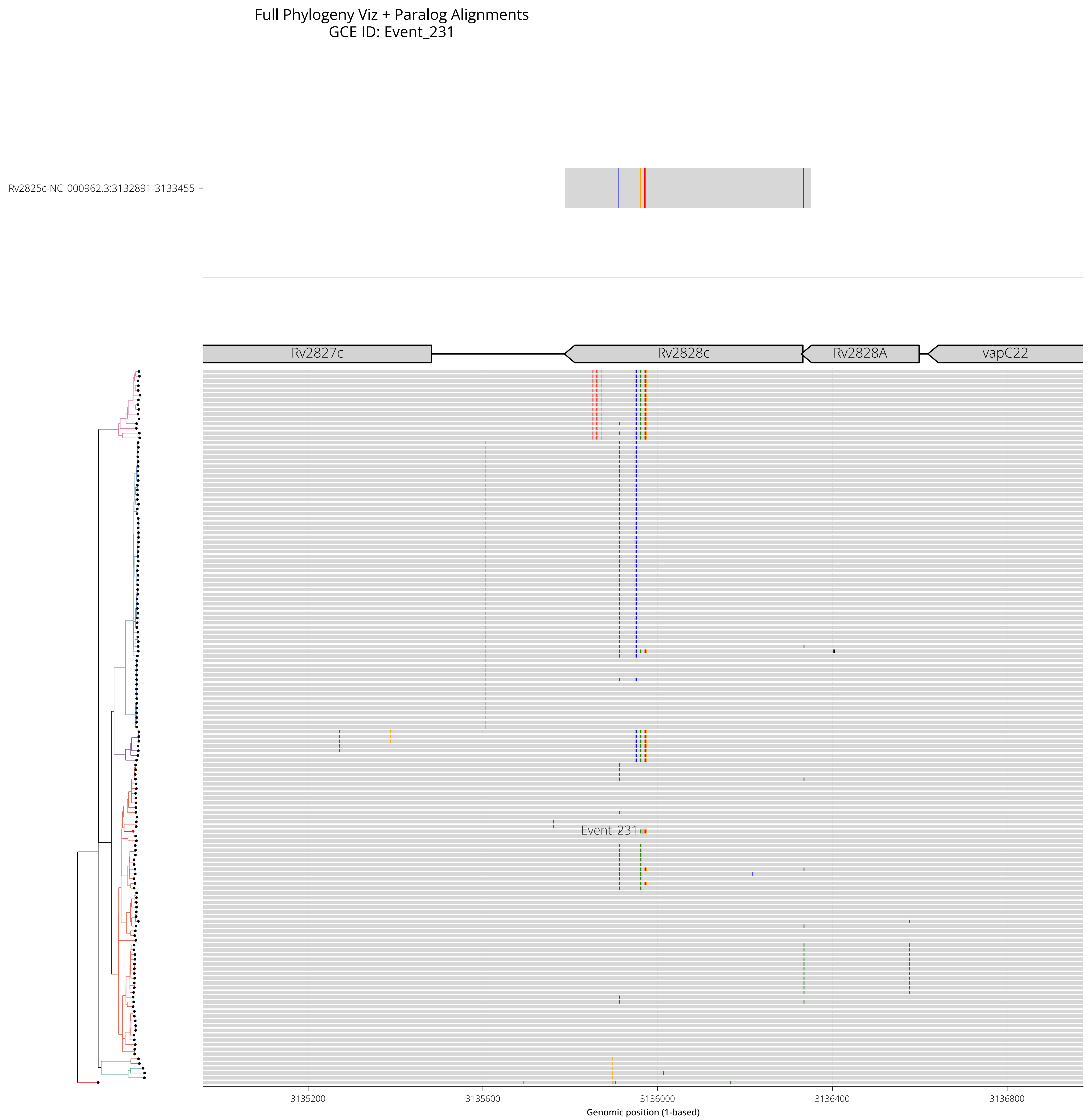

Full Phylogeny Viz + Paralog Alignments
GCE ID: Event_231
Rv2825c-NC_000962.3:3132891-3133455
Rv2827c
Rv2828c
Rv2828A
vapC22
Event_231
3135200
3135600
3136000
3136400
3136800
Genomic position (1-based)

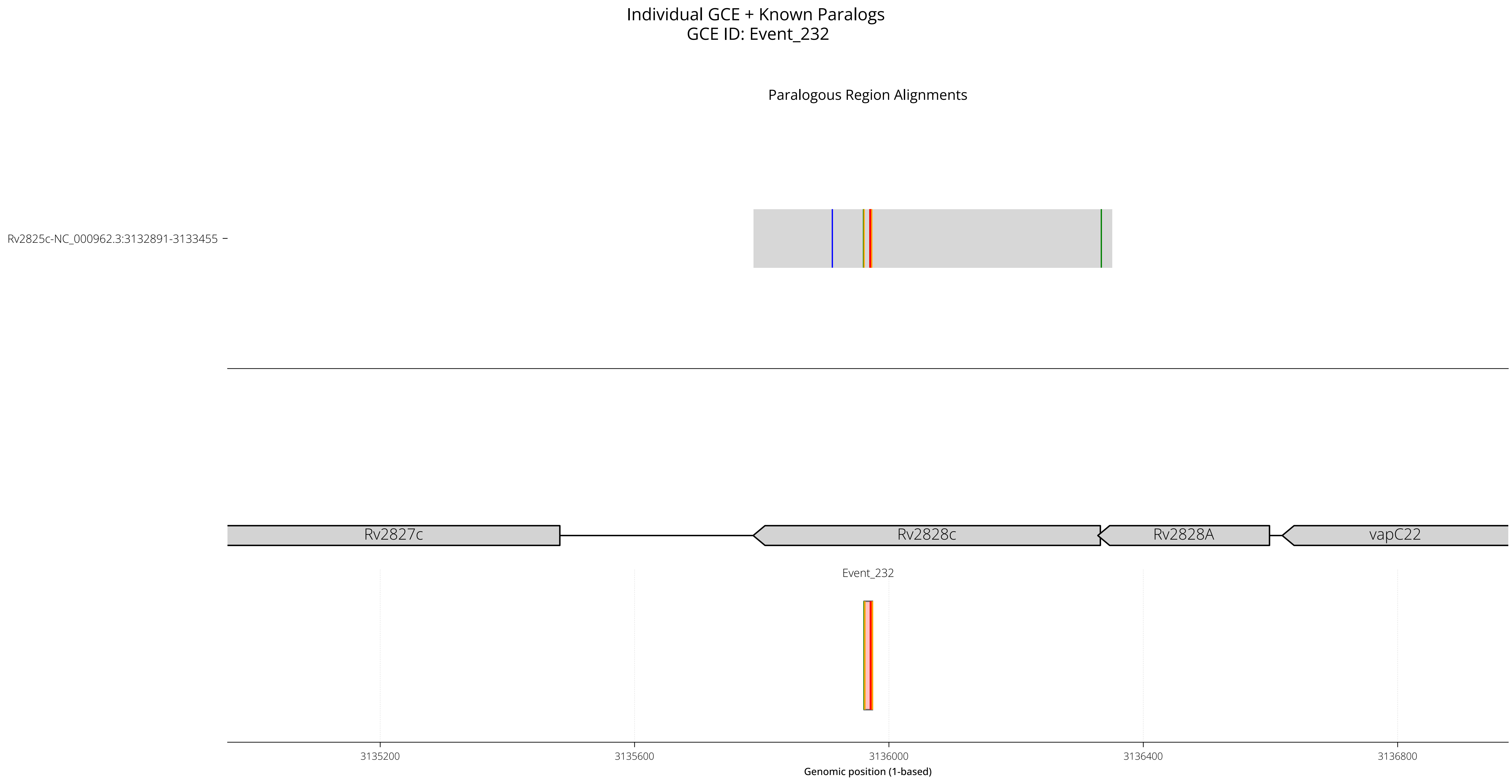
Individual GCE + Known Paralogs
GCE ID: Event_232
Paralogous Region Alignments
Rv2825c-NC_000962.3:3132891-3133455
Rv2827c
Rv2828c
Rv2828A
vapC22
Event_232
3135200
3135600
3136000
3136400
3136800
Genomic position (1-based)

Circos Viz - GCE to Known Paralogs Locations
GCE ID: Event_232

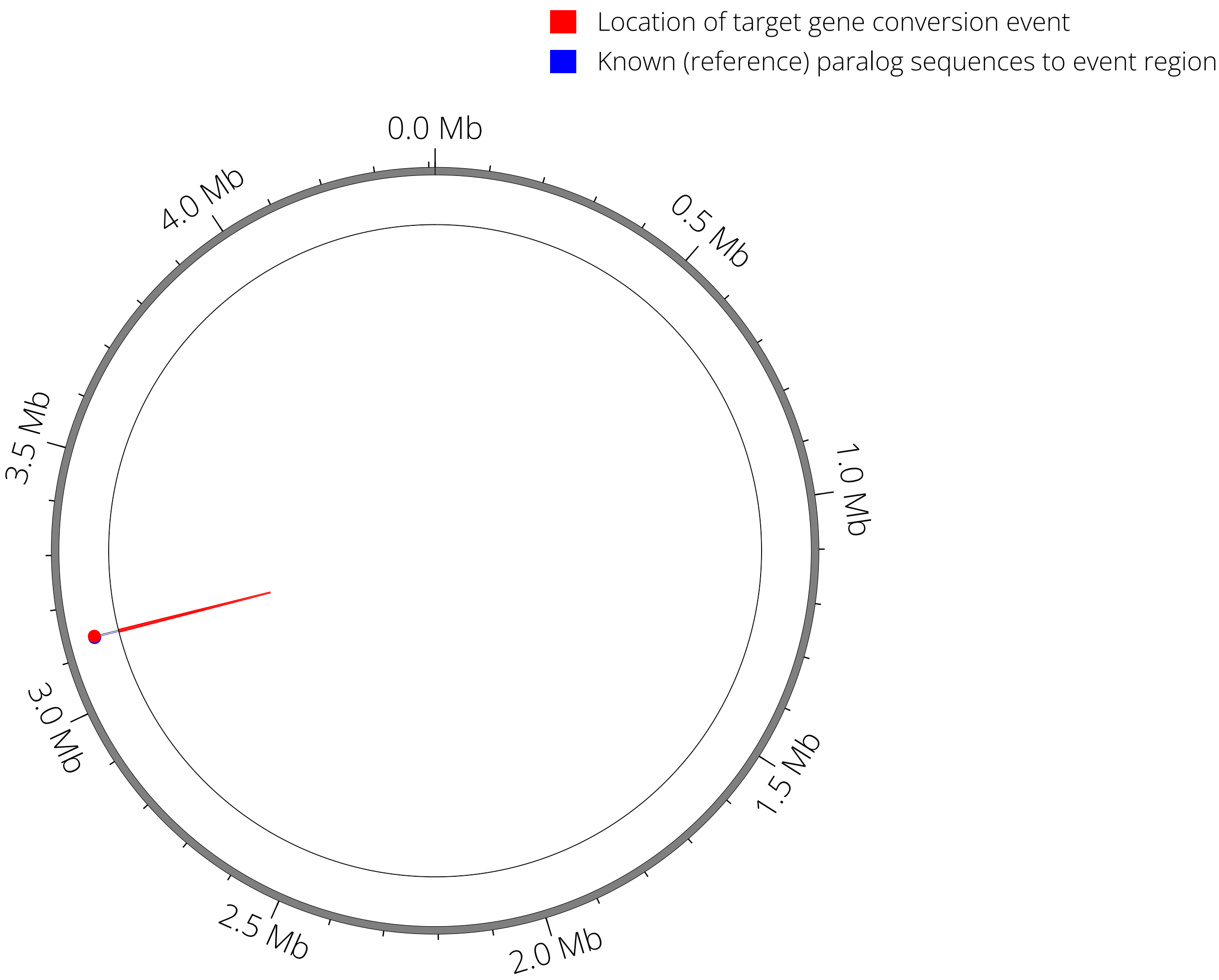

### Event_232 - 6 SNPs - k-mer analysis relative to paralogs

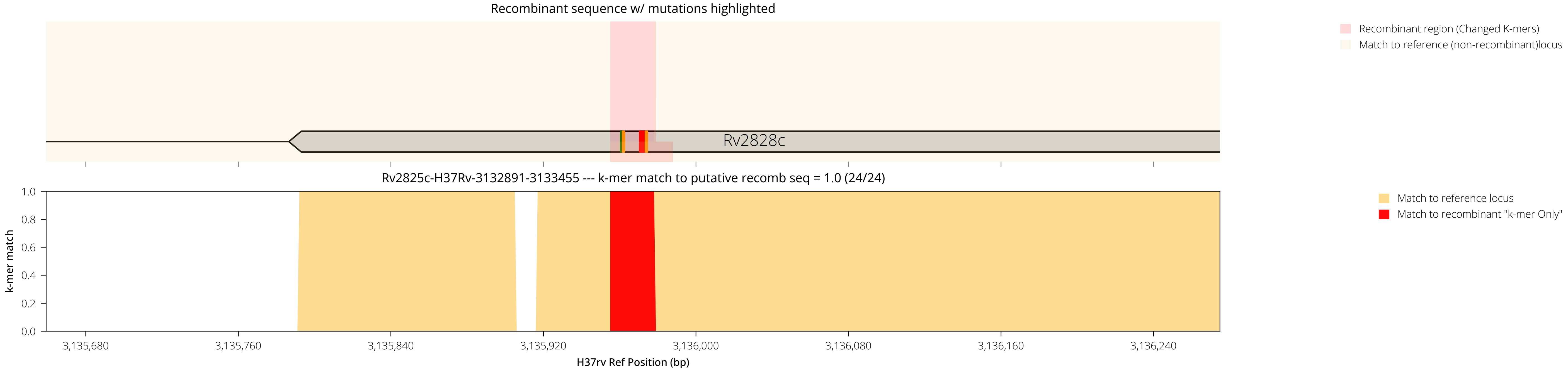

Sub-phylogeny Viz + Paralog Alignments
GCE ID: Event_232

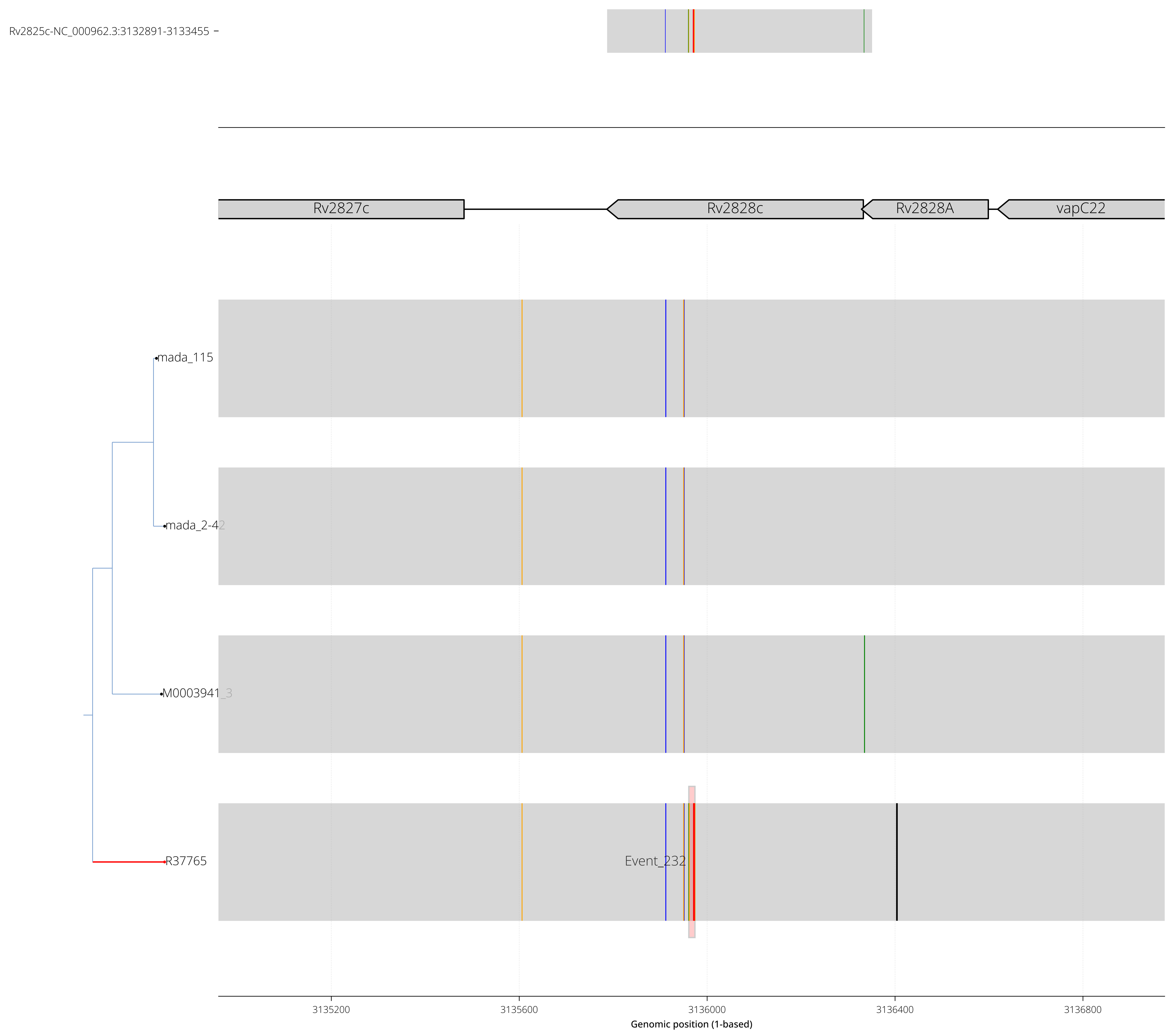

Full Phylogeny Viz + Paralog Alignments
GCE ID: Event_232

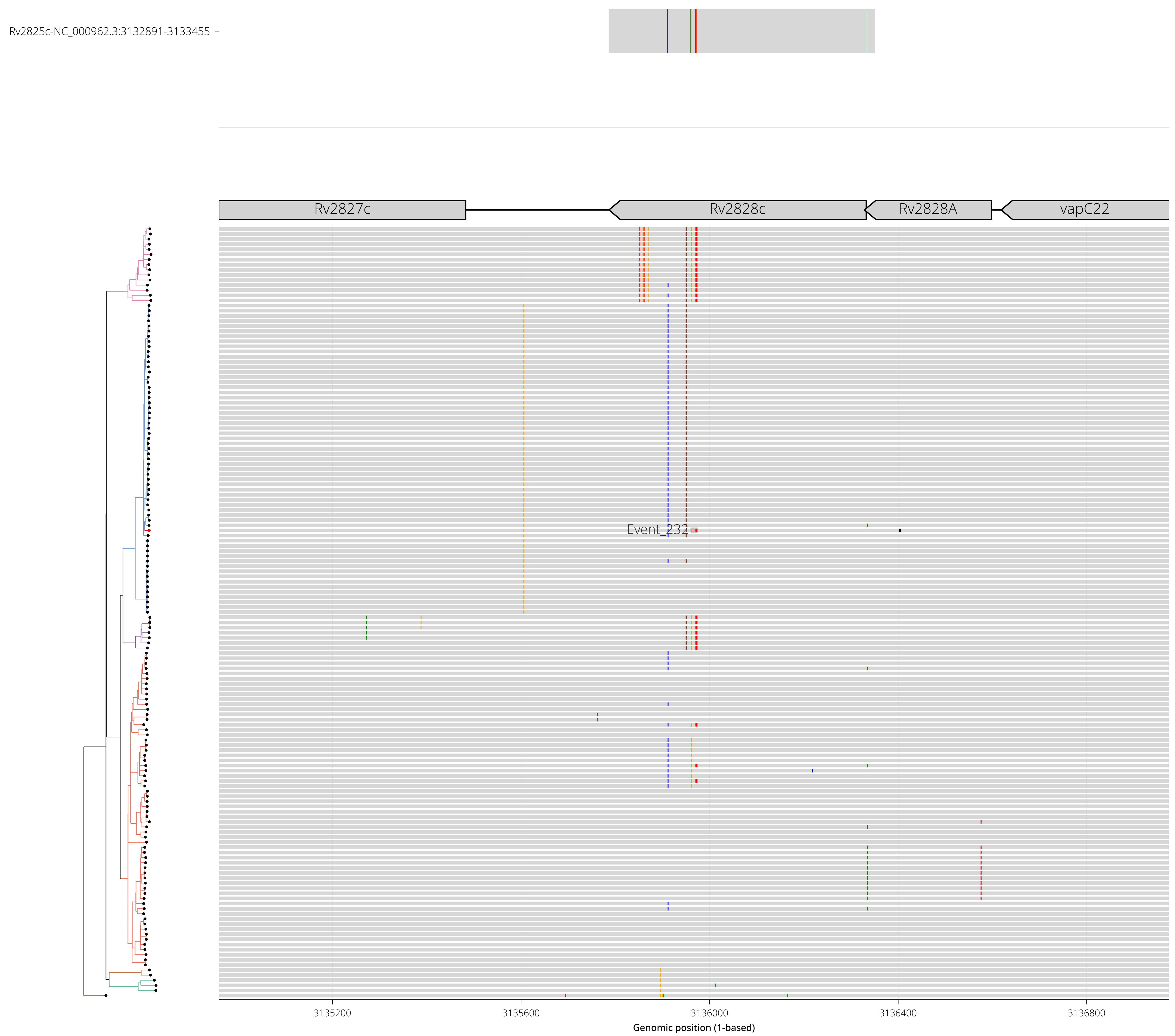

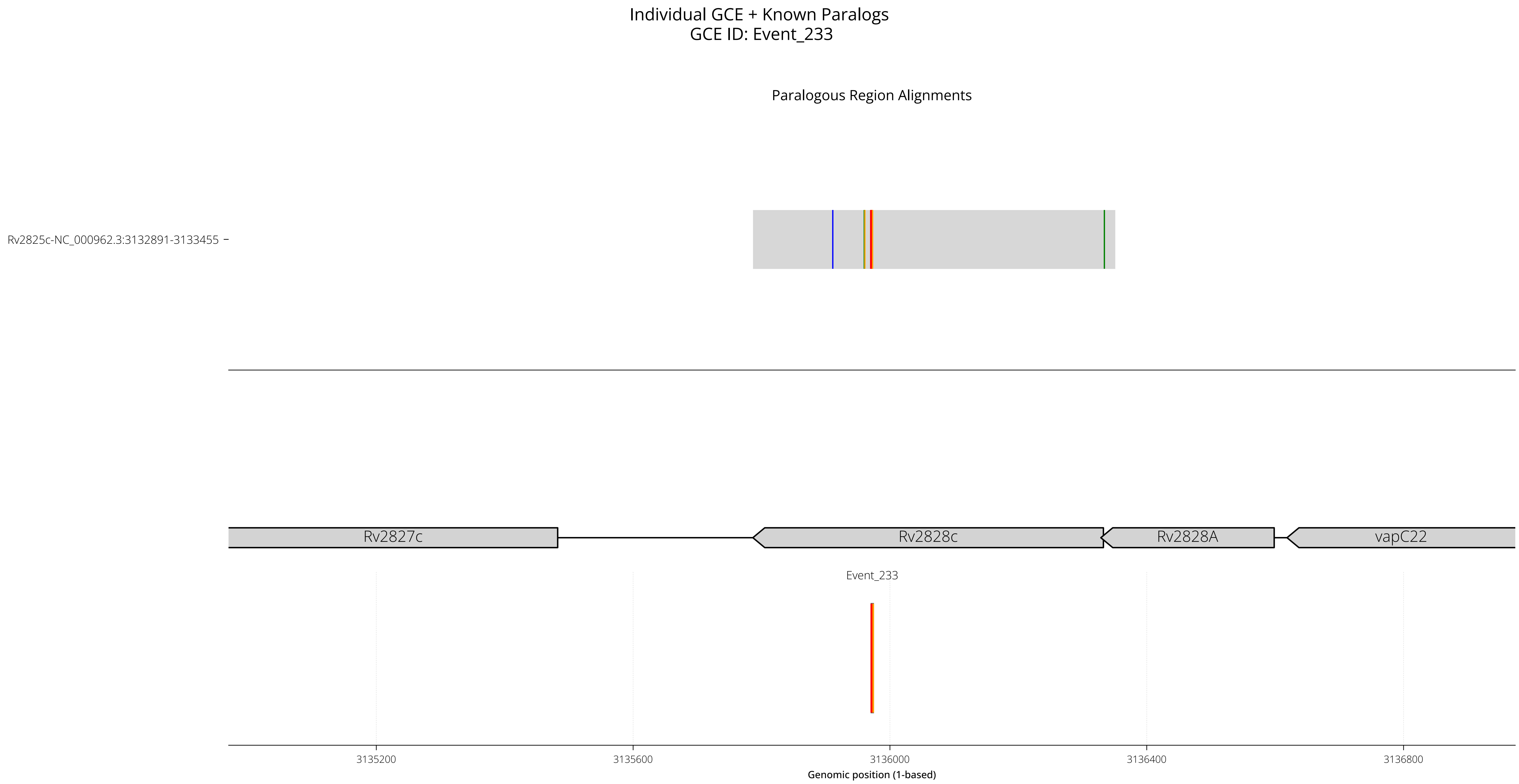
Individual GCE + Known Paralogs
GCE ID: Event_233
Paralogous Region Alignments
Rv2825c-NC_000962.3:3132891-3133455
Rv2827c
Rv2828c
Rv2828A
vapC22
Event_233
3135200
3135600
3136000
3136400
3136800
Genomic position (1-based)

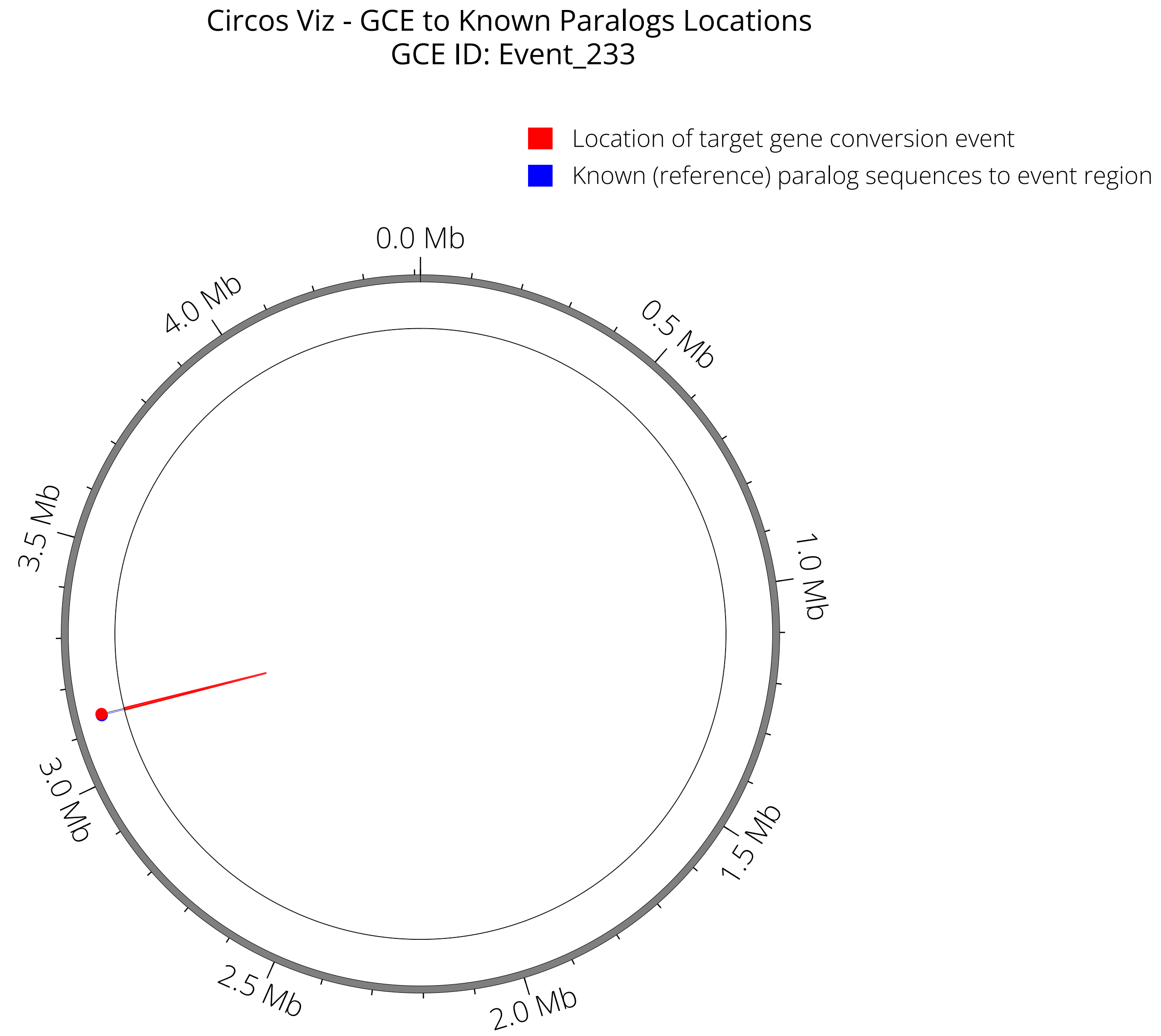

Circos Viz - GCE to Known Paralogs Locations
GCE ID: Event_233
Location of target gene conversion event
Known (reference) paralog sequences to event region
0.0 Mb
0.5 Mb
1.0 Mb
1.5 Mb
2.0 Mb
2.5 Mb
3.0 Mb
3.5 Mb
4.0 Mb

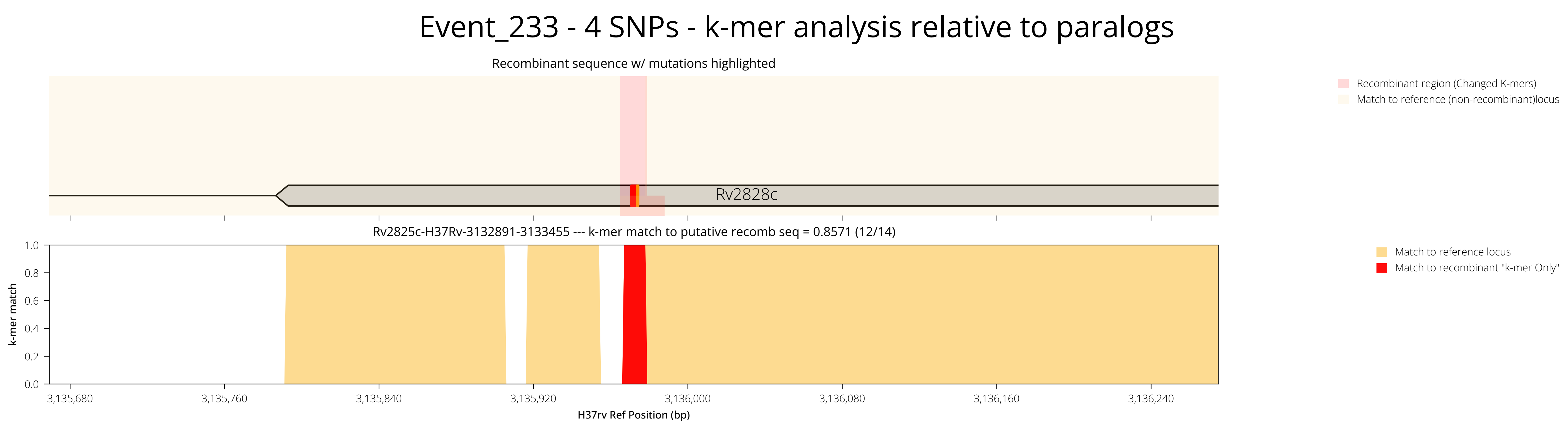
Event_233 - 4 SNPs - k-mer analysis relative to paralogs
Recombinant sequence w/ mutations highlighted
Rv2828c
Recombinant region (Changed K-mers)
Match to reference (non-recombinant)locus
Rv2825c-H37Rv-3132891-3133455 --- k-mer match to putative recomb seq = 0.8571 (12/14)
k-mer match
1.0
0.8
0.6
0.4
0.2
0.0
3,135,680
3,135,760
3,135,840
3,135,920
3,136,000
3,136,080
3,136,160
3,136,240
H37rv Ref Position (bp)
Match to reference locus
Match to recombinant "k-mer Only"

Sub-phylogeny Viz + Paralog Alignments
GCE ID: Event_233

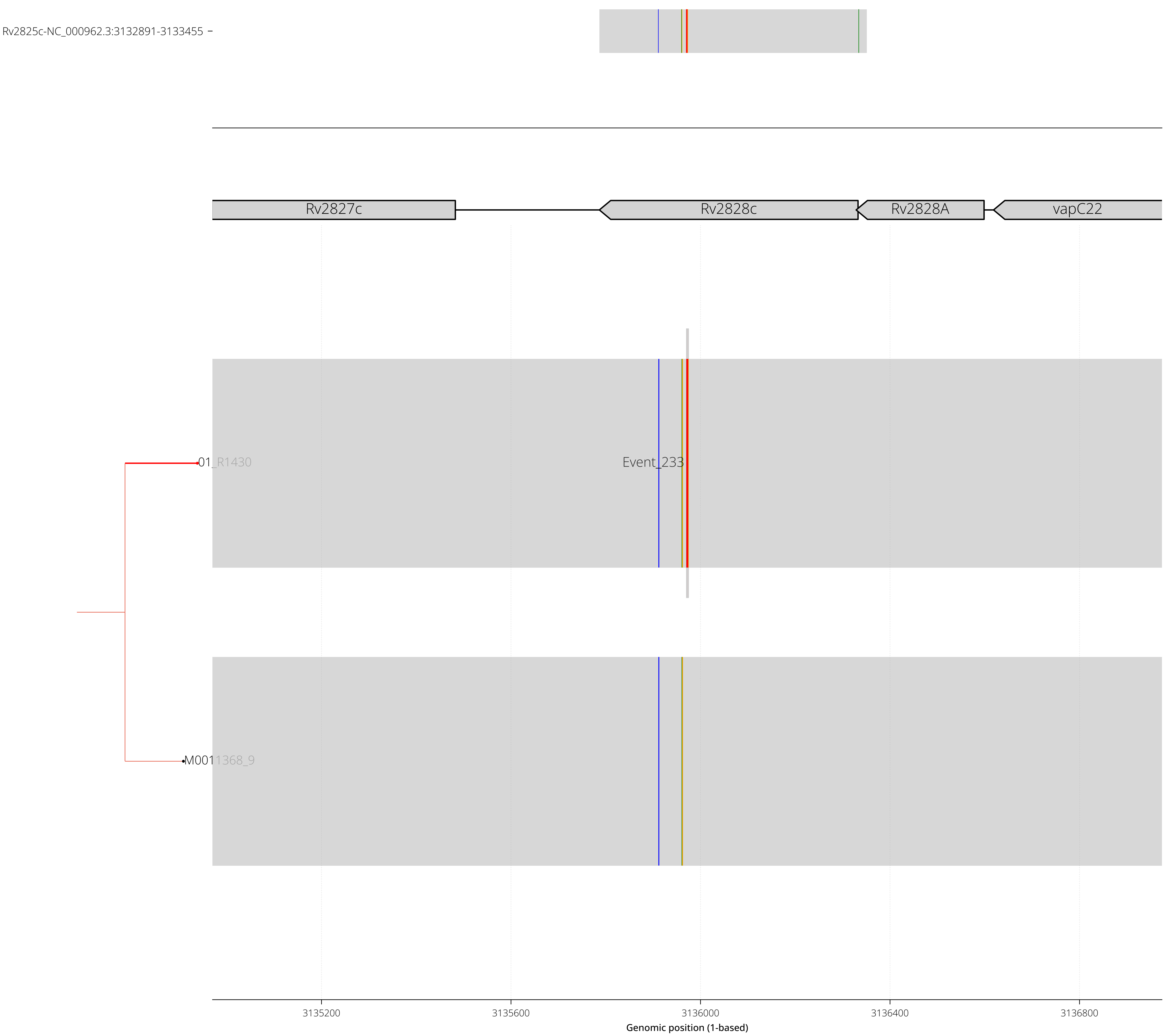

Full Phylogeny Viz + Paralog Alignments
GCE ID: Event_233

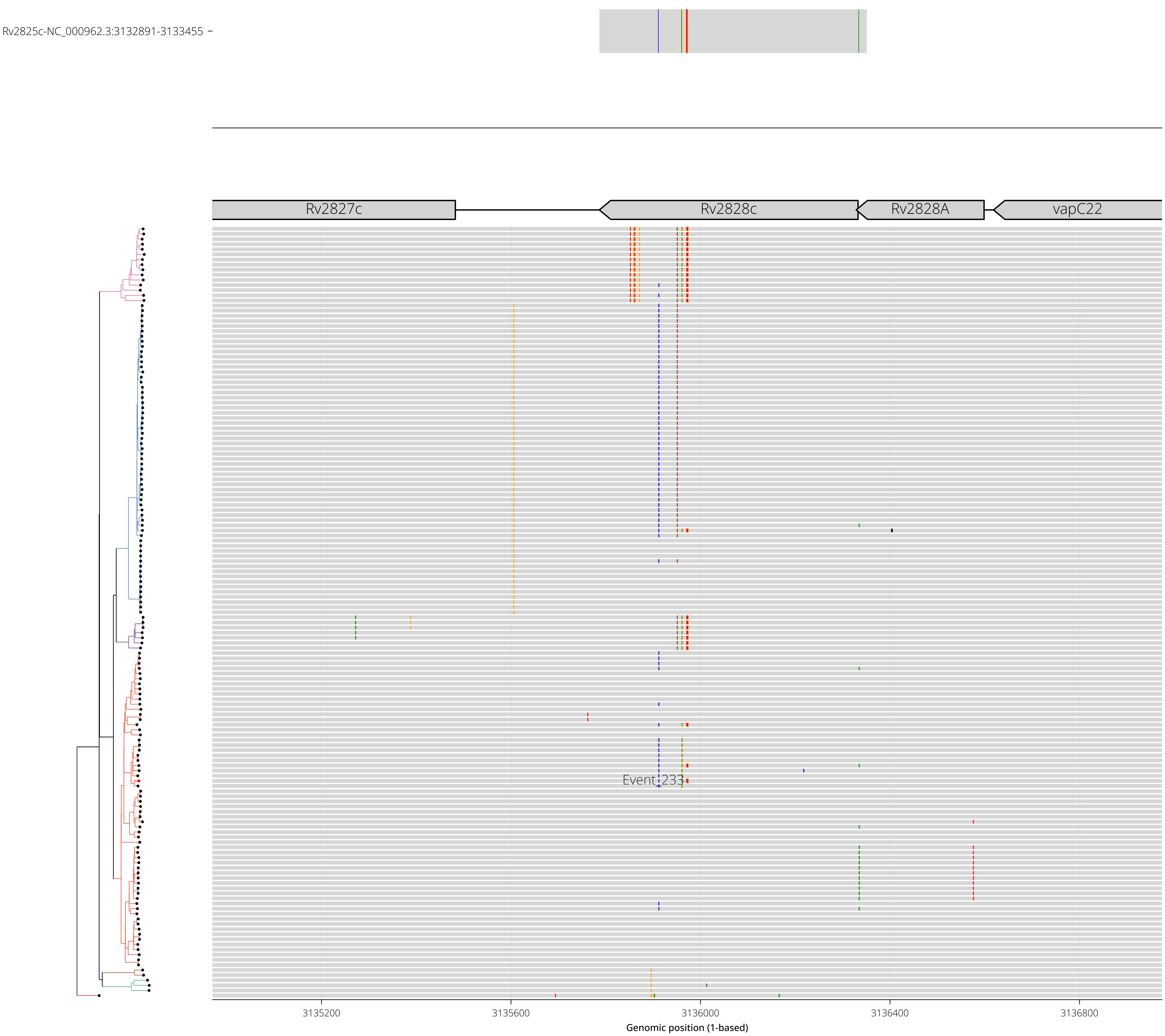

#### Individual GCE + Known Paralogs
#### GCE ID: Event_234

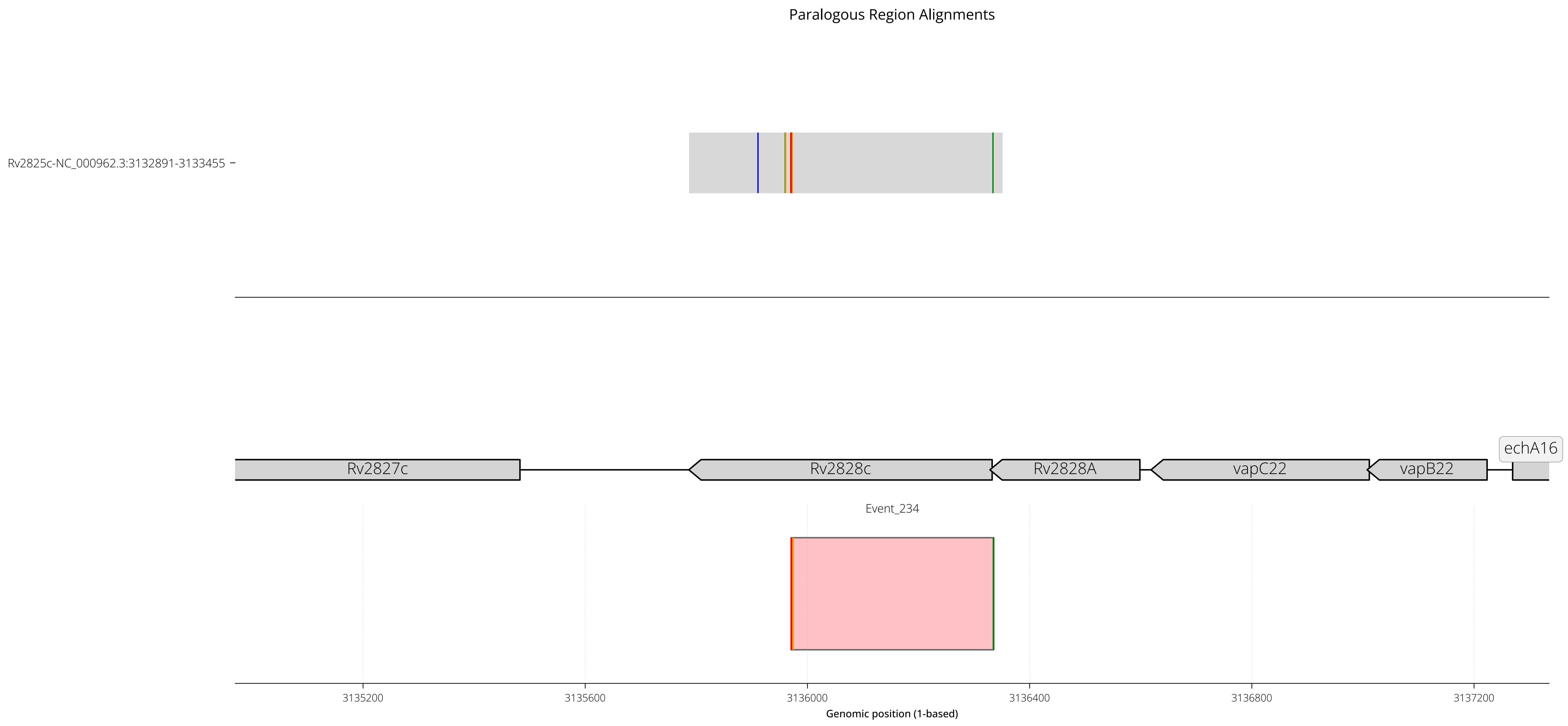

Circos Viz - GCE to Known Paralogs Locations
GCE ID: Event_234

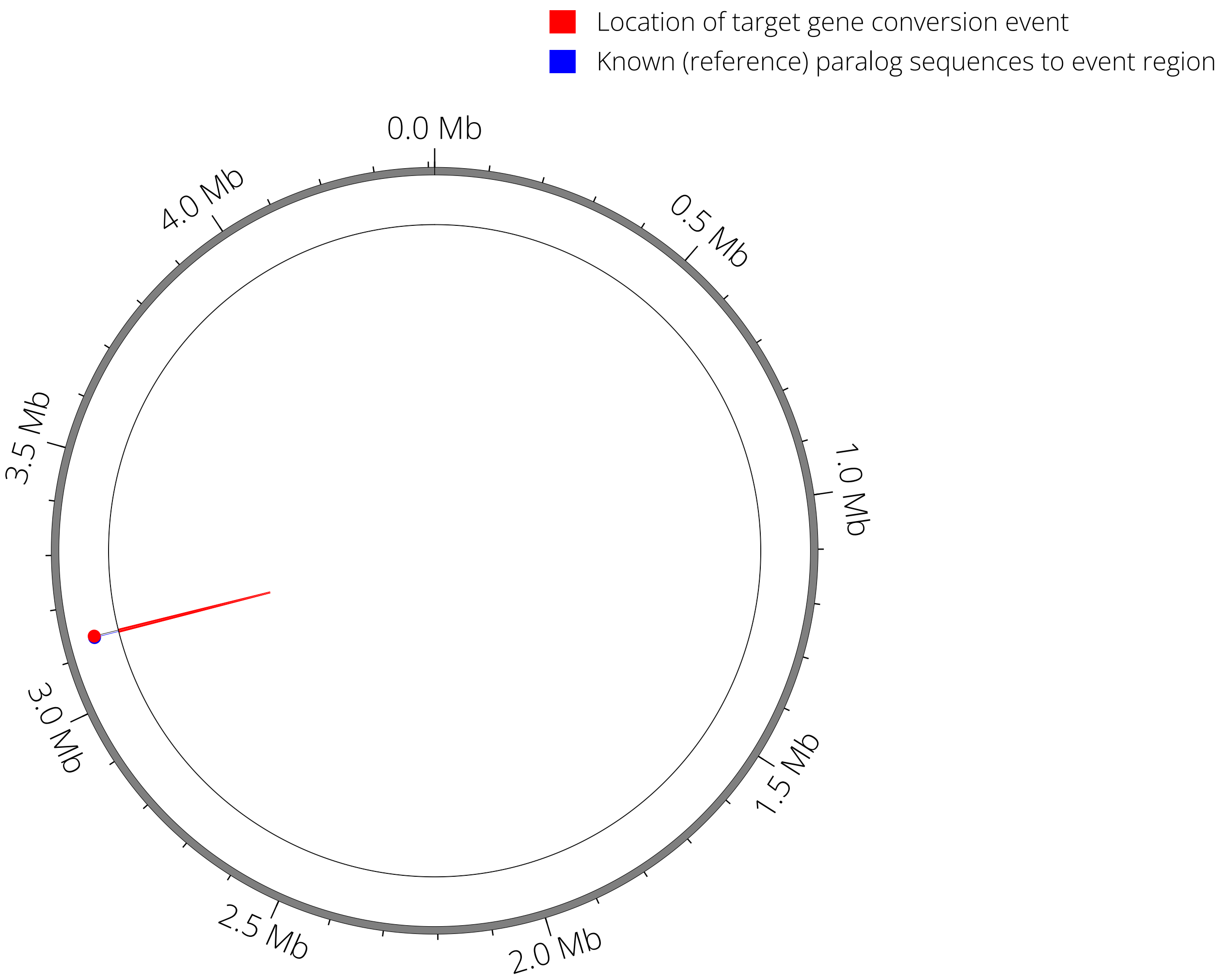

### Event_234 - 5 SNPs - k-mer analysis relative to paralogs

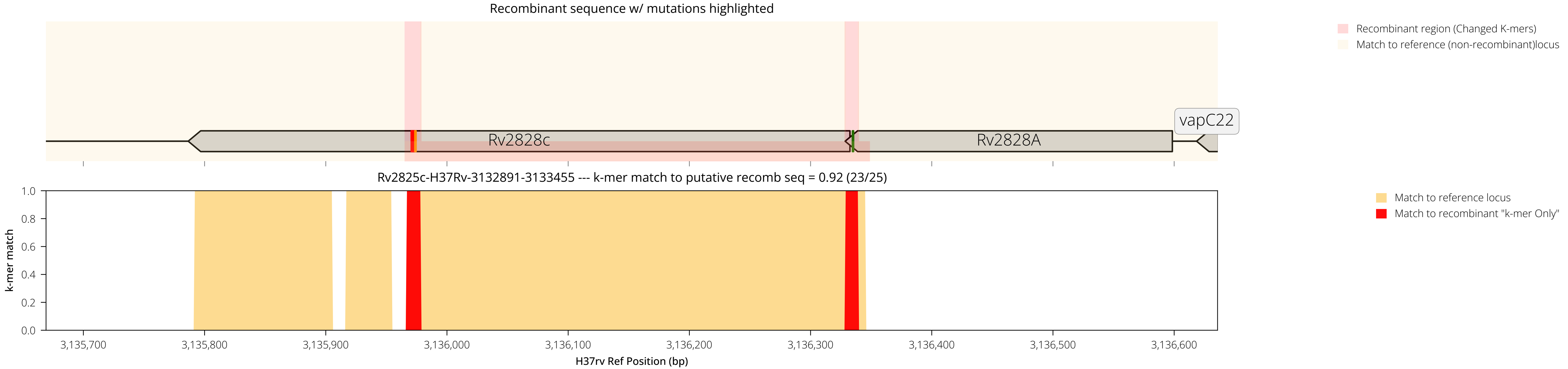

Sub-phylogeny Viz + Paralog Alignments
GCE ID: Event_234

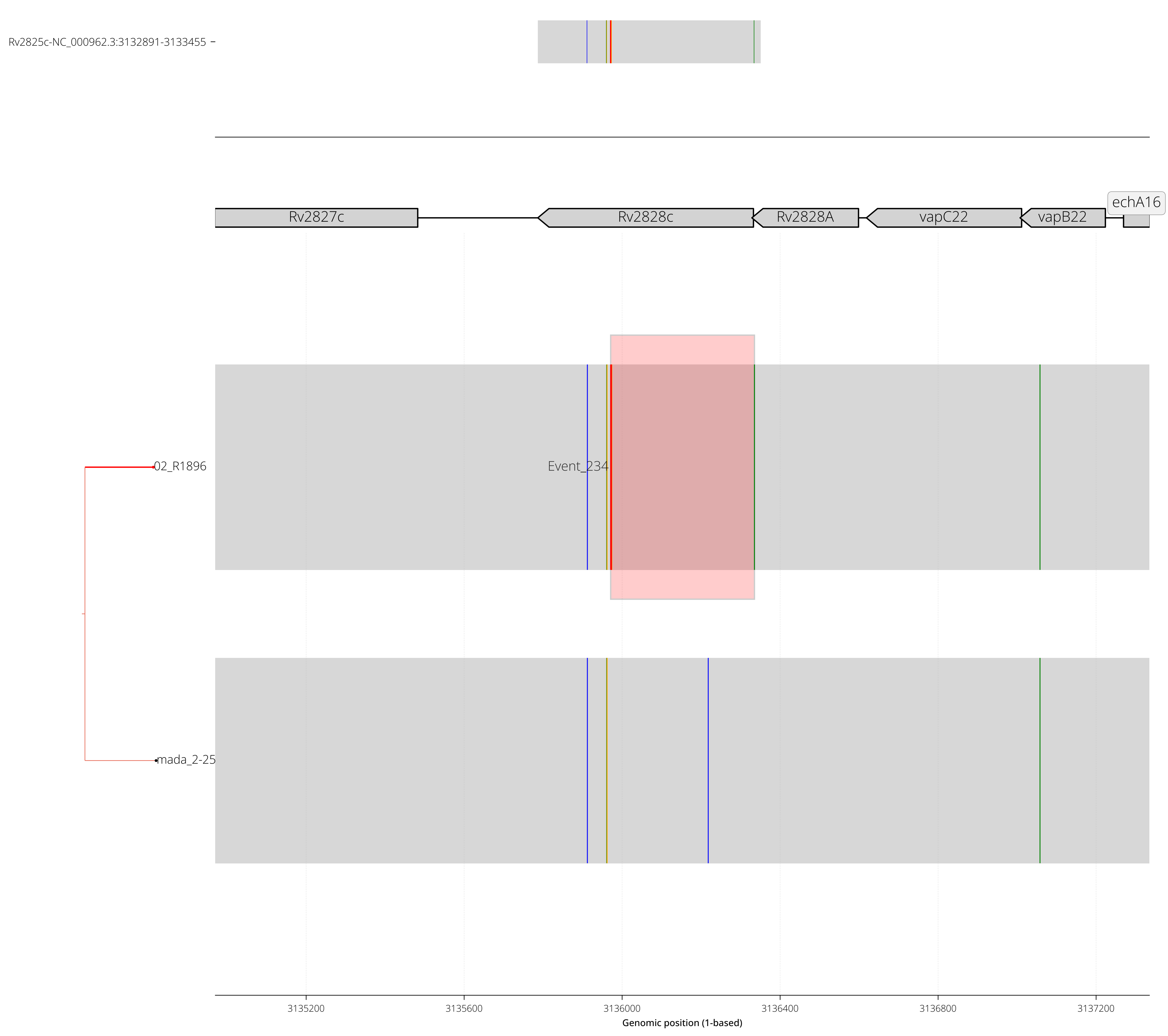

### Full Phylogeny Viz + Paralog Alignments
GCE ID: Event_234

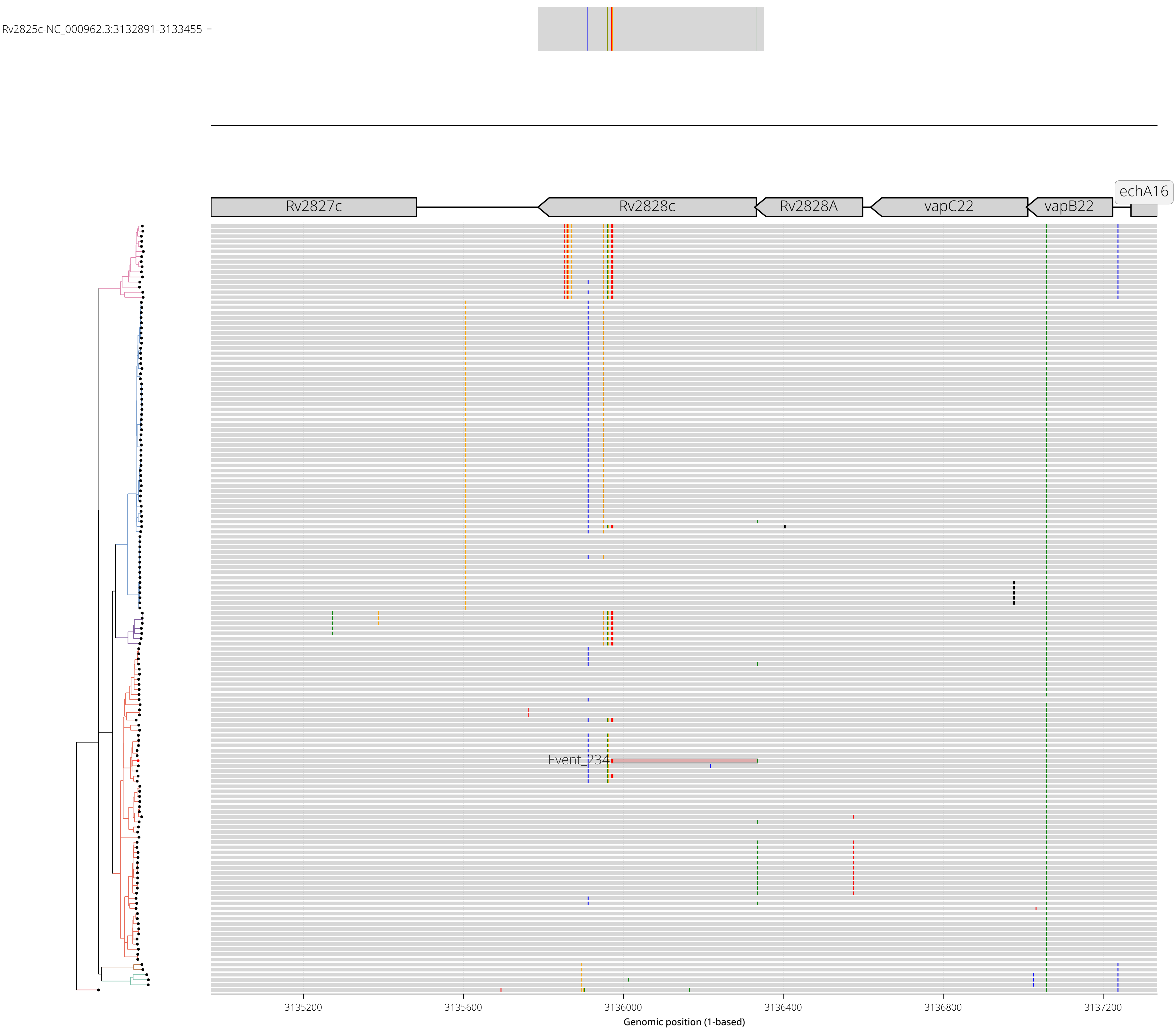

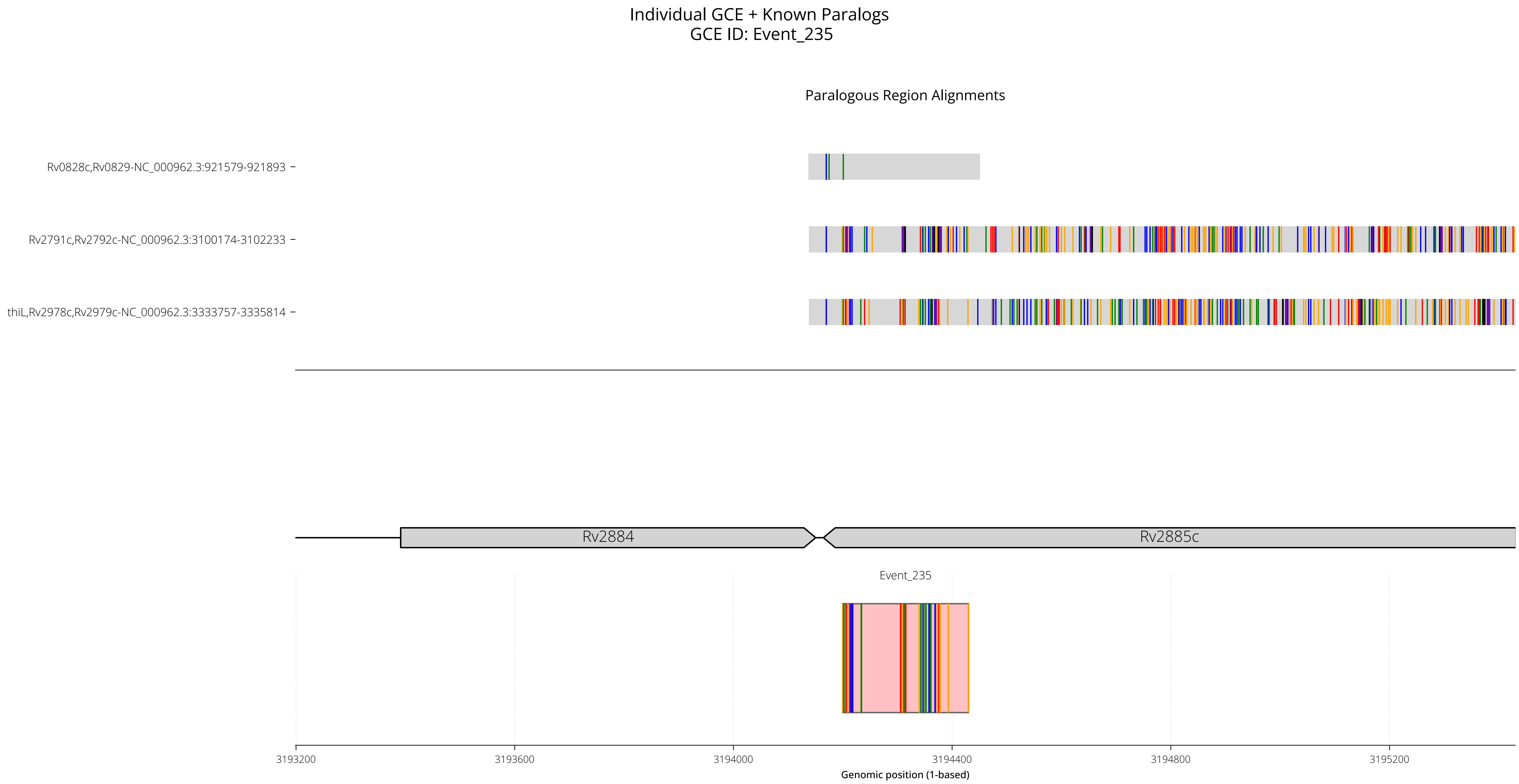

Individual GCE + Known Paralogs
GCE ID: Event_235
Paralogous Region Alignments
Rv0828c,Rv0829-NC_000962.3:921579-921893
Rv2791c,Rv2792c-NC_000962.3:3100174-3102233
thiL,Rv2978c,Rv2979c-NC_000962.3:3333757-3335814
Rv2884
Rv2885c
Event_235
3193200
3193600
3194000
3194400
3194800
3195200
Genomic position (1-based)

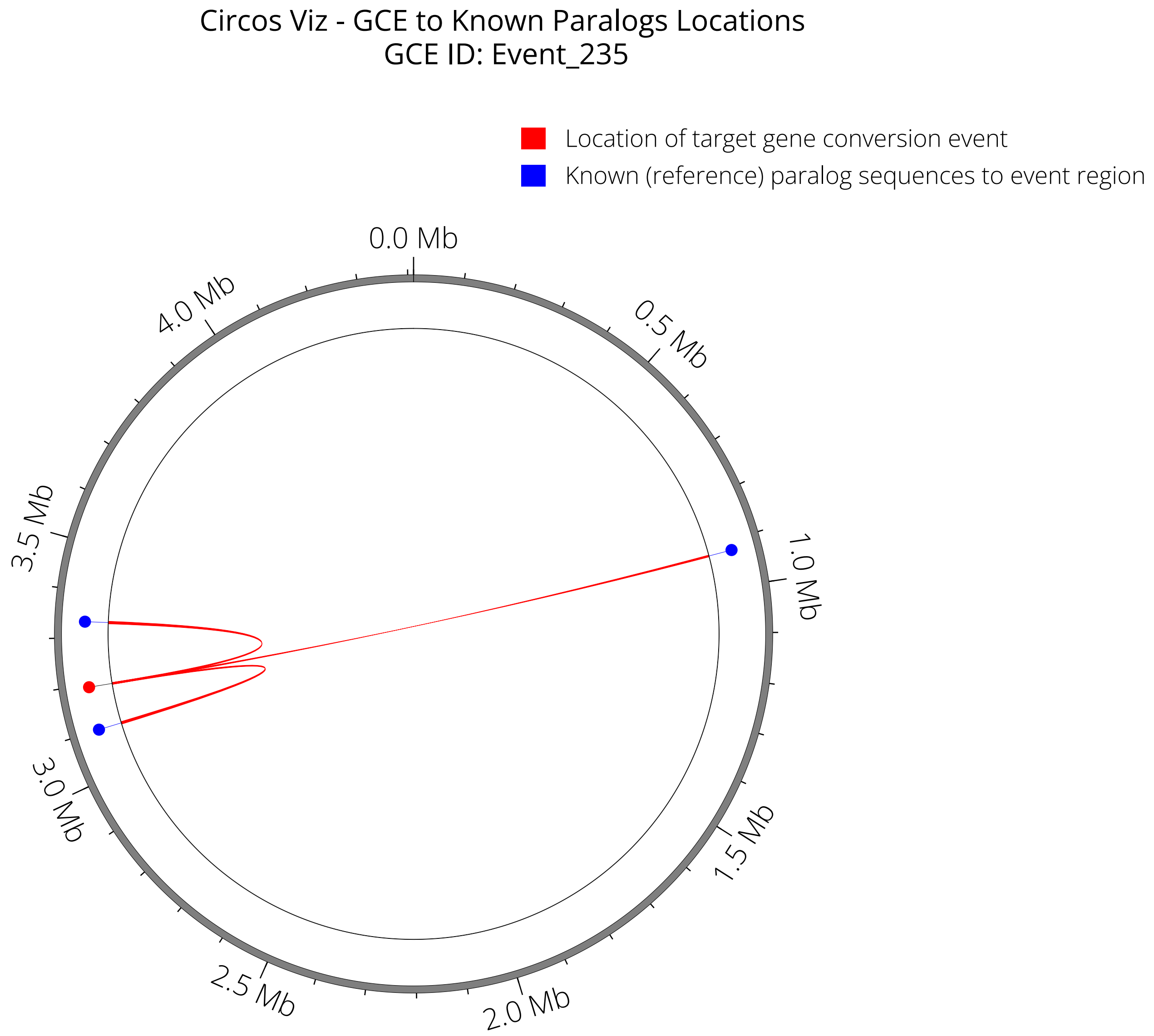
Circos Viz - GCE to Known Paralogs Locations
GCE ID: Event_235
Location of target gene conversion event
Known (reference) paralog sequences to event region
0.0 Mb
0.5 Mb
1.0 Mb
1.5 Mb
2.0 Mb
2.5 Mb
3.0 Mb
3.5 Mb
4.0 Mb

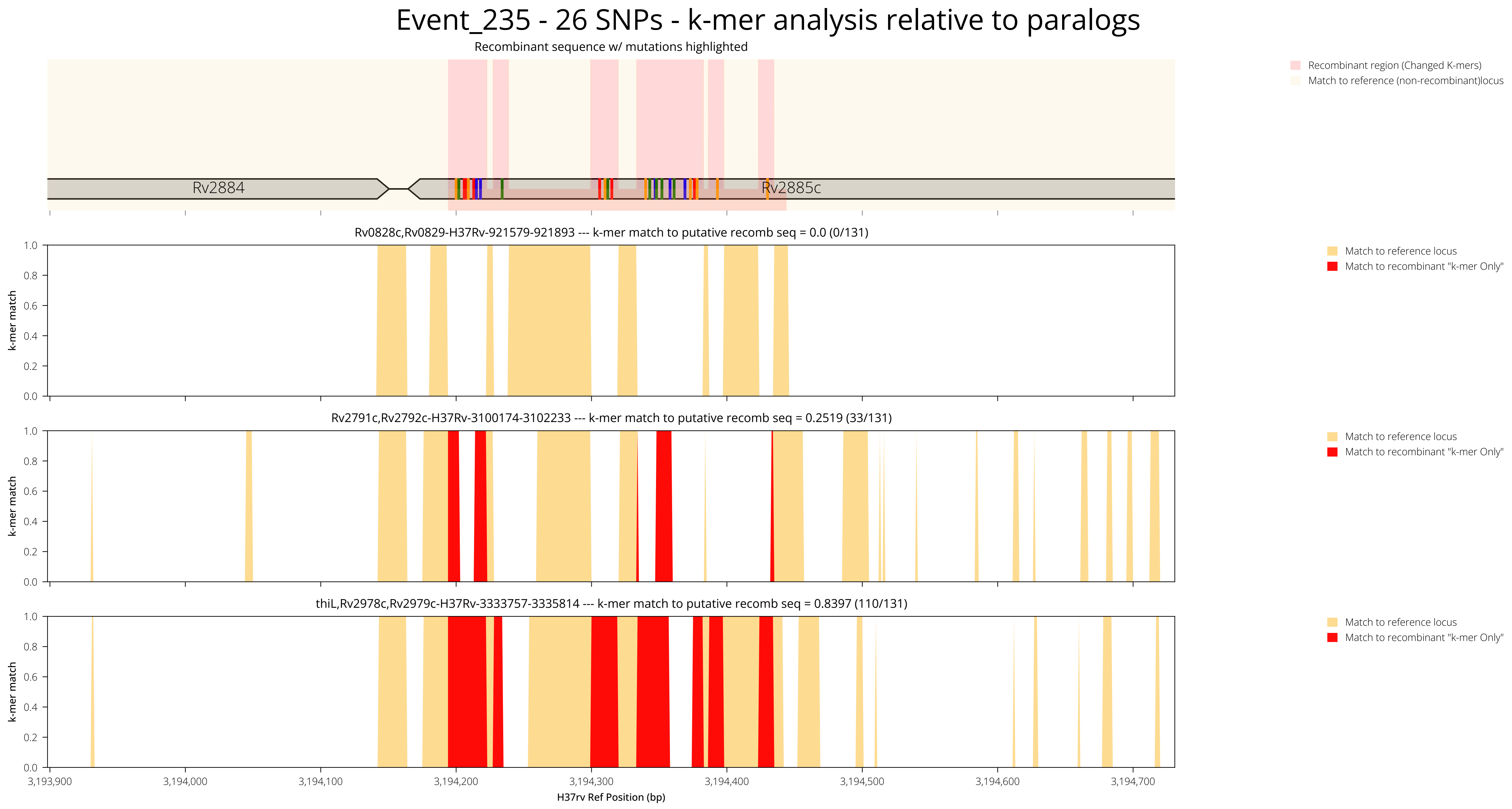

Event_235 - 26 SNPs - k-mer analysis relative to paralogs
Recombinant sequence w/ mutations highlighted
Recombinant region (Changed K-mers)
Match to reference (non-recombinant)locus
Rv2884
Rv2885c
Rv0828c,Rv0829-H37Rv-921579-921893 --- k-mer match to putative recomb seq = 0.0 (0/131)
Rv2791c,Rv2792c-H37Rv-3100174-3102233 --- k-mer match to putative recomb seq = 0.2519 (33/131)
thiL,Rv2978c,Rv2979c-H37Rv-3333757-3335814 --- k-mer match to putative recomb seq = 0.8397 (110/131)
Match to reference locus
Match to recombinant "k-mer Only"
k-mer match
1.0
0.8
0.6
0.4
0.2
0.0
3,193,900
3,194,000
3,194,100
3,194,200
3,194,300
3,194,400
3,194,500
3,194,600
3,194,700
H37rv Ref Position (bp)

Sub-phylogeny Viz + Paralog Alignments
GCE ID: Event_235

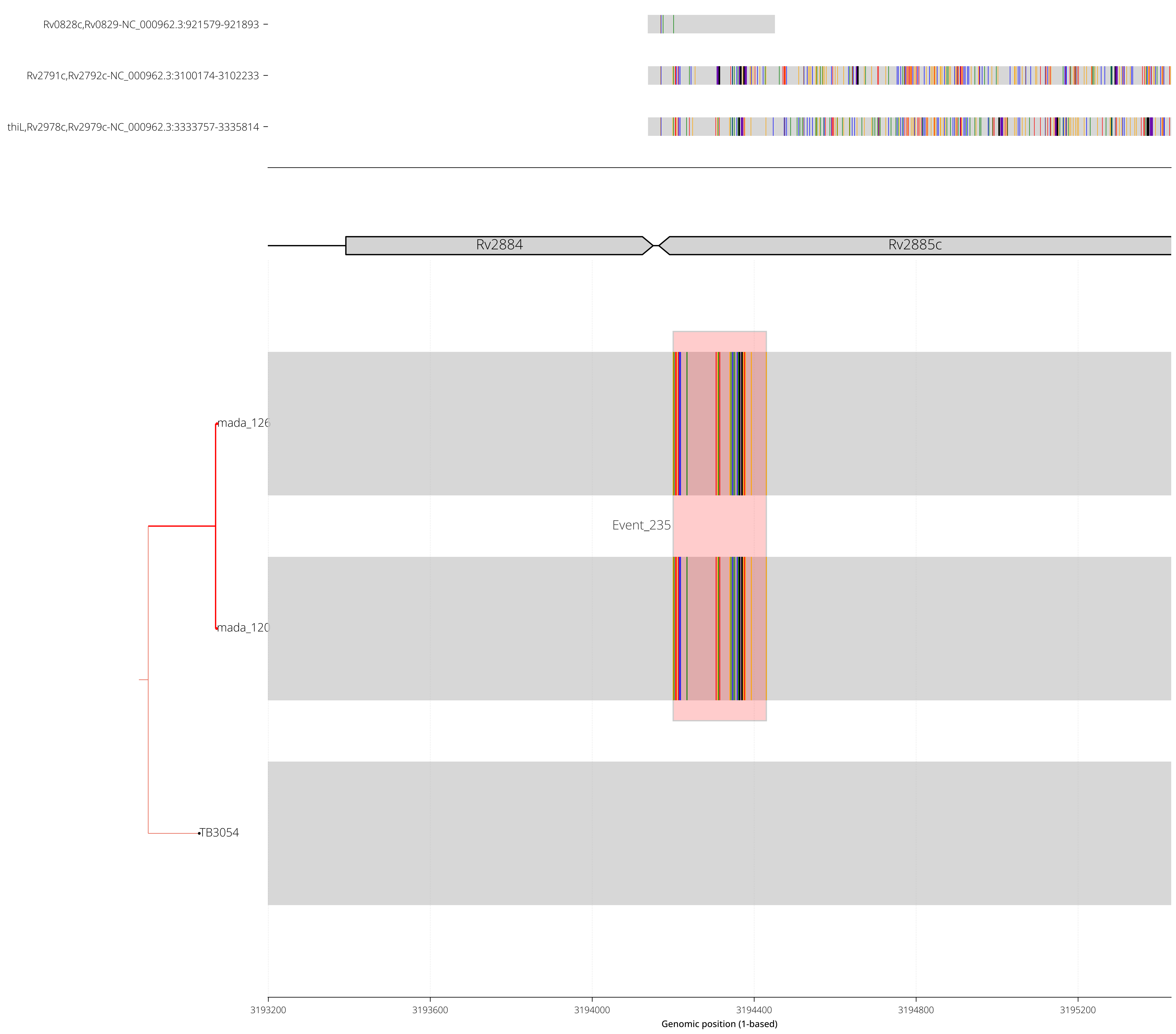

Full Phylogeny Viz + Paralog Alignments
GCE ID: Event_235

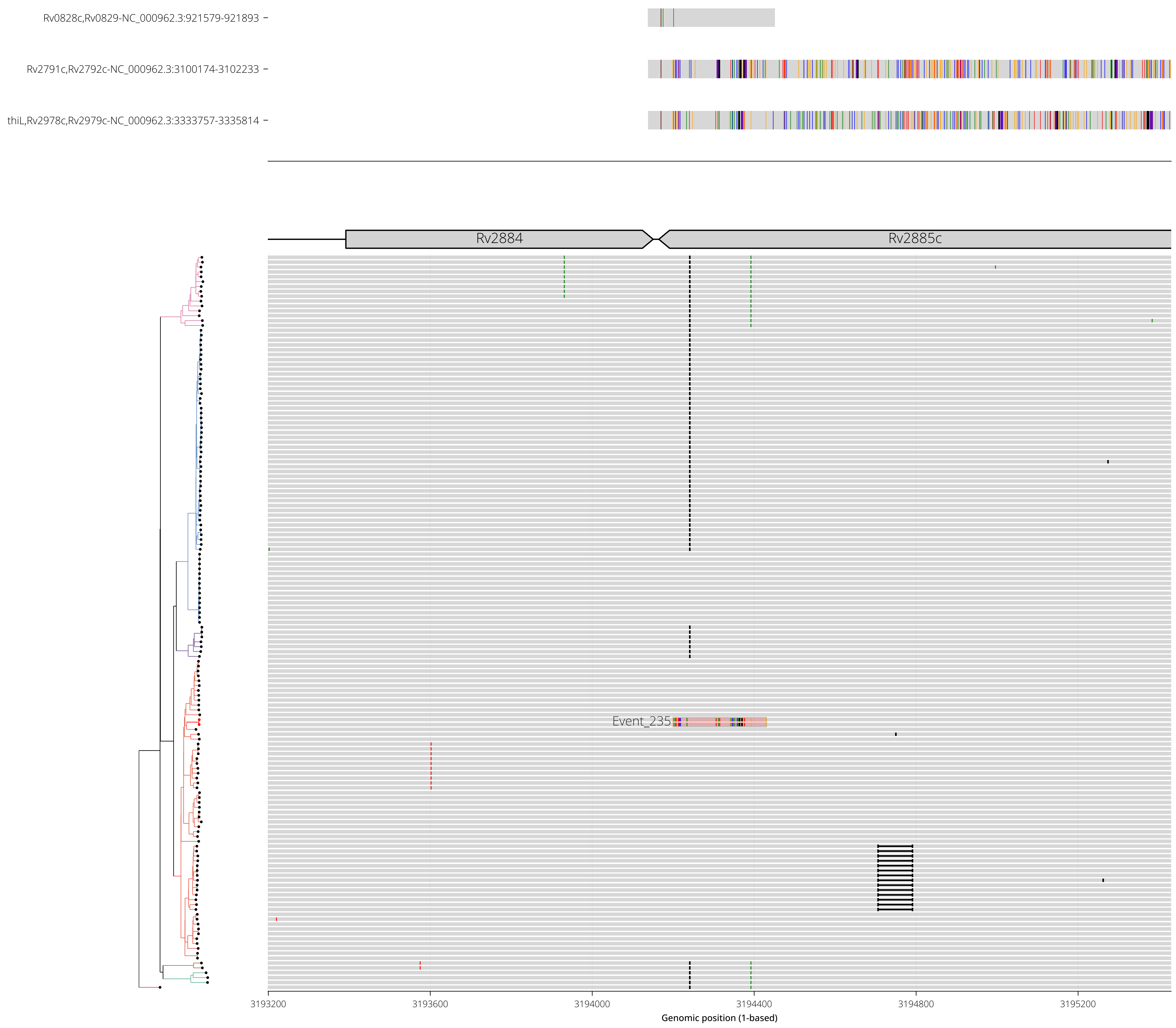

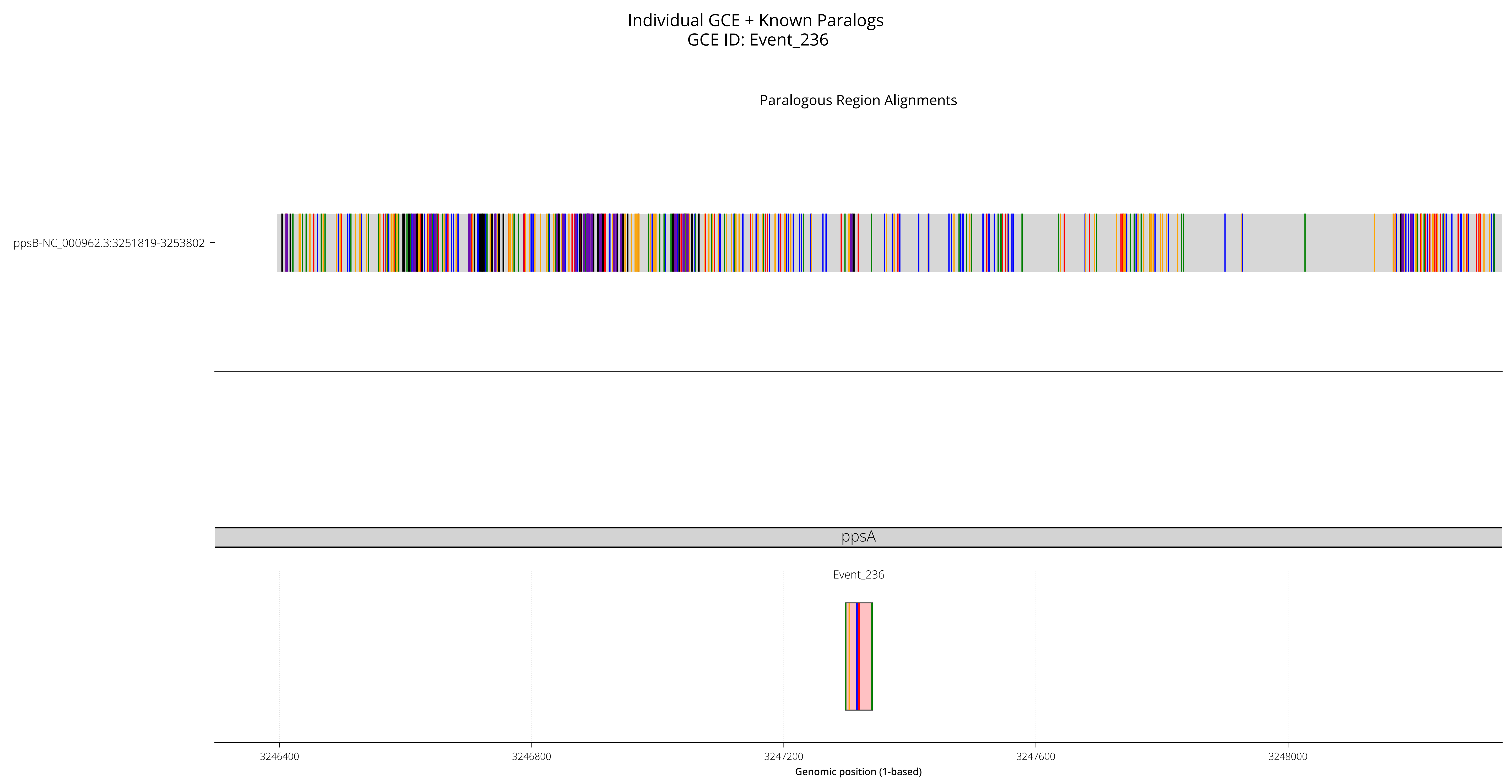
Individual GCE + Known Paralogs
GCE ID: Event_236
Paralogous Region Alignments
ppsB-NC_000962.3:3251819-3253802
ppsA
Event_236
3246400
3246800
3247200
3247600
3248000
Genomic position (1-based)

Circos Viz - GCE to Known Paralogs Locations
GCE ID: Event_236

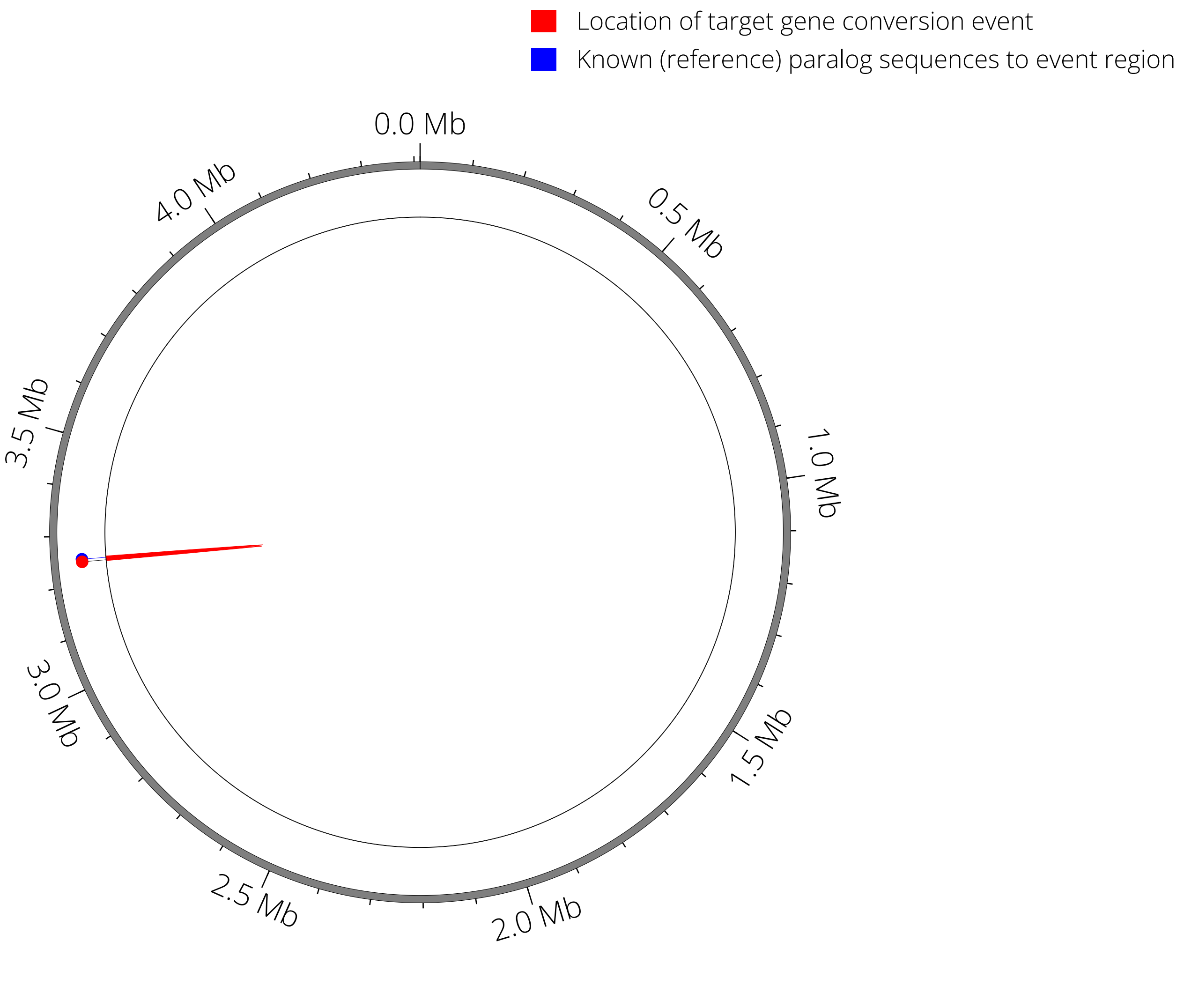

### Event_236 - 5 SNPs - k-mer analysis relative to paralogs

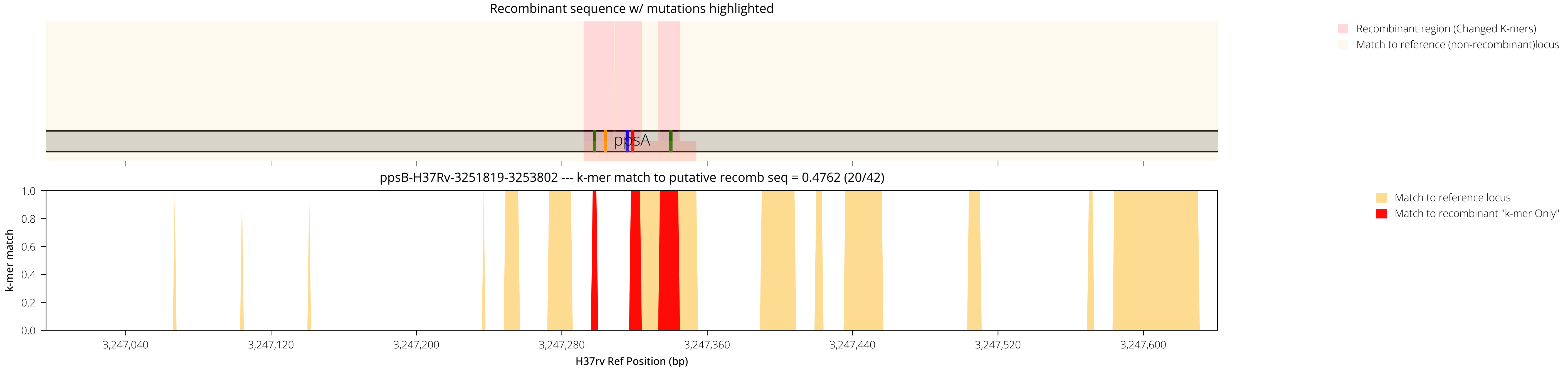

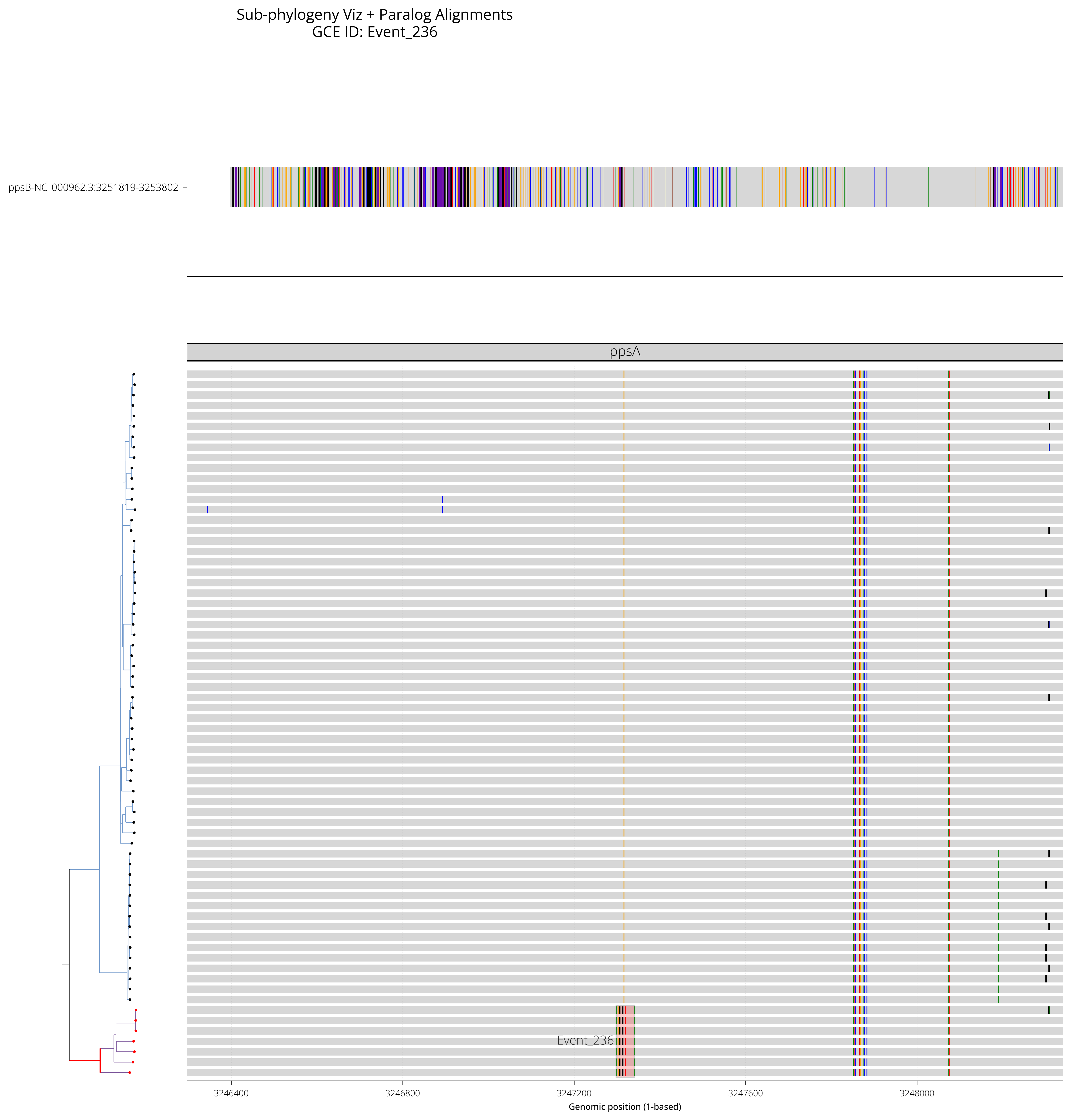

Sub-phylogeny Viz + Paralog Alignments
GCE ID: Event_236
ppsB-NC_000962.3:3251819-3253802
ppsA
Event_236
3246400
3246800
3247200
3247600
3248000
Genomic position (1-based)

Full Phylogeny Viz + Paralog Alignments
GCE ID: Event_236

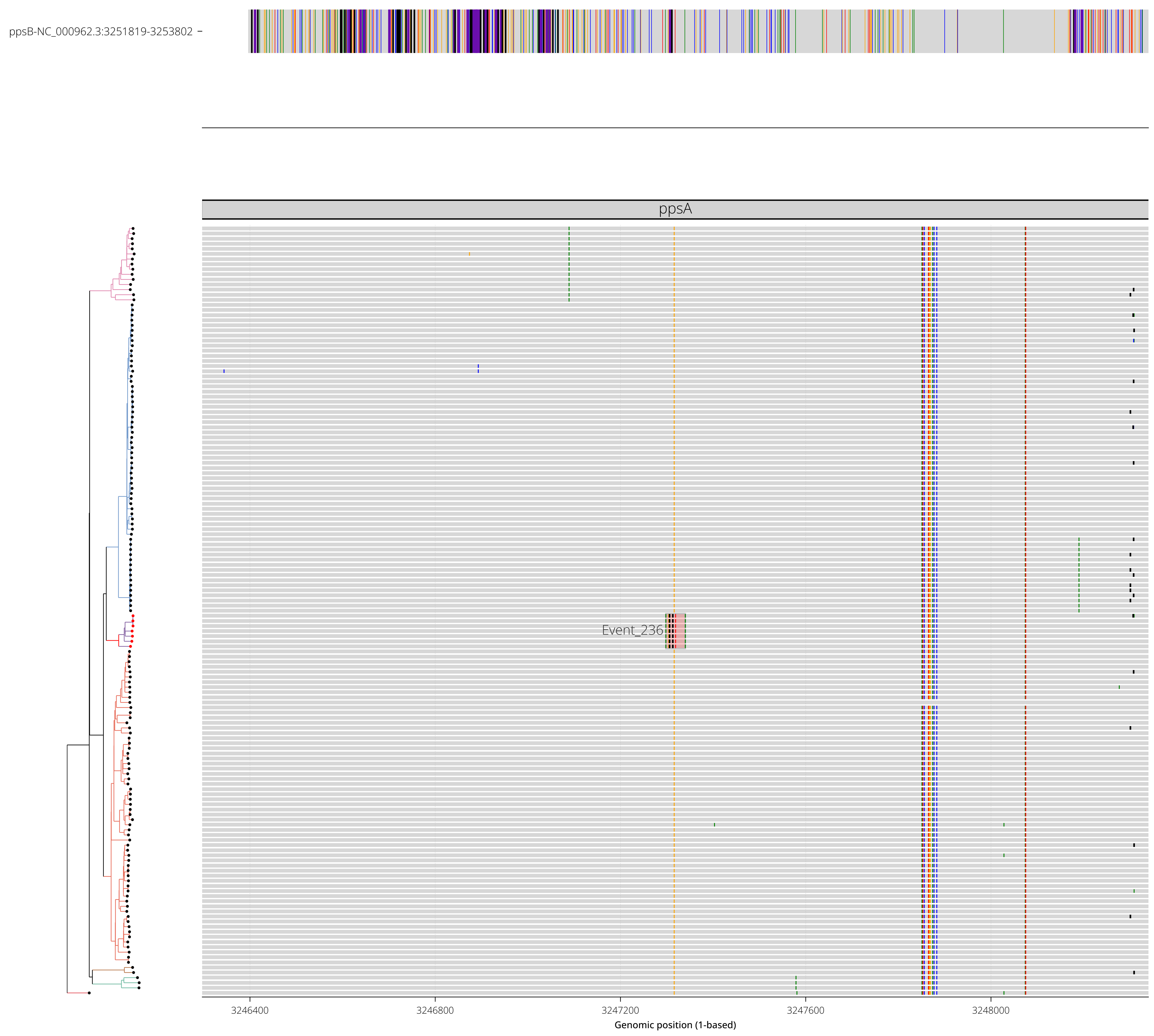

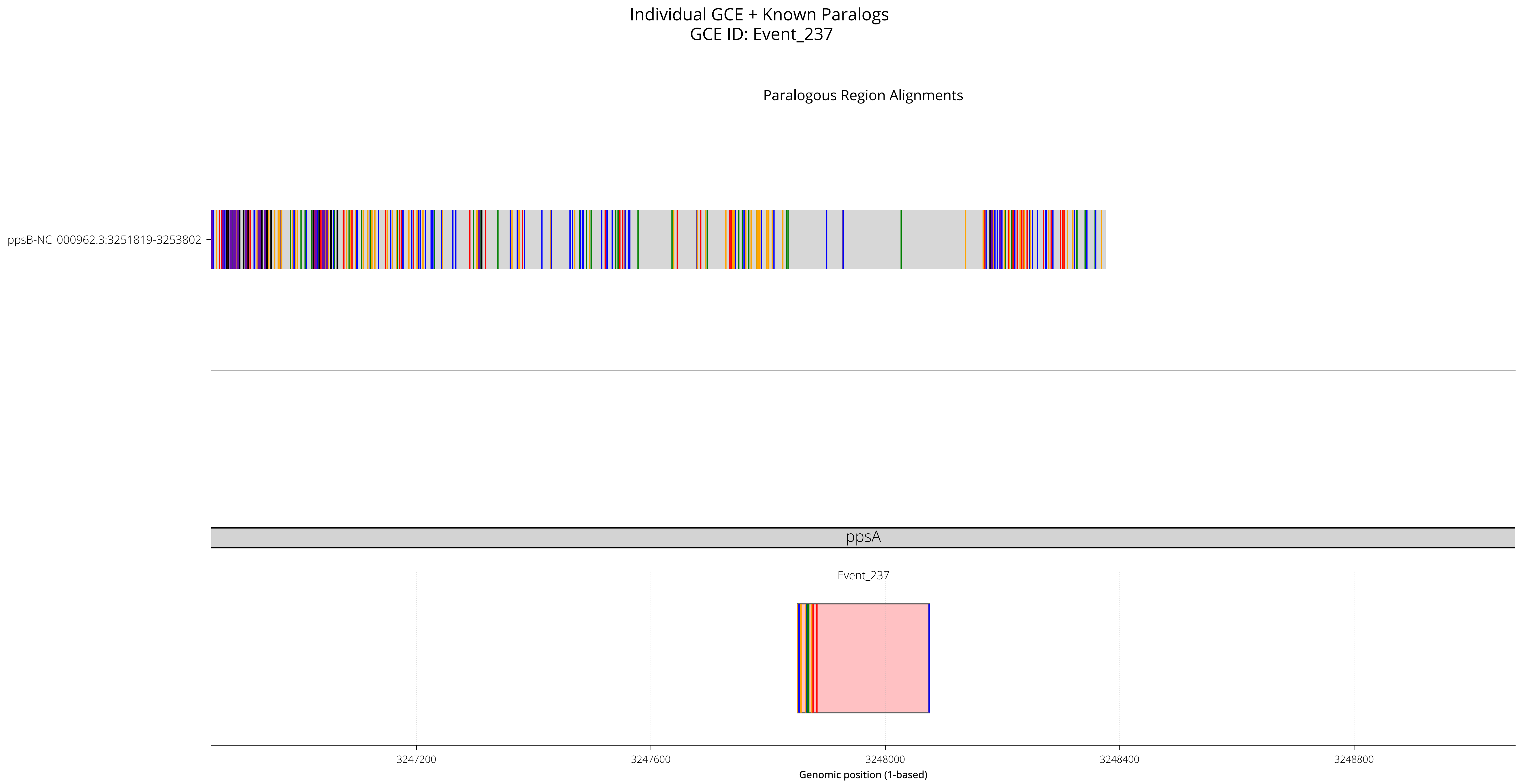
Individual GCE + Known Paralogs
GCE ID: Event_237
Paralogous Region Alignments
ppsB-NC_000962.3:3251819-3253802
ppsA
Event_237
3247200
3247600
3248000
3248400
3248800
Genomic position (1-based)

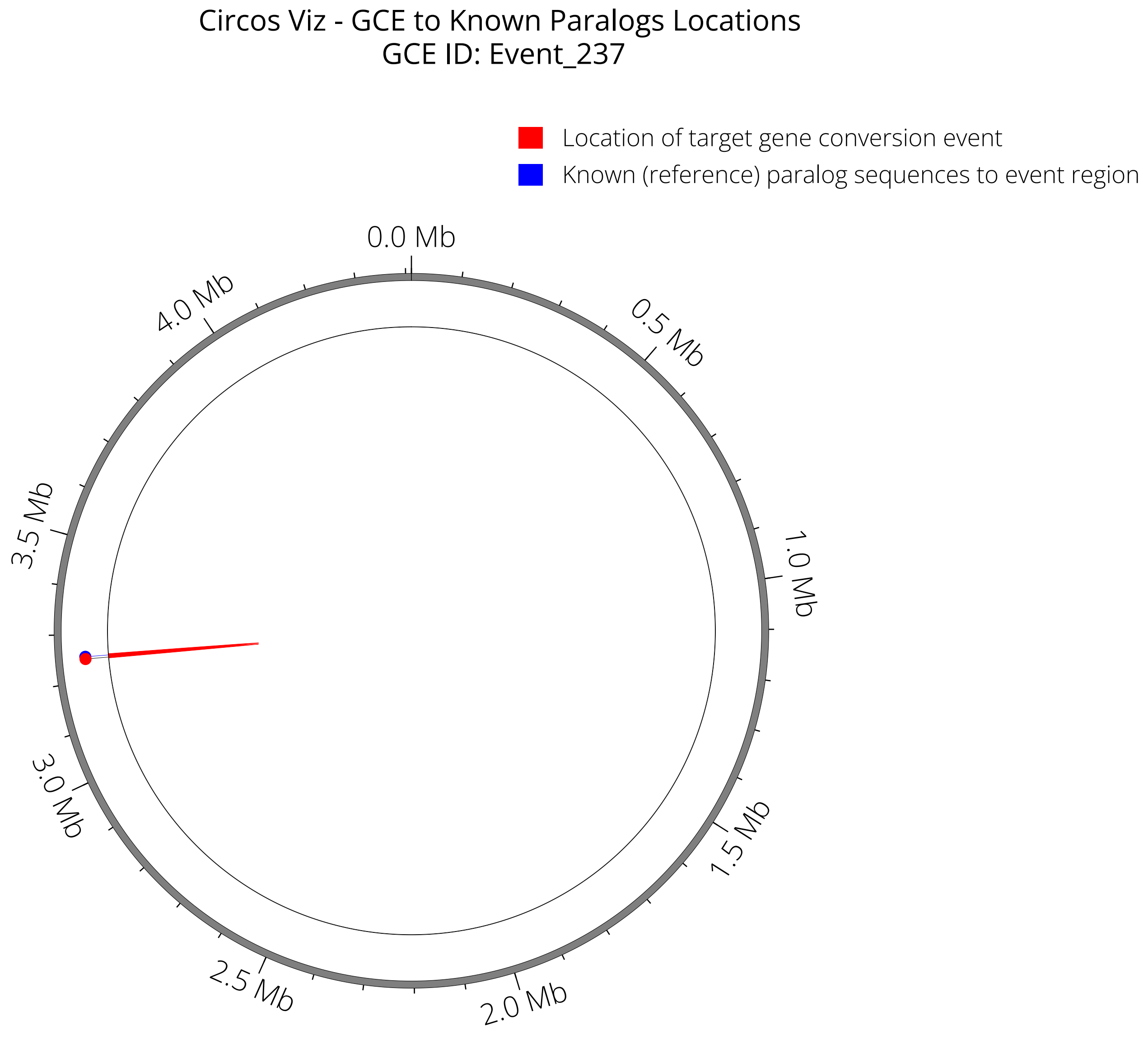
Circos Viz - GCE to Known Paralogs Locations
GCE ID: Event_237
Location of target gene conversion event
Known (reference) paralog sequences to event region
0.0 Mb
0.5 Mb
1.0 Mb
1.5 Mb
2.0 Mb
2.5 Mb
3.0 Mb
3.5 Mb
4.0 Mb

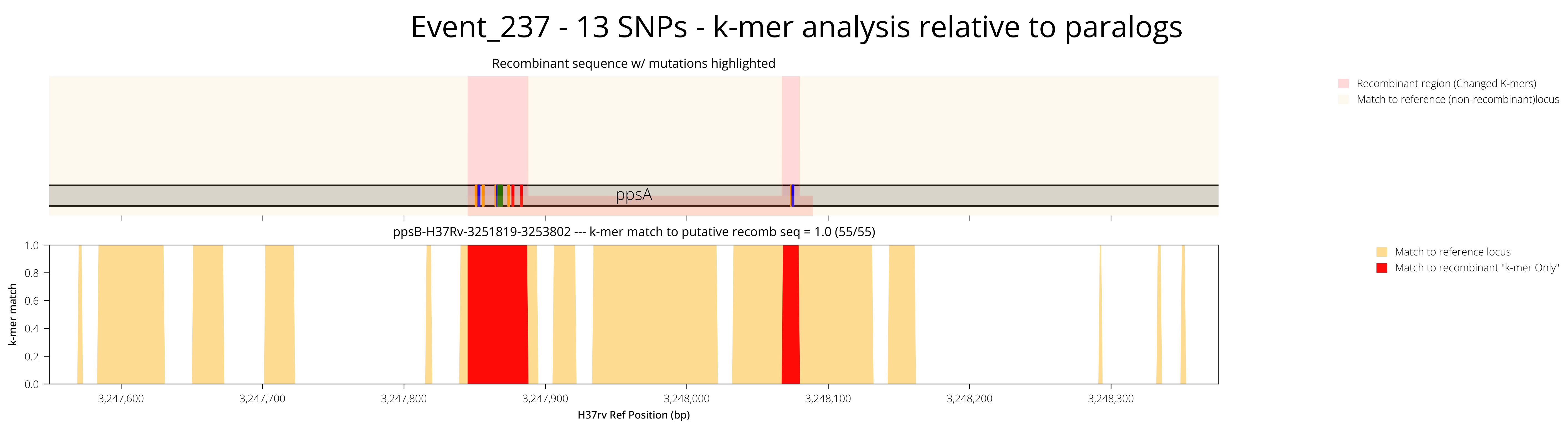

Event_237 - 13 SNPs - k-mer analysis relative to paralogs
Recombinant sequence w/ mutations highlighted
Recombinant region (Changed K-mers)
Match to reference (non-recombinant)locus
ppsA
ppsB-H37Rv-3251819-3253802 --- k-mer match to putative recomb seq = 1.0 (55/55)
Match to reference locus
Match to recombinant "k-mer Only"
k-mer match
1.0
0.8
0.6
0.4
0.2
0.0
3,247,600
3,247,700
3,247,800
3,247,900
3,248,000
3,248,100
3,248,200
3,248,300
H37rv Ref Position (bp)

Sub-phylogeny Viz + Paralog Alignments
GCE ID: Event_237

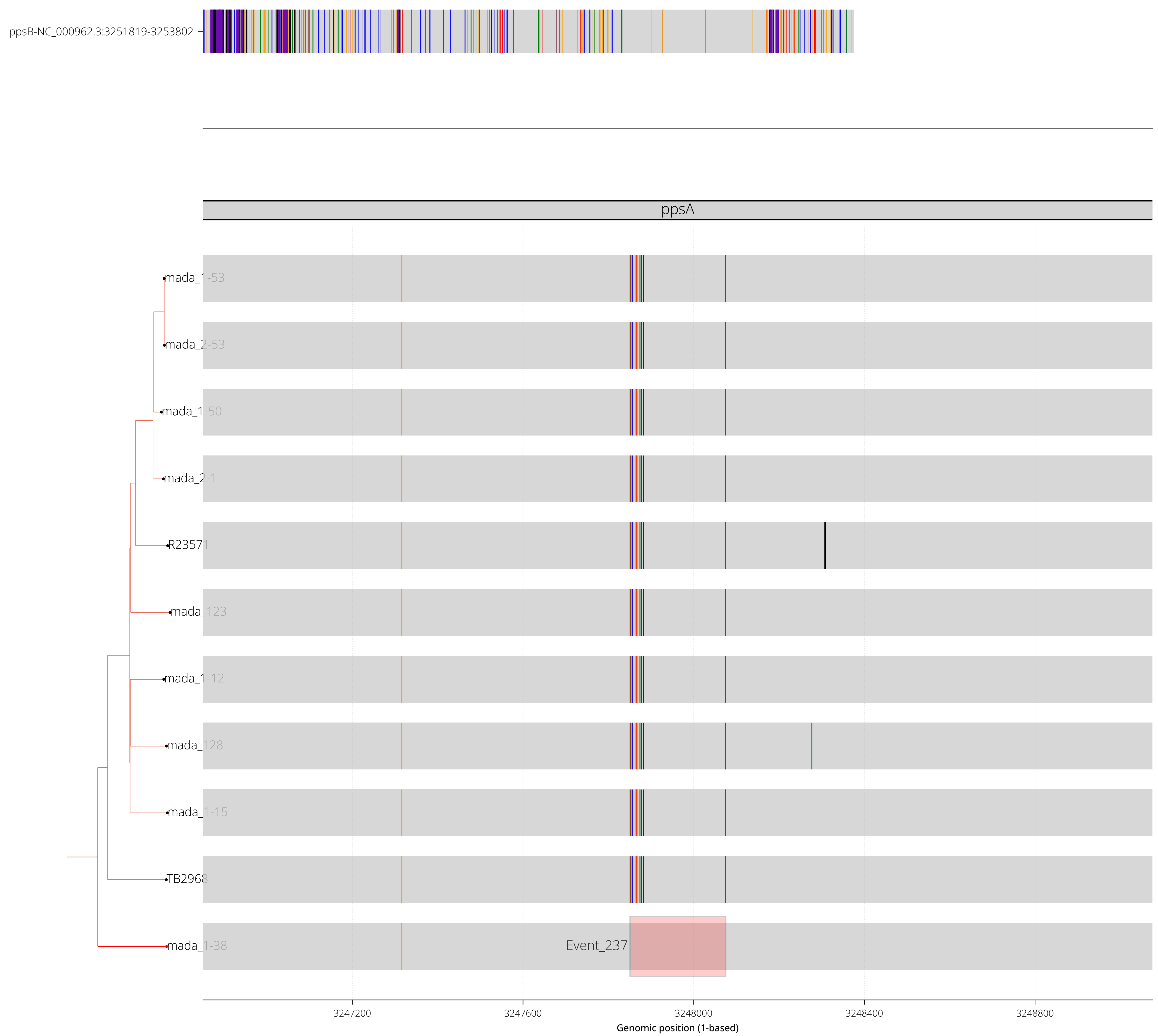

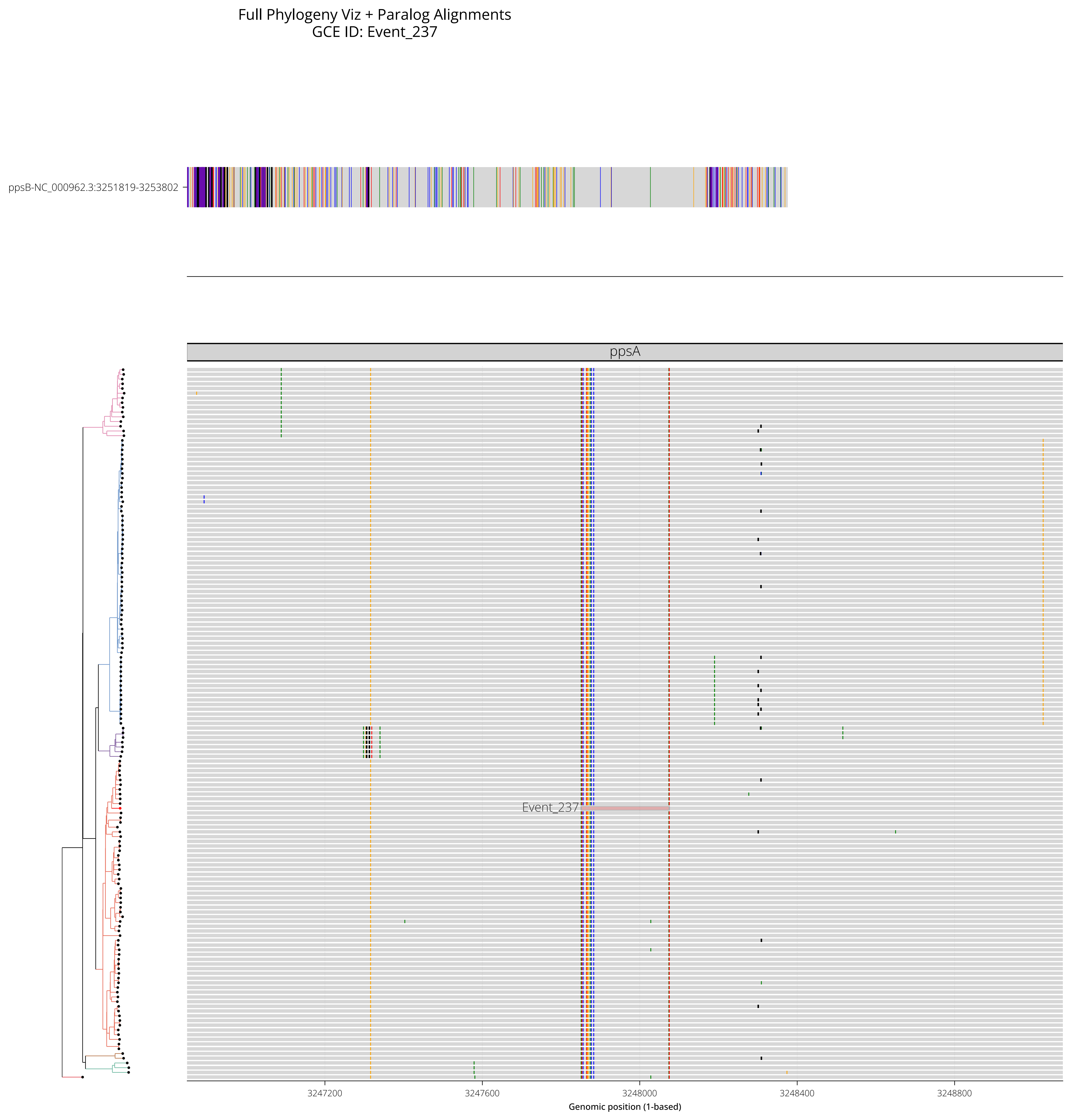
Full Phylogeny Viz + Paralog Alignments
GCE ID: Event_237
ppsB-NC_000962.3:3251819-3253802
ppsA
Event_237
3247200
3247600
3248000
3248400
3248800
Genomic position (1-based)

Individual GCE + Known Paralogs
GCE ID: Event_238

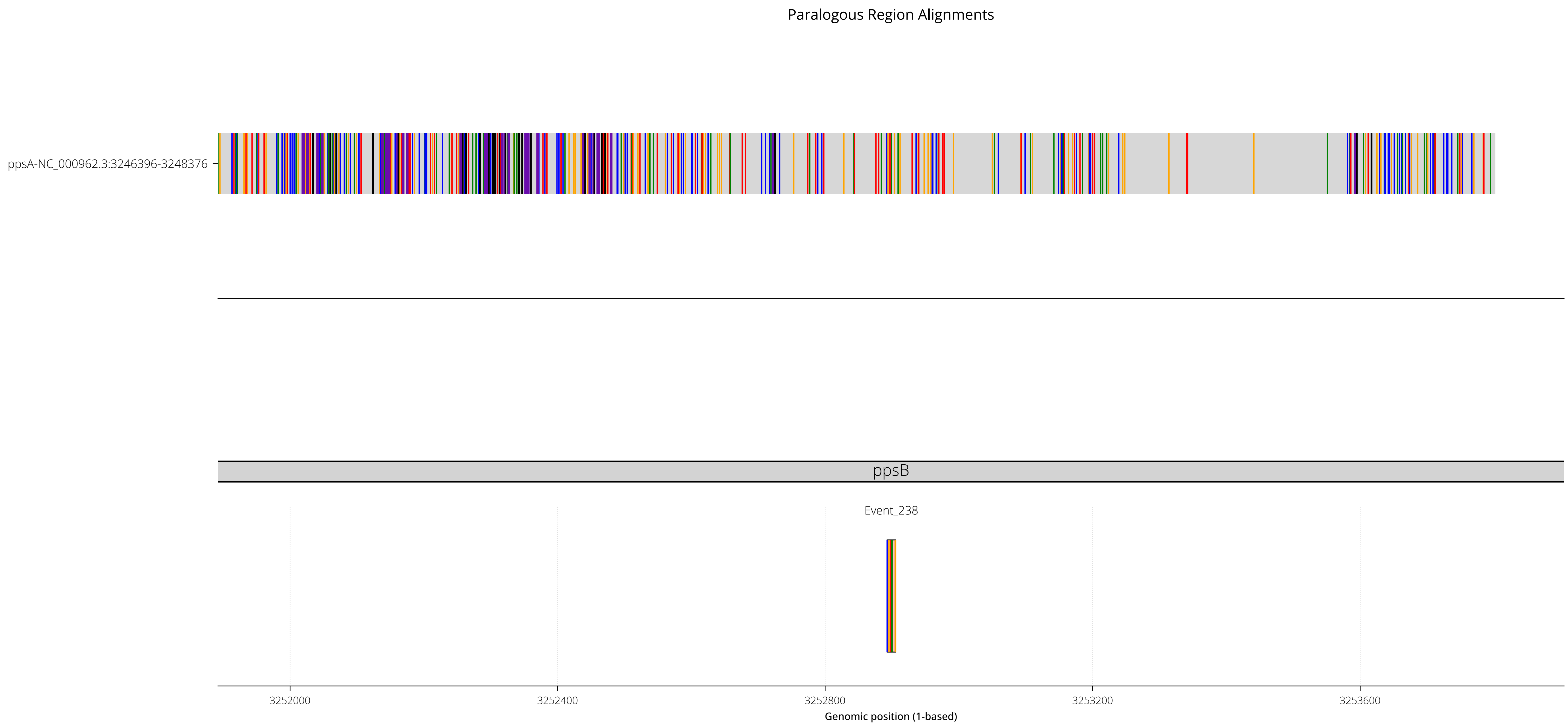

Circos Viz - GCE to Known Paralogs Locations
GCE ID: Event_238

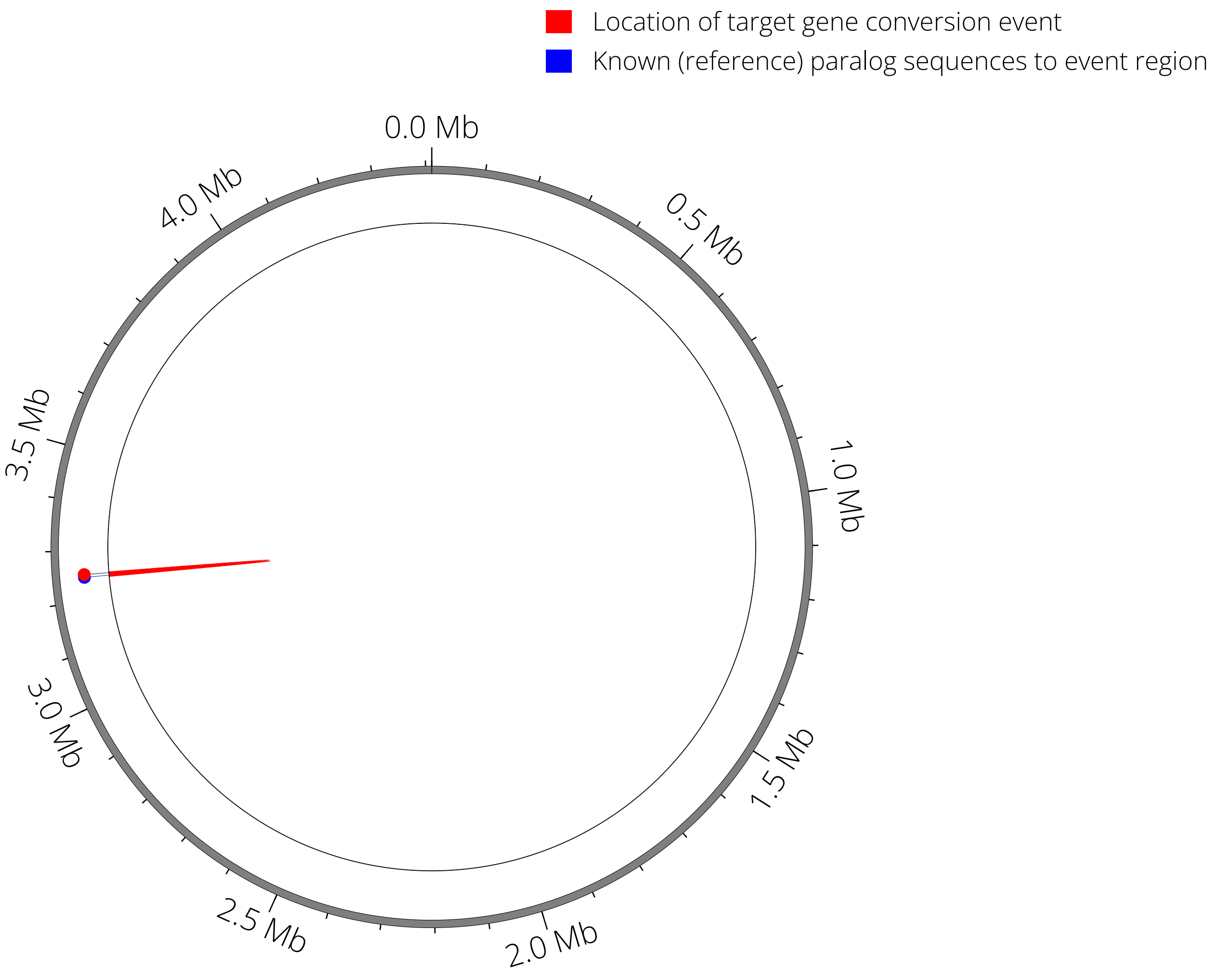

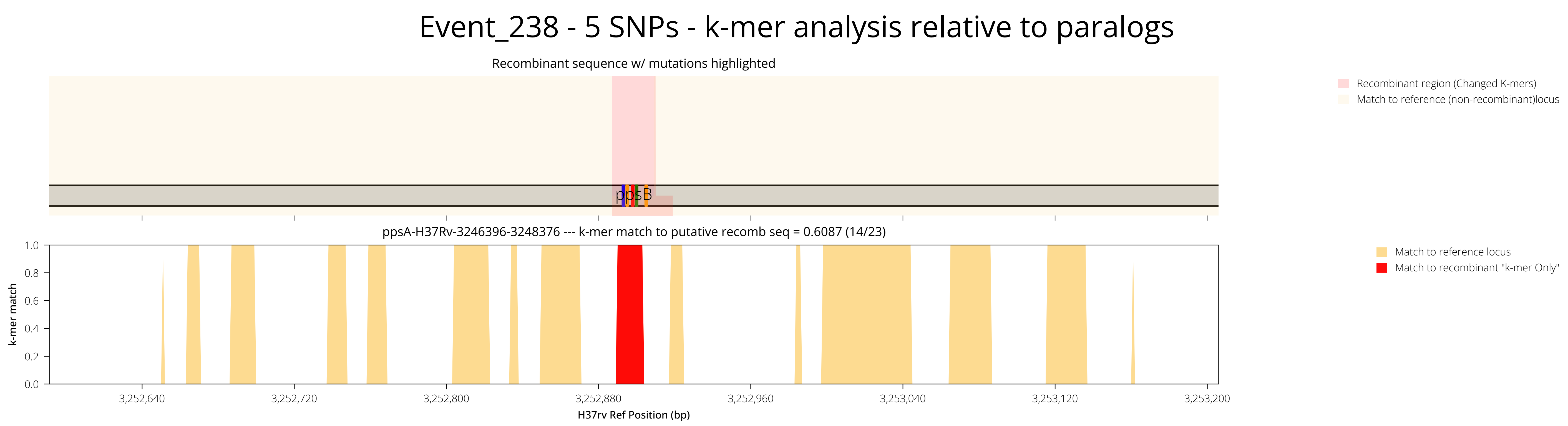

Event_238 - 5 SNPs - k-mer analysis relative to paralogs
Recombinant sequence w/ mutations highlighted
Recombinant region (Changed K-mers)
Match to reference (non-recombinant)locus
ppsB
ppsA-H37Rv-3246396-3248376 --- k-mer match to putative recomb seq = 0.6087 (14/23)
Match to reference locus
Match to recombinant "k-mer Only"
k-mer match
1.0
0.8
0.6
0.4
0.2
0.0
3,252,640
3,252,720
3,252,800
3,252,880
3,252,960
3,253,040
3,253,120
3,253,200
H37rv Ref Position (bp)

Sub-phylogeny Viz + Paralog Alignments
GCE ID: Event_238

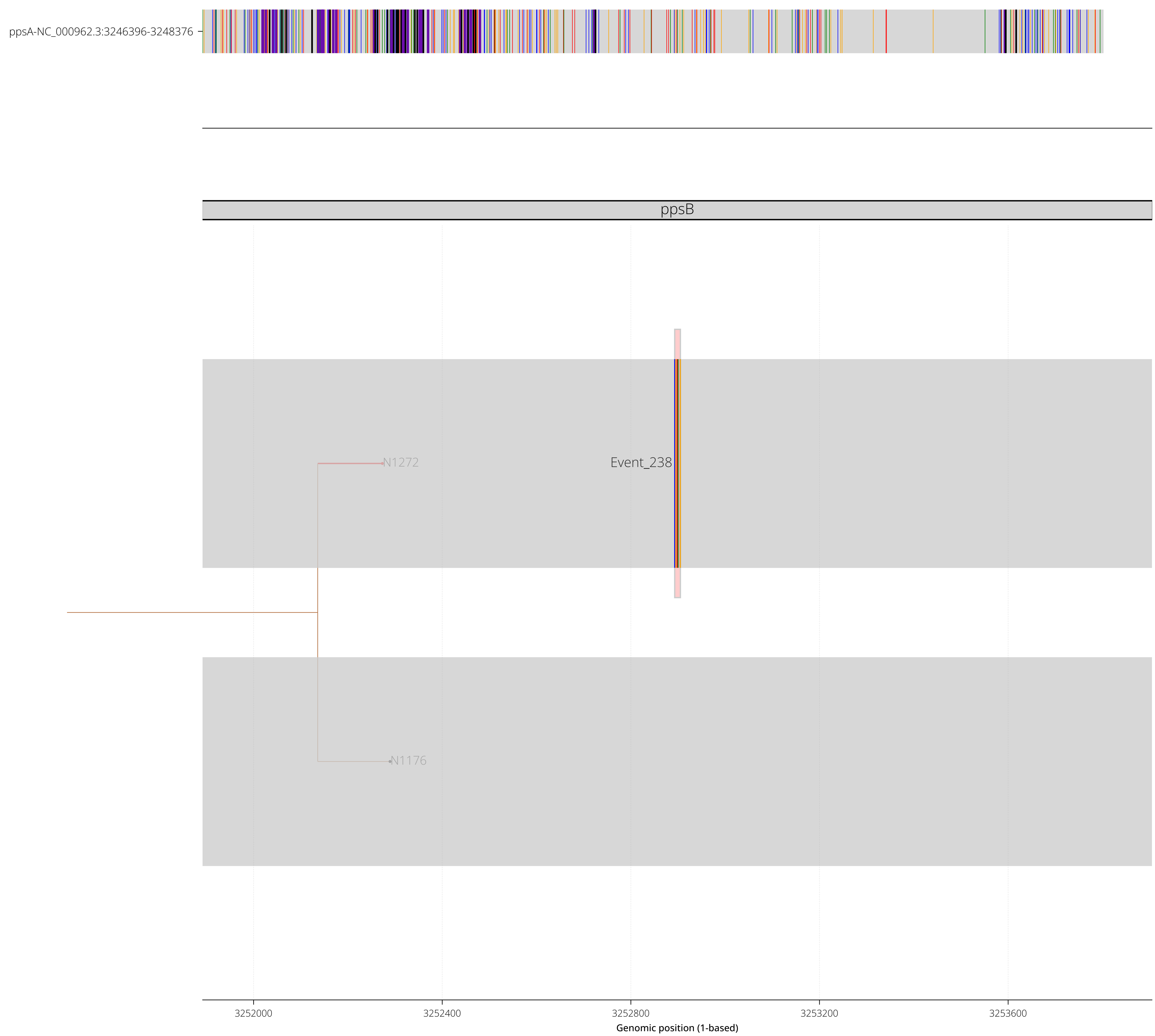

Full Phylogeny Viz + Paralog Alignments
GCE ID: Event_238

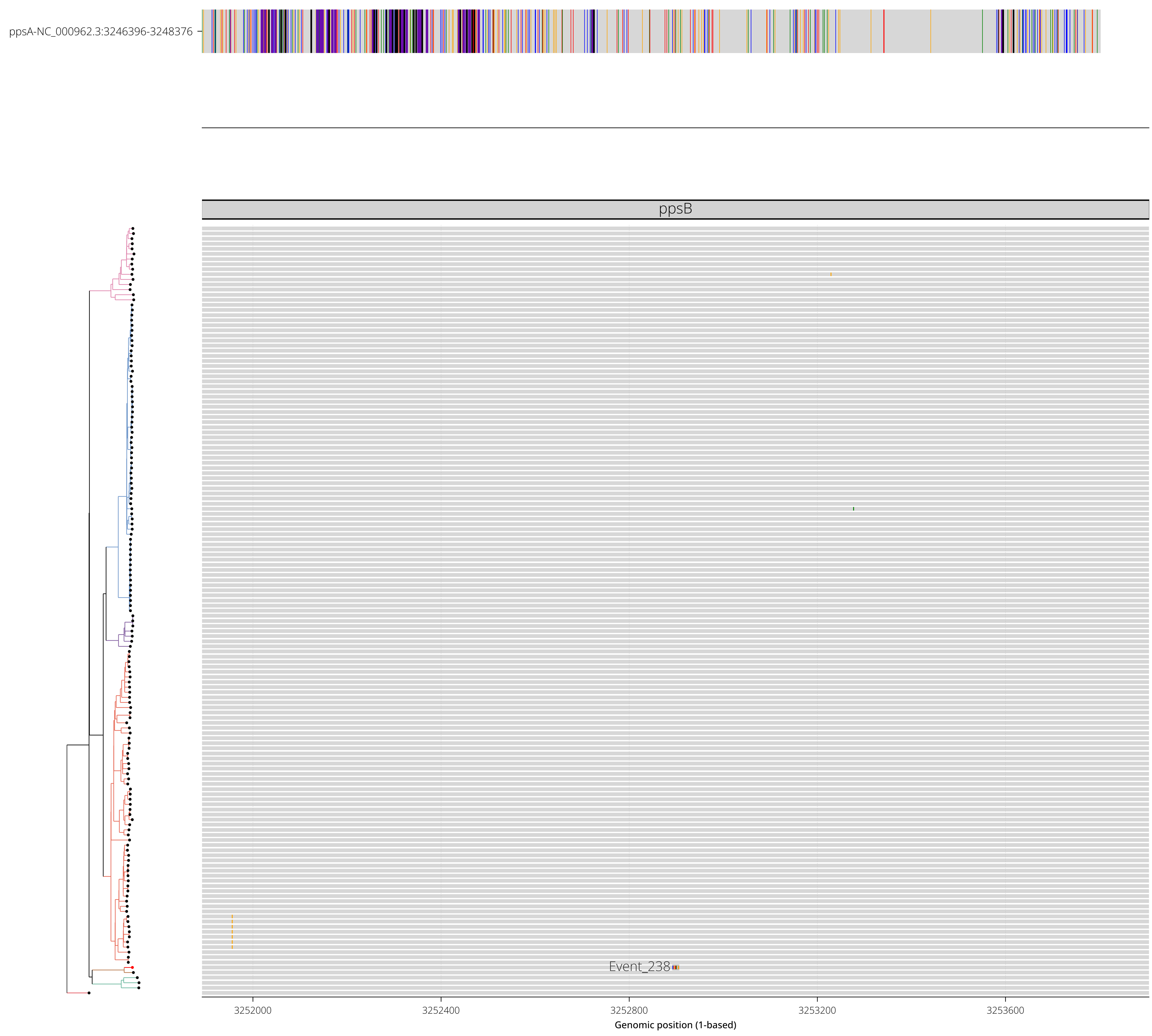

Individual GCE + Known Paralogs
GCE ID: Event_239

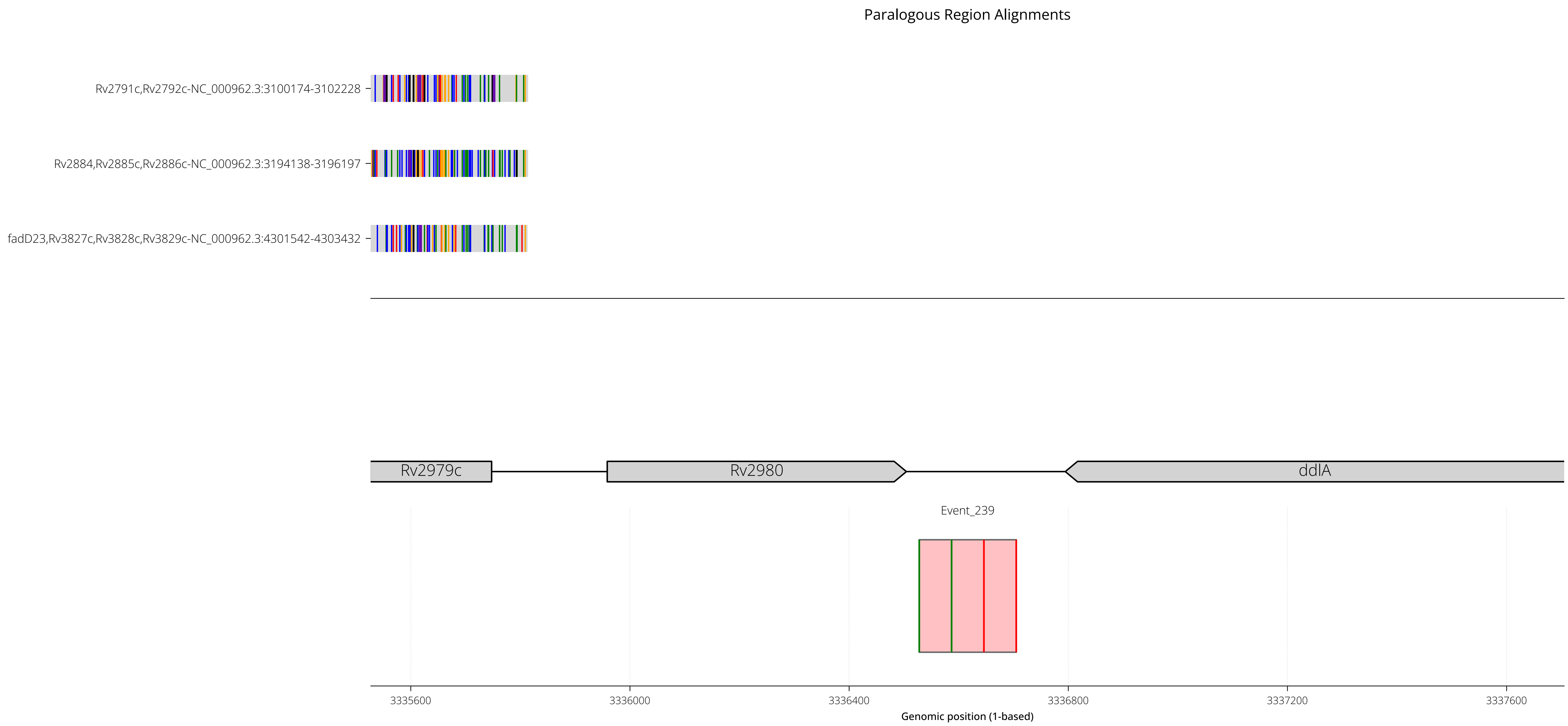

Sub-phylogeny Viz + Paralog Alignments
GCE ID: Event_239

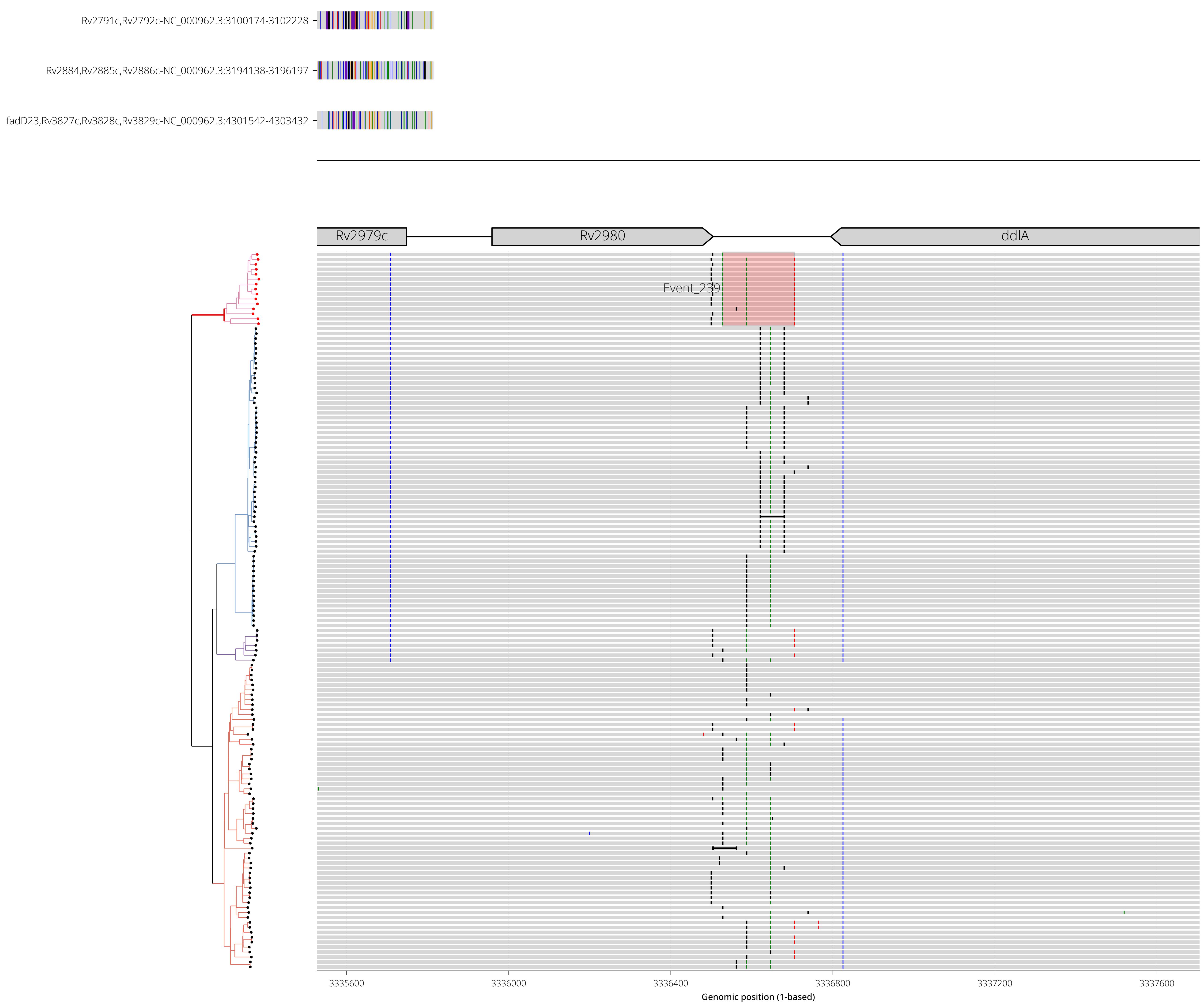

Full Phylogeny Viz + Paralog Alignments
GCE ID: Event_239

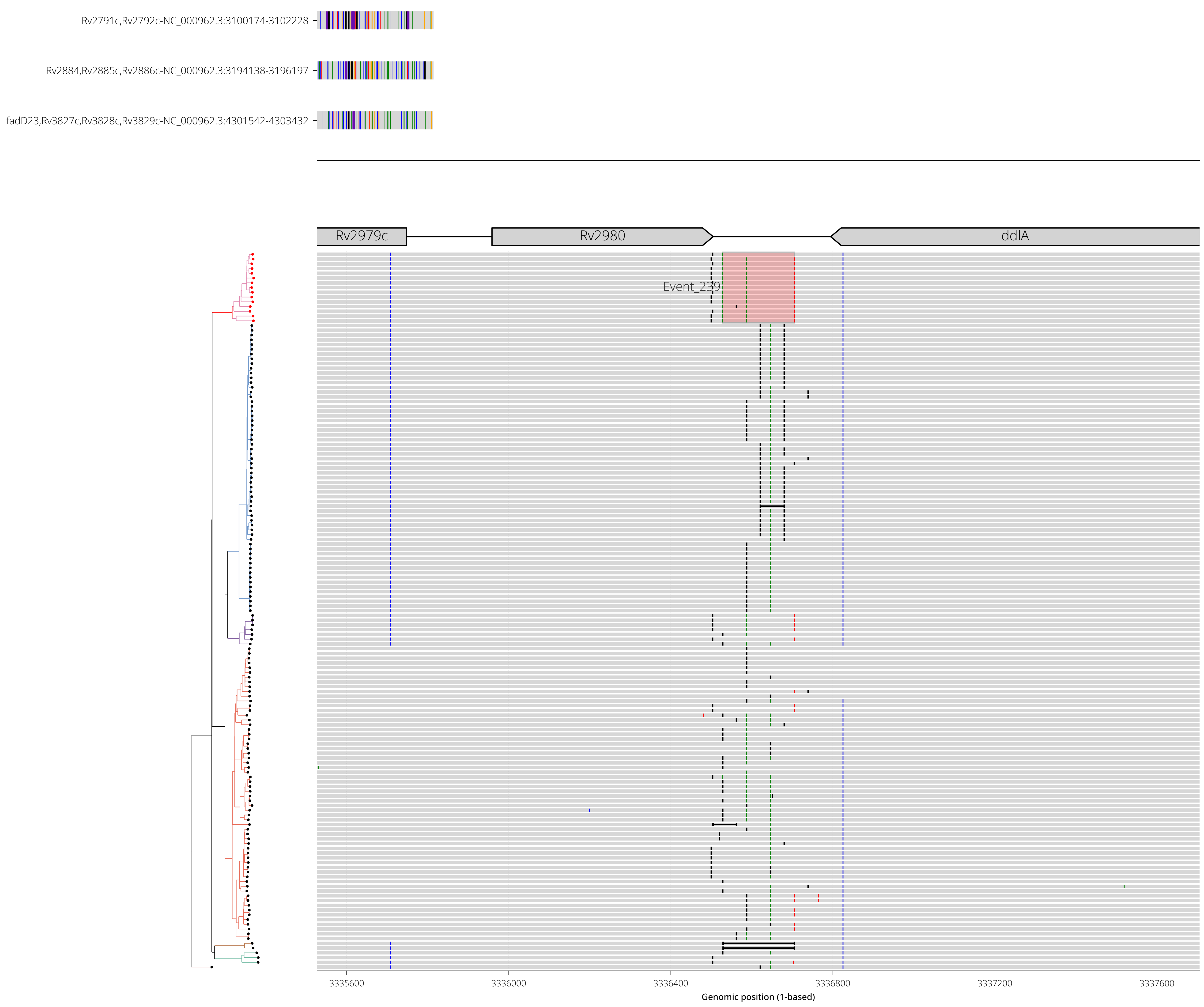

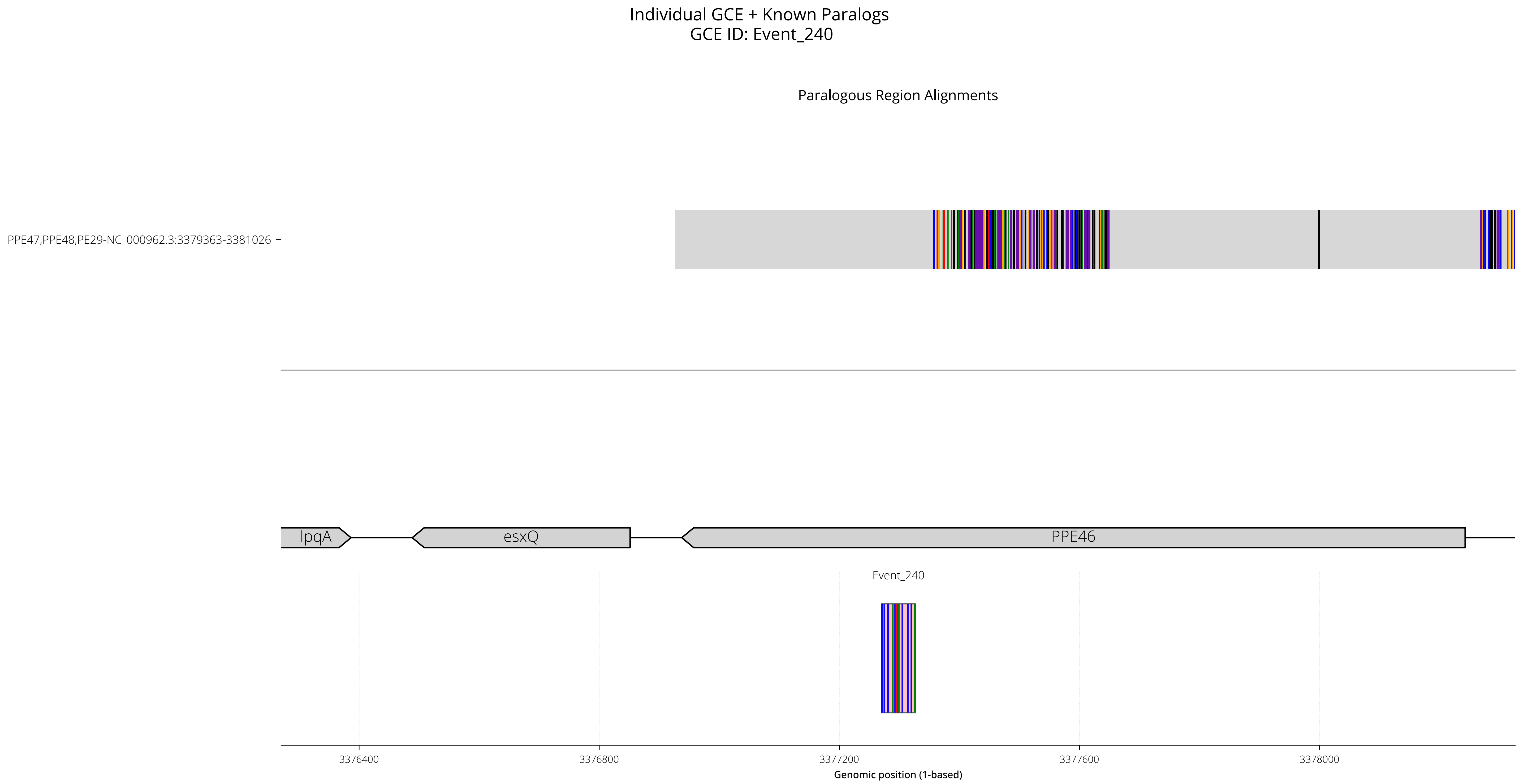

Individual GCE + Known Paralogs
GCE ID: Event_240
Paralogous Region Alignments
PPE47,PPE48,PE29-NC_000962.3:3379363-3381026
lpqA
esxQ
PPE46
Event_240
3376400
3376800
3377200
3377600
3378000
Genomic position (1-based)

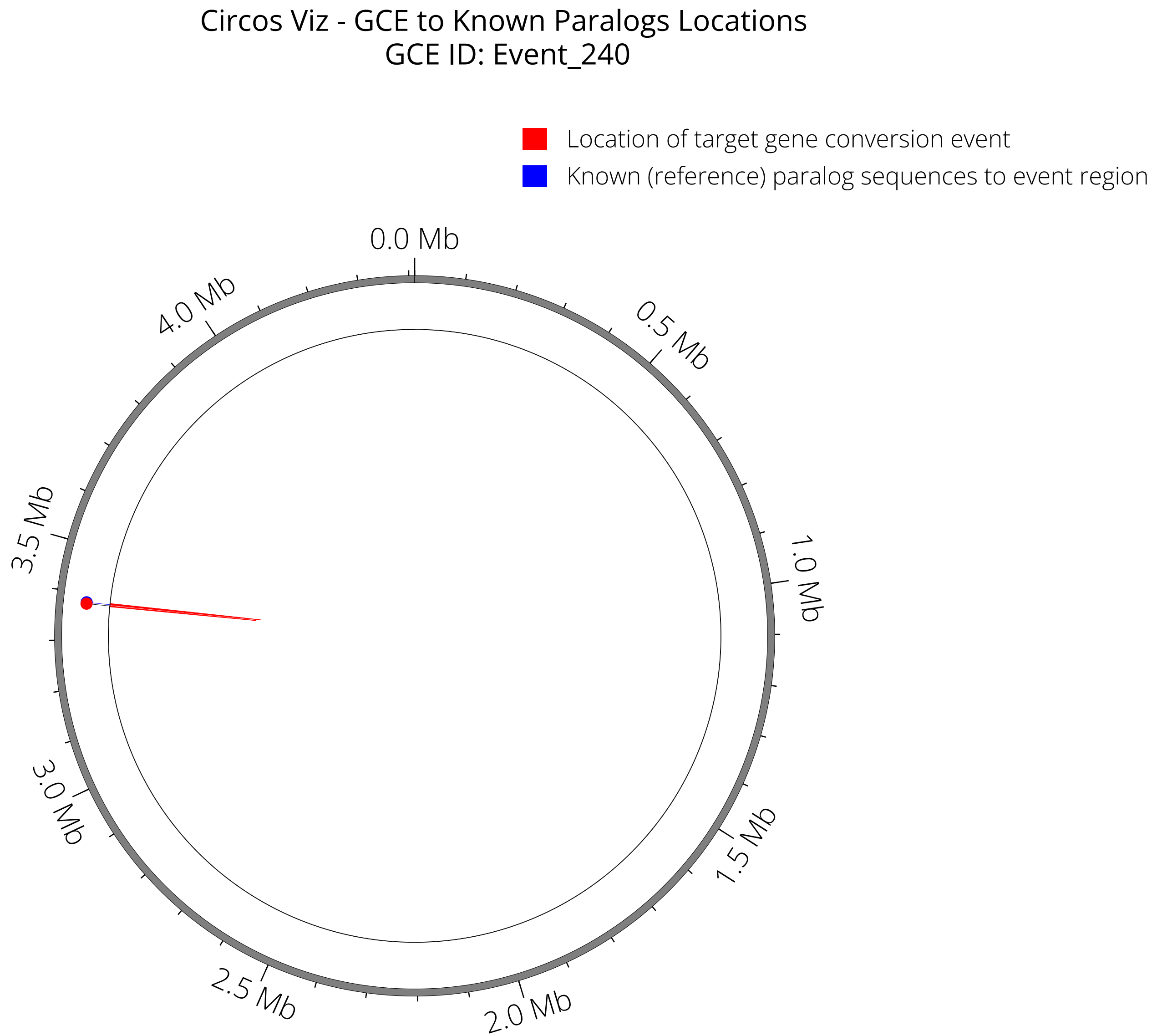

Circos Viz - GCE to Known Paralogs Locations
GCE ID: Event_240
Location of target gene conversion event
Known (reference) paralog sequences to event region
0.0 Mb
0.5 Mb
1.0 Mb
1.5 Mb
2.0 Mb
2.5 Mb
3.0 Mb
3.5 Mb
4.0 Mb

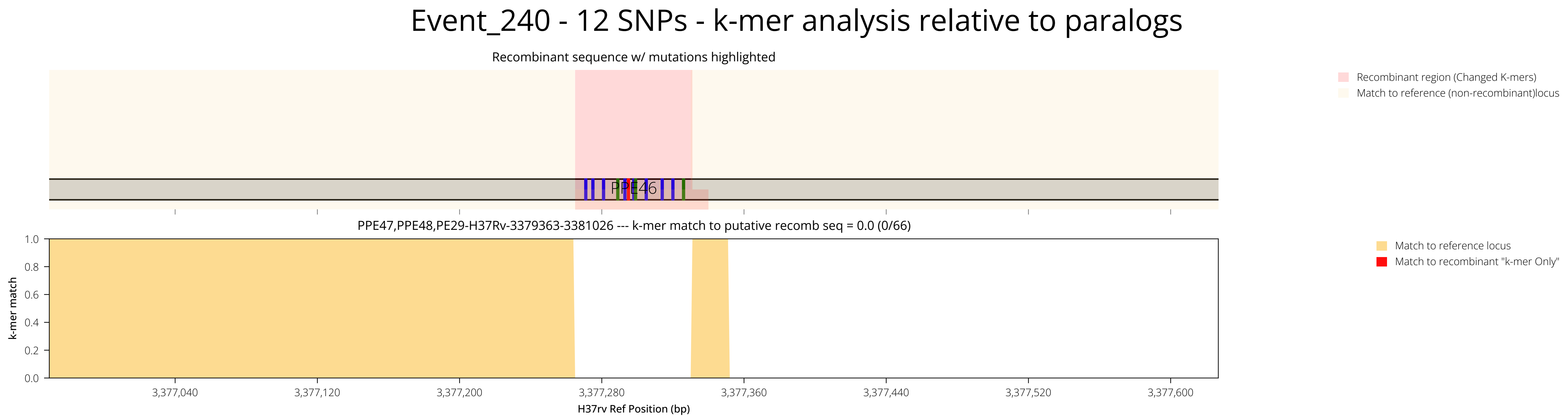

Event_240 - 12 SNPs - k-mer analysis relative to paralogs
Recombinant sequence w/ mutations highlighted
Recombinant region (Changed K-mers)
Match to reference (non-recombinant)locus
PPE46
PPE47,PPE48,PE29-H37Rv-3379363-3381026 --- k-mer match to putative recomb seq = 0.0 (0/66)
Match to reference locus
Match to recombinant "k-mer Only"
k-mer match
1.0
0.8
0.6
0.4
0.2
0.0
3,377,040
3,377,120
3,377,200
3,377,280
3,377,360
3,377,440
3,377,520
3,377,600
H37rv Ref Position (bp)

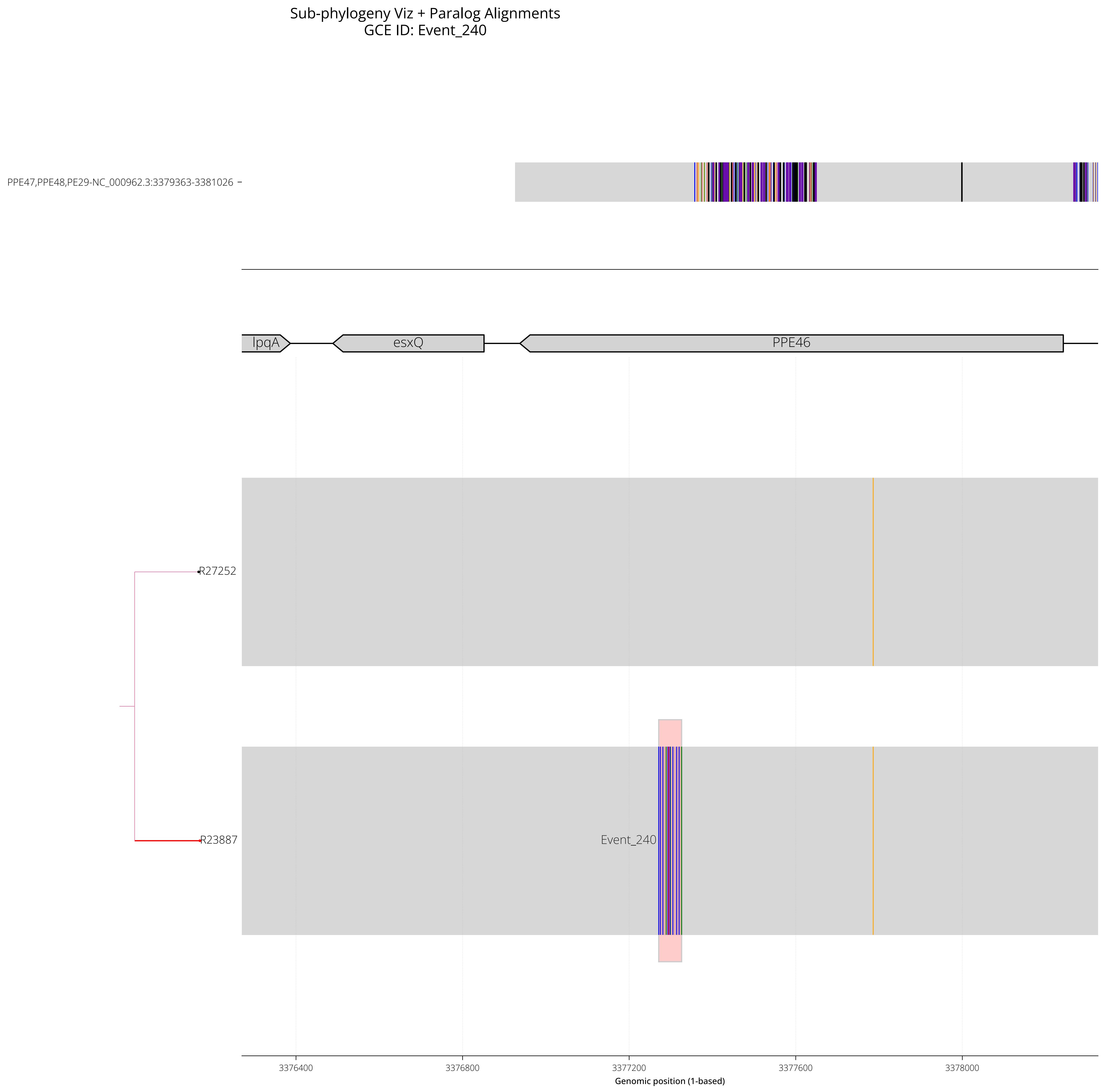

Sub-phylogeny Viz + Paralog Alignments
GCE ID: Event_240
PPE47,PPE48,PE29-NC_000962.3:3379363-3381026
lpqA
esxQ
PPE46
R27252
R23887
Event_240
3376400
3376800
3377200
3377600
3378000
Genomic position (1-based)

Full Phylogeny Viz + Paralog Alignments
GCE ID: Event_240

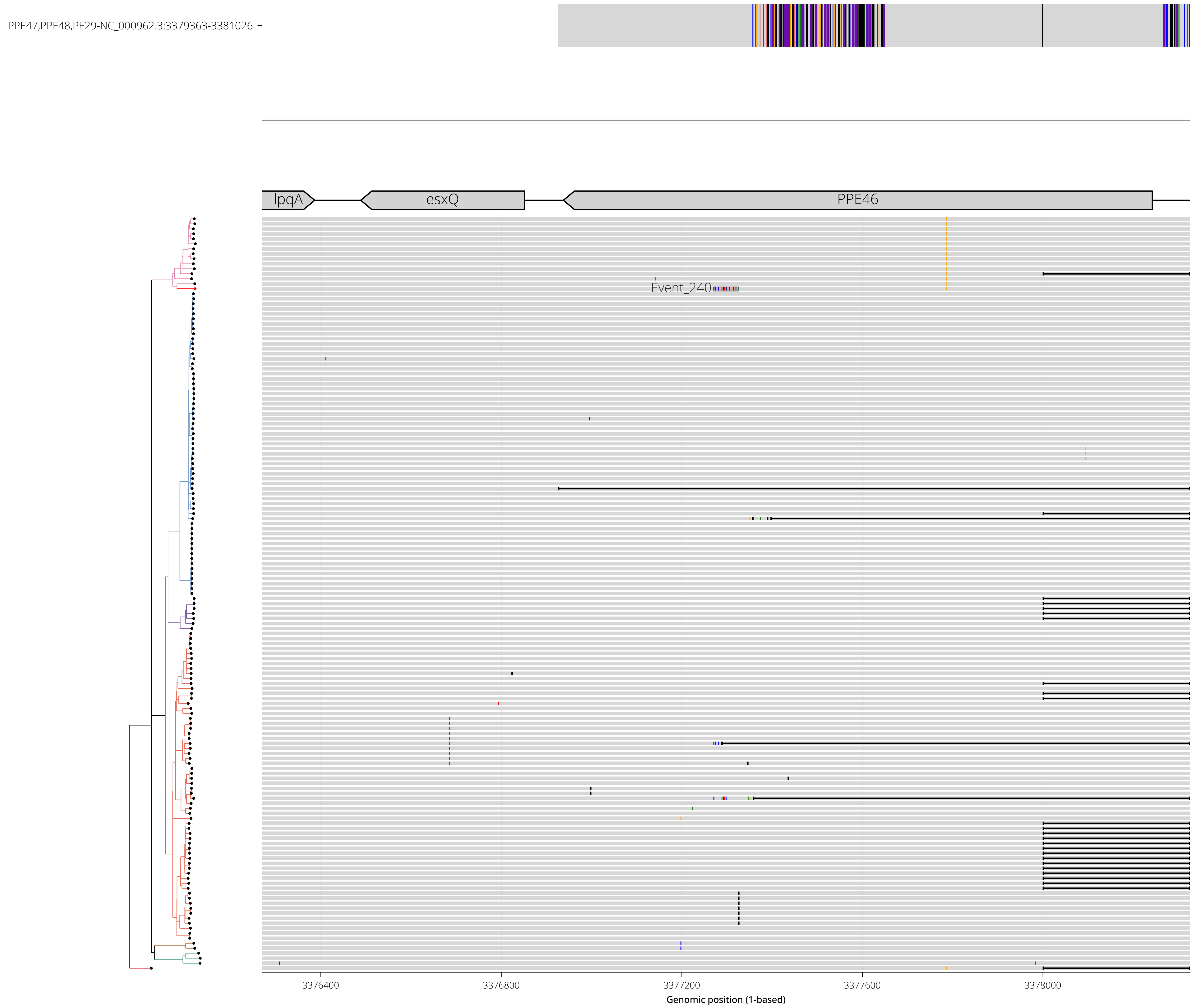

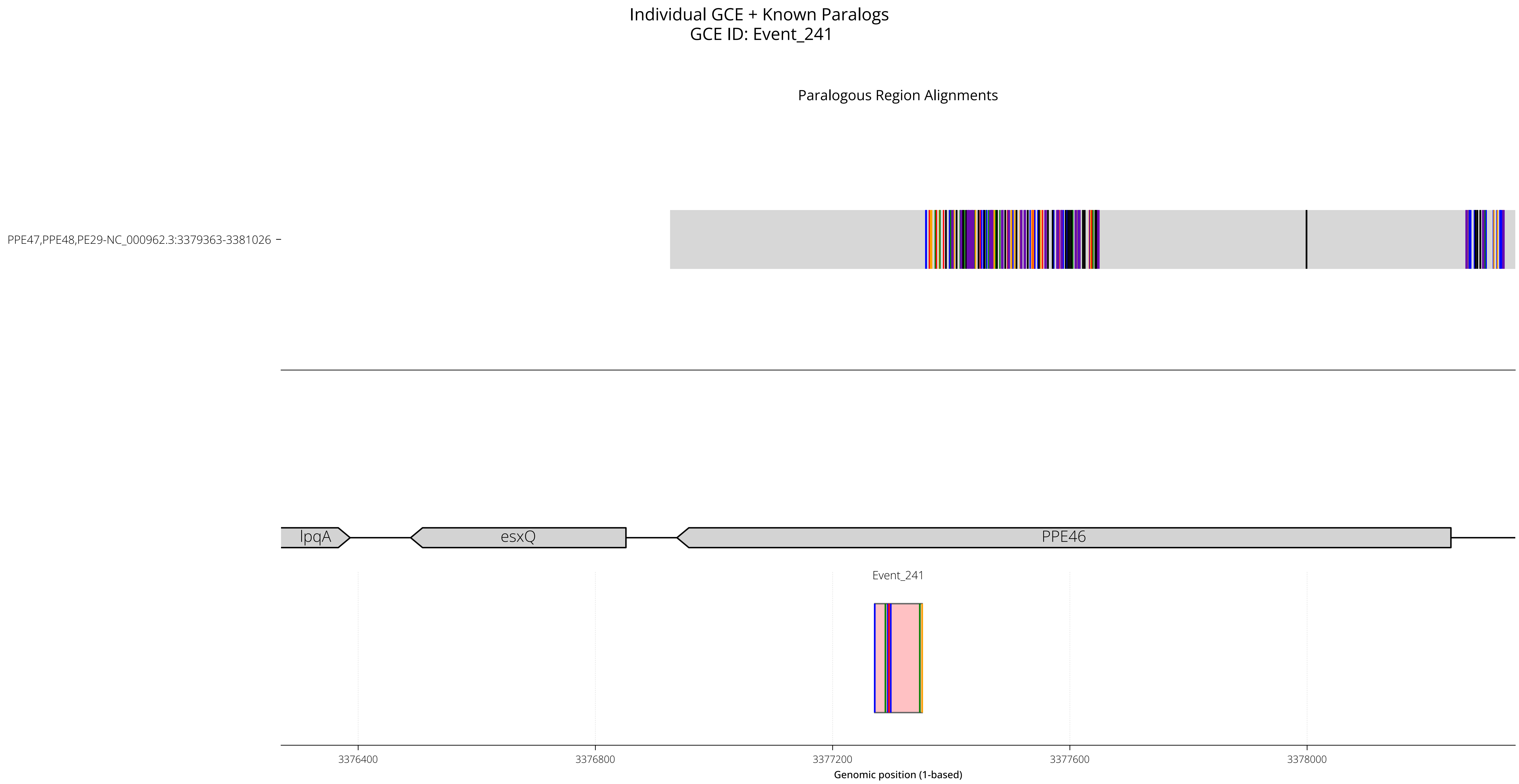

Individual GCE + Known Paralogs
GCE ID: Event_241
Paralogous Region Alignments
PPE47,PPE48,PE29-NC_000962.3:3379363-3381026
lpqA
esxQ
PPE46
Event_241
3376400
3376800
3377200
3377600
3378000
Genomic position (1-based)

Circos Viz - GCE to Known Paralogs Locations
GCE ID: Event_241

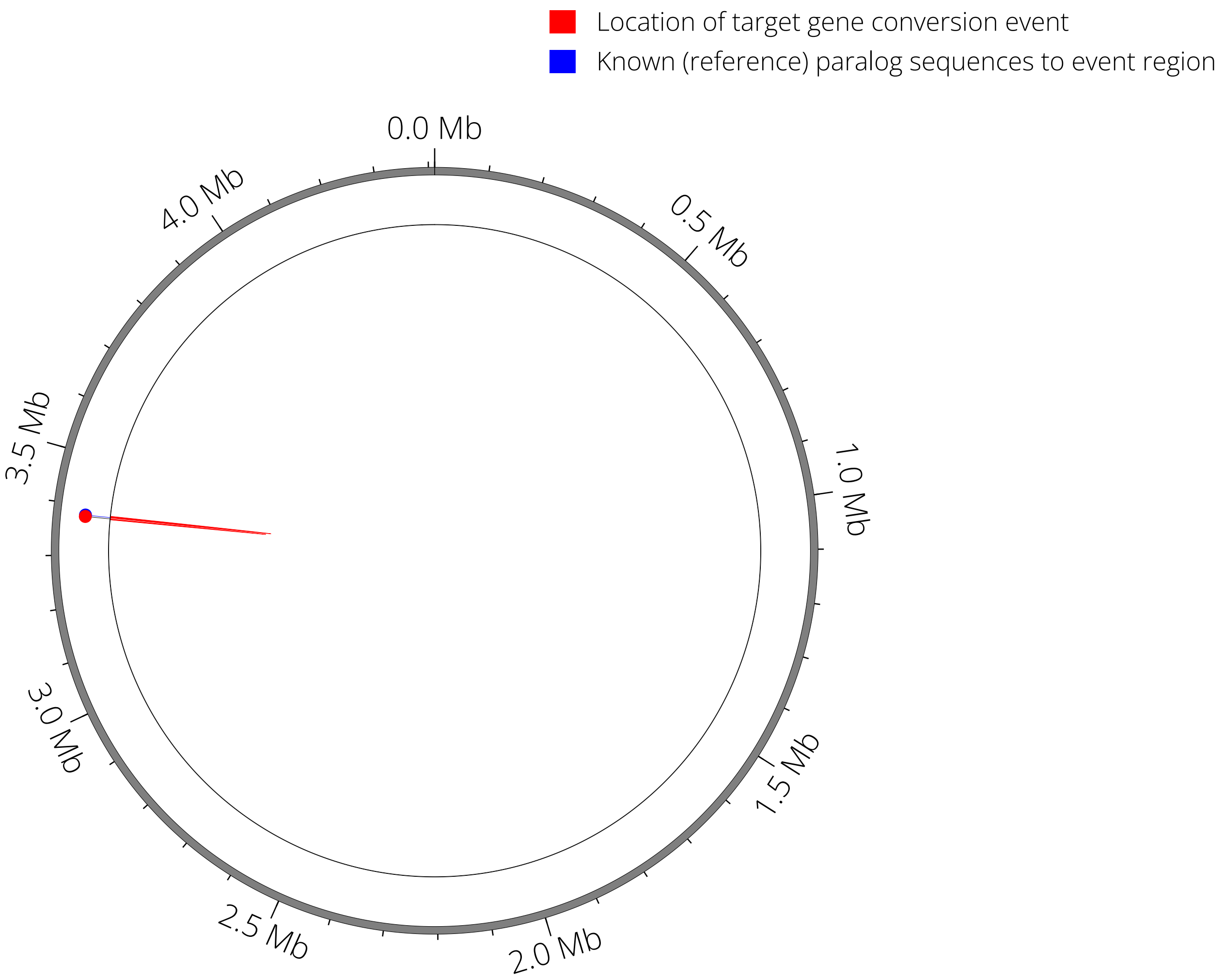

### Event_241 - 7 SNPs - k-mer analysis relative to paralogs

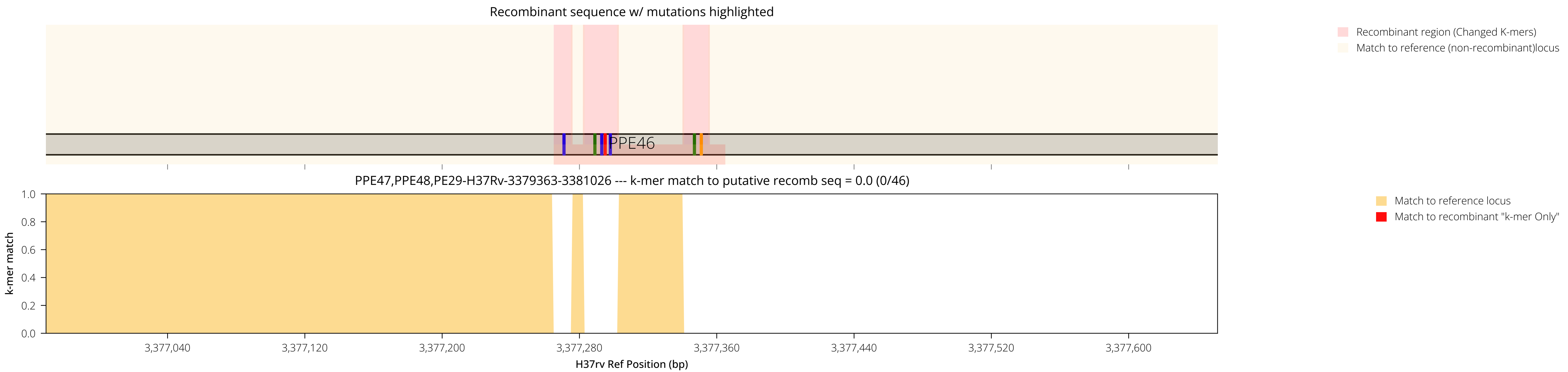

Sub-phylogeny Viz + Paralog Alignments
GCE ID: Event_241

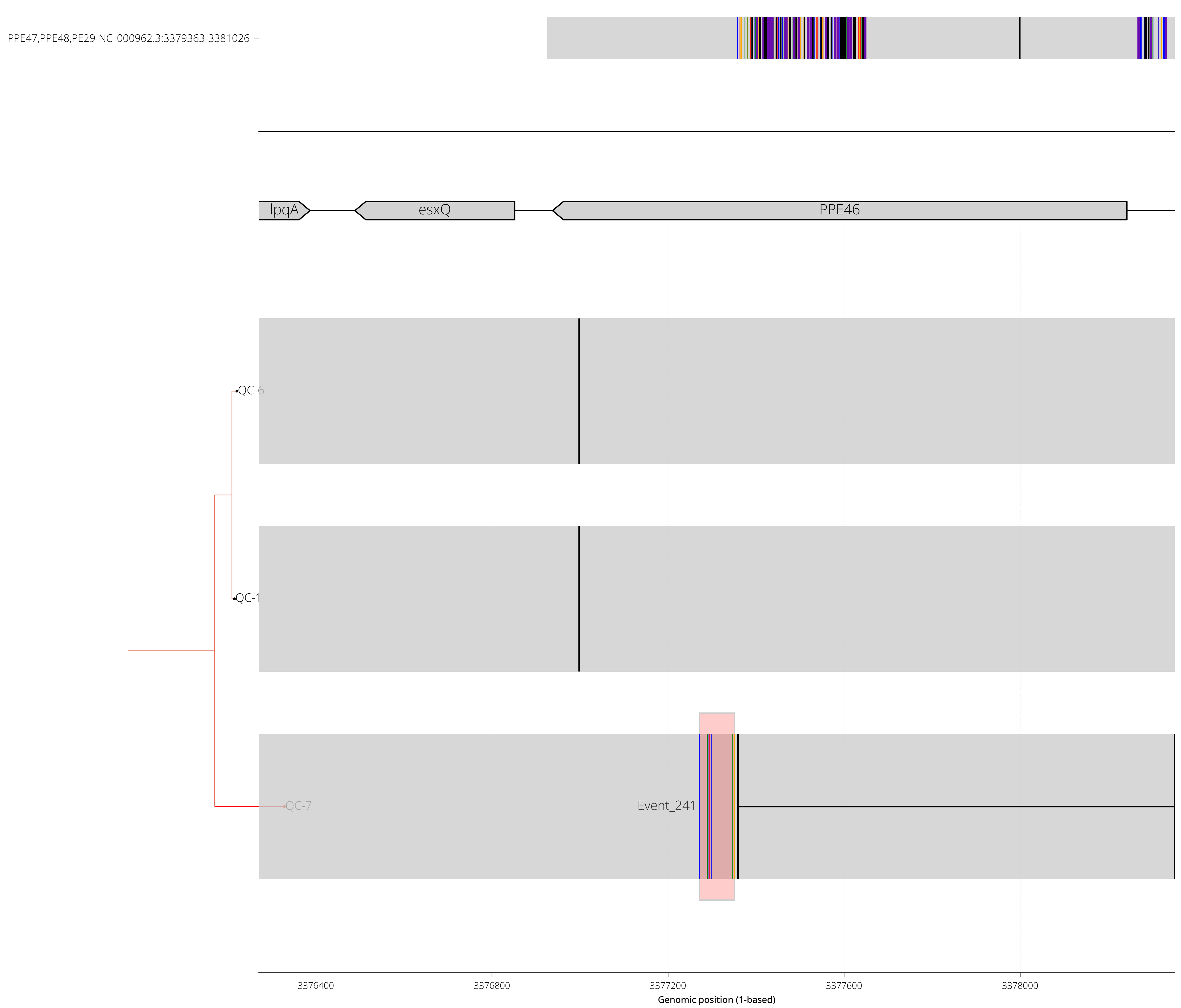

Full Phylogeny Viz + Paralog Alignments
GCE ID: Event_241

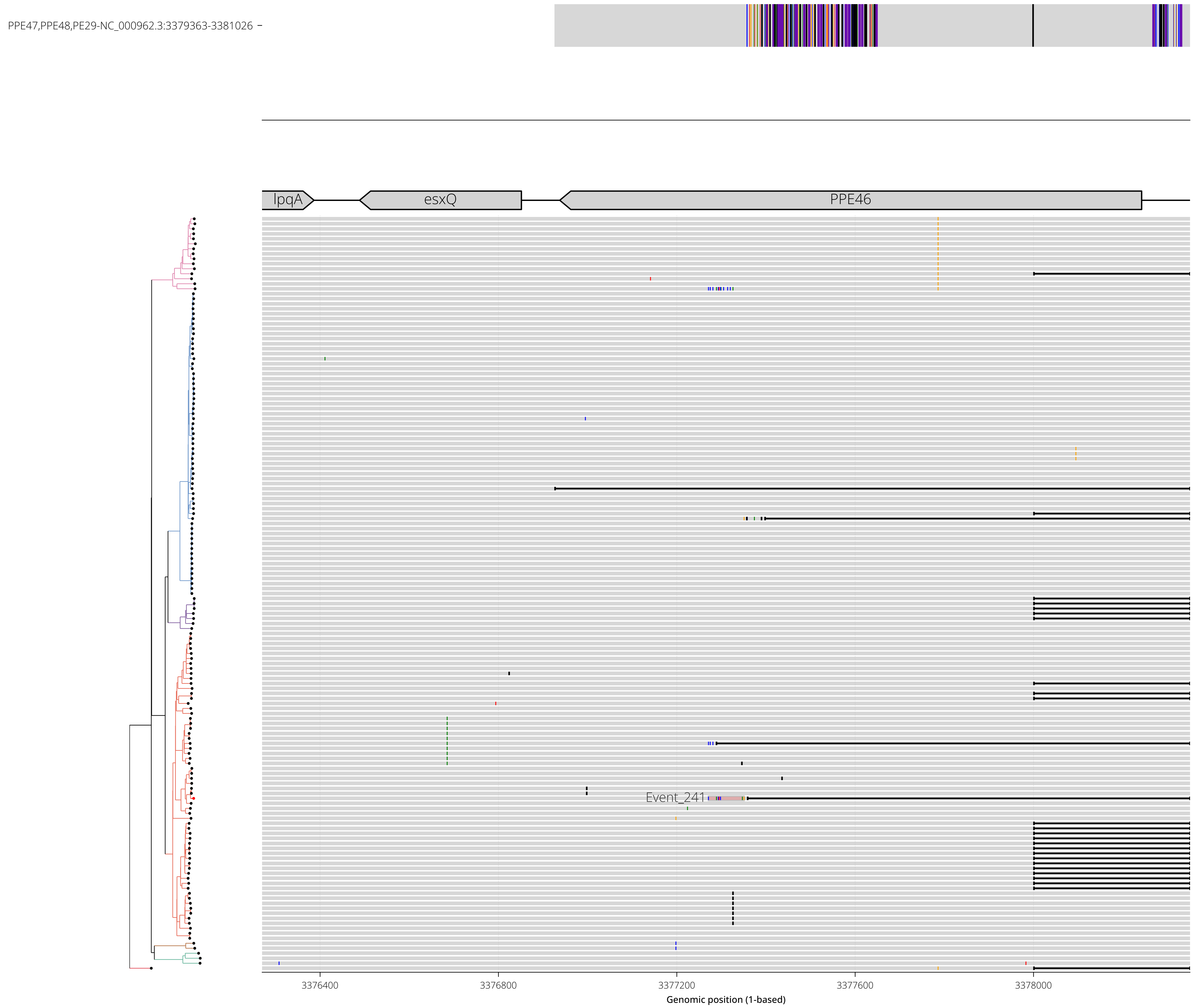

Individual GCE + Known Paralogs
GCE ID: Event_242

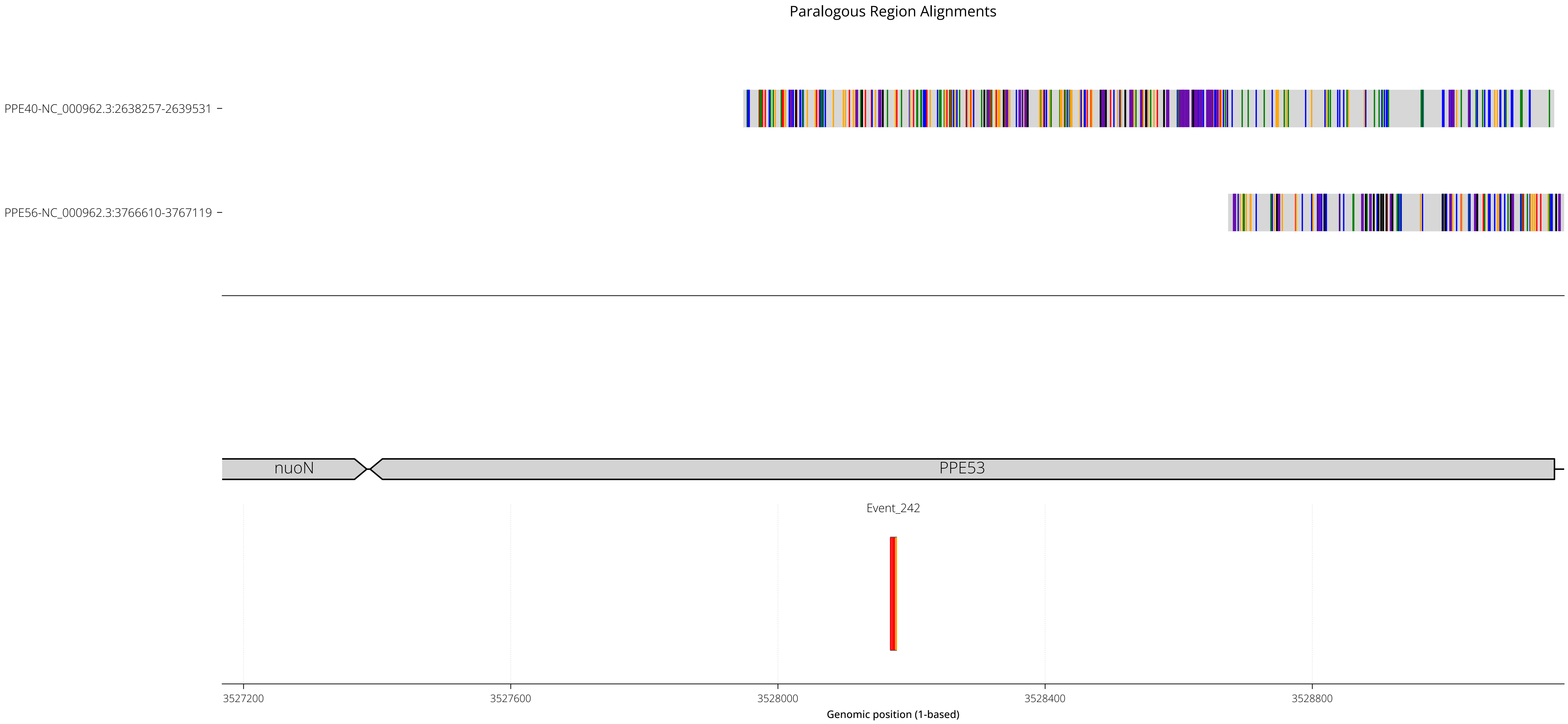

Circos Viz - GCE to Known Paralogs Locations
GCE ID: Event_242

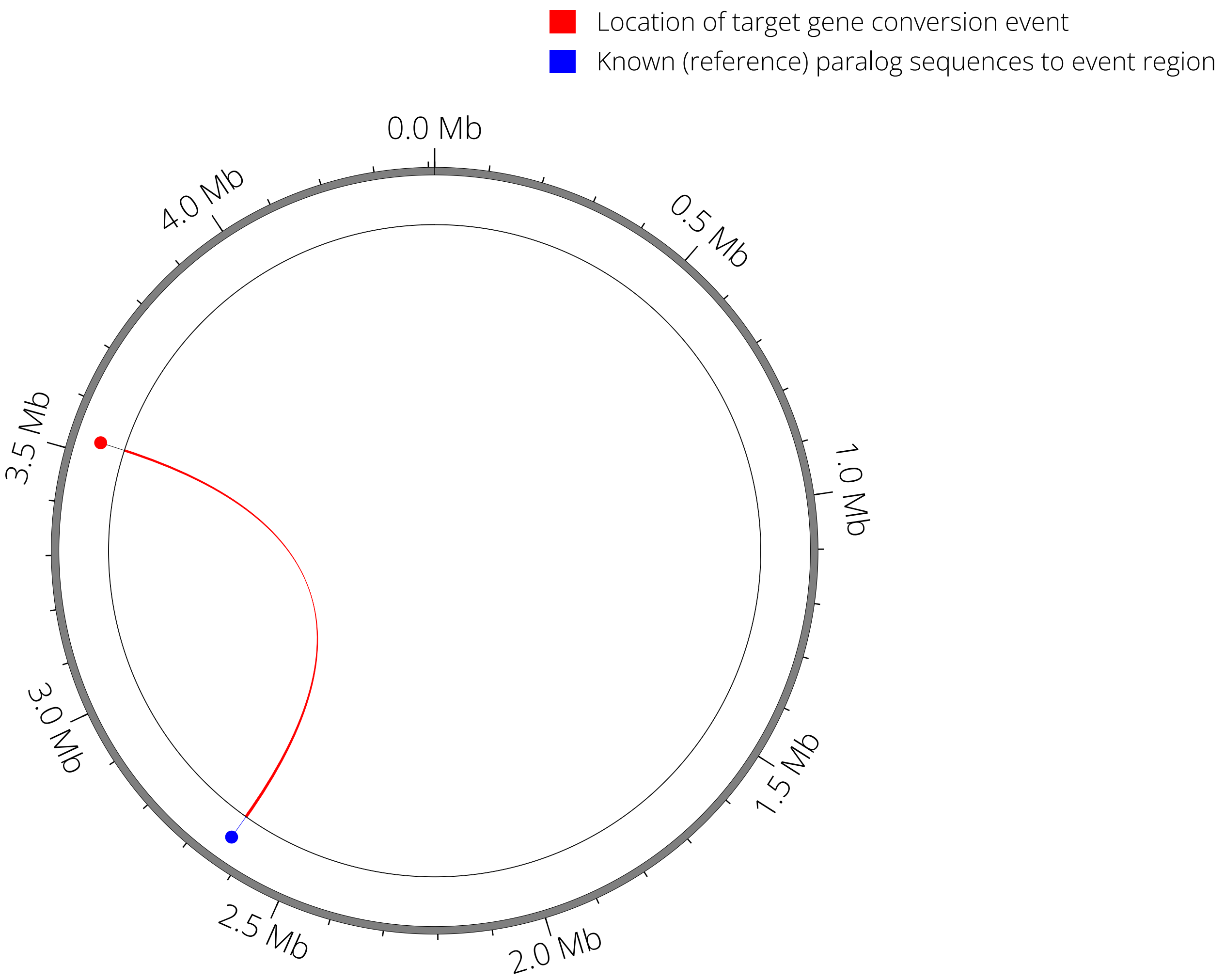

### Event_242 - 5 SNPs - k-mer analysis relative to paralogs

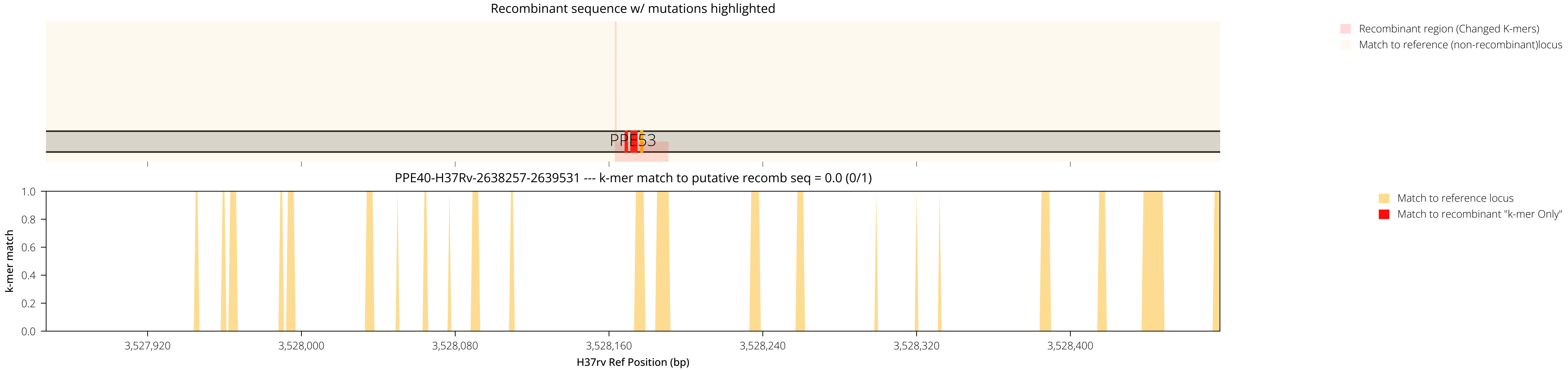

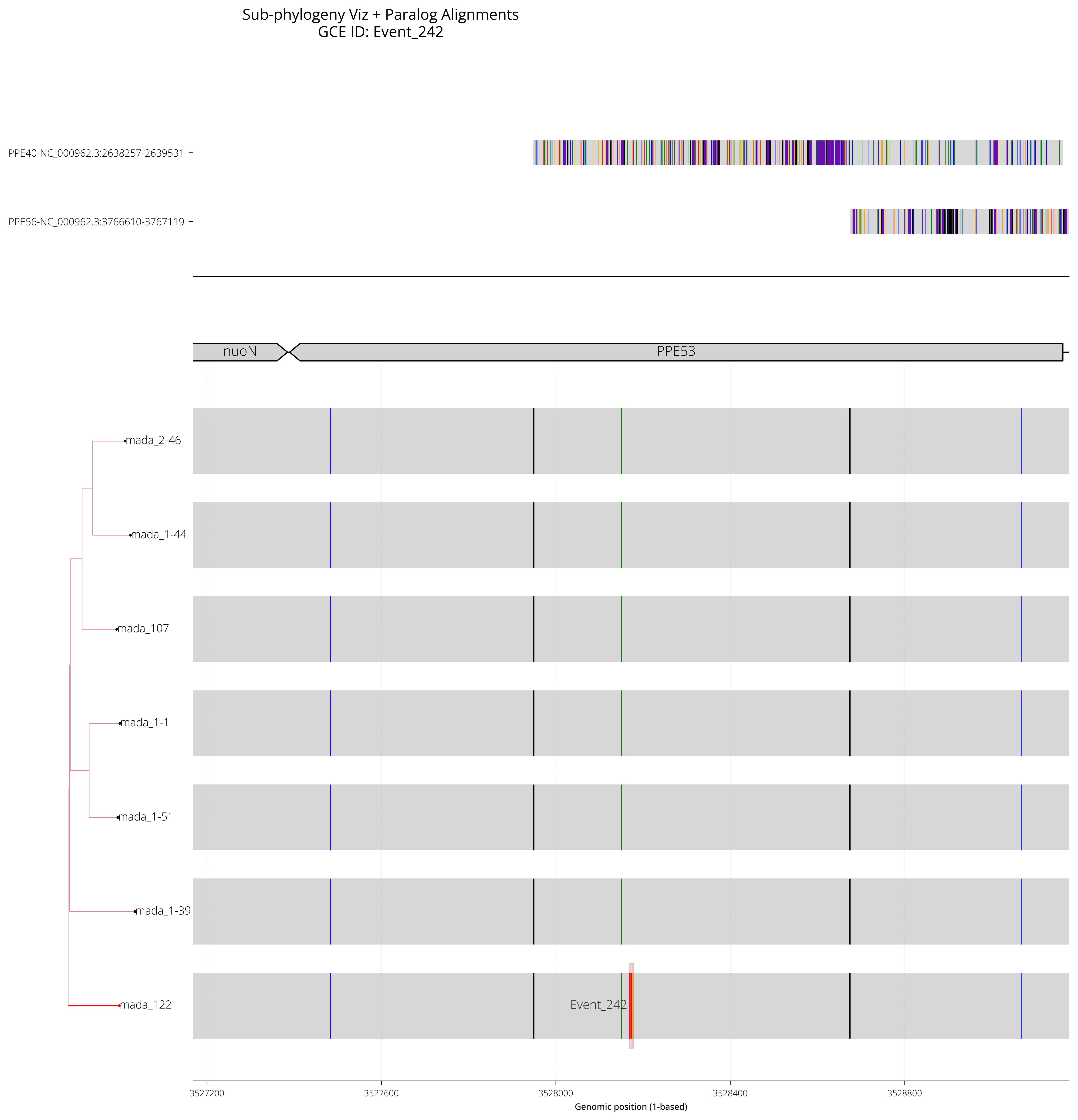

Sub-phylogeny Viz + Paralog Alignments
GCE ID: Event_242
PPE40-NC_000962.3:2638257-2639531
PPE56-NC_000962.3:3766610-3767119
nuoN
PPE53
mada_2-46
mada_1-44
mada_107
mada_1-1
mada_1-51
mada_1-39
mada_122
Event_242
3527200
3527600
3528000
3528400
3528800
Genomic position (1-based)

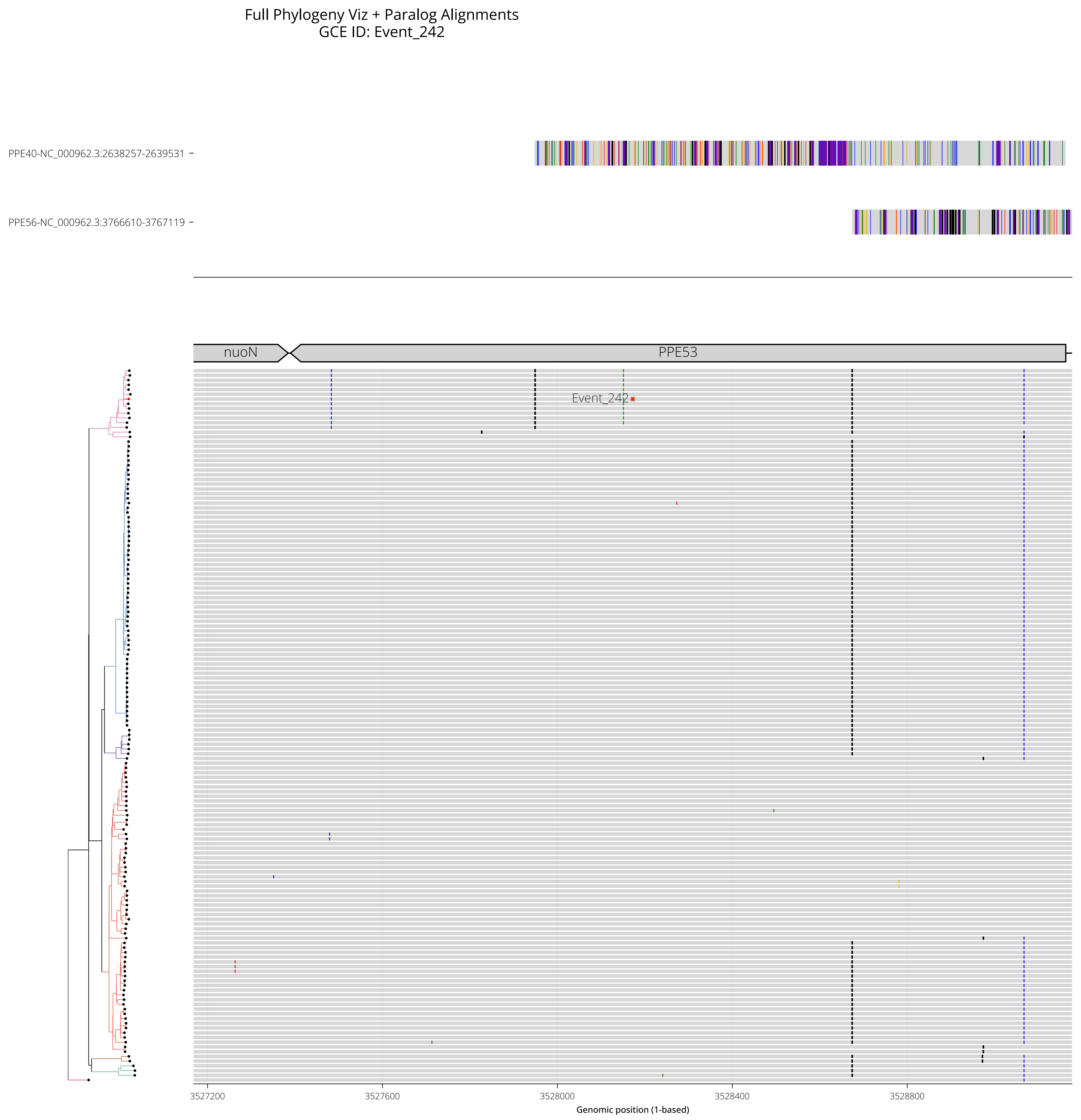

Full Phylogeny Viz + Paralog Alignments
GCE ID: Event_242
PPE40-NC_000962.3:2638257-2639531
PPE56-NC_000962.3:3766610-3767119
nuoN
PPE53
Event_242
3527200
3527600
3528000
3528400
3528800
Genomic position (1-based)

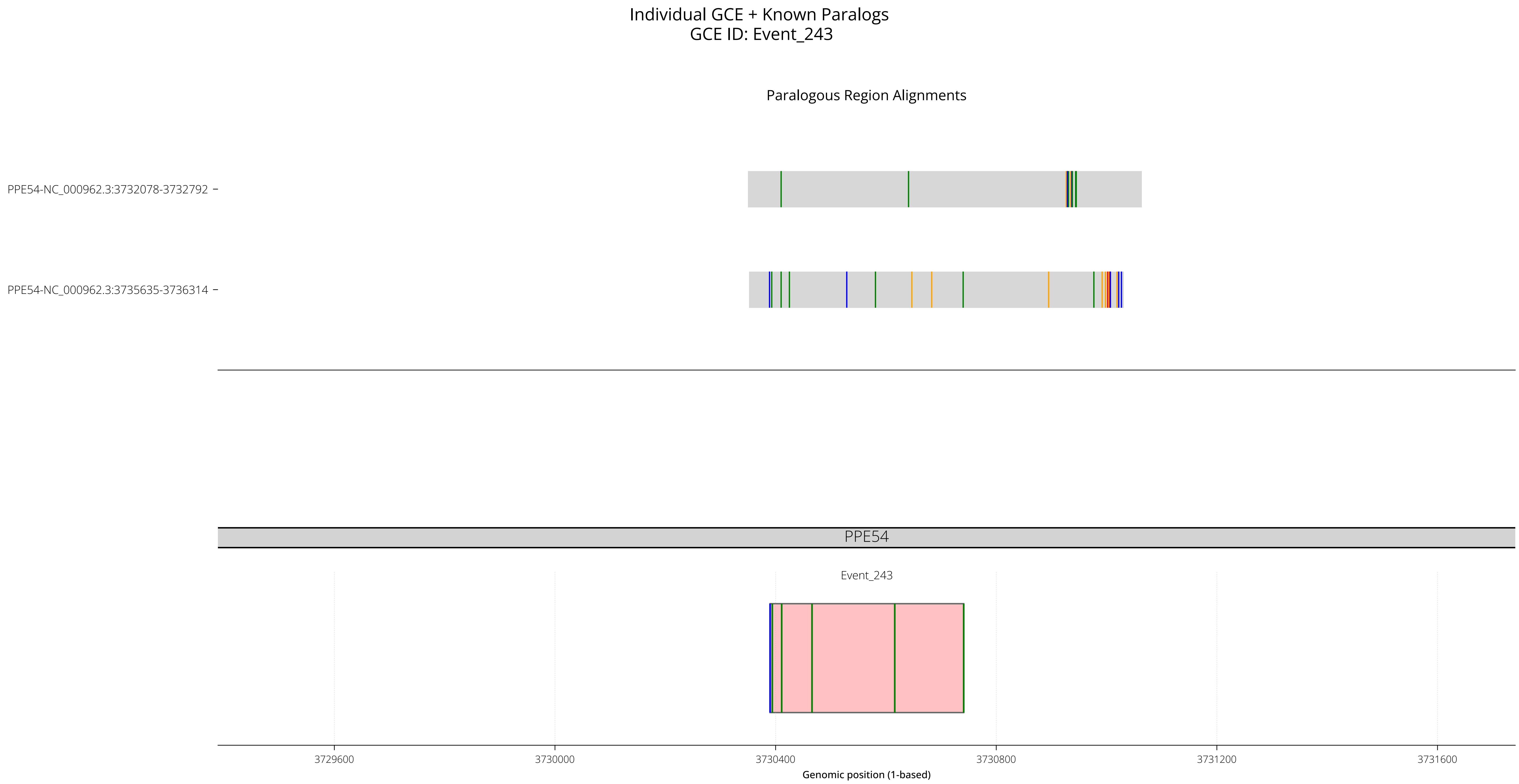

Individual GCE + Known Paralogs
GCE ID: Event_243
Paralogous Region Alignments
PPE54-NC_000962.3:3732078-3732792
PPE54-NC_000962.3:3735635-3736314
PPE54
Event_243
3729600
3730000
3730400
3730800
3731200
3731600
Genomic position (1-based)

Circos Viz - GCE to Known Paralogs Locations
GCE ID: Event_243

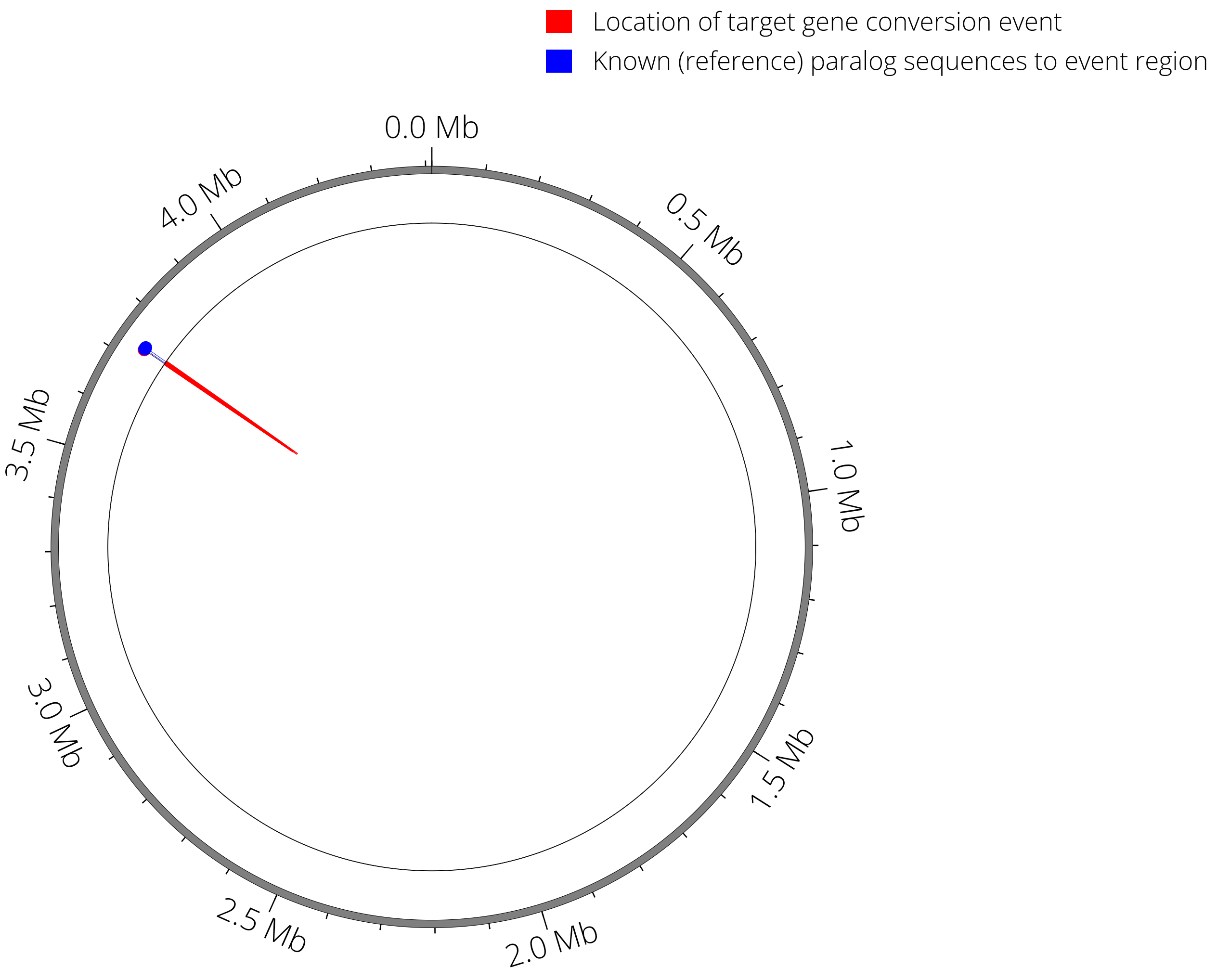

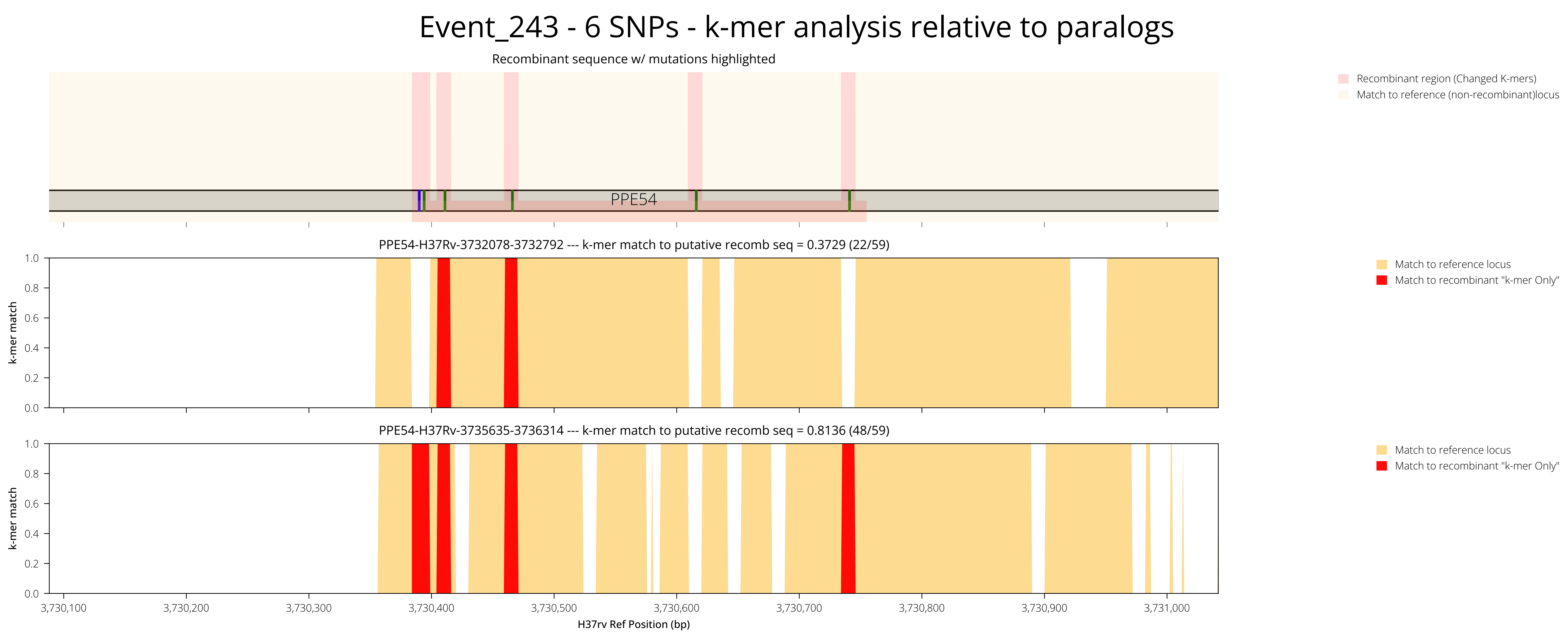

Event_243 - 6 SNPs - k-mer analysis relative to paralogs
Recombinant sequence w/ mutations highlighted
Recombinant region (Changed K-mers)
Match to reference (non-recombinant)locus
PPE54
PPE54-H37Rv-3732078-3732792 --- k-mer match to putative recomb seq = 0.3729 (22/59)
Match to reference locus
Match to recombinant "k-mer Only"
k-mer match
PPE54-H37Rv-3735635-3736314 --- k-mer match to putative recomb seq = 0.8136 (48/59)
Match to reference locus
Match to recombinant "k-mer Only"
k-mer match
1.0
0.8
0.6
0.4
0.2
0.0
3,730,100
3,730,200
3,730,300
3,730,400
3,730,500
3,730,600
3,730,700
3,730,800
3,730,900
3,731,000
H37rv Ref Position (bp)

Sub-phylogeny Viz + Paralog Alignments
GCE ID: Event_243

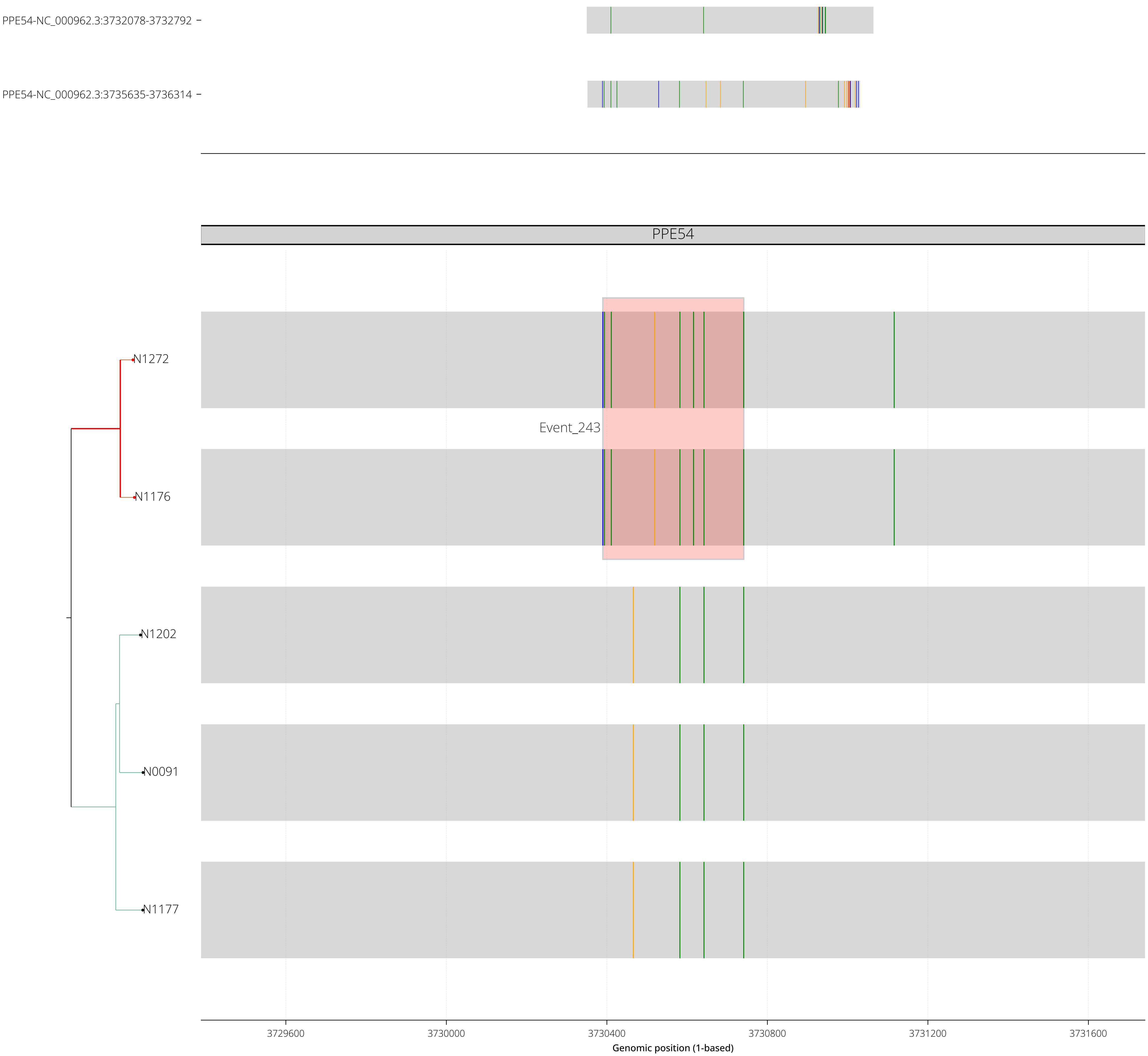

Full Phylogeny Viz + Paralog Alignments
GCE ID: Event_243

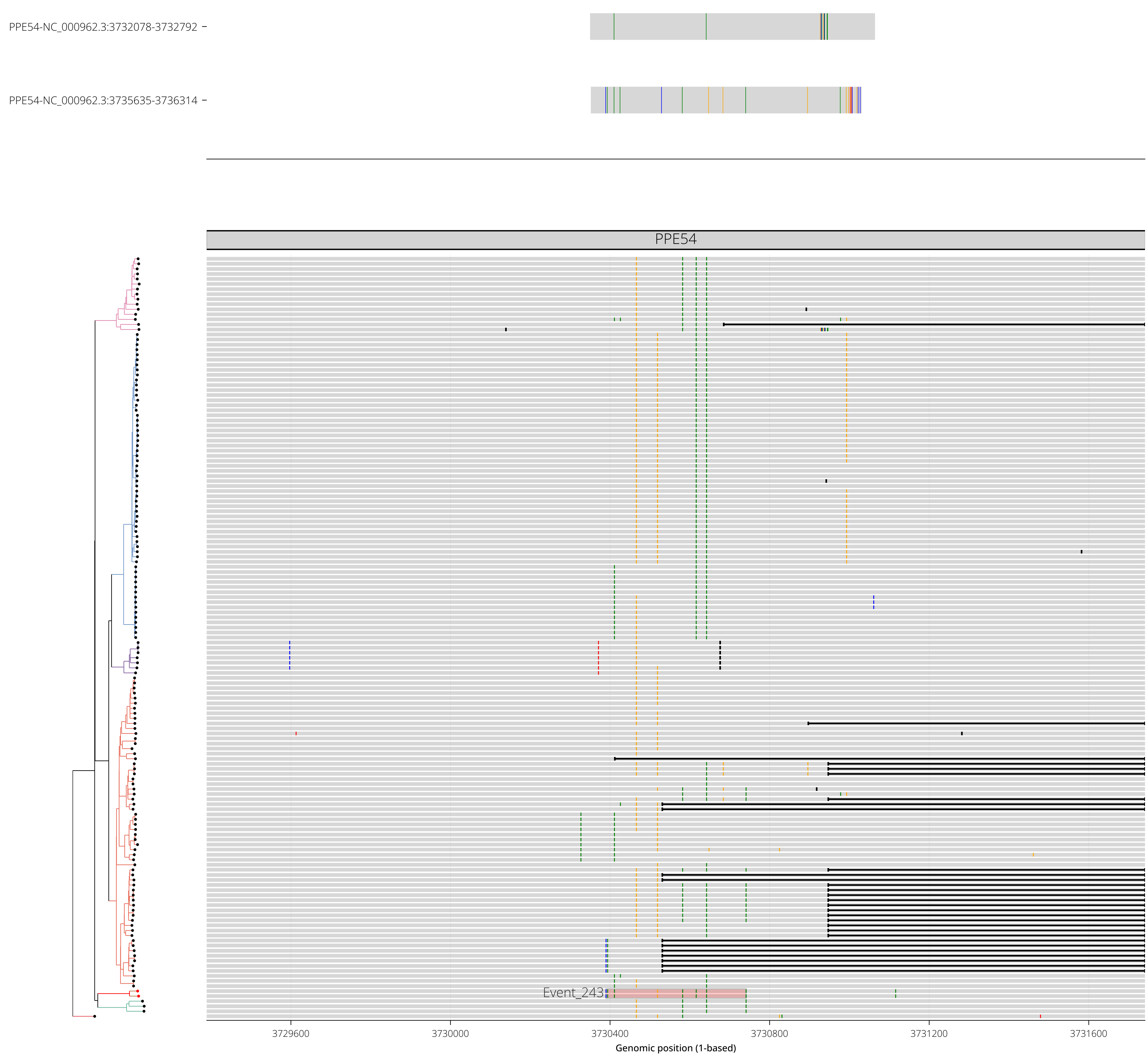

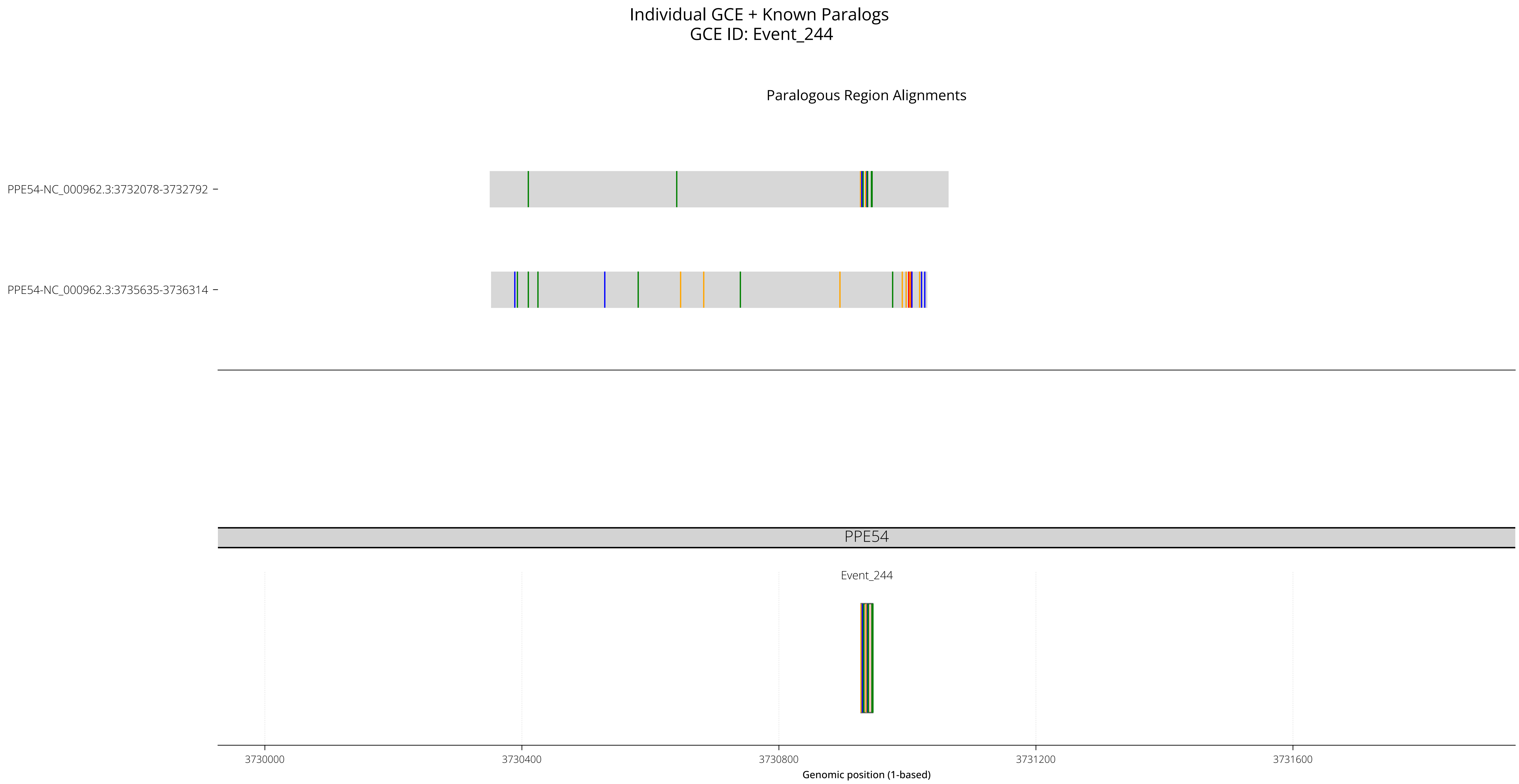

Individual GCE + Known Paralogs
GCE ID: Event_244
Paralogous Region Alignments
PPE54-NC_000962.3:3732078-3732792
PPE54-NC_000962.3:3735635-3736314
PPE54
Event_244
3730000
3730400
3730800
3731200
3731600
Genomic position (1-based)

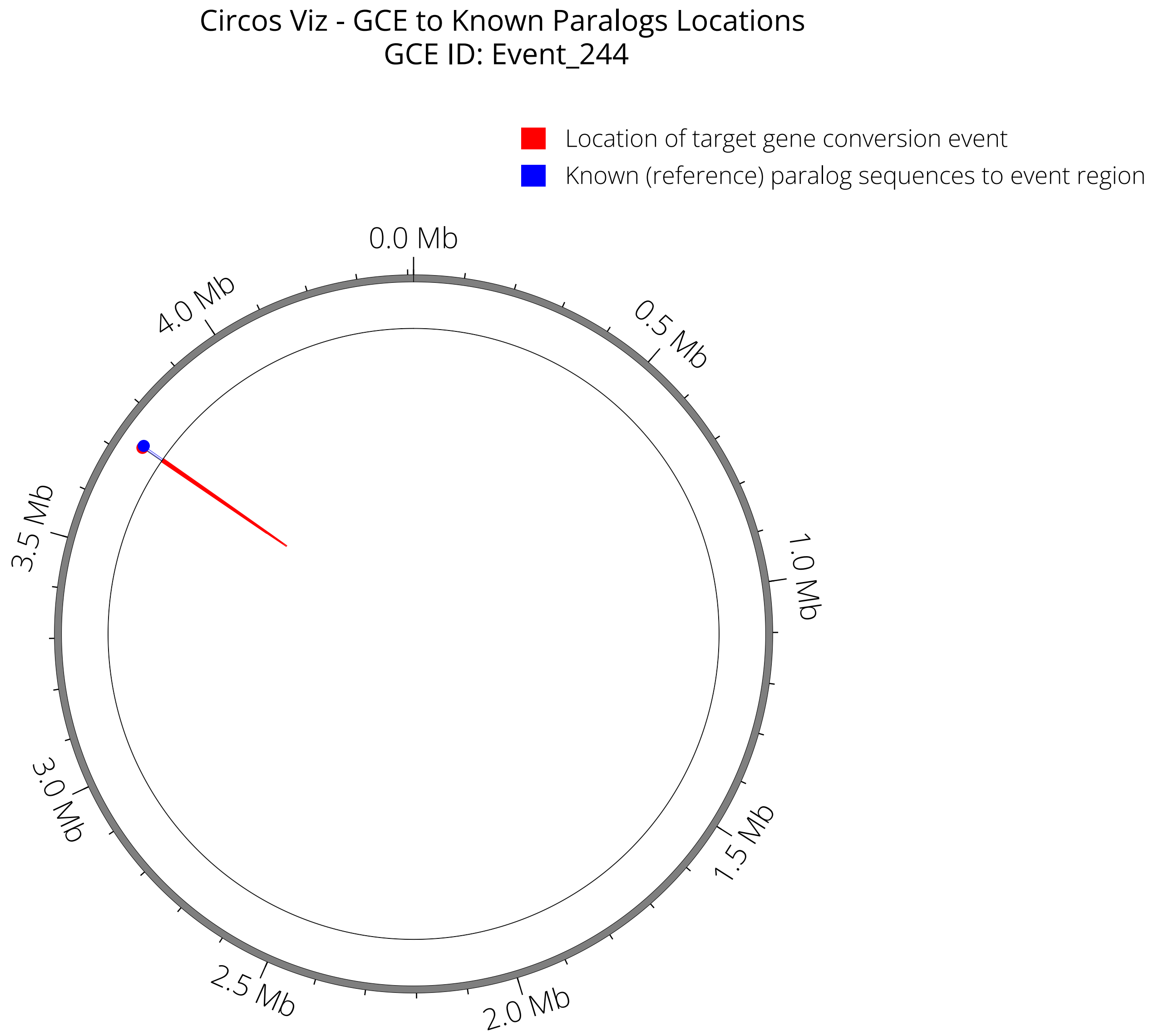

Circos Viz - GCE to Known Paralogs Locations
GCE ID: Event_244
Location of target gene conversion event
Known (reference) paralog sequences to event region
0.0 Mb
0.5 Mb
1.0 Mb
1.5 Mb
2.0 Mb
2.5 Mb
3.0 Mb
3.5 Mb
4.0 Mb

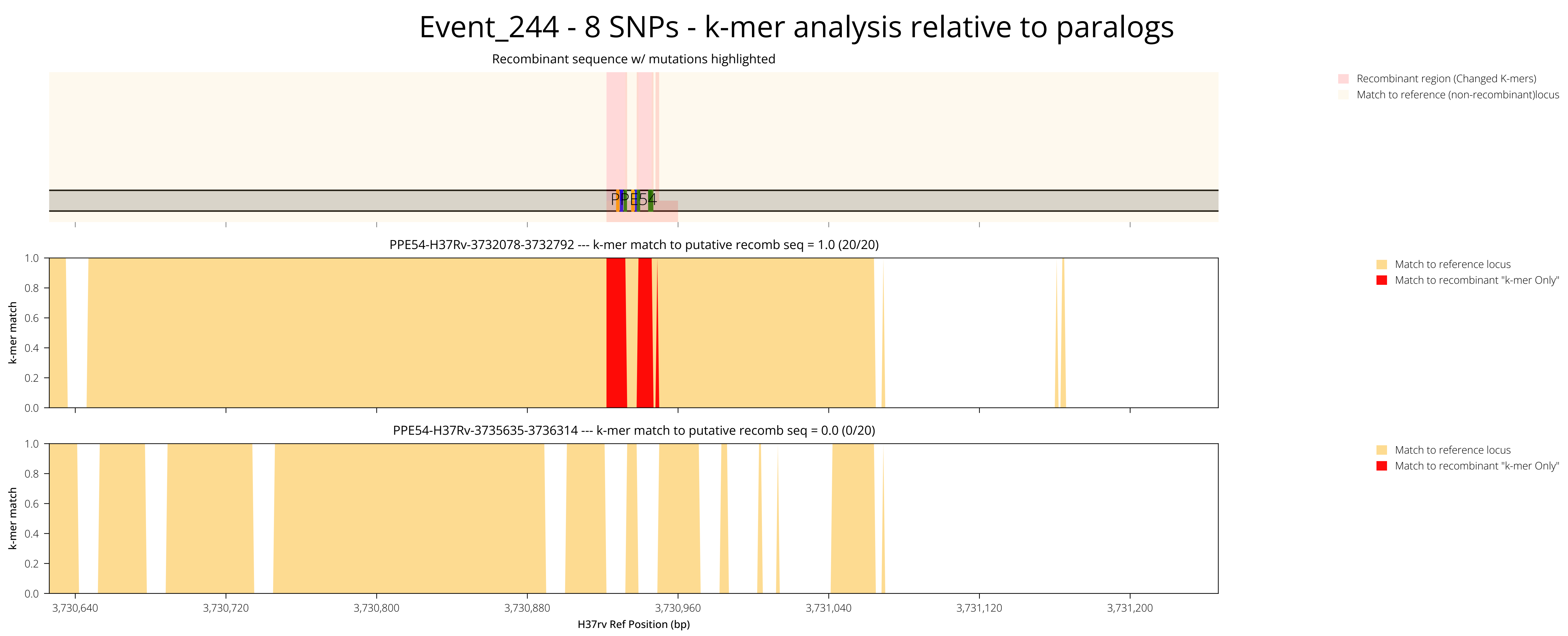

Event_244 - 8 SNPs - k-mer analysis relative to paralogs
Recombinant sequence w/ mutations highlighted
Recombinant region (Changed K-mers)
Match to reference (non-recombinant)locus
PPE54
PPE54-H37Rv-3732078-3732792 --- k-mer match to putative recomb seq = 1.0 (20/20)
Match to reference locus
Match to recombinant "k-mer Only"
k-mer match
1.0
0.8
0.6
0.4
0.2
0.0
PPE54-H37Rv-3735635-3736314 --- k-mer match to putative recomb seq = 0.0 (0/20)
Match to reference locus
Match to recombinant "k-mer Only"
k-mer match
3,730,640
3,730,720
3,730,800
3,730,880
3,730,960
3,731,040
3,731,120
3,731,200
H37rv Ref Position (bp)

Sub-phylogeny Viz + Paralog Alignments
GCE ID: Event_244

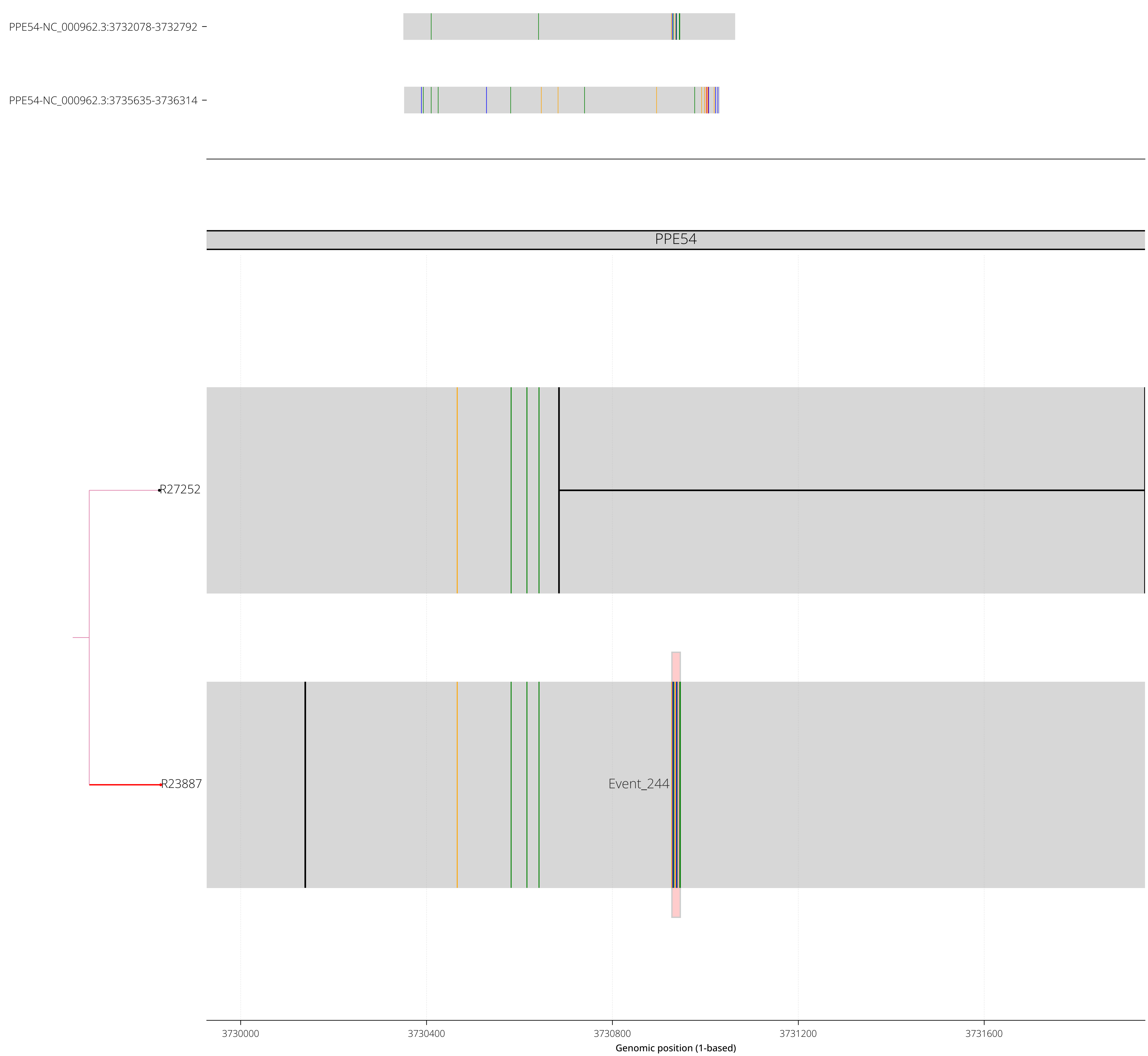

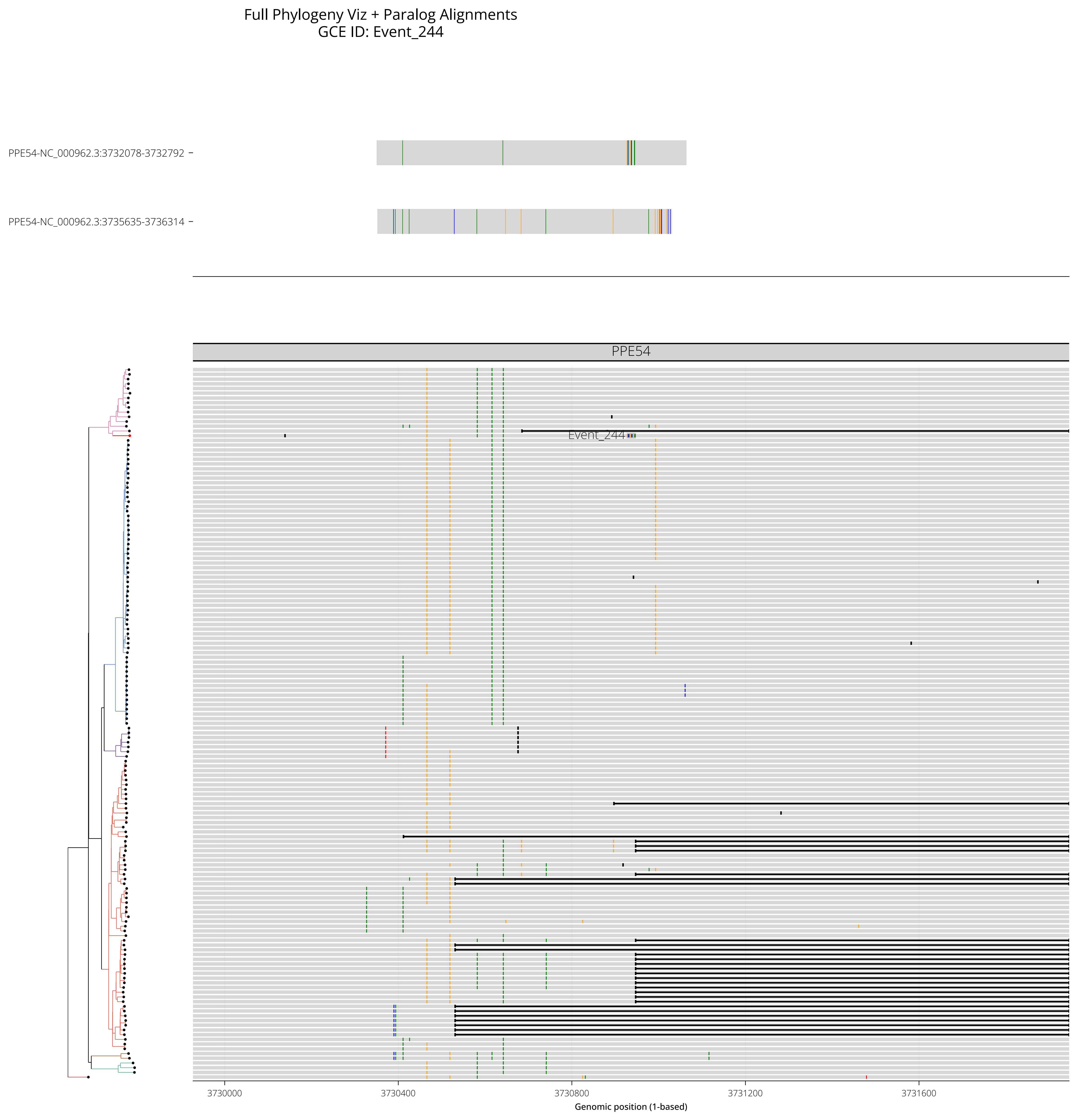

Full Phylogeny Viz + Paralog Alignments
GCE ID: Event_244
PPE54-NC_000962.3:3732078-3732792
PPE54-NC_000962.3:3735635-3736314
PPE54
Event_244
3730000
3730400
3730800
3731200
3731600
Genomic position (1-based)

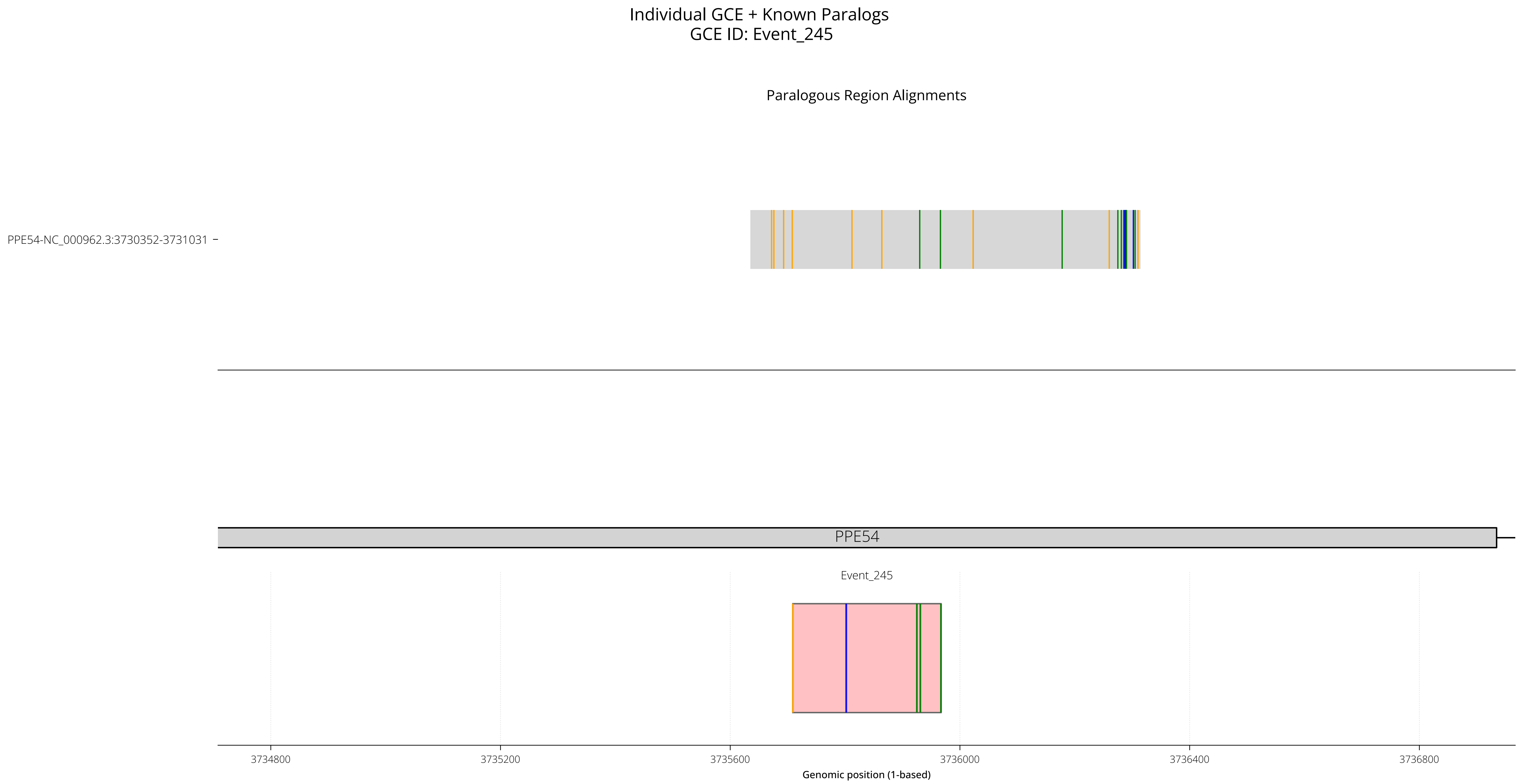

Individual GCE + Known Paralogs
GCE ID: Event_245
Paralogous Region Alignments
PPE54-NC_000962.3:3730352-3731031
PPE54
Event_245
3734800
3735200
3735600
3736000
3736400
3736800
Genomic position (1-based)

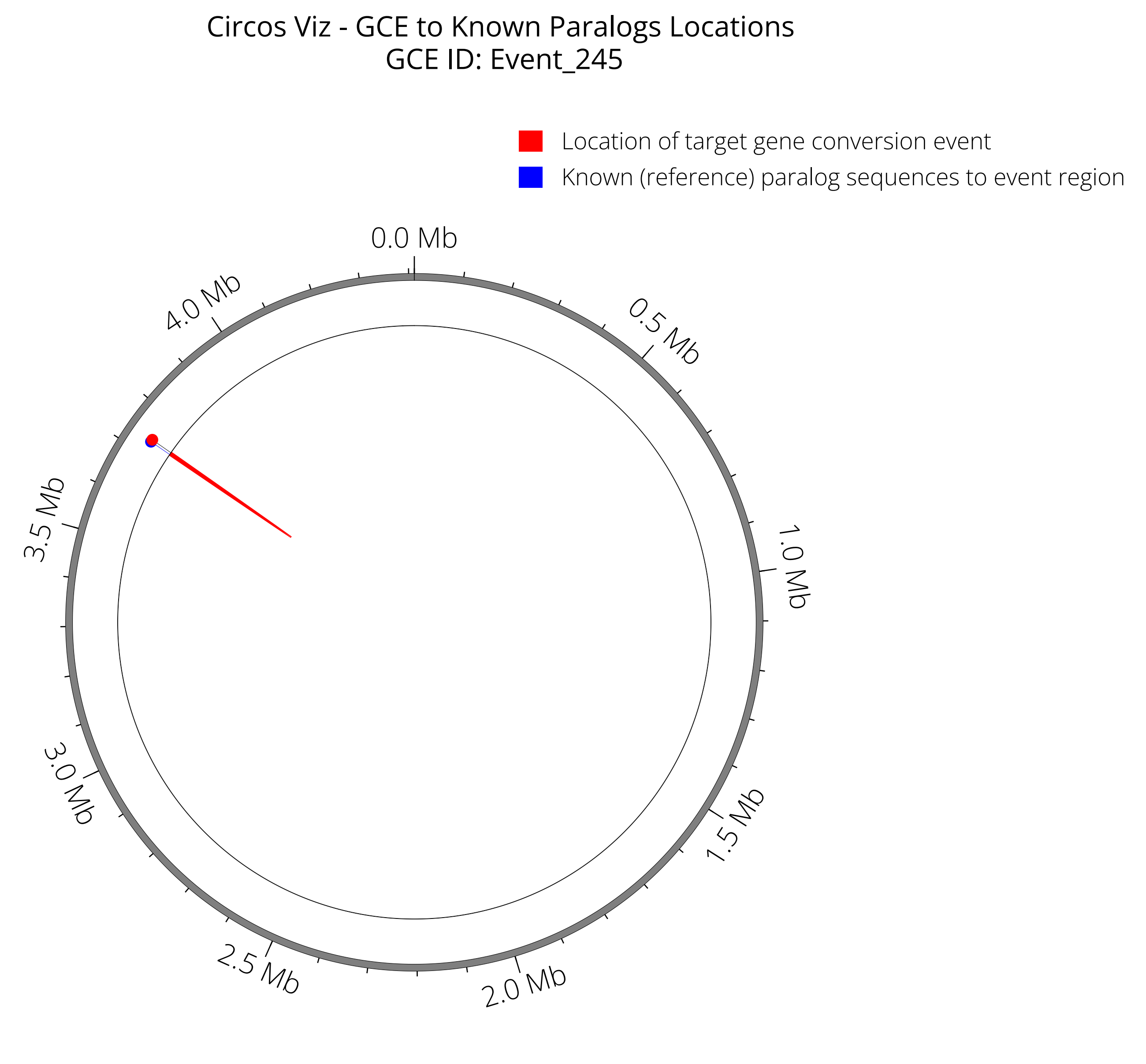
Circos Viz - GCE to Known Paralogs Locations
GCE ID: Event_245
Location of target gene conversion event
Known (reference) paralog sequences to event region
0.0 Mb
0.5 Mb
1.0 Mb
1.5 Mb
2.0 Mb
2.5 Mb
3.0 Mb
3.5 Mb
4.0 Mb

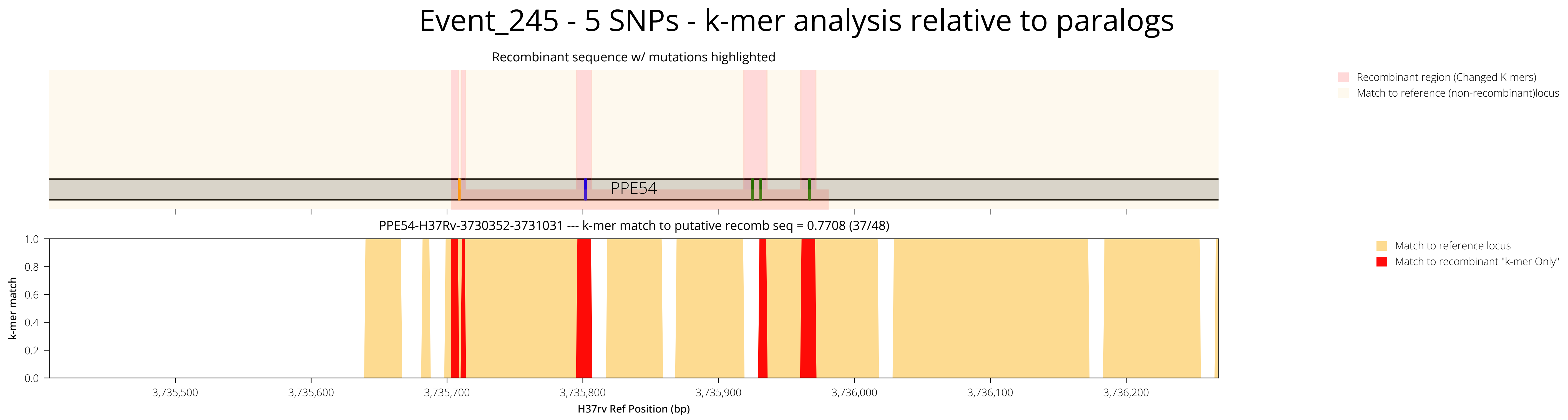

Event_245 - 5 SNPs - k-mer analysis relative to paralogs
Recombinant sequence w/ mutations highlighted
PPE54
Recombinant region (Changed K-mers)
Match to reference (non-recombinant)locus
PPE54-H37Rv-3730352-3731031 --- k-mer match to putative recomb seq = 0.7708 (37/48)
k-mer match
1.0
0.8
0.6
0.4
0.2
0.0
3,735,500
3,735,600
3,735,700
3,735,800
3,735,900
3,736,000
3,736,100
3,736,200
H37rv Ref Position (bp)
Match to reference locus
Match to recombinant "k-mer Only"

Sub-phylogeny Viz + Paralog Alignments
GCE ID: Event_245

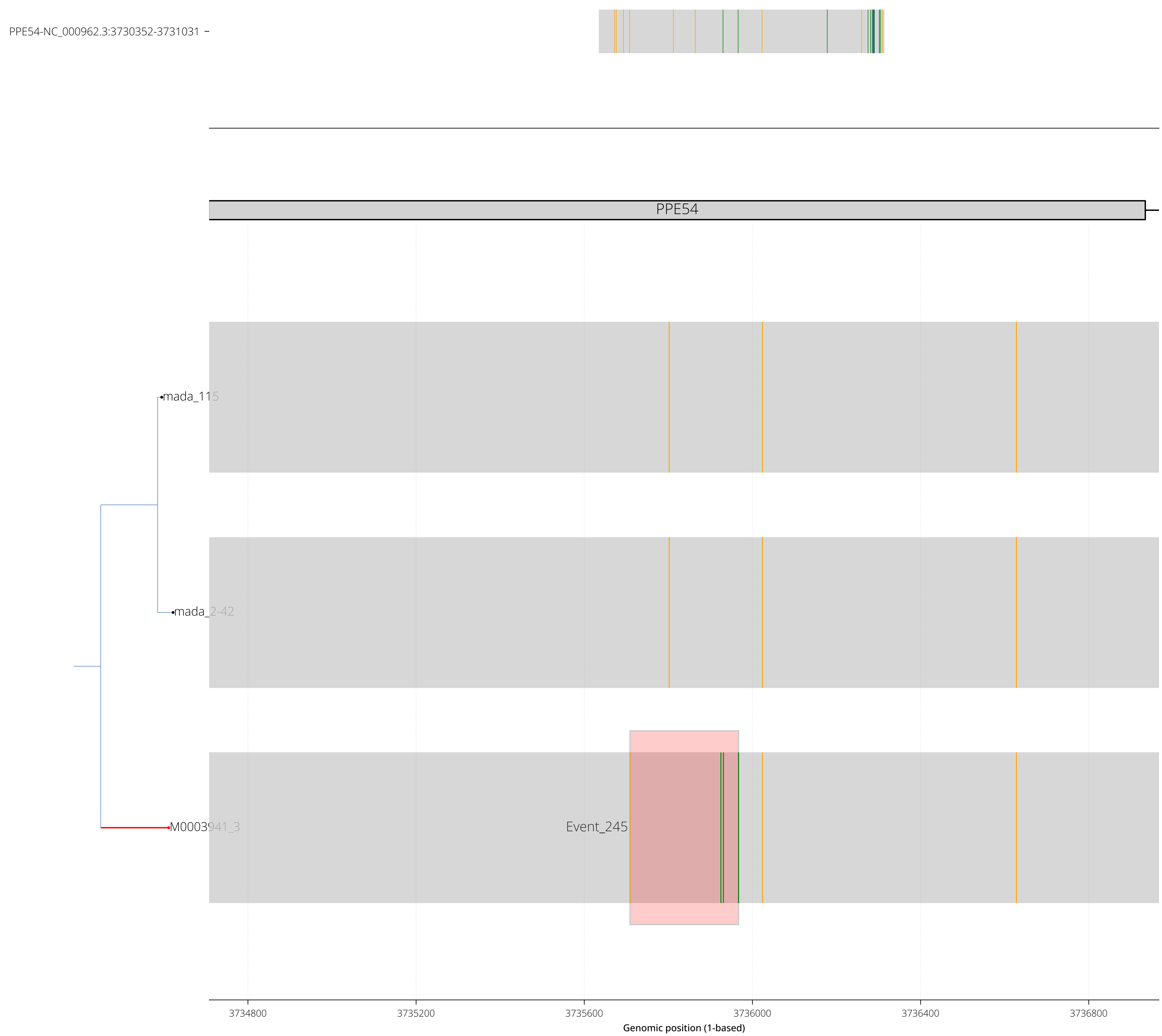

Full Phylogeny Viz + Paralog Alignments
GCE ID: Event_245

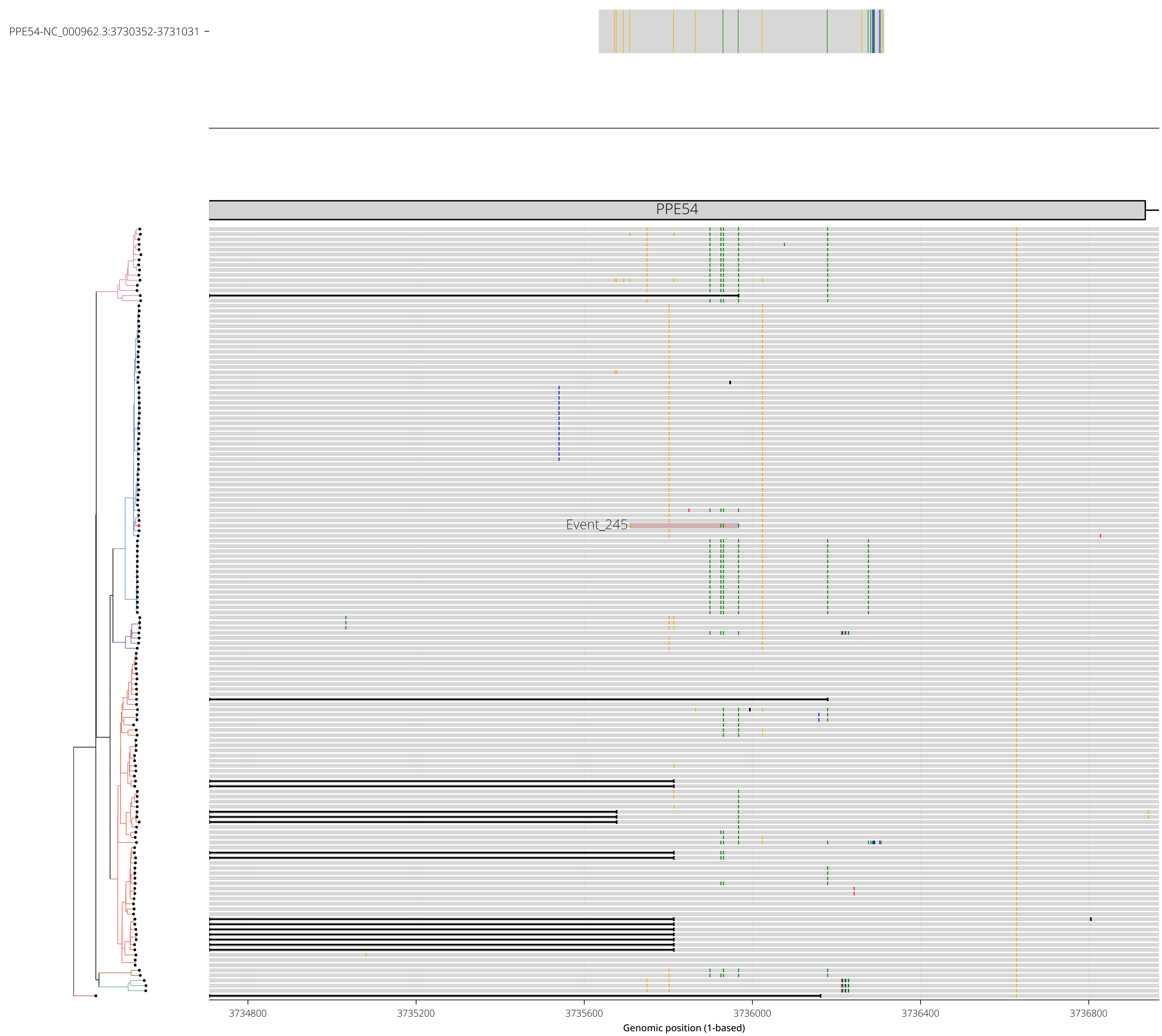

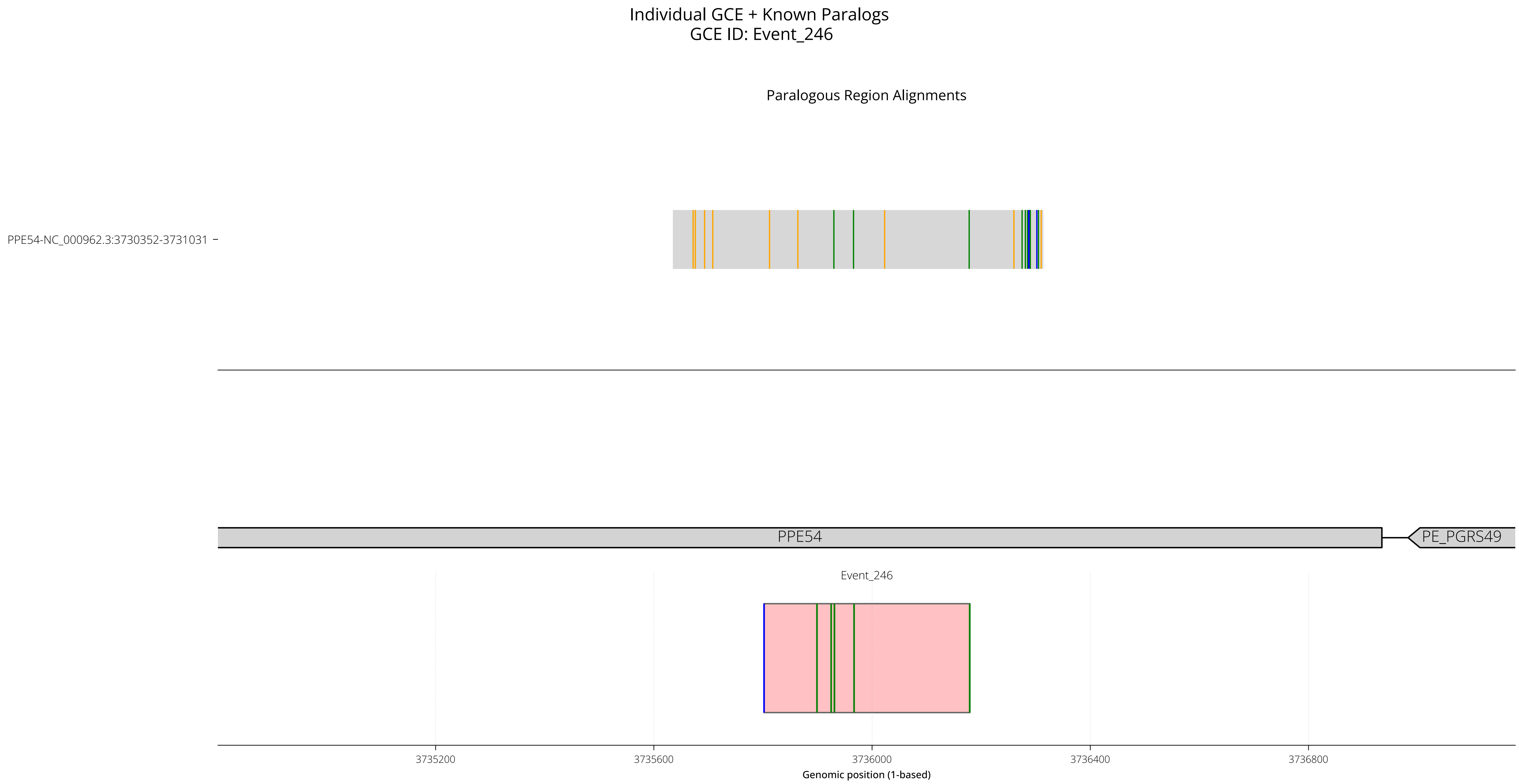

Individual GCE + Known Paralogs
GCE ID: Event_246
Paralogous Region Alignments
PPE54-NC_000962.3:3730352-3731031
PPE54
PE_PGRS49
Event_246
3735200
3735600
3736000
3736400
3736800
Genomic position (1-based)

Circos Viz - GCE to Known Paralogs Locations
GCE ID: Event_246

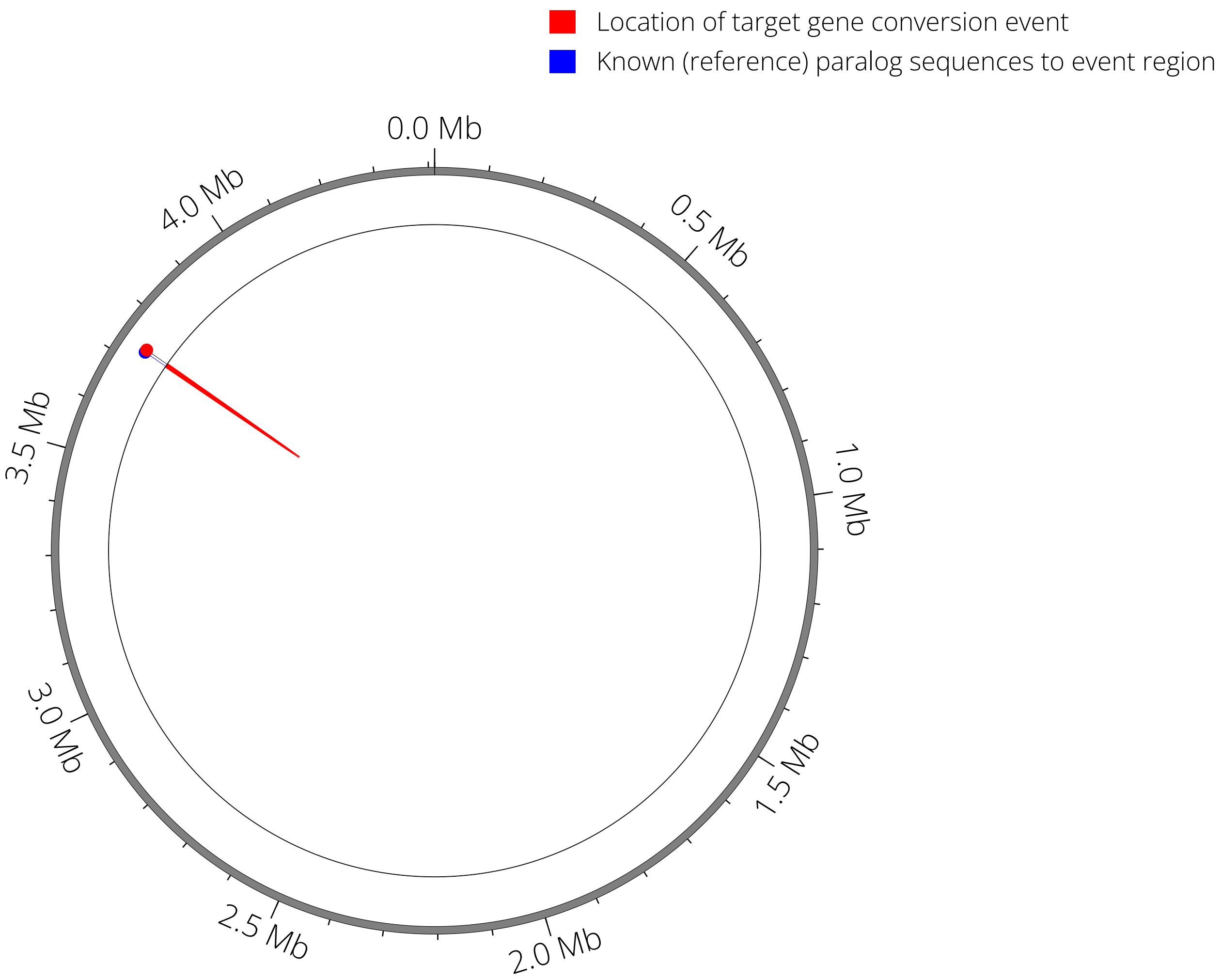

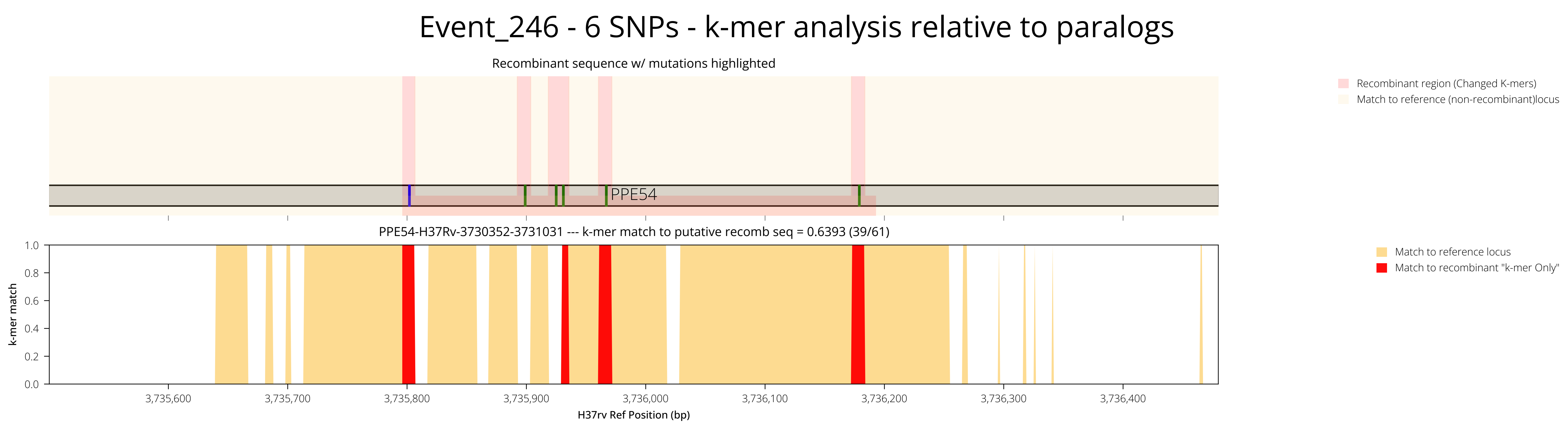
Event_246 - 6 SNPs - k-mer analysis relative to paralogs
Recombinant sequence w/ mutations highlighted
PPE54
Recombinant region (Changed K-mers)
Match to reference (non-recombinant)locus
PPE54-H37Rv-3730352-3731031 --- k-mer match to putative recomb seq = 0.6393 (39/61)
k-mer match
1.0
0.8
0.6
0.4
0.2
0.0
3,735,600
3,735,700
3,735,800
3,735,900
3,736,000
3,736,100
3,736,200
3,736,300
3,736,400
H37rv Ref Position (bp)
Match to reference locus
Match to recombinant "k-mer Only"

#### Sub-phylogeny Viz + Paralog Alignments
#### GCE ID: Event_246

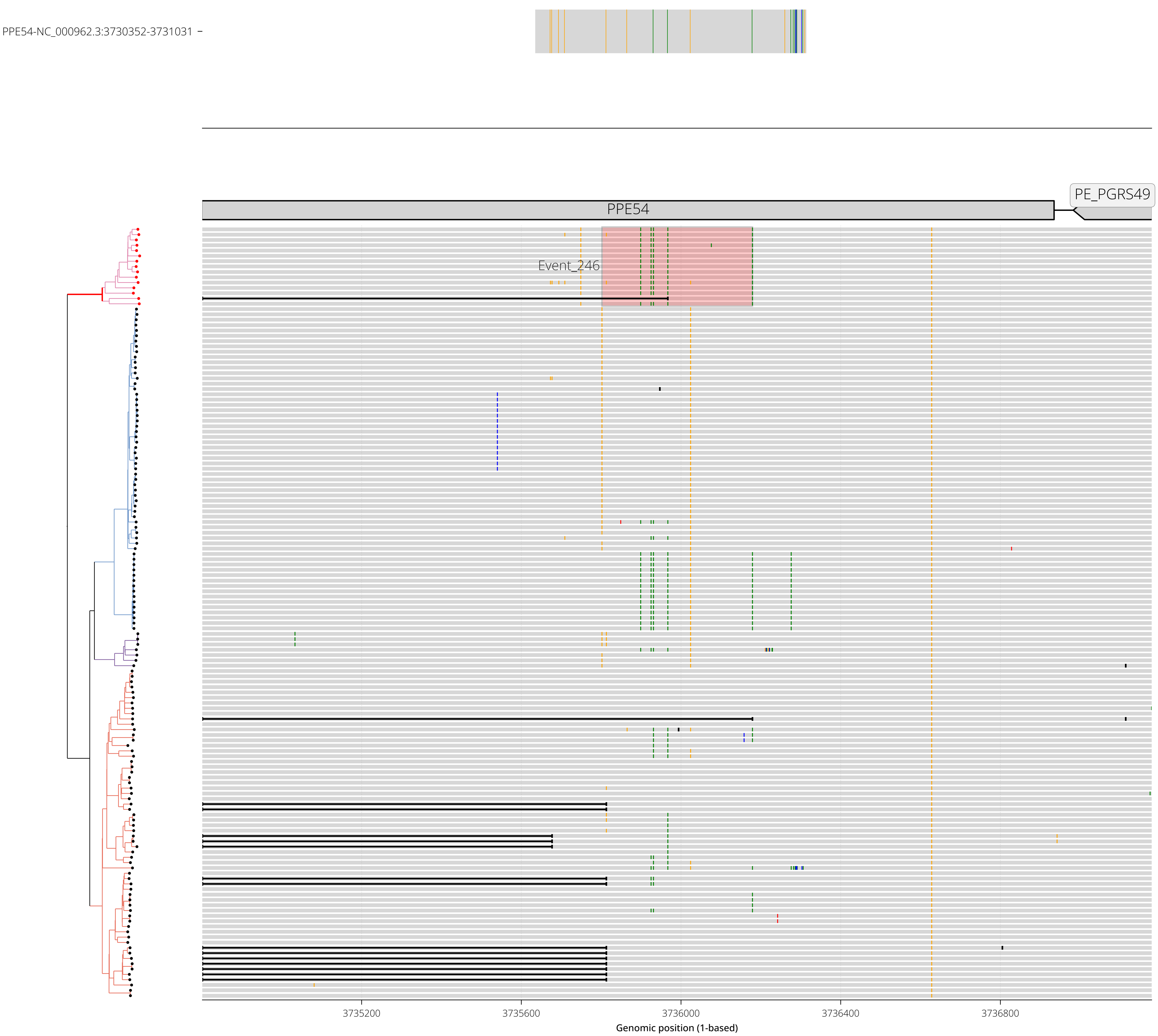

Full Phylogeny Viz + Paralog Alignments
GCE ID: Event_246

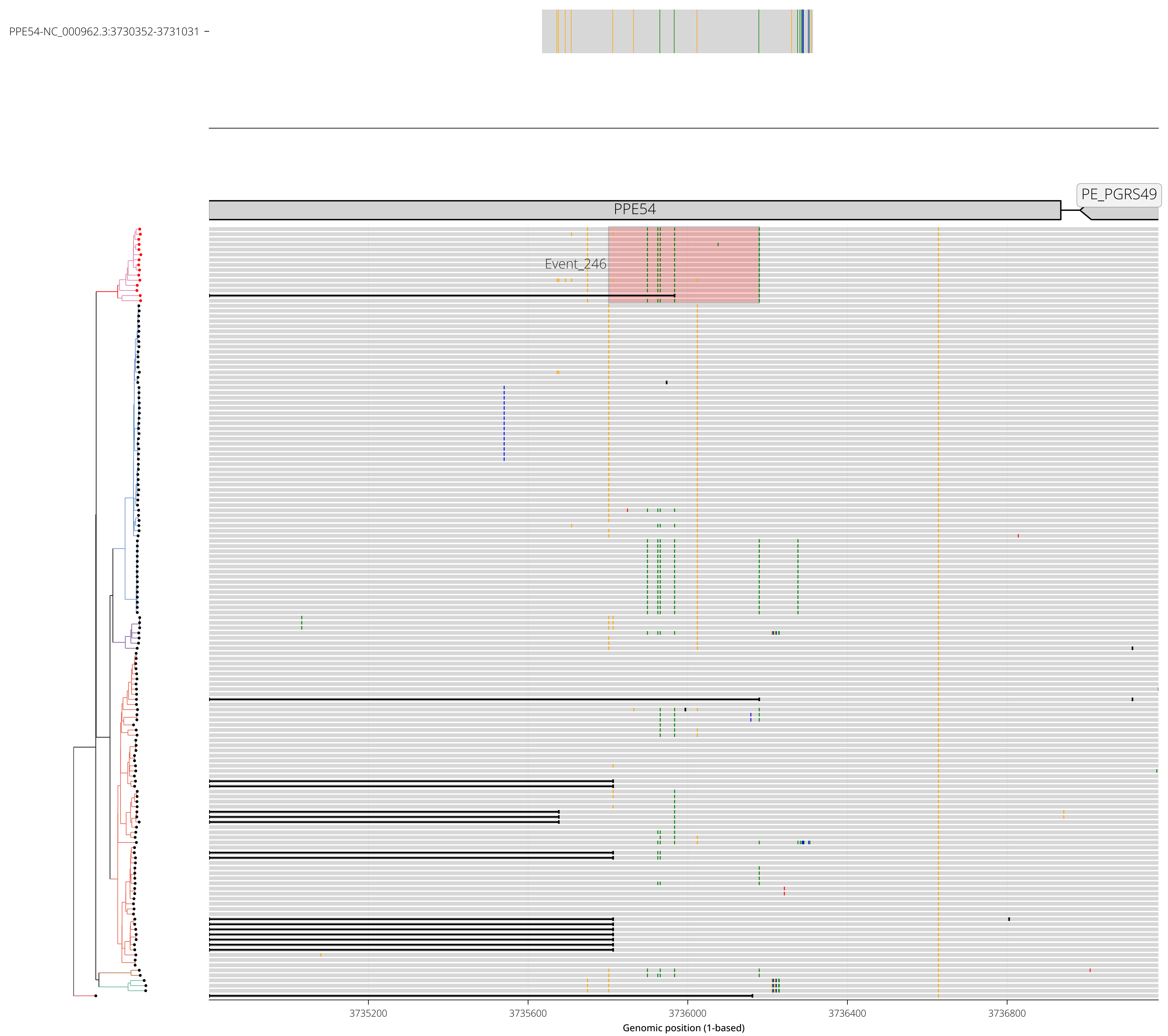

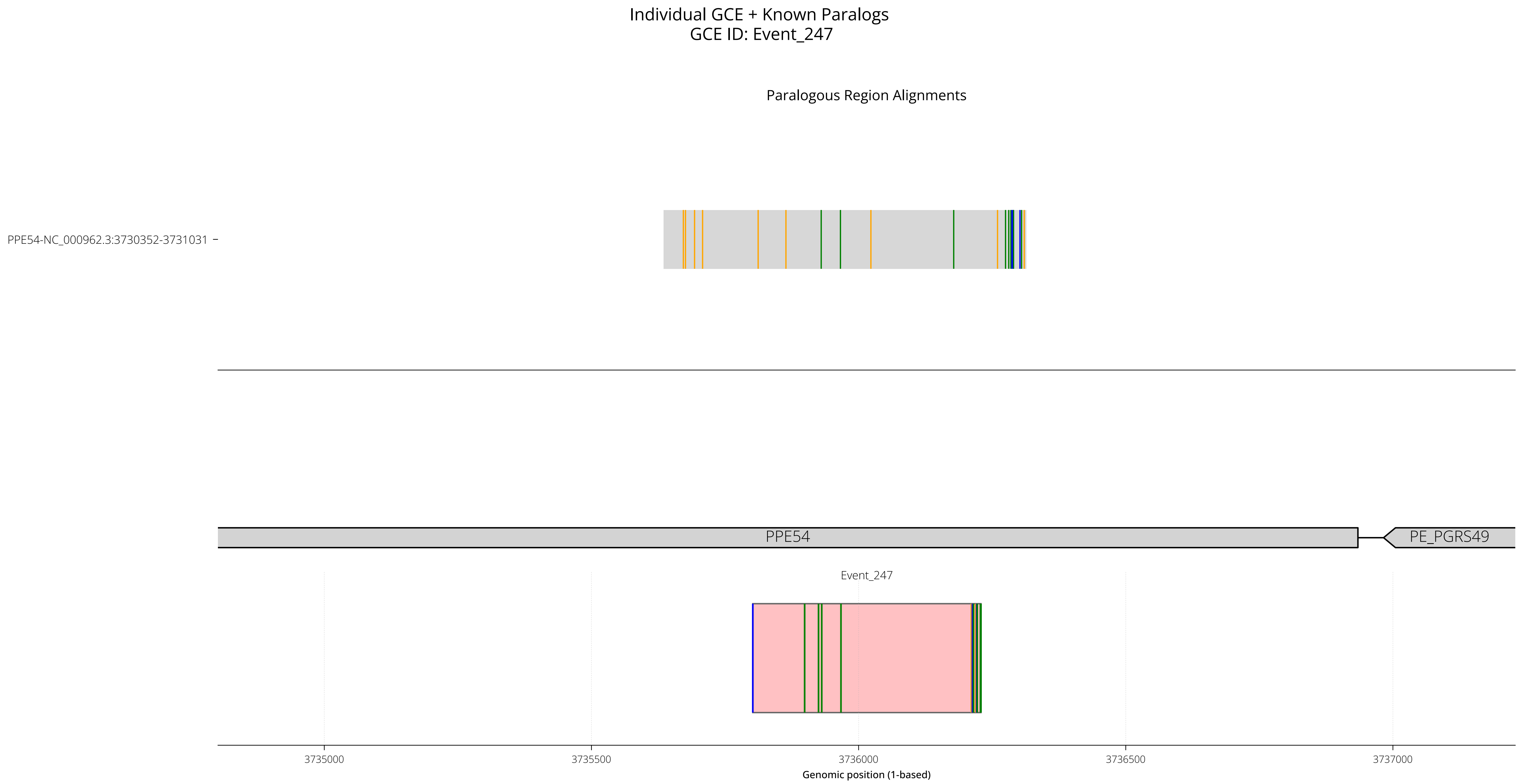
Individual GCE + Known Paralogs
GCE ID: Event_247
Paralogous Region Alignments
PPE54-NC_000962.3:3730352-3731031
PPE54
PE_PGRS49
Event_247
3735000
3735500
3736000
3736500
3737000
Genomic position (1-based)

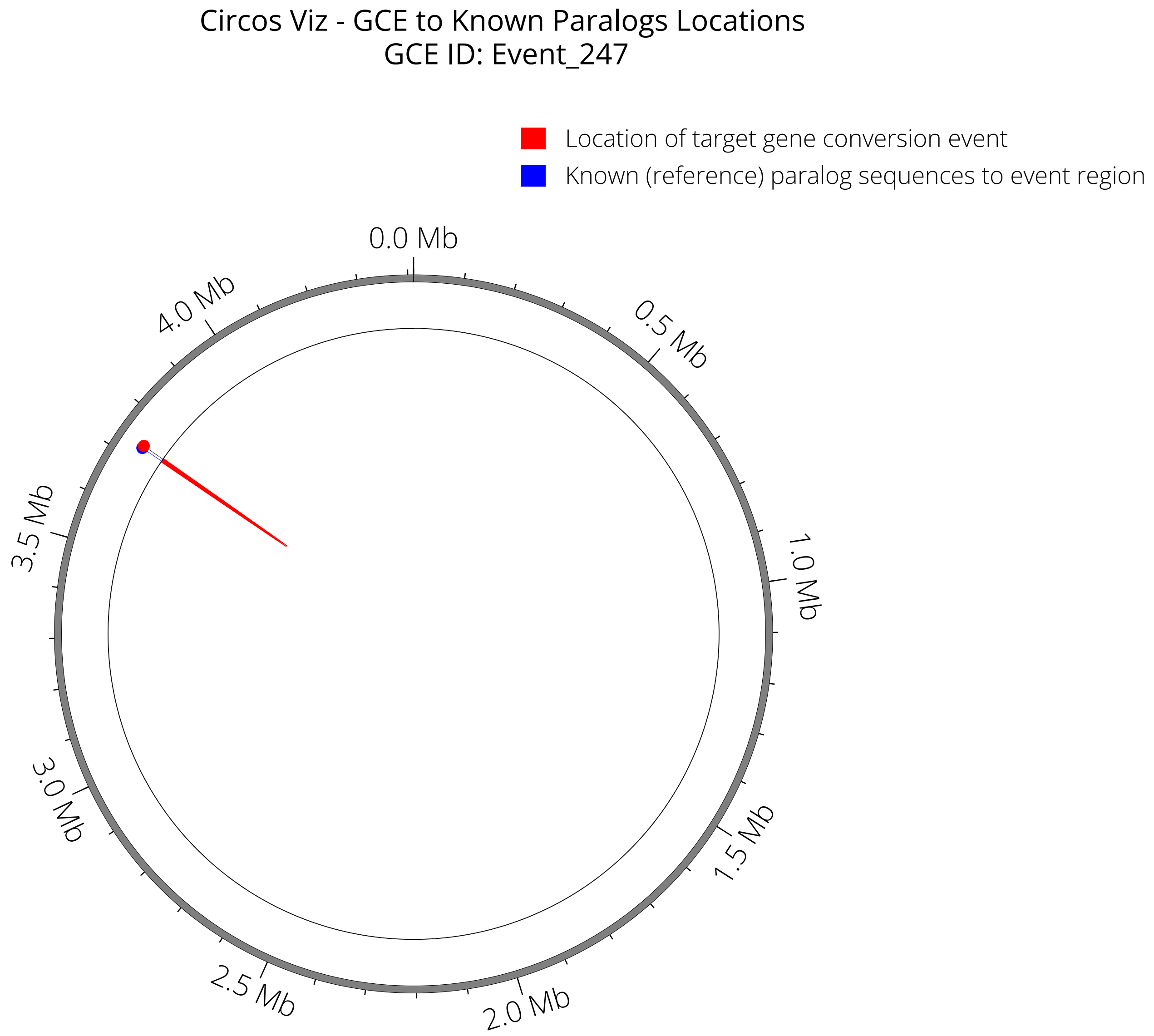

Circos Viz - GCE to Known Paralogs Locations
GCE ID: Event_247
Location of target gene conversion event
Known (reference) paralog sequences to event region
0.0 Mb
0.5 Mb
1.0 Mb
1.5 Mb
2.0 Mb
2.5 Mb
3.0 Mb
3.5 Mb
4.0 Mb

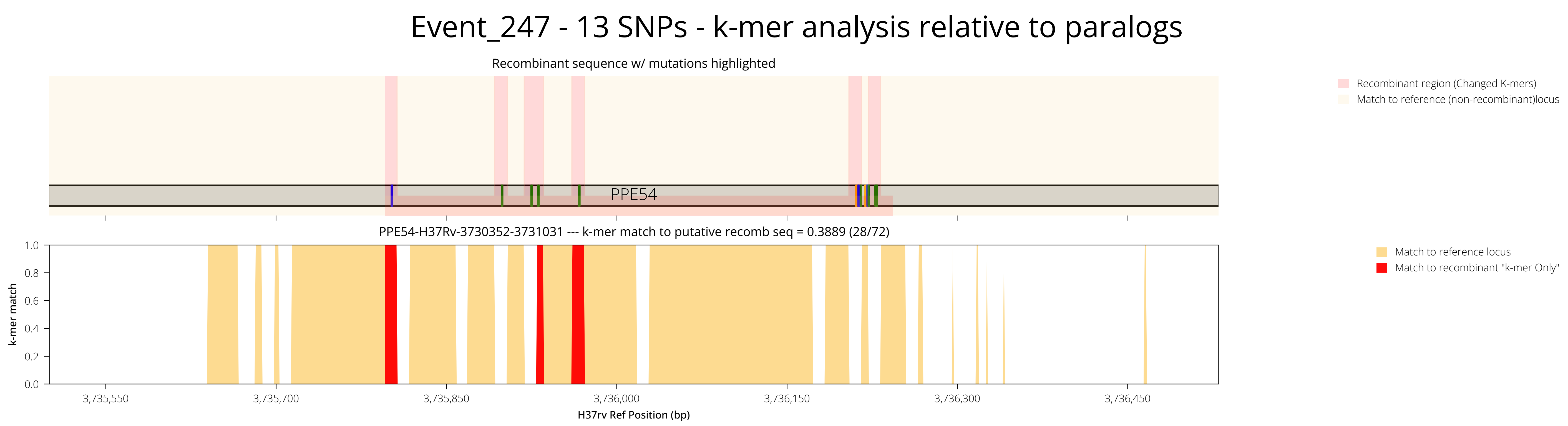

Event_247 - 13 SNPs - k-mer analysis relative to paralogs
Recombinant sequence w/ mutations highlighted
PPE54
Recombinant region (Changed K-mers)
Match to reference (non-recombinant)locus
PPE54-H37Rv-3730352-3731031 --- k-mer match to putative recomb seq = 0.3889 (28/72)
k-mer match
1.0
0.8
0.6
0.4
0.2
0.0
3,735,550
3,735,700
3,735,850
3,736,000
3,736,150
3,736,300
3,736,450
H37rv Ref Position (bp)
Match to reference locus
Match to recombinant "k-mer Only"

Sub-phylogeny Viz + Paralog Alignments
GCE ID: Event_247

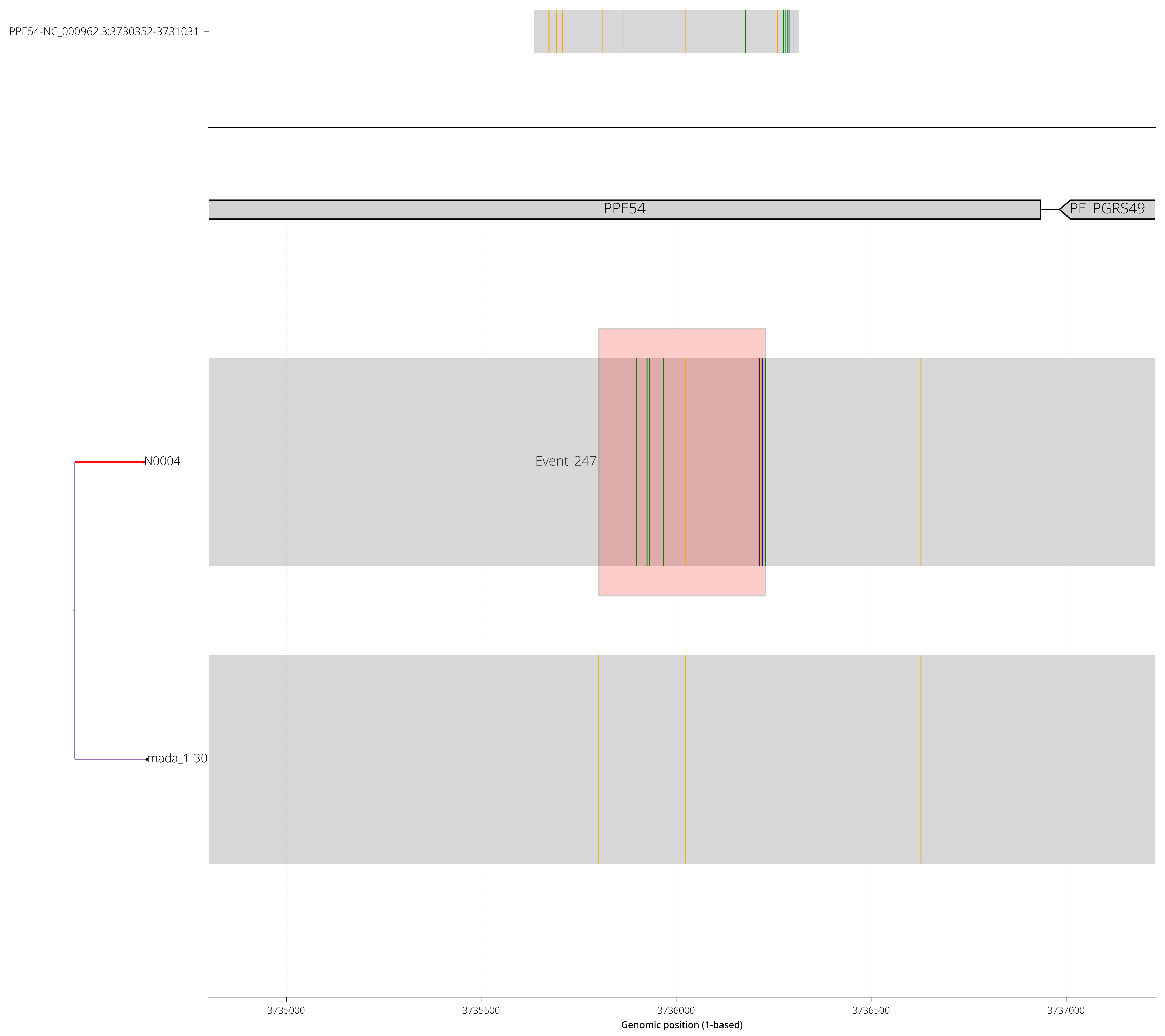

Full Phylogeny Viz + Paralog Alignments
GCE ID: Event_247

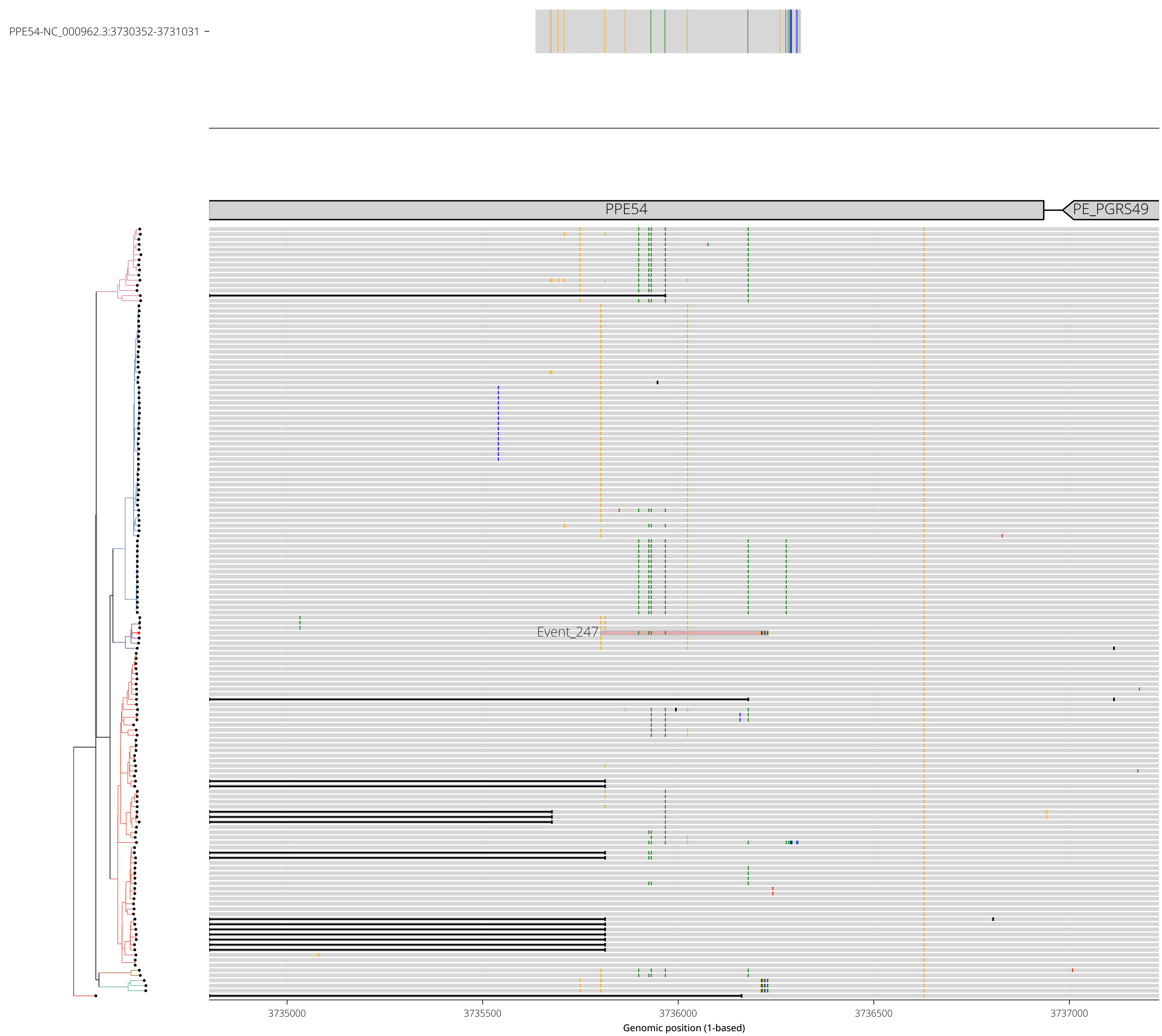

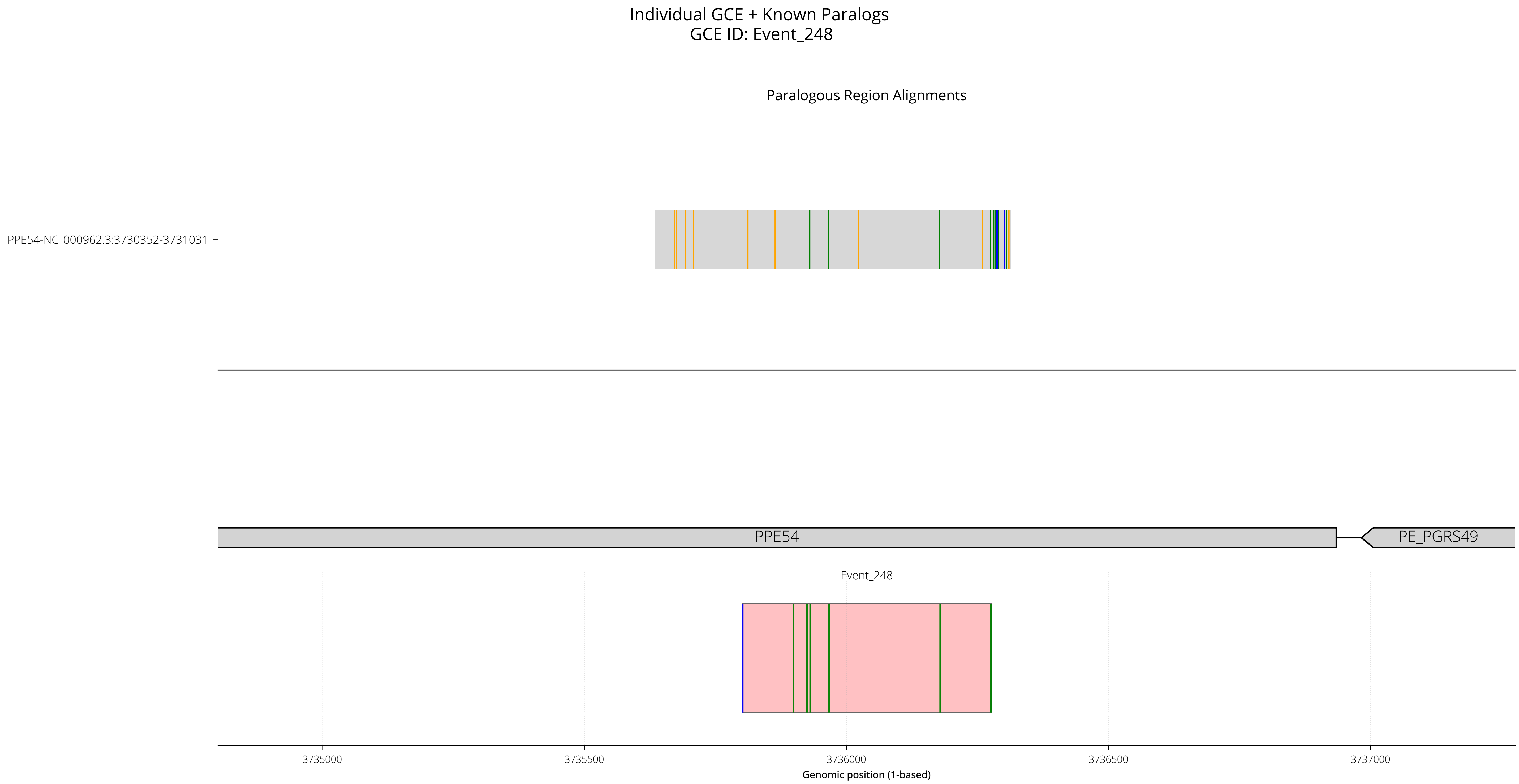

Individual GCE + Known Paralogs
GCE ID: Event_248
Paralogous Region Alignments
PPE54-NC_000962.3:3730352-3731031
PPE54
PE_PGRS49
Event_248
3735000
3735500
3736000
3736500
3737000
Genomic position (1-based)

### Circos Viz - GCE to Known Paralogs Locations
GCE ID: Event_248

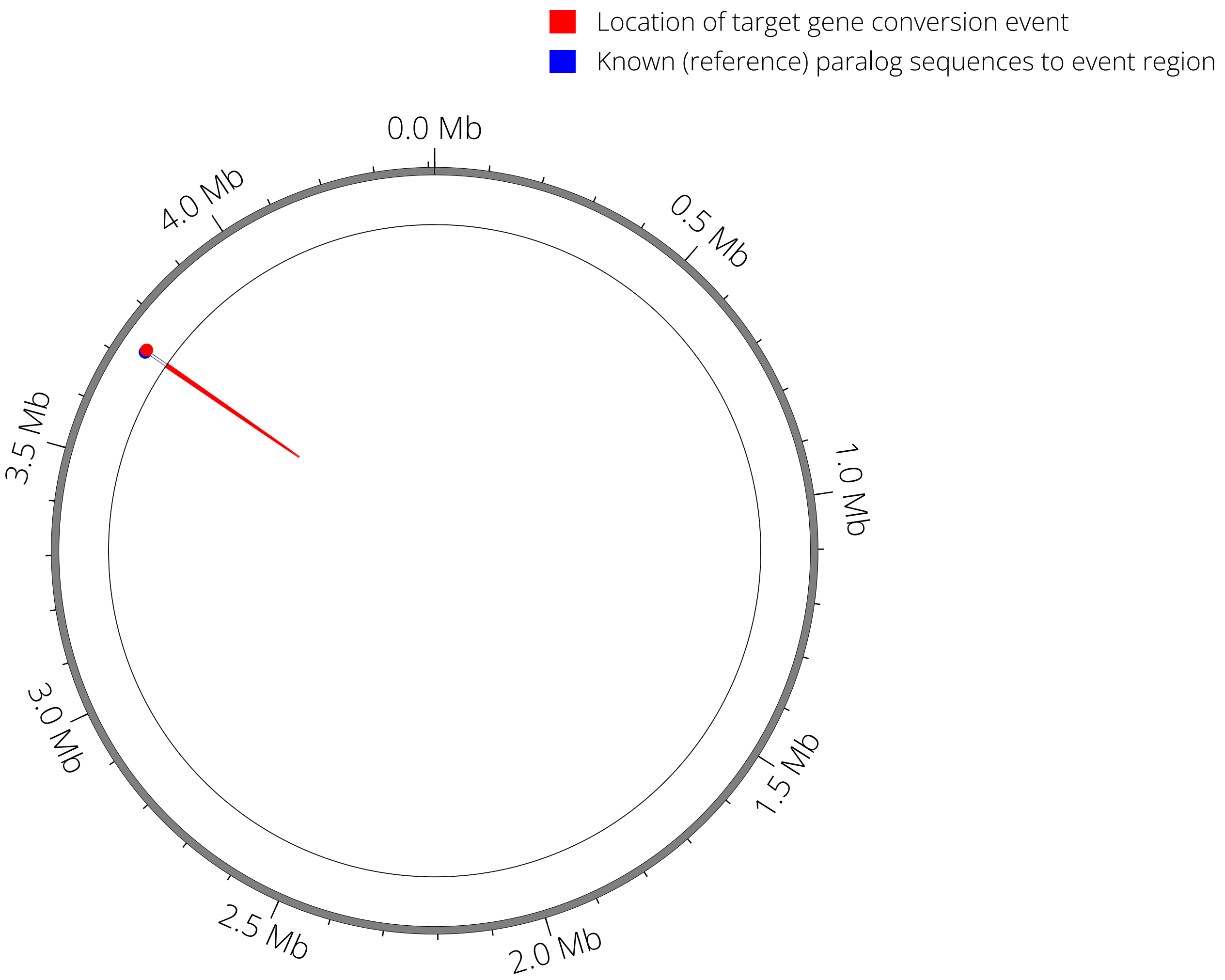

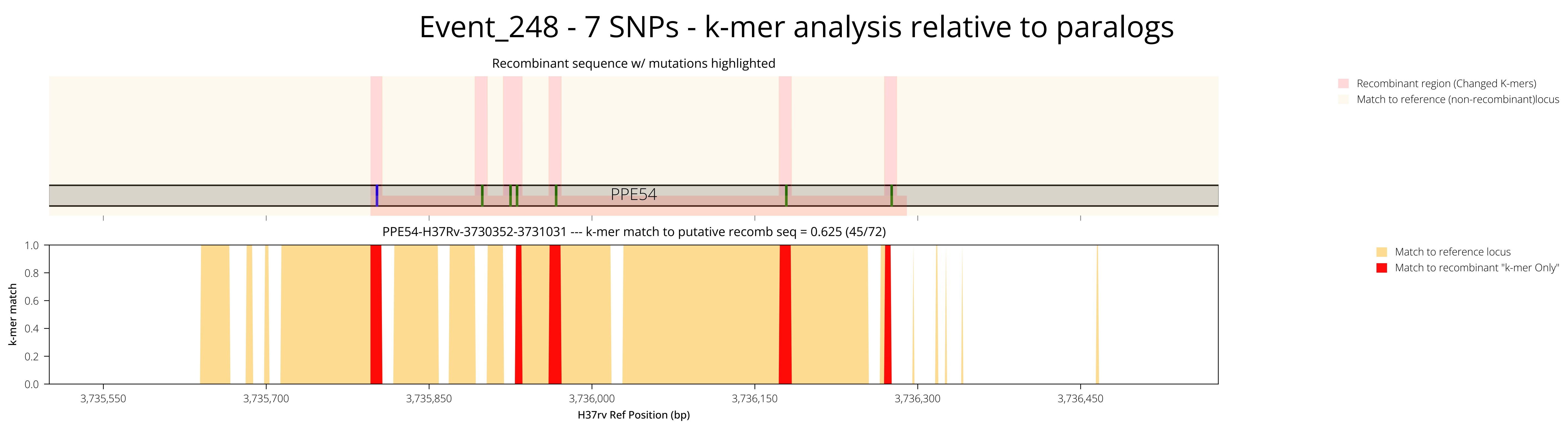

Event_248 - 7 SNPs - k-mer analysis relative to paralogs
Recombinant sequence w/ mutations highlighted
Recombinant region (Changed K-mers)
Match to reference (non-recombinant)locus
PPE54
PPE54-H37Rv-3730352-3731031 --- k-mer match to putative recomb seq = 0.625 (45/72)
Match to reference locus
Match to recombinant "k-mer Only"
k-mer match
1.0
0.8
0.6
0.4
0.2
0.0
3,735,550
3,735,700
3,735,850
3,736,000
3,736,150
3,736,300
3,736,450
H37rv Ref Position (bp)

Sub-phylogeny Viz + Paralog Alignments
GCE ID: Event_248

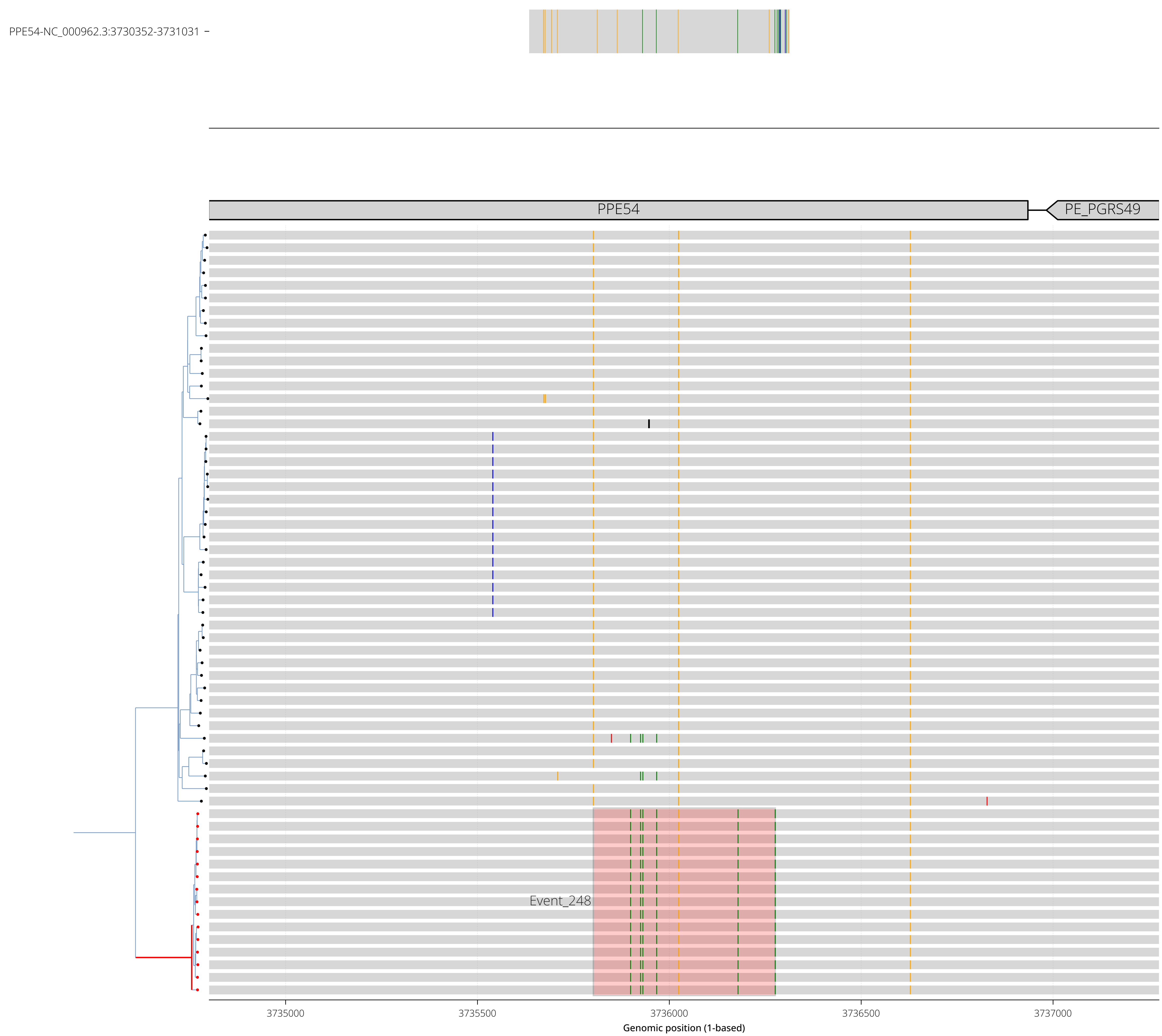

Full Phylogeny Viz + Paralog Alignments
GCE ID: Event_248

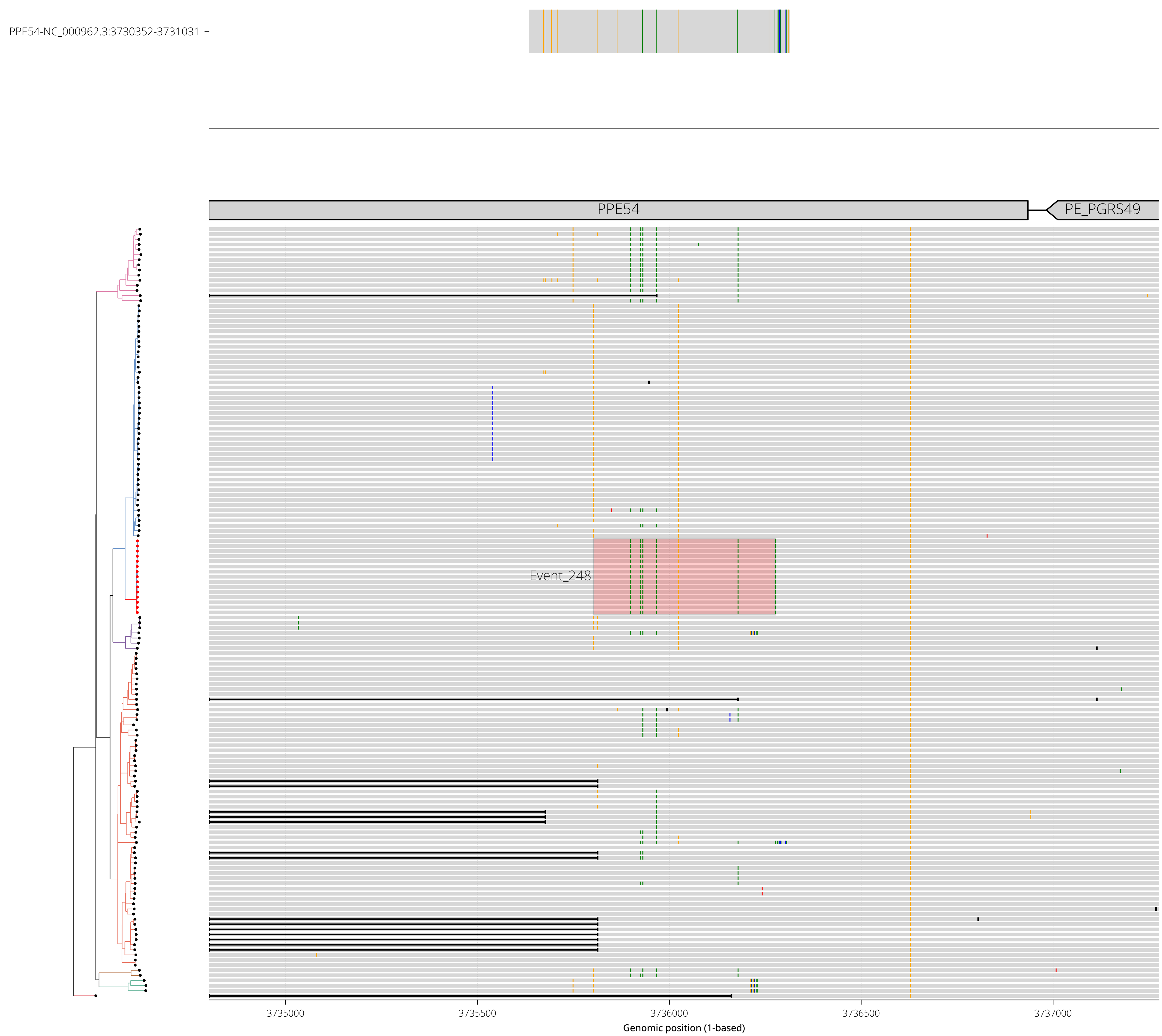

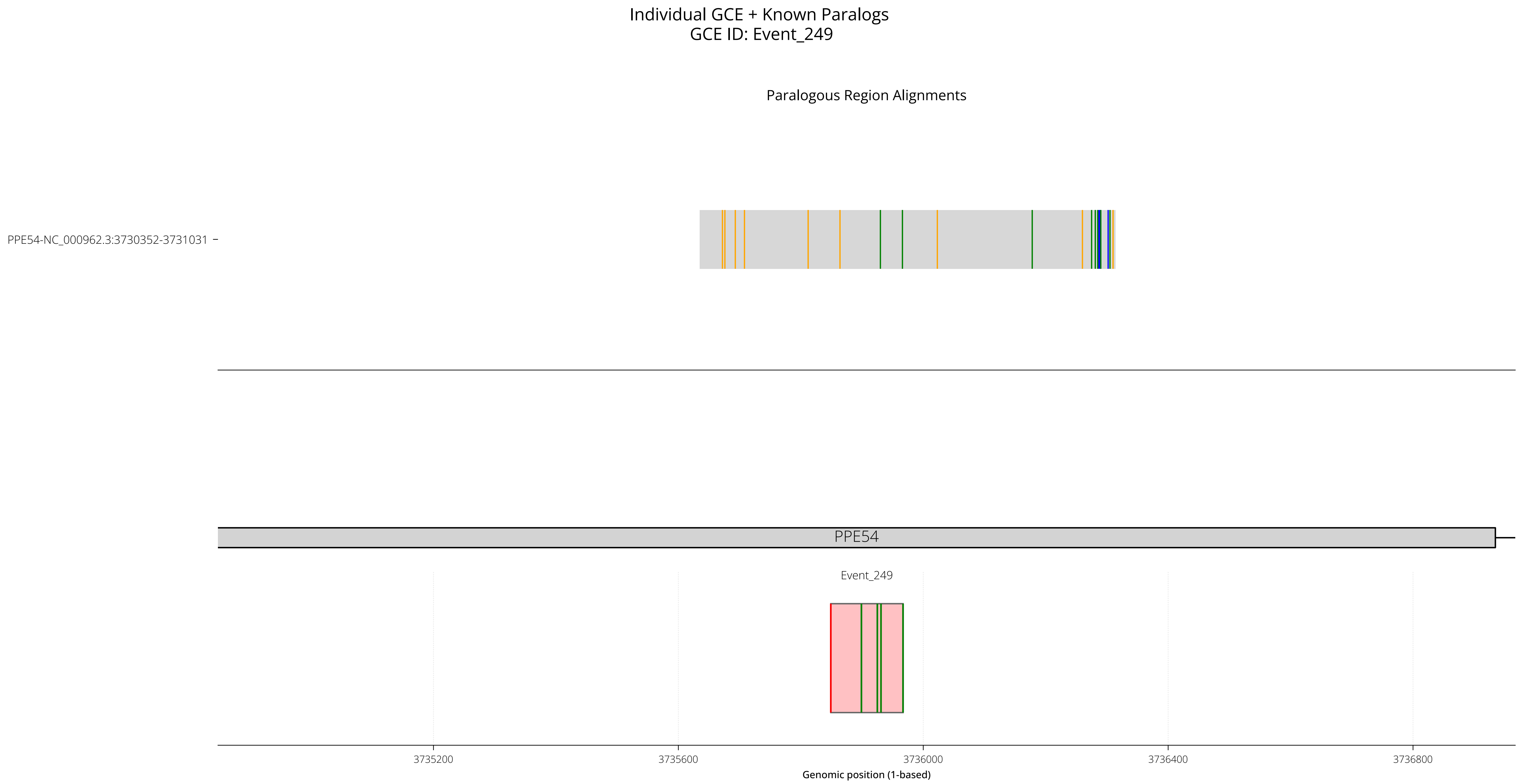

Individual GCE + Known Paralogs
GCE ID: Event_249
Paralogous Region Alignments
PPE54-NC_000962.3:3730352-3731031
PPE54
Event_249
3735200
3735600
3736000
3736400
3736800
Genomic position (1-based)

Circos Viz - GCE to Known Paralogs Locations
GCE ID: Event_249

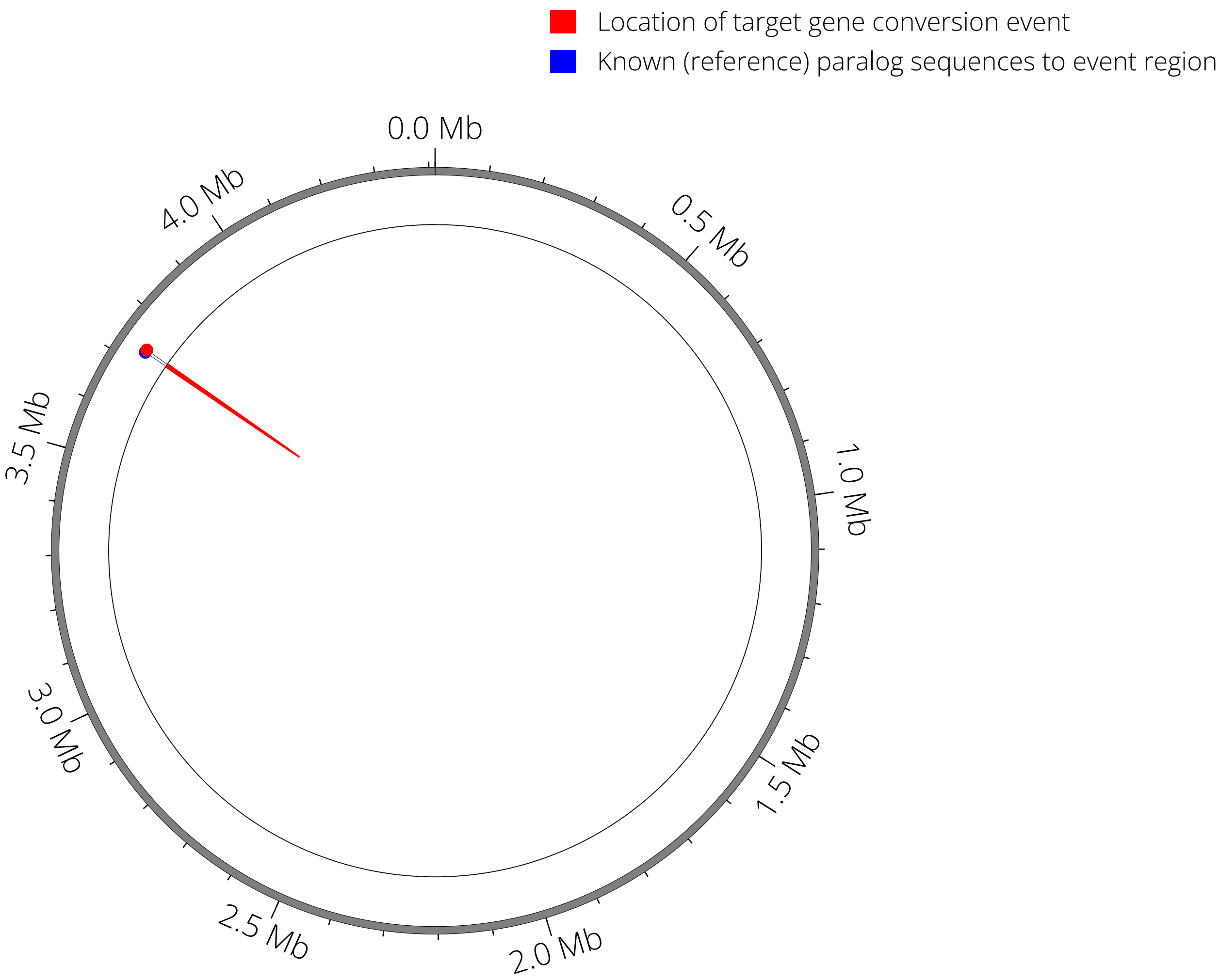

### Event_249 - 5 SNPs - k-mer analysis relative to paralogs

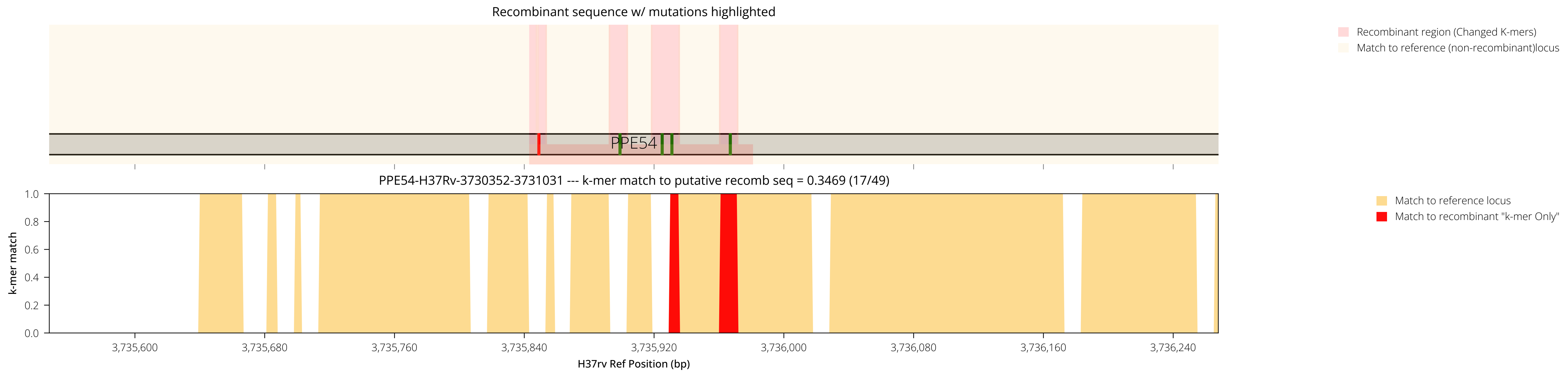

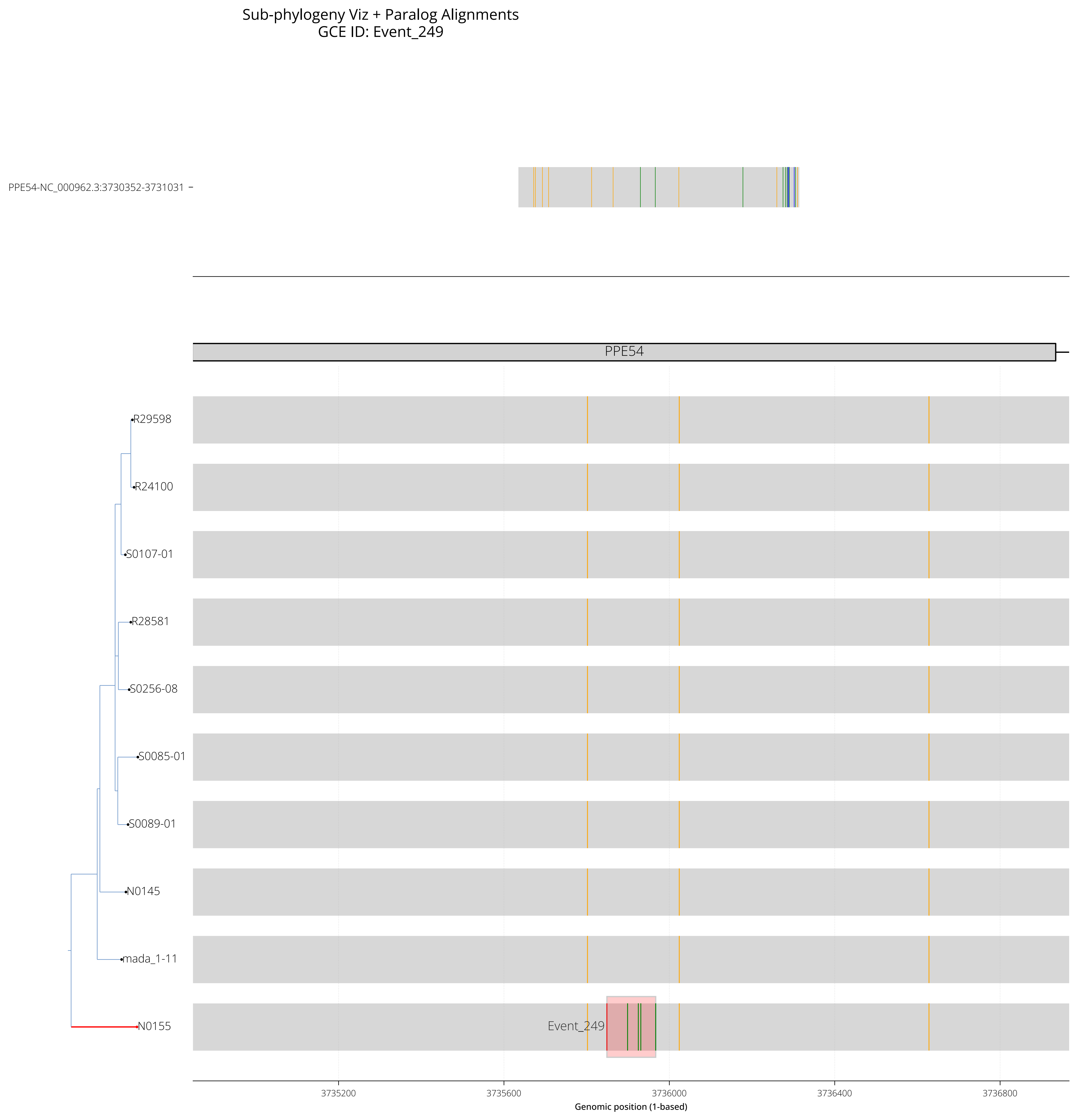
Sub-phylogeny Viz + Paralog Alignments
GCE ID: Event_249
PPE54-NC_000962.3:3730352-3731031
PPE54
R29598
R24100
S0107-01
R28581
S0256-08
S0085-01
S0089-01
N0145
mada_1-11
N0155
Event_249
3735200
3735600
3736000
3736400
3736800
Genomic position (1-based)

Full Phylogeny Viz + Paralog Alignments
GCE ID: Event_249

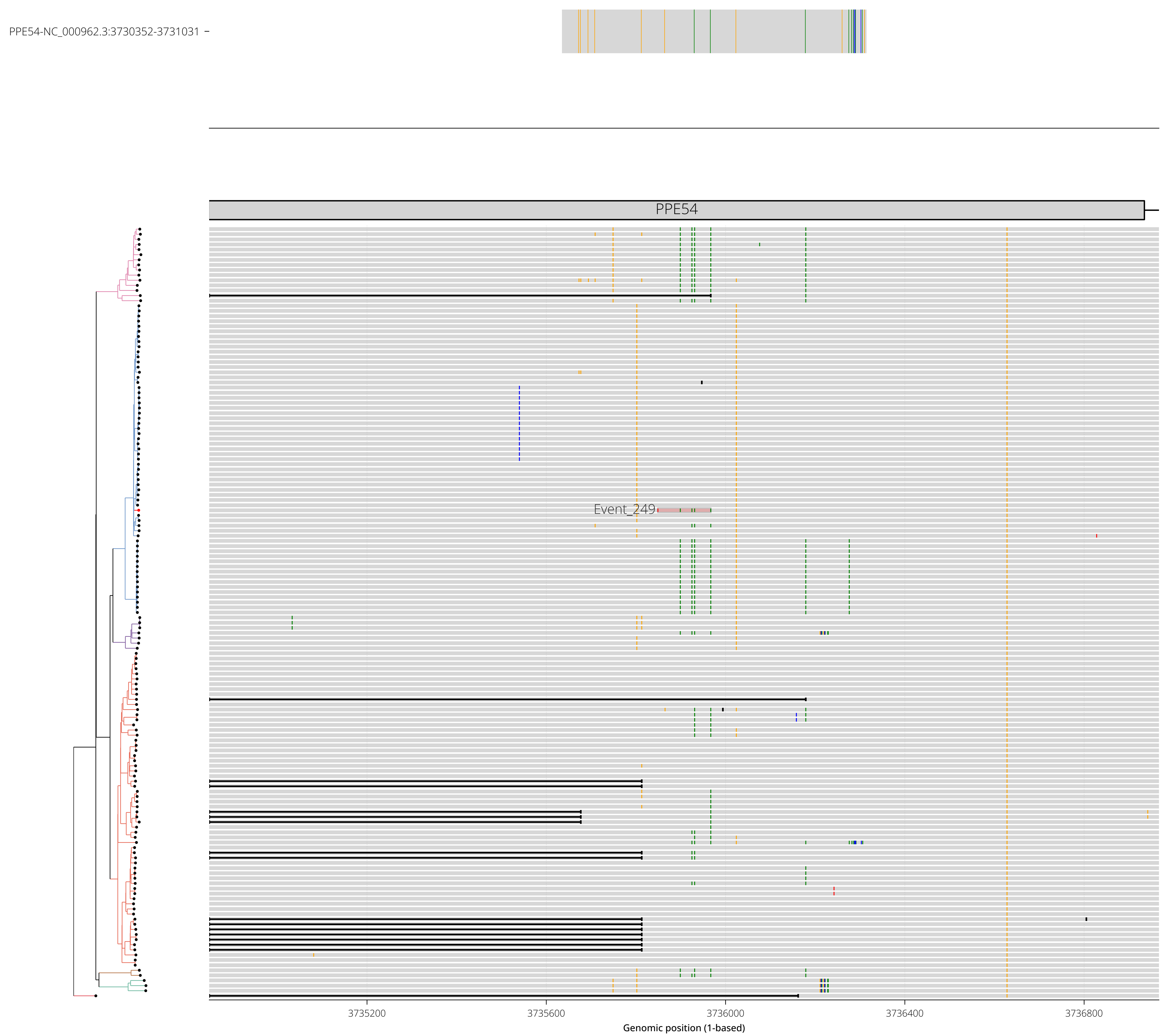

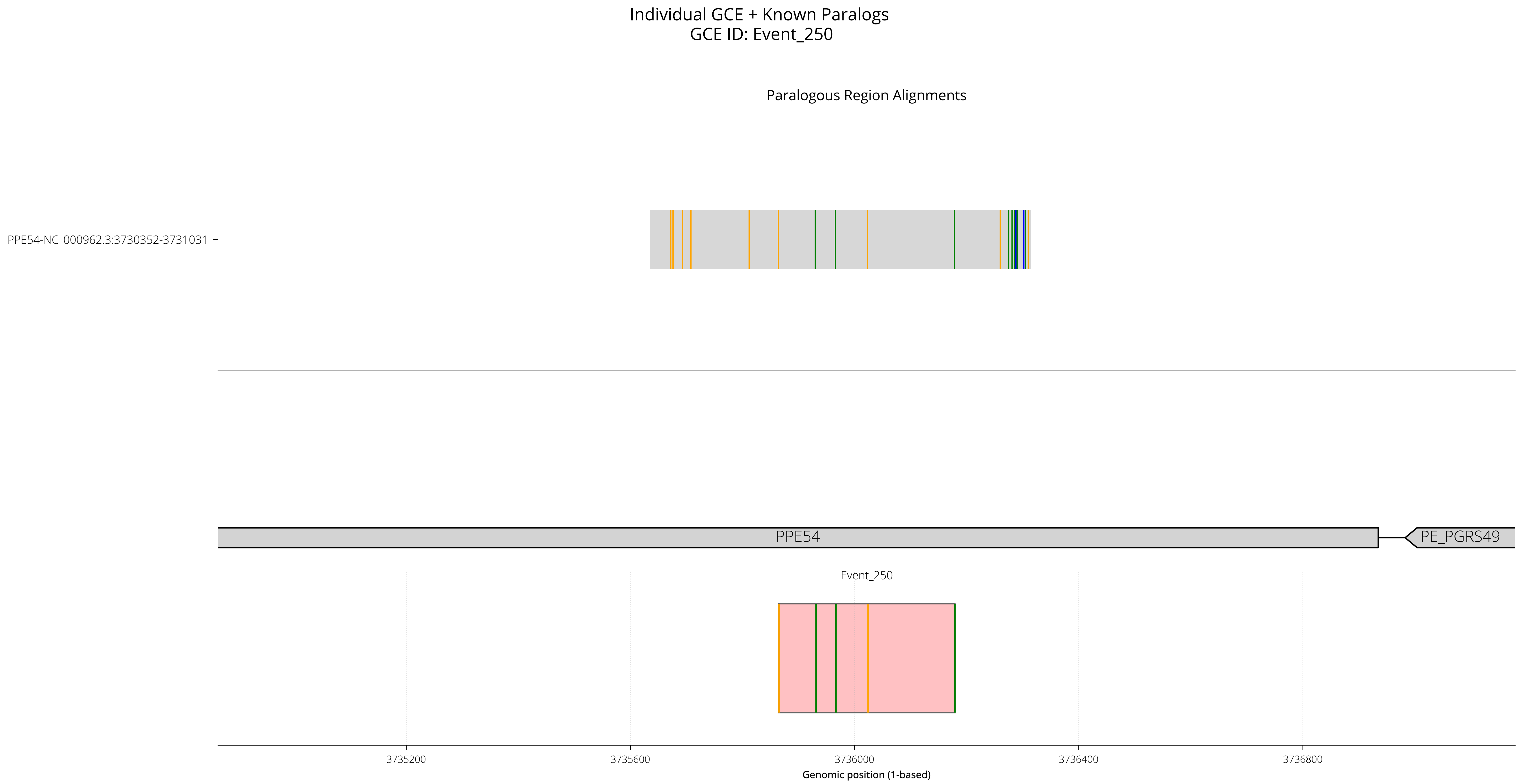

Individual GCE + Known Paralogs
GCE ID: Event_250
Paralogous Region Alignments
PPE54-NC_000962.3:3730352-3731031
PPE54
PE_PGRS49
Event_250
3735200
3735600
3736000
3736400
3736800
Genomic position (1-based)

Circos Viz - GCE to Known Paralogs Locations
GCE ID: Event_250

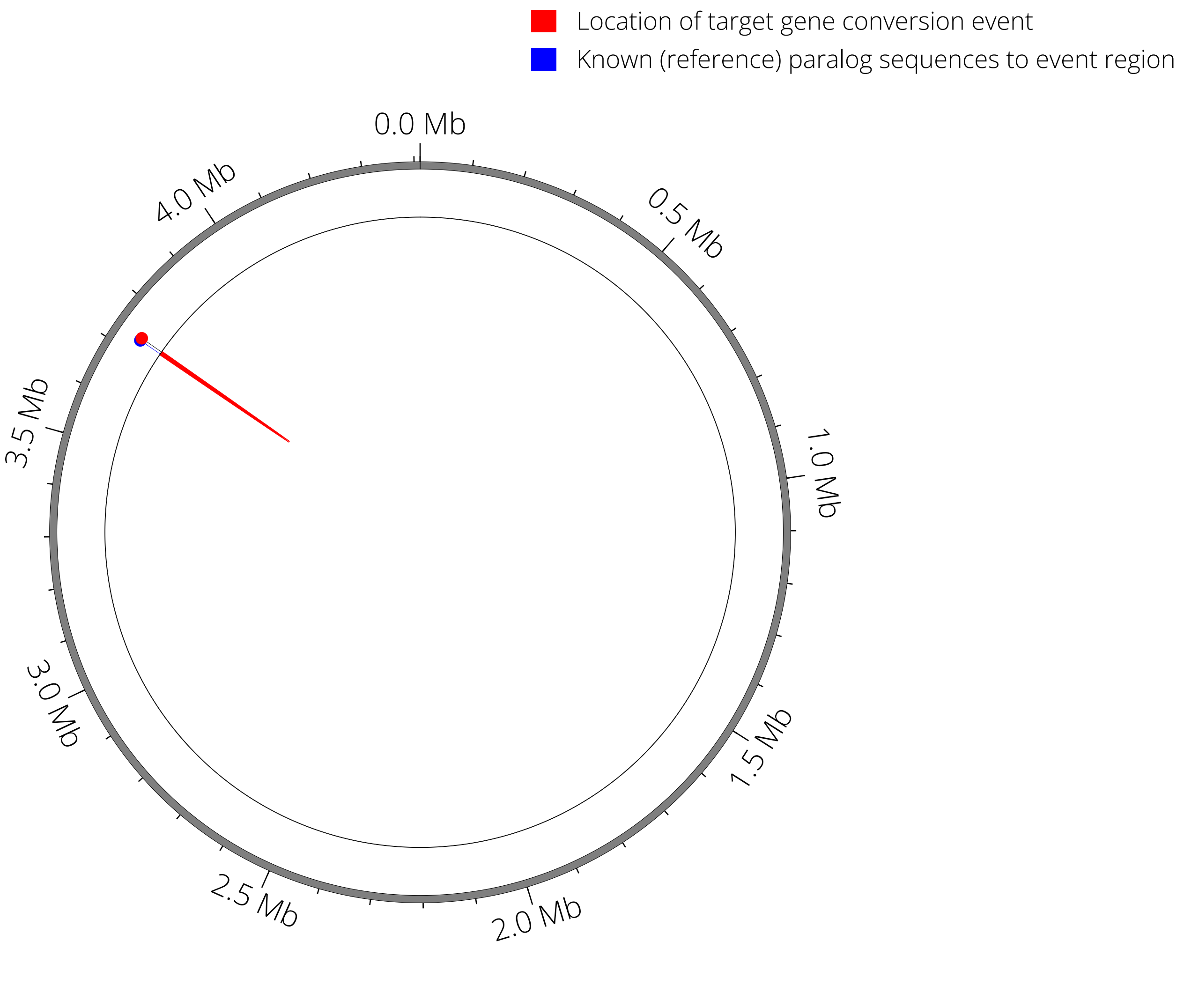

### Event_250 - 5 SNPs - k-mer analysis relative to paralogs

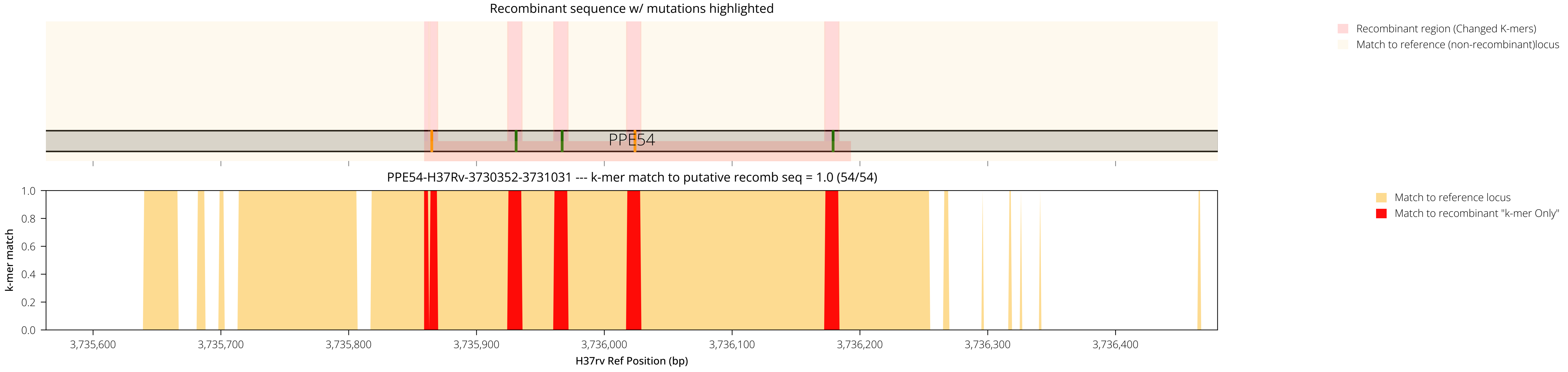

Sub-phylogeny Viz + Paralog Alignments
GCE ID: Event_250

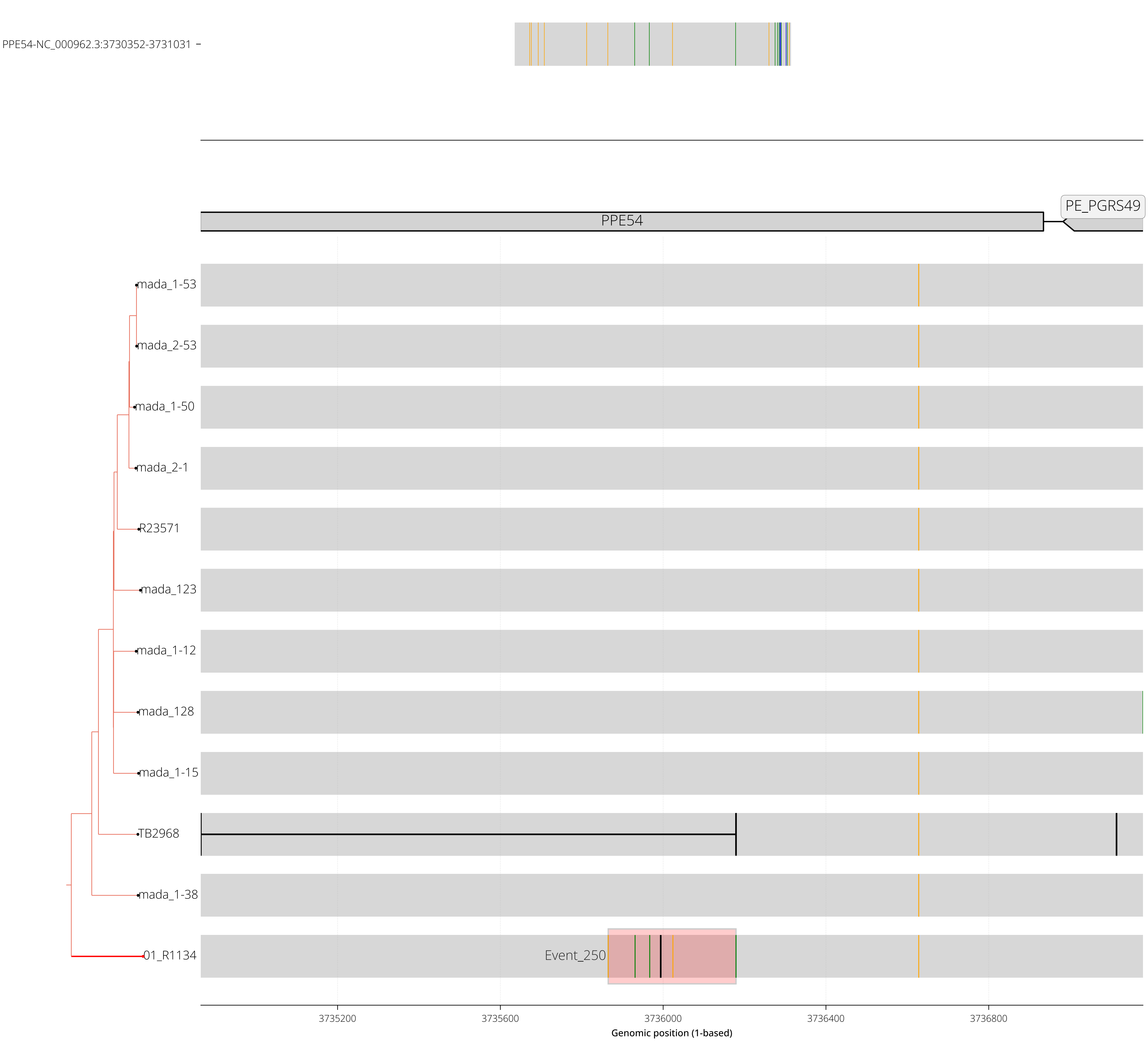

Full Phylogeny Viz + Paralog Alignments
GCE ID: Event_250

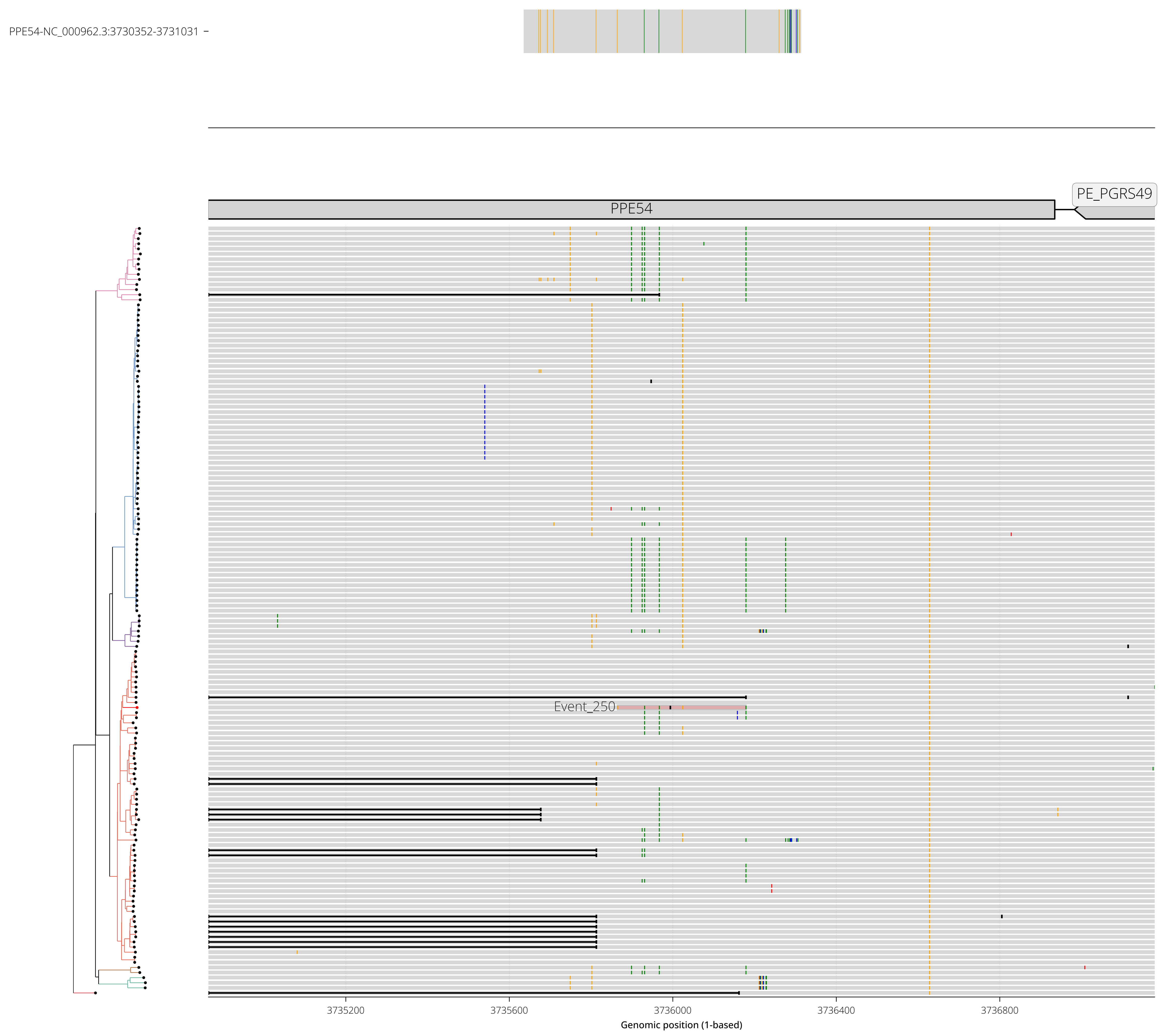

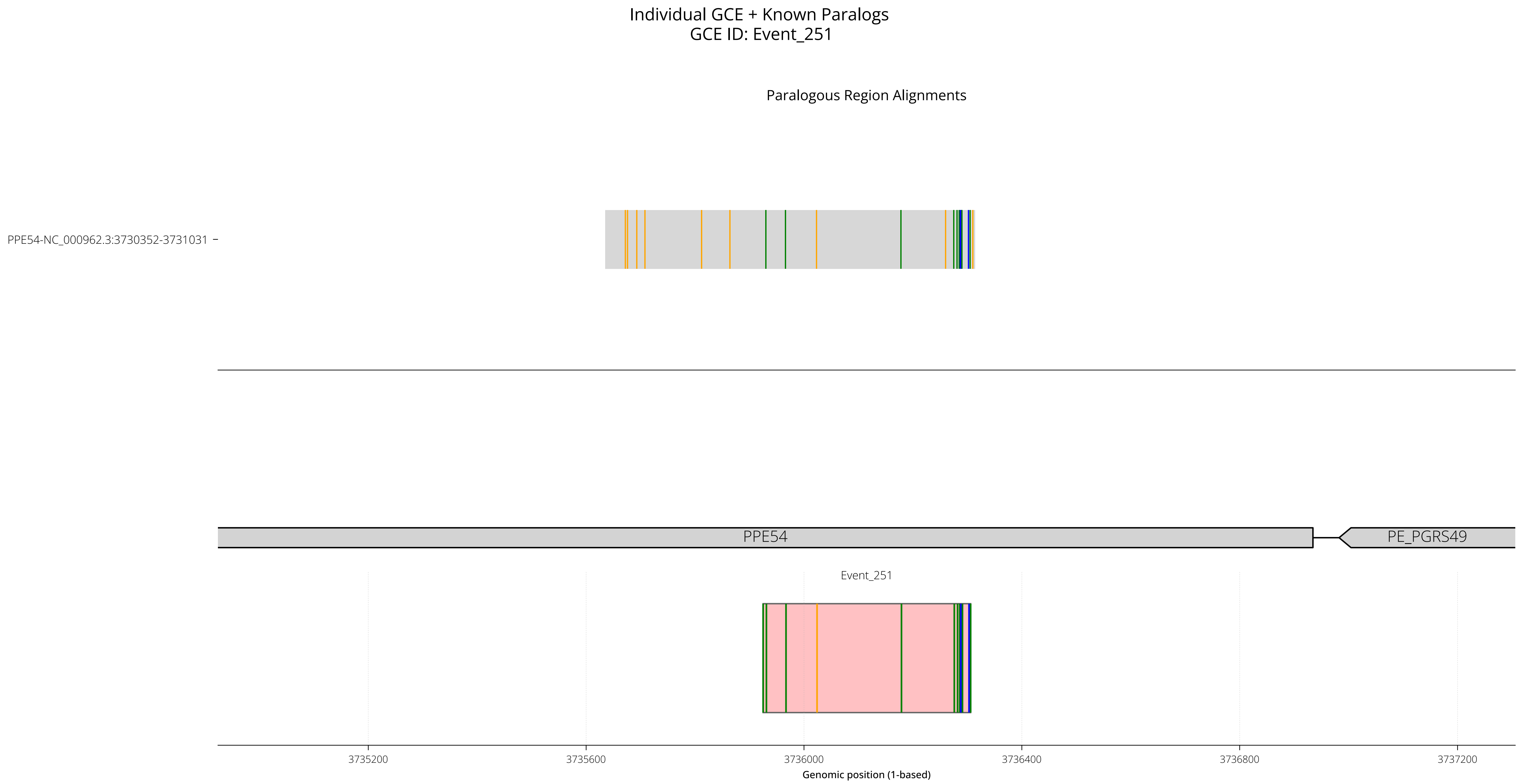

Individual GCE + Known Paralogs
GCE ID: Event_251
Paralogous Region Alignments
PPE54-NC_000962.3:3730352-3731031
PPE54
PE_PGRS49
Event_251
3735200
3735600
3736000
3736400
3736800
3737200
Genomic position (1-based)

Circos Viz - GCE to Known Paralogs Locations
GCE ID: Event_251

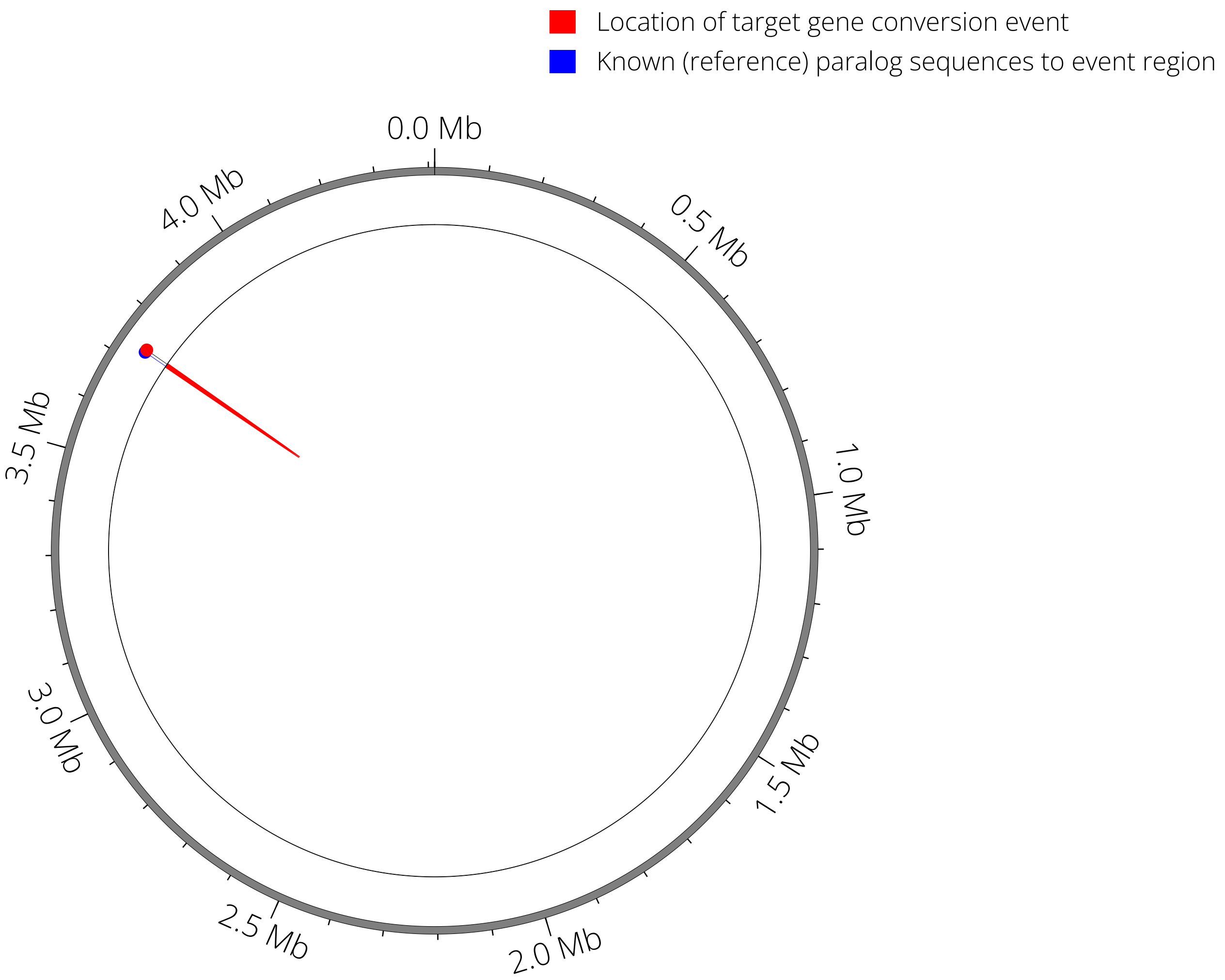

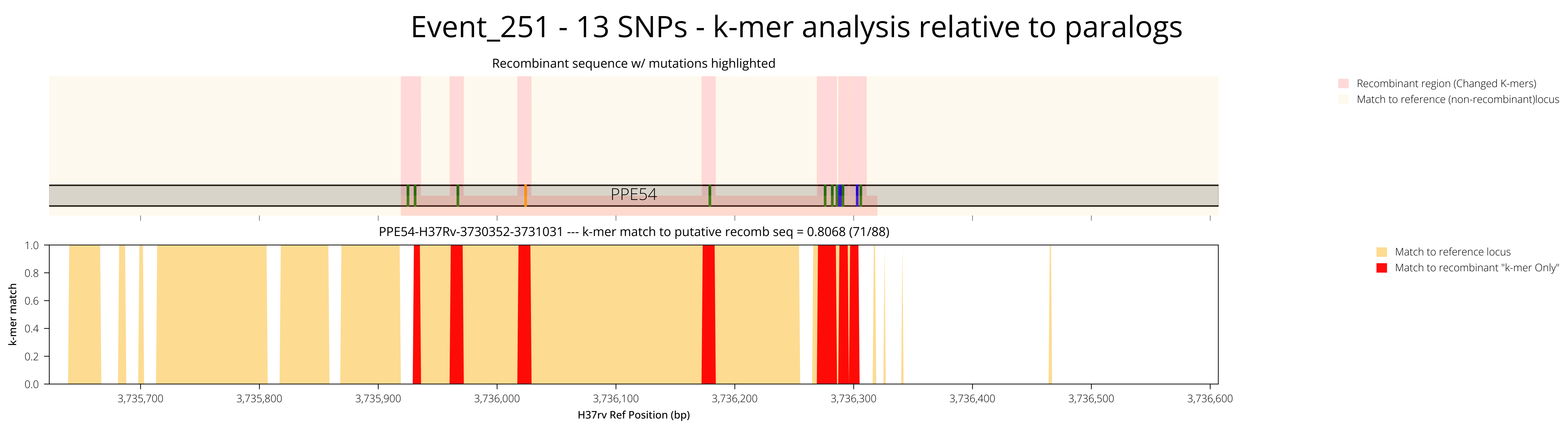

Event_251 - 13 SNPs - k-mer analysis relative to paralogs
Recombinant sequence w/ mutations highlighted
PPE54
Recombinant region (Changed K-mers)
Match to reference (non-recombinant)locus
PPE54-H37Rv-3730352-3731031 --- k-mer match to putative recomb seq = 0.8068 (71/88)
Match to reference locus
Match to recombinant "k-mer Only"
k-mer match
1.0
0.8
0.6
0.4
0.2
0.0
3,735,700
3,735,800
3,735,900
3,736,000
3,736,100
3,736,200
3,736,300
3,736,400
3,736,500
3,736,600
H37rv Ref Position (bp)

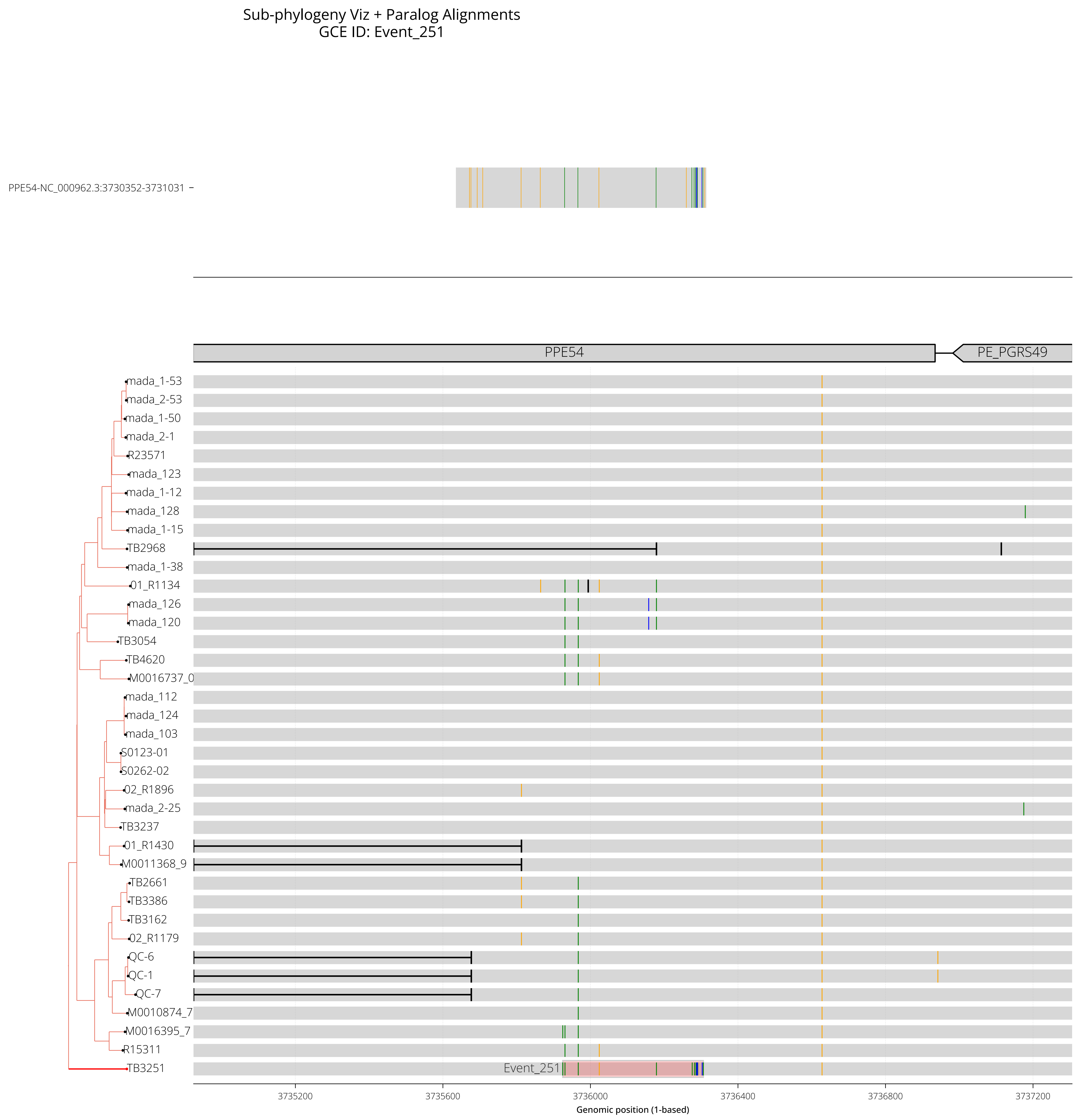

Sub-phylogeny Viz + Paralog Alignments
GCE ID: Event_251
PPE54-NC_000962.3:3730352-3731031
PPE54
PE_PGRS49
mada_1-53
mada_2-53
mada_1-50
mada_2-1
R23571
mada_123
mada_1-12
mada_128
mada_1-15
TB2968
mada_1-38
01_R1134
mada_126
mada_120
TB3054
TB4620
M0016737_0
mada_112
mada_124
mada_103
S0123-01
S0262-02
02_R1896
mada_2-25
TB3237
01_R1430
M0011368_9
TB2661
TB3386
TB3162
02_R1179
QC-6
QC-1
QC-7
M0010874_7
M0016395_7
R15311
TB3251
Event_251
3735200
3735600
3736000
3736400
3736800
3737200
Genomic position (1-based)

Full Phylogeny Viz + Paralog Alignments
GCE ID: Event_251

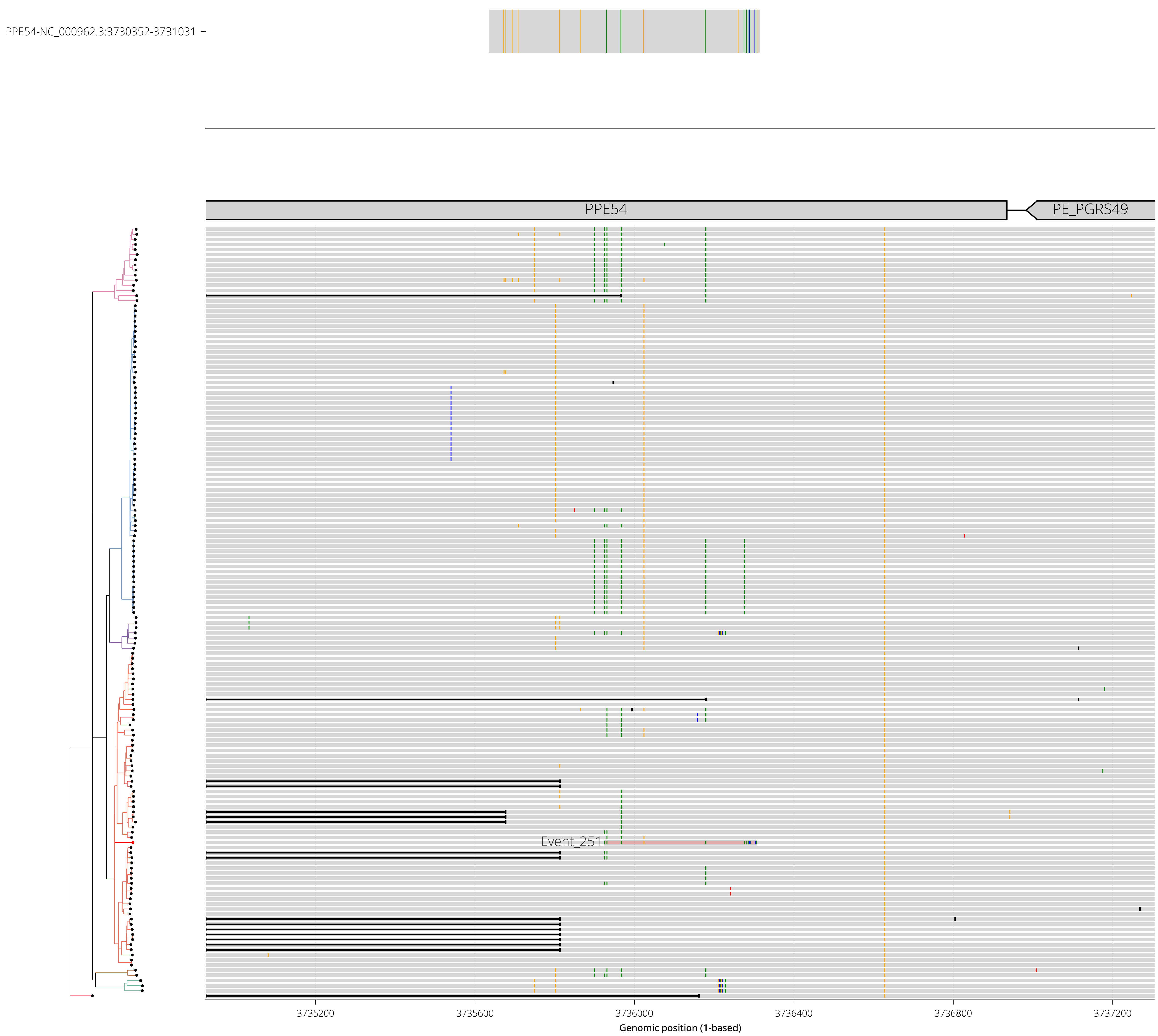

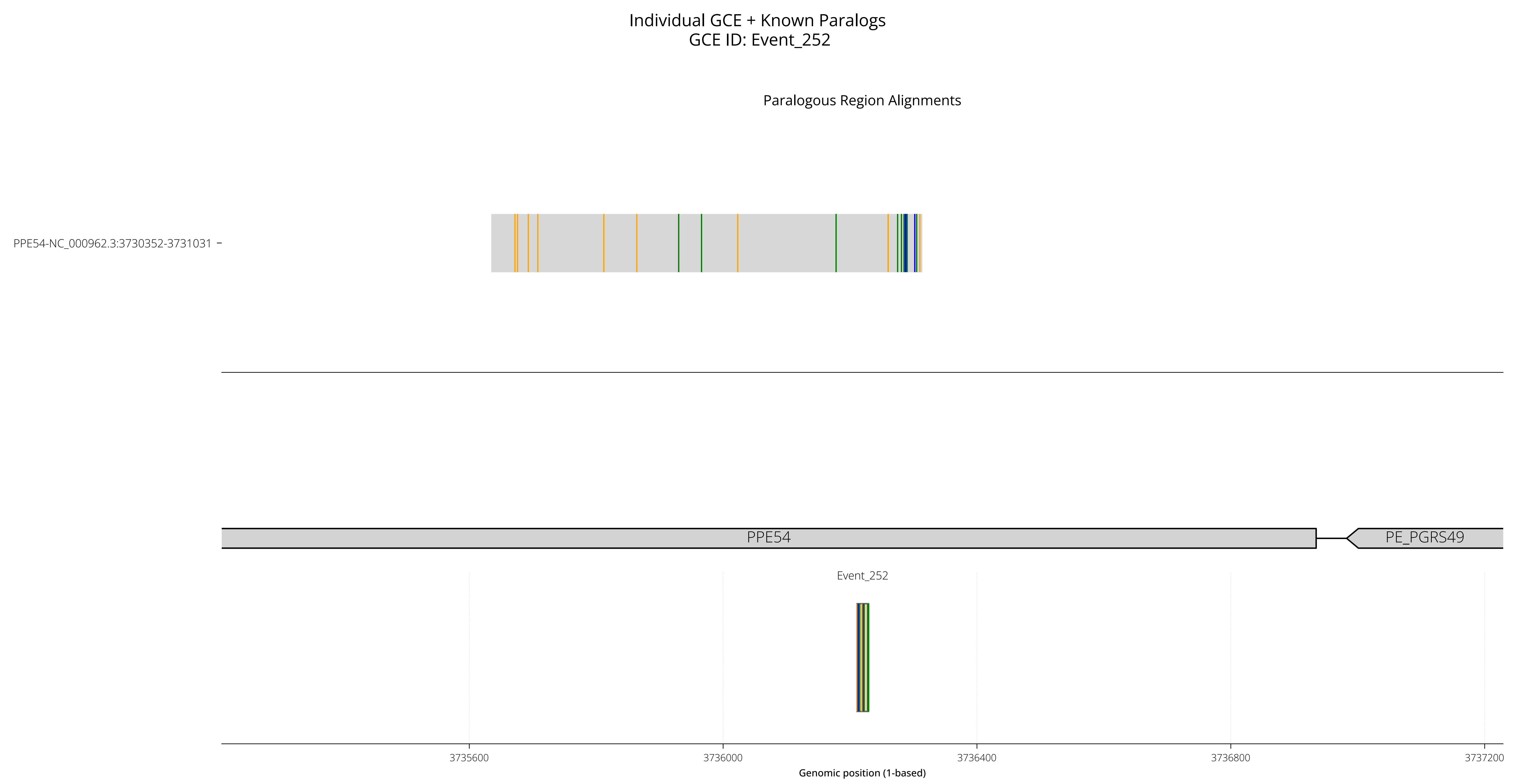
Individual GCE + Known Paralogs
GCE ID: Event_252
Paralogous Region Alignments
PPE54-NC_000962.3:3730352-3731031
PPE54
PE_PGRS49
Event_252
3735600
3736000
3736400
3736800
3737200
Genomic position (1-based)

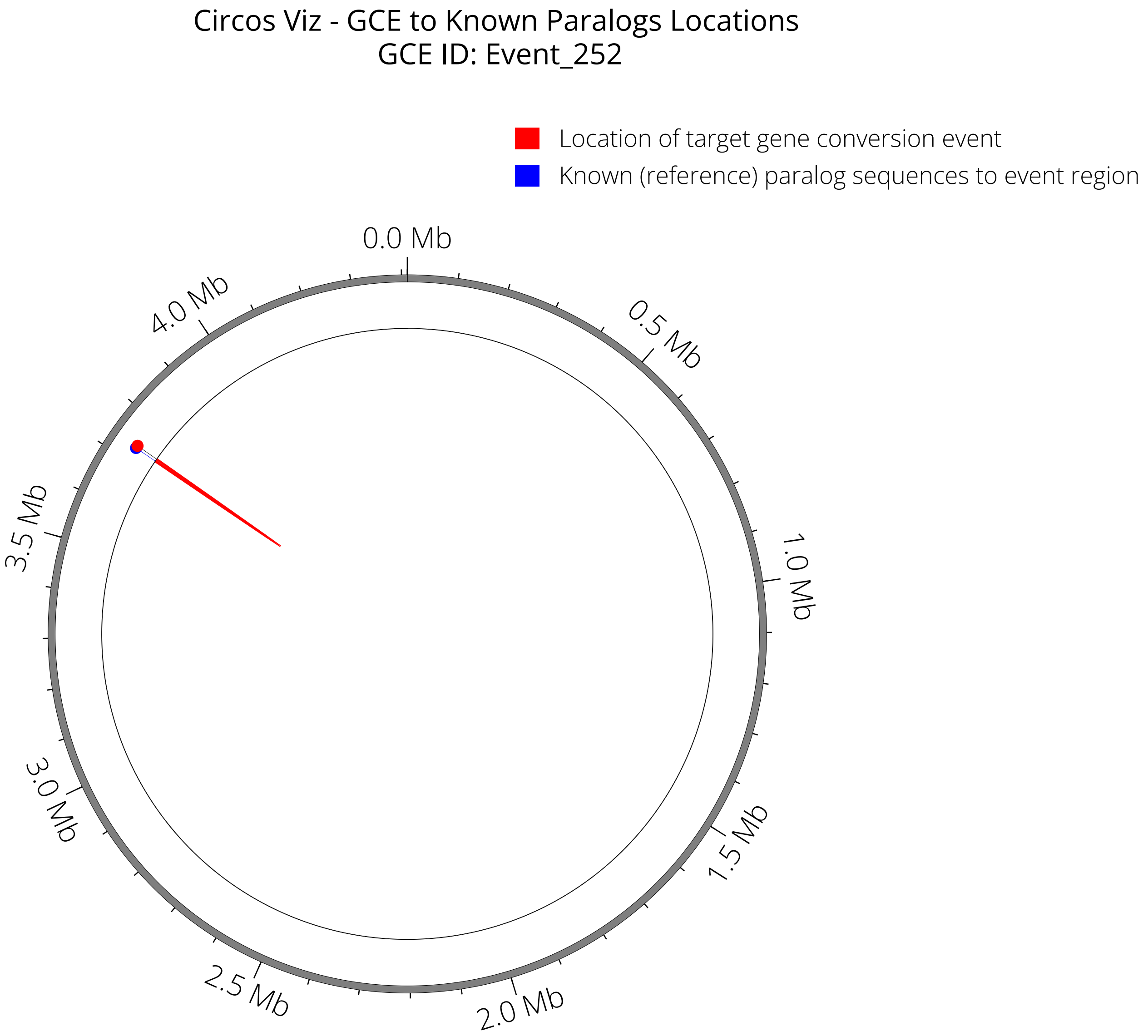
Circos Viz - GCE to Known Paralogs Locations
GCE ID: Event_252
Location of target gene conversion event
Known (reference) paralog sequences to event region
0.0 Mb
0.5 Mb
1.0 Mb
1.5 Mb
2.0 Mb
2.5 Mb
3.0 Mb
3.5 Mb
4.0 Mb

### Event_252 - 8 SNPs - k-mer analysis relative to paralogs

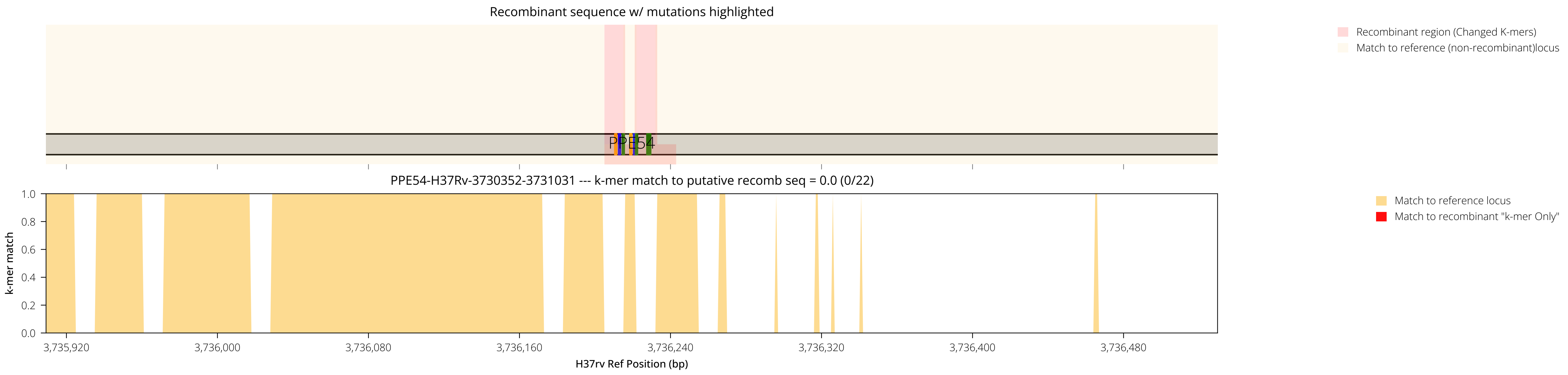

Sub-phylogeny Viz + Paralog Alignments
GCE ID: Event_252

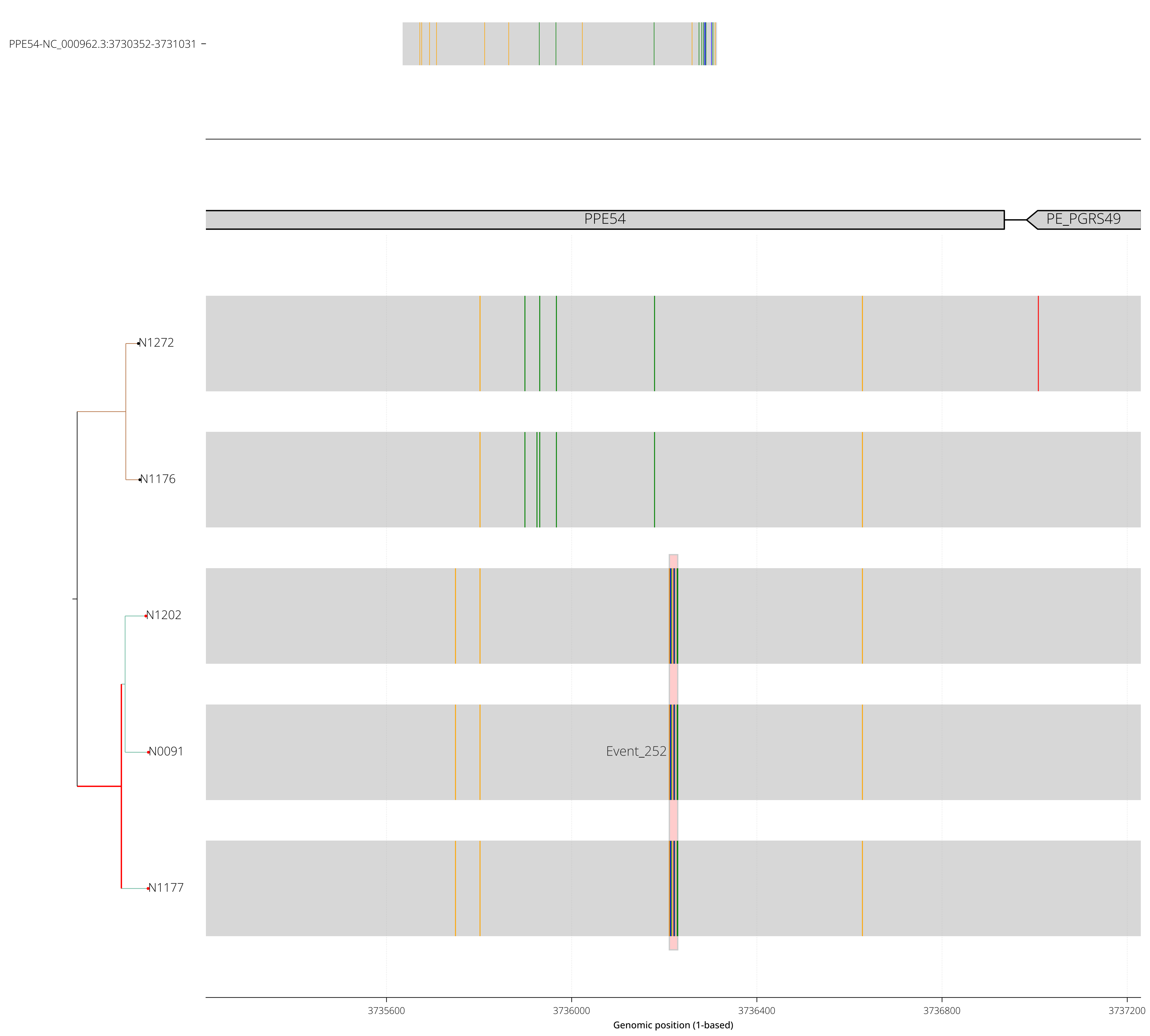

Full Phylogeny Viz + Paralog Alignments
GCE ID: Event_252

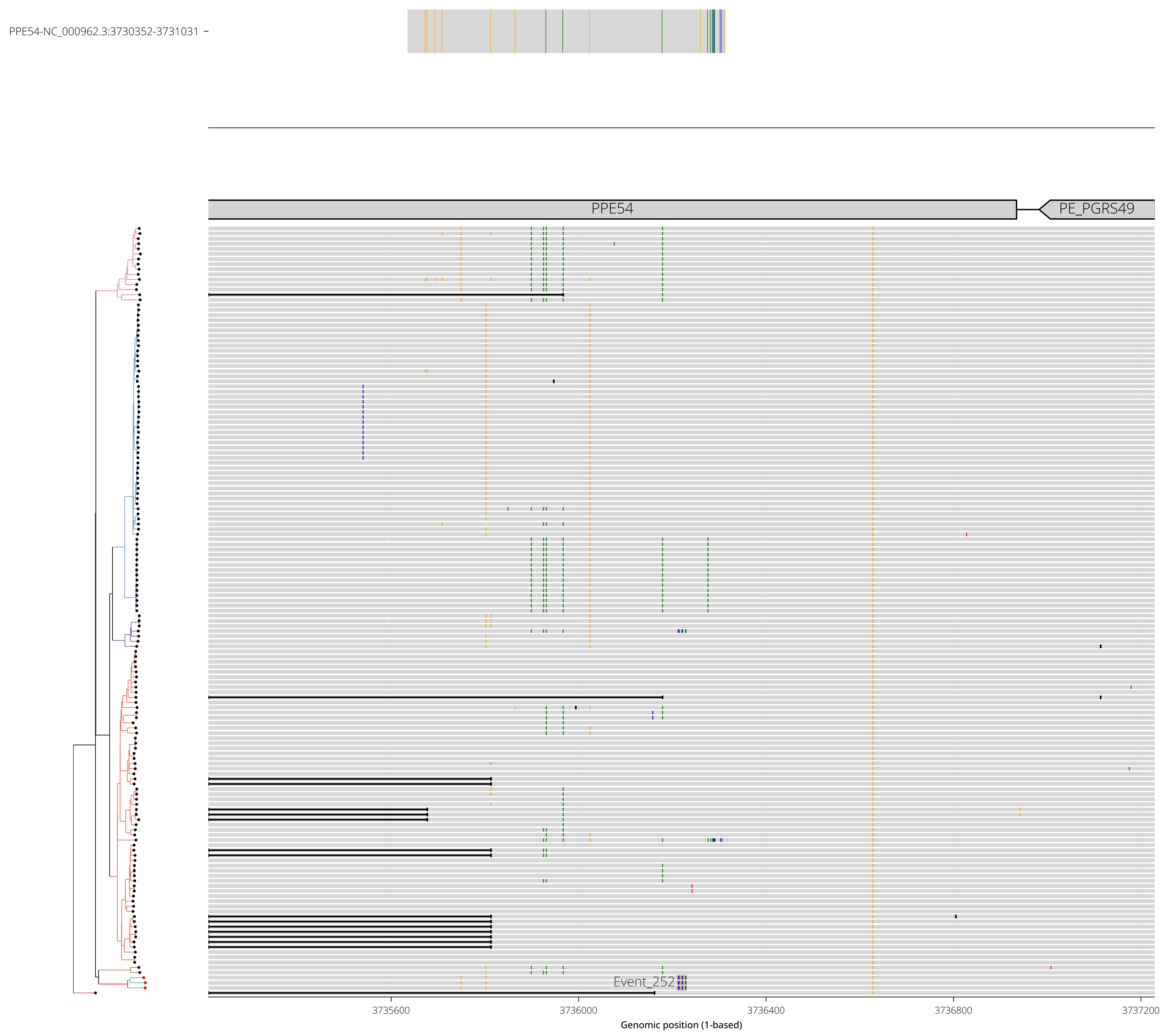

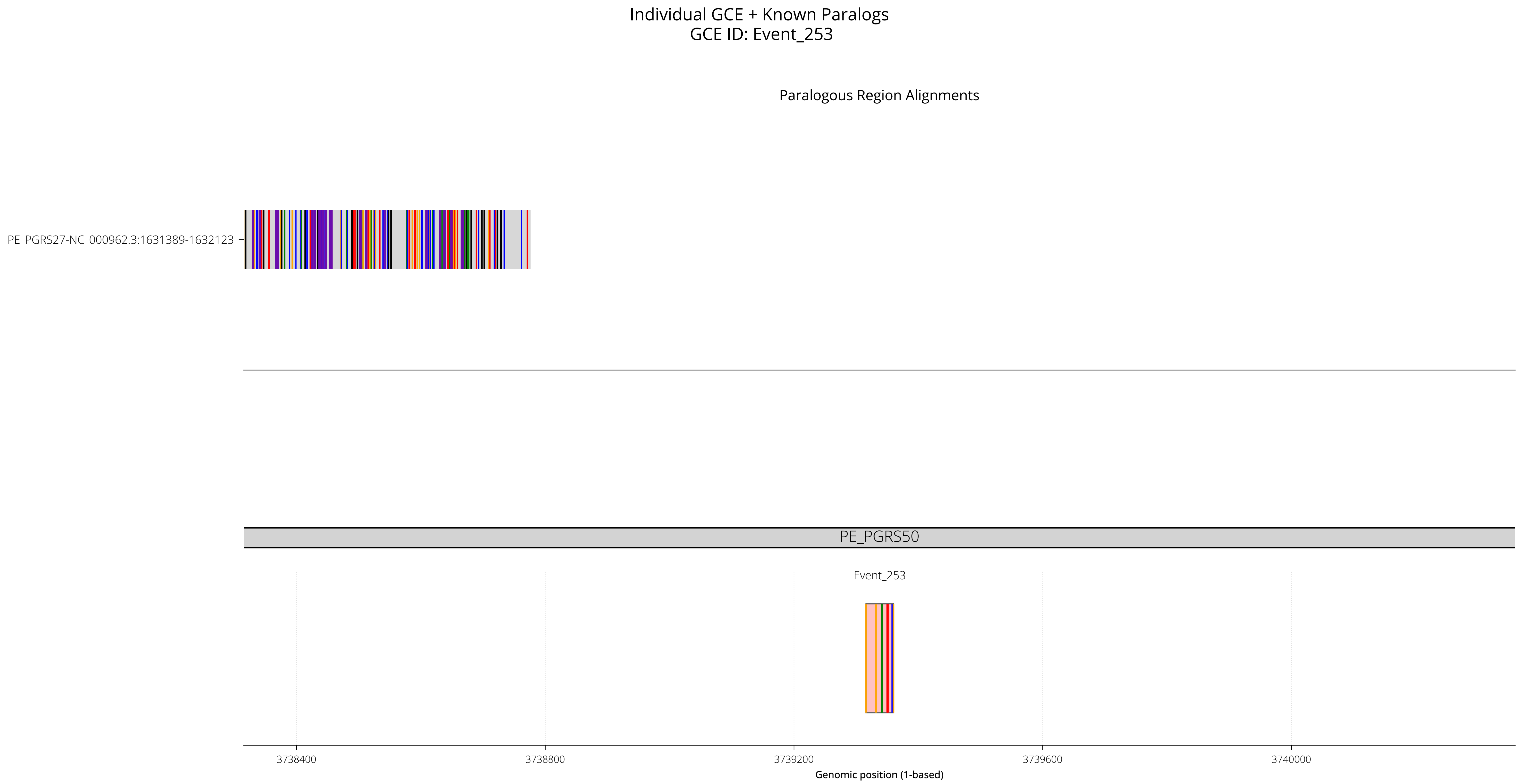

Individual GCE + Known Paralogs
GCE ID: Event_253
Paralogous Region Alignments
PE_PGRS27-NC_000962.3:1631389-1632123
PE_PGRS50
Event_253
3738400
3738800
3739200
3739600
3740000
Genomic position (1-based)

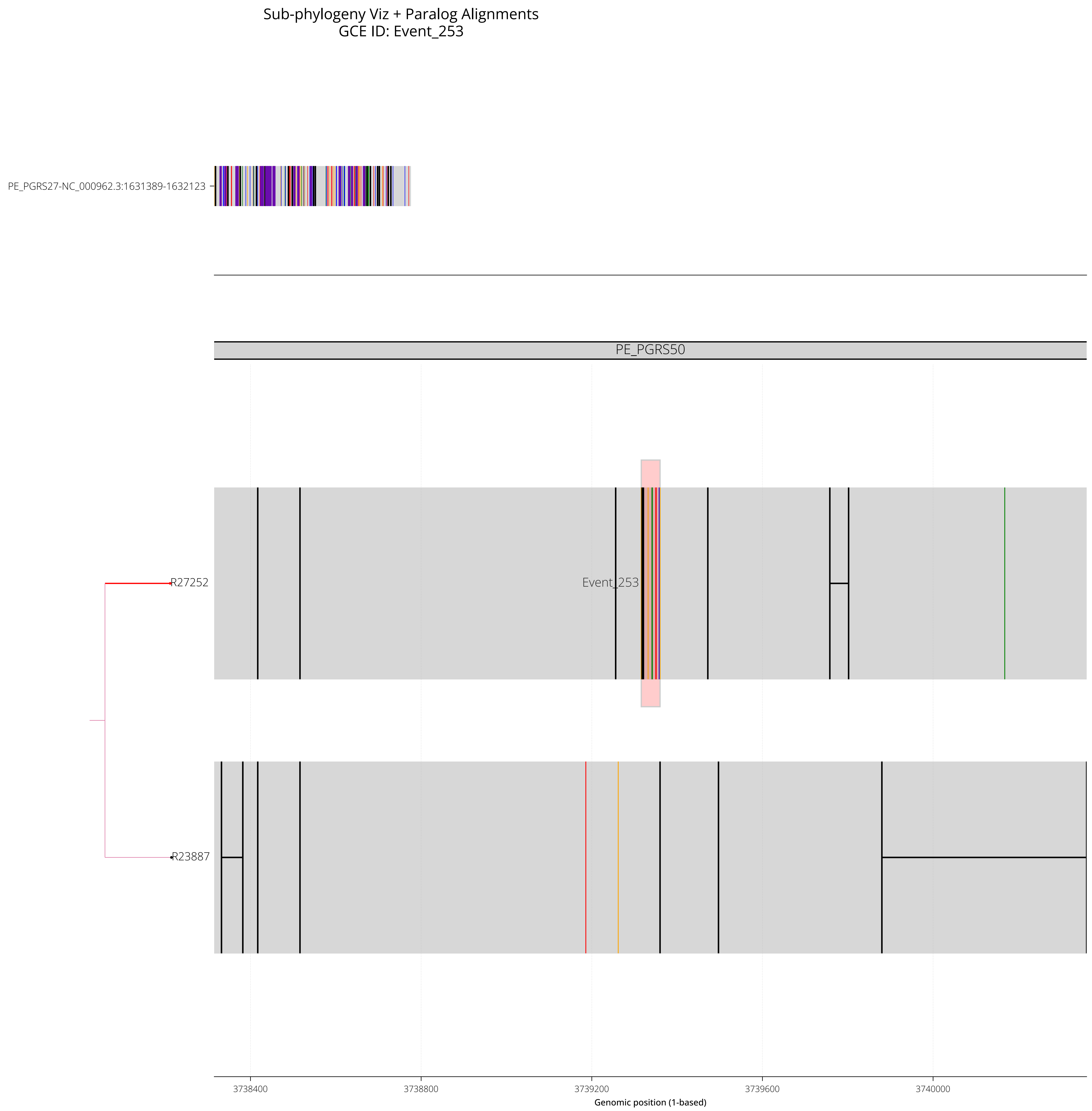

Sub-phylogeny Viz + Paralog Alignments
GCE ID: Event_253
PE_PGRS27-NC_000962.3:1631389-1632123
PE_PGRS50
R27252
Event_253
R23887
3738400
3738800
3739200
3739600
3740000
Genomic position (1-based)

Full Phylogeny Viz + Paralog Alignments
GCE ID: Event_253

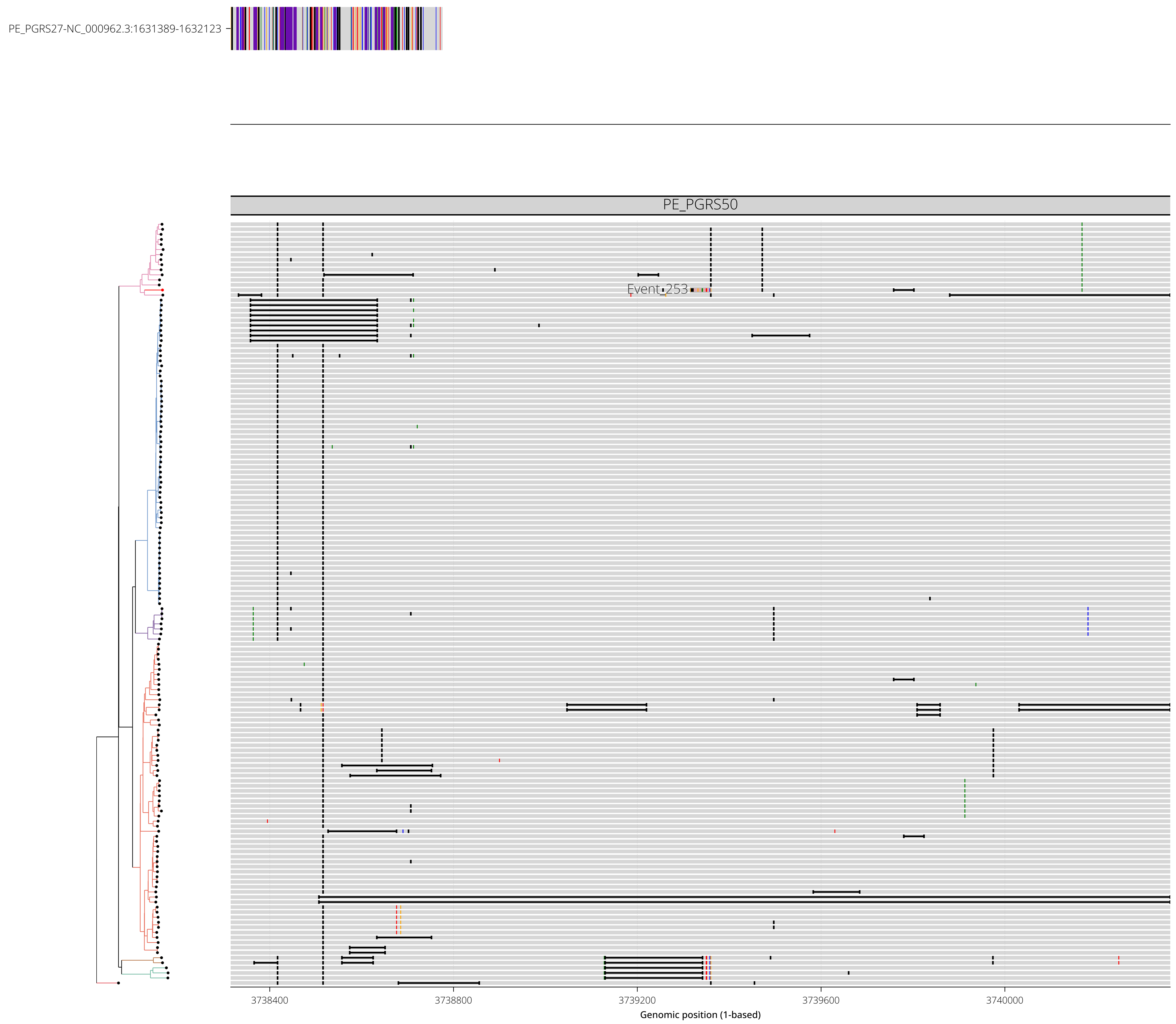

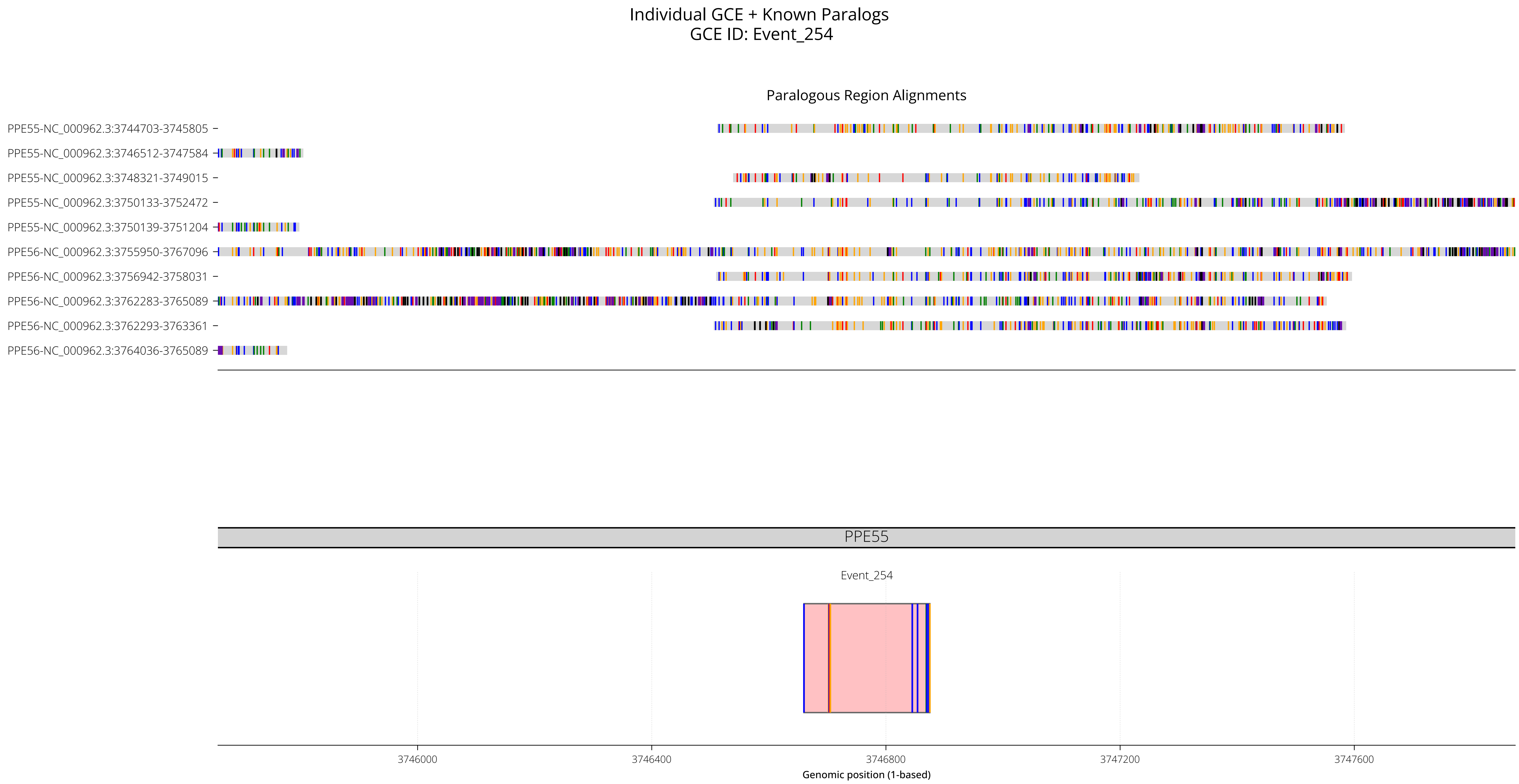
Individual GCE + Known Paralogs
GCE ID: Event_254
Paralogous Region Alignments
PPE55-NC_000962.3:3744703-3745805
PPE55-NC_000962.3:3746512-3747584
PPE55-NC_000962.3:3748321-3749015
PPE55-NC_000962.3:3750133-3752472
PPE55-NC_000962.3:3750139-3751204
PPE56-NC_000962.3:3755950-3767096
PPE56-NC_000962.3:3756942-3758031
PPE56-NC_000962.3:3762283-3765089
PPE56-NC_000962.3:3762293-3763361
PPE56-NC_000962.3:3764036-3765089
PPE55
Event_254
3746000
3746400
3746800
3747200
3747600
Genomic position (1-based)

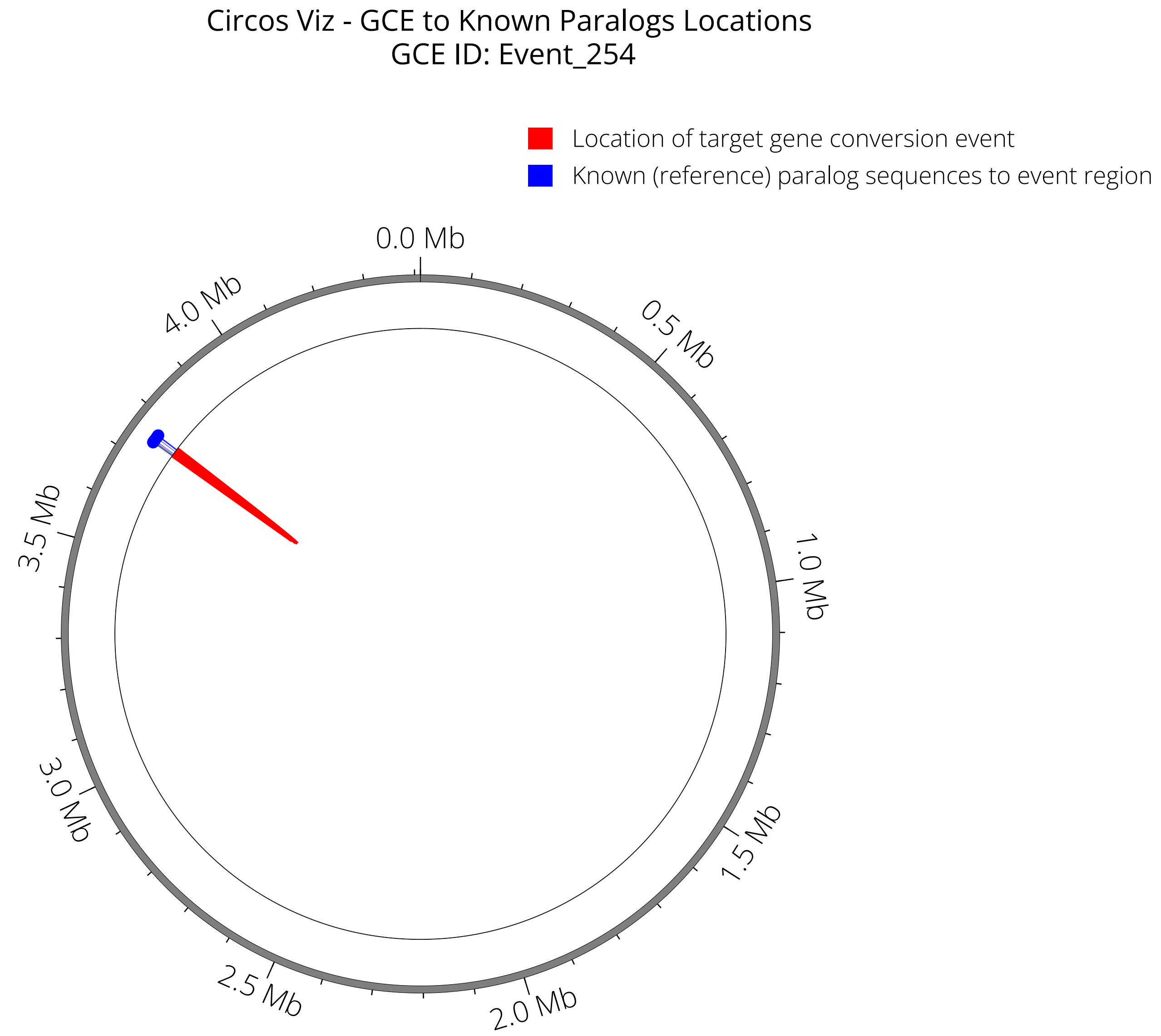
Circos Viz - GCE to Known Paralogs Locations
GCE ID: Event_254
Location of target gene conversion event
Known (reference) paralog sequences to event region
0.0 Mb
0.5 Mb
1.0 Mb
1.5 Mb
2.0 Mb
2.5 Mb
3.0 Mb
3.5 Mb
4.0 Mb

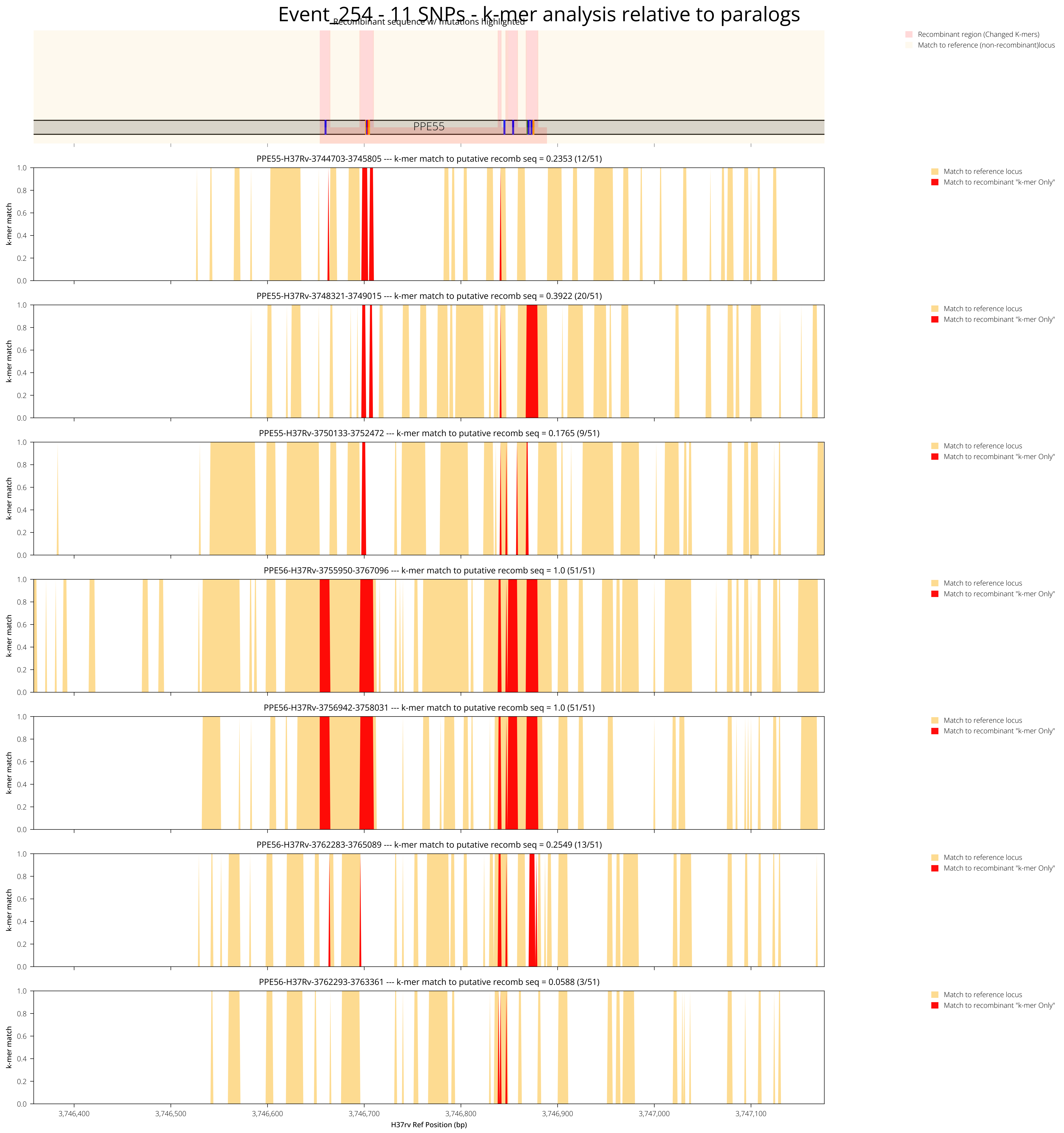
Event_254 - 11 SNPs - k-mer analysis relative to paralogs
Recombinant sequence w/ mutations highlighted
PPE55
Recombinant region (Changed K-mers)
Match to reference (non-recombinant)locus
PPE55-H37Rv-3744703-3745805 --- k-mer match to putative recomb seq = 0.2353 (12/51)
PPE55-H37Rv-3748321-3749015 --- k-mer match to putative recomb seq = 0.3922 (20/51)
PPE55-H37Rv-3750133-3752472 --- k-mer match to putative recomb seq = 0.1765 (9/51)
PPE56-H37Rv-3755950-3767096 --- k-mer match to putative recomb seq = 1.0 (51/51)
PPE56-H37Rv-3756942-3758031 --- k-mer match to putative recomb seq = 1.0 (51/51)
PPE56-H37Rv-3762283-3765089 --- k-mer match to putative recomb seq = 0.2549 (13/51)
PPE56-H37Rv-3762293-3763361 --- k-mer match to putative recomb seq = 0.0588 (3/51)
Match to reference locus
Match to recombinant "k-mer Only"
k-mer match
H37rv Ref Position (bp)
3,746,400
3,746,500
3,746,600
3,746,700
3,746,800
3,746,900
3,747,000
3,747,100

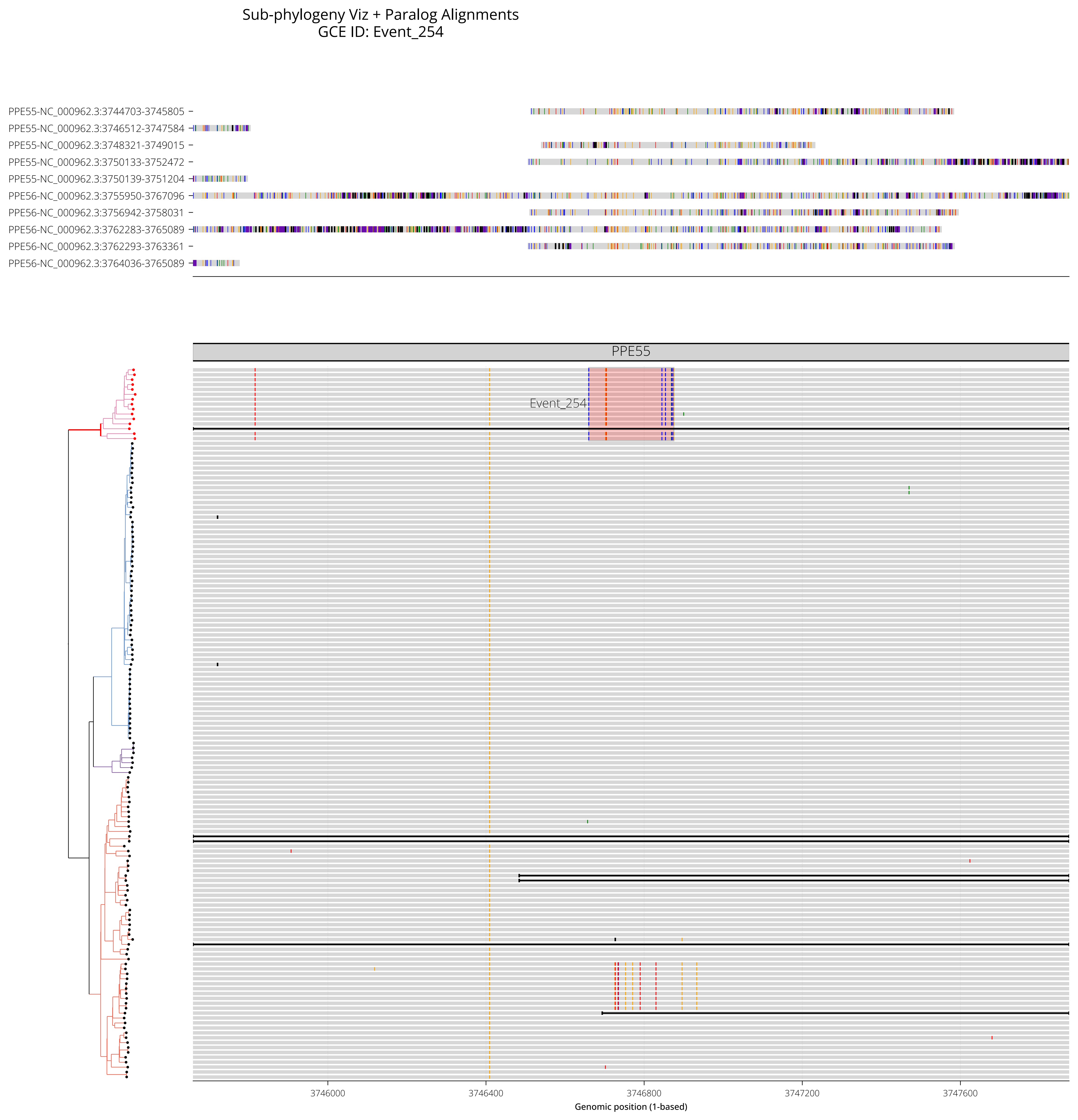

Sub-phylogeny Viz + Paralog Alignments
GCE ID: Event_254
PPE55-NC_000962.3:3744703-3745805
PPE55-NC_000962.3:3746512-3747584
PPE55-NC_000962.3:3748321-3749015
PPE55-NC_000962.3:3750133-3752472
PPE55-NC_000962.3:3750139-3751204
PPE56-NC_000962.3:3755950-3767096
PPE56-NC_000962.3:3756942-3758031
PPE56-NC_000962.3:3762283-3765089
PPE56-NC_000962.3:3762293-3763361
PPE56-NC_000962.3:3764036-3765089
PPE55
Event_254
3746000
3746400
3746800
3747200
3747600
Genomic position (1-based)

Full Phylogeny Viz + Paralog Alignments
GCE ID: Event_254

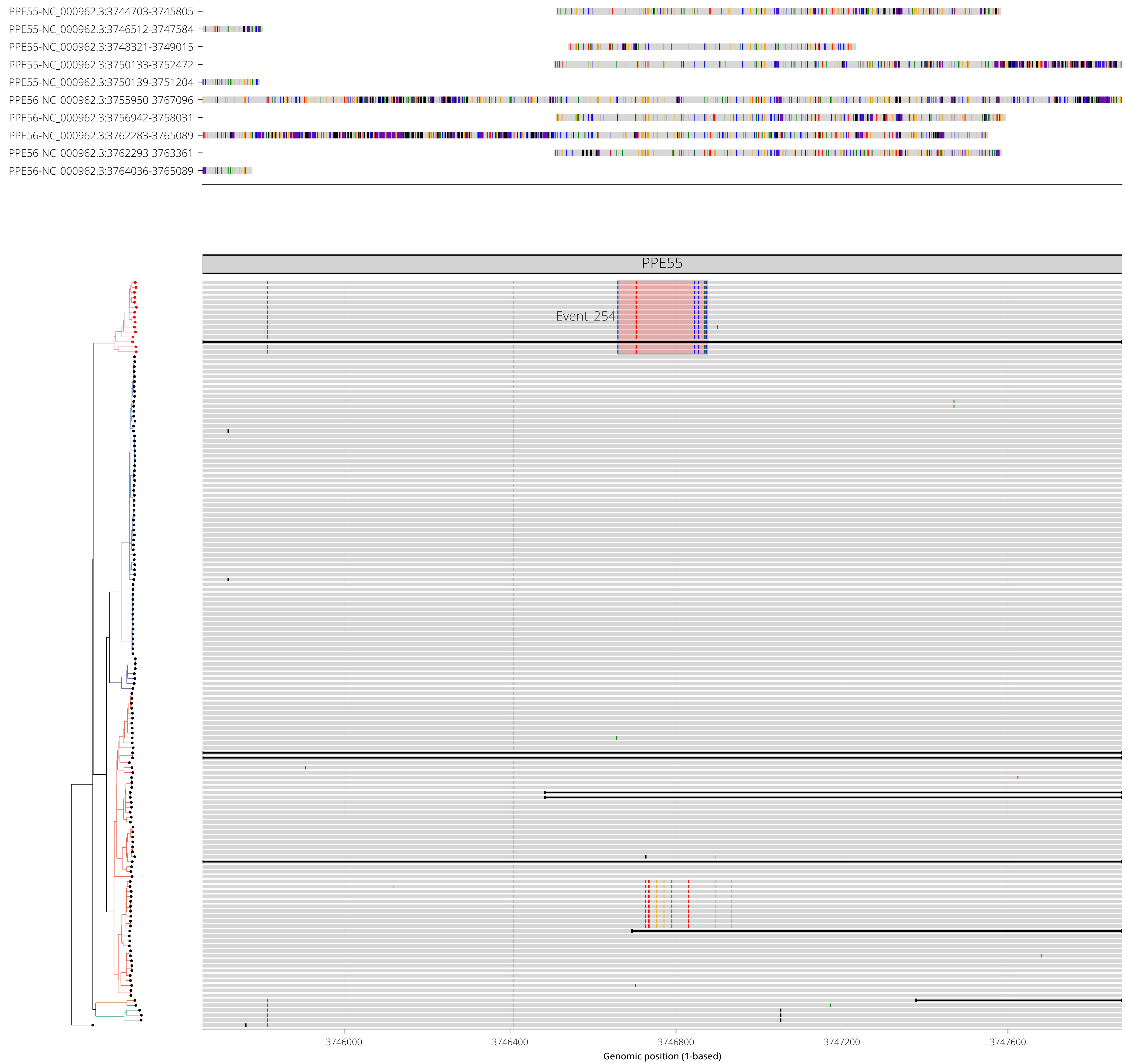

Individual GCE + Known Paralogs
GCE ID: Event_255

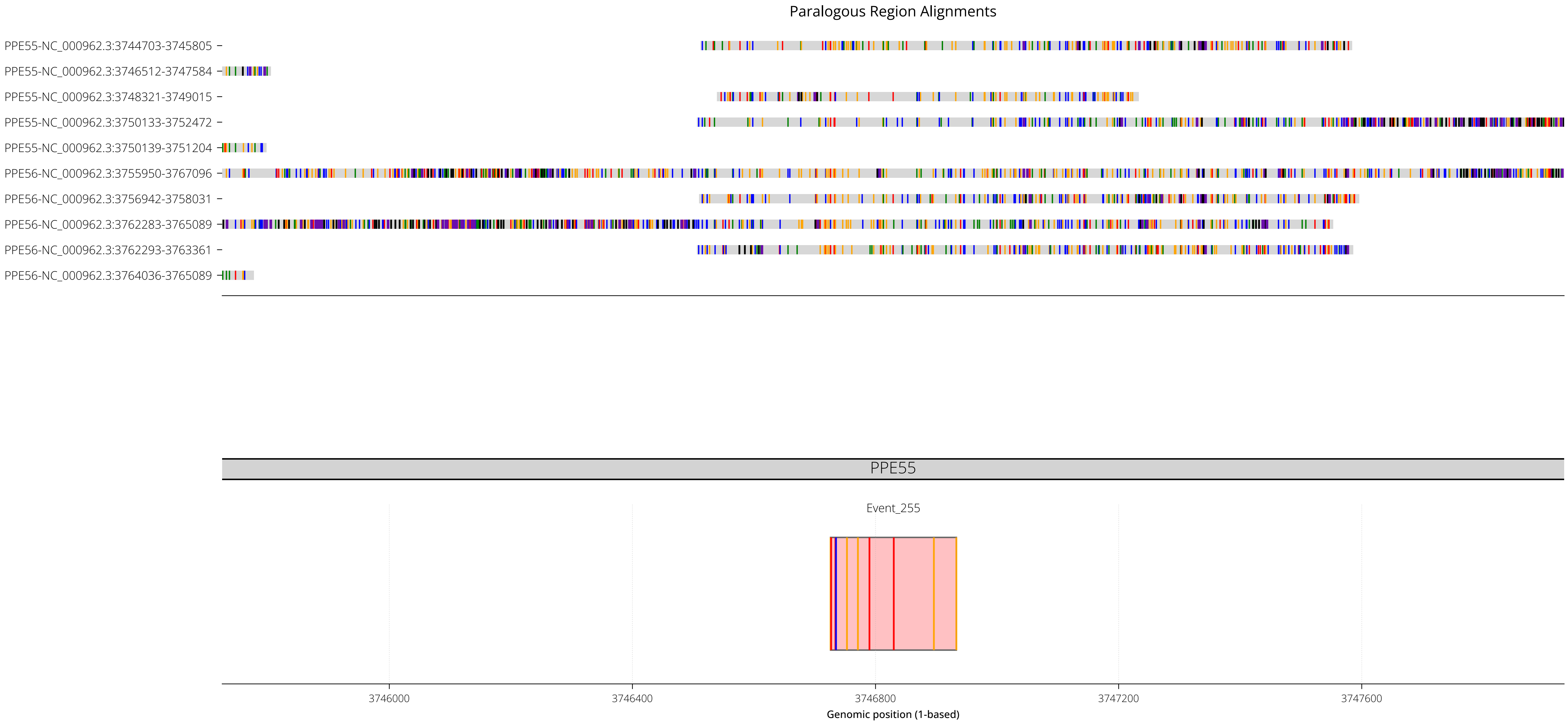

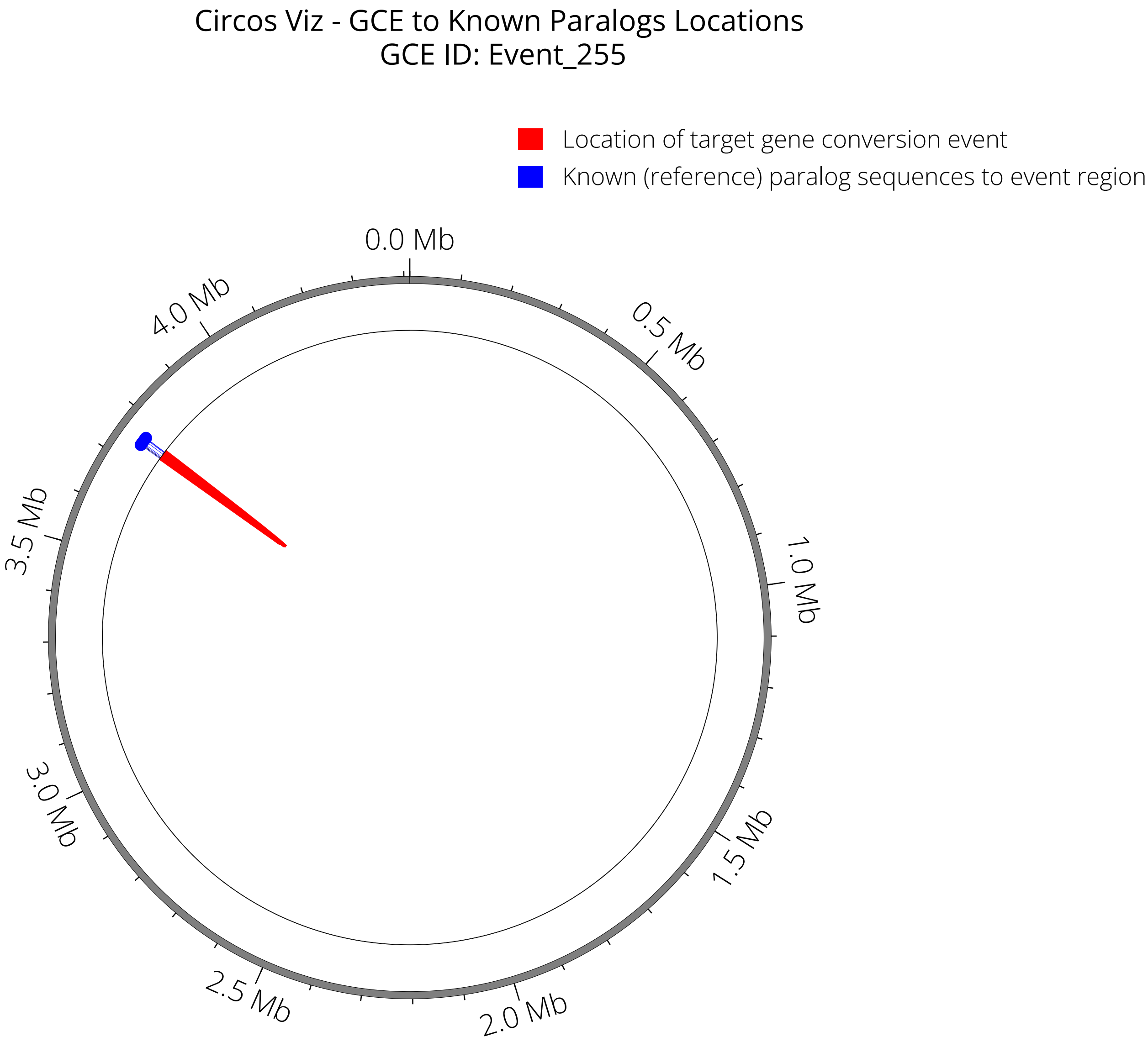

Circos Viz - GCE to Known Paralogs Locations
GCE ID: Event_255
Location of target gene conversion event
Known (reference) paralog sequences to event region
0.0 Mb
0.5 Mb
1.0 Mb
1.5 Mb
2.0 Mb
2.5 Mb
3.0 Mb
3.5 Mb
4.0 Mb

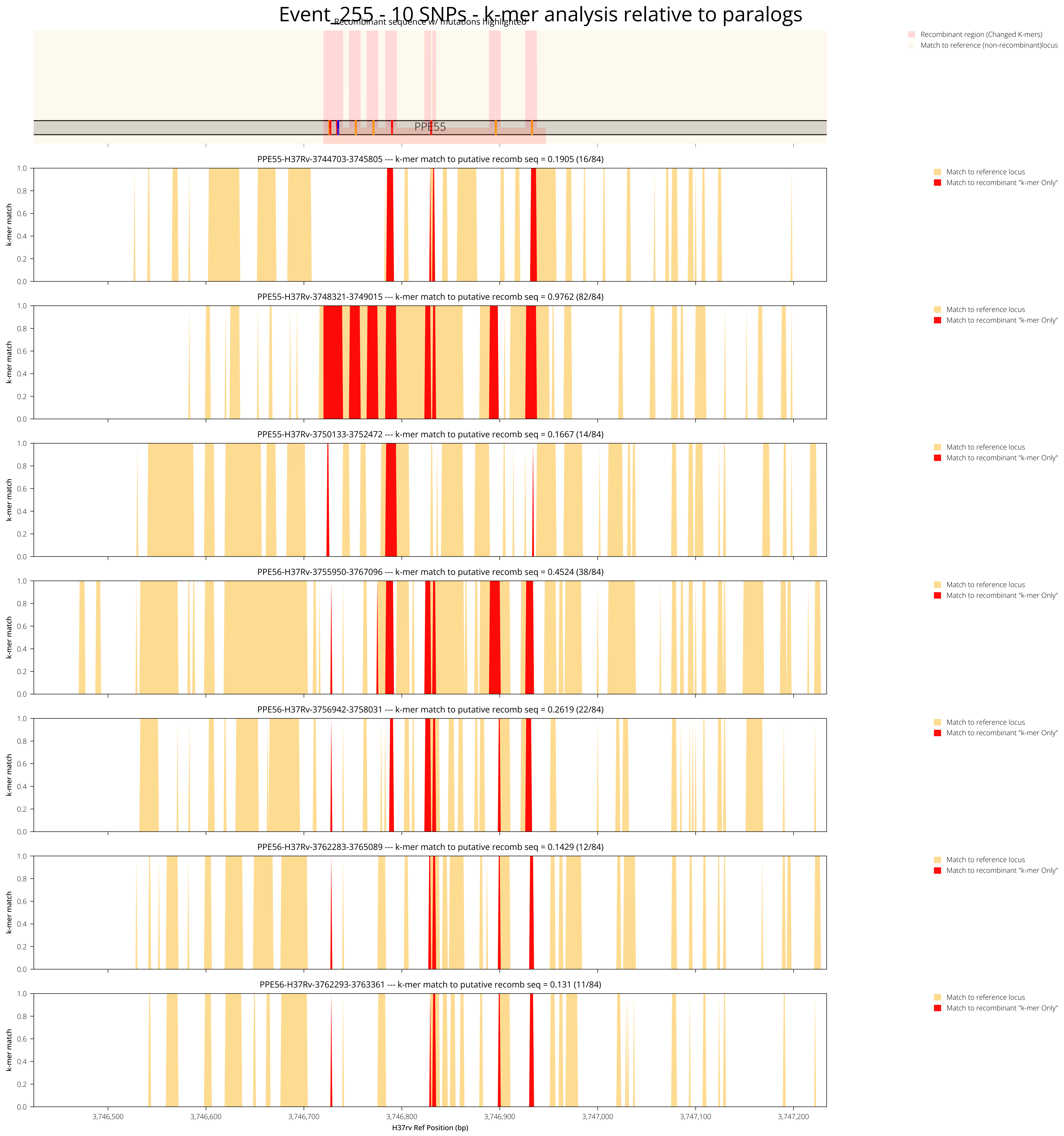
Event_255 - 10 SNPs - k-mer analysis relative to paralogs
Recombinant sequence w/ mutations highlighted
Recombinant region (Changed K-mers)
Match to reference (non-recombinant)locus
PPE55
PPE55-H37Rv-3744703-3745805 --- k-mer match to putative recomb seq = 0.1905 (16/84)
PPE55-H37Rv-3748321-3749015 --- k-mer match to putative recomb seq = 0.9762 (82/84)
PPE55-H37Rv-3750133-3752472 --- k-mer match to putative recomb seq = 0.1667 (14/84)
PPE56-H37Rv-3755950-3767096 --- k-mer match to putative recomb seq = 0.4524 (38/84)
PPE56-H37Rv-3756942-3758031 --- k-mer match to putative recomb seq = 0.2619 (22/84)
PPE56-H37Rv-3762283-3765089 --- k-mer match to putative recomb seq = 0.1429 (12/84)
PPE56-H37Rv-3762293-3763361 --- k-mer match to putative recomb seq = 0.131 (11/84)
Match to reference locus
Match to recombinant "k-mer Only"
k-mer match
0.0
0.2
0.4
0.6
0.8
1.0
3,746,500
3,746,600
3,746,700
3,746,800
3,746,900
3,747,000
3,747,100
3,747,200
H37rv Ref Position (bp)

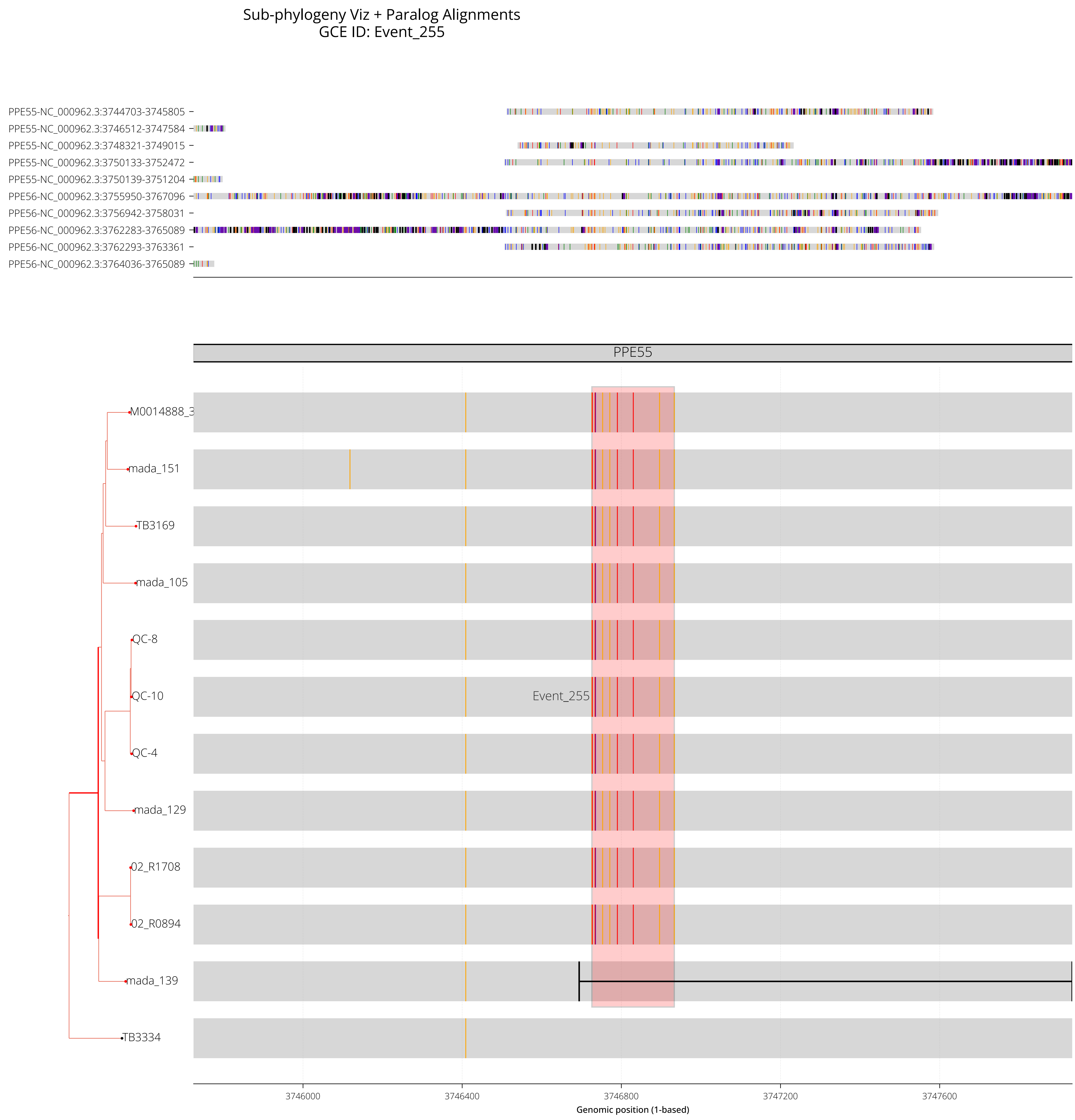

Sub-phylogeny Viz + Paralog Alignments
GCE ID: Event_255
PPE55-NC_000962.3:3744703-3745805
PPE55-NC_000962.3:3746512-3747584
PPE55-NC_000962.3:3748321-3749015
PPE55-NC_000962.3:3750133-3752472
PPE55-NC_000962.3:3750139-3751204
PPE56-NC_000962.3:3755950-3767096
PPE56-NC_000962.3:3756942-3758031
PPE56-NC_000962.3:3762283-3765089
PPE56-NC_000962.3:3762293-3763361
PPE56-NC_000962.3:3764036-3765089
PPE55
M0014888_3
mada_151
TB3169
mada_105
QC-8
QC-10
Event_255
QC-4
mada_129
02_R1708
02_R0894
mada_139
TB3334
3746000
3746400
3746800
3747200
3747600
Genomic position (1-based)

Full Phylogeny Viz + Paralog Alignments
GCE ID: Event_255

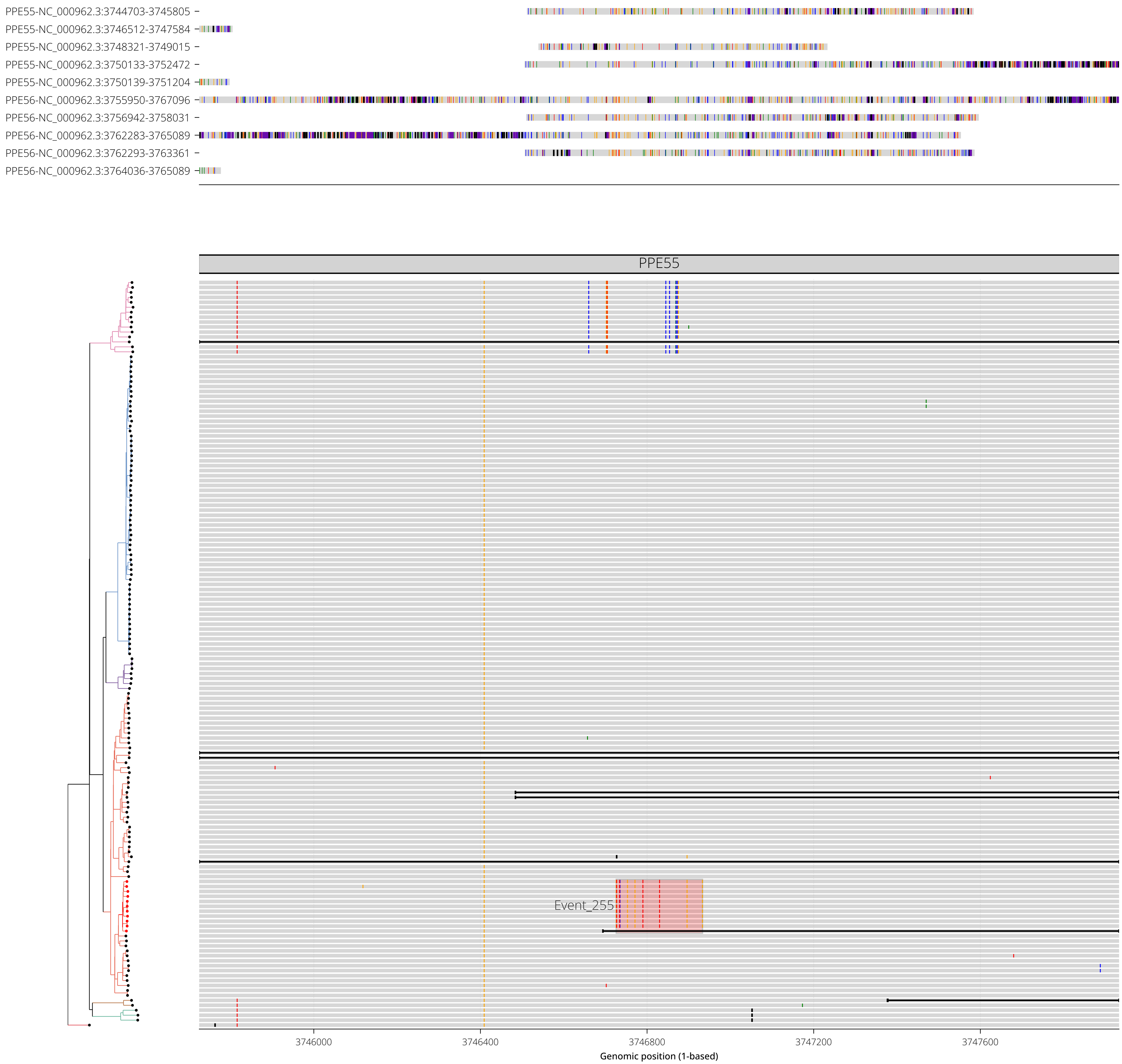

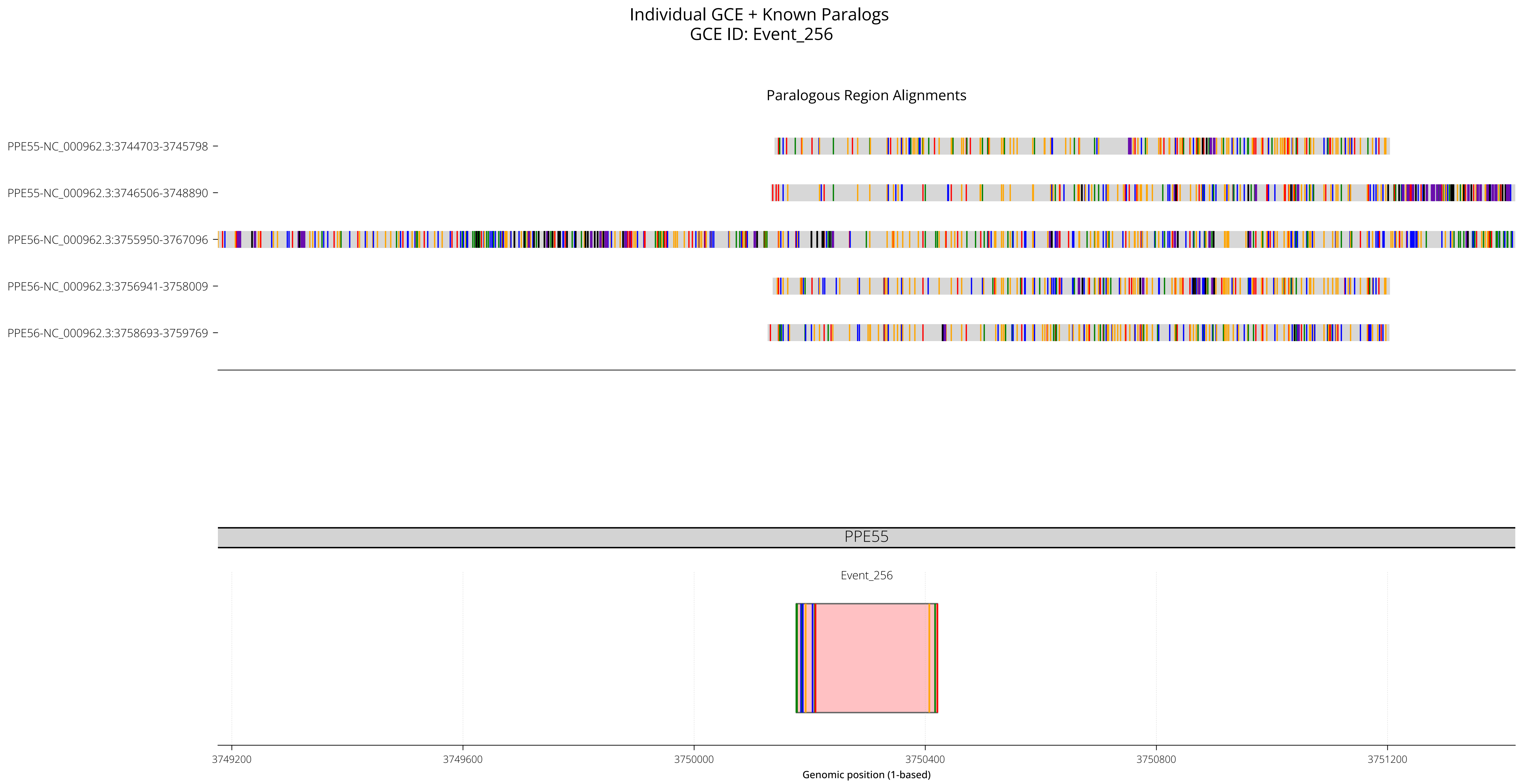
Individual GCE + Known Paralogs
GCE ID: Event_256
Paralogous Region Alignments
PPE55-NC_000962.3:3744703-3745798
PPE55-NC_000962.3:3746506-3748890
PPE56-NC_000962.3:3755950-3767096
PPE56-NC_000962.3:3756941-3758009
PPE56-NC_000962.3:3758693-3759769
PPE55
Event_256
3749200
3749600
3750000
3750400
3750800
3751200
Genomic position (1-based)

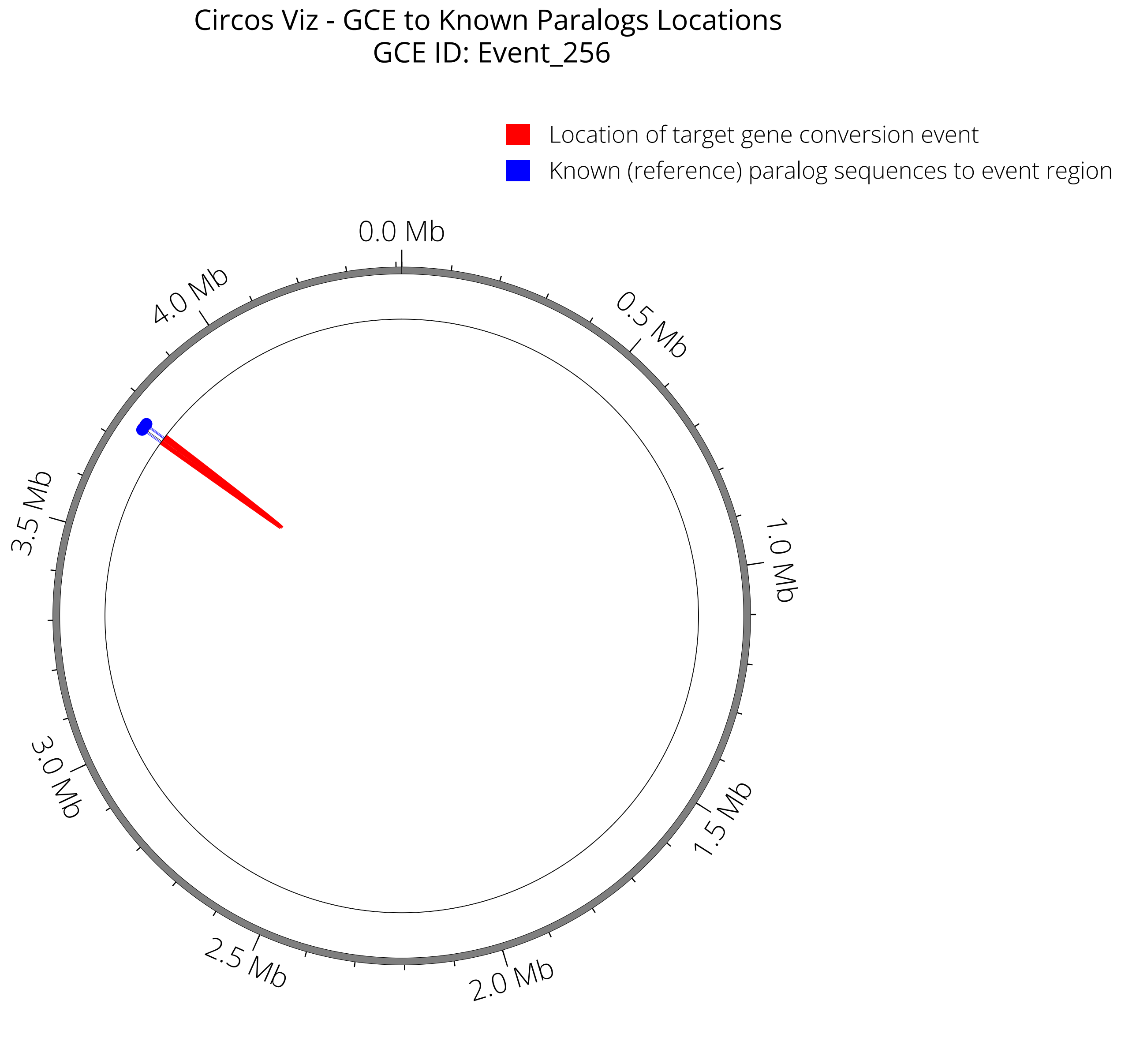

Circos Viz - GCE to Known Paralogs Locations
GCE ID: Event_256
Location of target gene conversion event
Known (reference) paralog sequences to event region
0.0 Mb
0.5 Mb
1.0 Mb
1.5 Mb
2.0 Mb
2.5 Mb
3.0 Mb
3.5 Mb
4.0 Mb

### Event_256 - 12 SNPs - k-mer analysis relative to paralogs

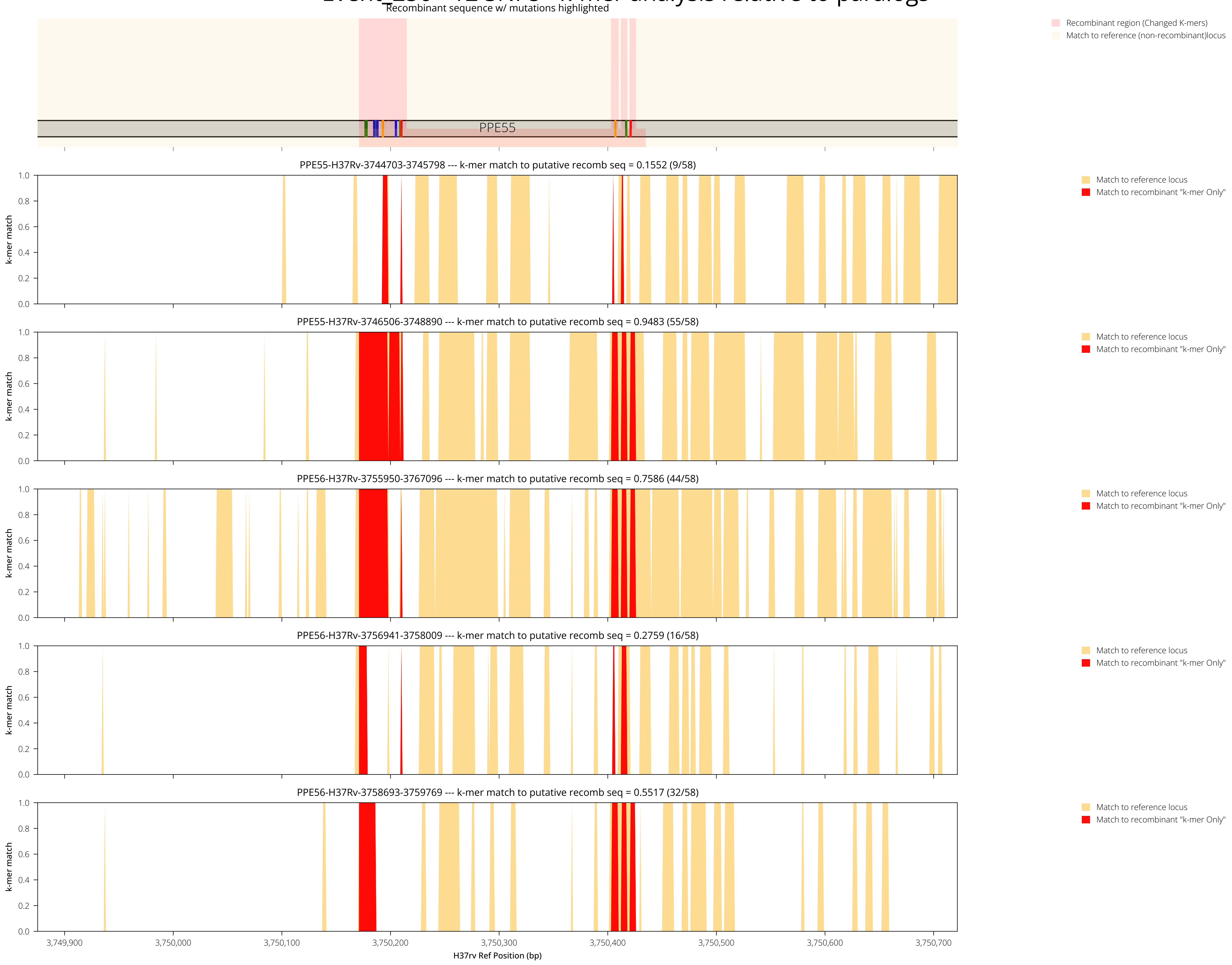

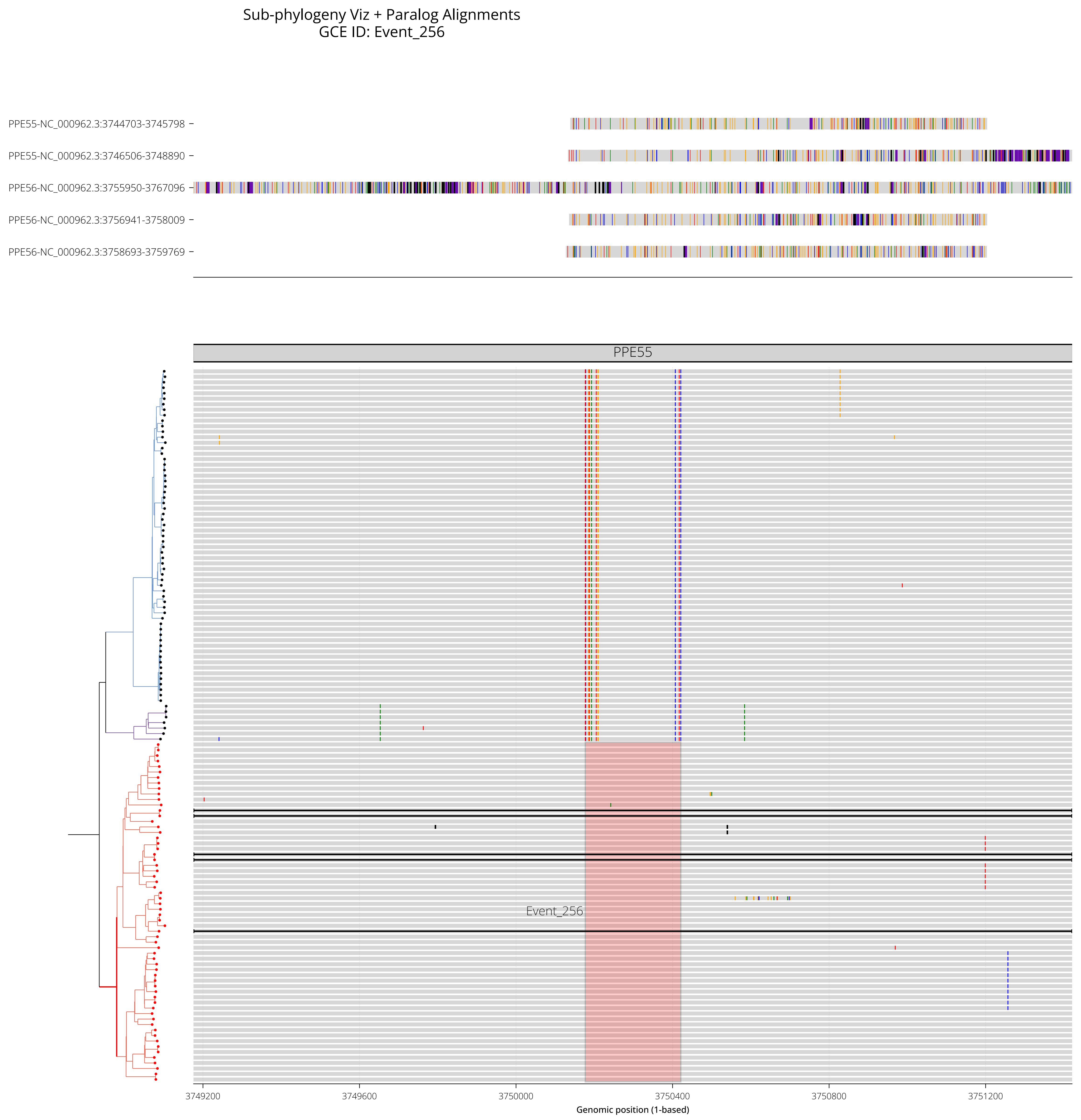

Sub-phylogeny Viz + Paralog Alignments
GCE ID: Event_256
PPE55-NC_000962.3:3744703-3745798
PPE55-NC_000962.3:3746506-3748890
PPE56-NC_000962.3:3755950-3767096
PPE56-NC_000962.3:3756941-3758009
PPE56-NC_000962.3:3758693-3759769
PPE55
Event_256
3749200
3749600
3750000
3750400
3750800
3751200
Genomic position (1-based)

Full Phylogeny Viz + Paralog Alignments
GCE ID: Event_256

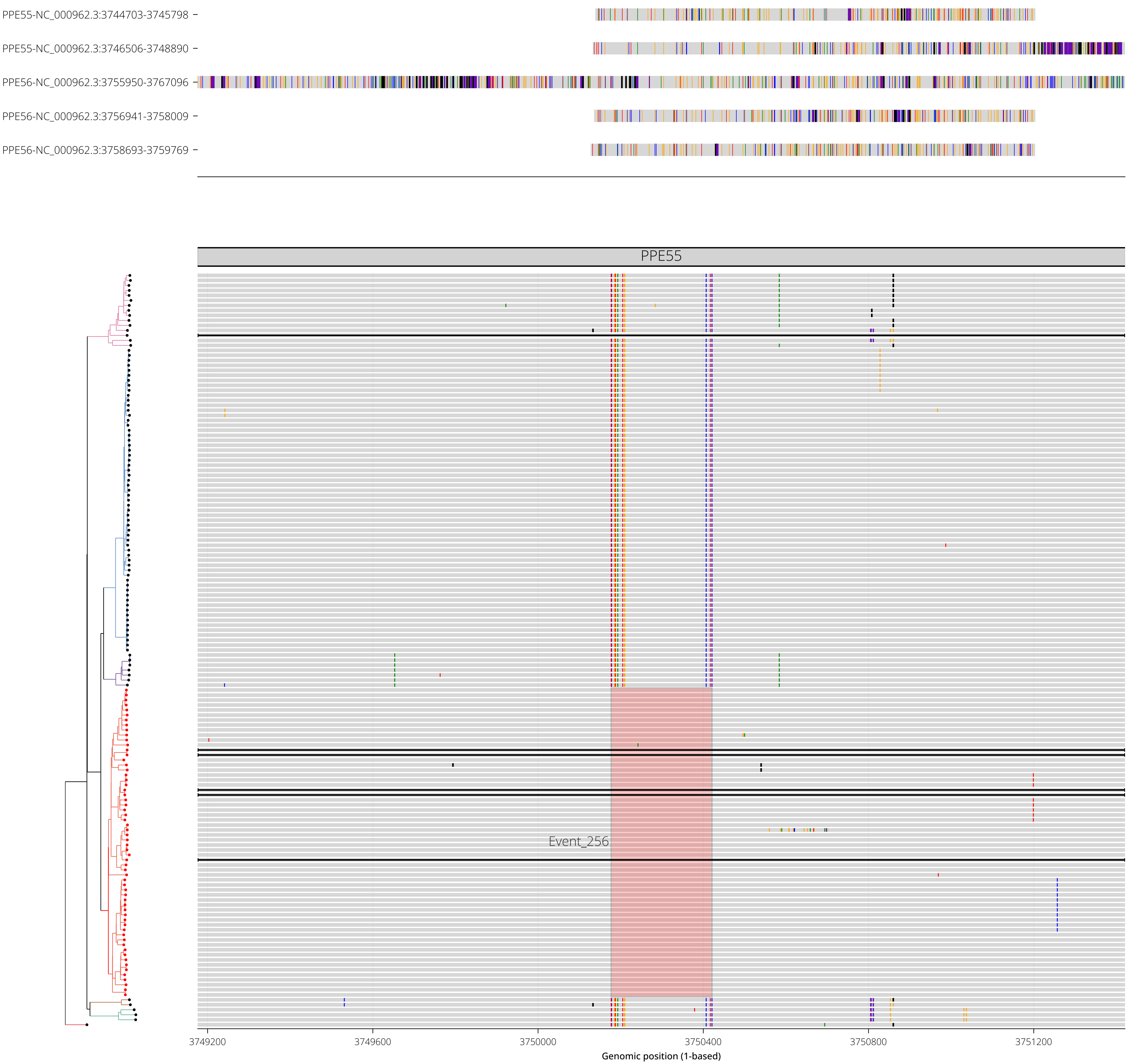

Individual GCE + Known Paralogs
GCE ID: Event_257

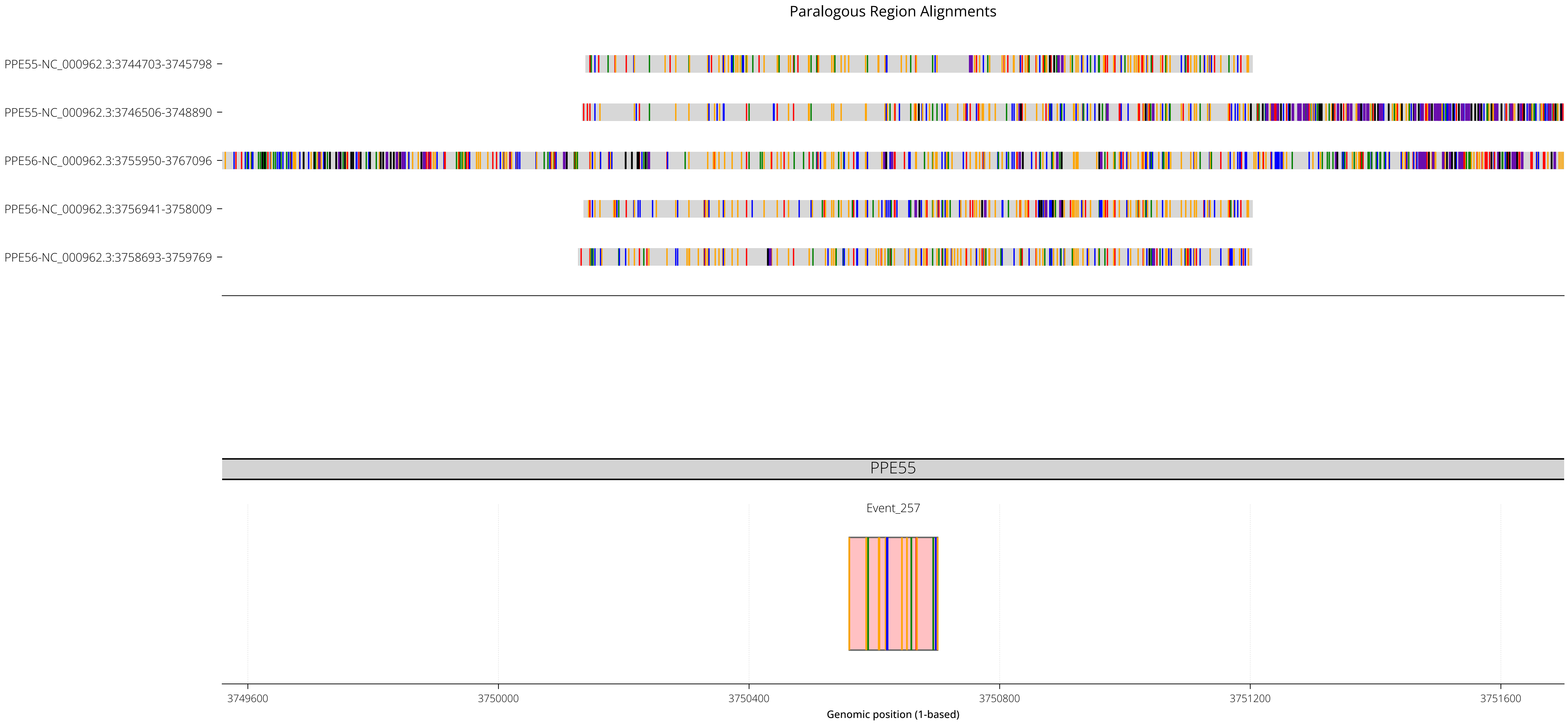

Circos Viz - GCE to Known Paralogs Locations
GCE ID: Event_257

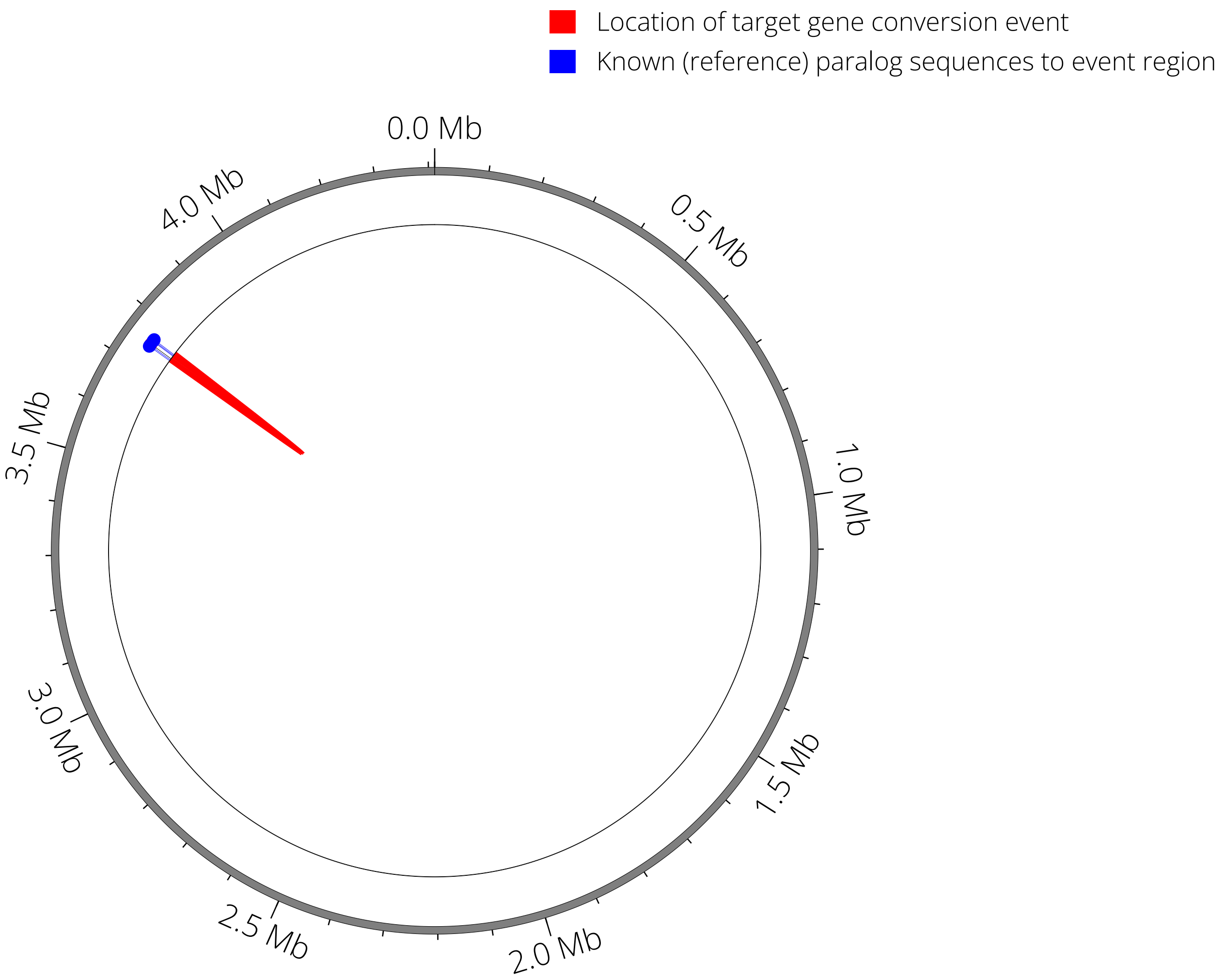

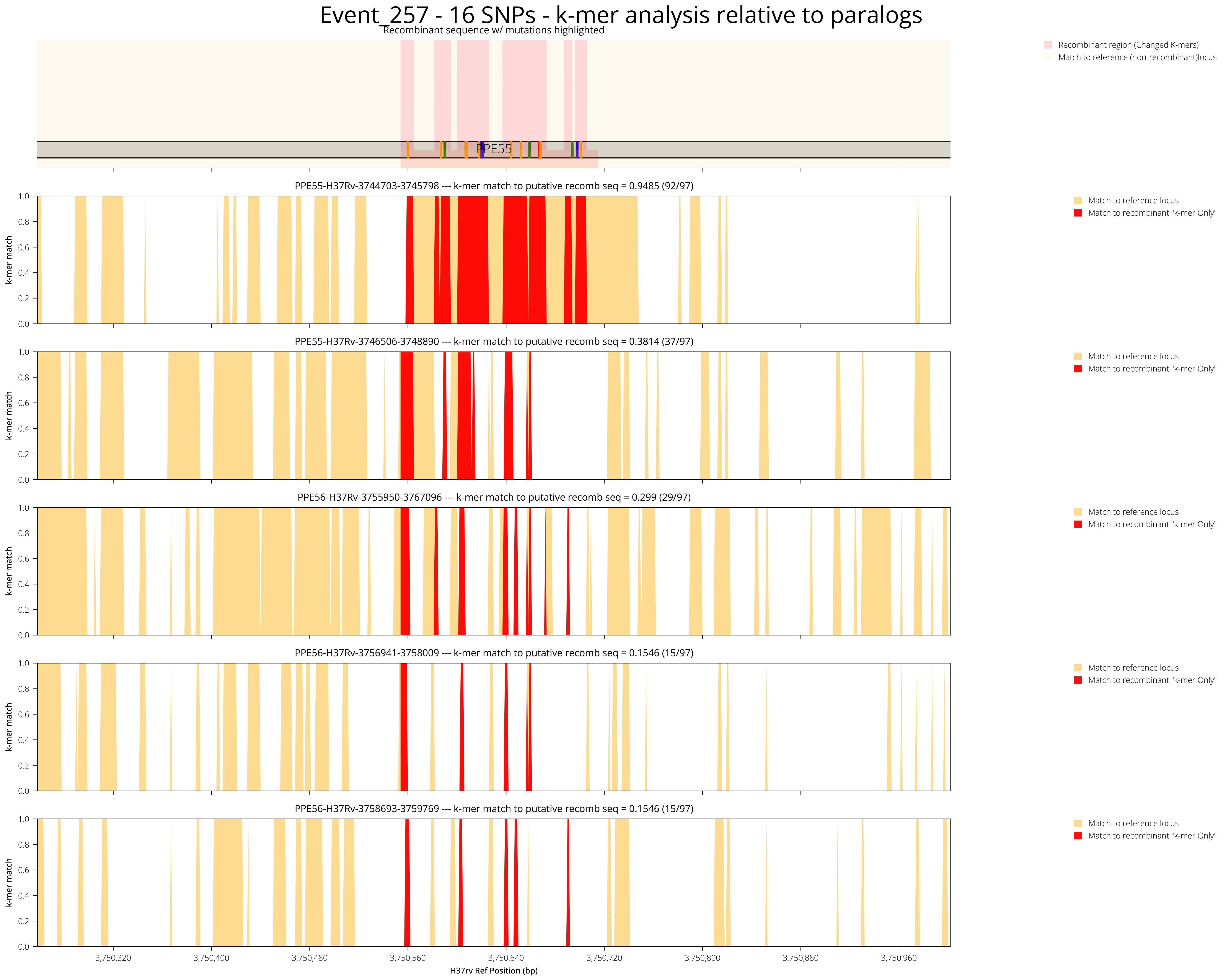

Event_257 - 16 SNPs - k-mer analysis relative to paralogs
Recombinant sequence w/ mutations highlighted
Recombinant region (Changed K-mers)
Match to reference (non-recombinant)locus
PPE55
PPE55-H37Rv-3744703-3745798 --- k-mer match to putative recomb seq = 0.9485 (92/97)
PPE55-H37Rv-3746506-3748890 --- k-mer match to putative recomb seq = 0.3814 (37/97)
PPE56-H37Rv-3755950-3767096 --- k-mer match to putative recomb seq = 0.299 (29/97)
PPE56-H37Rv-3756941-3758009 --- k-mer match to putative recomb seq = 0.1546 (15/97)
PPE56-H37Rv-3758693-3759769 --- k-mer match to putative recomb seq = 0.1546 (15/97)
Match to reference locus
Match to recombinant "k-mer Only"
k-mer match
1.0
0.8
0.6
0.4
0.2
0.0
3,750,320
3,750,400
3,750,480
3,750,560
3,750,640
3,750,720
3,750,800
3,750,880
3,750,960
H37rv Ref Position (bp)

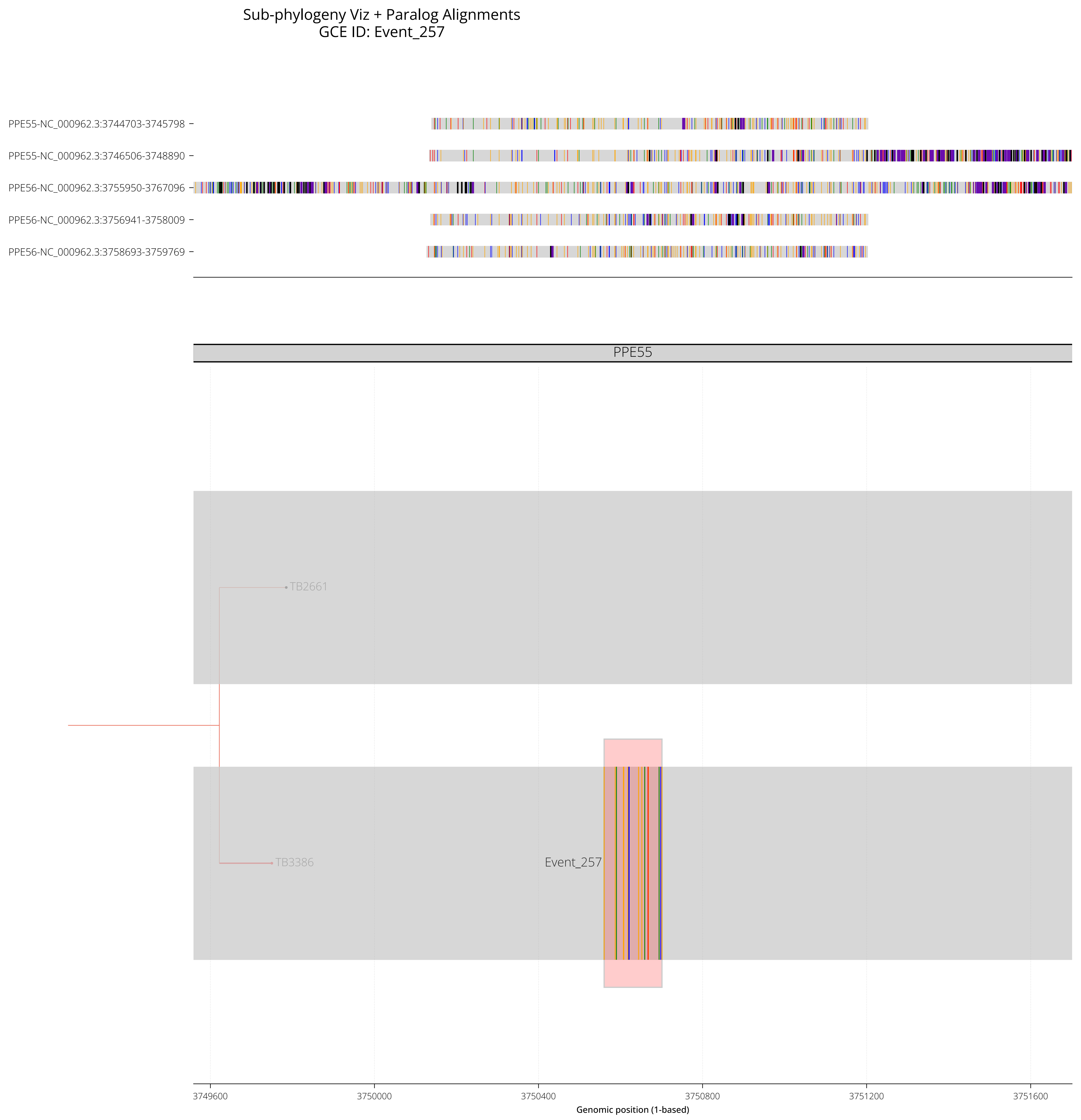

Sub-phylogeny Viz + Paralog Alignments
GCE ID: Event_257
PPE55-NC_000962.3:3744703-3745798
PPE55-NC_000962.3:3746506-3748890
PPE56-NC_000962.3:3755950-3767096
PPE56-NC_000962.3:3756941-3758009
PPE56-NC_000962.3:3758693-3759769
PPE55
TB2661
TB3386
Event_257
3749600
3750000
3750400
3750800
3751200
3751600
Genomic position (1-based)

Full Phylogeny Viz + Paralog Alignments
GCE ID: Event_257

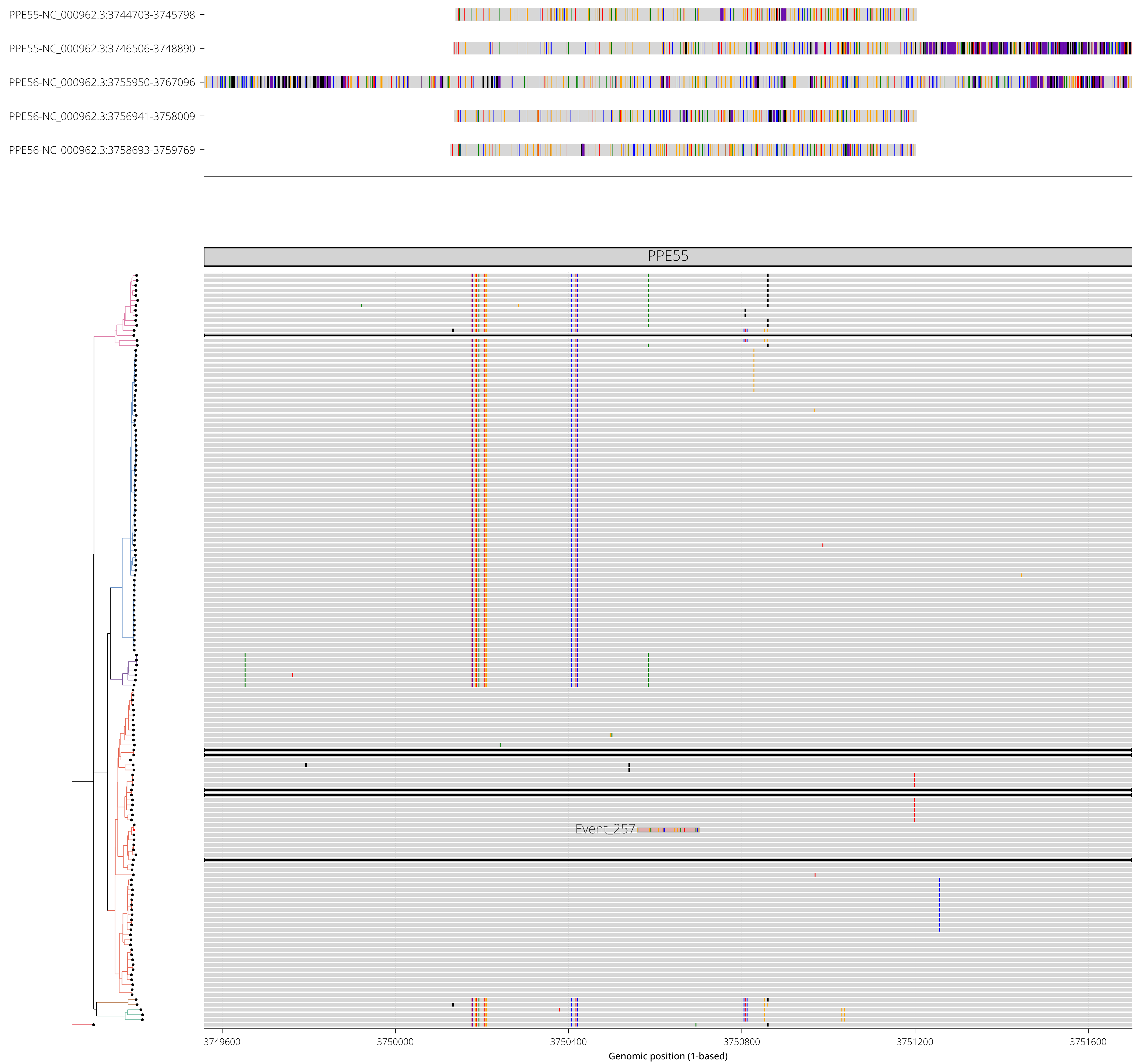

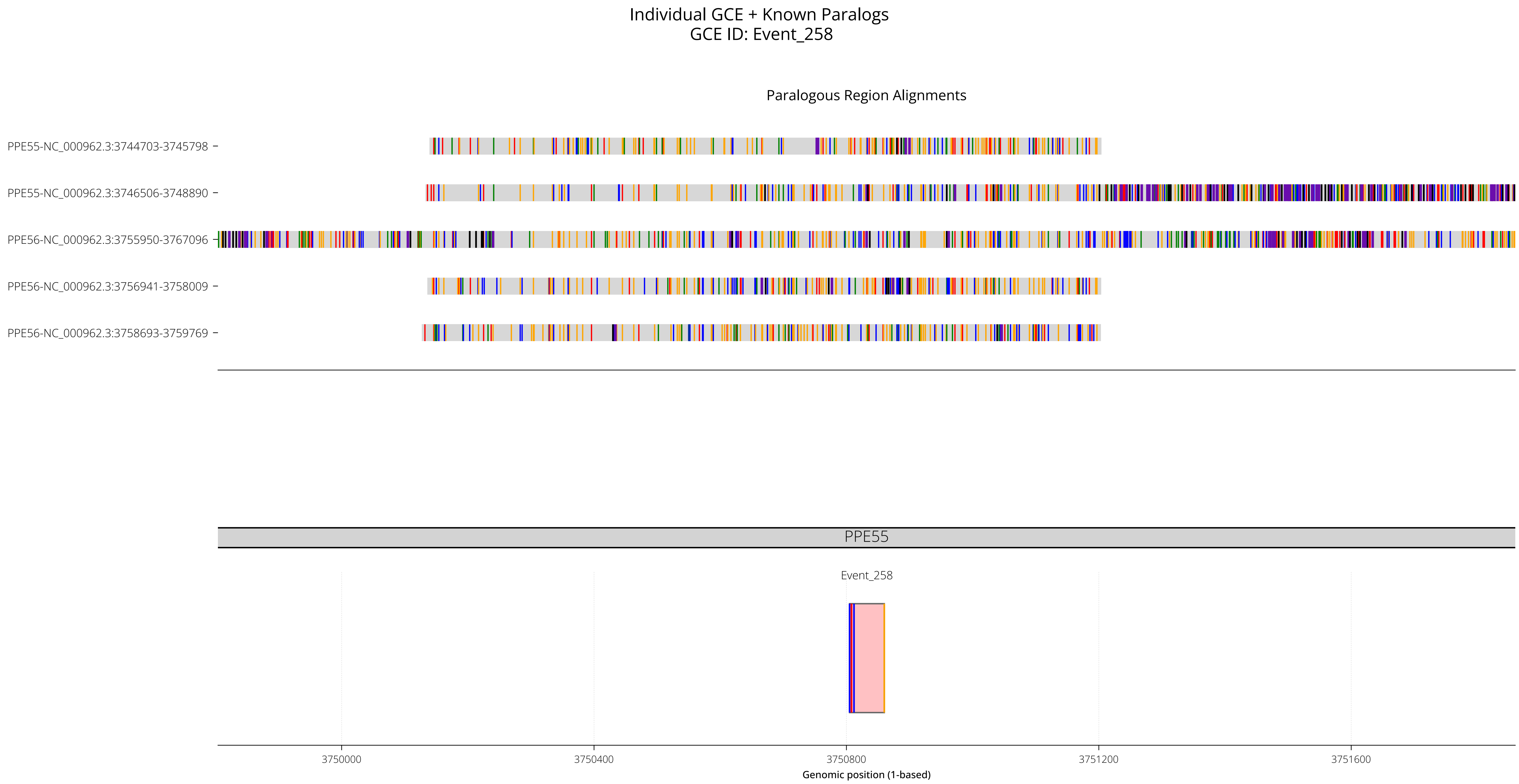
Individual GCE + Known Paralogs
GCE ID: Event_258
Paralogous Region Alignments
PPE55-NC_000962.3:3744703-3745798
PPE55-NC_000962.3:3746506-3748890
PPE56-NC_000962.3:3755950-3767096
PPE56-NC_000962.3:3756941-3758009
PPE56-NC_000962.3:3758693-3759769
PPE55
Event_258
3750000
3750400
3750800
3751200
3751600
Genomic position (1-based)

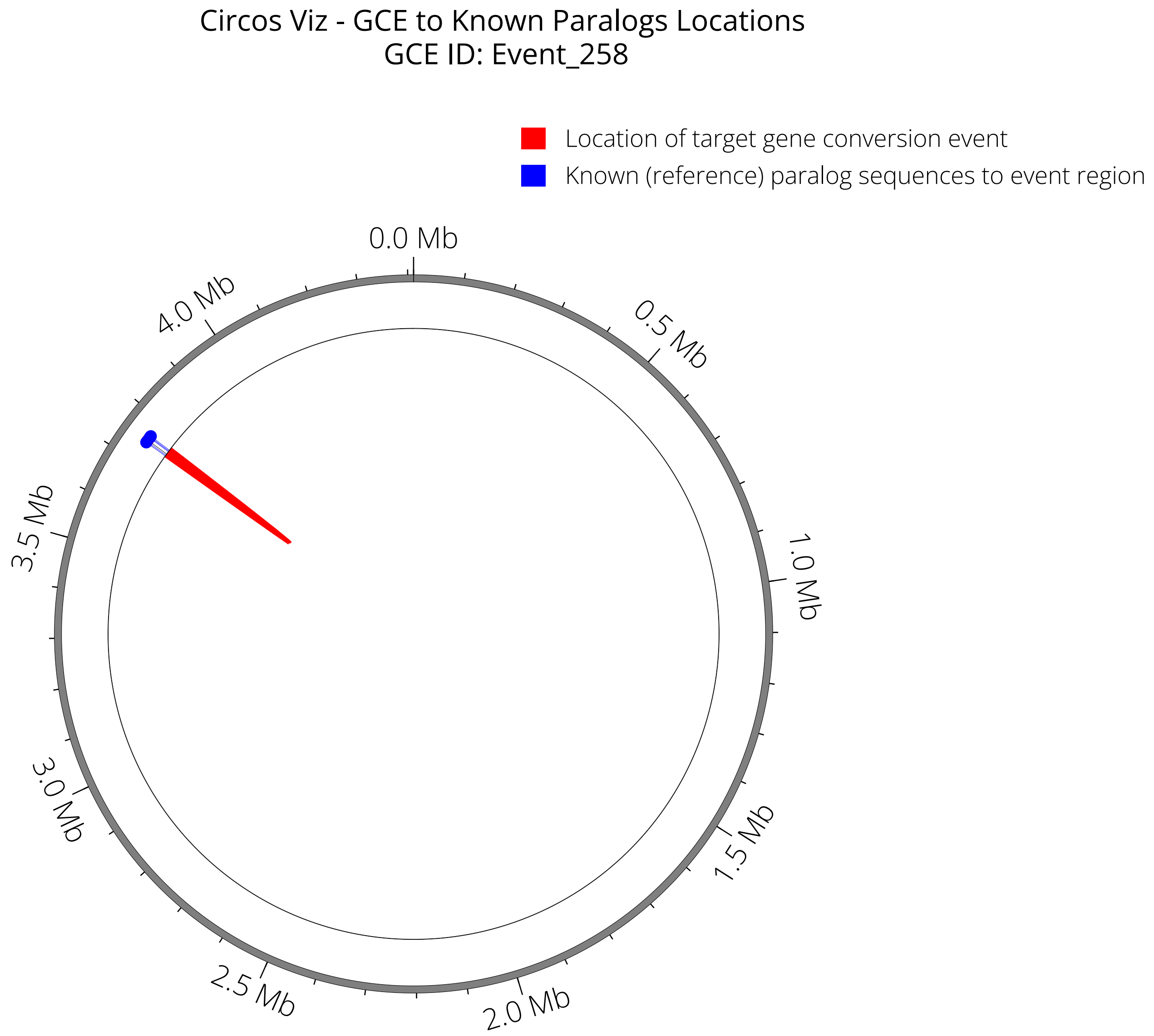

Circos Viz - GCE to Known Paralogs Locations
GCE ID: Event_258
Location of target gene conversion event
Known (reference) paralog sequences to event region
0.0 Mb
0.5 Mb
1.0 Mb
1.5 Mb
2.0 Mb
2.5 Mb
3.0 Mb
3.5 Mb
4.0 Mb

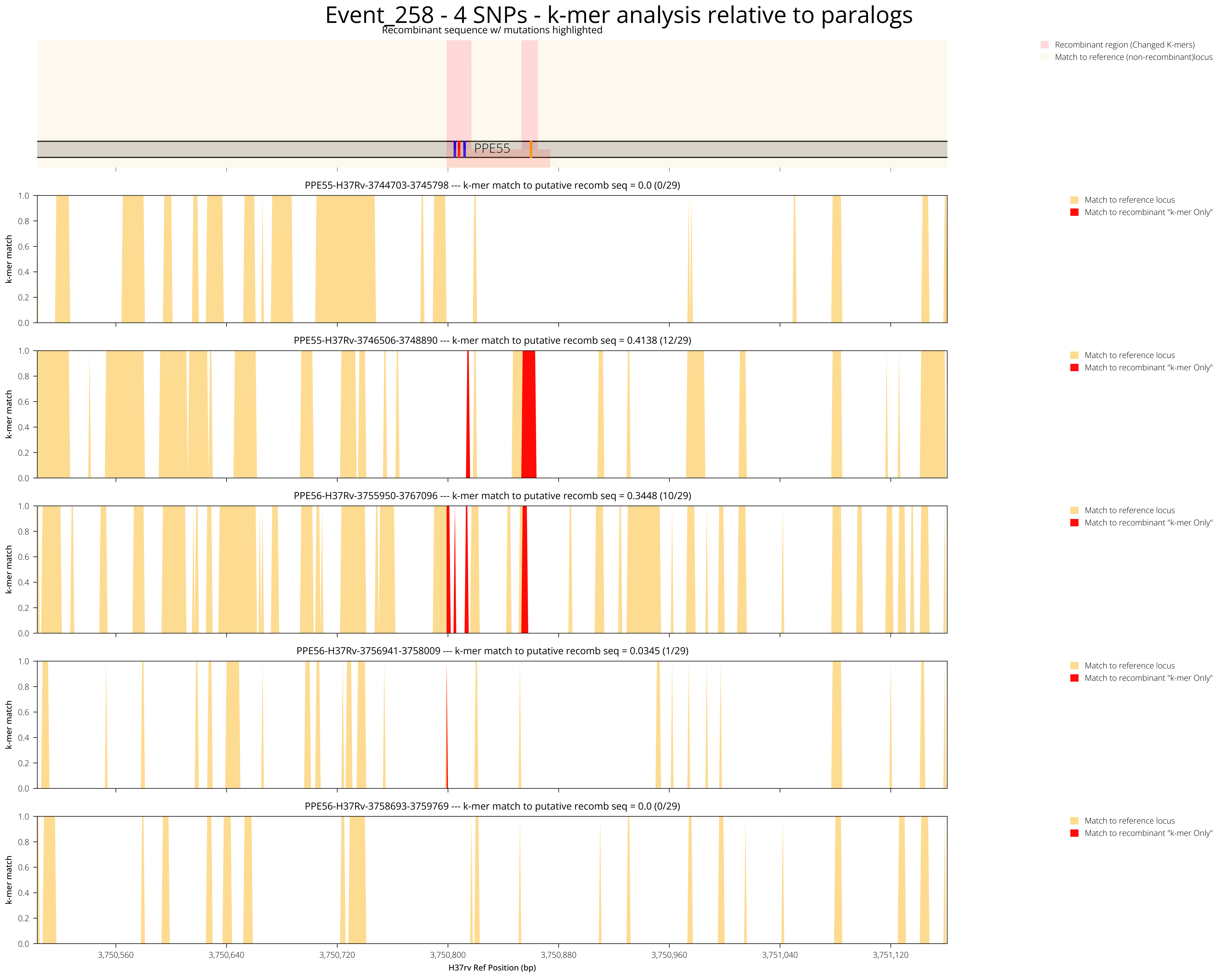
Event_258 - 4 SNPs - k-mer analysis relative to paralogs
Recombinant sequence w/ mutations highlighted
Recombinant region (Changed K-mers)
Match to reference (non-recombinant)locus
PPE55
PPE55-H37Rv-3744703-3745798 --- k-mer match to putative recomb seq = 0.0 (0/29)
PPE55-H37Rv-3746506-3748890 --- k-mer match to putative recomb seq = 0.4138 (12/29)
PPE56-H37Rv-3755950-3767096 --- k-mer match to putative recomb seq = 0.3448 (10/29)
PPE56-H37Rv-3756941-3758009 --- k-mer match to putative recomb seq = 0.0345 (1/29)
PPE56-H37Rv-3758693-3759769 --- k-mer match to putative recomb seq = 0.0 (0/29)
Match to reference locus
Match to recombinant "k-mer Only"
k-mer match
1.0
0.8
0.6
0.4
0.2
0.0
3,750,560
3,750,640
3,750,720
3,750,800
3,750,880
3,750,960
3,751,040
3,751,120
H37rv Ref Position (bp)

Sub-phylogeny Viz + Paralog Alignments
GCE ID: Event_258

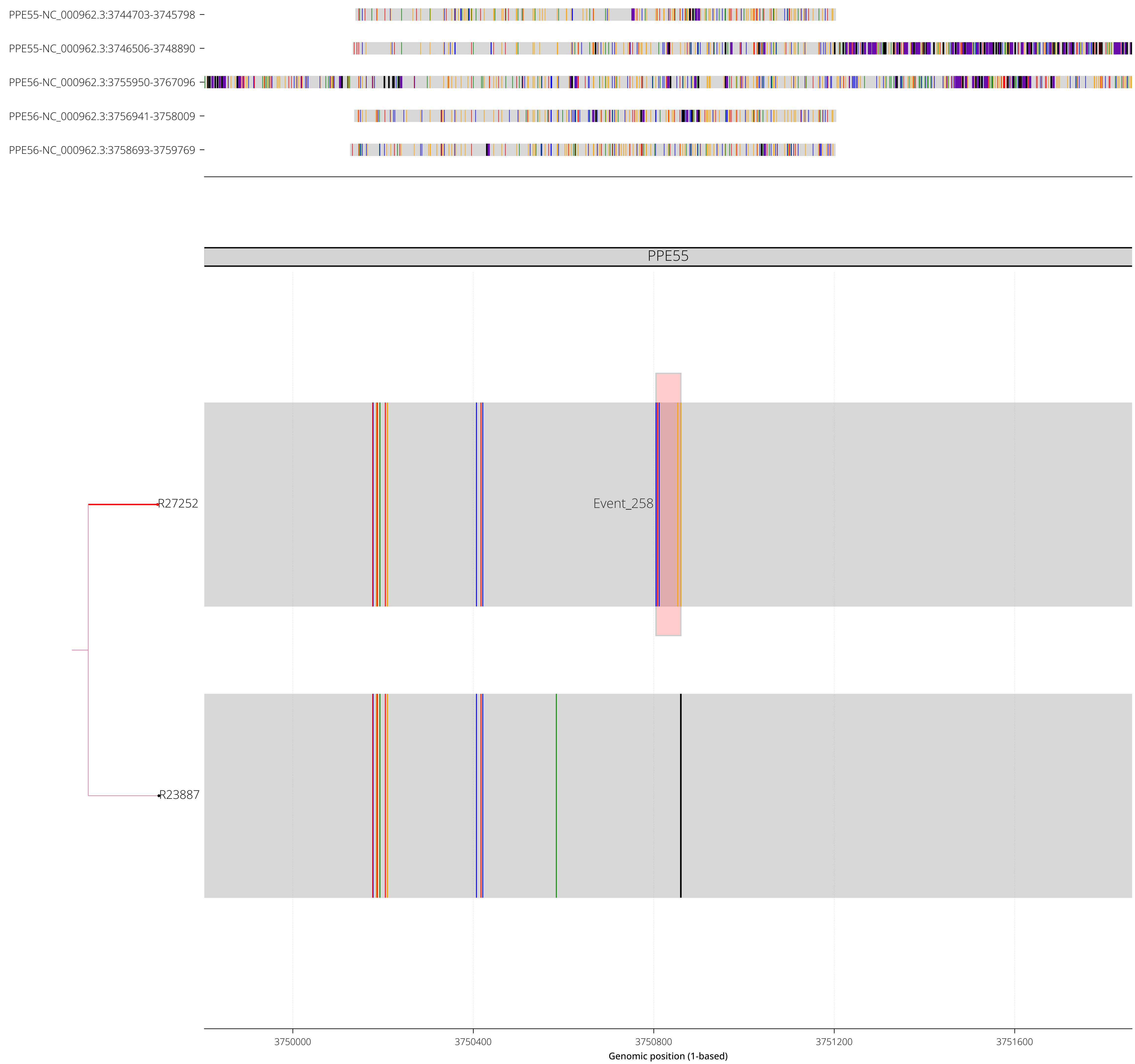

Full Phylogeny Viz + Paralog Alignments
GCE ID: Event_258

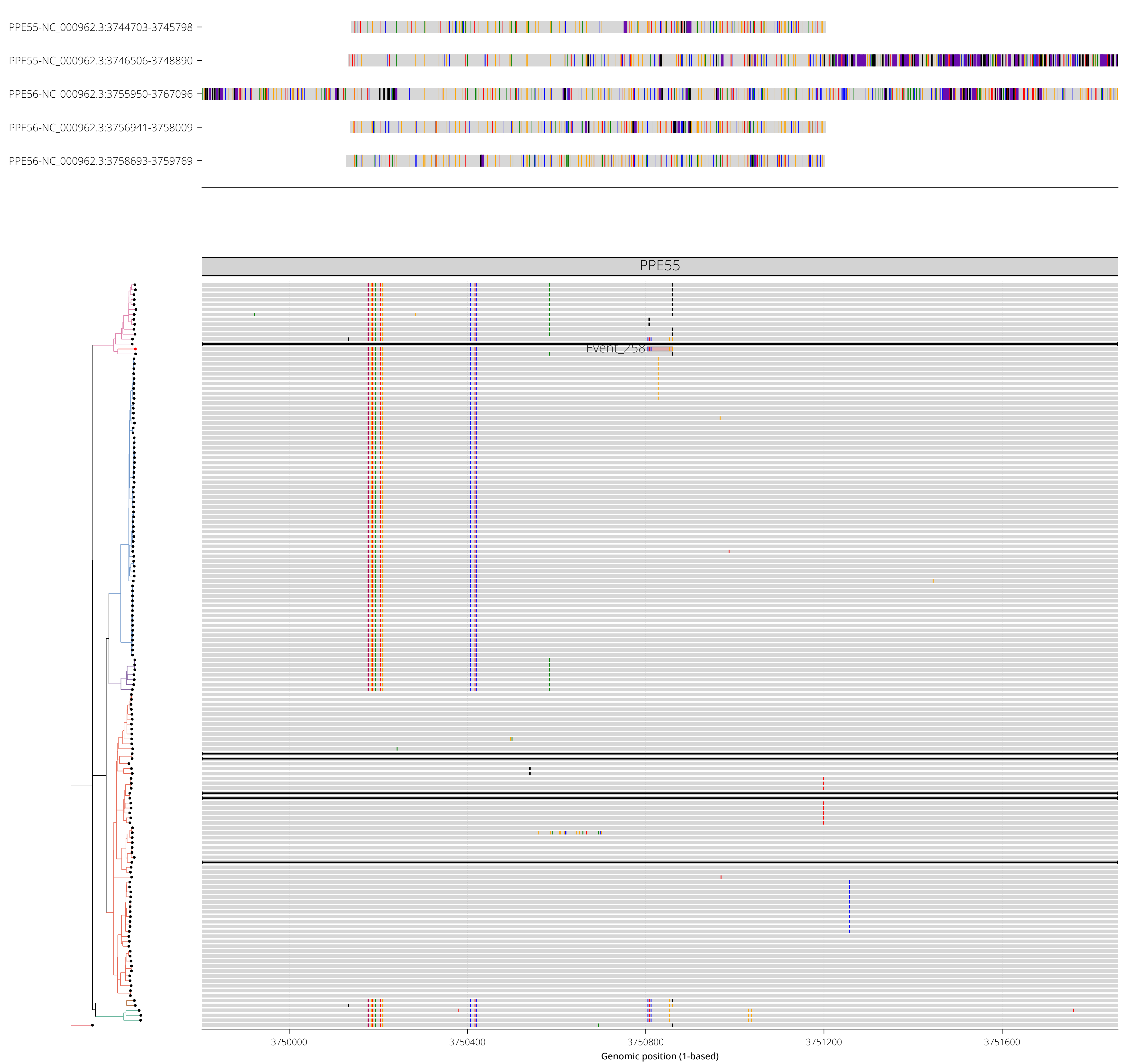

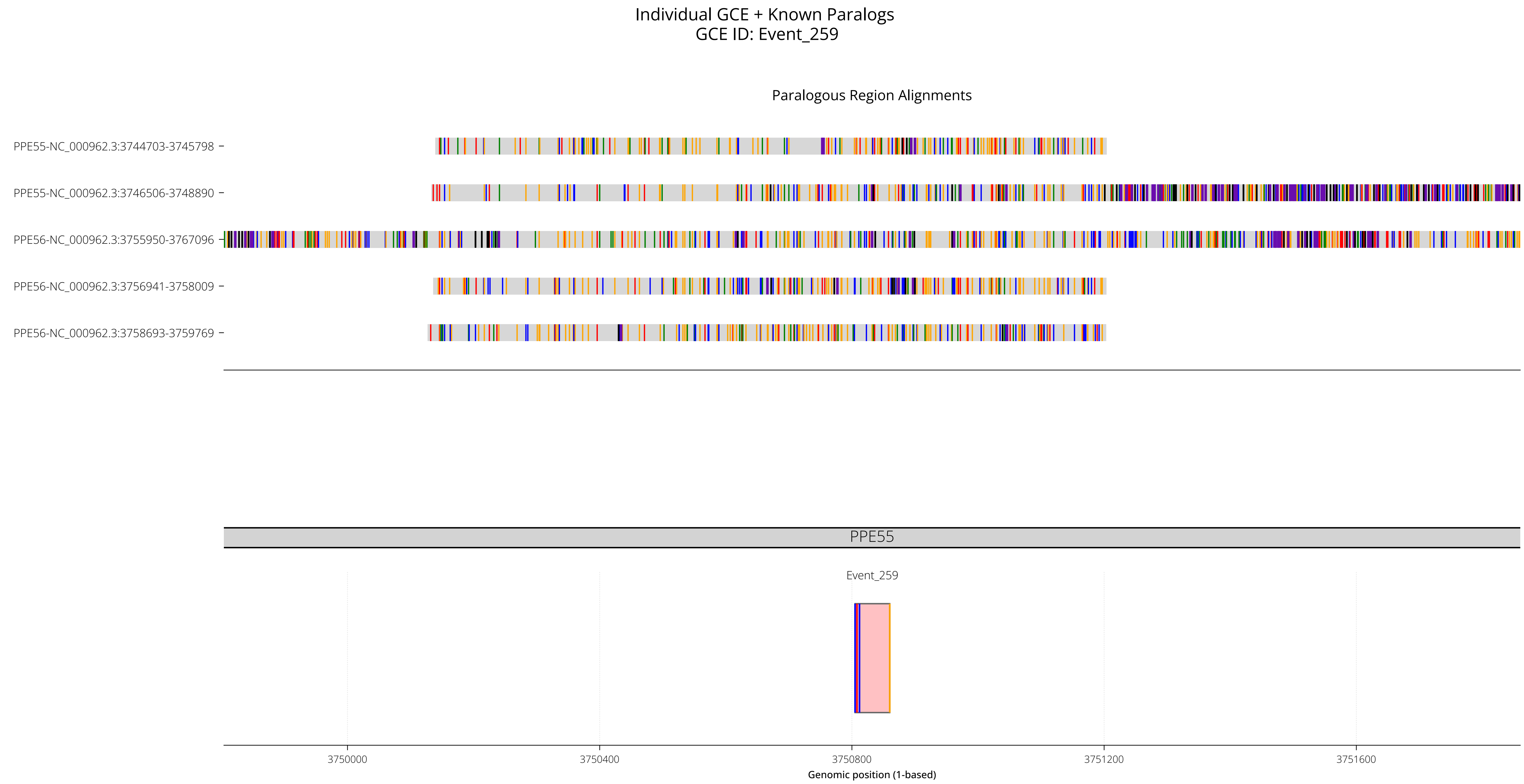
Individual GCE + Known Paralogs
GCE ID: Event_259
Paralogous Region Alignments
PPE55-NC_000962.3:3744703-3745798
PPE55-NC_000962.3:3746506-3748890
PPE56-NC_000962.3:3755950-3767096
PPE56-NC_000962.3:3756941-3758009
PPE56-NC_000962.3:3758693-3759769
PPE55
Event_259
3750000
3750400
3750800
3751200
3751600
Genomic position (1-based)

Circos Viz - GCE to Known Paralogs Locations
GCE ID: Event_259

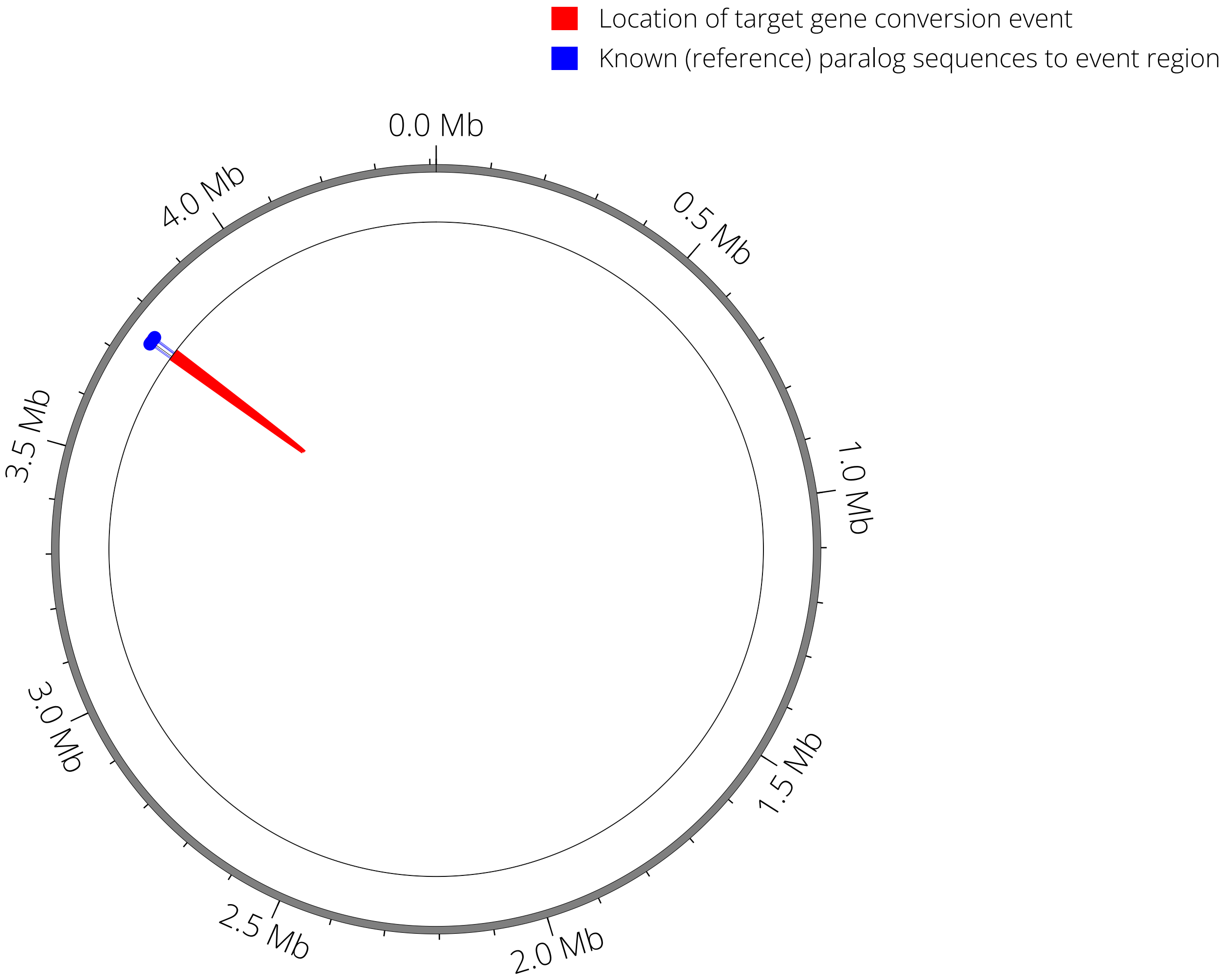

### Event_259 - 4 SNPs - k-mer analysis relative to paralogs

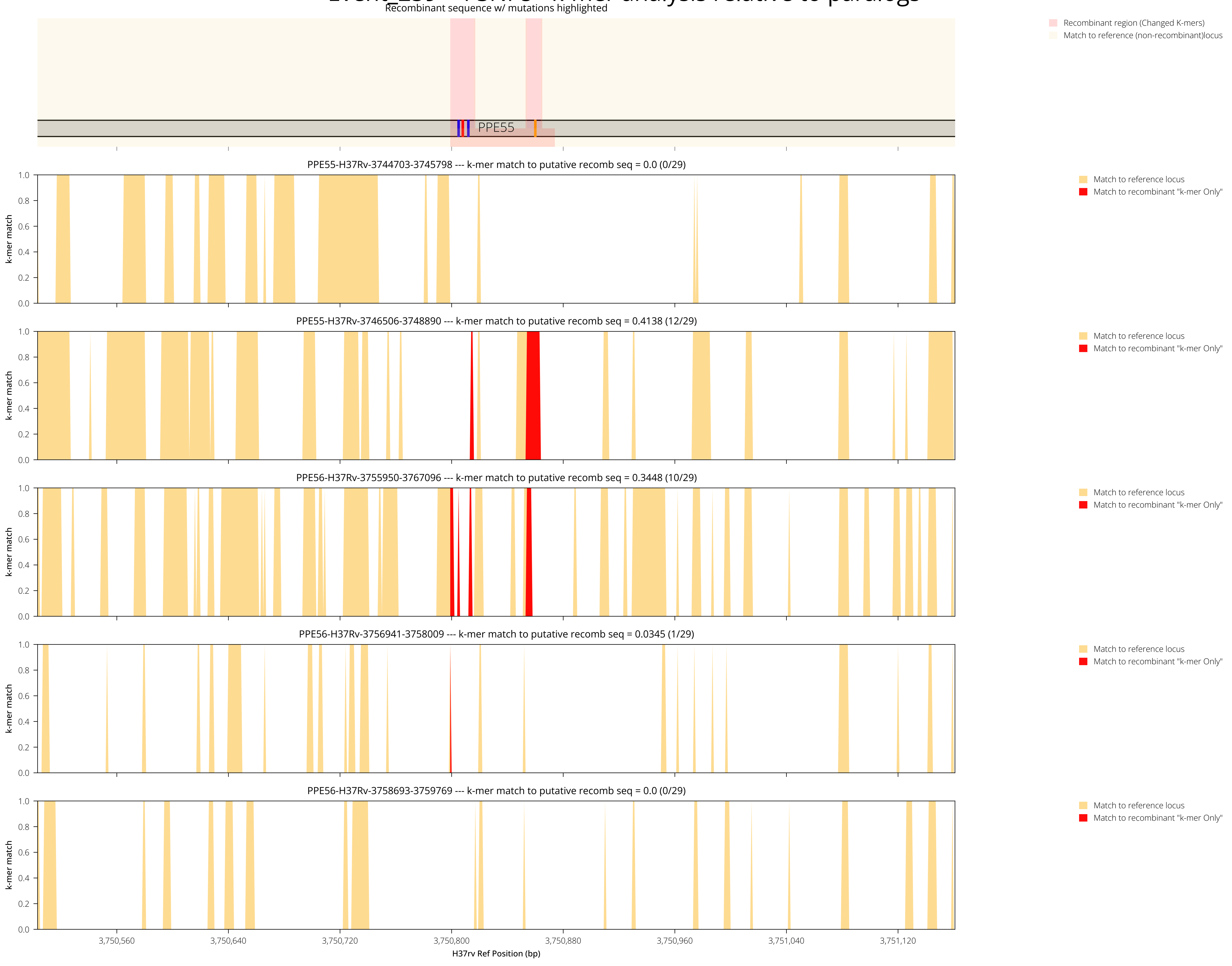

Sub-phylogeny Viz + Paralog Alignments
GCE ID: Event_259

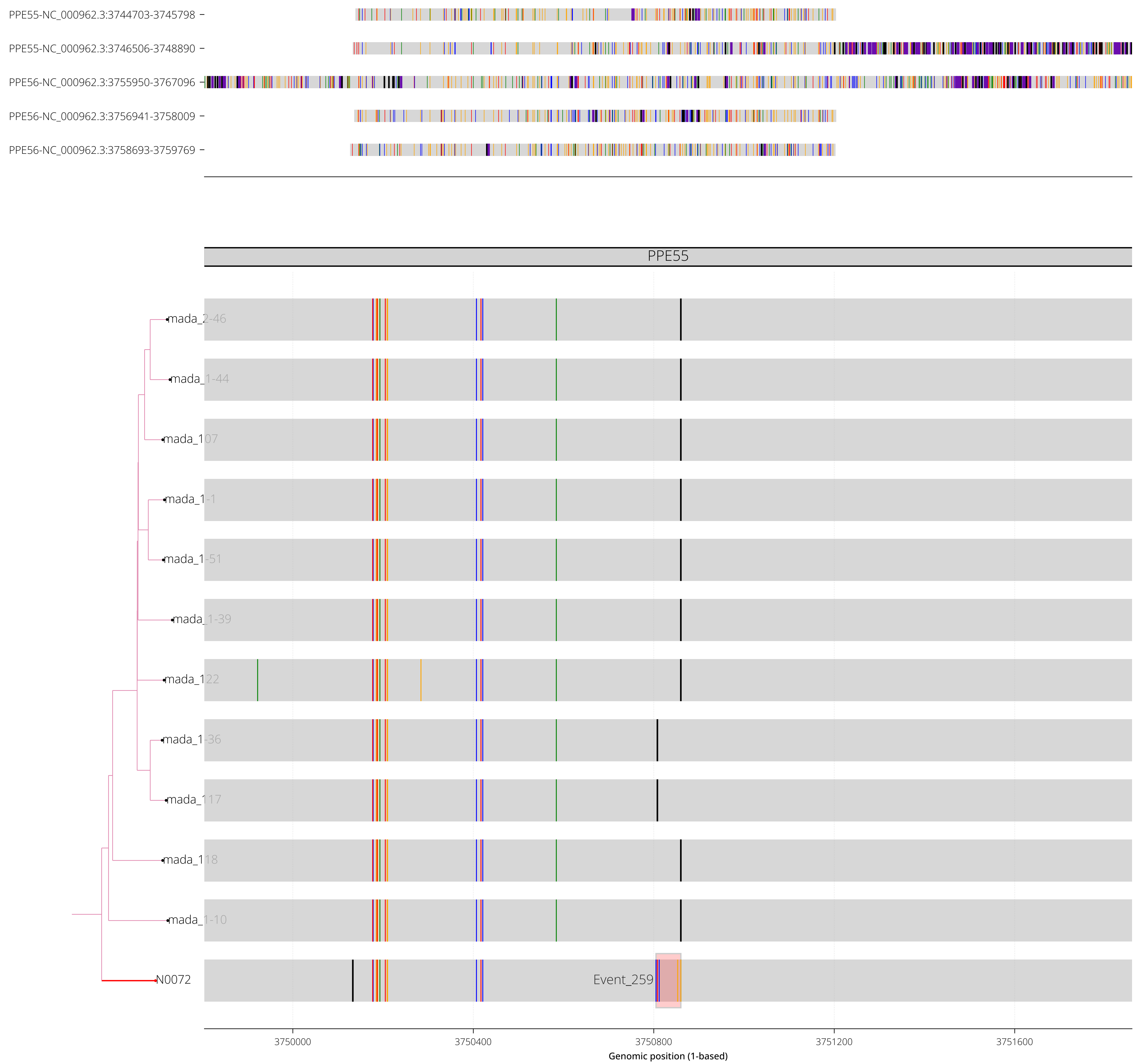

Full Phylogeny Viz + Paralog Alignments
GCE ID: Event_259

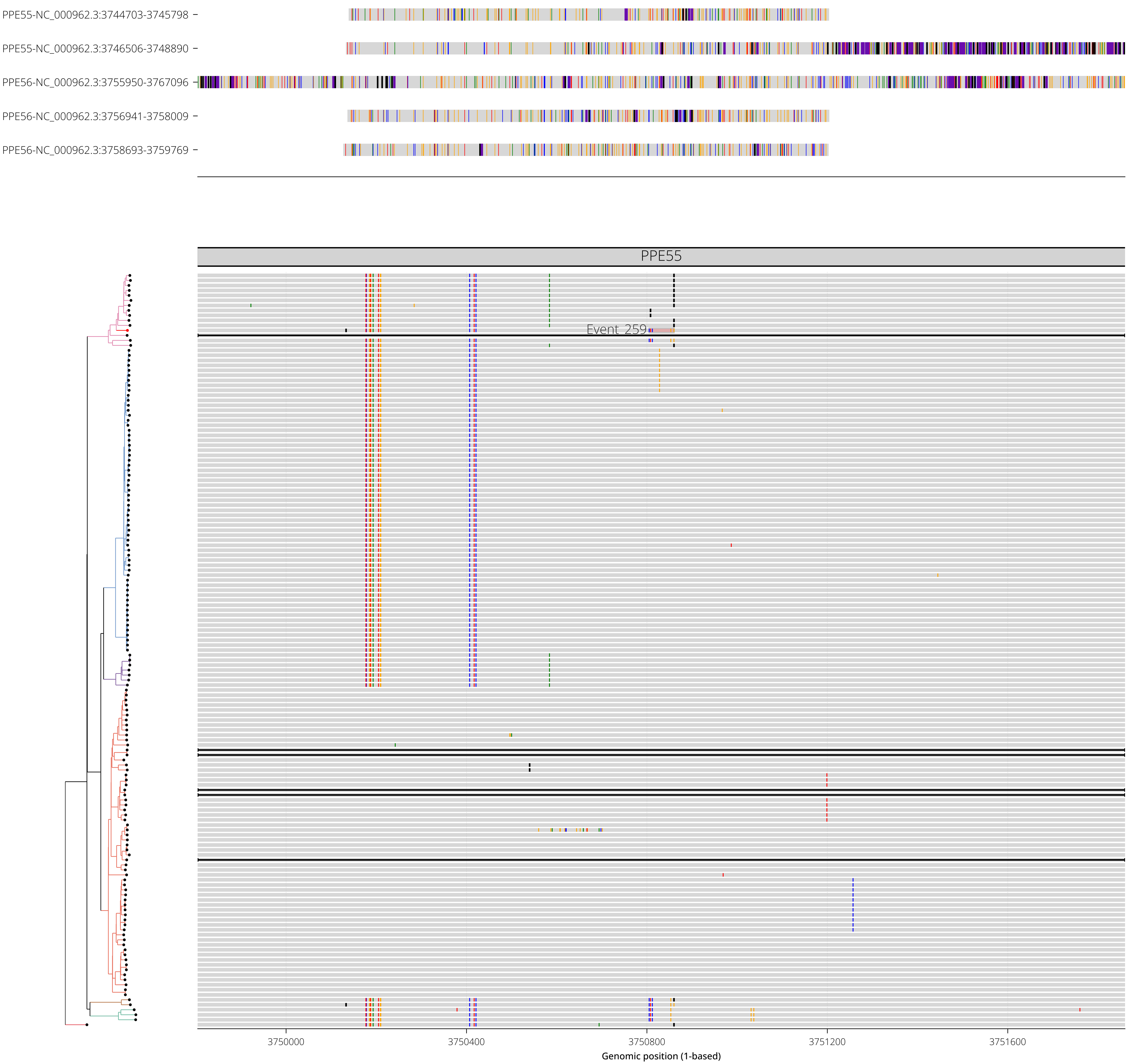

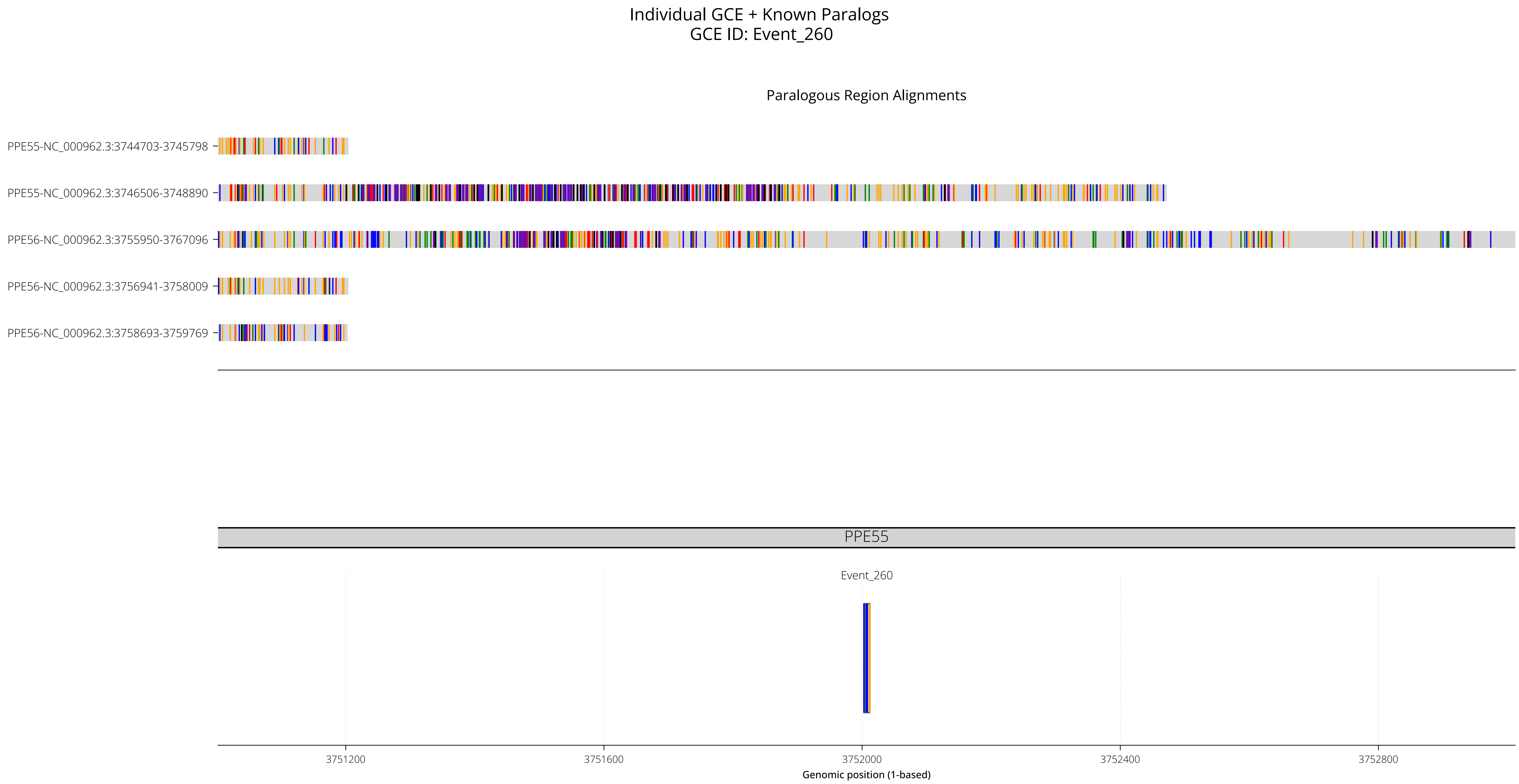

Individual GCE + Known Paralogs
GCE ID: Event_260
Paralogous Region Alignments
PPE55-NC_000962.3:3744703-3745798
PPE55-NC_000962.3:3746506-3748890
PPE56-NC_000962.3:3755950-3767096
PPE56-NC_000962.3:3756941-3758009
PPE56-NC_000962.3:3758693-3759769
PPE55
Event_260
3751200
3751600
3752000
3752400
3752800
Genomic position (1-based)

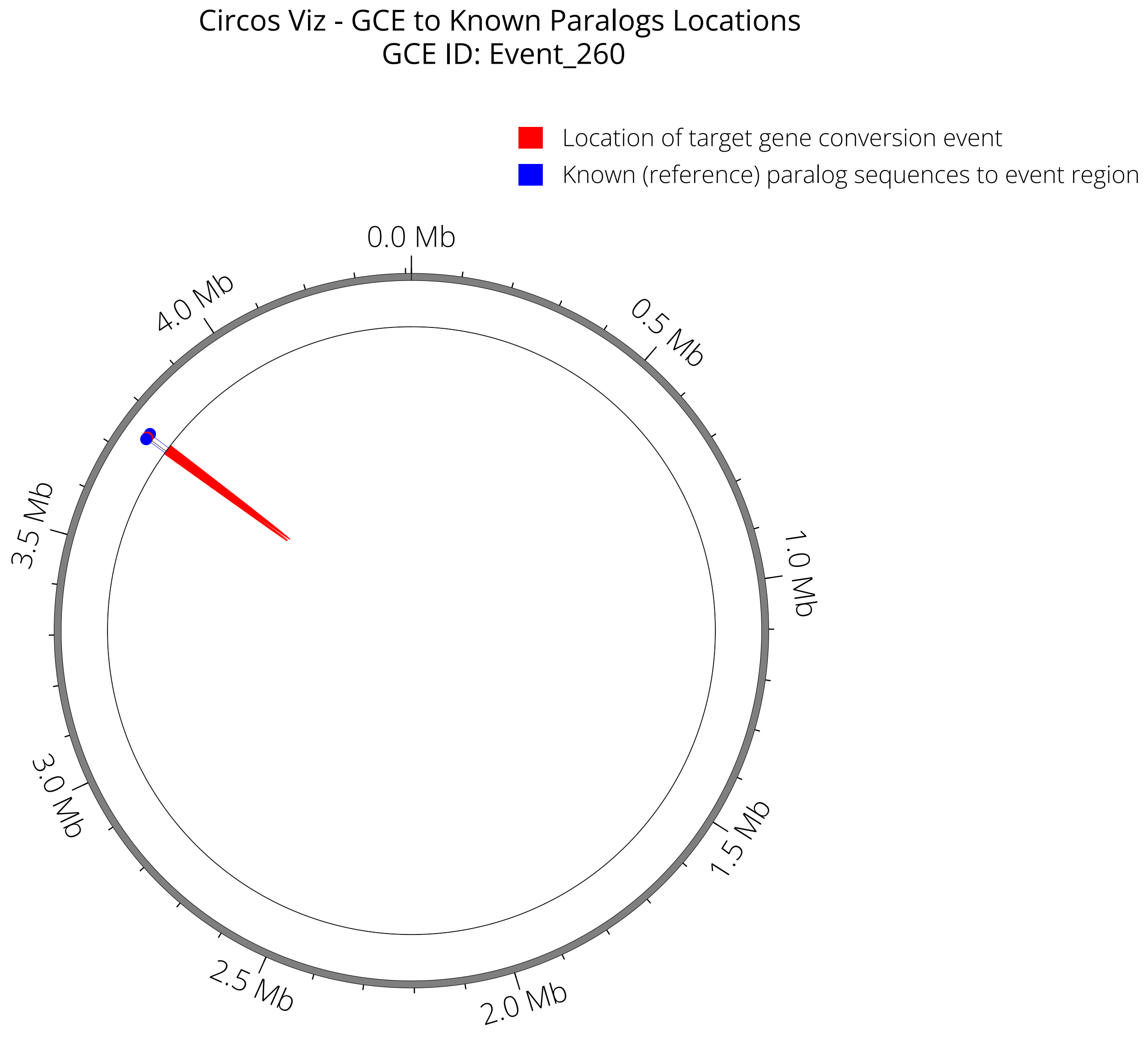
Circos Viz - GCE to Known Paralogs Locations
GCE ID: Event_260
Location of target gene conversion event
Known (reference) paralog sequences to event region
0.0 Mb
0.5 Mb
1.0 Mb
1.5 Mb
2.0 Mb
2.5 Mb
3.0 Mb
3.5 Mb
4.0 Mb

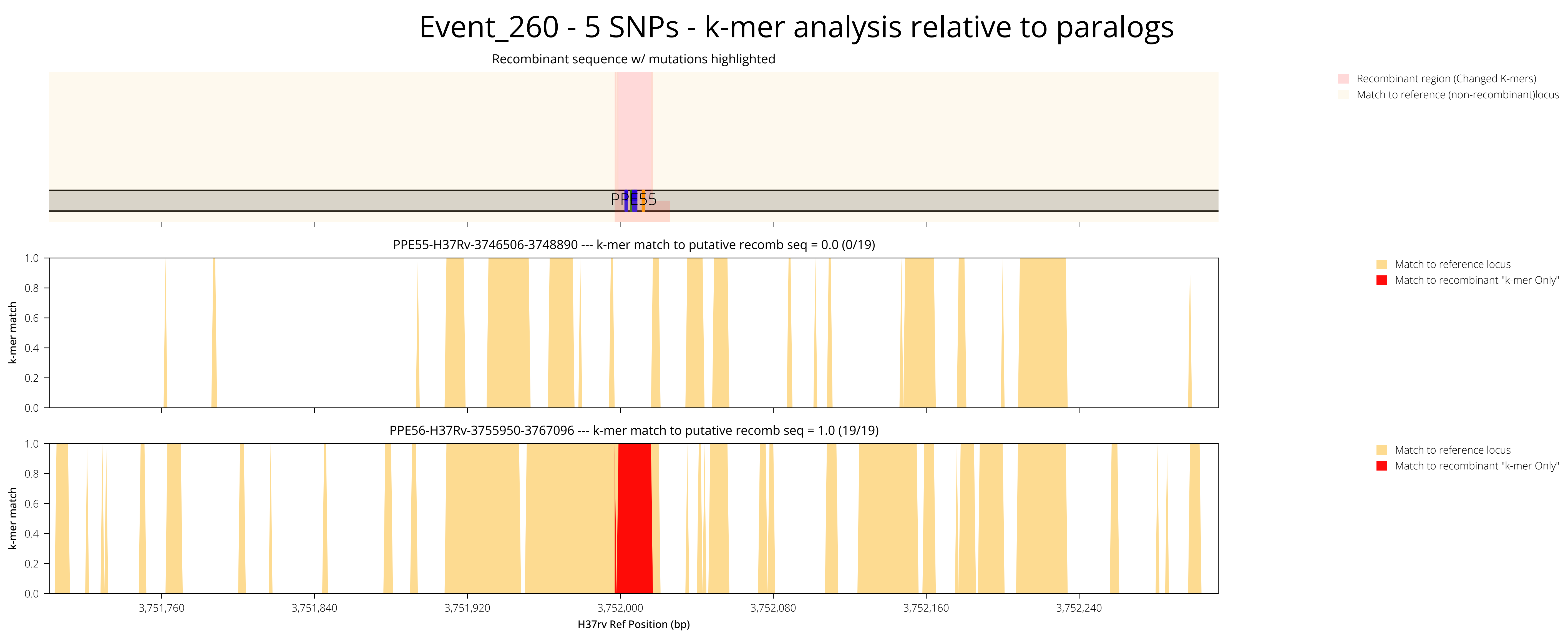

Event_260 - 5 SNPs - k-mer analysis relative to paralogs
Recombinant sequence w/ mutations highlighted
Recombinant region (Changed K-mers)
Match to reference (non-recombinant)locus
PPE55
PPE55-H37Rv-3746506-3748890 --- k-mer match to putative recomb seq = 0.0 (0/19)
Match to reference locus
Match to recombinant "k-mer Only"
k-mer match
1.0
0.8
0.6
0.4
0.2
0.0
PPE56-H37Rv-3755950-3767096 --- k-mer match to putative recomb seq = 1.0 (19/19)
Match to reference locus
Match to recombinant "k-mer Only"
k-mer match
1.0
0.8
0.6
0.4
0.2
0.0
3,751,760
3,751,840
3,751,920
3,752,000
3,752,080
3,752,160
3,752,240
H37rv Ref Position (bp)

Sub-phylogeny Viz + Paralog Alignments
GCE ID: Event_260

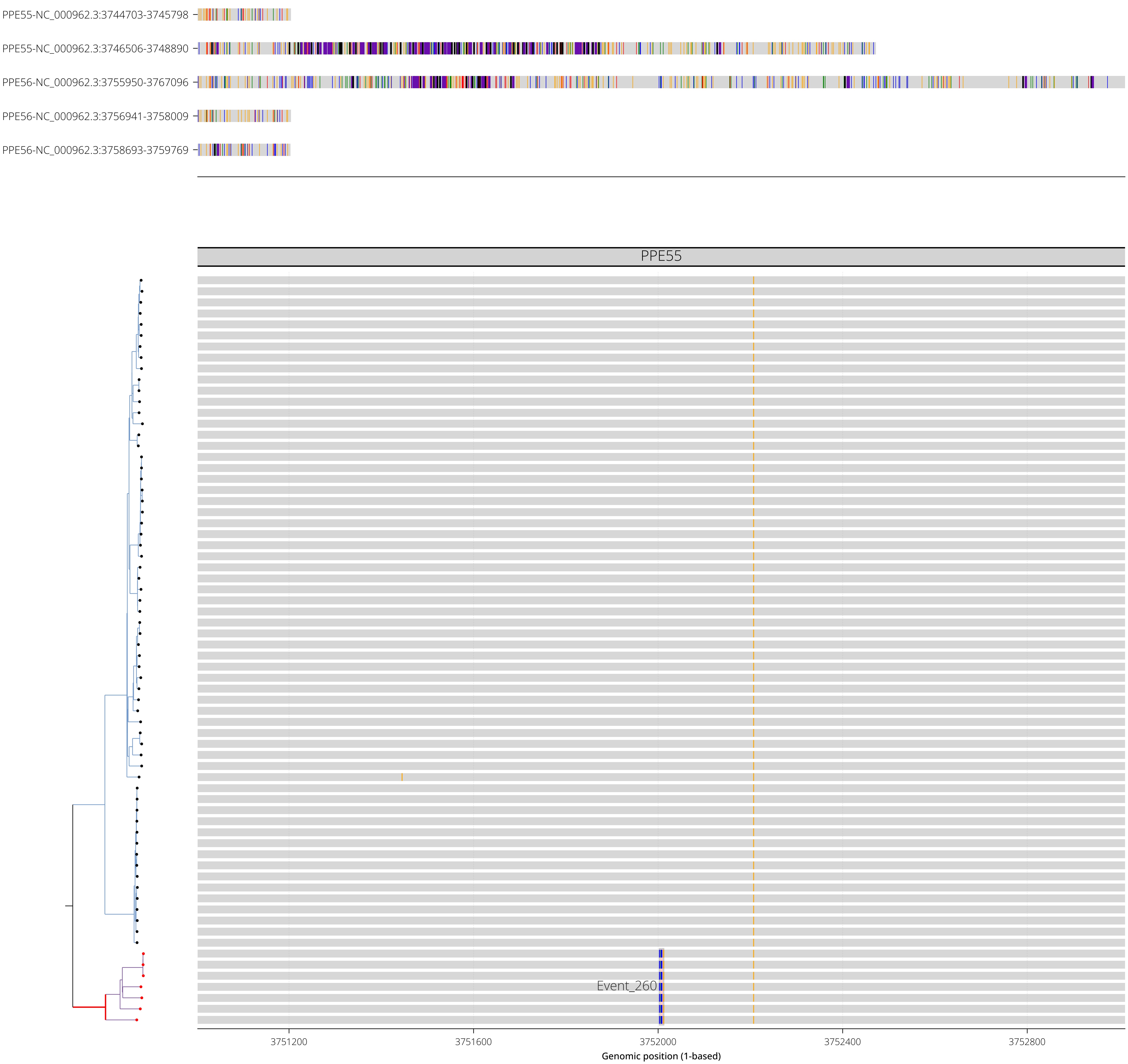

Full Phylogeny Viz + Paralog Alignments
GCE ID: Event_260

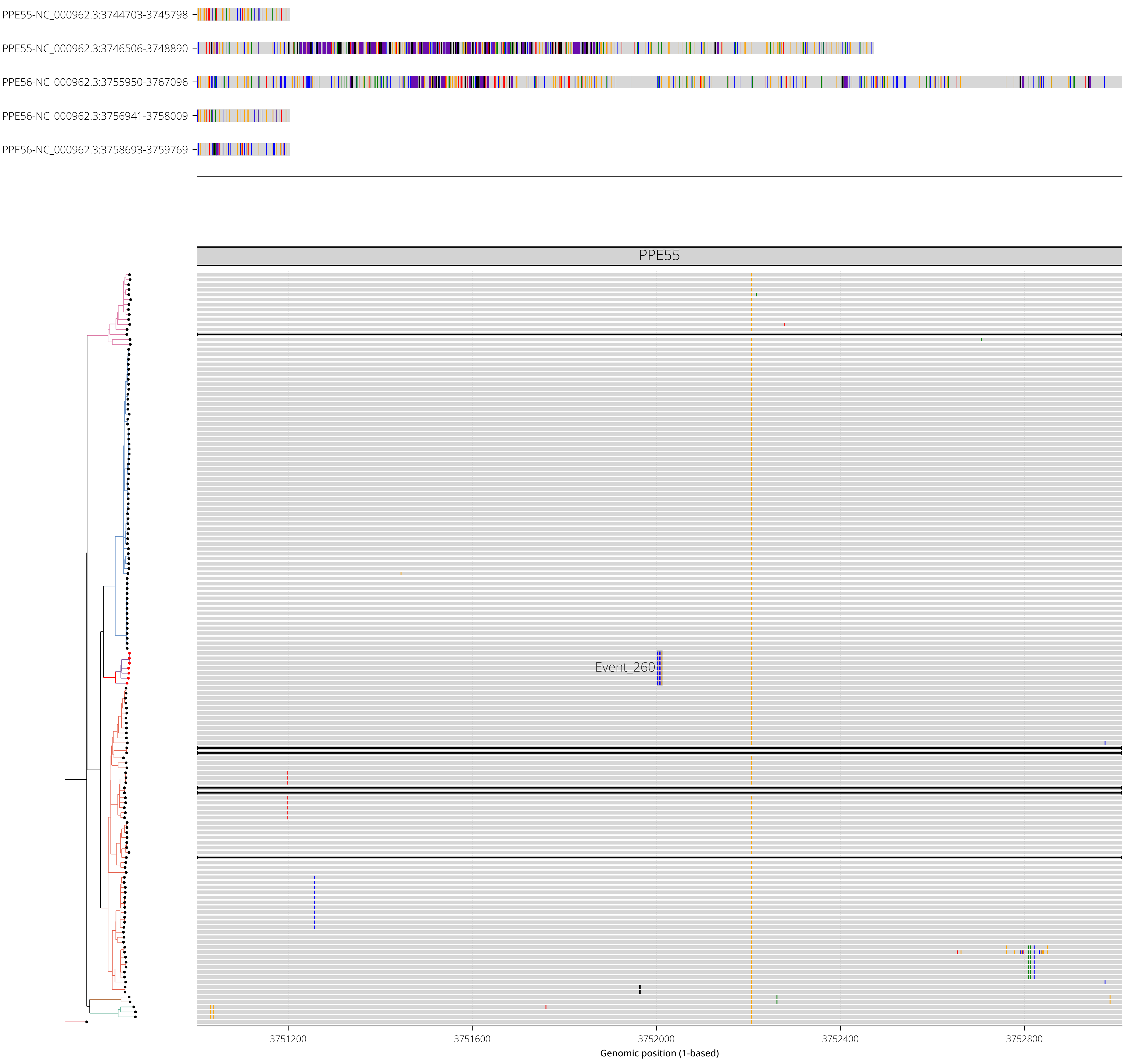

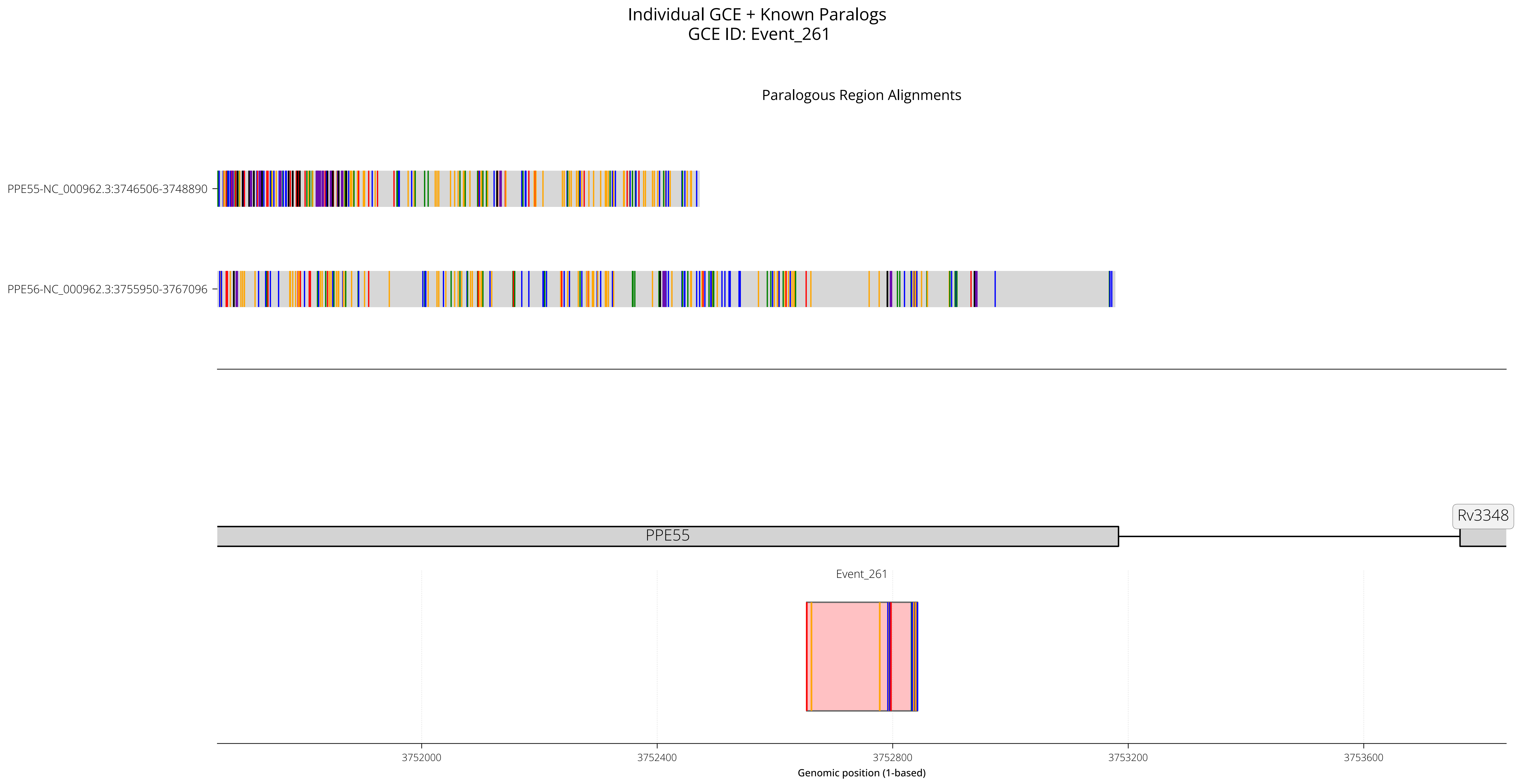

Individual GCE + Known Paralogs
GCE ID: Event_261
Paralogous Region Alignments
PPE55-NC_000962.3:3746506-3748890
PPE56-NC_000962.3:3755950-3767096
Rv3348
PPE55
Event_261
3752000
3752400
3752800
3753200
3753600
Genomic position (1-based)

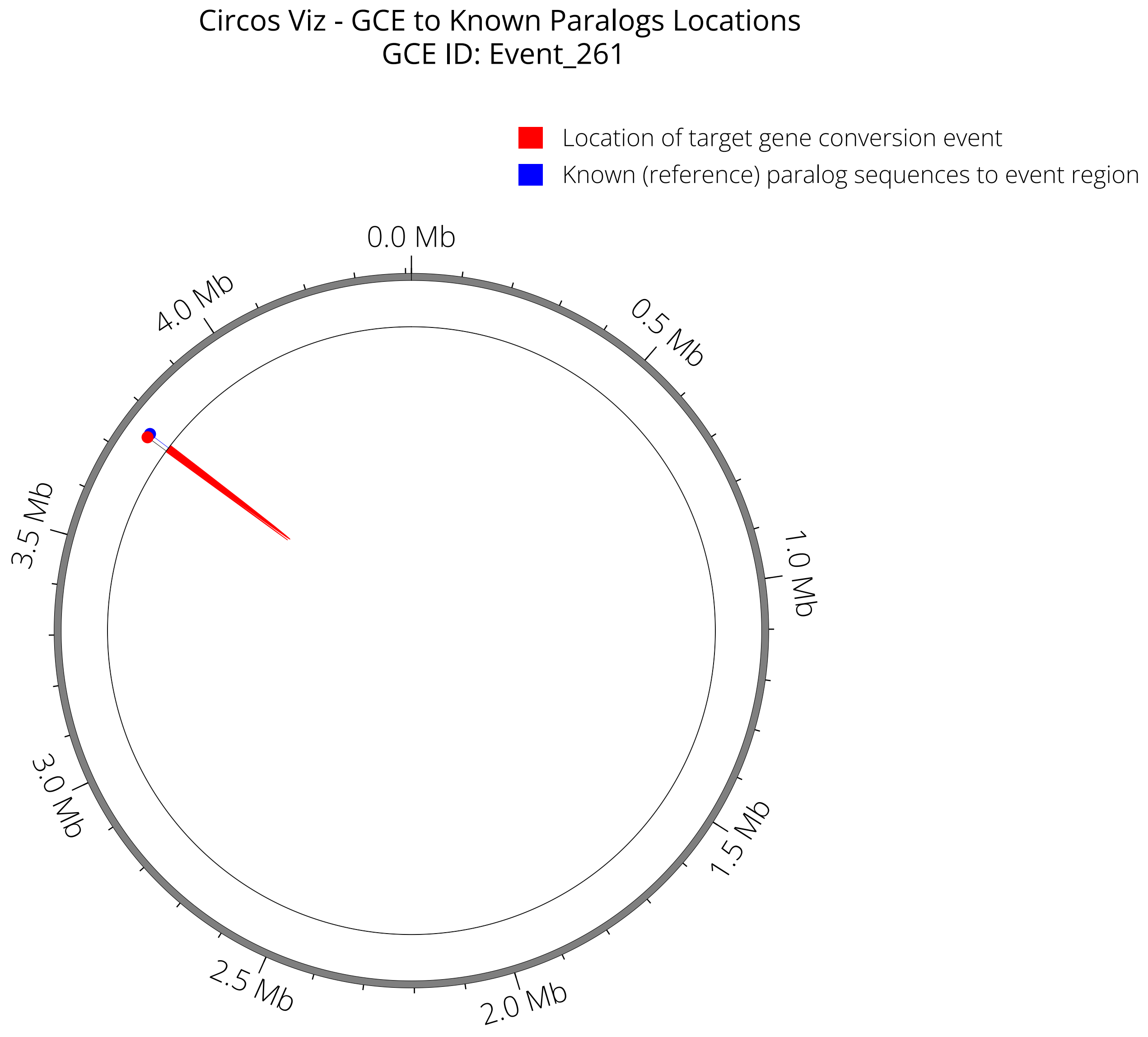
Circos Viz - GCE to Known Paralogs Locations
GCE ID: Event_261
Location of target gene conversion event
Known (reference) paralog sequences to event region
0.0 Mb
0.5 Mb
1.0 Mb
1.5 Mb
2.0 Mb
2.5 Mb
3.0 Mb
3.5 Mb
4.0 Mb

### Event_261 - 14 SNPs - k-mer analysis relative to paralogs

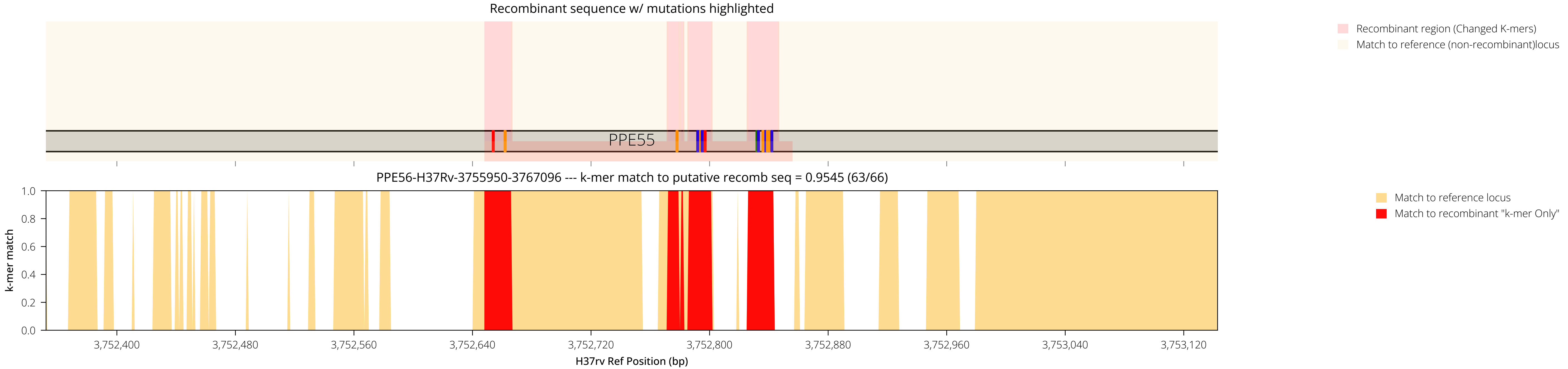

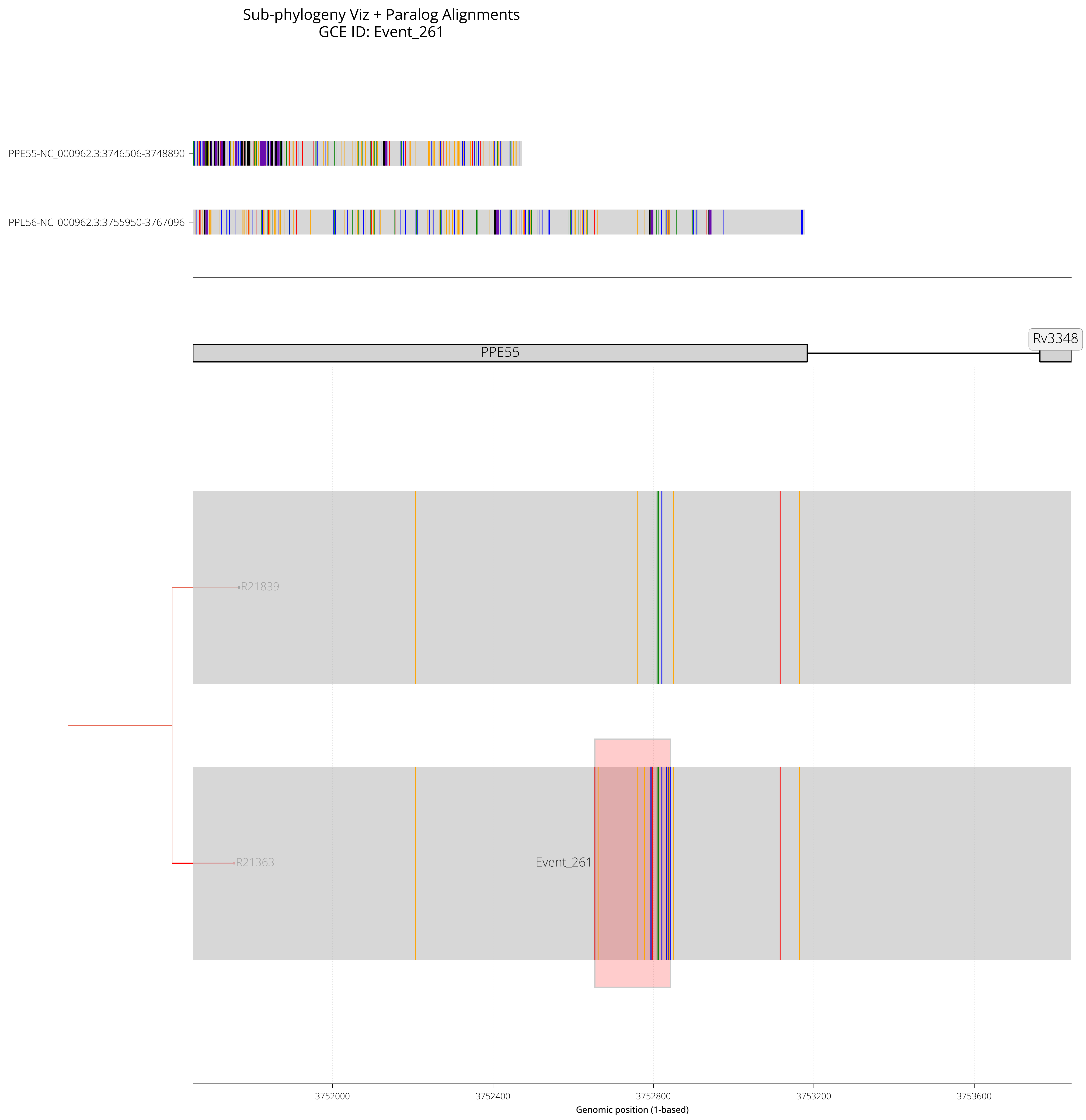
Sub-phylogeny Viz + Paralog Alignments
GCE ID: Event_261
PPE55-NC_000962.3:3746506-3748890
PPE56-NC_000962.3:3755950-3767096
PPE55
Rv3348
R21839
R21363
Event_261
3752000
3752400
3752800
3753200
3753600
Genomic position (1-based)

Full Phylogeny Viz + Paralog Alignments
GCE ID: Event_261

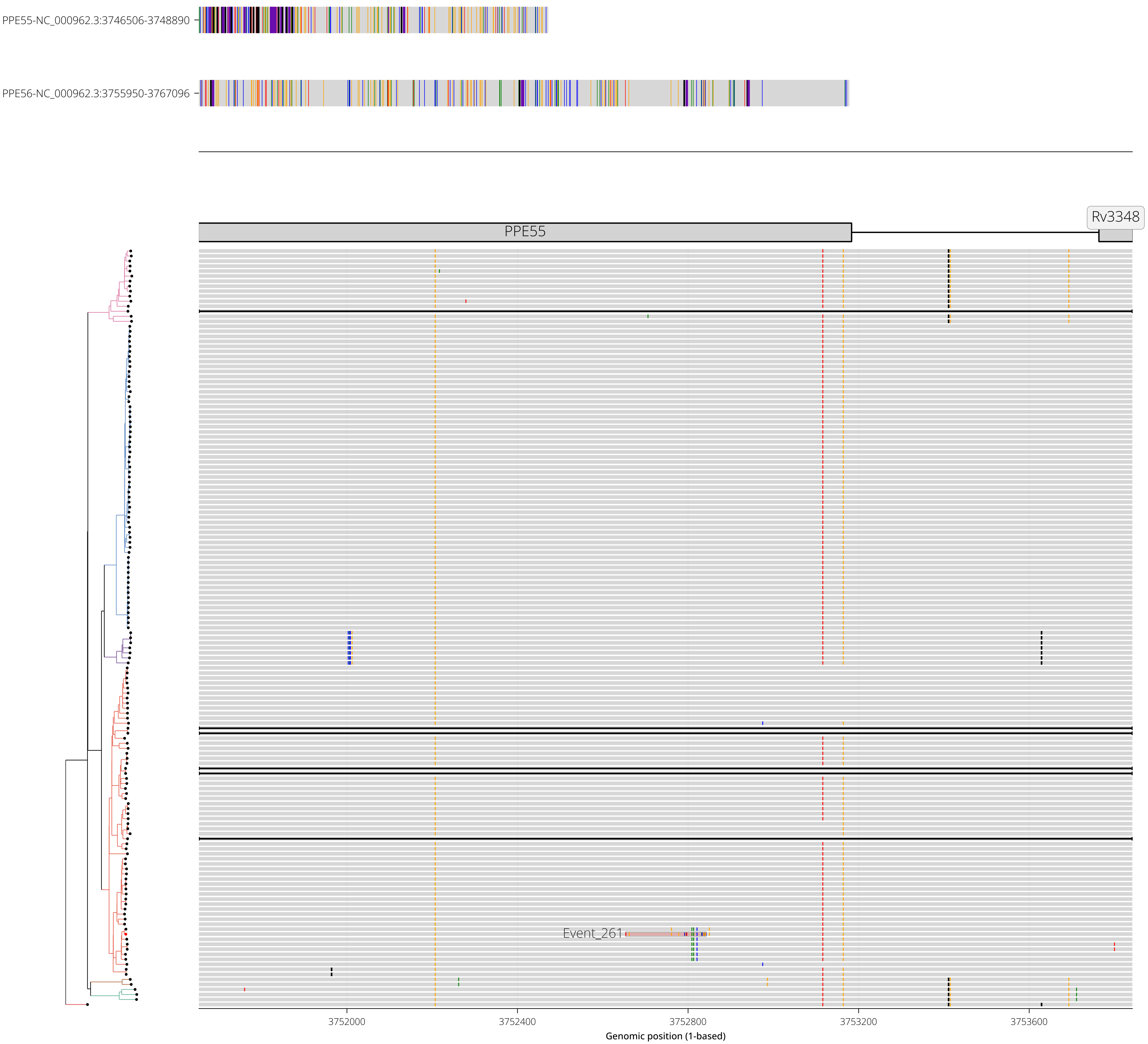

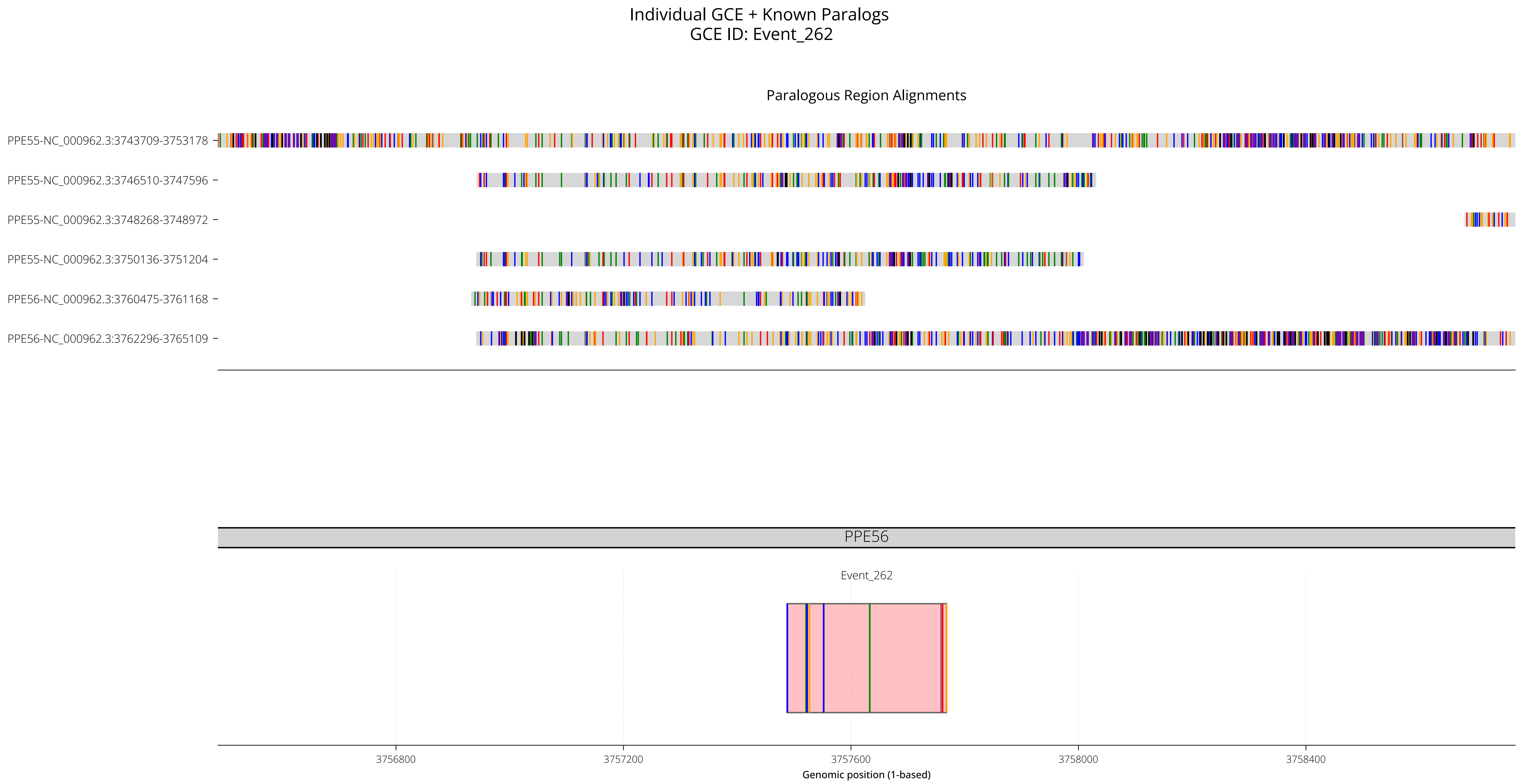

Individual GCE + Known Paralogs
GCE ID: Event_262
Paralogous Region Alignments
PPE55-NC_000962.3:3743709-3753178
PPE55-NC_000962.3:3746510-3747596
PPE55-NC_000962.3:3748268-3748972
PPE55-NC_000962.3:3750136-3751204
PPE56-NC_000962.3:3760475-3761168
PPE56-NC_000962.3:3762296-3765109
PPE56
Event_262
3756800
3757200
3757600
3758000
3758400
Genomic position (1-based)

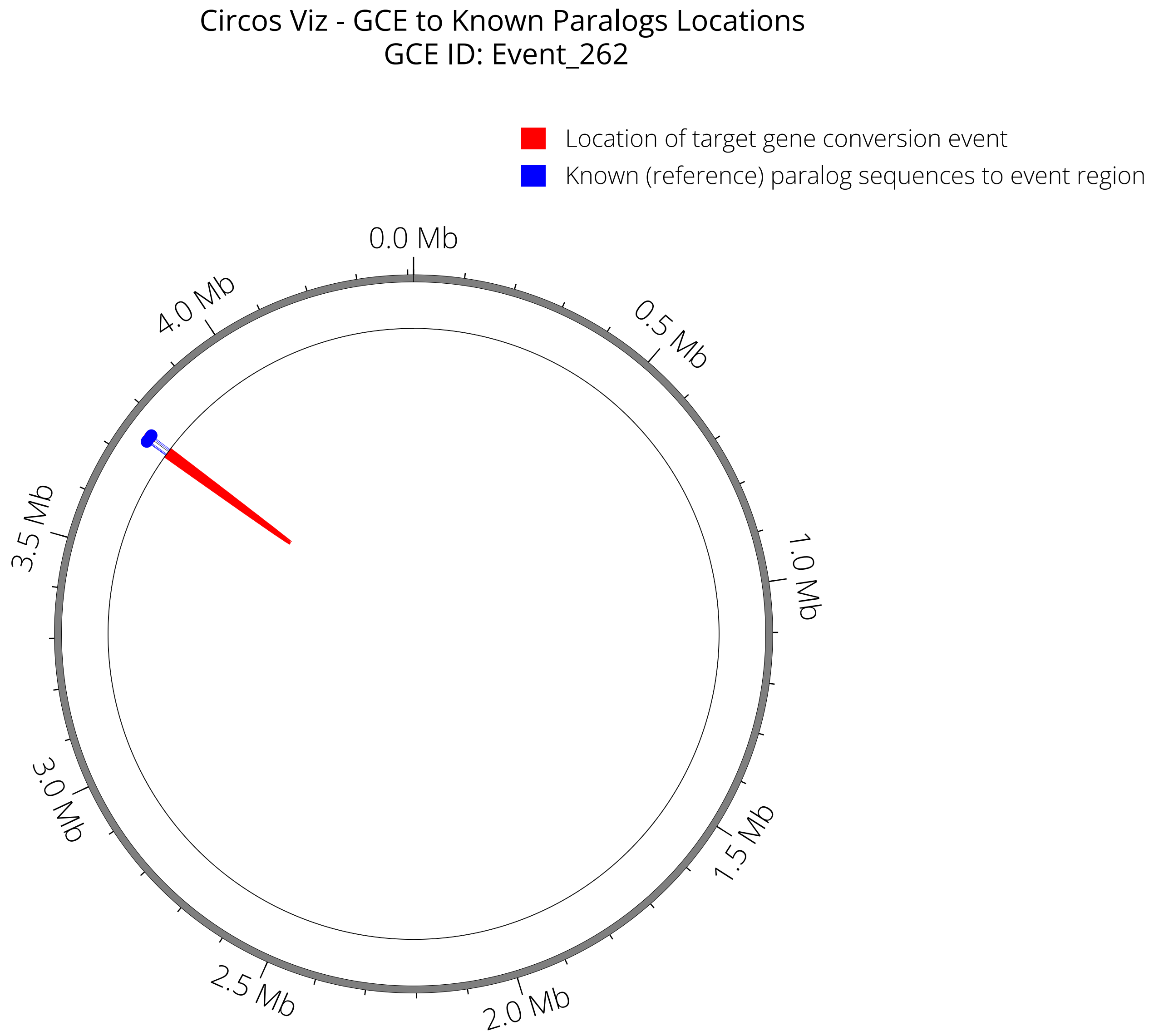
Circos Viz - GCE to Known Paralogs Locations
GCE ID: Event_262
Location of target gene conversion event
Known (reference) paralog sequences to event region
0.0 Mb
0.5 Mb
1.0 Mb
1.5 Mb
2.0 Mb
2.5 Mb
3.0 Mb
3.5 Mb
4.0 Mb

### Event_262 - 10 SNPs - k-mer analysis relative to paralogs

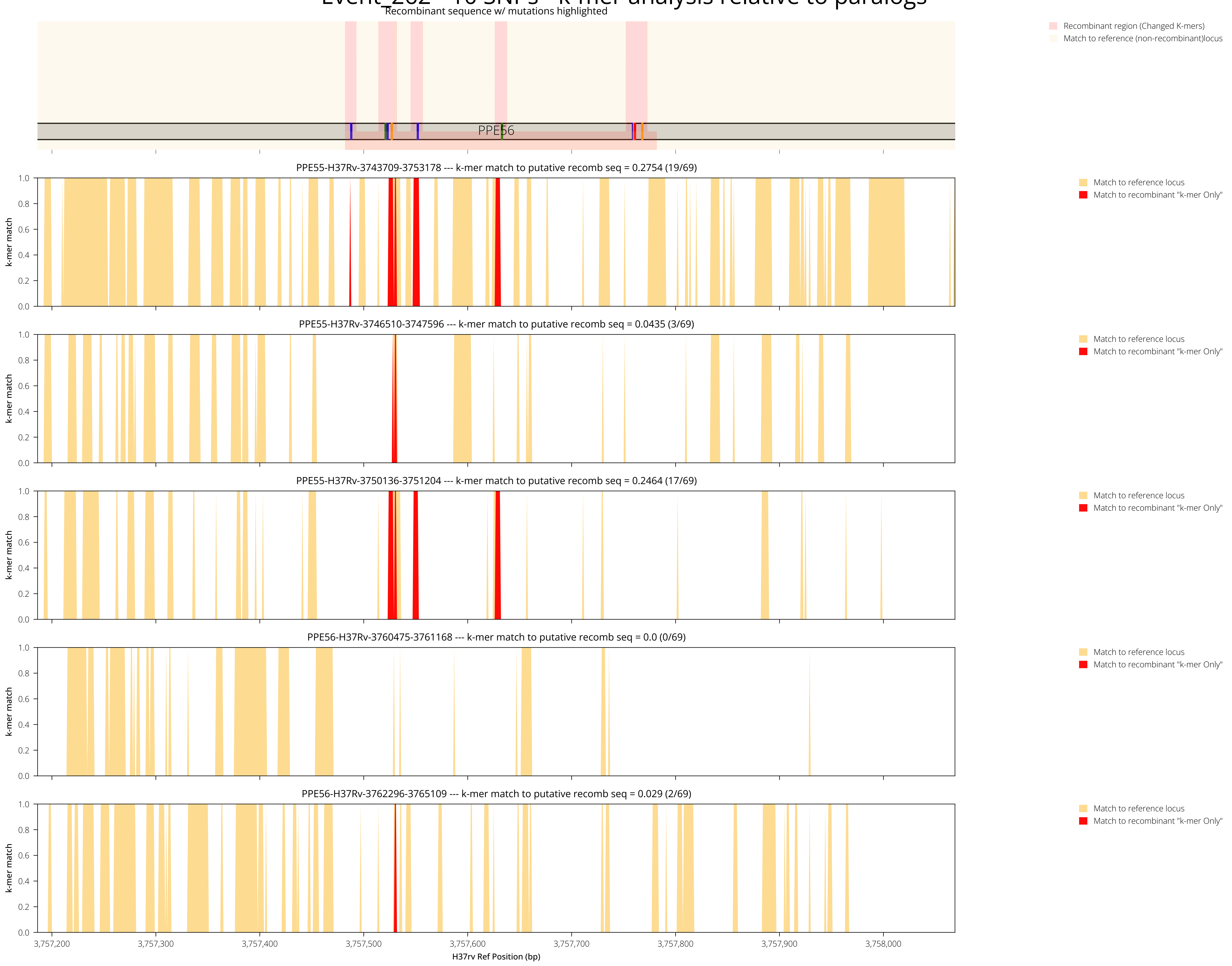

Sub-phylogeny Viz + Paralog Alignments
GCE ID: Event_262

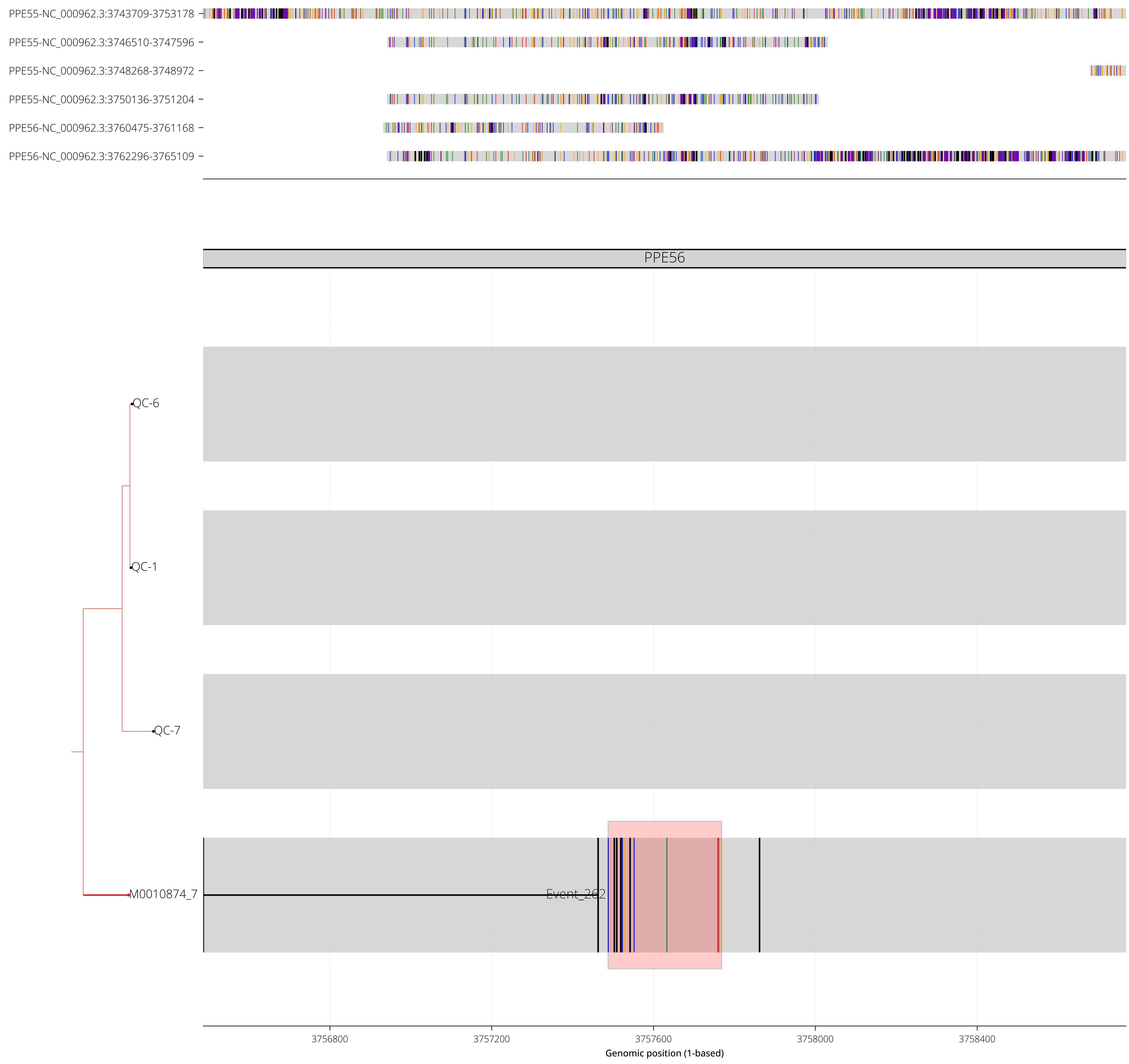

### Full Phylogeny Viz + Paralog Alignments
GCE ID: Event_262

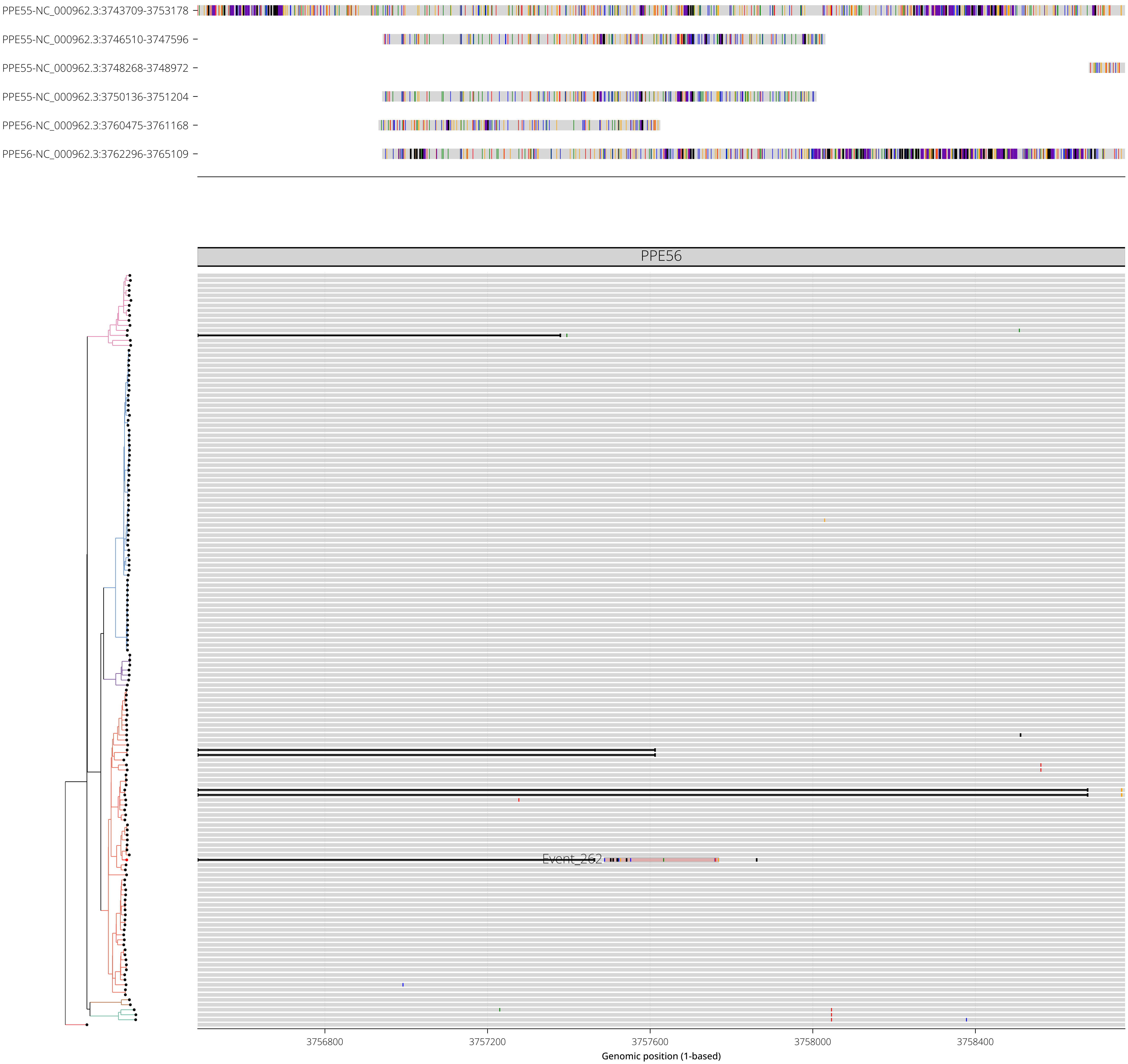

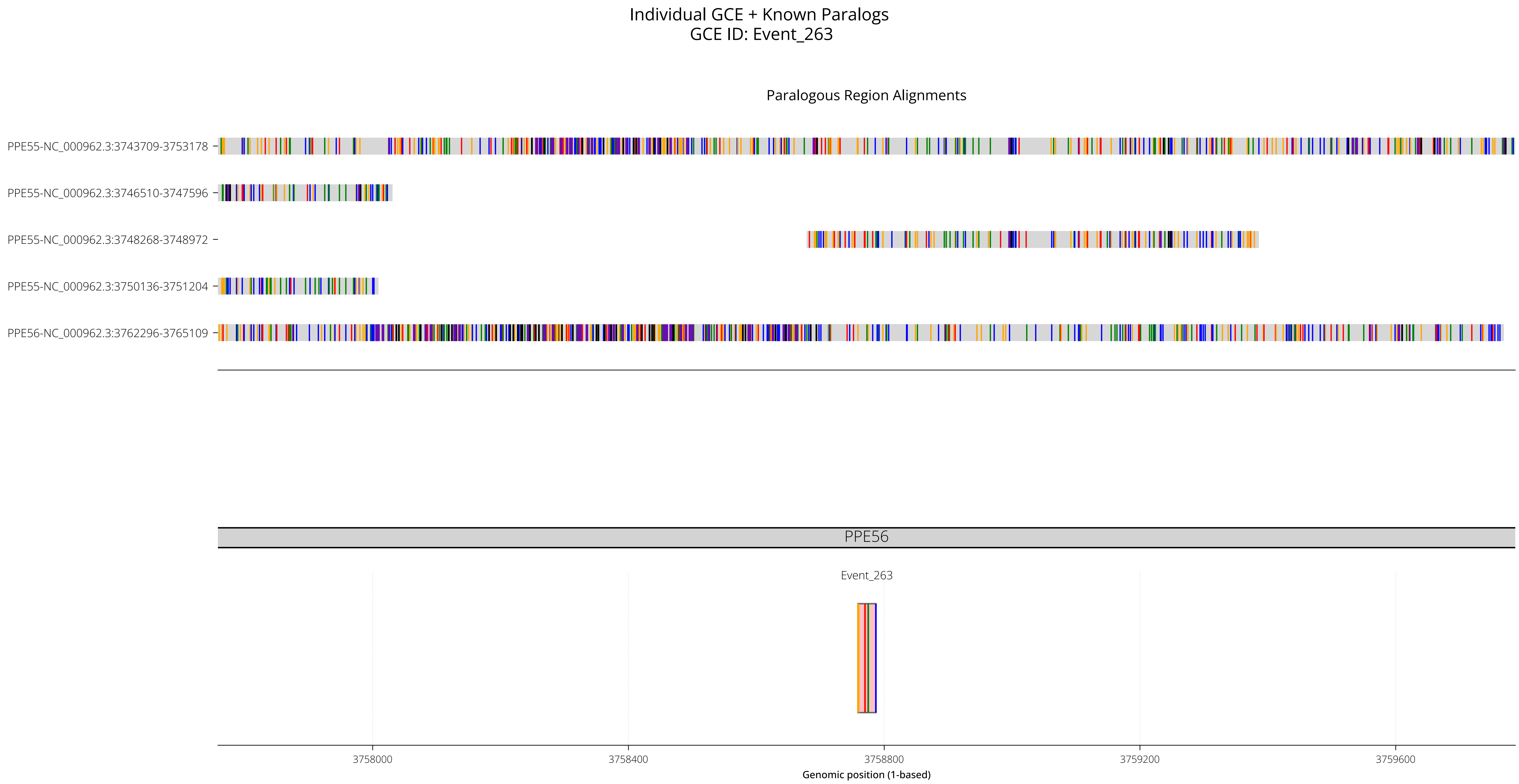
Individual GCE + Known Paralogs
GCE ID: Event_263
Paralogous Region Alignments
PPE55-NC_000962.3:3743709-3753178
PPE55-NC_000962.3:3746510-3747596
PPE55-NC_000962.3:3748268-3748972
PPE55-NC_000962.3:3750136-3751204
PPE56-NC_000962.3:3762296-3765109
PPE56
Event_263
3758000
3758400
3758800
3759200
3759600
Genomic position (1-based)

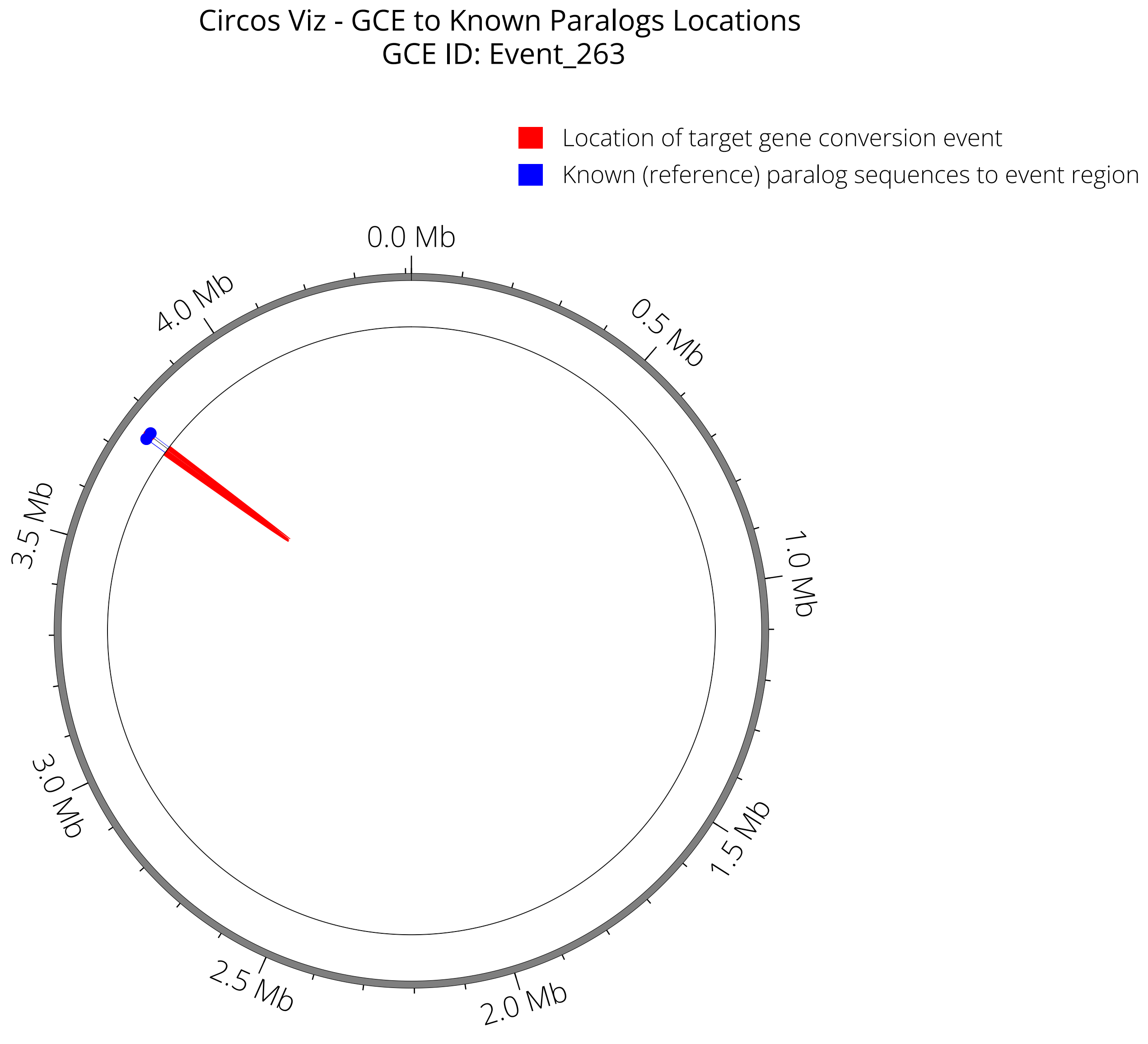

Circos Viz - GCE to Known Paralogs Locations
GCE ID: Event_263
Location of target gene conversion event
Known (reference) paralog sequences to event region
0.0 Mb
0.5 Mb
1.0 Mb
1.5 Mb
2.0 Mb
2.5 Mb
3.0 Mb
3.5 Mb
4.0 Mb

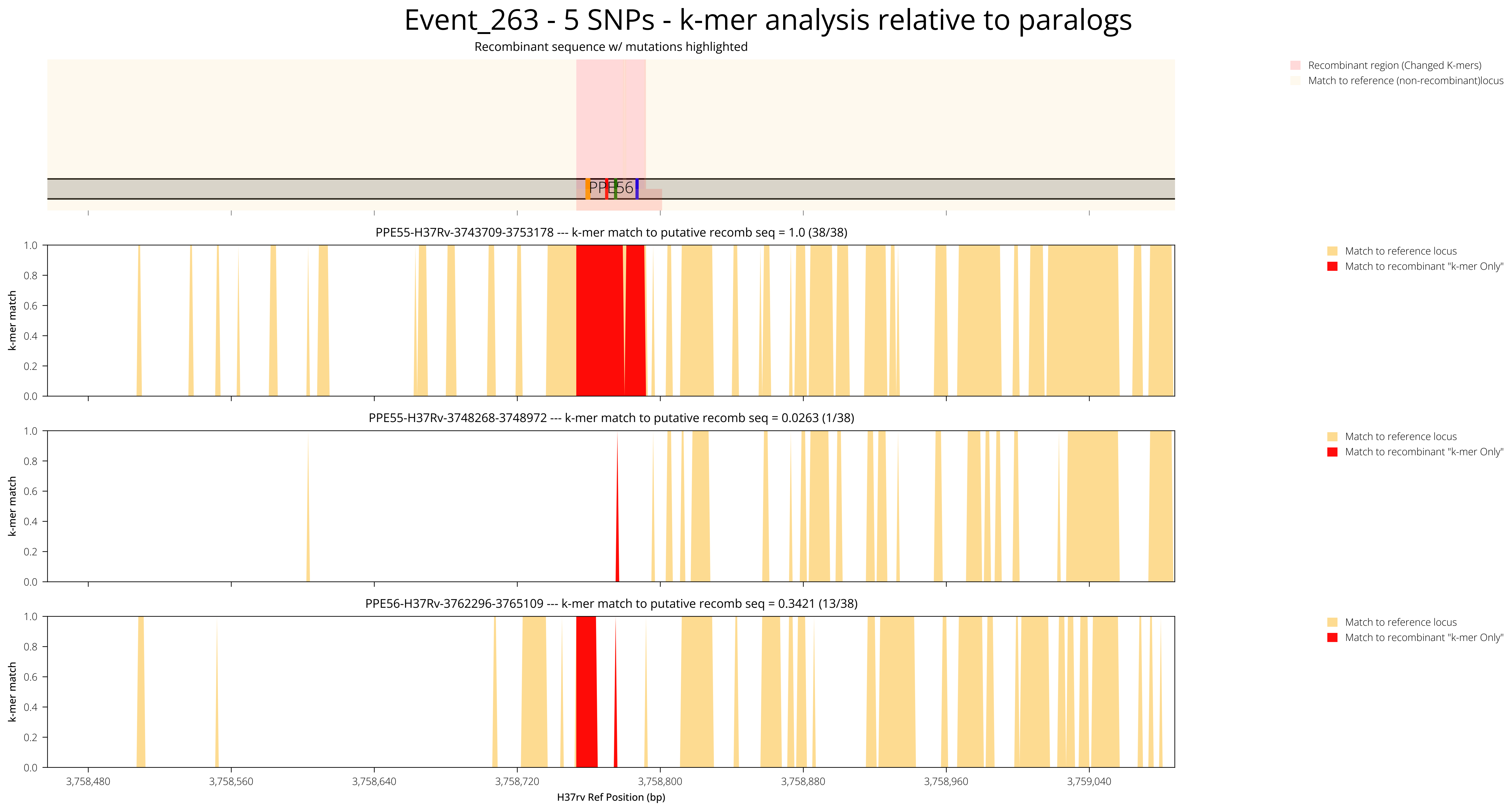

Event_263 - 5 SNPs - k-mer analysis relative to paralogs
Recombinant sequence w/ mutations highlighted
Recombinant region (Changed K-mers)
Match to reference (non-recombinant)locus
PPE56
PPE55-H37Rv-3743709-3753178 --- k-mer match to putative recomb seq = 1.0 (38/38)
PPE55-H37Rv-3748268-3748972 --- k-mer match to putative recomb seq = 0.0263 (1/38)
PPE56-H37Rv-3762296-3765109 --- k-mer match to putative recomb seq = 0.3421 (13/38)
Match to reference locus
Match to recombinant "k-mer Only"
k-mer match
0.0
0.2
0.4
0.6
0.8
1.0
3,758,480
3,758,560
3,758,640
3,758,720
3,758,800
3,758,880
3,758,960
3,759,040
H37rv Ref Position (bp)

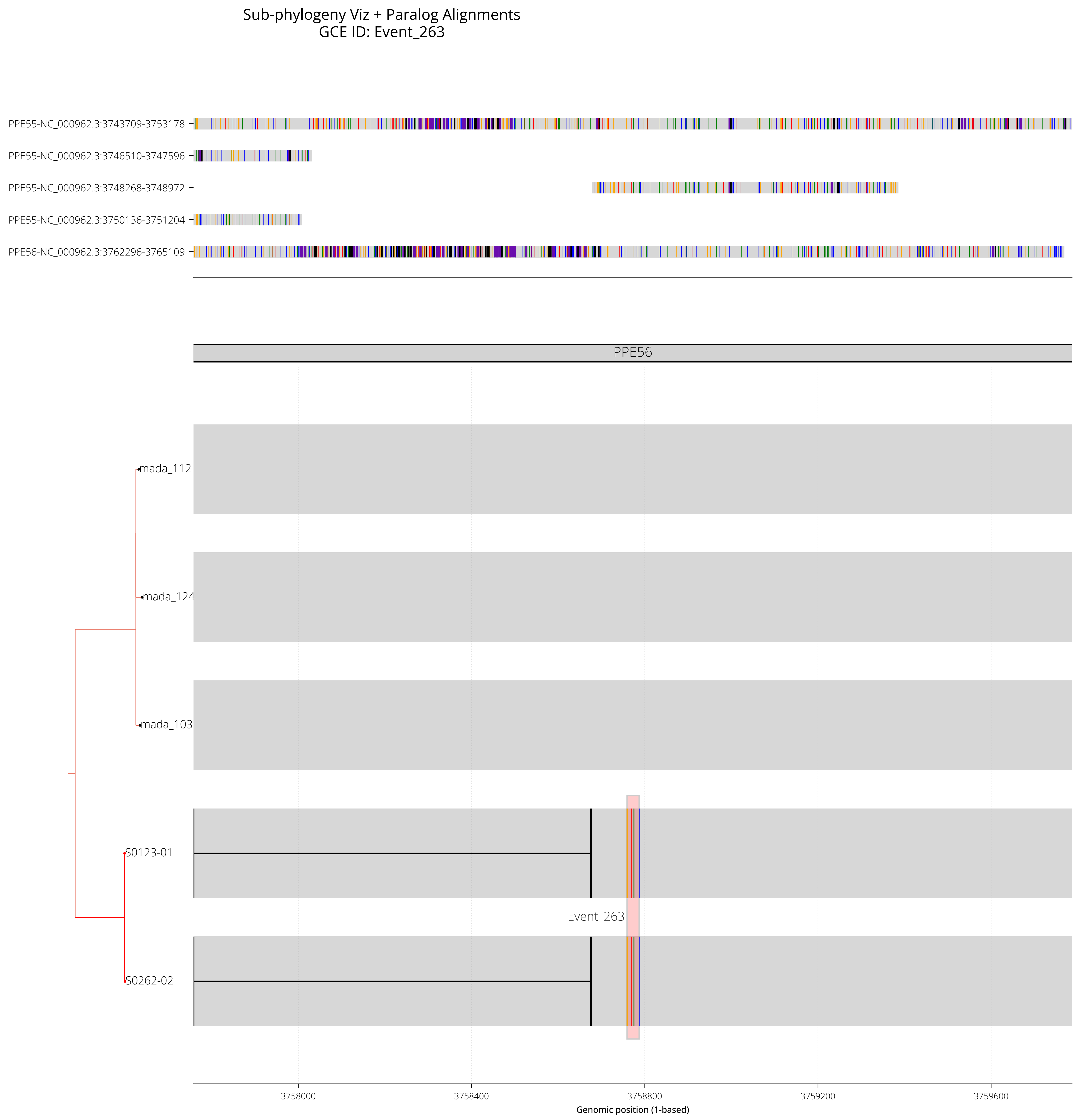

Sub-phylogeny Viz + Paralog Alignments
GCE ID: Event_263
PPE55-NC_000962.3:3743709-3753178
PPE55-NC_000962.3:3746510-3747596
PPE55-NC_000962.3:3748268-3748972
PPE55-NC_000962.3:3750136-3751204
PPE56-NC_000962.3:3762296-3765109
PPE56
mada_112
mada_124
mada_103
S0123-01
S0262-02
Event_263
3758000
3758400
3758800
3759200
3759600
Genomic position (1-based)

Full Phylogeny Viz + Paralog Alignments
GCE ID: Event_263

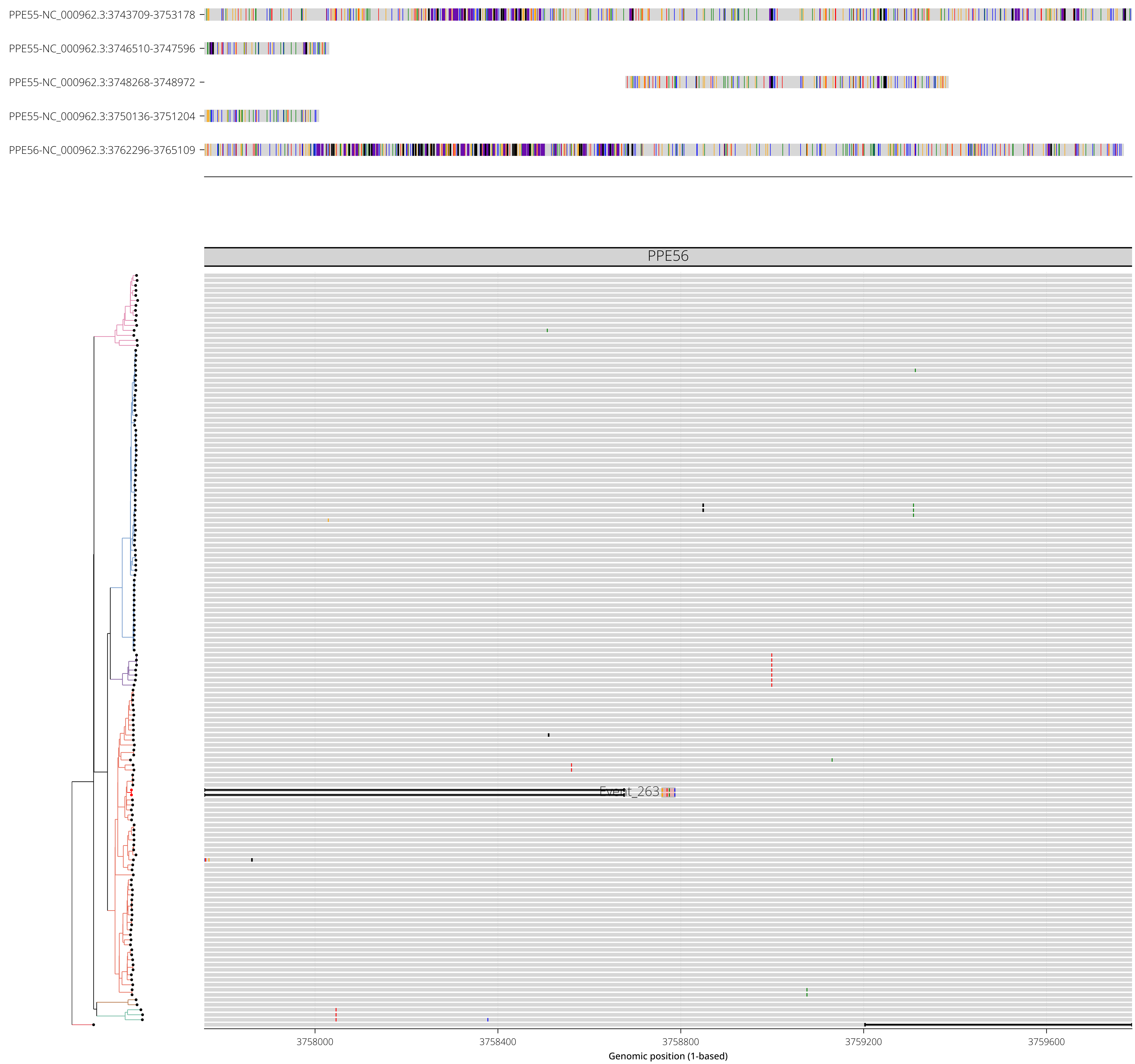

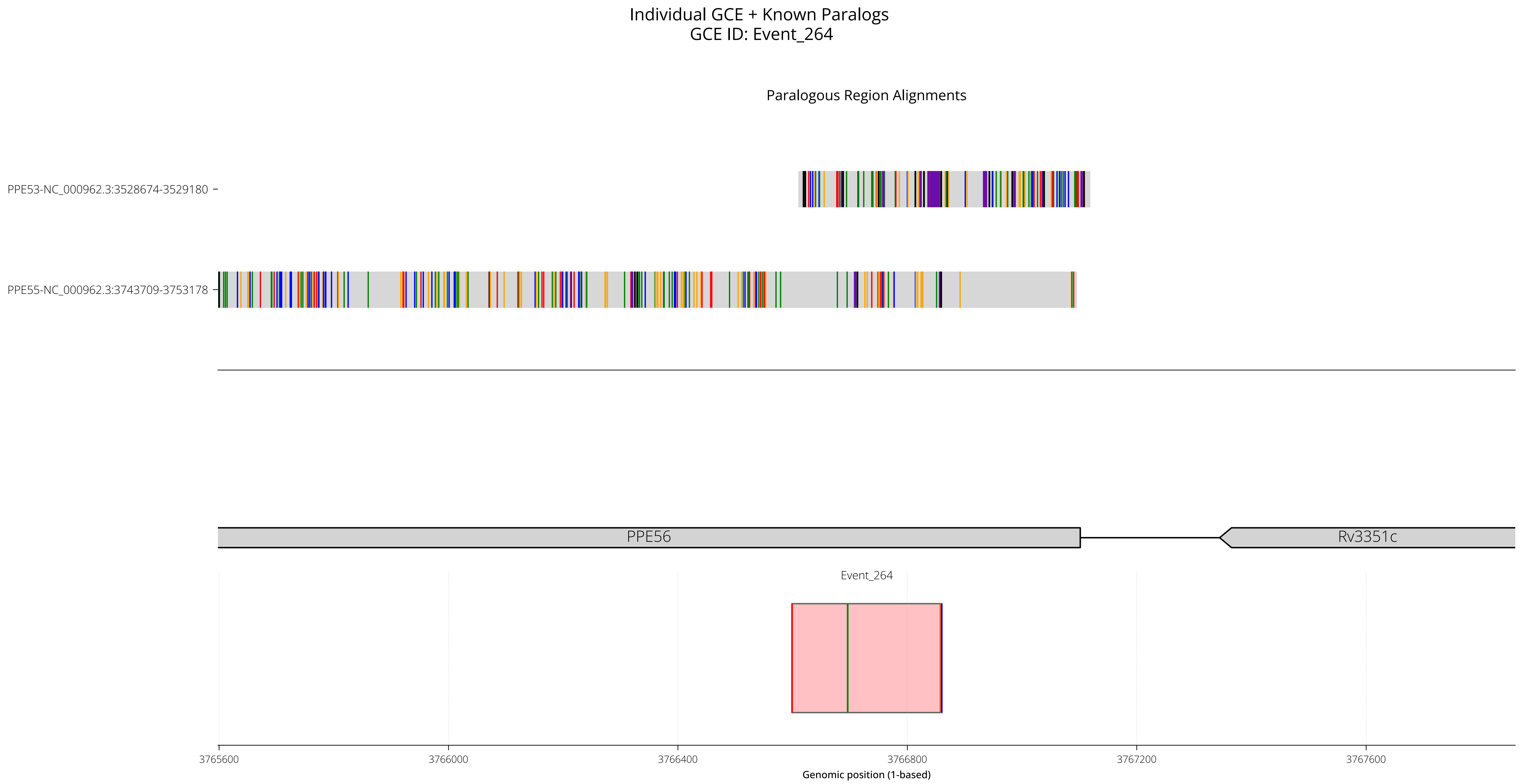
Individual GCE + Known Paralogs
GCE ID: Event_264
Paralogous Region Alignments
PPE53-NC_000962.3:3528674-3529180
PPE55-NC_000962.3:3743709-3753178
PPE56
Rv3351c
Event_264
3765600
3766000
3766400
3766800
3767200
3767600
Genomic position (1-based)

Circos Viz - GCE to Known Paralogs Locations
GCE ID: Event_264

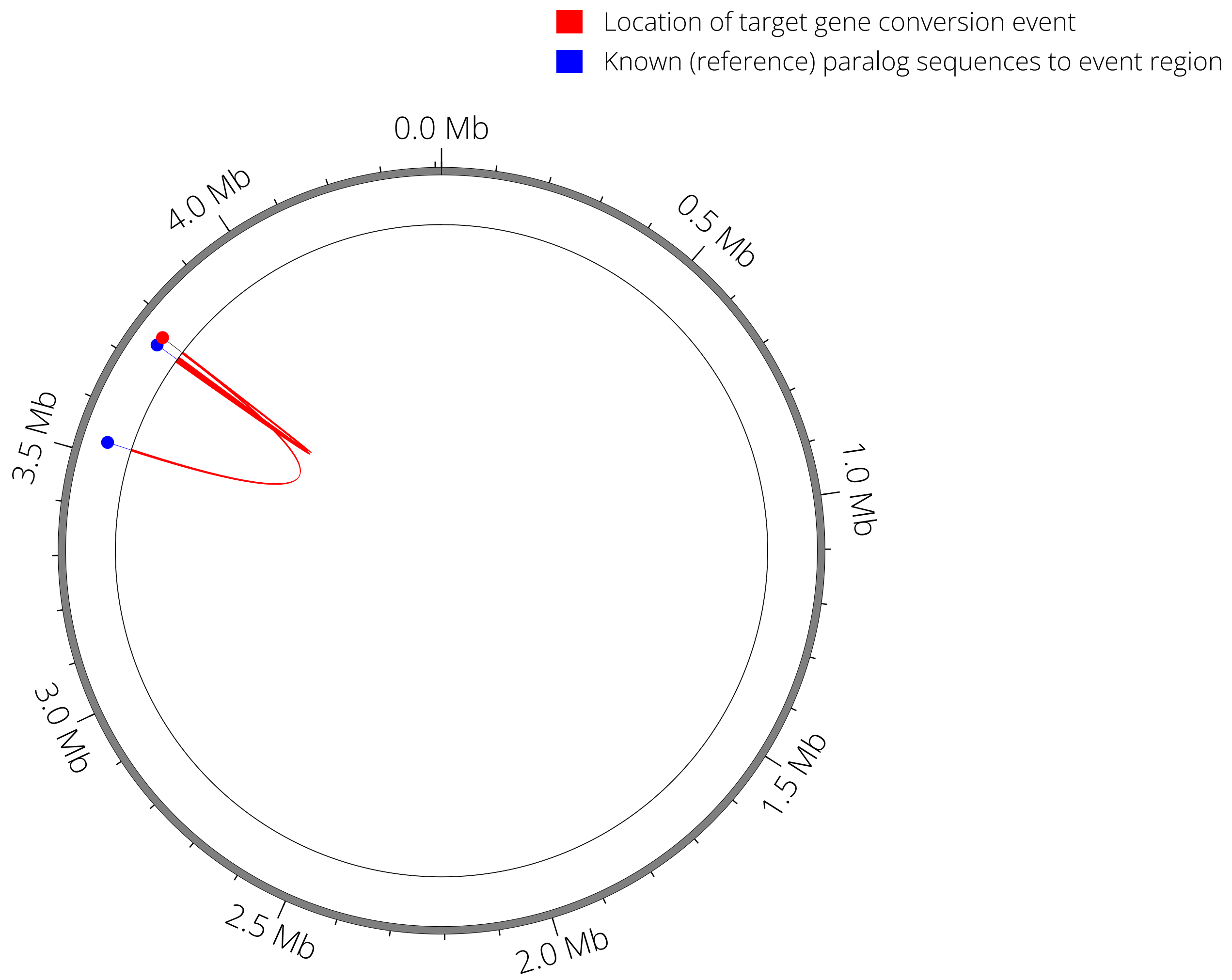

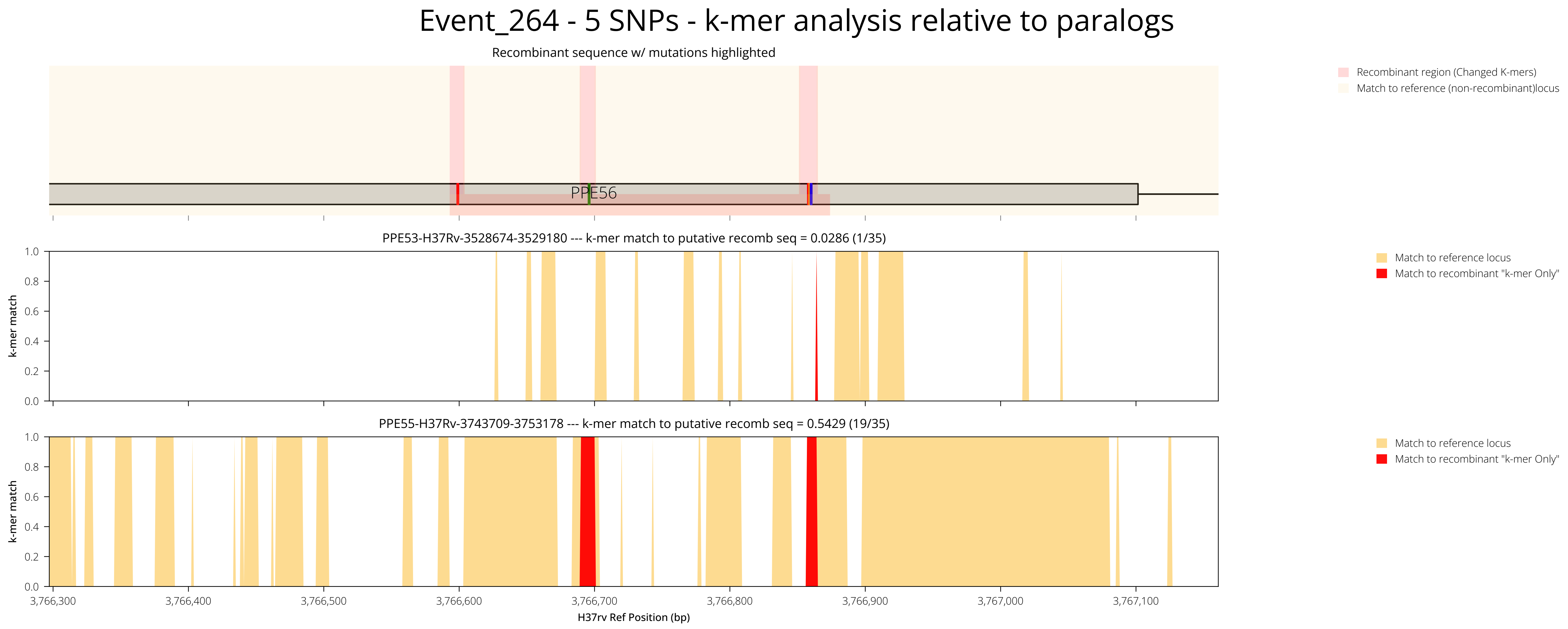
Event_264 - 5 SNPs - k-mer analysis relative to paralogs
Recombinant sequence w/ mutations highlighted
PPE56
Recombinant region (Changed K-mers)
Match to reference (non-recombinant)locus
PPE53-H37Rv-3528674-3529180 --- k-mer match to putative recomb seq = 0.0286 (1/35)
Match to reference locus
Match to recombinant "k-mer Only"
PPE55-H37Rv-3743709-3753178 --- k-mer match to putative recomb seq = 0.5429 (19/35)
Match to reference locus
Match to recombinant "k-mer Only"
k-mer match
1.0
0.8
0.6
0.4
0.2
0.0
3,766,300
3,766,400
3,766,500
3,766,600
3,766,700
3,766,800
3,766,900
3,767,000
3,767,100
H37rv Ref Position (bp)

Sub-phylogeny Viz + Paralog Alignments
GCE ID: Event_264

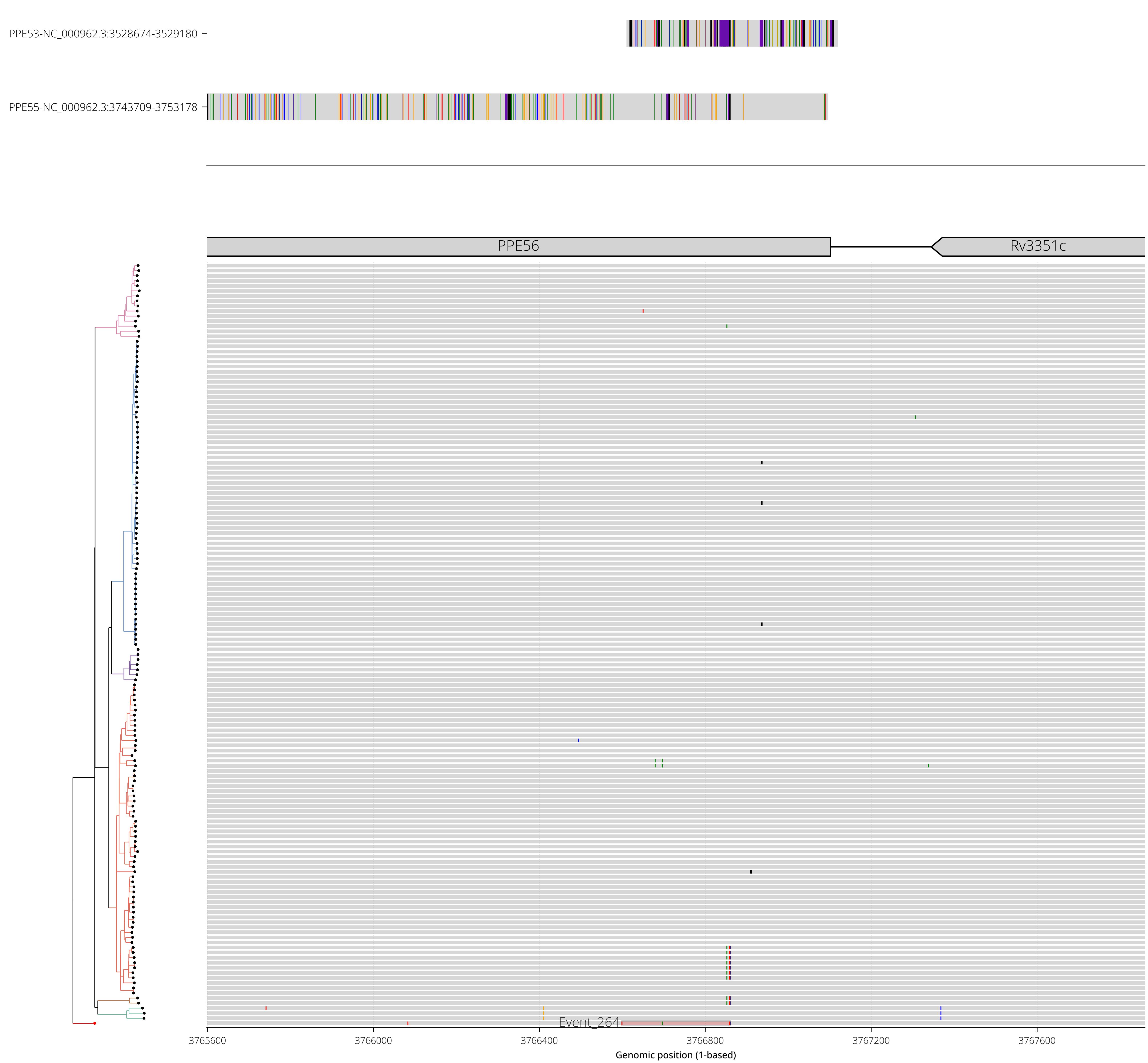

Full Phylogeny Viz + Paralog Alignments
GCE ID: Event_264

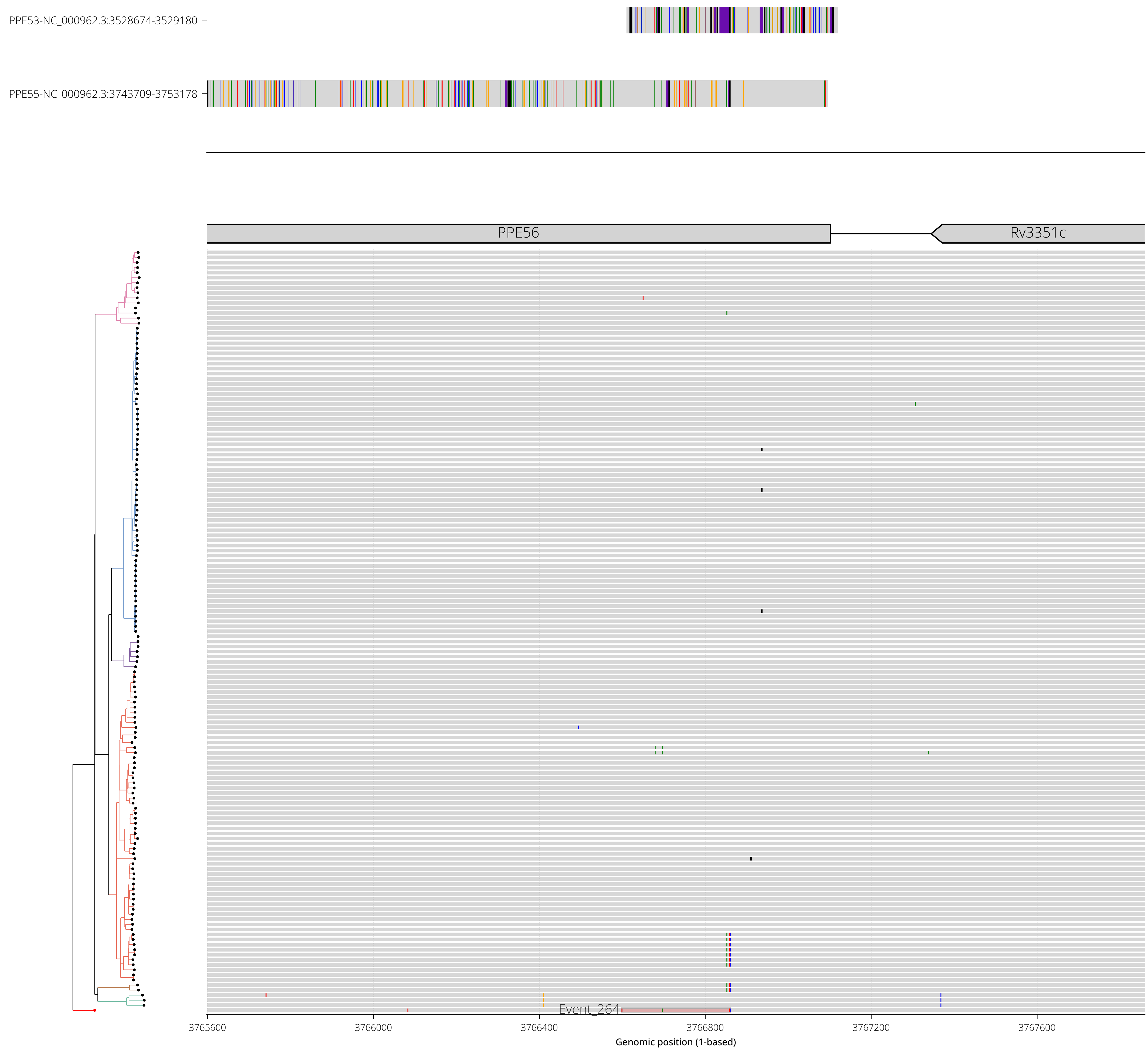

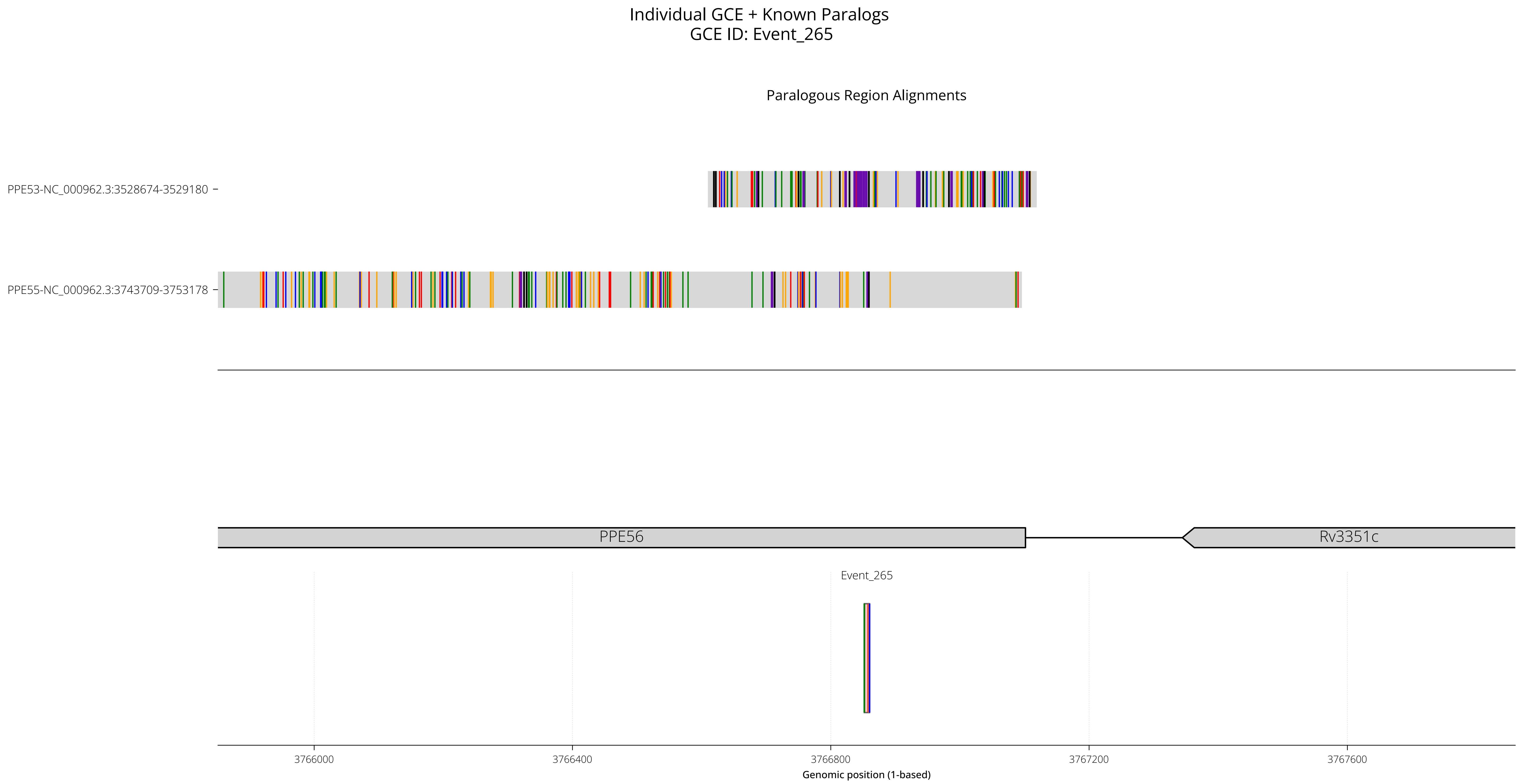

Individual GCE + Known Paralogs
GCE ID: Event_265
Paralogous Region Alignments
PPE53-NC_000962.3:3528674-3529180
PPE55-NC_000962.3:3743709-3753178
PPE56
Rv3351c
Event_265
3766000
3766400
3766800
3767200
3767600
Genomic position (1-based)

Circos Viz - GCE to Known Paralogs Locations
GCE ID: Event_265

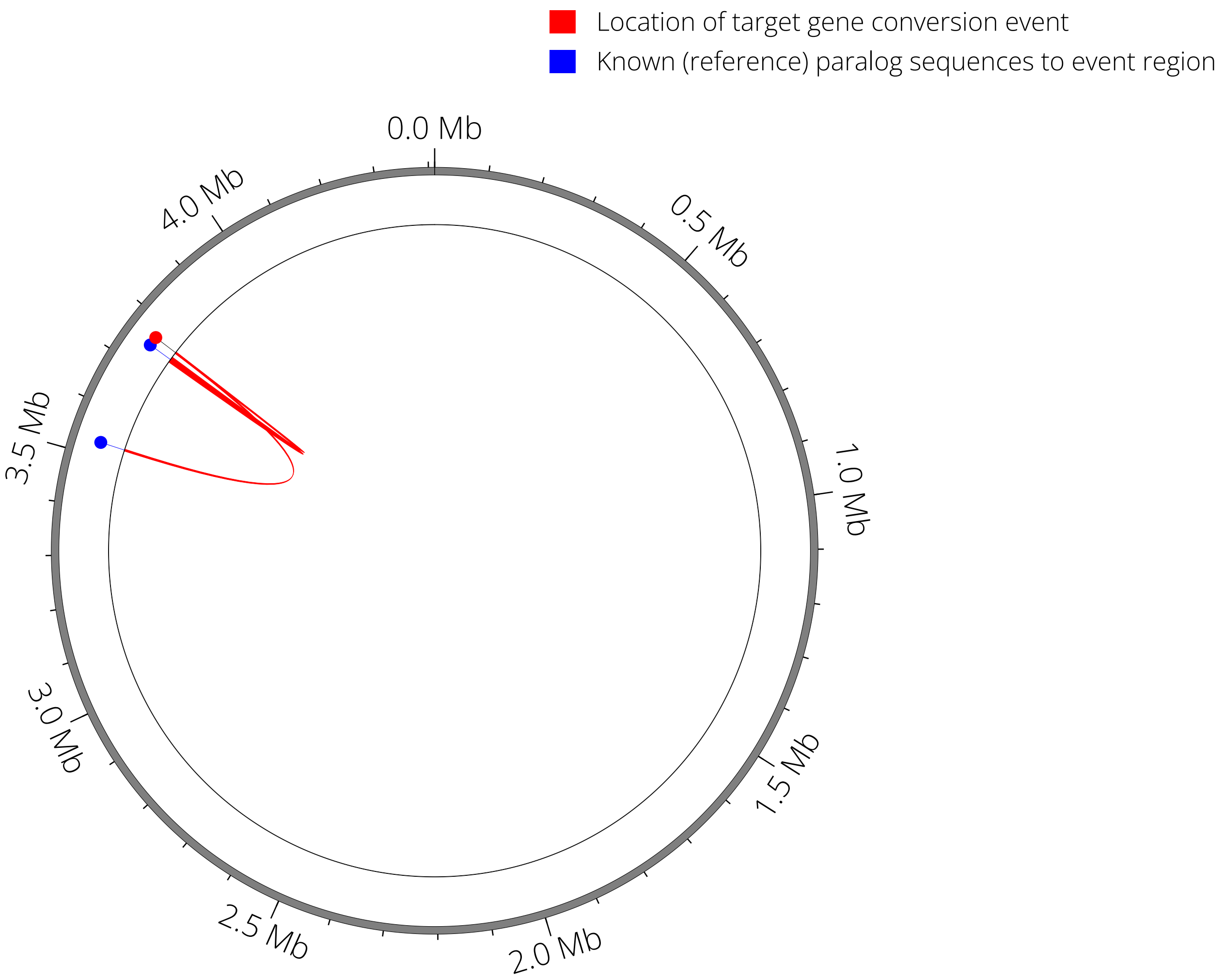

### Event_265 - 4 SNPs - k-mer analysis relative to paralogs

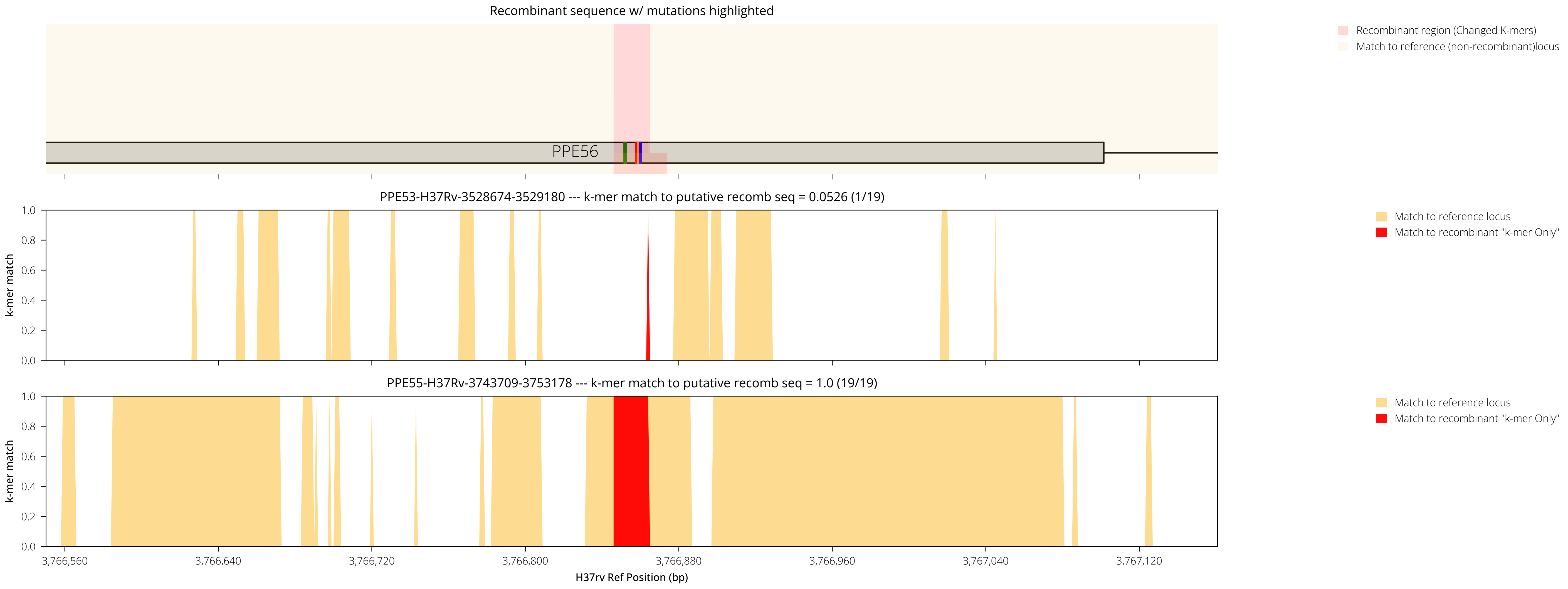

Sub-phylogeny Viz + Paralog Alignments
GCE ID: Event_265

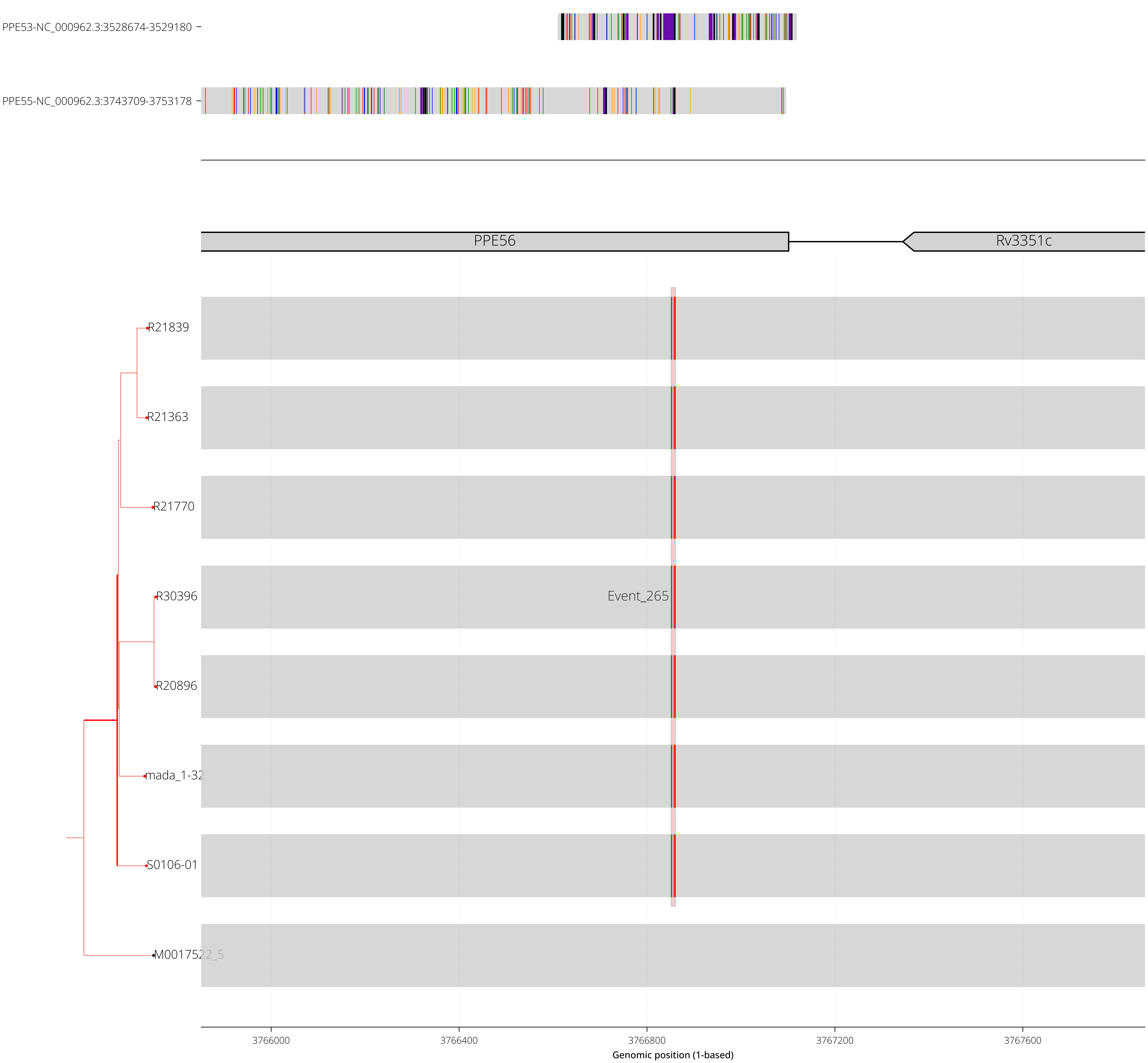

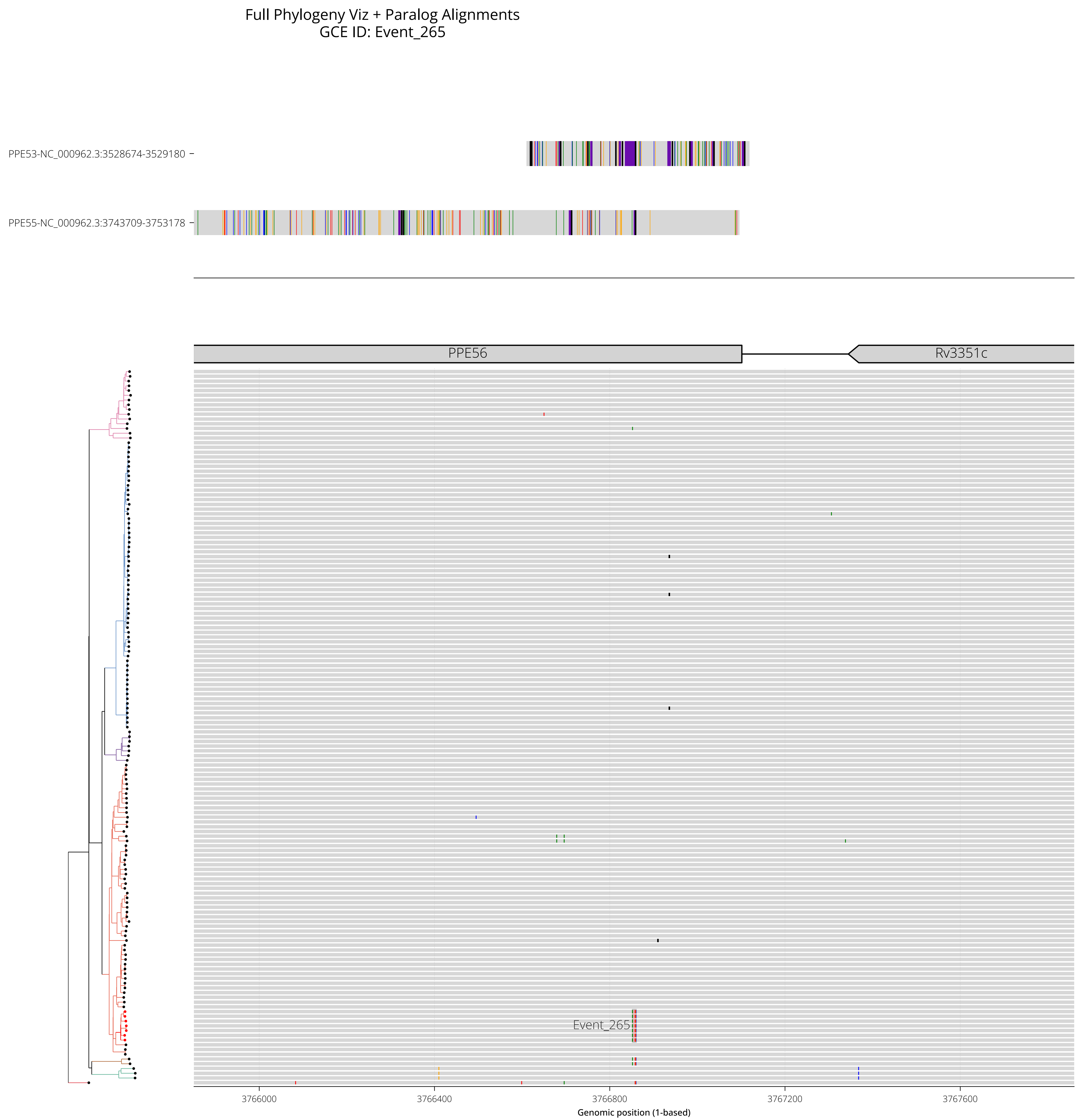

Full Phylogeny Viz + Paralog Alignments
GCE ID: Event_265
PPE53-NC_000962.3:3528674-3529180
PPE55-NC_000962.3:3743709-3753178
PPE56
Rv3351c
Event_265
3766000
3766400
3766800
3767200
3767600
Genomic position (1-based)

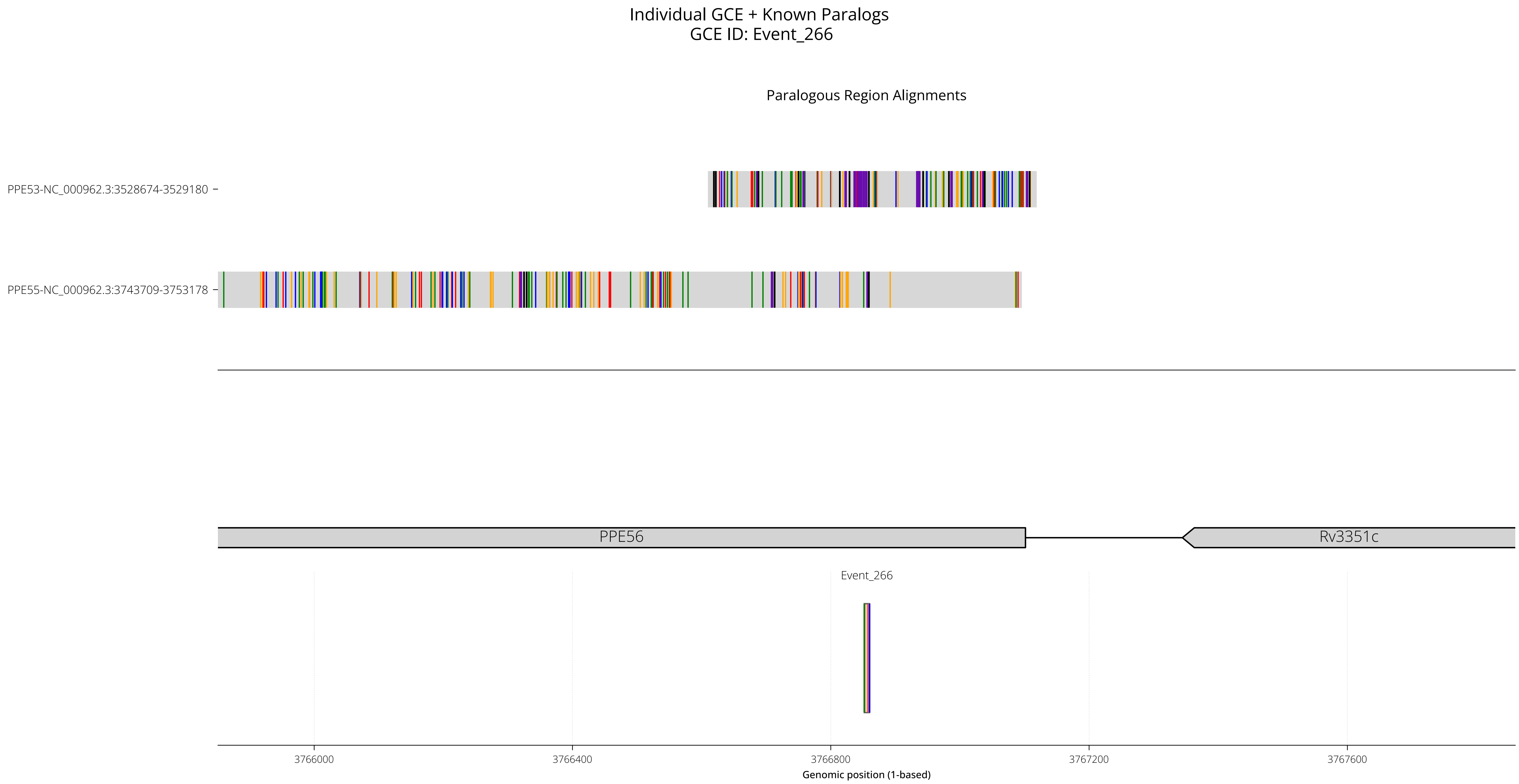
Individual GCE + Known Paralogs
GCE ID: Event_266
Paralogous Region Alignments
PPE53-NC_000962.3:3528674-3529180
PPE55-NC_000962.3:3743709-3753178
PPE56
Rv3351c
Event_266
3766000
3766400
3766800
3767200
3767600
Genomic position (1-based)

Circos Viz - GCE to Known Paralogs Locations
GCE ID: Event_266

Location of target gene conversion event
Known (reference) paralog sequences to event region

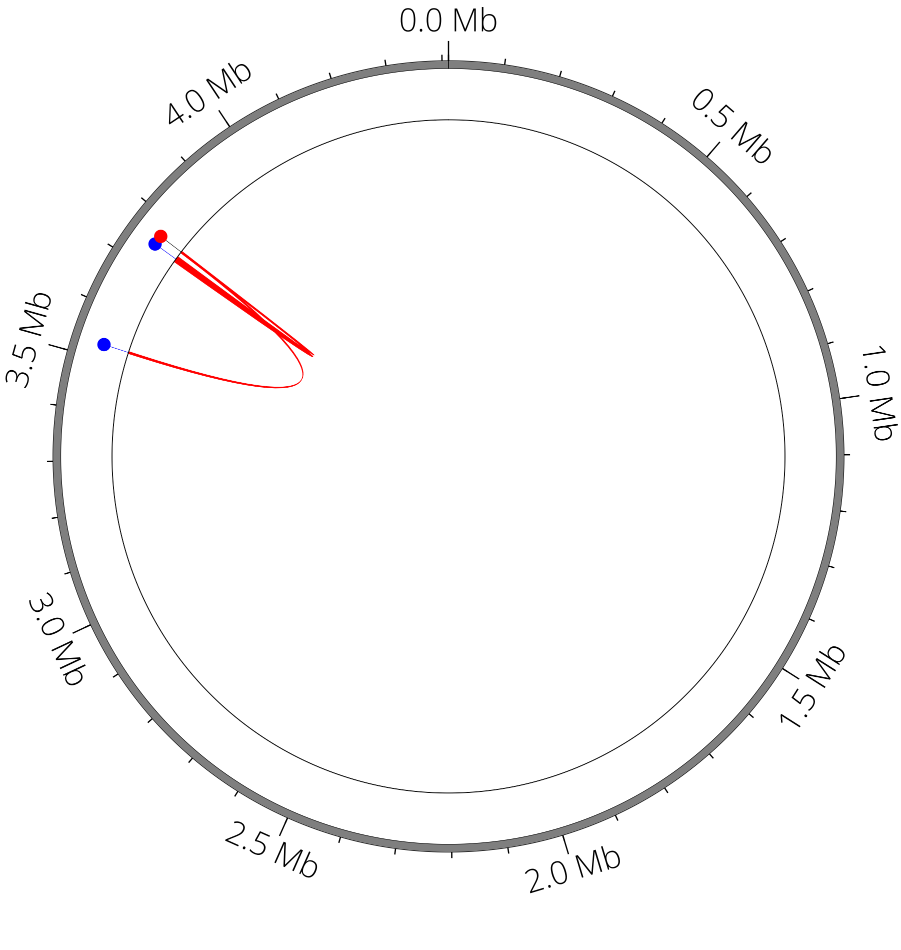

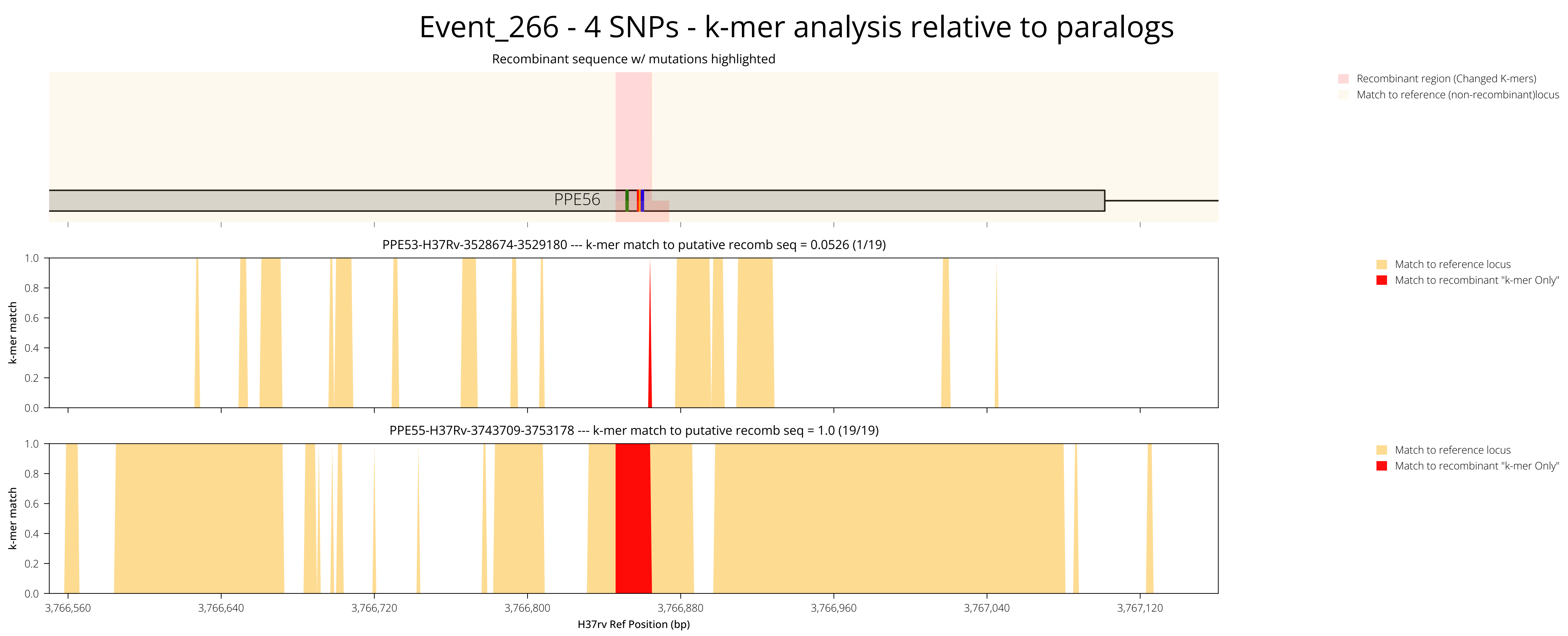
Event_266 - 4 SNPs - k-mer analysis relative to paralogs
Recombinant sequence w/ mutations highlighted
Recombinant region (Changed K-mers)
Match to reference (non-recombinant)locus
PPE56
PPE53-H37Rv-3528674-3529180 --- k-mer match to putative recomb seq = 0.0526 (1/19)
Match to reference locus
Match to recombinant "k-mer Only"
k-mer match
PPE55-H37Rv-3743709-3753178 --- k-mer match to putative recomb seq = 1.0 (19/19)
Match to reference locus
Match to recombinant "k-mer Only"
k-mer match
3,766,560
3,766,640
3,766,720
3,766,800
3,766,880
3,766,960
3,767,040
3,767,120
H37rv Ref Position (bp)

Sub-phylogeny Viz + Paralog Alignments
GCE ID: Event_266

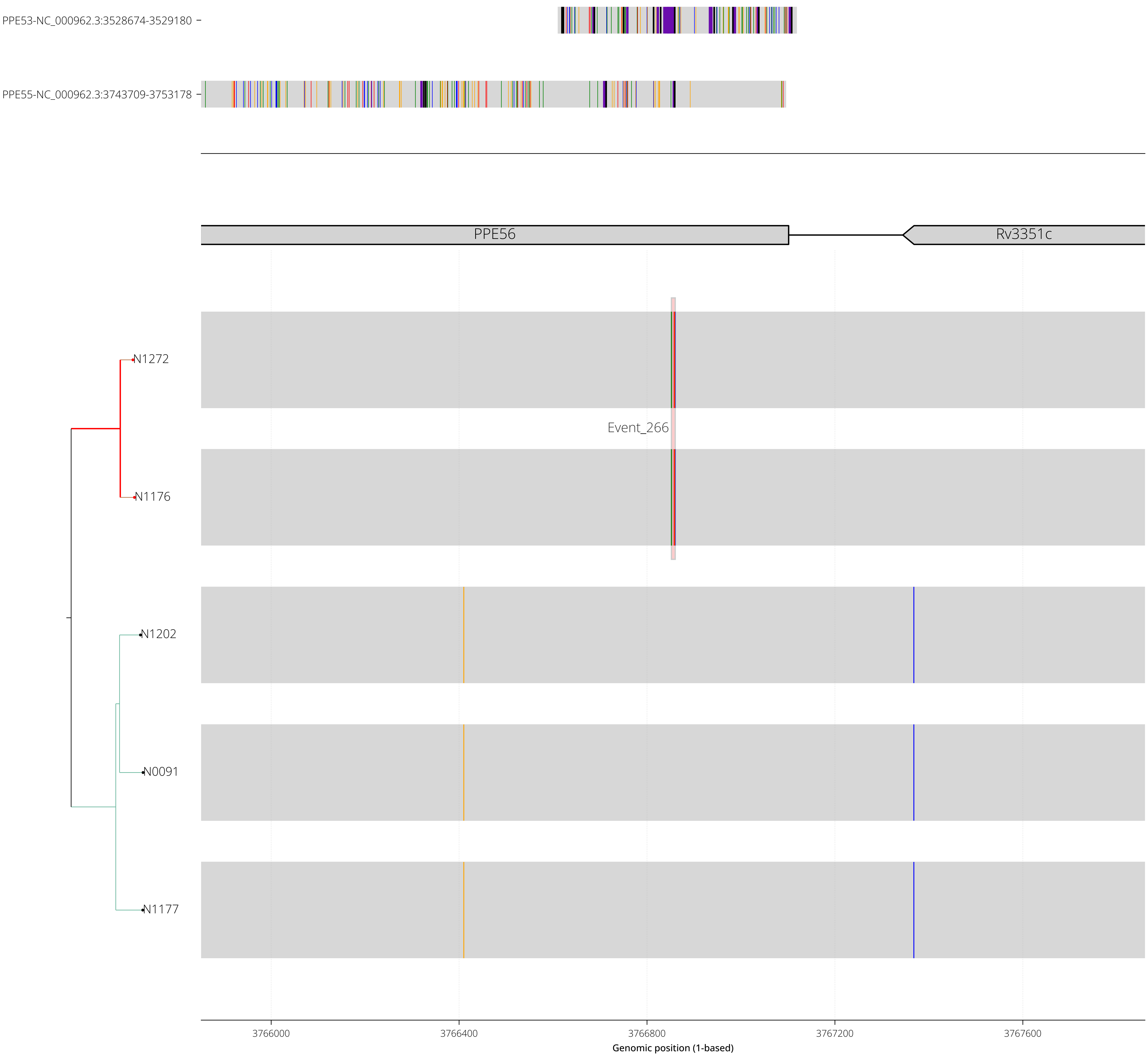

Full Phylogeny Viz + Paralog Alignments
GCE ID: Event_266

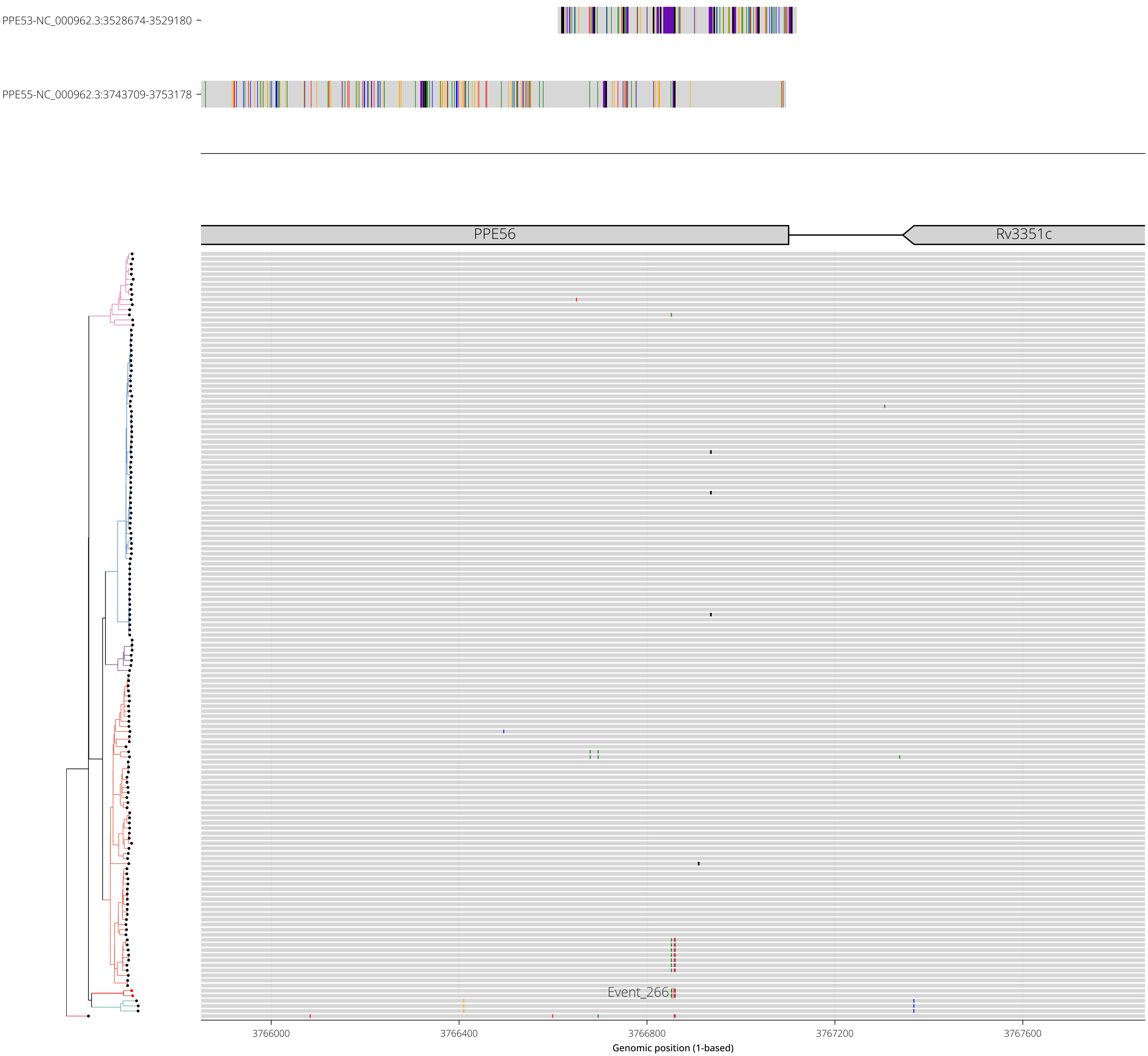

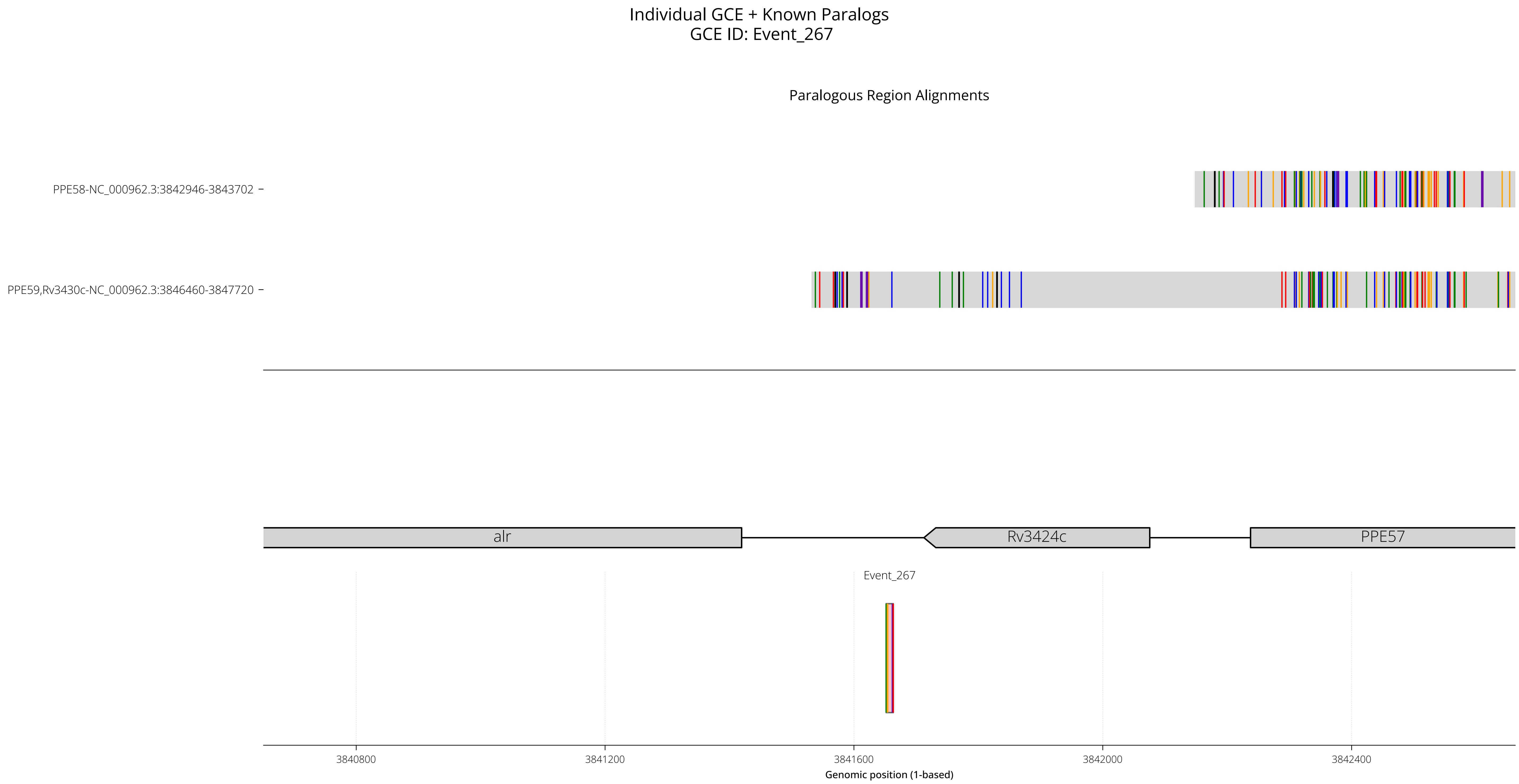

Individual GCE + Known Paralogs
GCE ID: Event_267
Paralogous Region Alignments
PPE58-NC_000962.3:3842946-3843702
PPE59,Rv3430c-NC_000962.3:3846460-3847720
alr
Rv3424c
PPE57
Event_267
3840800
3841200
3841600
3842000
3842400
Genomic position (1-based)

Circos Viz - GCE to Known Paralogs Locations
GCE ID: Event_267

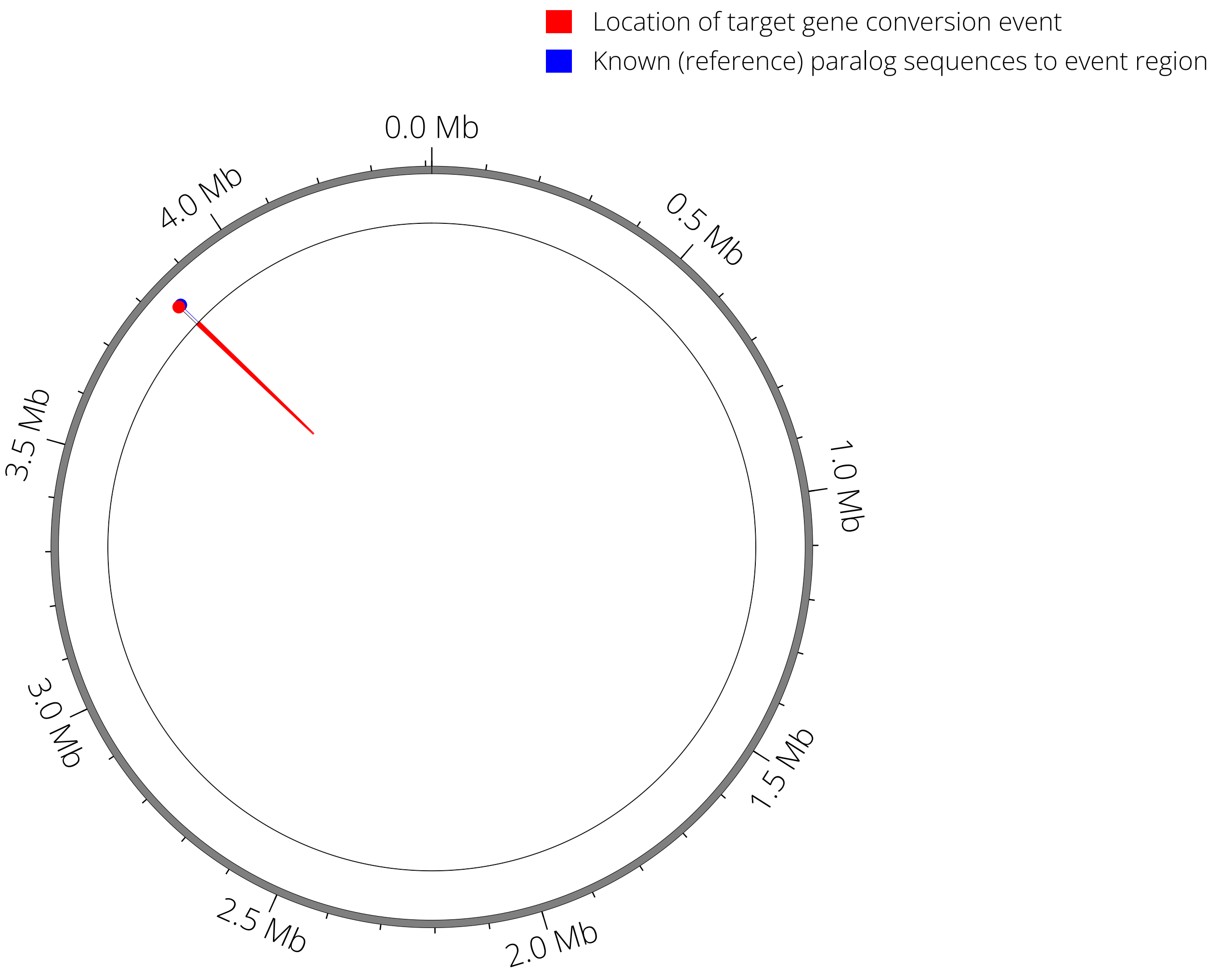

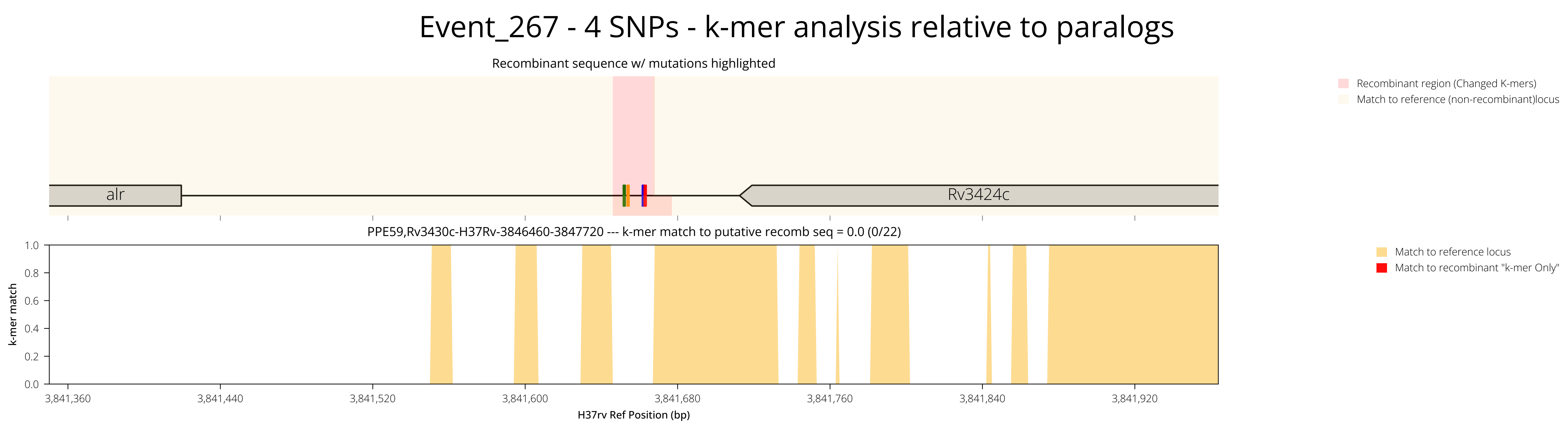
Event_267 - 4 SNPs - k-mer analysis relative to paralogs
Recombinant sequence w/ mutations highlighted
alr
Rv3424c
Recombinant region (Changed K-mers)
Match to reference (non-recombinant)locus
PPE59,Rv3430c-H37Rv-3846460-3847720 --- k-mer match to putative recomb seq = 0.0 (0/22)
1.0
0.8
0.6
0.4
0.2
0.0
k-mer match
3,841,360
3,841,440
3,841,520
3,841,600
3,841,680
3,841,760
3,841,840
3,841,920
H37rv Ref Position (bp)
Match to reference locus
Match to recombinant "k-mer Only"

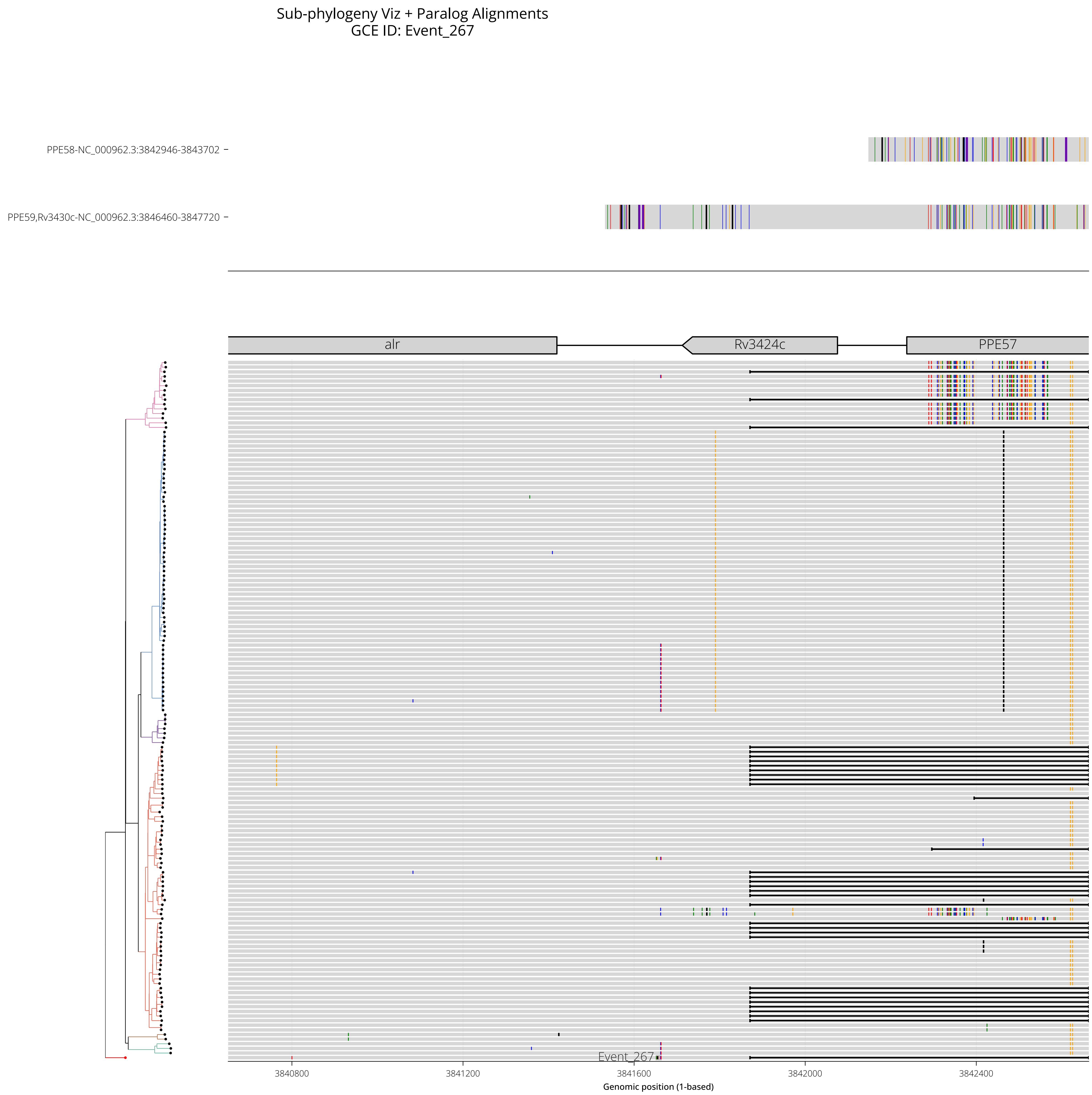

Sub-phylogeny Viz + Paralog Alignments
GCE ID: Event_267
PPE58-NC_000962.3:3842946-3843702
PPE59,Rv3430c-NC_000962.3:3846460-3847720
alr
Rv3424c
PPE57
Event_267
3840800
3841200
3841600
3842000
3842400
Genomic position (1-based)

Full Phylogeny Viz + Paralog Alignments
GCE ID: Event_267

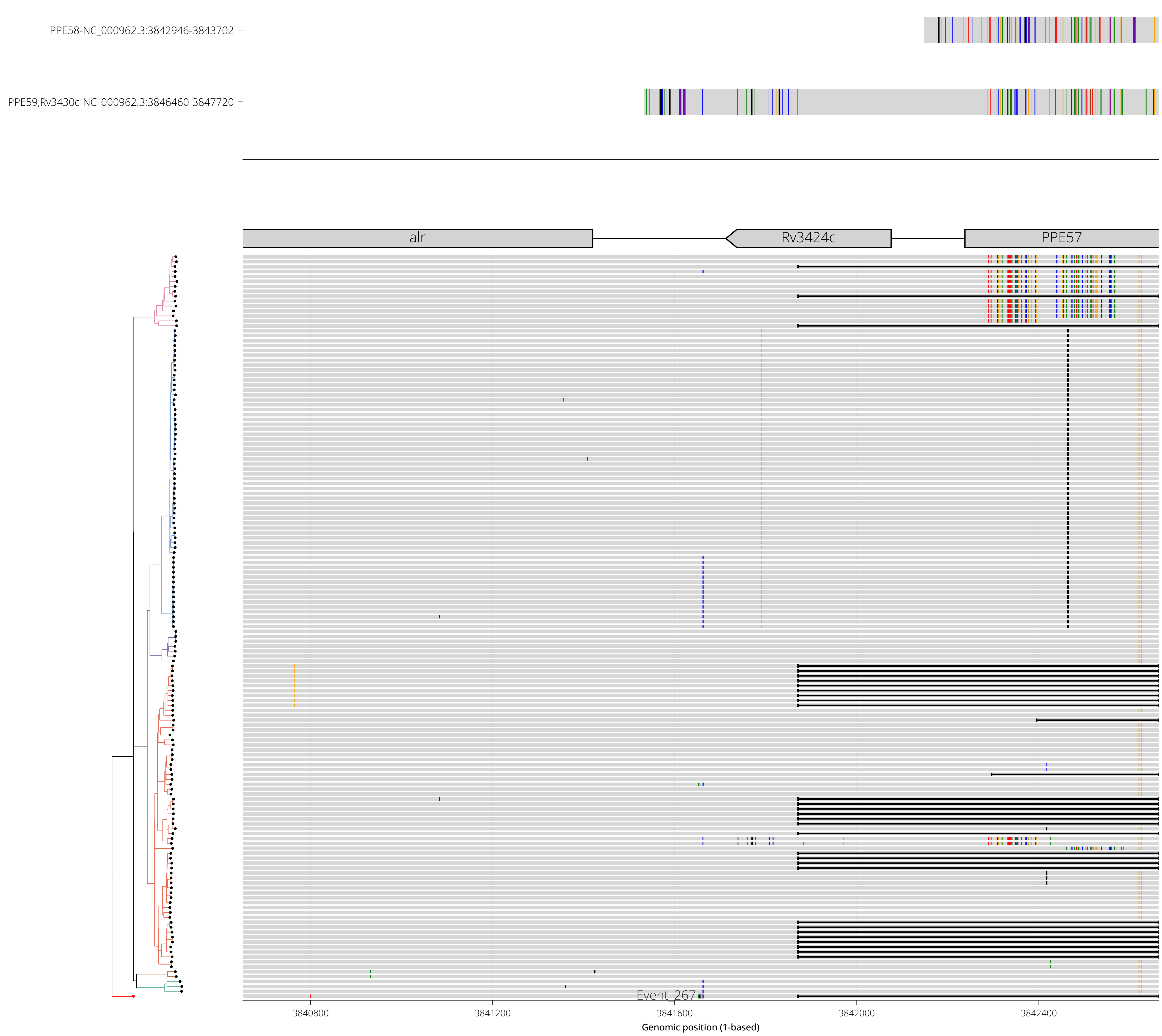

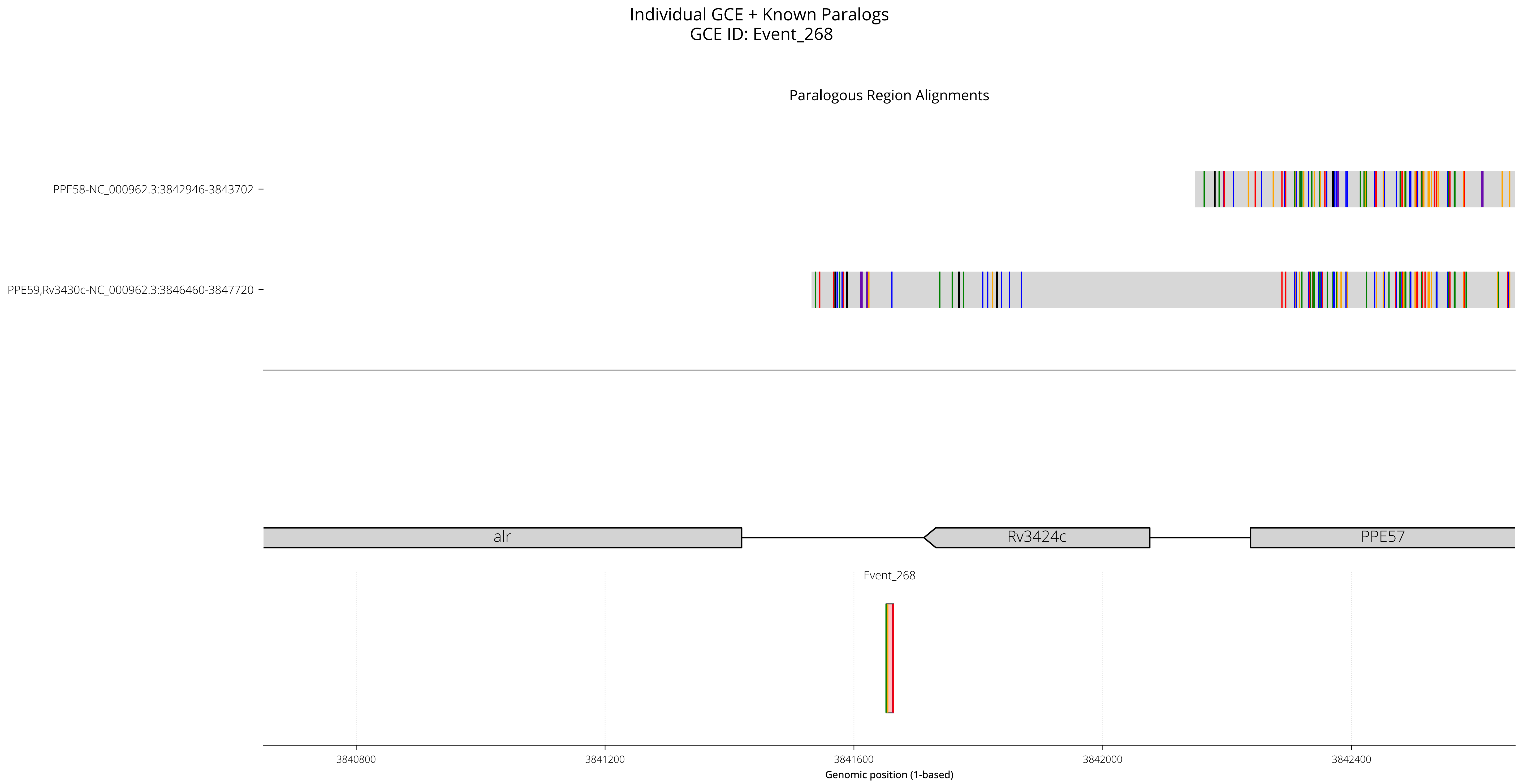

Individual GCE + Known Paralogs
GCE ID: Event_268
Paralogous Region Alignments
PPE58-NC_000962.3:3842946-3843702
PPE59,Rv3430c-NC_000962.3:3846460-3847720
alr
Rv3424c
PPE57
Event_268
3840800
3841200
3841600
3842000
3842400
Genomic position (1-based)

Circos Viz - GCE to Known Paralogs Locations
GCE ID: Event_268

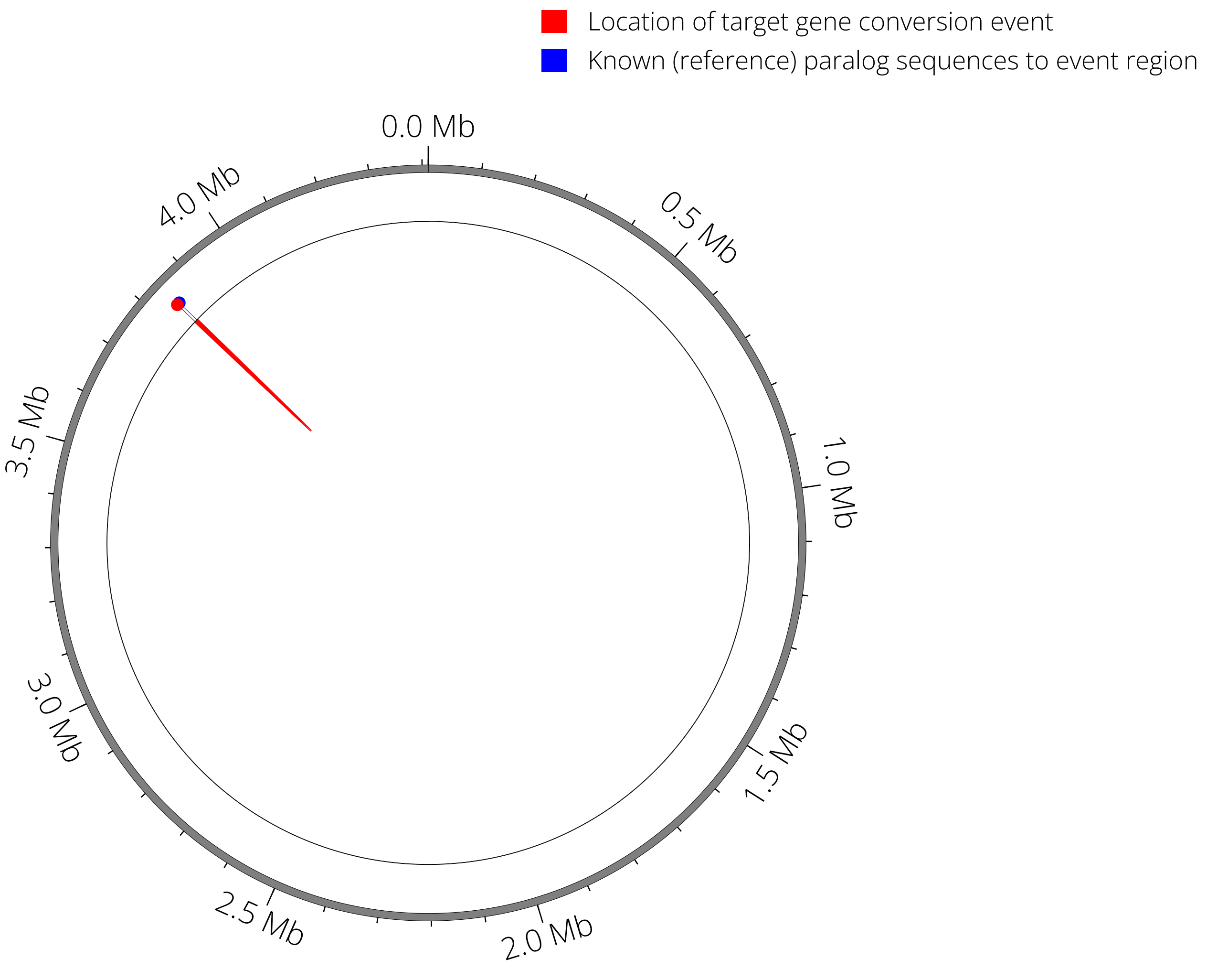

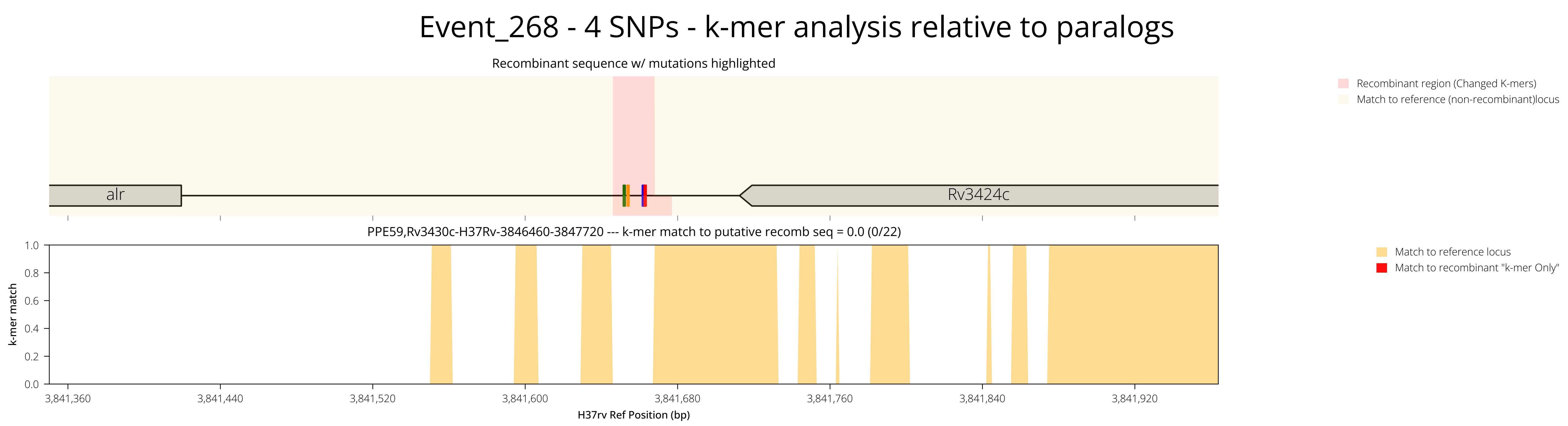

Event_268 - 4 SNPs - k-mer analysis relative to paralogs
Recombinant sequence w/ mutations highlighted
Recombinant region (Changed K-mers)
Match to reference (non-recombinant)locus
alr
Rv3424c
PPE59,Rv3430c-H37Rv-3846460-3847720 --- k-mer match to putative recomb seq = 0.0 (0/22)
Match to reference locus
Match to recombinant "k-mer Only"
k-mer match
1.0
0.8
0.6
0.4
0.2
0.0
3,841,360
3,841,440
3,841,520
3,841,600
3,841,680
3,841,760
3,841,840
3,841,920
H37rv Ref Position (bp)

Sub-phylogeny Viz + Paralog Alignments
GCE ID: Event_268

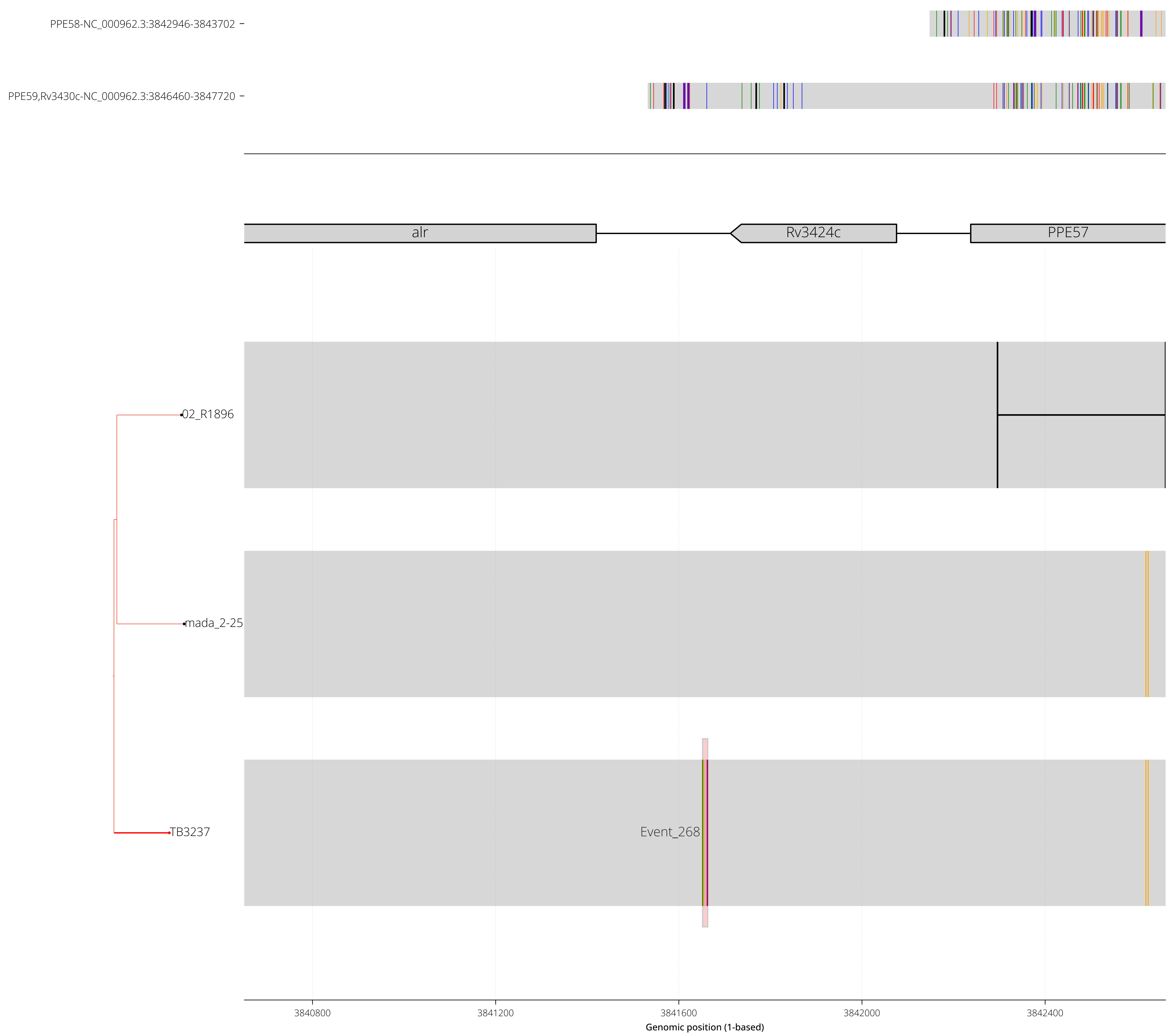

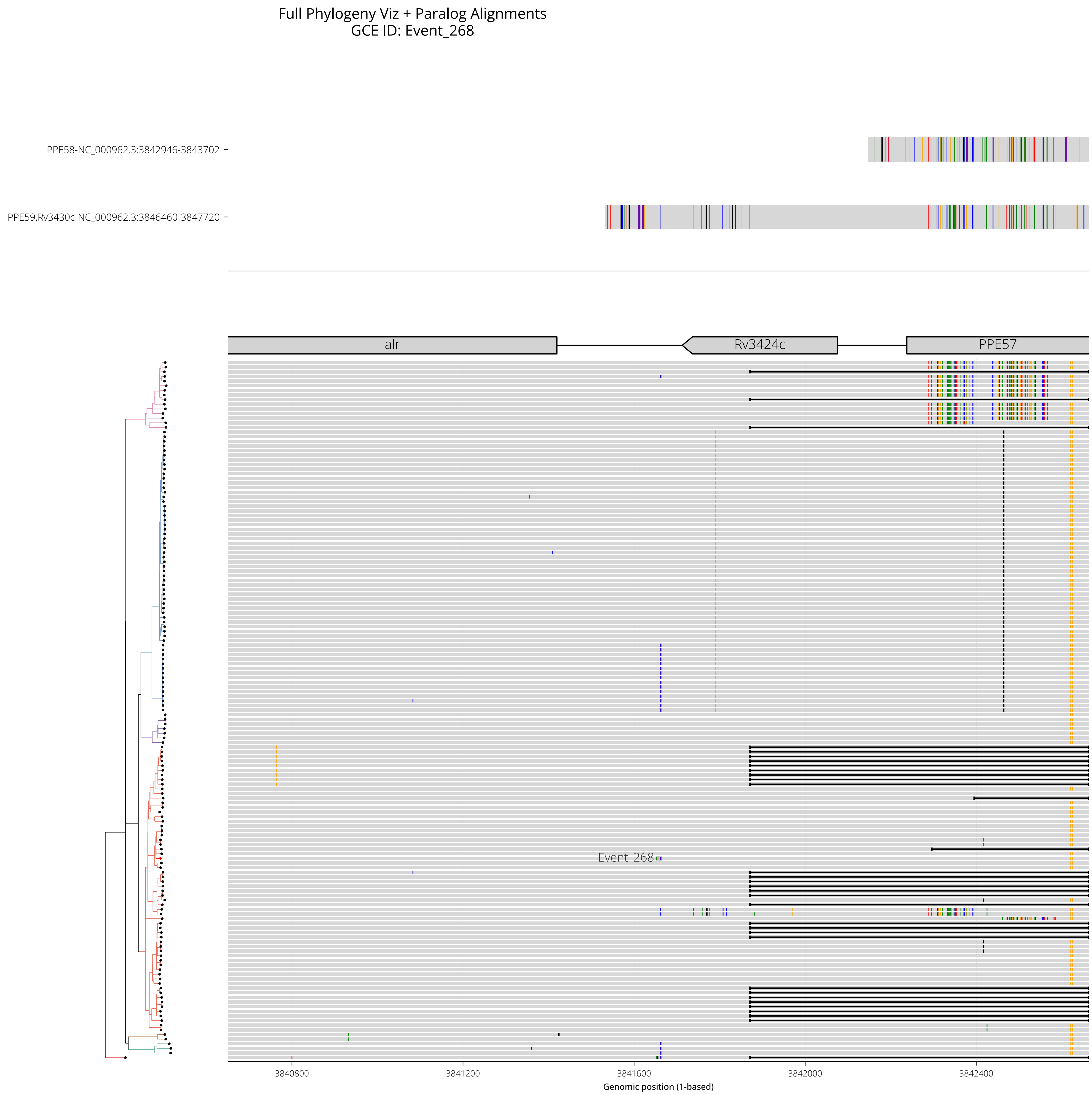

Full Phylogeny Viz + Paralog Alignments
GCE ID: Event_268
PPE58-NC_000962.3:3842946-3843702
PPE59,Rv3430c-NC_000962.3:3846460-3847720
alr
Rv3424c
PPE57
Event_268
3840800
3841200
3841600
3842000
3842400
Genomic position (1-based)

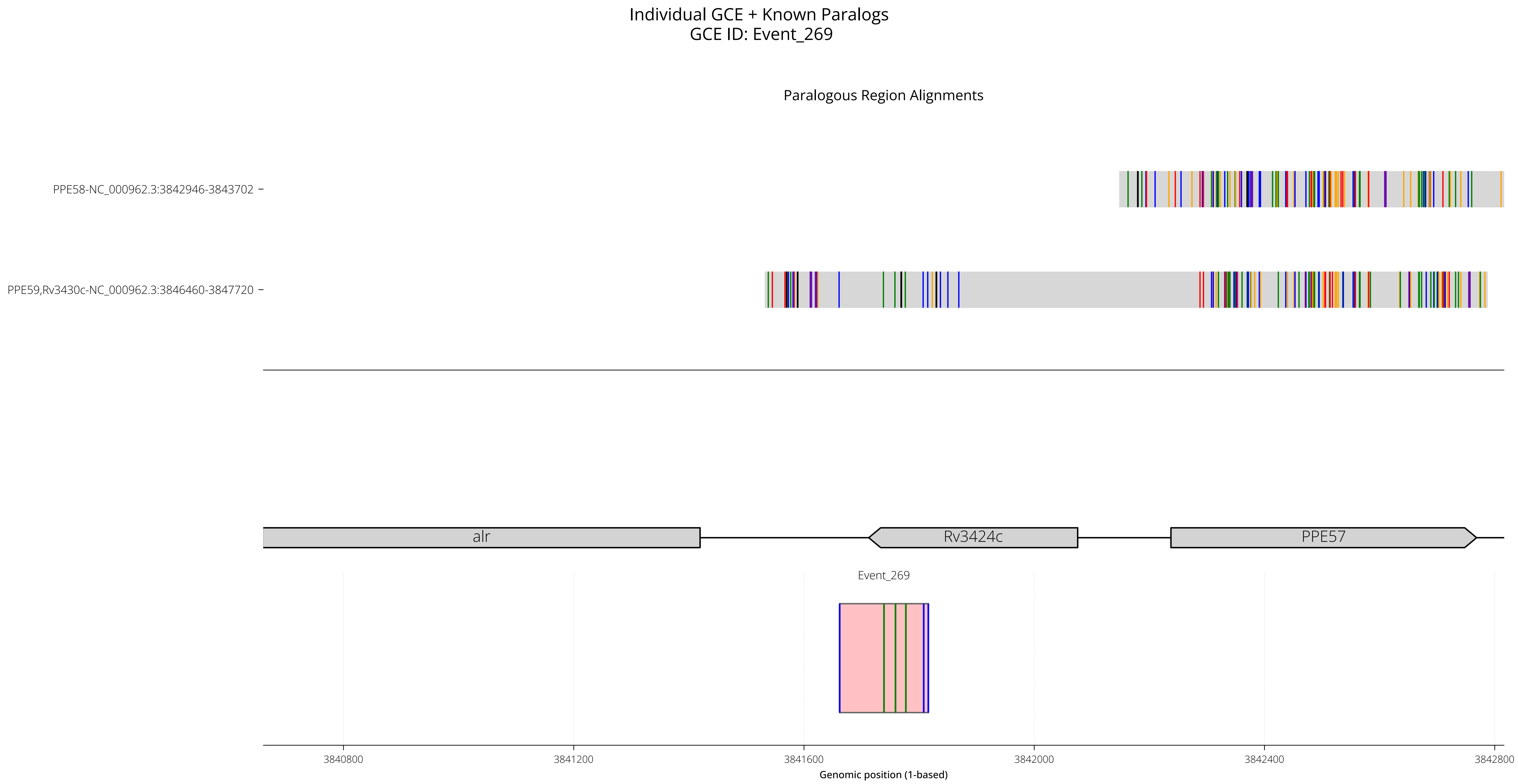
Individual GCE + Known Paralogs
GCE ID: Event_269
Paralogous Region Alignments
PPE58-NC_000962.3:3842946-3843702
PPE59,Rv3430c-NC_000962.3:3846460-3847720
alr
Rv3424c
PPE57
Event_269
3840800
3841200
3841600
3842000
3842400
3842800
Genomic position (1-based)

### Circos Viz - GCE to Known Paralogs Locations
GCE ID: Event_269

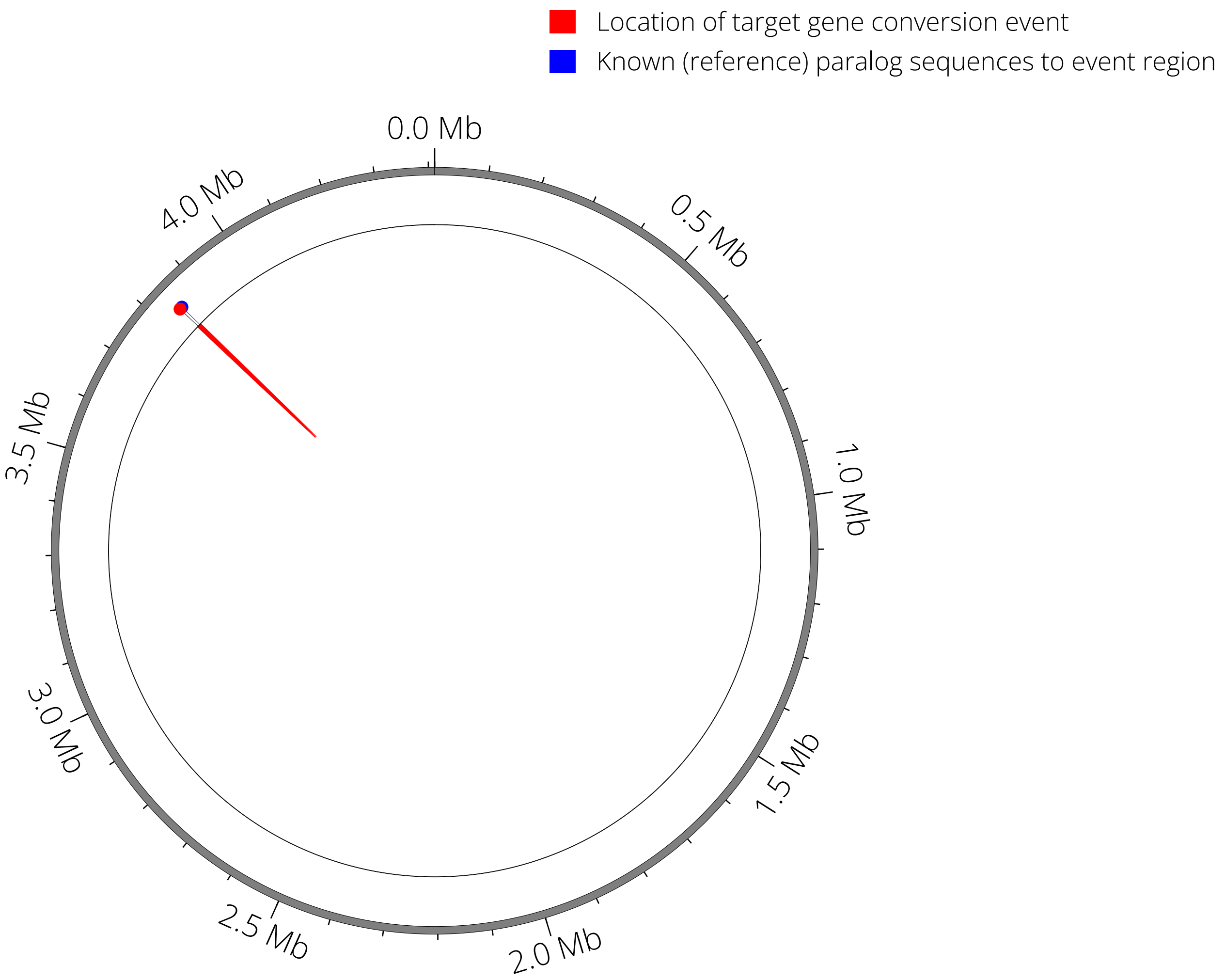

### Event_269 - 6 SNPs - k-mer analysis relative to paralogs

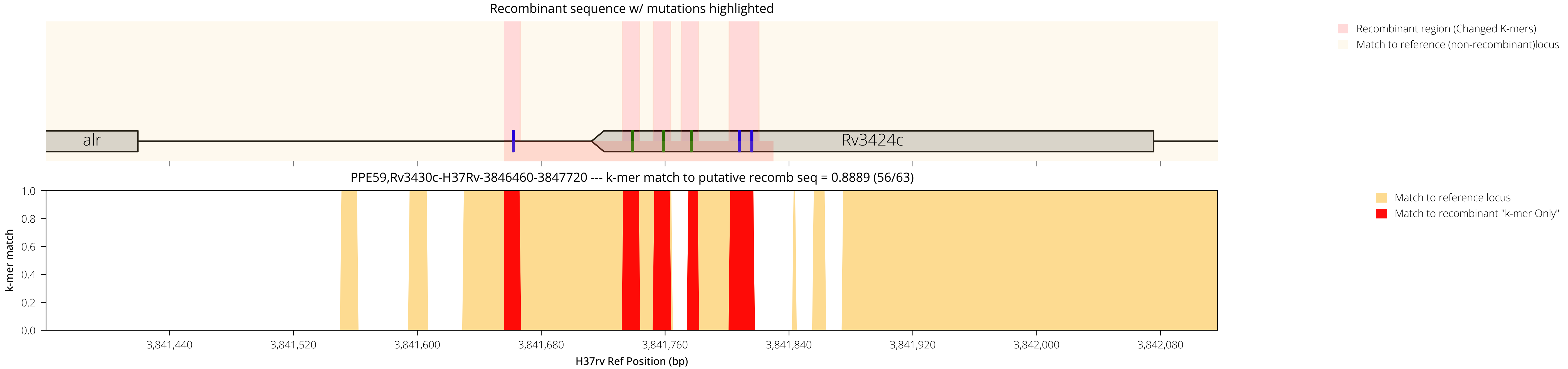

Sub-phylogeny Viz + Paralog Alignments
GCE ID: Event_269

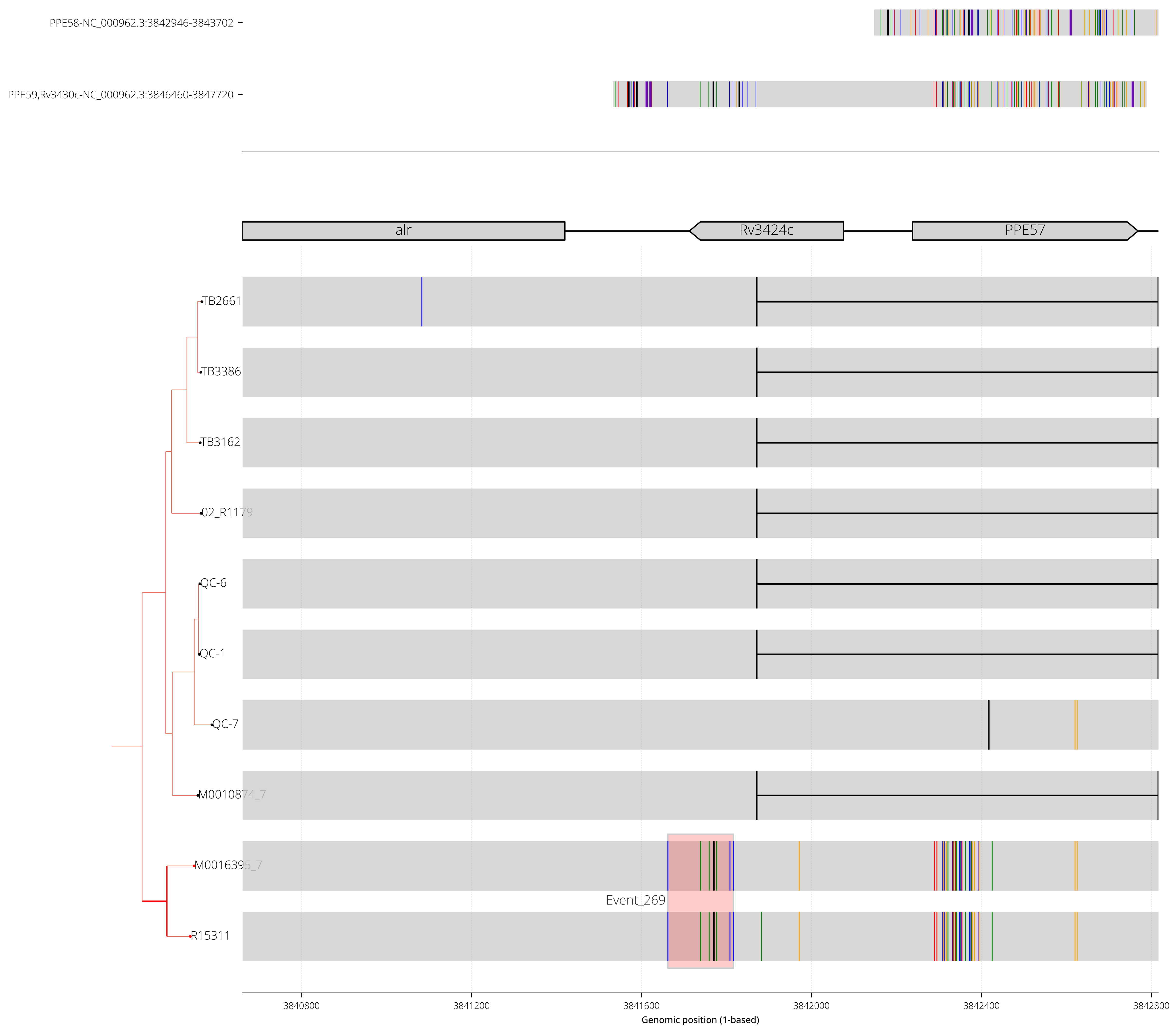

Full Phylogeny Viz + Paralog Alignments
GCE ID: Event_269

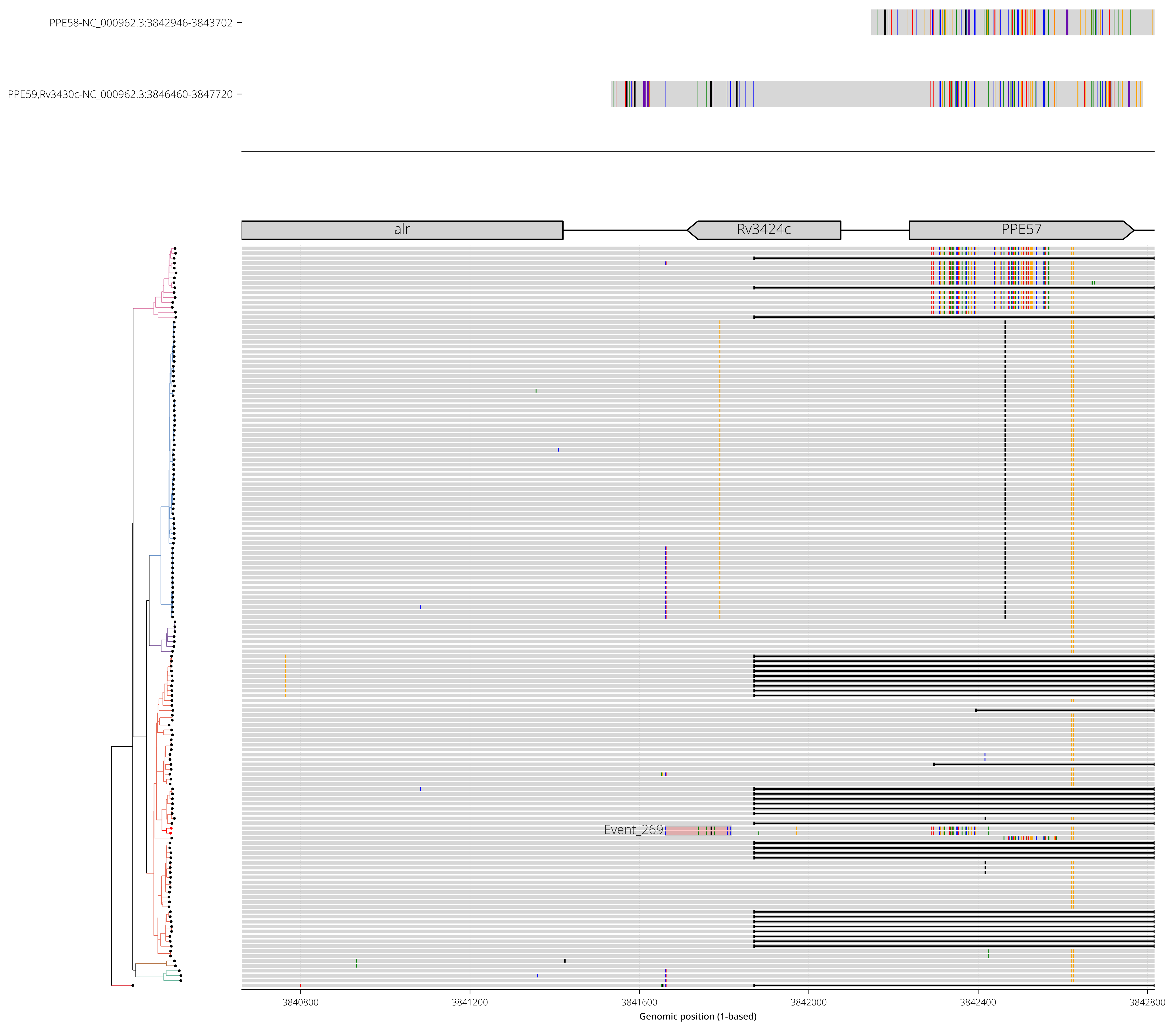

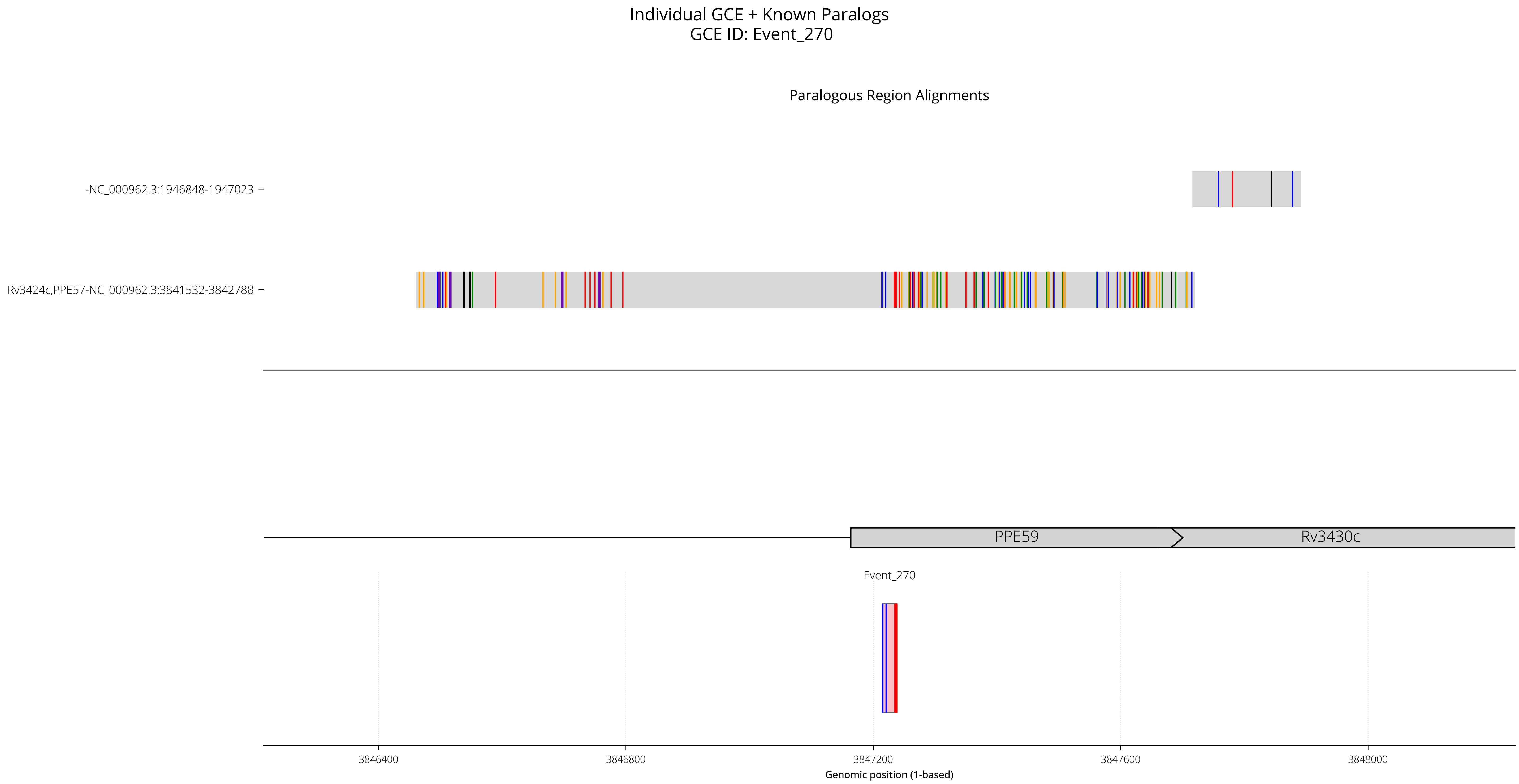
Individual GCE + Known Paralogs
GCE ID: Event_270
Paralogous Region Alignments
-NC_000962.3:1946848-1947023
Rv3424c,PPE57-NC_000962.3:3841532-3842788
PPE59
Rv3430c
Event_270
3846400
3846800
3847200
3847600
3848000
Genomic position (1-based)

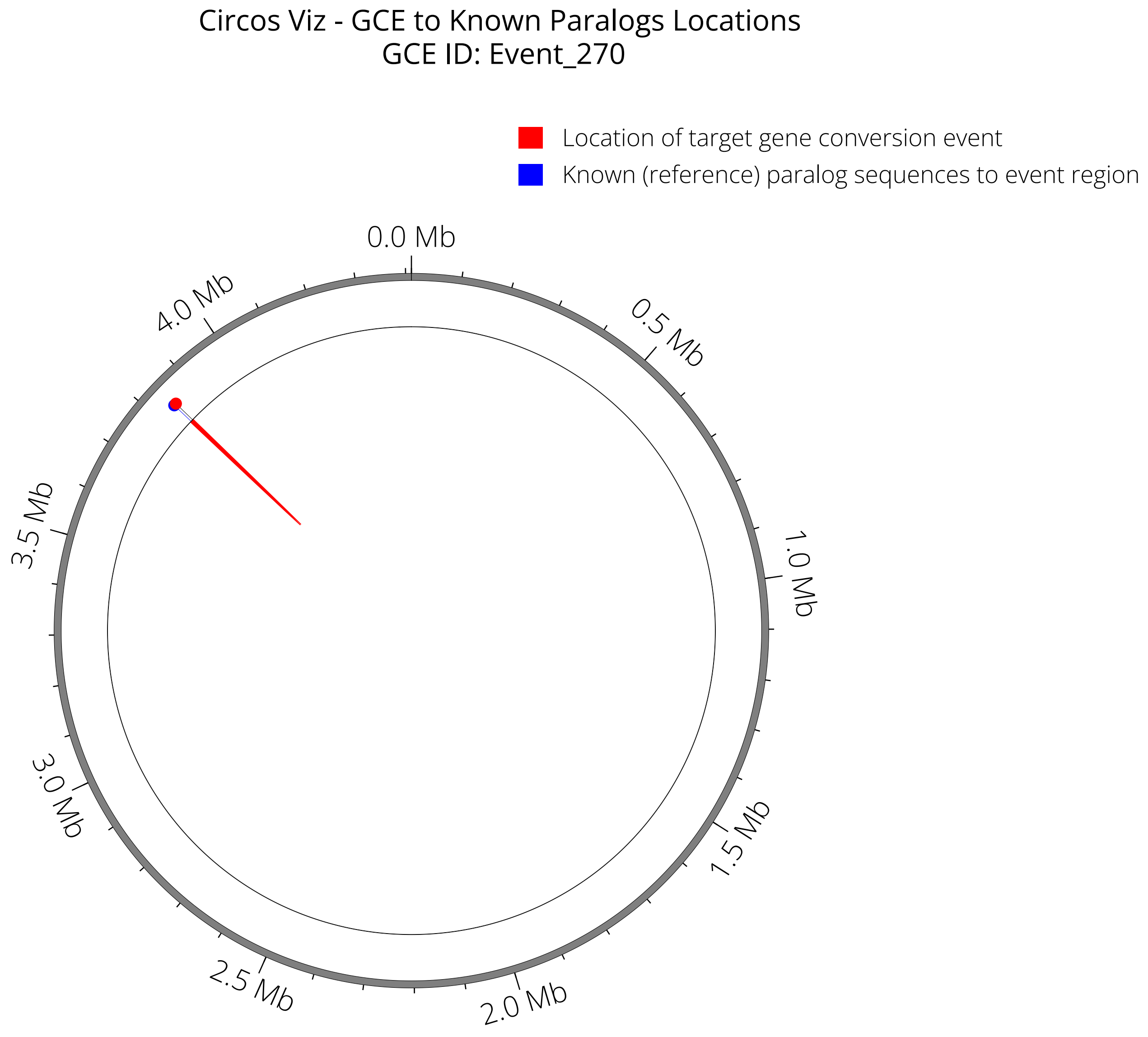
Circos Viz - GCE to Known Paralogs Locations
GCE ID: Event_270
Location of target gene conversion event
Known (reference) paralog sequences to event region
0.0 Mb
0.5 Mb
1.0 Mb
1.5 Mb
2.0 Mb
2.5 Mb
3.0 Mb
3.5 Mb
4.0 Mb

### Event_270 - 5 SNPs - k-mer analysis relative to paralogs

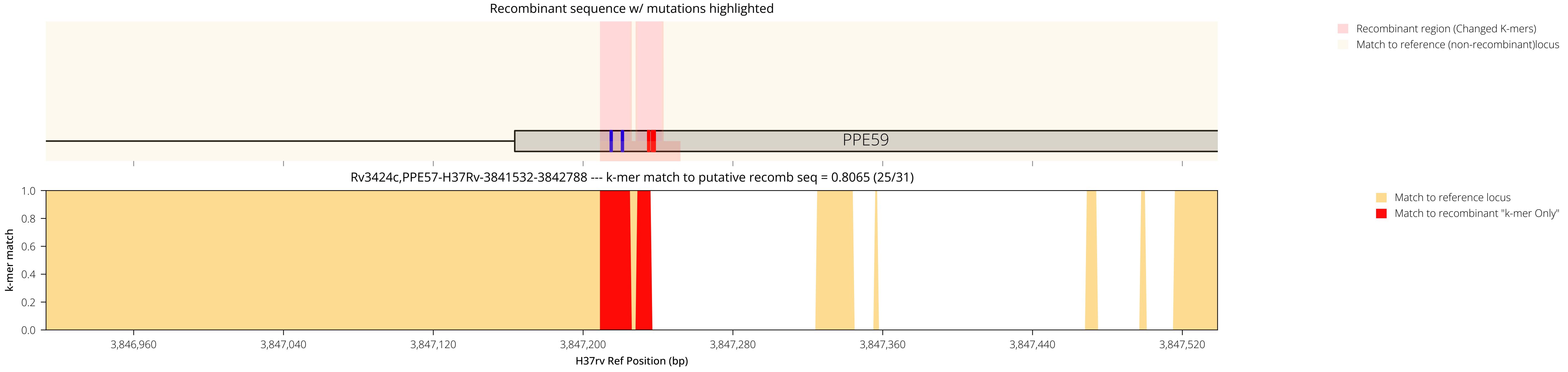

Sub-phylogeny Viz + Paralog Alignments
GCE ID: Event_270

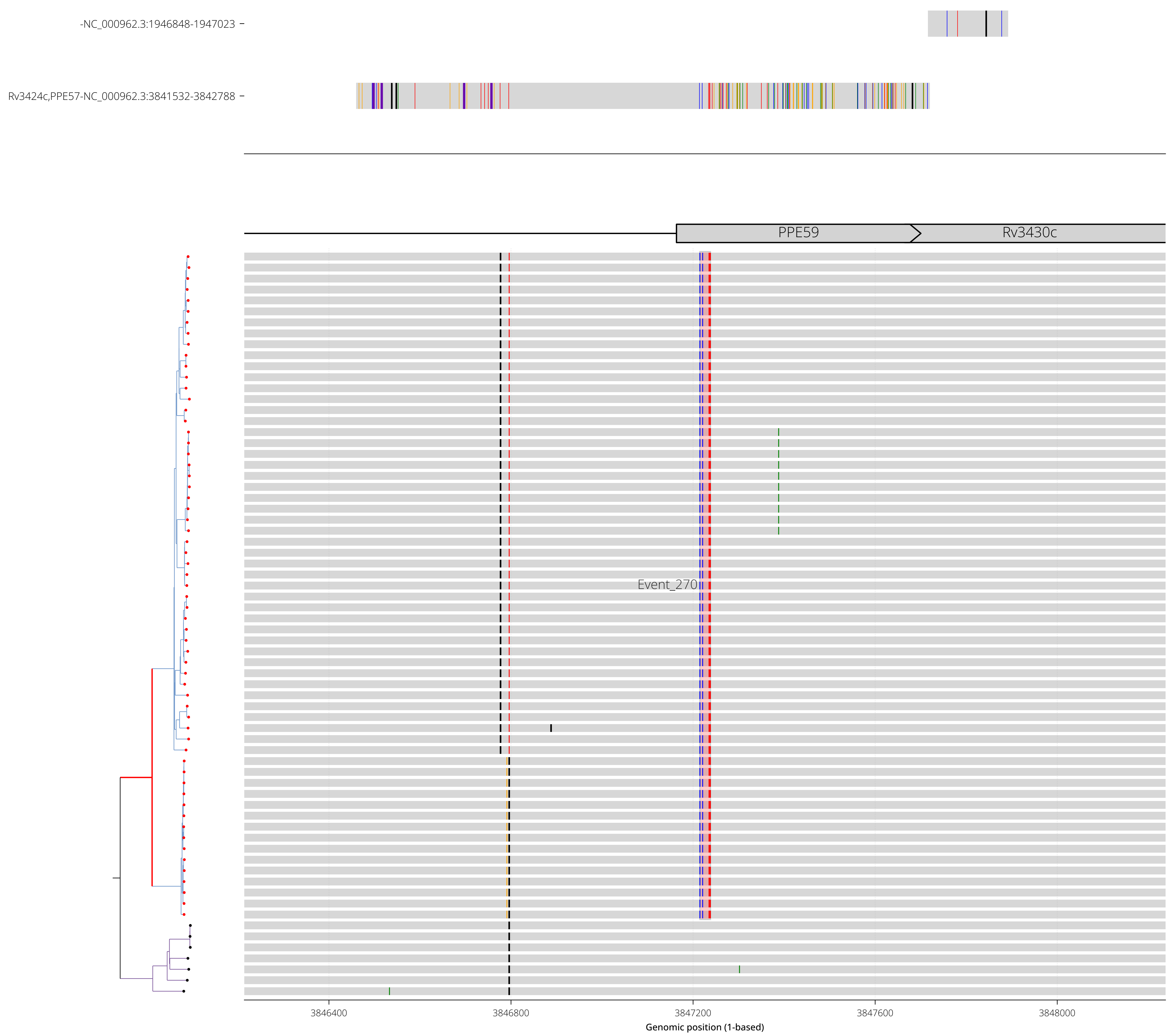

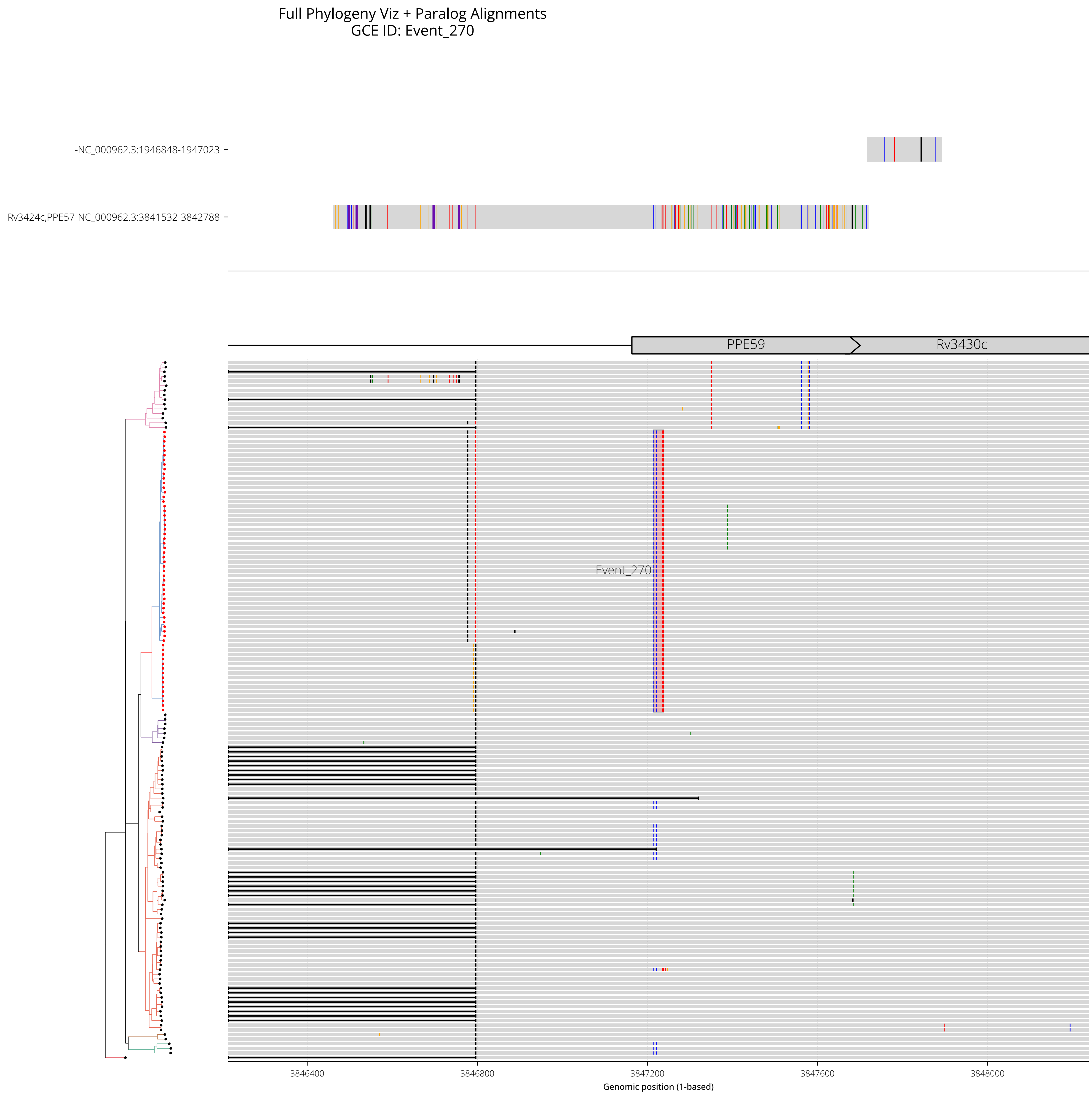

Full Phylogeny Viz + Paralog Alignments
GCE ID: Event_270
-NC_000962.3:1946848-1947023
Rv3424c,PPE57-NC_000962.3:3841532-3842788
PPE59
Rv3430c
Event_270
3846400
3846800
3847200
3847600
3848000
Genomic position (1-based)

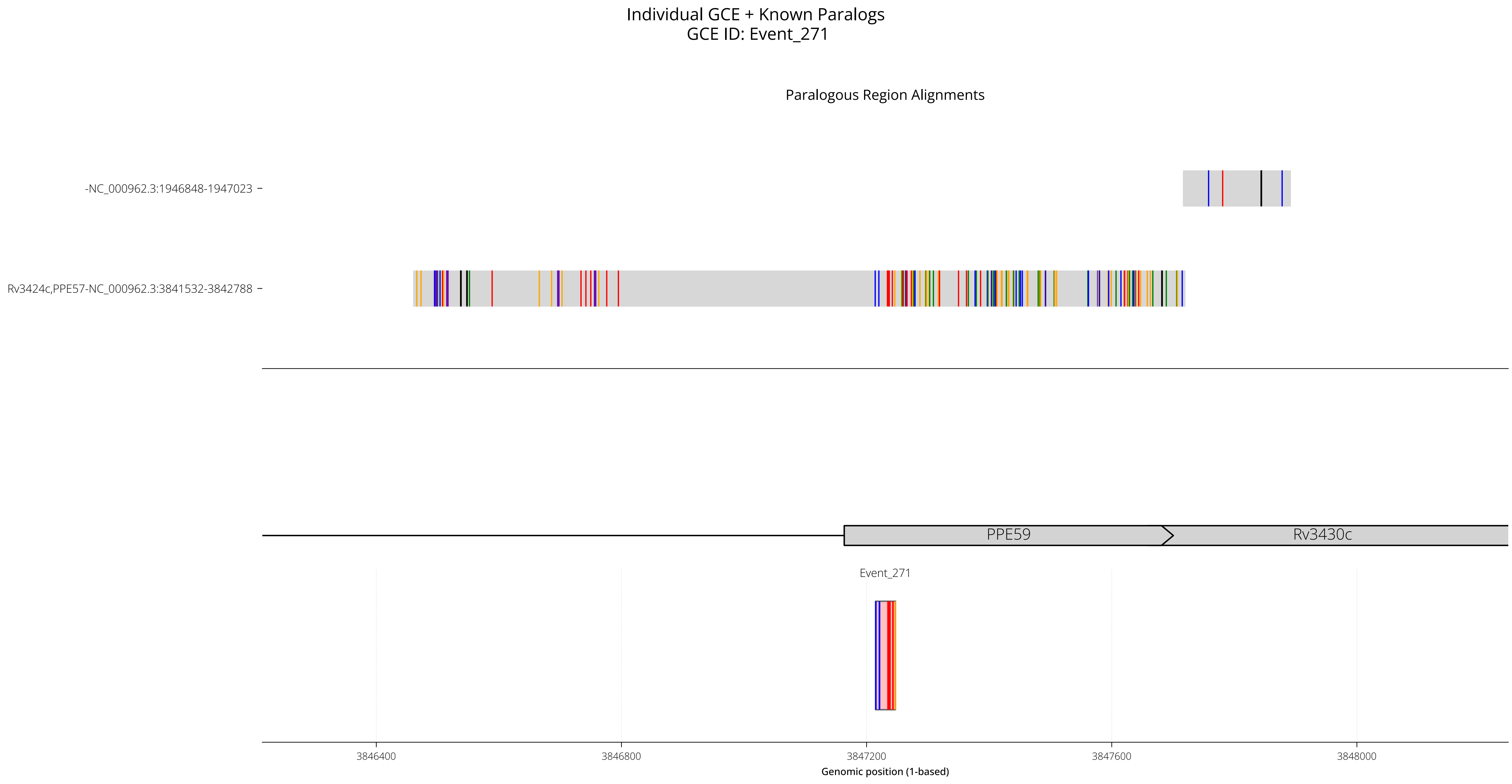
Individual GCE + Known Paralogs
GCE ID: Event_271
Paralogous Region Alignments
-NC_000962.3:1946848-1947023
Rv3424c,PPE57-NC_000962.3:3841532-3842788
PPE59
Rv3430c
Event_271
3846400
3846800
3847200
3847600
3848000
Genomic position (1-based)

Circos Viz - GCE to Known Paralogs Locations
GCE ID: Event_271

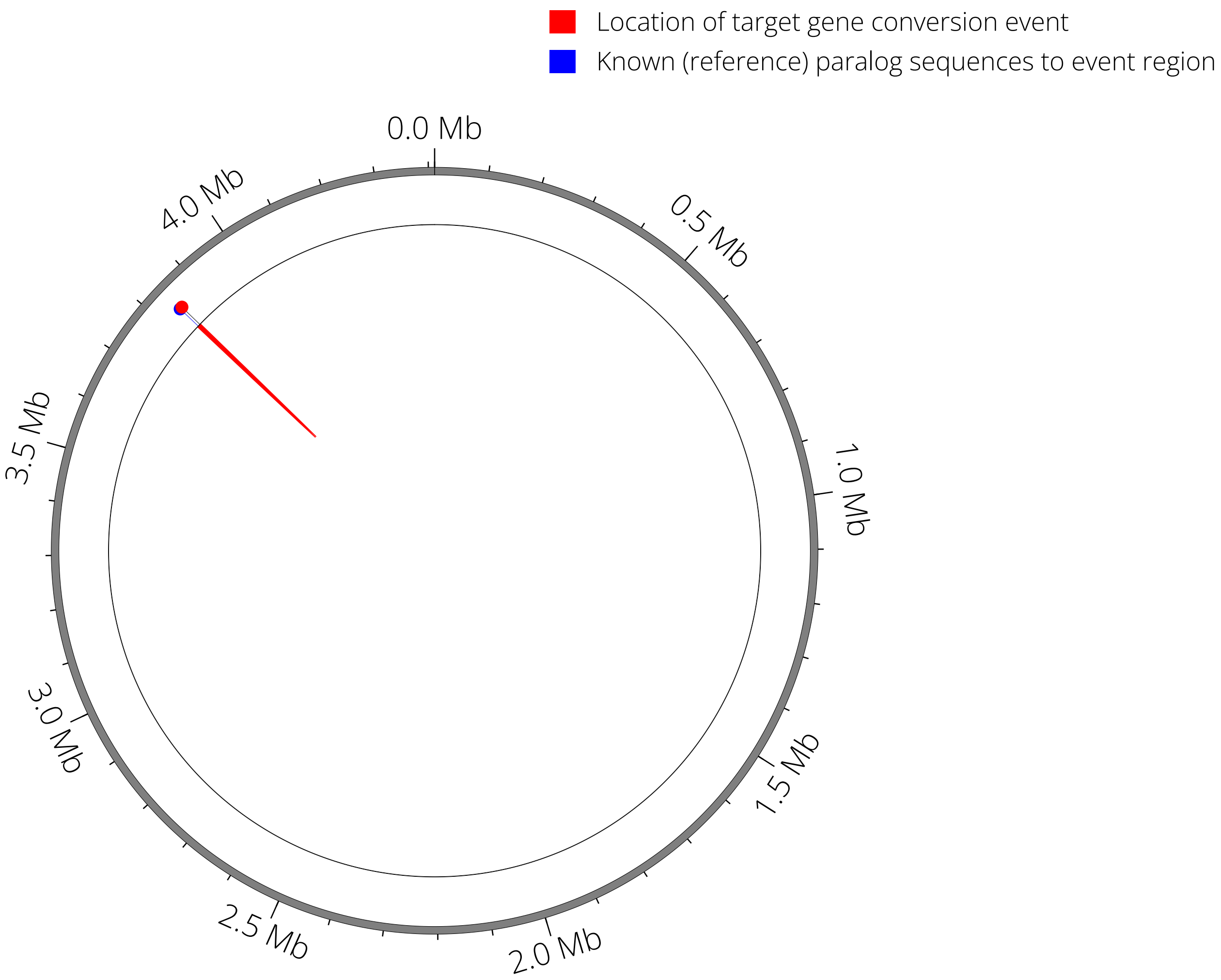

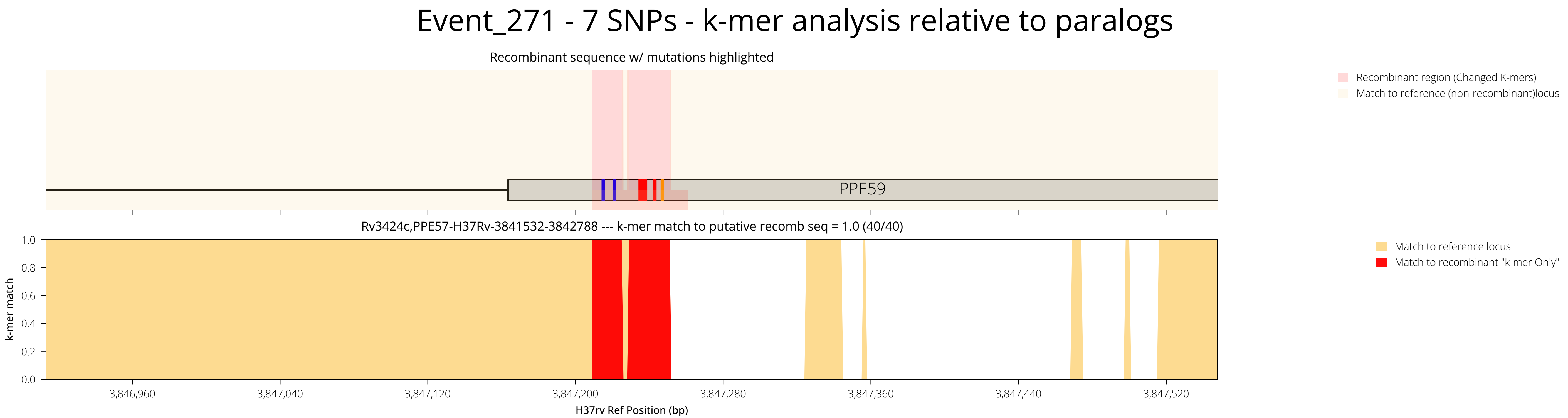
Event_271 - 7 SNPs - k-mer analysis relative to paralogs
Recombinant sequence w/ mutations highlighted
PPE59
Recombinant region (Changed K-mers)
Match to reference (non-recombinant)locus
Rv3424c,PPE57-H37Rv-3841532-3842788 --- k-mer match to putative recomb seq = 1.0 (40/40)
Match to reference locus
Match to recombinant "k-mer Only"
k-mer match
1.0
0.8
0.6
0.4
0.2
0.0
3,846,960
3,847,040
3,847,120
3,847,200
3,847,280
3,847,360
3,847,440
3,847,520
H37rv Ref Position (bp)

Sub-phylogeny Viz + Paralog Alignments
GCE ID: Event_271

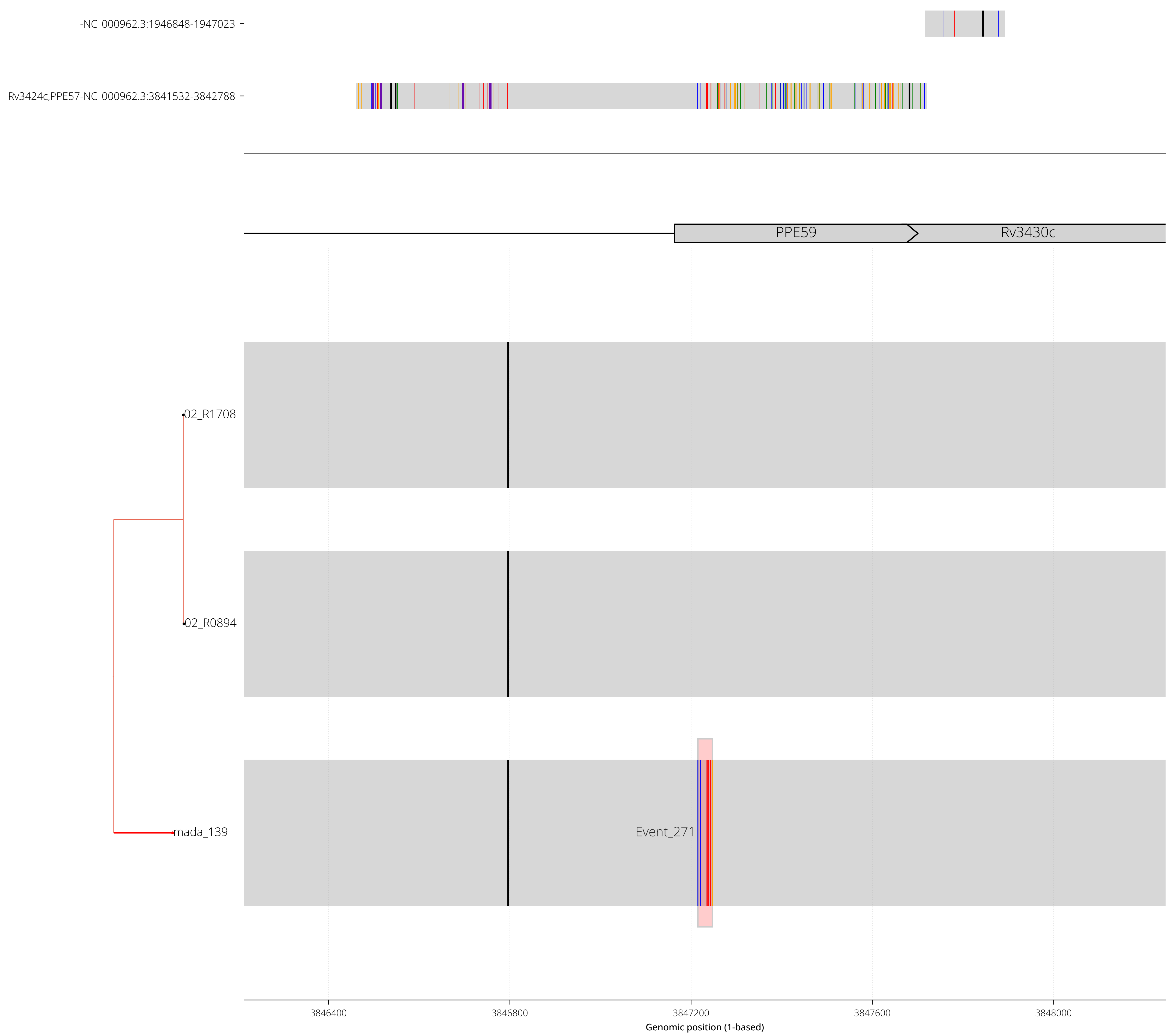

Full Phylogeny Viz + Paralog Alignments
GCE ID: Event_271

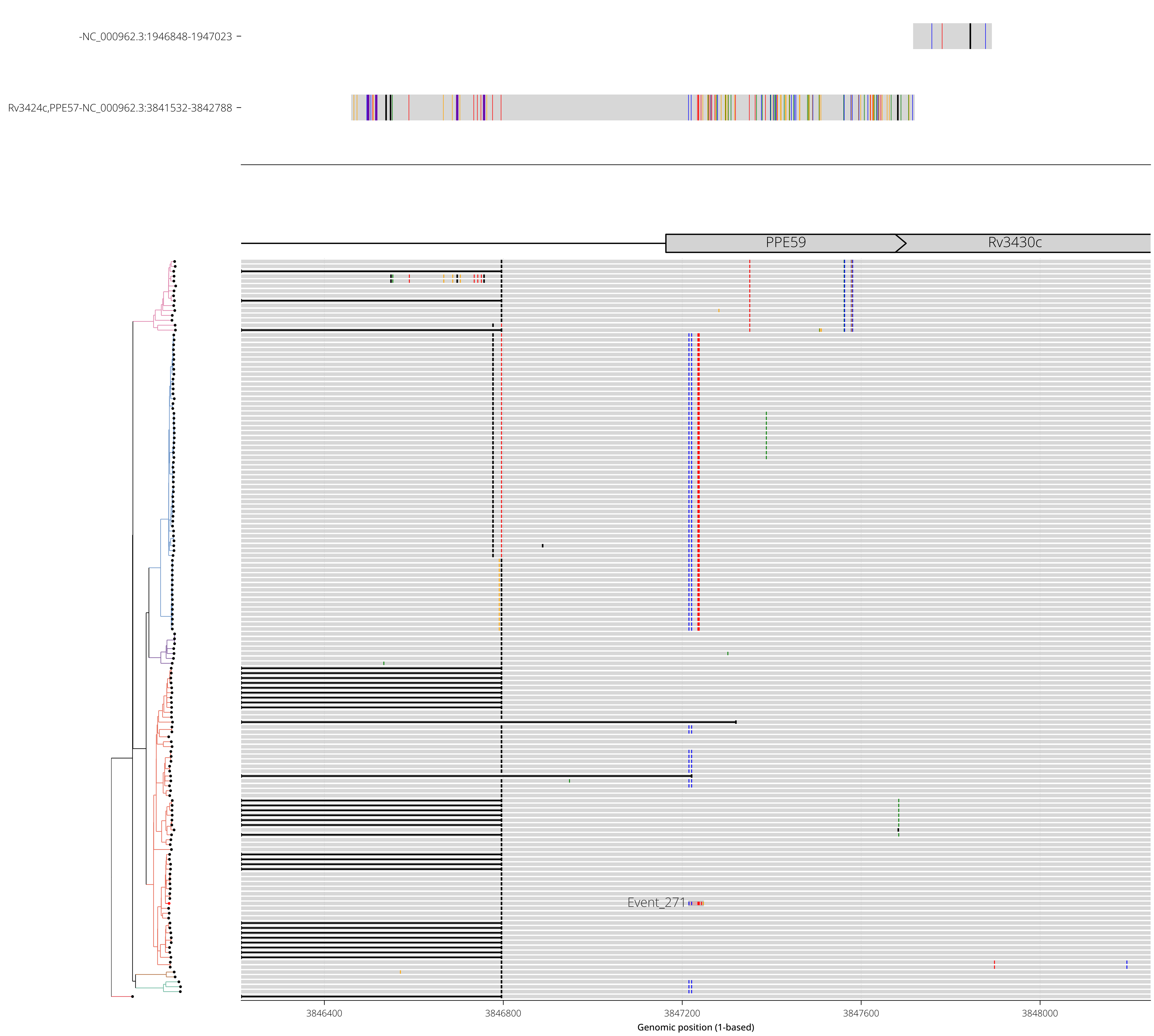

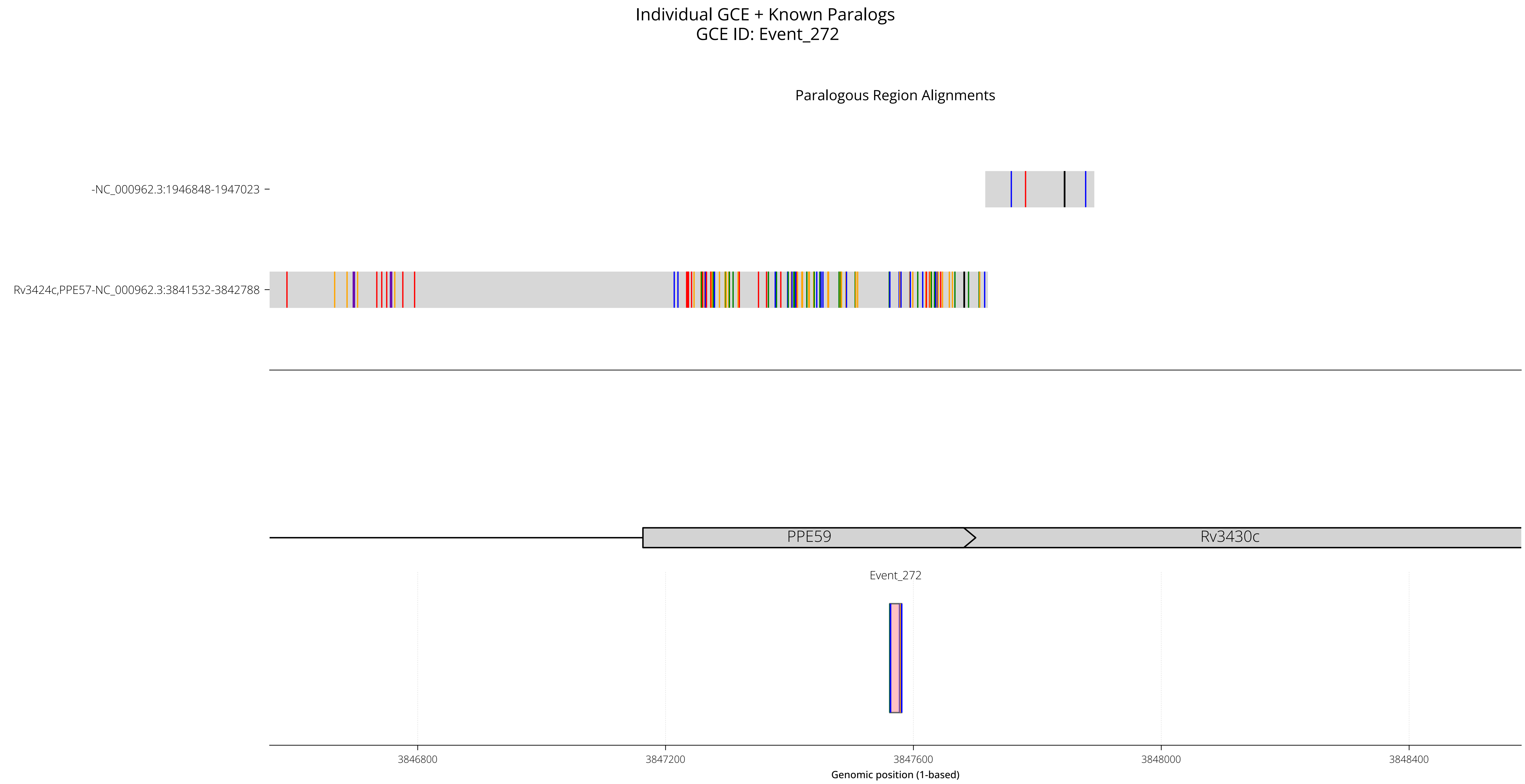
Individual GCE + Known Paralogs
GCE ID: Event_272
Paralogous Region Alignments
-NC_000962.3:1946848-1947023
Rv3424c,PPE57-NC_000962.3:3841532-3842788
PPE59
Rv3430c
Event_272
3846800
3847200
3847600
3848000
3848400
Genomic position (1-based)

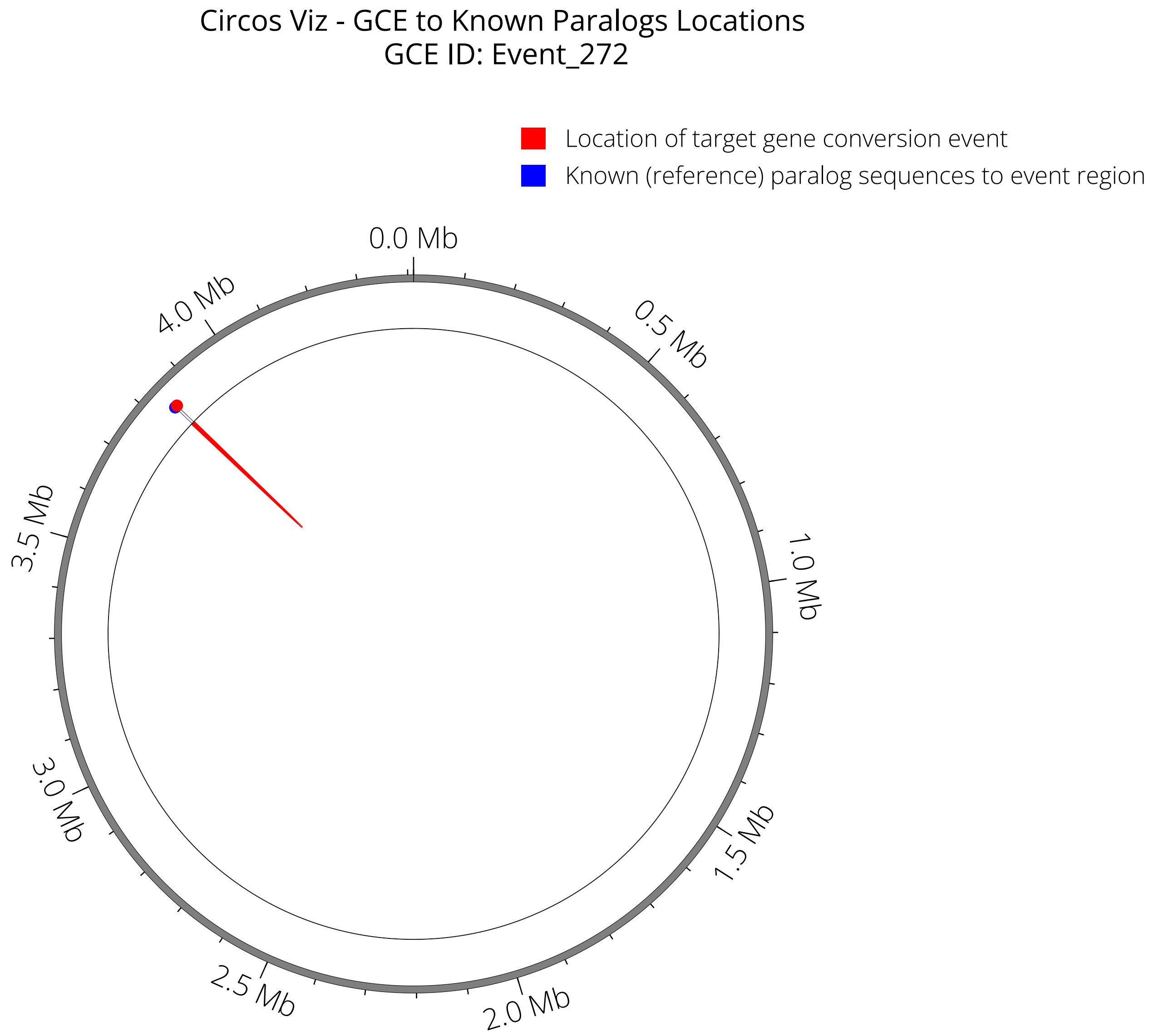

Circos Viz - GCE to Known Paralogs Locations
GCE ID: Event_272
Location of target gene conversion event
Known (reference) paralog sequences to event region
0.0 Mb
0.5 Mb
1.0 Mb
1.5 Mb
2.0 Mb
2.5 Mb
3.0 Mb
3.5 Mb
4.0 Mb

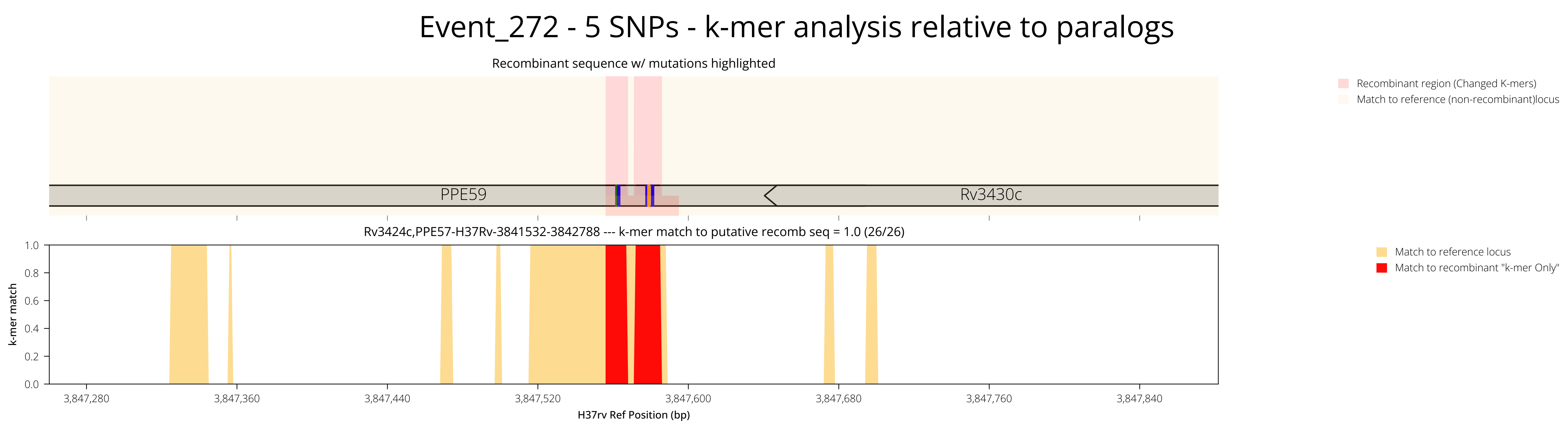
Event_272 - 5 SNPs - k-mer analysis relative to paralogs
Recombinant sequence w/ mutations highlighted
PPE59
Rv3430c
Recombinant region (Changed K-mers)
Match to reference (non-recombinant)locus
Rv3424c,PPE57-H37Rv-3841532-3842788 --- k-mer match to putative recomb seq = 1.0 (26/26)
k-mer match
1.0
0.8
0.6
0.4
0.2
0.0
3,847,280
3,847,360
3,847,440
3,847,520
3,847,600
3,847,680
3,847,760
3,847,840
H37rv Ref Position (bp)
Match to reference locus
Match to recombinant "k-mer Only"

Sub-phylogeny Viz + Paralog Alignments
GCE ID: Event_272

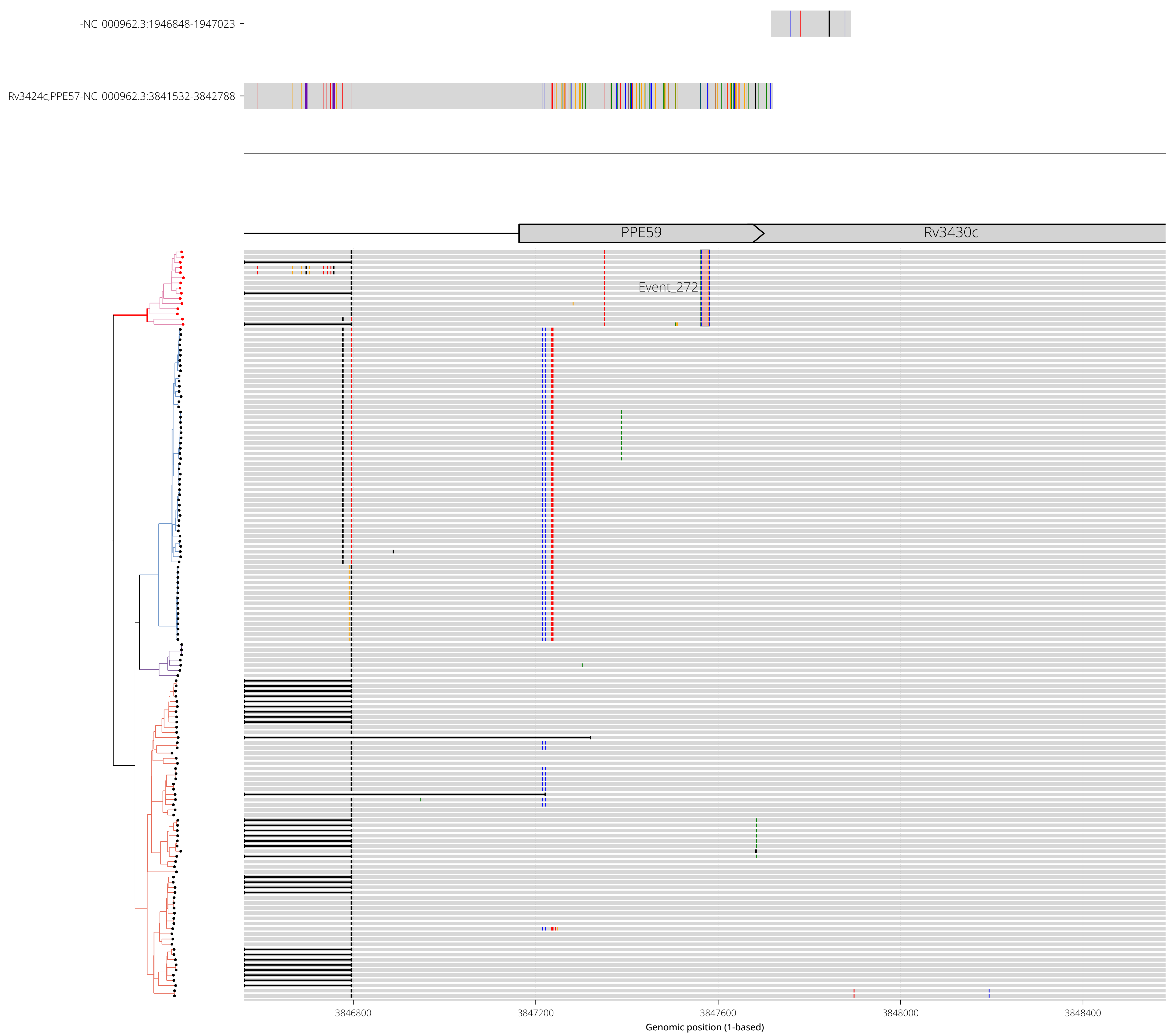

### Full Phylogeny Viz + Paralog Alignments
GCE ID: Event_272

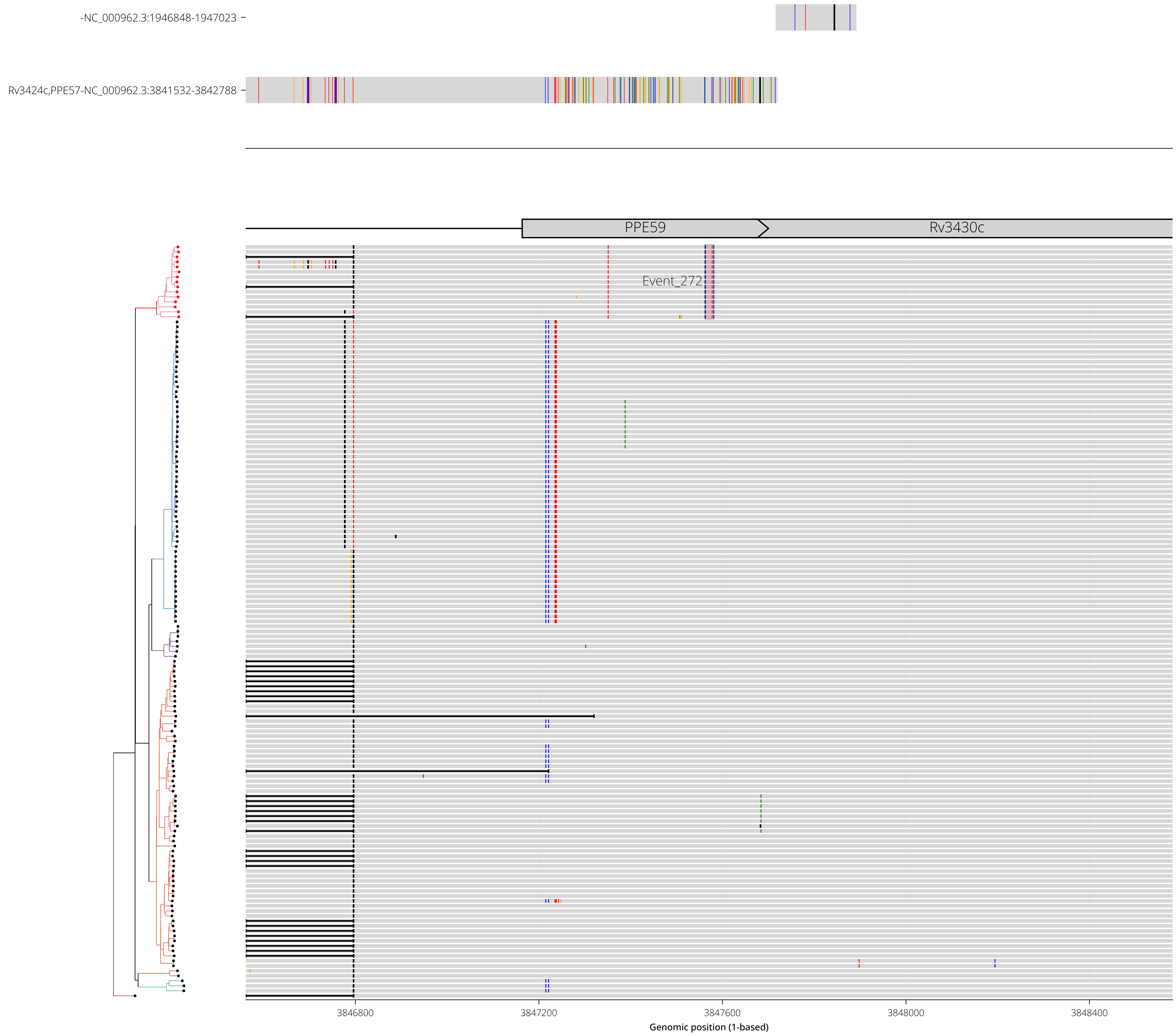

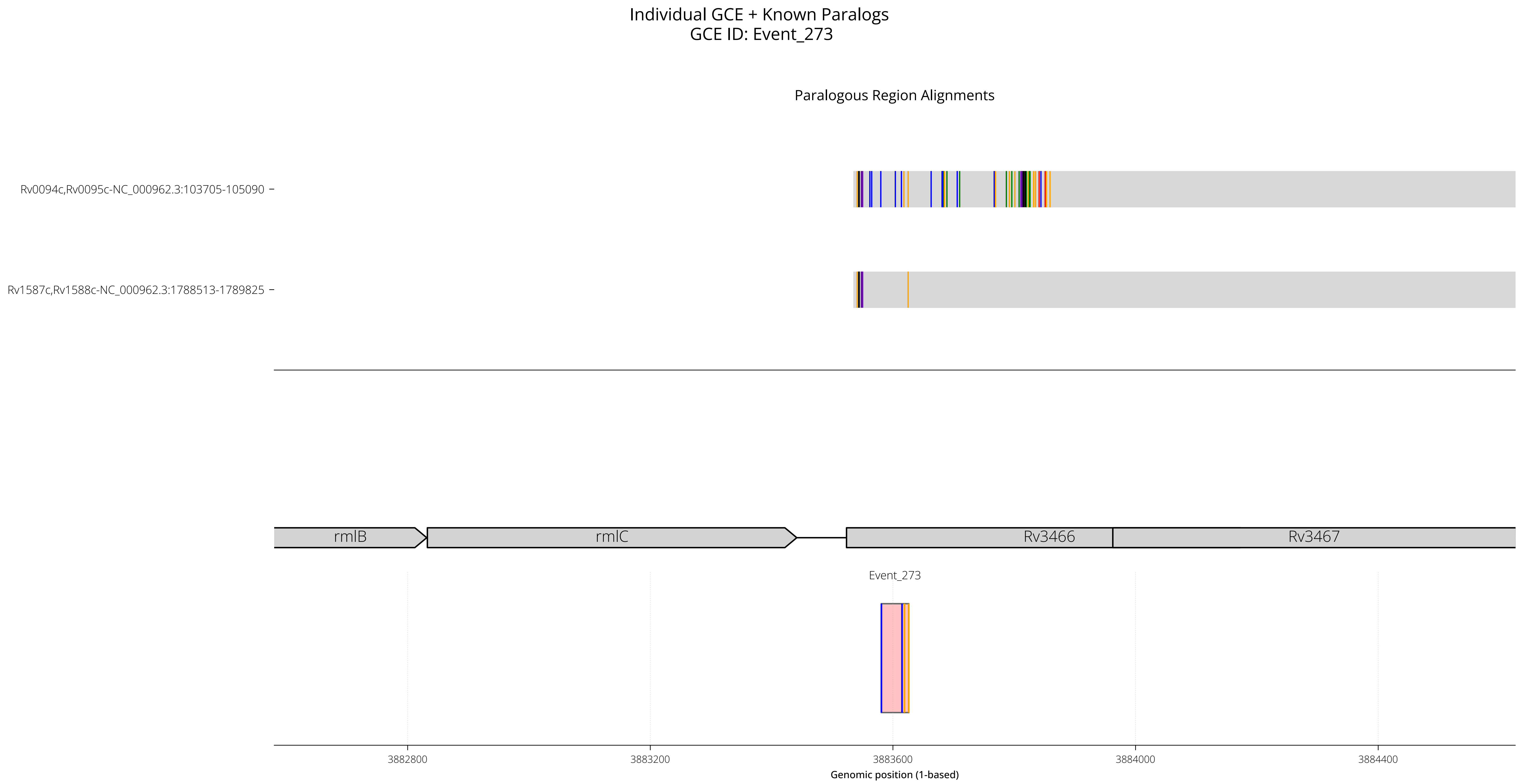
Individual GCE + Known Paralogs
GCE ID: Event_273
Paralogous Region Alignments
Rv0094c,Rv0095c-NC_000962.3:103705-105090
Rv1587c,Rv1588c-NC_000962.3:1788513-1789825
rmlB
rmlC
Rv3466
Rv3467
Event_273
3882800
3883200
3883600
3884000
3884400
Genomic position (1-based)

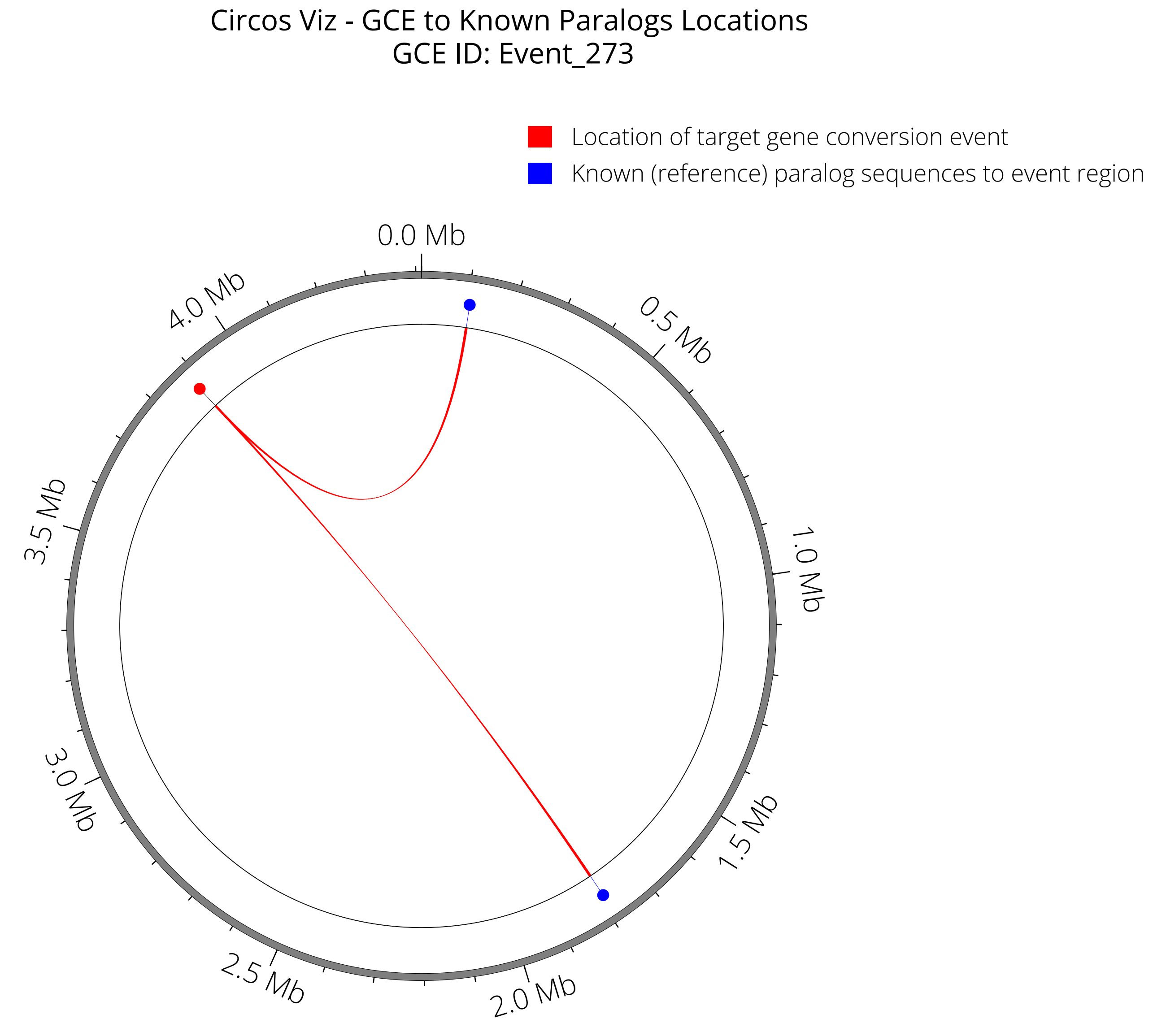
Circos Viz - GCE to Known Paralogs Locations
GCE ID: Event_273
Location of target gene conversion event
Known (reference) paralog sequences to event region
0.0 Mb
0.5 Mb
1.0 Mb
1.5 Mb
2.0 Mb
2.5 Mb
3.0 Mb
3.5 Mb
4.0 Mb

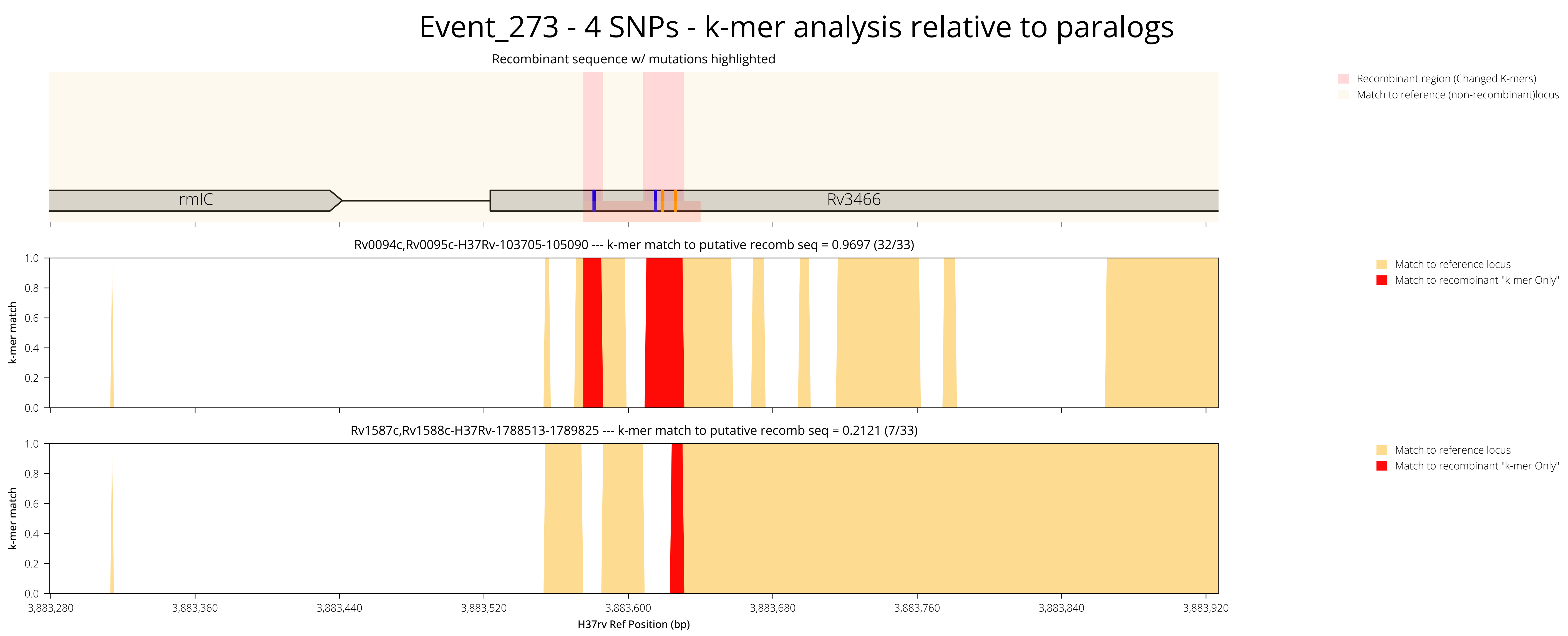

Event_273 - 4 SNPs - k-mer analysis relative to paralogs
Recombinant sequence w/ mutations highlighted
rmlC
Rv3466
Recombinant region (Changed K-mers)
Match to reference (non-recombinant)locus
Rv0094c,Rv0095c-H37Rv-103705-105090 --- k-mer match to putative recomb seq = 0.9697 (32/33)
Match to reference locus
Match to recombinant "k-mer Only"
Rv1587c,Rv1588c-H37Rv-1788513-1789825 --- k-mer match to putative recomb seq = 0.2121 (7/33)
Match to reference locus
Match to recombinant "k-mer Only"
k-mer match
1.0
0.8
0.6
0.4
0.2
0.0
3,883,280
3,883,360
3,883,440
3,883,520
3,883,600
3,883,680
3,883,760
3,883,840
3,883,920
H37rv Ref Position (bp)

Sub-phylogeny Viz + Paralog Alignments
GCE ID: Event_273

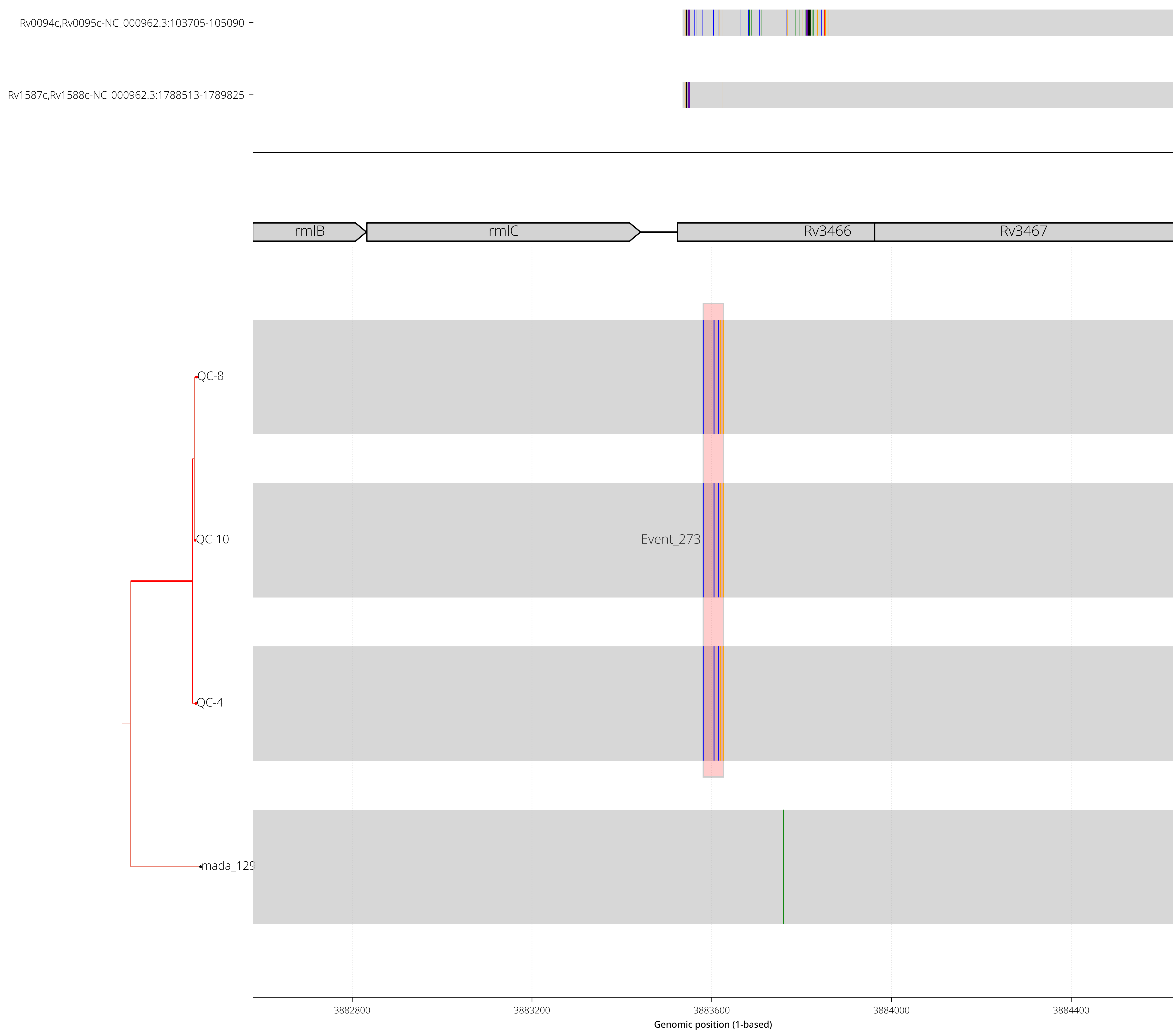

Full Phylogeny Viz + Paralog Alignments
GCE ID: Event_273

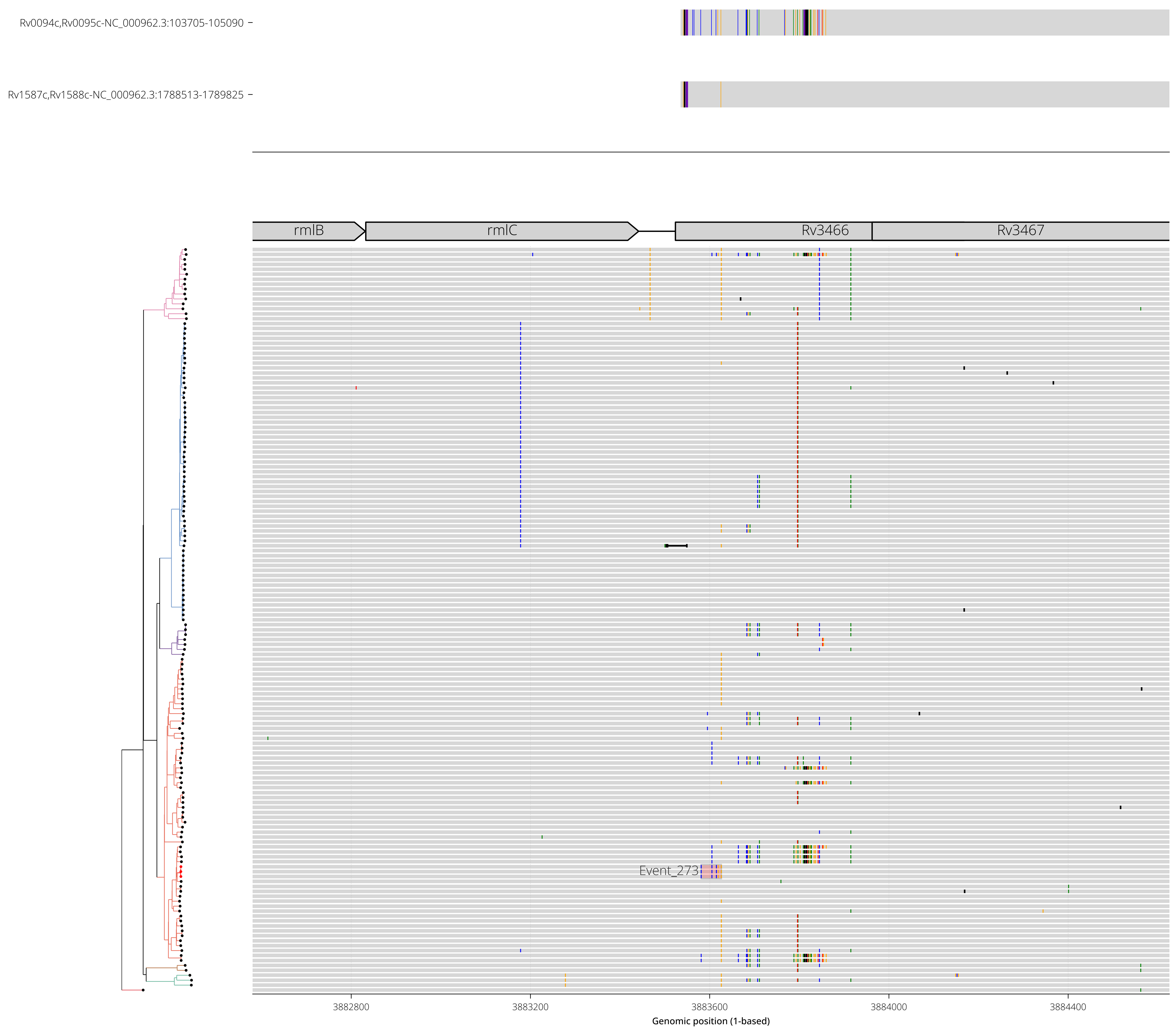

Individual GCE + Known Paralogs
GCE ID: Event_274

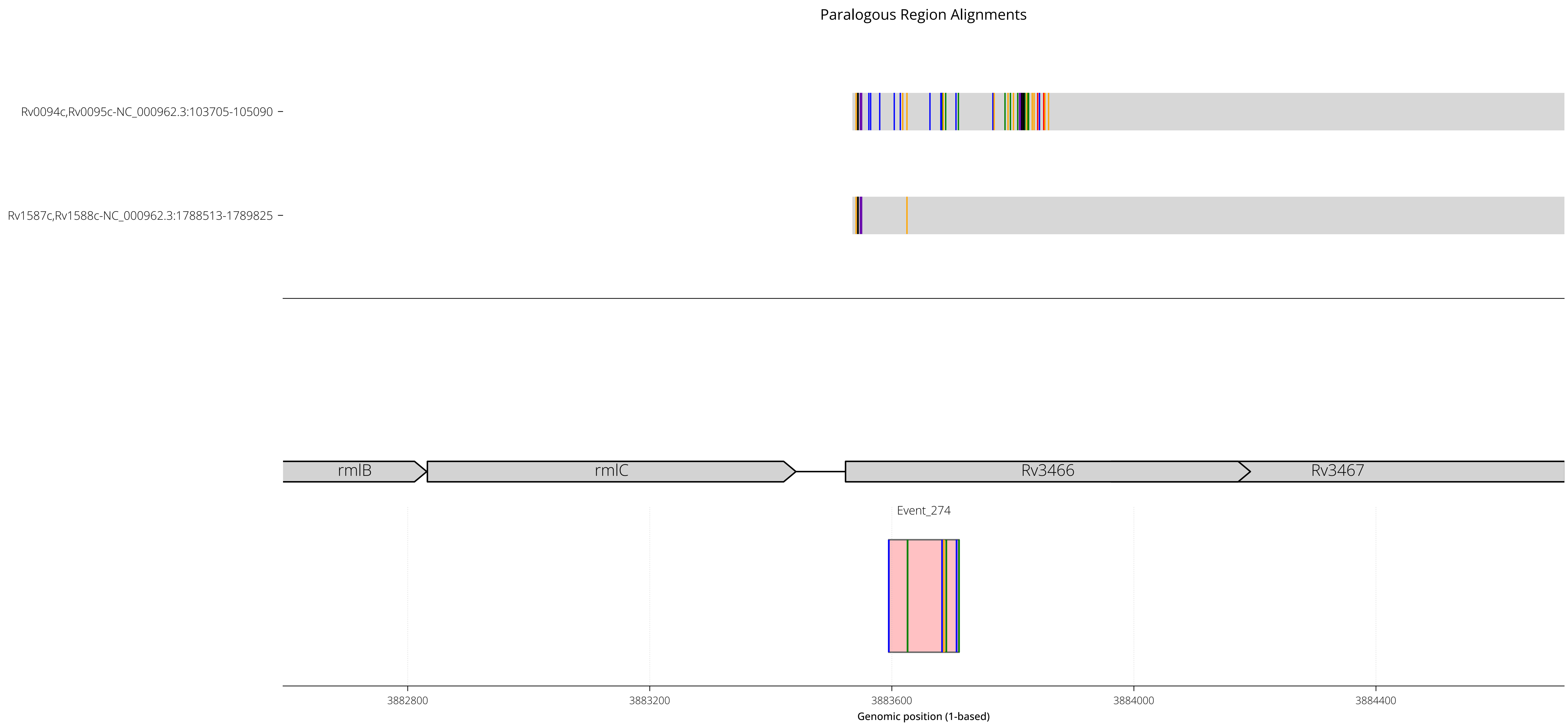

### Circos Viz - GCE to Known Paralogs Locations
### GCE ID: Event_274

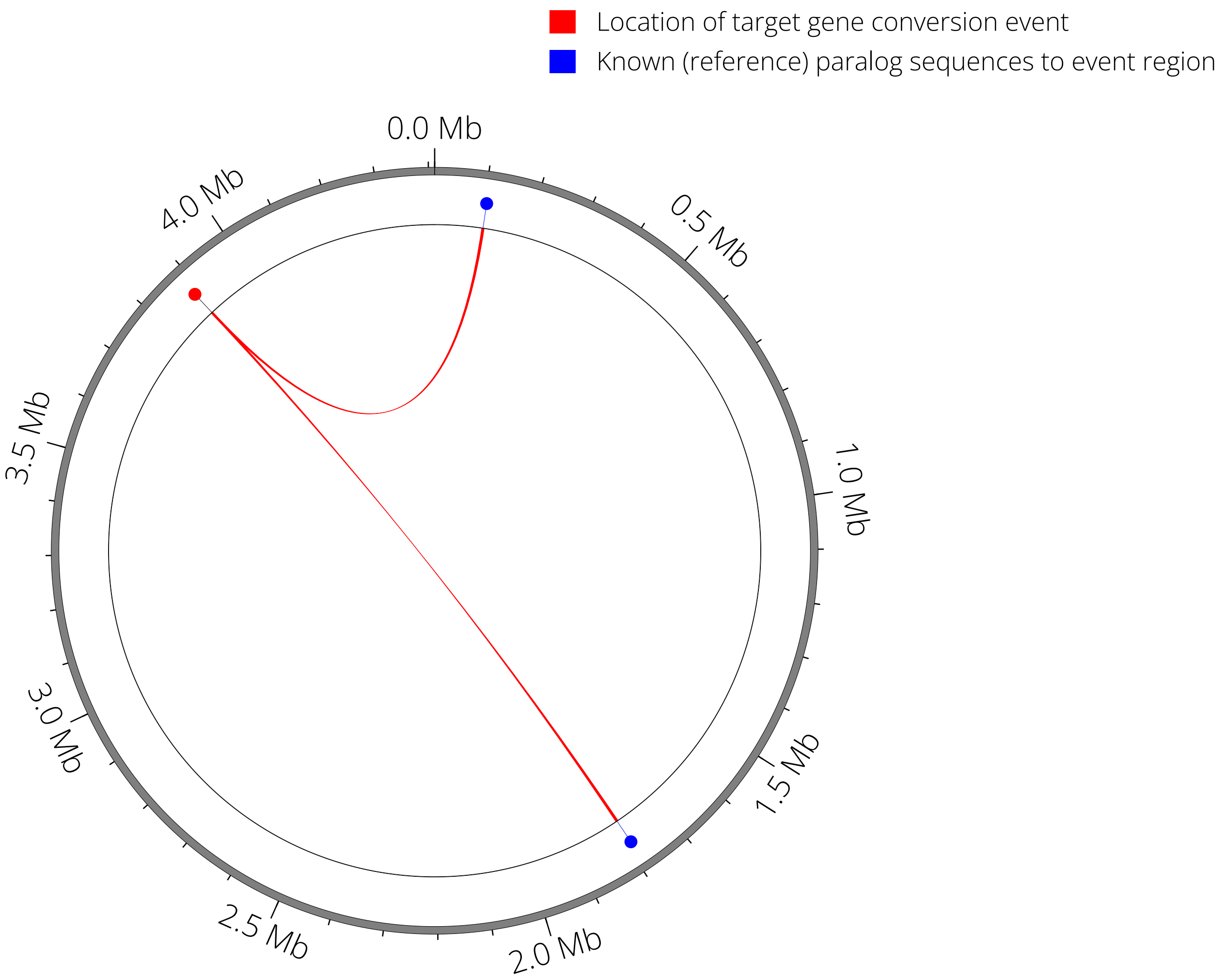

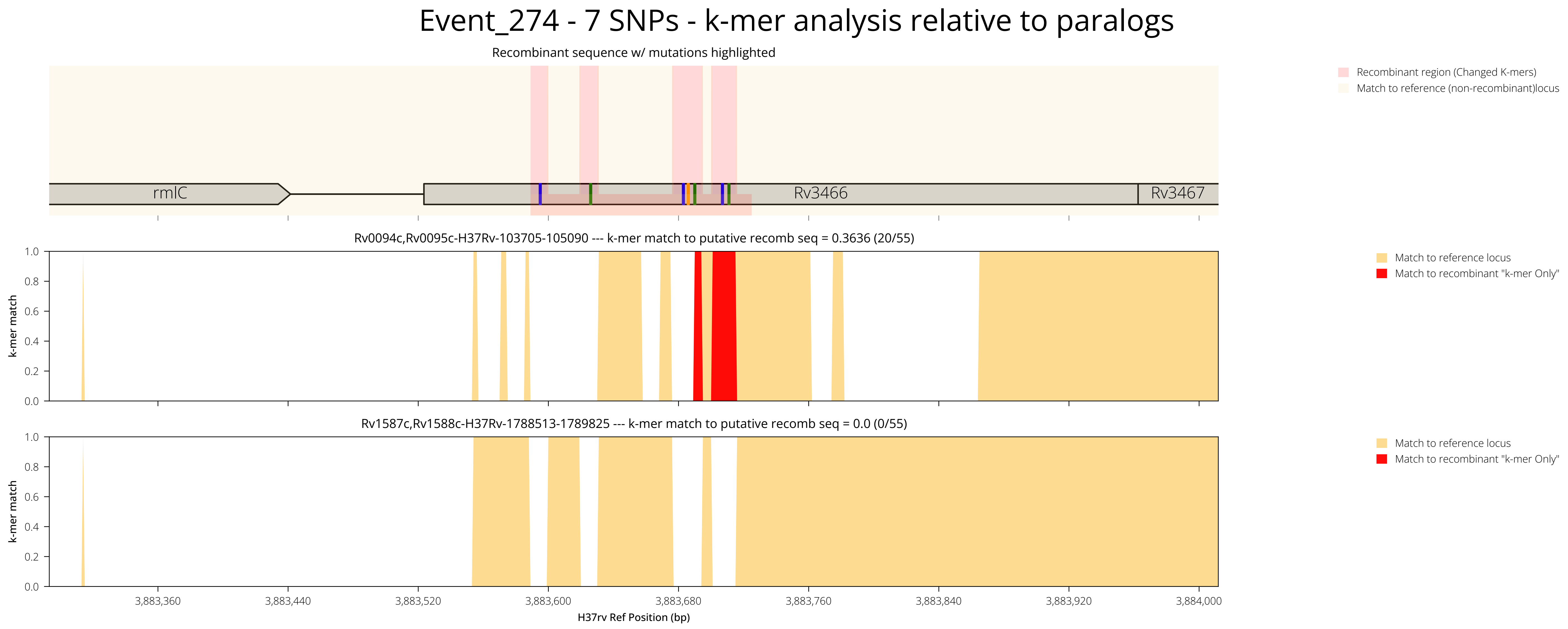

Event_274 - 7 SNPs - k-mer analysis relative to paralogs
Recombinant sequence w/ mutations highlighted
Recombinant region (Changed K-mers)
Match to reference (non-recombinant)locus
rmlC
Rv3466
Rv3467
Rv0094c,Rv0095c-H37Rv-103705-105090 --- k-mer match to putative recomb seq = 0.3636 (20/55)
Match to reference locus
Match to recombinant "k-mer Only"
Rv1587c,Rv1588c-H37Rv-1788513-1789825 --- k-mer match to putative recomb seq = 0.0 (0/55)
Match to reference locus
Match to recombinant "k-mer Only"
k-mer match
1.0
0.8
0.6
0.4
0.2
0.0
3,883,360
3,883,440
3,883,520
3,883,600
3,883,680
3,883,760
3,883,840
3,883,920
3,884,000
H37rv Ref Position (bp)

Sub-phylogeny Viz + Paralog Alignments
GCE ID: Event_274

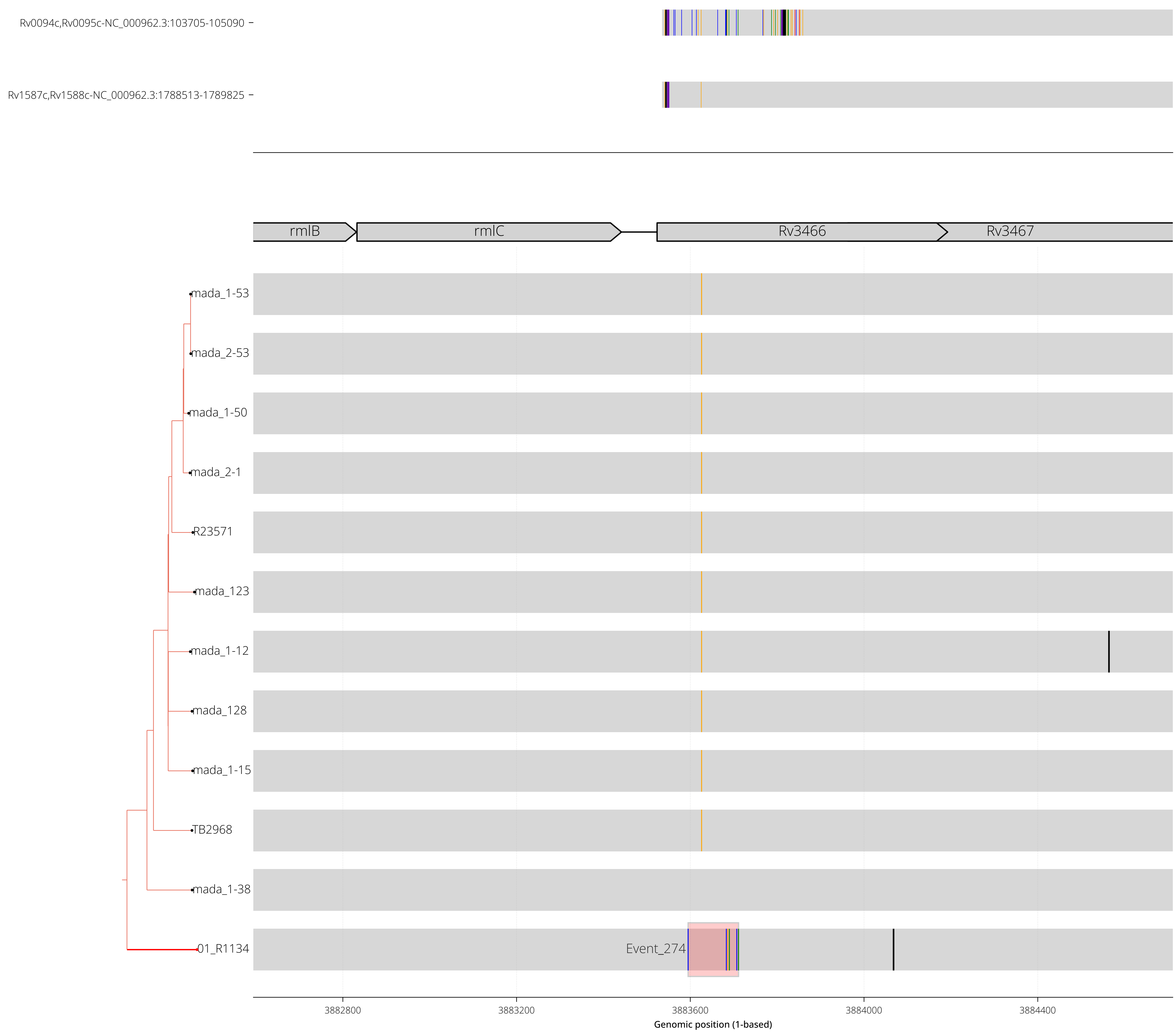

Full Phylogeny Viz + Paralog Alignments
GCE ID: Event_274

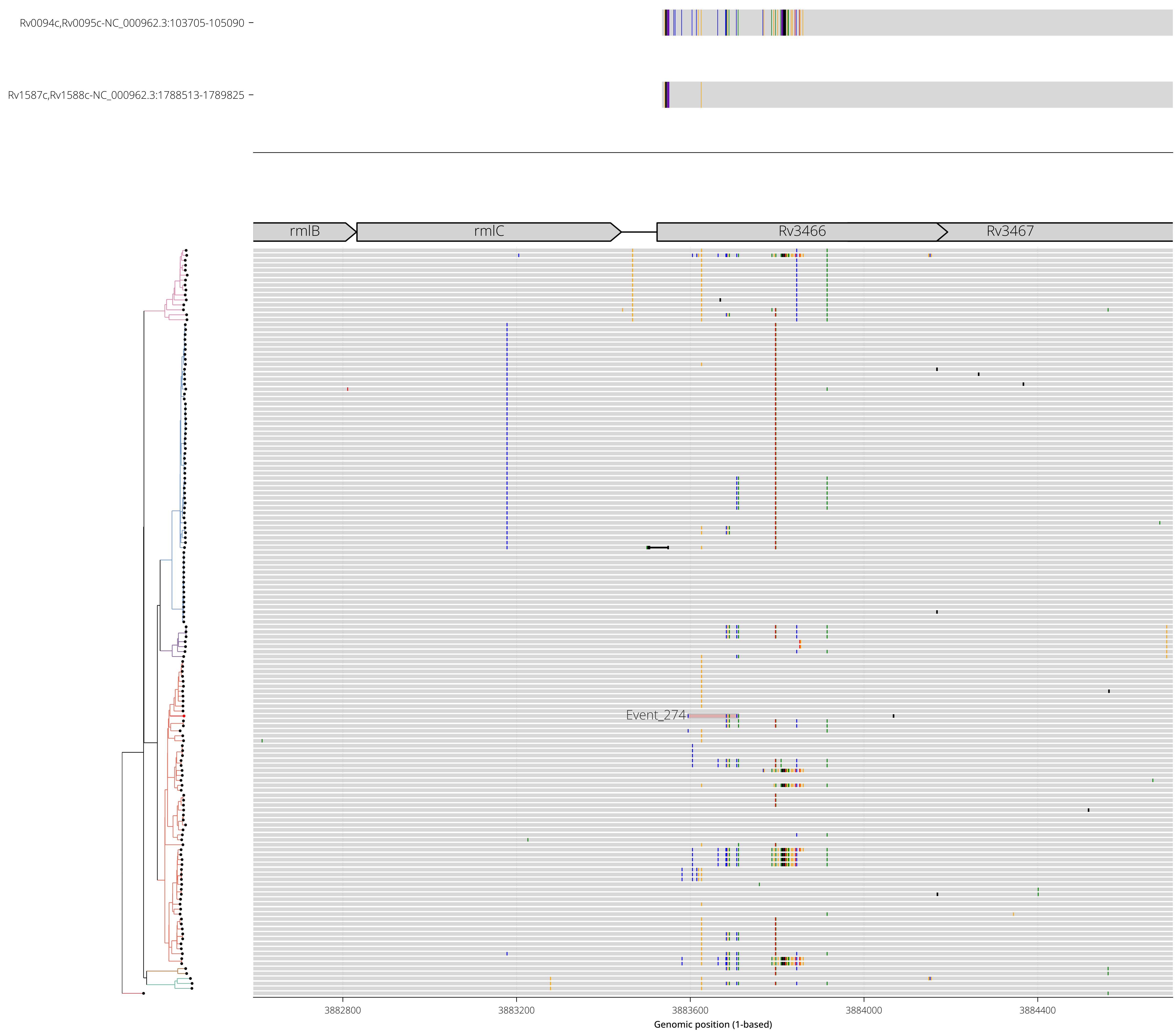

Individual GCE + Known Paralogs
GCE ID: Event_275

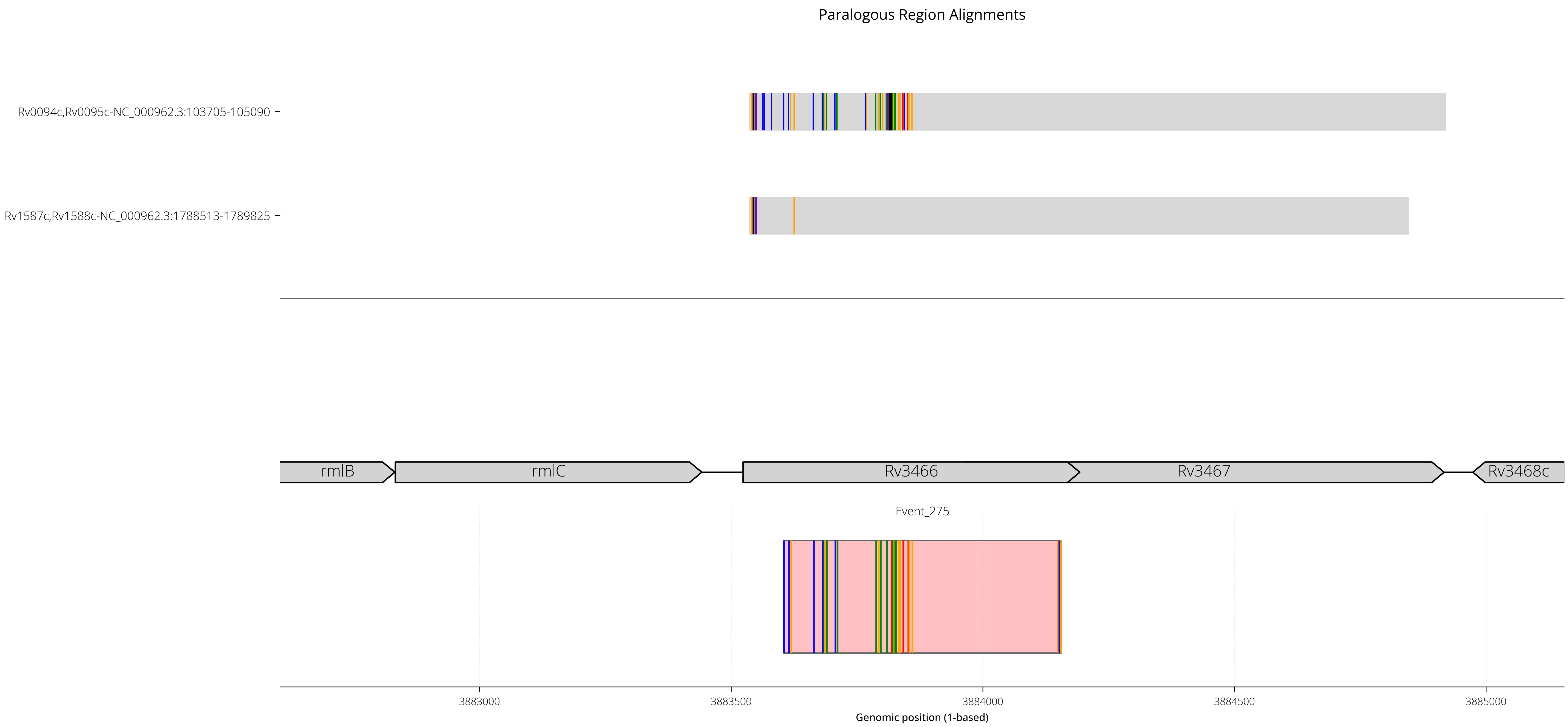

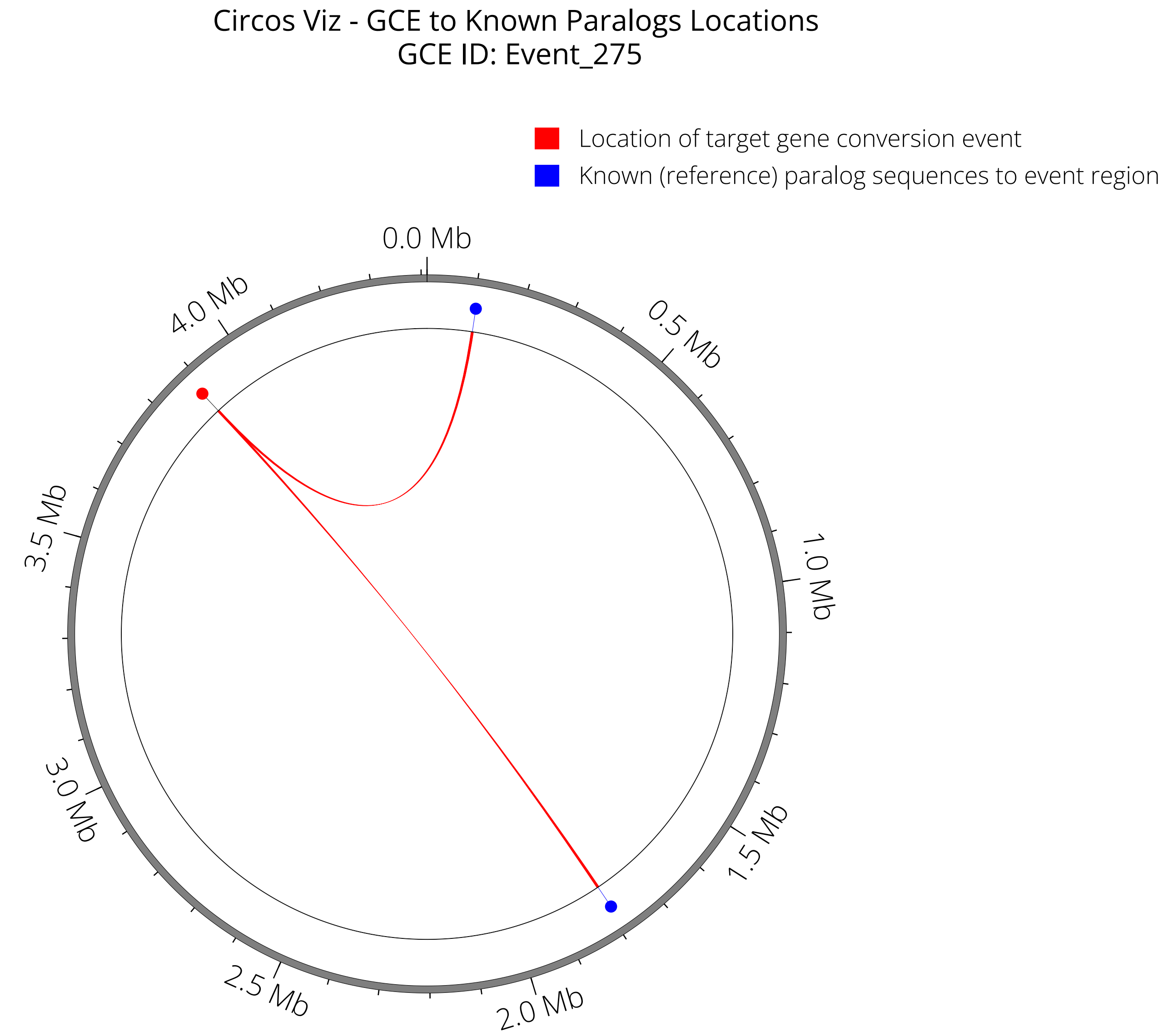
Circos Viz - GCE to Known Paralogs Locations
GCE ID: Event_275
Location of target gene conversion event
Known (reference) paralog sequences to event region
0.0 Mb
0.5 Mb
1.0 Mb
1.5 Mb
2.0 Mb
2.5 Mb
3.0 Mb
3.5 Mb
4.0 Mb

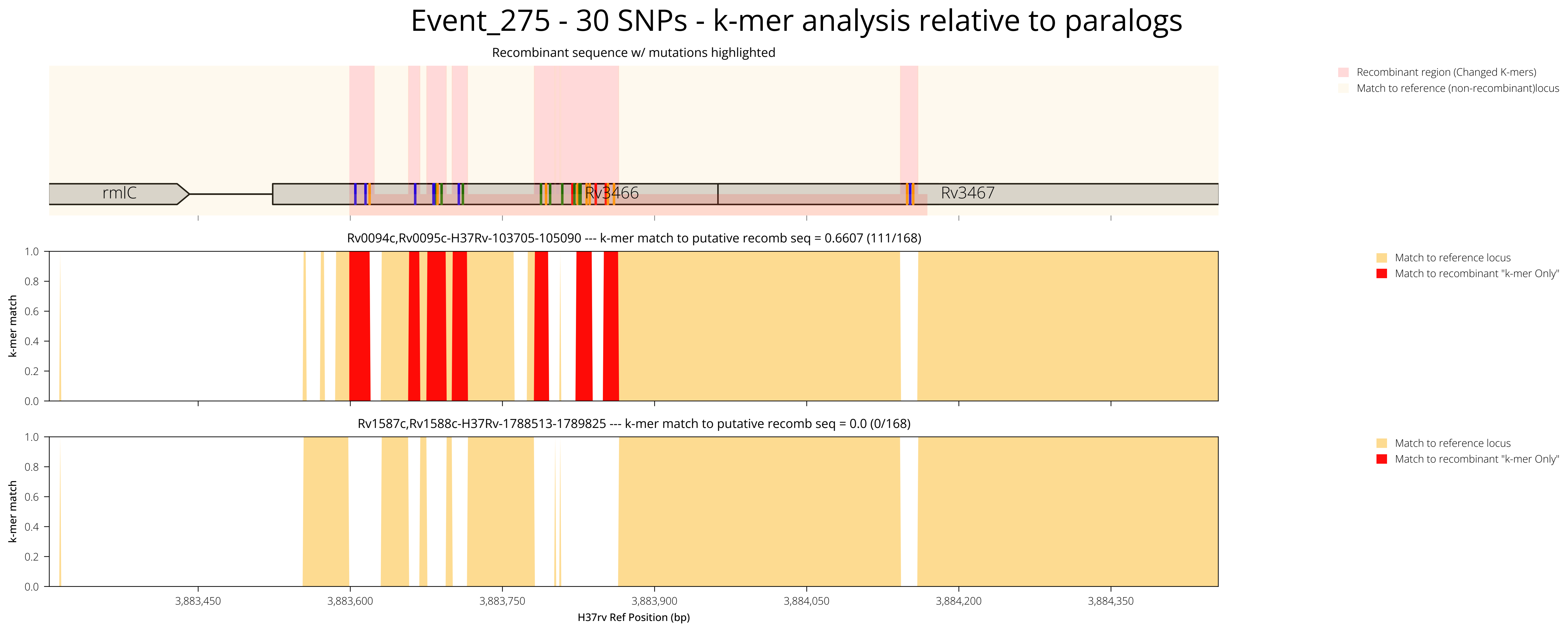

Event_275 - 30 SNPs - k-mer analysis relative to paralogs
Recombinant sequence w/ mutations highlighted
Recombinant region (Changed K-mers)
Match to reference (non-recombinant)locus
rmlC
Rv3466
Rv3467
Rv0094c,Rv0095c-H37Rv-103705-105090 --- k-mer match to putative recomb seq = 0.6607 (111/168)
Match to reference locus
Match to recombinant "k-mer Only"
k-mer match
Rv1587c,Rv1588c-H37Rv-1788513-1789825 --- k-mer match to putative recomb seq = 0.0 (0/168)
Match to reference locus
Match to recombinant "k-mer Only"
k-mer match
3,883,450
3,883,600
3,883,750
3,883,900
3,884,050
3,884,200
3,884,350
H37rv Ref Position (bp)

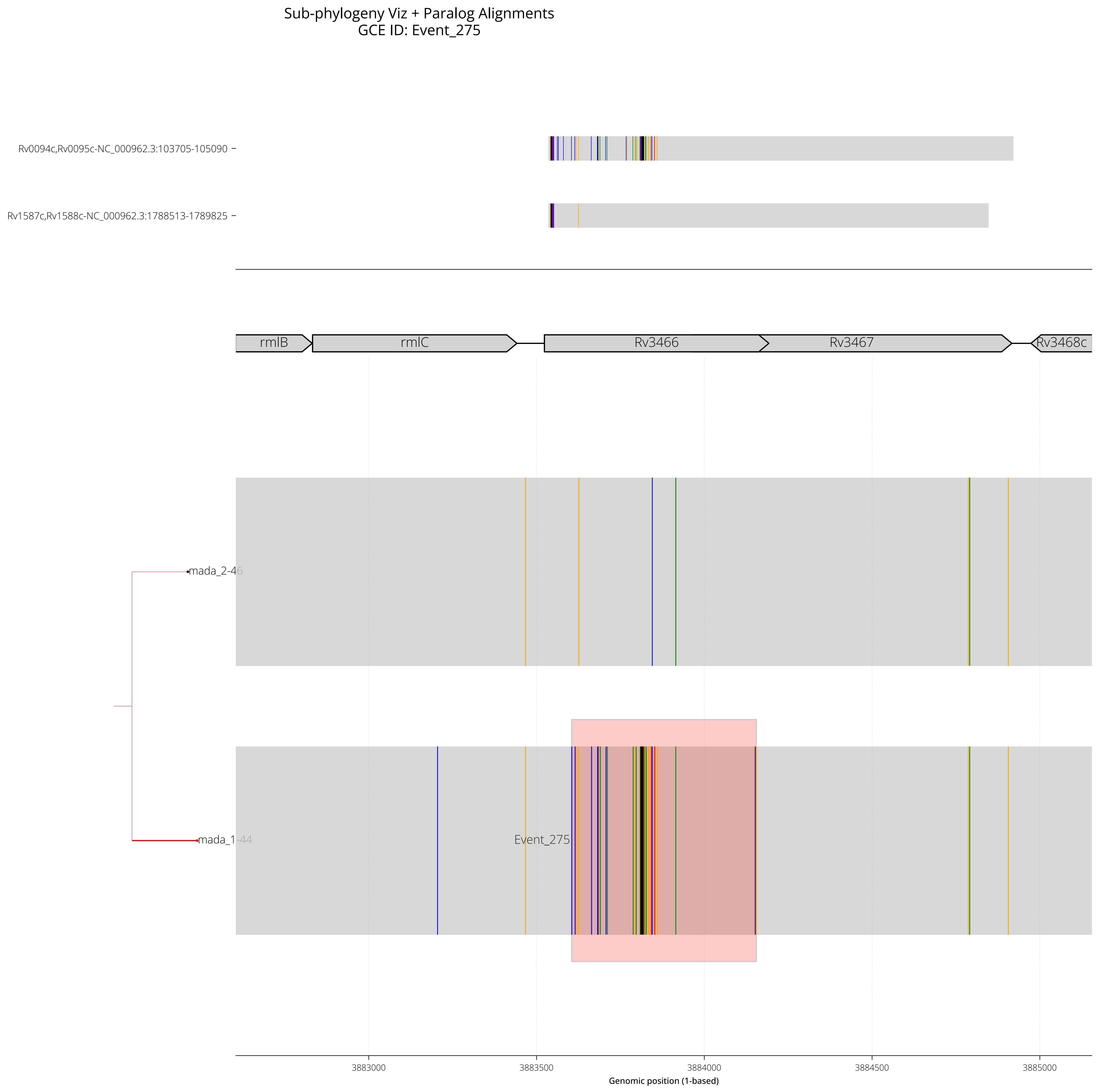

Sub-phylogeny Viz + Paralog Alignments
GCE ID: Event_275
Rv0094c,Rv0095c-NC_000962.3:103705-105090
Rv1587c,Rv1588c-NC_000962.3:1788513-1789825
rmlB
rmlC
Rv3466
Rv3467
Rv3468c
mada_2-46
mada_1-44
Event_275
3883000
3883500
3884000
3884500
3885000
Genomic position (1-based)

Full Phylogeny Viz + Paralog Alignments
GCE ID: Event_275

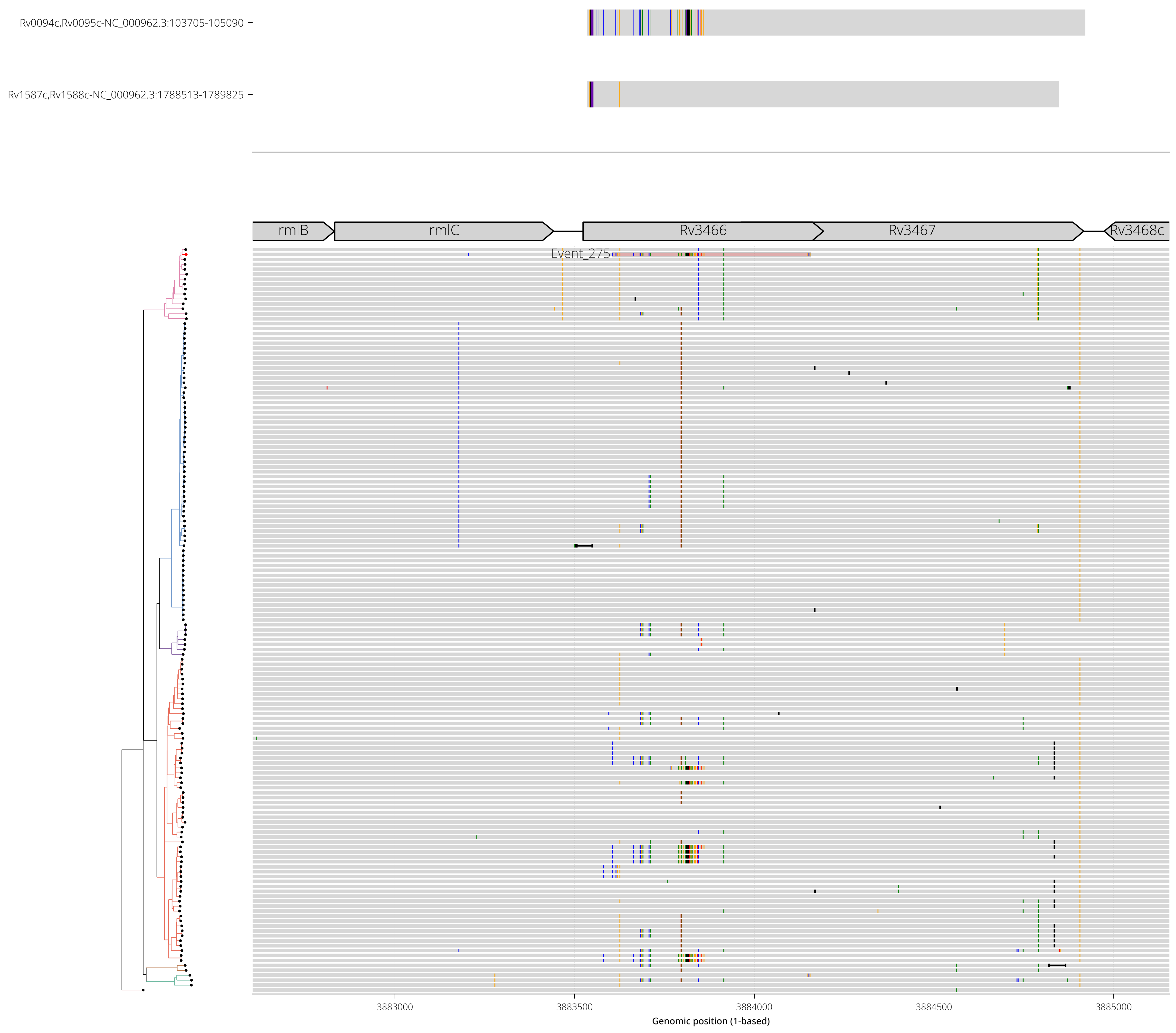

Individual GCE + Known Paralogs
GCE ID: Event_276

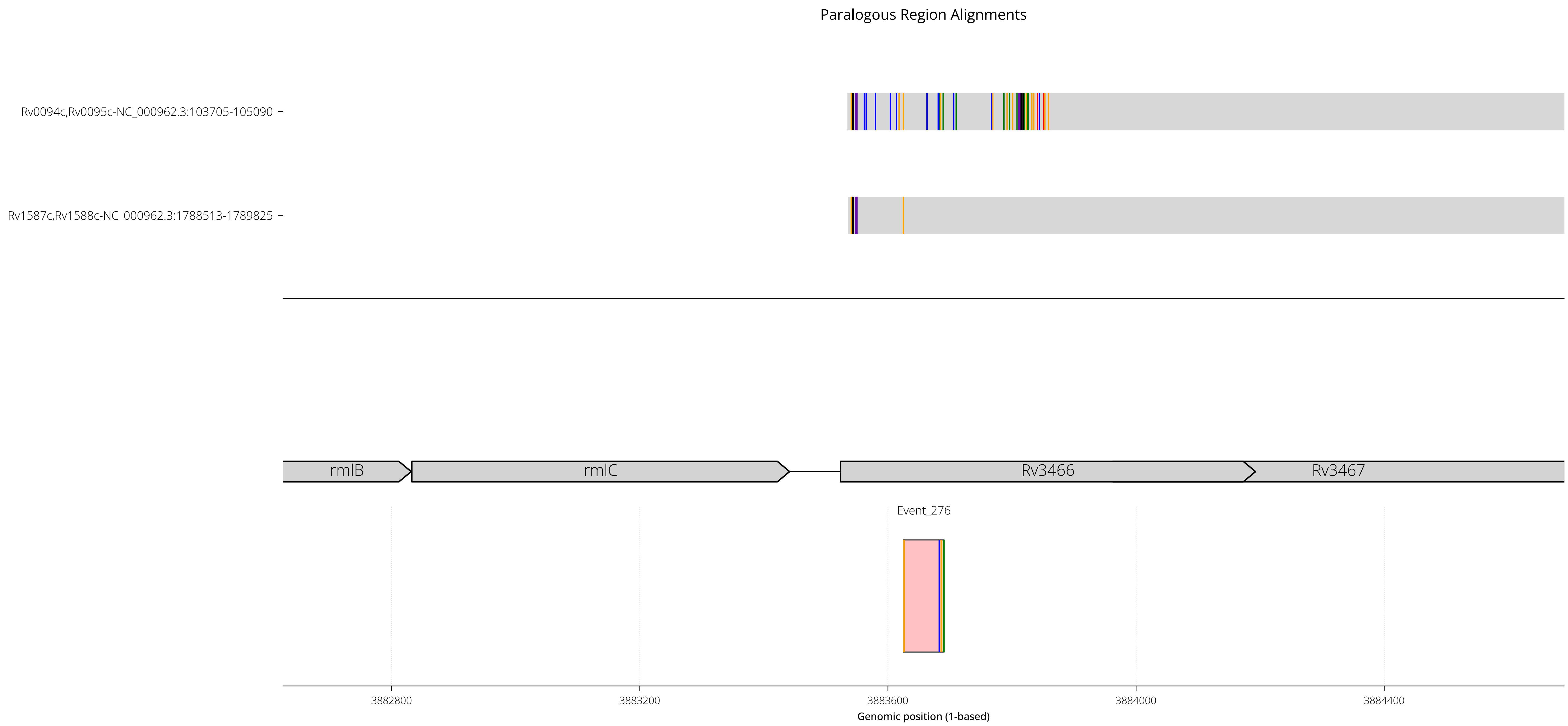

Circos Viz - GCE to Known Paralogs Locations
GCE ID: Event_276

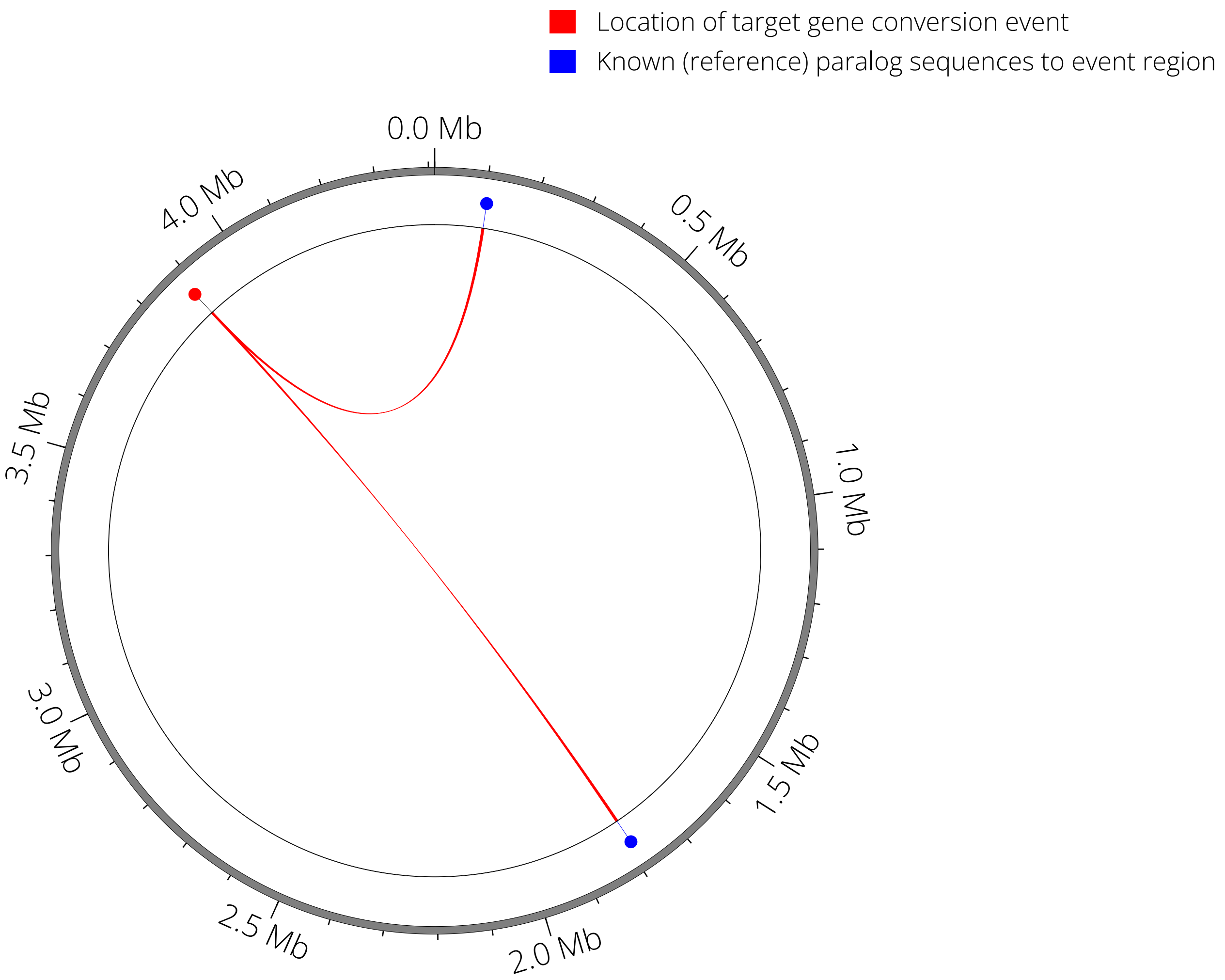

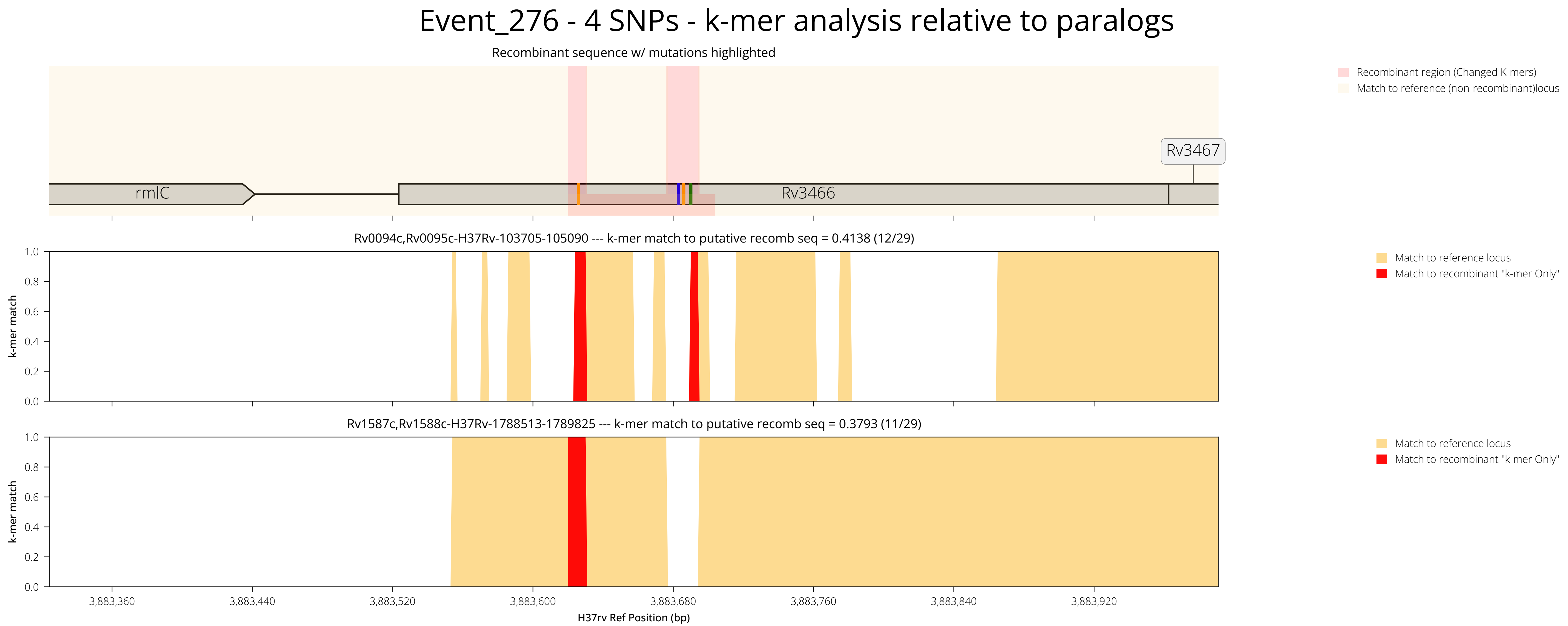

Event_276 - 4 SNPs - k-mer analysis relative to paralogs
Recombinant sequence w/ mutations highlighted
Recombinant region (Changed K-mers)
Match to reference (non-recombinant)locus
rmlC
Rv3466
Rv3467
Rv0094c,Rv0095c-H37Rv-103705-105090 --- k-mer match to putative recomb seq = 0.4138 (12/29)
Match to reference locus
Match to recombinant "k-mer Only"
k-mer match
1.0
0.8
0.6
0.4
0.2
0.0
Rv1587c,Rv1588c-H37Rv-1788513-1789825 --- k-mer match to putative recomb seq = 0.3793 (11/29)
Match to reference locus
Match to recombinant "k-mer Only"
k-mer match
3,883,360
3,883,440
3,883,520
3,883,600
3,883,680
3,883,760
3,883,840
3,883,920
H37rv Ref Position (bp)

Sub-phylogeny Viz + Paralog Alignments
GCE ID: Event_276

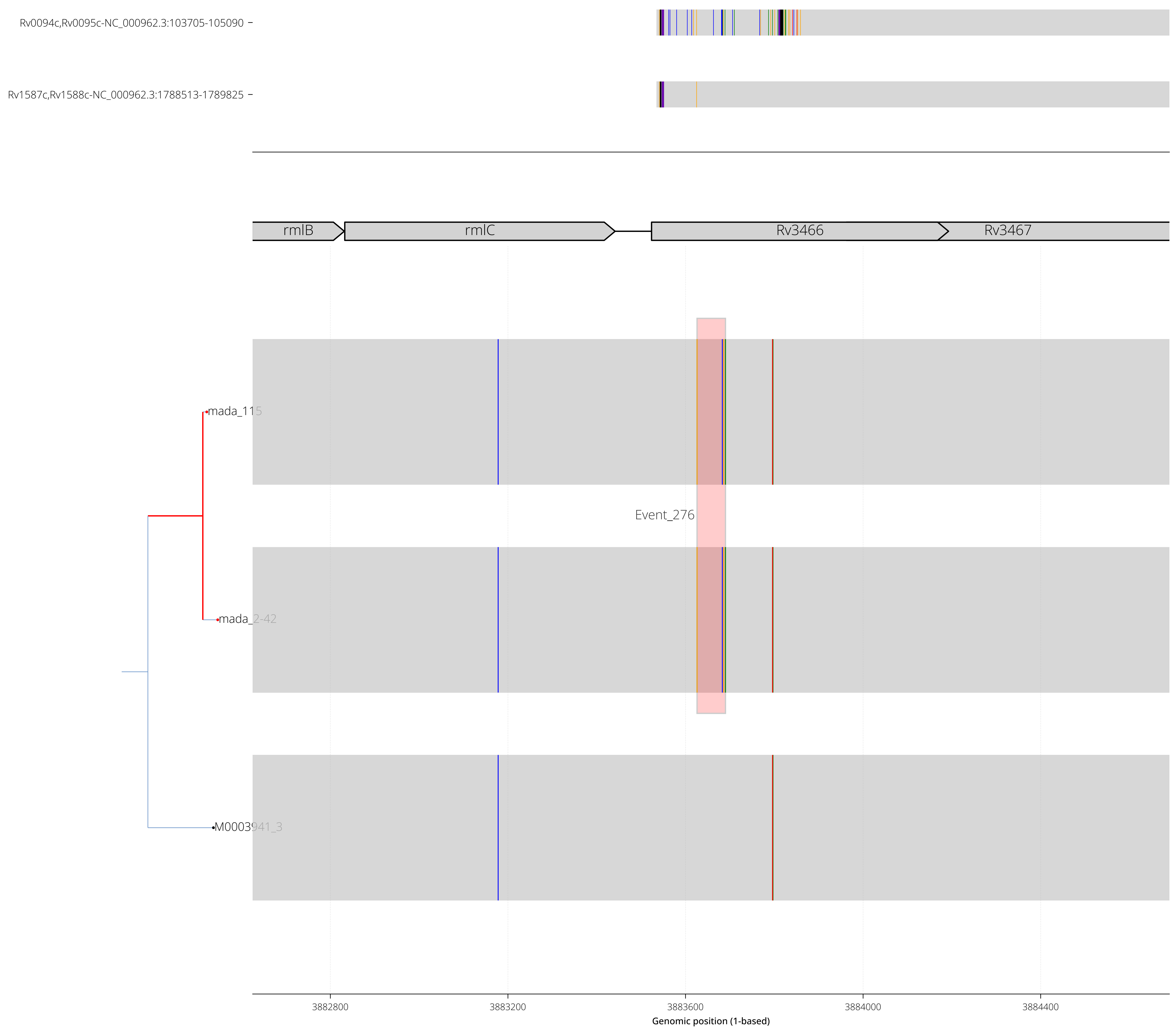

Full Phylogeny Viz + Paralog Alignments
GCE ID: Event_276

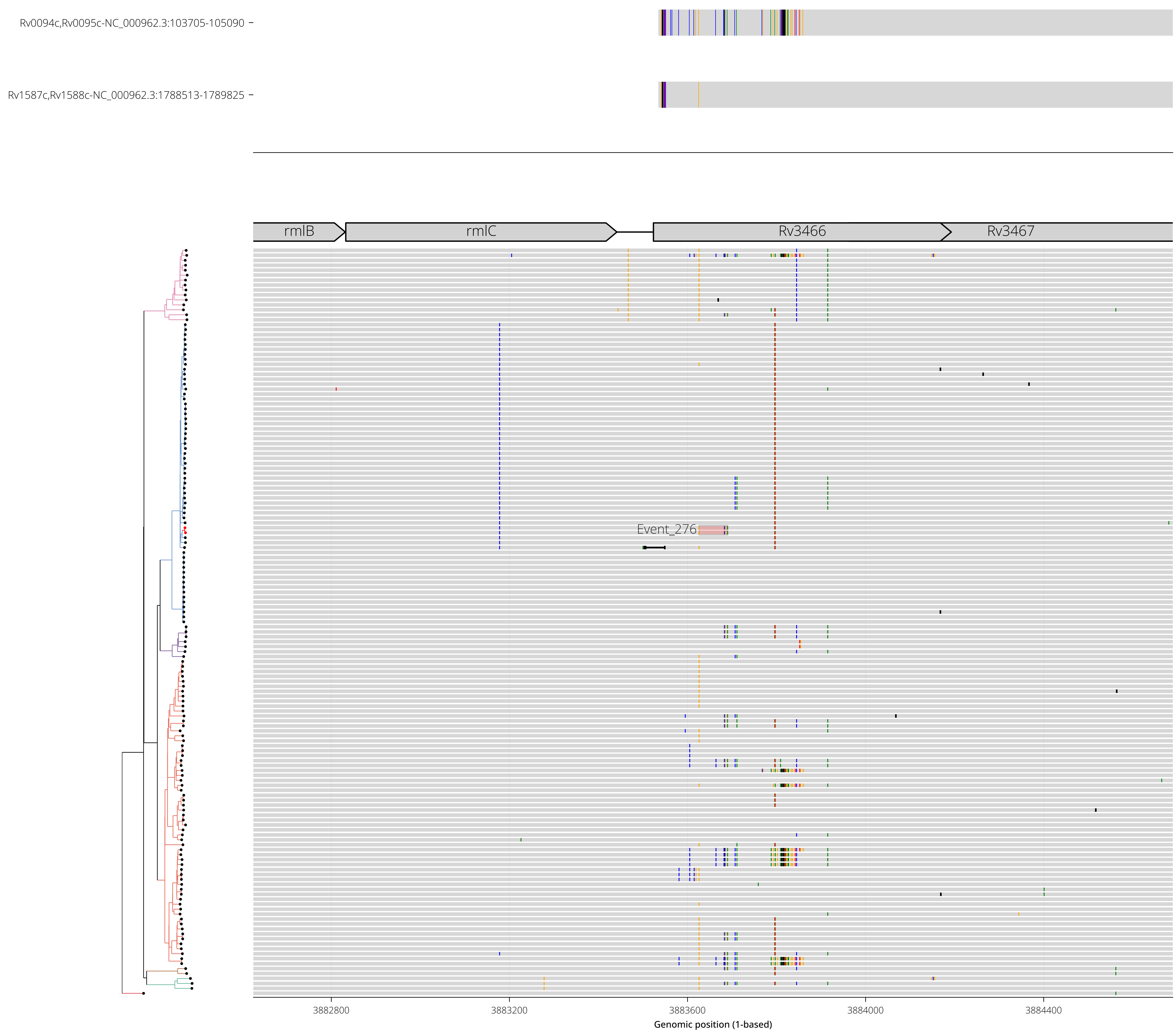

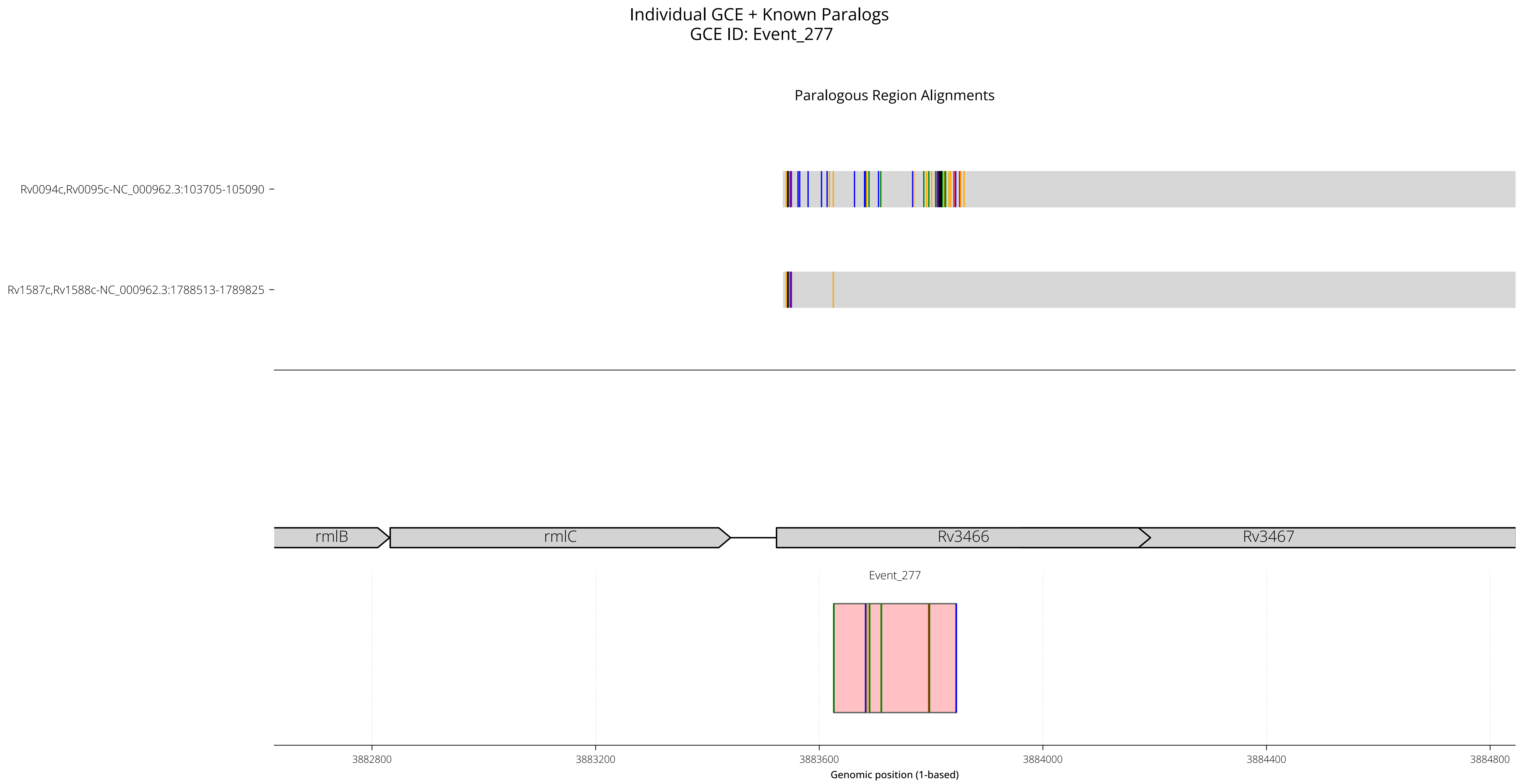
Individual GCE + Known Paralogs
GCE ID: Event_277
Paralogous Region Alignments
Rv0094c,Rv0095c-NC_000962.3:103705-105090
Rv1587c,Rv1588c-NC_000962.3:1788513-1789825
rmlB
rmlC
Rv3466
Rv3467
Event_277
3882800
3883200
3883600
3884000
3884400
3884800
Genomic position (1-based)

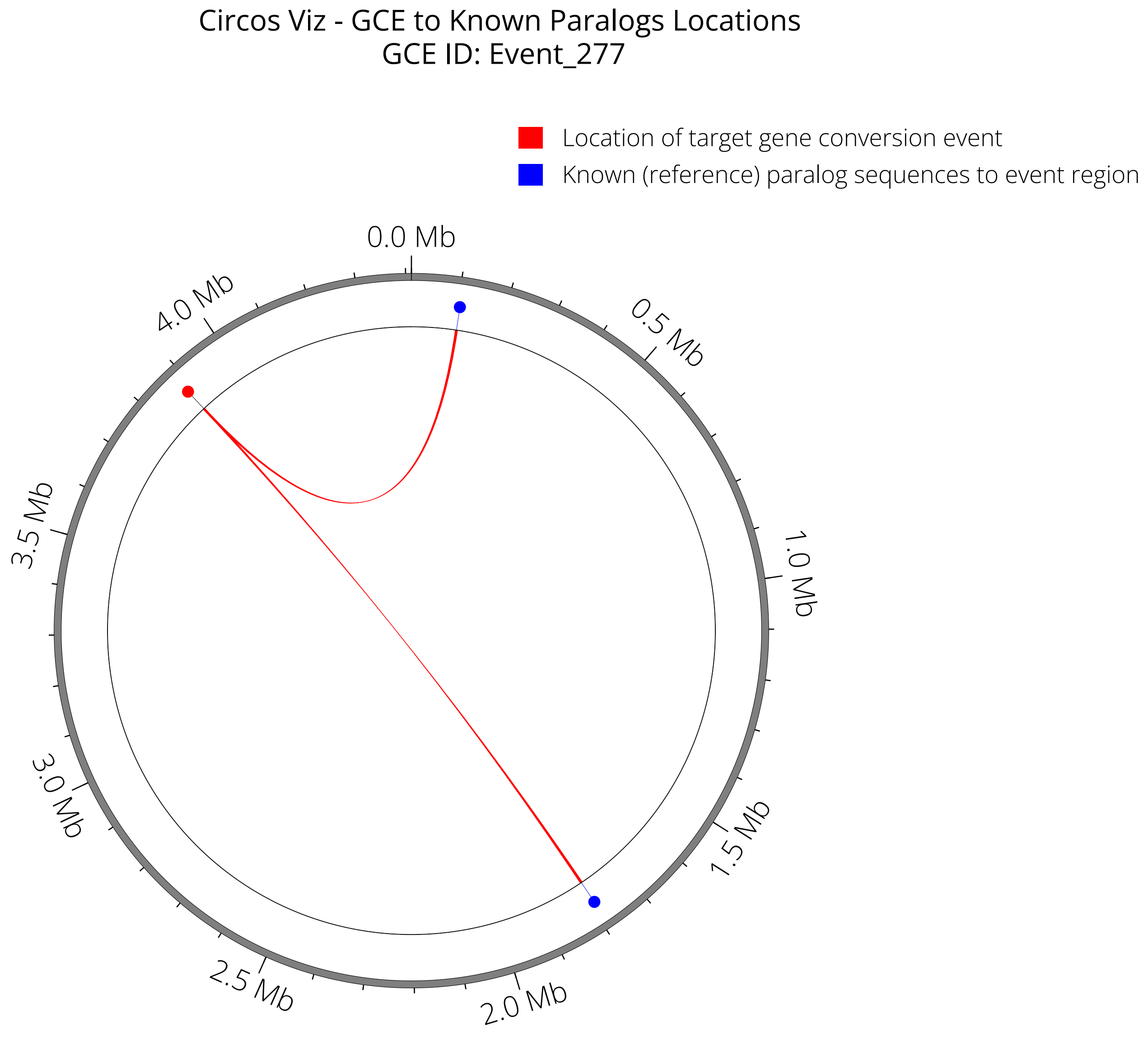
Circos Viz - GCE to Known Paralogs Locations
GCE ID: Event_277
Location of target gene conversion event
Known (reference) paralog sequences to event region
0.0 Mb
0.5 Mb
1.0 Mb
1.5 Mb
2.0 Mb
2.5 Mb
3.0 Mb
3.5 Mb
4.0 Mb

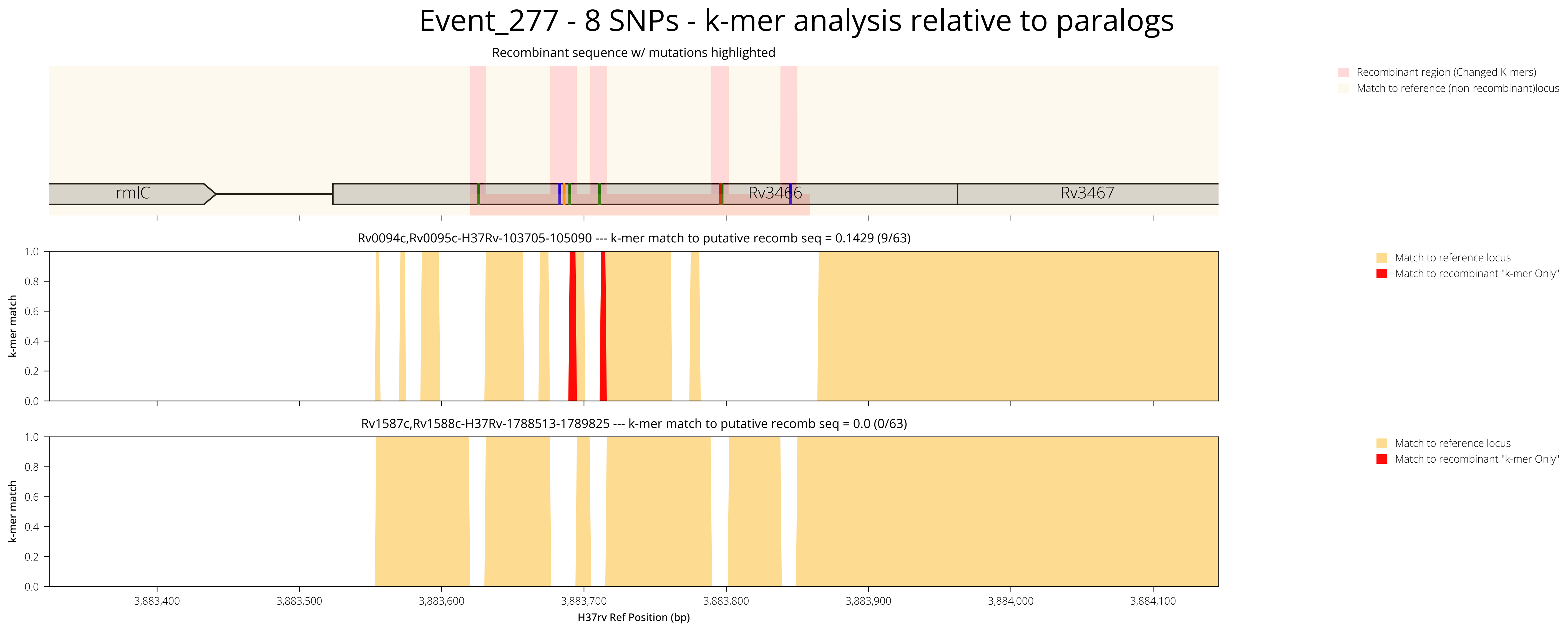

Event_277 - 8 SNPs - k-mer analysis relative to paralogs
Recombinant sequence w/ mutations highlighted
rmlC
Rv3466
Rv3467
Recombinant region (Changed K-mers)
Match to reference (non-recombinant)locus
Rv0094c,Rv0095c-H37Rv-103705-105090 --- k-mer match to putative recomb seq = 0.1429 (9/63)
Match to reference locus
Match to recombinant "k-mer Only"
Rv1587c,Rv1588c-H37Rv-1788513-1789825 --- k-mer match to putative recomb seq = 0.0 (0/63)
Match to reference locus
Match to recombinant "k-mer Only"
k-mer match
1.0
0.8
0.6
0.4
0.2
0.0
3,883,400
3,883,500
3,883,600
3,883,700
3,883,800
3,883,900
3,884,000
3,884,100
H37rv Ref Position (bp)

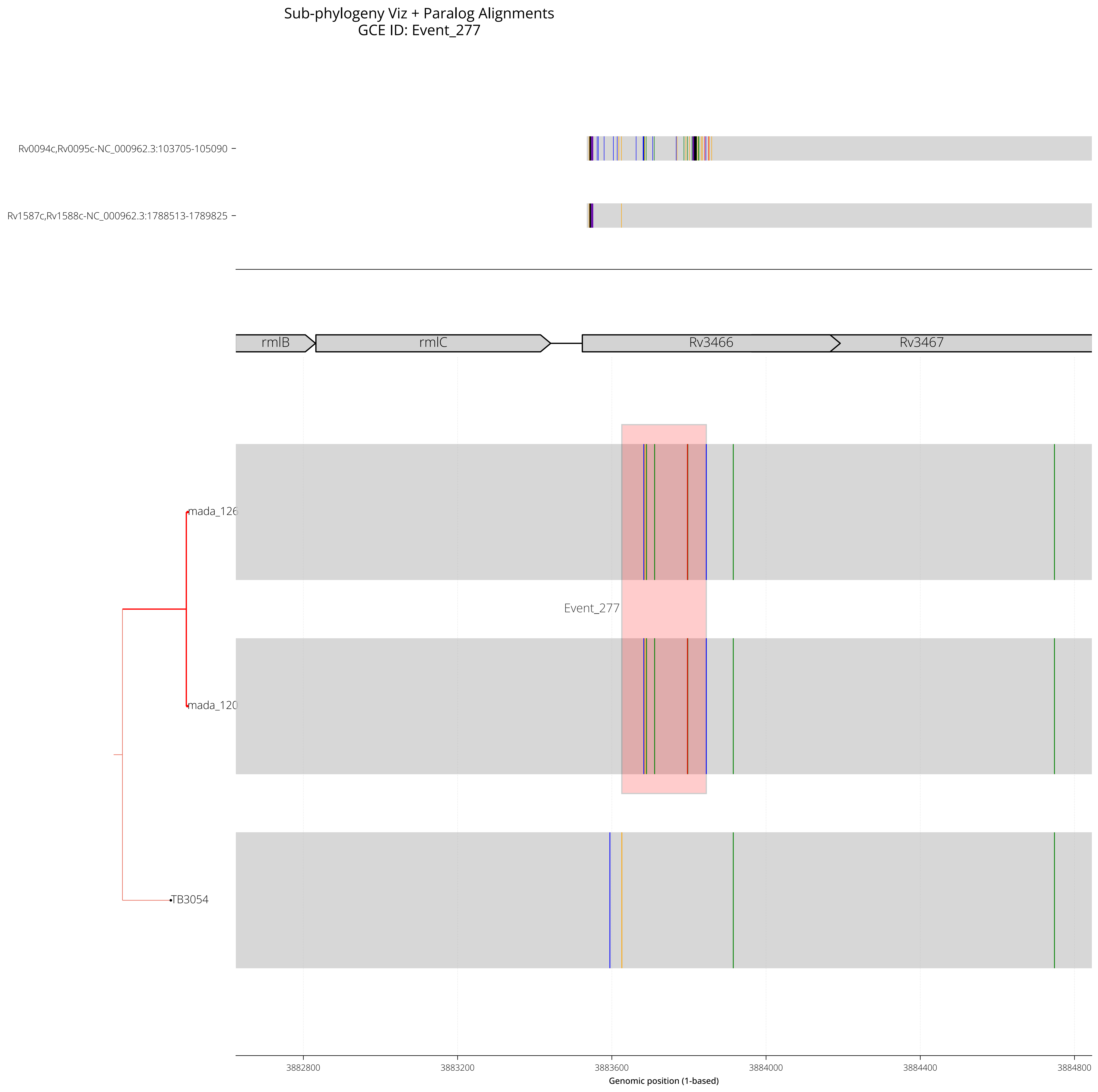

Sub-phylogeny Viz + Paralog Alignments
GCE ID: Event_277
Rv0094c,Rv0095c-NC_000962.3:103705-105090
Rv1587c,Rv1588c-NC_000962.3:1788513-1789825
rmlB
rmlC
Rv3466
Rv3467
mada_126
Event_277
mada_120
TB3054
3882800
3883200
3883600
3884000
3884400
3884800
Genomic position (1-based)

Full Phylogeny Viz + Paralog Alignments
GCE ID: Event_277

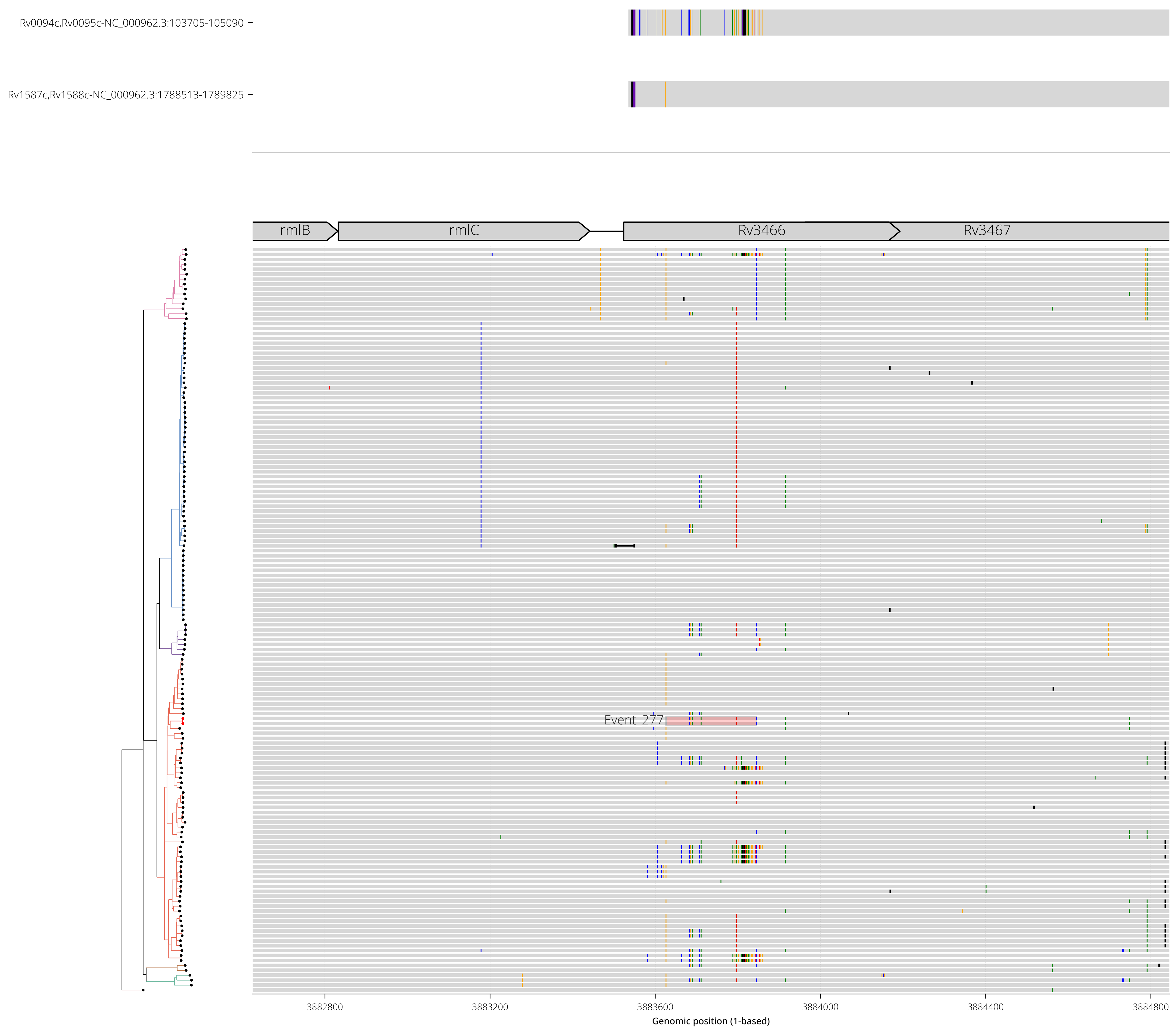

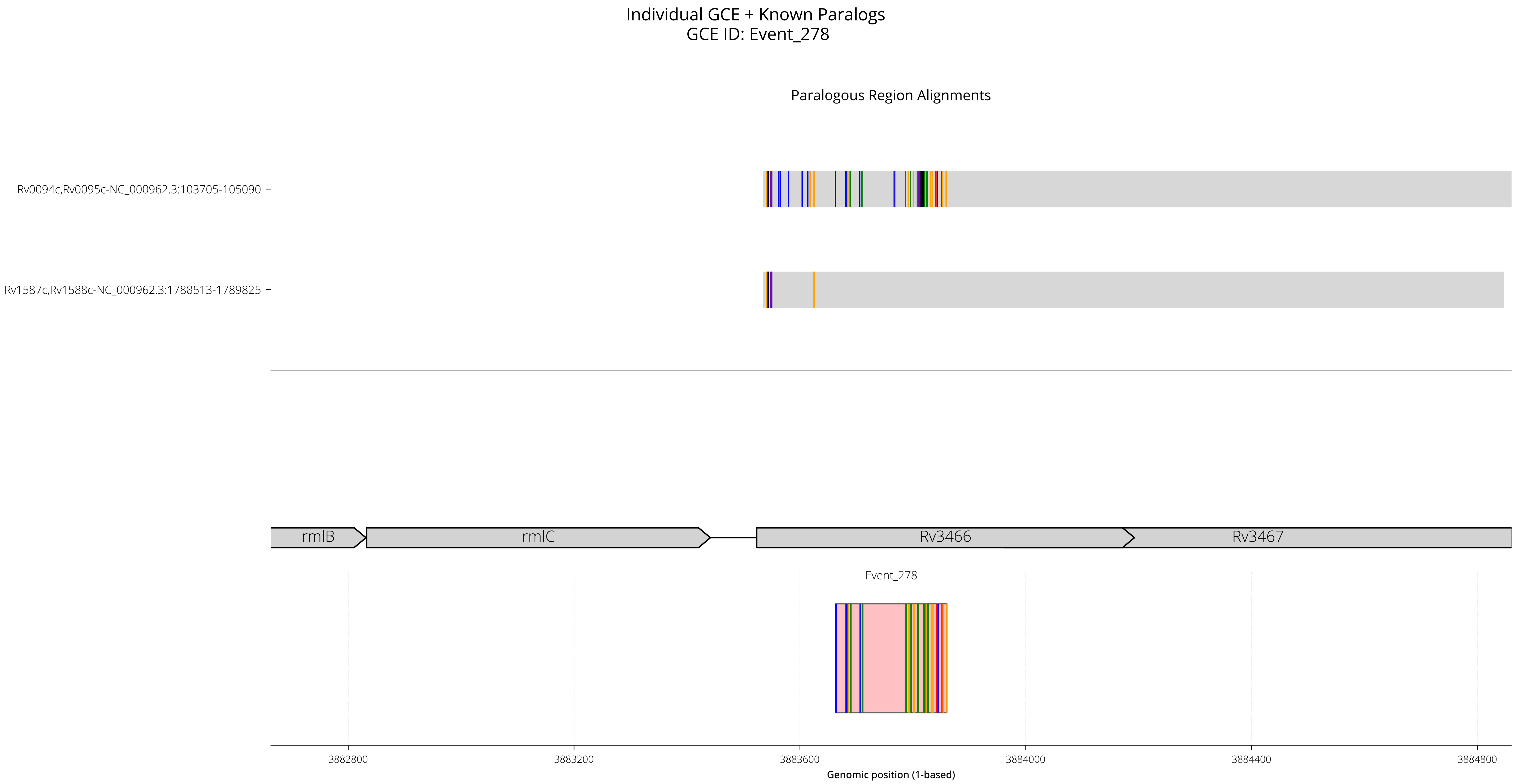

Individual GCE + Known Paralogs
GCE ID: Event_278
Paralogous Region Alignments
Rv0094c,Rv0095c-NC_000962.3:103705-105090
Rv1587c,Rv1588c-NC_000962.3:1788513-1789825
rmlB
rmlC
Rv3466
Rv3467
Event_278
3882800
3883200
3883600
3884000
3884400
3884800
Genomic position (1-based)

Circos Viz - GCE to Known Paralogs Locations
GCE ID: Event_278

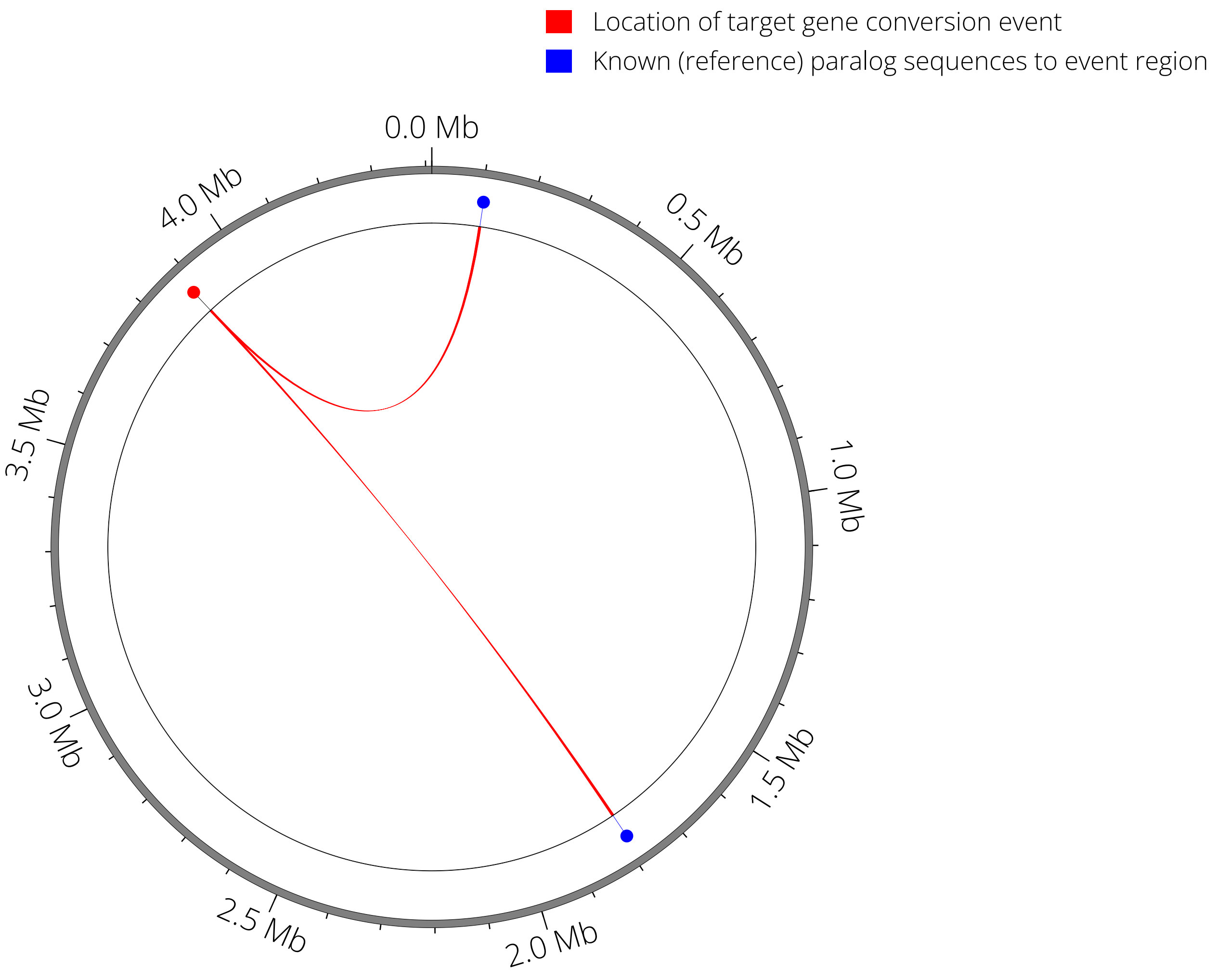

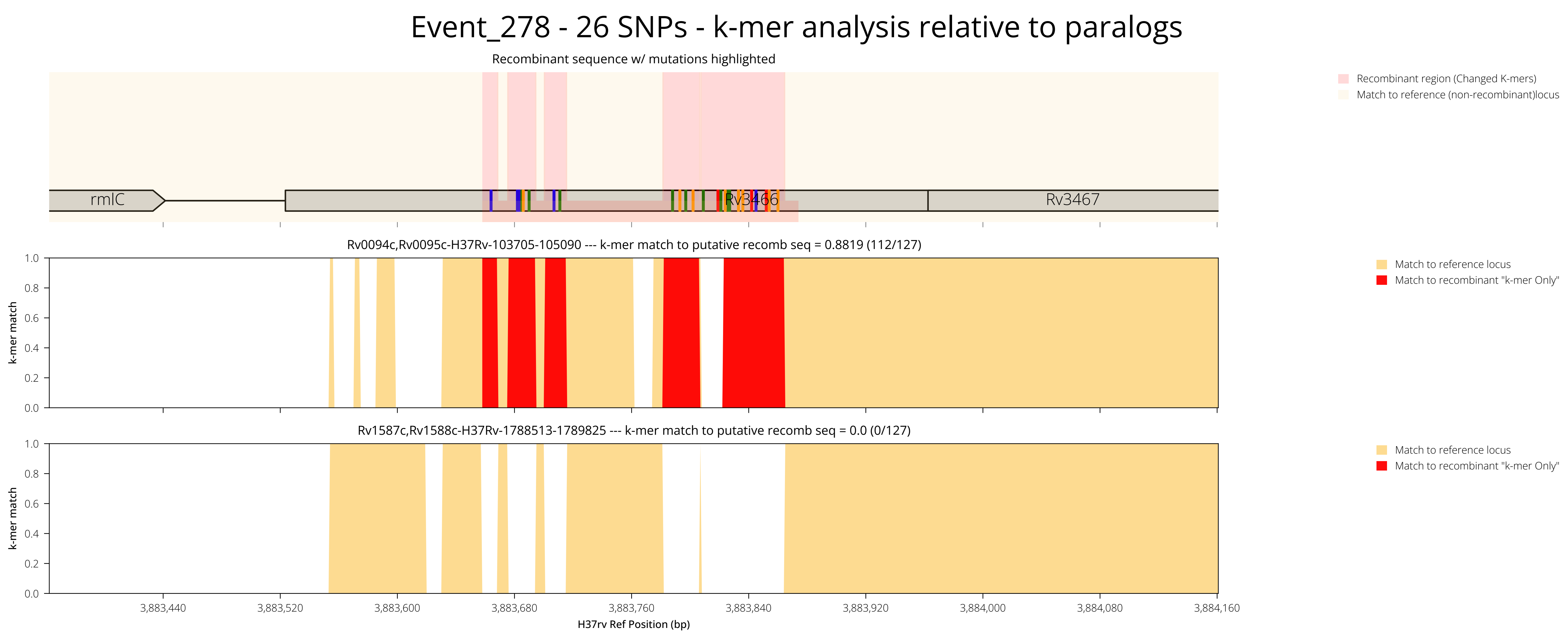

Event_278 - 26 SNPs - k-mer analysis relative to paralogs
Recombinant sequence w/ mutations highlighted
Recombinant region (Changed K-mers)
Match to reference (non-recombinant)locus
rmlC
Rv3466
Rv3467
Rv0094c,Rv0095c-H37Rv-103705-105090 --- k-mer match to putative recomb seq = 0.8819 (112/127)
Match to reference locus
Match to recombinant "k-mer Only"
k-mer match
1.0
0.8
0.6
0.4
0.2
0.0
Rv1587c,Rv1588c-H37Rv-1788513-1789825 --- k-mer match to putative recomb seq = 0.0 (0/127)
Match to reference locus
Match to recombinant "k-mer Only"
k-mer match
1.0
0.8
0.6
0.4
0.2
0.0
3,883,440
3,883,520
3,883,600
3,883,680
3,883,760
3,883,840
3,883,920
3,884,000
3,884,080
3,884,160
H37rv Ref Position (bp)

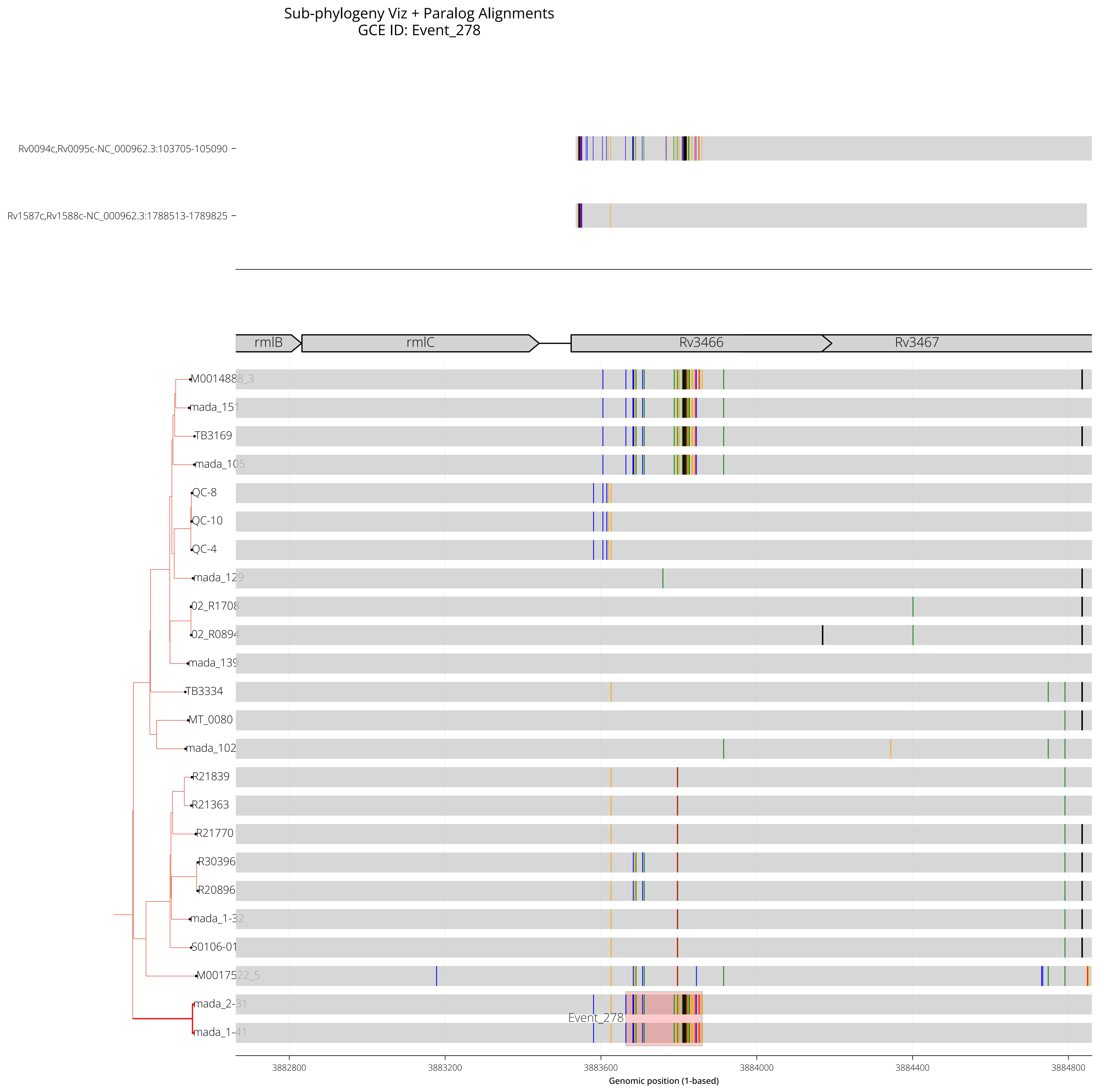

Sub-phylogeny Viz + Paralog Alignments
GCE ID: Event_278
Rv0094c,Rv0095c-NC_000962.3:103705-105090
Rv1587c,Rv1588c-NC_000962.3:1788513-1789825
rmlB
rmlC
Rv3466
Rv3467
M0014888_3
mada_151
TB3169
mada_105
QC-8
QC-10
QC-4
mada_129
02_R1708
02_R0894
mada_139
TB3334
MT_0080
mada_102
R21839
R21363
R21770
R30396
R20896
mada_1-32
S0106-01
M0017522_5
mada_2-31
mada_1-41
Event_278
3882800
3883200
3883600
3884000
3884400
3884800
Genomic position (1-based)

Full Phylogeny Viz + Paralog Alignments
GCE ID: Event_278

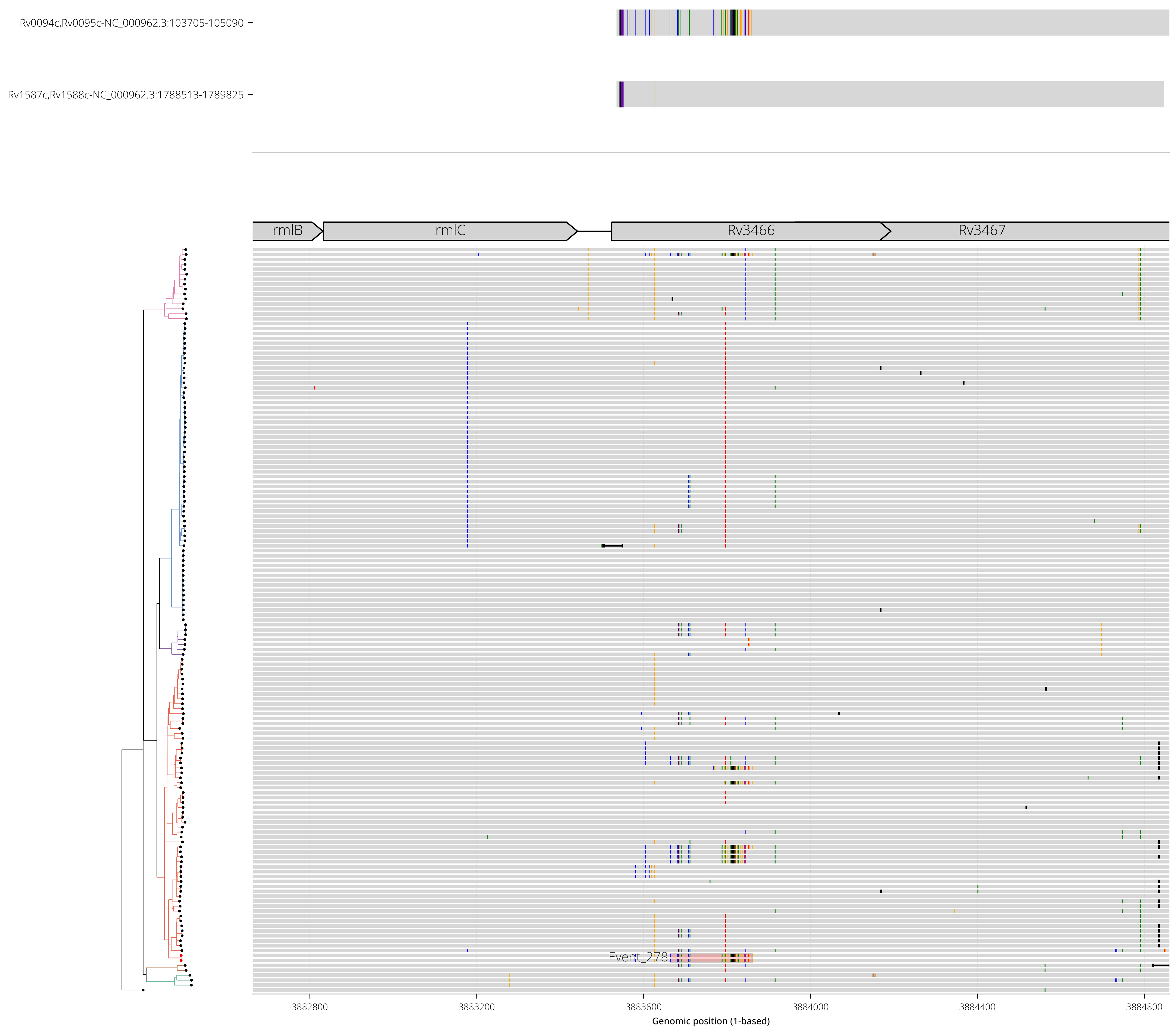

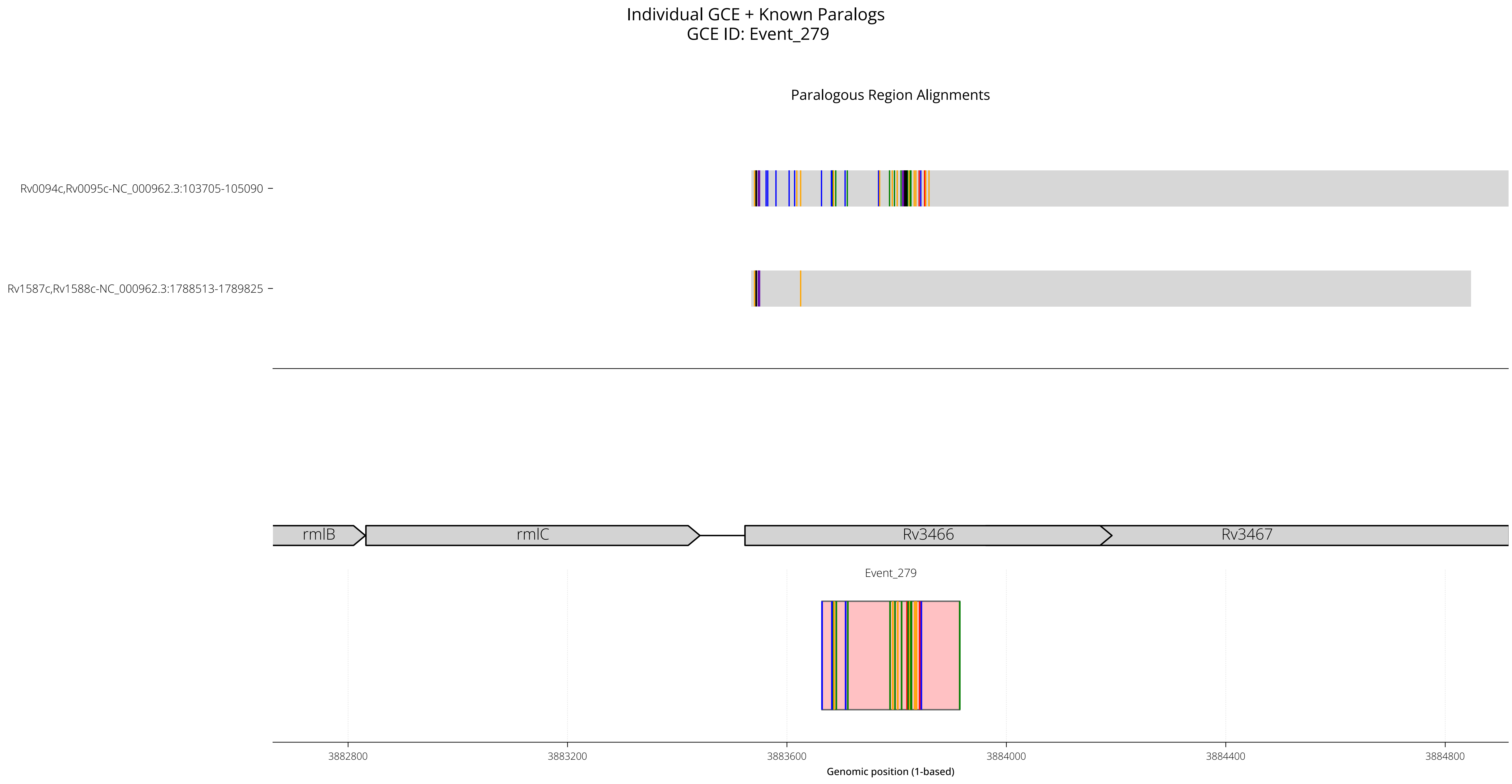

Individual GCE + Known Paralogs
GCE ID: Event_279
Paralogous Region Alignments
Rv0094c,Rv0095c-NC_000962.3:103705-105090
Rv1587c,Rv1588c-NC_000962.3:1788513-1789825
rmlB
rmlC
Rv3466
Rv3467
Event_279
3882800
3883200
3883600
3884000
3884400
3884800
Genomic position (1-based)

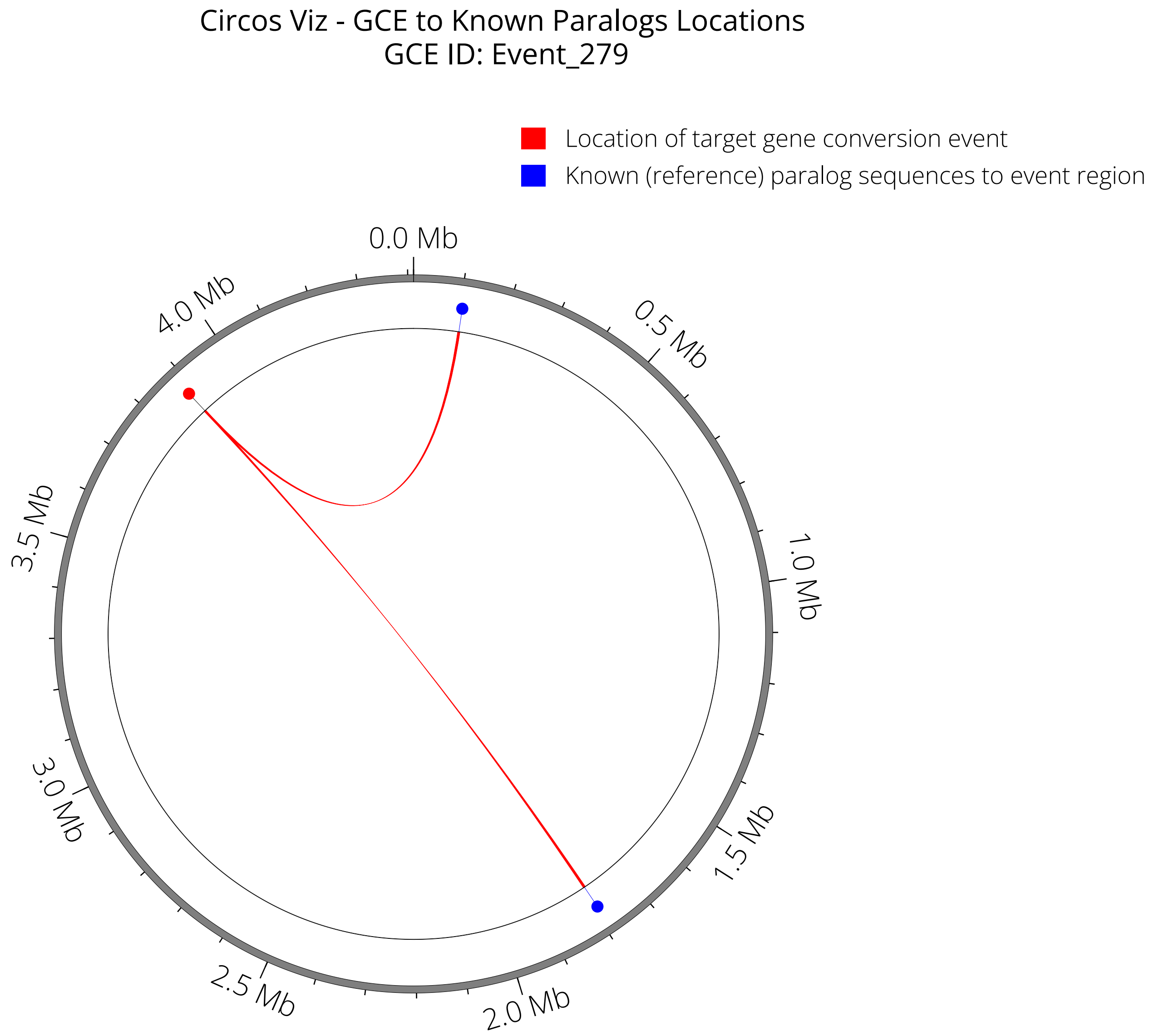
Circos Viz - GCE to Known Paralogs Locations
GCE ID: Event_279
Location of target gene conversion event
Known (reference) paralog sequences to event region
0.0 Mb
0.5 Mb
1.0 Mb
1.5 Mb
2.0 Mb
2.5 Mb
3.0 Mb
3.5 Mb
4.0 Mb

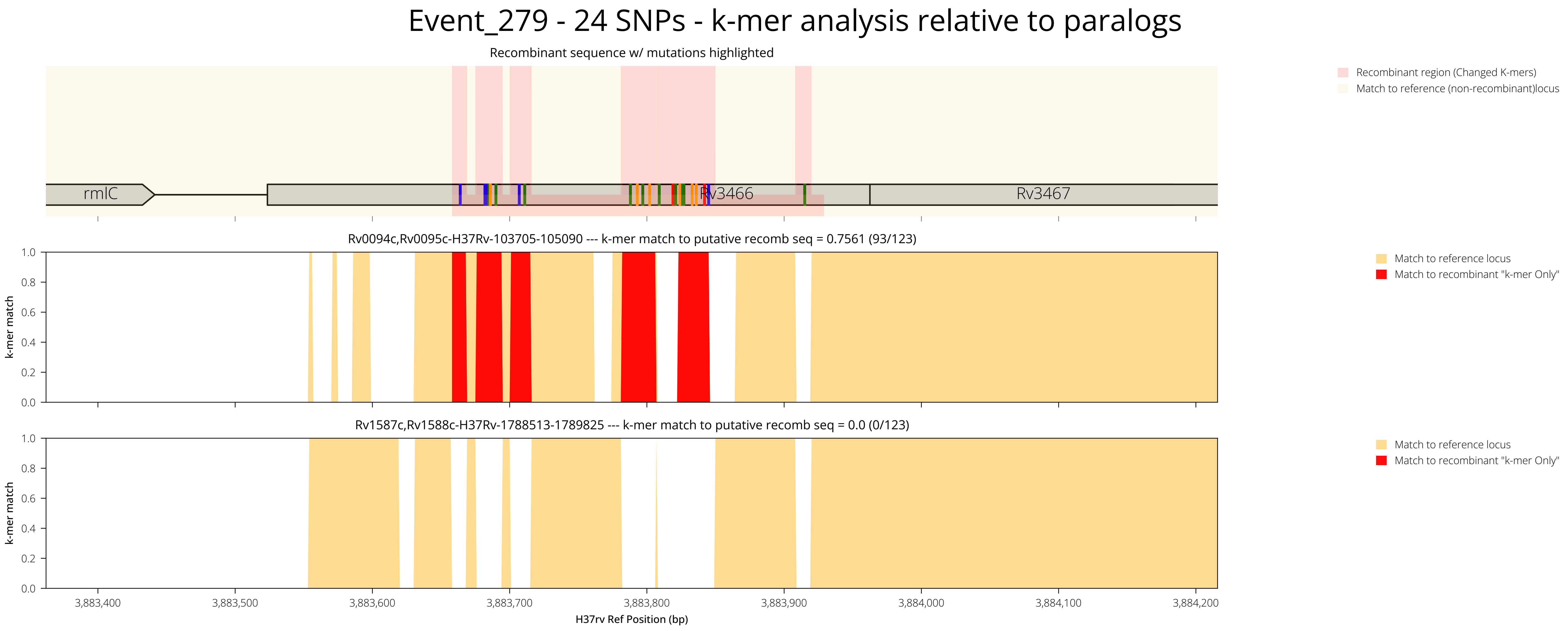

Event_279 - 24 SNPs - k-mer analysis relative to paralogs
Recombinant sequence w/ mutations highlighted
Recombinant region (Changed K-mers)
Match to reference (non-recombinant)locus
rmlC
Rv3466
Rv3467
Rv0094c,Rv0095c-H37Rv-103705-105090 --- k-mer match to putative recomb seq = 0.7561 (93/123)
Match to reference locus
Match to recombinant "k-mer Only"
k-mer match
1.0
0.8
0.6
0.4
0.2
0.0
Rv1587c,Rv1588c-H37Rv-1788513-1789825 --- k-mer match to putative recomb seq = 0.0 (0/123)
Match to reference locus
Match to recombinant "k-mer Only"
k-mer match
1.0
0.8
0.6
0.4
0.2
0.0
3,883,400
3,883,500
3,883,600
3,883,700
3,883,800
3,883,900
3,884,000
3,884,100
3,884,200
H37rv Ref Position (bp)

Sub-phylogeny Viz + Paralog Alignments
GCE ID: Event_279

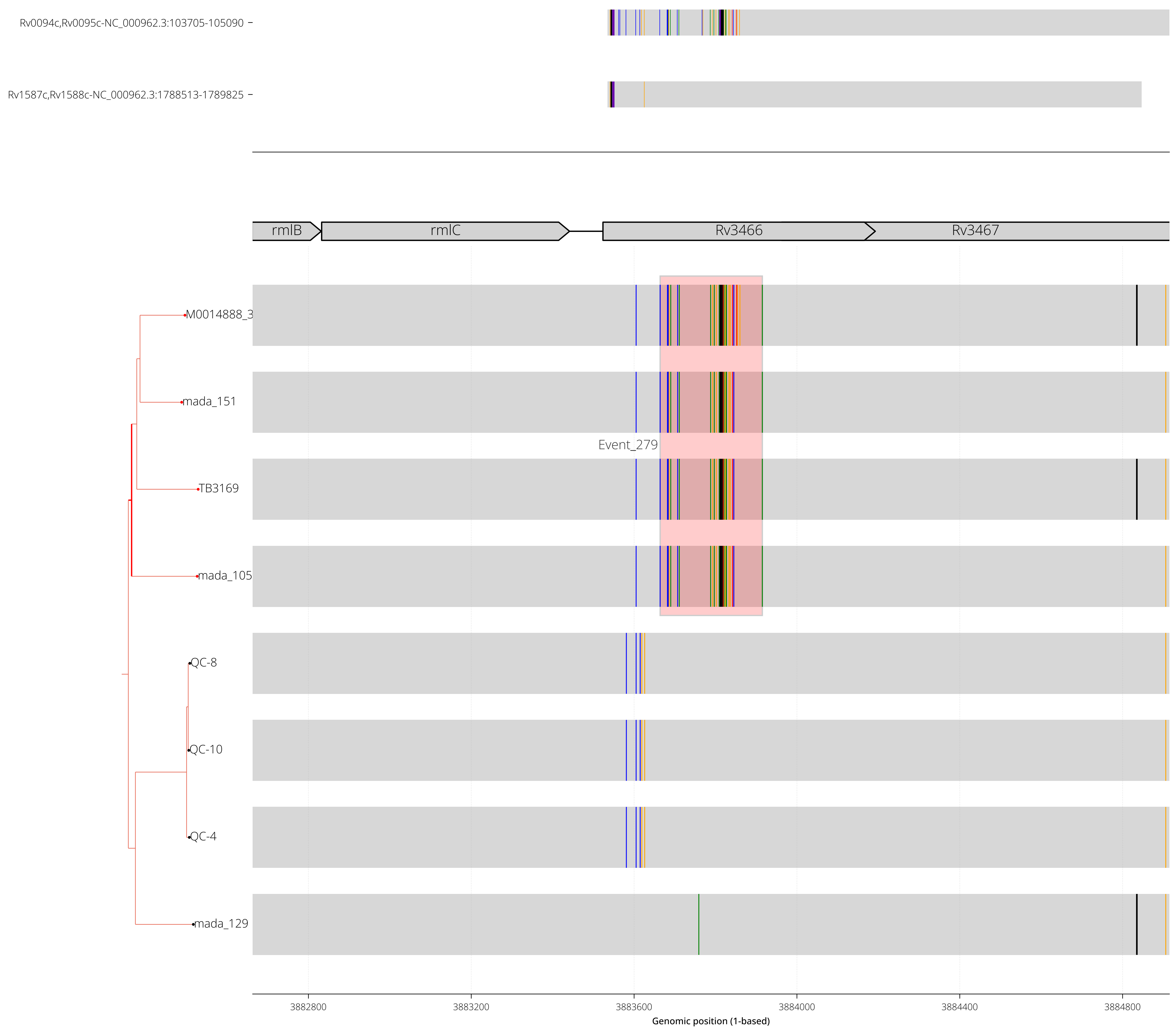

Full Phylogeny Viz + Paralog Alignments
GCE ID: Event_279

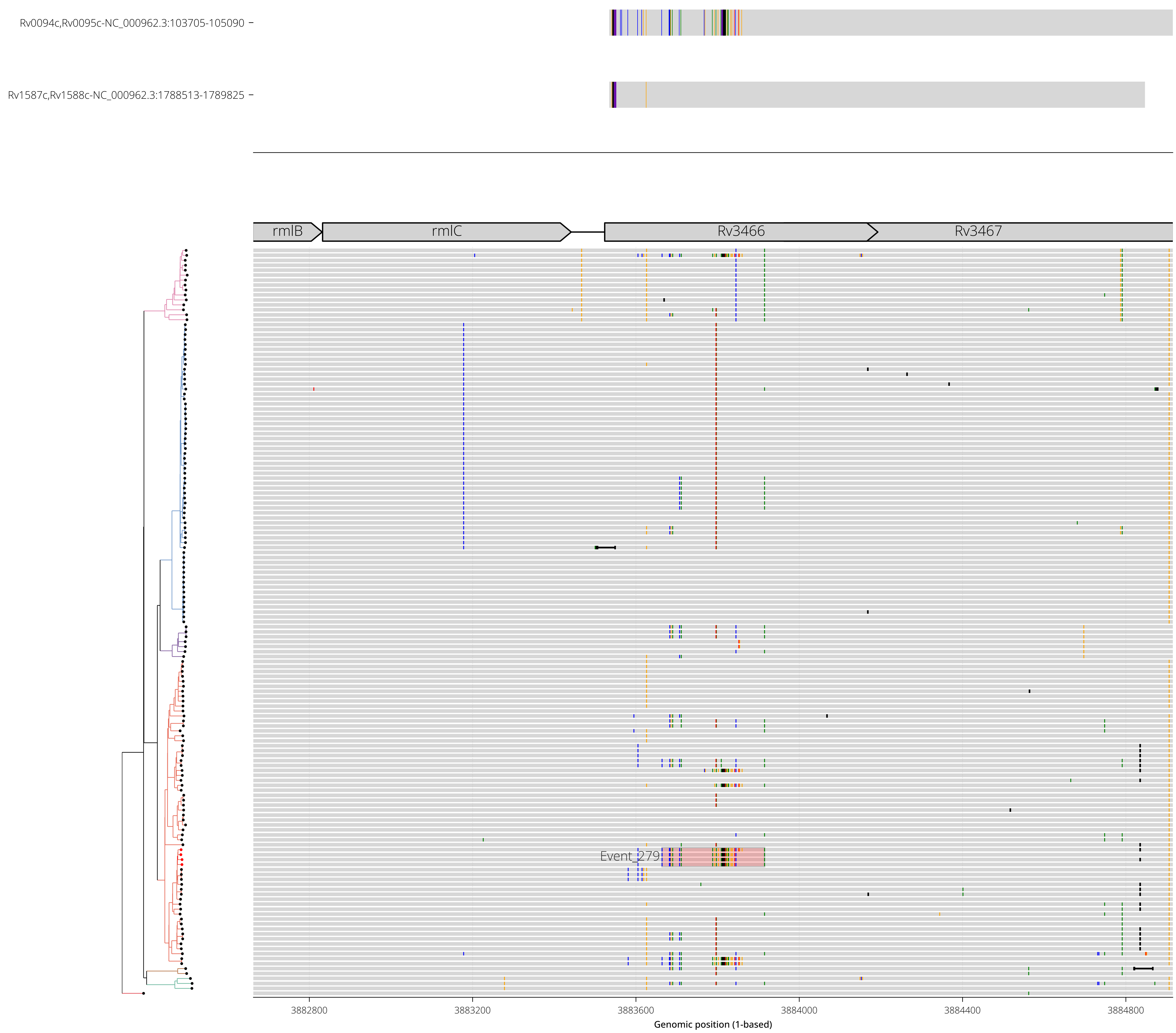

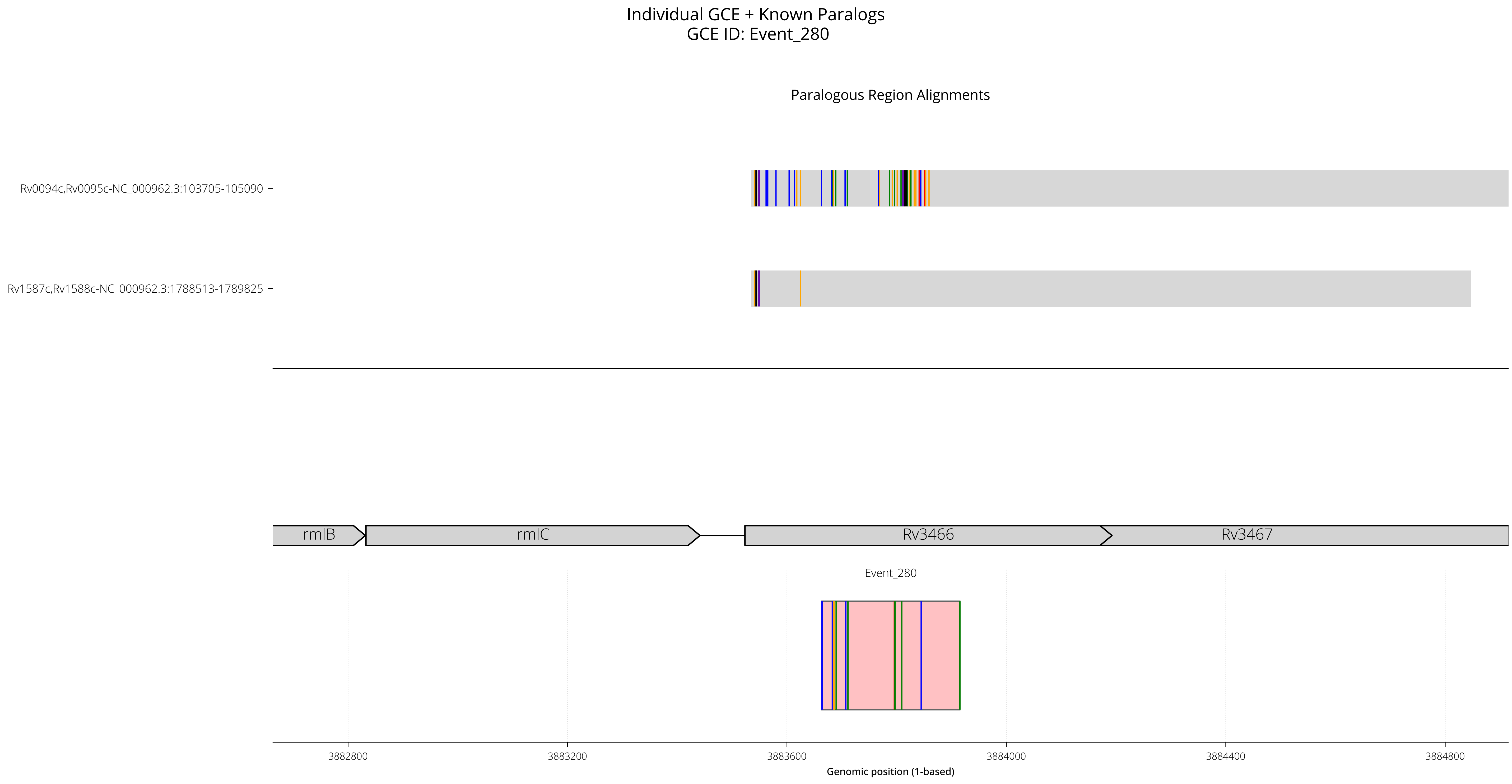

Individual GCE + Known Paralogs
GCE ID: Event_280
Paralogous Region Alignments
Rv0094c,Rv0095c-NC_000962.3:103705-105090
Rv1587c,Rv1588c-NC_000962.3:1788513-1789825
rmlB
rmlC
Rv3466
Rv3467
Event_280
3882800
3883200
3883600
3884000
3884400
3884800
Genomic position (1-based)

Circos Viz - GCE to Known Paralogs Locations
GCE ID: Event_280

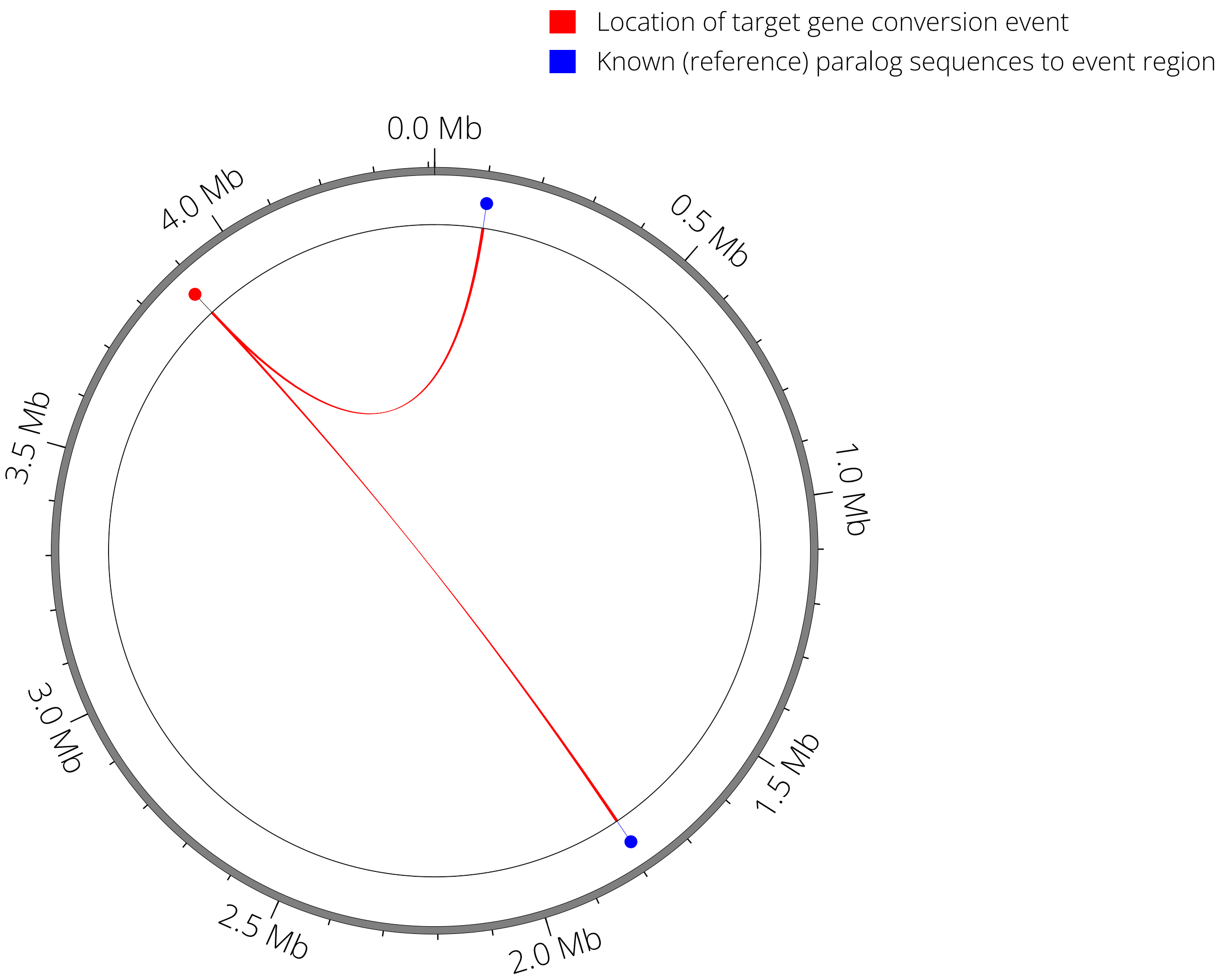

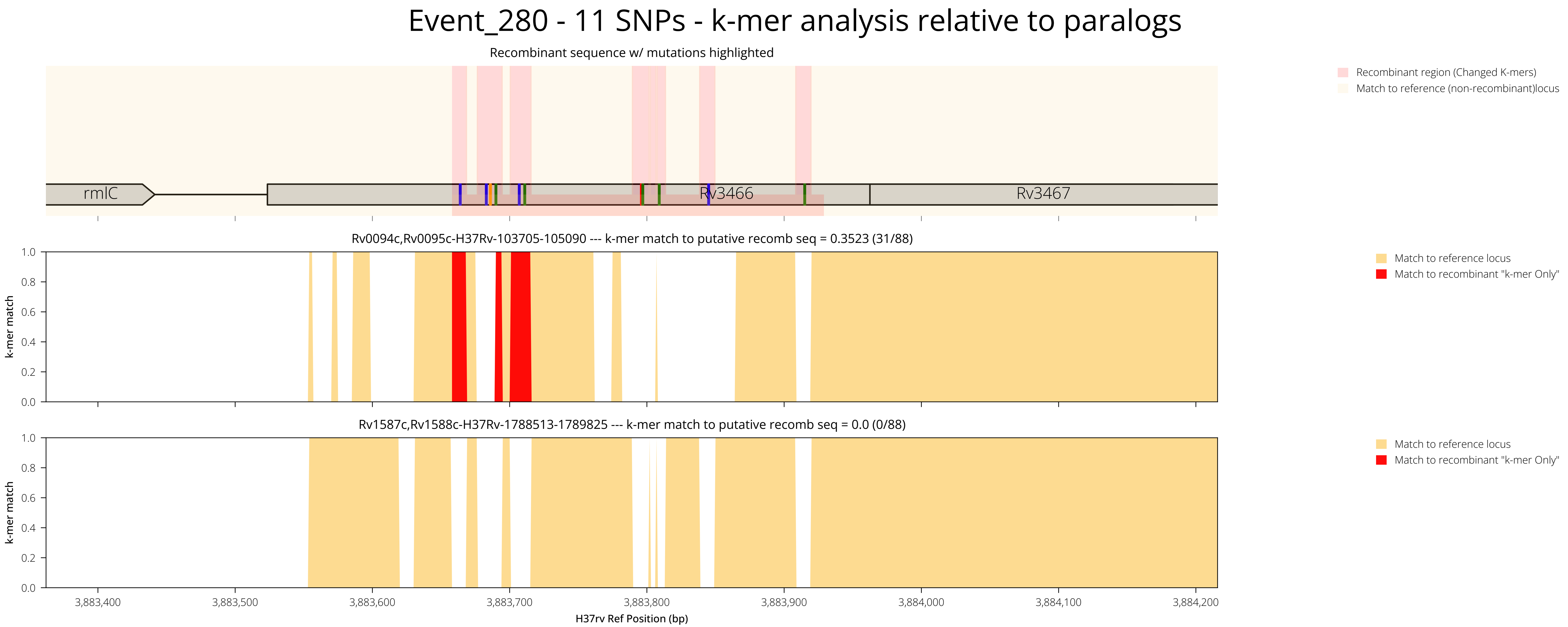

Event_280 - 11 SNPs - k-mer analysis relative to paralogs
Recombinant sequence w/ mutations highlighted
rmlC
Rv3466
Rv3467
Recombinant region (Changed K-mers)
Match to reference (non-recombinant)locus
Rv0094c,Rv0095c-H37Rv-103705-105090 --- k-mer match to putative recomb seq = 0.3523 (31/88)
Match to reference locus
Match to recombinant "k-mer Only"
k-mer match
1.0
0.8
0.6
0.4
0.2
0.0
Rv1587c,Rv1588c-H37Rv-1788513-1789825 --- k-mer match to putative recomb seq = 0.0 (0/88)
3,883,400
3,883,500
3,883,600
3,883,700
3,883,800
3,883,900
3,884,000
3,884,100
3,884,200
H37rv Ref Position (bp)

Sub-phylogeny Viz + Paralog Alignments
GCE ID: Event_280

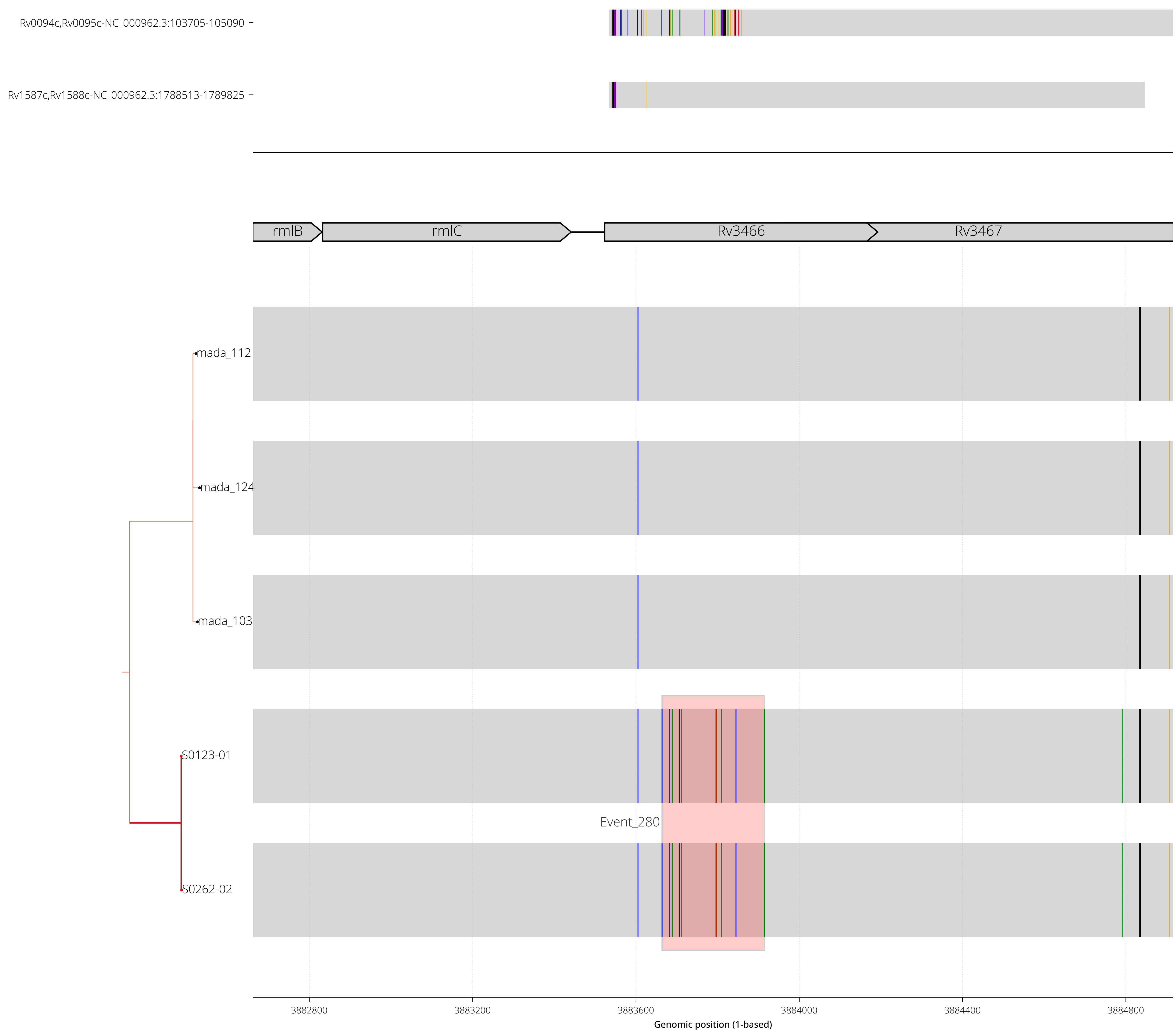

Full Phylogeny Viz + Paralog Alignments
GCE ID: Event_280

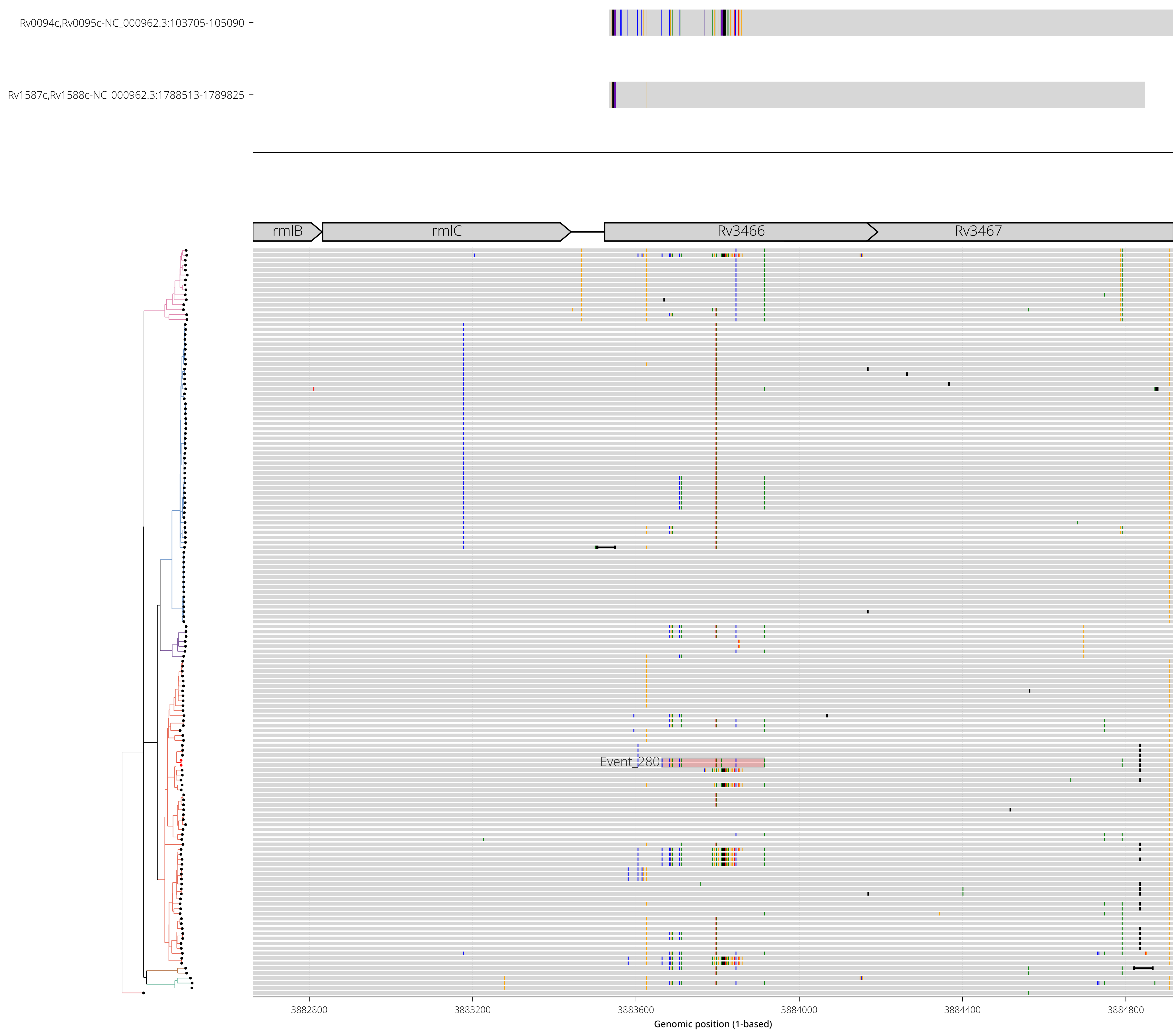

Individual GCE + Known Paralogs
GCE ID: Event_281

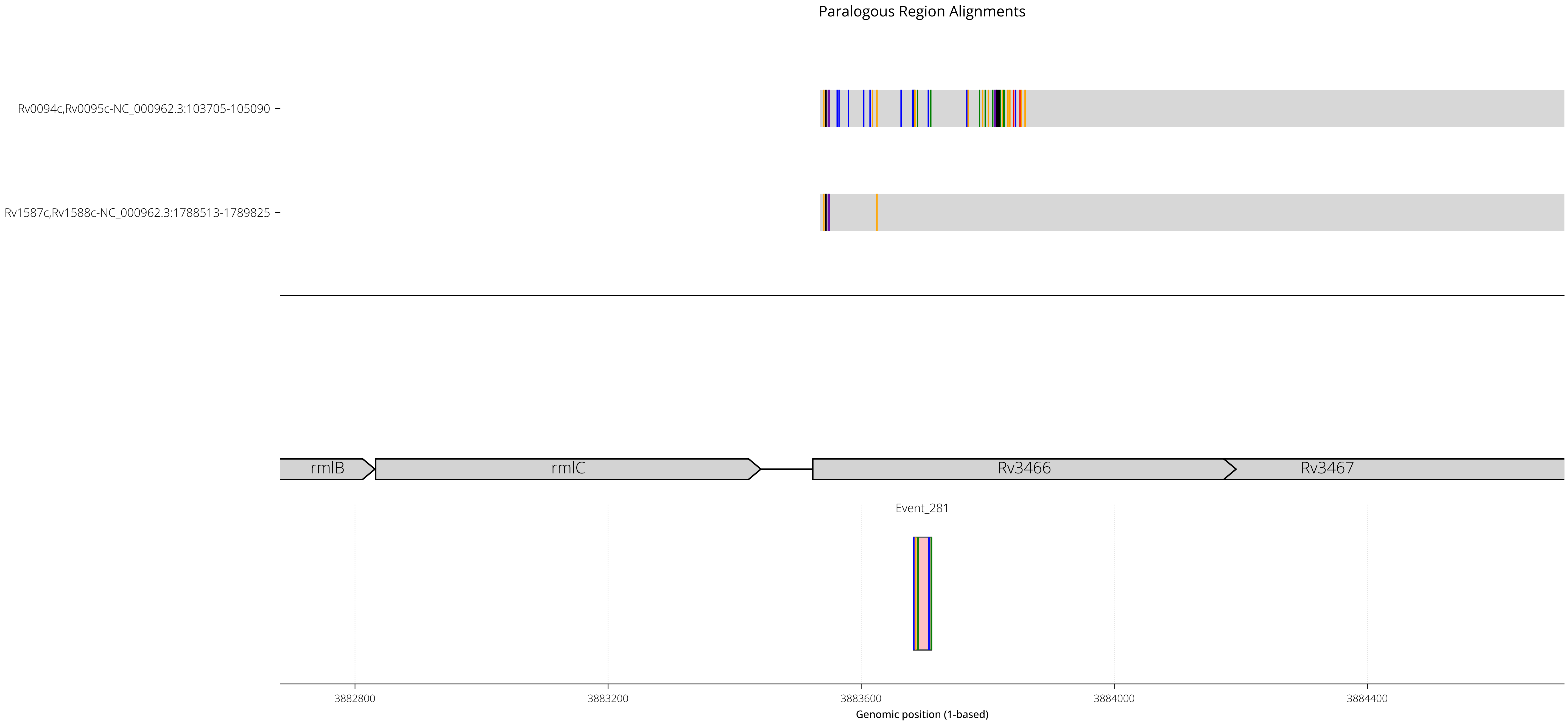

Circos Viz - GCE to Known Paralogs Locations
GCE ID: Event_281

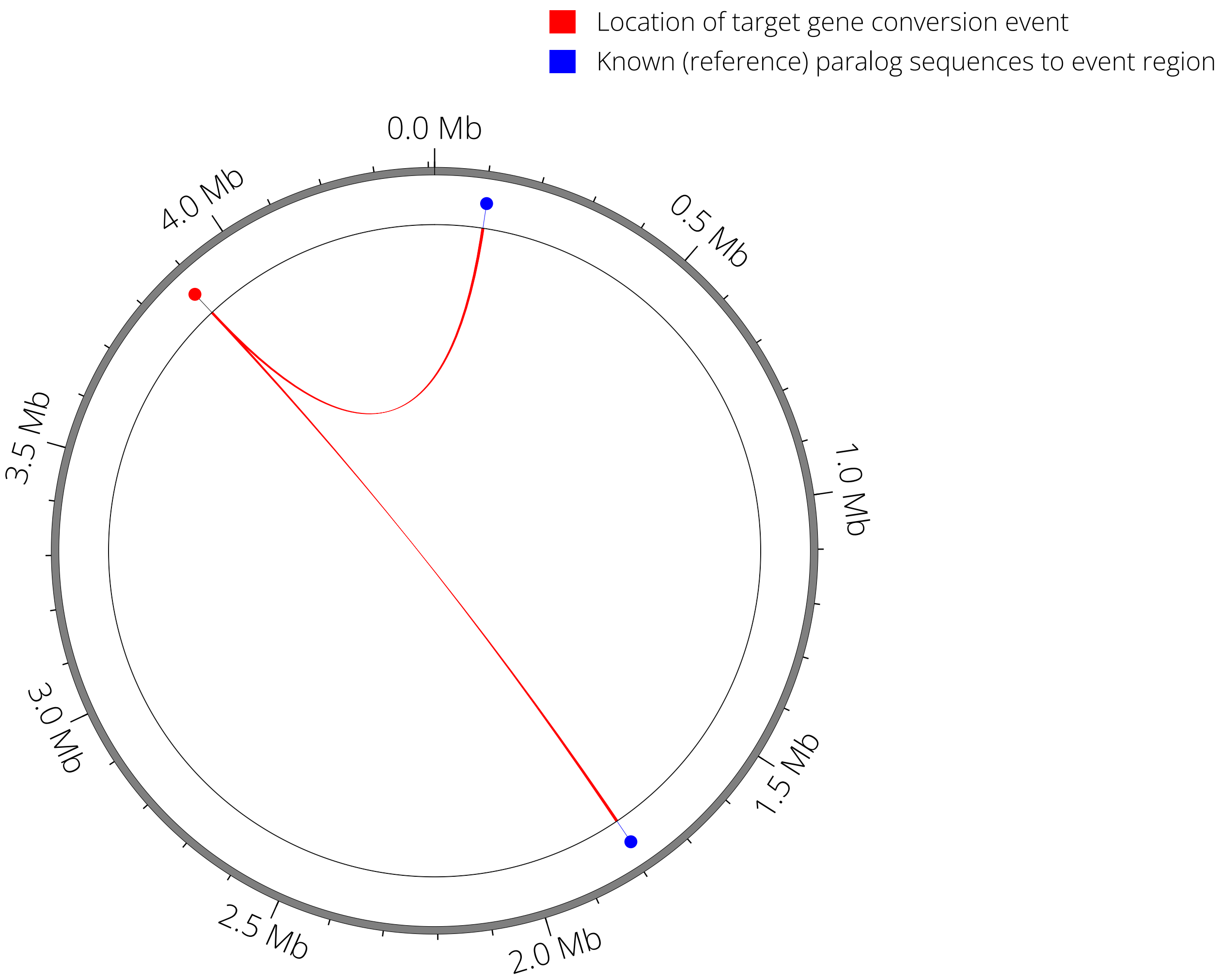

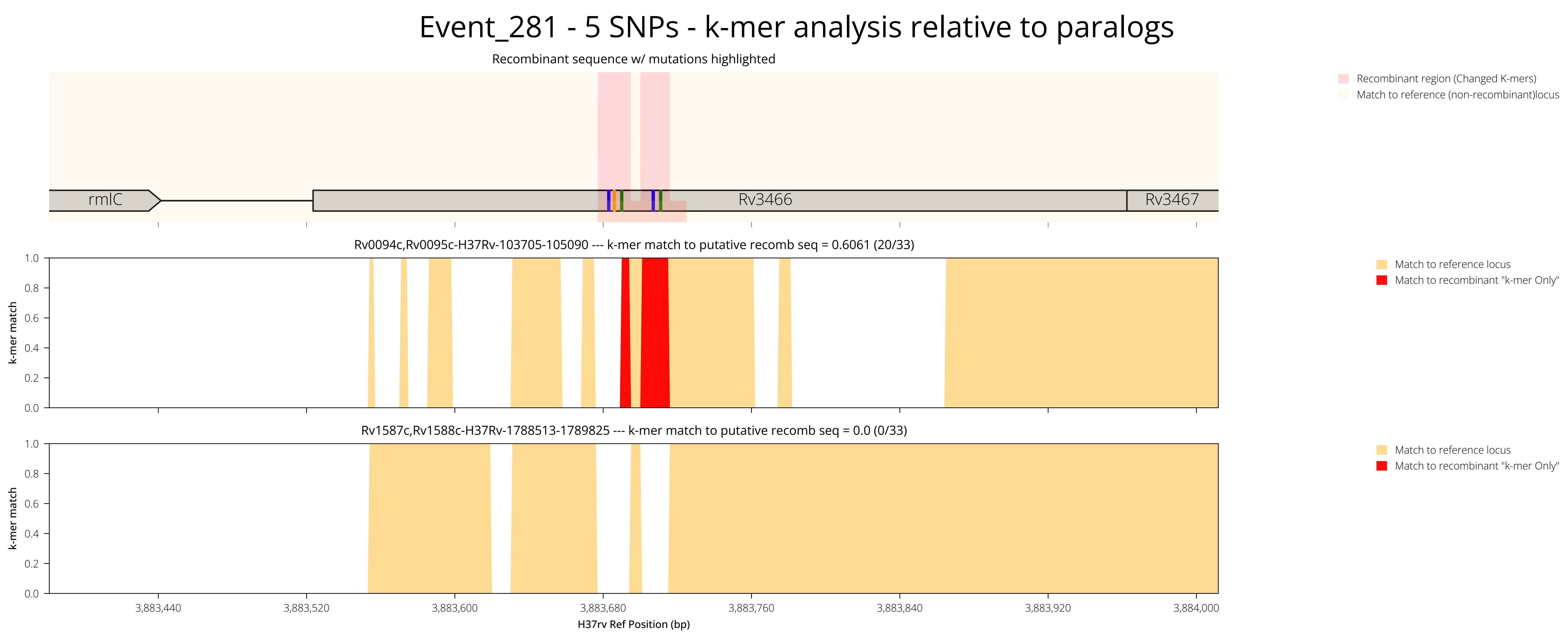
Event_281 - 5 SNPs - k-mer analysis relative to paralogs
Recombinant sequence w/ mutations highlighted
Recombinant region (Changed K-mers)
Match to reference (non-recombinant)locus
rmlC
Rv3466
Rv3467
Rv0094c,Rv0095c-H37Rv-103705-105090 --- k-mer match to putative recomb seq = 0.6061 (20/33)
Match to reference locus
Match to recombinant "k-mer Only"
k-mer match
1.0
0.8
0.6
0.4
0.2
0.0
Rv1587c,Rv1588c-H37Rv-1788513-1789825 --- k-mer match to putative recomb seq = 0.0 (0/33)
3,883,440
3,883,520
3,883,600
3,883,680
3,883,760
3,883,840
3,883,920
3,884,000
H37rv Ref Position (bp)

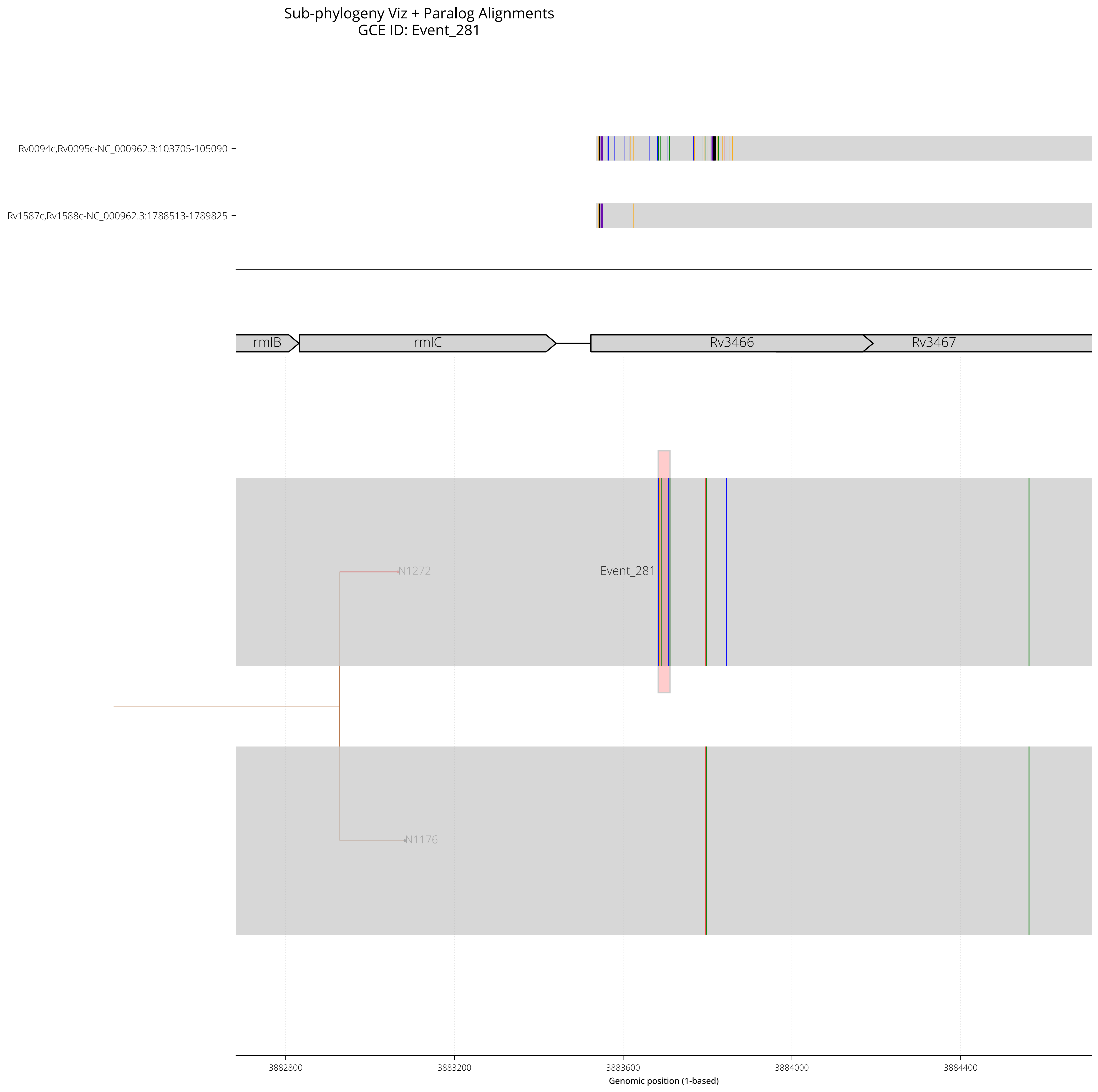
Sub-phylogeny Viz + Paralog Alignments
GCE ID: Event_281
Rv0094c,Rv0095c-NC_000962.3:103705-105090
Rv1587c,Rv1588c-NC_000962.3:1788513-1789825
rmlB
rmlC
Rv3466
Rv3467
N1272
Event_281
N1176
3882800
3883200
3883600
3884000
3884400
Genomic position (1-based)

Full Phylogeny Viz + Paralog Alignments
GCE ID: Event_281

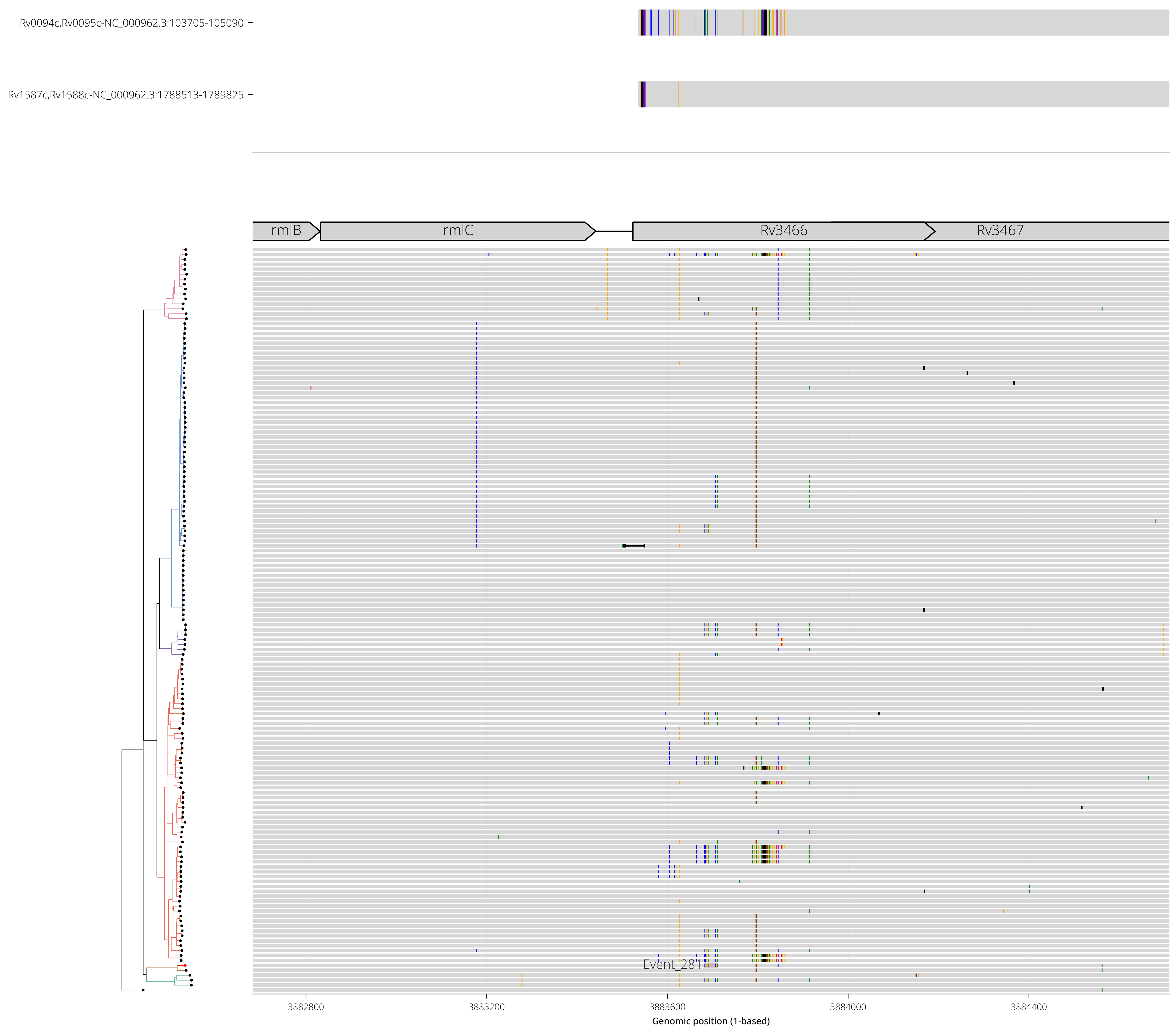

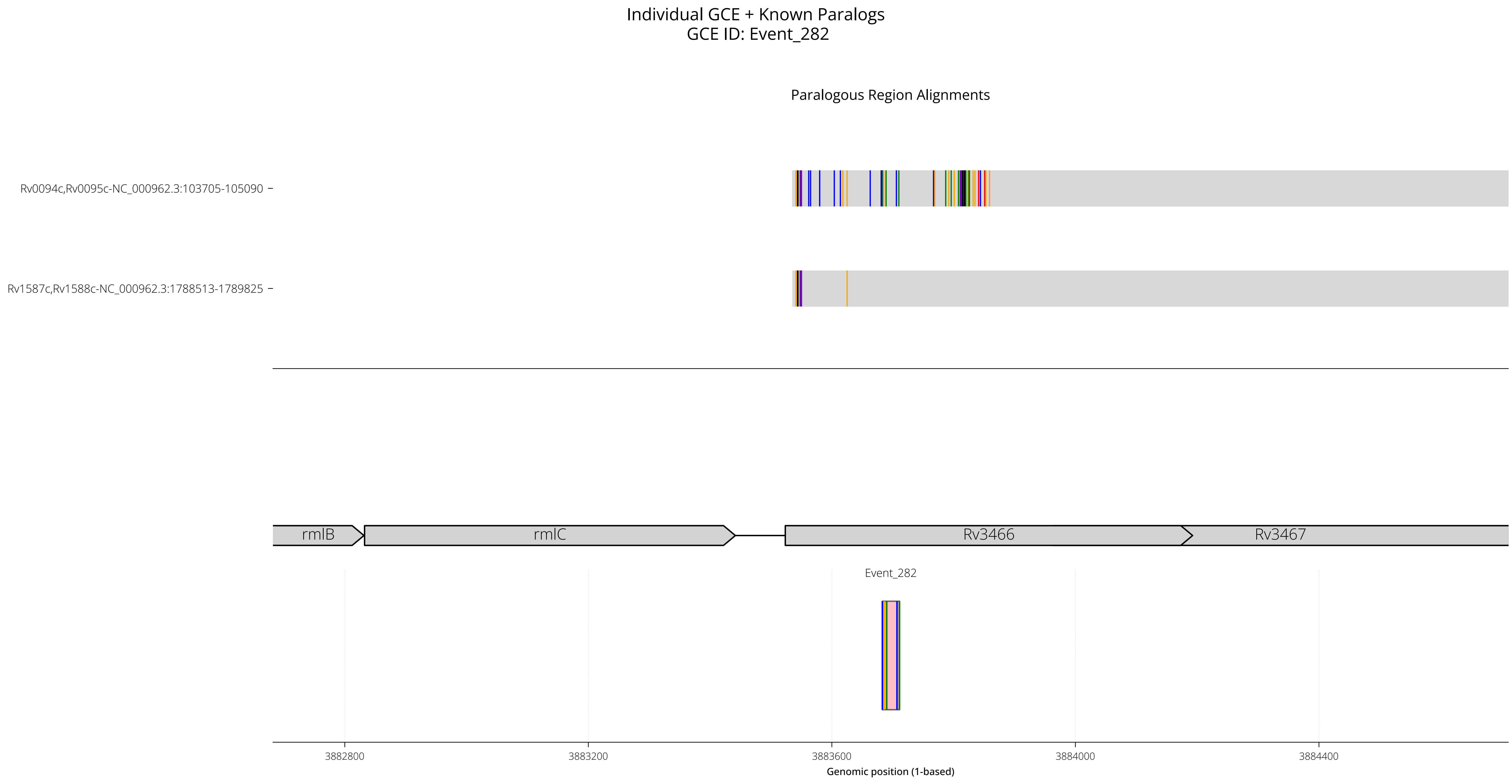
Individual GCE + Known Paralogs
GCE ID: Event_282
Paralogous Region Alignments
Rv0094c,Rv0095c-NC_000962.3:103705-105090
Rv1587c,Rv1588c-NC_000962.3:1788513-1789825
rmlB
rmlC
Rv3466
Rv3467
Event_282
3882800
3883200
3883600
3884000
3884400
Genomic position (1-based)

Circos Viz - GCE to Known Paralogs Locations
GCE ID: Event_282

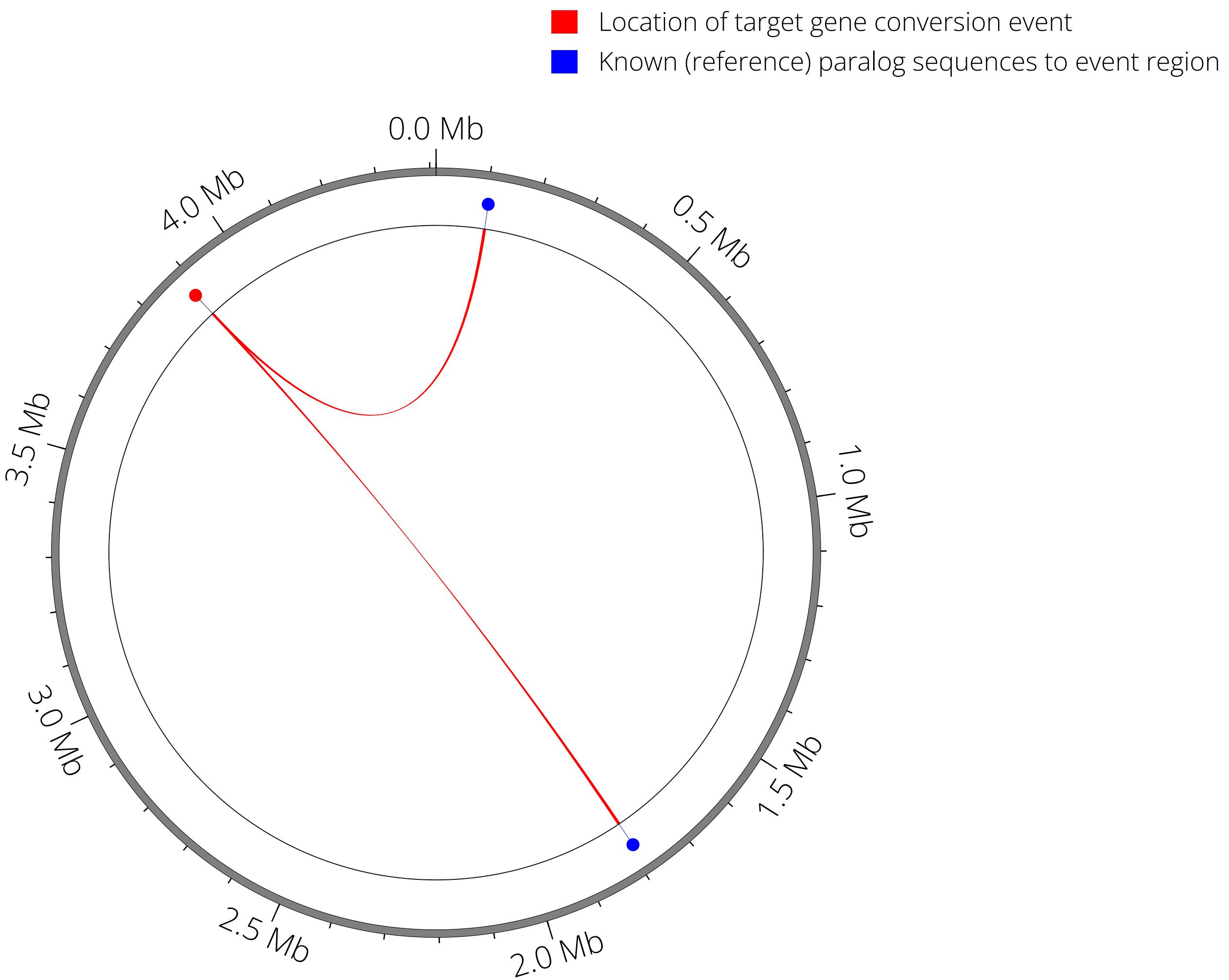

### Event_282 - 5 SNPs - k-mer analysis relative to paralogs

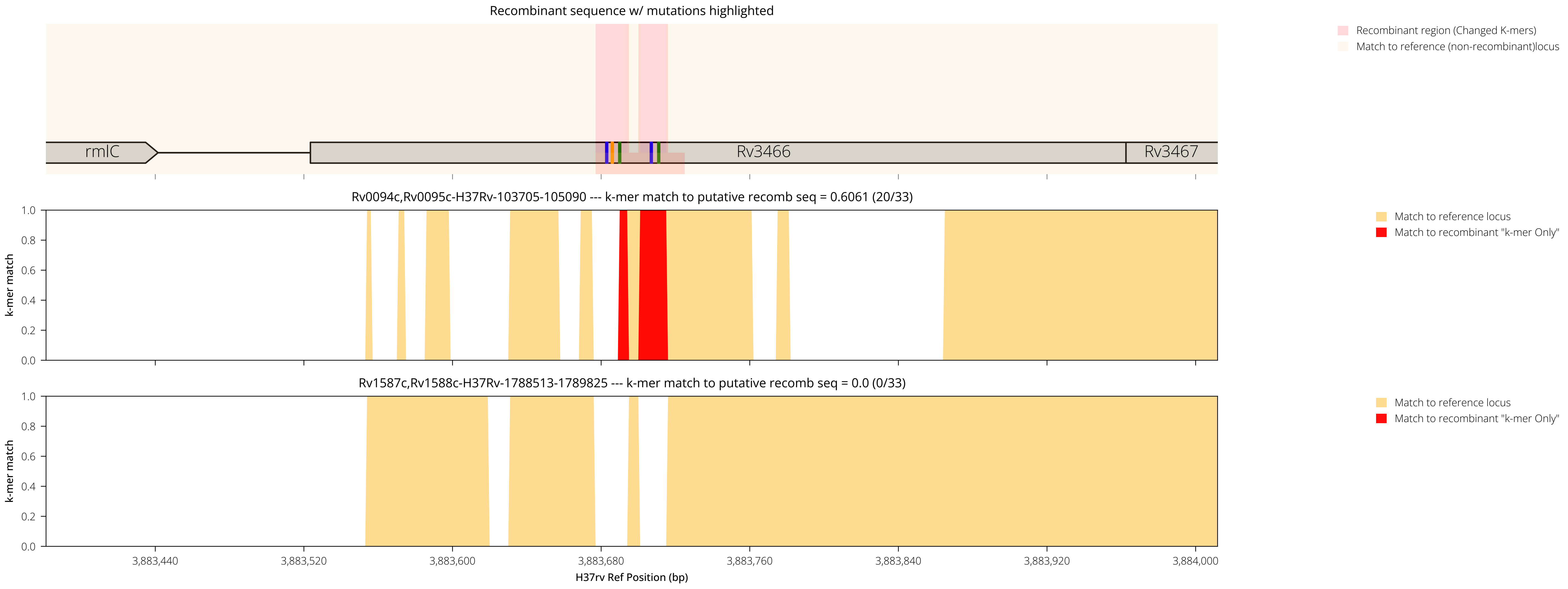

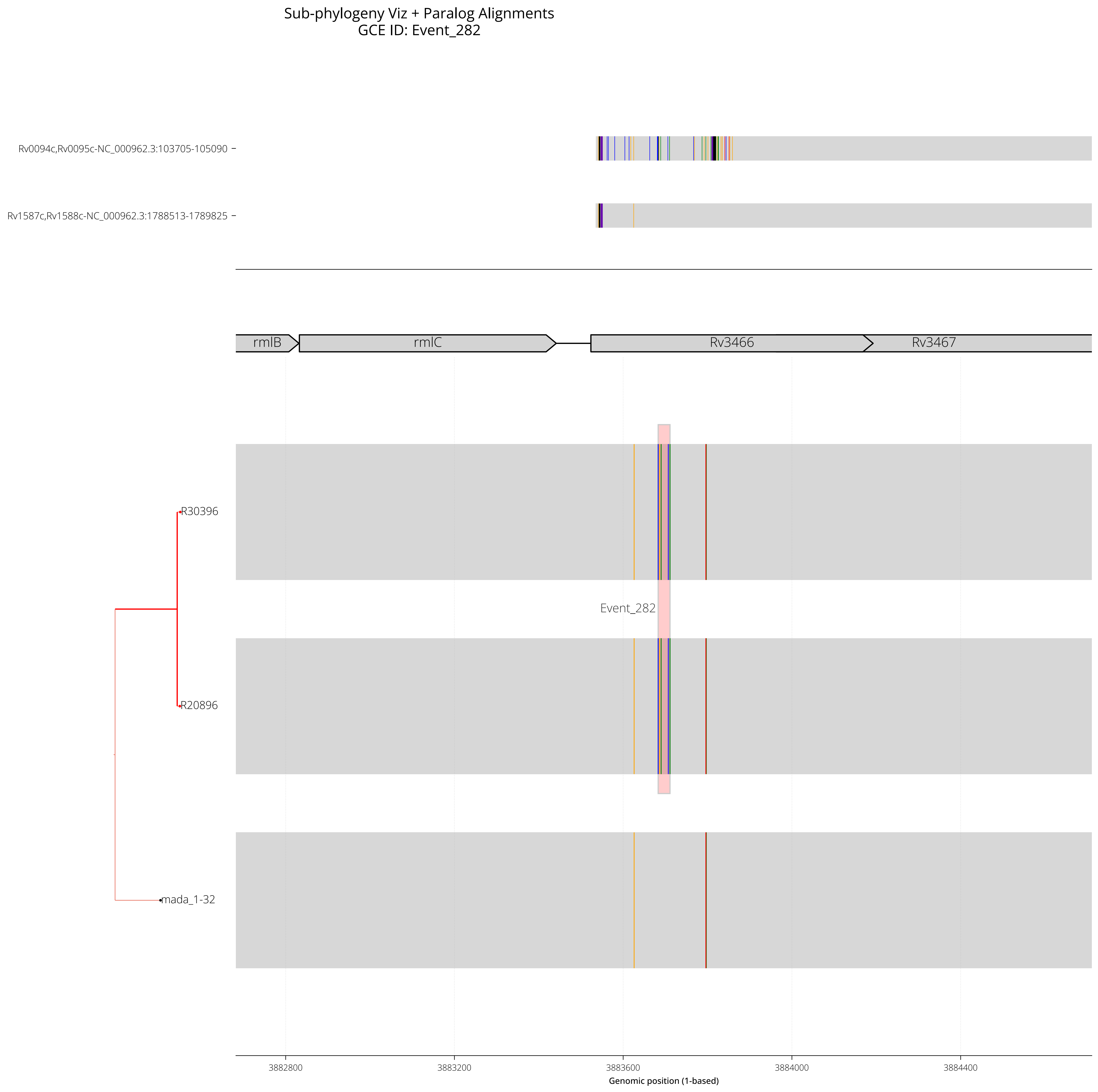
Sub-phylogeny Viz + Paralog Alignments
GCE ID: Event_282
Rv0094c,Rv0095c-NC_000962.3:103705-105090
Rv1587c,Rv1588c-NC_000962.3:1788513-1789825
rmlB
rmlC
Rv3466
Rv3467
R30396
Event_282
R20896
mada_1-32
3882800
3883200
3883600
3884000
3884400
Genomic position (1-based)

Full Phylogeny Viz + Paralog Alignments
GCE ID: Event_282

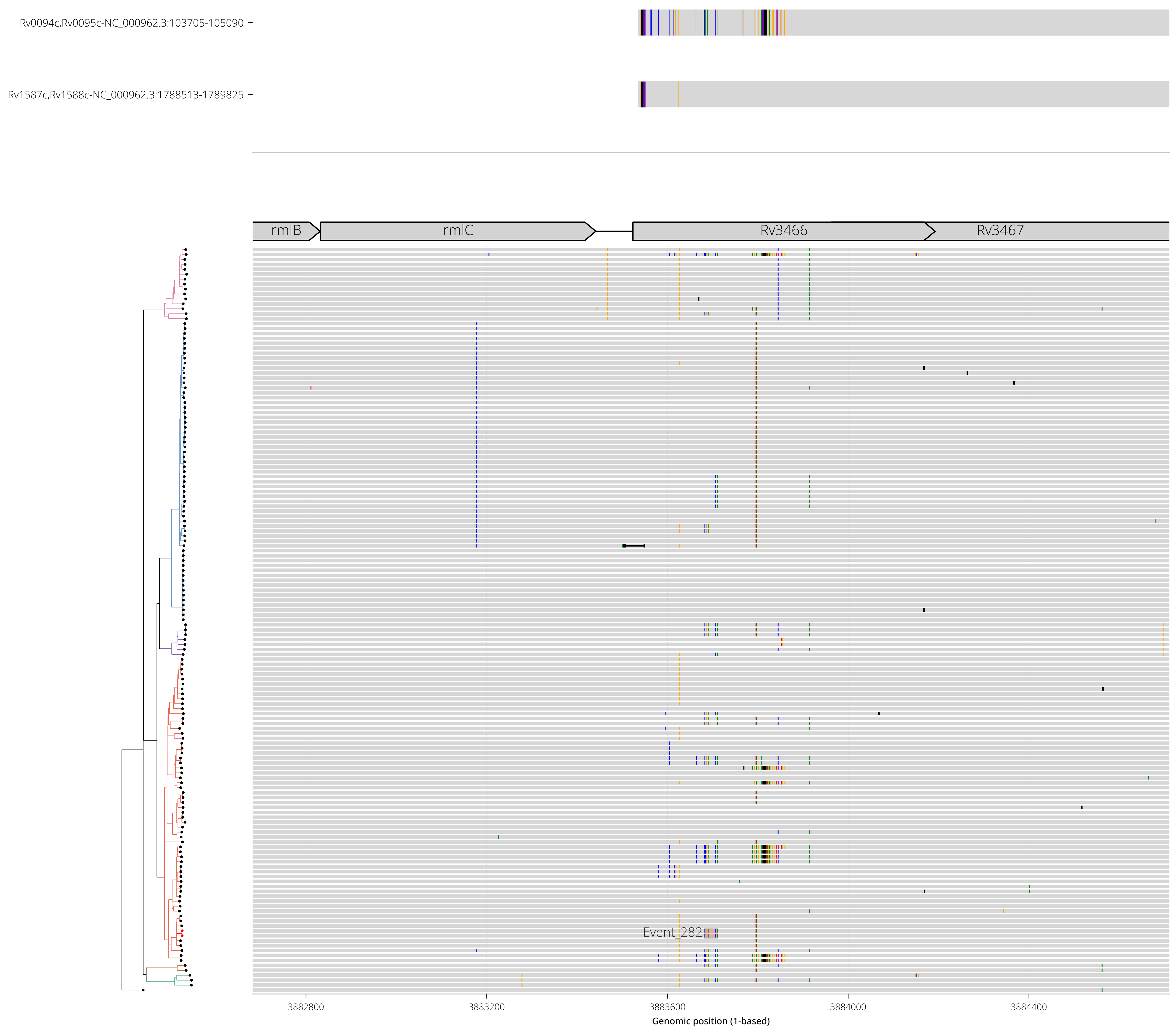

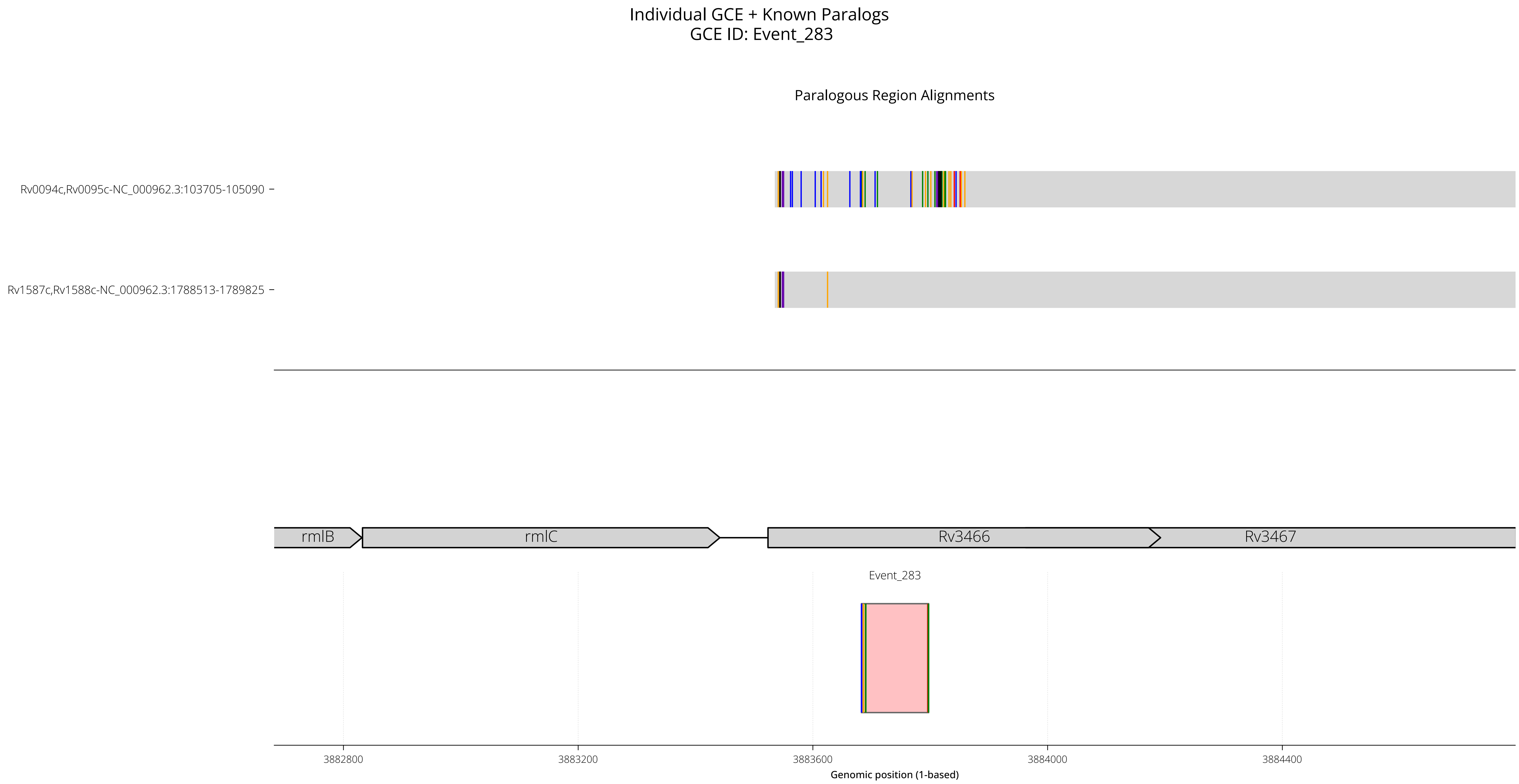
Individual GCE + Known Paralogs
GCE ID: Event_283
Paralogous Region Alignments
Rv0094c,Rv0095c-NC_000962.3:103705-105090
Rv1587c,Rv1588c-NC_000962.3:1788513-1789825
rmlB
rmlC
Rv3466
Rv3467
Event_283
3882800
3883200
3883600
3884000
3884400
Genomic position (1-based)

Circos Viz - GCE to Known Paralogs Locations
GCE ID: Event_283

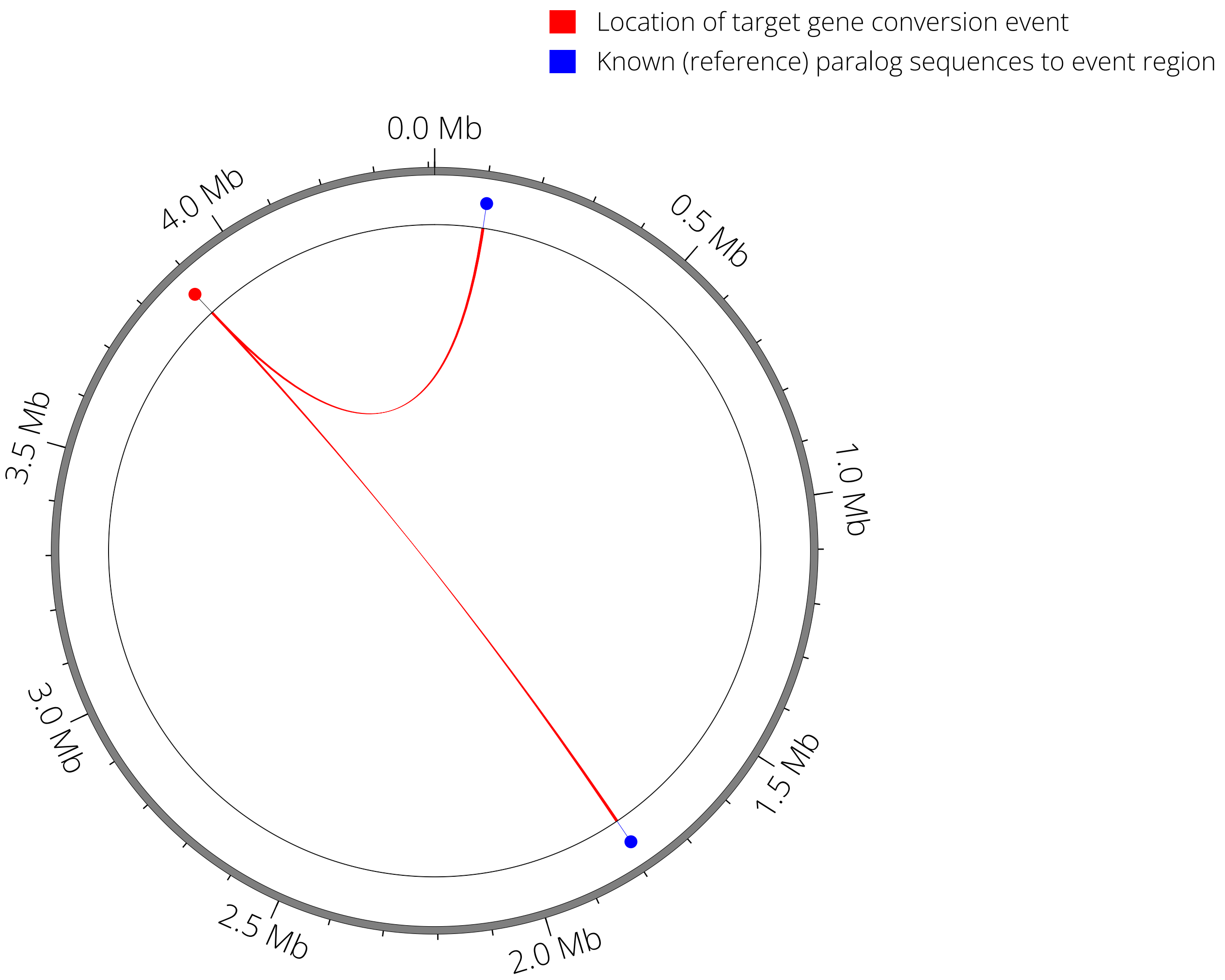

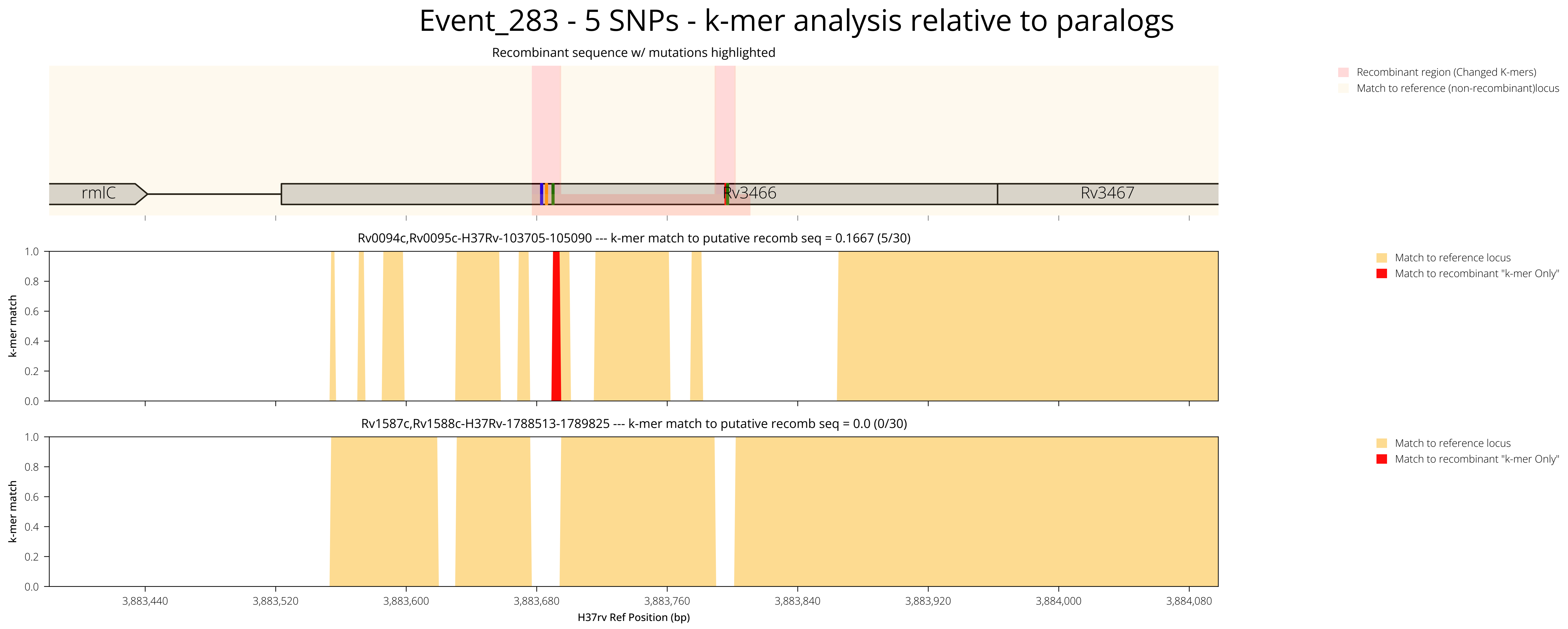
Event_283 - 5 SNPs - k-mer analysis relative to paralogs
Recombinant sequence w/ mutations highlighted
Recombinant region (Changed K-mers)
Match to reference (non-recombinant)locus
rmlC
Rv3466
Rv3467
Rv0094c,Rv0095c-H37Rv-103705-105090 --- k-mer match to putative recomb seq = 0.1667 (5/30)
Match to reference locus
Match to recombinant "k-mer Only"
k-mer match
1.0
0.8
0.6
0.4
0.2
0.0
Rv1587c,Rv1588c-H37Rv-1788513-1789825 --- k-mer match to putative recomb seq = 0.0 (0/30)
Match to reference locus
Match to recombinant "k-mer Only"
k-mer match
3,883,440
3,883,520
3,883,600
3,883,680
3,883,760
3,883,840
3,883,920
3,884,000
3,884,080
H37rv Ref Position (bp)

Sub-phylogeny Viz + Paralog Alignments
GCE ID: Event_283

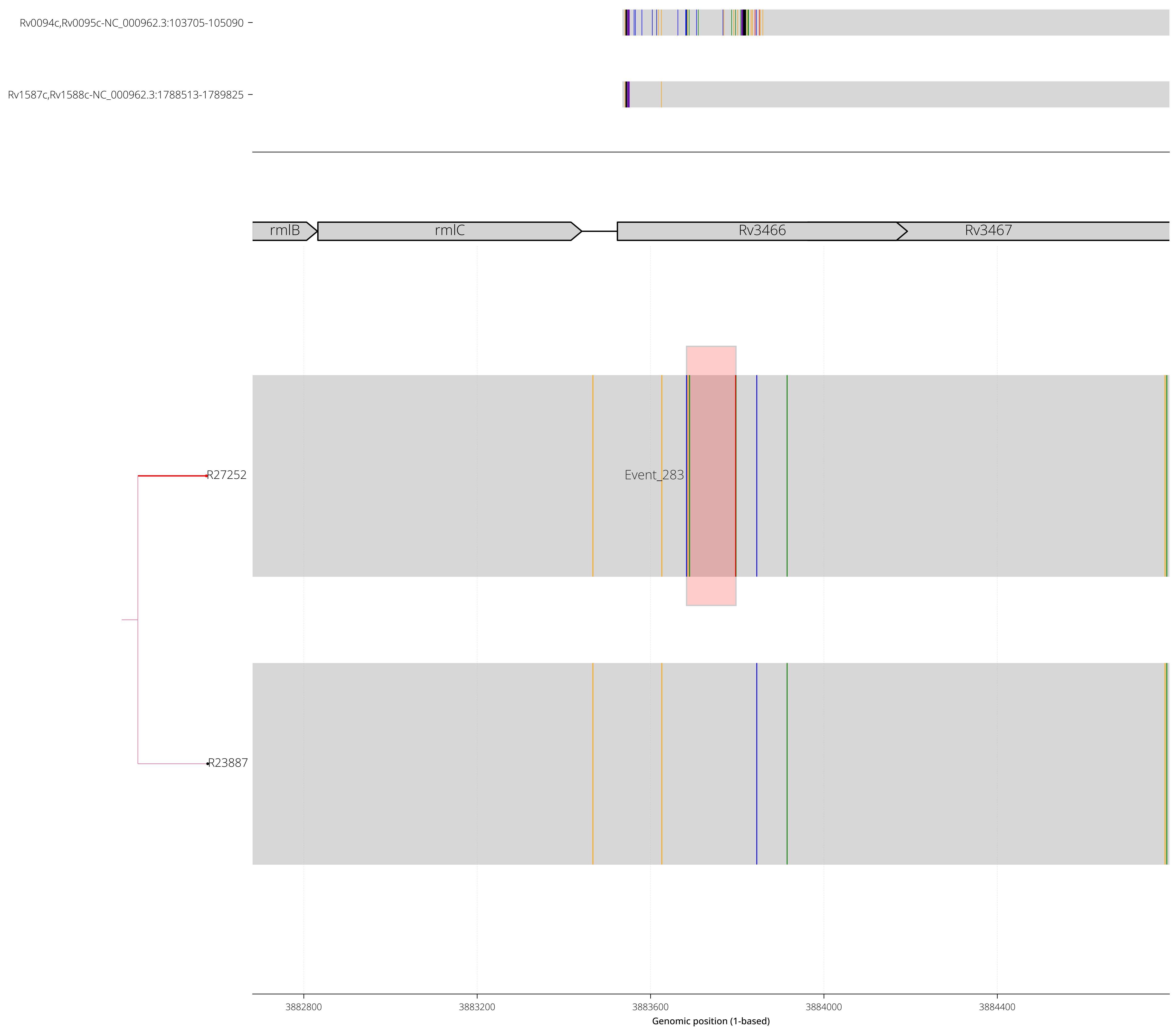

Full Phylogeny Viz + Paralog Alignments
GCE ID: Event_283

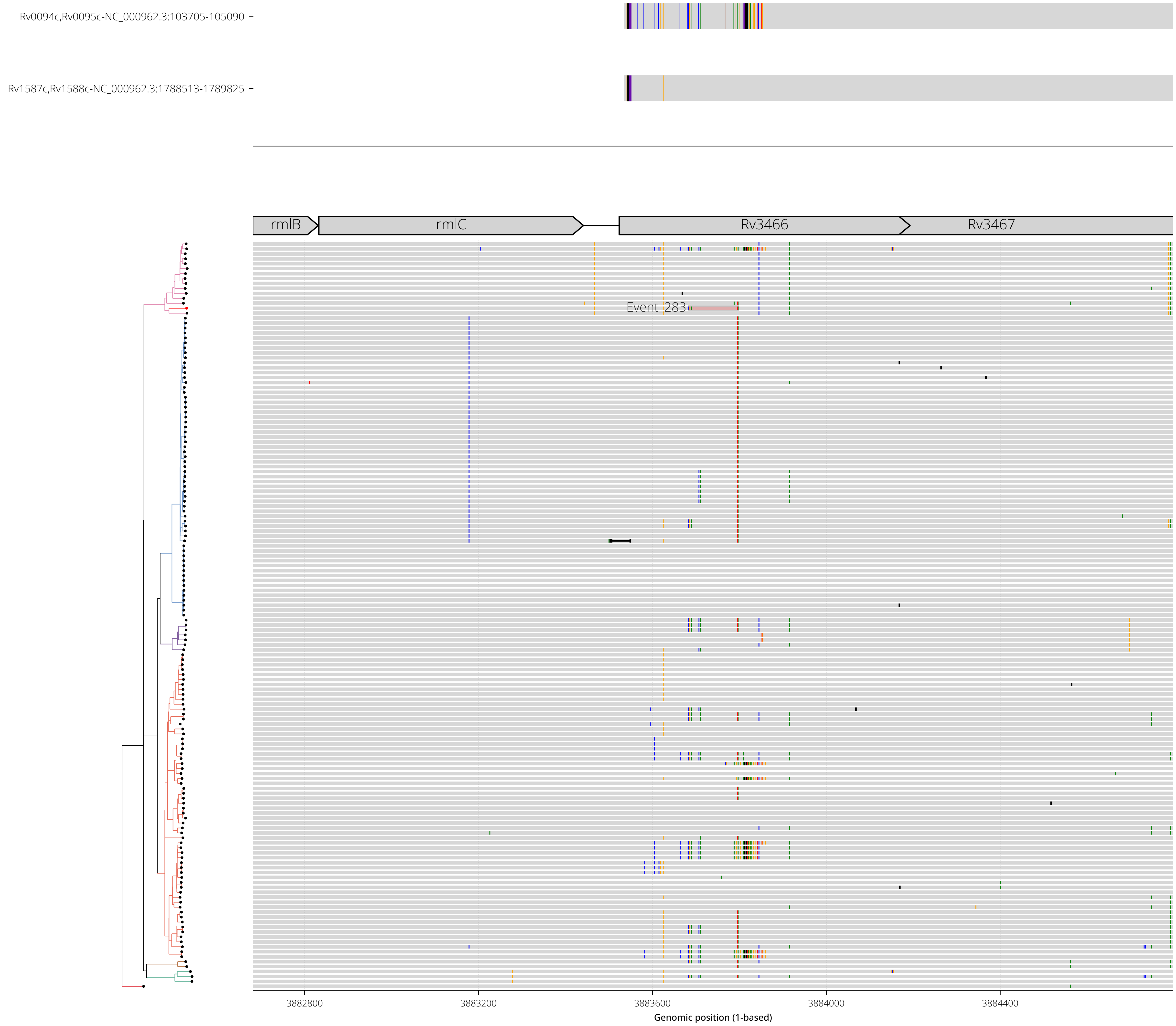

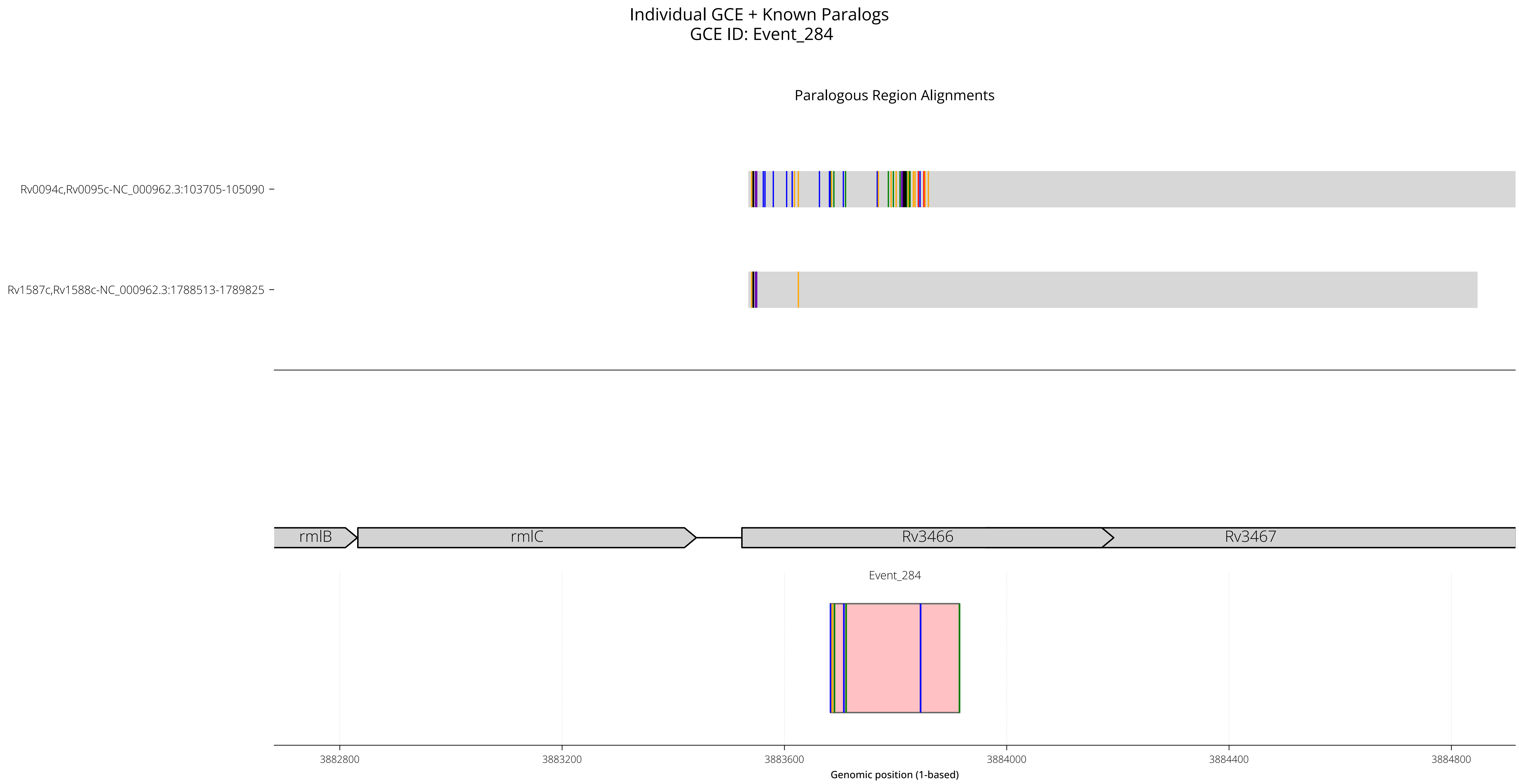
Individual GCE + Known Paralogs
GCE ID: Event_284
Paralogous Region Alignments
Rv0094c,Rv0095c-NC_000962.3:103705-105090
Rv1587c,Rv1588c-NC_000962.3:1788513-1789825
rmlB
rmlC
Rv3466
Rv3467
Event_284
3882800
3883200
3883600
3884000
3884400
3884800
Genomic position (1-based)

Circos Viz - GCE to Known Paralogs Locations
GCE ID: Event_284

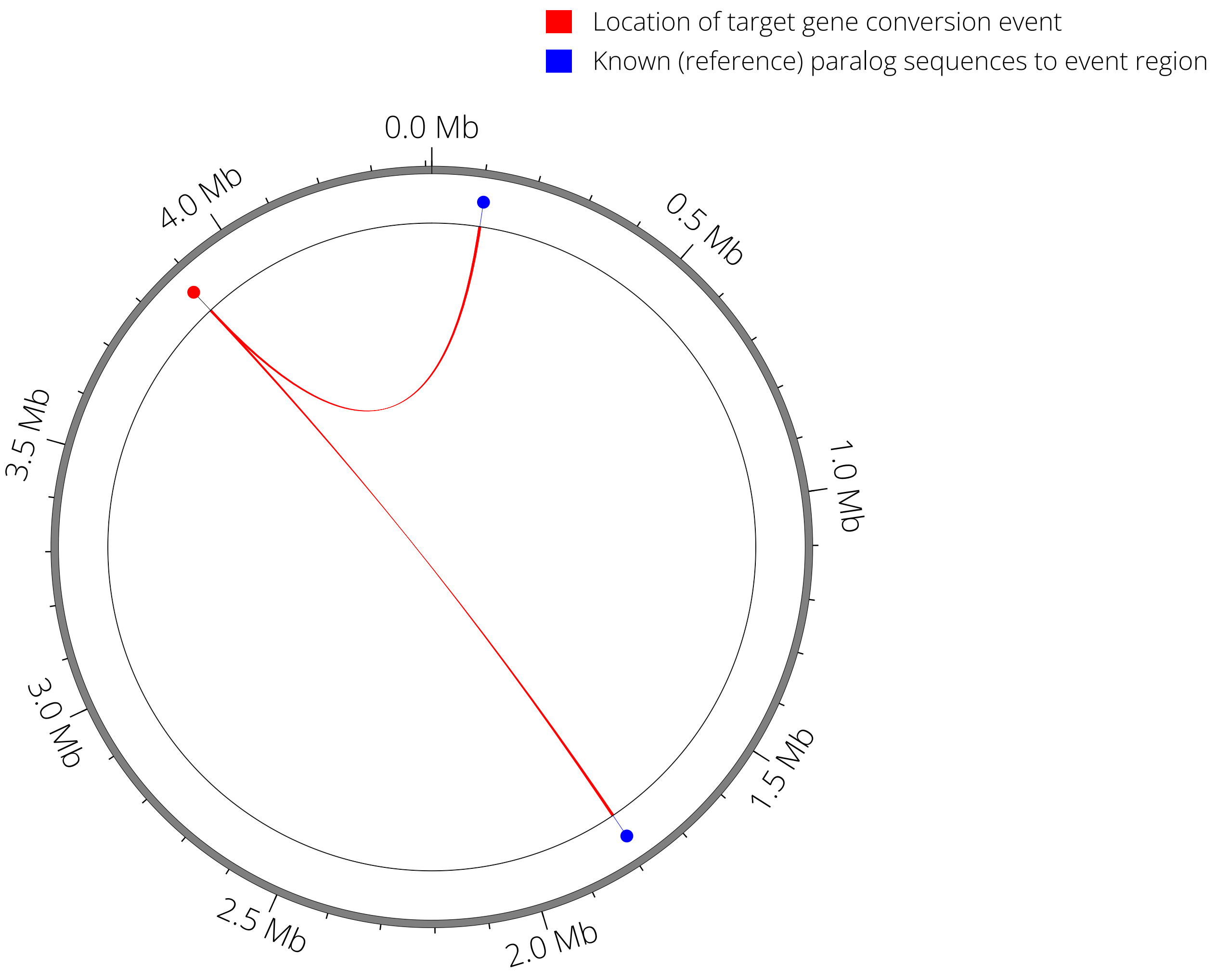

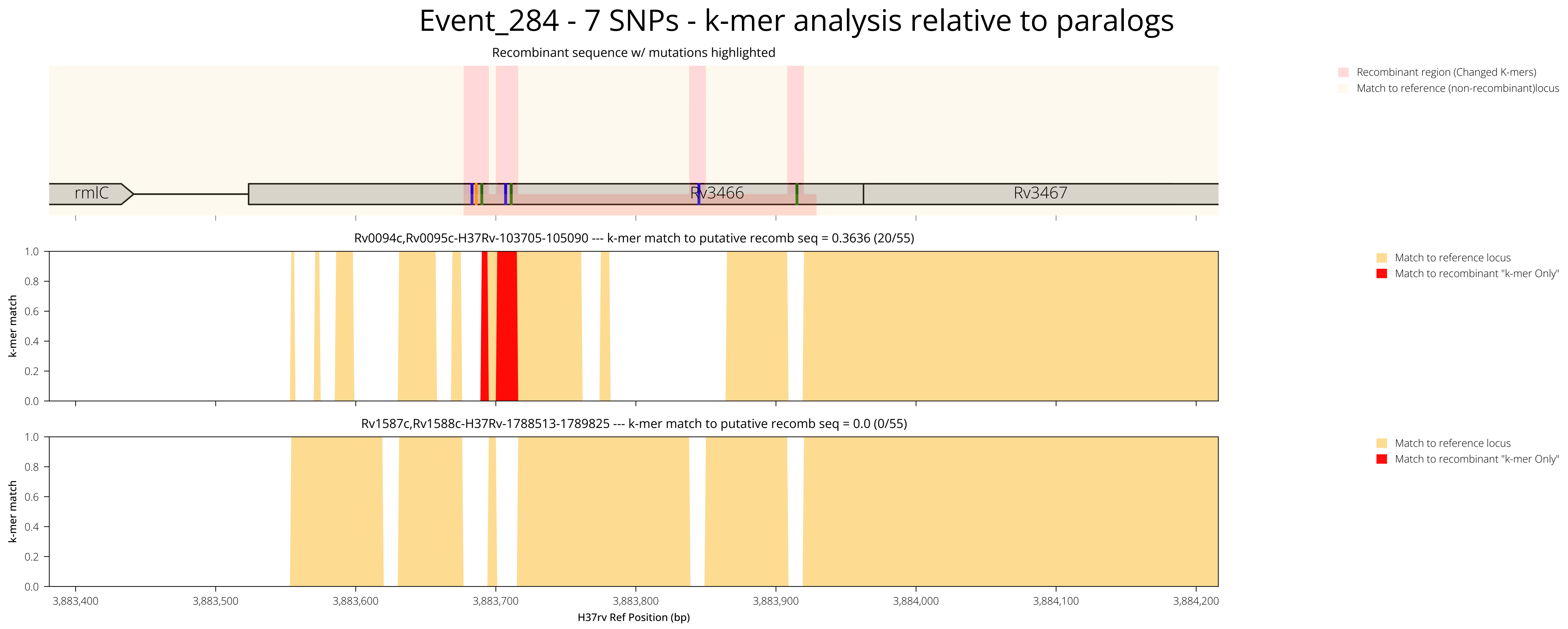
Event_284 - 7 SNPs - k-mer analysis relative to paralogs
Recombinant sequence w/ mutations highlighted
Recombinant region (Changed K-mers)
Match to reference (non-recombinant)locus
rmlC
Rv3466
Rv3467
Rv0094c,Rv0095c-H37Rv-103705-105090 --- k-mer match to putative recomb seq = 0.3636 (20/55)
Match to reference locus
Match to recombinant "k-mer Only"
k-mer match
1.0
0.8
0.6
0.4
0.2
0.0
Rv1587c,Rv1588c-H37Rv-1788513-1789825 --- k-mer match to putative recomb seq = 0.0 (0/55)
Match to reference locus
Match to recombinant "k-mer Only"
k-mer match
1.0
0.8
0.6
0.4
0.2
0.0
3,883,400
3,883,500
3,883,600
3,883,700
3,883,800
3,883,900
3,884,000
3,884,100
3,884,200
H37rv Ref Position (bp)

Sub-phylogeny Viz + Paralog Alignments
GCE ID: Event_284

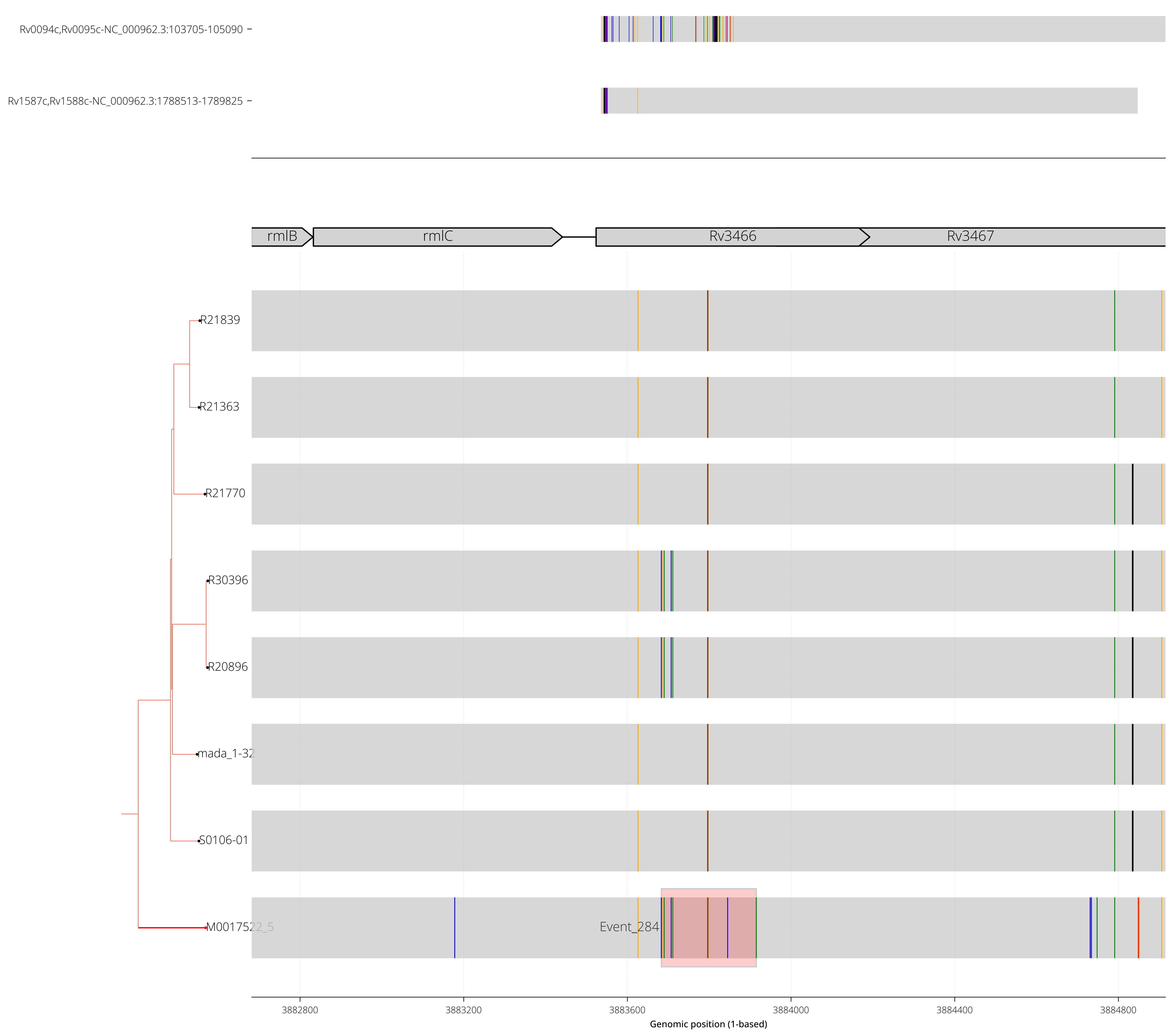

### Full Phylogeny Viz + Paralog Alignments
GCE ID: Event_284

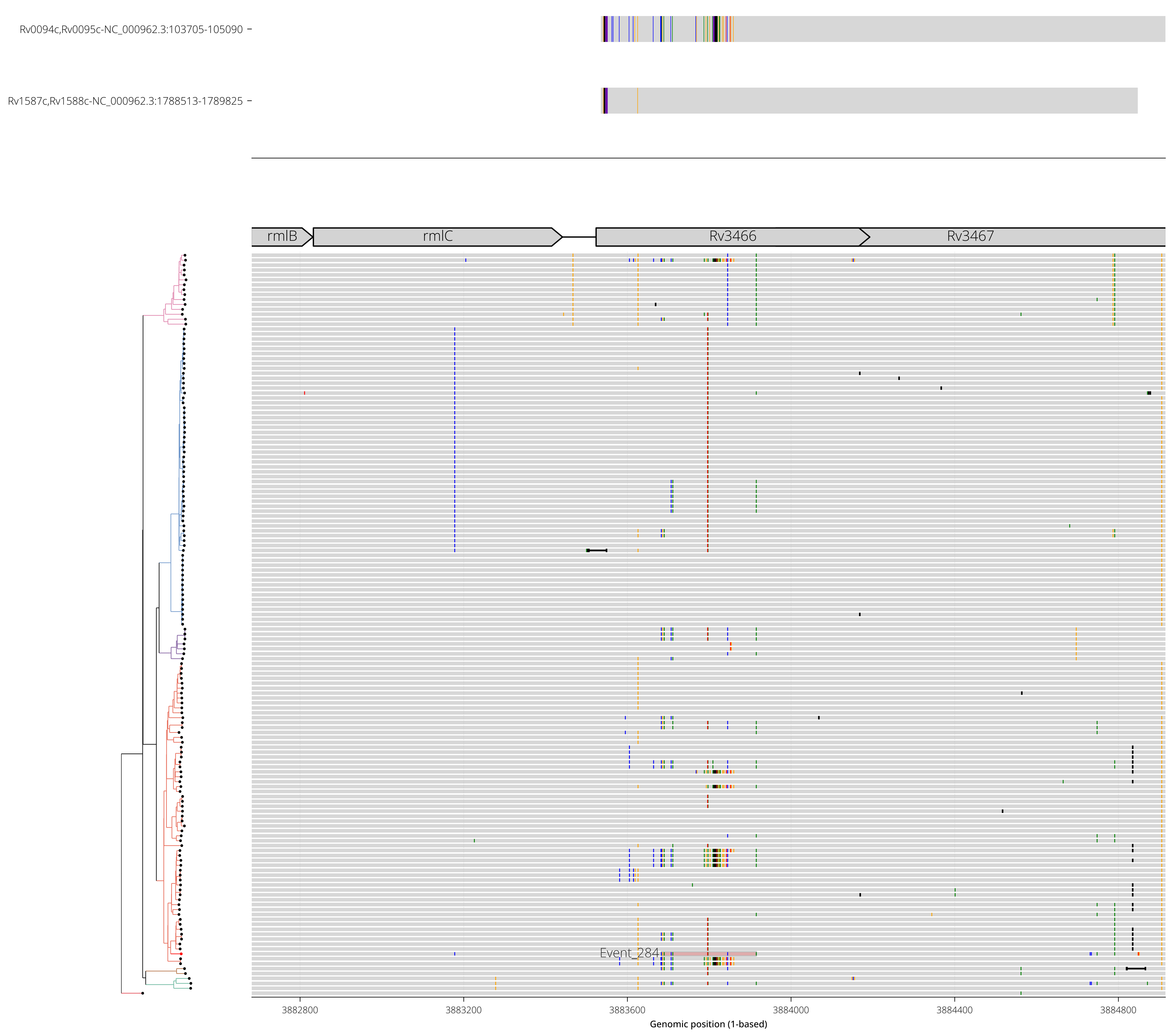

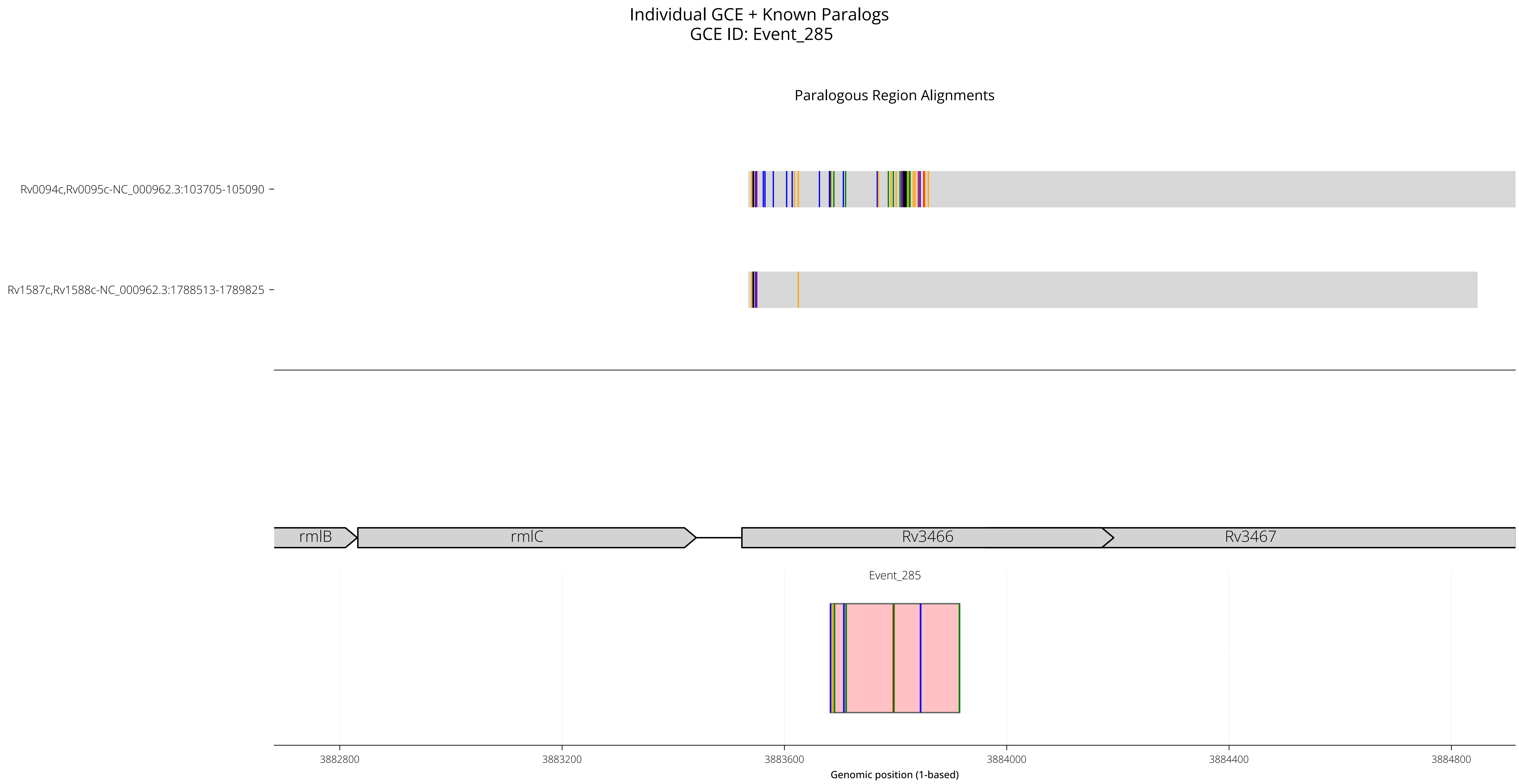

Individual GCE + Known Paralogs
GCE ID: Event_285
Paralogous Region Alignments
Rv0094c,Rv0095c-NC_000962.3:103705-105090
Rv1587c,Rv1588c-NC_000962.3:1788513-1789825
rmlB
rmlC
Rv3466
Rv3467
Event_285
3882800
3883200
3883600
3884000
3884400
3884800
Genomic position (1-based)

Circos Viz - GCE to Known Paralogs Locations
GCE ID: Event_285

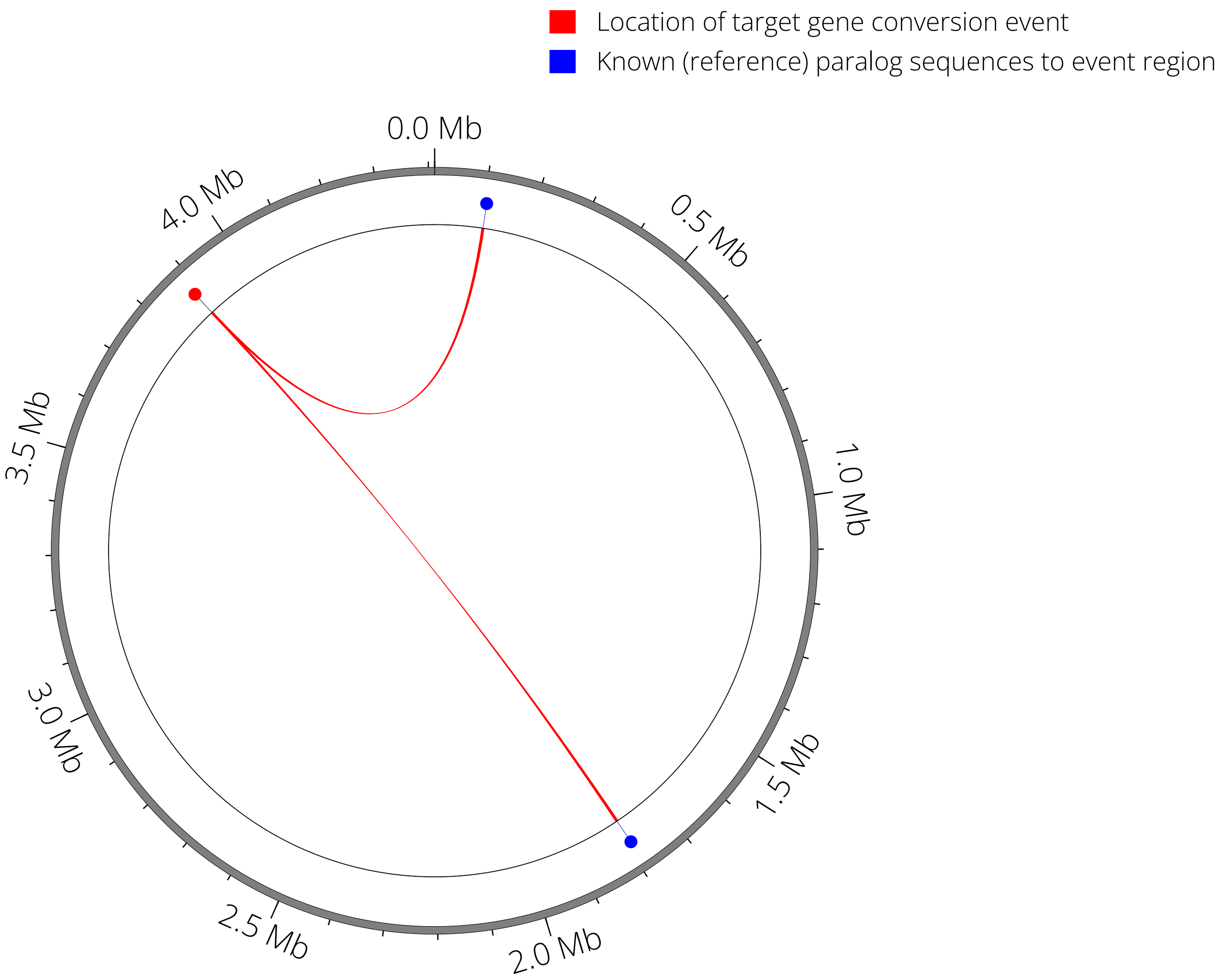

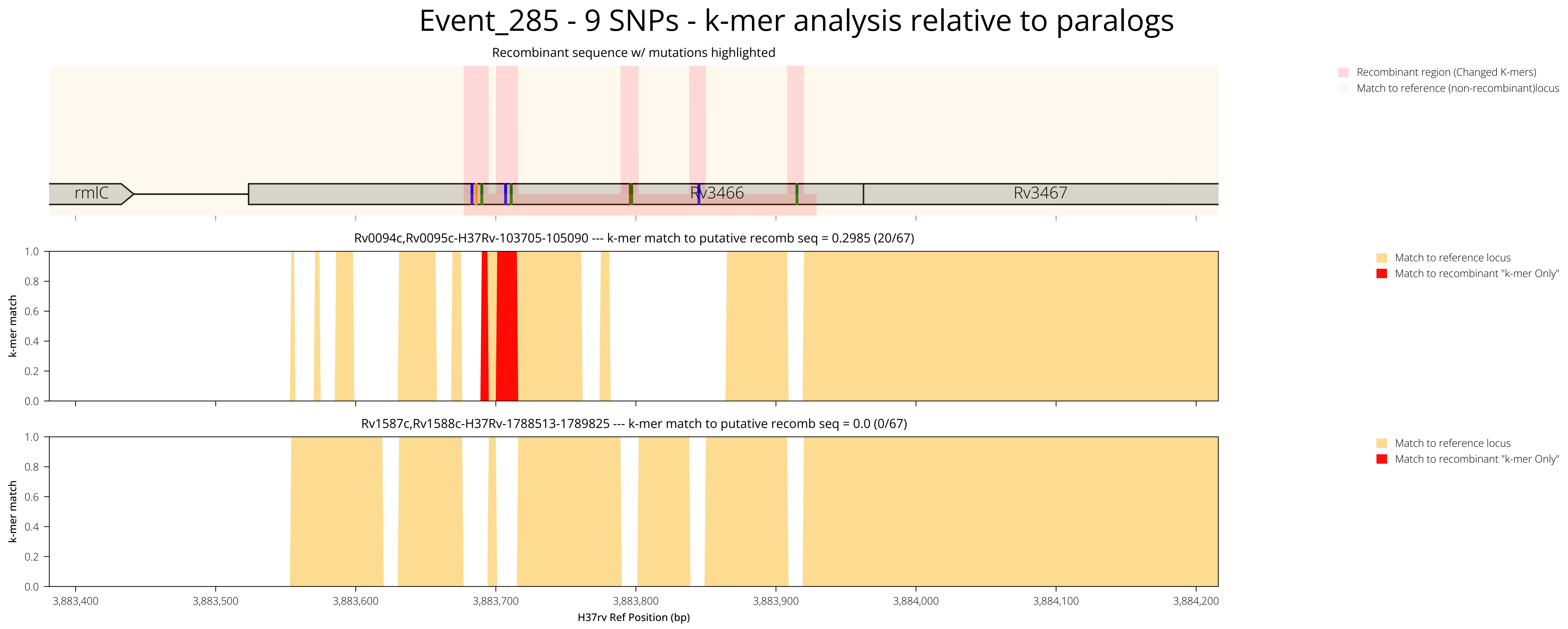
Event_285 - 9 SNPs - k-mer analysis relative to paralogs
Recombinant sequence w/ mutations highlighted
rmlC
Rv3466
Rv3467
Recombinant region (Changed K-mers)
Match to reference (non-recombinant)locus
Rv0094c,Rv0095c-H37Rv-103705-105090 --- k-mer match to putative recomb seq = 0.2985 (20/67)
Match to reference locus
Match to recombinant "k-mer Only"
Rv1587c,Rv1588c-H37Rv-1788513-1789825 --- k-mer match to putative recomb seq = 0.0 (0/67)
Match to reference locus
Match to recombinant "k-mer Only"
k-mer match
1.0
0.8
0.6
0.4
0.2
0.0
3,883,400
3,883,500
3,883,600
3,883,700
3,883,800
3,883,900
3,884,000
3,884,100
3,884,200
H37rv Ref Position (bp)

Sub-phylogeny Viz + Paralog Alignments
GCE ID: Event_285

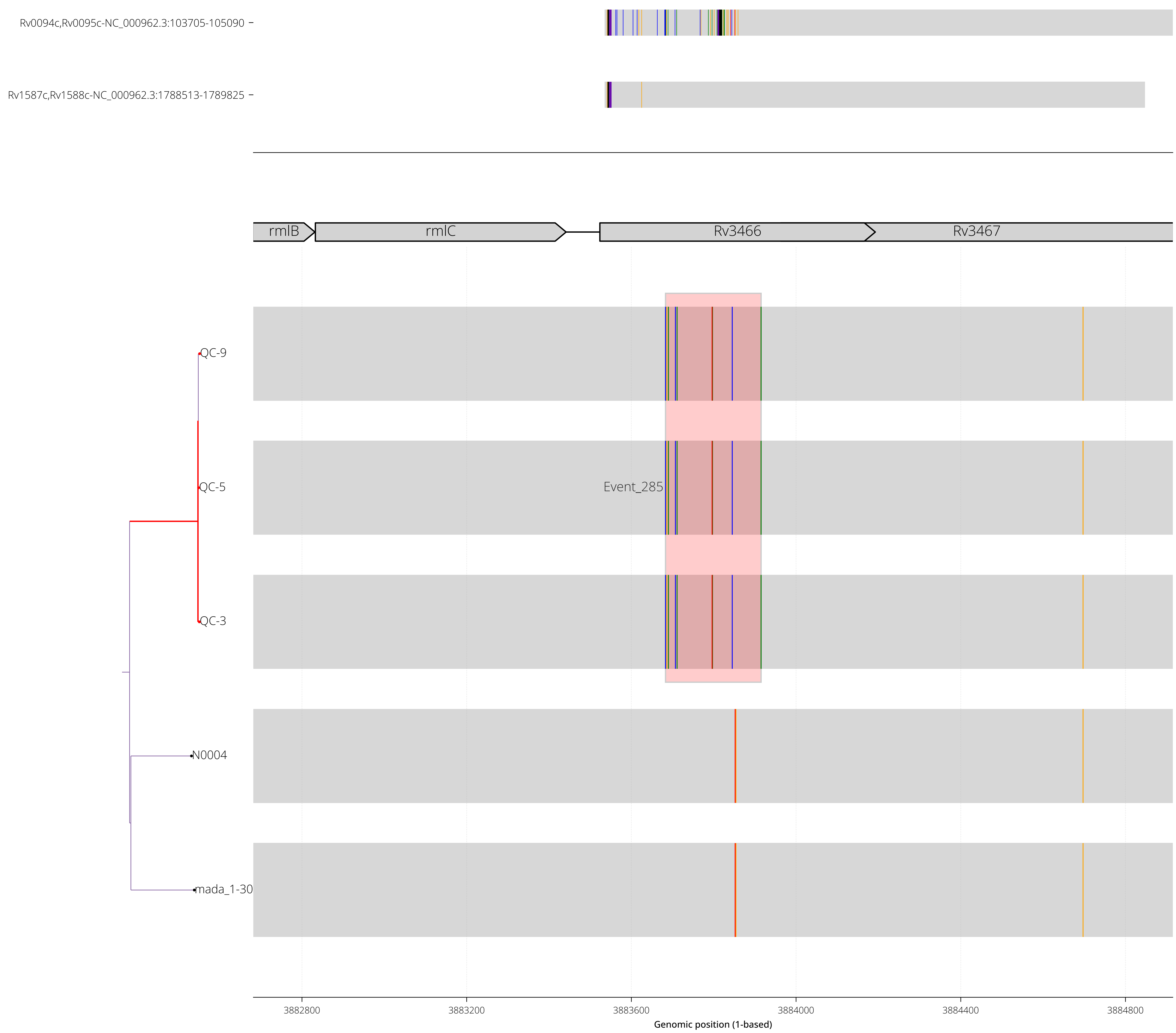

### Full Phylogeny Viz + Paralog Alignments
GCE ID: Event_285

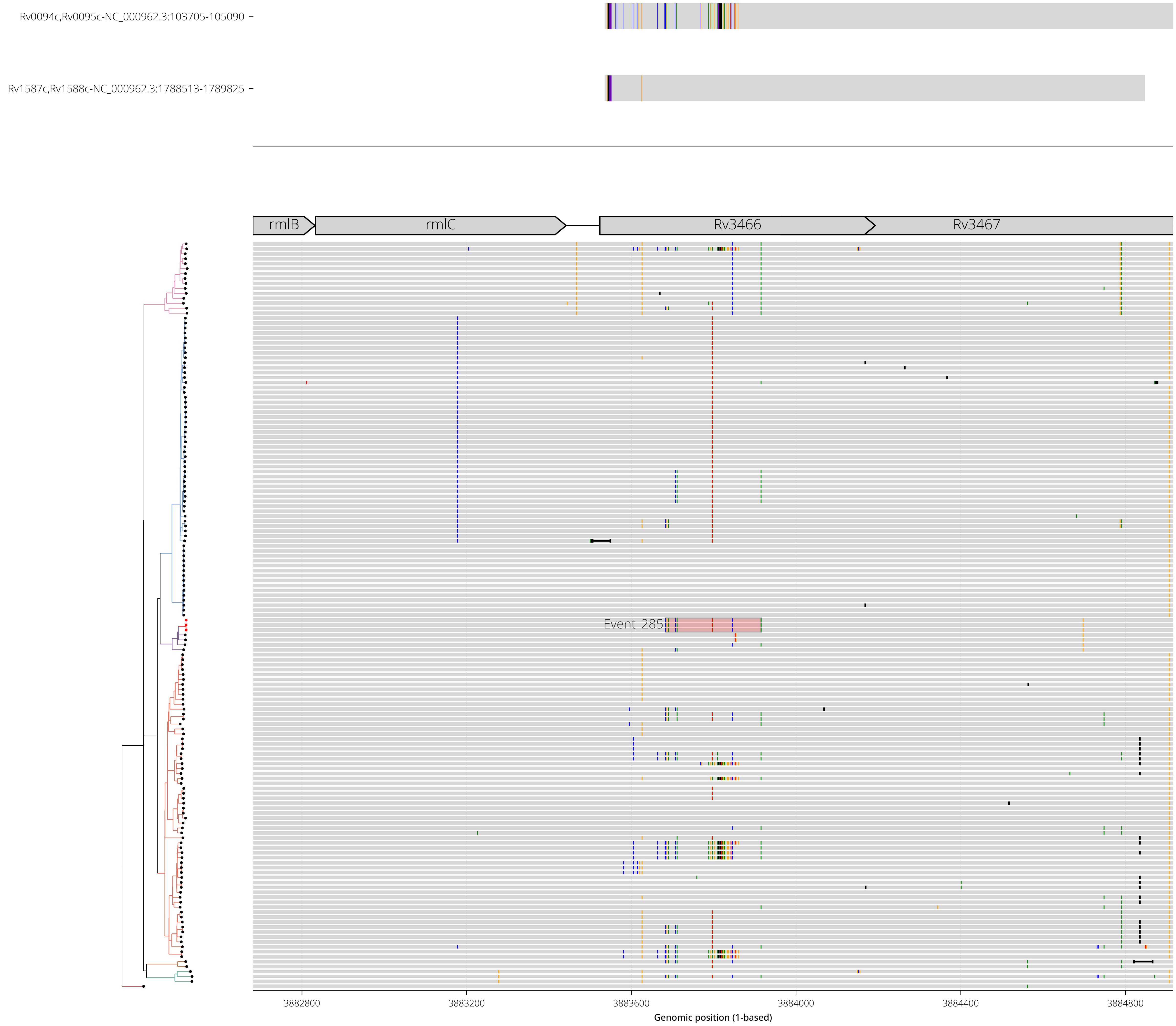

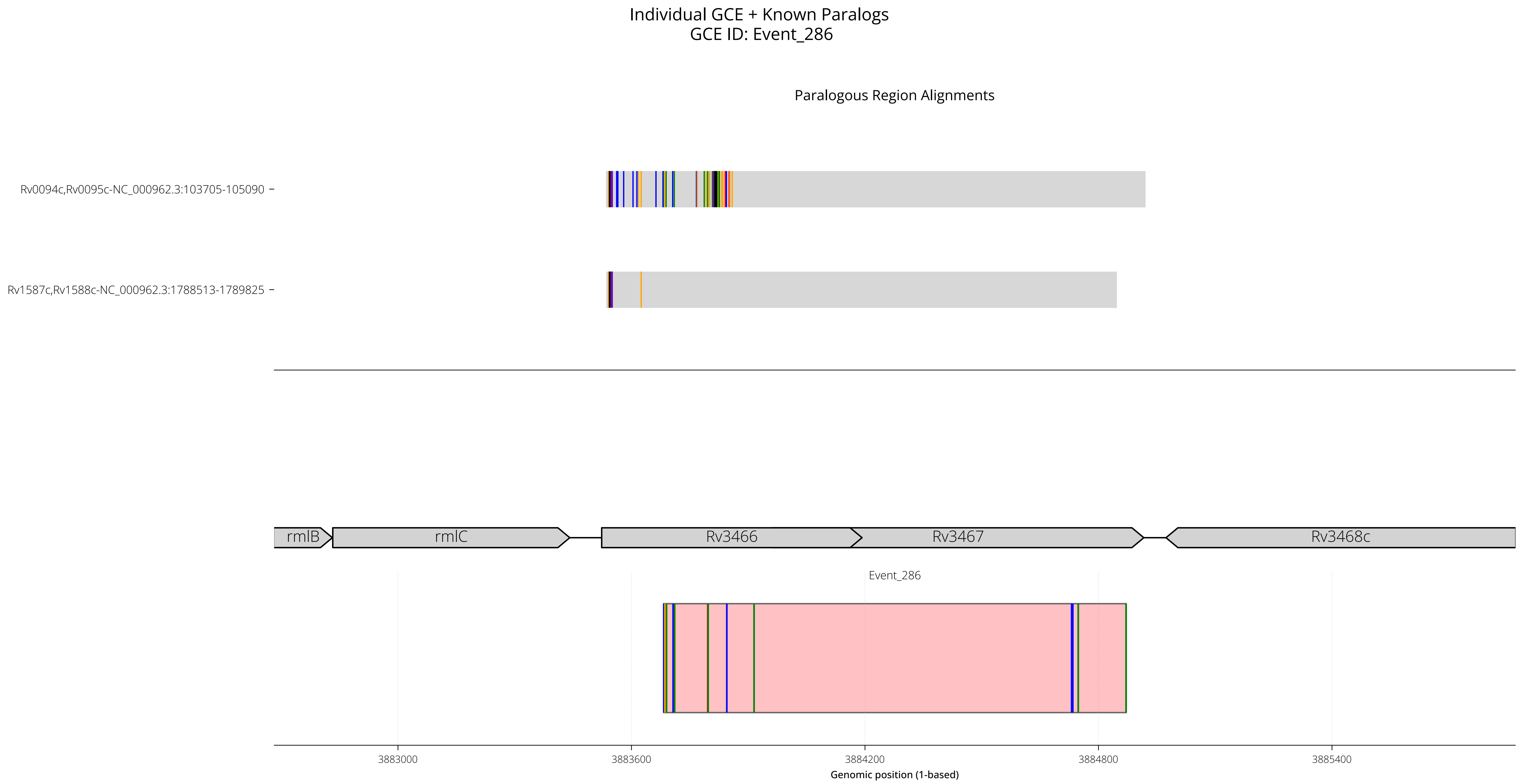

Individual GCE + Known Paralogs
GCE ID: Event_286
Paralogous Region Alignments
Rv0094c,Rv0095c-NC_000962.3:103705-105090
Rv1587c,Rv1588c-NC_000962.3:1788513-1789825
rmlB
rmlC
Rv3466
Rv3467
Rv3468c
Event_286
3883000
3883600
3884200
3884800
3885400
Genomic position (1-based)

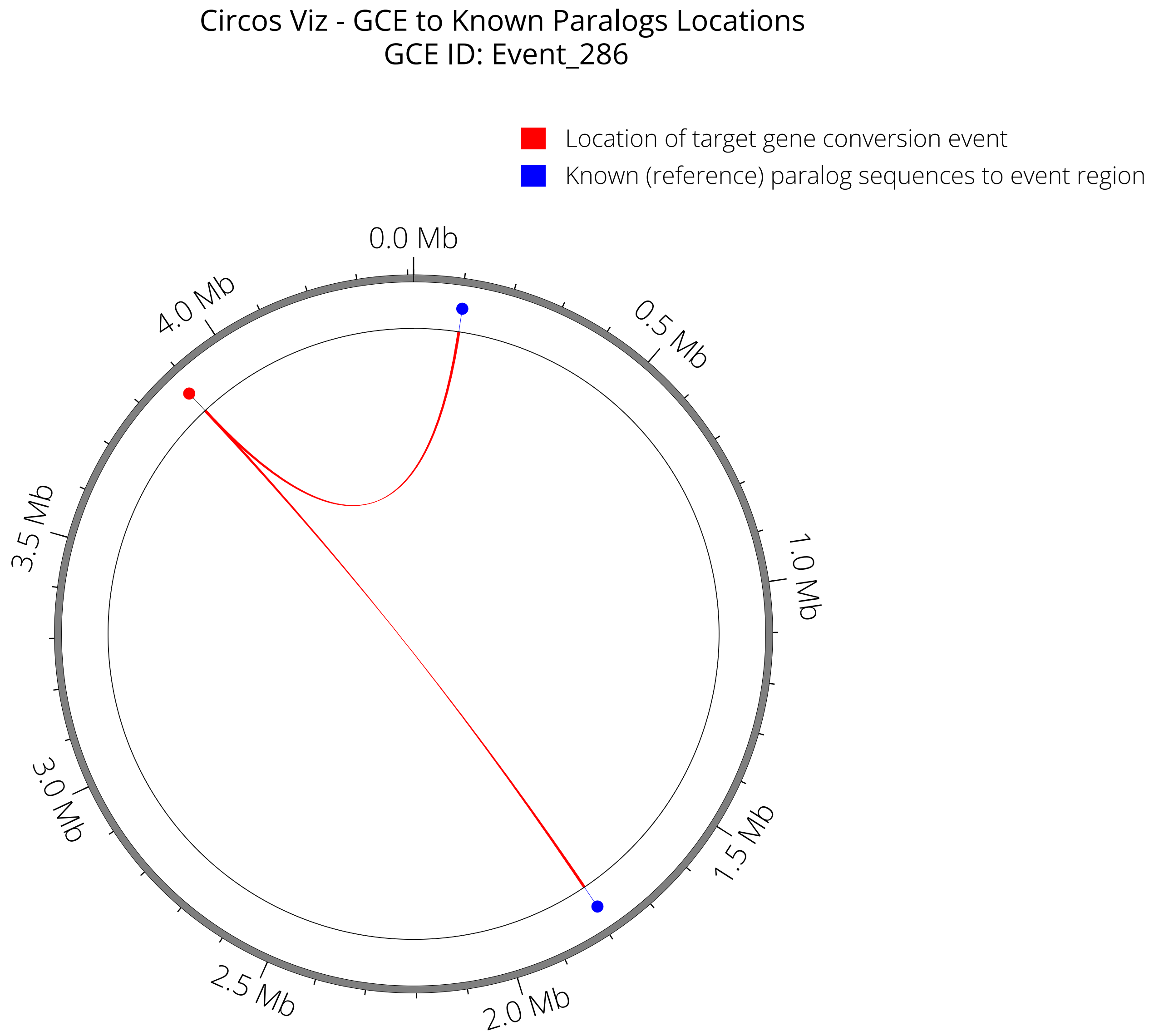

Circos Viz - GCE to Known Paralogs Locations
GCE ID: Event_286
Location of target gene conversion event
Known (reference) paralog sequences to event region
0.0 Mb
0.5 Mb
1.0 Mb
1.5 Mb
2.0 Mb
2.5 Mb
3.0 Mb
3.5 Mb
4.0 Mb

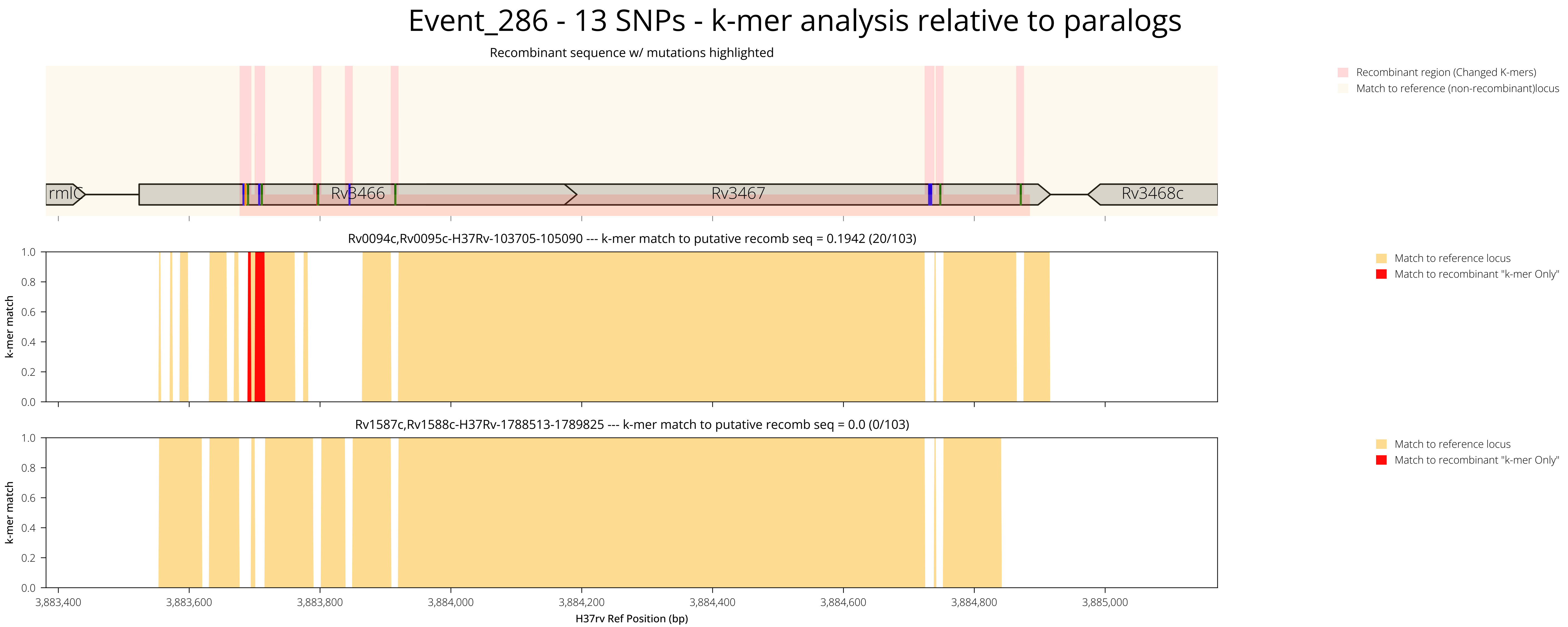

Event_286 - 13 SNPs - k-mer analysis relative to paralogs
Recombinant sequence w/ mutations highlighted
Recombinant region (Changed K-mers)
Match to reference (non-recombinant)locus
rmlC
Rv3466
Rv3467
Rv3468c
Rv0094c,Rv0095c-H37Rv-103705-105090 --- k-mer match to putative recomb seq = 0.1942 (20/103)
Match to reference locus
Match to recombinant "k-mer Only"
k-mer match
Rv1587c,Rv1588c-H37Rv-1788513-1789825 --- k-mer match to putative recomb seq = 0.0 (0/103)
Match to reference locus
Match to recombinant "k-mer Only"
k-mer match
3,883,400
3,883,600
3,883,800
3,884,000
3,884,200
3,884,400
3,884,600
3,884,800
3,885,000
H37rv Ref Position (bp)

Sub-phylogeny Viz + Paralog Alignments
GCE ID: Event_286

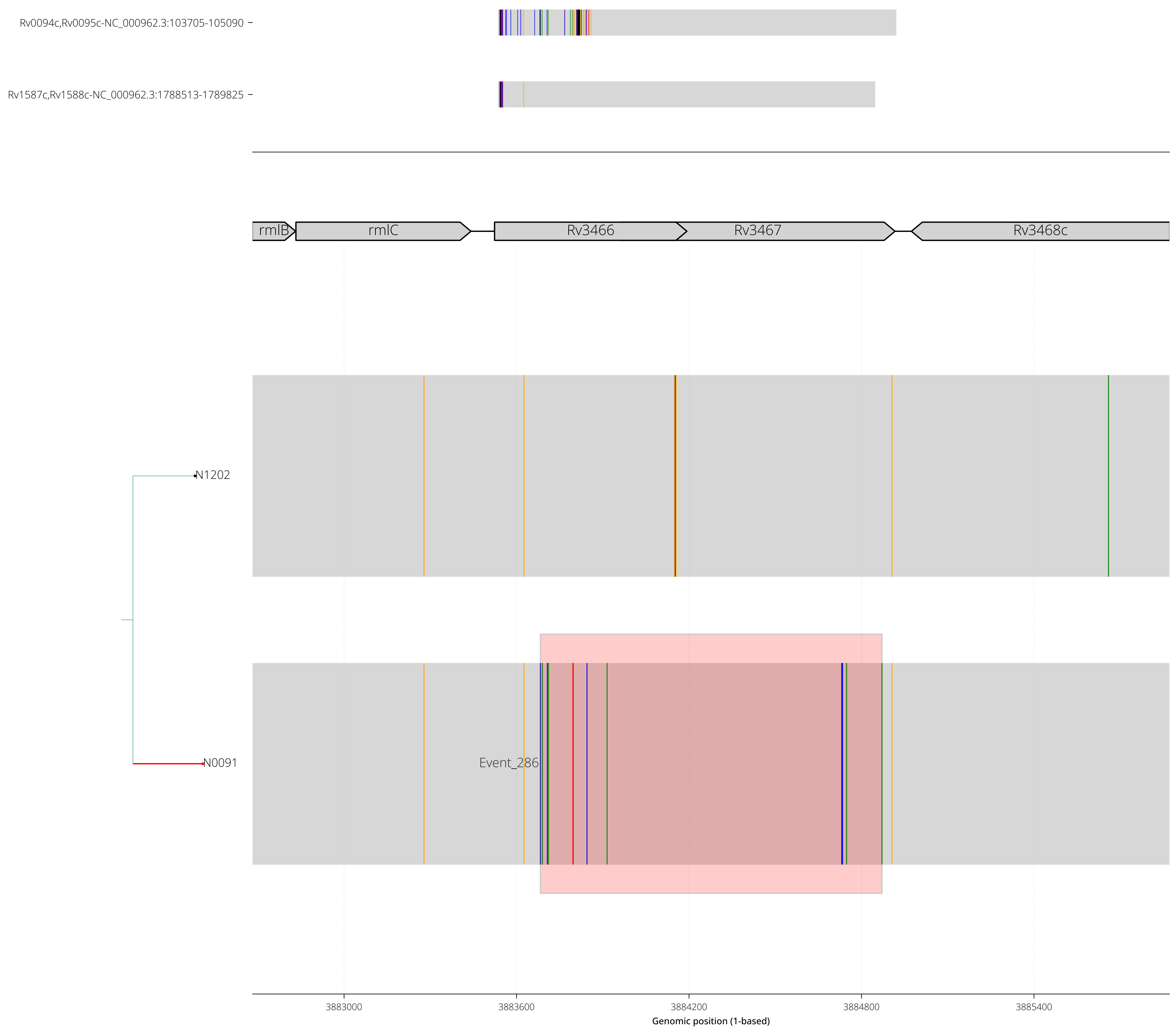

Full Phylogeny Viz + Paralog Alignments
GCE ID: Event_286

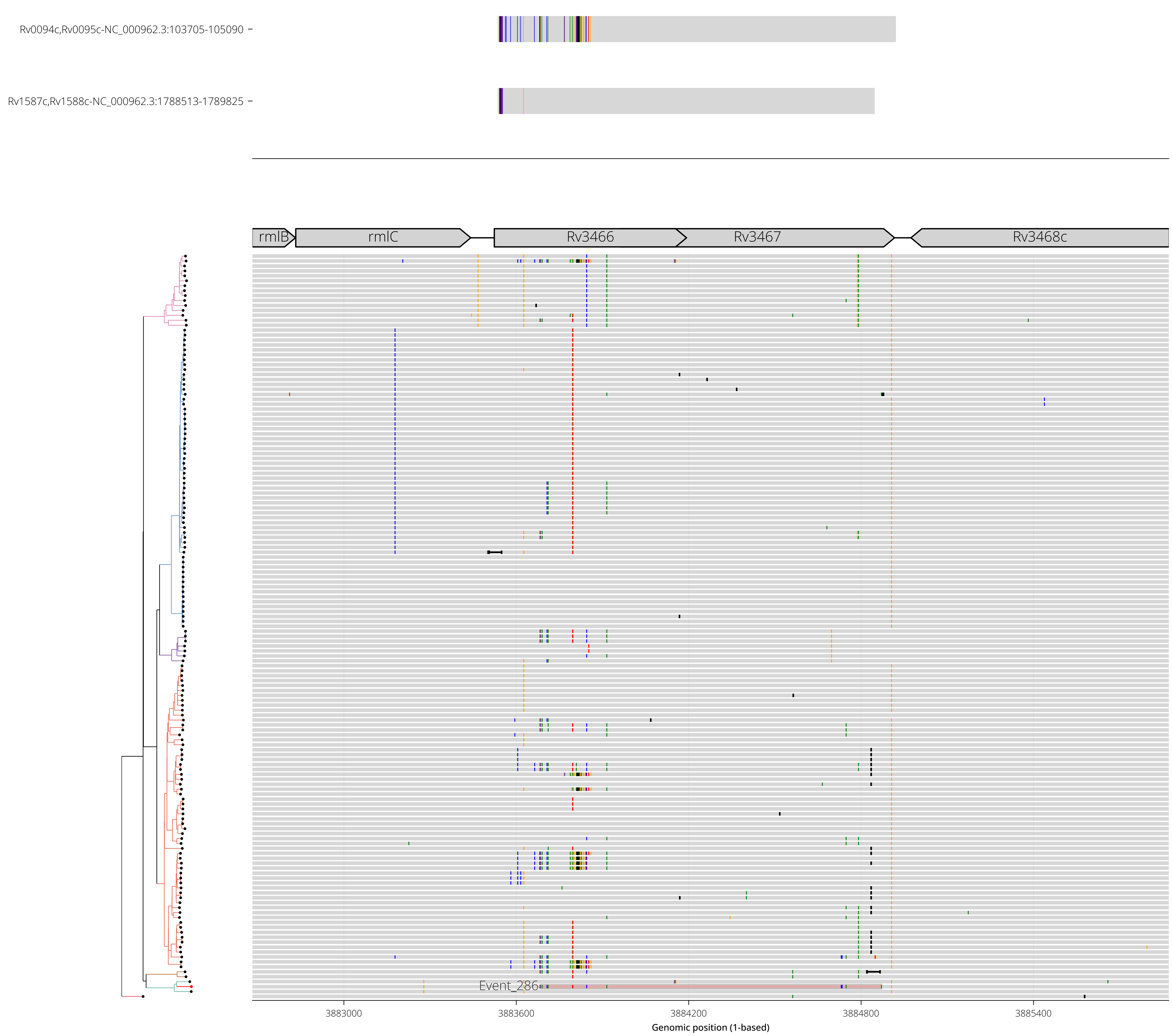

Individual GCE + Known Paralogs
GCE ID: Event_287

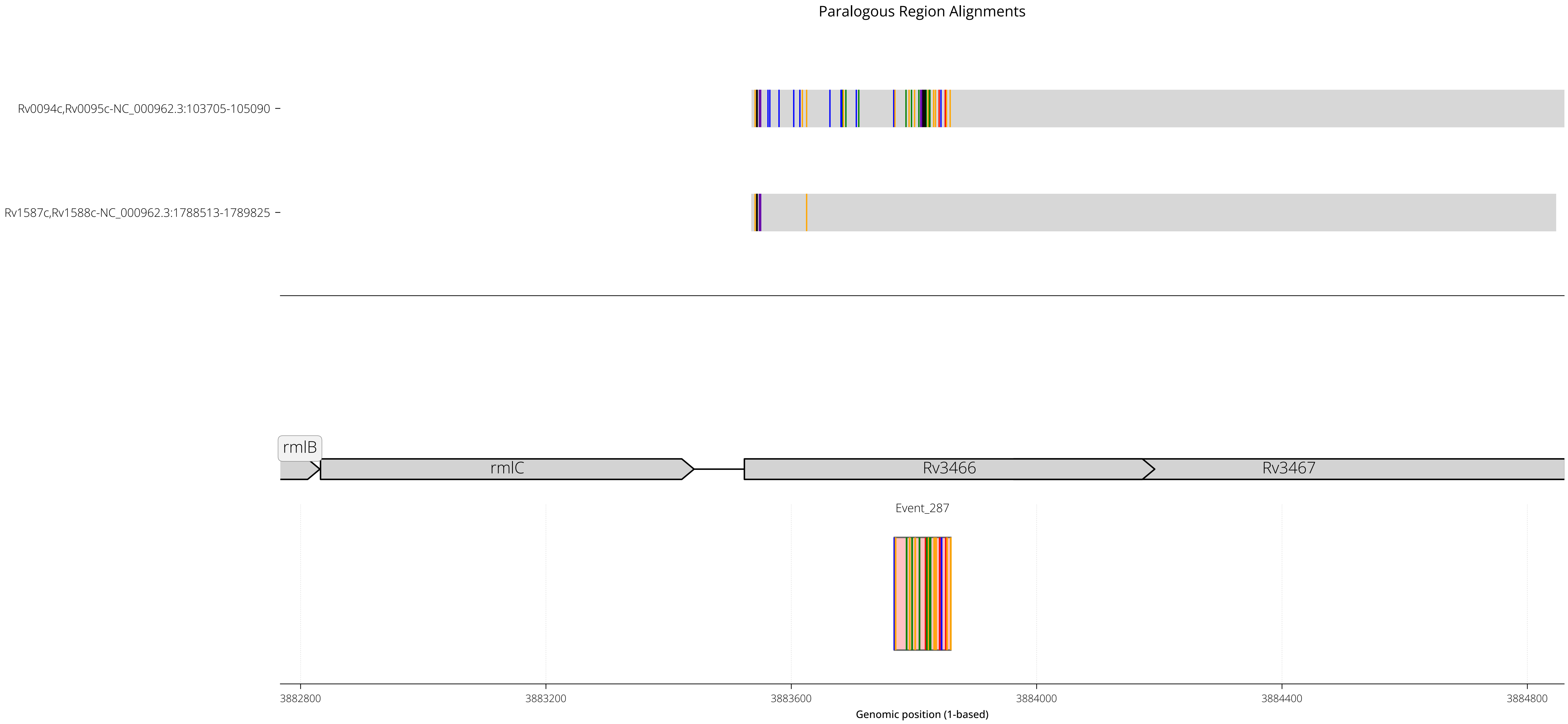

Circos Viz - GCE to Known Paralogs Locations
GCE ID: Event_287

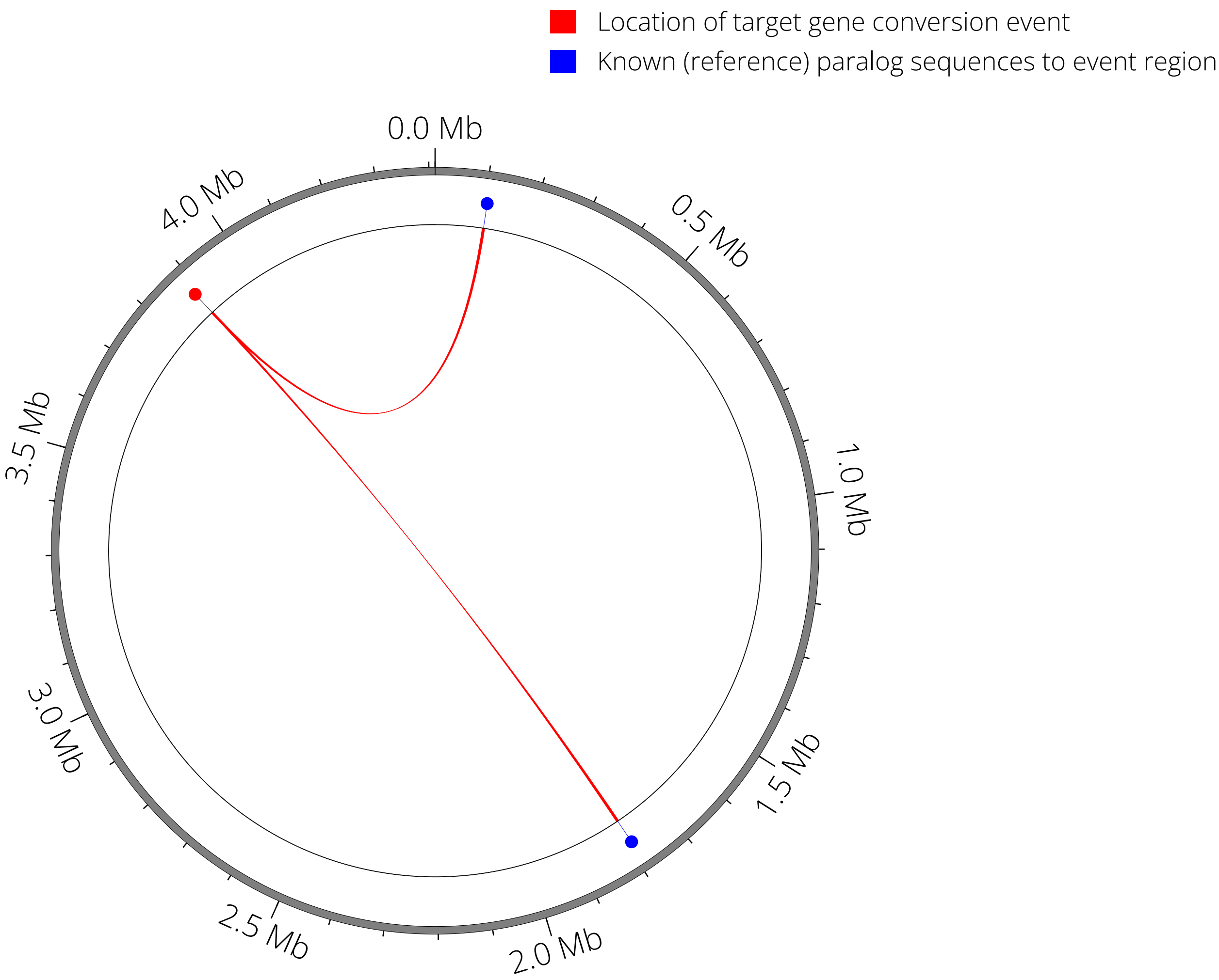

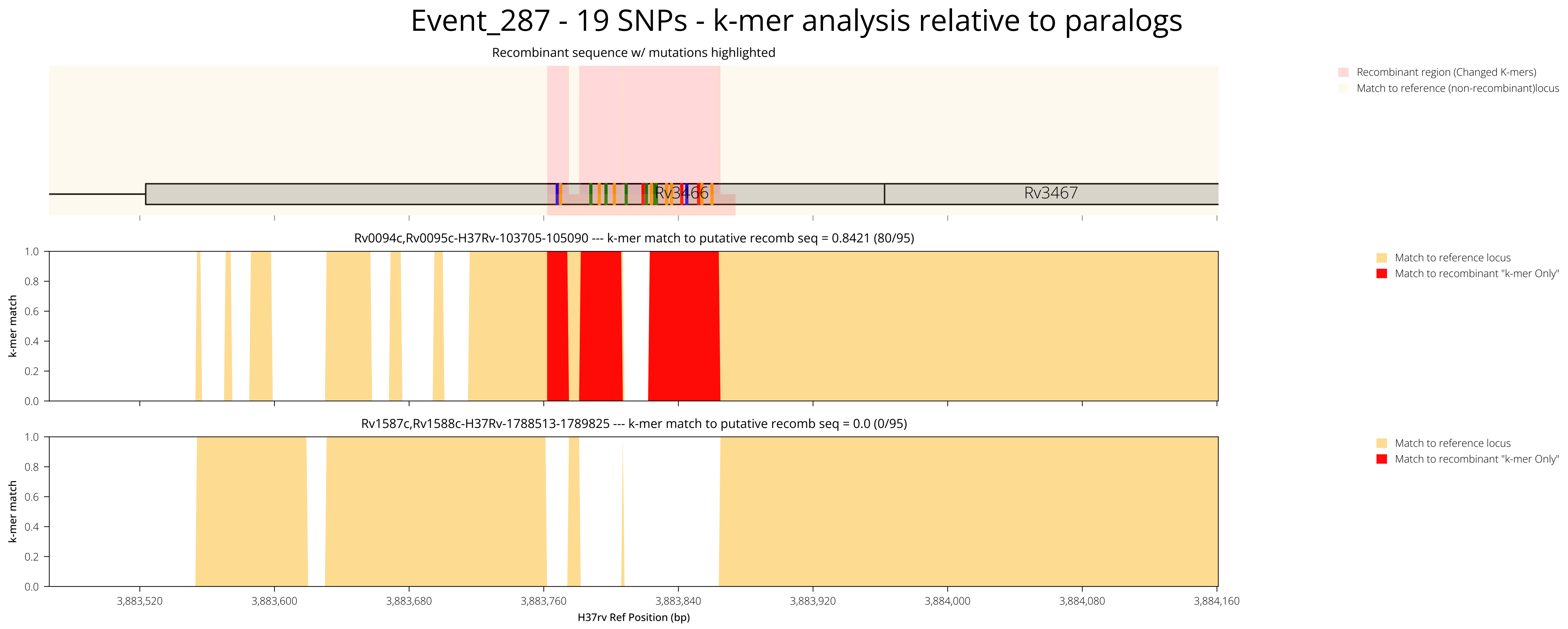
Event_287 - 19 SNPs - k-mer analysis relative to paralogs
Recombinant sequence w/ mutations highlighted
Recombinant region (Changed K-mers)
Match to reference (non-recombinant)locus
Rv3466
Rv3467
Rv0094c,Rv0095c-H37Rv-103705-105090 --- k-mer match to putative recomb seq = 0.8421 (80/95)
Match to reference locus
Match to recombinant "k-mer Only"
k-mer match
1.0
0.8
0.6
0.4
0.2
0.0
Rv1587c,Rv1588c-H37Rv-1788513-1789825 --- k-mer match to putative recomb seq = 0.0 (0/95)
Match to reference locus
Match to recombinant "k-mer Only"
k-mer match
1.0
0.8
0.6
0.4
0.2
0.0
3,883,520
3,883,600
3,883,680
3,883,760
3,883,840
3,883,920
3,884,000
3,884,080
3,884,160
H37rv Ref Position (bp)

Sub-phylogeny Viz + Paralog Alignments
GCE ID: Event_287

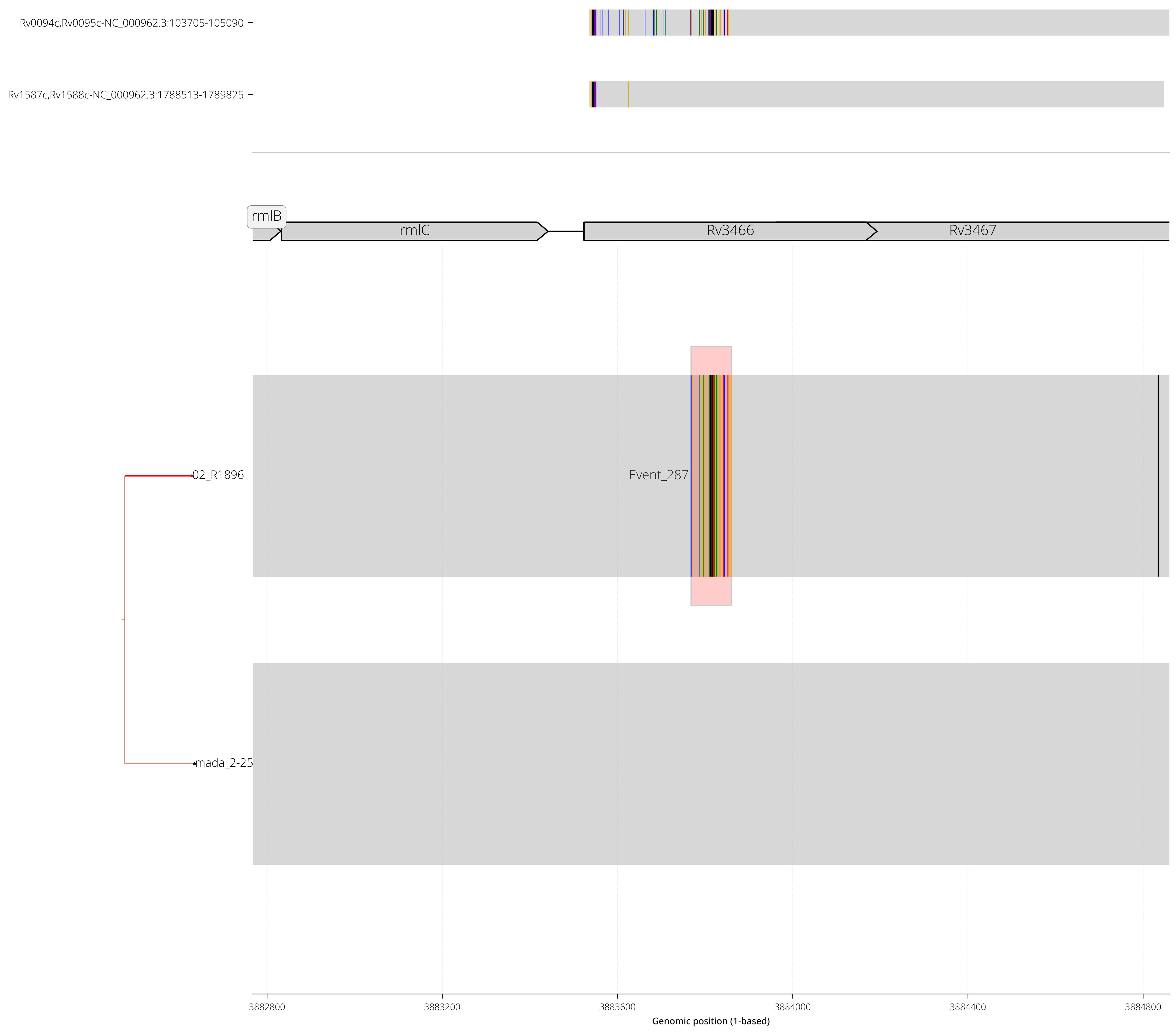

Full Phylogeny Viz + Paralog Alignments
GCE ID: Event_287

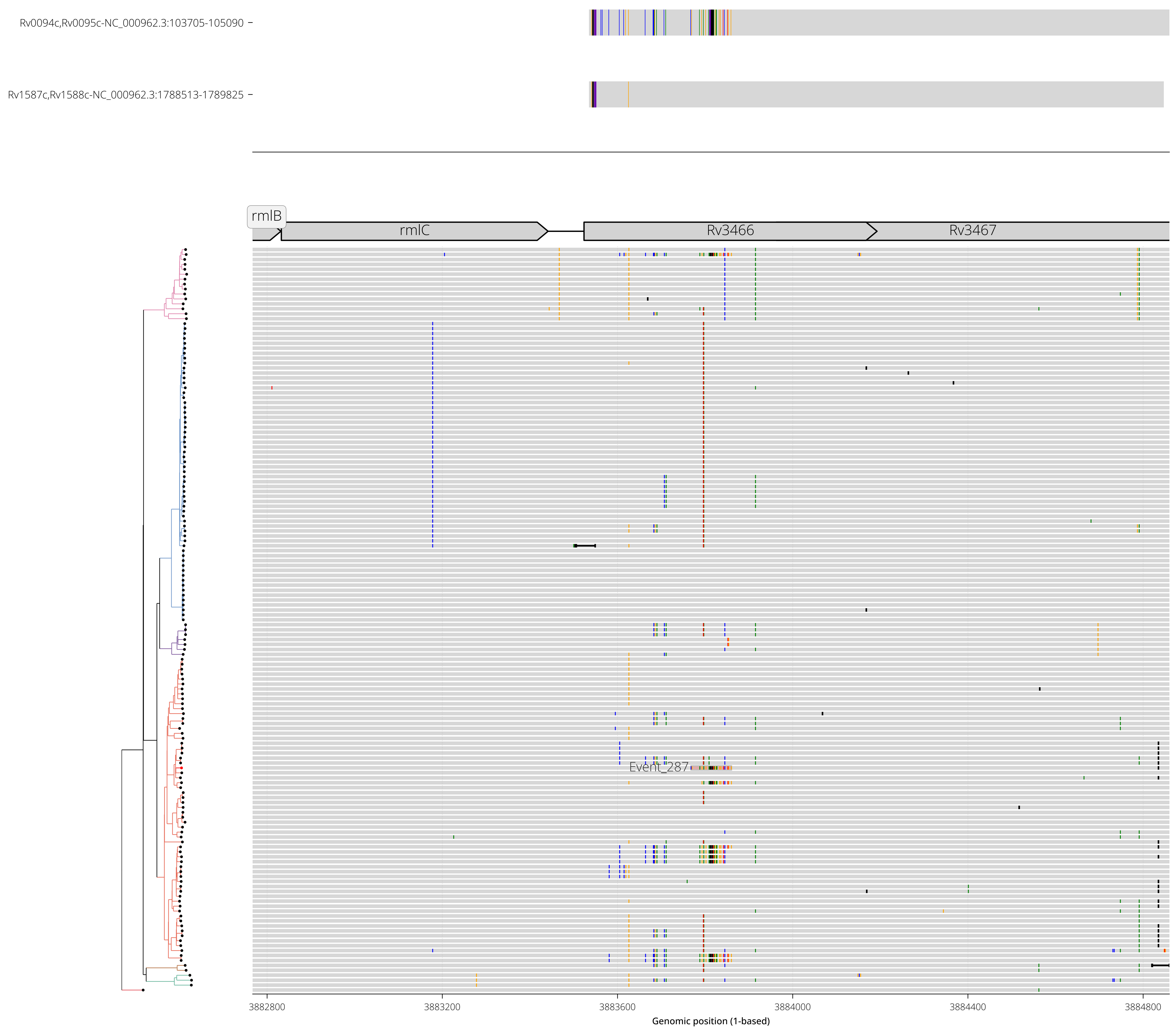

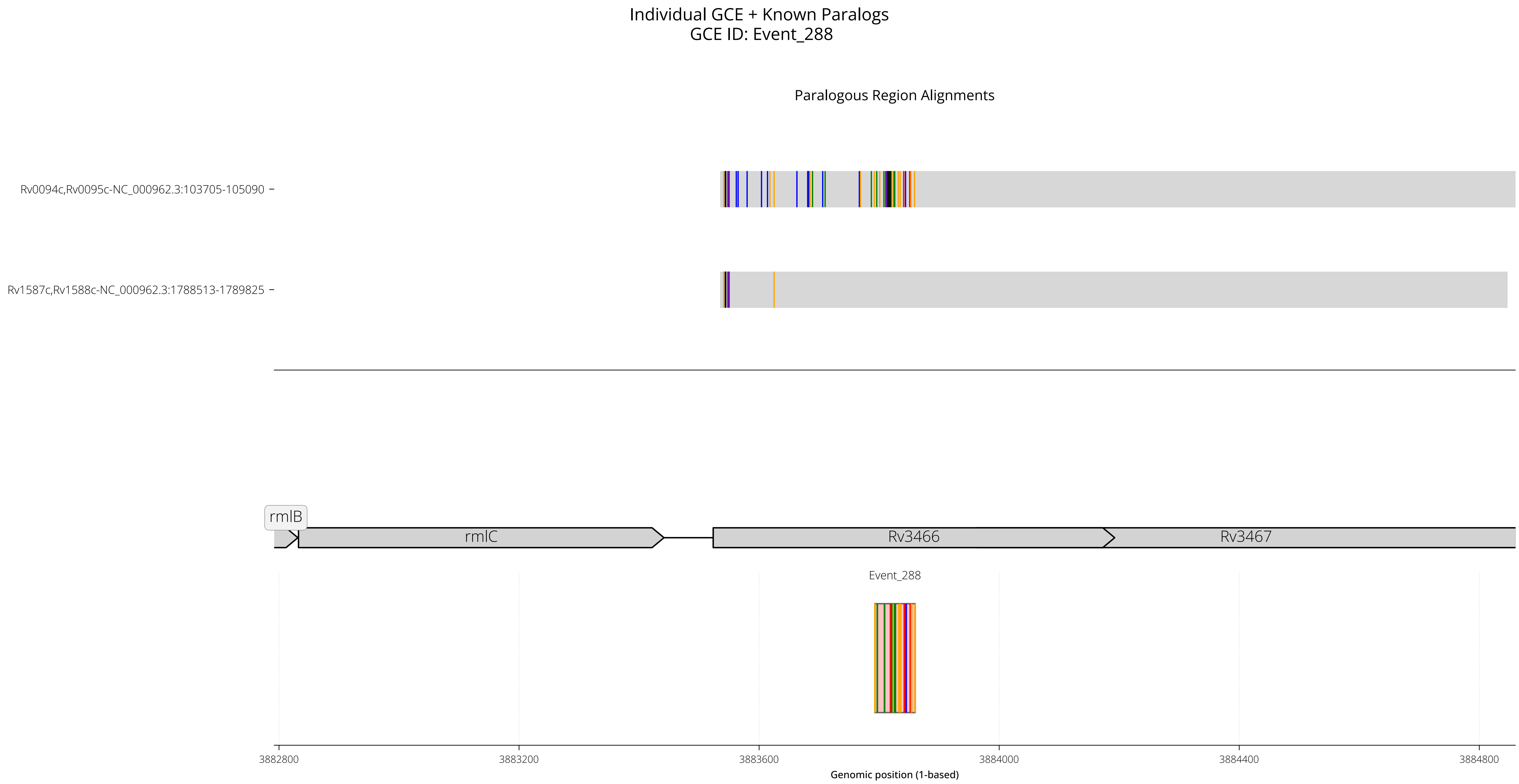

Individual GCE + Known Paralogs
GCE ID: Event_288
Paralogous Region Alignments
Rv0094c,Rv0095c-NC_000962.3:103705-105090
Rv1587c,Rv1588c-NC_000962.3:1788513-1789825
rmlB
rmlC
Rv3466
Rv3467
Event_288
3882800
3883200
3883600
3884000
3884400
3884800
Genomic position (1-based)

Circos Viz - GCE to Known Paralogs Locations
GCE ID: Event_288

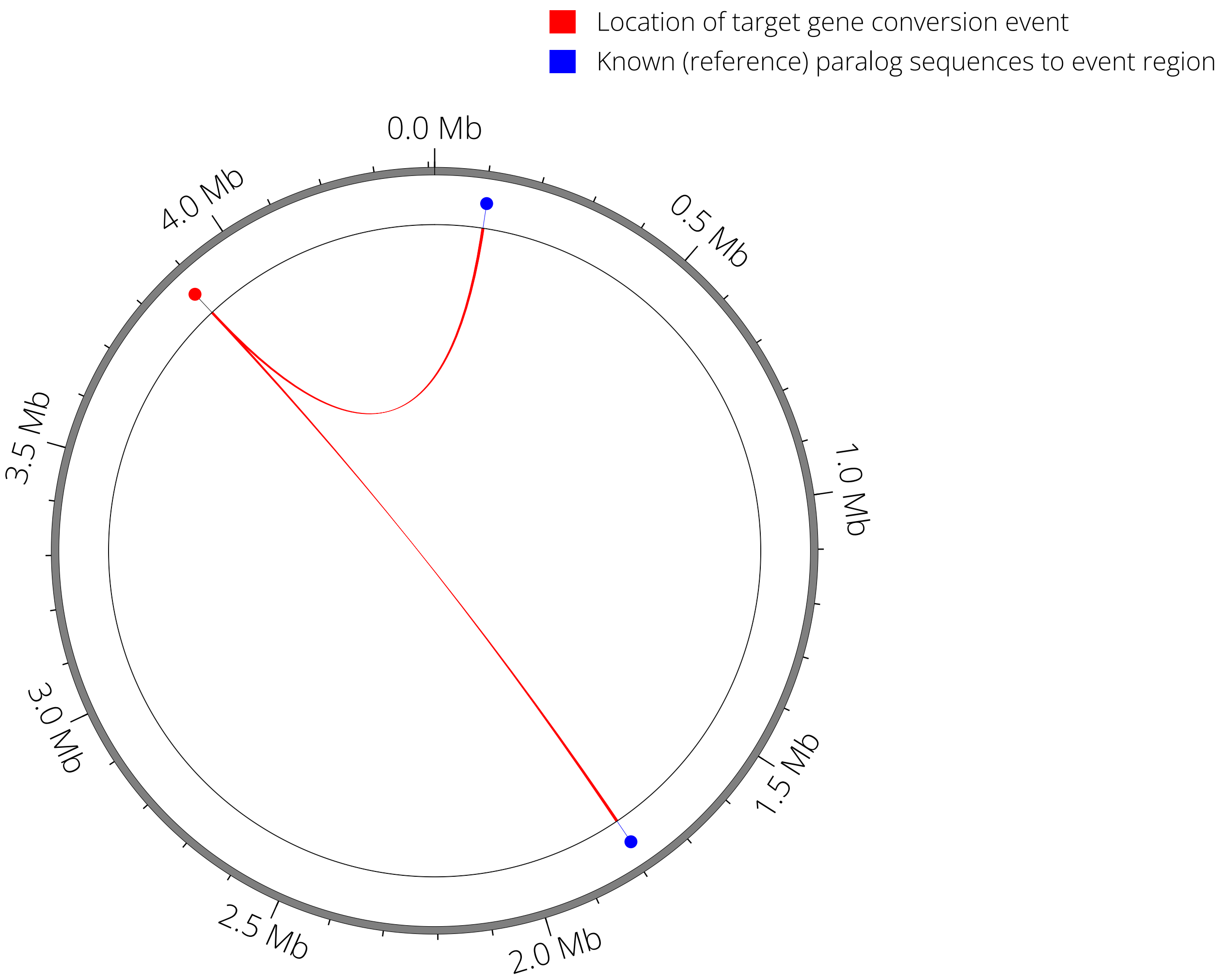

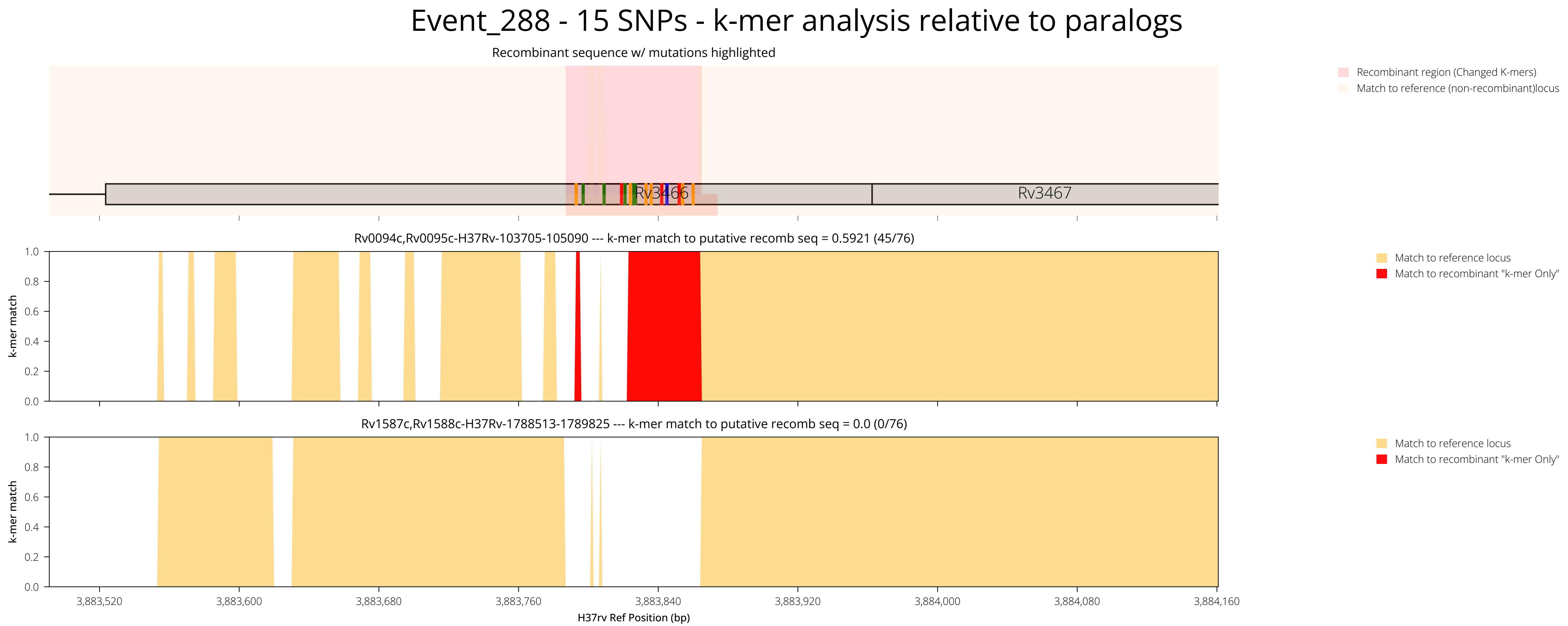

Event_288 - 15 SNPs - k-mer analysis relative to paralogs
Recombinant sequence w/ mutations highlighted
Recombinant region (Changed K-mers)
Match to reference (non-recombinant)locus
Rv3466
Rv3467
Rv0094c,Rv0095c-H37Rv-103705-105090 --- k-mer match to putative recomb seq = 0.5921 (45/76)
Match to reference locus
Match to recombinant "k-mer Only"
k-mer match
Rv1587c,Rv1588c-H37Rv-1788513-1789825 --- k-mer match to putative recomb seq = 0.0 (0/76)
Match to reference locus
Match to recombinant "k-mer Only"
k-mer match
3,883,520
3,883,600
3,883,680
3,883,760
3,883,840
3,883,920
3,884,000
3,884,080
3,884,160
H37rv Ref Position (bp)

Sub-phylogeny Viz + Paralog Alignments
GCE ID: Event_288

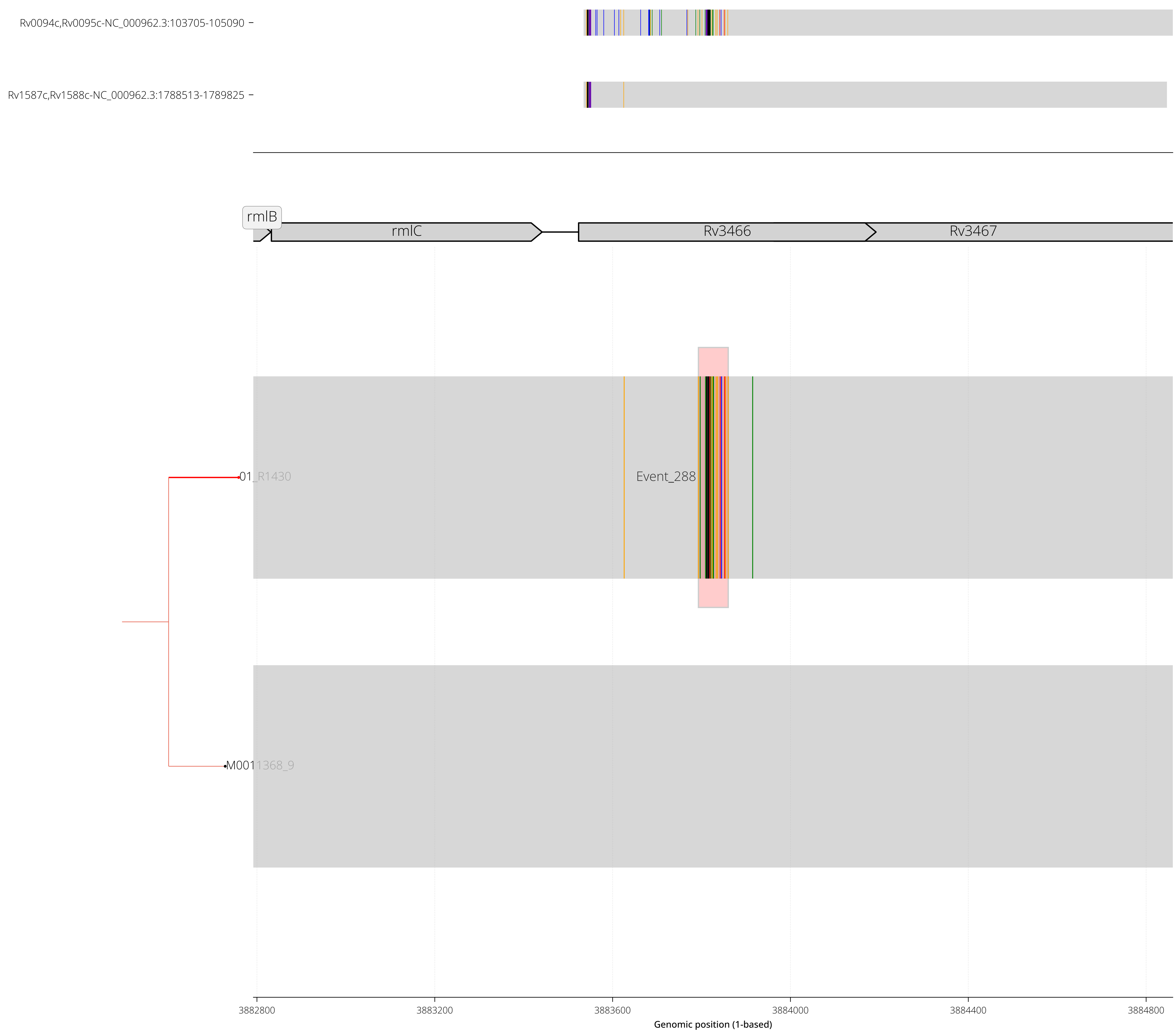

Full Phylogeny Viz + Paralog Alignments
GCE ID: Event_288

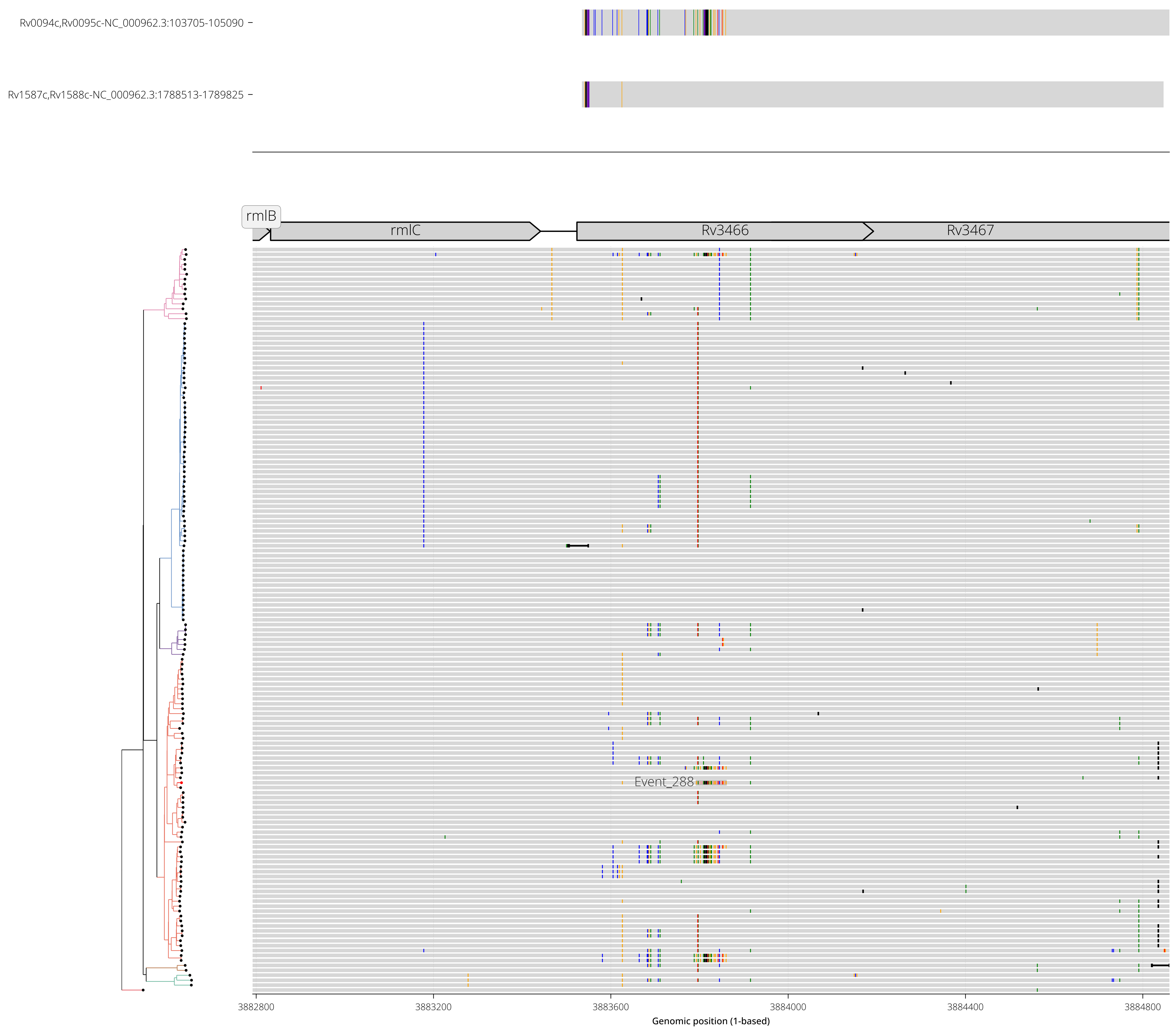

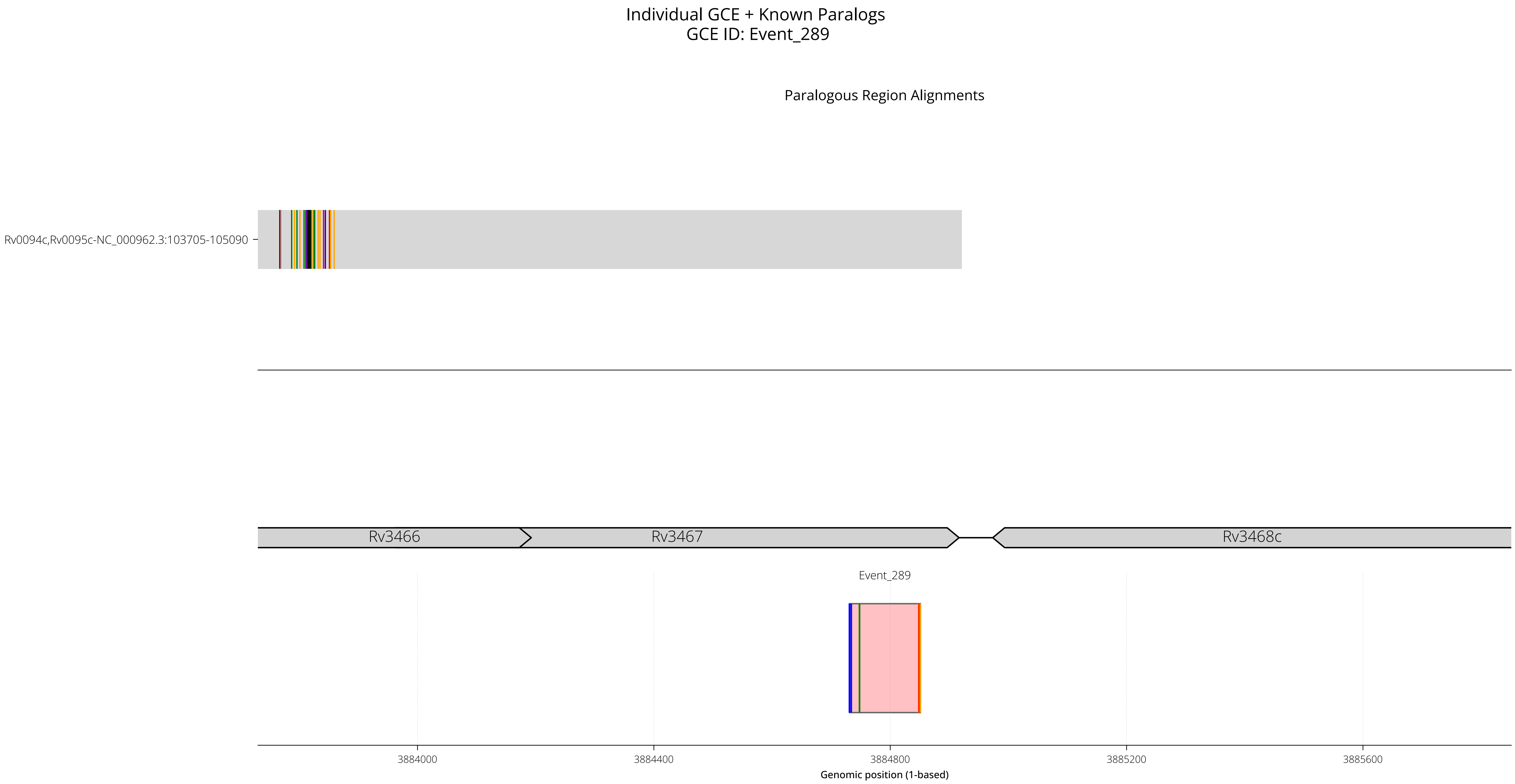

Individual GCE + Known Paralogs
GCE ID: Event_289
Paralogous Region Alignments
Rv0094c,Rv0095c-NC_000962.3:103705-105090
Rv3466
Rv3467
Rv3468c
Event_289
3884000
3884400
3884800
3885200
3885600
Genomic position (1-based)

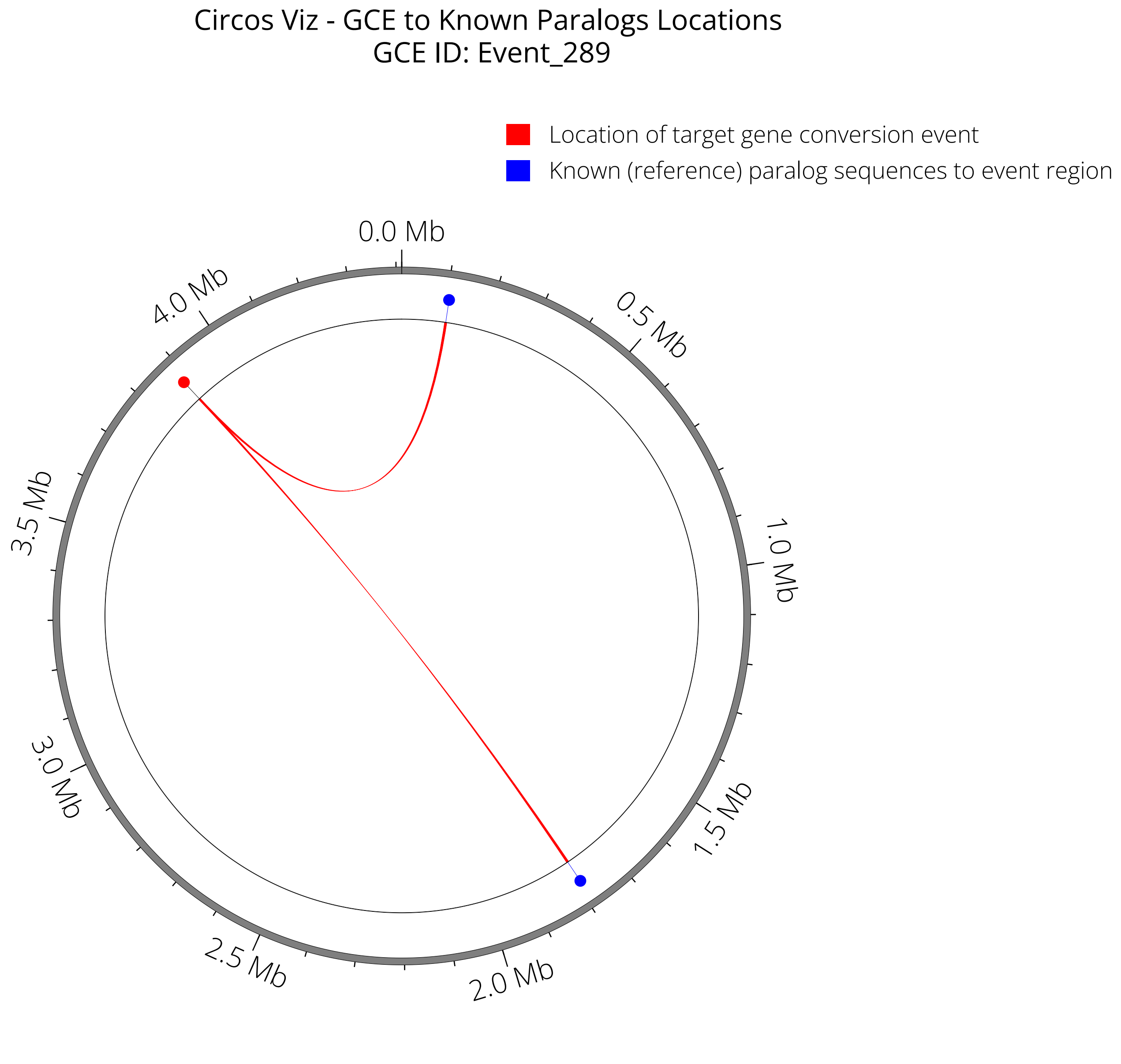
Circos Viz - GCE to Known Paralogs Locations
GCE ID: Event_289
Location of target gene conversion event
Known (reference) paralog sequences to event region
0.0 Mb
0.5 Mb
1.0 Mb
1.5 Mb
2.0 Mb
2.5 Mb
3.0 Mb
3.5 Mb
4.0 Mb

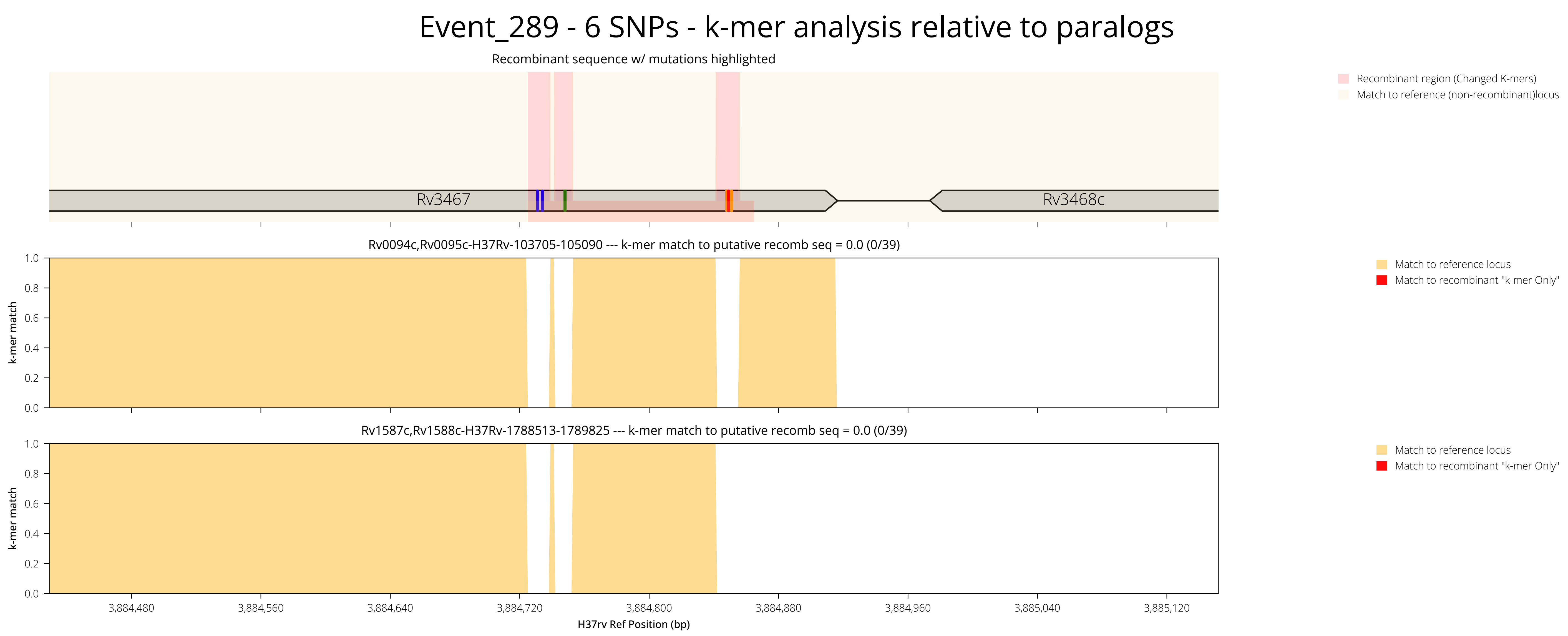
Event_289 - 6 SNPs - k-mer analysis relative to paralogs
Recombinant sequence w/ mutations highlighted
Rv3467
Rv3468c
Recombinant region (Changed K-mers)
Match to reference (non-recombinant)locus
Rv0094c,Rv0095c-H37Rv-103705-105090 --- k-mer match to putative recomb seq = 0.0 (0/39)
Match to reference locus
Match to recombinant "k-mer Only"
Rv1587c,Rv1588c-H37Rv-1788513-1789825 --- k-mer match to putative recomb seq = 0.0 (0/39)
Match to reference locus
Match to recombinant "k-mer Only"
k-mer match
1.0
0.8
0.6
0.4
0.2
0.0
3,884,480
3,884,560
3,884,640
3,884,720
3,884,800
3,884,880
3,884,960
3,885,040
3,885,120
H37rv Ref Position (bp)

Sub-phylogeny Viz + Paralog Alignments
GCE ID: Event_289

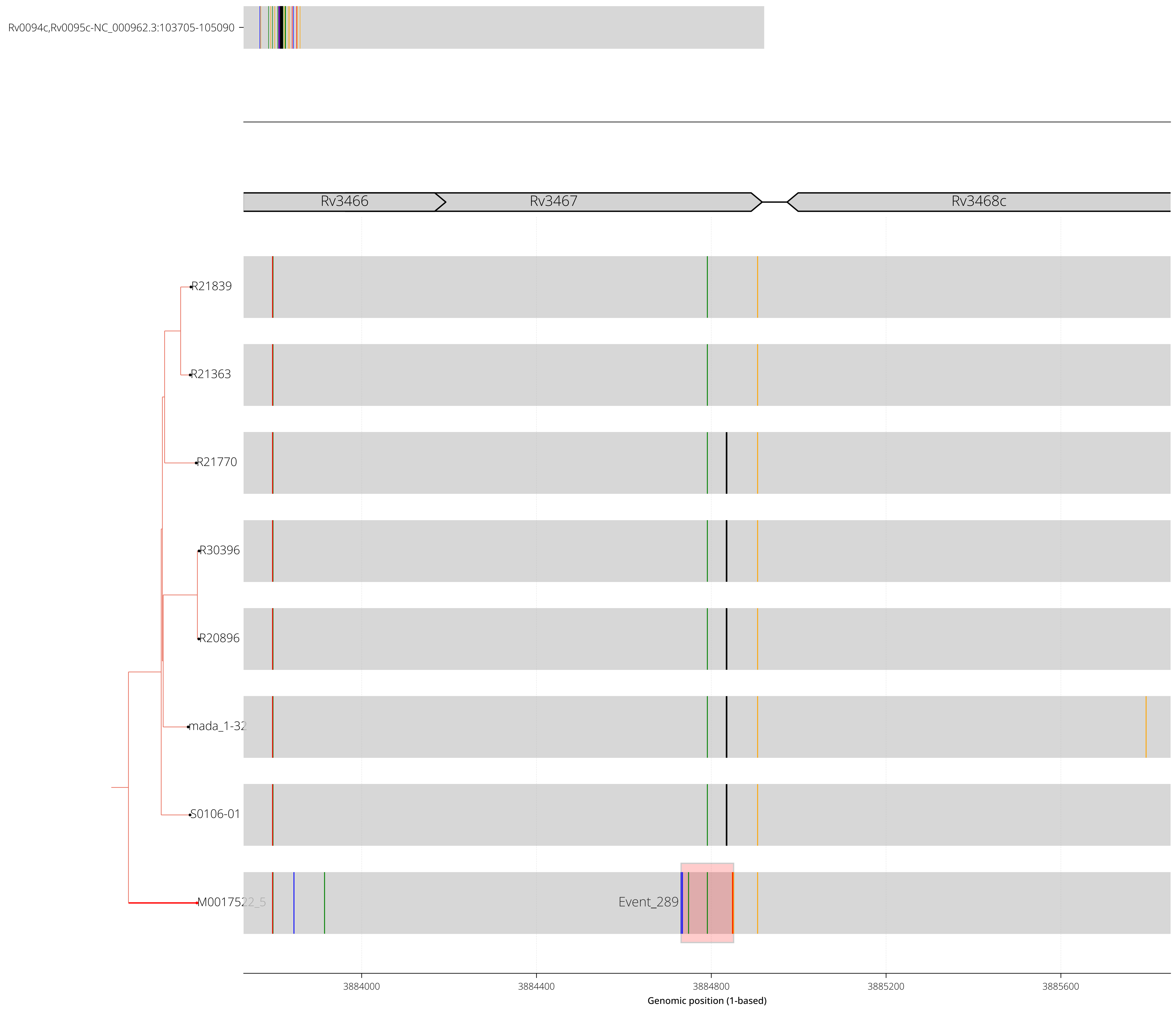

Full Phylogeny Viz + Paralog Alignments
GCE ID: Event_289

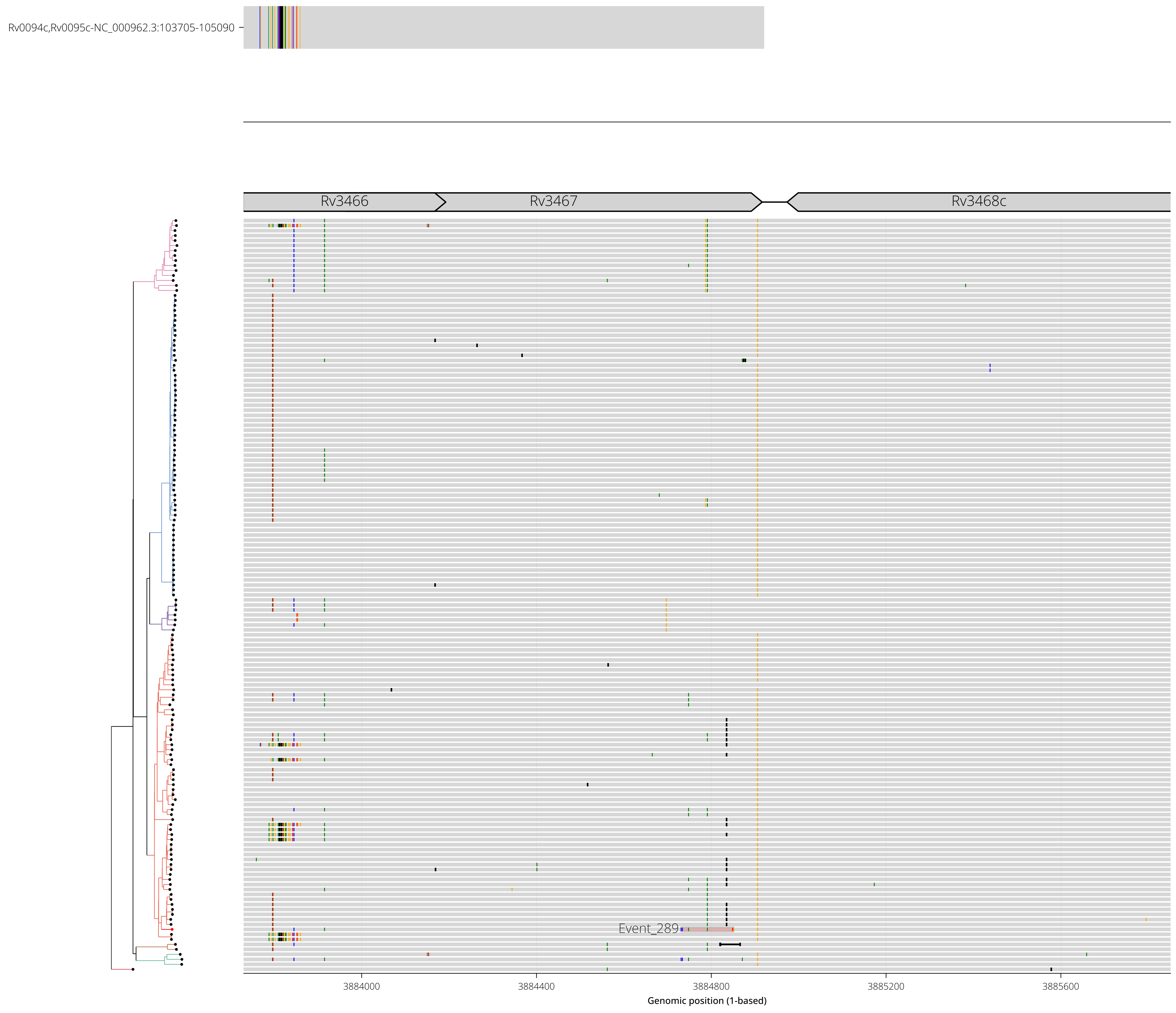

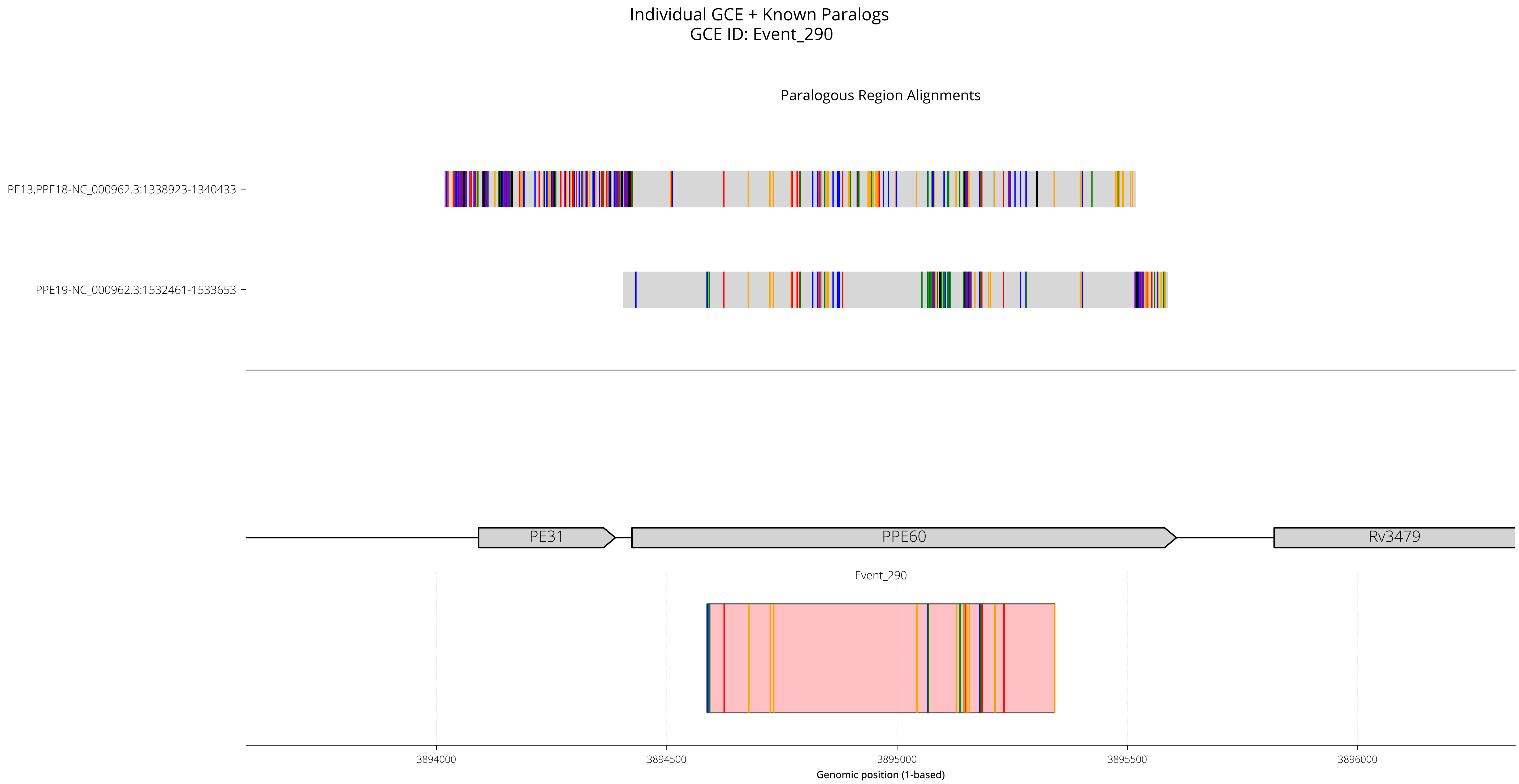
Individual GCE + Known Paralogs
GCE ID: Event_290
Paralogous Region Alignments
PE13,PPE18-NC_000962.3:1338923-1340433
PPE19-NC_000962.3:1532461-1533653
PE31
PPE60
Rv3479
Event_290
3894000
3894500
3895000
3895500
3896000
Genomic position (1-based)

Circos Viz - GCE to Known Paralogs Locations
GCE ID: Event_290

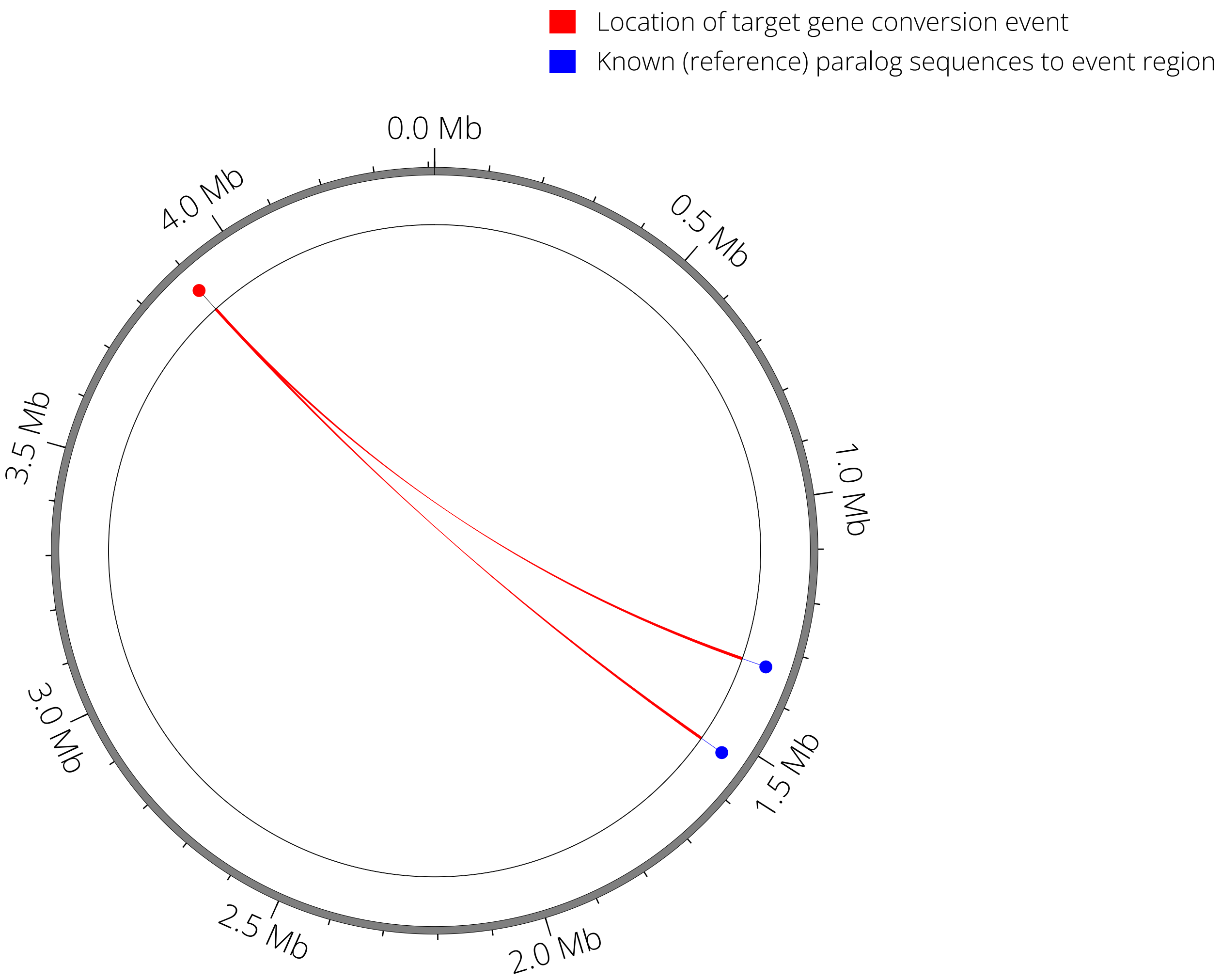

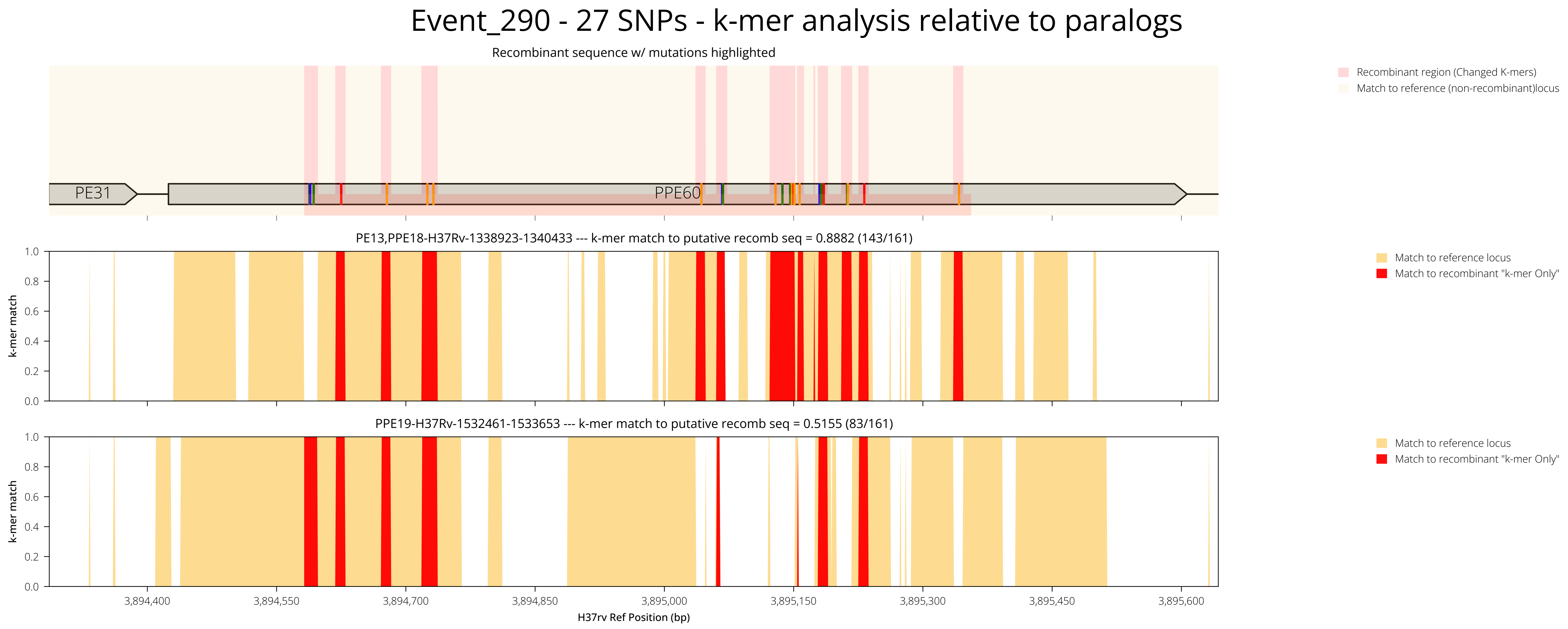

Event_290 - 27 SNPs - k-mer analysis relative to paralogs
Recombinant sequence w/ mutations highlighted
Recombinant region (Changed K-mers)
Match to reference (non-recombinant)locus
PE31
PPE60
PE13,PPE18-H37Rv-1338923-1340433 --- k-mer match to putative recomb seq = 0.8882 (143/161)
Match to reference locus
Match to recombinant "k-mer Only"
k-mer match
PPE19-H37Rv-1532461-1533653 --- k-mer match to putative recomb seq = 0.5155 (83/161)
Match to reference locus
Match to recombinant "k-mer Only"
k-mer match
1.0
0.8
0.6
0.4
0.2
0.0
3,894,400
3,894,550
3,894,700
3,894,850
3,895,000
3,895,150
3,895,300
3,895,450
3,895,600
H37rv Ref Position (bp)

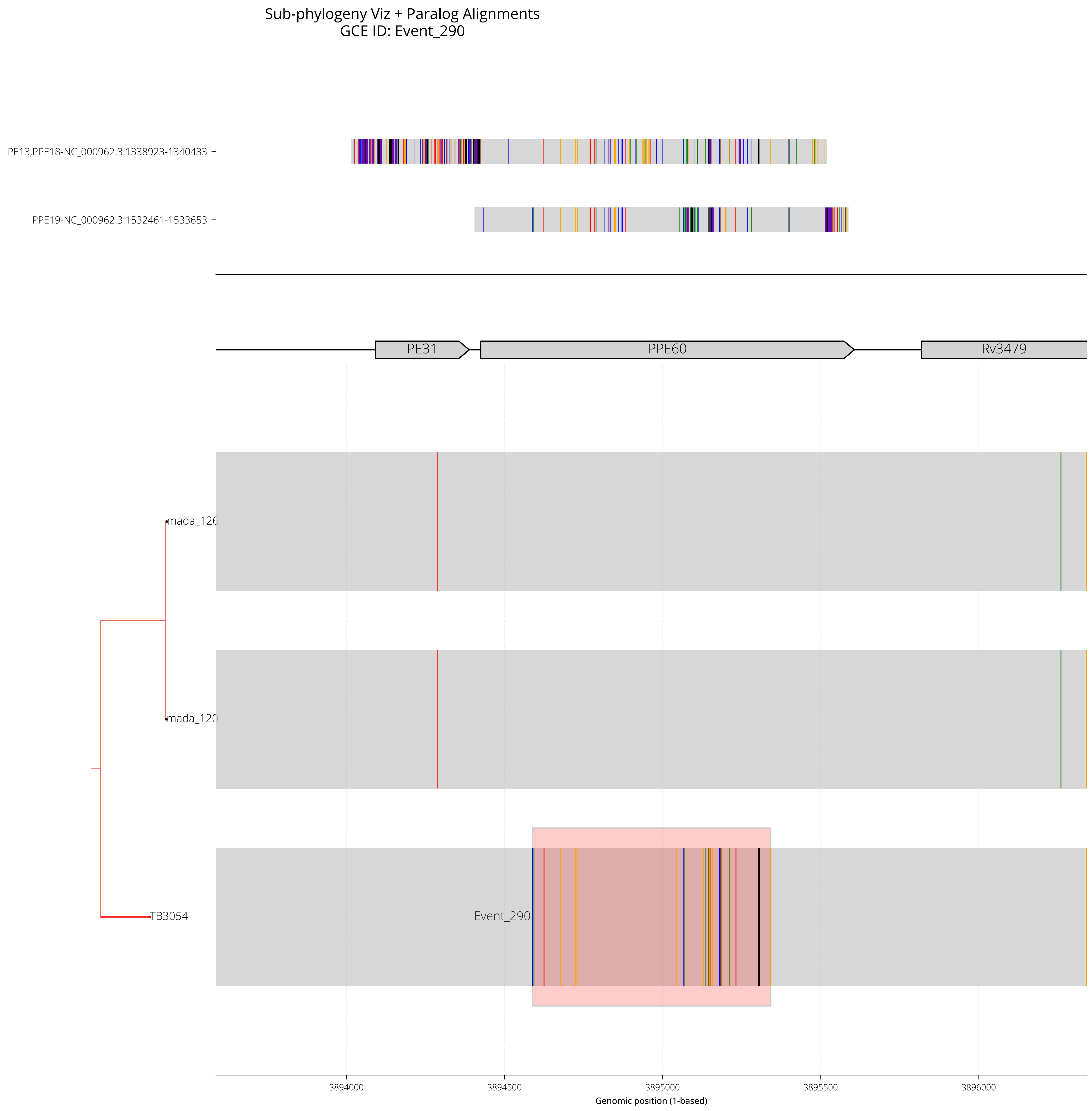

Sub-phylogeny Viz + Paralog Alignments
GCE ID: Event_290
PE13,PPE18-NC_000962.3:1338923-1340433
PPE19-NC_000962.3:1532461-1533653
PE31
PPE60
Rv3479
mada_126
mada_120
TB3054
Event_290
3894000
3894500
3895000
3895500
3896000
Genomic position (1-based)

Full Phylogeny Viz + Paralog Alignments
GCE ID: Event_290

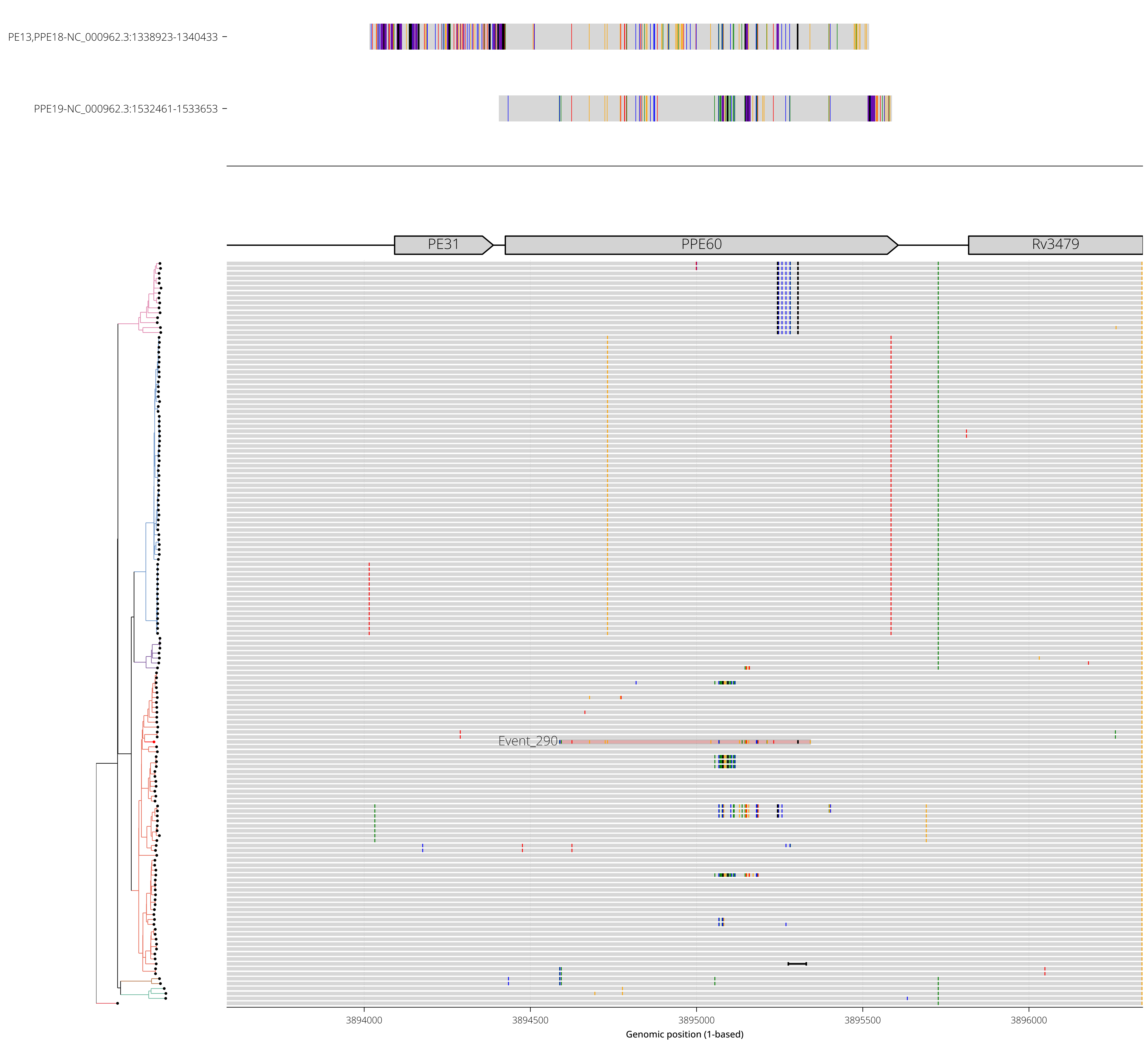

Individual GCE + Known Paralogs
GCE ID: Event_291

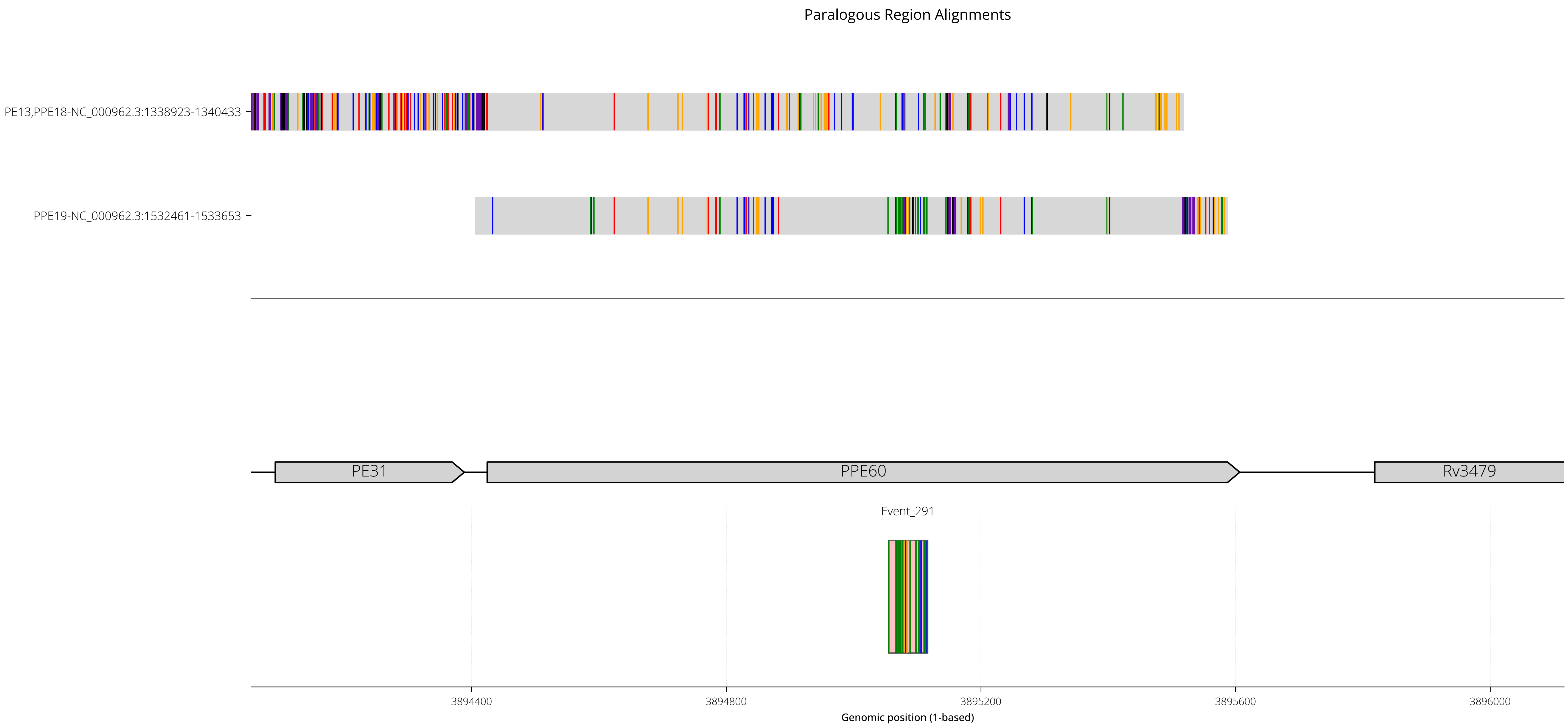

Circos Viz - GCE to Known Paralogs Locations
GCE ID: Event_291

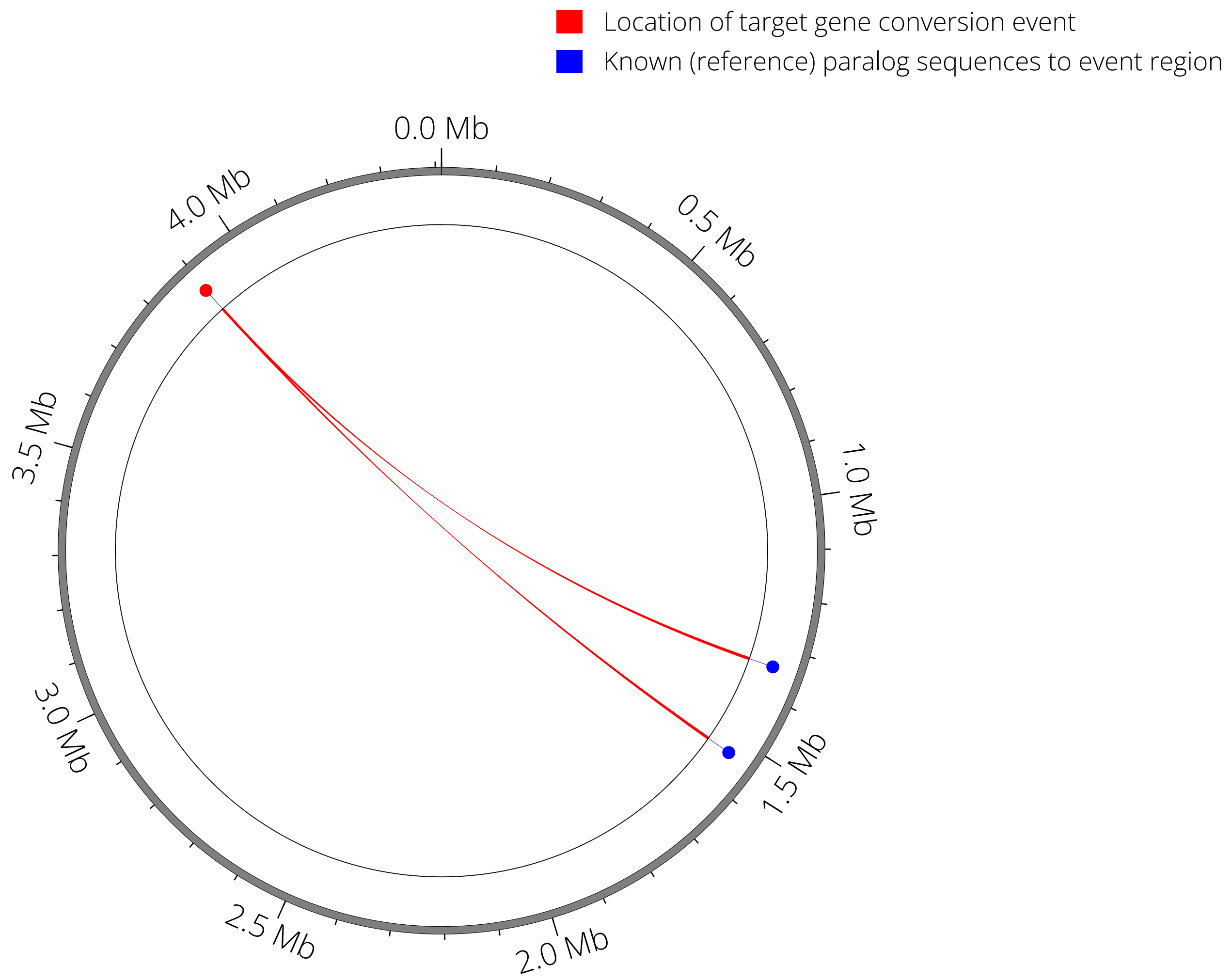

### Event_291 - 20 SNPs - k-mer analysis relative to paralogs

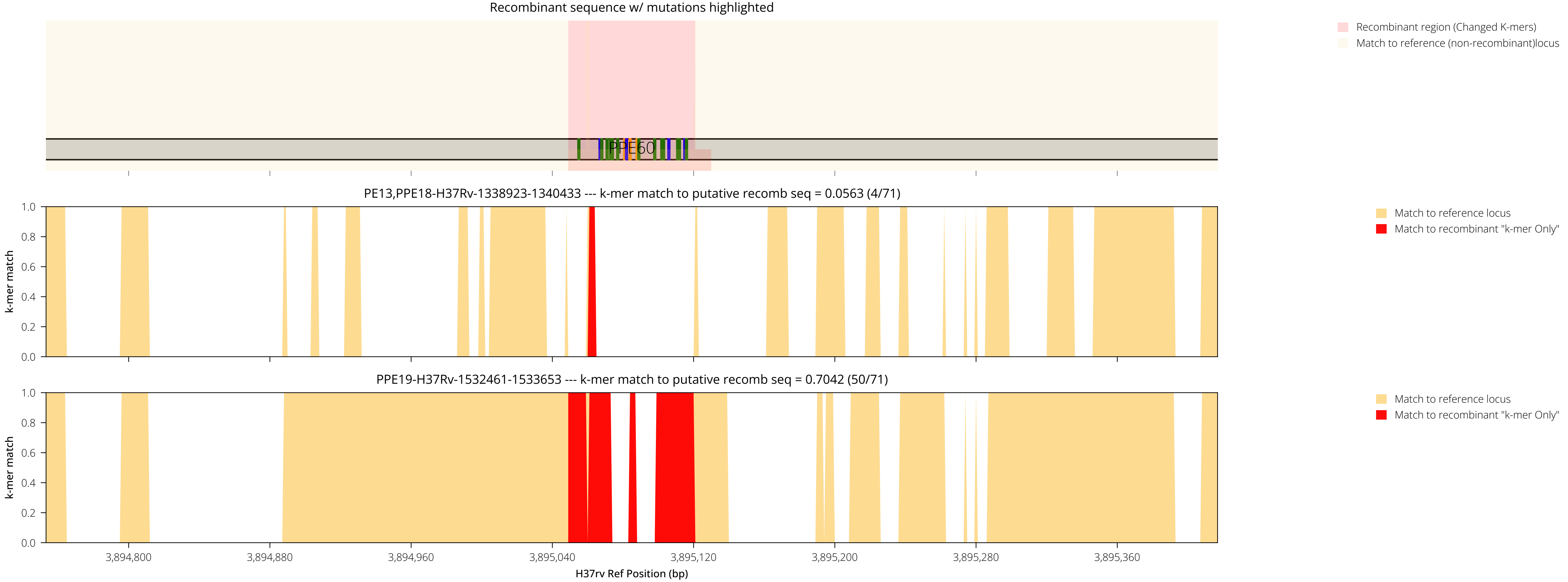

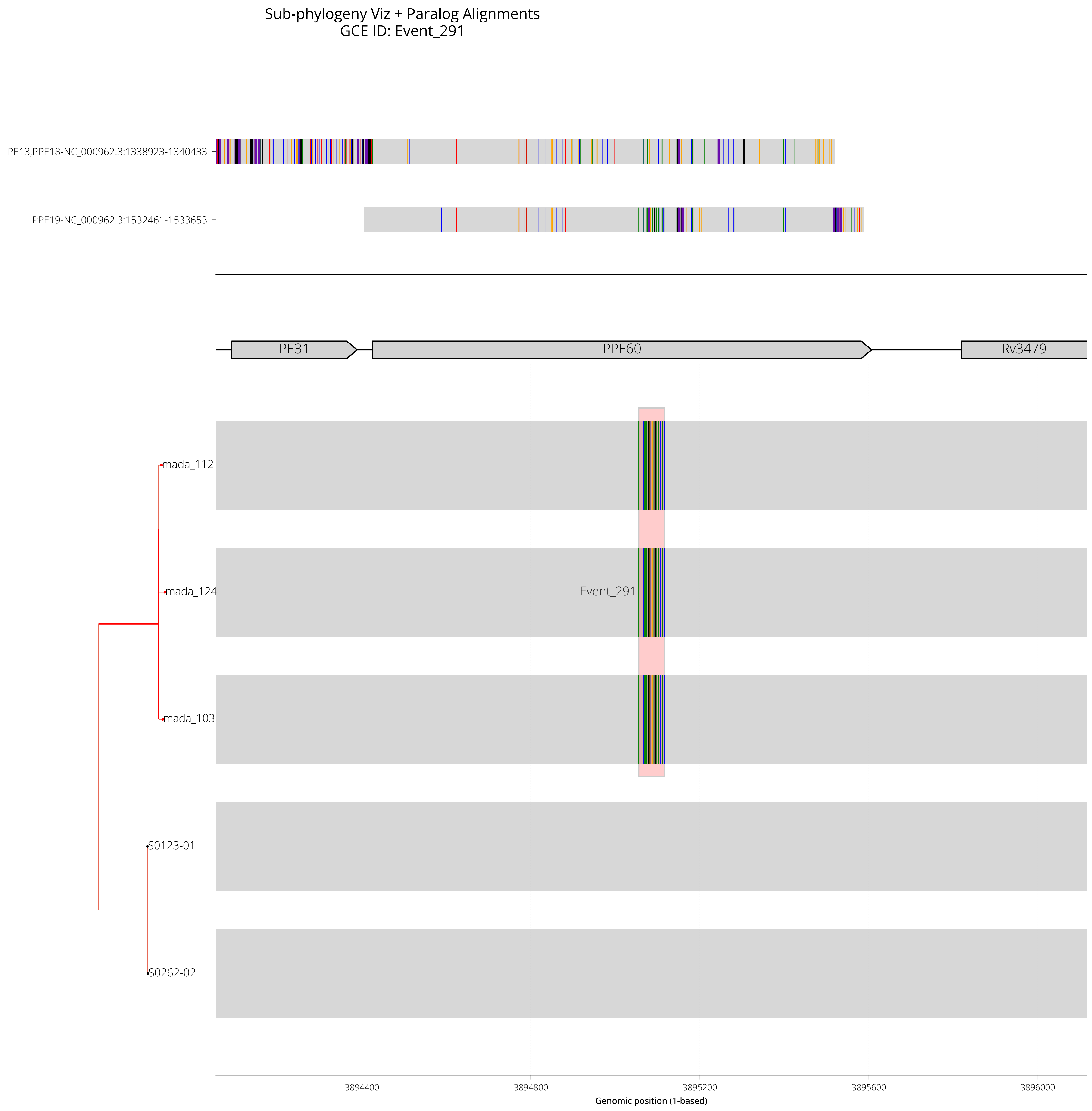

Sub-phylogeny Viz + Paralog Alignments
GCE ID: Event_291
PE13,PPE18-NC_000962.3:1338923-1340433
PPE19-NC_000962.3:1532461-1533653
PE31
PPE60
Rv3479
mada_112
mada_124
mada_103
S0123-01
S0262-02
Event_291
3894400
3894800
3895200
3895600
3896000
Genomic position (1-based)

Full Phylogeny Viz + Paralog Alignments
GCE ID: Event_291

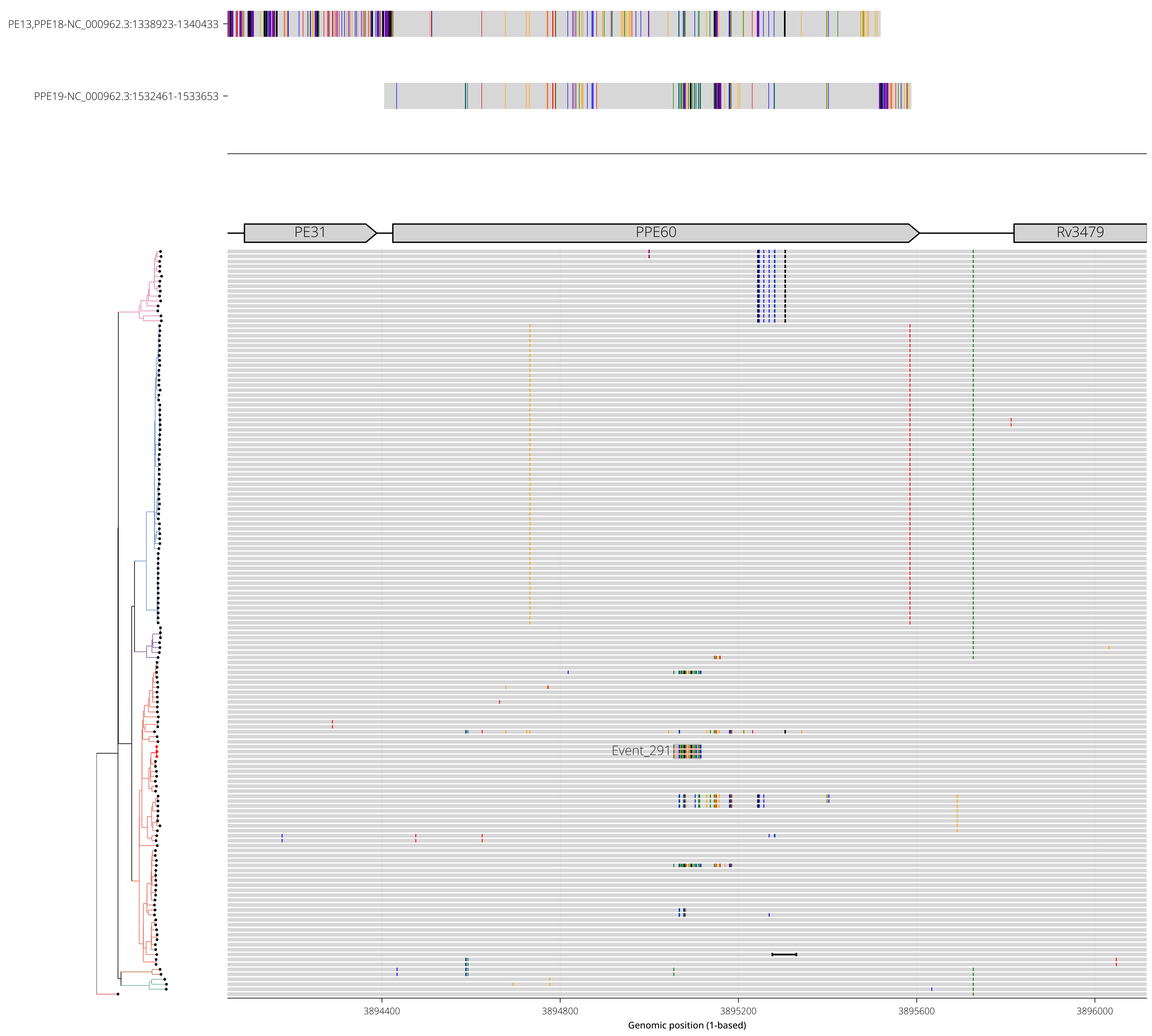

Individual GCE + Known Paralogs
GCE ID: Event_292

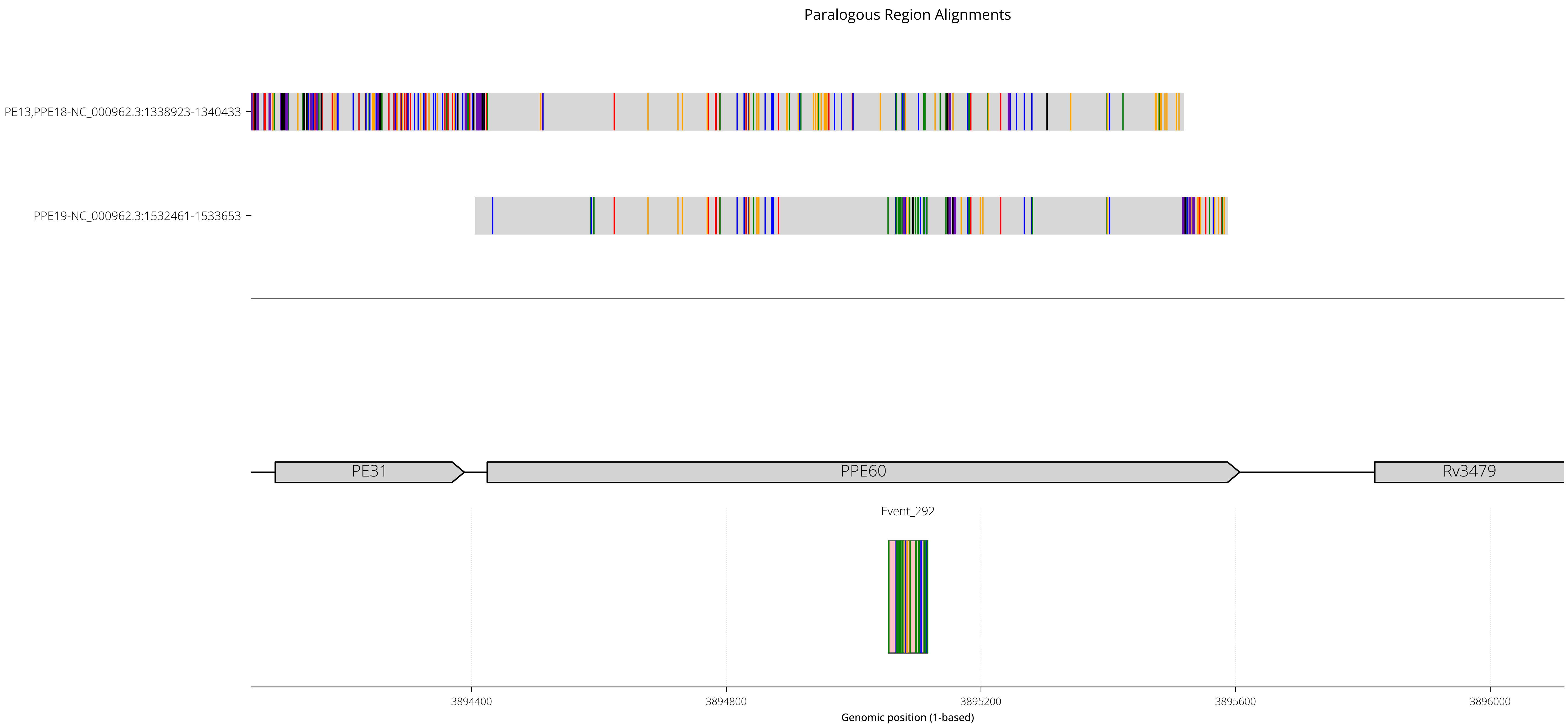

Circos Viz - GCE to Known Paralogs Locations
GCE ID: Event_292

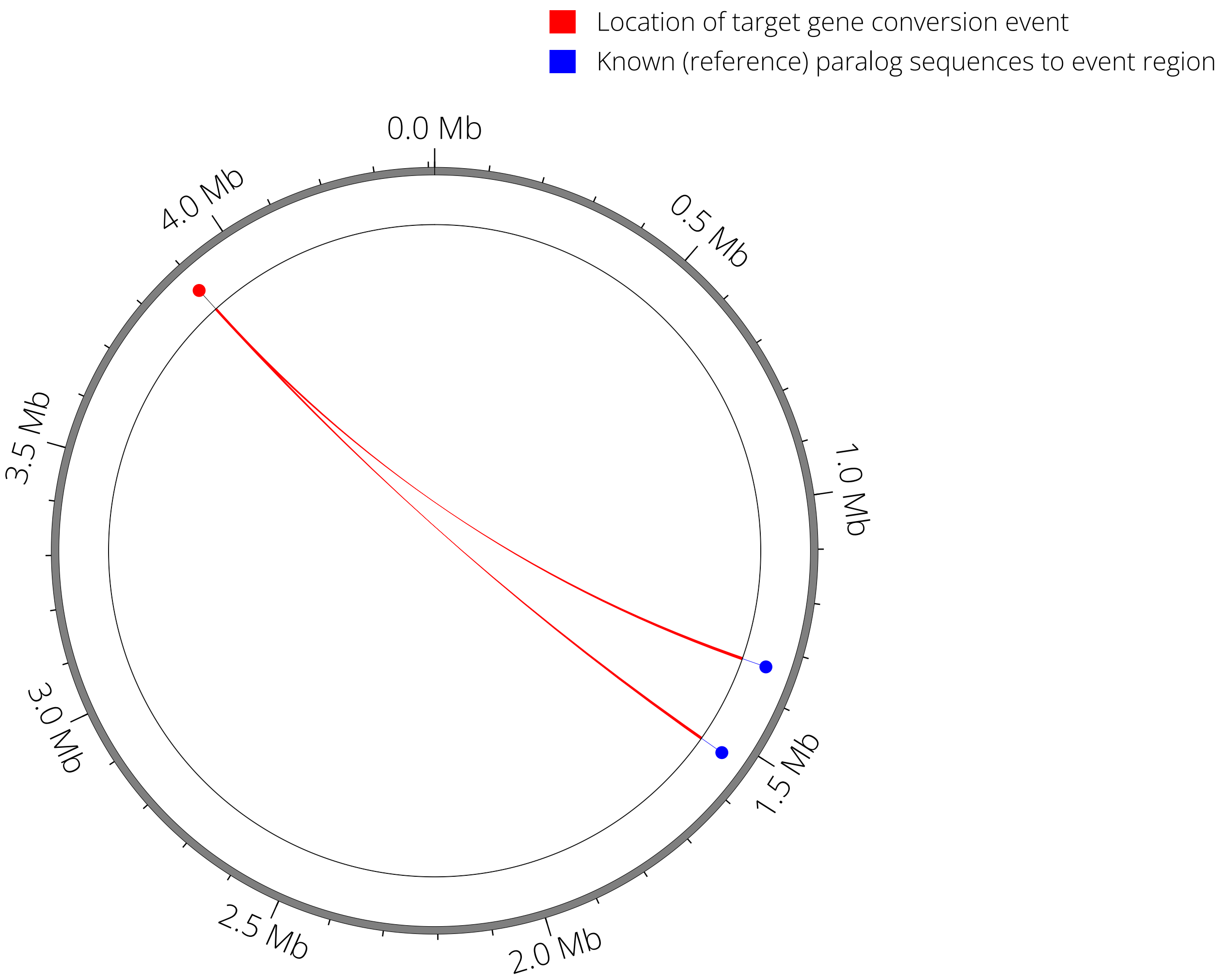

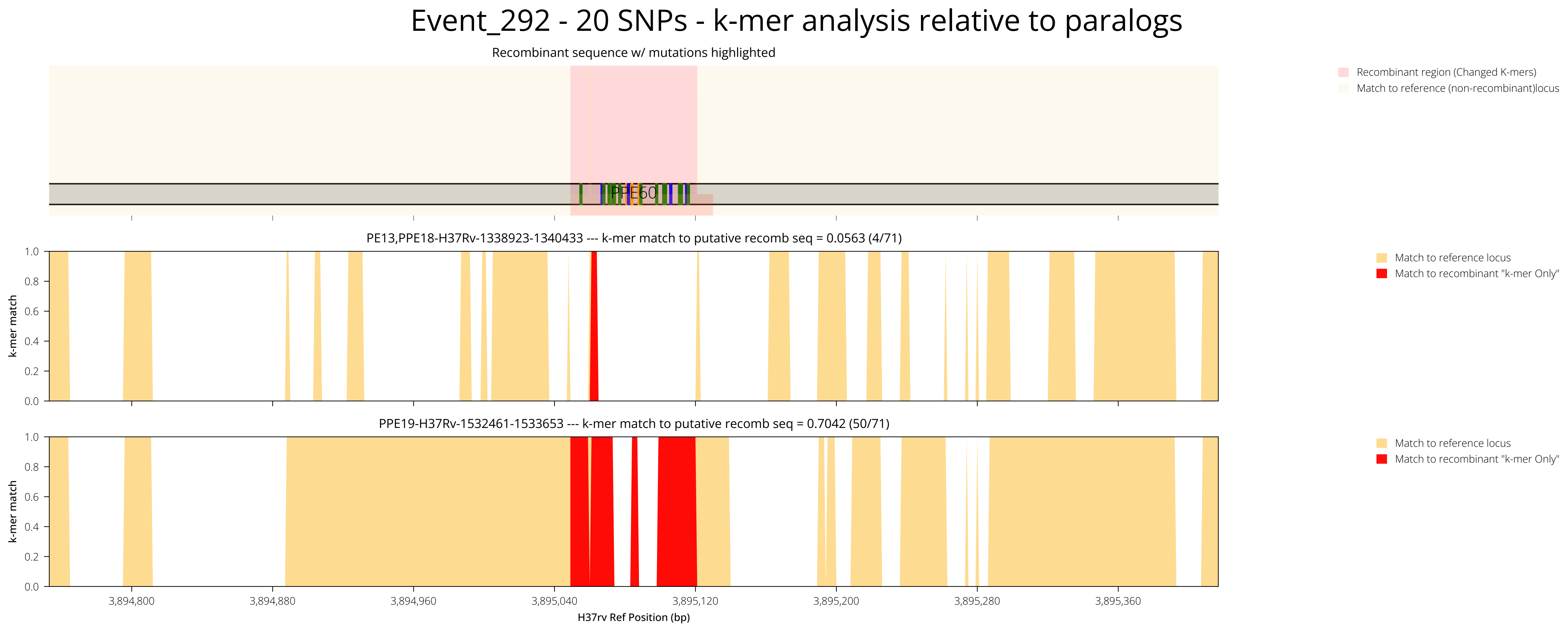

Event_292 - 20 SNPs - k-mer analysis relative to paralogs
Recombinant sequence w/ mutations highlighted
Recombinant region (Changed K-mers)
Match to reference (non-recombinant)locus
PPE60
PE13,PPE18-H37Rv-1338923-1340433 --- k-mer match to putative recomb seq = 0.0563 (4/71)
Match to reference locus
Match to recombinant "k-mer Only"
PPE19-H37Rv-1532461-1533653 --- k-mer match to putative recomb seq = 0.7042 (50/71)
Match to reference locus
Match to recombinant "k-mer Only"
k-mer match
1.0
0.8
0.6
0.4
0.2
0.0
3,894,800
3,894,880
3,894,960
3,895,040
3,895,120
3,895,200
3,895,280
3,895,360
H37rv Ref Position (bp)

Sub-phylogeny Viz + Paralog Alignments
GCE ID: Event_292

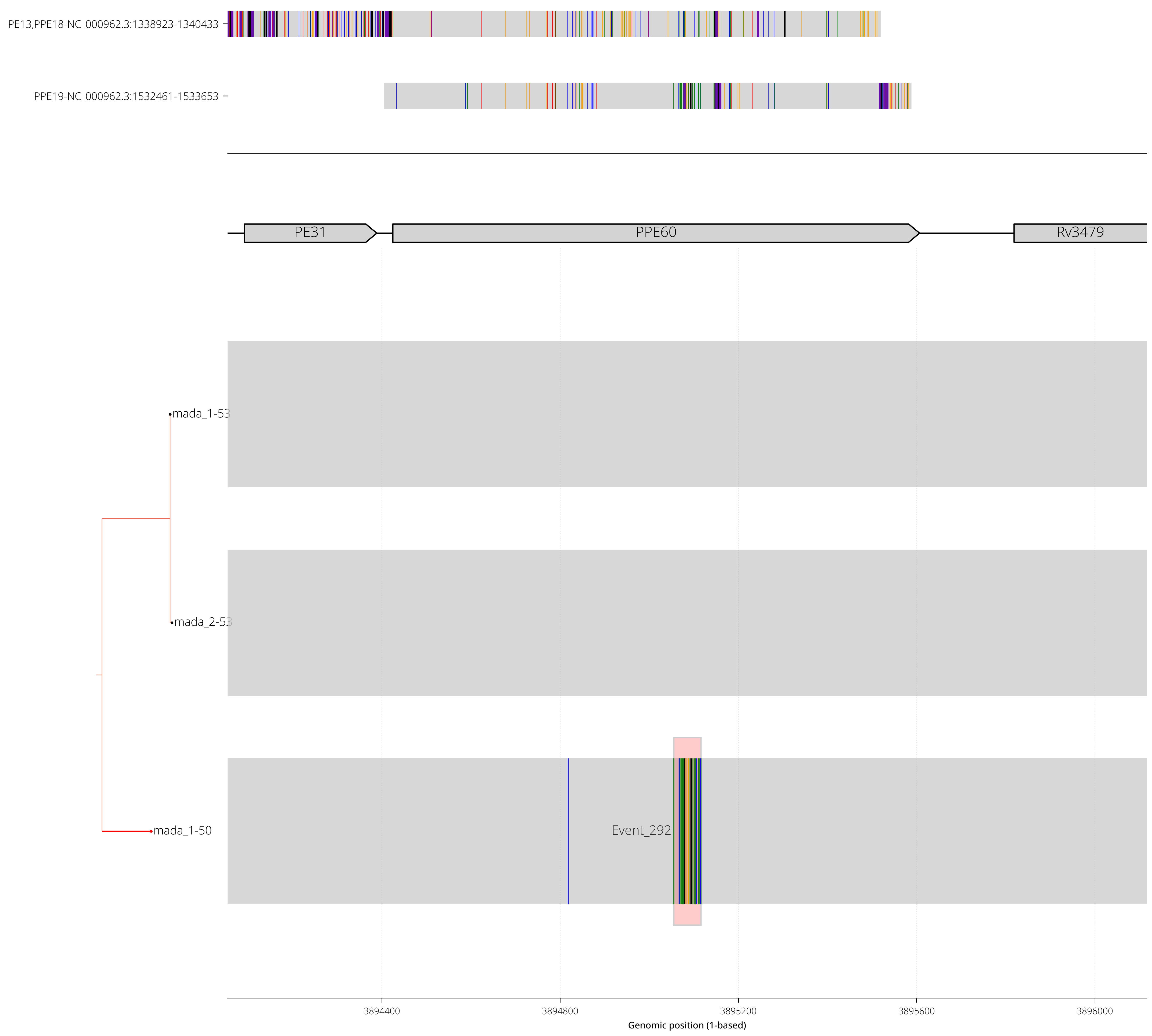

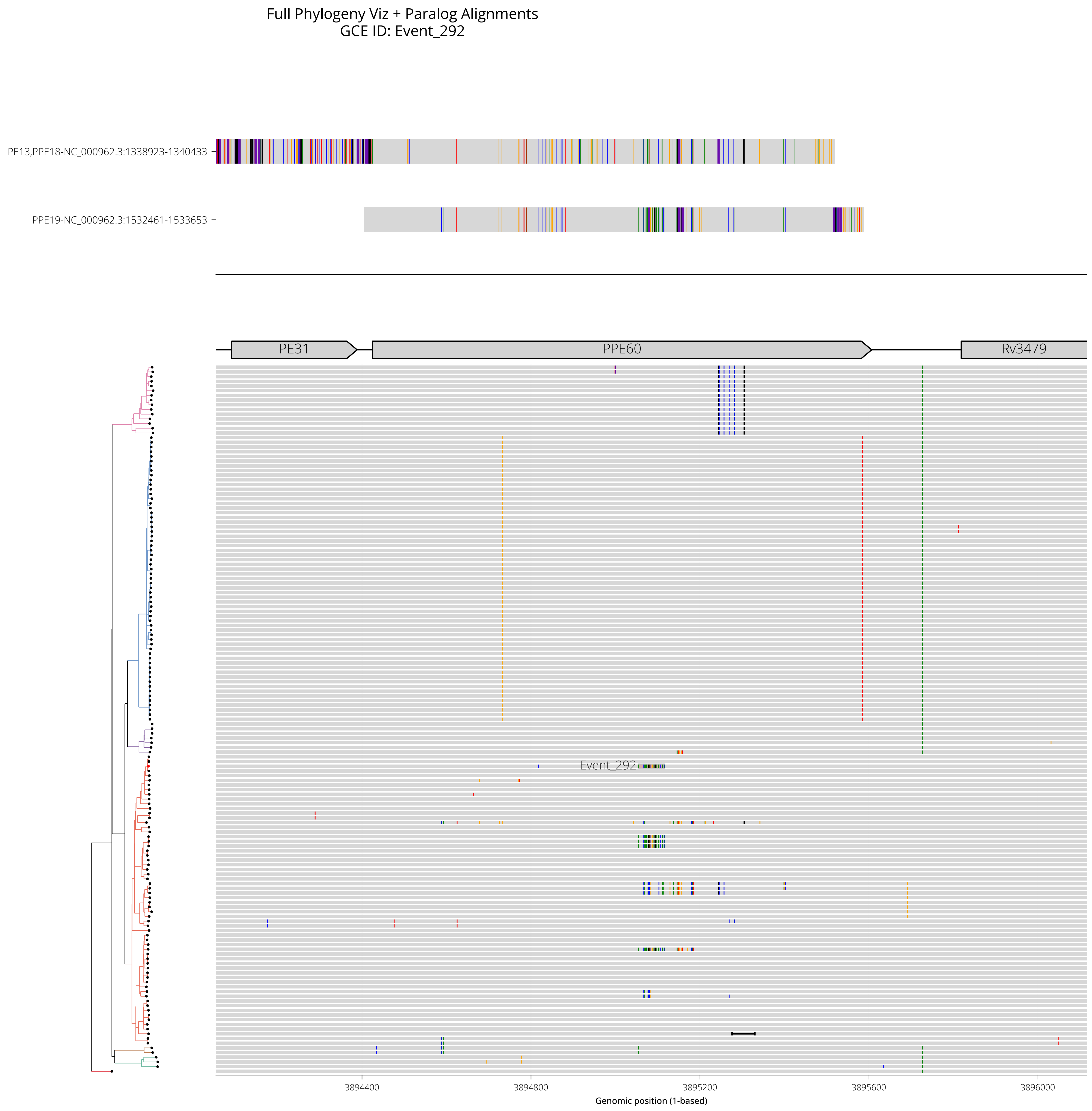

Full Phylogeny Viz + Paralog Alignments
GCE ID: Event_292
PE13,PPE18-NC_000962.3:1338923-1340433
PPE19-NC_000962.3:1532461-1533653
PE31
PPE60
Rv3479
Event_292
3894400
3894800
3895200
3895600
3896000
Genomic position (1-based)

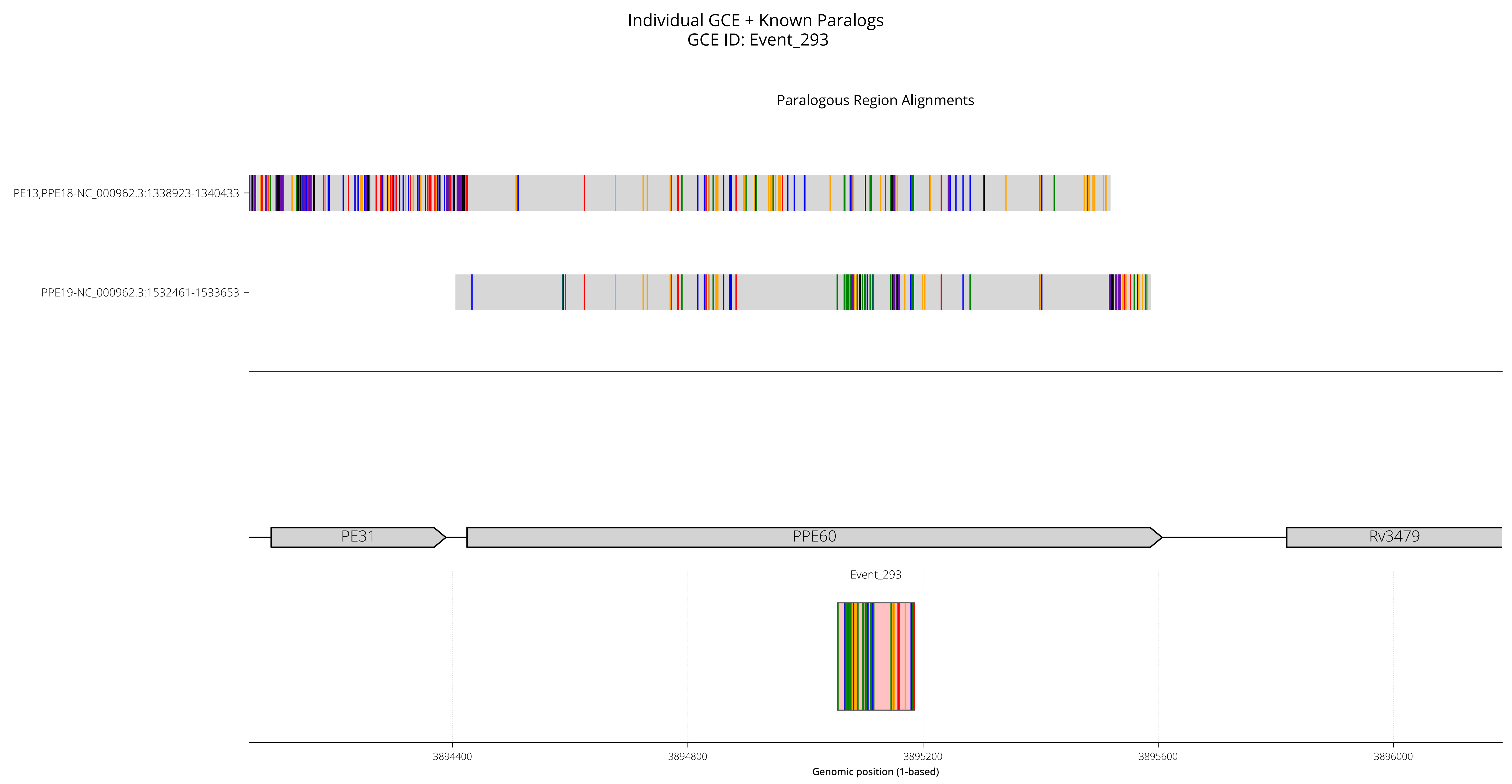

Individual GCE + Known Paralogs
GCE ID: Event_293
Paralogous Region Alignments
PE13,PPE18-NC_000962.3:1338923-1340433
PPE19-NC_000962.3:1532461-1533653
PE31
PPE60
Rv3479
Event_293
3894400
3894800
3895200
3895600
3896000
Genomic position (1-based)

Circos Viz - GCE to Known Paralogs Locations
GCE ID: Event_293

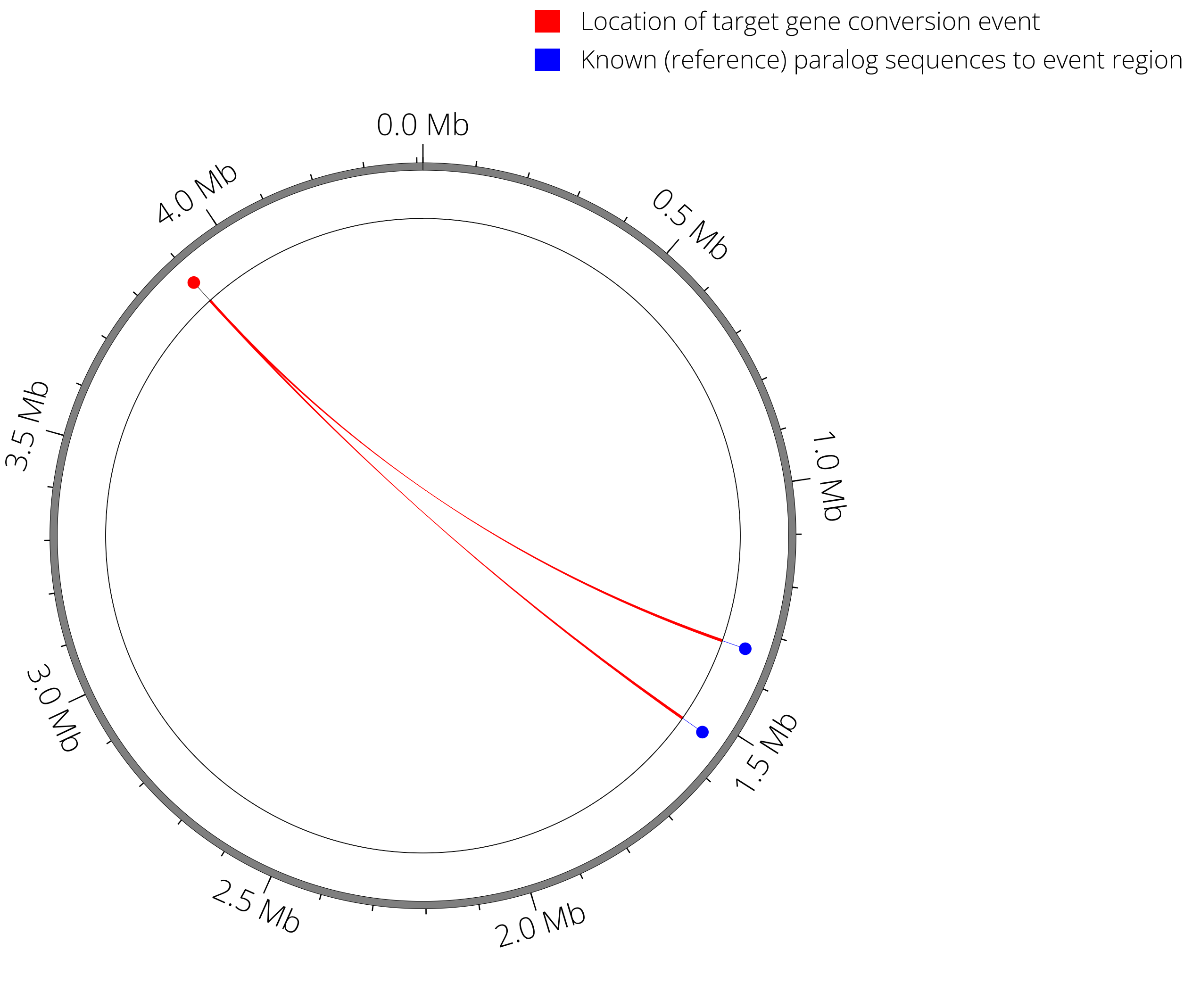

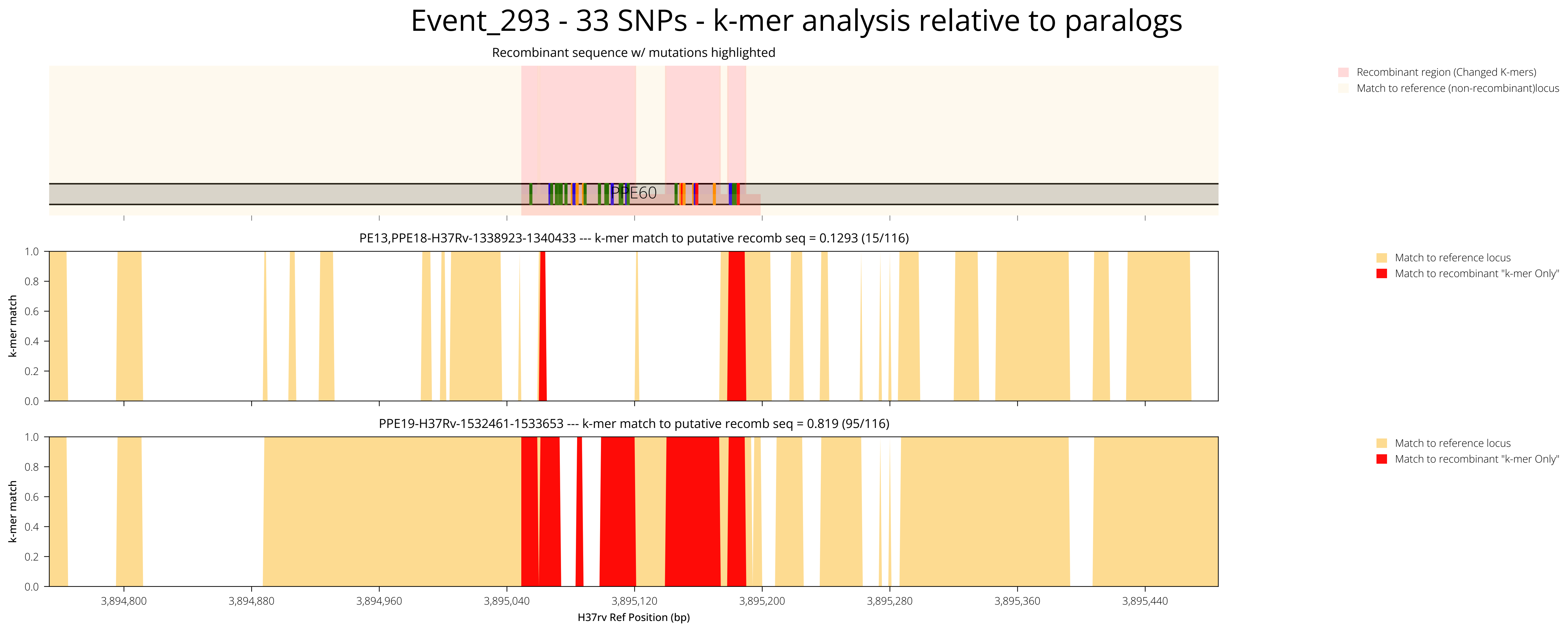

Event_293 - 33 SNPs - k-mer analysis relative to paralogs
Recombinant sequence w/ mutations highlighted
Recombinant region (Changed K-mers)
Match to reference (non-recombinant)locus
PPE60
PE13,PPE18-H37Rv-1338923-1340433 --- k-mer match to putative recomb seq = 0.1293 (15/116)
Match to reference locus
Match to recombinant "k-mer Only"
k-mer match
1.0
0.8
0.6
0.4
0.2
0.0
PPE19-H37Rv-1532461-1533653 --- k-mer match to putative recomb seq = 0.819 (95/116)
Match to reference locus
Match to recombinant "k-mer Only"
k-mer match
3,894,800
3,894,880
3,894,960
3,895,040
3,895,120
3,895,200
3,895,280
3,895,360
3,895,440
H37rv Ref Position (bp)

Sub-phylogeny Viz + Paralog Alignments
GCE ID: Event_293

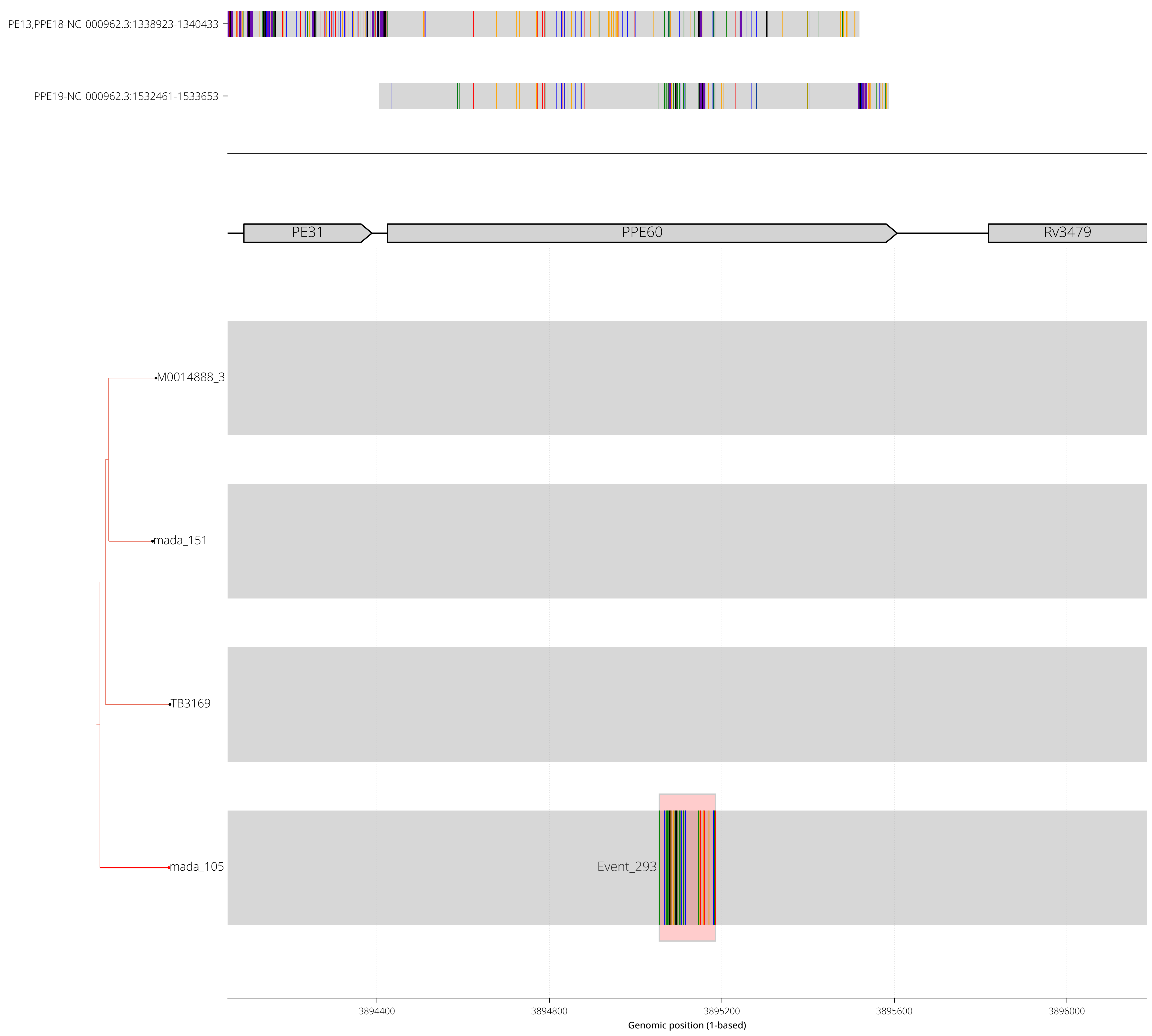

Full Phylogeny Viz + Paralog Alignments
GCE ID: Event_293

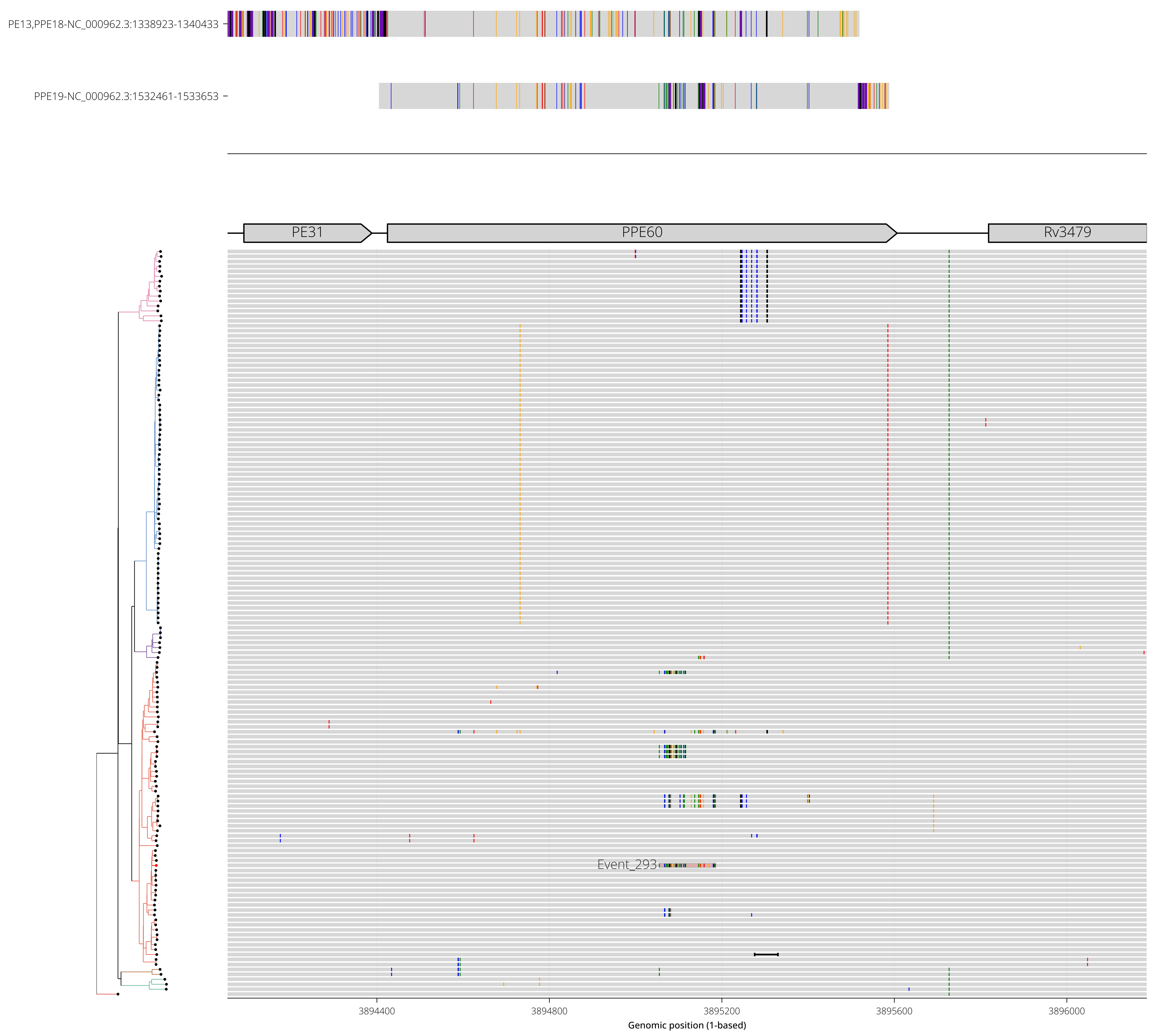

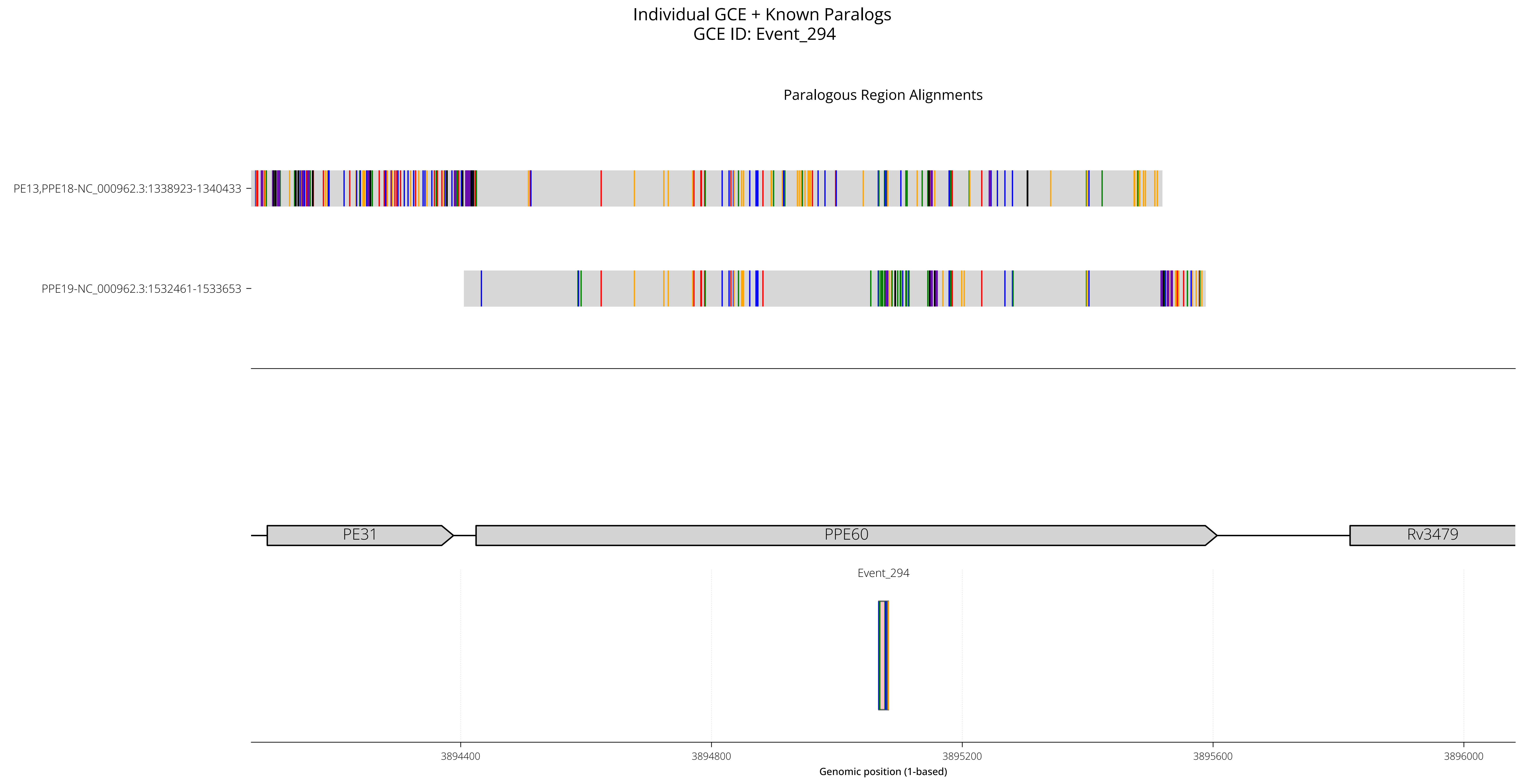

Individual GCE + Known Paralogs
GCE ID: Event_294
Paralogous Region Alignments
PE13,PPE18-NC_000962.3:1338923-1340433
PPE19-NC_000962.3:1532461-1533653
PE31
PPE60
Rv3479
Event_294
3894400
3894800
3895200
3895600
3896000
Genomic position (1-based)

### Circos Viz - GCE to Known Paralogs Locations
GCE ID: Event_294

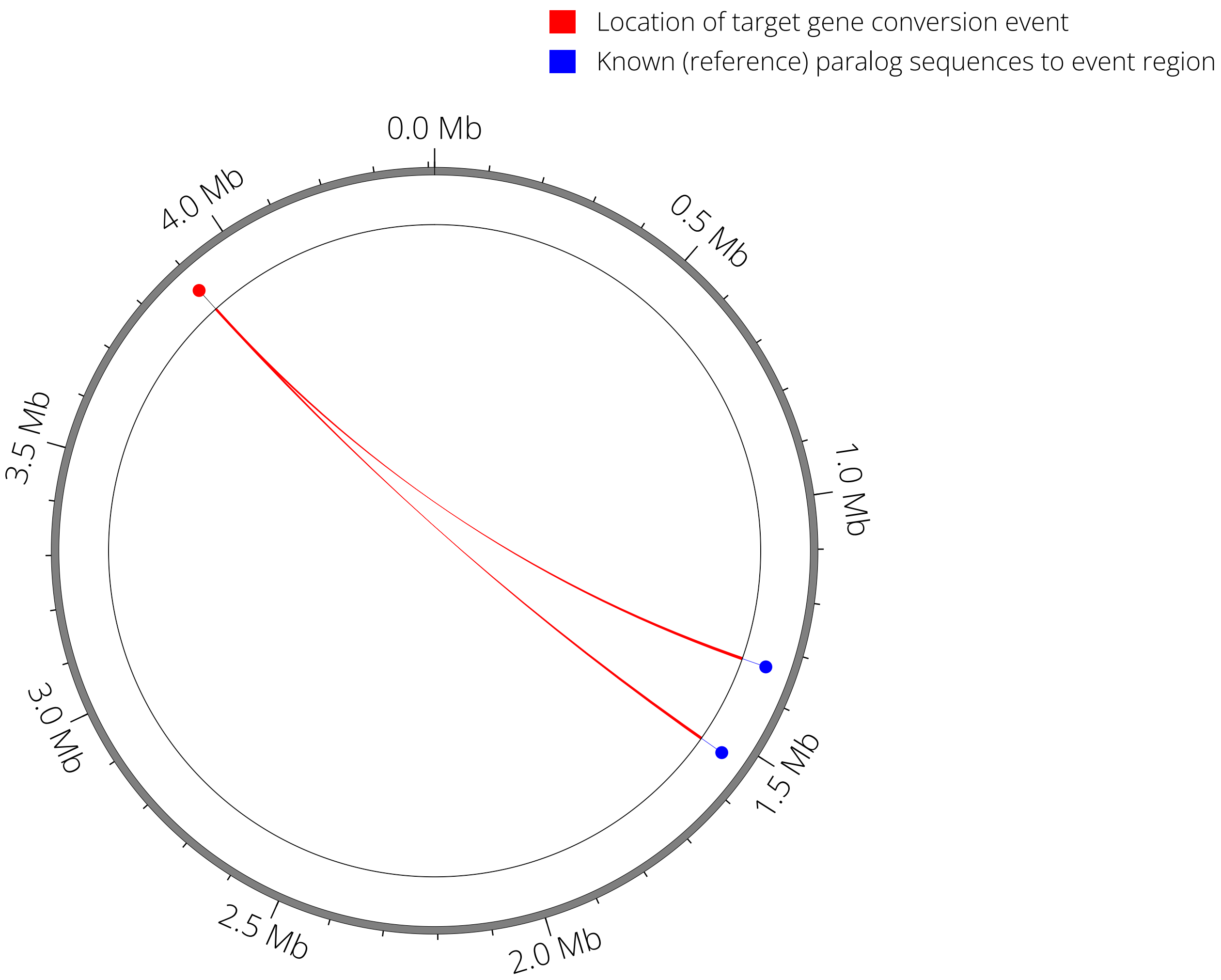

### Event_294 - 7 SNPs - k-mer analysis relative to paralogs

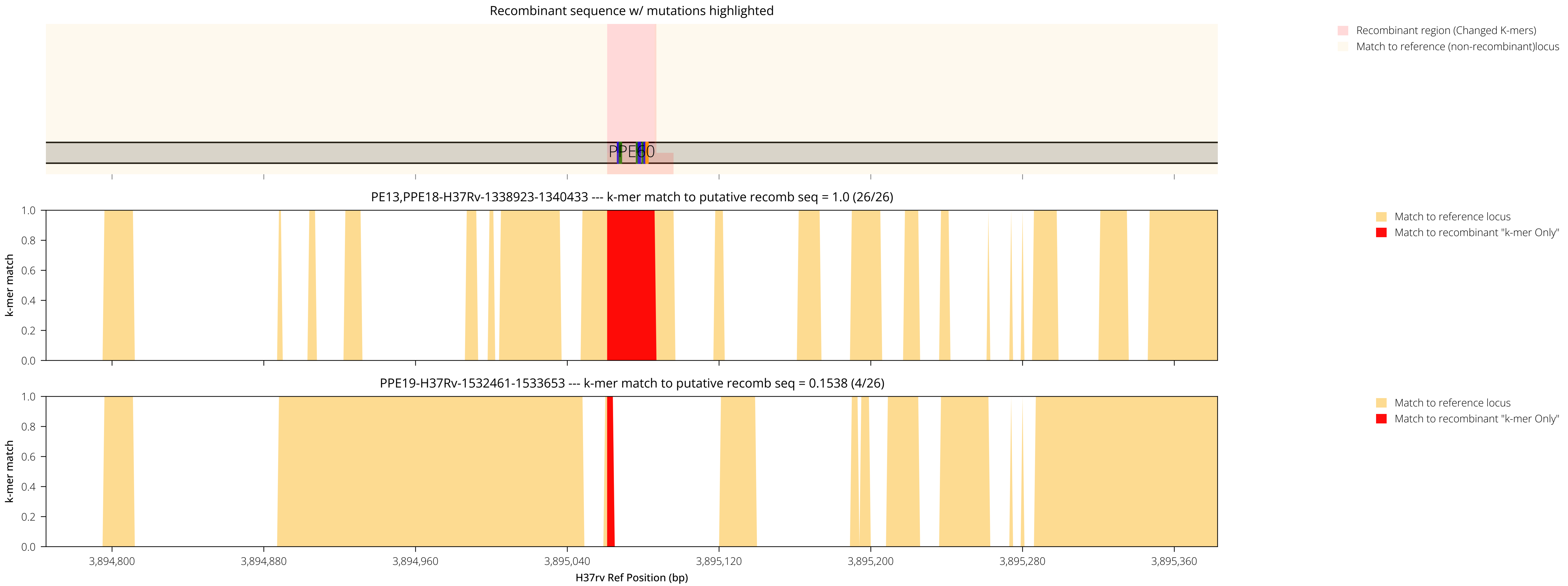

Sub-phylogeny Viz + Paralog Alignments
GCE ID: Event_294

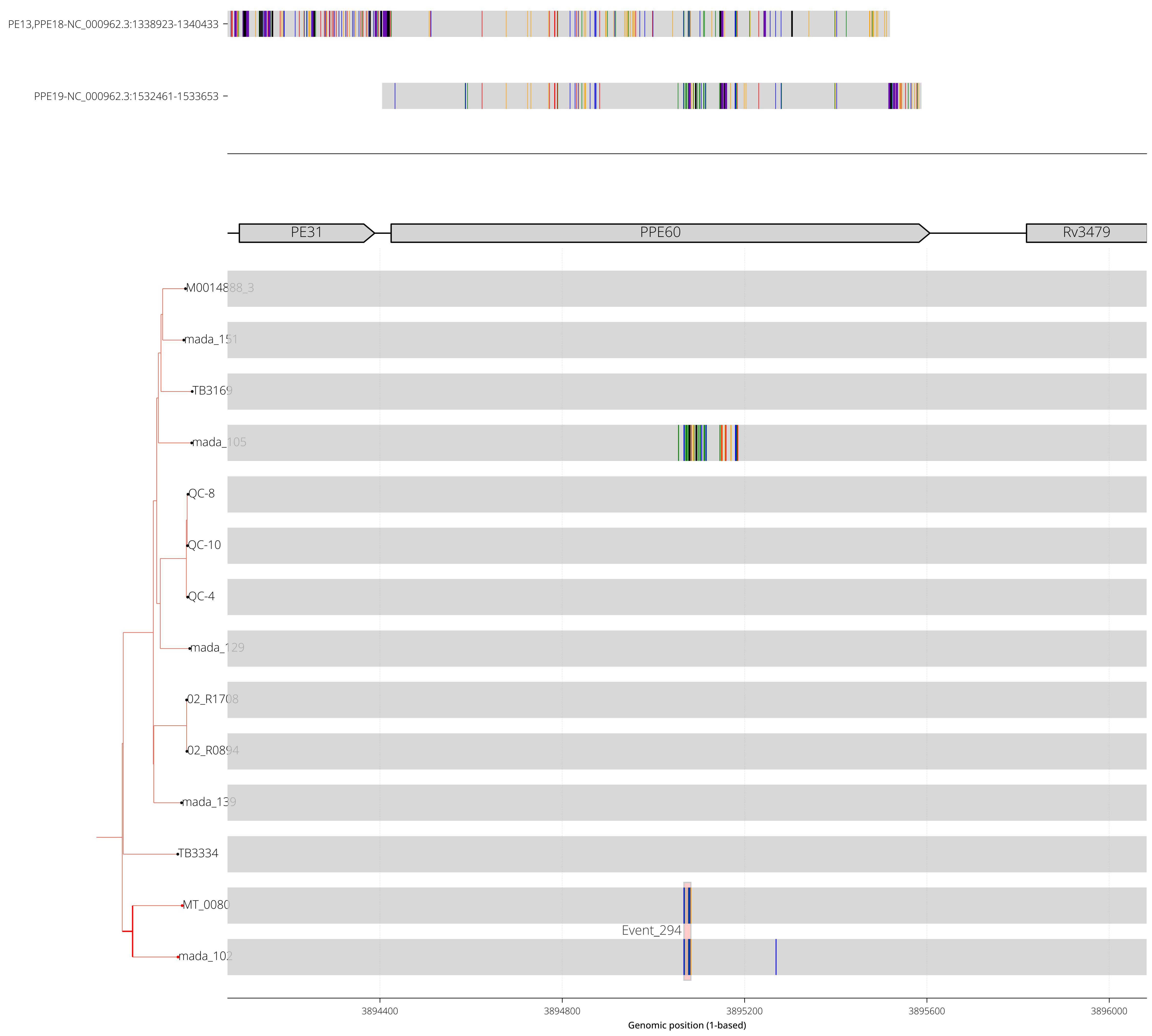

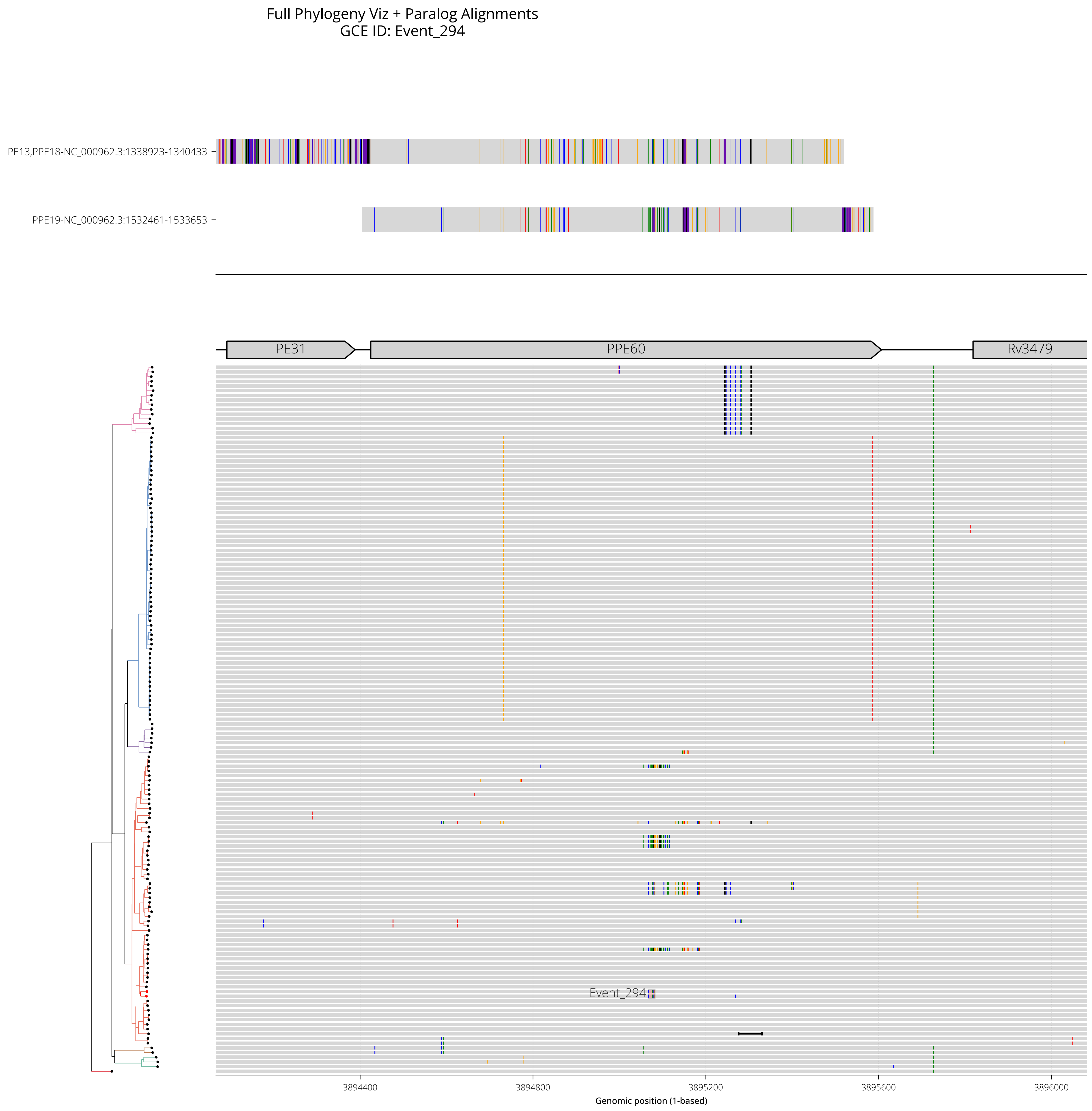

Full Phylogeny Viz + Paralog Alignments
GCE ID: Event_294
PE13,PPE18-NC_000962.3:1338923-1340433
PPE19-NC_000962.3:1532461-1533653
PE31
PPE60
Rv3479
Event_294
3894400
3894800
3895200
3895600
3896000
Genomic position (1-based)

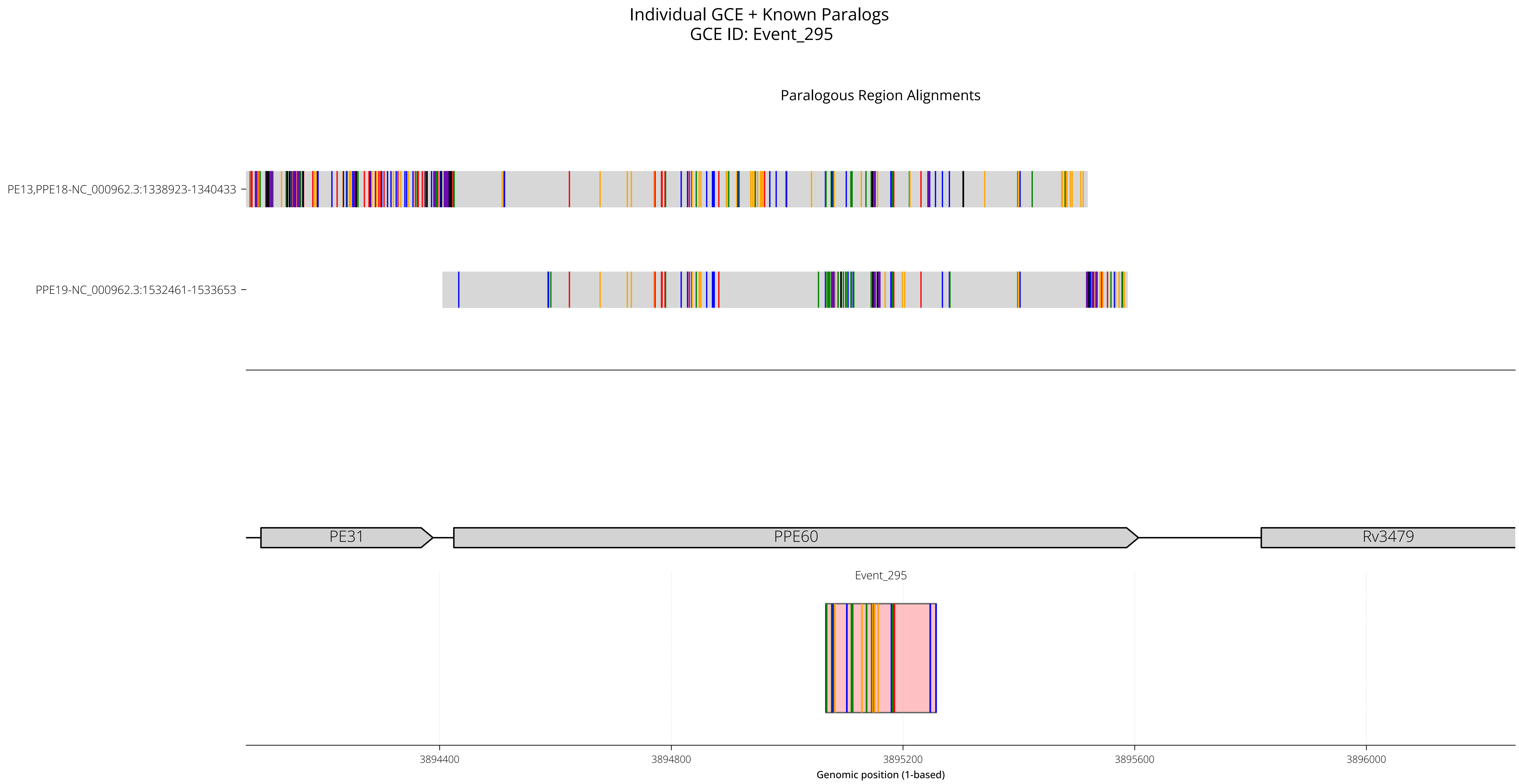
Individual GCE + Known Paralogs
GCE ID: Event_295
Paralogous Region Alignments
PE13,PPE18-NC_000962.3:1338923-1340433
PPE19-NC_000962.3:1532461-1533653
PE31
PPE60
Rv3479
Event_295
3894400
3894800
3895200
3895600
3896000
Genomic position (1-based)

Circos Viz - GCE to Known Paralogs Locations
GCE ID: Event_295

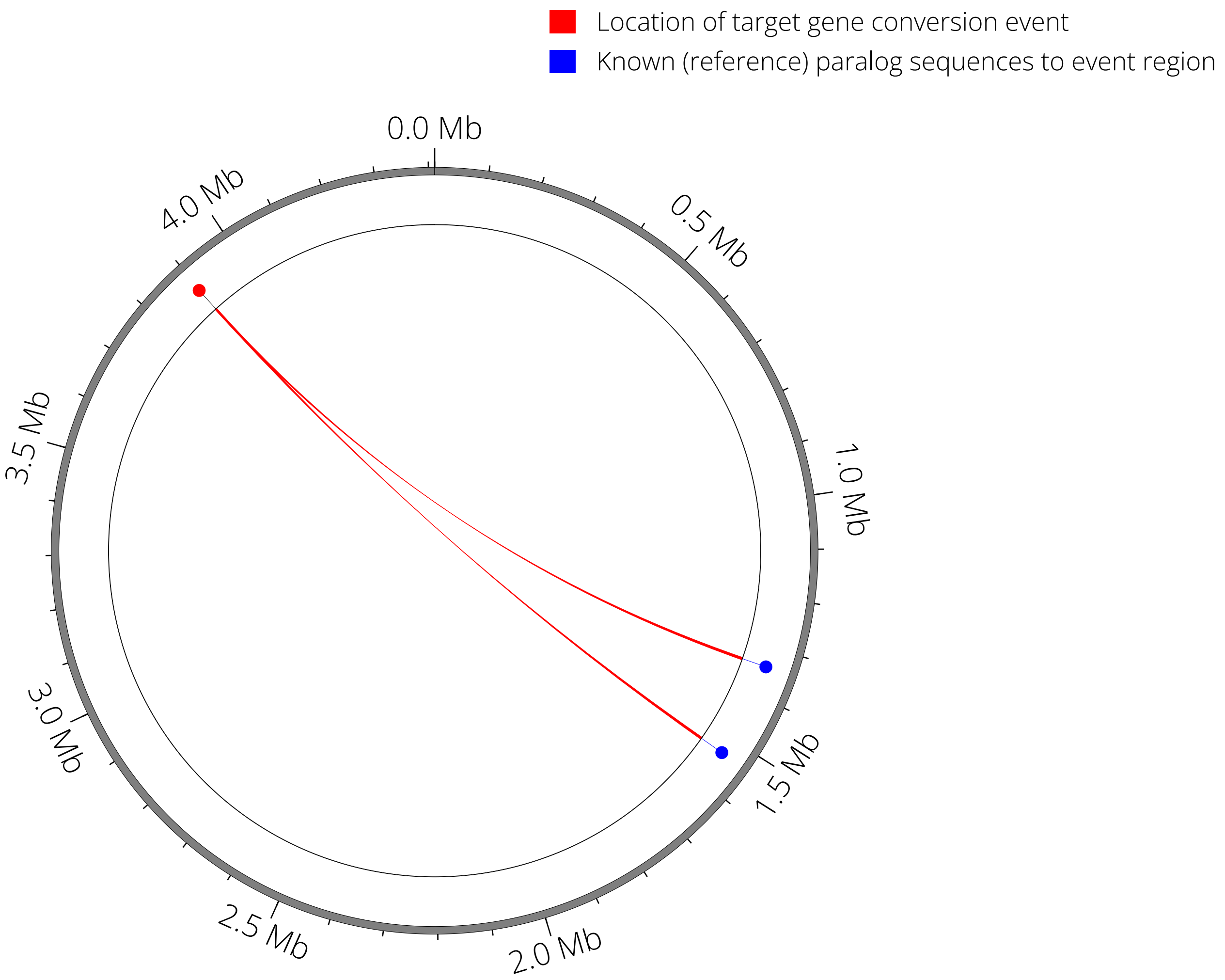

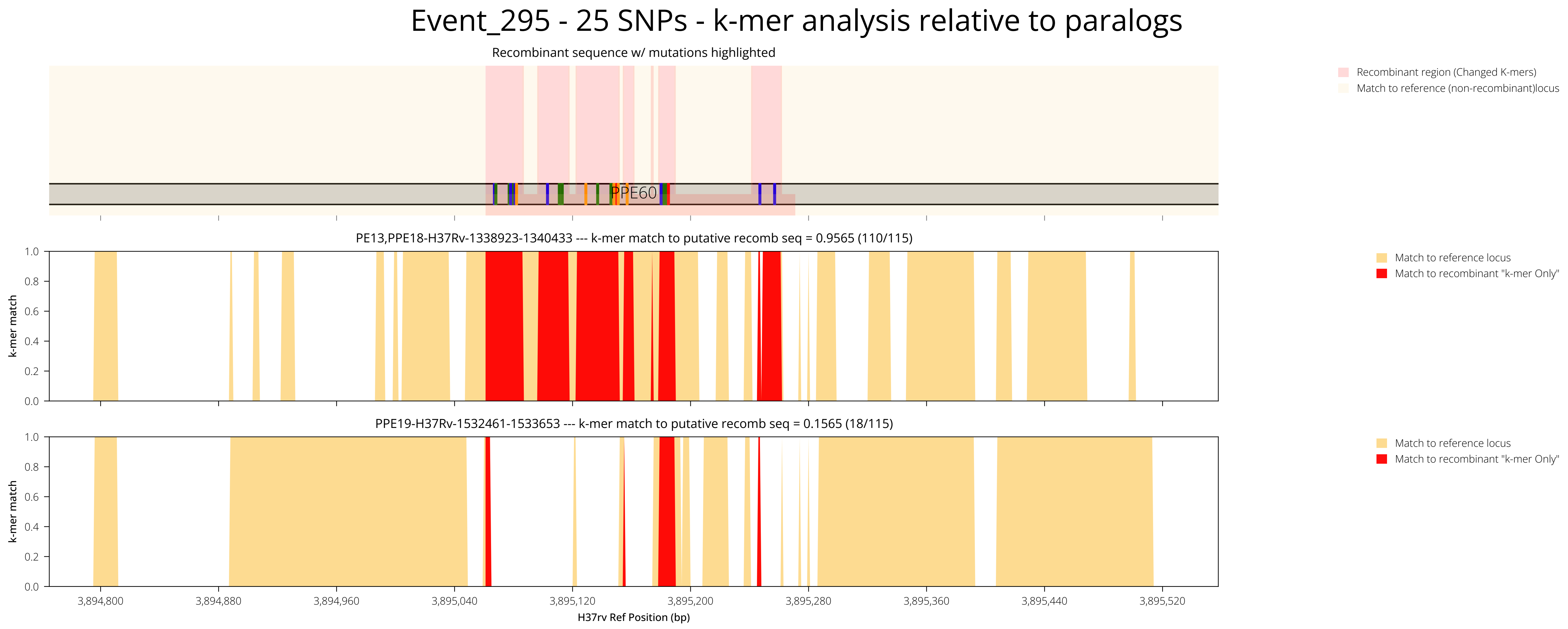

Event_295 - 25 SNPs - k-mer analysis relative to paralogs
Recombinant sequence w/ mutations highlighted
PPE60
Recombinant region (Changed K-mers)
Match to reference (non-recombinant)locus
PE13,PPE18-H37Rv-1338923-1340433 --- k-mer match to putative recomb seq = 0.9565 (110/115)
Match to reference locus
Match to recombinant "k-mer Only"
PPE19-H37Rv-1532461-1533653 --- k-mer match to putative recomb seq = 0.1565 (18/115)
Match to reference locus
Match to recombinant "k-mer Only"
k-mer match
1.0
0.8
0.6
0.4
0.2
0.0
3,894,800
3,894,880
3,894,960
3,895,040
3,895,120
3,895,200
3,895,280
3,895,360
3,895,440
3,895,520
H37rv Ref Position (bp)

Sub-phylogeny Viz + Paralog Alignments
GCE ID: Event_295

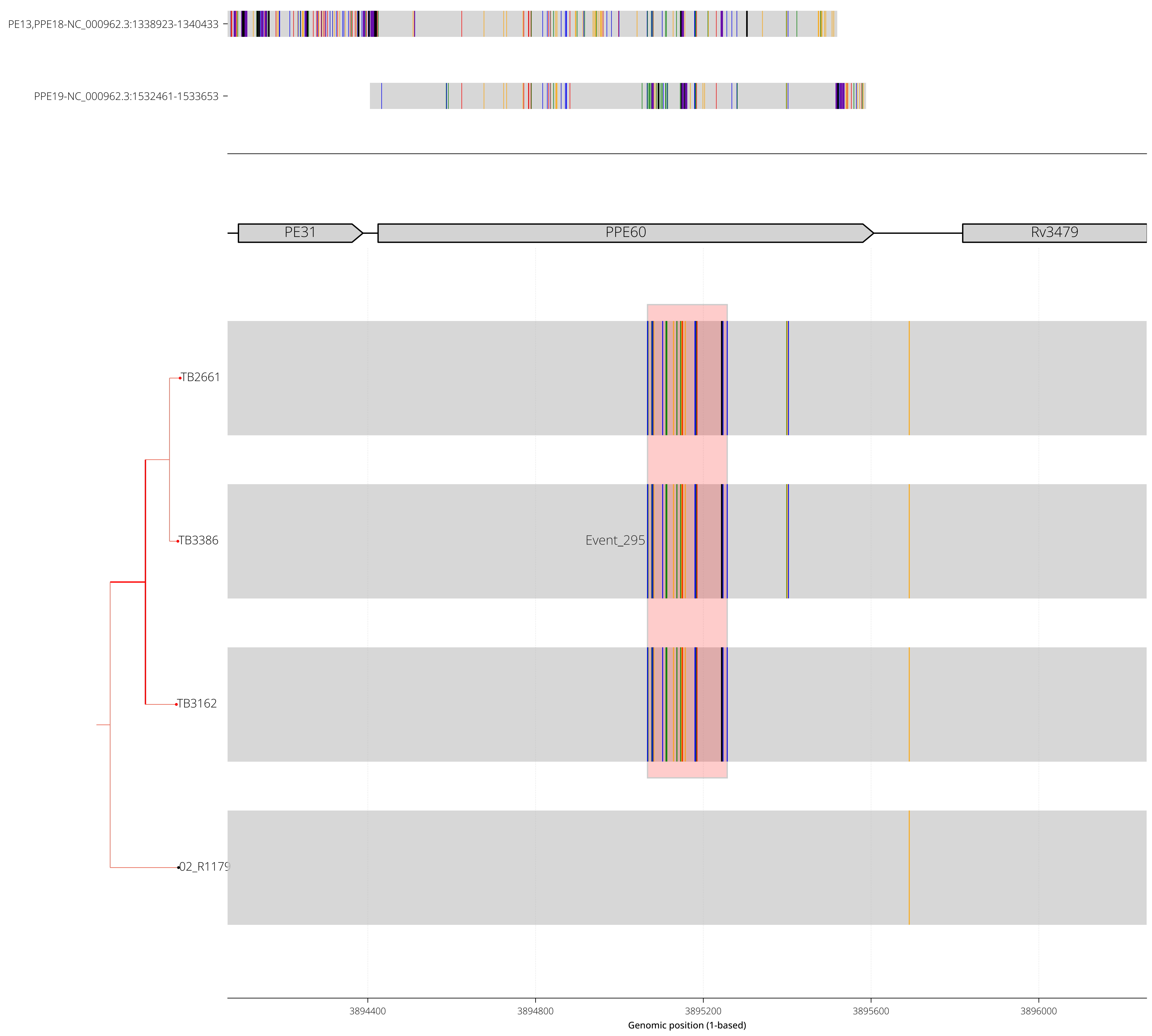

Full Phylogeny Viz + Paralog Alignments
GCE ID: Event_295

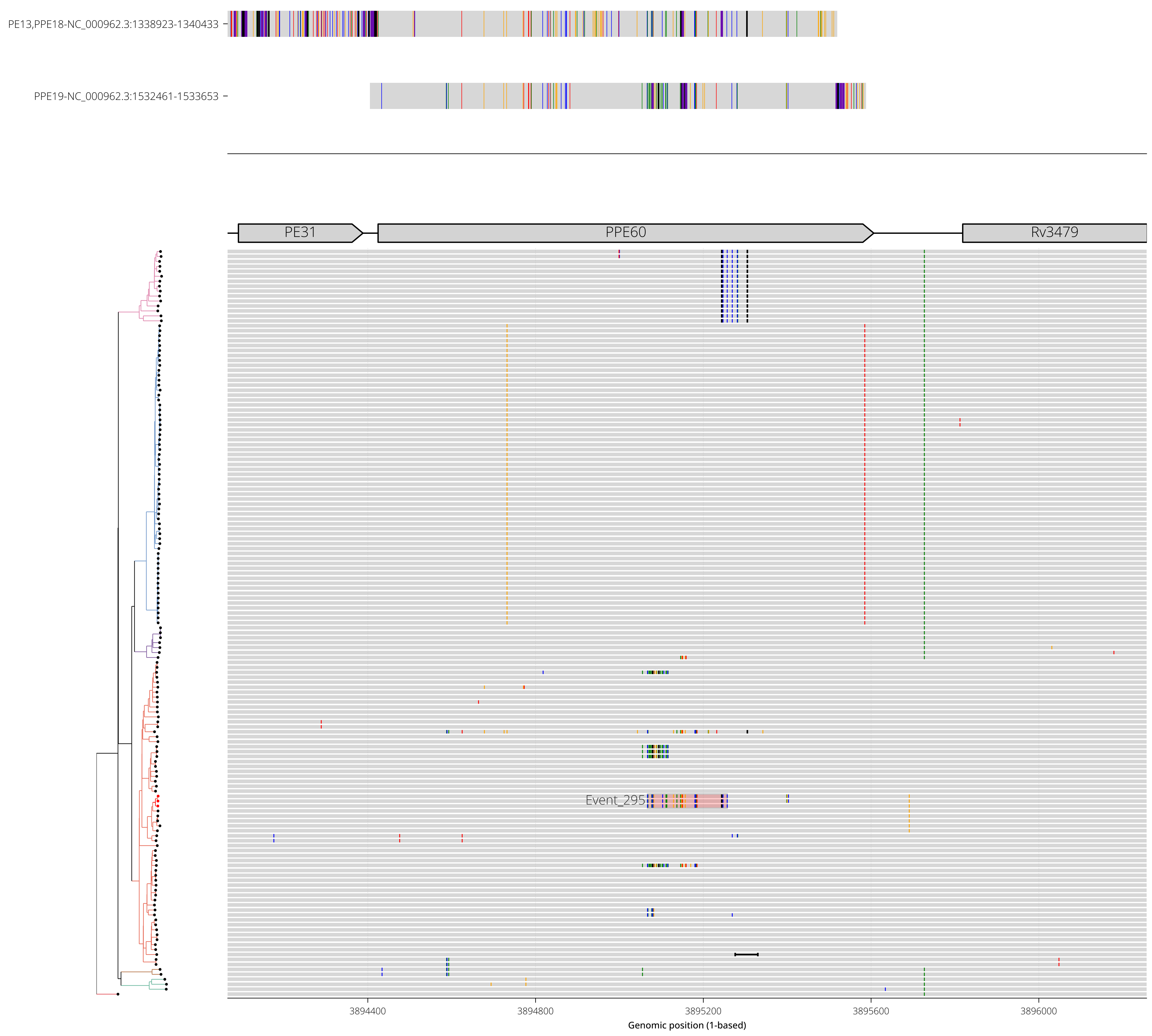

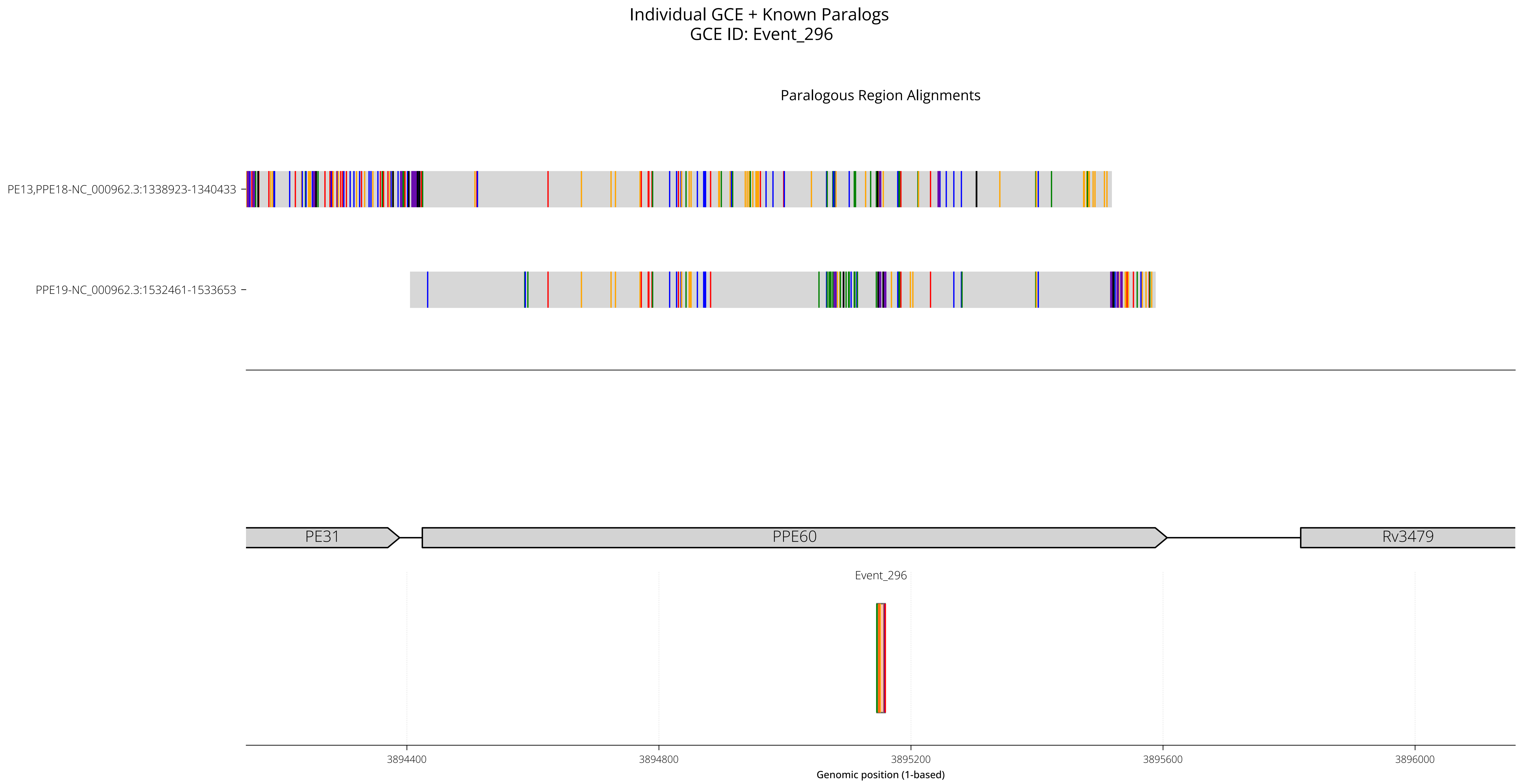

Individual GCE + Known Paralogs
GCE ID: Event_296
Paralogous Region Alignments
PE13,PPE18-NC_000962.3:1338923-1340433
PPE19-NC_000962.3:1532461-1533653
PE31
PPE60
Rv3479
Event_296
3894400
3894800
3895200
3895600
3896000
Genomic position (1-based)

Circos Viz - GCE to Known Paralogs Locations
GCE ID: Event_296

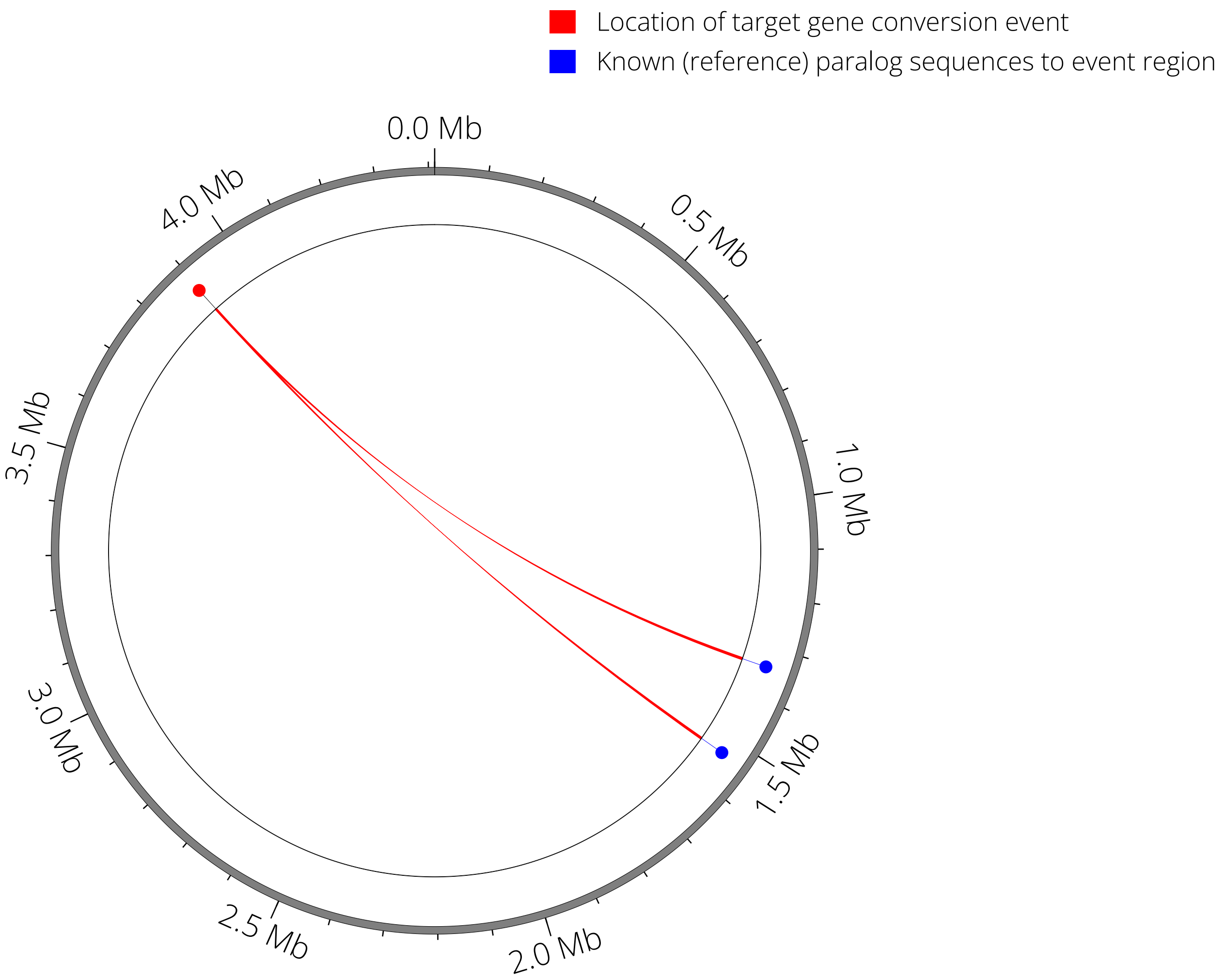

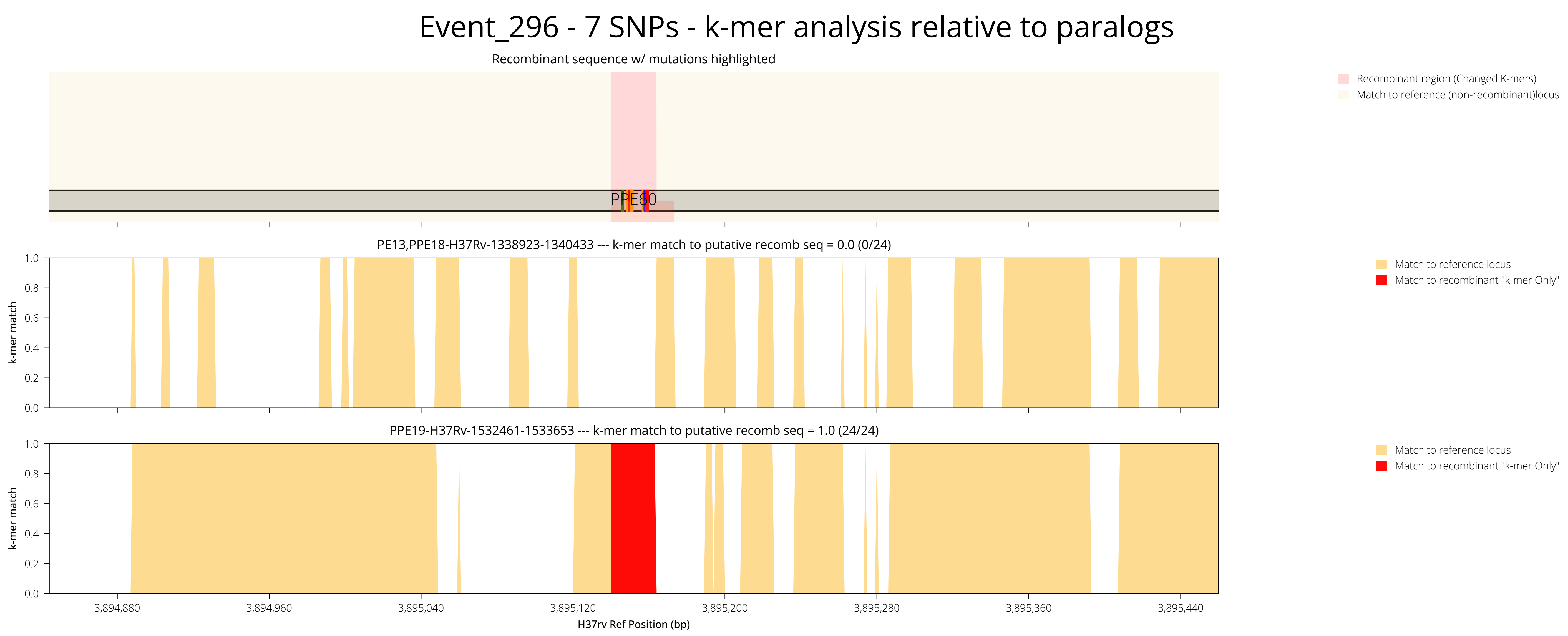
Event_296 - 7 SNPs - k-mer analysis relative to paralogs
Recombinant sequence w/ mutations highlighted
Recombinant region (Changed K-mers)
Match to reference (non-recombinant)locus
PPE60
PE13,PPE18-H37Rv-1338923-1340433 --- k-mer match to putative recomb seq = 0.0 (0/24)
Match to reference locus
Match to recombinant "k-mer Only"
PPE19-H37Rv-1532461-1533653 --- k-mer match to putative recomb seq = 1.0 (24/24)
Match to reference locus
Match to recombinant "k-mer Only"
k-mer match
1.0
0.8
0.6
0.4
0.2
0.0
3,894,880
3,894,960
3,895,040
3,895,120
3,895,200
3,895,280
3,895,360
3,895,440
H37rv Ref Position (bp)

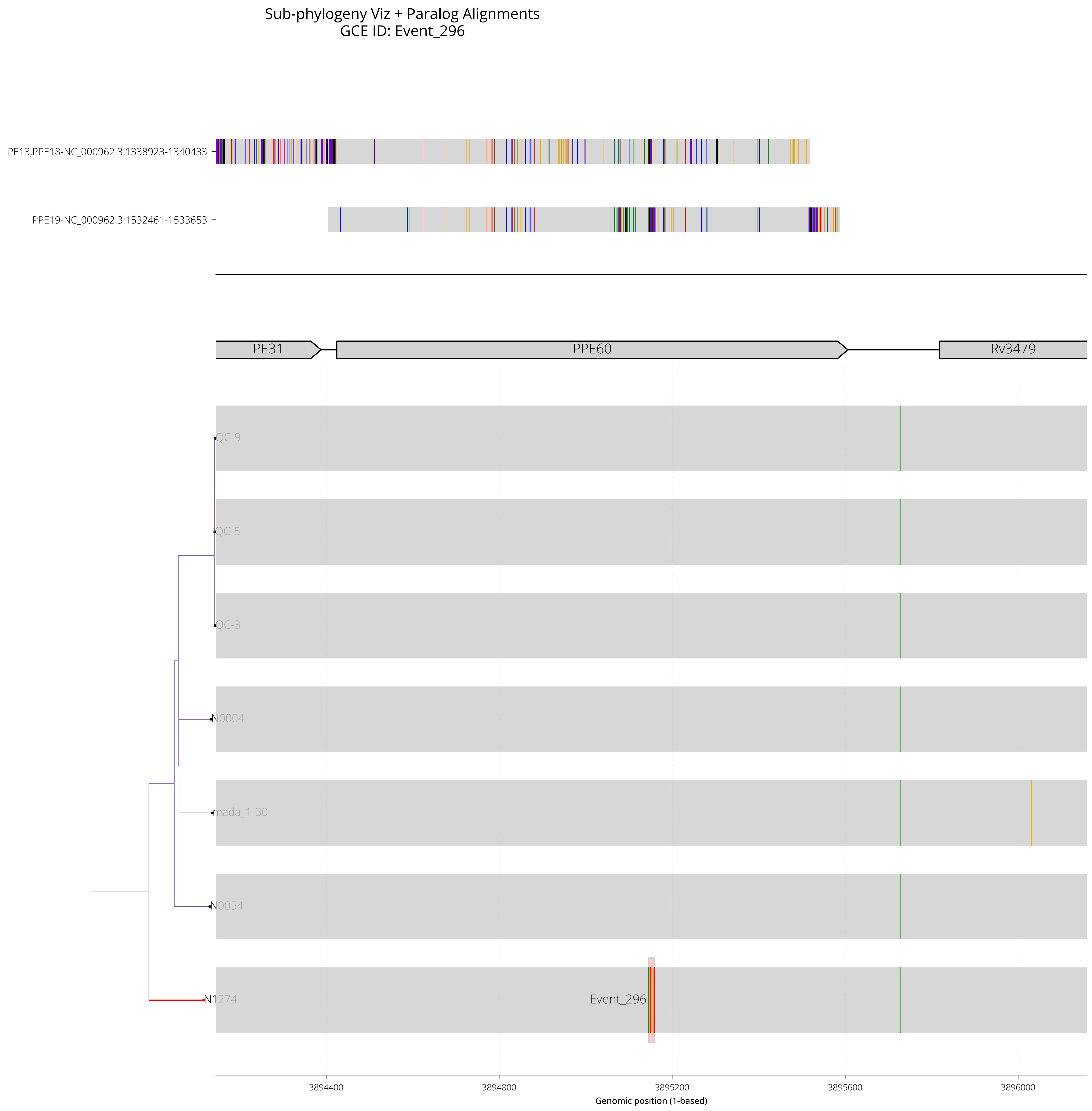

Sub-phylogeny Viz + Paralog Alignments
GCE ID: Event_296
PE13,PPE18-NC_000962.3:1338923-1340433
PPE19-NC_000962.3:1532461-1533653
PE31
PPE60
Rv3479
QC-9
QC-5
QC-3
N0004
mada_1-30
N0054
N1274
Event_296
3894400
3894800
3895200
3895600
3896000
Genomic position (1-based)

Full Phylogeny Viz + Paralog Alignments
GCE ID: Event_296

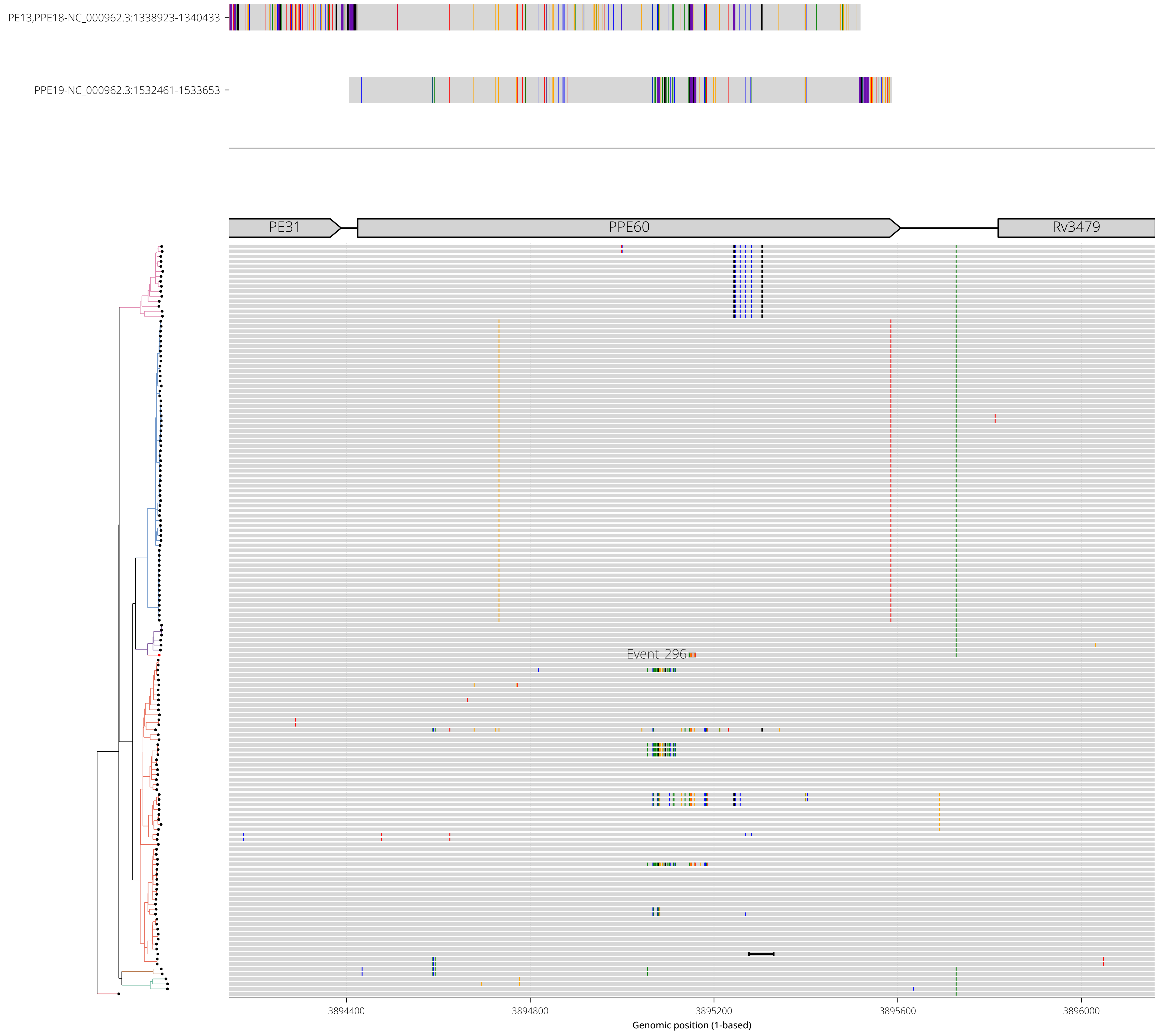

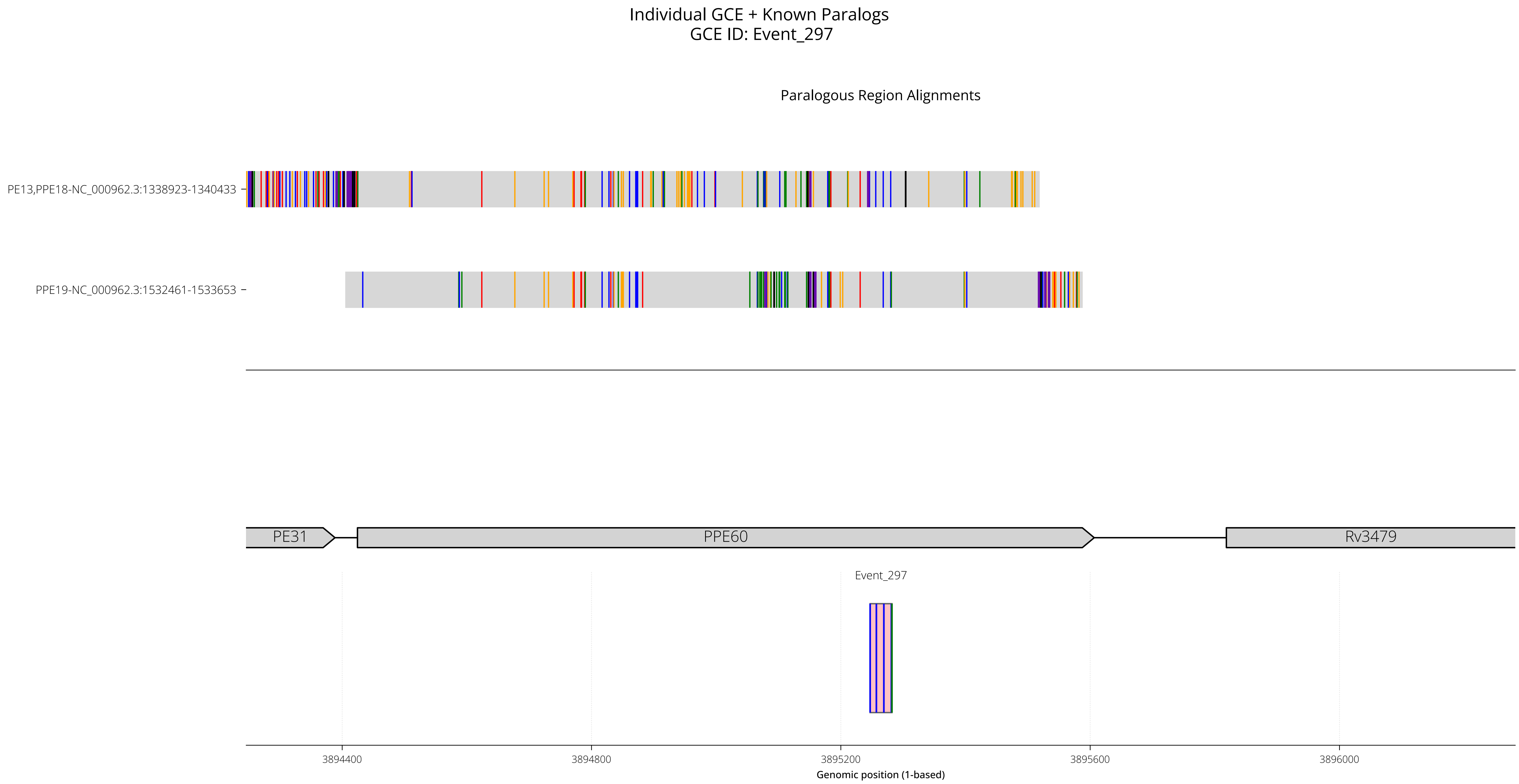

Individual GCE + Known Paralogs
GCE ID: Event_297
Paralogous Region Alignments
PE13,PPE18-NC_000962.3:1338923-1340433
PPE19-NC_000962.3:1532461-1533653
PE31
PPE60
Rv3479
Event_297
3894400
3894800
3895200
3895600
3896000
Genomic position (1-based)

Circos Viz - GCE to Known Paralogs Locations
GCE ID: Event_297

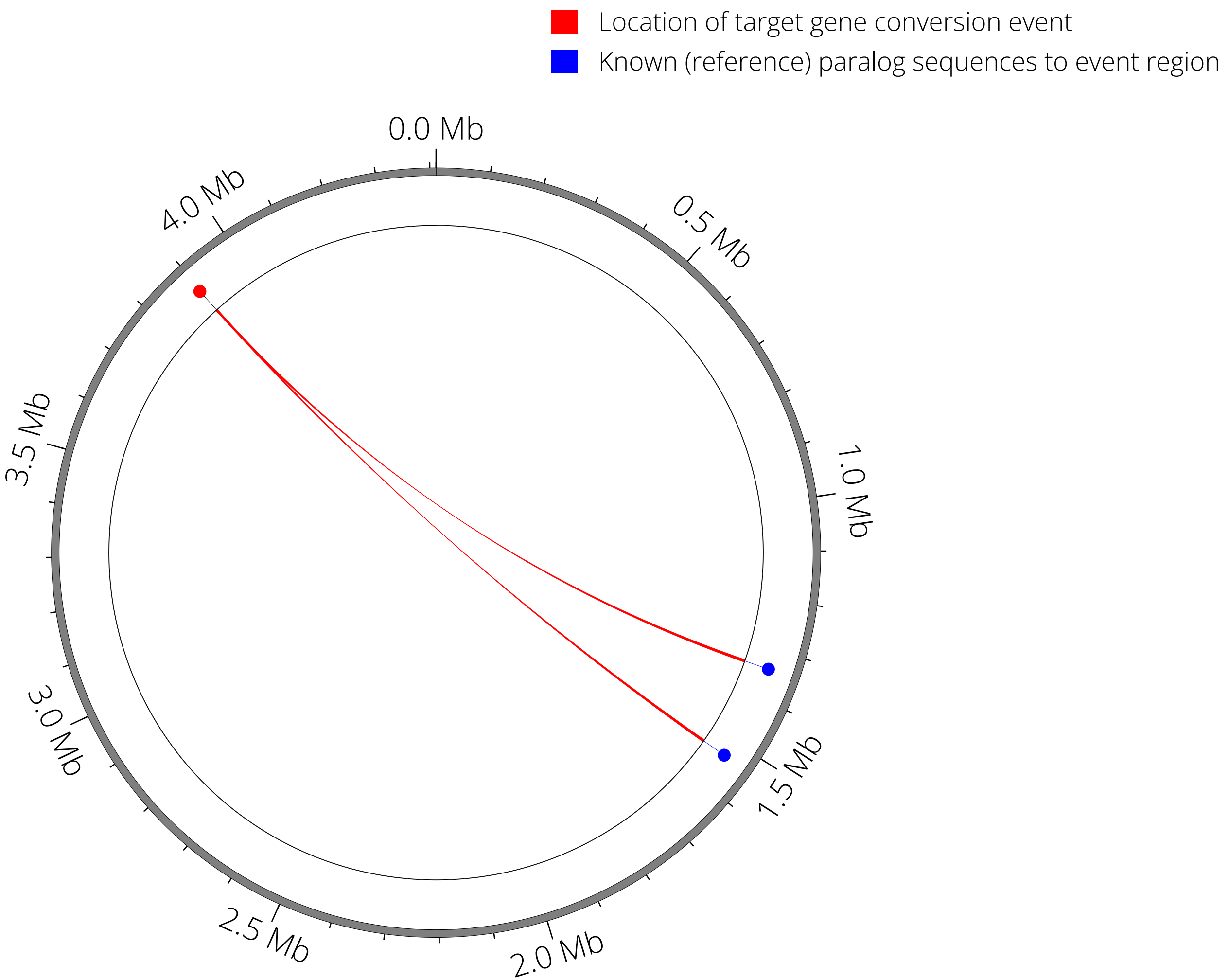

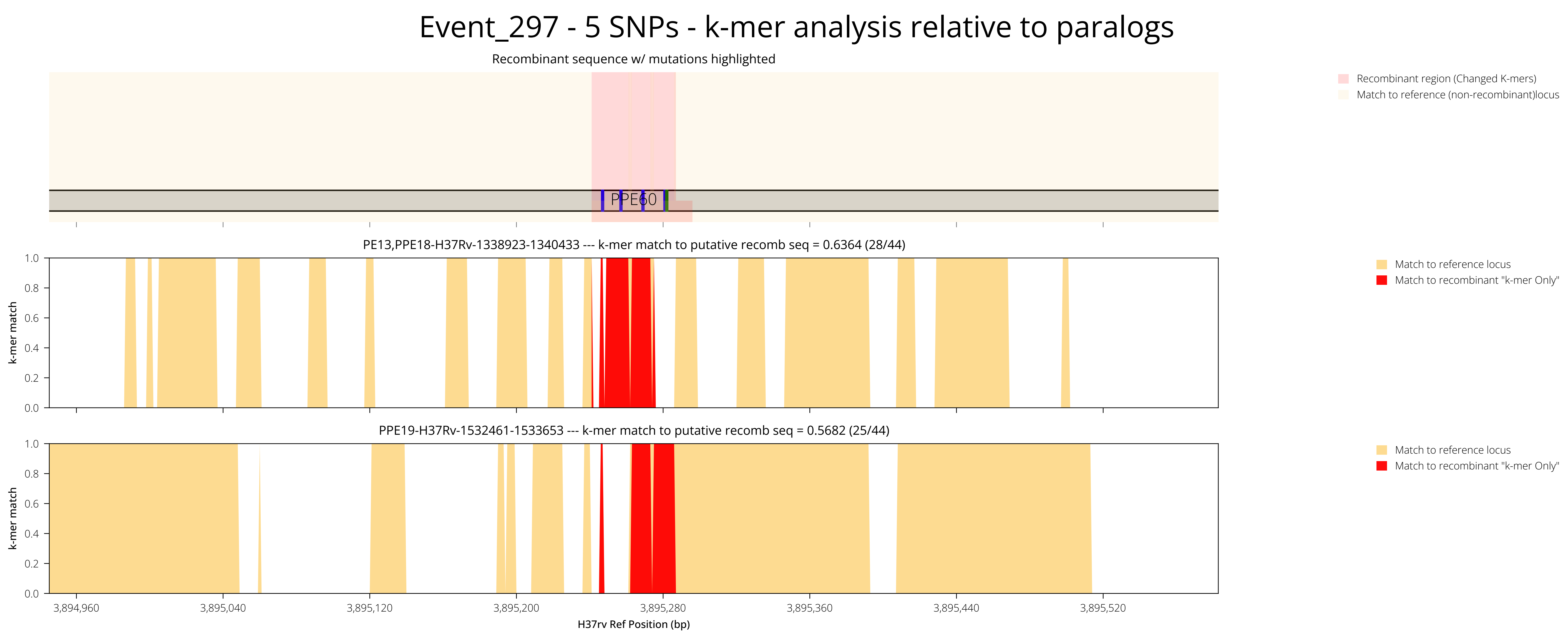
Event_297 - 5 SNPs - k-mer analysis relative to paralogs
Recombinant sequence w/ mutations highlighted
Recombinant region (Changed K-mers)
Match to reference (non-recombinant)locus
PPE60
PE13,PPE18-H37Rv-1338923-1340433 --- k-mer match to putative recomb seq = 0.6364 (28/44)
Match to reference locus
Match to recombinant "k-mer Only"
k-mer match
1.0
0.8
0.6
0.4
0.2
0.0
PPE19-H37Rv-1532461-1533653 --- k-mer match to putative recomb seq = 0.5682 (25/44)
Match to reference locus
Match to recombinant "k-mer Only"
k-mer match
1.0
0.8
0.6
0.4
0.2
0.0
3,894,960
3,895,040
3,895,120
3,895,200
3,895,280
3,895,360
3,895,440
3,895,520
H37rv Ref Position (bp)

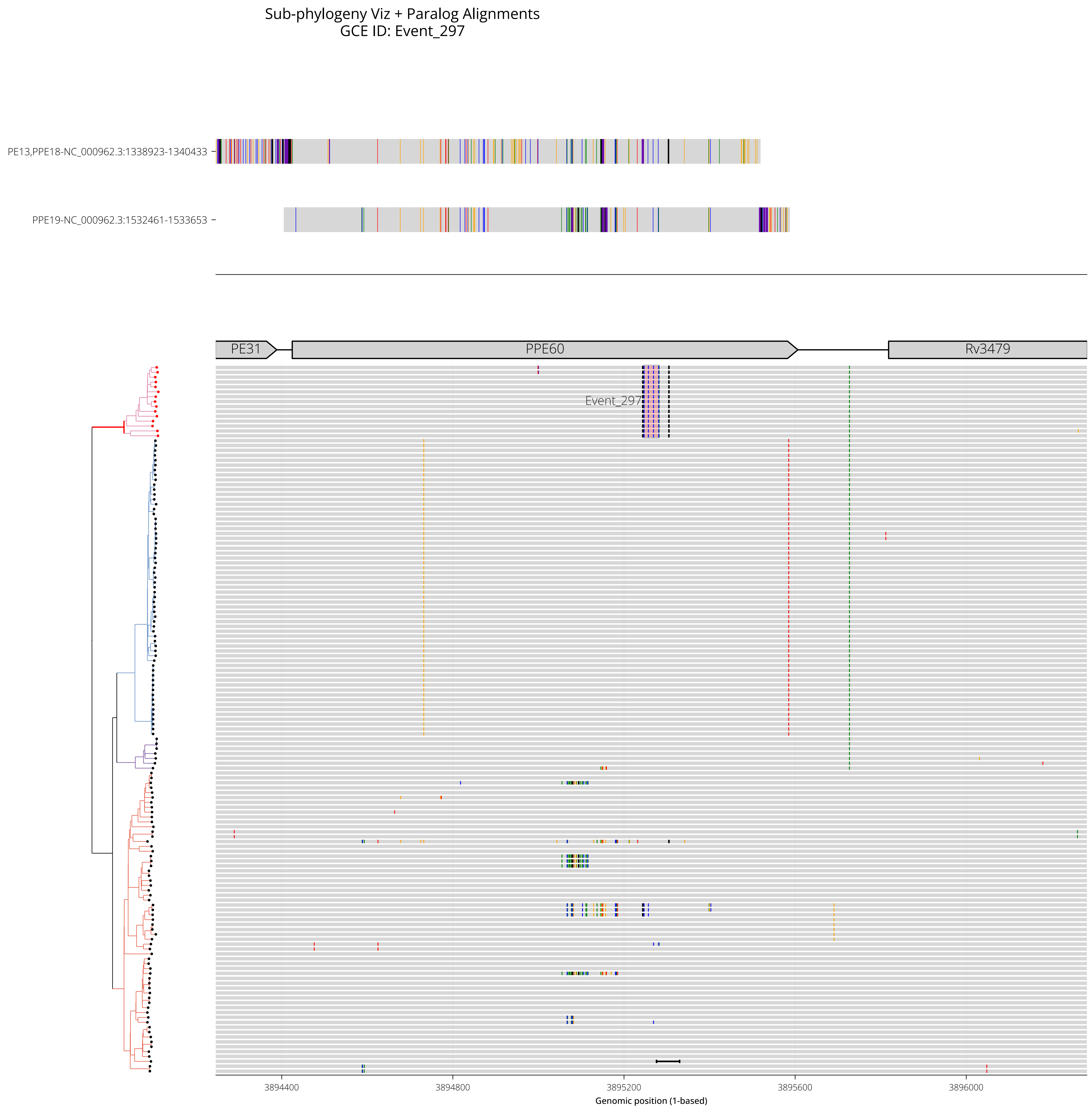

Sub-phylogeny Viz + Paralog Alignments
GCE ID: Event_297
PE13,PPE18-NC_000962.3:1338923-1340433
PPE19-NC_000962.3:1532461-1533653
PE31
PPE60
Rv3479
Event_297
3894400
3894800
3895200
3895600
3896000
Genomic position (1-based)

Full Phylogeny Viz + Paralog Alignments
GCE ID: Event_297

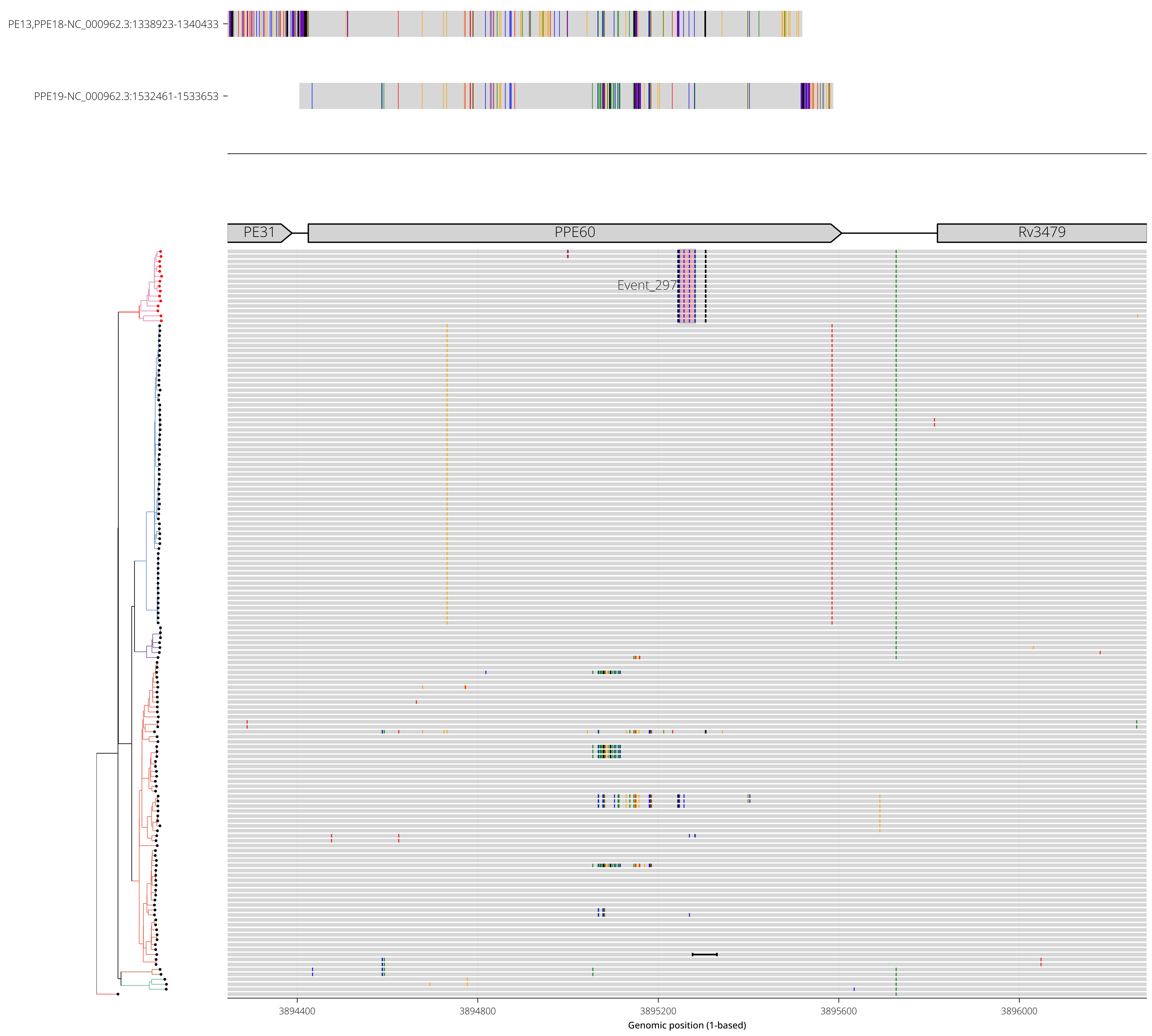

Individual GCE + Known Paralogs
GCE ID: Event_298

Paralogous Region Alignments

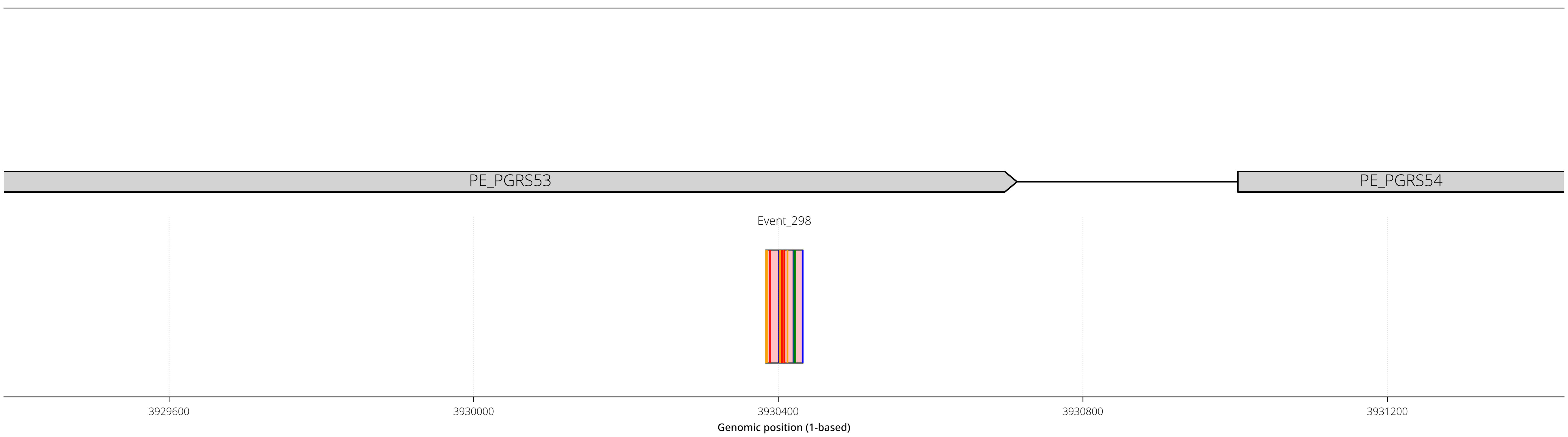

Sub-phylogeny Viz + Paralog Alignments
GCE ID: Event_298

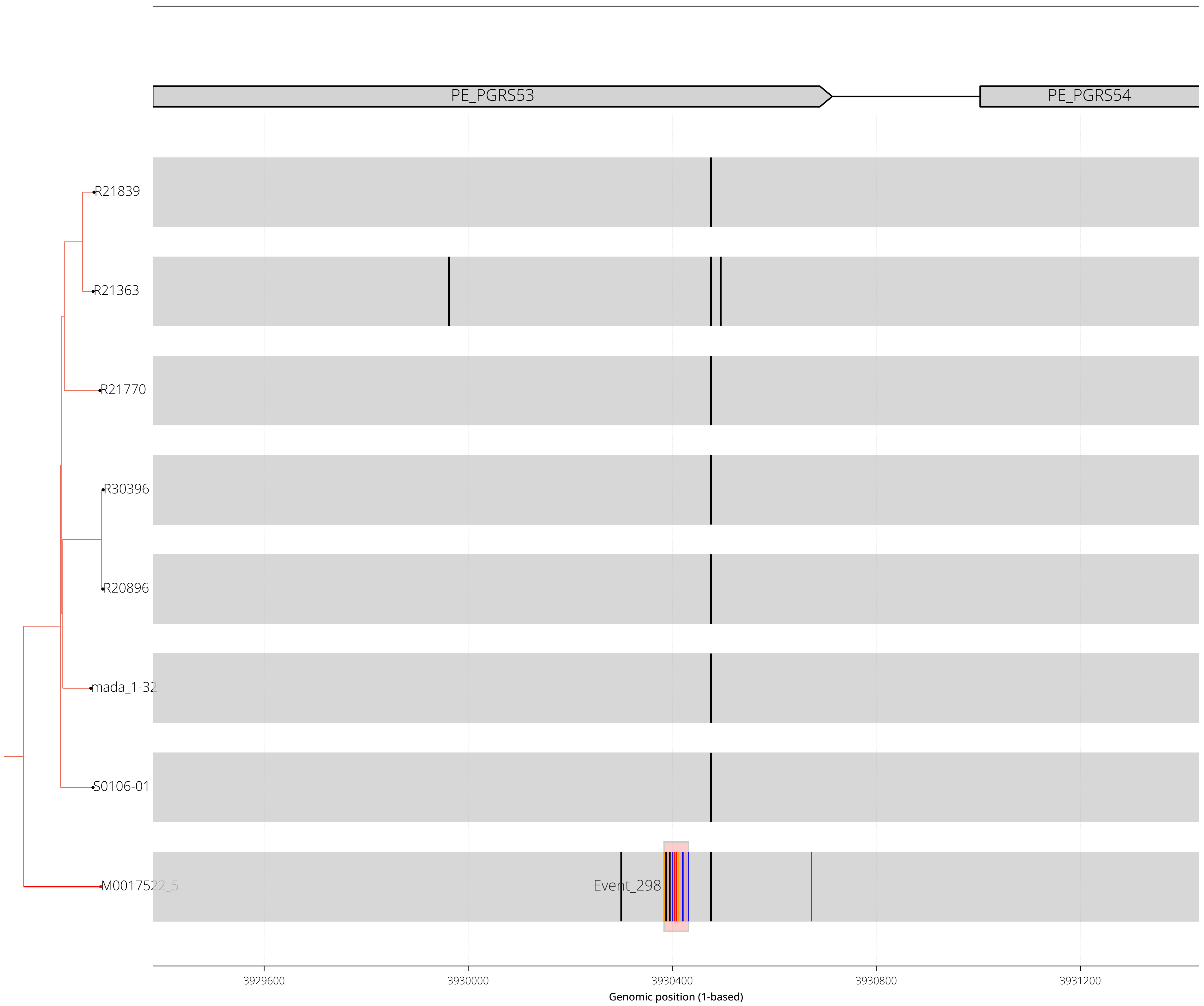

Full Phylogeny Viz + Paralog Alignments
GCE ID: Event_298

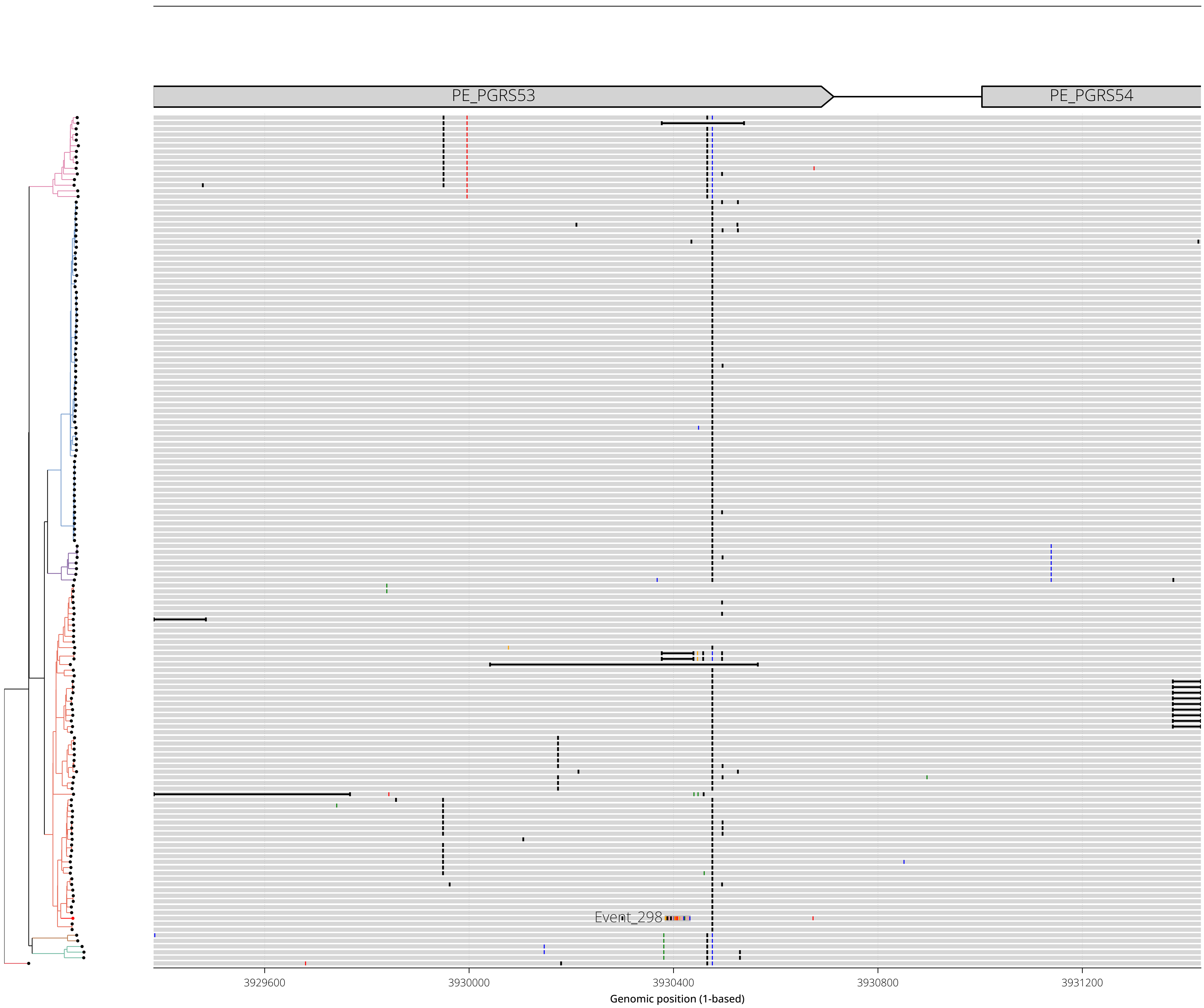

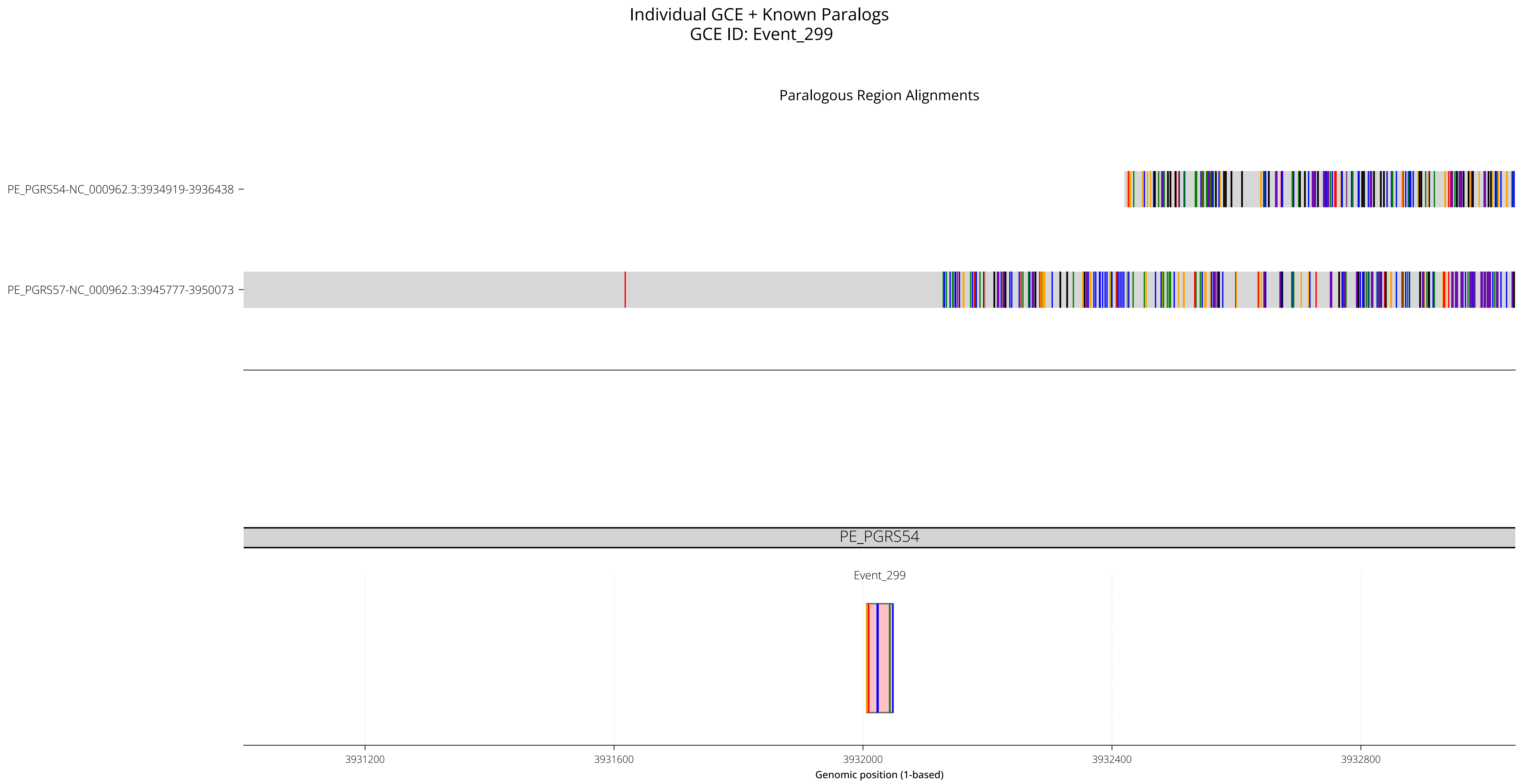

Individual GCE + Known Paralogs
GCE ID: Event_299
Paralogous Region Alignments
PE_PGRS54-NC_000962.3:3934919-3936438
PE_PGRS57-NC_000962.3:3945777-3950073
PE_PGRS54
Event_299
3931200
3931600
3932000
3932400
3932800
Genomic position (1-based)

Circos Viz - GCE to Known Paralogs Locations
GCE ID: Event_299

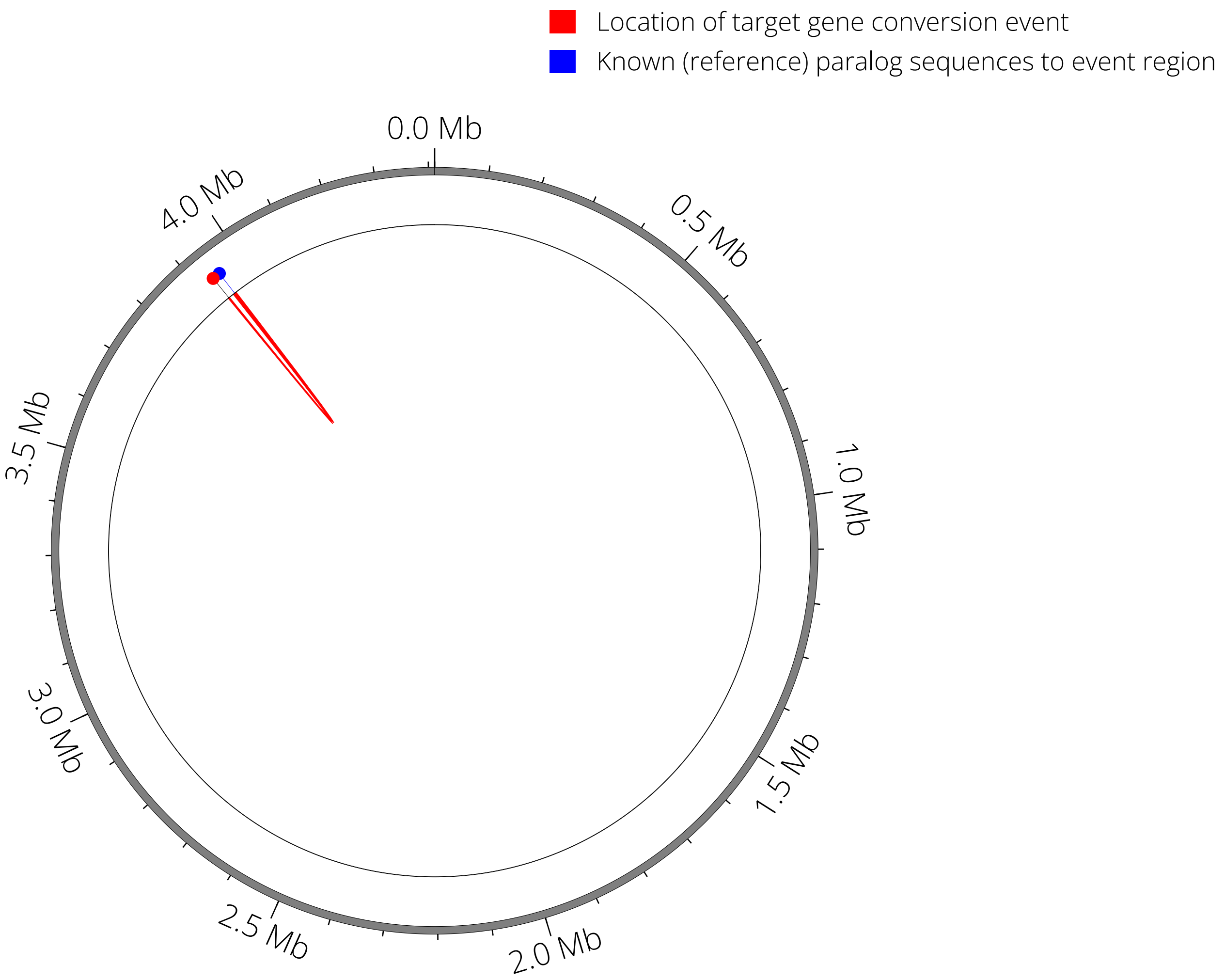

### Event_299 - 6 SNPs - k-mer analysis relative to paralogs

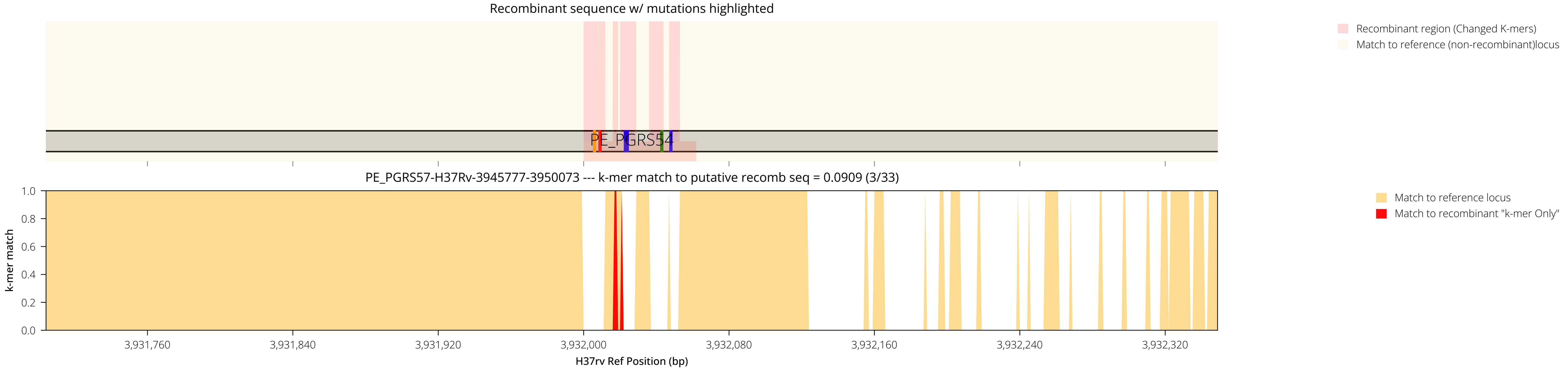

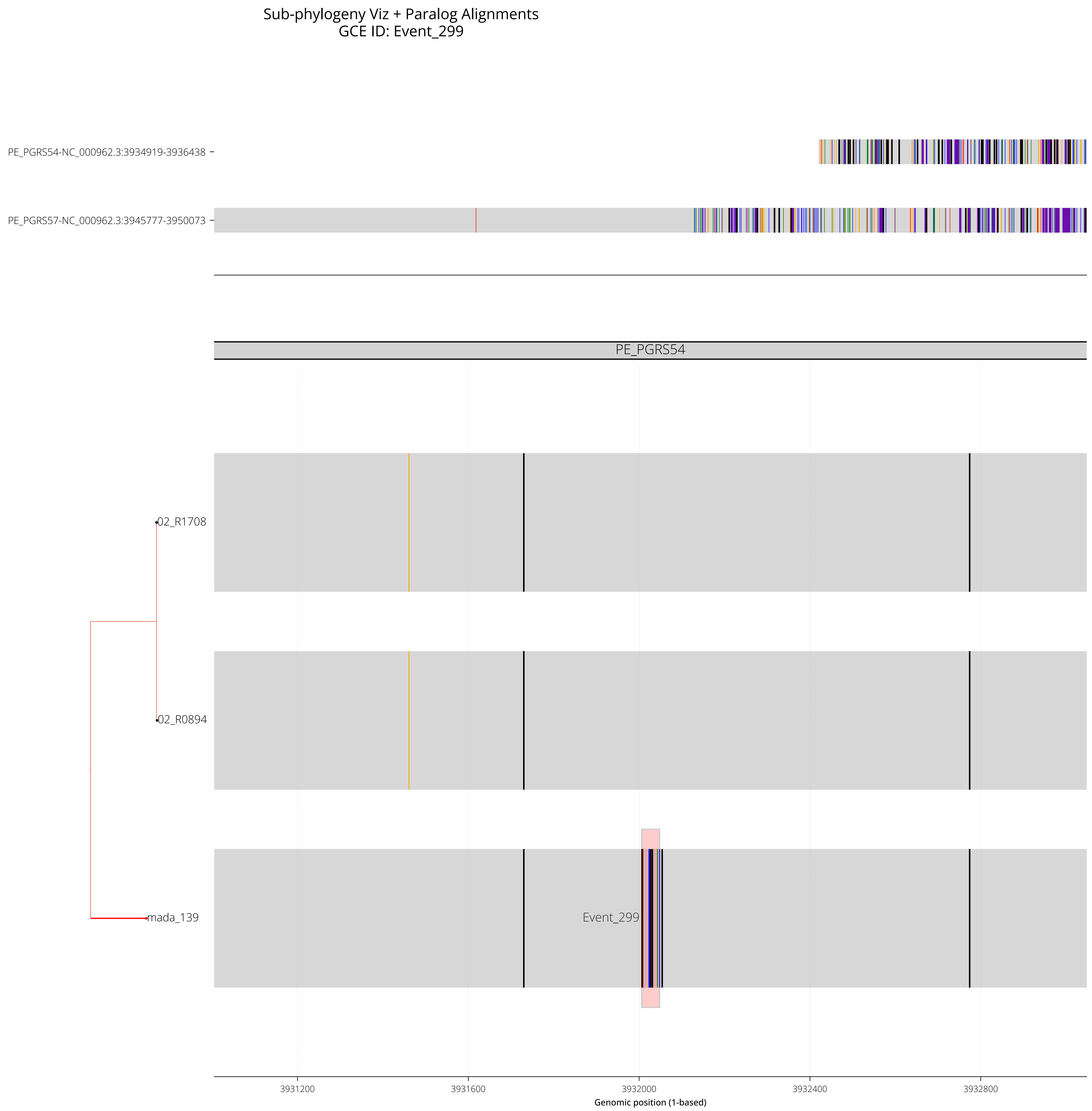
Sub-phylogeny Viz + Paralog Alignments
GCE ID: Event_299
PE_PGRS54-NC_000962.3:3934919-3936438
PE_PGRS57-NC_000962.3:3945777-3950073
PE_PGRS54
02_R1708
02_R0894
mada_139
Event_299
3931200
3931600
3932000
3932400
3932800
Genomic position (1-based)

Full Phylogeny Viz + Paralog Alignments
GCE ID: Event_299

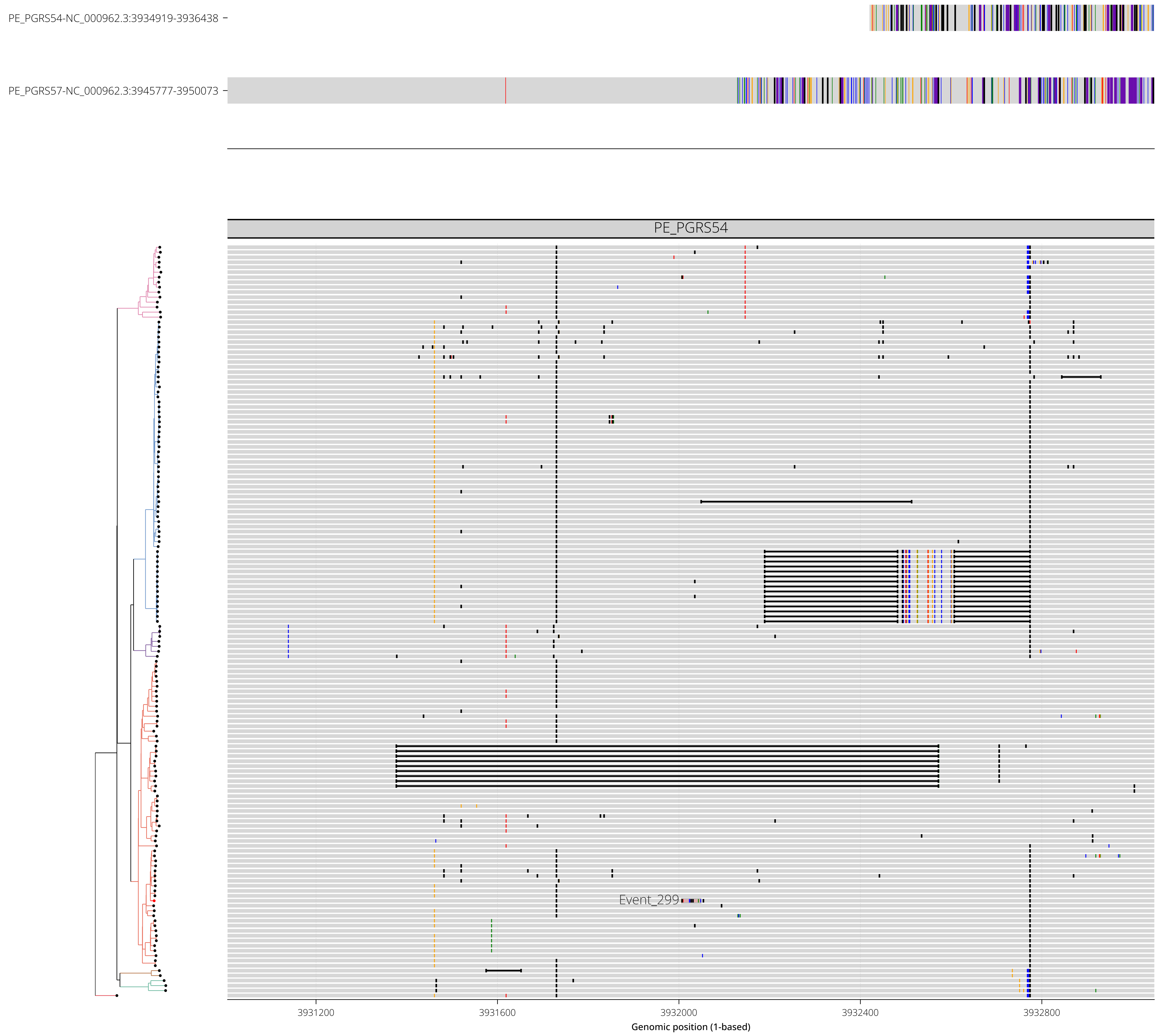

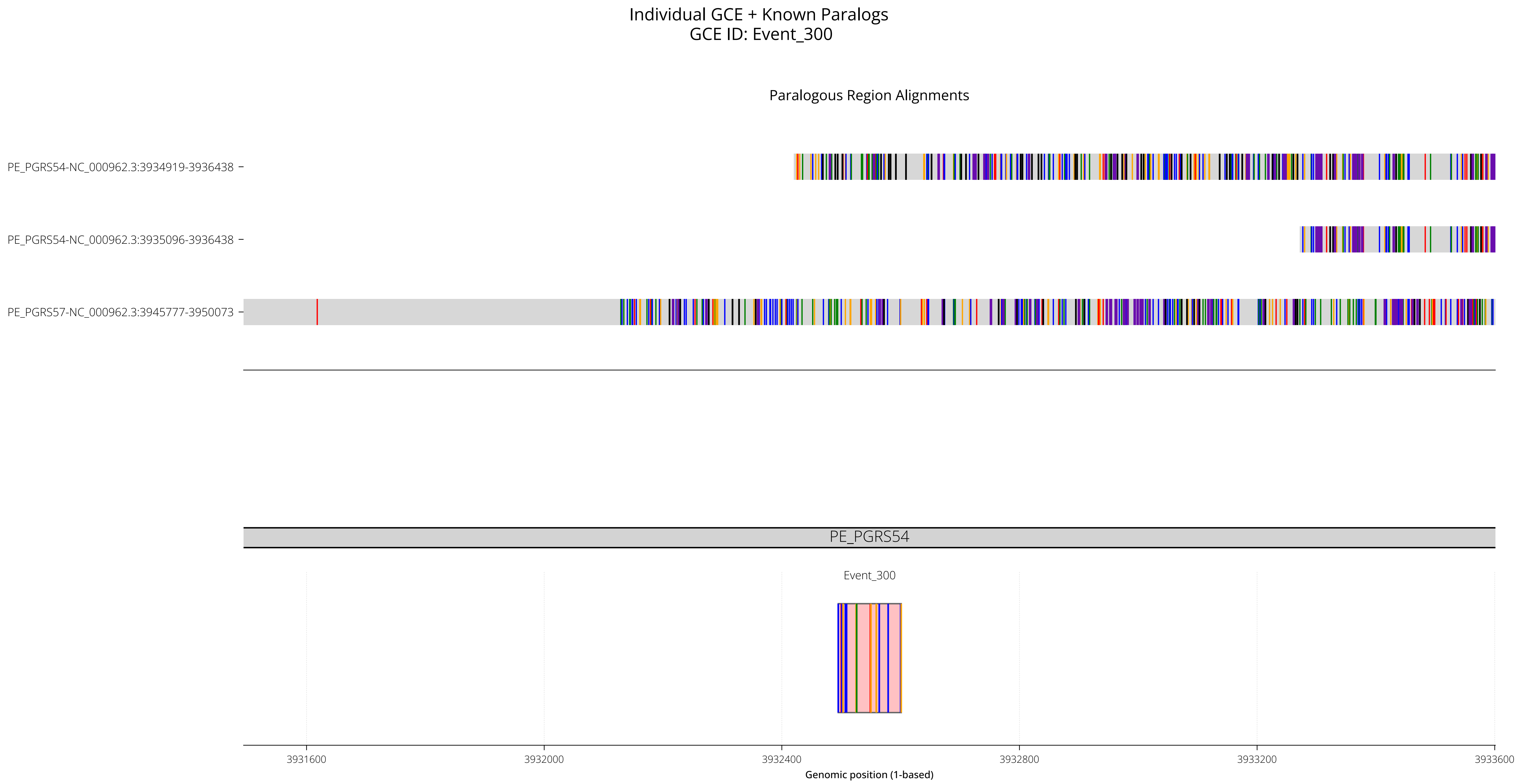

Individual GCE + Known Paralogs
GCE ID: Event_300
Paralogous Region Alignments
PE_PGRS54-NC_000962.3:3934919-3936438
PE_PGRS54-NC_000962.3:3935096-3936438
PE_PGRS57-NC_000962.3:3945777-3950073
PE_PGRS54
Event_300
3931600
3932000
3932400
3932800
3933200
3933600
Genomic position (1-based)

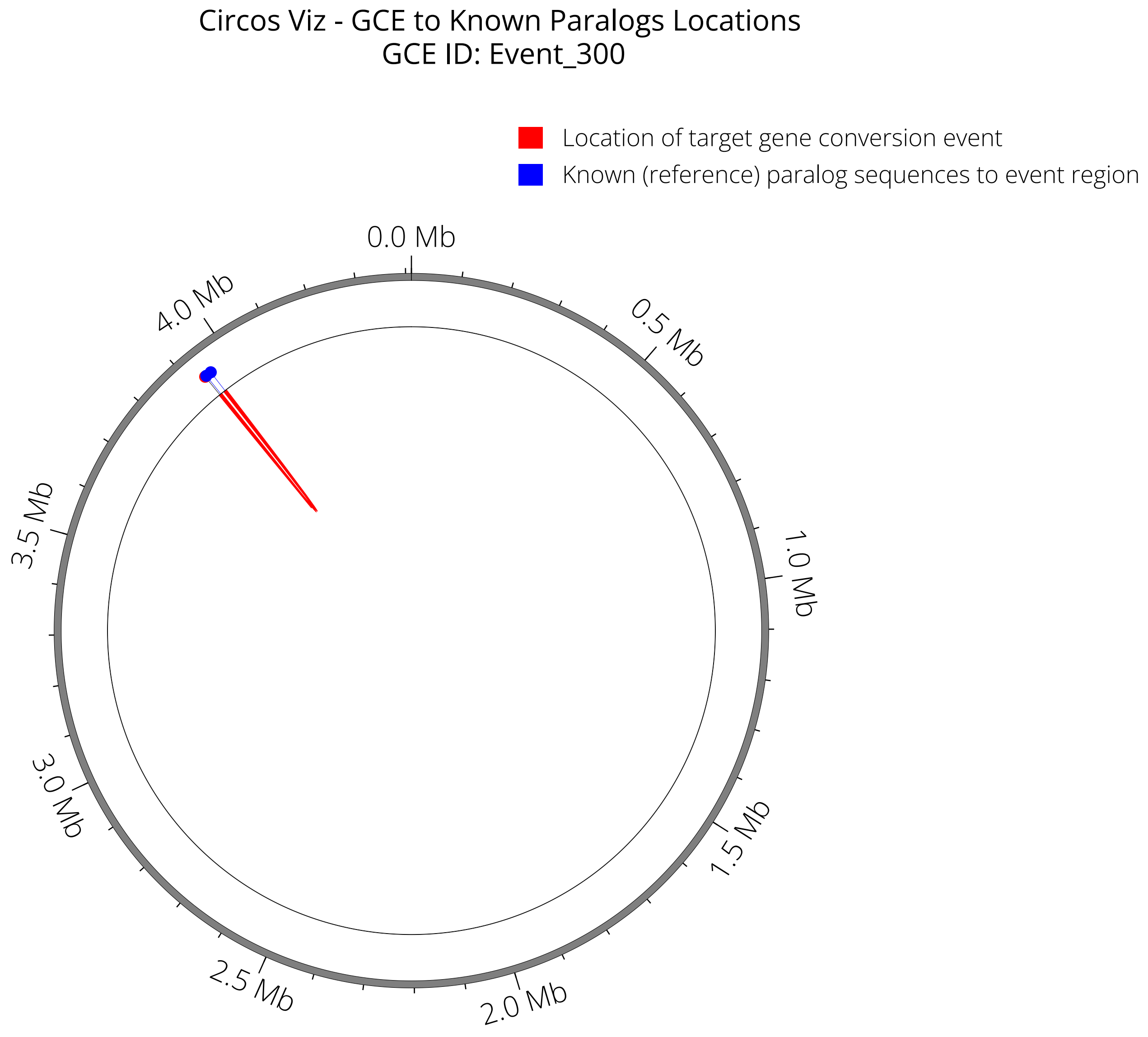

Circos Viz - GCE to Known Paralogs Locations
GCE ID: Event_300
Location of target gene conversion event
Known (reference) paralog sequences to event region
0.0 Mb
0.5 Mb
1.0 Mb
1.5 Mb
2.0 Mb
2.5 Mb
3.0 Mb
3.5 Mb
4.0 Mb

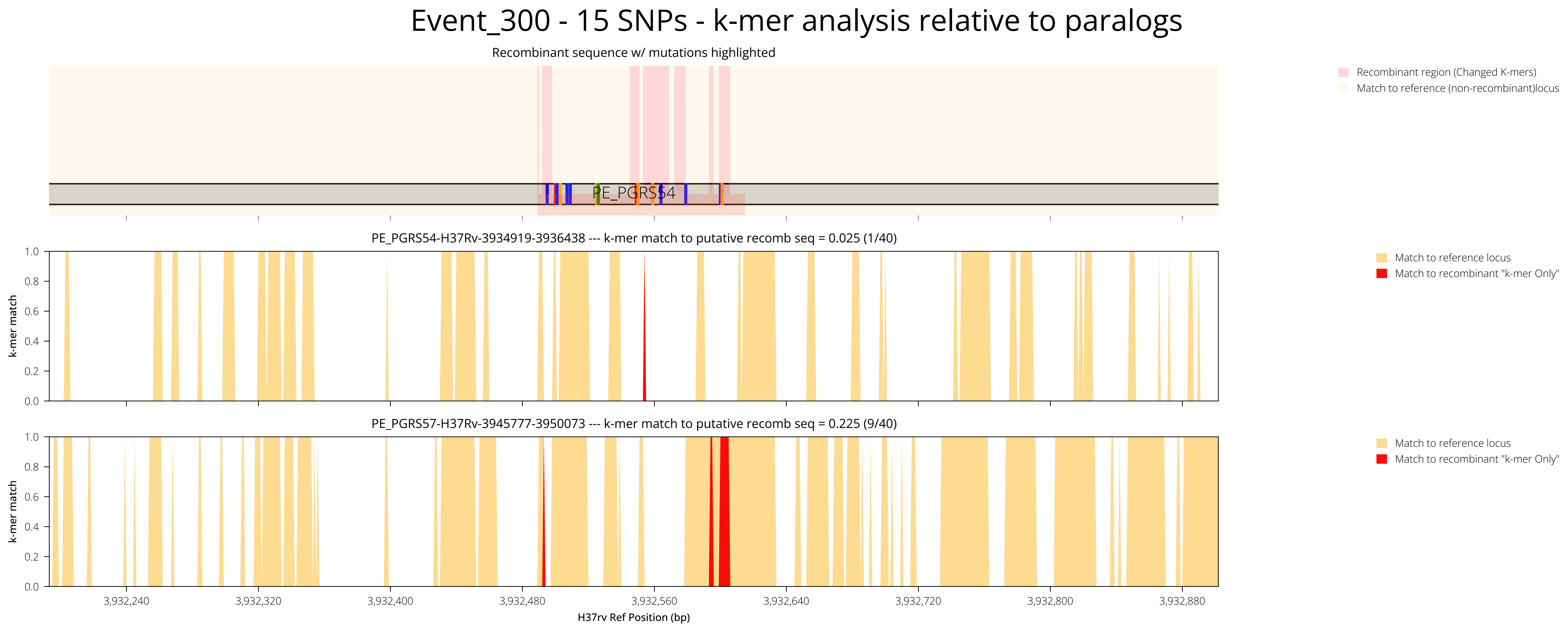
Event_300 - 15 SNPs - k-mer analysis relative to paralogs
Recombinant sequence w/ mutations highlighted
PE_PGRS54
Recombinant region (Changed K-mers)
Match to reference (non-recombinant)locus
PE_PGRS54-H37Rv-3934919-3936438 --- k-mer match to putative recomb seq = 0.025 (1/40)
k-mer match
1.0
0.8
0.6
0.4
0.2
0.0
Match to reference locus
Match to recombinant "k-mer Only"
PE_PGRS57-H37Rv-3945777-3950073 --- k-mer match to putative recomb seq = 0.225 (9/40)
k-mer match
1.0
0.8
0.6
0.4
0.2
0.0
Match to reference locus
Match to recombinant "k-mer Only"
3,932,240
3,932,320
3,932,400
3,932,480
3,932,560
3,932,640
3,932,720
3,932,800
3,932,880
H37rv Ref Position (bp)

Sub-phylogeny Viz + Paralog Alignments
GCE ID: Event_300

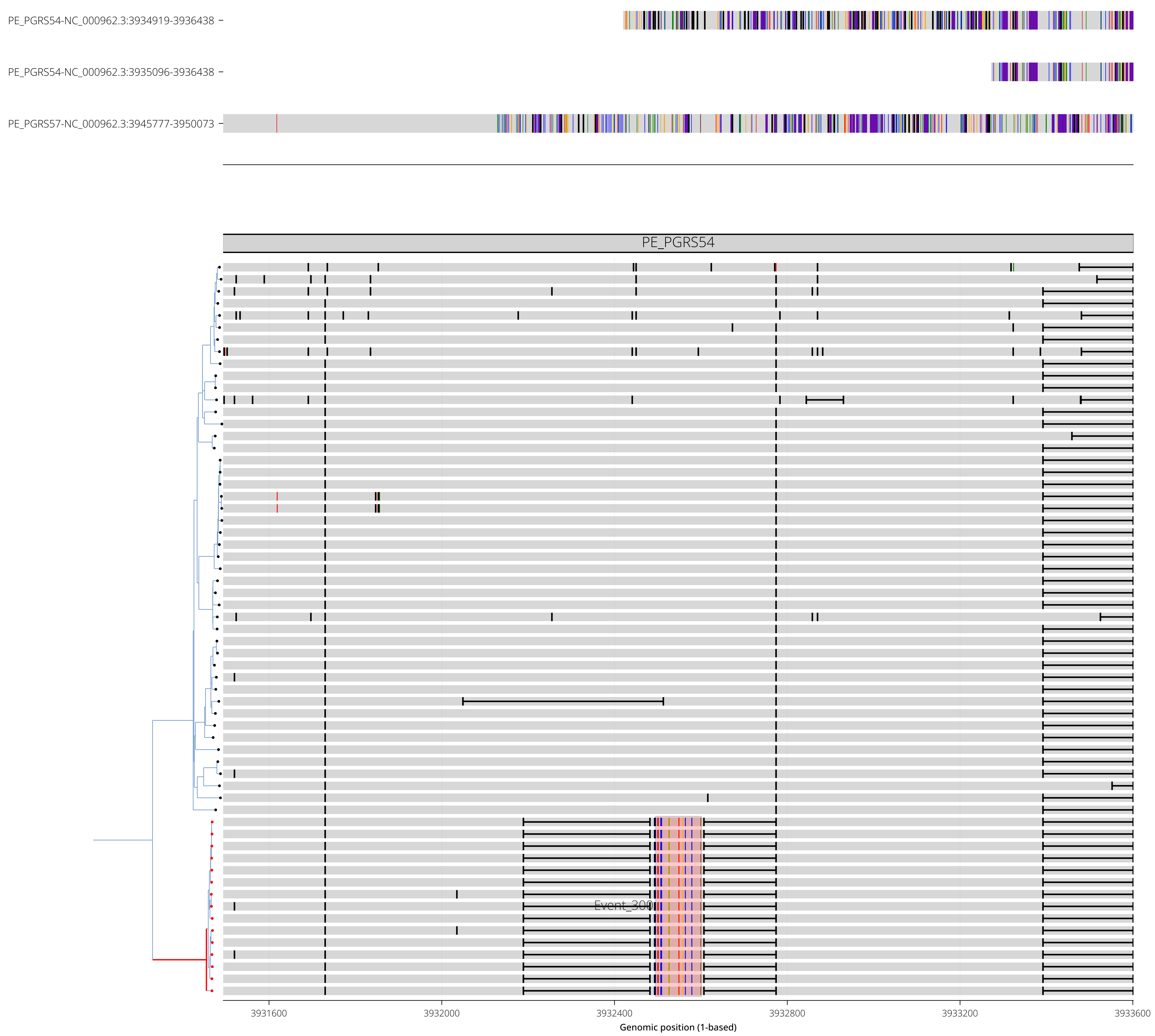

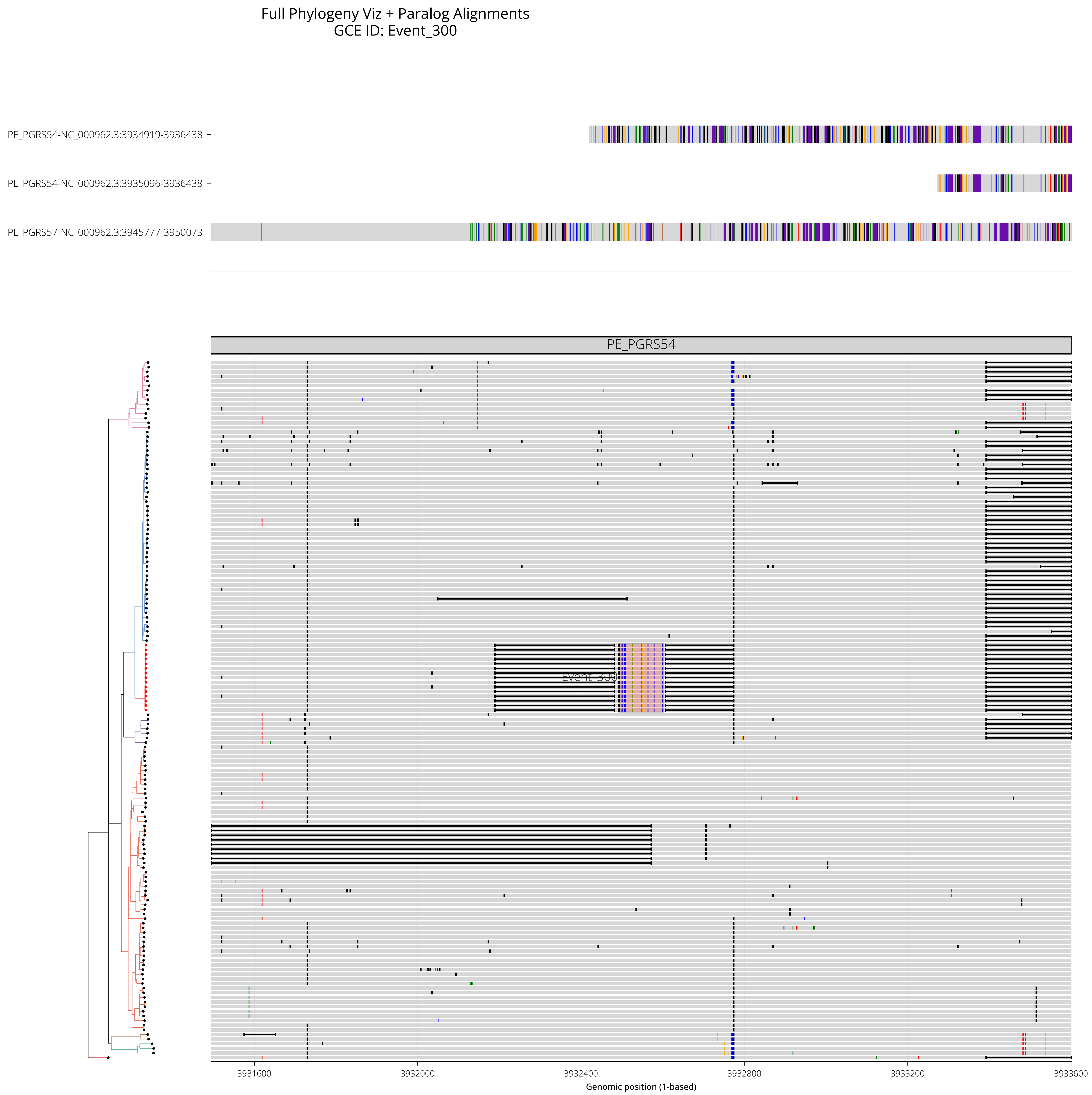
Full Phylogeny Viz + Paralog Alignments
GCE ID: Event_300
PE_PGRS54-NC_000962.3:3934919-3936438
PE_PGRS54-NC_000962.3:3935096-3936438
PE_PGRS57-NC_000962.3:3945777-3950073
PE_PGRS54
Event_300
3931600
3932000
3932400
3932800
3933200
3933600
Genomic position (1-based)

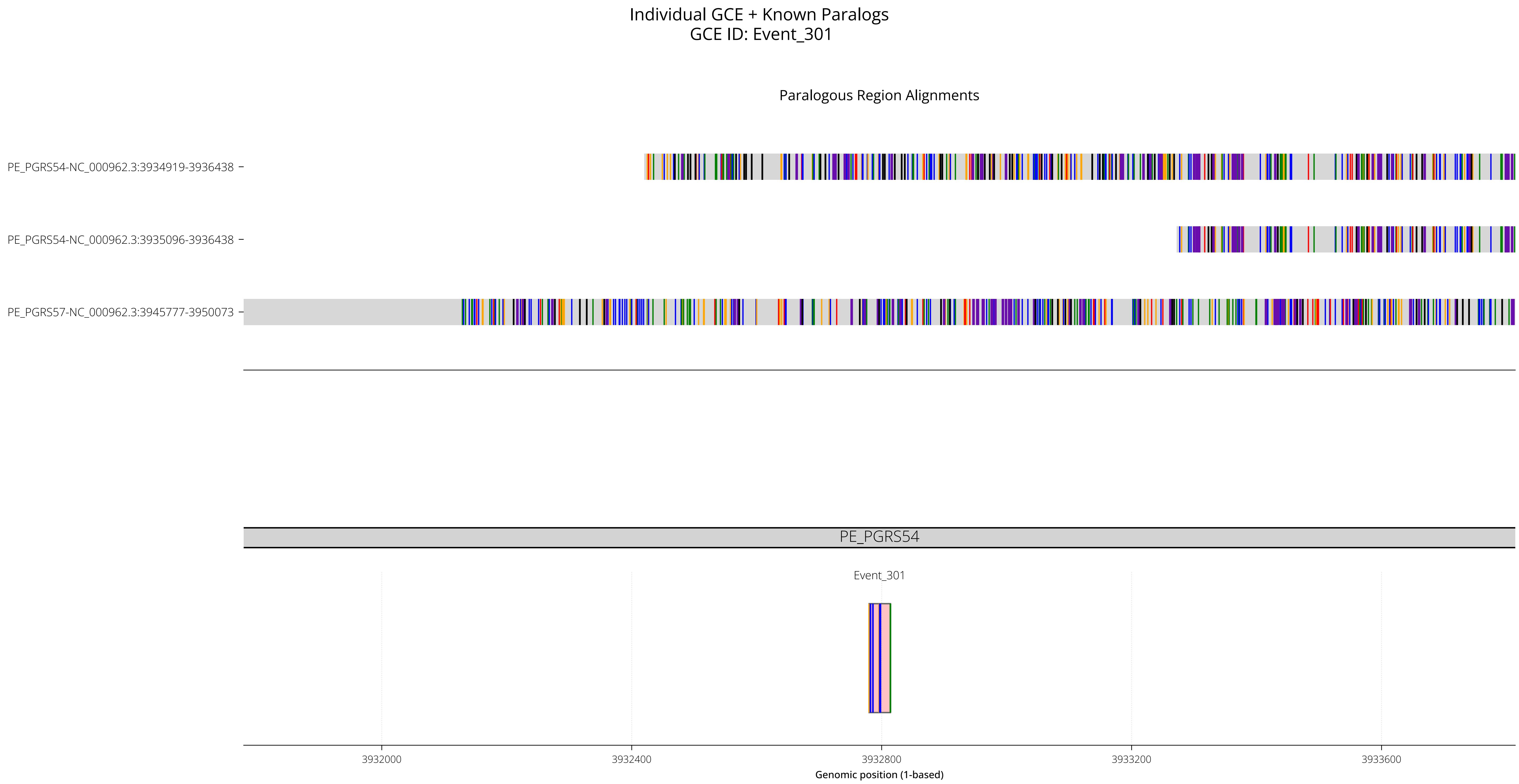
Individual GCE + Known Paralogs
GCE ID: Event_301
Paralogous Region Alignments
PE_PGRS54-NC_000962.3:3934919-3936438
PE_PGRS54-NC_000962.3:3935096-3936438
PE_PGRS57-NC_000962.3:3945777-3950073
PE_PGRS54
Event_301
3932000
3932400
3932800
3933200
3933600
Genomic position (1-based)

Circos Viz - GCE to Known Paralogs Locations
GCE ID: Event_301

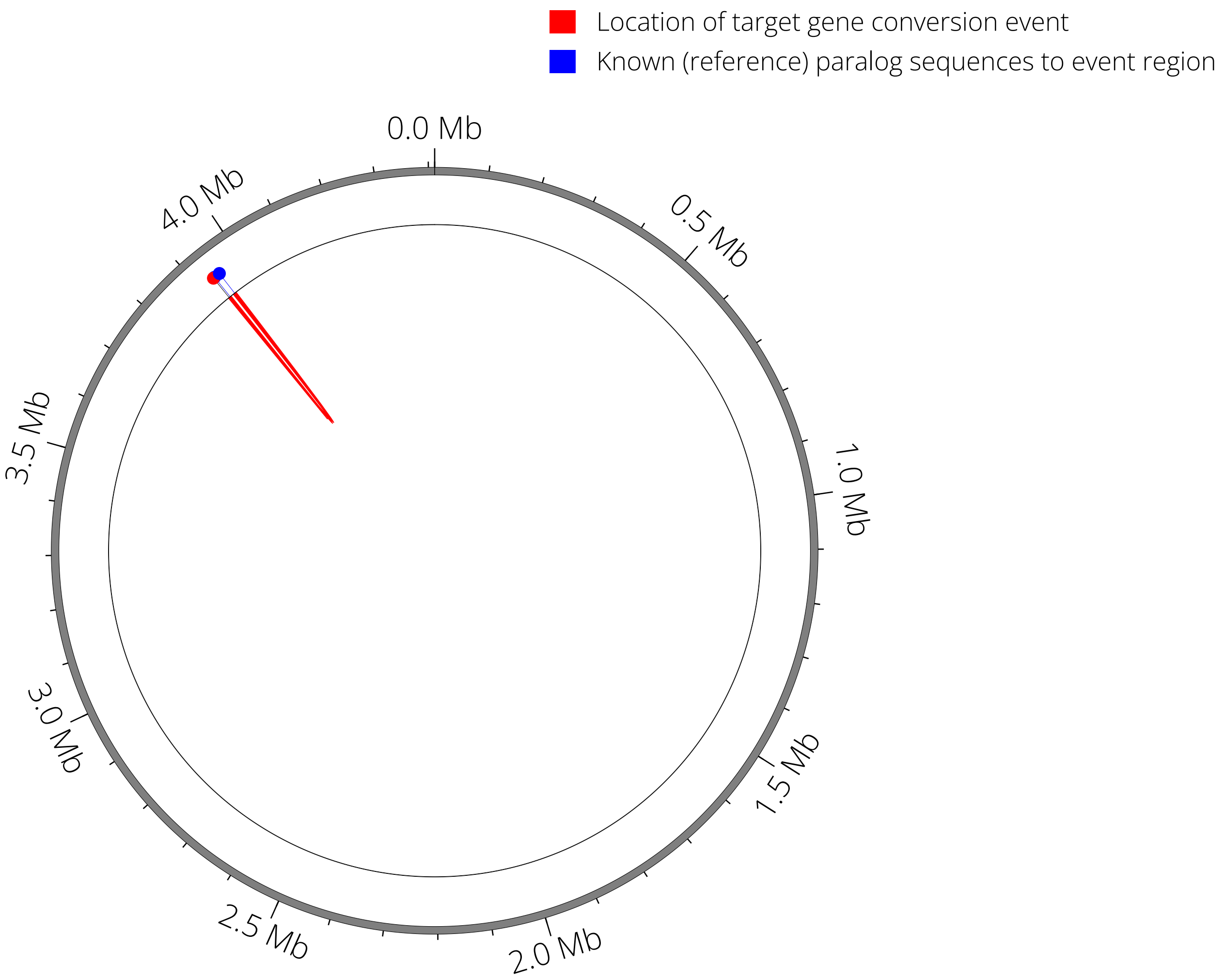

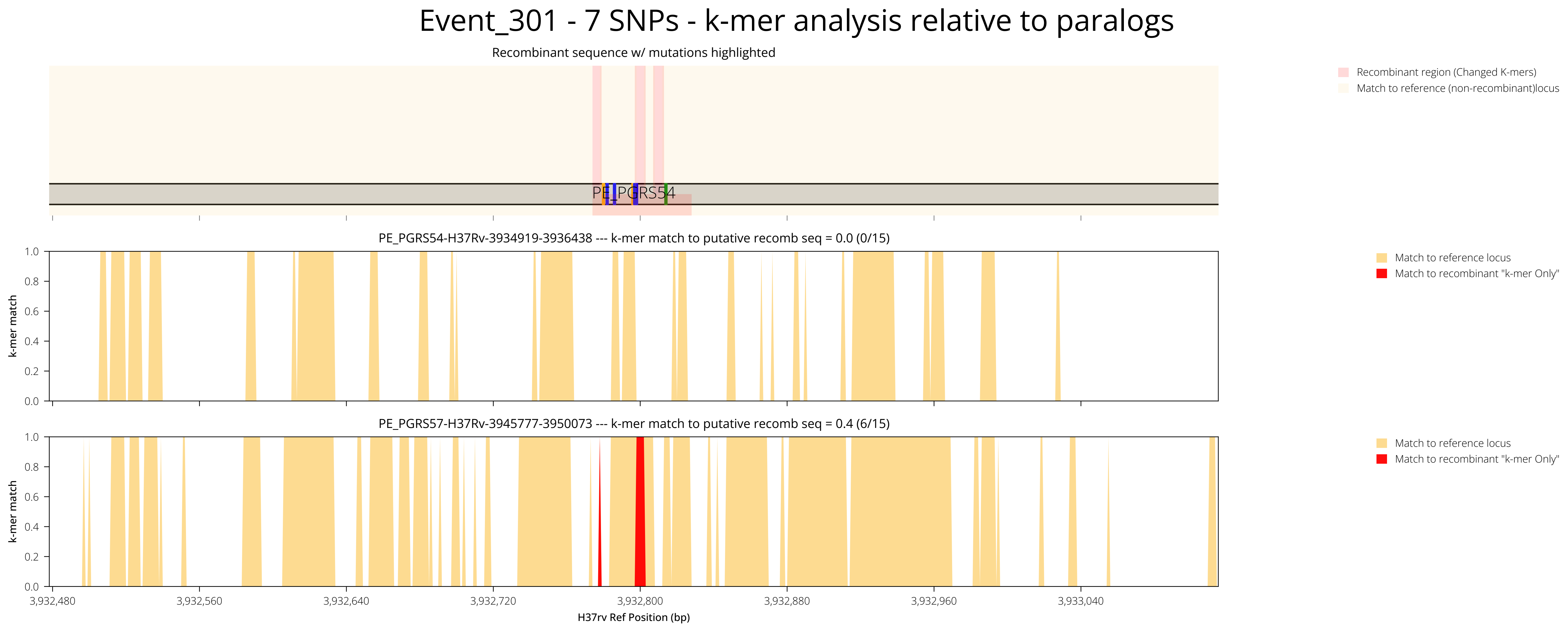

Event_301 - 7 SNPs - k-mer analysis relative to paralogs
Recombinant sequence w/ mutations highlighted
PE_PGRS54
Recombinant region (Changed K-mers)
Match to reference (non-recombinant)locus
PE_PGRS54-H37Rv-3934919-3936438 --- k-mer match to putative recomb seq = 0.0 (0/15)
PE_PGRS57-H37Rv-3945777-3950073 --- k-mer match to putative recomb seq = 0.4 (6/15)
k-mer match
Match to reference locus
Match to recombinant "k-mer Only"
1.0
0.8
0.6
0.4
0.2
0.0
3,932,480
3,932,560
3,932,640
3,932,720
3,932,800
3,932,880
3,932,960
3,933,040
H37rv Ref Position (bp)

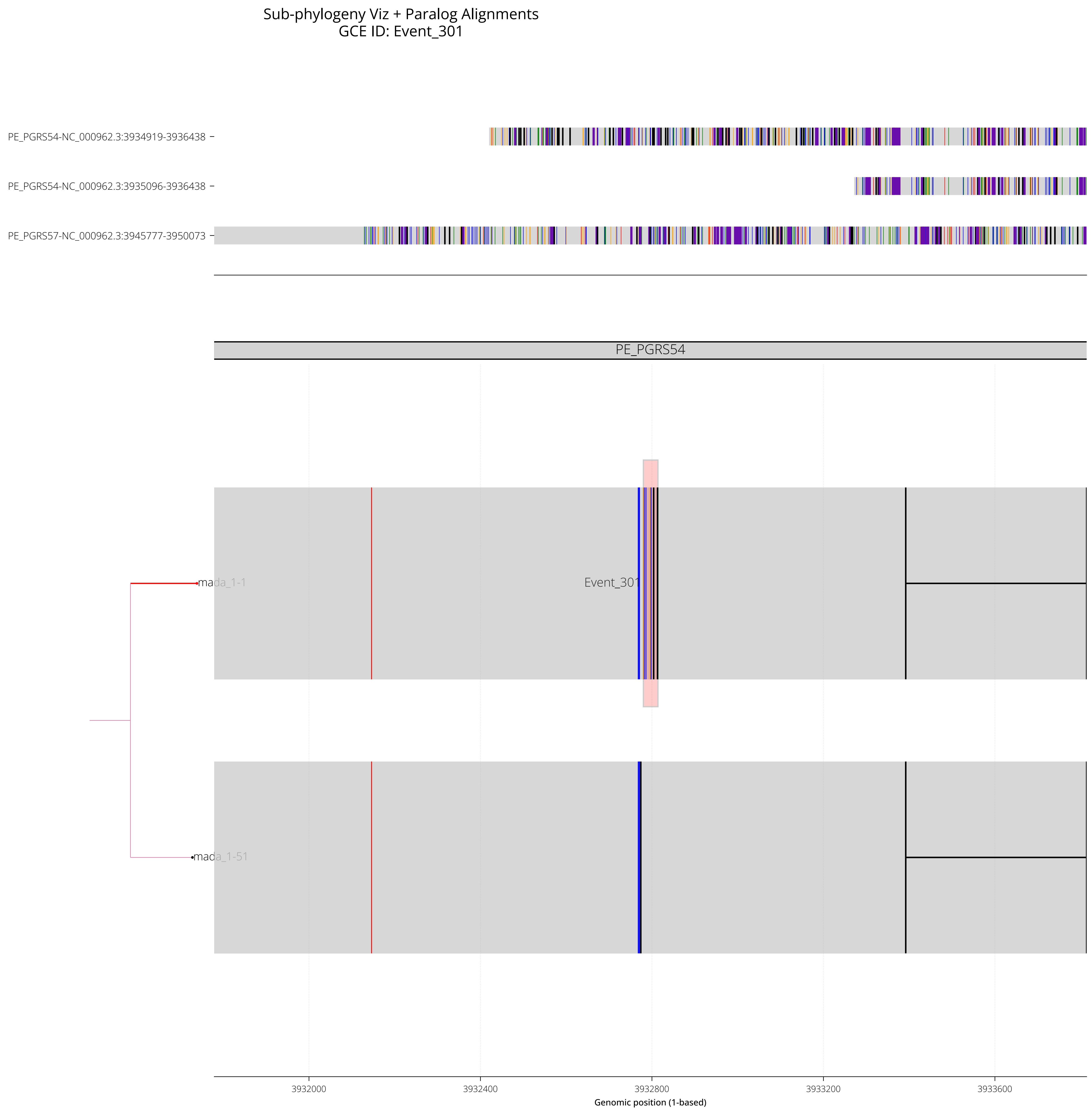

Sub-phylogeny Viz + Paralog Alignments
GCE ID: Event_301
PE_PGRS54-NC_000962.3:3934919-3936438
PE_PGRS54-NC_000962.3:3935096-3936438
PE_PGRS57-NC_000962.3:3945777-3950073
PE_PGRS54
mada_1-1
Event_301
mada_1-51
3932000
3932400
3932800
3933200
3933600
Genomic position (1-based)

Full Phylogeny Viz + Paralog Alignments
GCE ID: Event_301

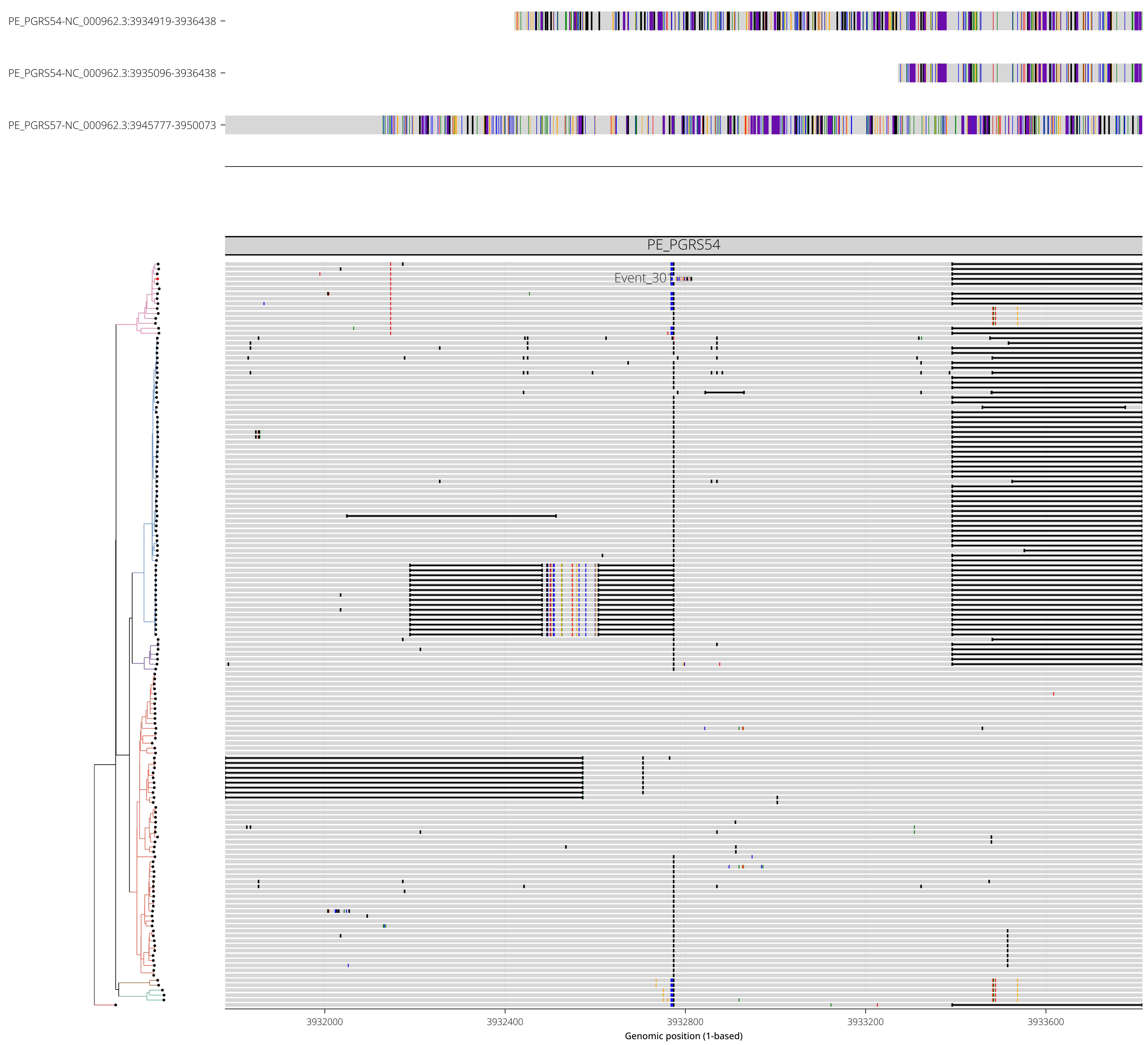

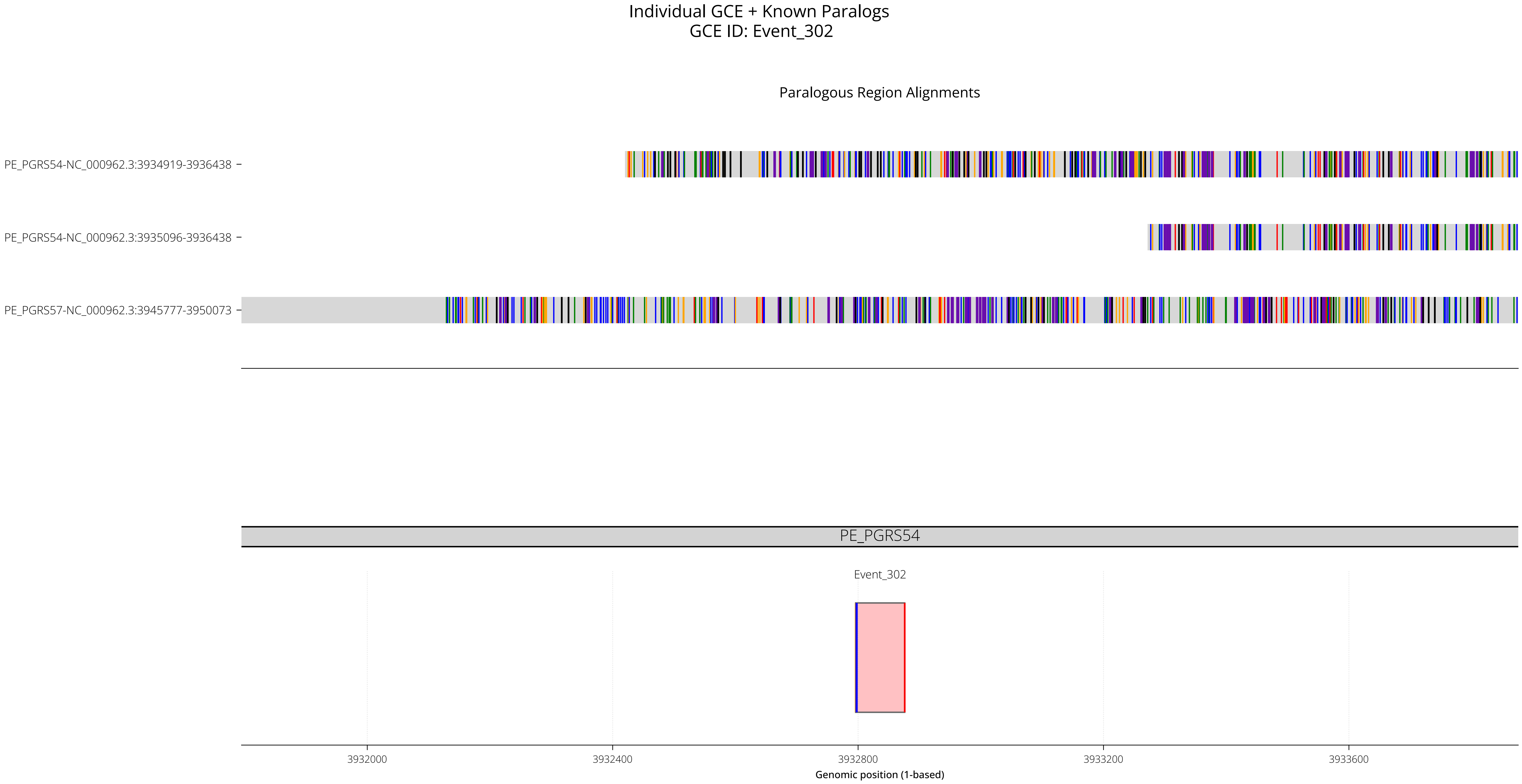

Individual GCE + Known Paralogs
GCE ID: Event_302
Paralogous Region Alignments
PE_PGRS54-NC_000962.3:3934919-3936438
PE_PGRS54-NC_000962.3:3935096-3936438
PE_PGRS57-NC_000962.3:3945777-3950073
PE_PGRS54
Event_302
3932000
3932400
3932800
3933200
3933600
Genomic position (1-based)

Circos Viz - GCE to Known Paralogs Locations
GCE ID: Event_302

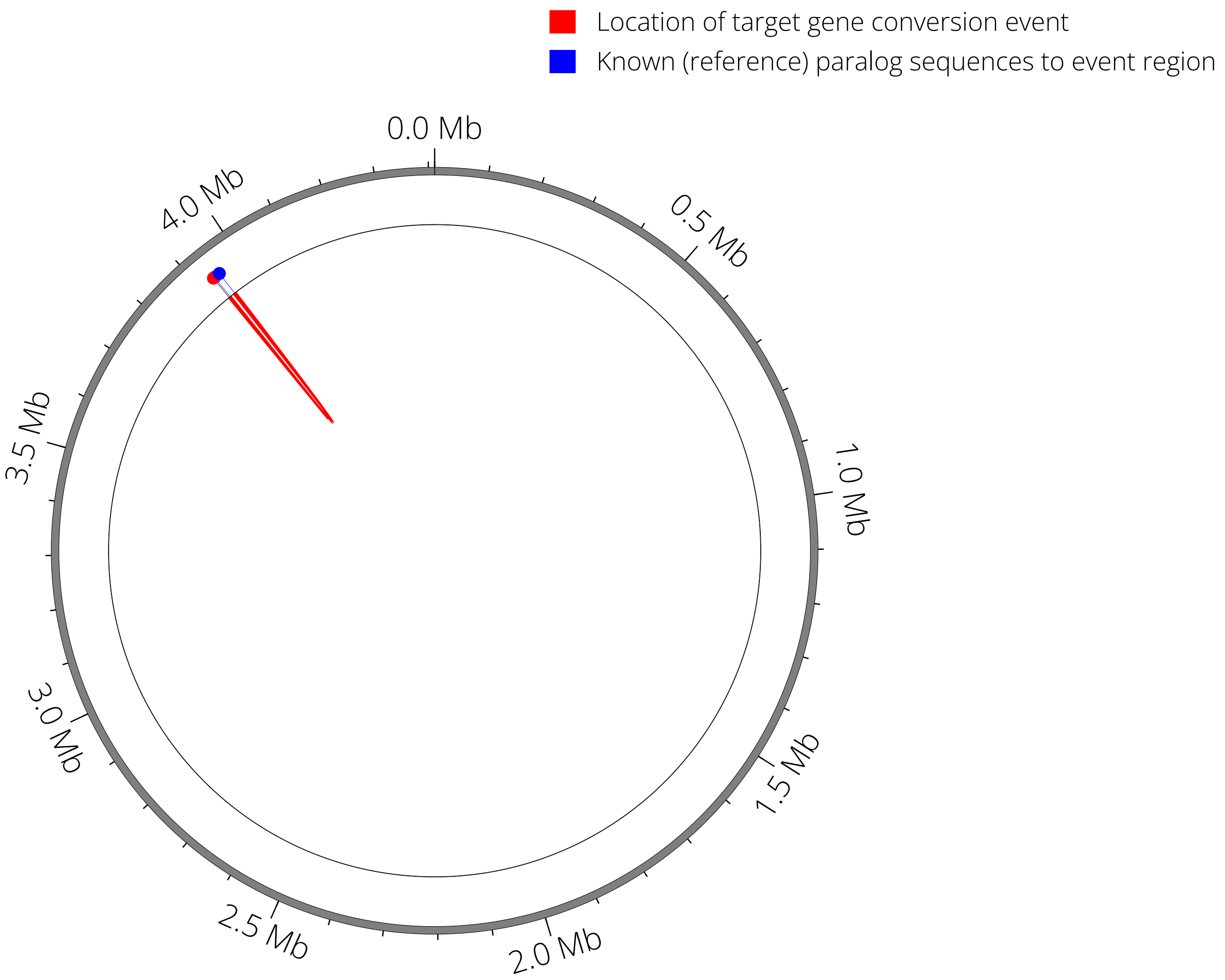

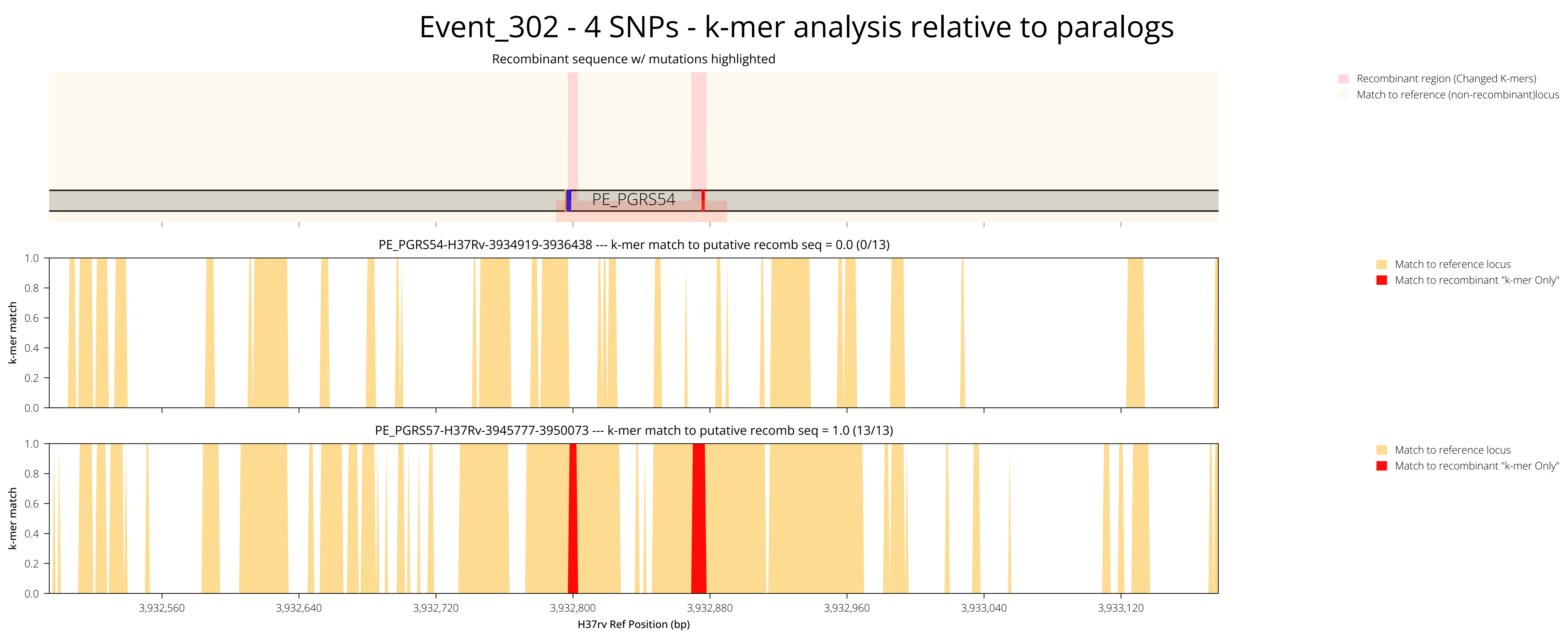
Event_302 - 4 SNPs - k-mer analysis relative to paralogs
Recombinant sequence w/ mutations highlighted
PE_PGRS54
Recombinant region (Changed K-mers)
Match to reference (non-recombinant)locus
PE_PGRS54-H37Rv-3934919-3936438 --- k-mer match to putative recomb seq = 0.0 (0/13)
k-mer match
1.0
0.8
0.6
0.4
0.2
0.0
Match to reference locus
Match to recombinant "k-mer Only"
PE_PGRS57-H37Rv-3945777-3950073 --- k-mer match to putative recomb seq = 1.0 (13/13)
k-mer match
1.0
0.8
0.6
0.4
0.2
0.0
Match to reference locus
Match to recombinant "k-mer Only"
3,932,560
3,932,640
3,932,720
3,932,800
3,932,880
3,932,960
3,933,040
3,933,120
H37rv Ref Position (bp)

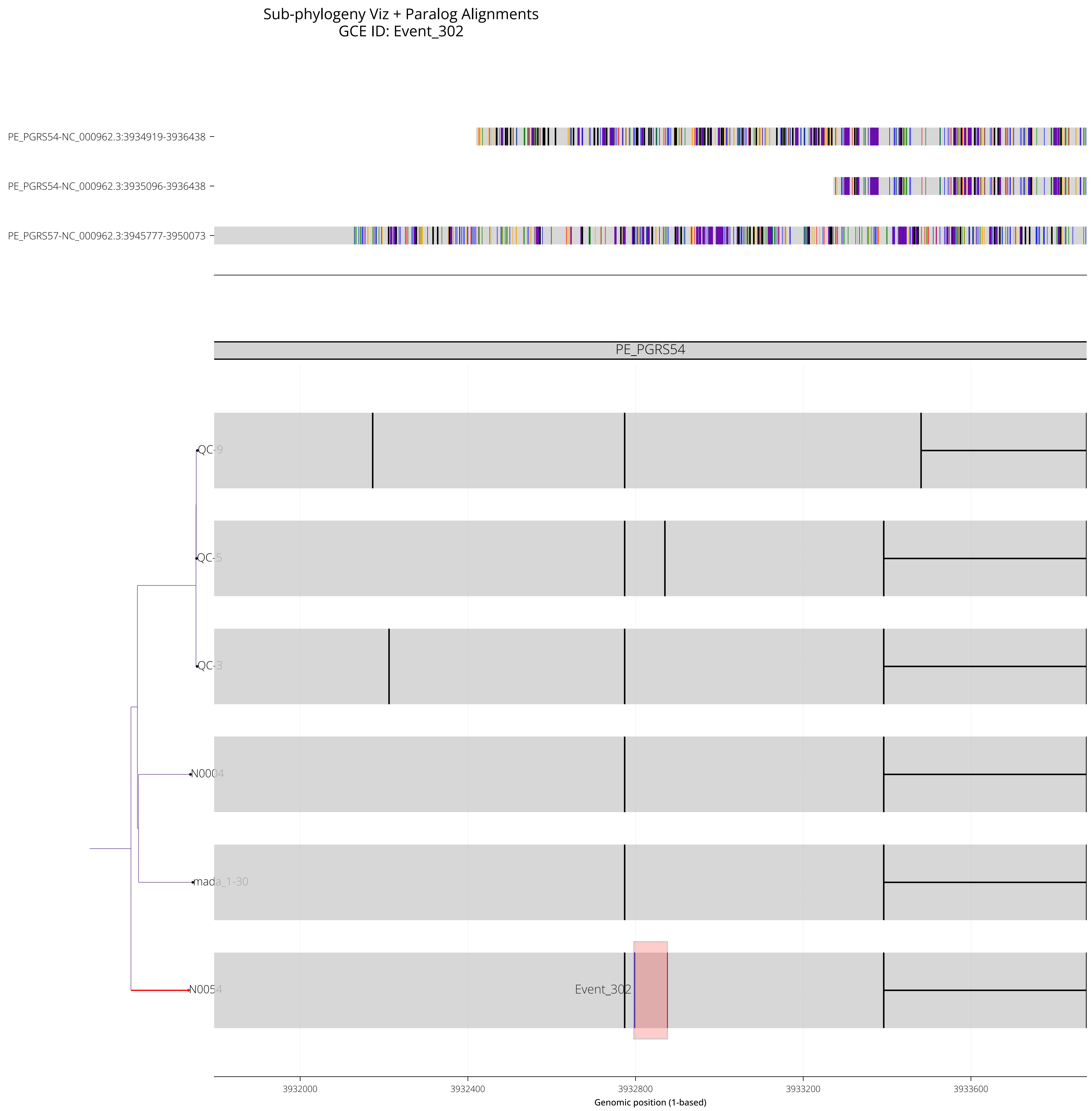

Sub-phylogeny Viz + Paralog Alignments
GCE ID: Event_302
PE_PGRS54-NC_000962.3:3934919-3936438
PE_PGRS54-NC_000962.3:3935096-3936438
PE_PGRS57-NC_000962.3:3945777-3950073
PE_PGRS54
QC-9
QC-5
QC-3
N0004
mada_1-30
N0054
Event_302
3932000
3932400
3932800
3933200
3933600
Genomic position (1-based)

Full Phylogeny Viz + Paralog Alignments
GCE ID: Event_302

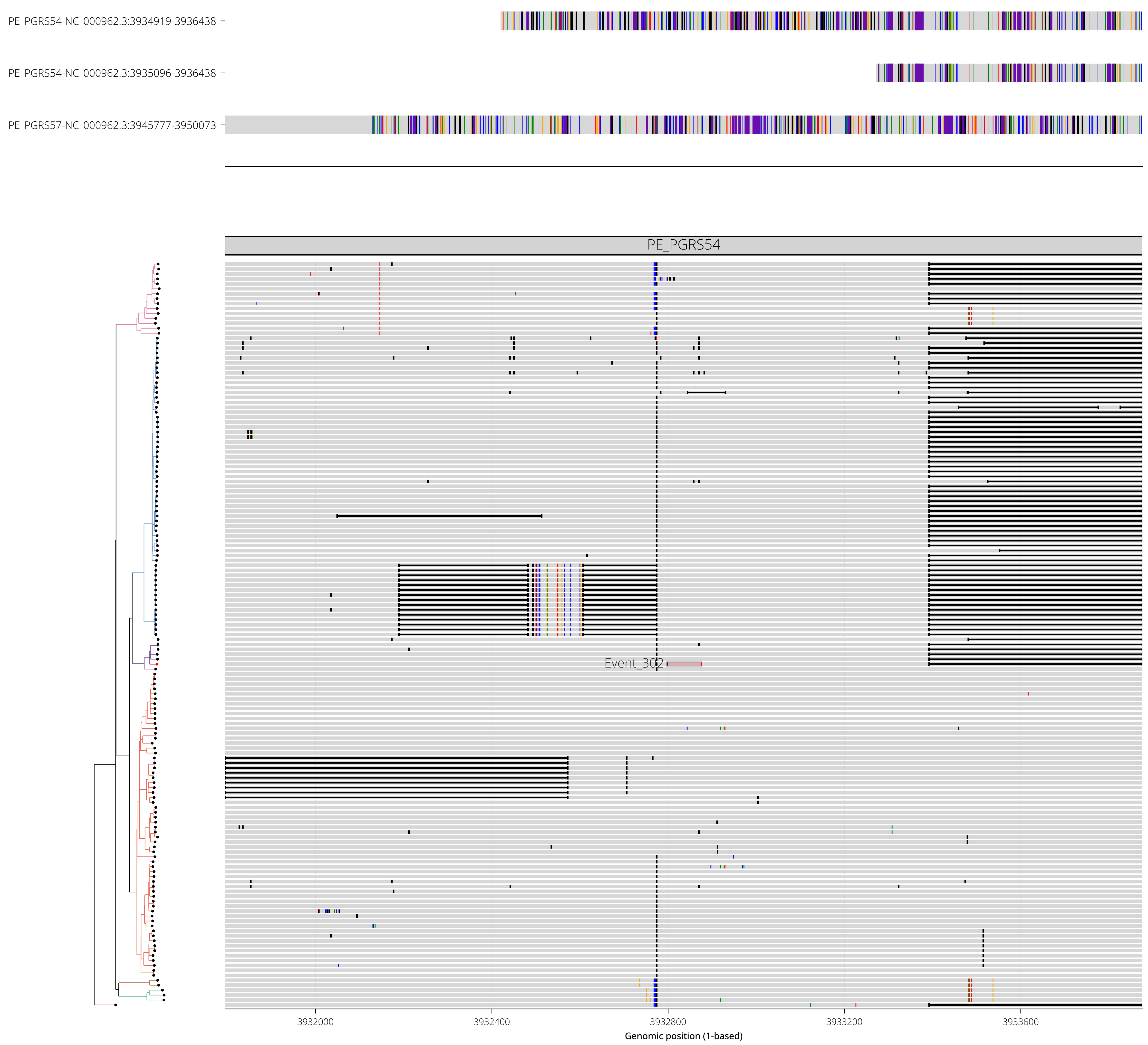

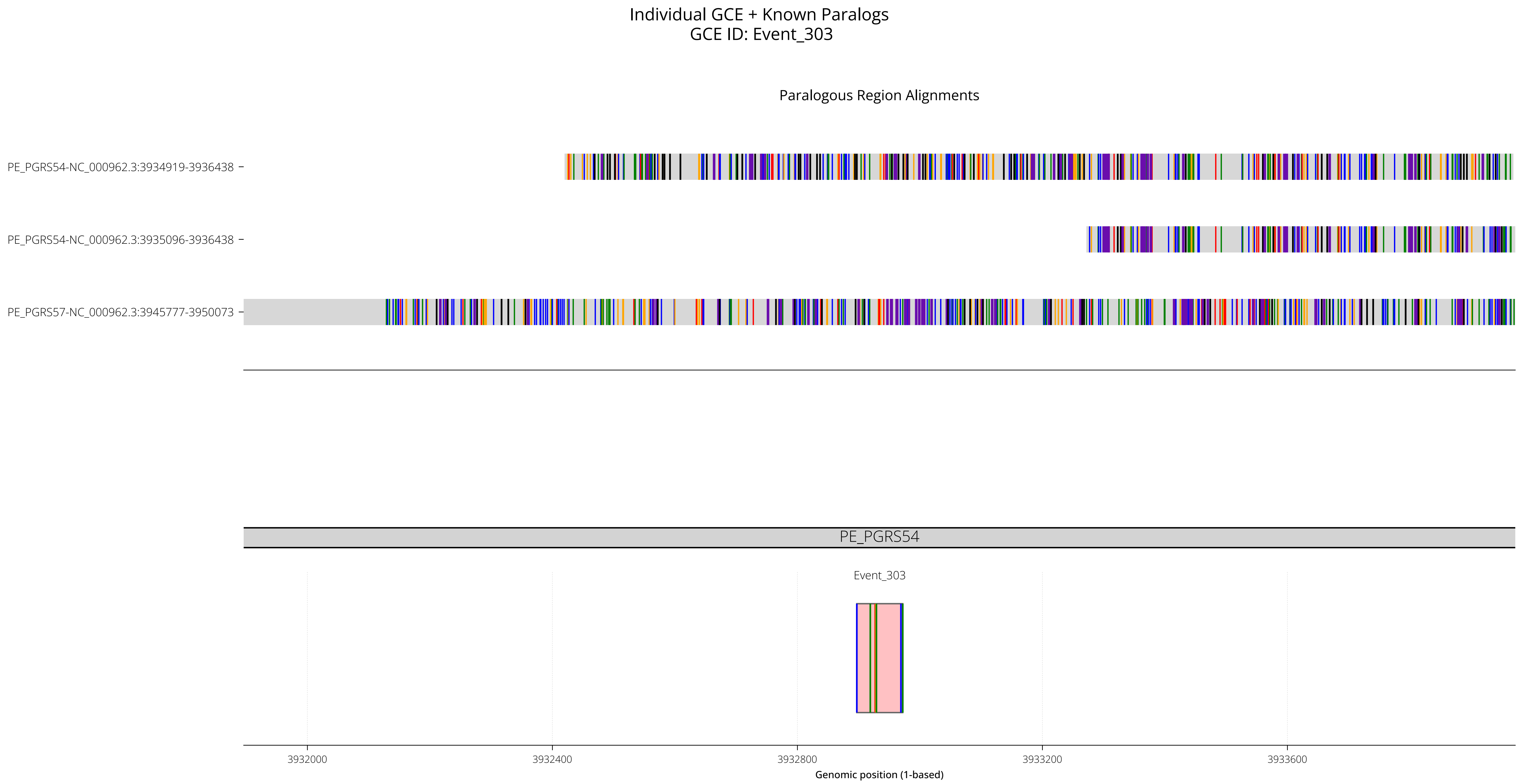
Individual GCE + Known Paralogs
GCE ID: Event_303
Paralogous Region Alignments
PE_PGRS54-NC_000962.3:3934919-3936438
PE_PGRS54-NC_000962.3:3935096-3936438
PE_PGRS57-NC_000962.3:3945777-3950073
PE_PGRS54
Event_303
3932000
3932400
3932800
3933200
3933600
Genomic position (1-based)

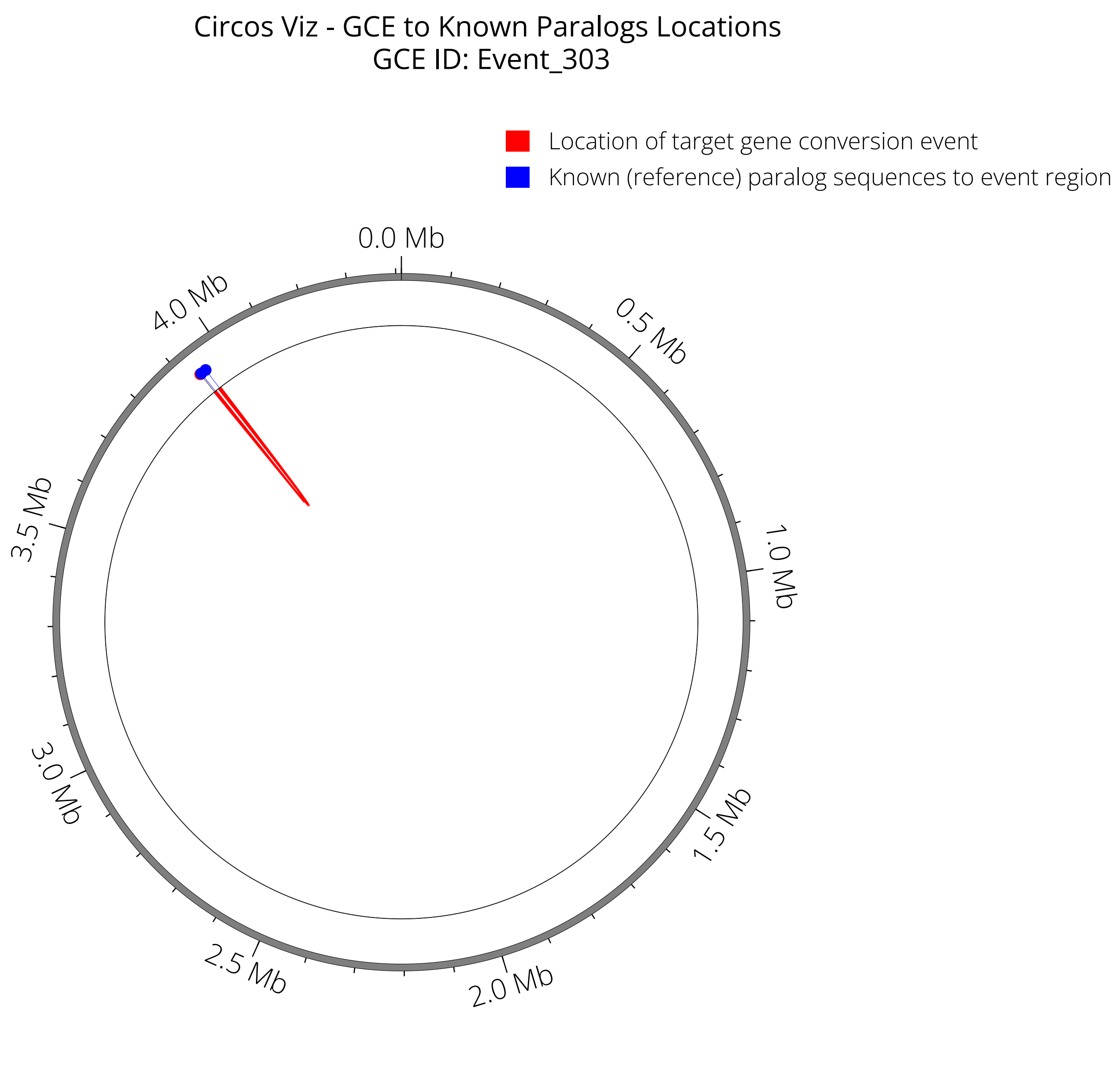
Circos Viz - GCE to Known Paralogs Locations
GCE ID: Event_303
Location of target gene conversion event
Known (reference) paralog sequences to event region
0.0 Mb
0.5 Mb
1.0 Mb
1.5 Mb
2.0 Mb
2.5 Mb
3.0 Mb
3.5 Mb
4.0 Mb

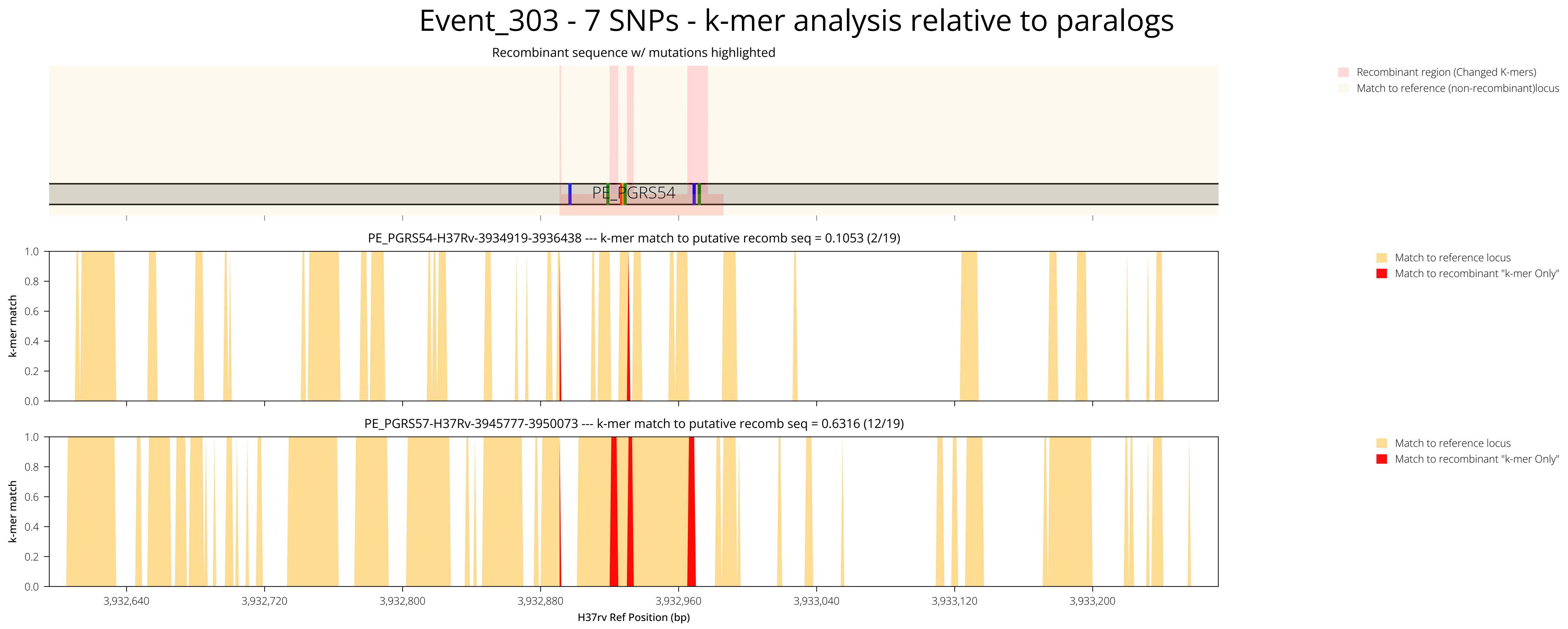

Event_303 - 7 SNPs - k-mer analysis relative to paralogs
Recombinant sequence w/ mutations highlighted
PE_PGRS54
Recombinant region (Changed K-mers)
Match to reference (non-recombinant)locus
PE_PGRS54-H37Rv-3934919-3936438 --- k-mer match to putative recomb seq = 0.1053 (2/19)
Match to reference locus
Match to recombinant "k-mer Only"
k-mer match
1.0
0.8
0.6
0.4
0.2
0.0
PE_PGRS57-H37Rv-3945777-3950073 --- k-mer match to putative recomb seq = 0.6316 (12/19)
3,932,640
3,932,720
3,932,800
3,932,880
3,932,960
3,933,040
3,933,120
3,933,200
H37rv Ref Position (bp)

Sub-phylogeny Viz + Paralog Alignments
GCE ID: Event_303

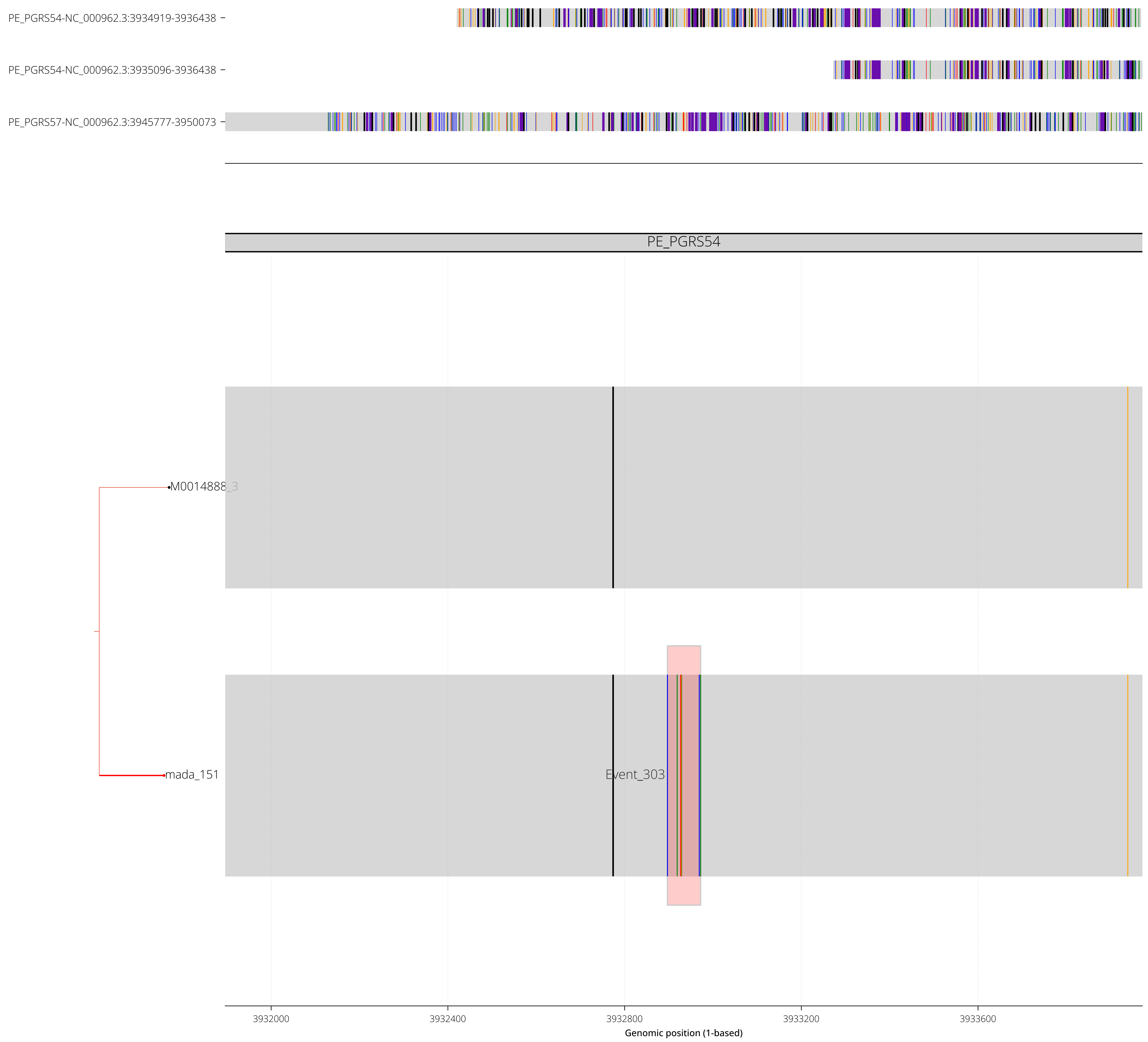

Full Phylogeny Viz + Paralog Alignments
GCE ID: Event_303

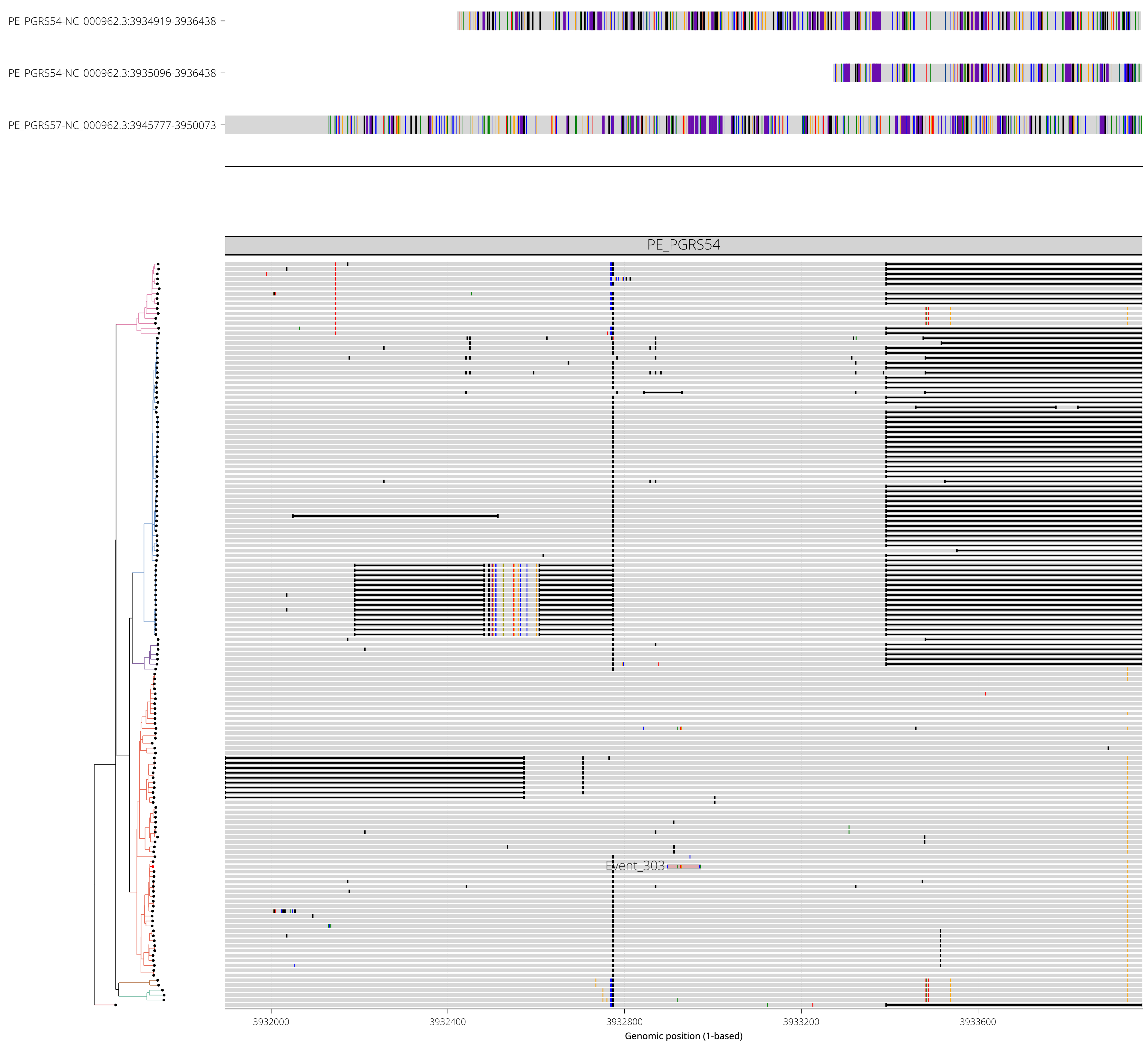

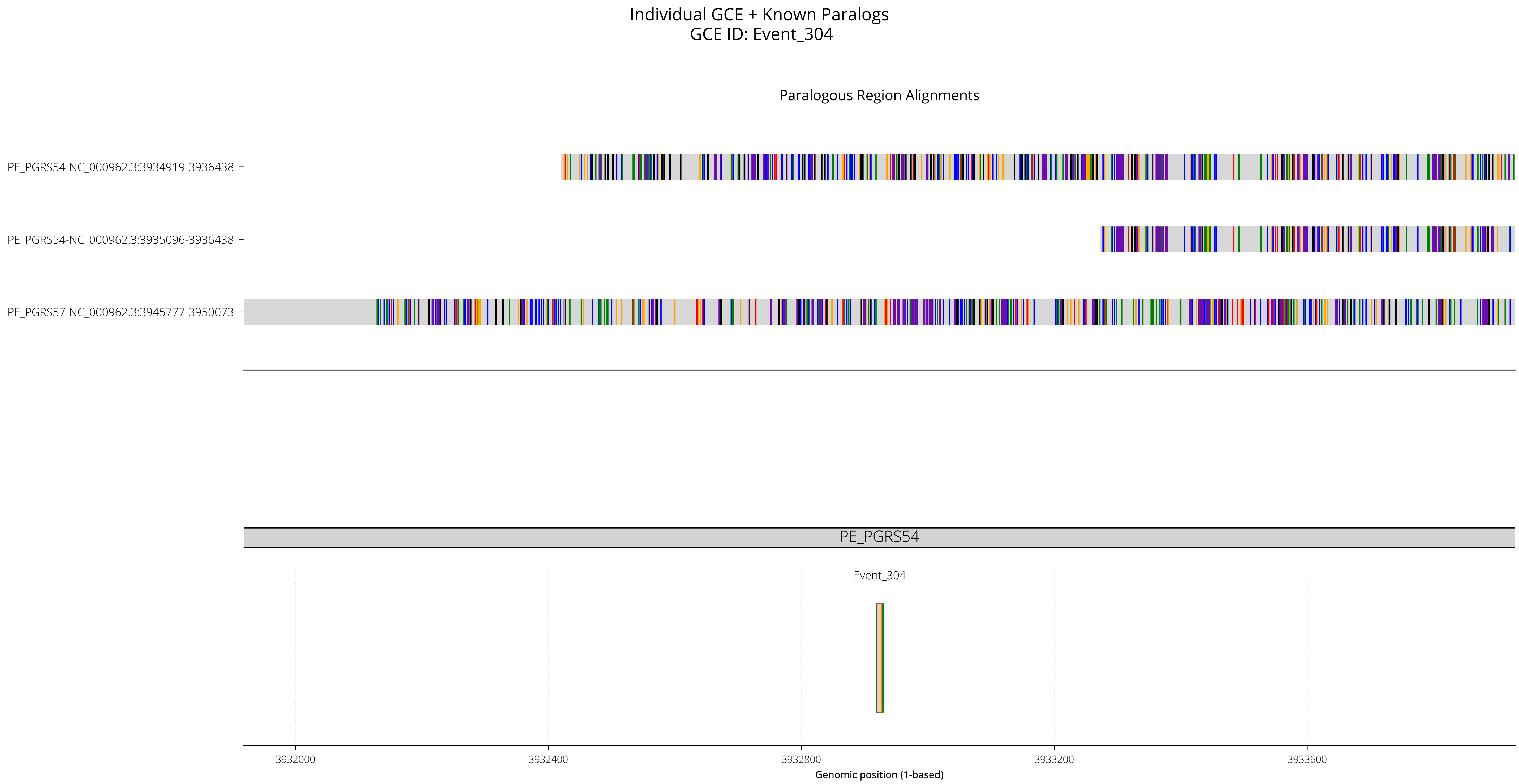

Individual GCE + Known Paralogs
GCE ID: Event_304
Paralogous Region Alignments
PE_PGRS54-NC_000962.3:3934919-3936438
PE_PGRS54-NC_000962.3:3935096-3936438
PE_PGRS57-NC_000962.3:3945777-3950073
PE_PGRS54
Event_304
3932000
3932400
3932800
3933200
3933600
Genomic position (1-based)

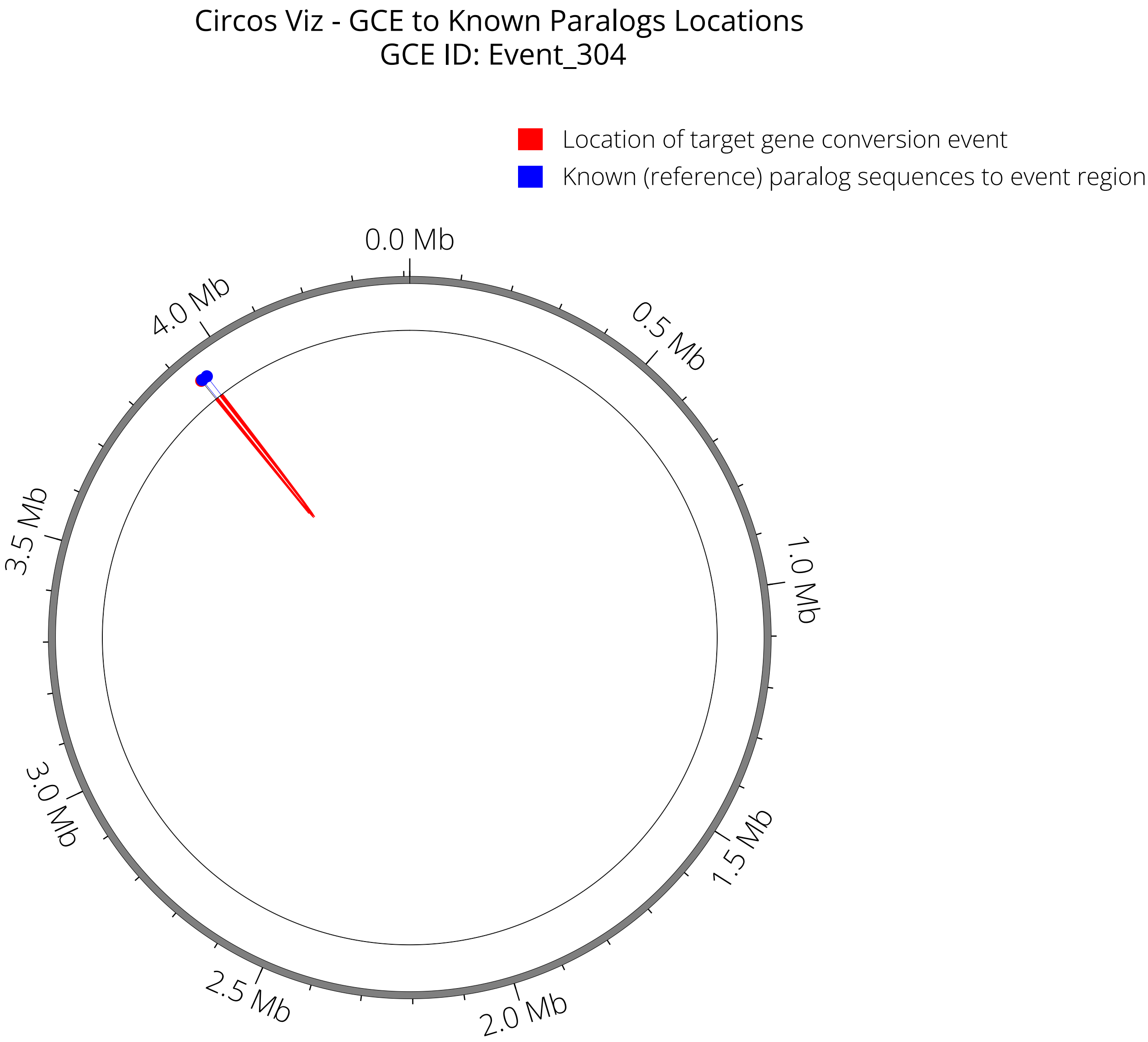
Circos Viz - GCE to Known Paralogs Locations
GCE ID: Event_304
Location of target gene conversion event
Known (reference) paralog sequences to event region
0.0 Mb
0.5 Mb
1.0 Mb
1.5 Mb
2.0 Mb
2.5 Mb
3.0 Mb
3.5 Mb
4.0 Mb

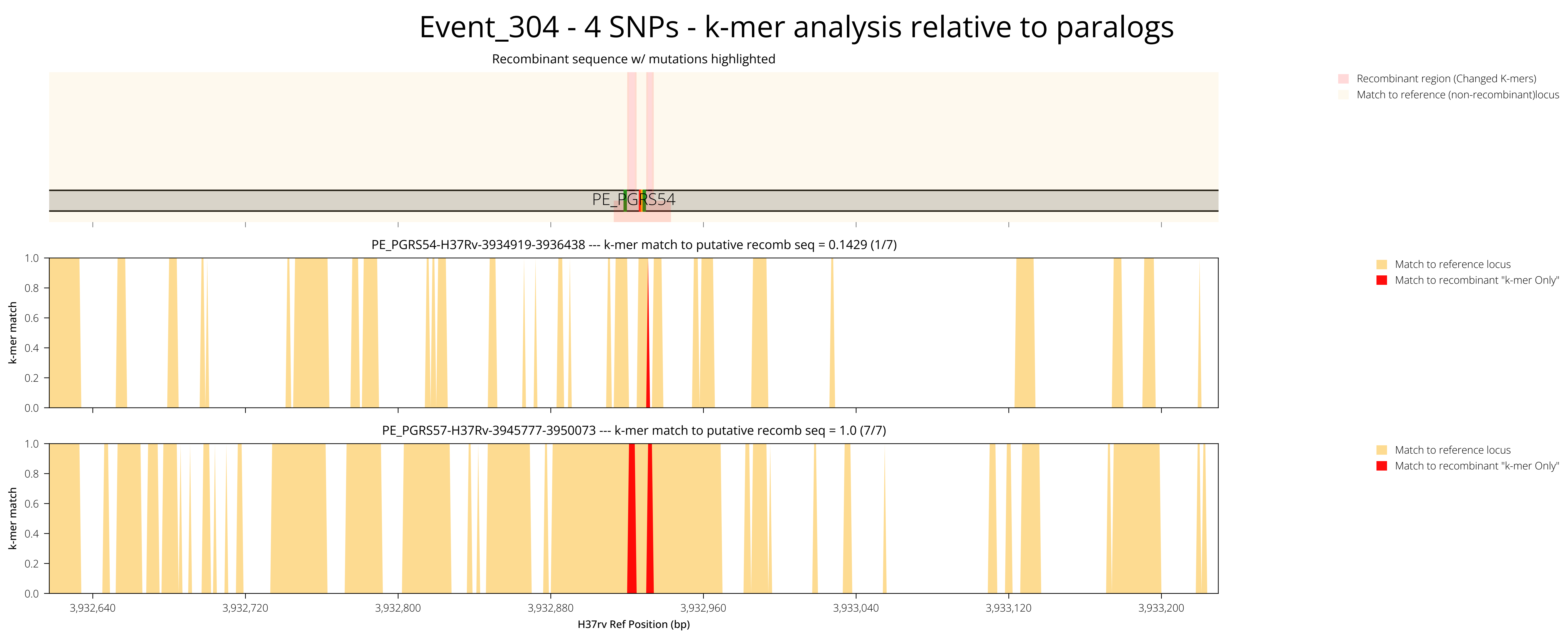

Event_304 - 4 SNPs - k-mer analysis relative to paralogs
Recombinant sequence w/ mutations highlighted
Recombinant region (Changed K-mers)
Match to reference (non-recombinant)locus
PE_PGRS54
PE_PGRS54-H37Rv-3934919-3936438 --- k-mer match to putative recomb seq = 0.1429 (1/7)
Match to reference locus
Match to recombinant "k-mer Only"
k-mer match
1.0
0.8
0.6
0.4
0.2
0.0
PE_PGRS57-H37Rv-3945777-3950073 --- k-mer match to putative recomb seq = 1.0 (7/7)
Match to reference locus
Match to recombinant "k-mer Only"
k-mer match
1.0
0.8
0.6
0.4
0.2
0.0
3,932,640
3,932,720
3,932,800
3,932,880
3,932,960
3,933,040
3,933,120
3,933,200
H37rv Ref Position (bp)

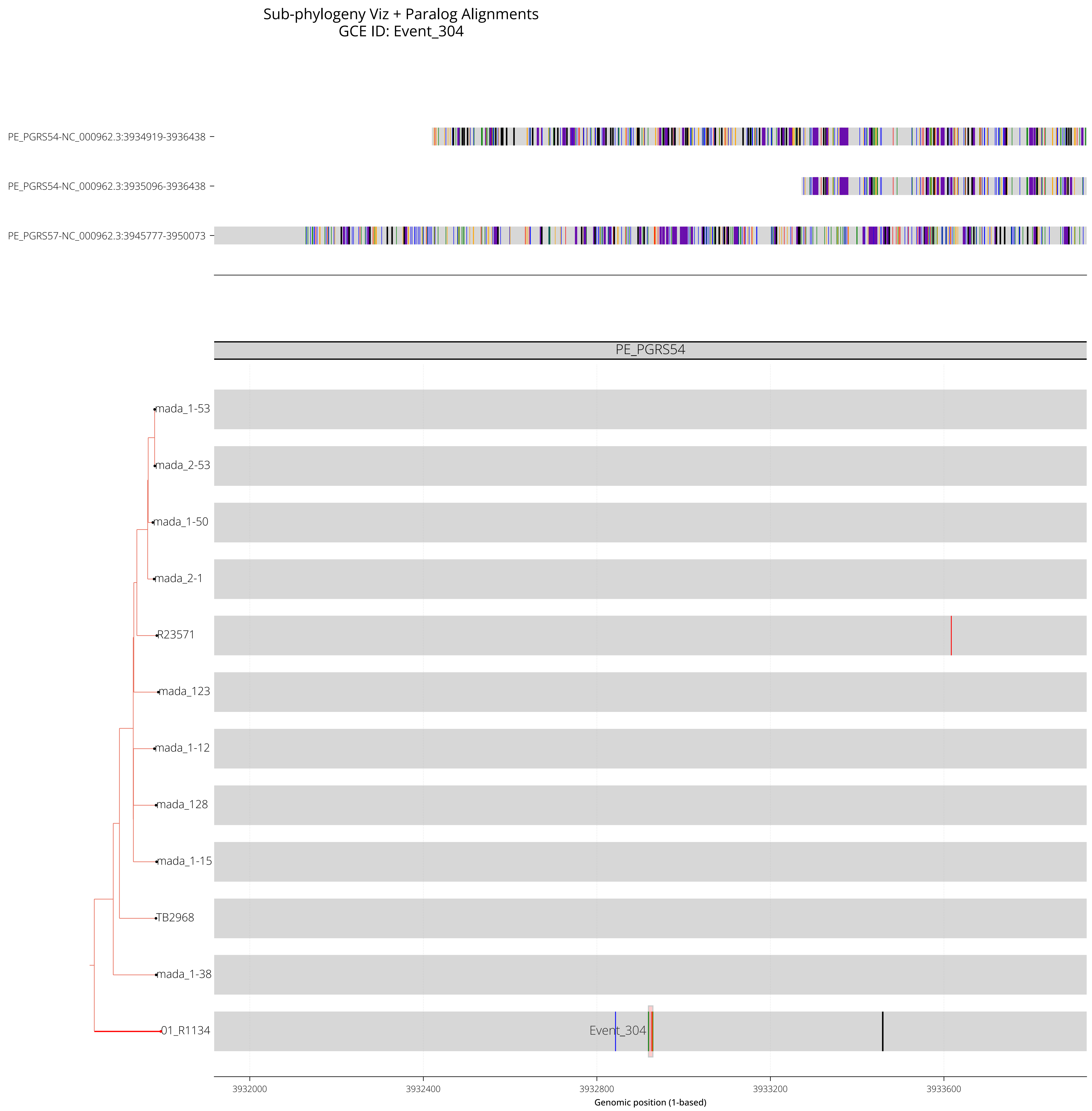

Sub-phylogeny Viz + Paralog Alignments
GCE ID: Event_304
PE_PGRS54-NC_000962.3:3934919-3936438
PE_PGRS54-NC_000962.3:3935096-3936438
PE_PGRS57-NC_000962.3:3945777-3950073
PE_PGRS54
mada_1-53
mada_2-53
mada_1-50
mada_2-1
R23571
mada_123
mada_1-12
mada_128
mada_1-15
TB2968
mada_1-38
01_R1134
Event_304
3932000
3932400
3932800
3933200
3933600
Genomic position (1-based)

Full Phylogeny Viz + Paralog Alignments
GCE ID: Event_304

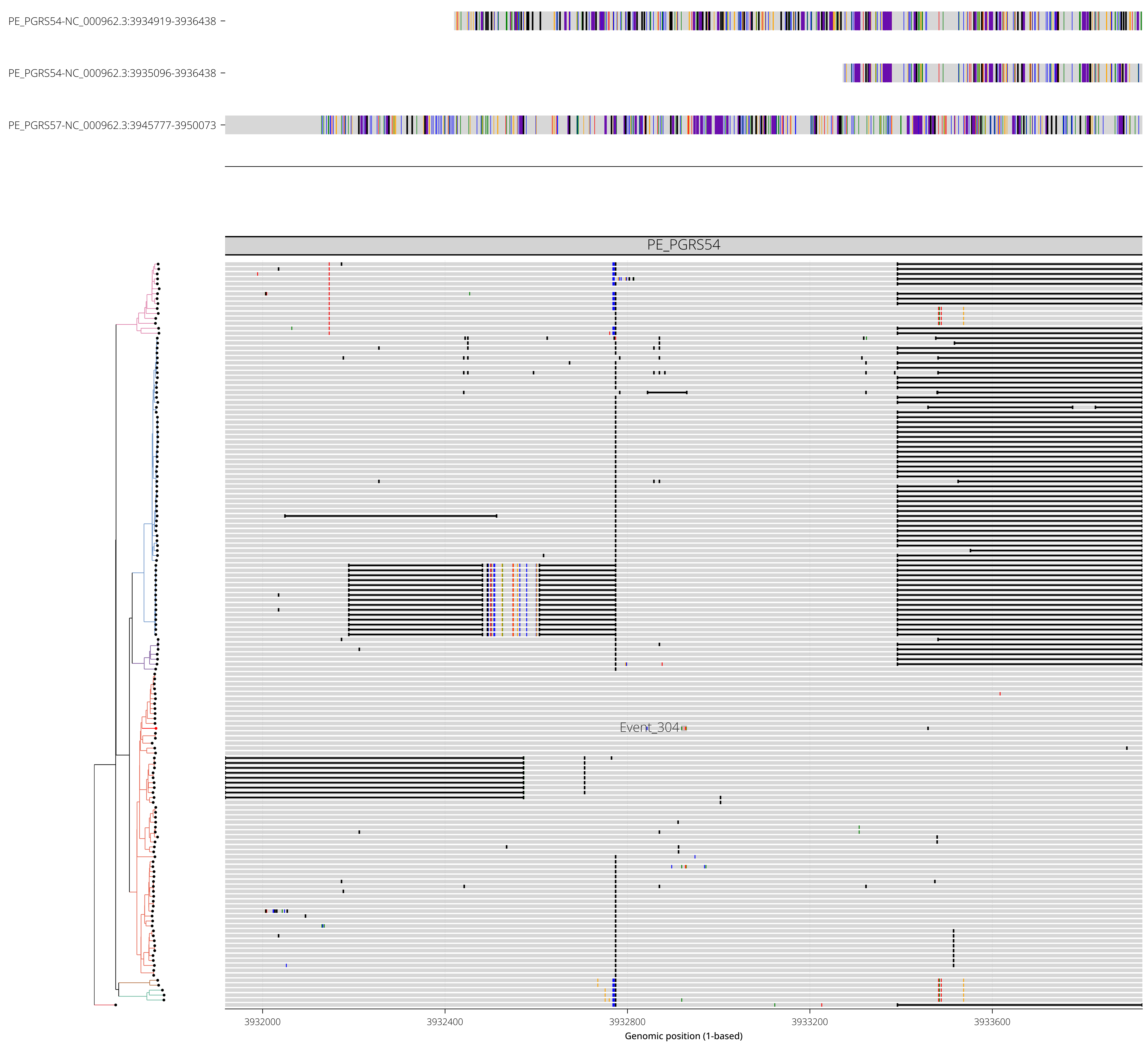

Individual GCE + Known Paralogs
GCE ID: Event_305

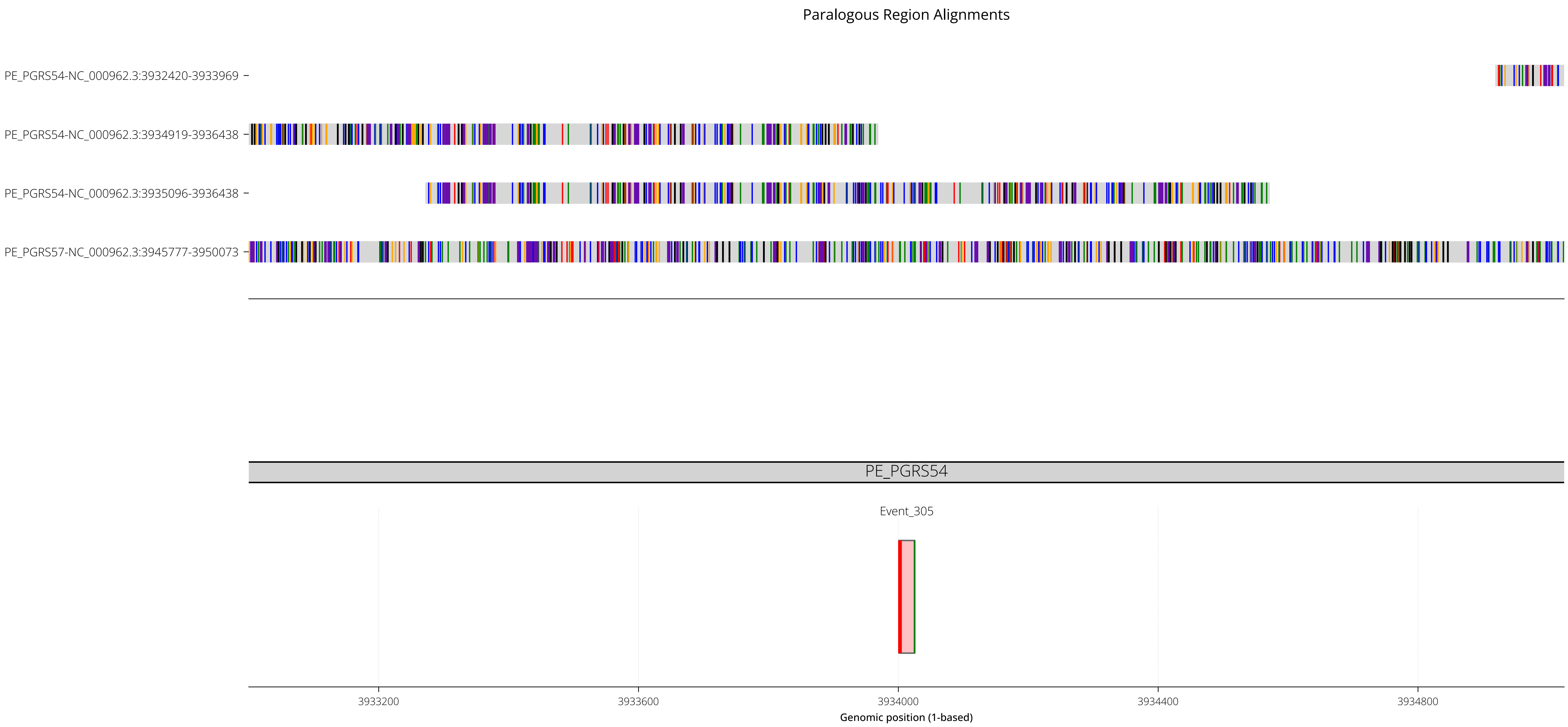

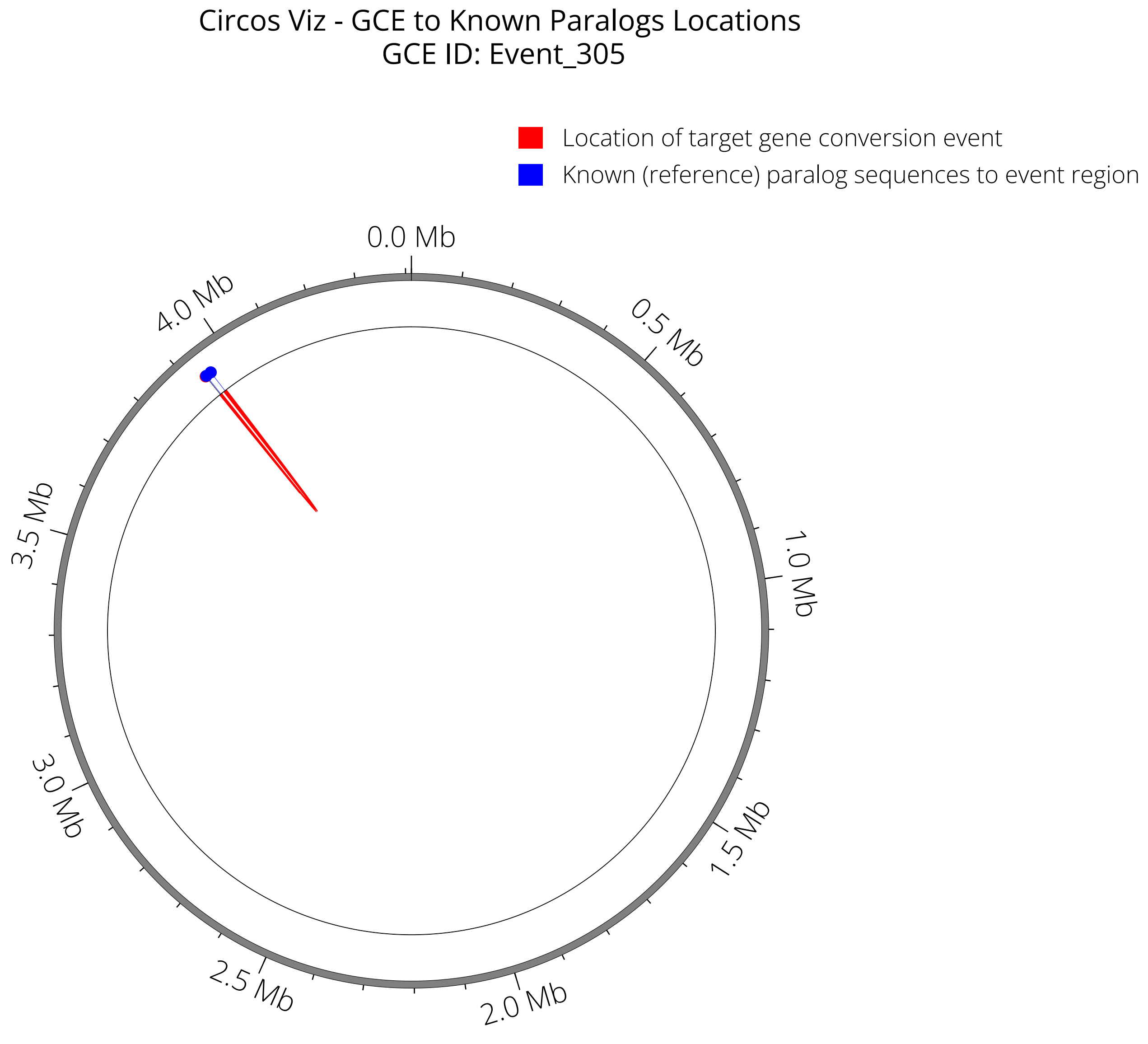
Circos Viz - GCE to Known Paralogs Locations
GCE ID: Event_305
Location of target gene conversion event
Known (reference) paralog sequences to event region
0.0 Mb
0.5 Mb
1.0 Mb
1.5 Mb
2.0 Mb
2.5 Mb
3.0 Mb
3.5 Mb
4.0 Mb

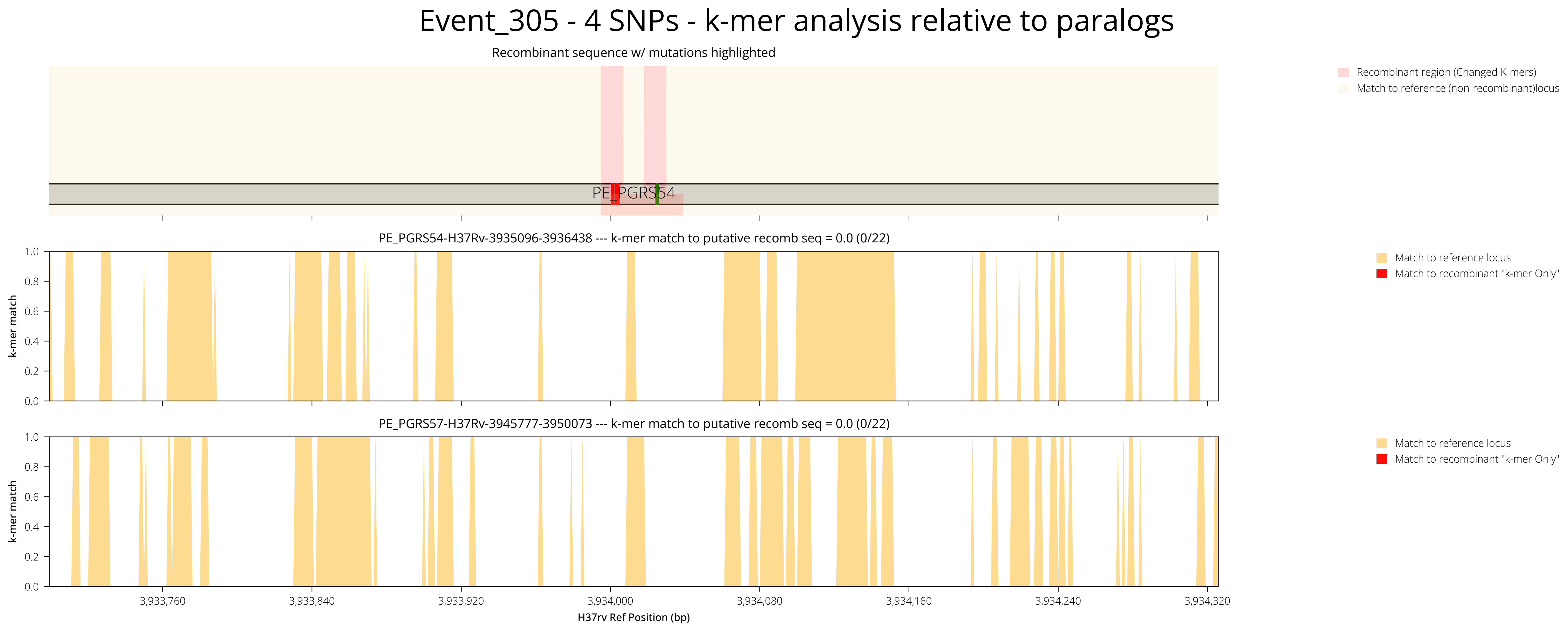

Event_305 - 4 SNPs - k-mer analysis relative to paralogs
Recombinant sequence w/ mutations highlighted
Recombinant region (Changed K-mers)
Match to reference (non-recombinant)locus
PE_PGRS54
PE_PGRS54-H37Rv-3935096-3936438 --- k-mer match to putative recomb seq = 0.0 (0/22)
Match to reference locus
Match to recombinant "k-mer Only"
k-mer match
1.0
0.8
0.6
0.4
0.2
0.0
PE_PGRS57-H37Rv-3945777-3950073 --- k-mer match to putative recomb seq = 0.0 (0/22)
Match to reference locus
Match to recombinant "k-mer Only"
k-mer match
1.0
0.8
0.6
0.4
0.2
0.0
3,933,760
3,933,840
3,933,920
3,934,000
3,934,080
3,934,160
3,934,240
3,934,320
H37rv Ref Position (bp)

Sub-phylogeny Viz + Paralog Alignments
GCE ID: Event_305

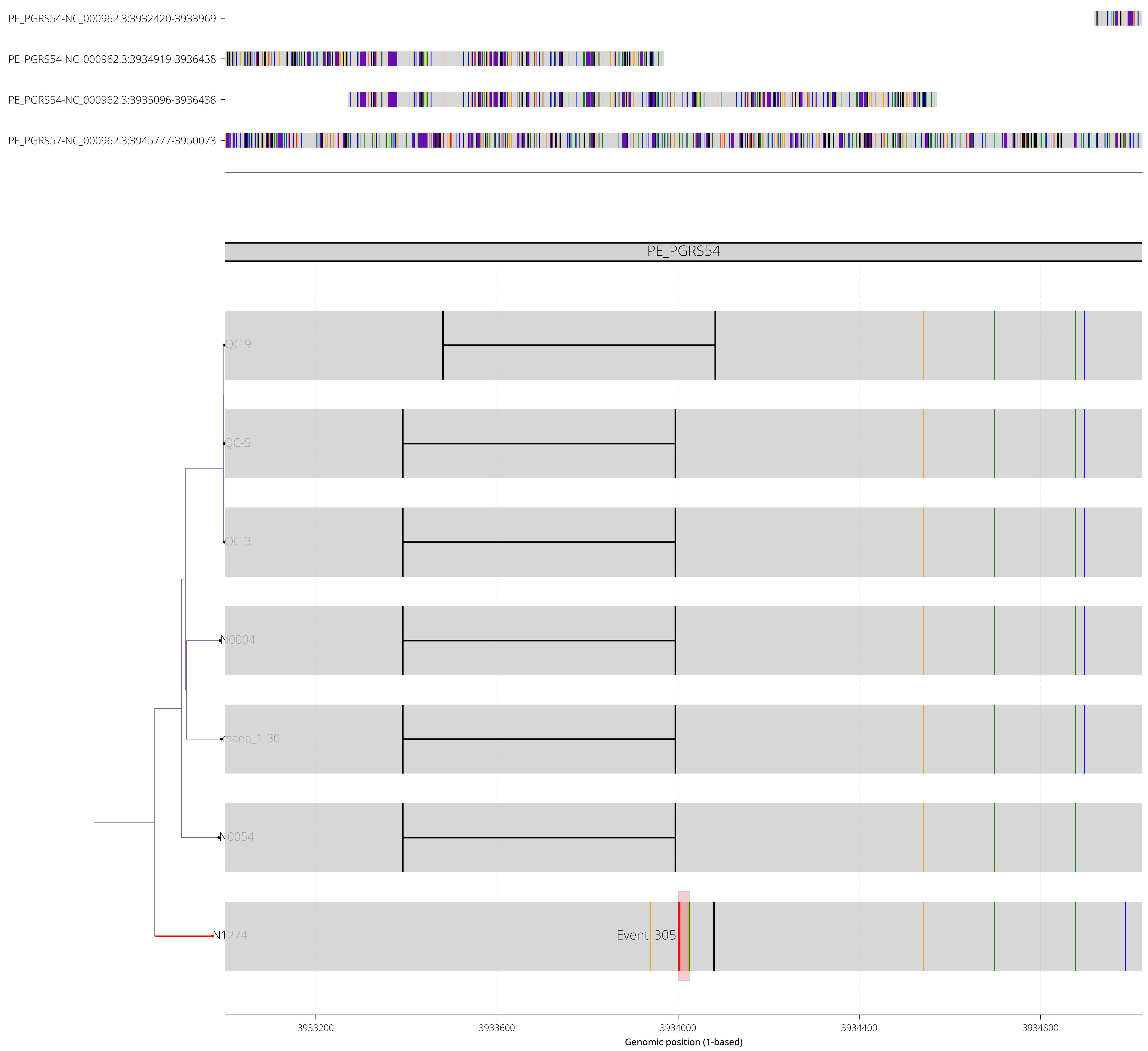

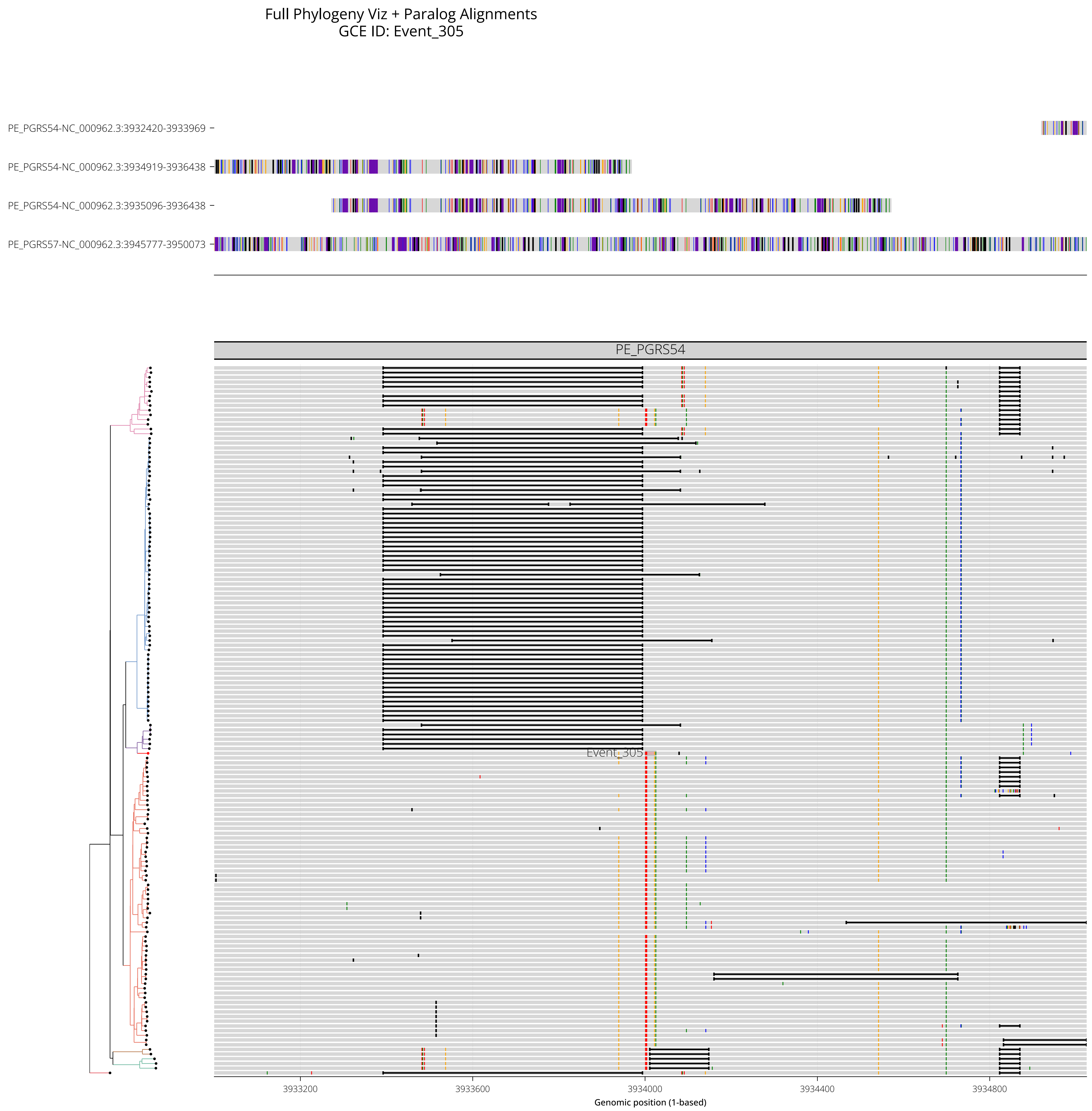

Full Phylogeny Viz + Paralog Alignments
GCE ID: Event_305
PE_PGRS54-NC_000962.3:3932420-3933969
PE_PGRS54-NC_000962.3:3934919-3936438
PE_PGRS54-NC_000962.3:3935096-3936438
PE_PGRS57-NC_000962.3:3945777-3950073
PE_PGRS54
Event_305
3933200
3933600
3934000
3934400
3934800
Genomic position (1-based)

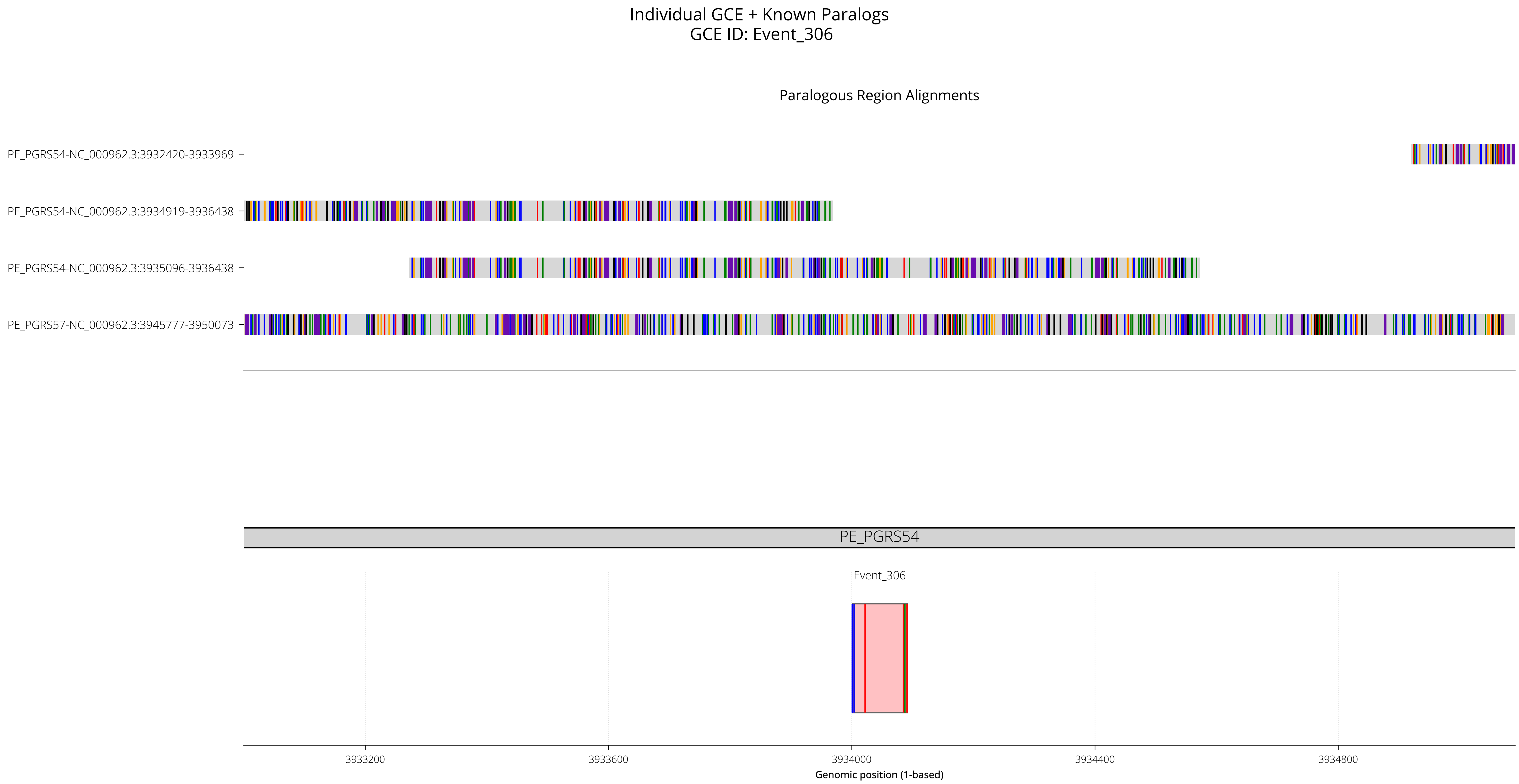
Individual GCE + Known Paralogs
GCE ID: Event_306
Paralogous Region Alignments
PE_PGRS54-NC_000962.3:3932420-3933969
PE_PGRS54-NC_000962.3:3934919-3936438
PE_PGRS54-NC_000962.3:3935096-3936438
PE_PGRS57-NC_000962.3:3945777-3950073
PE_PGRS54
Event_306
3933200
3933600
3934000
3934400
3934800
Genomic position (1-based)

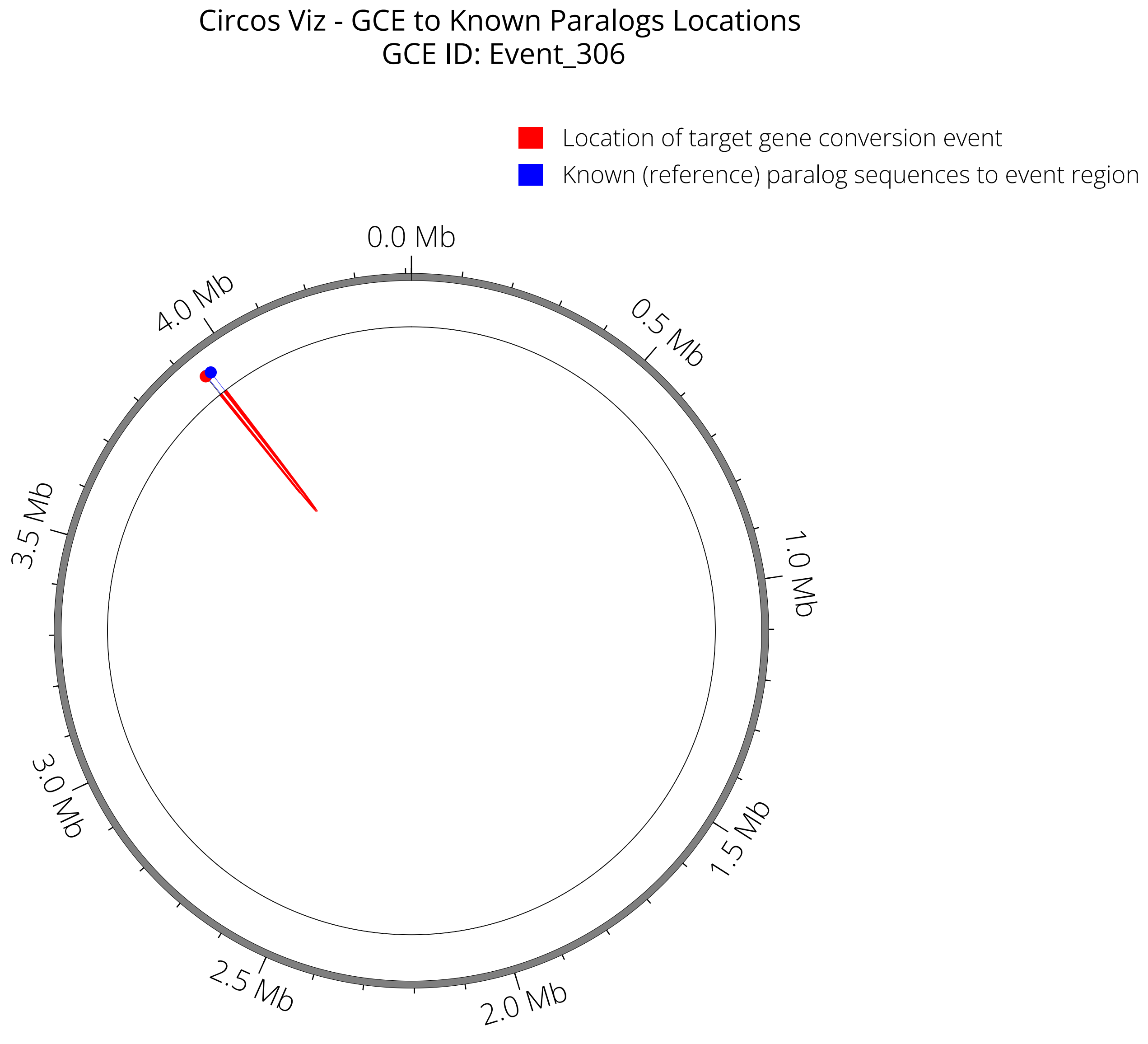

Circos Viz - GCE to Known Paralogs Locations
GCE ID: Event_306
Location of target gene conversion event
Known (reference) paralog sequences to event region
0.0 Mb
0.5 Mb
1.0 Mb
1.5 Mb
2.0 Mb
2.5 Mb
3.0 Mb
3.5 Mb
4.0 Mb

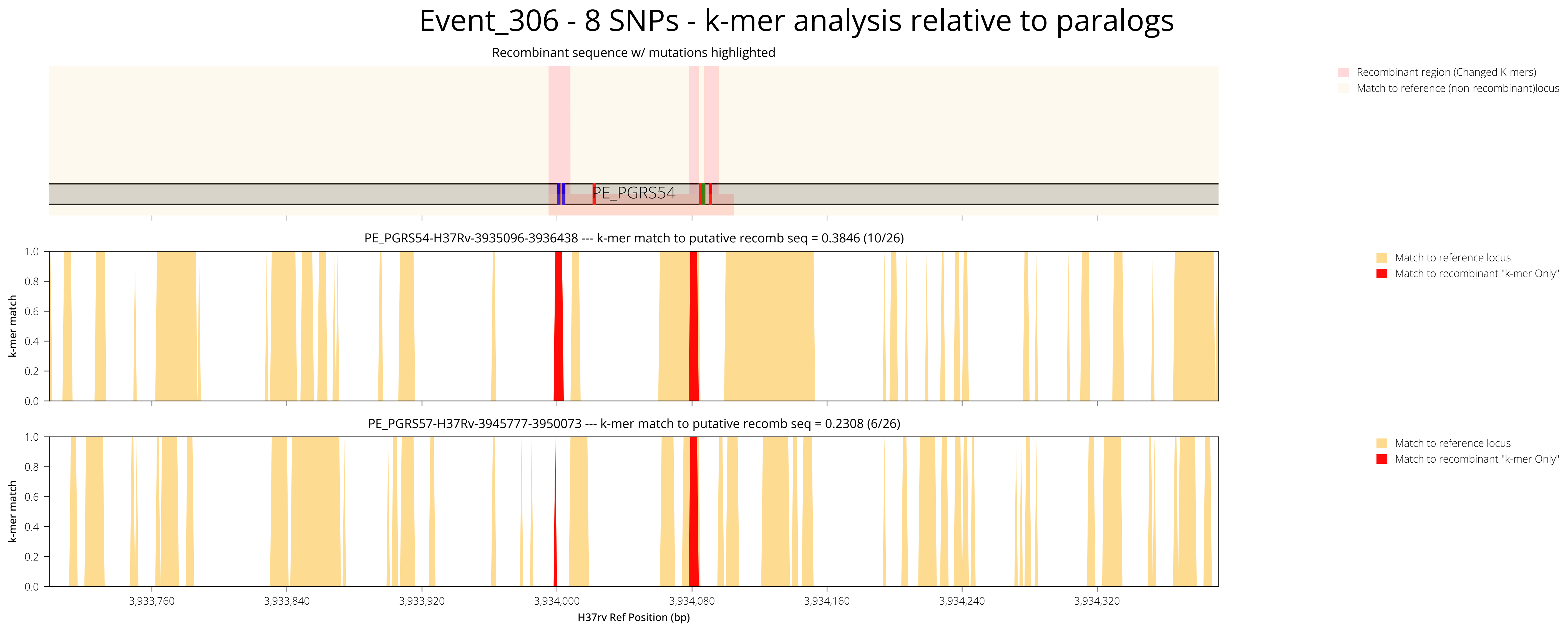
Event_306 - 8 SNPs - k-mer analysis relative to paralogs
Recombinant sequence w/ mutations highlighted
PE_PGRS54
Recombinant region (Changed K-mers)
Match to reference (non-recombinant)locus
PE_PGRS54-H37Rv-3935096-3936438 --- k-mer match to putative recomb seq = 0.3846 (10/26)
k-mer match
1.0
0.8
0.6
0.4
0.2
0.0
Match to reference locus
Match to recombinant "k-mer Only"
PE_PGRS57-H37Rv-3945777-3950073 --- k-mer match to putative recomb seq = 0.2308 (6/26)
k-mer match
Match to reference locus
Match to recombinant "k-mer Only"
3,933,760
3,933,840
3,933,920
3,934,000
3,934,080
3,934,160
3,934,240
3,934,320
H37rv Ref Position (bp)

Sub-phylogeny Viz + Paralog Alignments
GCE ID: Event_306

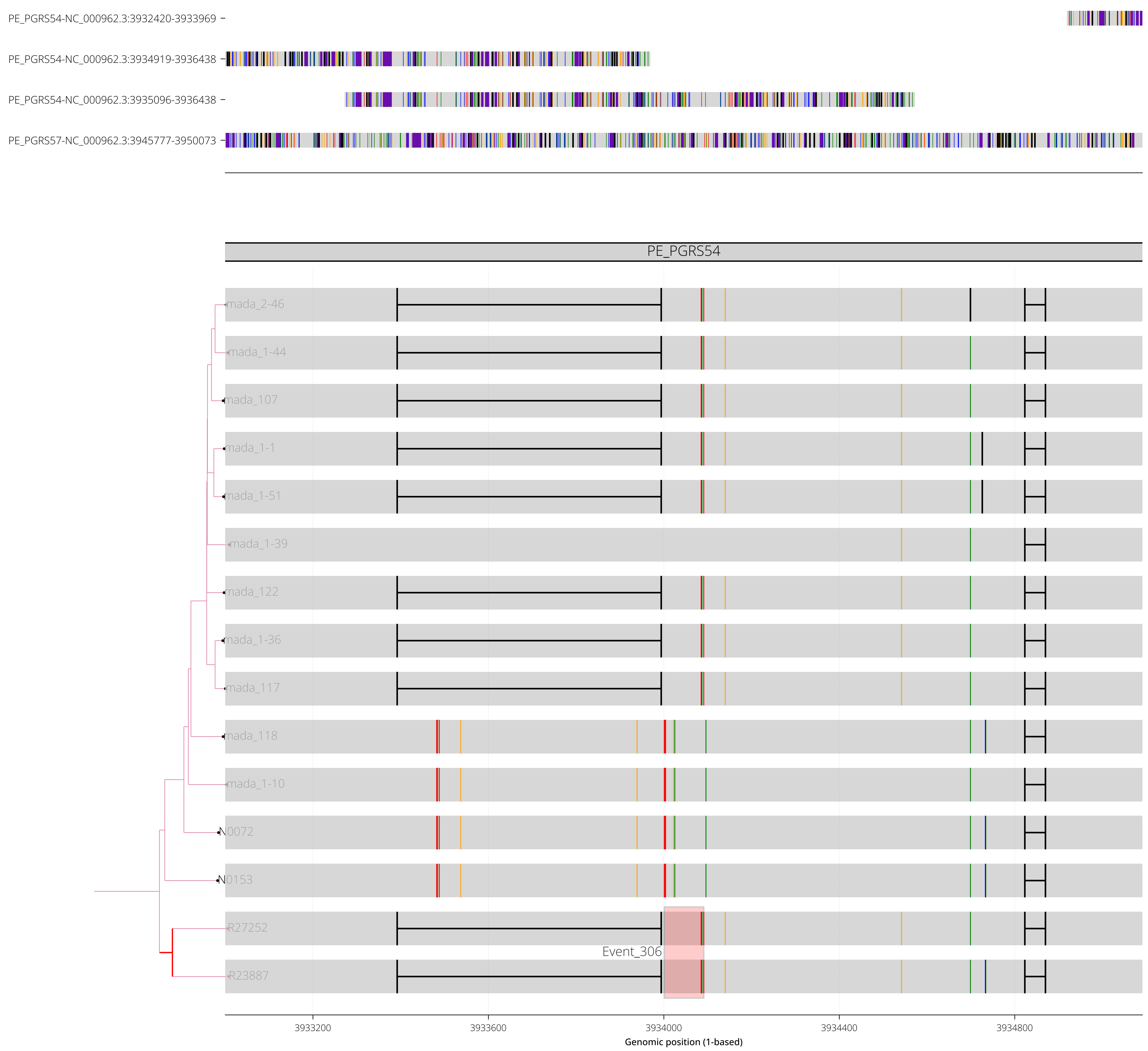

Full Phylogeny Viz + Paralog Alignments
GCE ID: Event_306

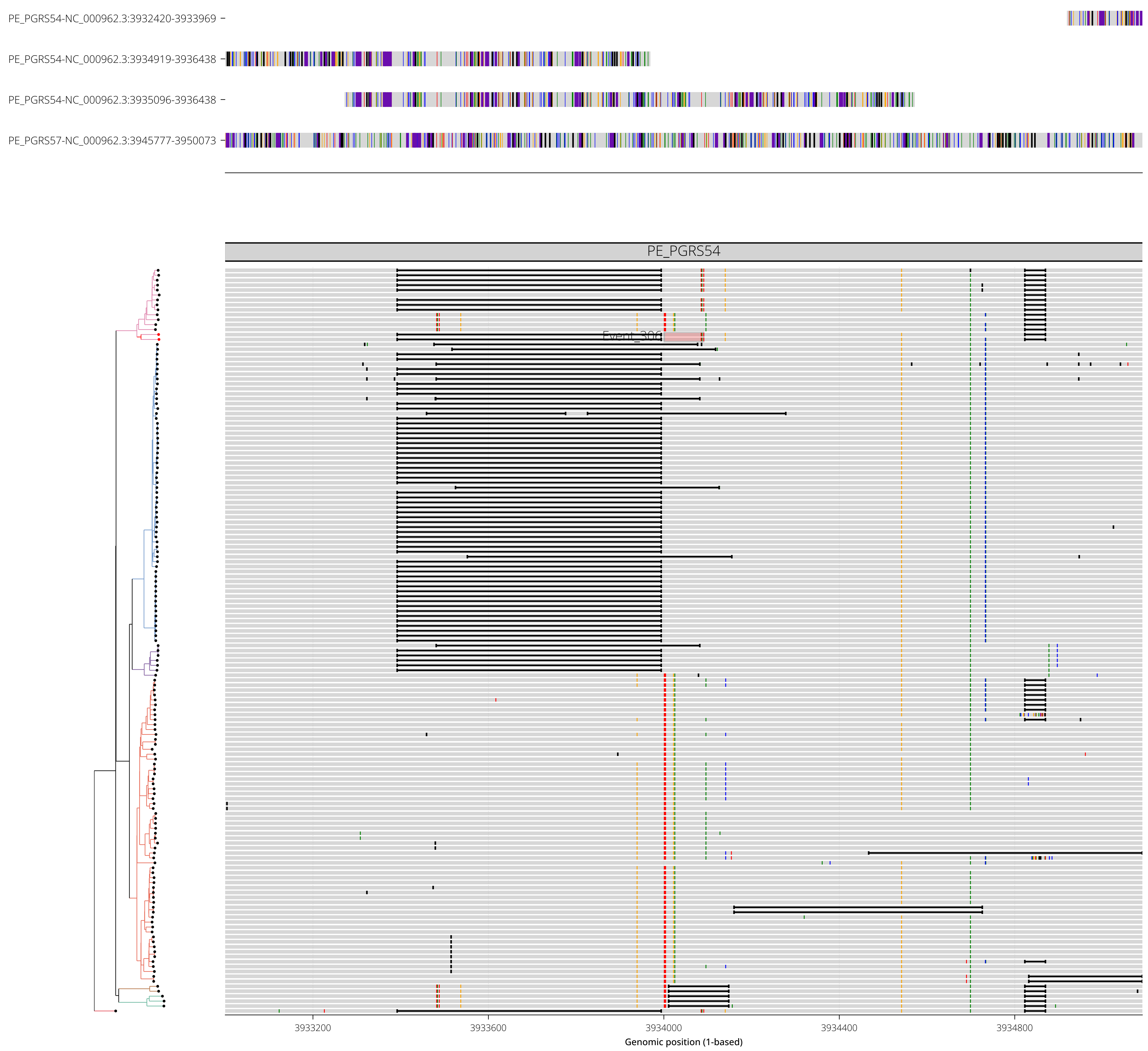

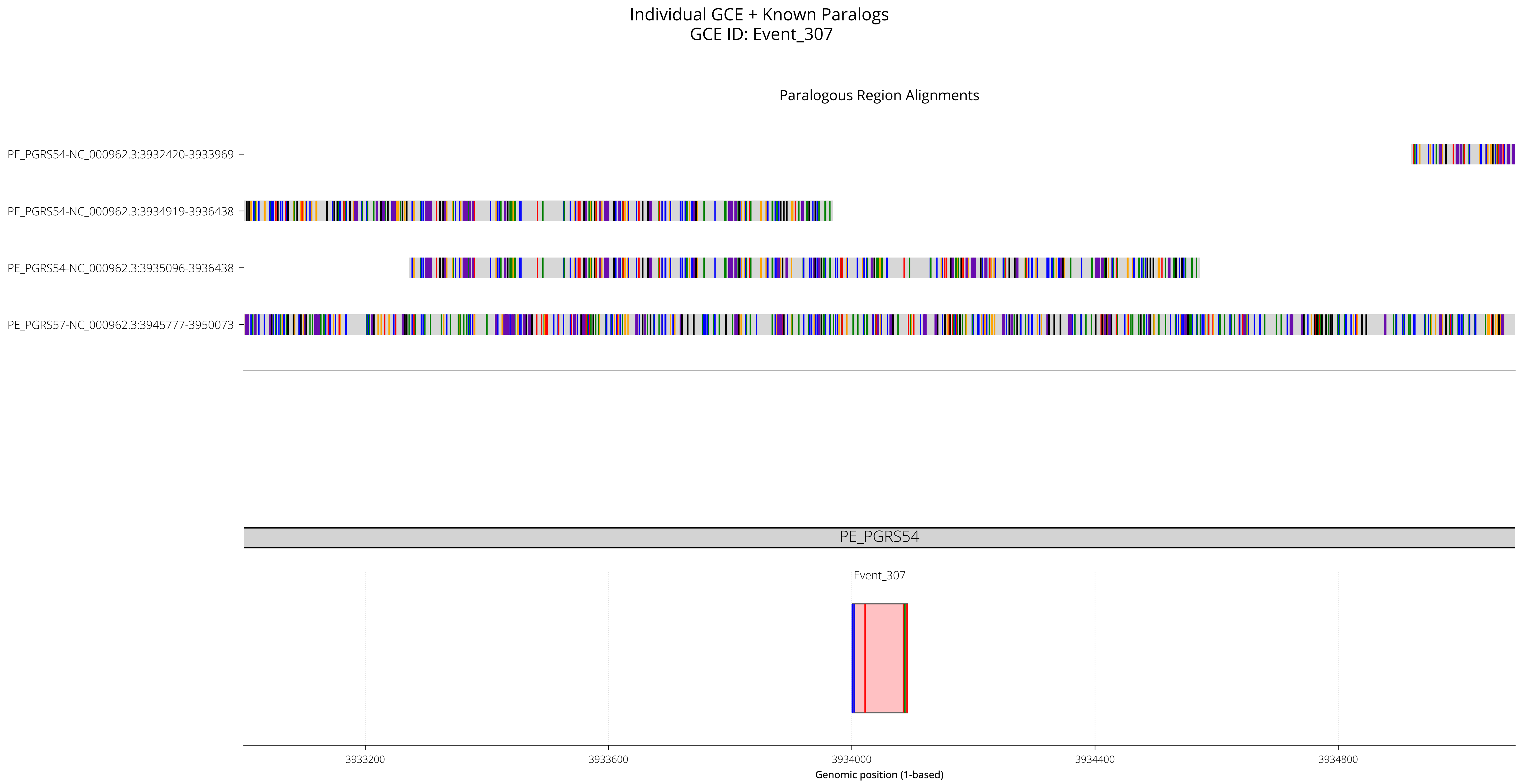
Individual GCE + Known Paralogs
GCE ID: Event_307
Paralogous Region Alignments
PE_PGRS54-NC_000962.3:3932420-3933969
PE_PGRS54-NC_000962.3:3934919-3936438
PE_PGRS54-NC_000962.3:3935096-3936438
PE_PGRS57-NC_000962.3:3945777-3950073
PE_PGRS54
Event_307
3933200
3933600
3934000
3934400
3934800
Genomic position (1-based)

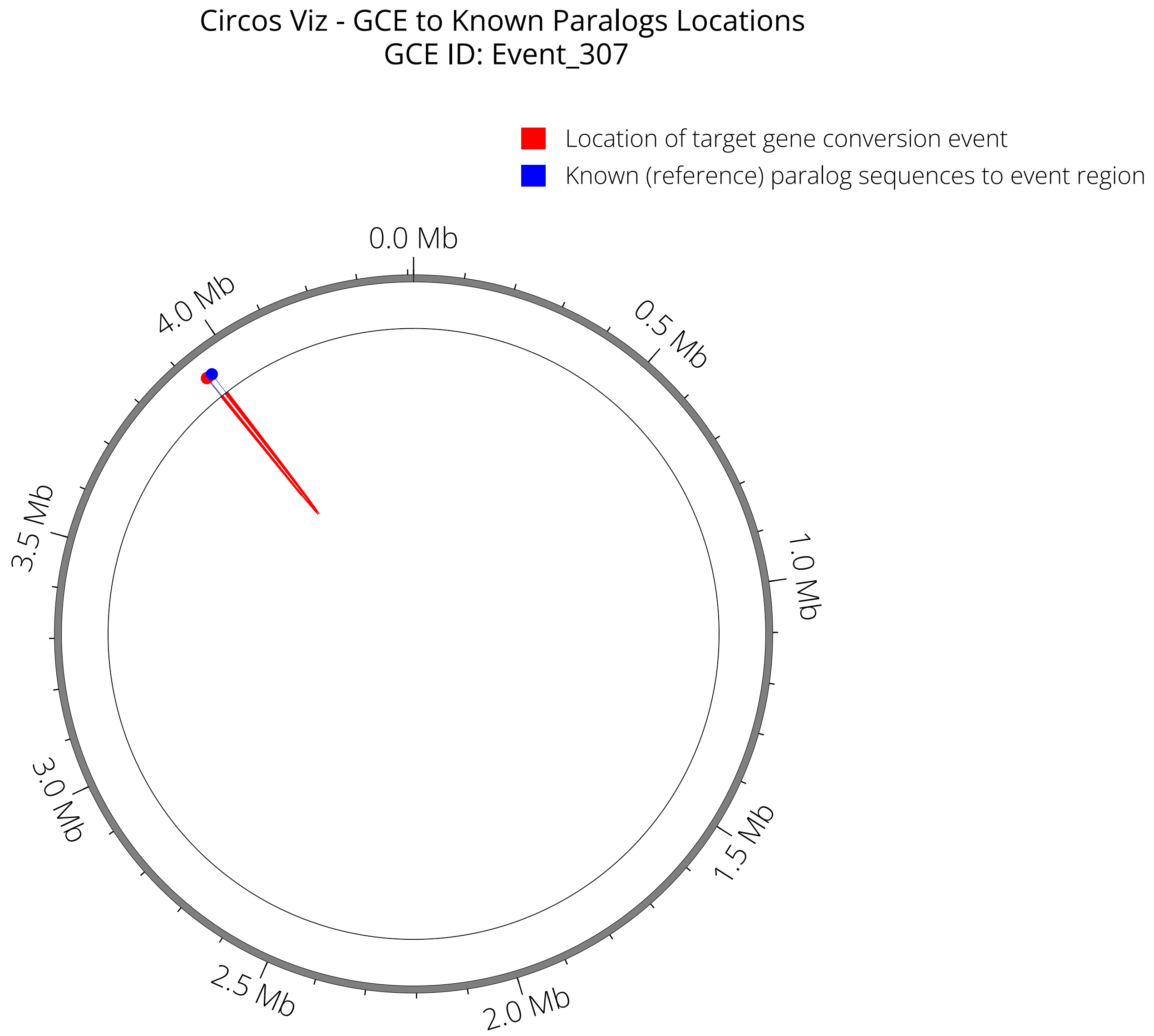
Circos Viz - GCE to Known Paralogs Locations
GCE ID: Event_307
Location of target gene conversion event
Known (reference) paralog sequences to event region
0.0 Mb
0.5 Mb
1.0 Mb
1.5 Mb
2.0 Mb
2.5 Mb
3.0 Mb
3.5 Mb
4.0 Mb

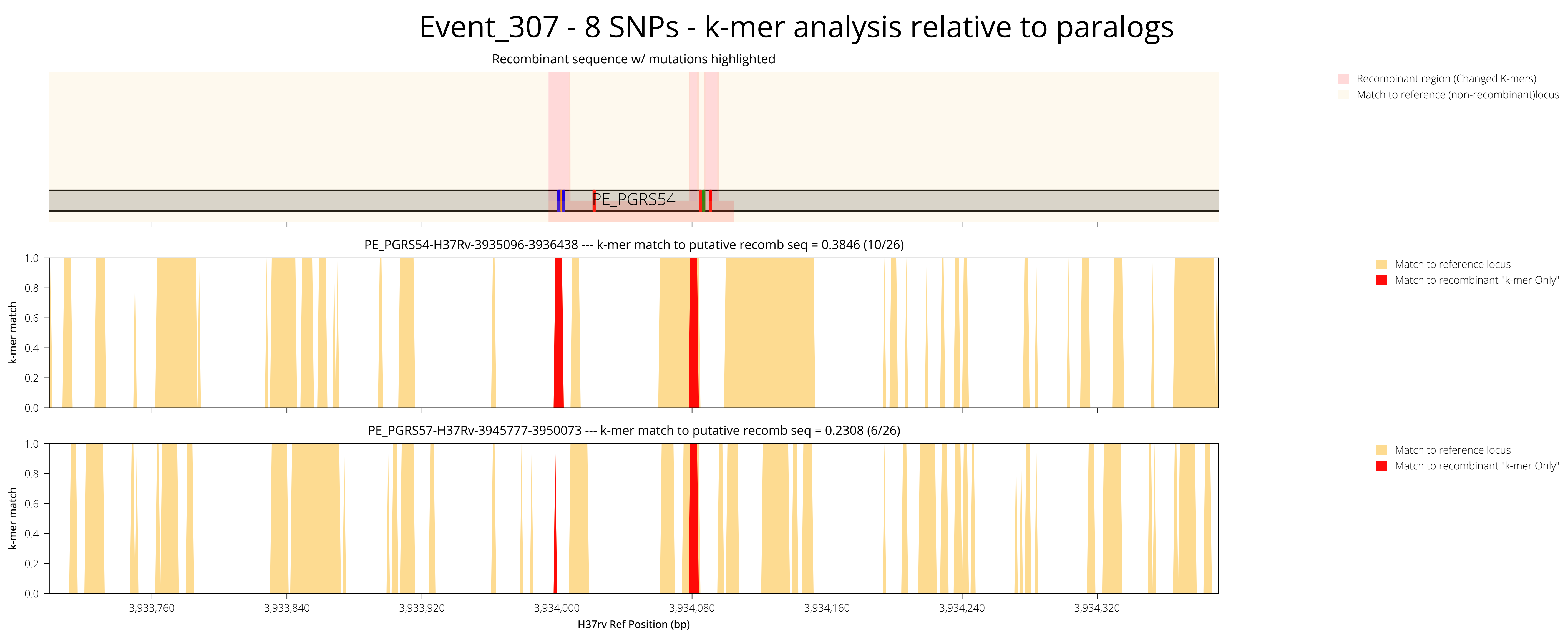

Event_307 - 8 SNPs - k-mer analysis relative to paralogs
Recombinant sequence w/ mutations highlighted
PE_PGRS54
Recombinant region (Changed K-mers)
Match to reference (non-recombinant)locus
PE_PGRS54-H37Rv-3935096-3936438 --- k-mer match to putative recomb seq = 0.3846 (10/26)
Match to reference locus
Match to recombinant "k-mer Only"
PE_PGRS57-H37Rv-3945777-3950073 --- k-mer match to putative recomb seq = 0.2308 (6/26)
k-mer match
0.0
0.2
0.4
0.6
0.8
1.0
3,933,760
3,933,840
3,933,920
3,934,000
3,934,080
3,934,160
3,934,240
3,934,320
H37rv Ref Position (bp)

Sub-phylogeny Viz + Paralog Alignments
GCE ID: Event_307

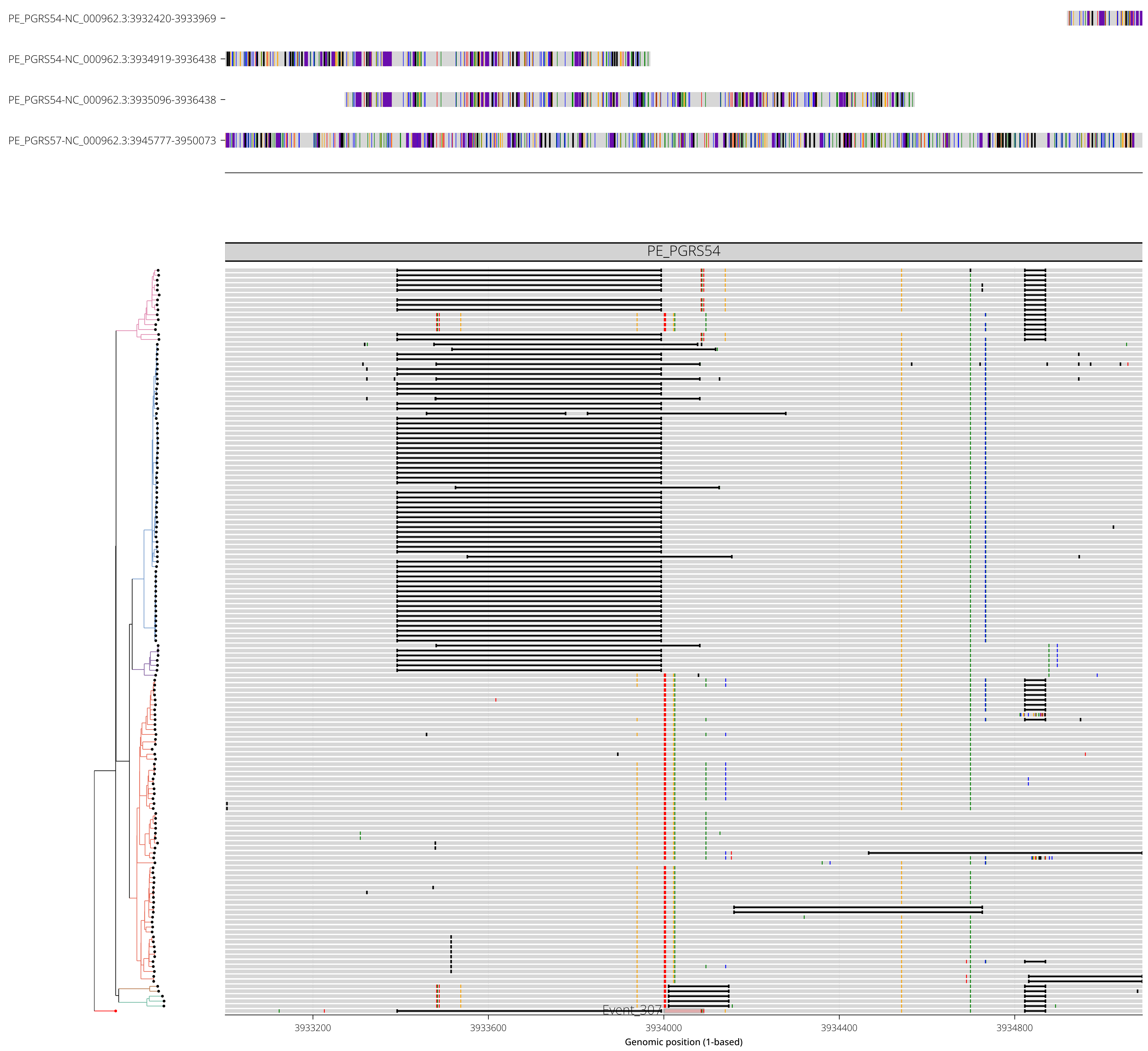

Full Phylogeny Viz + Paralog Alignments
GCE ID: Event_307

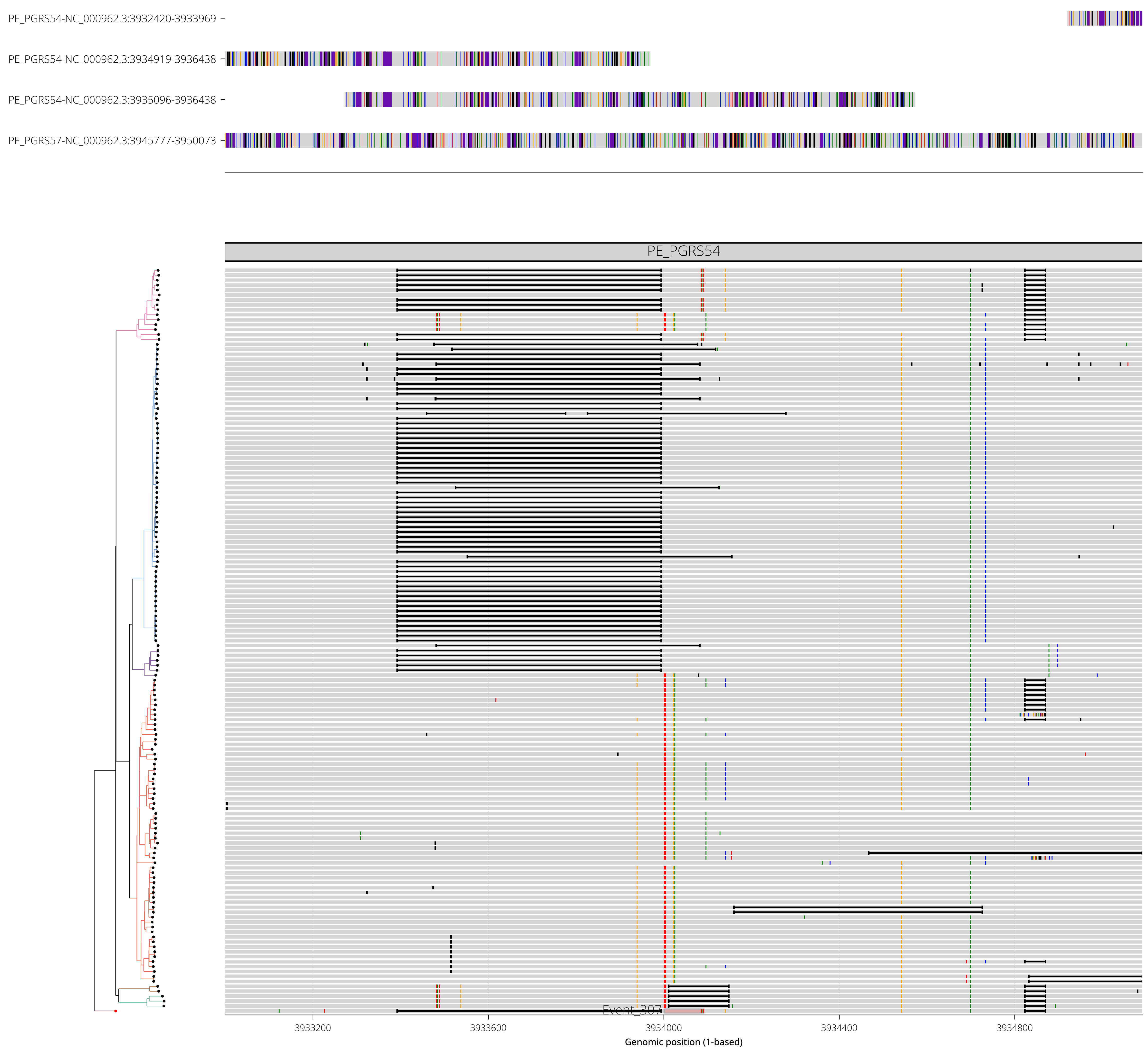

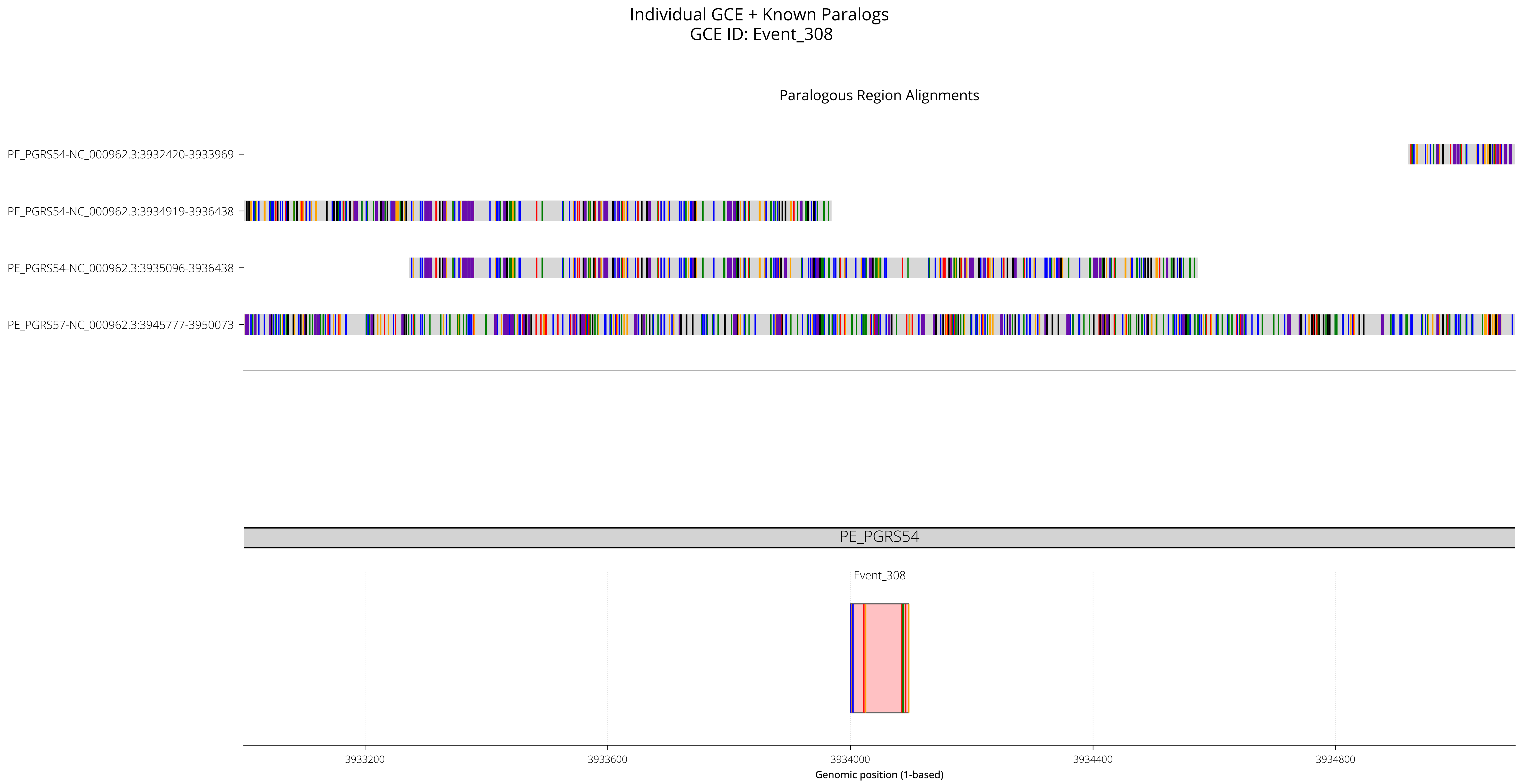

Individual GCE + Known Paralogs
GCE ID: Event_308
Paralogous Region Alignments
PE_PGRS54-NC_000962.3:3932420-3933969
PE_PGRS54-NC_000962.3:3934919-3936438
PE_PGRS54-NC_000962.3:3935096-3936438
PE_PGRS57-NC_000962.3:3945777-3950073
PE_PGRS54
Event_308
3933200
3933600
3934000
3934400
3934800
Genomic position (1-based)

Circos Viz - GCE to Known Paralogs Locations
GCE ID: Event_308

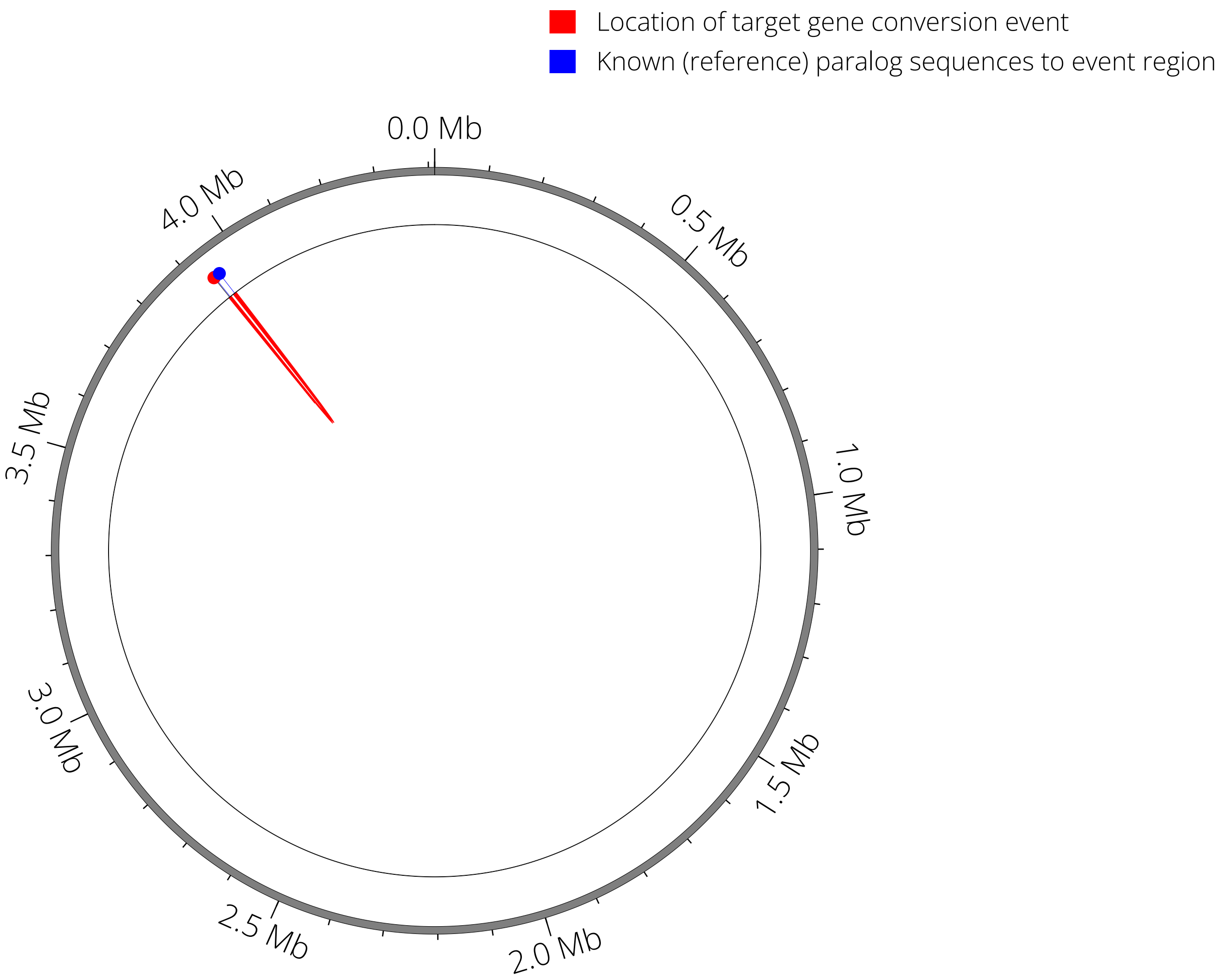

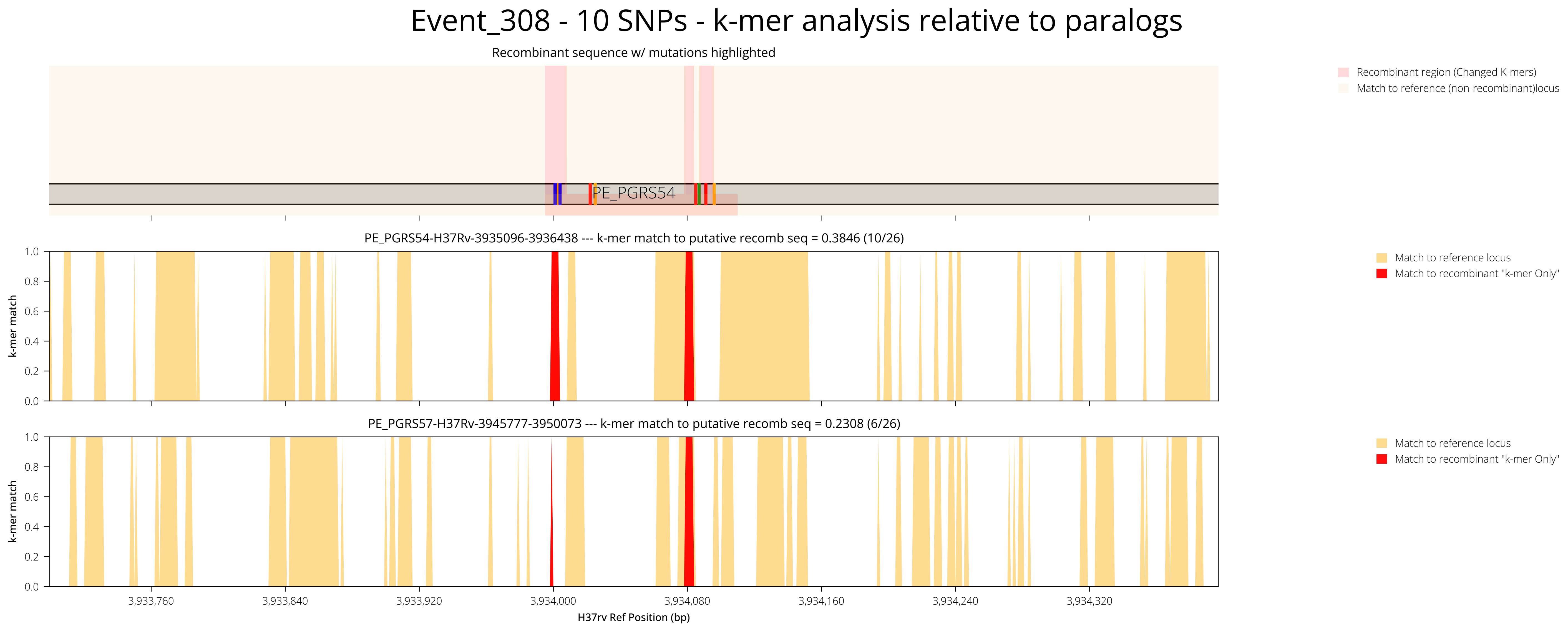
Event_308 - 10 SNPs - k-mer analysis relative to paralogs
Recombinant sequence w/ mutations highlighted
Recombinant region (Changed K-mers)
Match to reference (non-recombinant)locus
PE_PGRS54
PE_PGRS54-H37Rv-3935096-3936438 --- k-mer match to putative recomb seq = 0.3846 (10/26)
Match to reference locus
Match to recombinant "k-mer Only"
k-mer match
1.0
0.8
0.6
0.4
0.2
0.0
PE_PGRS57-H37Rv-3945777-3950073 --- k-mer match to putative recomb seq = 0.2308 (6/26)
Match to reference locus
Match to recombinant "k-mer Only"
k-mer match
1.0
0.8
0.6
0.4
0.2
0.0
3,933,760
3,933,840
3,933,920
3,934,000
3,934,080
3,934,160
3,934,240
3,934,320
H37rv Ref Position (bp)

Sub-phylogeny Viz + Paralog Alignments
GCE ID: Event_308

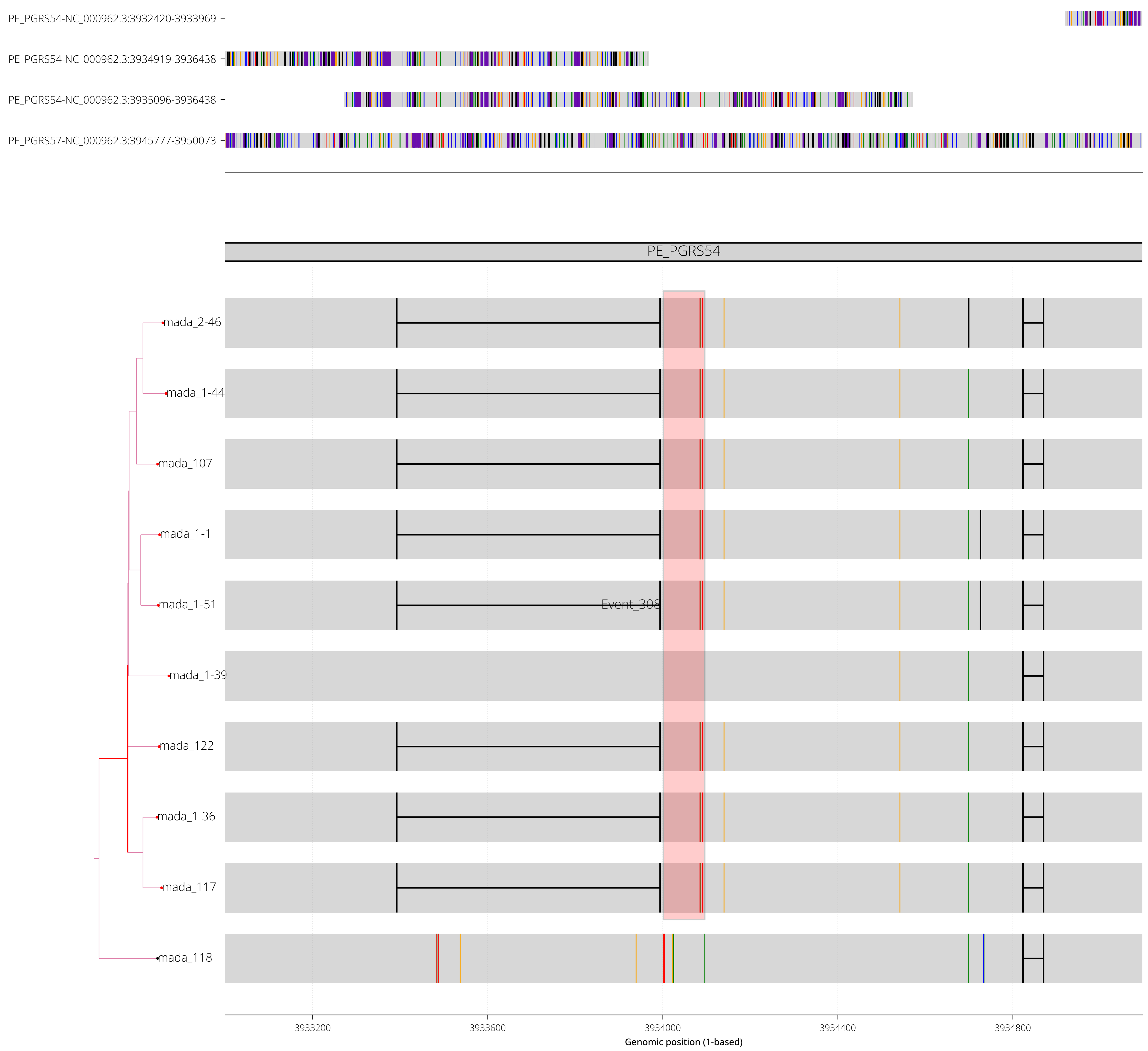

Full Phylogeny Viz + Paralog Alignments
GCE ID: Event_308

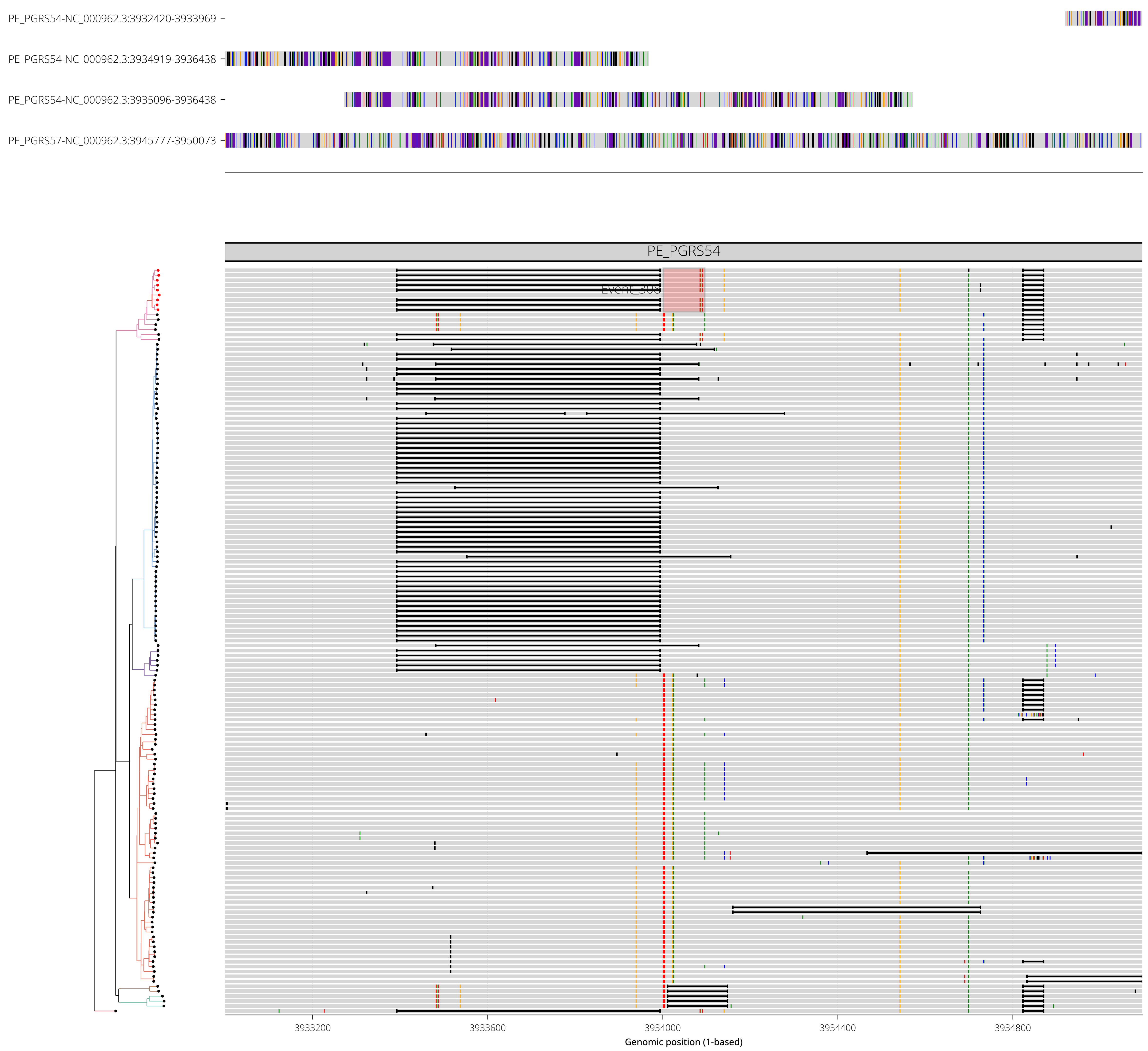

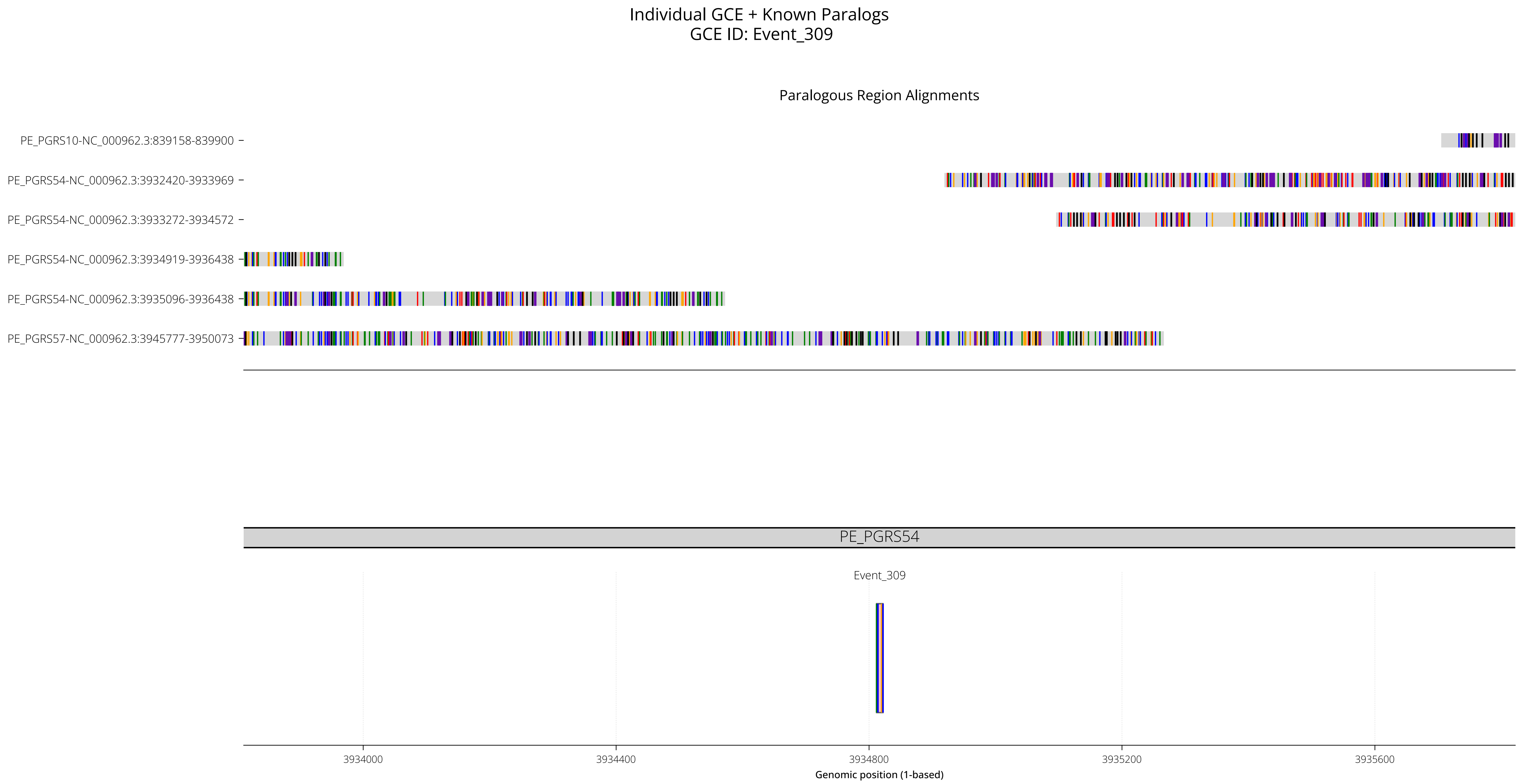

Individual GCE + Known Paralogs
GCE ID: Event_309
Paralogous Region Alignments
PE_PGRS10-NC_000962.3:839158-839900
PE_PGRS54-NC_000962.3:3932420-3933969
PE_PGRS54-NC_000962.3:3933272-3934572
PE_PGRS54-NC_000962.3:3934919-3936438
PE_PGRS54-NC_000962.3:3935096-3936438
PE_PGRS57-NC_000962.3:3945777-3950073
PE_PGRS54
Event_309
3934000
3934400
3934800
3935200
3935600
Genomic position (1-based)

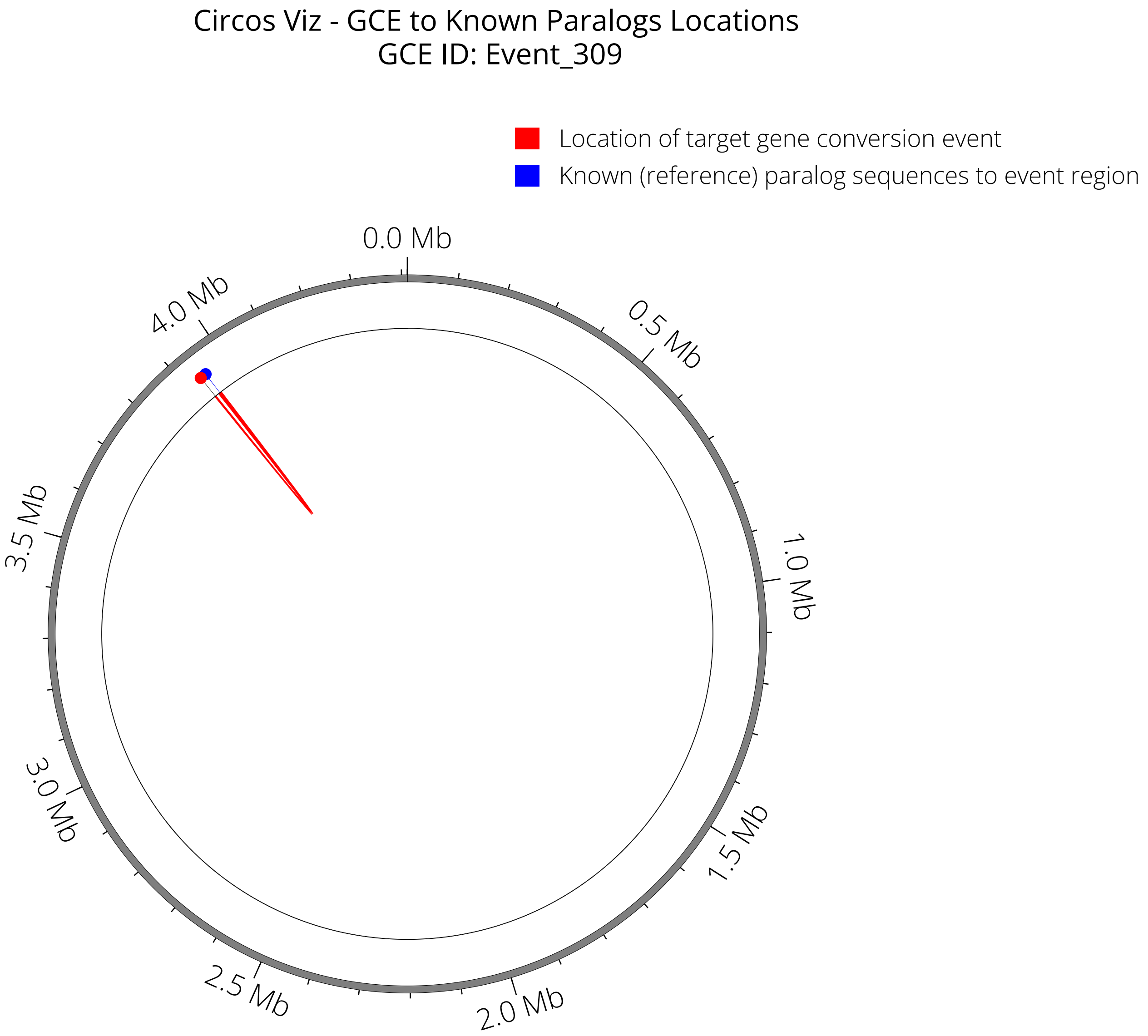
Circos Viz - GCE to Known Paralogs Locations
GCE ID: Event_309
Location of target gene conversion event
Known (reference) paralog sequences to event region
0.0 Mb
0.5 Mb
1.0 Mb
1.5 Mb
2.0 Mb
2.5 Mb
3.0 Mb
3.5 Mb
4.0 Mb

### Event_309 - 5 SNPs - k-mer analysis relative to paralogs

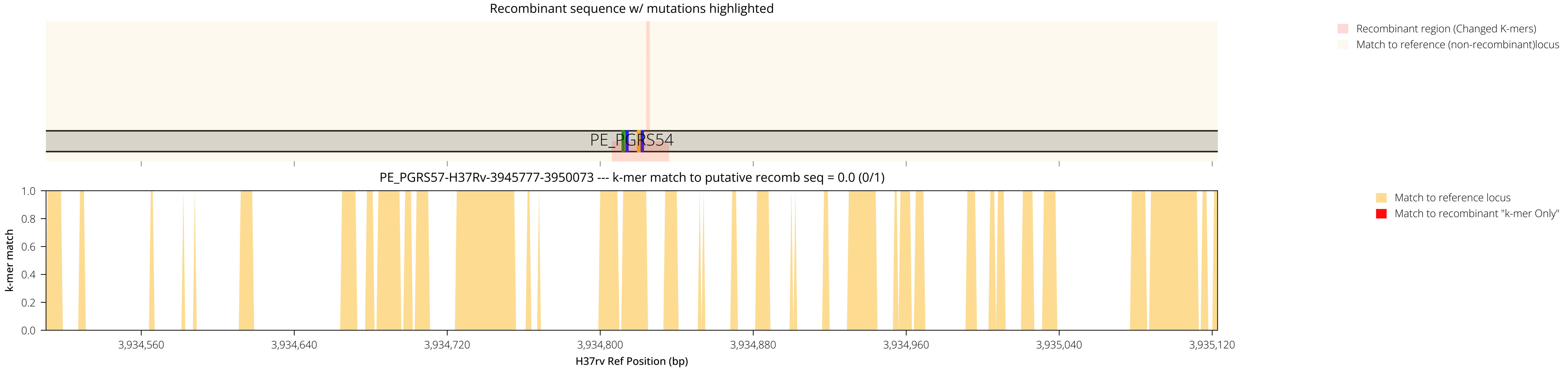

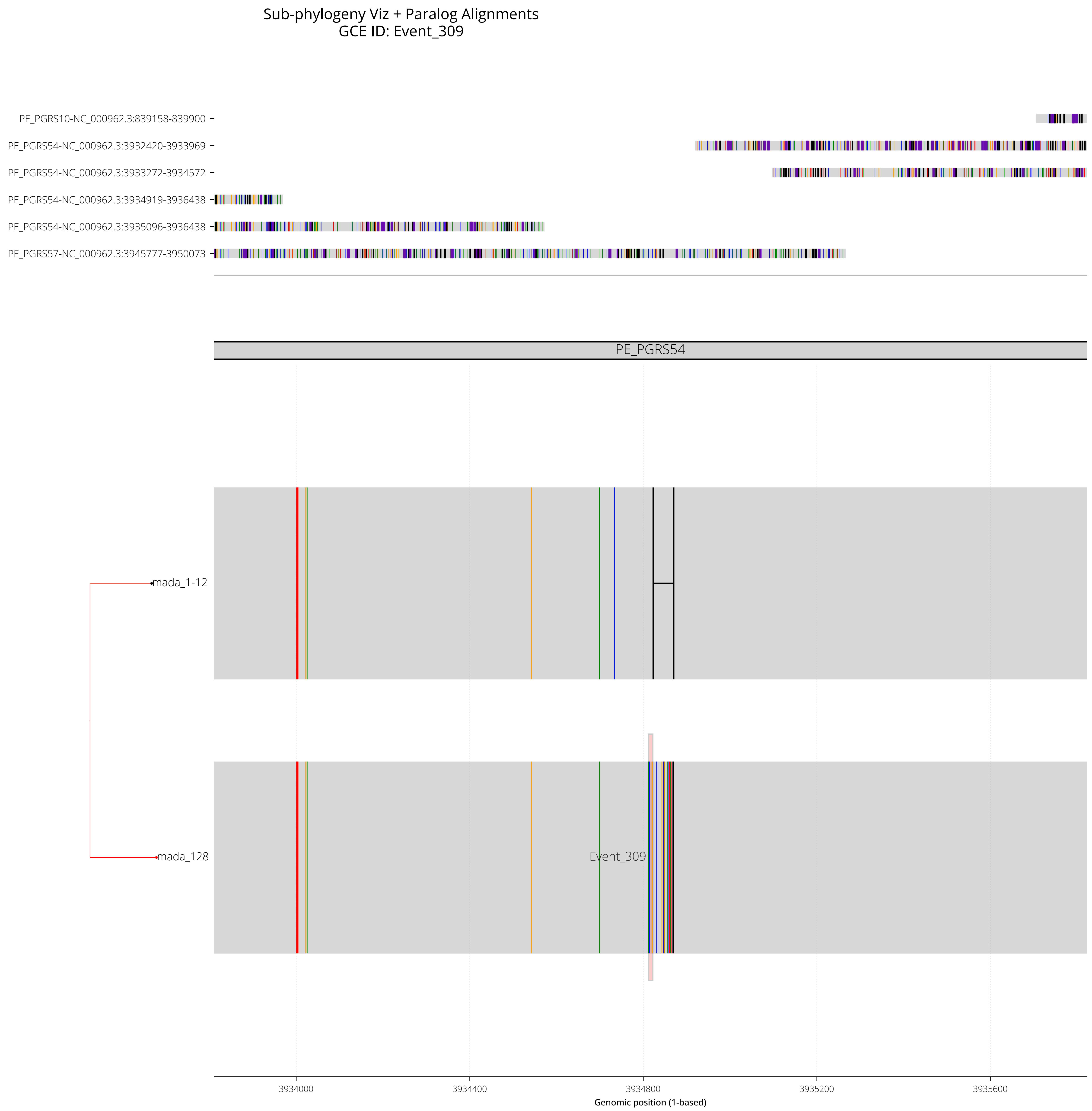

Sub-phylogeny Viz + Paralog Alignments
GCE ID: Event_309
PE_PGRS10-NC_000962.3:839158-839900
PE_PGRS54-NC_000962.3:3932420-3933969
PE_PGRS54-NC_000962.3:3933272-3934572
PE_PGRS54-NC_000962.3:3934919-3936438
PE_PGRS54-NC_000962.3:3935096-3936438
PE_PGRS57-NC_000962.3:3945777-3950073
PE_PGRS54
mada_1-12
mada_128
Event_309
3934000
3934400
3934800
3935200
3935600
Genomic position (1-based)

Full Phylogeny Viz + Paralog Alignments
GCE ID: Event_309

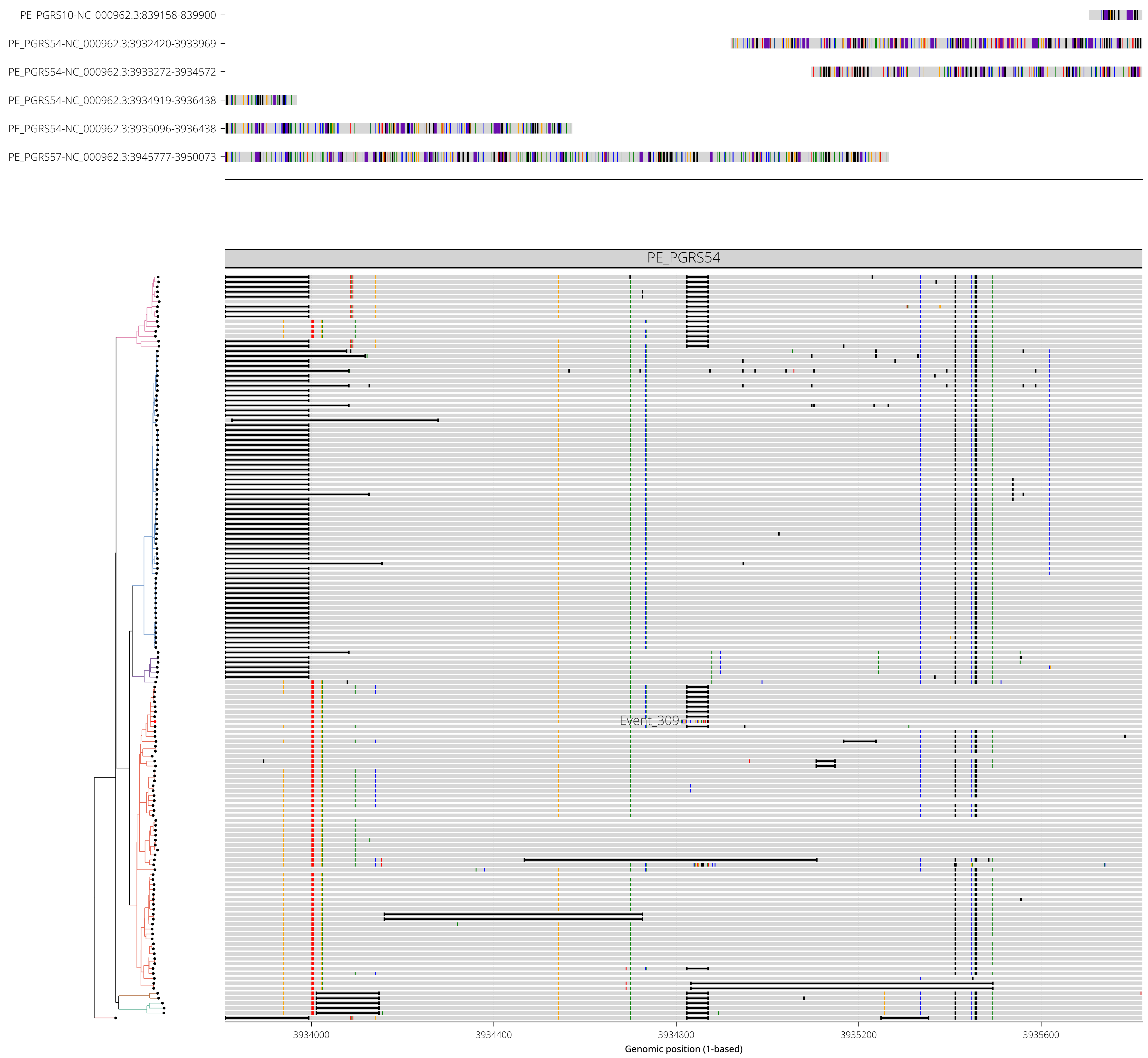

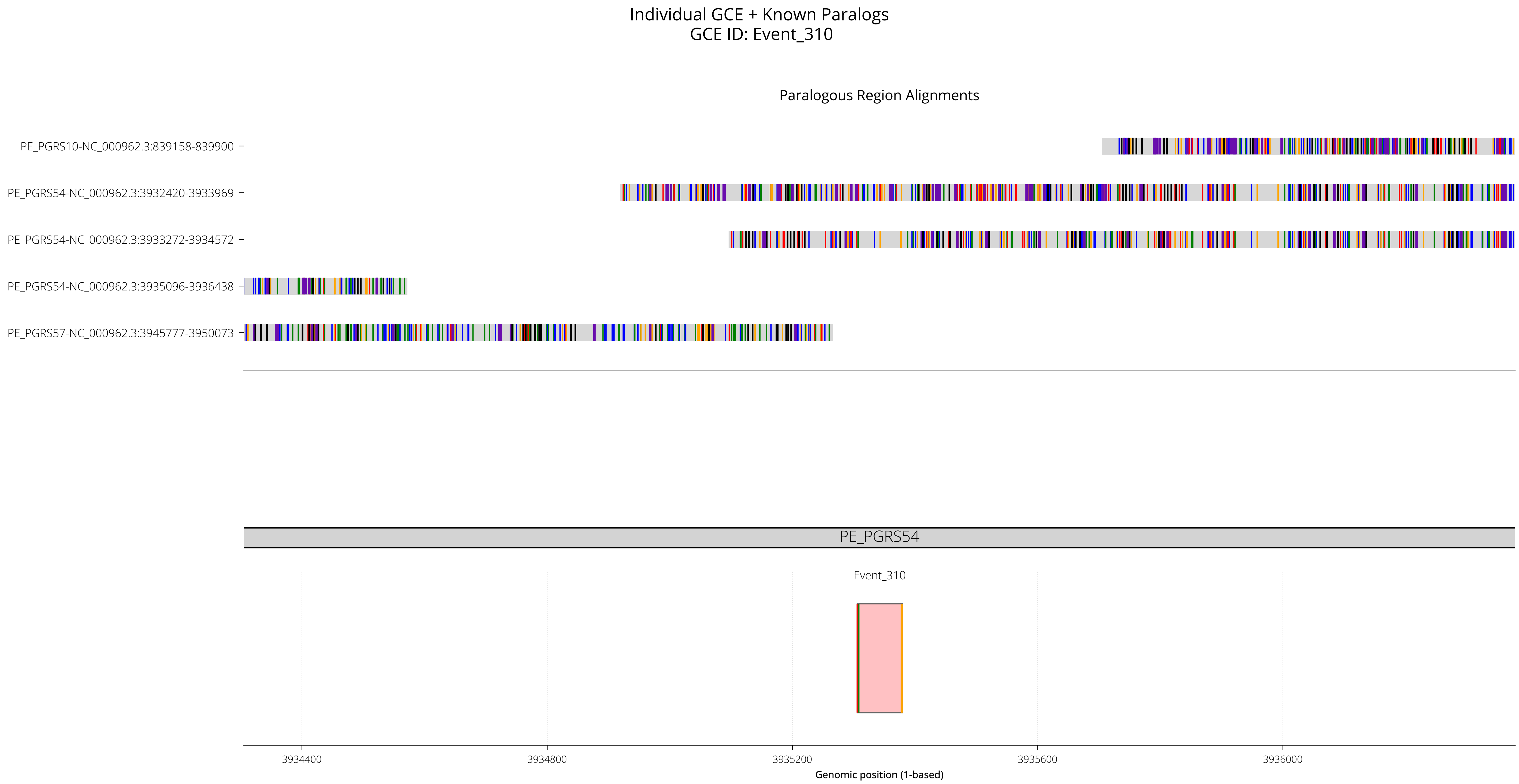

Individual GCE + Known Paralogs
GCE ID: Event_310
Paralogous Region Alignments
PE_PGRS10-NC_000962.3:839158-839900
PE_PGRS54-NC_000962.3:3932420-3933969
PE_PGRS54-NC_000962.3:3933272-3934572
PE_PGRS54-NC_000962.3:3935096-3936438
PE_PGRS57-NC_000962.3:3945777-3950073
PE_PGRS54
Event_310
3934400
3934800
3935200
3935600
3936000
Genomic position (1-based)

### Circos Viz - GCE to Known Paralogs Locations
GCE ID: Event_310

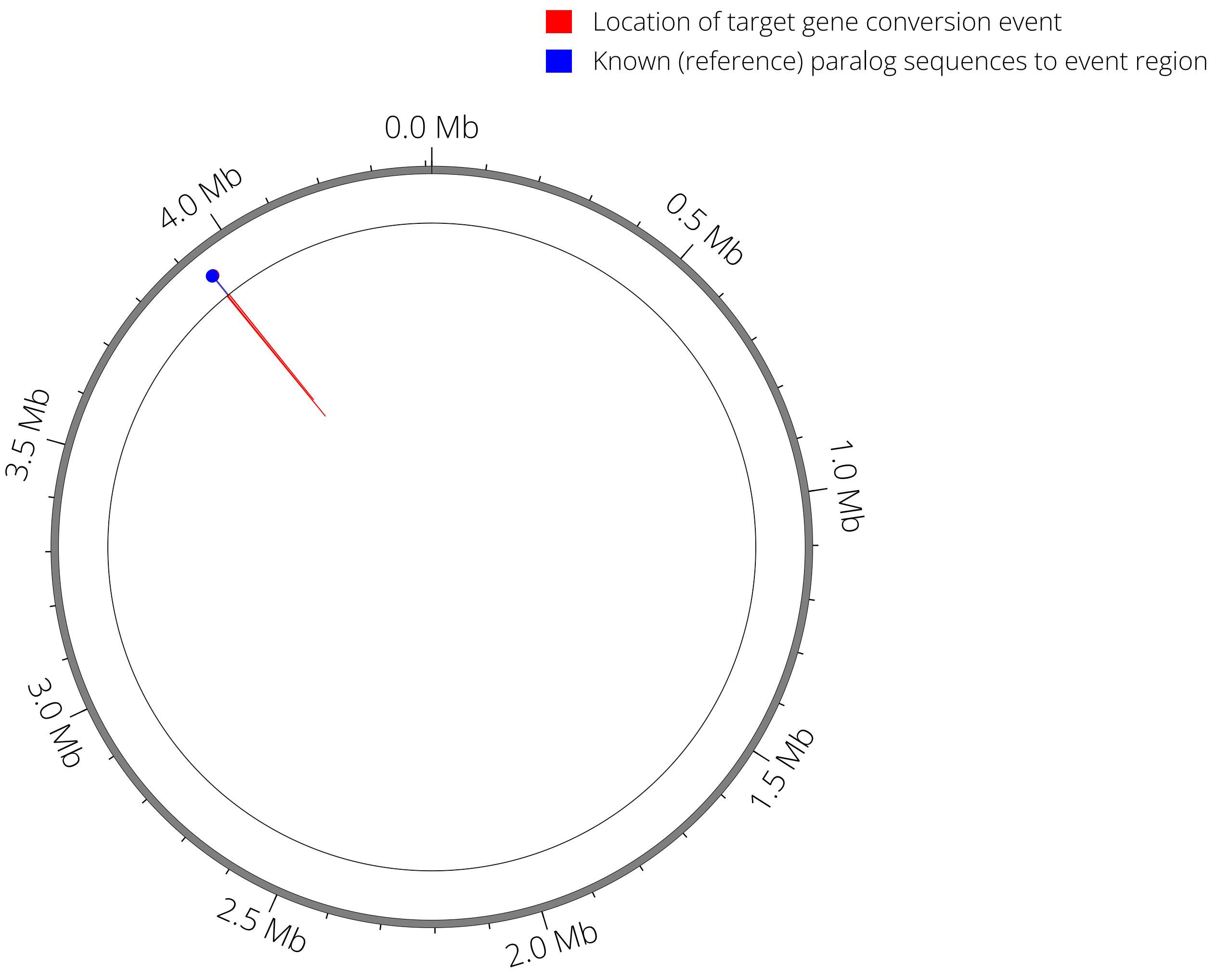

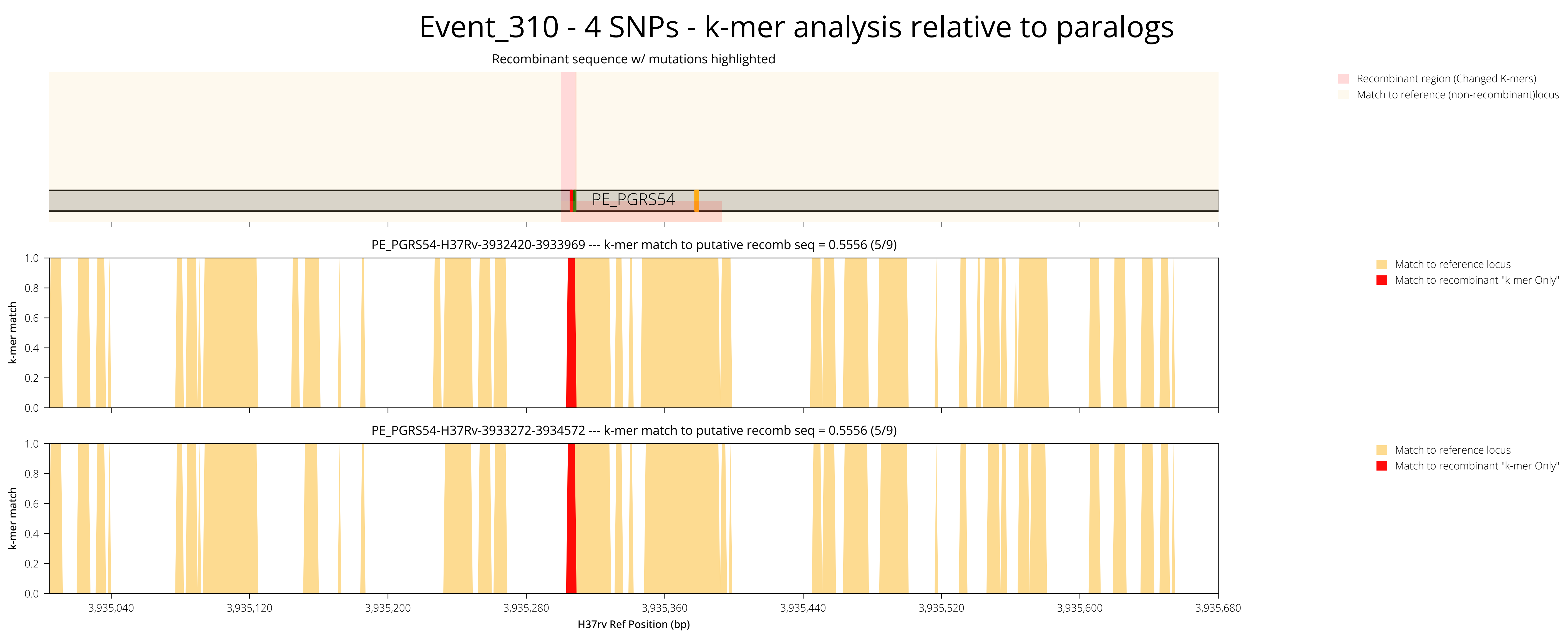
Event_310 - 4 SNPs - k-mer analysis relative to paralogs
Recombinant sequence w/ mutations highlighted
PE_PGRS54
Recombinant region (Changed K-mers)
Match to reference (non-recombinant)locus
PE_PGRS54-H37Rv-3932420-3933969 --- k-mer match to putative recomb seq = 0.5556 (5/9)
Match to reference locus
Match to recombinant "k-mer Only"
PE_PGRS54-H37Rv-3933272-3934572 --- k-mer match to putative recomb seq = 0.5556 (5/9)
Match to reference locus
Match to recombinant "k-mer Only"
k-mer match
1.0
0.8
0.6
0.4
0.2
0.0
3,935,040
3,935,120
3,935,200
3,935,280
3,935,360
3,935,440
3,935,520
3,935,600
3,935,680
H37rv Ref Position (bp)

Sub-phylogeny Viz + Paralog Alignments
GCE ID: Event_310

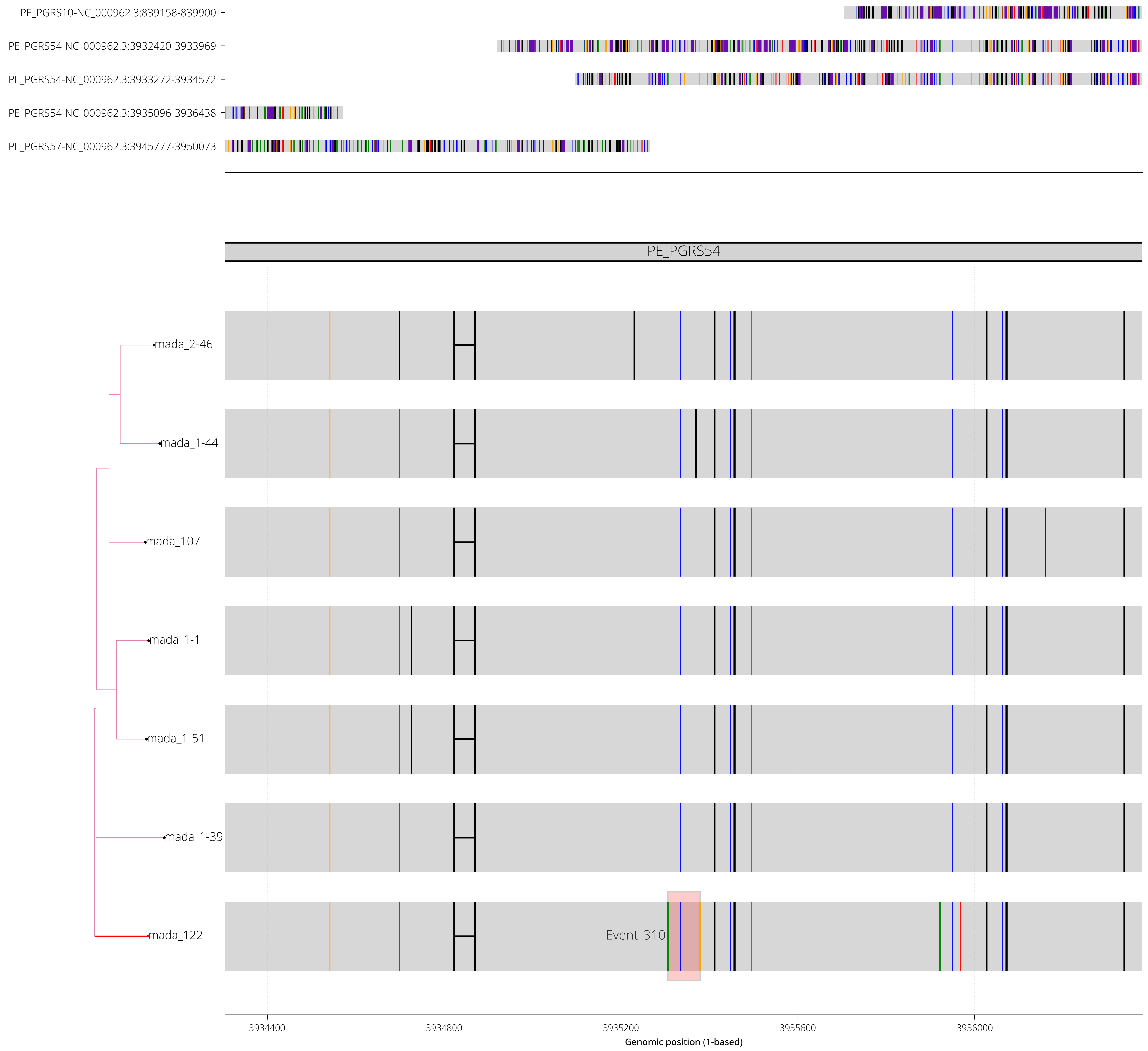

Full Phylogeny Viz + Paralog Alignments
GCE ID: Event_310

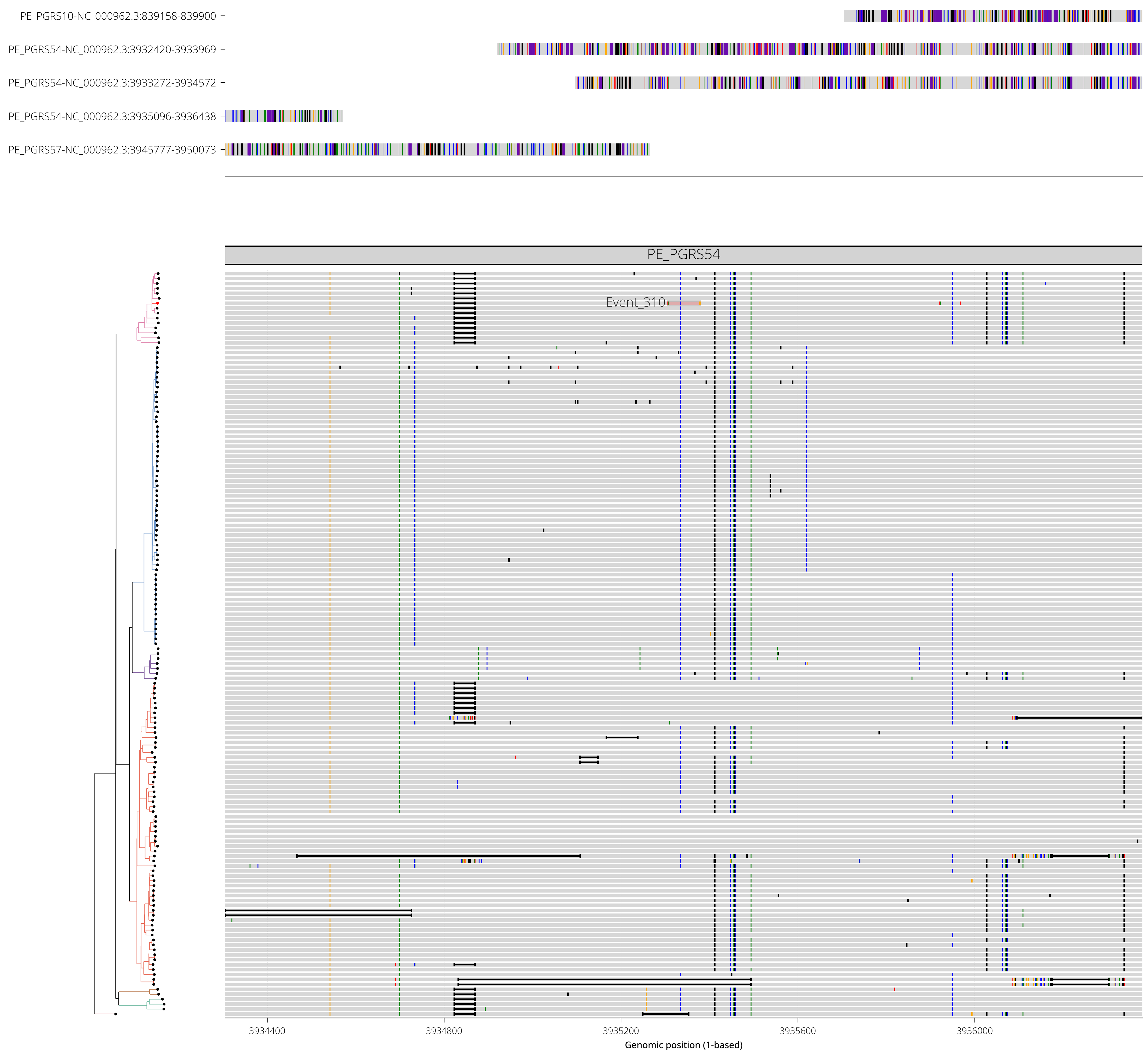

Individual GCE + Known Paralogs
GCE ID: Event_311

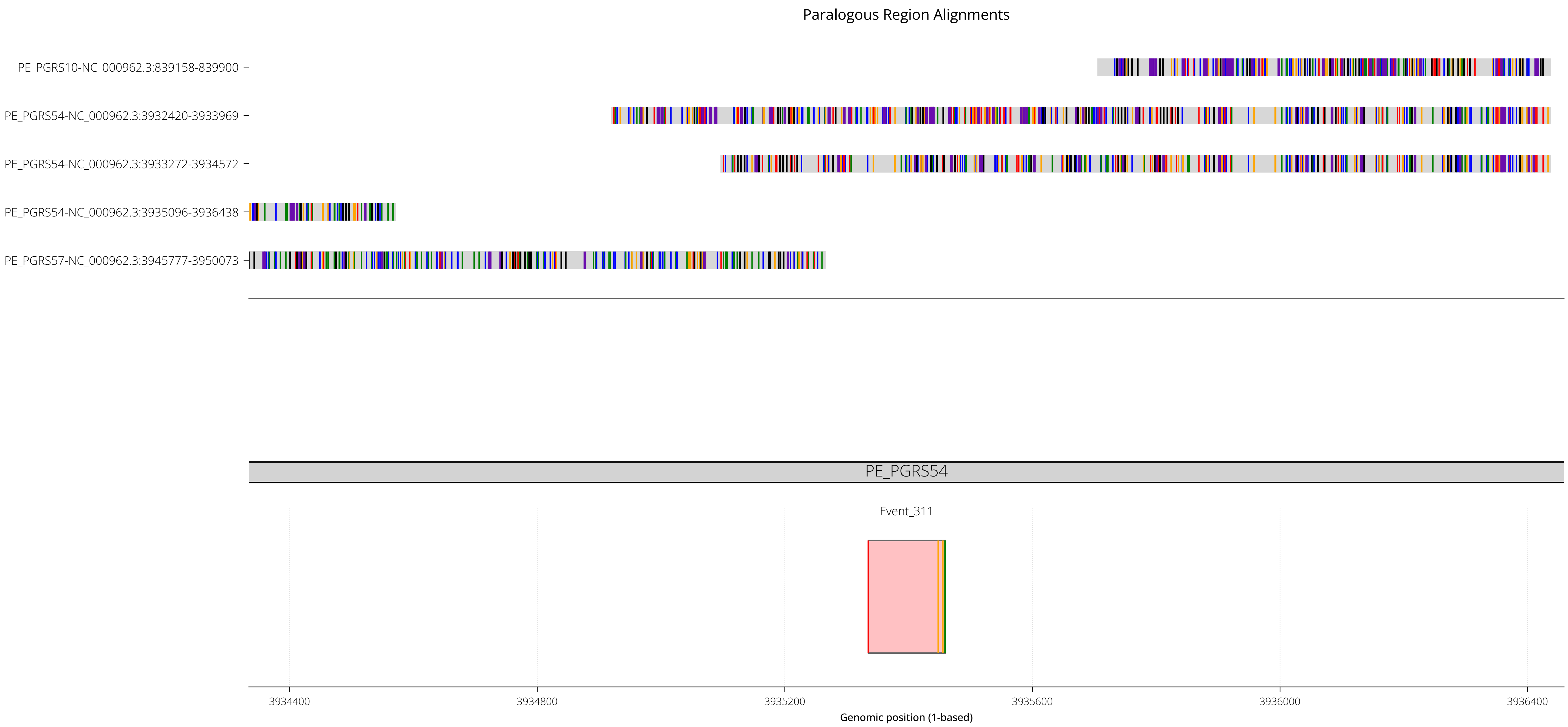

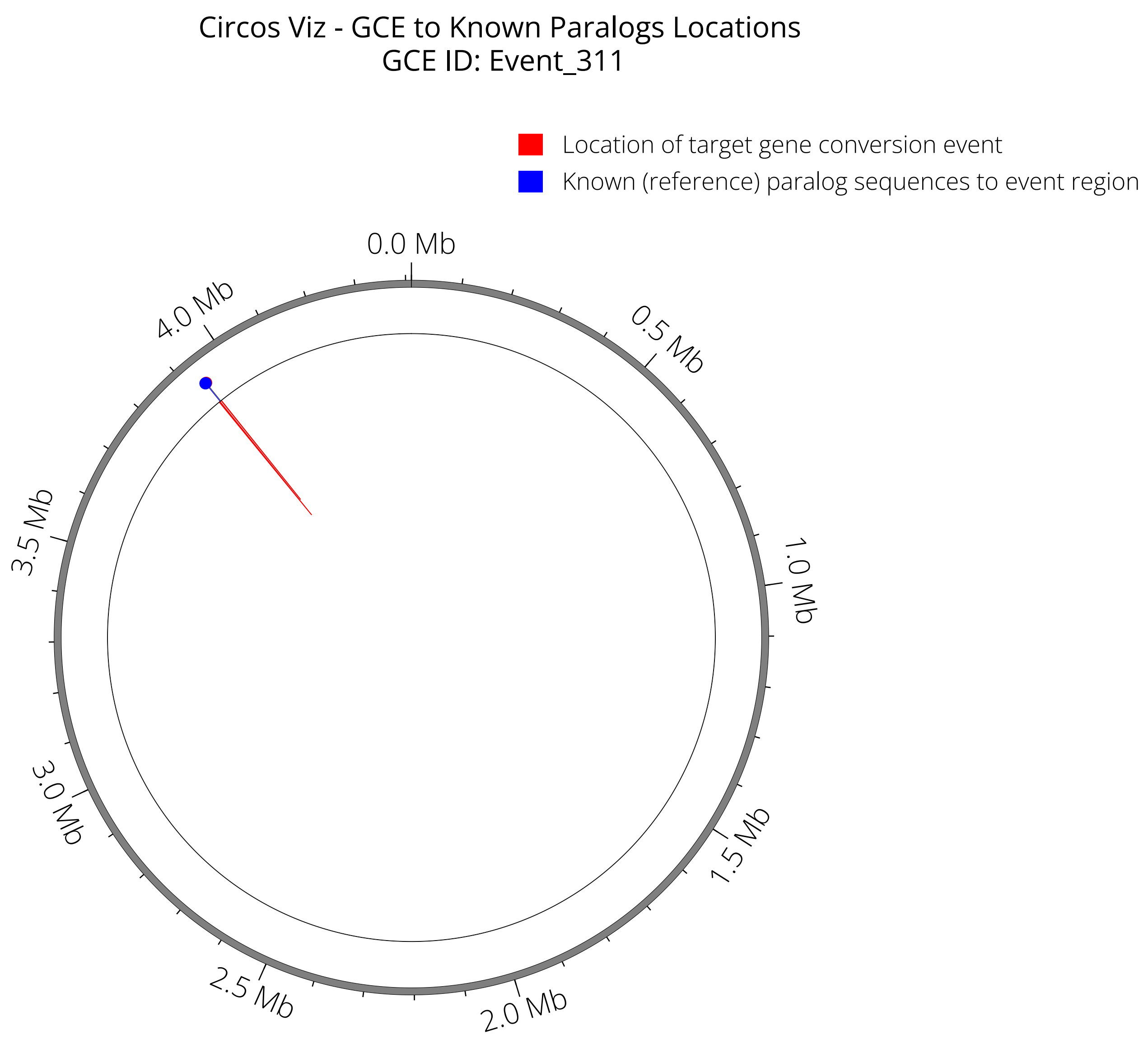
Circos Viz - GCE to Known Paralogs Locations
GCE ID: Event_311
Location of target gene conversion event
Known (reference) paralog sequences to event region
0.0 Mb
0.5 Mb
1.0 Mb
1.5 Mb
2.0 Mb
2.5 Mb
3.0 Mb
3.5 Mb
4.0 Mb

### Event_311 - 4 SNPs - k-mer analysis relative to paralogs

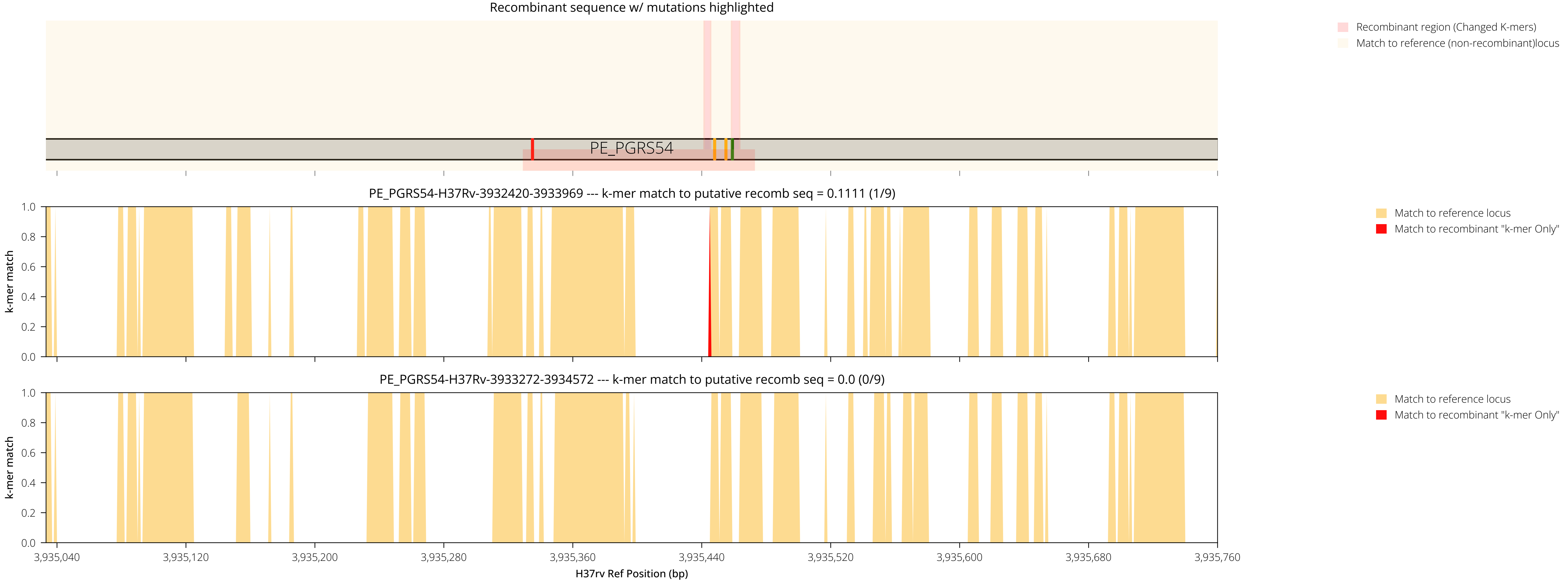

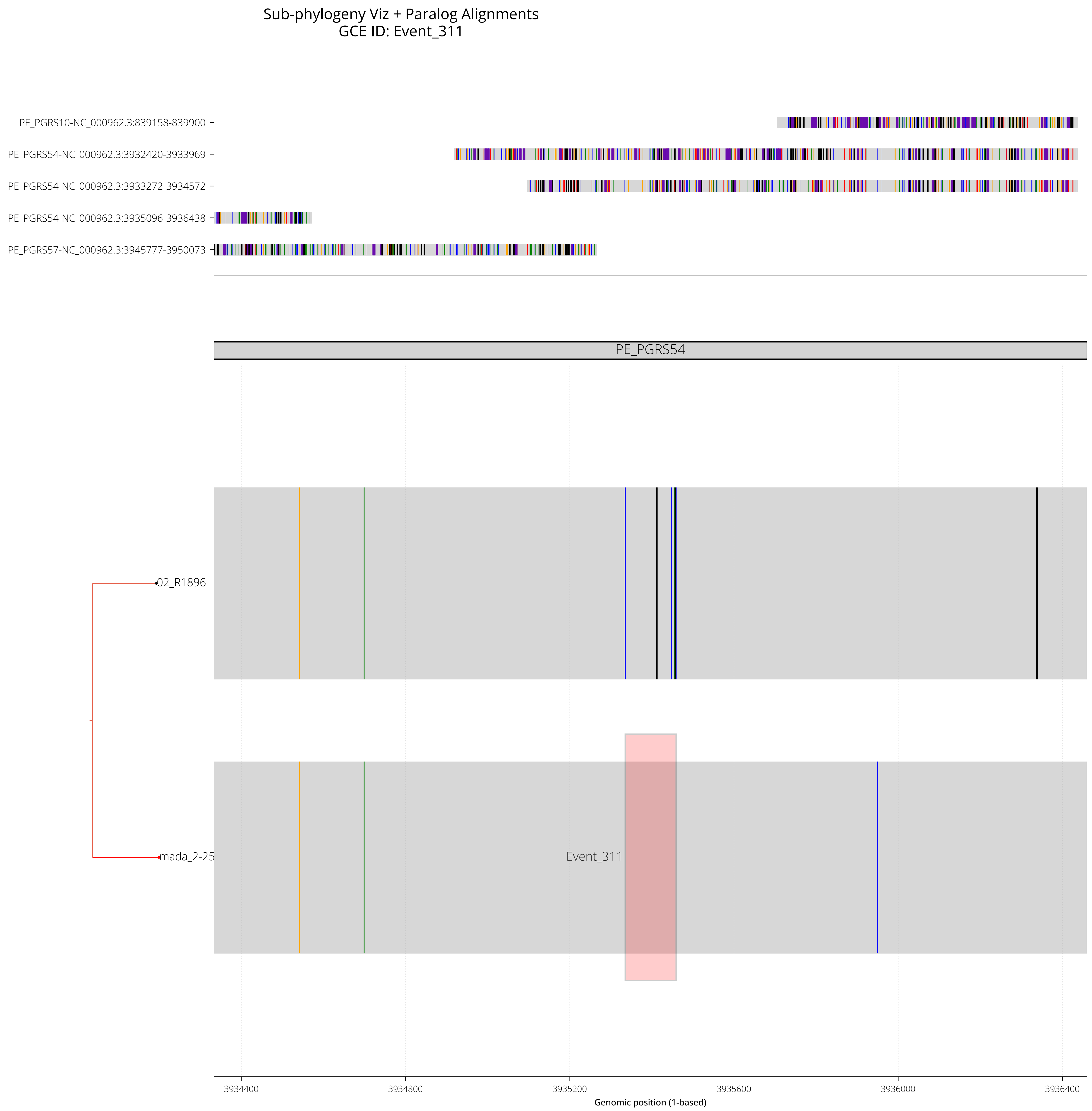
Sub-phylogeny Viz + Paralog Alignments
GCE ID: Event_311
PE_PGRS10-NC_000962.3:839158-839900
PE_PGRS54-NC_000962.3:3932420-3933969
PE_PGRS54-NC_000962.3:3933272-3934572
PE_PGRS54-NC_000962.3:3935096-3936438
PE_PGRS57-NC_000962.3:3945777-3950073
PE_PGRS54
02_R1896
mada_2-25
Event_311
3934400
3934800
3935200
3935600
3936000
3936400
Genomic position (1-based)

Full Phylogeny Viz + Paralog Alignments
GCE ID: Event_311

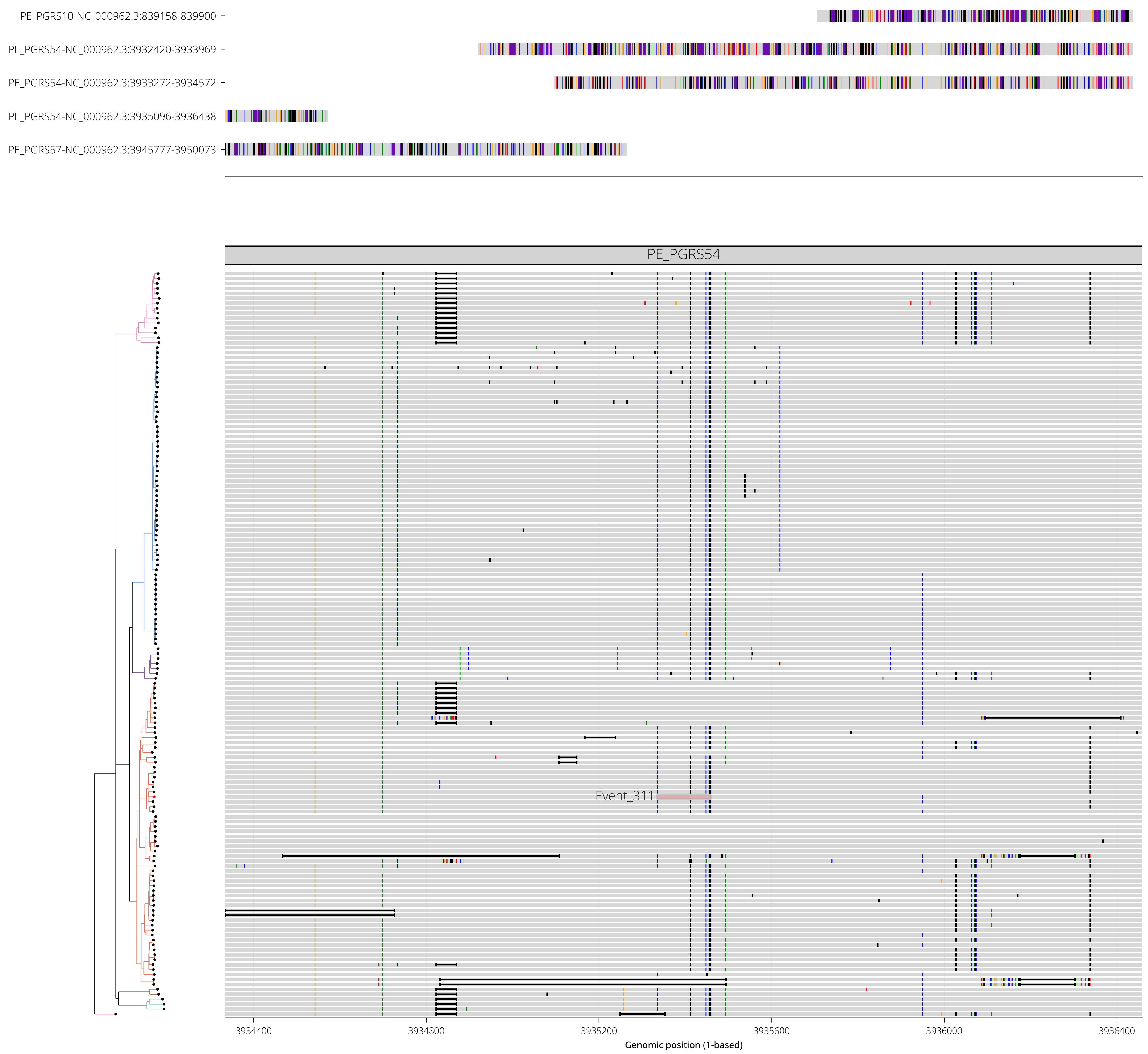

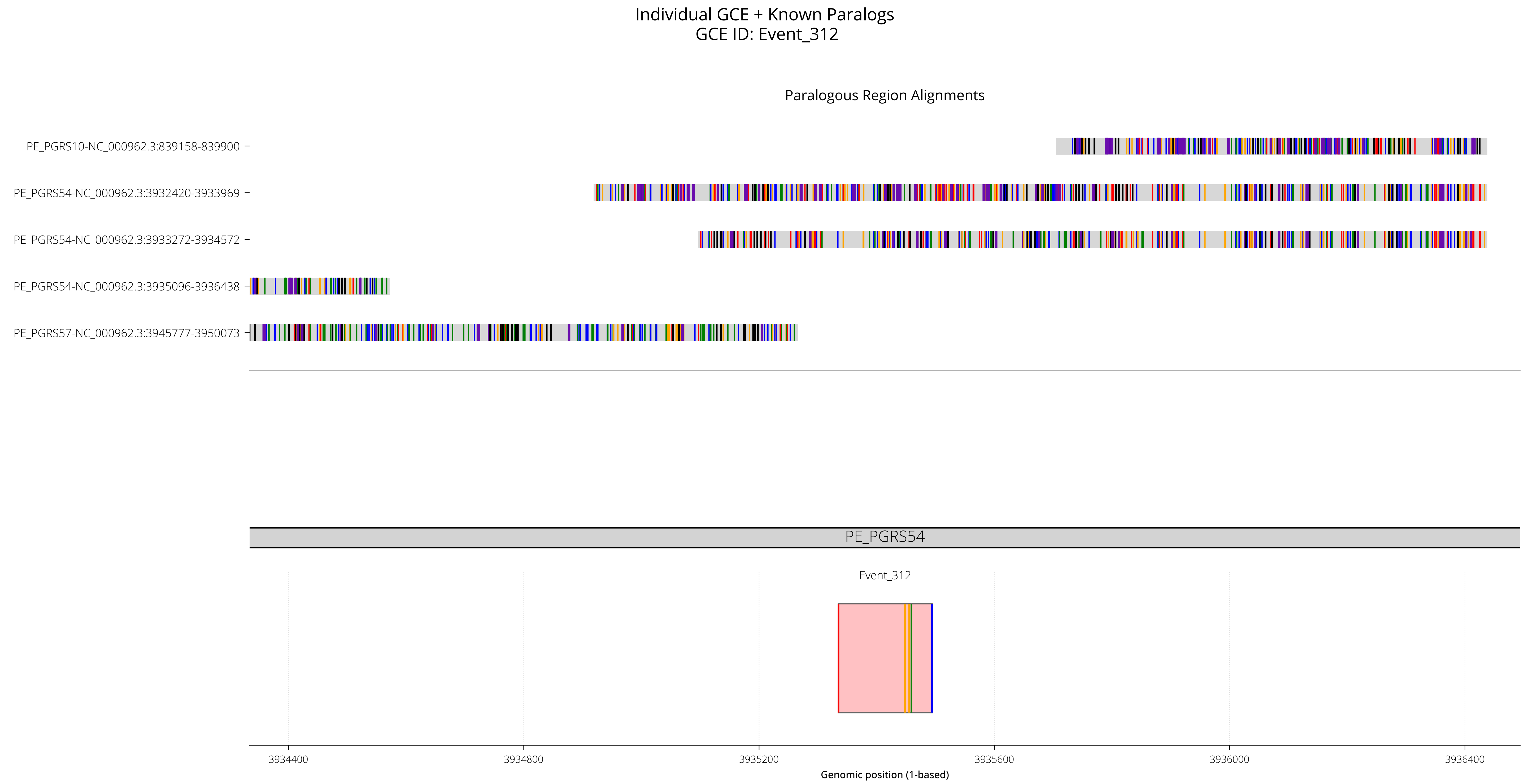

Individual GCE + Known Paralogs
GCE ID: Event_312
Paralogous Region Alignments
PE_PGRS10-NC_000962.3:839158-839900
PE_PGRS54-NC_000962.3:3932420-3933969
PE_PGRS54-NC_000962.3:3933272-3934572
PE_PGRS54-NC_000962.3:3935096-3936438
PE_PGRS57-NC_000962.3:3945777-3950073
PE_PGRS54
Event_312
3934400
3934800
3935200
3935600
3936000
3936400
Genomic position (1-based)

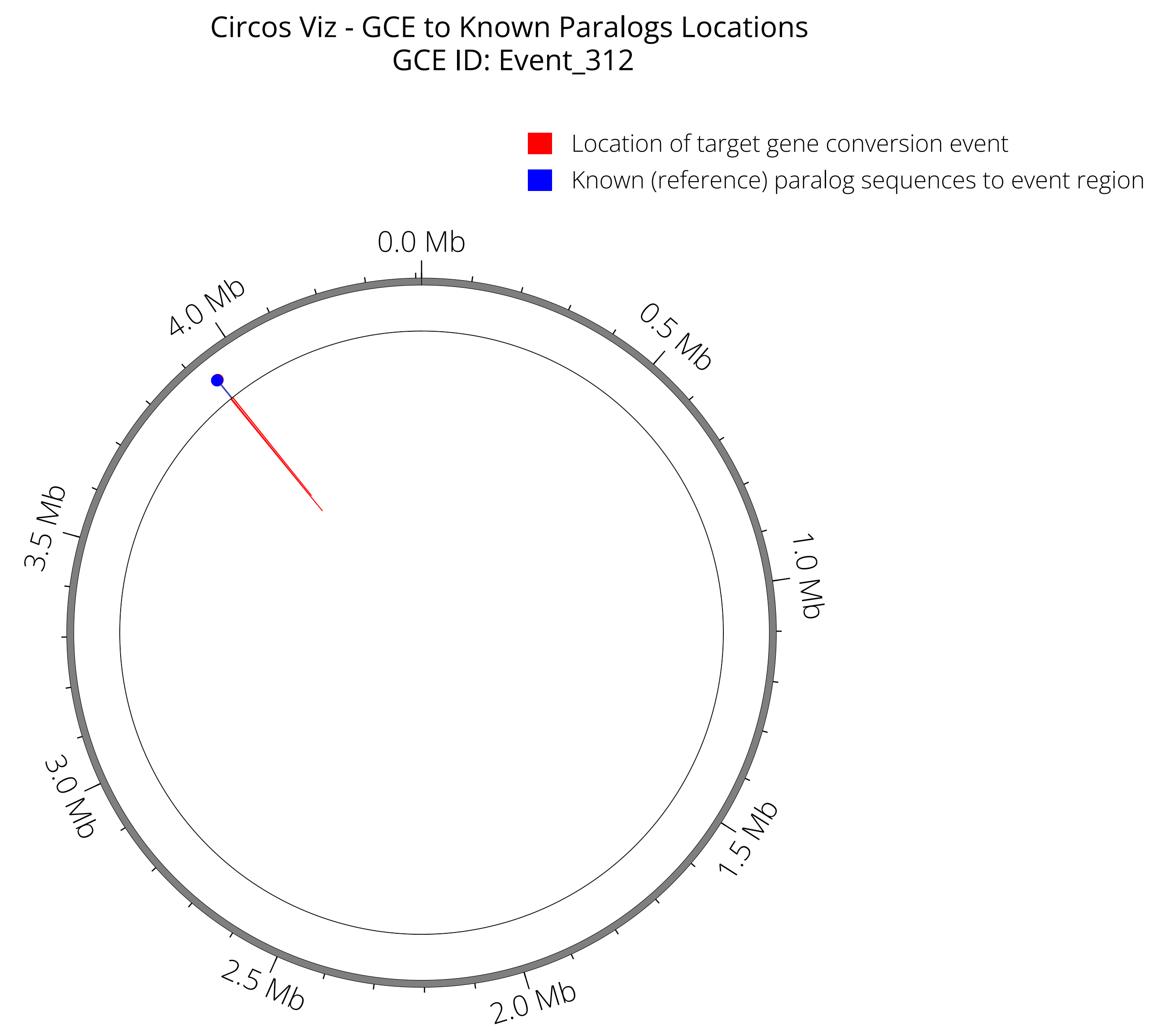
Circos Viz - GCE to Known Paralogs Locations
GCE ID: Event_312
Location of target gene conversion event
Known (reference) paralog sequences to event region
0.0 Mb
0.5 Mb
1.0 Mb
1.5 Mb
2.0 Mb
2.5 Mb
3.0 Mb
3.5 Mb
4.0 Mb

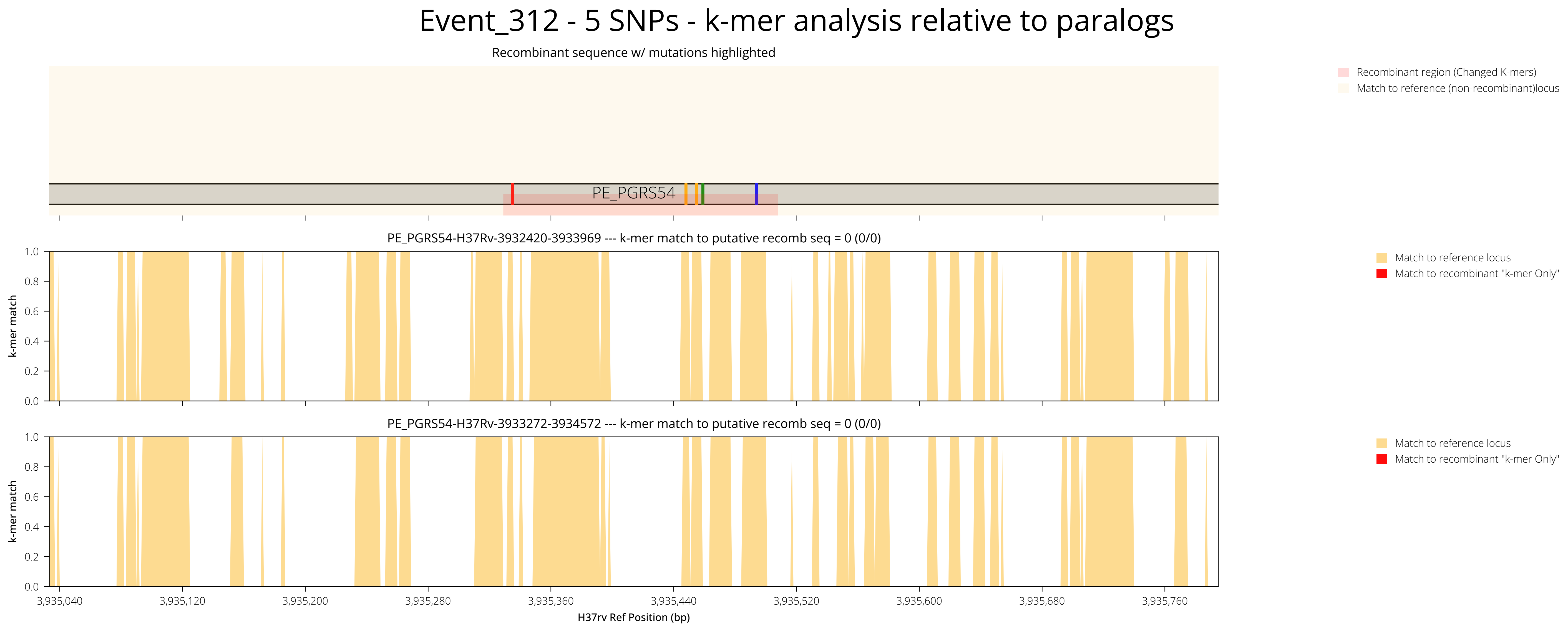

Event_312 - 5 SNPs - k-mer analysis relative to paralogs
Recombinant sequence w/ mutations highlighted
Recombinant region (Changed K-mers)
Match to reference (non-recombinant)locus
PE_PGRS54
PE_PGRS54-H37Rv-3932420-3933969 --- k-mer match to putative recomb seq = 0 (0/0)
Match to reference locus
Match to recombinant "k-mer Only"
k-mer match
1.0
0.8
0.6
0.4
0.2
0.0
PE_PGRS54-H37Rv-3933272-3934572 --- k-mer match to putative recomb seq = 0 (0/0)
Match to reference locus
Match to recombinant "k-mer Only"
k-mer match
1.0
0.8
0.6
0.4
0.2
0.0
3,935,040
3,935,120
3,935,200
3,935,280
3,935,360
3,935,440
3,935,520
3,935,600
3,935,680
3,935,760
H37rv Ref Position (bp)

Sub-phylogeny Viz + Paralog Alignments
GCE ID: Event_312

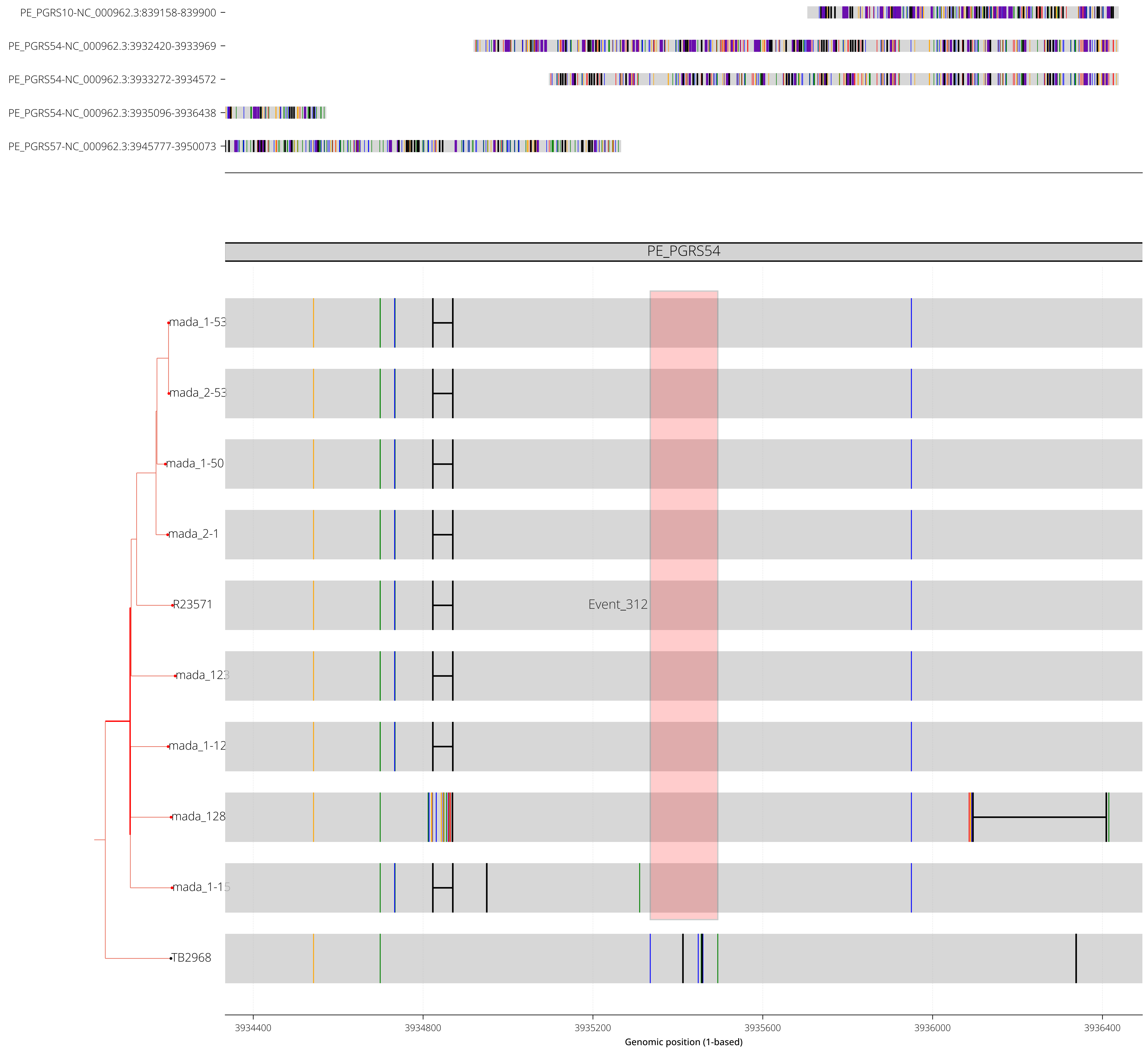

Full Phylogeny Viz + Paralog Alignments
GCE ID: Event_312

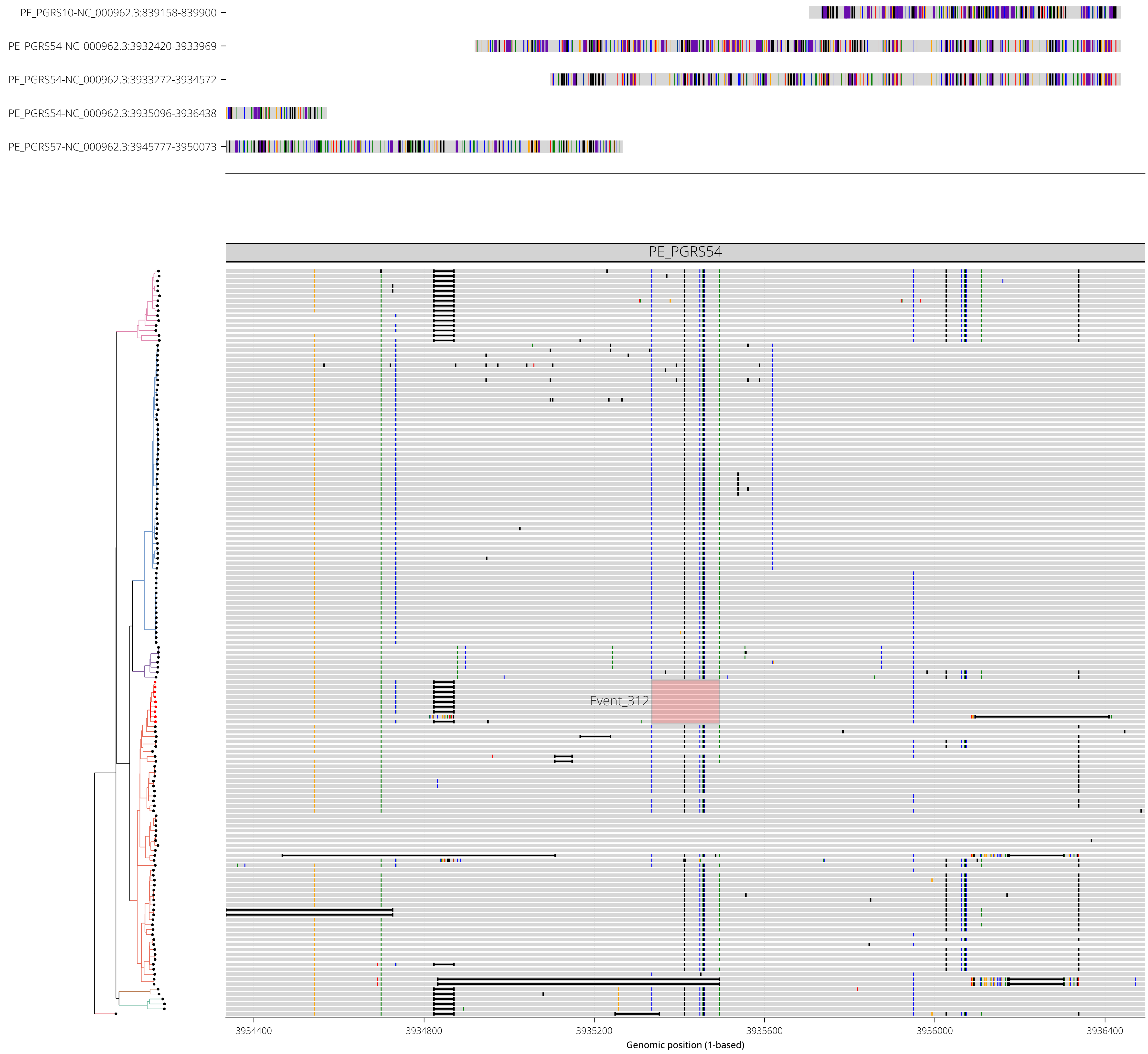

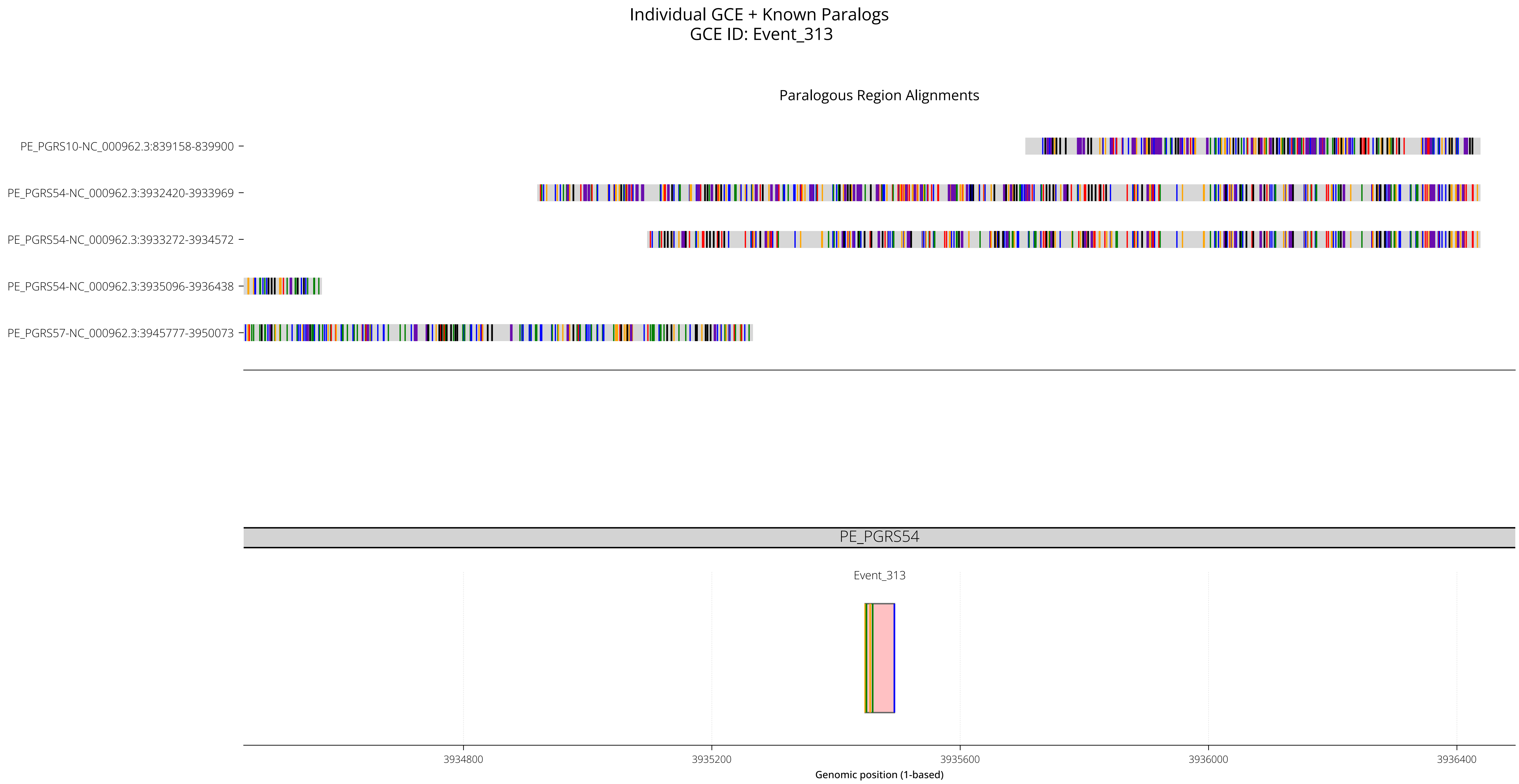

Individual GCE + Known Paralogs
GCE ID: Event_313
Paralogous Region Alignments
PE_PGRS10-NC_000962.3:839158-839900
PE_PGRS54-NC_000962.3:3932420-3933969
PE_PGRS54-NC_000962.3:3933272-3934572
PE_PGRS54-NC_000962.3:3935096-3936438
PE_PGRS57-NC_000962.3:3945777-3950073
PE_PGRS54
Event_313
3934800
3935200
3935600
3936000
3936400
Genomic position (1-based)

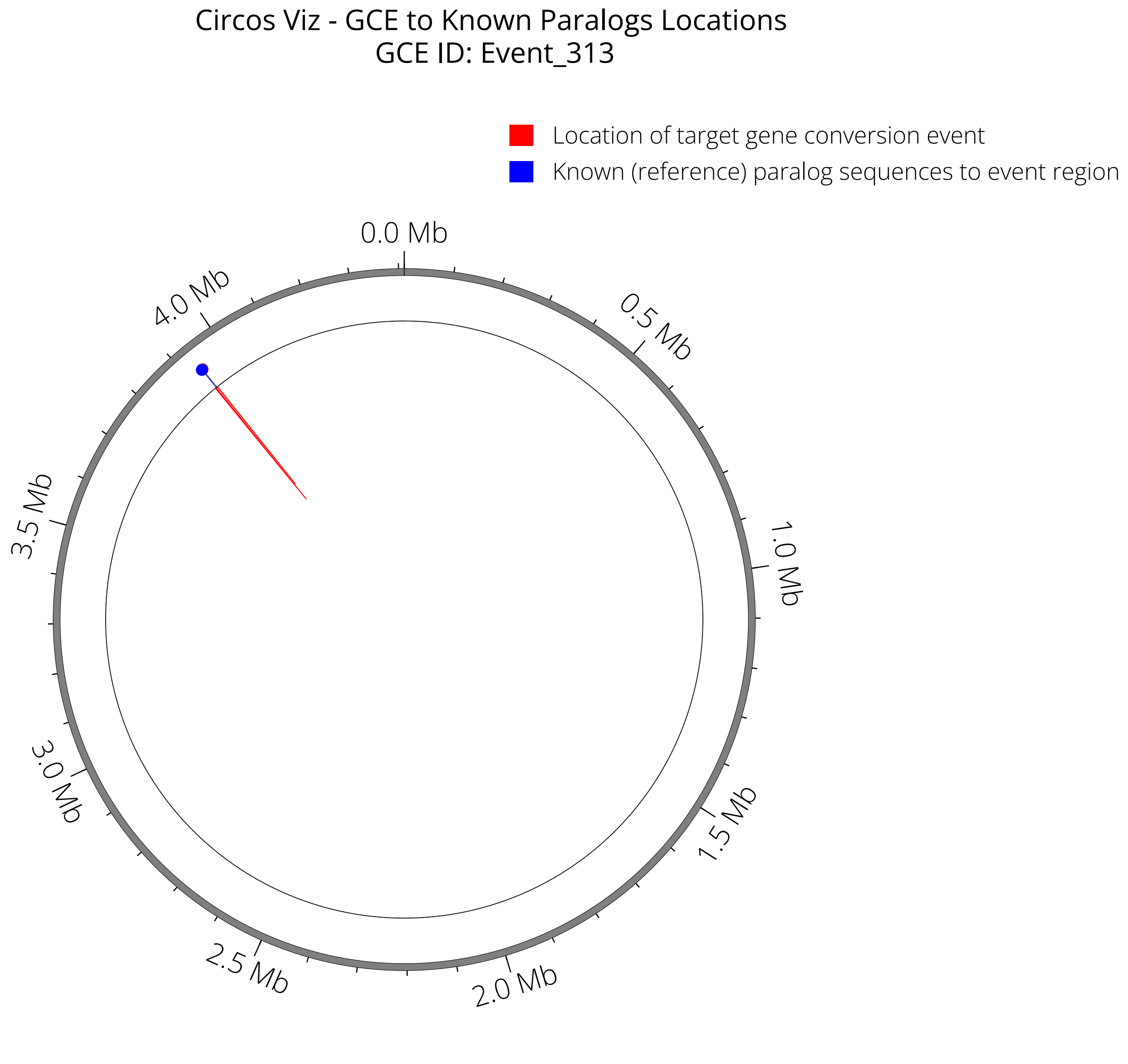
Circos Viz - GCE to Known Paralogs Locations
GCE ID: Event_313
Location of target gene conversion event
Known (reference) paralog sequences to event region
0.0 Mb
0.5 Mb
1.0 Mb
1.5 Mb
2.0 Mb
2.5 Mb
3.0 Mb
3.5 Mb
4.0 Mb

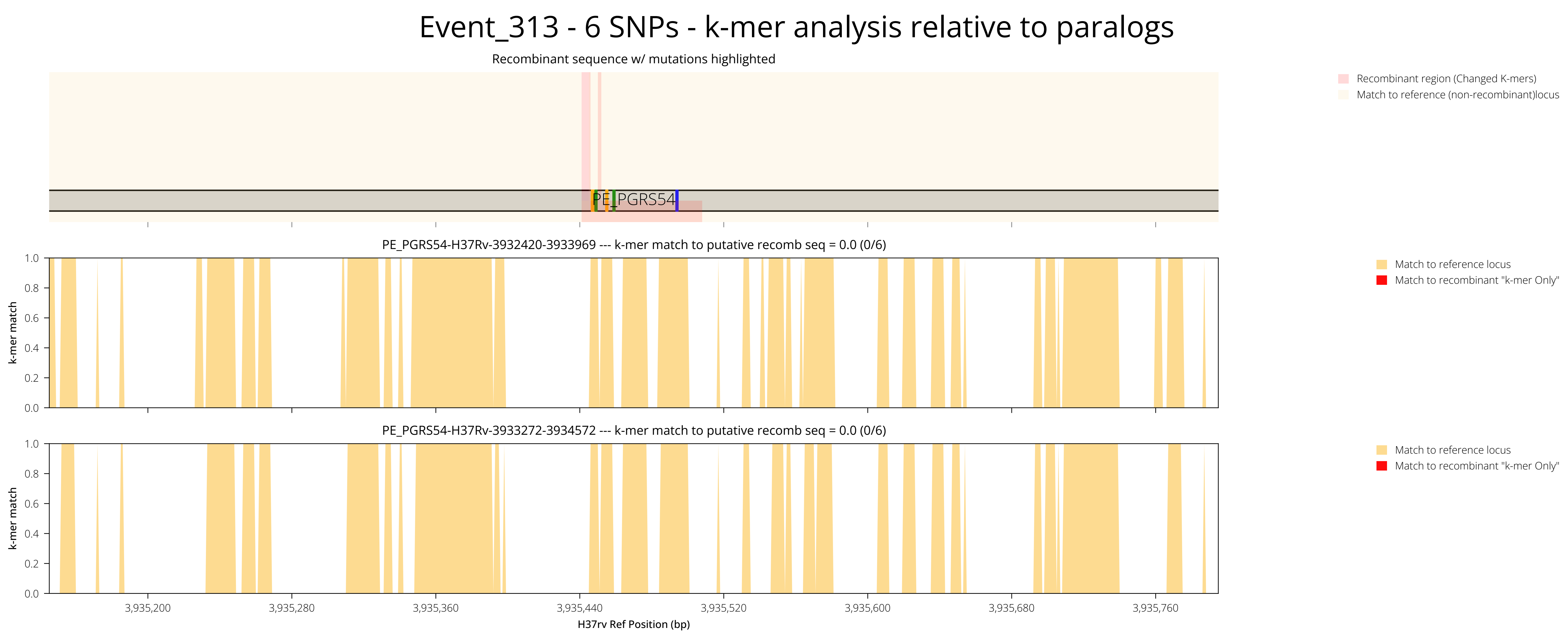
Event_313 - 6 SNPs - k-mer analysis relative to paralogs
Recombinant sequence w/ mutations highlighted
PE_PGRS54
Recombinant region (Changed K-mers)
Match to reference (non-recombinant)locus
PE_PGRS54-H37Rv-3932420-3933969 --- k-mer match to putative recomb seq = 0.0 (0/6)
k-mer match
1.0
0.8
0.6
0.4
0.2
0.0
Match to reference locus
Match to recombinant "k-mer Only"
PE_PGRS54-H37Rv-3933272-3934572 --- k-mer match to putative recomb seq = 0.0 (0/6)
k-mer match
1.0
0.8
0.6
0.4
0.2
0.0
Match to reference locus
Match to recombinant "k-mer Only"
3,935,200
3,935,280
3,935,360
3,935,440
3,935,520
3,935,600
3,935,680
3,935,760
H37rv Ref Position (bp)

Sub-phylogeny Viz + Paralog Alignments
GCE ID: Event_313

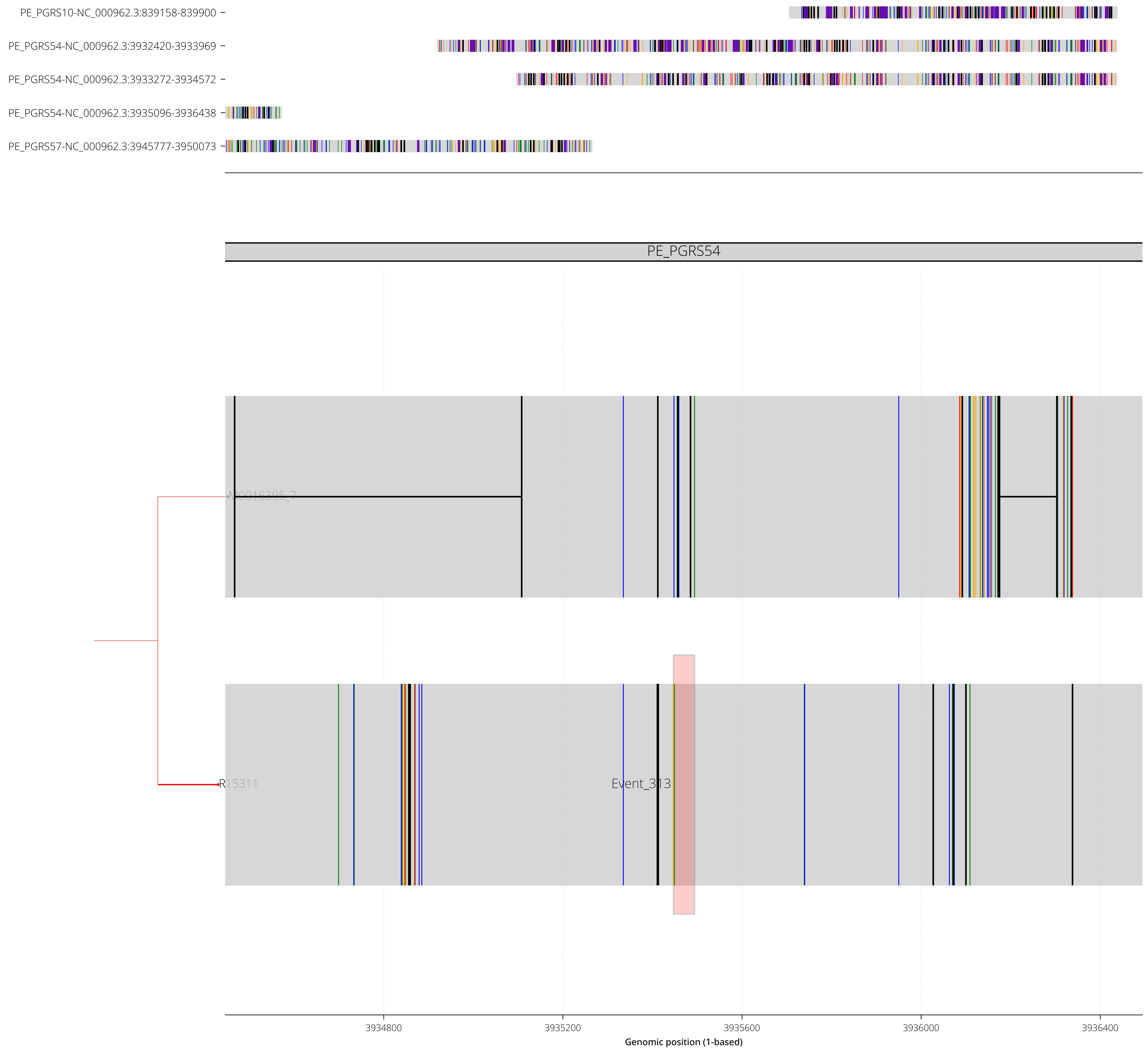

Full Phylogeny Viz + Paralog Alignments
GCE ID: Event_313

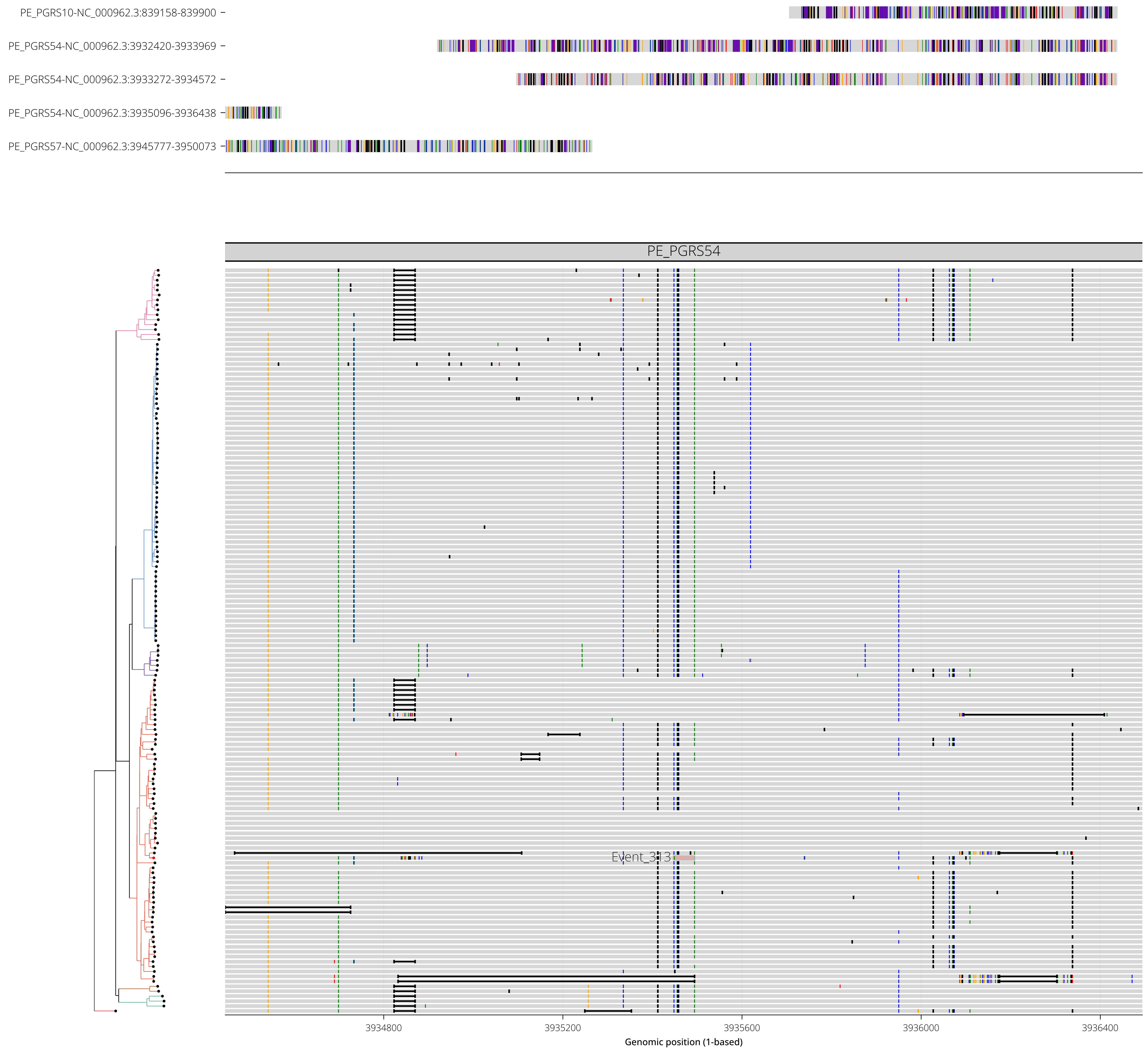

Individual GCE + Known Paralogs
GCE ID: Event_314

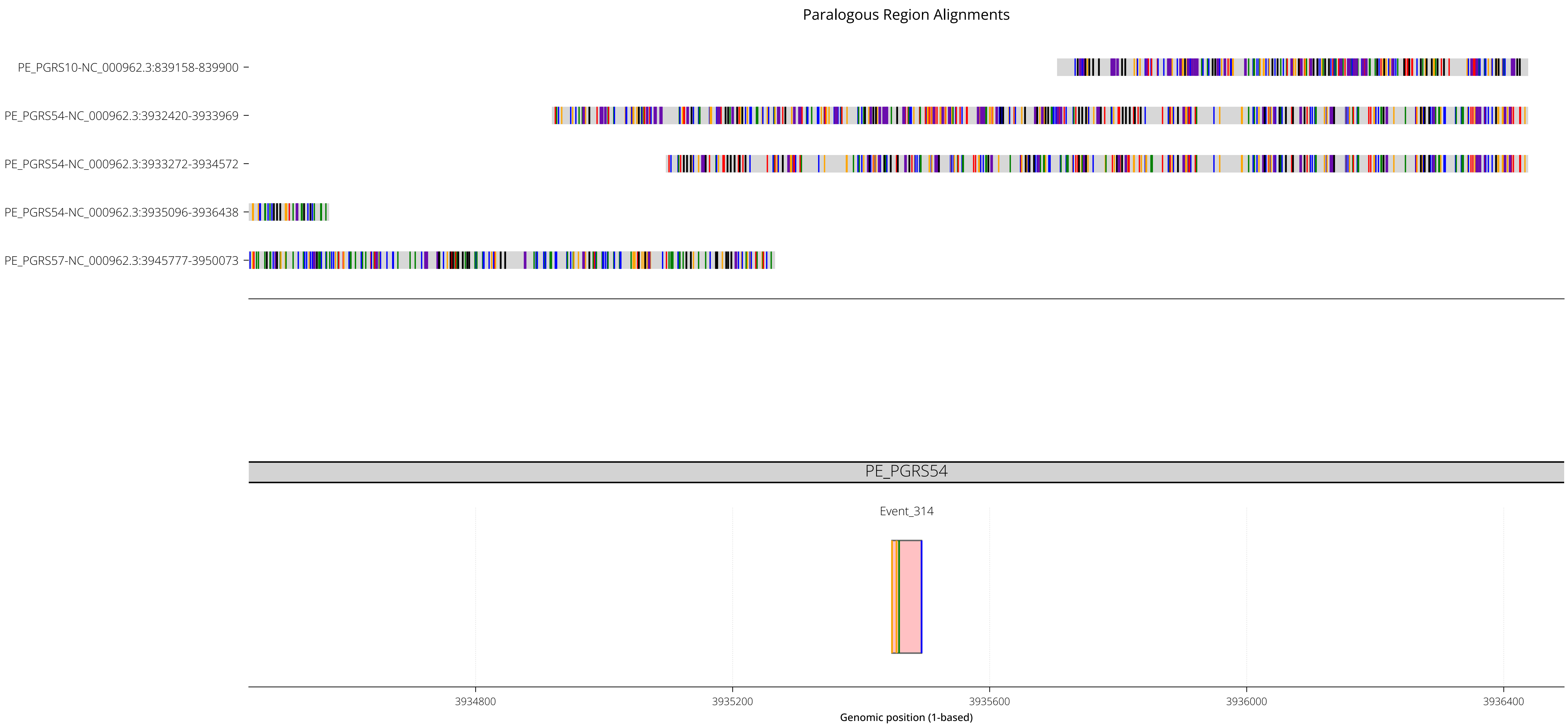

Circos Viz - GCE to Known Paralogs Locations
GCE ID: Event_314

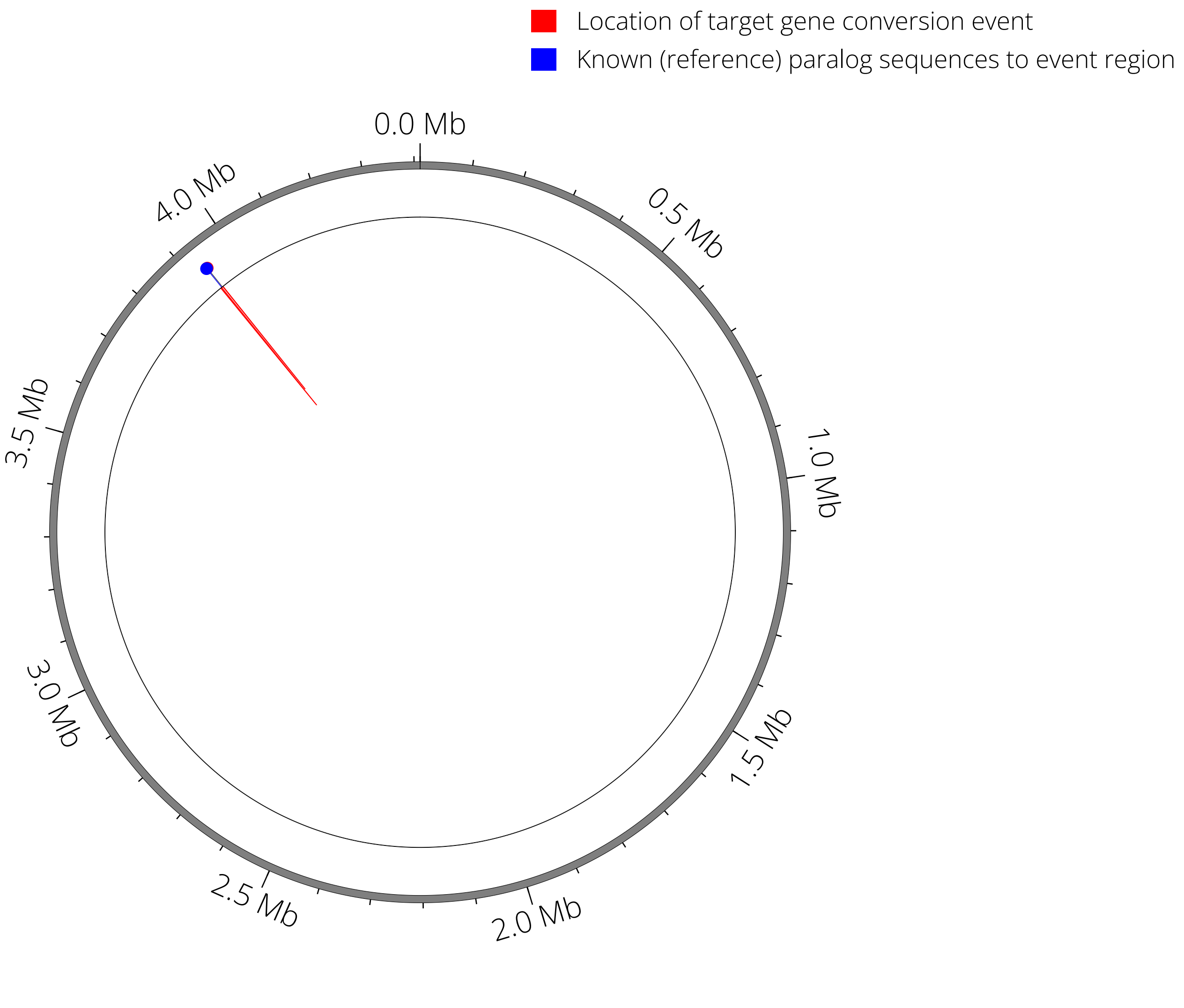

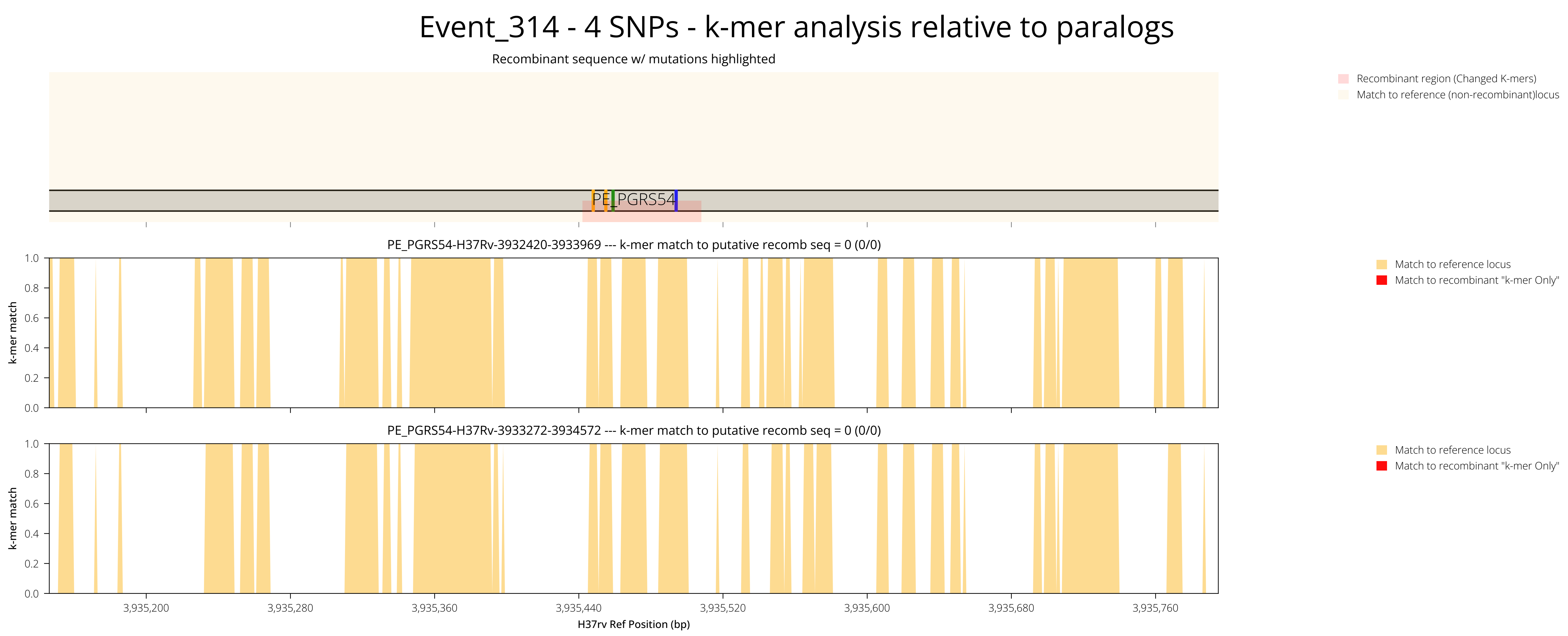

Event_314 - 4 SNPs - k-mer analysis relative to paralogs
Recombinant sequence w/ mutations highlighted
PE_PGRS54
Recombinant region (Changed K-mers)
Match to reference (non-recombinant)locus
PE_PGRS54-H37Rv-3932420-3933969 --- k-mer match to putative recomb seq = 0 (0/0)
PE_PGRS54-H37Rv-3933272-3934572 --- k-mer match to putative recomb seq = 0 (0/0)
k-mer match
1.0
0.8
0.6
0.4
0.2
0.0
Match to reference locus
Match to recombinant "k-mer Only"
3,935,200
3,935,280
3,935,360
3,935,440
3,935,520
3,935,600
3,935,680
3,935,760
H37rv Ref Position (bp)

Sub-phylogeny Viz + Paralog Alignments
GCE ID: Event_314

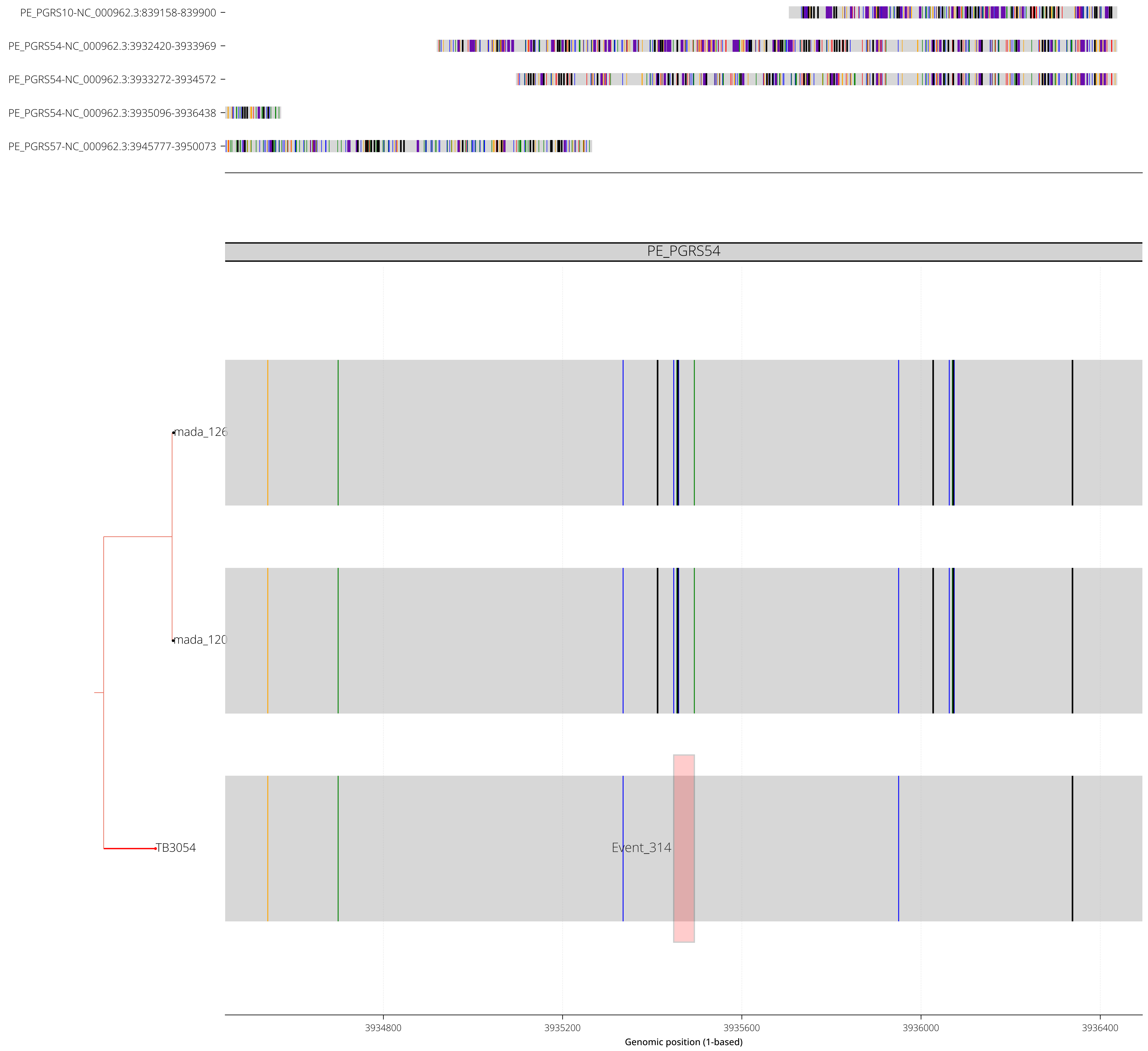

Full Phylogeny Viz + Paralog Alignments
GCE ID: Event_314

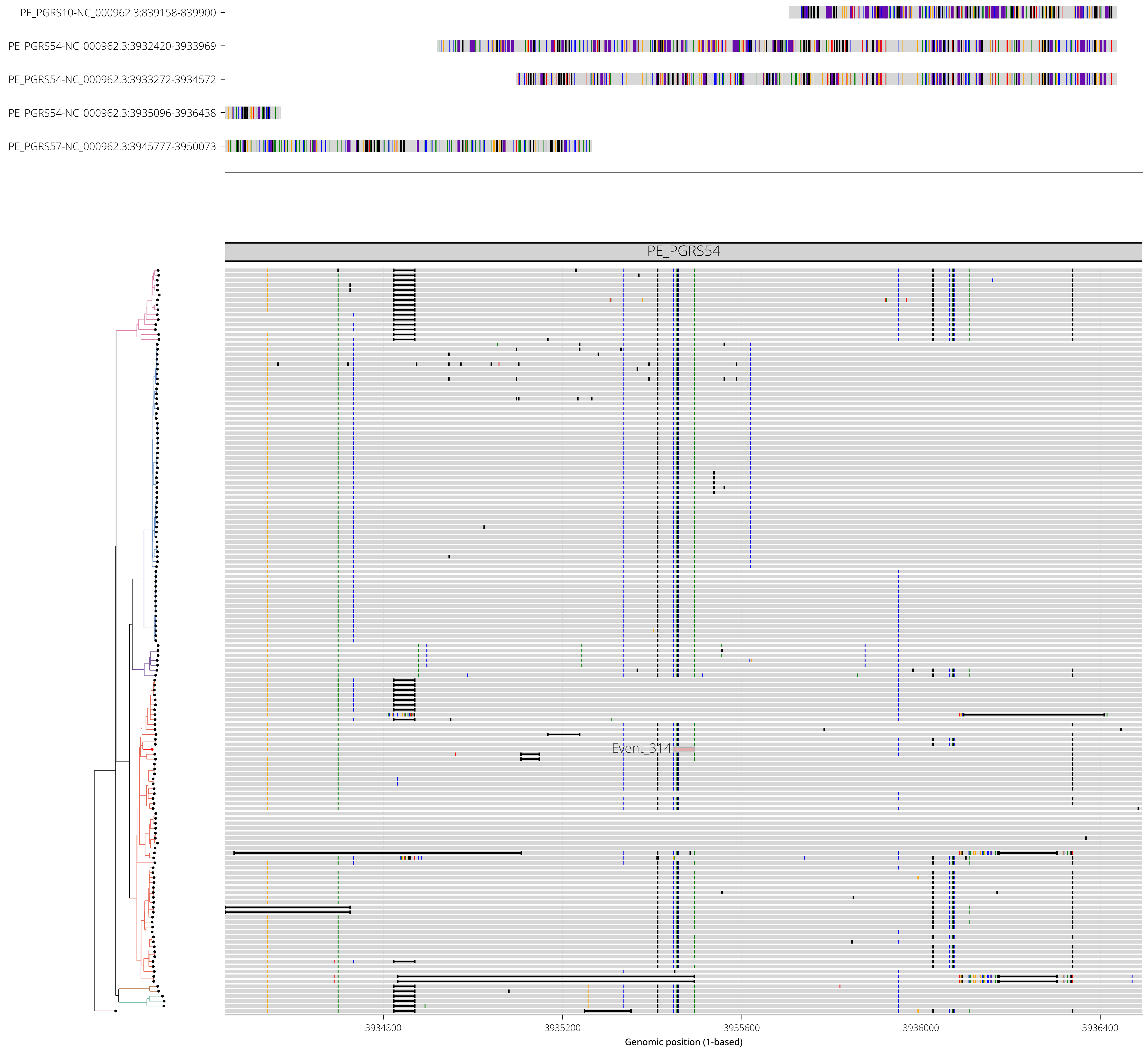

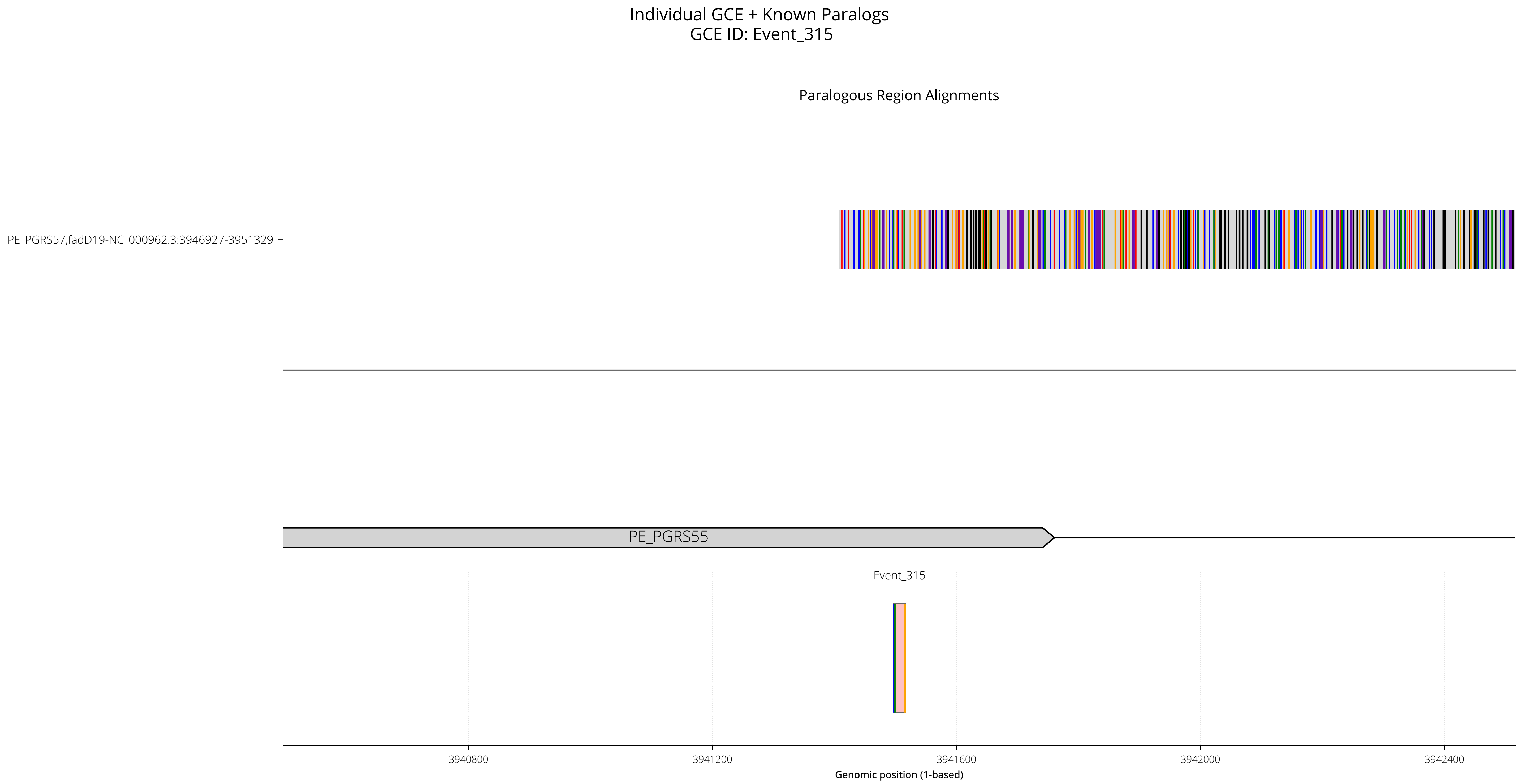
Individual GCE + Known Paralogs
GCE ID: Event_315
Paralogous Region Alignments
PE_PGRS57,fadD19-NC_000962.3:3946927-3951329
PE_PGRS55
Event_315
3940800
3941200
3941600
3942000
3942400
Genomic position (1-based)

Circos Viz - GCE to Known Paralogs Locations
GCE ID: Event_315

Location of target gene conversion event
Known (reference) paralog sequences to event region

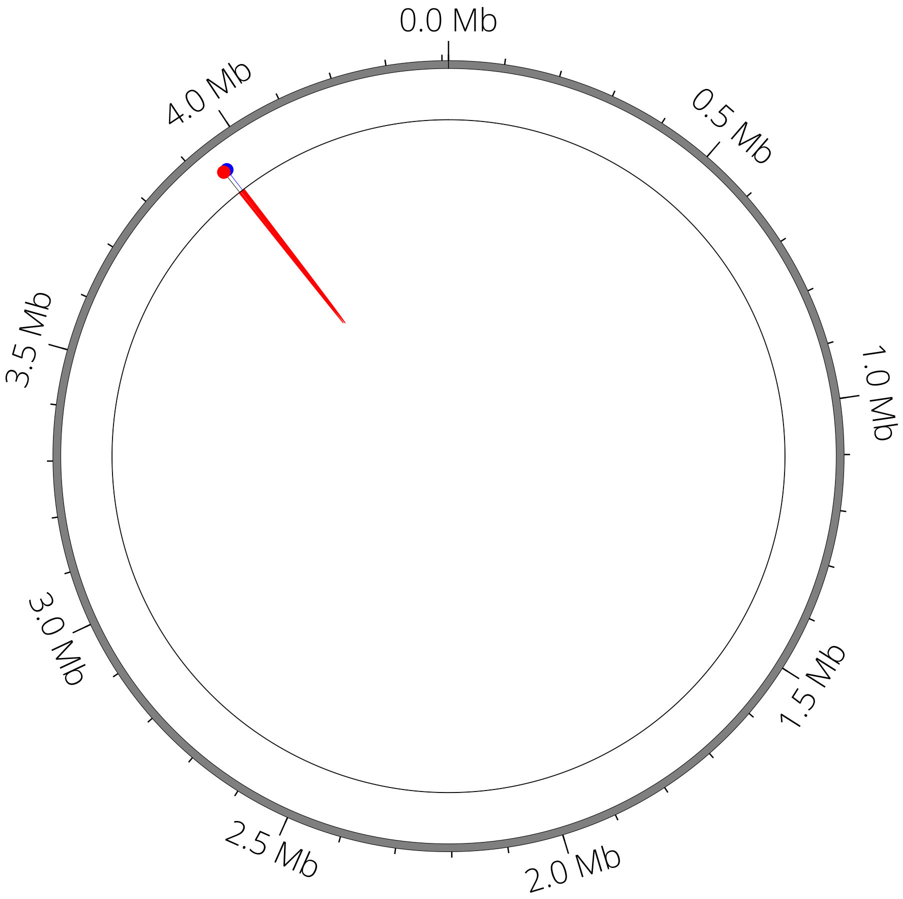

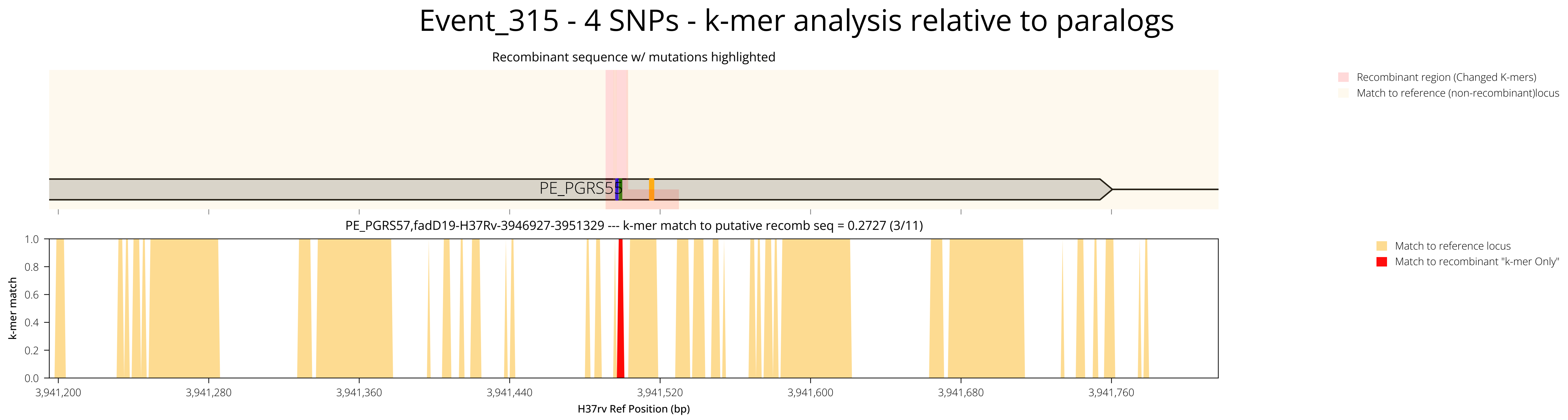

Event_315 - 4 SNPs - k-mer analysis relative to paralogs
Recombinant sequence w/ mutations highlighted
Recombinant region (Changed K-mers)
Match to reference (non-recombinant)locus
PE_PGRS55
PE_PGRS57,fadD19-H37Rv-3946927-3951329 --- k-mer match to putative recomb seq = 0.2727 (3/11)
Match to reference locus
Match to recombinant "k-mer Only"
k-mer match
1.0
0.8
0.6
0.4
0.2
0.0
3,941,200
3,941,280
3,941,360
3,941,440
3,941,520
3,941,600
3,941,680
3,941,760
H37rv Ref Position (bp)

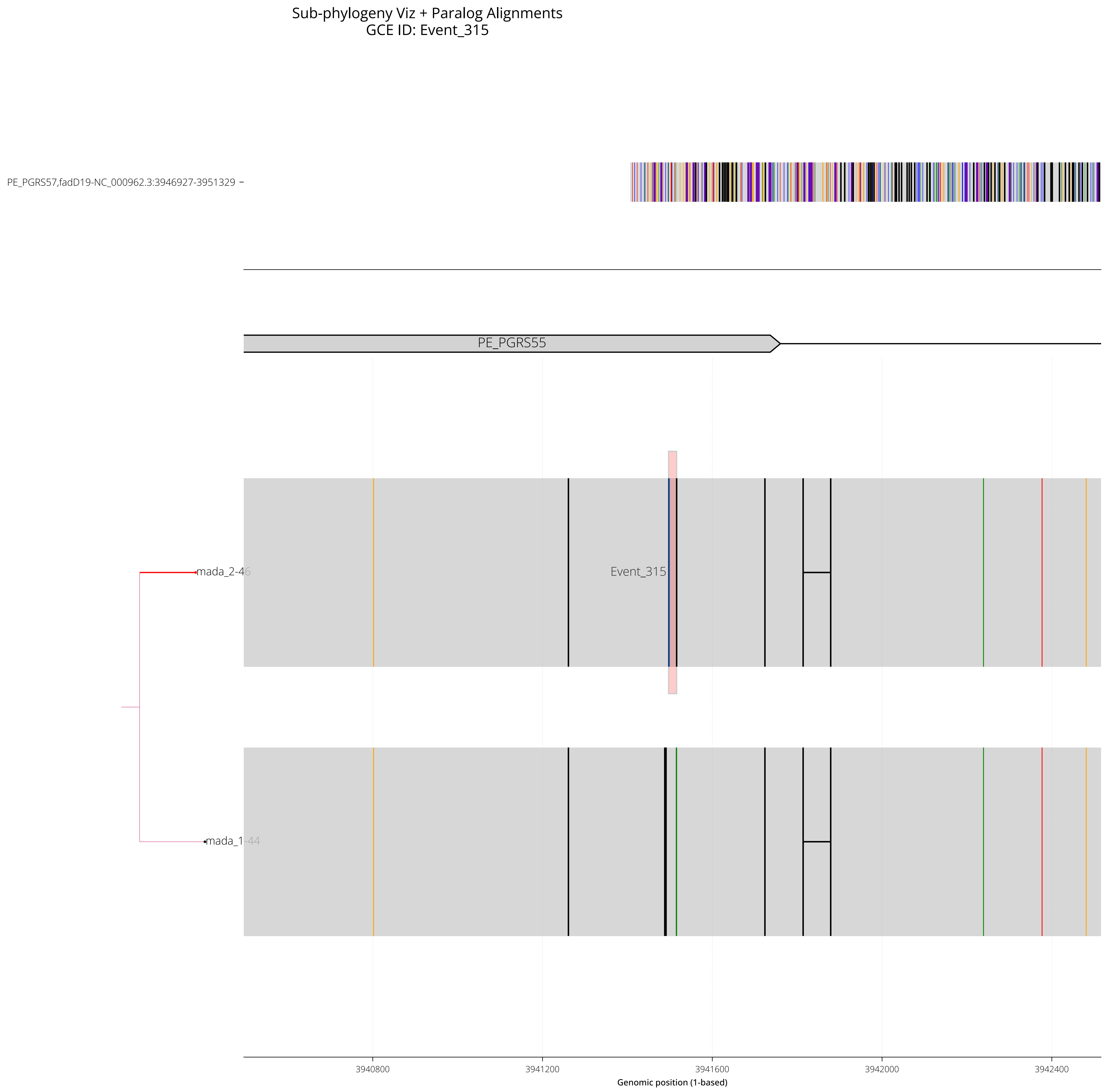

Sub-phylogeny Viz + Paralog Alignments
GCE ID: Event_315
PE_PGRS57,fadD19-NC_000962.3:3946927-3951329
PE_PGRS55
mada_2-46
Event_315
mada_1-44
3940800
3941200
3941600
3942000
3942400
Genomic position (1-based)

### Full Phylogeny Viz + Paralog Alignments
GCE ID: Event_315

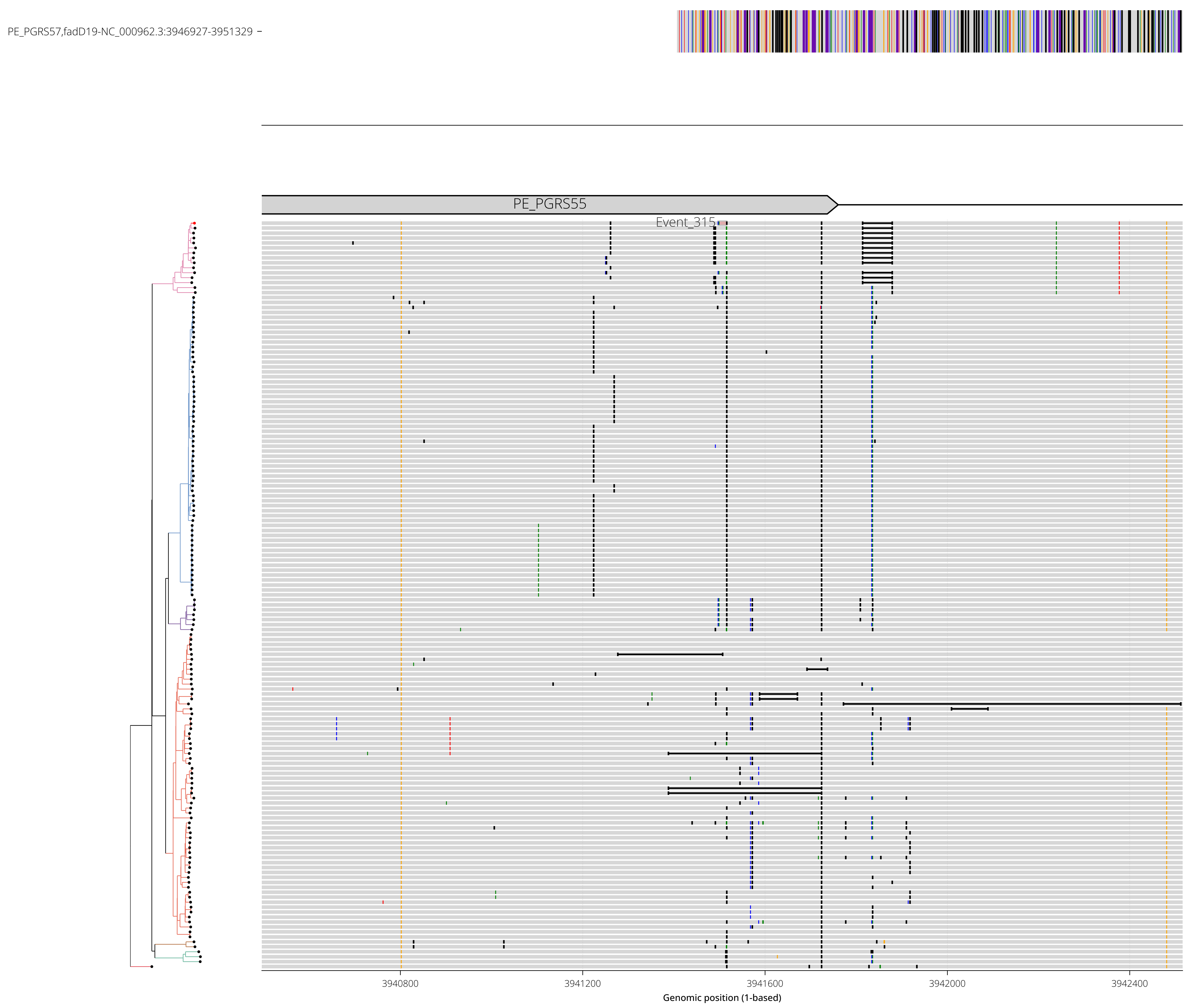

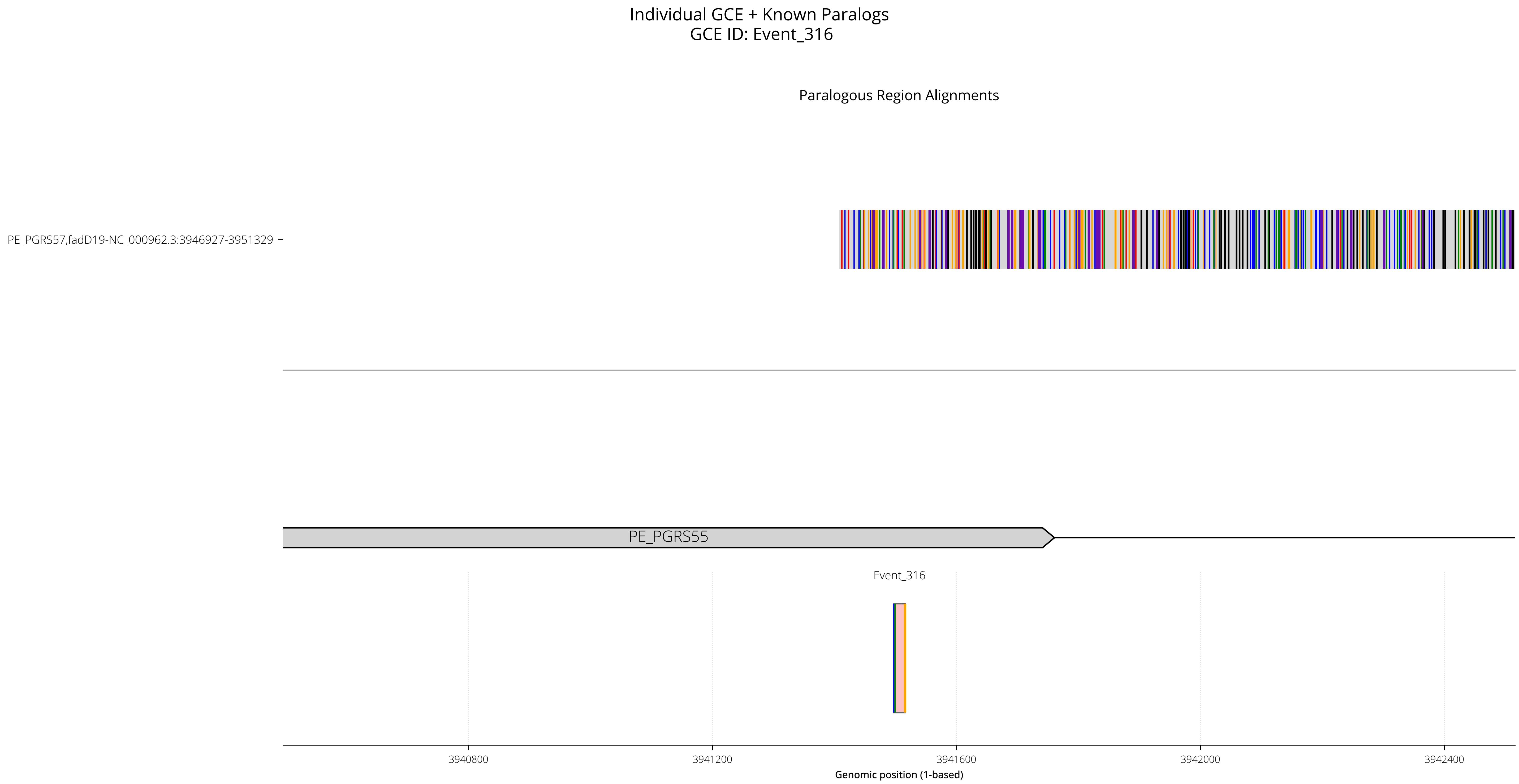

Individual GCE + Known Paralogs
GCE ID: Event_316
Paralogous Region Alignments
PE_PGRS57,fadD19-NC_000962.3:3946927-3951329
PE_PGRS55
Event_316
3940800
3941200
3941600
3942000
3942400
Genomic position (1-based)

Circos Viz - GCE to Known Paralogs Locations
GCE ID: Event_316

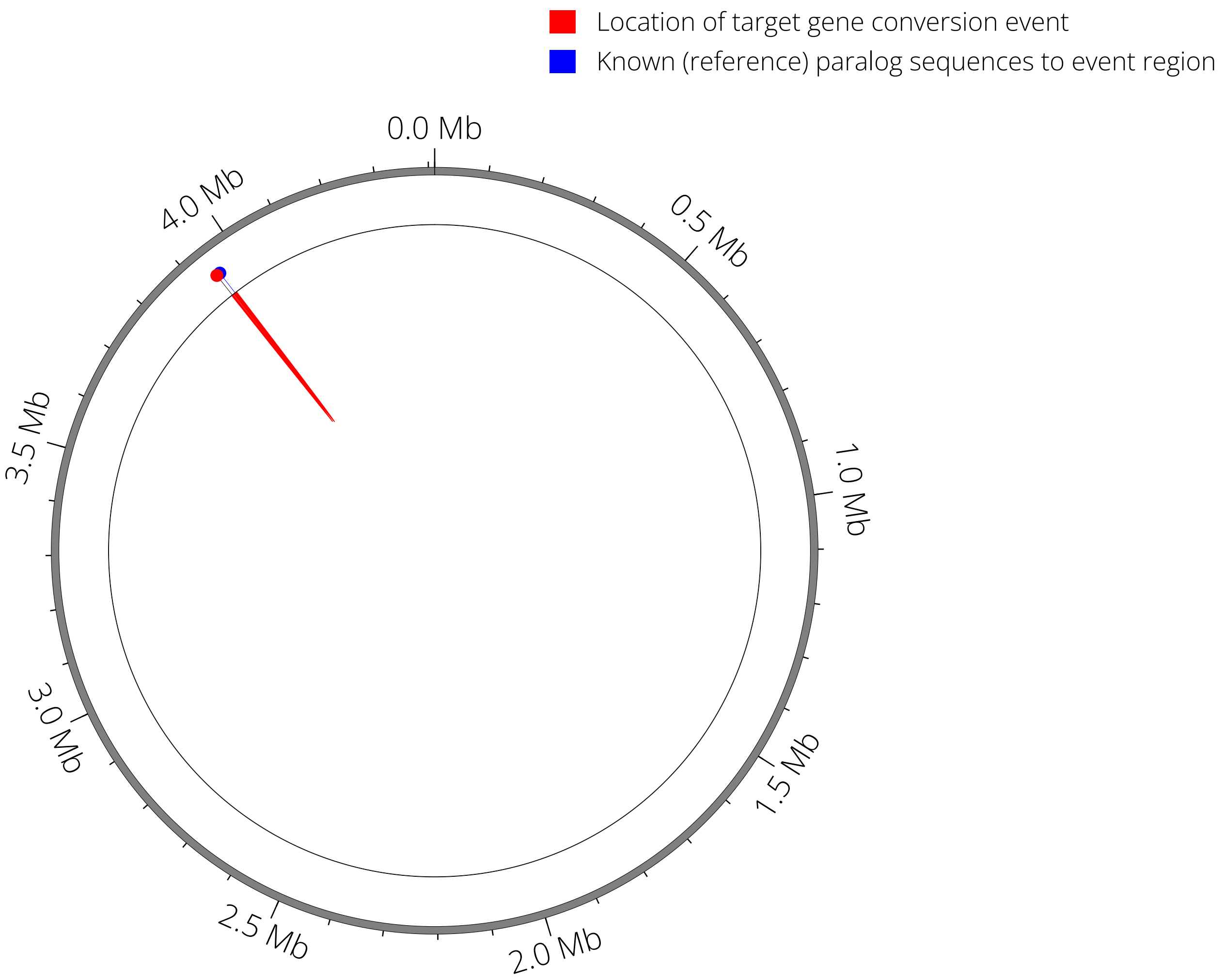

### Event_316 - 4 SNPs - k-mer analysis relative to paralogs

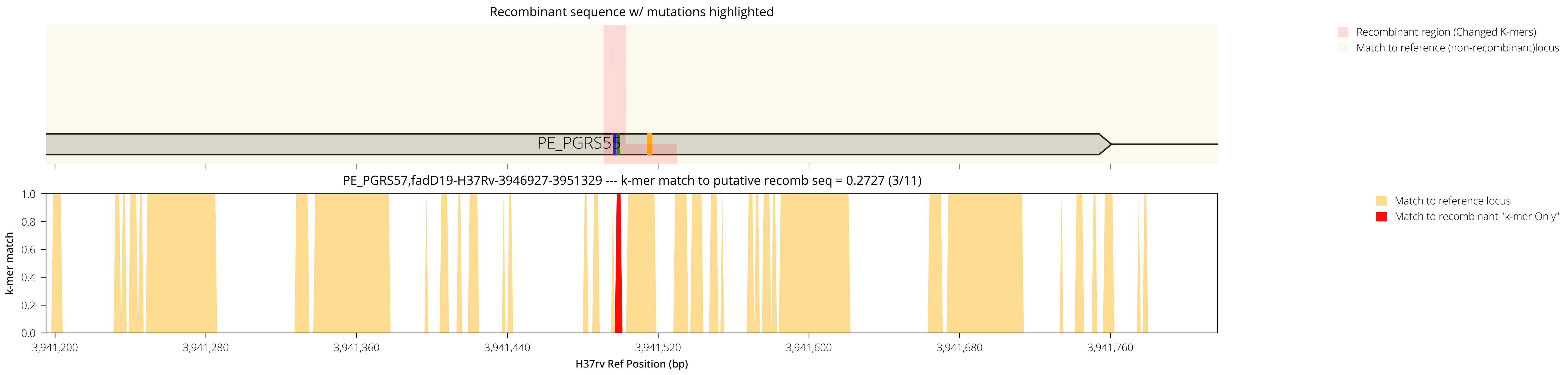

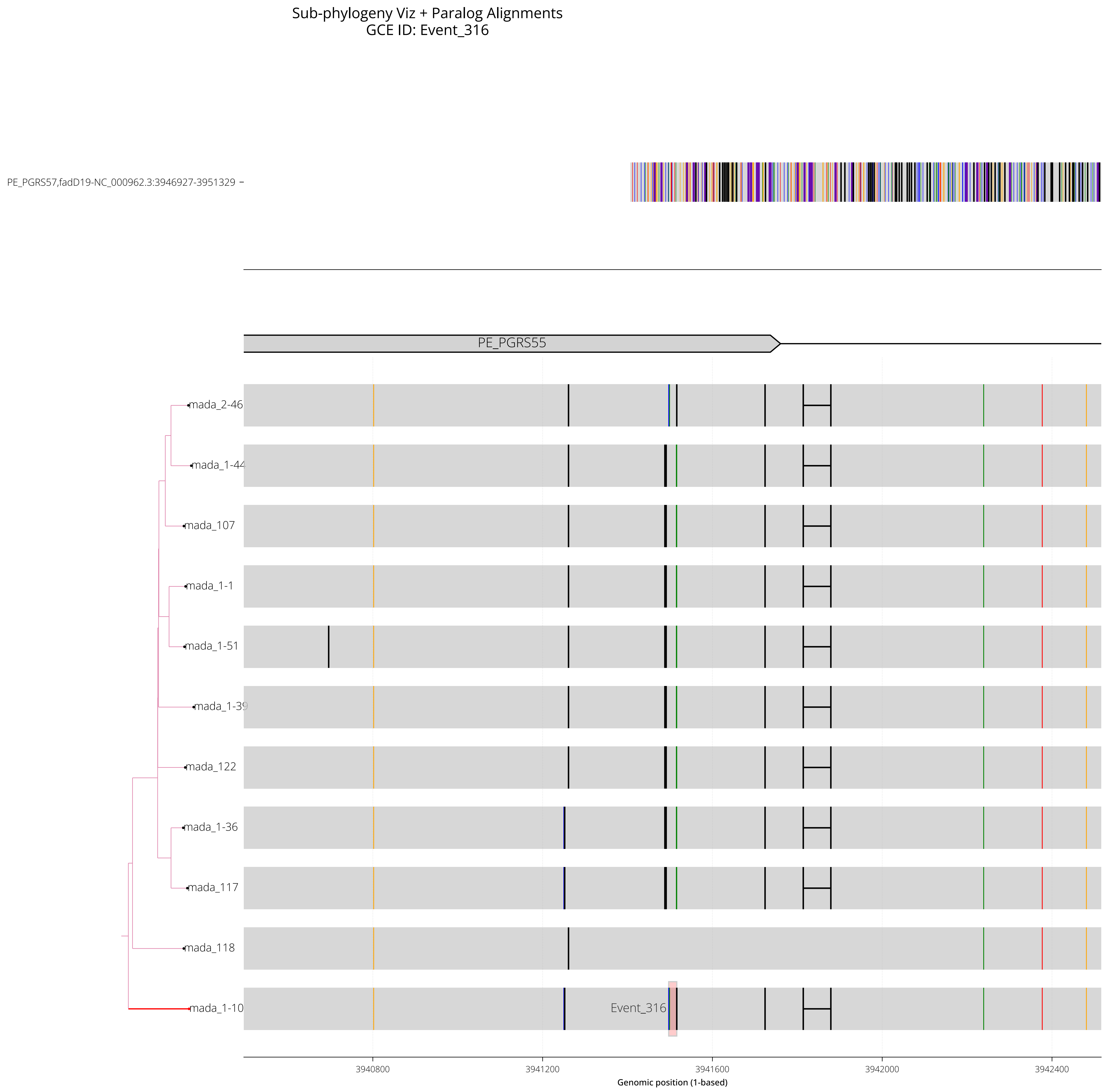
Sub-phylogeny Viz + Paralog Alignments
GCE ID: Event_316
PE_PGRS57,fadD19-NC_000962.3:3946927-3951329
PE_PGRS55
mada_2-46
mada_1-44
mada_107
mada_1-1
mada_1-51
mada_1-39
mada_122
mada_1-36
mada_117
mada_118
mada_1-10
Event_316
3940800
3941200
3941600
3942000
3942400
Genomic position (1-based)

### Full Phylogeny Viz + Paralog Alignments
GCE ID: Event_316

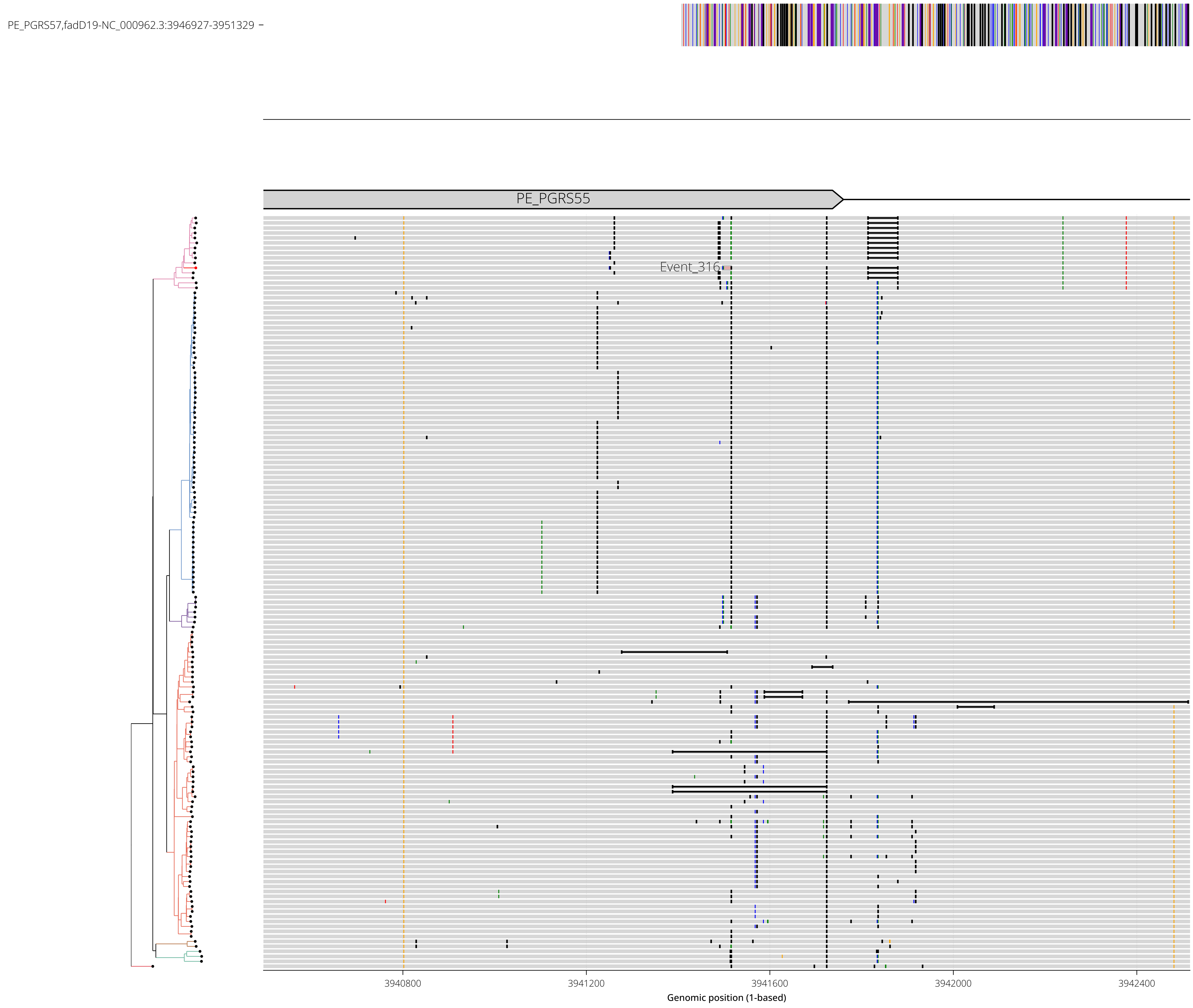

Individual GCE + Known Paralogs
GCE ID: Event_317

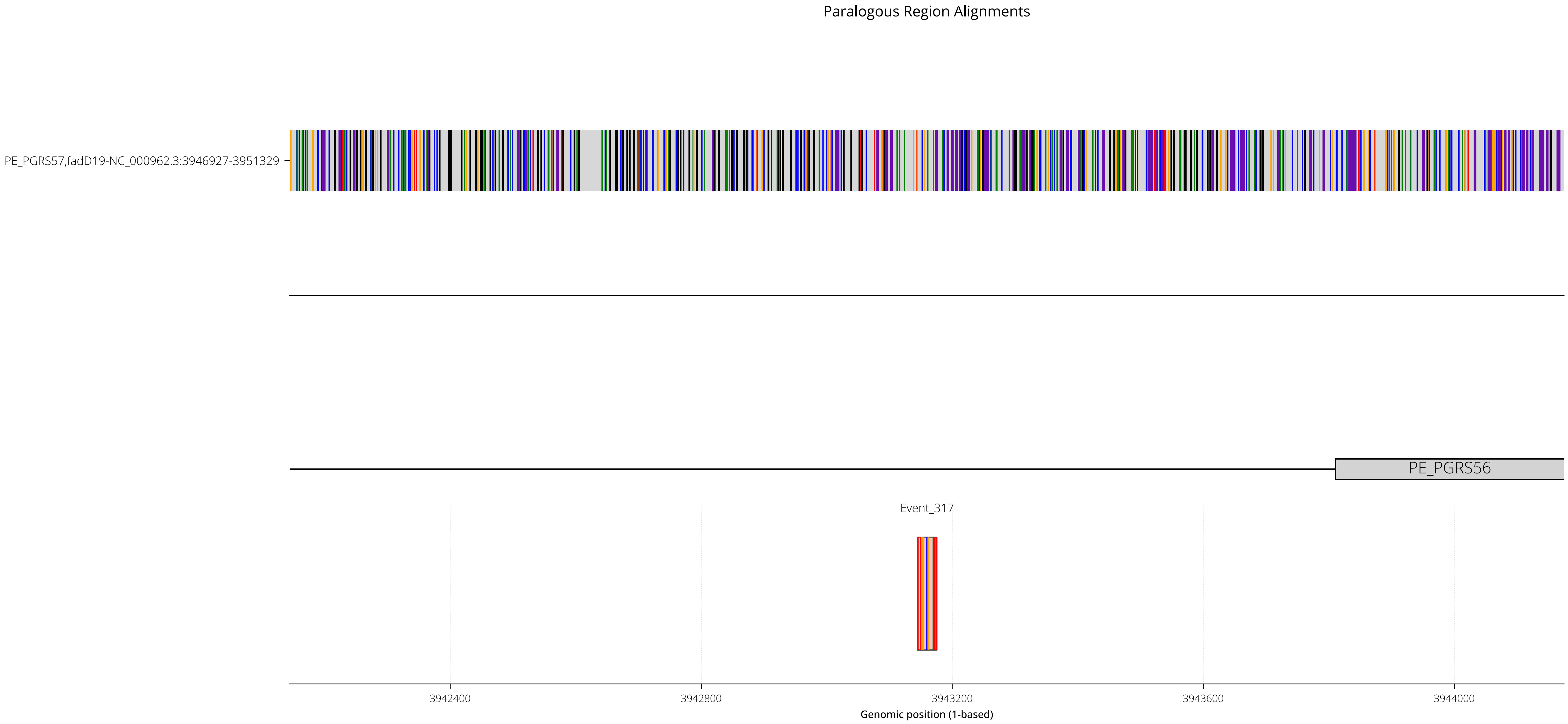

Circos Viz - GCE to Known Paralogs Locations
GCE ID: Event_317

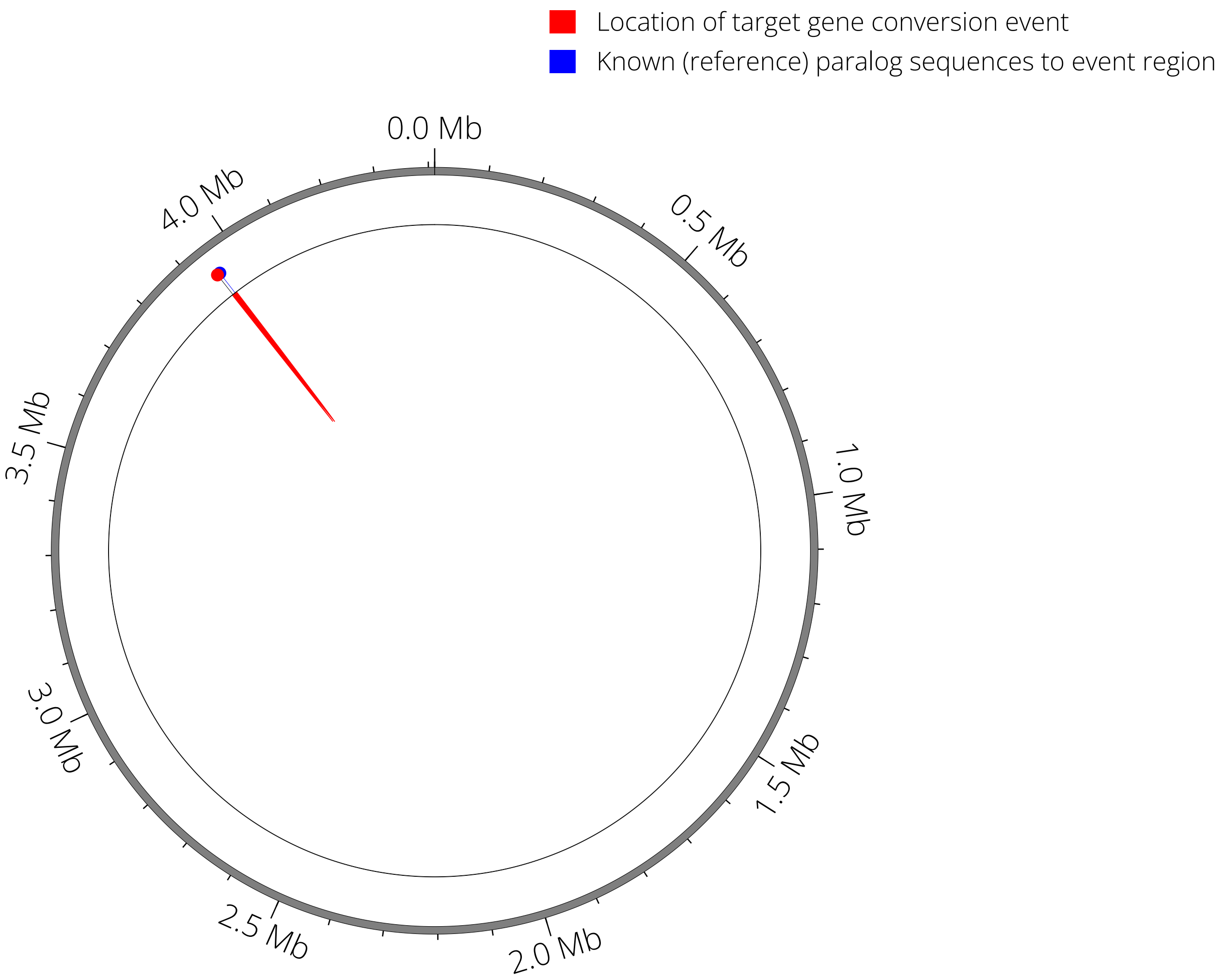

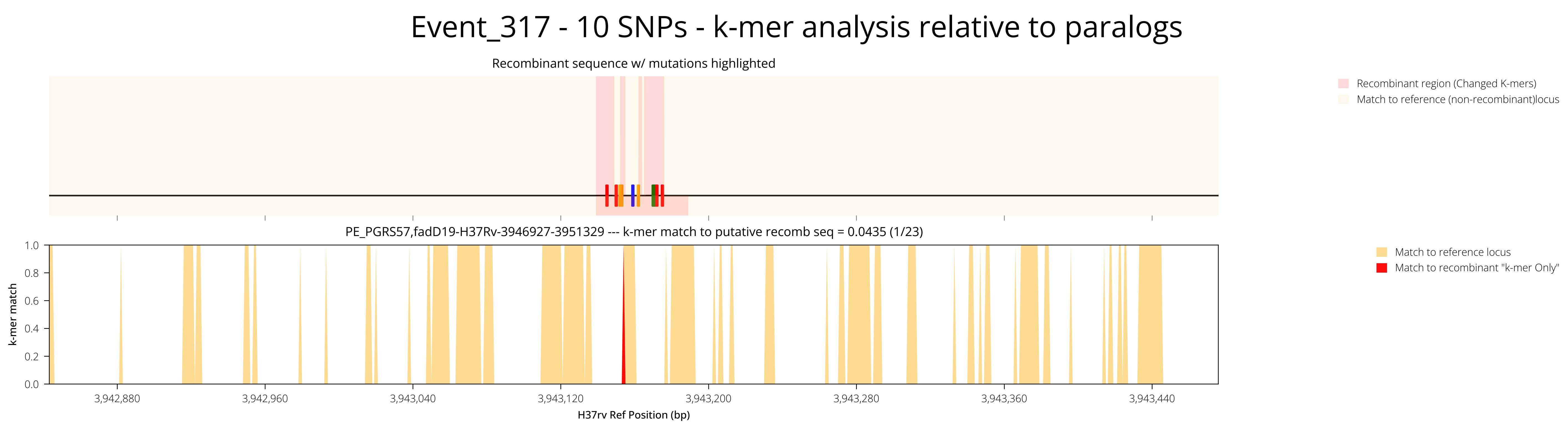

Event_317 - 10 SNPs - k-mer analysis relative to paralogs
Recombinant sequence w/ mutations highlighted
Recombinant region (Changed K-mers)
Match to reference (non-recombinant)locus
PE_PGRS57,fadD19-H37Rv-3946927-3951329 --- k-mer match to putative recomb seq = 0.0435 (1/23)
Match to reference locus
Match to recombinant "k-mer Only"
k-mer match
1.0
0.8
0.6
0.4
0.2
0.0
3,942,880
3,942,960
3,943,040
3,943,120
3,943,200
3,943,280
3,943,360
3,943,440
H37rv Ref Position (bp)

Sub-phylogeny Viz + Paralog Alignments
GCE ID: Event_317

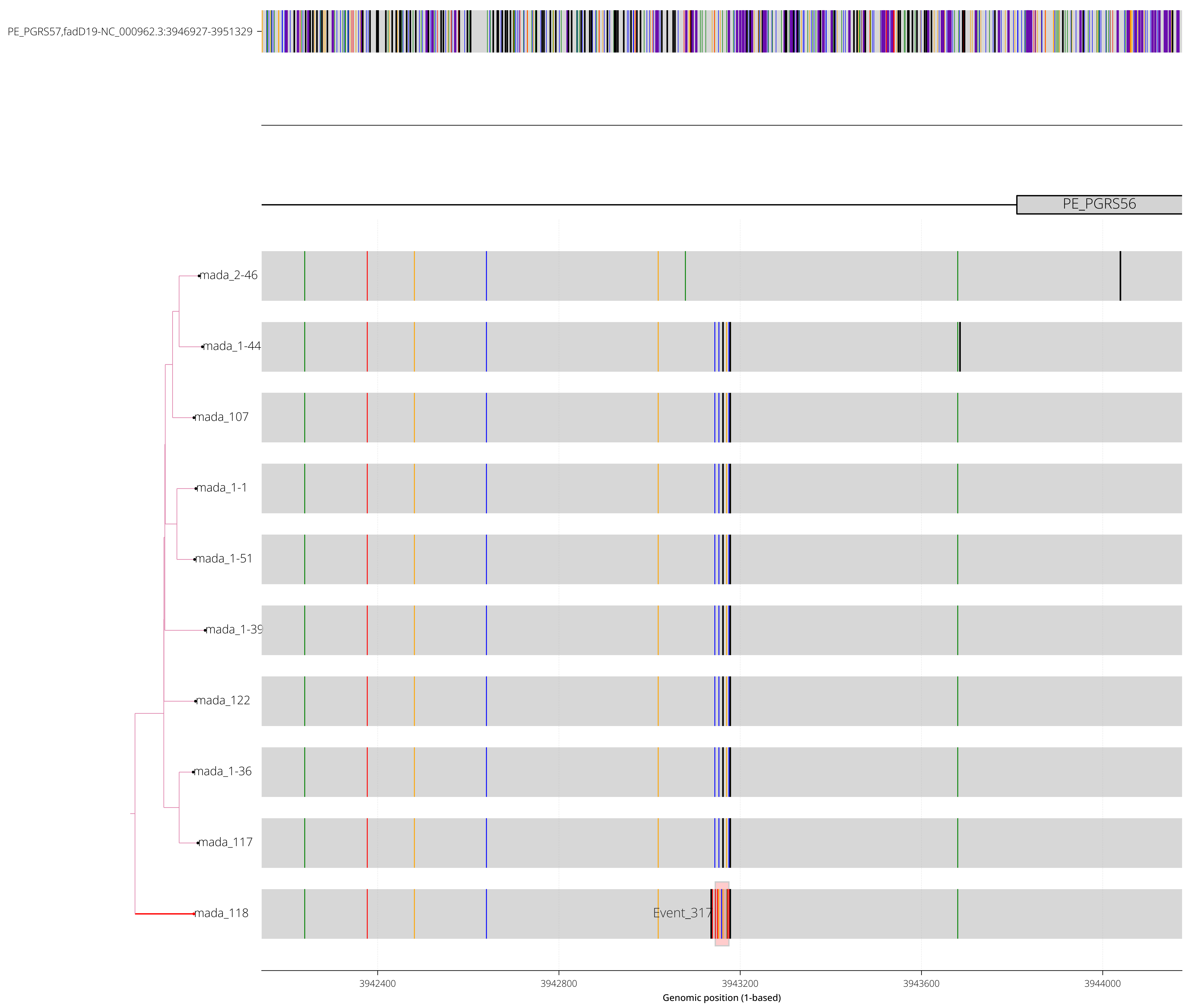

Full Phylogeny Viz + Paralog Alignments
GCE ID: Event_317

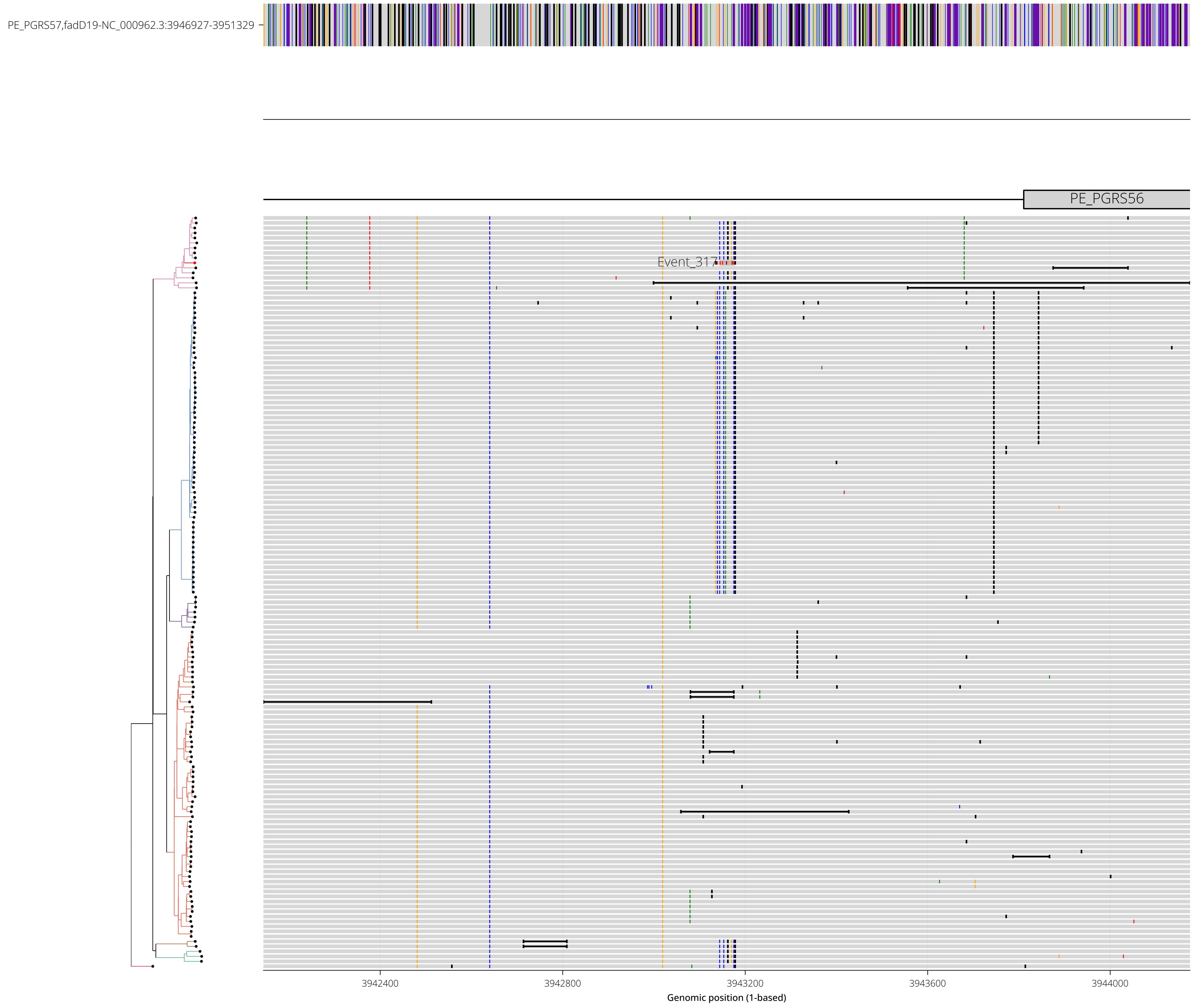

Individual GCE + Known Paralogs
GCE ID: Event_318

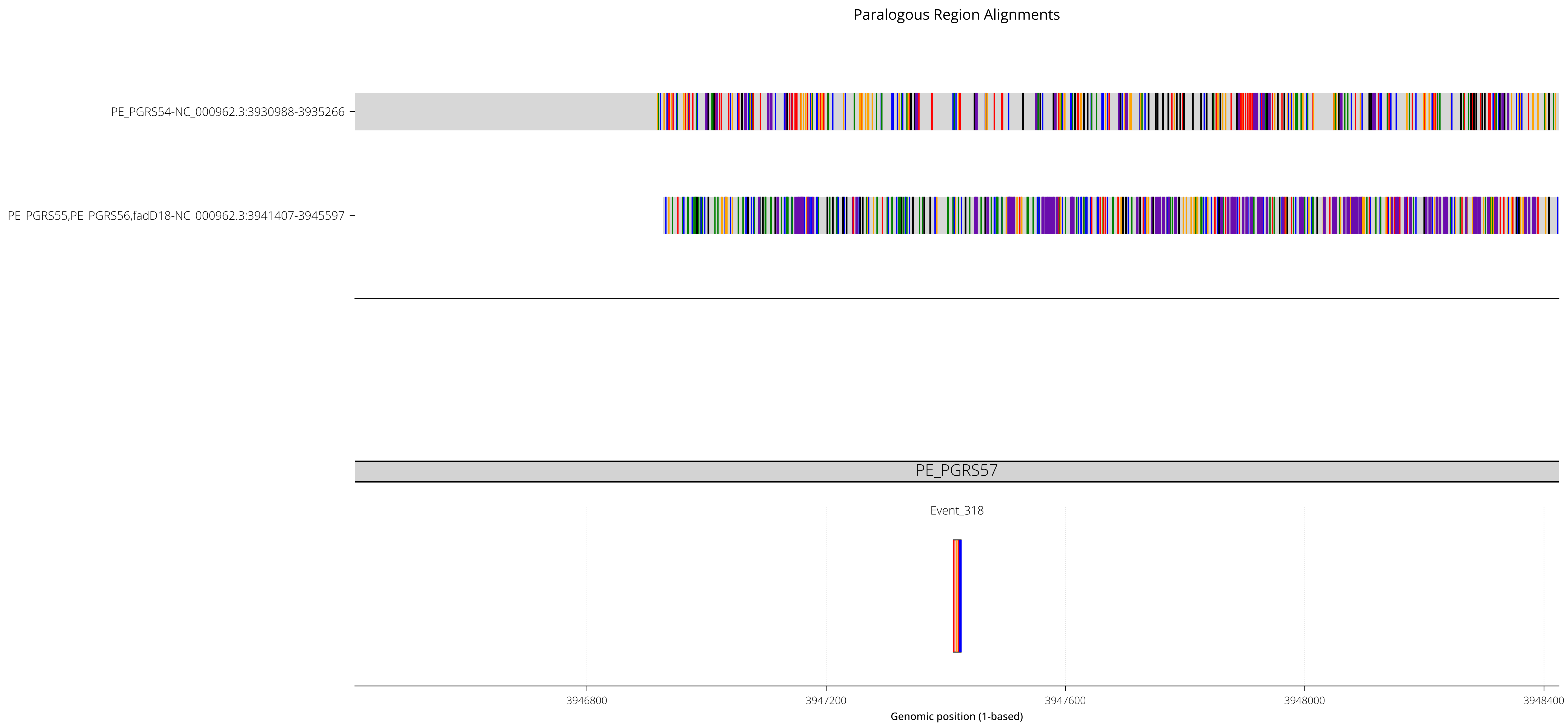

### Circos Viz - GCE to Known Paralogs Locations
GCE ID: Event_318

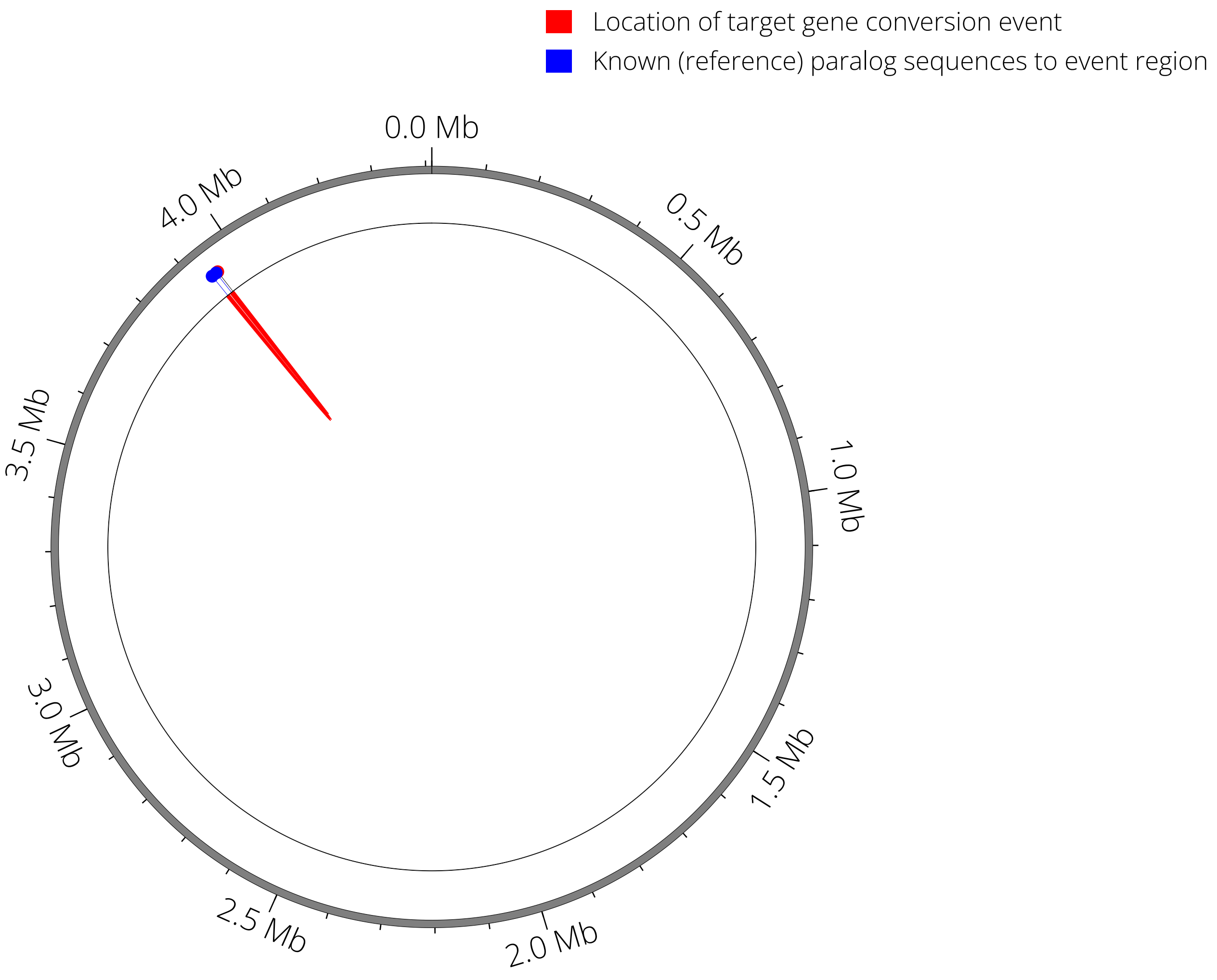

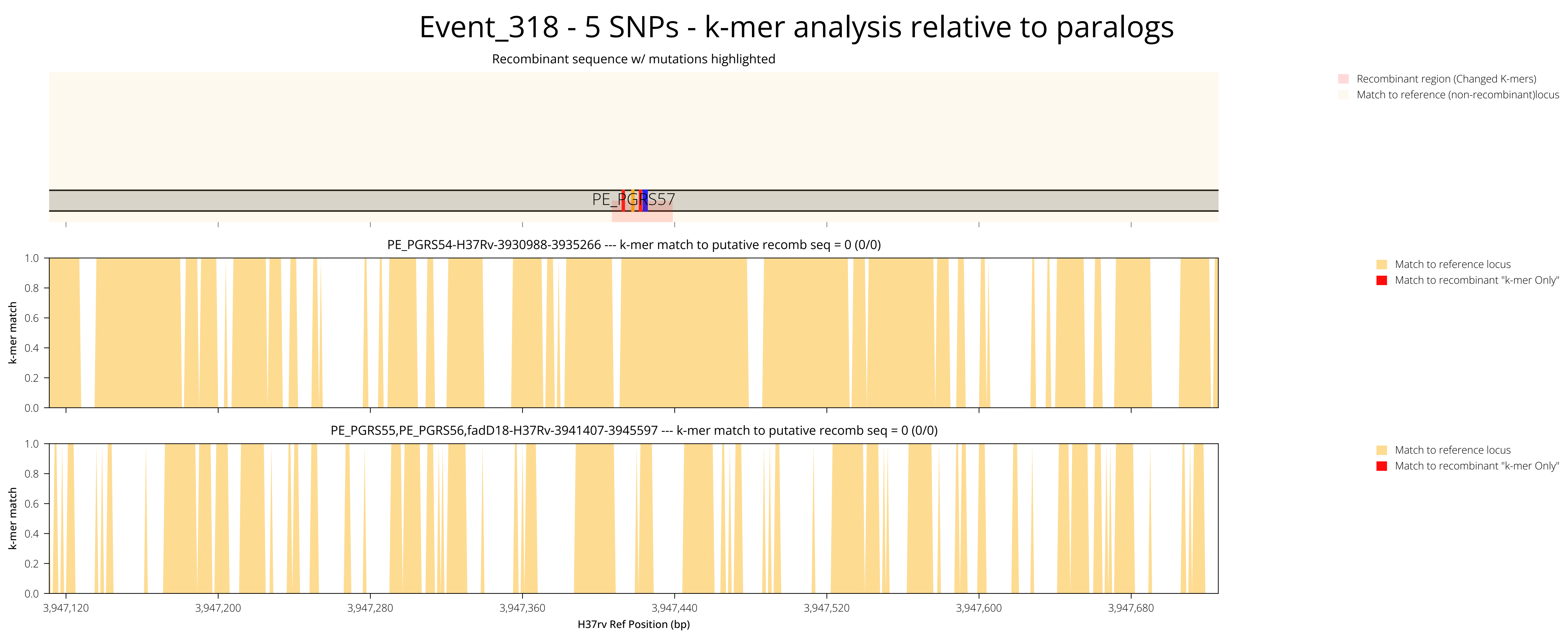

Event_318 - 5 SNPs - k-mer analysis relative to paralogs
Recombinant sequence w/ mutations highlighted
Recombinant region (Changed K-mers)
Match to reference (non-recombinant)locus
PE_PGRS57
PE_PGRS54-H37Rv-3930988-3935266 --- k-mer match to putative recomb seq = 0 (0/0)
Match to reference locus
Match to recombinant "k-mer Only"
PE_PGRS55,PE_PGRS56,fadD18-H37Rv-3941407-3945597 --- k-mer match to putative recomb seq = 0 (0/0)
Match to reference locus
Match to recombinant "k-mer Only"
k-mer match
1.0
0.8
0.6
0.4
0.2
0.0
3,947,120
3,947,200
3,947,280
3,947,360
3,947,440
3,947,520
3,947,600
3,947,680
H37rv Ref Position (bp)

Sub-phylogeny Viz + Paralog Alignments
GCE ID: Event_318

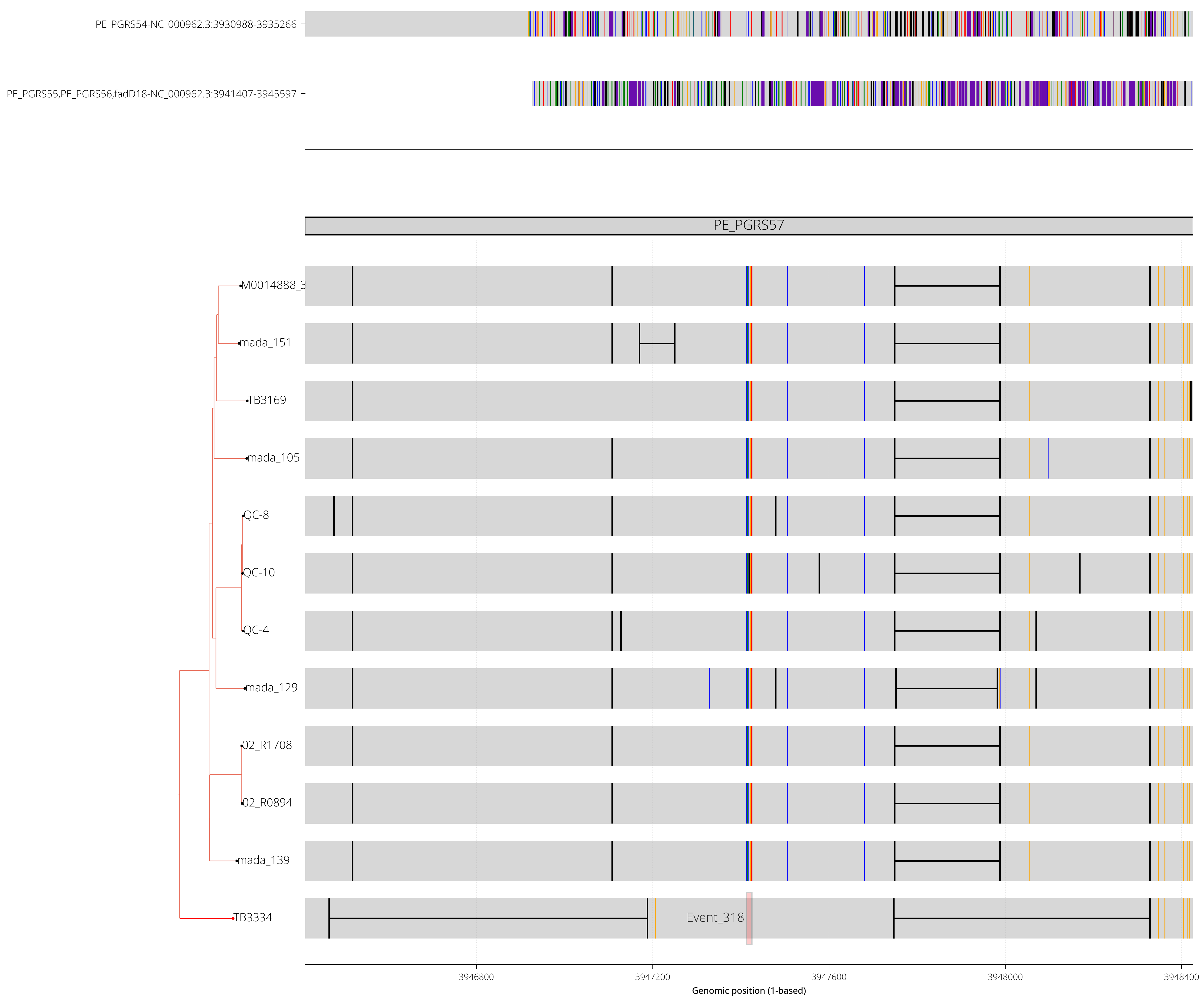

Full Phylogeny Viz + Paralog Alignments
GCE ID: Event_318

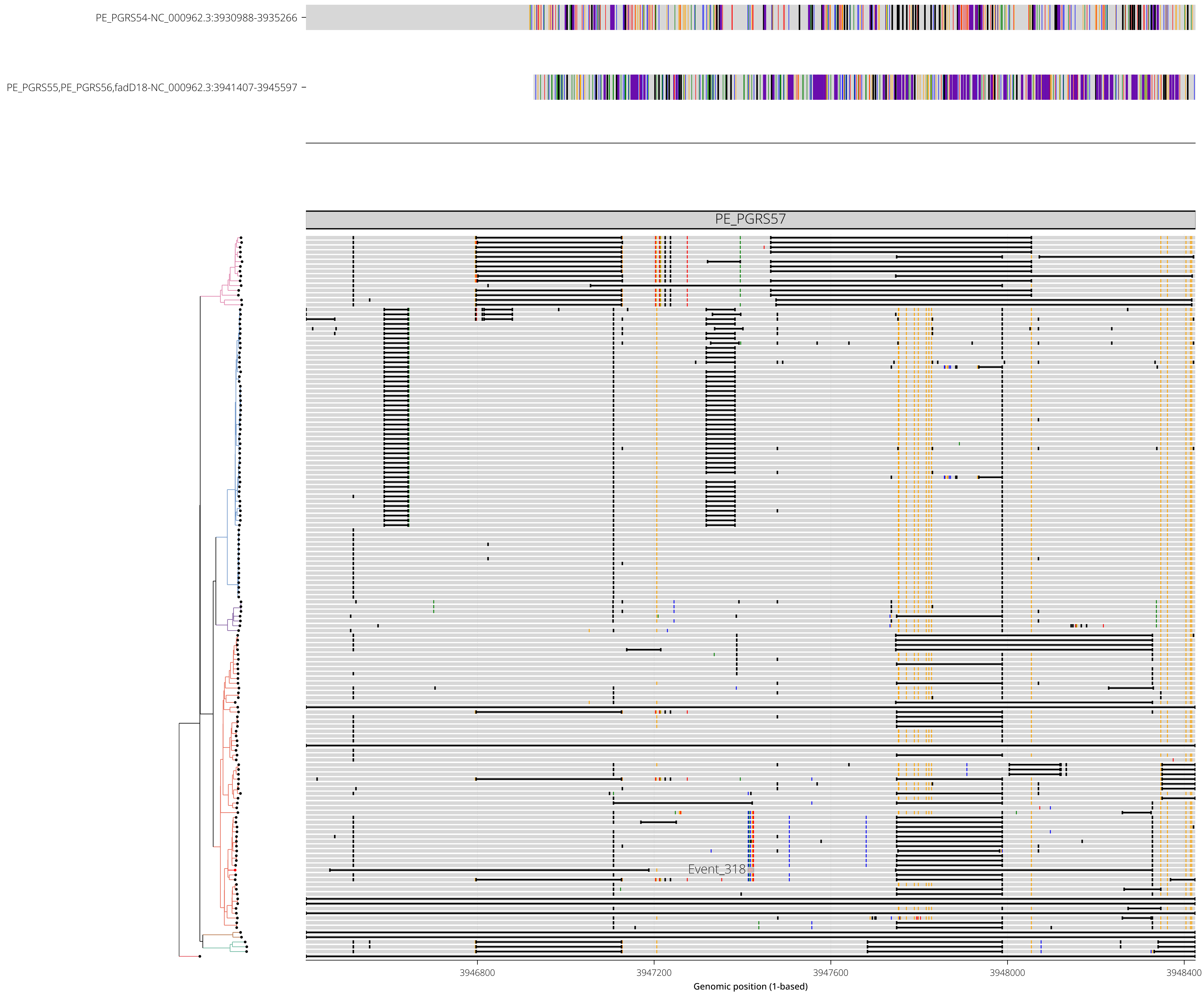

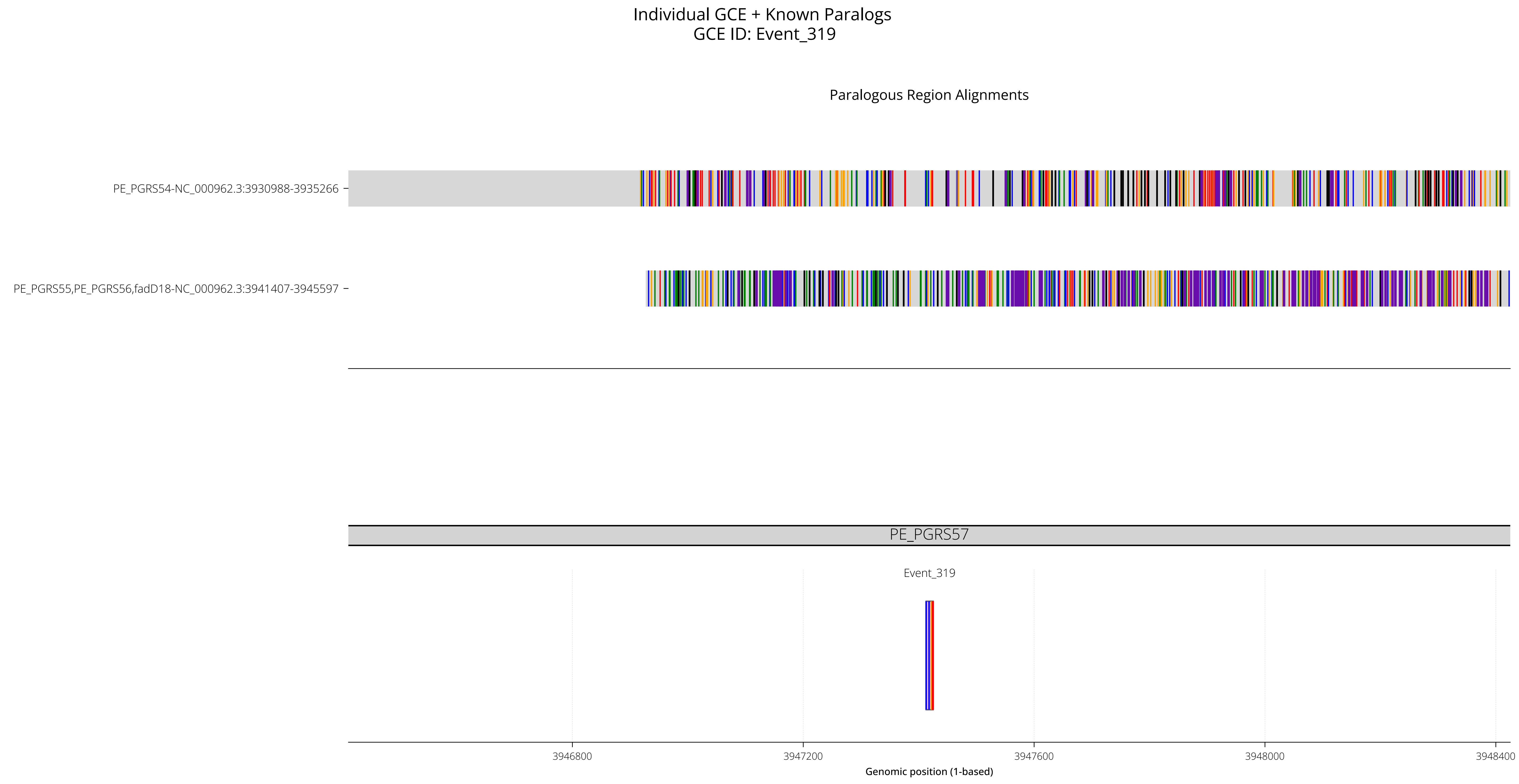

Individual GCE + Known Paralogs
GCE ID: Event_319
Paralogous Region Alignments
PE_PGRS54-NC_000962.3:3930988-3935266
PE_PGRS55,PE_PGRS56,fadD18-NC_000962.3:3941407-3945597
PE_PGRS57
Event_319
3946800
3947200
3947600
3948000
3948400
Genomic position (1-based)

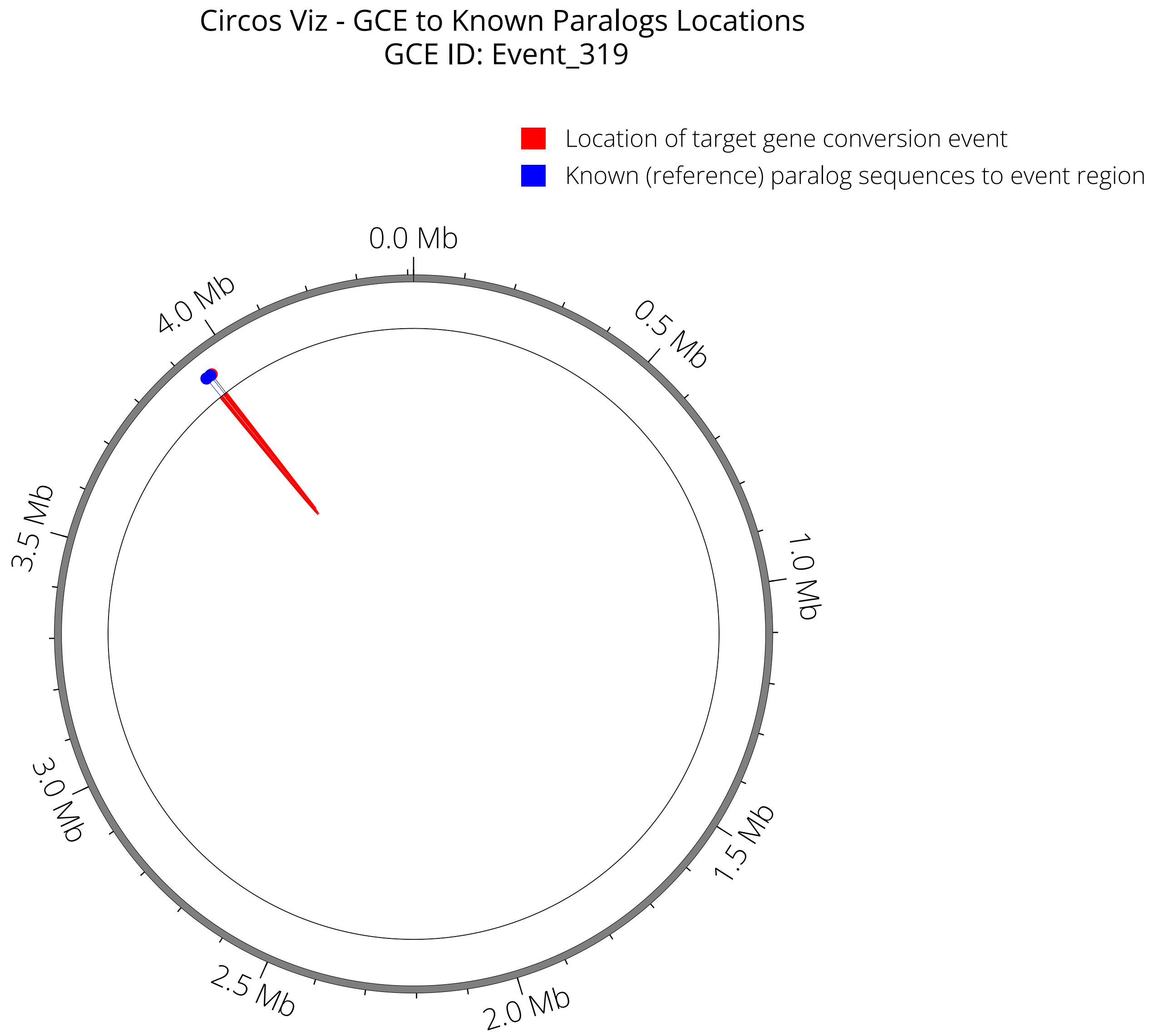

Circos Viz - GCE to Known Paralogs Locations
GCE ID: Event_319
Location of target gene conversion event
Known (reference) paralog sequences to event region
0.0 Mb
0.5 Mb
1.0 Mb
1.5 Mb
2.0 Mb
2.5 Mb
3.0 Mb
3.5 Mb
4.0 Mb

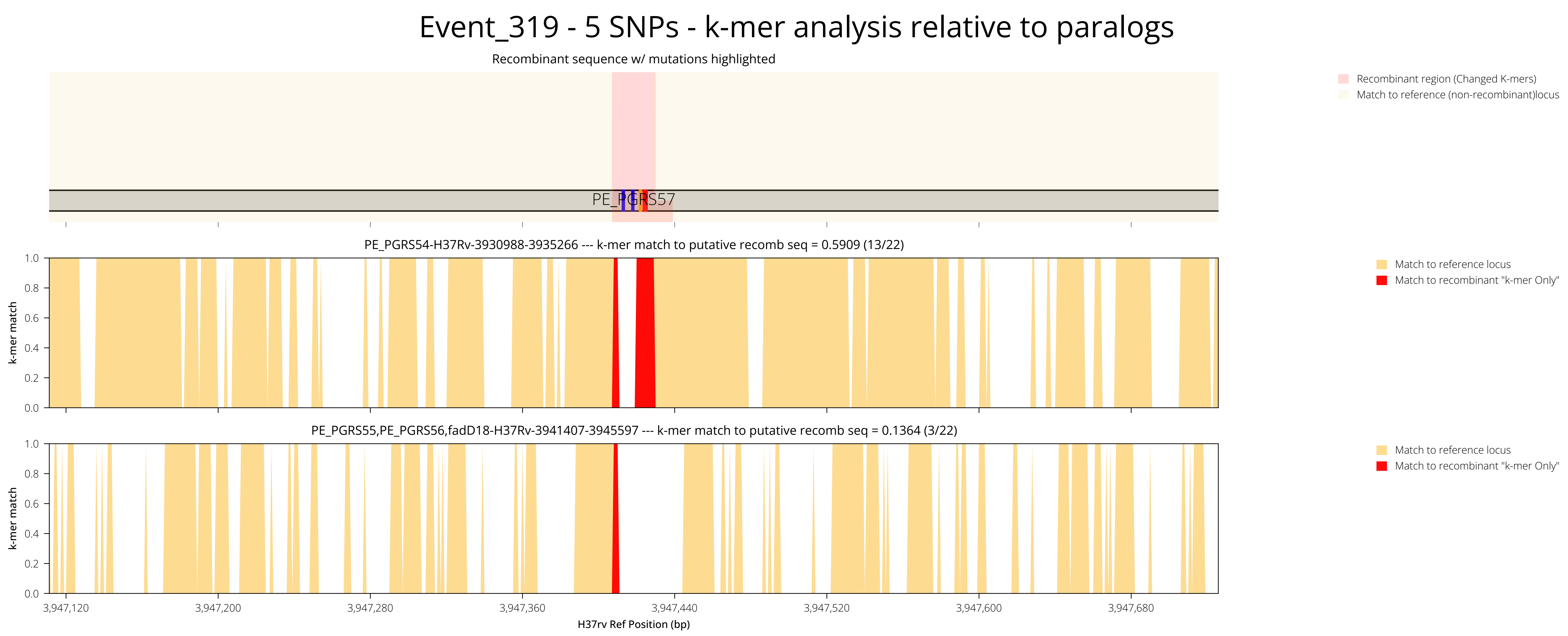

Event_319 - 5 SNPs - k-mer analysis relative to paralogs
Recombinant sequence w/ mutations highlighted
PE_PGRS57
Recombinant region (Changed K-mers)
Match to reference (non-recombinant)locus
PE_PGRS54-H37Rv-3930988-3935266 --- k-mer match to putative recomb seq = 0.5909 (13/22)
Match to reference locus
Match to recombinant "k-mer Only"
PE_PGRS55,PE_PGRS56,fadD18-H37Rv-3941407-3945597 --- k-mer match to putative recomb seq = 0.1364 (3/22)
Match to reference locus
Match to recombinant "k-mer Only"
k-mer match
1.0
0.8
0.6
0.4
0.2
0.0
3,947,120
3,947,200
3,947,280
3,947,360
3,947,440
3,947,520
3,947,600
3,947,680
H37rv Ref Position (bp)

Sub-phylogeny Viz + Paralog Alignments
GCE ID: Event_319

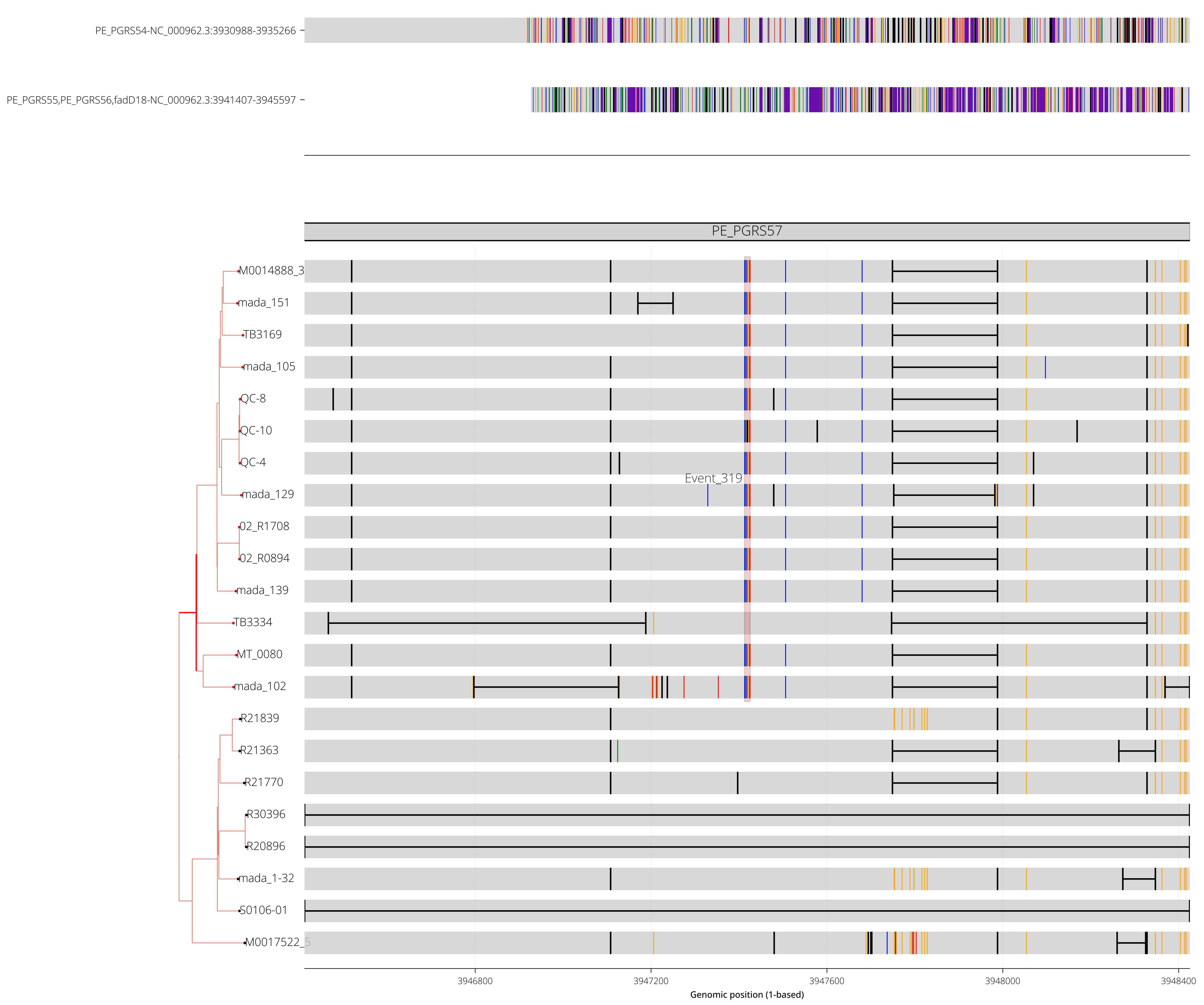

Full Phylogeny Viz + Paralog Alignments
GCE ID: Event_319

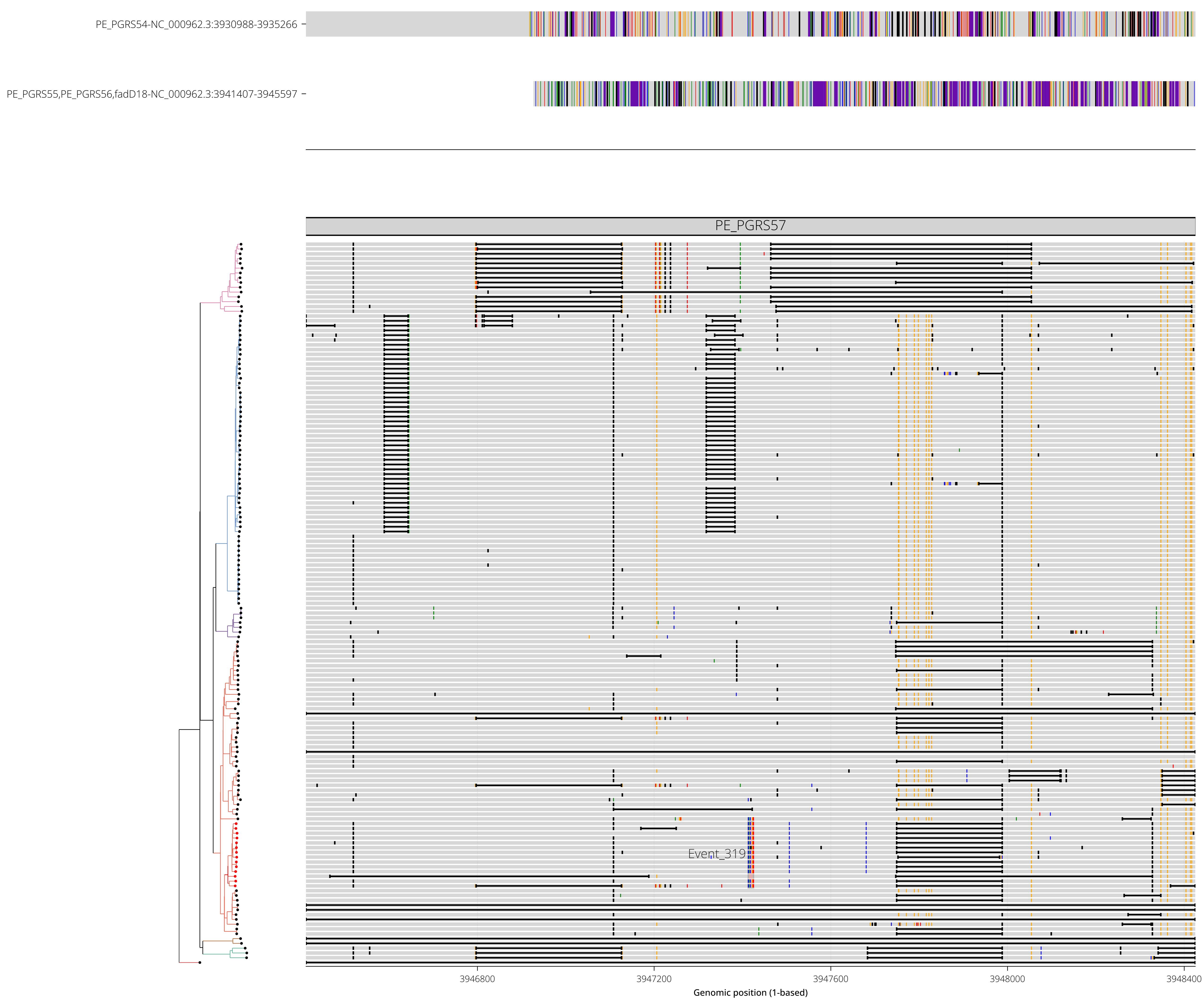

Individual GCE + Known Paralogs
GCE ID: Event_320

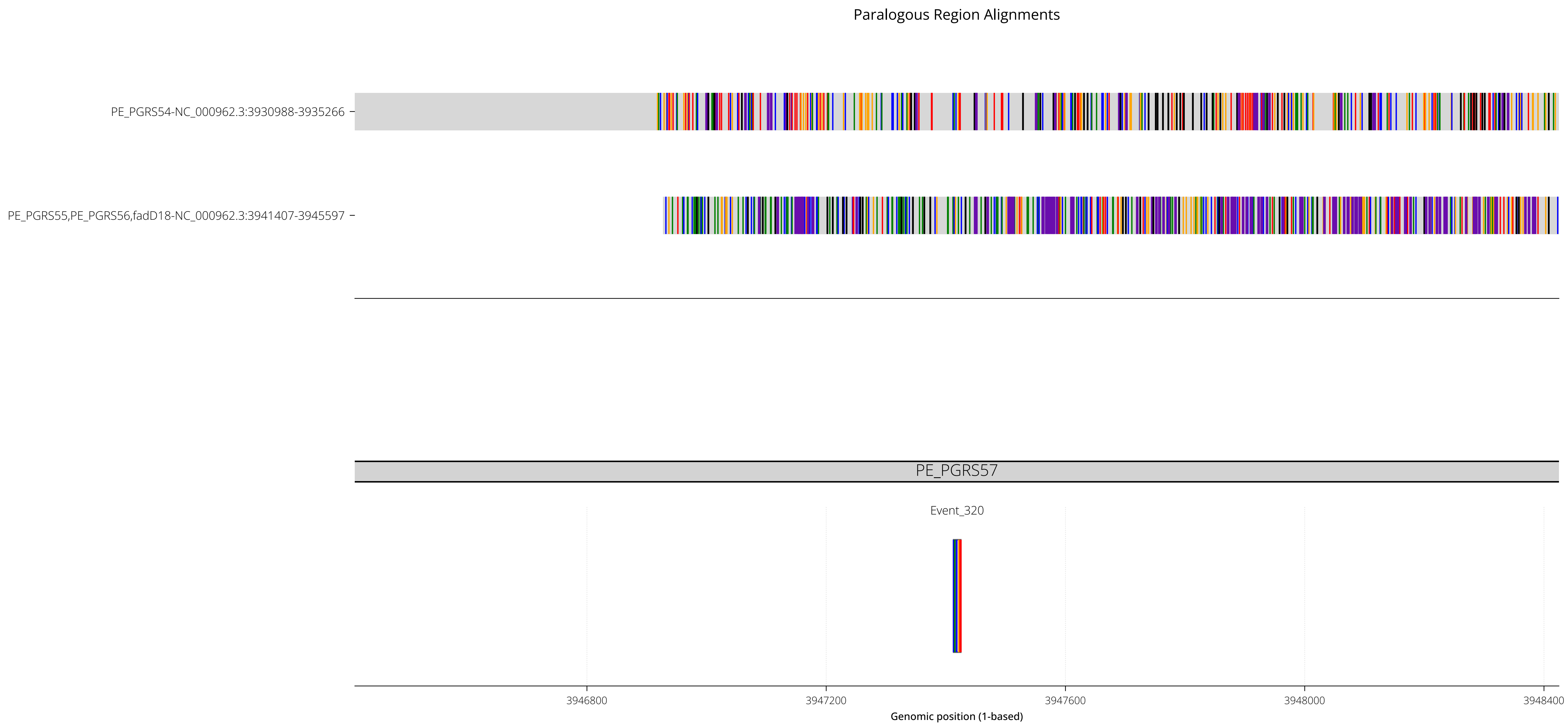

Circos Viz - GCE to Known Paralogs Locations
GCE ID: Event_320

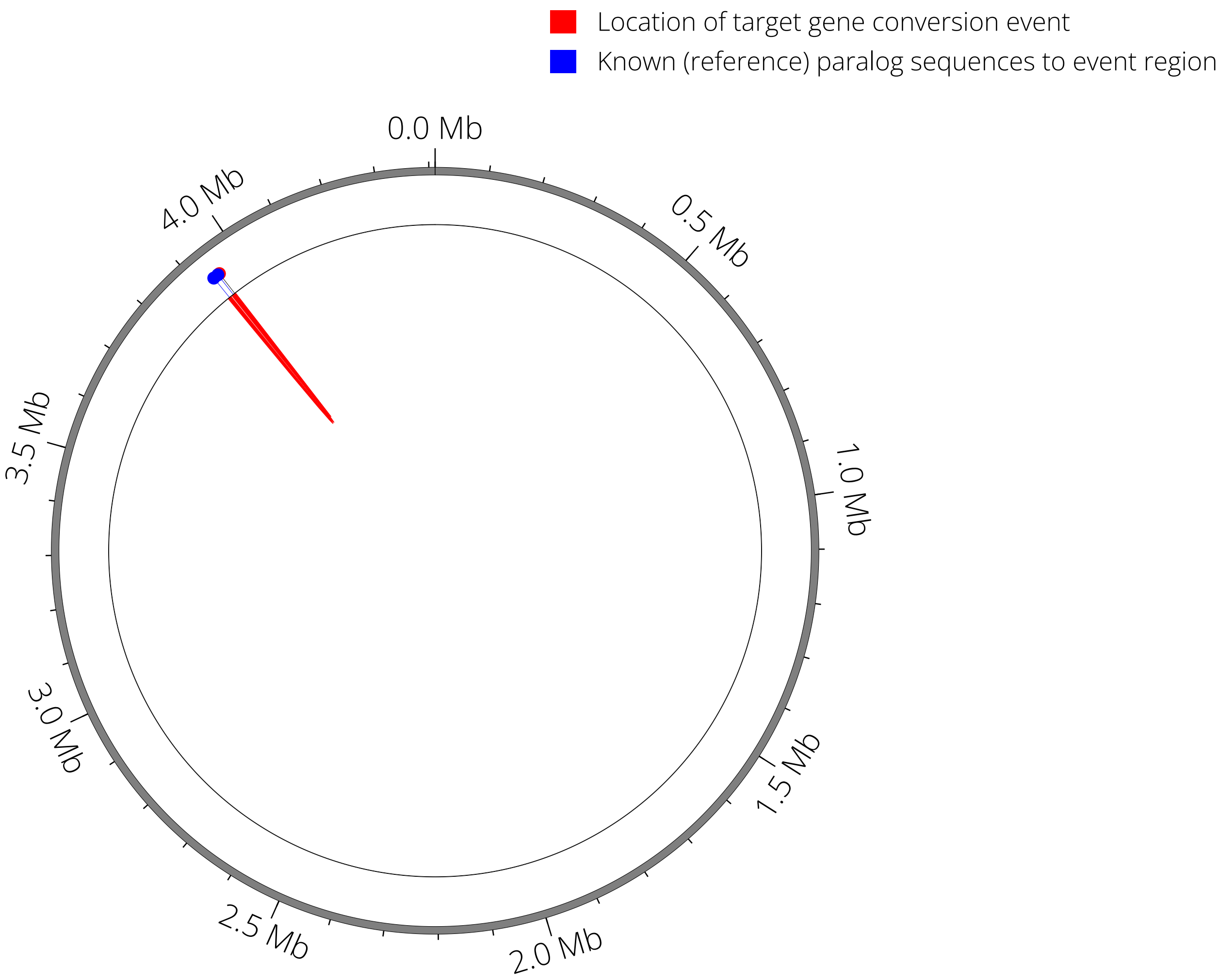

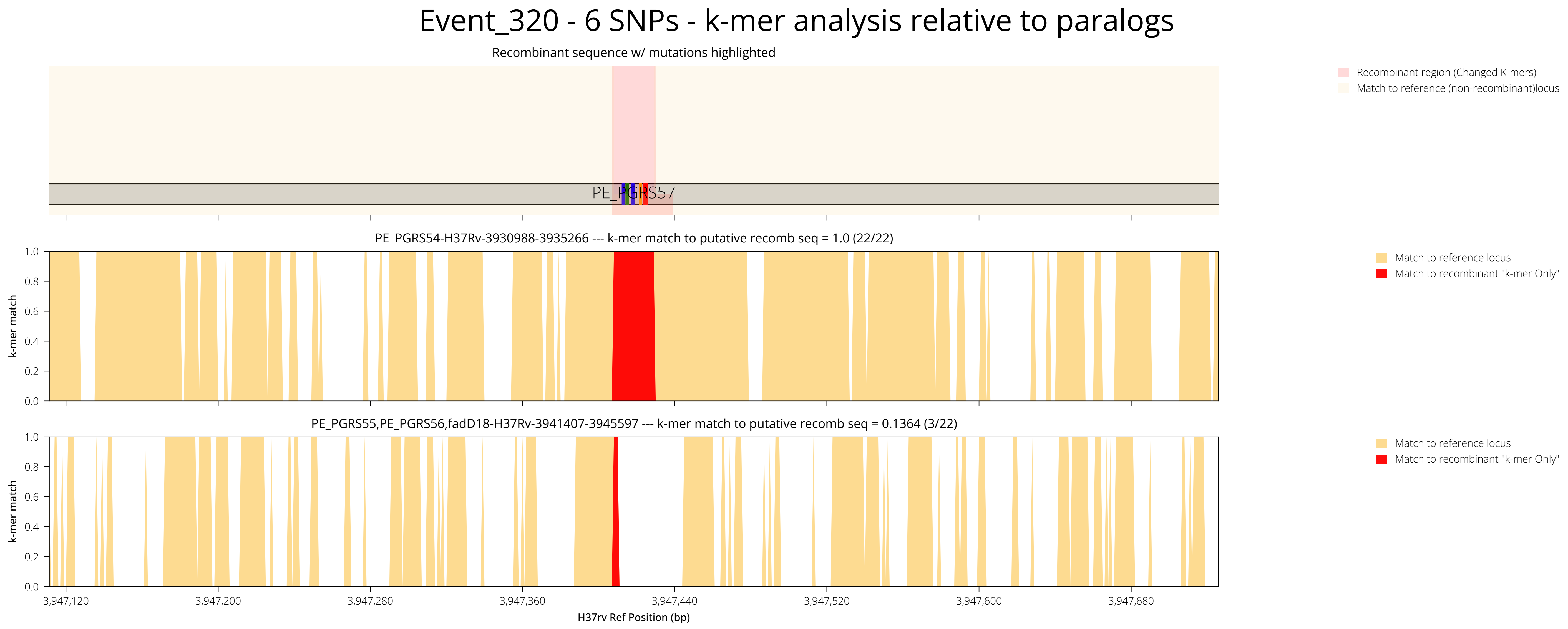

Event_320 - 6 SNPs - k-mer analysis relative to paralogs
Recombinant sequence w/ mutations highlighted
PE_PGRS57
Recombinant region (Changed K-mers)
Match to reference (non-recombinant)locus
PE_PGRS54-H37Rv-3930988-3935266 --- k-mer match to putative recomb seq = 1.0 (22/22)
k-mer match
Match to reference locus
Match to recombinant "k-mer Only"
PE_PGRS55,PE_PGRS56,fadD18-H37Rv-3941407-3945597 --- k-mer match to putative recomb seq = 0.1364 (3/22)
k-mer match
Match to reference locus
Match to recombinant "k-mer Only"
1.0
0.8
0.6
0.4
0.2
0.0
3,947,120
3,947,200
3,947,280
3,947,360
3,947,440
3,947,520
3,947,600
3,947,680
H37rv Ref Position (bp)

Sub-phylogeny Viz + Paralog Alignments
GCE ID: Event_320

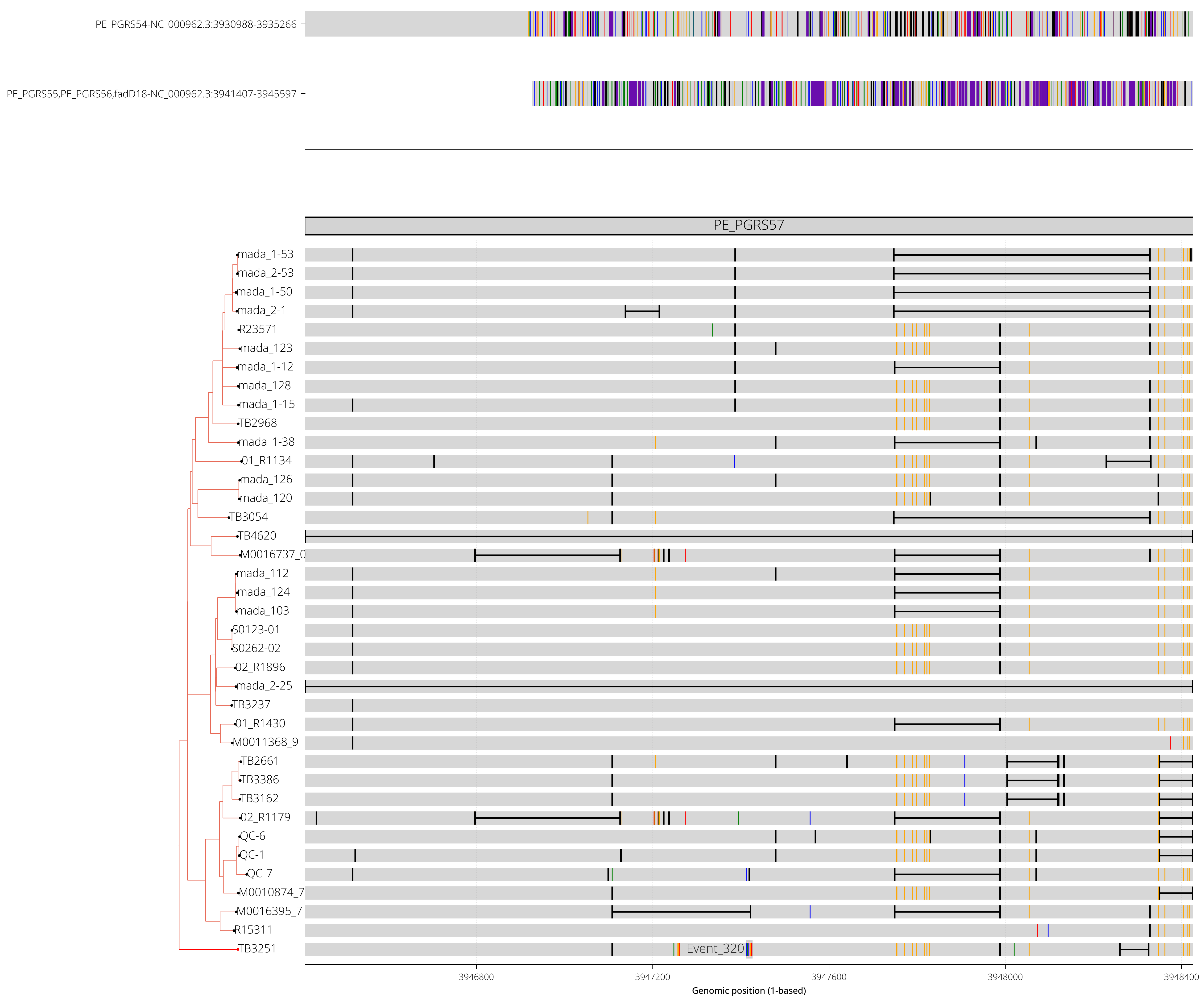

Full Phylogeny Viz + Paralog Alignments
GCE ID: Event_320

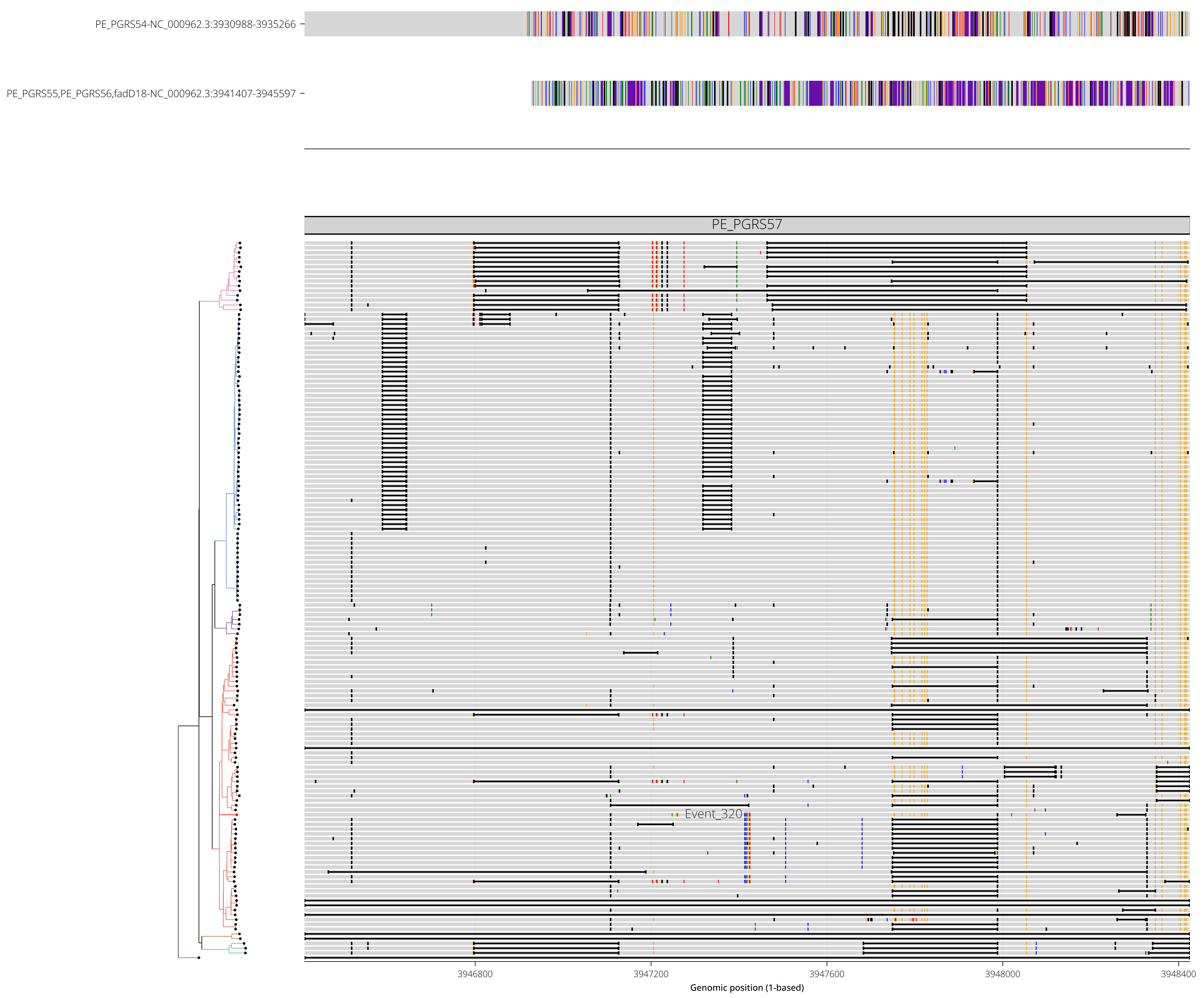

Individual GCE + Known Paralogs
GCE ID: Event_321

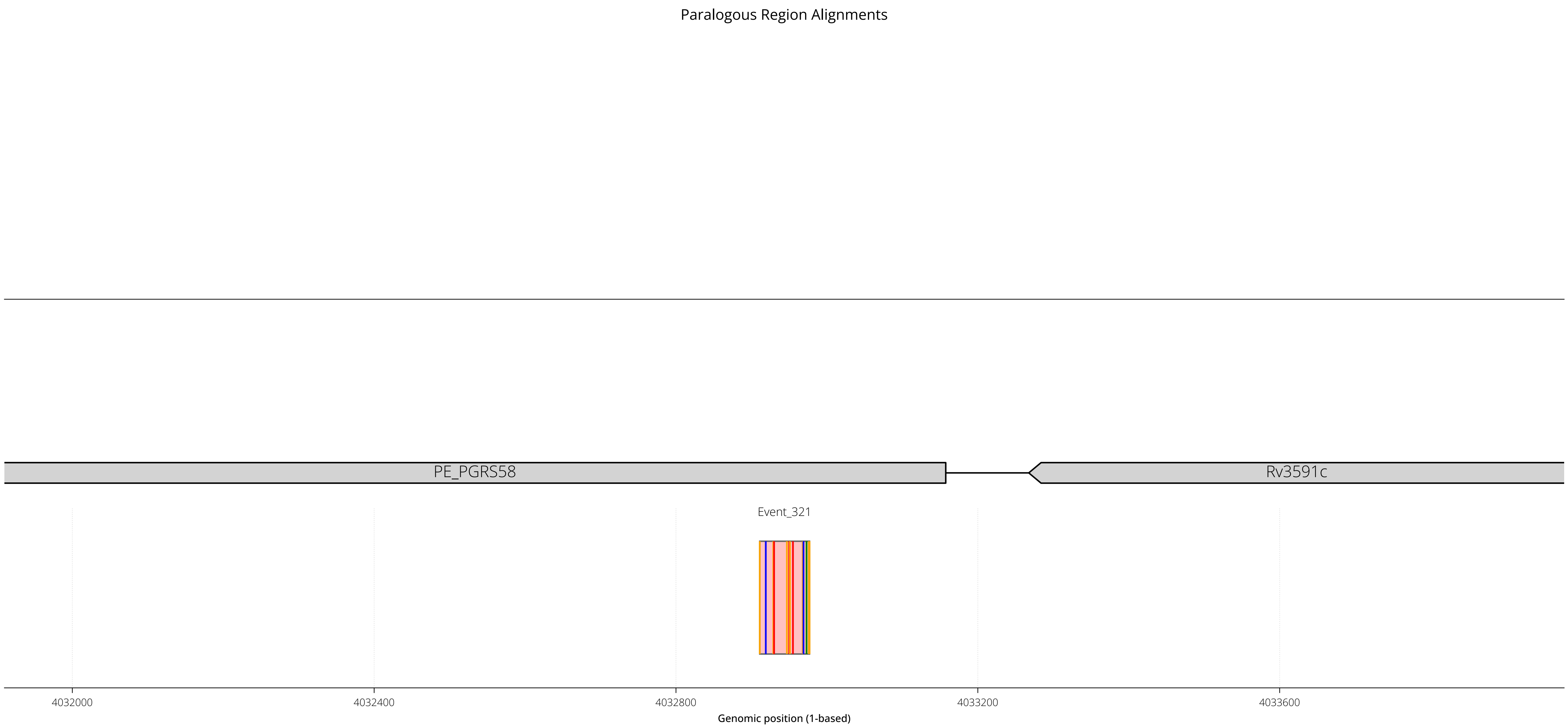

Sub-phylogeny Viz + Paralog Alignments
GCE ID: Event_321

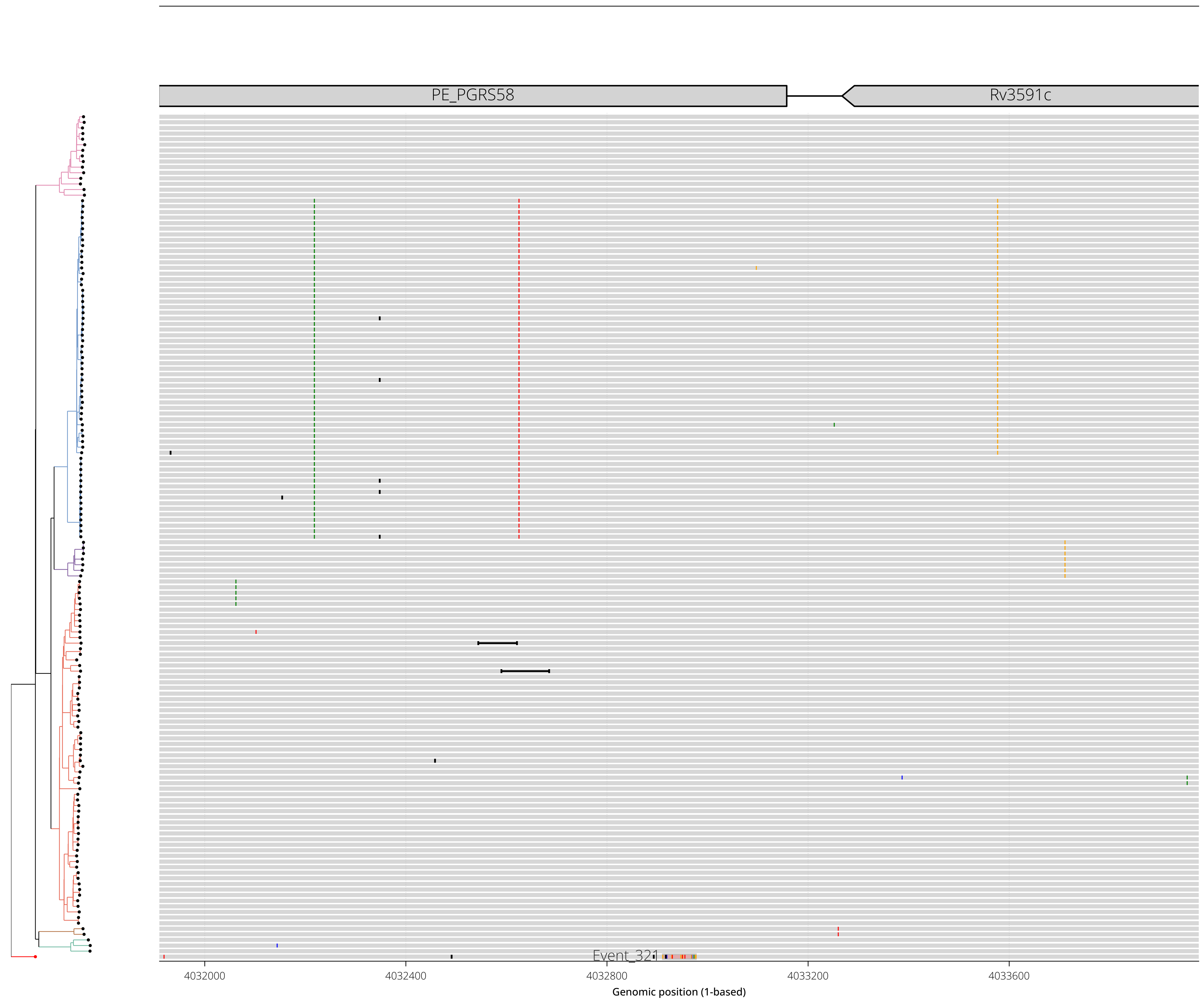

Full Phylogeny Viz + Paralog Alignments
GCE ID: Event_321

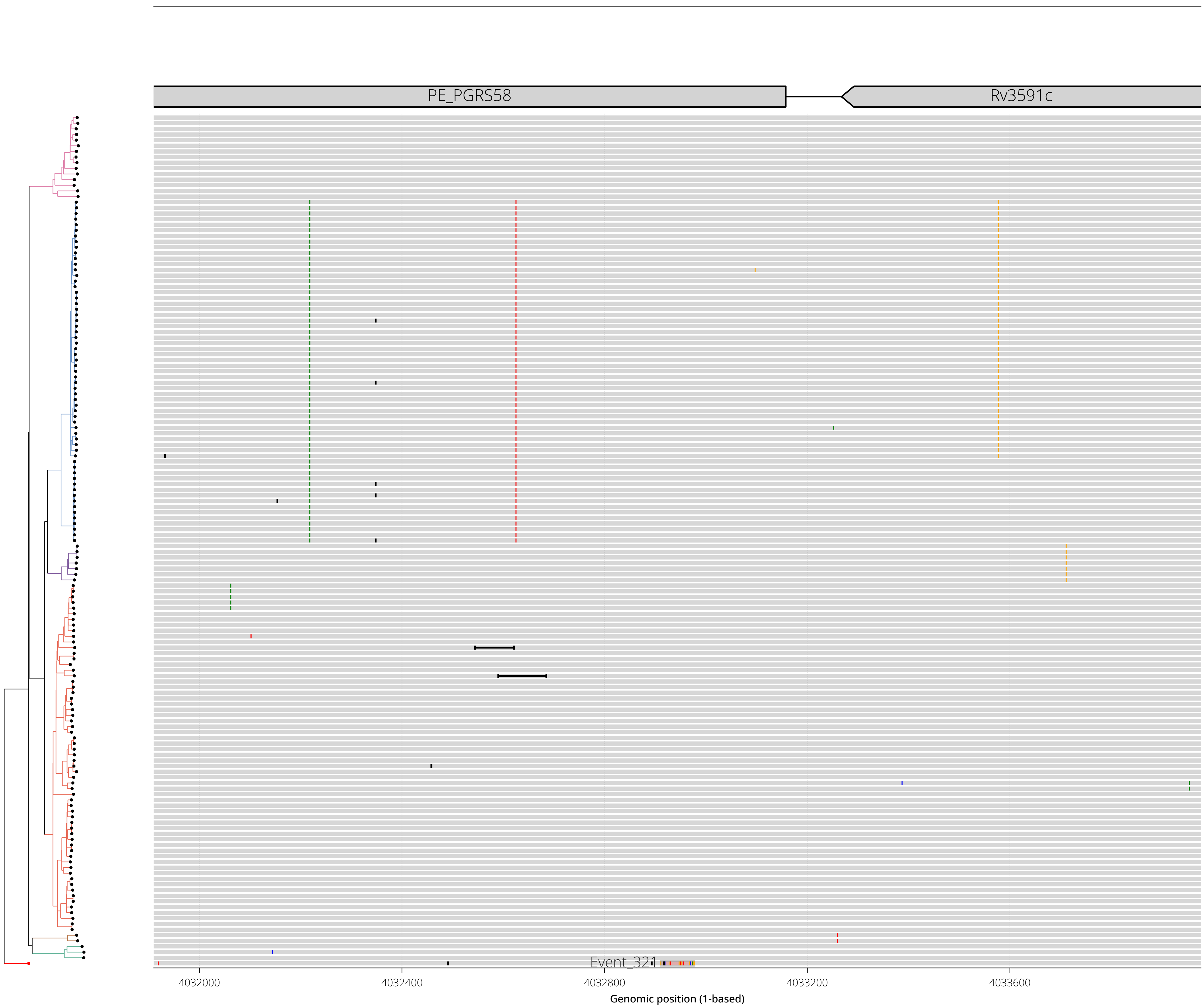

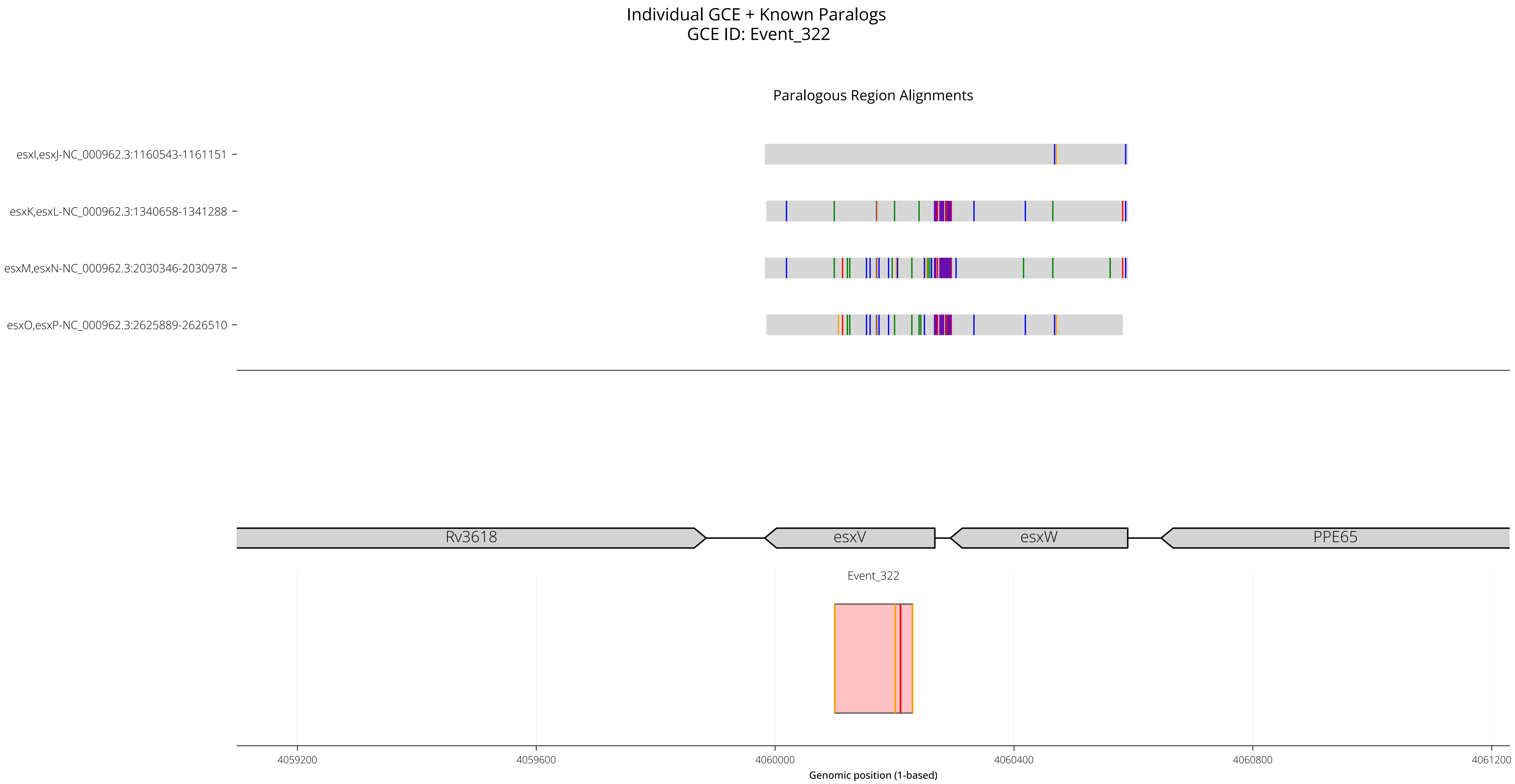

Individual GCE + Known Paralogs
GCE ID: Event_322
Paralogous Region Alignments
esxI,esxJ-NC_000962.3:1160543-1161151
esxK,esxL-NC_000962.3:1340658-1341288
esxM,esxN-NC_000962.3:2030346-2030978
esxO,esxP-NC_000962.3:2625889-2626510
Rv3618
esxV
esxW
PPE65
Event_322
4059200
4059600
4060000
4060400
4060800
4061200
Genomic position (1-based)

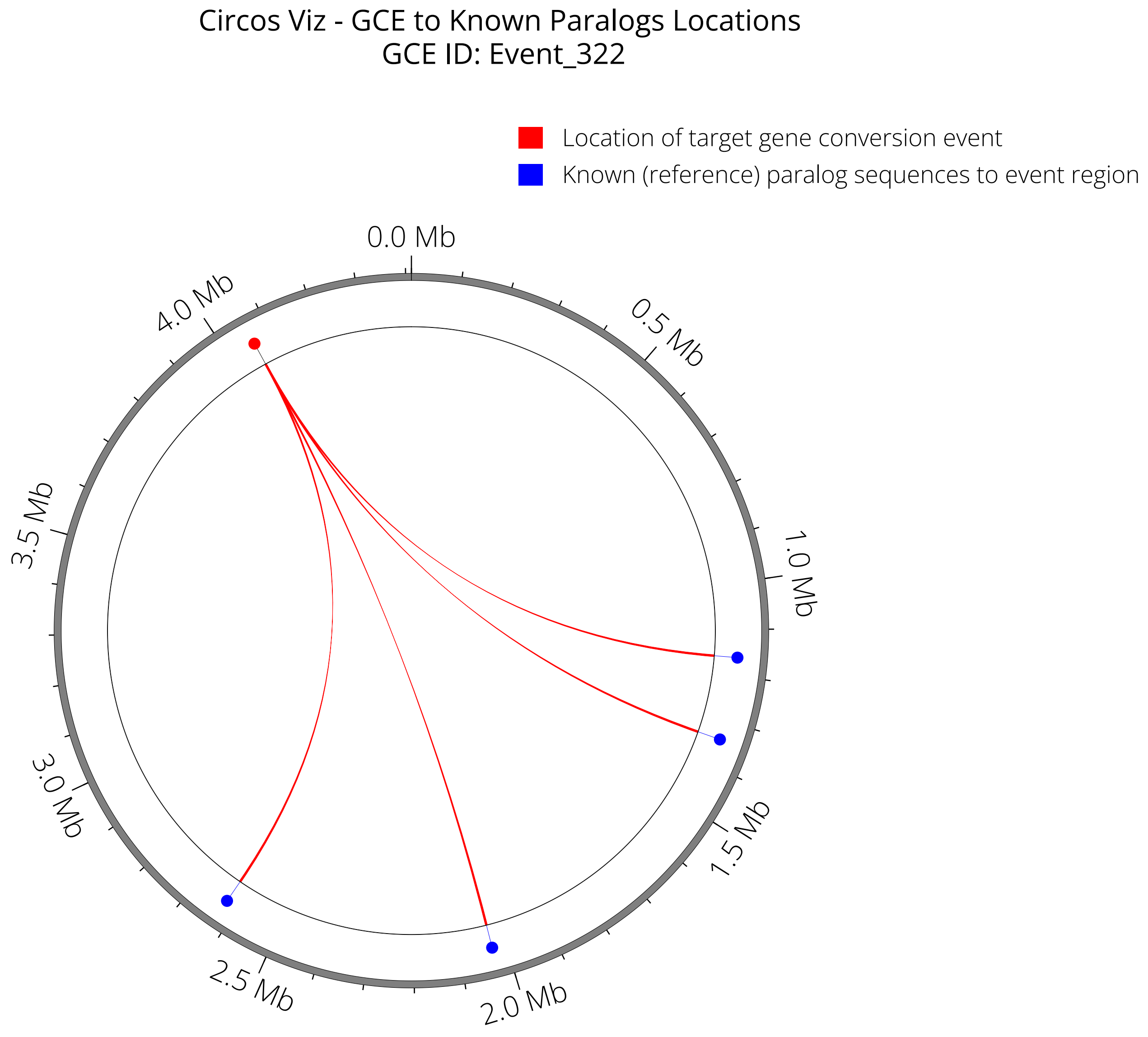
Circos Viz - GCE to Known Paralogs Locations
GCE ID: Event_322
Location of target gene conversion event
Known (reference) paralog sequences to event region
0.0 Mb
0.5 Mb
1.0 Mb
1.5 Mb
2.0 Mb
2.5 Mb
3.0 Mb
3.5 Mb
4.0 Mb

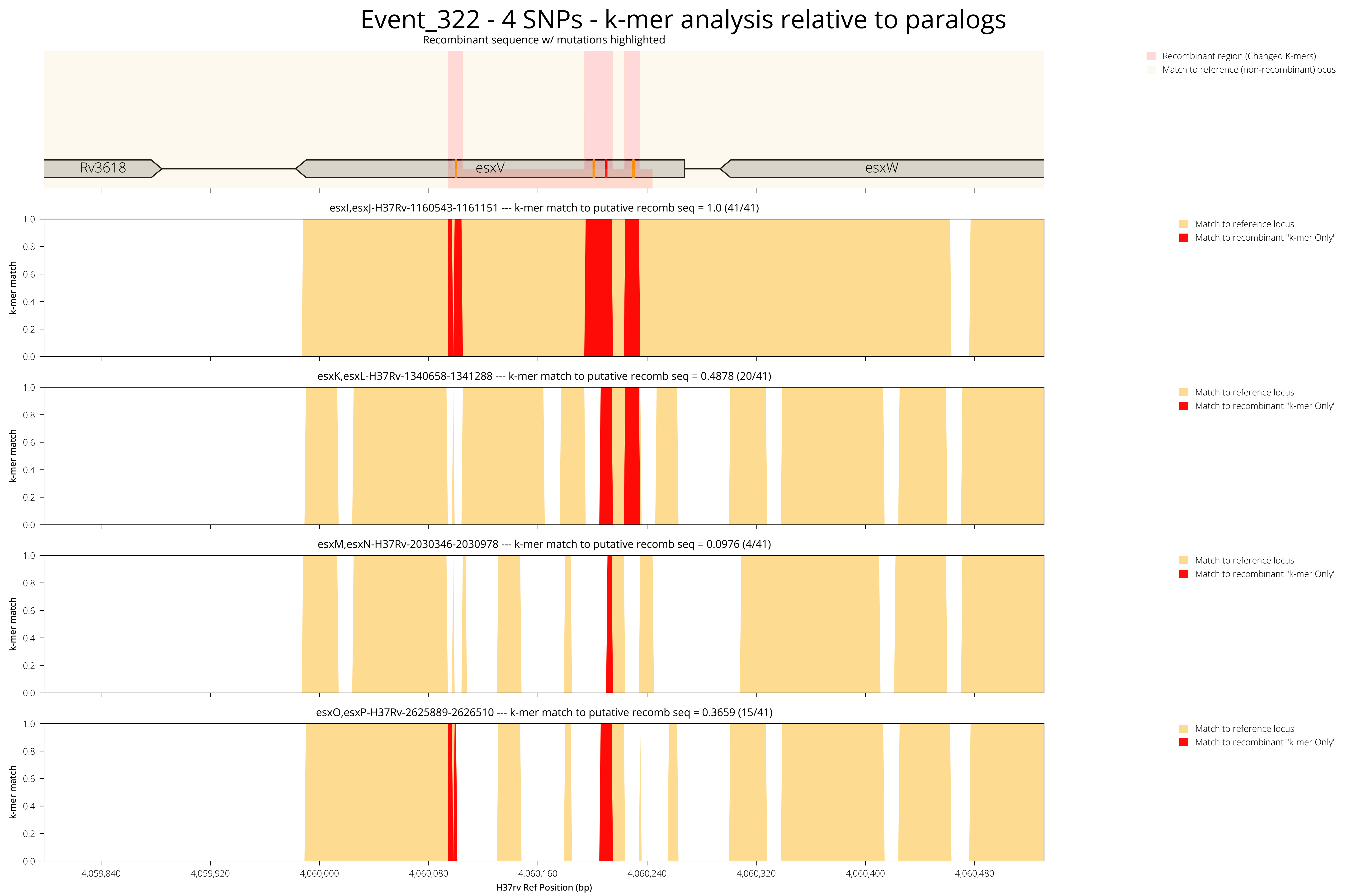

Event_322 - 4 SNPs - k-mer analysis relative to paralogs
Recombinant sequence w/ mutations highlighted
Recombinant region (Changed K-mers)
Match to reference (non-recombinant)locus
Rv3618
esxV
esxW
esxI,esxJ-H37Rv-1160543-1161151 --- k-mer match to putative recomb seq = 1.0 (41/41)
esxK,esxL-H37Rv-1340658-1341288 --- k-mer match to putative recomb seq = 0.4878 (20/41)
esxM,esxN-H37Rv-2030346-2030978 --- k-mer match to putative recomb seq = 0.0976 (4/41)
esxO,esxP-H37Rv-2625889-2626510 --- k-mer match to putative recomb seq = 0.3659 (15/41)
Match to reference locus
Match to recombinant "k-mer Only"
k-mer match
0.0
0.2
0.4
0.6
0.8
1.0
4,059,840
4,059,920
4,060,000
4,060,080
4,060,160
4,060,240
4,060,320
4,060,400
4,060,480
H37rv Ref Position (bp)

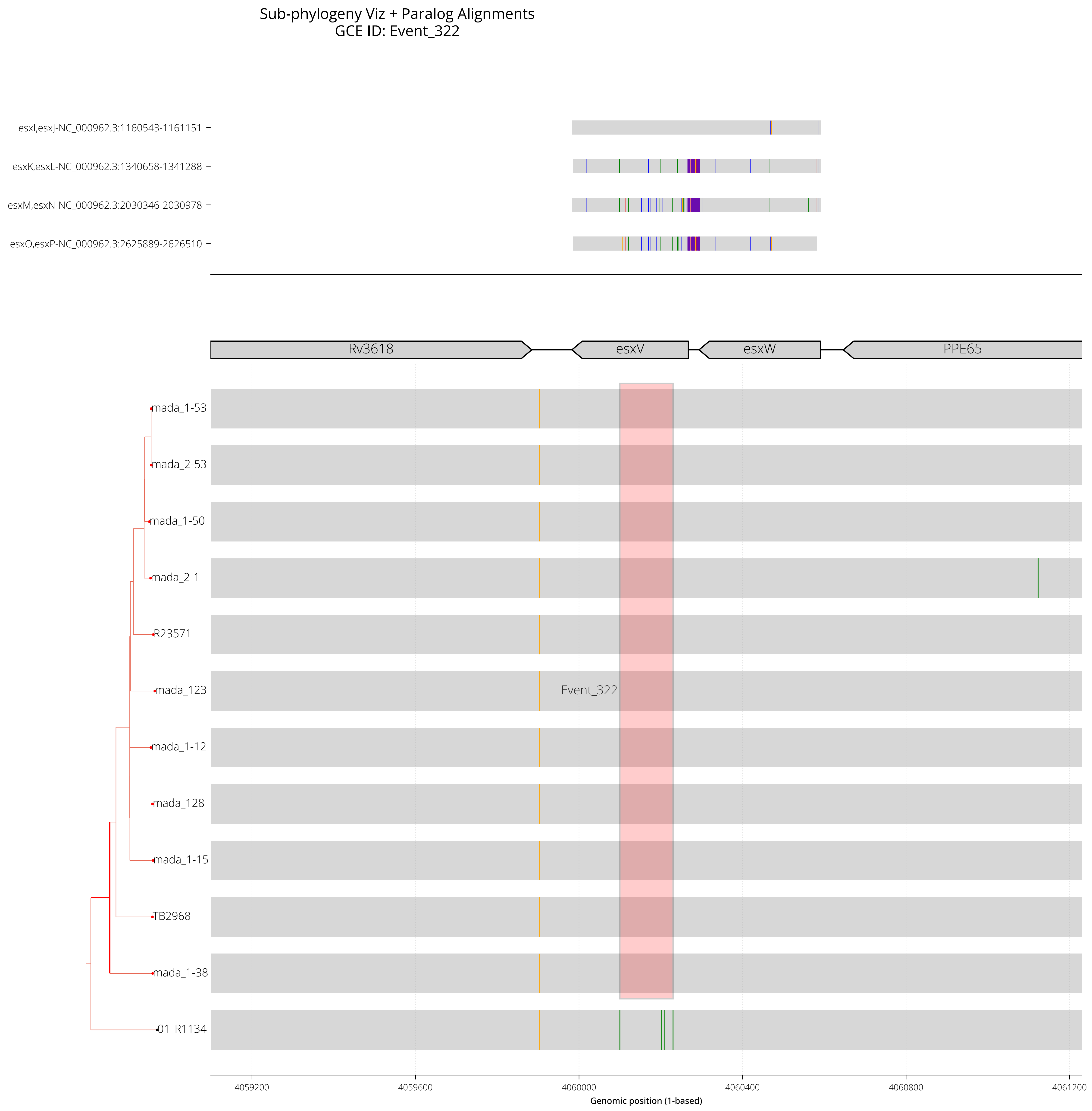

Sub-phylogeny Viz + Paralog Alignments
GCE ID: Event_322
esxI,esxJ-NC_000962.3:1160543-1161151
esxK,esxL-NC_000962.3:1340658-1341288
esxM,esxN-NC_000962.3:2030346-2030978
esxO,esxP-NC_000962.3:2625889-2626510
Rv3618
esxV
esxW
PPE65
mada_1-53
mada_2-53
mada_1-50
mada_2-1
R23571
mada_123
Event_322
mada_1-12
mada_128
mada_1-15
TB2968
mada_1-38
01_R1134
4059200
4059600
4060000
4060400
4060800
4061200
Genomic position (1-based)

Full Phylogeny Viz + Paralog Alignments
GCE ID: Event_322

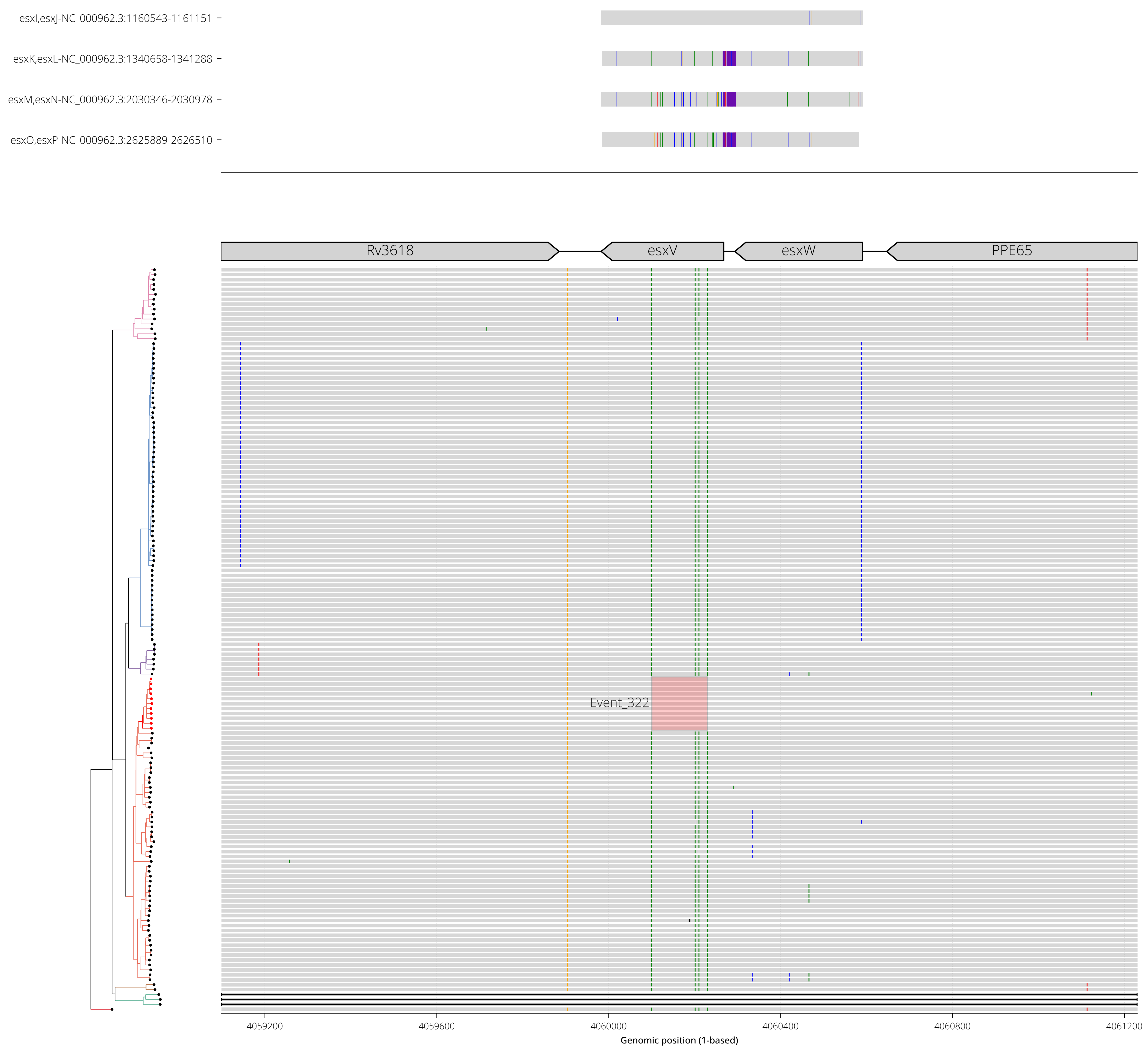

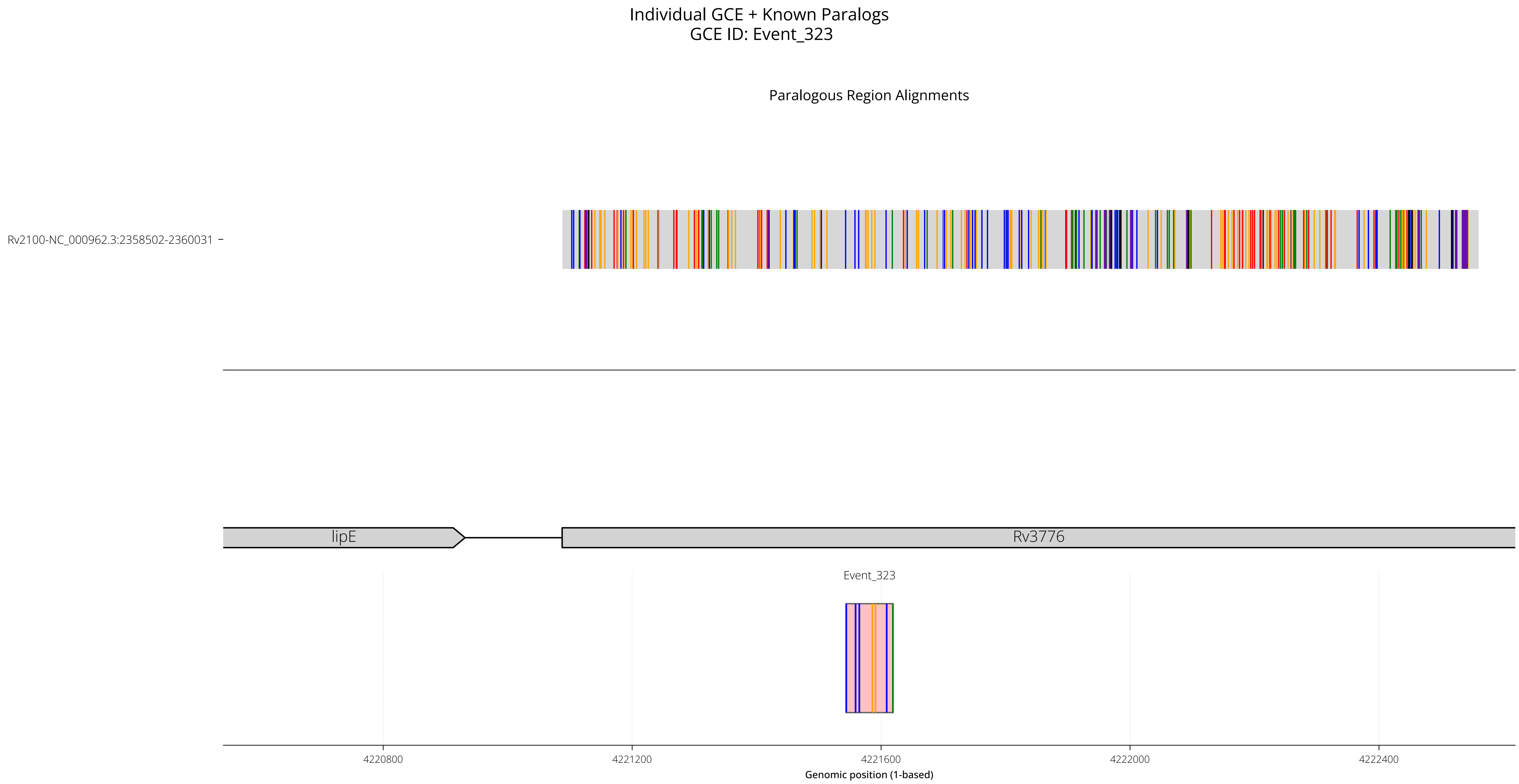

Individual GCE + Known Paralogs
GCE ID: Event_323
Paralogous Region Alignments
Rv2100-NC_000962.3:2358502-2360031
lipE
Rv3776
Event_323
4220800
4221200
4221600
4222000
4222400
Genomic position (1-based)

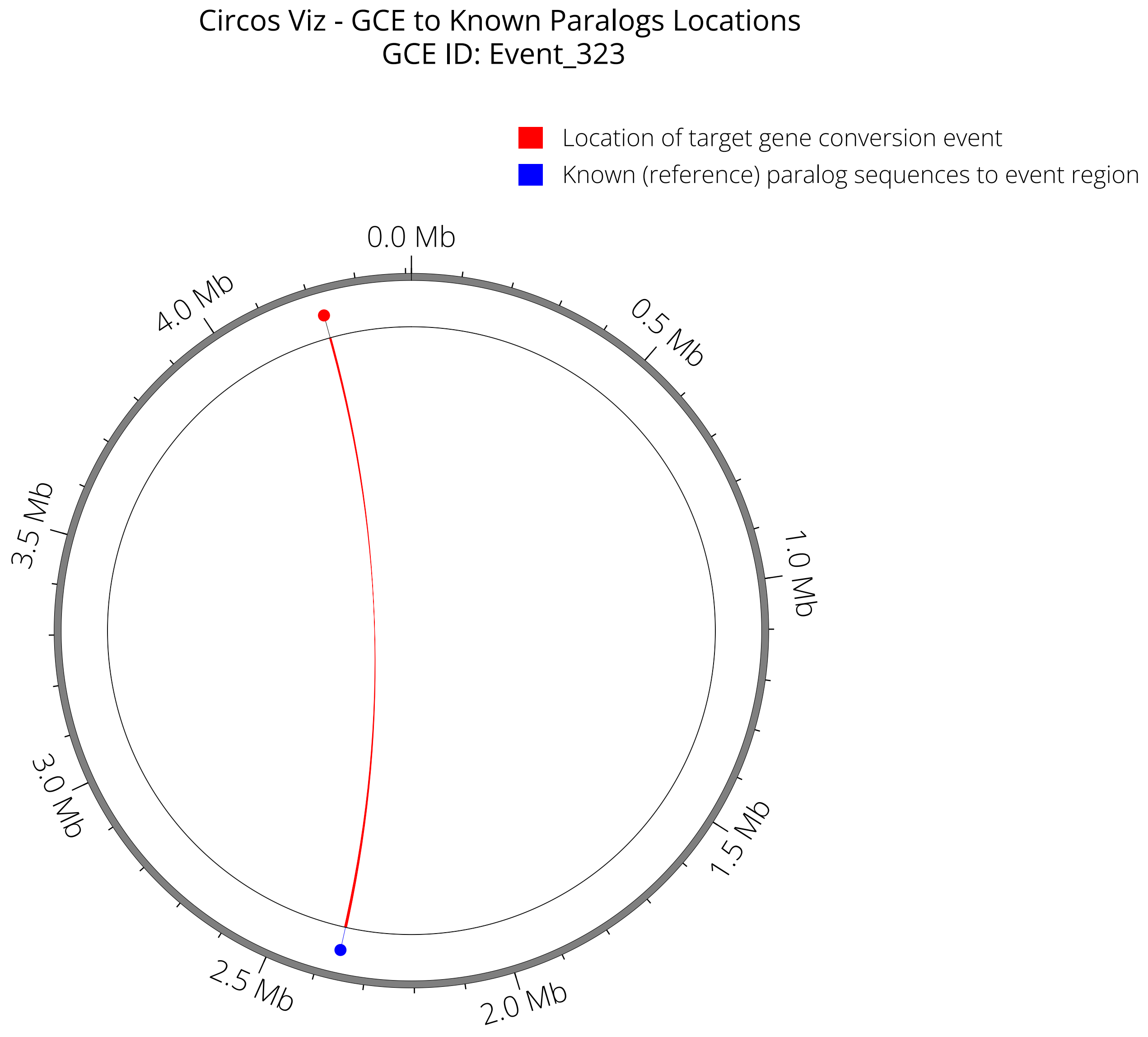
Circos Viz - GCE to Known Paralogs Locations
GCE ID: Event_323
Location of target gene conversion event
Known (reference) paralog sequences to event region
0.0 Mb
0.5 Mb
1.0 Mb
1.5 Mb
2.0 Mb
2.5 Mb
3.0 Mb
3.5 Mb
4.0 Mb

### Event_323 - 7 SNPs - k-mer analysis relative to paralogs

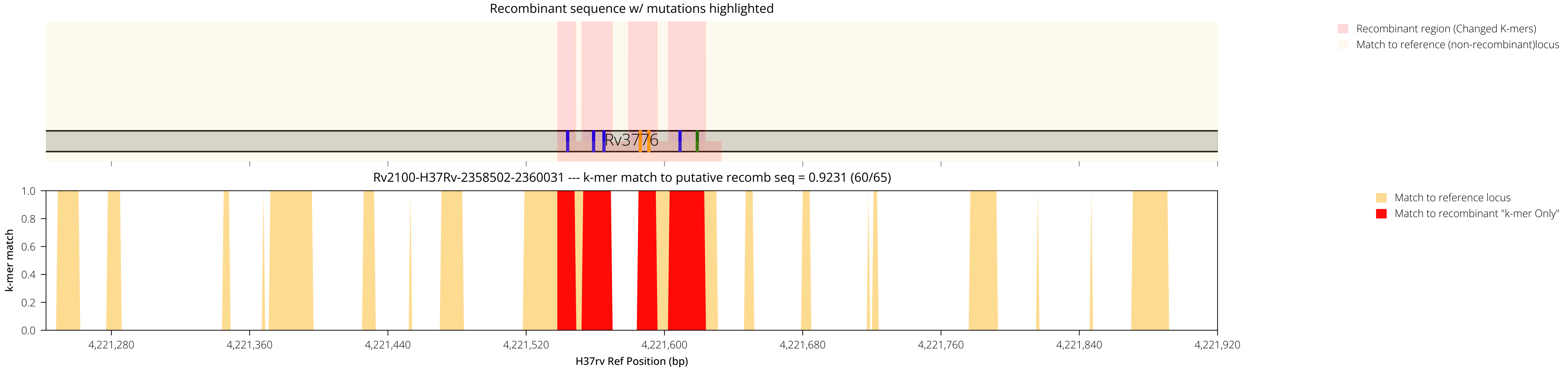

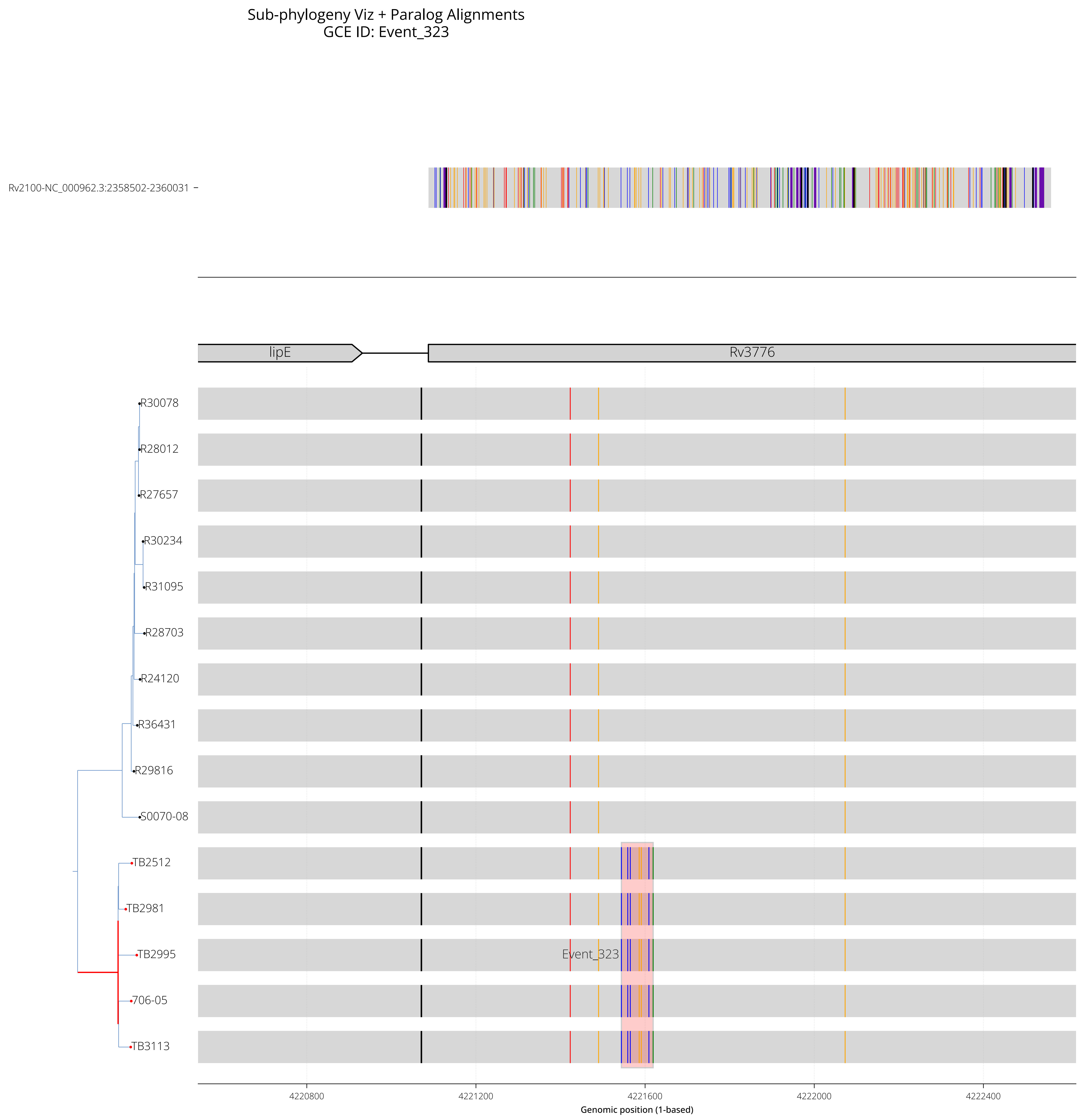

Sub-phylogeny Viz + Paralog Alignments
GCE ID: Event_323
Rv2100-NC_000962.3:2358502-2360031
lipE
Rv3776
R30078
R28012
R27657
R30234
R31095
R28703
R24120
R36431
R29816
S0070-08
TB2512
TB2981
TB2995
706-05
TB3113
Event_323
4220800
4221200
4221600
4222000
4222400
Genomic position (1-based)

Full Phylogeny Viz + Paralog Alignments
GCE ID: Event_323

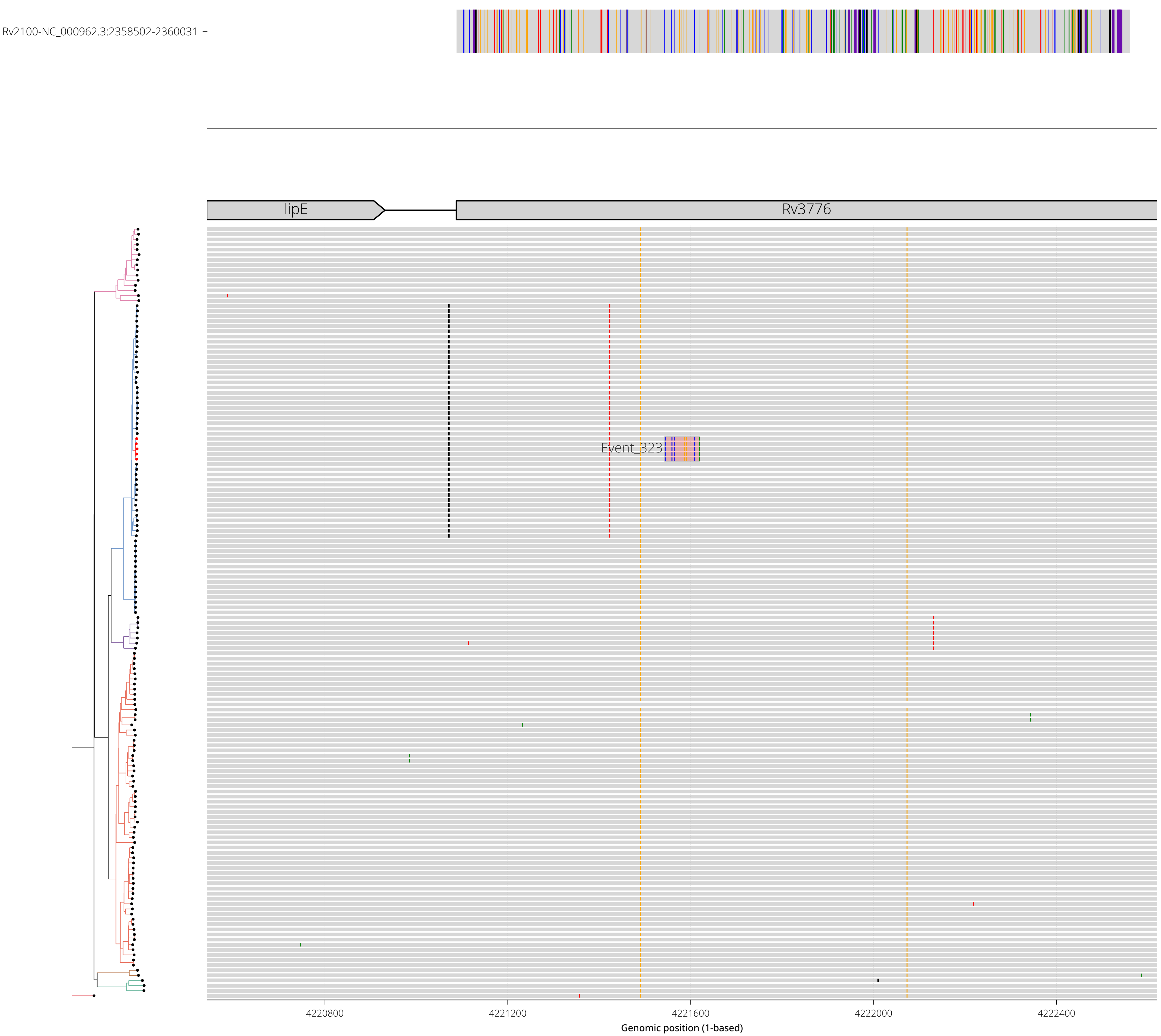

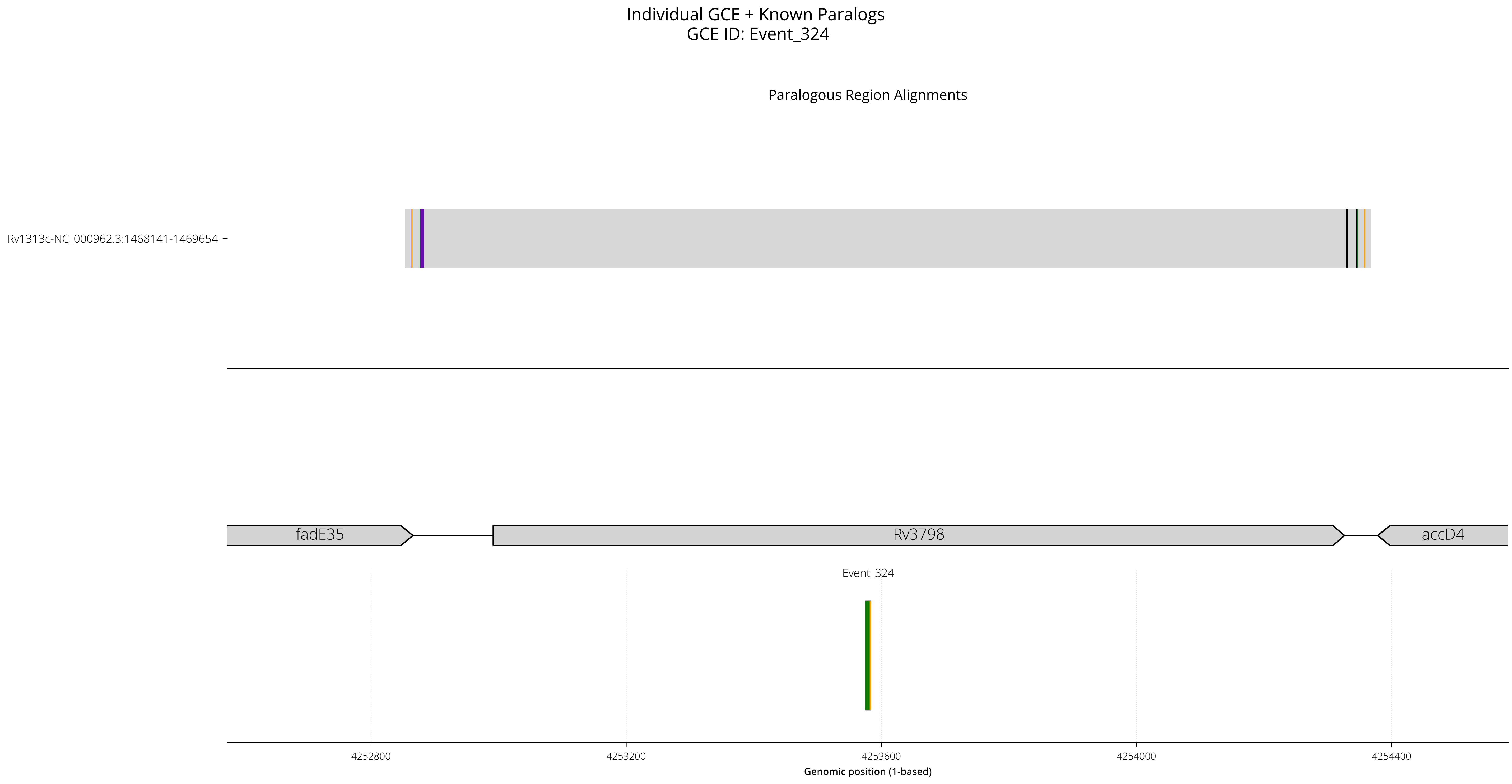

Individual GCE + Known Paralogs
GCE ID: Event_324
Paralogous Region Alignments
Rv1313c-NC_000962.3:1468141-1469654
fadE35
Rv3798
accD4
Event_324
4252800
4253200
4253600
4254000
4254400
Genomic position (1-based)

Circos Viz - GCE to Known Paralogs Locations
GCE ID: Event_324

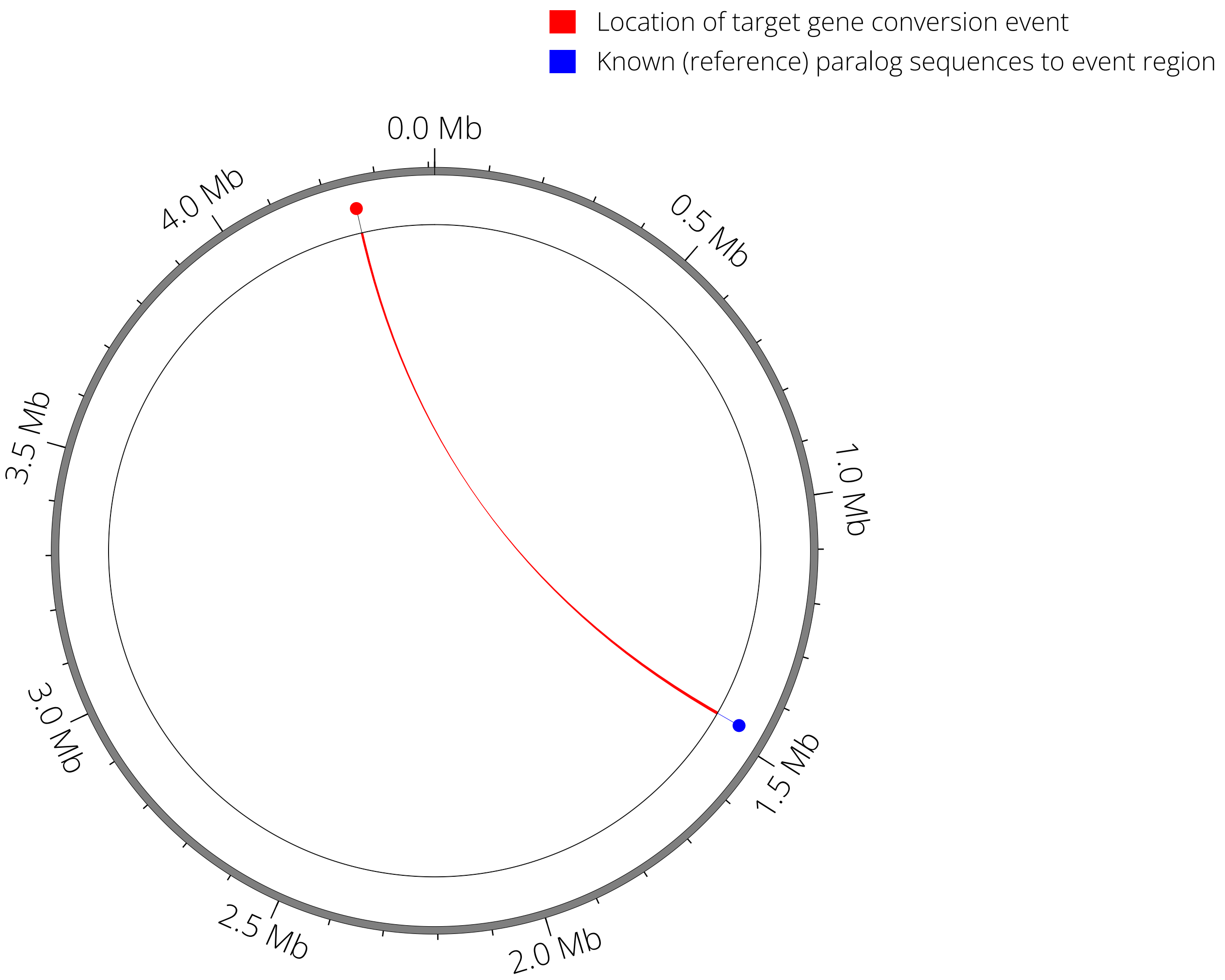

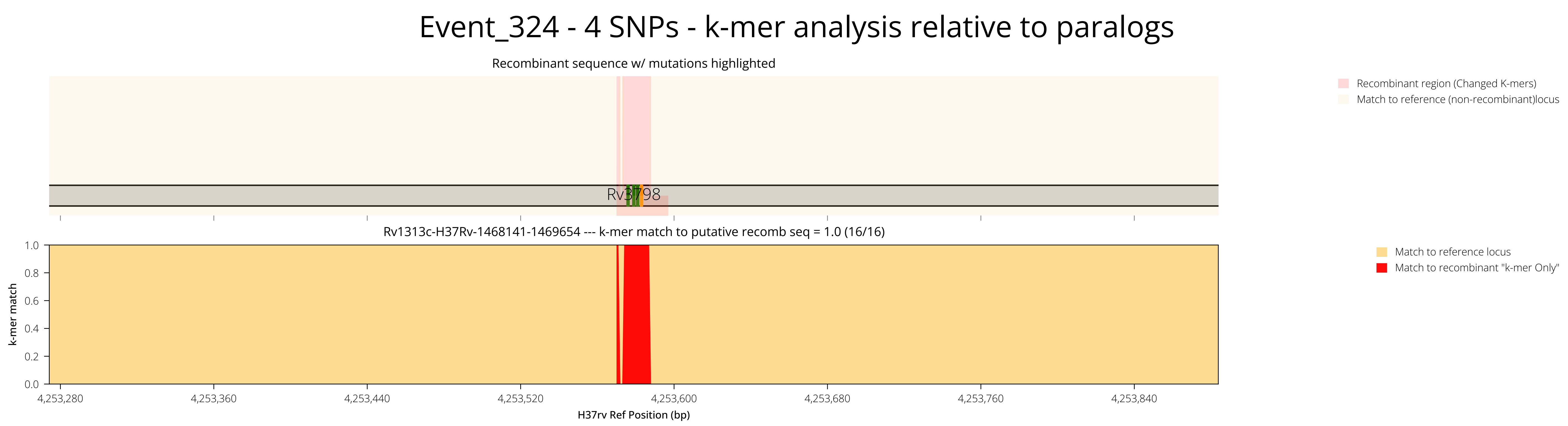
Event_324 - 4 SNPs - k-mer analysis relative to paralogs
Recombinant sequence w/ mutations highlighted
Recombinant region (Changed K-mers)
Match to reference (non-recombinant)locus
Rv3798
Rv1313c-H37Rv-1468141-1469654 --- k-mer match to putative recomb seq = 1.0 (16/16)
Match to reference locus
Match to recombinant "k-mer Only"
k-mer match
1.0
0.8
0.6
0.4
0.2
0.0
4,253,280
4,253,360
4,253,440
4,253,520
4,253,600
4,253,680
4,253,760
4,253,840
H37rv Ref Position (bp)

Sub-phylogeny Viz + Paralog Alignments
GCE ID: Event_324

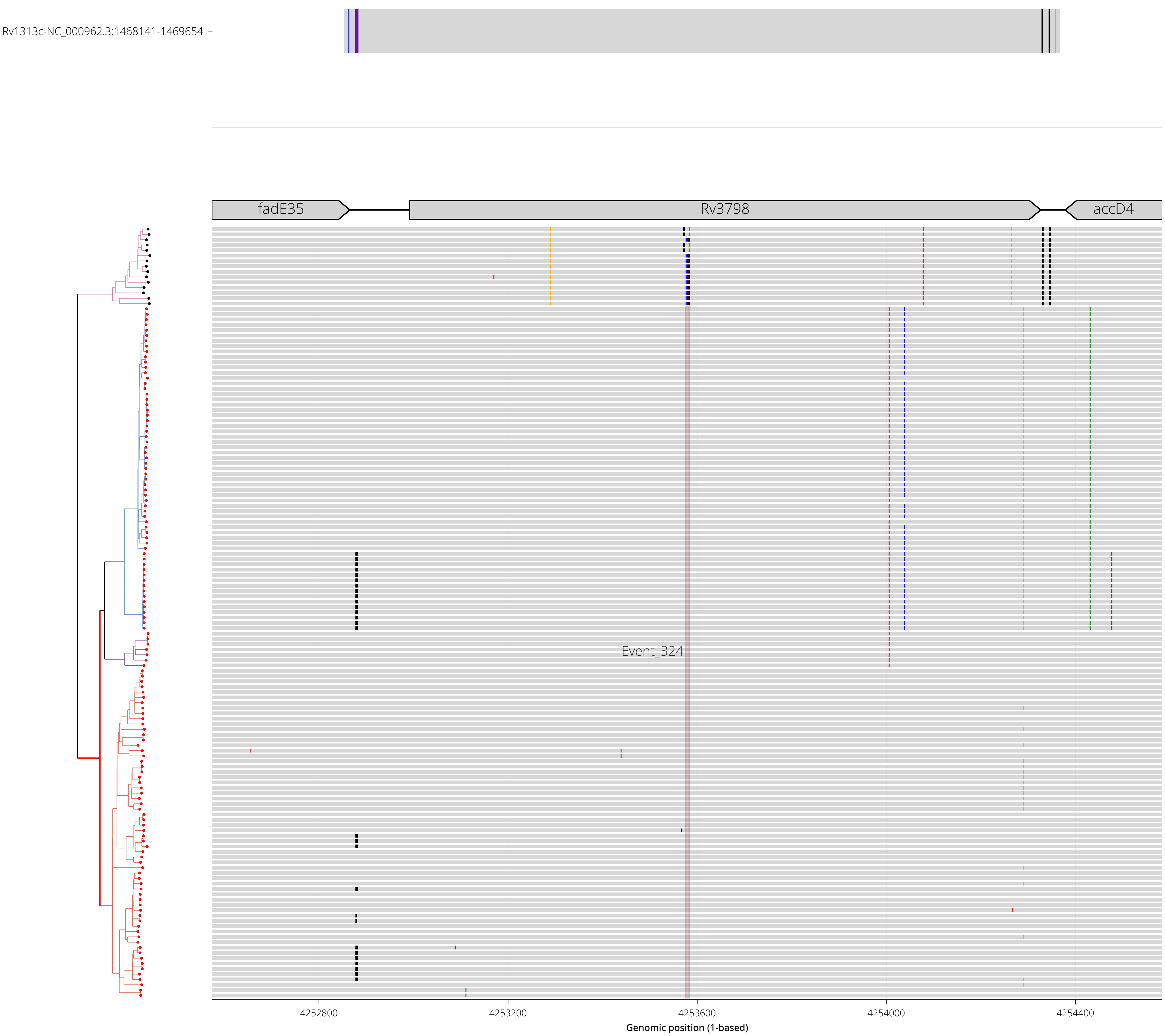

Full Phylogeny Viz + Paralog Alignments
GCE ID: Event_324

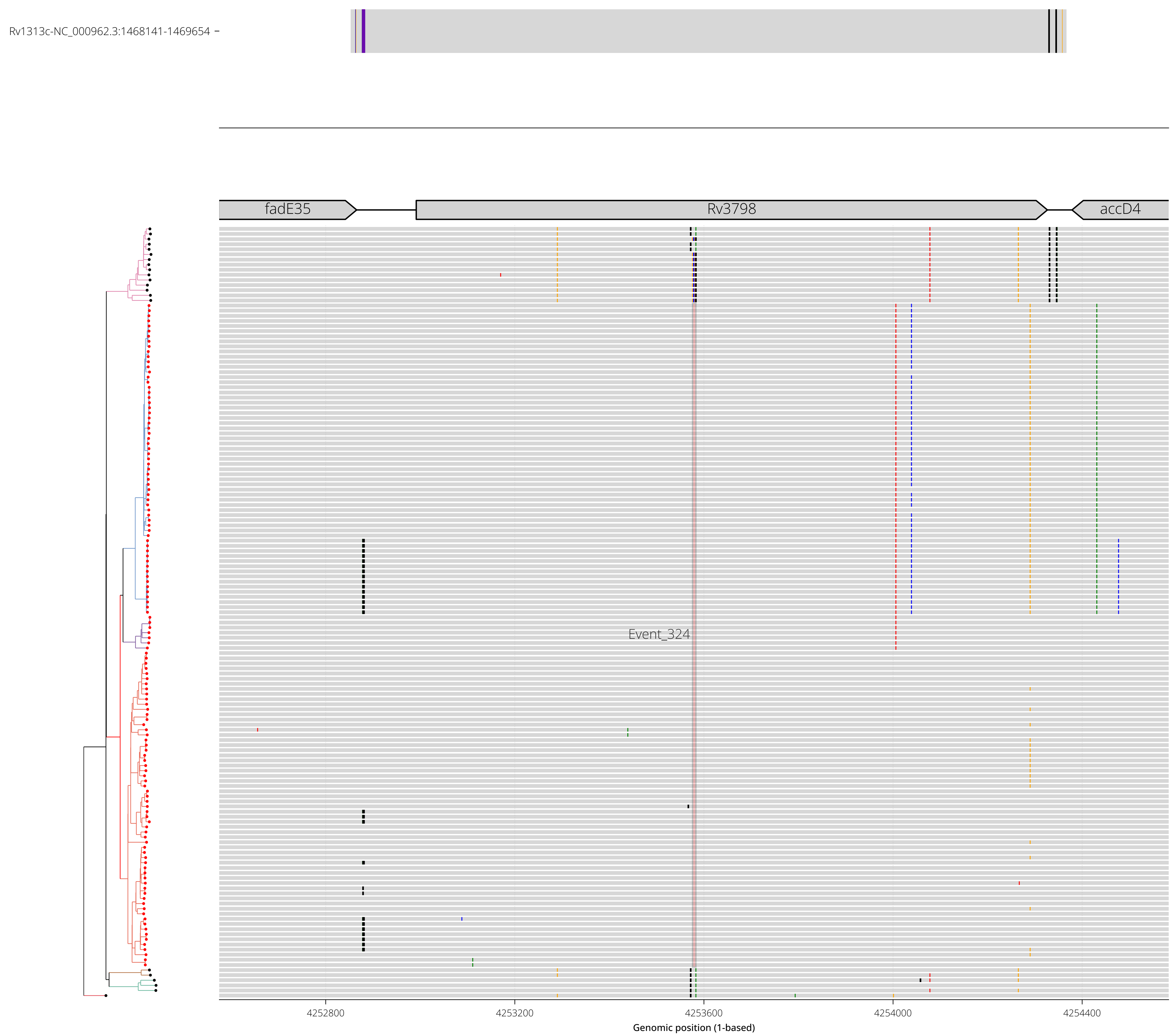
