## Supplemental Data File 14 for "Gene conversion is a key driver of diversity hotspots in *M. tuberculosis* antigens and virulence-associated loci"

### SI-14 — Per-antigen visualization of mutation frequency, epitope mapping, and gene conversion events

#### Overview

This file contains one visualization for the gene of a target antigen. Each visualization summarizes the missense mutational frequency across all codon positions of the target gene(s), alongside the available T cell epitope mapping data and the location of each detected gene conversion event. All gene annotations and coordinates shown are relative to the H37Rv reference [NC_000962.3].

#### What is shown

Each page corresponds to a genomic region encoding T cell antigen(s). Summary metadata and statistics for the region are shown in the header. Below the metadata header, from top to bottom, each page contains the following tracks:

- Gene annotations: H37Rv annotated genes for the region, drawn at the top.
- Missense mutation event frequency: per-codon missense mutation event frequency, drawn as a vertical line per codon. This frequency is based on the number of individual mutation events inferred on the phylogeny, not a count of mutations observed across genomes.
- Epitope density: a summary bar where each position is colored from white (low) to red (high) by epitope density.
- Epitope mapping results: Overlapping peptides assayed for T cell reactivity are drawn at its mapped position within the genome. Positive (reactive) T-cell epitopes are colored dark red and non-reactive peptides are colored grey.
- Gene conversion events: the recombination tract and mutations of all overlapping putative gene conversion events. Missense (non-synonymous) mutations are colored red and synonymous (silent) mutations are colored blue.

#### Figure key

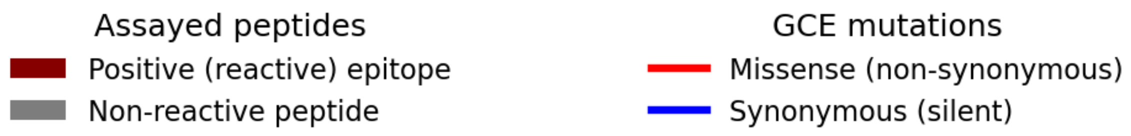

Gene(s): PPE18
In Paralogous Region (PR)? : [ True]
### of Gene Conversion Events (GCE) detected: 8
Missense Mutation Events Detected (Total): 61
Missense Mutation Events Detected (In Epitopes): 44
Total # of assayed peptides: 83
### Positive T-cell epitopes: 27

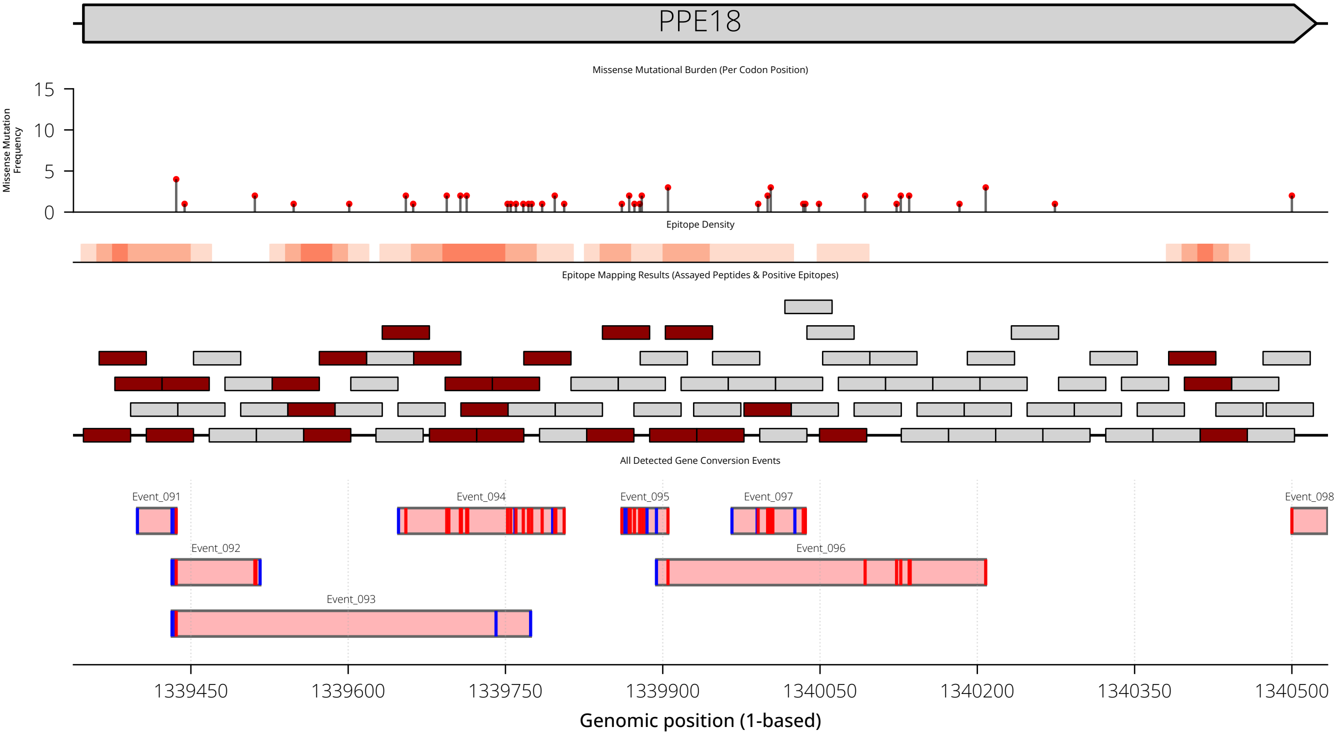

Gene(s): PPE19
In Paralogous Region (PR)? : [ True]
### of Gene Conversion Events (GCE) detected: 5
Missense Mutation Events Detected (Total): 61
Missense Mutation Events Detected (In Epitopes): 9
Total # of assayed peptides: 37
### Positive T-cell epitopes: 16

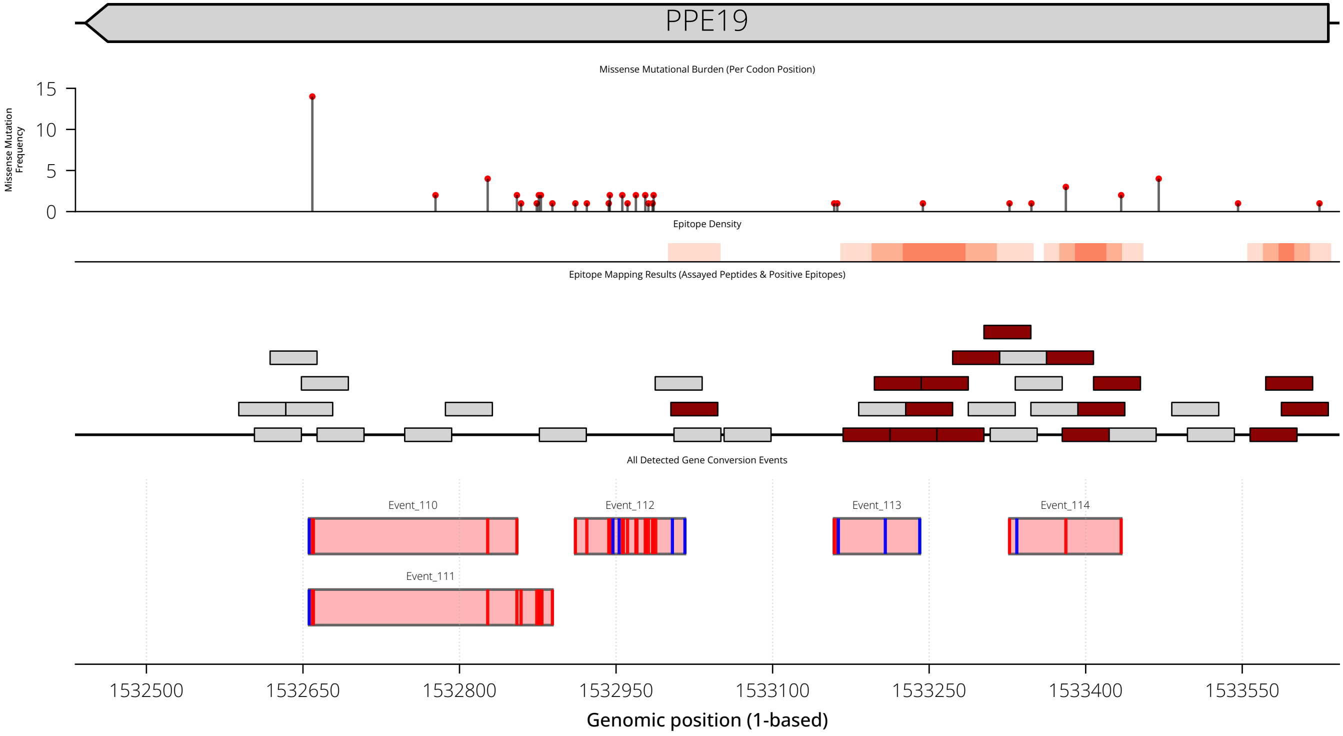

Gene(s): PPE60
In Paralogous Region (PR)? : [ True]
### of Gene Conversion Events (GCE) detected: 8
Missense Mutation Events Detected (Total): 141
Missense Mutation Events Detected (In Epitopes): 24
Total # of assayed peptides: 20
### Positive T-cell epitopes: 5

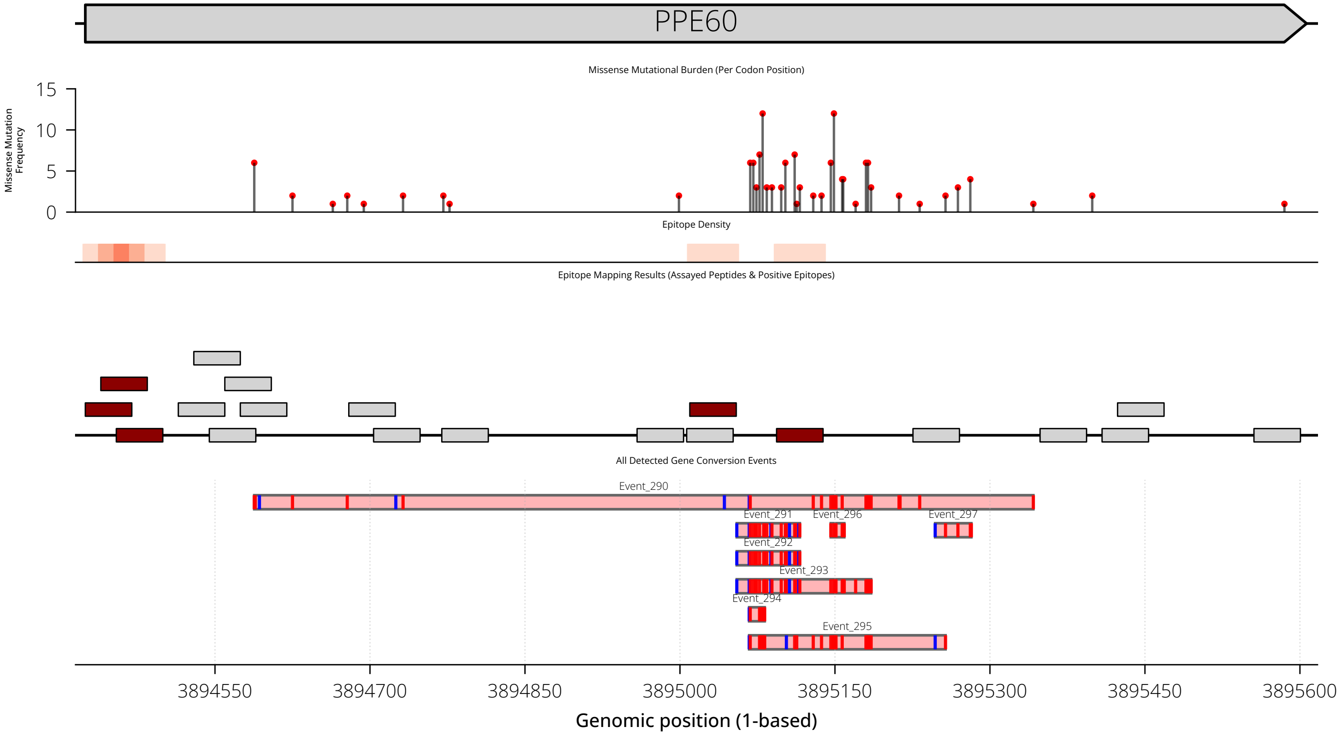

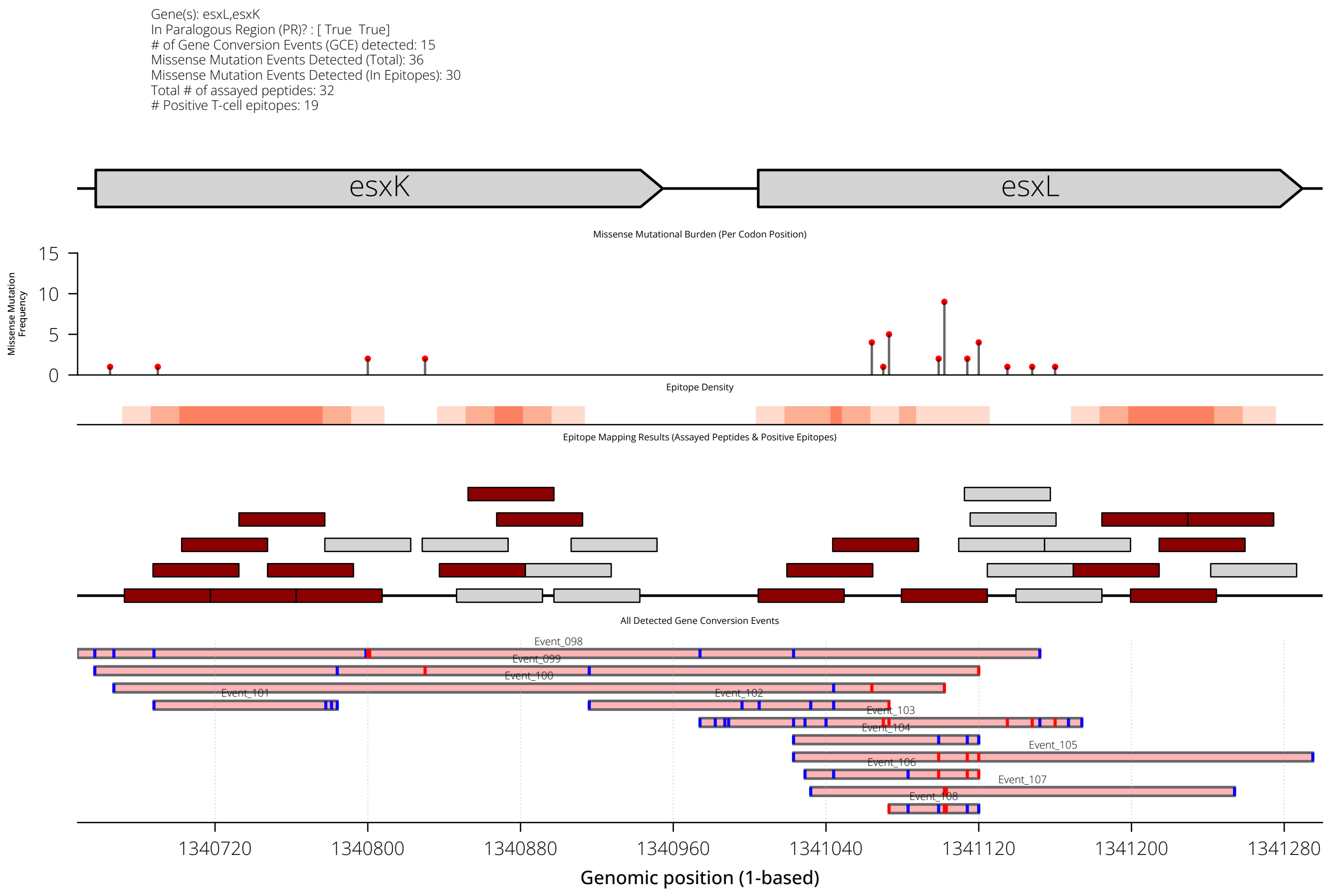
Gene(s): esxL,esxK
In Paralogous Region (PR)? : [ True  True]
### of Gene Conversion Events (GCE) detected: 15
Missense Mutation Events Detected (Total): 36
Missense Mutation Events Detected (In Epitopes): 30
Total # of assayed peptides: 32
### Positive T-cell epitopes: 19
esxK
esxL
Missense Mutational Burden (Per Codon Position)
Missense Mutation Frequency
15
10
5
0
Epitope Density
Epitope Mapping Results (Assayed Peptides & Positive Epitopes)
All Detected Gene Conversion Events
Event_098
Event_099
Event_100
Event_101
Event_102
Event_103
Event_104
Event_105
Event_106
Event_107
Event_108
1340720
1340800
1340880
1340960
1341040
1341120
1341200
1341280
Genomic position (1-based)

Gene(s): esxO,esxP
In Paralogous Region (PR)? : [ True  True]
### of Gene Conversion Events (GCE) detected: 14
Missense Mutation Events Detected (Total): 27
Missense Mutation Events Detected (In Epitopes): 5
Total # of assayed peptides: 26
### Positive T-cell epitopes: 19

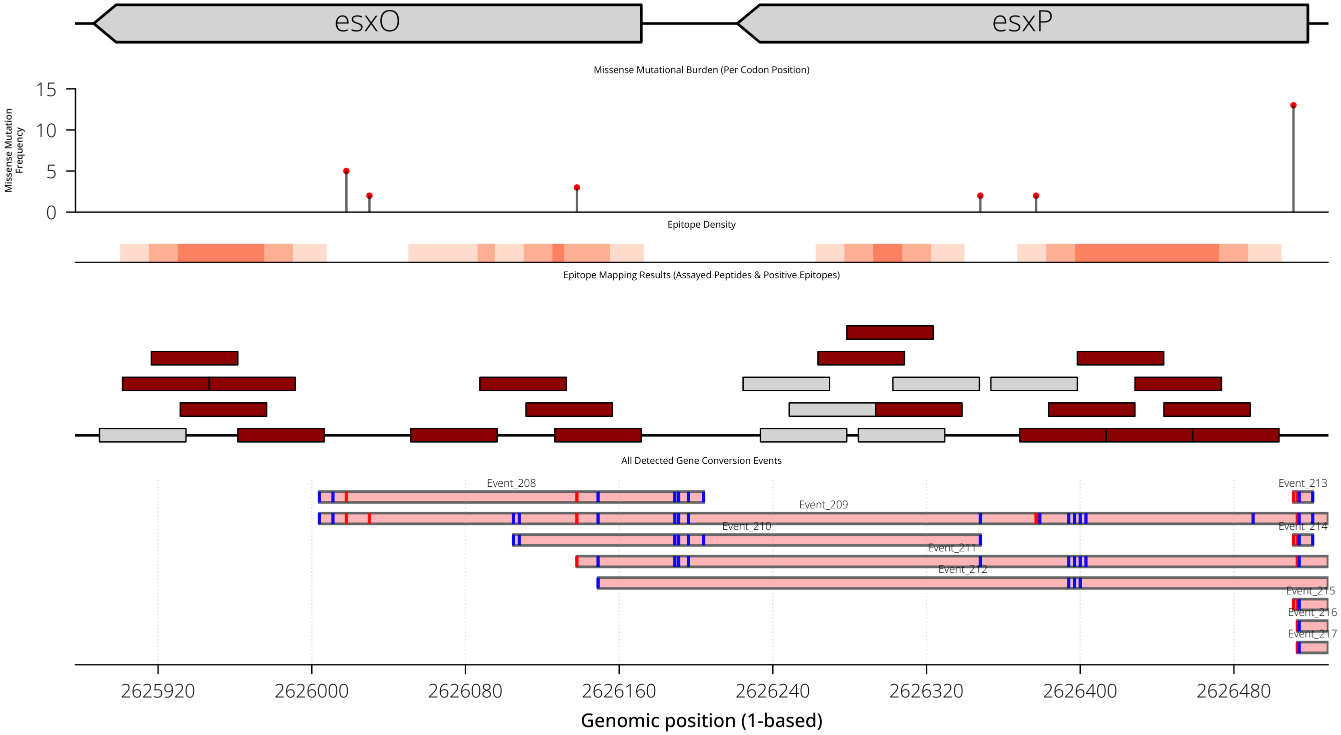

Gene(s): esxM,esxN
In Paralogous Region (PR)? : [ True  True]
### of Gene Conversion Events (GCE) detected: 6
Missense Mutation Events Detected (Total): 11
Missense Mutation Events Detected (In Epitopes): 1
Total # of assayed peptides: 25
### Positive T-cell epitopes: 18

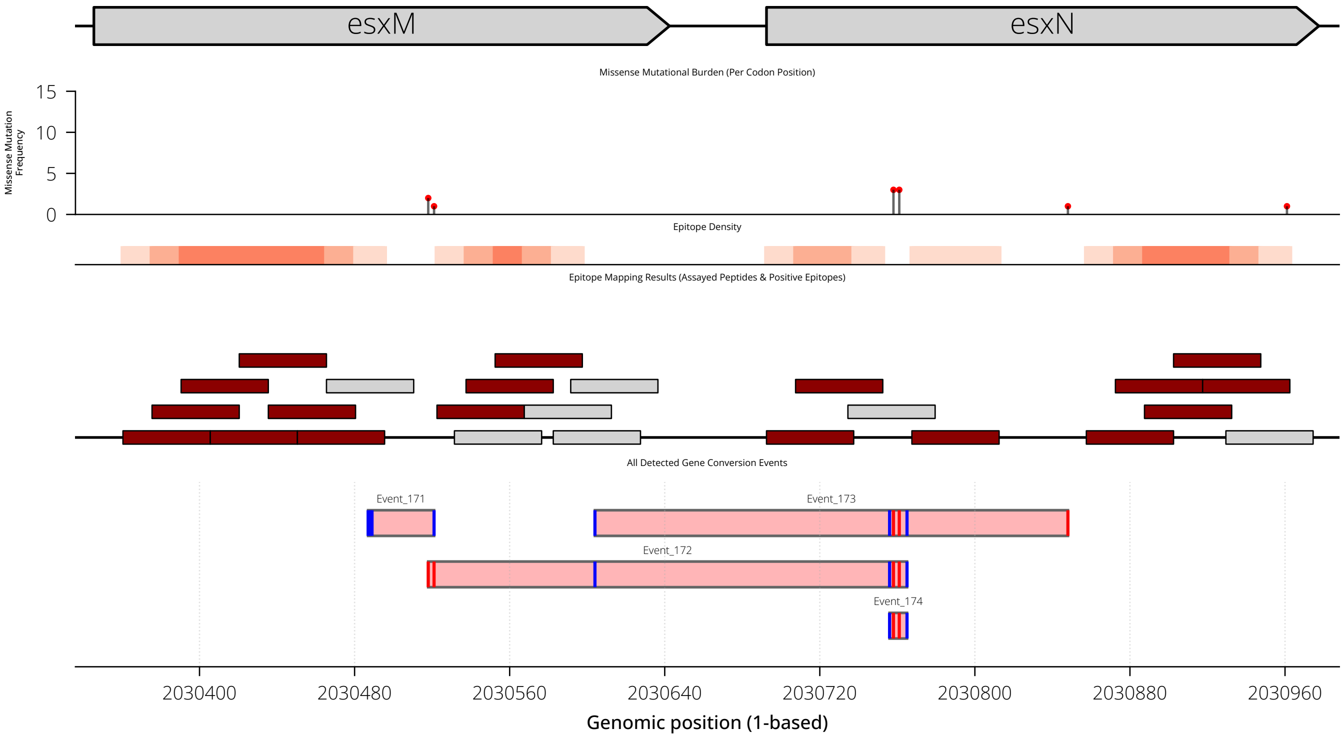

Gene(s): pepA
In Paralogous Region (PR)? : [False]
### of Gene Conversion Events (GCE) detected: 0
Missense Mutation Events Detected (Total): 5
Missense Mutation Events Detected (In Epitopes): 0
Total # of assayed peptides: 72
### Positive T-cell epitopes: 3

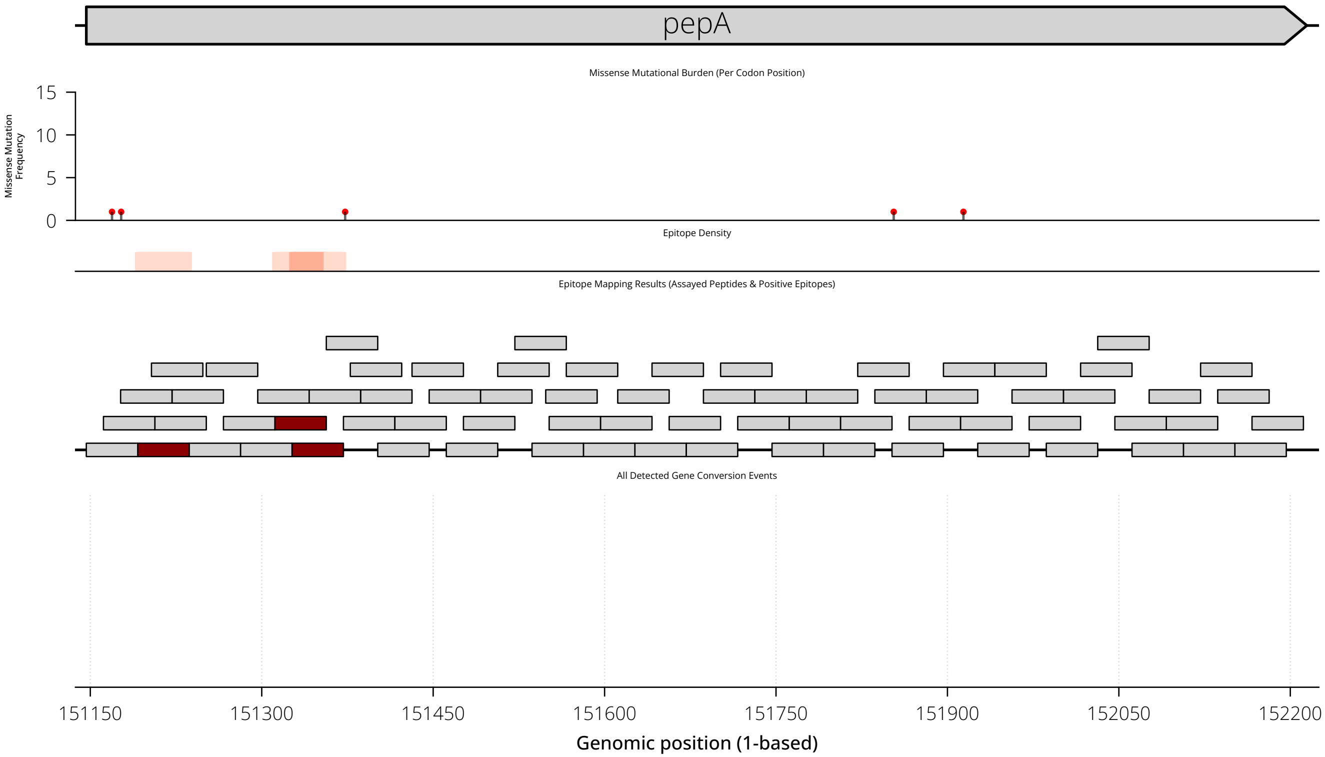

Gene(s): esxA,esxB
In Paralogous Region (PR)? : [False False]
### of Gene Conversion Events (GCE) detected: 0
Missense Mutation Events Detected (Total): 3
Missense Mutation Events Detected (In Epitopes): 3
Total # of assayed peptides: 45
### Positive T-cell epitopes: 40

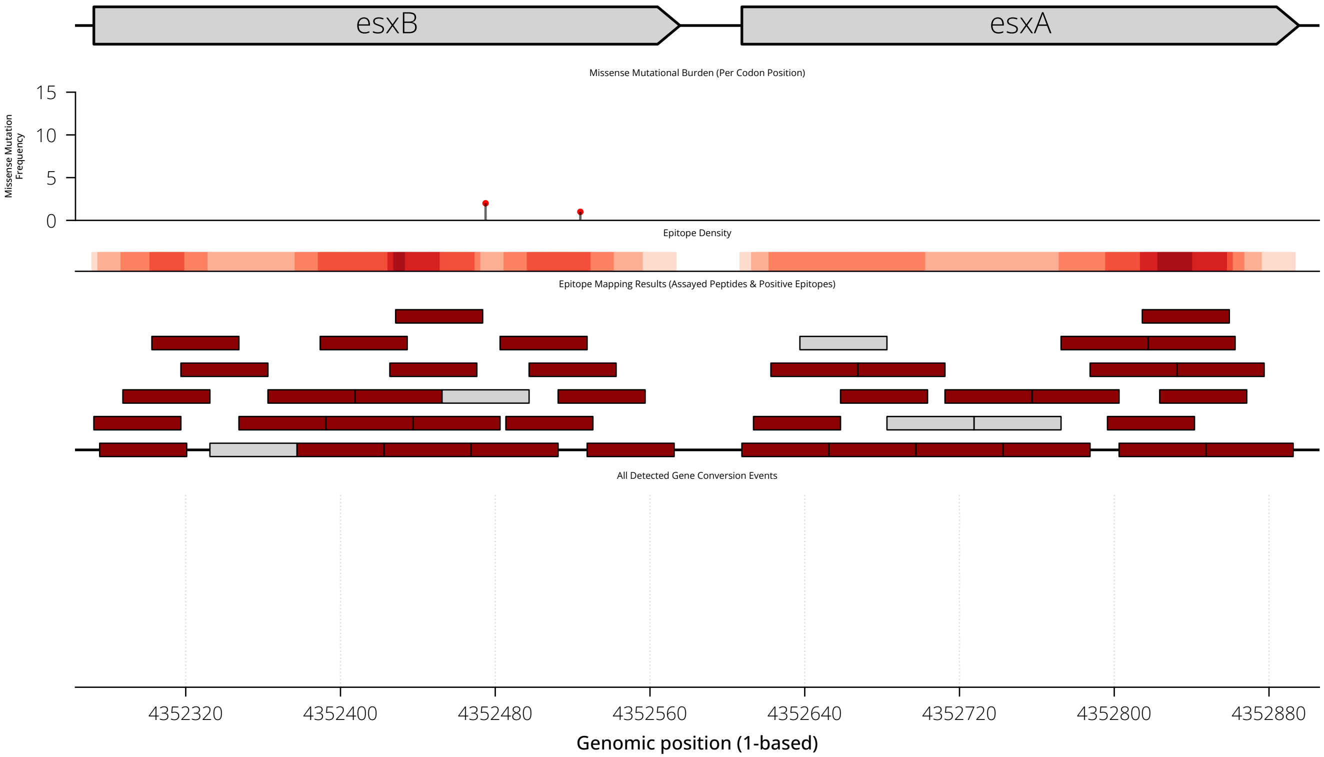

Gene(s): Rv0010c
In Paralogous Region (PR)? : [False]
### of Gene Conversion Events (GCE) detected: 0
Missense Mutation Events Detected (Total): 9
Missense Mutation Events Detected (In Epitopes): 2
Total # of assayed peptides: 6
### Positive T-cell epitopes: 2

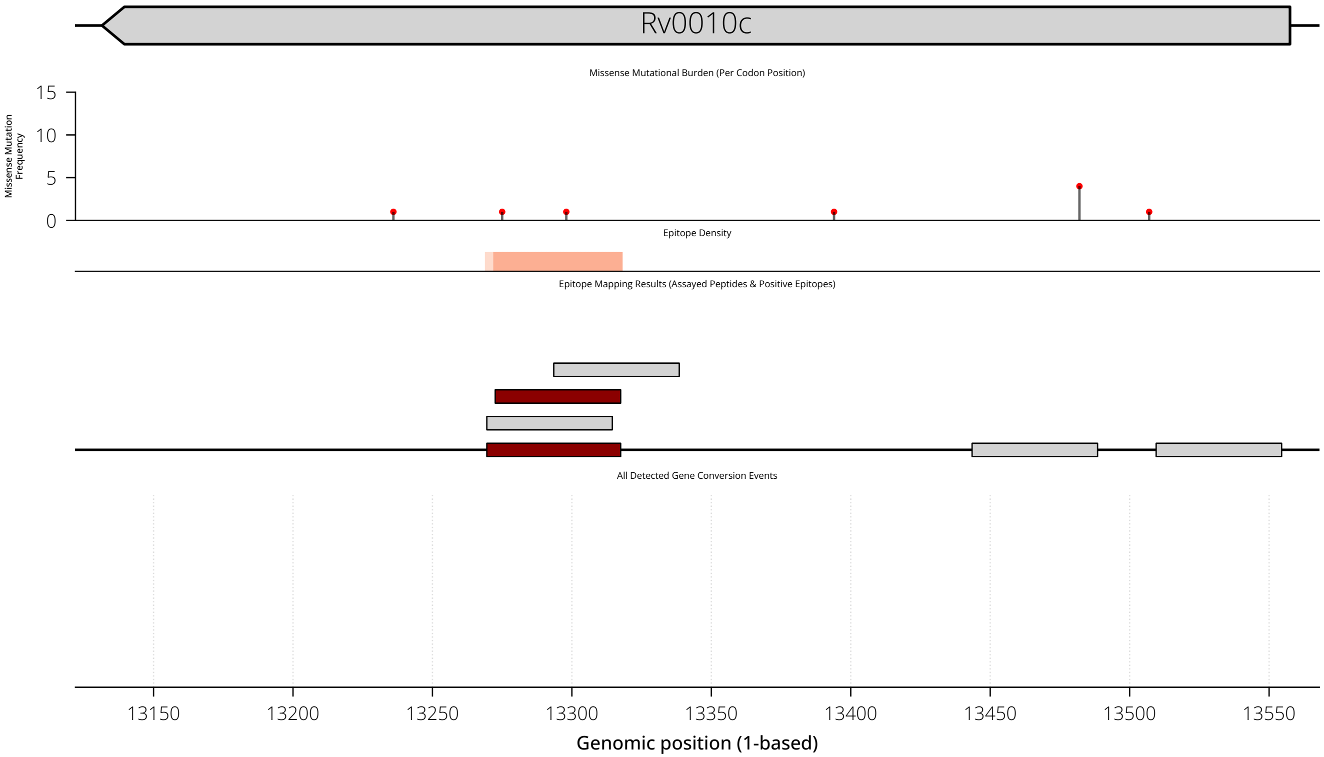

Gene(s): fbpA
In Paralogous Region (PR)? : [False]
### of Gene Conversion Events (GCE) detected: 0
Missense Mutation Events Detected (Total): 3
Missense Mutation Events Detected (In Epitopes): 1
Total # of assayed peptides: 78
### Positive T-cell epitopes: 13

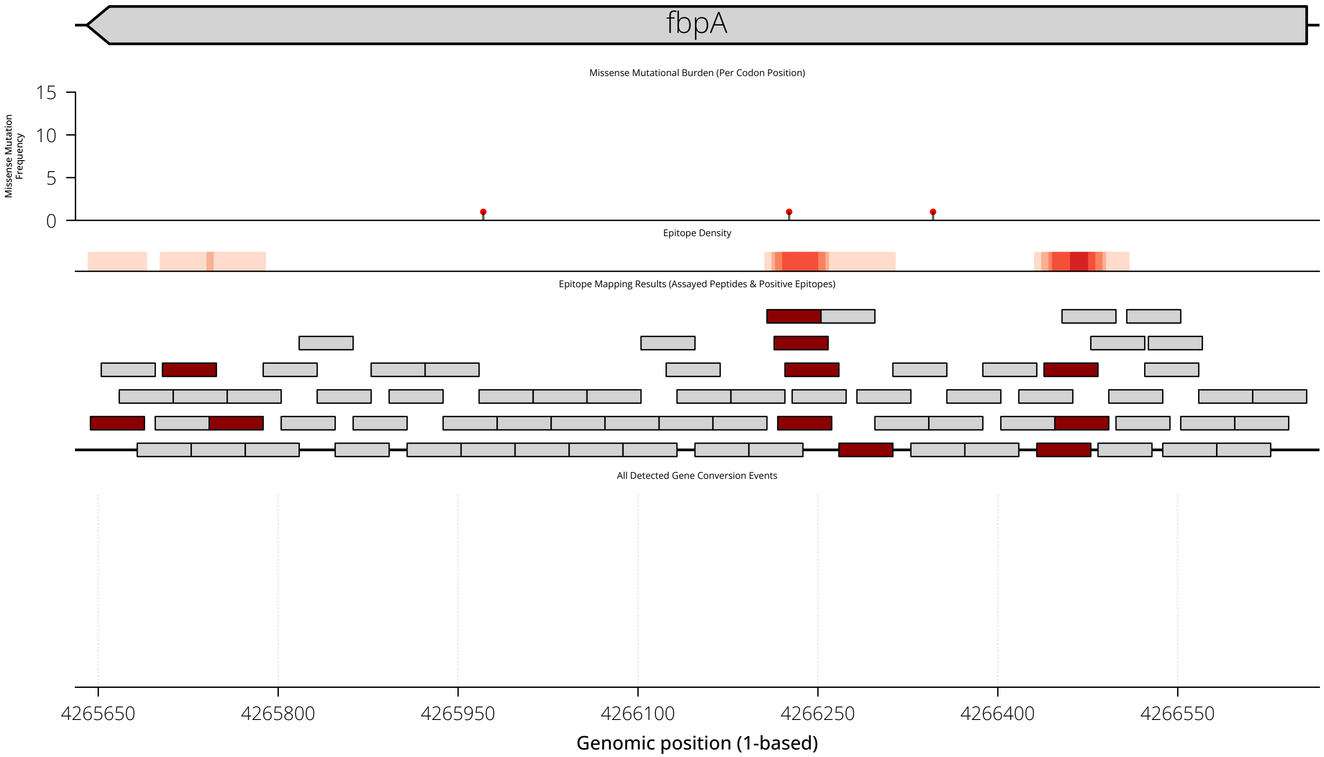

Gene(s): fbpB
In Paralogous Region (PR)? : [False]
### of Gene Conversion Events (GCE) detected: 0
Missense Mutation Events Detected (Total): 3
Missense Mutation Events Detected (In Epitopes): 2
Total # of assayed peptides: 72
### Positive T-cell epitopes: 16

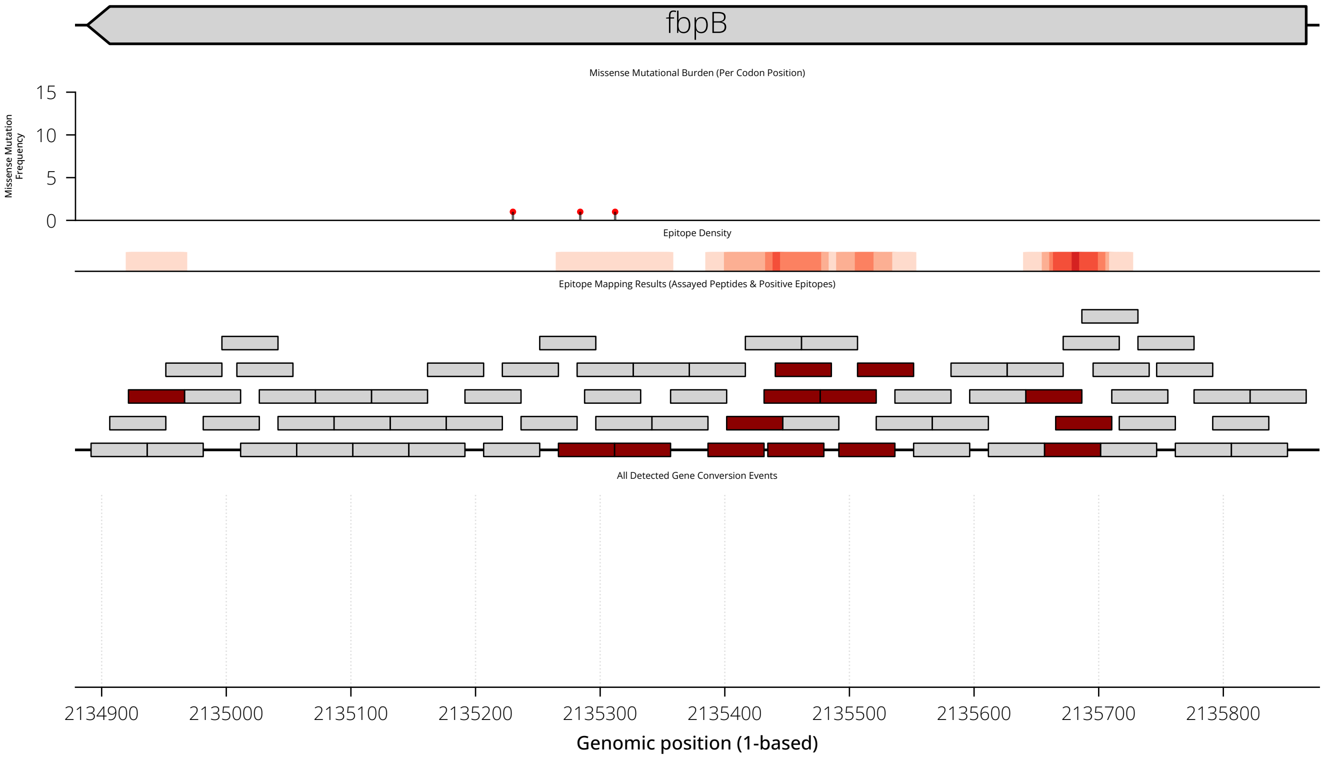
