## Supplemental Appendix for "Gene conversion is a key driver of diversity hotspots in *M. tuberculosis* antigens and virulence-associated loci"

**This PDF file includes:**

Overview of all supplementary materials and data
Supplemental Text
Figures S1 to S16
Tables S1 to S16

#### Supplementary Materials Overview

**Supplementary Information**: Supplemental Text, Figures (S1-16), and Tables (S1-16)

**Dataset S1**: Isolate metadata, genome assemblies, and sequencing accessions for all data used.

**Dataset S2**: Catalog of non-unique (high-homology or repetitive) sequence content annotated across the H37Rv reference genome.

**Dataset S3**: Genome-wide nucleotide diversity estimates and complete variant catalogs per genome assembly.

**Dataset S4**: Mutation characteristics of paralogous regions compared to the rest of the genome.

**Dataset S5**: Catalog of detected gene conversion events (N=324) along with associated phylogenetic assignments, event-to-paralog mapping results, and quality-control metrics.

**Dataset S6**: Distribution of gene conversion events across paralogous regions and paralog networks in the H37Rv genome.

**Dataset S7**: Supporting data for gene conversion analyses of TGEN-937-SR short-read WGS dataset and TBP-22 long-read (PacBio HiFi) resequencing dataset.

**Dataset S8**: Curated T cell epitope mapping data from Lindestam et al. (2016) and Panda et al. (2024), and supporting analysis results.

**Dataset S9**: *In silico* HLA class II binding predictions (netMHCpanII) and mutational effect analyses of PPE18 gene conversion events.

**Dataset S10:** Phylogenies used in this study for the Mtb151CI, TBP-22CI, and TGEN-937CI datasets (Nexus format).

**Dataset S11**: Visualization of each detected nucleotide diversity hotspot with all detected variation shown in phylogenetic context.

**Dataset S12**: Visualization of paralogous regions with all detected gene conversion events shown in both genomic and phylogenetic context.

**Dataset S13**: Event-level visualizations of inferred gene conversion tracts, including variant patterns, phylogenetic context, and paralog sequence comparisons.

**Dataset S14**: Visualization of per-codon mutation frequency and T cell epitope mapping coverage across selected antigens.

**Dataset S15:** Assembly quality control results from NucFlag pipeline and summary depth and base quality statistics.

#### Supplemental Results

##### Identification of non-unique (repetitive and homologous) sequence content in the Mtb genome

We systematically identified repetitive and homologous sequence features in the *Mycobacterium tuberculosis* H37Rv genome (NC_000962.3) based on four non-exclusive categories: **paralogous regions (PRs), local repeat regions (LRRs), low-complexity regions (LCRs), and low mappability regions (LMRs)**. PRs were defined as distinct genomic segments with detectable homology elsewhere in the genome. Local repeat regions reflect cases where consecutive sequence elements share homology. Low-complexity regions are sequences enriched for short tandem motifs, including imperfect repeats. Low-mappability regions correspond to sequences with non-unique k-mer content (using a k-mer size of 50 bp and allowing up to 4 mismatches).

Across the 4.4 Mb H37Rv genome, the four categories had the following coverage: PRs 257 kb (5.8%), LRRs 150 kb (3.4%), LCRs 127 kb (2.9%), and LMRs 189 kb (4.3%). The union of all four categories spanned 447 kb (10.1% of the genome), representing all regions associated with repetitive and/or high-homology sequences. A summary of region categories and their definitions is provided in **Table S5**, and their genomic distribution is shown in **Figure S2**. Notably, these categories frequently overlapped within the genome, with many paralogous regions also containing local repeats, low-complexity motifs, or non-unique sequence content.

##### *In silico* analysis of effect of HLA binding of Gene Conversion events in PPE18 on HLA binding of european populations

As described in the main text (**Figure S15**, **Table S15**), we evaluated the predicted effects of gene conversion events in PPE18 on HLA-II binding using NetMHCpanII (1). We extended this analysis to specifically assess the impact of *PPE18* gene-conversion mutations on binding of the subset of HLA-II alleles most common in the context of European host populations (**Figure S16, Table S16**) Of the seven GCEs mutating PPE18, three (Events_091–093) introduced no amino acid changes in regions predicted to bind any of the tested HLA alleles. In contrast, the largest event (Event-94), which introduced 17 missense mutations, resulted in both predicted gains and losses of strong-binding interactions across multiple HLA alleles (10 gains, 4 losses). The remaining three events (Events 95–97) were each associated exclusively with gains in predicted HLA binding. These results indicate that some gene conversion events in PPE18 have the potential to alter the antigenic landscape by modifying HLA binding affinity.

#### Supplemental Methods

##### Hybrid genome assembly using long and short-read whole genome sequencing data

All genome assemblies in this study were generated using the same core hybrid assembly approach, consisting of two stages: (1) de novo assembly from the long-read data, with iterative long-read polishing performed internally by the assembler, followed by (2) polishing of the resulting assembly with the matched paired-end Illumina data to correct residual base-level errors. Because the underlying isolates were sequenced with three distinct long-read data types, this core strategy was implemented as three pipeline variants, each tailored to one long-read data type:

- **PBclr_LR_Flye_I3_SR_Pilon** — PacBio CLR (subread) data
- **PBccs_LR_Flye_I3_SR_Pilon** — PacBio CCS (HiFi) read data
- **ONT_LR_FlyeI3M_SR_Pilon_PolyPolish** — Oxford Nanopore (R9.4.1) read data

The specific pipeline applied to each isolate is recorded in **Table S1** and **Dataset S1**. The Mtb151 primary dataset includes isolates processed with all three pipeline variants, whereas the isolates of the TBP-22-LR were processed exclusively with the PBccs pipeline.

**Hybrid Assembly Pipeline 1 - PBclr_LR_Flye_I3_SR_Pilon (PacBio CLR/subreads)**

PacBio CLR (Continuous Long Read) subreads were assembled de novo with Flye (v2.6; `--pacbio-raw`, `--genome-size 5m`, `--asm-coverage 200`, `--iterations 3`), which includes three rounds of internal long-read polishing(2). The assembly origin was set to the start of *dnaA* using Circlator `fixstart` (v1.5.1) with the H37Rv *dnaA* sequence, and contigs were renamed by sample and length-filtered to retain those greater than 100 kb. The assembly was then polished with the matched paired-end Illumina data: reads were adapter- and quality-trimmed with Trimmomatic (Snakemake wrapper v0.38.0; `ILLUMINACLIP` with a custom adapter set, `SLIDINGWINDOW:4:20`, `MINLEN:75`), aligned to the draft assembly with BWA-MEM (v0.7.17), coordinate-sorted with SAMtools (v1.9), and duplicate-marked with Picard MarkDuplicates (v2.21.4; `REMOVE_DUPLICATES=true`), followed by a single round of polishing with Pilon (v1.23; `--fix snps,indels`)(3–5). Final assemblies were annotated with Bakta (v4.8)(6). The full Snakemake pipeline code and configuration files are found in 2.Mtb.Generate.HybridAsm.PBccs.smk in the manuscript GitHub repo.

**Hybrid Assembly Pipeline 2 - PBccs_LR_Flye_I3_SR_Pilon (PacBio CCS/HiFi)**

PacBio CCS reads were assembled *de novo* with Flye (v2.9.2; `--pacbio-hifi`, `--genome-size 4.4m`, `--asm-coverage 200`, `--iterations 3`), which includes three rounds of internal long-read polishing(2). Because HiFi reads have high intrinsic base accuracy, no additional long-read polishing step was applied. The assembly origin was set to the start of *dnaA* using Circlator `fixstart` (v1.5.1) with the H37Rv *dnaA* sequence, and contigs were renamed by sample and length-filtered to retain those greater than 100 kb. The assembly was then polished with the matched paired-end Illumina data: reads were adapter- and quality-trimmed with Trimmomatic (Snakemake wrapper v0.38.0; `ILLUMINACLIP` with a custom adapter set, `SLIDINGWINDOW:4:20`, `MINLEN:75`), aligned to the draft assembly with BWA-MEM (v0.7.17), coordinate-sorted with SAMtools (v1.9), and duplicate-marked with Picard MarkDuplicates (v2.21.4; `REMOVE_DUPLICATES=true`), followed by a single round of polishing with Pilon (v1.23; `--fix snps,indels`)(3–5). Final assemblies were annotated with Bakta (v4.8)(6). The full Snakemake pipeline code and configuration files are found in 2.Mtb.Generate.HybridAsm.PBccs.smk in the manuscript GitHub repo.

##### Hybrid Assembly Pipeline 3 - ONT_LR_FlyeI3M_SR_Pilon_PolyPolish (Oxford Nanopore R9.4.1)

Nanopore R9.4.1 reads were assembled de novo with Flye (v2.6; `--nano-raw`, `--genome-size 4.4m`, `--asm-coverage 200`, `--iterations 3`), which includes three rounds of internal long-read polishing (denoted "I3")(2). To address the higher base-calling error rate of this chemistry, the assembly underwent an additional long-read polishing step with Medaka (v1.5.0; model `r941_min_high_g303`, denoted "M"). Contigs were renamed by sample and length-filtered to retain those greater than 100 kb, and the assembly origin was set to the start of *dnaA* using Circlator `fixstart` (v1.5.1) with the H37Rv *dnaA* sequence. The assembly was then polished with the matched paired-end Illumina data: reads were adapter- and quality-trimmed with Trimmomatic (Snakemake wrapper v0.38.0; `ILLUMINACLIP` with a custom adapter set, `SLIDINGWINDOW:4:20`, `MINLEN:75`), aligned with BWA-MEM (v0.7.17), coordinate-sorted with SAMtools (v1.9), and duplicate-marked with Picard MarkDuplicates (v2.21.4; `REMOVE_DUPLICATES=true`), followed by polishing with Pilon (v1.23; `--fix snps,indels`)(3–5). A further short-read polishing step was then applied with PolyPolish (v0.5.0), for which trimmed Illumina reads were aligned in all-alignments mode (`bwa mem -a`) and processed with `polypolish_insert_filter` before polishing. Final assemblies were annotated with Bakta (v4.8)(6). The full Snakemake pipeline code and configuration files are found in 3.Mtb.Generate.HybridAsm.ONT.smk in the manuscript GitHub repo.

#### Defining non-unique (repetitive and homologous) sequence content of the H37Rv genome

We sought to identify repetitive and homologous sequence content within the *Mycobacterium tuberculosis* H37Rv reference genome (NC_000962.3). We defined four non-exclusive categories of non-unique sequence: (i) paralogous regions (PRs), (ii) local repeat regions (LRRs), (iii) low-complexity regions (LCRs), and (iv) low mappability regions (LMRs). PRs and LRRs were identified from a genome-wide intra-genomic homology map; LCRs and LMRs were identified independently using dedicated tools, as described below. The genomic ranges for all four categories are provided as separate annotated tables in **Dataset S2**. **Table S5** specifies the cumulative size of each category and the union of all categories relative to the H37Rv reference genome. The Python notebooks for related processing and analysis can be found in the [Analysis/1_Reference_Preprocessing]() directory of the manuscript GitHub repository.

**Detection of paralogous and local repeat regions via an intra-genomic homology map**

To generate a genome-wide homology map, we performed a self-alignment of the H37Rv genome against itself using minimap2 (v2.26) with the parameters `--MD -DP -k19 -w19 --cs`, outputting results in both PAF and SAM formats (as described in the minimap2 Cookbook, https://github.com/lh3/minimap2/blob/master/cookbook.md)(7). This approach detects sequence homology across the entire genome independent of predefined gene boundaries, and does not impose a fixed sequence-identity threshold; reported alignments are instead determined by the internal alignment-scoring heuristics of minimap2. Each detected homologous alignment was then classified based on whether its two aligned segments overlap one another in genomic coordinates: alignments between overlapping segments were classified as local repeat regions (LRRs), reflecting tandem or locally proximal repeats, and alignments between distinct, non-overlapping segments were classified as paralogous regions (PRs). For each alignment, we additionally inferred variants between the two aligned segments using paftools.js (v2.26) `call -q 0 -l 100 -L 100`. In total, 200 distinct PRs and 78 distinct LRRs were identified across the H37Rv reference genome (**Dataset S2**, tables "PR - ParalogousRegions" and "LRR - LocalRepeatRegions").

**Detection of low-complexity regions (LCRs)**

Low-complexity regions (LCRs) were identified by running Longdust (v1.2) on the H37Rv genome sequence (NC_000962.3) with the parameters `-k6 -w1000 -t 0.5`(8). Longdust scans nucleotide sequences for intervals enriched in short tandem repeat motifs, including imperfect or approximate repeats, and all reported intervals were retained as low-complexity regions. The annotated regions are provided in **Dataset S2** (table "LCR - LowComplexityRegions").

**Detection of low mappability regions (LMRs)**

Regions with low pileup-mappability were identified using Pupmapper software (v0.2.0, https://github.com/maxgmarin/pupmapper), run with the `pupmapper all` command and the parameters `-k 50 -e 4` on the H37Rv genome sequence (NC_000962.3). Pupmapper integrates Genmap-based k-mer uniqueness profiles (Genmap v1.3; k-mer size 50 bp, up to 4 mismatches) with per-base pileup mappability statistics. All genomic positions with a non-perfect pileup mappability score (< 1) were designated as low pileup-mappability regions. The annotated regions are provided in **Dataset S2** (table "LMR - LowPileupMappability").

#### H37Rv reference genome and annotations

The H37Rv genome (NCBI accession NC_000962.3) and its annotations were used as the reference for all analyses. Functional category annotations for all H37Rv genes were downloaded from Release 3 (2018-06-05) of MycoBrowser (https://mycobrowser.epfl.ch/releases).

#### Variant detection and CDS effect annotation

Genetic variants relative to the H37Rv reference were inferred for each complete genome assembly using minimap2 (v2.24) and its companion script paftools.js(7). Each assembly was aligned to H37Rv with the asm10 preset and base-level alignment enabled via the --cs tag (minimap2 -cx asm10 --cs), and alignments were filtered to retain only primary, uniquely mapped records (mapping quality = 60). Filtered alignments were sorted by reference coordinate and passed to paftools.js call to call variants, requiring a minimum alignment length of 1000 bp for both variant calling (-L 1000) and coverage assessment (-l 1000), with the H37Rv sequence supplied (-f) to emit VCF output. Variant calls from all assemblies were processed uniformly and annotated using Biopython (v1.87) to infer coding-sequence consequences relative to standard H37Rv gene annotations. All genome alignment, variant calling, and processing steps are implemented in the Mtb.WGA.Core.V3.smk Snakemake workflow available in the manuscript repository.

#### Manual inspection of genome assemblies, variant calls, and read alignments in nucleotide diversity hotspots

Variant calls for each genome were derived from whole-genome assembly-to-reference alignments generated using Minimap2. To assess the accuracy of these calls within regions of elevated nucleotide diversity, we performed systematic manual inspection of all 37 identified diversity hotspots across all 151 genome assemblies using the Integrative Genomics Viewer (IGV)(9).

For each hotspot, inspection was conducted by simultaneously visualizing two complementary alignment tracks: (1) the whole-genome assembly-to-reference alignment, which formed the basis of variant calling, and (2) the underlying long-read alignment pileup relative to the reference genome. In cases where multiple closely spaced variants were observed within a single isolate, we paid particular attention to the assembly-to-reference alignment to verify that the broader synteny and genomic structure of the assembled sequence were preserved relative to the reference. confirming that the local alignment containing the variants of interest was not indicative of a misassembly or structural artefact. This side-by-side approach allowed direct comparison between the called variants and the raw sequencing data that was used as the basis for the genome assembly.

During inspection, we evaluated each hotspot for two primary indicators of potential assembly or variant-calling error: visual disagreement between the assembly alignment and the long-read pileup, and uneven or interrupted read coverage. Variant calls were considered well-supported when the assembly-to-reference alignment and the long-read pileup were concordant and coverage was uniform across the region.

To enable independent verification of these inspections, we have made all alignment data publicly available through an interactive browser-based viewer built with igv.js. The viewer allows any sample in the dataset to be selected and the alignments at each of the 37 nucleotide diversity hotspots to be examined directly: https://maxgmarin.github.io/mtb-geneconv-dataviz/NucDivHotspots_Viewer/

##### Comparing sequencing depth and base quality in paralogous regions versus the rest of the genome

To compare read support in paralogous regions (PRs) with the rest of the genome, we summarized sequencing depth and base quality for each sample's long-read alignments to the H37Rv reference. Per-base depth was computed genome-wide with mosdepth (v0.3.14)(10). The resulting per-base depth profile was intersected with the PR coordinate ranges and its complement (bedtools intersect, v2.31.1) to obtain pooled mean and median depth across PR and non-PR positions, respectively(11). Mean base quality was obtained with samtools coverage (v1.20), run genome-wide and on the reads aligning to PRs, using the **meanbaseq** field. Per-sample depth and base-quality statistics are provided in **Dataset S15** (sheets "DepthStats_PR_vs_NonPR" and "BaseQual_Stats_PRvsGenomeWide"). Data processing steps are implemented in the AsmAndWGS.QC.smk snakemake workflow in the manuscript GitHub repository.

##### Systematic detection of misassemblies with NucFlag

To assess the accuracy of each genome assembly independently of the reference, we screened all 151 assemblies for misassemblies using NucFlag (v1.0.0), a reference-free method that identifies regions where an assembly is inconsistent with the long reads from which it was built. For each isolate, the sample's own long reads were realigned to its own assembly with minimap2 (v2.28), using the platform-appropriate preset and the --MD and --eqx flags required for NucFlag's per-base mismatch pileups. Alignments were sorted and indexed with SAMtools (v1.20). Next, samtools view -F 2308 was used to filter to primary alignments only  prior to misassembly calling.

Misassemblies were then called with `nucflag call`, which flags positions where a substantial fraction of the aligned reads disagree with the assembled sequence and classifies each flagged region into one of NucFlag's misassembly types. To avoid spurious calls at contig ends arising from circular-genome linearization, the first and last 3 bp of each contig were masked from calling. Each assembly was additionally assigned a consensus quality value with nucflag qv.

To relate flagged regions to the reference genome and to our regions of interest, each sample's diversity-hotspot windows and paralogous regions were lifted from H37Rv coordinates onto that sample's assembly with `paftools.js` (packaged with Minimap2 v2.28). The lifted over misassembly calls were intersected against these lifted regions using the `nucflag status` command (in both per-region length and count modes). Flagged regions were then lifted back to H37Rv coordinates and annotated with their overlap of diversity hotspots and paralogous regions.

Finally, all misassembly calls and all assembly-derived SNP calls were compiled in H37Rv coordinates and intersected to determine how many variant calls fell within a flagged region, genome-wide and within diversity hotspots. The complete per-flag inventory, the per-isolate summary, and the SNP-level overlap annotations are provided in Dataset S15 (sheets "NucFlag - All Misassemblies Detected," "NucFlag -

Summary Stats Per Genome," and "AllAsm_SNPs_AnnoByNucFlagCalls"). The full pipeline is available in the manuscript GitHub repository.

#### Mapping of gene conversion events to donor paralogs

To identify the candidate donor paralog(s) underlying each putative GCE, we compared each inferred recombination tract to all paralogous sequences in the *M. tuberculosis* genome using a k-mer–based similarity approach. We first extracted all overlapping 11-mers spanning each recombination tract (a sliding window of length 11, step size 1) and compared them to the 11-mer content of all reference paralogous sequences of the affected locus. Each 11-mer was reduced to its canonical form (the lexicographically smaller of the k-mer and its reverse complement), so that each sequence was represented as a set of unique canonical k-mers. Similarity between a recombination tract and each candidate paralog was quantified using the Jaccard containment metric, defined as the proportion of recombination tract 11-mers found in the target paralogous sequence. Containment scores range from 0 to 1, with higher values indicating greater sequence similarity. GCEs were classified as mapped (mGCEs) to a paralog sequence if the Jaccard containment was ≥ 0.5. If no paralog was found with a Jaccard containment over 0.5, the GCE was classified as unmapped. For each mapped GCE, the paralogous region(s) with the highest containment score were classified as potential donor sequences. The per-event paralog-mapping results, including the k-mer containment scores for each event against each candidate paralog, are provided in **Dataset S5**. The event to paralog comparison implementation is available in the manuscript GitHub repository (gcutils/kmer.py and gcutils/eventtoparalogcomparison.py).

#### Paralog Network Construction and Gene Conversion Event Distribution Analysis

To characterize the genomic distribution of detected gene conversion events (GCEs), we constructed a genome-wide paralog network graph using the *M. tuberculosis* H37Rv intra-genomic homology map generated with minimap2(7). Each paralogous region (PR) with sequence homology to at least one other locus was represented as a node, and edges were drawn between nodes sharing a homologous segment. This produced an undirected graph in which connected components reflect paralog networks. Each connected component of the graph was assigned a unique paralog network identifier (PN-ID), grouping the 200 PRs into 71 discrete paralog networks. Of the 71 networks, 24 had at least one detected GCE. These 24 paralog networks were classified by GCE burden into three tiers: low (1–4 detected GCEs), intermediate (5–19), and high (≥ 20). All supporting results for this analysis are provided in **Dataset S6**, including detailed statistics for every annotated paralogous region and summary GCE statistics for each of the paralog networks these regions assemble into.

To quantify how unevenly detected GCEs were distributed across paralog networks, we computed the Gini coefficient of putative GCE counts across all 71 networks. The Gini coefficient was calculated as

$G = \frac{2\sum_{i=1}^{n} i \cdot x_i}{n\sum_{i=1}^{n} x_i} - \frac{n+1}{n}$ where $x_i$ are the per-network putative GCE counts sorted in ascending order and n=71. A value of G = 0 indicates perfect equality across networks and G=1 indicates maximal concentration in a single network.

#### Analysis of phylogenetic distribution of gene conversion events

For every branch of the dataset phylogeny, Gubbins inferred all gene conversion events (GCEs), SNPs inside recombinations (SNPs falling within an inferred recombination tract; equivalently, GCE-associated SNPs), and SNPs outside recombinations (SNPs inferred to have occurred independently, not as part of a putative recombination event signature) as part of its standard output. We first calculated the phylogeny-wide GCE rate as the total number of inferred GCEs per SNP outside recombinations, summed across all branches, yielding 0.0137 GCEs per SNP outside recombinations (equivalently, 1.37 GCEs per 100 SNPs outside recombinations). To additionally assess the relative contribution of gene conversion at the substitution level, we quantified the ratio of SNPs inside recombinations to SNPs outside recombinations, which was 0.124. The per-branch Gubbins statistics and inferred rates described here are provided for all branches of the Mtb151 phylogeny in the "Gubbins - Per Branch Stats" table of **Dataset S5**.

**Lineage-specific variation in gene conversion rate.** To evaluate whether gene conversion rates varied across lineages, each node of the Gubbins phylogeny was assigned a lineage label. A node was labeled with a given lineage if all of its descendant tips belonged to that lineage, and was otherwise labeled Ancestral. Each branch of the phylogeny then inherited the lineage label of its child node. GCE counts and SNP-outside-recombination counts were then summed across all branches assigned to each of the seven lineages tested (L1–L6 and L8); Ancestral branches were excluded.

For each lineage, the observed GCE count was compared to the count expected under the phylogeny-wide rate using a two-sided Poisson test, where the expected count was the phylogeny-wide GCE rate (0.0137 GCEs per SNP outside recombinations) multiplied by that lineage's total SNP-outside-recombination count, thereby accounting for differences in evolutionary depth across lineages. Two-sided p-values were computed by doubling the smaller one-tailed tail probability. p-values were corrected for multiple comparisons across the seven tested lineages using the Benjamini–Hochberg false discovery rate (FDR). Lineages with $q < 0.05$ were considered to have a significantly altered gene conversion rate. The direction of each significant deviation was determined from the observed versus expected counts. The per-lineage gene conversion rates and Poisson test results are reported in **Table 3.** Poisson tests were performed using SciPy (scipy.stats.poisson, v1.13.0) and Benjamini–Hochberg FDR correction using statsmodels (statsmodels.stats.multitest.fdrcorrection, v0.14.1)(12, 13).

**Identification of gene conversion on lineage-ancestral branches.** To characterize gene conversion events inherited by entire lineages, we examined the ancestral branch of each lineage represented by two or more isolates (L1–L6). For each lineage, we identified the internal node corresponding to the most recent common ancestor of all its isolates (L1: Node_147; L2: Node_131; L3: Node_71; L4: Node_65; L5: Node_1; L6: Node_3) and extracted all putative GCEs inferred by Gubbins to have occurred on the branch leading to that node. All node identifiers referenced here correspond to internal node names in the final Mtb151 phylogeny generated by Gubbins, provided in Newick format in **Dataset S10**.

Lineage 8 was excluded because it is represented by a single isolate and thus has no internal ancestral branch. All GCEs detected on these six lineage-ancestral branches, together with their affected loci and the number of SNPs in each recombination tract, are listed in **Table S9**.

#### Gene-level enrichment analysis across functional categories of H37Rv genome

To test whether specific functional categories were enriched for a given set of genes, we used a one-sided Fisher's exact test applied at the gene level. Each annotated gene in the *M. tuberculosis* H37Rv genome was assigned to one of 13 functional categories (based on the Mycobrowser H37Rv annotation). For each category, the 2×2 contingency table compared the number of genes in the set and not in the set within the category against the corresponding counts across all remaining annotated genes (universe = 4,079 genes). Raw p-values were corrected for multiple comparisons across the 13 categories using the Bonferroni method, and categories with an adjusted p-value (p_adj) < 0.05 were considered significantly enriched. Tests were performed using SciPy (scipy.stats.fisher_exact, v1.13.0)(12). This framework was applied to two gene sets: (1) genes overlapping identified nucleotide diversity hotspots, and (2) genes mutated by detected gene conversion events. Results are reported in **Table S4** and **Table 1,** respectively.

#### Assessing correlation between gene conversion frequency and nucleotide diversity

For our primary dataset of 151 Mtb genomes, we recorded the number of overlapping GCEs and the window's nucleotide diversity for each 1-kb window of the H37Rv reference genome overlapping a paralogous region (PR) (see Methods "Nucleotide diversity analysis and diversity hotspot identification"). The association between these two quantities was summarized using the Spearman rank correlation coefficient (ρ; scipy.stats.spearmanr, SciPy v1.13.0)(12). The per-window GCE counts and nucleotide diversity values for all 1-kb windows of the H37Rv genome are provided in the "GCE_And_NucDiv_Stats_Per1kb" table of **Dataset S5.**

#### Comparison of mutational characteristics of paralogous regions versus the rest of the genome

**Inter-SNP distance analysis.**

To assess whether substitutions cluster more tightly within paralogous regions (PRs) than elsewhere in the genome, we compared inter-SNP distances between PR and non-PR genomic regions. For every genome in the Mtb151 dataset, each SNP variant was annotated with (i) the distance to the nearest neighboring SNP within that same genome and (ii) whether it fell within an identified paralogous region (**Dataset S2**). The distribution of inter-SNP distances for SNPs located in PRs was then compared to that of SNPs in non-PRs using a two-sided Mann-Whitney U test (scipy.stats.mannwhitneyu, SciPy v1.13.0). The unit of observation was the individual SNP, and no multiple-comparison correction was applied as this constituted a single comparison. The inter-SNP distances for every SNP underlying this comparison are provided in the "SNPs.With_Inter_SNP_Distances" table of **Dataset S4**.

**Test of enrichment for mutations that match existing variants in paralogs.**

To assess whether substitutions within a paralogous region (PR) preferentially reproduce sequence variants already present in its paralogs, we tested each PR for enrichment of paralog-matching substitutions using a one-sided hypergeometric test. Substitution events were taken from the Gubbins ancestral-state reconstruction, which infers, for every substitution in the dataset, the number of independent mutational events and the phylogenetic branch on which each occurred (**Dataset S5**); this ensures that recurrent substitutions are counted as distinct events rather than as a single shared polymorphism. For each PR, every inferred substitution event was classified as paralog-matching if its identity exactly matched a variant observed at the homologous position in a paralog.

For each PR, the hypergeometric test compared the observed number of paralog-matching substitution events to that expected by chance, where the population comprised all possible point substitutions in the region (region length × 3, assuming each of the three alternative bases is equally likely at every position), the number of success states was the count of possible substitutions that would match a paralog variant, and the number of draws was the total number of observed substitution events in the region. The test was applied to all 157 PRs containing at least one observed substitution event (of 200 PRs total) using SciPy (scipy.stats.hypergeom, v1.13.0)(12). Raw p-values were Bonferroni-corrected for the number of tested regions, and PRs with an adjusted p-value (p_adj) $< 0.01$ were considered significantly enriched. Effect size was quantified as the fold-enrichment of observed over expected paralog-matching substitutions. Per-region test results are provided in the "PerPR.Paralog_SNP_Enrichment" table of **Dataset S4**.

**Mutational spectrum analysis.**

For each genome in the Mtb151 dataset, every substitution variant was classified by its trinucleotide context, defined by the reference bases immediately 5′ and 3′ of the substituted position together with the base change. Following the standard single-base-substitution (SBS-96) convention, each substitution was represented relative to the pyrimidine base of the mutated pair, giving six substitution classes (C>A, C>G, C>T, T>A, T>C, T>G); combined with the 16 possible 5′/3′ flanking-base combinations, this produced 96 trinucleotide substitution classes. Substitutions were tallied separately for those occurring within paralogous regions (PRs) and those in the rest of the genome (non-PRs).

To account for differences in trinucleotide sequence composition between PR and non-PR regions, the substitution count in each of the 96 classes was normalized by the background frequency of the corresponding reference trinucleotide within that region type. Each background-normalized spectrum was then converted to a relative-frequency distribution by L1 normalization (dividing each class by the sum across all 96 classes) so that each spectrum summed to one. The per-class difference (PR minus non-PR L1-normalized frequency) was used to summarize divergence between the two spectra. The full 96-class count and spectrum table is provided in the "SBS_Signatures.PRvsNonPR.Counts" table of **Dataset S4**.

To test whether individual substitution classes differed in frequency between PRs and the rest of the genome, we compared the two L1-normalized spectra for each of the six substitution classes. For a given class, the 16 trinucleotide-context frequencies in PRs were compared to the corresponding 16 frequencies in non-PR regions using a two-sided Mann-Whitney U test (scipy.stats.mannwhitneyu, SciPy v1.13.0); the unit of observation was the trinucleotide context ($n = 16$ per region type). Raw p-values were Bonferroni-corrected across the six classes, and classes with an adjusted p-value (p_adj) $< 0.05$ were considered significantly different. Effect sizes were quantified as the absolute rank-biserial correlation. Statistical test results for all six classes are provided in the "SBS_Signatures.PRvsNonPR.MannWhitneyU.Results" table of **Dataset S4**.

#### Regression analysis of sequence features with gene conversion frequency across all paralogous regions

To evaluate whether general sequence features predict gene conversion frequency between paralogous regions, we modeled the number of mapped gene conversion events (mGCEs) detected for each paralog pair using a generalized linear modeling framework. Analyses were restricted to paralog pairs that were non-perfect repeats of each other (n = 284 pairs). Non-perfect repeats were defined as any homology alignment between paralogous regions that did not have 100% sequence identity.

Because gene conversion counts are discrete and overdispersed, we fit a generalized linear model assuming a negative binomial distribution (dispersion parameter fixed at $\alpha = 1.0$) with a log link function. The response variable was the number of mapped gene conversion events detected between each paralog pair. Predictor variables included: 1) Sequence divergence, quantified as the number of SNPs per kilobase of aligned paralogous sequence. 2) Genomic distance between paralogs (kilobases). 3) Paralog copy number, measured as the number of overlapping paralogous alignments. 4) Local GC content, calculated as the percentage of bases that are either G or C within the defined region. All predictors were included simultaneously in the model, and an intercept term was added. Robust (HC0) standard errors were used to account for potential heteroskedasticity. Model coefficients were exponentiated to obtain incidence rate ratios (IRRs), representing the multiplicative change in expected gene conversion counts per unit increase in each predictor. The per-paralog-pair input data (event counts and all predictor values) used for this model are provided in **Dataset S6**. Statistical significance for the regression was assessed using Wald tests, and results are reported in **Table S7**. Model fitting and statistical inference were performed using the `statsmodels (v0.14.1)` Python package.

To summarize overall model fit, we computed McFadden's pseudo-$R^2$, defined as $1 - \frac{\ln \hat{L}_{full}}{\ln \hat{L}_{null}}$, where $\hat{L}_{full}$ is the maximized likelihood of the fitted model and $\hat{L}_{null}$ is the maximized likelihood of the intercept-only null model(14). This metric quantifies the relative improvement in fit of the evaluated model over a null model containing no predictors, with values ranging from 0 (no improvement) to 1 (perfect fit).

In addition to the multivariable model, we assessed the univariate association between each sequence feature and mGCE frequency using Spearman's rank correlation, computed across all 284 paralog pairs (**Figure S9, Table S8**). Spearman correlations were computed using SciPy (scipy.stats.spearmanr, v1.13.0).

The full implementation of this analysis is available in the manuscript GitHub repository in the following python notebook: [251211.2.B.Mtb151.GCE.EventFreq_Vs_SeqFeatures.V1.ipynb]()

#### Curation of validated T cell (CD4+/MHC-II) epitopes from two published studies

We compiled $CD4^+$ T cell epitope-mapping data from two large-scale peptide screening studies, each of which deposited their complete assay datasets in the Immune Epitope Database (IEDB). The first dataset, from Lindestam et al. 2016 (IEDB accession: 1031648), assayed 693 unique 15-mer peptides derived from 88 *Mtb* proteins, identifying 207 unique positive epitope sequences. This study screened peptides with IFN-γ ELISPOT assays using PBMCs from 63 latently infected individuals in South Africa, targeting known immunogenic proteins and their homologs. The second dataset, from Panda et al. 2024 (IEDB accession: 1042336), assayed 18,325 unique 15-mer peptides spanning the entire H37Rv proteome (with bias toward known antigens). This study screened peptides with IFN-γ ELISPOT assays of PBMCs from 21 patients with active tuberculosis in Peru, identifying 153 unique positive epitope sequences.

We merged the results of these two studies into a combined dataset (referred to as LPM), representing the union of all assayed peptides and their reactivity results (Positive epitope or non-reactive peptide). LPM contains 18,371 unique assayed peptides spanning 3,840 annotated protein-coding genes of H37Rv genome, with a total of 307 unique positive CD4+ epitope sequences (**Dataset S8**). All assayed peptide sequences were harmonized to H37Rv genomic coordinates using reference protein sequences, excluding any without a match to the reference proteome. Epitopes were considered experimentally validated if they were positive in at least one human $CD4^+$ T cell assay (IFN-γ ELISPOT) and matched exactly a H37Rv reference protein sequence. Proteins were classified as high-confidence antigens if they contained ≥2 experimentally validated epitopes in the LPM dataset. This resulted in 53 high-confidence antigens containing a total of 226 validated epitope sequences (**Dataset S8**). Each annotated protein in the H37Rv reference (including Antigens) were also classified based on their overlap with annotated paralogous regions (PR).

The full implementation of this analysis is available in the manuscript GitHub repository in the following python notebook: 250807.A.MtbEpitopeInfo.ProcessIEDB.EpitopeInfo.Part1.V1.ipynb

#### Analysis of mutational patterns among T cell antigens and epitopes

**Quantification of mutational burden across gene classes.** Non-synonymous mutational burden was quantified for every annotated coding sequence (CDS) in the H37Rv reference genome. Starting from the Gubbins ancestral-state reconstruction of all substitution events (see "Analysis of phylogenetic distribution of gene conversion events"), each inferred substitution was mapped onto the H37Rv reference protein sequence and its codon consequence determined using custom Python functions built on Biopython (v1.87). Substitutions producing any amino acid change were classified as non-synonymous events. Because mutational events were taken from the ancestral reconstruction, each corresponds to an independent mutational event on a specific phylogenetic branch, so recurrent changes at the same site are counted separately. For each CDS, mutational burden was calculated as the number of non-synonymous events divided by the CDS length in base pairs and multiplied by 300, yielding missense events per 100 per 100 amino acids. Epitope-region burden was computed identically but counting only non-synonymous events overlapping a validated T cell epitope and then normalized by total CDS length. All calculated supporting annotations and statistics per protein-coding gene are provided in the "Stats Per CDS Gene" table of **Dataset S8**.

Each protein coding gene was then assigned to one of four classes by combining its antigen classification (high-confidence antigen vs. non-antigen) with its genomic context (overlapping a paralogous region or not): PR-antigen, unique antigen, PR-non-antigen, and unique non-antigen. Missense mutational burden was compared between classes using two-sided Mann-Whitney U tests with Bonferroni corrections for multiple hypothesis testing. All six pairwise comparisons between the four classes of genes were performed and comparisons with an adjusted p-value (p_adj) $< 0.05$ were considered significant (**Table S13**). Epitope-region burden was compared between PR-antigens and unique antigens using a separate two-sided Mann-Whitney U test (**Table S14**). Tests were performed using SciPy (scipy.stats.mannwhitneyu, v1.13.0).

The implementation of this analysis is available in the manuscript GitHub repository in the following python notebook: 250808.C.MtbAntigenicVariation.Part1.InitialAnalysis.V1.ipynb

**Test for enrichment of gene conversion events within antigen epitopes of paralogous regions.** To test whether gene conversion events (GCEs) preferentially introduced missense mutations into T cell epitope sequences within paralogous regions (PRs), we compared the observed number of GCEs producing such mutations to that expected under a null of length-proportional distribution across PR sequence, using a one-sided hypergeometric test. The population comprised all base pairs of PR sequence in the H37Rv reference genome (257,094 bp), of which 4,395 bp (1.71%) overlapped a validated T cell epitope. Treating the 295 GCEs located within PRs as draws from this sequence, we asked whether the observed number introducing at least one missense mutation within a validated epitope (25) exceeded the number expected if GCEs were distributed in proportion to sequence length (5.05). Effect size was quantified as the fold-enrichment of observed over expected. The test was performed using SciPy (scipy.stats.hypergeom, v1.13.0).

**Comparison of missense mutation rate between epitope and non-epitope regions of PPE18.** For each of the 392 amino acid positions in PPE18, we counted the number of inferred missense mutation events (from the Gubbins ancestral-state reconstruction; see "Analysis of phylogenetic distribution of gene conversion events") and recorded whether the position overlapped a validated T cell epitope. Missense mutation rate was modeled as a function of epitope overlap using a Poisson generalized linear model (per-position missense event count ~ epitope overlap), fit with statsmodels (v0.14.1). Poisson was appropriate as the fitted model was not overdispersed (dispersion = 0.74). The "PPE18.PerAAPos.DetectedEpitopesVsMutFreq.V1" table found in **Dataset S8 has the supporting input data for this analysis, defining the mutation frequency and epitope classification for each codon of PPE18.**

**Association between T cell responsiveness and missense mutation burden in PPE18.** Using peptide-level T cell response-frequency data for PPE18 from Lindestam et al. 2016 (assayed in 63 latently infected individuals), we tested whether the missense mutation burden of a peptide was associated with the number of donors responding to it. The analysis was restricted to assayed peptides overlapping a validated positive epitope (n = 50 peptides, **Dataset S8**). The count of missense mutation events per peptide was modeled as a function of the number of responding donors using a negative binomial generalized linear model (dispersion parameter fixed at $\alpha = 1.0$), fit with statsmodels (v0.14.1). A negative binomial distribution was used because the corresponding Poisson model was overdispersed (dispersion = 2.2). The rate ratio (exponentiated coefficient) with its 95% confidence interval quantified the association, and model fit was summarized with McFadden's pseudo-$R^2$.

#### HLA-II binding prediction of PPE18 protein sequences mutated by gene conversion

HLA-II binding was predicted using NetMHCIIpan-EL (v4.1; IEDB, https://tools.iedb.org/mhcii/) for the reference PPE18 sequence and for each mutant PPE18 sequence produced by a PPE18 gene conversion event (Event-091 through Event-097)(1). The exact mutated sequences are provided in the **Dataset S9**. For each sequence, all possible 15-mer peptides were scored across a reference panel of 27 HLA-II alleles (**Dataset S9**). A peptide was classified as a Strong Binder (SB) if its predicted eluted-ligand percentile rank was ≤ 1%. For each gene conversion event and allele, we counted binding "gains" (peptide positions changing from non-SB in the reference sequence to SB in the mutant) and "losses" (SB to non-SB). Per-event and per-allele counts are provided in the "Stats_PerGCE_PerAllele" sheet of **Dataset S9**. To test whether gains and losses occurred at unequal frequencies, all SB-status changes were pooled across the seven events and 27 alleles (n = 66 changes) and compared to a null of equal gains and losses using an exact two-sided binomial test ($p_0 = 0.5$; scipy.stats.binomtest, SciPy v1.13.0). To assess population-specific patterns, the analysis was repeated using the seven most frequent HLA-II alleles in European populations (HLA-DRB1*01:01, *03:01, *04:01, *07:01, *11:01, *13:02, *15:01); results are reported in the Supplementary Results. Full per-peptide binding predictions for every input sequence are provided as individual sheets in **Dataset S9**.

#### Validation of gene conversion events in an independent dataset of clinical *Mtb* isolates

##### Part 1: Identification of suspected gene conversion events from short-read WGS dataset

We screened an independent short-read WGS dataset for gene conversion signatures. This dataset, referred to as TGEN-937-SR, comprises publicly available short-read WGS from 937 clinical *M. tuberculosis* isolates for which genomic DNA was archived at the Translational Genomics Research Institute (TGen). Sample metadata and SRA run accessions for all 937 isolates are provided in Dataset S1 ("TGEN-937-SR").

Variants were called conservatively from each isolate's short-read WGS data as follows. Paired-end short reads were adapter and quality trimmed with Trimmomatic (Snakemake wrapper v0.38.0; ILLUMINACLIP with a custom adapter set, SLIDINGWINDOW:4:20, MINLEN:75). Trimmed reads were then aligned to the H37Rv reference genome (NC_000962.3) with BWA-MEM (v0.7.17). Alignments were then coordinate-sorted with samtools (v1.9), and duplicate-marked with Picard MarkDuplicates (v2.21.4; REMOVE_DUPLICATES=true). Variants were called with Pilon (v1.23) with the --variant flag. To filter the variants further, only SNP variants called PASS by Pilon and with a mean mapping quality (MQ) score ≥40 were retained for downstream analysis.

A concatenated whole-genome SNP alignment of the retained SNPs (PASS, MQ ≥40) was constructed and used to build an initial phylogeny with FastTree (v2.1.10; -nt -gtr). Suspected gene conversion signatures were then identified from this alignment using Gubbins (v3.1.4) with default parameters, providing the FastTree phylogeny as the starting tree. Each recombination tract reported by Gubbins was treated as a suspected gene conversion event. This yielded 27 suspected gene conversion events across the 937 isolates. The complete set of events, with genomic coordinates, overlapping genes, SNP counts, and phylogenetic placement, is provided in **Table S10** and **Dataset S7**. Of these 27 events, 24 (89%) occurred in genes that also harbored gene conversion events in the primary analysis of the Mtb151 dataset.

##### Part 2: Resequencing of TGEN isolates with PacBio HiFi sequencing

Of the 27 suspected events, 12 occurred in isolates for which archived genomic DNA was available in sufficient quantity (at least 1 μg) for long-read library preparation. For each of these 12 events, we resequenced two isolates: one predicted to carry the recombinant sequence and a phylogenetically matched control predicted to lack the event-defining variants. Two isolates each served in two events (as the verification and control), so the 12 events corresponded to 22 distinct resequenced isolates rather than 24. The mapping between the 12 events and the 22 resequenced isolates (including verification/control roles and short-read run accessions) is provided in the "TBP-22-LR - GCE_To_IsolateIDs" and "TBP-22-LR - IsolateIDs_To_GCE" tables of **Dataset S7**.

Long-read sequencing libraries were generated using the SMRTbell Express Template Prep Kit 2.0 (Pacific Biosciences, Menlo Park, CA), following the manufacturer's protocol. PacBio HiFi sequencing was performed on the Sequel II platform, and HiFi reads were generated using the Circular Consensus Sequencing (CCS) mode in SMRT Link (v13; Pacific Biosciences). Complete genome assemblies were then generated from the resequencing data using the same PacBio HiFi hybrid assembly approach described previously. Snakemake pipeline code and configuration files are available at 2.Mtb.Generate.HybridAsm.PBccs.smk in the manuscript GitHub repository. Assembly sequence statistics for the 22 resequenced isolates are provided in the "TBP-22-LR – Asm Stats" table of **Dataset S7**.

#### Part 3: Evaluation of gene conversion signatures by PacBio sequencing.

For the 22 resequenced isolates, we generated complete genome assemblies (above) and then applied the same gene conversion analysis pipeline used for the primary Mtb151 complete genomes. Alignment to the H37Rv reference, variant calling, and gene conversion detection with Gubbins were performed identically to the primary Mtb151 analysis (Gubbins v3.2.1, extensive-search mode, --min-snps 4, --min-window-size 25, --max-window-size 1000; recombination-detection pipeline Snakemake_Workflows/2_CoreAnalysis_SMK/Mtb.WGA.Core.V3.smk). This produced an independent set of gene conversion events called directly from the complete genome assemblies and phylogeny of these 22 isolates. All 84 detected gene conversion events are provided in the "TBP-22-LR – GCEs" table of **Dataset S7**.

Validation of each suspected gene conversion event was then assessed by directly comparing the two independently generated event sets in H37Rv reference coordinates: the suspected events detected in the short-read screen (TGEN-937-SR) and the events detected in the long-read assemblies of the same isolates (TBP-22-LR). For each short-read-detected event, we compared (i) the H37Rv genomic coordinates of the Gubbins-called recombination tracts between the two datasets to determine whether an overlapping event was recovered in the long-read data, and (ii) the coordinates and allele identities of the individual SNPs comprising each event, to determine whether the same substitutions were independently called from the long-read assembly. An event was considered validated when the long-read data recovered a coordinate-overlapping recombination tract that contained the same SNPs.

We assessed concordance at two levels. At the event level, all 12 suspected short-read events were recovered as overlapping gene conversion events in the long-read assemblies (precision = 12/12; 100%). At the individual-SNP level, all 86 SNPs defining the 12 short-read events were called at the identical reference position and allele in the corresponding long-read assembly (86/86; 100% concordance). Per-event results, including the number of SNPs resolved by short-read and by long-read WGS and their agreement, are provided in the "TBP-22-LR – Validation Results" table of **Dataset S7**.

We additionally quantified the sensitivity (recall) of short-read WGS for gene conversion detection across this dataset of 22 Mtb isolates. The long-read analysis of the isolates recovered substantially more events than the short-read screen. In addition to the 12 events also detected with short reads, 72 further gene conversion events were identified from the long-read data that had not been detected by short-read WGS (84 events total). The 12 short-read-detected events therefore represent a recall of 12/84 events (14.3%), consistent with the expected loss of sensitivity when detecting recombination in paralogous regions from short-read data.

#### Supplemental Figures & Tables

Figure S1

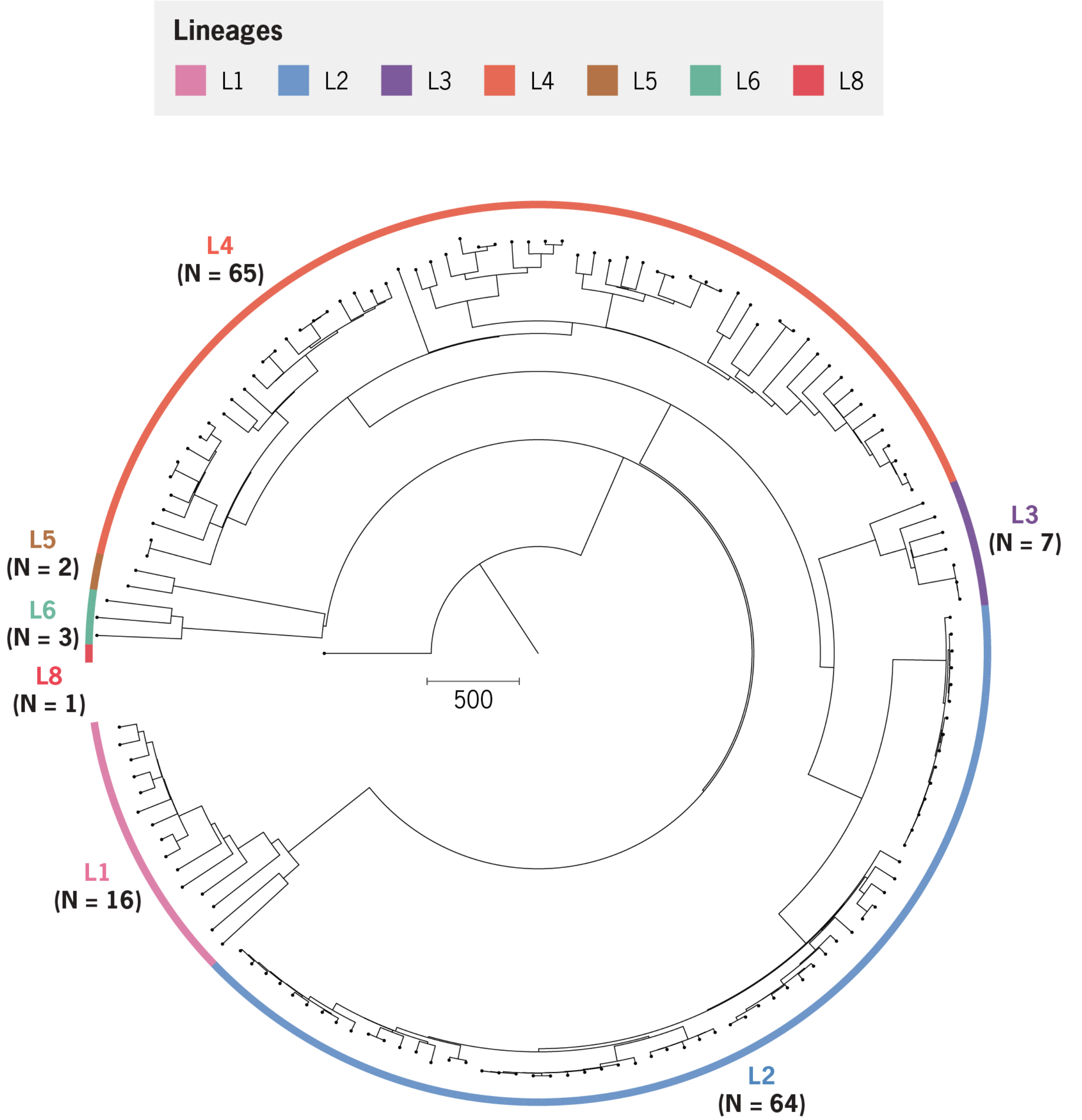


**Figure S1.** Maximum likelihood phylogeny of 151 *Mtb* isolates with complete genome assemblies. All assemblies were generated through *de novo* assembly and polishing with long-read whole genome sequencing data (Oxford Nanopore & PacBio), followed by polishing with short-read whole genome sequencing data.

#### Figure S2

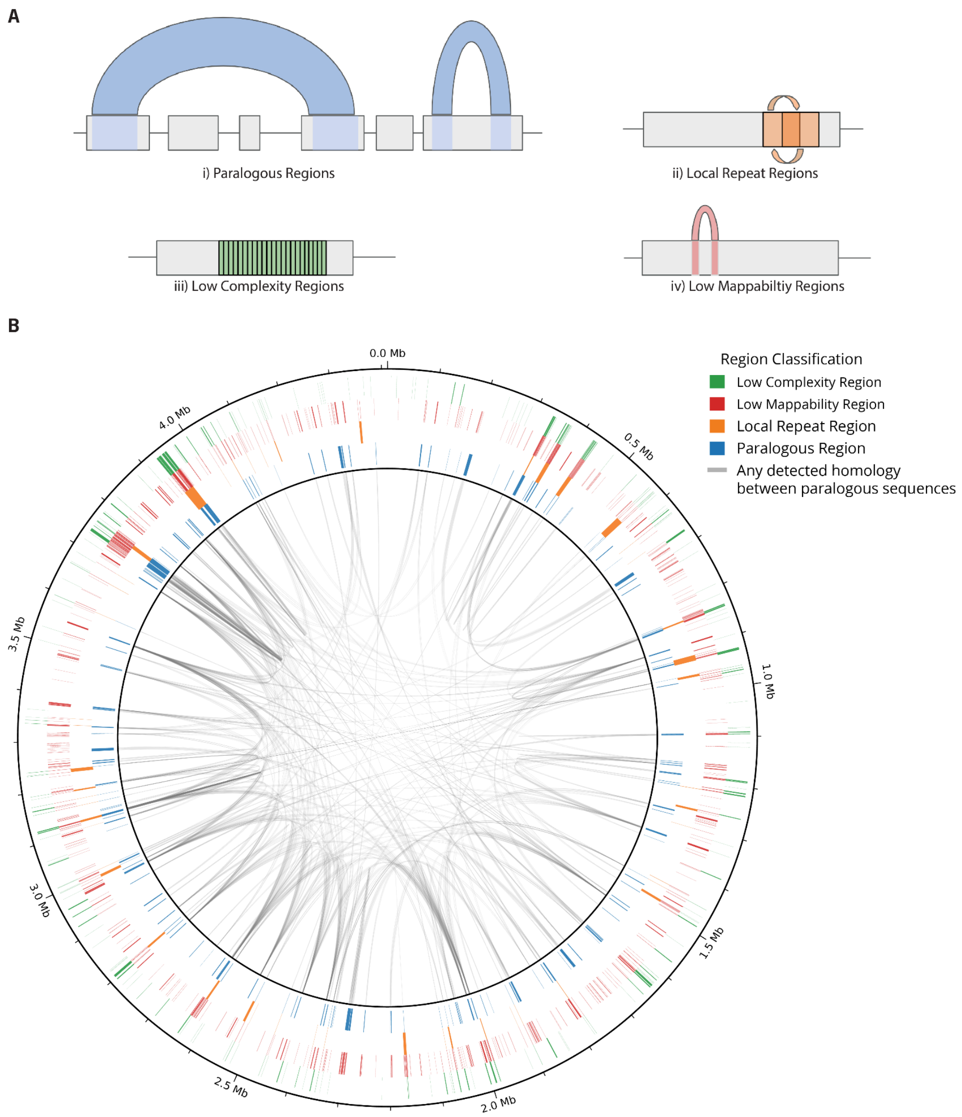


**Figure S2. Distribution of repetitive and high-homology regions in the *M. tuberculosis* H37Rv genome.** (**A)** Cartoon diagrams illustrate the four categories detected: paralogous regions (PRs), defined as distinct genomic segments with detectable homology; local repeat regions (LRRs), tandem overlapping repeats with detectable homology; low-complexity regions (LCRs), enriched for short tandem motifs; and low pileup-mappability regions (LMRs), sequences with non-unique k-mer content. **(B)** The genomic distribution of these categories (PRs, blue; LRRs, orange; LCRs, green; LowPmap, red), with inner links marking pairwise homology between paralogous loci. Many genomic regions fall into multiple categories, highlighting that non-unique sequence content in H37Rv is often simultaneously paralogous, repetitive, and low complexity.

**Figure S3**

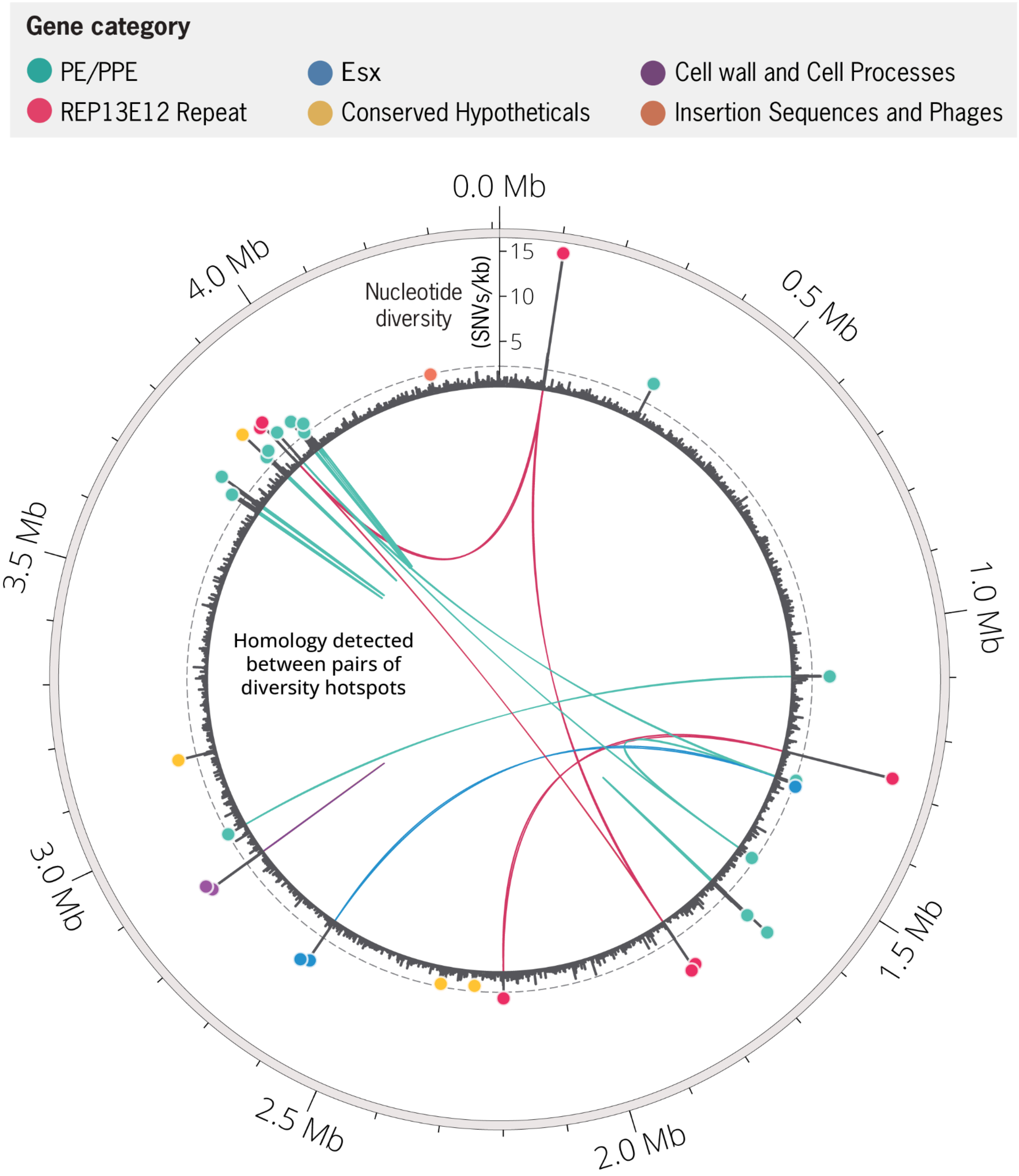


**Figure S3. Highlighting pattern of homology between identified nucleotide diversity hotspots**
Circular visualization of genome-wide nucleotide diversity (π) across non-overlapping 1-kb windows in the H37Rv reference genome. Nucleotide diversity hotspots are indicated as colored points according to the functional category of the associated gene(s). Links within the center of the plot denote sequence homology between diversity hotspots. In total, 31 of the 37 identified hotspots are paralogous to at least one other hotspot, forming an interconnected network of homologous loci distributed across the genome.

### Figure S4

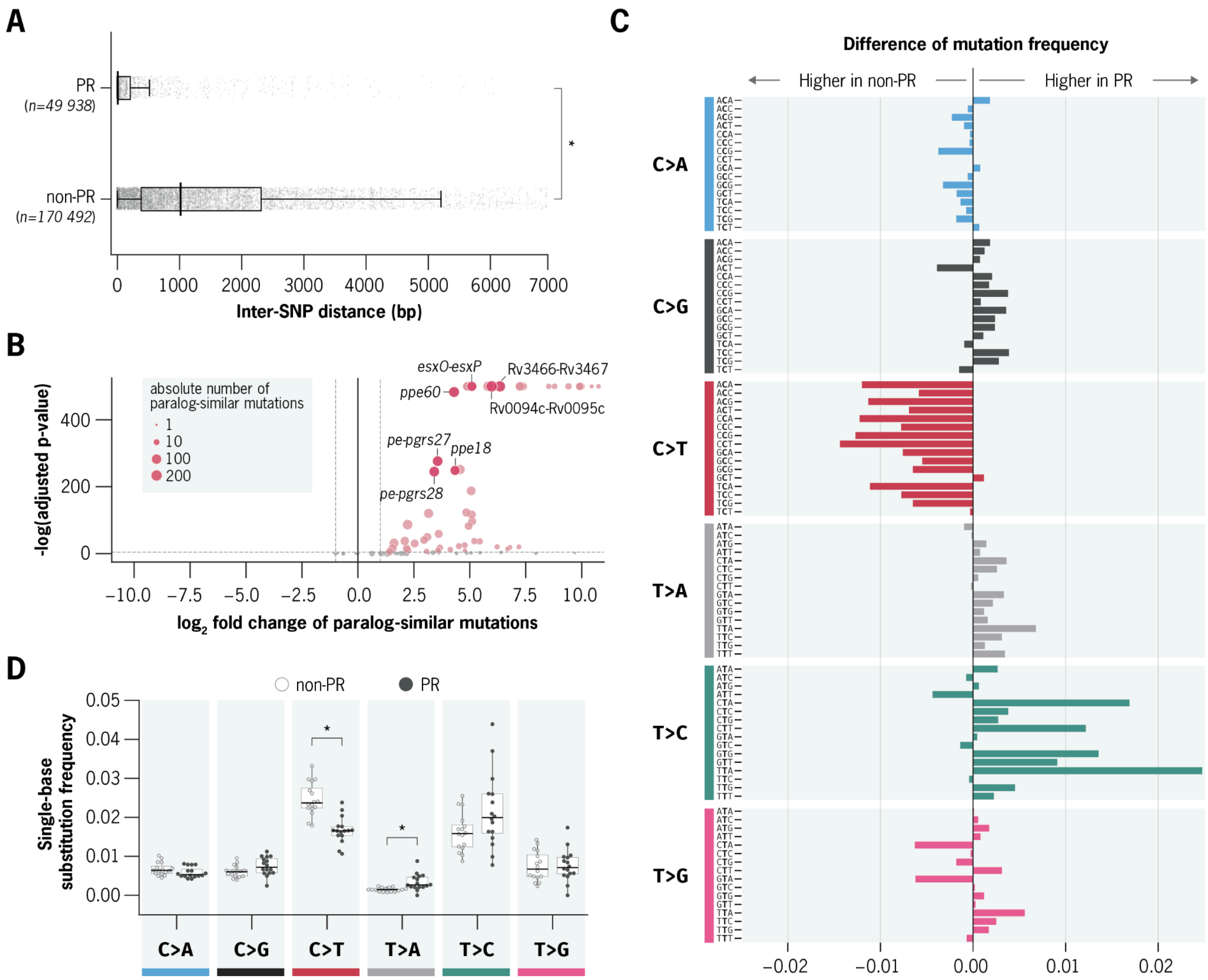


**Figure S4. Paralogous regions of the *Mtb* genome harbor distinct mutational characteristics suggestive of gene conversion. a,** Distribution of inter-SNP distances in paralogous regions (PRs, top) and non-PR regions (bottom) across all substitutions detected relative to the H37Rv reference genome. For each substitution, inter-SNP distance was defined as the distance to the nearest co-occurring substitution within the same genome. Mutations in PRs were significantly more clustered, with a median inter-SNP distance of 7 bp compared to 768 bp in non-PR regions (Wilcoxon rank-sum test, $p < 4e\text{-}8$). **b,** Quantification of paralog-similar mutations across PRs. Paralog-similar mutations were defined as substitutions that exactly match known sequence differences between paralogs in the reference genome. Each point represents one PR, with circle size indicating the absolute number of paralog-similar mutations. PRs highlighted in red are significantly enriched for paralog-similar mutations after multiple hypothesis correction (FDR < 0.05, Poisson test). **c,** Relative frequencies of substitution type across all trinucleotide contexts in PRs and non-PR regions. Boxplots summarize the distribution of normalized frequencies of each trinucleotide context within each substitution type. PRs exhibit a distinct mutational spectrum, including a depletion of C>T and an enrichment of T>C substitutions compared to non-PR regions. **d,** Visualization of the difference in trinucleotide substitution frequencies between PRs and non-PRs. Bars represent the Δ mutation rate (PR minus non-PR) for each substitution type.

#### Figure S5

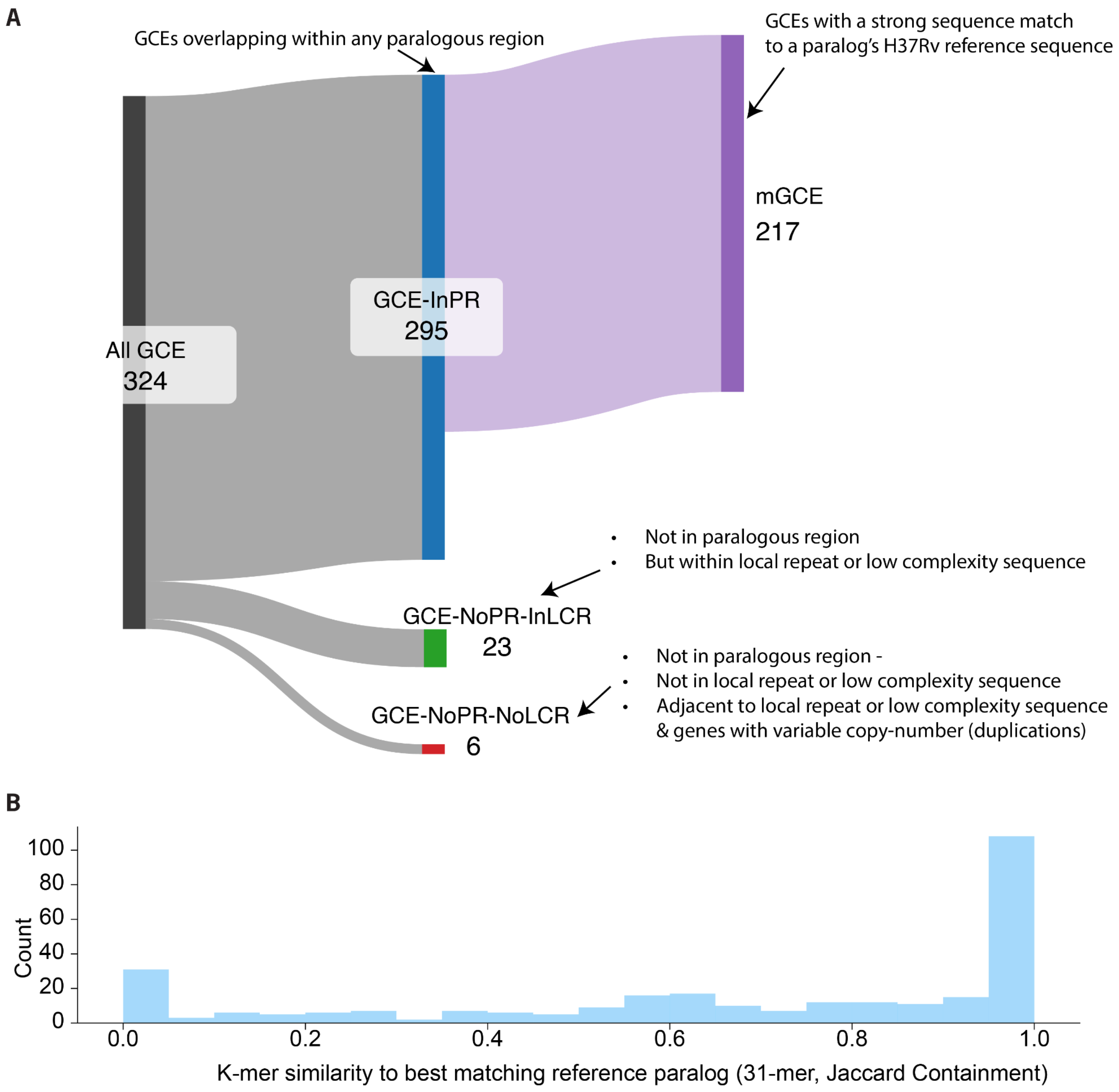


**Figure S5. Breakdown of detected GCEs by overlap with non-unique features. a**, Sankey diagram summarizing the classification of detected GCEs by overlap with paralogous regions, mapping status, and local repeat sequences. **b**, Histogram of k-mer similarity values between GCEs and their best-matching reference paralog.

#### Figure S6

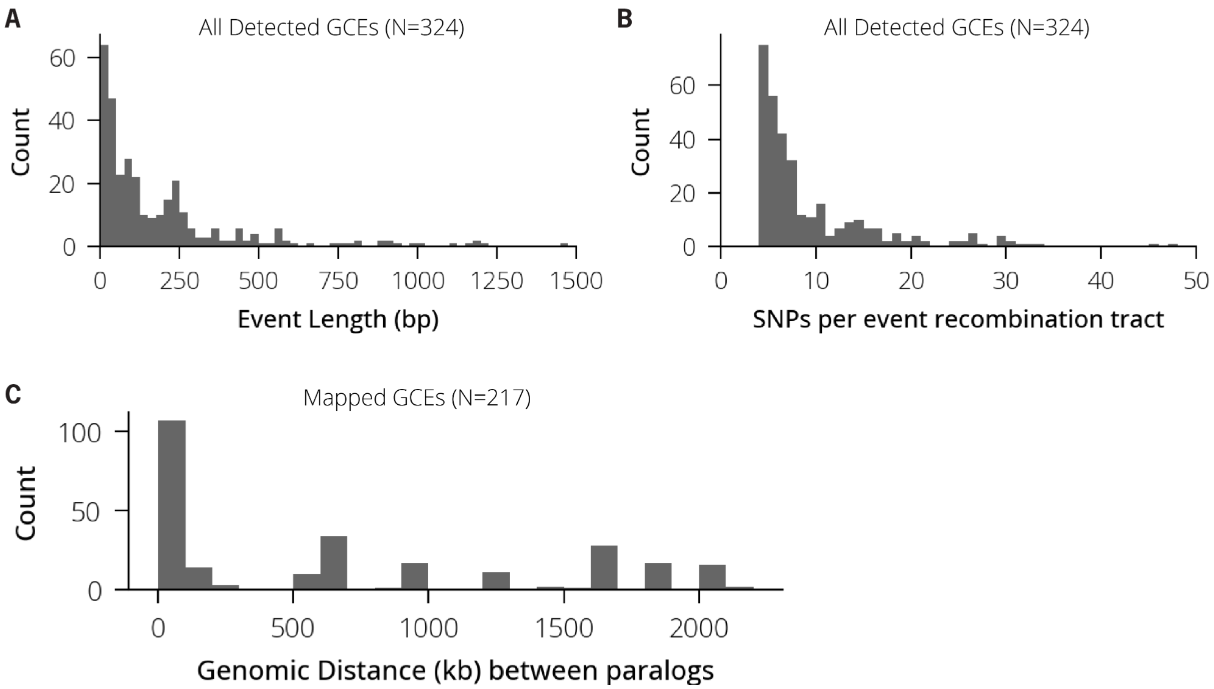


**Figure S6. Overview of general characteristics of detected gene conversion events. a)** Distribution of recombination tract lengths associated with detected GCEs. **b)** Distribution of number of substitutions associated with detected GCE. **c)** The distribution of genomic distances between the inferred donor paralog sequences and mapped GCEs.

**Figure S7**

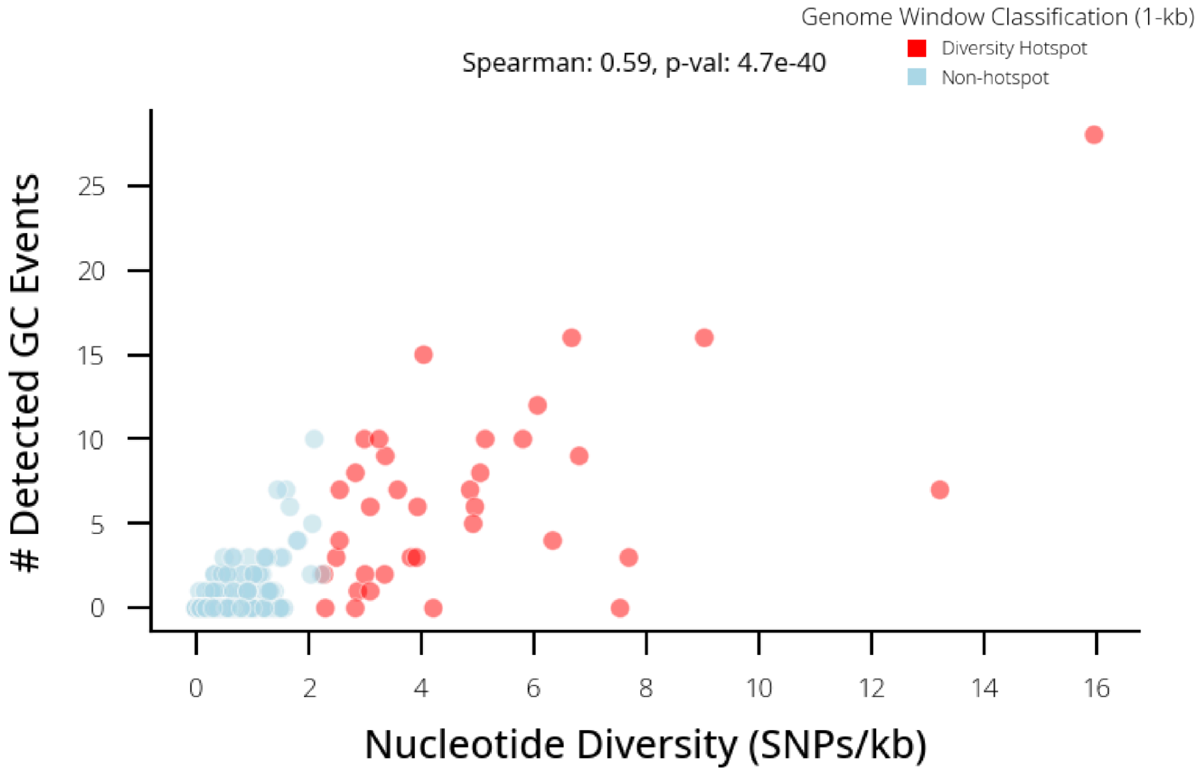


**Figure S7. Scatterplot showing nucleotide diversity versus detected GCEs per 1-kb window overlapping paralogous regions of the H37Rv genome.** Windows classified as diversity hotspots are highlighted in red. Spearman's rank correlation: $\rho = 0.59$, $p = 4.7 \times 10^{-40}$.

**Figure S8**

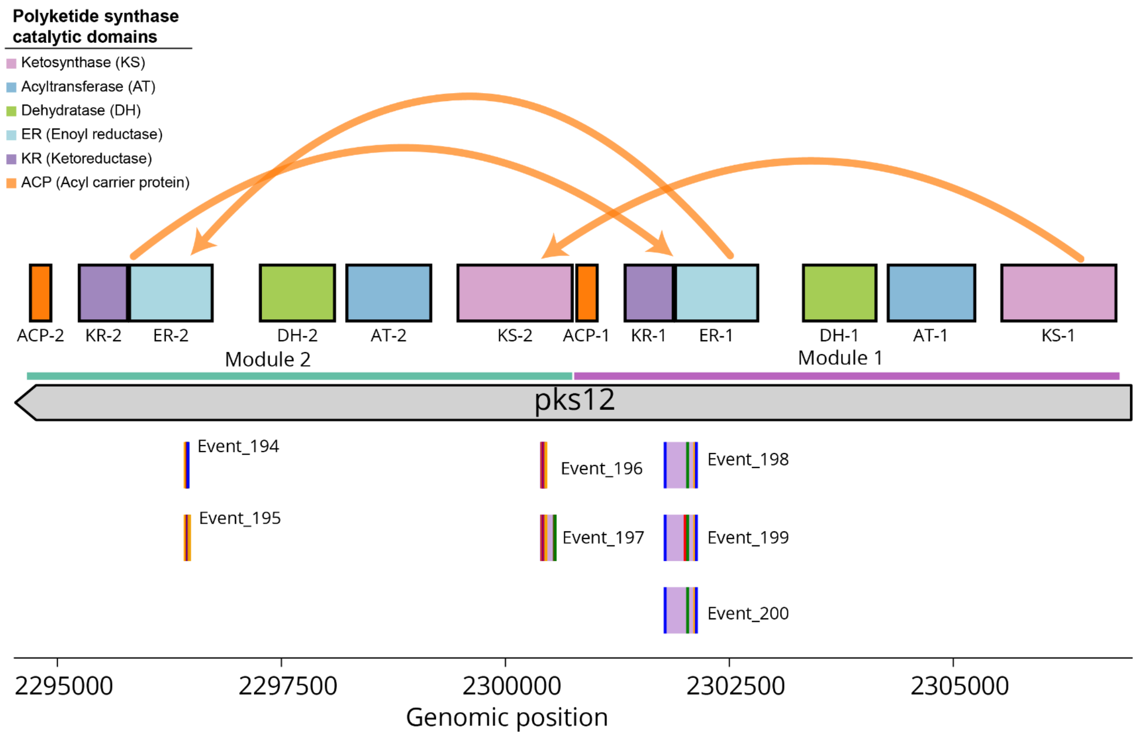


**Figure S8. Overview of intragenic gene conversion events detected in pks12**. Top: Schematic representation of the pks12 locus showing gene structure and annotated protein domains. pks12 comprises two modular units containing six homologous polyketide synthase catalytic domains. Bottom: Detected intragenic gene conversion events (n = 7) shown with associated substitution variants.

Figure S9

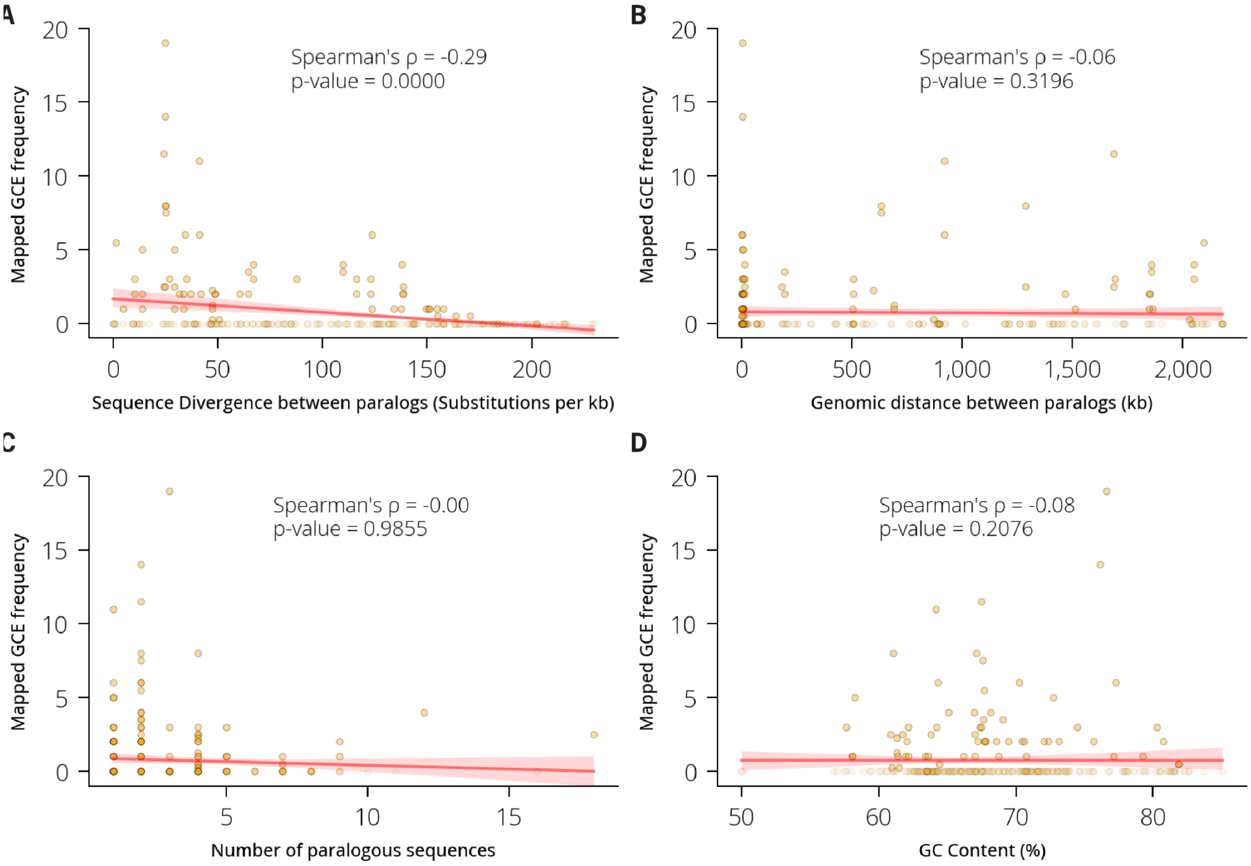


**Figure S9. Relationship between gene conversion frequency and general sequence characteristics**. (A–D) Scatterplots showing associations between mapped gene conversion event frequency (mGCEs) and general sequence or genomic characteristics across all evaluated paralog pairs (n = 284). Spearman's rank correlation coefficient (ρ) is shown for each comparison.

**Figure S10**

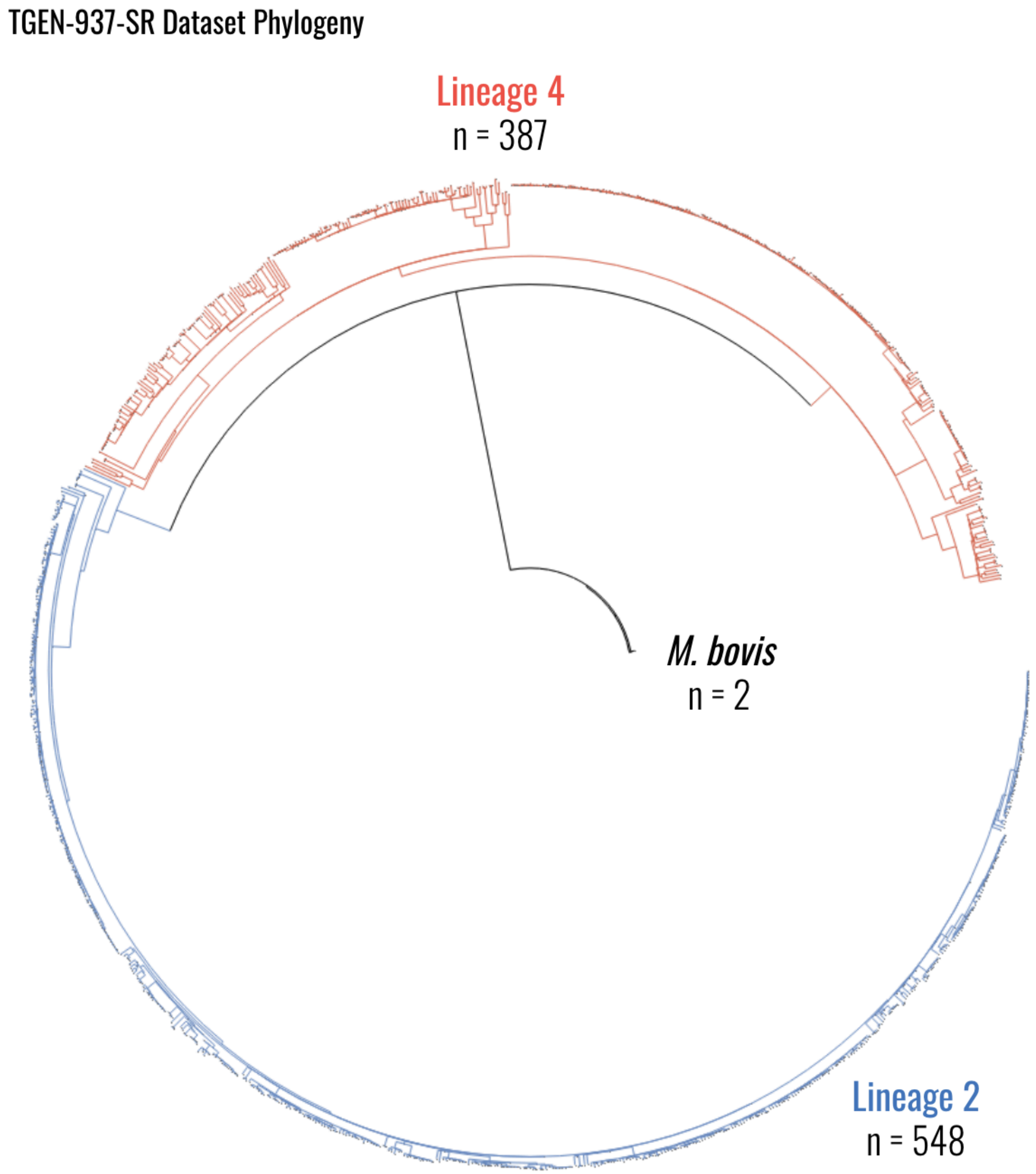


**Figure S10.** Maximum-likelihood phylogeny of 937 clinical isolates sequenced with short-read whole-genome sequencing (TGENSR dataset). The dataset comprises *M. tuberculosis* isolates from lineage 2 (n = 548) and lineage 4 (n = 387), with two additional isolates identified as Mycobacterium bovis.

**Figure S11**

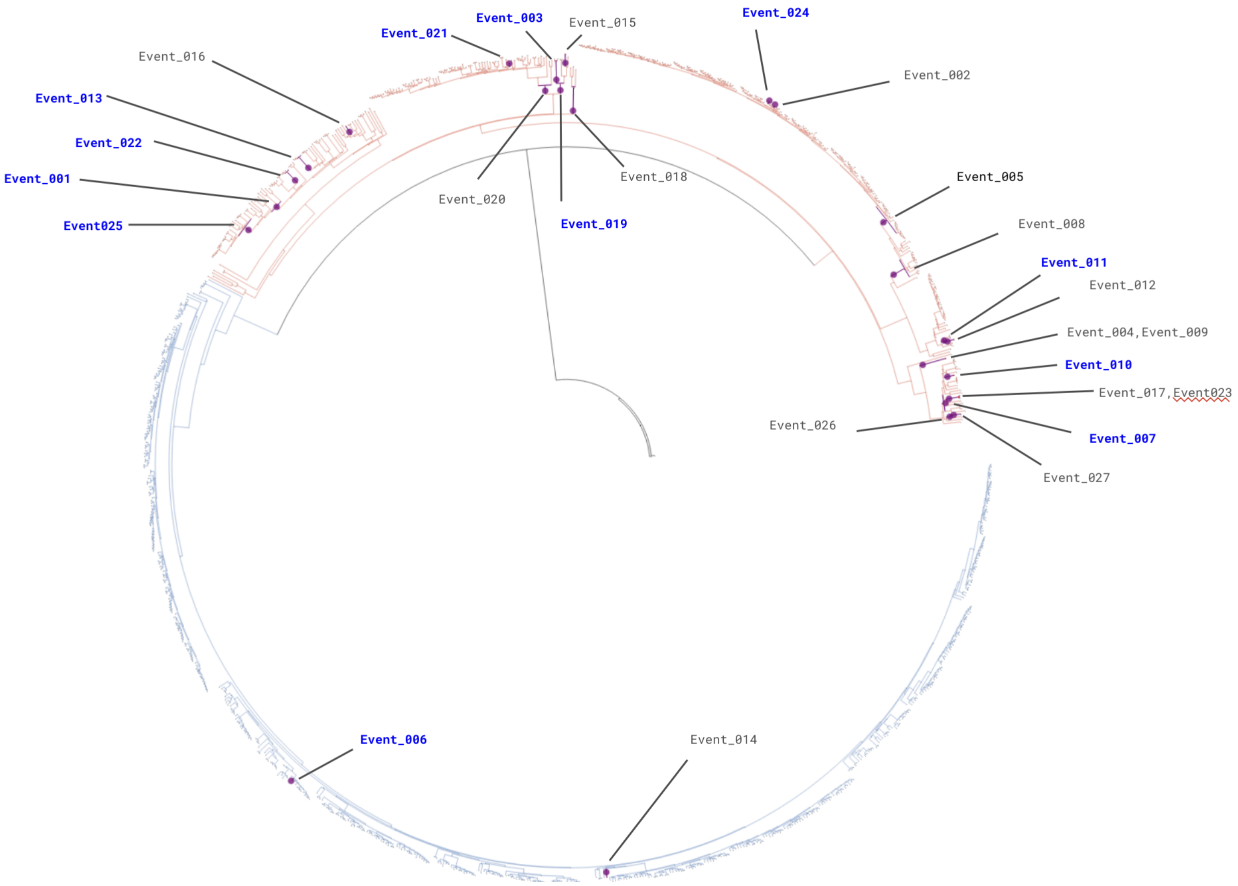


**Figure S11. Phylogenetic distribution of suspected gene conversion events detected in the TGEN-937-SR dataset**. Phylogenetic branches inferred to harbor suspected gene conversion events are highlighted in purple. Event labels shown in blue denote loci selected for long-read resequencing and validation.

#### Figure S12

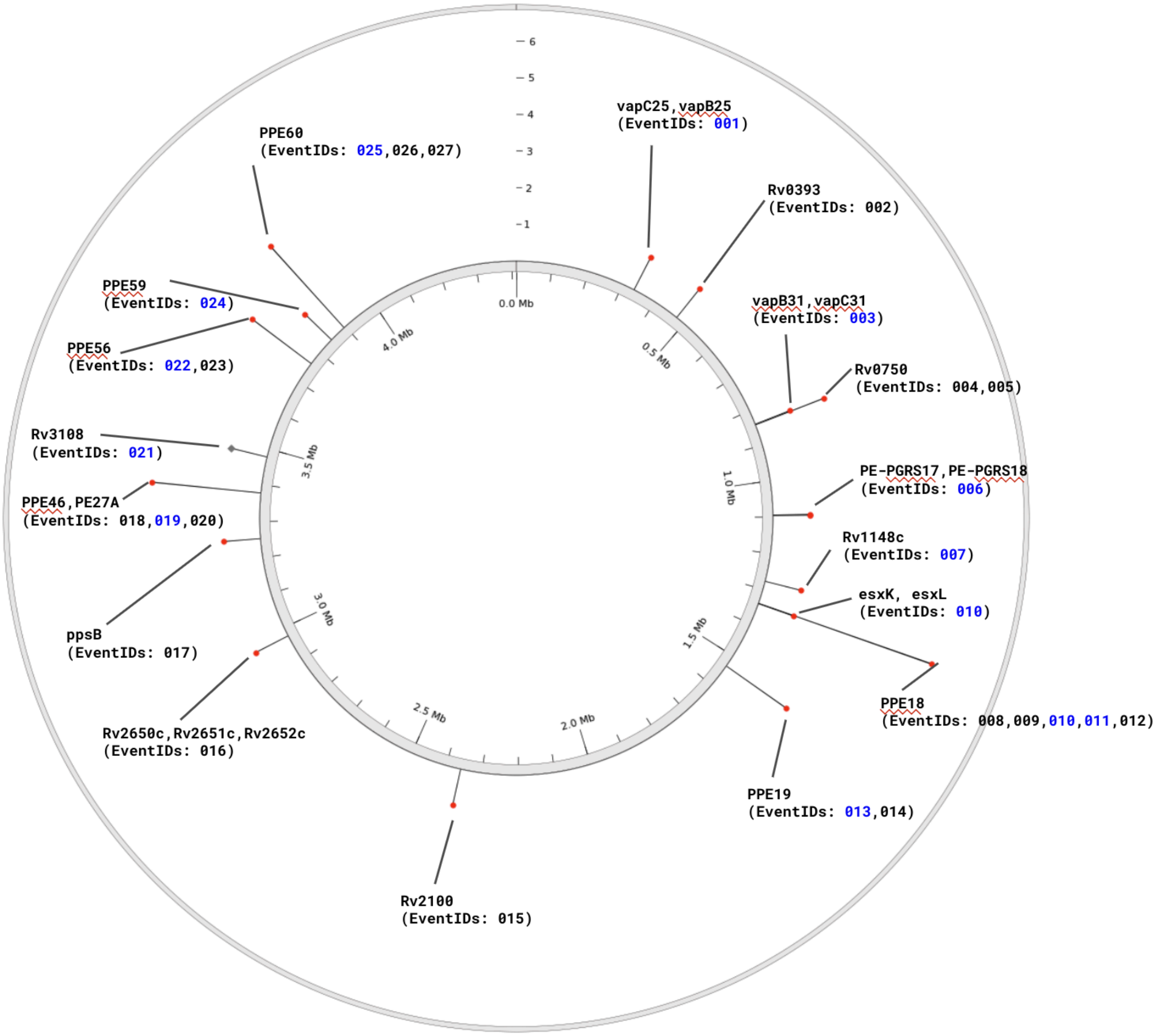


**Figure S12. Genomic distribution of suspected gene conversion events detected in the TGEN-937-SR dataset.** Visualization of the genomic locations of all detected suspected gene conversion events in the TGEN-937-SR dataset. Events selected for long-read resequencing are highlighted in blue.

#### Figure S13

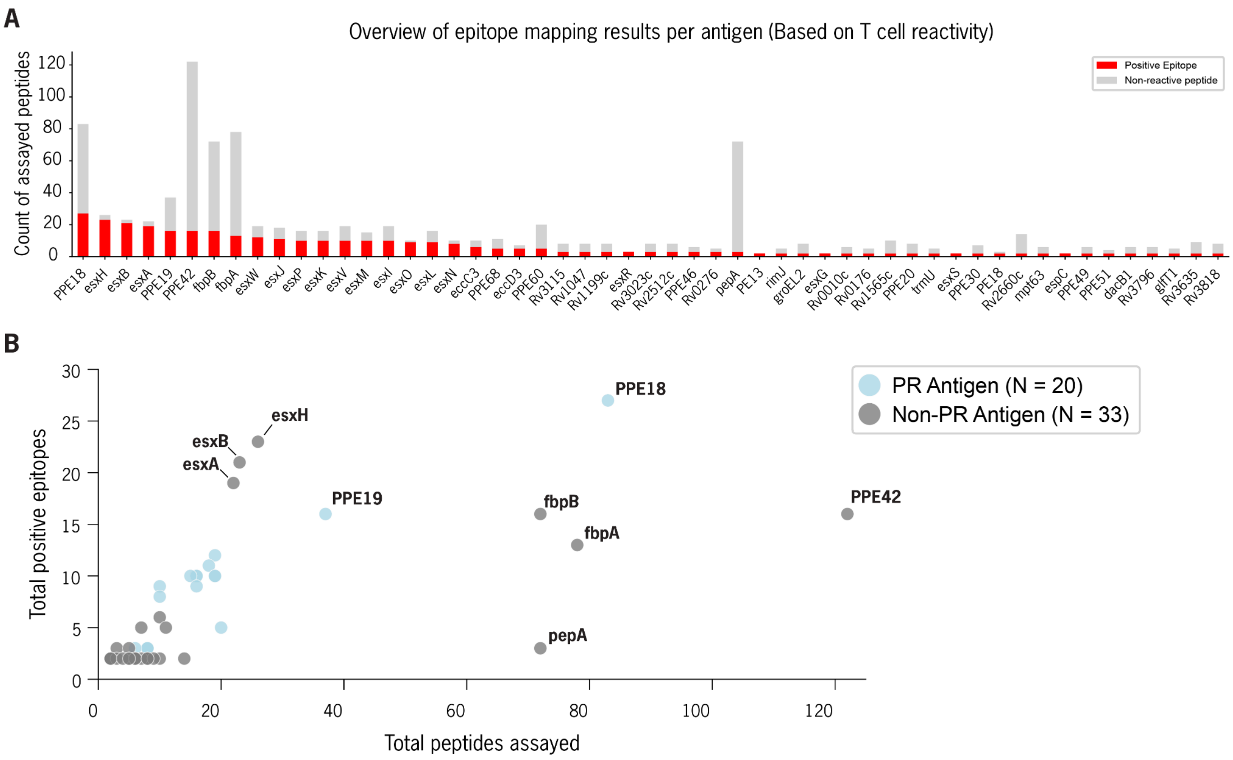


**Figure S13. Overview of identified epitopes and assayed peptides per T cell antigen.** a, Stacked bar plot showing the number of assayed peptides for each antigen (≥2 positive epitopes) in the curated epitope mapping dataset. Total bar height represents the total number of peptides assayed per antigen. Grey segments indicate non-reactive peptides, while red segments indicate reactive peptides (positive epitopes). b, Scatter plot of the same 53 T cell antigens, showing the relationship between the total number of peptides assayed (x-axis) and the total number of positive epitopes identified (y-axis).

**Figure S14**

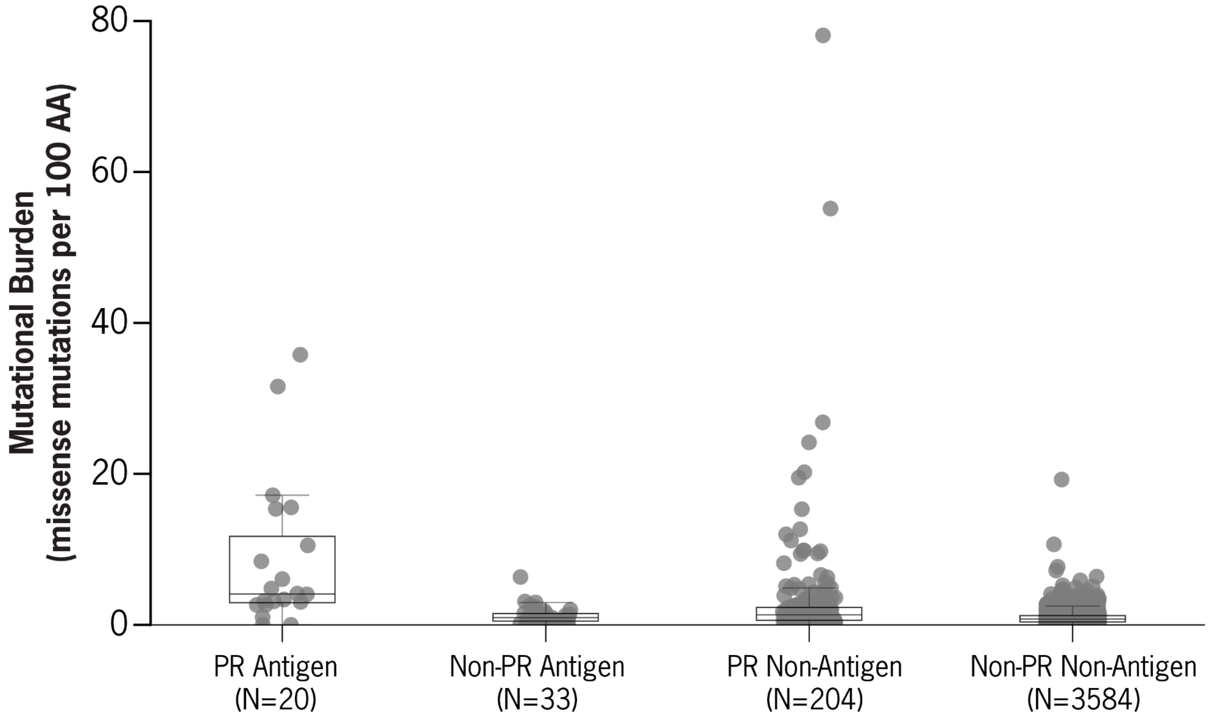


**Figure S14. Mutational burden across antigens and non-antigens stratified by genomic context**. Boxplot with overlaid scatter points showing normalized non-synonymous mutational burden (non-synonymous mutations per 100 amino acids) for four categories: PR-Antigen, Unique-Antigen, NonAntigen-PR, and NonAntigen-Unique. The number of coding sequences (CDSs) in each category is indicated on the x-axis. Box boundaries represent the IQR, horizontal lines denote the median, and whiskers extend to 1.5× IQR. Statistical comparisons between all category pairs are provided in **Table S12**.

**Figure S15**

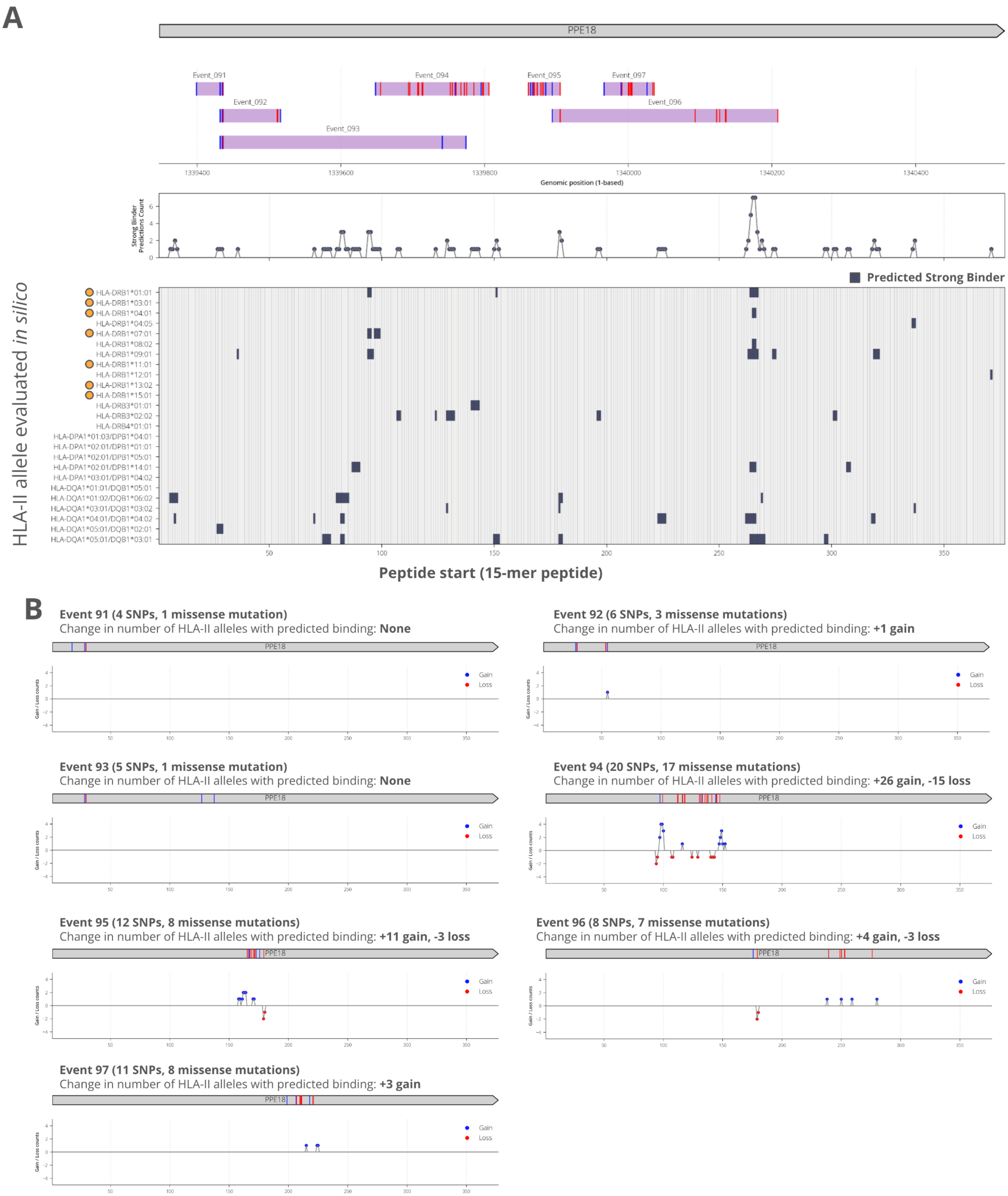
A
PPE18
Event_091
Event_092
Event_093
Event_094
Event_095
Event_096
Event_097
1339400
1339600
1339800
1340000
1340200
1340400
Genomic position (1-based)
Strong Binder Predictions Count
Predicted Strong Binder
HLA-II allele evaluated in silico
HLA-DRB1*01:01
HLA-DRB1*03:01
HLA-DRB1*04:01
HLA-DRB1*04:05
HLA-DRB1*07:01
HLA-DRB1*08:02
HLA-DRB1*09:01
HLA-DRB1*11:01
HLA-DRB1*12:01
HLA-DRB1*13:02
HLA-DRB1*15:01
HLA-DRB3*01:01
HLA-DRB3*02:02
HLA-DRB4*01:01
HLA-DPA1*01:03/DPB1*04:01
HLA-DPA1*02:01/DPB1*01:01
HLA-DPA1*02:01/DPB1*05:01
HLA-DPA1*02:01/DPB1*14:01
HLA-DPA1*03:01/DPB1*04:02
HLA-DQA1*01:01/DQB1*05:01
HLA-DQA1*01:02/DQB1*06:02
HLA-DQA1*03:01/DQB1*03:02
HLA-DQA1*04:01/DQB1*04:02
HLA-DQA1*05:01/DQB1*02:01
HLA-DQA1*05:01/DQB1*03:01
Peptide start (15-mer peptide)
B
Event 91 (4 SNPs, 1 missense mutation)
Change in number of HLA-II alleles with predicted binding: None
Event 92 (6 SNPs, 3 missense mutations)
Change in number of HLA-II alleles with predicted binding: +1 gain
Event 93 (5 SNPs, 1 missense mutation)
Change in number of HLA-II alleles with predicted binding: None
Event 94 (20 SNPs, 17 missense mutations)
Change in number of HLA-II alleles with predicted binding: +26 gain, -15 loss
Event 95 (12 SNPs, 8 missense mutations)
Change in number of HLA-II alleles with predicted binding: +11 gain, -3 loss
Event 96 (8 SNPs, 7 missense mutations)
Change in number of HLA-II alleles with predicted binding: +4 gain, -3 loss
Event 97 (11 SNPs, 8 missense mutations)
Change in number of HLA-II alleles with predicted binding: +3 gain
Gain
Loss
Gain / Loss counts

**Figure S15. in silico analysis of the effect of PPE18 gene conversion events on HLA-II binding for 27 common HLA-II alleles**. **A.** Heatmap of whether netMHCpanII predicted a strong binding interaction between each HLA-II allele from the IEDB reference HLA-II set (N= 27) for each 15-mer peptide of the PPE18 reference (H37Rv) protein sequence. Orange dots next highlight the most common HLA-II alleles across European populations (N = 7). **B.** For each gene conversion event in PPE18, the number of alleles with a predicted strong binding interaction with a 15-mer peptide sequence was plotted along the protein length of PPE18 before and after mutation by each gene conversion event.

#### Figure S16

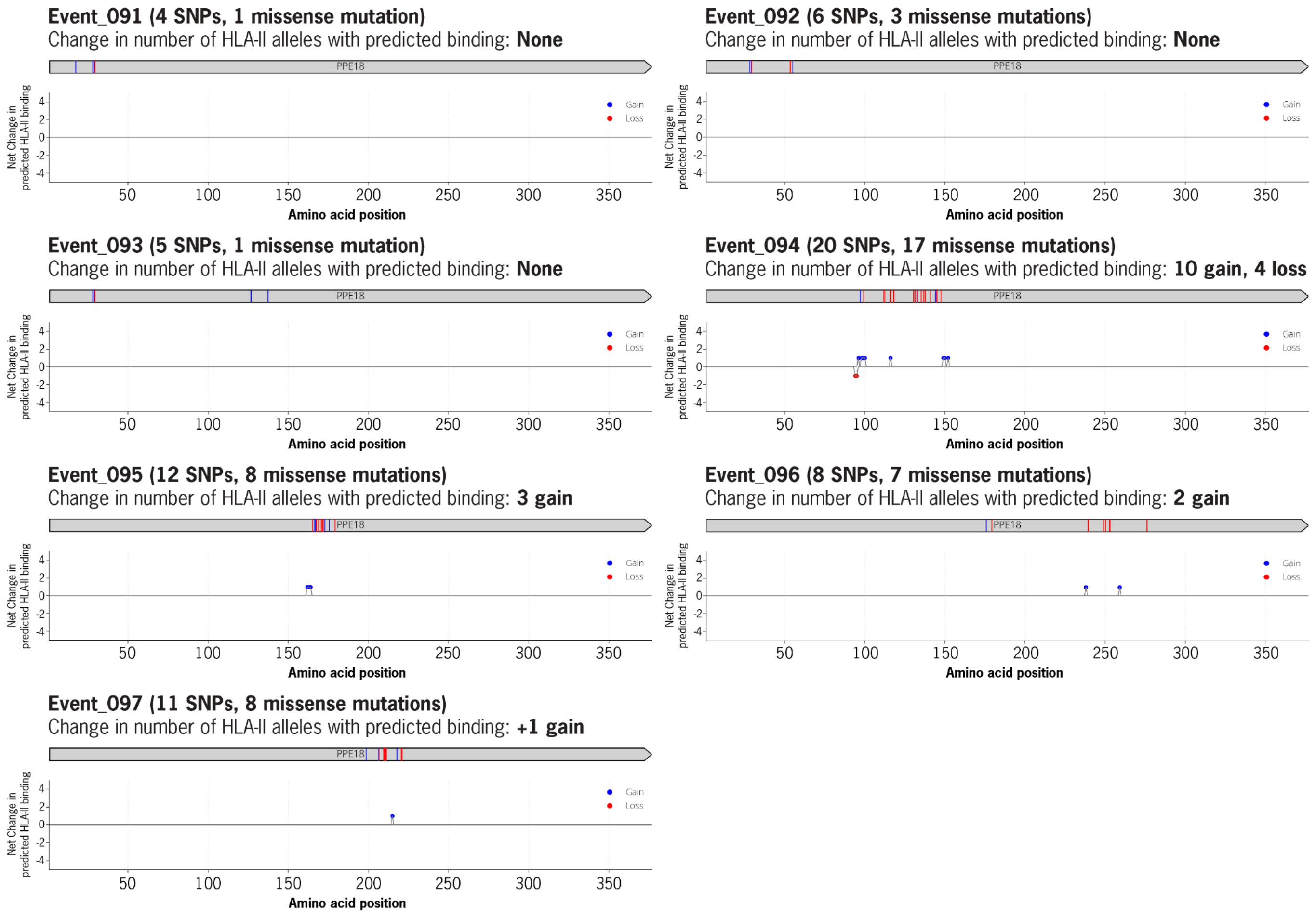


**Figure S16. *in silico* analysis of the effect of PPE18 gene conversion events on HLA-II binding for 7 top HLA-II alleles in European populations**. For each gene conversion event in PPE18, the number of alleles with a predicted strong binding interaction with a 15-mer peptide sequence was plotted along the protein length of PPE18 before and after mutation by each gene conversion event.

#### Table S1

| Dataset Label | N | Long Read Platform & Read Type | Assembly Pipeline Label | Reference |
|---|---|---|---|---|
| Farhat_Peru_2019 | 13 | PacBio (RS II) - CLR Subreads | PBclr_LR_Flye_I3_SR_Pilon | Marin et al. 2022 PMID: 35020793 |
| ChinerOms_2019 | 12 | PacBio (RS II) - CLR Subreads | PBclr_LR_Flye_I3_SR_Pilon | Chiner-Oms et al. 2019 PMID: 31488832 |
| Ngabonziza_2020 | 1 | PacBio (RS II) - CLR Subreads | PBclr_LR_Flye_I3_SR_Pilon | Ngabonziza et al. 2020 PMID: 32518235 |
| Lee_2020 | 1 | PacBio (RS II) - CLR Subreads | PBclr_LR_Flye_I3_SR_Pilon | Lee et al. 2020 PMID: 32014110 |
| TBPortals_2020 | 21 | PacBio (Sequel II) - CLR Subreads | PBclr_LR_Flye_I3_SR_Pilon | Marin et al. 2022 PMID: 35020793 |
| TRUST_PB_Set1 | 8 | PacBio (Sequel II) - CCS Reads | PBccs_LR_Flye_I3_SR_Pilon | Marin et al. 2025 PMID: 40341387 |
| Hall2022 | 78 | Oxford Nanopore - R9.4.1 Reads | ONT_LR_FlyeI3M_SR_Pilon_PolyPolish | Hall et al. 2023 PMID: 36549315 |
| Peker2021 | 17 | Oxford Nanopore - R9.4.1 Reads | ONT_LR_FlyeI3M_SR_Pilon_PolyPolish | Peker et al. 2021 PMID: 34825880 |

**Table S1. Composition of the *Mtb151* genome assembly dataset.** The 151 complete *M. tuberculosis* genome assemblies analyzed in this study were generated in Marin et al. 2025 (PMID: 40341387) using platform-appropriate hybrid long- and short-read assembly pipelines. Isolates were aggregated from previously published or publicly archived collections, listed here by dataset label with the number of isolates contributed (N), the long-read sequencing platform and read type, the assembly pipeline applied, and the source reference. All 151 isolates additionally had matched Illumina short-read WGS, used for short-read polishing in every pipeline. Per-isolate sequencing run accessions and metadata are provided in **Dataset S1**. CLR, continuous long reads (PacBio subread data); CCS, circular consensus sequencing (PacBio HiFi reads).

**Table S2**

| Stat | Median | Interquartile range | Min to Max range |
|---|---|---|---|
| BUSCO Completeness Score | 99.4 | 99.3 - 99.5 | 98.6 - 99.6 |
| Chromosome size | 4,413 kb | 4,407 - 4,421 kb | 4,380 - 4,439 |
| # of predicted CDSs<br><br>(*De novo* annotation via Bakta) | 4074 CDSs | 4065 - 4088 | 4020 - 4135 |
| GC Content | 65.6% | 65.6 - 65.6% | 65.6 - 65.6% |

**Table S2. Distribution of characteristics for dataset of 151 complete Mtb genome assemblies**

**Table S3**

| # | Coordinates (kb) | Affected gene(s) | $\pi$ (SNPs/kb) | Gene Categories |
|---|---|---|---|---|
| 1 | 103 - 104 | Rv0094c, Rv0095c | 4.05 | REP13E12 repeat region |
| 2 | 104 - 105 | | 15.96 | |
| 3 | 105 -106 | | 3.37 | |
| 4 | 338 - 339 | PE_PGRS4 | 5.14 | PE/PPE protein families |
| 5 | 1,095 - 1,096 | PE_PGRS18 | 3.10 | PE/PPE protein families |
| 6 | 1,096 - 1,097 | | 4.88 | |
| 7 | 1,276 - 1,277 | Rv1148c | 13.22 | REP13E12 repeat region |
| 8 | 1,340 - 1,341 | PPE18, esxK, esxL | 2.84 | PE/PPE protein families & esx |
| 9 | 1,341 - 1,342 | | 3.00 | |
| 10 | 1,533 - 1,534 | PPE19 | 2.50 | PE/PPE protein families |
| 11 | 1,634 - 1,635 | PE_PGRS27 | 9.04 | PE/PPE protein families |
| 12 | 1,637 - 1,638 | PE_PGRS28 | 6.07 | PE/PPE protein families |
| 13 | 1,638 - 1,639 | | 3.26 | |
| 14 | 1,788 - 1,789 | Rv1587c,Rv1588c | 2.28 | REP13E12 repeat region |
| 15 | 1,789 - 1,790 | | 6.81 | |
| 16 | 2,196 - 2,197 | Rv1945 | 3.59 | REP13E12 repeat region |
| 17 | 2,262 - 2,263 | Rv2015c | 2.30 | conserved hypotheticals |
| 18 | 2,338 - 2,339 | Rv2082 | 2.56 | conserved hypotheticals |
| 19 | 2,339 - 2,340 | Rv2082 | 2.41 | |
| 20 | 2,626 - 2,627 | esxO,esxP | 5.81 | esx |
| 21 | 2,867 - 2,868 | lppA,lppB | 7.70 | cell wall and cell processes |
| 22 | 2,944 - 2,945 | PE_PGRS45 | 2.88 | PE/PPE protein families |
| 23 | 3,135 - 3,136 | Rv2827c,Rv2828c | 4.96 | conserved hypotheticals |
| 24 | 3,730 - 3,731 | PPE54, PPE55 | 3.36 | PE/PPE protein families |
| 25 | 3,732 - 3,733 | | 4.22 | |
| 26 | 3,735 - 3,736 | | 2.56 | |
| 27 | 3,746 - 3,747 | | 3.01 | |
| 28 | 3,750 - 3,751 | | 6.34 | |
| 29 | 3,842 - 3,843 | Rv3424c, PPE57 | 7.54 | conserved hypotheticals & PE/PPE |
| 30 | 3,847 - 3,848 | PPE59 | 3.83 | PE/PPE protein families |
| 31 | 3,883 - 3,884 | Rv3466,Rv3467 | 6.68 | REP13E12 repeat region |
| 32 | 3,895 - 3,896 | PPE60 | 5.06 | PE/PPE protein families |
| 33 | 3,932 - 3,933 | PE_PGRS54 | 3.94 | PE/PPE protein families |
| 34 | 3,934 - 3,935 | | 4.93 | |
| 35 | 3,943 - 3,944 | PE_PGRS56 | 3.10 | PE/PPE protein families |
| 36 | 3,947 - 3,948 | PE_PGRS57 | 3.92 | PE/ PPE protein families |
| 37 | 4,254 - 4,255 | Rv3798 | 2.84 | Insertion sequences |

**Table S3. Summary of identified hotspots of nucleotide diversity (1-kb windows)**

#### Table S4

| Gene Category | # genes overlapping diversity hotspots | Total genes in H37Rv genome | OR | p_adj (Bonferroni) |
|---|---|---|---|---|
| PE & PPE proteins | 15 | 168 | 20.1 | $1.1 \times 10^{-11}$ |
| REP13E12 Repeat Regions | 8 | 14 | 207.1 | $3.6 \times 10^{-13}$ |
| ESX proteins | 4 | 23 | 28.3 | $4.1 \times 10^{-4}$ |
| conserved hypotheticals | 4 | 1042 | 0.4 | 1.000 |
| lipid metabolism | 0 | 272 | - | 1.000 |
| insertion sequences and phages | 1 | 133 | 0.9 | 1.000 |
| cell wall and cell processes | 2 | 749 | 0.3 | 1.000 |
| Virulence, detoxification, adaptation | 0 | 239 | - | 1.000 |
| unknown | 0 | 15 | - | 1.000 |
| intermediary metabolism and respiration | 0 | 936 | - | 1.000 |
| information pathways | 0 | 242 | - | 1.000 |
| regulatory proteins | 0 | 198 | - | 1.000 |
| stable RNAs | 0 | 48 | - | 1.000 |

**Table S4. Enrichment of nucleotide diversity hotspots across gene functional categories.** Enrichment was tested at the gene level using a one-sided Fisher's exact test (genes overlapping diversity hotspot vs. all other annotated genes), with p-values Bonferroni-corrected across 13 categories. The odds ratio (OR) and Bonferroni-adjusted p-value (p_adj) are reported for each test. Categories with $p_{adj} < 0.05$ were considered enriched.

**Table S5**

| **Category** | **Total size (bp)** | **Percent of total genome** | **Definition** |
|---|---|---|---|
| Paralogous Regions (PR) | 257 kb | 5.8% | Paralogous regions identified by within genome homology map generated by **`minimap2`** |
| Local Repeat Regions (LRR) | 150 kb | 3.4% | Local Repeat Regions identified by within genome homology map generated by **`minimap2`** |
| Low-Complexity Regions (LCR) | 127 kb | 2.9% | Low complexity sequences identified by **`Longdust`** |
| Low Mappability Regions (LMR) | 189 kb | 4.3% | Regions with non-unique 50 bp k-mers (≤4 mismatches) identified by **`Genmap`** |
| Union of Non-unique Regions | 447 kb | 10.1% | Union of the genomic regions of all categories |

**Table S5. Breakdown of repetitive and high-homology regions detected in the H37Rv genome**

#### Table S6

| DNA Repair Pathway | # Genes | Genes (RvID; protein) | Loss-of-function variant summary |
|---|---|---|---|
| Homologous recombination: end resection and RecA loading | 5 | Rv3202c (AdnA); Rv3201c (AdnB); Rv0003 (RecF); Rv2362c (RecO); Rv3715c (RecR) | None detected across all 151 genomes |
| Homologous recombination: resolution | 5 | Rv2593c (RuvA); Rv2592c (RuvB); Rv2973c (RecG); Rv2594c (RuvC); Rv2554c (RuvX) | None detected across all 151 genomes |
| Homologous recombination: strand exchange | 3 | Rv2737c (RecA); Rv0054 (SSBa); Rv2478c (SSBb) | None detected across all 151 genomes |
| AP endonucleases | 2 | Rv0670 (End); Rv0427c (XthA) | None detected across all 151 genomes |
| Base excision repair – DNA glycosylases | 10 | Rv2924c (Fpg); Rv0944 (Fpg2); Rv3589 (MutY); Rv2976c (Ung); Rv1259 (UdgB); Rv1210 (TagA); Rv1317c (AlkA); Rv2464c (Nei1); Rv3297 (Nei2); Rv3674c (Nth) | None detected across all 151 genomes |
| DNA ligases | 3 | Rv3104c (LigA); Rv3062 (LigB); Rv3731 (LigC) | A single isolate (N1202, Lineage 6) was found to have a nonsense mutation (p.Trp435*) in ligB (Rv3062). |
| DNA polymerases | 7 | Rv1629 (PolA); Rv1547 (DnaE1); Rv3370c (DnaE2); Rv1537 (DinB1); Rv3056 (DinB2); Rv3730c (PolD1); Rv0269c (PolD2) | None detected across all 151 genomes |
| Mismatch repair | 1 | Rv1321 (NucS) | None detected across all 151 genomes |
| Non-homologous end joining | 2 | Rv0937c (Ku); Rv0938 (LigD) | None detected across all 151 genomes |
| Nucleotide excision repair | 7 | Rv1638 (UvrA); Rv1633 (UvrB); Rv1420 (UvrC); Rv1020 (Mfd); Rv2191 (Cho); Rv0949 (UvrD1); Rv3198c (UvrD2) | None detected across all 151 genomes |
| Nucleotide pool sanitization enzymes | 6 | Rv2985 (MutT1); Rv1160 (MutT2); Rv0413 (MutT3); Rv3908 (MutT4); Rv2697c (Dut); Rv1021 (MazG) | None detected across all 151 genomes |
| Other proteins | 4 | Rv1696 (RecN); Rv2736c (RecX); Rv3585 (RadA); Rv2694c (RecG) | A single isolate (R21770, Lineage 4.1.1.3) was found to have a 128 bp deletion (4,026,829–4,026,956) causing a frameshift (p.Gln129fs) in radA (Rv3585) |
| Ribonucleotide excision | 2 | Rv2228c (RNaseH1); Rv2902 (RNaseH2) | None detected across all 151 genomes |
| Single-strand annealing pathway | 3 | Rv0630c (RecB); Rv0631c (RecC); Rv0629c (RecD) | None detected across all 151 genomes |

**Table S6. Predicted loss-of-function variants across *M. tuberculosis* DNA repair genes.** Genes are grouped by repair pathway (14 categories). Across all 60 genes, all 151 genomes were screened for predicted loss-of-function (nonsense/frameshift) variants. Only two LoF events were found: ligB (Rv3062; p.Trp435*) in one isolate (N1202, Lineage 6) and radA (Rv3585) with a 128 bp deletion (4,026,829–4,026,956) causing p.Gln129fs in one isolate (R21770, Lineage 4.1.1.3). Protein changes follow HGVS notation.

#### Table S7

| Negative binomial regression of gene conversion frequency between paralogous regions | | | | |
|---|---|---|---|---|
| **Predictor** | **Coefficient (log IRR)** | **IRR** | **95% CI (IRR)** | **p-value** |
| Sequence divergence (SNPs per kb) | −0.0171 | **0.983** | 0.978 – 0.988 | **$2.9 \times 10^{-10}$** |
| Genomic distance (kb) | −0.0002 | 0.9998 | 0.9994 – 1.0002 | 0.264 |
| Paralog copy number | 0.0153 | 1.015 | 0.880 – 1.171 | 0.834 |
| GC content % | 0.0193 | 1.020 | 0.973 – 1.068 | 0.417 |
| **Model Summary Statistics** | | | | |
| **McFadden pseudo-$R^2$: 0.1377**<br>**Sample Size (N) = 284 pairs of paralogous regions** | | | | |

**Table S7. Negative binomial regression of gene conversion frequency between paralogous regions.** A generalized linear model with a negative binomial distribution and log link was used to model the number of mapped gene conversion events (mGCEs) detected between non-perfect repeat paralog pairs (n = 284). Predictor variables included sequence divergence (SNPs per kb of aligned paralogous sequence), genomic distance between paralogs (kb), paralog copy number (number of overlapping paralogous alignments), and GC content percent. The dispersion parameter was fixed at $\alpha = 1.0$, and model fit is reported as McFadden's pseudo-$R^2$. Reported values are incidence rate ratios (IRRs) with 95% confidence intervals, calculated using robust (HC0) standard errors. IRRs represent the multiplicative change in the expected number of mGCEs per unit increase in each predictor.

#### Table S8

| Predictor Variable | Spearman's ρ | p-value |
|---|---|---|
| Sequence divergence (substitutions/kb) | −0.287 | $8.4 \times 10^{-7}$ |
| Genomic distance (kb) | −0.059 | 0.320 |
| Paralog copy number | −0.001 | 0.985 |
| GC content % | −0.075 | 0.208 |

**Table S8. Univariate correlations between paralog sequence features and mapped gene conversion frequency.** Spearman's rank correlation between each sequence feature and the number of mapped gene conversion events (mGCEs) per paralog pair (**Dataset S6**), computed across all non-perfect-repeat paralog pairs (n = 284). ρ, Spearman's rank correlation coefficient.

**Table S9**

| MTBC Lineage | # GC Events on ancestral branch | GC Event ID | Affected loci | # SNPs associated with recombination tract |
|---|---|---|---|---|
| Lineage 1 | 9 | Event-013 | Rv0094c | *6* |
| | | Event-076 | *pe-pgrs18* | *7* |
| | | Event-124 | *pe-pgrs27* | *7* |
| | | Event-229 | Rv2828c | *14* |
| | | Event-246 | *ppe54* | *6* |
| | | Event-254 | *ppe55* | *11* |
| | | Event-272 | *ppe59* | *5* |
| | | Event-297 | *ppe60* | *5* |
| | | Event-239 | Intergenic | 4 |
| Lineage 2 | 4 | Event-023 | Rv0095c | *25* |
| | | Event-132 | *pe-pgrs27* | *5* |
| | | Event-220 | *lppA, lppB* | *10* |
| | | Event-270 | *ppe59* | *5* |
| Lineage 3 | 6 | Event-105 | *esxL* | *5* |
| | | Event-152 | *pe-pgrs28* | *5* |
| | | Event-230 | Rv2828c | *8* |
| | | Event-236 | *ppsA* | *5* |
| | | Event-260 | *ppe55* | *5* |
| | | Event-225 | Intergenic | 4 |
| Lineage 4 | 2 | Event-125 | *pe-pgrs27* | *4* |
| | | Event-256 | *ppe55* | *12* |
| Lineage 5 | 6 | Event-010 | Rv0094c-Rv0095c | *31* |
| | | Event-053 | *pe-pgrs6* | *5* |
| | | Event-192 | Rv1945 | *14* |
| | | Event-204 | Rv2082 | *6* |
| | | Event-243 | *ppe54* | *6* |
| | | Event-266 | *ppe56* | *4* |
| Lineage 6 | 5 | Event-005 | Rv0094c | *7* |
| | | Event-050 | Rv0397 | *10* |
| | | Event-057 | *pe-pgrs10* | *7* |
| | | Event-122 | *pe-pgrs27* | *9* |
| | | Event-252 | *ppe54* | *8* |

**Table S9. All gene conversion events detected on ancestral branches of each lineage within the phylogeny.** For each lineage represented by two or more isolates (L1–L6), all detected gene conversion events inferred to have occurred on its ancestral branch (the branch leading to the lineage's most recent common ancestor; Methods) are listed. Each event is annotated by the overlapping loci and the number of SNPs in the associated recombination tract.

**Table S10**

| TGENSR EventID | Overlapping Genes | # of SNPs within recombination event signature | Lineage | Within a paralogous region? | In a region with detected GC events in primary analysis? |
|---|---|---|---|---|---|
| Event_001 | vapC25,vapB25 | 8 | L4 | Yes | Yes |
| Event_002 | Rv0393 | 6 | L4 | Yes | Yes |
| Event_003 | vapB31,vapC31 | 14 | L4 | Yes | Yes |
| Event_004 | Rv0750 | 9 | L4 | Yes | Yes |
| Event_005 | Rv0750 | 7 | L4 | Yes | Yes |
| Event_006 | PE_PGRS18 | 7 | L2 | Yes | Yes |
| Event_007 | Rv1148c | 5 | L4 | Yes | Yes |
| Event_008 | PPE18 | 5 | L4 | Yes | Yes |
| Event_009 | PPE18 | 8 | L4 | Yes | Yes |
| Event_010 | PPE18, esxK, esxL | 5 | L4 | Yes | Yes |
| Event_011 | PPE18 | 7 | L4 | Yes | Yes |
| Event_012 | PPE18 | 7 | L4 | Yes | Yes |
| Event_013 | PPE19 | 6 | L4 | Yes | Yes |
| Event_014 | PPE19 | 4 | L2 | Yes | Yes |
| Event_015 | Rv2100 | 4 | L4 | Yes | No |
| Event_016 | Rv2651c | 5 | L4 | Yes | No |
| Event_017 | ppsB | 6 | L4 | Yes | Yes |
| Event_018 | esxQ, PPE46 | 8 | L4 | Yes | Yes |
| Event_019 | PPE46 | 8 | L4 | Yes | Yes |
| Event_020 | PPE46 | 5 | L4 | Yes | Yes |
| Event_021 | Rv3108 | 5 | L4 | No | No |
| Event_022 | PPE56 | 7 | L4 | Yes | Yes |
| Event_023 | PPE56, Rv3351c | 7 | L4 | Yes | Yes |
| Event_024 | PPE59, Rv3430c | 8 | L4 | Yes | Yes |
| Event_025 | PPE60 | 6 | L4 | Yes | Yes |
| Event_026 | PPE60 | 5 | L4 | Yes | Yes |
| Event_027 | PPE60 | 5 | L4 | Yes | Yes |

**Table S10. Putative gene conversion events detected in TGEN-SR dataset.**

**Table S11**

| TGEN-SR EventID | Overlapping Genes | In a Paralogous Region? | Supported by PacBio WGS? | SNPs resolved by SR-WGS | SNPs resolved by PacBio (LR-WGS) | SR-SNPs agree with PacBio-SNPs |
|---|---|---|---|---|---|---|
| Event-001 | vapC25, vapB25 | Yes | Yes | 8 | 19 | 8/8 (100%) |
| Event-003 | vapB31, vapC31 | Yes | Yes | 14 | 21 | 14/14 (100%) |
| Event-006 | PE_PGRS18 | Yes | Yes | 7 | 7 | 7/7 (100%) |
| Event-007 | Rv1148c | Yes | Yes | 5 | 42 | 5/5 (100%) |
| Event-010 | PPE18, esxK, esxL | Yes | Yes | 5 | 13 | 5/5 (100%) |
| Event-011 | PPE18 | Yes | Yes | 7 | 8 | 7/7 (100%) |
| Event-013 | PPE19 | Yes | Yes | 6 | 9 | 6/6 (100%) |
| Event-019 | PPE46 | Yes | Yes | 8 | 13 | 8/8 (100%) |
| Event-021 | Rv3108 | No | Yes | 5 | 5 | 5/5 (100%) |
| Event-022 | PPE56 | Yes | Yes | 7 | 16 | 7/7 (100%) |
| Event-024 | PPE59 | Yes | Yes | 8 | 29 | 8/8 (100%) |
| Event-025 | PPE60 | Yes | Yes | 6 | 17 | 6/6 (100%) |

**Table S11. Validation of suspected gene conversion events by PacBio HiFi long-read resequencing.** *SR-SNPs agree with PacBio-SNPs* gives the number and percentage of short-read-detected SNPs called at the identical reference position and allele in the long-read assembly. Across all 12 events, all 86 short-read-detected SNPs were confirmed by long-read data (86/86), while long-read assemblies resolved additional SNPs within these events (199 total).

**Table S12**

| Antigen | Protein Length (aa) | # Positive Epitopes | # Negative Peptides | # Peptides Assayed | Region Type | Vaccine candidates containing antigen of interest |
|---|---|---|---|---|---|---|
| pepA | 356 | 3 | 69 | 72 | Non-PR | M72/AS01E vaccine candidate |
| esxH | 97 | 23 | 3 | 26 | Non-PR | Aeras402, HyVac4 vaccine candidate |
| PPE18 | 392 | 27 | 56 | 83 | PR | M72/AS01E vaccine candidate |
| fbpB (Ag85B) | 326 | 16 | 56 | 72 | Non-PR | Aeras402, H1, HyVac4 vaccine candidate |
| PPE42 | 581 | 16 | 106 | 122 | Non-PR | ID93 vaccine candidate |
| Rv2660c | 76 | 2 | 12 | 14 | Non-PR | H56 vaccine candidate |
| esxV | 95 | 10 | 9 | 19 | PR | ID93 vaccine candidate |
| esxW | 99 | 12 | 7 | 19 | PR | ID93 vaccine candidate |
| fbpA (Ag85A) | 339 | 13 | 65 | 78 | Non-PR | MVA85A, Aeras402, Ad85A vaccine candidate |
| esxB (CFP10) | 101 | 21 | 2 | 23 | Non-PR | IGRA diagnostic antigen |
| esxA (ESAT-6) | 96 | 19 | 3 | 22 | Non-PR | IGRA diagnostic antigen;<br>H1, H56 vaccine candidate |

**Table S12. T cell Epitope mapping statistics for IGRA diagnostic antigens and vaccine targets (N=11) within the merged dataset of T cell antigens.** The above 11 proteins are included in our dataset of 53 high-confidence T cell antigens that contain multiple experimentally validated $CD4^{+}$ T cell epitopes. Many are among the most extensively studied antigens in TB research, including IGRA diagnostic targets (EsxA, EsxB) and antigens incorporated into the following candidate vaccines: M72/AS01E, Aeras402, HyVac4, H1, H56, ID93, MVA85A, and Ad85A.

#### Table S13

| Pairwise Comparison | # of CDSs | | Normalized Mutational Burden | | U | \|r\| | p | p_adj (Bonferroni) |
|---|---|---|---|---|---|---|---|---|
| | $n_1$ | $n_2$ | $Median_1$ | $Median_2$ | | | | |
| NonPR-NonAg vs NonPR-Ag | 3584 | 33 | 0.76 | 0.96 | 51356 | 0.13 | 0.192 | 1.000 |
| NonPR-NonAg vs PR-NonAg | 3584 | 204 | 0.76 | 1.33 | 244506 | 0.33 | $1.4 \times 10^{-15}$ | $8.5 \times 10^{-15}$ |
| NonPR-NonAg vs PR-Ag | 3584 | 20 | 0.76 | 4.09 | 8272 | 0.77 | $2.6 \times 10^{-9}$ | $1.6 \times 10^{-8}$ |
| NonPR-Ag vs PR-NonAg | 33 | 204 | 0.96 | 1.33 | 2670 | 0.21 | 0.056 | 0.339 |
| NonPR-Ag vs PR-Ag | 33 | 20 | 0.96 | 4.09 | 92 | 0.72 | $1.3 \times 10^{-5}$ | $7.8 \times 10^{-5}$ |
| PR-NonAg vs PR-Ag | 204 | 20 | 1.33 | 4.09 | 899 | 0.56 | $3.6 \times 10^{-5}$ | $2.2 \times 10^{-4}$ |

**Table S13. Pairwise comparisons of length-normalized non-synonymous mutation burden across antigen and non-antigen protein groups stratified by genomic context.** Each pair of groups was compared using a two-sided Mann-Whitney U test on per-gene non-synonymous mutation burden (non-synonymous events per 100 amino acids). The unit of observation was each annotated protein in the H37Rv genome. Groups: PR-Ag (paralogous-region antigens), PR-NonAg (paralogous-region non-antigens), NonPR-Ag (unique-region antigens), NonPR-NonAg (unique-region non-antigens). p-values were Bonferroni-corrected across the six pairwise comparisons. Comparisons with $p\_adj < 0.05$ were considered significant. Effect size is the absolute rank-biserial correlation (|r|). Group medians indicate the direction of each difference. U, Mann-Whitney U statistic; p_adj, Bonferroni-adjusted p-value.

**Table S14**

| Group | Antigen Classification | Genomic context | # of CDSs | **Epitope Mutational Burden** (Detected non-synonymous mutation events in a validated T cell epitope per 100 coding residues) | |
|---|---|---|---|---|---|
| | | | | **Median** | **Interquartile Range** |
| PR-Ag | Antigen | PR Encoded | 20 | 1.49 | 0.42 - 3.15 |
| NonPR-Ag | Antigen | Non-PR Encoded | 33 | 0.08 | 0.00 - 0.33 |

**Table S14. Mutational burden within validated T cell epitopes, compared between antigens encoded in paralogous versus non-paralogous regions.** Per-antigen epitope mutational burden was compared between paralogous-region antigens (PR-Ag, n = 20) and unique-region antigens (NonPR-Ag, n = 33) using a two-sided Mann-Whitney U test (unit of observation = antigen). Median and interquartile range are shown per group. $U = 145$, $p = 5.0 \times 10^{-4}$; effect size (absolute rank-biserial correlation) = 0.56.

**Table S15**

| GC EventID | Mapped to Paralog? | # of SNPs | # of non-synonymous mutations | Change in predicted HLA-binding interactions (27 alleles) | |
|---|---|---|---|---|---|
| | | | | # gained HLA-binding interactions | # loss HLA-binding interactions |
| Event-091 | Yes | 4 | 1 | 0 | 0 |
| Event-092 | Yes | 6 | 3 | +1 | 0 |
| Event-093 | Yes | 5 | 1 | 0 | 0 |
| Event-094 | Yes | 20 | 17 | +26 | -15 |
| Event-095 | Yes | 12 | 8 | +11 | -3 |
| Event-096 | Yes | 8 | 7 | +4 | -3 |
| Event-097 | Yes | 11 | 8 | +3 | 0 |

**Table S15. Net effect of PPE18 gene conversion events on predicted HLA-II binding interactions for 27 common HLA-II alleles.**

**Table S16**

| GC EventID | # of SNPs | # of non-synonymous mutations | Change predicted HLA-binding interactions (7 common alleles in european populations) | |
|---|---|---|---|---|
| | | | # gained HLA-binding interactions | # loss HLA-binding interactions |
| Event-091 | 4 | 1 | 0 | 0 |
| Event-092 | 6 | 3 | 0 | 0 |
| Event-093 | 5 | 1 | 0 | 0 |
| Event-094 | 20 | 17 | +10 | -4 |
| Event-095 | 12 | 8 | +3 | 0 |
| Event-096 | 8 | 7 | +2 | 0 |
| Event-097 | 11 | 8 | +1 | 0 |

**Table S16. Net effect of PPE18 GC events on predicted HLA-II binding interactions for 7 common European HLA-II alleles.**
